# Reversible m^6^Am methylation of snRNA by FTO controls morphine reward and tolerance without altering analgesia

**DOI:** 10.64898/2026.08.06.743062

**Authors:** Shan Liu, Ayma F. Malik, Raymond Chien, Jianheng Liu, Luke S. Nicholson, Jin Xu, Kimia Didehvar, Vipin Rai, Valerie P. Le Rouzic, Arlene Martínez-Rivera, Konstantinos Boulias, Bing Wang, Anjali M. Rajadhyaksha, Yuan-Xian Tao, Eric Lieberman Greer, Samie R. Jaffrey, Ying-Xian Pan

**Affiliations:** Department of Anesthesiology, New Jersey Medical School, Newark, NJ 07103, USA; Rutgers Addiction Research Center, Brain Health Institute, Rutgers Health, Piscataway, NJ 08854, USA; Department of Pharmacology, Weill Cornell Medical College, Cornell University, New York, NY 10065, USA; Department of Neurology, Memorial Sloan-Kettering Cancer Center, New York, NY, 10065, USA; Division of Pediatric Neurology, Department of Pediatrics, Weill Cornell Medicine, New York, NY 10065, USA; Center for Substance Abuse Research and Department of Neural Sciences, Lewis Katz School of Medicine at Temple University, Philadelphia, PA 19140, USA; Feil Family Brain and Mind Research Institute, Weill Cornell Medicine, New York, NY 10065, USA; Department of Pediatrics, Washington University School of Medicine, St. Louis, MO, 63110, USA

**Author notes:** Correspondence (S.R.J.), (Y.X.P.). These authors contributed equally (Shan Liu, Ayma F. Malik, Raymond Chien, Jianheng Liu, Luke S. Nicholson).

## Abstract

Mu opioids, such as morphine, are effective analgesics, but their reward and tolerance drive opioid use disorder. A major goal is to achieve analgesia without these harmful effects. Here we show that morphine reward and tolerance require the RNA demethylase FTO. Genetic depletion and pharmacologic inhibition of FTO each reduced morphine reward, measured by conditioned-place preference, and reduced antinociceptive tolerance to morphine and fentanyl, without altering analgesia. Although FTO is known to erase m^6^A on mRNA, we found no effect of FTO depletion on m^6^A sites, but markedly increased levels of m^6^Am on snRNA. The effects of FTO depletion were suppressed in mice that cannot make m^6^Am, supporting the role of m^6^Am in morphine reward and tolerance. We show that FTO depletion regulates a gene expression network linked to morphine signaling. FTO inhibitors may therefore provide useful adjuvants to mu opioids in pain management and treatment of opioid use disorder.

## INTRODUCTION

Mu opioids, including natural derivatives like morphine and synthetic analogs such as fentanyl, remain the standard for managing moderate to severe pain in clinical settings, despite associated side effects like tolerance, reward and addiction. However, the misuse or illegal use of these mu opioids often leads to opioid use disorder, a primary driver of the global opioid epidemic and rising rates of overdose deaths^1–4^. A major goal is to develop pain treatment strategies in which the analgesic effects of morphine are retained while the risk of opioid use disorder is minimized.

Mu opioids produce analgesia, tolerance, and reward through the mu opioid receptors (MORs). These effects diverge downstream of MORs, at the level of signaling pathways, cell types, and circuits. Reward and addiction depend on the nucleus accumbens (NAc), ventral tegmental area (VTA), and prefrontal cortex (mPFC)^5–9^. Tolerance has been linked to N-methyl-D-aspartate (NMDA) receptors^10^, nitric oxide signaling^11,12^, P-glycoprotein^13^, cAMP signaling^14,15^, alternatively spliced *Oprm1*^16^, MOR/DOR heterodimerization^17,18^, protein kinase C (PKC) pathways^19,20^, β-arrestin signaling^21,22^, PDGFβ^23^, and MOR trafficking^24^. It remains challenging to target these pathways to block tolerance and reward while preserving analgesia.

Fat mass and obesity-associated protein (FTO), also known as alpha-ketoglutarate-dependent dioxygenase, is an RNA demethylase that has been shown to affect diverse neuronal signaling mechanisms^25–28^. FTO is abundantly expressed in both the central and peripheral nervous systems^29–31^ and can influence gene expression by modulating mRNA stability, alternative splicing, translation, and epigenetic regulation^27,28,32–35^. More significantly, FTO has been implicated in various neuronal functions, including locomotion^36^, stress response^37^, and the sensation of neuropathic pain^31^, as well as in several neuropsychiatric disorders such as Alzheimer’s disease, Parkinson’s disease, epilepsy, anxiety, and depression^38–40^.

Early studies showed that the mechanism of FTO involves demethylating *N*^6^-methyladenosine (m^6^A)^25,26^, a modified nucleotide enriched in mRNA. The effects of FTO have primarily been linked to demethylation of m^6^A in specific transcripts^41^. FTO has been reported to target specific mRNAs, maintaining m^6^A sites in these transcripts at low levels. Upon FTO depletion, m^6^A levels rise, thus causing mRNA instability or other effects on these mRNAs^41^. More recent studies have shown that FTO can also demethylate *N*^6^,2’-O-dimethyladenosine (m^6^Am) with a catalytic efficiency of up to 100 times greater than that for m^6^A^27,28^. m^6^Am is located at the first transcribed nucleotide in both mRNA and small nuclear RNA (snRNA), but FTO primarily demethylates m^6^Am in snRNA, leading to the formation of 2’-O-methyladenosine in snRNA. By regulating these epitranscriptomic pathways, FTO can influence diverse neuronal functions.

Here we assessed the role of FTO in the analgesic, reward, and tolerance effects of morphine. Using either genomic deletion of *Fto* or pharmacologic inhibition of FTO we found that FTO is required to establish morphine reward and antinociceptive tolerance of morphine. However, depleting FTO does not alter morphine analgesia. We found that depleting *Fto* in the nucleus accumbens shell (NAcSh) reduced reward but not tolerance, while depleting *Fto* in the dorsal root ganglion (DRG) reduced tolerance but not reward. To determine how FTO produces these effects, we mapped m^6^A in control and FTO-depleted NAcSh transcriptome. We saw no change in m^6^A stoichiometry across 20,139 detected m^6^A sites. Instead, we found that FTO depletion led to a marked increase in m^6^Am in snRNA, as well as altered gene expression patterns in genes linked to morphine signaling pathway. To determine whether the effects of FTO depletion were due to increased m^6^Am levels, we examined the effects of FTO depletion in mice lacking m^6^Am due to depletion of PCIF1, the enzyme that synthesizes m^6^Am. In these mice, m^6^Am levels cannot increase, and FTO depletion was no longer able to suppress morphine reward and tolerance. Overall, these data point to m^6^Am levels in snRNA as a key regulator of morphine reward and tolerance and suggest that FTO inhibitors may be useful adjuvants to mu opioid in pain management and opioid use disorder treatment.

## RESULTS

### Targeting FTO reduces morphine-induced conditioned place preference

To investigate the role of FTO in opioid reward, we assessed the acquisition of morphine conditioned place preference (CPP) in *Fto*^+/-^ mice using a 6-day CPP paradigm (**Figure 1A**). *Fto* homozygous knockout (KO) mice were excluded from these studies owing to their increased postnatal mortality and developmental delays^42^. *Fto*^+/-^mice exhibited a significant reduction in morphine CPP scores relative to wild-type C57BL/6 (WT-B6) mice (**Figure 1B**), indicating that partial loss of *Fto* is sufficient to impair the acquisition of morphine CPP. Morphine analgesia assessed using the hot plate assay was unaffected in *Fto*^+/-^ mice (**Figure 1C**), demonstrating that reducing *Fto* selectively modulates the rewarding properties of morphine without altering its antinociceptive potency and efficacy.

**Figure 1.**
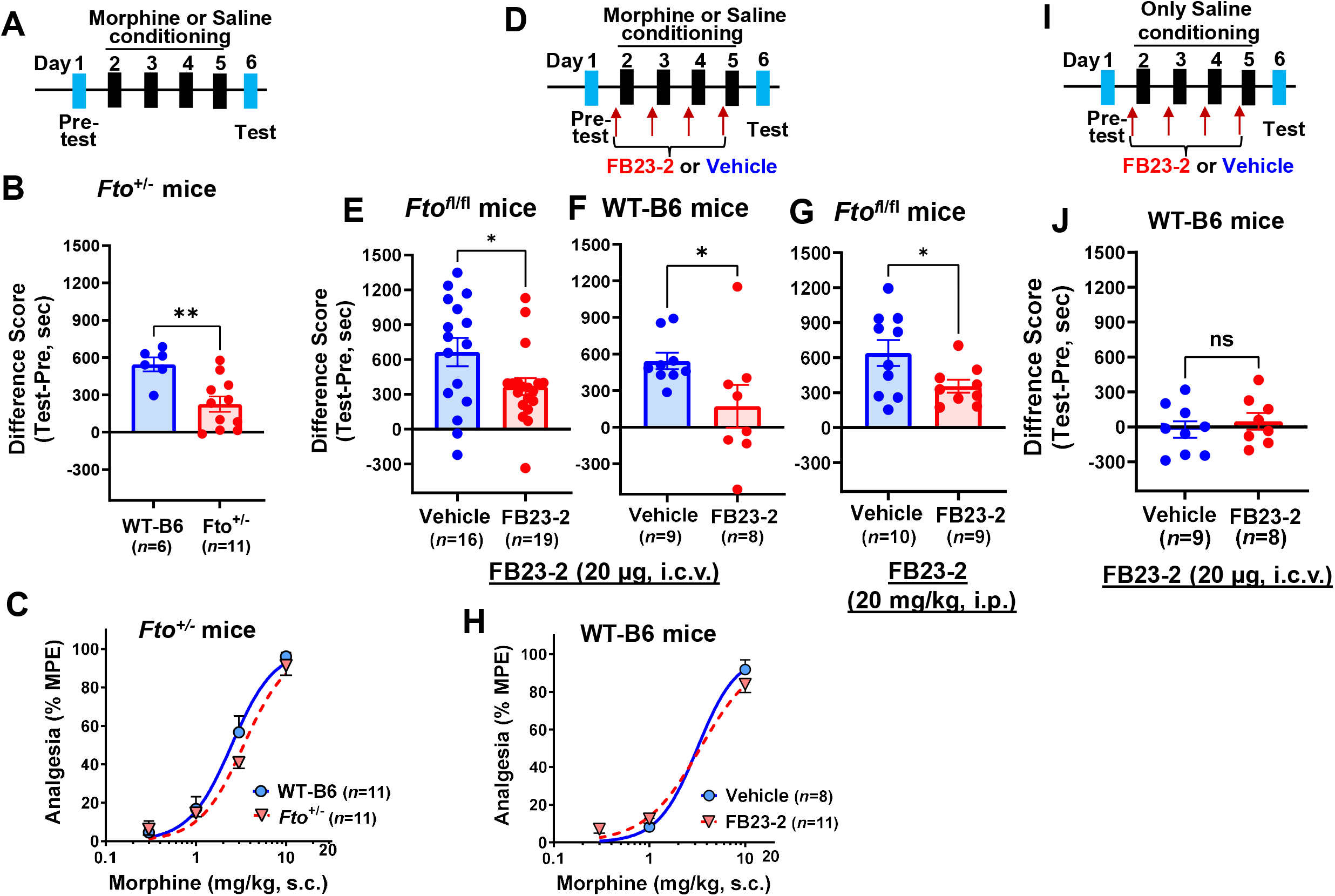
Targeting FTO reduced morphine reward. (A) Schematic of morphine CPP in male Fto heterozygous (*Fto*^+/-^) mice. CPP was performed using a 3-chamber apparatus (Med Associates) with a 6-day protocol described in the Methods. After pre-test without any injection on Day 1, mice were i.p. injected with saline (morning) or morphine (10 mg/kg, afternoon) and conditioned for 30 min on Days 2 – 5, followed by post-test without any injection on Day 6. (B) Reduced morphine CPP in *Fto*^+/-^ mice. The numbers of mice used in WT-B6 and *Fto*^+/-^ mice were 6 and 11, respectively. Preference (Difference Score) was calculated as difference between the times spent in the morphine-paired chamber minus time spent in the saline-paired chamber on the post-test day (day 6) and the pre-test day (day 1). **: *p* < 0.01, two-tailed Student *t*-test. (C) No change of morphine analgesia in *Fto*^+/-^ mice. Morphine analgesic dose-response was measured using a Hot-plate assay and quantified as the percentage of maximum possible effect (% MPE) as described in the Methods. The ED_50_ values were calculated using non-linear regression analysis (GraphPad Prism 10). The ED_50_ in WT-B6 mice, 2.48 mg/kg, 95% confident intervals (CIs), 1.78 – 3.46. The ED_50_ in *Fto*^+/-^ mice, 3.38 mg/kg, 95% CIs, 2.70 – 4.24. There was no significant difference in the ED_50_ values between WT-B6 and *Fto*^+/-^ mice, as determined by an extra sum-of-square F-test (*p* = 0.1230). (D) Schematic of morphine CPP in mice treated with FB23-2 for E, F & G. Morphine CPP was performed as (A) except for FB23-2 or vehicle treatment. **(E & F)** Reduced morphine CPP by i.c.v. administration of FB23-2 in *Fto*^fl/fl^ **(E)** or WT-B6 mice **(F)**. FB23-2 (20µg) or Vehicle (DMSO) was i.c.v. administered 2 hours before morphine conditioning for 4 consecutive days (Day 2 – Day 5) in *Fto*^fl/fl^ (E) or WT-B6 mice (F). *: *p* < 0.05, two-tailed Student *t*-test. **(G)** Reduced morphine CPP by i.p. administration of FB23-2 in *Fto*^fl/fl^ mice. 20 mg/kg of FB23-2 or DMSO was i.p. administered 2 hours before morphine conditioning for 4 consecutive days (Day 2 – Day 5) in *Fto*^fl/fl^ mice. *: *p* < 0.05, two-tailed Student *t*-test. **(H)** No change of morphine analgesia in WT-B6 mice. Morphine analgesic dose-response was measured using a Hot-plate assay and quantified as the percentage of maximum possible effect (% MPE) as described in the Methods. The ED_50_ values were calculated using non-linear regression analysis (GraphPad Prism 10). The ED_50_ in WT-B6 mice treated i.c.v. FB23-2, 3.45 mg/kg, 95% confident intervals (CIs), 2.58 – 4.61. The ED_50_ in WT-B6 mice treated i.c.v. Vehicle, 3.38 mg/kg, 95% confident intervals (CIs), 2.07 – 5.52. There was no significant difference in the ED_50_ values between WT-B6 mice treated with FB23-2 and Vehicle, as determined by the extra sum-of-square F-test (*p* = 0.9249). **(I)** Schematic of CPP in WT-B6 mice treated with i.c.v. FB23-2. Saline conditioning was performed twice per day for 4 consecutive days (Day 2 – Day 5). 20µg of FB23-2 or Vehicle (DMSO) was i.c.v. administered 2 hours before second saline conditioning. **(J)** No effect of FB23-2 on saline CPP in WT-B6 mice. n.s.: No statistical significance, two-tailed Student *t*-test.

To extend these findings pharmacologically, we examined the effect of the selective, potent FTO inhibitor FB23-2^43^ on morphine CPP acquisition in both *Fto*^fl/fl^ and WT-B6 mice. Vehicle (DMSO) or 20 µg of FB23-2 was intracerebroventricularly (i.c.v.) administered, two hours prior to each morphine conditioning session over four consecutive days (Days 2–5) (**Figure 1D**). FB23-2 significantly attenuated the acquisition of morphine CPP in both *Fto*^fl/fl^ and WT-B6 mice (**Figures 1E, F**). FB23-2 administered alone, in the absence of morphine, produced no CPP (**Figures 1I, 1J**), confirming that the observed attenuation reflects suppression of morphine’s rewarding properties rather than a non-specific effect on CPP behavior.

We next assessed morphine analgesia following i.c.v. administration of FB23-2. Morphine analgesia, evaluated using the hot plate assay, was similarly unaffected (**Figure 1H**), reinforcing the idea that pharmacological inhibition of FTO selectively attenuates the acquisition of morphine CPP without compromising its analgesia potency and efficacy.

Since FB23-2 has been reported to penetrate the blood-brain barrier following systemic administration^44^, we next asked whether intraperitoneal (i.p.) delivery would recapitulate the suppression of morphine CPP acquisition observed with i.c.v. administration, thereby establishing a more tractable route of inhibition. *Fto*^fl/fl^ mice received FB23-2 (20 mg/kg, i.p.) two hours prior to morphine conditioning over four consecutive days (Days 2–5). Consistent with the i.c.v. findings, systemic FB23-2 administration significantly attenuated the acquisition of morphine CPP (**Figure 1G**), demonstrating that peripheral delivery of an FTO inhibitor is sufficient to suppress morphine CPP, further supporting the feasibility of targeting FTO pharmacologically to mitigate morphine reward.

### Selective depletion of FTO in the NAcSh reduces morphine reward

Having established that both genetic reduction and pharmacological inhibition of FTO attenuate the acquisition of morphine CPP, we next sought to identify the brain region through which FTO acts to modulate morphine reward. The nucleus accumbens shell (NAcSh) is a key node of the mesolimbic reward circuitry and has been directly implicated in the rewarding properties of opioids^45–47^.

To test whether FTO within the NAcSh is required for morphine CPP, *Fto* was selectively depleted in this region by bilateral stereotaxic microinjection of adenovirus expressing Cre recombinase/GFP under the control of the cytomegalovirus (CMV) promoter (AAV5-CMV-Cre/GFP; hereafter AAV-Cre) into *Fto*^fl/fl^ mice, with AAV5-CMV-GFP (AAV-GFP) serving as the control. The viral transduction and efficiency of FTO depletion were examined by GFP imaging combined with RNAScope *in situ* hybridization using *Fto*-specific probes in the same brain sections. GFP imaging revealed robust expression of Cre/GFP and GFP transduced by AAV-Cre or AAV-GFP, respectively, in the NAcSh (**Figure 2A**), confirming transduction by both AAV vectors. RNAScope in situ hybridization showed that over 95% of *Fto* transcripts were eliminated in the NAcSh of AAV-Cre-injected mice, with no detectable effect on *Fto* mRNA levels in AAV-GFP-injected controls (**Figure 2A**).

**Figure 2.**
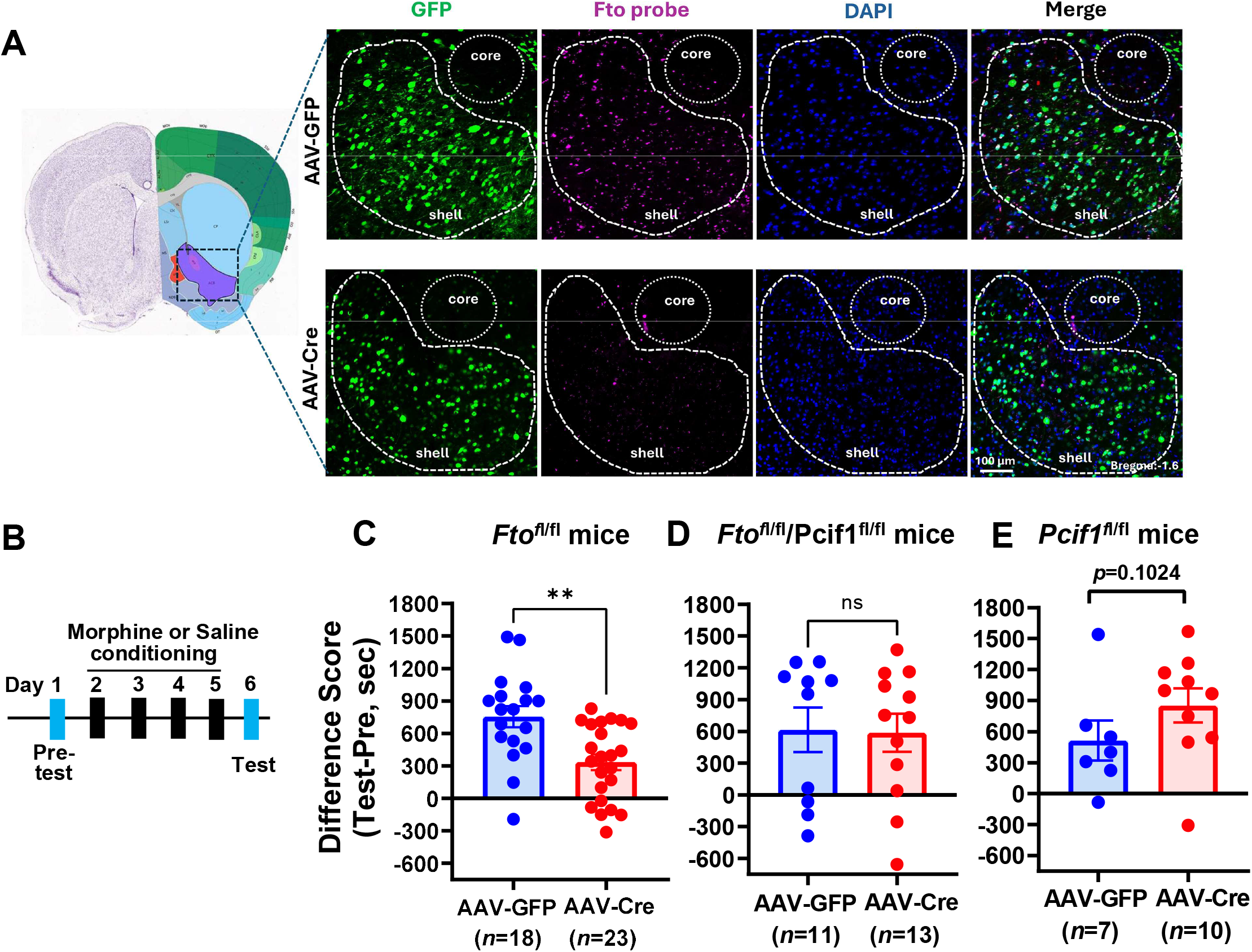
Targeting FTO in the NAcSh reduced morphine reward. **(A)** Depletion of Fto in the NAcSh using microinjected AAV-Cre/GFP. RNAscope with Fto probe and Fto/GFP imaging was performed on brain sections from *Fto*^fl/fl^ mice microinjected with AAV-Cre/GFP or AAV-GFP in the NAcSh after morphine CPP study, as described in the Methods. AAV-Cre: AAV5-CMV-Cre/GFP; AAV-GFP: AAV5-CMV-GFP; core: NAc core; shell: NAc shell (NAcSh). **(B)** Schematic of CPP. **(C)** Reduced morphine CPP by disrupting Fto in the NAcSh via AAV-Cre in *Fto*^fl/fl^ mice. *Fto*^fl/fl^ mice were bilaterally microinjected in the NAcSh with AAV-Cre/GFP or AAV-GFP. Morphine CPP was performed as shown in Figure 1A five weeks post-injection. **: *p* < 0.01, two-tailed Student *t*-test. **(D)** No significant changes in morphine CPP by disrupting both Fto and Pcif1 in the NAcSh in *Fto*^fl/fl^;*Pcif1*^fl/fl^ mice. *Fto*^fl/fl^;*Pcif1*^fl/fl^ mice were microinjected in the NAcSh with AAV-Cre/GFP or AAV-GFP. Morphine CPP was performed as shown in Figure 2B five weeks post-injection. n.s.: No statistical significance, two-tailed Student *t*-test. **(E)** No significant changes in morphine CPP by disrupting Pcif1 in the NAcSh in *Pcif1*^fl/fl^ mice. *Pcif1*^fl/fl^ mice were microinjected in the NAcSh with AAV-Cre/GFP or AAV-GFP. Morphine CPP was performed as shown in Figure 2B five weeks post-injection. *p* = 0.1024, no statistical significance, two-tailed Student *t*-test.

We assessed morphine CPP in mice five weeks after viral transduction. Mice underwent morphine CPP acquisition using the 6-day paradigm described above (**Figure 2B**). *Fto*^fl/fl^ mice receiving NAcSh microinjections of AAV-Cre exhibited significantly lower morphine CPP scores than AAV-GFP-injected controls (**Figure 2C**). Taken together, these results demonstrate that selective depletion of *Fto* within the NAcSh is sufficient to impair the acquisition of morphine-induced CPP, identifying the NAcSh as a key locus through which FTO regulates morphine reward.

### FTO inhibitors markedly reduce morphine antinociceptive tolerance

Having demonstrated that FTO modulates morphine reward, we next asked whether FTO also governs the development of morphine antinociceptive tolerance. Antinociceptive tolerance was induced in WT-B6 mice by subcutaneous (s.c.) administration of morphine (10 mg/kg, twice daily) over five days, a well-established protocol for producing robust antinociceptive tolerance^16^. FB23-2 (20 mg/kg, i.p.) or vehicle (DMSO) was administered two hours prior to the first morphine injection each day throughout the five-day period. Analgesia was assessed using the radiant heat tail-flick assay through morphine dose-response curves on Days 1 and 5 and following the second morphine dose (10 mg/kg) on Days 3 and 4. Morphine antinociceptive tolerance was readily apparent in vehicle-treated mice by Days 4 and 5 (**Figures 3A**). In contrast, FB23-2-treated mice exhibited significantly attenuated morphine antinociceptive tolerance, as reflected in the time course (**Figure 3A**) and dose-response curves, which had a smaller rightward shift in ED_50_ between Days 1 and 5 relative to vehicle-treated controls (**Figure 3B** & **Table 1**).

**Figure 3.**
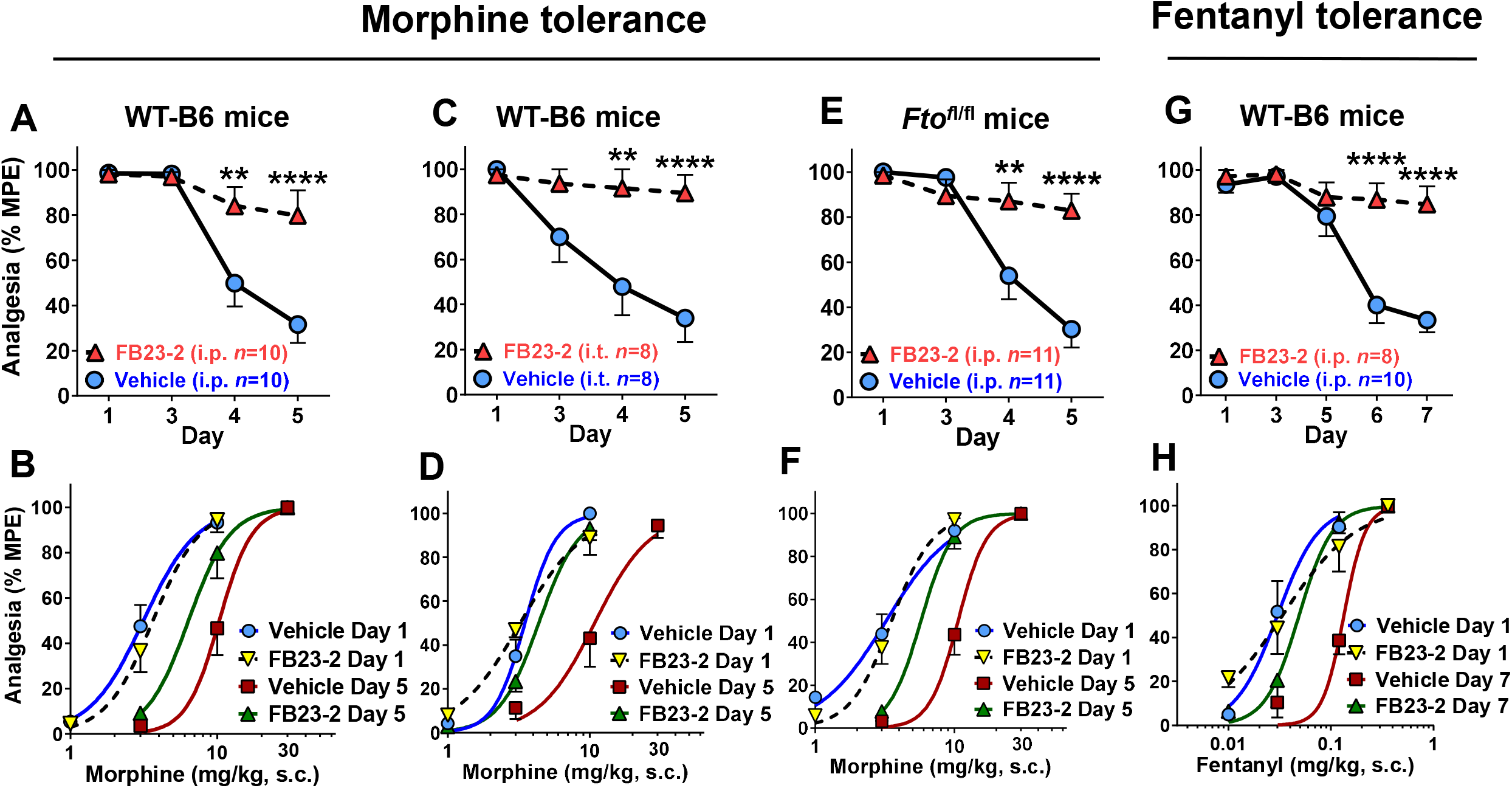
Targeting FTO reduced mu opioid nociceptive tolerance (A &. **B)** Reduced morphine tolerance by i.p. administration of FB23-2 in WT-B6 mice. FB23-2 (20 mg/kg) and DMSO (vehicle) were i.p. administered daily 2 hours before the first morphine dosing for 5 days in WT-B6 mice. Morphine analgesia was measured by using a radiant heat tail-flick assay and quantified as the percentage of maximum possible effect (% MPE) as described in the Methods. Morphine dose-response curves (1, 3,10 and 30 mg/kg, s.c.) were measured on Days 1 and 5 with single dose (10 mg/kg, s.c.) on Days 2, 3 and 4. The ED_50_ values were calculated using non-linear regression analysis (GraphPad Prism 10) and listed on Table 1. A: Time course; B: Dose-response curve. **: *p* < 0.01; ****: *p* < 0.0001, 2-way ANOVA with Bonferroni’s *post hoc* test. **(C & D)** Reduced morphine tolerance by i.t. administration of FB23-2 in WT-B6 mice. FB23-2 (20 µg) and DMSO (vehicle) were i.t. administered daily 2 hours before the first morphine dose for 5 days in WT-B6. Morphine analgesia was measured and quantified as above. C: Time course; D: Dose-response curve. The ED_50_ values were listed on Table 1. **: *p* < 0.01; ****: *p* < 0.0001, 2-way ANOVA with Bonferroni’s *post hoc* test. **(E & F)** Reduced morphine tolerance by i.p. administration of FB23-2 in *Fto*^fl/fl^ mice. FB23-2 (20 mg/kg) and DMSO (vehicle) were i.p. administered daily 2 hours before the first morphine dose for 5 days in *Fto*^fl/fl^ mice. Morphine analgesia was measured and quantified as A. E: Time course; F: Dose-response curve. The ED_50_ values were listed on Table 1. **: *p* < 0.01; ****: *p* < 0.0001, 2-way ANOVA with Bonferroni’s *post hoc* test. **(G & H)** Reduced fentanyl tolerance by i.p. administration of FB23-2 in WT-B6 mice. FB23-2 (20 mg/kg) and DMSO (vehicle) were i.p. administered daily for 7 days in WT-B6 mice. Fentanyl tolerance was induced by an s.c. implanted osmotic pump (Alzet Model: 1007D; 1 mg/kg/day/mouse for 7 days). Tolerance was assessed by cumulative dose-response curve (0.01, 0.04, 0.1 and 0.3 mg/kg) on Days 1 and 7, and by a fixed dose on Days 3, 5 or 6 (0.12 mg/kg, s.c.). Fentanyl analgesia was measured and quantified as A. G: Time course; H: Dose-response curve. The ED_50_ values were listed on Table 1. ****: *p* < 0.0001, 2-way ANOVA with Bonferroni’s *post hoc* test.

**Table 1.** Mu opioid analgesic dose-responses in tolerance studies.

| Mouse model | Drug or AAV | Route | # of mice | Morphine ED<sub>50</sub> (mg/kg, s.c.)<sup>A</sup> |  | ED<sub>50</sub> shift |
| --- | --- | --- | --- | --- | --- | --- |
|  |  |  |  | Day 1 | Day 5 |  |
| WT-B6 | FB23-2 | i.p. | 10 | 3.7 (2.9-4.5) | 6.4 (4.9-8.3)<sup>B,C</sup> | 1.7 |
|  | Vehicle | i.p. | 10 | 3.2 (2.6-3.8) | 10.3 (8.8-12.1)<sup>B</sup> | 3.3 |
| WT-B6 | FB23-2 | i.t. | 8 | 3.2 (2.7-3.9) | 4.4 (3.6-5.4)<sup>C</sup> | 1.3 |
|  | Vehicle | i.t. | 8 | 3.5 (2.9-4.3) | 10.8 (8.0-14.6)<sup>B</sup> | 3.1 |
| *Fto*<sup>fl/fl</sup> | FB23-2 | i.p. | 11 | 3.5 (3.0-4.1) | 5.7 (4.8-6.9)<sup>B,C</sup> | 1.6 |
|  | Vehicle | i.p. | 11 | 3.2 (2.5-4.2) | 10.6 (9.1-12.4)<sup>B</sup> | 3.3 |
| *Fto*<sup>fl/fl</sup> | NAcSh-Cre |  | 10 | 3.6 (2.7-4.7) | 10.8 (9.0-12.9) | 3.0 |
|  | NAcSh-GFP |  | 9 | 4.1 (3.7-7.0) | 10.9 (8.5-14.9) | 2.7 |
| *Fto*<sup>fl/fl</sup>;AvCreERT2 | Tamoxifen | i.p. | 12 | 3.8 (2.9-4.5) | 5.1 (4.1-5.5) | 1.3 |
|  | Vehicle | i.p. | 14 | 3.5 (2.7-4.4) | 10.6 (8.4-13.3) | 3.0 |
| *Fto*<sup>fl/fl</sup>;Pcif1<sup>fl/fl</sup>;AvCreERT2 | Tamoxifen | i.p. | 8 | 4.3 (3.5-6.2) | 11.6 (9.3-14.3) | 2.7 |
|  | Vehicle | i.p. | 6 | 3.4 (1.9-4.9) | 10.6 (8.8-12.5) | 3.1 |
|  |  |  |  | Fentanyl ED<sub>50</sub> (μg/kg, s.c.)<sup>A</sup> |  | ED<sub>50</sub> shift |
|  |  |  |  | Day 1 | Day 7 |  |
| WT-B6 | FB23-2 | i.p. | 8 | 33.6 (22-51) | 48.0 (36-66)<sup>C</sup> | 1.5 |
|  | Vehicle | i.p. | 10 | 30.1 (22-42) | 133.9 (109-164)<sup>B</sup> | 4.0 |

<sup>A</sup>: ED<sub>50</sub> values with 95% confidence intervals (Cis) were determined by nonlinear regression analysis (Prism). ED<sub>50</sub> values with nonoverlapping 95% Cis were considered significantly different<sup>16</sup>. <sup>B</sup>: *p*<0.05, compared with Day 1; <sup>C</sup>: *p*<0.05, compared with Vehicle.

We next examined the effect of FB23-2 on morphine tolerance in WT B6 mice through intrathecal (i.t.) administration. 20 µg of FB23-2 or vehicle (DMSO) was administered i.t. two hours prior to the first morphine injection each day for five days. Similar to i.p. administration, Intrathecal FB23-2 significantly attenuated the development of antinociceptive tolerance on Days 4 and 5, as reflected in the time course (**Figure 3C**) and dose-response curves with a much smaller rightward shift in ED_50_ between Days 1 and 5 relative to vehicle-treated controls (**Figure 3D** & **Table 1**).

To confirm that the effect of FB23-2 on morphine antinociceptive tolerance is not background-dependent, we conducted an identical morphine tolerance study in WT B6-like *Fto*^fl/fl^ mice. Systemic FB23-2 (20 mg/kg, i.p.) similarly attenuated morphine antinociceptive tolerance on Days 4 and 5 in these mice (**Figures 3E**), with a minimal rightward shift in ED_50_ between Days 1 and 5 relative to vehicle-treated controls (**Figure 3F** & **Table 1**), consistent with the findings in WT-B6 mice.

Fentanyl is a commonly used opioid analgesics in clinic and displays distinct pharmacological profiles that set it apart from opium-derived morphine^48^. We next asked whether FTO inhibition extends to fentanyl antinociceptive tolerance. Fentanyl antinociceptive tolerance was induced by continuous s.c. delivery via an osmotic pump (Alzet Model 1007D; 1 mg/kg/day for 7 days). Analgesia was assessed using the radiant heat tail-flick assay through fentanyl dose-response curves on Days 1 and 7 and following a single fentanyl challenge (0.12 mg/kg) on Days 3, 5, and 6. FB23-2 (20 mg/kg, i.p.) or vehicle was co-administered once daily throughout the 7-day period. FB23-2 significantly attenuated the development of fentanyl antinociceptive tolerance on Days 6 and 7, as reflected in the time course (**Figure 3G**) and dose-response curves, with a reduced rightward shift in ED_50_ between Days 1 and 7 relative to vehicle-treated controls (**Figure 3H** & **Table 1**).

Taken together, these findings demonstrate that FTO is required for the development of antinociceptive tolerance to both morphine and fentanyl, and that pharmacological inhibition of FTO with FB23-2 markedly attenuates this process across multiple routes of administration and mu opioid compounds.

### Targeting FTO in the DRG attenuates morphine antinociceptive tolerance

The DRG has been established as a critical site for the involvement of opioid antinociceptive tolerance^49–63^. Having shown that systemic and intrathecal FTO inhibition attenuates the development of morphine antinociceptive tolerance, we next sought to determine whether *Fto* expression in DRG sensory neurons specifically accounts for this effect. To achieve inducible, DRG-restricted *Fto* depletion, we used *Fto*^fl/fl^;AvCreERT2 mice, generated by crossing *Fto*^fl/fl^ mice with Advillin-CreERT2 (AvCreERT2) mice in which CreERT2 expression is driven by the Advillin promoter and is therefore restricted to DRG sensory neurons^64^. Following i.p. administration of tamoxifen (75 mg/kg) for 10 consecutive days, *Fto* transcript levels were markedly depleted in the DRG (**Figure 4A**). Morphine antinociceptive tolerance was then assessed using the 5-day paradigm described above. Tamoxifen-treated *Fto*^fl/fl^;AvCreERT2 mice exhibited markedly attenuated morphine antinociceptive tolerance relative to vehicle-treated controls (**Figures 4B, C**), demonstrating that *Fto* expression in DRG sensory neurons is required for the full development of morphine antinociceptive tolerance. On the other hand, Fto depletion by tamoxifen in *Fto*^fl/fl^;AvCreERT2 mice had no effect on morphine CPP (**Figure 4D**), demonstrating that *Fto* in the DRG selectively modulates morphine tolerance without impacting its rewarding properties.

**Figure 4.**
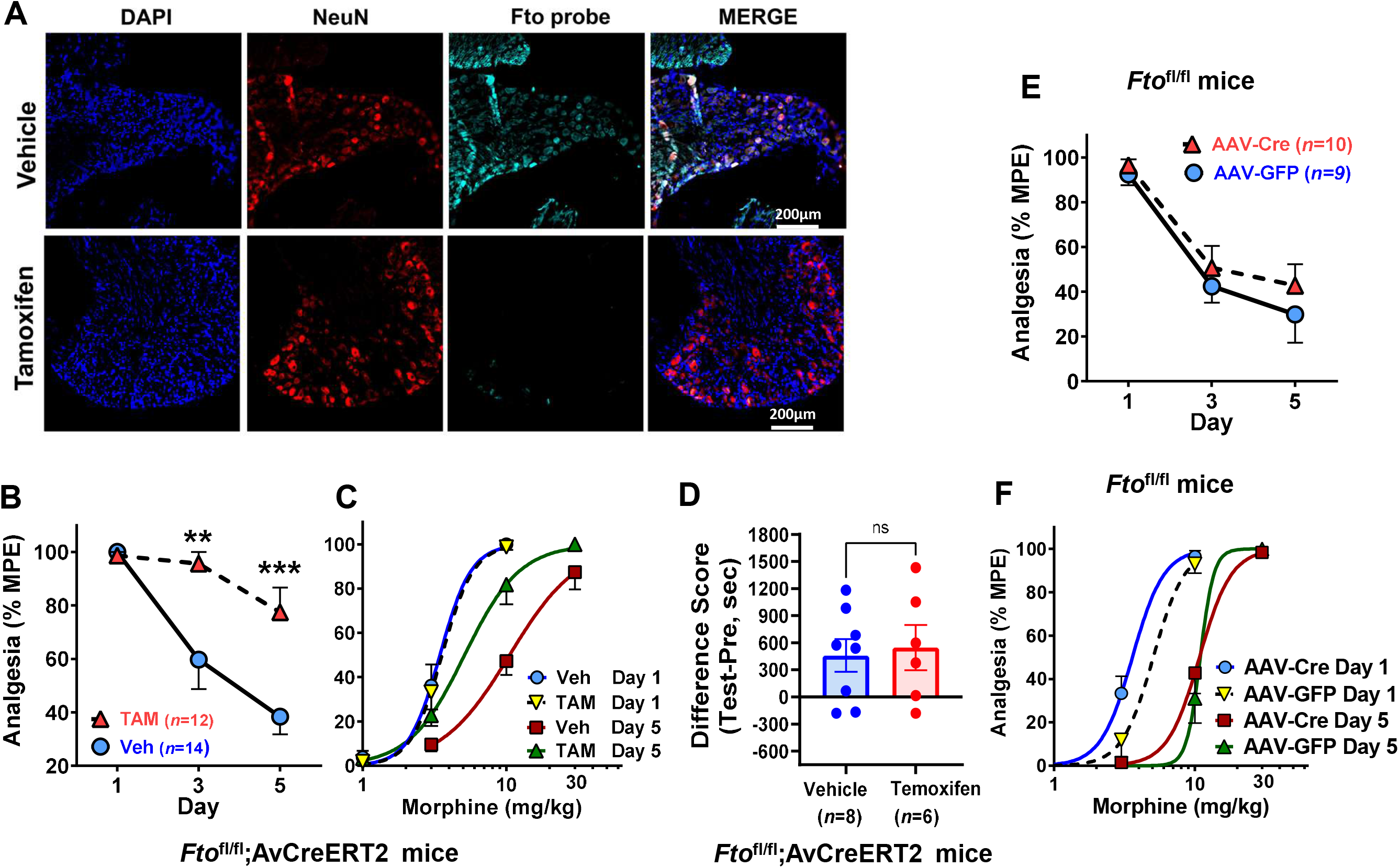
Targeting FTO in the DRG reduced morphine nociceptive tolerance. **(A)** Depletion of Fto in the DRG sensory neurons using tamoxifen in *Fto*^fl/fl^;AvCreERT2 mice. Tamoxifen (75 mg/kg) or Corn oil (Vehicle) was i.p. administered daily in *Fto*^fl/fl^;AvCreERT2 mice for 10 days. RNAScope combined with IHC with anti-NeuN antibody was performed on the DRG sections from perfused mice (4% PFA) after morphine tolerance study, as described in the Methods. **(B & C)** Depleting Fto in the DRG sensory neurons attenuated morphine tolerance in *Fto*^fl/fl^;AvCreERT2 mice. Morphine analgesia was measured by using a radiant heat tail-flick assay and quantified as the percentage of maximum possible effect (% MPE) as described in the Methods. Morphine dose-response curves (1, 3,10 and 30 mg/kg, s.c.) were measured on Days 1 and 5 with single dose (10 mg/kg, s.c.) on Day 3. The ED_50_ values were calculated using non-linear regression analysis (GraphPad Prism 10) and listed on Table 1. B: Time course; C: Dose-response curve. **: *p* < 0.01; ***: *p* < 0.001, 2-way ANOVA with Bonferroni’s *post hoc* test. **(D)** Depleting Fto in the DRG sensory neurons had no effect on morphine CPP in *Fto*^fl/fl^;AvCreERT2 mice. Morphine CPP was performed in *Fto*^fl/fl^;AvCreERT2 mice treated with tamoxifen or vehicle same as Figure 1A. Preference (Difference Score) was calculated as difference between the times spent in the morphine-paired chamber minus time spent in the saline-paired chamber on the post-test day (day 6) and the pre-test day (day 1). n.s: no statistical significance, two-tailed Student *t*-test. **(E & F)** Depleting Fto in the NAcSh had no significant effect on morphine tolerance in *Fto*^fl/fl^ mice. Morphine tolerance in *Fto*^fl/fl^ mice microinjected with AAV-Cre or AAV-GFP was performed same as B (E, time course) & C (F, dose-response curve). No statistical significance, 2-way ANOVA with Bonferroni’s *post hoc* test.

The NAc also involves morphine tolerance in addition to morphine reward^65–68^. The results above, combined with the previous NAcSh-specific depletion data, raised the question of whether the NAcSh, the primary locus governing FTO-dependent morphine reward, also contributes to morphine antinociceptive tolerance. To address this directly, morphine antinociceptive tolerance was assessed in *Fto*^fl/fl^ mice following bilateral NAcSh microinjection of AAV-Cre or AAV-GFP. *Fto*^fl/fl^ mice receiving NAcSh injections of AAV-Cre exhibited a morphine tolerance profile indistinguishable from that of AAV-GFP-injected controls, both in terms of the time course and the rightward shift in ED_50_ between Days 1 and 5 (**Figures 4E, F**). These findings indicate that *Fto* within the NAcSh does not contribute to morphine antinociceptive tolerance. Instead, FTO in the DRG contributes to the development of morphine antinociceptive tolerance.

### FTO depletion selectively increases snRNA m^6^Am levels in the NAcSh transcriptome

Our finding that FTO depletion in the NAcSh impairs morphine-induced CPP suggests that RNA modification levels markedly influence the rewarding properties of morphine. The canonical substrate of FTO is m^6^A, which FTO demethylates in specific transcripts, thereby maintaining those mRNAs in a hypomethylated state^41^. Loss of FTO is therefore expected to result in elevated m^6^A stoichiometry at regulated sites, with downstream consequences including enhanced mRNA decay through m^6^A-dependent degradation pathways. To determine whether this mechanism accounts for the effects of FTO depletion on morphine reward, we mapped m^6^A sites across the NAc transcriptome using Oxford Nanopore direct RNA sequencing of poly(A)+ RNA^69–76^. NAc tissue was harvested five weeks after bilateral microinjection of AAV-GFP or AAV-Cre into *Fto*^fl/fl^ mice. Four biological replicates of each condition were sequenced, and m^6^A stoichiometry values showed high concordance across replicates at sites covered by a minimum of 25 independent reads per replicate (**Figure S1A, B**, **Table S1**). To maximize the number of sites meeting this coverage threshold, reads from the four replicates were merged for each condition, yielding ∼990,000 primary-mapped reads. m^6^A stoichiometry was then compared between conditions at all sites with a minimum coverage of 25 independent reads and a minimum stoichiometry of 0.05.

Comparison of m^6^A stoichiometries in AAV-GFP and AAV-Cre revealed a high similarity in m^6^A stoichiometry across all detected sites between FTO-depleted and control NAcSh samples. This contrasts with FTO studies in leukemia where specific m^6^A sites in the *MYC* mRNA show marked increases in m6A levels after FTO depletion^77^. This indicates that FTO depletion produces no specific m^6^A sites that have clear increases in m^6^A levels in the NAcSh transcriptome (**Figure 5A**, **B**).

**Figure 5.**
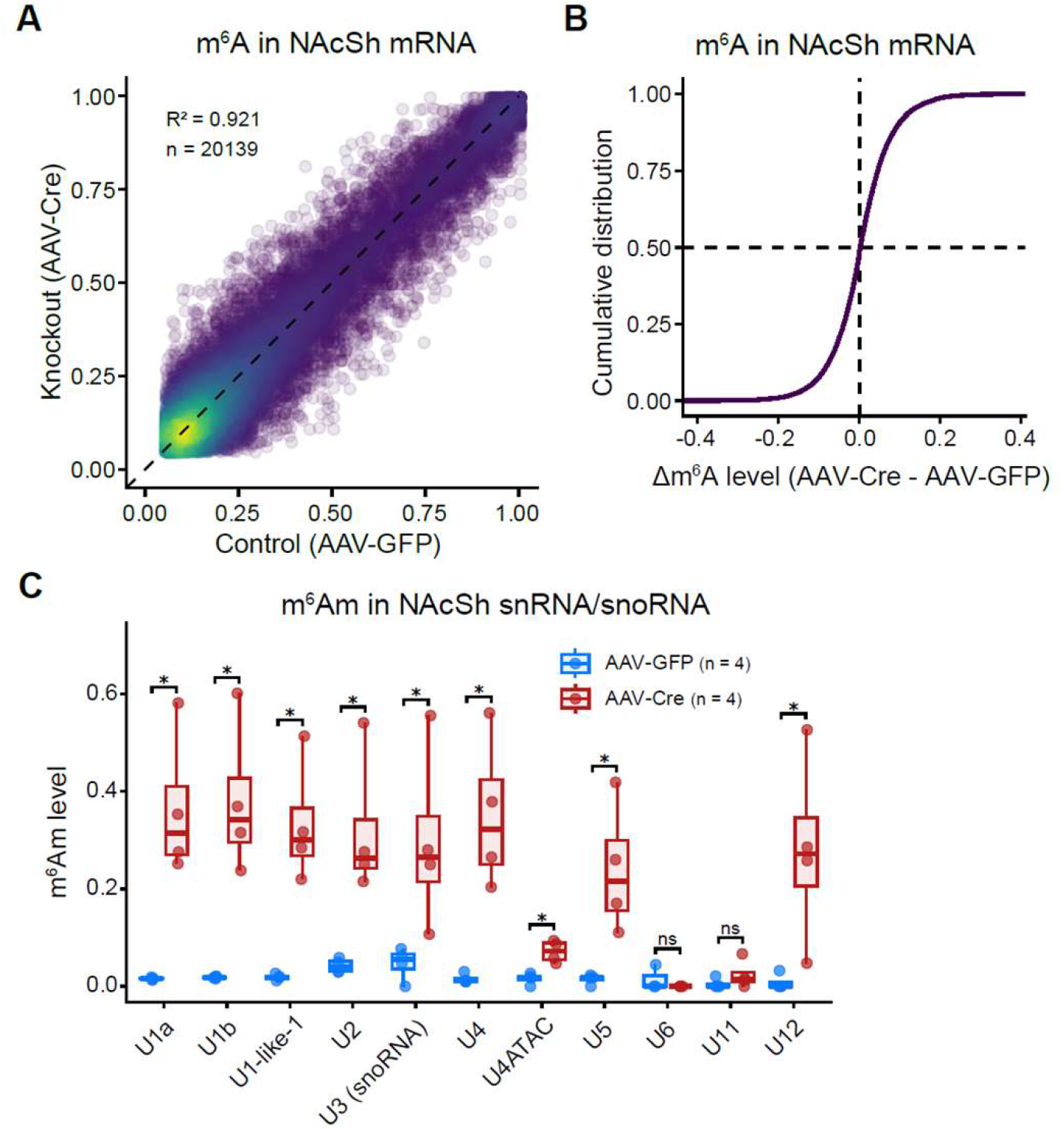
m6A and m6Am profiling in the Ftofl/fl NAcSh upon FTO depletion. **(A)** Scatter plot comparing site-specific m^6^A stoichiometries between AAV-Cre and AAV-GFP treated *Fto^fl/fl^* mice NAcSh tissue (n = 20,139 shared sites, ≥25 reads). Each condition consists of four biological replicates merged for stoichiometry calculation. Pearson’s coefficient of determination is given by R^2^ = 0.921. **(B)** Cumulative distribution plot of m^6^A differences between AAV-Cre and AAV-GFP treated samples (Δm^6^A = Cre – GFP). **(C)** Boxplots of m^6^Am stoichiometry in different snRNA and snoRNA transcripts. For each condition, four biological replicates were shown. Compared with AAV-GFP control NAcSh, AAV-Cre NAcSh show a robust increase in m^6^Am at most snRNA and snoRNA (U3). Only transcripts with ≥25 read coverage for each replicate are shown. Statistical significance determined by a Wilcoxon rank-sum test for each snRNA/snoRNA type (ns, non-significant, \**p* < 0.05).

Since m^6^A stoichiometry was unaffected by FTO depletion, we next examined whether FTO instead regulates levels of m^6^Am, the other established FTO substrate. m^6^Am occupies the first transcribed nucleotide position in both mRNAs and snRNAs. To analyze m^6^Am, we previously developed CROWN-seq to specifically capture and quantify the stoichiometry at the first nucleotide^78^. In CROWN-seq, unmethylated 2’-*O*-methyladenosine (Am) is chemically deaminated into 2’-*O*-methylinosine (Im), which appears in sequencing data as G. However, m^6^Am resists deamination and therefore appears as A. Thus, the percent of transcripts that contain A at the first transcribed position indicates the stoichiometry of m^6^Am.

CROWN-seq analysis revealed that m^6^Am levels in snRNA were markedly elevated in FTO-depleted NAcSh relative to controls (**Figures 5C, S1C**, **Table S2**). Analysis at the level of individual snRNA species revealed particularly prominent increases in m^6^Am in U1, U2, U4, U5, and U12 snRNA, along with the U3 snoRNA, with mean m^6^Am stoichiometries rising from a range of 0.008-0.042 under control conditions to 0.240-0.381 following FTO depletion (**Figure 5C**, **Table S2**) . The maximum increase for any given replicate was found to be +0.580 for the U1b snRNA. Notably, this data demonstrates that FTO in the NAcSh very efficiently demethylates m^6^Am in snRNAs to near-zero levels in basal conditions, and that, upon conditional knockout of FTO, these levels can rise substantially.

Taken together, these data demonstrate that FTO depletion in the NAcSh does not alter m^6^A stoichiometry at any detectable site in the transcriptome but instead produces widespread increases in m^6^Am levels in snRNAs, most prominently in U1. These findings suggest that the mechanism of FTO inhibition does not involve the canonical m^6^A pathway but instead points to the less explored m^6^Am-snRNA axis as the molecular basis for FTO’s role in morphine reward.

### Targeting *Pcif1* abolishes the effect of FTO depletion on morphine reward

To determine whether the increase in m^6^Am upon FTO depletion is causally required for the suppression of morphine reward, we employed a genetic epistasis strategy using *Fto*^fl/fl^;*Pcif1*^fl/fl^ mice. PCIF1 is the methyltransferase responsible for m^6^Am synthesis; accordingly, *Pcif1* depletion abolishes m^6^Am and prevents any FTO depletion-dependent increase in this modification. If FTO depletion mediates its effects on morphine reward by elevating m^6^Am, FTO depletion should no longer have an effect in *Fto*^fl/fl^;*Pcif1*^fl/fl^ mice when m6Am synthesis is blocked by *Pcif1* deletion.

To test this prediction, both *Fto* and *Pcif1* were simultaneously depleted in the NAcSh of *Fto*^fl/fl^;*Pcif1*^fl/fl^ mice by bilateral AAV-Cre microinjection, with AAV-GFP serving as the control. Successful elimination of both *Fto* and *Pcif1* transcripts was confirmed by RNAScope in situ hybridization in AAV-Cre-injected mice relative to AAV-GFP-injected controls (**Figure S2**). Five weeks after injection, mice underwent morphine CPP acquisition using the 6-day paradigm described above. In AAV-GFP-injected *Fto*^fl/fl^;*Pcif1*^fl/fl^ mice, morphine induced robust CPP (**Figure 2D**). However, in contrast to *Fto* depletion alone which impaired morphine-induced CPP, depletion of *Fto* along with *Pcif1* restored morphine-induced CPP acquisition (**Figure 2D**). These results demonstrate that the suppression of morphine CPP by FTO depletion requires the capacity to elevate m^6^Am, establishing increased m^6^Am in snRNA as the causal mediator of FTO’s effect on morphine reward.

Since increased m^6^Am caused by FTO depletion reduces morphine-induced CPP, we next asked if reduced m^6^Am would enhance the acquisition of morphine-induced CPP. Although m^6^Am is already low in snRNA (see **Figure 5C**), we nevertheless measured morphine CPP acquisition in *Pcif1*^fl/fl^ mice following NAcSh microinjection of AAV-Cre or AAV-GFP. NAcSh-specific depletion of *Pcif1* led to increased CPP scores compared to the AAV-GFP group, but this did not achieve statistical significance (p = 0.1024) (**Figure 2E**). Nevertheless, the trend toward increased CPP scores raises the possibility that endogenous m^6^Am levels in the NAcSh are inversely correlated with morphine reward as measured by CPP.

### Targeting *Pcif1* abolishes the effect of FTO depletion on morphine antinociceptive tolerance

Having established that elevated m^6^Am in snRNA mediates the suppression of morphine reward by FTO depletion in the NAcSh, we next asked whether the same mechanism accounts for the attenuation of morphine antinociceptive tolerance produced by FTO depletion in the DRG. To test this, a parallel genetic epistasis approach was employed in *Fto*^fl/fl^;*Pcif1*^fl/fl^;Advillin-CreERT2 mice, enabling inducible, DRG-restricted co-depletion of both *Fto* and *Pcif1* by tamoxifen administration. Successful elimination of both *Fto* and *Pcif1* transcripts in the DRG was confirmed by RNAScope in situ hybridization in tamoxifen-treated mice relative to vehicle-treated controls (**Figure S3**).

Morphine antinociceptive tolerance was then assessed using the 5-day paradigm described above. Vehicle-treated controls developed robust morphine antinociceptive tolerance, as expected. In contrast to *Fto* depletion, co-depletion of both *Fto* and *Pcif1* in the DRG was associated with restored morphine antinociceptive tolerance. Morphine antinociceptive tolerance in *Fto* and *Pcif1* depleted mice achieved levels comparable to those of vehicle-treated controls (**Figures 6A, B**). These results demonstrate that the attenuation of morphine antinociceptive tolerance produced by DRG-specific FTO depletion is mediated through its capacity to elevate m^6^Am, establishing increased m^6^Am as a causal determinant of both morphine reward and antinociceptive tolerance.

**Figure 6.**
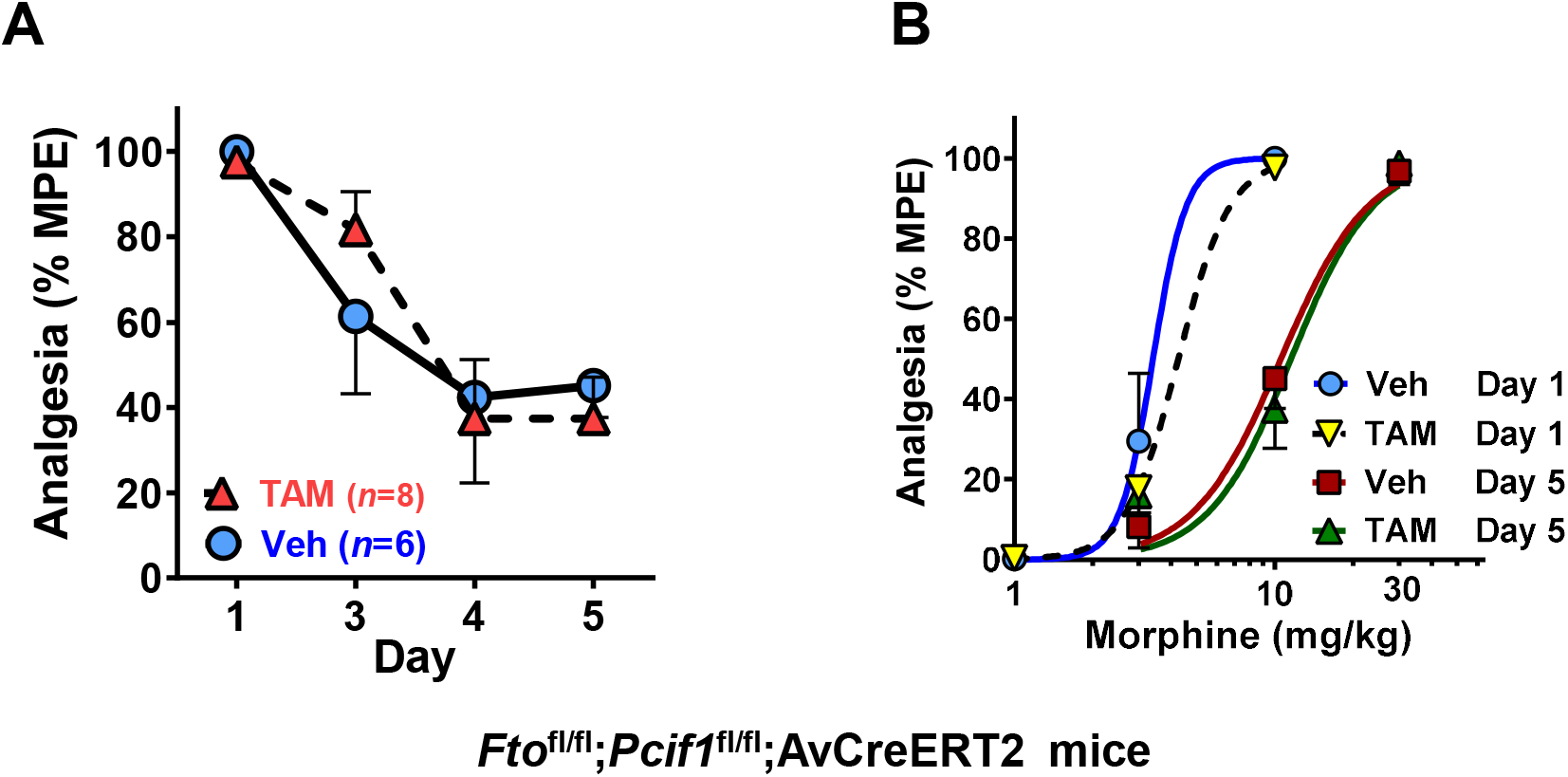
Depletion of FTO along with PCIF1 in the DRG restores morphine nociceptive tolerance (A &. **B)** Morphine tolerance was performed in *Fto*^fl/fl^;*Pcif1*^fl/fl^;AvCreERT2 mice after treatment of tamoxifen or vehicle same as Figures 4B/4C. The ED_50_ values were calculated using non-linear regression analysis (GraphPad Prism 10) and listed on Table 1. A: Time course; B: Dose-response curve.

### Impact of FTO depletion on alternative splicing

We next asked how increased m^6^Am could affect gene expression related to morphine reward and tolerance. Our data, as well as previous studies^27^, show that FTO primarily regulates m^6^Am on snRNA, not mRNA. However, the function of the m^6^Am form of snRNA compared to the more abundant Am form of snRNA is not known. The canonical function of snRNAs is in pre-mRNA splicing, where they direct intron recognition and removal as core components of the spliceosome^79^. However, U1 snRNA serves an additional, splicing-independent function to suppress premature cleavage and polyadenylation at cryptic intronic sites, thereby sustaining transcriptional elongation through intronic regions and ensuring production of full-length transcripts^80,81^. Elevated m^6^Am in U1 snRNA and other snRNA species following FTO depletion could therefore perturb either spliceosome fidelity and telescripting activity, with broad consequences for gene expression in the NAcSh.

To distinguish transcriptomic changes strictly attributable to FTO depletion from those induced by opioid exposure, we performed RNA-seq on NAc tissue harvested from *Fto*^fl/fl^ mice that received bilateral AAV-GFP or AAV-Cre microinjections and underwent the CPP paradigm without morphine exposure.

We first examined the impact of Fto depletion on splicing. We performed alternative splicing analysis with rMATS^82^ (**Figure S4**, **Table S3**). The major splicing change in Fto-depleted NAc is exon skipping, although other types of alternative splicing were also found (**Figure 7A**). In total, we observed 552 significant exon skipping events across 483 genes (FDR <0.1), as well as other forms of alternative splicing. Alternative splicing was seen across diverse types of genes, without a clear enrichment in specific Gene Ontology pathways (**Table S3**).

**Figure 7.**
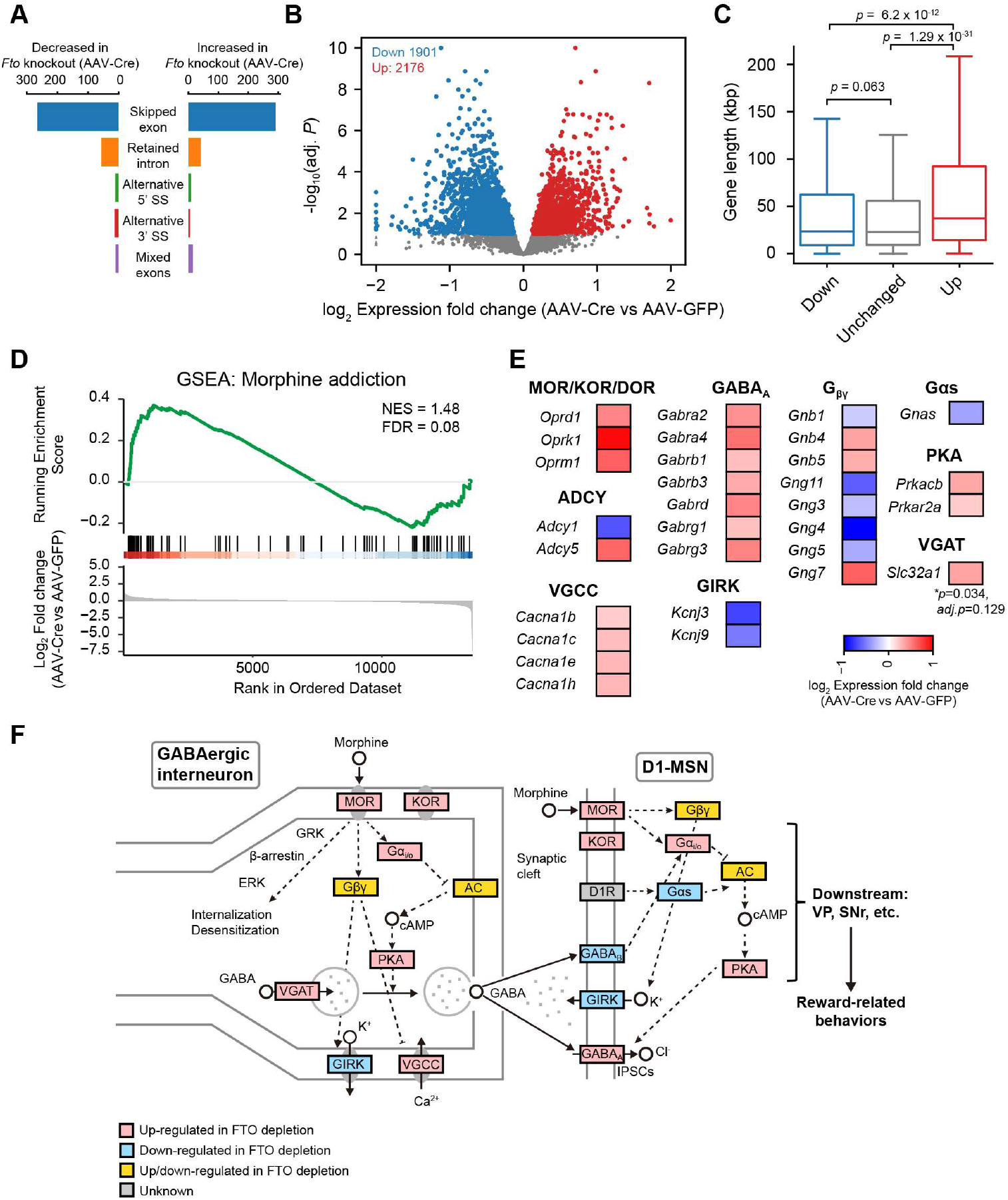
The transcriptomic changes in Ftofl/fl NAcSh upon FTO depletion. **(A)** The number of alternative splicing events found between AAV-Cre (Fto knockout) and AAV-GFP (control). The corresponding volcano plots can be found in **Figure S4**. **(B)** A volcano plot showing genes with significant differential expression upon FTO depletion. Genes with at least 100 reads on average across four biological replicates were analyzed. Genes with adjusted P-value <0.1 were considered to have significant expression change. **(C)** Genes that exhibit up-regulated gene expression upon FTO depletion show longer gene length. The classifications of the genes correspond to **(B)**. P-values, Student’s t-test. **(D)** GSEA enrichment of the genes related to morphine addiction pathway (mmu05032). **(E)** Shown are heatmaps demonstrating the genes in the core components in the morphine addiction pathway with significant differential expression (adjusted P-value < 0.1). **(F)** A schematic diagram showing the impact on different components in the morphine addiction pathway upon FTO depletion. MOR, mu opioid receptor; KOR, kappa opioid receptor; GIRK, G-protein-gated inwardly rectifying K^+^ channels; VGCC, voltage-gated Ca^2+^ channel; D1-MSN, D1 receptor–expressing medium spiny neuron; GABA_A_, γ-Aminobutyric acid sub-type A receptor; VGAT, vesicular inhibitory amino acid transporter; VP, the ventral pallidum; SNr, the substantia Nigra pars reticulata.

We next asked whether FTO depletion affects U1 snRNA-dependent transcriptional elongation. U1 snRNA promotes elongation through introns and 3’UTRs, which preferentially enhances expression of long genes^80^. We performed differential gene expression analysis between control and FTO-depleted NAcSh. We found 2,176 upregulated genes and 1,901 downregulated genes (adjusted *p* < 0.1) (**Figure 7B**, **Table S4**). The upregulated genes had significantly longer gene bodies than the downregulated and unchanged genes (median 37.6 kbp versus 23.5 and 23.2 kbp) (**Figure 7C**). This length bias indicates that elevated m^6^Am promotes U1-dependent elongation of long genes. FTO depletion raises snRNA m^6^Am, so FTO normally opposes this elongation and limits expression of long genes.

### Impact of FTO depletion on genes in morphine addiction pathways

We next asked whether the transcriptomic changes produced by FTO depletion converge on genes that control morphine reward. We performed Gene Set Enrichment Analysis (GSEA) of the differentially expressed genes against KEGG pathways. Many signaling pathways were altered, and the morphine addiction pathway was significantly enriched among the genes changed in the FTO-depleted NAcSh (**Figure 7D**, **Table S5**).

Many of these genes were altered in a coordinated direction that opposes MOR-driven reward signaling in the GABAergic interneuron/D1-MSN circuit of the NAc shell, the circuit through which morphine produces reward^45,83,84^ (**Figure 7E, F**). MOR normally suppresses presynaptic GABA release by inhibiting adenylyl cyclase/cAMP/PKA signaling through Gαi/o and by regulating voltage-gated Ca^2+^ channels (VGCCs) and G-protein-gated inwardly rectifying K+ channels (GIRKs) through Gβγ, which disinhibits D1-MSNs and promotes reward^85,86^. FTO depletion reversed this gene expression signature at multiple nodes. It lowered Ca^2+^/calmodulin-stimulated *Adcy1* and raised Ca^2+^-inhibited *Adcy5* and PKA subunits (*Prkacb*, *Prkar2a*), which may lead to restoration of cAMP/PKA signaling that MOR normally controls^88^. It raised VGCC subunits (*Cacna1b*, *Cacna1c*, *Cacna1h*) and lowered GIRK subunits (*Kcnj3*, *Kcnj9*), which would neutralize the G_βγ_-mediated effects of MOR on these channels^85^. It raised GABAA receptor subunits (*Gabrb1*, *Gabrb2*, *Gabrb3*, *Gabrd*, *Gabrg1*, *Gabrg3*) in D1-MSNs, which would strengthen the inhibitory currents that MOR activation reduces^84,87^. FTO-depletion led to increased expression of opioid receptor genes, most notably *Oprk1* (KOR), which would increase dynorphin/KOR anti-reward tone that counteracts MOR-driven reward^89–91^, as well as *Oprm1* and *Oprd1*.

Together, these changes suggest a coordinated shift in the NAc shell transcriptome. Rather than acting on a single effector, FTO depletion alters many genes across the same circuit in directions that oppose MOR signaling, and this network-level shift may explain some of the reduction in morphine reward.

## DISCUSSION

Morphine and other mu opioids remain among the most effective analgesics in pain management, yet their clinical use is limited by tolerance, reward and addiction, the properties that drive the escalation underlying opioid use disorder. A long-standing goal has been to separate these harmful properties from analgesia, so that pain can be relieved without promoting opioid misuse. Here we describe a mechanism that achieves this separation, in which elevated m^6^Am in snRNA suppresses morphine reward and tolerance while preserving analgesia. We uncovered this mechanism using genetic depletion of FTO and pharmacological FTO inhibition, which both reduced morphine reward and antinociceptive tolerance without altering morphine analgesia. Because FTO is known mainly as a demethylase of m^6^A in mRNA, we expected these effects would be associated with increased levels of m^6^A on specific mRNAs. Instead, we found that FTO depletion leads to increases in m^6^Am in snRNA, pointing to m^6^Am as a novel regulator of morphine reward and tolerance. Overall, our work shows that raising m^6^Am in snRNA reduces the reward and tolerance produced by opioids while sparing their analgesic action, and establishes the control of RNA modifications in snRNA, and thus of snRNA function, as a new approach to separating the therapeutic and harmful effects of opioids.

To find the RNA substrate of FTO, we mapped m^6^A and m^6^Am across the NAcSh transcriptome using direct RNA sequencing and CROWN-seq, respectively. These methods provide highly quantitative measurements of m^6^A and m^6^Am stoichiometry. Our analysis of 20,139 m^6^A sites showed no clear increase in stoichiometry in any m^6^A site in any mRNA after FTO depletion. FTO depletion instead led to marked increases in m6Am stoichiometry in snRNA. To test whether this rise in m^6^Am was responsible for the behavioral effects, we used a genetic strategy, reasoning that if elevated m^6^Am mediates the effects of FTO loss, then removing the m^6^Am biosynthetic enzyme PCIF1 should mitigate the effects of FTO depletion. When we co-depleted *Fto* and *Pcif1*, FTO depletion no longer suppressed morphine reward and tolerance. This epistasis experiment supports the conclusion that elevated m^6^Am mediates the beneficial effects of FTO depletion.

How could a modification on snRNA control morphine reward and tolerance? snRNAs are the core components of the spliceosome, where they direct the recognition and removal of introns, and U1 snRNA has a further role in promoting transcriptional elongation through introns and 3’UTRs^80^. A modification that alters snRNA function could therefore reshape gene expression broadly, through both splicing and elongation. Our data supports this idea. FTO depletion changed alternative splicing at thousands of sites across the transcriptome, with exon skipping as the predominant change, and it also shifted gene expression in a way that favored longer genes, consistent with an effect on U1-dependent elongation. These changes were not confined to a few transcripts but were spread across the transcriptome, and the resulting set of altered genes was enriched for components of the morphine addiction pathway. We therefore propose that elevated m^6^Am in snRNA reduces morphine reward and tolerance by altering snRNA function, which remodels splicing and elongation across many genes and, through this broad change, shifts the expression of morphine-pathway genes.

An interesting feature of this mechanism is that it operates in two separate areas for the two behaviors. When we depleted *Fto* in NAcSh, morphine reward was reduced but tolerance was intact, whereas depletion in the DRG reduced tolerance but left reward unchanged. Reward and tolerance are therefore controlled by FTO at anatomically distinct sites, a central reward circuit for one and a peripheral sensory ganglion for the other, suggesting that the role of FTO in regulating the actions of mu opioids is region-specific.

These findings point to FTO as a therapeutic target for reducing the harmful effects of opioids. An FTO inhibitor given alongside an opioid could lessen the reward and tolerance while preserving pain relief. The reduction of fentanyl tolerance by the FTO inhibitor suggests that this benefit would extend across mu opioids rather than being specific to morphine. A therapy that inhibits FTO would also be expected to have an acceptable safety profile. Global loss of FTO early in development is poorly tolerated in mice^42,92^, but deletion restricted to adult animals has minimal pathologic effects^93^. The manipulations used in this study were both adult-onset and confined to specific tissues. Small-molecule FTO inhibitors have been used in animal cancer studies without severe toxicity^43^. FTO inhibitors may therefore represent a new class of adjuncts to mu opioids in pain management and opioid use disorder treatment.

### Limitations of the study

Our transcriptomic and m^6^A/m^6^Am mapping studies used bulk NAc tissue, which cannot resolve changes at single cell level. Future work using single cell-RNAseq or spatial transcriptomics will be needed to decipher the molecular mechanisms at the level of individual cells or neurons. While this study focused on the NAcSh and DRG, several other brain regions, such as the VTA, mPFC, and PAG, are also involved in opioid reward or tolerance. We plan to explore the role of FTO in these regions in future studies.

## STAR*METHODS

### KEY RESOURCES TABLE

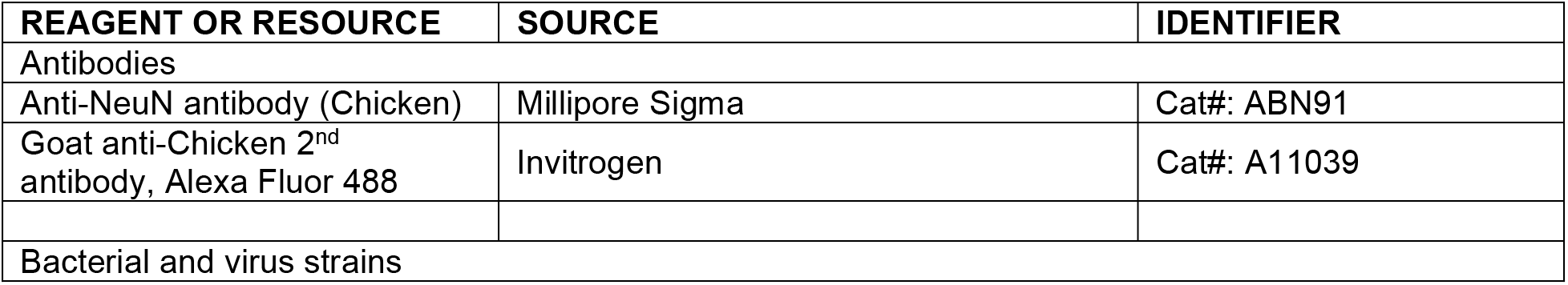
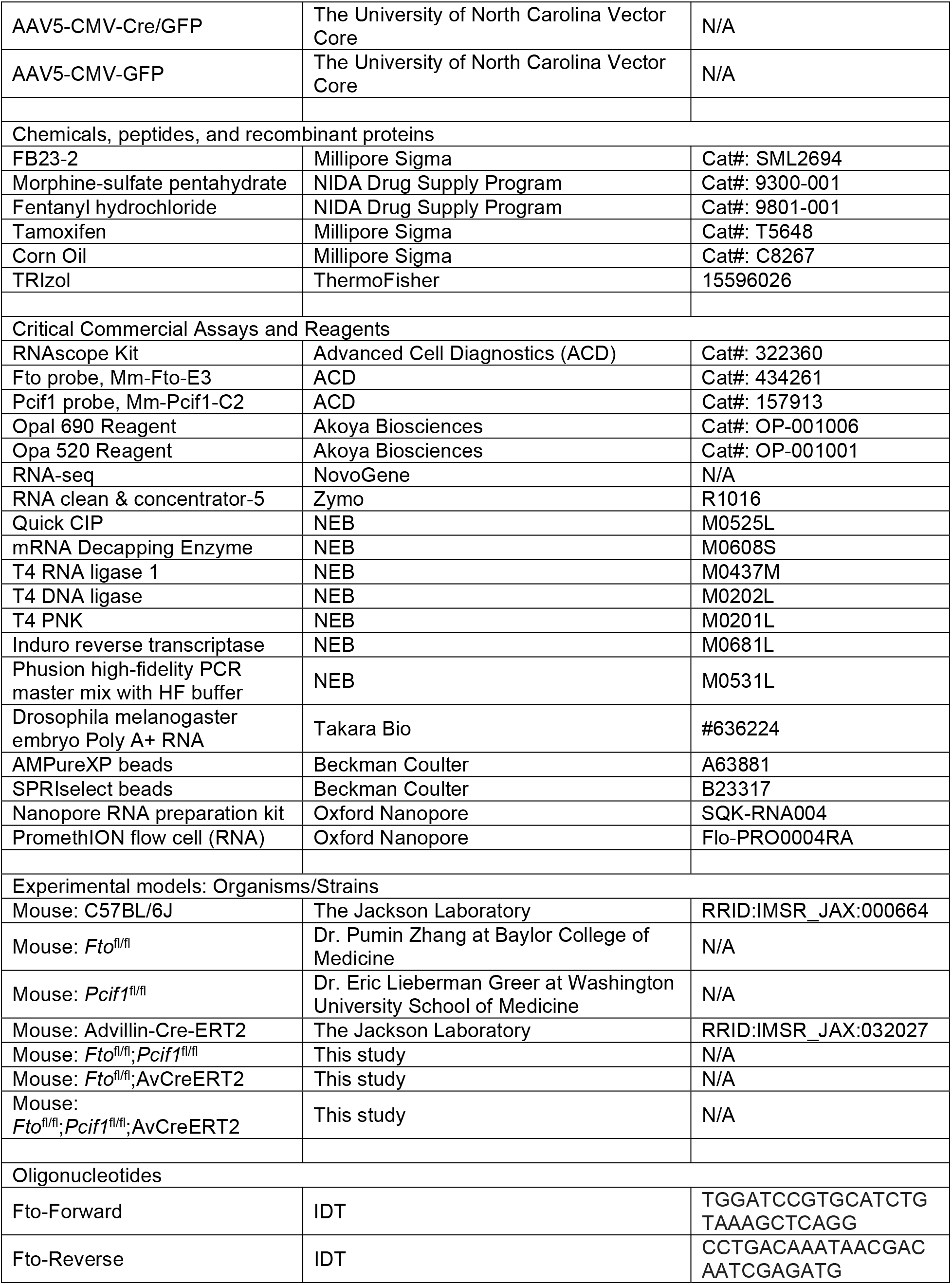
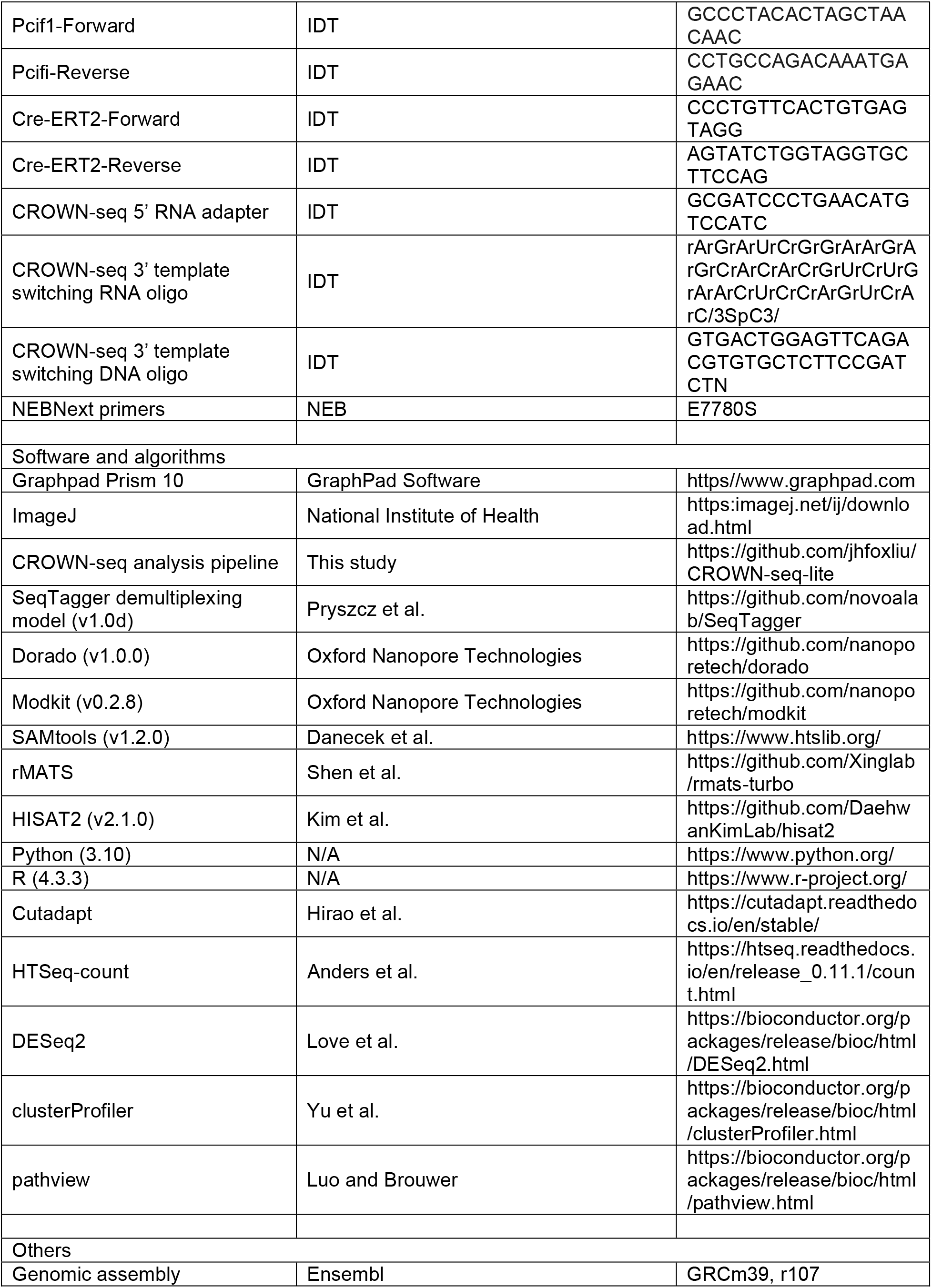

## EXPERIMENTAL MODEL AND STUDY PARTICIPANT DETAILS

### Animals

C57BL/6J (B6) (stock#: 000664) and Advillin-Cre-ERT2 mouse (Stock #: 032027) were obtained from Jackson Laboratory. *Fto*^fl/fl^ mice in B6 background were obtained from Dr. Pumin Zhang, Baylor College of Medicine. *Pcif1*^fl/fl^ mice in B6 background were obtained from Dr. Eric Lieberman Greer, Washington University School of Medicine. We bred *Fto*^fl/fl^ mice with *Pcif1*^fl/fl^ mice to generate *Fto*^fl/fl^;*Pcif1*^fl/fl^ mice and bred *Fto*^fl/fl^ mice with Advillin-Cre-ERT2 (AvCreERT2) mice to produce *Fto*^fl/fl^;AvCreERT2 mice. *Fto*^fl/fl^;*Pcif1*^fl/fl^ mice were bred with AvCreERT2 mice to produce *Fto*^fl/fl^;*Pcif1*^fl/fl^;AvCreERT2 mice. Adult (9 – 16 weeks-old) male mice were used in all experiments. All mice were housed in groups of five, maintained on a 12-h light/dark cycle and given *ad libitum* access to food and water. All animal studies were approved by the Institutional Animal Care and Use Committee of Rutgers New Jersey medical School.

## METHODS

### Drug administration

FB23-2 (Millipore Sigma) or DMSO (Vehicle control) was administered by i.c.v. (20 μg) or i.p. (20 mg/kg) once a day for 4 or 5 days followed by morphine (s.c.) injection two hours later. *Fto*^fl/fl^;AvCreERT2 or *Fto*^fl/fl^;*Pcif1*^fl/fl^;AvCreERT2 mice were administered with tamoxifen (75 mg/kg, i.p., once per day for consecutive 10 days) prepared in corn oil or with corn oil as vehicle control.

### Opioid analgesia

Analgesia was determined 30 min after subcutaneous (s.c.) injection using a radiant-heat tail-flick assay or hot-plate assay, with a maximal latency of 10 sec or 30 sec, respectively, to minimize tissue damage, as previously described^94,95^. Results were calculated as the percentage of maximum possible effect (% MPE) [(latency after drug – baseline latency)/(10 – baseline latency)*100]. Morphine doses used in cumulative dose-response studies were 0.3, 1, 3, 10 or 30 mg/kg. ED_50_ values were determined using nonlinear regression analysis (GraphPad Prism 10).

### Opioid tolerance

Morphine tolerance was induced by twice-daily injections with morphine (10 mg/kg, s.c.) for 5 days. Morphine analgesia was examined on Days 1 and 5 by dose-response curves. Fentanyl tolerance was induced by an s.c. implanted osmotic pump (Alzet Model: 1007D; 1 mg/kg/day/mouse for 7 days). Tolerance was assessed by cumulative dose-response curve (Morphine doses: 1, 3, 10 and 30 mg/kg; Fentanyl doses: 0.01, 0.04, 0.1 and 0.3 mg/kg) on Days 1 and 5 or 7 with radiant heat tail-flick assay, by which the ED_50_ values were determined, as well as analgesic time courses with a fixed dose on Days 3, 4, 5 or 6 (morphine, 10 mg/kg, s.c.; fentanyl, 0.12 mg/kg, s.c.).

### Conditioned place preference (CPP)

CPP was performed using a three-chamber apparatus (Med Associates) with a 6-day paradigm. On day 1, mice were placed in the central chamber for 1 min of habituation with the sliding doors closed, followed by a 20 min preconditioning period of free exploration in the three chambers. Time spent in each chamber was recorded. On days 2 – 5, mice were intraperitoneally (i.p.) injected with saline (morning) or morphine (10 mg/kg, afternoon), and confined to either the black or the white chamber for 20 min. On day 6, mice were placed for a 1 min habituation in the central chamber and followed by a 20 min free exploration in the three chambers. Time spent in each chamber was recorded. Preference is defined as difference between the times spent in the morphine-paired side of the chamber on the test day (Day 6) as compared to the preconditioning day (Day 1).

### Stereotaxic microinjection of AAV vectors

AAV5-CMV-GFP/Cre (AAV-Cre) and AAV5-CMV-GFP (AAV-GFP, Control) with >5.6 x 10^12^ titers were obtained from the Vector Core at the University of North Carolina. AAVs were bilaterally microinjected into the NAc shell (NAcSh, Coordinates (in mm): AP +1.6, DV -4.55, ML ±0.55.) using a robot stereotaxic system (Neurostar). Mice were recovered for 4-5 weeks after microinjection to allow sufficient Cre expression before behavioral tests.

### RNAscope and immunohistochemistry (IHC)

Mice were anaesthetized with Ketamine/Xylazine and transcardially perfused with PBS followed by 4% paraformaldehyde. Brain and DRG were dissected, post-fixed overnight, incubated in a sucrose gradient (20-30%), embedded in OCT medium (Tissue-Tek) and cryosectioned with 10 µm sections. RNAscope was performed on sections using RNAscope Kit (ACD, Catalog #: 322360) with Fto probe (Advanced Cell Diagnostics (ACD), Catalog #: 434261) and/or Pcif1 probe (ACD, Catalog #: 157913), as well as Opal fluorophores (Akoya Biosciences), in ACD HybEZ™ II Hybridization System, following the manufactural protocol. The DRG sections were then used in IHC with an anti-NeuN antibody (Millipore Sigma) and a Goat anti-Chicken 2^nd^ antibody conjugated with Alexa Flour 488 following RNAscope procedure. The sections were then imaged for Fto or Pcif1/GFP in brain sections and for Fto or Pcif1/NeuN with a Nikon Ti2 HCA inverted fluorescent microscope.

### Statistics

All mice were randomized and assigned to groups. Some, but not all, experiments were performed under blind conditions. All statistical analysis was carried out using GraphPad Prism 10. A student *t*-test, 1-way ANOVA or 2-way ANOVA was performed with *post hoc* Bonferroni’s multiple comparisons test as described in the figure legend. Data represented the means ± SEM. Statistical significance was set at *p* < 0.05.

### Differential gene expression analysis via RNA-seq

RNA-seq were performed by NovoGene. The sequencing reads were first trimmed by Cutadapt to remove adapter sequences^96^. The trimmed reads were then mapped to reference genome (GRCm39, Ensembl r107) via HISAT2 (key parameters: --fr unstranded^97^. To count the number of reads mapped to each genes in the reference, HTSeq-count was used^98^. Differential expression was performed with DESeq2 based on the read counts^99^. Only protein coding genes with at least 100 reads on average across all samples were considered. To analyze the gene lengths, for each gene, we selected the longest isoform (including all exons and introns) based on the Ensembl gene annotations.

### Alternative splicing analysis

Alternative splicing analysis was performed with rMATS (v4.3)^82^. The BAM files mapped by HISAT2 were used. Key parameters: -t paired, --readLength 150, --library-type fr-unstranded --variable-read-length.

### GSEA analysis

GSEA analyses were performed with gseGO (nPerm = 10000, minGSSize = 3, maxGSSize = 800, pvalueCutoff = 0.1, pAdjustMethod = “fdr” ) and gseKEGG (minGSSize = 10, maxGSSize = 800, pvalueCutoff = 0.1, pAdjustMethod = “fdr”) from clusterProfiler^100^.

### Nanopore direct RNA library preparation and sequencing

RNA sequencing libraries were prepared from total RNA using the nanopore direct RNA sequencing kit (ONT, SQK-RNA004) according to the manufacturer’s instructions, with some notable adjustments. Samples were multiplexed following the SeqTagger method from Pryszcz et al. 2025^73^ using the custom RT adapters for the b96_RNA004 model. In brief, 500 ng total RNA was barcoded with custom RT adapters via ligation with T4 DNA ligase (NEB, M0202L); next, samples were reverse transcribed with Induro Reverse Transcriptase (NEB, M0681L), cleaned-up with 1.8X AMPureXP beads (Beckman, A63881), and pooled together. The remaining library preparation steps were followed according to ONT kit instructions. Samples were loaded onto a PromethION flow cell (ONT, Flo-PRO0004RA) and sequenced on a PromethION for 48 hours, acquiring 6.19 Gigabases and 5.87 M reads with an N50 read length of 1.66 kb.

### Nanopore data analysis

The raw POD5 sequencing data files were basecalled with Dorado v1.0.0 with the rna004_130bps_sup@5.2.0 m^6^A_DRACH modification model. The POD5 files were also used to generate a demultiplexed pod5.demux.tsv.gz file following the SeqTagger protocol with model b96_RNA004^73^. Basecalled BAM files were mapped to the GRCm39 mouse genome using the Dorado aligner with the parameter --mm2-opts “-ax splice -k 14”. Aligned reads were demultiplexed with the SeqTagger bam_split_by_barcode.py script, and then sorted and indexed with SAMtools v1.2.20. For increased read coverage, replicates were merged with SAMtools merge. Modification bed tables (mod.bed) were compiled using Modkit v0.2.8 with the pileup command and the parameter --filter-threshold 0.99.

### CROWN-seq experiments

The manuscript of modified CROWN-seq is under submission at the time of submission of this study. Briefly, to study snRNA m^6^Am stoichiometries, we used a modified protocol of CROWN-seq^78^, in which 2’-*O-*methyladenosine (Am) is chemically converted into 2’*O*-methylinosine (Im) and *N*6,2’*O*-dimethyladenosine (m^6^Am) remains unconverted, allowing for m^6^Am quantification. To enrich snRNA, short RNAs (17-200 nt) were isolated from total RNA using RCC-5 (Zymo Research, Cat#: R1016), and 100 ng of the small RNA fraction was used as input. Samples were chemically converted following the GLORI v2.0^101^ conversion conditions. After clean-up, 10 ng of Drosophila melanogaster embryo Poly A+ RNA (Takara Bio, Cat#: 636224) was spiked-in for QC. For precise capture of capped RNA species, samples were sequentially dephosphorylated with Quick CIP (NEB, Cat#: M0525L), decapped with mRNA Decapping Enzyme (NEB, Cat#: M0608S), 5’ ligated with a 5’ adapter (rCrCrUrArCrArCrGrArCrGrCrUrCrUrUrCrCrGrArUrCrUrNrNrNrNrNrNrNrNrNrNrNrUrUr A) and T4 RNA ligase 1 (NEB, Cat#: M0437M), and end-repaired with T4 PNK (NEB, Cat#: M0201L). Library preparation was completed by 3’ adapter incorporation and cDNA synthesis via a template switching reaction with Induro reverse transcriptase (NEB, Cat#: M0681L) and indexing PCR with Phusion High-Fidelity PCR Master Mix with HF Buffer (NEB, Cat#: M0531L) and NEBNext primers (NEB, Cat#: E7780S). The amplified libraries were purified and size selected sequentially with AMPureXP (Beckman Coulter, Cat#: A63881) and SPRIselect (Beckman Coulter, Cat#: B23317) beads to remove short amplicons without inserted sequences. Libraries were pooled and sequenced by NovaSeqX by Weill Cornell Genomics Resources Core Facility.

### CROWN-seq analysis

Quantifying snRNA methylation in mice is challenging because there are many copies of the same snRNA sequence in the mouse genome. As a result, many reads originating from snRNA fail to uniquely map to a genomic locus via genomic mapping or a transcript isoform via transcriptomic mapping. To overcome this issue, we performed a snRNA-type-specific analysis. In this analysis, we focused on the type of snRNA the reads mapped to. To do so, we first mapped the reads to all possible snRNA sequences via Bowtie2. We then grouped the reads based on a customized dictionary which maps different snRNA isoforms to their type of snRNA. After assigning the reads to a specific snRNA type, we calculate the non-conversion rates of the first nucleotides.

## Supplementary Tables

Table S1. m^6^A stoichiometry quantified by Nanopore sequencing

Table S2. m^6^Am stoichiometry quantified by CROWN-seq

Table S3. Alternative splicing

Table S4. Differential gene expression

Table S5. GSEA analysis

## Supplemental Figures

**Figure S1. m6A stoichiometry comparisons across biological replicates at each site (A)** and (**B)** Pairwise scatter plots comparing per-site m^6^A stoichiometry calculations between four biological replicates (≥25 reads) of AAV-GFP treated (**A**) or AAV-Cre treated (**B**) *Fto*^fl/fl^ mice. Pearson’s coefficient of determination (R^2^) and number of shared sites (n) are reported in each plot.

**Figure S2. Depletion of both Fto and Pcif1 in the NAcSh using microinjected AAV-Cre/GFP** RNAscope with Fto probe/Pcifi1 probe and Fto/Pcif1/GFP imaging was performed on brain sections from *Fto^fl/fl^*;*Pcif1*^fl/fl^ mice microinjected with AAV-Cre/GFP or AAV-GFP in the NAcSh after morphine CPP study, as described in the Methods. AAV-Cre: AAV5-CMV-Cre/GFP; AAV-GFP: AAV5-CMV-GFP; core: NAc core; shell: NAc shell (NAcSh).

**Figure S3. Depletion of both Fto and Pcif1 in the DRG sensory neurons using tamoxifen** Tamoxifen (75 mg/kg) or Corn oil (Vehicle) was i.p. administered daily in *Fto*^fl/fl^;*Pcif1*^fl/fl^;AvCreERT2 mice for 10 days. RNAScope combined with IHC with anti-NeuN antibody was performed on the DRG sections from perfused mice (4% PFA) after morphine tolerance study, as described in the Methods.

**Figure S4. Alternative splicing in mice NAcSh upon FTO depletion (A)** - **(E)**, volcano plots showing the significant alternative splicing events (FDR < 0.1, |Δinclusion level| > 0.1 ) in different categories.

**Figure S5. KEGG pathway analysis on mice NAcSh upon FTO depletion** Shown are top 17 KEGG pathways demonstrating significant enrichment in GSEA analysis. Pathways activated upon FTO depletion are shown on the left, while pathways suppressed upon FTO depletion are shown on the right.

## Data and Code Availability

All sequencing data generated from this study can be accessed from NCBI Gene Expression Omnibus (GEO):

- <u>GSE334880</u> for nanopore direct RNA sequencing data (reviewer token: mpozqeokxnihvqj);
- <u>GSE334881</u> for CROWN-seq data (reviewer token: wdituecqvrmdlcp) ;
- <u>GSE335428</u> for NovoGene RNA-seq data (reviewer token: epehwmeihpipnsn). The CROWN-seq analysis pipeline is available on GitHub: https://github.com/jhfoxliu/CROWN-seq-lite

## Acknowledgements

We thank Dr. Pumin Zhang at Baylor College of Medicine to provide *Fto*^fl/fl^ mice. This work is supported by grants from the National Institutes of Health (R01DA09544 to Y.X.P., and S.R.J, DA060222 to Y.-X.P., R01DA037755 and RM1HG011563 to S.R.J., R01DK142785 to Y.-X.T., R01AI151215 to E.L.G., R01DA054368 and R01DA053261 to A.M.R., F31 DA060029 to A.F.M. and T32 DA055569 to R.C); and grants from the Department of Anesthesiology, New Jersey Medical School and Brain Health Institute, Rutgers.

## AUTHOR CONTRIBUTIONS

Conceptualization, Y.-X.P., S.R.J., A.M.R., Y.-X.T. and E.L.G.; methodology, L.S., A.F.M., R.C., J.L., L.S.N., J.X., K.D., V.P., V.P.L., A.M.R., K.B. and B.W.; supervision, Y.-X.P., S.R.J., A.M.R., Y.-X.T. and E.L.G.; validation, L.S., A.F.M., R.C., J.L., L.S.N., J.X., K.D. and V.P.; investigation, L.S., A.F.M., R.C., J.L. and L.S.N.; writing, Y.-X.P., S.R.J., L.S., A.F.M., R.C., J.L. and L.S.N.

