## Supplementary figures and images for "Reversible m^6^Am methylation of snRNA by FTO controls morphine reward and tolerance without altering analgesia"

### Figure S1; Figure S2; Figure S3; Figure S4; Figure S5

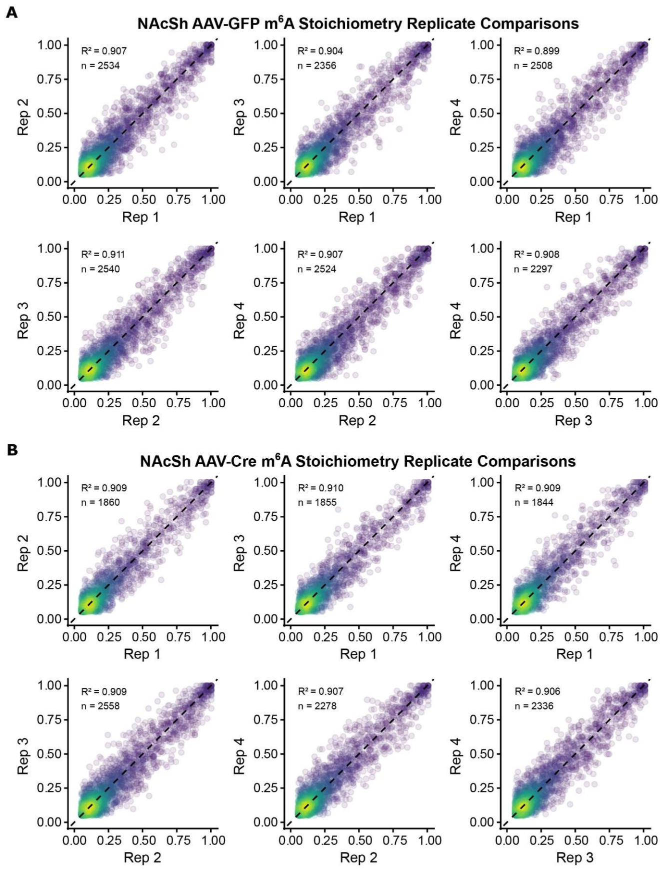

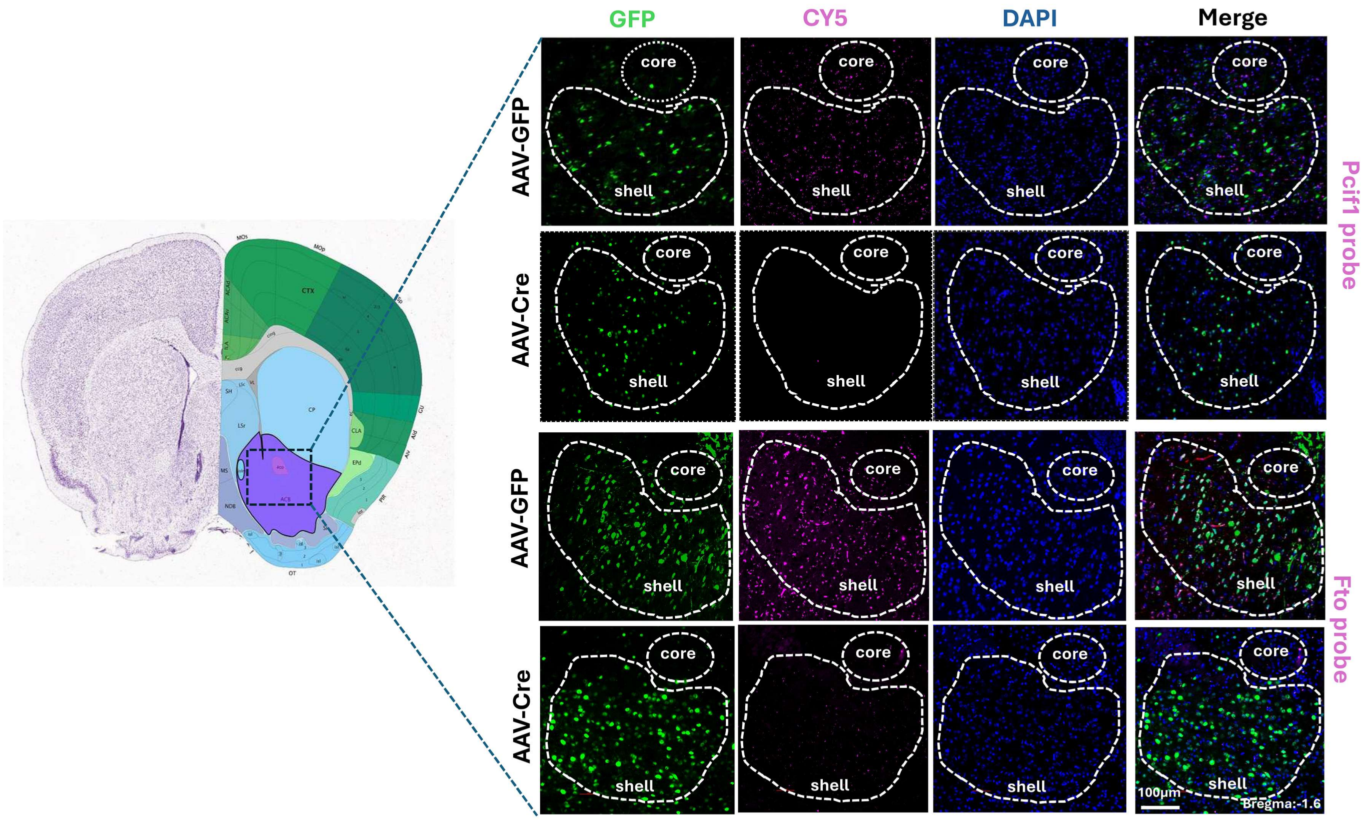

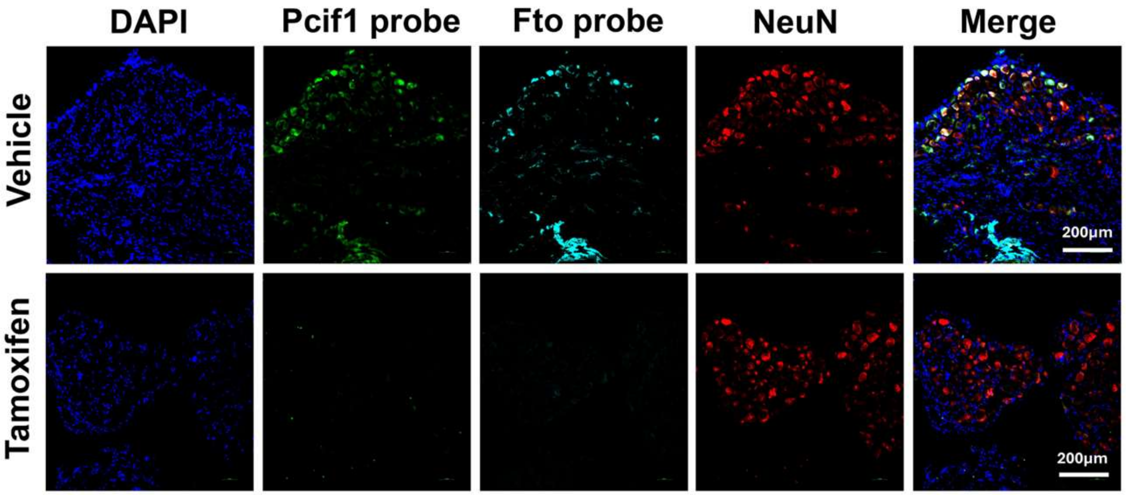

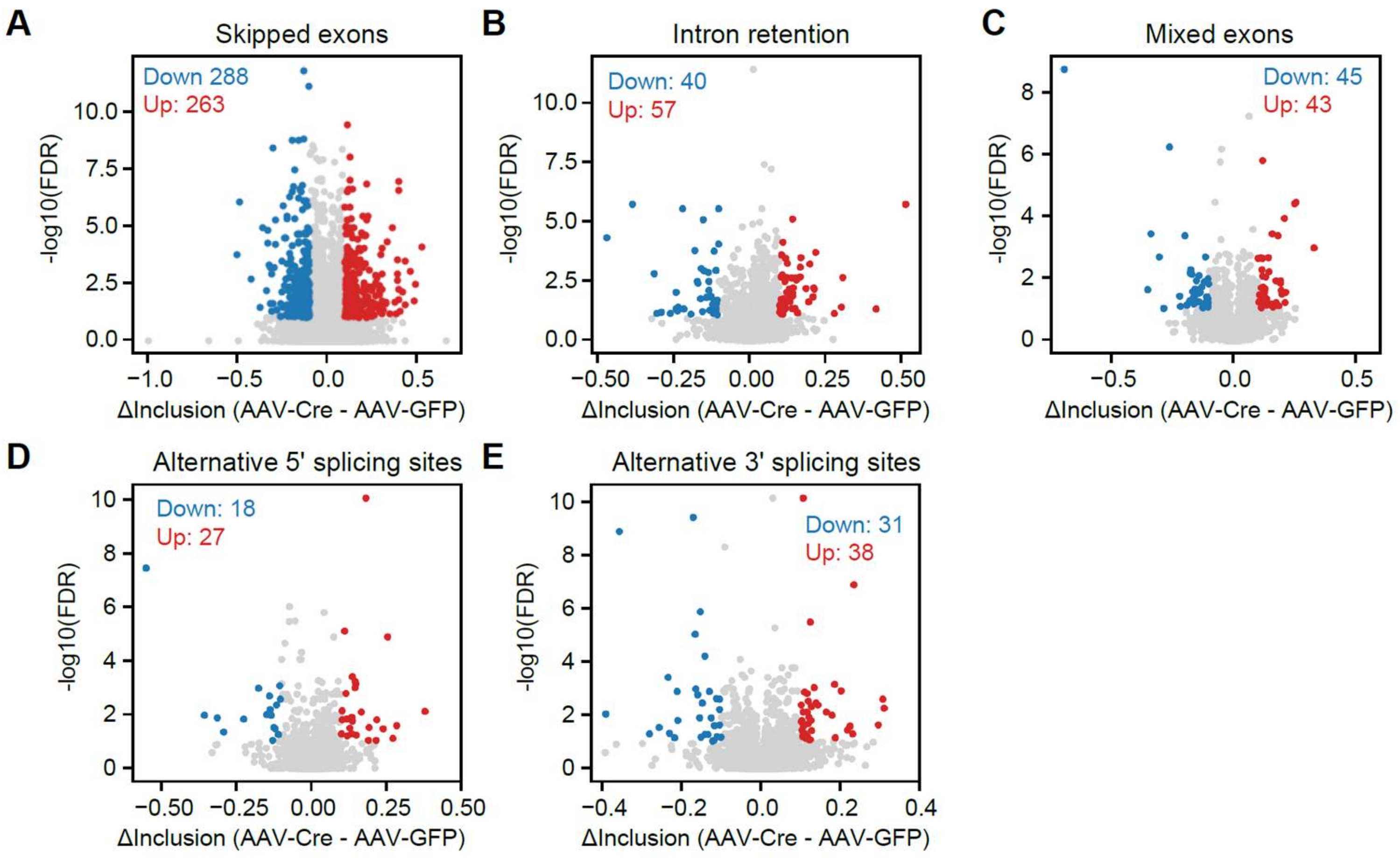

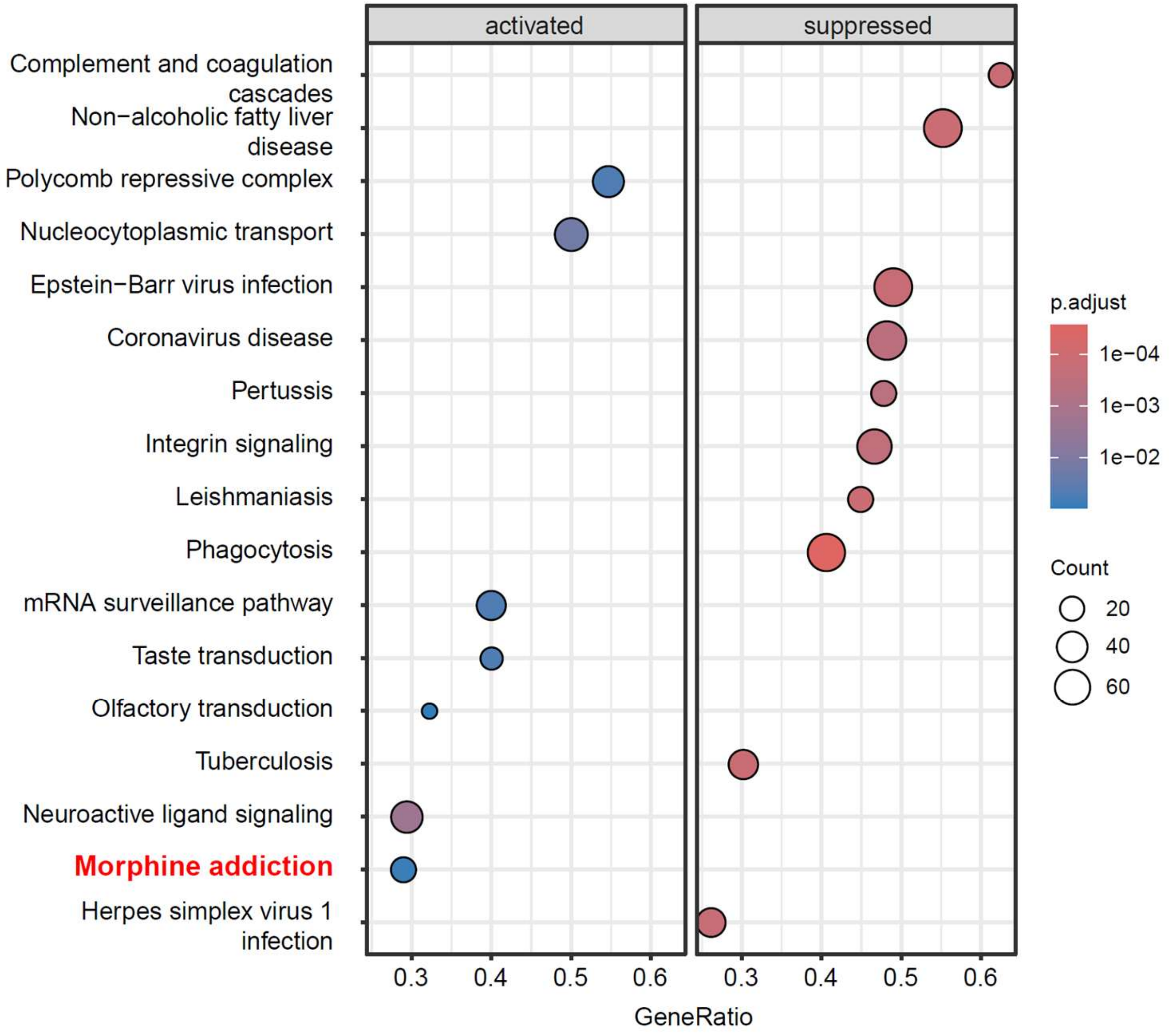
