## Supplementary material for "Reversible m^6^Am methylation of snRNA by FTO controls morphine reward and tolerance without altering analgesia": Table S2

Table S2. m6Am_methylation

| sample | condition | replicate | A | T | C | G | coverage | m6Am_leve | snRNA |
|---|---|---|---|---|---|---|---|---|---|
| NAc_Cre_1 | AAV-Cre | 1 | 3 | 0 | 0 | 4 | 7 | 0.4285714 | U7 |
| NAc_Cre_1 | AAV-Cre | 1 | 0 | 0 | 0 | 23 | 23 | 0 | U6 |
| NAc_Cre_1 | AAV-Cre | 1 | 79 | 0 | 1 | 225 | 304 | 0.2598684 | U5 |
| NAc_Cre_1 | AAV-Cre | 1 | 6 | 0 | 0 | 58 | 64 | 0.09375 | U4ATAC |
| NAc_Cre_1 | AAV-Cre | 1 | 35 | 0 | 0 | 97 | 132 | 0.2651515 | U4 |
| NAc_Cre_1 | AAV-Cre | 1 | 90 | 1 | 0 | 269 | 359 | 0.2506964 | U2 |
| NAc_Cre_1 | AAV-Cre | 1 | 132 | 0 | 0 | 332 | 464 | 0.2844828 | U1-like-1 |
| NAc_Cre_1 | AAV-Cre | 1 | 282 | 2 | 1 | 482 | 764 | 0.3691099 | U1b |
| NAc_Cre_1 | AAV-Cre | 1 | 617 | 5 | 2 | 1130 | 1747 | 0.3531769 | U1a |
| NAc_Cre_1 | AAV-Cre | 1 | 6 | 0 | 0 | 15 | 21 | 0.2857143 | U12 |
| NAc_Cre_1 | AAV-Cre | 1 | 2 | 0 | 0 | 151 | 153 | 0.0130719 | U11 |
| NAc_Cre_1 | AAV-Cre | 1 | 0 | 2 | 0 | 0 | 0 | NA | NA |
| NAc_Cre_1 | AAV-Cre | 1 | 3 | 0 | 0 | 25 | 28 | 0.1071429 | U3 |
| NAc_Cre_1 | AAV-Cre | 1 | 0 | 0 | 0 | 0 | 0 | NA | NA |
| NAc_Cre_1 | AAV-Cre | 1 | 0 | 0 | 0 | 0 | 0 | NA | NA |
| NAc_Cre_1 | AAV-Cre | 1 | 10 | 0 | 0 | 27 | 37 | 0.2702703 | NA |
| NAc_Cre_1 | AAV-Cre | 1 | 9 | 0 | 0 | 13 | 22 | 0.4090909 | NA |
| NAc_Cre_1 | AAV-Cre | 1 | 2 | 0 | 0 | 0 | 2 | 1 | NA |
| NAc_Cre_1 | AAV-Cre | 1 | 0 | 0 | 0 | 4 | 4 | 0 | NA |
| NAc_Cre_1 | AAV-Cre | 1 | 3 | 12 | 0 | 3 | 6 | 0.5 | NA |
| NAc_Cre_1 | AAV-Cre | 1 | 0 | 0 | 0 | 0 | 0 | NA | NA |
| NAc_Cre_2 | AAV-Cre | 2 | 0 | 0 | 0 | 1 | 1 | 0 | U7 |
| NAc_Cre_2 | AAV-Cre | 2 | 0 | 0 | 0 | 15 | 15 | 0 | U6 |
| NAc_Cre_2 | AAV-Cre | 2 | 29 | 3 | 0 | 233 | 262 | 0.110687 | U5 |
| NAc_Cre_2 | AAV-Cre | 2 | 8 | 0 | 0 | 130 | 138 | 0.057971 | U4ATAC |
| NAc_Cre_2 | AAV-Cre | 2 | 22 | 0 | 0 | 86 | 108 | 0.2037037 | U4 |
| NAc_Cre_2 | AAV-Cre | 2 | 62 | 0 | 0 | 226 | 288 | 0.2152778 | U2 |

| | | | | | | | | | |
|---|---|---|---|---|---|---|---|---|---|
| NAc_Cre_2 | AAV-Cre | 2 | 91 | 0 | 0 | 323 | 414 | 0.2198068 | U1-like-1 |
| NAc_Cre_2 | AAV-Cre | 2 | 145 | 2 | 0 | 465 | 610 | 0.2377049 | U1b |
| NAc_Cre_2 | AAV-Cre | 2 | 404 | 2 | 3 | 1200 | 1604 | 0.2518703 | U1a |
| NAc_Cre_2 | AAV-Cre | 2 | 1 | 0 | 0 | 20 | 21 | 0.047619 | U12 |
| NAc_Cre_2 | AAV-Cre | 2 | 0 | 0 | 0 | 126 | 126 | 0 | U11 |
| NAc_Cre_2 | AAV-Cre | 2 | 0 | 2 | 0 | 0 | 0 | NA | NA |
| NAc_Cre_2 | AAV-Cre | 2 | 7 | 0 | 0 | 18 | 25 | 0.28 | U3 |
| NAc_Cre_2 | AAV-Cre | 2 | 0 | 0 | 0 | 0 | 0 | NA | NA |
| NAc_Cre_2 | AAV-Cre | 2 | 0 | 0 | 0 | 0 | 0 | NA | NA |
| NAc_Cre_2 | AAV-Cre | 2 | 8 | 0 | 0 | 21 | 29 | 0.2758621 | NA |
| NAc_Cre_2 | AAV-Cre | 2 | 1 | 0 | 0 | 9 | 10 | 0.1 | NA |
| NAc_Cre_2 | AAV-Cre | 2 | 0 | 0 | 0 | 12 | 12 | 0 | NA |
| NAc_Cre_2 | AAV-Cre | 2 | 0 | 0 | 0 | 2 | 2 | 0 | NA |
| NAc_Cre_2 | AAV-Cre | 2 | 2 | 0 | 1 | 3 | 5 | 0.4 | NA |
| NAc_Cre_2 | AAV-Cre | 2 | 0 | 0 | 0 | 0 | 0 | NA | NA |
| NAc_Cre_3 | AAV-Cre | 3 | 0 | 0 | 0 | 2 | 2 | 0 | U7 |
| NAc_Cre_3 | AAV-Cre | 3 | 0 | 0 | 0 | 21 | 21 | 0 | U6 |
| NAc_Cre_3 | AAV-Cre | 3 | 47 | 1 | 0 | 229 | 276 | 0.1702899 | U5 |
| NAc_Cre_3 | AAV-Cre | 3 | 11 | 1 | 0 | 223 | 234 | 0.0470085 | U4ATAC |
| NAc_Cre_3 | AAV-Cre | 3 | 50 | 0 | 0 | 82 | 132 | 0.3787879 | U4 |
| NAc_Cre_3 | AAV-Cre | 3 | 87 | 0 | 0 | 228 | 315 | 0.2761905 | U2 |
| NAc_Cre_3 | AAV-Cre | 3 | 146 | 2 | 1 | 315 | 461 | 0.3167028 | U1-like-1 |
| NAc_Cre_3 | AAV-Cre | 3 | 235 | 2 | 1 | 510 | 745 | 0.3154362 | U1b |
| NAc_Cre_3 | AAV-Cre | 3 | 594 | 7 | 3 | 1559 | 2153 | 0.2758941 | U1a |
| NAc_Cre_3 | AAV-Cre | 3 | 8 | 0 | 0 | 23 | 31 | 0.2580645 | U12 |
| NAc_Cre_3 | AAV-Cre | 3 | 2 | 0 | 0 | 119 | 121 | 0.0165289 | U11 |
| NAc_Cre_3 | AAV-Cre | 3 | 0 | 0 | 0 | 0 | 0 | NA | NA |
| NAc_Cre_3 | AAV-Cre | 3 | 5 | 0 | 0 | 15 | 20 | 0.25 | U3 |

| | | | | | | | | | |
|---|---|---|---|---|---|---|---|---|---|
| NAc_Cre_3 | AAV-Cre | 3 | 0 | 0 | 0 | 0 | 0 | NA | NA |
| NAc_Cre_3 | AAV-Cre | 3 | 0 | 0 | 0 | 0 | 0 | NA | NA |
| NAc_Cre_3 | AAV-Cre | 3 | 15 | 0 | 0 | 19 | 34 | 0.4411765 | NA |
| NAc_Cre_3 | AAV-Cre | 3 | 5 | 0 | 0 | 14 | 19 | 0.2631579 | NA |
| NAc_Cre_3 | AAV-Cre | 3 | 0 | 0 | 0 | 1 | 1 | 0 | NA |
| NAc_Cre_3 | AAV-Cre | 3 | 0 | 0 | 0 | 1 | 1 | 0 | NA |
| NAc_Cre_3 | AAV-Cre | 3 | 5 | 5 | 0 | 7 | 12 | 0.4166667 | NA |
| NAc_Cre_3 | AAV-Cre | 3 | 0 | 0 | 0 | 0 | 0 | NA | NA |
| NAc_Cre_4 | AAV-Cre | 4 | 2 | 0 | 0 | 0 | 2 | 1 | U7 |
| NAc_Cre_4 | AAV-Cre | 4 | 0 | 0 | 0 | 1 | 1 | 0 | U6 |
| NAc_Cre_4 | AAV-Cre | 4 | 72 | 0 | 0 | 100 | 172 | 0.4186047 | U5 |
| NAc_Cre_4 | AAV-Cre | 4 | 22 | 1 | 0 | 230 | 252 | 0.0873016 | U4ATAC |
| NAc_Cre_4 | AAV-Cre | 4 | 46 | 0 | 1 | 36 | 82 | 0.5609756 | U4 |
| NAc_Cre_4 | AAV-Cre | 4 | 113 | 1 | 0 | 96 | 209 | 0.5406699 | U2 |
| NAc_Cre_4 | AAV-Cre | 4 | 172 | 0 | 1 | 163 | 335 | 0.5134328 | U1-like-1 |
| NAc_Cre_4 | AAV-Cre | 4 | 205 | 1 | 0 | 136 | 341 | 0.601173 | U1b |
| NAc_Cre_4 | AAV-Cre | 4 | 611 | 3 | 3 | 440 | 1051 | 0.5813511 | U1a |
| NAc_Cre_4 | AAV-Cre | 4 | 10 | 0 | 0 | 9 | 19 | 0.5263158 | U12 |
| NAc_Cre_4 | AAV-Cre | 4 | 4 | 0 | 0 | 56 | 60 | 0.0666667 | U11 |
| NAc_Cre_4 | AAV-Cre | 4 | 0 | 2 | 0 | 0 | 0 | NA | NA |
| NAc_Cre_4 | AAV-Cre | 4 | 10 | 0 | 0 | 8 | 18 | 0.5555556 | U3 |
| NAc_Cre_4 | AAV-Cre | 4 | 0 | 0 | 0 | 0 | 0 | NA | NA |
| NAc_Cre_4 | AAV-Cre | 4 | 0 | 0 | 0 | 0 | 0 | NA | NA |
| NAc_Cre_4 | AAV-Cre | 4 | 7 | 0 | 0 | 8 | 15 | 0.4666667 | NA |
| NAc_Cre_4 | AAV-Cre | 4 | 9 | 0 | 0 | 6 | 15 | 0.6 | NA |
| NAc_Cre_4 | AAV-Cre | 4 | 3 | 0 | 0 | 1 | 4 | 0.75 | NA |
| NAc_Cre_4 | AAV-Cre | 4 | 1 | 0 | 0 | 1 | 2 | 0.5 | NA |
| NAc_Cre_4 | AAV-Cre | 4 | 4 | 6 | 0 | 0 | 4 | 1 | NA |

| | | | | | | | | | |
|---|---|---|---|---|---|---|---|---|---|
| NAc_Cre_4 | AAV-Cre | 4 | 0 | 0 | 0 | 0 | 0 | NA | NA |
| NAc_GFP_1 | AAV-GFP | 1 | 0 | 0 | 0 | 5 | 5 | 0 | U7 |
| NAc_GFP_1 | AAV-GFP | 1 | 2 | 0 | 0 | 43 | 45 | 0.0444444 | U6 |
| NAc_GFP_1 | AAV-GFP | 1 | 9 | 0 | 0 | 384 | 393 | 0.0229008 | U5 |
| NAc_GFP_1 | AAV-GFP | 1 | 4 | 3 | 0 | 196 | 200 | 0.02 | U4ATAC |
| NAc_GFP_1 | AAV-GFP | 1 | 2 | 0 | 0 | 158 | 160 | 0.0125 | U4 |
| NAc_GFP_1 | AAV-GFP | 1 | 14 | 6 | 0 | 439 | 453 | 0.0309051 | U2 |
| NAc_GFP_1 | AAV-GFP | 1 | 9 | 0 | 0 | 752 | 761 | 0.0118265 | U1-like-1 |
| NAc_GFP_1 | AAV-GFP | 1 | 17 | 1 | 0 | 945 | 962 | 0.0176715 | U1b |
| NAc_GFP_1 | AAV-GFP | 1 | 39 | 5 | 0 | 2388 | 2427 | 0.0160692 | U1a |
| NAc_GFP_1 | AAV-GFP | 1 | 2 | 0 | 0 | 60 | 62 | 0.0322581 | U12 |
| NAc_GFP_1 | AAV-GFP | 1 | 0 | 0 | 0 | 180 | 180 | 0 | U11 |
| NAc_GFP_1 | AAV-GFP | 1 | 0 | 1 | 0 | 0 | 0 | NA | NA |
| NAc_GFP_1 | AAV-GFP | 1 | 0 | 0 | 0 | 37 | 37 | 0 | U3 |
| NAc_GFP_1 | AAV-GFP | 1 | 0 | 0 | 0 | 0 | 0 | NA | NA |
| NAc_GFP_1 | AAV-GFP | 1 | 0 | 0 | 0 | 1 | 1 | 0 | NA |
| NAc_GFP_1 | AAV-GFP | 1 | 1 | 0 | 0 | 33 | 34 | 0.0294118 | NA |
| NAc_GFP_1 | AAV-GFP | 1 | 1 | 1 | 0 | 29 | 30 | 0.0333333 | NA |
| NAc_GFP_1 | AAV-GFP | 1 | 0 | 0 | 0 | 7 | 7 | 0 | NA |
| NAc_GFP_1 | AAV-GFP | 1 | 0 | 0 | 0 | 1 | 1 | 0 | NA |
| NAc_GFP_1 | AAV-GFP | 1 | 0 | 5 | 0 | 12 | 12 | 0 | NA |
| NAc_GFP_1 | AAV-GFP | 1 | 0 | 0 | 0 | 0 | 0 | NA | NA |
| NAc_GFP_2 | AAV-GFP | 2 | 1 | 0 | 0 | 5 | 6 | 0.1666667 | U7 |
| NAc_GFP_2 | AAV-GFP | 2 | 0 | 0 | 0 | 11 | 11 | 0 | U6 |
| NAc_GFP_2 | AAV-GFP | 2 | 0 | 0 | 0 | 223 | 223 | 0 | U5 |
| NAc_GFP_2 | AAV-GFP | 2 | 0 | 0 | 0 | 76 | 76 | 0 | U4ATAC |
| NAc_GFP_2 | AAV-GFP | 2 | 1 | 0 | 0 | 106 | 107 | 0.0093458 | U4 |
| NAc_GFP_2 | AAV-GFP | 2 | 7 | 3 | 0 | 232 | 239 | 0.0292887 | U2 |

| | | | | | | | | | |
|---|---|---|---|---|---|---|---|---|---|
| NAc_GFP_2 | AAV-GFP | 2 | 8 | 1 | 0 | 392 | 400 | 0.02 | U1-like-1 |
| NAc_GFP_2 | AAV-GFP | 2 | 7 | 0 | 0 | 437 | 444 | 0.0157658 | U1b |
| NAc_GFP_2 | AAV-GFP | 2 | 22 | 1 | 1 | 1148 | 1170 | 0.0188034 | U1a |
| NAc_GFP_2 | AAV-GFP | 2 | 0 | 1 | 0 | 17 | 17 | 0 | U12 |
| NAc_GFP_2 | AAV-GFP | 2 | 2 | 0 | 0 | 93 | 95 | 0.0210526 | U11 |
| NAc_GFP_2 | AAV-GFP | 2 | 0 | 0 | 0 | 0 | 0 | NA | NA |
| NAc_GFP_2 | AAV-GFP | 2 | 1 | 0 | 0 | 20 | 21 | 0.047619 | U3 |
| NAc_GFP_2 | AAV-GFP | 2 | 0 | 0 | 0 | 0 | 0 | NA | NA |
| NAc_GFP_2 | AAV-GFP | 2 | 0 | 0 | 0 | 2 | 2 | 0 | NA |
| NAc_GFP_2 | AAV-GFP | 2 | 0 | 0 | 0 | 33 | 33 | 0 | NA |
| NAc_GFP_2 | AAV-GFP | 2 | 0 | 0 | 0 | 26 | 26 | 0 | NA |
| NAc_GFP_2 | AAV-GFP | 2 | 0 | 0 | 0 | 5 | 5 | 0 | NA |
| NAc_GFP_2 | AAV-GFP | 2 | 0 | 0 | 0 | 1 | 1 | 0 | NA |
| NAc_GFP_2 | AAV-GFP | 2 | 0 | 3 | 0 | 5 | 5 | 0 | NA |
| NAc_GFP_2 | AAV-GFP | 2 | 0 | 0 | 0 | 0 | 0 | NA | NA |
| NAc_GFP_3 | AAV-GFP | 3 | 0 | 0 | 0 | 1 | 1 | 0 | U7 |
| NAc_GFP_3 | AAV-GFP | 3 | 1 | 0 | 0 | 4 | 5 | 0.2 | U6 |
| NAc_GFP_3 | AAV-GFP | 3 | 3 | 0 | 0 | 160 | 163 | 0.0184049 | U5 |
| NAc_GFP_3 | AAV-GFP | 3 | 2 | 0 | 0 | 73 | 75 | 0.0266667 | U4ATAC |
| NAc_GFP_3 | AAV-GFP | 3 | 1 | 0 | 0 | 93 | 94 | 0.0106383 | U4 |
| NAc_GFP_3 | AAV-GFP | 3 | 11 | 0 | 0 | 212 | 223 | 0.0493274 | U2 |
| NAc_GFP_3 | AAV-GFP | 3 | 8 | 0 | 0 | 293 | 301 | 0.0265781 | U1-like-1 |
| NAc_GFP_3 | AAV-GFP | 3 | 8 | 1 | 0 | 372 | 380 | 0.0210526 | U1b |
| NAc_GFP_3 | AAV-GFP | 3 | 12 | 3 | 0 | 931 | 943 | 0.0127253 | U1a |
| NAc_GFP_3 | AAV-GFP | 3 | 0 | 0 | 0 | 17 | 17 | 0 | U12 |
| NAc_GFP_3 | AAV-GFP | 3 | 0 | 0 | 0 | 58 | 58 | 0 | U11 |
| NAc_GFP_3 | AAV-GFP | 3 | 0 | 1 | 0 | 0 | 0 | NA | NA |
| NAc_GFP_3 | AAV-GFP | 3 | 1 | 0 | 0 | 12 | 13 | 0.0769231 | U3 |

| | | | | | | | | | |
|---|---|---|---|---|---|---|---|---|---|
| NAc_GFP_3 | AAV-GFP | 3 | 0 | 0 | 0 | 0 | 0 | NA | NA |
| NAc_GFP_3 | AAV-GFP | 3 | 0 | 0 | 0 | 0 | 0 | NA | NA |
| NAc_GFP_3 | AAV-GFP | 3 | 0 | 0 | 0 | 19 | 19 | 0 | NA |
| NAc_GFP_3 | AAV-GFP | 3 | 0 | 0 | 0 | 14 | 14 | 0 | NA |
| NAc_GFP_3 | AAV-GFP | 3 | 0 | 0 | 0 | 2 | 2 | 0 | NA |
| NAc_GFP_3 | AAV-GFP | 3 | 0 | 0 | 0 | 2 | 2 | 0 | NA |
| NAc_GFP_3 | AAV-GFP | 3 | 0 | 1 | 0 | 4 | 4 | 0 | NA |
| NAc_GFP_3 | AAV-GFP | 3 | 0 | 0 | 0 | 0 | 0 | NA | NA |
| NAc_GFP_4 | AAV-GFP | 4 | 0 | 0 | 0 | 0 | 0 | NA | U7 |
| NAc_GFP_4 | AAV-GFP | 4 | 0 | 0 | 0 | 11 | 11 | 0 | U6 |
| NAc_GFP_4 | AAV-GFP | 4 | 4 | 0 | 0 | 228 | 232 | 0.0172414 | U5 |
| NAc_GFP_4 | AAV-GFP | 4 | 3 | 0 | 0 | 169 | 172 | 0.0174419 | U4ATAC |
| NAc_GFP_4 | AAV-GFP | 4 | 4 | 0 | 0 | 129 | 133 | 0.0300752 | U4 |
| NAc_GFP_4 | AAV-GFP | 4 | 14 | 0 | 0 | 224 | 238 | 0.0588235 | U2 |
| NAc_GFP_4 | AAV-GFP | 4 | 6 | 0 | 0 | 330 | 336 | 0.0178571 | U1-like-1 |
| NAc_GFP_4 | AAV-GFP | 4 | 10 | 0 | 0 | 521 | 531 | 0.0188324 | U1b |
| NAc_GFP_4 | AAV-GFP | 4 | 24 | 1 | 0 | 1447 | 1471 | 0.0163154 | U1a |
| NAc_GFP_4 | AAV-GFP | 4 | 0 | 0 | 0 | 17 | 17 | 0 | U12 |
| NAc_GFP_4 | AAV-GFP | 4 | 0 | 0 | 0 | 86 | 86 | 0 | U11 |
| NAc_GFP_4 | AAV-GFP | 4 | 0 | 4 | 0 | 0 | 0 | NA | NA |
| NAc_GFP_4 | AAV-GFP | 4 | 1 | 0 | 0 | 15 | 16 | 0.0625 | U3 |
| NAc_GFP_4 | AAV-GFP | 4 | 0 | 0 | 0 | 0 | 0 | NA | NA |
| NAc_GFP_4 | AAV-GFP | 4 | 0 | 0 | 0 | 0 | 0 | NA | NA |
| NAc_GFP_4 | AAV-GFP | 4 | 0 | 0 | 0 | 28 | 28 | 0 | NA |
| NAc_GFP_4 | AAV-GFP | 4 | 0 | 0 | 0 | 17 | 17 | 0 | NA |
| NAc_GFP_4 | AAV-GFP | 4 | 0 | 0 | 0 | 6 | 6 | 0 | NA |
| NAc_GFP_4 | AAV-GFP | 4 | 0 | 0 | 0 | 0 | 0 | NA | NA |
| NAc_GFP_4 | AAV-GFP | 4 | 0 | 4 | 0 | 5 | 5 | 0 | NA |

| | | | | | | | | | |
|---|---|---|---|---|---|---|---|---|---|
| NAc_GFP_4 | AAV-GFP | 4 | 0 | 0 | 0 | 0 | 0 | NA | NA |
