## Supplementary material for "Reversible m^6^Am methylation of snRNA by FTO controls morphine reward and tolerance without altering analgesia": Table S3

Table S3. Alternative_splicing_Legend

**Alternative splicing analysis: Fto KO vs Control**

| | |
|---|---|
| Sample 1, IncLevel1 | Fto fl/fl |
| Sample 2, IncLevel2 | Control |
| IncLevelDifference | IncLevel1 - IncLevel2 |

Data sheets:

| | |
|---|---|
| A5SS | Alternative 5' splicing |
| A3SS | Alternative 3' splicing |
| SE | Skipped exon |
| RI | Intron retention |
| MXE | Mutually exclusive exons |

### Table S3. Alternative_splicing_A5SS

| GeneID | geneSymbol | chr | strand | longExonS | longExonE | shortES | shortEE | flankingES | flankingEE | IJC_SAMPLE_1 | SJC_SAMPLE_1 | IJC_SAMPLE_2 | SJC_SAMPLE_2 | IncFormL | SkipForm | PValue | FDR | IncLevel1 | IncLevel2 | IncLevelDifference |
|---|---|---|---|---|---|---|---|---|---|---|---|---|---|---|---|---|---|---|---|---|
| ENSMUSG00000024299 | Adamts10 | chr17 | + | 33747653 | 33748000 | 33747653 | 33747802 | 33749106 | 33749263 | 103,103,133,109 | 0,0,1,0 | 93,103,119,110 | 12,14,4,12 | 347 | 149 | 3.06E-14 | 8.72E-11 | 1.0,1.0,0.9 | 0.769,0.76 | 0.182 |
| ENSMUSG00000022335 | Zfat | chr15 | - | 67982292 | 67982411 | 67982325 | 67982411 | 67976804 | 67976928 | 17,11,23,7 | 20,16,32,5 | 7,2,7,1 | 13,34,8,33 | 182 | 149 | 0.00059 | 0.01624 | 0.41,0.36, | 0.306,0.04 | 0.22 |
| ENSMUSG00000096995 | 2810029C07R | chr12 | - | 111540306 | 111540826 | 111540379 | 111540826 | 111540011 | 111540174 | 38,36,34,42 | 4,1,0,0 | 19,32,19,14 | 4,3,3,4 | 222 | 149 | 0.00022 | 0.00844 | 0.864,0.96 | 0.761,0.87 | 0.169 |
| ENSMUSG00000022948 | Setd4 | chr16 | - | 93387143 | 93388191 | 93387761 | 93388191 | 93386798 | 93386973 | 188,155,218,172 | 17,13,16,9 | 135,102,98,109 | 23,21,29,25 | 767 | 149 | 4.63E-08 | 1.32E-05 | 0.682,0.69 | 0.533,0.48 | 0.255 |
| ENSMUSG00000104674 | Gm42756 | chr9 | + | 109373893 | 109374245 | 109373893 | 109374230 | 109383035 | 109383509 | 19,9,19,24 | 6,6,0,2 | 14,12,22,14 | 17,3,7,18 | 164 | 149 | 0.00686 | 0.09169 | 0.742,0.57 | 0.428,0.78 | 0.217 |
| ENSMUSG00000028797 | Tmem234 | chr4 | + | 129495199 | 129495418 | 129495199 | 129495326 | 129495987 | 129496054 | 77,73,85,59 | 24,0,17,11 | 41,30,43,37 | 15,11,13,24 | 241 | 149 | 0.00145 | 0.03151 | 0.665,1.0, | 0.628,0.62 | 0.193 |
| ENSMUSG00000026005 | Rpe | chr1 | + | 66740775 | 66740807 | 66740775 | 66740795 | 66745604 | 66745684 | 31,20,78,41 | 26,34,46,10 | 19,18,1,12 | 24,41,33,17 | 161 | 149 | 0.00114 | 0.0264 | 0.525,0.35 | 0.423,0.28 | 0.286 |
| ENSMUSG00000032827 | Ppp1r9a | chr6 | + | 5154008 | 5154318 | 5154008 | 5154207 | 5156120 | 5156240 | 51,50,34,31 | 14,16,32,0 | 14,21,6,18 | 16,17,22,25 | 260 | 149 | 0.00019 | 0.00768 | 0.676,0.64 | 0.334,0.41 | 0.38 |
| ENSMUSG00000021288 | Klc1 | chr12 | + | 111755758 | 111755920 | 111755758 | 111755893 | 111760082 | 111760133 | 1,594,120,022,611,180 | 713,487,748,678 | 157,711,311,032,963 | 818,643,957,704 | 176 | 149 | 0.00018 | 0.00735 | 0.654,0.67 | 0.62,0.598 | 0.103 |
| ENSMUSG00000032413 | Rasa2 | chr9 | - | 96448068 | 96450540 | 96450416 | 96450540 | 96442750 | 96442834 | 98,65,114,90 | 0,5,0,1 | 88,83,91,76 | 4,4,9,2 | 2497 | 149 | 0.00558 | 0.07901 | 1.0,0.437, | 0.568,0.55 | 0.272 |
| ENSMUSG00000026867 | Gapvd1 | chr2 | - | 34607264 | 34607394 | 34607327 | 34607394 | 34605216 | 34605342 | 241,207,331,228 | 41,33,23,35 | 150,170,135,158 | 42,30,59,56 | 212 | 149 | 2.71E-06 | 0.00041 | 0.805,0.81 | 0.715,0.79 | 0.139 |
| ENSMUSG00000027748 | Trpc4 | chr3 | + | 54173504 | 54173841 | 54173504 | 54173788 | 54187283 | 54187423 | 46,32,101,25 | 0,2,3,1 | 66,20,42,26 | 5,5,10,4 | 202 | 149 | 7.31E-06 | 0.00074 | 1.0,0.922, | 0.907,0.74 | 0.149 |
| ENSMUSG00000055553 | Kxd1 | chr8 | - | 70972521 | 70972711 | 70972586 | 70972711 | 70968044 | 70968091 | 12,12,2,7 | 17,34,42,25 | 4,1,1,3 | 12,40,35,24 | 214 | 149 | 0.00059 | 0.01624 | 0.33,0.197 | 0.188,0.01 | 0.104 |
| ENSMUSG00000086370 | Ftx | chrX | - | 102656628 | 102656801 | 102656649 | 102656801 | 102652909 | 102653022 | 52,59,39,39 | 1,0,0,0 | 67,62,33,35 | 6,9,1,8 | 170 | 149 | 2.17E-08 | 7.73E-06 | 0.979,1.0, | 0.907,0.85 | 0.113 |
| ENSMUSG00000024830 | Rps6kb2 | chr19 | - | 4208801 | 4208918 | 4208827 | 4208918 | 4208608 | 4208716 | 40,66,95,66 | 21,19,22,15 | 33,32,51,48 | 23,14,23,36 | 175 | 149 | 0.00307 | 0.05296 | 0.619,0.74 | 0.55,0.661 | 0.136 |
| ENSMUSG00000036792 | Mbd5 | chr2 | + | 49162052 | 49163069 | 49162052 | 49162379 | 49164574 | 49164786 | 395,423,456,434 | 33,32,52,55 | 227,297,239,377 | 38,54,53,49 | 839 | 149 | 0.00042 | 0.01285 | 0.68,0.701 | 0.515,0.49 | 0.136 |
| ENSMUSG00000063810 | Alms1 | chr6 | + | 85585967 | 85587514 | 85585967 | 85586107 | 85587799 | 85587918 | 16,23,30,22 | 7,11,10,6 | 25,22,2,3 | 18,27,5,15 | 1556 | 149 | 1.08E-05 | 0.00099 | 0.18,0.167 | 0.117,0.07 | 0.146 |
| ENSMUSG00000068329 | Htra2 | chr6 | - | 83029981 | 83030051 | 83030018 | 83030051 | 83029453 | 83029523 | 21,30,37,22 | 10,11,10,4 | 38,18,9,13 | 24,11,9,9 | 186 | 149 | 0.00694 | 0.09185 | 0.627,0.68 | 0.559,0.56 | 0.192 |
| ENSMUSG00000032637 | Atxn2l | chr7 | - | 126091929 | 126092173 | 126091983 | 126092173 | 126091615 | 126091780 | 114,92,109,79 | 31,27,40,29 | 61,57,66,93 | 44,33,46,37 | 203 | 149 | 0.00069 | 0.01811 | 0.73,0.714 | 0.504,0.55 | 0.138 |
| ENSMUSG00000005682 | Pan2 | chr10 | + | 128139963 | 128140362 | 128139963 | 128140258 | 128143933 | 128144103 | 132,134,137,137 | 3,19,6,1 | 69,127,101,107 | 21,10,9,15 | 253 | 149 | 0.00055 | 0.01553 | 0.963,0.80 | 0.659,0.88 | 0.118 |
| ENSMUSG00000043923 | Ccdc84 | chr9 | - | 44324739 | 44324830 | 44324767 | 44324830 | 44324433 | 44324498 | 90,61,90,88 | 1,1,4,6 | 65,39,45,62 | 7,12,8,4 | 177 | 149 | 2.20E-05 | 0.00168 | 0.987,0.98 | 0.887,0.73 | 0.117 |
| ENSMUSG00000019817 | Plagl1 | chr10 | + | 12966575 | 12966768 | 12966575 | 12966764 | 12981598 | 12981637 | 36,21,44,25 | 1,0,7,0 | 29,10,37,28 | 0,6,7,8 | 153 | 149 | 0.0037 | 0.05884 | 0.972,1.0, | 1.0,0.619, | 0.151 |
| ENSMUSG00000038354 | Ankrd35 | chr3 | + | 96585448 | 96585583 | 96585448 | 96585579 | 96586495 | 96586584 | 8,8,8,11 | 51,48,25,36 | 3,4,6,1 | 31,44,33,47 | 153 | 149 | 0.0032 | 0.05387 | 0.133,0.14 | 0.086,0.08 | 0.101 |
| ENSMUSG00000086003 | B230206L02R | chr11 | + | 94029312 | 94030060 | 94029312 | 94029489 | 94046281 | 94046323 | 55,40,57,27 | 61,17,20,29 | 4,29,23,28 | 31,37,28,25 | 720 | 149 | 0.00156 | 0.0333 | 0.157,0.32 | 0.026,0.14 | 0.129 |
| ENSMUSG00000047153 | Khnyn | chr14 | + | 56131448 | 56131616 | 56131448 | 56131556 | 56131743 | 56131845 | 8,5,19,29 | 33,32,53,40 | 2,3,5,0 | 25,35,33,24 | 209 | 149 | 5.22E-06 | 0.00059 | 0.147,0.1, | 0.054,0.05 | 0.146 |
| ENSMUSG00000021733 | Slc4a7 | chr14 | - | 7729849 | 7730038 | 7729888 | 7730038 | 7723679 | 7723795 | 51,35,41,55 | 5,0,7,7 | 33,47,23,41 | 6,13,4,13 | 188 | 149 | 0.00402 | 0.06208 | 0.89,1.0,0 | 0.813,0.74 | 0.122 |
| ENSMUSG00000020831 | 0610010K14R | chr11 | - | 70126822 | 70127037 | 70126849 | 70127037 | 70126033 | 70126290 | 19,5,18,4 | 8,10,18,8 | 10,5,3,3 | 27,12,15,20 | 176 | 149 | 0.00172 | 0.03507 | 0.668,0.29 | 0.239,0.26 | 0.24 |
| ENSMUSG00000022702 | Hira | chr16 | + | 18714652 | 18714796 | 18714652 | 18714677 | 18715203 | 18715314 | 86,75,50,47 | 23,12,38,10 | 76,121,93,59 | 1,18,20,0 | 268 | 149 | 0.00055 | 0.01554 | 0.675,0.77 | 0.977,0.78 | -0.223 |
| ENSMUSG00000023026 | Dip2b | chr15 | + | 100052914 | 100053037 | 100052914 | 100053034 | 100055050 | 100055248 | 6,2,1,16 | 126,84,127,81 | 14,18,2,15 | 70,68,66,46 | 152 | 149 | 5.64E-05 | 0.00277 | 0.045,0.02 | 0.164,0.20 | -0.101 |
| ENSMUSG00000022335 | Zfat | chr15 | - | 68084441 | 68084693 | 68084462 | 68084693 | 68058899 | 68059085 | 36,30,36,13 | 4,5,1,6 | 16,21,23,15 | 0,2,0,0 | 170 | 149 | 3.54E-05 | 0.0021 | 0.887,0.84 | 1.0,0.902, | -0.138 |
| ENSMUSG00000042050 | Dync2li1 | chr12 | - | 116189584 | 116189757 | 116189595 | 116189757 | 116188506 | 116188669 | 77,59,50,94 | 18,11,31,19 | 70,70,63,68 | 6,6,6,12 | 160 | 149 | 0.00034 | 0.01111 | 0.799,0.83 | 0.916,0.91 | -0.131 |
| ENSMUSG00000039233 | Tbce | chr13 | - | 14178117 | 14178281 | 14178127 | 14178281 | 14175694 | 14175763 | 89,52,51,90 | 185,116,237,152 | 60,64,64,94 | 70,112,120,114 | 159 | 149 | 0.00337 | 0.05611 | 0.311,0.29 | 0.445,0.34 | -0.108 |
| ENSMUSG00000030428 | Ttyh1 | chr7 | + | 4125475 | 4125650 | 4125475 | 4125646 | 4127618 | 4127730 | 57,66,107,63 | 8,6,12,21 | 65,55,58,41 | 3,4,2,0 | 153 | 149 | 9.25E-06 | 0.00088 | 0.874,0.91 | 0.955,0.93 | -0.105 |
| ENSMUSG00000073705 | Cenps | chr4 | - | 149216715 | 149216839 | 149216734 | 149216839 | 149216095 | 149216129 | 59,35,30,39 | 6,2,8,9 | 67,32,53,33 | 5,1,0,1 | 168 | 149 | 0.0001 | 0.00456 | 0.897,0.93 | 0.922,0.96 | -0.114 |
| ENSMUSG00000063626 | Unc5d | chr8 | - | 29214248 | 29214434 | 29214287 | 29214434 | 29209644 | 29210022 | 20,12,28,10 | 12,13,18,23 | 33,18,27,24 | 4,10,12,1 | 188 | 149 | 0.00046 | 0.01373 | 0.569,0.42 | 0.867,0.58 | -0.312 |
| ENSMUSG00000092060 | Bend4 | chr5 | - | 67557403 | 67557644 | 67557474 | 67557644 | 67549490 | 67555793 | 18,28,17,23 | 6,3,0,4 | 18,16,30,21 | 0,0,3,0 | 220 | 149 | 0.00015 | 0.00648 | 0.67,0.863 | 1.0,1.0,0.8 | -0.136 |
| ENSMUSG00000038733 | Wdr26 | chr1 | - | 181036567 | 181036672 | 181036615 | 181036672 | 181030588 | 181030789 | 41,31,42,19 | 2,6,14,8 | 30,35,20,32 | 3,0,2,0 | 197 | 149 | 1.22E-05 | 0.00109 | 0.939,0.79 | 0.883,1.0, | -0.174 |
| ENSMUSG00000024187 | Fam234a | chr17 | - | 26444885 | 26444997 | 26444901 | 26444997 | 26439254 | 26439554 | 24,13,14,13 | 2,0,6,5 | 50,18,8,26 | 0,3,0,2 | 165 | 149 | 0.00172 | 0.03507 | 0.916,1.0, | 1.0,0.844, | -0.118 |
| ENSMUSG00000029916 | Agk | chr6 | + | 40329286 | 40331741 | 40329286 | 40329366 | 40331947 | 40332027 | 63,42,47,80 | 4,3,13,5 | 59,49,65,69 | 0,0,1,0 | 2524 | 149 | 2.55E-11 | 3.63E-08 | 0.482,0.45 | 1.0,1.0,0.7 | -0.549 |
| ENSMUSG00000040213 | Kyat3 | chr3 | + | 142412449 | 142412556 | 142412449 | 142412552 | 142424136 | 142424195 | 5,2,5,1 | 50,27,79,33 | 4,5,9,13 | 36,38,19,39 | 153 | 149 | 0.00034 | 0.01111 | 0.089,0.06 | 0.098,0.11 | -0.133 |
| ENSMUSG00000032280 | Tle3 | chr9 | + | 61301889 | 61301994 | 61301889 | 61301964 | 61309137 | 61309339 | 3,0,5,6 | 10,25,10,12 | 27,3,16,11 | 22,6,13,1 | 179 | 149 | 0.00032 | 0.01074 | 0.2,0.0,0.2 | 0.505,0.29 | -0.355 |
| ENSMUSG00000030815 | Phkg2 | chr7 | + | 127176706 | 127176882 | 127176706 | 127176860 | 127178838 | 127178904 | 35,30,35,34 | 8,0,11,7 | 45,25,28,36 | 8,0,0,0 | 171 | 149 | 0.00142 | 0.03101 | 0.792,1.0, | 0.831,1.0, | -0.124 |
| ENSMUSG00000036180 | Gatad2a | chr8 | - | 70369209 | 70369350 | 70369218 | 70369350 | 70368933 | 70369098 | 0,0,4,12 | 51,48,53,41 | 11,5,8,12 | 63,19,17,40 | 158 | 149 | 0.00029 | 0.01003 | 0.0,0.0,0.0 | 0.141,0.19 | -0.146 |
| ENSMUSG00000090086 | AI480526 | chr5 | - | 123276013 | 123276129 | 123276017 | 123276129 | 123272394 | 123272493 | 6,3,8,1 | 24,14,30,18 | 11,4,7,3 | 12,2,14,6 | 153 | 149 | 0.00244 | 0.04502 | 0.196,0.17 | 0.472,0.66 | -0.29 |
| ENSMUSG00000070643 | Sox13 | chr1 | - | 133312150 | 133312411 | 133312194 | 133312411 | 133310040 | 133311662 | 1,1,6,2 | 16,20,31,19 | 8,5,7,5 | 22,34,12,19 | 193 | 149 | 0.00749 | 0.0952 | 0.046,0.03 | 0.219,0.10 | -0.128 |

### Table S3. Alternative_splicing_A3SS

| GeneID | geneSymbol | chr | strand | longExonS | longExonE | shortES | shortEE | flankingES | flankingEE | IJC_SAMPLE_1 | SJC_SAMPLE_1 | IJC_SAMPLE_2 | SJC_SAMPLE_2 | IncFormL | SkipForm | PValue | FDR | IncLevel1 | IncLevel2 | IncLevelDifference |
|---|---|---|---|---|---|---|---|---|---|---|---|---|---|---|---|---|---|---|---|---|
| ENSMUSG00000022552 | Sharpin | chr15 | - | 76231863 | 76232007 | 76231863 | 76231989 | 76232095 | 76232204 | 34,27,42,29 | 17,1,4,2 | 33,41,27,23 | 2,0,0,2 | 167 | 149 | 4.30E-05 | 0.00258 | 0.641,0.9 | 0.936,1.0, | -0.104 |
| ENSMUSG00000029563 | Foxp2 | chr6 | + | 15377868 | 15378087 | 15377949 | 15378087 | 15376687 | 15376825 | 50,32,52,65 | 22,18,18,16 | 13,21,8,26 | 0,1,0,0 | 230 | 149 | 1.07E-12 | 1.26E-09 | 0.596,0.5 | 1.0,0.932, | -0.356 |
| ENSMUSG00000029563 | Foxp2 | chr6 | + | 15377886 | 15378087 | 15377949 | 15378087 | 15376687 | 15376825 | 149,90,132,136 | 22,18,18,16 | 26,64,37,71 | 0,1,0,0 | 212 | 149 | 2.45E-13 | 3.86E-10 | 0.826,0.7 | 1.0,0.978, | -0.17 |
| ENSMUSG00000047193 | Dync2h1 | chr9 | - | 7111424 | 7111592 | 7111424 | 7111571 | 7112047 | 7112150 | 1,11,0,1 | 33,44,57,51 | 7,2,11,13 | 16,27,24,49 | 170 | 149 | 7.10E-05 | 0.00367 | 0.026,0.1 | 0.277,0.0 | -0.148 |
| ENSMUSG00000075470 | Alg10b | chr15 | + | 90111526 | 90117674 | 90111675 | 90117674 | 90109859 | 90110057 | 226,173,212,167 | 26,15,49,26 | 166,155,152,143 | 0,6,9,12 | 298 | 149 | 1.96E-07 | 6.17E-05 | 0.813,0.8 | 1.0,0.928, | -0.141 |
| ENSMUSG00000085517 | Gm12963 | chr4 | - | 129921759 | 129922139 | 129921759 | 129921948 | 129931605 | 129931675 | 16,24,32,16 | 6,2,7,0 | 25,19,15,10 | 0,0,0,2 | 340 | 149 | 2.66E-05 | 0.00177 | 0.539,0.8 | 1.0,1.0,1. | -0.16 |
| ENSMUSG00000054752 | Fsd1l | chr4 | + | 53693983 | 53694084 | 53693986 | 53694084 | 53686387 | 53686517 | 10,4,0,3 | 33,32,52,25 | 12,11,18,9 | 19,17,40,32 | 152 | 149 | 1.66E-05 | 0.00136 | 0.229,0.1 | 0.382,0.3 | -0.212 |
| ENSMUSG00000055723 | Rras2 | chr7 | - | 113649535 | 113650933 | 113649535 | 113649654 | 113657774 | 113657883 | 17,36,115,40 | 41,23,37,53 | 48,53,108,69 | 46,11,22,35 | 1428 | 149 | 0.00437 | 0.06657 | 0.041,0.1 | 0.098,0.3 | -0.111 |
| ENSMUSG00000037270 | 4932438A13Ri | chr3 | + | 36991137 | 36991317 | 36991140 | 36991317 | 36985146 | 36985284 | 29,24,31,40 | 32,23,29,26 | 32,42,17,37 | 7,27,7,9 | 152 | 149 | 0.00058 | 0.01637 | 0.47,0.50 | 0.818,0.6 | -0.21 |
| ENSMUSG00000038095 | Sbno1 | chr5 | - | 124548132 | 124548445 | 124548132 | 124548442 | 124552461 | 124552590 | 0,42,45,0 | 29,30,19,20 | 41,56,35,30 | 25,28,8,10 | 152 | 149 | 0.00026 | 0.00947 | 0.0,0.578, | 0.617,0.6 | -0.39 |
| ENSMUSG00000025144 | Cenpx | chr11 | - | 120602528 | 120602721 | 120602528 | 120602582 | 120604518 | 120604564 | 55,43,50,63 | 96,71,77,58 | 62,66,47,47 | 50,55,39,33 | 288 | 149 | 0.00043 | 0.01326 | 0.229,0.2 | 0.391,0.3 | -0.126 |
| ENSMUSG00000025932 | Eya1 | chr1 | - | 14344288 | 14344434 | 14344288 | 14344367 | 14344757 | 14344827 | 13,22,21,2 | 8,8,21,2 | 18,30,14,20 | 6,0,9,1 | 216 | 149 | 0.00309 | 0.05378 | 0.529,0.6 | 0.674,1.0, | -0.281 |
| ENSMUSG00000026288 | Inpp5d | chr1 | + | 87642759 | 87643035 | 87642998 | 87643035 | 87640914 | 87641011 | 85,93,120,89 | 7,9,7,4 | 93,73,94,86 | 2,0,2,2 | 388 | 149 | 4.03E-05 | 0.00251 | 0.823,0.7 | 0.947,1.0, | -0.113 |
| ENSMUSG00000026269 | Rnpepl1 | chr1 | + | 92846627 | 92846825 | 92846659 | 92846825 | 92845739 | 92845848 | 32,21,42,32 | 9,8,5,1 | 33,39,25,16 | 0,0,0,1 | 181 | 149 | 2.53E-08 | 9.22E-06 | 0.745,0.6 | 1.0,1.0,1. | -0.166 |
| ENSMUSG00000039988 | Ankrd13c | chr3 | + | 157678636 | 157680931 | 157680864 | 157680931 | 157667914 | 157668019 | 24,22,27,22 | 7,7,0,0 | 41,31,38,31 | 2,0,1,0 | 2377 | 149 | 0.00276 | 0.05055 | 0.177,0.1 | 0.562,1.0, | -0.231 |
| ENSMUSG00000033439 | Trmt13 | chr3 | - | 116376142 | 116376532 | 116376142 | 116376373 | 116378844 | 116378919 | 38,39,43,47 | 37,48,43,48 | 30,46,63,51 | 24,26,43,21 | 308 | 149 | 0.0032 | 0.05444 | 0.332,0.2 | 0.377,0.4 | -0.133 |
| ENSMUSG00000034928 | Rnf44 | chr13 | - | 54831773 | 54831963 | 54831773 | 54831902 | 54832150 | 54832301 | 140,105,219,95 | 37,14,15,7 | 100,96,112,74 | 0,0,19,0 | 210 | 149 | 0.00014 | 0.00621 | 0.729,0.8 | 1.0,1.0,0. | -0.104 |
| ENSMUSG00000044037 | Als2cl | chr9 | + | 110713082 | 110713214 | 110713085 | 110713214 | 110711781 | 110711891 | 32,23,20,19 | 8,2,7,5 | 26,14,48,46 | 5,1,3,0 | 152 | 149 | 0.00102 | 0.02484 | 0.797,0.9 | 0.836,0.9 | -0.117 |
| ENSMUSG00000044550 | Tceal3 | chrX | + | 135567808 | 135567939 | 135567872 | 135567939 | 135494950 | 135494999 | 29,33,35,23 | 11,13,35,23 | 27,31,27,44 | 0,14,6,9 | 213 | 149 | 0.00131 | 0.03003 | 0.648,0.6 | 1.0,0.608, | -0.257 |
| ENSMUSG00000038070 | Cntln | chr4 | + | 84981414 | 84981592 | 84981417 | 84981592 | 84967840 | 84968373 | 29,32,22,40 | 8,3,7,4 | 13,18,31,22 | 1,0,0,4 | 152 | 149 | 0.00094 | 0.02346 | 0.78,0.91 | 0.927,1.0, | -0.104 |
| ENSMUSG00000014418 | Hps5 | chr7 | - | 46427444 | 46427623 | 46427444 | 46427524 | 46428495 | 46428584 | 40,39,57,59 | 5,6,12,6 | 69,59,50,37 | 4,4,1,4 | 248 | 149 | 0.0045 | 0.06744 | 0.828,0.7 | 0.912,0.8 | -0.102 |
| ENSMUSG00000023191 | P3h3 | chr6 | - | 124827880 | 124827948 | 124827880 | 124827946 | 124828041 | 124828094 | 92,82,53,69 | 21,19,23,20 | 35,68,59,57 | 5,3,9,5 | 151 | 149 | 1.65E-05 | 0.00136 | 0.812,0.8 | 0.874,0.9 | -0.131 |
| ENSMUSG00000029638 | Glcci1 | chr6 | + | 8573181 | 8573298 | 8573184 | 8573298 | 8558499 | 8558586 | 27,49,59,49 | 105,94,131,68 | 32,54,42,54 | 41,51,78,61 | 152 | 149 | 0.00794 | 0.09701 | 0.201,0.3 | 0.433,0.5 | -0.123 |
| ENSMUSG00000015149 | Sirt2 | chr7 | + | 28487341 | 28488085 | 28487344 | 28488085 | 28487007 | 28487078 | 26,11,14,15 | 0,0,21,0 | 52,11,35,38 | 0,0,0,0 | 152 | 149 | 0.00488 | 0.07064 | 1.0,1.0,0. | 1.0,1.0,1. | -0.151 |
| ENSMUSG00000032816 | Igdcc4 | chr9 | + | 65034036 | 65034210 | 65034039 | 65034210 | 65032566 | 65032704 | 29,36,45,22 | 55,37,64,31 | 74,37,49,44 | 52,29,33,38 | 152 | 149 | 0.00044 | 0.0133 | 0.341,0.4 | 0.582,0.5 | -0.154 |
| ENSMUSG00000029238 | Clock | chr5 | - | 76377204 | 76377410 | 76377204 | 76377407 | 76378011 | 76378206 | 108,75,115,55 | 55,47,71,56 | 56,97,60,92 | 31,21,23,44 | 152 | 149 | 0.0073 | 0.09278 | 0.658,0.6 | 0.639,0.8 | -0.119 |
| ENSMUSG00000000759 | Tubgcp3 | chr8 | - | 12698620 | 12699822 | 12698620 | 12698787 | 12700153 | 12700222 | 285,253,358,320 | 6,4,14,13 | 333,229,280,256 | 0,3,1,0 | 1184 | 149 | 1.98E-09 | 1.34E-06 | 0.857,0.8 | 1.0,0.906, | -0.153 |
| ENSMUSG00000031520 | Vegfc | chr8 | + | 54612327 | 54612518 | 54612448 | 54612518 | 54609992 | 54610194 | 6,12,12,11 | 1,0,5,5 | 30,22,17,40 | 0,0,2,0 | 270 | 149 | 2.26E-06 | 0.00038 | 0.768,1.0, | 1.0,1.0,0. | -0.234 |
| ENSMUSG00000032323 | Cyp11a1 | chr9 | + | 57925476 | 57926779 | 57926575 | 57926779 | 57923556 | 57923712 | 47,20,54,43 | 0,1,10,0 | 49,34,50,54 | 3,0,0,0 | 1248 | 149 | 0.00323 | 0.05471 | 1.0,0.705, | 0.661,1.0, | -0.141 |
| ENSMUSG00000001034 | Mapk7 | chr11 | - | 61383729 | 61384011 | 61383729 | 61383895 | 61384349 | 61385092 | 14,8,7,5 | 18,13,14,18 | 11,11,22,20 | 12,4,21,13 | 265 | 149 | 0.00545 | 0.07518 | 0.304,0.2 | 0.34,0.60 | -0.217 |
| ENSMUSG00000023051 | Tarbp2 | chr15 | + | 102429555 | 102429648 | 102429572 | 102429648 | 102427556 | 102427726 | 39,35,42,37 | 12,10,20,5 | 24,22,36,45 | 1,4,2,2 | 166 | 149 | 1.00E-05 | 0.00106 | 0.745,0.7 | 0.956,0.8 | -0.164 |
| ENSMUSG00000036036 | Zfp57 | chr17 | + | 37316941 | 37317086 | 37316950 | 37317086 | 37315688 | 37315795 | 15,24,21,30 | 0,3,0,0 | 17,14,23,13 | 6,0,10,0 | 158 | 149 | 5.54E-05 | 0.00312 | 1.0,0.883, | 0.728,1.0, | 0.118 |
| ENSMUSG00000023972 | Ptk7 | chr17 | - | 46878850 | 46879955 | 46878850 | 46879002 | 46882528 | 46882609 | 101,56,121,87 | 74,21,29,31 | 36,40,34,27 | 25,36,24,27 | 1102 | 149 | 0.00041 | 0.01251 | 0.156,0.2 | 0.163,0.1 | 0.121 |
| ENSMUSG00000041199 | Rpusd1 | chr17 | + | 25947154 | 25947365 | 25947176 | 25947365 | 25946690 | 25946821 | 102,65,103,66 | 214,107,228,165 | 44,59,30,26 | 151,167,119,157 | 171 | 149 | 1.75E-05 | 0.0014 | 0.293,0.3 | 0.202,0.2 | 0.109 |
| ENSMUSG00000036158 | Prickle1 | chr15 | - | 93410330 | 93410517 | 93410330 | 93410514 | 93417144 | 93417404 | 7,20,22,24 | 2,5,11,6 | 9,13,10,12 | 18,4,17,18 | 152 | 149 | 0.00094 | 0.02353 | 0.774,0.7 | 0.329,0.7 | 0.295 |
| ENSMUSG00000004267 | Eno2 | chr6 | - | 124745200 | 124745683 | 124745200 | 124745297 | 124746211 | 124746513 | 354,350,430,267 | 18,0,23,9 | 215,219,243,261 | 22,11,14,23 | 535 | 149 | 0.0006 | 0.01653 | 0.846,1.0, | 0.731,0.8 | 0.103 |
| ENSMUSG00000025006 | Sorbs1 | chr19 | - | 40287906 | 40288097 | 40287906 | 40287965 | 40288686 | 40288746 | 330,270,292,220 | 22,6,27,16 | 142,216,169,167 | 41,36,24,8 | 281 | 149 | 0.00019 | 0.00778 | 0.888,0.9 | 0.647,0.7 | 0.116 |
| ENSMUSG00000025006 | Sorbs1 | chr19 | - | 40287906 | 40288097 | 40287906 | 40287965 | 40300112 | 40300307 | 315,259,304,228 | 160,153,237,195 | 138,225,170,173 | 212,172,201,160 | 281 | 149 | 0.00107 | 0.02573 | 0.511,0.4 | 0.257,0.4 | 0.108 |
| ENSMUSG00000028758 | Kif17 | chr4 | + | 138016548 | 138016635 | 138016551 | 138016635 | 138015239 | 138015796 | 51,60,74,43 | 82,51,92,49 | 19,11,19,44 | 42,76,36,87 | 152 | 149 | 0.0003 | 0.01045 | 0.379,0.5 | 0.307,0.1 | 0.179 |
| ENSMUSG00000085517 | Gm12963 | chr4 | - | 129921759 | 129922139 | 129921759 | 129921851 | 129931605 | 129931675 | 22,29,53,26 | 0,0,0,2 | 33,29,23,20 | 6,2,1,1 | 437 | 149 | 8.30E-05 | 0.00418 | 1.0,1.0,1. | 0.652,0.8 | 0.143 |
| ENSMUSG00000057897 | Camk2b | chr11 | - | 5932720 | 5932761 | 5932720 | 5932758 | 5938100 | 5938143 | 36,24,52,51 | 12,15,28,23 | 41,25,21,11 | 26,24,61,8 | 152 | 149 | 0.00537 | 0.07453 | 0.746,0.6 | 0.607,0.5 | 0.187 |
| ENSMUSG00000028456 | Unc13b | chr4 | + | 43249456 | 43249581 | 43249459 | 43249581 | 43245467 | 43245596 | 33,12,35,34 | 7,0,13,2 | 31,12,25,24 | 17,17,23,10 | 152 | 149 | 4.25E-05 | 0.00258 | 0.822,1.0, | 0.641,0.4 | 0.306 |
| ENSMUSG00000060862 | Zbtb40 | chr4 | - | 136715858 | 136716076 | 136715858 | 136716073 | 136718808 | 136718973 | 8,9,6,0 | 22,22,23,18 | 2,2,1,1 | 18,29,22,19 | 152 | 149 | 0.00058 | 0.01637 | 0.263,0.2 | 0.098,0.0 | 0.125 |
| ENSMUSG00000034729 | Mrps10 | chr17 | + | 47683500 | 47686034 | 47685897 | 47686034 | 47683351 | 47683416 | 656,578,797,529 | 45,53,63,45 | 485,382,418,456 | 56,66,46,56 | 2546 | 149 | 0.00021 | 0.0079 | 0.46,0.39, | 0.336,0.2 | 0.106 |
| ENSMUSG00000037355 | Uvssa | chr5 | + | 33568202 | 33568386 | 33568306 | 33568386 | 33566770 | 33566893 | 30,20,19,40 | 0,1,0,3 | 16,20,16,22 | 4,0,1,4 | 253 | 149 | 0.00295 | 0.05237 | 1.0,0.922, | 0.702,1.0, | 0.11 |
| ENSMUSG00000041375 | Ccdc9 | chr7 | - | 16018195 | 16018338 | 16018195 | 16018287 | 16018419 | 16018471 | 25,17,41,22 | 0,0,0,0 | 20,36,13,10 | 2,2,5,2 | 200 | 149 | 5.40E-06 | 0.00073 | 1.0,1.0,1. | 0.882,0.9 | 0.185 |
| ENSMUSG00000023707 | Ogfod2 | chr5 | + | 124252127 | 124252708 | 124252450 | 124252708 | 124251491 | 124251591 | 263,179,199,141 | 15,2,13,12 | 181,147,140,151 | 13,10,15,32 | 472 | 149 | 0.00583 | 0.0792 | 0.847,0.9 | 0.815,0.8 | 0.112 |
| ENSMUSG00000034973 | Dop1a | chr9 | + | 86394610 | 86395232 | 86394613 | 86395232 | 86389556 | 86389790 | 81,68,97,60 | 26,14,14,23 | 56,38,34,33 | 26,18,20,15 | 152 | 149 | 0.00201 | 0.04029 | 0.753,0.8 | 0.679,0.6 | 0.127 |
| ENSMUSG00000024963 | Dnajc4 | chr19 | - | 6969526 | 6969689 | 6969526 | 6969640 | 6969916 | 6969940 | 46,22,43,26 | 45,20,42,34 | 27,32,13,8 | 70,51,43,33 | 198 | 149 | 1.50E-05 | 0.00129 | 0.435,0.4 | 0.225,0.3 | 0.201 |
| ENSMUSG00000119994 | | chr10 | + | 128337248 | 128337558 | 128337503 | 128337558 | 128335177 | 128335395 | 74,93,103,111 | 39,6,19,15 | 112,58,69,59 | 34,36,28,42 | 404 | 149 | 0.0029 | 0.05201 | 0.412,0.8 | 0.549,0.3 | 0.231 |
| ENSMUSG00000023764 | Sfi1 | chr11 | - | 3083071 | 3085542 | 3083071 | 3083207 | 3085653 | 3085830 | 985,763,970,885 | 0,2,0,0 | 536,517,514,632 | 2,10,5,4 | 2484 | 149 | 6.37E-09 | 3.35E-06 | 1.0,0.958, | 0.941,0.7 | 0.124 |
| ENSMUSG00000051323 | Pcdh19 | chrX | - | 132526017 | 132526190 | 132526017 | 132526187 | 132533643 | 132533702 | 13,25,21,18 | 21,15,43,24 | 12,4,5,5 | 29,22,25,15 | 152 | 149 | 0.00114 | 0.02674 | 0.378,0.6 | 0.289,0.1 | 0.224 |
| ENSMUSG00000040370 | Etfrf1 | chr6 | + | 145160247 | 145160997 | 145160909 | 145160997 | 145156859 | 145157202 | 261,316,357,377 | 11,5,1,5 | 161,306,170,207 | 16,12,7,9 | 811 | 149 | 6.81E-05 | 0.00362 | 0.813,0.9 | 0.649,0.8 | 0.138 |
| ENSMUSG00000030922 | Lyrm1 | chr7 | + | 119508893 | 119509185 | 119509026 | 119509185 | 119495514 | 119495795 | 66,57,72,55 | 5,2,0,1 | 47,34,57,37 | 5,3,3,6 | 282 | 149 | 0.00428 | 0.06609 | 0.875,0.9 | 0.832,0.8 | 0.104 |
| ENSMUSG00000033767 | Tmem131l | chr3 | - | 83807633 | 83807725 | 83807633 | 83807722 | 83808652 | 83808773 | 96,76,90,76 | 6,12,7,4 | 39,32,48,49 | 10,6,10,11 | 152 | 149 | 0.00067 | 0.01787 | 0.94,0.86 | 0.793,0.8 | 0.101 |
| ENSMUSG00000035762 | Tmem161b | chr13 | + | 84430829 | 84430981 | 84430867 | 84430981 | 84420289 | 84420446 | 92,91,113,80 | 16,2,7,2 | 71,56,55,69 | 13,13,15,3 | 187 | 149 | 0.00181 | 0.03732 | 0.821,0.9 | 0.813,0.7 | 0.103 |
| ENSMUSG00000029165 | Agbl5 | chr5 | + | 31062361 | 31062499 | 31062452 | 31062499 | 31062046 | 31062159 | 41,23,26,34 | 65,46,42,53 | 12,10,10,8 | 35,40,37,36 | 240 | 149 | 9.85E-05 | 0.00471 | 0.281,0.2 | 0.176,0.1 | 0.127 |
| ENSMUSG00000032280 | Tle3 | chr9 | + | 61309134 | 61309339 | 61309137 | 61309339 | 61301889 | 61301994 | 10,19,20,29 | 0,0,0,0 | 28,30,27,19 | 16,1,8,8 | 152 | 149 | 1.61E-10 | 1.27E-07 | 1.0,1.0,1. | 0.632,0.9 | 0.233 |
| ENSMUSG00000043923 | Ccdc84 | chr9 | - | 44324213 | 44324340 | 44324213 | 44324319 | 44324433 | 44324498 | 16,19,11,13 | 4,20,20,29 | 5,5,5,10 | 35,40,25,28 | 170 | 149 | 0.00012 | 0.00562 | 0.778,0.4 | 0.111,0.0 | 0.31 |
| ENSMUSG00000057788 | Ddx49 | chr8 | - | 70747336 | 70747485 | 70747336 | 70747411 | 70748016 | 70748114 | 45,20,34,29 | 100,79,64,91 | 10,14,11,41 | 83,99,100,122 | 223 | 149 | 8.65E-05 | 0.00428 | 0.231,0.1 | 0.075,0.0 | 0.101 |
| ENSMUSG00000093803 | Ppp2r3d | chr9 | + | 124202110 | 124204759 | 124202202 | 124204759 | 124201794 | 124201907 | 15,22,33,19 | 2,4,0,0 | 13,7,7,21 | 8,0,5,4 | 241 | 149 | 0.00189 | 0.03848 | 0.823,0.7 | 0.501,1.0, | 0.217 |
| ENSMUSG00000033419 | Snap91 | chr9 | - | 86663622 | 86663772 | 86663622 | 86663757 | 86665352 | 86665487 | 214,178,240,183 | 652,480,924,542 | 108,97,99,74 | 679,555,617,634 | 164 | 149 | 2.73E-14 | 7.12E-11 | 0.23,0.25 | 0.126,0.1 | 0.106 |
| ENSMUSG00000056228 | Cars2 | chr8 | - | 11580294 | 11580427 | 11580294 | 11580424 | 11587271 | 11587355 | 14,20,11,21 | 55,44,44,36 | 1,12,10,0 | 47,40,49,41 | 152 | 149 | 0.0002 | 0.00782 | 0.2,0.308, | 0.02,0.22 | 0.164 |
| ENSMUSG00000097277 | 2900076A07Ri | chr7 | + | 81178904 | 81179275 | 81179094 | 81179275 | 81178534 | 81178688 | 123,86,103,90 | 51,35,49,46 | 73,53,50,51 | 64,23,62,34 | 339 | 149 | 0.00669 | 0.088 | 0.515,0.5 | 0.334,0.5 | 0.12 |
| ENSMUSG00000004562 | Arhgef40 | chr14 | + | 52242056 | 52242199 | 52242068 | 52242199 | 52241429 | 52241573 | 11,1,8,4 | 39,16,15,33 | 0,3,0,2 | 39,26,35,29 | 161 | 149 | 8.26E-06 | 0.00095 | 0.207,0.0 | 0.0,0.096, | 0.134 |
| ENSMUSG00000042185 | Nfrkb | chr9 | + | 31308089 | 31308433 | 31308331 | 31308433 | 31307618 | 31307793 | 190,202,256,130 | 26,15,11,18 | 113,103,118,105 | 18,24,12,24 | 391 | 149 | 0.00438 | 0.06657 | 0.736,0.8 | 0.705,0.6 | 0.116 |
| ENSMUSG00000004110 | Cacna1e | chr1 | - | 154362243 | 154363879 | 154362243 | 154362453 | 154364530 | 154364584 | 419,304,440,259 | 66,80,102,22 | 263,235,171,189 | 75,44,60,91 | 1575 | 149 | 0.00683 | 0.08885 | 0.375,0.2 | 0.249,0.3 | 0.124 |
| ENSMUSG00000057649 | Brd9 | chr13 | + | 74090749 | 74090925 | 74090809 | 74090925 | 74090062 | 74090173 | 149,150,242,184 | 75,63,119,76 | 131,121,150,97 | 95,88,80,101 | 209 | 149 | 0.00086 | 0.02191 | 0.586,0.6 | 0.496,0.4 | 0.118 |
| ENSMUSG00000063445 | Nmral1 | chr16 | - | 4529180 | 4529544 | 4529180 | 4529467 | 4531503 | 4531694 | 49,21,37,19 | 69,56,96,113 | 18,16,11,9 | 96,89,102,71 | 226 | 149 | 2.25E-05 | 0.00159 | 0.319,0.1 | 0.11,0.10 | 0.115 |

### Table S3. Alternative_splicing_SE

| GeneID | geneSym | chr | strand | exonStart_ | exonEnd | upstreamE | upstreamE | downstrea | downstrea | IJC_SAMPLE_1 | SJC_SAMPLE_1 | IJC_SAMPLE_2 | SJC_SAMPLE_2 | IncFormL | SkipForm | PValue | FDR | IncLevel1 | IncLevel2 | IncLevelDifference |
|---|---|---|---|---|---|---|---|---|---|---|---|---|---|---|---|---|---|---|---|---|
| ENSMUSG00000085396 | Firre | chrX | - | 49695839 | 49695981 | 49694884 | 49695039 | 49697354 | 49697481 | 25,16,8,0 | 0,44,0,32 | 0,0,0,3 | 37,14,39,34 | 291 | 149 | 4.34E-07 | 8.31E-05 | 1.0,0.157, | 0.0,0.0,0. | 0.528 |
| ENSMUSG00000032076 | Cadm1 | chr9 | + | 47748014 | 47748068 | 47725070 | 47725243 | 47759464 | 47759596 | 12,11,8,16 | 0,220,318,0 | 4,0,9,7 | 175,176,147,132 | 203 | 149 | 5.76E-05 | 0.00331 | 1.0,0.035, | 0.017,0.0, | 0.489 |
| ENSMUSG00000066735 | Vkorc1l1 | chr5 | + | 130005829 | 130006033 | 129971014 | 129971206 | 130007359 | 130007469 | 7,6,9,8 | 0,0,0,156 | 8,4,10,6 | 84,180,0,129 | 353 | 149 | 0.00059 | 0.0191 | 1.0,1.0,1. | 0.039,0.0 | 0.488 |
| ENSMUSG00000025006 | Sorbs1 | chr19 | - | 40329128 | 40329194 | 40325429 | 40325479 | 40332799 | 40332883 | 44,41,21,20 | 32,0,30,0 | 12,19,6,16 | 20,42,26,29 | 215 | 149 | 1.11E-05 | 0.00095 | 0.488,1.0, | 0.294,0.2 | 0.467 |
| ENSMUSG00000032567 | Aste1 | chr9 | + | 105279916 | 105279975 | 105278673 | 105278885 | 105280594 | 105280790 | 10,22,23,39 | 2,8,4,0 | 3,5,26,9 | 7,7,15,10 | 208 | 149 | 2.76E-06 | 0.00034 | 0.782,0.6 | 0.235,0.3 | 0.433 |
| ENSMUSG00000028344 | Invs | chr4 | + | 48391304 | 48391414 | 48389958 | 48390139 | 48392583 | 48392693 | 22,28,28,4 | 12,4,5,0 | 8,1,5,17 | 6,10,5,10 | 259 | 149 | 0.00096 | 0.02672 | 0.513,0.8 | 0.434,0.0 | 0.432 |
| ENSMUSG00000022565 | Plec | chr15 | - | 76078758 | 76078773 | 76078540 | 76078630 | 76078950 | 76079012 | 199,171,259,143 | 49,29,59,15 | 0,128,0,69 | 36,25,20,15 | 164 | 149 | 0.00018 | 0.00791 | 0.787,0.8 | 0.0,0.823, | 0.424 |
| ENSMUSG00000057541 | Pus7 | chr5 | - | 23968331 | 23968349 | 23967360 | 23967472 | 23973765 | 23973910 | 52,35,74,35 | 12,14,13,0 | 47,29,0,1 | 21,0,23,45 | 167 | 149 | 0.00242 | 0.05326 | 0.795,0.6 | 0.666,1.0, | 0.409 |
| ENSMUSG00000053453 | Thoc7 | chr14 | + | 8512131 | 8512209 | 8508547 | 8508578 | 8515106 | 8515224 | 59,30,52,39 | 93,0,14,7 | 15,41,28,42 | 53,40,46,54 | 227 | 149 | 0.00012 | 0.00587 | 0.294,1.0, | 0.157,0.4 | 0.402 |
| ENSMUSG00000074219 | Gm10644 | chr8 | - | 84661487 | 84661699 | 84658957 | 84661235 | 84662360 | 84662457 | 27,55,20,16 | 0,0,0,0 | 35,19,32,50 | 15,14,11,0 | 361 | 149 | 8.54E-11 | 1.01E-07 | 1.0,1.0,1. | 0.491,0.3 | 0.401 |
| ENSMUSG00000049606 | Zfp644 | chr5 | - | 106843002 | 106843565 | 106814650 | 106814711 | 106844099 | 106844412 | 92,56,66,52 | 0,15,0,32 | 49,19,29,53 | 17,18,17,28 | 712 | 149 | 0.00049 | 0.01654 | 1.0,0.439, | 0.376,0.1 | 0.397 |
| ENSMUSG00000041219 | Arhgap11 | chr2 | - | 113665122 | 113665261 | 113661890 | 113664807 | 113667210 | 113667316 | 8,21,39,17 | 0,0,0,0 | 31,7,21,8 | 4,4,2,10 | 288 | 149 | 2.72E-10 | 2.58E-07 | 1.0,1.0,1. | 0.8,0.475, | 0.397 |
| ENSMUSG00000025089 | Gfra1 | chr19 | - | 58381004 | 58381019 | 58288673 | 58289010 | 58440400 | 58440484 | 0,200,223,0 | 56,58,46,28 | 0,0,0,0 | 79,32,42,44 | 164 | 149 | 2.25E-06 | 0.0003 | 0.0,0.758, | 0.0,0.0,0. | 0.393 |
| ENSMUSG00000085396 | Firre | chrX | - | 49687010 | 49687165 | 49684337 | 49684492 | 49694884 | 49695039 | 67,73,76,68 | 3,0,0,0 | 12,47,0,83 | 11,0,17,0 | 304 | 149 | 3.82E-05 | 0.00244 | 0.916,1.0, | 0.348,1.0, | 0.392 |
| ENSMUSG00000097881 | Celrr | chr1 | + | 121045037 | 121045826 | 121026767 | 121026816 | 121047586 | 121047738 | 38,52,125,27 | 0,2,0,4 | 46,65,42,34 | 10,5,9,17 | 938 | 149 | 1.57E-05 | 0.00124 | 1.0,0.805, | 0.422,0.6 | 0.39 |
| ENSMUSG00000026211 | Obsl1 | chr1 | - | 75468871 | 75468986 | 75468001 | 75468101 | 75469149 | 75469275 | 10,20,14,15 | 20,0,7,0 | 14,7,0,4 | 0,24,14,28 | 264 | 149 | 0.00362 | 0.07102 | 0.22,1.0,0 | 1.0,0.141, | 0.384 |
| ENSMUSG00000027455 | Nsfl1c | chr2 | + | 151344374 | 151344380 | 151342634 | 151342709 | 151344933 | 151345099 | 30,30,25,490 | 0,54,72,70 | 12,22,21,2 | 67,43,34,17 | 155 | 149 | 0.00075 | 0.0225 | 1.0,0.348, | 0.147,0.3 | 0.379 |
| ENSMUSG00000056602 | Fry | chr5 | + | 150384853 | 150384862 | 150381209 | 150381380 | 150389708 | 150389906 | 36,48,429,33 | 105,72,82,56 | 15,0,0,18 | 88,84,91,61 | 158 | 149 | 3.41E-08 | 1.09E-05 | 0.244,0.3 | 0.138,0.0, | 0.365 |
| ENSMUSG00000022607 | Ptk2 | chr15 | - | 73236889 | 73237044 | 73214981 | 73215208 | 73263607 | 73263729 | 37,24,19,30 | 7,6,0,12 | 0,41,30,15 | 15,12,14,11 | 304 | 149 | 0.00347 | 0.06901 | 0.722,0.6 | 0.0,0.626, | 0.349 |
| ENSMUSG00000024998 | Plce1 | chr19 | + | 38736853 | 38736895 | 38734209 | 38734339 | 38737735 | 38737857 | 10,20,24,41 | 25,19,13,21 | 6,6,3,14 | 54,30,23,38 | 191 | 149 | 2.12E-07 | 4.77E-05 | 0.238,0.4 | 0.08,0.13 | 0.338 |
| ENSMUSG00000071042 | Rasgrp3 | chr17 | + | 75795811 | 75795944 | 75772480 | 75772655 | 75798752 | 75798785 | 14,23,36,10 | 0,4,8,4 | 30,3,35,24 | 18,3,27,13 | 282 | 149 | 0.00077 | 0.02278 | 1.0,0.752, | 0.468,0.3 | 0.327 |
| ENSMUSG00000028649 | Macf1 | chr4 | - | 123257849 | 123257867 | 123254702 | 123254820 | 123259065 | 123259140 | 810,631,821,725 | 154,130,222,191 | 8,46,547,542 | 141,92,119,191 | 167 | 149 | 0.00084 | 0.02449 | 0.824,0.8 | 0.048,0.3 | 0.324 |
| ENSMUSG00000034593 | Myo5a | chr9 | + | 75094803 | 75094812 | 75093176 | 75093404 | 75097181 | 75097282 | 42,0,21,61 | 15,62,35,37 | 0,0,27,0 | 35,32,31,20 | 158 | 149 | 0.002 | 0.04602 | 0.725,0.0, | 0.0,0.0,0. | 0.311 |
| ENSMUSG00000042632 | Pla2g6 | chr15 | - | 79211834 | 79211914 | 79201960 | 79202210 | 79212409 | 79212453 | 22,16,32,30 | 29,9,0,27 | 7,8,20,20 | 32,28,17,26 | 229 | 149 | 0.00345 | 0.06868 | 0.33,0.53 | 0.125,0.1 | 0.309 |
| ENSMUSG00000032076 | Cadm1 | chr9 | + | 47740674 | 47740707 | 47725070 | 47725243 | 47759464 | 47759596 | 160,50,158,194 | 0,220,318,0 | 117,106,74,49 | 175,176,147,132 | 182 | 149 | 0.00013 | 0.00604 | 1.0,0.157, | 0.354,0.3 | 0.309 |
| ENSMUSG00000022340 | Sybu | chr15 | - | 44611704 | 44611794 | 44582183 | 44582381 | 44650984 | 44651459 | 21,30,18,11 | 0,0,0,4 | 11,19,11,26 | 6,0,10,19 | 239 | 149 | 1.52E-05 | 0.0012 | 1.0,1.0,1. | 0.533,1.0, | 0.308 |
| ENSMUSG00000047996 | Prrg1 | chrX | - | 77526031 | 77526067 | 77493218 | 77496778 | 77527356 | 77527517 | 5,5,12,10 | 5,6,15,13 | 0,3,3,6 | 22,11,21,36 | 185 | 149 | 5.03E-07 | 9.22E-05 | 0.446,0.4 | 0.0,0.18,0 | 0.306 |
| ENSMUSG00000037369 | Kdm6a | chrX | + | 18113213 | 18113348 | 18112435 | 18112655 | 18113805 | 18113961 | 54,41,42,43 | 42,13,11,0 | 26,25,18,27 | 24,23,24,16 | 284 | 149 | 0.00122 | 0.03164 | 0.403,0.6 | 0.362,0.3 | 0.304 |
| ENSMUSG00000040929 | Rfx3 | chr19 | - | 27900607 | 27900666 | 27878178 | 27878303 | 27960253 | 27960346 | 33,23,46,19 | 0,1,0,10 | 13,13,13,8 | 10,0,10,11 | 208 | 149 | 0.00056 | 0.01848 | 1.0,0.943, | 0.482,1.0, | 0.303 |
| ENSMUSG00000031391 | L1cam | chrX | - | 72910627 | 72910642 | 72909660 | 72909766 | 72913334 | 72913511 | 311,287,468,226 | 5,0,1,0 | 79,136,165,0 | 12,7,4,2 | 164 | 149 | 0.00309 | 0.06325 | 0.983,1.0, | 0.857,0.9 | 0.301 |
| ENSMUSG00000040690 | Col16a1 | chr4 | + | 129943119 | 129943219 | 129941667 | 129941876 | 129943425 | 129943526 | 14,10,20,17 | 13,17,18,6 | 4,8,5,6 | 12,45,23,26 | 249 | 149 | 1.11E-06 | 0.00017 | 0.392,0.2 | 0.166,0.0 | 0.296 |
| ENSMUSG00000031791 | Tmem38a | chr8 | + | 73335063 | 73335116 | 73333803 | 73333988 | 73339705 | 73340019 | 20,22,33,11 | 7,3,0,2 | 14,6,19,3 | 6,17,4,2 | 202 | 149 | 0.00196 | 0.04523 | 0.678,0.8 | 0.633,0.2 | 0.295 |
| ENSMUSG00000034799 | Unc13a | chr8 | - | 72108146 | 72108152 | 72107776 | 72107848 | 72108306 | 72108385 | 15,344,206,55 | 2,0,2,0 | 63,0,18,33 | 2,7,1,9 | 155 | 149 | 7.07E-05 | 0.00386 | 0.878,1.0, | 0.968,0.0, | 0.294 |
| ENSMUSG00000031367 | Ap1s2 | chrX | + | 162714358 | 162714367 | 162700701 | 162700839 | 162715128 | 162716662 | 57,52,40,39 | 0,17,12,7 | 16,32,14,25 | 23,9,28,11 | 158 | 149 | 0.00026 | 0.01016 | 1.0,0.743, | 0.396,0.7 | 0.293 |
| ENSMUSG00000006498 | Ptbp1 | chr10 | + | 79695950 | 79696028 | 79695614 | 79695789 | 79696657 | 79696803 | 51,69,63,64 | 39,0,39,0 | 55,37,38,49 | 37,23,46,37 | 227 | 149 | 0.00014 | 0.00643 | 0.462,1.0, | 0.494,0.5 | 0.288 |
| ENSMUSG00000085396 | Firre | chrX | - | 49692247 | 49692402 | 49689620 | 49689775 | 49697563 | 49697691 | 27,20,32,62 | 2,5,9,0 | 15,15,15,7 | 8,3,6,8 | 304 | 149 | 0.00144 | 0.03618 | 0.869,0.6 | 0.479,0.7 | 0.282 |
| ENSMUSG00000005871 | Apc | chr18 | + | 34394057 | 34394210 | 34354036 | 34354329 | 34399110 | 34399275 | 25,40,50,38 | 10,0,0,22 | 29,13,35,43 | 15,7,25,20 | 302 | 149 | 0.00101 | 0.02763 | 0.552,1.0, | 0.488,0.4 | 0.28 |
| ENSMUSG00000031644 | Nek1 | chr8 | + | 61503061 | 61503193 | 61502817 | 61502981 | 61507553 | 61507656 | 177,132,158,132 | 51,5,83,32 | 72,145,124,135 | 61,70,115,109 | 281 | 149 | 2.78E-05 | 0.00193 | 0.648,0.9 | 0.385,0.5 | 0.275 |
| ENSMUSG00000053453 | Thoc7 | chr14 | + | 8512131 | 8512303 | 8508550 | 8508578 | 8515106 | 8515197 | 116,110,163,90 | 93,0,14,7 | 72,106,120,111 | 53,40,46,54 | 321 | 149 | 0.00031 | 0.01185 | 0.367,1.0, | 0.387,0.5 | 0.273 |
| ENSMUSG00000040374 | Pex2 | chr3 | - | 5635546 | 5635689 | 5630192 | 5630258 | 5641097 | 5641188 | 16,18,41,22 | 0,0,3,0 | 16,12,3,20 | 6,4,1,0 | 292 | 149 | 5.39E-05 | 0.00314 | 1.0,1.0,0. | 0.576,0.6 | 0.272 |
| ENSMUSG00000052384 | Nrros | chr16 | - | 31981055 | 31982215 | 31961642 | 31963823 | 31984197 | 31984294 | 48,49,65,57 | 0,0,1,2 | 64,43,56,53 | 4,1,2,12 | 1309 | 149 | 1.81E-05 | 0.00138 | 1.0,1.0,0. | 0.646,0.8 | 0.268 |
| ENSMUSG00000025892 | Gria4 | chr9 | - | 4432772 | 4432887 | 4427029 | 4427144 | 4456004 | 4456252 | 52,45,74,60 | 24,11,0,0 | 43,41,13,55 | 9,23,22,13 | 264 | 149 | 0.00142 | 0.03573 | 0.55,0.69 | 0.729,0.5 | 0.266 |
| ENSMUSG00000057738 | Sptan1 | chr2 | + | 29904169 | 29904184 | 29903687 | 29903850 | 29905600 | 29905732 | 210,791,718,562,091 | 26,34,36,16 | 1391,0,359,1716 | 26,29,30,44 | 164 | 149 | 0.00091 | 0.02586 | 0.987,0.9 | 0.98,0.0,0 | 0.263 |
| ENSMUSG00000020385 | Clk4 | chr11 | + | 51159595 | 51159673 | 51158991 | 51159058 | 51161363 | 51161422 | 19,23,26,22 | 7,3,11,16 | 8,12,16,8 | 13,7,16,12 | 227 | 149 | 0.00425 | 0.07957 | 0.64,0.83 | 0.288,0.5 | 0.26 |
| ENSMUSG00000027598 | Itch | chr2 | + | 154999693 | 154999895 | 154980115 | 154980192 | 155005312 | 155005403 | 289,207,295,173 | 137,25,0,57 | 167,213,178,170 | 98,79,78,135 | 351 | 149 | 9.48E-05 | 0.00483 | 0.472,0.7 | 0.42,0.53 | 0.255 |
| ENSMUSG00000061904 | Slc25a3 | chr10 | - | 90958068 | 90958193 | 90955388 | 90955568 | 90959419 | 90959568 | 24,31,29,24 | 1,0,0,5 | 8,8,11,35 | 10,2,2,2 | 274 | 149 | 0.00216 | 0.04874 | 0.929,1.0, | 0.303,0.6 | 0.252 |
| ENSMUSG00000022701 | Ccdc191 | chr16 | + | 43718452 | 43718594 | 43710162 | 43710401 | 43725708 | 43725846 | 46,14,23,42 | 1,0,4,3 | 23,23,37,16 | 6,9,2,10 | 291 | 149 | 0.00066 | 0.02032 | 0.959,1.0, | 0.662,0.5 | 0.25 |
| ENSMUSG00000031093 | Dock11 | chrX | + | 35339569 | 35339587 | 35333500 | 35333639 | 35339751 | 35339880 | 5,9,15,19 | 13,9,18,5 | 7,7,6,4 | 12,24,25,13 | 167 | 149 | 0.00467 | 0.08532 | 0.255,0.4 | 0.342,0.2 | 0.247 |
| ENSMUSG00000052707 | Tnrc6a | chr7 | + | 122751386 | 122751408 | 122723890 | 122723938 | 122761716 | 122761886 | 28,50,68,25 | 0,15,7,0 | 29,21,11,26 | 7,7,5,23 | 171 | 149 | 0.00014 | 0.00645 | 1.0,0.744, | 0.783,0.7 | 0.245 |
| ENSMUSG00000056870 | Gulp1 | chr1 | + | 44829855 | 44829866 | 44827607 | 44827702 | 44834109 | 44835985 | 16,13,5,6 | 14,9,28,21 | 0,5,0,3 | 14,13,20,11 | 160 | 149 | 0.00129 | 0.03328 | 0.516,0.5 | 0.0,0.264, | 0.244 |
| ENSMUSG00000058174 | Gm5148 | chr3 | - | 37776447 | 37777130 | 37770270 | 37770381 | 37778351 | 37778480 | 135,122,152,112 | 10,3,14,5 | 60,76,44,68 | 17,18,5,6 | 832 | 149 | 0.0009 | 0.02572 | 0.707,0.8 | 0.387,0.4 | 0.237 |
| ENSMUSG00000034813 | Grip1 | chr10 | + | 119821378 | 119821534 | 119814342 | 119814512 | 119822236 | 119822389 | 89,57,100,68 | 11,0,2,10 | 47,27,45,54 | 9,5,18,18 | 305 | 149 | 9.70E-06 | 0.00086 | 0.798,1.0, | 0.718,0.7 | 0.235 |
| ENSMUSG00000022607 | Ptk2 | chr15 | - | 73236889 | 73237044 | 73231874 | 73232186 | 73266496 | 73266593 | 25,20,66,30 | 0,0,3,0 | 14,34,19,22 | 14,9,0,0 | 304 | 149 | 8.72E-05 | 0.00453 | 1.0,1.0,0. | 0.329,0.6 | 0.234 |
| ENSMUSG00000112117 | Rmst | chr10 | - | 91939567 | 91939774 | 91919327 | 91919944 | 91945500 | 91945595 | 10,18,23,28 | 2,3,0,2 | 9,27,34,16 | 3,10,3,10 | 356 | 149 | 0.0044 | 0.08155 | 0.677,0.7 | 0.557,0.5 | 0.233 |
| ENSMUSG00000022607 | Ptk2 | chr15 | - | 73231874 | 73232186 | 73214981 | 73215208 | 73263607 | 73263729 | 61,54,111,36 | 7,6,0,12 | 58,33,57,29 | 15,12,14,11 | 461 | 149 | 0.00239 | 0.05281 | 0.738,0.7 | 0.556,0.4 | 0.23 |
| ENSMUSG00000032076 | Cadm1 | chr9 | + | 47730031 | 47730115 | 47725070 | 47725243 | 47759464 | 47759596 | 201,180,205,191 | 0,220,318,0 | 231,124,164,245 | 175,176,147,132 | 233 | 149 | 0.00036 | 0.01308 | 1.0,0.343, | 0.458,0.3 | 0.227 |
| ENSMUSG00000025949 | Pikfyve | chr1 | + | 65231692 | 65231725 | 65231303 | 65231453 | 65234814 | 65234933 | 48,14,56,44 | 6,0,1,10 | 30,42,26,19 | 2,20,10,16 | 182 | 149 | 0.00062 | 0.01963 | 0.868,1.0, | 0.925,0.6 | 0.225 |
| ENSMUSG00000050545 | Fam228b | chr12 | - | 4811858 | 4811987 | 4798263 | 4798351 | 4812326 | 4812407 | 10,19,3,9 | 11,14,11,29 | 3,1,0,0 | 17,14,10,25 | 278 | 149 | 8.79E-09 | 3.49E-06 | 0.328,0.4 | 0.086,0.0 | 0.224 |
| ENSMUSG00000025006 | Sorbs1 | chr19 | - | 40328488 | 40328590 | 40325429 | 40325479 | 40332799 | 40332883 | 218,187,207,192 | 32,0,30,0 | 68,135,118,97 | 20,42,26,29 | 251 | 149 | 1.29E-10 | 1.44E-07 | 0.802,1.0, | 0.669,0.6 | 0.222 |
| ENSMUSG00000008496 | Pou2f2 | chr7 | - | 24814895 | 24815174 | 24799562 | 24799668 | 24815462 | 24815645 | 30,27,45,21 | 0,0,1,2 | 7,23,22,37 | 7,0,4,1 | 428 | 149 | 0.00036 | 0.0131 | 1.0,1.0,0. | 0.258,1.0, | 0.221 |
| ENSMUSG00000040990 | Sh3kbp1 | chrX | + | 158607470 | 158607545 | 158599572 | 158599702 | 158611369 | 158611575 | 153,106,159,91 | 8,0,10,46 | 106,63,67,69 | 66,0,58,33 | 224 | 149 | 0.00406 | 0.07714 | 0.927,1.0, | 0.517,1.0, | 0.219 |
| ENSMUSG00000026767 | Mindy3 | chr2 | - | 12402303 | 12402377 | 12391419 | 12391490 | 12423969 | 12424099 | 56,0,41,38 | 29,25,36,39 | 0,15,13,0 | 35,22,34,20 | 223 | 149 | 0.00568 | 0.09764 | 0.563,0.0, | 0.0,0.313, | 0.218 |
| ENSMUSG00000086587 | Gm11837 | chr4 | + | 14947438 | 14947545 | 14942199 | 14942275 | 14952620 | 14952905 | 40,33,34,48 | 0,0,0,0 | 41,21,32,29 | 1,0,12,13 | 256 | 149 | 1.56E-08 | 5.35E-06 | 1.0,1.0,1. | 0.96,1.0,0 | 0.217 |
| ENSMUSG00000021809 | Nudt13 | chr14 | + | 20345155 | 20345334 | 20344781 | 20344860 | 20354020 | 20354160 | 66,60,83,70 | 0,9,23,12 | 74,42,45,49 | 16,25,23,12 | 328 | 149 | 0.00349 | 0.06935 | 1.0,0.752, | 0.678,0.4 | 0.217 |
| ENSMUSG00000026017 | Carf | chr1 | + | 60141785 | 60141943 | 60137405 | 60137466 | 60144469 | 60144593 | 32,31,55,27 | 22,17,41,14 | 19,11,6,5 | 23,7,21,42 | 307 | 149 | 0.00014 | 0.00652 | 0.414,0.4 | 0.286,0.4 | 0.216 |
| ENSMUSG00000004952 | Rasa4 | chr5 | + | 136130798 | 136130911 | 136130097 | 136130543 | 136131088 | 136131149 | 28,28,54,19 | 2,0,1,4 | 19,21,11,32 | 11,0,5,9 | 262 | 149 | 0.00065 | 0.02018 | 0.888,1.0, | 0.496,1.0, | 0.216 |
| ENSMUSG00000086370 | Ftx | chrX | - | 102635408 | 102635512 | 102616105 | 102616167 | 102639536 | 102639613 | 38,25,11,11 | 0,0,0,0 | 42,21,6,26 | 3,5,1,5 | 253 | 149 | 9.77E-08 | 2.43E-05 | 1.0,1.0,1. | 0.892,0.7 | 0.216 |
| ENSMUSG00000075020 | Mir670hg | chr2 | - | 93997188 | 93997335 | 93990980 | 93991553 | 94014911 | 94015006 | 34,13,24,19 | 33,37,11,17 | 11,5,9,4 | 25,16,27,37 | 296 | 149 | 9.75E-05 | 0.00494 | 0.342,0.1 | 0.181,0.1 | 0.215 |
| ENSMUSG00000073609 | D2hgdh | chr1 | + | 93760317 | 93760511 | 93757478 | 93757618 | 93763008 | 93763177 | 127,70,136,72 | 7,0,0,0 | 94,53,31,53 | 0,21,3,10 | 343 | 149 | 4.00E-06 | 0.00044 | 0.887,1.0, | 1.0,0.523, | 0.212 |
| ENSMUSG00000056938 | Acbd4 | chr11 | + | 102994060 | 102994195 | 102993864 | 102993971 | 102994351 | 102994470 | 84,53,102,63 | 2,11,0,37 | 135,116,74,86 | 55,31,33,40 | 284 | 149 | 0.00056 | 0.01845 | 0.957,0.7 | 0.563,0.6 | 0.212 |
| ENSMUSG00000035919 | Bbs9 | chr9 | + | 22559315 | 22559330 | 22557259 | 22557364 | 22566520 | 22566661 | 96,52,64,71 | 3,6,2,13 | 17,21,8,26 | 4,5,8,8 | 164 | 149 | 0.00029 | 0.01137 | 0.967,0.8 | 0.794,0.7 | 0.211 |
| ENSMUSG00000045962 | Wnk1 | chr6 | - | 119906786 | 119906828 | 119904085 | 119904195 | 119911216 | 119911411 | 198,157,293,193 | 194,63,159,153 | 121,105,128,123 | 216,144,185,193 | 191 | 149 | 1.06E-07 | 2.60E-05 | 0.443,0.6 | 0.304,0.3 | 0.21 |
| ENSMUSG00000066442 | Mthfs | chr9 | + | 89097261 | 89097523 | 89093209 | 89093364 | 89121894 | 89122168 | 114,54,104,44 | 0,0,0,0 | 78,49,22,49 | 0,12,5,1 | 411 | 149 | 9.45E-08 | 2.38E-05 | 1.0,1.0,1. | 1.0,0.597, | 0.21 |

| | | | | | | | | | | | | | | | | | | | | |
|---|---|---|---|---|---|---|---|---|---|---|---|---|---|---|---|---|---|---|---|---|
| ENSMUSG00000085316 | D330050 | chr2 | + | 116734522 | 116734703 | 116730697 | 116730812 | 116741201 | 116741317 | 39,11,39,14 | 0,0,4,0 | 16,5,26,9 | 1,1,0,6 | 330 | 149 | 0.00413 | 0.07804 | 1.0,1.0,0.8 | 0.878,0.69 | 0.21 |
| ENSMUSG00000000131 | Xpo6 | chr7 | - | 125748675 | 125748709 | 125748405 | 125748532 | 125750395 | 125752451 | 23,27,31,26 | 0,0,3,0 | 6,9,30,32 | 8,1,6,0 | 183 | 149 | 0.00013 | 0.00611 | 1.0,1.0,0.8 | 0.379,0.88 | 0.208 |
| ENSMUSG00000028163 | Nfkb1 | chr3 | - | 135344900 | 135344917 | 135332355 | 135332504 | 135361260 | 135361359 | 15,0,15,13 | 21,42,41,20 | 9,0,0,1 | 43,22,24,47 | 166 | 149 | 8.47E-05 | 0.00442 | 0.391,0.0, | 0.158,0.0, | 0.207 |
| ENSMUSG00000113701 | B230303A | chr13 | - | 16091134 | 16091232 | 15988132 | 15990573 | 16197849 | 16197945 | 19,17,32,21 | 27,10,33,27 | 8,0,16,30 | 34,32,31,41 | 247 | 149 | 0.00072 | 0.02165 | 0.298,0.50 | 0.124,0.0, | 0.206 |
| ENSMUSG00000010538 | Tsacc | chr3 | - | 88194404 | 88194524 | 88190063 | 88190291 | 88202697 | 88202757 | 33,7,23,20 | 0,0,0,0 | 18,12,17,19 | 3,2,0,6 | 269 | 149 | 4.96E-07 | 9.13E-05 | 1.0,1.0,1.0 | 0.769,0.76 | 0.206 |
| ENSMUSG00000022376 | Adcy8 | chr15 | - | 64616689 | 64616779 | 64609238 | 64609411 | 64618745 | 64618947 | 35,20,55,29 | 0,3,5,0 | 38,27,42,26 | 8,7,6,11 | 239 | 149 | 5.29E-05 | 0.0031 | 1.0,0.806, | 0.748,0.70 | 0.204 |
| ENSMUSG00000113701 | B230303A | chr13 | - | 16147822 | 16147933 | 16119336 | 16119925 | 16197849 | 16198008 | 15,20,15,9 | 29,7,31,18 | 4,3,7,0 | 22,13,11,10 | 260 | 149 | 0.00459 | 0.08426 | 0.229,0.62 | 0.094,0.11 | 0.203 |
| ENSMUSG00000033021 | Gmppa | chr1 | + | 75412789 | 75412956 | 75412586 | 75412655 | 75413421 | 75413482 | 196,171,194,158 | 0,40,74,67 | 125,130,102,120 | 76,49,51,67 | 316 | 149 | 0.00107 | 0.02877 | 1.0,0.668, | 0.437,0.55 | 0.203 |
| ENSMUSG00000087056 | Gm14004 | chr2 | - | 124973860 | 124974024 | 124971568 | 124971661 | 124976154 | 124976247 | 62,25,46,52 | 34,23,36,27 | 37,6,4,14 | 21,36,19,22 | 313 | 149 | 0.00063 | 0.01978 | 0.465,0.34 | 0.456,0.07 | 0.202 |
| ENSMUSG00000035126 | Dnai4 | chr4 | - | 102923456 | 102923683 | 102917074 | 102917232 | 102929845 | 102930040 | 72,62,111,60 | 3,0,7,8 | 52,39,27,51 | 6,9,8,7 | 376 | 149 | 0.00103 | 0.02806 | 0.905,1.0, | 0.774,0.63 | 0.199 |
| ENSMUSG00000015968 | Cacna1d | chr14 | - | 29811252 | 29811336 | 29804908 | 29804953 | 29817274 | 29817385 | 16,23,26,30 | 0,0,0,0 | 22,8,27,8 | 4,0,7,2 | 233 | 149 | 9.74E-08 | 2.43E-05 | 1.0,1.0,1.0 | 0.779,1.0, | 0.198 |
| ENSMUSG00000119994 | | chr10 | + | 128337248 | 128337558 | 128334812 | 128335395 | 128338277 | 128338836 | 87,93,112,113 | 21,0,0,0 | 126,64,77,60 | 10,4,20,14 | 459 | 149 | 0.00018 | 0.00791 | 0.574,1.0, | 0.804,0.83 | 0.198 |
| ENSMUSG00000052151 | Plpp2 | chr10 | - | 79366710 | 79366815 | 79366331 | 79366606 | 79369457 | 79369594 | 13,17,25,23 | 1,0,0,0 | 32,7,18,14 | 3,4,4,0 | 254 | 149 | 6.64E-06 | 0.00065 | 0.884,1.0, | 0.862,0.50 | 0.198 |
| ENSMUSG00000025316 | Banp | chr8 | + | 122747257 | 122747383 | 122734491 | 122734530 | 122750741 | 122750885 | 122,102,137,86 | 43,47,72,45 | 109,81,69,78 | 112,65,82,91 | 275 | 149 | 8.58E-09 | 3.43E-06 | 0.606,0.54 | 0.345,0.40 | 0.196 |
| ENSMUSG00000038816 | Ctnnal1 | chr4 | - | 56838004 | 56838047 | 56837731 | 56837902 | 56838965 | 56839055 | 38,33,76,34 | 6,9,12,7 | 30,44,29,42 | 19,10,27,19 | 192 | 149 | 0.00057 | 0.01866 | 0.831,0.74 | 0.551,0.77 | 0.195 |
| ENSMUSG00000087095 | Emx2os | chr19 | - | 59418431 | 59418531 | 59413535 | 59417693 | 59421234 | 59421464 | 19,20,22,21 | 0,3,0,0 | 22,25,21,31 | 3,3,3,14 | 249 | 149 | 3.73E-06 | 0.00042 | 1.0,0.8,1.0 | 0.814,0.83 | 0.194 |
| ENSMUSG00000054280 | Prr14l | chr5 | - | 33009225 | 33009342 | 32992959 | 32993032 | 33011436 | 33011600 | 55,58,66,49 | 22,8,0,0 | 36,46,22,46 | 13,13,9,8 | 266 | 149 | 0.00263 | 0.05664 | 0.583,0.80 | 0.608,0.66 | 0.193 |
| ENSMUSG00000031441 | Atp11a | chr8 | + | 12911639 | 12911724 | 12909313 | 12909479 | 12913758 | 12913862 | 70,49,96,77 | 7,0,3,0 | 65,68,44,57 | 13,5,10,19 | 234 | 149 | 7.18E-08 | 1.98E-05 | 0.864,1.0, | 0.761,0.89 | 0.192 |
| ENSMUSG00000096401 | Gm21811 | chr8 | + | 20723149 | 20723317 | 20721120 | 20721240 | 20727081 | 20727266 | 30,38,63,46 | 30,23,36,38 | 20,34,17,23 | 39,40,48,47 | 317 | 149 | 3.80E-06 | 0.00043 | 0.32,0.437 | 0.194,0.28 | 0.191 |
| ENSMUSG00000040723 | Rcsd1 | chr1 | - | 165491134 | 165491224 | 165486941 | 165487013 | 165492517 | 165492619 | 23,27,38,24 | 0,3,0,0 | 39,15,21,24 | 10,0,13,2 | 239 | 149 | 4.64E-06 | 0.0005 | 1.0,0.849, | 0.709,1.0, | 0.189 |
| ENSMUSG00000054099 | Slc25a40 | chr5 | + | 8474796 | 8474887 | 8472872 | 8473039 | 8477195 | 8477267 | 19,27,15,38 | 18,26,10,22 | 9,8,12,9 | 13,22,27,11 | 240 | 149 | 0.00074 | 0.02224 | 0.396,0.39 | 0.301,0.18 | 0.187 |
| ENSMUSG00000051736 | Fam229b | chr10 | - | 38996221 | 38996392 | 38994863 | 38995030 | 38998172 | 38998331 | 71,88,65,49 | 18,0,5,13 | 47,41,79,61 | 15,12,21,23 | 320 | 149 | 0.00211 | 0.04796 | 0.647,1.0, | 0.593,0.61 | 0.186 |
| ENSMUSG00000035293 | G2e3 | chr12 | + | 51412037 | 51412342 | 51410199 | 51410329 | 51414557 | 51414739 | 52,63,71,72 | 0,0,0,3 | 55,38,42,75 | 9,6,0,6 | 454 | 149 | 6.39E-07 | 0.00011 | 1.0,1.0,1.0 | 0.667,0.67 | 0.185 |
| ENSMUSG00000075415 | Fnbp1 | chr2 | - | 30942996 | 30943149 | 30930416 | 30930526 | 30944035 | 30944230 | 49,44,60,46 | 6,1,5,0 | 50,38,23,33 | 9,5,6,7 | 302 | 149 | 0.00089 | 0.02541 | 0.801,0.95 | 0.733,0.78 | 0.184 |
| ENSMUSG00000028760 | Eif4g3 | chr4 | + | 137811567 | 137811681 | 137810203 | 137810299 | 137814843 | 137814876 | 48,68,69,40 | 52,32,24,29 | 22,27,37,32 | 37,29,37,51 | 263 | 149 | 0.00137 | 0.03475 | 0.343,0.54 | 0.252,0.34 | 0.182 |
| ENSMUSG00000027677 | Ttc14 | chr3 | + | 33862004 | 33862097 | 33860964 | 33861082 | 33862834 | 33863826 | 48,60,57,61 | 0,0,0,1 | 13,27,24,27 | 6,0,0,8 | 242 | 149 | 2.09E-06 | 0.00028 | 1.0,1.0,1.0 | 0.572,1.0, | 0.182 |
| ENSMUSG00000034442 | Trmt5 | chr12 | - | 73332677 | 73332749 | 73331465 | 73332049 | 73333457 | 73333484 | 38,20,18,12 | 22,19,17,24 | 7,9,4,15 | 27,19,17,17 | 221 | 149 | 0.00564 | 0.09731 | 0.538,0.41 | 0.149,0.24 | 0.18 |
| ENSMUSG00000025766 | D3Ertd751 | chr3 | + | 41699130 | 41699247 | 41697045 | 41697124 | 41700982 | 41701163 | 28,11,22,15 | 22,8,37,19 | 0,12,5,12 | 17,23,25,12 | 266 | 149 | 0.00474 | 0.08634 | 0.416,0.43 | 0.0,0.226, | 0.18 |
| ENSMUSG00000058897 | Col25a1 | chr3 | + | 130366605 | 130366686 | 130363957 | 130363984 | 130369023 | 130369059 | 7,10,15,2 | 5,27,54,31 | 0,0,7,0 | 31,42,27,36 | 230 | 149 | 1.42E-05 | 0.00114 | 0.476,0.19 | 0.0,0.0,0.1 | 0.18 |
| ENSMUSG00000034334 | Fam151b | chr13 | - | 92610470 | 92610636 | 92586340 | 92586760 | 92614335 | 92614461 | 69,44,78,82 | 10,2,11,7 | 28,31,46,57 | 6,9,15,12 | 315 | 149 | 0.00164 | 0.03942 | 0.765,0.91 | 0.688,0.62 | 0.176 |
| ENSMUSG00000103428 | Gm20754 | chr3 | + | 73146629 | 73146743 | 72973683 | 72973915 | 73466781 | 73466859 | 50,23,50,47 | 0,0,0,0 | 19,28,22,14 | 3,0,5,2 | 263 | 149 | 1.36E-05 | 0.00111 | 1.0,1.0,1.0 | 0.782,1.0, | 0.176 |
| ENSMUSG00000033295 | Ptprf | chr4 | - | 118109272 | 118109290 | 118106095 | 118106206 | 118114685 | 118114874 | 106,54,110,143 | 58,61,67,57 | 71,61,53,26 | 101,53,47,69 | 167 | 149 | 0.00289 | 0.06044 | 0.62,0.441 | 0.385,0.50 | 0.175 |
| ENSMUSG00000071550 | Cfap44 | chr16 | + | 44259359 | 44259534 | 44257489 | 44257686 | 44269392 | 44269570 | 21,3,35,5 | 23,11,23,14 | 1,1,3,2 | 12,28,16,6 | 324 | 149 | 4.79E-07 | 8.93E-05 | 0.296,0.11 | 0.037,0.01 | 0.174 |
| ENSMUSG00000028184 | Adgrl2 | chr3 | - | 148526915 | 148527044 | 148521221 | 148523564 | 148528590 | 148528687 | 214,179,166,200 | 0,35,46,49 | 66,114,42,113 | 23,39,25,26 | 278 | 149 | 0.00156 | 0.03815 | 1.0,0.733, | 0.606,0.61 | 0.172 |
| ENSMUSG00000073633 | Fbxo36 | chr1 | + | 84825600 | 84825799 | 84817573 | 84817689 | 84858812 | 84858921 | 16,7,17,16 | 12,24,24,27 | 7,5,4,0 | 28,28,30,43 | 348 | 149 | 1.54E-07 | 3.58E-05 | 0.363,0.11 | 0.097,0.07 | 0.172 |
| ENSMUSG00000036377 | Cracd | chr5 | + | 76988634 | 76988770 | 76970312 | 76970484 | 76996663 | 76996828 | 84,61,133,65 | 23,11,0,2 | 87,46,97,47 | 11,23,14,19 | 285 | 149 | 0.00433 | 0.08075 | 0.656,0.74 | 0.805,0.51 | 0.17 |
| ENSMUSG00000028681 | Ptch2 | chr4 | + | 116968711 | 116968849 | 116968321 | 116968602 | 116971208 | 116971351 | 34,26,52,35 | 0,0,0,0 | 32,10,18,23 | 0,6,0,2 | 287 | 149 | 0.00248 | 0.05419 | 1.0,1.0,1.0 | 1.0,0.464, | 0.17 |
| ENSMUSG00000016763 | Scube1 | chr15 | - | 83535855 | 83535945 | 83522904 | 83523021 | 83538369 | 83538486 | 28,20,62,22 | 11,1,0,6 | 44,26,75,36 | 17,9,26,12 | 239 | 149 | 0.00403 | 0.07687 | 0.613,0.92 | 0.617,0.64 | 0.17 |
| ENSMUSG00000040929 | Rfx3 | chr19 | - | 27900607 | 27900666 | 27878178 | 27878303 | 27988371 | 27988566 | 47,30,54,36 | 16,15,17,12 | 26,23,27,18 | 20,11,17,23 | 208 | 149 | 0.00484 | 0.08757 | 0.678,0.58 | 0.482,0.6, | 0.168 |
| ENSMUSG00000087177 | E130307A | chr10 | - | 39529707 | 39529980 | 39521551 | 39521804 | 39530517 | 39530560 | 99,60,105,97 | 3,2,0,11 | 74,63,38,54 | 6,5,12,6 | 422 | 149 | 0.00281 | 0.0593 | 0.921,0.91 | 0.813,0.81 | 0.168 |
| ENSMUSG00000034902 | Pip5k1c | chr10 | + | 81152525 | 81152794 | 81151712 | 81151790 | 81153159 | 81153237 | 70,73,87,42 | 36,30,80,25 | 36,18,53,39 | 53,44,48,41 | 418 | 149 | 0.00014 | 0.00647 | 0.409,0.46 | 0.195,0.12 | 0.168 |
| ENSMUSG00000086363 | A330102I | chr13 | + | 29202290 | 29202528 | 29200692 | 29200862 | 29207158 | 29207195 | 29,27,45,16 | 14,14,18,24 | 20,22,15,19 | 27,23,29,23 | 387 | 149 | 0.00461 | 0.0845 | 0.444,0.42 | 0.222,0.26 | 0.166 |
| ENSMUSG00000028865 | Cd164l2 | chr4 | + | 132949479 | 132949524 | 132949015 | 132949087 | 132950913 | 132951058 | 33,24,65,35 | 3,0,0,0 | 12,25,29,19 | 13,0,3,1 | 194 | 149 | 0.00014 | 0.00644 | 0.894,1.0, | 0.415,1.0, | 0.165 |
| ENSMUSG00000029452 | Tmem116 | chr5 | + | 121602861 | 121602925 | 121601799 | 121601896 | 121605916 | 121606048 | 29,21,21,14 | 0,0,0,0 | 22,30,11,16 | 0,4,1,7 | 213 | 149 | 2.31E-06 | 0.0003 | 1.0,1.0,1.0 | 1.0,0.84,0 | 0.165 |
| ENSMUSG00000045482 | Trrap | chr5 | + | 144746960 | 144747014 | 144744132 | 144744201 | 144748704 | 144748824 | 95,71,73,50 | 35,32,24,23 | 56,54,39,24 | 35,41,25,30 | 203 | 149 | 0.00074 | 0.02231 | 0.666,0.62 | 0.54,0.492 | 0.164 |
| ENSMUSG00000060798 | Intu | chr3 | + | 40618689 | 40618775 | 40608135 | 40608683 | 40627000 | 40627204 | 34,22,49,33 | 0,4,0,0 | 40,38,39,37 | 6,16,10,0 | 235 | 149 | 4.87E-06 | 0.00051 | 1.0,0.777, | 0.809,0.60 | 0.164 |
| ENSMUSG00000045659 | Plekha7 | chr7 | - | 115747487 | 115747568 | 115744404 | 115744473 | 115753867 | 115753921 | 26,12,19,20 | 26,16,39,25 | 10,12,11,2 | 40,17,64,13 | 230 | 149 | 0.00046 | 0.01576 | 0.393,0.32 | 0.139,0.31 | 0.164 |
| ENSMUSG00000027012 | Dync1i2 | chr2 | + | 71058170 | 71058188 | 71049644 | 71049762 | 71058943 | 71059034 | 333,235,236,162 | 271,227,408,288 | 0,186,158,101 | 370,247,284,224 | 167 | 149 | 8.31E-05 | 0.00435 | 0.523,0.48 | 0.0,0.402, | 0.164 |
| ENSMUSG00000028906 | Epb41 | chr4 | - | 131664325 | 131664775 | 131663049 | 131663178 | 131684953 | 131685130 | 571,466,508,421 | 33,25,76,58 | 252,325,266,338 | 58,66,67,40 | 599 | 149 | 0.00038 | 0.01358 | 0.811,0.82 | 0.519,0.55 | 0.164 |
| ENSMUSG00000032411 | Tfdp2 | chr9 | + | 96172632 | 96172680 | 96169653 | 96169775 | 96177017 | 96177180 | 127,90,126,100 | 1,18,22,11 | 80,83,61,80 | 29,27,12,33 | 197 | 149 | 5.22E-06 | 0.00053 | 0.99,0.791 | 0.676,0.69 | 0.163 |
| ENSMUSG00000084885 | 3010001F | chrX | + | 151168640 | 151168748 | 151152837 | 151153018 | 151199286 | 151199694 | 32,20,16,20 | 0,0,0,0 | 11,21,27,23 | 3,6,0,0 | 257 | 149 | 0.00036 | 0.01317 | 1.0,1.0,1.0 | 0.68,0.67, | 0.162 |
| ENSMUSG00000096929 | A330023F | chr1 | + | 194699782 | 194699859 | 194658682 | 194659108 | 194700908 | 194701248 | 69,84,85,49 | 14,13,20,20 | 50,40,63,67 | 25,19,32,33 | 226 | 149 | 0.00045 | 0.01553 | 0.765,0.81 | 0.569,0.58 | 0.161 |
| ENSMUSG00000028657 | Ppt1 | chr4 | + | 122748172 | 122748271 | 122742225 | 122742328 | 122751302 | 122752871 | 27,8,57,13 | 2,1,0,0 | 13,15,6,17 | 4,0,5,0 | 248 | 149 | 0.00186 | 0.04361 | 0.89,0.828 | 0.661,1.0, | 0.159 |
| ENSMUSG00000042790 | Rnf214 | chr9 | - | 45816067 | 45816217 | 45811043 | 45811554 | 45818035 | 45818159 | 60,35,45,25 | 0,1,0,0 | 43,14,16,18 | 2,1,0,8 | 299 | 149 | 3.40E-05 | 0.00223 | 1.0,0.946, | 0.915,0.87 | 0.157 |
| ENSMUSG00000040648 | Ppip5k2 | chr1 | - | 97648830 | 97649004 | 97647558 | 97647631 | 97651394 | 97651559 | 153,134,145,119 | 18,15,6,32 | 59,91,63,117 | 26,23,16,21 | 323 | 149 | 0.00312 | 0.06382 | 0.797,0.80 | 0.511,0.64 | 0.157 |
| ENSMUSG00000044252 | Osbpl1a | chr18 | - | 12974240 | 12974403 | 12952561 | 12952718 | 12995163 | 12995297 | 716,537,959,582 | 102,0,219,108 | 759,618,644,719 | 230,136,242,156 | 312 | 149 | 1.67E-05 | 0.00129 | 0.77,1.0,0 | 0.612,0.68 | 0.155 |
| ENSMUSG00000025812 | Pard3 | chr8 | + | 128137209 | 128137254 | 128136036 | 128136188 | 128142050 | 128142278 | 155,81,82,100 | 16,4,23,6 | 43,60,31,33 | 13,21,9,9 | 194 | 149 | 0.00013 | 0.00615 | 0.882,0.94 | 0.718,0.68 | 0.153 |
| ENSMUSG00000023007 | Prpf40b | chr15 | + | 99199667 | 99199865 | 99193121 | 99193154 | 99201163 | 99201244 | 12,30,27,22 | 50,27,31,29 | 11,7,10,7 | 34,33,51,66 | 347 | 149 | 1.05E-06 | 0.00016 | 0.093,0.32 | 0.122,0.08 | 0.152 |
| ENSMUSG00000021140 | Pcnx | chr12 | + | 81975015 | 81975148 | 81964438 | 81966154 | 81993704 | 81993795 | 117,68,150,82 | 39,44,51,50 | 53,64,60,49 | 44,51,46,52 | 282 | 149 | 0.00064 | 0.02001 | 0.613,0.45 | 0.389,0.39 | 0.152 |
| ENSMUSG00000028556 | Dock7 | chr4 | - | 98881883 | 98881973 | 98880605 | 98880695 | 98889269 | 98889439 | 63,30,53,79 | 6,0,0,0 | 40,28,35,38 | 18,0,0,11 | 239 | 149 | 0.00022 | 0.00911 | 0.867,1.0, | 0.581,1.0, | 0.151 |
| ENSMUSG00000023044 | Csad | chr15 | - | 102110878 | 102111009 | 102097321 | 102097397 | 102112130 | 102112151 | 40,33,38,31 | 0,0,0,0 | 31,30,37,35 | 9,4,1,0 | 280 | 149 | 1.25E-06 | 0.00019 | 1.0,1.0,1.0 | 0.647,0.8, | 0.15 |
| ENSMUSG00000021301 | Hecw1 | chr13 | - | 14486273 | 14486375 | 14481367 | 14481518 | 14490643 | 14491751 | 61,38,59,22 | 5,0,0,0 | 16,51,39,42 | 5,1,5,5 | 251 | 149 | 0.0005 | 0.01701 | 0.879,1.0, | 0.655,0.96 | 0.15 |
| ENSMUSG00000022607 | Ptk2 | chr15 | - | 73231874 | 73232186 | 73202697 | 73202864 | 73236889 | 73237044 | 45,44,80,26 | 0,1,2,3 | 37,11,32,12 | 6,0,3,3 | 461 | 149 | 0.00508 | 0.09048 | 1.0,0.934, | 0.666,1.0, | 0.149 |
| ENSMUSG00000026743 | Mllt10 | chr2 | + | 18130944 | 18131040 | 18114592 | 18114702 | 18151587 | 18151683 | 73,46,91,51 | 0,0,6,4 | 30,43,38,46 | 13,14,1,0 | 245 | 149 | 0.00294 | 0.06107 | 1.0,1.0,0.9 | 0.584,0.65 | 0.149 |
| ENSMUSG00000061306 | Slc38a10 | chr11 | - | 119999292 | 119999379 | 119997259 | 119997436 | 120000257 | 120000362 | 80,92,67,52 | 40,41,55,19 | 58,34,40,41 | 46,42,29,43 | 236 | 149 | 0.00456 | 0.08398 | 0.558,0.58 | 0.443,0.33 | 0.148 |
| ENSMUSG00000071267 | Zfp942 | chr17 | - | 22160706 | 22160831 | 22149346 | 22149473 | 22172868 | 22173034 | 36,64,62,55 | 8,0,9,3 | 23,21,41,32 | 4,4,13,8 | 274 | 149 | 0.00457 | 0.08411 | 0.71,1.0,0 | 0.758,0.74 | 0.148 |
| ENSMUSG00000021271 | Zfp839 | chr12 | + | 110827229 | 110827457 | 110821433 | 110822306 | 110830397 | 110830490 | 57,45,61,48 | 18,16,17,15 | 34,44,34,39 | 12,30,22,27 | 377 | 149 | 0.00398 | 0.07616 | 0.556,0.52 | 0.528,0.36 | 0.147 |
| ENSMUSG00000041670 | Rims1 | chr1 | - | 22496120 | 22496183 | 22467674 | 22467787 | 22497711 | 22497837 | 86,65,130,92 | 41,49,85,69 | 71,74,71,75 | 113,51,88,113 | 212 | 149 | 0.00069 | 0.02113 | 0.596,0.48 | 0.306,0.50 | 0.147 |
| ENSMUSG00000004018 | Fancl | chr11 | + | 26353321 | 26353378 | 26349648 | 26349709 | 26357547 | 26357648 | 42,46,49,56 | 0,0,0,0 | 20,26,48,17 | 5,0,0,6 | 206 | 149 | 5.92E-05 | 0.00338 | 1.0,1.0,1.0 | 0.743,1.0, | 0.146 |
| ENSMUSG00000030922 | Lyrm1 | chr7 | + | 119499333 | 119499531 | 119495089 | 119495113 | 119509026 | 119509185 | 48,43,46,20 | 0,0,8,2 | 25,41,2,35 | 3,14,0,10 | 347 | 149 | 0.0048 | 0.08704 | 1.0,1.0,0.7 | 0.782,0.55 | 0.146 |
| ENSMUSG00000049295 | Zfp219 | chr14 | - | 52257647 | 52257831 | 52247426 | 52247515 | 52257979 | 52258143 | 11,6,13,14 | 27,27,27,14 | 2,0,4,0 | 26,35,14,17 | 333 | 149 | 3.45E-08 | 1.09E-05 | 0.154,0.09 | 0.033,0.0, | 0.146 |
| ENSMUSG00000040433 | Zbtb38 | chr9 | - | 96600262 | 96600362 | 96564822 | 96571082 | 96613344 | 96613728 | 83,89,150,57 | 14,11,32,4 | 72,44,59,49 | 22,8,18,22 | 249 | 149 | 0.00396 | 0.07598 | 0.78,0.829 | 0.662,0.76 | 0.145 |
| ENSMUSG00000033671 | Cep350 | chr1 | - | 155762881 | 155763004 | 155761652 | 155761852 | 155770339 | 155770343 | 108,123,119,122 | 20,26,18,23 | 81,65,37,74 | 30,20,17,23 | 272 | 149 | 0.00054 | 0.01793 | 0.747,0.72 | 0.597,0.64 | 0.144 |
| ENSMUSG00000073468 | Sft2d1 | chr17 | + | 8536681 | 8536703 | 8529970 | 8530117 | 8537675 | 8537762 | 353,302,379,298 | 129,102,127,118 | 315,256,255,183 | 205,129,171,164 | 171 | 149 | 1.47E-07 | 3.44E-05 | 0.705,0.72 | 0.572,0.63 | 0.143 |

| | | | | | | | | | | | | | | | | | | | | |
|---|---|---|---|---|---|---|---|---|---|---|---|---|---|---|---|---|---|---|---|---|
| ENSMUSG00000029538 | Srsf9 | chr5 | + | 115466405 | 115466763 | 115465453 | 115465558 | 115468556 | 115468717 | 9,327,741,069,770 | 120,77,117,101 | 706,669,625,579 | 167,120,133,146 | 507 | 149 | 2.10E-10 | 2.17E-07 | 0.695,0.7 | 0.554,0.6 | 0.142 |
| ENSMUSG00000087174 | 5530601H | chrX | - | 104113216 | 104113305 | 104112069 | 104112234 | 104113605 | 104113693 | 42,58,56,42 | 2,7,4,5 | 30,38,24,26 | 12,11,5,2 | 238 | 149 | 0.00423 | 0.07938 | 0.929,0.8 | 0.61,0.68 | 0.142 |
| ENSMUSG00000043987 | Cep164 | chr9 | - | 45693249 | 45693546 | 45690987 | 45691134 | 45696180 | 45696327 | 172,110,173,112 | 1,1,5,0 | 103,88,87,78 | 8,8,1,9 | 446 | 149 | 1.77E-06 | 0.00024 | 0.983,0.9 | 0.811,0.7 | 0.142 |
| ENSMUSG00000091474 | 2610021A | chr7 | + | 41252859 | 41252958 | 41248653 | 41248824 | 41261099 | 41261343 | 67,55,115,49 | 55,52,89,42 | 39,43,38,45 | 54,55,64,100 | 248 | 149 | 1.28E-05 | 0.00106 | 0.423,0.3 | 0.303,0.3 | 0.141 |
| ENSMUSG00000026987 | Baz2b | chr2 | - | 59790164 | 59790449 | 59788775 | 59788836 | 59792364 | 59792974 | 176,189,227,167 | 25,24,6,17 | 123,144,113,122 | 24,17,26,29 | 434 | 149 | 0.00187 | 0.04371 | 0.707,0.7 | 0.638,0.7 | 0.141 |
| ENSMUSG00000041769 | Ppp2r2d | chr7 | + | 138466749 | 138466846 | 138462565 | 138462638 | 138470138 | 138470236 | 25,26,23,8 | 1,0,0,3 | 9,11,23,21 | 4,1,5,3 | 246 | 149 | 0.00115 | 0.03036 | 0.938,1.0, | 0.577,0.8 | 0.141 |
| ENSMUSG00000007888 | Crlf1 | chr8 | + | 70951246 | 70951528 | 70945807 | 70946050 | 70951742 | 70951872 | 25,1,55,16 | 0,0,3,0 | 71,68,36,34 | 2,0,5,6 | 431 | 149 | 0.00136 | 0.0345 | 1.0,1.0,0. | 0.925,1.0, | 0.141 |
| ENSMUSG00000025949 | Pikfyve | chr1 | + | 65231689 | 65231725 | 65231303 | 65231453 | 65234814 | 65234933 | 53,35,68,38 | 6,0,1,10 | 44,51,37,43 | 2,20,10,16 | 185 | 149 | 0.00135 | 0.03434 | 0.877,1.0, | 0.947,0.6 | 0.14 |
| ENSMUSG00000085396 | Firre | chrX | - | 49695839 | 49695981 | 49694884 | 49695039 | 49697347 | 49697481 | 21,7,0,0 | 18,16,41,26 | 0,0,0,0 | 31,3,6,12 | 291 | 149 | 3.64E-06 | 0.00042 | 0.374,0.1 | 0.0,0.0,0. | 0.139 |
| ENSMUSG00000047342 | Zfp286 | chr11 | - | 62675704 | 62675835 | 62674497 | 62674587 | 62678784 | 62678870 | 78,53,48,58 | 0,0,3,0 | 51,64,61,42 | 8,7,11,0 | 280 | 149 | 9.14E-08 | 2.32E-05 | 1.0,1.0,0. | 0.772,0.8 | 0.137 |
| ENSMUSG00000040003 | Magi2 | chr5 | + | 20748582 | 20748624 | 20739335 | 20739518 | 20755204 | 20755296 | 71,56,74,65 | 2,9,3,7 | 44,49,32,44 | 4,20,7,12 | 191 | 149 | 0.00023 | 0.00935 | 0.965,0.8 | 0.896,0.6 | 0.137 |
| ENSMUSG00000043079 | Synpo | chr18 | - | 60742982 | 60743073 | 60733277 | 60737562 | 60757242 | 60757308 | 342,215,335,168 | 129,106,232,102 | 207,205,152,138 | 181,149,142,166 | 240 | 149 | 5.63E-05 | 0.00325 | 0.622,0.5 | 0.415,0.4 | 0.136 |
| ENSMUSG00000054715 | Zscan22 | chr7 | + | 12637527 | 12638012 | 12631762 | 12631907 | 12640160 | 12641464 | 68,46,53,58 | 34,21,47,15 | 41,32,26,36 | 30,46,30,25 | 634 | 149 | 0.0044 | 0.08153 | 0.32,0.34, | 0.243,0.1 | 0.135 |
| ENSMUSG00000033530 | Ttc7b | chr12 | - | 100306755 | 100306806 | 100298837 | 100298978 | 100314194 | 100314292 | 456,364,418,292 | 170,122,256,166 | 341,274,248,246 | 329,151,247,184 | 200 | 149 | 0.00019 | 0.00824 | 0.666,0.6 | 0.436,0.5 | 0.133 |
| ENSMUSG00000097040 | 2610316D | chr3 | - | 45241766 | 45241851 | 45236429 | 45236599 | 45281500 | 45281597 | 10,18,28,7 | 72,50,40,95 | 0,3,2,2 | 42,52,39,57 | 234 | 149 | 1.15E-08 | 4.48E-06 | 0.081,0.1 | 0.0,0.035, | 0.133 |
| ENSMUSG00000100826 | Snhg14 | chr7 | - | 59328606 | 59328753 | 59326140 | 59326232 | 59331151 | 59331298 | 240,211,306,186 | 0,11,0,0 | 139,101,47,144 | 5,0,23,5 | 296 | 149 | 0.00017 | 0.00743 | 1.0,0.906, | 0.933,1.0, | 0.133 |
| ENSMUSG00000031371 | Haus7 | chrX | - | 72492416 | 72492508 | 72491797 | 72491956 | 72496604 | 72496666 | 26,20,16,40 | 4,0,0,0 | 27,16,19,29 | 5,2,1,6 | 241 | 149 | 0.00042 | 0.01481 | 0.801,1.0, | 0.77,0.83 | 0.132 |
| ENSMUSG00000039449 | Prpf18 | chr2 | - | 4652907 | 4652934 | 4650379 | 4650484 | 4653051 | 4653129 | 380,211,353,253 | 72,47,76,68 | 150,246,144,165 | 106,67,62,68 | 176 | 149 | 3.29E-05 | 0.00218 | 0.817,0.7 | 0.545,0.7 | 0.132 |
| ENSMUSG00000018900 | Slc22a5 | chr11 | - | 53774486 | 53774590 | 53766834 | 53766989 | 53781972 | 53782486 | 52,28,46,41 | 0,2,0,0 | 39,42,41,28 | 4,3,4,5 | 253 | 149 | 2.46E-06 | 0.00032 | 1.0,0.892, | 0.852,0.8 | 0.131 |
| ENSMUSG00000059921 | Unc5c | chr3 | + | 141492617 | 141492674 | 141476778 | 141476943 | 141494631 | 141494823 | 18,15,21,12 | 31,35,55,28 | 7,5,5,8 | 35,31,37,35 | 206 | 149 | 0.00028 | 0.01082 | 0.296,0.2 | 0.126,0.1 | 0.131 |
| ENSMUSG00000021619 | Atg10 | chr13 | - | 91188968 | 91189107 | 91170688 | 91170786 | 91356436 | 91356552 | 33,28,53,51 | 0,0,0,0 | 39,32,49,54 | 3,3,5,2 | 288 | 149 | 4.56E-06 | 0.00049 | 1.0,1.0,1. | 0.871,0.8 | 0.129 |
| ENSMUSG00000021760 | Gpx8 | chr13 | - | 113181965 | 113182227 | 113179292 | 113179833 | 113182705 | 113182944 | 105,66,130,77 | 1,6,4,0 | 98,81,113,102 | 5,15,9,8 | 411 | 149 | 0.00036 | 0.01318 | 0.974,0.8, | 0.877,0.6 | 0.129 |
| ENSMUSG00000028550 | Atg4c | chr4 | + | 99116792 | 99116871 | 99112650 | 99112787 | 99123303 | 99123423 | 106,72,115,90 | 2,0,0,0 | 77,105,86,73 | 16,12,0,8 | 228 | 149 | 7.48E-11 | 9.58E-08 | 0.972,1.0, | 0.759,0.8 | 0.127 |
| ENSMUSG00000000827 | Tpd52l2 | chr2 | + | 181153366 | 181153408 | 181149958 | 181150060 | 181154831 | 181154849 | 208,183,210,164 | 52,59,61,67 | 145,115,118,98 | 96,68,51,53 | 191 | 149 | 1.78E-05 | 0.00136 | 0.757,0.7 | 0.541,0.5 | 0.127 |
| ENSMUSG00000034573 | Ptpn13 | chr5 | + | 103649232 | 103649778 | 103640088 | 103640176 | 103664193 | 103664289 | 237,165,207,208 | 5,13,11,6 | 132,181,121,142 | 10,17,12,13 | 695 | 149 | 0.00269 | 0.05745 | 0.91,0.73 | 0.739,0.6 | 0.126 |
| ENSMUSG00000023764 | Sfi1 | chr11 | - | 3085282 | 3085542 | 3083071 | 3083207 | 3085653 | 3085830 | 164,123,157,155 | 0,2,0,0 | 65,101,109,66 | 2,10,5,4 | 409 | 149 | 2.87E-09 | 1.42E-06 | 1.0,0.957, | 0.922,0.7 | 0.126 |
| ENSMUSG00000025728 | Pigq | chr17 | - | 26153067 | 26153247 | 26150435 | 26150547 | 26153731 | 26153852 | 23,15,56,27 | 0,0,4,0 | 17,13,33,26 | 0,1,2,7 | 329 | 149 | 0.00251 | 0.0546 | 1.0,1.0,0. | 1.0,0.855, | 0.125 |
| ENSMUSG00000037773 | Pced1a | chr2 | - | 130265331 | 130265472 | 130264732 | 130264812 | 130266412 | 130266607 | 36,42,67,19 | 5,0,1,0 | 50,36,37,23 | 1,2,7,6 | 290 | 149 | 0.00485 | 0.08766 | 0.787,1.0, | 0.963,0.9 | 0.125 |
| ENSMUSG00000045962 | Wnk1 | chr6 | - | 119930731 | 119931193 | 119928858 | 119928956 | 119933031 | 119933157 | 412,319,487,339 | 85,66,105,97 | 291,274,227,222 | 129,75,102,83 | 611 | 149 | 3.53E-05 | 0.00229 | 0.542,0.5 | 0.355,0.4 | 0.125 |
| ENSMUSG00000036712 | Cyld | chr8 | + | 89444917 | 89444926 | 89436518 | 89436624 | 89445925 | 89446024 | 13,11,258,0 | 319,270,312,216 | 0,2,0,0 | 193,171,152,171 | 158 | 149 | 7.34E-08 | 2.01E-05 | 0.037,0.0 | 0.0,0.011, | 0.125 |
| ENSMUSG00000039765 | Cc2d2a | chr5 | + | 43874278 | 43874293 | 43873095 | 43873187 | 43875894 | 43876062 | 14,0,0,12 | 44,35,38,29 | 0,0,0,0 | 26,42,36,40 | 164 | 149 | 3.91E-08 | 1.19E-05 | 0.224,0.0, | 0.0,0.0,0. | 0.124 |
| ENSMUSG00000110195 | Pde2a | chr7 | + | 101110686 | 101110748 | 101101334 | 101101640 | 101130577 | 101130667 | 110,106,104,74 | 0,0,0,1 | 62,107,83,70 | 1,12,7,16 | 211 | 149 | 4.52E-12 | 8.96E-09 | 1.0,1.0,1. | 0.978,0.8 | 0.123 |
| ENSMUSG00000033526 | Ppip5k1 | chr2 | - | 121153674 | 121153857 | 121152170 | 121152273 | 121157332 | 121157395 | 113,64,78,86 | 91,78,101,72 | 45,40,44,52 | 92,62,108,105 | 332 | 149 | 1.83E-06 | 0.00025 | 0.358,0.2 | 0.18,0.22 | 0.123 |
| ENSMUSG00000027799 | Nbea | chr3 | - | 55901468 | 55901477 | 55899754 | 55899935 | 55907907 | 55908061 | 0,103,0,0 | 131,100,250,95 | 0,0,0,0 | 117,78,85,71 | 158 | 149 | 0.00138 | 0.0348 | 0.0,0.493, | 0.0,0.0,0. | 0.123 |
| ENSMUSG00000026159 | Agfg1 | chr1 | + | 82869180 | 82869228 | 82864095 | 82864188 | 82871144 | 82871303 | 248,185,217,189 | 34,21,39,29 | 165,154,130,119 | 56,44,33,39 | 197 | 149 | 7.25E-07 | 0.00012 | 0.847,0.8 | 0.69,0.72 | 0.123 |
| ENSMUSG00000026975 | Dph7 | chr2 | + | 24853467 | 24853555 | 24852434 | 24852822 | 24855571 | 24855666 | 33,23,23,39 | 0,0,0,0 | 18,30,16,16 | 11,0,0,0 | 237 | 149 | 0.00324 | 0.06572 | 1.0,1.0,1. | 0.507,1.0, | 0.123 |
| ENSMUSG00000037519 | Ppfia1 | chr7 | - | 144056603 | 144056633 | 144053988 | 144054140 | 144058639 | 144058871 | 160,135,157,149 | 30,31,38,22 | 88,92,90,62 | 39,24,37,28 | 179 | 149 | 0.00012 | 0.00565 | 0.816,0.7 | 0.653,0.7 | 0.123 |
| ENSMUSG00000032463 | Faim | chr9 | + | 98872967 | 98873024 | 98868426 | 98868565 | 98874153 | 98874286 | 141,125,201,126 | 48,45,71,35 | 178,112,137,164 | 98,65,66,105 | 206 | 149 | 3.94E-05 | 0.00249 | 0.68,0.66 | 0.568,0.5 | 0.122 |
| ENSMUSG00000041617 | Ccdc74a | chr16 | + | 17466658 | 17466824 | 17465930 | 17465990 | 17467858 | 17467938 | 44,34,100,38 | 0,4,0,0 | 84,72,47,34 | 12,5,8,1 | 315 | 149 | 4.07E-06 | 0.00045 | 1.0,0.801, | 0.768,0.8 | 0.121 |
| ENSMUSG00000032641 | Gpr19 | chr6 | - | 134854179 | 134854297 | 134846119 | 134847445 | 134864592 | 134864748 | 23,18,40,10 | 43,51,41,37 | 0,11,13,8 | 78,28,59,38 | 267 | 149 | 2.17E-05 | 0.00158 | 0.23,0.16 | 0.0,0.18,0 | 0.121 |
| ENSMUSG00000026643 | Nmt2 | chr2 | + | 3306433 | 3306476 | 3305848 | 3305984 | 3310503 | 3310648 | 201,172,266,171 | 87,51,128,93 | 174,177,202,194 | 120,130,160,124 | 192 | 149 | 9.83E-05 | 0.00496 | 0.642,0.7 | 0.529,0.5 | 0.121 |
| ENSMUSG00000054162 | Spock3 | chr8 | + | 63566467 | 63566476 | 63404833 | 63405022 | 63566576 | 63566622 | 0,27,0,0 | 42,27,46,46 | 0,0,0,0 | 28,24,48,60 | 158 | 149 | 0.00182 | 0.04285 | 0.0,0.485, | 0.0,0.0,0. | 0.121 |
| ENSMUSG00000051736 | Fam229b | chr10 | - | 38996253 | 38996392 | 38994799 | 38995030 | 38998172 | 38998328 | 120,138,116,68 | 18,0,5,13 | 81,89,106,100 | 15,12,21,23 | 288 | 149 | 0.00109 | 0.02908 | 0.775,1.0, | 0.736,0.7 | 0.121 |
| ENSMUSG00000059439 | Bcas3 | chr11 | + | 85434753 | 85434798 | 85422618 | 85422769 | 85445015 | 85445140 | 177,141,232,112 | 55,53,87,57 | 89,114,88,89 | 84,65,46,55 | 194 | 149 | 0.0006 | 0.01918 | 0.712,0.6 | 0.449,0.5 | 0.121 |
| ENSMUSG00000027012 | Dync1i2 | chr2 | + | 71063992 | 71064052 | 71058943 | 71059034 | 71066258 | 71066374 | 1,267,102,213,401,050 | 123,80,151,113 | 664,712,682,711 | 200,147,137,146 | 209 | 149 | 0 | 0 | 0.88,0.90 | 0.703,0.7 | 0.12 |
| ENSMUSG00000062115 | Rai1 | chr11 | + | 60031066 | 60031196 | 59995838 | 59996220 | 60075921 | 60076257 | 221,193,195,158 | 11,18,10,10 | 118,150,114,152 | 18,24,21,20 | 279 | 149 | 5.14E-06 | 0.00053 | 0.915,0.8 | 0.778,0.7 | 0.12 |
| ENSMUSG00000034006 | Slc66a2 | chr18 | + | 80315676 | 80315730 | 80306634 | 80306768 | 80326490 | 80326707 | 10,24,45,34 | 131,77,87,51 | 1,10,7,18 | 66,56,60,70 | 203 | 149 | 0.00127 | 0.03292 | 0.053,0.1 | 0.011,0.1 | 0.119 |
| ENSMUSG00000028759 | Hp1bp3 | chr4 | + | 137948836 | 137949029 | 137944578 | 137944605 | 137949399 | 137949499 | 229,172,216,160 | 120,79,164,112 | 117,124,110,119 | 121,130,107,112 | 342 | 149 | 7.63E-05 | 0.00408 | 0.454,0.4 | 0.296,0.2 | 0.119 |
| ENSMUSG00000004110 | Cacna1e | chr1 | - | 154291806 | 154291903 | 154289721 | 154289827 | 154293695 | 154293792 | 195,183,276,119 | 9,2,13,11 | 133,151,100,109 | 31,2,18,21 | 246 | 149 | 2.89E-05 | 0.00198 | 0.929,0.9 | 0.722,0.9 | 0.119 |
| ENSMUSG00000032878 | Ccdc85a | chr11 | - | 28342853 | 28342952 | 28338843 | 28338948 | 28346045 | 28346165 | 193,154,358,196 | 32,41,60,49 | 149,106,172,139 | 59,41,54,50 | 248 | 149 | 6.88E-05 | 0.00379 | 0.784,0.6 | 0.603,0.6 | 0.118 |
| ENSMUSG00000024524 | Gnal | chr18 | + | 67266795 | 67267041 | 67221405 | 67221944 | 67267283 | 67267356 | 3,481,271,230,452,190 | 146,82,237,90 | 1,199,211,810,441,560 | 193,99,166,155 | 395 | 149 | 4.50E-08 | 1.33E-05 | 0.9,0.926, | 0.701,0.8 | 0.118 |
| ENSMUSG00000028809 | Srrm1 | chr4 | - | 135072506 | 135072788 | 135071838 | 135071980 | 135074045 | 135074168 | 511,529,667,380 | 326,313,400,226 | 268,269,254,238 | 356,261,247,247 | 431 | 149 | 2.41E-10 | 2.34E-07 | 0.351,0.3 | 0.207,0.2 | 0.118 |
| ENSMUSG00000033545 | Znrf1 | chr8 | + | 112333378 | 112333480 | 112264974 | 112265098 | 112335924 | 112335996 | 11,28,31,26 | 0,0,0,0 | 28,7,15,17 | 0,0,0,9 | 251 | 149 | 0.00553 | 0.09584 | 1.0,1.0,1. | 1.0,1.0,1. | 0.118 |
| ENSMUSG00000042557 | Sin3a | chr9 | + | 56988378 | 56988486 | 56979323 | 56979347 | 56993864 | 56994085 | 33,27,11,23 | 0,0,0,0 | 13,35,16,13 | 0,6,3,0 | 257 | 149 | 0.00029 | 0.01114 | 1.0,1.0,1. | 1.0,0.772, | 0.118 |
| ENSMUSG00000024462 | Gabbr1 | chr17 | + | 37360544 | 37360689 | 37359469 | 37359498 | 37361890 | 37362019 | 74,88,95,97 | 3,1,7,0 | 67,68,44,47 | 7,0,5,12 | 294 | 149 | 0.00159 | 0.03861 | 0.926,0.9 | 0.829,1.0, | 0.117 |
| ENSMUSG00000040690 | Col16a1 | chr4 | + | 129943398 | 129943532 | 129941737 | 129941876 | 129945447 | 129945522 | 20,24,30,13 | 0,0,0,0 | 27,32,23,37 | 3,0,5,0 | 283 | 149 | 0.00513 | 0.09091 | 1.0,1.0,1. | 0.826,1.0, | 0.117 |
| ENSMUSG00000058402 | Zfp420 | chr7 | + | 29560440 | 29560493 | 29559403 | 29559518 | 29572316 | 29572358 | 77,45,79,34 | 5,0,0,2 | 43,15,50,46 | 8,2,4,7 | 202 | 149 | 5.85E-05 | 0.00335 | 0.919,1.0, | 0.799,0.8 | 0.117 |
| ENSMUSG00000020831 | 0610010K | chr11 | - | 70126521 | 70126626 | 70126031 | 70126290 | 70126849 | 70127037 | 156,97,126,113 | 8,10,18,8 | 120,116,94,61 | 27,12,15,20 | 254 | 149 | 0.00145 | 0.0363 | 0.92,0.85 | 0.723,0.8 | 0.117 |
| ENSMUSG00000035126 | Dnai4 | chr4 | - | 102923456 | 102923683 | 102919492 | 102919644 | 102929845 | 102930040 | 96,95,158,78 | 2,3,0,1 | 63,73,44,58 | 9,0,3,6 | 376 | 149 | 6.57E-05 | 0.00366 | 0.95,0.92 | 0.735,1.0, | 0.116 |
| ENSMUSG00000052151 | Plpp2 | chr10 | - | 79366710 | 79366862 | 79366331 | 79366606 | 79369457 | 79369630 | 33,36,41,40 | 1,0,0,0 | 81,23,34,31 | 3,4,4,0 | 301 | 149 | 0.00012 | 0.0058 | 0.942,1.0, | 0.93,0.74, | 0.116 |
| ENSMUSG00000031174 | Rpgr | chrX | - | 10028840 | 10028929 | 10024610 | 10025088 | 10030960 | 10031018 | 100,101,67,84 | 3,6,9,11 | 43,77,58,59 | 10,16,13,7 | 238 | 149 | 0.00134 | 0.03417 | 0.954,0.9 | 0.729,0.7 | 0.115 |
| ENSMUSG00000038056 | Kmt2c | chr5 | - | 25535793 | 25535898 | 25534444 | 25534573 | 25543410 | 25543579 | 126,107,166,112 | 6,5,13,19 | 93,101,59,55 | 10,13,12,18 | 254 | 149 | 0.00499 | 0.08923 | 0.925,0.9 | 0.845,0.8 | 0.115 |
| ENSMUSG00000040481 | Bptf | chr11 | - | 106977537 | 106977723 | 106973338 | 106973526 | 106986541 | 106986751 | 183,126,166,118 | 3,0,4,0 | 124,117,91,64 | 2,12,10,4 | 335 | 149 | 5.23E-07 | 9.48E-05 | 0.964,1.0, | 0.965,0.8 | 0.114 |
| ENSMUSG00000060961 | Slc4a4 | chr5 | + | 89380622 | 89380719 | 89376684 | 89376846 | 89382655 | 89382720 | 630,379,418,429 | 204,191,292,238 | 256,397,224,337 | 255,220,235,232 | 246 | 149 | 0.00417 | 0.0786 | 0.652,0.5 | 0.378,0.5 | 0.113 |
| ENSMUSG00000038170 | Pde4dip | chr3 | - | 97614187 | 97614373 | 97610482 | 97610734 | 97617536 | 97617760 | 382,285,412,272 | 9,5,25,15 | 207,264,230,172 | 22,12,28,28 | 335 | 149 | 3.46E-06 | 0.00041 | 0.95,0.96 | 0.807,0.9 | 0.113 |
| ENSMUSG00000026596 | Abl2 | chr1 | + | 156448408 | 156448471 | 156443492 | 156443628 | 156450054 | 156450225 | 58,18,32,43 | 0,0,0,0 | 36,57,23,17 | 1,0,6,2 | 212 | 149 | 0.00298 | 0.06169 | 1.0,1.0,1. | 0.962,1.0, | 0.113 |
| ENSMUSG00000038879 | Nipal2 | chr15 | - | 34582866 | 34582955 | 34578853 | 34578917 | 34600160 | 34600257 | 48,42,48,35 | 0,0,2,0 | 36,29,16,43 | 2,2,5,0 | 238 | 149 | 0.00231 | 0.05161 | 1.0,1.0,0. | 0.918,0.9 | 0.113 |
| ENSMUSG00000049971 | Glt1d1 | chr5 | + | 127721382 | 127721531 | 127709325 | 127709415 | 127727746 | 127727852 | 284,247,243,221 | 0,0,7,0 | 170,221,163,153 | 9,17,24,4 | 298 | 149 | 9.31E-14 | 3.69E-10 | 1.0,1.0,0. | 0.904,0.8 | 0.113 |
| ENSMUSG00000033216 | Eefsec | chr6 | - | 88332464 | 88332629 | 88274558 | 88275218 | 88423187 | 88423466 | 71,66,65,54 | 0,0,0,0 | 71,58,65,57 | 0,7,10,0 | 314 | 149 | 7.48E-07 | 0.00013 | 1.0,1.0,1. | 1.0,0.797, | 0.112 |
| ENSMUSG00000023330 | Dtwd1 | chr2 | + | 126000329 | 126000473 | 125996460 | 125996780 | 126001689 | 126001948 | 177,130,177,125 | 4,0,0,0 | 101,78,72,104 | 17,11,1,0 | 293 | 149 | 1.38E-08 | 4.95E-06 | 0.957,1.0, | 0.751,0.7 | 0.112 |
| ENSMUSG00000021969 | Zdhhc20 | chr14 | - | 58075047 | 58075124 | 58070159 | 58074103 | 58076546 | 58076662 | 266,253,310,263 | 40,54,45,40 | 152,172,117,158 | 46,34,37,63 | 226 | 149 | 8.14E-05 | 0.00428 | 0.814,0.7 | 0.685,0.7 | 0.112 |
| ENSMUSG00000022668 | Gtpbp8 | chr16 | - | 44564100 | 44564231 | 44562866 | 44562966 | 44566361 | 44566646 | 74,49,91,91 | 3,0,0,1 | 46,29,77,40 | 6,7,1,0 | 280 | 149 | 0.00013 | 0.00628 | 0.929,1.0, | 0.803,0.6 | 0.111 |
| ENSMUSG00000021693 | Kif2a | chr13 | - | 107106211 | 107106325 | 107105145 | 107105296 | 107111002 | 107111070 | 340,287,487,405 | 20,16,27,10 | 300,208,349,273 | 39,35,37,35 | 263 | 149 | 3.88E-10 | 3.12E-07 | 0.906,0.9 | 0.813,0.7 | 0.111 |
| ENSMUSG00000025006 | Sorbs1 | chr19 | - | 40298024 | 40298777 | 40288686 | 40288746 | 40300112 | 40300307 | 130,612,381,133,927 | 25,0,30,15 | 790,810,612,805 | 52,40,21,13 | 902 | 149 | 1.31E-06 | 0.00019 | 0.896,1.0, | 0.715,0.7 | 0.111 |

| | | | | | | | | | | | | | | | | | | | | |
|---|---|---|---|---|---|---|---|---|---|---|---|---|---|---|---|---|---|---|---|---|
| ENSMUSG00000074892 | B3galt5 | chr16 | + | 96098391 | 96098563 | 96074810 | 96075097 | 96114798 | 96114975 | 45,31,51,37 | 91,37,78,82 | 10,40,13,20 | 86,68,78,90 | 321 | 149 | 1.63E-05 | 0.00127 | 0.187,0.2 | 0.051,0.2 | 0.11 |
| ENSMUSG00000029475 | Kdm2b | chr5 | - | 123019699 | 123019813 | 123019082 | 123019121 | 123020026 | 123020287 | 73,71,88,60 | 10,1,4,9 | 69,66,62,51 | 19,13,8,6 | 263 | 149 | 0.00511 | 0.09074 | 0.805,0.9 | 0.673,0.7 | 0.11 |
| ENSMUSG00000012126 | Ubxn11 | chr4 | + | 133843341 | 133843468 | 133836870 | 133836969 | 133850702 | 133850914 | 45,37,66,55 | 0,0,0,3 | 36,20,31,26 | 0,8,2,0 | 276 | 149 | 0.00183 | 0.04303 | 1.0,1.0,1. | 1.0,0.574, | 0.11 |
| ENSMUSG00000035268 | Pkig | chr2 | + | 163501069 | 163501155 | 163500334 | 163500412 | 163563047 | 163563118 | 11,1,1,8 | 21,3,28,18 | 1,6,1,0 | 39,19,15,15 | 235 | 149 | 0.00266 | 0.05713 | 0.249,0.1 | 0.016,0.1 | 0.11 |
| ENSMUSG00000078773 | Rad54b | chr4 | + | 11612617 | 11612818 | 11612371 | 11612438 | 11615442 | 11615805 | 18,23,17,10 | 1,0,0,0 | 22,25,27,19 | 5,0,3,0 | 350 | 149 | 0.00132 | 0.03384 | 0.885,1.0, | 0.652,1.0, | 0.11 |
| ENSMUSG00000019817 | Plagl1 | chr10 | + | 12967518 | 12967556 | 12966575 | 12966768 | 12981598 | 12981637 | 7,0,0,13 | 34,18,44,24 | 0,0,0,0 | 29,10,37,28 | 187 | 149 | 1.21E-06 | 0.00018 | 0.141,0.0, | 0.0,0.0,0. | 0.11 |
| ENSMUSG00000020776 | Fbf1 | chr11 | - | 116057958 | 116058054 | 116056657 | 116056738 | 116058387 | 116058974 | 26,19,30,26 | 4,0,0,0 | 22,19,25,32 | 4,2,0,7 | 245 | 149 | 0.00104 | 0.02812 | 0.798,1.0, | 0.77,0.85 | 0.11 |
| ENSMUSG00000055923 | Aasdh | chr5 | - | 77025373 | 77025505 | 77023781 | 77024269 | 77026250 | 77026373 | 63,50,21,39 | 0,0,0,0 | 23,49,36,43 | 3,3,3,0 | 281 | 149 | 0.00135 | 0.03424 | 1.0,1.0,1. | 0.803,0.8 | 0.109 |
| ENSMUSG00000033216 | Eefsec | chr6 | - | 88353185 | 88353393 | 88274558 | 88275218 | 88423187 | 88423484 | 95,92,79,107 | 0,0,0,0 | 82,89,63,47 | 0,7,10,0 | 357 | 149 | 3.97E-07 | 7.74E-05 | 1.0,1.0,1. | 1.0,0.841, | 0.109 |
| ENSMUSG00000021870 | Slmap | chr14 | - | 26180564 | 26180615 | 26163729 | 26163789 | 26181058 | 26181109 | 272,195,157,230 | 0,41,0,26 | 86,159,77,151 | 27,0,25,26 | 200 | 149 | 0.00393 | 0.07571 | 1.0,0.78,1 | 0.704,1.0, | 0.109 |
| ENSMUSG00000056310 | Tyw1 | chr5 | + | 130291659 | 130291782 | 130285875 | 130286034 | 130295823 | 130295925 | 76,51,79,45 | 7,0,0,0 | 70,39,14,40 | 0,5,0,14 | 272 | 149 | 0.00459 | 0.08426 | 0.856,1.0, | 1.0,0.81,1 | 0.109 |
| ENSMUSG00000032340 | Neo1 | chr9 | - | 58809365 | 58809398 | 58806447 | 58806516 | 58810196 | 58810391 | 192,187,223,195 | 97,63,101,90 | 163,119,109,129 | 104,64,92,99 | 182 | 149 | 0.00037 | 0.01334 | 0.618,0.7 | 0.562,0.6 | 0.109 |
| ENSMUSG00000031137 | Fgf13 | chrX | - | 58296158 | 58296326 | 58177236 | 58177310 | 58612499 | 58612832 | 227,159,214,179 | 45,4,21,15 | 145,147,160,122 | 32,21,30,21 | 317 | 149 | 0.00402 | 0.07665 | 0.703,0.9 | 0.68,0.76 | 0.109 |
| ENSMUSG00000075289 | Carns1 | chr19 | - | 4219526 | 4219807 | 4214322 | 4216563 | 4219897 | 4220070 | 110,58,145,91 | 0,2,0,4 | 93,101,71,71 | 12,1,4,6 | 430 | 149 | 6.15E-05 | 0.00347 | 1.0,0.909, | 0.729,0.9 | 0.108 |
| ENSMUSG00000035954 | Dock4 | chr12 | + | 40844595 | 40844622 | 40840074 | 40840160 | 40844766 | 40844862 | 214,200,205,173 | 33,25,28,31 | 131,179,122,126 | 44,34,33,50 | 176 | 149 | 3.11E-05 | 0.00209 | 0.846,0.8 | 0.716,0.8 | 0.108 |
| ENSMUSG00000054226 | Tprkb | chr6 | + | 85896898 | 85897006 | 85888888 | 85888962 | 85898404 | 85898493 | 51,17,20,17 | 0,0,0,0 | 25,44,32,25 | 6,0,3,0 | 257 | 149 | 0.00162 | 0.03905 | 1.0,1.0,1. | 0.707,1.0, | 0.108 |
| ENSMUSG00000062110 | Scfd2 | chr5 | - | 74618965 | 74619303 | 74558314 | 74558564 | 74623220 | 74623396 | 46,56,47,41 | 61,40,85,64 | 17,18,18,12 | 45,45,47,73 | 487 | 149 | 2.42E-06 | 0.00031 | 0.187,0.3, | 0.104,0.1 | 0.108 |
| ENSMUSG00000053768 | Chchd3 | chr6 | - | 32870326 | 32870341 | 32869226 | 32869310 | 32945135 | 32945253 | 159,0,0,0 | 189,148,167,133 | 0,0,0,0 | 217,157,137,123 | 164 | 149 | 0.00049 | 0.01671 | 0.433,0.0, | 0.0,0.0,0. | 0.108 |
| ENSMUSG00000034730 | Adgrb1 | chr15 | + | 74449243 | 74449342 | 74447687 | 74447773 | 74452513 | 74452585 | 32,27,49,28 | 71,81,107,86 | 20,9,15,9 | 80,96,79,116 | 248 | 149 | 3.61E-08 | 1.11E-05 | 0.213,0.1 | 0.131,0.0 | 0.107 |
| ENSMUSG00000041351 | Rap1gap | chr4 | + | 137451056 | 137451134 | 137449136 | 137449268 | 137452002 | 137452112 | 2,691,187,119,802,160 | 676,444,647,662 | 92,815,667,721,284 | 566,439,464,480 | 227 | 149 | 0.00075 | 0.02235 | 0.723,0.7 | 0.518,0.7 | 0.107 |
| ENSMUSG00000030189 | Ybx3 | chr6 | - | 131352936 | 131353143 | 131347285 | 131347377 | 131356320 | 131356443 | 278,226,443,194 | 119,98,131,95 | 293,208,201,213 | 141,162,142,125 | 356 | 149 | 0.00044 | 0.01529 | 0.494,0.4 | 0.465,0.3 | 0.107 |
| ENSMUSG00000110027 | C030029 | chr7 | + | 135870642 | 135870676 | 135870060 | 135870205 | 135877802 | 135878260 | 3,0,12,2 | 26,7,14,26 | 3,0,2,0 | 40,20,21,34 | 183 | 149 | 0.00082 | 0.02392 | 0.086,0.0, | 0.058,0.0, | 0.107 |
| ENSMUSG00000036249 | Rbm43 | chr2 | - | 51822411 | 51822531 | 51819671 | 51819861 | 51824843 | 51825019 | 63,30,52,71 | 0,5,0,3 | 43,28,34,44 | 8,8,3,0 | 269 | 149 | 0.00413 | 0.07804 | 1.0,0.769, | 0.749,0.6 | 0.107 |
| ENSMUSG00000022372 | Sla | chr15 | - | 66684377 | 66684646 | 66673279 | 66673377 | 66703418 | 66703495 | 45,25,150,36 | 0,0,0,0 | 197,72,130,113 | 8,0,14,4 | 418 | 149 | 8.48E-08 | 2.20E-05 | 1.0,1.0,1. | 0.898,1.0, | 0.106 |
| ENSMUSG00000062785 | Kcnc3 | chr7 | + | 44248229 | 44248357 | 44247807 | 44247999 | 44250292 | 44250352 | 31,22,56,21 | 63,53,97,50 | 3,21,40,10 | 85,71,115,65 | 277 | 149 | 8.87E-06 | 0.00081 | 0.209,0.1 | 0.019,0.1 | 0.106 |
| ENSMUSG00000034602 | Mon2 | chr10 | - | 122845817 | 122845835 | 122845407 | 122845536 | 122846398 | 122846542 | 215,202,217,188 | 23,32,23,19 | 158,178,86,165 | 47,29,43,21 | 167 | 149 | 0.00057 | 0.01866 | 0.893,0.8 | 0.75,0.84 | 0.106 |
| ENSMUSG00000025384 | Faap100 | chr11 | - | 120269153 | 120269272 | 120266995 | 120268487 | 120269367 | 120269535 | 42,39,29,34 | 0,0,0,2 | 20,22,28,24 | 3,1,0,4 | 268 | 149 | 0.00238 | 0.05271 | 1.0,1.0,1. | 0.788,0.9 | 0.106 |
| ENSMUSG00000022377 | Asap1 | chr15 | - | 64025692 | 64025701 | 64024678 | 64024779 | 64030766 | 64030853 | 0,4,0,5 | 6,28,28,11 | 0,0,0,0 | 23,20,18,26 | 158 | 149 | 0.00269 | 0.05754 | 0.0,0.119, | 0.0,0.0,0. | 0.105 |
| ENSMUSG00000069227 | Gprin1 | chr13 | - | 54889821 | 54889921 | 54884485 | 54888426 | 54897382 | 54897482 | 99,71,106,57 | 142,97,218,93 | 72,14,88,60 | 232,107,218,109 | 249 | 149 | 0.0001 | 0.00513 | 0.294,0.3 | 0.157,0.0 | 0.105 |
| ENSMUSG00000047656 | Trpt1 | chr19 | + | 6975559 | 6975734 | 6974063 | 6974145 | 6975876 | 6975933 | 142,106,160,125 | 6,2,10,4 | 119,119,130,126 | 13,17,7,14 | 324 | 149 | 0.00027 | 0.01055 | 0.916,0.9 | 0.808,0.7 | 0.105 |
| ENSMUSG00000027244 | Atg13 | chr2 | - | 91512370 | 91512481 | 91511903 | 91512040 | 91515013 | 91515104 | 375,309,426,287 | 41,1,0,10 | 255,270,248,200 | 30,19,21,34 | 260 | 149 | 2.91E-08 | 9.47E-06 | 0.84,0.99 | 0.83,0.89 | 0.104 |
| ENSMUSG00000041215 | Yeats2 | chr16 | + | 20026373 | 20026543 | 20024833 | 20024995 | 20027168 | 20027327 | 144,171,182,110 | 2,8,8,1 | 138,96,76,97 | 14,8,6,9 | 319 | 149 | 0.00013 | 0.00611 | 0.971,0.9 | 0.822,0.8 | 0.104 |
| ENSMUSG00000035258 | Abi3bp | chr16 | + | 56471073 | 56471133 | 56467387 | 56467462 | 56472499 | 56472580 | 7,5,45,8 | 10,13,68,25 | 26,7,27,20 | 92,24,128,68 | 209 | 149 | 0.00422 | 0.07929 | 0.333,0.2 | 0.168,0.1 | 0.103 |
| ENSMUSG00000039458 | Mtmr12 | chr15 | + | 12238014 | 12238105 | 12233939 | 12234012 | 12245069 | 12245138 | 79,50,98,56 | 0,1,0,0 | 48,70,43,75 | 6,0,10,0 | 240 | 149 | 6.57E-06 | 0.00064 | 1.0,0.969, | 0.832,1.0, | 0.103 |
| ENSMUSG00000011382 | Dhdh | chr7 | - | 45125028 | 45125176 | 45120410 | 45124799 | 45128432 | 45128557 | 176,85,131,161 | 4,2,6,0 | 78,68,67,101 | 5,10,7,4 | 297 | 149 | 7.77E-05 | 0.00413 | 0.957,0.9 | 0.887,0.7 | 0.103 |
| ENSMUSG00000028759 | Hp1bp3 | chr4 | + | 137948836 | 137949034 | 137944441 | 137944605 | 137949399 | 137949499 | 806,591,694,677 | 120,79,164,112 | 457,505,398,394 | 121,130,107,112 | 347 | 149 | 4.12E-05 | 0.00257 | 0.743,0.7 | 0.619,0.6 | 0.103 |
| ENSMUSG00000042590 | Ipo11 | chr13 | - | 106994006 | 106994033 | 106993736 | 106993816 | 106997368 | 106997445 | 145,111,180,125 | 34,28,31,29 | 86,92,136,70 | 41,28,36,36 | 176 | 149 | 0.0023 | 0.05129 | 0.783,0.7 | 0.64,0.73 | 0.102 |
| ENSMUSG00000025006 | Sorbs1 | chr19 | - | 40298024 | 40298777 | 40287906 | 40287965 | 40300112 | 40300307 | 135,712,451,179,960 | 160,153,237,195 | 807,860,648,840 | 212,172,201,160 | 902 | 149 | 0.00296 | 0.0615 | 0.584,0.5 | 0.386,0.4 | 0.102 |
| ENSMUSG00000048249 | Crebrf | chr17 | + | 26955789 | 26955957 | 26934628 | 26934748 | 26961015 | 26961933 | 41,28,33,15 | 2,0,0,0 | 32,30,25,15 | 0,6,3,0 | 317 | 149 | 0.00092 | 0.02592 | 0.906,1.0, | 1.0,0.702, | 0.102 |
| ENSMUSG00000021621 | Zcchc9 | chr13 | - | 91946941 | 91947010 | 91944651 | 91945373 | 91948729 | 91948822 | 159,92,126,125 | 4,1,1,0 | 110,100,73,105 | 4,18,6,8 | 218 | 149 | 2.91E-09 | 1.43E-06 | 0.964,0.9 | 0.949,0.7 | 0.101 |
| ENSMUSG00000055717 | Slain1 | chr14 | + | 103921669 | 103921819 | 103894368 | 103894508 | 103923104 | 103923254 | 252,202,288,181 | 87,51,117,72 | 244,182,164,178 | 123,67,103,110 | 299 | 149 | 0.00479 | 0.08699 | 0.591,0.6 | 0.497,0.5 | 0.101 |
| ENSMUSG00000033767 | Tmem131 | chr3 | - | 83843303 | 83843373 | 83842126 | 83842244 | 83844728 | 83844899 | 82,74,56,97 | 0,3,0,0 | 40,48,38,55 | 4,7,3,2 | 219 | 149 | 3.41E-06 | 0.00041 | 1.0,0.944, | 0.872,0.8 | 0.101 |
| ENSMUSG00000058498 | Rnf207 | chr4 | - | 152400059 | 152400135 | 152399848 | 152399974 | 152400207 | 152400289 | 81,61,140,70 | 9,9,0,6 | 65,80,70,86 | 0,2,0,0 | 225 | 149 | 4.12E-07 | 7.93E-05 | 0.856,0.8 | 1.0,0.964, | -0.101 |
| ENSMUSG00000000085 | Scmh1 | chr4 | + | 120319209 | 120319352 | 120313481 | 120313706 | 120320271 | 120320281 | 308,180,260,208 | 23,0,19,19 | 224,204,162,198 | 0,0,0,0 | 292 | 149 | 1.33E-15 | 7.27E-12 | 0.872,1.0, | 1.0,1.0,1. | -0.101 |
| ENSMUSG00000098789 | Jmjd7 | chr2 | + | 119862055 | 119862132 | 119861296 | 119861353 | 119862321 | 119862481 | 31,17,40,25 | 0,0,7,4 | 26,17,13,12 | 0,0,0,0 | 226 | 149 | 5.34E-05 | 0.00312 | 1.0,1.0,0. | 1.0,1.0,1. | -0.101 |
| ENSMUSG00000025138 | Sirt7 | chr11 | - | 120509974 | 120510116 | 120509245 | 120509875 | 120510508 | 120510593 | 28,28,19,23 | 62,43,69,63 | 38,31,34,18 | 64,46,42,18 | 291 | 149 | 0.00462 | 0.08465 | 0.188,0.2 | 0.233,0.2 | -0.101 |
| ENSMUSG00000020821 | Kif1c | chr11 | + | 70593137 | 70593377 | 70591386 | 70591535 | 70593597 | 70593731 | 275,227,301,221 | 42,38,51,33 | 336,258,252,244 | 53,9,18,23 | 329 | 149 | 0.0007 | 0.02135 | 0.748,0.7 | 0.742,0.9 | -0.101 |
| ENSMUSG00000042104 | Uggt2 | chr14 | - | 119240237 | 119240398 | 119235526 | 119235625 | 119243975 | 119244129 | 25,26,34,24 | 0,0,11,0 | 17,30,22,26 | 0,0,0,0 | 310 | 149 | 0.00086 | 0.02491 | 1.0,1.0,0. | 1.0,1.0,1. | -0.101 |
| ENSMUSG00000023826 | Prkn | chr17 | + | 11286015 | 11286179 | 11059270 | 11059376 | 11456348 | 11456589 | 73,60,76,63 | 7,9,0,0 | 96,58,36,57 | 0,0,0,0 | 313 | 149 | 2.49E-06 | 0.00032 | 0.832,0.7 | 1.0,1.0,1. | -0.102 |
| ENSMUSG00000109857 | Gm53058 | chr16 | + | 13281736 | 13281801 | 13265766 | 13266056 | 13284690 | 13285094 | 17,32,7,29 | 6,5,0,0 | 8,35,26,30 | 0,3,0,0 | 214 | 149 | 0.00064 | 0.01998 | 0.664,0.8 | 1.0,0.89,1 | -0.102 |
| ENSMUSG00000032855 | Pkd1 | chr17 | + | 24783552 | 24783722 | 24783309 | 24783381 | 24783984 | 24784656 | 252,201,262,147 | 20,51,26,27 | 217,163,132,155 | 20,12,11,7 | 319 | 149 | 0.00231 | 0.05161 | 0.855,0.6 | 0.835,0.8 | -0.103 |
| ENSMUSG00000028573 | Fggy | chr4 | + | 95688793 | 95688901 | 95657662 | 95657766 | 95700344 | 95700406 | 60,53,73,61 | 15,0,0,13 | 67,53,90,77 | 5,0,0,2 | 257 | 149 | 0.0006 | 0.01934 | 0.699,1.0, | 0.886,1.0, | -0.103 |
| ENSMUSG00000057897 | Camk2b | chr11 | - | 5931069 | 5931114 | 5929672 | 5929721 | 5932500 | 5932572 | 3,246,267,431,792,660 | 700,710,736,633 | 2,441,264,924,102,410 | 294,393,218,272 | 194 | 149 | 0 | 0 | 0.781,0.7 | 0.864,0.8 | -0.103 |
| ENSMUSG00000029635 | Cdk8 | chr5 | + | 146229448 | 146229577 | 146222917 | 146223049 | 146231972 | 146232042 | 113,110,153,91 | 19,5,13,9 | 104,103,102,84 | 0,7,5,0 | 278 | 149 | 6.12E-05 | 0.00347 | 0.761,0.9 | 1.0,0.887, | -0.103 |
| ENSMUSG00000034275 | Igsf9b | chr9 | + | 27253806 | 27253928 | 27244373 | 27246029 | 27257065 | 27268842 | 126,97,100,125 | 20,20,13,0 | 99,81,114,123 | 5,0,15,0 | 271 | 149 | 0.00349 | 0.06936 | 0.776,0.7 | 0.916,1.0, | -0.103 |
| ENSMUSG00000037935 | Smarce1 | chr11 | - | 99115863 | 99115968 | 99111480 | 99111561 | 99119100 | 99119144 | 228,155,256,222 | 125,90,209,97 | 215,233,216,235 | 95,97,75,77 | 254 | 149 | 0.00107 | 0.02885 | 0.517,0.5 | 0.57,0.58 | -0.103 |
| ENSMUSG00000052512 | Nav2 | chr7 | + | 49234891 | 49234992 | 49233797 | 49233960 | 49237454 | 49237550 | 144,96,131,117 | 42,42,63,44 | 148,132,125,121 | 29,26,40,35 | 250 | 149 | 0.00453 | 0.08357 | 0.671,0.5 | 0.753,0.7 | -0.104 |
| ENSMUSG00000021288 | Klc1 | chr12 | + | 111760082 | 111760133 | 111755758 | 111755893 | 111761981 | 111762112 | 171,119,200,155 | 91,63,69,94 | 196,136,228,170 | 37,60,72,57 | 200 | 149 | 0.00361 | 0.07091 | 0.583,0.5 | 0.798,0.6 | -0.104 |
| ENSMUSG00000031731 | Ap1g1 | chr8 | + | 110557244 | 110557253 | 110556230 | 110556307 | 110559348 | 110559444 | 0,0,0,0 | 66,78,93,64 | 0,45,0,0 | 65,59,50,52 | 158 | 149 | 0.0002 | 0.00852 | 0.0,0.0,0. | 0.0,0.418, | -0.104 |
| ENSMUSG00000032123 | Dpagt1 | chr9 | + | 44238848 | 44238969 | 44237847 | 44238502 | 44239249 | 44239463 | 116,61,54,85 | 11,10,4,7 | 95,77,71,101 | 2,4,2,2 | 270 | 149 | 8.00E-05 | 0.00422 | 0.853,0.7 | 0.963,0.9 | -0.104 |
| ENSMUSG00000022721 | Trmt2a | chr16 | + | 18070787 | 18070884 | 18070265 | 18070341 | 18070963 | 18070976 | 116,72,90,85 | 11,6,15,3 | 93,99,73,65 | 1,6,0,0 | 246 | 149 | 4.31E-06 | 0.00047 | 0.865,0.8 | 0.983,0.9 | -0.105 |
| ENSMUSG00000034574 | Daam1 | chr12 | + | 72005520 | 72005547 | 71999225 | 71999333 | 72006579 | 72006686 | 109,51,87,67 | 18,21,19,23 | 67,51,82,96 | 12,11,2,15 | 176 | 149 | 0.00533 | 0.09342 | 0.837,0.6 | 0.825,0.7 | -0.105 |
| ENSMUSG00000021071 | Trim9 | chr12 | - | 70314007 | 70314040 | 70301811 | 70302000 | 70314835 | 70314974 | 411,221,380,196 | 144,98,200,122 | 388,273,283,265 | 105,88,106,57 | 182 | 149 | 0.00016 | 0.00715 | 0.7,0.649, | 0.752,0.7 | -0.105 |
| ENSMUSG00000035069 | Oma1 | chr4 | + | 103186034 | 103186163 | 103182489 | 103182597 | 103188501 | 103188576 | 58,81,155,56 | 7,7,25,4 | 79,77,62,76 | 11,0,0,2 | 278 | 149 | 0.00057 | 0.01854 | 0.816,0.8 | 0.794,1.0, | -0.105 |
| ENSMUSG00000027285 | Haus2 | chr2 | + | 120442818 | 120442911 | 120439863 | 120440026 | 120446335 | 120446468 | 62,77,90,54 | 9,3,11,1 | 67,71,59,44 | 0,0,1,0 | 242 | 149 | 5.12E-08 | 1.47E-05 | 0.809,0.9 | 1.0,1.0,0. | -0.105 |
| ENSMUSG00000022629 | Kif21a | chr15 | - | 90833208 | 90833247 | 90827967 | 90828104 | 90836913 | 90837021 | 15,40,31,33 | 121,113,148,165 | 37,18,40,37 | 76,63,89,73 | 188 | 149 | 6.62E-05 | 0.00367 | 0.089,0.2 | 0.278,0.1 | -0.106 |
| ENSMUSG00000021268 | Meg3 | chr12 | + | 109514446 | 109514548 | 109512312 | 109512425 | 109515634 | 109515806 | 78,72,112,108 | 18,7,8,1 | 51,55,72,86 | 0,0,0,7 | 251 | 149 | 0.00021 | 0.00886 | 0.72,0.85 | 1.0,1.0,1. | -0.106 |
| ENSMUSG00000040183 | Ankrd6 | chr4 | - | 32874206 | 32874370 | 32860321 | 32860588 | 32950746 | 32950841 | 12,12,3,26 | 100,55,56,82 | 23,29,28,8 | 35,38,59,86 | 313 | 149 | 0.00054 | 0.01779 | 0.054,0.0 | 0.238,0.2 | -0.106 |
| ENSMUSG00000070883 | Ccdc173 | chr2 | - | 69606473 | 69606613 | 69592065 | 69592230 | 69608403 | 69608560 | 47,43,52,42 | 11,1,8,3 | 13,30,54,34 | 2,0,0,1 | 289 | 149 | 1.21E-05 | 0.00101 | 0.688,0.9 | 0.77,1.0,1 | -0.106 |
| ENSMUSG00000041757 | Plekha6 | chr1 | + | 133203549 | 133203603 | 133201556 | 133202112 | 133207110 | 133207191 | 64,56,55,43 | 0,14,6,7 | 42,45,29,34 | 5,0,0,0 | 203 | 149 | 0.00016 | 0.0072 | 1.0,0.746, | 0.86,1.0,1 | -0.106 |
| ENSMUSG00000029729 | Zkscan1 | chr5 | + | 138092253 | 138092401 | 138083345 | 138083493 | 138098851 | 138102753 | 34,27,35,31 | 0,0,13,0 | 23,20,30,28 | 0,0,0,0 | 297 | 149 | 0.00073 | 0.0221 | 1.0,1.0,0. | 1.0,1.0,1. | -0.106 |
| ENSMUSG00000026956 | Uap1l1 | chr2 | - | 25253255 | 25253396 | 25252674 | 25252860 | 25253817 | 25254011 | 128,85,170,110 | 18,17,6,9 | 192,127,137,124 | 6,0,6,10 | 290 | 149 | 0.00034 | 0.01265 | 0.785,0.7 | 0.943,1.0, | -0.106 |
| ENSMUSG00000071753 | Cdr1os | chrX | + | 60226853 | 60229780 | 60178326 | 60178402 | 60247301 | 60247391 | 4,389,393,350,624,280 | 74,59,74,69 | 1,706,333,734,903,750 | 15,23,33,22 | 3076 | 149 | 5.98E-08 | 1.68E-05 | 0.742,0.7 | 0.846,0.8 | -0.106 |
| ENSMUSG00000040247 | Tbc1d10c | chr19 | - | 4235350 | 4235505 | 4233409 | 4235072 | 4237940 | 4238079 | 40,35,34,43 | 4,2,3,0 | 33,15,10,12 | 0,0,0,0 | 304 | 149 | 0.00099 | 0.02727 | 0.831,0.8 | 1.0,1.0,1. | -0.107 |

| | | | | | | | | | | | | | | | | | | | | |
|---|---|---|---|---|---|---|---|---|---|---|---|---|---|---|---|---|---|---|---|---|
| ENSMUSG00000052551 | Adarb2 | chr13 | + | 8609139 | 8609226 | 8252901 | 8253356 | 8619702 | 8620610 | 71,49,137,66 | 1,6,10,9 | 74,51,95,94 | 1,0,1,0 | 236 | 149 | 3.53E-08 | 1.10E-05 | 0.978,0.8 | 0.979,1.0, | -0.107 |
| ENSMUSG00000038569 | Rad9b | chr5 | - | 122489623 | 122489773 | 122489321 | 122489436 | 122490606 | 122490677 | 70,65,88,65 | 4,3,11,3 | 72,52,41,41 | 0,0,1,0 | 299 | 149 | 2.73E-07 | 5.76E-05 | 0.897,0.9 | 1.0,1.0,0. | -0.107 |
| ENSMUSG00000031608 | Galnt7 | chr8 | - | 57998362 | 57998442 | 57995545 | 57995728 | 57998702 | 57998833 | 34,16,33,43 | 0,0,16,0 | 18,29,16,21 | 0,0,0,0 | 229 | 149 | 0.00057 | 0.01852 | 1.0,1.0,0. | 1.0,1.0,1. | -0.107 |
| ENSMUSG00000029775 | Klhdc10 | chr6 | + | 30427707 | 30427794 | 30401894 | 30402149 | 30439642 | 30439864 | 60,23,42,33 | 190,139,173,128 | 50,49,46,61 | 91,114,103,104 | 236 | 149 | 7.85E-07 | 0.00013 | 0.166,0.0 | 0.258,0.2 | -0.107 |
| ENSMUSG00000012114 | Med15 | chr16 | - | 17481240 | 17481360 | 17473541 | 17473669 | 17489408 | 17489537 | 61,35,57,66 | 70,65,139,68 | 82,73,66,46 | 65,69,54,50 | 269 | 149 | 0.00172 | 0.04093 | 0.326,0.2 | 0.411,0.3 | -0.108 |
| ENSMUSG00000032525 | Nktr | chr9 | + | 121567453 | 121569141 | 121560543 | 121560631 | 121570187 | 121570217 | 12,529,921,487,912 | 140,184,232,144 | 10,339,669,071,073 | 112,84,104,83 | 1837 | 149 | 7.58E-05 | 0.00406 | 0.42,0.30 | 0.428,0.4 | -0.108 |
| ENSMUSG00000014592 | Camta1 | chr4 | - | 151670871 | 151670939 | 151537740 | 151537876 | 151914874 | 151914993 | 135,77,105,101 | 0,8,19,12 | 92,100,122,94 | 0,4,0,0 | 217 | 149 | 2.11E-09 | 1.21E-06 | 1.0,0.869, | 1.0,0.945, | -0.108 |
| ENSMUSG00000058174 | Gm5148 | chr3 | - | 37776447 | 37777130 | 37768474 | 37769284 | 37778351 | 37778480 | 135,122,152,112 | 17,0,4,4 | 60,76,43,71 | 0,1,2,0 | 832 | 149 | 0.00015 | 0.00667 | 0.587,1.0, | 1.0,0.932, | -0.108 |
| ENSMUSG00000020212 | Mdm1 | chr10 | + | 117983885 | 117984020 | 117982499 | 117982876 | 117986693 | 117986861 | 60,17,27,34 | 7,1,5,1 | 19,10,21,47 | 0,1,0,0 | 284 | 149 | 4.17E-06 | 0.00046 | 0.818,0.8 | 1.0,0.84,1 | -0.109 |
| ENSMUSG00000021706 | Zfyve16 | chr13 | - | 92653038 | 92653200 | 92650294 | 92650437 | 92655974 | 92656071 | 71,88,70,64 | 10,3,7,12 | 82,69,59,57 | 0,3,4,3 | 311 | 149 | 0.001 | 0.02733 | 0.773,0.9 | 1.0,0.917, | -0.11 |
| ENSMUSG00000030279 | C2cd5 | chr6 | - | 143004871 | 143004937 | 142995850 | 142995965 | 143007478 | 143007587 | 144,120,139,122 | 199,162,186,166 | 78,126,107,166 | 72,117,85,130 | 215 | 149 | 3.69E-05 | 0.00237 | 0.334,0.3 | 0.429,0.4 | -0.11 |
| ENSMUSG00000025144 | Cenpx | chr11 | - | 120603265 | 120603538 | 120602528 | 120602582 | 120604518 | 120604558 | 55,41,66,75 | 96,71,77,58 | 66,63,50,63 | 50,55,39,33 | 422 | 149 | 0.00137 | 0.03476 | 0.168,0.1 | 0.318,0.2 | -0.11 |
| ENSMUSG00000006728 | Cdk4 | chr10 | + | 126900088 | 126900322 | 126899402 | 126899628 | 126900453 | 126900499 | 939,639,990,705 | 215,125,162,98 | 865,642,664,678 | 63,84,56,80 | 383 | 149 | 2.65E-07 | 5.62E-05 | 0.63,0.66 | 0.842,0.7 | -0.111 |
| ENSMUSG00000086370 | Ftx | chrX | - | 102658670 | 102658798 | 102652909 | 102653022 | 102667152 | 102667269 | 33,45,51,32 | 4,0,7,1 | 48,67,19,26 | 0,0,0,0 | 277 | 149 | 1.24E-05 | 0.00104 | 0.816,1.0, | 1.0,1.0,1. | -0.111 |
| ENSMUSG00000024074 | Crim1 | chr17 | + | 78610415 | 78610537 | 78545170 | 78545344 | 78620552 | 78620617 | 125,80,91,105 | 16,19,32,25 | 117,150,119,96 | 12,13,15,18 | 271 | 149 | 0.00329 | 0.06636 | 0.811,0.6 | 0.843,0.8 | -0.112 |
| ENSMUSG00000040473 | Cfap69 | chr5 | - | 5675749 | 5675927 | 5671919 | 5672043 | 5676008 | 5676158 | 87,117,136,122 | 9,14,22,5 | 60,75,97,92 | 0,0,10,5 | 327 | 149 | 0.00249 | 0.05444 | 0.815,0.7 | 1.0,1.0,0. | -0.112 |
| ENSMUSG00000026977 | Marchf7 | chr2 | + | 60075555 | 60075604 | 60073900 | 60074014 | 60078233 | 60078301 | 164,91,97,101 | 12,25,56,23 | 48,143,152,101 | 20,1,3,17 | 198 | 149 | 0.00354 | 0.06997 | 0.911,0.7 | 0.644,0.9 | -0.112 |
| ENSMUSG00000036555 | Iqce | chr5 | - | 140678548 | 140678683 | 140677354 | 140677425 | 140679169 | 140679298 | 88,63,56,59 | 92,72,102,56 | 58,67,64,71 | 51,52,37,50 | 284 | 149 | 0.00173 | 0.04125 | 0.334,0.3 | 0.374,0.4 | -0.113 |
| ENSMUSG00000055897 | Ppp4r1l-p | chr2 | - | 173431580 | 173431741 | 173431297 | 173431467 | 173433909 | 173434014 | 26,37,60,42 | 7,0,8,0 | 29,21,34,30 | 2,0,0,0 | 310 | 149 | 9.24E-05 | 0.00476 | 0.641,1.0, | 0.875,1.0, | -0.113 |
| ENSMUSG00000006958 | Chrd | chr16 | + | 20553655 | 20553719 | 20553435 | 20553523 | 20553866 | 20554008 | 0,2,0,0 | 16,12,45,11 | 9,0,4,7 | 25,15,27,14 | 213 | 149 | 1.34E-06 | 0.0002 | 0.0,0.104, | 0.201,0.0, | -0.113 |
| ENSMUSG00000022378 | Cyrib | chr15 | - | 63829433 | 63829516 | 63821854 | 63821976 | 63842778 | 63842817 | 182,128,213,151 | 25,28,46,26 | 145,151,127,157 | 15,11,6,15 | 232 | 149 | 4.69E-06 | 0.0005 | 0.824,0.7 | 0.861,0.8 | -0.114 |
| ENSMUSG00000046138 | 9930021J | chr19 | - | 29712515 | 29712605 | 29700789 | 29700905 | 29720932 | 29721012 | 109,140,136,124 | 32,53,87,38 | 91,118,122,113 | 31,24,25,21 | 239 | 149 | 0.00443 | 0.08187 | 0.68,0.62 | 0.647,0.7 | -0.114 |
| ENSMUSG00000014592 | Camta1 | chr4 | - | 151876875 | 151876959 | 151537740 | 151537876 | 151914874 | 151914993 | 124,83,133,73 | 0,8,19,12 | 106,94,94,112 | 0,4,0,0 | 233 | 149 | 2.15E-09 | 1.22E-06 | 1.0,0.869, | 1.0,0.938, | -0.114 |
| ENSMUSG00000046897 | Zfp740 | chr15 | + | 102113663 | 102113740 | 102113005 | 102113521 | 102116205 | 102116355 | 69,36,48,77 | 99,75,122,96 | 54,66,73,82 | 59,84,84,58 | 226 | 149 | 0.00264 | 0.05692 | 0.315,0.2 | 0.376,0.3 | -0.114 |
| ENSMUSG00000053580 | Tanc2 | chr11 | + | 105649747 | 105649858 | 105604207 | 105604279 | 105667631 | 105667723 | 85,71,101,75 | 37,15,22,15 | 63,53,70,82 | 11,7,8,11 | 260 | 149 | 0.00527 | 0.0926 | 0.568,0.7 | 0.766,0.8 | -0.114 |
| ENSMUSG00000038930 | Rccd1 | chr7 | - | 79971024 | 79971091 | 79969839 | 79970929 | 79973497 | 79973873 | 110,84,76,58 | 23,18,32,9 | 50,101,70,87 | 10,8,10,5 | 216 | 149 | 0.001 | 0.02733 | 0.767,0.7 | 0.775,0.8 | -0.114 |
| ENSMUSG00000022307 | Oxr1 | chr15 | + | 41712065 | 41712146 | 41694353 | 41694525 | 41713854 | 41713980 | 261,191,497,229 | 367,266,356,260 | 289,287,386,313 | 187,200,247,251 | 230 | 149 | 0.00023 | 0.00933 | 0.315,0.3 | 0.5,0.482, | -0.115 |
| ENSMUSG00000032534 | Cep63 | chr9 | - | 102500754 | 102500792 | 102498894 | 102498927 | 102503102 | 102503180 | 29,28,52,16 | 8,3,11,3 | 22,40,22,20 | 0,2,3,2 | 187 | 149 | 0.00155 | 0.03807 | 0.743,0.8 | 1.0,0.941, | -0.115 |
| ENSMUSG00000032468 | Armc8 | chr9 | - | 99384564 | 99384651 | 99381345 | 99381438 | 99402539 | 99402635 | 101,65,115,84 | 20,10,38,20 | 28,99,59,60 | 9,6,4,3 | 236 | 149 | 0.00119 | 0.03114 | 0.761,0.8 | 0.663,0.9 | -0.115 |
| ENSMUSG00000026074 | Map4k4 | chr1 | + | 40042916 | 40043147 | 40040589 | 40040751 | 40043966 | 40044073 | 310,172,274,174 | 56,38,59,46 | 223,150,159,201 | 14,35,17,26 | 380 | 149 | 0.00413 | 0.07808 | 0.685,0.6 | 0.862,0.6 | -0.115 |
| ENSMUSG00000020300 | Cpeb4 | chr11 | + | 31868800 | 31868824 | 31858824 | 31858875 | 31870408 | 31870582 | 114,79,153,114 | 38,16,36,27 | 72,87,65,78 | 23,15,0,0 | 173 | 149 | 0.00058 | 0.01891 | 0.721,0.8 | 0.729,0.8 | -0.115 |
| ENSMUSG00000024069 | Slc30a6 | chr17 | + | 74725614 | 74725662 | 74722614 | 74722717 | 74730011 | 74730661 | 14,13,34,21 | 4,5,4,0 | 10,20,16,16 | 2,0,1,0 | 197 | 149 | 0.00037 | 0.01329 | 0.726,0.6 | 0.791,1.0, | -0.115 |
| ENSMUSG00000066892 | Fbxl12 | chr9 | - | 20553319 | 20553392 | 20549081 | 20550494 | 20555868 | 20556051 | 27,25,35,39 | 3,0,11,0 | 34,24,19,28 | 0,0,0,0 | 222 | 149 | 8.74E-06 | 0.0008 | 0.858,1.0, | 1.0,1.0,1. | -0.115 |
| ENSMUSG00000097709 | 2810429I( | chr13 | + | 3529521 | 3529574 | 3528214 | 3528390 | 3529908 | 3530001 | 0,1,0,0 | 38,28,27,30 | 0,0,9,10 | 6,9,16,30 | 202 | 149 | 8.27E-08 | 2.20E-05 | 0.0,0.026, | 0.0,0.0,0. | -0.116 |
| ENSMUSG00000034621 | Gpatch8 | chr11 | - | 102422138 | 102422156 | 102418919 | 102418992 | 102429107 | 102429182 | 30,1,16,14 | 124,74,149,89 | 29,27,11,27 | 47,111,63,98 | 167 | 149 | 0.00087 | 0.02493 | 0.178,0.0 | 0.355,0.1 | -0.116 |
| ENSMUSG00000031153 | Gripap1 | chrX | + | 7660498 | 7660525 | 7658514 | 7658576 | 7665528 | 7665667 | 164,121,206,107 | 267,215,294,134 | 103,150,102,177 | 134,151,113,92 | 176 | 149 | 0.0033 | 0.06647 | 0.342,0.3 | 0.394,0.4 | -0.116 |
| ENSMUSG00000014602 | Kif1a | chr1 | - | 92993273 | 92993300 | 92991870 | 92992004 | 92993768 | 92993911 | 11,148,778,781,237 | 10,167,541,417,662 | 121,893,411,311,146 | 789,583,634,516 | 176 | 149 | 0.00382 | 0.07409 | 0.481,0.4 | 0.567,0.5 | -0.116 |
| ENSMUSG00000030970 | Ctbp2 | chr7 | - | 132627441 | 132627601 | 132600919 | 132601074 | 132683485 | 132683589 | 91,58,88,91 | 14,12,22,23 | 115,86,104,110 | 14,5,16,14 | 309 | 149 | 0.00512 | 0.09088 | 0.758,0.7, | 0.798,0.8 | -0.116 |
| ENSMUSG00000044807 | Zfp354c | chr11 | - | 50708115 | 50708186 | 50708019 | 50708038 | 50708635 | 50708762 | 10,8,19,2 | 130,108,123,75 | 26,20,5,25 | 53,79,70,56 | 220 | 149 | 1.56E-06 | 0.00022 | 0.05,0.04 | 0.249,0.1 | -0.116 |
| ENSMUSG00000036555 | Iqce | chr5 | - | 140654776 | 140654860 | 140651762 | 140651963 | 140656221 | 140656322 | 5,7,14,15 | 80,51,69,40 | 12,12,25,10 | 47,44,26,22 | 233 | 149 | 0.00157 | 0.03828 | 0.038,0.0 | 0.14,0.14 | -0.117 |
| ENSMUSG00000025925 | Terf1 | chr1 | + | 15886109 | 15886196 | 15883264 | 15883386 | 15889157 | 15889307 | 93,47,66,85 | 9,7,16,8 | 31,58,74,55 | 0,7,1,3 | 236 | 149 | 0.00087 | 0.02497 | 0.867,0.8 | 1.0,0.84,0 | -0.117 |
| ENSMUSG00000060176 | Kif27 | chr13 | - | 58491679 | 58492638 | 58485356 | 58485500 | 58502378 | 58502676 | 235,182,158,223 | 9,7,2,6 | 96,150,104,97 | 0,2,1,1 | 1108 | 149 | 0.00017 | 0.00757 | 0.778,0.7 | 1.0,0.91,0 | -0.117 |
| ENSMUSG00000057176 | Ccdc189 | chr7 | - | 127184316 | 127184433 | 127184164 | 127184225 | 127184625 | 127184696 | 22,23,67,37 | 6,1,0,7 | 44,32,27,28 | 0,4,0,0 | 266 | 149 | 0.00127 | 0.03288 | 0.673,0.9 | 1.0,0.818, | -0.117 |
| ENSMUSG00000032567 | Aste1 | chr9 | + | 105273996 | 105275057 | 105273743 | 105273904 | 105278674 | 105278885 | 260,103,262,179 | 6,3,10,6 | 211,133,247,203 | 4,1,5,0 | 1210 | 149 | 0.00503 | 0.08971 | 0.842,0.8 | 0.867,0.9 | -0.117 |
| ENSMUSG00000035293 | G2e3 | chr12 | + | 51410199 | 51410329 | 51409968 | 51410093 | 51414557 | 51414739 | 29,14,22,12 | 0,0,6,4 | 30,21,14,24 | 1,0,0,3 | 279 | 149 | 0.00569 | 0.09773 | 1.0,1.0,0. | 0.941,1.0, | -0.118 |
| ENSMUSG00000035954 | Dock4 | chr12 | + | 40888439 | 40888553 | 40886628 | 40886701 | 40891235 | 40891297 | 161,80,171,154 | 156,152,146,143 | 90,146,104,141 | 90,80,71,72 | 263 | 149 | 0.00393 | 0.07574 | 0.369,0.2 | 0.362,0.5 | -0.118 |
| ENSMUSG00000032560 | Dnajc13 | chr9 | - | 104087067 | 104087082 | 104086503 | 104086586 | 104089959 | 104090103 | 0,0,0,0 | 28,29,29,35 | 0,11,0,5 | 22,17,19,39 | 164 | 149 | 5.64E-06 | 0.00056 | 0.0,0.0,0. | 0.0,0.37,0 | -0.118 |
| ENSMUSG00000016382 | Pls3 | chrX | - | 74843322 | 74843349 | 74843000 | 74843096 | 74844107 | 74844250 | 234,170,398,217 | 154,105,192,119 | 298,202,400,223 | 135,89,88,62 | 176 | 149 | 8.94E-05 | 0.00464 | 0.563,0.5 | 0.651,0.6 | -0.118 |
| ENSMUSG00000026792 | Lrsam1 | chr2 | - | 32845192 | 32845261 | 32844213 | 32844298 | 32849528 | 32849625 | 27,31,27,25 | 5,0,8,0 | 48,32,28,39 | 0,1,0,0 | 218 | 149 | 1.05E-05 | 0.00091 | 0.787,1.0, | 1.0,0.956, | -0.118 |
| ENSMUSG00000057110 | Cntrl | chr2 | + | 35023969 | 35024123 | 35019632 | 35019827 | 35025479 | 35025700 | 20,39,92,47 | 6,0,16,2 | 32,72,51,41 | 1,2,0,3 | 303 | 149 | 0.00195 | 0.04497 | 0.621,1.0, | 0.94,0.94 | -0.119 |
| ENSMUSG00000074796 | Slc4a11 | chr2 | - | 130538567 | 130538675 | 130537960 | 130538023 | 130539278 | 130539389 | 58,105,85,90 | 15,5,5,4 | 28,72,12,56 | 0,3,0,0 | 257 | 149 | 8.42E-06 | 0.00077 | 0.692,0.9 | 1.0,0.933, | -0.12 |
| ENSMUSG00000029575 | Mmab | chr5 | - | 114575110 | 114575183 | 114574770 | 114574868 | 114575878 | 114575936 | 125,79,160,93 | 20,8,0,16 | 116,95,99,82 | 4,0,0,0 | 222 | 149 | 1.19E-09 | 7.57E-07 | 0.808,0.8 | 0.951,1.0, | -0.12 |
| ENSMUSG00000037514 | Pank2 | chr2 | + | 131124511 | 131124688 | 131116104 | 131116183 | 131138192 | 131138228 | 42,29,40,36 | 3,2,5,0 | 50,41,28,34 | 0,0,0,0 | 326 | 149 | 9.40E-05 | 0.00481 | 0.865,0.8 | 1.0,1.0,1. | -0.12 |
| ENSMUSG00000035713 | Usp35 | chr7 | - | 96971255 | 96971385 | 96970766 | 96970868 | 96973414 | 96973547 | 88,75,140,53 | 15,11,4,11 | 104,74,98,67 | 5,7,0,3 | 279 | 149 | 0.00099 | 0.02723 | 0.758,0.7 | 0.917,0.8 | -0.12 |
| ENSMUSG00000047495 | Dlgap2 | chr8 | + | 14229582 | 14229681 | 14145874 | 14146141 | 14307971 | 14308026 | 160,142,207,129 | 24,12,31,15 | 180,119,167,143 | 10,0,13,0 | 248 | 149 | 8.51E-08 | 2.20E-05 | 0.8,0.877, | 0.915,1.0, | -0.121 |
| ENSMUSG00000063142 | Kcnma1 | chr14 | - | 23356792 | 23356821 | 23348761 | 23350110 | 23359107 | 23359332 | 31,12,20,16 | 62,51,87,52 | 39,10,35,36 | 59,28,59,47 | 178 | 149 | 0.00159 | 0.03861 | 0.295,0.1 | 0.356,0.2 | -0.121 |
| ENSMUSG00000038784 | Cnot4 | chr6 | - | 35001062 | 35001275 | 34999005 | 35000367 | 35027996 | 35028494 | 133,121,219,152 | 18,19,17,21 | 157,84,132,97 | 8,4,3,9 | 362 | 149 | 0.00022 | 0.00919 | 0.753,0.7 | 0.89,0.89 | -0.121 |
| ENSMUSG00000039153 | Runx2 | chr17 | - | 44949399 | 44949592 | 44806873 | 44808130 | 44950551 | 44950713 | 11,1,38,22 | 6,0,7,0 | 64,12,58,18 | 0,3,0,0 | 342 | 149 | 0.00032 | 0.01219 | 0.444,1.0, | 1.0,0.635, | -0.122 |
| ENSMUSG00000000827 | Tpd52l2 | chr2 | + | 181152296 | 181152323 | 181149958 | 181150060 | 181153366 | 181153408 | 183,177,155,176 | 143,130,138,111 | 173,176,143,145 | 104,73,69,50 | 176 | 149 | 7.87E-05 | 0.00417 | 0.52,0.53 | 0.585,0.6 | -0.122 |
| ENSMUSG00000037940 | Inpp4b | chr8 | + | 82571084 | 82571135 | 82497361 | 82497553 | 82583320 | 82583400 | 39,47,53,31 | 13,5,7,7 | 10,35,19,16 | 1,2,1,1 | 200 | 149 | 0.00061 | 0.0195 | 0.691,0.8 | 0.882,0.9 | -0.122 |
| ENSMUSG00000002342 | Tmem161 | chr8 | + | 70633585 | 70633699 | 70633355 | 70633518 | 70633798 | 70633862 | 73,54,36,39 | 5,2,7,6 | 67,60,44,64 | 1,0,2,2 | 263 | 149 | 5.46E-05 | 0.00317 | 0.892,0.9 | 0.974,1.0, | -0.122 |
| ENSMUSG00000041471 | Shld2 | chr14 | - | 33981581 | 33981689 | 33971133 | 33971328 | 34030996 | 34031051 | 19,27,27,23 | 3,0,3,4 | 11,13,16,27 | 0,1,0,0 | 257 | 149 | 0.00013 | 0.00633 | 0.786,1.0, | 1.0,0.883, | -0.122 |
| ENSMUSG00000031626 | Sorbs2 | chr8 | + | 46194523 | 46194616 | 46189189 | 46189239 | 46198456 | 46198483 | 171,161,297,139 | 74,81,95,65 | 147,211,162,166 | 34,71,36,36 | 242 | 149 | 0.00031 | 0.01168 | 0.587,0.5 | 0.727,0.6 | -0.122 |
| ENSMUSG00000027752 | Exosc8 | chr3 | - | 54641523 | 54641587 | 54641078 | 54641152 | 54641716 | 54641734 | 253,97,203,166 | 32,33,47,19 | 169,196,137,152 | 23,1,10,14 | 213 | 149 | 2.02E-05 | 0.0015 | 0.847,0.6 | 0.837,0.9 | -0.123 |
| ENSMUSG00000046897 | Zfp740 | chr15 | + | 102113663 | 102113740 | 102113005 | 102113318 | 102116205 | 102116355 | 29,37,11,43 | 129,88,115,98 | 42,25,47,51 | 100,78,64,50 | 226 | 149 | 0.0012 | 0.03123 | 0.129,0.2 | 0.217,0.1 | -0.123 |
| ENSMUSG00000020536 | Llgl1 | chr11 | + | 60597577 | 60597632 | 60597263 | 60597399 | 60597784 | 60597939 | 136,101,139,141 | 27,34,55,12 | 156,106,147,103 | 16,10,17,10 | 204 | 149 | 0.00036 | 0.01318 | 0.786,0.6 | 0.877,0.8 | -0.123 |
| ENSMUSG00000021692 | Dimt1 | chr13 | + | 107089941 | 107090006 | 107088823 | 107088916 | 107093601 | 107093708 | 103,69,104,52 | 2,21,22,14 | 74,41,51,67 | 0,7,2,5 | 214 | 149 | 0.00076 | 0.02265 | 0.973,0.6 | 1.0,0.803, | -0.124 |
| ENSMUSG00000015377 | Dennd6b | chr15 | - | 89072661 | 89072724 | 89071534 | 89071657 | 89072813 | 89072893 | 138,141,169,122 | 26,17,25,18 | 85,88,101,111 | 1,2,7,6 | 212 | 149 | 6.48E-08 | 1.81E-05 | 0.789,0.8 | 0.984,0.9 | -0.124 |
| ENSMUSG00000022201 | Zfr | chr15 | + | 12149613 | 12149805 | 12146234 | 12146482 | 12150258 | 12150550 | 1,242,107,911,061,300 | 190,168,498,98 | 88,410,249,571,125 | 166,58,66,72 | 341 | 149 | 0.0053 | 0.09304 | 0.741,0.7 | 0.699,0.8 | -0.124 |
| ENSMUSG00000026566 | Mpzl1 | chr1 | - | 165429318 | 165429433 | 165421222 | 165421246 | 165432178 | 165432311 | 22,23,29,13 | 50,47,63,57 | 32,27,26,49 | 47,52,29,43 | 264 | 149 | 0.00102 | 0.02786 | 0.199,0.2 | 0.278,0.2 | -0.124 |
| ENSMUSG00000109006 | B230209E | chr7 | - | 61254263 | 61254448 | 61184823 | 61185236 | 61264907 | 61265075 | 437,384,633,316 | 79,110,112,81 | 277,207,325,246 | 53,39,45,0 | 334 | 149 | 0.00209 | 0.04759 | 0.712,0.6 | 0.7,0.703, | -0.124 |
| ENSMUSG00000021071 | Trim9 | chr12 | - | 70314007 | 70314037 | 70301811 | 70302000 | 70314835 | 70314974 | 255,174,274,113 | 144,98,200,122 | 291,175,214,171 | 105,88,106,57 | 179 | 149 | 0.00046 | 0.01569 | 0.596,0.5 | 0.698,0.6 | -0.125 |
| ENSMUSG00000060147 | Serpinb6a | chr13 | - | 34180225 | 34180334 | 34115454 | 34115629 | 34186448 | 34186750 | 38,28,18,14 | 5,7,0,4 | 24,15,23,40 | 0,2,2,0 | 258 | 149 | 0.00029 | 0.01129 | 0.814,0.6 | 1.0,0.812, | -0.125 |
| ENSMUSG00000045672 | Col27a1 | chr4 | + | 63211437 | 63211491 | 63209759 | 63209813 | 63211706 | 63211760 | 19,35,22,21 | 7,6,7,1 | 6,12,22,34 | 2,0,0,2 | 203 | 149 | 0.00021 | 0.00868 | 0.666,0.8 | 0.688,1.0, | -0.125 |

| | | | | | | | | | | | | | | | | | | | | |
|---|---|---|---|---|---|---|---|---|---|---|---|---|---|---|---|---|---|---|---|---|
| ENSMUSG00000027259 | Adal | chr2 | + | 120978721 | 120978803 | 120973629 | 120973690 | 120980700 | 120980827 | 33,23,42,19 | 8,0,8,0 | 31,11,21,12 | 0,0,0,0 | 231 | 149 | 1.02E-06 | 0.00016 | 0.727,1.0, | 1.0,1.0,1. | -0.125 |
| ENSMUSG00000019986 | Ahi1 | chr10 | + | 20946186 | 20946442 | 20934051 | 20934149 | 20948441 | 20948500 | 7,528,111,317,831 | 827,660,903,865 | 749,704,797,846 | 428,575,287,361 | 405 | 149 | 9.10E-05 | 0.0047 | 0.251,0.3 | 0.392,0.3 | -0.125 |
| ENSMUSG00000027200 | Sema6d | chr2 | + | 124502624 | 124502681 | 124502163 | 124502264 | 124504123 | 124504279 | 36,43,69,47 | 56,40,67,38 | 84,37,79,68 | 53,24,37,51 | 206 | 149 | 0.00402 | 0.07664 | 0.317,0.4 | 0.534,0.5 | -0.126 |
| ENSMUSG00000039671 | Zmynd8 | chr2 | - | 165726098 | 165726159 | 165716796 | 165717695 | 165726527 | 165726794 | 3,6,15,1 | 41,41,60,15 | 13,7,12,8 | 25,32,28,22 | 210 | 149 | 0.0027 | 0.0576 | 0.049,0.0 | 0.27,0.13 | -0.126 |
| ENSMUSG00000039842 | Mcph1 | chr8 | + | 18838254 | 18838492 | 18738997 | 18739075 | 18851421 | 18853205 | 72,62,88,60 | 0,7,13,0 | 68,57,66,65 | 0,0,0,0 | 387 | 149 | 2.13E-07 | 4.77E-05 | 1.0,0.773, | 1.0,1.0,1. | -0.126 |
| ENSMUSG00000030096 | Slc6a6 | chr6 | + | 91718158 | 91718283 | 91717929 | 91718033 | 91721897 | 91722010 | 375,223,309,293 | 76,58,125,69 | 296,316,268,289 | 51,44,25,47 | 274 | 149 | 1.63E-05 | 0.00127 | 0.728,0.6 | 0.759,0.7 | -0.126 |
| ENSMUSG00000110080 | Gm6145 | chr14 | - | 105865947 | 105866015 | 105862842 | 105864927 | 105867921 | 105868248 | 27,13,23,19 | 0,0,7,5 | 17,23,18,18 | 0,0,0,1 | 217 | 149 | 2.16E-05 | 0.00157 | 1.0,1.0,0. | 1.0,1.0,1. | -0.127 |
| ENSMUSG00000028525 | Pde4b | chr4 | + | 102278745 | 102279017 | 102112426 | 102112653 | 102344407 | 102344602 | 70,67,120,62 | 126,75,139,84 | 106,73,55,58 | 92,29,70,34 | 421 | 149 | 0.00246 | 0.05385 | 0.164,0.2 | 0.29,0.47 | -0.128 |
| ENSMUSG00000024174 | Pot1b | chr17 | - | 56002026 | 56002317 | 55999721 | 55999877 | 56005456 | 56005587 | 72,67,68,56 | 8,6,3,0 | 60,47,76,46 | 0,1,0,0 | 440 | 149 | 6.63E-07 | 0.00012 | 0.753,0.7 | 1.0,0.941, | -0.128 |
| ENSMUSG00000039375 | Wdr17 | chr8 | - | 55156868 | 55156974 | 55146091 | 55146275 | 55177322 | 55177423 | 46,23,9,38 | 1,0,4,4 | 14,35,11,26 | 1,0,0,0 | 255 | 149 | 0.00155 | 0.03809 | 0.964,1.0, | 0.891,1.0, | -0.128 |
| ENSMUSG00000039934 | Gsap | chr5 | + | 21427387 | 21427437 | 21426223 | 21426293 | 21431241 | 21431346 | 30,19,14,12 | 3,6,2,3 | 9,27,26,12 | 1,0,0,2 | 199 | 149 | 9.15E-05 | 0.00472 | 0.882,0.7 | 0.871,1.0, | -0.129 |
| ENSMUSG00000084885 | 3010001F | chrX | + | 151168640 | 151168748 | 151152837 | 151153018 | 151186308 | 151186473 | 32,22,20,21 | 3,0,10,1 | 11,22,27,23 | 0,0,3,0 | 257 | 149 | 0.00051 | 0.01715 | 0.861,1.0, | 1.0,1.0,0. | -0.129 |
| ENSMUSG00000003226 | Ranbp2 | chr10 | + | 58303211 | 58303295 | 58302936 | 58303116 | 58306448 | 58306584 | 197,171,324,160 | 22,19,20,18 | 98,169,134,202 | 0,0,1,1 | 233 | 149 | 0 | 0 | 0.851,0.8 | 1.0,1.0,0. | -0.129 |
| ENSMUSG00000037989 | Wnk2 | chr13 | - | 49203959 | 49204058 | 49197849 | 49197894 | 49206034 | 49206249 | 87,54,93,41 | 88,44,89,88 | 96,66,78,84 | 55,61,54,42 | 248 | 149 | 0.00468 | 0.08559 | 0.373,0.4 | 0.512,0.3 | -0.129 |
| ENSMUSG00000019338 | Zfp687 | chr3 | - | 94921143 | 94921239 | 94919150 | 94919787 | 94922630 | 94922759 | 3,15,14,16 | 128,95,186,107 | 26,23,29,41 | 85,87,62,83 | 245 | 149 | 4.47E-13 | 1.50E-09 | 0.014,0.0 | 0.157,0.1 | -0.13 |
| ENSMUSG00000024050 | Wiz | chr17 | - | 32597813 | 32598092 | 32586797 | 32587121 | 32606546 | 32606791 | 27,13,24,23 | 41,35,43,38 | 32,16,38,34 | 29,16,22,36 | 428 | 149 | 0.00036 | 0.01311 | 0.187,0.1 | 0.278,0.2 | -0.13 |
| ENSMUSG00000032212 | Sltm | chr9 | + | 70451251 | 70451339 | 70450035 | 70450363 | 70479430 | 70479458 | 63,57,143,82 | 6,5,17,12 | 70,65,74,78 | 2,0,2,0 | 237 | 149 | 1.17E-09 | 7.57E-07 | 0.868,0.8 | 0.957,1.0, | -0.13 |
| ENSMUSG00000022556 | Hsf1 | chr15 | + | 76384330 | 76384396 | 76384147 | 76384253 | 76384466 | 76384536 | 129,100,155,133 | 50,34,38,33 | 138,133,136,132 | 40,28,18,0 | 215 | 149 | 0.00087 | 0.02495 | 0.641,0.6 | 0.705,0.7 | -0.131 |
| ENSMUSG00000023051 | Tarbp2 | chr15 | + | 102428909 | 102429012 | 102427160 | 102427726 | 102429572 | 102429648 | 91,104,78,68 | 12,10,20,5 | 83,68,78,55 | 1,4,2,2 | 252 | 149 | 2.81E-06 | 0.00035 | 0.818,0.8 | 0.98,0.91, | -0.131 |
| ENSMUSG00000024982 | Zdhhc6 | chr19 | - | 55290989 | 55291036 | 55286732 | 55287320 | 55291122 | 55291268 | 153,111,142,171 | 40,60,76,47 | 161,198,143,120 | 52,38,24,15 | 196 | 149 | 0.00091 | 0.02572 | 0.744,0.5 | 0.702,0.7 | -0.132 |
| ENSMUSG00000056724 | Nbeal2 | chr9 | - | 110472779 | 110472862 | 110471452 | 110471538 | 110473076 | 110473195 | 38,17,31,34 | 4,5,1,3 | 20,44,29,5 | 0,0,2,0 | 232 | 149 | 6.14E-05 | 0.00347 | 0.859,0.6 | 1.0,1.0,0. | -0.132 |
| ENSMUSG00000031376 | Atp2b3 | chrX | + | 72577490 | 72577532 | 72573902 | 72574028 | 72578961 | 72579126 | 219,167,275,159 | 183,166,202,146 | 174,217,228,266 | 121,146,82,100 | 191 | 149 | 0.00033 | 0.01225 | 0.483,0.4 | 0.529,0.5 | -0.132 |
| ENSMUSG00000032872 | Cyb5r4 | chr9 | + | 86931154 | 86931277 | 86924832 | 86924926 | 86937817 | 86937958 | 247,149,171,111 | 15,13,26,7 | 126,185,162,153 | 2,0,0,0 | 272 | 149 | 2.22E-16 | 1.38E-12 | 0.9,0.863, | 0.972,1.0, | -0.132 |
| ENSMUSG00000042439 | Zfp532 | chr18 | + | 65777263 | 65777440 | 65758208 | 65758405 | 65789421 | 65789703 | 20,17,29,24 | 4,3,6,0 | 16,19,27,22 | 1,1,2,0 | 326 | 149 | 0.0026 | 0.05616 | 0.696,0.7 | 0.88,0.89 | -0.133 |
| ENSMUSG00000029478 | Ncor2 | chr5 | - | 125230670 | 125230716 | 125196545 | 125196673 | 125256060 | 125256244 | 35,18,15,12 | 0,0,13,0 | 25,26,26,45 | 0,0,0,0 | 195 | 149 | 0.0042 | 0.07905 | 1.0,1.0,0. | 1.0,1.0,1. | -0.133 |
| ENSMUSG00000024943 | Smc5 | chr19 | + | 23192516 | 23192563 | 23192008 | 23192104 | 23195768 | 23195894 | 95,48,79,100 | 7,21,8,23 | 49,48,58,68 | 1,2,3,7 | 194 | 149 | 0.0002 | 0.00852 | 0.912,0.6 | 0.974,0.9 | -0.135 |
| ENSMUSG00000020821 | Kif1c | chr11 | + | 70593271 | 70593377 | 70591373 | 70591535 | 70593597 | 70593731 | 145,130,192,133 | 42,38,51,33 | 238,179,154,145 | 53,9,18,23 | 255 | 149 | 0.00011 | 0.00525 | 0.669,0.6 | 0.724,0.9 | -0.135 |
| ENSMUSG00000033852 | Gm28042 | chr2 | + | 119861982 | 119862132 | 119861296 | 119861353 | 119862321 | 119862481 | 27,21,40,20 | 0,0,7,4 | 20,22,17,19 | 0,0,0,0 | 299 | 149 | 1.61E-05 | 0.00126 | 1.0,1.0,0. | 1.0,1.0,1. | -0.136 |
| ENSMUSG00000026017 | Carf | chr1 | + | 60147241 | 60147359 | 60144469 | 60144593 | 60148485 | 60148608 | 24,26,21,33 | 0,0,14,0 | 29,36,34,40 | 0,0,0,0 | 267 | 149 | 0.00308 | 0.06325 | 1.0,1.0,0. | 1.0,1.0,1. | -0.136 |
| ENSMUSG00000020821 | Kif1c | chr11 | + | 70593257 | 70593377 | 70591475 | 70591535 | 70593597 | 70593731 | 134,132,161,128 | 42,38,51,33 | 228,165,122,128 | 53,9,18,23 | 269 | 149 | 0.00032 | 0.012 | 0.639,0.6 | 0.704,0.9 | -0.136 |
| ENSMUSG00000074527 | Gm14296 | chr2 | - | 176610459 | 176610520 | 176594470 | 176594653 | 176610725 | 176610852 | 68,102,125,40 | 7,19,26,21 | 75,55,71,62 | 12,0,4,10 | 210 | 149 | 0.00441 | 0.08167 | 0.873,0.7 | 0.816,1.0, | -0.136 |
| ENSMUSG00000086968 | 4933431E | chr3 | - | 107801904 | 107801972 | 107801366 | 107801544 | 107802176 | 107802308 | 0,4,29,3 | 109,78,162,109 | 59,30,17,23 | 110,81,120,91 | 217 | 149 | 1.52E-10 | 1.65E-07 | 0.0,0.034, | 0.269,0.2 | -0.137 |
| ENSMUSG00000027751 | Supt20 | chr3 | + | 54614508 | 54614562 | 54614280 | 54614397 | 54615677 | 54615817 | 136,127,145,206 | 33,72,91,26 | 116,103,152,126 | 26,18,19,19 | 203 | 149 | 0.00509 | 0.09053 | 0.752,0.5 | 0.766,0.8 | -0.137 |
| ENSMUSG00000026074 | Map4k4 | chr1 | + | 40042922 | 40043147 | 40040589 | 40040751 | 40043966 | 40044070 | 249,136,192,140 | 56,38,59,46 | 191,125,121,167 | 14,35,17,26 | 374 | 149 | 0.00232 | 0.05174 | 0.639,0.5 | 0.845,0.5 | -0.137 |
| ENSMUSG00000054256 | Msi1 | chr5 | + | 115585188 | 115585245 | 115583453 | 115583534 | 115588273 | 115588342 | 85,91,158,122 | 23,19,44,27 | 127,111,115,122 | 17,0,13,18 | 206 | 149 | 9.36E-06 | 0.00085 | 0.728,0.7 | 0.844,1.0, | -0.137 |
| ENSMUSG00000028906 | Epb41 | chr4 | - | 131682853 | 131682889 | 131682038 | 131682467 | 131684953 | 131685130 | 10,8,22,17 | 93,68,97,58 | 25,26,10,18 | 40,55,34,42 | 185 | 149 | 2.92E-05 | 0.00199 | 0.08,0.08 | 0.335,0.2 | -0.137 |
| ENSMUSG00000024082 | Ndufaf7 | chr17 | + | 79251262 | 79251373 | 79247023 | 79247134 | 79253724 | 79253811 | 32,27,22,25 | 0,1,12,0 | 22,12,11,14 | 0,0,0,0 | 260 | 149 | 0.00041 | 0.01431 | 1.0,0.939, | 1.0,1.0,1. | -0.137 |
| ENSMUSG00000068250 | Amn1 | chr6 | - | 149084891 | 149085036 | 149072334 | 149072548 | 149090103 | 149090193 | 188,98,173,120 | 31,33,26,26 | 149,93,143,102 | 37,1,0,19 | 294 | 149 | 0.00236 | 0.05246 | 0.755,0.6 | 0.671,0.9 | -0.138 |
| ENSMUSG00000034579 | Pla2g3 | chr11 | + | 3440142 | 3440277 | 3438989 | 3439894 | 3440806 | 3441090 | 52,7,14,25 | 2,1,1,4 | 33,20,63,29 | 0,0,3,0 | 284 | 149 | 0.00513 | 0.09091 | 0.932,0.7 | 1.0,1.0,0. | -0.138 |
| ENSMUSG00000026527 | Rgs7 | chr1 | - | 174894182 | 174894260 | 174886652 | 174887360 | 174905754 | 174905844 | 181,102,170,107 | 73,55,85,100 | 145,106,161,128 | 33,42,48,45 | 227 | 149 | 0.00086 | 0.02491 | 0.619,0.5 | 0.743,0.6 | -0.139 |
| ENSMUSG00000030733 | Sh2b1 | chr7 | - | 126067121 | 126067221 | 126066165 | 126066945 | 126067591 | 126067813 | 47,36,63,42 | 134,137,162,100 | 84,57,86,82 | 106,91,99,106 | 249 | 149 | 4.18E-09 | 1.92E-06 | 0.173,0.1 | 0.322,0.2 | -0.139 |
| ENSMUSG00000051113 | Fam71e1 | chr7 | + | 44146829 | 44147057 | 44146338 | 44146457 | 44149573 | 44149790 | 37,15,47,17 | 0,0,13,3 | 27,20,13,19 | 0,0,1,0 | 377 | 149 | 1.40E-05 | 0.00113 | 1.0,1.0,0. | 1.0,1.0,0. | -0.139 |
| ENSMUSG00000071337 | Tia1 | chr6 | + | 86396082 | 86396115 | 86395859 | 86395914 | 86397305 | 86397393 | 90,68,69,59 | 123,185,183,123 | 48,71,77,71 | 62,89,57,97 | 182 | 149 | 0.00026 | 0.01016 | 0.375,0.2 | 0.388,0.3 | -0.14 |
| ENSMUSG00000026566 | Mpzl1 | chr1 | - | 165429318 | 165429424 | 165419808 | 165421246 | 165432178 | 165432311 | 51,50,68,49 | 50,47,63,57 | 91,60,57,96 | 47,52,29,43 | 255 | 149 | 0.00107 | 0.02885 | 0.373,0.3 | 0.531,0.4 | -0.14 |
| ENSMUSG00000049658 | Bdp1 | chr13 | - | 100220648 | 100220703 | 100215166 | 100215307 | 100225860 | 100225955 | 56,43,80,36 | 17,8,5,6 | 36,31,34,54 | 0,0,3,4 | 204 | 149 | 7.26E-05 | 0.00394 | 0.706,0.7 | 1.0,1.0,0. | -0.14 |
| ENSMUSG00000052917 | Senp7 | chr16 | + | 55893085 | 55893247 | 55877887 | 55877987 | 55895839 | 55895941 | 31,26,27,28 | 1,0,13,0 | 16,12,5,17 | 0,0,0,0 | 311 | 149 | 0.00026 | 0.01017 | 0.937,1.0, | 1.0,1.0,1. | -0.141 |
| ENSMUSG00000040430 | Pitpnc1 | chr11 | - | 107107503 | 107107622 | 107098717 | 107103484 | 107117056 | 107117120 | 149,105,166,123 | 261,168,172,205 | 163,127,143,150 | 126,124,81,133 | 268 | 149 | 2.64E-06 | 0.00033 | 0.241,0.2 | 0.418,0.3 | -0.141 |
| ENSMUSG00000045983 | Eif4g1 | chr16 | + | 20492348 | 20492405 | 20491498 | 20491626 | 20492977 | 20493064 | 79,75,93,32 | 4,2,25,19 | 65,55,74,98 | 8,0,1,7 | 206 | 149 | 0.0047 | 0.08578 | 0.935,0.9 | 0.855,1.0, | -0.142 |
| ENSMUSG00000033467 | Crlf2 | chr5 | - | 109703372 | 109703493 | 109703138 | 109703223 | 109703884 | 109704047 | 89,73,95,72 | 21,16,12,26 | 77,68,68,67 | 9,6,15,0 | 270 | 149 | 0.00415 | 0.07826 | 0.7,0.716, | 0.825,0.8 | -0.142 |
| ENSMUSG00000029122 | Evc | chr5 | - | 37472105 | 37472171 | 37471908 | 37471915 | 37473668 | 37473767 | 2,2,0,5 | 23,17,12,10 | 9,4,6,11 | 21,13,24,11 | 215 | 149 | 0.003 | 0.06202 | 0.057,0.0 | 0.229,0.1 | -0.143 |
| ENSMUSG00000051669 | AU02109 | chr16 | - | 5035661 | 5035812 | 5034605 | 5034723 | 5038159 | 5038309 | 18,31,13,27 | 12,0,0,0 | 44,26,23,17 | 0,0,0,0 | 300 | 149 | 0.00512 | 0.09082 | 0.427,1.0, | 1.0,1.0,1. | -0.143 |
| ENSMUSG00000035356 | Nfkbiz | chr16 | - | 55642130 | 55642478 | 55639929 | 55640069 | 55656683 | 55656755 | 20,15,45,22 | 1,1,13,0 | 53,22,48,23 | 2,0,0,1 | 497 | 149 | 0.00016 | 0.0071 | 0.857,0.8 | 0.888,1.0, | -0.144 |
| ENSMUSG00000018774 | Cd68 | chr11 | - | 69556328 | 69556762 | 69556072 | 69556192 | 69556845 | 69556979 | 456,236,306,304 | 88,49,71,87 | 400,296,190,273 | 48,40,16,45 | 583 | 149 | 7.50E-05 | 0.00404 | 0.57,0.55 | 0.68,0.65 | -0.144 |
| ENSMUSG00000031938 | 4931406C | chr9 | - | 15205981 | 15206055 | 15205727 | 15205829 | 15212662 | 15212809 | 42,16,50,18 | 5,0,0,9 | 45,24,32,29 | 0,0,0,0 | 223 | 149 | 1.02E-05 | 0.0009 | 0.849,1.0, | 1.0,1.0,1. | -0.145 |
| ENSMUSG00000031258 | Xkrx | chrX | - | 133057961 | 133058230 | 133049791 | 133052045 | 133062324 | 133062677 | 72,21,33,22 | 2,4,1,2 | 23,43,13,31 | 0,2,0,0 | 418 | 149 | 0.00093 | 0.02603 | 0.928,0.6 | 1.0,0.885, | -0.146 |
| ENSMUSG00000027882 | Stxbp3 | chr3 | - | 108717192 | 108717317 | 108712553 | 108712649 | 108720193 | 108720284 | 160,104,157,193 | 16,13,34,23 | 138,187,178,147 | 2,3,10,6 | 274 | 149 | 3.90E-10 | 3.12E-07 | 0.845,0.8 | 0.974,0.9 | -0.147 |
| ENSMUSG00000026867 | Gapvd1 | chr2 | - | 34589837 | 34589918 | 34585497 | 34585620 | 34590774 | 34590862 | 197,184,126,150 | 85,97,123,81 | 150,197,147,132 | 72,35,39,53 | 230 | 149 | 0.00181 | 0.04263 | 0.6,0.551, | 0.574,0.7 | -0.147 |
| ENSMUSG00000035601 | Trmt10b | chr4 | + | 45299746 | 45299897 | 45297154 | 45297195 | 45300416 | 45300627 | 1,0,0,0 | 90,47,85,95 | 21,1,2,0 | 8,52,57,48 | 300 | 149 | 0.00541 | 0.09463 | 0.005,0.0, | 0.566,0.0 | -0.147 |
| ENSMUSG00000018774 | Cd68 | chr11 | - | 69556301 | 69556762 | 69556072 | 69556192 | 69556845 | 69556888 | 389,208,248,262 | 88,49,71,87 | 319,262,151,243 | 48,40,16,45 | 610 | 149 | 7.42E-05 | 0.00401 | 0.519,0.5 | 0.619,0.6 | -0.147 |
| ENSMUSG00000003812 | Dnase2a | chr8 | + | 85635885 | 85635964 | 85635613 | 85635791 | 85636400 | 85636593 | 13,10,31,20 | 0,8,3,0 | 28,23,22,32 | 0,0,0,2 | 228 | 149 | 0.00069 | 0.02124 | 1.0,0.45,0 | 1.0,1.0,1. | -0.148 |
| ENSMUSG00000014837 | 4931428F | chr8 | - | 106015400 | 106015797 | 106011887 | 106012501 | 106016035 | 106016160 | 246,118,237,165 | 45,14,37,35 | 233,186,219,152 | 20,13,14,16 | 546 | 149 | 3.37E-05 | 0.00222 | 0.599,0.6 | 0.761,0.7 | -0.148 |
| ENSMUSG00000042323 | Pbrm1 | chr14 | + | 30835805 | 30835961 | 30829065 | 30829267 | 30838969 | 30839040 | 74,56,72,51 | 92,98,120,102 | 58,63,66,92 | 34,70,68,57 | 305 | 149 | 3.29E-05 | 0.00218 | 0.282,0.2 | 0.455,0.3 | -0.15 |
| ENSMUSG00000006728 | Cdk4 | chr10 | + | 126900070 | 126900322 | 126899472 | 126899628 | 126900453 | 126900589 | 441,346,433,453 | 215,125,162,98 | 498,377,362,314 | 63,84,56,80 | 401 | 149 | 0.00015 | 0.00676 | 0.433,0.5 | 0.746,0.6 | -0.15 |
| ENSMUSG00000024830 | Rps6kb2 | chr19 | - | 4209401 | 4209459 | 4209225 | 4209326 | 4211066 | 4211214 | 89,79,95,98 | 41,60,52,47 | 76,101,98,108 | 36,22,28,22 | 207 | 149 | 0.00039 | 0.014 | 0.61,0.48 | 0.603,0.7 | -0.15 |
| ENSMUSG00000028653 | Trit1 | chr4 | + | 122942944 | 122943057 | 122938283 | 122938429 | 122943269 | 122943347 | 37,26,31,45 | 4,11,9,3 | 21,30,51,25 | 4,0,0,3 | 262 | 149 | 0.0016 | 0.03874 | 0.84,0.57 | 0.749,1.0, | -0.151 |
| ENSMUSG00000028545 | Bend5 | chr4 | + | 111305702 | 111305851 | 111290283 | 111290750 | 111311281 | 111311495 | 109,57,34,74 | 10,11,5,2 | 68,77,70,89 | 4,0,0,0 | 298 | 149 | 8.70E-08 | 2.23E-05 | 0.845,0.7 | 0.895,1.0, | -0.152 |
| ENSMUSG00000079657 | Rab26 | chr17 | - | 24748887 | 24748944 | 24748714 | 24748791 | 24749364 | 24749417 | 62,29,86,44 | 8,6,4,7 | 69,54,72,60 | 0,0,0,0 | 206 | 149 | 2.33E-10 | 2.31E-07 | 0.849,0.7 | 1.0,1.0,1. | -0.153 |
| ENSMUSG00000041890 | Git2 | chr5 | - | 114875408 | 114875498 | 114871921 | 114872004 | 114877093 | 114877342 | 31,28,42,39 | 59,49,81,47 | 77,28,46,50 | 42,37,36,44 | 239 | 149 | 0.00033 | 0.01225 | 0.247,0.2 | 0.533,0.3 | -0.154 |
| ENSMUSG00000026288 | Inpp5d | chr1 | + | 87642759 | 87642815 | 87640914 | 87641011 | 87642998 | 87643035 | 23,24,45,36 | 7,9,7,4 | 19,25,32,21 | 2,0,2,2 | 205 | 149 | 0.0004 | 0.01423 | 0.705,0.6 | 0.873,1.0, | -0.155 |
| ENSMUSG00000019961 | Tmpo | chr10 | - | 90988489 | 90988600 | 90987570 | 90987659 | 90994609 | 90994707 | 55,46,44,38 | 119,135,219,141 | 66,72,79,77 | 107,87,125,70 | 260 | 149 | 8.52E-10 | 5.81E-07 | 0.209,0.1 | 0.261,0.3 | -0.157 |
| ENSMUSG00000021240 | Abcd4 | chr12 | - | 84652873 | 84653082 | 84652121 | 84652158 | 84655343 | 84655433 | 91,54,63,92 | 13,18,16,15 | 62,106,60,92 | 10,4,6,8 | 358 | 149 | 0.00143 | 0.03583 | 0.744,0.5 | 0.721,0.9 | -0.158 |
| ENSMUSG00000039542 | Ncam1 | chr9 | - | 49453187 | 49453202 | 49446508 | 49446550 | 49454310 | 49454330 | 23,13,30,23 | 16,3,7,1 | 22,23,14,25 | 2,4,0,0 | 164 | 149 | 0.00067 | 0.0206 | 0.566,0.7 | 0.909,0.8 | -0.159 |
| ENSMUSG00000023044 | Csad | chr15 | - | 102087215 | 102087267 | 102085753 | 102086221 | 102087395 | 102087595 | 62,70,79,86 | 27,31,40,30 | 75,62,84,91 | 30,14,16,3 | 201 | 149 | 0.00067 | 0.02053 | 0.63,0.62 | 0.65,0.76 | -0.16 |
| ENSMUSG00000059326 | Csf2ra | chr19 | - | 61215709 | 61215803 | 61215467 | 61215622 | 61216776 | 61216856 | 18,26,30,18 | 2,3,3,11 | 28,27,25,25 | 2,0,0,4 | 243 | 149 | 0.00277 | 0.05866 | 0.847,0.8 | 0.896,1.0, | -0.16 |

| | | | | | | | | | | | | | | | | | | | |
|---|---|---|---|---|---|---|---|---|---|---|---|---|---|---|---|---|---|---|---|
| ENSMUSG00000053046 | Brsk2 | chr7 | + | 141554578 | 141554626 | 141552656 | 141552751 | 141556211 | 141556412 | 6,3,2,5 | 53,57,96,77 | 26,16,43,24 | 117,61,97,52 | 197 | 149 | 5.75E-13 | 1.75E-09 | 0.079,0.0 | 0.144,0.1 | -0.16 |
| ENSMUSG00000090063 | Dlx6os1 | chr6 | - | 6822297 | 6822323 | 6820542 | 6820771 | 6823742 | 6824567 | 27,32,35,15 | 3,12,9,0 | 9,14,6,16 | 0,0,0,0 | 175 | 149 | 7.92E-08 | 2.15E-05 | 0.885,0.6 | 1.0,1.0,1. | -0.163 |
| ENSMUSG00000029769 | Ccdc136 | chr6 | + | 29417930 | 29418152 | 29417070 | 29417577 | 29426623 | 29426952 | 44,47,44,20 | 3,4,11,0 | 45,29,45,43 | 0,0,1,0 | 371 | 149 | 2.18E-07 | 4.82E-05 | 0.855,0.8 | 1.0,1.0,0. | -0.163 |
| ENSMUSG00000033902 | Mapkbp1 | chr2 | + | 119843781 | 119843799 | 119843392 | 119843575 | 119844099 | 119844260 | 146,113,149,83 | 56,60,80,54 | 130,131,124,94 | 33,33,18,26 | 167 | 149 | 2.36E-07 | 5.16E-05 | 0.699,0.6 | 0.779,0.7 | -0.163 |
| ENSMUSG00000032340 | Neo1 | chr9 | - | 58791752 | 58791911 | 58787874 | 58788138 | 58795683 | 58795776 | 146,125,189,157 | 253,140,203,155 | 255,137,142,193 | 93,101,122,97 | 308 | 149 | 1.40E-05 | 0.00113 | 0.218,0.3 | 0.57,0.39 | -0.164 |
| ENSMUSG00000028020 | Glrb | chr3 | - | 80816093 | 80816245 | 80769416 | 80769428 | 80819274 | 80819425 | 14,31,19,15 | 2,5,2,3 | 23,21,16,22 | 0,2,1,0 | 301 | 149 | 0.00014 | 0.00645 | 0.776,0.7 | 1.0,0.839, | -0.165 |
| ENSMUSG00000040473 | Cfap69 | chr5 | - | 5694423 | 5694500 | 5690128 | 5690227 | 5696935 | 5697045 | 56,33,92,38 | 3,12,14,17 | 44,38,55,30 | 3,6,3,0 | 226 | 149 | 0.00158 | 0.03846 | 0.925,0.6 | 0.906,0.8 | -0.165 |
| ENSMUSG00000043531 | Sorcs1 | chr19 | - | 50141197 | 50141358 | 50131738 | 50132585 | 50141491 | 50141597 | 2,2,5,7 | 42,25,15,38 | 9,14,19,15 | 13,18,34,26 | 310 | 149 | 8.39E-08 | 2.20E-05 | 0.022,0.0 | 0.25,0.27 | -0.168 |
| ENSMUSG00000108713 | Gm33027 | chr7 | - | 129004149 | 129004388 | 129001719 | 129003017 | 129004713 | 129005049 | 21,23,17,16 | 3,3,0,4 | 32,8,12,34 | 4,0,0,0 | 388 | 149 | 0.00298 | 0.06167 | 0.729,0.7 | 0.754,1.0, | -0.168 |
| ENSMUSG00000068250 | Amn1 | chr6 | - | 149086489 | 149086622 | 149072330 | 149072548 | 149090103 | 149090210 | 81,85,107,77 | 31,33,26,26 | 70,83,73,66 | 37,1,0,19 | 282 | 149 | 0.0046 | 0.08443 | 0.58,0.57 | 0.5,0.978, | -0.168 |
| ENSMUSG00000097589 | Dleu2 | chr14 | - | 61862668 | 61862811 | 61840479 | 61841255 | 61869885 | 61869932 | 26,18,53,37 | 3,14,0,0 | 32,22,44,59 | 0,0,0,4 | 292 | 149 | 0.0039 | 0.07537 | 0.816,0.3 | 1.0,1.0,1. | -0.168 |
| ENSMUSG00000058174 | Gm5148 | chr3 | - | 37770270 | 37770381 | 37768338 | 37769284 | 37778351 | 37778559 | 30,13,27,26 | 17,0,4,4 | 23,33,15,13 | 0,1,2,0 | 260 | 149 | 1.75E-05 | 0.00134 | 0.503,1.0, | 1.0,0.95,0 | -0.169 |
| ENSMUSG00000021713 | Ppwd1 | chr13 | - | 104350033 | 104350223 | 104346121 | 104346303 | 104353596 | 104353787 | 159,73,134,106 | 12,24,31,21 | 98,86,125,137 | 6,8,8,9 | 339 | 149 | 0.0001 | 0.00524 | 0.853,0.5 | 0.878,0.8 | -0.169 |
| ENSMUSG00000028789 | Azin2 | chr4 | - | 128853576 | 128853750 | 128844392 | 128844565 | 128855876 | 128855994 | 135,108,103,134 | 27,19,22,19 | 116,140,114,71 | 19,0,12,0 | 323 | 149 | 5.79E-05 | 0.00332 | 0.698,0.7 | 0.738,1.0, | -0.17 |
| ENSMUSG00000030768 | Disp1 | chr1 | - | 182955059 | 182955187 | 182916918 | 182917440 | 182984515 | 182984603 | 39,33,13,31 | 5,0,8,4 | 16,12,22,33 | 0,2,0,0 | 277 | 149 | 2.07E-05 | 0.00153 | 0.808,1.0, | 1.0,0.763, | -0.17 |
| ENSMUSG00000118604 | Gm53013 | chrX | + | 60165617 | 60165765 | 60162947 | 60163030 | 60178326 | 60178402 | 32,58,73,41 | 9,5,11,15 | 34,34,43,33 | 2,6,0,2 | 297 | 149 | 0.00217 | 0.04898 | 0.641,0.8 | 0.895,0.7 | -0.171 |
| ENSMUSG00000045503 | Sys1 | chr2 | + | 164309067 | 164309247 | 164305220 | 164305288 | 164314057 | 164314242 | 60,57,101,90 | 15,19,10,10 | 51,69,66,46 | 0,0,13,4 | 329 | 149 | 0.00261 | 0.05631 | 0.644,0.5 | 1.0,1.0,0. | -0.173 |
| ENSMUSG00000034518 | Hmgxb4 | chr8 | + | 75726626 | 75726741 | 75726440 | 75726519 | 75727277 | 75728212 | 30,40,34,25 | 58,34,57,22 | 43,30,39,25 | 18,22,29,13 | 264 | 149 | 0.0012 | 0.03123 | 0.226,0.3 | 0.574,0.4 | -0.173 |
| ENSMUSG00000028948 | Nol9 | chr4 | + | 152130404 | 152130492 | 152130151 | 152130287 | 152131003 | 152131101 | 86,63,93,69 | 13,12,31,32 | 76,75,75,42 | 7,16,1,3 | 237 | 149 | 0.00023 | 0.00952 | 0.806,0.7 | 0.872,0.7 | -0.174 |
| ENSMUSG00000004677 | Myo9b | chr8 | + | 71789519 | 71789609 | 71786956 | 71787038 | 71795405 | 71795528 | 10,12,41,19 | 0,7,0,14 | 19,17,10,10 | 0,0,0,3 | 239 | 149 | 0.00092 | 0.02592 | 1.0,0.517, | 1.0,1.0,1. | -0.175 |
| ENSMUSG00000048603 | Gm9828 | chr13 | + | 98455581 | 98455653 | 98453582 | 98454103 | 98457710 | 98457966 | 16,36,45,26 | 7,0,0,8 | 8,11,11,5 | 0,0,0,0 | 221 | 149 | 1.06E-05 | 0.00092 | 0.606,1.0, | 1.0,1.0,1. | -0.177 |
| ENSMUSG00000086363 | A330102I | chr13 | + | 29200692 | 29200862 | 29198240 | 29198517 | 29214140 | 29214224 | 34,25,36,34 | 0,10,4,3 | 20,31,37,20 | 0,0,2,0 | 319 | 149 | 9.99E-06 | 0.00088 | 1.0,0.539, | 1.0,1.0,0. | -0.177 |
| ENSMUSG00000056938 | Acbd4 | chr11 | + | 102994060 | 102994195 | 102993507 | 102993922 | 102994351 | 102994470 | 67,47,106,56 | 22,19,17,24 | 113,90,62,80 | 13,10,8,13 | 284 | 149 | 0.00025 | 0.00985 | 0.615,0.5 | 0.82,0.82 | -0.179 |
| ENSMUSG00000025144 | Cenpx | chr11 | - | 120602669 | 120602721 | 120602528 | 120602582 | 120604518 | 120604564 | 45,24,46,52 | 96,71,77,58 | 58,66,45,43 | 50,55,39,33 | 201 | 149 | 2.77E-05 | 0.00192 | 0.258,0.2, | 0.462,0.4 | -0.18 |
| ENSMUSG00000037134 | Prmt9 | chr8 | + | 78289077 | 78289145 | 78287435 | 78287603 | 78290604 | 78290711 | 73,54,27,44 | 11,0,12,16 | 29,55,48,20 | 0,0,5,1 | 217 | 149 | 4.74E-05 | 0.00287 | 0.82,1.0,0 | 1.0,1.0,0. | -0.18 |
| ENSMUSG00000020674 | Pxdn | chr12 | + | 30049497 | 30049692 | 30049163 | 30049272 | 30049890 | 30050048 | 112,70,88,66 | 59,96,106,85 | 116,91,63,66 | 44,59,24,33 | 344 | 149 | 0.00012 | 0.0059 | 0.451,0.2 | 0.533,0.4 | -0.181 |
| ENSMUSG00000031109 | Enox2 | chrX | - | 48375721 | 48375781 | 48257517 | 48257661 | 48376977 | 48377089 | 30,18,43,15 | 7,5,3,10 | 19,21,22,36 | 0,1,7,0 | 209 | 149 | 0.00512 | 0.09088 | 0.753,0.7 | 1.0,0.937, | -0.182 |
| ENSMUSG00000028028 | Alpk1 | chr3 | - | 127471334 | 127471488 | 127466968 | 127467131 | 127473009 | 127475101 | 19,19,32,24 | 3,2,6,2 | 33,15,28,32 | 0,0,0,2 | 303 | 149 | 5.15E-06 | 0.00053 | 0.757,0.8 | 1.0,1.0,1. | -0.182 |
| ENSMUSG00000097993 | Ptprv | chr1 | - | 135055774 | 135055851 | 135054753 | 135054820 | 135056854 | 135057488 | 53,28,27,31 | 39,0,20,7 | 19,37,11,23 | 0,0,10,0 | 226 | 149 | 0.00493 | 0.08842 | 0.473,1.0, | 1.0,1.0,0. | -0.183 |
| ENSMUSG00000022770 | Dlg1 | chr16 | + | 31673485 | 31673506 | 31672660 | 31672702 | 31676364 | 31676466 | 23,18,24,11 | 7,4,8,0 | 16,18,19,13 | 0,0,0,0 | 170 | 149 | 1.15E-08 | 4.48E-06 | 0.742,0.7 | 1.0,1.0,1. | -0.184 |
| ENSMUSG00000053046 | Brsk2 | chr7 | + | 141552656 | 141552751 | 141552369 | 141552459 | 141556211 | 141556666 | 205,169,246,180 | 144,100,190,101 | 288,183,273,207 | 89,59,62,76 | 244 | 149 | 2.04E-11 | 3.29E-08 | 0.465,0.5 | 0.664,0.6 | -0.184 |
| ENSMUSG00000051910 | Sox6 | chr7 | - | 115178250 | 115178373 | 115149244 | 115149394 | 115179799 | 115179879 | 5,1,1,12 | 29,15,30,6 | 10,18,11,13 | 15,22,8,11 | 272 | 149 | 0.00098 | 0.02703 | 0.086,0.0 | 0.268,0.3 | -0.184 |
| ENSMUSG00000029004 | Kmt2e | chr5 | + | 23705448 | 23705574 | 23705253 | 23705342 | 23705724 | 23705810 | 396,282,513,326 | 252,195,228,259 | 320,345,437,318 | 98,105,107,102 | 275 | 149 | 1.62E-10 | 1.73E-07 | 0.46,0.43 | 0.639,0.6 | -0.186 |
| ENSMUSG00000034269 | Setd5 | chr6 | + | 113086384 | 113086577 | 113081895 | 113082082 | 113086845 | 113086915 | 73,65,77,46 | 30,33,67,32 | 93,60,55,60 | 25,14,10,30 | 342 | 149 | 0.00168 | 0.04019 | 0.515,0.4 | 0.618,0.6 | -0.186 |
| ENSMUSG00000022668 | Gtpbp8 | chr16 | - | 44565645 | 44565856 | 44564170 | 44564231 | 44566361 | 44566672 | 88,69,73,61 | 26,9,6,13 | 86,77,78,64 | 5,0,13,0 | 360 | 149 | 0.00033 | 0.01225 | 0.583,0.7 | 0.877,1.0, | -0.188 |
| ENSMUSG00000042423 | Fbrs | chr7 | + | 127086031 | 127086061 | 127082092 | 127082393 | 127086436 | 127086508 | 13,14,19,13 | 8,1,7,6 | 31,16,32,24 | 1,6,2,0 | 179 | 149 | 0.00121 | 0.03164 | 0.575,0.9 | 0.963,0.6 | -0.188 |
| ENSMUSG00000049493 | Pls1 | chr9 | - | 95677814 | 95677919 | 95668985 | 95669149 | 95727223 | 95727339 | 68,57,61,7 | 2,0,12,6 | 94,59,47,135 | 0,0,2,6 | 254 | 149 | 0.00397 | 0.07603 | 0.952,1.0, | 1.0,1.0,0. | -0.189 |
| ENSMUSG00000087177 | E130307 | chr10 | - | 39516375 | 39516507 | 39508945 | 39509012 | 39557468 | 39557545 | 17,16,32,22 | 6,2,8,3 | 24,23,19,13 | 0,1,4,0 | 281 | 149 | 0.00108 | 0.02895 | 0.6,0.809, | 1.0,0.924, | -0.189 |
| ENSMUSG00000003812 | Dnase2a | chr8 | + | 85636154 | 85636319 | 85635613 | 85635791 | 85636400 | 85636554 | 14,8,13,19 | 0,8,3,0 | 35,23,30,13 | 0,0,0,2 | 314 | 149 | 0.0003 | 0.0115 | 1.0,0.322, | 1.0,1.0,1. | -0.19 |
| ENSMUSG00000058174 | Gm5148 | chr3 | - | 37770270 | 37770487 | 37768346 | 37769284 | 37778351 | 37778478 | 34,18,28,30 | 17,0,4,4 | 14,21,21,12 | 0,1,2,0 | 366 | 149 | 9.94E-05 | 0.00501 | 0.449,1.0, | 1.0,0.895, | -0.191 |
| ENSMUSG00000041633 | Kctd12b | chrX | - | 152478619 | 152478674 | 152468149 | 152472693 | 152479080 | 152479276 | 13,31,21,15 | 0,7,6,15 | 11,10,26,21 | 0,2,0,2 | 204 | 149 | 0.00011 | 0.00543 | 1.0,0.764, | 1.0,0.785, | -0.191 |
| ENSMUSG00000048720 | Tbc1d12 | chr19 | + | 38887312 | 38887430 | 38884430 | 38884513 | 38896215 | 38896296 | 34,26,39,27 | 25,8,28,13 | 23,30,34,33 | 6,12,7,4 | 267 | 149 | 0.00428 | 0.07998 | 0.431,0.6 | 0.681,0.5 | -0.191 |
| ENSMUSG00000030822 | Prr14 | chr7 | + | 127071077 | 127071149 | 127070156 | 127070291 | 127071269 | 127071438 | 41,38,44,29 | 10,23,14,20 | 38,65,32,49 | 10,10,4,7 | 221 | 149 | 0.00079 | 0.02322 | 0.734,0.5 | 0.719,0.8 | -0.192 |
| ENSMUSG00000021090 | Lrrc9 | chr12 | + | 72555744 | 72555816 | 72553059 | 72553200 | 72557109 | 72557339 | 19,21,18,11 | 8,9,0,0 | 14,18,9,33 | 0,0,0,0 | 221 | 149 | 3.74E-07 | 7.41E-05 | 0.616,0.6 | 1.0,1.0,1. | -0.193 |
| ENSMUSG00000044712 | Slc38a6 | chr12 | + | 73390442 | 73390522 | 73388480 | 73388534 | 73391566 | 73391667 | 50,65,96,72 | 21,11,44,12 | 42,45,25,52 | 4,7,2,0 | 229 | 149 | 2.62E-05 | 0.00183 | 0.608,0.7 | 0.872,0.8 | -0.196 |
| ENSMUSG00000021619 | Atg10 | chr13 | - | 91170688 | 91170786 | 91085411 | 91085509 | 91302335 | 91302431 | 59,57,92,64 | 11,9,26,1 | 40,66,48,72 | 0,0,0,0 | 247 | 149 | 6.03E-13 | 1.75E-09 | 0.764,0.7 | 1.0,1.0,1. | -0.197 |
| ENSMUSG00000001039 | B9d1 | chr11 | + | 61399897 | 61399994 | 61397169 | 61397238 | 61400295 | 61400358 | 55,36,44,32 | 5,8,17,0 | 51,40,45,44 | 0,0,0,0 | 246 | 149 | 3.61E-10 | 3.03E-07 | 0.869,0.7 | 1.0,1.0,1. | -0.197 |
| ENSMUSG00000027810 | Eif2a | chr3 | + | 58447010 | 58447067 | 58444954 | 58445029 | 58448460 | 58448560 | 74,52,69,76 | 68,49,87,36 | 61,62,49,56 | 19,16,33,19 | 206 | 149 | 0.00061 | 0.0195 | 0.44,0.43 | 0.699,0.7 | -0.198 |
| ENSMUSG00000041617 | Ccdc74a | chr16 | + | 17466658 | 17466824 | 17465930 | 17465990 | 17468230 | 17468602 | 44,34,100,38 | 18,18,17,9 | 86,72,47,31 | 0,24,0,9 | 315 | 149 | 0.00467 | 0.08532 | 0.536,0.4 | 1.0,0.587, | -0.199 |
| ENSMUSG00000062209 | Erbb4 | chr1 | - | 68329597 | 68329642 | 68329313 | 68329476 | 68337445 | 68337477 | 25,18,32,14 | 14,0,8,12 | 13,23,18,14 | 5,0,1,0 | 194 | 149 | 0.0019 | 0.04423 | 0.578,1.0, | 0.666,1.0, | -0.199 |
| ENSMUSG00000026956 | Uap1l1 | chr2 | - | 25253323 | 25253396 | 25252674 | 25252860 | 25253817 | 25253869 | 38,22,51,35 | 18,17,6,9 | 56,36,49,37 | 6,0,6,10 | 222 | 149 | 0.00426 | 0.0797 | 0.586,0.4 | 0.862,1.0, | -0.199 |
| ENSMUSG00000111375 | Btbd8 | chr5 | + | 107609303 | 107609421 | 107599854 | 107600051 | 107618573 | 107618663 | 25,14,37,19 | 2,10,5,1 | 18,21,18,34 | 0,0,2,0 | 267 | 149 | 4.21E-06 | 0.00046 | 0.875,0.4 | 1.0,1.0,0. | -0.2 |
| ENSMUSG00000032305 | Fam219b | chr9 | + | 57446528 | 57446577 | 57446093 | 57446171 | 57447937 | 57450467 | 32,26,8,16 | 49,22,37,26 | 41,45,34,43 | 33,23,36,24 | 198 | 149 | 0.00194 | 0.04486 | 0.33,0.47 | 0.483,0.5 | -0.203 |
| ENSMUSG00000079109 | Pms2 | chr5 | + | 143858842 | 143858918 | 143856396 | 143856426 | 143860294 | 143860450 | 37,33,47,32 | 4,7,17,11 | 24,19,20,20 | 0,0,2,2 | 225 | 149 | 1.81E-06 | 0.00025 | 0.86,0.75 | 1.0,1.0,0. | -0.204 |
| ENSMUSG00000041189 | Chrnb1 | chr11 | - | 69677706 | 69677930 | 69676532 | 69676705 | 69683524 | 69683734 | 10,26,31,26 | 2,0,8,1 | 16,8,18,15 | 0,0,0,0 | 373 | 149 | 4.50E-06 | 0.00049 | 0.666,1.0, | 1.0,1.0,1. | -0.204 |
| ENSMUSG00000020181 | Nav3 | chr10 | - | 109702791 | 109702860 | 109699612 | 109699752 | 109716395 | 109716579 | 5,7,24,1 | 15,15,46,21 | 26,13,14,18 | 32,14,21,13 | 218 | 149 | 0.00152 | 0.03756 | 0.186,0.2 | 0.357,0.3 | -0.205 |
| ENSMUSG00000000782 | Tcf7 | chr11 | - | 52149679 | 52149772 | 52148510 | 52148630 | 52151357 | 52151448 | 22,13,3,14 | 30,36,61,31 | 5,34,6,17 | 12,11,9,15 | 242 | 149 | 0.00301 | 0.06218 | 0.311,0.1 | 0.204,0.6 | -0.206 |
| ENSMUSG00000109849 | Gm45645 | chr14 | - | 27574756 | 27574906 | 27560121 | 27561704 | 27591752 | 27591861 | 75,86,74,40 | 12,21,8,3 | 24,44,8,37 | 0,0,0,1 | 299 | 149 | 7.47E-10 | 5.26E-07 | 0.757,0.6 | 1.0,1.0,1. | -0.208 |
| ENSMUSG00000086968 | 4933431E | chr3 | - | 107801904 | 107801972 | 107801366 | 107801531 | 107802958 | 107803082 | 0,1,10,0 | 10,20,48,9 | 17,6,0,15 | 25,15,18,12 | 217 | 149 | 0.00123 | 0.03195 | 0.0,0.033, | 0.318,0.2 | -0.209 |
| ENSMUSG00000074829 | 2010315B | chr9 | - | 124074305 | 124074380 | 124057987 | 124058114 | 124075189 | 124075301 | 27,21,36,19 | 12,8,3,6 | 10,15,19,37 | 0,0,3,5 | 224 | 149 | 0.00405 | 0.07714 | 0.599,0.6 | 1.0,1.0,0. | -0.209 |
| ENSMUSG00000073434 | Wdr90 | chr17 | - | 26078992 | 26079107 | 26078831 | 26078899 | 26079345 | 26079516 | 51,33,37,15 | 22,12,8,15 | 44,29,32,49 | 9,4,5,8 | 264 | 149 | 0.00333 | 0.0671 | 0.567,0.6 | 0.734,0.8 | -0.21 |
| ENSMUSG00000020271 | Fbxw11 | chr11 | + | 32658187 | 32658250 | 32602747 | 32602904 | 32661771 | 32661997 | 52,55,76,89 | 32,23,49,18 | 65,49,73,59 | 0,18,0,23 | 212 | 149 | 0.00036 | 0.01306 | 0.533,0.6 | 1.0,0.657, | -0.21 |
| ENSMUSG00000022668 | Gtpbp8 | chr16 | - | 44565757 | 44565856 | 44564100 | 44564231 | 44566361 | 44566726 | 52,40,43,56 | 26,9,6,13 | 87,61,63,59 | 5,0,13,0 | 248 | 149 | 1.11E-05 | 0.00095 | 0.546,0.7 | 0.913,1.0, | -0.212 |
| ENSMUSG00000043531 | Sorcs1 | chr19 | - | 50141197 | 50141358 | 50139404 | 50139789 | 50141491 | 50141597 | 9,7,20,14 | 58,50,74,55 | 19,26,32,16 | 10,48,31,32 | 310 | 149 | 8.89E-07 | 0.00014 | 0.069,0.0 | 0.477,0.2 | -0.213 |
| ENSMUSG00000016624 | Phf21b | chr15 | - | 84678096 | 84678119 | 84675983 | 84676142 | 84679280 | 84679335 | 24,26,15,21 | 4,7,15,16 | 22,15,9,16 | 0,2,1,6 | 172 | 149 | 0.00281 | 0.0593 | 0.839,0.7 | 1.0,0.867, | -0.213 |
| ENSMUSG00000022971 | Ifnar2 | chr16 | + | 91196105 | 91196233 | 91190644 | 91190801 | 91200597 | 91202477 | 40,25,35,52 | 13,7,13,11 | 33,44,31,47 | 1,4,4,5 | 277 | 149 | 9.32E-05 | 0.00479 | 0.623,0.6 | 0.947,0.8 | -0.213 |
| ENSMUSG00000046962 | Zbtb21 | chr16 | - | 97757957 | 97758100 | 97748634 | 97754293 | 97763613 | 97763809 | 83,46,100,96 | 28,25,67,40 | 71,89,63,52 | 11,11,11,17 | 292 | 149 | 2.45E-05 | 0.00175 | 0.602,0.4 | 0.767,0.8 | -0.215 |
| ENSMUSG00000052387 | Trpm3 | chr19 | + | 22877497 | 22877527 | 22875033 | 22875174 | 22878409 | 22878699 | 37,33,78,73 | 7,36,47,27 | 66,121,76,63 | 23,11,11,4 | 179 | 149 | 0.00015 | 0.00675 | 0.815,0.4 | 0.705,0.9 | -0.217 |
| ENSMUSG00000027200 | Sema6d | chr2 | + | 124502624 | 124502681 | 124502163 | 124502264 | 124505940 | 124509689 | 37,13,35,42 | 73,60,61,48 | 46,36,48,47 | 10,54,48,42 | 206 | 149 | 0.00514 | 0.09099 | 0.268,0.1 | 0.769,0.3 | -0.219 |
| ENSMUSG00000059897 | Zfp930 | chr8 | + | 69679323 | 69679381 | 69678994 | 69679121 | 69680494 | 69681632 | 46,63,34,60 | 15,26,30,14 | 17,47,59,58 | 0,3,13,16 | 207 | 149 | 0.00265 | 0.05703 | 0.688,0.6 | 1.0,0.919, | -0.22 |
| ENSMUSG00000029863 | Casp2 | chr6 | + | 42253628 | 42253719 | 42250850 | 42250979 | 42256710 | 42256831 | 41,36,54,18 | 13,4,10,5 | 25,30,30,46 | 0,0,3,0 | 240 | 149 | 8.51E-09 | 3.43E-06 | 0.662,0.8 | 1.0,1.0,0. | -0.222 |
| ENSMUSG00000105265 | Sox2ot | chr3 | + | 34562641 | 34562735 | 34251309 | 34251378 | 34665847 | 34666032 | 22,38,20,21 | 11,4,13,5 | 30,11,21,8 | 0,5,1,0 | 243 | 149 | 0.00138 | 0.03495 | 0.551,0.8 | 1.0,0.574, | -0.223 |
| ENSMUSG00000038578 | Susd1 | chr4 | - | 59369530 | 59369640 | 59365721 | 59365914 | 59379598 | 59379785 | 39,36,25,37 | 19,16,11,14 | 34,42,57,26 | 0,6,11,9 | 259 | 149 | 0.00113 | 0.02985 | 0.541,0.5 | 1.0,0.801, | -0.225 |
| ENSMUSG00000028174 | Rpe65 | chr3 | + | 159305953 | 159306036 | 159304811 | 159304880 | 159307154 | 159307305 | 59,36,33,17 | 5,9,9,4 | 20,20,16,23 | 0,0,0,1 | 232 | 149 | 1.26E-08 | 4.77E-06 | 0.883,0.7 | 1.0,1.0,1. | -0.225 |

| | | | | | | | | | | | | | | | | | | | | |
|---|---|---|---|---|---|---|---|---|---|---|---|---|---|---|---|---|---|---|---|---|
| ENSMUSG00000059890 | Ube4a | chr9 | - | 44874661 | 44874730 | 44871276 | 44871437 | 44876782 | 44876823 | 2,5,1,6 | 21,26,36,22 | 8,7,12,6 | 14,13,16,7 | 218 | 149 | 3.70E-06 | 0.00042 | 0.061,0.1 | 0.281,0.2 | -0.226 |
| ENSMUSG00000097881 | Celrr | chr1 | + | 121045037 | 121045826 | 121015141 | 121015285 | 121047586 | 121047738 | 48,53,133,28 | 4,0,0,6 | 50,60,47,46 | 0,0,0,0 | 938 | 149 | 3.33E-05 | 0.0022 | 0.656,1.0, | 1.0,1.0,1. | -0.229 |
| ENSMUSG00000032567 | Aste1 | chr9 | + | 105274823 | 105275057 | 105273743 | 105273904 | 105278674 | 105279342 | 49,14,24,41 | 6,3,10,6 | 48,26,52,19 | 4,1,5,0 | 383 | 149 | 0.00401 | 0.07664 | 0.761,0.6 | 0.824,0.9 | -0.23 |
| ENSMUSG00000054715 | Zscan22 | chr7 | + | 12637527 | 12638012 | 12631767 | 12631804 | 12640160 | 12641464 | 66,38,46,49 | 0,2,3,13 | 34,23,26,31 | 0,0,0,0 | 634 | 149 | 1.39E-07 | 3.28E-05 | 1.0,0.817, | 1.0,1.0,1. | -0.232 |
| ENSMUSG00000020598 | Nrcam | chr12 | + | 44630449 | 44630575 | 44624927 | 44624963 | 44631617 | 44631770 | 67,31,59,55 | 26,10,31,13 | 78,59,66,50 | 21,5,7,0 | 275 | 149 | 7.08E-05 | 0.00386 | 0.583,0.6 | 0.668,0.8 | -0.239 |
| ENSMUSG00000045813 | Gm9801 | chr7 | - | 61767470 | 61767650 | 61764393 | 61764524 | 61768642 | 61768791 | 30,42,40,28 | 6,5,8,5 | 32,8,23,26 | 0,0,0,2 | 329 | 149 | 1.32E-07 | 3.14E-05 | 0.694,0.7 | 1.0,1.0,1. | -0.24 |
| ENSMUSG00000051727 | Kctd14 | chr7 | + | 97104063 | 97104184 | 97102410 | 97102669 | 97106800 | 97108760 | 9,4,1,7 | 19,19,23,7 | 19,25,3,10 | 15,6,7,10 | 270 | 149 | 0.00113 | 0.02988 | 0.207,0.1 | 0.411,0.6 | -0.241 |
| ENSMUSG00000027522 | Stx16 | chr2 | + | 173926438 | 173926450 | 173918714 | 173918982 | 173932413 | 173932521 | 25,34,27,20 | 18,18,20,30 | 15,15,13,92 | 0,10,6,14 | 161 | 149 | 0.00159 | 0.03861 | 0.562,0.6 | 1.0,0.581, | -0.243 |
| ENSMUSG00000053046 | Brsk2 | chr7 | + | 141554578 | 141554626 | 141552656 | 141552751 | 141555799 | 141555884 | 12,11,2,3 | 33,46,73,46 | 51,30,26,25 | 58,27,54,46 | 197 | 149 | 2.32E-09 | 1.24E-06 | 0.216,0.1 | 0.399,0.4 | -0.245 |
| ENSMUSG00000046962 | Zbtb21 | chr16 | - | 97757957 | 97758094 | 97753729 | 97754293 | 97763613 | 97763801 | 25,24,47,64 | 28,25,67,40 | 44,39,39,22 | 11,11,11,17 | 286 | 149 | 0.00019 | 0.00798 | 0.317,0.3 | 0.676,0.6 | -0.251 |
| ENSMUSG00000039607 | Rbms3 | chr9 | - | 116507714 | 116507765 | 116465436 | 116465547 | 116510198 | 116510295 | 6,5,4,4 | 30,50,44,44 | 9,7,5,26 | 17,24,9,13 | 200 | 149 | 2.88E-06 | 0.00035 | 0.13,0.06 | 0.283,0.1 | -0.257 |
| ENSMUSG00000020598 | Nrcam | chr12 | + | 44624927 | 44624963 | 44623386 | 44623572 | 44630449 | 44630575 | 21,9,25,20 | 19,13,18,13 | 25,18,39,24 | 6,14,16,0 | 185 | 149 | 0.00489 | 0.08802 | 0.471,0.3 | 0.77,0.50 | -0.258 |
| ENSMUSG00000058498 | Rnf207 | chr4 | - | 152398492 | 152398565 | 152398335 | 152398404 | 152399498 | 152399545 | 38,9,3,20 | 0,4,5,1 | 14,13,17,37 | 0,0,0,4 | 222 | 149 | 0.00479 | 0.08695 | 1.0,0.602, | 1.0,1.0,1. | -0.26 |
| ENSMUSG00000057715 | A830018L | chr1 | + | 12003767 | 12003815 | 11900038 | 11900115 | 12021186 | 12021280 | 27,31,34,23 | 25,0,22,13 | 33,32,21,37 | 0,15,0,0 | 197 | 149 | 0.00089 | 0.02541 | 0.45,1.0,0 | 1.0,0.617, | -0.264 |
| ENSMUSG00000024317 | Rnf138 | chr18 | + | 21157465 | 21157573 | 21153944 | 21154056 | 21159142 | 21160592 | 58,49,39,66 | 52,46,84,51 | 39,63,50,30 | 0,34,19,23 | 257 | 149 | 0.00208 | 0.04738 | 0.393,0.3 | 1.0,0.518, | -0.284 |
| ENSMUSG00000020042 | Btbd11 | chr10 | + | 85467156 | 85467292 | 85465378 | 85467045 | 85469547 | 85469726 | 10,28,43,38 | 12,15,2,7 | 41,30,39,39 | 15,0,0,0 | 285 | 149 | 0.00215 | 0.04866 | 0.303,0.4 | 0.588,1.0, | -0.284 |
| ENSMUSG00000041623 | D11Wsu4 | chr11 | + | 113576015 | 113576097 | 113575237 | 113575636 | 113578599 | 113579039 | 18,16,41,25 | 5,10,3,9 | 15,9,9,22 | 1,0,0,0 | 231 | 149 | 1.63E-08 | 5.52E-06 | 0.699,0.5 | 0.906,1.0, | -0.29 |
| ENSMUSG00000015790 | Surf1 | chr2 | - | 26805966 | 26806018 | 26805607 | 26805781 | 26806268 | 26806312 | 164,109,192,176 | 94,107,178,132 | 213,132,143,186 | 106,96,0,0 | 201 | 149 | 3.03E-07 | 6.33E-05 | 0.564,0.4 | 0.598,0.5 | -0.292 |
| ENSMUSG00000044566 | Cage1 | chr13 | - | 38216340 | 38216600 | 38212078 | 38212166 | 38220763 | 38220899 | 18,14,51,26 | 12,6,0,0 | 21,13,10,15 | 0,0,0,0 | 409 | 149 | 6.43E-06 | 0.00063 | 0.353,0.4 | 1.0,1.0,1. | -0.297 |
| ENSMUSG00000021619 | Atg10 | chr13 | - | 91188968 | 91189107 | 91085411 | 91085509 | 91302335 | 91302431 | 50,36,43,46 | 11,9,26,1 | 50,50,32,60 | 0,0,0,0 | 288 | 149 | 1.50E-12 | 3.85E-09 | 0.702,0.6 | 1.0,1.0,1. | -0.301 |
| ENSMUSG00000069631 | Strada | chr11 | - | 106071780 | 106071809 | 106064504 | 106064607 | 106077926 | 106078006 | 14,3,18,18 | 33,15,18,27 | 16,30,21,16 | 11,29,9,4 | 178 | 149 | 0.0009 | 0.02567 | 0.262,0.1 | 0.549,0.4 | -0.306 |
| ENSMUSG00000037108 | Zcwpw1 | chr5 | + | 137798247 | 137798405 | 137797450 | 137797863 | 137799235 | 137799323 | 40,42,65,50 | 12,40,13,19 | 41,58,51,50 | 10,7,0,0 | 307 | 149 | 1.61E-05 | 0.00126 | 0.618,0.3 | 0.666,0.8 | -0.31 |
| ENSMUSG00000031109 | Enox2 | chrX | - | 48375732 | 48375781 | 48257517 | 48257661 | 48376977 | 48377089 | 25,5,32,6 | 7,5,3,10 | 13,9,27,24 | 0,1,7,0 | 198 | 149 | 0.00264 | 0.05688 | 0.729,0.4 | 1.0,0.871, | -0.314 |
| ENSMUSG00000072591 | Fzd10os | chr5 | - | 128671271 | 128672828 | 128662335 | 128662470 | 128677600 | 128677751 | 34,48,59,39 | 11,2,3,2 | 33,40,49,46 | 4,0,0,0 | 1706 | 149 | 0.00092 | 0.02595 | 0.213,0.6 | 0.419,1.0, | -0.317 |
| ENSMUSG00000028559 | Osbpl9 | chr4 | - | 108948857 | 108948896 | 108944591 | 108944682 | 108955699 | 108955750 | 45,22,28,18 | 0,12,0,88 | 22,37,34,24 | 0,0,0,0 | 188 | 149 | 6.25E-06 | 0.00062 | 1.0,0.592, | 1.0,1.0,1. | -0.317 |
| ENSMUSG00000054457 | 9430021M | chr2 | + | 162507728 | 162507944 | 162503082 | 162503346 | 162508308 | 162508583 | 25,23,58,19 | 19,7,12,9 | 70,12,55,54 | 0,3,8,0 | 365 | 149 | 5.03E-06 | 0.00052 | 0.349,0.5 | 1.0,0.62,0 | -0.327 |
| ENSMUSG00000032409 | Atr | chr9 | + | 95787790 | 95787911 | 95785715 | 95785831 | 95789348 | 95789486 | 30,27,11,36 | 7,10,14,9 | 14,21,19,27 | 2,1,1,0 | 270 | 149 | 2.41E-07 | 5.23E-05 | 0.703,0.5 | 0.794,0.9 | -0.334 |
| ENSMUSG00000014329 | Bicc1 | chr10 | - | 70779199 | 70779365 | 70776829 | 70776869 | 70781054 | 70781211 | 24,15,28,11 | 0,13,9,3 | 8,47,21,50 | 0,0,0,2 | 315 | 149 | 4.87E-08 | 1.41E-05 | 1.0,0.353, | 1.0,1.0,1. | -0.335 |
| ENSMUSG00000037110 | Ralgapa2 | chr2 | - | 146199007 | 146199066 | 146195090 | 146195199 | 146199876 | 146200005 | 12,22,15,22 | 36,15,35,19 | 20,16,23,30 | 14,7,7,2 | 208 | 149 | 0.00015 | 0.0068 | 0.193,0.5 | 0.506,0.6 | -0.338 |
| ENSMUSG00000086968 | 4933431E | chr3 | - | 107801904 | 107801972 | 107801366 | 107801531 | 107802176 | 107802308 | 0,0,27,0 | 30,38,49,22 | 46,33,8,30 | 25,20,30,22 | 217 | 149 | 4.00E-08 | 1.20E-05 | 0.0,0.0,0. | 0.558,0.5 | -0.363 |
| ENSMUSG00000067786 | Nnat | chr2 | + | 157403132 | 157403208 | 157402059 | 157402455 | 157403461 | 157404158 | 27,9,12,15 | 11,29,10,21 | 30,14,15,8 | 0,19,4,0 | 225 | 149 | 0.00135 | 0.03423 | 0.619,0.1 | 1.0,0.328, | -0.372 |
| ENSMUSG00000002279 | Lmf1 | chr17 | + | 25807649 | 25807716 | 25804516 | 25804826 | 25831269 | 25831418 | 33,10,16,15 | 9,24,33,10 | 21,7,8,20 | 0,0,8,0 | 216 | 149 | 2.81E-05 | 0.00195 | 0.717,0.2 | 1.0,1.0,0. | -0.427 |
| ENSMUSG00000060149 | BC00205 | chr17 | + | 17191161 | 17191288 | 17171810 | 17171970 | 17191432 | 17191493 | 24,12,6,41 | 2,28,35,16 | 4,8,25,8 | 1,0,0,0 | 276 | 149 | 1.39E-09 | 8.65E-07 | 0.866,0.1 | 0.683,1.0, | -0.491 |
| ENSMUSG00000078877 | Gm14295 | chr2 | + | 176493973 | 176494028 | 176490404 | 176490511 | 176499154 | 176499281 | 0,295,0,197 | 7,12,21,4 | 214,117,310,262 | 0,0,12,1 | 204 | 149 | 1.22E-06 | 0.00018 | 0.0,0.947, | 1.0,1.0,0. | -0.506 |

### Table S3. Alternative_splicing_RI

| GeneID | geneSym | chr | strand | riExonStar | riExonEnd | upstreamE | upstreamE | downstrea | downstrea | IJC_SAMPLE_1 | SJC_SAMPLE_1 | IJC_SAMPLE_2 | SJC_SAMPLE_2 | IncFormL | SkipForm | PValue | FDR | IncLevel1 | IncLevel2 | IncLevelDifference |
|---|---|---|---|---|---|---|---|---|---|---|---|---|---|---|---|---|---|---|---|---|
| ENSMUSG00000096727 | Psmb9 | chr17 | - | 34402588 | 34403387 | 34402588 | 34402718 | 34403255 | 34403387 | 2,4,2,2 | 7,13,27,9 | 15,8,5,16 | 25,11,13,3 | 686 | 149 | 0.00014 | 0.00368 | 0.058,0.0 | 0.115,0.1 | -0.17 |
| ENSMUSG00000005899 | Smpd4 | chr16 | + | 17443583 | 17444402 | 17443583 | 17443726 | 17444293 | 17444402 | 164,106,168,178 | 2,6,5,9 | 186,78,101,168 | 0,0,3,0 | 716 | 149 | 1.92E-06 | 0.0002 | 0.945,0.7 | 1.0,1.0,0.8 | -0.116 |
| ENSMUSG00000062906 | Hdac10 | chr15 | - | 89011796 | 89012221 | 89011796 | 89011894 | 89012124 | 89012221 | 23,6,16,4 | 28,40,58,45 | 26,16,23,10 | 27,16,45,28 | 379 | 149 | 0.00187 | 0.02188 | 0.244,0.0 | 0.275,0.2 | -0.104 |
| ENSMUSG00000097709 | 2810429I | chr13 | + | 3528242 | 3530001 | 3528242 | 3528390 | 3529908 | 3530001 | 44,34,44,40 | 38,28,27,30 | 26,49,60,61 | 6,9,16,30 | 1667 | 149 | 2.89E-05 | 0.00133 | 0.094,0.0 | 0.279,0.3 | -0.147 |
| ENSMUSG00000029716 | Tfr2 | chr5 | + | 137581666 | 137582328 | 137581666 | 137581894 | 137582187 | 137582328 | 25,7,17,40 | 4,6,11,4 | 33,21,32,47 | 5,4,3,0 | 442 | 149 | 0.01194 | 0.07996 | 0.678,0.2 | 0.69,0.63 | -0.259 |
| ENSMUSG00000045903 | Npas4 | chr19 | - | 5037342 | 5038376 | 5037342 | 5037478 | 5038102 | 5038376 | 68,108,169,42 | 9,10,7,0 | 82,57,59,141 | 0,1,2,5 | 773 | 149 | 0.00798 | 0.05973 | 0.593,0.6 | 1.0,0.917, | -0.13 |
| ENSMUSG00000037262 | Kin | chr2 | + | 10094923 | 10095174 | 10094923 | 10094967 | 10095051 | 10095174 | 32,25,22,29 | 77,70,74,100 | 31,20,36,46 | 56,55,37,48 | 233 | 149 | 0.00102 | 0.01505 | 0.21,0.18 | 0.261,0.1 | -0.125 |
| ENSMUSG00000092060 | Bend4 | chr5 | - | 67549490 | 67557644 | 67549490 | 67555793 | 67557474 | 67557644 | 42,52,56,81 | 6,3,0,4 | 49,40,41,61 | 0,0,3,0 | 1830 | 149 | 0.0006 | 0.01055 | 0.363,0.5 | 1.0,1.0,0. | -0.239 |
| ENSMUSG00000000561 | Wdr77 | chr3 | + | 105873080 | 105873741 | 105873080 | 105873172 | 105873672 | 105873741 | 431,348,383,336 | 11,10,16,14 | 209,286,223,316 | 3,0,3,0 | 649 | 149 | 7.97E-09 | 3.17E-06 | 0.9,0.889, | 0.941,1.0, | -0.101 |
| ENSMUSG00000028057 | Rit1 | chr3 | + | 88633301 | 88633778 | 88633301 | 88633486 | 88633586 | 88633778 | 37,29,12,19 | 0,2,7,1 | 25,21,36,2 | 0,0,0,0 | 249 | 149 | 0.00018 | 0.00438 | 1.0,0.897, | 1.0,1.0,1. | -0.169 |
| ENSMUSG00000060373 | Hnrnpc | chr14 | - | 52335471 | 52336001 | 52335471 | 52335497 | 52335772 | 52336001 | 190,136,174,155 | 28,4,31,10 | 121,114,111,119 | 1,4,8,2 | 424 | 149 | 0.00013 | 0.00359 | 0.705,0.9 | 0.977,0.9 | -0.133 |
| ENSMUSG00000054199 | Gon4l | chr3 | + | 88805411 | 88806381 | 88805411 | 88805667 | 88806196 | 88806381 | 29,5,33,26 | 0,3,6,0 | 9,20,16,14 | 2,0,0,1 | 678 | 149 | 0.00738 | 0.05628 | 1.0,0.268, | 0.497,1.0, | -0.109 |
| ENSMUSG00000038225 | Primpol | chr8 | - | 47045627 | 47046738 | 47045627 | 47045790 | 47046504 | 47046738 | 16,20,12,9 | 35,20,19,9 | 7,16,13,32 | 2,13,12,19 | 863 | 149 | 0.01276 | 0.0839 | 0.073,0.1 | 0.377,0.1 | -0.118 |
| ENSMUSG00000050211 | Pla2g4e | chr2 | - | 120016827 | 120017719 | 120016827 | 120016891 | 120017676 | 120017719 | 12,10,17,5 | 6,13,10,4 | 15,16,27,35 | 7,4,6,14 | 934 | 149 | 0.00924 | 0.06651 | 0.242,0.1 | 0.255,0.3 | -0.154 |
| ENSMUSG00000091476 | Catspere2 | chr1 | + | 177968974 | 177969156 | 177968974 | 177969008 | 177969076 | 177969156 | 15,16,45,10 | 6,5,16,11 | 9,17,5,10 | 4,0,0,0 | 217 | 149 | 4.13E-05 | 0.0017 | 0.632,0.6 | 0.607,1.0, | -0.311 |
| ENSMUSG00000026269 | Rnpepl1 | chr1 | + | 92845739 | 92846825 | 92845739 | 92845848 | 92846659 | 92846825 | 225,125,245,168 | 9,8,5,1 | 176,170,124,142 | 0,0,0,1 | 960 | 149 | 2.94E-08 | 8.75E-06 | 0.795,0.7 | 1.0,1.0,1. | -0.152 |
| ENSMUSG00000043065 | Spice1 | chr16 | + | 44190242 | 44190741 | 44190242 | 44190382 | 44190600 | 44190741 | 30,30,13,23 | 70,44,66,21 | 40,27,41,23 | 35,49,25,24 | 367 | 149 | 0.00577 | 0.04835 | 0.148,0.2 | 0.317,0.1 | -0.108 |
| ENSMUSG00000071337 | Tia1 | chr6 | + | 86395859 | 86397393 | 86395859 | 86395914 | 86397305 | 86397393 | 2,068,167,723,161,850 | 123,185,183,123 | 1,690,164,918,341,820 | 62,89,57,97 | 1540 | 149 | 3.38E-05 | 0.00147 | 0.619,0.4 | 0.725,0.6 | -0.135 |
| ENSMUSG00000028845 | Tekt2 | chr4 | - | 126218022 | 126218512 | 126218022 | 126218228 | 126218386 | 126218512 | 33,25,54,43 | 5,4,9,5 | 33,29,15,30 | 1,0,0,0 | 307 | 149 | 7.65E-09 | 3.17E-06 | 0.762,0.7 | 0.941,1.0, | -0.219 |
| ENSMUSG00000005882 | Uqcc1 | chr2 | - | 155752276 | 155753756 | 155752276 | 155752387 | 155753660 | 155753756 | 78,93,122,88 | 42,32,33,36 | 117,136,101,71 | 25,22,12,22 | 1422 | 149 | 0.00129 | 0.01741 | 0.163,0.2 | 0.329,0.3 | -0.141 |
| ENSMUSG00000034928 | Rnf44 | chr13 | - | 54831773 | 54832301 | 54831773 | 54831902 | 54832150 | 54832301 | 172,154,186,159 | 37,14,15,7 | 94,112,113,122 | 0,0,19,0 | 397 | 149 | 0.00046 | 0.00886 | 0.636,0.8 | 1.0,1.0,1. | -0.133 |
| ENSMUSG00000053158 | Fes | chr7 | - | 80032817 | 80033154 | 80032817 | 80032937 | 80033016 | 80033154 | 2,0,2,2 | 13,17,33,13 | 4,2,8,4 | 21,18,9,12 | 228 | 149 | 0.00115 | 0.01618 | 0.091,0.0 | 0.111,0.0 | -0.126 |
| ENSMUSG00000026107 | Nabp1 | chr1 | - | 51516636 | 51517558 | 51516636 | 51516775 | 51516983 | 51517558 | 46,29,24,16 | 9,14,9,5 | 45,18,26,52 | 4,0,7,7 | 357 | 149 | 0.00832 | 0.062 | 0.681,0.4 | 0.824,1.0, | -0.236 |
| ENSMUSG00000028948 | Nol9 | chr4 | + | 152130151 | 152131101 | 152130151 | 152130287 | 152131003 | 152131101 | 139,90,130,106 | 13,12,31,32 | 91,74,75,141 | 7,16,1,3 | 865 | 149 | 0.00663 | 0.05204 | 0.648,0.5 | 0.691,0.4 | -0.239 |
| ENSMUSG00000055235 | Wdr86 | chr5 | - | 24917847 | 24920659 | 24917847 | 24917951 | 24920523 | 24920659 | 21,11,32,14 | 0,1,6,7 | 6,33,25,13 | 0,0,1,0 | 2721 | 149 | 3.12E-07 | 5.12E-05 | 1.0,0.376, | 1.0,1.0,0. | -0.469 |
| ENSMUSG00000010803 | Gabra1 | chr11 | - | 42072992 | 42073757 | 42072992 | 42073216 | 42073700 | 42073757 | 325,265,375,350 | 13,13,5,19 | 274,249,388,344 | 5,1,5,1 | 633 | 149 | 7.48E-07 | 9.91E-05 | 0.855,0.8 | 0.928,0.9 | -0.101 |
| ENSMUSG00000021868 | Ppif | chr14 | + | 25696427 | 25698765 | 25696427 | 25696516 | 25698689 | 25698765 | 657,489,729,520 | 10,16,28,18 | 466,390,426,431 | 10,3,1,8 | 2322 | 149 | 0.00152 | 0.01948 | 0.808,0.6 | 0.749,0.8 | -0.159 |
| ENSMUSG00000028064 | Sema4a | chr3 | - | 88358651 | 88359093 | 88358651 | 88358804 | 88358994 | 88359093 | 73,42,44,29 | 10,18,19,12 | 39,23,37,57 | 1,8,0,10 | 339 | 149 | 0.00496 | 0.04402 | 0.762,0.5 | 0.945,0.5 | -0.233 |
| ENSMUSG00000023191 | P3h3 | chr6 | - | 124827880 | 124828094 | 124827880 | 124827946 | 124828041 | 124828094 | 59,77,113,105 | 21,19,23,20 | 65,72,77,78 | 5,3,9,5 | 244 | 149 | 1.66E-06 | 0.00017 | 0.632,0.7 | 0.888,0.9 | -0.178 |
| ENSMUSG00000038538 | Ubn2 | chr6 | + | 38464042 | 38468866 | 38464042 | 38464085 | 38467318 | 38468866 | 539,465,477,398 | 99,67,85,86 | 449,466,235,419 | 29,35,51,41 | 3382 | 149 | 0.00211 | 0.02383 | 0.193,0.2 | 0.406,0.3 | -0.115 |
| ENSMUSG00000031756 | Cenpn | chr8 | + | 117662932 | 117664013 | 117662932 | 117662990 | 117663900 | 117664013 | 21,30,32,25 | 4,2,1,3 | 26,26,20,10 | 0,0,0,0 | 1059 | 149 | 3.29E-09 | 1.96E-06 | 0.425,0.6 | 1.0,1.0,1. | -0.384 |
| ENSMUSG00000030871 | Ears2 | chr7 | - | 121643524 | 121643889 | 121643524 | 121643655 | 121643735 | 121643889 | 3,8,10,15 | 50,58,28,35 | 10,15,11,16 | 33,27,14,34 | 229 | 149 | 0.00373 | 0.03519 | 0.038,0.0 | 0.165,0.2 | -0.119 |
| ENSMUSG00000120042 | | chr7 | + | 78532372 | 78533464 | 78532372 | 78532484 | 78532684 | 78533464 | 27,41,18,13 | 2,5,10,3 | 2,17,18,18 | 1,0,0,1 | 349 | 149 | 1.87E-05 | 0.00101 | 0.852,0.7 | 0.461,1.0, | -0.158 |
| ENSMUSG00000034813 | Grip1 | chr10 | + | 119821378 | 119822389 | 119821378 | 119821534 | 119822236 | 119822389 | 18,28,36,19 | 42,29,52,35 | 39,33,43,17 | 20,18,27,20 | 851 | 149 | 2.58E-05 | 0.00123 | 0.07,0.14 | 0.255,0.2 | -0.109 |
| ENSMUSG00000112117 | Rmst | chr10 | - | 91969084 | 91970178 | 91969084 | 91969349 | 91970103 | 91970178 | 30,24,50,16 | 3,7,5,7 | 51,57,43,45 | 0,6,0,8 | 903 | 149 | 0.01215 | 0.08106 | 0.623,0.3 | 1.0,0.611, | -0.303 |
| ENSMUSG00000097277 | 2900076A | chr7 | + | 81178534 | 81179275 | 81178534 | 81178688 | 81178904 | 81179275 | 114,88,93,87 | 10,8,10,17 | 80,51,117,77 | 5,4,6,2 | 365 | 149 | 0.00307 | 0.0303 | 0.823,0.8 | 0.867,0.8 | -0.106 |
| ENSMUSG00000025484 | Bet1l | chr7 | - | 140434677 | 140435078 | 140434677 | 140434850 | 140434986 | 140435078 | 20,18,40,20 | 3,7,11,4 | 16,17,10,15 | 6,0,0,0 | 285 | 149 | 0.00655 | 0.05195 | 0.777,0.5 | 0.582,1.0, | -0.214 |
| ENSMUSG00000079434 | Neu2 | chr1 | + | 87522271 | 87522811 | 87522271 | 87522339 | 87522599 | 87522811 | 43,43,68,50 | 8,2,6,3 | 51,38,29,57 | 5,0,2,0 | 409 | 149 | 0.01545 | 0.09542 | 0.662,0.8 | 0.788,1.0, | -0.104 |
| ENSMUSG00000021572 | Cep72 | chr13 | - | 74197014 | 74198483 | 74197014 | 74197153 | 74198178 | 74198483 | 63,50,28,58 | 11,9,4,5 | 16,20,21,45 | 1,4,0,0 | 1174 | 149 | 0.01019 | 0.07134 | 0.421,0.4 | 0.67,0.38 | -0.289 |
| ENSMUSG00000021363 | Mak | chr13 | - | 41195494 | 41199806 | 41195494 | 41195661 | 41199494 | 41199806 | 264,193,354,246 | 10,13,12,21 | 130,134,157,157 | 4,2,4,4 | 3982 | 149 | 0.01376 | 0.08781 | 0.497,0.3 | 0.549,0.7 | -0.192 |
| ENSMUSG00000022550 | Adck5 | chr15 | + | 76478352 | 76478647 | 76478352 | 76478484 | 76478563 | 76478647 | 74,86,141,75 | 54,37,69,47 | 47,52,68,44 | 45,39,47,55 | 228 | 149 | 0.00843 | 0.06251 | 0.472,0.6 | 0.406,0.4 | 0.114 |
| ENSMUSG00000022556 | Hsf1 | chr15 | + | 76384330 | 76384536 | 76384330 | 76384396 | 76384466 | 76384536 | 177,157,151,137 | 75,65,99,77 | 84,89,69,101 | 92,74,98,83 | 219 | 149 | 4.61E-06 | 0.00037 | 0.616,0.6 | 0.383,0.4 | 0.171 |
| ENSMUSG00000022621 | Rabl2 | chr15 | - | 89468112 | 89468593 | 89468112 | 89468210 | 89468481 | 89468593 | 337,308,246,253 | 55,48,69,43 | 180,214,252,240 | 70,71,51,63 | 420 | 149 | 0.0089 | 0.06471 | 0.685,0.6 | 0.477,0.5 | 0.102 |
| ENSMUSG00000053137 | Mapk11 | chr15 | - | 89029298 | 89029649 | 89029298 | 89029378 | 89029577 | 89029649 | 108,81,202,68 | 20,20,48,22 | 169,76,122,134 | 42,36,55,65 | 348 | 149 | 0.00643 | 0.05123 | 0.698,0.6 | 0.633,0.4 | 0.12 |
| ENSMUSG00000061740 | Cyp2d22 | chr15 | - | 82257311 | 82258034 | 82257311 | 82257488 | 82257873 | 82258034 | 212,171,222,154 | 111,82,88,114 | 129,123,122,138 | 144,113,93,133 | 534 | 149 | 4.91E-06 | 0.00038 | 0.348,0.3 | 0.2,0.233, | 0.119 |
| ENSMUSG00000097494 | 4933406C | chr12 | - | 33003196 | 33003804 | 33003196 | 33003370 | 33003503 | 33003804 | 28,18,31,32 | 1,1,0,0 | 24,10,28,16 | 3,0,1,4 | 282 | 149 | 0.00162 | 0.02035 | 0.937,0.9 | 0.809,1.0, | 0.104 |
| ENSMUSG00000032085 | Tagln | chr9 | - | 45842130 | 45842775 | 45842130 | 45842233 | 45842597 | 45842775 | 63,51,60,64 | 46,24,64,51 | 37,44,38,30 | 34,95,67,40 | 513 | 149 | 0.00013 | 0.00363 | 0.285,0.3 | 0.24,0.11 | 0.117 |
| ENSMUSG00000026048 | Ercc5 | chr1 | + | 44212497 | 44213191 | 44212497 | 44212739 | 44213071 | 44213191 | 126,96,137,126 | 61,62,69,96 | 92,79,46,74 | 98,76,75,68 | 481 | 149 | 9.68E-06 | 0.00064 | 0.39,0.32 | 0.225,0.2 | 0.126 |
| ENSMUSG00000120369 | | chr2 | - | 26527179 | 26529483 | 26527179 | 26527645 | 26528052 | 26529483 | 172,115,161,134 | 37,29,45,40 | 100,86,92,73 | 28,37,41,32 | 556 | 149 | 0.00999 | 0.07021 | 0.555,0.5 | 0.489,0.3 | 0.101 |
| ENSMUSG00000028453 | Fancg | chr4 | - | 43003226 | 43003932 | 43003226 | 43003344 | 43003782 | 43003932 | 172,133,127,134 | 100,38,51,70 | 52,55,62,59 | 44,51,49,35 | 587 | 149 | 0.00058 | 0.01046 | 0.304,0.4 | 0.231,0.2 | 0.125 |
| ENSMUSG00000037326 | Capn15 | chr17 | - | 26179038 | 26179537 | 26179038 | 26179205 | 26179310 | 26179537 | 201,196,236,197 | 8,7,17,2 | 158,160,146,119 | 37,22,11,7 | 254 | 149 | 3.11E-06 | 0.00028 | 0.936,0.9 | 0.715,0.8 | 0.108 |
| ENSMUSG00000033728 | Lrrc14 | chr15 | + | 76597066 | 76598185 | 76597066 | 76597420 | 76598015 | 76598185 | 342,213,368,203 | 20,3,8,1 | 282,217,232,212 | 17,13,12,12 | 744 | 149 | 0.00223 | 0.02454 | 0.774,0.9 | 0.769,0.7 | 0.118 |
| ENSMUSG00000020460 | Rps27a | chr11 | - | 29496702 | 29497260 | 29496702 | 29496788 | 29497205 | 29497260 | 153,112,207,145 | 370,0,1371,0 | 147,108,156,125 | 435,925,192,211,786 | 566 | 149 | 3.14E-09 | 1.96E-06 | 0.098,1.0, | 0.009,0.0 | 0.515 |
| ENSMUSG00000068744 | Psrc1 | chr3 | + | 108291551 | 108291747 | 108291551 | 108291601 | 108291689 | 108291747 | 7,4,40,6 | 8,0,17,0 | 15,24,27,9 | 27,23,26,19 | 237 | 149 | 0.00693 | 0.05388 | 0.355,1.0, | 0.259,0.3 | 0.418 |
| ENSMUSG00000042286 | Stab1 | chr14 | - | 30861882 | 30862224 | 30861882 | 30862032 | 30862104 | 30862224 | 23,15,44,18 | 32,31,67,60 | 19,13,14,8 | 67,51,49,39 | 221 | 149 | 0.00156 | 0.01993 | 0.326,0.2 | 0.161,0.1 | 0.114 |
| ENSMUSG00000028582 | Cc2d1b | chr4 | + | 108483497 | 108483787 | 108483497 | 108483573 | 108483676 | 108483787 | 65,76,114,67 | 86,57,70,86 | 33,46,44,73 | 69,94,69,99 | 252 | 149 | 0.00095 | 0.0146 | 0.309,0.4 | 0.22,0.22 | 0.134 |
| ENSMUSG00000039382 | Wdr45 | chrX | + | 7589553 | 7589992 | 7589553 | 7589603 | 7589803 | 7589992 | 96,60,101,60 | 4,0,6,0 | 67,41,70,49 | 5,11,10,9 | 349 | 149 | 2.30E-06 | 0.00022 | 0.911,1.0, | 0.851,0.6 | 0.219 |
| ENSMUSG00000038828 | Tmem21 | chr5 | + | 31030030 | 31030488 | 31030030 | 31030112 | 31030386 | 31030488 | 471,392,432,355 | 166,92,150,142 | 235,220,237,311 | 129,118,142,104 | 423 | 149 | 0.00376 | 0.03531 | 0.5,0.6,0. | 0.391,0.3 | 0.101 |
| ENSMUSG00000028041 | Adam15 | chr3 | - | 89250694 | 89251284 | 89250694 | 89250890 | 89250979 | 89251284 | 243,249,269,255 | 165,138,193,155 | 158,149,229,186 | 216,194,181,162 | 238 | 149 | 6.65E-05 | 0.0024 | 0.48,0.53, | 0.314,0.3 | 0.121 |
| ENSMUSG00000033191 | Tie1 | chr4 | - | 118336058 | 118337100 | 118336058 | 118336349 | 118336809 | 118337100 | 70,37,51,25 | 16,7,26,16 | 31,29,36,47 | 43,39,23,35 | 609 | 149 | 0.00033 | 0.00688 | 0.517,0.5 | 0.15,0.15 | 0.213 |
| ENSMUSG00000036073 | Galt | chr4 | + | 41756512 | 41756995 | 41756512 | 41756811 | 41756938 | 41756995 | 274,164,184,186 | 98,71,134,67 | 152,120,121,151 | 151,79,130,87 | 276 | 149 | 0.001 | 0.01487 | 0.601,0.5 | 0.352,0.4 | 0.14 |
| ENSMUSG00000026200 | Glb1l | chr1 | - | 75185330 | 75185866 | 75185330 | 75185481 | 75185699 | 75185866 | 45,65,81,24 | 2,9,11,3 | 51,36,59,49 | 23,11,15,11 | 367 | 149 | 0.00037 | 0.00744 | 0.901,0.7 | 0.474,0.5 | 0.214 |
| ENSMUSG00000024561 | Mbd1 | chr18 | + | 74410371 | 74410649 | 74410371 | 74410500 | 74410587 | 74410649 | 67,98,91,103 | 170,122,152,143 | 36,30,61,48 | 139,153,117,130 | 236 | 149 | 4.91E-05 | 0.0019 | 0.199,0.3 | 0.141,0.1 | 0.109 |
| ENSMUSG00000034729 | Mrps10 | chr17 | + | 47683351 | 47683570 | 47683351 | 47683416 | 47683497 | 47683570 | 39,29,31,39 | 68,51,46,48 | 25,15,18,15 | 54,50,45,98 | 230 | 149 | 6.81E-05 | 0.00243 | 0.271,0.2 | 0.231,0.1 | 0.125 |
| ENSMUSG00000030083 | Abtb1 | chr6 | - | 88817691 | 88817882 | 88817691 | 88817747 | 88817854 | 88817882 | 147,80,152,82 | 26,13,28,13 | 69,85,81,133 | 34,22,36,30 | 256 | 149 | 0.00104 | 0.01514 | 0.767,0.7 | 0.542,0.6 | 0.143 |
| ENSMUSG00000030689 | Ino80e | chr7 | - | 126456397 | 126456959 | 126456397 | 126456463 | 126456847 | 126456959 | 138,135,132,116 | 0,0,0,2 | 73,97,89,115 | 3,9,6,2 | 533 | 149 | 2.62E-08 | 8.53E-06 | 1.0,1.0,1. | 0.872,0.7 | 0.143 |
| ENSMUSG00000019518 | Ap4m1 | chr5 | + | 138176184 | 138176442 | 138176184 | 138176235 | 138176330 | 138176442 | 65,70,91,67 | 39,62,61,31 | 50,31,15,43 | 38,40,32,45 | 244 | 149 | 0.00649 | 0.05155 | 0.504,0.4 | 0.446,0.3 | 0.15 |
| ENSMUSG00000028809 | Srrm1 | chr4 | - | 135071809 | 135074168 | 135071809 | 135071980 | 135074045 | 135074168 | 6,577,610,680,755,370 | 326,313,400,226 | 3,580,363,234,813,810 | 356,261,247,247 | 2214 | 149 | 5.70E-07 | 7.84E-05 | 0.576,0.5 | 0.404,0.4 | 0.112 |
| ENSMUSG00000026860 | Sh3glb2 | chr2 | - | 30235710 | 30236465 | 30235710 | 30235951 | 30236364 | 30236465 | 599,469,581,485 | 53,21,61,72 | 316,372,387,371 | 56,71,67,63 | 562 | 149 | 8.21E-05 | 0.00263 | 0.75,0.85 | 0.599,0.5 | 0.142 |
| ENSMUSG00000015790 | Surf1 | chr2 | - | 26805607 | 26806018 | 26805607 | 26805761 | 26805966 | 26806018 | 123,146,219,127 | 4,3,47,22 | 77,117,102,98 | 23,21,17,19 | 354 | 149 | 0.00621 | 0.05008 | 0.928,0.9 | 0.585,0.7 | 0.141 |
| ENSMUSG00000014873 | Surf2 | chr2 | + | 26807439 | 26809067 | 26807439 | 26807543 | 26808940 | 26809067 | 835,755,961,818 | 38,25,22,40 | 644,503,548,784 | 40,41,48,32 | 1546 | 149 | 0.00195 | 0.02252 | 0.679,0.7 | 0.608,0.5 | 0.13 |
| ENSMUSG00000044906 | 4930503L | chr18 | - | 70600405 | 70601588 | 70600405 | 70600556 | 70600829 | 70601588 | 165,112,133,150 | 14,17,23,8 | 81,78,108,122 | 15,9,24,32 | 422 | 149 | 0.01362 | 0.08731 | 0.806,0.6 | 0.656,0.7 | 0.112 |
| ENSMUSG00000053253 | Ndfip2 | chr14 | + | 105535436 | 105542270 | 105535436 | 105535561 | 105542164 | 105542270 | 548,581,817,600 | 0,0,0,0 | 610,547,523,695 | 0,0,2,6 | 6752 | 149 | 0.00254 | 0.02657 | 1.0,1.0,1. | 1.0,1.0,0. | 0.107 |
| ENSMUSG00000086290 | Snhg12 | chr4 | + | 132035986 | 132037214 | 132035986 | 132036252 | 132037148 | 132037214 | 517,357,478,406 | 5,18,29,0 | 335,361,204,335 | 29,17,13,14 | 1045 | 149 | 0.00235 | 0.02514 | 0.936,0.7 | 0.622,0.7 | 0.135 |
| ENSMUSG00000026193 | Fn1 | chr1 | - | 71636439 | 71637622 | 71636439 | 71636546 | 71637532 | 71637622 | 322,208,467,413 | 6,4,12,0 | 259,478,471,320 | 14,9,14,12 | 1135 | 149 | 0.00477 | 0.04319 | 0.876,0.8 | 0.708,0.8 | 0.102 |
| ENSMUSG00000078700 | D030028 | chr11 | + | 96835068 | 96835919 | 96835068 | 96835176 | 96835670 | 96835919 | 66,37,61,24 | 0,9,4,8 | 17,14,24,43 | 13,11,7,8 | 643 | 149 | 0.0048 | 0.04337 | 1.0,0.488, | 0.233,0.2 | 0.304 |

| | | | | | | | | | | | | | | | | | | | |
|---|---|---|---|---|---|---|---|---|---|---|---|---|---|---|---|---|---|---|---|
| ENSMUSG00000097772 | 5430416N | chr5 | - | 100568692 | 100569900 | 100568692 | 100568831 | 100569826 | 100569900 | 144,151,208,183 | 49,27,38,24 | 153,99,134,132 | 48,49,50,45 | 1144 | 149 | 0.00068 | 0.01145 | 0.277,0.42 | 0.293,0.20 | 0.144 |
| ENSMUSG00000033416 | Gucd1 | chr10 | - | 75345425 | 75345957 | 75345425 | 75345667 | 75345865 | 75345957 | 47,28,36,44 | 26,23,34,40 | 29,27,26,23 | 28,43,37,31 | 347 | 149 | 0.00958 | 0.06812 | 0.437,0.34 | 0.308,0.21 | 0.105 |
| ENSMUSG00000042099 | Kank3 | chr17 | + | 34041259 | 34041891 | 34041259 | 34041349 | 34041648 | 34041891 | 225,146,283,162 | 73,36,55,40 | 145,152,223,193 | 66,81,58,78 | 448 | 149 | 0.00394 | 0.03683 | 0.506,0.57 | 0.422,0.38 | 0.117 |
| ENSMUSG00000091625 | Lsm5 | chr6 | - | 56679990 | 56680394 | 56679990 | 56680018 | 56680302 | 56680394 | 27,81,89,57 | 0,1,2,0 | 23,37,33,51 | 6,8,1,0 | 433 | 149 | 1.03E-05 | 0.00067 | 1.0,0.965, | 0.569,0.61 | 0.201 |
| ENSMUSG00000041477 | Dcp1b | chr6 | + | 119191730 | 119194940 | 119191730 | 119192567 | 119194763 | 119194940 | 1,135,107,414,481,250 | 32,30,35,50 | 790,819,812,900 | 39,46,58,33 | 2345 | 149 | 0.00037 | 0.00744 | 0.693,0.69 | 0.563,0.53 | 0.132 |
| ENSMUSG00000054708 | Ankrd24 | chr10 | + | 81475861 | 81476646 | 81475861 | 81475969 | 81476601 | 81476646 | 340,340,329,398 | 7,5,11,14 | 227,238,264,369 | 5,17,27,13 | 781 | 149 | 0.00445 | 0.04079 | 0.903,0.92 | 0.896,0.72 | 0.102 |
| ENSMUSG00000041777 | Cir1 | chr2 | - | 73136233 | 73136685 | 73136233 | 73136282 | 73136620 | 73136685 | 128,142,146,127 | 71,132,123,63 | 79,48,63,91 | 123,43,66,128 | 487 | 149 | 6.47E-05 | 0.00236 | 0.355,0.24 | 0.164,0.25 | 0.106 |
| ENSMUSG00000002963 | Pnkp | chr7 | + | 44507586 | 44508173 | 44507586 | 44507633 | 44507876 | 44508173 | 133,105,90,88 | 62,34,57,32 | 68,61,73,109 | 54,60,63,80 | 392 | 149 | 0.00011 | 0.00323 | 0.449,0.54 | 0.324,0.27 | 0.156 |
| ENSMUSG00000002043 | Trappc6a | chr7 | + | 19248346 | 19249015 | 19248346 | 19248464 | 19248949 | 19249015 | 68,55,56,24 | 2,1,1,0 | 36,26,20,24 | 0,3,2,4 | 634 | 149 | 0.00246 | 0.02607 | 0.889,0.92 | 1.0,0.671, | 0.197 |
| ENSMUSG00000036067 | Slc2a6 | chr2 | - | 26913099 | 26913655 | 26913099 | 26913261 | 26913546 | 26913655 | 89,92,186,88 | 34,28,62,22 | 99,59,62,101 | 53,34,64,52 | 434 | 149 | 6.29E-05 | 0.00232 | 0.473,0.53 | 0.391,0.37 | 0.169 |
| ENSMUSG00000062031 | Pgghg | chr7 | + | 140524863 | 140525262 | 140524863 | 140524995 | 140525150 | 140525262 | 23,12,15,10 | 5,7,12,8 | 2,2,8,17 | 19,8,8,35 | 304 | 149 | 7.16E-05 | 0.00245 | 0.693,0.45 | 0.049,0.10 | 0.308 |
| ENSMUSG00000025504 | Eps8l2 | chr7 | + | 140937154 | 140937604 | 140937154 | 140937229 | 140937455 | 140937604 | 16,19,26,9 | 4,7,0,2 | 5,18,14,13 | 10,4,14,4 | 375 | 149 | 0.01257 | 0.08298 | 0.614,0.51 | 0.166,0.64 | 0.28 |
| ENSMUSG00000025473 | Adam8 | chr7 | - | 139567091 | 139567650 | 139567091 | 139567181 | 139567472 | 139567650 | 91,83,113,46 | 1,0,3,3 | 69,36,64,48 | 3,4,4,3 | 440 | 149 | 0.007 | 0.05432 | 0.969,1.0, | 0.886,0.75 | 0.102 |
| ENSMUSG00000035773 | Kiss1r | chr10 | + | 79752804 | 79755385 | 79752804 | 79754751 | 79755260 | 79755385 | 56,49,60,55 | 13,25,11,22 | 44,23,29,47 | 24,30,26,14 | 658 | 149 | 0.01116 | 0.07664 | 0.494,0.30 | 0.293,0.14 | 0.16 |
| ENSMUSG00000097391 | Mirg | chr12 | + | 109697675 | 109698346 | 109697675 | 109697784 | 109698150 | 109698346 | 47,77,54,66 | 23,19,17,18 | 20,17,35,52 | 11,15,17,30 | 515 | 149 | 0.00642 | 0.05123 | 0.372,0.54 | 0.345,0.24 | 0.152 |
| ENSMUSG00000018442 | Derl2 | chr11 | - | 70904268 | 70906626 | 70904268 | 70904464 | 70906552 | 70906626 | 196,166,289,158 | 0,0,0,1 | 158,145,112,170 | 1,6,3,0 | 2237 | 149 | 1.64E-05 | 0.00093 | 1.0,1.0,1.0 | 0.913,0.61 | 0.168 |
| ENSMUSG00000014782 | Plekhg4 | chr8 | + | 106103380 | 106103767 | 106103380 | 106103497 | 106103668 | 106103767 | 31,33,33,27 | 26,19,20,8 | 11,7,4,19 | 14,19,4,23 | 320 | 149 | 0.00113 | 0.01614 | 0.357,0.44 | 0.268,0.14 | 0.21 |
| ENSMUSG00000092203 | 1110038E | chr17 | - | 35171090 | 35171444 | 35171090 | 35171149 | 35171327 | 35171444 | 71,77,75,72 | 66,53,66,34 | 61,27,37,46 | 81,46,55,64 | 327 | 149 | 7.83E-05 | 0.00259 | 0.329,0.39 | 0.255,0.21 | 0.153 |
| ENSMUSG00000039308 | Ndst2 | chr14 | - | 20774528 | 20774888 | 20774528 | 20774631 | 20774778 | 20774888 | 160,127,114,94 | 46,34,42,30 | 69,55,69,75 | 29,39,37,25 | 296 | 149 | 0.01397 | 0.08796 | 0.636,0.65 | 0.545,0.41 | 0.108 |
| ENSMUSG00000032579 | Hemk1 | chr9 | - | 107213770 | 107214577 | 107213770 | 107213862 | 107214199 | 107214577 | 152,129,136,109 | 33,26,28,23 | 70,109,82,81 | 19,51,30,35 | 486 | 149 | 0.00033 | 0.00693 | 0.585,0.60 | 0.53,0.396 | 0.145 |
| ENSMUSG00000019312 | Grb7 | chr11 | + | 98344006 | 98344419 | 98344006 | 98344117 | 98344320 | 98344419 | 15,14,39,10 | 20,18,14,13 | 4,10,2,7 | 9,19,26,16 | 352 | 149 | 0.00035 | 0.00712 | 0.241,0.24 | 0.158,0.18 | 0.187 |

### Table S3. Alternative_splicing_MXE

| GeneID | geneSymbol | chr | strand | 1stExonSt | 1stExonEn | 2ndExonS | 2ndExonE | upstreamE | upstreamE | downstrea | downstrea | ID.1 | IJC_SAMPLE_1 | SJC_SAMPLE_1 | IJC_SAMPLE_2 | SJC_SAMPLE_2 | IncFormL | SkipForm | PValue | FDR | IncLevel1 | IncLevel2 | IncLevelDifference |
|---|---|---|---|---|---|---|---|---|---|---|---|---|---|---|---|---|---|---|---|---|---|---|---|
| ENSMUSG00000100826 | Snhg14 | chr7 | - | 59055222 | 59055340 | 59057088 | 59057207 | 59054792 | 59054838 | 59058529 | 59058571 | 4722 | 3,2,2,11 | 73,27,43,0 | 33,1,12,5 | 0,0,1,0 | 268 | 267 | 6.58E-13 | 1.87E-09 | 0.039,0.0 | 1.0,1.0,0. | -0.693 |
| ENSMUSG00000026883 | Dab2ip | chr2 | + | 35533899 | 35534003 | 35551503 | 35551637 | 35448277 | 35448624 | 35597659 | 35597813 | 3706 | 22,10,32,16 | 128,126,151,68 | 104,64,100,52 | ############ | 253 | 283 | 5.42E-10 | 6.17E-07 | 0.161,0.0 | 0.505,0.3 | -0.263 |
| ENSMUSG00000097277 | 2900076A07Ri | chr7 | + | 81178904 | 81179069 | 81179094 | 81179275 | 81178534 | 81178688 | 81180958 | 81181250 | 5414 | 28,15,22,27 | 123,84,144,105 | 8,5,9,2 | 147,60,139,98 | 314 | 330 | 2.10E-09 | 1.71E-06 | 0.193,0.1 | 0.054,0.0 | 0.12 |
| ENSMUSG00000085218 | BB218582 | chr2 | - | 1.06E+08 | 1.06E+08 | 1.06E+08 | 1.06E+08 | 1.06E+08 | 1.06E+08 | 1.06E+08 | 1.06E+08 | 3156 | 0,2,6,4 | 25,17,39,29 | 0,0,0,0 | 10,17,10,13 | 259 | 983 | 7.38E-08 | 3.81E-05 | 0.0,0.309 | 0.0,0.0,0. | 0.255 |
| ENSMUSG00000085218 | BB218582 | chr2 | - | 1.06E+08 | 1.06E+08 | 1.06E+08 | 1.06E+08 | 1.06E+08 | 1.06E+08 | 1.06E+08 | 1.06E+08 | 3157 | 0,2,6,4 | 25,17,43,29 | 0,0,0,0 | 10,17,10,13 | 259 | 986 | 8.85E-08 | 4.19E-05 | 0.0,0.309 | 0.0,0.0,0. | 0.25 |
| ENSMUSG00000022763 | Aifm3 | chr16 | + | 17324093 | 17324198 | 17324762 | 17324783 | 17322781 | 17322880 | 17324990 | 17325349 | 452 | 39,44,81,41 | 3,4,2,4 | 40,42,72,42 | 9,12,15,21 | 254 | 170 | 3.10E-07 | 0.00013 | 0.897,0.8 | 0.748,0.7 | 0.208 |
| ENSMUSG00000097040 | 2610316D01Ri | chr3 | - | 45241766 | 45241854 | 45261531 | 45262093 | 45236429 | 45236599 | 45281500 | 45281597 | 2340 | 35,10,33,30 | 7,13,10,2 | 22,15,7,24 | 0,0,2,0 | 711 | 237 | 1.09E-06 | 0.00039 | 0.625,0.2 | 1.0,1.0,0. | -0.338 |
| ENSMUSG00000040374 | Pex2 | chr3 | - | 5630192 | 5630258 | 5635546 | 5635689 | 5628679 | 5628789 | 5641097 | 5641210 | 1769 | 24,35,77,29 | 0,0,3,0 | 38,34,6,29 | 6,12,1,0 | 292 | 215 | 1.19E-06 | 0.0004 | 1.0,1.0,0. | 0.823,0.6 | 0.159 |
| ENSMUSG00000026017 | Carf | chr1 | + | 60147241 | 60147359 | 60148485 | 60148608 | 60144469 | 60144593 | 60164009 | 60164070 | 3159 | 16,6,17,7 | 52,58,110,50 | 20,18,18,28 | 40,46,26,52 | 267 | 272 | 1.50E-06 | 0.00045 | 0.239,0.0 | 0.337,0.2 | -0.199 |
| ENSMUSG00000039652 | Cpeb3 | chr19 | - | 37103731 | 37103788 | 37116530 | 37116621 | 37065819 | 37065987 | 37151366 | 37152385 | 4099 | 104,89,109,72 | 2,11,6,16 | 72,41,51,47 | 14,20,14,22 | 240 | 206 | 1.43E-06 | 0.00045 | 0.978,0.8 | 0.815,0.6 | 0.182 |
| ENSMUSG00000021458 | Aopep | chr13 | + | 63208848 | 63209008 | 63215853 | 63216099 | 63180836 | 63181003 | 63304359 | 63304549 | 3887 | 27,31,32,18 | 11,17,19,13 | 6,8,7,15 | 27,25,11,19 | 309 | 395 | 4.31E-06 | 0.00111 | 0.758,0.7 | 0.221,0.2 | 0.33 |
| ENSMUSG00000097040 | 2610316D01Ri | chr3 | - | 45241766 | 45241887 | 45261531 | 45262093 | 45236429 | 45236599 | 45281500 | 45281597 | 2343 | 35,10,33,30 | 8,13,10,2 | 22,15,7,24 | 0,0,2,1 | 711 | 270 | 1.20E-05 | 0.0022 | 0.624,0.2 | 1.0,1.0,0. | -0.304 |
| ENSMUSG00000026765 | Lypd6b | chr2 | + | 49730099 | 49730155 | 49753822 | 49753918 | 49677742 | 49677785 | 49820936 | 49820996 | 5268 | 0,0,0,0 | 38,28,37,19 | 9,0,2,1 | 33,22,12,25 | 205 | 245 | 1.19E-05 | 0.0022 | 0.0,0.0,0. | 0.246,0.0 | -0.115 |
| ENSMUSG00000034164 | Emid1 | chr11 | - | 5059543 | 5059634 | 5060689 | 5060774 | 5056264 | 5056969 | 5066879 | 5066924 | 974 | 30,17,101,21 | 1,0,7,5 | 55,52,66,58 | 23,14,21,8 | 234 | 240 | 1.44E-05 | 0.00234 | 0.969,1.0 | 0.71,0.79 | 0.143 |
| ENSMUSG00000026885 | Ttll11 | chr2 | - | 35780283 | 35780379 | 35792581 | 35793157 | 35779276 | 35779392 | 35830685 | 35830818 | 3712 | 623,492,637,407 | 93,57,67,55 | 447,340,381,410 | 101,83,82,65 | 725 | 245 | 1.42E-05 | 0.00234 | 0.694,0.7 | 0.599,0.5 | 0.111 |
| ENSMUSG00000031433 | Rbm41 | chrX | - | 1.39E+08 | 1.39E+08 | 1.39E+08 | 1.39E+08 | 1.39E+08 | 1.39E+08 | 1.39E+08 | 1.39E+08 | 3462 | 11,1,4,9 | 81,50,76,71 | 0,2,3,0 | 82,43,89,65 | 221 | 553 | 1.62E-05 | 0.00243 | 0.254,0.0 | 0.0,0.104 | 0.119 |
| ENSMUSG00000026825 | Dnm1 | chr2 | - | 32223164 | 32223216 | 32224693 | 32224832 | 32217972 | 32218059 | 32225787 | 32225855 | 3522 | 2,196,182,122,071,820 | 8,175,011,018,586 | 1,602,160,211,801,210 | ############ | 288 | 201 | 1.62E-05 | 0.00243 | 0.652,0.7 | 0.581,0.6 | 0.102 |
| ENSMUSG00000002688 | Prkd1 | chr12 | - | 50413084 | 50413246 | 50430182 | 50430289 | 50412396 | 50412495 | 50431925 | 50431998 | 714 | 23,40,40,20 | 78,103,84,72 | 53,38,35,36 | 68,49,38,64 | 256 | 311 | 4.68E-05 | 0.00566 | 0.264,0.3 | 0.486,0.4 | -0.176 |
| ENSMUSG00000022668 | Gtpbp8 | chr16 | - | 44562866 | 44562966 | 44564100 | 44564231 | 44560391 | 44560510 | 44566361 | 44566646 | 467 | 22,9,6,11 | 64,32,80,58 | 6,0,10,0 | 76,57,74,42 | 280 | 249 | 5.06E-05 | 0.00576 | 0.234,0.2 | 0.066,0.0 | 0.117 |
| ENSMUSG00000039842 | Mcph1 | chr8 | + | 18679561 | 18679651 | 18681531 | 18682659 | 18677133 | 18677256 | 18691580 | 18691681 | 3797 | 38,21,40,32 | 442,314,362,299 | 12,11,6,15 | ############ | 239 | 1277 | 6.33E-05 | 0.00654 | 0.315,0.2 | 0.217,0.1 | 0.145 |
| ENSMUSG00000057715 | A830018L16Ri | chr1 | + | 12003767 | 12003815 | 12021186 | 12021280 | 11900038 | 11900115 | 12042269 | 12042326 | 2713 | 6,5,12,9 | 75,26,70,53 | 10,10,10,15 | 29,28,21,31 | 197 | 243 | 7.23E-05 | 0.00734 | 0.09,0.19 | 0.298,0.3 | -0.179 |
| ENSMUSG00000055897 | Ppp4r1l-ps | chr2 | - | 1.73E+08 | 1.73E+08 | 1.73E+08 | 1.73E+08 | 1.73E+08 | 1.73E+08 | 1.74E+08 | 1.74E+08 | 2226 | 4,1,0,3 | 35,29,39,37 | 4,12,2,14 | 33,21,23,41 | 243 | 296 | 8.25E-05 | 0.00795 | 0.122,0.0 | 0.129,0.4 | -0.169 |
| ENSMUSG00000034902 | Pip5k1c | chr10 | + | 81151712 | 81151790 | 81152525 | 81152794 | 81150825 | 81150952 | 81153159 | 81153237 | 5632 | 67,54,116,45 | 70,73,85,42 | 77,65,81,67 | 38,18,53,39 | 227 | 418 | 9.64E-05 | 0.00884 | 0.638,0.5 | 0.789,0.8 | -0.14 |
| ENSMUSG00000021326 | Trim27 | chr13 | + | 21374266 | 21374294 | 21374762 | 21374856 | 21374024 | 21374140 | 21375389 | 21375416 | 3112 | 52,35,30,42 | 6,2,0,7 | 32,37,28,33 | 10,13,6,10 | 177 | 243 | 0.0001 | 0.00925 | 0.922,0.9 | 0.815,0.7 | 0.12 |
| ENSMUSG00000026836 | Acvr1 | chr2 | - | 58390495 | 58390569 | 58406035 | 58406312 | 58369676 | 58369940 | 58427789 | 58427885 | 3788 | 109,83,124,75 | 74,41,68,48 | 110,60,78,50 | 96,58,80,61 | 426 | 223 | 0.00011 | 0.00973 | 0.435,0.5 | 0.375,0.3 | 0.131 |
| ENSMUSG00000024948 | Map4k2 | chr19 | + | 6393133 | 6393206 | 6393284 | 6393416 | 6392773 | 6392816 | 6393490 | 6393553 | 56 | 90,46,90,72 | 243,188,237,218 | 88,86,102,97 | ############ | 222 | 281 | 0.00014 | 0.01113 | 0.319,0.2 | 0.378,0.3 | -0.105 |
| ENSMUSG00000046876 | Atxn1 | chr13 | - | 45851296 | 45851421 | 45888590 | 45888654 | 45720048 | 45722056 | 45948984 | 45949113 | 5521 | 38,47,56,30 | 124,92,139,72 | 45,25,48,45 | 60,43,66,43 | 213 | 274 | 0.00017 | 0.01268 | 0.283,0.3 | 0.491,0.4 | -0.152 |
| ENSMUSG00000037463 | Fbxo27 | chr7 | + | 28394430 | 28394526 | 28396064 | 28396200 | 28394155 | 28394267 | 28397663 | 28398760 | 2253 | 49,37,60,41 | 32,38,27,35 | 43,20,34,32 | 62,25,60,47 | 245 | 285 | 0.00018 | 0.01268 | 0.64,0.53 | 0.447,0.4 | 0.175 |
| ENSMUSG00000034543 | Morc2a | chr11 | + | 3600307 | 3600447 | 3608671 | 3608725 | 3599493 | 3600146 | 3611828 | 3611863 | 6038 | 37,16,24,21 | 53,17,34,20 | 4,9,15,7 | 25,25,29,30 | 289 | 203 | 0.00019 | 0.01285 | 0.329,0.3 | 0.101,0.2 | 0.193 |
| ENSMUSG00000079657 | Rab26 | chr17 | - | 24749028 | 24749094 | 24749364 | 24749417 | 24748887 | 24748944 | 24749616 | 24749683 | 3420 | 107,68,100,63 | 133,75,111,73 | 120,112,99,109 | 81,76,92,64 | 202 | 215 | 0.00022 | 0.01432 | 0.461,0.4 | 0.612,0.6 | -0.12 |
| ENSMUSG00000028760 | Eif4g3 | chr4 | + | 1.38E+08 | 1.38E+08 | 1.38E+08 | 1.38E+08 | 1.38E+08 | 1.38E+08 | 1.38E+08 | 1.38E+08 | 1806 | 60,76,72,60 | 43,33,34,23 | 28,36,43,32 | 40,31,31,49 | 263 | 182 | 0.00026 | 0.01593 | 0.491,0.6 | 0.326,0.4 | 0.193 |
| ENSMUSG00000021969 | Zdhhc20 | chr14 | - | 58102977 | 58103098 | 58111349 | 58111453 | 58095996 | 58096066 | 58115889 | 58115916 | 4780 | 148,153,199,175 | 225,187,225,172 | 146,149,156,191 | ############ | 253 | 270 | 0.00027 | 0.0163 | 0.412,0.4 | 0.571,0.5 | -0.102 |
| ENSMUSG00000044712 | Slc38a6 | chr12 | + | 73340472 | 73340573 | 73356918 | 73356971 | 73338933 | 73339007 | 73365750 | 73365790 | 720 | 33,10,43,13 | 0,0,0,0 | 18,20,18,18 | 3,4,2,0 | 250 | 202 | 0.00038 | 0.02054 | 1.0,1.0,1. | 0.829,0.8 | 0.123 |
| ENSMUSG00000030518 | Fam189a1 | chr7 | - | 64469512 | 64469647 | 64505856 | 64505967 | 64436437 | 64436545 | 64633128 | 64633185 | 4453 | 37,33,44,37 | 107,102,154,115 | 67,60,51,43 | 121,81,69,117 | 260 | 284 | 0.00039 | 0.02095 | 0.274,0.2 | 0.377,0.4 | -0.131 |
| ENSMUSG00000027203 | Dut | chr2 | + | 1.25E+08 | 1.25E+08 | 1.25E+08 | 1.25E+08 | 1.25E+08 | 1.25E+08 | 1.25E+08 | 1.25E+08 | 3301 | 134,100,173,105 | 46,25,37,30 | 118,102,83,95 | 65,41,49,34 | 241 | 194 | 0.00043 | 0.02163 | 0.701,0.7 | 0.594,0.6 | 0.115 |
| ENSMUSG00000034949 | Zfr2 | chr10 | + | 81084817 | 81084912 | 81084998 | 81085094 | 81084223 | 81084362 | 81085518 | 81085630 | 2290 | 121,129,140,136 | 51,45,73,53 | 96,59,70,105 | 72,42,75,54 | 244 | 245 | 0.00047 | 0.02353 | 0.704,0.7 | 0.572,0.5 | 0.13 |
| ENSMUSG00000029575 | Mmab | chr5 | - | 1.15E+08 | 1.15E+08 | 1.15E+08 | 1.15E+08 | 1.15E+08 | 1.15E+08 | 1.15E+08 | 1.15E+08 | 1436 | 48,31,48,36 | 84,50,64,56 | 51,40,44,29 | 52,39,28,21 | 222 | 247 | 0.0005 | 0.024 | 0.389,0.4 | 0.522,0.5 | -0.157 |
| ENSMUSG00000097040 | 2610316D01Ri | chr3 | - | 45241766 | 45241851 | 45261531 | 45262093 | 45236429 | 45236599 | 45281500 | 45281597 | 2337 | 35,10,33,30 | 10,18,28,7 | 22,15,7,24 | 0,3,2,2 | 711 | 234 | 0.00056 | 0.02553 | 0.535,0.1 | 1.0,0.622 | -0.35 |
| ENSMUSG00000026767 | Mindy3 | chr2 | - | 12400989 | 12401069 | 12402303 | 12402377 | 12391419 | 12391490 | 12423969 | 12424099 | 2283 | 56,0,41,38 | 55,40,130,59 | 0,15,13,0 | 76,51,79,80 | 223 | 229 | 0.00058 | 0.02598 | 0.511,0.0 | 0.0,0.232 | 0.194 |
| ENSMUSG00000024330 | Col11a2 | chr17 | + | 34264744 | 34264822 | 34266071 | 34266251 | 34263792 | 34263984 | 34268592 | 34268652 | 195 | 38,25,43,25 | 59,24,64,33 | 46,22,45,50 | 29,20,36,30 | 227 | 329 | 0.00061 | 0.02696 | 0.483,0.6 | 0.697,0.6 | -0.14 |
| ENSMUSG00000020100 | Slc29a3 | chr10 | - | 60559542 | 60559769 | 60566348 | 60566431 | 60549804 | 60552269 | 60586164 | 60586463 | 4072 | 17,27,20,17 | 17,13,21,18 | 71,54,21,34 | 15,6,18,14 | 232 | 376 | 0.00067 | 0.02855 | 0.618,0.7 | 0.885,0.9 | -0.168 |
| ENSMUSG00000032913 | Lrig2 | chr3 | - | 1.04E+08 | 1.04E+08 | 1.04E+08 | 1.04E+08 | 1.04E+08 | 1.04E+08 | 1.04E+08 | 1.04E+08 | 1697 | 17,27,35,36 | 88,77,103,81 | 37,32,28,37 | 65,50,54,44 | 218 | 313 | 0.0007 | 0.02923 | 0.217,0.3 | 0.45,0.47 | -0.158 |
| ENSMUSG00000037935 | Smarce1 | chr11 | - | 99119100 | 99119144 | 99119322 | 99119374 | 99115546 | 99115968 | 99121227 | 99121271 | 5869 | 9,10,20,11 | 50,37,37,34 | 2,9,6,3 | 43,39,34,53 | 201 | 193 | 0.00074 | 0.03008 | 0.147,0.2 | 0.043,0.1 | 0.128 |
| ENSMUSG00000040543 | Pitpnm3 | chr11 | - | 72003082 | 72003190 | 72010912 | 72011008 | 71983353 | 71983401 | 72026479 | 72026604 | 3325 | 92,45,85,88 | 75,73,85,85 | 121,91,145,104 | 77,45,85,78 | 245 | 257 | 0.00079 | 0.031 | 0.563,0.3 | 0.622,0.6 | -0.135 |
| ENSMUSG00000018372 | Cep95 | chr11 | + | 1.07E+08 | 1.07E+08 | 1.07E+08 | 1.07E+08 | 1.07E+08 | 1.07E+08 | 1.07E+08 | 1.07E+08 | 4966 | 62,47,54,55 | 58,41,64,47 | 6,17,10,28 | 29,27,40,35 | 265 | 278 | 0.00078 | 0.031 | 0.529,0.5 | 0.178,0.3 | 0.214 |
| ENSMUSG00000052605 | Isoc2b | chr7 | - | 4852448 | 4852566 | 4854391 | 4854532 | 4847957 | 4848364 | 4869059 | 4869192 | 2945 | 92,48,94,56 | 47,52,56,24 | 107,84,84,88 | 23,31,31,29 | 290 | 267 | 0.00089 | 0.03358 | 0.643,0.4 | 0.811,0.7 | -0.146 |
| ENSMUSG00000015112 | Slc25a13 | chr6 | - | 6180970 | 6181113 | 6191263 | 6191317 | 6152407 | 6152523 | 6216995 | 6217118 | 3804 | 20,8,17,22 | 23,17,35,36 | 6,5,11,8 | 53,22,48,16 | 203 | 292 | 0.00095 | 0.03501 | 0.556,0.4 | 0.14,0.24 | 0.197 |
| ENSMUSG00000029202 | Pds5a | chr5 | - | 65787261 | 65787398 | 65789055 | 65789124 | 65785341 | 65785465 | 65791514 | 65791673 | 5398 | 92,53,84,50 | 124,107,173,104 | 49,35,45,49 | ############ | 218 | 286 | 0.00104 | 0.03765 | 0.493,0.3 | 0.31,0.29 | 0.106 |
| ENSMUSG00000036086 | Zranb3 | chr1 | - | 1.28E+08 | 1.28E+08 | 1.28E+08 | 1.28E+08 | 1.28E+08 | 1.28E+08 | 1.28E+08 | 1.28E+08 | 2781 | 9,7,6,5 | 41,20,23,16 | 8,14,8,7 | 14,13,10,12 | 235 | 321 | 0.00112 | 0.03983 | 0.231,0.3 | 0.438,0.5 | -0.22 |
| ENSMUSG00000047888 | Tnrc6b | chr15 | + | 80668939 | 80669007 | 80742881 | 80742969 | 80643740 | 80643831 | 80748585 | 80748607 | 596 | 14,8,16,17 | 83,100,102,61 | 17,25,15,15 | 58,57,59,45 | 217 | 237 | 0.00123 | 0.04273 | 0.156,0.0 | 0.242,0.3 | -0.109 |
| ENSMUSG00000030782 | Tgfb1i1 | chr7 | + | 1.28E+08 | 1.28E+08 | 1.28E+08 | 1.28E+08 | 1.28E+08 | 1.28E+08 | 1.28E+08 | 1.28E+08 | 2400 | 26,18,33,25 | 27,17,45,16 | 55,48,83,36 | 19,26,38,24 | 292 | 237 | 0.00132 | 0.0448 | 0.439,0.4 | 0.701,0.6 | -0.164 |
| ENSMUSG00000021870 | Slmap | chr14 | - | 26180564 | 26180615 | 26181058 | 26181109 | 26174692 | 26174755 | 26181196 | 26181365 | 2265 | 164,130,133,142 | 167,108,111,131 | 67,131,107,121 | 46,86,40,76 | 200 | 200 | 0.00136 | 0.04542 | 0.495,0.5 | 0.593,0.6 | -0.108 |
| ENSMUSG00000049411 | Tmem241 | chr18 | - | 12246451 | 12246529 | 12251460 | 12251507 | 12237155 | 12237237 | 12254437 | 12254537 | 1104 | 5,5,11,4 | 15,14,52,29 | 19,5,14,18 | 31,15,22,27 | 196 | 227 | 0.00149 | 0.04828 | 0.279,0.2 | 0.415,0.2 | -0.162 |
| ENSMUSG00000028232 | Tmem68 | chr4 | - | 3569630 | 3569752 | 3570320 | 3570372 | 3569361 | 3569463 | 3574703 | 3574804 | 2185 | 25,24,26,24 | 8,3,16,2 | 23,19,33,28 | 4,1,4,0 | 201 | 271 | 0.00149 | 0.04828 | 0.808,0.9 | 0.886,0.9 | -0.103 |
| ENSMUSG00000025184 | R3hcc1l | chr19 | + | 42564464 | 42564640 | 42570358 | 42570429 | 42550990 | 42552780 | 42571263 | 42571359 | 4593 | 91,55,154,96 | 35,19,44,37 | 67,98,59,53 | 52,54,43,21 | 325 | 220 | 0.00149 | 0.04828 | 0.638,0.6 | 0.466,0.5 | 0.128 |
| ENSMUSG00000033257 | Ttll4 | chr1 | + | 74729118 | 74729265 | 74734866 | 74734969 | 74728481 | 74728616 | 74735563 | 74735774 | 2354 | 65,60,80,35 | 58,31,40,41 | 58,63,63,56 | 25,17,25,20 | 296 | 252 | 0.00153 | 0.04892 | 0.488,0.6 | 0.664,0.7 | -0.162 |
| ENSMUSG00000026361 | Cdc73 | chr1 | - | 1.44E+08 | 1.44E+08 | 1.44E+08 | 1.44E+08 | 1.44E+08 | 1.44E+08 | 1.44E+08 | 1.44E+08 | 3066 | 302,231,279,260 | 92,89,133,62 | 153,179,147,154 | 111,68,87,94 | 366 | 248 | 0.00153 | 0.04892 | 0.69,0.63 | 0.483,0.6 | 0.118 |
| ENSMUSG00000006498 | Ptbp1 | chr10 | + | 79695950 | 79696028 | 79696657 | 79696803 | 79695614 | 79695789 | 79697847 | 79697881 | 2269 | 34,27,22,33 | 113,59,119,57 | 23,20,16,19 | 130,102,98,117 | 227 | 295 | 0.00161 | 0.04983 | 0.281,0.3 | 0.187,0.2 | 0.135 |
| ENSMUSG00000032507 | Fbxl2 | chr9 | - | 1.14E+08 | 1.14E+08 | 1.14E+08 | 1.14E+08 | 1.14E+08 | 1.14E+08 | 1.14E+08 | 1.14E+08 | 4236 | 186,120,132,116 | 54,57,70,51 | 102,107,101,56 | 59,90,56,65 | 276 | 224 | 0.00159 | 0.04983 | 0.737,0.6 | 0.584,0.4 | 0.135 |
| ENSMUSG00000031378 | Abcd1 | chrX | + | 72777614 | 72777760 | 72778067 | 72778213 | 72774399 | 72774494 | 72780395 | 72780480 | 1059 | 26,12,15,15 | 39,53,24,30 | 45,25,36,41 | 47,36,35,27 | 295 | 295 | 0.00193 | 0.05746 | 0.4,0.185 | 0.489,0.4 | -0.177 |
| ENSMUSG00000031536 | Polb | chr8 | - | 23135060 | 23135110 | 23138197 | 23138272 | 23132320 | 23132372 | 23140504 | 23140571 | 5282 | 139,72,97,74 | 104,72,80,85 | 89,85,95,117 | 59,9,65,53 | 224 | 199 | 0.00199 | 0.05904 | 0.543,0.4 | 0.573,0.8 | -0.181 |
| ENSMUSG00000055897 | Ppp4r1l-ps | chr2 | - | 1.73E+08 | 1.73E+08 | 1.73E+08 | 1.73E+08 | 1.73E+08 | 1.73E+08 | 1.74E+08 | 1.74E+08 | 2212 | 4,1,0,3 | 11,10,43,10 | 4,12,2,14 | 14,13,18,26 | 243 | 247 | 0.0021 | 0.06107 | 0.27,0.09 | 0.225,0.4 | -0.142 |
| ENSMUSG00000026349 | Ccnt2 | chr1 | + | 1.28E+08 | 1.28E+08 | 1.28E+08 | 1.28E+08 | 1.28E+08 | 1.28E+08 | 1.28E+08 | 1.28E+08 | 2583 | 63,70,137,52 | 63,36,61,42 | 78,68,50,51 | 26,23,11,25 | 210 | 212 | 0.00215 | 0.06131 | 0.502,0.6 | 0.752,0.7 | -0.145 |
| ENSMUSG00000032087 | Dscaml1 | chr9 | + | 45572430 | 45572577 | 45579286 | 45579565 | 45361779 | 45361926 | 45581293 | 45581569 | 5718 | 97,63,110,51 | 124,101,228,98 | 79,50,83,47 | ############ | 296 | 428 | 0.00214 | 0.06131 | 0.531,0.4 | 0.351,0.3 | 0.103 |
| ENSMUSG00000034574 | Daam1 | chr12 | + | 71942500 | 71942720 | 71962472 | 71962562 | 71877851 | 71877970 | 71983332 | 71983404 | 726 | 135,95,161,137 | 88,70,141,72 | 126,148,121,115 | 68,47,50,59 | 369 | 239 | 0.0022 | 0.06212 | 0.498,0.4 | 0.545,0.6 | -0.111 |
| ENSMUSG00000032411 | Tfdp2 | chr9 | + | 96105146 | 96105251 | 96169653 | 96169775 | 96084183 | 96084241 | 96177017 | 96177180 | 5675 | 46,55,50,33 | 1,17,21,6 | 53,33,37,34 | 23,27,10,29 | 254 | 271 | 0.00228 | 0.06389 | 0.98,0.77 | 0.711,0.5 | 0.174 |
| ENSMUSG00000017692 | Rhbdl3 | chr11 | + | 80193489 | 80193513 | 80210383 | 80210542 | 80191737 | 80192097 | 80214184 | 80214409 | 5941 | 27,28,18,27 | 163,122,170,134 | 43,22,22,19 | 124,78,79,76 | 173 | 308 | 0.00232 | 0.06448 | 0.228,0.2 | 0.382,0.3 | -0.103 |
| ENSMUSG00000042208 | Sanbr | chr11 | - | 23570228 | 23570467 | 23572435 | 23572529 | 23567137 | 23567196 | 23574935 | 23575122 | 1838 | 53,42,78,77 | 172,151,340,201 | 66,42,58,71 | ############ | 243 | 388 | 0.00238 | 0.06529 | 0.33,0.30 | 0.442,0.3 | -0.103 |
| ENSMUSG00000030882 | Dnhd1 | chr7 | + | 1.05E+08 | 1.05E+08 | 1.05E+08 | 1.05E+08 | 1.05E+08 | 1.05E+08 | 1.05E+08 | 1.05E+08 | 4487 | 18,24,24,34 | 4,2,0,2 | 15,28,14,23 | 8,0,10,8 | 404 | 287 | 0.00247 | 0.06667 | 0.762,0.8 | 0.571,1.0 | 0.21 |
| ENSMUSG00000035126 | Dnai4 | chr4 | - | 1.03E+08 | 1.03E+08 | 1.03E+08 | 1.03E+08 | 1.03E+08 | 1.03E+08 | 1.03E+08 | 1.03E+08 | 1616 | 72,62,111,60 | 37,52,51,36 | 52,39,27,51 | 56,38,28,45 | 376 | 301 | 0.00253 | 0.06795 | 0.609,0.4 | 0.426,0.4 | 0.129 |
| ENSMUSG00000028886 | Eya3 | chr4 | + | 1.32E+08 | 1.32E+08 | 1.32E+08 | 1.32E+08 | 1.32E+08 | 1.32E+08 | 1.32E+08 | 1.32E+08 | 5444 | 52,30,73,62 | 60,56,62,58 | 31,26,18,25 | 65,37,37,66 | 287 | 235 | 0.00267 | 0.07048 | 0.415,0.3 | 0.281,0.3 | 0.127 |
| ENSMUSG00000001576 | Ergic1 | chr17 | + | 26848874 | 26849503 | 26850899 | 26851000 | 26848528 | 26848653 | 26853390 | 26853495 | 6058 | 17,28,29,19 | 0,3,0,1 | 24,15,10,17 | 7,3,0,1 | 778 | 250 | 0.00277 | 0.07177 | 1.0,0.75,1 | 0.524,0.6 | 0.156 |
| ENSMUSG00000074776 | Gm10754 | chr10 | - | 97643349 | 97643437 | 97758813 | 97758904 | 97588367 | 97588474 | 97802850 | 97802952 | 2454 | 61,57,55,39 | 91,68,54,58 | 11,26,24,24 | 28,60,38,60 | 240 | 237 | 0.00283 | 0.07245 | 0.398,0.4 | 0.28,0.3,( | 0.126 |
| ENSMUSG00000063889 | Crem | chr18 | - | 3276693 | 3276729 | 3287903 | 3288092 | 3273421 | 3273578 | 3295039 | 3295182 | 1047 | 11,15,29,6 | 15,13,29,14 | 7,3,15,2 | 18,22,17,25 | 338 | 185 | 0.00289 | 0.07296 | 0.286,0.3 | 0.175,0.0 | 0.151 |
| ENSMUSG00000048720 | Tbc1d12 | chr19 | + | 38871102 | 38871218 | 38884430 | 38884513 | 38854059 | 38854183 | 38896215 | 38896296 | 4563 | 24,29,46,31 | 21,4,21,9 | 37,30,21,28 | 5,5,3,4 | 265 | 232 | 0.00297 | 0.07305 | 0.5,0.864 | 0.866,0.8 | -0.163 |
| ENSMUSG00000073016 | Uprt | chrX | + | 1.04E+08 | 1.04E+08 | 1.04E+08 | 1.04E+08 | 1.04E+08 | 1.04E+08 | 1.04E+08 | 1.04E+08 | 1127 | 8,2,6,11 | 47,38,24,21 | 7,12,13,6 | 16,18,15,15 | 192 | 219 | 0.00327 | 0.07774 | 0.163,0.0 | 0.333,0.4 | -0.19 |
| ENSMUSG00000029647 | Pan3 | chr5 | + | 1.47E+08 | 1.47E+08 | 1.47E+08 | 1.47E+08 | 1.47E+08 | 1.47E+08 | 1.47E+08 | 1.47E+08 | 5970 | 42,42,60,47 | 99,93,147,88 | 50,57,52,62 | 58,97,81,89 | 297 | 397 | 0.00341 | 0.07974 | 0.362,0.3 | 0.535,0.4 | -0.103 |
| ENSMUSG00000032017 | Grik4 | chr9 | - | 42502645 | 42502849 | 42509149 | 42509257 | 42477594 | 42477708 | 42533354 | 42533459 | 5077 | 85,42,70,58 | 98,46,139,109 | 40,31,25,20 | 76,87,74,59 | 257 | 353 | 0.00338 | 0.07974 | 0.544,0.5 | 0.42,0.32 | 0.137 |
| ENSMUSG00000023079 | Gtf2ird1 | chr5 | - | 1.34E+08 | 1.34E+08 | 1.34E+08 | 1.34E+08 | 1.34E+08 | 1.34E+08 | 1.34E+08 | 1.34E+08 | 3221 | 103,76,67,73 | 40,29,33,58 | 105,76,85,48 | 17,33,23,13 | 342 | 342 | 0.0035 | 0.08123 | 0.72,0.72 | 0.861,0.6 | -0.115 |
| ENSMUSG00000053111 | Fank1 | chr7 | + | 1.33E+08 | 1.33E+08 | 1.33E+08 | 1.33E+08 | 1.33E+08 | 1.33E+08 | 1.33E+08 | 1.33E+08 | 4759 | 23,14,54,28 | 4,9,11,8 | 27,22,14,16 | 16,15,8,15 | 327 | 274 | 0.00353 | 0.08127 | 0.828,0.5 | 0.586,0.5 | 0.185 |
| ENSMUSG00000023923 | Tbc1d5 | chr17 | - | 51291632 | 51291702 | 51450386 | 51450449 | 51275217 | 51275326 | 51486127 | 51486196 | 346 | 54,47,50,52 | 91,45,75,63 | 47,39,51,75 | 47,31,45,45 | 212 | 219 | 0.0036 | 0.08231 | 0.38,0.51 | 0.508,0.5 | -0.12 |
| ENSMUSG00000012609 | Ttll5 | chr12 | + | 85980055 | 85980448 | 85986099 | 85986284 | 85979498 | 85979582 | 86003339 | 86003494 | 1257 | 196,171,249,186 | 67,54,144,54 | 253,159,239,168 | 128,85,128,150 | 542 | 334 | 0.00375 | 0.08468 | 0.643,0.6 | 0.549,0.5 | 0.118 |
| ENSMUSG00000026767 | Mindy3 | chr2 | - | 12402303 | 12402377 | 12414811 | 12414891 | 12391419 | 12391490 | 12423969 | 12424259 | 2285 | 51,74,78,45 | 56,0,41,38 | 69,40,53,51 | 0,15,13,0 | 229 | 223 | 0.00384 | 0.08569 | 0.47,1.0,( | 1.0,0.722 | -0.217 |
| ENSMUSG00000036377 | Cracd | chr5 | + | 76907432 | 76907578 | 76970312 | 76970484 | 76805529 | 76805700 | 76988634 | 76988728 | 1689 | 119,116,119,75 | 92,76,126,62 | 190,127,132,114 | 83,51,109,57 | 295 | 321 | 0.00421 | 0.09171 | 0.585,0.6 | 0.714,0.7 | -0.103 |
| ENSMUSG00000040929 | Rfx3 | chr19 | - | 27790977 | 27791090 | 27792884 | 27793006 | 27782542 | 27784308 | 27803432 | 27803552 | 77 | 70,78,142,65 | 39,65,55,73 | 48,36,53,66 | 79,76,53,58 | 271 | 262 | 0.00431 | 0.09355 | 0.634,0.5 | 0.37,0.31 | 0.162 |
| ENSMUSG00000086370 | Ftx | chrX | - | 1.03E+08 | 1.03E+08 | 1.03E+08 | 1.03E+08 | 1.03E+08 | 1.03E+08 | 1.03E+08 | 1.03E+08 | 3734 | 634,742,895,605 | 65,51,45,52 | 843,863,402,602 | 22,21,38,22 | 1652 | 211 | 0.00436 | 0.09385 | 0.555,0.6 | 0.83,0.84 | -0.126 |

| | | | | | | | | | | | | | | | | | | | | | | |
|---|---|---|---|---|---|---|---|---|---|---|---|---|---|---|---|---|---|---|---|---|---|---|
| ENSMUSG00000032741 | Tpcn1 | chr5 | - | 1.21E+08 | 1.21E+08 | 1.21E+08 | 1.21E+08 | 1.21E+08 | 1.21E+08 | 1.21E+08 | 1.21E+08 | 1813 | 74,65,79,56 | 92,71,92,58 | 46,33,40,27 | 61,61,73,65 | 203 | 222 | 0.00463 | 0.09801 | 0.468,0.5 | 0.452,0.3 | 0.114 |
| ENSMUSG00000045915 | Ccdc42 | chr11 | + | 68481701 | 68481899 | 68484979 | 68485201 | 68479002 | 68479107 | 68485356 | 68485515 | 5940 | 5,25,14,12 | 24,11,37,12 | 13,14,9,17 | 5,11,4,4 | 347 | 371 | 0.00476 | 0.0985 | 0.182,0.7 | 0.735,0.5 | -0.285 |
