## Supplementary material for "Reversible m^6^Am methylation of snRNA by FTO controls morphine reward and tolerance without altering analgesia": Table S4

Table S4. Differential_expression

| gene name | baseMear | log2FoldCl | lfcSE | stat | pvalue | padj | gene length (kbp) |
|---|---|---|---|---|---|---|---|
| 0610005C13Rik | 41.1018 | 0.365236 | 0.35027 | 1.042727 | 0.29707 | 0.50725 | 7.539 |
| 0610009B22Rik | 723.849 | -0.16377 | 0.10155 | -1.61263 | 0.10682 | 0.27117 | 3.489 |
| 0610009E02Rik | 80.808 | 0.141283 | 0.23108 | 0.611414 | 0.54093 | 0.72658 | 13.701 |
| 0610009L18Rik | 172.755 | 0.338857 | 0.17113 | 1.980128 | 0.04769 | 0.1606 | 2.533 |
| 0610010K14Rik | 57.9821 | 0.199963 | 0.23224 | 0.861022 | 0.38923 | 0.59834 | 2.709 |
| 0610012G03Rik | 1145.17 | -0.16941 | 0.08541 | -1.9835 | 0.04731 | 0.1597 | 1.445 |
| 0610030E20Rik | 1223.91 | 0.165752 | 0.08482 | 1.954245 | 0.05067 | 0.16741 | 5.832 |
| 0610038B21Rik | 25.9842 | -0.02747 | 0.42348 | -0.06487 | 0.94828 | 0.97427 | 6.843 |
| 0610039K10Rik | 13.333 | -0.20965 | 0.5578 | -0.37584 | 0.70703 | 0.84193 | 0.952 |
| 0610040B10Rik | 67.7066 | 0.033173 | 0.24364 | 0.136155 | 0.8917 | 0.94605 | 3.458 |
| 0610040F04Rik | 36.4083 | 0.085729 | 0.33187 | 0.258324 | 0.79616 | 0.89487 | 83.896 |
| 0610040J01Rik | 116.781 | -0.01298 | 0.17161 | -0.07561 | 0.93973 | 0.96985 | 87.263 |
| 0610043K17Rik | 61.5642 | -0.13012 | 0.23899 | -0.54446 | 0.58612 | 0.76022 | 45.399 |
| 0710001A04Rik | 16.8998 | 0.335773 | 0.4281 | 0.784336 | 0.43284 | 0.63894 | 5.378 |
| 1010001B22Rik | 13.7948 | -0.13804 | 0.49221 | -0.28046 | 0.77913 | 0.88492 | 0.888 |
| 1110002E22Rik | 53.7761 | -0.86717 | 0.35504 | -2.44248 | 0.01459 | 0.07302 | 16.455 |
| 1110002J07Rik | 71.8459 | -0.20643 | 0.26866 | -0.76835 | 0.44228 | 0.64682 | 7.77 |
| 1110002L01Rik | 250.228 | 0.145255 | 0.13417 | 1.08259 | 0.27899 | 0.48891 | 23.676 |
| 1110003F10Rik | 64.5085 | 0.884902 | 0.27439 | 3.225021 | 0.00126 | 0.01476 | 0.767 |
| 1110004F10Rik | 4037.76 | -0.04855 | 0.06772 | -0.71701 | 0.47337 | 0.67166 | 65.814 |
| 1110006O24Rik | 13.1489 | 0.392768 | 0.61052 | 0.643336 | 0.52001 | 0.70963 | 0.775 |
| 1110015O18Rik | 60.9022 | 0.838659 | 0.28172 | 2.97691 | 0.00291 | 0.02572 | 17.584 |
| 1110017D15Rik | 86.7411 | 0.227425 | 0.24634 | 0.923204 | 0.3559 | 0.56636 | 12.325 |
| 1110018N20Rik | 245.515 | 0.354367 | 0.14706 | 2.409716 | 0.01596 | 0.07759 | 4.107 |
| 1110019D14Rik | 115.898 | 0.045083 | 0.17449 | 0.258366 | 0.79612 | 0.89487 | 172.863 |
| 1110020A21Rik | 30.3372 | -0.89322 | 0.375 | -2.38193 | 0.01722 | 0.08153 | 40.529 |
| 1110025M09Rik | 34.7815 | -0.03774 | 0.29388 | -0.12842 | 0.89782 | 0.94882 | 0.855 |
| 1110032A03Rik | 898.929 | -0.1404 | 0.09653 | -1.45437 | 0.14584 | 0.32872 | 12.693 |
| 1110032F04Rik | 354.475 | -0.54023 | 0.19873 | -2.71836 | 0.00656 | 0.04367 | 2.578 |
| 1110035H17Rik | 25.449 | 0.275149 | 0.34463 | 0.798396 | 0.42464 | 0.63135 | 1.241 |
| 1110038B12Rik | 264.022 | -0.22321 | 0.1767 | -1.26321 | 0.20651 | 0.40586 | 4.264 |
| 1110038F14Rik | 668.216 | -0.00343 | 0.0846 | -0.04051 | 0.96768 | 0.98368 | 2.474 |
| 1110046J04Rik | 71.722 | 0.026925 | 0.20304 | 0.132608 | 0.8945 | 0.94736 | 24.625 |

| | | | | | | | |
|---|---|---|---|---|---|---|---|
| 1110051M20Rik | 2720.1 | 0.111586 | 0.07326 | 1.523241 | 0.1277 | 0.3042 | 169.637 |
| 1110059E24Rik | 557.076 | 0.256335 | 0.13233 | 1.937022 | 0.05274 | 0.17185 | 71.775 |
| 1110059G10Rik | 659.665 | 0.144467 | 0.11093 | 1.302268 | 0.19282 | 0.38957 | 5.912 |
| 1110065P20Rik | 458.982 | -0.35719 | 0.10096 | -3.53785 | 0.0004 | 0.00692 | 2.117 |
| 1190007I07Rik | 352.772 | 0.239623 | 0.1338 | 1.790876 | 0.07331 | 0.2132 | 3.378 |
| 1300002E11Rik | 607.659 | 0.229071 | 0.14156 | 1.618143 | 0.10563 | 0.26915 | 14.723 |
| 1300014J16Rik | 100.572 | 0.20351 | 0.2025 | 1.005008 | 0.31489 | 0.52545 | 2.796 |
| 1500002C15Rik | 87.1181 | 0.041237 | 0.20169 | 0.204462 | 0.83799 | 0.91664 | 2.183 |
| 1500004A13Rik | 2349.23 | 0.017582 | 0.07824 | 0.224707 | 0.82221 | 0.90785 | 49.635 |
| 1500009L16Rik | 390.055 | 0.090045 | 0.12627 | 0.713116 | 0.47577 | 0.67391 | 39.898 |
| 1500011B03Rik | 3480.4 | 0.061485 | 0.07519 | 0.817744 | 0.4135 | 0.6216 | 15.273 |
| 1500015A07Rik | 298.46 | 0.385932 | 0.11896 | 3.244339 | 0.00118 | 0.01413 | 9.541 |
| 1500015L24Rik | 47.3819 | -0.41213 | 0.36509 | -1.12883 | 0.25897 | 0.46721 | 120.028 |
| 1500026H17Rik | 109.037 | -0.14518 | 0.19423 | -0.74748 | 0.45478 | 0.65702 | 14.529 |
| 1500035N22Rik | 70.6658 | 0.621281 | 0.24015 | 2.587043 | 0.00968 | 0.05614 | 12.336 |
| 1520401A03Rik | 18.9049 | -0.15117 | 0.43066 | -0.35101 | 0.72558 | 0.85388 | 18.296 |
| 1600002D24Rik | 13.1688 | 1.371502 | 0.58737 | 2.334994 | 0.01954 | 0.08818 | 172.537 |
| 1600010M07Rik | 18.1626 | -0.24895 | 0.49717 | -0.50073 | 0.61656 | 0.78174 | 152.827 |
| 1600012H06Rik | 776.818 | -0.00683 | 0.09921 | -0.06888 | 0.94508 | 0.97268 | 16.387 |
| 1600014C10Rik | 1174.52 | -0.15809 | 0.078 | -2.02687 | 0.04268 | 0.14888 | 14.351 |
| 1600020E01Rik | 579.063 | 0.297174 | 0.12319 | 2.412338 | 0.01585 | 0.07724 | 56.22 |
| 1600023N17Rik | 25.3906 | 0.30797 | 0.34048 | 0.904531 | 0.36571 | 0.57626 | 1.007 |
| 1600029O15Rik | 55.8044 | -0.13519 | 0.28523 | -0.47398 | 0.63552 | 0.79462 | 5.912 |
| 1700001G11Rik | 14.5767 | 0.919548 | 0.45834 | 2.006275 | 0.04483 | 0.15385 | 1.808 |
| 1700001G17Rik | 14.6363 | 0.43149 | 0.51836 | 0.832406 | 0.40518 | 0.61393 | 1.264 |
| 1700001K19Rik | 209.532 | 0.407381 | 0.17913 | 2.274234 | 0.02295 | 0.09807 | 14.932 |
| 1700001K23Rik | 34.3092 | -0.31231 | 0.36632 | -0.85256 | 0.39391 | 0.60303 | 16.67 |
| 1700001L05Rik | 164.453 | -0.41583 | 0.16498 | -2.52051 | 0.01172 | 0.0637 | 13.436 |
| 1700001L19Rik | 330.981 | -0.41884 | 0.17808 | -2.35204 | 0.01867 | 0.08566 | 16.811 |
| 1700001O22Rik | 177.408 | -0.04256 | 0.19969 | -0.21314 | 0.83122 | 0.91335 | 8.893 |
| 1700001P01Rik | 27.5311 | 0.353928 | 0.34261 | 1.03302 | 0.30159 | 0.51133 | 4.438 |
| 1700003D09Rik | 367.298 | 0.885215 | 0.26806 | 3.302338 | 0.00096 | 0.01244 | 7.566 |
| 1700003E16Rik | 767.496 | -0.14859 | 0.10655 | -1.3945 | 0.16317 | 0.35173 | 6.542 |
| 1700003F12Rik | 111.3 | -0.64017 | 0.28909 | -2.21445 | 0.0268 | 0.1092 | 2.919 |

| | | | | | | | |
|---|---|---|---|---|---|---|---|
| 1700003M07Rik | 723.01 | 0.17549 | 0.09966 | 1.760939 | 0.07825 | 0.22144 | 4.765 |
| 1700007E05Rik | 14.0929 | 0.0443 | 0.54114 | 0.081865 | 0.93475 | 0.96717 | 2.628 |
| 1700007F19Rik | 29.8325 | 0.672132 | 0.49956 | 1.345443 | 0.17848 | 0.37102 | 22.101 |
| 1700007J10Rik | 231.984 | 0.52312 | 0.17778 | 2.94259 | 0.00325 | 0.02753 | 14.238 |
| 1700007K13Rik | 58.4637 | 0.256311 | 0.29649 | 0.864491 | 0.38732 | 0.59665 | 4.332 |
| 1700007L15Rik | 14.095 | 0.327386 | 0.52693 | 0.62131 | 0.5344 | 0.72154 | 7.512 |
| 1700008A23Rik | 15.1907 | 0.8203 | 0.57868 | 1.417531 | 0.15633 | 0.34312 | 0.691 |
| 1700008B11Rik | 14.0829 | 0.565903 | 0.47425 | 1.193266 | 0.23277 | 0.43754 | 0.634 |
| 1700008J07Rik | 53.5779 | 0.245788 | 0.2887 | 0.851361 | 0.39457 | 0.60364 | 2.432 |
| 1700008O03Rik | 81.8691 | -0.01919 | 0.22931 | -0.08371 | 0.93329 | 0.96643 | 25.778 |
| 1700010I14Rik | 61.088 | 0.755364 | 0.23567 | 3.205113 | 0.00135 | 0.01537 | 23.617 |
| 1700012D14Rik | 81.6258 | 0.47387 | 0.23851 | 1.986772 | 0.04695 | 0.15894 | 39.641 |
| 1700013F07Rik | 68.294 | 0.462657 | 0.25593 | 1.807769 | 0.07064 | 0.20776 | 10.574 |
| 1700014D04Rik | 57.8853 | 0.415757 | 0.22465 | 1.85071 | 0.06421 | 0.19501 | 5.911 |
| 1700016A09Rik | 23.0482 | -0.23647 | 0.51659 | -0.45775 | 0.64713 | 0.80359 | 0.835 |
| 1700016K19Rik | 25.1122 | -1.14585 | 0.49005 | -2.33823 | 0.01938 | 0.08767 | 3.658 |
| 1700016P03Rik | 454.213 | -0.12436 | 0.17317 | -0.71815 | 0.47267 | 0.6709 | 5.08 |
| 1700017B05Rik | 445.758 | -0.11136 | 0.12153 | -0.91638 | 0.35947 | 0.56983 | 9.496 |
| 1700018G05Rik | 32.1835 | 0.695456 | 0.31382 | 2.21608 | 0.02669 | 0.10891 | 0.723 |
| 1700018L02Rik | 86.8366 | 0.160918 | 0.24705 | 0.651364 | 0.51481 | 0.70524 | 2.485 |
| 1700019A02Rik | 23.2272 | 0.597085 | 0.35066 | 1.702749 | 0.08862 | 0.24052 | 29.06 |
| 1700019D03Rik | 1052.11 | -0.18316 | 0.10581 | -1.731 | 0.08345 | 0.23096 | 97.856 |
| 1700019N19Rik | 21.9458 | 1.041802 | 0.41813 | 2.491547 | 0.01272 | 0.06724 | 8.659 |
| 1700020A23Rik | 23.523 | 0.401575 | 0.39126 | 1.026357 | 0.30472 | 0.51453 | 0.816 |
| 1700020L24Rik | 13.4055 | -0.78972 | 0.5151 | -1.53314 | 0.12524 | 0.30024 | 3.557 |
| 1700022I11Rik | 15.9435 | 0.256806 | 0.56362 | 0.455634 | 0.64865 | 0.80422 | 14.037 |
| 1700022N22Rik | 77.6228 | -0.35688 | 0.20645 | -1.72866 | 0.08387 | 0.23177 | 5.245 |
| 1700023B13Rik | 31.4774 | 0.581524 | 0.48709 | 1.193884 | 0.23252 | 0.43737 | 1.046 |
| 1700023H06Rik | 19.5171 | 0.060211 | 0.42784 | 0.140733 | 0.88808 | 0.94406 | 1.301 |
| 1700025F24Rik | 24.7908 | 0.008891 | 0.36718 | 0.024214 | 0.98068 | 0.98986 | 13.421 |
| 1700025G04Rik | 4192.99 | -0.21606 | 0.10995 | -1.9651 | 0.0494 | 0.1646 | 237.723 |
| 1700027A07Rik | 17.7055 | -0.06353 | 0.52386 | -0.12128 | 0.90347 | 0.95134 | 1.695 |
| 1700027H10Rik | 25.961 | 0.557742 | 0.4527 | 1.232023 | 0.21794 | 0.41996 | 22.727 |
| 1700028E10Rik | 49.1918 | -0.10967 | 0.28827 | -0.38042 | 0.70363 | 0.83999 | 63.444 |

| | | | | | | | |
|---|---|---|---|---|---|---|---|
| 1700028J19Rik | 61.773 | -0.00464 | 0.219 | -0.02118 | 0.9831 | 0.99078 | 16.885 |
| 1700028K03Rik | 100.131 | 0.349588 | 0.21897 | 1.596538 | 0.11037 | 0.27695 | 16.834 |
| 1700028P14Rik | 18.4679 | -1.08589 | 0.56302 | -1.92868 | 0.05377 | 0.17403 | 94.06 |
| 1700029I15Rik | 98.7717 | 0.260497 | 0.19189 | 1.357548 | 0.17461 | 0.36621 | 3.171 |
| 1700029J07Rik | 421.605 | 0.525333 | 0.13761 | 3.817657 | 0.00013 | 0.0033 | 21.835 |
| 1700030C10Rik | 18.9939 | -0.21548 | 0.5652 | -0.38124 | 0.70303 | 0.83971 | 11.399 |
| 1700030C14Rik | 21.8709 | 0.702319 | 0.43332 | 1.620786 | 0.10506 | 0.26817 | 1.449 |
| 1700030J22Rik | 384.477 | 0.216655 | 0.12969 | 1.670538 | 0.09481 | 0.25135 | 9.366 |
| 1700030K09Rik | 604.969 | 0.291577 | 0.09998 | 2.916459 | 0.00354 | 0.02897 | 17.122 |
| 1700034H15Rik | 233.296 | 0.421625 | 0.13058 | 3.228966 | 0.00124 | 0.01463 | 21.398 |
| 1700037C18Rik | 136.147 | -0.26431 | 0.15284 | -1.72936 | 0.08375 | 0.23153 | 13.511 |
| 1700037H04Rik | 2162.69 | 0.485464 | 0.17456 | 2.78101 | 0.00542 | 0.03831 | 13.758 |
| 1700040D17Rik | 89.6739 | 0.537606 | 0.20297 | 2.648693 | 0.00808 | 0.04991 | 3.525 |
| 1700041B01Rik | 47.4446 | 0.38101 | 0.3589 | 1.061597 | 0.28842 | 0.49847 | 5.33 |
| 1700041G16Rik | 17.4804 | -0.01095 | 0.45808 | -0.0239 | 0.98093 | 0.99 | 3.772 |
| 1700047K16Rik | 26.274 | 0.209941 | 0.41401 | 0.507092 | 0.61209 | 0.77898 | 1.452 |
| 1700047M11Rik | 658.781 | -0.29561 | 0.16715 | -1.76852 | 0.07697 | 0.21936 | 15.385 |
| 1700048M11Rik | 45.6863 | 0.242299 | 0.2769 | 0.875055 | 0.38154 | 0.59096 | 3.753 |
| 1700048O20Rik | 316.064 | 0.440731 | 0.2616 | 1.684781 | 0.09203 | 0.24693 | 9.794 |
| 1700052K11Rik | 225.92 | -0.04454 | 0.13656 | -0.32618 | 0.74429 | 0.86539 | 3.708 |
| 1700055D18Rik | 17.4328 | 0.209579 | 0.43108 | 0.486176 | 0.62684 | 0.78888 | 3.72 |
| 1700056N10Rik | 123.935 | 0.364036 | 0.16015 | 2.273097 | 0.02302 | 0.09815 | 1.592 |
| 1700058P15Rik | 26.854 | 0.216364 | 0.36468 | 0.593295 | 0.55298 | 0.73584 | 1.248 |
| 1700060J05Rik | 26.321 | 0.18789 | 0.34131 | 0.550493 | 0.58198 | 0.75769 | 1.715 |
| 1700063D05Rik | 68.3997 | -1.019 | 0.40227 | -2.53309 | 0.01131 | 0.06214 | 25.545 |
| 1700065J18Rik | 20.1183 | 1.278119 | 0.39555 | 3.231224 | 0.00123 | 0.01456 | 1.08 |
| 1700066M21Rik | 1005.19 | 0.027156 | 0.09641 | 0.28168 | 0.77819 | 0.88423 | 7.804 |
| 1700067K01Rik | 48.2699 | 0.449202 | 0.29814 | 1.506701 | 0.13189 | 0.31073 | 3.269 |
| 1700071M16Rik | 114.737 | 0.363673 | 0.24311 | 1.495896 | 0.13468 | 0.31449 | 4.103 |
| 1700080N15Rik | 40.5858 | -0.15113 | 0.26635 | -0.56742 | 0.57043 | 0.74969 | 9.079 |
| 1700081N11Rik | 15.102 | -0.14482 | 0.48796 | -0.29679 | 0.76663 | 0.87802 | 1.034 |
| 1700084E18Rik | 41.6011 | -0.28378 | 0.25506 | -1.1126 | 0.26588 | 0.47503 | 1.429 |
| 1700086L19Rik | 1615.85 | 0.25049 | 0.08787 | 2.850702 | 0.00436 | 0.03338 | 12.153 |
| 1700086O06Rik | 212.534 | 0.142005 | 0.16521 | 0.859528 | 0.39005 | 0.59915 | 12.261 |

| | | | | | | | |
|---|---|---|---|---|---|---|---|
| 1700086P04Rik | 18.1096 | -0.49736 | 0.50456 | -0.98572 | 0.32427 | 0.53539 | 7.897 |
| 1700087I21Rik | 757.525 | 0.025608 | 0.18131 | 0.141239 | 0.88768 | 0.94402 | 102.961 |
| 1700088E04Rik | 83.7717 | 0.210047 | 0.19379 | 1.083878 | 0.27842 | 0.48828 | 11.905 |
| 1700093J21Rik | 37.3537 | -0.06549 | 0.32476 | -0.20167 | 0.84018 | 0.918 | 6.395 |
| 1700093K21Rik | 33.6313 | -0.04025 | 0.37932 | -0.10611 | 0.9155 | 0.95671 | 4.953 |
| 1700094D03Rik | 174.242 | 0.547648 | 0.14285 | 3.833814 | 0.00013 | 0.00313 | 5.552 |
| 1700095A13Rik | 12.9345 | 1.464831 | 0.64013 | 2.288336 | 0.02212 | 0.09584 | 32.583 |
| 1700096K18Rik | 77.7679 | 0.11721 | 0.21459 | 0.546216 | 0.58492 | 0.75957 | 1.454 |
| 1700101I11Rik | 39.087 | -0.13501 | 0.27258 | -0.4953 | 0.62039 | 0.78411 | 1.871 |
| 1700102H20Rik | 14.8424 | -0.01746 | 0.42816 | -0.04078 | 0.96747 | 0.98365 | 2.388 |
| 1700102P08Rik | 84.4984 | 0.088864 | 0.24071 | 0.36917 | 0.712 | 0.845 | 18.13 |
| 1700105P06Rik | 29.7458 | 0.195924 | 0.35355 | 0.554169 | 0.57946 | 0.75605 | 1.836 |
| 1700109H08Rik | 315.849 | 0.215795 | 0.12244 | 1.762406 | 0.078 | 0.22104 | 12.791 |
| 1700109K24Rik | 382.395 | 0.052784 | 0.14676 | 0.359658 | 0.7191 | 0.84995 | 11.917 |
| 1700110C19Rik | 178.773 | 0.181488 | 0.15893 | 1.141903 | 0.25349 | 0.46087 | 9.426 |
| 1700110K17Rik | 22.3303 | -0.04711 | 0.43556 | -0.10815 | 0.91387 | 0.95639 | 20.139 |
| 1700111E14Rik | 12.55 | 0.008555 | 0.46937 | 0.018227 | 0.98546 | 0.99203 | 48.737 |
| 1700112D23Rik | 34.5633 | -0.30013 | 0.34317 | -0.87459 | 0.3818 | 0.59125 | 78.498 |
| 1700112J16Rik | 49.5588 | 0.033056 | 0.32906 | 0.100458 | 0.91998 | 0.95948 | 18.455 |
| 1700113A16Rik | 43.0576 | -0.31415 | 0.35653 | -0.88113 | 0.37825 | 0.58818 | 6.24 |
| 1700120C14Rik | 46.402 | 0.239155 | 0.30159 | 0.792976 | 0.42779 | 0.6344 | 12.809 |
| 1700121C08Rik | 16.8451 | -0.01542 | 0.45214 | -0.0341 | 0.9728 | 0.98639 | 25.892 |
| 1700123K08Rik | 13.4915 | 0.836755 | 0.69327 | 1.206976 | 0.22744 | 0.43163 | 2.873 |
| 1700123M08Rik | 123.555 | -0.14986 | 0.15597 | -0.96087 | 0.33662 | 0.54718 | 28.089 |
| 1700123O20Rik | 557.894 | -0.34824 | 0.1153 | -3.02025 | 0.00253 | 0.02352 | 4.526 |
| 1700123O21Rik | 20.5703 | -0.11159 | 0.43489 | -0.25659 | 0.79749 | 0.8956 | 22.912 |
| 1700124L16Rik | 54.724 | 0.061643 | 0.2644 | 0.233139 | 0.81565 | 0.90517 | 1.11 |
| 1700125G02Rik | 190.824 | -0.58336 | 0.20515 | -2.84365 | 0.00446 | 0.03381 | 3.895 |
| 1700125H03Rik | 136.49 | 0.560409 | 0.21915 | 2.557197 | 0.01055 | 0.0596 | 7.277 |
| 1700126G02Rik | 71.1754 | -0.33269 | 0.1938 | -1.71665 | 0.08604 | 0.23562 | 20.052 |
| 1810010H24Rik | 135.645 | -0.40643 | 0.16328 | -2.48916 | 0.0128 | 0.06755 | 4.763 |
| 1810010K12Rik | 13.265 | -0.71658 | 0.58081 | -1.23376 | 0.21729 | 0.41935 | 3.17 |
| 1810012K08Rik | 79.0817 | -0.08911 | 0.2307 | -0.38626 | 0.6993 | 0.83742 | 0.998 |
| 1810013L24Rik | 3395.14 | 0.072775 | 0.05467 | 1.331241 | 0.18311 | 0.37714 | 28.823 |

| | | | | | | | |
|---|---|---|---|---|---|---|---|
| 1810014B01Rik | 236.402 | 0.068581 | 0.14997 | 0.457306 | 0.64745 | 0.80377 | 9.284 |
| 1810020O05Rik | 331.214 | 1.208235 | 0.44124 | 2.738281 | 0.00618 | 0.04206 | 47.212 |
| 1810021B22Rik | 157.189 | -0.27677 | 0.18333 | -1.50968 | 0.13113 | 0.30963 | 4.74 |
| 1810024B03Rik | 45.6618 | 0.091766 | 0.26623 | 0.344685 | 0.73033 | 0.85705 | 21.926 |
| 1810030O07Rik | 1366.89 | 0.24435 | 0.0759 | 3.219562 | 0.00128 | 0.01487 | 18.79 |
| 1810034E14Rik | 56.8201 | -0.14387 | 0.22078 | -0.65166 | 0.51462 | 0.70511 | 23.216 |
| 1810037I17Rik | 1779.95 | -0.19055 | 0.08257 | -2.30772 | 0.02101 | 0.0926 | 1.982 |
| 1810044D09Rik | 15.1054 | 0.219979 | 0.42686 | 0.515345 | 0.60631 | 0.77467 | 0.815 |
| 1810055G02Rik | 564.316 | -0.22914 | 0.12959 | -1.7682 | 0.07703 | 0.21941 | 9.589 |
| 1810058I24Rik | 318.128 | -0.18417 | 0.12199 | -1.50969 | 0.13112 | 0.30963 | 10.843 |
| 1810059H22Rik | 51.6702 | -0.24583 | 0.28896 | -0.85074 | 0.39491 | 0.6039 | 16.375 |
| 1810062O18Rik | 106.135 | 0.188754 | 0.19028 | 0.991969 | 0.32121 | 0.53206 | 24.609 |
| 2010001A14Rik | 155.846 | -0.26422 | 0.18276 | -1.44571 | 0.14826 | 0.33188 | 5.993 |
| 2010007H06Rik | 64.0226 | 0.038358 | 0.26708 | 0.143621 | 0.8858 | 0.9432 | 5.698 |
| 2010106C02Rik | 14.8119 | 0.25365 | 0.50862 | 0.498703 | 0.61799 | 0.78287 | 2.015 |
| 2010109A12Rik | 22.3119 | 0.866051 | 0.5989 | 1.446071 | 0.14816 | 0.33179 | 6.958 |
| 2010204K13Rik | 547.947 | 0.269678 | 0.15665 | 1.721508 | 0.08516 | 0.23404 | 33.327 |
| 2010315B03Rik | 1013.4 | 0.367887 | 0.12805 | 2.872898 | 0.00407 | 0.03184 | 20.893 |
| 2010320M18Rik | 137.119 | -0.49486 | 0.1943 | -2.54688 | 0.01087 | 0.06069 | 1.072 |
| 2010320O07Rik | 12.9436 | -0.81687 | 0.57985 | -1.40877 | 0.1589 | 0.34639 | 19.392 |
| 2200002D01Rik | 22.5791 | 0.439058 | 0.3897 | 1.126661 | 0.25989 | 0.46839 | 1.906 |
| 2210016F16Rik | 351.677 | -0.34283 | 0.12011 | -2.85422 | 0.00431 | 0.03314 | 5.409 |
| 2210016L21Rik | 3229.69 | -0.09961 | 0.05954 | -1.67282 | 0.09436 | 0.25065 | 7.626 |
| 2210406H18Rik | 37.8935 | 0.372187 | 0.31776 | 1.171282 | 0.24149 | 0.44749 | 1.108 |
| 2210406O10Rik | 91.9916 | -0.40812 | 0.19084 | -2.13856 | 0.03247 | 0.12479 | 17.243 |
| 2210408F21Rik | 927.716 | -0.09087 | 0.10349 | -0.87809 | 0.37989 | 0.58968 | 149.392 |
| 2210408I21Rik | 398.966 | 0.100292 | 0.11296 | 0.88784 | 0.37463 | 0.58497 | 478.245 |
| 2210412B16Rik | 51.8317 | 0.329322 | 0.28807 | 1.143214 | 0.25295 | 0.46033 | 0.498 |
| 2210418O10Rik | 14.9623 | 0.654911 | 0.47725 | 1.372254 | 0.16998 | 0.3604 | 13.76 |
| 2300009A05Rik | 277.141 | 0.213831 | 0.17085 | 1.251605 | 0.21071 | 0.41079 | 4.8 |
| 2310001H17Rik | 30.817 | -0.22741 | 0.36395 | -0.62486 | 0.53206 | 0.71958 | 30.314 |
| 2310002D06Rik | 19.1647 | 0.42583 | 0.40199 | 1.059316 | 0.28946 | 0.49973 | 10.764 |
| 2310002F09Rik | 20.3741 | -0.60202 | 0.59436 | -1.0129 | 0.31111 | 0.52135 | 102.75 |
| 2310009B15Rik | 205.44 | 0.098403 | 0.15403 | 0.638846 | 0.52292 | 0.71143 | 4.87 |

| | | | | | | | |
|---|---|---|---|---|---|---|---|
| 2310010J17Rik | 77.8966 | -0.4329 | 0.228 | -1.8987 | 0.0576 | 0.18222 | 5.866 |
| 2310011J03Rik | 1043.38 | -0.10097 | 0.10606 | -0.95196 | 0.34112 | 0.55206 | 2.284 |
| 2310014F06Rik | 41.985 | 0.217283 | 0.32133 | 0.676193 | 0.49892 | 0.6922 | 67.522 |
| 2310015A10Rik | 451.007 | 0.060865 | 0.12494 | 0.487161 | 0.62614 | 0.78844 | 12.321 |
| 2310016G11Rik | 16.1854 | -0.66443 | 0.44311 | -1.49948 | 0.13375 | 0.31339 | 10.029 |
| 2310022A10Rik | 952.822 | 0.05458 | 0.11371 | 0.479989 | 0.63123 | 0.79177 | 28.867 |
| 2310022B05Rik | 2031.88 | 0.092496 | 0.10391 | 0.890187 | 0.37337 | 0.58357 | 27.614 |
| 2310026I22Rik | 14.1709 | 0.894348 | 0.55836 | 1.601748 | 0.10921 | 0.275 | 23.404 |
| 2310026L22Rik | 30.8917 | -0.13377 | 0.36218 | -0.36935 | 0.71187 | 0.845 | 0.476 |
| 2310030G06Rik | 113.953 | 0.385247 | 0.2746 | 1.402919 | 0.16064 | 0.34889 | 6.846 |
| 2310033P09Rik | 406.022 | -0.01225 | 0.10404 | -0.11772 | 0.90629 | 0.95309 | 2.418 |
| 2310034G01Rik | 68.301 | 0.000478 | 0.26446 | 0.001809 | 0.99856 | 0.99919 | 2.102 |
| 2310039H08Rik | 75.854 | -0.10228 | 0.24669 | -0.41459 | 0.67844 | 0.82401 | 0.762 |
| 2310040G07Rik | 22.9304 | -0.52568 | 0.42883 | -1.22586 | 0.22025 | 0.42283 | 19.391 |
| 2310040G24Rik | 16.4872 | -0.78045 | 0.4326 | -1.80408 | 0.07122 | 0.20889 | 5.013 |
| 2310043M15Rik | 118.45 | 0.753556 | 0.24627 | 3.05987 | 0.00221 | 0.02151 | 2.933 |
| 2310043P16Rik | 26.7419 | 0.131449 | 0.3708 | 0.3545 | 0.72296 | 0.85223 | 3.881 |
| 2310057M21Rik | 1277.83 | -0.02623 | 0.07914 | -0.33144 | 0.74031 | 0.863 | 24.54 |
| 2310058D17Rik | 304.331 | 0.032773 | 0.1669 | 0.196365 | 0.84432 | 0.92016 | 12.334 |
| 2310061I04Rik | 1708.26 | 0.005287 | 0.08476 | 0.062373 | 0.95027 | 0.97539 | 4.755 |
| 2310074N15Rik | 17.1484 | 0.68512 | 0.42667 | 1.605754 | 0.10833 | 0.27367 | 0.582 |
| 2310075C17Rik | 64.7759 | 0.66539 | 0.25683 | 2.590785 | 0.00958 | 0.05571 | 0.809 |
| 2410002F23Rik | 1661.27 | 0.231759 | 0.09178 | 2.525072 | 0.01157 | 0.06306 | 5.598 |
| 2410004B18Rik | 948.063 | -0.21645 | 0.07607 | -2.84538 | 0.00444 | 0.03369 | 6.816 |
| 2410004I01Rik | 18.5123 | -0.65514 | 0.51143 | -1.281 | 0.20019 | 0.39817 | 6.088 |
| 2410004P03Rik | 37.0356 | -0.51652 | 0.46872 | -1.10198 | 0.27047 | 0.47998 | 8.409 |
| 2410006H16Rik | 419.721 | -0.11554 | 0.16466 | -0.7017 | 0.48287 | 0.67933 | 2.076 |
| 2410018L13Rik | 43.6359 | 0.414579 | 0.33799 | 1.226606 | 0.21997 | 0.42245 | 56.288 |
| 2410021H03Rik | 151.746 | 0.073104 | 0.17186 | 0.425361 | 0.67057 | 0.81845 | 1.847 |
| 2410022M11Rik | 157.958 | 0.31892 | 0.16731 | 1.906137 | 0.05663 | 0.18018 | 1.911 |
| 2410080I02Rik | 15.7986 | 0.623627 | 0.47639 | 1.309055 | 0.19052 | 0.3867 | 0.818 |
| 2410124H12Rik | 15.0005 | 1.761773 | 0.51572 | 3.416123 | 0.00064 | 0.00932 | 26.478 |
| 2500002B13Rik | 82.1436 | 0.1078 | 0.22147 | 0.486753 | 0.62643 | 0.78864 | 23.425 |
| 2500004C02Rik | 357.252 | 0.42578 | 0.12987 | 3.278412 | 0.00104 | 0.01319 | 4.957 |

| | | | | | | | |
|---|---|---|---|---|---|---|---|
| 2510002D24Rik | 423.805 | 0.088554 | 0.09303 | 0.951925 | 0.34114 | 0.55206 | 3.583 |
| 2510009E07Rik | 4171.41 | 0.368617 | 0.10589 | 3.481122 | 0.0005 | 0.00791 | 45.621 |
| 2510016D11Rik | 70.5182 | 0.3075 | 0.27 | 1.138892 | 0.25475 | 0.46228 | 0.985 |
| 2510017J16Rik | 112.176 | 0.349728 | 0.20738 | 1.686379 | 0.09172 | 0.24633 | 1.263 |
| 2510039O18Rik | 1112.98 | -0.04259 | 0.07838 | -0.54346 | 0.58681 | 0.7605 | 6.42 |
| 2600002D14Rik | 15.0669 | 0.262293 | 0.54509 | 0.481193 | 0.63038 | 0.79109 | 0.292 |
| 2600006K01Rik | 43.4685 | -0.39047 | 0.29327 | -1.33143 | 0.18305 | 0.37709 | 1.595 |
| 2600014E21Rik | 132.553 | -1.55669 | 0.95141 | -1.63619 | 0.1018 | 0.26328 | 178.978 |
| 2610002M06Rik | 3370.64 | 0.220445 | 0.07678 | 2.871221 | 0.00409 | 0.03197 | 33.584 |
| 2610005L07Rik | 875.17 | 0.133086 | 0.09865 | 1.349027 | 0.17733 | 0.36948 | 39.033 |
| 2610008E11Rik | 1955.3 | 0.375059 | 0.17443 | 2.150156 | 0.03154 | 0.12226 | 33.227 |
| 2610011E03Rik | 14.4811 | -0.23574 | 0.49198 | -0.47916 | 0.63182 | 0.79212 | 2.235 |
| 2610017A05Rik | 18.294 | 0.900171 | 0.41575 | 2.165177 | 0.03037 | 0.11902 | 1.663 |
| 2610020C07Rik | 123.095 | 0.784875 | 0.16153 | 4.859138 | 1.18E-06 | 0.00012 | 42.876 |
| 2610021A01Rik | 1073.71 | 0.227764 | 0.15215 | 1.496928 | 0.13441 | 0.31425 | 29.304 |
| 2610027K06Rik | 36.7865 | 0.086606 | 0.36072 | 0.240091 | 0.81026 | 0.90233 | 40.64 |
| 2610028D06Rik | 135.355 | 0.152162 | 0.17257 | 0.881743 | 0.37792 | 0.588 | 1.713 |
| 2610028E06Rik | 163.503 | 0.531567 | 0.15566 | 3.414845 | 0.00064 | 0.00933 | 34.567 |
| 2610035D17Rik | 607.43 | -0.05203 | 0.11113 | -0.46819 | 0.63965 | 0.79795 | 158.02 |
| 2610035F20Rik | 67.1612 | -0.81014 | 0.25638 | -3.15994 | 0.00158 | 0.01723 | 8.156 |
| 2610037D02Rik | 351.297 | -0.08725 | 0.12406 | -0.7033 | 0.48187 | 0.67858 | 149.677 |
| 2610044O15Rik8 | 1014.09 | 0.384138 | 0.17487 | 2.196651 | 0.02805 | 0.11271 | 18.053 |
| 2610203C22Rik | 118.464 | -0.76643 | 0.36474 | -2.10132 | 0.03561 | 0.13257 | 70.702 |
| 2610306M01Rik | 168.292 | -0.22796 | 0.16123 | -1.41387 | 0.1574 | 0.34463 | 1.042 |
| 2610316D01Rik | 448.678 | 0.439637 | 0.11587 | 3.794144 | 0.00015 | 0.0035 | 98.102 |
| 2610318N02Rik | 65.1013 | -0.28722 | 0.24828 | -1.15688 | 0.24732 | 0.45418 | 11.77 |
| 2610507B11Rik | 15739.2 | 0.005208 | 0.05499 | 0.094713 | 0.92454 | 0.96202 | 28.872 |
| 2610507I01Rik | 956.326 | -0.02492 | 0.08599 | -0.28985 | 0.77193 | 0.88117 | 5.573 |
| 2700029L08Rik | 108.45 | -0.23715 | 0.21717 | -1.09199 | 0.27484 | 0.48444 | 1.465 |
| 2700033N17Rik | 888.23 | 0.568273 | 0.14279 | 3.979833 | 6.90E-05 | 0.00207 | 1.257 |
| 2700038G22Rik | 68.3922 | -0.10829 | 0.28584 | -0.37883 | 0.70481 | 0.84105 | 4.506 |
| 2700046A07Rik | 52.9791 | -0.66129 | 0.31527 | -2.09753 | 0.03595 | 0.13342 | 4.674 |
| 2700046G09Rik | 66.059 | -0.07802 | 0.225 | -0.34676 | 0.72877 | 0.8556 | 2.115 |
| 2700049A03Rik | 523.167 | 0.200662 | 0.09257 | 2.167757 | 0.03018 | 0.11868 | 106.456 |

| | | | | | | | |
|---|---|---|---|---|---|---|---|
| 2700052C19Rik | 235.536 | 0.110643 | 0.14294 | 0.774077 | 0.43889 | 0.64412 | 0.335 |
| 2700054A10Rik | 105.971 | 0.125984 | 0.2228 | 0.565463 | 0.57176 | 0.75056 | 186.339 |
| 2700062C07Rik | 576.119 | 0.27891 | 0.10408 | 2.679882 | 0.00736 | 0.04718 | 6.949 |
| 2700069I18Rik | 24.8314 | -1.22613 | 0.45425 | -2.69922 | 0.00695 | 0.04524 | 43.256 |
| 2700078F05Rik | 16.7371 | -0.75645 | 0.51458 | -1.47003 | 0.14155 | 0.32333 | 3.244 |
| 2700097O09Rik | 795.681 | 0.103785 | 0.10206 | 1.016912 | 0.3092 | 0.51908 | 34.45 |
| 2700099C18Rik | 296.54 | 0.032138 | 0.11863 | 0.270902 | 0.78647 | 0.88899 | 25.106 |
| 2810001G20Rik | 428.437 | 0.109061 | 0.11972 | 0.910982 | 0.36231 | 0.57285 | 82.477 |
| 2810002D19Rik | 192.89 | 0.149068 | 0.13784 | 1.081479 | 0.27948 | 0.48948 | 16.615 |
| 2810004N23Rik | 1121.65 | 0.091361 | 0.0685 | 1.333799 | 0.18227 | 0.37605 | 26.176 |
| 2810013P06Rik | 895.714 | -0.05258 | 0.09754 | -0.53911 | 0.58981 | 0.76287 | 2.194 |
| 2810021J22Rik | 625.628 | 0.234157 | 0.10867 | 2.154705 | 0.03118 | 0.12124 | 16.073 |
| 2810025M15Rik | 471.065 | -0.68799 | 0.16588 | -4.14763 | 3.36E-05 | 0.00124 | 7.893 |
| 2810029C07Rik | 372.639 | 0.164396 | 0.11529 | 1.425882 | 0.1539 | 0.3397 | 2.133 |
| 2810032G03Rik | 442.306 | 0.095012 | 0.09751 | 0.974373 | 0.32987 | 0.54112 | 40.762 |
| 2810039B14Rik | 414.038 | -0.07877 | 0.13528 | -0.58228 | 0.56038 | 0.74173 | 11.515 |
| 2810049E08Rik | 124.362 | -0.06749 | 0.20728 | -0.32562 | 0.74471 | 0.86574 | 37.503 |
| 2810402E24Rik | 110.4 | 0.025802 | 0.18612 | 0.138635 | 0.88974 | 0.9448 | 3.049 |
| 2810403D21Rik | 47.4986 | -0.38783 | 0.30235 | -1.28272 | 0.19959 | 0.39758 | 141.107 |
| 2810405F15Rik | 17.8481 | 0.801897 | 0.5077 | 1.579461 | 0.11423 | 0.28303 | 0.932 |
| 2810405F17Rik | 77.4927 | 0.152401 | 0.21504 | 0.708721 | 0.4785 | 0.67612 | 1.18 |
| 2810407A14Rik | 54.368 | 0.313087 | 0.34293 | 0.912971 | 0.36126 | 0.57155 | 55.219 |
| 2810408A11Rik | 16.9644 | -0.22507 | 0.49768 | -0.45224 | 0.6511 | 0.8058 | 3.636 |
| 2810410L24Rik | 407.888 | -0.04241 | 0.1552 | -0.27327 | 0.78465 | 0.88833 | 4.589 |
| 2810414N06Rik | 73.6033 | -0.18957 | 0.25563 | -0.74158 | 0.45834 | 0.65953 | 6.105 |
| 2810428J06Rik | 64.8698 | 0.040033 | 0.24191 | 0.165486 | 0.86856 | 0.9334 | 2.696 |
| 2810429I04Rik | 506.958 | 0.222289 | 0.12166 | 1.827058 | 0.06769 | 0.20167 | 24.191 |
| 2810442N19Rik | 23.422 | -0.60602 | 0.5579 | -1.08625 | 0.27737 | 0.48702 | 11.526 |
| 2810454H06Rik | 28.756 | -0.4174 | 0.34422 | -1.21259 | 0.22528 | 0.4293 | 3.148 |
| 2810455O05Rik | 1059.47 | 0.259348 | 0.10075 | 2.574059 | 0.01005 | 0.05765 | 2.233 |
| 2810459M11Rik | 349.039 | -0.5322 | 0.17255 | -3.0843 | 0.00204 | 0.02034 | 9.594 |
| 2900005J15Rik | 110.054 | -0.06977 | 0.22648 | -0.30804 | 0.75805 | 0.87261 | 9.179 |
| 2900022M07Rik | 26.6172 | -0.12815 | 0.37724 | -0.33969 | 0.73409 | 0.85925 | 1.017 |
| 2900026A02Rik | 1792.28 | -0.32597 | 0.13469 | -2.4201 | 0.01552 | 0.07614 | 134.914 |

| | | | | | | | |
|---|---|---|---|---|---|---|---|
| 2900027M19Rik | 44.4358 | -1.19128 | 0.34766 | -3.4266 | 0.00061 | 0.00909 | 4.448 |
| 2900037B21Rik | 98.3231 | -0.1915 | 0.18419 | -1.03974 | 0.29846 | 0.50856 | 0.337 |
| 2900041M22Rik | 92.131 | -0.26457 | 0.28481 | -0.92892 | 0.35293 | 0.56319 | 2.611 |
| 2900042K21Rik | 18.3218 | 0.615096 | 0.41841 | 1.470089 | 0.14154 | 0.32333 | 0.653 |
| 2900045O20Rik | 26.6004 | -0.35646 | 0.38618 | -0.92304 | 0.35599 | 0.56636 | 13.054 |
| 2900052L18Rik | 322.949 | 0.032252 | 0.14343 | 0.224865 | 0.82208 | 0.90785 | 5.251 |
| 2900052N01Rik | 2726.17 | 0.212401 | 0.10733 | 1.97895 | 0.04782 | 0.16088 | 13.891 |
| 2900060N12Rik | 280.607 | 0.190307 | 0.13976 | 1.361675 | 0.1733 | 0.36452 | 35.463 |
| 2900064F13Rik | 288.262 | 0.162832 | 0.15552 | 1.047007 | 0.2951 | 0.5055 | 1.008 |
| 2900064K03Rik | 27.9532 | -0.59943 | 0.34707 | -1.72711 | 0.08415 | 0.23231 | 3.113 |
| 2900072N19Rik | 33.6631 | -0.12284 | 0.31897 | -0.38512 | 0.70014 | 0.83799 | 4.381 |
| 2900076A07Rik | 396.101 | 0.116837 | 0.11271 | 1.036581 | 0.29993 | 0.51006 | 7.991 |
| 2900078I11Rik | 44.3007 | 0.238567 | 0.30878 | 0.772608 | 0.43975 | 0.64503 | 0.999 |
| 2900079G21Rik | 81.8883 | -0.4334 | 0.30167 | -1.43668 | 0.15081 | 0.33567 | 114.728 |
| 2900089D17Rik | 107.582 | -0.14725 | 0.16986 | -0.86692 | 0.38599 | 0.59548 | 14.497 |
| 2900092N22Rik | 16.1128 | -0.85913 | 0.68205 | -1.25962 | 0.20781 | 0.40738 | 1.409 |
| 2900093K20Rik | 62.388 | 0.06541 | 0.24291 | 0.269282 | 0.78771 | 0.88973 | 0.602 |
| 3000002C10Rik | 331.555 | 0.376862 | 0.13078 | 2.881567 | 0.00396 | 0.03123 | 4.764 |
| 3010001F23Rik | 113.553 | -0.04647 | 0.21288 | -0.21828 | 0.82721 | 0.91108 | 48.126 |
| 3110001I22Rik | 74.2507 | -0.17553 | 0.25282 | -0.69428 | 0.48751 | 0.68284 | 6.46 |
| 3110009E18Rik | 70.0477 | 0.211717 | 0.25492 | 0.830526 | 0.40624 | 0.61467 | 67.003 |
| 3110021N24Rik | 23.5974 | -0.1281 | 0.38041 | -0.33673 | 0.73632 | 0.86061 | 62.256 |
| 3110039M20Rik | 115.709 | 0.273975 | 0.17859 | 1.534065 | 0.12501 | 0.30012 | 17.709 |
| 3110040N11Rik | 309.862 | -0.1543 | 0.13864 | -1.113 | 0.26571 | 0.4749 | 7.297 |
| 3110045C21Rik | 24.0122 | -0.21705 | 0.41879 | -0.51828 | 0.60426 | 0.7733 | 3.559 |
| 3110053B16Rik | 123.086 | -0.19274 | 0.17922 | -1.07545 | 0.28217 | 0.49237 | 47.138 |
| 3110056K07Rik | 639.224 | 0.301781 | 0.10047 | 3.003573 | 0.00267 | 0.02437 | 24.218 |
| 3110067C02Rik | 62.1692 | 0.592062 | 0.25334 | 2.33703 | 0.01944 | 0.08783 | 2.431 |
| 3110070M22Rik | 22.5556 | -0.20216 | 0.40019 | -0.50518 | 0.61343 | 0.77991 | 1.133 |
| 3110080E11Rik | 50.0411 | -0.53938 | 0.31772 | -1.69764 | 0.08958 | 0.24255 | 3.477 |
| 3110082I17Rik | 290.543 | -0.29419 | 0.14393 | -2.04391 | 0.04096 | 0.14491 | 100.789 |
| 3110082J24Rik | 25.6356 | -0.22361 | 0.3911 | -0.57176 | 0.56749 | 0.7475 | 2.503 |
| 3110083C13Rik | 24.3846 | -0.68979 | 0.34772 | -1.98372 | 0.04729 | 0.15968 | 5.823 |
| 3110099E03Rik | 55.7627 | 0.016851 | 0.55913 | 0.030137 | 0.97596 | 0.98762 | 34.232 |

| | | | | | | | |
|---|---|---|---|---|---|---|---|
| 3222401L13Rik | 116.169 | -0.24939 | 0.1817 | -1.37253 | 0.1699 | 0.3603 | 6.292 |
| 3300002I08Rik | 106.297 | 0.275589 | 0.23982 | 1.149155 | 0.25049 | 0.45785 | 51.831 |
| 3300002P13Rik | 1062.91 | -0.57538 | 0.18811 | -3.05869 | 0.00222 | 0.02156 | 1.781 |
| 3632454L22Rik | 122.03 | 0.875195 | 0.24453 | 3.579043 | 0.00034 | 0.00617 | 37.603 |
| 3830406C13Rik | 674.559 | 0.004058 | 0.08724 | 0.04651 | 0.9629 | 0.98198 | 19.059 |
| 3830408C21Rik | 326.735 | 0.350843 | 0.15185 | 2.310512 | 0.02086 | 0.09212 | 32.945 |
| 3930402G23Rik | 17.097 | 0.215765 | 0.58327 | 0.369925 | 0.71144 | 0.84471 | 4.27 |
| 4631405J19Rik | 23.297 | 0.229704 | 0.36868 | 0.62304 | 0.53326 | 0.7205 | 16.777 |
| 4632404H12Rik | 1321.82 | 0.260669 | 0.1272 | 2.049322 | 0.04043 | 0.14368 | 5.79 |
| 4632411P08Rik | 31.217 | 0.073955 | 0.39241 | 0.188463 | 0.85051 | 0.92407 | 11.79 |
| 4632415L05Rik | 1894.11 | -0.14277 | 0.086 | -1.66004 | 0.09691 | 0.25501 | 6.16 |
| 4632418H02Rik | 67.9725 | -0.05178 | 0.24937 | -0.20766 | 0.83549 | 0.91568 | 2.721 |
| 4632427E13Rik | 206.792 | -0.08941 | 0.17082 | -0.5234 | 0.6007 | 0.77104 | 7.345 |
| 4632428C04Rik | 70.3368 | 0.12623 | 0.23854 | 0.529175 | 0.59668 | 0.76858 | 12.806 |
| 4633401B06Rik | 34.0567 | 0.325285 | 0.32321 | 1.006411 | 0.31422 | 0.52468 | 1.073 |
| 4731419I09Rik | 20.0085 | -0.47782 | 0.39469 | -1.21064 | 0.22603 | 0.43008 | 9.982 |
| 4732414G09Rik | 22.5835 | 0.305069 | 0.42218 | 0.722604 | 0.46992 | 0.66875 | 2.986 |
| 4732416N19Rik | 60.2538 | -0.03412 | 0.23686 | -0.14406 | 0.88546 | 0.9431 | 3.565 |
| 4732419C18Rik | 40.9291 | -1.24967 | 0.48033 | -2.60168 | 0.00928 | 0.05454 | 10.096 |
| 4732440D04Rik | 353.457 | 0.179768 | 0.11396 | 1.577467 | 0.11469 | 0.28365 | 51.165 |
| 4732463B04Rik | 28.0472 | -0.24515 | 0.33366 | -0.73472 | 0.46251 | 0.66283 | 14.339 |
| 4732471J01Rik | 533.177 | 0.0289 | 0.10188 | 0.283659 | 0.77667 | 0.88366 | 198.256 |
| 4732487G21Rik | 125.351 | -0.08899 | 0.16277 | -0.54675 | 0.58455 | 0.75928 | 15.246 |
| 4732491K20Rik | 99.3201 | 0.16034 | 0.22678 | 0.707016 | 0.47956 | 0.67719 | 17.451 |
| 4732496C06Rik | 271.435 | 0.276719 | 0.14436 | 1.916848 | 0.05526 | 0.17718 | 2.5 |
| 4833404L02Rik | 15.8265 | 0.21614 | 0.43002 | 0.502633 | 0.61522 | 0.78109 | 0.908 |
| 4833408A19Rik | 15.8523 | -0.26523 | 0.54687 | -0.48499 | 0.62768 | 0.78931 | 0.995 |
| 4833411C07Rik | 34.8171 | -0.85299 | 0.39544 | -2.15706 | 0.031 | 0.12075 | 23.761 |
| 4833412C15Rik | 24.1264 | 0.18548 | 0.45658 | 0.406238 | 0.68457 | 0.82814 | 2.071 |
| 4833412K13Rik | 170.737 | 0.590663 | 0.15861 | 3.724039 | 0.0002 | 0.00421 | 1.479 |
| 4833413G10Rik | 206.09 | 0.152807 | 0.16486 | 0.926871 | 0.35399 | 0.56422 | 1.946 |
| 4833415N18Rik | 279.947 | 0.252833 | 0.1864 | 1.356409 | 0.17497 | 0.36659 | 3.082 |
| 4833418N02Rik | 151.938 | 0.509329 | 0.1891 | 2.693366 | 0.00707 | 0.04577 | 7.935 |
| 4833420G17Rik | 1777.34 | 0.060951 | 0.12366 | 0.492883 | 0.6221 | 0.7853 | 23.353 |

| | | | | | | | |
|---|---|---|---|---|---|---|---|
| 4833422C13Rik | 581.762 | 0.222722 | 0.14545 | 1.53122 | 0.12572 | 0.30095 | 40.208 |
| 4833422M21Rik | 68.9174 | 0.268058 | 0.28631 | 0.936249 | 0.34915 | 0.55955 | 22.815 |
| 4833438C02Rik | 49.0133 | -0.17455 | 0.32173 | -0.54254 | 0.58745 | 0.76091 | 2.056 |
| 4833439L19Rik | 5737.28 | -0.11069 | 0.06419 | -1.7244 | 0.08464 | 0.23318 | 14.218 |
| 4833445I07Rik | 187.294 | -0.06488 | 0.15712 | -0.41293 | 0.67965 | 0.82491 | 2.189 |
| 4921504A21Rik | 79.8981 | 0.039491 | 0.19662 | 0.200856 | 0.84081 | 0.91835 | 24.353 |
| 4921507P07Rik | 51.9944 | -0.05723 | 0.27293 | -0.2097 | 0.8339 | 0.9148 | 23.331 |
| 4921513H07Rik | 38.0336 | -0.60228 | 0.29728 | -2.02596 | 0.04277 | 0.14905 | 2.238 |
| 4921514A10Rik | 58.8772 | 0.258259 | 0.24055 | 1.073629 | 0.28299 | 0.49316 | 2.428 |
| 4921515G04Rik | 17.5652 | 0.04022 | 0.50169 | 0.080168 | 0.9361 | 0.96792 | 4.695 |
| 4921522P10Rik | 44.1188 | 0.368853 | 0.35132 | 1.049914 | 0.29376 | 0.5038 | 2.928 |
| 4921524J17Rik | 443.09 | 0.28672 | 0.12964 | 2.211633 | 0.02699 | 0.10973 | 24.075 |
| 4921525O09Rik | 23.5642 | 1.147051 | 0.46229 | 2.481241 | 0.01309 | 0.0685 | 27.421 |
| 4921531C22Rik | 117.701 | -0.05644 | 0.17064 | -0.33075 | 0.74083 | 0.86313 | 3.021 |
| 4921536K21Rik | 29.9179 | 0.253513 | 0.35041 | 0.723471 | 0.46939 | 0.6683 | 9.011 |
| 4921539H07Rik | 14.8722 | -0.46647 | 0.6171 | -0.75591 | 0.4497 | 0.65278 | 50.564 |
| 4930403P22Rik | 12.9198 | 0.014347 | 0.51425 | 0.0279 | 0.97774 | 0.98807 | 0.713 |
| 4930404I05Rik | 106.13 | 0.259846 | 0.17367 | 1.496166 | 0.13461 | 0.31449 | 5.5 |
| 4930404N11Rik | 17.2826 | 0.410649 | 0.51991 | 0.78984 | 0.42962 | 0.63602 | 2.75 |
| 4930405A21Rik | 57.1979 | 0.480465 | 0.28604 | 1.679727 | 0.09301 | 0.24834 | 6.37 |
| 4930405J17Rik | 14.6314 | -0.05554 | 0.62846 | -0.08837 | 0.92958 | 0.96459 | 2.599 |
| 4930408O17Rik | 16.9946 | -0.63866 | 0.39996 | -1.5968 | 0.11031 | 0.27686 | 14.165 |
| 4930412C18Rik | 70.702 | -0.07213 | 0.25521 | -0.28264 | 0.77746 | 0.88413 | 52.83 |
| 4930412F12Rik | 21.3874 | 0.476711 | 0.42035 | 1.134091 | 0.25676 | 0.46474 | 1.471 |
| 4930412M03Rik | 63.8447 | 0.104652 | 0.24016 | 0.435766 | 0.66301 | 0.81385 | 3.041 |
| 4930413G21Rik | 276.362 | 0.027379 | 0.12965 | 0.211168 | 0.83276 | 0.91425 | 0.993 |
| 4930414N06Rik | 15.49 | 0.934892 | 0.50472 | 1.852314 | 0.06398 | 0.19465 | 24.178 |
| 4930417O13Rik | 171.591 | 0.651166 | 0.24696 | 2.636695 | 0.00837 | 0.05104 | 25.253 |
| 4930419G24Rik | 15.0185 | -0.78356 | 0.44651 | -1.75486 | 0.07928 | 0.2235 | 63.831 |
| 4930422M22Rik | 205.208 | 0.362302 | 0.16125 | 2.246769 | 0.02465 | 0.103 | 5.669 |
| 4930425K24Rik | 20.4836 | -0.12303 | 0.59984 | -0.20511 | 0.83749 | 0.91642 | 2.143 |
| 4930426D05Rik | 270.065 | -1.24092 | 0.58055 | -2.13749 | 0.03256 | 0.12503 | 4.658 |
| 4930426I24Rik | 32.4521 | 0.285135 | 0.32461 | 0.878382 | 0.37974 | 0.58952 | 1.232 |
| 4930426L09Rik | 23.5448 | 0.150993 | 0.47057 | 0.320876 | 0.7483 | 0.86814 | 2.419 |

| | | | | | | | |
|---|---|---|---|---|---|---|---|
| 4930428E07Rik | 17.9528 | -0.54811 | 0.49933 | -1.09769 | 0.27234 | 0.48159 | 61.714 |
| 4930428N03Rik | 16.9299 | -1.2196 | 0.70827 | -1.72193 | 0.08508 | 0.23389 | 121.32 |
| 4930429F24Rik | 271.864 | 0.584493 | 0.24844 | 2.352659 | 0.01864 | 0.08557 | 9.637 |
| 4930429H19Rik | 32.3624 | -0.09738 | 0.31888 | -0.3054 | 0.76006 | 0.87418 | 4.814 |
| 4930430F08Rik | 439.494 | 0.129801 | 0.16242 | 0.799195 | 0.42418 | 0.63094 | 18.159 |
| 4930431P19Rik | 130.432 | -0.04563 | 0.20039 | -0.22769 | 0.81989 | 0.90734 | 3.009 |
| 4930432B10Rik | 32.5497 | 0.452342 | 0.3068 | 1.474409 | 0.14037 | 0.3219 | 36.432 |
| 4930435N07Rik | 14.203 | -0.96604 | 0.62908 | -1.53563 | 0.12463 | 0.29958 | 2.127 |
| 4930444F02Rik | 20.5727 | 0.663202 | 0.45132 | 1.469472 | 0.1417 | 0.3235 | 35.16 |
| 4930444P10Rik | 31.464 | 0.608979 | 0.29686 | 2.051423 | 0.04023 | 0.1433 | 27.347 |
| 4930447C04Rik | 806.786 | -0.19283 | 0.22037 | -0.87503 | 0.38156 | 0.59096 | 59.666 |
| 4930447F24Rik | 139.013 | 0.00158 | 0.19203 | 0.008228 | 0.99343 | 0.99627 | 4.85 |
| 4930447M23Rik | 580.537 | 0.551055 | 0.10966 | 5.025229 | 5.03E-07 | 6.37E-05 | 7.938 |
| 4930447N08Rik | 14.8775 | -0.54016 | 0.66595 | -0.81111 | 0.4173 | 0.62473 | 40.094 |
| 4930451E10Rik | 27.6605 | 0.675572 | 0.38052 | 1.775406 | 0.07583 | 0.21743 | 9.37 |
| 4930452G13Rik | 27.8423 | 0.200597 | 0.42222 | 0.475104 | 0.63471 | 0.79438 | 11.545 |
| 4930453N24Rik | 1159.48 | -0.05539 | 0.08053 | -0.6879 | 0.49152 | 0.68643 | 19.411 |
| 4930455C13Rik | 16.9568 | -0.62436 | 0.56068 | -1.11356 | 0.26547 | 0.47463 | 22.97 |
| 4930455M05Rik | 22.7294 | 0.53599 | 0.37917 | 1.413571 | 0.15749 | 0.34464 | 4.054 |
| 4930456G14Rik | 61.4343 | 0.29478 | 0.38119 | 0.773318 | 0.43933 | 0.64469 | 1.33 |
| 4930458D05Rik | 42.5572 | 0.884137 | 0.28213 | 3.133797 | 0.00173 | 0.01828 | 6.371 |
| 4930461G14Rik | 39.3232 | 0.193788 | 0.39363 | 0.492306 | 0.6225 | 0.78561 | 14.536 |
| 4930466K18Rik | 158.889 | 0.230234 | 0.16037 | 1.435633 | 0.15111 | 0.33587 | 0.906 |
| 4930470B04Rik | 26.511 | 0.10481 | 0.37206 | 0.281704 | 0.77817 | 0.88423 | 1.926 |
| 4930471C04Rik | 31.2475 | -0.55827 | 0.33151 | -1.68405 | 0.09217 | 0.24718 | 4.941 |
| 4930473A02Rik | 108.004 | 0.390716 | 0.19867 | 1.966631 | 0.04923 | 0.16423 | 19.959 |
| 4930473H19Rik | 32.3018 | 0.661307 | 0.33857 | 1.953227 | 0.05079 | 0.16767 | 8.142 |
| 4930477O15Rik | 19.3331 | 0.296778 | 0.54821 | 0.541359 | 0.58826 | 0.76163 | 1.089 |
| 4930478K11Rik | 22.4546 | 0.295354 | 0.34254 | 0.862235 | 0.38856 | 0.59781 | 14.252 |
| 4930479D17Rik | 61.0733 | 0.150617 | 0.26424 | 0.570007 | 0.56867 | 0.74839 | 8.186 |
| 4930480K15Rik | 36.0778 | -0.37976 | 0.37525 | -1.01202 | 0.31153 | 0.52179 | 5.934 |
| 4930481A15Rik | 81.375 | 0.012833 | 0.27901 | 0.045995 | 0.96331 | 0.98208 | 16.108 |
| 4930481B07Rik | 51.6512 | 0.267522 | 0.35819 | 0.746869 | 0.45514 | 0.65722 | 4.514 |
| 4930483K19Rik | 13.0538 | 0.118913 | 0.64362 | 0.184757 | 0.85342 | 0.92567 | 16.356 |

| | | | | | | | |
|---|---|---|---|---|---|---|---|
| 4930483P17Rik | 116.453 | -0.51996 | 0.23523 | -2.21044 | 0.02707 | 0.10988 | 2.393 |
| 4930488L21Rik | 210.862 | -0.06518 | 0.1808 | -0.36051 | 0.71846 | 0.84959 | 63.576 |
| 4930500F10Rik | 27.5512 | 0.67349 | 0.37814 | 1.781047 | 0.0749 | 0.21572 | 1.454 |
| 4930503L19Rik | 1064 | 0.258526 | 0.11619 | 2.225013 | 0.02608 | 0.10718 | 20.298 |
| 4930505A04Rik | 33.2057 | 0.032565 | 0.46313 | 0.070316 | 0.94394 | 0.97194 | 45.822 |
| 4930506C21Rik | 270.515 | 0.348927 | 0.14588 | 2.391947 | 0.01676 | 0.08012 | 17.749 |
| 4930507D05Rik | 26.0169 | 0.528518 | 0.39314 | 1.344345 | 0.17884 | 0.37148 | 1.865 |
| 4930509G22Rik | 27.241 | 0.482297 | 0.35475 | 1.359547 | 0.17397 | 0.36537 | 14.117 |
| 4930509J09Rik | 14.4192 | 0.152663 | 0.70729 | 0.215841 | 0.82911 | 0.91246 | 130.446 |
| 4930511M06Rik | 21.326 | -0.96037 | 0.51906 | -1.85019 | 0.06429 | 0.19515 | 178.953 |
| 4930512B01Rik | 62.5987 | -0.13663 | 0.26584 | -0.51397 | 0.60727 | 0.77542 | 50.192 |
| 4930513N10Rik | 15.6075 | -0.09707 | 0.4782 | -0.20298 | 0.83915 | 0.91722 | 14.985 |
| 4930516B21Rik | 396.463 | 0.321073 | 0.14063 | 2.283056 | 0.02243 | 0.09666 | 4.553 |
| 4930517E14Rik | 18.7888 | 0.183239 | 0.46438 | 0.394587 | 0.69315 | 0.83358 | 2.972 |
| 4930517G19Rik | 24.0876 | 0.068285 | 0.38198 | 0.178767 | 0.85812 | 0.92854 | 1.676 |
| 4930517O19Rik | 13.8792 | 0.128962 | 0.5206 | 0.247717 | 0.80435 | 0.89895 | 32.198 |
| 4930518I15Rik | 125.841 | 0.08088 | 0.17617 | 0.459108 | 0.64616 | 0.80293 | 1.597 |
| 4930519K11Rik | 48.2659 | 0.077085 | 0.2525 | 0.305288 | 0.76015 | 0.87418 | 4.492 |
| 4930520O04Rik | 41.0023 | -0.20149 | 0.2883 | -0.6989 | 0.48462 | 0.68089 | 8.927 |
| 4930522L14Rik | 51.0379 | 0.125474 | 0.33628 | 0.373128 | 0.70905 | 0.84326 | 15.897 |
| 4930523C07Rik | 52.3068 | -0.70174 | 0.249 | -2.81828 | 0.00483 | 0.03547 | 35.827 |
| 4930524O07Rik | 34.3241 | -0.04516 | 0.3704 | -0.12191 | 0.90297 | 0.95118 | 3.636 |
| 4930525C09Rik | 22.3643 | 0.775428 | 0.42429 | 1.827605 | 0.06761 | 0.20161 | 38.931 |
| 4930525G20Rik | 141.038 | 0.270452 | 0.17897 | 1.511179 | 0.13074 | 0.30908 | 34.392 |
| 4930528D03Rik | 16.9068 | 0.80121 | 0.40593 | 1.973769 | 0.04841 | 0.16228 | 5.556 |
| 4930532G15Rik | 13.4089 | 0.800983 | 0.46129 | 1.736381 | 0.0825 | 0.22904 | 3.405 |
| 4930532I03Rik | 26.1387 | 0.270616 | 0.41965 | 0.644855 | 0.51902 | 0.70872 | 47.286 |
| 4930537H20Rik | 19.0941 | 0.700123 | 0.43121 | 1.623606 | 0.10446 | 0.26741 | 2.499 |
| 4930538E20Rik | 63.6566 | 0.057692 | 0.24544 | 0.235054 | 0.81417 | 0.90456 | 12.297 |
| 4930539J05Rik | 87.2599 | 0.130672 | 0.2007 | 0.651069 | 0.515 | 0.70535 | 2.498 |
| 4930545L23Rik | 29.3297 | -0.16184 | 0.44589 | -0.36296 | 0.71664 | 0.84856 | 46.044 |
| 4930547M16Rik | 31.2811 | 0.271921 | 0.40831 | 0.665969 | 0.50543 | 0.69703 | 8.623 |
| 4930549G23Rik | 48.5397 | 0.443074 | 0.31735 | 1.396169 | 0.16266 | 0.35109 | 31.673 |
| 4930550C14Rik | 13.9462 | -0.15236 | 0.54506 | -0.27952 | 0.77984 | 0.88529 | 32.078 |

| | | | | | | | |
|---|---|---|---|---|---|---|---|
| 4930553P18Rik | 36.2786 | -0.3601 | 0.30607 | -1.17653 | 0.23938 | 0.44538 | 4.976 |
| 4930555F03Rik | 17.2789 | -0.17119 | 0.49867 | -0.3433 | 0.73138 | 0.85734 | 178.109 |
| 4930556H04Rik | 15.5332 | -0.5041 | 0.41912 | -1.20276 | 0.22907 | 0.43319 | 1.675 |
| 4930556I23Rik | 352.16 | 0.37944 | 0.13751 | 2.7593 | 0.00579 | 0.04027 | 1.114 |
| 4930556M19Rik | 13.3647 | -0.79821 | 0.50196 | -1.5902 | 0.11179 | 0.27929 | 75.298 |
| 4930556N13Rik | 17.306 | -0.07704 | 0.4379 | -0.17593 | 0.86035 | 0.92937 | 29.792 |
| 4930557K07Rik | 105.369 | 0.408965 | 0.20505 | 1.994498 | 0.0461 | 0.15671 | 39.679 |
| 4930562C15Rik | 94.3711 | 0.326702 | 0.191 | 1.71045 | 0.08718 | 0.23781 | 32.271 |
| 4930563E22Rik | 115.356 | 0.052404 | 0.173 | 0.302906 | 0.76196 | 0.87518 | 3.307 |
| 4930565N06Rik | 50.0991 | -0.16693 | 0.32726 | -0.51007 | 0.61 | 0.77739 | 16.671 |
| 4930567K12Rik | 19.0463 | 0.526725 | 0.45128 | 1.167183 | 0.24314 | 0.44937 | 27.306 |
| 4930568G15Rik | 18.7655 | 0.657295 | 0.40279 | 1.631858 | 0.10271 | 0.26462 | 27.905 |
| 4930570G19Rik | 984.743 | 0.503448 | 0.13929 | 3.614323 | 0.0003 | 0.00565 | 73.253 |
| 4930572G02Rik | 23.268 | 0.291367 | 0.60132 | 0.484547 | 0.628 | 0.78952 | 33.898 |
| 4930577N17Rik | 24.2901 | -0.38568 | 0.4044 | -0.9537 | 0.34023 | 0.55103 | 1.625 |
| 4930578G10Rik | 37.8753 | -0.73066 | 0.40889 | -1.78693 | 0.07395 | 0.21427 | 110.704 |
| 4930578M01Rik | 280.446 | 0.128923 | 0.13151 | 0.980364 | 0.32691 | 0.53825 | 13.099 |
| 4930579G18Rik | 394.09 | -0.01434 | 0.13411 | -0.10696 | 0.91482 | 0.95656 | 4.097 |
| 4930579G24Rik | 53.67 | -0.27699 | 0.32551 | -0.85093 | 0.39481 | 0.60387 | 3.742 |
| 4930579K19Rik | 33.0482 | 0.573144 | 0.32152 | 1.7826 | 0.07465 | 0.21534 | 1.548 |
| 4930581F22Rik | 258.039 | 0.066834 | 0.12414 | 0.538395 | 0.5903 | 0.76324 | 14.195 |
| 4930588G17Rik | 20.7018 | -0.1036 | 0.41666 | -0.24864 | 0.80364 | 0.89865 | 5.607 |
| 4930590J08Rik | 169.293 | 0.020836 | 0.19563 | 0.106508 | 0.91518 | 0.95656 | 47.917 |
| 4930593A02Rik | 21.5665 | 0.610649 | 0.54584 | 1.118731 | 0.26326 | 0.47208 | 104.743 |
| 4930594M22Rik | 31.1341 | -0.34321 | 0.31201 | -1.09998 | 0.27134 | 0.48066 | 24.44 |
| 4930598N05Rik | 54.0785 | 1.357616 | 0.38011 | 3.57162 | 0.00035 | 0.00628 | 73.9 |
| 4931403E22Rik | 55.6381 | 0.547377 | 0.28054 | 1.951167 | 0.05104 | 0.16815 | 55.107 |
| 4931406C07Rik | 1480.76 | -0.05378 | 0.07771 | -0.69208 | 0.48889 | 0.68402 | 23.112 |
| 4931408D14Rik | 31.2658 | -0.08575 | 0.32661 | -0.26255 | 0.7929 | 0.89266 | 5.777 |
| 4931413K12Rik | 59.1883 | -0.13624 | 0.23048 | -0.59111 | 0.55445 | 0.73705 | 5.107 |
| 4931414P19Rik | 149.423 | -0.29474 | 0.15152 | -1.94528 | 0.05174 | 0.1697 | 22.331 |
| 4931415C17Rik | 195.252 | 0.358415 | 0.17103 | 2.095575 | 0.03612 | 0.13382 | 7.044 |
| 4931422A03Rik | 101.216 | 0.322195 | 0.23404 | 1.376655 | 0.16862 | 0.35867 | 61.419 |
| 4931428F04Rik | 749.211 | -0.15006 | 0.1011 | -1.48429 | 0.13773 | 0.31857 | 9.456 |

| | | | | | | | |
|---|---|---|---|---|---|---|---|
| 4931429L15Rik | 27.0447 | 0.614905 | 0.41477 | 1.482522 | 0.1382 | 0.31934 | 16.626 |
| 4931440P22Rik | 53.8575 | -0.13559 | 0.26996 | -0.50224 | 0.6155 | 0.78122 | 22.123 |
| 4932422M17Rik | 119.69 | 0.281561 | 0.21229 | 1.326311 | 0.18474 | 0.37937 | 3.346 |
| 4932438A13Rik | 7694.22 | 0.184913 | 0.14431 | 1.281392 | 0.20006 | 0.39817 | 189.93 |
| 4932438H23Rik | 24.9998 | 0.272533 | 0.34947 | 0.779852 | 0.43548 | 0.64111 | 41.188 |
| 4932441J04Rik | 125.098 | -0.36978 | 0.21732 | -1.70157 | 0.08884 | 0.24096 | 150.12 |
| 4932442E05Rik | 58.7905 | -0.23565 | 0.31273 | -0.75351 | 0.45114 | 0.65374 | 2.717 |
| 4932443L11Rik | 33.9383 | 0.066939 | 0.29423 | 0.227507 | 0.82003 | 0.90734 | 44.361 |
| 4932702P03Rik | 95.4071 | 0.643017 | 0.23064 | 2.787928 | 0.0053 | 0.03768 | 8.68 |
| 4933400F21Rik | 17.2006 | -0.237 | 0.46155 | -0.51348 | 0.60762 | 0.77569 | 33.38 |
| 4933402D24Rik | 69.5743 | -0.03752 | 0.21359 | -0.17567 | 0.86055 | 0.92938 | 14.361 |
| 4933403J19Rik | 22.4414 | -0.38475 | 0.45964 | -0.83707 | 0.40255 | 0.61125 | 7.456 |
| 4933404O12Rik | 678.499 | 0.009563 | 0.08965 | 0.106678 | 0.91504 | 0.95656 | 18.005 |
| 4933406B15Rik | 40.4059 | 0.372996 | 0.3369 | 1.107129 | 0.26824 | 0.47751 | 14.13 |
| 4933406C10Rik | 157.378 | 0.179944 | 0.15787 | 1.139838 | 0.25435 | 0.46172 | 34.456 |
| 4933406I18Rik | 57.8027 | -0.22277 | 0.27534 | -0.80908 | 0.41847 | 0.62566 | 99.683 |
| 4933406P04Rik | 19.4736 | -0.13732 | 0.40521 | -0.33889 | 0.73469 | 0.85968 | 1.827 |
| 4933407E24Rik | 13.4732 | 1.049702 | 0.48429 | 2.167516 | 0.0302 | 0.11869 | 5.456 |
| 4933407I05Rik | 19.4853 | 0.293304 | 0.4664 | 0.628871 | 0.52943 | 0.71727 | 12.577 |
| 4933407K13Rik | 469.819 | 0.265589 | 0.16282 | 1.631184 | 0.10285 | 0.26481 | 57.81 |
| 4933407L21Rik | 275.043 | 0.918316 | 0.18176 | 5.052252 | 4.37E-07 | 5.71E-05 | 12.143 |
| 4933409K07Rik | 78.3638 | -0.21937 | 0.21161 | -1.03668 | 0.29988 | 0.51004 | 7.553 |
| 4933412E12Rik | 137.026 | -0.28233 | 0.17345 | -1.62775 | 0.10358 | 0.26613 | 13.53 |
| 4933412L11Rik | 31.2138 | 0.436165 | 0.31074 | 1.403628 | 0.16043 | 0.3486 | 1.044 |
| 4933412O06Rik | 88.8253 | -0.0437 | 0.21405 | -0.20417 | 0.83822 | 0.91679 | 20.967 |
| 4933413J09Rik | 21.25 | -0.02819 | 0.4911 | -0.0574 | 0.95423 | 0.97748 | 43.883 |
| 4933415A04Rik | 13.1426 | 0.16655 | 0.53975 | 0.308568 | 0.75765 | 0.87241 | 2.009 |
| 4933416E14Rik | 14.5143 | 0.081286 | 0.46806 | 0.173666 | 0.86213 | 0.93002 | 1.938 |
| 4933416I08Rik | 716.576 | 0.17816 | 0.11802 | 1.50953 | 0.13116 | 0.30963 | 34.379 |
| 4933417D19Rik | 95.2572 | 0.211155 | 0.21236 | 0.994337 | 0.32006 | 0.53067 | 7.269 |
| 4933421O10Rik | 641.927 | 0.099044 | 0.09252 | 1.070533 | 0.28438 | 0.49449 | 4.182 |
| 4933423P22Rik | 45.3611 | 0.500763 | 0.2764 | 1.811745 | 0.07003 | 0.20664 | 9.19 |
| 4933424G05Rik | 37.3085 | 0.264277 | 0.33768 | 0.782623 | 0.43385 | 0.6396 | 58.993 |
| 4933424G06Rik | 42.9727 | 0.717198 | 0.29764 | 2.409633 | 0.01597 | 0.07759 | 47.805 |

| | | | | | | | |
|---|---|---|---|---|---|---|---|
| 4933426B08Rik | 38.6454 | 0.269042 | 0.33783 | 0.796372 | 0.42582 | 0.63232 | 10.021 |
| 4933427G23Rik | 31.4104 | 0.142196 | 0.31387 | 0.453039 | 0.65052 | 0.80552 | 16.28 |
| 4933427I04Rik | 38.1726 | -0.12265 | 0.29952 | -0.4095 | 0.68218 | 0.82633 | 3.487 |
| 4933427I22Rik | 26.2026 | -0.57514 | 0.36301 | -1.58434 | 0.11312 | 0.2815 | 17.409 |
| 4933428G20Rik | 198.173 | -0.32113 | 0.20507 | -1.56593 | 0.11736 | 0.28827 | 9.558 |
| 4933430I17Rik | 23.9574 | -0.10167 | 0.41949 | -0.24236 | 0.8085 | 0.90156 | 22.625 |
| 4933431C10Rik | 71.5763 | 0.354653 | 0.22565 | 1.571722 | 0.11601 | 0.28589 | 1.266 |
| 4933431E20Rik | 2771.26 | -0.14916 | 0.10364 | -1.43929 | 0.15007 | 0.33457 | 7.364 |
| 4933431G14Rik | 69.0431 | 0.166797 | 0.27228 | 0.612604 | 0.54014 | 0.72606 | 2.985 |
| 4933431K14Rik | 74.7697 | 0.294162 | 0.26975 | 1.090515 | 0.27549 | 0.4847 | 1.352 |
| 4933432I03Rik | 25.0518 | 0.134725 | 0.43192 | 0.311918 | 0.7551 | 0.87123 | 46.392 |
| 4933432I09Rik | 84.4516 | -0.5115 | 0.26126 | -1.95784 | 0.05025 | 0.1666 | 6.042 |
| 4933432K03Rik | 19.3808 | -0.77836 | 0.52904 | -1.47125 | 0.14122 | 0.32297 | 140.491 |
| 4933433G15Rik | 33.9691 | 0.949014 | 0.31761 | 2.988032 | 0.00281 | 0.02514 | 7.526 |
| 4933433G19Rik | 153.597 | 0.059133 | 0.2064 | 0.2865 | 0.77449 | 0.88245 | 35.59 |
| 4933434E20Rik | 538.868 | 0.171452 | 0.12945 | 1.324468 | 0.18535 | 0.38013 | 11.706 |
| 4933439C10Rik | 484.464 | 0.153088 | 0.12468 | 1.227866 | 0.2195 | 0.42192 | 7.986 |
| 4933439K11Rik | 29.1854 | 0.193358 | 0.33309 | 0.5805 | 0.56158 | 0.7427 | 19.997 |
| 4933440N22Rik | 18.5403 | 0.091653 | 0.51528 | 0.177871 | 0.85882 | 0.92876 | 1.953 |
| 5031415H12Rik | 45.5274 | -0.43261 | 0.32883 | -1.31561 | 0.1883 | 0.38384 | 10.254 |
| 5031425E22Rik | 965.23 | 0.131204 | 0.07623 | 1.721107 | 0.08523 | 0.23406 | 52.1 |
| 5031434O11Rik | 303.928 | -0.08804 | 0.1484 | -0.59328 | 0.553 | 0.73584 | 7.083 |
| 5031439G07Rik | 8751.87 | -0.13276 | 0.08123 | -1.63446 | 0.10216 | 0.26377 | 44.616 |
| 5033404E19Rik | 21.1813 | 0.296006 | 0.44037 | 0.672172 | 0.50147 | 0.69396 | 10.874 |
| 5033417F24Rik | 100.313 | 0.164934 | 0.1784 | 0.924542 | 0.3552 | 0.56553 | 2.387 |
| 5033430I15Rik | 14.1506 | 0.538663 | 0.5526 | 0.974776 | 0.32967 | 0.54093 | 2.246 |
| 5330406M23Rik | 114.237 | 0.151079 | 0.23358 | 0.646809 | 0.51776 | 0.70756 | 2.094 |
| 5330413P13Rik | 98.8656 | 0.527001 | 0.28017 | 1.881036 | 0.05997 | 0.18671 | 27.678 |
| 5330416C01Rik | 13.0702 | -1.28434 | 0.61705 | -2.08141 | 0.0374 | 0.13676 | 13.914 |
| 5330425B07Rik | 17.7981 | 1.377699 | 0.53977 | 2.552385 | 0.0107 | 0.06008 | 1.027 |
| 5330431K02Rik | 20.0533 | 0.207307 | 0.46737 | 0.443562 | 0.65736 | 0.80979 | 1.963 |
| 5330432J10Rik | 15.5705 | -0.2433 | 0.56881 | -0.42773 | 0.66885 | 0.81737 | 4.233 |
| 5330434G04Rik | 770.365 | -0.15936 | 0.11089 | -1.43709 | 0.15069 | 0.33561 | 50.337 |
| 5330437M03Rik | 62.9474 | 1.4045 | 0.36404 | 3.858074 | 0.00011 | 0.00294 | 4.083 |

| | | | | | | | |
|---|---|---|---|---|---|---|---|
| 5330438D12Rik | 251.885 | 0.264524 | 0.16787 | 1.575772 | 0.11508 | 0.28437 | 106.919 |
| 5330438I03Rik | 53.4908 | 0.20481 | 0.25069 | 0.816989 | 0.41393 | 0.622 | 2.845 |
| 5330439A09Rik | 13.1647 | 0.852286 | 0.53167 | 1.603023 | 0.10893 | 0.27443 | 32.438 |
| 5330439K02Rik | 98.9405 | -0.23763 | 0.19584 | -1.21337 | 0.22499 | 0.42882 | 8.952 |
| 5430400D12Rik | 73.1126 | 0.423741 | 0.21454 | 1.975155 | 0.04825 | 0.16192 | 1.799 |
| 5430402O13Rik | 83.7074 | 0.330826 | 0.20642 | 1.602661 | 0.10901 | 0.27456 | 57.757 |
| 5430403G16Rik | 105.499 | 0.313803 | 0.29078 | 1.079177 | 0.28051 | 0.49031 | 16.497 |
| 5430405H02Rik | 200.625 | 0.223982 | 0.1466 | 1.527842 | 0.12655 | 0.30235 | 19.882 |
| 5430414B19Rik | 21.7592 | -0.0887 | 0.38521 | -0.23026 | 0.81789 | 0.90637 | 0.75 |
| 5430416N02Rik | 337.356 | 0.157878 | 0.10844 | 1.455874 | 0.14543 | 0.32825 | 9.071 |
| 5430431A17Rik | 38.604 | -0.31436 | 0.33767 | -0.93096 | 0.35187 | 0.56223 | 12.64 |
| 5430434F05Rik | 20.1164 | 0.074709 | 0.44529 | 0.167778 | 0.86676 | 0.9324 | 3.553 |
| 5530401A14Rik | 37.633 | 0.470789 | 0.38962 | 1.208338 | 0.22692 | 0.43112 | 33.91 |
| 5530601H04Rik | 1051.48 | 0.297171 | 0.12044 | 2.467363 | 0.01361 | 0.07022 | 42.413 |
| 5730405O15Rik | 82.7675 | 0.216302 | 0.19742 | 1.095646 | 0.27323 | 0.48255 | 3.548 |
| 5730409E04Rik | 3363.78 | -0.13297 | 0.06968 | -1.90819 | 0.05637 | 0.17956 | 4.554 |
| 5730414N17Rik | 42.3891 | 0.787388 | 0.35168 | 2.238926 | 0.02516 | 0.10439 | 1.439 |
| 5730455P16Rik | 3862.93 | 0.053892 | 0.07991 | 0.674425 | 0.50004 | 0.69306 | 17.543 |
| 5730471H19Rik | 24.9521 | -0.11444 | 0.39021 | -0.29328 | 0.7693 | 0.87965 | 37.686 |
| 5730480H06Rik | 645.145 | 0.116488 | 0.10813 | 1.077287 | 0.28135 | 0.49119 | 86.343 |
| 5730507A11Rik | 20.4209 | -0.44388 | 0.44135 | -1.00574 | 0.31454 | 0.52507 | 3.371 |
| 5730521K06Rik | 65.101 | 0.370415 | 0.25342 | 1.461643 | 0.14384 | 0.32621 | 1.608 |
| 5730522E02Rik | 236.157 | -0.15179 | 0.1384 | -1.09679 | 0.27274 | 0.48188 | 608.001 |
| 5830408C22Rik | 176.736 | -0.71029 | 0.53139 | -1.33668 | 0.18133 | 0.3748 | 9.405 |
| 5830417I10Rik | 890.482 | 0.246381 | 0.09155 | 2.691319 | 0.00712 | 0.04598 | 50.372 |
| 5830418P13Rik | 16.3502 | 0.899631 | 0.40209 | 2.237373 | 0.02526 | 0.10462 | 37.683 |
| 5830432E09Rik | 29.1518 | 0.862737 | 0.43397 | 1.988025 | 0.04681 | 0.15868 | 16.384 |
| 5830444B04Rik | 174.689 | 0.602559 | 0.17161 | 3.511272 | 0.00045 | 0.00735 | 22.942 |
| 5830448L01Rik | 86.3475 | 0.224623 | 0.21957 | 1.023029 | 0.30629 | 0.51635 | 1.321 |
| 5830454E08Rik | 27.4438 | 0.357105 | 0.44698 | 0.798924 | 0.42433 | 0.63107 | 0.744 |
| 5830462I19Rik | 36.2102 | 0.022314 | 0.35365 | 0.063098 | 0.94969 | 0.97495 | 3.383 |
| 5830468F06Rik | 34.9357 | 0.67649 | 0.34059 | 1.986215 | 0.04701 | 0.15907 | 3.819 |
| 5830487J09Rik | 81.1453 | 0.488281 | 0.25539 | 1.911941 | 0.05588 | 0.17867 | 2.448 |
| 5930403N24Rik | 12.8686 | 0.304533 | 0.5385 | 0.565523 | 0.57172 | 0.75056 | 11.914 |

| | | | | | | | |
|---|---|---|---|---|---|---|---|
| 5930409G06Rik | 219.828 | 0.897165 | 0.21387 | 4.194904 | 2.73E-05 | 0.00108 | 1.544 |
| 5930420M18Rik | 188.823 | 0.321606 | 0.1351 | 2.380443 | 0.01729 | 0.08154 | 2.281 |
| 5930430L01Rik | 163.43 | -0.12122 | 0.16828 | -0.72037 | 0.4713 | 0.67008 | 5.168 |
| 5S_rRNA | 433.185 | 0.104171 | 0.1243 | 0.838067 | 0.40199 | 0.6106 | 0.138 |
| 6030400A10Rik | 28.4296 | 0.058789 | 0.33503 | 0.175471 | 0.86071 | 0.92947 | 1.212 |
| 6030407O03Rik | 21.5576 | 0.027139 | 0.40091 | 0.067694 | 0.94603 | 0.973 | 246.379 |
| 6030442K20Rik | 52.4329 | 0.470979 | 0.27637 | 1.704169 | 0.08835 | 0.24015 | 10.632 |
| 6030443J06Rik | 85.1756 | 0.160866 | 0.22833 | 0.704527 | 0.4811 | 0.6782 | 257.46 |
| 6030458C11Rik | 2489.52 | 0.141349 | 0.08252 | 1.712959 | 0.08672 | 0.23706 | 16.473 |
| 6230400D17Rik | 263.062 | 0.20837 | 0.11982 | 1.739051 | 0.08203 | 0.22823 | 15.796 |
| 6330403K07Rik | 11684.6 | 0.144859 | 0.09499 | 1.525068 | 0.12724 | 0.30346 | 1.573 |
| 6330403L08Rik | 847.283 | -0.26808 | 0.16994 | -1.57751 | 0.11468 | 0.28365 | 5.522 |
| 6330408M09Rik | 32.2927 | 0.449923 | 0.33102 | 1.359222 | 0.17408 | 0.36555 | 1.133 |
| 6330409D20Rik | 34.6612 | -0.13682 | 0.30213 | -0.45286 | 0.65065 | 0.80552 | 8.397 |
| 6330415B21Rik | 21.8753 | 0.29943 | 0.46302 | 0.646689 | 0.51783 | 0.70757 | 2.2 |
| 6330415G19Rik | 28.6418 | 0.087866 | 0.32933 | 0.266802 | 0.78962 | 0.89082 | 14.493 |
| 6330418K02Rik | 21.8505 | 0.100796 | 0.34836 | 0.289346 | 0.77232 | 0.88141 | 2.622 |
| 6330537M06Rik | 65.691 | 0.303278 | 0.22299 | 1.360074 | 0.17381 | 0.36524 | 2.184 |
| 6330549D23Rik | 104.093 | 0.169717 | 0.20409 | 0.831592 | 0.40564 | 0.61433 | 23.162 |
| 6330562C20Rik | 15.0411 | -0.08896 | 0.50563 | -0.17593 | 0.86035 | 0.92937 | 8.06 |
| 6330576A10Rik | 12.9221 | -0.30987 | 0.52576 | -0.58937 | 0.55561 | 0.73792 | 20.103 |
| 6430503K07Rik | 23.5017 | 0.079186 | 0.33786 | 0.234374 | 0.8147 | 0.90481 | 0.984 |
| 6430511E19Rik | 248.53 | 0.273959 | 0.17325 | 1.581262 | 0.11382 | 0.28245 | 3.835 |
| 6430548M08Rik | 14474 | -0.4814 | 0.12902 | -3.73112 | 0.00019 | 0.00417 | 51.155 |
| 6430550D23Rik | 81.517 | -0.11189 | 0.25204 | -0.44393 | 0.65709 | 0.80956 | 34.699 |
| 6430553K19Rik | 33.975 | -0.97303 | 0.35054 | -2.77577 | 0.00551 | 0.0388 | 36.428 |
| 6430562O15Rik | 93.0101 | 0.335856 | 0.19953 | 1.683258 | 0.09233 | 0.2474 | 15.953 |
| 6430571L13Rik | 274.279 | -0.28481 | 0.18216 | -1.56347 | 0.11794 | 0.28927 | 11.821 |
| 6430573P05Rik | 89.2826 | -0.59346 | 0.29316 | -2.02439 | 0.04293 | 0.14943 | 2.676 |
| 6430584L05Rik | 261.168 | 0.898512 | 0.14278 | 6.292966 | 3.11E-10 | 2.27E-07 | 15.399 |
| 6430590A07Rik | 283.898 | -0.00328 | 0.15702 | -0.02091 | 0.98331 | 0.99092 | 8.297 |
| 6430628N08Rik | 17.6967 | 1.658022 | 0.49804 | 3.329073 | 0.00087 | 0.01163 | 8.563 |
| 6430710M23Rik | 53.0862 | -0.2336 | 0.26534 | -0.88037 | 0.37866 | 0.5884 | 1.191 |
| 6530401F13Rik | 25.0208 | 1.254182 | 0.36273 | 3.457592 | 0.00055 | 0.0084 | 24.523 |

| | | | | | | | |
|---|---|---|---|---|---|---|---|
| 6530402F18Rik | 623.256 | -0.28188 | 0.14989 | -1.88055 | 0.06003 | 0.1868 | 7.9 |
| 6530403H02Rik | 270.427 | 0.428429 | 0.17536 | 2.443174 | 0.01456 | 0.07292 | 282.655 |
| 6530409C15Rik | 79.7735 | 0.18848 | 0.18839 | 1.000492 | 0.31707 | 0.52744 | 45.308 |
| 6530411M01Rik | 16.0622 | -0.5893 | 0.40184 | -1.46651 | 0.14251 | 0.32436 | 20.118 |
| 6720427I07Rik | 372.785 | -0.00283 | 0.13636 | -0.02072 | 0.98347 | 0.99095 | 13.596 |
| 6720483E21Rik | 22.4063 | 0.868074 | 0.39573 | 2.193614 | 0.02826 | 0.11321 | 1.824 |
| 6720489N17Rik | 159.756 | 0.072722 | 0.21101 | 0.344639 | 0.73037 | 0.85705 | 21.17 |
| 6820402A03Rik | 24.4708 | -0.39237 | 0.38179 | -1.0277 | 0.30409 | 0.51398 | 2.654 |
| 6820408C15Rik | 153.779 | 0.522447 | 0.16086 | 3.247891 | 0.00116 | 0.01403 | 28.744 |
| 6820431F20Rik | 975.983 | 0.137092 | 0.10986 | 1.247893 | 0.21207 | 0.41271 | 36.434 |
| 7630403G23Rik | 13.8907 | 0.413782 | 0.55498 | 0.745574 | 0.45593 | 0.65758 | 16.651 |
| 7SK | 747.485 | 0.161402 | 0.10522 | 1.533878 | 0.12506 | 0.30014 | 0.371 |
| 8030445P17Rik | 29.944 | 0.169181 | 0.43365 | 0.390133 | 0.69644 | 0.83541 | 2.741 |
| 8030451O07Rik | 17.1421 | -0.08234 | 0.65723 | -0.12529 | 0.90029 | 0.94992 | 7.945 |
| 8030453O22Rik | 85.5282 | -0.58808 | 0.27546 | -2.1349 | 0.03277 | 0.12566 | 16.974 |
| 8030456M14Rik | 36.2436 | 0.147778 | 0.38508 | 0.383757 | 0.70116 | 0.83856 | 1.019 |
| 8030462N17Rik | 3339.47 | 0.130754 | 0.08045 | 1.625373 | 0.10408 | 0.2667 | 82.994 |
| 8430406P12Rik | 62.5081 | 0.321971 | 0.25697 | 1.252951 | 0.21022 | 0.4102 | 1.07 |
| 8430422M14Rik | 306.475 | 0.208691 | 0.16144 | 1.29269 | 0.19612 | 0.3937 | 0.822 |
| 8430426J06Rik | 63.5388 | 0.807703 | 0.26483 | 3.049864 | 0.00229 | 0.02199 | 70.214 |
| 8430429K09Rik | 271.207 | 0.051185 | 0.13405 | 0.381851 | 0.70257 | 0.8396 | 27.467 |
| 8430432A02Rik | 22.6846 | 0.155942 | 0.46886 | 0.332596 | 0.73944 | 0.86264 | 1.248 |
| 9030025P20Rik | 183.69 | -0.2563 | 0.23462 | -1.0924 | 0.27466 | 0.4843 | 13.105 |
| 9030407P20Rik | 157.541 | -0.25707 | 0.18209 | -1.41178 | 0.15802 | 0.34527 | 4.135 |
| 9030612E09Rik | 32.5196 | -0.6885 | 0.33324 | -2.06611 | 0.03882 | 0.14042 | 1.864 |
| 9030624G23Rik | 57.331 | 0.192832 | 0.27225 | 0.708299 | 0.47876 | 0.67635 | 54.045 |
| 9130002K18Rik | 39.7911 | 0.57117 | 0.41401 | 1.379594 | 0.16771 | 0.35762 | 3.126 |
| 9130015G15Rik | 15.5691 | 0.12753 | 0.48332 | 0.263864 | 0.79188 | 0.89181 | 37.486 |
| 9130019O22Rik | 148.3 | -0.1464 | 0.17857 | -0.81984 | 0.41231 | 0.62038 | 4.907 |
| 9130019P16Rik | 77.6836 | 0.337092 | 0.23171 | 1.454817 | 0.14572 | 0.32865 | 160.541 |
| 9130023H24Rik | 133.719 | -0.31787 | 0.18074 | -1.75866 | 0.07863 | 0.22224 | 3.574 |
| 9130024F11Rik | 38.5406 | -0.87588 | 0.45555 | -1.92269 | 0.05452 | 0.17559 | 78.673 |
| 9130213A22Rik | 13.1051 | -0.90262 | 0.48432 | -1.86369 | 0.06237 | 0.19151 | 2.536 |
| 9130221H12Rik | 69.447 | -0.02978 | 0.22854 | -0.13031 | 0.89632 | 0.94789 | 1.577 |

| | | | | | | | |
|---|---|---|---|---|---|---|---|
| 9130230N09Rik | 29.4026 | 0.295408 | 0.40265 | 0.733663 | 0.46315 | 0.66313 | 2.334 |
| 9130401M01Rik | 313.751 | 0.264436 | 0.11674 | 2.265114 | 0.02351 | 0.09951 | 12.067 |
| 9130604C24Rik | 24.5652 | 0.219913 | 0.38287 | 0.574385 | 0.56571 | 0.74579 | 0.956 |
| 9230009I02Rik | 12.9174 | 0.164917 | 0.52384 | 0.314821 | 0.7529 | 0.87023 | 6.747 |
| 9230112E08Rik | 867.126 | 0.041248 | 0.09043 | 0.456115 | 0.64831 | 0.80408 | 3.352 |
| 9230112J17Rik | 38.5654 | -0.11965 | 0.33634 | -0.35574 | 0.72204 | 0.85169 | 21.936 |
| 9230114K14Rik | 138.197 | 0.238717 | 0.19625 | 1.216402 | 0.22383 | 0.42709 | 88.38 |
| 9230116N13Rik | 16.8799 | 0.348701 | 0.43398 | 0.803493 | 0.42169 | 0.62866 | 3.604 |
| 9230117E06Rik | 25.7964 | -0.0318 | 0.53091 | -0.0599 | 0.95224 | 0.97658 | 6.202 |
| 9330020H09Rik | 37.0858 | 0.188672 | 0.28433 | 0.663576 | 0.50696 | 0.69814 | 2.26 |
| 9330102E08Rik | 1261.87 | -0.13641 | 0.09274 | -1.4708 | 0.14134 | 0.32311 | 14.1 |
| 9330104G04Rik | 177.215 | -0.11865 | 0.16163 | -0.73406 | 0.46291 | 0.66297 | 3.54 |
| 9330111N05Rik | 43.8465 | -0.36965 | 0.29933 | -1.23492 | 0.21686 | 0.41876 | 165.977 |
| 9330117O12Rik | 65.9851 | -0.60404 | 0.25189 | -2.39803 | 0.01648 | 0.07926 | 28.127 |
| 9330118I20Rik | 58.991 | 0.228389 | 0.30698 | 0.743978 | 0.45689 | 0.65827 | 9.132 |
| 9330121K16Rik | 1510.92 | -0.11916 | 0.20372 | -0.58493 | 0.55859 | 0.73995 | 3.886 |
| 9330133O14Rik | 178.005 | 0.098283 | 0.14658 | 0.670521 | 0.50253 | 0.69511 | 2.851 |
| 9330136K24Rik | 48.8776 | 0.410364 | 0.25831 | 1.588653 | 0.11214 | 0.27968 | 7.685 |
| 9330151L19Rik | 646.795 | 0.247658 | 0.0853 | 2.903271 | 0.00369 | 0.02987 | 2.657 |
| 9330154J02Rik | 63.4618 | 0.161436 | 0.29518 | 0.546902 | 0.58445 | 0.75925 | 24.083 |
| 9330158H04Rik | 31.4022 | -0.13582 | 0.36647 | -0.3706 | 0.71093 | 0.84441 | 180.572 |
| 9330159F19Rik | 5092.57 | 0.287855 | 0.12431 | 2.315618 | 0.02058 | 0.09128 | 19.174 |
| 9330159M07Rik | 60.0154 | -0.46573 | 0.24661 | -1.88852 | 0.05896 | 0.1848 | 18.635 |
| 9330160F10Rik | 44.5976 | -0.24072 | 0.25771 | -0.93407 | 0.35027 | 0.5609 | 2.766 |
| 9330161L09Rik | 16.257 | -0.05333 | 0.43328 | -0.12309 | 0.90203 | 0.95085 | 1.155 |
| 9330162012Rik | 168.781 | 0.398905 | 0.16176 | 2.466028 | 0.01366 | 0.07036 | 4.017 |
| 9330162G02Rik | 2457.33 | 0.343611 | 0.17706 | 1.940641 | 0.0523 | 0.17088 | 30.942 |
| 9330171B17Rik | 22.1776 | -0.45511 | 0.38534 | -1.18107 | 0.23757 | 0.44294 | 2.075 |
| 9330175M20Rik | 20.1355 | 0.180819 | 0.36945 | 0.489428 | 0.62454 | 0.78719 | 115.394 |
| 9330179D12Rik | 30.1107 | -0.17791 | 0.38171 | -0.4661 | 0.64115 | 0.79912 | 63.035 |
| 9330185C12Rik | 14.5843 | 0.503197 | 0.57919 | 0.868791 | 0.38496 | 0.59437 | 72.75 |
| 9330188P03Rik | 141.269 | -0.26041 | 0.23679 | -1.09975 | 0.27144 | 0.4808 | 42.6 |
| 9330198I05Rik | 60.3874 | -0.10068 | 0.24265 | -0.41492 | 0.6782 | 0.82377 | 3.699 |
| 9330198N18Rik | 14.1353 | 0.737673 | 0.48295 | 1.527425 | 0.12666 | 0.30249 | 59.556 |

| | | | | | | | |
|---|---|---|---|---|---|---|---|
| 9330199G10Rik | 21.8427 | -0.86905 | 0.4714 | -1.84356 | 0.06525 | 0.19707 | 41.521 |
| 9430015G10Rik | 1228.35 | 0.165369 | 0.07336 | 2.254095 | 0.02419 | 0.10158 | 17.284 |
| 9430021M05Rik | 352.003 | -0.38632 | 0.15897 | -2.4302 | 0.01509 | 0.07475 | 14.043 |
| 9430025C20Rik | 64.793 | 0.575823 | 0.29006 | 1.9852 | 0.04712 | 0.15929 | 2.117 |
| 9430034N14Rik | 32.4661 | -0.57221 | 0.35273 | -1.62223 | 0.10475 | 0.26777 | 3.785 |
| 9430037G07Rik | 98.877 | 0.262536 | 0.17808 | 1.474231 | 0.14042 | 0.3219 | 4.192 |
| 9430038I01Rik | 98.3055 | -0.38922 | 0.28851 | -1.34909 | 0.17731 | 0.36948 | 35.189 |
| 9430041J12Rik | 355.138 | 0.172389 | 0.11752 | 1.466881 | 0.14241 | 0.32434 | 46.604 |
| 9430053O09Rik | 24.6645 | -0.19926 | 0.37417 | -0.53255 | 0.59435 | 0.76649 | 2.78 |
| 9430060I03Rik | 18.4088 | 0.947149 | 0.55177 | 1.716552 | 0.08606 | 0.23563 | 17.065 |
| 9430064I24Rik | 21.9135 | 0.310304 | 0.40976 | 0.757281 | 0.44888 | 0.65217 | 2.669 |
| 9430065F17Rik | 23.3438 | -0.17967 | 0.44305 | -0.40553 | 0.68509 | 0.82843 | 10.976 |
| 9430081H08Rik | 23.4679 | -0.30167 | 0.44132 | -0.68357 | 0.49425 | 0.68845 | 0.83 |
| 9430085M18Rik | 35.8487 | -0.32316 | 0.46374 | -0.69684 | 0.4859 | 0.68204 | 2.897 |
| 9430091E24Rik | 263.57 | 0.284982 | 0.11773 | 2.420733 | 0.01549 | 0.07606 | 16.595 |
| 9430092D12Rik | 23.3237 | 0.2183 | 0.40775 | 0.535374 | 0.59239 | 0.76489 | 1.274 |
| 9530018F02Rik | 16.1316 | -0.15091 | 0.45582 | -0.33107 | 0.74059 | 0.86302 | 2.599 |
| 9530026P05Rik | 87.61 | -0.62232 | 0.25253 | -2.46431 | 0.01373 | 0.07049 | 332.661 |
| 9530036O11Rik | 21.0821 | -0.34361 | 0.46024 | -0.7466 | 0.4553 | 0.65729 | 475.722 |
| 9530046B11Rik | 72.423 | 0.534385 | 0.23043 | 2.319052 | 0.02039 | 0.09068 | 3.506 |
| 9530051G07Rik | 58.3418 | 0.55923 | 0.227 | 2.463548 | 0.01376 | 0.07056 | 35.287 |
| 9530052C20Rik | 17.1798 | 0.38309 | 0.42431 | 0.902858 | 0.3666 | 0.57697 | 10.427 |
| 9530052E02Rik | 96.0028 | 0.481698 | 0.18543 | 2.597793 | 0.00938 | 0.05502 | 46.692 |
| 9530053A07Rik | 17.4623 | 0.439585 | 0.55306 | 0.794819 | 0.42672 | 0.63327 | 35.346 |
| 9530056E24Rik | 27.262 | -0.01306 | 0.31876 | -0.04099 | 0.96731 | 0.98357 | 3.29 |
| 9530056K15Rik | 30.7973 | -0.57932 | 0.31322 | -1.84956 | 0.06438 | 0.19522 | 5.875 |
| 9530059O14Rik | 704.943 | 0.083366 | 0.12716 | 0.655622 | 0.51207 | 0.70326 | 107.098 |
| 9530062K07Rik | 83.2896 | 0.301157 | 0.31754 | 0.948406 | 0.34292 | 0.55368 | 8.586 |
| 9530068E07Rik | 3418.66 | 0.038494 | 0.07511 | 0.512538 | 0.60827 | 0.77621 | 12.301 |
| 9530077C05Rik | 397.296 | -0.06053 | 0.11768 | -0.51434 | 0.60701 | 0.77527 | 33.169 |
| 9530080O11Rik | 44.0378 | 0.5067 | 0.28769 | 1.761245 | 0.0782 | 0.22138 | 7.945 |
| 9530082P21Rik | 94.9672 | -0.05715 | 0.24386 | -0.23434 | 0.81472 | 0.90481 | 10.832 |
| 9530085L11Rik | 265.751 | 0.365958 | 0.22633 | 1.616934 | 0.10589 | 0.26954 | 2.115 |
| 9630001P10Rik | 201.864 | 0.549495 | 0.18196 | 3.019942 | 0.00253 | 0.02353 | 6.771 |

| | | | | | | | |
|---|---|---|---|---|---|---|---|
| 9630009A06Rik | 22.3138 | -0.44641 | 0.34986 | -1.27595 | 0.20197 | 0.40049 | 1.579 |
| 9630010A21Rik | 86.6174 | 0.019458 | 0.26653 | 0.073008 | 0.9418 | 0.97095 | 2.703 |
| 9630013A20Rik | 142.548 | -0.08766 | 0.19171 | -0.45725 | 0.64749 | 0.80377 | 17.786 |
| 9630013K17Rik | 73.132 | 0.054283 | 0.20212 | 0.268566 | 0.78826 | 0.89 | 3.815 |
| 9630014M24Rik | 38.3302 | -1.21554 | 0.39268 | -3.09547 | 0.00197 | 0.01989 | 187.512 |
| 9630015K15Rik | 67.9874 | 0.58582 | 0.24294 | 2.411365 | 0.01589 | 0.07735 | 11.292 |
| 9630028B13Rik | 19.9478 | -0.60757 | 0.39075 | -1.55487 | 0.11998 | 0.29247 | 12.465 |
| 9630041A04Rik | 29.7031 | 1.451968 | 0.43219 | 3.359581 | 0.00078 | 0.01076 | 81.388 |
| 9630050E16Rik | 12.6386 | -1.34207 | 0.69116 | -1.94177 | 0.05216 | 0.17057 | 7.441 |
| 9830144P21Rik | 369.78 | 0.272843 | 0.15289 | 1.784612 | 0.07432 | 0.21492 | 24.622 |
| 9830166K06Rik | 68.9052 | 0.292475 | 0.27946 | 1.046558 | 0.2953 | 0.50577 | 11.537 |
| 9930004E17Rik | 21.7112 | 0.003514 | 0.39513 | 0.008894 | 0.9929 | 0.99593 | 0.563 |
| 9930012K11Rik | 61.4408 | -0.29664 | 0.24725 | -1.19979 | 0.23022 | 0.43456 | 5.098 |
| 9930014A18Rik | 42.665 | 0.090869 | 0.3146 | 0.288836 | 0.77271 | 0.88166 | 9.523 |
| 9930017N22Rik | 128.473 | 0.597443 | 0.24206 | 2.468202 | 0.01358 | 0.07019 | 3.745 |
| 9930021J03Rik | 3380.84 | 0.255419 | 0.11344 | 2.251539 | 0.02435 | 0.10207 | 91.588 |
| 9930104L06Rik | 1267.35 | 0.046913 | 0.09882 | 0.474708 | 0.63499 | 0.79438 | 7.609 |
| 9930111J21Rik1 | 24.4581 | -0.11096 | 0.44432 | -0.24973 | 0.80279 | 0.89815 | 33.249 |
| 9930111J21Rik2 | 88.5855 | 0.281447 | 0.31475 | 0.894184 | 0.37122 | 0.58145 | 35.369 |
| 9930120I10Rik | 58.8942 | 0.13258 | 0.24652 | 0.537814 | 0.59071 | 0.76358 | 0.865 |
| A030001D20Rik | 237.721 | -0.41599 | 0.14316 | -2.9057 | 0.00366 | 0.02973 | 4.766 |
| A030005L19Rik | 24.276 | 0.205179 | 0.43723 | 0.469275 | 0.63887 | 0.79738 | 0.806 |
| A130006I12Rik | 26.6957 | 0.185477 | 0.38504 | 0.481708 | 0.63001 | 0.79082 | 0.416 |
| A130010J15Rik | 204.634 | -0.00292 | 0.15955 | -0.01828 | 0.98541 | 0.99203 | 9.787 |
| A130014A01Rik | 172.504 | -0.06522 | 0.17385 | -0.37518 | 0.70752 | 0.84212 | 2.493 |
| A230001M10Rik | 151.531 | -0.59336 | 0.20941 | -2.83341 | 0.00461 | 0.03446 | 182.735 |
| A230004M16Rik | 21.3125 | 0.201369 | 0.452 | 0.445511 | 0.65595 | 0.80893 | 262.776 |
| A230005M16Rik | 28.8523 | -0.46772 | 0.36554 | -1.27951 | 0.20072 | 0.39886 | 3.867 |
| A230006K03Rik | 819.155 | 0.34178 | 0.16155 | 2.115679 | 0.03437 | 0.1296 | 183.726 |
| A230009B12Rik | 61.4036 | -0.29728 | 0.25007 | -1.18881 | 0.23452 | 0.43973 | 36.842 |
| A230020J21Rik | 14.0173 | 0.558514 | 0.50839 | 1.098596 | 0.27194 | 0.48136 | 4.432 |
| A230028O05Rik | 18.3865 | -0.35793 | 0.37368 | -0.95786 | 0.33814 | 0.54899 | 9.686 |
| A230051N06Rik | 44.558 | 0.057299 | 0.26472 | 0.216446 | 0.82864 | 0.91212 | 1.116 |
| A230052G05Rik | 26.3364 | -1.17737 | 0.38487 | -3.05915 | 0.00222 | 0.02155 | 2.302 |

| | | | | | | | |
|---|---|---|---|---|---|---|---|
| A230056P14Rik | 1017.87 | 0.007169 | 0.1025 | 0.069942 | 0.94424 | 0.97202 | 18.358 |
| A230057D06Rik | 1851.65 | 0.620048 | 0.20529 | 3.020409 | 0.00252 | 0.02352 | 224.172 |
| A230072C01Rik | 601.287 | 0.154973 | 0.09039 | 1.714426 | 0.08645 | 0.23648 | 25.705 |
| A230072E10Rik | 76.7374 | 0.504093 | 0.26932 | 1.871697 | 0.06125 | 0.18912 | 52.873 |
| A230077H06Rik | 512.797 | -0.61241 | 0.24038 | -2.54764 | 0.01085 | 0.06059 | 82.462 |
| A230081H15Rik | 37.3834 | -0.48333 | 0.34394 | -1.40529 | 0.15994 | 0.34784 | 2.929 |
| A230083G16Rik | 146.795 | 0.362772 | 0.16047 | 2.260655 | 0.02378 | 0.10039 | 8.982 |
| A230085B16Rik | 16.8914 | -0.07996 | 0.4751 | -0.16829 | 0.86635 | 0.93232 | 1.856 |
| A230087F16Rik | 104.649 | 0.949724 | 0.24 | 3.957246 | 7.58E-05 | 0.00221 | 9.227 |
| A230103J11Rik | 124.83 | 0.664163 | 0.17052 | 3.894978 | 9.82E-05 | 0.00266 | 8.017 |
| A230103L15Rik | 459.646 | 0.31886 | 0.39218 | 0.813041 | 0.41619 | 0.62376 | 3.872 |
| A230107N01Rik | 93.9567 | 0.257188 | 0.18553 | 1.386265 | 0.16567 | 0.35498 | 2.595 |
| A2ml1 | 176.773 | -0.69352 | 0.1893 | -3.66362 | 0.00025 | 0.00496 | 41.788 |
| A330008L17Rik | 218.269 | 0.766046 | 0.1488 | 5.14799 | 2.63E-07 | 3.78E-05 | 316.797 |
| A330009N23Rik | 163.863 | -0.13699 | 0.16545 | -0.82799 | 0.40768 | 0.61541 | 4.242 |
| A330015K06Rik | 196.965 | -0.33873 | 0.15983 | -2.11938 | 0.03406 | 0.12885 | 169.63 |
| A330023F24Rik | 6052.86 | 0.154689 | 0.11961 | 1.293247 | 0.19593 | 0.39362 | 61.559 |
| A330032B11Rik | 22.9414 | 0.845958 | 0.38032 | 2.224339 | 0.02613 | 0.10726 | 22.698 |
| A330032P22Rik | 16.3499 | -0.98971 | 0.6081 | -1.62754 | 0.10362 | 0.26619 | 13.829 |
| A330033J07Rik | 14.7088 | 0.33948 | 0.49388 | 0.687375 | 0.49185 | 0.68654 | 218.393 |
| A330041J22Rik | 30.1777 | -0.22178 | 0.39958 | -0.55504 | 0.57887 | 0.75561 | 5.794 |
| A330048O09Rik | 65.9356 | -0.48086 | 0.26965 | -1.78323 | 0.07455 | 0.21518 | 1.505 |
| A330049N07Rik | 105.481 | -0.68136 | 0.24751 | -2.75288 | 0.00591 | 0.04085 | 113.407 |
| A330069E16Rik | 36.62 | 0.848159 | 0.29852 | 2.841187 | 0.00449 | 0.03392 | 17.026 |
| A330070K13Rik | 22.6724 | -0.46499 | 0.552 | -0.84238 | 0.39958 | 0.6084 | 5.854 |
| A330074H02Rik | 321.866 | 0.137501 | 0.13961 | 0.984867 | 0.32469 | 0.53586 | 1.039 |
| A330074K22Rik | 260.22 | -0.38541 | 0.14585 | -2.64248 | 0.00823 | 0.05049 | 23.797 |
| A330076C08Rik | 46.4006 | -0.08054 | 0.34197 | -0.23551 | 0.81382 | 0.90437 | 30.728 |
| A330076H08Rik | 677.581 | 0.480549 | 0.16015 | 3.000659 | 0.00269 | 0.02452 | 52.222 |
| A330084C13Rik | 33.092 | -0.34483 | 0.33888 | -1.01756 | 0.30889 | 0.51869 | 4.244 |
| A330093E20Rik | 67.4883 | -0.21645 | 0.22372 | -0.96749 | 0.3333 | 0.54421 | 361.774 |
| A330094K24Rik | 111.661 | 0.15583 | 0.29504 | 0.528162 | 0.59739 | 0.7692 | 3.299 |
| A330102I10Rik | 985.961 | 0.221933 | 0.09119 | 2.433839 | 0.01494 | 0.07415 | 27.399 |
| A3galt2 | 44.9756 | -0.04514 | 0.29307 | -0.15404 | 0.87758 | 0.93882 | 13.935 |

| | | | | | | | |
|---|---|---|---|---|---|---|---|
| A430005L14Rik | 371.909 | -0.12197 | 0.17336 | -0.70356 | 0.48171 | 0.67858 | 4.689 |
| A430010J10Rik | 22.8054 | 0.793306 | 0.40959 | 1.936822 | 0.05277 | 0.17188 | 52.754 |
| A430027C01Rik | 20.9813 | 0.335087 | 0.37416 | 0.895576 | 0.37048 | 0.58072 | 1.644 |
| A430027H14Rik | 44.5245 | 0.604119 | 0.33792 | 1.787762 | 0.07381 | 0.21412 | 1.467 |
| A430033K04Rik | 724.118 | 0.263705 | 0.11072 | 2.381724 | 0.01723 | 0.08153 | 29.556 |
| A430035B10Rik | 147.603 | 0.063362 | 0.18851 | 0.336114 | 0.73678 | 0.86096 | 21.636 |
| A430057M04Rik | 14.0208 | -0.64769 | 0.46175 | -1.40269 | 0.16071 | 0.34889 | 9.354 |
| A430073D23Rik | 27.5523 | 0.42375 | 0.33056 | 1.281905 | 0.19988 | 0.39801 | 98.641 |
| A430090L17Rik | 21.761 | 0.591526 | 0.43135 | 1.371334 | 0.17027 | 0.3606 | 27.245 |
| A430103D13Rik | 107.617 | -0.13809 | 0.21855 | -0.63186 | 0.52748 | 0.71566 | 0.857 |
| A430105J06Rik | 25.5419 | 0.537944 | 0.39331 | 1.367733 | 0.1714 | 0.3618 | 3.526 |
| A430108G06Rik | 82.1748 | 0.314792 | 0.19031 | 1.654077 | 0.09811 | 0.25712 | 2.626 |
| A430110C17Rik | 19.8073 | -0.49413 | 0.39961 | -1.23654 | 0.21626 | 0.41816 | 4.348 |
| A430110L20Rik | 144.795 | 0.382571 | 0.1666 | 2.296366 | 0.02165 | 0.09443 | 2.513 |
| A4galt | 99.5395 | -0.45954 | 0.23536 | -1.95249 | 0.05088 | 0.16778 | 25.053 |
| A530010L16Rik | 12.9445 | -0.0933 | 0.5043 | -0.18502 | 0.85322 | 0.92555 | 2.977 |
| A530016L24Rik | 42.3759 | -0.35021 | 0.34061 | -1.02817 | 0.30387 | 0.51369 | 10.48 |
| A530017D24Rik | 520.957 | 0.004383 | 0.11004 | 0.039828 | 0.96823 | 0.98402 | 3.491 |
| A530041M06Rik | 22.4504 | -0.23779 | 0.47777 | -0.49771 | 0.61869 | 0.78312 | 2.288 |
| A530058N18Rik | 110.111 | -0.30181 | 0.2281 | -1.32314 | 0.18579 | 0.38054 | 54.762 |
| A530058O07Rik | 14.924 | 0.214893 | 0.57858 | 0.371417 | 0.71033 | 0.84412 | 3.364 |
| A530064N14Rik | 18.4825 | 0.936189 | 0.61624 | 1.519194 | 0.12871 | 0.30583 | 1.492 |
| A530072M11Rik | 36.7309 | -0.24618 | 0.34212 | -0.71957 | 0.47179 | 0.67032 | 102.162 |
| A530083I20Rik | 29.8841 | 0.443629 | 0.3715 | 1.194152 | 0.23242 | 0.43733 | 18.452 |
| A630001G21Rik | 60.6276 | -0.17948 | 0.27319 | -0.65696 | 0.51121 | 0.70233 | 61.561 |
| A630023P12Rik | 18.0223 | -1.09639 | 0.54966 | -1.99466 | 0.04608 | 0.15668 | 11.118 |
| A630036G19Rik | 30.3768 | -0.27506 | 0.45902 | -0.59923 | 0.54902 | 0.73289 | 4.08 |
| A630052C17Rik | 286.603 | 0.251163 | 0.12837 | 1.956583 | 0.0504 | 0.16686 | 2.65 |
| A630066F11Rik | 24.5285 | 0.012403 | 0.40284 | 0.030789 | 0.97544 | 0.9876 | 1.31 |
| A630072L19Rik | 19.6876 | 0.341253 | 0.40816 | 0.836075 | 0.40311 | 0.6118 | 8.134 |
| A630089N07Rik | 113.895 | 0.541152 | 0.29609 | 1.827631 | 0.0676 | 0.20161 | 81.141 |
| A730009L09Rik | 26.2932 | -0.38263 | 0.45746 | -0.83643 | 0.40291 | 0.61163 | 8.768 |
| A730011C13Rik | 50.0275 | 0.114982 | 0.32721 | 0.351401 | 0.72529 | 0.85364 | 2.941 |
| A730017L22Rik | 471.089 | 0.19912 | 0.10308 | 1.931684 | 0.0534 | 0.1731 | 33.338 |

| | | | | | | | |
|---|---|---|---|---|---|---|---|
| A730018C14Rik | 53.309 | 0.939154 | 0.36426 | 2.578232 | 0.00993 | 0.05718 | 12.19 |
| A730020E08Rik | 173.512 | 0.181923 | 0.13803 | 1.317985 | 0.18751 | 0.38287 | 8.518 |
| A730020M07Rik | 56.022 | 0.052122 | 0.25152 | 0.207225 | 0.83583 | 0.91581 | 8.55 |
| A730035I17Rik | 16.1647 | -0.94854 | 0.63868 | -1.48516 | 0.1375 | 0.31822 | 2.864 |
| A730036I17Rik | 25.8185 | -0.40205 | 0.42587 | -0.94407 | 0.34514 | 0.55606 | 16.636 |
| A730043L09Rik | 13.7358 | -0.6967 | 0.6298 | -1.10621 | 0.26863 | 0.47791 | 1.505 |
| A730046J19Rik | 35.4309 | 1.649197 | 0.55109 | 2.992595 | 0.00277 | 0.02494 | 12.58 |
| A730056A06Rik | 109.555 | -0.21083 | 0.19114 | -1.10299 | 0.27003 | 0.4796 | 61.772 |
| A730060N03Rik | 116.828 | 0.235856 | 0.20593 | 1.145316 | 0.25208 | 0.4594 | 24.437 |
| A730061H03Rik | 20.6258 | 1.424234 | 0.44883 | 3.173219 | 0.00151 | 0.01664 | 1.312 |
| A730063M14Rik | 111.161 | 0.150124 | 0.22828 | 0.657642 | 0.51077 | 0.70199 | 4.422 |
| A730081D07Rik | 80.0098 | -0.13643 | 0.23524 | -0.57999 | 0.56192 | 0.74286 | 44.738 |
| A730085K08Rik | 28.3766 | 0.497572 | 0.36038 | 1.38069 | 0.16737 | 0.35707 | 18.227 |
| A730090N16Rik | 42.0043 | 0.764914 | 0.33762 | 2.265585 | 0.02348 | 0.09941 | 24.654 |
| A730091E23Rik | 50.9028 | -0.18988 | 0.31466 | -0.60345 | 0.54621 | 0.73071 | 3.825 |
| A830005F24Rik | 41.0145 | 0.062807 | 0.2777 | 0.22617 | 0.82107 | 0.90748 | 1.29 |
| A830008E24Rik | 56.3383 | -0.38888 | 0.29619 | -1.31296 | 0.1892 | 0.38511 | 80.355 |
| A830009L08Rik | 29.7667 | -1.14572 | 0.35991 | -3.18336 | 0.00146 | 0.01621 | 19.143 |
| A830011K09Rik | 115.987 | -0.10133 | 0.1641 | -0.61746 | 0.53693 | 0.72386 | 6.488 |
| A830018L16Rik | 4819.06 | 0.156132 | 0.10726 | 1.455662 | 0.14549 | 0.32834 | 561.797 |
| A830019P07Rik | 43.4295 | -0.98039 | 0.35909 | -2.73024 | 0.00633 | 0.04265 | 89.362 |
| A830031A19Rik | 46.0041 | -0.30269 | 0.31494 | -0.96111 | 0.3365 | 0.54716 | 26.1 |
| A830035O19Rik | 56.7337 | 0.003269 | 0.25107 | 0.013021 | 0.98961 | 0.99424 | 1.898 |
| A830036E02Rik | 3156.94 | 0.356894 | 0.17555 | 2.033018 | 0.04205 | 0.14748 | 8.542 |
| A830052D11Rik | 30.7464 | -0.29428 | 0.35632 | -0.82591 | 0.40886 | 0.61666 | 19.226 |
| A830073O21Rik | 560.634 | 0.570577 | 0.10873 | 5.247534 | 1.54E-07 | 2.51E-05 | 3.096 |
| A830082K12Rik | 431.791 | -0.99267 | 0.23306 | -4.25938 | 2.05E-05 | 0.0009 | 81.375 |
| A830082N09Rik | 1433.47 | 0.06358 | 0.09743 | 0.652566 | 0.51404 | 0.7047 | 7.84 |
| A830092H15Rik | 85.5201 | 0.647741 | 0.29077 | 2.227684 | 0.0259 | 0.10662 | 2.228 |
| A930001C03Rik | 39.8709 | 0.000582 | 0.29002 | 0.002006 | 0.9984 | 0.99913 | 9.971 |
| A930001M01Rik | 44.9857 | 0.075019 | 0.30368 | 0.247033 | 0.80488 | 0.89917 | 21.747 |
| A930002C04Rik | 64.4934 | 0.847706 | 0.27659 | 3.064901 | 0.00218 | 0.02131 | 4.729 |
| A930003O13Rik | 78.6042 | -0.10749 | 0.31171 | -0.34483 | 0.73022 | 0.85698 | 7.998 |
| A930004D18Rik | 80.877 | -0.35678 | 0.21045 | -1.6953 | 0.09002 | 0.24339 | 12.545 |

| | | | | | | | |
|---|---|---|---|---|---|---|---|
| A930004J17Rik | 802.913 | 0.219233 | 0.1434 | 1.528841 | 0.1263 | 0.30205 | 4.61 |
| A930005G22Rik | 37.6022 | -0.1289 | 0.30418 | -0.42375 | 0.67175 | 0.8194 | 1.904 |
| A930005H10Rik | 118.279 | -0.17312 | 0.1718 | -1.00771 | 0.31359 | 0.52413 | 6.569 |
| A930006K02Rik | 14.8005 | 0.261317 | 0.51314 | 0.509251 | 0.61058 | 0.77784 | 5.104 |
| A930007A09Rik | 74.4679 | -0.02072 | 0.21521 | -0.09626 | 0.92331 | 0.9614 | 3.972 |
| A930007I19Rik | 224.784 | 0.266009 | 0.14432 | 1.843159 | 0.06531 | 0.19714 | 58.423 |
| A930009A15Rik | 48.868 | 0.079184 | 0.29028 | 0.27279 | 0.78501 | 0.88852 | 12.463 |
| A930012L18Rik | 39.8375 | 0.03171 | 0.33681 | 0.094146 | 0.92499 | 0.96217 | 14.607 |
| A930015D03Rik | 657.518 | 0.188648 | 0.12515 | 1.507422 | 0.1317 | 0.31047 | 43.991 |
| A930016O22Rik | 16.6937 | 0.074019 | 0.48853 | 0.151515 | 0.87957 | 0.93973 | 4.917 |
| A930017K11Rik | 28.4919 | -1.03319 | 0.4176 | -2.4741 | 0.01336 | 0.06933 | 6.179 |
| A930017M01Rik | 523.62 | 0.169559 | 0.12696 | 1.335511 | 0.18171 | 0.37535 | 132.462 |
| A930019D19Rik | 16.7605 | 0.970818 | 0.43827 | 2.215116 | 0.02675 | 0.10907 | 7.875 |
| A930024E05Rik | 88.7729 | -0.10342 | 0.19765 | -0.52326 | 0.60079 | 0.7711 | 8.992 |
| A930027P06Rik | 41.4034 | 0.786079 | 0.32497 | 2.418941 | 0.01557 | 0.07631 | 2.975 |
| A930028N01Rik | 113.204 | 0.249172 | 0.2007 | 1.241526 | 0.21441 | 0.4158 | 1.585 |
| A930028O11Rik | 19.8855 | 0.990674 | 0.49543 | 1.999644 | 0.04554 | 0.15533 | 1.108 |
| A930029G22Rik | 23.8925 | -0.59312 | 0.40244 | -1.47383 | 0.14053 | 0.32203 | 22.645 |
| A930030B08Rik | 28.8947 | 0.58015 | 0.37047 | 1.566001 | 0.11735 | 0.28827 | 3.965 |
| A930033H14Rik | 58.6958 | -0.36668 | 0.3064 | -1.19674 | 0.23141 | 0.43599 | 3.337 |
| A930036I15Rik | 123.582 | -0.10441 | 0.23677 | -0.44099 | 0.65922 | 0.8112 | 1.045 |
| A930037H05Rik | 80.7917 | -0.08831 | 0.19697 | -0.44832 | 0.65392 | 0.8079 | 42.511 |
| A930040O22Rik | 44.8799 | 0.25363 | 0.26312 | 0.963948 | 0.33507 | 0.5458 | 0.973 |
| AA386476 | 35.7951 | 0.363529 | 0.36825 | 0.987178 | 0.32356 | 0.5346 | 9.849 |
| AA387200 | 24.8859 | 0.631002 | 0.3539 | 1.783014 | 0.07458 | 0.21525 | 1.954 |
| AA387883 | 195.053 | -0.38353 | 0.19813 | -1.93571 | 0.0529 | 0.1721 | 18.029 |
| AA388235 | 98.6916 | 0.009141 | 0.17037 | 0.053652 | 0.95721 | 0.97902 | 1.284 |
| AA413626 | 21.7468 | 0.129112 | 0.4036 | 0.319903 | 0.74904 | 0.86842 | 0.494 |
| AA414768 | 342.746 | -0.07006 | 0.13751 | -0.5095 | 0.6104 | 0.77775 | 1.257 |
| AA465934 | 130.554 | 0.02126 | 0.16393 | 0.129692 | 0.89681 | 0.94826 | 3.015 |
| AA474408 | 115.936 | 0.439607 | 0.24298 | 1.809243 | 0.07041 | 0.20732 | 3.339 |
| AA914427 | 36.8143 | 0.367429 | 0.3296 | 1.114757 | 0.26495 | 0.47401 | 1.177 |
| AA986860 | 156.908 | 0.210531 | 0.17709 | 1.188829 | 0.23451 | 0.43973 | 12.647 |
| AB041806 | 45.0913 | 0.305251 | 0.25149 | 1.213751 | 0.22484 | 0.42858 | 2.517 |

| | | | | | | | |
|---|---|---|---|---|---|---|---|
| AB124611 | 29.1987 | -0.06504 | 0.34431 | -0.18889 | 0.85018 | 0.92383 | 19.158 |
| AF357399 | 15.652 | 0.392885 | 0.42877 | 0.916303 | 0.35951 | 0.56984 | 0.125 |
| AI115009 | 21.0559 | -0.62922 | 0.43802 | -1.4365 | 0.15086 | 0.33575 | 14.52 |
| AI182371 | 23.1504 | 0.103267 | 0.40066 | 0.257743 | 0.79661 | 0.89517 | 19.683 |
| AI197445 | 167.335 | 0.886803 | 0.21817 | 4.064784 | 4.81E-05 | 0.0016 | 12.916 |
| AI413582 | 326.603 | -0.57388 | 0.1376 | -4.17049 | 3.04E-05 | 0.00117 | 2.184 |
| AI429214 | 100.54 | -0.05156 | 0.21307 | -0.24198 | 0.80879 | 0.90165 | 1.93 |
| AI467606 | 54.2792 | -0.55806 | 0.30607 | -1.82335 | 0.06825 | 0.20283 | 2.628 |
| AI480526 | 1073.13 | 0.187179 | 0.11443 | 1.635749 | 0.10189 | 0.26347 | 7.944 |
| AI504432 | 2662.99 | -0.03958 | 0.16132 | -0.24533 | 0.8062 | 0.90032 | 14.844 |
| AI506816 | 123.03 | -0.08999 | 0.22871 | -0.39346 | 0.69398 | 0.83408 | 14.391 |
| AI593442 | 6036.57 | 0.186523 | 0.10341 | 1.803677 | 0.07128 | 0.20897 | 6.739 |
| AI597479 | 2233.69 | 0.098154 | 0.07144 | 1.373973 | 0.16945 | 0.35966 | 17.238 |
| AI606473 | 92.4008 | 1.297335 | 0.41569 | 3.120898 | 0.0018 | 0.01883 | 3.434 |
| AI661453 | 29.8942 | -0.64265 | 0.38416 | -1.67285 | 0.09436 | 0.25065 | 34.024 |
| AI837181 | 1506.65 | -0.32802 | 0.0869 | -3.77481 | 0.00016 | 0.0037 | 2.17 |
| AI839979 | 90.0335 | 0.004068 | 0.2603 | 0.01563 | 0.98753 | 0.99316 | 14.652 |
| AI849053 | 50.1311 | -0.40173 | 0.23898 | -1.68101 | 0.09276 | 0.24807 | 5.494 |
| AI854703 | 1175.99 | -0.04651 | 0.07603 | -0.61172 | 0.54073 | 0.72648 | 6.926 |
| AI987944 | 411.295 | 0.155054 | 0.25501 | 0.608026 | 0.54317 | 0.72866 | 20.457 |
| AK157302 | 29.5895 | 0.101981 | 0.3105 | 0.328444 | 0.74258 | 0.86446 | 1.924 |
| AU020206 | 144.777 | -0.70365 | 0.23034 | -3.05477 | 0.00225 | 0.02176 | 13.084 |
| AU021092 | 80.19 | -0.1174 | 0.19657 | -0.59728 | 0.55032 | 0.73386 | 10.477 |
| AU022252 | 1962.23 | -0.04475 | 0.07624 | -0.58692 | 0.55726 | 0.73902 | 29.143 |
| AU022754 | 91.9476 | -0.8427 | 0.31072 | -2.71213 | 0.00669 | 0.04414 | 13.457 |
| AU023762 | 97.6274 | 0.040486 | 0.17854 | 0.226761 | 0.82061 | 0.90737 | 115.038 |
| AU040320 | 3632.2 | 0.095094 | 0.08494 | 1.119487 | 0.26293 | 0.47176 | 116.527 |
| AU040972 | 19.9311 | -0.16662 | 0.46244 | -0.36031 | 0.71862 | 0.84962 | 2.313 |
| AU041133 | 217.917 | 0.367045 | 0.26973 | 1.360767 | 0.17359 | 0.36493 | 25.053 |
| AV039307 | 73.6206 | 0.354759 | 0.23575 | 1.504785 | 0.13238 | 0.31143 | 16.064 |
| AV356131 | 79.1315 | 0.314987 | 0.2174 | 1.448883 | 0.14737 | 0.33065 | 0.785 |
| AW011738 | 775.207 | 0.241311 | 0.0957 | 2.521617 | 0.01168 | 0.06353 | 7.294 |
| AW046200 | 217.818 | 0.305877 | 0.14171 | 2.158442 | 0.03089 | 0.12052 | 11.713 |
| AW047730 | 195.102 | -0.57897 | 0.16206 | -3.57264 | 0.00035 | 0.00627 | 2.859 |

| | | | | | | | |
|---|---|---|---|---|---|---|---|
| AW112010 | 36.0034 | -0.78596 | 0.34645 | -2.26863 | 0.02329 | 0.09892 | 11.689 |
| AW146154 | 307.355 | 0.691376 | 0.25701 | 2.690065 | 0.00714 | 0.04613 | 21.017 |
| AW209491 | 822.478 | -0.04292 | 0.08162 | -0.52589 | 0.59896 | 0.76993 | 8.943 |
| AW495222 | 33.3103 | 0.0154 | 0.34623 | 0.04448 | 0.96452 | 0.98273 | 12.994 |
| AW549877 | 4799 | 0.059238 | 0.06208 | 0.954227 | 0.33997 | 0.55078 | 13.71 |
| AW551984 | 352.266 | 1.430528 | 0.47816 | 2.991737 | 0.00277 | 0.02495 | 17.008 |
| AW554918 | 729.565 | 0.167707 | 0.115 | 1.458335 | 0.14475 | 0.32747 | 298.323 |
| AW822252 | 639.377 | 0.823399 | 0.21298 | 3.866008 | 0.00011 | 0.00287 | 20.919 |
| AY036118 | 385.23 | -0.02294 | 0.16896 | -0.13577 | 0.89201 | 0.94613 | 0.725 |
| Aaas | 530.256 | -0.12965 | 0.09352 | -1.38637 | 0.16563 | 0.35495 | 12.52 |
| Aacs | 1252.75 | -0.23659 | 0.11534 | -2.05118 | 0.04025 | 0.14335 | 41.597 |
| Aagab | 2500.68 | -0.01525 | 0.09088 | -0.16781 | 0.86674 | 0.9324 | 41.929 |
| Aak1 | 26315.5 | 0.033621 | 0.05442 | 0.617841 | 0.53668 | 0.72362 | 153.707 |
| Aamdc | 741.03 | -0.05685 | 0.08345 | -0.68129 | 0.49569 | 0.68961 | 29.167 |
| Aamp | 3582.71 | -0.01704 | 0.08356 | -0.20393 | 0.83841 | 0.91681 | 4.899 |
| Aanat | 55.419 | 0.557059 | 0.29915 | 1.862129 | 0.06258 | 0.19198 | 11.249 |
| Aar2 | 1880.99 | 0.047144 | 0.0821 | 0.574234 | 0.56581 | 0.74583 | 21.389 |
| Aard | 132.727 | -0.42186 | 0.17869 | -2.36082 | 0.01823 | 0.08438 | 5.616 |
| Aars | 6639.65 | -0.16513 | 0.06026 | -2.74011 | 0.00614 | 0.04191 | 24.521 |
| Aars2 | 1357.27 | -0.11307 | 0.09647 | -1.17203 | 0.24118 | 0.44734 | 14.003 |
| Aarsd1 | 59.8911 | 0.338741 | 0.24198 | 1.399887 | 0.16155 | 0.34979 | 10.777 |
| Aasdh | 1249.2 | 0.337252 | 0.09776 | 3.44995 | 0.00056 | 0.00857 | 31.856 |
| Aasdhppt | 1512.98 | 0.176049 | 0.08524 | 2.065343 | 0.03889 | 0.14058 | 14.679 |
| Aass | 372.174 | -0.24029 | 0.13248 | -1.81379 | 0.06971 | 0.20594 | 60.814 |
| Aatf | 551.067 | -0.0875 | 0.10807 | -0.80967 | 0.41813 | 0.62541 | 90.668 |
| Aatk | 11771.5 | -0.18188 | 0.08218 | -2.21306 | 0.02689 | 0.10946 | 39.855 |
| Abat | 25955.2 | 0.343553 | 0.14233 | 2.41372 | 0.01579 | 0.0771 | 108.14 |
| Abca1 | 2997.53 | -0.05713 | 0.12547 | -0.45536 | 0.64885 | 0.80429 | 129.109 |
| Abca17 | 68.7427 | 0.444177 | 0.24402 | 1.820277 | 0.06872 | 0.20383 | 86.81 |
| Abca2 | 16100.6 | -0.15445 | 0.07552 | -2.04523 | 0.04083 | 0.14468 | 19.838 |
| Abca3 | 3828.51 | 0.070233 | 0.06358 | 1.1046 | 0.26933 | 0.47881 | 58.252 |
| Abca4 | 405 | -0.45201 | 0.14172 | -3.18943 | 0.00143 | 0.016 | 135.681 |
| Abca5 | 3166.56 | -0.00448 | 0.07555 | -0.05935 | 0.95267 | 0.97677 | 68.348 |
| Abca6 | 115.974 | -0.47581 | 0.17446 | -2.72734 | 0.00638 | 0.04287 | 74.957 |

| | | | | | | | |
|---|---|---|---|---|---|---|---|
| Abca7 | 1405.18 | -0.47501 | 0.15989 | -2.97089 | 0.00297 | 0.026 | 19.079 |
| Abca8a | 890.791 | -0.29898 | 0.12851 | -2.32652 | 0.01999 | 0.08956 | 70.345 |
| Abca8b | 1787.99 | 0.300351 | 0.09334 | 3.217976 | 0.00129 | 0.01492 | 63.656 |
| Abca9 | 888.991 | -0.5332 | 0.13375 | -3.98659 | 6.70E-05 | 0.00204 | 67.448 |
| Abcb10 | 1464.72 | 0.07524 | 0.075 | 1.003179 | 0.31577 | 0.52635 | 30.664 |
| Abcb1a | 1826.68 | -0.50401 | 0.06922 | -7.28094 | 3.31E-13 | 1.35E-09 | 88.499 |
| Abcb1b | 151.289 | -0.48625 | 0.21815 | -2.22894 | 0.02582 | 0.10638 | 68.169 |
| Abcb4 | 498.308 | -0.21199 | 0.1011 | -2.09685 | 0.03601 | 0.13358 | 65.515 |
| Abcb6 | 44.9989 | 0.233291 | 0.28718 | 0.812354 | 0.41659 | 0.62421 | 8.676 |
| Abcb7 | 3059.34 | 0.202969 | 0.08747 | 2.320441 | 0.02032 | 0.09037 | 133.2 |
| Abcb8 | 1902.42 | 0.190088 | 0.07528 | 2.525096 | 0.01157 | 0.06306 | 16.392 |
| Abcb9 | 2303.54 | 0.060619 | 0.09935 | 0.610177 | 0.54174 | 0.72732 | 34.269 |
| Abcc1 | 1375.17 | 0.002229 | 0.0895 | 0.024906 | 0.98013 | 0.98945 | 114.18 |
| Abcc10 | 1225.01 | 0.376833 | 0.09928 | 3.795659 | 0.00015 | 0.00349 | 25.132 |
| Abcc12 | 188.322 | -0.19217 | 0.31042 | -0.61906 | 0.53588 | 0.72278 | 98.427 |
| Abcc3 | 111.939 | -0.37011 | 0.19709 | -1.87785 | 0.0604 | 0.18755 | 49.703 |
| Abcc4 | 873.165 | -0.40322 | 0.13157 | -3.06458 | 0.00218 | 0.02131 | 223.528 |
| Abcc5 | 16630.3 | 0.110737 | 0.07206 | 1.536748 | 0.12436 | 0.2992 | 95.092 |
| Abcc6 | 50.5051 | -0.06823 | 0.30271 | -0.22539 | 0.82167 | 0.90766 | 62.748 |
| Abcc8 | 1262.02 | 0.202263 | 0.11821 | 1.710999 | 0.08708 | 0.23761 | 75.511 |
| Abcc9 | 590.111 | -0.35665 | 0.11546 | -3.08891 | 0.00201 | 0.02016 | 114.454 |
| Abcd1 | 779.89 | -0.02849 | 0.09167 | -0.31082 | 0.75594 | 0.87157 | 21.938 |
| Abcd2 | 2073.01 | -0.10752 | 0.11688 | -0.91988 | 0.35764 | 0.56794 | 45.929 |
| Abcd3 | 4030.83 | -0.01786 | 0.0587 | -0.3043 | 0.7609 | 0.8747 | 56.529 |
| Abcd4 | 1056.84 | 0.067555 | 0.12263 | 0.550897 | 0.5817 | 0.75754 | 15.95 |
| Abce1 | 4357.74 | 0.136353 | 0.0818 | 1.666848 | 0.09554 | 0.25277 | 28.279 |
| Abcf1 | 4475.71 | -0.01703 | 0.05473 | -0.31115 | 0.75568 | 0.87155 | 12.943 |
| Abcf2 | 4013.8 | 0.015337 | 0.06061 | 0.253024 | 0.80025 | 0.89682 | 12.123 |
| Abcf3 | 2988.62 | 0.002143 | 0.07129 | 0.030067 | 0.97601 | 0.98762 | 12.803 |
| Abcg1 | 4110.28 | 0.177951 | 0.09407 | 1.891696 | 0.05853 | 0.18401 | 60.314 |
| Abcg2 | 793.952 | -0.33458 | 0.11012 | -3.03828 | 0.00238 | 0.02255 | 111.154 |
| Abcg4 | 2992.86 | 0.068311 | 0.06149 | 1.110899 | 0.26661 | 0.47589 | 15.428 |
| Abhd1 | 16.5512 | 0.534417 | 0.47008 | 1.136874 | 0.25559 | 0.46331 | 5.026 |
| Abhd10 | 816.167 | 0.117691 | 0.09093 | 1.294359 | 0.19554 | 0.39318 | 13.231 |

| | | | | | | | |
|---|---|---|---|---|---|---|---|
| Abhd11 | 269.654 | -0.28417 | 0.12968 | -2.19138 | 0.02842 | 0.1137 | 3.024 |
| Abhd11os | 138.607 | -0.04726 | 0.19092 | -0.24754 | 0.80449 | 0.89905 | 1.259 |
| Abhd12 | 10008.4 | -0.13352 | 0.06965 | -1.91702 | 0.05523 | 0.17717 | 72.249 |
| Abhd13 | 2283.86 | 0.042779 | 0.08351 | 0.512246 | 0.60848 | 0.77621 | 14.449 |
| Abhd14a | 1135.46 | 0.279192 | 0.08231 | 3.392028 | 0.00069 | 0.00993 | 7.628 |
| Abhd14b | 380.025 | 0.09729 | 0.15854 | 0.613675 | 0.53943 | 0.72589 | 5.465 |
| Abhd15 | 46.9643 | -0.08967 | 0.2735 | -0.32788 | 0.74301 | 0.86466 | 23.487 |
| Abhd16a | 2729.25 | -0.12329 | 0.06126 | -2.01273 | 0.04414 | 0.15229 | 13.725 |
| Abhd17a | 3587.46 | -0.23989 | 0.09602 | -2.49838 | 0.01248 | 0.06631 | 6.688 |
| Abhd17b | 1812.36 | 0.111049 | 0.07958 | 1.395436 | 0.16288 | 0.35138 | 32.453 |
| Abhd17c | 1507.38 | 0.213064 | 0.093 | 2.290917 | 0.02197 | 0.09543 | 42.538 |
| Abhd18 | 1222.71 | 0.125011 | 0.11183 | 1.117826 | 0.26364 | 0.47241 | 91.169 |
| Abhd2 | 7789.76 | 0.085145 | 0.07674 | 1.109521 | 0.26721 | 0.47653 | 92.31 |
| Abhd3 | 2219.66 | -0.15539 | 0.09157 | -1.6969 | 0.08971 | 0.2428 | 64.68 |
| Abhd4 | 2747.81 | -0.30201 | 0.13194 | -2.28897 | 0.02208 | 0.09575 | 16.45 |
| Abhd5 | 575.09 | -0.04309 | 0.09091 | -0.47395 | 0.63553 | 0.79462 | 29.917 |
| Abhd6 | 2103.11 | -0.0291 | 0.07277 | -0.39986 | 0.68926 | 0.83062 | 53.862 |
| Abhd8 | 7755.47 | -0.06517 | 0.07445 | -0.87534 | 0.38139 | 0.59084 | 6.951 |
| Abi1 | 6299.26 | 0.058743 | 0.07279 | 0.806974 | 0.41968 | 0.62696 | 100.169 |
| Abi2 | 10517.8 | 0.177165 | 0.05381 | 3.29242 | 0.00099 | 0.01277 | 71.54 |
| Abi3 | 22.4719 | -0.45448 | 0.36853 | -1.23325 | 0.21748 | 0.41946 | 12.403 |
| Abi3bp | 3131.39 | -1.42524 | 0.62143 | -2.29347 | 0.02182 | 0.095 | 212.258 |
| Abitram | 1455.06 | 0.162727 | 0.07074 | 2.300402 | 0.02143 | 0.09376 | 7.257 |
| Abl1 | 3612.78 | -0.05714 | 0.0637 | -0.89695 | 0.36975 | 0.58022 | 115.852 |
| Abl2 | 7674.72 | 0.147119 | 0.08324 | 1.767345 | 0.07717 | 0.21961 | 90.783 |
| Ablim1 | 4940.33 | -0.29278 | 0.09977 | -2.93445 | 0.00334 | 0.02791 | 282.187 |
| Ablim2 | 8713.49 | 0.529392 | 0.17847 | 2.966352 | 0.00301 | 0.02623 | 127.094 |
| Ablim3 | 1241.93 | 0.182152 | 0.17441 | 1.044376 | 0.29631 | 0.50667 | 112.433 |
| Abr | 22804.9 | -0.09474 | 0.0703 | -1.3477 | 0.17776 | 0.37003 | 206.825 |
| Abracl | 183.661 | 0.025433 | 0.15214 | 0.167167 | 0.86724 | 0.93259 | 12.029 |
| Abraxas1 | 316.291 | 0.144472 | 0.13641 | 1.059123 | 0.28954 | 0.49978 | 15.774 |
| Abraxas2 | 1215.47 | 0.164269 | 0.06647 | 2.471495 | 0.01345 | 0.0697 | 25.887 |
| Abt1 | 659.368 | -0.07666 | 0.09931 | -0.77191 | 0.44017 | 0.64519 | 5.506 |
| Abtb1 | 1469.56 | 0.075795 | 0.09082 | 0.834555 | 0.40397 | 0.61269 | 6.071 |

| | | | | | | | |
|---|---|---|---|---|---|---|---|
| Abtb2 | 582.752 | -0.20447 | 0.13399 | -1.52601 | 0.12701 | 0.30318 | 152.114 |
| Acaa1a | 3155.02 | -0.06696 | 0.08733 | -0.76673 | 0.44324 | 0.64777 | 10.624 |
| Acaa1b | 21.5133 | -0.01202 | 0.39145 | -0.03071 | 0.9755 | 0.98761 | 9.071 |
| Acaa2 | 1034.11 | -0.15499 | 0.10401 | -1.49017 | 0.13618 | 0.31645 | 27.012 |
| Acaca | 3928.87 | -0.24452 | 0.08733 | -2.7999 | 0.00511 | 0.03683 | 271.98 |
| Acacb | 601.255 | -0.40618 | 0.15024 | -2.70356 | 0.00686 | 0.04485 | 104.227 |
| Acad10 | 308.208 | 0.016442 | 0.13404 | 0.122668 | 0.90237 | 0.95108 | 39.489 |
| Acad11 | 722.321 | 0.010385 | 0.11058 | 0.093916 | 0.92518 | 0.96225 | 64.349 |
| Acad12 | 182.654 | -0.19985 | 0.13769 | -1.45145 | 0.14665 | 0.32981 | 22.164 |
| Acad8 | 1414.11 | -0.00416 | 0.07633 | -0.05453 | 0.95651 | 0.97874 | 25.432 |
| Acad9 | 1289.07 | -0.0598 | 0.07479 | -0.79957 | 0.42396 | 0.63068 | 26.875 |
| Acadl | 1810.76 | -0.19583 | 0.08567 | -2.28584 | 0.02226 | 0.09619 | 32.439 |
| Acadm | 2109.56 | 0.001483 | 0.06626 | 0.022386 | 0.98214 | 0.99059 | 22.276 |
| Acads | 332.353 | 0.016728 | 0.14735 | 0.11352 | 0.90962 | 0.95456 | 9.048 |
| Acadsb | 3313.3 | 0.111352 | 0.06263 | 1.777936 | 0.07541 | 0.21667 | 38.344 |
| Acadvl | 1072.07 | -0.09495 | 0.0779 | -1.21877 | 0.22293 | 0.4261 | 5.229 |
| Acan | 258.392 | -0.22433 | 0.15692 | -1.42953 | 0.15285 | 0.33835 | 61.617 |
| Acap2 | 10530.4 | 0.089617 | 0.06542 | 1.369942 | 0.1707 | 0.36103 | 108.834 |
| Acap3 | 4828.22 | -0.01332 | 0.07989 | -0.16672 | 0.86759 | 0.9329 | 15.43 |
| Acat1 | 4186.41 | -0.01719 | 0.06594 | -0.26076 | 0.79428 | 0.89351 | 29.861 |
| Acat2 | 1620.19 | -0.02611 | 0.09265 | -0.28178 | 0.77811 | 0.88423 | 17.858 |
| Acat3 | 166.605 | -0.22641 | 0.20538 | -1.1024 | 0.27029 | 0.47985 | 16.788 |
| Acbd3 | 2267.6 | 0.067797 | 0.06672 | 1.016074 | 0.30959 | 0.51958 | 28.162 |
| Acbd4 | 1216.83 | -0.09606 | 0.07301 | -1.31564 | 0.1883 | 0.38384 | 10.519 |
| Acbd5 | 4830.04 | 0.085608 | 0.09287 | 0.921782 | 0.35664 | 0.56705 | 47.392 |
| Acbd6 | 1721.03 | -0.06001 | 0.07782 | -0.77119 | 0.44059 | 0.64547 | 133.211 |
| Accs | 232.396 | -0.09377 | 0.15314 | -0.61231 | 0.54033 | 0.72614 | 16.477 |
| Accsl | 15.5856 | -0.37097 | 0.55709 | -0.6659 | 0.50547 | 0.69703 | 13.797 |
| Acd | 928.779 | 0.055004 | 0.08901 | 0.617984 | 0.53659 | 0.72354 | 5.243 |
| Ace | 399.656 | -0.40131 | 0.17929 | -2.2384 | 0.0252 | 0.10447 | 22.02 |
| Ace2 | 102.458 | -0.20809 | 0.1687 | -1.23352 | 0.21738 | 0.4194 | 49.089 |
| Acer2 | 644.617 | -0.29073 | 0.14127 | -2.05804 | 0.03959 | 0.14209 | 60.427 |
| Acer3 | 1257.89 | -0.06338 | 0.07186 | -0.88204 | 0.37776 | 0.58784 | 114.82 |
| Ache | 3775.76 | 0.3349 | 0.11675 | 2.868565 | 0.00412 | 0.03208 | 6.948 |

| | | | | | | | |
|---|---|---|---|---|---|---|---|
| Acin1 | 8009.25 | -0.03533 | 0.07 | -0.5047 | 0.61377 | 0.78 | 44.771 |
| Ackr1 | 1134.8 | -0.1247 | 0.1019 | -1.22371 | 0.22106 | 0.42394 | 1.865 |
| Ackr2 | 108.687 | 0.713524 | 0.23211 | 3.07407 | 0.00211 | 0.02088 | 12.72 |
| Ackr3 | 422.936 | -0.15853 | 0.15229 | -1.041 | 0.29788 | 0.50794 | 12.772 |
| Ackr4 | 24.0963 | 0.05625 | 0.3814 | 0.147482 | 0.88275 | 0.9416 | 42.777 |
| Acly | 10317.5 | 0.138567 | 0.07927 | 1.747986 | 0.08047 | 0.22574 | 51.648 |
| Acmsd | 24.913 | 0.770245 | 0.41379 | 1.861444 | 0.06268 | 0.19214 | 38.566 |
| Aco1 | 1357.16 | -0.08513 | 0.07605 | -1.11933 | 0.263 | 0.47183 | 55.258 |
| Aco2 | 16993.8 | -0.09202 | 0.06191 | -1.48619 | 0.13723 | 0.31787 | 42.825 |
| Acot1 | 294.929 | -0.61698 | 0.1218 | -5.06537 | 4.08E-07 | 5.47E-05 | 8.882 |
| Acot11 | 1715.2 | -0.37422 | 0.09037 | -4.14078 | 3.46E-05 | 0.00126 | 60.444 |
| Acot12 | 50.9641 | -0.18504 | 0.29256 | -0.63249 | 0.52706 | 0.71515 | 44.637 |
| Acot13 | 1436.41 | -0.19111 | 0.0799 | -2.39186 | 0.01676 | 0.08012 | 13.593 |
| Acot2 | 212.476 | -0.35287 | 0.22312 | -1.58154 | 0.11375 | 0.28233 | 6.013 |
| Acot3 | 43.8422 | -1.06188 | 0.29287 | -3.62577 | 0.00029 | 0.00547 | 8.259 |
| Acot4 | 179.455 | -0.31716 | 0.21759 | -1.4576 | 0.14495 | 0.32781 | 10.223 |
| Acot5 | 47.1238 | -1.41053 | 0.40003 | -3.52605 | 0.00042 | 0.00714 | 6.696 |
| Acot6 | 357.787 | -0.0652 | 0.11168 | -0.58378 | 0.55937 | 0.74078 | 10.696 |
| Acot7 | 6623.35 | -0.29918 | 0.0965 | -3.10033 | 0.00193 | 0.01969 | 93.722 |
| Acot8 | 535.882 | -0.06553 | 0.10029 | -0.6534 | 0.5135 | 0.70439 | 12.118 |
| Acot9 | 551.41 | -0.19951 | 0.13199 | -1.51155 | 0.13065 | 0.30899 | 35.212 |
| Acox1 | 11081 | -0.02818 | 0.07256 | -0.38831 | 0.69778 | 0.83624 | 27.158 |
| Acox3 | 1237.81 | -0.22026 | 0.08146 | -2.70388 | 0.00685 | 0.04485 | 32.313 |
| Acp1 | 1937.63 | -0.07991 | 0.0668 | -1.19623 | 0.2316 | 0.43628 | 18.264 |
| Acp2 | 2904.86 | -0.03285 | 0.06047 | -0.54321 | 0.58699 | 0.76058 | 11.214 |
| Acp6 | 1335.31 | 0.255927 | 0.09433 | 2.713039 | 0.00667 | 0.04408 | 18.523 |
| Acp7 | 104.04 | -1.27295 | 0.41547 | -3.06389 | 0.00218 | 0.02131 | 24.057 |
| Acr | 90.0592 | 0.207662 | 0.22484 | 0.923596 | 0.3557 | 0.56614 | 6.26 |
| Acrbp | 522.965 | 0.36807 | 0.13428 | 2.741124 | 0.00612 | 0.04184 | 13.579 |
| Acsbg1 | 6996.7 | -0.31537 | 0.08987 | -3.50932 | 0.00045 | 0.00739 | 56.994 |
| Acsbg3 | 54.3207 | 0.126804 | 0.37489 | 0.338243 | 0.73518 | 0.85989 | 13.428 |
| Acsf2 | 774.908 | -0.24691 | 0.13523 | -1.82577 | 0.06788 | 0.20204 | 44.77 |
| Acsf3 | 1003.05 | 0.265702 | 0.10343 | 2.56892 | 0.0102 | 0.05832 | 42.395 |
| Acsl1 | 2566.8 | -0.09514 | 0.06985 | -1.3621 | 0.17317 | 0.3644 | 65.015 |

| | | | | | | | |
|---|---|---|---|---|---|---|---|
| Acsl3 | 10329.1 | -0.18131 | 0.0808 | -2.24407 | 0.02483 | 0.10351 | 49.919 |
| Acsl4 | 4044.42 | -0.12994 | 0.13233 | -0.98192 | 0.32614 | 0.53752 | 72.543 |
| Acsl5 | 1263.7 | -0.27842 | 0.111 | -2.50821 | 0.01213 | 0.06506 | 45.783 |
| Acsl6 | 10102.4 | -0.16123 | 0.07571 | -2.12949 | 0.03321 | 0.12659 | 60.959 |
| Acsm3 | 28.009 | -0.60577 | 0.36581 | -1.65596 | 0.09773 | 0.2564 | 26.591 |
| Acsm5 | 88.0642 | 0.046403 | 0.19424 | 0.238898 | 0.81118 | 0.90277 | 26.089 |
| Acss1 | 1299.25 | -0.22439 | 0.0954 | -2.35208 | 0.01867 | 0.08566 | 50.396 |
| Acss2 | 1721.73 | -0.21488 | 0.10089 | -2.12981 | 0.03319 | 0.12657 | 67.777 |
| Acss2os | 24.6 | -0.05931 | 0.36991 | -0.16035 | 0.87261 | 0.93573 | 9.891 |
| Acss3 | 66.8932 | -0.2923 | 0.45203 | -0.64663 | 0.51787 | 0.70758 | 190.152 |
| Acta1 | 162.334 | -0.19815 | 0.16323 | -1.21392 | 0.22478 | 0.42853 | 2.983 |
| Acta2 | 245.008 | -0.53018 | 0.24013 | -2.20784 | 0.02726 | 0.1103 | 14.501 |
| Actb | 30219.4 | -0.04814 | 0.08081 | -0.59571 | 0.55137 | 0.73445 | 3.64 |
| Actg1 | 17682.8 | -0.04073 | 0.09477 | -0.42979 | 0.66735 | 0.8162 | 2.853 |
| Actl10 | 159.361 | -0.6285 | 0.2375 | -2.64627 | 0.00814 | 0.05009 | 1.501 |
| Actl6a | 562.22 | 0.041107 | 0.10064 | 0.408468 | 0.68293 | 0.8267 | 20.676 |
| Actl6b | 4181.6 | 0.169717 | 0.08518 | 1.992438 | 0.04632 | 0.1573 | 16.066 |
| Actn1 | 5845.34 | 0.359101 | 0.11223 | 3.199824 | 0.00138 | 0.01557 | 92.825 |
| Actn2 | 3061.14 | 0.474797 | 0.16718 | 2.840075 | 0.00451 | 0.034 | 71.33 |
| Actn3 | 49.4004 | -0.85763 | 0.26303 | -3.26063 | 0.00111 | 0.0137 | 16.694 |
| Actn4 | 5405.31 | -0.17343 | 0.07407 | -2.34155 | 0.0192 | 0.08705 | 69.093 |
| Actr10 | 5097.68 | -0.0333 | 0.06241 | -0.53361 | 0.59361 | 0.76589 | 26.862 |
| Actr1a | 7762.83 | -0.07907 | 0.07267 | -1.08805 | 0.27657 | 0.48588 | 18.938 |
| Actr1b | 10521.2 | -0.11347 | 0.05516 | -2.05704 | 0.03968 | 0.14226 | 16.309 |
| Actr2 | 17430.8 | 0.092892 | 0.06709 | 1.384507 | 0.1662 | 0.3559 | 50.61 |
| Actr3 | 10304.7 | 0.10694 | 0.07074 | 1.511746 | 0.1306 | 0.30894 | 42.823 |
| Actr3-ps | 192.795 | 0.206332 | 0.13476 | 1.531111 | 0.12574 | 0.30098 | 2.347 |
| Actr3b | 4325.06 | 0.063922 | 0.06148 | 1.039722 | 0.29847 | 0.50856 | 90.692 |
| Actr5 | 625.699 | 0.260279 | 0.09618 | 2.706172 | 0.00681 | 0.04467 | 14.324 |
| Actr6 | 898.715 | 0.134586 | 0.09965 | 1.350633 | 0.17681 | 0.36894 | 20.325 |
| Actr8 | 2742.38 | 0.125835 | 0.06662 | 1.888876 | 0.05891 | 0.18475 | 28.018 |
| Actrt3 | 30.7347 | -0.37969 | 0.32111 | -1.18241 | 0.23704 | 0.44215 | 2.867 |
| Acvr1 | 1032.64 | -0.23348 | 0.11309 | -2.06448 | 0.03897 | 0.14064 | 178.514 |
| Acvr1b | 7097.07 | -0.06155 | 0.05649 | -1.08953 | 0.27592 | 0.48511 | 39.618 |

| | | | | | | | |
|---|---|---|---|---|---|---|---|
| Acvr1c | 1421.55 | 0.148482 | 0.29597 | 0.501684 | 0.61589 | 0.78132 | 90.443 |
| Acvr2a | 4280.18 | 0.506947 | 0.1081 | 4.689486 | 2.74E-06 | 0.00022 | 89.161 |
| Acvr2b | 631.424 | 0.308195 | 0.09872 | 3.121909 | 0.0018 | 0.01879 | 32.878 |
| Acvrl1 | 668.035 | -0.29849 | 0.12971 | -2.30122 | 0.02138 | 0.09364 | 16.815 |
| Acy1 | 742.183 | 0.428198 | 0.18314 | 2.338074 | 0.01938 | 0.08767 | 5.339 |
| Acy3 | 75.0421 | -1.04436 | 0.27597 | -3.78432 | 0.00015 | 0.00361 | 3.438 |
| Acyp1 | 431.769 | -0.24356 | 0.10968 | -2.22062 | 0.02638 | 0.10794 | 16.041 |
| Acyp2 | 583.205 | 0.043636 | 0.09845 | 0.443232 | 0.6576 | 0.81004 | 143.597 |
| Ada | 110.491 | -0.11272 | 0.25812 | -0.43669 | 0.66234 | 0.81342 | 23.656 |
| Adal | 1039.52 | 0.126162 | 0.08591 | 1.468508 | 0.14197 | 0.3238 | 16.253 |
| Adam10 | 4179.67 | 0.086331 | 0.0664 | 1.300193 | 0.19353 | 0.39052 | 101.233 |
| Adam11 | 4843.44 | -0.28759 | 0.10486 | -2.74247 | 0.0061 | 0.04175 | 18.824 |
| Adam12 | 779.66 | 0.668777 | 0.17401 | 3.843311 | 0.00012 | 0.00305 | 348.948 |
| Adam15 | 2543.79 | -0.07273 | 0.06201 | -1.17295 | 0.24082 | 0.44705 | 11.455 |
| Adam17 | 1226.62 | -0.10852 | 0.07836 | -1.38486 | 0.1661 | 0.35575 | 50.124 |
| Adam18 | 32.3822 | -0.43714 | 0.37699 | -1.15956 | 0.24623 | 0.45299 | 72.51 |
| Adam19 | 560.48 | -0.32927 | 0.14583 | -2.25786 | 0.02395 | 0.10083 | 91.352 |
| Adam1a | 517.597 | 0.460734 | 0.10517 | 4.381057 | 1.18E-05 | 0.00063 | 3.496 |
| Adam1b | 37.1375 | 0.437473 | 0.31366 | 1.394742 | 0.16309 | 0.35165 | 3.338 |
| Adam21 | 66.9008 | 0.516517 | 0.26541 | 1.946081 | 0.05165 | 0.16944 | 9.891 |
| Adam22 | 12982 | 0.070293 | 0.07021 | 1.00124 | 0.31671 | 0.52722 | 295.809 |
| Adam23 | 11357.8 | 0.024534 | 0.07583 | 0.323556 | 0.74627 | 0.86683 | 150.386 |
| Adam32 | 16.4679 | 0.09293 | 0.49003 | 0.189641 | 0.84959 | 0.92352 | 112.665 |
| Adam33 | 84.1928 | -1.36407 | 0.33572 | -4.06314 | 4.84E-05 | 0.0016 | 13.224 |
| Adam4 | 47.1934 | 0.212863 | 0.29407 | 0.723861 | 0.46915 | 0.66805 | 2.504 |
| Adam7 | 50.1948 | 0.64583 | 0.30277 | 2.133049 | 0.03292 | 0.12609 | 36.406 |
| Adam8 | 334.442 | -0.29228 | 0.20061 | -1.45697 | 0.14513 | 0.32787 | 13.631 |
| Adam9 | 3155.69 | -0.0662 | 0.07629 | -0.86767 | 0.38557 | 0.59501 | 67.317 |
| Adamts1 | 465.569 | -0.62971 | 0.17388 | -3.62154 | 0.00029 | 0.00553 | 9.287 |
| Adamts10 | 1369.61 | -0.33361 | 0.12517 | -2.66525 | 0.00769 | 0.04845 | 29.579 |
| Adamts13 | 14.9821 | -1.73355 | 0.54595 | -3.1753 | 0.0015 | 0.01655 | 36.213 |
| Adamts14 | 43.3797 | -1.25676 | 0.30537 | -4.11553 | 3.86E-05 | 0.00137 | 76.327 |
| Adamts15 | 226.644 | -0.75721 | 0.2009 | -3.76909 | 0.00016 | 0.00377 | 23.298 |
| Adamts16 | 161.006 | 0.334428 | 0.22051 | 1.516607 | 0.12937 | 0.30688 | 114.01 |

| | | | | | | | |
|---|---|---|---|---|---|---|---|
| Adamts17 | 251.727 | 0.110952 | 0.13355 | 0.830803 | 0.40608 | 0.61461 | 313.437 |
| Adamts18 | 66.7567 | 1.449823 | 0.38803 | 3.736328 | 0.00019 | 0.00412 | 151.613 |
| Adamts19 | 90.5601 | -0.49104 | 0.65061 | -0.75475 | 0.4504 | 0.65308 | 217.394 |
| Adamts2 | 503.607 | -0.67419 | 0.24033 | -2.80523 | 0.00503 | 0.03641 | 205.49 |
| Adamts20 | 925.656 | -0.27134 | 0.16842 | -1.61107 | 0.10717 | 0.27174 | 195.256 |
| Adamts3 | 2858.54 | 0.62096 | 0.18517 | 3.353471 | 0.0008 | 0.01096 | 206.248 |
| Adamts4 | 1203.96 | -0.65793 | 0.14994 | -4.38787 | 1.14E-05 | 0.00061 | 10.217 |
| Adamts5 | 169.564 | 0.353178 | 0.16895 | 2.090383 | 0.03658 | 0.1348 | 45.656 |
| Adamts6 | 640.248 | 0.291814 | 0.14672 | 1.988937 | 0.04671 | 0.15842 | 208.861 |
| Adamts7 | 26.3015 | -0.76725 | 0.41987 | -1.82735 | 0.06765 | 0.20161 | 45.003 |
| Adamts8 | 105.188 | -0.09738 | 0.25922 | -0.37566 | 0.70717 | 0.842 | 21.277 |
| Adamts9 | 264.185 | 0.344403 | 0.13229 | 2.6033 | 0.00923 | 0.05434 | 170.794 |
| Adamtsl1 | 447.287 | -0.42143 | 0.21969 | -1.91825 | 0.05508 | 0.17684 | 914.214 |
| Adamtsl2 | 171.337 | -0.26568 | 0.23536 | -1.12881 | 0.25898 | 0.46721 | 29.603 |
| Adamtsl3 | 147.075 | -1.11836 | 0.32258 | -3.46698 | 0.00053 | 0.00822 | 278.757 |
| Adamtsl4 | 306.472 | -0.87288 | 0.17668 | -4.94058 | 7.79E-07 | 9.01E-05 | 11.718 |
| Adamtsl5 | 265.885 | -0.13395 | 0.17835 | -0.75105 | 0.45262 | 0.65504 | 9.303 |
| Adap1 | 5346.29 | -0.34759 | 0.08937 | -3.88924 | 0.0001 | 0.0027 | 53.747 |
| Adap2 | 329.102 | -0.17006 | 0.13612 | -1.24936 | 0.21153 | 0.41203 | 24.854 |
| Adap2os | 34.4503 | -0.32854 | 0.31791 | -1.03341 | 0.30141 | 0.5111 | 18.769 |
| Adar | 7819.03 | 0.080278 | 0.06295 | 1.275349 | 0.20219 | 0.40058 | 38.425 |
| Adarb1 | 14419.7 | -0.11906 | 0.06369 | -1.86936 | 0.06157 | 0.18978 | 127.545 |
| Adarb2 | 1255.28 | -0.22498 | 0.1374 | -1.6374 | 0.10155 | 0.26294 | 565.882 |
| Adat1 | 787.571 | 0.20458 | 0.09194 | 2.225248 | 0.02606 | 0.10714 | 25.395 |
| Adat2 | 337.653 | 0.231709 | 0.10432 | 2.221067 | 0.02635 | 0.10784 | 10.483 |
| Adck1 | 1209.13 | 0.107891 | 0.09266 | 1.16434 | 0.24429 | 0.45076 | 101.171 |
| Adck2 | 818.795 | 0.115999 | 0.09606 | 1.207617 | 0.22719 | 0.43135 | 14.897 |
| Adck5 | 782.5 | 0.271261 | 0.08854 | 3.063814 | 0.00219 | 0.02131 | 19.459 |
| Adcy1 | 21142.5 | -0.67056 | 0.27787 | -2.41324 | 0.01581 | 0.07716 | 115.018 |
| Adcy10 | 39.258 | -0.09593 | 0.33412 | -0.28711 | 0.77403 | 0.88228 | 91.592 |
| Adcy2 | 4946.64 | -0.30295 | 0.12556 | -2.4128 | 0.01583 | 0.07722 | 379.499 |
| Adcy3 | 2187.1 | 0.209749 | 0.08073 | 2.598185 | 0.00937 | 0.05499 | 80.423 |
| Adcy4 | 211.271 | -0.30853 | 0.16252 | -1.89837 | 0.05765 | 0.1823 | 15.039 |
| Adcy5 | 45184 | 0.599296 | 0.18608 | 3.220579 | 0.00128 | 0.01485 | 150.862 |

| | | | | | | | |
|---|---|---|---|---|---|---|---|
| Adcy6 | 2117.5 | -0.40266 | 0.16171 | -2.48993 | 0.01278 | 0.06744 | 20.104 |
| Adcy7 | 560.676 | 0.044001 | 0.10893 | 0.403949 | 0.68625 | 0.82887 | 57.56 |
| Adcy8 | 448.255 | -0.38206 | 0.19213 | -1.98854 | 0.04675 | 0.15854 | 225.213 |
| Adcy9 | 3984.86 | 0.018639 | 0.09029 | 0.206435 | 0.83645 | 0.91608 | 132.97 |
| Adcyap1 | 707.745 | -1.59418 | 0.5993 | -2.66006 | 0.00781 | 0.04892 | 7.521 |
| Adcyap1r1 | 12237.8 | -0.03282 | 0.07473 | -0.43913 | 0.66057 | 0.81233 | 49.474 |
| Add1 | 19650.3 | -0.1103 | 0.05091 | -2.1666 | 0.03027 | 0.11875 | 58.645 |
| Add2 | 20235.5 | 0.171314 | 0.07962 | 2.15163 | 0.03143 | 0.12193 | 95.729 |
| Add3 | 7383.59 | -0.0584 | 0.06781 | -0.86111 | 0.38918 | 0.59831 | 106.957 |
| Adgb | 12.6548 | -0.22111 | 0.51468 | -0.42961 | 0.66748 | 0.81624 | 136.624 |
| Adgra1 | 1466.9 | -0.41235 | 0.17576 | -2.34607 | 0.01897 | 0.08652 | 43.915 |
| Adgra2 | 1196.3 | -0.10811 | 0.11619 | -0.9304 | 0.35217 | 0.56253 | 37.854 |
| Adgra3 | 973.788 | -0.14929 | 0.12984 | -1.14981 | 0.25022 | 0.45761 | 99.051 |
| Adgrb1 | 24406.2 | 0.055436 | 0.07461 | 0.743046 | 0.45745 | 0.65867 | 73.271 |
| Adgrb2 | 18277.4 | 0.078631 | 0.06236 | 1.261008 | 0.20731 | 0.40683 | 37.764 |
| Adgrb3 | 5266.22 | 0.284215 | 0.09756 | 2.913343 | 0.00358 | 0.02918 | 762.232 |
| Adgrd1 | 96.5818 | -0.27495 | 0.33103 | -0.83059 | 0.40621 | 0.61466 | 107.85 |
| Adgre1 | 243.986 | -0.54843 | 0.18602 | -2.94827 | 0.0032 | 0.02719 | 124.837 |
| Adgre5 | 283.941 | -0.25068 | 0.15135 | -1.65627 | 0.09767 | 0.25637 | 18.076 |
| Adgrf2 | 81.8923 | -0.02472 | 0.23148 | -0.1068 | 0.91495 | 0.95656 | 46.05 |
| Adgrf4 | 101.568 | 0.283585 | 0.21506 | 1.318626 | 0.18729 | 0.38255 | 35.394 |
| Adgrf5 | 2323.37 | -0.33498 | 0.08684 | -3.85735 | 0.00011 | 0.00294 | 99.107 |
| Adgrg1 | 3572.18 | -0.02109 | 0.11019 | -0.19144 | 0.84818 | 0.92288 | 39.467 |
| Adgrg2 | 67.5379 | 0.030899 | 0.30162 | 0.102444 | 0.9184 | 0.95861 | 107.381 |
| Adgrg3 | 42.7893 | 0.233757 | 0.30241 | 0.772972 | 0.43954 | 0.6449 | 27.559 |
| Adgrg6 | 468.697 | -0.6201 | 0.17783 | -3.48715 | 0.00049 | 0.00779 | 143.077 |
| Adgrl1 | 19456.4 | -0.03682 | 0.06544 | -0.56258 | 0.57372 | 0.75227 | 41.85 |
| Adgrl2 | 3818.71 | 0.661201 | 0.18984 | 3.482859 | 0.0005 | 0.00788 | 174.973 |
| Adgrl3 | 4395.56 | 0.183333 | 0.0775 | 2.365479 | 0.01801 | 0.08366 | 804.996 |
| Adgrl4 | 1415.92 | -0.26448 | 0.12246 | -2.15975 | 0.03079 | 0.12022 | 107.2 |
| Adgrv1 | 153.257 | 0.238347 | 0.16784 | 1.420052 | 0.15559 | 0.34222 | 538.087 |
| Adh5 | 1721.79 | -0.12989 | 0.09275 | -1.40049 | 0.16137 | 0.3496 | 12.407 |
| Adhfe1 | 752.966 | -0.15634 | 0.11988 | -1.30417 | 0.19218 | 0.38882 | 32.726 |
| Adi1 | 1364.39 | -0.07084 | 0.08682 | -0.81593 | 0.41454 | 0.62254 | 6.945 |

| | | | | | | | |
|---|---|---|---|---|---|---|---|
| Adipor1 | 3055.29 | -0.07638 | 0.07072 | -1.08002 | 0.28013 | 0.49017 | 17.974 |
| Adipor2 | 4948.42 | -0.4654 | 0.08426 | -5.52308 | 3.33E-08 | 8.59E-06 | 64.555 |
| Adk | 2116.29 | -0.13828 | 0.09416 | -1.46854 | 0.14196 | 0.3238 | 395.996 |
| Adm | 56.609 | -0.72938 | 0.2509 | -2.90711 | 0.00365 | 0.02963 | 2.16 |
| Adnp | 239.447 | 0.01789 | 0.16013 | 0.111719 | 0.91105 | 0.95504 | 26.127 |
| Adnp2 | 1621.31 | 0.107358 | 0.0646 | 1.661795 | 0.09655 | 0.25453 | 25.172 |
| Ado | 2685.68 | -0.23252 | 0.07356 | -3.16085 | 0.00157 | 0.01719 | 4.558 |
| Adora1 | 5406.25 | -0.12035 | 0.08153 | -1.47616 | 0.1399 | 0.3212 | 36.209 |
| Adora2a | 6628.92 | 0.808555 | 0.37103 | 2.179211 | 0.02932 | 0.11624 | 17.908 |
| Adora2b | 122.602 | -0.29462 | 0.21741 | -1.35512 | 0.17538 | 0.36715 | 17.47 |
| Adora3 | 20.418 | -0.02396 | 0.57867 | -0.04141 | 0.96697 | 0.98345 | 38.069 |
| Adpgk | 949.388 | -0.14482 | 0.08089 | -1.79031 | 0.0734 | 0.21336 | 32.495 |
| Adprh | 2843.89 | -0.17724 | 0.07011 | -2.5281 | 0.01147 | 0.06276 | 8.674 |
| Adprm | 715.619 | 0.249548 | 0.10431 | 2.39234 | 0.01674 | 0.08007 | 14.69 |
| Adprs | 831.591 | 0.035131 | 0.08971 | 0.391631 | 0.69533 | 0.83491 | 5.657 |
| Adra1a | 708.664 | -0.00163 | 0.14435 | -0.01128 | 0.991 | 0.99495 | 135.918 |
| Adra1b | 296.925 | -1.07694 | 0.21978 | -4.89997 | 9.59E-07 | 0.0001 | 126.605 |
| Adra1d | 103.381 | -1.1076 | 0.29241 | -3.78778 | 0.00015 | 0.00357 | 16.434 |
| Adra2a | 1293.04 | -1.05182 | 0.48418 | -2.1724 | 0.02983 | 0.11778 | 4.832 |
| Adra2b | 207.828 | 0.567183 | 0.27775 | 2.04206 | 0.04115 | 0.14536 | 4.014 |
| Adra2c | 1992.35 | 0.475277 | 0.15876 | 2.993735 | 0.00276 | 0.0249 | 3.445 |
| Adrb1 | 1005.86 | -0.13892 | 0.09707 | -1.43116 | 0.15238 | 0.33764 | 10.915 |
| Adrb2 | 87.3662 | -0.42686 | 0.24463 | -1.74496 | 0.08099 | 0.22645 | 2.144 |
| Adrb3 | 33.4091 | -0.4553 | 0.35766 | -1.27302 | 0.20301 | 0.40164 | 24.841 |
| Adrm1 | 2332.22 | 0.031448 | 0.06987 | 0.450074 | 0.65266 | 0.80683 | 4.802 |
| Adsl | 1143.59 | 0.188514 | 0.09141 | 2.06227 | 0.03918 | 0.14101 | 22.457 |
| Adss | 4319.93 | 0.056083 | 0.08883 | 0.631367 | 0.5278 | 0.71577 | 33.55 |
| Adssl1 | 645.83 | -0.71669 | 0.1303 | -5.50019 | 3.79E-08 | 9.54E-06 | 21.316 |
| Aebp1 | 990.71 | -1.07481 | 0.41133 | -2.613 | 0.00898 | 0.05332 | 10.142 |
| Aebp2 | 5256.2 | 0.218974 | 0.0831 | 2.635056 | 0.00841 | 0.05116 | 55.81 |
| Aen | 1011.93 | -0.06052 | 0.14084 | -0.42972 | 0.6674 | 0.8162 | 15.356 |
| Afap1 | 1251.14 | -0.1711 | 0.09153 | -1.86934 | 0.06158 | 0.18978 | 110.605 |
| Afap1l1 | 483.785 | -0.26721 | 0.11223 | -2.38094 | 0.01727 | 0.08154 | 56.442 |
| Afap1l2 | 497.081 | -0.18777 | 0.13087 | -1.43485 | 0.15133 | 0.3362 | 95.868 |

| | | | | | | | |
|---|---|---|---|---|---|---|---|
| Afdn | 6166.64 | 0.108245 | 0.06707 | 1.613897 | 0.10655 | 0.27076 | 145.612 |
| Aff1 | 2716.65 | -0.0874 | 0.09216 | -0.94839 | 0.34293 | 0.55368 | 162.949 |
| Aff2 | 330.788 | 0.111787 | 0.10979 | 1.018179 | 0.30859 | 0.51837 | 507.744 |
| Aff3 | 2495.42 | -0.11402 | 0.08163 | -1.39684 | 0.16246 | 0.35091 | 487.63 |
| Aff4 | 14376.8 | 0.228783 | 0.05794 | 3.948768 | 7.86E-05 | 0.00227 | 70.998 |
| Afg1l | 735.006 | -0.10094 | 0.10681 | -0.94504 | 0.34464 | 0.55549 | 165.981 |
| Afg3l1 | 2138.14 | -0.02449 | 0.10677 | -0.22939 | 0.81857 | 0.90663 | 26.014 |
| Afg3l2 | 4461.56 | 0.053275 | 0.06853 | 0.777407 | 0.43692 | 0.64258 | 44.409 |
| Afmid | 34.9349 | -0.11402 | 0.36186 | -0.31509 | 0.75269 | 0.87017 | 13.985 |
| Aftph | 4759.35 | 0.175549 | 0.076 | 2.309894 | 0.02089 | 0.09217 | 56.506 |
| Aga | 234.437 | -0.57886 | 0.13849 | -4.17988 | 2.92E-05 | 0.00113 | 11.695 |
| Agap1 | 9851.26 | -0.05782 | 0.06804 | -0.8499 | 0.39538 | 0.60438 | 442.812 |
| Agap2 | 52291 | 0.061661 | 0.07292 | 0.845585 | 0.39778 | 0.60678 | 17.886 |
| Agap3 | 9336.37 | 0.004362 | 0.07119 | 0.061265 | 0.95115 | 0.97595 | 49.871 |
| Agbl2 | 341.265 | 0.78398 | 0.23922 | 3.277212 | 0.00105 | 0.01322 | 51.711 |
| Agbl3 | 485.186 | 0.27574 | 0.1014 | 2.719441 | 0.00654 | 0.0436 | 79.028 |
| Agbl4 | 407.821 | -0.199 | 0.14182 | -1.40316 | 0.16057 | 0.34883 | 1266.664 |
| Agbl5 | 1222.3 | 0.35727 | 0.07473 | 4.780679 | 1.75E-06 | 0.00016 | 18.272 |
| Ager | 36.0801 | -0.19881 | 0.2992 | -0.66446 | 0.50639 | 0.69789 | 3.075 |
| Agfg1 | 5043.82 | 0.075088 | 0.05597 | 1.341518 | 0.17975 | 0.37289 | 61.7 |
| Agfg2 | 2753.44 | 0.137282 | 0.11848 | 1.158726 | 0.24657 | 0.45341 | 34.264 |
| Aggf1 | 3323.21 | -0.0608 | 0.06482 | -0.93809 | 0.3482 | 0.55865 | 24.67 |
| Agk | 1602.56 | 0.168752 | 0.09099 | 1.854555 | 0.06366 | 0.19397 | 71.591 |
| Agl | 2736.04 | -0.16396 | 0.0876 | -1.87166 | 0.06125 | 0.18912 | 68.168 |
| Agmat | 16.5009 | 0.09859 | 0.4621 | 0.213353 | 0.83105 | 0.91334 | 12.592 |
| Agmo | 401.737 | -0.26238 | 0.16557 | -1.58474 | 0.11302 | 0.28134 | 340.562 |
| Ago1 | 5117.88 | 0.092445 | 0.06481 | 1.426309 | 0.15378 | 0.3396 | 33.572 |
| Ago2 | 9869.25 | 0.302905 | 0.07159 | 4.231313 | 2.32E-05 | 0.00097 | 89.092 |
| Ago3 | 5255.25 | 0.08105 | 0.0859 | 0.943569 | 0.34539 | 0.55616 | 97.853 |
| Ago4 | 1047.03 | 0.145663 | 0.12504 | 1.164905 | 0.24406 | 0.45042 | 43.932 |
| Agpat1 | 4562.81 | -0.12872 | 0.05912 | -2.17708 | 0.02947 | 0.11675 | 9.188 |
| Agpat2 | 41.0047 | -0.23742 | 0.38931 | -0.60985 | 0.54196 | 0.72752 | 11.361 |
| Agpat3 | 11791.5 | -0.14542 | 0.05513 | -2.63753 | 0.00835 | 0.05095 | 83.312 |
| Agpat4 | 1584.76 | -0.41898 | 0.12107 | -3.46056 | 0.00054 | 0.00836 | 100.942 |

| | | | | | | | |
|---|---|---|---|---|---|---|---|
| Agpat5 | 3101.63 | 0.015514 | 0.06231 | 0.249006 | 0.80336 | 0.89848 | 45.085 |
| Agps | 2604.31 | 0.103553 | 0.06844 | 1.512951 | 0.13029 | 0.30843 | 99.174 |
| Agrn | 11021.7 | 0.143543 | 0.08742 | 1.641944 | 0.1006 | 0.26113 | 32.199 |
| Agrp | 52.5972 | 0.10878 | 0.25919 | 0.419686 | 0.67471 | 0.82139 | 70.696 |
| Agt | 254.138 | 0.674276 | 0.3052 | 2.209317 | 0.02715 | 0.11008 | 13.173 |
| Agtpbp1 | 8754.58 | -0.17219 | 0.0725 | -2.375 | 0.01755 | 0.0823 | 139.486 |
| Agtr1a | 29.7463 | 0.60827 | 0.34695 | 1.753177 | 0.07957 | 0.22403 | 46.427 |
| Agtr2 | 49.7547 | 0.380375 | 0.322 | 1.181302 | 0.23748 | 0.44288 | 4.621 |
| Agtrap | 487.539 | -0.46655 | 0.12058 | -3.86925 | 0.00011 | 0.00285 | 10.971 |
| Ahctf1 | 4745.91 | 0.35676 | 0.07898 | 4.517124 | 6.27E-06 | 0.00039 | 58.787 |
| Ahcy | 1068.29 | 0.15549 | 0.10577 | 1.470028 | 0.14155 | 0.32333 | 15.188 |
| Ahcyl | 149.395 | -0.10751 | 0.16121 | -0.66693 | 0.50482 | 0.6967 | 2.093 |
| Ahcyl1 | 22488.4 | -0.11455 | 0.06976 | -1.642 | 0.10059 | 0.26113 | 33.443 |
| Ahcyl2 | 9050.57 | -0.02569 | 0.05897 | -0.43562 | 0.66311 | 0.81387 | 144.3 |
| Ahdc1 | 2886.63 | 0.102236 | 0.07084 | 1.443225 | 0.14896 | 0.3329 | 66.851 |
| Ahi1 | 14271.7 | 0.415584 | 0.07544 | 5.509023 | 3.61E-08 | 9.19E-06 | 127.883 |
| Ahnak | 4949.38 | -0.23048 | 0.10757 | -2.14262 | 0.03214 | 0.12384 | 87.631 |
| Ahnak2 | 49.465 | -1.2314 | 0.31332 | -3.93011 | 8.49E-05 | 0.00241 | 27.65 |
| Ahr | 466.817 | -0.0042 | 0.10623 | -0.03955 | 0.96845 | 0.98409 | 37.065 |
| Ahrr | 45.4513 | -0.77497 | 0.32652 | -2.37339 | 0.01763 | 0.0825 | 81.214 |
| Ahsa1 | 2815.08 | -0.10733 | 0.07022 | -1.52838 | 0.12642 | 0.30219 | 7.52 |
| Ahsa2 | 906.501 | 0.149423 | 0.13208 | 1.131286 | 0.25793 | 0.46603 | 10.149 |
| Aida | 2059.17 | 0.076803 | 0.07498 | 1.0243 | 0.30569 | 0.51566 | 28.198 |
| Aif1 | 208.79 | -0.19558 | 0.20765 | -0.94189 | 0.34625 | 0.55684 | 5.078 |
| Aif1l | 232.648 | -0.52696 | 0.1557 | -3.38452 | 0.00071 | 0.0101 | 23.304 |
| Aifm1 | 1714.29 | 0.067874 | 0.07282 | 0.932066 | 0.3513 | 0.56182 | 38.62 |
| Aifm2 | 542.898 | -0.09958 | 0.10327 | -0.96424 | 0.33493 | 0.54565 | 23.998 |
| Aifm3 | 4339.59 | -0.39602 | 0.13517 | -2.92971 | 0.00339 | 0.02816 | 17.875 |
| Aig1 | 1482.84 | -0.0641 | 0.07595 | -0.84389 | 0.39873 | 0.60773 | 221.927 |
| Aim2 | 312.295 | 0.203899 | 0.15802 | 1.290332 | 0.19694 | 0.39476 | 115.162 |
| Aimp1 | 1574.05 | 0.047018 | 0.07343 | 0.640344 | 0.52195 | 0.71105 | 23.89 |
| Aimp2 | 391.122 | 0.014191 | 0.10551 | 0.1345 | 0.89301 | 0.94632 | 7.144 |
| Aip | 2715.57 | -0.1223 | 0.07216 | -1.69479 | 0.09012 | 0.24355 | 11.413 |
| Aire | 20.3055 | -0.5536 | 0.41699 | -1.3276 | 0.18431 | 0.37877 | 13.589 |

| | | | | | | | |
|---|---|---|---|---|---|---|---|
| Airn | 75.1961 | -0.26662 | 0.32584 | -0.81824 | 0.41322 | 0.62139 | 118.826 |
| Ajap1 | 2483.45 | 0.209301 | 0.07467 | 2.803146 | 0.00506 | 0.03657 | 109.591 |
| Ajm1 | 5707.84 | 0.164067 | 0.07662 | 2.141373 | 0.03224 | 0.12412 | 6.342 |
| Ajuba | 68.935 | -0.56046 | 0.24105 | -2.3251 | 0.02007 | 0.08969 | 10.087 |
| Ak1 | 3629.67 | -0.2484 | 0.06641 | -3.74036 | 0.00018 | 0.00406 | 13.301 |
| Ak2 | 1102.93 | 0.106636 | 0.07813 | 1.364866 | 0.1723 | 0.36312 | 19.572 |
| Ak3 | 4323.84 | -0.19347 | 0.06931 | -2.79148 | 0.00525 | 0.03741 | 27.129 |
| Ak3l2-ps | 19.7986 | -0.37383 | 0.50357 | -0.74236 | 0.45787 | 0.65913 | 0.672 |
| Ak4 | 1345.86 | -0.22279 | 0.13556 | -1.64344 | 0.10029 | 0.26056 | 47.719 |
| Ak5 | 3742.01 | -0.40764 | 0.16178 | -2.5198 | 0.01174 | 0.06375 | 205.546 |
| Ak6 | 267.944 | -0.19727 | 0.12545 | -1.57244 | 0.11585 | 0.2857 | 15.381 |
| Ak7 | 67.6646 | 1.424859 | 0.29174 | 4.883948 | 1.04E-06 | 0.00011 | 76.472 |
| Ak8 | 40.0547 | 0.073885 | 0.3099 | 0.238419 | 0.81156 | 0.90289 | 113.002 |
| Ak9 | 23.5468 | 0.57603 | 0.40606 | 1.418587 | 0.15602 | 0.3427 | 130.555 |
| Akain1 | 206.858 | -0.28188 | 0.19044 | -1.4801 | 0.13885 | 0.32009 | 49.908 |
| Akap1 | 2781.97 | 0.107082 | 0.06057 | 1.76798 | 0.07706 | 0.21943 | 33.795 |
| Akap10 | 1611.95 | 0.016972 | 0.08769 | 0.193537 | 0.84654 | 0.92184 | 58.946 |
| Akap11 | 17775.9 | 0.303202 | 0.08766 | 3.458759 | 0.00054 | 0.00839 | 44.563 |
| Akap12 | 1355.7 | -0.36878 | 0.15321 | -2.40703 | 0.01608 | 0.07798 | 93.091 |
| Akap13 | 3321.07 | -0.23065 | 0.1181 | -1.953 | 0.05082 | 0.1677 | 299.076 |
| Akap14 | 49.7641 | 0.456104 | 0.28437 | 1.603887 | 0.10874 | 0.27415 | 18.145 |
| Akap17b | 1617.51 | 0.345432 | 0.14062 | 2.456517 | 0.01403 | 0.07132 | 37.082 |
| Akap5 | 13685.9 | 0.79305 | 0.19479 | 4.071339 | 4.67E-05 | 0.00156 | 9.263 |
| Akap6 | 14475.2 | 0.109362 | 0.06647 | 1.645201 | 0.09993 | 0.25998 | 456.217 |
| Akap7 | 1084.72 | -0.08268 | 0.09588 | -0.86226 | 0.38855 | 0.59781 | 138.781 |
| Akap8 | 9015.25 | 0.169389 | 0.08408 | 2.014646 | 0.04394 | 0.15174 | 17.567 |
| Akap8l | 9520.51 | 0.182794 | 0.11303 | 1.617152 | 0.10585 | 0.26949 | 29.184 |
| Akap9 | 10214.4 | 0.525389 | 0.32157 | 1.633839 | 0.10229 | 0.26399 | 153.257 |
| Akip1 | 90.3497 | -0.35326 | 0.21013 | -1.68116 | 0.09273 | 0.24803 | 8.5 |
| Akirin1 | 3008.27 | -0.09127 | 0.0698 | -1.30771 | 0.19097 | 0.38722 | 15.787 |
| Akirin2 | 2463.21 | -0.08911 | 0.09216 | -0.96692 | 0.33359 | 0.54445 | 15.972 |
| Akna | 202.097 | -0.15735 | 0.15595 | -1.009 | 0.31297 | 0.5235 | 36.23 |
| Akr1a1 | 8573.11 | -0.17716 | 0.08306 | -2.13298 | 0.03293 | 0.12609 | 15.171 |
| Akr1b10 | 423.827 | -0.21902 | 0.11714 | -1.86968 | 0.06153 | 0.18974 | 12.733 |

| | | | | | | | |
|---|---|---|---|---|---|---|---|
| Akr1b3 | 1276.31 | 0.069698 | 0.08829 | 0.789418 | 0.42987 | 0.63616 | 15.045 |
| Akr1b8 | 22.4192 | -0.94496 | 0.38734 | -2.43965 | 0.0147 | 0.07327 | 14.345 |
| Akr1c13 | 14.1413 | -0.02723 | 0.54358 | -0.05009 | 0.96005 | 0.98055 | 14.447 |
| Akr1c14 | 65.4223 | -0.38651 | 0.2902 | -1.33188 | 0.1829 | 0.37689 | 41.678 |
| Akr1e1 | 1066.06 | -0.04328 | 0.08295 | -0.52178 | 0.60183 | 0.77182 | 18.425 |
| Akr7a5 | 569.651 | -0.12331 | 0.10965 | -1.12463 | 0.26075 | 0.46932 | 7.683 |
| Akt1 | 6157.62 | -0.02528 | 0.0825 | -0.30648 | 0.75924 | 0.87354 | 21.064 |
| Akt1s1 | 1064.66 | -0.19759 | 0.09444 | -2.09214 | 0.03643 | 0.13444 | 6.431 |
| Akt2 | 5286.7 | 0.035872 | 0.08839 | 0.405839 | 0.68486 | 0.82838 | 49.275 |
| Akt2-ps | 124.798 | -0.19759 | 0.1992 | -0.99192 | 0.32124 | 0.53206 | 1.449 |
| Akt3 | 7267.65 | -0.06498 | 0.06755 | -0.96199 | 0.33606 | 0.54683 | 238.131 |
| Aktip | 4179.32 | -0.09589 | 0.07868 | -1.21875 | 0.22294 | 0.4261 | 88.193 |
| Alad | 645.381 | 0.132387 | 0.10605 | 1.248327 | 0.21191 | 0.41257 | 10.75 |
| Alas1 | 2230.37 | -0.04026 | 0.09128 | -0.44104 | 0.65919 | 0.8112 | 15.2 |
| Alas2 | 186.318 | -0.39893 | 0.23661 | -1.68602 | 0.09179 | 0.24642 | 23.264 |
| Alcam | 18801.7 | 0.477057 | 0.13208 | 3.611924 | 0.0003 | 0.00568 | 205.079 |
| Aldh16a1 | 231.065 | -0.15296 | 0.15252 | -1.00285 | 0.31593 | 0.52651 | 13.901 |
| Aldh18a1 | 1065.92 | -0.13663 | 0.10213 | -1.33779 | 0.18096 | 0.37434 | 38.207 |
| Aldh1a1 | 2334.78 | -0.88183 | 0.12824 | -6.87635 | 6.14E-12 | 1.04E-08 | 150.751 |
| Aldh1a2 | 393.311 | -1.29132 | 0.47933 | -2.69399 | 0.00706 | 0.04574 | 80.455 |
| Aldh1a3 | 33.4808 | -0.43047 | 0.3143 | -1.36962 | 0.1708 | 0.36117 | 36.628 |
| Aldh1a7 | 61.3526 | -1.10865 | 0.3767 | -2.94303 | 0.00325 | 0.02752 | 34.61 |
| Aldh1b1 | 505.893 | 0.108135 | 0.12531 | 0.862939 | 0.38817 | 0.59748 | 5.583 |
| Aldh1l1 | 2345.85 | -0.34582 | 0.10118 | -3.41773 | 0.00063 | 0.00928 | 113.777 |
| Aldh1l2 | 184.154 | -0.40205 | 0.18686 | -2.15161 | 0.03143 | 0.12193 | 46.691 |
| Aldh2 | 4111.11 | -0.10703 | 0.08017 | -1.33495 | 0.18189 | 0.37561 | 27.798 |
| Aldh3a2 | 2288.11 | -0.12416 | 0.08359 | -1.48547 | 0.13742 | 0.3181 | 44.048 |
| Aldh3b1 | 213.321 | -0.34685 | 0.16576 | -2.0925 | 0.03639 | 0.13442 | 16.318 |
| Aldh3b2 | 26.319 | -0.50212 | 0.45109 | -1.11313 | 0.26565 | 0.47488 | 9.319 |
| Aldh4a1 | 770.666 | -0.05112 | 0.11043 | -0.46288 | 0.64345 | 0.80108 | 26.825 |
| Aldh5a1 | 5569.64 | 0.164596 | 0.08623 | 1.908871 | 0.05628 | 0.17947 | 30.083 |
| Aldh6a1 | 3498.05 | -0.0796 | 0.06797 | -1.17114 | 0.24154 | 0.44756 | 20.288 |
| Aldh7a1 | 4247.88 | -0.15897 | 0.08666 | -1.83442 | 0.06659 | 0.19977 | 63.265 |
| Aldh9a1 | 1581.47 | -0.27757 | 0.09794 | -2.83403 | 0.0046 | 0.03443 | 18.542 |

| | | | | | | | |
|---|---|---|---|---|---|---|---|
| Aldoa | 32271.3 | -0.17629 | 0.05455 | -3.23171 | 0.00123 | 0.01456 | 8.319 |
| Aldob | 81.4262 | 0.114927 | 0.21792 | 0.52737 | 0.59794 | 0.76947 | 13.552 |
| Aldoc | 30546.6 | -0.31187 | 0.09202 | -3.38918 | 0.0007 | 0.00999 | 4.814 |
| Alg1 | 919.73 | 0.261419 | 0.08137 | 3.212626 | 0.00132 | 0.01511 | 11.292 |
| Alg10b | 3595.61 | 0.201444 | 0.07061 | 2.852865 | 0.00433 | 0.0332 | 9.161 |
| Alg11 | 1786.05 | 0.05082 | 0.0831 | 0.611558 | 0.54083 | 0.72656 | 10.907 |
| Alg12 | 588.384 | 0.185188 | 0.09683 | 1.912434 | 0.05582 | 0.17857 | 14.075 |
| Alg13 | 451.95 | 0.038889 | 0.12316 | 0.315751 | 0.75219 | 0.86997 | 56.647 |
| Alg14 | 441.928 | -0.16652 | 0.11316 | -1.47163 | 0.14112 | 0.32296 | 71.322 |
| Alg2 | 5783.11 | -0.05601 | 0.07179 | -0.78025 | 0.43525 | 0.641 | 9.267 |
| Alg3 | 359.541 | 0.010777 | 0.1252 | 0.086074 | 0.93141 | 0.96532 | 6.362 |
| Alg5 | 615.739 | 0.128494 | 0.10241 | 1.254699 | 0.20959 | 0.40977 | 15.78 |
| Alg6 | 623.322 | 0.092476 | 0.1161 | 0.79651 | 0.42574 | 0.63229 | 47.797 |
| Alg8 | 387.107 | 0.135362 | 0.11288 | 1.199198 | 0.23045 | 0.43483 | 20.58 |
| Alg9 | 607.33 | 0.039364 | 0.09194 | 0.428145 | 0.66855 | 0.81715 | 68.524 |
| Alk | 829.213 | 0.0877 | 0.11853 | 0.739908 | 0.45936 | 0.65991 | 735.656 |
| Alkal2 | 46.9773 | -1.17152 | 0.39857 | -2.93935 | 0.00329 | 0.02768 | 9.533 |
| Alkbh1 | 935.206 | 0.156828 | 0.08434 | 1.859519 | 0.06295 | 0.19263 | 18.178 |
| Alkbh2 | 141.436 | 0.028955 | 0.17462 | 0.16582 | 0.8683 | 0.93337 | 4.293 |
| Alkbh3 | 419.056 | 0.159418 | 0.13176 | 1.209923 | 0.22631 | 0.43032 | 30.258 |
| Alkbh3os1 | 69.4517 | -0.05741 | 0.24218 | -0.23707 | 0.8126 | 0.90362 | 3.542 |
| Alkbh4 | 240.993 | -0.28432 | 0.1438 | -1.97714 | 0.04803 | 0.16139 | 5.47 |
| Alkbh5 | 3609.5 | -0.09899 | 0.06031 | -1.64131 | 0.10073 | 0.2614 | 22.132 |
| Alkbh6 | 224.175 | -0.08248 | 0.14638 | -0.56344 | 0.57313 | 0.75183 | 5.588 |
| Alkbh7 | 387.464 | -0.01508 | 0.10497 | -0.14365 | 0.88577 | 0.9432 | 1.984 |
| Alkbh8 | 2780.95 | 0.065163 | 0.08369 | 0.778574 | 0.43623 | 0.64189 | 56.015 |
| Alms1 | 681.038 | -0.05924 | 0.08304 | -0.71339 | 0.4756 | 0.67371 | 115.223 |
| Alms1-ps1 | 27.7754 | -0.03243 | 0.47845 | -0.06779 | 0.94595 | 0.973 | 34.602 |
| Alms1-ps2 | 36.4682 | 0.749172 | 0.29987 | 2.4983 | 0.01248 | 0.06631 | 29.665 |
| Alox12 | 23.5282 | -0.2391 | 0.35566 | -0.67229 | 0.5014 | 0.69395 | 13.897 |
| Alox12b | 46.7254 | -0.65441 | 0.40031 | -1.63476 | 0.1021 | 0.26371 | 12.804 |
| Alox5 | 38.4824 | -0.6128 | 0.31687 | -1.9339 | 0.05312 | 0.17246 | 51.102 |
| Alox5ap | 30.2599 | -0.88273 | 0.36382 | -2.42629 | 0.01525 | 0.07527 | 23.387 |
| Alox8 | 131.32 | -0.15793 | 0.19382 | -0.81482 | 0.41518 | 0.62301 | 13.912 |

| | | | | | | | |
|---|---|---|---|---|---|---|---|
| Aloxe3 | 222.199 | -0.02808 | 0.19859 | -0.1414 | 0.88755 | 0.94401 | 23.22 |
| Alpk1 | 315.191 | -0.29482 | 0.14176 | -2.07966 | 0.03756 | 0.1372 | 110.218 |
| Alpk3 | 35.063 | 0.080752 | 0.32291 | 0.250078 | 0.80253 | 0.89814 | 48.013 |
| Alpl | 251.698 | -0.10319 | 0.18878 | -0.54662 | 0.58464 | 0.75936 | 54.652 |
| Als2 | 2200.45 | -0.14021 | 0.07755 | -1.80788 | 0.07063 | 0.20776 | 74.306 |
| Als2cl | 606.632 | 0.080579 | 0.09497 | 0.848449 | 0.39619 | 0.60534 | 20.661 |
| Alx3 | 34.3663 | -1.09744 | 0.4648 | -2.36111 | 0.01822 | 0.08435 | 10.746 |
| Alx4 | 77.8056 | -1.24811 | 0.38786 | -3.21799 | 0.00129 | 0.01492 | 38.956 |
| Alyref | 1186.82 | -0.07845 | 0.10901 | -0.71959 | 0.47178 | 0.67032 | 6.245 |
| Alyref2 | 28.8952 | -0.32787 | 0.36367 | -0.90157 | 0.36729 | 0.57768 | 1.273 |
| Amacr | 368.484 | -0.07729 | 0.13443 | -0.57496 | 0.56532 | 0.74547 | 14.871 |
| Ambra1 | 3509.36 | 0.089465 | 0.06132 | 1.458963 | 0.14458 | 0.32724 | 188.716 |
| Amd-ps3 | 24.1636 | 0.045151 | 0.32421 | 0.139267 | 0.88924 | 0.94457 | 1.005 |
| Amd-ps4 | 38.252 | 0.376755 | 0.38132 | 0.988021 | 0.32314 | 0.53418 | 1.003 |
| Amd1 | 6619.63 | 0.101211 | 0.07859 | 1.287902 | 0.19778 | 0.39558 | 14.731 |
| Amd2 | 83.4798 | -0.17389 | 0.22708 | -0.76575 | 0.44382 | 0.6481 | 3.218 |
| Amdhd2 | 705.656 | 0.16564 | 0.1181 | 1.402596 | 0.16074 | 0.34889 | 7.934 |
| Amer1 | 534.821 | -0.04905 | 0.0938 | -0.52289 | 0.60105 | 0.77133 | 24.555 |
| Amer2 | 6048.62 | 0.061283 | 0.06599 | 0.928718 | 0.35304 | 0.56331 | 14.939 |
| Amer3 | 1373.36 | 0.189898 | 0.09127 | 2.080694 | 0.03746 | 0.13695 | 11.288 |
| Amfr | 8456.5 | 0.033255 | 0.08714 | 0.381635 | 0.70273 | 0.8396 | 41.255 |
| Amh | 117.412 | 0.585447 | 0.18721 | 3.127148 | 0.00177 | 0.01859 | 2.406 |
| Amhr2 | 48.5246 | 0.084685 | 0.26839 | 0.315525 | 0.75236 | 0.86997 | 9.267 |
| Amigo1 | 4712.52 | -0.21964 | 0.07778 | -2.82395 | 0.00474 | 0.03514 | 5.952 |
| Amigo2 | 487.335 | 0.181734 | 0.19575 | 0.928399 | 0.3532 | 0.56348 | 3.163 |
| Amigo3 | 270.152 | 0.305399 | 0.1495 | 2.042751 | 0.04108 | 0.14519 | 2.543 |
| Ammecr1 | 309.589 | 0.01347 | 0.14049 | 0.095875 | 0.92362 | 0.96155 | 115.583 |
| Ammecr1l | 997.155 | 0.049307 | 0.09042 | 0.545309 | 0.58554 | 0.75994 | 22.918 |
| Amn | 129.189 | -0.30228 | 0.17821 | -1.69619 | 0.08985 | 0.24306 | 5.332 |
| Amn1 | 910.454 | 0.109751 | 0.08427 | 1.302315 | 0.19281 | 0.38957 | 31.566 |
| Amot | 1851.53 | -0.02308 | 0.08402 | -0.27474 | 0.78352 | 0.88753 | 58.757 |
| Amotl1 | 3830.61 | 0.337525 | 0.08849 | 3.814395 | 0.00014 | 0.00332 | 103.091 |
| Amotl2 | 1119.89 | -0.38274 | 0.08409 | -4.55174 | 5.32E-06 | 0.00035 | 16.747 |
| Ampd2 | 2784.45 | 0.056716 | 0.09183 | 0.617603 | 0.53684 | 0.72378 | 12.59 |

| | | | | | | | |
|---|---|---|---|---|---|---|---|
| Ampd3 | 1849.34 | 0.116312 | 0.09561 | 1.216512 | 0.22379 | 0.42705 | 44.2 |
| Amph | 6817.95 | -0.27768 | 0.10784 | -2.5749 | 0.01003 | 0.05756 | 202.717 |
| Amt | 1882.24 | 0.098036 | 0.07275 | 1.347521 | 0.17781 | 0.37008 | 5.45 |
| Amy1 | 1461.95 | 1.038919 | 0.16317 | 6.367024 | 1.93E-10 | 1.64E-07 | 50.99 |
| Amz1 | 918.642 | 0.544782 | 0.16102 | 3.38341 | 0.00072 | 0.01013 | 37.313 |
| Amz2 | 2421.17 | 0.129617 | 0.06228 | 2.08104 | 0.03743 | 0.13686 | 26.868 |
| Anapc1 | 5312.56 | 0.087431 | 0.08666 | 1.008925 | 0.31301 | 0.5235 | 77.288 |
| Anapc10 | 457.941 | 0.152592 | 0.12063 | 1.264953 | 0.20589 | 0.40514 | 67.24 |
| Anapc11 | 1640.1 | -0.15094 | 0.08075 | -1.86932 | 0.06158 | 0.18978 | 9.778 |
| Anapc13 | 412.975 | 0.024328 | 0.10769 | 0.225915 | 0.82127 | 0.90759 | 8.13 |
| Anapc15 | 206.922 | -0.10607 | 0.14286 | -0.74252 | 0.45777 | 0.65903 | 38.135 |
| Anapc16 | 2392.03 | 0.08266 | 0.07579 | 1.090618 | 0.27544 | 0.4847 | 15.727 |
| Anapc2 | 3408.61 | -0.05356 | 0.07486 | -0.71553 | 0.47428 | 0.67238 | 13.438 |
| Anapc4 | 2737.59 | 0.118949 | 0.07343 | 1.619827 | 0.10527 | 0.26856 | 33.786 |
| Anapc5 | 9591.87 | 0.131615 | 0.05085 | 2.588516 | 0.00964 | 0.05596 | 33.881 |
| Anapc7 | 1506.34 | 0.005962 | 0.06811 | 0.087535 | 0.93025 | 0.96485 | 23.22 |
| Ang | 52.9353 | -0.15997 | 0.26811 | -0.59667 | 0.55073 | 0.73421 | 10.86 |
| Angel1 | 383.165 | -0.38198 | 0.17782 | -2.14807 | 0.03171 | 0.12267 | 25.959 |
| Angel2 | 3448.18 | 0.145886 | 0.06925 | 2.106515 | 0.03516 | 0.13143 | 21.851 |
| Angpt1 | 161.229 | -0.24008 | 0.25541 | -0.93997 | 0.34724 | 0.55778 | 252.255 |
| Angpt2 | 78.1356 | -0.89769 | 0.30428 | -2.95017 | 0.00318 | 0.02711 | 51.3 |
| Angptl1 | 21.2446 | -0.08036 | 0.44209 | -0.18177 | 0.85576 | 0.92708 | 22.517 |
| Angptl2 | 156.367 | -0.3216 | 0.18099 | -1.77696 | 0.07557 | 0.217 | 31.649 |
| Angptl3 | 19.8541 | -0.07783 | 0.43008 | -0.18095 | 0.8564 | 0.92757 | 15.158 |
| Angptl4 | 348.406 | -0.79261 | 0.46895 | -1.69019 | 0.09099 | 0.24505 | 8.081 |
| Angptl6 | 171.133 | 0.188661 | 0.19187 | 0.983256 | 0.32548 | 0.53678 | 5.92 |
| Angptl7 | 71.6526 | -0.0884 | 0.288 | -0.30694 | 0.75889 | 0.87333 | 5.278 |
| Angptl8 | 21.9646 | 0.118653 | 0.40554 | 0.29258 | 0.76984 | 0.87985 | 1.838 |
| Ank | 8620.8 | -0.04765 | 0.07137 | -0.6676 | 0.50439 | 0.69662 | 128.233 |
| Ank1 | 3229.43 | -0.17621 | 0.09385 | -1.87751 | 0.06045 | 0.18757 | 175.654 |
| Ank2 | 37119.5 | -0.1131 | 0.08194 | -1.38033 | 0.16748 | 0.35726 | 577.739 |
| Ank3 | 29632.6 | 0.183658 | 0.06608 | 2.779282 | 0.00545 | 0.03847 | 628.666 |
| Ankdd1a | 236.301 | 0.992134 | 0.23867 | 4.156922 | 3.23E-05 | 0.00122 | 31.724 |
| Ankdd1b | 15.5683 | -0.47121 | 0.5495 | -0.85751 | 0.39116 | 0.60033 | 55.134 |

| | | | | | | | |
|---|---|---|---|---|---|---|---|
| Ankef1 | 77.0706 | 0.418483 | 0.31732 | 1.318795 | 0.18724 | 0.3825 | 60.182 |
| Ankfn1 | 204.063 | 0.398698 | 0.21918 | 1.819038 | 0.06891 | 0.20415 | 451.044 |
| Ankfy1 | 3641.78 | 0.024016 | 0.07028 | 0.341723 | 0.73256 | 0.85819 | 82.141 |
| Ankhd1 | 3377.93 | 0.083786 | 0.08841 | 0.94769 | 0.34329 | 0.55397 | 98.927 |
| Ankib1 | 3396.17 | 0.081031 | 0.06257 | 1.294961 | 0.19533 | 0.3929 | 112.926 |
| Ankk1 | 544.379 | 1.213133 | 0.24653 | 4.920786 | 8.62E-07 | 9.55E-05 | 11.828 |
| Ankle1 | 20.9171 | -0.00528 | 0.45636 | -0.01158 | 0.99076 | 0.99495 | 3.895 |
| Ankle2 | 2515.25 | 0.200959 | 0.07543 | 2.664271 | 0.00772 | 0.04854 | 25.648 |
| Ankmy1 | 17.091 | 0.198424 | 0.50963 | 0.389349 | 0.69702 | 0.8358 | 43.104 |
| Ankmy2 | 1975.94 | 0.151401 | 0.08124 | 1.863558 | 0.06238 | 0.19154 | 40.178 |
| Ankra2 | 1427.59 | 0.09796 | 0.07354 | 1.332085 | 0.18283 | 0.37683 | 11.681 |
| Ankrd10 | 5368.44 | 0.245832 | 0.0826 | 2.976101 | 0.00292 | 0.02574 | 24.175 |
| Ankrd11 | 13135.9 | 0.059765 | 0.05736 | 1.041972 | 0.29742 | 0.50755 | 158.456 |
| Ankrd12 | 5132.96 | 0.398054 | 0.33323 | 1.194516 | 0.23228 | 0.43719 | 111.392 |
| Ankrd13a | 3216.06 | -0.1643 | 0.07815 | -2.10235 | 0.03552 | 0.1324 | 31.524 |
| Ankrd13b | 4247.12 | 0.156794 | 0.08867 | 1.7683 | 0.07701 | 0.2194 | 19.194 |
| Ankrd13c | 4304.86 | 0.142952 | 0.06514 | 2.194498 | 0.0282 | 0.11309 | 60.796 |
| Ankrd13d | 2367.36 | 0.086794 | 0.08423 | 1.030429 | 0.30281 | 0.51232 | 12.958 |
| Ankrd16 | 2525.21 | 0.149944 | 0.10136 | 1.479307 | 0.13906 | 0.3203 | 12.454 |
| Ankrd17 | 9541.86 | 0.179999 | 0.06821 | 2.638716 | 0.00832 | 0.05084 | 139.412 |
| Ankrd2 | 19.0041 | 0.440565 | 0.41767 | 1.05482 | 0.29151 | 0.50128 | 9.111 |
| Ankrd23 | 28.7157 | 0.103817 | 0.34638 | 0.299718 | 0.76439 | 0.87659 | 6.696 |
| Ankrd24 | 939.846 | -2.03E-06 | 0.12538 | -1.62E-05 | 0.99999 | 0.99999 | 19.071 |
| Ankrd26 | 1350.18 | 0.267306 | 0.1009 | 2.649165 | 0.00807 | 0.04987 | 60.919 |
| Ankrd27 | 1722.64 | -0.10157 | 0.09206 | -1.10338 | 0.26986 | 0.47946 | 52.983 |
| Ankrd28 | 4884.54 | -0.03503 | 0.08492 | -0.41252 | 0.67996 | 0.82501 | 131.884 |
| Ankrd29 | 807.692 | -0.30538 | 0.13105 | -2.33034 | 0.01979 | 0.0889 | 53.436 |
| Ankrd31 | 15.4208 | -0.68015 | 0.65805 | -1.03358 | 0.30133 | 0.51105 | 161.768 |
| Ankrd33b | 6103.65 | 0.115294 | 0.10027 | 1.149882 | 0.25019 | 0.45761 | 76.249 |
| Ankrd34a | 1945.65 | -0.03398 | 0.07282 | -0.46654 | 0.64083 | 0.79893 | 3.14 |
| Ankrd34b | 1384.62 | 0.250114 | 0.12205 | 2.049349 | 0.04043 | 0.14368 | 15.763 |
| Ankrd34c | 391.173 | -1.03365 | 0.23094 | -4.47584 | 7.61E-06 | 0.00045 | 13.231 |
| Ankrd35 | 693.954 | -0.00565 | 0.16191 | -0.03487 | 0.97218 | 0.98602 | 20.902 |
| Ankrd37 | 139.687 | 0.159213 | 0.15058 | 1.057314 | 0.29037 | 0.50071 | 8.806 |

| | | | | | | | |
|---|---|---|---|---|---|---|---|
| Ankrd39 | 335.134 | -0.01078 | 0.10259 | -0.10503 | 0.91635 | 0.95722 | 9.939 |
| Ankrd40 | 6872.82 | -0.18246 | 0.08073 | -2.2601 | 0.02382 | 0.10044 | 13.841 |
| Ankrd40cl | 12.4357 | -0.11165 | 0.51628 | -0.21625 | 0.82879 | 0.91221 | 4.381 |
| Ankrd42 | 649.406 | -0.08034 | 0.12848 | -0.62535 | 0.53174 | 0.71934 | 55.42 |
| Ankrd44 | 528.433 | 0.028762 | 0.09497 | 0.302865 | 0.76199 | 0.87518 | 281.048 |
| Ankrd45 | 4379.49 | 0.028657 | 0.08628 | 0.332145 | 0.73978 | 0.86273 | 27.808 |
| Ankrd46 | 8242.98 | -0.04797 | 0.06609 | -0.72584 | 0.46793 | 0.66697 | 19.153 |
| Ankrd49 | 479.887 | 0.011255 | 0.10114 | 0.111277 | 0.9114 | 0.95526 | 5.239 |
| Ankrd50 | 3494.13 | -0.13618 | 0.072 | -1.89138 | 0.05857 | 0.18403 | 35.586 |
| Ankrd52 | 4373.23 | -0.00284 | 0.07267 | -0.03907 | 0.96883 | 0.98419 | 31.59 |
| Ankrd53 | 225.865 | 0.296401 | 0.14501 | 2.04399 | 0.04095 | 0.14491 | 5.683 |
| Ankrd54 | 575.764 | -0.07965 | 0.09617 | -0.82824 | 0.40753 | 0.61541 | 9.8 |
| Ankrd55 | 134.31 | 0.313164 | 0.21269 | 1.47239 | 0.14092 | 0.32265 | 95.552 |
| Ankrd6 | 2734.2 | 0.331717 | 0.1131 | 2.933071 | 0.00336 | 0.028 | 146.807 |
| Ankrd61 | 176.338 | 0.348774 | 0.15792 | 2.208596 | 0.0272 | 0.11019 | 6.945 |
| Ankrd63 | 6834.32 | 0.533762 | 0.17266 | 3.091379 | 0.00199 | 0.02005 | 4.861 |
| Ankrd9 | 280.632 | -0.40721 | 0.13219 | -3.08044 | 0.00207 | 0.02052 | 3.688 |
| Anks1 | 2058.73 | 0.050517 | 0.07583 | 0.666167 | 0.5053 | 0.69699 | 153.47 |
| Anks1b | 18456 | 0.224139 | 0.06194 | 3.61864 | 0.0003 | 0.00559 | 1099.792 |
| Anks3 | 3137.34 | 0.258367 | 0.07028 | 3.676232 | 0.00024 | 0.00478 | 22.77 |
| Anks4b | 32.2739 | 1.229127 | 0.33727 | 3.644387 | 0.00027 | 0.00518 | 11.729 |
| Anks6 | 781.841 | 0.080196 | 0.08342 | 0.961336 | 0.33638 | 0.54715 | 41.759 |
| Ankub1 | 204.803 | -0.34981 | 0.23665 | -1.47816 | 0.13937 | 0.32067 | 35.407 |
| Ankzf1 | 1567.92 | 0.140996 | 0.08241 | 1.710974 | 0.08709 | 0.23761 | 7.237 |
| Anln | 2031.57 | -0.68348 | 0.1824 | -3.74723 | 0.00018 | 0.004 | 57.177 |
| Ano1 | 115.689 | -0.20158 | 0.20552 | -0.98084 | 0.32667 | 0.538 | 163.426 |
| Ano10 | 537.118 | -0.17841 | 0.1331 | -1.34042 | 0.18011 | 0.37337 | 118.55 |
| Ano2 | 622.083 | 1.142484 | 0.43881 | 2.603625 | 0.00922 | 0.05431 | 349.708 |
| Ano3 | 9262.33 | 0.657131 | 0.15033 | 4.371194 | 1.24E-05 | 0.00065 | 295.723 |
| Ano4 | 1028.82 | -0.12573 | 0.09232 | -1.36195 | 0.17321 | 0.36445 | 395.769 |
| Ano5 | 175.118 | 0.299691 | 0.19765 | 1.516237 | 0.12946 | 0.30701 | 87.681 |
| Ano6 | 1459.09 | -0.00284 | 0.0676 | -0.04208 | 0.96643 | 0.98322 | 183.909 |
| Ano7 | 44.6049 | 1.151059 | 0.28325 | 4.063685 | 4.83E-05 | 0.0016 | 30.374 |
| Ano8 | 4266.05 | 0.077886 | 0.07641 | 1.019253 | 0.30808 | 0.51789 | 9.945 |

| | | | | | | | |
|---|---|---|---|---|---|---|---|
| Anp32a | 8436.51 | 0.063237 | 0.0713 | 0.886957 | 0.3751 | 0.58522 | 37.52 |
| Anp32b | 3190.45 | 0.089361 | 0.08991 | 0.993895 | 0.32027 | 0.53099 | 21.756 |
| Anp32e | 6391.84 | -0.12185 | 0.07479 | -1.62928 | 0.10325 | 0.26552 | 18.145 |
| Anpep | 177.321 | -1.01458 | 0.21539 | -4.71042 | 2.47E-06 | 0.00021 | 39.257 |
| Antkmt | 1211.89 | -0.04148 | 0.08451 | -0.4909 | 0.6235 | 0.78618 | 2.822 |
| Antxr1 | 953.361 | -0.43401 | 0.19347 | -2.24324 | 0.02488 | 0.10359 | 201.969 |
| Antxr2 | 64.7994 | -0.80137 | 0.29419 | -2.72402 | 0.00645 | 0.04314 | 148.261 |
| Anxa1 | 53.4045 | -1.53795 | 0.53209 | -2.89038 | 0.00385 | 0.03065 | 17.517 |
| Anxa11 | 675.971 | -0.53562 | 0.16565 | -3.23347 | 0.00122 | 0.01452 | 44.649 |
| Anxa11os | 15.8525 | 0.179467 | 0.4913 | 0.365288 | 0.7149 | 0.84719 | 13.49 |
| Anxa13 | 15.0436 | 0.767873 | 0.44916 | 1.70956 | 0.08735 | 0.23805 | 56.227 |
| Anxa2 | 395.566 | -0.74498 | 0.20181 | -3.69157 | 0.00022 | 0.00458 | 38.176 |
| Anxa3 | 638.315 | -0.37982 | 0.10866 | -3.49539 | 0.00047 | 0.00765 | 52.628 |
| Anxa4 | 173.291 | -0.79623 | 0.1836 | -4.3368 | 1.45E-05 | 0.00072 | 56.745 |
| Anxa5 | 2138.43 | -0.48834 | 0.1009 | -4.84007 | 1.30E-06 | 0.00013 | 26.972 |
| Anxa6 | 3870.32 | -0.16033 | 0.06755 | -2.37331 | 0.01763 | 0.0825 | 54.338 |
| Anxa7 | 3087.84 | -0.09532 | 0.08222 | -1.15932 | 0.24632 | 0.45308 | 24.874 |
| Anxa9 | 65.5797 | 0.050599 | 0.25171 | 0.201018 | 0.84068 | 0.91835 | 11.081 |
| Aoah | 14.9171 | -2 | 0.63563 | -4.18648 | 2.83E-05 | 0.00111 | 242.505 |
| Aoc2 | 545.994 | 0.54396 | 0.1214 | 4.480667 | 7.44E-06 | 0.00045 | 4.64 |
| Aoc3 | 29.702 | -0.14797 | 0.31391 | -0.47136 | 0.63739 | 0.79601 | 11.334 |
| Aopep | 1435.25 | -0.1004 | 0.10112 | -0.99289 | 0.32076 | 0.53154 | 361.204 |
| Aox1 | 390.798 | -0.49338 | 0.14004 | -3.52314 | 0.00043 | 0.00721 | 76.483 |
| Aox3 | 163.369 | -0.7965 | 0.31703 | -2.51237 | 0.01199 | 0.06465 | 87.569 |
| Aox4 | 34.2809 | -0.96296 | 0.42487 | -2.26646 | 0.02342 | 0.09929 | 58.201 |
| Ap1ar | 4759.47 | 0.256379 | 0.10101 | 2.538275 | 0.01114 | 0.06152 | 30.517 |
| Ap1b1 | 7512.46 | -0.02562 | 0.05687 | -0.45055 | 0.65231 | 0.8066 | 55.968 |
| Ap1g1 | 3653.45 | 0.047508 | 0.05308 | 0.895008 | 0.37078 | 0.58094 | 85.651 |
| Ap1g2 | 571.722 | 0.282362 | 0.0952 | 2.966045 | 0.00302 | 0.02624 | 8.019 |
| Ap1m1 | 3074.58 | 0.039601 | 0.06283 | 0.63033 | 0.52848 | 0.71632 | 17.368 |
| Ap1m2 | 38.0675 | -0.59338 | 0.33695 | -1.76105 | 0.07823 | 0.22144 | 18.063 |
| Ap1s1 | 10280.4 | 0.113096 | 0.09192 | 1.230373 | 0.21856 | 0.42065 | 11.143 |
| Ap1s2 | 3696.09 | 0.36629 | 0.11845 | 3.092251 | 0.00199 | 0.02005 | 24.65 |
| Ap1s3 | 161.844 | -0.36151 | 0.22903 | -1.57846 | 0.11446 | 0.28336 | 65.097 |

| | | | | | | | |
|---|---|---|---|---|---|---|---|
| Ap2a1 | 7063.22 | 0.025702 | 0.05639 | 0.455831 | 0.64851 | 0.80419 | 29.124 |
| Ap2a2 | 9188.96 | -0.11679 | 0.08329 | -1.40217 | 0.16086 | 0.34901 | 70.839 |
| Ap2b1 | 16526.3 | -0.0445 | 0.06608 | -0.67346 | 0.50065 | 0.69348 | 106.012 |
| Ap2m1 | 16221.8 | -0.22736 | 0.06501 | -3.49727 | 0.00047 | 0.00761 | 9.432 |
| Ap2m1-ps | 2274.39 | -0.06893 | 0.15962 | -0.43181 | 0.66588 | 0.81557 | 1.308 |
| Ap2s1 | 2827.7 | -0.18181 | 0.09147 | -1.98752 | 0.04686 | 0.15876 | 10.885 |
| Ap3b1 | 2943.34 | 0.02681 | 0.06017 | 0.445541 | 0.65593 | 0.80893 | 207.358 |
| Ap3b2 | 6449.36 | 0.005271 | 0.07197 | 0.073231 | 0.94162 | 0.97087 | 33.527 |
| Ap3d1 | 9418.18 | 0.048968 | 0.06099 | 0.802842 | 0.42207 | 0.62886 | 35.309 |
| Ap3m1 | 2604.79 | 0.038076 | 0.06262 | 0.608063 | 0.54315 | 0.72866 | 21.067 |
| Ap3m1-ps | 12.7154 | 0.643444 | 0.60634 | 1.061191 | 0.2886 | 0.49868 | 1.258 |
| Ap3m2 | 6006.57 | -0.15426 | 0.05617 | -2.74617 | 0.00603 | 0.04142 | 18.269 |
| Ap3s1 | 2509.47 | -0.04554 | 0.07848 | -0.5803 | 0.56171 | 0.74283 | 48.951 |
| Ap3s1-ps1 | 111.177 | 0.298433 | 0.30145 | 0.989999 | 0.32217 | 0.53314 | 0.579 |
| Ap3s2 | 3591.44 | -0.12837 | 0.08237 | -1.5585 | 0.11912 | 0.29131 | 45.325 |
| Ap4b1 | 17.1408 | -0.16894 | 0.50857 | -0.33218 | 0.73975 | 0.86273 | 12.506 |
| Ap4e1 | 1288.12 | 0.178338 | 0.08685 | 2.053397 | 0.04003 | 0.14298 | 59.193 |
| Ap4m1 | 718.766 | 0.262357 | 0.09766 | 2.686369 | 0.00722 | 0.04651 | 8.428 |
| Ap4s1 | 1240.38 | 0.063984 | 0.08766 | 0.729882 | 0.46546 | 0.66495 | 53.754 |
| Ap5b1 | 131.808 | -0.36234 | 0.18927 | -1.91448 | 0.05556 | 0.17789 | 3.237 |
| Ap5m1 | 1649.94 | 0.059926 | 0.09867 | 0.60734 | 0.54363 | 0.72894 | 28.138 |
| Ap5s1 | 692.652 | -0.07697 | 0.12694 | -0.60638 | 0.54426 | 0.72945 | 6.437 |
| Ap5z1 | 1192.73 | -0.04184 | 0.07075 | -0.59133 | 0.5543 | 0.73694 | 14.767 |
| Apaf1 | 950.817 | 0.169155 | 0.10535 | 1.605676 | 0.10835 | 0.27367 | 93.46 |
| Apba1 | 7251.67 | -0.45397 | 0.13042 | -3.48081 | 0.0005 | 0.00791 | 190.71 |
| Apba2 | 5586.63 | -0.20597 | 0.08343 | -2.46885 | 0.01355 | 0.07011 | 252.173 |
| Apba3 | 659.119 | -0.0148 | 0.10443 | -0.1417 | 0.88732 | 0.94396 | 6.287 |
| Apbb1 | 12841.6 | -0.28861 | 0.09454 | -3.0527 | 0.00227 | 0.02186 | 23.171 |
| Apbb1ip | 191.267 | -0.10049 | 0.16831 | -0.59703 | 0.55049 | 0.73403 | 101.56 |
| Apbb2 | 2318.21 | -0.24952 | 0.08743 | -2.85405 | 0.00432 | 0.03314 | 320.082 |
| Apbb3 | 2328.56 | 0.173162 | 0.08065 | 2.147202 | 0.03178 | 0.12282 | 8.213 |
| Apc | 16930.6 | -0.2242 | 0.14577 | -1.53804 | 0.12404 | 0.29865 | 101.629 |
| Apc2 | 8130.64 | 0.099988 | 0.05137 | 1.946285 | 0.05162 | 0.16941 | 22.287 |
| Apcdd1 | 1120.54 | -0.55271 | 0.11592 | -4.76816 | 1.86E-06 | 0.00017 | 38.949 |

| | | | | | | | |
|---|---|---|---|---|---|---|---|
| Apeh | 1156.76 | 0.232685 | 0.09842 | 2.36419 | 0.01807 | 0.08383 | 9.194 |
| Apex1 | 321.519 | 0.122475 | 0.12055 | 1.015927 | 0.30966 | 0.51965 | 2.172 |
| Apex2 | 395.097 | 0.15109 | 0.13138 | 1.15006 | 0.25012 | 0.45759 | 70.35 |
| Aph1a | 1618.63 | -0.09668 | 0.08472 | -1.14118 | 0.2538 | 0.46112 | 4.32 |
| Aph1b | 2152.77 | -0.1138 | 0.11736 | -0.96968 | 0.33221 | 0.5431 | 20.289 |
| Aph1c | 484.037 | -0.02929 | 0.11255 | -0.26022 | 0.7947 | 0.89379 | 19.733 |
| Api5 | 4395.06 | 0.088762 | 0.06315 | 1.405582 | 0.15985 | 0.34778 | 26.455 |
| Apip | 439.362 | -0.10973 | 0.10607 | -1.03453 | 0.30089 | 0.51075 | 18.97 |
| Aplf | 292.796 | 0.085904 | 0.15331 | 0.560337 | 0.57525 | 0.7533 | 43.77 |
| Apln | 1176.51 | -0.54236 | 0.11767 | -4.60899 | 4.05E-06 | 0.00029 | 9.708 |
| Aplp1 | 42917.7 | -0.08555 | 0.07222 | -1.18458 | 0.23618 | 0.44147 | 10.554 |
| Aplp2 | 29382.4 | -0.09744 | 0.05676 | -1.71664 | 0.08604 | 0.23562 | 62.259 |
| Apmap | 1946.53 | -0.09263 | 0.06582 | -1.4073 | 0.15934 | 0.34704 | 25.488 |
| Apobec1 | 94.8556 | -0.65149 | 0.22425 | -2.90518 | 0.00367 | 0.02974 | 24.653 |
| Apobec2 | 21.6572 | -0.0728 | 0.36461 | -0.19966 | 0.84175 | 0.91879 | 13.7 |
| Apobec3 | 181.604 | -0.44831 | 0.15879 | -2.82337 | 0.00475 | 0.03518 | 24.248 |
| Apobr | 41.354 | -0.64375 | 0.29333 | -2.19465 | 0.02819 | 0.11309 | 4.171 |
| Apoc1 | 20.8251 | -0.44237 | 0.6308 | -0.70128 | 0.48313 | 0.67955 | 3.18 |
| Apod | 7116.51 | -1.00858 | 0.16489 | -6.11655 | 9.56E-10 | 5.13E-07 | 18.617 |
| Apoe | 60435.1 | -0.24086 | 0.13986 | -1.72212 | 0.08505 | 0.23388 | 3.08 |
| Apol8 | 264.404 | 0.322 | 0.15718 | 2.048662 | 0.0405 | 0.14378 | 9.206 |
| Apold1 | 299.155 | -0.35479 | 0.26035 | -1.36274 | 0.17297 | 0.36423 | 5.119 |
| Apom | 14.1901 | 0.451998 | 0.57216 | 0.789982 | 0.42954 | 0.63601 | 3.054 |
| Apoo | 1281.64 | 0.069846 | 0.07253 | 0.963023 | 0.33554 | 0.54638 | 49.977 |
| Apoo-ps | 59.5479 | -0.82485 | 0.38206 | -2.15894 | 0.03085 | 0.12042 | 0.586 |
| Apool | 180.044 | 0.066966 | 0.16012 | 0.418221 | 0.67579 | 0.82201 | 61.422 |
| App | 61870.8 | -0.10933 | 0.05345 | -2.04532 | 0.04082 | 0.14468 | 224.082 |
| Appbp2 | 5212.01 | 0.03011 | 0.05498 | 0.547664 | 0.58392 | 0.75891 | 47.869 |
| Appbp2os | 33.2384 | 0.168389 | 0.4387 | 0.383837 | 0.7011 | 0.83856 | 3.53 |
| Appl1 | 6439.96 | 0.116102 | 0.09184 | 1.264119 | 0.20619 | 0.40534 | 52.245 |
| Appl2 | 4605.42 | -0.19882 | 0.08008 | -2.48288 | 0.01303 | 0.06831 | 48.706 |
| Aprt | 525.031 | -0.15525 | 0.10567 | -1.46925 | 0.14177 | 0.32357 | 2.275 |
| Aptx | 1556.05 | -0.0214 | 0.08623 | -0.24824 | 0.80395 | 0.89875 | 20.813 |
| Aqp1 | 23.0056 | -0.75052 | 0.41534 | -1.807 | 0.07076 | 0.20791 | 12.124 |

| | | | | | | | |
|---|---|---|---|---|---|---|---|
| Aqp11 | 559.413 | -0.69856 | 0.1337 | -5.22499 | 1.74E-07 | 2.75E-05 | 14.284 |
| Aqp4 | 6163.13 | -0.45492 | 0.12771 | -3.56211 | 0.00037 | 0.00646 | 21.487 |
| Aqp6 | 14.0546 | 0.768052 | 0.59298 | 1.295243 | 0.19524 | 0.39279 | 4.078 |
| Aqp9 | 67.1641 | -0.32224 | 0.23901 | -1.34821 | 0.17759 | 0.36981 | 58.024 |
| Aqr | 2720.53 | 0.264984 | 0.07224 | 3.667899 | 0.00024 | 0.00489 | 85.855 |
| Ar | 1365.86 | -0.11754 | 0.07854 | -1.49656 | 0.13451 | 0.31436 | 174.447 |
| Araf | 8805.87 | 0.144718 | 0.06201 | 2.333812 | 0.01961 | 0.08838 | 62.706 |
| Arap1 | 851.871 | -0.18203 | 0.097 | -1.87668 | 0.06056 | 0.18784 | 64.52 |
| Arap2 | 3246.08 | -0.21965 | 0.12073 | -1.81935 | 0.06886 | 0.20404 | 163.715 |
| Arap3 | 527.844 | -0.04372 | 0.13406 | -0.32615 | 0.74431 | 0.86539 | 26.342 |
| Arc | 3146.77 | -0.63698 | 0.28986 | -2.19754 | 0.02798 | 0.11257 | 3.488 |
| Arcn1 | 5816.02 | -0.0001 | 0.05522 | -0.00188 | 0.9985 | 0.99919 | 26.282 |
| Arel1 | 12804.9 | 0.491386 | 0.15498 | 3.17065 | 0.00152 | 0.01675 | 52.753 |
| Arf1 | 15980 | -0.00983 | 0.06958 | -0.14124 | 0.88768 | 0.94402 | 16.859 |
| Arf2 | 2055.3 | 0.056488 | 0.08531 | 0.662158 | 0.50787 | 0.6992 | 18.599 |
| Arf3 | 63636.2 | 0.041108 | 0.07979 | 0.515167 | 0.60644 | 0.77468 | 26.581 |
| Arf4 | 4490.62 | 0.179046 | 0.1095 | 1.63519 | 0.10201 | 0.26371 | 27.011 |
| Arf5 | 2759.68 | -0.13823 | 0.07137 | -1.93675 | 0.05278 | 0.17188 | 3.043 |
| Arf6 | 867.86 | -0.17128 | 0.0773 | -2.21571 | 0.02671 | 0.10896 | 1.634 |
| Arfgap1 | 4333.51 | 0.023151 | 0.0634 | 0.365139 | 0.71501 | 0.84727 | 15.302 |
| Arfgap2 | 1629.8 | 0.108198 | 0.07552 | 1.432756 | 0.15193 | 0.33702 | 11.958 |
| Arfgap3 | 768.229 | -0.38945 | 0.11682 | -3.3338 | 0.00086 | 0.0115 | 50.509 |
| Arfgef1 | 5955.95 | 0.099736 | 0.09048 | 1.102234 | 0.27036 | 0.47993 | 95.1 |
| Arfgef2 | 5162.95 | 0.037547 | 0.05482 | 0.684943 | 0.49338 | 0.6878 | 92.465 |
| Arfgef3 | 8127.68 | 0.206998 | 0.08363 | 2.475027 | 0.01332 | 0.06927 | 162.111 |
| Arfip1 | 109.327 | -0.01689 | 0.20611 | -0.08194 | 0.9347 | 0.96717 | 86.533 |
| Arfip2 | 3763.67 | 0.007607 | 0.06979 | 0.109004 | 0.9132 | 0.95612 | 6.214 |
| Arfrp1 | 1702.77 | 0.151386 | 0.07828 | 1.933844 | 0.05313 | 0.17246 | 7.715 |
| Arg2 | 168.427 | 0.253679 | 0.22052 | 1.150344 | 0.25 | 0.45741 | 25.525 |
| Arglu1 | 14385.1 | 0.378813 | 0.08761 | 4.323742 | 1.53E-05 | 0.00075 | 25.447 |
| Arhgap1 | 2581.14 | -0.01245 | 0.08686 | -0.14334 | 0.88602 | 0.94329 | 22.467 |
| Arhgap10 | 1653.42 | 0.40224 | 0.12797 | 3.143223 | 0.00167 | 0.0179 | 267.588 |
| Arhgap11a | 279.81 | -0.28094 | 0.16442 | -1.70866 | 0.08751 | 0.23839 | 17.17 |
| Arhgap12 | 2320.41 | 0.213691 | 0.08724 | 2.449598 | 0.0143 | 0.07211 | 111.672 |

| | | | | | | | |
|---|---|---|---|---|---|---|---|
| Arhgap15 | 274.897 | -0.60336 | 0.53785 | -1.12179 | 0.26195 | 0.4707 | 647.13 |
| Arhgap17 | 1316.24 | 0.259903 | 0.07868 | 3.303273 | 0.00096 | 0.01242 | 90.698 |
| Arhgap18 | 1212.67 | 0.650471 | 0.16643 | 3.908452 | 9.29E-05 | 0.00255 | 165.228 |
| Arhgap19 | 535.12 | 0.305843 | 0.17891 | 1.709469 | 0.08736 | 0.23805 | 35.46 |
| Arhgap20 | 5411.18 | 0.182555 | 0.07782 | 2.345981 | 0.01898 | 0.08652 | 88.52 |
| Arhgap20os | 62.6317 | 0.161911 | 0.23027 | 0.703143 | 0.48197 | 0.67862 | 43.073 |
| Arhgap21 | 11333.2 | -0.02792 | 0.07832 | -0.35654 | 0.72144 | 0.85135 | 120.963 |
| Arhgap22 | 128.683 | -0.4786 | 0.21121 | -2.26605 | 0.02345 | 0.09933 | 155.909 |
| Arhgap23 | 4587.92 | -0.4597 | 0.11573 | -3.97211 | 7.12E-05 | 0.00212 | 86.87 |
| Arhgap24 | 359.89 | -0.03933 | 0.14061 | -0.2797 | 0.77971 | 0.88519 | 416.547 |
| Arhgap25 | 106.553 | -0.67776 | 0.25912 | -2.61561 | 0.00891 | 0.0531 | 74.715 |
| Arhgap26 | 7546.7 | 0.300224 | 0.09849 | 3.048171 | 0.0023 | 0.02206 | 433.817 |
| Arhgap27 | 790.588 | 0.10742 | 0.11087 | 0.968915 | 0.33259 | 0.54345 | 32.196 |
| Arhgap27os3 | 27.4962 | 0.022835 | 0.34154 | 0.066859 | 0.94669 | 0.9732 | 5.372 |
| Arhgap28 | 52.5684 | -0.51809 | 0.28367 | -1.82638 | 0.06779 | 0.20188 | 161.408 |
| Arhgap29 | 1606.49 | -0.69728 | 0.14097 | -4.94613 | 7.57E-07 | 8.82E-05 | 64.213 |
| Arhgap30 | 204.201 | -0.38788 | 0.15073 | -2.57337 | 0.01007 | 0.05775 | 21.345 |
| Arhgap31 | 1668.57 | -0.44205 | 0.08762 | -5.04509 | 4.53E-07 | 5.85E-05 | 114.935 |
| Arhgap32 | 22267.4 | 0.156334 | 0.07841 | 1.993707 | 0.04618 | 0.15693 | 152.311 |
| Arhgap33 | 11261.7 | 0.353673 | 0.11665 | 3.031853 | 0.00243 | 0.02289 | 12.835 |
| Arhgap33os | 15.0749 | 1.018303 | 0.56914 | 1.789198 | 0.07358 | 0.2136 | 7.072 |
| Arhgap35 | 9542.16 | -0.08138 | 0.07181 | -1.13317 | 0.25714 | 0.46513 | 121.275 |
| Arhgap36 | 132.571 | 1.77332 | 0.65124 | 2.722969 | 0.00647 | 0.04327 | 36.3 |
| Arhgap39 | 3633.13 | -0.05678 | 0.09436 | -0.60178 | 0.54732 | 0.73143 | 94.186 |
| Arhgap4 | 208.108 | 0.174481 | 0.13924 | 1.253087 | 0.21017 | 0.4102 | 30.429 |
| Arhgap40 | 102.74 | 0.613918 | 0.27895 | 2.200812 | 0.02775 | 0.1119 | 37.967 |
| Arhgap42 | 574.812 | -0.39308 | 0.1249 | -3.14725 | 0.00165 | 0.0177 | 244.777 |
| Arhgap44 | 7006.19 | -0.27009 | 0.09512 | -2.83954 | 0.00452 | 0.03403 | 160.923 |
| Arhgap45 | 376.708 | -0.66719 | 0.13543 | -4.92642 | 8.38E-07 | 9.38E-05 | 14.82 |
| Arhgap5 | 6568.91 | 0.171803 | 0.19238 | 0.893021 | 0.37185 | 0.58216 | 68.004 |
| Arhgap6 | 1756.6 | 0.91208 | 0.20811 | 4.382745 | 1.17E-05 | 0.00062 | 509.337 |
| Arhgap9 | 32.247 | -0.4371 | 0.33231 | -1.31536 | 0.18839 | 0.38397 | 7.98 |
| Arhgdia | 8713.8 | -0.21032 | 0.07315 | -2.87533 | 0.00404 | 0.03169 | 3.521 |
| Arhgdib | 188.171 | -0.16709 | 0.16203 | -1.03127 | 0.30241 | 0.51199 | 18.245 |

| | | | | | | | |
|---|---|---|---|---|---|---|---|
| Arhgdig | 880.434 | -0.49001 | 0.10686 | -4.58562 | 4.53E-06 | 0.00031 | 8.604 |
| Arhgef1 | 3607.35 | 0.08644 | 0.1051 | 0.822496 | 0.41079 | 0.61876 | 23.683 |
| Arhgef10 | 3005.39 | -0.42945 | 0.11156 | -3.84947 | 0.00012 | 0.00301 | 89.423 |
| Arhgef10l | 2056.25 | 0.109519 | 0.10862 | 1.008276 | 0.31332 | 0.52381 | 151.528 |
| Arhgef11 | 3548.05 | -0.11925 | 0.05892 | -2.02414 | 0.04296 | 0.1495 | 120.476 |
| Arhgef12 | 14573.9 | 0.033065 | 0.06029 | 0.548445 | 0.58339 | 0.75845 | 143.397 |
| Arhgef15 | 1299.75 | 0.334644 | 0.13373 | 2.502338 | 0.01234 | 0.06589 | 14.326 |
| Arhgef16 | 22.2506 | -0.40483 | 0.40344 | -1.00345 | 0.31564 | 0.52626 | 23.191 |
| Arhgef17 | 8324.31 | 0.188351 | 0.08542 | 2.204933 | 0.02746 | 0.11092 | 62.356 |
| Arhgef18 | 4039.53 | 0.269056 | 0.11263 | 2.388913 | 0.0169 | 0.08054 | 103.187 |
| Arhgef19 | 962.503 | -0.07427 | 0.09428 | -0.78779 | 0.43082 | 0.63702 | 18.066 |
| Arhgef2 | 9349.77 | 0.023109 | 0.07249 | 0.318791 | 0.74989 | 0.8687 | 42.087 |
| Arhgef25 | 5415.72 | -0.48554 | 0.13109 | -3.70397 | 0.00021 | 0.00444 | 7.559 |
| Arhgef26 | 1070.54 | -0.13053 | 0.11829 | -1.10349 | 0.26982 | 0.47946 | 123.878 |
| Arhgef28 | 2106.42 | -0.69824 | 0.44543 | -1.56756 | 0.11698 | 0.28768 | 306.971 |
| Arhgef3 | 3973.52 | 0.103017 | 0.06717 | 1.533669 | 0.12511 | 0.30014 | 289.013 |
| Arhgef33 | 121.89 | 0.781079 | 0.18812 | 4.151958 | 3.30E-05 | 0.00123 | 92.675 |
| Arhgef37 | 260.557 | -0.67521 | 0.17728 | -3.80864 | 0.00014 | 0.00336 | 44.938 |
| Arhgef39 | 103.074 | 0.331003 | 0.28331 | 1.168342 | 0.24267 | 0.44883 | 3.554 |
| Arhgef4 | 5004.76 | -0.11553 | 0.07525 | -1.5352 | 0.12474 | 0.29973 | 135.122 |
| Arhgef40 | 1252.71 | 0.038291 | 0.10115 | 0.378547 | 0.70502 | 0.84115 | 21.533 |
| Arhgef5 | 223.626 | -0.39972 | 0.1875 | -2.13183 | 0.03302 | 0.12617 | 23.739 |
| Arhgef6 | 731.66 | -0.22186 | 0.10404 | -2.13248 | 0.03297 | 0.12614 | 107.245 |
| Arhgef7 | 7352.35 | -0.05921 | 0.08614 | -0.68738 | 0.49184 | 0.68654 | 107.499 |
| Arhgef9 | 26951.1 | 0.233827 | 0.10238 | 2.283952 | 0.02237 | 0.09652 | 147.922 |
| Arid1a | 8368.23 | 0.157248 | 0.06489 | 2.423241 | 0.01538 | 0.0757 | 77.762 |
| Arid1b | 5245.22 | -0.02599 | 0.05705 | -0.45564 | 0.64865 | 0.80422 | 353.325 |
| Arid2 | 2656.37 | 0.041618 | 0.07063 | 0.58928 | 0.55567 | 0.73795 | 117.475 |
| Arid3a | 474.3 | -0.05788 | 0.1049 | -0.55181 | 0.58108 | 0.75714 | 27.976 |
| Arid3b | 502.449 | 0.106024 | 0.12645 | 0.838499 | 0.40175 | 0.61037 | 46.441 |
| Arid4a | 2825.22 | 0.216557 | 0.12751 | 1.698385 | 0.08944 | 0.24227 | 82.603 |
| Arid4b | 3639.41 | 0.278202 | 0.18162 | 1.531822 | 0.12557 | 0.30073 | 136.372 |
| Arid5a | 459.351 | -0.43294 | 0.12572 | -3.44362 | 0.00057 | 0.00871 | 16.297 |
| Arid5b | 2660.92 | -0.10874 | 0.11462 | -0.94871 | 0.34277 | 0.55356 | 186.221 |

| | | | | | | | |
|---|---|---|---|---|---|---|---|
| Arih1 | 6211.2 | 0.106659 | 0.06096 | 1.749692 | 0.08017 | 0.22519 | 98.361 |
| Arih2 | 2889.35 | 0.133564 | 0.07225 | 1.848668 | 0.06451 | 0.19551 | 46.445 |
| Arl1 | 5334.02 | -0.07999 | 0.07164 | -1.11653 | 0.2642 | 0.47298 | 13.237 |
| Arl10 | 465.989 | -0.08778 | 0.12225 | -0.71802 | 0.47274 | 0.67097 | 6.114 |
| Arl11 | 18.6169 | -0.46927 | 0.50285 | -0.93321 | 0.35071 | 0.56136 | 2.184 |
| Arl13b | 694.115 | 0.144826 | 0.1307 | 1.108101 | 0.26782 | 0.47712 | 53.356 |
| Arl14ep | 2795.14 | 0.269269 | 0.08616 | 3.125099 | 0.00178 | 0.01865 | 11.898 |
| Arl15 | 2241.07 | -0.04475 | 0.09527 | -0.4697 | 0.63857 | 0.7971 | 362.957 |
| Arl16 | 754.418 | 0.050105 | 0.09377 | 0.534333 | 0.59311 | 0.76548 | 2.742 |
| Arl2 | 1282.91 | 0.166136 | 0.08414 | 1.974609 | 0.04831 | 0.16205 | 7.175 |
| Arl2bp | 4768.93 | -0.13218 | 0.06339 | -2.08522 | 0.03705 | 0.13608 | 7.826 |
| Arl3 | 3375.08 | -0.07654 | 0.07138 | -1.07238 | 0.28355 | 0.49363 | 42.103 |
| Arl4a | 2428.91 | 0.430934 | 0.11056 | 3.897743 | 9.71E-05 | 0.00263 | 32.579 |
| Arl4c | 1821.04 | -0.0882 | 0.08511 | -1.03626 | 0.30008 | 0.51023 | 29.097 |
| Arl4d | 1333.95 | -0.09382 | 0.15897 | -0.59016 | 0.55508 | 0.7376 | 2.292 |
| Arl5a | 4400.77 | 0.063296 | 0.07748 | 0.816953 | 0.41396 | 0.622 | 26.951 |
| Arl5b | 1499.24 | -0.06931 | 0.09611 | -0.72123 | 0.47077 | 0.66942 | 33.062 |
| Arl5c | 164.748 | 0.336696 | 0.15691 | 2.145763 | 0.03189 | 0.12317 | 6.604 |
| Arl6 | 1056.84 | 0.287335 | 0.11185 | 2.568902 | 0.0102 | 0.05832 | 26.443 |
| Arl6ip1 | 13341.5 | -0.11764 | 0.09339 | -1.25973 | 0.20777 | 0.40734 | 10.772 |
| Arl6ip4 | 1256.5 | 0.01868 | 0.07955 | 0.234827 | 0.81434 | 0.90466 | 2.108 |
| Arl6ip5 | 3189.08 | -0.11434 | 0.06866 | -1.6654 | 0.09583 | 0.25327 | 22.627 |
| Arl6ip6 | 615.016 | 0.081349 | 0.10749 | 0.756824 | 0.44916 | 0.65243 | 27.495 |
| Arl8a | 5841.22 | -0.20462 | 0.06809 | -3.00507 | 0.00266 | 0.0243 | 9.446 |
| Arl8b | 8624.59 | -0.07531 | 0.05108 | -1.47434 | 0.14039 | 0.3219 | 42.18 |
| Arl9 | 14.9717 | 0.626614 | 0.46887 | 1.336429 | 0.18141 | 0.37488 | 6.552 |
| Armc1 | 4112.42 | 0.07322 | 0.07181 | 1.019595 | 0.30792 | 0.51771 | 31.664 |
| Armc10 | 744.177 | -0.03461 | 0.08015 | -0.4318 | 0.66589 | 0.81557 | 16.882 |
| Armc2 | 505.413 | 0.196901 | 0.09729 | 2.023795 | 0.04299 | 0.14957 | 103.453 |
| Armc3 | 15.8007 | -0.50343 | 0.41937 | -1.20043 | 0.22997 | 0.43425 | 110.94 |
| Armc5 | 560.399 | 0.013587 | 0.09651 | 0.140781 | 0.88804 | 0.94406 | 7.759 |
| Armc6 | 885.156 | 0.02528 | 0.08249 | 0.306444 | 0.75927 | 0.87354 | 14.295 |
| Armc7 | 307.798 | 0.067108 | 0.1409 | 0.476289 | 0.63387 | 0.79385 | 14.801 |
| Armc8 | 4152.29 | 0.22361 | 0.09409 | 2.376469 | 0.01748 | 0.08211 | 90.528 |

| | | | | | | | |
|---|---|---|---|---|---|---|---|
| Armc9 | 1771.31 | 0.163332 | 0.08031 | 2.033786 | 0.04197 | 0.14726 | 123.505 |
| Armcx1 | 3028.15 | -0.02663 | 0.10496 | -0.25372 | 0.79971 | 0.89651 | 3.955 |
| Armcx2 | 3830.41 | -0.05041 | 0.07149 | -0.7051 | 0.48075 | 0.67798 | 5.077 |
| Armcx3 | 3049.18 | 0.005246 | 0.07767 | 0.06754 | 0.94615 | 0.97302 | 4.861 |
| Armcx4 | 3979.97 | 0.024877 | 0.0973 | 0.255676 | 0.7982 | 0.89581 | 10.239 |
| Armcx5 | 674.885 | 0.024366 | 0.10177 | 0.239412 | 0.81079 | 0.90258 | 4.59 |
| Armcx6 | 216.917 | 0.307446 | 0.17507 | 1.756098 | 0.07907 | 0.22311 | 2.957 |
| Armh3 | 945.967 | 0.011518 | 0.10164 | 0.113317 | 0.90978 | 0.9546 | 181.125 |
| Armh4 | 1381.99 | -0.4363 | 0.12405 | -3.51725 | 0.00044 | 0.00729 | 107.432 |
| Armt1 | 1393.86 | 0.17035 | 0.08886 | 1.917063 | 0.05523 | 0.17717 | 22.675 |
| Arnt | 3191.48 | 0.060193 | 0.0992 | 0.60678 | 0.544 | 0.72924 | 62.853 |
| Arnt2 | 14082 | -0.07264 | 0.07146 | -1.0165 | 0.30939 | 0.51936 | 163.899 |
| Arntl | 1172 | 0.15031 | 0.09363 | 1.60534 | 0.10842 | 0.27367 | 106.662 |
| Arntl2 | 781.518 | -0.02791 | 0.11597 | -0.24062 | 0.80985 | 0.90197 | 37.475 |
| Arpc1a | 6665.4 | 0.188359 | 0.10134 | 1.858596 | 0.06308 | 0.19285 | 24.932 |
| Arpc1b | 648.785 | -0.5027 | 0.1453 | -3.45967 | 0.00054 | 0.00838 | 16.491 |
| Arpc2 | 6975.96 | 0.002832 | 0.07187 | 0.039399 | 0.96857 | 0.9841 | 32.126 |
| Arpc3 | 1976.07 | -0.29295 | 0.10123 | -2.89383 | 0.00381 | 0.03043 | 22.307 |
| Arpc4 | 3217.83 | -0.24867 | 0.09866 | -2.52042 | 0.01172 | 0.0637 | 12.334 |
| Arpc5 | 4423.06 | -0.21269 | 0.11484 | -1.85202 | 0.06402 | 0.19475 | 9.056 |
| Arpc5l | 3382.6 | -0.03652 | 0.06603 | -0.5531 | 0.5802 | 0.75652 | 10.53 |
| Arpin | 1072.24 | 0.067641 | 0.10451 | 0.647241 | 0.51748 | 0.70733 | 9.999 |
| Arpp19 | 16421.4 | 0.411405 | 0.13869 | 2.96641 | 0.00301 | 0.02623 | 22.7 |
| Arpp21 | 43665.2 | 0.644344 | 0.16242 | 3.967156 | 7.27E-05 | 0.00215 | 170.848 |
| Arr3 | 78.7821 | 0.603453 | 0.22327 | 2.702773 | 0.00688 | 0.04491 | 12.997 |
| Arrb1 | 19733.1 | -0.03432 | 0.06954 | -0.49361 | 0.62159 | 0.78484 | 71.306 |
| Arrb2 | 1547.98 | -0.11372 | 0.08287 | -1.37232 | 0.16996 | 0.3604 | 8.194 |
| Arrdc1 | 234.17 | -0.17697 | 0.12814 | -1.381 | 0.16728 | 0.35697 | 9.901 |
| Arrdc2 | 1353.91 | -0.26382 | 0.13416 | -1.96641 | 0.04925 | 0.16429 | 4.592 |
| Arrdc3 | 3874.37 | -0.01702 | 0.10964 | -0.15522 | 0.87664 | 0.93831 | 12.659 |
| Arrdc4 | 510.767 | 0.342043 | 0.10635 | 3.216311 | 0.0013 | 0.01497 | 12.247 |
| Arsa | 1180.13 | -0.08411 | 0.08431 | -0.99764 | 0.31845 | 0.52908 | 4.95 |
| Arsb | 4136.59 | -0.18834 | 0.09002 | -2.09224 | 0.03642 | 0.13444 | 171.387 |
| Arsg | 724.472 | -0.54164 | 0.12528 | -4.32361 | 1.53E-05 | 0.00075 | 99.957 |

| | | | | | | | |
|---|---|---|---|---|---|---|---|
| Arsi | 32.0057 | -1.37956 | 0.36984 | -3.73011 | 0.00019 | 0.00418 | 6.782 |
| Arsj | 95.5782 | -0.47672 | 0.35828 | -1.33057 | 0.18333 | 0.3774 | 76.692 |
| Arsk | 520.565 | 0.010906 | 0.11306 | 0.096468 | 0.92315 | 0.96135 | 38.239 |
| Art3 | 129.926 | 0.08678 | 0.19605 | 0.442636 | 0.65803 | 0.81018 | 82.802 |
| Artn | 30.4908 | -0.41914 | 0.4242 | -0.98807 | 0.32312 | 0.53418 | 3.602 |
| Arv1 | 404.938 | 0.220945 | 0.11993 | 1.842261 | 0.06544 | 0.19735 | 11.985 |
| Arvcf | 1483.32 | 0.061343 | 0.09204 | 0.666476 | 0.50511 | 0.69681 | 59.781 |
| Arx | 830.472 | -0.24071 | 0.11592 | -2.07657 | 0.03784 | 0.13791 | 11.913 |
| Arxes1 | 904.237 | 0.206513 | 0.10868 | 1.900276 | 0.0574 | 0.18181 | 1.58 |
| Arxes2 | 1475.97 | 0.235499 | 0.0869 | 2.710092 | 0.00673 | 0.04429 | 1.532 |
| As3mt | 645.101 | -0.24437 | 0.10991 | -2.22329 | 0.0262 | 0.10748 | 33.642 |
| Asah1 | 2896.35 | 0.005232 | 0.07141 | 0.07327 | 0.94159 | 0.97087 | 34.577 |
| Asah2 | 970.474 | 0.160927 | 0.12493 | 1.288118 | 0.1977 | 0.39553 | 125.615 |
| Asap1 | 4073.91 | -0.01741 | 0.06906 | -0.25208 | 0.80098 | 0.89715 | 296.063 |
| Asap2 | 1847.61 | -0.10964 | 0.09233 | -1.18742 | 0.23506 | 0.44032 | 279.713 |
| Asap3 | 314.026 | -0.03244 | 0.11083 | -0.29269 | 0.76976 | 0.87983 | 38.852 |
| Asb1 | 2246.61 | -0.08753 | 0.07979 | -1.0969 | 0.27268 | 0.48183 | 19.046 |
| Asb11 | 341.19 | 0.592939 | 0.23602 | 2.512229 | 0.012 | 0.06466 | 22.177 |
| Asb13 | 2404.22 | -0.19841 | 0.08093 | -2.45163 | 0.01422 | 0.07191 | 19.791 |
| Asb14 | 47.0409 | 0.17145 | 0.2607 | 0.65766 | 0.51076 | 0.70199 | 20.702 |
| Asb16 | 115.285 | -0.08602 | 0.20584 | -0.41788 | 0.67603 | 0.82226 | 10.72 |
| Asb18 | 194.1 | 0.143071 | 0.20603 | 0.694417 | 0.48742 | 0.68276 | 64.447 |
| Asb2 | 96.6319 | -0.35968 | 0.20126 | -1.78717 | 0.07391 | 0.21422 | 34.86 |
| Asb3 | 811.38 | 0.207463 | 0.08167 | 2.540326 | 0.01107 | 0.06133 | 217.289 |
| Asb4 | 34.9304 | -0.35564 | 0.45562 | -0.78056 | 0.43506 | 0.64087 | 49.637 |
| Asb5 | 35.7295 | 0.52382 | 0.31819 | 1.64624 | 0.09971 | 0.25969 | 58.296 |
| Asb6 | 1167.04 | -0.07574 | 0.07745 | -0.97791 | 0.32812 | 0.53939 | 7.237 |
| Asb7 | 989.6 | 0.036338 | 0.09702 | 0.374544 | 0.708 | 0.84235 | 45.032 |
| Asb8 | 1445.83 | -0.10554 | 0.06865 | -1.53743 | 0.12419 | 0.29891 | 30.989 |
| Ascc1 | 587.561 | 0.020761 | 0.11953 | 0.173682 | 0.86212 | 0.93002 | 97.184 |
| Ascc2 | 2288.75 | 0.219724 | 0.06989 | 3.143948 | 0.00167 | 0.01787 | 47.953 |
| Ascc3 | 1294.17 | 0.290207 | 0.11407 | 2.544196 | 0.01095 | 0.06094 | 258.817 |
| Ascl1 | 233.967 | -0.85484 | 0.22127 | -3.8633 | 0.00011 | 0.0029 | 2.842 |
| Asf1a | 293.857 | -0.06998 | 0.14133 | -0.49512 | 0.62052 | 0.78421 | 12.469 |

| | | | | | | | |
|---|---|---|---|---|---|---|---|
| Asf1b | 31.1017 | -0.47001 | 0.35704 | -1.3164 | 0.18804 | 0.38365 | 14.691 |
| Asgr1 | 31.4656 | -1.01613 | 0.39192 | -2.59273 | 0.00952 | 0.05554 | 3.81 |
| Ash1l | 10264.2 | 0.165633 | 0.07597 | 2.180334 | 0.02923 | 0.116 | 128.754 |
| Ash2l | 2553.68 | 0.063719 | 0.0825 | 0.772372 | 0.43989 | 0.64503 | 31.699 |
| Asic1 | 5133.8 | 0.202985 | 0.09749 | 2.082077 | 0.03734 | 0.13666 | 30.763 |
| Asic2 | 1700.93 | -0.21561 | 0.1261 | -1.70982 | 0.0873 | 0.23803 | 1088.289 |
| Asic3 | 63.4526 | 0.21838 | 0.239 | 0.913706 | 0.36087 | 0.57112 | 4.444 |
| Asic4 | 1986.01 | 0.492073 | 0.17909 | 2.747629 | 0.006 | 0.0413 | 23.908 |
| Asl | 633.81 | 0.091285 | 0.11402 | 0.800616 | 0.42335 | 0.63018 | 17.99 |
| Asns | 3841.29 | -0.05614 | 0.07379 | -0.76072 | 0.44682 | 0.65034 | 18.086 |
| Asnsd1 | 44.3183 | 0.257248 | 0.28478 | 0.903325 | 0.36635 | 0.57688 | 8.136 |
| Aspa | 600.449 | -0.51351 | 0.17322 | -2.96452 | 0.00303 | 0.02631 | 24.605 |
| Aspdh | 150.039 | -0.1352 | 0.18153 | -0.7448 | 0.4564 | 0.65798 | 2.368 |
| Aspg | 156.042 | -1.27952 | 0.24722 | -5.17556 | 2.27E-07 | 3.32E-05 | 20.881 |
| Asph | 4582.06 | -0.19251 | 0.08211 | -2.34444 | 0.01906 | 0.08668 | 221.276 |
| Asphd1 | 1167.71 | 0.055409 | 0.07179 | 0.771784 | 0.44024 | 0.64523 | 4.016 |
| Asphd2 | 4441.43 | 0.148246 | 0.07815 | 1.89686 | 0.05785 | 0.18276 | 10.513 |
| Aspm | 56.2682 | -0.42793 | 0.35594 | -1.20224 | 0.22927 | 0.43337 | 39.32 |
| Aspn | 41.0953 | -0.04855 | 0.34801 | -0.13951 | 0.88905 | 0.94457 | 23.123 |
| Asprv1 | 64.5329 | 0.610142 | 0.21182 | 2.880431 | 0.00397 | 0.03132 | 1.547 |
| Aspscr1 | 816.302 | -0.06384 | 0.0874 | -0.7304 | 0.46515 | 0.66476 | 36.475 |
| Asrgl1 | 9674.07 | -0.22396 | 0.08238 | -2.71877 | 0.00655 | 0.04365 | 25.769 |
| Ass1 | 1577.76 | 0.323467 | 0.16463 | 1.964777 | 0.04944 | 0.16463 | 50.466 |
| Aste1 | 305.104 | -0.07319 | 0.13697 | -0.53434 | 0.5931 | 0.76548 | 16.838 |
| Astn1 | 12738 | 0.031121 | 0.0695 | 0.447806 | 0.65429 | 0.80821 | 329.509 |
| Astn2 | 1170.22 | -0.41261 | 0.12616 | -3.27047 | 0.00107 | 0.01341 | 1023.809 |
| Asxl1 | 3144.51 | 0.140963 | 0.06572 | 2.144791 | 0.03197 | 0.12342 | 58.179 |
| Asxl2 | 2843.06 | 0.248521 | 0.08584 | 2.895262 | 0.00379 | 0.03032 | 79.996 |
| Asxl3 | 770.094 | -0.56118 | 0.21463 | -2.61466 | 0.00893 | 0.05322 | 185.982 |
| Atad1 | 5162.95 | 0.042965 | 0.06142 | 0.699545 | 0.48421 | 0.68059 | 68.149 |
| Atad2 | 465.996 | -0.09476 | 0.14459 | -0.65533 | 0.51225 | 0.70333 | 41.039 |
| Atad2b | 1811.12 | 0.153567 | 0.08827 | 1.739795 | 0.0819 | 0.22809 | 130.042 |
| Atad3a | 1807.03 | 0.010751 | 0.0719 | 0.149529 | 0.88114 | 0.94072 | 20.453 |
| Atad3aos | 41.4482 | 0.251102 | 0.27617 | 0.90922 | 0.36323 | 0.57357 | 8.123 |

| | | | | | | | |
|---|---|---|---|---|---|---|---|
| Atad5 | 410.028 | 0.237837 | 0.11658 | 2.040085 | 0.04134 | 0.14589 | 46.395 |
| Atat1 | 1584.49 | 0.161882 | 0.08249 | 1.962394 | 0.04972 | 0.16521 | 12.481 |
| Atcay | 7271.57 | -0.12447 | 0.06094 | -2.04231 | 0.04112 | 0.14532 | 26.326 |
| Atcayos | 182.357 | -0.27373 | 0.15498 | -1.76627 | 0.07735 | 0.21999 | 16.269 |
| Ate1 | 3841.05 | 0.092779 | 0.0741 | 1.25203 | 0.21056 | 0.41056 | 128.877 |
| Atf1 | 643.583 | -0.00298 | 0.1151 | -0.02588 | 0.97935 | 0.9889 | 33.426 |
| Atf2 | 5828.71 | 0.166273 | 0.06339 | 2.622841 | 0.00872 | 0.05239 | 76.131 |
| Atf3 | 51.0633 | -0.50372 | 0.29565 | -1.70379 | 0.08842 | 0.24021 | 47.744 |
| Atf4 | 3646.62 | -0.37924 | 0.08168 | -4.64282 | 3.44E-06 | 0.00026 | 2.358 |
| Atf5 | 559.923 | -0.74618 | 0.10888 | -6.85298 | 7.23E-12 | 1.13E-08 | 4.403 |
| Atf6 | 4996.56 | 0.128347 | 0.08824 | 1.454439 | 0.14582 | 0.32872 | 163.098 |
| Atf6b | 3309.93 | -0.09703 | 0.0903 | -1.07453 | 0.28259 | 0.49279 | 7.929 |
| Atf7 | 1735.29 | -0.14293 | 0.07276 | -1.96441 | 0.04948 | 0.16471 | 99.519 |
| Atf7ip | 4808.8 | 0.123849 | 0.05704 | 2.171211 | 0.02992 | 0.11809 | 104.696 |
| Atg10 | 328.566 | -0.03208 | 0.1216 | -0.2638 | 0.79193 | 0.89181 | 288.613 |
| Atg101 | 856.696 | -0.15329 | 0.08351 | -1.83554 | 0.06643 | 0.19938 | 6.674 |
| Atg12 | 3593.03 | 0.076887 | 0.07843 | 0.98026 | 0.32696 | 0.53826 | 11.437 |
| Atg13 | 4831.93 | 0.100367 | 0.06818 | 1.47207 | 0.141 | 0.32275 | 35.959 |
| Atg14 | 1074.27 | 0.261712 | 0.09696 | 2.699179 | 0.00695 | 0.04524 | 29.757 |
| Atg16l1 | 3640.43 | 0.139567 | 0.06089 | 2.292278 | 0.02189 | 0.09522 | 36.559 |
| Atg16l2 | 1338.77 | 0.258738 | 0.08797 | 2.94129 | 0.00327 | 0.02757 | 39.218 |
| Atg2a | 3446.82 | 0.323524 | 0.08323 | 3.886882 | 0.0001 | 0.00272 | 20.668 |
| Atg2b | 3305.54 | 0.156351 | 0.06174 | 2.532329 | 0.01133 | 0.0622 | 69.076 |
| Atg3 | 2006.44 | -0.19144 | 0.08764 | -2.18432 | 0.02894 | 0.11507 | 29.754 |
| Atg4a | 403.118 | -0.03785 | 0.12279 | -0.30825 | 0.7579 | 0.87258 | 89.611 |
| Atg4b | 2784.61 | 0.056421 | 0.06373 | 0.885289 | 0.376 | 0.5861 | 39.111 |
| Atg4c | 1914.09 | 0.017169 | 0.0621 | 0.27646 | 0.78219 | 0.88669 | 65.854 |
| Atg4d | 1170.49 | -0.02599 | 0.09524 | -0.27286 | 0.78496 | 0.88852 | 12.48 |
| Atg5 | 982.9 | 0.065809 | 0.08194 | 0.803182 | 0.42187 | 0.6287 | 95.934 |
| Atg7 | 981.891 | -0.26139 | 0.08671 | -3.01437 | 0.00258 | 0.02385 | 217.518 |
| Atg9a | 2791.41 | 0.045543 | 0.06899 | 0.660125 | 0.50917 | 0.70067 | 11.337 |
| Atg9b | 233.802 | 0.334468 | 0.18853 | 1.774056 | 0.07605 | 0.21774 | 7.963 |
| Atic | 1480.18 | -0.0401 | 0.0749 | -0.5354 | 0.59237 | 0.76489 | 22.482 |
| Atl1 | 5239.39 | 0.122045 | 0.07927 | 1.539697 | 0.12363 | 0.29802 | 73.804 |

| | | | | | | | |
|---|---|---|---|---|---|---|---|
| Atl2 | 4931.9 | 0.167487 | 0.07668 | 2.184272 | 0.02894 | 0.11507 | 47.734 |
| Atl3 | 3731.01 | 0.013087 | 0.07105 | 0.184205 | 0.85385 | 0.92589 | 45.635 |
| Atm | 1246.34 | 0.032354 | 0.13029 | 0.248318 | 0.80389 | 0.89875 | 97.592 |
| Atmin | 3237.61 | 0.118091 | 0.06984 | 1.690898 | 0.09086 | 0.24474 | 17.053 |
| Atn1 | 462.665 | -0.13071 | 0.15147 | -0.86293 | 0.38818 | 0.59748 | 13.981 |
| Atoh7 | 32.6755 | 0.314151 | 0.35294 | 0.890092 | 0.37342 | 0.58357 | 1.629 |
| Atoh8 | 125.114 | -0.1344 | 0.16684 | -0.80556 | 0.4205 | 0.62772 | 29.401 |
| Atox1 | 681.092 | -0.35732 | 0.11861 | -3.0126 | 0.00259 | 0.02395 | 14.599 |
| Atp10a | 1176.38 | -0.45446 | 0.07834 | -5.80147 | 6.57E-09 | 2.16E-06 | 173.255 |
| Atp10b | 55.9091 | -0.28457 | 0.24948 | -1.14064 | 0.25402 | 0.46137 | 112.409 |
| Atp10d | 339.435 | 0.097306 | 0.14346 | 0.67827 | 0.4976 | 0.691 | 95.447 |
| Atp11a | 6569.69 | 0.228577 | 0.08257 | 2.768322 | 0.00563 | 0.03947 | 111.715 |
| Atp11b | 5567.64 | -0.34602 | 0.12533 | -2.76083 | 0.00577 | 0.04014 | 102.171 |
| Atp11c | 1479.37 | 0.460757 | 0.16195 | 2.845103 | 0.00444 | 0.0337 | 367.655 |
| Atp13a1 | 2428.99 | 0.103911 | 0.06235 | 1.666527 | 0.09561 | 0.25289 | 16.587 |
| Atp13a2 | 5723.83 | -0.03301 | 0.06683 | -0.49396 | 0.62133 | 0.78471 | 20.458 |
| Atp13a3 | 4809.25 | 0.152224 | 0.06873 | 2.214727 | 0.02678 | 0.10915 | 93.553 |
| Atp13a4 | 435.537 | -0.00311 | 0.19852 | -0.01569 | 0.98748 | 0.99316 | 149.012 |
| Atp13a5 | 582.137 | -0.36061 | 0.10074 | -3.57973 | 0.00034 | 0.00616 | 146.948 |
| Atp1a1 | 25984.6 | -0.02248 | 0.07491 | -0.30016 | 0.76406 | 0.8765 | 28.466 |
| Atp1a2 | 50030.6 | -0.39222 | 0.07351 | -5.33542 | 9.53E-08 | 1.66E-05 | 26.356 |
| Atp1a3 | 95320 | -0.14506 | 0.057 | -2.54467 | 0.01094 | 0.06094 | 27.792 |
| Atp1b1 | 57630.9 | -0.20572 | 0.07189 | -2.8617 | 0.00421 | 0.03257 | 21.247 |
| Atp1b2 | 15783 | -0.38401 | 0.09616 | -3.99355 | 6.51E-05 | 0.002 | 6.207 |
| Atp1b3 | 4536.08 | -0.14198 | 0.06761 | -2.10014 | 0.03572 | 0.13287 | 31.788 |
| Atp23 | 381.631 | 0.22282 | 0.12257 | 1.817887 | 0.06908 | 0.20453 | 32.929 |
| Atp2a1 | 12.431 | -0.06283 | 0.54345 | -0.11562 | 0.90796 | 0.95386 | 17.251 |
| Atp2a2 | 40349.5 | -0.00777 | 0.06565 | -0.11829 | 0.90584 | 0.95277 | 48.713 |
| Atp2a3 | 290.767 | -0.55216 | 0.17709 | -3.11788 | 0.00182 | 0.01894 | 31.876 |
| Atp2b1 | 43936.5 | 0.182034 | 0.10886 | 1.672212 | 0.09448 | 0.25084 | 111.738 |
| Atp2b2 | 31635.7 | 0.02332 | 0.09229 | 0.252671 | 0.80052 | 0.89692 | 298.783 |
| Atp2b3 | 5762.03 | 0.08197 | 0.0681 | 1.20371 | 0.2287 | 0.43274 | 67.92 |
| Atp2b4 | 12171.2 | -0.15298 | 0.16161 | -0.94664 | 0.34382 | 0.55451 | 101.585 |
| Atp2c1 | 7742.3 | 0.008438 | 0.0607 | 0.139001 | 0.88945 | 0.94464 | 123.781 |

| | | | | | | | |
|---|---|---|---|---|---|---|---|
| Atp2c2 | 75.8891 | 0.329211 | 0.22143 | 1.486717 | 0.13709 | 0.31781 | 57.709 |
| Atp4a | 17.2271 | -0.12647 | 0.46087 | -0.27441 | 0.78377 | 0.88762 | 13.326 |
| Atp5a1 | 34760.5 | -0.01676 | 0.06507 | -0.25758 | 0.79673 | 0.89517 | 9.141 |
| Atp5b | 47221.5 | -0.14467 | 0.05721 | -2.52895 | 0.01144 | 0.06262 | 7.119 |
| Atp5c1 | 7296.74 | -0.10317 | 0.06614 | -1.55985 | 0.1188 | 0.29084 | 24.495 |
| Atp5d | 5136.53 | -0.18179 | 0.08515 | -2.13504 | 0.03276 | 0.12565 | 7.187 |
| Atp5e | 2361.69 | -0.23182 | 0.09234 | -2.51054 | 0.01205 | 0.06482 | 3.034 |
| Atp5g1 | 4403.8 | -0.17684 | 0.08248 | -2.14417 | 0.03202 | 0.12355 | 6.819 |
| Atp5g2 | 1985.19 | -0.11741 | 0.09553 | -1.22904 | 0.21906 | 0.42125 | 9.489 |
| Atp5g3 | 8084.11 | -0.14944 | 0.06296 | -2.37346 | 0.01762 | 0.0825 | 2.88 |
| Atp5h | 5581.86 | -0.1431 | 0.06472 | -2.21108 | 0.02703 | 0.10985 | 4.274 |
| Atp5j | 4629.35 | -0.25978 | 0.06941 | -3.7425 | 0.00018 | 0.00405 | 7.76 |
| Atp5j2 | 3640.22 | -0.21205 | 0.08106 | -2.61608 | 0.00889 | 0.05308 | 8.365 |
| Atp5k | 1355.15 | -0.33032 | 0.1683 | -1.96264 | 0.04969 | 0.16518 | 1.205 |
| Atp5l | 1158.55 | -0.04206 | 0.11051 | -0.38062 | 0.70348 | 0.83993 | 8.186 |
| Atp5md | 3939.28 | -0.13465 | 0.08127 | -1.65682 | 0.09756 | 0.25626 | 7.184 |
| Atp5mpl | 2550.76 | -0.25448 | 0.0988 | -2.5756 | 0.01001 | 0.05751 | 5.603 |
| Atp5o | 161.006 | -0.2134 | 0.24904 | -0.85692 | 0.39149 | 0.60055 | 242.585 |
| Atp5pb | 7968.59 | -0.11339 | 0.06168 | -1.83841 | 0.066 | 0.19847 | 17.402 |
| Atp5pb-ps | 45.0468 | -0.14054 | 0.26632 | -0.52771 | 0.5977 | 0.76926 | 0.695 |
| Atp6ap1 | 12809.2 | -0.16025 | 0.06342 | -2.52698 | 0.0115 | 0.06287 | 7.625 |
| Atp6ap1l | 431.666 | -0.84305 | 0.1748 | -4.82286 | 1.42E-06 | 0.00013 | 21.923 |
| Atp6ap2 | 11073 | -0.02822 | 0.06937 | -0.40681 | 0.68414 | 0.82778 | 29.249 |
| Atp6v0a1 | 22169.6 | -0.0022 | 0.06689 | -0.03286 | 0.97379 | 0.98681 | 54.268 |
| Atp6v0a2 | 4219.45 | 0.086096 | 0.07229 | 1.191001 | 0.23365 | 0.43864 | 34.879 |
| Atp6v0b | 4275.88 | -0.10977 | 0.07224 | -1.51952 | 0.12863 | 0.30583 | 3.008 |
| Atp6v0c | 13225.5 | -0.17824 | 0.08491 | -2.09921 | 0.0358 | 0.13304 | 5.837 |
| Atp6v0c-ps2 | 119.218 | -0.25285 | 0.17922 | -1.41087 | 0.15828 | 0.34558 | 0.465 |
| Atp6v0d1 | 6878.17 | -0.18365 | 0.064 | -2.86953 | 0.00411 | 0.03204 | 41.583 |
| Atp6v0e | 1576.86 | -0.08974 | 0.08094 | -1.10874 | 0.26754 | 0.47692 | 36.417 |
| Atp6v0e2 | 10346.8 | -0.08185 | 0.0641 | -1.27687 | 0.20165 | 0.40013 | 4.187 |
| Atp6v1a | 19934.2 | -0.06925 | 0.07831 | -0.88441 | 0.37647 | 0.58653 | 54.304 |
| Atp6v1b1 | 42.8981 | 0.196547 | 0.39962 | 0.491831 | 0.62284 | 0.78588 | 15.866 |
| Atp6v1b2 | 29469.7 | -0.09987 | 0.06844 | -1.45914 | 0.14453 | 0.32723 | 25.066 |

| | | | | | | | |
|---|---|---|---|---|---|---|---|
| Atp6v1c1 | 9343.56 | -0.0499 | 0.05942 | -0.83972 | 0.40106 | 0.60978 | 30.514 |
| Atp6v1c2 | 181.35 | 1.710609 | 0.59295 | 2.88491 | 0.00392 | 0.03099 | 44.639 |
| Atp6v1d | 9317.15 | 0.051941 | 0.07019 | 0.740021 | 0.45929 | 0.65991 | 20.914 |
| Atp6v1e1 | 11381.9 | -0.08625 | 0.06196 | -1.39214 | 0.16388 | 0.35265 | 28.489 |
| Atp6v1f | 2350.17 | -0.20704 | 0.06347 | -3.26196 | 0.00111 | 0.01368 | 2.795 |
| Atp6v1g1 | 2943.51 | -0.19639 | 0.07916 | -2.48092 | 0.0131 | 0.06854 | 5.979 |
| Atp6v1g2 | 13119 | -0.21493 | 0.063 | -3.41151 | 0.00065 | 0.00942 | 5.108 |
| Atp6v1h | 7212.75 | 0.047797 | 0.07027 | 0.680207 | 0.49637 | 0.68999 | 92.512 |
| Atp7a | 495.715 | -0.01402 | 0.09414 | -0.14897 | 0.88158 | 0.94099 | 97.651 |
| Atp7b | 175.11 | -0.19879 | 0.1902 | -1.04519 | 0.29594 | 0.50629 | 67.521 |
| Atp8a1 | 14157.1 | 0.028549 | 0.06628 | 0.430716 | 0.66667 | 0.8158 | 229.295 |
| Atp8a2 | 3504.19 | -0.07101 | 0.08234 | -0.86235 | 0.3885 | 0.5978 | 558.64 |
| Atp8b1 | 91.7204 | -0.85991 | 0.27513 | -3.12542 | 0.00178 | 0.01864 | 132.301 |
| Atp8b2 | 5247.33 | 0.223641 | 0.09481 | 2.358952 | 0.01833 | 0.08467 | 24.028 |
| Atp8b3 | 16.8875 | 0.007699 | 0.42251 | 0.018221 | 0.98546 | 0.99203 | 19.541 |
| Atp8b5 | 23.2788 | 0.295171 | 0.39687 | 0.74374 | 0.45703 | 0.65829 | 106.675 |
| Atp9a | 13655.4 | 0.011778 | 0.06267 | 0.187927 | 0.85093 | 0.9241 | 107.972 |
| Atp9b | 3442.43 | 0.001626 | 0.06277 | 0.025909 | 0.97933 | 0.9889 | 199.92 |
| Atpaf1 | 2527.77 | -0.08944 | 0.07217 | -1.23925 | 0.21525 | 0.41673 | 37.844 |
| Atpaf2 | 896.75 | -0.0333 | 0.07611 | -0.43752 | 0.66173 | 0.81292 | 17.832 |
| Atpif1 | 249.433 | -0.17468 | 0.21512 | -0.81203 | 0.41677 | 0.62432 | 3.105 |
| Atpsckmt | 1239.71 | 0.149073 | 0.08506 | 1.752525 | 0.07968 | 0.22422 | 19.376 |
| Atr | 899.793 | 0.226827 | 0.16946 | 1.338563 | 0.18071 | 0.37405 | 94.185 |
| Atraid | 1442.3 | -0.02701 | 0.09077 | -0.29751 | 0.76608 | 0.8779 | 6.312 |
| Atrip | 1209.27 | 0.053781 | 0.07688 | 0.699554 | 0.48421 | 0.68059 | 16.192 |
| Atrn | 12557.2 | 0.068405 | 0.05372 | 1.273334 | 0.2029 | 0.40154 | 123.839 |
| Atrnl1 | 4566.63 | -0.43594 | 0.1191 | -3.66032 | 0.00025 | 0.00497 | 522.31 |
| Atrx | 11215.3 | 0.371377 | 0.32276 | 1.150639 | 0.24988 | 0.45723 | 131.789 |
| Atxn1 | 5468.31 | 0.153893 | 0.06874 | 2.23871 | 0.02517 | 0.10441 | 415.254 |
| Atxn10 | 8429.68 | -0.06695 | 0.07736 | -0.86546 | 0.38679 | 0.5961 | 126.968 |
| Atxn1l | 3383.75 | -0.05619 | 0.07651 | -0.73443 | 0.46269 | 0.66283 | 11.289 |
| Atxn2 | 4524.57 | -0.02663 | 0.06641 | -0.40107 | 0.68837 | 0.83003 | 105.157 |
| Atxn2l | 9271.61 | 0.051673 | 0.09891 | 0.522424 | 0.60138 | 0.77152 | 11.73 |
| Atxn3 | 2530.51 | 0.223441 | 0.08319 | 2.685881 | 0.00723 | 0.04653 | 39.346 |

| | | | | | | | |
|---|---|---|---|---|---|---|---|
| Atxn7 | 2334.52 | 0.251501 | 0.08976 | 2.801889 | 0.00508 | 0.03666 | 145.863 |
| Atxn7l1 | 1481.01 | 0.128686 | 0.07966 | 1.61543 | 0.10622 | 0.27017 | 225.493 |
| Atxn7l1os1 | 12.5429 | 0.814848 | 0.62653 | 1.300577 | 0.1934 | 0.39029 | 34.861 |
| Atxn7l1os2 | 32.8504 | 0.436612 | 0.31278 | 1.395888 | 0.16275 | 0.3512 | 36.518 |
| Atxn7l2 | 1361.26 | 0.05779 | 0.09986 | 0.578695 | 0.5628 | 0.74354 | 8.713 |
| Atxn7l3 | 9449.67 | -0.29405 | 0.08404 | -3.49883 | 0.00047 | 0.00759 | 7.332 |
| Atxn7l3b | 7442.81 | -0.07435 | 0.06265 | -1.18687 | 0.23528 | 0.44047 | 3.572 |
| Auh | 2202.38 | -0.25596 | 0.06918 | -3.70015 | 0.00022 | 0.00448 | 94.563 |
| Aup1 | 1457.49 | 0.008201 | 0.06469 | 0.126759 | 0.89913 | 0.94943 | 3.288 |
| Aurka | 98.4866 | -0.14771 | 0.17117 | -0.86289 | 0.3882 | 0.59748 | 14.346 |
| Aurkaip1 | 1588.86 | -0.1109 | 0.0885 | -1.25302 | 0.2102 | 0.4102 | 1.859 |
| Aurkb | 15.3108 | -0.79896 | 0.55721 | -1.43386 | 0.15161 | 0.33658 | 6.018 |
| Auts2 | 4903.48 | 0.317513 | 0.10678 | 2.973581 | 0.00294 | 0.02585 | 1106.013 |
| Aven | 362.228 | -0.11821 | 0.13469 | -0.87766 | 0.38013 | 0.58974 | 141.61 |
| Avil | 59.4757 | -0.16368 | 0.24294 | -0.67374 | 0.50047 | 0.69333 | 20.286 |
| Avl9 | 3995.23 | 0.208488 | 0.06801 | 3.065379 | 0.00217 | 0.02128 | 47.014 |
| Avpi1 | 578.341 | -0.21346 | 0.10444 | -2.04391 | 0.04096 | 0.14491 | 5.787 |
| Avpr1a | 33.8223 | 0.832406 | 0.49589 | 1.678603 | 0.09323 | 0.24855 | 4.954 |
| Avpr2 | 15.8334 | 0.52705 | 0.49527 | 1.06416 | 0.28726 | 0.49736 | 3.401 |
| Axdnd1 | 27.473 | -0.18435 | 0.34431 | -0.53543 | 0.59235 | 0.76489 | 97.651 |
| Axin1 | 1412.54 | 0.114084 | 0.07164 | 1.592467 | 0.11128 | 0.2786 | 57.124 |
| Axin2 | 1219.74 | -0.09758 | 0.09189 | -1.06188 | 0.28829 | 0.49843 | 30.435 |
| Axl | 2290.11 | -0.39648 | 0.10268 | -3.8612 | 0.00011 | 0.00291 | 31.433 |
| Azi2 | 4162.68 | 0.222734 | 0.08153 | 2.73209 | 0.00629 | 0.04246 | 29.296 |
| Azin1 | 9870.08 | -0.06537 | 0.05585 | -1.17049 | 0.2418 | 0.4478 | 31.84 |
| Azin2 | 1569.1 | 0.09161 | 0.09635 | 0.950852 | 0.34168 | 0.55259 | 32.21 |
| B130021K23Rik | 76.6044 | 0.354893 | 0.24621 | 1.441412 | 0.14947 | 0.33376 | 2.593 |
| B130024G19Rik | 22.5571 | -0.87591 | 0.37371 | -2.34381 | 0.01909 | 0.08673 | 125.063 |
| B130034C11Rik | 125.152 | 0.234029 | 0.17971 | 1.30223 | 0.19284 | 0.38957 | 17.147 |
| B130046B21Rik | 53.9428 | -0.19415 | 0.26699 | -0.72717 | 0.46712 | 0.66622 | 9.054 |
| B130055M24Rik | 426.373 | 0.325534 | 0.1182 | 2.754178 | 0.00588 | 0.04073 | 11.726 |
| B230110C06Rik | 28.4353 | 0.211453 | 0.45043 | 0.469446 | 0.63875 | 0.79727 | 14.823 |
| B230206H07Rik | 72.9016 | -0.45769 | 0.33309 | -1.37405 | 0.16943 | 0.35966 | 6.051 |
| B230206I08Rik | 61.6708 | 0.183655 | 0.31317 | 0.586444 | 0.55758 | 0.73923 | 3.191 |

| | | | | | | | |
|---|---|---|---|---|---|---|---|
| B230206L02Rik | 85.0591 | 0.126921 | 0.24229 | 0.523845 | 0.60039 | 0.77095 | 22.362 |
| B230208B08Rik | 201.922 | 0.038148 | 0.14023 | 0.272032 | 0.7856 | 0.88863 | 3.655 |
| B230209E15Rik | 1894 | 0.385603 | 0.10752 | 3.58639 | 0.00034 | 0.0061 | 85.457 |
| B230216N24Rik | 197.895 | -0.24502 | 0.22815 | -1.07392 | 0.28286 | 0.4931 | 26.567 |
| B230217C12Rik | 1983.35 | -0.27357 | 0.09045 | -3.02448 | 0.00249 | 0.02331 | 3.32 |
| B230217J21Rik | 25.6099 | 0.992625 | 0.45069 | 2.202446 | 0.02763 | 0.11152 | 25.676 |
| B230217O12Rik | 226.672 | 0.039513 | 0.16476 | 0.239828 | 0.81046 | 0.90251 | 37.679 |
| B230219D22Rik | 7067.4 | 0.169883 | 0.06759 | 2.513421 | 0.01196 | 0.0645 | 10.377 |
| B230303A05Rik | 383.252 | 0.238382 | 0.13907 | 1.714105 | 0.08651 | 0.2366 | 209.884 |
| B230303O12Rik | 83.3904 | 0.265833 | 0.25128 | 1.057904 | 0.2901 | 0.50029 | 6.19 |
| B230307C23Rik | 240.513 | -0.03834 | 0.15039 | -0.25494 | 0.79877 | 0.89614 | 34.487 |
| B230311B06Rik | 29.2834 | -0.05102 | 0.38581 | -0.13225 | 0.89479 | 0.94736 | 4.569 |
| B230312C02Rik | 23.0719 | 0.279544 | 0.47743 | 0.58552 | 0.5582 | 0.73975 | 15.395 |
| B230317F23Rik | 56.7083 | 0.157245 | 0.2468 | 0.637134 | 0.52404 | 0.71242 | 2.472 |
| B230319C09Rik | 42.9366 | 0.142817 | 0.29231 | 0.488574 | 0.62514 | 0.78752 | 6.568 |
| B230334C09Rik | 11629.4 | 0.379754 | 0.12314 | 3.083851 | 0.00204 | 0.02036 | 8.927 |
| B230354K17Rik | 774.074 | 0.001924 | 0.08625 | 0.022304 | 0.98221 | 0.99059 | 8.947 |
| B230362B09Rik | 15.343 | -0.29307 | 0.5164 | -0.56753 | 0.57035 | 0.74968 | 1.621 |
| B230364G03Rik | 53.9761 | -0.86834 | 0.46145 | -1.88176 | 0.05987 | 0.18643 | 2.086 |
| B230369F24Rik | 380.412 | 0.355051 | 0.12069 | 2.941757 | 0.00326 | 0.02757 | 40.953 |
| B230377A18Rik | 17.7407 | -0.43601 | 0.38933 | -1.1199 | 0.26276 | 0.47157 | 1.364 |
| B2m | 2860.71 | -0.6532 | 0.13243 | -4.93239 | 8.12E-07 | 9.20E-05 | 5.398 |
| B330016D10Rik | 149.993 | -0.09057 | 0.16479 | -0.54958 | 0.58261 | 0.758 | 1.704 |
| B3galnt1 | 1494.41 | -0.14726 | 0.09402 | -1.56631 | 0.11728 | 0.28825 | 24.803 |
| B3galnt2 | 1281.81 | 0.121327 | 0.08367 | 1.450078 | 0.14704 | 0.3302 | 44.635 |
| B3galt1 | 2407.16 | 0.323936 | 0.11215 | 2.888358 | 0.00387 | 0.03079 | 556.819 |
| B3galt2 | 1537.43 | -0.42443 | 0.36199 | -1.17249 | 0.241 | 0.44713 | 13.951 |
| B3galt4 | 134.514 | -0.14425 | 0.17444 | -0.82697 | 0.40826 | 0.61607 | 1.58 |
| B3galt5 | 5125.92 | 0.10894 | 0.0911 | 1.195871 | 0.23175 | 0.43651 | 84.059 |
| B3galt6 | 575.334 | 0.123557 | 0.09304 | 1.32806 | 0.18416 | 0.37857 | 3.184 |
| B3gat1 | 9438.43 | -0.45988 | 0.14889 | -3.08871 | 0.00201 | 0.02016 | 29.374 |
| B3gat2 | 289.947 | 0.171061 | 0.11897 | 1.437822 | 0.15048 | 0.33528 | 87.406 |
| B3gat3 | 2660.05 | -0.2964 | 0.08429 | -3.51643 | 0.00044 | 0.00729 | 6.863 |
| B3glct | 1074.2 | -0.17222 | 0.13595 | -1.26676 | 0.20524 | 0.40426 | 84.37 |

| | | | | | | | |
|---|---|---|---|---|---|---|---|
| B3gnt2 | 70.4117 | 0.283817 | 0.21865 | 1.298041 | 0.19427 | 0.39147 | 101.224 |
| B3gnt3 | 74.2202 | -1.00279 | 0.30744 | -3.2618 | 0.00111 | 0.01368 | 11.034 |
| B3gnt5 | 39.6721 | -0.10982 | 0.42707 | -0.25715 | 0.79706 | 0.89539 | 12.546 |
| B3gnt6 | 23.8912 | 0.025847 | 0.43291 | 0.059706 | 0.95239 | 0.97658 | 7.065 |
| B3gnt7 | 19.0339 | 0.319089 | 0.43441 | 0.734529 | 0.46263 | 0.66283 | 4.474 |
| B3gnt8 | 83.4827 | -0.2196 | 0.23972 | -0.91608 | 0.35963 | 0.56994 | 8.447 |
| B3gnt9 | 114.033 | -0.58702 | 0.18974 | -3.09376 | 0.00198 | 0.01998 | 2.516 |
| B3gntl1 | 367.79 | 0.215691 | 0.12425 | 1.735934 | 0.08258 | 0.2292 | 56.957 |
| B430010I23Rik | 19.4663 | 0.671951 | 0.39222 | 1.713196 | 0.08668 | 0.237 | 6.749 |
| B430203G13Rik | 41.3315 | 0.429199 | 0.31289 | 1.371737 | 0.17015 | 0.36052 | 1.375 |
| B430212C06Rik | 14.6077 | -0.90756 | 0.49656 | -1.82771 | 0.06759 | 0.20161 | 22.895 |
| B430218F22Rik | 13.0857 | -0.23536 | 0.55614 | -0.4232 | 0.67215 | 0.81974 | 1.495 |
| B430319F04Rik | 42.4126 | 0.005352 | 0.28306 | 0.018908 | 0.98491 | 0.99174 | 2.669 |
| B430319G15Rik | 18.2047 | 0.034663 | 0.58959 | 0.058791 | 0.95312 | 0.97691 | 4.069 |
| B4galnt1 | 3117.99 | -0.01569 | 0.05771 | -0.27185 | 0.78574 | 0.88863 | 7.106 |
| B4galnt2 | 20.9888 | -0.468 | 0.38337 | -1.22073 | 0.22219 | 0.42538 | 48.949 |
| B4galnt3 | 23.9154 | -0.72669 | 0.45529 | -1.59609 | 0.11047 | 0.27711 | 91.487 |
| B4galnt4 | 3596.91 | 0.161127 | 0.06638 | 2.427334 | 0.01521 | 0.07514 | 11.297 |
| B4galt1 | 478.743 | -0.16438 | 0.16574 | -0.99183 | 0.32128 | 0.53209 | 49.404 |
| B4galt2 | 2161.07 | -0.0889 | 0.09055 | -0.98177 | 0.32621 | 0.53752 | 14.228 |
| B4galt3 | 1246.79 | 0.036214 | 0.07118 | 0.50879 | 0.6109 | 0.77809 | 6.572 |
| B4galt4 | 1000.69 | -0.04208 | 0.0902 | -0.4665 | 0.64086 | 0.79893 | 26.786 |
| B4galt5 | 2241.31 | -0.37577 | 0.10892 | -3.45001 | 0.00056 | 0.00857 | 50.74 |
| B4galt6 | 8339.3 | 0.006852 | 0.04953 | 0.138337 | 0.88997 | 0.94486 | 61.806 |
| B4galt7 | 779.19 | -0.08184 | 0.08999 | -0.90945 | 0.36311 | 0.57346 | 10.548 |
| B630019K06Rik | 1733.05 | 0.317761 | 0.10207 | 3.113238 | 0.00185 | 0.0191 | 2.583 |
| B830012L14Rik | 258.902 | 0.078217 | 0.2009 | 0.389328 | 0.69703 | 0.8358 | 38.279 |
| B830017H08Rik | 74.1265 | 0.001665 | 0.20995 | 0.007931 | 0.99367 | 0.99633 | 1.888 |
| B830042I05Rik | 61.8242 | 0.479064 | 0.26884 | 1.781969 | 0.07475 | 0.21547 | 5.848 |
| B930059L03Rik | 45.2614 | -0.04251 | 0.26449 | -0.16073 | 0.87231 | 0.93551 | 2.189 |
| B930095G15Rik | 3958.89 | -0.57001 | 0.12282 | -4.64092 | 3.47E-06 | 0.00026 | 5.266 |
| B9d1 | 612.332 | 0.005676 | 0.09322 | 0.060892 | 0.95145 | 0.97617 | 7.788 |
| B9d1os | 37.0269 | -0.34738 | 0.30975 | -1.12149 | 0.26208 | 0.47085 | 0.674 |
| B9d2 | 616.363 | 0.132658 | 0.09739 | 1.362137 | 0.17315 | 0.3644 | 5.779 |

| | | | | | | | |
|---|---|---|---|---|---|---|---|
| BB031773 | 95.3191 | 0.53836 | 0.18642 | 2.88796 | 0.00388 | 0.0308 | 8.057 |
| BB187690 | 33.8763 | -0.18279 | 0.40992 | -0.44592 | 0.65566 | 0.80879 | 1.176 |
| BB218582 | 109.039 | 0.501813 | 0.18473 | 2.716494 | 0.0066 | 0.04379 | 7.496 |
| BB557941 | 12.3662 | -0.86006 | 0.50654 | -1.69791 | 0.08952 | 0.24244 | 54.295 |
| BC002059 | 352.172 | 0.44416 | 0.1864 | 2.382833 | 0.01718 | 0.08153 | 22.625 |
| BC003965 | 1116.18 | -0.09162 | 0.08685 | -1.05494 | 0.29145 | 0.50126 | 3.109 |
| BC004004 | 1797.39 | -0.03107 | 0.07937 | -0.39148 | 0.69544 | 0.83491 | 45.278 |
| BC005537 | 12430.9 | -0.06422 | 0.06143 | -1.04545 | 0.29581 | 0.50616 | 14.541 |
| BC005624 | 3165.58 | -0.02235 | 0.05634 | -0.3967 | 0.69159 | 0.83252 | 10.025 |
| BC006965 | 47.8126 | 0.016195 | 0.29799 | 0.054348 | 0.95666 | 0.97879 | 118.05 |
| BC018473 | 14.0191 | -0.6948 | 0.56249 | -1.23521 | 0.21675 | 0.41862 | 7.207 |
| BC022960 | 191.728 | -0.08844 | 0.22592 | -0.39149 | 0.69544 | 0.83491 | 1.815 |
| BC024063 | 151.24 | 0.680802 | 0.51687 | 1.317158 | 0.18779 | 0.38324 | 27.864 |
| BC024139 | 33.5744 | -0.62195 | 0.40713 | -1.52766 | 0.1266 | 0.30242 | 7.08 |
| BC024978 | 625.69 | -0.04892 | 0.0937 | -0.52211 | 0.60159 | 0.77164 | 14.406 |
| BC025920 | 258.229 | 0.151731 | 0.18218 | 0.832879 | 0.40491 | 0.61384 | 14.434 |
| BC026513 | 13.7065 | 0.455087 | 0.50064 | 0.909005 | 0.36335 | 0.5737 | 2.156 |
| BC028528 | 40.4267 | -0.24278 | 0.33405 | -0.72678 | 0.46736 | 0.66644 | 8.052 |
| BC029722 | 1056.84 | -0.18067 | 0.08615 | -2.09716 | 0.03598 | 0.13352 | 1.697 |
| BC030343 | 301.706 | 0.245369 | 0.14347 | 1.7102 | 0.08723 | 0.23787 | 2.473 |
| BC030500 | 340.019 | 0.042495 | 0.1287 | 0.330184 | 0.74126 | 0.86333 | 2.544 |
| BC031181 | 4208.88 | -0.18515 | 0.07055 | -2.62452 | 0.00868 | 0.05225 | 4.285 |
| BC034090 | 496.301 | -0.35238 | 0.11921 | -2.95599 | 0.00312 | 0.02673 | 31.974 |
| BC035044 | 25.968 | -0.27787 | 0.35292 | -0.78734 | 0.43108 | 0.63717 | 42.037 |
| BC035947 | 76.4373 | -0.05476 | 0.37317 | -0.14674 | 0.88334 | 0.94179 | 15.133 |
| BC037032 | 57.7291 | 0.123095 | 0.25492 | 0.48288 | 0.62918 | 0.79031 | 7.292 |
| BC037039 | 21.9861 | 0.243208 | 0.36522 | 0.665919 | 0.50546 | 0.69703 | 2.205 |
| BC037704 | 57.2659 | 0.007867 | 0.24028 | 0.032739 | 0.97388 | 0.98686 | 2.044 |
| BC043934 | 38.374 | -0.13456 | 0.34681 | -0.388 | 0.69802 | 0.83637 | 2.571 |
| BC046251 | 25.4554 | -0.56269 | 0.42189 | -1.33374 | 0.18229 | 0.37605 | 36.815 |
| BC046401 | 165.946 | 0.232047 | 0.14561 | 1.59358 | 0.11103 | 0.27814 | 1.351 |
| BC048644 | 19.6191 | -0.00655 | 0.39628 | -0.01652 | 0.98682 | 0.99281 | 11.858 |
| BC049352 | 99.2513 | 0.548293 | 0.2432 | 2.254532 | 0.02416 | 0.10151 | 68.014 |
| BC049715 | 46.9655 | -0.4797 | 0.25983 | -1.84624 | 0.06486 | 0.19618 | 13.037 |

| | | | | | | | |
|---|---|---|---|---|---|---|---|
| BC051142 | 201.837 | 0.201754 | 0.17909 | 1.126567 | 0.25993 | 0.46842 | 61.915 |
| BC051226 | 131.653 | 0.035781 | 0.17249 | 0.207441 | 0.83567 | 0.91572 | 2.192 |
| BC051408 | 30.0542 | 0.484116 | 0.32257 | 1.500824 | 0.1334 | 0.31306 | 14.987 |
| BC055308 | 53.2779 | 0.282907 | 0.22614 | 1.250999 | 0.21093 | 0.41098 | 2.538 |
| BC055324 | 88.1711 | 0.330451 | 0.19093 | 1.730709 | 0.0835 | 0.23102 | 48.804 |
| BC055402 | 20.1647 | -0.69517 | 0.42407 | -1.63929 | 0.10115 | 0.26232 | 14.349 |
| BC060293 | 14.4577 | -0.04249 | 0.48108 | -0.08832 | 0.92963 | 0.96459 | 1.104 |
| BC064078 | 118.384 | -0.01755 | 0.24628 | -0.07125 | 0.9432 | 0.97165 | 15.134 |
| BC065397 | 209.701 | 0.210369 | 0.15844 | 1.327767 | 0.18426 | 0.37873 | 61.541 |
| BC106175 | 20.6229 | 0.386698 | 0.5196 | 0.744229 | 0.45674 | 0.65819 | 3.284 |
| BC106179 | 20.5903 | -0.55468 | 0.44604 | -1.24354 | 0.21367 | 0.41501 | 4.829 |
| BE692007 | 19.1904 | -1.24979 | 0.51955 | -2.40554 | 0.01615 | 0.07815 | 14.533 |
| Baalc | 6104.94 | -0.41586 | 0.12427 | -3.34634 | 0.00082 | 0.01116 | 75.524 |
| Babam1 | 2092 | -0.02548 | 0.05756 | -0.44272 | 0.65797 | 0.81018 | 7.759 |
| Babam2 | 497.759 | -0.21185 | 0.12373 | -1.71217 | 0.08687 | 0.23726 | 387.279 |
| Bace1 | 3312.51 | -0.02856 | 0.0621 | -0.45995 | 0.64555 | 0.80247 | 25.82 |
| Bace2 | 134.944 | -0.5833 | 0.18725 | -3.11513 | 0.00184 | 0.01906 | 86.195 |
| Bach1 | 1122.35 | -0.28131 | 0.11334 | -2.48201 | 0.01306 | 0.06843 | 34.402 |
| Bach2 | 441.413 | 0.202853 | 0.11463 | 1.769642 | 0.07679 | 0.21916 | 347.305 |
| Bach2os | 22.2493 | 0.175454 | 0.36845 | 0.476189 | 0.63394 | 0.79385 | 12.029 |
| Bad | 112.322 | -0.13621 | 0.19523 | -0.69769 | 0.48537 | 0.68158 | 10.039 |
| Bag1 | 4578.51 | -0.19188 | 0.07041 | -2.72524 | 0.00643 | 0.04301 | 11.897 |
| Bag2 | 263.098 | -0.18422 | 0.1502 | -1.22648 | 0.22002 | 0.42246 | 12.312 |
| Bag3 | 320.36 | -0.55466 | 0.12719 | -4.361 | 1.29E-05 | 0.00067 | 23.366 |
| Bag4 | 2706.88 | 0.129424 | 0.0926 | 1.397592 | 0.16224 | 0.35062 | 20.75 |
| Bag5 | 2658.67 | 0.256875 | 0.11111 | 2.311901 | 0.02078 | 0.09188 | 3.77 |
| Bag6 | 10297.7 | 0.080321 | 0.06243 | 1.28647 | 0.19828 | 0.39603 | 12.145 |
| Bahcc1 | 2097.15 | 0.089339 | 0.06623 | 1.348964 | 0.17735 | 0.36949 | 59.336 |
| Bahd1 | 2450.13 | -0.01741 | 0.07468 | -0.23316 | 0.81563 | 0.90517 | 24.152 |
| Baiap2 | 21876.9 | 0.406811 | 0.12044 | 3.377737 | 0.00073 | 0.01031 | 64.02 |
| Baiap2l1 | 74.7038 | -0.07236 | 0.25094 | -0.28837 | 0.77306 | 0.88187 | 93.587 |
| Baiap2l2 | 129.205 | -0.42928 | 0.20598 | -2.08404 | 0.03716 | 0.13625 | 27.343 |
| Baiap3 | 2438.17 | 1.131132 | 0.28585 | 3.957062 | 7.59E-05 | 0.00221 | 13.706 |
| Bak1 | 306.759 | -0.19517 | 0.13124 | -1.48716 | 0.13697 | 0.31768 | 9.2 |

| | | | | | | | |
|---|---|---|---|---|---|---|---|
| Bambi | 151.907 | -0.3787 | 0.22391 | -1.69127 | 0.09078 | 0.24468 | 8.482 |
| Bambi-ps1 | 18.6205 | -0.14823 | 0.42018 | -0.35279 | 0.72425 | 0.85296 | 0.592 |
| Banf1 | 1038.66 | 0.008597 | 0.08894 | 0.096662 | 0.92299 | 0.96131 | 2.531 |
| Bank1 | 24.4155 | 0.387952 | 0.42994 | 0.902344 | 0.36687 | 0.57726 | 272.704 |
| Banp | 1417.05 | 0.035328 | 0.095 | 0.371871 | 0.70999 | 0.84387 | 79.509 |
| Bap1 | 6072.96 | -0.07994 | 0.07177 | -1.11385 | 0.26534 | 0.4745 | 8.495 |
| Bard1 | 47.5032 | -0.08477 | 0.28975 | -0.29255 | 0.76987 | 0.87985 | 75.649 |
| Barx2 | 101.051 | -0.10031 | 0.2114 | -0.47451 | 0.63514 | 0.79438 | 67.419 |
| Basp1 | 13379.8 | -0.65048 | 0.18292 | -3.55603 | 0.00038 | 0.00656 | 50.488 |
| Batf3 | 38.4109 | -0.44745 | 0.31096 | -1.43893 | 0.15017 | 0.33476 | 11.099 |
| Bax | 486.723 | -0.17169 | 0.13575 | -1.26474 | 0.20596 | 0.40522 | 5.202 |
| Baz1a | 354.425 | 0.656688 | 0.19619 | 3.347124 | 0.00082 | 0.01114 | 121.36 |
| Baz1b | 10906.2 | 0.174409 | 0.06226 | 2.801429 | 0.00509 | 0.03668 | 58.866 |
| Baz2a | 4969.02 | 0.047515 | 0.07647 | 0.621387 | 0.53435 | 0.72154 | 37.727 |
| Baz2b | 2488.63 | -0.02456 | 0.09785 | -0.25095 | 0.80185 | 0.89768 | 310.477 |
| Bbc3 | 255.695 | -0.02809 | 0.13583 | -0.20682 | 0.83615 | 0.91592 | 9.813 |
| Bbip1 | 1872.98 | 0.099333 | 0.09668 | 1.027489 | 0.30419 | 0.51405 | 14.767 |
| Bbln | 1206.74 | -0.46733 | 0.12435 | -3.75813 | 0.00017 | 0.00389 | 4.842 |
| Bbof1 | 535.659 | 0.033609 | 0.12058 | 0.27873 | 0.78045 | 0.88564 | 33.817 |
| Bbox1 | 43.2131 | -0.25847 | 0.37209 | -0.69464 | 0.48728 | 0.68271 | 51.864 |
| Bbs1 | 3077.11 | 0.187727 | 0.05974 | 3.14244 | 0.00168 | 0.01794 | 19.751 |
| Bbs10 | 218.362 | 0.197372 | 0.15771 | 1.251479 | 0.21076 | 0.4108 | 3.049 |
| Bbs12 | 255.49 | 0.36868 | 0.13965 | 2.640014 | 0.00829 | 0.05072 | 8.9 |
| Bbs2 | 2157.06 | 0.314837 | 0.11069 | 2.844286 | 0.00445 | 0.03375 | 30.975 |
| Bbs4 | 1838.63 | 0.121772 | 0.0906 | 1.344088 | 0.17892 | 0.37156 | 31.519 |
| Bbs5 | 955.269 | 0.200131 | 0.09755 | 2.051488 | 0.04022 | 0.1433 | 20.401 |
| Bbs7 | 1170.05 | 0.33699 | 0.08481 | 3.97325 | 7.09E-05 | 0.00212 | 40.336 |
| Bbs9 | 746.653 | 0.51684 | 0.14764 | 3.500574 | 0.00046 | 0.00757 | 412.566 |
| Bbx | 2403.65 | 0.160168 | 0.08765 | 1.827378 | 0.06764 | 0.20161 | 240.547 |
| Bcam | 638.621 | -0.34087 | 0.10173 | -3.35065 | 0.00081 | 0.01103 | 14.886 |
| Bcan | 12723.3 | -0.02544 | 0.07537 | -0.33752 | 0.73572 | 0.86012 | 12.7 |
| Bcap29 | 2199.52 | 0.159013 | 0.1129 | 1.408496 | 0.15898 | 0.34641 | 43.692 |
| Bcap31 | 1907.87 | -0.25609 | 0.07942 | -3.22446 | 0.00126 | 0.01476 | 29.998 |
| Bcar1 | 2310 | -0.02999 | 0.06847 | -0.438 | 0.66139 | 0.81269 | 33.336 |

| | | | | | | | |
|---|---|---|---|---|---|---|---|
| Bcar3 | 520.802 | -0.28542 | 0.10777 | -2.6484 | 0.00809 | 0.04992 | 236.056 |
| Bcas1 | 4448.97 | -0.82624 | 0.13813 | -5.98171 | 2.21E-09 | 1.02E-06 | 80.855 |
| Bcas1os1 | 12.8315 | -0.63616 | 0.60299 | -1.05501 | 0.29142 | 0.50126 | 1.995 |
| Bcas1os2 | 66.8506 | -0.13474 | 0.265 | -0.50844 | 0.61114 | 0.77825 | 24.075 |
| Bcas2 | 2824.94 | 0.000514 | 0.06775 | 0.007588 | 0.99395 | 0.99634 | 7.512 |
| Bcas3 | 1774.82 | 0.104567 | 0.07796 | 1.34131 | 0.17982 | 0.37296 | 472.892 |
| Bcas3os1 | 15.5737 | 0.581781 | 0.46978 | 1.238415 | 0.21556 | 0.41717 | 17.545 |
| Bcat1 | 4715.34 | -0.47258 | 0.11214 | -4.21408 | 2.51E-05 | 0.00103 | 82.35 |
| Bcat2 | 689.798 | 0.001704 | 0.11122 | 0.015318 | 0.98778 | 0.99316 | 19.559 |
| Bccip | 2471.21 | 0.174986 | 0.09133 | 1.915934 | 0.05537 | 0.17747 | 11.813 |
| Bcdin3d | 387.817 | 0.217532 | 0.111 | 1.959745 | 0.05003 | 0.16605 | 4.652 |
| Bche | 211.61 | -0.58844 | 0.20453 | -2.87708 | 0.00401 | 0.03157 | 72.608 |
| Bckdha | 451.91 | 0.024593 | 0.11711 | 0.210005 | 0.83366 | 0.91471 | 29.036 |
| Bckdhb | 822.578 | -0.09581 | 0.10172 | -0.94193 | 0.34623 | 0.55684 | 199.096 |
| Bckdk | 1093.18 | -0.0279 | 0.07796 | -0.3579 | 0.72042 | 0.85081 | 6.14 |
| Bcl10 | 657.985 | -0.13825 | 0.0825 | -1.67587 | 0.09376 | 0.24964 | 11.553 |
| Bcl11a | 4197.59 | -0.08874 | 0.09075 | -0.97793 | 0.32811 | 0.53939 | 96.068 |
| Bcl11b | 12113.6 | 0.247882 | 0.08867 | 2.795641 | 0.00518 | 0.03711 | 93.2 |
| Bcl2 | 1247.18 | 0.261201 | 0.0801 | 3.260887 | 0.00111 | 0.0137 | 176.097 |
| Bcl2l1 | 4485.68 | -0.16393 | 0.09302 | -1.76227 | 0.07802 | 0.22104 | 51.061 |
| Bcl2l11 | 227.418 | -0.47267 | 0.15546 | -3.04048 | 0.00236 | 0.02243 | 36.51 |
| Bcl2l12 | 38.5655 | -0.61479 | 0.32325 | -1.90188 | 0.05719 | 0.18129 | 7.491 |
| Bcl2l13 | 2279.47 | -0.11364 | 0.07128 | -1.59428 | 0.11087 | 0.27788 | 56.631 |
| Bcl2l15 | 41.4648 | -0.15814 | 0.27724 | -0.57043 | 0.56838 | 0.7483 | 22.018 |
| Bcl2l2 | 6286.85 | -0.2241 | 0.08199 | -2.73332 | 0.00627 | 0.04238 | 4.858 |
| Bcl3 | 16.4597 | -0.49261 | 0.53371 | -0.92301 | 0.356 | 0.56636 | 14.309 |
| Bcl6 | 1756.11 | -0.39851 | 0.13098 | -3.04261 | 0.00235 | 0.02234 | 23.801 |
| Bcl6b | 63.4354 | -0.45826 | 0.26895 | -1.7039 | 0.0884 | 0.24019 | 5.671 |
| Bcl7a | 4520.57 | -0.0919 | 0.06771 | -1.3573 | 0.17469 | 0.36622 | 31.159 |
| Bcl7b | 1787.33 | -0.05996 | 0.08059 | -0.74405 | 0.45685 | 0.65826 | 13.573 |
| Bcl7c | 455.263 | -0.50321 | 0.13563 | -3.71023 | 0.00021 | 0.00437 | 47.478 |
| Bcl9 | 2024.29 | -0.0583 | 0.08759 | -0.66558 | 0.50568 | 0.69719 | 94.256 |
| Bcl9l | 3246.35 | 0.249439 | 0.09345 | 2.669128 | 0.0076 | 0.04816 | 29.072 |
| Bclaf1 | 9872.31 | 0.232704 | 0.13028 | 1.786221 | 0.07406 | 0.21445 | 32.145 |

| | | | | | | | |
|---|---|---|---|---|---|---|---|
| Bclaf3 | 625.673 | 0.245881 | 0.13156 | 1.868967 | 0.06163 | 0.18982 | 66.394 |
| Bco2 | 168.055 | -0.29007 | 0.16614 | -1.74589 | 0.08083 | 0.22626 | 22.095 |
| Bcor | 1451.81 | -0.08332 | 0.07809 | -1.06695 | 0.28599 | 0.49604 | 123.616 |
| Bcorl1 | 1391.93 | 0.163936 | 0.07737 | 2.118815 | 0.03411 | 0.12896 | 66.692 |
| Bcr | 19413.4 | 0.51938 | 0.15458 | 3.359848 | 0.00078 | 0.01076 | 124.33 |
| Bcs1l | 811.883 | -0.01503 | 0.09205 | -0.16328 | 0.8703 | 0.93448 | 4.155 |
| Bdh1 | 1382.72 | -0.16222 | 0.07617 | -2.1297 | 0.0332 | 0.12658 | 36.622 |
| Bdh2 | 42.1805 | -0.75751 | 0.34295 | -2.20884 | 0.02719 | 0.11015 | 23.206 |
| Bdkrb2 | 62.4671 | -0.99793 | 0.28124 | -3.54833 | 0.00039 | 0.00669 | 32.012 |
| Bdnf | 422.156 | -1.2116 | 0.52282 | -2.31743 | 0.02048 | 0.09101 | 52.308 |
| Bdp1 | 2697.97 | 0.148836 | 0.10519 | 1.41495 | 0.15708 | 0.34408 | 86.077 |
| Bean1 | 240.847 | -0.37564 | 0.17458 | -2.15163 | 0.03143 | 0.12193 | 48.681 |
| Becn1 | 2980.25 | -0.02221 | 0.06157 | -0.3608 | 0.71825 | 0.84954 | 16.335 |
| Begain | 6189.04 | 0.33385 | 0.08299 | 4.022585 | 5.76E-05 | 0.00183 | 36.031 |
| Bend3 | 245.729 | -0.15957 | 0.15661 | -1.01891 | 0.30825 | 0.51812 | 36.566 |
| Bend4 | 762.218 | -0.04802 | 0.09941 | -0.48307 | 0.62905 | 0.79023 | 36.164 |
| Bend5 | 245.659 | -0.11534 | 0.20631 | -0.55907 | 0.57612 | 0.75395 | 45.293 |
| Bend6 | 4030.66 | -0.10805 | 0.07331 | -1.47395 | 0.1405 | 0.32203 | 55.765 |
| Bend7 | 281.838 | 0.540997 | 0.19327 | 2.799206 | 0.00512 | 0.0369 | 84.312 |
| Best1 | 25.4859 | 0.091735 | 0.33677 | 0.272397 | 0.78532 | 0.88859 | 16.46 |
| Bet1 | 1023.85 | 0.047085 | 0.08022 | 0.586958 | 0.55723 | 0.73902 | 10.074 |
| Bet1l | 526.734 | -0.04248 | 0.09807 | -0.43312 | 0.66493 | 0.81503 | 4.184 |
| Bex1 | 1310.9 | 0.101296 | 0.0792 | 1.27899 | 0.2009 | 0.39907 | 1.542 |
| Bex2 | 9096.87 | 0.018966 | 0.06659 | 0.284837 | 0.77577 | 0.88326 | 1.673 |
| Bex3 | 5046.24 | 0.102407 | 0.09222 | 1.110464 | 0.2668 | 0.47601 | 1.726 |
| Bex4 | 837.528 | 0.300272 | 0.09866 | 3.043516 | 0.00234 | 0.0223 | 1.448 |
| Bfar | 1404.23 | 0.01945 | 0.07617 | 0.255351 | 0.79845 | 0.89601 | 31.755 |
| Bfsp1 | 21.2202 | 0.067341 | 0.51766 | 0.130086 | 0.8965 | 0.94798 | 36.646 |
| Bfsp2 | 109.28 | -0.52506 | 0.24209 | -2.16881 | 0.0301 | 0.11854 | 55.497 |
| Bgn | 622.326 | -0.83809 | 0.20617 | -4.06498 | 4.80E-05 | 0.0016 | 12.332 |
| Bhlha9 | 18.5673 | -1.20166 | 0.48929 | -2.45592 | 0.01405 | 0.07136 | 1.207 |
| Bhlhb9 | 5102.3 | 0.400491 | 0.09172 | 4.366301 | 1.26E-05 | 0.00066 | 93.488 |
| Bhlhe22 | 792.375 | -1.20164 | 0.6241 | -1.92538 | 0.05418 | 0.17506 | 3.344 |
| Bhlhe40 | 2247.37 | -0.29386 | 0.10774 | -2.72741 | 0.00638 | 0.04287 | 6.297 |

| | | | | | | | |
|---|---|---|---|---|---|---|---|
| Bhlhe41 | 2103.04 | 0.25331 | 0.09847 | 2.572446 | 0.0101 | 0.05786 | 7.316 |
| Bhmt2 | 12.4206 | 0.239018 | 0.52493 | 0.455329 | 0.64887 | 0.80429 | 18.583 |
| Bicc1 | 327.447 | -0.45197 | 0.19684 | -2.29615 | 0.02167 | 0.09443 | 236.869 |
| Bicd1 | 2841.68 | 0.083718 | 0.1586 | 0.527864 | 0.59759 | 0.76924 | 154.444 |
| Bicd2 | 4684.94 | -0.12537 | 0.07105 | -1.76469 | 0.07762 | 0.22047 | 45.442 |
| Bicdl1 | 2455.12 | -0.08984 | 0.08398 | -1.0698 | 0.28471 | 0.49463 | 83.447 |
| Bicra | 1156.8 | 0.104374 | 0.07913 | 1.318948 | 0.18719 | 0.38244 | 77.25 |
| Bicral | 4183.95 | 0.122474 | 0.06842 | 1.789993 | 0.07346 | 0.21344 | 60.363 |
| Bid | 255.207 | -0.28656 | 0.14609 | -1.96157 | 0.04981 | 0.16545 | 24.924 |
| Bin1 | 8280.45 | -0.21681 | 0.09534 | -2.27416 | 0.02296 | 0.09807 | 58.508 |
| Bin2 | 154.777 | -0.32806 | 0.20462 | -1.60324 | 0.10888 | 0.27434 | 28.477 |
| Bin3 | 389.141 | 0.000506 | 0.09779 | 0.005174 | 0.99587 | 0.99773 | 38.102 |
| Birc2 | 2225.15 | 0.164364 | 0.07297 | 2.252596 | 0.02428 | 0.10183 | 18.838 |
| Birc3 | 115.48 | -0.43475 | 0.16274 | -2.67142 | 0.00755 | 0.04795 | 24.488 |
| Birc5 | 31.6164 | -0.41971 | 0.31381 | -1.33747 | 0.18107 | 0.37448 | 6.493 |
| Birc6 | 8803.01 | 0.129974 | 0.0728 | 1.785275 | 0.07422 | 0.2148 | 175.062 |
| Birc7 | 12.4613 | 0.360226 | 0.57185 | 0.629934 | 0.52874 | 0.71654 | 4.988 |
| Bivm | 1240.03 | 0.151176 | 0.08699 | 1.737914 | 0.08223 | 0.22863 | 25.814 |
| Blcap | 4864.41 | 0.257833 | 0.08418 | 3.062692 | 0.00219 | 0.02136 | 14.913 |
| Blm | 249.711 | 0.008604 | 0.14138 | 0.060858 | 0.95147 | 0.97617 | 80.387 |
| Blmh | 2656.22 | 0.136231 | 0.07077 | 1.925099 | 0.05422 | 0.1751 | 62.571 |
| Blnk | 224.015 | -0.69286 | 0.27842 | -2.48857 | 0.01283 | 0.06761 | 65.608 |
| Bloc1s1 | 19.1479 | 0.613646 | 0.43097 | 1.423886 | 0.15448 | 0.34053 | 6.154 |
| Bloc1s2 | 1098.63 | 0.029353 | 0.07638 | 0.384304 | 0.70075 | 0.83841 | 7.201 |
| Bloc1s3 | 419.859 | -0.39158 | 0.10552 | -3.71079 | 0.00021 | 0.00437 | 3.882 |
| Bloc1s4 | 325.161 | -0.05274 | 0.13067 | -0.40363 | 0.68648 | 0.82889 | 1.273 |
| Bloc1s5 | 550.028 | 0.040908 | 0.13218 | 0.309484 | 0.75695 | 0.87207 | 42.268 |
| Bloc1s6 | 1218.83 | 0.040375 | 0.07425 | 0.543792 | 0.58658 | 0.76038 | 10.973 |
| Blvra | 532.023 | -0.07464 | 0.09771 | -0.76385 | 0.44495 | 0.64896 | 26.42 |
| Blvrb | 178.619 | -0.32669 | 0.15202 | -2.14895 | 0.03164 | 0.12253 | 18.167 |
| Blzf1 | 1297.51 | 0.064188 | 0.07891 | 0.813395 | 0.41599 | 0.62364 | 17.69 |
| Bmerb1 | 4727.93 | -0.10302 | 0.05409 | -1.90461 | 0.05683 | 0.18045 | 114.897 |
| Bmf | 189.25 | -0.52232 | 0.19056 | -2.74098 | 0.00613 | 0.04185 | 20.931 |
| Bmi1 | 2861.66 | 0.163849 | 0.0805 | 2.035382 | 0.04181 | 0.14713 | 9.612 |

| | | | | | | | |
|---|---|---|---|---|---|---|---|
| Bmp1 | 2895.55 | -0.2104 | 0.10855 | -1.93832 | 0.05258 | 0.17154 | 45.677 |
| Bmp2 | 237.596 | 0.28364 | 0.24253 | 1.169522 | 0.24219 | 0.4482 | 10.727 |
| Bmp2k | 870.085 | -0.3249 | 0.10862 | -2.99124 | 0.00278 | 0.02498 | 94.179 |
| Bmp3 | 388.914 | -0.94821 | 0.65383 | -1.45025 | 0.14699 | 0.33019 | 29.982 |
| Bmp4 | 109.402 | -1.01391 | 0.33237 | -3.05053 | 0.00228 | 0.02196 | 7.15 |
| Bmp5 | 36.1345 | -0.98281 | 0.43085 | -2.28108 | 0.02254 | 0.09695 | 124.947 |
| Bmp6 | 737.414 | -0.92249 | 0.19909 | -4.63358 | 3.59E-06 | 0.00027 | 155.196 |
| Bmp7 | 612.039 | -0.33764 | 0.20805 | -1.62292 | 0.10461 | 0.26751 | 72.31 |
| Bmp8b | 15.2086 | -0.24643 | 0.46013 | -0.53556 | 0.59226 | 0.76487 | 26.935 |
| Bmper | 446.138 | 0.019765 | 0.13803 | 0.143188 | 0.88614 | 0.94329 | 262.125 |
| Bmpr1a | 3129.45 | 0.072528 | 0.06489 | 1.117711 | 0.26369 | 0.47245 | 92.608 |
| Bmpr1b | 1821.69 | -0.25528 | 0.12037 | -2.12083 | 0.03394 | 0.12863 | 332.29 |
| Bmpr2 | 10463.4 | 0.02275 | 0.0825 | 0.275773 | 0.78272 | 0.88707 | 115.615 |
| Bms1 | 2420.4 | 0.064451 | 0.09094 | 0.708753 | 0.47848 | 0.67612 | 36.094 |
| Bmt2 | 1761.21 | 0.158615 | 0.08909 | 1.780387 | 0.07501 | 0.21594 | 52.294 |
| Bmx | 66.8137 | -0.27813 | 0.37308 | -0.74551 | 0.45597 | 0.65759 | 65.352 |
| Bmyc | 89.2245 | -0.12497 | 0.19822 | -0.63048 | 0.52838 | 0.71627 | 0.983 |
| Bnc2 | 33.0544 | -0.60774 | 0.476 | -1.27678 | 0.20168 | 0.40014 | 400.181 |
| Bnip1 | 958.666 | 0.18875 | 0.0802 | 2.35335 | 0.01861 | 0.08547 | 11.5 |
| Bnip2 | 1720.6 | -0.0215 | 0.07905 | -0.27196 | 0.78566 | 0.88863 | 18.852 |
| Bnip3 | 1932.05 | -0.05022 | 0.08658 | -0.58005 | 0.56188 | 0.74286 | 18.684 |
| Bnip3l | 5442.27 | -0.08482 | 0.06454 | -1.31423 | 0.18877 | 0.38449 | 23.639 |
| Bnip3l-ps | 255.061 | -0.02059 | 0.12201 | -0.16872 | 0.86602 | 0.93214 | 1.132 |
| Bnip5 | 312.481 | -0.10001 | 0.12605 | -0.79343 | 0.42753 | 0.63412 | 18.933 |
| Bnipl | 33.5757 | 0.473064 | 0.32344 | 1.462603 | 0.14358 | 0.32574 | 9.923 |
| Boc | 448.637 | -0.81869 | 0.20771 | -3.94152 | 8.10E-05 | 0.00232 | 73.849 |
| Bod1 | 2448.62 | -0.01137 | 0.08388 | -0.1355 | 0.89222 | 0.94617 | 6.705 |
| Bod1l | 5024.68 | 0.40759 | 0.27275 | 1.49435 | 0.13508 | 0.31502 | 56.778 |
| Bok | 1451.99 | -0.58513 | 0.14332 | -4.08261 | 4.45E-05 | 0.00151 | 10.105 |
| Bola1 | 285.256 | -0.20224 | 0.12695 | -1.59312 | 0.11113 | 0.27833 | 23.122 |
| Bola2 | 690.133 | -0.21363 | 0.10376 | -2.05889 | 0.03951 | 0.14182 | 4.398 |
| Bola3 | 607.601 | -0.066 | 0.12113 | -0.54482 | 0.58588 | 0.76012 | 10.99 |
| Bop1 | 1561.08 | -0.09583 | 0.08836 | -1.08451 | 0.27814 | 0.48804 | 24.289 |
| Bora | 272.084 | 0.330972 | 0.17198 | 1.924497 | 0.05429 | 0.17522 | 28.319 |

| | | | | | | | |
|---|---|---|---|---|---|---|---|
| Borcs5 | 852.555 | -0.0579 | 0.07431 | -0.77917 | 0.43588 | 0.64152 | 71.678 |
| Borcs6 | 431.07 | -0.05286 | 0.12434 | -0.42511 | 0.67076 | 0.8186 | 1.862 |
| Borcs7 | 641.921 | -0.00487 | 0.1125 | -0.04327 | 0.96549 | 0.9828 | 13.477 |
| Borcs8 | 711.893 | 0.025755 | 0.08701 | 0.296012 | 0.76722 | 0.8783 | 7.821 |
| Bpgm | 2255.7 | -0.14126 | 0.07863 | -1.79651 | 0.07241 | 0.21144 | 29.407 |
| Bphl | 859.373 | 0.075875 | 0.11057 | 0.686196 | 0.49259 | 0.68717 | 36.478 |
| Bpnt1 | 2507.67 | -0.0198 | 0.05751 | -0.34424 | 0.73067 | 0.85728 | 25.629 |
| Bpnt2 | 8754.48 | 0.189936 | 0.0789 | 2.407181 | 0.01608 | 0.07797 | 30.872 |
| Bptf | 8343.92 | 0.181945 | 0.05994 | 3.035635 | 0.0024 | 0.02269 | 99.047 |
| Braf | 8631.11 | 0.139254 | 0.09769 | 1.425527 | 0.154 | 0.33978 | 122.227 |
| Brap | 3045.34 | -0.12215 | 0.06742 | -1.81173 | 0.07003 | 0.20664 | 26.694 |
| Brat1 | 1069.16 | -0.01106 | 0.07799 | -0.14187 | 0.88718 | 0.94391 | 14.369 |
| Brca1 | 153.216 | 0.378888 | 0.1395 | 2.715953 | 0.00661 | 0.04382 | 63.192 |
| Brca2 | 484.103 | 0.185003 | 0.1491 | 1.240814 | 0.21467 | 0.41608 | 47.7 |
| Brcc3 | 1640.91 | 0.184313 | 0.13094 | 1.407658 | 0.15923 | 0.34685 | 37.374 |
| Brd1 | 3314.51 | 0.132192 | 0.0657 | 2.011961 | 0.04422 | 0.15247 | 47.2 |
| Brd2 | 7692.13 | -0.11496 | 0.0634 | -1.8131 | 0.06982 | 0.20619 | 10.612 |
| Brd3 | 3296.15 | 0.08561 | 0.05919 | 1.446346 | 0.14808 | 0.33176 | 62.084 |
| Brd3os | 831.707 | 0.11528 | 0.08295 | 1.389727 | 0.16461 | 0.35361 | 1.898 |
| Brd4 | 5680.78 | 0.079488 | 0.05871 | 1.353989 | 0.17574 | 0.36754 | 88.449 |
| Brd7 | 2271.03 | 0.161256 | 0.06815 | 2.366024 | 0.01798 | 0.08358 | 31.156 |
| Brd8 | 3467.14 | 0.190021 | 0.08862 | 2.144151 | 0.03202 | 0.12355 | 25.987 |
| Brd8dc | 139.491 | 0.476463 | 0.19479 | 2.446047 | 0.01444 | 0.07264 | 17.623 |
| Brd9 | 3682.98 | -0.1118 | 0.08874 | -1.25988 | 0.20771 | 0.40727 | 23.085 |
| Brdt | 827.168 | -0.04304 | 0.09411 | -0.45732 | 0.64744 | 0.80377 | 55.9 |
| Brf1 | 2022.71 | 0.00248 | 0.06995 | 0.035456 | 0.97172 | 0.98579 | 40.62 |
| Brf2 | 546.364 | 0.013286 | 0.09509 | 0.139716 | 0.88888 | 0.94449 | 5.317 |
| Bri3 | 662.163 | -0.09109 | 0.11588 | -0.78608 | 0.43182 | 0.63787 | 202.321 |
| Bri3bp | 5076.28 | -0.25303 | 0.09391 | -2.69432 | 0.00705 | 0.04573 | 18.803 |
| Brinp1 | 7866.55 | -0.13175 | 0.06861 | -1.92037 | 0.05481 | 0.17623 | 192.884 |
| Brinp2 | 4571.74 | -0.31249 | 0.11787 | -2.65107 | 0.00802 | 0.04967 | 111.058 |
| Brinp3 | 936.394 | -0.27248 | 0.12554 | -2.17036 | 0.02998 | 0.11822 | 407.713 |
| Brip1os | 540.702 | -0.42695 | 0.10122 | -4.21816 | 2.46E-05 | 0.00101 | 103.133 |
| Brix1 | 964.795 | 0.121119 | 0.10428 | 1.161497 | 0.24544 | 0.45204 | 11.169 |

| | | | | | | | |
|---|---|---|---|---|---|---|---|
| Brk1 | 4696.23 | -0.09943 | 0.05955 | -1.66972 | 0.09497 | 0.25165 | 12.18 |
| Brme1 | 121.208 | -0.33703 | 0.19131 | -1.7617 | 0.07812 | 0.2212 | 24.566 |
| Brms1 | 20.6816 | -0.6358 | 0.44051 | -1.44333 | 0.14893 | 0.33287 | 8.564 |
| Brms1l | 3597.58 | 0.066723 | 0.0669 | 0.997419 | 0.31856 | 0.52914 | 33.413 |
| Brox | 3752.52 | 0.088005 | 0.06447 | 1.365094 | 0.17222 | 0.36301 | 20.898 |
| Brpf1 | 1621.63 | -0.06117 | 0.07609 | -0.80391 | 0.42145 | 0.6284 | 17.724 |
| Brpf3 | 2484.37 | 0.196251 | 0.08482 | 2.313668 | 0.02069 | 0.09163 | 38.86 |
| Brsk1 | 9437.16 | -0.3541 | 0.0995 | -3.55894 | 0.00037 | 0.00651 | 25.394 |
| Brsk2 | 6769.65 | -0.16711 | 0.09067 | -1.84306 | 0.06532 | 0.19714 | 54.497 |
| Brwd1 | 7099.96 | 0.161994 | 0.07676 | 2.110279 | 0.03483 | 0.13081 | 90.435 |
| Brwd3 | 1374.63 | 0.300366 | 0.16115 | 1.863852 | 0.06234 | 0.1915 | 97.357 |
| Bscl2 | 1910.49 | 0.008394 | 0.0684 | 0.122731 | 0.90232 | 0.95108 | 11.217 |
| Bsdc1 | 2696.39 | 0.04427 | 0.07218 | 0.613296 | 0.53968 | 0.72589 | 26.918 |
| Bsg | 11682.3 | -0.24401 | 0.0694 | -3.51621 | 0.00044 | 0.00729 | 7.479 |
| Bsn | 20385.6 | -0.26495 | 0.11129 | -2.38075 | 0.01728 | 0.08154 | 94.363 |
| Bspry | 61.7972 | 0.822706 | 0.25969 | 3.167999 | 0.00153 | 0.01689 | 17.246 |
| Bst2 | 124.076 | -0.90904 | 0.25589 | -3.55241 | 0.00038 | 0.00662 | 3.202 |
| Btaf1 | 3586.57 | 0.26334 | 0.10344 | 2.545833 | 0.0109 | 0.06082 | 86.674 |
| Btbd1 | 6943.74 | 0.014222 | 0.05074 | 0.280298 | 0.77925 | 0.88501 | 37.358 |
| Btbd10 | 3436.27 | -0.21676 | 0.10169 | -2.13153 | 0.03305 | 0.12624 | 53.767 |
| Btbd11 | 814.525 | -0.07906 | 0.13548 | -0.58352 | 0.55954 | 0.74096 | 273.479 |
| Btbd16 | 51.6835 | -0.1008 | 0.30113 | -0.33476 | 0.73781 | 0.86163 | 51.831 |
| Btbd17 | 667.17 | -0.21247 | 0.12603 | -1.68581 | 0.09183 | 0.24647 | 4.729 |
| Btbd19 | 411.319 | 0.369015 | 0.13458 | 2.742064 | 0.00611 | 0.04178 | 6.507 |
| Btbd2 | 2855.86 | 0.033324 | 0.08217 | 0.405541 | 0.68508 | 0.82843 | 13.878 |
| Btbd3 | 9652.46 | -0.32312 | 0.09887 | -3.26796 | 0.00108 | 0.01349 | 332.728 |
| Btbd6 | 1752.9 | -0.01053 | 0.10486 | -0.10045 | 0.91999 | 0.95948 | 2.464 |
| Btbd7 | 2174.26 | -0.01977 | 0.06238 | -0.31688 | 0.75133 | 0.86954 | 97.675 |
| Btbd8 | 3368.37 | 0.158037 | 0.09866 | 1.601763 | 0.10921 | 0.275 | 73.211 |
| Btbd9 | 2392.3 | 0.057235 | 0.08033 | 0.712484 | 0.47616 | 0.67418 | 360.965 |
| Btc | 28.2137 | -0.28004 | 0.4016 | -0.69732 | 0.4856 | 0.68176 | 45.734 |
| Btd | 309.969 | -0.13135 | 0.12165 | -1.07974 | 0.28026 | 0.49019 | 27.552 |
| Btf3 | 2221.59 | -0.01008 | 0.0739 | -0.13634 | 0.89155 | 0.94599 | 7.111 |
| Btf3-ps1 | 124.975 | -0.33387 | 0.24467 | -1.36462 | 0.17237 | 0.36321 | 0.489 |

| | | | | | | | |
|---|---|---|---|---|---|---|---|
| Btf3-ps18 | 13.7442 | 0.706154 | 0.4766 | 1.481655 | 0.13843 | 0.31961 | 2.524 |
| Btf3l4 | 2177.17 | -0.04727 | 0.07901 | -0.59823 | 0.54969 | 0.73326 | 19.32 |
| Btf3l4b | 313.232 | -0.13111 | 0.10462 | -1.25319 | 0.21014 | 0.41017 | 2.161 |
| Btg1 | 2333.53 | 0.252905 | 0.10998 | 2.299604 | 0.02147 | 0.09392 | 5.804 |
| Btg2 | 672.984 | -0.64822 | 0.18876 | -3.434 | 0.00059 | 0.0089 | 3.951 |
| Btg3 | 989.486 | 0.339232 | 0.12719 | 2.667064 | 0.00765 | 0.04834 | 44.556 |
| Btk | 50.5921 | -0.09558 | 0.30302 | -0.31543 | 0.75243 | 0.86997 | 41.235 |
| Btla | 46.6434 | -0.02942 | 0.38197 | -0.07702 | 0.93861 | 0.96942 | 33.356 |
| Btn2a2 | 28.939 | 0.145761 | 0.34237 | 0.425735 | 0.6703 | 0.81832 | 11.182 |
| Btrc | 5171.89 | 0.049497 | 0.08863 | 0.558463 | 0.57653 | 0.75404 | 166.28 |
| Bub1 | 14.8493 | -0.00133 | 0.49425 | -0.0027 | 0.99785 | 0.99888 | 30.75 |
| Bub1b | 109.97 | -0.8767 | 0.38468 | -2.27905 | 0.02266 | 0.09728 | 43.381 |
| Bub3 | 2615.09 | -0.03618 | 0.07788 | -0.46461 | 0.64221 | 0.80012 | 11.674 |
| Bud13 | 451.21 | -0.01865 | 0.10151 | -0.18377 | 0.85419 | 0.9262 | 16.103 |
| Bud23 | 1159.94 | 0.118181 | 0.07063 | 1.673324 | 0.09426 | 0.25054 | 12.003 |
| Bud31 | 1109.31 | -0.04074 | 0.09727 | -0.4188 | 0.67536 | 0.82188 | 7.717 |
| Bves | 138.651 | 0.2888 | 0.15802 | 1.82756 | 0.06762 | 0.20161 | 36.708 |
| Bvht | 81.7989 | -0.21041 | 0.23617 | -0.89092 | 0.37297 | 0.58339 | 33.032 |
| Bysl | 1039.73 | 0.009899 | 0.08788 | 0.112639 | 0.91032 | 0.95475 | 12.162 |
| Bzw1 | 8258.66 | -0.09488 | 0.07668 | -1.23733 | 0.21596 | 0.41775 | 14.456 |
| Bzw2 | 1608.72 | 0.132598 | 0.07828 | 1.693952 | 0.09027 | 0.24379 | 66.246 |
| C030004G16Rik | 73.6877 | 0.046936 | 0.2807 | 0.167208 | 0.86721 | 0.93259 | 6.825 |
| C030005K06Rik | 42.8533 | 0.442917 | 0.30274 | 1.463043 | 0.14346 | 0.32563 | 11.115 |
| C030006K11Rik | 575.293 | -0.10297 | 0.09992 | -1.03045 | 0.3028 | 0.51232 | 2.381 |
| C030006N10Rik | 22.8351 | 0.593535 | 0.40415 | 1.468615 | 0.14194 | 0.3238 | 0.517 |
| C030013G03Rik | 20.0635 | -0.71297 | 0.524 | -1.36063 | 0.17363 | 0.36499 | 12.769 |
| C030015A19Rik | 272.997 | 0.036197 | 0.1362 | 0.265773 | 0.79041 | 0.89129 | 2.551 |
| C030017B01Rik | 2928.23 | 0.393152 | 0.14699 | 2.674598 | 0.00748 | 0.04763 | 3.241 |
| C030018K13Rik | 165.715 | -0.30964 | 0.19949 | -1.55215 | 0.12063 | 0.29349 | 6.96 |
| C030029H02Rik | 160.354 | -0.19505 | 0.24234 | -0.80485 | 0.42091 | 0.62796 | 50.201 |
| C030034I22Rik | 193.468 | 0.29254 | 0.15353 | 1.90542 | 0.05673 | 0.18033 | 4.278 |
| C030037D09Rik | 340.362 | -0.40698 | 0.12897 | -3.15566 | 0.0016 | 0.01735 | 10.633 |
| C030047K22Rik | 16.2632 | -0.66458 | 0.42739 | -1.55499 | 0.11995 | 0.29246 | 121.091 |
| C130012C08Rik | 81.906 | 0.806625 | 0.27524 | 2.930666 | 0.00338 | 0.02813 | 3.665 |

| | | | | | | | |
|---|---|---|---|---|---|---|---|
| C130013H08Rik | 40.976 | -0.14307 | 0.28085 | -0.50941 | 0.61046 | 0.77778 | 3.692 |
| C130023A14Rik | 160.555 | 0.067879 | 0.1915 | 0.354455 | 0.723 | 0.85223 | 3.087 |
| C130026L21Rik | 253.55 | -0.09389 | 0.13878 | -0.67656 | 0.49868 | 0.69208 | 23.511 |
| C130036L24Rik | 249.979 | 0.454521 | 0.16725 | 2.71769 | 0.00657 | 0.04371 | 14.218 |
| C130046K22Rik | 225.663 | 0.033658 | 0.13545 | 0.248486 | 0.80376 | 0.89873 | 26.233 |
| C130050O18Rik | 28.7136 | 0.454767 | 0.30906 | 1.471436 | 0.14117 | 0.32297 | 10.344 |
| C130051F05Rik | 14.0596 | 0.057849 | 0.47008 | 0.123062 | 0.90206 | 0.95085 | 3.838 |
| C130060C02Rik | 36.1783 | 0.287074 | 0.32778 | 0.875816 | 0.38113 | 0.59057 | 25.622 |
| C130071C03Rik | 3612 | 0.031772 | 0.0877 | 0.36228 | 0.71714 | 0.84899 | 237.209 |
| C130073E24Rik | 147.641 | 0.24095 | 0.20482 | 1.176376 | 0.23944 | 0.44541 | 173.707 |
| C130074G19Rik | 1235.33 | -0.84115 | 0.14187 | -5.92908 | 3.05E-09 | 1.32E-06 | 11.293 |
| C130083M11Rik | 248.161 | 0.339184 | 0.16249 | 2.087387 | 0.03685 | 0.13545 | 16.451 |
| C130089K02Rik | 25.8112 | -0.15927 | 0.40558 | -0.39269 | 0.69455 | 0.8343 | 6.96 |
| C1d | 1734.41 | 0.086281 | 0.09694 | 0.890061 | 0.37343 | 0.58357 | 11.598 |
| C1galt1 | 1406.05 | -0.00724 | 0.10942 | -0.06614 | 0.94727 | 0.97353 | 30.846 |
| C1galt1c1 | 872.715 | 0.02817 | 0.08005 | 0.351908 | 0.72491 | 0.85334 | 4.303 |
| C1qa | 1382.03 | -0.64613 | 0.15312 | -4.21974 | 2.45E-05 | 0.00101 | 2.887 |
| C1qb | 1369.49 | -0.56868 | 0.16432 | -3.46071 | 0.00054 | 0.00836 | 6.059 |
| C1qbp | 1765.35 | 0.014161 | 0.07496 | 0.188909 | 0.85016 | 0.92383 | 5.191 |
| C1qc | 1123.67 | -0.55131 | 0.16733 | -3.2948 | 0.00098 | 0.01267 | 3.262 |
| C1ql1 | 316.933 | -0.63395 | 0.1953 | -3.24606 | 0.00117 | 0.0141 | 7.425 |
| C1ql2 | 22.5256 | 0.932914 | 0.64179 | 1.453621 | 0.14605 | 0.329 | 2.592 |
| C1ql3 | 760.289 | -1.11658 | 0.68772 | -1.62359 | 0.10446 | 0.26741 | 8.35 |
| C1qtnf1 | 70.3337 | -0.9966 | 0.30973 | -3.21763 | 0.00129 | 0.01493 | 21.761 |
| C1qtnf12 | 557.861 | 0.341715 | 0.15235 | 2.242915 | 0.0249 | 0.10366 | 4.312 |
| C1qtnf2 | 28.7254 | -0.8588 | 0.3575 | -2.40222 | 0.0163 | 0.07862 | 17.25 |
| C1qtnf3 | 16.3044 | 0.338896 | 0.56593 | 0.598832 | 0.54929 | 0.73299 | 27.819 |
| C1qtnf4 | 5120.15 | -0.14311 | 0.10408 | -1.37506 | 0.16911 | 0.35927 | 4.666 |
| C1qtnf6 | 74.1921 | -0.11393 | 0.29928 | -0.38068 | 0.70344 | 0.83993 | 8.071 |
| C1qtnf7 | 49.6825 | -0.89627 | 0.32483 | -2.7592 | 0.00579 | 0.04027 | 103.265 |
| C1qtnf9 | 21.7555 | 0.047029 | 0.44311 | 0.106133 | 0.91548 | 0.95671 | 12.736 |
| C1ra | 47.092 | -1.52026 | 0.38742 | -3.92403 | 8.71E-05 | 0.00245 | 11.039 |
| C1s1 | 46.1708 | -1.32806 | 0.39968 | -3.32283 | 0.00089 | 0.0118 | 12.015 |
| C1s2 | 22.6496 | -0.2433 | 0.37924 | -0.64154 | 0.52117 | 0.71037 | 11.461 |

| | | | | | | | |
|---|---|---|---|---|---|---|---|
| C2 | 28.5027 | -0.09395 | 0.3242 | -0.2898 | 0.77197 | 0.88117 | 35.662 |
| C230012O17Rik | 69.6984 | -0.1723 | 0.22728 | -0.75808 | 0.4484 | 0.6519 | 6.66 |
| C230013L11Rik | 118.442 | 0.077438 | 0.205 | 0.377741 | 0.70562 | 0.84142 | 3.026 |
| C230014O12Rik | 103.265 | 1.270974 | 0.2912 | 4.364671 | 1.27E-05 | 0.00066 | 89.447 |
| C230031I18Rik | 13.779 | -0.8851 | 0.68182 | -1.29815 | 0.19424 | 0.39143 | 3.698 |
| C230034O21Rik | 53.7153 | -0.89686 | 0.4188 | -2.14151 | 0.03223 | 0.12411 | 13.69 |
| C230035I16Rik | 44.5419 | -0.39401 | 0.27248 | -1.44603 | 0.14817 | 0.33179 | 3.064 |
| C230037L18Rik | 140.539 | 0.100835 | 0.18122 | 0.556419 | 0.57792 | 0.75477 | 8.612 |
| C230038L03Rik | 13.9711 | -0.14132 | 0.57704 | -0.24491 | 0.80653 | 0.90042 | 45.061 |
| C230057M02Rik | 1438.78 | -0.18135 | 0.12365 | -1.46662 | 0.14248 | 0.32436 | 27.449 |
| C230062I16Rik | 142.396 | -0.23698 | 0.19778 | -1.19817 | 0.23085 | 0.43527 | 16.732 |
| C230086J09Rik | 56.2115 | -0.90706 | 0.32562 | -2.78563 | 0.00534 | 0.03789 | 4.033 |
| C230096K16Rik | 35.5438 | -0.625 | 0.363 | -1.72177 | 0.08511 | 0.23394 | 1.722 |
| C2cd2 | 831.267 | -0.21275 | 0.0891 | -2.38763 | 0.01696 | 0.0807 | 107.39 |
| C2cd2l | 9082.8 | 0.312025 | 0.11054 | 2.822626 | 0.00476 | 0.03524 | 11.049 |
| C2cd3 | 2106.17 | 0.17613 | 0.09402 | 1.873356 | 0.06102 | 0.1886 | 97.92 |
| C2cd4c | 528.635 | 0.017304 | 0.16683 | 0.103719 | 0.91739 | 0.95781 | 7.172 |
| C2cd4d | 42.5935 | 0.50398 | 0.30665 | 1.643488 | 0.10028 | 0.26056 | 2.129 |
| C2cd5 | 6908.71 | 0.324347 | 0.11104 | 2.920879 | 0.00349 | 0.02869 | 89.222 |
| C330011M18Rik | 96.2778 | -0.24946 | 0.21763 | -1.14624 | 0.2517 | 0.45907 | 2.052 |
| C330013E15Rik | 30.2209 | -0.41949 | 0.41064 | -1.02155 | 0.30699 | 0.5168 | 1.09 |
| C330018D20Rik | 671.43 | -0.02247 | 0.10021 | -0.22428 | 0.82254 | 0.90798 | 19.536 |
| C3ar1 | 106.9 | -0.85318 | 0.22812 | -3.7401 | 0.00018 | 0.00406 | 9.024 |
| C430049B03Rik | 18.4891 | -0.33189 | 0.47337 | -0.70113 | 0.48322 | 0.67959 | 2.889 |
| C4a | 279.751 | 0.625234 | 0.16132 | 3.875791 | 0.00011 | 0.0028 | 14.372 |
| C4b | 2584.59 | -0.81989 | 0.34416 | -2.38228 | 0.01721 | 0.08153 | 15.503 |
| C530005A16Rik | 158.31 | 0.205702 | 0.15223 | 1.351268 | 0.17661 | 0.36865 | 7.898 |
| C530025M09Rik | 65.1191 | 0.309877 | 0.31139 | 0.995156 | 0.31966 | 0.53036 | 3.304 |
| C530044C16Rik | 14.2896 | -0.59203 | 0.425 | -1.39302 | 0.16361 | 0.35238 | 38.919 |
| C5ar1 | 42.6701 | -0.37573 | 0.33441 | -1.12357 | 0.26119 | 0.4698 | 12.798 |
| C5ar2 | 45.0058 | -0.76098 | 0.29549 | -2.57534 | 0.01001 | 0.05752 | 9.57 |
| C630031E19Rik | 49.5691 | 0.082697 | 0.26162 | 0.316098 | 0.75193 | 0.86994 | 10.818 |
| C630043F03Rik | 394.665 | 0.151482 | 0.11522 | 1.314738 | 0.1886 | 0.38428 | 14.109 |
| C630044B11Rik | 24.442 | -0.40968 | 0.44961 | -0.9112 | 0.36219 | 0.57276 | 2.211 |

| | | | | | | | |
|---|---|---|---|---|---|---|---|
| C7 | 75.3633 | 0.243122 | 0.24321 | 0.999625 | 0.31749 | 0.52783 | 74.979 |
| C730002L08Rik | 270.89 | -1.14823 | 0.73113 | -1.57048 | 0.1163 | 0.28642 | 141.787 |
| C730034F03Rik | 71.8522 | 0.336392 | 0.24553 | 1.370061 | 0.17067 | 0.36101 | 1.28 |
| C77080 | 328.653 | -0.23442 | 0.12422 | -1.88715 | 0.05914 | 0.18512 | 41.827 |
| C78197 | 77.1704 | 0.341829 | 0.2419 | 1.413112 | 0.15762 | 0.34467 | 3.961 |
| C78859 | 5204.37 | 0.442639 | 0.16555 | 2.673669 | 0.0075 | 0.04773 | 5.943 |
| C79130 | 63.6505 | -0.05279 | 0.23003 | -0.22951 | 0.81848 | 0.90658 | 4.296 |
| C87436 | 584.39 | 0.09321 | 0.11303 | 0.82464 | 0.40958 | 0.61733 | 35.127 |
| C8g | 145.181 | 0.280599 | 0.14737 | 1.904012 | 0.05691 | 0.18066 | 3.069 |
| C920006O11Rik | 171.858 | 0.289427 | 0.15939 | 1.815822 | 0.0694 | 0.20525 | 3 |
| C920021L13Rik | 283.887 | 0.092043 | 0.13305 | 0.691788 | 0.48907 | 0.68423 | 17.586 |
| C9orf72 | 1328.23 | 0.149015 | 0.08824 | 1.688663 | 0.09128 | 0.24554 | 34.891 |
| CK137956 | 18.6136 | -0.48157 | 0.39085 | -1.23211 | 0.21791 | 0.41996 | 43.36 |
| CN725425 | 21.7507 | -0.15371 | 0.47696 | -0.32227 | 0.74725 | 0.86757 | 61.401 |
| Caap1 | 551.62 | 0.106042 | 0.10597 | 1.000636 | 0.317 | 0.52743 | 56.716 |
| Cab39 | 9392.98 | 0.004529 | 0.06672 | 0.067887 | 0.94588 | 0.973 | 58.136 |
| Cab39l | 1359.01 | -0.22229 | 0.09934 | -2.23768 | 0.02524 | 0.10458 | 144.793 |
| Cabcoco1 | 67.0019 | 0.845607 | 0.27813 | 3.040361 | 0.00236 | 0.02243 | 110.944 |
| Cabin1 | 5052.64 | 0.1593 | 0.07673 | 2.076095 | 0.03789 | 0.13802 | 118.23 |
| Cables1 | 908.215 | 0.086163 | 0.12229 | 0.704604 | 0.48106 | 0.6782 | 106.411 |
| Cables2 | 1257.07 | -0.16209 | 0.0932 | -1.7392 | 0.082 | 0.22822 | 14.957 |
| Cabp1 | 7125.59 | -0.73298 | 0.14974 | -4.89494 | 9.83E-07 | 0.0001 | 25.693 |
| Cabp4 | 106.65 | 0.011694 | 0.20348 | 0.057467 | 0.95417 | 0.97748 | 8.611 |
| Cabp7 | 945.461 | -0.26041 | 0.08324 | -3.12861 | 0.00176 | 0.01853 | 9.887 |
| Cabyr | 295.2 | -0.30562 | 0.13978 | -2.18641 | 0.02879 | 0.11465 | 14.939 |
| Cacfd1 | 1932.79 | -0.0885 | 0.07254 | -1.21999 | 0.22247 | 0.42568 | 11.164 |
| Cachd1 | 982.385 | 0.03772 | 0.10648 | 0.354243 | 0.72316 | 0.85223 | 252.546 |
| Cacna1a | 7458.33 | 0.063411 | 0.08259 | 0.767799 | 0.44261 | 0.64716 | 301.608 |
| Cacna1b | 8333.82 | 0.198342 | 0.06138 | 3.231305 | 0.00123 | 0.01456 | 159.266 |
| Cacna1c | 9028.99 | 0.256269 | 0.08432 | 3.03938 | 0.00237 | 0.02249 | 609.651 |
| Cacna1d | 4285.12 | 0.028066 | 0.08601 | 0.326292 | 0.7442 | 0.86539 | 451.517 |
| Cacna1e | 29822 | 0.280338 | 0.07532 | 3.722129 | 0.0002 | 0.00423 | 493.771 |
| Cacna1f | 61.0846 | 0.202669 | 0.23398 | 0.866196 | 0.38638 | 0.59577 | 28.114 |
| Cacna1g | 3840.05 | -0.66859 | 0.34552 | -1.93506 | 0.05298 | 0.17222 | 65.808 |

| | | | | | | | |
|---|---|---|---|---|---|---|---|
| Cacna1h | 7126.76 | 0.275391 | 0.09612 | 2.865219 | 0.00417 | 0.03233 | 59.499 |
| Cacna1i | 7806.07 | -0.09931 | 0.05422 | -1.83144 | 0.06704 | 0.20046 | 111.042 |
| Cacna1s | 27.9263 | -1.12527 | 0.41127 | -2.73607 | 0.00622 | 0.04216 | 67.073 |
| Cacna2d1 | 10437.7 | -0.14461 | 0.16035 | -0.90187 | 0.36713 | 0.57748 | 439.821 |
| Cacna2d2 | 6443 | 0.416745 | 0.13954 | 2.986663 | 0.00282 | 0.02517 | 129.732 |
| Cacna2d3 | 8070.2 | 0.662973 | 0.26996 | 2.455831 | 0.01406 | 0.07136 | 816.922 |
| Cacna2d4 | 103.111 | -0.31647 | 0.23927 | -1.32265 | 0.18595 | 0.38075 | 115.882 |
| Cacnb1 | 6398.07 | 0.175296 | 0.06121 | 2.863904 | 0.00418 | 0.03243 | 21.527 |
| Cacnb2 | 3899.03 | 0.168909 | 0.10104 | 1.671771 | 0.09457 | 0.25104 | 384.821 |
| Cacnb3 | 6966.72 | -0.29501 | 0.09865 | -2.99042 | 0.00279 | 0.02503 | 13.69 |
| Cacnb4 | 10483.4 | 0.332806 | 0.1392 | 2.390788 | 0.01681 | 0.08026 | 248.512 |
| Cacng1 | 13.145 | 0.652438 | 0.60152 | 1.084646 | 0.27808 | 0.48804 | 13.305 |
| Cacng2 | 2521.64 | -0.11496 | 0.10438 | -1.10133 | 0.27075 | 0.48026 | 128.281 |
| Cacng3 | 2718.97 | 0.203617 | 0.0852 | 2.390001 | 0.01685 | 0.08037 | 98.902 |
| Cacng4 | 2927.63 | 0.477825 | 0.17646 | 2.707887 | 0.00677 | 0.0445 | 62.201 |
| Cacng5 | 1273.38 | -0.06526 | 0.09564 | -0.68232 | 0.49504 | 0.68917 | 40.451 |
| Cacng6 | 37.8981 | -1.03397 | 0.48233 | -2.14368 | 0.03206 | 0.12363 | 11.473 |
| Cacng7 | 8217.51 | -0.51061 | 0.10618 | -4.80908 | 1.52E-06 | 0.00014 | 35.267 |
| Cacng8 | 2456.01 | -0.13235 | 0.10571 | -1.25208 | 0.21054 | 0.41056 | 24.966 |
| Cactin | 1371.91 | -0.08569 | 0.08133 | -1.05365 | 0.29204 | 0.50186 | 5.14 |
| Cacul1 | 3619.89 | 0.13283 | 0.07085 | 1.874892 | 0.06081 | 0.18829 | 56.33 |
| Cacybp | 3321.35 | 0.170172 | 0.07676 | 2.216819 | 0.02664 | 0.10874 | 10.509 |
| Cad | 1362.69 | 0.03345 | 0.08285 | 0.40372 | 0.68642 | 0.82888 | 23.7 |
| Cadm1 | 8322.25 | 0.244067 | 0.08199 | 2.97688 | 0.00291 | 0.02572 | 327.943 |
| Cadm2 | 10429.8 | 0.041554 | 0.18616 | 0.223217 | 0.82337 | 0.90849 | 965.49 |
| Cadm3 | 17394.6 | -0.19911 | 0.06887 | -2.89109 | 0.00384 | 0.03059 | 34.437 |
| Cadm4 | 9633.67 | -0.32965 | 0.07704 | -4.27887 | 1.88E-05 | 0.00085 | 22.517 |
| Cadps | 12618.9 | -0.39394 | 0.12396 | -3.17799 | 0.00148 | 0.01643 | 450.517 |
| Cadps2 | 2238.17 | -0.65421 | 0.35701 | -1.83247 | 0.06688 | 0.20022 | 576.649 |
| Cage1 | 212.651 | 0.376602 | 0.17764 | 2.120077 | 0.034 | 0.12875 | 31.018 |
| Calb1 | 3993.5 | 0.07966 | 0.13009 | 0.612336 | 0.54032 | 0.72614 | 26.801 |
| Calb2 | 749.423 | -0.84118 | 0.19805 | -4.24722 | 2.16E-05 | 0.00092 | 30.709 |
| Calca | 17.2619 | -1.16421 | 0.56259 | -2.06939 | 0.03851 | 0.13975 | 4.88 |
| Calcoco1 | 4210.85 | 0.052054 | 0.0648 | 0.803301 | 0.4218 | 0.6287 | 15.402 |

| | | | | | | | |
|---|---|---|---|---|---|---|---|
| Calcr | 52.3987 | 2 | 0.96184 | 3.128476 | 0.00176 | 0.01853 | 79.035 |
| Calcrl | 332.465 | -0.04596 | 0.11843 | -0.3881 | 0.69795 | 0.83634 | 94.786 |
| Cald1 | 1507.33 | -0.43568 | 0.14845 | -2.93486 | 0.00334 | 0.0279 | 176.974 |
| Calhm2 | 158.431 | -0.17041 | 0.22029 | -0.77355 | 0.4392 | 0.64453 | 32.942 |
| Calhm5 | 247.144 | -0.03309 | 0.1968 | -0.16814 | 0.86647 | 0.93232 | 8.705 |
| Calm1 | 126138 | -0.0903 | 0.05313 | -1.69957 | 0.08921 | 0.24185 | 10.38 |
| Calm2 | 67593.2 | -0.07125 | 0.06116 | -1.16494 | 0.24404 | 0.45042 | 13.524 |
| Calm3 | 55374.6 | -0.35862 | 0.0664 | -5.40078 | 6.64E-08 | 1.29E-05 | 8.736 |
| Caln1 | 34546.8 | 0.370592 | 0.12528 | 2.958014 | 0.0031 | 0.02662 | 477.958 |
| Calr | 15964.5 | -0.0989 | 0.09252 | -1.06895 | 0.28509 | 0.49488 | 5.085 |
| Calr3 | 94.0325 | -0.12163 | 0.23179 | -0.52474 | 0.59976 | 0.77047 | 19.695 |
| Calr4 | 27.7277 | -0.43752 | 0.31912 | -1.37102 | 0.17037 | 0.36066 | 20.087 |
| Calu | 4157.57 | -0.16823 | 0.06983 | -2.40913 | 0.01599 | 0.07767 | 29.042 |
| Caly | 4981.4 | 0.168834 | 0.08773 | 1.924435 | 0.0543 | 0.17522 | 12.669 |
| Camk1 | 2102.81 | -0.49362 | 0.168 | -2.93816 | 0.0033 | 0.02774 | 9.861 |
| Camk1d | 7248.64 | 0.021216 | 0.07961 | 0.266487 | 0.78986 | 0.89088 | 421.059 |
| Camk1g | 1926.47 | 0.939539 | 0.19854 | 4.732226 | 2.22E-06 | 0.00019 | 23.953 |
| Camk2a | 108327 | -0.2273 | 0.08731 | -2.60326 | 0.00923 | 0.05434 | 62.535 |
| Camk2b | 50343.6 | 0.134493 | 0.08459 | 1.589949 | 0.11185 | 0.27929 | 96.719 |
| Camk2d | 5866.49 | -0.49024 | 0.19141 | -2.56124 | 0.01043 | 0.05907 | 250.025 |
| Camk2g | 7249.27 | -0.26979 | 0.09063 | -2.97676 | 0.00291 | 0.02572 | 59.214 |
| Camk2n1 | 46988.2 | -0.24324 | 0.08114 | -2.99786 | 0.00272 | 0.02464 | 5.81 |
| Camk2n2 | 2164.83 | -0.81638 | 0.16754 | -4.87272 | 1.10E-06 | 0.00011 | 2.073 |
| Camk4 | 3727.42 | 0.484236 | 0.18777 | 2.57889 | 0.00991 | 0.05714 | 256.298 |
| Camkk1 | 4722.47 | 0.106554 | 0.08988 | 1.185478 | 0.23583 | 0.44108 | 23.066 |
| Camkk2 | 11921.7 | -0.06735 | 0.06244 | -1.07859 | 0.28077 | 0.4905 | 48.24 |
| Camkmt | 431.748 | 0.295201 | 0.10386 | 2.842273 | 0.00448 | 0.03387 | 368.037 |
| Camkv | 34796.7 | 0.310989 | 0.11711 | 2.655485 | 0.00792 | 0.04931 | 14.018 |
| Caml | 1363.28 | -0.05763 | 0.07292 | -0.79021 | 0.42941 | 0.63598 | 9.407 |
| Camsap1 | 6393.34 | 0.170724 | 0.06246 | 2.733434 | 0.00627 | 0.04238 | 56.445 |
| Camsap2 | 9397.06 | 0.100015 | 0.05565 | 1.797227 | 0.0723 | 0.21123 | 77.982 |
| Camsap3 | 939.118 | 0.156438 | 0.07996 | 1.95647 | 0.05041 | 0.16686 | 21.783 |
| Camta1 | 13860.5 | 0.087301 | 0.09531 | 0.915943 | 0.3597 | 0.56997 | 944.555 |
| Camta2 | 9080.56 | -0.19812 | 0.06186 | -3.20283 | 0.00136 | 0.01545 | 18.643 |

| | | | | | | | |
|---|---|---|---|---|---|---|---|
| Cand1 | 7355.1 | 0.018696 | 0.06611 | 0.282789 | 0.77734 | 0.88407 | 40.801 |
| Cand2 | 212.837 | 0.198973 | 0.14103 | 1.410833 | 0.15829 | 0.34558 | 31.02 |
| Cant1 | 2721.96 | 0.053813 | 0.06967 | 0.772415 | 0.43987 | 0.64503 | 12.798 |
| Canx | 18974.6 | 0.051048 | 0.05934 | 0.860237 | 0.38966 | 0.59887 | 31.713 |
| Cap1 | 11647.8 | -0.01087 | 0.06316 | -0.17211 | 0.86335 | 0.93092 | 27.01 |
| Cap2 | 9653.44 | 0.162591 | 0.06958 | 2.336823 | 0.01945 | 0.08786 | 148.434 |
| Capg | 62.5061 | -1.06898 | 0.35486 | -3.01241 | 0.00259 | 0.02395 | 18.593 |
| Capn1 | 834.786 | -0.23181 | 0.10046 | -2.30746 | 0.02103 | 0.09263 | 27.355 |
| Capn10 | 1033.4 | 0.114388 | 0.0814 | 1.405283 | 0.15994 | 0.34784 | 13.566 |
| Capn11 | 66.7448 | -0.77444 | 1.30617 | -0.59291 | 0.55324 | 0.73607 | 29.122 |
| Capn12 | 32.5639 | 0.196343 | 0.4018 | 0.488656 | 0.62509 | 0.78752 | 12.142 |
| Capn15 | 1944.8 | 0.06602 | 0.07595 | 0.869297 | 0.38468 | 0.59416 | 27.433 |
| Capn2 | 3443.98 | -0.19403 | 0.08321 | -2.33189 | 0.01971 | 0.08869 | 50.349 |
| Capn3 | 56.9627 | -0.36967 | 0.30043 | -1.23048 | 0.21852 | 0.42062 | 41.347 |
| Capn5 | 1643.89 | -0.15243 | 0.11514 | -1.32389 | 0.18554 | 0.38034 | 56.716 |
| Capn6 | 119.911 | 0.267512 | 0.25405 | 1.052998 | 0.29234 | 0.50216 | 25.184 |
| Capn7 | 2243.89 | 0.176039 | 0.0773 | 2.277336 | 0.02277 | 0.09762 | 35.349 |
| Capns1 | 3738.28 | -0.14165 | 0.07338 | -1.93042 | 0.05356 | 0.17347 | 11.876 |
| Capns2 | 25.2107 | -0.13262 | 0.3823 | -0.3469 | 0.72866 | 0.85554 | 1.005 |
| Caprin1 | 12781.1 | 0.122414 | 0.06499 | 1.883594 | 0.05962 | 0.18591 | 34.709 |
| Caprin2 | 585.416 | 0.119932 | 0.11497 | 1.043193 | 0.29686 | 0.50702 | 53.746 |
| Caps2 | 15.0406 | 0.110049 | 0.52442 | 0.209848 | 0.83379 | 0.91475 | 52.935 |
| Capsl | 35.7272 | -1.32052 | 0.41355 | -3.19312 | 0.00141 | 0.01585 | 30.011 |
| Capza1 | 1472.16 | 0.029175 | 0.08698 | 0.335423 | 0.73731 | 0.86132 | 41.727 |
| Capza1-ps1 | 134.939 | -0.01564 | 0.18385 | -0.08507 | 0.93221 | 0.96589 | 2.859 |
| Capza2 | 10949.1 | 0.01461 | 0.05521 | 0.264634 | 0.79129 | 0.89159 | 30.739 |
| Capzb | 7065.81 | -0.15287 | 0.07639 | -2.00119 | 0.04537 | 0.15497 | 98.92 |
| Car10 | 1574.35 | -0.96345 | 0.29909 | -3.2213 | 0.00128 | 0.01485 | 503.722 |
| Car11 | 10650.4 | 0.293904 | 0.11396 | 2.578936 | 0.00991 | 0.05714 | 4.84 |
| Car12 | 1787.55 | 0.813548 | 0.17525 | 4.642268 | 3.45E-06 | 0.00026 | 53.16 |
| Car13 | 82.9274 | -1.3862 | 0.47011 | -2.94865 | 0.00319 | 0.02718 | 21.276 |
| Car14 | 368.012 | -0.82888 | 0.12299 | -6.7396 | 1.59E-11 | 2.16E-08 | 6.924 |
| Car15 | 51.1178 | -0.09143 | 0.29556 | -0.30934 | 0.75707 | 0.87207 | 3.089 |
| Car2 | 5069.19 | -0.30873 | 0.14465 | -2.13428 | 0.03282 | 0.1258 | 14.498 |

| | | | | | | | |
|---|---|---|---|---|---|---|---|
| Car3 | 26.0959 | -1.668 | 0.44118 | -3.78072 | 0.00016 | 0.00365 | 9.012 |
| Car4 | 456.197 | -0.58578 | 0.14553 | -4.02506 | 5.70E-05 | 0.00181 | 8.259 |
| Car5b | 226.485 | -0.04591 | 0.163 | -0.28165 | 0.77821 | 0.88423 | 51.176 |
| Car7 | 733.121 | -0.30283 | 0.13195 | -2.29496 | 0.02174 | 0.09467 | 15.655 |
| Car8 | 334.471 | 0.253389 | 0.16176 | 1.56649 | 0.11723 | 0.28823 | 95.68 |
| Car9 | 21.8522 | 0.495301 | 0.4615 | 1.073233 | 0.28317 | 0.49334 | 6.764 |
| Card10 | 288.758 | -0.48997 | 0.18139 | -2.70121 | 0.00691 | 0.04501 | 27.905 |
| Card14 | 19.0863 | -0.41005 | 0.57714 | -0.71049 | 0.4774 | 0.67527 | 37.608 |
| Card19 | 201.756 | -0.14033 | 0.12721 | -1.10315 | 0.26996 | 0.47957 | 13.088 |
| Card6 | 126.283 | -0.36929 | 0.1546 | -2.38865 | 0.01691 | 0.08058 | 12.559 |
| Card9 | 186.483 | 0.204278 | 0.18878 | 1.082076 | 0.27922 | 0.48914 | 8.743 |
| Carf | 1531.72 | 0.462723 | 0.1069 | 4.328434 | 1.50E-05 | 0.00074 | 55.707 |
| Carhsp1 | 5067.62 | 0.184813 | 0.13047 | 1.416543 | 0.15662 | 0.3435 | 13.576 |
| Carlr | 33.4285 | -0.52736 | 0.34082 | -1.54731 | 0.12179 | 0.29526 | 2.574 |
| Carm1 | 2141.43 | 0.112556 | 0.07262 | 1.550021 | 0.12114 | 0.29433 | 45.73 |
| Carmil1 | 1235.48 | 0.068954 | 0.07651 | 0.901221 | 0.36747 | 0.57793 | 268.452 |
| Carmil2 | 712.826 | -0.13939 | 0.09996 | -1.39445 | 0.16318 | 0.35173 | 11.914 |
| Carmil3 | 3104 | 0.380849 | 0.11926 | 3.193433 | 0.00141 | 0.01584 | 17.622 |
| Carmn | 29.7541 | -0.16182 | 0.35785 | -0.45222 | 0.65111 | 0.8058 | 15.913 |
| Carnmt1 | 579.974 | 0.024229 | 0.1158 | 0.209228 | 0.83427 | 0.91498 | 36.437 |
| Carns1 | 303.105 | -0.45894 | 0.12969 | -3.53887 | 0.0004 | 0.00691 | 11.156 |
| Cars | 1980.24 | -0.15066 | 0.08772 | -1.71743 | 0.0859 | 0.23546 | 42.861 |
| Cars2 | 813.845 | 0.133333 | 0.10379 | 1.28463 | 0.19892 | 0.39679 | 36.807 |
| Cartpt | 1736.49 | 0.733698 | 0.2051 | 3.57734 | 0.00035 | 0.0062 | 2.603 |
| Casc1 | 77.3309 | -0.38584 | 0.19943 | -1.93477 | 0.05302 | 0.17223 | 36.172 |
| Casc3 | 3555.56 | -0.0445 | 0.06955 | -0.63984 | 0.52228 | 0.71129 | 28.91 |
| Casd1 | 4175.75 | 0.227569 | 0.07563 | 3.009059 | 0.00262 | 0.0241 | 42.517 |
| Cask | 4127.67 | 0.009338 | 0.06447 | 0.144855 | 0.88483 | 0.94273 | 334.288 |
| Caskin1 | 7540.35 | 0.133439 | 0.07721 | 1.728196 | 0.08395 | 0.23195 | 20.071 |
| Caskin2 | 1284.74 | -0.11986 | 0.07576 | -1.58212 | 0.11362 | 0.28217 | 14.457 |
| Casp1 | 50.4618 | -0.31771 | 0.32522 | -0.9769 | 0.32862 | 0.53979 | 8.783 |
| Casp12 | 47.8818 | -0.85829 | 0.30624 | -2.80266 | 0.00507 | 0.0366 | 27.603 |
| Casp16 | 33.3898 | 0.649375 | 0.31592 | 2.055489 | 0.03983 | 0.14259 | 4.74 |
| Casp2 | 788.924 | 0.100752 | 0.09362 | 1.076128 | 0.28187 | 0.49192 | 17.524 |

| | | | | | | | |
|---|---|---|---|---|---|---|---|
| Casp3 | 424.421 | -0.21877 | 0.15587 | -1.40357 | 0.16045 | 0.3486 | 22.399 |
| Casp4 | 23.1253 | -0.83013 | 0.52331 | -1.58631 | 0.11267 | 0.28066 | 27.956 |
| Casp6 | 215.565 | -0.0616 | 0.13231 | -0.4656 | 0.6415 | 0.79934 | 12.679 |
| Casp7 | 150.456 | 0.200283 | 0.17107 | 1.17078 | 0.24169 | 0.4477 | 45.216 |
| Casp8 | 100.949 | -0.14768 | 0.17454 | -0.84613 | 0.39748 | 0.60658 | 52.13 |
| Casp8ap2 | 882.953 | 0.33665 | 0.23105 | 1.457059 | 0.1451 | 0.32787 | 37.815 |
| Casp9 | 910.569 | -0.14652 | 0.08431 | -1.73789 | 0.08223 | 0.22863 | 22.365 |
| Casq1 | 48.505 | 0.07782 | 0.28756 | 0.270619 | 0.78668 | 0.88907 | 9.975 |
| Casq2 | 112.028 | -0.05348 | 0.24627 | -0.21718 | 0.82807 | 0.9118 | 60.1 |
| Cass4 | 54.7047 | 0.418043 | 0.26648 | 1.568754 | 0.11671 | 0.28707 | 39.964 |
| Cast | 526.722 | -0.35797 | 0.14996 | -2.3872 | 0.01698 | 0.08078 | 116.443 |
| Castor1 | 173.516 | -0.03801 | 0.19094 | -0.19904 | 0.84223 | 0.91879 | 4.185 |
| Castor2 | 8407.72 | -0.10903 | 0.06778 | -1.60857 | 0.10771 | 0.27261 | 44.633 |
| Casz1 | 481.86 | 0.395488 | 0.13352 | 2.96212 | 0.00306 | 0.02638 | 150.461 |
| Cat | 1554.47 | -0.37468 | 0.08221 | -4.55762 | 5.17E-06 | 0.00035 | 31.312 |
| Catip | 74.8978 | -0.56725 | 0.24086 | -2.35508 | 0.01852 | 0.08529 | 7.214 |
| Catsper2 | 367.464 | 0.053654 | 0.11774 | 0.455692 | 0.64861 | 0.80422 | 21.162 |
| Catsperd | 72.3143 | 0.290759 | 0.28083 | 1.035343 | 0.30051 | 0.51055 | 36.314 |
| Catspere2 | 278.909 | 0.688053 | 0.11823 | 5.819423 | 5.91E-09 | 2.02E-06 | 189.283 |
| Catsperg1 | 111.474 | 0.370807 | 0.21501 | 1.724607 | 0.0846 | 0.23316 | 32.715 |
| Catsperz | 26.6372 | -0.89838 | 0.39712 | -2.26224 | 0.02368 | 0.10007 | 3.01 |
| Cav1 | 525.415 | -0.46948 | 0.12327 | -3.80845 | 0.00014 | 0.00336 | 35.118 |
| Cav2 | 879.456 | -0.45322 | 0.12403 | -3.65408 | 0.00026 | 0.00505 | 7.931 |
| Cavin1 | 921.321 | -0.45702 | 0.15734 | -2.90459 | 0.00368 | 0.02977 | 14.155 |
| Cavin2 | 663.411 | -0.30166 | 0.14819 | -2.03558 | 0.04179 | 0.14708 | 13.835 |
| Cavin3 | 256.553 | -0.35391 | 0.13674 | -2.58824 | 0.00965 | 0.05599 | 2.218 |
| Cavin4 | 76.7861 | -0.19632 | 0.27803 | -0.70611 | 0.48012 | 0.67761 | 9.989 |
| Cbarp | 10039 | -0.02606 | 0.07406 | -0.35182 | 0.72497 | 0.85337 | 10.385 |
| Cbfa2t2 | 2673.75 | 0.018536 | 0.07238 | 0.256105 | 0.79787 | 0.89568 | 102.876 |
| Cbfa2t2-ps1 | 110.187 | -0.03546 | 0.17977 | -0.19726 | 0.84362 | 0.91962 | 0.73 |
| Cbfa2t3 | 1260.49 | 0.046461 | 0.09773 | 0.475391 | 0.63451 | 0.79427 | 73.969 |
| Cbfb | 857.567 | -0.14299 | 0.12194 | -1.1726 | 0.24096 | 0.4471 | 47.316 |
| Cbl | 7471.11 | 0.15051 | 0.0598 | 2.516771 | 0.01184 | 0.06404 | 91.074 |
| Cblb | 1195.69 | -0.04786 | 0.0716 | -0.66843 | 0.50386 | 0.69618 | 176.824 |

| | | | | | | | |
|---|---|---|---|---|---|---|---|
| Cbll1 | 873.437 | 0.049314 | 0.08021 | 0.614823 | 0.53867 | 0.72535 | 14.788 |
| Cbln1 | 115.122 | -0.61539 | 0.33812 | -1.82005 | 0.06875 | 0.20383 | 4.205 |
| Cbln2 | 619.96 | -0.95762 | 0.21632 | -4.42682 | 9.56E-06 | 0.00054 | 7.174 |
| Cbln3 | 18.8991 | 0.071564 | 0.4069 | 0.175878 | 0.86039 | 0.92937 | 5.48 |
| Cbln4 | 345.143 | 0.666997 | 0.27538 | 2.422123 | 0.01543 | 0.07584 | 7.234 |
| Cbr1 | 1596.01 | 0.127404 | 0.07625 | 1.670918 | 0.09474 | 0.25135 | 4.653 |
| Cbr2 | 38.6915 | -0.6632 | 0.2756 | -2.4064 | 0.01611 | 0.07804 | 2.626 |
| Cbr3 | 867.825 | 0.606239 | 0.22573 | 2.685711 | 0.00724 | 0.04654 | 7.776 |
| Cbr4 | 784.747 | 0.217461 | 0.10529 | 2.06543 | 0.03888 | 0.14058 | 18.961 |
| Cbs | 1889.87 | -0.08408 | 0.13624 | -0.61716 | 0.53713 | 0.72389 | 28.345 |
| Cbwd1 | 633.447 | -0.04413 | 0.09736 | -0.4533 | 0.65033 | 0.80552 | 41.695 |
| Cbx1 | 2606.58 | -0.07238 | 0.07395 | -0.97876 | 0.3277 | 0.53908 | 19.514 |
| Cbx2 | 280.865 | 0.219337 | 0.15326 | 1.431105 | 0.1524 | 0.33764 | 8.309 |
| Cbx3 | 3673.26 | 0.05907 | 0.08821 | 0.669628 | 0.5031 | 0.69545 | 13.345 |
| Cbx4 | 2841.96 | 0.124091 | 0.08229 | 1.508 | 0.13155 | 0.31028 | 8.649 |
| Cbx5 | 13286.7 | 0.01779 | 0.05676 | 0.313404 | 0.75397 | 0.87067 | 48.273 |
| Cbx6 | 21203.8 | -0.07533 | 0.05602 | -1.34477 | 0.1787 | 0.37128 | 10.793 |
| Cbx7 | 3492.68 | 0.038445 | 0.06893 | 0.557739 | 0.57702 | 0.75436 | 55.313 |
| Cbx8 | 633.57 | -0.08393 | 0.11828 | -0.7096 | 0.47795 | 0.67573 | 4.665 |
| Cby1 | 1733.35 | -0.15415 | 0.08042 | -1.91686 | 0.05526 | 0.17718 | 8.462 |
| Cc2d1a | 2386.21 | 0.067982 | 0.07582 | 0.896651 | 0.36991 | 0.58027 | 15.109 |
| Cc2d1b | 1464.87 | 0.094753 | 0.10354 | 0.915156 | 0.36011 | 0.57053 | 14.184 |
| Cc2d2a | 1416.81 | 0.046213 | 0.10334 | 0.447198 | 0.65473 | 0.80841 | 78.627 |
| Cc2d2b | 29.5613 | -0.08338 | 0.36815 | -0.22648 | 0.82083 | 0.90745 | 78.991 |
| Ccar1 | 4081.26 | 0.246511 | 0.1073 | 2.297423 | 0.02159 | 0.09434 | 48.359 |
| Ccar2 | 4476.46 | 0.081489 | 0.0621 | 1.312257 | 0.18943 | 0.38536 | 15.648 |
| Ccbe1 | 133.484 | 0.316288 | 0.25059 | 1.262172 | 0.20689 | 0.40632 | 257.44 |
| Ccdc102a | 55.7938 | -0.33312 | 0.27742 | -1.20078 | 0.22983 | 0.43407 | 15.23 |
| Ccdc103 | 97.0013 | -0.21482 | 0.18816 | -1.14168 | 0.25359 | 0.46097 | 4.013 |
| Ccdc105 | 33.3926 | 0.191981 | 0.36096 | 0.531859 | 0.59482 | 0.76686 | 6.14 |
| Ccdc106 | 483.325 | 0.3274 | 0.09784 | 3.346382 | 0.00082 | 0.01116 | 40.233 |
| Ccdc107 | 579.353 | 0.027715 | 0.09633 | 0.287699 | 0.77358 | 0.88191 | 3.022 |
| Ccdc110 | 56.823 | 0.53889 | 0.29434 | 1.830864 | 0.06712 | 0.20061 | 9.527 |
| Ccdc112 | 743.296 | 0.456665 | 0.10811 | 4.224034 | 2.40E-05 | 0.00099 | 29.778 |

| | | | | | | | |
|---|---|---|---|---|---|---|---|
| Ccdc113 | 87.234 | 0.410624 | 0.20489 | 2.004087 | 0.04506 | 0.15432 | 24.806 |
| Ccdc115 | 712.965 | 0.084679 | 0.09044 | 0.936325 | 0.34911 | 0.55954 | 3.003 |
| Ccdc116 | 89.6688 | 0.504105 | 0.23357 | 2.158291 | 0.03091 | 0.12055 | 8.166 |
| Ccdc117 | 1372.48 | -0.07488 | 0.1089 | -0.6876 | 0.4917 | 0.68654 | 13.301 |
| Ccdc12 | 919.176 | -0.16044 | 0.08277 | -1.93828 | 0.05259 | 0.17154 | 55.104 |
| Ccdc120 | 576.472 | 0.123581 | 0.10703 | 1.154652 | 0.24823 | 0.45532 | 19.192 |
| Ccdc122 | 80.1517 | 0.051302 | 0.24233 | 0.211706 | 0.83234 | 0.91402 | 75.486 |
| Ccdc124 | 2231.38 | -0.43664 | 0.08426 | -5.18183 | 2.20E-07 | 3.27E-05 | 5.709 |
| Ccdc125 | 120.639 | 0.009137 | 0.1799 | 0.050791 | 0.95949 | 0.98032 | 27.524 |
| Ccdc126 | 679.82 | 0.117164 | 0.09149 | 1.280583 | 0.20034 | 0.39838 | 22.309 |
| Ccdc127 | 3380.9 | -0.01781 | 0.06572 | -0.27103 | 0.78637 | 0.88899 | 15.475 |
| Ccdc13 | 265.294 | 0.042757 | 0.12543 | 0.340885 | 0.73319 | 0.85857 | 41.835 |
| Ccdc130 | 452.987 | 0.220246 | 0.11683 | 1.885236 | 0.0594 | 0.18544 | 12.586 |
| Ccdc134 | 1010.37 | 0.048756 | 0.08308 | 0.586857 | 0.5573 | 0.73902 | 14.282 |
| Ccdc136 | 2461.55 | -0.38327 | 0.09884 | -3.87764 | 0.00011 | 0.00279 | 30.68 |
| Ccdc137 | 1498.01 | 0.08688 | 0.07173 | 1.211124 | 0.22585 | 0.42991 | 6.244 |
| Ccdc138 | 206.116 | 0.084356 | 0.16655 | 0.506478 | 0.61252 | 0.77934 | 78.297 |
| Ccdc14 | 257.424 | 0.216316 | 0.13067 | 1.655424 | 0.09784 | 0.25657 | 34.587 |
| Ccdc141 | 1790.78 | -0.20586 | 0.10984 | -1.87412 | 0.06091 | 0.18839 | 160.735 |
| Ccdc142os | 28.1622 | 0.068293 | 0.35802 | 0.190752 | 0.84872 | 0.92308 | 2.447 |
| Ccdc146 | 18.7378 | -0.29613 | 0.37552 | -0.7886 | 0.43035 | 0.63672 | 131.717 |
| Ccdc148 | 711.65 | 0.072703 | 0.09621 | 0.755663 | 0.44985 | 0.65289 | 279.946 |
| Ccdc149 | 2958.38 | 0.089255 | 0.05645 | 1.581022 | 0.11387 | 0.28252 | 96.871 |
| Ccdc15 | 130.495 | 0.048677 | 0.17049 | 0.285509 | 0.77525 | 0.88294 | 72.598 |
| Ccdc150 | 19.4435 | -0.05917 | 0.3768 | -0.15704 | 0.87522 | 0.93783 | 118.045 |
| Ccdc152 | 34.2354 | -0.05739 | 0.34677 | -0.16549 | 0.86856 | 0.9334 | 23.457 |
| Ccdc153 | 13.3378 | 2 | 0.58973 | 3.811763 | 0.00014 | 0.00334 | 6.631 |
| Ccdc157 | 1578.34 | -0.13504 | 0.08168 | -1.65323 | 0.09828 | 0.25734 | 19.171 |
| Ccdc158 | 30.6279 | -0.57431 | 0.3419 | -1.67975 | 0.09301 | 0.24834 | 67.318 |
| Ccdc159 | 640.297 | 0.211728 | 0.12051 | 1.756944 | 0.07893 | 0.22283 | 8.402 |
| Ccdc160 | 58.3004 | -0.05114 | 0.27151 | -0.18834 | 0.85061 | 0.92407 | 8.269 |
| Ccdc162 | 49.9095 | -0.10397 | 0.27483 | -0.3783 | 0.70521 | 0.84118 | 177.789 |
| Ccdc163 | 162.83 | -0.32336 | 0.2091 | -1.54646 | 0.12199 | 0.29544 | 42.698 |
| Ccdc166 | 243.745 | 0.002694 | 0.17818 | 0.01512 | 0.98794 | 0.99319 | 2.584 |

| | | | | | | | |
|---|---|---|---|---|---|---|---|
| Ccdc167 | 273.264 | 0.039611 | 0.14639 | 0.270579 | 0.78671 | 0.88907 | 29.688 |
| Ccdc169 | 19.4426 | 0.447802 | 0.4105 | 1.090874 | 0.27533 | 0.48468 | 35.597 |
| Ccdc17 | 72.876 | -0.07185 | 0.20027 | -0.35878 | 0.71976 | 0.85033 | 3.613 |
| Ccdc170 | 13.9532 | 0.175533 | 0.60107 | 0.292032 | 0.77026 | 0.87996 | 79.73 |
| Ccdc171 | 175.235 | 0.127618 | 0.16031 | 0.796071 | 0.42599 | 0.63253 | 339.126 |
| Ccdc173 | 155.004 | 0.199961 | 0.18146 | 1.101944 | 0.27049 | 0.47998 | 31.543 |
| Ccdc174 | 1195.15 | 0.230374 | 0.09824 | 2.344964 | 0.01903 | 0.0866 | 21.791 |
| Ccdc177 | 611.339 | -0.23209 | 0.13976 | -1.66059 | 0.09679 | 0.25492 | 6.645 |
| Ccdc18 | 129.516 | 0.208984 | 0.26017 | 0.803272 | 0.42182 | 0.6287 | 100.754 |
| Ccdc181 | 1010.85 | 0.138186 | 0.13207 | 1.046338 | 0.29541 | 0.50588 | 12.263 |
| Ccdc183 | 67.5107 | 0.324952 | 0.24955 | 1.302174 | 0.19286 | 0.38957 | 9.044 |
| Ccdc184 | 588.11 | -0.23317 | 0.10698 | -2.17967 | 0.02928 | 0.11612 | 2.977 |
| Ccdc186 | 2843.57 | 0.157164 | 0.09037 | 1.739131 | 0.08201 | 0.22822 | 34.71 |
| Ccdc187 | 5619.47 | 0.956023 | 0.3015 | 3.170888 | 0.00152 | 0.01675 | 51.089 |
| Ccdc188 | 157.987 | -0.26326 | 0.1827 | -1.44094 | 0.1496 | 0.33394 | 2.47 |
| Ccdc189 | 216.699 | 0.274794 | 0.13406 | 2.049839 | 0.04038 | 0.14365 | 6.213 |
| Ccdc190 | 151.139 | -0.57784 | 0.15298 | -3.77729 | 0.00016 | 0.00368 | 6.006 |
| Ccdc191 | 408.876 | 0.08918 | 0.14708 | 0.606359 | 0.54428 | 0.72945 | 74.515 |
| Ccdc24 | 15.946 | -0.49062 | 0.45892 | -1.06907 | 0.28504 | 0.49486 | 6.034 |
| Ccdc25 | 1706.91 | -0.03955 | 0.08549 | -0.46269 | 0.64359 | 0.80116 | 29.306 |
| Ccdc27 | 43.1986 | 0.008621 | 0.29171 | 0.029552 | 0.97642 | 0.98767 | 16.039 |
| Ccdc28a | 1025.25 | 0.056514 | 0.07186 | 0.786469 | 0.43159 | 0.63765 | 21.323 |
| Ccdc28b | 820.574 | 0.053385 | 0.08868 | 0.602001 | 0.54717 | 0.7314 | 4.674 |
| Ccdc3 | 251.774 | -1.23729 | 0.47906 | -2.58274 | 0.0098 | 0.05673 | 93.103 |
| Ccdc30 | 1202.58 | 0.295767 | 0.09115 | 3.244823 | 0.00118 | 0.01412 | 93.056 |
| Ccdc32 | 1688.85 | -0.30218 | 0.07996 | -3.77912 | 0.00016 | 0.00367 | 11.615 |
| Ccdc33 | 19.2823 | 0.331254 | 0.37421 | 0.8852 | 0.37605 | 0.58613 | 90.147 |
| Ccdc34 | 469.477 | -0.15235 | 0.1209 | -1.26017 | 0.20761 | 0.40716 | 155.544 |
| Ccdc38 | 44.4765 | 0.726227 | 0.30506 | 2.38057 | 0.01729 | 0.08154 | 43.696 |
| Ccdc39 | 1533.83 | 0.288288 | 0.18628 | 1.547612 | 0.12172 | 0.29516 | 31.949 |
| Ccdc40 | 106.695 | -0.08662 | 0.18567 | -0.46651 | 0.64085 | 0.79893 | 36.667 |
| Ccdc42 | 47.7942 | 0.630338 | 0.2505 | 2.516284 | 0.01186 | 0.06409 | 10.946 |
| Ccdc43 | 1075.26 | 0.042346 | 0.08091 | 0.52338 | 0.60071 | 0.77104 | 13.097 |
| Ccdc47 | 5864.51 | 0.077031 | 0.07072 | 1.08924 | 0.27605 | 0.48521 | 18.937 |

| | | | | | | | |
|---|---|---|---|---|---|---|---|
| Ccdc50 | 3674.76 | -0.06225 | 0.09328 | -0.66733 | 0.50456 | 0.6967 | 63.35 |
| Ccdc51 | 261.753 | -0.08945 | 0.13193 | -0.67803 | 0.49775 | 0.6911 | 9.997 |
| Ccdc57 | 691.825 | 0.106092 | 0.10385 | 1.021631 | 0.30696 | 0.5168 | 106.344 |
| Ccdc58 | 523.815 | 0.105301 | 0.1022 | 1.030293 | 0.30287 | 0.51234 | 20.436 |
| Ccdc59 | 1311.34 | 0.061749 | 0.07798 | 0.791872 | 0.42844 | 0.6351 | 6.767 |
| Ccdc6 | 3787.02 | 0.200179 | 0.07897 | 2.53483 | 0.01125 | 0.06195 | 96.08 |
| Ccdc60 | 36.8794 | 0.306256 | 0.27318 | 1.121086 | 0.26225 | 0.47111 | 164.345 |
| Ccdc61 | 437.968 | -0.18502 | 0.09105 | -2.03199 | 0.04215 | 0.14774 | 19.533 |
| Ccdc62 | 294.916 | 0.134575 | 0.17813 | 0.755478 | 0.44996 | 0.65294 | 39.217 |
| Ccdc65 | 184.303 | -0.3775 | 0.18406 | -2.05093 | 0.04027 | 0.14337 | 15.13 |
| Ccdc66 | 960.967 | 0.350339 | 0.18084 | 1.937255 | 0.05271 | 0.17179 | 27.371 |
| Ccdc69 | 18.0769 | 0.380051 | 0.49434 | 0.768806 | 0.44201 | 0.64653 | 28.401 |
| Ccdc71 | 1026.79 | -0.04821 | 0.09076 | -0.53115 | 0.59532 | 0.76727 | 5.412 |
| Ccdc71l | 890.672 | -0.18273 | 0.09651 | -1.89332 | 0.05832 | 0.18358 | 4.24 |
| Ccdc73 | 985.012 | 0.310787 | 0.16167 | 1.922349 | 0.05456 | 0.17569 | 150.1 |
| Ccdc74a | 839.48 | -0.23103 | 0.1157 | -1.99681 | 0.04585 | 0.15609 | 4.269 |
| Ccdc77 | 457.838 | 0.119708 | 0.09879 | 1.211793 | 0.22559 | 0.42964 | 40.052 |
| Ccdc78 | 45.8947 | 0.079389 | 0.29931 | 0.265236 | 0.79083 | 0.89136 | 3.934 |
| Ccdc7b | 64.8228 | 0.160749 | 0.29523 | 0.544496 | 0.5861 | 0.76022 | 116.599 |
| Ccdc8 | 236.675 | -0.20622 | 0.13874 | -1.48636 | 0.13718 | 0.31781 | 4.809 |
| Ccdc80 | 256.272 | -0.82723 | 0.23292 | -3.55152 | 0.00038 | 0.00663 | 34.676 |
| Ccdc81 | 84.4625 | -0.13322 | 0.23827 | -0.55911 | 0.57609 | 0.75395 | 37.482 |
| Ccdc82 | 1615.9 | 0.129041 | 0.12051 | 1.070783 | 0.28427 | 0.49437 | 46.332 |
| Ccdc84 | 778.438 | 0.083984 | 0.10285 | 0.816554 | 0.41418 | 0.62213 | 8.411 |
| Ccdc85a | 2820.74 | -0.00958 | 0.09238 | -0.10369 | 0.91741 | 0.95781 | 198.64 |
| Ccdc85b | 3129.42 | -0.08744 | 0.09539 | -0.91671 | 0.35929 | 0.56968 | 4.715 |
| Ccdc85c | 4401.88 | 0.176636 | 0.09021 | 1.95806 | 0.05022 | 0.16657 | 71.824 |
| Ccdc86 | 303.157 | 0.132535 | 0.14615 | 0.906813 | 0.36451 | 0.57517 | 7.786 |
| Ccdc87 | 68.9513 | -0.32189 | 0.22995 | -1.39982 | 0.16157 | 0.34979 | 3.163 |
| Ccdc88a | 4149.15 | 0.194475 | 0.31137 | 0.624584 | 0.53224 | 0.71973 | 137.151 |
| Ccdc88b | 290.911 | -0.03712 | 0.11941 | -0.31082 | 0.75594 | 0.87157 | 13.589 |
| Ccdc88c | 4252.27 | 0.433355 | 0.09434 | 4.593387 | 4.36E-06 | 0.00031 | 117.534 |
| Ccdc89 | 29.5987 | 0.45502 | 0.35577 | 1.278989 | 0.2009 | 0.39907 | 2.093 |
| Ccdc9 | 2628.06 | -0.02502 | 0.08561 | -0.29225 | 0.7701 | 0.87991 | 12.754 |

| | | | | | | | |
|---|---|---|---|---|---|---|---|
| Ccdc90b | 442.726 | 0.221438 | 0.12331 | 1.795792 | 0.07253 | 0.21168 | 21.146 |
| Ccdc91 | 1265.45 | -0.11258 | 0.0767 | -1.46777 | 0.14217 | 0.32405 | 156.785 |
| Ccdc92 | 2854.94 | -0.01645 | 0.07042 | -0.23363 | 0.81527 | 0.90512 | 28.007 |
| Ccdc92b | 1756.24 | 0.059772 | 0.08176 | 0.73103 | 0.46476 | 0.66445 | 21.912 |
| Ccdc93 | 2378.81 | 0.056819 | 0.11902 | 0.477388 | 0.63309 | 0.79325 | 75.412 |
| Ccdc96 | 95.5964 | -0.33731 | 0.21748 | -1.551 | 0.1209 | 0.29397 | 3.585 |
| Ccdc97 | 2333.82 | 0.009328 | 0.07639 | 0.122108 | 0.90281 | 0.95118 | 7.983 |
| Ccdc9b | 579.964 | -0.27812 | 0.12373 | -2.24786 | 0.02459 | 0.10276 | 8.504 |
| Ccer2 | 26.7971 | -0.29682 | 0.45702 | -0.64947 | 0.51603 | 0.70609 | 2.795 |
| Cchcr1 | 300.602 | -0.29 | 0.17601 | -1.64767 | 0.09942 | 0.25912 | 13.916 |
| Cck | 2962.31 | -1.13759 | 0.4727 | -2.4066 | 0.0161 | 0.07804 | 5.871 |
| Cckbr | 686.543 | -0.39666 | 0.1743 | -2.27571 | 0.02286 | 0.09783 | 45.168 |
| Ccl17 | 41.3732 | -0.13409 | 0.26794 | -0.50044 | 0.61676 | 0.7819 | 1.584 |
| Ccl19 | 35.7335 | -0.23124 | 0.39967 | -0.57858 | 0.56288 | 0.74359 | 2.053 |
| Ccl2 | 12.5854 | -1.34949 | 0.93296 | -1.44646 | 0.14805 | 0.33174 | 1.883 |
| Ccl21a | 18.5177 | -0.25235 | 0.44547 | -0.5665 | 0.57106 | 0.75015 | 1.134 |
| Ccl21b | 19.3294 | -1.54062 | 0.63168 | -2.43893 | 0.01473 | 0.07338 | 1.133 |
| Ccl25 | 708.095 | 0.164323 | 0.10422 | 1.576622 | 0.11488 | 0.28399 | 34.811 |
| Ccl27a | 2443.49 | 0.01511 | 0.08793 | 0.171843 | 0.86356 | 0.93094 | 4.781 |
| Ccl27b | 14.2879 | -1.82062 | 0.79645 | -2.28592 | 0.02226 | 0.09619 | 0.748 |
| Ccl28 | 175.487 | 0.034026 | 0.22134 | 0.153731 | 0.87782 | 0.93893 | 30.541 |
| Ccl5 | 14.5875 | -0.75769 | 0.51197 | -1.47995 | 0.13889 | 0.32011 | 4.741 |
| Ccl6 | 54.1247 | -0.36724 | 0.33664 | -1.0909 | 0.27532 | 0.48468 | 5.206 |
| Ccl9 | 48.8819 | -0.47227 | 0.29094 | -1.62328 | 0.10453 | 0.26745 | 5.718 |
| Ccm2 | 3728.75 | 0.184009 | 0.12214 | 1.506553 | 0.13193 | 0.31074 | 49.858 |
| Ccm2l | 316.482 | -0.15754 | 0.12384 | -1.27208 | 0.20335 | 0.40179 | 15.781 |
| Ccn1 | 323.593 | -0.2377 | 0.1572 | -1.51204 | 0.13052 | 0.30883 | 3.006 |
| Ccn2 | 727.414 | -0.88572 | 0.1933 | -4.58217 | 4.60E-06 | 0.00032 | 3.242 |
| Ccn3 | 3231.13 | -1.42752 | 0.56766 | -2.51473 | 0.01191 | 0.06432 | 8.338 |
| Ccn4 | 45.3946 | -1.82631 | 0.53841 | -3.39208 | 0.00069 | 0.00993 | 31.882 |
| Ccn6 | 15.9087 | -0.32867 | 0.54438 | -0.60376 | 0.546 | 0.73068 | 12.825 |
| Ccna1 | 13.4368 | -0.50768 | 0.48686 | -1.04278 | 0.29705 | 0.50725 | 10.033 |
| Ccna2 | 150.45 | 0.07801 | 0.16659 | 0.468266 | 0.63959 | 0.79795 | 7.286 |
| Ccnb1 | 18.3462 | -0.41372 | 0.48033 | -0.86134 | 0.38905 | 0.5982 | 7.921 |

| | | | | | | | |
|---|---|---|---|---|---|---|---|
| Ccnb2 | 63.9929 | 0.067856 | 0.20539 | 0.33037 | 0.74112 | 0.86331 | 13.856 |
| Ccnb2-ps | 260.21 | 0.297727 | 0.14178 | 2.099983 | 0.03573 | 0.13288 | 2.296 |
| Ccnc | 1874.05 | 0.125526 | 0.08716 | 1.440109 | 0.14984 | 0.33427 | 32.222 |
| Ccnd1 | 1101.63 | -0.61913 | 0.2241 | -2.76272 | 0.00573 | 0.03997 | 9.995 |
| Ccnd2 | 2850.18 | -0.29503 | 0.16904 | -1.74533 | 0.08093 | 0.22645 | 27.032 |
| Ccnd3 | 745.891 | -0.15533 | 0.11332 | -1.37075 | 0.17045 | 0.36077 | 94.641 |
| Ccndbp1 | 2278.03 | 0.078933 | 0.07924 | 0.996062 | 0.31922 | 0.5298 | 8.502 |
| Ccne1 | 576.162 | -0.66364 | 0.15686 | -4.23088 | 2.33E-05 | 0.00097 | 9.551 |
| Ccne2 | 405.023 | 0.388879 | 0.12528 | 3.103975 | 0.00191 | 0.01951 | 13.429 |
| Ccnf | 256.012 | -0.05617 | 0.16313 | -0.34434 | 0.73059 | 0.85727 | 29.287 |
| Ccng1 | 3532.52 | -0.05219 | 0.11133 | -0.46875 | 0.63925 | 0.79775 | 6.76 |
| Ccng2 | 3402.34 | 0.066769 | 0.09112 | 0.732716 | 0.46373 | 0.66349 | 8.975 |
| Ccnh | 2873.06 | 0.216184 | 0.0964 | 2.242678 | 0.02492 | 0.1037 | 34.062 |
| Ccni | 11288.6 | -0.39137 | 0.08465 | -4.62318 | 3.78E-06 | 0.00028 | 24.563 |
| Ccnj | 624.362 | 0.126191 | 0.09526 | 1.324637 | 0.18529 | 0.38008 | 17.294 |
| Ccnjl | 184.089 | 0.371307 | 0.15793 | 2.351103 | 0.01872 | 0.0858 | 58.214 |
| Ccnk | 1728.69 | 0.203345 | 0.08363 | 2.431534 | 0.01504 | 0.07453 | 23.781 |
| Ccnl1 | 3233.42 | 0.341872 | 0.08005 | 4.270954 | 1.95E-05 | 0.00087 | 12.099 |
| Ccnl2 | 8162.65 | 0.241561 | 0.08923 | 2.707214 | 0.00679 | 0.04456 | 12.055 |
| Ccno | 93.8133 | 0.350015 | 0.32511 | 1.076604 | 0.28166 | 0.49168 | 2.976 |
| Ccnq | 384.465 | -0.00651 | 0.12903 | -0.05044 | 0.95977 | 0.98045 | 1.224 |
| Ccnt1 | 2702.02 | 0.063453 | 0.07547 | 0.840754 | 0.40049 | 0.60926 | 32.235 |
| Ccnt2 | 7996.16 | 0.15769 | 0.07503 | 2.101595 | 0.03559 | 0.13252 | 33.898 |
| Ccny | 7974.17 | 0.013284 | 0.0611 | 0.217396 | 0.8279 | 0.91172 | 137.851 |
| Ccnyl1 | 1376.31 | 0.380154 | 0.1394 | 2.727109 | 0.00639 | 0.04288 | 35.19 |
| Ccp110 | 4247.87 | -0.08562 | 0.12552 | -0.68214 | 0.49515 | 0.68919 | 24.473 |
| Ccpg1 | 125.437 | 0.070869 | 0.19228 | 0.368581 | 0.71244 | 0.84516 | 30.912 |
| Ccpg1os | 167.62 | 0.100916 | 0.16189 | 0.623361 | 0.53305 | 0.72034 | 5.661 |
| Ccr10 | 27.3406 | -0.05439 | 0.3127 | -0.17394 | 0.86192 | 0.92994 | 2.447 |
| Ccr2 | 22.4841 | -0.91247 | 0.58331 | -1.56429 | 0.11775 | 0.28887 | 11.608 |
| Ccr5 | 271.934 | -0.30976 | 0.17644 | -1.75562 | 0.07915 | 0.22325 | 26.157 |
| Ccr9 | 17.196 | -0.39533 | 0.51638 | -0.76558 | 0.44393 | 0.6481 | 105.892 |
| Ccrl2 | 24.3793 | -1.28606 | 0.44575 | -2.88514 | 0.00391 | 0.03099 | 3.034 |
| Ccs | 773.509 | -0.06879 | 0.09445 | -0.72827 | 0.46645 | 0.66569 | 13.967 |

| | | | | | | | |
|---|---|---|---|---|---|---|---|
| Ccsap | 3759.45 | 0.178802 | 0.09147 | 1.954794 | 0.05061 | 0.16736 | 19.379 |
| Ccser1 | 483.151 | 0.179157 | 0.13563 | 1.320927 | 0.18653 | 0.38147 | 1202.542 |
| Ccser2 | 6122.58 | 0.104011 | 0.07168 | 1.451143 | 0.14674 | 0.32983 | 93.842 |
| Cct2 | 5713.76 | 0.04458 | 0.06688 | 0.666522 | 0.50508 | 0.69681 | 12.814 |
| Cct3 | 4863.93 | 0.060976 | 0.07108 | 0.857811 | 0.391 | 0.60016 | 24.652 |
| Cct4 | 4417.94 | 0.031393 | 0.06789 | 0.462407 | 0.64379 | 0.80129 | 13.262 |
| Cct5 | 5322.83 | -0.01839 | 0.07293 | -0.2521 | 0.80096 | 0.89715 | 11.005 |
| Cct6a | 6315.35 | 0.022207 | 0.06941 | 0.31996 | 0.749 | 0.86841 | 11.151 |
| Cct7 | 6422.93 | 0.018099 | 0.0719 | 0.251724 | 0.80125 | 0.8972 | 16.962 |
| Cct8 | 6682.57 | 0.046945 | 0.06544 | 0.717348 | 0.47316 | 0.67151 | 12.548 |
| Cct8l1 | 29.5591 | 0.302427 | 0.31386 | 0.96356 | 0.33527 | 0.54607 | 1.961 |
| Ccz1 | 1290.51 | -0.01484 | 0.07456 | -0.19909 | 0.8422 | 0.91879 | 26.969 |
| Cd101 | 41.5031 | -0.39273 | 0.28262 | -1.38959 | 0.16465 | 0.35367 | 36.028 |
| Cd109 | 79.2925 | -1.11538 | 0.3736 | -2.9855 | 0.00283 | 0.02525 | 100.708 |
| Cd14 | 119.181 | -0.47444 | 0.26846 | -1.76727 | 0.07718 | 0.21961 | 1.732 |
| Cd151 | 728.388 | -0.61227 | 0.09798 | -6.24903 | 4.13E-10 | 2.72E-07 | 4.082 |
| Cd160 | 45.8051 | -0.12713 | 0.28126 | -0.45199 | 0.65128 | 0.80589 | 30.589 |
| Cd163 | 518.948 | 0.258816 | 0.20638 | 1.254088 | 0.20981 | 0.40989 | 25.872 |
| Cd164 | 3929.45 | -0.13402 | 0.06039 | -2.21934 | 0.02646 | 0.10823 | 11.635 |
| Cd164l2 | 136.938 | -0.27951 | 0.17092 | -1.63537 | 0.10197 | 0.26365 | 12.425 |
| Cd180 | 45.0103 | -0.54516 | 0.27441 | -1.98665 | 0.04696 | 0.15896 | 46.072 |
| Cd1d1 | 93.5071 | -0.15063 | 0.18547 | -0.81213 | 0.41672 | 0.62431 | 3.608 |
| Cd200 | 3961.26 | -0.01904 | 0.09036 | -0.21075 | 0.83308 | 0.91456 | 26.919 |
| Cd200r1 | 15.484 | -0.40347 | 0.51441 | -0.78435 | 0.43284 | 0.63894 | 29.243 |
| Cd226 | 27.6246 | -0.27855 | 0.4293 | -0.64884 | 0.51644 | 0.70643 | 95.263 |
| Cd248 | 103.341 | -0.63128 | 0.19479 | -3.24079 | 0.00119 | 0.01424 | 2.605 |
| Cd24a | 336.544 | -0.61749 | 0.28102 | -2.19728 | 0.028 | 0.11262 | 5.982 |
| Cd27 | 70.0801 | 0.246185 | 0.23678 | 1.039721 | 0.29847 | 0.50856 | 4.389 |
| Cd274 | 74.6868 | -0.34246 | 0.24065 | -1.42309 | 0.15471 | 0.34078 | 20.641 |
| Cd276 | 538.983 | 0.080777 | 0.11337 | 0.712512 | 0.47615 | 0.67418 | 31.14 |
| Cd2ap | 2210.97 | 0.164167 | 0.15405 | 1.065685 | 0.28657 | 0.4965 | 83.715 |
| Cd2bp2 | 3875.94 | -0.02209 | 0.07135 | -0.30968 | 0.75681 | 0.87201 | 4.391 |
| Cd300a | 63.1918 | -0.39936 | 0.25889 | -1.54261 | 0.12293 | 0.29696 | 14.614 |
| Cd300c2 | 123.247 | -0.5689 | 0.24673 | -2.3058 | 0.02112 | 0.0929 | 5.16 |

| | | | | | | | |
|---|---|---|---|---|---|---|---|
| Cd300lf | 25.3901 | -1.57882 | 0.38549 | -4.09559 | 4.21E-05 | 0.00145 | 17.779 |
| Cd300lg | 23.0225 | -0.32766 | 0.43602 | -0.75147 | 0.45237 | 0.65477 | 14.112 |
| Cd302 | 580.651 | 0.018035 | 0.09426 | 0.19133 | 0.84827 | 0.92288 | 32.496 |
| Cd320 | 729.45 | 0.11571 | 0.10517 | 1.100271 | 0.27121 | 0.48061 | 6.684 |
| Cd33 | 525.682 | -0.33012 | 0.10932 | -3.01978 | 0.00253 | 0.02353 | 20.213 |
| Cd34 | 670.337 | -0.33161 | 0.11771 | -2.81722 | 0.00484 | 0.03553 | 22.461 |
| Cd36 | 17.8249 | -0.39456 | 0.49208 | -0.80181 | 0.42266 | 0.62942 | 107.112 |
| Cd37 | 157.482 | -0.36893 | 0.21118 | -1.74697 | 0.08064 | 0.22601 | 5.484 |
| Cd38 | 154.23 | -0.30355 | 0.23677 | -1.28202 | 0.19984 | 0.39799 | 43.823 |
| Cd3eap | 671.163 | -0.02367 | 0.08781 | -0.26961 | 0.78746 | 0.88953 | 3.47 |
| Cd4 | 1024.26 | 0.661766 | 0.43473 | 1.522256 | 0.12794 | 0.30475 | 23.53 |
| Cd40 | 12.5302 | -0.19389 | 0.47129 | -0.41139 | 0.68078 | 0.82523 | 17.322 |
| Cd44 | 115.377 | -0.79621 | 0.36826 | -2.1621 | 0.03061 | 0.11973 | 90.525 |
| Cd46 | 433.01 | 0.221723 | 0.19176 | 1.156258 | 0.24758 | 0.45457 | 55.424 |
| Cd47 | 8474.39 | 0.111395 | 0.06775 | 1.644293 | 0.10012 | 0.26033 | 114.478 |
| Cd48 | 27.808 | -0.65113 | 0.47939 | -1.35825 | 0.17438 | 0.36601 | 23.25 |
| Cd52 | 39.8239 | -0.95697 | 0.41423 | -2.31024 | 0.02087 | 0.09216 | 12.635 |
| Cd53 | 348.771 | -0.48315 | 0.14454 | -3.3426 | 0.00083 | 0.01126 | 30.229 |
| Cd55 | 150.257 | -0.11225 | 0.1714 | -0.6549 | 0.51253 | 0.70349 | 23.718 |
| Cd59a | 258.052 | -0.45542 | 0.15742 | -2.89292 | 0.00382 | 0.0305 | 19.554 |
| Cd59b | 50.8989 | -0.4297 | 0.31317 | -1.37209 | 0.17004 | 0.3604 | 21.339 |
| Cd6 | 26.6165 | 0.01739 | 0.43503 | 0.039974 | 0.96811 | 0.984 | 40.718 |
| Cd63 | 1990.82 | -0.51193 | 0.13082 | -3.91327 | 9.11E-05 | 0.00253 | 11.834 |
| Cd63-ps | 115.742 | -0.69473 | 0.19477 | -3.56696 | 0.00036 | 0.00637 | 0.875 |
| Cd68 | 401.967 | 0.060332 | 0.16078 | 0.375236 | 0.70749 | 0.84212 | 1.941 |
| Cd72 | 25.8586 | 0.270348 | 0.41172 | 0.656634 | 0.51142 | 0.70252 | 8.167 |
| Cd74 | 262.275 | -1.65248 | 0.5625 | -2.93776 | 0.00331 | 0.02776 | 8.805 |
| Cd79a | 17.6496 | -0.54448 | 0.51293 | -1.06151 | 0.28846 | 0.49847 | 4.817 |
| Cd80 | 87.1101 | 0.006482 | 0.19656 | 0.032977 | 0.97369 | 0.98681 | 40.775 |
| Cd81 | 6506.82 | -0.30829 | 0.09183 | -3.35725 | 0.00079 | 0.01084 | 15.196 |
| Cd82 | 667.814 | -0.63688 | 0.14774 | -4.31082 | 1.63E-05 | 0.00078 | 44.03 |
| Cd83 | 307.947 | -0.19622 | 0.13491 | -1.45452 | 0.1458 | 0.32872 | 18.358 |
| Cd84 | 151.056 | -0.53538 | 0.25453 | -2.1034 | 0.03543 | 0.13224 | 51.021 |
| Cd86 | 109.429 | -0.80674 | 0.18559 | -4.347 | 1.38E-05 | 0.0007 | 62.213 |

| | | | | | | | |
|---|---|---|---|---|---|---|---|
| Cd9 | 916.238 | -0.48951 | 0.16461 | -2.97372 | 0.00294 | 0.02585 | 34.526 |
| Cd93 | 378.65 | -0.26621 | 0.15385 | -1.73032 | 0.08357 | 0.23115 | 6.924 |
| Cd99l2 | 4362.23 | -0.05528 | 0.07936 | -0.69656 | 0.48608 | 0.68219 | 72.79 |
| Cda | 35.323 | -0.28273 | 0.38563 | -0.73316 | 0.46346 | 0.66324 | 29.569 |
| Cdadc1 | 3429.36 | 0.140987 | 0.06975 | 2.021258 | 0.04325 | 0.15025 | 38.572 |
| Cdan1 | 1187.92 | -0.02767 | 0.08157 | -0.33918 | 0.73447 | 0.85958 | 133.975 |
| Cdc123 | 3207.2 | 0.003371 | 0.06241 | 0.05401 | 0.95693 | 0.97897 | 50.871 |
| Cdc14a | 270.635 | -0.01926 | 0.14036 | -0.13719 | 0.89088 | 0.94557 | 156.193 |
| Cdc14b | 851.011 | 0.277812 | 0.10744 | 2.585743 | 0.00972 | 0.0563 | 86.023 |
| Cdc16 | 2297.67 | 0.033574 | 0.07074 | 0.4746 | 0.63507 | 0.79438 | 24.263 |
| Cdc20 | 73.0757 | -0.33196 | 0.25543 | -1.29963 | 0.19373 | 0.39071 | 4.452 |
| Cdc23 | 2409.74 | 0.154429 | 0.08079 | 1.911476 | 0.05594 | 0.17876 | 20.785 |
| Cdc25a | 293.024 | -0.39582 | 0.16783 | -2.35849 | 0.01835 | 0.08475 | 18.317 |
| Cdc25b | 642.213 | 0.011279 | 0.11342 | 0.099447 | 0.92078 | 0.95978 | 11.549 |
| Cdc25c | 27.9245 | -1.2492 | 0.55268 | -2.26028 | 0.0238 | 0.10042 | 18.541 |
| Cdc26 | 1278.35 | -0.07291 | 0.0871 | -0.83704 | 0.40257 | 0.61125 | 24.989 |
| Cdc27 | 3200.38 | 0.05986 | 0.07359 | 0.813447 | 0.41596 | 0.62364 | 47.876 |
| Cdc34 | 1089.14 | -0.22652 | 0.09844 | -2.30112 | 0.02138 | 0.09364 | 6.204 |
| Cdc37 | 4633.27 | -0.18371 | 0.0648 | -2.83524 | 0.00458 | 0.03434 | 16.761 |
| Cdc37l1 | 9675.68 | -0.18471 | 0.07996 | -2.31004 | 0.02089 | 0.09217 | 36.33 |
| Cdc40 | 2591.83 | 0.031784 | 0.06141 | 0.517536 | 0.60478 | 0.7735 | 51.691 |
| Cdc42 | 11708.8 | -0.06773 | 0.05824 | -1.163 | 0.24483 | 0.45139 | 38.025 |
| Cdc42bpa | 15898.8 | -0.01243 | 0.08403 | -0.14788 | 0.88244 | 0.9416 | 205.132 |
| Cdc42bpb | 6847.25 | -0.00322 | 0.0558 | -0.05778 | 0.95393 | 0.97742 | 84.743 |
| Cdc42bpg | 584.675 | 0.13375 | 0.10515 | 1.272038 | 0.20336 | 0.40179 | 19.197 |
| Cdc42ep1 | 441.476 | -0.47649 | 0.17908 | -2.66071 | 0.0078 | 0.04888 | 8.274 |
| Cdc42ep2 | 280.829 | -0.5913 | 0.19932 | -2.96654 | 0.00301 | 0.02623 | 9.181 |
| Cdc42ep3 | 614.935 | 0.389013 | 0.18586 | 2.093026 | 0.03635 | 0.13438 | 21.365 |
| Cdc42ep4 | 1258.3 | -0.28742 | 0.09335 | -3.07896 | 0.00208 | 0.02061 | 25.032 |
| Cdc42ep5 | 21.1699 | -0.69098 | 0.41092 | -1.68153 | 0.09266 | 0.24797 | 13.601 |
| Cdc42se1 | 2498.59 | -0.35274 | 0.11053 | -3.19141 | 0.00142 | 0.01592 | 7.678 |
| Cdc42se2 | 7816.1 | -0.06524 | 0.0751 | -0.86874 | 0.38499 | 0.59437 | 70.22 |
| Cdc45 | 93.9582 | -0.34085 | 0.18591 | -1.8334 | 0.06674 | 0.2001 | 31.541 |
| Cdc5l | 1889.92 | 0.103306 | 0.08011 | 1.289632 | 0.19718 | 0.39502 | 41.854 |

| | | | | | | | |
|---|---|---|---|---|---|---|---|
| Cdc6 | 37.1679 | -0.04911 | 0.33875 | -0.14498 | 0.88473 | 0.94271 | 16.14 |
| Cdc7 | 1420.41 | 0.302512 | 0.10109 | 2.992357 | 0.00277 | 0.02495 | 20.111 |
| Cdc73 | 3504.27 | 0.21388 | 0.10547 | 2.027854 | 0.04258 | 0.14866 | 104.094 |
| Cdca2 | 25.7031 | -0.18535 | 0.34547 | -0.53652 | 0.5916 | 0.76416 | 39.511 |
| Cdca3 | 24.7255 | -0.86313 | 0.37752 | -2.28634 | 0.02223 | 0.0961 | 4.155 |
| Cdca4 | 302.821 | -0.08812 | 0.12547 | -0.7023 | 0.48249 | 0.67901 | 9.195 |
| Cdca7 | 254.179 | -0.61284 | 0.23865 | -2.56798 | 0.01023 | 0.05841 | 10.735 |
| Cdca7l | 43.2883 | -0.64088 | 0.43024 | -1.48957 | 0.13634 | 0.3167 | 74.418 |
| Cdca8 | 83.1578 | -0.11796 | 0.19019 | -0.62021 | 0.53512 | 0.72204 | 20.847 |
| Cdh1 | 95.1359 | -1.56443 | 0.69632 | -2.24672 | 0.02466 | 0.103 | 66.896 |
| Cdh10 | 1948.18 | 0.071709 | 0.08233 | 0.870965 | 0.38377 | 0.59313 | 195.29 |
| Cdh11 | 2280.24 | 0.225801 | 0.10301 | 2.192036 | 0.02838 | 0.11359 | 153.548 |
| Cdh12 | 429.772 | -0.37541 | 0.22213 | -1.69002 | 0.09102 | 0.24507 | 1140.269 |
| Cdh13 | 2875.91 | 0.032028 | 0.0946 | 0.338566 | 0.73494 | 0.85984 | 1041.189 |
| Cdh15 | 159.305 | 0.298123 | 0.23137 | 1.288525 | 0.19756 | 0.39543 | 19.432 |
| Cdh18 | 598.602 | 0.203676 | 0.11113 | 1.832698 | 0.06685 | 0.20018 | 925.397 |
| Cdh19 | 1145.53 | -0.08801 | 0.33753 | -0.26075 | 0.79429 | 0.89351 | 89.259 |
| Cdh2 | 5840.16 | 0.080167 | 0.07433 | 1.078466 | 0.28083 | 0.49056 | 220.37 |
| Cdh20 | 739.245 | -0.34504 | 0.13706 | -2.51732 | 0.01183 | 0.06399 | 226.953 |
| Cdh22 | 1839.92 | -0.03008 | 0.09034 | -0.33295 | 0.73917 | 0.86247 | 123.347 |
| Cdh23 | 123.896 | -0.46088 | 0.19577 | -2.35415 | 0.01857 | 0.08539 | 393.743 |
| Cdh24 | 699.304 | -0.44518 | 0.13563 | -3.2822 | 0.00103 | 0.01305 | 10.134 |
| Cdh26 | 13.2334 | 0.263005 | 0.6156 | 0.427231 | 0.66921 | 0.81761 | 56.836 |
| Cdh4 | 2023.65 | 0.30806 | 0.11417 | 2.698177 | 0.00697 | 0.0453 | 456.943 |
| Cdh5 | 541.136 | -0.5997 | 0.14919 | -4.01984 | 5.82E-05 | 0.00184 | 42.887 |
| Cdh6 | 1163.5 | -0.62168 | 0.362 | -1.71736 | 0.08591 | 0.23546 | 144.975 |
| Cdh7 | 1862.05 | 0.848945 | 0.18706 | 4.538373 | 5.67E-06 | 0.00036 | 157.727 |
| Cdh8 | 1970.18 | 0.732973 | 0.15753 | 4.652885 | 3.27E-06 | 0.00025 | 392.001 |
| Cdh9 | 927.502 | -0.31462 | 0.17872 | -1.76041 | 0.07834 | 0.22159 | 128.339 |
| Cdhr1 | 682.152 | -1.02752 | 0.39549 | -2.5981 | 0.00937 | 0.05499 | 20.491 |
| Cdhr3 | 170.098 | -0.05144 | 0.24123 | -0.21326 | 0.83112 | 0.91334 | 59.08 |
| Cdin1 | 503.906 | 0.215909 | 0.10478 | 2.060607 | 0.03934 | 0.14138 | 197.053 |
| Cdip1 | 8324.12 | -0.06682 | 0.06091 | -1.09707 | 0.27261 | 0.48182 | 39.945 |
| Cdipt | 4005.81 | -0.20254 | 0.08433 | -2.40184 | 0.01631 | 0.07867 | 4.588 |

| | | | | | | | |
|---|---|---|---|---|---|---|---|
| Cdiptos | 209.025 | -0.11941 | 0.18326 | -0.65161 | 0.51465 | 0.70511 | 4.377 |
| Cdk1 | 16.9965 | -0.92385 | 0.61299 | -1.50712 | 0.13178 | 0.31058 | 17.793 |
| Cdk10 | 1984.3 | 0.056309 | 0.09798 | 0.57472 | 0.56548 | 0.74563 | 7.409 |
| Cdk11b | 3708.85 | -0.03444 | 0.05811 | -0.59264 | 0.55342 | 0.73626 | 25.085 |
| Cdk12 | 1746.66 | 0.014825 | 0.08031 | 0.184595 | 0.85355 | 0.92571 | 75.446 |
| Cdk13 | 3666.6 | 0.126419 | 0.07572 | 1.669569 | 0.095 | 0.2517 | 94.783 |
| Cdk14 | 5385.33 | -0.7162 | 0.19339 | -3.70345 | 0.00021 | 0.00444 | 616.922 |
| Cdk15 | 18.5205 | 0.082288 | 0.36937 | 0.222778 | 0.82371 | 0.90858 | 96.088 |
| Cdk16 | 8478.49 | -0.37977 | 0.08231 | -4.61374 | 3.95E-06 | 0.00028 | 11.927 |
| Cdk17 | 16594.2 | 0.412576 | 0.11792 | 3.498793 | 0.00047 | 0.00759 | 106.197 |
| Cdk18 | 1441.23 | -0.93315 | 0.34854 | -2.6773 | 0.00742 | 0.04738 | 27.448 |
| Cdk19 | 5014.23 | -0.00618 | 0.08277 | -0.07469 | 0.94046 | 0.97035 | 144.255 |
| Cdk2 | 141.431 | -0.31485 | 0.19007 | -1.65645 | 0.09763 | 0.25633 | 7.093 |
| Cdk20 | 535.87 | 0.17667 | 0.08622 | 2.049002 | 0.04046 | 0.14371 | 9.46 |
| Cdk2ap1 | 1035.67 | -0.21634 | 0.12825 | -1.68678 | 0.09165 | 0.24619 | 17.666 |
| Cdk2ap2 | 615.402 | -0.02866 | 0.09071 | -0.31597 | 0.75202 | 0.86994 | 1.838 |
| Cdk3 | 81.1431 | -0.12445 | 0.23992 | -0.51874 | 0.60395 | 0.7733 | 4.285 |
| Cdk4 | 1842.97 | 0.117691 | 0.08329 | 1.412965 | 0.15767 | 0.34472 | 4.387 |
| Cdk5 | 3224.99 | -0.0238 | 0.06082 | -0.39131 | 0.69557 | 0.83501 | 5.29 |
| Cdk5r1 | 14794.3 | -0.00847 | 0.06497 | -0.13042 | 0.89623 | 0.94785 | 4.162 |
| Cdk5r2 | 9360.01 | -0.39121 | 0.09442 | -4.14335 | 3.42E-05 | 0.00125 | 2.799 |
| Cdk5rap1 | 420.909 | 0.300555 | 0.11403 | 2.635835 | 0.00839 | 0.05112 | 37.631 |
| Cdk5rap2 | 1203.35 | -0.16599 | 0.12049 | -1.3776 | 0.16833 | 0.35831 | 193.588 |
| Cdk5rap3 | 1053.96 | -0.13117 | 0.10538 | -1.24478 | 0.21321 | 0.41433 | 9.071 |
| Cdk6 | 180.014 | -0.35913 | 0.20904 | -1.71802 | 0.08579 | 0.23534 | 189.524 |
| Cdk7 | 1124.1 | 0.095325 | 0.08514 | 1.119591 | 0.26289 | 0.47176 | 28.309 |
| Cdk8 | 2006.37 | 0.213886 | 0.07185 | 2.976681 | 0.00291 | 0.02572 | 71.645 |
| Cdk9 | 3528.78 | 0.121644 | 0.07853 | 1.54893 | 0.1214 | 0.29466 | 7.293 |
| Cdkal1 | 515.197 | 0.01994 | 0.0889 | 0.224286 | 0.82253 | 0.90798 | 663.929 |
| Cdkl1 | 541.567 | 0.297261 | 0.1091 | 2.724603 | 0.00644 | 0.04308 | 44.42 |
| Cdkl2 | 2242.44 | 0.106594 | 0.06378 | 1.671239 | 0.09467 | 0.25125 | 37.81 |
| Cdkl3 | 602.698 | 0.207203 | 0.10208 | 2.029755 | 0.04238 | 0.14813 | 85.564 |
| Cdkl4 | 494.201 | -0.31136 | 0.19726 | -1.57845 | 0.11446 | 0.28336 | 54.264 |
| Cdkl5 | 11303.1 | 0.12834 | 0.08203 | 1.56448 | 0.1177 | 0.28886 | 222.782 |

| | | | | | | | |
|---|---|---|---|---|---|---|---|
| Cdkn1a | 1916.97 | -0.36159 | 0.13815 | -2.61736 | 0.00886 | 0.05295 | 9.752 |
| Cdkn1b | 2807.22 | -0.16504 | 0.06489 | -2.54343 | 0.01098 | 0.061 | 5.113 |
| Cdkn1c | 55.7502 | -0.33453 | 0.26242 | -1.27478 | 0.20239 | 0.40077 | 2.712 |
| Cdkn2aip | 823.872 | 0.186681 | 0.0933 | 2.000772 | 0.04542 | 0.1551 | 4.589 |
| Cdkn2aipnl | 1665.52 | 0.055284 | 0.07315 | 0.755791 | 0.44977 | 0.65282 | 9.674 |
| Cdkn2b | 17.6517 | -0.3498 | 0.43469 | -0.80471 | 0.42099 | 0.62803 | 4.741 |
| Cdkn2c | 93.7484 | -0.70268 | 0.18606 | -3.77662 | 0.00016 | 0.00369 | 6.314 |
| Cdkn2d | 1504.2 | -0.06286 | 0.10392 | -0.60493 | 0.54523 | 0.73012 | 2.998 |
| Cdkn3 | 14.1601 | 0.335674 | 0.63278 | 0.530474 | 0.59578 | 0.76757 | 11.129 |
| Cdnf | 89.3195 | 0.064558 | 0.25836 | 0.249874 | 0.80268 | 0.89815 | 13.347 |
| Cdo1 | 2145.94 | 0.335725 | 0.11931 | 2.81398 | 0.00489 | 0.03572 | 15.203 |
| Cdon | 817.016 | 0.138553 | 0.08423 | 1.644995 | 0.09997 | 0.26006 | 86.525 |
| Cdpf1 | 274.177 | -0.06242 | 0.11855 | -0.52653 | 0.59852 | 0.76969 | 4.726 |
| Cdr1os | 9027.18 | 0.527101 | 0.29395 | 1.793144 | 0.07295 | 0.21243 | 87.408 |
| Cdr2 | 518.461 | -0.26365 | 0.089 | -2.96246 | 0.00305 | 0.02637 | 25.277 |
| Cdr2l | 1120.92 | -0.25043 | 0.11817 | -2.11925 | 0.03407 | 0.12887 | 14.217 |
| Cds1 | 3076.49 | -0.25942 | 0.08679 | -2.98893 | 0.0028 | 0.02512 | 58.729 |
| Cds2 | 34327.2 | -0.05481 | 0.05935 | -0.92361 | 0.35569 | 0.56614 | 48.903 |
| Cdt1 | 33.9185 | -0.5244 | 0.38299 | -1.36923 | 0.17093 | 0.36136 | 5.54 |
| Cdv3 | 3993.21 | -0.06154 | 0.06267 | -0.98205 | 0.32607 | 0.53752 | 12.747 |
| Cdv3-ps | 13.0324 | -0.28039 | 0.47335 | -0.59235 | 0.55362 | 0.73632 | 1.535 |
| Cdyl | 413.265 | -0.06899 | 0.11236 | -0.61402 | 0.5392 | 0.72582 | 214.231 |
| Cdyl2 | 257.74 | -0.18943 | 0.12217 | -1.55052 | 0.12102 | 0.29412 | 158.592 |
| Ceacam1 | 169.663 | -0.08002 | 0.17123 | -0.46734 | 0.64026 | 0.79859 | 15.897 |
| Ceacam16 | 27.8188 | -0.2051 | 0.34561 | -0.59343 | 0.55289 | 0.7358 | 9.203 |
| Ceacam2 | 51.5333 | -0.52929 | 0.37576 | -1.40857 | 0.15896 | 0.34641 | 23.816 |
| Cebpa | 492.135 | -0.40765 | 0.15282 | -2.6676 | 0.00764 | 0.0483 | 2.636 |
| Cebpb | 133.731 | -0.38088 | 0.18548 | -2.0535 | 0.04002 | 0.14298 | 1.504 |
| Cebpd | 277.619 | -0.26246 | 0.21704 | -1.2093 | 0.22655 | 0.4307 | 3.746 |
| Cebpg | 2034.84 | -0.10813 | 0.07668 | -1.41016 | 0.15849 | 0.34586 | 10.152 |
| Cebpz | 1925.61 | 0.176529 | 0.11099 | 1.590436 | 0.11174 | 0.27929 | 18.061 |
| Cebpzos | 95.1139 | 0.213821 | 0.17396 | 1.229172 | 0.21901 | 0.42125 | 3.807 |
| Cecr2 | 228.132 | -0.47118 | 0.15735 | -2.99451 | 0.00275 | 0.02487 | 104.822 |
| Cela1 | 21.3892 | -0.66759 | 0.42263 | -1.57963 | 0.11419 | 0.283 | 13.501 |

| | | | | | | | |
|---|---|---|---|---|---|---|---|
| Celf1 | 16483.7 | 0.273834 | 0.08574 | 3.193934 | 0.0014 | 0.01582 | 79.116 |
| Celf2 | 32603.5 | 0.181657 | 0.08105 | 2.241373 | 0.025 | 0.10396 | 969.87 |
| Celf3 | 8314.25 | 0.406803 | 0.08274 | 4.916397 | 8.82E-07 | 9.71E-05 | 13.904 |
| Celf4 | 29307.3 | 0.041644 | 0.07526 | 0.553321 | 0.58004 | 0.75648 | 276.526 |
| Celf5 | 14843 | 0.346248 | 0.08581 | 4.035037 | 5.46E-05 | 0.00176 | 23.483 |
| Celf6 | 1126.68 | 0.120402 | 0.09782 | 1.230789 | 0.2184 | 0.42055 | 29.376 |
| Celrr | 187.592 | 0.008967 | 0.19801 | 0.045287 | 0.96388 | 0.98239 | 91.055 |
| Celsr1 | 455.094 | -0.22227 | 0.13139 | -1.69167 | 0.09071 | 0.24468 | 135.275 |
| Celsr2 | 12286 | -0.05918 | 0.08327 | -0.71071 | 0.47727 | 0.67518 | 25.217 |
| Celsr3 | 9249.82 | 0.226119 | 0.08258 | 2.738137 | 0.00618 | 0.04207 | 26.65 |
| Cemip | 498.056 | -0.35479 | 0.16454 | -2.15621 | 0.03107 | 0.12095 | 153.646 |
| Cemip2 | 985.61 | -0.02362 | 0.09308 | -0.2538 | 0.79965 | 0.89651 | 80.019 |
| Cend1 | 9557.23 | -0.20543 | 0.06051 | -3.39496 | 0.00069 | 0.00985 | 3.046 |
| Cenpa | 95.678 | -0.87139 | 0.29176 | -2.98667 | 0.00282 | 0.02517 | 8.054 |
| Cenpb | 6385.53 | -0.35487 | 0.09867 | -3.5966 | 0.00032 | 0.00595 | 4.886 |
| Cenpc1 | 1529.44 | 0.280067 | 0.13248 | 2.114078 | 0.03451 | 0.13002 | 53.56 |
| Cenpe | 90.3849 | 0.439319 | 0.31234 | 1.406547 | 0.15956 | 0.34742 | 61.075 |
| Cenpf | 43.1891 | -0.47056 | 0.53969 | -0.8719 | 0.38326 | 0.59252 | 47.481 |
| Cenph | 46.3415 | 0.257501 | 0.39252 | 0.656013 | 0.51182 | 0.70298 | 16.226 |
| Cenpi | 17.2668 | -0.23828 | 0.43305 | -0.55024 | 0.58216 | 0.75777 | 54.556 |
| Cenpj | 258.637 | 0.327298 | 0.15215 | 2.15119 | 0.03146 | 0.12204 | 48.665 |
| Cenpk | 43.0371 | 0.167802 | 0.33388 | 0.502574 | 0.61526 | 0.78109 | 23.782 |
| Cenpl | 72.9882 | -0.18529 | 0.19893 | -0.93139 | 0.35165 | 0.56204 | 16.012 |
| Cenpm | 53.8127 | -0.9165 | 0.35998 | -2.54596 | 0.0109 | 0.06082 | 10.97 |
| Cenpn | 90.3293 | 0.188659 | 0.19237 | 0.980715 | 0.32673 | 0.53806 | 19.778 |
| Cenpo | 216.213 | -0.08481 | 0.13355 | -0.63507 | 0.52538 | 0.71367 | 38.291 |
| Cenpp | 39.5556 | -0.14421 | 0.32819 | -0.43943 | 0.66035 | 0.81225 | 188.763 |
| Cenpq | 178.29 | -0.12054 | 0.15515 | -0.77692 | 0.43721 | 0.64268 | 11.997 |
| Cenps | 93.0483 | -0.32467 | 0.21068 | -1.54107 | 0.1233 | 0.29747 | 10.509 |
| Cenpt | 816.55 | 0.014586 | 0.10541 | 0.138372 | 0.88995 | 0.94486 | 8.606 |
| Cenpu | 93.4868 | -0.18306 | 0.2119 | -0.86392 | 0.38763 | 0.59697 | 27.98 |
| Cenpv | 952.017 | 0.14972 | 0.0894 | 1.674657 | 0.094 | 0.25005 | 14.316 |
| Cenpw | 102.214 | -0.22154 | 0.20026 | -1.10626 | 0.26861 | 0.47791 | 5.992 |
| Cenpx | 366.514 | -0.06319 | 0.12682 | -0.49822 | 0.61833 | 0.7831 | 2.797 |

| | | | | | | | |
|---|---|---|---|---|---|---|---|
| Cep104 | 2820.59 | 0.058752 | 0.06628 | 0.88648 | 0.37536 | 0.58555 | 33.539 |
| Cep112 | 192.206 | 0.107056 | 0.16649 | 0.643037 | 0.5202 | 0.70963 | 435.424 |
| Cep120 | 3391.58 | 0.095243 | 0.07893 | 1.206742 | 0.22753 | 0.43172 | 62.886 |
| Cep126 | 1061.57 | 0.421449 | 0.13776 | 3.059222 | 0.00222 | 0.02155 | 57.834 |
| Cep128 | 334.679 | 0.076085 | 0.11128 | 0.683746 | 0.49414 | 0.68845 | 385.918 |
| Cep131 | 1345.44 | -0.00839 | 0.07264 | -0.11545 | 0.90808 | 0.95395 | 22.398 |
| Cep135 | 1016.82 | 0.299943 | 0.11877 | 2.525337 | 0.01156 | 0.06306 | 57.769 |
| Cep152 | 346.187 | 0.07683 | 0.12659 | 0.606928 | 0.5439 | 0.72924 | 62.026 |
| Cep162 | 1060.17 | 0.245319 | 0.12006 | 2.043255 | 0.04103 | 0.14504 | 65.96 |
| Cep164 | 1695.63 | 0.186797 | 0.06379 | 2.928282 | 0.00341 | 0.02822 | 61.746 |
| Cep170 | 4342.72 | 0.072714 | 0.12063 | 0.602802 | 0.54664 | 0.73092 | 80.415 |
| Cep170b | 8177.74 | -0.26225 | 0.10677 | -2.4562 | 0.01404 | 0.07134 | 26.138 |
| Cep19 | 2532.78 | -0.04175 | 0.07184 | -0.58118 | 0.56112 | 0.74218 | 8.27 |
| Cep192 | 683.34 | -0.00756 | 0.11137 | -0.06793 | 0.94584 | 0.973 | 85.065 |
| Cep20 | 1586.17 | 0.085115 | 0.09412 | 0.904308 | 0.36583 | 0.57628 | 18.162 |
| Cep250 | 2776.81 | 0.084025 | 0.085 | 0.988564 | 0.32288 | 0.53387 | 42.443 |
| Cep290 | 947.577 | 0.454262 | 0.38545 | 1.178535 | 0.23858 | 0.44422 | 87.283 |
| Cep295 | 1322.72 | 0.286895 | 0.12048 | 2.381323 | 0.01725 | 0.08154 | 40.874 |
| Cep295nl | 27.2103 | 0.674831 | 0.37901 | 1.78051 | 0.07499 | 0.21591 | 10.141 |
| Cep350 | 4259.1 | 0.197644 | 0.07554 | 2.616393 | 0.00889 | 0.05306 | 128.292 |
| Cep41 | 327.296 | -0.0371 | 0.11287 | -0.32873 | 0.74236 | 0.8643 | 40.293 |
| Cep43 | 1099.75 | 0.068631 | 0.08244 | 0.832532 | 0.40511 | 0.61387 | 31.304 |
| Cep44 | 720.514 | 0.101928 | 0.10101 | 1.009042 | 0.31295 | 0.5235 | 19.526 |
| Cep55 | 15.48 | -0.70039 | 0.49179 | -1.42417 | 0.1544 | 0.34039 | 19.413 |
| Cep57 | 1083.65 | 0.270708 | 0.09969 | 2.715379 | 0.00662 | 0.04386 | 19.316 |
| Cep57l1 | 344.363 | 0.219276 | 0.12406 | 1.767474 | 0.07715 | 0.21958 | 91.032 |
| Cep63 | 2901.21 | 0.549336 | 0.08865 | 6.196711 | 5.77E-10 | 3.46E-07 | 41.947 |
| Cep68 | 1632.69 | 0.038807 | 0.07845 | 0.49467 | 0.62083 | 0.78437 | 22.393 |
| Cep70 | 1286.49 | 0.290176 | 0.12603 | 2.302528 | 0.02131 | 0.09342 | 57.038 |
| Cep72 | 132.904 | 0.07567 | 0.21027 | 0.359864 | 0.71895 | 0.8499 | 25.8 |
| Cep76 | 434.06 | 0.108709 | 0.10322 | 1.053156 | 0.29227 | 0.50207 | 23.537 |
| Cep78 | 692.668 | 0.068256 | 0.08701 | 0.78445 | 0.43278 | 0.63893 | 29.217 |
| Cep83 | 1848.24 | 0.219643 | 0.09526 | 2.305774 | 0.02112 | 0.0929 | 102.2 |
| Cep83os | 1369.69 | 0.044623 | 0.12795 | 0.348743 | 0.72728 | 0.85443 | 17.576 |

| | | | | | | | |
|---|---|---|---|---|---|---|---|
| Cep85 | 1316.24 | 0.006049 | 0.07005 | 0.086359 | 0.93118 | 0.96532 | 57.255 |
| Cep85l | 1292.35 | 0.137826 | 0.0914 | 1.507968 | 0.13156 | 0.31028 | 106.505 |
| Cep89 | 1161.04 | 0.103518 | 0.09459 | 1.094395 | 0.27378 | 0.48317 | 41.655 |
| Cep95 | 819.437 | 0.163555 | 0.11634 | 1.405878 | 0.15976 | 0.34774 | 30.679 |
| Cep97 | 2177.11 | 0.076411 | 0.07306 | 1.045872 | 0.29562 | 0.50608 | 34.968 |
| Cept1 | 1678.51 | 0.070558 | 0.07991 | 0.882961 | 0.37726 | 0.58738 | 45.543 |
| Cercam | 294.202 | -0.21943 | 0.13868 | -1.5823 | 0.11358 | 0.28217 | 13.677 |
| Cerk | 3266.09 | -0.35704 | 0.09817 | -3.63676 | 0.00028 | 0.0053 | 47.014 |
| Cerkl | 18.0865 | -1.40891 | 0.55926 | -2.51923 | 0.01176 | 0.06378 | 126.243 |
| Cerox1 | 1635.87 | 0.275286 | 0.10033 | 2.743905 | 0.00607 | 0.04164 | 4.721 |
| Cers1 | 2493.42 | 0.016386 | 0.07324 | 0.223739 | 0.82296 | 0.90818 | 15.818 |
| Cers2 | 2642.65 | -0.35702 | 0.09404 | -3.79635 | 0.00015 | 0.00348 | 8.809 |
| Cers3 | 19.9667 | 0.254762 | 0.48237 | 0.528147 | 0.5974 | 0.7692 | 80.188 |
| Cers4 | 3464.59 | -0.01128 | 0.10828 | -0.10422 | 0.917 | 0.95768 | 38.655 |
| Cers5 | 2365.1 | 0.13986 | 0.07015 | 1.993748 | 0.04618 | 0.15693 | 38.005 |
| Cers6 | 3429.15 | 0.044954 | 0.10083 | 0.445843 | 0.65571 | 0.80879 | 252.842 |
| Cert1 | 5070.81 | 0.057523 | 0.0784 | 0.733671 | 0.46315 | 0.66313 | 97.55 |
| Ces5a | 35.9196 | -0.09791 | 0.3081 | -0.31778 | 0.75065 | 0.86924 | 36.767 |
| Cetn2 | 1349.51 | 0.131544 | 0.07308 | 1.800006 | 0.07186 | 0.21034 | 4.88 |
| Cetn3 | 1916.56 | 0.00287 | 0.10951 | 0.026212 | 0.97909 | 0.98889 | 14.079 |
| Cetn4 | 430.784 | 0.481015 | 0.13132 | 3.662949 | 0.00025 | 0.00496 | 4.96 |
| Cfap100 | 183.277 | 0.225705 | 0.16472 | 1.370198 | 0.17063 | 0.361 | 25.319 |
| Cfap126 | 29.9398 | -0.49429 | 0.35306 | -1.40003 | 0.1615 | 0.34978 | 13.05 |
| Cfap157 | 144.102 | 0.26855 | 0.16505 | 1.627108 | 0.10371 | 0.26623 | 7.048 |
| Cfap161 | 17.9289 | 0.798111 | 0.42484 | 1.878606 | 0.0603 | 0.18734 | 20.78 |
| Cfap20 | 2016.96 | 0.225241 | 0.10408 | 2.164116 | 0.03046 | 0.11925 | 14.621 |
| Cfap206 | 86.5545 | 0.701155 | 0.24876 | 2.818596 | 0.00482 | 0.03546 | 41.648 |
| Cfap20dc | 617.338 | 0.017108 | 0.10001 | 0.17106 | 0.86418 | 0.93125 | 235.049 |
| Cfap221 | 13.1287 | 0.624341 | 0.47636 | 1.310656 | 0.18997 | 0.38603 | 73.894 |
| Cfap298 | 311.465 | 0.080926 | 0.12726 | 0.6359 | 0.52484 | 0.71318 | 9.306 |
| Cfap300 | 521.543 | 0.100403 | 0.13243 | 0.758142 | 0.44837 | 0.65189 | 21.152 |
| Cfap36 | 3516.37 | 0.172151 | 0.11299 | 1.523624 | 0.1276 | 0.30407 | 25.878 |
| Cfap410 | 1397.02 | 0.150685 | 0.06824 | 2.208043 | 0.02724 | 0.11026 | 8.382 |
| Cfap418 | 1675.97 | 0.150719 | 0.08013 | 1.880858 | 0.05999 | 0.18675 | 24.928 |

| | | | | | | | |
|---|---|---|---|---|---|---|---|
| Cfap43 | 109.952 | -0.23783 | 0.21599 | -1.1011 | 0.27085 | 0.48026 | 182.431 |
| Cfap44 | 412.257 | 0.620026 | 0.13225 | 4.68813 | 2.76E-06 | 0.00022 | 87.633 |
| Cfap45 | 110.946 | -0.08248 | 0.18542 | -0.44484 | 0.65644 | 0.8092 | 25.07 |
| Cfap46 | 189.038 | 0.529666 | 0.1913 | 2.768796 | 0.00563 | 0.03945 | 82.867 |
| Cfap47 | 63.0727 | 0.543842 | 0.2969 | 1.831724 | 0.06699 | 0.20038 | 250.727 |
| Cfap54 | 446.142 | 0.203345 | 0.0895 | 2.271994 | 0.02309 | 0.09837 | 306 |
| Cfap57 | 32.6594 | 0.859983 | 0.30248 | 2.843099 | 0.00447 | 0.03383 | 66.227 |
| Cfap61 | 92.5971 | 0.298337 | 0.18967 | 1.572955 | 0.11573 | 0.28553 | 280.256 |
| Cfap65 | 258.808 | 0.167491 | 0.15792 | 1.060587 | 0.28888 | 0.49907 | 33.529 |
| Cfap69 | 1341.14 | 0.257299 | 0.145 | 1.77443 | 0.07599 | 0.21768 | 84.962 |
| Cfap70 | 46.127 | 0.013553 | 0.35157 | 0.03855 | 0.96925 | 0.98441 | 58.034 |
| Cfap74 | 1307.03 | 0.382148 | 0.10399 | 3.674966 | 0.00024 | 0.00479 | 57.634 |
| Cfap91 | 34.8503 | 0.795908 | 0.34347 | 2.317245 | 0.02049 | 0.09101 | 44.39 |
| Cfap97 | 1734.66 | 0.244295 | 0.10564 | 2.312619 | 0.02074 | 0.09176 | 43.82 |
| Cfap97d1 | 16.4808 | -0.10065 | 0.5031 | -0.20007 | 0.84143 | 0.91872 | 7.986 |
| Cfap97d2 | 48.8856 | -0.2541 | 0.27643 | -0.91923 | 0.35798 | 0.56813 | 37.526 |
| Cfap99 | 89.2761 | 0.187122 | 0.29789 | 0.628156 | 0.5299 | 0.71752 | 38.728 |
| Cfdp1 | 2438.69 | 0.080737 | 0.07653 | 1.054972 | 0.29144 | 0.50126 | 85.801 |
| Cfh | 1816.72 | -0.48109 | 0.17687 | -2.7201 | 0.00653 | 0.04353 | 99.057 |
| Cfl1 | 12508.6 | -0.29694 | 0.06753 | -4.39685 | 1.10E-05 | 0.0006 | 4.747 |
| Cfl2 | 3073.49 | -0.07037 | 0.07602 | -0.92571 | 0.3546 | 0.565 | 4.069 |
| Cflar | 1296.82 | -0.45899 | 0.15192 | -3.02137 | 0.00252 | 0.02346 | 47.377 |
| Cfp | 999.936 | 0.243966 | 0.10191 | 2.393925 | 0.01667 | 0.0799 | 6.102 |
| Cftr | 184.014 | 0.68335 | 0.22682 | 3.012703 | 0.00259 | 0.02395 | 152.082 |
| Cgas | 86.3308 | -0.02798 | 0.23249 | -0.12034 | 0.90421 | 0.95164 | 12.712 |
| Cggbp1 | 3524.53 | 0.15285 | 0.0583 | 2.621865 | 0.00875 | 0.05244 | 7.512 |
| Cgn | 500.327 | -0.34966 | 0.18976 | -1.84258 | 0.06539 | 0.19729 | 26.451 |
| Cgnl1 | 1307.79 | -0.15068 | 0.11914 | -1.26472 | 0.20597 | 0.40522 | 145.094 |
| Cgref1 | 842.641 | -0.23855 | 0.12043 | -1.9809 | 0.0476 | 0.16041 | 12.449 |
| Cgrrf1 | 812.905 | -0.05356 | 0.08853 | -0.60506 | 0.54514 | 0.73012 | 22.069 |
| Ch25h | 14.9938 | -0.30997 | 0.53728 | -0.57691 | 0.564 | 0.74426 | 1.387 |
| Chac1 | 195.908 | -0.3988 | 0.15836 | -2.51834 | 0.01179 | 0.06387 | 3.153 |
| Chac2 | 394.202 | 0.004336 | 0.13065 | 0.033189 | 0.97352 | 0.98679 | 9.644 |
| Chad | 42.5256 | -0.31475 | 0.30729 | -1.0243 | 0.3057 | 0.51566 | 4.081 |

| | | | | | | | |
|---|---|---|---|---|---|---|---|
| Chadl | 476.967 | -0.03145 | 0.09763 | -0.32213 | 0.74735 | 0.86759 | 18.992 |
| Chaf1a | 183.777 | 0.072468 | 0.14494 | 0.500004 | 0.61707 | 0.78204 | 31.851 |
| Champ1 | 1782.79 | 0.031281 | 0.06918 | 0.452157 | 0.65116 | 0.8058 | 11.999 |
| Chaserr | 1007.74 | 0.114528 | 0.09228 | 1.241157 | 0.21455 | 0.41591 | 18.595 |
| Chat | 646.221 | 0.973873 | 0.37727 | 2.581348 | 0.00984 | 0.05688 | 57.787 |
| Chchd1 | 792.136 | 0.014179 | 0.08144 | 0.174108 | 0.86178 | 0.92991 | 1.42 |
| Chchd10 | 288.657 | -0.20133 | 0.25833 | -0.77936 | 0.43576 | 0.64139 | 4.618 |
| Chchd2 | 6752.34 | -0.06297 | 0.09115 | -0.69079 | 0.4897 | 0.68476 | 6.315 |
| Chchd3 | 1845.04 | 0.017684 | 0.09231 | 0.191568 | 0.84808 | 0.92287 | 269.285 |
| Chchd4 | 1522.51 | 0.144509 | 0.08769 | 1.648002 | 0.09935 | 0.25904 | 11.375 |
| Chchd5 | 262.219 | 0.130521 | 0.15279 | 0.854278 | 0.39295 | 0.6022 | 4.435 |
| Chchd6 | 1462.81 | -0.00865 | 0.11292 | -0.07658 | 0.93896 | 0.96951 | 212.507 |
| Chchd7 | 1136.53 | 0.119704 | 0.10683 | 1.120512 | 0.2625 | 0.4713 | 12.159 |
| Chd1 | 1767.44 | 0.16809 | 0.09476 | 1.773937 | 0.07607 | 0.21776 | 67.644 |
| Chd1l | 545.533 | 0.002549 | 0.09959 | 0.025595 | 0.97958 | 0.98899 | 49.462 |
| Chd2 | 4922.3 | 0.171272 | 0.06835 | 2.505996 | 0.01221 | 0.06535 | 115.193 |
| Chd3 | 44947.6 | -0.15986 | 0.10206 | -1.56626 | 0.11729 | 0.28825 | 26.134 |
| Chd4 | 14181.6 | 0.107869 | 0.09072 | 1.188969 | 0.23445 | 0.4397 | 34.611 |
| Chd5 | 14898.7 | -0.20623 | 0.12274 | -1.68012 | 0.09293 | 0.24829 | 51.544 |
| Chd6 | 6548.33 | 0.112208 | 0.06614 | 1.696579 | 0.08978 | 0.2429 | 162.098 |
| Chd7 | 1451.87 | 0.010411 | 0.11149 | 0.093382 | 0.9256 | 0.96248 | 177.254 |
| Chd8 | 4733.33 | 0.098982 | 0.06301 | 1.570941 | 0.1162 | 0.28619 | 59.63 |
| Chd9 | 6580.87 | 0.12195 | 0.09877 | 1.234708 | 0.21694 | 0.4188 | 226.165 |
| Chdh | 365.467 | -0.52691 | 0.22312 | -2.3616 | 0.0182 | 0.0843 | 31.505 |
| Chek1 | 113.075 | -0.03941 | 0.16048 | -0.24555 | 0.80603 | 0.90018 | 18.584 |
| Chek2 | 61.0095 | -0.5661 | 0.23251 | -2.43476 | 0.0149 | 0.07405 | 34.167 |
| Cherp | 2694.92 | 0.127002 | 0.0718 | 1.768737 | 0.07694 | 0.21936 | 14.738 |
| Chfr | 1860.19 | 0.151223 | 0.07271 | 2.079729 | 0.03755 | 0.1372 | 36.131 |
| Chga | 6800.18 | -0.40841 | 0.11547 | -3.53702 | 0.0004 | 0.00692 | 10.06 |
| Chgb | 14094.4 | -0.45847 | 0.11385 | -4.02693 | 5.65E-05 | 0.00181 | 13.802 |
| Chia1 | 317.825 | 1.058129 | 0.46924 | 2.254985 | 0.02413 | 0.10143 | 18.892 |
| Chic1 | 2924.03 | 0.226308 | 0.08372 | 2.703092 | 0.00687 | 0.04488 | 39.617 |
| Chic2 | 858.452 | -0.02738 | 0.07716 | -0.35483 | 0.72272 | 0.85219 | 46.787 |
| Chid1 | 3103.89 | 0.078987 | 0.07357 | 1.073646 | 0.28298 | 0.49316 | 46.722 |

| | | | | | | | |
|---|---|---|---|---|---|---|---|
| Chil1 | 491.004 | -1.09468 | 0.615 | -1.77996 | 0.07508 | 0.21608 | 8.006 |
| Chil5 | 34.7118 | 0.172059 | 0.48956 | 0.351457 | 0.72525 | 0.85364 | 15.896 |
| Chit1 | 19.4292 | 0.848702 | 0.64222 | 1.321503 | 0.18633 | 0.38123 | 40.299 |
| Chka | 5953.43 | 0.13922 | 0.08829 | 1.576874 | 0.11482 | 0.28391 | 42.597 |
| Chkb | 23.9853 | -0.04037 | 0.43906 | -0.09195 | 0.92674 | 0.9631 | 3.549 |
| Chl1 | 4855.96 | 0.017127 | 0.17076 | 0.100294 | 0.92011 | 0.95953 | 239.626 |
| Chm | 2727.62 | 0.259403 | 0.14857 | 1.746007 | 0.08081 | 0.22624 | 144.925 |
| Chml | 541.755 | 0.044759 | 0.18696 | 0.239404 | 0.81079 | 0.90258 | 10.665 |
| Chmp1a | 3604.58 | -0.1455 | 0.06119 | -2.37774 | 0.01742 | 0.08192 | 8.5 |
| Chmp1b | 802.637 | -0.02931 | 0.09754 | -0.3005 | 0.7638 | 0.87635 | 2.524 |
| Chmp2a | 2517.13 | 0.077074 | 0.07915 | 0.973719 | 0.3302 | 0.54139 | 2.798 |
| Chmp2b | 2019.94 | 0.10201 | 0.0675 | 1.511285 | 0.13072 | 0.30907 | 23.599 |
| Chmp3 | 4006.54 | 0.046245 | 0.08043 | 0.574991 | 0.5653 | 0.74547 | 38.813 |
| Chmp4b | 6816.19 | -0.12832 | 0.07058 | -1.81805 | 0.06906 | 0.20451 | 43.081 |
| Chmp4c | 157.623 | 0.119903 | 0.14473 | 0.828452 | 0.40741 | 0.61538 | 61.95 |
| Chmp5 | 3054.49 | 0.046184 | 0.07807 | 0.591566 | 0.55414 | 0.73681 | 16.897 |
| Chmp6 | 470.217 | -0.20845 | 0.1071 | -1.94634 | 0.05161 | 0.16941 | 6.108 |
| Chmp7 | 3931.53 | -0.04625 | 0.06263 | -0.73838 | 0.46028 | 0.66087 | 15.542 |
| Chn1 | 17065 | -0.08265 | 0.0719 | -1.14949 | 0.25036 | 0.45774 | 164.687 |
| Chn1os3 | 362.382 | 0.377366 | 0.12294 | 3.069456 | 0.00214 | 0.0211 | 19.487 |
| Chn2 | 3134.77 | 0.573203 | 0.16205 | 3.537298 | 0.0004 | 0.00692 | 262.257 |
| Chodl | 82.2247 | 0.716211 | 0.23677 | 3.02491 | 0.00249 | 0.02329 | 20.786 |
| Chordc1 | 5275.48 | 0.183819 | 0.09105 | 2.018988 | 0.04349 | 0.15084 | 25.318 |
| Chp1 | 5978.13 | -0.19433 | 0.06438 | -3.01835 | 0.00254 | 0.0236 | 39.331 |
| Chp2 | 108.696 | -0.36018 | 0.2557 | -1.40865 | 0.15894 | 0.34639 | 3.329 |
| Chpf | 2687.48 | 0.085087 | 0.08291 | 1.026315 | 0.30474 | 0.51453 | 4.739 |
| Chpf2 | 2085.85 | -0.00506 | 0.06672 | -0.07582 | 0.93956 | 0.96977 | 7.816 |
| Chpt1 | 2951.93 | 0.041284 | 0.12887 | 0.320345 | 0.74871 | 0.86822 | 51.329 |
| Chrac1 | 468.723 | 0.119068 | 0.117 | 1.017638 | 0.30885 | 0.51867 | 7.163 |
| Chrd | 1273.89 | -0.65081 | 0.31902 | -2.04004 | 0.04135 | 0.14589 | 9.258 |
| Chrdl1 | 524.216 | -0.56406 | 0.14815 | -3.80735 | 0.00014 | 0.00337 | 108.589 |
| Chrdl2 | 40.4518 | 1.089922 | 0.30435 | 3.581111 | 0.00034 | 0.00615 | 28.557 |
| Chrm1 | 6027.71 | -0.05289 | 0.10541 | -0.50173 | 0.61586 | 0.78132 | 19.799 |
| Chrm2 | 672.117 | -0.09571 | 0.18469 | -0.51821 | 0.60431 | 0.7733 | 140.331 |

| | | | | | | | |
|---|---|---|---|---|---|---|---|
| Chrm3 | 904.352 | -0.59631 | 0.19438 | -3.06767 | 0.00216 | 0.02119 | 485.362 |
| Chrm4 | 2327.2 | 0.640011 | 0.17701 | 3.615723 | 0.0003 | 0.00563 | 1.44 |
| Chrm5 | 48.8793 | 0.714365 | 0.39225 | 1.821175 | 0.06858 | 0.20357 | 1.599 |
| Chrna1 | 18.3897 | -0.66462 | 0.47554 | -1.39762 | 0.16223 | 0.35062 | 17.124 |
| Chrna10 | 28.8406 | -0.22936 | 0.41416 | -0.5538 | 0.57971 | 0.75628 | 5.563 |
| Chrna1os | 18.0473 | -0.72206 | 0.44167 | -1.63483 | 0.10208 | 0.26371 | 4.393 |
| Chrna2 | 112.269 | 0.476359 | 0.22134 | 2.152195 | 0.03138 | 0.12185 | 17.91 |
| Chrna3 | 21.8704 | 0.229858 | 0.47189 | 0.487105 | 0.62618 | 0.78844 | 16.452 |
| Chrna4 | 1355.64 | -0.42128 | 0.16127 | -2.61224 | 0.009 | 0.05339 | 25.167 |
| Chrna5 | 25.3659 | -0.69369 | 0.34101 | -2.03422 | 0.04193 | 0.1472 | 26.9 |
| Chrna7 | 569.254 | -0.04254 | 0.16462 | -0.25839 | 0.79611 | 0.89487 | 113.878 |
| Chrnb1 | 199.474 | -0.07657 | 0.1666 | -0.45959 | 0.64581 | 0.80274 | 11.908 |
| Chrnb2 | 3187.58 | 0.066359 | 0.06742 | 0.984309 | 0.32496 | 0.53614 | 18.438 |
| Chrnb4 | 13.6528 | 0.184363 | 0.47288 | 0.389871 | 0.69663 | 0.8356 | 20.626 |
| Chrne | 19.1088 | -0.013 | 0.40671 | -0.03197 | 0.97449 | 0.98728 | 4.334 |
| Chst1 | 11844 | -0.14758 | 0.10222 | -1.44377 | 0.1488 | 0.33268 | 15.544 |
| Chst10 | 3990.54 | 0.241518 | 0.0902 | 2.677729 | 0.00741 | 0.04733 | 34.295 |
| Chst11 | 3665.03 | 0.055733 | 0.06996 | 0.796593 | 0.42569 | 0.63229 | 210.403 |
| Chst12 | 483.576 | -0.12524 | 0.10341 | -1.21109 | 0.22586 | 0.42991 | 20.175 |
| Chst14 | 64.5922 | -0.2551 | 0.26347 | -0.96821 | 0.33294 | 0.54382 | 2.09 |
| Chst15 | 4592.87 | 0.489834 | 0.16626 | 2.946227 | 0.00322 | 0.02732 | 81.449 |
| Chst2 | 7422.07 | -0.03892 | 0.09513 | -0.40909 | 0.68247 | 0.82653 | 7.431 |
| Chst3 | 159.813 | -0.64308 | 0.2052 | -3.13391 | 0.00172 | 0.01828 | 37.729 |
| Chst5 | 35.8075 | -0.13777 | 0.31442 | -0.43818 | 0.66126 | 0.81269 | 21.312 |
| Chst7 | 62.3684 | -0.09827 | 0.30399 | -0.32328 | 0.74648 | 0.86693 | 37.962 |
| Chst8 | 148.216 | -1.09009 | 0.27358 | -3.98446 | 6.76E-05 | 0.00205 | 138.244 |
| Chsy1 | 684.873 | 0.102427 | 0.09935 | 1.030947 | 0.30257 | 0.51217 | 64.284 |
| Chsy3 | 573.508 | 0.424088 | 0.1222 | 3.470369 | 0.00052 | 0.00814 | 360.599 |
| Chtf18 | 68.7643 | -0.17105 | 0.20216 | -0.84609 | 0.3975 | 0.60658 | 8.494 |
| Chtf8 | 37.7831 | 0.419855 | 0.27884 | 1.505726 | 0.13214 | 0.31099 | 9.739 |
| Chtop | 7284.16 | 0.049325 | 0.07554 | 0.652975 | 0.51377 | 0.70456 | 10.543 |
| Chuk | 3917.93 | 0.253402 | 0.09087 | 2.788633 | 0.00529 | 0.03766 | 34.146 |
| Churc1 | 208.832 | 0.32277 | 0.15437 | 2.090911 | 0.03654 | 0.1347 | 17.643 |
| Ciao1 | 2277.84 | 0.055496 | 0.07816 | 0.710006 | 0.4777 | 0.67551 | 6.879 |

| | | | | | | | |
|---|---|---|---|---|---|---|---|
| Ciao2a | 875.656 | -0.03102 | 0.09983 | -0.31069 | 0.75604 | 0.87157 | 12.345 |
| Ciao2b | 517.201 | -0.29577 | 0.09695 | -3.05079 | 0.00228 | 0.02196 | 2.128 |
| Ciao3 | 1241.95 | 0.079755 | 0.07418 | 1.075104 | 0.28233 | 0.49247 | 9.557 |
| Ciapin1 | 2995.23 | -0.09233 | 0.08716 | -1.05924 | 0.28949 | 0.49974 | 18.555 |
| Ciart | 889.207 | 0.394859 | 0.11855 | 3.330705 | 0.00087 | 0.01158 | 3.749 |
| Cib1 | 351.223 | -0.45825 | 0.14592 | -3.14042 | 0.00169 | 0.01803 | 5.667 |
| Cib2 | 431.007 | -0.34384 | 0.16935 | -2.03038 | 0.04232 | 0.14806 | 15.425 |
| Cibar1 | 1674.48 | 0.065662 | 0.07332 | 0.895578 | 0.37048 | 0.58072 | 18.607 |
| Cibar2 | 18.9086 | -0.44563 | 0.39338 | -1.13282 | 0.25729 | 0.46522 | 11.07 |
| Cic | 8402.47 | -0.02623 | 0.07733 | -0.33914 | 0.73451 | 0.85958 | 26.456 |
| Cidea | 88.8142 | -0.36898 | 0.22078 | -1.67128 | 0.09467 | 0.25125 | 24.182 |
| Ciita | 300.719 | -0.23905 | 0.13287 | -1.79909 | 0.072 | 0.21067 | 48.36 |
| Cilk1 | 1340.26 | -0.24001 | 0.09931 | -2.41684 | 0.01566 | 0.07658 | 62.916 |
| Cilp2 | 61.8916 | -0.3335 | 0.26725 | -1.24789 | 0.21207 | 0.41271 | 7.319 |
| Cinp | 1766.45 | 0.062047 | 0.07625 | 0.81368 | 0.41583 | 0.62362 | 16.536 |
| Cip2a | 192.452 | 0.008839 | 0.15176 | 0.058245 | 0.95355 | 0.97719 | 25.525 |
| Cipc | 5634.57 | 0.122821 | 0.0767 | 1.601318 | 0.10931 | 0.27513 | 18.32 |
| Cir1 | 833.872 | 0.138205 | 0.1212 | 1.140314 | 0.25416 | 0.46153 | 29.597 |
| Cirbp | 3012.42 | -0.10258 | 0.10484 | -0.97845 | 0.32785 | 0.53925 | 6.802 |
| Cisd1 | 2263.5 | -0.11567 | 0.06877 | -1.68206 | 0.09256 | 0.24776 | 14.475 |
| Cisd2 | 2914.1 | 0.105789 | 0.06957 | 1.520645 | 0.12835 | 0.30535 | 17.514 |
| Cisd3 | 396.636 | -0.16759 | 0.10832 | -1.54712 | 0.12183 | 0.2953 | 2.796 |
| Cish | 188.639 | 0.172274 | 0.24531 | 0.702276 | 0.48251 | 0.67901 | 6.759 |
| Cit | 7222.69 | 0.178125 | 0.09924 | 1.794882 | 0.07267 | 0.21196 | 163.67 |
| Cited1 | 39.4139 | -0.32804 | 0.27153 | -1.20813 | 0.227 | 0.43117 | 4.801 |
| Cited2 | 2224.28 | 0.098764 | 0.10017 | 0.985932 | 0.32417 | 0.53531 | 2.457 |
| Cited4 | 242.643 | 0.077456 | 0.2041 | 0.3795 | 0.70432 | 0.84056 | 1.249 |
| Ciz1 | 2476.75 | 0.03869 | 0.0787 | 0.491578 | 0.62302 | 0.78589 | 28.644 |
| Ckap2 | 26.4119 | -0.08528 | 0.38765 | -0.22 | 0.82587 | 0.91013 | 17.66 |
| Ckap2l | 27.8508 | -0.58886 | 0.50314 | -1.17036 | 0.24186 | 0.44782 | 29.003 |
| Ckap4 | 648.636 | -0.68015 | 0.15827 | -4.29733 | 1.73E-05 | 0.0008 | 7.758 |
| Ckap5 | 8047.83 | 0.076366 | 0.07472 | 1.022061 | 0.30675 | 0.51671 | 93.903 |
| Ckb | 49145.1 | 0.008982 | 0.10074 | 0.089159 | 0.92896 | 0.96429 | 2.978 |
| Cklf | 363.496 | 0.120348 | 0.10746 | 1.119959 | 0.26273 | 0.47156 | 14.078 |

| | | | | | | | |
|---|---|---|---|---|---|---|---|
| Ckmt1 | 5755.26 | -0.02407 | 0.07706 | -0.31234 | 0.75478 | 0.87116 | 6.024 |
| Ckmt2 | 22.4332 | 0.673616 | 0.44497 | 1.51385 | 0.13006 | 0.30796 | 23.499 |
| Cks1b | 63.3774 | -0.43228 | 0.31708 | -1.36333 | 0.17278 | 0.36397 | 2.912 |
| Cks2 | 37.2564 | -0.48405 | 0.30984 | -1.56226 | 0.11823 | 0.28972 | 5.429 |
| Clasp1 | 6372.21 | 0.008768 | 0.06773 | 0.129456 | 0.897 | 0.94835 | 223.621 |
| Clasp2 | 14315.3 | 0.052751 | 0.05631 | 0.936814 | 0.34885 | 0.55931 | 178.21 |
| Clasrp | 3959.84 | 0.071928 | 0.10528 | 0.683178 | 0.49449 | 0.68865 | 23.452 |
| Clba1 | 442.618 | 0.019346 | 0.15908 | 0.121618 | 0.9032 | 0.95127 | 7.316 |
| Clcc1 | 1210.54 | 0.121596 | 0.07688 | 1.581534 | 0.11376 | 0.28233 | 24.928 |
| Clcf1 | 110.844 | -0.509 | 0.19863 | -2.56256 | 0.01039 | 0.05891 | 9.253 |
| Clcn1 | 275.83 | -0.12659 | 0.14326 | -0.88366 | 0.37688 | 0.58697 | 29.072 |
| Clcn2 | 2714.11 | 0.036041 | 0.0954 | 0.377774 | 0.7056 | 0.84142 | 14.783 |
| Clcn3 | 10573 | 0.000976 | 0.05183 | 0.018822 | 0.98498 | 0.99174 | 72.912 |
| Clcn4 | 8702.46 | -0.05698 | 0.0676 | -0.84289 | 0.39929 | 0.60812 | 18.543 |
| Clcn5 | 703.085 | 0.230013 | 0.08932 | 2.575084 | 0.01002 | 0.05754 | 165.549 |
| Clcn6 | 4960.63 | -0.05507 | 0.05815 | -0.94702 | 0.34363 | 0.55429 | 34.563 |
| Clcn7 | 2943.59 | 0.003214 | 0.07232 | 0.044445 | 0.96455 | 0.98273 | 28.714 |
| Cldn1 | 122.461 | 0.255564 | 0.26777 | 0.954411 | 0.33988 | 0.55075 | 15.2 |
| Cldn10 | 1096.34 | -0.38895 | 0.1311 | -2.96679 | 0.00301 | 0.02623 | 87.582 |
| Cldn11 | 6335.55 | -0.60311 | 0.1278 | -4.71908 | 2.37E-06 | 0.0002 | 14.405 |
| Cldn12 | 2864.32 | 0.229586 | 0.10282 | 2.232974 | 0.02555 | 0.10545 | 25.422 |
| Cldn14 | 34.7515 | -1.05966 | 0.34021 | -3.11477 | 0.00184 | 0.01906 | 93.778 |
| Cldn15 | 27.9585 | 0.085889 | 0.41961 | 0.204689 | 0.83782 | 0.91655 | 9.243 |
| Cldn19 | 24.9047 | -1.00826 | 0.44419 | -2.26986 | 0.02322 | 0.09876 | 7.025 |
| Cldn20 | 25.4049 | 0.10543 | 0.38996 | 0.270362 | 0.78688 | 0.88907 | 0.656 |
| Cldn34c1 | 60.7773 | 0.012856 | 0.26112 | 0.049235 | 0.96073 | 0.98085 | 85.301 |
| Cldn5 | 833.303 | -0.72407 | 0.14523 | -4.98565 | 6.18E-07 | 7.64E-05 | 1.416 |
| Cldnd1 | 2866.66 | -0.12369 | 0.0774 | -1.59797 | 0.11005 | 0.2764 | 6.342 |
| Clec10a | 23.1835 | -0.19931 | 0.45655 | -0.43657 | 0.66243 | 0.81348 | 14.638 |
| Clec11a | 130.833 | -0.71564 | 0.2174 | -3.29186 | 0.001 | 0.01278 | 4.216 |
| Clec12a | 148.175 | 0.90177 | 0.30782 | 2.929495 | 0.0034 | 0.02816 | 22.613 |
| Clec12b | 14.3973 | 1.514304 | 0.64128 | 2.361363 | 0.01821 | 0.08431 | 10.36 |
| Clec14a | 116.618 | -0.83718 | 0.27124 | -3.08655 | 0.00202 | 0.02025 | 4.571 |
| Clec16a | 4516.63 | -0.00257 | 0.06176 | -0.04162 | 0.9668 | 0.9834 | 199.54 |

| | | | | | | | |
|---|---|---|---|---|---|---|---|
| Clec18a | 13.5799 | -0.90492 | 0.60441 | -1.49719 | 0.13434 | 0.31425 | 33.507 |
| Clec1a | 33.343 | -0.71342 | 0.43549 | -1.6382 | 0.10138 | 0.26262 | 27.229 |
| Clec2d | 149.371 | -1.18039 | 0.21477 | -5.49609 | 3.88E-08 | 9.54E-06 | 5.92 |
| Clec2l | 1225.71 | -0.45144 | 0.10833 | -4.1673 | 3.08E-05 | 0.00118 | 17.799 |
| Clec3b | 95.3496 | -1.35369 | 0.40921 | -3.30807 | 0.00094 | 0.01226 | 6.487 |
| Clec4a1 | 17.4725 | -0.47166 | 0.58211 | -0.81027 | 0.41779 | 0.62518 | 12.772 |
| Clec4a2 | 47.6419 | -0.23035 | 0.32318 | -0.71275 | 0.476 | 0.67416 | 37.572 |
| Clec4a3 | 51.3257 | -0.59496 | 0.30677 | -1.93943 | 0.05245 | 0.17131 | 17.361 |
| Clec5a | 62.7483 | -0.22835 | 0.24828 | -0.91976 | 0.3577 | 0.56794 | 10.928 |
| Clec7a | 27.4991 | -1.68324 | 0.61922 | -2.71833 | 0.00656 | 0.04367 | 11.189 |
| Clgn | 759.061 | -0.32024 | 0.14168 | -2.26029 | 0.0238 | 0.10042 | 38.686 |
| Clic1 | 305.039 | -0.51667 | 0.15845 | -3.26085 | 0.00111 | 0.0137 | 8.784 |
| Clic3 | 144.012 | 0.219741 | 0.15448 | 1.422496 | 0.15488 | 0.34109 | 1.939 |
| Clic4 | 5168.91 | -0.54992 | 0.10515 | -5.22984 | 1.70E-07 | 2.70E-05 | 58.846 |
| Clic5 | 583.435 | -0.19664 | 0.1171 | -1.67928 | 0.0931 | 0.2484 | 145.401 |
| Clic6 | 1536.55 | 0.778756 | 0.32705 | 2.381166 | 0.01726 | 0.08154 | 55.508 |
| Clint1 | 3321.4 | 0.032066 | 0.06479 | 0.494906 | 0.62067 | 0.78424 | 58.662 |
| Clip1 | 8049.24 | 0.067517 | 0.07631 | 0.884754 | 0.37629 | 0.58637 | 106.824 |
| Clip2 | 5726.88 | -0.36259 | 0.11839 | -3.06262 | 0.00219 | 0.02136 | 63.052 |
| Clip3 | 13241 | -0.20926 | 0.09049 | -2.31245 | 0.02075 | 0.09178 | 16.64 |
| Clip4 | 5897.37 | 0.461928 | 0.16209 | 2.849773 | 0.00438 | 0.03343 | 95.801 |
| Clk1 | 11126.5 | 0.265067 | 0.09899 | 2.677747 | 0.00741 | 0.04733 | 13.878 |
| Clk2 | 2127.68 | 0.139381 | 0.07833 | 1.779363 | 0.07518 | 0.21627 | 12.127 |
| Clk3 | 3060.98 | -0.06027 | 0.07711 | -0.78163 | 0.43443 | 0.64022 | 15.237 |
| Clk4 | 5564.56 | 0.180067 | 0.08757 | 2.056156 | 0.03977 | 0.14241 | 20.037 |
| Clmn | 7306.9 | -0.07296 | 0.0916 | -0.79646 | 0.42576 | 0.63229 | 101.96 |
| Clmp | 1315.38 | -0.17485 | 0.10513 | -1.66316 | 0.09628 | 0.25407 | 99.358 |
| Cln3 | 703.86 | 0.031906 | 0.0978 | 0.326242 | 0.74424 | 0.86539 | 14.611 |
| Cln5 | 815.207 | -0.03198 | 0.08192 | -0.39037 | 0.69627 | 0.83541 | 7.413 |
| Cln6 | 285.052 | -0.26293 | 0.12258 | -2.14497 | 0.03195 | 0.12339 | 13.222 |
| Cln8 | 1461.71 | -0.17826 | 0.07165 | -2.48795 | 0.01285 | 0.06768 | 20.386 |
| Clnk | 22.068 | -0.61442 | 0.43802 | -1.4027 | 0.1607 | 0.34889 | 170.351 |
| Clns1a | 1905.34 | -0.09352 | 0.09268 | -1.00914 | 0.31291 | 0.5235 | 24.163 |
| Clock | 5376.47 | 0.188672 | 0.1007 | 1.873637 | 0.06098 | 0.18857 | 94.925 |

| | | | | | | | |
|---|---|---|---|---|---|---|---|
| Clp1 | 304.817 | 0.039436 | 0.1102 | 0.357846 | 0.72046 | 0.85081 | 5.247 |
| Clpb | 3024.15 | 0.060317 | 0.06885 | 0.876054 | 0.381 | 0.59053 | 131.874 |
| Clpp | 1303.75 | -0.14914 | 0.0932 | -1.60027 | 0.10954 | 0.27546 | 5.884 |
| Clptm1 | 6548.61 | -0.0474 | 0.06741 | -0.70321 | 0.48193 | 0.67861 | 33.463 |
| Clptm1l | 4136.79 | 0.054131 | 0.07869 | 0.68792 | 0.4915 | 0.68643 | 16.6 |
| Clpx | 1654.09 | 0.053709 | 0.08749 | 0.61391 | 0.53927 | 0.72582 | 36.399 |
| Clspn | 102.437 | 0.202731 | 0.29154 | 0.695375 | 0.48682 | 0.6826 | 36.969 |
| Clstn1 | 46538.7 | -0.08124 | 0.06227 | -1.30476 | 0.19198 | 0.38879 | 62.432 |
| Clstn2 | 2560.25 | -0.29677 | 0.15304 | -1.93913 | 0.05249 | 0.17137 | 588.787 |
| Clstn3 | 20245.9 | -0.00623 | 0.06332 | -0.0984 | 0.92161 | 0.96044 | 34.214 |
| Clta | 5277.1 | -0.08388 | 0.06293 | -1.33302 | 0.18252 | 0.37638 | 28.395 |
| Cltb | 7582.1 | -0.18459 | 0.06062 | -3.0453 | 0.00232 | 0.0222 | 18.944 |
| Cltc | 31314 | 0.018401 | 0.06604 | 0.278626 | 0.78053 | 0.88564 | 63.215 |
| Clu | 31859.7 | -0.33969 | 0.13424 | -2.53051 | 0.01139 | 0.06243 | 13.065 |
| Cluap1 | 518.646 | 0.001322 | 0.10968 | 0.012051 | 0.99039 | 0.99473 | 32.347 |
| Cluh | 5488.61 | 0.062896 | 0.07091 | 0.88693 | 0.37512 | 0.58522 | 21.353 |
| Clvs1 | 816.408 | 0.373818 | 0.09556 | 3.911747 | 9.16E-05 | 0.00254 | 182.399 |
| Clvs2 | 1247.9 | -0.22436 | 0.1074 | -2.08894 | 0.03671 | 0.13513 | 112.484 |
| Clybl | 607.765 | -0.16315 | 0.12414 | -1.31421 | 0.18877 | 0.38449 | 220.531 |
| Cmah | 19.0212 | -1.44531 | 0.60925 | -2.37228 | 0.01768 | 0.08263 | 149.974 |
| Cmas | 3673.62 | -0.13037 | 0.07228 | -1.80363 | 0.07129 | 0.20897 | 19.029 |
| Cmbl | 201.602 | -0.53313 | 0.15274 | -3.49038 | 0.00048 | 0.00774 | 24.731 |
| Cmc1 | 537.265 | -0.01443 | 0.11492 | -0.12556 | 0.90008 | 0.94992 | 85.656 |
| Cmc2 | 435.747 | -0.13994 | 0.09312 | -1.5028 | 0.13289 | 0.31233 | 32.771 |
| Cmc4 | 207.307 | 0.267322 | 0.13328 | 2.005761 | 0.04488 | 0.15394 | 11.739 |
| Cmip | 6978.9 | -0.15639 | 0.10103 | -1.54798 | 0.12163 | 0.29505 | 202.367 |
| Cmklr1 | 153.183 | -0.34349 | 0.18715 | -1.83536 | 0.06645 | 0.19944 | 38.073 |
| Cmpk1 | 3230.69 | -0.02927 | 0.07037 | -0.41592 | 0.67747 | 0.82322 | 27.906 |
| Cmpk2 | 1249.19 | -0.03354 | 0.07678 | -0.43685 | 0.66222 | 0.81337 | 10.634 |
| Cmss1 | 337.605 | 0.068786 | 0.14973 | 0.459397 | 0.64595 | 0.80278 | 304.865 |
| Cmtm3 | 426.303 | -0.12553 | 0.17167 | -0.73124 | 0.46463 | 0.66431 | 8.263 |
| Cmtm4 | 8501.18 | -0.01256 | 0.08749 | -0.14359 | 0.88582 | 0.9432 | 47.617 |
| Cmtm5 | 1083.84 | -0.45464 | 0.09189 | -4.9479 | 7.50E-07 | 8.82E-05 | 3.014 |
| Cmtm6 | 1286.82 | -0.19562 | 0.08448 | -2.31547 | 0.02059 | 0.09128 | 18.229 |

| | | | | | | | |
|---|---|---|---|---|---|---|---|
| Cmtm7 | 53.0522 | -0.4836 | 0.26959 | -1.79386 | 0.07284 | 0.21231 | 25.021 |
| Cmtm8 | 37.6056 | -0.44521 | 0.29738 | -1.49709 | 0.13437 | 0.31425 | 54.812 |
| Cmtr1 | 151.83 | 0.075378 | 0.20304 | 0.371238 | 0.71046 | 0.84419 | 45.385 |
| Cmtr2 | 342.669 | 0.314985 | 0.1054 | 2.988471 | 0.0028 | 0.02513 | 8.822 |
| Cmya5 | 383.235 | 0.144781 | 0.1341 | 1.079625 | 0.28031 | 0.49019 | 104.012 |
| Cnbd2 | 136.988 | -0.05969 | 0.16851 | -0.3542 | 0.72319 | 0.85223 | 63.34 |
| Cnbp | 10703.4 | 0.089312 | 0.075 | 1.190888 | 0.2337 | 0.43869 | 8.492 |
| Cndp2 | 1998.9 | -0.11915 | 0.10312 | -1.15542 | 0.24792 | 0.45503 | 18.238 |
| Cnep1r1 | 1510.71 | 0.131175 | 0.07805 | 1.680739 | 0.09281 | 0.24813 | 16.342 |
| Cnga2 | 21.5987 | 0.337027 | 0.41665 | 0.808897 | 0.41857 | 0.62571 | 18.37 |
| Cnga4 | 46.4797 | 0.000976 | 0.23956 | 0.004073 | 0.99675 | 0.99821 | 4.175 |
| Cngb1 | 18.2488 | -0.39915 | 0.48124 | -0.82941 | 0.40687 | 0.61498 | 67.541 |
| Cnih1 | 2092.81 | 0.045956 | 0.07964 | 0.577057 | 0.5639 | 0.74426 | 12.845 |
| Cnih2 | 7427.59 | -0.12967 | 0.09628 | -1.34681 | 0.17804 | 0.3703 | 5.654 |
| Cnih3 | 1892.2 | -0.8792 | 0.38438 | -2.2873 | 0.02218 | 0.096 | 108.014 |
| Cnih4 | 2118.29 | -0.16119 | 0.07837 | -2.05681 | 0.0397 | 0.14229 | 24.302 |
| Cnksr1 | 22.1854 | 0.137652 | 0.46893 | 0.293548 | 0.7691 | 0.87961 | 10.359 |
| Cnksr2 | 13405.5 | 0.203484 | 0.08015 | 2.538635 | 0.01113 | 0.06149 | 221.859 |
| Cnksr3 | 284.527 | -0.42864 | 0.19022 | -2.25342 | 0.02423 | 0.10168 | 93.175 |
| Cnmd | 34.1885 | 0.556939 | 0.33572 | 1.65893 | 0.09713 | 0.2554 | 24.481 |
| Cnn1 | 21.27 | -1.08938 | 0.53538 | -2.03478 | 0.04187 | 0.1472 | 10.415 |
| Cnn2 | 191.54 | -0.37961 | 0.14857 | -2.55516 | 0.01061 | 0.05981 | 7.479 |
| Cnn3 | 2178.92 | -0.38859 | 0.10888 | -3.56895 | 0.00036 | 0.00633 | 31.711 |
| Cnnm1 | 5533.78 | 0.157597 | 0.05679 | 2.774964 | 0.00552 | 0.03887 | 56.775 |
| Cnnm2 | 1772.92 | -0.1026 | 0.06526 | -1.57224 | 0.11589 | 0.28576 | 118.597 |
| Cnnm3 | 1803.02 | 0.110482 | 0.07945 | 1.390626 | 0.16434 | 0.35329 | 16.371 |
| Cnnm4 | 653.622 | 0.040758 | 0.08436 | 0.48315 | 0.62899 | 0.79023 | 37.145 |
| Cnot1 | 6980.94 | 0.18058 | 0.05961 | 3.029596 | 0.00245 | 0.023 | 88.014 |
| Cnot10 | 3606.02 | 0.126925 | 0.06483 | 1.95777 | 0.05026 | 0.1666 | 54.307 |
| Cnot11 | 1469 | 0.08408 | 0.0806 | 1.043177 | 0.29687 | 0.50702 | 11.898 |
| Cnot2 | 2976.55 | 0.18122 | 0.06299 | 2.87717 | 0.00401 | 0.03157 | 96.351 |
| Cnot3 | 2470.69 | 0.148477 | 0.06417 | 2.313835 | 0.02068 | 0.09161 | 15.842 |
| Cnot4 | 1735.02 | 0.166042 | 0.07379 | 2.250347 | 0.02443 | 0.10224 | 111.66 |
| Cnot6 | 4545.12 | 0.236803 | 0.08134 | 2.911351 | 0.0036 | 0.02933 | 41.221 |

| | | | | | | | |
|---|---|---|---|---|---|---|---|
| Cnot6l | 4020.46 | 0.24281 | 0.0824 | 2.94656 | 0.00321 | 0.0273 | 93.839 |
| Cnot7 | 4872.24 | 0.235246 | 0.08528 | 2.758415 | 0.00581 | 0.04034 | 23.308 |
| Cnot8 | 2272.77 | 0.090924 | 0.07819 | 1.16284 | 0.24489 | 0.45147 | 14.442 |
| Cnot9 | 1402.23 | -0.058 | 0.07964 | -0.72832 | 0.46642 | 0.66569 | 24.785 |
| Cnp | 10768 | -0.52698 | 0.11765 | -4.4791 | 7.50E-06 | 0.00045 | 16.826 |
| Cnppd1 | 1306.28 | 0.052608 | 0.07882 | 0.66748 | 0.50447 | 0.69668 | 8.158 |
| Cnpy2 | 926.822 | -0.06118 | 0.08549 | -0.71562 | 0.47422 | 0.67238 | 5.095 |
| Cnpy3 | 1727.29 | -0.22241 | 0.08569 | -2.59539 | 0.00945 | 0.05528 | 16.51 |
| Cnpy4 | 524.713 | -0.21152 | 0.09387 | -2.25345 | 0.02423 | 0.10168 | 6.434 |
| Cnr1 | 8300.95 | -0.59926 | 0.29562 | -2.02713 | 0.04265 | 0.14884 | 24.239 |
| Cnr2 | 20.3807 | -0.68664 | 0.47807 | -1.43627 | 0.15093 | 0.33582 | 24.814 |
| Cnrip1 | 1171.41 | -0.19951 | 0.09712 | -2.05422 | 0.03995 | 0.14286 | 27.786 |
| Cnst | 3513.36 | 0.152636 | 0.08365 | 1.824693 | 0.06805 | 0.20237 | 81.109 |
| Cntd1 | 37.366 | 0.278451 | 0.36589 | 0.761023 | 0.44664 | 0.65022 | 17.306 |
| Cntfr | 3269.81 | -0.24639 | 0.06367 | -3.8696 | 0.00011 | 0.00285 | 39.592 |
| Cntln | 633.354 | 0.307578 | 0.17234 | 1.784706 | 0.07431 | 0.21491 | 247.613 |
| Cntn1 | 17539.4 | 0.033938 | 0.07632 | 0.444712 | 0.65653 | 0.80921 | 290.803 |
| Cntn2 | 9642.61 | -0.50294 | 0.13996 | -3.59359 | 0.00033 | 0.00599 | 33.83 |
| Cntn3 | 1477.55 | 0.491108 | 0.14215 | 3.454927 | 0.00055 | 0.00846 | 410.447 |
| Cntn4 | 1808.22 | -0.3243 | 0.15298 | -2.11997 | 0.03401 | 0.12876 | 1021.651 |
| Cntn5 | 508.698 | -0.02084 | 0.16732 | -0.12453 | 0.9009 | 0.95027 | 1243.885 |
| Cntn6 | 992.907 | 0.05409 | 0.12096 | 0.447164 | 0.65476 | 0.80841 | 370.617 |
| Cntnap1 | 15404.8 | -0.04088 | 0.07927 | -0.51568 | 0.60608 | 0.77461 | 20.202 |
| Cntnap2 | 7230.75 | 0.387369 | 0.12164 | 3.184482 | 0.00145 | 0.01616 | 2244.857 |
| Cntnap3 | 1236.82 | 0.804647 | 0.1267 | 6.35097 | 2.14E-10 | 1.68E-07 | 167.774 |
| Cntnap4 | 690.445 | -0.16992 | 0.11185 | -1.51924 | 0.1287 | 0.30583 | 312.675 |
| Cntnap5a | 2830.94 | 0.185481 | 0.11115 | 1.668736 | 0.09517 | 0.25201 | 902.568 |
| Cntnap5b | 602.658 | 0.140839 | 0.13607 | 1.035016 | 0.30066 | 0.51068 | 713.178 |
| Cntnap5c | 608.56 | -0.14378 | 0.11948 | -1.2034 | 0.22882 | 0.43288 | 640.786 |
| Cntrl | 1038.39 | -0.12418 | 0.12425 | -0.99942 | 0.31759 | 0.52796 | 69.331 |
| Cntrob | 1281.55 | 0.116631 | 0.08903 | 1.310038 | 0.19018 | 0.38629 | 24.289 |
| Coa3 | 1655.75 | -0.20753 | 0.07594 | -2.73264 | 0.00628 | 0.04244 | 1.147 |
| Coa4 | 93.7911 | -0.3314 | 0.22374 | -1.48121 | 0.13855 | 0.31964 | 3.299 |
| Coa5 | 5289.64 | 0.004162 | 0.07994 | 0.052065 | 0.95848 | 0.97972 | 13.02 |

| | | | | | | | |
|---|---|---|---|---|---|---|---|
| Coa6 | 507.882 | 0.004708 | 0.0981 | 0.047994 | 0.96172 | 0.98141 | 2.919 |
| Coa7 | 755.27 | 0.134144 | 0.10527 | 1.274328 | 0.20255 | 0.40104 | 13.405 |
| Coa8 | 565.669 | -0.04188 | 0.10035 | -0.41731 | 0.67645 | 0.82252 | 41.793 |
| Coasy | 957.386 | -0.03556 | 0.09865 | -0.36042 | 0.71853 | 0.84962 | 4.055 |
| Cobl | 2298.89 | 0.340651 | 0.08245 | 4.131361 | 3.61E-05 | 0.0013 | 228.353 |
| Cobll1 | 522.572 | -0.41039 | 0.13187 | -3.11218 | 0.00186 | 0.01912 | 151.065 |
| Coch | 570.763 | -0.47916 | 0.16695 | -2.87017 | 0.0041 | 0.03204 | 12.431 |
| Cog1 | 1949.44 | 0.066003 | 0.0706 | 0.934839 | 0.34987 | 0.56045 | 17.886 |
| Cog2 | 1468.75 | 0.036522 | 0.09047 | 0.403691 | 0.68644 | 0.82888 | 31.242 |
| Cog3 | 2245.24 | 0.029154 | 0.07151 | 0.407713 | 0.68348 | 0.82718 | 52.209 |
| Cog4 | 1969.37 | -0.02167 | 0.07975 | -0.27167 | 0.78587 | 0.88867 | 35.628 |
| Cog5 | 1475.8 | 0.227144 | 0.09068 | 2.504804 | 0.01225 | 0.06552 | 282.762 |
| Cog6 | 2660.8 | 0.116071 | 0.06557 | 1.770091 | 0.07671 | 0.21907 | 35.363 |
| Cog7 | 2082.95 | 0.063129 | 0.07801 | 0.809287 | 0.41835 | 0.62561 | 102.349 |
| Cog8 | 1051.83 | -0.10585 | 0.08032 | -1.31776 | 0.18758 | 0.38298 | 10.401 |
| Coil | 1051.19 | 0.106664 | 0.07429 | 1.435808 | 0.15106 | 0.33587 | 21.362 |
| Col11a1 | 2752.37 | 0.749107 | 0.19405 | 3.860344 | 0.00011 | 0.00292 | 190.179 |
| Col11a2 | 1733.6 | 0.034898 | 0.13978 | 0.24967 | 0.80284 | 0.89815 | 27.249 |
| Col12a1 | 408.981 | -0.32993 | 0.2047 | -1.61178 | 0.10701 | 0.27147 | 119.841 |
| Col13a1 | 65.2184 | -0.81875 | 0.2754 | -2.97293 | 0.00295 | 0.02588 | 140.873 |
| Col14a1 | 85.2608 | 0.173536 | 0.25189 | 0.688946 | 0.49086 | 0.68583 | 213.054 |
| Col15a1 | 134.715 | -0.27441 | 0.23593 | -1.16309 | 0.24479 | 0.45139 | 105.007 |
| Col16a1 | 981.537 | -0.1253 | 0.10708 | -1.17025 | 0.2419 | 0.44786 | 51.444 |
| Col17a1 | 106.649 | 0.952913 | 0.27111 | 3.514811 | 0.00044 | 0.0073 | 45.751 |
| Col18a1 | 324.436 | -0.31759 | 0.16785 | -1.89212 | 0.05847 | 0.18391 | 114.371 |
| Col19a1 | 562.704 | -0.48493 | 0.201 | -2.41256 | 0.01584 | 0.07722 | 325.583 |
| Col1a1 | 322.246 | -0.63583 | 0.20522 | -3.0983 | 0.00195 | 0.01974 | 16.819 |
| Col1a2 | 857.36 | -0.49996 | 0.32935 | -1.51802 | 0.12901 | 0.30621 | 36.731 |
| Col20a1 | 362.028 | -0.18986 | 0.19538 | -0.97173 | 0.33119 | 0.54208 | 31.829 |
| Col22a1 | 61.5261 | -1.03486 | 0.32877 | -3.14762 | 0.00165 | 0.01769 | 238.433 |
| Col23a1 | 336.998 | -0.48033 | 0.17256 | -2.78347 | 0.00538 | 0.03807 | 293.999 |
| Col24a1 | 299.818 | -0.77367 | 0.26349 | -2.93628 | 0.00332 | 0.02779 | 259.534 |
| Col25a1 | 3340.08 | 0.22107 | 0.09397 | 2.352445 | 0.01865 | 0.0856 | 468.377 |
| Col26a1 | 159.632 | -0.2294 | 0.18439 | -1.24416 | 0.21344 | 0.41472 | 141.451 |

| | | | | | | | |
|---|---|---|---|---|---|---|---|
| Col27a1 | 530.9 | 0.118734 | 0.15869 | 0.748233 | 0.45432 | 0.65643 | 120.988 |
| Col28a1 | 30.375 | 0.854991 | 0.36843 | 2.320651 | 0.02031 | 0.09035 | 194.81 |
| Col2a1 | 58.0122 | 0.017401 | 0.34146 | 0.05096 | 0.95936 | 0.98028 | 29.094 |
| Col3a1 | 98.7513 | -0.97706 | 0.35322 | -2.76612 | 0.00567 | 0.03967 | 38.169 |
| Col4a1 | 2325.57 | -0.33758 | 0.11398 | -2.9617 | 0.00306 | 0.02639 | 114.404 |
| Col4a2 | 3069.25 | -0.12051 | 0.09467 | -1.27298 | 0.20303 | 0.40164 | 136.483 |
| Col4a3 | 27.1313 | -0.46427 | 0.47983 | -0.96757 | 0.33326 | 0.54421 | 135.139 |
| Col4a4 | 44.1531 | -0.11452 | 0.32659 | -0.35065 | 0.72585 | 0.85388 | 138.427 |
| Col4a5 | 243.514 | -0.43975 | 0.22462 | -1.95777 | 0.05026 | 0.1666 | 213.85 |
| Col4a6 | 112.578 | -0.56934 | 0.29926 | -1.90253 | 0.0571 | 0.18105 | 308.674 |
| Col5a1 | 494.998 | 0.327243 | 0.12688 | 2.579235 | 0.0099 | 0.05714 | 153.09 |
| Col5a2 | 240.176 | -0.69494 | 0.24489 | -2.83783 | 0.00454 | 0.03411 | 128.962 |
| Col5a3 | 240.472 | -0.15033 | 0.21466 | -0.70031 | 0.48374 | 0.68008 | 45.018 |
| Col6a1 | 11463.7 | 0.536018 | 0.09912 | 5.407612 | 6.39E-08 | 1.26E-05 | 17.377 |
| Col6a2 | 1066.36 | -0.03681 | 0.10894 | -0.33792 | 0.73543 | 0.85989 | 27.869 |
| Col6a3 | 263.286 | -0.25721 | 0.16741 | -1.53636 | 0.12445 | 0.29933 | 78.049 |
| Col6a4 | 286.927 | 1.277321 | 0.62477 | 2.044465 | 0.04091 | 0.14484 | 107.33 |
| Col6a5 | 104.017 | 0.984806 | 0.21 | 4.689565 | 2.74E-06 | 0.00022 | 104.566 |
| Col6a6 | 19.8492 | 0.01162 | 0.4035 | 0.028798 | 0.97703 | 0.98796 | 140.352 |
| Col7a1 | 411.471 | 0.184903 | 0.21912 | 0.843832 | 0.39876 | 0.60773 | 31.29 |
| Col8a1 | 35.1316 | -0.18335 | 0.41169 | -0.44536 | 0.65606 | 0.80893 | 130.48 |
| Col8a2 | 40.39 | -0.51482 | 0.58591 | -0.87868 | 0.37957 | 0.58932 | 27.538 |
| Col9a1 | 16.8873 | -0.62292 | 0.49703 | -1.25328 | 0.21011 | 0.41015 | 75.075 |
| Col9a2 | 351.227 | -0.35586 | 0.24372 | -1.46014 | 0.14425 | 0.32697 | 15.938 |
| Col9a3 | 983.158 | 0.533419 | 0.12359 | 4.31617 | 1.59E-05 | 0.00077 | 24.4 |
| Colca2 | 37.8114 | -0.21601 | 0.29365 | -0.73559 | 0.46198 | 0.66235 | 9.394 |
| Colec11 | 20.3198 | -0.59807 | 0.41152 | -1.45331 | 0.14614 | 0.32908 | 29.205 |
| Colec12 | 668.114 | -0.73646 | 0.22249 | -3.31013 | 0.00093 | 0.01221 | 175.05 |
| Colgalt1 | 2226.71 | 0.175656 | 0.10031 | 1.751084 | 0.07993 | 0.22477 | 13.914 |
| Colgalt2 | 345.445 | -0.37324 | 0.13498 | -2.76515 | 0.00569 | 0.03976 | 110.866 |
| Commd1 | 122.21 | 0.157197 | 0.24234 | 0.648667 | 0.51655 | 0.70653 | 86.247 |
| Commd10 | 988.033 | 0.047076 | 0.08379 | 0.561846 | 0.57422 | 0.75253 | 275.469 |
| Commd2 | 594.344 | -0.00306 | 0.08392 | -0.03641 | 0.97096 | 0.98532 | 7.367 |
| Commd3 | 1745.39 | -0.01769 | 0.0727 | -0.24333 | 0.80775 | 0.90112 | 3.848 |

| | | | | | | | |
|---|---|---|---|---|---|---|---|
| Commd4 | 1862.49 | 0.0639 | 0.07435 | 0.85947 | 0.39008 | 0.59916 | 3.297 |
| Commd6 | 1292.23 | -0.06566 | 0.0736 | -0.89213 | 0.37232 | 0.58269 | 7.706 |
| Commd7 | 1225.24 | 0.030002 | 0.07159 | 0.419071 | 0.67516 | 0.82179 | 15.849 |
| Commd8 | 2069.62 | -0.00082 | 0.06246 | -0.01313 | 0.98952 | 0.99424 | 11.615 |
| Commd9 | 784.148 | -0.0633 | 0.09951 | -0.63606 | 0.52474 | 0.71313 | 15.4 |
| Comp | 19.5306 | -0.35183 | 0.51837 | -0.67874 | 0.4973 | 0.69077 | 8.509 |
| Comt | 2784.2 | 0.104448 | 0.08831 | 1.182774 | 0.2369 | 0.44201 | 19.967 |
| Comtd1 | 29.4233 | -0.3358 | 0.32556 | -1.03148 | 0.30231 | 0.51199 | 3.123 |
| Cop1 | 2498.96 | -0.00379 | 0.12397 | -0.03058 | 0.9756 | 0.98762 | 115.321 |
| Copa | 9547.44 | -0.12339 | 0.05703 | -2.16356 | 0.0305 | 0.11939 | 39.802 |
| Copb1 | 3240.43 | 0.139112 | 0.06128 | 2.26993 | 0.02321 | 0.09876 | 39.153 |
| Copb2 | 4279.18 | -0.10245 | 0.08026 | -1.27651 | 0.20177 | 0.40021 | 24.662 |
| Cope | 2820.27 | -0.09964 | 0.06823 | -1.46041 | 0.14418 | 0.32683 | 10.476 |
| Copg1 | 12435.1 | 0.1191 | 0.07406 | 1.608253 | 0.10778 | 0.27268 | 25.782 |
| Copg2 | 4190.57 | 0.169892 | 0.08568 | 1.982851 | 0.04738 | 0.15989 | 126.177 |
| Coprs | 568.056 | -0.31491 | 0.12068 | -2.60936 | 0.00907 | 0.05373 | 5.488 |
| Cops2 | 3944.29 | 0.357823 | 0.14254 | 2.510341 | 0.01206 | 0.06482 | 28.836 |
| Cops3 | 1619.1 | 0.025324 | 0.06383 | 0.396757 | 0.69155 | 0.83252 | 22.044 |
| Cops4 | 2638.7 | 0.020011 | 0.07541 | 0.265356 | 0.79074 | 0.8913 | 29.495 |
| Cops5 | 2681.42 | -0.09057 | 0.07042 | -1.28612 | 0.1984 | 0.3961 | 13.568 |
| Cops6 | 3216.37 | -0.17424 | 0.07711 | -2.25961 | 0.02385 | 0.10049 | 3.576 |
| Cops7a | 3901.26 | -0.07332 | 0.07058 | -1.03881 | 0.29889 | 0.50895 | 7.126 |
| Cops7b | 998.383 | 0.1145 | 0.0985 | 1.162384 | 0.24508 | 0.45169 | 26.472 |
| Cops8 | 3407.57 | 0.048093 | 0.07057 | 0.681519 | 0.49554 | 0.68945 | 10.361 |
| Cops9 | 1846.04 | -0.29648 | 0.07956 | -3.72665 | 0.00019 | 0.0042 | 4.841 |
| Copz1 | 2745.28 | 0.045497 | 0.07204 | 0.63153 | 0.52769 | 0.71577 | 27.155 |
| Copz2 | 155.054 | -0.45782 | 0.1847 | -2.47867 | 0.01319 | 0.06875 | 11.334 |
| Coq10a | 1255.75 | -0.19279 | 0.07558 | -2.55094 | 0.01074 | 0.06024 | 5.898 |
| Coq10b | 877.892 | 0.242222 | 0.09901 | 2.446498 | 0.01443 | 0.0726 | 19.933 |
| Coq2 | 2958.46 | 0.076678 | 0.06032 | 1.271192 | 0.20366 | 0.40223 | 20.418 |
| Coq3 | 656.103 | -0.06463 | 0.11174 | -0.5784 | 0.56299 | 0.74365 | 32.49 |
| Coq4 | 452.44 | -0.00279 | 0.11864 | -0.02351 | 0.98124 | 0.99018 | 10.443 |
| Coq5 | 1372.93 | 0.049858 | 0.0793 | 0.628711 | 0.52954 | 0.71736 | 17.307 |
| Coq6 | 719.918 | 0.018074 | 0.08794 | 0.205531 | 0.83716 | 0.91632 | 12.14 |

| | | | | | | | |
|---|---|---|---|---|---|---|---|
| Coq7 | 746.657 | -0.10063 | 0.09025 | -1.11503 | 0.26484 | 0.47384 | 23.698 |
| Coq8a | 1619.12 | -0.06653 | 0.09572 | -0.69502 | 0.48705 | 0.68268 | 34.365 |
| Coq8b | 690.906 | 0.02021 | 0.08659 | 0.233407 | 0.81545 | 0.90514 | 24.928 |
| Coq9 | 2195.42 | 0.026129 | 0.06586 | 0.396754 | 0.69155 | 0.83252 | 16.575 |
| Corin | 19.3416 | 0.165177 | 0.36963 | 0.446868 | 0.65497 | 0.80851 | 204.449 |
| Coro1a | 1991.2 | -0.32619 | 0.11582 | -2.81632 | 0.00486 | 0.03557 | 8.015 |
| Coro1b | 1854.73 | -0.10939 | 0.0828 | -1.32117 | 0.18645 | 0.38139 | 5.417 |
| Coro1c | 3465.03 | -0.07894 | 0.06295 | -1.25402 | 0.20984 | 0.4099 | 66.323 |
| Coro2a | 638.113 | -0.5306 | 0.11068 | -4.79385 | 1.64E-06 | 0.00015 | 65.266 |
| Coro2b | 7126.83 | 0.114738 | 0.07604 | 1.508847 | 0.13134 | 0.30993 | 117.553 |
| Coro6 | 1810.31 | -0.48782 | 0.20868 | -2.33771 | 0.0194 | 0.08771 | 8.074 |
| Coro7 | 4086.2 | 0.065381 | 0.06472 | 1.010157 | 0.31242 | 0.52285 | 53.645 |
| Cort | 151.238 | -1.19461 | 0.30883 | -3.86819 | 0.00011 | 0.00285 | 1.73 |
| Cotl1 | 2093.74 | -0.01598 | 0.10984 | -0.14547 | 0.88434 | 0.94244 | 31.323 |
| Cox10 | 830.096 | -0.19175 | 0.08049 | -2.38225 | 0.01721 | 0.08153 | 116.842 |
| Cox11 | 427.022 | -0.15637 | 0.14442 | -1.08276 | 0.27892 | 0.48886 | 6.759 |
| Cox14 | 2112.88 | -0.15777 | 0.0759 | -2.07853 | 0.03766 | 0.13735 | 3.886 |
| Cox15 | 1424.81 | 0.093942 | 0.08819 | 1.065205 | 0.28678 | 0.4967 | 19.747 |
| Cox16 | 43.7523 | 0.061254 | 0.28959 | 0.211519 | 0.83248 | 0.91405 | 126.102 |
| Cox17 | 456.632 | 0.080189 | 0.13079 | 0.613131 | 0.53979 | 0.72592 | 5.773 |
| Cox18 | 694.123 | 0.298773 | 0.09334 | 3.200795 | 0.00137 | 0.01554 | 9.278 |
| Cox19 | 1528.46 | 0.007526 | 0.08532 | 0.088207 | 0.92971 | 0.96459 | 15.23 |
| Cox20 | 55.5258 | -0.25527 | 0.3131 | -0.81528 | 0.41491 | 0.62289 | 3.564 |
| Cox4i1 | 8467.96 | -0.25725 | 0.08869 | -2.90045 | 0.00373 | 0.03002 | 5.986 |
| Cox4i2 | 70.4654 | 0.319625 | 0.2378 | 1.34407 | 0.17893 | 0.37156 | 10.865 |
| Cox5a | 1772.37 | -0.23118 | 0.104 | -2.22285 | 0.02623 | 0.10754 | 11.153 |
| Cox5b | 6111.21 | -0.24429 | 0.08048 | -3.03527 | 0.0024 | 0.02271 | 2.233 |
| Cox5b-ps | 528.714 | -0.30426 | 0.10212 | -2.97959 | 0.00289 | 0.0256 | 0.39 |
| Cox6a1 | 8285.49 | -0.38684 | 0.09003 | -4.29699 | 1.73E-05 | 0.0008 | 3.34 |
| Cox6a2 | 190.269 | -0.38351 | 0.15388 | -2.49219 | 0.0127 | 0.0672 | 0.953 |
| Cox6b1 | 4638.75 | -0.27142 | 0.07412 | -3.66196 | 0.00025 | 0.00496 | 9.291 |
| Cox6b2 | 117.887 | 0.002821 | 0.21212 | 0.013299 | 0.98939 | 0.99417 | 1.303 |
| Cox6c | 7203.95 | -0.30119 | 0.09288 | -3.24289 | 0.00118 | 0.01416 | 12.361 |
| Cox7a1 | 46.4836 | -0.3359 | 0.26396 | -1.27253 | 0.20319 | 0.40179 | 1.935 |

| | | | | | | | |
|---|---|---|---|---|---|---|---|
| Cox7a2 | 3475.08 | -0.22772 | 0.08069 | -2.82229 | 0.00477 | 0.03524 | 4.518 |
| Cox7a2l | 4515.42 | 0.039435 | 0.08698 | 0.453382 | 0.65027 | 0.80549 | 15.413 |
| Cox7b | 4395.43 | -0.18989 | 0.06953 | -2.73112 | 0.00631 | 0.04257 | 6.751 |
| Cox7c | 5848.03 | -0.25136 | 0.07596 | -3.30901 | 0.00094 | 0.01223 | 2.089 |
| Cox7c-ps1 | 35.823 | 0.426445 | 0.36156 | 1.179462 | 0.23821 | 0.44369 | 0.372 |
| Cox8a | 8225.8 | -0.38343 | 0.09696 | -3.95436 | 7.67E-05 | 0.00223 | 2.464 |
| Cp | 1140.02 | -0.82593 | 0.18043 | -4.57763 | 4.70E-06 | 0.00032 | 52.092 |
| Cpa2 | 48.1743 | 0.476971 | 0.28843 | 1.653698 | 0.09819 | 0.25724 | 22.895 |
| Cpa4 | 22.3915 | 0.404126 | 0.56264 | 0.718269 | 0.47259 | 0.6709 | 24.05 |
| Cpa6 | 17.2019 | -1.68888 | 0.69388 | -2.43394 | 0.01494 | 0.07415 | 395.226 |
| Cpd | 4978.69 | -0.1705 | 0.09227 | -1.84778 | 0.06463 | 0.19571 | 68.595 |
| Cpe | 96704.3 | -0.33097 | 0.09071 | -3.64865 | 0.00026 | 0.00513 | 100.513 |
| Cpeb1 | 2293.19 | -0.55712 | 0.14584 | -3.81999 | 0.00013 | 0.00328 | 108.44 |
| Cpeb1os1 | 123.258 | -0.39453 | 0.23574 | -1.6736 | 0.09421 | 0.25054 | 2.112 |
| Cpeb2 | 5506.78 | 0.145107 | 0.07219 | 2.009985 | 0.04443 | 0.15302 | 56.555 |
| Cpeb3 | 5024.03 | 0.048425 | 0.06339 | 0.763884 | 0.44494 | 0.64896 | 187.31 |
| Cpeb4 | 8128.08 | 0.090303 | 0.07676 | 1.176378 | 0.23944 | 0.44541 | 63.424 |
| Cped1 | 277.879 | -0.92819 | 0.28818 | -3.22083 | 0.00128 | 0.01485 | 270.489 |
| Cplane1 | 4987.82 | 0.207171 | 0.05611 | 3.691982 | 0.00022 | 0.00458 | 102.053 |
| Cplane2 | 134.482 | -0.12352 | 0.1703 | -0.72531 | 0.46826 | 0.66717 | 12.801 |
| Cplx1 | 26734.1 | -0.41049 | 0.0873 | -4.70233 | 2.57E-06 | 0.00021 | 31.471 |
| Cplx2 | 63487.3 | -0.012 | 0.11269 | -0.10649 | 0.91519 | 0.95656 | 12.569 |
| Cplx3 | 461.885 | -1.0892 | 0.25873 | -4.20985 | 2.56E-05 | 0.00104 | 6.29 |
| Cpm | 776.137 | -0.45488 | 0.1375 | -3.30828 | 0.00094 | 0.01226 | 57.853 |
| Cpn1 | 36.5612 | 0.45699 | 0.32881 | 1.389812 | 0.16459 | 0.3536 | 30.25 |
| Cpne1 | 12.7678 | -0.25393 | 0.46299 | -0.54846 | 0.58338 | 0.75845 | 40.124 |
| Cpne2 | 1427.22 | 0.470566 | 0.16491 | 2.853405 | 0.00433 | 0.03317 | 37.542 |
| Cpne3 | 1122.21 | -0.03032 | 0.08216 | -0.36904 | 0.7121 | 0.845 | 50.855 |
| Cpne4 | 2083.92 | -0.29874 | 0.1541 | -1.93855 | 0.05256 | 0.17152 | 487.259 |
| Cpne5 | 9839.6 | 0.593611 | 0.18227 | 3.256713 | 0.00113 | 0.01378 | 82.612 |
| Cpne6 | 6417.01 | 0.019367 | 0.11755 | 0.164758 | 0.86913 | 0.93357 | 6.987 |
| Cpne7 | 3850.06 | -0.69493 | 0.37364 | -1.8599 | 0.0629 | 0.19257 | 17.809 |
| Cpne8 | 801.712 | -0.14242 | 0.18078 | -0.7878 | 0.43081 | 0.63702 | 191.951 |
| Cpne9 | 474.272 | -0.68481 | 0.18467 | -3.70834 | 0.00021 | 0.00438 | 23.321 |

| | | | | | | | |
|---|---|---|---|---|---|---|---|
| Cpox | 2047 | -0.12071 | 0.08643 | -1.39663 | 0.16252 | 0.35091 | 47.345 |
| Cpped1 | 1842.48 | 0.009148 | 0.08511 | 0.107482 | 0.91441 | 0.95656 | 105.725 |
| Cpq | 392.92 | -0.28745 | 0.13854 | -2.07491 | 0.03799 | 0.1383 | 511.424 |
| Cps1 | 12.9067 | 0.284665 | 0.66663 | 0.427021 | 0.66936 | 0.81764 | 108.234 |
| Cpsf1 | 3232.34 | 0.158747 | 0.0741 | 2.142307 | 0.03217 | 0.12391 | 11.789 |
| Cpsf2 | 2424.87 | 0.048754 | 0.06621 | 0.73637 | 0.46151 | 0.66207 | 30.437 |
| Cpsf3 | 2929.94 | 0.326258 | 0.10624 | 3.070867 | 0.00213 | 0.02104 | 29.666 |
| Cpsf4 | 1277.63 | -0.20248 | 0.1126 | -1.79826 | 0.07214 | 0.21084 | 14.829 |
| Cpsf4l | 325.92 | 0.336029 | 0.13153 | 2.55482 | 0.01062 | 0.05984 | 11.846 |
| Cpsf6 | 8151.9 | 0.256078 | 0.07111 | 3.601028 | 0.00032 | 0.00587 | 35.343 |
| Cpsf7 | 4549.21 | 0.123467 | 0.05859 | 2.107348 | 0.03509 | 0.13139 | 22.388 |
| Cpt1a | 1842.01 | -0.39081 | 0.08148 | -4.79656 | 1.61E-06 | 0.00015 | 63.4 |
| Cpt1b | 14.3136 | 0.194799 | 0.47659 | 0.408731 | 0.68274 | 0.82662 | 9.459 |
| Cpt1c | 3944.95 | 0.170626 | 0.06552 | 2.604383 | 0.0092 | 0.05422 | 15.48 |
| Cpt2 | 304.108 | -0.29975 | 0.13181 | -2.27413 | 0.02296 | 0.09807 | 19.63 |
| Cptp | 645.728 | 0.009634 | 0.12278 | 0.078472 | 0.93745 | 0.96877 | 4.718 |
| Cpxm1 | 278.898 | -0.03964 | 0.13902 | -0.28514 | 0.77554 | 0.88307 | 6.8 |
| Cpxm2 | 108.454 | -0.07788 | 0.24891 | -0.31289 | 0.75437 | 0.87077 | 122.053 |
| Cr1l | 1395.86 | -0.05618 | 0.09258 | -0.60687 | 0.54394 | 0.72924 | 34.205 |
| Cr2 | 58.3376 | -0.79957 | 0.3077 | -2.59853 | 0.00936 | 0.05497 | 39.906 |
| Crabp1 | 100.596 | 1.166788 | 0.32867 | 3.549997 | 0.00039 | 0.00665 | 8.363 |
| Crabp2 | 122.013 | -0.81435 | 0.32456 | -2.50908 | 0.0121 | 0.06496 | 4.711 |
| Cracd | 2831.92 | -0.11854 | 0.11428 | -1.03725 | 0.29962 | 0.50972 | 217.043 |
| Cracdl | 19010.1 | 0.045123 | 0.07708 | 0.58537 | 0.5583 | 0.73975 | 108.409 |
| Cracr2a | 191.081 | 0.073834 | 0.14995 | 0.492384 | 0.62245 | 0.78561 | 112.911 |
| Cracr2b | 50.4134 | -0.0731 | 0.24102 | -0.30327 | 0.76168 | 0.87511 | 5.52 |
| Cradd | 193.726 | 0.315273 | 0.15739 | 2.003094 | 0.04517 | 0.15456 | 149.388 |
| Cramp1 | 2462.92 | 0.046253 | 0.06376 | 0.725402 | 0.46821 | 0.66717 | 54.563 |
| Crat | 2278.61 | 0.013378 | 0.06499 | 0.205854 | 0.8369 | 0.91619 | 15.343 |
| Crb1 | 96.6543 | -0.15881 | 0.21656 | -0.73331 | 0.46337 | 0.66317 | 180.045 |
| Crb2 | 176.886 | -0.38825 | 0.18821 | -2.0629 | 0.03912 | 0.14095 | 22.855 |
| Crb3 | 47.8624 | -0.12232 | 0.24716 | -0.4949 | 0.62067 | 0.78424 | 6.814 |
| Crbn | 2596.13 | 0.064451 | 0.07666 | 0.840764 | 0.40048 | 0.60926 | 19.877 |
| Crcp | 1169.86 | 0.040933 | 0.08435 | 0.485261 | 0.62749 | 0.78925 | 31.5 |

| | | | | | | | |
|---|---|---|---|---|---|---|---|
| Creb1 | 3537.46 | 0.114718 | 0.09995 | 1.147714 | 0.25109 | 0.45847 | 71.904 |
| Creb3 | 1297.72 | -0.12918 | 0.0699 | -1.84826 | 0.06457 | 0.1956 | 4.729 |
| Creb3l1 | 101.885 | -0.38912 | 0.17753 | -2.19186 | 0.02839 | 0.1136 | 42.175 |
| Creb3l2 | 1187.39 | -0.3962 | 0.09646 | -4.10755 | 4.00E-05 | 0.0014 | 114.892 |
| Creb3l3 | 15.5651 | 0.087816 | 0.44139 | 0.198954 | 0.8423 | 0.91879 | 14.551 |
| Creb3l4 | 28.9417 | 0.59518 | 0.3251 | 1.830738 | 0.06714 | 0.20064 | 6.013 |
| Creb5 | 201.894 | -0.48877 | 0.16531 | -2.95671 | 0.00311 | 0.02669 | 413.107 |
| Crebbp | 5117.07 | 0.162397 | 0.06583 | 2.46708 | 0.01362 | 0.07023 | 132.67 |
| Crebl2 | 2648.19 | 0.088749 | 0.10485 | 0.846417 | 0.39732 | 0.60651 | 28.778 |
| Crebrf | 4099.86 | 0.160768 | 0.07413 | 2.168686 | 0.03011 | 0.11854 | 60.986 |
| Crebzf | 3958.14 | 0.005058 | 0.06262 | 0.080773 | 0.93562 | 0.96776 | 5.266 |
| Creg1 | 1060.21 | -0.16051 | 0.08737 | -1.83712 | 0.06619 | 0.19883 | 11.563 |
| Creg2 | 4667.62 | -0.23071 | 0.09825 | -2.34821 | 0.01886 | 0.0862 | 33.069 |
| Creld1 | 3969.49 | 0.031879 | 0.07507 | 0.424633 | 0.6711 | 0.81882 | 10.047 |
| Creld2 | 869.228 | 0.107876 | 0.11539 | 0.934865 | 0.34986 | 0.56045 | 7.038 |
| Crem | 645.806 | 0.028682 | 0.12407 | 0.231181 | 0.81717 | 0.90593 | 71.701 |
| Crh | 179.36 | -0.29523 | 0.30222 | -0.97688 | 0.32863 | 0.53979 | 1.996 |
| Crhbp | 588.459 | -0.96272 | 0.3408 | -2.82487 | 0.00473 | 0.03509 | 13.554 |
| Crhr1 | 490.384 | 0.144778 | 0.14164 | 1.022183 | 0.30669 | 0.51671 | 42.669 |
| Crhr2 | 97.8045 | -0.12543 | 0.24201 | -0.51828 | 0.60426 | 0.7733 | 42.968 |
| Crim1 | 3119.65 | 0.031119 | 0.07135 | 0.436175 | 0.66271 | 0.81368 | 176.345 |
| Crip1 | 76.7593 | -0.38439 | 0.22334 | -1.72109 | 0.08523 | 0.23406 | 7.564 |
| Crip2 | 3431.98 | -0.04616 | 0.0874 | -0.52821 | 0.59735 | 0.7692 | 5.271 |
| Crip3 | 292.651 | 0.429259 | 0.13816 | 3.106899 | 0.00189 | 0.01936 | 2.851 |
| Cript | 1939.66 | -0.02668 | 0.07712 | -0.34594 | 0.72939 | 0.8561 | 10.261 |
| Crispld1 | 329.944 | 0.081371 | 0.1711 | 0.475584 | 0.63437 | 0.79419 | 39.3 |
| Crispld2 | 137.927 | -0.9226 | 0.29873 | -3.08846 | 0.00201 | 0.02016 | 60.356 |
| Crk | 7504.93 | 0.054693 | 0.05644 | 0.969033 | 0.33253 | 0.54344 | 27.65 |
| Crkl | 3218.09 | -0.07227 | 0.07505 | -0.9629 | 0.3356 | 0.54639 | 35.448 |
| Crlf1 | 103.814 | -0.72093 | 0.37103 | -1.94307 | 0.05201 | 0.17024 | 10.924 |
| Crlf2 | 661.568 | 0.093144 | 0.0863 | 1.079287 | 0.28046 | 0.4903 | 4.285 |
| Crlf3 | 834.966 | -0.08725 | 0.08076 | -1.08031 | 0.28001 | 0.49005 | 34.499 |
| Crls1 | 1032.32 | -0.01508 | 0.09372 | -0.16095 | 0.87214 | 0.93543 | 20.12 |
| Crmp1 | 6559.8 | -0.18202 | 0.11071 | -1.6442 | 0.10013 | 0.26034 | 50.194 |

| | | | | | | | |
|---|---|---|---|---|---|---|---|
| Crnde | 26.6774 | -0.89541 | 0.57578 | -1.55512 | 0.11992 | 0.29246 | 31.589 |
| Crnkl1 | 1103.28 | 0.13441 | 0.10101 | 1.330594 | 0.18332 | 0.3774 | 17.536 |
| Crocc | 8252.57 | 0.566221 | 0.15252 | 3.7125 | 0.00021 | 0.00435 | 43.914 |
| Crot | 2356.72 | -0.11799 | 0.09428 | -1.25142 | 0.21078 | 0.4108 | 31.292 |
| Crppa | 277.618 | -0.05922 | 0.13245 | -0.44715 | 0.65477 | 0.80841 | 308.054 |
| Crtac1 | 3262.55 | -0.75083 | 0.33967 | -2.21049 | 0.02707 | 0.10988 | 163.499 |
| Crtap | 166.71 | -0.31231 | 0.17815 | -1.7531 | 0.07958 | 0.22404 | 15.542 |
| Crtc1 | 12000.5 | 0.262982 | 0.08509 | 3.090497 | 0.002 | 0.0201 | 57.225 |
| Crtc2 | 1608.73 | 0.116536 | 0.06226 | 1.871895 | 0.06122 | 0.18908 | 9.963 |
| Crtc3 | 1192.69 | -0.08975 | 0.07595 | -1.18171 | 0.23732 | 0.44263 | 102.251 |
| Cry1 | 1389.03 | 0.197784 | 0.11729 | 1.68625 | 0.09175 | 0.24635 | 53.365 |
| Cry2 | 6883.4 | 0.043155 | 0.06977 | 0.618553 | 0.53621 | 0.72318 | 30.398 |
| Cryab | 2862.26 | -0.66336 | 0.15594 | -4.25387 | 2.10E-05 | 0.00091 | 5.312 |
| Cryba4 | 25.3344 | -0.97229 | 0.35514 | -2.73774 | 0.00619 | 0.04207 | 6.026 |
| Crybb1 | 212.647 | -0.2987 | 0.12235 | -2.44131 | 0.01463 | 0.07308 | 13.771 |
| Crybb3 | 58.9367 | -0.38808 | 0.24107 | -1.60985 | 0.10743 | 0.27217 | 5.746 |
| Crybg1 | 112.767 | -0.35955 | 0.22456 | -1.60109 | 0.10936 | 0.2752 | 198.218 |
| Crybg2 | 61.3369 | -0.07446 | 0.32703 | -0.22769 | 0.81989 | 0.90734 | 31.69 |
| Crybg3 | 1797.52 | 0.509986 | 0.14911 | 3.420109 | 0.00063 | 0.00924 | 110.205 |
| Cryl1 | 373.541 | -0.63339 | 0.17295 | -3.66228 | 0.00025 | 0.00496 | 123.537 |
| Crym | 8580.88 | 1.027773 | 0.29228 | 3.516354 | 0.00044 | 0.00729 | 15.732 |
| Cryz | 992.414 | 0.224429 | 0.1419 | 1.581626 | 0.11374 | 0.28233 | 26.472 |
| Cryzl1 | 2397.5 | 0.114106 | 0.07632 | 1.495192 | 0.13486 | 0.31476 | 39.654 |
| Cryzl2 | 237.301 | -0.06192 | 0.18501 | -0.33469 | 0.73786 | 0.86163 | 34.062 |
| Cs | 10054.4 | -0.11206 | 0.05725 | -1.95753 | 0.05029 | 0.16664 | 24.746 |
| Csad | 1294.81 | 0.130072 | 0.09078 | 1.43288 | 0.15189 | 0.33702 | 27.726 |
| Csdc2 | 4718.14 | -0.19339 | 0.066 | -2.93029 | 0.00339 | 0.02815 | 14.183 |
| Csde1 | 10581.1 | -0.02481 | 0.07398 | -0.33533 | 0.73738 | 0.86132 | 37.761 |
| Cse1l | 3038.27 | 0.108974 | 0.07697 | 1.41583 | 0.15683 | 0.3437 | 40.35 |
| Csf1 | 990.735 | -0.5267 | 0.13268 | -3.96985 | 7.19E-05 | 0.00213 | 19.422 |
| Csf1r | 4116.75 | -0.28341 | 0.12834 | -2.20817 | 0.02723 | 0.11025 | 31.552 |
| Csf2ra | 1669.74 | 0.104303 | 0.14066 | 0.741507 | 0.45839 | 0.65954 | 4.473 |
| Csf2rb | 50.4014 | -0.80308 | 0.28758 | -2.79254 | 0.00523 | 0.03734 | 28.096 |
| Csf2rb2 | 33.6483 | -0.5047 | 0.39675 | -1.27208 | 0.20335 | 0.40179 | 23.215 |

| | | | | | | | |
|---|---|---|---|---|---|---|---|
| Csf3r | 326.905 | -0.2998 | 0.15491 | -1.93532 | 0.05295 | 0.17218 | 19.891 |
| Csgalnact1 | 1527.72 | 0.252001 | 0.0906 | 2.781346 | 0.00541 | 0.03829 | 378.366 |
| Csgalnact2 | 1178.81 | 0.21154 | 0.08172 | 2.588559 | 0.00964 | 0.05596 | 31.689 |
| Csk | 1106.21 | -0.1368 | 0.0822 | -1.66435 | 0.09604 | 0.25372 | 26.986 |
| Csmd1 | 5016.59 | 0.101102 | 0.07177 | 1.408691 | 0.15893 | 0.34639 | 1643.066 |
| Csmd2 | 3823.53 | -0.08539 | 0.11451 | -0.74571 | 0.45584 | 0.65758 | 579.8 |
| Csmd3 | 2670.72 | 0.305348 | 0.13598 | 2.245526 | 0.02473 | 0.10325 | 1211.427 |
| Csnk1a1 | 18683.9 | 0.155373 | 0.07761 | 2.001909 | 0.04529 | 0.15478 | 34.788 |
| Csnk1d | 6649.67 | 0.017577 | 0.07264 | 0.241973 | 0.8088 | 0.90165 | 32.341 |
| Csnk1e | 3600.63 | -0.11944 | 0.08218 | -1.45335 | 0.14613 | 0.32908 | 37.712 |
| Csnk1g1 | 2605.41 | 0.227678 | 0.09417 | 2.417765 | 0.01562 | 0.0765 | 136.092 |
| Csnk1g2 | 5880.91 | -0.18072 | 0.07469 | -2.41973 | 0.01553 | 0.0762 | 17.912 |
| Csnk1g3 | 3057.07 | 0.055272 | 0.07322 | 0.75482 | 0.45036 | 0.65308 | 93.427 |
| Csnk2a1 | 5617.59 | 0.043163 | 0.05857 | 0.736969 | 0.46114 | 0.66177 | 55.014 |
| Csnk2a1-ps3 | 832.285 | -0.29527 | 0.13641 | -2.16463 | 0.03042 | 0.11912 | 2.633 |
| Csnk2a2 | 3156.15 | 0.048004 | 0.07515 | 0.638801 | 0.52295 | 0.71143 | 43.944 |
| Csnk2b | 2034.21 | -0.15023 | 0.07046 | -2.13198 | 0.03301 | 0.12614 | 5.858 |
| Cspg4 | 942.992 | -0.26116 | 0.10399 | -2.51136 | 0.01203 | 0.0647 | 34.838 |
| Cspg4b | 133.888 | 0.254295 | 0.21099 | 1.205236 | 0.22811 | 0.43202 | 86.553 |
| Cspg5 | 11450.5 | -0.07534 | 0.08235 | -0.91485 | 0.36027 | 0.57061 | 18.794 |
| Cspp1 | 2646.8 | 0.228928 | 0.08182 | 2.797989 | 0.00514 | 0.03696 | 98.782 |
| Csrnp1 | 440.332 | 0.052128 | 0.12156 | 0.428827 | 0.66805 | 0.81679 | 13.496 |
| Csrnp2 | 1299.07 | 0.012214 | 0.071 | 0.17202 | 0.86342 | 0.93092 | 15.919 |
| Csrnp3 | 6030.43 | 0.236449 | 0.08218 | 2.877302 | 0.00401 | 0.03157 | 185.78 |
| Csrp1 | 9628.55 | -0.48729 | 0.10105 | -4.8221 | 1.42E-06 | 0.00013 | 32.172 |
| Csrp2 | 168.629 | 0.311339 | 0.22518 | 1.382643 | 0.16677 | 0.35646 | 19.834 |
| Cst3 | 37803.3 | -0.40184 | 0.1206 | -3.332 | 0.00086 | 0.01155 | 3.971 |
| Cst6 | 118.024 | 0.241562 | 0.18227 | 1.325281 | 0.18508 | 0.37981 | 4.87 |
| Cstad | 45.7296 | 0.141141 | 0.3519 | 0.401081 | 0.68836 | 0.83003 | 13.904 |
| Cstb | 580.464 | -0.30263 | 0.10345 | -2.92527 | 0.00344 | 0.02843 | 1.954 |
| Cstf1 | 1008.62 | 0.02836 | 0.08109 | 0.349735 | 0.72654 | 0.85418 | 11.768 |
| Cstf2 | 6281.89 | 0.123762 | 0.08133 | 1.521686 | 0.12809 | 0.30498 | 27.633 |
| Cstf2t | 4911.48 | -0.02044 | 0.0716 | -0.28541 | 0.77533 | 0.88298 | 4.233 |
| Cstf3 | 2385.41 | 0.123511 | 0.07695 | 1.605028 | 0.10849 | 0.27376 | 74.907 |

| | | | | | | | |
|---|---|---|---|---|---|---|---|
| Cstl1 | 20.4699 | -0.35422 | 0.59432 | -0.59601 | 0.55117 | 0.73445 | 5.076 |
| Ctbp1 | 8614.62 | -0.0828 | 0.08346 | -0.99208 | 0.32116 | 0.53202 | 27.272 |
| Ctbp2 | 450.744 | -0.08965 | 0.10569 | -0.84824 | 0.39631 | 0.60539 | 136.792 |
| Ctbs | 414.956 | -0.04968 | 0.11411 | -0.43535 | 0.66331 | 0.81392 | 16.055 |
| Ctc1 | 1666.19 | 0.072443 | 0.0711 | 1.018823 | 0.30829 | 0.51815 | 20.563 |
| Ctcf | 3829.16 | 0.048168 | 0.06726 | 0.716136 | 0.47391 | 0.67223 | 46.355 |
| Ctcfl | 12.9869 | 1.364798 | 0.53612 | 2.545706 | 0.01091 | 0.06082 | 25.917 |
| Ctdnep1 | 1599.83 | -0.01018 | 0.0775 | -0.13131 | 0.89553 | 0.94746 | 9.446 |
| Ctdp1 | 706.837 | -0.09379 | 0.08093 | -1.15895 | 0.24648 | 0.45328 | 61.737 |
| Ctdsp1 | 1222.63 | -0.31861 | 0.0889 | -3.58374 | 0.00034 | 0.00613 | 5.777 |
| Ctdsp2 | 4671.65 | -0.07124 | 0.05841 | -1.21953 | 0.22264 | 0.42587 | 21.259 |
| Ctdsp2-ps | 112.33 | -0.02501 | 0.18292 | -0.13671 | 0.89126 | 0.94585 | 0.811 |
| Ctdspl | 2545.69 | 0.182773 | 0.08435 | 2.166924 | 0.03024 | 0.11875 | 117.546 |
| Ctdspl2 | 1474.76 | 0.158553 | 0.08606 | 1.842438 | 0.06541 | 0.19731 | 57.642 |
| Ctf1 | 59.7386 | -0.23497 | 0.23871 | -0.98436 | 0.32494 | 0.53614 | 5.457 |
| Ctf2 | 13.6732 | -0.43563 | 0.57375 | -0.75926 | 0.4477 | 0.65129 | 14.061 |
| Cth | 271.648 | 0.123518 | 0.16146 | 0.765026 | 0.44426 | 0.64827 | 30.83 |
| Cthrc1 | 283.667 | -0.96496 | 0.16359 | -5.89857 | 3.67E-09 | 1.53E-06 | 10.19 |
| Ctif | 2714.78 | -0.24384 | 0.07987 | -3.05292 | 0.00227 | 0.02185 | 266.331 |
| Ctla2a | 55.8158 | -0.49591 | 0.24806 | -1.99914 | 0.04559 | 0.15547 | 2.471 |
| Ctnna1 | 3333.13 | -0.09063 | 0.07312 | -1.23945 | 0.21518 | 0.41662 | 135.921 |
| Ctnna2 | 5243.93 | 0.059807 | 0.06813 | 0.877776 | 0.38007 | 0.58974 | 1098.063 |
| Ctnna3 | 123.892 | 0.376265 | 0.19088 | 1.971239 | 0.0487 | 0.16299 | 1573.57 |
| Ctnnal1 | 1510.38 | -0.07489 | 0.07658 | -0.97793 | 0.32811 | 0.53939 | 54.254 |
| Ctnnb1 | 13436.9 | -0.05372 | 0.07085 | -0.75818 | 0.44834 | 0.65189 | 31.292 |
| Ctnnbip1 | 886.121 | -0.17416 | 0.10535 | -1.65306 | 0.09832 | 0.25734 | 48.202 |
| Ctnnbl1 | 1589.44 | 0.077032 | 0.10779 | 0.714675 | 0.47481 | 0.67287 | 154.214 |
| Ctnnd1 | 405.638 | -0.31736 | 0.12646 | -2.50953 | 0.01209 | 0.0649 | 50.695 |
| Ctnnd2 | 8261.02 | -0.22206 | 0.09007 | -2.4654 | 0.01369 | 0.07038 | 856.749 |
| Ctns | 986.951 | 0.092599 | 0.098 | 0.944876 | 0.34472 | 0.55557 | 15.447 |
| Ctps | 1536.87 | -0.03716 | 0.07037 | -0.5281 | 0.59743 | 0.7692 | 30.409 |
| Ctps2 | 1566.61 | -0.12928 | 0.10163 | -1.27208 | 0.20334 | 0.40179 | 131.271 |
| Ctr9 | 3215.42 | 0.05659 | 0.07183 | 0.787847 | 0.43079 | 0.63702 | 27.427 |
| Ctrl | 44.1196 | 0.069039 | 0.32598 | 0.211786 | 0.83227 | 0.91402 | 1.86 |

| | | | | | | | |
|---|---|---|---|---|---|---|---|
| Ctsa | 3289.42 | -0.1354 | 0.06502 | -2.08257 | 0.03729 | 0.13654 | 8.16 |
| Ctsb | 20375.3 | -0.19009 | 0.07673 | -2.4773 | 0.01324 | 0.06894 | 23.462 |
| Ctsc | 434.485 | -0.5128 | 0.14063 | -3.6465 | 0.00027 | 0.00516 | 32.804 |
| Ctsd | 4353.78 | -0.38509 | 0.12379 | -3.1108 | 0.00187 | 0.01917 | 12.128 |
| Ctsf | 2796.23 | -0.04708 | 0.06247 | -0.75364 | 0.45107 | 0.65372 | 5.789 |
| Ctsh | 479.253 | -0.61793 | 0.16027 | -3.85549 | 0.00012 | 0.00295 | 21.938 |
| Ctsk | 119.448 | -0.43228 | 0.19885 | -2.17387 | 0.02972 | 0.11741 | 10.107 |
| Ctsl | 3403.45 | -0.33245 | 0.09296 | -3.57637 | 0.00035 | 0.00622 | 11.554 |
| Ctso | 1861.71 | -0.20597 | 0.09697 | -2.12398 | 0.03367 | 0.12789 | 24.125 |
| Ctss | 2720.55 | -0.5547 | 0.18005 | -3.08085 | 0.00206 | 0.02052 | 29.618 |
| Ctsw | 32.4956 | 0.09764 | 0.41818 | 0.23349 | 0.81538 | 0.90512 | 3.465 |
| Ctsz | 1552.9 | -0.40334 | 0.11126 | -3.6253 | 0.00029 | 0.00547 | 11.547 |
| Cttn | 5570.24 | -0.01343 | 0.064 | -0.20984 | 0.83379 | 0.91475 | 35.277 |
| Cttnbp2 | 10034.6 | 0.158469 | 0.10743 | 1.475088 | 0.14019 | 0.32155 | 148.366 |
| Cttnbp2nl | 1702.5 | 0.014746 | 0.07111 | 0.207373 | 0.83572 | 0.91573 | 51.232 |
| Ctu1 | 374.653 | 0.169768 | 0.12893 | 1.316769 | 0.18792 | 0.38343 | 6.283 |
| Ctu2 | 488.077 | 0.079901 | 0.10692 | 0.747319 | 0.45487 | 0.65702 | 7.996 |
| Ctxn1 | 9717.47 | 0.023535 | 0.07721 | 0.304834 | 0.76049 | 0.87444 | 1.615 |
| Ctxn2 | 352.107 | 0.208999 | 0.11491 | 1.818785 | 0.06894 | 0.20424 | 11.209 |
| Ctxn3 | 130.155 | -1.28895 | 0.28388 | -4.5405 | 5.61E-06 | 0.00036 | 9.618 |
| Cubn | 33.0238 | -0.22632 | 0.49495 | -0.45726 | 0.64748 | 0.80377 | 215.476 |
| Cuedc1 | 1546.5 | -0.24971 | 0.08862 | -2.81789 | 0.00483 | 0.0355 | 96.083 |
| Cuedc2 | 3779.98 | -0.24275 | 0.08227 | -2.95056 | 0.00317 | 0.02709 | 8.891 |
| Cul1 | 5302.66 | 0.121568 | 0.06318 | 1.924244 | 0.05432 | 0.17524 | 72.742 |
| Cul2 | 2738.02 | 0.158432 | 0.08303 | 1.908161 | 0.05637 | 0.17956 | 53.39 |
| Cul3 | 6584.81 | 0.123177 | 0.085 | 1.449147 | 0.1473 | 0.33053 | 75.558 |
| Cul4a | 3634.28 | 0.068192 | 0.06668 | 1.022709 | 0.30645 | 0.51646 | 42.32 |
| Cul4b | 2264.47 | 0.160165 | 0.08146 | 1.966194 | 0.04928 | 0.16434 | 42.923 |
| Cul5 | 4260.02 | 0.234667 | 0.08587 | 2.732946 | 0.00628 | 0.04241 | 55.433 |
| Cul7 | 1674.82 | 0.179375 | 0.10141 | 1.768789 | 0.07693 | 0.21936 | 14.028 |
| Cul9 | 5041.24 | 0.136133 | 0.06562 | 2.074702 | 0.03801 | 0.13835 | 45.817 |
| Cuta | 968.424 | -0.20447 | 0.08704 | -2.34928 | 0.01881 | 0.08608 | 5.751 |
| Cutal | 25.9059 | -0.4268 | 0.34225 | -1.24707 | 0.21237 | 0.41311 | 17.737 |
| Cutc | 282.491 | 0.211205 | 0.13011 | 1.623306 | 0.10452 | 0.26745 | 15.643 |

| | | | | | | | |
|---|---|---|---|---|---|---|---|
| Cux1 | 5102.99 | 0.017558 | 0.06436 | 0.272818 | 0.78499 | 0.88852 | 319.356 |
| Cux2 | 2775.92 | 0.303926 | 0.12977 | 2.342082 | 0.01918 | 0.087 | 193.737 |
| Cwc15 | 2117.52 | -0.05403 | 0.07861 | -0.68736 | 0.49186 | 0.68654 | 9.961 |
| Cwc22 | 1117.28 | 0.307577 | 0.13218 | 2.326935 | 0.01997 | 0.08948 | 65.217 |
| Cwc25 | 936.332 | -0.15369 | 0.08156 | -1.88442 | 0.05951 | 0.18562 | 21.076 |
| Cwc27 | 421.961 | -0.10142 | 0.17497 | -0.57964 | 0.56216 | 0.74299 | 186.003 |
| Cwf19l1 | 777.677 | 0.146844 | 0.09173 | 1.600857 | 0.10941 | 0.27521 | 27.24 |
| Cwf19l2 | 922.654 | 0.22162 | 0.16189 | 1.368983 | 0.171 | 0.36141 | 75.645 |
| Cwh43 | 147.598 | -0.9725 | 0.26082 | -3.72859 | 0.00019 | 0.0042 | 47.361 |
| Cx3cl1 | 26099.1 | 0.131136 | 0.08097 | 1.619658 | 0.10531 | 0.26859 | 10.419 |
| Cx3cr1 | 1728.39 | -0.18298 | 0.12008 | -1.52382 | 0.12755 | 0.30403 | 168.264 |
| Cxadr | 2314.83 | 0.217294 | 0.09492 | 2.289122 | 0.02207 | 0.09575 | 58.286 |
| Cxcl12 | 2403.24 | -0.48091 | 0.09712 | -4.95154 | 7.36E-07 | 8.82E-05 | 12.833 |
| Cxcl13 | 19.3502 | -0.44184 | 0.50657 | -0.87222 | 0.38309 | 0.5923 | 4.118 |
| Cxcl14 | 6092.11 | -0.26696 | 0.09495 | -2.81162 | 0.00493 | 0.0359 | 7.909 |
| Cxcl16 | 83.1438 | -0.70261 | 0.26623 | -2.63908 | 0.00831 | 0.05081 | 6.002 |
| Cxcl5 | 34.667 | -1.93788 | 0.57608 | -3.36394 | 0.00077 | 0.01068 | 2.265 |
| Cxcr4 | 38.1899 | -0.75787 | 0.44851 | -1.68976 | 0.09107 | 0.24511 | 4.095 |
| Cxxc1 | 2779.94 | 0.088384 | 0.07978 | 1.107838 | 0.26793 | 0.47725 | 5.373 |
| Cxxc4 | 2008.19 | 0.253742 | 0.117 | 2.168742 | 0.0301 | 0.11854 | 25.678 |
| Cxxc5 | 1776.61 | -0.68711 | 0.11751 | -5.84721 | 5.00E-09 | 1.82E-06 | 32.292 |
| Cyb561 | 2426.08 | -0.22954 | 0.10443 | -2.19806 | 0.02794 | 0.11249 | 19.635 |
| Cyb561a3 | 420.633 | 0.00981 | 0.10602 | 0.092528 | 0.92628 | 0.96294 | 18.79 |
| Cyb561d1 | 1257.93 | -0.06793 | 0.07021 | -0.96744 | 0.33332 | 0.54421 | 5.526 |
| Cyb561d2 | 392.874 | -0.19639 | 0.11726 | -1.67482 | 0.09397 | 0.25 | 3.187 |
| Cyb5a | 1656.09 | -0.01841 | 0.09217 | -0.19974 | 0.84168 | 0.91879 | 41.168 |
| Cyb5b | 8354.56 | -0.01298 | 0.05033 | -0.2579 | 0.79648 | 0.89514 | 36.832 |
| Cyb5d1 | 1229.83 | 0.078754 | 0.08981 | 0.876893 | 0.38054 | 0.59002 | 3.707 |
| Cyb5d2 | 663.445 | -0.03065 | 0.10359 | -0.2959 | 0.76731 | 0.87835 | 18.911 |
| Cyb5r1 | 730.54 | -0.21889 | 0.08523 | -2.56821 | 0.01022 | 0.05839 | 6.182 |
| Cyb5r2 | 21.7399 | -0.48658 | 0.38154 | -1.27532 | 0.2022 | 0.40058 | 9.578 |
| Cyb5r3 | 2989.65 | -0.0808 | 0.11468 | -0.70458 | 0.48107 | 0.6782 | 19.099 |
| Cyb5r4 | 1767.67 | -0.06505 | 0.07589 | -0.8572 | 0.39133 | 0.6004 | 55.761 |
| Cyb5rl | 188.969 | -0.00368 | 0.16006 | -0.02299 | 0.98166 | 0.9904 | 21.281 |

| | | | | | | | |
|---|---|---|---|---|---|---|---|
| Cyba | 182.616 | -0.41227 | 0.20107 | -2.0504 | 0.04033 | 0.1435 | 8.155 |
| Cybb | 64.894 | -1.52254 | 0.36753 | -4.14264 | 3.43E-05 | 0.00125 | 52.52 |
| Cybc1 | 938.593 | -0.01985 | 0.10543 | -0.18824 | 0.85069 | 0.92407 | 6.735 |
| Cybrd1 | 112.861 | -0.91021 | 0.23285 | -3.90901 | 9.27E-05 | 0.00255 | 25.004 |
| Cyc1 | 4823.59 | -0.20334 | 0.05468 | -3.71879 | 0.0002 | 0.00427 | 2.738 |
| Cycs | 4114.23 | -0.13729 | 0.07511 | -1.82792 | 0.06756 | 0.20161 | 3.976 |
| Cyfip1 | 4119.8 | -0.09852 | 0.07564 | -1.30244 | 0.19277 | 0.38957 | 90.858 |
| Cyfip2 | 34272.4 | -0.01773 | 0.05969 | -0.29705 | 0.76643 | 0.8779 | 119.01 |
| Cygb | 2011.5 | -0.08051 | 0.12781 | -0.62993 | 0.52874 | 0.71654 | 8.719 |
| Cyhr1 | 6584.99 | -0.04367 | 0.06832 | -0.63913 | 0.52274 | 0.71143 | 16.723 |
| Cyld | 13813.7 | 0.516931 | 0.15144 | 3.413415 | 0.00064 | 0.00937 | 54.918 |
| Cyp11a1 | 152.974 | -0.30245 | 0.24719 | -1.22354 | 0.22113 | 0.42403 | 20.613 |
| Cyp19a1 | 16.3032 | 0.369802 | 0.43401 | 0.852052 | 0.39419 | 0.60332 | 102.174 |
| Cyp1b1 | 304.136 | -1.59241 | 0.43151 | -3.69034 | 0.00022 | 0.00459 | 13.525 |
| Cyp20a1 | 534.706 | -0.41722 | 0.11417 | -3.65439 | 0.00026 | 0.00505 | 44.738 |
| Cyp26b1 | 151.975 | -0.10156 | 0.3449 | -0.29445 | 0.76841 | 0.87901 | 22.495 |
| Cyp27a1 | 208.957 | -0.84274 | 0.22933 | -3.67483 | 0.00024 | 0.00479 | 24.319 |
| Cyp27b1 | 81.9221 | 0.846392 | 0.21144 | 4.00301 | 6.25E-05 | 0.00194 | 4.757 |
| Cyp2a5 | 24.7032 | 0.370565 | 0.49985 | 0.741347 | 0.45848 | 0.65959 | 8.244 |
| Cyp2c23 | 35.1458 | 1.048561 | 0.38517 | 2.722302 | 0.00648 | 0.04333 | 24.187 |
| Cyp2c52-ps | 18.737 | 0.81992 | 0.40386 | 2.030192 | 0.04234 | 0.14806 | 26.643 |
| Cyp2d22 | 1619 | -0.02952 | 0.09202 | -0.32079 | 0.74837 | 0.86814 | 9.734 |
| Cyp2e1 | 14.1431 | -0.09803 | 0.6187 | -0.15845 | 0.87411 | 0.93694 | 11.249 |
| Cyp2j12 | 111.893 | -0.60045 | 0.21215 | -2.83034 | 0.00465 | 0.03467 | 41.835 |
| Cyp2j6 | 951.222 | -0.12539 | 0.09691 | -1.29387 | 0.19571 | 0.39339 | 37.524 |
| Cyp2j9 | 1263.47 | -0.17223 | 0.10759 | -1.60083 | 0.10942 | 0.27521 | 23.15 |
| Cyp2r1 | 19.0552 | 0.085989 | 0.51858 | 0.165817 | 0.8683 | 0.93337 | 13.291 |
| Cyp2s1 | 395.886 | -0.04437 | 0.1844 | -0.24063 | 0.80984 | 0.90197 | 14.439 |
| Cyp2t4 | 133.916 | 0.166121 | 0.18596 | 0.893312 | 0.37169 | 0.58205 | 4.955 |
| Cyp2u1 | 488.073 | -0.04378 | 0.08833 | -0.49559 | 0.62018 | 0.78389 | 14.787 |
| Cyp39a1 | 87.0911 | -0.3013 | 0.23042 | -1.3076 | 0.19101 | 0.38726 | 84.007 |
| Cyp3a13 | 57.5541 | -0.57859 | 0.39447 | -1.46677 | 0.14244 | 0.32435 | 28.688 |
| Cyp46a1 | 8418.49 | 0.008375 | 0.07494 | 0.111756 | 0.91102 | 0.95504 | 27.854 |
| Cyp4f13 | 698.68 | -0.32525 | 0.10149 | -3.2048 | 0.00135 | 0.01537 | 22.715 |

| | | | | | | | |
|---|---|---|---|---|---|---|---|
| Cyp4f14 | 383.59 | -0.11988 | 0.13201 | -0.90812 | 0.36382 | 0.57434 | 12.273 |
| Cyp4f15 | 1209.85 | 0.13828 | 0.12904 | 1.071572 | 0.28391 | 0.49409 | 17.726 |
| Cyp4f16 | 258.462 | -0.54818 | 0.17996 | -3.04606 | 0.00232 | 0.02218 | 15.241 |
| Cyp4f17 | 186.686 | -0.1526 | 0.17674 | -0.86342 | 0.38791 | 0.59725 | 24.094 |
| Cyp4f18 | 44.4447 | -0.36399 | 0.28291 | -1.28659 | 0.19824 | 0.39602 | 21.145 |
| Cyp4f41-ps | 36.852 | 0.220101 | 0.29502 | 0.746049 | 0.45564 | 0.65749 | 14.772 |
| Cyp4v3 | 335.536 | -0.53179 | 0.12475 | -4.26272 | 2.02E-05 | 0.00089 | 28.273 |
| Cyp4x1 | 518.011 | -0.29323 | 0.15675 | -1.87062 | 0.0614 | 0.18951 | 27.959 |
| Cyp4x1os | 18.7117 | 0.493209 | 0.39729 | 1.241428 | 0.21445 | 0.4158 | 4.033 |
| Cyp51 | 2641.61 | -0.21184 | 0.07594 | -2.78951 | 0.00528 | 0.03757 | 23.602 |
| Cyp7b1 | 503.449 | -0.05084 | 0.10995 | -0.46237 | 0.64381 | 0.80129 | 171.389 |
| Cyp8b1 | 13.3326 | 1.232223 | 0.51729 | 2.382055 | 0.01722 | 0.08153 | 1.95 |
| Cyren | 147.605 | 0.035201 | 0.2375 | 0.148218 | 0.88217 | 0.94146 | 6.29 |
| Cyria | 13488.7 | 0.32131 | 0.08816 | 3.644634 | 0.00027 | 0.00518 | 118.827 |
| Cyrib | 7670.63 | 0.189906 | 0.08235 | 2.306192 | 0.0211 | 0.09285 | 131.382 |
| Cys1 | 454.148 | -0.40431 | 0.19094 | -2.11749 | 0.03422 | 0.12921 | 15.981 |
| Cysltr1 | 38.4566 | -0.44618 | 0.33474 | -1.33291 | 0.18256 | 0.37638 | 29.334 |
| Cysltr2 | 18.0018 | -0.09266 | 0.4204 | -0.2204 | 0.82556 | 0.90998 | 23.512 |
| Cysrt1 | 272.151 | -0.11162 | 0.12096 | -0.9228 | 0.35611 | 0.56647 | 3.635 |
| Cystm1 | 1128.04 | -0.12178 | 0.08946 | -1.36129 | 0.17342 | 0.36474 | 60.578 |
| Cyth1 | 4726.16 | -0.01663 | 0.07257 | -0.22918 | 0.81873 | 0.90676 | 116.574 |
| Cyth2 | 2273.9 | -0.19234 | 0.13584 | -1.416 | 0.15678 | 0.34363 | 7.945 |
| Cyth3 | 1813.81 | -0.1203 | 0.0854 | -1.40865 | 0.15894 | 0.34639 | 87.804 |
| Cyth4 | 599.155 | -0.35689 | 0.10778 | -3.31119 | 0.00093 | 0.01219 | 24.973 |
| Cytl1 | 35.3664 | -1.06401 | 1.08629 | -0.97949 | 0.32734 | 0.53868 | 4.302 |
| Cyyr1 | 275.463 | -0.59892 | 0.16974 | -3.52834 | 0.00042 | 0.00709 | 131.865 |
| Czib | 1991.61 | 0.092723 | 0.06973 | 1.329722 | 0.18361 | 0.37774 | 9.572 |
| D030028A08Rik | 343.892 | 0.211834 | 0.13051 | 1.623087 | 0.10457 | 0.26745 | 48.791 |
| D030034A15Rik | 15.7314 | -1.16916 | 0.50365 | -2.32136 | 0.02027 | 0.0903 | 3.953 |
| D030047H15Rik | 1183.08 | 0.493649 | 0.10695 | 4.615895 | 3.91E-06 | 0.00028 | 10.587 |
| D030055H07Rik | 42.7781 | 0.402398 | 0.29664 | 1.356515 | 0.17494 | 0.36656 | 11.674 |
| D030056L22Rik | 1004.04 | -0.08321 | 0.09139 | -0.9105 | 0.36256 | 0.57294 | 5.236 |
| D030068K23Rik | 25.7806 | -0.19124 | 0.34062 | -0.56144 | 0.57449 | 0.75265 | 198.59 |
| D11Wsu47e | 478.077 | 0.138391 | 0.09686 | 1.428829 | 0.15305 | 0.3386 | 10.236 |

| | | | | | | | |
|---|---|---|---|---|---|---|---|
| D130007C19Rik | 46.4736 | -0.03351 | 0.2662 | -0.12588 | 0.89983 | 0.94981 | 1.506 |
| D130017N08Rik | 1142.98 | 0.035271 | 0.10149 | 0.347538 | 0.72819 | 0.85513 | 10.238 |
| D130020L05Rik | 159.355 | 0.341625 | 0.16316 | 2.093852 | 0.03627 | 0.13417 | 6.477 |
| D130040H23Rik | 760.721 | 0.162971 | 0.13877 | 1.174419 | 0.24023 | 0.44624 | 43.128 |
| D130043K22Rik | 2254.94 | -0.01177 | 0.06316 | -0.18642 | 0.85212 | 0.92477 | 56.136 |
| D130058E05Rik | 17.2909 | 0.445206 | 0.5549 | 0.802313 | 0.42237 | 0.62917 | 3.973 |
| D16Ertd472e | 570.224 | -0.36486 | 0.17986 | -2.02858 | 0.0425 | 0.14845 | 36.371 |
| D17H6S53E | 548.146 | -0.13503 | 0.08995 | -1.50108 | 0.13334 | 0.31294 | 2.483 |
| D230022J07Rik | 15.9452 | -0.02417 | 0.43737 | -0.05526 | 0.95593 | 0.97839 | 33.529 |
| D230049E03Rik | 36.369 | 0.217795 | 0.36532 | 0.596176 | 0.55106 | 0.73441 | 4.251 |
| D2hgdh | 628.607 | 0.137949 | 0.11947 | 1.154702 | 0.24821 | 0.45532 | 27.44 |
| D330023K18Rik | 175.971 | 0.04614 | 0.17781 | 0.259487 | 0.79526 | 0.89438 | 5.864 |
| D330041H03Rik | 240.595 | 0.2942 | 0.13958 | 2.107697 | 0.03506 | 0.13131 | 9.894 |
| D330050G23Rik | 227.569 | 0.282455 | 0.17823 | 1.58481 | 0.11301 | 0.28134 | 14.463 |
| D330050I16Rik | 19.7677 | 0.177104 | 0.43358 | 0.40847 | 0.68293 | 0.8267 | 1.266 |
| D3Ertd751e | 738.157 | 0.009985 | 0.21707 | 0.046 | 0.96331 | 0.98208 | 60.71 |
| D430013B06Rik | 91.1115 | 0.259789 | 0.19453 | 1.335441 | 0.18173 | 0.37536 | 1.872 |
| D430018E03Rik | 105.784 | 0.417292 | 0.1837 | 2.271591 | 0.02311 | 0.09843 | 4.297 |
| D430019H16Rik | 16592.2 | 0.284408 | 0.11685 | 2.433869 | 0.01494 | 0.07415 | 39.24 |
| D430020J02Rik | 192.481 | 0.459169 | 0.16738 | 2.74329 | 0.00608 | 0.04167 | 3.264 |
| D430036J16Rik | 658.483 | 0.955054 | 0.34398 | 2.776446 | 0.0055 | 0.03875 | 15.099 |
| D430040D24Rik | 30.6778 | -0.50902 | 0.34076 | -1.49377 | 0.13523 | 0.31507 | 7.913 |
| D430041D05Rik | 6026.95 | 0.044456 | 0.07787 | 0.570888 | 0.56808 | 0.74799 | 267.941 |
| D530018E20Rik | 18.6824 | 0.11519 | 0.43582 | 0.264305 | 0.79155 | 0.89163 | 0.958 |
| D5Ertd579e | 6645.34 | 0.13705 | 0.06023 | 2.275603 | 0.02287 | 0.09783 | 95.54 |
| D5Ertd605e | 14.9911 | 0.682555 | 0.47045 | 1.45085 | 0.14682 | 0.32993 | 4.45 |
| D630003M21Rik | 336.399 | 0.181724 | 0.11406 | 1.593291 | 0.11109 | 0.27827 | 46.69 |
| D630023F18Rik | 84.3962 | -0.03688 | 0.21052 | -0.1752 | 0.86093 | 0.92962 | 17.93 |
| D630024D03Rik | 20.6239 | 0.263279 | 0.43736 | 0.601977 | 0.54719 | 0.7314 | 35.384 |
| D630030B08Rik | 19.6143 | 0.26969 | 0.43636 | 0.61804 | 0.53655 | 0.72354 | 3.479 |
| D630033A02Rik | 102.913 | -0.18381 | 0.21267 | -0.86429 | 0.38743 | 0.5967 | 1.482 |
| D630033O11Rik | 154.536 | -0.44802 | 0.16154 | -2.77352 | 0.00555 | 0.03899 | 36.478 |
| D630039A03Rik | 25.7249 | -0.27215 | 0.41835 | -0.65054 | 0.51534 | 0.70568 | 7.815 |
| D630044L22Rik | 28.998 | 0.563237 | 0.36914 | 1.525796 | 0.12706 | 0.30324 | 12.141 |

| | | | | | | | |
|---|---|---|---|---|---|---|---|
| D630045J12Rik | 3213.17 | 0.123647 | 0.0724 | 1.707746 | 0.08768 | 0.23866 | 130.836 |
| D6Ertd474e | 15.2403 | -0.83815 | 0.45745 | -1.83222 | 0.06692 | 0.20024 | 51.385 |
| D6Wsu163e | 1417.05 | 0.024021 | 0.08622 | 0.278611 | 0.78054 | 0.88564 | 36.006 |
| D730003I15Rik | 101.082 | -0.26368 | 0.20947 | -1.25875 | 0.20812 | 0.40775 | 1.737 |
| D730045A05Rik | 28.1316 | -0.01097 | 0.39122 | -0.02805 | 0.97762 | 0.98803 | 2.182 |
| D7Bwg0826e | 16.7103 | -0.06488 | 0.47976 | -0.13524 | 0.89242 | 0.94617 | 3.795 |
| D7Ertd443e | 492.211 | 0.088703 | 0.10875 | 0.815687 | 0.41468 | 0.62268 | 235.382 |
| D830024N08Rik | 98.3788 | 0.257669 | 0.22553 | 1.142508 | 0.25324 | 0.46062 | 9.876 |
| D830025C05Rik | 13.1246 | -0.63682 | 0.48222 | -1.32059 | 0.18664 | 0.38159 | 2.196 |
| D830030K20Rik | 24.9808 | -0.48652 | 0.3638 | -1.33733 | 0.18112 | 0.3745 | 25.864 |
| D830035M03Rik | 20.7041 | 0.045984 | 0.41659 | 0.11038 | 0.91211 | 0.9557 | 2.016 |
| D830036C21Rik | 138.932 | 0.365218 | 0.21252 | 1.718514 | 0.0857 | 0.23512 | 22.399 |
| D830050J10Rik | 26.2066 | -0.03606 | 0.37697 | -0.09567 | 0.92378 | 0.96165 | 1.813 |
| D8Ertd738e | 1259.59 | -0.30241 | 0.08242 | -3.66923 | 0.00024 | 0.00487 | 3.482 |
| D930007J09Rik | 37.1336 | 0.36384 | 0.33128 | 1.098294 | 0.27208 | 0.48148 | 2.459 |
| D930015M05Rik | 181.893 | 0.332678 | 0.15604 | 2.132018 | 0.03301 | 0.12614 | 23.417 |
| D930019O06Rik | 70.5435 | -1.31886 | 0.45757 | -2.8823 | 0.00395 | 0.03118 | 7.207 |
| D930020B18Rik | 36.4853 | 0.12264 | 0.35116 | 0.34924 | 0.72691 | 0.85431 | 52.328 |
| D930028M14Rik | 17.7976 | 0.286866 | 0.45937 | 0.624478 | 0.53231 | 0.71978 | 4.209 |
| D930030I03Rik | 30.2169 | -0.28774 | 0.35375 | -0.8134 | 0.41599 | 0.62364 | 1.562 |
| D930036K23Rik | 79.5614 | -0.13726 | 0.24873 | -0.55185 | 0.58105 | 0.75714 | 3.964 |
| D930048N14Rik | 310.979 | -0.33716 | 0.19004 | -1.77416 | 0.07604 | 0.21772 | 6.728 |
| Daam1 | 3655.66 | -0.06699 | 0.08353 | -0.80201 | 0.42255 | 0.62935 | 161.256 |
| Daam2 | 9233.67 | -0.34274 | 0.09939 | -3.44851 | 0.00056 | 0.0086 | 108.322 |
| Dab1 | 7123.7 | -0.14261 | 0.09261 | -1.53999 | 0.12356 | 0.2979 | 1125.486 |
| Dab2 | 407.263 | -0.55171 | 0.19612 | -2.81318 | 0.00491 | 0.03576 | 140.925 |
| Dab2ip | 4687.56 | -0.29922 | 0.08515 | -3.51389 | 0.00044 | 0.00731 | 172.729 |
| Dach1 | 1132.61 | 0.807068 | 0.21887 | 3.687389 | 0.00023 | 0.00463 | 382.913 |
| Dach2 | 31.3418 | 0.094846 | 0.44114 | 0.215002 | 0.82977 | 0.91274 | 538.877 |
| Dact1 | 154.586 | -0.57835 | 0.22698 | -2.54809 | 0.01083 | 0.06059 | 10.224 |
| Dact2 | 1435.35 | 0.404949 | 0.13577 | 2.982634 | 0.00286 | 0.02542 | 9.253 |
| Dact3 | 4820.51 | -0.49125 | 0.12545 | -3.91603 | 9.00E-05 | 0.00251 | 12.146 |
| Dad1 | 1982.16 | -0.12994 | 0.07873 | -1.65049 | 0.09884 | 0.25828 | 18.626 |
| Dag1 | 5799.67 | 0.055916 | 0.09482 | 0.589706 | 0.55539 | 0.73781 | 59.325 |

| | | | | | | | |
|---|---|---|---|---|---|---|---|
| Dagla | 3379.03 | -0.00807 | 0.07499 | -0.10755 | 0.91435 | 0.95656 | 59.613 |
| Daglb | 1151.72 | -0.2517 | 0.0814 | -3.09202 | 0.00199 | 0.02005 | 41.359 |
| Dalir | 96.2269 | 0.338878 | 0.21118 | 1.604687 | 0.10856 | 0.27385 | 2.17 |
| Dalrd3 | 1839.88 | -0.04737 | 0.07455 | -0.63536 | 0.52519 | 0.71355 | 2.889 |
| Dancr | 99.2435 | -0.11705 | 0.21762 | -0.53788 | 0.59066 | 0.76357 | 2.373 |
| Dand5 | 296.682 | -0.16446 | 0.15509 | -1.06044 | 0.28895 | 0.4991 | 17.006 |
| Dao | 13.1582 | 0.950813 | 0.50358 | 1.88812 | 0.05901 | 0.18494 | 21.98 |
| Dap | 211.888 | -0.1682 | 0.13943 | -1.20634 | 0.22769 | 0.43174 | 50.028 |
| Dap3 | 2996.65 | -0.02826 | 0.06586 | -0.42902 | 0.66791 | 0.81667 | 30.379 |
| Dapk1 | 10026.6 | 0.228659 | 0.08606 | 2.657065 | 0.00788 | 0.04922 | 161.245 |
| Dapk2 | 43.7393 | -0.68914 | 0.30013 | -2.29613 | 0.02167 | 0.09443 | 114.02 |
| Dapk3 | 1196.21 | -0.02761 | 0.09415 | -0.2933 | 0.76929 | 0.87965 | 10.193 |
| Dapl1 | 15.1732 | -1.11887 | 0.62796 | -1.78175 | 0.07479 | 0.21551 | 20.368 |
| Dapp1 | 361.864 | -0.33558 | 0.13693 | -2.45078 | 0.01425 | 0.07201 | 50.539 |
| Dars | 2370.34 | -0.03951 | 0.06651 | -0.59409 | 0.55245 | 0.7354 | 53.662 |
| Dars2 | 740.779 | -0.09155 | 0.11681 | -0.78376 | 0.43318 | 0.63921 | 30.058 |
| Daxx | 1163.2 | -0.1168 | 0.08959 | -1.30375 | 0.19232 | 0.38895 | 6.177 |
| Dazap1 | 2241.31 | 0.027805 | 0.06928 | 0.401334 | 0.68817 | 0.83002 | 26.923 |
| Dazap2 | 4914.03 | -0.18797 | 0.08412 | -2.23458 | 0.02544 | 0.1052 | 5.413 |
| Dazl | 303.407 | 0.994785 | 0.16509 | 6.02563 | 1.68E-09 | 8.18E-07 | 14.242 |
| Dbf4 | 109.895 | -0.23849 | 0.19252 | -1.23881 | 0.21542 | 0.41696 | 25.744 |
| Dbi | 3353.39 | -0.25427 | 0.10902 | -2.33235 | 0.01968 | 0.08867 | 7.799 |
| Dbil5 | 20.8435 | 0.085892 | 0.42827 | 0.200557 | 0.84105 | 0.91851 | 1.059 |
| Dbn1 | 12583.3 | 0.213265 | 0.06881 | 3.09919 | 0.00194 | 0.01971 | 14.683 |
| Dbndd1 | 917.635 | 0.105103 | 0.08858 | 1.186515 | 0.23542 | 0.44058 | 10.746 |
| Dbndd2 | 3809.26 | -0.0926 | 0.0935 | -0.99038 | 0.32199 | 0.53296 | 7.214 |
| Dbnl | 2916.41 | -7.14E-05 | 0.06123 | -0.00117 | 0.99907 | 0.99966 | 12.475 |
| Dbp | 2802.7 | 0.558441 | 0.1453 | 3.843487 | 0.00012 | 0.00305 | 5.116 |
| Dbpht2 | 4008.94 | 0.143858 | 0.10508 | 1.369096 | 0.17097 | 0.36141 | 5.341 |
| Dbr1 | 585.751 | 0.165795 | 0.09013 | 1.839513 | 0.06584 | 0.19821 | 8.703 |
| Dbt | 1352.54 | -0.02945 | 0.08073 | -0.36476 | 0.71529 | 0.84755 | 36.912 |
| Dbx2 | 539.39 | -0.04888 | 0.16057 | -0.30442 | 0.7608 | 0.87469 | 32.398 |
| Dcaf1 | 2284.95 | 0.154149 | 0.0796 | 1.936584 | 0.0528 | 0.17189 | 59.119 |
| Dcaf10 | 1189.04 | 0.178664 | 0.07524 | 2.374741 | 0.01756 | 0.08233 | 37.659 |

| | | | | | | | |
|---|---|---|---|---|---|---|---|
| Dcaf11 | 3228.32 | -0.02362 | 0.06283 | -0.37587 | 0.70702 | 0.84193 | 10.06 |
| Dcaf12 | 2383.3 | -0.14324 | 0.07743 | -1.84986 | 0.06433 | 0.19522 | 23.59 |
| Dcaf12l1 | 2294.58 | 0.183716 | 0.08947 | 2.053487 | 0.04003 | 0.14298 | 3.628 |
| Dcaf12l2 | 42.6495 | 0.167497 | 0.36537 | 0.458431 | 0.64664 | 0.80324 | 2.885 |
| Dcaf13 | 2346.09 | 0.366116 | 0.10869 | 3.368589 | 0.00076 | 0.01056 | 33.992 |
| Dcaf15 | 1210.78 | 0.338812 | 0.08876 | 3.817174 | 0.00013 | 0.0033 | 7.697 |
| Dcaf17 | 1247.99 | 0.274967 | 0.09095 | 3.023187 | 0.0025 | 0.02336 | 43.815 |
| Dcaf4 | 1022.52 | 0.247046 | 0.13364 | 1.848622 | 0.06451 | 0.19551 | 21.455 |
| Dcaf5 | 3768.96 | -0.01411 | 0.05891 | -0.23954 | 0.81068 | 0.90256 | 100.754 |
| Dcaf6 | 3532.54 | 0.077951 | 0.07338 | 1.06222 | 0.28814 | 0.49829 | 131.778 |
| Dcaf7 | 10637 | 0.001239 | 0.07447 | 0.016634 | 0.98673 | 0.99277 | 22.453 |
| Dcaf8 | 8634.24 | -0.07494 | 0.06669 | -1.12357 | 0.2612 | 0.4698 | 48.922 |
| Dcakd | 1158.39 | -0.17218 | 0.12295 | -1.4004 | 0.16139 | 0.34961 | 34.287 |
| Dcbld1 | 663.303 | 0.533415 | 0.16943 | 3.148275 | 0.00164 | 0.01766 | 87.76 |
| Dcbld2 | 1293.92 | -0.35292 | 0.09466 | -3.72818 | 0.00019 | 0.0042 | 61.285 |
| Dcc | 2292.68 | 0.53636 | 0.1532 | 3.501036 | 0.00046 | 0.00757 | 1097.436 |
| Dcdc2a | 431.147 | -0.18514 | 0.15992 | -1.15768 | 0.24699 | 0.45383 | 154.703 |
| Dcdc2b | 421.389 | -0.08739 | 0.15643 | -0.55864 | 0.57641 | 0.75396 | 5.927 |
| Dcdc5 | 30.403 | -0.43956 | 0.35878 | -1.22516 | 0.22051 | 0.42317 | 402.823 |
| Dchs1 | 1328.38 | 0.215249 | 0.09189 | 2.342418 | 0.01916 | 0.08697 | 34.665 |
| Dchs2 | 846.495 | 0.71701 | 0.21242 | 3.375513 | 0.00074 | 0.01036 | 229.262 |
| Dck | 1265.11 | 0.056711 | 0.08029 | 0.706331 | 0.47998 | 0.67751 | 18.286 |
| Dclk1 | 46133.9 | -0.08764 | 0.0903 | -0.97056 | 0.33176 | 0.54275 | 296.705 |
| Dclk2 | 6175.35 | 0.156108 | 0.07917 | 1.971826 | 0.04863 | 0.16288 | 134.702 |
| Dclk3 | 6743.31 | 0.597697 | 0.17465 | 3.42231 | 0.00062 | 0.0092 | 50.038 |
| Dclre1a | 376.594 | -0.06495 | 0.129 | -0.50346 | 0.61464 | 0.78066 | 19.077 |
| Dclre1b | 363.597 | -0.0934 | 0.1443 | -0.64724 | 0.51747 | 0.70733 | 8.84 |
| Dclre1c | 730.002 | 0.017363 | 0.12588 | 0.137933 | 0.89029 | 0.94514 | 40 |
| Dcn | 506.277 | -1.05933 | 0.51142 | -2.07133 | 0.03833 | 0.13927 | 38.535 |
| Dcp1a | 1453.9 | 0.106486 | 0.07126 | 1.494345 | 0.13509 | 0.31502 | 47.529 |
| Dcp1b | 1431.92 | 0.197773 | 0.07235 | 2.733727 | 0.00626 | 0.04235 | 46.364 |
| Dcp2 | 4939.35 | 0.115064 | 0.06428 | 1.790165 | 0.07343 | 0.21339 | 44.468 |
| Dcps | 358.755 | -0.05011 | 0.09937 | -0.50428 | 0.61406 | 0.78027 | 51.654 |
| Dcst1 | 240.263 | 0.06447 | 0.15152 | 0.425473 | 0.67049 | 0.81841 | 15.035 |

| | | | | | | | |
|---|---|---|---|---|---|---|---|
| Dcst2 | 48.0349 | -0.68672 | 0.28537 | -2.40646 | 0.01611 | 0.07804 | 15.575 |
| Dct | 52.0484 | -2 | 0.58343 | -3.53857 | 0.0004 | 0.00691 | 39.455 |
| Dctd | 506.329 | 0.261553 | 0.10453 | 2.502079 | 0.01235 | 0.06592 | 54.142 |
| Dctn1 | 19580.2 | 0.077156 | 0.05417 | 1.424296 | 0.15436 | 0.34035 | 34.198 |
| Dctn2 | 7002.54 | -0.08662 | 0.07422 | -1.16707 | 0.24318 | 0.44942 | 15.583 |
| Dctn3 | 2667.65 | -0.1829 | 0.08367 | -2.18608 | 0.02881 | 0.11468 | 8.373 |
| Dctn4 | 4466.91 | -0.02519 | 0.05432 | -0.46371 | 0.64286 | 0.80073 | 32.582 |
| Dctn5 | 3307.77 | -0.06306 | 0.05984 | -1.05392 | 0.29192 | 0.5018 | 16.004 |
| Dctn6 | 2648.51 | 0.079196 | 0.07469 | 1.060379 | 0.28897 | 0.4991 | 18.377 |
| Dctpp1 | 206.71 | -0.1519 | 0.17056 | -0.89059 | 0.37315 | 0.58352 | 3.751 |
| Dcun1d1 | 2780.49 | 0.203645 | 0.10305 | 1.976115 | 0.04814 | 0.16167 | 45.341 |
| Dcun1d2 | 2507.13 | 0.175347 | 0.06722 | 2.608672 | 0.00909 | 0.05381 | 32.169 |
| Dcun1d3 | 1257.58 | -0.14137 | 0.11502 | -1.22905 | 0.21905 | 0.42125 | 43.503 |
| Dcun1d4 | 6840.95 | 0.103813 | 0.06065 | 1.711634 | 0.08696 | 0.2374 | 79.795 |
| Dcun1d5 | 2182.15 | 0.142243 | 0.06369 | 2.233424 | 0.02552 | 0.10539 | 23.686 |
| Dcx | 2599.46 | 0.082849 | 0.1474 | 0.562054 | 0.57408 | 0.7525 | 77.47 |
| Dcxr | 159.772 | -0.39614 | 0.21554 | -1.83792 | 0.06607 | 0.19859 | 1.883 |
| Dda1 | 2739.03 | 0.066612 | 0.07935 | 0.839425 | 0.40123 | 0.6099 | 6.9 |
| Ddah1 | 5786.7 | -0.24013 | 0.08895 | -2.69978 | 0.00694 | 0.04519 | 135.603 |
| Ddah2 | 433.513 | -0.60736 | 0.18046 | -3.3657 | 0.00076 | 0.01064 | 3.061 |
| Ddb1 | 11325.3 | 0.047047 | 0.06487 | 0.725298 | 0.46827 | 0.66717 | 24.493 |
| Ddb2 | 388.356 | 0.015743 | 0.10707 | 0.147039 | 0.8831 | 0.94166 | 25.411 |
| Ddc | 657.713 | 0.192312 | 0.1467 | 1.310887 | 0.1899 | 0.38596 | 84.044 |
| Ddhd1 | 4123.87 | -0.1548 | 0.06039 | -2.56353 | 0.01036 | 0.05879 | 69.677 |
| Ddhd2 | 3749.65 | 0.092318 | 0.06411 | 1.440103 | 0.14984 | 0.33427 | 29.252 |
| Ddi2 | 4207.18 | 0.088547 | 0.06003 | 1.475113 | 0.14018 | 0.32155 | 45.871 |
| Ddias | 34.5095 | 0.17048 | 0.34627 | 0.492337 | 0.62248 | 0.78561 | 16.723 |
| Ddit3 | 602.501 | 0.008767 | 0.0984 | 0.089095 | 0.92901 | 0.9643 | 5.515 |
| Ddit4 | 4488.1 | 0.028312 | 0.16564 | 0.170929 | 0.86428 | 0.93125 | 2.166 |
| Ddit4l | 326.393 | -0.12751 | 0.16787 | -0.75957 | 0.44751 | 0.65115 | 6.722 |
| Ddn | 47214.3 | 0.001113 | 0.0812 | 0.013713 | 0.98906 | 0.994 | 5.714 |
| Ddo | 374.935 | -0.3583 | 0.14318 | -2.5025 | 0.01233 | 0.06587 | 51.837 |
| Ddost | 3294.84 | 0.025595 | 0.07988 | 0.320438 | 0.74864 | 0.86819 | 7.899 |
| Ddr1 | 2985.24 | -0.24589 | 0.07378 | -3.33303 | 0.00086 | 0.01153 | 23.055 |

| | | | | | | | |
|---|---|---|---|---|---|---|---|
| Ddr2 | 550.106 | -0.6986 | 0.21381 | -3.26738 | 0.00109 | 0.01349 | 138.456 |
| Ddrgk1 | 2781.13 | -0.09266 | 0.08587 | -1.07912 | 0.28054 | 0.49031 | 10.7 |
| Ddt | 12.7408 | -0.37693 | 0.50247 | -0.75016 | 0.45316 | 0.65554 | 2.185 |
| Ddx1 | 5425.45 | -0.0579 | 0.06295 | -0.91972 | 0.35772 | 0.56794 | 32.241 |
| Ddx10 | 887.551 | 0.254244 | 0.10625 | 2.392883 | 0.01672 | 0.07998 | 149.419 |
| Ddx11 | 248.481 | 0.487059 | 0.18488 | 2.634447 | 0.00843 | 0.05116 | 28.655 |
| Ddx17 | 42214.4 | 0.153357 | 0.06094 | 2.516519 | 0.01185 | 0.06407 | 19.006 |
| Ddx18 | 934.035 | -0.01358 | 0.08661 | -0.15682 | 0.87538 | 0.93787 | 14.155 |
| Ddx19a | 1245.73 | 0.082176 | 0.07532 | 1.091034 | 0.27526 | 0.48468 | 22.805 |
| Ddx19b | 2041.89 | 0.009666 | 0.06789 | 0.142381 | 0.88678 | 0.94372 | 28.564 |
| Ddx20 | 831.88 | 0.203714 | 0.09264 | 2.198973 | 0.02788 | 0.11227 | 9.305 |
| Ddx21 | 1658.76 | 0.010454 | 0.06659 | 0.156981 | 0.87526 | 0.93783 | 22.031 |
| Ddx23 | 4699.79 | 0.042036 | 0.06328 | 0.664258 | 0.50653 | 0.69789 | 17.761 |
| Ddx24 | 7081.71 | -0.10281 | 0.06209 | -1.65599 | 0.09772 | 0.2564 | 17.849 |
| Ddx25 | 1936.09 | -0.0524 | 0.0756 | -0.69311 | 0.48824 | 0.68339 | 21.882 |
| Ddx27 | 1678.85 | 0.001892 | 0.08256 | 0.022912 | 0.98172 | 0.99042 | 19.755 |
| Ddx28 | 415.498 | 0.238445 | 0.10384 | 2.296195 | 0.02166 | 0.09443 | 2.262 |
| Ddx31 | 341.828 | 0.285755 | 0.14246 | 2.005815 | 0.04488 | 0.15394 | 65.166 |
| Ddx39a | 754.423 | 0.010645 | 0.09138 | 0.116498 | 0.90726 | 0.95354 | 11.716 |
| Ddx39b | 6825.98 | 0.045932 | 0.05952 | 0.771726 | 0.44028 | 0.64524 | 11.962 |
| Ddx3x | 12594.8 | 0.007148 | 0.09733 | 0.073441 | 0.94146 | 0.97087 | 13.083 |
| Ddx3y | 3387.3 | 0.309421 | 0.14477 | 2.137307 | 0.03257 | 0.12506 | 25.859 |
| Ddx4 | 226.542 | 0.302982 | 0.14236 | 2.128305 | 0.03331 | 0.12673 | 54.143 |
| Ddx41 | 1337.42 | 0.124808 | 0.07601 | 1.642001 | 0.10059 | 0.26113 | 6.249 |
| Ddx42 | 5889.1 | 0.05157 | 0.0774 | 0.666235 | 0.50526 | 0.69698 | 32.214 |
| Ddx46 | 5522.52 | 0.11874 | 0.08261 | 1.437399 | 0.1506 | 0.33547 | 46.23 |
| Ddx47 | 2261.97 | 0.096858 | 0.05951 | 1.627726 | 0.10358 | 0.26613 | 12.165 |
| Ddx49 | 808.786 | -0.00232 | 0.09785 | -0.02369 | 0.9811 | 0.99009 | 9.624 |
| Ddx5 | 60901.2 | 0.332132 | 0.08625 | 3.850743 | 0.00012 | 0.003 | 8.831 |
| Ddx50 | 3319.46 | 0.12587 | 0.07714 | 1.631637 | 0.10276 | 0.26467 | 35.324 |
| Ddx51 | 1132.63 | 0.155378 | 0.08867 | 1.752409 | 0.0797 | 0.22425 | 7.046 |
| Ddx52 | 1024.25 | 0.164711 | 0.07321 | 2.249802 | 0.02446 | 0.10237 | 21.027 |
| Ddx54 | 1306.69 | -0.03255 | 0.10808 | -0.30113 | 0.76332 | 0.876 | 15.854 |
| Ddx55 | 1975.4 | 0.199565 | 0.08649 | 2.307295 | 0.02104 | 0.09264 | 16.797 |

| | | | | | | | |
|---|---|---|---|---|---|---|---|
| Ddx56 | 1194.32 | -0.05645 | 0.08089 | -0.69787 | 0.48526 | 0.68147 | 8.854 |
| Ddx58 | 264.775 | -0.55926 | 0.16445 | -3.40084 | 0.00067 | 0.00971 | 36.056 |
| Ddx59 | 310.16 | 0.077785 | 0.14406 | 0.539948 | 0.58923 | 0.76226 | 24.888 |
| Ddx6 | 13578.2 | 0.029537 | 0.05076 | 0.581933 | 0.56061 | 0.74193 | 35.84 |
| Ddx60 | 49.4324 | -0.83487 | 0.33241 | -2.51159 | 0.01202 | 0.06468 | 110.158 |
| Deaf1 | 1646.43 | 0.012371 | 0.07345 | 0.168441 | 0.86624 | 0.93228 | 30.511 |
| Decr1 | 874.065 | -0.29058 | 0.08171 | -3.55598 | 0.00038 | 0.00656 | 28.268 |
| Decr2 | 1598.56 | -0.02343 | 0.06447 | -0.36341 | 0.7163 | 0.8483 | 9.13 |
| Dedd | 399.013 | 0.045441 | 0.13444 | 0.337992 | 0.73537 | 0.85989 | 13.187 |
| Dedd2 | 1247.66 | 0.096496 | 0.08648 | 1.115778 | 0.26452 | 0.47335 | 20.701 |
| Def6 | 56.4266 | -0.38532 | 0.28595 | -1.34753 | 0.17781 | 0.37008 | 20.831 |
| Def8 | 2688.56 | 0.075511 | 0.06972 | 1.08303 | 0.2788 | 0.48881 | 39.744 |
| Degs1 | 3194.67 | -0.03231 | 0.08568 | -0.3771 | 0.7061 | 0.8415 | 7.033 |
| Degs2 | 186.235 | -0.04551 | 0.14007 | -0.32492 | 0.74524 | 0.866 | 23.601 |
| Dek | 2837.86 | 0.340248 | 0.14758 | 2.305574 | 0.02113 | 0.09293 | 21.427 |
| Dele1 | 1980.79 | -0.02531 | 0.08556 | -0.29582 | 0.76737 | 0.87837 | 15.207 |
| Denn2b | 893.018 | -0.205 | 0.10658 | -1.92346 | 0.05442 | 0.17539 | 179.695 |
| Dennd10 | 1423.69 | 0.006539 | 0.0972 | 0.067281 | 0.94636 | 0.97302 | 24.643 |
| Dennd11 | 5928.68 | 0.282493 | 0.0848 | 3.331139 | 0.00086 | 0.01157 | 34.761 |
| Dennd1a | 2641.16 | 0.15562 | 0.06497 | 2.395327 | 0.01661 | 0.07972 | 488.4 |
| Dennd1b | 1013.4 | 0.293191 | 0.10757 | 2.725622 | 0.00642 | 0.04299 | 215.526 |
| Dennd1c | 106.949 | -0.00503 | 0.18886 | -0.02665 | 0.97874 | 0.98863 | 12.61 |
| Dennd2a | 1654.29 | -0.11971 | 0.08064 | -1.48446 | 0.13769 | 0.31853 | 95.49 |
| Dennd2c | 35.4943 | -0.54504 | 0.34106 | -1.5981 | 0.11002 | 0.27636 | 67.166 |
| Dennd2d | 16.2062 | 0.036065 | 0.44639 | 0.080791 | 0.93561 | 0.96776 | 20.626 |
| Dennd3 | 311.023 | -0.36096 | 0.14803 | -2.43841 | 0.01475 | 0.07343 | 59.683 |
| Dennd4a | 4210.27 | 0.062328 | 0.06157 | 1.012391 | 0.31135 | 0.52167 | 108.328 |
| Dennd4b | 2850.38 | 0.122803 | 0.08581 | 1.431081 | 0.15241 | 0.33764 | 15.485 |
| Dennd4c | 1654.61 | 0.191335 | 0.08279 | 2.311057 | 0.02083 | 0.09204 | 102.049 |
| Dennd5a | 15393.6 | -0.0873 | 0.07112 | -1.22756 | 0.21961 | 0.42204 | 66.691 |
| Dennd5b | 5048.37 | 0.056098 | 0.06572 | 0.853643 | 0.3933 | 0.60236 | 113.61 |
| Dennd6a | 1886.55 | 0.062229 | 0.06336 | 0.982204 | 0.326 | 0.53746 | 60.467 |
| Dennd6b | 3961.54 | 0.243508 | 0.09258 | 2.630167 | 0.00853 | 0.05162 | 14.284 |
| Denr | 1850.77 | -0.10594 | 0.08545 | -1.23979 | 0.21505 | 0.41642 | 21.661 |

| | | | | | | | |
|---|---|---|---|---|---|---|---|
| Depdc5 | 3334.28 | 0.097222 | 0.06976 | 1.393655 | 0.16342 | 0.35213 | 130.536 |
| Depdc7 | 49.5661 | -0.3671 | 0.30467 | -1.20489 | 0.22824 | 0.43223 | 21.095 |
| Depp1 | 63.38 | 0.213959 | 0.30072 | 0.711498 | 0.47678 | 0.67462 | 2.186 |
| Deptor | 1107.53 | -0.67934 | 0.13277 | -5.11682 | 3.11E-07 | 4.36E-05 | 146.955 |
| Dera | 121.485 | -0.22119 | 0.21979 | -1.00638 | 0.31423 | 0.52468 | 102.795 |
| Derl1 | 2928.56 | -0.09248 | 0.07169 | -1.28999 | 0.19705 | 0.39484 | 22.947 |
| Derl2 | 1648.7 | -0.00059 | 0.082 | -0.00718 | 0.99427 | 0.99657 | 12.678 |
| Derl3 | 110.035 | 0.116303 | 0.22424 | 0.518655 | 0.604 | 0.7733 | 2.529 |
| Des | 66.766 | -0.66773 | 0.25282 | -2.64116 | 0.00826 | 0.05062 | 8.251 |
| Desi1 | 4694.9 | -0.24614 | 0.09127 | -2.69686 | 0.007 | 0.04543 | 41.34 |
| Desi2 | 1807.13 | 0.242002 | 0.07045 | 3.434895 | 0.00059 | 0.00888 | 69.885 |
| Det1 | 289.915 | 0.062257 | 0.12631 | 0.492886 | 0.62209 | 0.7853 | 25.717 |
| Dexi | 1192.86 | -0.10343 | 0.06896 | -1.49981 | 0.13366 | 0.31339 | 40.524 |
| Dffa | 2129.32 | -0.12996 | 0.06401 | -2.03009 | 0.04235 | 0.14806 | 16.502 |
| Dffb | 577.837 | 0.046384 | 0.10764 | 0.430931 | 0.66652 | 0.8158 | 10.678 |
| Dgat1 | 1128.39 | 0.175507 | 0.0772 | 2.273268 | 0.02301 | 0.09815 | 9.939 |
| Dgat2 | 4637.13 | 0.245109 | 0.12226 | 2.004838 | 0.04498 | 0.15418 | 29.062 |
| Dgat2l6 | 149.542 | 0.317302 | 0.25274 | 1.25544 | 0.20932 | 0.40935 | 21.307 |
| Dgcr2 | 2932.45 | -0.00829 | 0.06605 | -0.12547 | 0.90015 | 0.94992 | 59.081 |
| Dgcr6 | 1373.64 | -0.15606 | 0.10054 | -1.55224 | 0.1206 | 0.29347 | 18.773 |
| Dgcr8 | 1691.72 | 0.229336 | 0.08152 | 2.813209 | 0.0049 | 0.03576 | 35.299 |
| Dgka | 607.31 | -0.24987 | 0.12086 | -2.06744 | 0.03869 | 0.14007 | 24.722 |
| Dgkb | 16233.3 | 0.415277 | 0.16984 | 2.445097 | 0.01448 | 0.07271 | 816.514 |
| Dgkd | 6733.7 | 0.091159 | 0.07552 | 1.20706 | 0.22741 | 0.43161 | 91.894 |
| Dgke | 4842.43 | -0.01311 | 0.06771 | -0.19363 | 0.84647 | 0.92184 | 31.672 |
| Dgkg | 2849.68 | -0.31703 | 0.17128 | -1.85097 | 0.06417 | 0.19501 | 188.761 |
| Dgkh | 10508.6 | 0.434123 | 0.11798 | 3.679673 | 0.00023 | 0.00475 | 174.027 |
| Dgki | 2810.09 | 0.144203 | 0.12009 | 1.200797 | 0.22983 | 0.43407 | 454.163 |
| Dgkk | 156.281 | 0.945828 | 0.36364 | 2.600977 | 0.0093 | 0.05459 | 124.193 |
| Dgkq | 1650.52 | 0.068482 | 0.0962 | 0.711886 | 0.47654 | 0.67447 | 22.98 |
| Dgkz | 14624 | -0.15467 | 0.09509 | -1.62661 | 0.10382 | 0.26629 | 43.041 |
| Dglucy | 362.656 | 0.166659 | 0.13052 | 1.276848 | 0.20166 | 0.40013 | 117.925 |
| Dguok | 982.116 | 0.286683 | 0.0888 | 3.228408 | 0.00124 | 0.01465 | 26.753 |
| Dhcr24 | 2662.83 | -0.39823 | 0.11091 | -3.59061 | 0.00033 | 0.00602 | 28.076 |

| | | | | | | | |
|---|---|---|---|---|---|---|---|
| Dhcr7 | 1432.36 | 0.110089 | 0.09496 | 1.159348 | 0.24631 | 0.45308 | 25.266 |
| Dhdds | 2755.86 | -0.00743 | 0.06395 | -0.11618 | 0.90751 | 0.95354 | 31.891 |
| Dhdh | 764.849 | 0.334583 | 0.10496 | 3.1878 | 0.00143 | 0.01602 | 17.819 |
| Dhfr | 230.408 | -0.1623 | 0.14668 | -1.10647 | 0.26852 | 0.47785 | 34.328 |
| Dhodh | 831.792 | 0.144679 | 0.10256 | 1.410737 | 0.15832 | 0.3456 | 17.331 |
| Dhps | 560.563 | -0.19374 | 0.09665 | -2.00451 | 0.04502 | 0.15421 | 3.406 |
| Dhrs1 | 1939.01 | -0.23659 | 0.10486 | -2.25637 | 0.02405 | 0.10113 | 6.671 |
| Dhrs11 | 142.23 | -0.19252 | 0.17214 | -1.11839 | 0.2634 | 0.47216 | 8.139 |
| Dhrs13 | 250.308 | 0.26138 | 0.14594 | 1.791017 | 0.07329 | 0.2132 | 5.587 |
| Dhrs13os | 20.294 | 0.482358 | 0.50839 | 0.948801 | 0.34272 | 0.55356 | 2.654 |
| Dhrs3 | 620.399 | -0.20055 | 0.1032 | -1.94329 | 0.05198 | 0.17024 | 35.383 |
| Dhrs4 | 551.006 | -0.05893 | 0.1066 | -0.55282 | 0.58039 | 0.75658 | 11.583 |
| Dhrs7 | 1323.07 | -0.13014 | 0.10991 | -1.18408 | 0.23638 | 0.44163 | 14.557 |
| Dhrs7b | 830.103 | -0.05314 | 0.0983 | -0.54057 | 0.58881 | 0.762 | 29.565 |
| Dhtkd1 | 728.584 | -0.00274 | 0.08729 | -0.03144 | 0.97492 | 0.98738 | 46.678 |
| Dhx15 | 6409.95 | 0.112753 | 0.05646 | 1.997133 | 0.04581 | 0.15602 | 40.312 |
| Dhx16 | 1538.1 | -0.00447 | 0.06394 | -0.06993 | 0.94425 | 0.97202 | 12.852 |
| Dhx29 | 1767.6 | 0.172498 | 0.09196 | 1.875721 | 0.06069 | 0.18808 | 41.979 |
| Dhx30 | 4930.41 | 0.128695 | 0.06738 | 1.910124 | 0.05612 | 0.1791 | 33.511 |
| Dhx32 | 1740.56 | 0.211332 | 0.11682 | 1.808988 | 0.07045 | 0.2074 | 61.785 |
| Dhx33 | 2177.28 | 0.101031 | 0.06364 | 1.587543 | 0.11239 | 0.28014 | 20.347 |
| Dhx34 | 681.728 | 0.002431 | 0.10549 | 0.023042 | 0.98162 | 0.9904 | 24.891 |
| Dhx35 | 1282.32 | 0.086031 | 0.08794 | 0.978332 | 0.32791 | 0.53926 | 63.408 |
| Dhx36 | 5403.44 | 0.250363 | 0.10071 | 2.485912 | 0.01292 | 0.06794 | 38.992 |
| Dhx37 | 813.099 | 0.044017 | 0.09973 | 0.441368 | 0.65895 | 0.81103 | 20.264 |
| Dhx38 | 2000.36 | 0.0596 | 0.06553 | 0.909547 | 0.36306 | 0.57343 | 17.851 |
| Dhx40 | 1743.53 | 0.242885 | 0.08038 | 3.021797 | 0.00251 | 0.02344 | 38.901 |
| Dhx57 | 2201.58 | 0.264901 | 0.08129 | 3.258692 | 0.00112 | 0.01374 | 52.168 |
| Dhx58 | 80.4832 | -0.12937 | 0.25753 | -0.50232 | 0.61544 | 0.78121 | 9.388 |
| Dhx8 | 2117.69 | 0.022312 | 0.07797 | 0.286155 | 0.77476 | 0.88253 | 34.44 |
| Dhx9 | 8046.59 | 0.134352 | 0.07136 | 1.882689 | 0.05974 | 0.18618 | 31.903 |
| Diablo | 362.242 | 0.162294 | 0.12794 | 1.26848 | 0.20463 | 0.40343 | 14.412 |
| Diaph1 | 3870.09 | 0.223499 | 0.1334 | 1.675434 | 0.09385 | 0.24984 | 91.876 |
| Diaph2 | 2881.37 | 0.444381 | 0.17505 | 2.538649 | 0.01113 | 0.06149 | 716.093 |

| | | | | | | | |
|---|---|---|---|---|---|---|---|
| Diaph3 | 92.1625 | -0.24718 | 0.24174 | -1.02252 | 0.30653 | 0.5165 | 485.869 |
| Dicer1 | 4589.37 | 0.145615 | 0.06525 | 2.231563 | 0.02564 | 0.10577 | 64.211 |
| Dido1 | 5009.31 | 0.162015 | 0.05373 | 3.01555 | 0.00257 | 0.02377 | 52.036 |
| Dimt1 | 1120.72 | 0.112599 | 0.08592 | 1.3105 | 0.19003 | 0.38608 | 13.098 |
| Dio2 | 8198.46 | -0.28047 | 0.19347 | -1.44969 | 0.14714 | 0.33033 | 14.487 |
| Dio3 | 53.5893 | 1.003501 | 0.38458 | 2.609316 | 0.00907 | 0.05373 | 2.03 |
| Dio3os | 21.6846 | 1.108612 | 0.52269 | 2.120971 | 0.03392 | 0.12861 | 2.792 |
| Dip2a | 4109.09 | -0.42105 | 0.1233 | -3.41495 | 0.00064 | 0.00933 | 85.863 |
| Dip2b | 5033.34 | -0.03277 | 0.06383 | -0.51344 | 0.60765 | 0.77569 | 180.81 |
| Dip2c | 5056.98 | 0.224468 | 0.0607 | 3.697896 | 0.00022 | 0.00451 | 392.401 |
| Dipk1a | 936.675 | 0.050951 | 0.10401 | 0.489863 | 0.62423 | 0.78686 | 79.033 |
| Dipk1b | 775.291 | -0.05142 | 0.09681 | -0.53112 | 0.59534 | 0.76727 | 8.041 |
| Dipk1c | 306.006 | -0.2227 | 0.13823 | -1.61103 | 0.10717 | 0.27174 | 21.201 |
| Dipk2a | 1718.5 | -0.24978 | 0.16566 | -1.50776 | 0.13162 | 0.31033 | 20.218 |
| Dipk2b | 62.7027 | -0.10508 | 0.27941 | -0.37606 | 0.70687 | 0.84193 | 46.517 |
| Diras1 | 5566.09 | -0.19504 | 0.0813 | -2.39906 | 0.01644 | 0.07912 | 6.074 |
| Diras2 | 9434.88 | -0.16826 | 0.05965 | -2.8206 | 0.00479 | 0.03532 | 26.9 |
| Dis3 | 978.072 | 0.11279 | 0.1455 | 0.77518 | 0.43823 | 0.64363 | 24.565 |
| Dis3l | 1474.09 | 0.075242 | 0.06772 | 1.111018 | 0.26656 | 0.47588 | 34.533 |
| Dis3l2 | 977.992 | -0.04633 | 0.0803 | -0.57693 | 0.56398 | 0.74426 | 346.288 |
| Disc1 | 169.006 | 0.163729 | 0.15511 | 1.055578 | 0.29116 | 0.50117 | 207.664 |
| Disp1 | 348.638 | 0.300248 | 0.11487 | 2.613733 | 0.00896 | 0.0533 | 135.257 |
| Disp2 | 12116.6 | -0.22701 | 0.1098 | -2.06745 | 0.03869 | 0.14007 | 31.575 |
| Disp3 | 2015.31 | -0.55905 | 0.14108 | -3.96275 | 7.41E-05 | 0.00218 | 47.702 |
| Dixdc1 | 5434.17 | 0.14201 | 0.08064 | 1.761001 | 0.07824 | 0.22144 | 76.766 |
| Dkc1 | 1372.29 | 0.184564 | 0.08663 | 2.130586 | 0.03312 | 0.12644 | 13.924 |
| Dkk2 | 13.7769 | -0.39152 | 0.49967 | -0.78355 | 0.4333 | 0.63924 | 95.013 |
| Dkk3 | 6283.38 | -0.0404 | 0.12604 | -0.32057 | 0.74854 | 0.86814 | 43.041 |
| Dkkl1 | 154.822 | -1.49929 | 0.28589 | -5.24421 | 1.57E-07 | 2.54E-05 | 4.629 |
| Dlat | 5616.3 | -0.07817 | 0.0878 | -0.89035 | 0.37328 | 0.58357 | 25.148 |
| Dlc1 | 2268.3 | -0.27639 | 0.08879 | -3.11283 | 0.00185 | 0.01911 | 385.393 |
| Dld | 5773.14 | 0.094109 | 0.08831 | 1.065623 | 0.28659 | 0.4965 | 20.177 |
| Dlec1 | 118.288 | 0.106225 | 0.17971 | 0.591099 | 0.55445 | 0.73705 | 45.769 |
| Dleu2 | 350.862 | -0.0535 | 0.1478 | -0.36197 | 0.71737 | 0.84903 | 79.558 |

| | | | | | | | |
|---|---|---|---|---|---|---|---|
| Dleu7 | 190.754 | -0.13748 | 0.13158 | -1.04486 | 0.29609 | 0.50641 | 16.817 |
| Dlg1 | 11681.8 | 0.246574 | 0.07916 | 3.114873 | 0.00184 | 0.01906 | 211.687 |
| Dlg2 | 22084.1 | 0.256449 | 0.07553 | 3.395191 | 0.00069 | 0.00985 | 1972.576 |
| Dlg3 | 9895.3 | 0.038933 | 0.06975 | 0.558182 | 0.57672 | 0.75408 | 50.689 |
| Dlg4 | 30720 | 0.069629 | 0.07163 | 0.972129 | 0.33099 | 0.54208 | 30.581 |
| Dlg5 | 2634.06 | 0.048568 | 0.0821 | 0.591593 | 0.55412 | 0.73681 | 111.968 |
| Dlgap1 | 13269.9 | -0.75331 | 0.19175 | -3.92867 | 8.54E-05 | 0.00242 | 852.341 |
| Dlgap2 | 3496.16 | 0.253357 | 0.09537 | 2.656601 | 0.00789 | 0.04923 | 751.816 |
| Dlgap3 | 13562.6 | -0.01173 | 0.09258 | -0.12668 | 0.89919 | 0.94945 | 67.819 |
| Dlgap4 | 11385.2 | 0.013031 | 0.06829 | 0.190814 | 0.84867 | 0.92307 | 150.659 |
| Dlgap5 | 16.0013 | -1.19595 | 0.58051 | -2.06018 | 0.03938 | 0.14147 | 30.629 |
| Dlk1 | 1626.48 | 1.71274 | 0.51395 | 3.332511 | 0.00086 | 0.01154 | 10.514 |
| Dlk2 | 1050.3 | 0.299007 | 0.10403 | 2.874158 | 0.00405 | 0.03175 | 5.851 |
| Dll1 | 586.804 | 0.237932 | 0.09426 | 2.524249 | 0.01159 | 0.06316 | 9.519 |
| Dll3 | 39.5136 | -0.12755 | 0.26876 | -0.4746 | 0.63507 | 0.79438 | 8.686 |
| Dll4 | 121.379 | 0.017357 | 0.17116 | 0.101408 | 0.91923 | 0.95914 | 10.179 |
| Dlst | 6263.15 | -0.05722 | 0.05483 | -1.04349 | 0.29672 | 0.50689 | 24.013 |
| Dlx1 | 696.041 | -0.9883 | 0.33212 | -2.97574 | 0.00292 | 0.02575 | 5.869 |
| Dlx1as | 745.562 | -0.07647 | 0.13164 | -0.58091 | 0.5613 | 0.74238 | 21.455 |
| Dlx2 | 387.161 | -0.2221 | 0.18424 | -1.20553 | 0.228 | 0.43189 | 3.347 |
| Dlx3 | 14.2579 | -0.85846 | 0.53092 | -1.61694 | 0.10589 | 0.26954 | 5.178 |
| Dlx4 | 14.7583 | -0.85986 | 0.5996 | -1.43407 | 0.15155 | 0.33655 | 5.817 |
| Dlx4os | 15.7368 | -0.87531 | 0.49297 | -1.77558 | 0.0758 | 0.21741 | 13.996 |
| Dlx5 | 738.379 | 0.445003 | 0.16554 | 2.688124 | 0.00719 | 0.04631 | 4.281 |
| Dlx6 | 970.29 | 0.580898 | 0.23662 | 2.454978 | 0.01409 | 0.07148 | 5.235 |
| Dlx6os1 | 2894.31 | 0.218056 | 0.1256 | 1.736162 | 0.08254 | 0.22912 | 121.893 |
| Dmac1 | 605.907 | -0.19319 | 0.1081 | -1.7871 | 0.07392 | 0.21422 | 0.952 |
| Dmac2 | 1785.77 | -0.00432 | 0.08361 | -0.05165 | 0.95881 | 0.97995 | 6.138 |
| Dmac2l | 745.389 | 0.055596 | 0.11865 | 0.468575 | 0.63937 | 0.79781 | 19.711 |
| Dmap1 | 974.679 | 0.078827 | 0.08562 | 0.920642 | 0.35724 | 0.56761 | 7.593 |
| Dmc1 | 121.236 | 0.020172 | 0.19895 | 0.101389 | 0.91924 | 0.95914 | 43.613 |
| Dmd | 3201.37 | 0.013049 | 0.13768 | 0.09478 | 0.92449 | 0.96202 | 2257.272 |
| Dmgdh | 22.5586 | -0.49298 | 0.38754 | -1.27208 | 0.20335 | 0.40179 | 78.401 |
| Dmkn | 180.899 | 0.559838 | 0.22205 | 2.521211 | 0.0117 | 0.06359 | 17.308 |

| | | | | | | | |
|---|---|---|---|---|---|---|---|
| Dmp1 | 379.661 | -0.77951 | 0.18731 | -4.16153 | 3.16E-05 | 0.0012 | 11.49 |
| Dmpk | 1981.33 | -0.03195 | 0.09411 | -0.33948 | 0.73425 | 0.85938 | 9.973 |
| Dmrta1 | 14.9108 | -0.01863 | 0.64314 | -0.02896 | 0.9769 | 0.98795 | 15.337 |
| Dmrta2 | 55.5772 | -0.05975 | 0.25953 | -0.23021 | 0.81793 | 0.90637 | 5.635 |
| Dmrta2os | 16.89 | -0.11317 | 0.51017 | -0.22183 | 0.82444 | 0.90912 | 2.083 |
| Dmrtb1 | 18.6889 | 1.541545 | 0.45295 | 3.403354 | 0.00067 | 0.00964 | 7.941 |
| Dmrtc1a | 173.686 | 0.120909 | 0.24531 | 0.492872 | 0.6221 | 0.7853 | 5.472 |
| Dmtf1 | 3836.39 | 0.247949 | 0.1088 | 2.278976 | 0.02267 | 0.09728 | 42.976 |
| Dmtn | 12349.5 | -0.03361 | 0.05535 | -0.60733 | 0.54363 | 0.72894 | 34.786 |
| Dmwd | 4161.29 | -0.03992 | 0.0673 | -0.59319 | 0.55305 | 0.73586 | 6.55 |
| Dmxl1 | 4599.33 | 0.224569 | 0.1413 | 1.589339 | 0.11198 | 0.2795 | 132.804 |
| Dmxl2 | 13319.3 | 0.045571 | 0.06567 | 0.693987 | 0.48769 | 0.68296 | 136.469 |
| Dna2 | 151.439 | 0.025159 | 0.16284 | 0.1545 | 0.87722 | 0.93864 | 27.16 |
| Dnaaf1 | 16.7528 | 1.262356 | 0.47554 | 2.6546 | 0.00794 | 0.04936 | 23.22 |
| Dnaaf10 | 546.823 | 0.296328 | 0.10226 | 2.897924 | 0.00376 | 0.03017 | 51.69 |
| Dnaaf2 | 29.0787 | 0.014421 | 0.37722 | 0.038229 | 0.96951 | 0.98458 | 9.343 |
| Dnaaf3 | 28.3166 | -0.06953 | 0.36937 | -0.18824 | 0.85069 | 0.92407 | 9.521 |
| Dnaaf5 | 347.656 | 0.151429 | 0.12577 | 1.203971 | 0.2286 | 0.43266 | 36.288 |
| Dnaaf8 | 225.658 | 0.113 | 0.16019 | 0.705417 | 0.48055 | 0.67789 | 14.737 |
| Dnaaf9 | 9204.27 | 0.004999 | 0.06487 | 0.077073 | 0.93857 | 0.96942 | 200.207 |
| Dnah1 | 665.631 | 0.536943 | 0.1469 | 3.655133 | 0.00026 | 0.00504 | 63.522 |
| Dnah10 | 118.057 | 0.237665 | 0.1968 | 1.207666 | 0.22718 | 0.43135 | 109.224 |
| Dnah11 | 68 | -2 | 0.97838 | -2.18501 | 0.02889 | 0.11492 | 321.062 |
| Dnah12 | 21.8466 | 0.84663 | 0.39059 | 2.167558 | 0.03019 | 0.11869 | 199.232 |
| Dnah14 | 58.3304 | -0.58626 | 0.30924 | -1.89582 | 0.05798 | 0.18305 | 239.216 |
| Dnah17 | 175.969 | 0.081024 | 0.16052 | 0.504748 | 0.61374 | 0.78 | 108.912 |
| Dnah2 | 525.538 | 0.738012 | 0.16434 | 4.490771 | 7.10E-06 | 0.00043 | 128.302 |
| Dnah3 | 52.9723 | 0.049536 | 0.32551 | 0.152178 | 0.87905 | 0.93947 | 172.61 |
| Dnah5 | 167.51 | -0.5784 | 0.32639 | -1.77211 | 0.07638 | 0.21848 | 268.301 |
| Dnah6 | 48.5501 | 0.954802 | 0.29533 | 3.232973 | 0.00123 | 0.01453 | 204.046 |
| Dnah7a | 67.5707 | -0.16957 | 0.22119 | -0.76663 | 0.4433 | 0.64781 | 309.779 |
| Dnah7b | 242.742 | -0.01439 | 0.19442 | -0.07403 | 0.94098 | 0.97075 | 307.236 |
| Dnah7c | 106.429 | -0.13176 | 0.19592 | -0.67253 | 0.50124 | 0.69395 | 381.885 |
| Dnah8 | 29.5523 | 0.20348 | 0.3587 | 0.567268 | 0.57053 | 0.74977 | 253.012 |

| | | | | | | | |
|---|---|---|---|---|---|---|---|
| Dnah9 | 136.671 | 0.13742 | 0.19747 | 0.695908 | 0.48649 | 0.68239 | 337.27 |
| Dnai1 | 141.938 | -0.04397 | 0.16644 | -0.26417 | 0.79165 | 0.89165 | 68.384 |
| Dnai2 | 111.167 | -0.39332 | 0.26813 | -1.46689 | 0.14241 | 0.32434 | 30.482 |
| Dnai4 | 880.285 | 0.286439 | 0.09683 | 2.958306 | 0.00309 | 0.02661 | 76.491 |
| Dnaja1 | 15021.3 | 0.105024 | 0.06313 | 1.663703 | 0.09617 | 0.25395 | 15 |
| Dnaja2 | 8825.03 | 0.049147 | 0.05857 | 0.83905 | 0.40144 | 0.60999 | 17.712 |
| Dnaja3 | 2632.03 | -0.01439 | 0.07373 | -0.19521 | 0.84523 | 0.9209 | 67.707 |
| Dnaja4 | 2014.16 | 0.101944 | 0.0809 | 1.260204 | 0.2076 | 0.40716 | 17.443 |
| Dnajb1 | 2361.32 | -0.06348 | 0.11532 | -0.55048 | 0.58199 | 0.75769 | 4.461 |
| Dnajb11 | 2301.36 | -0.16162 | 0.0935 | -1.72862 | 0.08388 | 0.23177 | 21.79 |
| Dnajb12 | 2097.22 | 0.010891 | 0.07718 | 0.141112 | 0.88778 | 0.94402 | 19.747 |
| Dnajb13 | 26.4684 | -0.49617 | 0.48619 | -1.02053 | 0.30748 | 0.51734 | 13.245 |
| Dnajb14 | 6957.78 | 0.025013 | 0.06653 | 0.375967 | 0.70694 | 0.84193 | 48.883 |
| Dnajb2 | 3728.34 | -0.10373 | 0.0848 | -1.22325 | 0.22124 | 0.42416 | 9.287 |
| Dnajb3 | 37.6766 | -0.06732 | 0.31434 | -0.21415 | 0.83043 | 0.91311 | 1.051 |
| Dnajb4 | 5162.3 | 0.333276 | 0.09992 | 3.335437 | 0.00085 | 0.01147 | 31.792 |
| Dnajb5 | 7316.64 | -0.14914 | 0.12929 | -1.15354 | 0.24869 | 0.45604 | 9.612 |
| Dnajb6 | 5635.24 | -0.06272 | 0.07291 | -0.86014 | 0.38971 | 0.59891 | 82.447 |
| Dnajb6-ps | 98.2229 | -0.18542 | 0.20634 | -0.89863 | 0.36885 | 0.5795 | 0.691 |
| Dnajb9 | 2227.31 | -0.05218 | 0.0813 | -0.64176 | 0.52103 | 0.71027 | 4.768 |
| Dnajc1 | 1725.5 | 0.019496 | 0.06707 | 0.290683 | 0.77129 | 0.88064 | 197.177 |
| Dnajc10 | 3935.26 | 0.082691 | 0.08656 | 0.955315 | 0.33942 | 0.55024 | 38.578 |
| Dnajc11 | 3312.04 | 0.01262 | 0.0604 | 0.20895 | 0.83449 | 0.91508 | 48.447 |
| Dnajc12 | 549.321 | 0.076987 | 0.11955 | 0.64396 | 0.5196 | 0.70925 | 28.134 |
| Dnajc13 | 3766.74 | 0.118462 | 0.06097 | 1.943108 | 0.052 | 0.17024 | 111.649 |
| Dnajc14 | 2368.06 | -0.15147 | 0.08461 | -1.79031 | 0.0734 | 0.21336 | 15.385 |
| Dnajc15 | 2994.86 | -0.03096 | 0.08606 | -0.35982 | 0.71898 | 0.8499 | 54.445 |
| Dnajc16 | 4268.28 | 0.030696 | 0.05617 | 0.54651 | 0.58472 | 0.75938 | 30.743 |
| Dnajc17 | 281.298 | 0.277796 | 0.11228 | 2.474186 | 0.01335 | 0.06933 | 36.296 |
| Dnajc18 | 5612.35 | -0.00945 | 0.06062 | -0.15594 | 0.87608 | 0.93819 | 37.312 |
| Dnajc19 | 729.113 | 0.012598 | 0.09365 | 0.134525 | 0.89299 | 0.94632 | 25.302 |
| Dnajc19-ps | 25.7004 | 0.052857 | 0.36591 | 0.144454 | 0.88514 | 0.94297 | 0.351 |
| Dnajc2 | 1685.11 | 0.357613 | 0.12514 | 2.857767 | 0.00427 | 0.03288 | 27.985 |
| Dnajc21 | 842.643 | -0.02759 | 0.13593 | -0.20299 | 0.83914 | 0.91722 | 23.761 |

| | | | | | | | |
|---|---|---|---|---|---|---|---|
| Dnajc24 | 408.261 | 0.024625 | 0.11629 | 0.211754 | 0.8323 | 0.91402 | 36.841 |
| Dnajc25 | 248.505 | -0.05267 | 0.13072 | -0.4029 | 0.68702 | 0.82926 | 30.512 |
| Dnajc27 | 7697.33 | -0.05799 | 0.06639 | -0.87347 | 0.38241 | 0.59192 | 28.024 |
| Dnajc28 | 762.87 | 0.052892 | 0.08742 | 0.605057 | 0.54514 | 0.73012 | 4.773 |
| Dnajc3 | 3661.58 | -0.12288 | 0.09287 | -1.3232 | 0.18577 | 0.38054 | 43.722 |
| Dnajc30 | 330.912 | -0.24166 | 0.14809 | -1.63185 | 0.10271 | 0.26462 | 1.661 |
| Dnajc4 | 434.41 | -0.06236 | 0.10946 | -0.56972 | 0.56887 | 0.7486 | 4.662 |
| Dnajc5 | 18359 | -0.03232 | 0.05797 | -0.55755 | 0.57715 | 0.75441 | 34.649 |
| Dnajc6 | 11018.7 | -0.15137 | 0.08217 | -1.84219 | 0.06545 | 0.19735 | 146.169 |
| Dnajc7 | 6428.9 | 0.322363 | 0.07779 | 4.143756 | 3.42E-05 | 0.00125 | 37.371 |
| Dnajc8 | 3915.71 | 0.012552 | 0.11473 | 0.109405 | 0.91288 | 0.95597 | 18.193 |
| Dnajc9 | 1412.53 | 0.032916 | 0.08924 | 0.368851 | 0.71224 | 0.84507 | 4.273 |
| Dnal1 | 8648.75 | 0.030324 | 0.06871 | 0.44131 | 0.65899 | 0.81103 | 33.133 |
| Dnal4 | 1639.26 | -0.01749 | 0.06793 | -0.25754 | 0.79676 | 0.89517 | 16.397 |
| Dnali1 | 21.5777 | -0.12477 | 0.46441 | -0.26868 | 0.78818 | 0.89 | 10.366 |
| Dnase1 | 316.319 | 0.268165 | 0.11203 | 2.393646 | 0.01668 | 0.07993 | 3.083 |
| Dnase1l1 | 63.481 | -0.22625 | 0.22773 | -0.99352 | 0.32046 | 0.53121 | 9.121 |
| Dnase1l2 | 462.97 | 0.296487 | 0.12165 | 2.437267 | 0.0148 | 0.07363 | 3.025 |
| Dnase2a | 227.684 | -0.32424 | 0.12704 | -2.55221 | 0.0107 | 0.06008 | 14.356 |
| Dnd1 | 104.868 | 0.356951 | 0.20118 | 1.774258 | 0.07602 | 0.21771 | 2.544 |
| Dner | 8772.59 | -0.41628 | 0.11768 | -3.53748 | 0.0004 | 0.00692 | 326.383 |
| Dnhd1 | 329.417 | -0.31822 | 0.18954 | -1.67891 | 0.09317 | 0.24849 | 70.973 |
| Dnlz | 1614.77 | -0.1211 | 0.08011 | -1.51174 | 0.1306 | 0.30894 | 3.99 |
| Dnm1 | 33269.6 | -0.16993 | 0.06464 | -2.62902 | 0.00856 | 0.05173 | 44.868 |
| Dnm1l | 10880.7 | 0.220123 | 0.10514 | 2.093715 | 0.03629 | 0.13419 | 46.73 |
| Dnm2 | 3549.28 | -0.12203 | 0.06469 | -1.88648 | 0.05923 | 0.18523 | 82.852 |
| Dnm3 | 8208.7 | 0.297266 | 0.12493 | 2.379398 | 0.01734 | 0.08167 | 495.582 |
| Dnm3os | 13.266 | -0.55493 | 0.64086 | -0.86591 | 0.38654 | 0.59587 | 7.928 |
| Dnmbp | 1104.74 | 0.289551 | 0.11804 | 2.452896 | 0.01417 | 0.07176 | 93.371 |
| Dnmt1 | 3133.9 | 0.01754 | 0.07159 | 0.245021 | 0.80644 | 0.90039 | 52.68 |
| Dnmt3a | 8849.57 | 0.22478 | 0.0895 | 2.511597 | 0.01202 | 0.06468 | 108.437 |
| Dnmt3aos | 41.2563 | 0.483083 | 0.27976 | 1.726762 | 0.08421 | 0.23239 | 2.952 |
| Dnmt3b | 44.3582 | -0.34137 | 0.27508 | -1.24095 | 0.21462 | 0.41602 | 38.281 |
| Dnpep | 1533.83 | 0.074217 | 0.09247 | 0.802594 | 0.42221 | 0.62898 | 10.095 |

| | | | | | | | |
|---|---|---|---|---|---|---|---|
| Dnph1 | 130.703 | 0.070368 | 0.16501 | 0.426452 | 0.66978 | 0.81795 | 2.914 |
| Dnttip1 | 617.364 | -0.04146 | 0.08758 | -0.47333 | 0.63597 | 0.79503 | 22.625 |
| Dnttip2 | 1177.67 | 0.119897 | 0.09147 | 1.310832 | 0.18991 | 0.38596 | 10.884 |
| Doc2a | 223.99 | -0.67897 | 0.27764 | -2.44547 | 0.01447 | 0.07271 | 5.29 |
| Doc2b | 5104.65 | 0.60611 | 0.1741 | 3.481472 | 0.0005 | 0.0079 | 27.084 |
| Doc2g | 126.877 | -0.44162 | 0.25111 | -1.75868 | 0.07863 | 0.22224 | 3.693 |
| Dock1 | 1736.49 | -0.21728 | 0.10246 | -2.1206 | 0.03396 | 0.12867 | 502.986 |
| Dock10 | 4092.37 | 0.414151 | 0.1527 | 2.712189 | 0.00668 | 0.04414 | 257.455 |
| Dock11 | 1059.79 | 0.488757 | 0.13356 | 3.659359 | 0.00025 | 0.00498 | 187.731 |
| Dock2 | 236.383 | -0.35766 | 0.1906 | -1.87652 | 0.06058 | 0.18785 | 497.905 |
| Dock3 | 14099.6 | -0.0303 | 0.06977 | -0.43424 | 0.66412 | 0.81447 | 339.085 |
| Dock4 | 6327.5 | 0.416699 | 0.0989 | 4.213545 | 2.51E-05 | 0.00103 | 400.923 |
| Dock5 | 847.57 | -0.2433 | 0.11587 | -2.09984 | 0.03574 | 0.13291 | 181.308 |
| Dock6 | 1191.3 | 0.119666 | 0.10533 | 1.136114 | 0.25591 | 0.46368 | 52.776 |
| Dock7 | 2502.4 | 0.281515 | 0.08731 | 3.224387 | 0.00126 | 0.01476 | 184.245 |
| Dock8 | 299.572 | -0.20937 | 0.17469 | -1.19856 | 0.2307 | 0.43503 | 202.899 |
| Dock9 | 7654.75 | -0.48589 | 0.11764 | -4.13018 | 3.62E-05 | 0.00131 | 255.792 |
| Dohh | 271.485 | -0.21286 | 0.1209 | -1.76067 | 0.07829 | 0.2215 | 3.925 |
| Dok1 | 52.8475 | -0.61724 | 0.24781 | -2.49078 | 0.01275 | 0.06731 | 2.538 |
| Dok2 | 75.5662 | 0.181291 | 0.26532 | 0.683283 | 0.49443 | 0.68865 | 12.46 |
| Dok3 | 894.196 | 0.262914 | 0.11844 | 2.219776 | 0.02643 | 0.10813 | 6.066 |
| Dok4 | 497.644 | -0.04152 | 0.09808 | -0.42331 | 0.67207 | 0.81969 | 12.503 |
| Dok5 | 581.897 | -0.22863 | 0.11563 | -1.97733 | 0.048 | 0.16134 | 147.963 |
| Dok6 | 7973.07 | 0.119651 | 0.14069 | 0.850466 | 0.39507 | 0.60407 | 477.105 |
| Dok7 | 875.79 | -0.00953 | 0.0942 | -0.10116 | 0.91942 | 0.95923 | 31.074 |
| Dolk | 797.589 | -0.21595 | 0.08347 | -2.58721 | 0.00968 | 0.05614 | 2.104 |
| Dolpp1 | 690.105 | 0.012002 | 0.1062 | 0.113006 | 0.91003 | 0.95466 | 8.276 |
| Donson | 1005.18 | 0.224199 | 0.08579 | 2.613461 | 0.00896 | 0.05331 | 11.963 |
| Dop1a | 3811.9 | 0.233041 | 0.06339 | 3.676495 | 0.00024 | 0.00478 | 88.77 |
| Dop1b | 4865.28 | 0.148825 | 0.07032 | 2.11649 | 0.0343 | 0.12942 | 98.687 |
| Dot1l | 6555.83 | 0.1396 | 0.1029 | 1.356679 | 0.17488 | 0.36656 | 40.256 |
| Dpagt1 | 626.064 | -0.09054 | 0.1186 | -0.76345 | 0.44519 | 0.64913 | 7.882 |
| Dpcd | 1855.81 | -0.23919 | 0.08401 | -2.84718 | 0.00441 | 0.03356 | 19.149 |
| Dpf1 | 2667.09 | 0.403756 | 0.07566 | 5.336513 | 9.48E-08 | 1.66E-05 | 14.293 |

| | | | | | | | |
|---|---|---|---|---|---|---|---|
| Dpf2 | 3611.14 | -0.00341 | 0.06047 | -0.05632 | 0.95509 | 0.97798 | 16.495 |
| Dpf3 | 193.317 | -0.12776 | 0.15431 | -0.82795 | 0.4077 | 0.61541 | 273.972 |
| Dph1 | 551.148 | 0.134232 | 0.11128 | 1.206242 | 0.22772 | 0.43174 | 13.599 |
| Dph2 | 553.453 | 0.149821 | 0.10487 | 1.428606 | 0.15312 | 0.33865 | 3.39 |
| Dph3 | 993.82 | -0.27391 | 0.10446 | -2.62211 | 0.00874 | 0.05244 | 5.164 |
| Dph5 | 395.123 | -0.04738 | 0.10509 | -0.45086 | 0.65209 | 0.80645 | 46.525 |
| Dph6 | 1252.32 | 0.379519 | 0.10113 | 3.752903 | 0.00017 | 0.00393 | 138.549 |
| Dph7 | 1040.48 | 0.163318 | 0.06867 | 2.378257 | 0.01739 | 0.08185 | 9.764 |
| Dpm1 | 866.497 | 0.259982 | 0.0844 | 3.080444 | 0.00207 | 0.02052 | 21.544 |
| Dpm2 | 1653.44 | 0.049139 | 0.07107 | 0.691384 | 0.48932 | 0.68444 | 2.722 |
| Dpm3 | 459.285 | 0.308968 | 0.14108 | 2.190018 | 0.02852 | 0.11395 | 7.722 |
| Dpp10 | 6843.21 | 0.349369 | 0.09722 | 3.593778 | 0.00033 | 0.00599 | 1524.577 |
| Dpp3 | 1567.53 | -0.03255 | 0.07399 | -0.43992 | 0.65999 | 0.81195 | 21.059 |
| Dpp4 | 113.306 | -1.1194 | 0.27766 | -4.03149 | 5.54E-05 | 0.00178 | 82.159 |
| Dpp6 | 11320.8 | 0.198284 | 0.08209 | 2.415338 | 0.01572 | 0.07681 | 910.303 |
| Dpp7 | 1061.64 | -0.07345 | 0.09129 | -0.80458 | 0.42106 | 0.62805 | 4.084 |
| Dpp8 | 10267 | 0.041536 | 0.05897 | 0.704406 | 0.48118 | 0.67822 | 50.238 |
| Dpp9 | 3209.45 | 0.03371 | 0.07027 | 0.47973 | 0.63142 | 0.79196 | 32.099 |
| Dpy19l1 | 4461.58 | -0.34093 | 0.13876 | -2.45695 | 0.01401 | 0.07132 | 91.365 |
| Dpy19l3 | 4603.15 | 0.570674 | 0.16602 | 3.43738 | 0.00059 | 0.00882 | 69.29 |
| Dpy19l4 | 3019.46 | 0.375083 | 0.09421 | 3.981329 | 6.85E-05 | 0.00207 | 60.823 |
| Dpy30 | 622.384 | -0.07765 | 0.09583 | -0.8103 | 0.41777 | 0.62518 | 24.471 |
| Dpyd | 295.971 | -0.2114 | 0.16825 | -1.25644 | 0.20896 | 0.40901 | 870.796 |
| Dpys | 15.1043 | -0.27106 | 0.54458 | -0.49775 | 0.61866 | 0.78312 | 88.984 |
| Dpysl2 | 8089.98 | -0.05546 | 0.05792 | -0.95758 | 0.33828 | 0.54905 | 65.825 |
| Dpysl3 | 1911.4 | -0.15405 | 0.12747 | -1.20851 | 0.22685 | 0.43103 | 117.303 |
| Dpysl4 | 2568.51 | 0.066513 | 0.07172 | 0.927378 | 0.35373 | 0.56399 | 16.704 |
| Dpysl5 | 2053.08 | 0.075381 | 0.09991 | 0.754459 | 0.45057 | 0.65319 | 87.812 |
| Dqx1 | 39.0218 | -0.0294 | 0.34774 | -0.08453 | 0.93263 | 0.96605 | 9.475 |
| Dr1 | 2054.79 | -0.01373 | 0.08091 | -0.16966 | 0.86528 | 0.93172 | 11.63 |
| Dram1 | 138.896 | 0.178764 | 0.18604 | 0.960904 | 0.3366 | 0.54718 | 56.277 |
| Dram2 | 1133.86 | 0.07138 | 0.10452 | 0.682946 | 0.49464 | 0.68876 | 28.093 |
| Drap1 | 4248.68 | -0.28514 | 0.0992 | -2.87436 | 0.00405 | 0.03175 | 2.175 |
| Draxin | 46.0129 | -0.88661 | 0.44076 | -2.01153 | 0.04427 | 0.15256 | 32.262 |

| | | | | | | | |
|---|---|---|---|---|---|---|---|
| Drc1 | 977.067 | 0.306472 | 0.12208 | 2.510431 | 0.01206 | 0.06482 | 37.654 |
| Drc3 | 362.32 | -0.08523 | 0.1115 | -0.76443 | 0.44461 | 0.64863 | 41.013 |
| Drc7 | 29.842 | -0.00949 | 0.43466 | -0.02183 | 0.98258 | 0.99061 | 23.039 |
| Drd1 | 9608.39 | 0.70625 | 0.31702 | 2.227747 | 0.0259 | 0.10662 | 4.523 |
| Drd2 | 4447.78 | 0.82512 | 0.3639 | 2.267416 | 0.02336 | 0.09912 | 67.551 |
| Drd3 | 550.396 | 0.898435 | 0.35884 | 2.503693 | 0.01229 | 0.06569 | 68.907 |
| Drd5 | 89.5045 | 0.219397 | 0.19883 | 1.10344 | 0.26984 | 0.47946 | 3.152 |
| Drg1 | 2828.69 | -0.04105 | 0.07526 | -0.54541 | 0.58547 | 0.75994 | 79.056 |
| Drg2 | 1844.1 | 0.005142 | 0.08003 | 0.064257 | 0.94877 | 0.97462 | 14.164 |
| Drosha | 5207.49 | 0.002797 | 0.05927 | 0.047198 | 0.96236 | 0.98172 | 110.477 |
| Drp2 | 4913.3 | 0.044494 | 0.06271 | 0.70946 | 0.47804 | 0.6758 | 52.023 |
| Dsc2 | 72.8118 | 0.973169 | 0.31101 | 3.129082 | 0.00175 | 0.01853 | 28.922 |
| Dsc3 | 26.5067 | 0.097635 | 0.36164 | 0.269976 | 0.78718 | 0.88931 | 41.422 |
| Dscam | 1506.56 | -0.42944 | 0.13669 | -3.14171 | 0.00168 | 0.01797 | 579.913 |
| Dscaml1 | 2675.62 | -0.27994 | 0.13939 | -2.0083 | 0.04461 | 0.15345 | 327.085 |
| Dse | 462.53 | 0.167509 | 0.14978 | 1.118357 | 0.26341 | 0.47216 | 56.323 |
| Dsel | 1181.1 | 0.339112 | 0.10002 | 3.390375 | 0.0007 | 0.00997 | 6.217 |
| Dsg1c | 390.894 | 0.828514 | 0.38406 | 2.157255 | 0.03099 | 0.12074 | 37.692 |
| Dsg2 | 252.028 | 0.354197 | 0.14367 | 2.4653 | 0.01369 | 0.07038 | 46.448 |
| Dsn1 | 184.235 | -0.01963 | 0.14856 | -0.13214 | 0.89487 | 0.94736 | 11.89 |
| Dsp | 17.5515 | -1.11782 | 0.42215 | -2.64793 | 0.0081 | 0.04993 | 47.284 |
| Dspp | 13.167 | -0.30621 | 0.53872 | -0.5684 | 0.56977 | 0.7492 | 9.416 |
| Dst | 26051.5 | 0.100382 | 0.07516 | 1.335586 | 0.18168 | 0.37534 | 400.437 |
| Dstn | 4755.06 | -0.25673 | 0.08878 | -2.89183 | 0.00383 | 0.03055 | 28.005 |
| Dstyk | 4886.27 | 0.233476 | 0.09143 | 2.553685 | 0.01066 | 0.05992 | 49.404 |
| Dtd1 | 1484.32 | -0.09915 | 0.0771 | -1.28599 | 0.19845 | 0.39615 | 168.862 |
| Dtd2 | 398.541 | -0.04981 | 0.12457 | -0.39989 | 0.68924 | 0.83062 | 18.19 |
| Dtl | 227.734 | -1.02792 | 0.44483 | -2.31082 | 0.02084 | 0.09208 | 38.189 |
| Dtna | 9110.05 | -0.03121 | 0.07618 | -0.40972 | 0.68201 | 0.82631 | 244.581 |
| Dtnb | 3784.24 | 0.217912 | 0.09069 | 2.402728 | 0.01627 | 0.07857 | 209.416 |
| Dtnbos | 113.23 | 0.096119 | 0.18237 | 0.527052 | 0.59816 | 0.76969 | 9.604 |
| Dtnbp1 | 2147.87 | -0.02011 | 0.10104 | -0.19903 | 0.84224 | 0.91879 | 80.073 |
| Dtwd1 | 354.019 | 0.329435 | 0.09968 | 3.305035 | 0.00095 | 0.01235 | 13.139 |
| Dtwd2 | 161.576 | -0.0526 | 0.16643 | -0.31607 | 0.75195 | 0.86994 | 59.457 |

| | | | | | | | |
|---|---|---|---|---|---|---|---|
| Dtx1 | 2944.42 | 0.098546 | 0.08133 | 1.211616 | 0.22566 | 0.42969 | 31.726 |
| Dtx2 | 456.541 | 0.137127 | 0.09647 | 1.421495 | 0.15517 | 0.34162 | 38.073 |
| Dtx3 | 9826.75 | 0.01298 | 0.06935 | 0.187179 | 0.85152 | 0.92445 | 5.351 |
| Dtx3l | 268.318 | -0.77169 | 0.20218 | -3.81692 | 0.00014 | 0.0033 | 12.641 |
| Dtx4 | 904.196 | -0.12841 | 0.11249 | -1.1415 | 0.25366 | 0.461 | 35.117 |
| Dtymk | 1220.68 | 0.006987 | 0.07935 | 0.088057 | 0.92983 | 0.96461 | 9.359 |
| Dubr | 490.311 | 0.297709 | 0.10759 | 2.767185 | 0.00565 | 0.03958 | 22.428 |
| Duox1 | 21.4438 | 0.214402 | 0.4532 | 0.473089 | 0.63615 | 0.79507 | 32.301 |
| Dus1l | 1278.61 | 0.13749 | 0.07041 | 1.952593 | 0.05087 | 0.16778 | 7.203 |
| Dus2 | 599.692 | -0.04204 | 0.092 | -0.45695 | 0.64771 | 0.80388 | 62.504 |
| Dus3l | 3000.45 | 0.069367 | 0.07373 | 0.940811 | 0.3468 | 0.55746 | 5.337 |
| Dus4l | 283.735 | 0.112723 | 0.13692 | 0.823285 | 0.41035 | 0.61822 | 14.777 |
| Dusp1 | 1009.59 | -0.24566 | 0.08635 | -2.84487 | 0.00444 | 0.0337 | 56.539 |
| Dusp10 | 168.85 | -0.57143 | 0.22373 | -2.55414 | 0.01065 | 0.05988 | 62.335 |
| Dusp11 | 7843.33 | 0.330819 | 0.07863 | 4.207306 | 2.58E-05 | 0.00105 | 19.4 |
| Dusp12 | 384.11 | 0.035056 | 0.1187 | 0.295335 | 0.76774 | 0.8786 | 12.043 |
| Dusp14 | 1149.84 | -0.14258 | 0.10395 | -1.37153 | 0.17021 | 0.36055 | 21.221 |
| Dusp15 | 856.012 | -0.28134 | 0.14434 | -1.94924 | 0.05127 | 0.16863 | 10.702 |
| Dusp16 | 527.425 | -0.21145 | 0.10557 | -2.00283 | 0.0452 | 0.15458 | 77.158 |
| Dusp18 | 1497.68 | 0.258194 | 0.09507 | 2.71574 | 0.00661 | 0.04384 | 6.057 |
| Dusp19 | 699.044 | 0.080172 | 0.08373 | 0.957551 | 0.33829 | 0.54905 | 15.317 |
| Dusp2 | 22.7861 | 0.026763 | 0.35038 | 0.076382 | 0.93912 | 0.96951 | 2.218 |
| Dusp22 | 1147.63 | -0.05149 | 0.07915 | -0.65056 | 0.51533 | 0.70568 | 51.233 |
| Dusp23 | 105.474 | -0.92061 | 0.24166 | -3.80948 | 0.00014 | 0.00336 | 2.199 |
| Dusp26 | 2314.36 | 0.020674 | 0.10385 | 0.19907 | 0.84221 | 0.91879 | 7.577 |
| Dusp27 | 21.305 | -0.69402 | 0.37015 | -1.87498 | 0.0608 | 0.18828 | 29.775 |
| Dusp28 | 502.58 | 0.182348 | 0.109 | 1.672898 | 0.09435 | 0.25065 | 1.455 |
| Dusp3 | 4930.44 | -0.39939 | 0.11841 | -3.37305 | 0.00074 | 0.01044 | 15.871 |
| Dusp4 | 409.265 | -0.51431 | 0.20179 | -2.54868 | 0.01081 | 0.06057 | 12.598 |
| Dusp5 | 272.455 | -0.23971 | 0.17626 | -1.35996 | 0.17384 | 0.36527 | 13.323 |
| Dusp6 | 1576.99 | -0.56728 | 0.1964 | -2.88835 | 0.00387 | 0.03079 | 4.259 |
| Dusp7 | 2202.94 | -0.38174 | 0.11196 | -3.40971 | 0.00065 | 0.00946 | 7.093 |
| Dusp8 | 7639.71 | 0.063785 | 0.07861 | 0.811389 | 0.41714 | 0.62461 | 16.354 |
| Dut | 920.477 | -0.07302 | 0.09231 | -0.79098 | 0.42896 | 0.63564 | 11.419 |

| | | | | | | | |
|---|---|---|---|---|---|---|---|
| Duxbl1 | 107.93 | 0.002749 | 0.24308 | 0.011308 | 0.99098 | 0.99495 | 11.512 |
| Dvl1 | 4213.97 | -0.02915 | 0.07886 | -0.36959 | 0.71169 | 0.84486 | 11.902 |
| Dvl2 | 1423.77 | 0.33773 | 0.09821 | 3.438738 | 0.00058 | 0.00882 | 11.707 |
| Dvl3 | 2732.04 | -0.04376 | 0.07452 | -0.58724 | 0.55704 | 0.739 | 17.029 |
| Dxo | 617.513 | 0.031038 | 0.10377 | 0.299093 | 0.76487 | 0.87704 | 2.215 |
| Dym | 2131.6 | 0.056111 | 0.08567 | 0.654926 | 0.51252 | 0.70349 | 268.184 |
| Dync1h1 | 34266.9 | 0.013522 | 0.05833 | 0.231792 | 0.8167 | 0.9056 | 65.494 |
| Dync1i1 | 3724.71 | -0.25805 | 0.10024 | -2.57432 | 0.01004 | 0.05762 | 302.401 |
| Dync1i2 | 8471.05 | 0.295307 | 0.07684 | 3.843076 | 0.00012 | 0.00305 | 51.598 |
| Dync1li1 | 4005.66 | 0.094465 | 0.06167 | 1.531883 | 0.12555 | 0.30073 | 35.513 |
| Dync1li2 | 16670 | 0.025629 | 0.05434 | 0.471629 | 0.63719 | 0.7959 | 25.368 |
| Dync2h1 | 2284.01 | 0.295647 | 0.16862 | 1.753322 | 0.07955 | 0.22403 | 255.944 |
| Dync2i1 | 1631.43 | 0.085364 | 0.13907 | 0.613804 | 0.53935 | 0.72587 | 56.761 |
| Dync2i2 | 1104.91 | 0.002844 | 0.10097 | 0.028165 | 0.97753 | 0.98803 | 17.336 |
| Dync2li1 | 690.931 | 0.19172 | 0.13257 | 1.446132 | 0.14814 | 0.33179 | 29.093 |
| Dynll1 | 7095.4 | -0.22538 | 0.08191 | -2.75168 | 0.00593 | 0.0409 | 3.89 |
| Dynll2 | 49661.2 | -0.06705 | 0.07584 | -0.88411 | 0.37664 | 0.58668 | 8.009 |
| Dynlrb1 | 4409.77 | -0.26199 | 0.0741 | -3.5358 | 0.00041 | 0.00694 | 13.745 |
| Dynlt1-ps1 | 106.375 | -0.38281 | 0.19322 | -1.98121 | 0.04757 | 0.16032 | 0.342 |
| Dynlt1a | 354.739 | 0.471583 | 0.10537 | 4.475574 | 7.62E-06 | 0.00045 | 11.161 |
| Dynlt1b | 223.397 | 0.00166 | 0.14697 | 0.011291 | 0.99099 | 0.99495 | 6.185 |
| Dynlt1c | 80.1924 | -0.20528 | 0.23125 | -0.8877 | 0.3747 | 0.58497 | 8.104 |
| Dynlt1f | 88.5397 | 0.202226 | 0.2619 | 0.772151 | 0.44002 | 0.6451 | 9.299 |
| Dynlt2b | 485.144 | 0.004942 | 0.13666 | 0.036162 | 0.97115 | 0.98546 | 9.398 |
| Dynlt3 | 5733.17 | 0.10817 | 0.09584 | 1.128619 | 0.25906 | 0.46729 | 8.737 |
| Dyrk1a | 3968.29 | -0.06811 | 0.05572 | -1.2225 | 0.22152 | 0.42444 | 125.508 |
| Dyrk1b | 1429.09 | 0.184514 | 0.09919 | 1.86014 | 0.06287 | 0.19256 | 7.826 |
| Dyrk2 | 1576.87 | -0.32454 | 0.19064 | -1.70235 | 0.08869 | 0.24064 | 14.607 |
| Dyrk3 | 179.589 | -0.39468 | 0.19565 | -2.01732 | 0.04366 | 0.15116 | 10.886 |
| Dysf | 205.461 | -0.73729 | 0.26283 | -2.80521 | 0.00503 | 0.03641 | 202.471 |
| Dzank1 | 15892 | 0.287918 | 0.08892 | 3.237878 | 0.0012 | 0.01436 | 56.858 |
| Dzip1 | 3979.27 | 0.118294 | 0.07272 | 1.626779 | 0.10378 | 0.26624 | 49.941 |
| Dzip1l | 2471.08 | 0.381164 | 0.08365 | 4.556522 | 5.20E-06 | 0.00035 | 39.761 |
| Dzip3 | 5441.4 | 0.3983 | 0.13596 | 2.929535 | 0.00339 | 0.02816 | 69.934 |

| | | | | | | | |
|---|---|---|---|---|---|---|---|
| E030013I19Rik | 26.9952 | -0.94984 | 0.44686 | -2.12559 | 0.03354 | 0.12748 | 11.37 |
| E030022I16Rik | 107.374 | 0.004842 | 0.22561 | 0.021464 | 0.98288 | 0.9907 | 2.766 |
| E030042O20Rik | 15.4852 | -0.44639 | 0.46116 | -0.96797 | 0.33306 | 0.54396 | 11.126 |
| E130006D01Rik | 24.6631 | -0.8771 | 0.40867 | -2.14624 | 0.03185 | 0.12307 | 28.099 |
| E130008D07Rik | 710.98 | 0.648236 | 0.15523 | 4.175877 | 2.97E-05 | 0.00115 | 20.494 |
| E130102H24Rik | 337.221 | 0.127667 | 0.11437 | 1.116282 | 0.2643 | 0.47305 | 10.212 |
| E130114P18Rik | 184.791 | -0.53221 | 0.15142 | -3.51488 | 0.00044 | 0.0073 | 209.946 |
| E130119H09Rik | 13.948 | 0.629633 | 0.51323 | 1.226814 | 0.21989 | 0.42242 | 5.136 |
| E130307A14Rik | 2220.18 | 0.244908 | 0.07862 | 3.11511 | 0.00184 | 0.01906 | 110.605 |
| E130308A19Rik | 892.325 | -0.15547 | 0.09053 | -1.71734 | 0.08592 | 0.23546 | 135.229 |
| E130309D02Rik | 616.762 | 0.131893 | 0.0973 | 1.355542 | 0.17524 | 0.36694 | 14.166 |
| E130311K13Rik | 380.33 | 0.142076 | 0.12585 | 1.128912 | 0.25893 | 0.46721 | 14.789 |
| E130317F20Rik | 130.459 | -0.1979 | 0.17997 | -1.09962 | 0.2715 | 0.48086 | 3.591 |
| E230001N04Rik | 52.0989 | 0.710442 | 0.2598 | 2.734609 | 0.00625 | 0.04229 | 2.683 |
| E230014E18Rik | 16.6221 | -0.84996 | 0.49756 | -1.70827 | 0.08759 | 0.23846 | 5.226 |
| E230015B07Rik | 13.2288 | 0.932884 | 0.50618 | 1.842996 | 0.06533 | 0.19714 | 9.042 |
| E230016M11Rik | 195.861 | 0.092905 | 0.17394 | 0.534117 | 0.59326 | 0.76557 | 44.056 |
| E230020A03Rik | 19.3133 | 0.142556 | 0.44378 | 0.32123 | 0.74804 | 0.86795 | 2.386 |
| E230020D15Rik | 132.987 | 0.173391 | 0.16248 | 1.067179 | 0.28589 | 0.49592 | 5.899 |
| E230025N22Rik | 26.1768 | 1.284824 | 0.48842 | 2.630556 | 0.00852 | 0.05159 | 11.003 |
| E230029C05Rik | 251.462 | 0.11763 | 0.13139 | 0.895286 | 0.37063 | 0.58084 | 88.074 |
| E2f1 | 415.773 | -0.28611 | 0.11696 | -2.44616 | 0.01444 | 0.07263 | 10.486 |
| E2f2 | 23.7118 | -0.85445 | 0.50525 | -1.69116 | 0.09081 | 0.24468 | 23.664 |
| E2f3 | 1226.55 | 0.071191 | 0.0772 | 0.922213 | 0.35642 | 0.56684 | 79.489 |
| E2f4 | 1318.71 | 0.014123 | 0.08305 | 0.170045 | 0.86498 | 0.93163 | 7.708 |
| E2f5 | 548.617 | 0.076318 | 0.10824 | 0.705101 | 0.48075 | 0.67798 | 27.669 |
| E2f6 | 2036.06 | 0.132205 | 0.08331 | 1.586981 | 0.11252 | 0.28042 | 28.78 |
| E2f7 | 18.2811 | -0.294 | 0.56552 | -0.51987 | 0.60316 | 0.77277 | 41.946 |
| E2f8 | 18.543 | -0.91417 | 0.44349 | -2.06133 | 0.03927 | 0.14122 | 15.168 |
| E330013P04Rik | 33.4909 | 0.007339 | 0.43155 | 0.017007 | 0.98643 | 0.99262 | 27.009 |
| E330020D12Rik | 37.3513 | -0.32439 | 0.40003 | -0.81092 | 0.41741 | 0.62485 | 9.409 |
| E330034L11Rik | 629.281 | -0.09168 | 0.11712 | -0.78276 | 0.43377 | 0.63959 | 1.742 |
| E330037G11Rik | 72.0885 | 0.318788 | 0.21694 | 1.469479 | 0.1417 | 0.3235 | 1.086 |
| E430018J23Rik | 219.111 | -0.0103 | 0.16325 | -0.06309 | 0.9497 | 0.97495 | 3.957 |

| | | | | | | | |
|---|---|---|---|---|---|---|---|
| E430024I08Rik | 1022.41 | 0.29881 | 0.19083 | 1.565817 | 0.11739 | 0.28827 | 30.509 |
| E430024P14Rik | 197.48 | 0.43566 | 0.16973 | 2.566821 | 0.01026 | 0.05849 | 65.321 |
| E4f1 | 1859.86 | 0.119067 | 0.07762 | 1.534073 | 0.12501 | 0.30012 | 26.536 |
| E530001F21Rik | 161.145 | 0.021989 | 0.19353 | 0.113625 | 0.90954 | 0.95456 | 0.73 |
| E530011L22Rik | 243.39 | -0.11082 | 0.15376 | -0.72077 | 0.47105 | 0.66978 | 5.282 |
| ENSMUSG0000009 | 9981.21 | -0.78879 | 0.45352 | -1.73926 | 0.08199 | 0.22822 | 7.501 |
| ENSMUSG0000009 | 62.5675 | -0.2753 | 0.26411 | -1.04237 | 0.29724 | 0.50736 | 1.414 |
| ENSMUSG0000009 | 133.198 | -0.17391 | 0.14868 | -1.1697 | 0.24212 | 0.4481 | 109.199 |
| ENSMUSG0000010 | 150.863 | -0.11031 | 0.1837 | -0.60047 | 0.54819 | 0.73227 | 1.946 |
| ENSMUSG0000010 | 49.7776 | -0.31717 | 0.27148 | -1.16828 | 0.24269 | 0.44884 | 34.973 |
| ENSMUSG0000010 | 1299.74 | 0.293715 | 0.09622 | 3.052473 | 0.00227 | 0.02186 | 52.93 |
| ENSMUSG0000011 | 147.208 | -0.31796 | 0.20928 | -1.51931 | 0.12868 | 0.30583 | 3.039 |
| ENSMUSG0000011 | 27.4766 | 0.62757 | 0.54017 | 1.161799 | 0.24532 | 0.45199 | 16.512 |
| ENSMUSG0000011 | 28.0575 | -0.47821 | 0.33729 | -1.41779 | 0.15625 | 0.34303 | 2.455 |
| ENSMUSG0000011 | 319.426 | 0.206103 | 0.13946 | 1.477842 | 0.13945 | 0.32071 | 0.953 |
| ENSMUSG0000011 | 1090.15 | 0.07889 | 0.15509 | 0.508659 | 0.61099 | 0.77816 | 3.704 |
| ENSMUSG0000011 | 235.87 | 0.312797 | 0.12733 | 2.456671 | 0.01402 | 0.07132 | 2.09 |
| ENSMUSG0000011 | 170.731 | 0.057671 | 0.18116 | 0.318337 | 0.75023 | 0.86885 | 1.757 |
| ENSMUSG0000011 | 12.6784 | 0.636188 | 0.52752 | 1.206007 | 0.22781 | 0.43174 | 12.516 |
| ENSMUSG0000011 | 18.7103 | 0.59393 | 0.4166 | 1.425662 | 0.15397 | 0.33978 | 11.327 |
| ENSMUSG0000011 | 44.1214 | 0.356369 | 0.33511 | 1.063441 | 0.28758 | 0.49771 | 1.078 |
| ENSMUSG0000011 | 28.2888 | -0.03885 | 0.30887 | -0.12578 | 0.89991 | 0.94981 | 1.263 |
| ENSMUSG0000011 | 664.019 | 0.006582 | 0.13112 | 0.050199 | 0.95996 | 0.98055 | 0.338 |
| ENSMUSG0000011 | 77.6811 | -0.05741 | 0.26006 | -0.22074 | 0.82529 | 0.90983 | 2.063 |
| ENSMUSG0000011 | 30.4716 | 1.692006 | 0.58685 | 2.883192 | 0.00394 | 0.0311 | 9.621 |
| ENSMUSG0000011 | 28.7223 | -0.23305 | 0.34732 | -0.67099 | 0.50222 | 0.69481 | 1.334 |
| ENSMUSG0000011 | 78.263 | 0.422683 | 0.26572 | 1.590685 | 0.11168 | 0.27927 | 1.229 |
| ENSMUSG0000011 | 29.9147 | -0.551 | 0.31573 | -1.74518 | 0.08095 | 0.22645 | 0.966 |
| ENSMUSG0000011 | 172.901 | -0.20635 | 0.21025 | -0.98142 | 0.32638 | 0.53762 | 3.61 |
| ENSMUSG0000011 | 115.195 | 0.189843 | 0.19086 | 0.994649 | 0.31991 | 0.53064 | 1.526 |
| ENSMUSG0000011 | 67.3446 | 0.520743 | 0.25233 | 2.063769 | 0.03904 | 0.14077 | 1.774 |
| ENSMUSG0000011 | 117.147 | 0.388747 | 0.20479 | 1.898266 | 0.05766 | 0.1823 | 54.215 |
| ENSMUSG0000011 | 13.3736 | 0.432265 | 0.57509 | 0.751651 | 0.45226 | 0.6547 | 1.422 |
| ENSMUSG0000011 | 29.2537 | 1.063647 | 0.3447 | 3.085676 | 0.00203 | 0.02026 | 1.285 |

| | | | | | | | |
|---|---|---|---|---|---|---|---|
| ENSMUSG0000011 | 61.7877 | 0.031193 | 0.22392 | 0.139307 | 0.88921 | 0.94457 | 0.608 |
| ENSMUSG0000011 | 39.3776 | 0.434788 | 0.29419 | 1.477906 | 0.13943 | 0.32071 | 2.526 |
| ENSMUSG0000011 | 793.688 | -0.09035 | 0.09811 | -0.92097 | 0.35707 | 0.56739 | 5.473 |
| ENSMUSG0000011 | 339.775 | 0.279968 | 0.11624 | 2.408486 | 0.01602 | 0.07776 | 2.009 |
| ENSMUSG0000012 | 80.4556 | 0.183252 | 0.19455 | 0.941913 | 0.34624 | 0.55684 | 22.717 |
| ENSMUSG0000012 | 26.0154 | 0.634723 | 0.37231 | 1.704834 | 0.08823 | 0.23994 | 1.391 |
| ENSMUSG0000012 | 104.709 | -0.14583 | 0.2163 | -0.67419 | 0.50019 | 0.69313 | 1.182 |
| ENSMUSG0000012 | 220.567 | -0.09138 | 0.17694 | -0.51644 | 0.60555 | 0.77423 | 3.334 |
| ENSMUSG0000012 | 99.8063 | -0.0304 | 0.18088 | -0.16807 | 0.86653 | 0.93232 | 2.284 |
| ENSMUSG0000012 | 86.8586 | 0.228134 | 0.24058 | 0.948268 | 0.34299 | 0.5537 | 1.528 |
| ENSMUSG0000012 | 22.8144 | -0.24187 | 0.3587 | -0.6743 | 0.50012 | 0.69308 | 1.562 |
| ENSMUSG0000012 | 36.2936 | 0.384166 | 0.31775 | 1.209004 | 0.22666 | 0.43083 | 12.622 |
| ENSMUSG0000012 | 20.2181 | 0.792732 | 0.44495 | 1.781631 | 0.07481 | 0.21551 | 5.831 |
| ENSMUSG0000012 | 116.674 | -0.04037 | 0.16887 | -0.23904 | 0.81107 | 0.90275 | 0.776 |
| ENSMUSG0000012 | 661.847 | 0.819381 | 0.2439 | 3.359462 | 0.00078 | 0.01076 | 11.632 |
| ENSMUSG0000012 | 144.158 | 0.674873 | 0.19227 | 3.510039 | 0.00045 | 0.00738 | 6.231 |
| ENSMUSG0000012 | 46.0581 | -0.6587 | 0.27969 | -2.35509 | 0.01852 | 0.08529 | 1.803 |
| ENSMUSG0000012 | 26.3057 | -0.1924 | 0.35964 | -0.53497 | 0.59267 | 0.76515 | 0.633 |
| ENSMUSG0000012 | 39.1985 | 0.167729 | 0.29951 | 0.560002 | 0.57548 | 0.75343 | 35.919 |
| ENSMUSG0000012 | 19.1526 | -0.20429 | 0.45128 | -0.4527 | 0.65077 | 0.80552 | 8.312 |
| ENSMUSG0000012 | 20.465 | 0.011459 | 0.43521 | 0.026329 | 0.97899 | 0.98884 | 1.594 |
| ENSMUSG0000012 | 20.8691 | -0.18922 | 0.38701 | -0.48893 | 0.62489 | 0.78745 | 0.959 |
| ENSMUSG0000012 | 15.37 | 0.586858 | 0.50155 | 1.170099 | 0.24196 | 0.44789 | 0.98 |
| ENSMUSG0000012 | 17.2036 | 1.056859 | 0.49193 | 2.148377 | 0.03168 | 0.12262 | 1.411 |
| ENSMUSG0000012 | 15.789 | -0.98914 | 0.48125 | -2.05534 | 0.03985 | 0.1426 | 1.53 |
| ENSMUSG0000012 | 20.0602 | -0.16268 | 0.40662 | -0.40008 | 0.6891 | 0.83056 | 2.247 |
| ENSMUSG0000012 | 21.4262 | -1.21522 | 0.57063 | -2.12962 | 0.0332 | 0.12658 | 43.798 |
| ENSMUSG0000012 | 28.0539 | -0.3726 | 0.3293 | -1.13148 | 0.25785 | 0.46595 | 19.098 |
| ENSMUSG0000012 | 748.702 | 0.037051 | 0.11337 | 0.326824 | 0.7438 | 0.86539 | 0.923 |
| ENSMUSG0000012 | 87.0471 | 0.013963 | 0.23262 | 0.060028 | 0.95213 | 0.97658 | 2.122 |
| ENSMUSG0000012 | 37.2313 | -0.02151 | 0.2979 | -0.07219 | 0.94245 | 0.97123 | 0.949 |
| ENSMUSG0000012 | 23.2131 | 0.23495 | 0.38833 | 0.60503 | 0.54516 | 0.73012 | 1.299 |
| ENSMUSG0000012 | 19.7638 | 0.142052 | 0.40519 | 0.350581 | 0.7259 | 0.85388 | 0.933 |
| ENSMUSG0000012 | 16.6894 | -0.07722 | 0.41252 | -0.18719 | 0.85151 | 0.92445 | 1.219 |

| | | | | | | | |
|---|---|---|---|---|---|---|---|
| ENSMUSG0000012 | 67.4434 | 0.12306 | 0.27585 | 0.446111 | 0.65552 | 0.80876 | 2.77 |
| ENSMUSG0000012 | 14.4003 | 0.32078 | 0.74158 | 0.432561 | 0.66533 | 0.81526 | 1.994 |
| ENSMUSG0000012 | 786.313 | 0.051877 | 0.10679 | 0.485779 | 0.62712 | 0.78903 | 1.577 |
| ENSMUSG0000012 | 14.2273 | 0.688938 | 0.45938 | 1.49972 | 0.13369 | 0.31339 | 8.457 |
| ENSMUSG0000012 | 117.268 | 0.183888 | 0.21482 | 0.856025 | 0.39198 | 0.60108 | 1.75 |
| ENSMUSG0000012 | 62.2139 | 0.54963 | 0.23418 | 2.347069 | 0.01892 | 0.08634 | 25.23 |
| ENSMUSG0000012 | 35.0326 | 0.012481 | 0.29015 | 0.043017 | 0.96569 | 0.9828 | 0.792 |
| ENSMUSG0000012 | 43.6784 | 0.218623 | 0.27399 | 0.797925 | 0.42491 | 0.6314 | 20.331 |
| ENSMUSG0000012 | 24.2152 | 0.277341 | 0.4043 | 0.685972 | 0.49273 | 0.68727 | 10.064 |
| ENSMUSG0000012 | 29.7037 | 0.047406 | 0.32171 | 0.147355 | 0.88285 | 0.94161 | 1.224 |
| ENSMUSG0000012 | 758.484 | -0.05018 | 0.09655 | -0.51979 | 0.60321 | 0.77279 | 1.377 |
| ENSMUSG0000012 | 15.9727 | 0.717173 | 0.49025 | 1.462886 | 0.1435 | 0.32566 | 0.934 |
| ENSMUSG0000012 | 15.1562 | 0.614117 | 0.44556 | 1.378317 | 0.16811 | 0.3581 | 0.822 |
| ENSMUSG0000012 | 13.2634 | 0.308866 | 0.51315 | 0.601901 | 0.54724 | 0.73142 | 15.81 |
| ENSMUSG0000012 | 37.3868 | 0.564336 | 0.30425 | 1.854872 | 0.06361 | 0.19392 | 3.184 |
| ENSMUSG0000012 | 48.7993 | -0.30507 | 0.27965 | -1.09091 | 0.27531 | 0.48468 | 0.945 |
| ENSMUSG0000012 | 226.64 | -0.56885 | 0.12491 | -4.55426 | 5.26E-06 | 0.00035 | 11.827 |
| ENSMUSG0000012 | 535.966 | 0.317689 | 0.11383 | 2.79096 | 0.00526 | 0.03745 | 12.98 |
| ENSMUSG0000012 | 182.829 | 0.007788 | 0.14223 | 0.054756 | 0.95633 | 0.97864 | 2.136 |
| ENSMUSG0000012 | 15.9794 | 0.487176 | 0.51844 | 0.939697 | 0.34737 | 0.55793 | 1.19 |
| ENSMUSG0000012 | 223.782 | 0.041899 | 0.16383 | 0.255744 | 0.79815 | 0.8958 | 1.09 |
| ENSMUSG0000012 | 23.1928 | 0.100106 | 0.34401 | 0.291002 | 0.77105 | 0.88041 | 0.424 |
| ENSMUSG0000012 | 907.706 | 0.242965 | 0.08548 | 2.842403 | 0.00448 | 0.03387 | 3.321 |
| ENSMUSG0000012 | 20.4048 | 0.180913 | 0.38461 | 0.470376 | 0.63809 | 0.79664 | 1.79 |
| ENSMUSG0000012 | 41.6261 | -0.5506 | 0.35234 | -1.5627 | 0.11812 | 0.28961 | 1.377 |
| ENSMUSG0000012 | 28.4936 | -0.43037 | 0.44251 | -0.97254 | 0.33078 | 0.54197 | 1.295 |
| ENSMUSG0000012 | 22.0863 | -0.28147 | 0.41498 | -0.67827 | 0.4976 | 0.691 | 1.257 |
| ENSMUSG0000012 | 15.9695 | 0.133196 | 0.57836 | 0.230299 | 0.81786 | 0.90637 | 0.671 |
| ENSMUSG0000012 | 101.626 | -0.28669 | 0.23227 | -1.23432 | 0.21709 | 0.41904 | 7.437 |
| ENSMUSG0000012 | 62.4576 | -0.24731 | 0.24679 | -1.00213 | 0.31628 | 0.52681 | 2.303 |
| ENSMUSG0000012 | 36.0432 | -0.90213 | 0.4652 | -1.93921 | 0.05248 | 0.17137 | 4.411 |
| ENSMUSG0000012 | 32.6689 | 0.531677 | 0.29591 | 1.796766 | 0.07237 | 0.21135 | 2.159 |
| ENSMUSG0000012 | 17.6142 | 0.751921 | 0.40315 | 1.865101 | 0.06217 | 0.19102 | 0.742 |
| ENSMUSG0000012 | 50.3908 | -0.03889 | 0.26983 | -0.14412 | 0.88541 | 0.9431 | 18.397 |

| | | | | | | | |
|---|---|---|---|---|---|---|---|
| ENSMUSG0000012 | 18.9482 | -0.26155 | 0.45265 | -0.57782 | 0.56338 | 0.74402 | 156.002 |
| ENSMUSG0000012 | 226.52 | -0.06908 | 0.16436 | -0.42031 | 0.67426 | 0.82113 | 0.68 |
| ENSMUSG0000012 | 23.4995 | 0.350407 | 0.45494 | 0.770219 | 0.44117 | 0.64576 | 9.944 |
| ENSMUSG0000012 | 218.418 | -0.05768 | 0.1768 | -0.32626 | 0.74423 | 0.86539 | 0.588 |
| ENSMUSG0000012 | 15.029 | -0.19221 | 0.48914 | -0.39295 | 0.69435 | 0.83419 | 1.096 |
| ENSMUSG0000012 | 17.543 | -1.28105 | 0.48774 | -2.62651 | 0.00863 | 0.05201 | 0.776 |
| ENSMUSG0000012 | 752.571 | 0.353945 | 0.17496 | 2.023063 | 0.04307 | 0.14978 | 42.745 |
| ENSMUSG0000012 | 657.288 | -0.12664 | 0.12129 | -1.04416 | 0.29641 | 0.50667 | 9.307 |
| ENSMUSG0000012 | 13.7697 | 1.518034 | 0.48746 | 3.114183 | 0.00184 | 0.01907 | 89.149 |
| ENSMUSG0000012 | 114.083 | 0.137401 | 0.21184 | 0.648594 | 0.5166 | 0.70655 | 0.717 |
| ENSMUSG0000012 | 14.0707 | 0.034341 | 0.49831 | 0.068915 | 0.94506 | 0.97268 | 1.087 |
| ENSMUSG0000012 | 118.282 | 0.550083 | 0.23096 | 2.38171 | 0.01723 | 0.08153 | 44.352 |
| ENSMUSG0000012 | 70.2656 | -0.05999 | 0.28139 | -0.2132 | 0.83117 | 0.91334 | 6.726 |
| ENSMUSG0000012 | 15.8805 | -0.03021 | 0.5113 | -0.05908 | 0.95289 | 0.97685 | 0.344 |
| ENSMUSG0000012 | 12.787 | -0.07987 | 0.55754 | -0.14325 | 0.88609 | 0.94329 | 1.453 |
| ENSMUSG0000012 | 44.3368 | 0.074642 | 0.25472 | 0.293031 | 0.7695 | 0.87972 | 0.563 |
| ENSMUSG0000012 | 49.6382 | 0.602212 | 0.28676 | 2.100079 | 0.03572 | 0.13287 | 0.496 |
| ENSMUSG0000012 | 27.2143 | 0.948897 | 0.45886 | 2.067923 | 0.03865 | 0.13997 | 9.904 |
| ENSMUSG0000012 | 53.6397 | 0.33366 | 0.25807 | 1.292919 | 0.19604 | 0.39362 | 1.334 |
| ENSMUSG0000012 | 123.394 | -0.20279 | 0.20585 | -0.98514 | 0.32455 | 0.53568 | 165.388 |
| ENSMUSG0000012 | 17.6556 | -0.16746 | 0.42365 | -0.39528 | 0.69264 | 0.83343 | 1.455 |
| ENSMUSG0000012 | 105.263 | 0.390406 | 0.17277 | 2.259643 | 0.02384 | 0.10049 | 1.266 |
| ENSMUSG0000012 | 171.329 | -0.01372 | 0.14129 | -0.0971 | 0.92264 | 0.96107 | 1.163 |
| ENSMUSG0000012 | 22.5653 | 0.085596 | 0.44506 | 0.192327 | 0.84749 | 0.92237 | 11.039 |
| ENSMUSG0000012 | 107.101 | 0.164233 | 0.19835 | 0.827983 | 0.40768 | 0.61541 | 1.478 |
| ENSMUSG0000012 | 227.969 | -0.06051 | 0.15497 | -0.39046 | 0.6962 | 0.83541 | 1.026 |
| ENSMUSG0000012 | 26.0885 | -0.59044 | 0.36868 | -1.6015 | 0.10927 | 0.27509 | 1.507 |
| ENSMUSG0000012 | 15.1009 | -0.29467 | 0.4497 | -0.65527 | 0.5123 | 0.70333 | 1.46 |
| ENSMUSG0000012 | 46.1573 | 1.168202 | 0.273 | 4.279062 | 1.88E-05 | 0.00085 | 7.912 |
| ENSMUSG0000012 | 54.7669 | 0.409341 | 0.30293 | 1.351264 | 0.17661 | 0.36865 | 0.536 |
| ENSMUSG0000012 | 19.8975 | 0.176557 | 0.40486 | 0.436098 | 0.66277 | 0.8137 | 0.929 |
| ENSMUSG0000012 | 36.9727 | -0.38178 | 0.46357 | -0.82355 | 0.4102 | 0.61804 | 1.193 |
| ENSMUSG0000012 | 19.0384 | 0.413818 | 0.44518 | 0.929562 | 0.3526 | 0.56294 | 17.87 |
| ENSMUSG0000012 | 35.8623 | -0.07685 | 0.33978 | -0.22619 | 0.82105 | 0.90748 | 1.902 |

| | | | | | | | |
|---|---|---|---|---|---|---|---|
| ENSMUSG0000012 | 28.4113 | 0.084236 | 0.46502 | 0.181147 | 0.85625 | 0.92746 | 1.265 |
| ENSMUSG0000012 | 66.3563 | -0.21545 | 0.22032 | -0.97791 | 0.32812 | 0.53939 | 2.044 |
| ENSMUSG0000012 | 36.8614 | -0.37198 | 0.32694 | -1.13776 | 0.25522 | 0.46289 | 1.08 |
| ENSMUSG0000012 | 242.394 | 0.235034 | 0.16987 | 1.38362 | 0.16647 | 0.356 | 3.116 |
| ENSMUSG0000012 | 22.2123 | 0.27785 | 0.39526 | 0.702958 | 0.48208 | 0.67869 | 0.73 |
| ENSMUSG0000012 | 21.6194 | 0.229569 | 0.44985 | 0.510325 | 0.60982 | 0.7773 | 0.631 |
| ENSMUSG0000012 | 63.287 | 0.105987 | 0.28788 | 0.36817 | 0.71275 | 0.84523 | 1.059 |
| ENSMUSG0000012 | 286.357 | -0.23758 | 0.17514 | -1.35654 | 0.17493 | 0.36656 | 0.378 |
| ENSMUSG0000012 | 38.9008 | 0.85468 | 0.36174 | 2.362708 | 0.01814 | 0.08412 | 3.003 |
| ENSMUSG0000012 | 50.3318 | 0.30456 | 0.2443 | 1.246659 | 0.21252 | 0.41333 | 13.476 |
| ENSMUSG0000012 | 18.0267 | 0.531712 | 0.40451 | 1.314454 | 0.18869 | 0.3844 | 0.888 |
| ENSMUSG0000012 | 23.111 | 0.245739 | 0.40096 | 0.612871 | 0.53996 | 0.726 | 27.65 |
| ENSMUSG0000012 | 57.0009 | -0.18866 | 0.27049 | -0.69746 | 0.48551 | 0.68173 | 1.184 |
| ENSMUSG0000012 | 15.7345 | 1.129915 | 0.64753 | 1.744967 | 0.08099 | 0.22645 | 0.379 |
| ENSMUSG0000012 | 19.5486 | 0.839072 | 0.44372 | 1.891013 | 0.05862 | 0.18415 | 1.37 |
| ENSMUSG0000012 | 27.4644 | 0.201913 | 0.32825 | 0.615124 | 0.53847 | 0.72517 | 1.575 |
| ENSMUSG0000012 | 15.0776 | 1.356377 | 0.61477 | 2.206298 | 0.02736 | 0.11067 | 23.887 |
| ENSMUSG0000012 | 19.8751 | -1.45661 | 0.45941 | -3.17058 | 0.00152 | 0.01675 | 2.866 |
| ENSMUSG0000012 | 29.8106 | -0.38415 | 0.31806 | -1.20779 | 0.22713 | 0.43132 | 0.473 |
| ENSMUSG0000012 | 64.463 | 0.172034 | 0.22749 | 0.756225 | 0.44951 | 0.65267 | 3.218 |
| ENSMUSG0000012 | 557.821 | 0.453657 | 0.12423 | 3.651772 | 0.00026 | 0.00509 | 3.614 |
| ENSMUSG0000012 | 15.3595 | -0.33797 | 0.44643 | -0.75705 | 0.44902 | 0.65228 | 0.544 |
| ENSMUSG0000012 | 51.3955 | 0.050126 | 0.34452 | 0.145496 | 0.88432 | 0.94244 | 1.194 |
| ENSMUSG0000012 | 14.4212 | 0.341245 | 0.48293 | 0.70661 | 0.47981 | 0.67732 | 1.472 |
| ENSMUSG0000012 | 67.0198 | -0.08351 | 0.23289 | -0.3586 | 0.71989 | 0.85044 | 1.235 |
| ENSMUSG0000012 | 113.352 | 1.022223 | 0.29041 | 3.519929 | 0.00043 | 0.00725 | 19.84 |
| ENSMUSG0000012 | 68.6727 | -0.90034 | 0.26574 | -3.38802 | 0.0007 | 0.01001 | 0.623 |
| ENSMUSG0000012 | 235.222 | 0.321131 | 0.20063 | 1.600631 | 0.10946 | 0.27529 | 0.322 |
| ENSMUSG0000012 | 21.4993 | -0.48649 | 0.42463 | -1.14567 | 0.25193 | 0.45929 | 1.344 |
| ENSMUSG0000012 | 42.0235 | 0.284525 | 0.33368 | 0.85268 | 0.39384 | 0.60297 | 52.081 |
| ENSMUSG0000012 | 26.4113 | -0.45284 | 0.59708 | -0.75842 | 0.4482 | 0.65174 | 1.001 |
| ENSMUSG0000012 | 82.0549 | 0.226094 | 0.2261 | 0.999964 | 0.31733 | 0.52765 | 1.117 |
| ENSMUSG0000012 | 14.485 | -0.26639 | 0.52943 | -0.50316 | 0.61485 | 0.78083 | 1.526 |
| ENSMUSG0000012 | 149.922 | -0.20065 | 0.14305 | -1.40268 | 0.16071 | 0.34889 | 2.186 |

| | | | | | | | |
|---|---|---|---|---|---|---|---|
| ENSMUSG0000012 | 62.2135 | -0.09464 | 0.26746 | -0.35385 | 0.72345 | 0.85232 | 5.382 |
| ENSMUSG0000012 | 30.8813 | 0.65385 | 0.30388 | 2.151658 | 0.03142 | 0.12193 | 0.688 |
| ENSMUSG0000012 | 79.8157 | -0.63219 | 0.26327 | -2.40131 | 0.01634 | 0.07875 | 1.49 |
| ENSMUSG0000012 | 23.6393 | 0.979377 | 0.39929 | 2.452785 | 0.01418 | 0.07176 | 9.122 |
| ENSMUSG0000012 | 33.516 | 0.590409 | 0.30895 | 1.91099 | 0.05601 | 0.17885 | 2.503 |
| ENSMUSG0000012 | 24.4546 | -0.07247 | 0.34934 | -0.20746 | 0.83565 | 0.91572 | 0.705 |
| ENSMUSG0000012 | 12.8999 | -0.45835 | 0.54457 | -0.84168 | 0.39997 | 0.60875 | 1.37 |
| ENSMUSG0000012 | 207.761 | 0.266588 | 0.18067 | 1.475561 | 0.14006 | 0.3214 | 3.324 |
| ENSMUSG0000012 | 27.8917 | -0.8924 | 0.43952 | -2.03041 | 0.04231 | 0.14806 | 5.966 |
| ENSMUSG0000012 | 29.5595 | -0.23205 | 0.36747 | -0.63147 | 0.52773 | 0.71577 | 0.62 |
| ENSMUSG0000012 | 17.8165 | 0.594086 | 0.40853 | 1.454209 | 0.14589 | 0.32878 | 9.01 |
| ENSMUSG0000012 | 64.6631 | -0.14016 | 0.25765 | -0.54401 | 0.58643 | 0.76037 | 0.424 |
| ENSMUSG0000012 | 22.6161 | -0.65929 | 0.50536 | -1.30459 | 0.19203 | 0.38879 | 1.262 |
| ENSMUSG0000012 | 16.7648 | 0.602643 | 0.41481 | 1.452825 | 0.14627 | 0.32924 | 3.079 |
| ENSMUSG0000012 | 36.6603 | 0.382317 | 0.38659 | 0.988955 | 0.32269 | 0.53366 | 0.725 |
| ENSMUSG0000012 | 70.4686 | 0.055712 | 0.24737 | 0.225216 | 0.82181 | 0.90776 | 1.15 |
| ENSMUSG0000012 | 19.0029 | 0.252872 | 0.4544 | 0.556493 | 0.57787 | 0.75477 | 61.952 |
| ENSMUSG0000012 | 15.5804 | 0.678259 | 0.5556 | 1.220774 | 0.22217 | 0.42538 | 0.332 |
| ENSMUSG0000012 | 12.8401 | -0.21561 | 0.70366 | -0.30641 | 0.75929 | 0.87354 | 25.974 |
| ENSMUSG0000012 | 13.0608 | -0.21609 | 0.50784 | -0.42551 | 0.67047 | 0.81841 | 2.177 |
| ENSMUSG0000012 | 40.1201 | 0.194178 | 0.33053 | 0.587472 | 0.55689 | 0.73889 | 1.637 |
| ENSMUSG0000012 | 31.2874 | -0.31523 | 0.40352 | -0.78118 | 0.43469 | 0.64042 | 2.135 |
| ENSMUSG0000012 | 28.8143 | 0.285888 | 0.39782 | 0.718639 | 0.47236 | 0.67066 | 0.515 |
| ENSMUSG0000012 | 113.343 | -0.0842 | 0.16712 | -0.50383 | 0.61438 | 0.78043 | 1.193 |
| ENSMUSG0000012 | 15.1961 | 0.226525 | 0.55643 | 0.407101 | 0.68393 | 0.82762 | 1.292 |
| ENSMUSG0000012 | 40.522 | 0.198835 | 0.27637 | 0.719456 | 0.47186 | 0.67032 | 1.124 |
| ENSMUSG0000012 | 51.7457 | -0.45761 | 0.33118 | -1.38175 | 0.16705 | 0.35691 | 0.802 |
| ENSMUSG0000012 | 109.953 | -0.17766 | 0.23055 | -0.77061 | 0.44094 | 0.64569 | 1.859 |
| ENSMUSG0000012 | 13.1154 | 0.048308 | 0.55157 | 0.087583 | 0.93021 | 0.96485 | 2.489 |
| ENSMUSG0000012 | 51.1649 | -0.72207 | 0.28884 | -2.49992 | 0.01242 | 0.06617 | 3.924 |
| ENSMUSG0000012 | 180.051 | -0.11118 | 0.26923 | -0.41294 | 0.67965 | 0.82491 | 1.675 |
| ENSMUSG0000012 | 18.1728 | -0.45313 | 0.52569 | -0.86197 | 0.38871 | 0.59793 | 0.167 |
| ENSMUSG0000012 | 89.9116 | 0.555862 | 0.34434 | 1.614269 | 0.10647 | 0.27067 | 26.608 |
| ENSMUSG0000012 | 45.8984 | 0.369041 | 0.33302 | 1.108159 | 0.26779 | 0.47712 | 0.796 |

| | | | | | | | |
|---|---|---|---|---|---|---|---|
| ENSMUSG0000012 | 13.5222 | 0.552124 | 0.53378 | 1.034366 | 0.30097 | 0.51075 | 1.364 |
| ENSMUSG0000012 | 49.2465 | -0.33412 | 0.32082 | -1.04145 | 0.29767 | 0.50774 | 40.842 |
| ENSMUSG0000012 | 134.895 | 0.039403 | 0.20442 | 0.192758 | 0.84715 | 0.92206 | 1.075 |
| ENSMUSG0000012 | 22.8109 | 0.036602 | 0.38494 | 0.095084 | 0.92425 | 0.96197 | 1.19 |
| ENSMUSG0000012 | 14.5801 | -0.53249 | 0.55049 | -0.9673 | 0.3334 | 0.54424 | 0.86 |
| ENSMUSG0000012 | 25.3286 | -0.14784 | 0.37437 | -0.39491 | 0.69291 | 0.83344 | 4.596 |
| ENSMUSG0000012 | 67.0161 | 0.28076 | 0.28824 | 0.974046 | 0.33003 | 0.54122 | 1.161 |
| ENSMUSG0000012 | 19.4035 | -0.53867 | 0.42236 | -1.2754 | 0.20217 | 0.40058 | 0.868 |
| ENSMUSG0000012 | 37.5701 | -0.20268 | 0.32294 | -0.6276 | 0.53027 | 0.71791 | 2.125 |
| ENSMUSG0000012 | 12.3611 | -0.0548 | 0.56097 | -0.0977 | 0.92217 | 0.96083 | 1.293 |
| ENSMUSG0000012 | 41.175 | 0.380984 | 0.29864 | 1.275712 | 0.20206 | 0.40058 | 0.672 |
| ENSMUSG0000012 | 45.0084 | -0.12577 | 0.26678 | -0.47144 | 0.63733 | 0.79598 | 0.237 |
| ENSMUSG0000012 | 16.2966 | 0.537198 | 0.51563 | 1.041835 | 0.29749 | 0.5076 | 1.358 |
| ENSMUSG0000012 | 23.5005 | 0.586499 | 0.36728 | 1.596852 | 0.1103 | 0.27686 | 1.771 |
| ENSMUSG0000012 | 144.636 | 0.035014 | 0.14986 | 0.233651 | 0.81526 | 0.90512 | 1.583 |
| ENSMUSG0000012 | 53.8446 | 0.060317 | 0.27965 | 0.215691 | 0.82923 | 0.91246 | 0.762 |
| ENSMUSG0000012 | 1816.94 | -0.32209 | 0.1076 | -2.9934 | 0.00276 | 0.02492 | 1.581 |
| ENSMUSG0000012 | 89.2999 | 0.560939 | 0.27835 | 2.01525 | 0.04388 | 0.15162 | 2.28 |
| ENSMUSG0000012 | 38.2628 | -0.15583 | 0.37211 | -0.41877 | 0.67538 | 0.82188 | 5.671 |
| ENSMUSG0000012 | 48.6009 | -0.3695 | 0.28501 | -1.29645 | 0.19482 | 0.39211 | 1.276 |
| ENSMUSG0000012 | 36.321 | 0.3067 | 0.43253 | 0.709087 | 0.47827 | 0.67599 | 0.633 |
| ENSMUSG0000012 | 28.4551 | -0.00095 | 0.34806 | -0.00274 | 0.99781 | 0.99888 | 2.172 |
| ENSMUSG0000012 | 25.7202 | -0.05965 | 0.35985 | -0.16575 | 0.86835 | 0.93337 | 1.379 |
| ENSMUSG0000012 | 20.7069 | -0.66999 | 0.40932 | -1.63686 | 0.10166 | 0.26311 | 1.741 |
| ENSMUSG0000012 | 25.6881 | -0.64232 | 0.59795 | -1.07421 | 0.28273 | 0.49296 | 0.634 |
| ENSMUSG0000012 | 30.4047 | 0.504066 | 0.31841 | 1.58308 | 0.1134 | 0.28194 | 23.289 |
| ENSMUSG0000012 | 39.4665 | 0.225075 | 0.29792 | 0.755489 | 0.44996 | 0.65294 | 0.898 |
| ENSMUSG0000012 | 13.7675 | 0.125599 | 0.44255 | 0.283806 | 0.77656 | 0.88361 | 17.294 |
| ENSMUSG0000012 | 102.691 | -0.05207 | 0.18079 | -0.28799 | 0.77335 | 0.88189 | 0.356 |
| ENSMUSG0000012 | 124.176 | -0.41026 | 0.30658 | -1.33816 | 0.18085 | 0.37423 | 5.023 |
| ENSMUSG0000012 | 14.3509 | -0.36972 | 0.54044 | -0.6841 | 0.49391 | 0.68836 | 0.954 |
| ENSMUSG0000012 | 130.265 | 0.19555 | 0.24911 | 0.784981 | 0.43246 | 0.63861 | 14.322 |
| ENSMUSG0000012 | 73.2153 | 0.344408 | 0.28843 | 1.194071 | 0.23245 | 0.43735 | 16.764 |
| ENSMUSG0000012 | 31.847 | -0.70963 | 0.565 | -1.25598 | 0.20912 | 0.40918 | 13.108 |

| | | | | | | | |
|---|---|---|---|---|---|---|---|
| ENSMUSG0000012 | 14.5356 | 0.063706 | 0.48268 | 0.131983 | 0.895 | 0.94738 | 0.129 |
| ENSMUSG0000012 | 166.277 | -0.01669 | 0.19896 | -0.08388 | 0.93315 | 0.96643 | 0.165 |
| ENSMUSG0000012 | 14.6271 | -0.33684 | 0.51795 | -0.65033 | 0.51548 | 0.70577 | 2.14 |
| ENSMUSG0000012 | 18.9454 | 1.058966 | 0.40794 | 2.595881 | 0.00943 | 0.05525 | 19.06 |
| ENSMUSG0000012 | 16.841 | -0.10671 | 0.42245 | -0.25261 | 0.80057 | 0.89692 | 0.398 |
| ENSMUSG0000012 | 16.4261 | -0.65097 | 0.42087 | -1.54672 | 0.12193 | 0.29538 | 0.155 |
| ENSMUSG0000012 | 97.3454 | 0.470195 | 0.26378 | 1.78253 | 0.07466 | 0.21534 | 1.566 |
| ENSMUSG0000012 | 17.5512 | 0.37888 | 0.63481 | 0.596838 | 0.55062 | 0.73411 | 34.25 |
| ENSMUSG0000012 | 13.9875 | -0.3431 | 0.47722 | -0.71896 | 0.47217 | 0.67052 | 1.943 |
| ENSMUSG0000012 | 43.3161 | 0.24925 | 0.24714 | 1.008529 | 0.3132 | 0.52364 | 2.528 |
| ENSMUSG0000012 | 26.7365 | 0.141325 | 0.31348 | 0.45082 | 0.65212 | 0.80645 | 2.06 |
| ENSMUSG0000012 | 83.5211 | 0.076809 | 0.22173 | 0.346413 | 0.72903 | 0.85579 | 0.143 |
| ENSMUSG0000012 | 16.2858 | 0.322964 | 0.40641 | 0.794684 | 0.4268 | 0.63329 | 0.815 |
| ENSMUSG0000012 | 29.4462 | -0.19806 | 0.3233 | -0.61261 | 0.54013 | 0.72606 | 0.096 |
| ENSMUSG0000012 | 20.8725 | -0.85534 | 0.40144 | -2.13071 | 0.03311 | 0.12642 | 0.793 |
| ENSMUSG0000012 | 436.698 | -0.0925 | 0.11568 | -0.79962 | 0.42393 | 0.63068 | 1.075 |
| ENSMUSG0000012 | 15.3136 | 0.184471 | 0.47594 | 0.387594 | 0.69832 | 0.83649 | 1.403 |
| ENSMUSG0000012 | 51.1147 | -0.23198 | 0.24536 | -0.94544 | 0.34443 | 0.55523 | 1.846 |
| ENSMUSG0000012 | 13.8902 | -0.7182 | 0.49433 | -1.45286 | 0.14626 | 0.32924 | 2.356 |
| ENSMUSG0000012 | 17.6294 | 0.298597 | 0.48087 | 0.620952 | 0.53463 | 0.7217 | 1.946 |
| ENSMUSG0000012 | 22.4716 | 0.366402 | 0.33538 | 1.092506 | 0.27461 | 0.4843 | 1.275 |
| ENSMUSG0000012 | 27.503 | -0.34892 | 0.31485 | -1.1082 | 0.26778 | 0.47712 | 0.579 |
| ENSMUSG0000012 | 128.205 | 0.028651 | 0.2078 | 0.137881 | 0.89033 | 0.94514 | 2.279 |
| ENSMUSG0000012 | 16.0654 | -0.74949 | 0.40276 | -1.8609 | 0.06276 | 0.19225 | 0.281 |
| ENSMUSG0000012 | 30.4646 | 0.486029 | 0.39833 | 1.220166 | 0.2224 | 0.42566 | 0.408 |
| ENSMUSG0000012 | 13.6317 | 0.013193 | 0.50844 | 0.025948 | 0.9793 | 0.9889 | 2.386 |
| ENSMUSG0000012 | 111.377 | 0.272241 | 0.18757 | 1.451386 | 0.14667 | 0.32981 | 1.788 |
| ENSMUSG0000012 | 44.0927 | 0.180545 | 0.31433 | 0.574384 | 0.56571 | 0.74579 | 10.8 |
| ENSMUSG0000012 | 12.871 | 0.151027 | 0.44397 | 0.340172 | 0.73373 | 0.85906 | 1.361 |
| ENSMUSG0000012 | 23.8503 | -0.8477 | 0.56443 | -1.50185 | 0.13314 | 0.31274 | 12.573 |
| ENSMUSG0000012 | 16.922 | -0.20286 | 0.43072 | -0.47098 | 0.63766 | 0.7962 | 1.266 |
| ENSMUSG0000012 | 61.7867 | 0.335744 | 0.27463 | 1.222556 | 0.2215 | 0.42444 | 0.926 |
| ENSMUSG0000012 | 57.0889 | -0.0259 | 0.23944 | -0.10815 | 0.91388 | 0.95639 | 2.008 |
| ENSMUSG0000012 | 69.6654 | -0.72338 | 0.40055 | -1.80598 | 0.07092 | 0.20828 | 8.268 |

| | | | | | | | |
|---|---|---|---|---|---|---|---|
| ENSMUSG0000012 | 20.4545 | 0.238284 | 0.48268 | 0.493669 | 0.62154 | 0.78483 | 1.993 |
| ENSMUSG0000012 | 16.3878 | 0.399756 | 0.52174 | 0.766197 | 0.44356 | 0.64792 | 5.811 |
| ENSMUSG0000012 | 17.7303 | -0.1833 | 0.52572 | -0.34867 | 0.72734 | 0.85443 | 1.404 |
| ENSMUSG0000012 | 179.11 | 0.499285 | 0.15855 | 3.14905 | 0.00164 | 0.01763 | 0.65 |
| ENSMUSG0000012 | 82.2902 | 0.499612 | 0.24115 | 2.071795 | 0.03828 | 0.13918 | 4.711 |
| ENSMUSG0000012 | 21.4826 | 0.050906 | 0.38514 | 0.132175 | 0.89485 | 0.94736 | 3.541 |
| ENSMUSG0000012 | 149.01 | 0.340749 | 0.19103 | 1.783716 | 0.07447 | 0.21507 | 1.793 |
| ENSMUSG0000012 | 58.9978 | 0.86094 | 0.29955 | 2.87412 | 0.00405 | 0.03175 | 4.33 |
| ENSMUSG0000012 | 28.9042 | -0.11103 | 0.41266 | -0.26905 | 0.78789 | 0.88982 | 1.922 |
| ENSMUSG0000012 | 14.6432 | 0.254665 | 0.45431 | 0.560554 | 0.5751 | 0.75324 | 1.85 |
| ENSMUSG0000012 | 156.652 | 0.278408 | 0.17855 | 1.559305 | 0.11892 | 0.29105 | 1.363 |
| ENSMUSG0000012 | 28.5014 | 0.50885 | 0.36001 | 1.413439 | 0.15753 | 0.34464 | 0.952 |
| ENSMUSG0000012 | 24.2326 | 0.526373 | 0.34848 | 1.510482 | 0.13092 | 0.30934 | 0.455 |
| ENSMUSG0000012 | 6695.57 | -0.04282 | 0.0913 | -0.46899 | 0.63908 | 0.79759 | 2.215 |
| ENSMUSG0000012 | 84.1606 | 0.034394 | 0.30413 | 0.113091 | 0.90996 | 0.95466 | 2.881 |
| ENSMUSG0000012 | 32.7485 | 0.369292 | 0.34901 | 1.058125 | 0.29 | 0.50018 | 1.934 |
| ENSMUSG0000012 | 328.606 | 0.292969 | 0.15537 | 1.885625 | 0.05935 | 0.1854 | 3.97 |
| ENSMUSG0000012 | 14.1037 | 0.398443 | 0.58103 | 0.685759 | 0.49287 | 0.68741 | 0.525 |
| ENSMUSG0000012 | 48.3118 | 0.011664 | 0.30059 | 0.038805 | 0.96905 | 0.98431 | 1.513 |
| ENSMUSG0000012 | 29.8508 | 0.21665 | 0.43203 | 0.501467 | 0.61604 | 0.78132 | 3.174 |
| ENSMUSG0000012 | 19.3835 | 0.626014 | 0.52504 | 1.192324 | 0.23313 | 0.43798 | 0.366 |
| ENSMUSG0000012 | 17.0508 | 0.815782 | 0.5433 | 1.501531 | 0.13322 | 0.31278 | 2.37 |
| ENSMUSG0000012 | 90.5613 | -0.56271 | 0.23709 | -2.37338 | 0.01763 | 0.0825 | 3.74 |
| ENSMUSG0000012 | 17.9865 | -0.5249 | 0.43513 | -1.20632 | 0.22769 | 0.43174 | 1.392 |
| ENSMUSG0000012 | 82.7854 | -0.00219 | 0.21837 | -0.01001 | 0.99201 | 0.99538 | 2.346 |
| ENSMUSG0000012 | 116.971 | -0.14851 | 0.21105 | -0.70365 | 0.48165 | 0.67858 | 1.447 |
| ENSMUSG0000012 | 162.28 | -0.14586 | 0.1551 | -0.94044 | 0.34699 | 0.55765 | 0.636 |
| ENSMUSG0000012 | 24.6046 | 0.424128 | 0.4013 | 1.056873 | 0.29057 | 0.50076 | 7.431 |
| ENSMUSG0000012 | 16.7404 | -0.08562 | 0.45523 | -0.18807 | 0.85082 | 0.9241 | 0.772 |
| ENSMUSG0000012 | 36.4142 | 1.528694 | 0.39087 | 3.911028 | 9.19E-05 | 0.00254 | 0.759 |
| ENSMUSG0000012 | 285.911 | 0.601118 | 0.17742 | 3.388075 | 0.0007 | 0.01001 | 1.798 |
| ENSMUSG0000012 | 288.245 | -0.86061 | 0.26458 | -3.2527 | 0.00114 | 0.01385 | 2.74 |
| ENSMUSG0000012 | 20.3353 | 0.173088 | 0.41644 | 0.415636 | 0.67768 | 0.82333 | 0.565 |
| ENSMUSG0000012 | 14.5061 | -0.3234 | 0.47122 | -0.6863 | 0.49252 | 0.68714 | 1.075 |

| | | | | | | | |
|---|---|---|---|---|---|---|---|
| ENSMUSG0000012 | 16.1346 | -0.44148 | 0.46484 | -0.94973 | 0.34225 | 0.553 | 2.025 |
| ENSMUSG0000012 | 16.9666 | 0.650682 | 0.45309 | 1.436107 | 0.15097 | 0.33585 | 0.377 |
| ENSMUSG0000012 | 31.6245 | 0.02368 | 0.45639 | 0.051886 | 0.95862 | 0.97982 | 1.112 |
| ENSMUSG0000012 | 113.311 | 0.143577 | 0.21912 | 0.65524 | 0.51231 | 0.70333 | 1.866 |
| ENSMUSG0000012 | 118.678 | 0.523368 | 0.19269 | 2.716112 | 0.00661 | 0.04381 | 2.218 |
| ENSMUSG0000012 | 83.8515 | 0.358232 | 0.21689 | 1.651684 | 0.0986 | 0.25787 | 1.166 |
| ENSMUSG0000012 | 14.2345 | 0.31821 | 0.46745 | 0.680741 | 0.49604 | 0.68983 | 0.448 |
| ENSMUSG0000012 | 21.1297 | -0.84765 | 0.44099 | -1.92217 | 0.05458 | 0.17572 | 1.557 |
| ENSMUSG0000012 | 31.5917 | -0.29651 | 0.36461 | -0.81323 | 0.41608 | 0.62368 | 0.477 |
| ENSMUSG0000012 | 797.416 | 0.554557 | 0.2212 | 2.507054 | 0.01217 | 0.06518 | 4.975 |
| ENSMUSG0000012 | 13.4609 | 0.58221 | 0.55682 | 1.045595 | 0.29575 | 0.50616 | 0.765 |
| ENSMUSG0000012 | 32.9216 | -0.09541 | 0.4104 | -0.23248 | 0.81617 | 0.9054 | 2.269 |
| ENSMUSG0000012 | 138.176 | -0.36036 | 0.17191 | -2.09626 | 0.03606 | 0.13369 | 0.334 |
| ENSMUSG0000012 | 44.5928 | 0.120595 | 0.28677 | 0.42053 | 0.6741 | 0.82108 | 2.305 |
| ENSMUSG0000012 | 159.703 | 0.025461 | 0.15959 | 0.159539 | 0.87324 | 0.93626 | 1.621 |
| ENSMUSG0000012 | 22.6118 | 0.602724 | 0.4639 | 1.299259 | 0.19386 | 0.39087 | 0.595 |
| ENSMUSG0000012 | 88.4013 | 0.283616 | 0.1952 | 1.452914 | 0.14625 | 0.32924 | 0.438 |
| ENSMUSG0000012 | 114.471 | 0.378759 | 0.1877 | 2.017856 | 0.04361 | 0.15104 | 0.534 |
| ENSMUSG0000012 | 35.7734 | -0.44962 | 0.33922 | -1.32547 | 0.18502 | 0.3798 | 3.221 |
| ENSMUSG0000012 | 36.8504 | -0.11071 | 0.32593 | -0.33968 | 0.7341 | 0.85925 | 0.604 |
| ENSMUSG0000012 | 741.981 | 0.003157 | 0.11495 | 0.027467 | 0.97809 | 0.98827 | 6.611 |
| ENSMUSG0000012 | 2142.74 | 0.137203 | 0.08801 | 1.559009 | 0.11899 | 0.29112 | 1.374 |
| ENSMUSG0000012 | 16.2536 | -0.21455 | 0.47904 | -0.44788 | 0.65424 | 0.80821 | 5.318 |
| ENSMUSG0000012 | 16.5907 | -0.12752 | 0.43931 | -0.29028 | 0.7716 | 0.88094 | 0.519 |
| ENSMUSG0000012 | 16.9653 | -0.02668 | 0.39914 | -0.06684 | 0.94671 | 0.9732 | 5.798 |
| ENSMUSG0000012 | 68.3009 | 0.073217 | 0.23338 | 0.313722 | 0.75373 | 0.87065 | 0.874 |
| ENSMUSG0000012 | 124.308 | 0.179459 | 0.20932 | 0.857357 | 0.39125 | 0.60036 | 2.116 |
| ENSMUSG0000012 | 40.1797 | 0.751846 | 0.42588 | 1.765401 | 0.0775 | 0.22023 | 0.511 |
| ENSMUSG0000012 | 39.5741 | -0.12072 | 0.27997 | -0.43119 | 0.66633 | 0.8158 | 0.324 |
| ENSMUSG0000012 | 22.867 | 0.537361 | 0.41296 | 1.301255 | 0.19317 | 0.39009 | 1.169 |
| ENSMUSG0000012 | 13.5673 | 0.034717 | 0.54113 | 0.064157 | 0.94885 | 0.97462 | 1.519 |
| ENSMUSG0000012 | 383.543 | -0.17899 | 0.13899 | -1.28782 | 0.19781 | 0.39558 | 2.093 |
| ENSMUSG0000012 | 82.4561 | -1.11665 | 0.24238 | -4.60692 | 4.09E-06 | 0.00029 | 15.94 |
| ENSMUSG0000012 | 8021.85 | -0.3177 | 0.10879 | -2.92046 | 0.0035 | 0.02872 | 1.796 |

| | | | | | | | |
|---|---|---|---|---|---|---|---|
| ENSMUSG0000012 | 27.1823 | 0.076844 | 0.39315 | 0.195458 | 0.84503 | 0.92074 | 0.156 |
| ENSMUSG0000012 | 32.9028 | 0.434511 | 0.35188 | 1.234845 | 0.21689 | 0.41877 | 2.434 |
| ENSMUSG0000012 | 22.9969 | -0.00881 | 0.33981 | -0.02594 | 0.97931 | 0.9889 | 0.585 |
| ENSMUSG0000012 | 15.7917 | -0.05003 | 0.46948 | -0.10657 | 0.91513 | 0.95656 | 0.833 |
| ENSMUSG0000012 | 8450.01 | 0.857735 | 0.32264 | 2.658455 | 0.00785 | 0.04908 | 3.093 |
| ENSMUSG0000012 | 24.5287 | -0.31106 | 0.40103 | -0.77566 | 0.43795 | 0.64331 | 0.309 |
| ENSMUSG0000012 | 1276.81 | 0.426964 | 0.1621 | 2.633906 | 0.00844 | 0.05121 | 1.821 |
| ENSMUSG0000012 | 32.4257 | -0.62623 | 0.33485 | -1.87017 | 0.06146 | 0.18957 | 8.532 |
| ENSMUSG0000012 | 22.8855 | -0.61672 | 0.41275 | -1.49418 | 0.13513 | 0.31502 | 1.509 |
| ENSMUSG0000012 | 17.3833 | -0.17217 | 0.41835 | -0.41154 | 0.68067 | 0.82518 | 0.637 |
| ENSMUSG0000012 | 236.862 | 0.545142 | 0.15884 | 3.431998 | 0.0006 | 0.00895 | 1.143 |
| ENSMUSG0000012 | 25.6712 | -0.08423 | 0.3988 | -0.21121 | 0.83272 | 0.91425 | 0.332 |
| ENSMUSG0000012 | 24.0885 | 0.625317 | 0.36361 | 1.719766 | 0.08547 | 0.23459 | 0.545 |
| ENSMUSG0000012 | 12.52 | -0.42757 | 0.46749 | -0.91462 | 0.36039 | 0.57074 | 1.482 |
| ENSMUSG0000012 | 12.5406 | -0.34509 | 0.52189 | -0.66125 | 0.50846 | 0.69973 | 28.819 |
| ENSMUSG0000012 | 45.0518 | -0.61824 | 0.50779 | -1.21751 | 0.22341 | 0.42664 | 16.638 |
| ENSMUSG0000012 | 16.9295 | 0.052469 | 0.4855 | 0.108071 | 0.91394 | 0.9564 | 0.957 |
| ENSMUSG0000012 | 26.8827 | -0.25958 | 0.38828 | -0.66855 | 0.50379 | 0.69618 | 1.426 |
| ENSMUSG0000012 | 303.794 | -0.19089 | 0.12077 | -1.58062 | 0.11397 | 0.28262 | 2.924 |
| ENSMUSG0000012 | 952.161 | 0.087119 | 0.09351 | 0.931619 | 0.35153 | 0.56204 | 0.829 |
| ENSMUSG0000012 | 19.9662 | -0.6858 | 0.54029 | -1.26933 | 0.20432 | 0.40307 | 2.17 |
| ENSMUSG0000012 | 14.7868 | -0.44129 | 0.60179 | -0.7333 | 0.46338 | 0.66317 | 0.736 |
| ENSMUSG0000012 | 171.706 | -0.25246 | 0.18718 | -1.34876 | 0.17741 | 0.36951 | 1.947 |
| ENSMUSG0000012 | 302.764 | -0.01789 | 0.16218 | -0.11028 | 0.91219 | 0.9557 | 1.643 |
| ENSMUSG0000012 | 12.6308 | -0.20531 | 0.70352 | -0.29183 | 0.77042 | 0.87998 | 1.086 |
| ENSMUSG0000012 | 23.0839 | 0.490005 | 0.42171 | 1.161939 | 0.24526 | 0.45194 | 1.376 |
| ENSMUSG0000012 | 21.3749 | 0.449297 | 0.5448 | 0.824708 | 0.40954 | 0.61732 | 0.842 |
| ENSMUSG0000012 | 13.4269 | 0.60939 | 0.53778 | 1.133165 | 0.25715 | 0.46513 | 4.281 |
| ENSMUSG0000012 | 13.0748 | 0.464954 | 0.56921 | 0.816843 | 0.41402 | 0.62205 | 1.067 |
| ENSMUSG0000012 | 244.954 | 0.327329 | 0.13204 | 2.479013 | 0.01317 | 0.06872 | 30.25 |
| ENSMUSG0000012 | 73.1019 | 0.244862 | 0.217 | 1.128382 | 0.25916 | 0.46734 | 4.919 |
| ENSMUSG0000207 | 31.1993 | 0.189042 | 0.33908 | 0.557521 | 0.57717 | 0.75441 | 0.124 |
| ENSMUSG0000207 | 69.0024 | -0.14959 | 0.20624 | -0.72531 | 0.46826 | 0.66717 | 0.155 |
| ENSMUSG0000207 | 19.9341 | 0.260885 | 0.4696 | 0.555548 | 0.57852 | 0.75535 | 0.114 |

| | | | | | | | |
|---|---|---|---|---|---|---|---|
| ENSMUSG0000207 | 45.8976 | 0.621563 | 0.2848 | 2.182418 | 0.02908 | 0.1155 | 0.123 |
| ENSMUSG0000207 | 100.859 | -0.28407 | 0.24216 | -1.17307 | 0.24077 | 0.44702 | 0.123 |
| ENSMUSG0000207 | 41.554 | 0.456388 | 0.27692 | 1.648092 | 0.09933 | 0.25904 | 0.076 |
| ENSMUSG0000207 | 42.0768 | 0.547347 | 0.34988 | 1.564372 | 0.11773 | 0.28887 | 0.135 |
| ENSMUSG0000207 | 20.2568 | 0.060105 | 0.45655 | 0.131649 | 0.89526 | 0.94744 | 0.054 |
| ENSMUSG0000207 | 15.0525 | 0.437715 | 0.54787 | 0.798935 | 0.42433 | 0.63107 | 0.124 |
| ENSMUSG0000207 | 80.8916 | -0.13972 | 0.3514 | -0.39761 | 0.69092 | 0.83217 | 0.057 |
| ENSMUSG0000207 | 28.2457 | -0.10347 | 0.40894 | -0.25302 | 0.80025 | 0.89682 | 0.088 |
| ENSMUSG0000207 | 12.5597 | 0.266302 | 0.52409 | 0.508122 | 0.61137 | 0.77842 | 0.118 |
| ENSMUSG0000207 | 43.4249 | 0.680374 | 0.31085 | 2.18876 | 0.02861 | 0.11419 | 0.135 |
| ENSMUSG0000207 | 26.7565 | 0.583987 | 0.4008 | 1.457053 | 0.1451 | 0.32787 | 0.121 |
| ENSMUSG0000207 | 26.2927 | -0.5731 | 0.42136 | -1.36011 | 0.1738 | 0.36524 | 0.091 |
| ENSMUSG0000207 | 409.225 | -0.42597 | 0.15981 | -2.66542 | 0.00769 | 0.04845 | 0.057 |
| ENSMUSG0000207 | 12.8874 | -0.0369 | 0.50236 | -0.07346 | 0.94144 | 0.97087 | 0.096 |
| ENSMUSG0000207 | 115.385 | 0.859747 | 0.22887 | 3.756483 | 0.00017 | 0.0039 | 0.095 |
| ENSMUSG0000207 | 17.1845 | -0.03657 | 0.49904 | -0.07328 | 0.94159 | 0.97087 | 0.091 |
| ENSMUSG0000207 | 18.327 | 1.087816 | 0.50328 | 2.161461 | 0.03066 | 0.11979 | 0.101 |
| ENSMUSG0000207 | 19.7223 | 0.056169 | 0.38623 | 0.145429 | 0.88437 | 0.94244 | 0.086 |
| ENSMUSG0000207 | 67.6708 | 0.04405 | 0.31444 | 0.140091 | 0.88859 | 0.9444 | 0.154 |
| Eaf1 | 1308.02 | -0.04012 | 0.08832 | -0.45422 | 0.64967 | 0.80496 | 15.46 |
| Eaf2 | 75.506 | 0.306458 | 0.25485 | 1.2025 | 0.22917 | 0.43326 | 82.12 |
| Eapp | 2125.79 | 0.032352 | 0.07511 | 0.430736 | 0.66666 | 0.8158 | 25.558 |
| Ears2 | 482.2 | 0.153746 | 0.09896 | 1.553638 | 0.12027 | 0.29298 | 30.051 |
| Ebag9 | 1130.75 | -0.00526 | 0.08301 | -0.06341 | 0.94944 | 0.97495 | 22.341 |
| Ebf1 | 235.197 | 0.333257 | 0.16309 | 2.043368 | 0.04102 | 0.14503 | 390.775 |
| Ebf2 | 13.7723 | -0.82998 | 0.69826 | -1.18863 | 0.23459 | 0.43974 | 197.627 |
| Ebf3 | 37.8895 | -0.79543 | 0.30439 | -2.61315 | 0.00897 | 0.05332 | 120.773 |
| Ebf4 | 534.192 | 0.267305 | 0.1744 | 1.532689 | 0.12535 | 0.30036 | 75.313 |
| Ebna1bp2 | 2325.46 | 0.074477 | 0.06143 | 1.212318 | 0.22539 | 0.42934 | 6.978 |
| Ebp | 456.434 | -0.15739 | 0.11429 | -1.37708 | 0.16849 | 0.35847 | 8.184 |
| Ebpl | 601.357 | -0.17538 | 0.09317 | -1.88244 | 0.05978 | 0.18623 | 28.698 |
| Ecd | 1653.93 | 0.017689 | 0.073 | 0.2423 | 0.80855 | 0.90156 | 28.27 |
| Ece1 | 2278.05 | -0.05438 | 0.07062 | -0.77002 | 0.44129 | 0.64588 | 102.993 |
| Ece2 | 276.168 | 0.190291 | 0.17916 | 1.06211 | 0.28819 | 0.49834 | 16.658 |

| | | | | | | | |
|---|---|---|---|---|---|---|---|
| Ecel1 | 1823.29 | 0.929349 | 0.30902 | 3.007453 | 0.00263 | 0.02416 | 8.867 |
| Ech1 | 1499.56 | -0.11105 | 0.09517 | -1.16687 | 0.24326 | 0.44952 | 7.031 |
| Echdc1 | 93.7231 | -0.34893 | 0.18445 | -1.89178 | 0.05852 | 0.184 | 34.304 |
| Echdc2 | 282.93 | 0.181285 | 0.14114 | 1.28444 | 0.19899 | 0.39689 | 13.872 |
| Echdc3 | 27.4218 | -0.29199 | 0.32341 | -0.90285 | 0.36661 | 0.57697 | 24.569 |
| Echs1 | 2004.75 | -0.06587 | 0.08462 | -0.77836 | 0.43636 | 0.64199 | 10.767 |
| Eci1 | 648.278 | -0.36453 | 0.12036 | -3.02871 | 0.00246 | 0.02306 | 12.708 |
| Eci2 | 1167.92 | -0.13221 | 0.08138 | -1.62451 | 0.10427 | 0.26711 | 49.349 |
| Ecm1 | 528.648 | 0.027926 | 0.12474 | 0.223883 | 0.82285 | 0.90817 | 5.423 |
| Ecm2 | 213.973 | -0.2417 | 0.19053 | -1.26857 | 0.20459 | 0.40342 | 27.98 |
| Ecpas | 7530.37 | 0.194563 | 0.06164 | 3.15658 | 0.0016 | 0.01734 | 113.839 |
| Ecrg4 | 22.9106 | -0.82565 | 0.41442 | -1.99229 | 0.04634 | 0.15732 | 11.977 |
| Ecscr | 47.7135 | -0.4785 | 0.28991 | -1.65049 | 0.09884 | 0.25828 | 9.271 |
| Ecsit | 966.937 | -0.05912 | 0.09367 | -0.63116 | 0.52794 | 0.71583 | 13.193 |
| Ect2 | 44.1847 | -0.19884 | 0.29322 | -0.67812 | 0.49769 | 0.69108 | 56.657 |
| Ect2l | 25.8591 | 0.771668 | 0.35074 | 2.20013 | 0.0278 | 0.11205 | 81.988 |
| Eda | 300.859 | 0.208384 | 0.1436 | 1.451119 | 0.14675 | 0.32983 | 425.157 |
| Eda2r | 40.0545 | -0.2286 | 0.53192 | -0.42976 | 0.66737 | 0.8162 | 43.377 |
| Edar | 22.0592 | -0.51354 | 0.38731 | -1.32594 | 0.18486 | 0.37959 | 74.866 |
| Edaradd | 12.3802 | -2 | 0.61757 | -3.60577 | 0.00031 | 0.00579 | 47.807 |
| Edc3 | 1438.17 | 0.165332 | 0.07693 | 2.149176 | 0.03162 | 0.12249 | 43.96 |
| Edc4 | 3478.09 | 0.166984 | 0.06781 | 2.462641 | 0.01379 | 0.07062 | 14.028 |
| Edem1 | 1427.62 | -0.20767 | 0.10795 | -1.92383 | 0.05438 | 0.17534 | 30.716 |
| Edem2 | 902.451 | -0.18888 | 0.09445 | -1.99981 | 0.04552 | 0.1553 | 27.799 |
| Edem3 | 3677.4 | 0.038262 | 0.0793 | 0.482516 | 0.62944 | 0.79054 | 66.681 |
| Edf1 | 2600.23 | -0.20429 | 0.08703 | -2.34746 | 0.0189 | 0.08627 | 4.236 |
| Edil3 | 4434.93 | -0.43421 | 0.13787 | -3.14953 | 0.00164 | 0.01761 | 501.752 |
| Edn1 | 157.232 | -0.3268 | 0.17424 | -1.87556 | 0.06072 | 0.1881 | 6.515 |
| Edn3 | 347.38 | -0.579 | 0.15182 | -3.81387 | 0.00014 | 0.00332 | 23.424 |
| Ednra | 144.085 | -0.45081 | 0.1473 | -3.06052 | 0.00221 | 0.02148 | 61.434 |
| Ednrb | 2664.47 | -0.1538 | 0.1391 | -1.10568 | 0.26887 | 0.47813 | 29.778 |
| Edrf1 | 1608.78 | 0.272108 | 0.06853 | 3.970545 | 7.17E-05 | 0.00213 | 35.429 |
| Eea1 | 3065.7 | 0.328996 | 0.1528 | 2.153158 | 0.03131 | 0.12162 | 104.869 |
| Eed | 959.802 | 0.142849 | 0.08756 | 1.63137 | 0.10281 | 0.26475 | 26.33 |

| | | | | | | | |
|---|---|---|---|---|---|---|---|
| Eef1a1 | 80225.3 | 0.101803 | 0.08984 | 1.133136 | 0.25716 | 0.46513 | 10.703 |
| Eef1a2 | 30925.2 | -0.21345 | 0.06285 | -3.39637 | 0.00068 | 0.00983 | 9.362 |
| Eef1akmt1 | 672.977 | -0.19861 | 0.09713 | -2.04474 | 0.04088 | 0.1448 | 22.016 |
| Eef1akmt2 | 830.032 | 0.296672 | 0.07815 | 3.796136 | 0.00015 | 0.00348 | 25.217 |
| Eef1akmt4 | 20.1449 | -0.09826 | 0.38414 | -0.25581 | 0.7981 | 0.8958 | 7.419 |
| Eef1aknmt | 841.363 | 0.138985 | 0.10514 | 1.321861 | 0.18621 | 0.38114 | 16.425 |
| Eef1b2 | 4462.62 | 0.008698 | 0.08397 | 0.10358 | 0.9175 | 0.95781 | 3.662 |
| Eef1d | 2791.36 | -0.0488 | 0.0696 | -0.70118 | 0.48319 | 0.67959 | 15.352 |
| Eef1e1 | 997.488 | -0.04427 | 0.08112 | -0.5458 | 0.5852 | 0.75979 | 14.852 |
| Eef1g | 9151.9 | 0.032074 | 0.06995 | 0.458524 | 0.64658 | 0.80321 | 11.439 |
| Eef2 | 44506.1 | -0.15468 | 0.0707 | -2.18784 | 0.02868 | 0.1143 | 5.868 |
| Eef2-ps2 | 241.581 | -0.11677 | 0.21412 | -0.54536 | 0.58551 | 0.75994 | 1.345 |
| Eef2k | 2012.86 | -0.0502 | 0.08861 | -0.56659 | 0.57099 | 0.75014 | 64.62 |
| Eef2kmt | 170.001 | 0.044079 | 0.15019 | 0.293496 | 0.76914 | 0.87961 | 11.832 |
| Eefsec | 434.724 | 0.0384 | 0.10232 | 0.375312 | 0.70743 | 0.84212 | 189.206 |
| Eepd1 | 563.125 | -0.28596 | 0.11957 | -2.39151 | 0.01678 | 0.08016 | 122.564 |
| Efcab1 | 367.775 | 0.753828 | 0.17034 | 4.425515 | 9.62E-06 | 0.00054 | 74.766 |
| Efcab10 | 20.9295 | 0.627494 | 0.37101 | 1.691296 | 0.09078 | 0.24468 | 6.414 |
| Efcab12 | 543.36 | 0.29549 | 0.1141 | 2.589641 | 0.00961 | 0.05584 | 27.684 |
| Efcab14 | 3484.4 | -0.10121 | 0.0668 | -1.51519 | 0.12972 | 0.30747 | 39.584 |
| Efcab15 | 89.339 | -0.16876 | 0.21433 | -0.78735 | 0.43107 | 0.63717 | 9.615 |
| Efcab2 | 899.797 | -0.20322 | 0.12693 | -1.60099 | 0.10938 | 0.2752 | 78.429 |
| Efcab5 | 222.813 | -0.07876 | 0.15345 | -0.51328 | 0.60776 | 0.77579 | 99.054 |
| Efcab6 | 285.359 | -0.57255 | 0.25724 | -2.22577 | 0.02603 | 0.10702 | 198.668 |
| Efcab7 | 128.734 | 0.706437 | 0.17325 | 4.07758 | 4.55E-05 | 0.00153 | 52.551 |
| Efcab9 | 31.2784 | 0.022153 | 0.39133 | 0.056609 | 0.95486 | 0.97783 | 9.331 |
| Efcc1 | 371.186 | 0.619533 | 0.18398 | 3.367391 | 0.00076 | 0.01059 | 25.04 |
| Efemp1 | 600.633 | -1.04464 | 0.23831 | -4.38351 | 1.17E-05 | 0.00062 | 73.54 |
| Efemp2 | 425.152 | -0.2602 | 0.12885 | -2.01951 | 0.04343 | 0.15068 | 8.564 |
| Efhb | 14.0658 | 1.175162 | 0.64934 | 1.809779 | 0.07033 | 0.20717 | 64.433 |
| Efhc1 | 44.8622 | 0.118289 | 0.29221 | 0.404811 | 0.68562 | 0.82872 | 39.216 |
| Efhc2 | 102.083 | 0.346705 | 0.1865 | 1.859054 | 0.06302 | 0.19274 | 187.32 |
| Efhd1 | 613.387 | -0.72417 | 0.15466 | -4.68247 | 2.83E-06 | 0.00023 | 46.477 |
| Efhd2 | 5624.02 | -0.64617 | 0.15254 | -4.23602 | 2.28E-05 | 0.00096 | 16.779 |

| | | | | | | | |
|---|---|---|---|---|---|---|---|
| Efl1 | 702.397 | 0.060432 | 0.07956 | 0.759603 | 0.44749 | 0.65115 | 129.239 |
| Efna1 | 100.699 | -0.51447 | 0.18408 | -2.79483 | 0.00519 | 0.03717 | 9.41 |
| Efna2 | 292.613 | -0.34969 | 0.13367 | -2.61603 | 0.0089 | 0.05308 | 10.529 |
| Efna3 | 958.984 | -0.22466 | 0.10541 | -2.1313 | 0.03306 | 0.12628 | 9.067 |
| Efna5 | 626.726 | -0.36145 | 0.13512 | -2.67505 | 0.00747 | 0.04761 | 277.134 |
| Efnb1 | 817.399 | -0.00488 | 0.12015 | -0.04061 | 0.96761 | 0.98368 | 12.862 |
| Efnb2 | 5759.25 | 0.314234 | 0.09746 | 3.224136 | 0.00126 | 0.01476 | 43.809 |
| Efnb3 | 6959.81 | 0.087639 | 0.1011 | 0.866851 | 0.38602 | 0.59548 | 6.114 |
| Efr3a | 4191.16 | -0.63539 | 0.31743 | -2.00163 | 0.04532 | 0.15486 | 86.783 |
| Efr3b | 13105.7 | -0.07145 | 0.05263 | -1.35766 | 0.17457 | 0.36618 | 76.362 |
| Efs | 407.623 | -0.2205 | 0.1124 | -1.96175 | 0.04979 | 0.16541 | 9.592 |
| Eftud2 | 4349.79 | 0.057241 | 0.06264 | 0.913864 | 0.36079 | 0.57103 | 42.513 |
| Egf | 161.984 | 0.173972 | 0.14694 | 1.183956 | 0.23643 | 0.44163 | 77.752 |
| Egfem1 | 341.403 | -0.34341 | 0.13199 | -2.60185 | 0.00927 | 0.05453 | 609.187 |
| Egfl6 | 86.9279 | -0.11142 | 0.29441 | -0.37846 | 0.70509 | 0.84115 | 62.714 |
| Egfl7 | 457.323 | -0.63275 | 0.16829 | -3.75988 | 0.00017 | 0.00387 | 13.107 |
| Egfl8 | 36.3619 | 0.372737 | 0.29358 | 1.269631 | 0.20422 | 0.40298 | 2.623 |
| Egflam | 143.082 | -0.38686 | 0.1838 | -2.10479 | 0.03531 | 0.13194 | 192.276 |
| Egfr | 1075.93 | 0.026306 | 0.10226 | 0.257231 | 0.797 | 0.89539 | 165.956 |
| Egln1 | 5203.45 | 0.127979 | 0.11539 | 1.109069 | 0.2674 | 0.47675 | 40.738 |
| Egln2 | 3816.19 | -0.06011 | 0.07897 | -0.76128 | 0.44649 | 0.65009 | 8.145 |
| Egln3 | 246.921 | -1.21668 | 0.2579 | -4.71769 | 2.39E-06 | 0.0002 | 24.88 |
| Egr1 | 6486.83 | -0.39869 | 0.11136 | -3.5801 | 0.00034 | 0.00616 | 5.162 |
| Egr2 | 106.032 | -1.19864 | 0.33074 | -3.62407 | 0.00029 | 0.00549 | 6.714 |
| Egr3 | 4958.81 | 0.224587 | 0.13168 | 1.705547 | 0.08809 | 0.23962 | 5.412 |
| Egr4 | 784.772 | -0.26786 | 0.14015 | -1.91132 | 0.05596 | 0.17878 | 2.469 |
| Ehbp1 | 2904.57 | 0.149921 | 0.09293 | 1.613275 | 0.10668 | 0.27095 | 336.465 |
| Ehbp1l1 | 1410.31 | 0.365234 | 0.10998 | 3.321046 | 0.0009 | 0.01186 | 18.942 |
| Ehd1 | 1919.95 | 0.144081 | 0.13163 | 1.094557 | 0.27371 | 0.48313 | 23.372 |
| Ehd2 | 402.442 | -0.25374 | 0.17667 | -1.43619 | 0.15095 | 0.33584 | 20.61 |
| Ehd3 | 3544.14 | -0.31069 | 0.08587 | -3.61833 | 0.0003 | 0.00559 | 27.961 |
| Ehd4 | 665.002 | -0.31764 | 0.12721 | -2.49708 | 0.01252 | 0.06645 | 65.432 |
| Ehhadh | 315.512 | 0.187634 | 0.14501 | 1.293979 | 0.19567 | 0.39335 | 26.521 |
| Ehmt1 | 2206.85 | 0.141782 | 0.07297 | 1.943084 | 0.05201 | 0.17024 | 129.687 |

| | | | | | | | |
|---|---|---|---|---|---|---|---|
| Ehmt2 | 9217.03 | 0.093342 | 0.06656 | 1.402267 | 0.16084 | 0.34901 | 15.584 |
| Ei24 | 3910.52 | -0.08146 | 0.08186 | -0.99509 | 0.31969 | 0.53037 | 18.235 |
| Eid1 | 4586.7 | -0.0242 | 0.0679 | -0.35636 | 0.72157 | 0.85138 | 1.691 |
| Eid2 | 1157.06 | 0.011501 | 0.09514 | 0.120886 | 0.90378 | 0.95146 | 1.392 |
| Eid2b | 1259.36 | 0.088124 | 0.08553 | 1.030383 | 0.30283 | 0.51232 | 1.751 |
| Eid3 | 32.4451 | 0.660534 | 0.30643 | 2.155592 | 0.03112 | 0.12109 | 1.305 |
| Eif1 | 14469.4 | 0.10249 | 0.09017 | 1.136655 | 0.25568 | 0.46344 | 2.212 |
| Eif1a | 2036.19 | 0.156892 | 0.07984 | 1.964971 | 0.04942 | 0.16462 | 18.756 |
| Eif1ad | 674.065 | -0.17989 | 0.08912 | -2.01855 | 0.04353 | 0.15094 | 4.786 |
| Eif1ax | 8168.18 | -0.11275 | 0.07701 | -1.46416 | 0.14315 | 0.32518 | 17.751 |
| Eif1b | 2397.34 | -0.05103 | 0.06855 | -0.74438 | 0.45665 | 0.65819 | 3.099 |
| Eif2a | 2064.48 | 0.240977 | 0.07526 | 3.201846 | 0.00137 | 0.01549 | 31.681 |
| Eif2ak1 | 2749.18 | 0.026447 | 0.06814 | 0.388156 | 0.6979 | 0.83633 | 37.527 |
| Eif2ak2 | 438.05 | -0.14426 | 0.12793 | -1.12772 | 0.25944 | 0.46775 | 30.01 |
| Eif2ak3 | 1199.63 | -0.19635 | 0.07962 | -2.46621 | 0.01366 | 0.07034 | 60.731 |
| Eif2ak4 | 1602.04 | 0.087328 | 0.08729 | 1.0004 | 0.31712 | 0.52747 | 86.617 |
| Eif2b1 | 980.462 | -0.02707 | 0.08537 | -0.31706 | 0.7512 | 0.86954 | 8.919 |
| Eif2b2 | 1207.93 | -0.09714 | 0.08232 | -1.18009 | 0.23796 | 0.44334 | 7.148 |
| Eif2b3 | 549.968 | -0.0539 | 0.09742 | -0.55322 | 0.58011 | 0.75651 | 67.905 |
| Eif2b4 | 1321.53 | -0.05589 | 0.07766 | -0.71963 | 0.47175 | 0.67032 | 5.873 |
| Eif2b5 | 2104.14 | -0.07611 | 0.07438 | -1.02322 | 0.3062 | 0.51631 | 10.507 |
| Eif2d | 1547.66 | 0.07619 | 0.07726 | 0.986186 | 0.32404 | 0.53514 | 34.478 |
| Eif2s1 | 1900.22 | -0.01204 | 0.07738 | -0.15555 | 0.87639 | 0.93824 | 25.192 |
| Eif2s2 | 2746.06 | 0.020258 | 0.10486 | 0.193196 | 0.84681 | 0.92198 | 21.526 |
| Eif2s3x | 3018.59 | -0.12847 | 0.06887 | -1.86539 | 0.06213 | 0.19095 | 24.156 |
| Eif2s3y | 3099.78 | 0.004724 | 0.13964 | 0.033832 | 0.97301 | 0.98656 | 18.305 |
| Eif3a | 10495.9 | 0.126575 | 0.10554 | 1.199364 | 0.23039 | 0.43483 | 29.542 |
| Eif3b | 4602.62 | -0.03177 | 0.07517 | -0.42255 | 0.67262 | 0.81992 | 24.033 |
| Eif3c | 7957.22 | 0.007895 | 0.06977 | 0.11315 | 0.90991 | 0.95466 | 19.957 |
| Eif3d | 857.205 | -0.0501 | 0.08966 | -0.55881 | 0.57629 | 0.75395 | 11.816 |
| Eif3e | 2905.19 | 0.177634 | 0.05732 | 3.099018 | 0.00194 | 0.01971 | 32.662 |
| Eif3f | 4096.26 | -0.0014 | 0.10266 | -0.01363 | 0.98912 | 0.994 | 8.535 |
| Eif3g | 1974.11 | -0.14691 | 0.07698 | -1.90834 | 0.05635 | 0.17956 | 4.275 |
| Eif3h | 4311.31 | -0.06247 | 0.07193 | -0.86851 | 0.38511 | 0.59448 | 78.966 |

| | | | | | | | |
|---|---|---|---|---|---|---|---|
| Eif3i | 2135.61 | -0.0947 | 0.08412 | -1.12587 | 0.26022 | 0.46875 | 8.689 |
| Eif3j1 | 1769.07 | 0.016748 | 0.0875 | 0.191405 | 0.84821 | 0.92288 | 28.053 |
| Eif3j2 | 47.797 | -0.65607 | 0.3536 | -1.85539 | 0.06354 | 0.19378 | 2.379 |
| Eif3k | 2280.61 | -0.04552 | 0.07737 | -0.58839 | 0.55627 | 0.73841 | 10.493 |
| Eif3l | 2645.73 | -0.04912 | 0.07029 | -0.69882 | 0.48466 | 0.68091 | 19.227 |
| Eif3m | 2121.45 | 0.131077 | 0.07573 | 1.730949 | 0.08346 | 0.23096 | 17.425 |
| Eif4a-ps4 | 1369.85 | 0.111967 | 0.11468 | 0.976372 | 0.32888 | 0.54005 | 1.218 |
| Eif4a1 | 8444.99 | 0.039551 | 0.06555 | 0.603351 | 0.54628 | 0.73071 | 5.488 |
| Eif4a2 | 32273.8 | 0.073485 | 0.07642 | 0.961573 | 0.33626 | 0.54704 | 6.693 |
| Eif4a3 | 1800.28 | -0.0439 | 0.09925 | -0.4423 | 0.65827 | 0.81038 | 11.727 |
| Eif4b | 10382.9 | 0.085656 | 0.06034 | 1.419437 | 0.15577 | 0.34238 | 23.401 |
| Eif4e | 4554.9 | 0.054334 | 0.08497 | 0.639442 | 0.52254 | 0.71132 | 33.518 |
| Eif4e2 | 2778.16 | -0.01237 | 0.06205 | -0.19941 | 0.84194 | 0.91879 | 26.575 |
| Eif4e3 | 674.749 | 0.182349 | 0.11307 | 1.612697 | 0.10681 | 0.27117 | 41.637 |
| Eif4ebp1 | 85.3465 | -0.61948 | 0.22017 | -2.81358 | 0.0049 | 0.03574 | 16.346 |
| Eif4ebp2 | 4654.25 | -0.05459 | 0.07024 | -0.77712 | 0.43709 | 0.64265 | 24.171 |
| Eif4ebp3 | 44.8174 | 0.323214 | 0.27029 | 1.195787 | 0.23178 | 0.43653 | 1.858 |
| Eif4enif1 | 2403.99 | 0.074001 | 0.06314 | 1.171982 | 0.2412 | 0.44734 | 42.197 |
| Eif4g1 | 15756.7 | -0.02723 | 0.0654 | -0.41628 | 0.6772 | 0.82314 | 24.572 |
| Eif4g2 | 31732.9 | -0.07067 | 0.05466 | -1.29307 | 0.19599 | 0.39362 | 15.281 |
| Eif4g3 | 11211.5 | -0.16737 | 0.08878 | -1.88526 | 0.05939 | 0.18544 | 215.487 |
| Eif4h | 14885.3 | -0.02844 | 0.0632 | -0.44999 | 0.65272 | 0.80685 | 19.77 |
| Eif5 | 12199.9 | 0.118351 | 0.0706 | 1.67637 | 0.09367 | 0.24948 | 8.737 |
| Eif5a | 9207.09 | -0.11908 | 0.08255 | -1.44244 | 0.14918 | 0.33328 | 5.245 |
| Eif5a2 | 3385.97 | -0.03091 | 0.06779 | -0.456 | 0.64839 | 0.80408 | 17.571 |
| Eif5b | 4144.3 | 0.066645 | 0.12243 | 0.544339 | 0.58621 | 0.76028 | 57.57 |
| Eif6 | 721.112 | -0.20176 | 0.10085 | -2.00058 | 0.04544 | 0.15512 | 7.094 |
| Eipr1 | 2449.23 | -0.09696 | 0.1007 | -0.96283 | 0.33563 | 0.54639 | 118.063 |
| Elac1 | 2290.78 | 0.151282 | 0.07095 | 2.132255 | 0.03299 | 0.12614 | 19.442 |
| Elac2 | 1699.29 | 0.024178 | 0.07368 | 0.328126 | 0.74282 | 0.86459 | 23.032 |
| Elapor1 | 3884.02 | 0.36709 | 0.09763 | 3.759941 | 0.00017 | 0.00387 | 80.843 |
| Elapor2 | 4197.08 | 0.641311 | 0.16989 | 3.774934 | 0.00016 | 0.0037 | 215.708 |
| Elavl1 | 4036.28 | 0.153963 | 0.06222 | 2.474628 | 0.01334 | 0.0693 | 40.032 |
| Elavl2 | 3741.85 | -0.18197 | 0.11683 | -1.55751 | 0.11935 | 0.29146 | 150.023 |

| | | | | | | | |
|---|---|---|---|---|---|---|---|
| Elavl3 | 10569.9 | -0.0296 | 0.0749 | -0.39525 | 0.69266 | 0.83343 | 37.019 |
| Elavl4 | 2927.09 | -0.61252 | 0.20308 | -3.01608 | 0.00256 | 0.02375 | 148.188 |
| Eldr | 22.9436 | 0.21266 | 0.34447 | 0.617357 | 0.537 | 0.72388 | 16.693 |
| Elf1 | 359.763 | -0.28037 | 0.14973 | -1.87246 | 0.06114 | 0.18892 | 101.301 |
| Elf2 | 1469.22 | -0.14 | 0.06808 | -2.05642 | 0.03974 | 0.14237 | 87.944 |
| Elf4 | 99.6823 | -0.51325 | 0.20946 | -2.45036 | 0.01427 | 0.07203 | 52.087 |
| Elf5 | 15.3246 | -0.40684 | 0.50055 | -0.8128 | 0.41633 | 0.62392 | 39.302 |
| Elfn1 | 1095.25 | -0.00597 | 0.13908 | -0.04295 | 0.96574 | 0.9828 | 66.78 |
| Elfn2 | 7879.51 | -0.12866 | 0.10541 | -1.22059 | 0.22224 | 0.42545 | 51.441 |
| Elk1 | 3528.72 | 0.199267 | 0.08216 | 2.425347 | 0.01529 | 0.07541 | 17.214 |
| Elk3 | 396.985 | 0.033553 | 0.14835 | 0.226184 | 0.82106 | 0.90748 | 63.722 |
| Elk4 | 1579.17 | -0.30853 | 0.11714 | -2.6338 | 0.00844 | 0.05121 | 25.006 |
| Ell | 1405.96 | 0.12847 | 0.10167 | 1.263654 | 0.20635 | 0.40562 | 53.402 |
| Ell2 | 1068.64 | -0.1233 | 0.09764 | -1.26277 | 0.20667 | 0.40605 | 65.608 |
| Elmo1 | 5637.51 | 0.245983 | 0.10081 | 2.440151 | 0.01468 | 0.07322 | 517.758 |
| Elmo2 | 6853.13 | -0.07048 | 0.06785 | -1.03885 | 0.29887 | 0.50895 | 38.449 |
| Elmo3 | 248.898 | 0.195183 | 0.17244 | 1.131875 | 0.25769 | 0.46576 | 5.16 |
| Elmod1 | 13100.5 | 0.41503 | 0.1094 | 3.793739 | 0.00015 | 0.0035 | 63.845 |
| Elmod2 | 1471.49 | 0.173718 | 0.10003 | 1.736685 | 0.08244 | 0.22898 | 19.855 |
| Elmod3 | 986.761 | -0.0158 | 0.07709 | -0.20501 | 0.83756 | 0.91642 | 32.492 |
| Eln | 463.51 | -0.69627 | 0.35991 | -1.93454 | 0.05305 | 0.17229 | 44.731 |
| Eloa | 1254.12 | -0.00414 | 0.08493 | -0.04879 | 0.96109 | 0.98111 | 18.396 |
| Elob | 2861.73 | -0.17583 | 0.07515 | -2.33982 | 0.01929 | 0.08737 | 4.399 |
| Eloc | 5147.55 | 0.025396 | 0.06669 | 0.380833 | 0.70333 | 0.83991 | 15.318 |
| Elof1 | 1078.54 | -0.22978 | 0.10069 | -2.2821 | 0.02248 | 0.09681 | 4.184 |
| Elovl1 | 1116.48 | -0.01921 | 0.07885 | -0.24365 | 0.8075 | 0.90088 | 4.861 |
| Elovl2 | 890.039 | -0.18514 | 0.13592 | -1.36218 | 0.17314 | 0.3644 | 38.025 |
| Elovl4 | 2286.87 | -0.21566 | 0.08206 | -2.62819 | 0.00858 | 0.05181 | 27.586 |
| Elovl5 | 4546.62 | -0.0401 | 0.07049 | -0.56891 | 0.56941 | 0.74898 | 67.156 |
| Elovl6 | 2503.76 | 0.024773 | 0.09903 | 0.250162 | 0.80246 | 0.89812 | 106.141 |
| Elovl7 | 566.796 | -0.50264 | 0.14226 | -3.53311 | 0.00041 | 0.00699 | 71.28 |
| Elp1 | 3831.23 | 0.047537 | 0.07881 | 0.603188 | 0.54638 | 0.73076 | 52.652 |
| Elp2 | 4166.94 | 0.022257 | 0.05421 | 0.410561 | 0.68139 | 0.82578 | 36.642 |
| Elp3 | 3216.32 | -0.0781 | 0.06417 | -1.21706 | 0.22358 | 0.42677 | 62.627 |

| | | | | | | | |
|---|---|---|---|---|---|---|---|
| Elp4 | 608.729 | -0.00242 | 0.09928 | -0.02442 | 0.98052 | 0.98974 | 203.538 |
| Elp5 | 1721.27 | -0.13987 | 0.08337 | -1.67774 | 0.0934 | 0.24883 | 14.296 |
| Elp6 | 414.326 | 0.060944 | 0.11163 | 0.545945 | 0.5851 | 0.75976 | 16.93 |
| Emb | 740.535 | 0.110886 | 0.12035 | 0.921386 | 0.35685 | 0.56713 | 65.88 |
| Emc1 | 3393.86 | -0.02869 | 0.05448 | -0.52655 | 0.5985 | 0.76969 | 26.144 |
| Emc10 | 6093.45 | -0.09071 | 0.06805 | -1.333 | 0.18253 | 0.37638 | 6.593 |
| Emc2 | 2079.76 | 0.082403 | 0.12867 | 0.640425 | 0.5219 | 0.71102 | 50.535 |
| Emc3 | 3002.52 | -0.10016 | 0.07989 | -1.25378 | 0.20992 | 0.40997 | 16.779 |
| Emc4 | 2467.38 | -0.10598 | 0.069 | -1.536 | 0.12454 | 0.29947 | 5.017 |
| Emc6 | 1141.52 | -0.10071 | 0.11147 | -0.90345 | 0.36629 | 0.57682 | 1.519 |
| Emc7 | 5231.61 | -0.06999 | 0.06844 | -1.02257 | 0.30651 | 0.5165 | 17.166 |
| Emc8 | 3630.23 | -0.11268 | 0.072 | -1.56515 | 0.11755 | 0.28858 | 14.66 |
| Emc9 | 1141.74 | 0.11189 | 0.11286 | 0.991401 | 0.32149 | 0.53233 | 3.787 |
| Emcn | 162.271 | 0.150675 | 0.20075 | 0.75058 | 0.45291 | 0.65531 | 91.119 |
| Emd | 1081.08 | -0.05795 | 0.09285 | -0.62414 | 0.53254 | 0.71989 | 6.862 |
| Eme2 | 246.393 | -0.00465 | 0.12278 | -0.03787 | 0.96979 | 0.98477 | 6.602 |
| Emg1 | 1279.28 | 0.076826 | 0.08462 | 0.907921 | 0.36392 | 0.57434 | 8.094 |
| Emid1 | 519.215 | -0.75297 | 0.36004 | -2.09133 | 0.0365 | 0.13459 | 45.993 |
| Emilin1 | 110.181 | -1.03711 | 0.33826 | -3.06603 | 0.00217 | 0.02127 | 7.876 |
| Emilin2 | 351.79 | -0.67861 | 0.17308 | -3.92088 | 8.82E-05 | 0.00247 | 59.807 |
| Eml1 | 2442.36 | 0.120072 | 0.07187 | 1.67063 | 0.09479 | 0.25135 | 168.661 |
| Eml2 | 6231.31 | -0.10419 | 0.08922 | -1.16777 | 0.2429 | 0.44905 | 30.062 |
| Eml3 | 447.507 | -0.32035 | 0.0987 | -3.2457 | 0.00117 | 0.01411 | 12.031 |
| Eml4 | 2317.11 | 0.129471 | 0.07465 | 1.734321 | 0.08286 | 0.22962 | 129.431 |
| Eml5 | 8046.46 | 0.182161 | 0.12324 | 1.478046 | 0.1394 | 0.3207 | 114.68 |
| Eml6 | 1864.38 | -0.03671 | 0.08933 | -0.41088 | 0.68116 | 0.82559 | 282.986 |
| Emp1 | 121.042 | -0.94143 | 0.32227 | -2.92124 | 0.00349 | 0.02868 | 20.629 |
| Emp2 | 929.599 | -0.19851 | 0.11125 | -1.78436 | 0.07437 | 0.21498 | 32.22 |
| Emp3 | 82.603 | -0.57167 | 0.24336 | -2.34911 | 0.01882 | 0.08608 | 3.382 |
| Emsy | 3958.89 | 0.176189 | 0.08708 | 2.023203 | 0.04305 | 0.14976 | 69.647 |
| Emx1 | 156.64 | -1.35001 | 0.35899 | -3.7606 | 0.00017 | 0.00387 | 17.025 |
| Emx2 | 434.164 | -0.33537 | 0.1396 | -2.40245 | 0.01629 | 0.0786 | 6.986 |
| Emx2os | 223.345 | -0.41344 | 0.14963 | -2.76311 | 0.00573 | 0.03996 | 33.532 |
| En2 | 28.1236 | -0.73664 | 0.39936 | -1.84457 | 0.0651 | 0.19674 | 6.473 |

| | | | | | | | |
|---|---|---|---|---|---|---|---|
| Enah | 12453.3 | 0.010152 | 0.05973 | 0.169967 | 0.86504 | 0.93163 | 123.607 |
| Enc1 | 22381.5 | -0.43944 | 0.28179 | -1.55944 | 0.11889 | 0.291 | 11.93 |
| Endod1 | 5326.84 | -0.41613 | 0.10721 | -3.88161 | 0.0001 | 0.00276 | 27.518 |
| Endog | 206.487 | -0.1822 | 0.13526 | -1.34705 | 0.17797 | 0.37028 | 2.577 |
| Endou | 71.0637 | -0.9109 | 0.30708 | -2.96631 | 0.00301 | 0.02623 | 20.325 |
| Endov | 1293.38 | -0.26872 | 0.09204 | -2.91956 | 0.00351 | 0.02878 | 20.091 |
| Eng | 883.152 | -0.3733 | 0.14015 | -2.66362 | 0.00773 | 0.0486 | 36.075 |
| Engase | 571.524 | -0.04265 | 0.09451 | -0.45124 | 0.65181 | 0.80618 | 12.381 |
| Enho | 1165.86 | -0.3326 | 0.09404 | -3.53681 | 0.0004 | 0.00692 | 2.181 |
| Enkd1 | 224.502 | 0.065174 | 0.14604 | 0.446275 | 0.6554 | 0.80876 | 4.56 |
| Enkur | 41.0417 | 0.402352 | 0.27253 | 1.476374 | 0.13984 | 0.32112 | 24.635 |
| Eno1 | 24366 | -0.13452 | 0.05945 | -2.26261 | 0.02366 | 0.10002 | 12.159 |
| Eno1b | 396.782 | -0.05387 | 0.12868 | -0.41864 | 0.67548 | 0.82192 | 3.078 |
| Eno2 | 40585 | 0.06497 | 0.07794 | 0.833551 | 0.40453 | 0.61341 | 9.621 |
| Eno3 | 330.742 | 0.083058 | 0.11469 | 0.724203 | 0.46894 | 0.66787 | 5.312 |
| Eno4 | 133.233 | 0.302617 | 0.18021 | 1.679261 | 0.0931 | 0.2484 | 27.997 |
| Enoph1 | 1746.43 | 0.02984 | 0.06684 | 0.446426 | 0.65529 | 0.80876 | 28.776 |
| Enox1 | 1812.81 | 0.008745 | 0.08484 | 0.103079 | 0.9179 | 0.95814 | 564.998 |
| Enox2 | 676.617 | 0.483921 | 0.19729 | 2.45289 | 0.01417 | 0.07176 | 278.553 |
| Enpep | 140.101 | -0.38446 | 0.2189 | -1.7563 | 0.07904 | 0.22308 | 63.546 |
| Enpp1 | 300.5 | -0.85481 | 0.13129 | -6.51091 | 7.47E-11 | 8.46E-08 | 74.246 |
| Enpp2 | 5890.49 | -0.64111 | 0.14197 | -4.51595 | 6.30E-06 | 0.00039 | 113.992 |
| Enpp3 | 19.3284 | -0.472 | 0.61173 | -0.77158 | 0.44036 | 0.64527 | 70.418 |
| Enpp4 | 2310.67 | -0.11549 | 0.08102 | -1.42557 | 0.15399 | 0.33978 | 9.502 |
| Enpp5 | 13301 | 0.134265 | 0.0546 | 2.459094 | 0.01393 | 0.07102 | 7.755 |
| Enpp6 | 245.868 | -0.46541 | 0.2123 | -2.19227 | 0.02836 | 0.11355 | 109.876 |
| Ensa | 11460.2 | -0.16475 | 0.07646 | -2.15462 | 0.03119 | 0.12124 | 7.11 |
| Entpd1 | 804.22 | -0.29406 | 0.12807 | -2.29617 | 0.02167 | 0.09443 | 129.237 |
| Entpd2 | 227.88 | -0.06962 | 0.22927 | -0.30364 | 0.7614 | 0.87503 | 5.448 |
| Entpd3 | 1106.3 | 0.371361 | 0.13987 | 2.655059 | 0.00793 | 0.04932 | 28.51 |
| Entpd5 | 1963.49 | -0.38749 | 0.09581 | -4.0442 | 5.25E-05 | 0.00171 | 35.173 |
| Entpd6 | 2353.96 | 0.089723 | 0.06817 | 1.316146 | 0.18812 | 0.38372 | 22.634 |
| Entpd7 | 567.657 | -0.07704 | 0.10255 | -0.75123 | 0.45252 | 0.65494 | 44.026 |
| Entr1 | 2783.68 | 0.062269 | 0.09356 | 0.66557 | 0.50569 | 0.69719 | 6.515 |

| | | | | | | | |
|---|---|---|---|---|---|---|---|
| Eny2 | 2968.84 | 0.170759 | 0.10863 | 1.571963 | 0.11596 | 0.28578 | 10.215 |
| Eogt | 2027.75 | 0.159598 | 0.12075 | 1.321768 | 0.18625 | 0.38115 | 39.159 |
| Eola1 | 286.016 | -0.14833 | 0.13837 | -1.07199 | 0.28373 | 0.49384 | 3.541 |
| Eomes | 22.2191 | -2 | 0.88194 | -6.33844 | 2.32E-10 | 1.75E-07 | 7.921 |
| Ep300 | 5171.82 | 0.105288 | 0.0548 | 1.921181 | 0.05471 | 0.17597 | 66.727 |
| Ep400 | 5620.78 | 0.119912 | 0.0759 | 1.579771 | 0.11416 | 0.28298 | 106.345 |
| Epas1 | 7765.23 | -0.30622 | 0.06093 | -5.02608 | 5.01E-07 | 6.37E-05 | 79.711 |
| Epb41 | 6909.59 | 0.364354 | 0.15371 | 2.370455 | 0.01777 | 0.08291 | 151.909 |
| Epb41l1 | 31711.5 | 0.037827 | 0.06592 | 0.573807 | 0.5661 | 0.74606 | 122.306 |
| Epb41l2 | 5266.57 | -0.3117 | 0.10284 | -3.03093 | 0.00244 | 0.02293 | 163.722 |
| Epb41l3 | 9760.67 | -0.11832 | 0.09151 | -1.29297 | 0.19602 | 0.39362 | 214.307 |
| Epb41l4a | 157.065 | 0.069792 | 0.1922 | 0.363125 | 0.71651 | 0.84846 | 210.64 |
| Epb41l4aos | 212.068 | 0.014687 | 0.13851 | 0.106038 | 0.91555 | 0.95671 | 1.162 |
| Epb41l4b | 2343.91 | 0.059513 | 0.09603 | 0.619711 | 0.53545 | 0.72244 | 151.466 |
| Epb41l5 | 539.972 | -0.02293 | 0.13169 | -0.17409 | 0.86179 | 0.92991 | 103.964 |
| Epc1 | 2676.54 | 0.163151 | 0.07158 | 2.279388 | 0.02264 | 0.09721 | 80.158 |
| Epc2 | 2411.1 | 0.093937 | 0.09757 | 0.962801 | 0.33565 | 0.54639 | 100.463 |
| Epdr1 | 5125.6 | -0.04267 | 0.05491 | -0.77701 | 0.43715 | 0.64268 | 28.222 |
| Epg5 | 3244.23 | 0.080372 | 0.07782 | 1.032756 | 0.30172 | 0.51136 | 96.549 |
| Epha1 | 31.412 | -0.55635 | 0.38696 | -1.43776 | 0.1505 | 0.33528 | 14.782 |
| Epha10 | 3370.56 | -0.64275 | 0.11942 | -5.38218 | 7.36E-08 | 1.40E-05 | 36.902 |
| Epha2 | 92.4029 | 0.26577 | 0.21059 | 1.262034 | 0.20694 | 0.40638 | 28.145 |
| Epha3 | 344.631 | -0.25232 | 0.20617 | -1.22388 | 0.221 | 0.4239 | 320.642 |
| Epha4 | 7427.53 | 0.386448 | 0.0825 | 4.684149 | 2.81E-06 | 0.00023 | 147.904 |
| Epha5 | 5208.9 | -0.20902 | 0.17169 | -1.2174 | 0.22345 | 0.42664 | 362.622 |
| Epha6 | 1085.07 | 0.300421 | 0.14201 | 2.115537 | 0.03438 | 0.1296 | 952.049 |
| Epha7 | 4496.14 | 0.30849 | 0.10841 | 2.84551 | 0.00443 | 0.03369 | 154.369 |
| Epha8 | 1180.62 | 0.861617 | 0.30358 | 2.838176 | 0.00454 | 0.0341 | 27.398 |
| Ephb1 | 2280.24 | 0.269217 | 0.12845 | 2.095911 | 0.03609 | 0.13377 | 432.566 |
| Ephb2 | 317.657 | 0.133495 | 0.14441 | 0.924434 | 0.35526 | 0.56556 | 188.45 |
| Ephb3 | 550.873 | -0.67272 | 0.20002 | -3.36333 | 0.00077 | 0.0107 | 18.551 |
| Ephb4 | 279.1 | -0.47113 | 0.13184 | -3.57363 | 0.00035 | 0.00625 | 28.561 |
| Ephb6 | 2050.11 | 0.237457 | 0.12463 | 1.905232 | 0.05675 | 0.18035 | 15.028 |
| Ephx1 | 1405.52 | 0.114472 | 0.17026 | 0.672347 | 0.50136 | 0.69395 | 44.695 |

| | | | | | | | |
|---|---|---|---|---|---|---|---|
| Ephx2 | 1197.68 | -0.1826 | 0.08548 | -2.13609 | 0.03267 | 0.12538 | 40.127 |
| Ephx4 | 1071.23 | -0.15586 | 0.07386 | -2.11007 | 0.03485 | 0.13081 | 27.3 |
| Epm2a | 1611.43 | 0.009039 | 0.08358 | 0.108149 | 0.91388 | 0.95639 | 116.241 |
| Epm2aip1 | 10353.9 | 0.089731 | 0.06652 | 1.348859 | 0.17738 | 0.36951 | 7.165 |
| Epn1 | 13018.5 | -0.27044 | 0.08649 | -3.12676 | 0.00177 | 0.01859 | 17.944 |
| Epn2 | 7046.76 | 0.11284 | 0.07037 | 1.60349 | 0.10883 | 0.27431 | 62.439 |
| Epn3 | 63.8329 | 0.189919 | 0.27349 | 0.694439 | 0.48741 | 0.68276 | 10.376 |
| Epop | 916.723 | -0.58581 | 0.17353 | -3.37588 | 0.00074 | 0.01036 | 2.314 |
| Epor | 285.769 | 0.235685 | 0.13143 | 1.793199 | 0.07294 | 0.21243 | 4.61 |
| Eprs | 5018.75 | 0.111402 | 0.08762 | 1.271453 | 0.20357 | 0.40212 | 65.317 |
| Eps15 | 9935.92 | -0.15375 | 0.08753 | -1.75665 | 0.07898 | 0.22295 | 107.55 |
| Eps15l1 | 2980.79 | 0.079733 | 0.08211 | 0.971074 | 0.33151 | 0.54247 | 80.462 |
| Eps8 | 1550.56 | -0.20641 | 0.08854 | -2.33122 | 0.01974 | 0.08875 | 177.632 |
| Eps8l1 | 595.966 | -0.2888 | 0.20303 | -1.42244 | 0.1549 | 0.34109 | 19.814 |
| Eps8l2 | 82.3434 | -0.3891 | 0.19938 | -1.95152 | 0.051 | 0.16804 | 24.141 |
| Eps8l3 | 19.2017 | 0.26597 | 0.506 | 0.525631 | 0.59914 | 0.77005 | 15.672 |
| Epsti1 | 36.7241 | -0.26822 | 0.28007 | -0.95772 | 0.33821 | 0.54905 | 98.419 |
| Epyc | 447.013 | 0.928904 | 0.41276 | 2.250485 | 0.02442 | 0.10223 | 38.387 |
| Eqtn | 75.8303 | -0.26922 | 0.2851 | -0.94429 | 0.34502 | 0.556 | 21.577 |
| Eral1 | 721.496 | 0.030532 | 0.09433 | 0.323668 | 0.74619 | 0.86678 | 7.008 |
| Erap1 | 511.632 | -0.18757 | 0.11931 | -1.57216 | 0.11591 | 0.28577 | 53.634 |
| Erbb2 | 122.995 | 0.072555 | 0.19129 | 0.379287 | 0.70447 | 0.8407 | 25.247 |
| Erbb3 | 737.341 | -0.56961 | 0.1217 | -4.68053 | 2.86E-06 | 0.00023 | 22.13 |
| Erbb4 | 5429.43 | 0.147194 | 0.12469 | 1.180453 | 0.23782 | 0.4432 | 1075.874 |
| Erbin | 6323.04 | 0.001896 | 0.11194 | 0.016939 | 0.98649 | 0.99262 | 101.728 |
| Erc1 | 2323.33 | 0.070948 | 0.08983 | 0.789791 | 0.42965 | 0.63602 | 277.372 |
| Erc2 | 9249.67 | 0.093179 | 0.08106 | 1.149474 | 0.25036 | 0.45774 | 856.11 |
| Ercc1 | 904.375 | 0.2082 | 0.0877 | 2.37389 | 0.0176 | 0.08244 | 11.747 |
| Ercc2 | 446.861 | 0.297995 | 0.10277 | 2.899695 | 0.00374 | 0.03006 | 13.685 |
| Ercc3 | 1206.69 | 0.084432 | 0.0727 | 1.161332 | 0.24551 | 0.45211 | 29.852 |
| Ercc4 | 1985.03 | 0.243638 | 0.12428 | 1.960435 | 0.04994 | 0.16584 | 40.934 |
| Ercc5 | 1527.56 | 0.177255 | 0.07529 | 2.354214 | 0.01856 | 0.08539 | 33.517 |
| Ercc6 | 2661.29 | 0.147848 | 0.07099 | 2.082749 | 0.03727 | 0.13651 | 67.47 |
| Ercc6l2 | 1394.95 | 0.225622 | 0.10703 | 2.108084 | 0.03502 | 0.13124 | 85.063 |

| | | | | | | | |
|---|---|---|---|---|---|---|---|
| Ercc8 | 456.596 | 0.003296 | 0.11105 | 0.029683 | 0.97632 | 0.98767 | 36.634 |
| Erdr1 | 469.556 | 0.669348 | 0.57384 | 1.166434 | 0.24344 | 0.44972 | 9.623 |
| Erf | 2431.21 | 0.143189 | 0.07255 | 1.973782 | 0.04841 | 0.16228 | 8.201 |
| Erg | 234.199 | -0.29655 | 0.13414 | -2.21076 | 0.02705 | 0.10988 | 227.425 |
| Erg28 | 1395.28 | -0.07678 | 0.07864 | -0.97634 | 0.3289 | 0.54005 | 9.103 |
| Ergic1 | 4295.04 | -0.0305 | 0.0632 | -0.48267 | 0.62933 | 0.79045 | 95.446 |
| Ergic2 | 1547.82 | 0.088308 | 0.08217 | 1.074671 | 0.28252 | 0.49272 | 33.296 |
| Ergic3 | 3448.72 | 0.0318 | 0.08085 | 0.393299 | 0.6941 | 0.83413 | 10.235 |
| Erh | 1402.09 | 0.032747 | 0.08881 | 0.36875 | 0.71231 | 0.84509 | 10.32 |
| Eri1 | 761.198 | 0.051507 | 0.09151 | 0.56284 | 0.57354 | 0.75223 | 30.944 |
| Eri2 | 409.087 | 0.079668 | 0.10895 | 0.731242 | 0.46463 | 0.66431 | 25.38 |
| Eri3 | 3833.59 | 0.020954 | 0.0733 | 0.285851 | 0.77499 | 0.88274 | 123.933 |
| Erich1 | 321.793 | 0.141271 | 0.1296 | 1.090013 | 0.27571 | 0.48494 | 62.741 |
| Erich2 | 33.7025 | 0.462142 | 0.30683 | 1.506186 | 0.13202 | 0.31082 | 32.066 |
| Erich3 | 3234.69 | 0.597339 | 0.14624 | 4.084584 | 4.42E-05 | 0.0015 | 103.932 |
| Erich5 | 38.5474 | -0.75024 | 0.42875 | -1.74983 | 0.08015 | 0.22519 | 20.588 |
| Erich6 | 136.484 | 0.07092 | 0.17832 | 0.397715 | 0.69084 | 0.83213 | 20.908 |
| Erlec1 | 1949.61 | -0.05095 | 0.10871 | -0.46863 | 0.63933 | 0.7978 | 23.562 |
| Erlin1 | 2261.61 | 0.529411 | 0.11319 | 4.677043 | 2.91E-06 | 0.00023 | 34.842 |
| Erlin2 | 3546.46 | 0.068189 | 0.07853 | 0.868361 | 0.3852 | 0.59452 | 17.068 |
| Ermap | 40.9307 | 0.410841 | 0.30341 | 1.354071 | 0.17571 | 0.36754 | 14.555 |
| Ermard | 347.211 | 0.072422 | 0.13396 | 0.540607 | 0.58878 | 0.762 | 48.837 |
| Ermn | 2436.08 | -0.38978 | 0.1916 | -2.03433 | 0.04192 | 0.1472 | 7.752 |
| Ermp1 | 3365.44 | -0.19358 | 0.05949 | -3.25411 | 0.00114 | 0.01382 | 40.202 |
| Ern1 | 874.644 | -0.10384 | 0.08951 | -1.16014 | 0.24599 | 0.45267 | 93.203 |
| Ero1a | 2695.24 | 0.121147 | 0.07434 | 1.629625 | 0.10318 | 0.26546 | 35.685 |
| Ero1b | 1034.78 | -0.19366 | 0.0952 | -2.03429 | 0.04192 | 0.1472 | 45.531 |
| Erp29 | 1952.88 | 0.033983 | 0.07578 | 0.448464 | 0.65382 | 0.80787 | 23.917 |
| Erp44 | 1732.19 | -0.01414 | 0.07425 | -0.19045 | 0.84895 | 0.92318 | 86.236 |
| Errfi1 | 1880.96 | -0.25895 | 0.11953 | -2.16636 | 0.03028 | 0.11878 | 14.974 |
| Esam | 623.475 | -0.2402 | 0.12681 | -1.89422 | 0.0582 | 0.18329 | 10.242 |
| Esco1 | 1242.78 | 0.438749 | 0.16829 | 2.607133 | 0.00913 | 0.05393 | 43.846 |
| Esco2 | 15.0456 | -0.58998 | 0.52173 | -1.13081 | 0.25813 | 0.46618 | 14.957 |
| Esd | 2430.47 | 0.033246 | 0.07157 | 0.4645 | 0.64229 | 0.80017 | 18.469 |

| | | | | | | | |
|---|---|---|---|---|---|---|---|
| Esf1 | 1349.8 | 0.348547 | 0.20881 | 1.669228 | 0.09507 | 0.25181 | 50.682 |
| Esm1 | 70.6017 | 0.261388 | 0.29345 | 0.890739 | 0.37307 | 0.58345 | 8.44 |
| Espl1 | 70.0218 | -0.36441 | 0.37876 | -0.96211 | 0.33599 | 0.54677 | 28.092 |
| Espn | 113.13 | 0.06497 | 0.21926 | 0.296318 | 0.76699 | 0.87819 | 32.041 |
| Esr1 | 154.1 | -0.26259 | 0.25899 | -1.01393 | 0.31062 | 0.52082 | 394.022 |
| Esr2 | 22.798 | 0.064222 | 0.34456 | 0.186389 | 0.85214 | 0.92477 | 56.841 |
| Esrra | 3447.75 | -0.20648 | 0.05803 | -3.55797 | 0.00037 | 0.00652 | 10.864 |
| Esrrb | 59.2211 | -0.47673 | 0.30386 | -1.56893 | 0.11667 | 0.28704 | 160.512 |
| Esrrg | 1444.29 | -0.16832 | 0.12093 | -1.39187 | 0.16396 | 0.35274 | 606.095 |
| Ess2 | 1117.93 | 0.097856 | 0.08449 | 1.158165 | 0.2468 | 0.45367 | 10.64 |
| Esyt1 | 589.254 | -0.48598 | 0.11097 | -4.37943 | 1.19E-05 | 0.00063 | 15.907 |
| Esyt2 | 3336.63 | 0.106149 | 0.08505 | 1.24802 | 0.21202 | 0.41271 | 109.855 |
| Esyt3 | 391.157 | 0.696037 | 0.18086 | 3.848444 | 0.00012 | 0.00302 | 48.595 |
| Etaa1 | 436.755 | 0.33256 | 0.1789 | 1.858898 | 0.06304 | 0.19278 | 15.12 |
| Etaa1os | 33.3014 | 0.333837 | 0.32276 | 1.034311 | 0.30099 | 0.51075 | 25.017 |
| Etf1 | 4436.47 | 0.078102 | 0.06819 | 1.145326 | 0.25207 | 0.4594 | 29.223 |
| Etfa | 1420.09 | -0.12399 | 0.0854 | -1.45191 | 0.14653 | 0.32967 | 57.736 |
| Etfb | 24.5718 | -0.41395 | 0.3552 | -1.1654 | 0.24386 | 0.45029 | 13.718 |
| Etfbkmt | 188.267 | 0.125345 | 0.20871 | 0.600559 | 0.54813 | 0.73223 | 11.323 |
| Etfdh | 1520.73 | -0.02273 | 0.11311 | -0.20094 | 0.84075 | 0.91835 | 25.713 |
| Etfrf1 | 1113.23 | 0.230996 | 0.15309 | 1.508903 | 0.13132 | 0.30993 | 5.806 |
| Ethe1 | 310.476 | -0.2508 | 0.10355 | -2.42193 | 0.01544 | 0.07587 | 21.383 |
| Etl4 | 3528.84 | -0.60295 | 0.18786 | -3.20952 | 0.00133 | 0.01522 | 900.934 |
| Etnk1 | 25561.1 | 0.159641 | 0.06346 | 2.515585 | 0.01188 | 0.0642 | 65.087 |
| Etnk2 | 75.4862 | -0.63115 | 0.30911 | -2.04183 | 0.04117 | 0.14541 | 16.765 |
| Etnppl | 2010.22 | -0.07974 | 0.18879 | -0.42238 | 0.67275 | 0.82003 | 20.074 |
| Etohd2 | 287.094 | 0.179367 | 0.11877 | 1.510188 | 0.131 | 0.30945 | 5.911 |
| Ets1 | 502.354 | -0.41551 | 0.14877 | -2.79293 | 0.00522 | 0.03732 | 121.6 |
| Ets2 | 7351.63 | 0.473623 | 0.15507 | 3.054205 | 0.00226 | 0.02178 | 19.154 |
| Etv1 | 2530.72 | -0.37679 | 0.14222 | -2.6493 | 0.00807 | 0.04987 | 91.105 |
| Etv3 | 2186.16 | 0.05118 | 0.06932 | 0.738336 | 0.46031 | 0.66087 | 14.75 |
| Etv4 | 325.883 | -0.08035 | 0.18807 | -0.42723 | 0.66921 | 0.81761 | 15.63 |
| Etv5 | 3433.07 | 0.004725 | 0.13354 | 0.035384 | 0.97177 | 0.9858 | 58.411 |
| Etv6 | 674.282 | -0.11418 | 0.08461 | -1.34946 | 0.17719 | 0.36946 | 234.459 |

| | | | | | | | |
|---|---|---|---|---|---|---|---|
| Eva1a | 499.438 | -0.34564 | 0.1213 | -2.84936 | 0.00438 | 0.03344 | 52.057 |
| Eva1b | 45.5147 | -0.35086 | 0.25481 | -1.37693 | 0.16853 | 0.35852 | 2.132 |
| Eva1c | 94.9508 | -0.54234 | 0.23228 | -2.33489 | 0.01955 | 0.08818 | 78.391 |
| Evc | 487.284 | -0.30257 | 0.10829 | -2.79402 | 0.00521 | 0.03723 | 47.797 |
| Evc2 | 541.445 | -0.29853 | 0.10533 | -2.83422 | 0.00459 | 0.03443 | 86.557 |
| Evi2a | 406.841 | -0.59356 | 0.20096 | -2.95359 | 0.00314 | 0.02688 | 4.05 |
| Evi5 | 3569.82 | 0.060812 | 0.07917 | 0.768093 | 0.44243 | 0.64699 | 130.313 |
| Evi5l | 3523.41 | -0.08089 | 0.07409 | -1.09168 | 0.27497 | 0.48452 | 44.691 |
| Evl | 4191.72 | 0.102437 | 0.07167 | 1.429333 | 0.15291 | 0.33835 | 133.797 |
| Evpl | 792.765 | 0.157747 | 0.11269 | 1.399777 | 0.16158 | 0.34979 | 17.519 |
| Ewsr1 | 14886.5 | 0.274261 | 0.07543 | 3.636095 | 0.00028 | 0.00531 | 29.578 |
| Exd1 | 135.088 | -0.09622 | 0.21092 | -0.45619 | 0.64825 | 0.80406 | 31.123 |
| Exd2 | 5910.85 | 0.010519 | 0.09461 | 0.111177 | 0.91148 | 0.95529 | 37.133 |
| Exo1 | 26.9463 | -0.20469 | 0.36865 | -0.55525 | 0.57872 | 0.75547 | 32.909 |
| Exo5 | 320.827 | 0.150317 | 0.11536 | 1.303069 | 0.19255 | 0.38927 | 3.822 |
| Exoc1 | 2971.17 | 0.076131 | 0.06592 | 1.154952 | 0.24811 | 0.45524 | 40.984 |
| Exoc2 | 2177.71 | -0.03943 | 0.10272 | -0.38388 | 0.70107 | 0.83856 | 160.175 |
| Exoc3 | 4880.08 | 0.083419 | 0.06047 | 1.379557 | 0.16772 | 0.35762 | 39.245 |
| Exoc3l | 140.712 | -0.40565 | 0.17219 | -2.35583 | 0.01848 | 0.08517 | 6.178 |
| Exoc3l4 | 39.6501 | -0.25127 | 0.31704 | -0.79255 | 0.42804 | 0.63465 | 14.662 |
| Exoc4 | 3722.01 | 0.026379 | 0.078 | 0.338185 | 0.73522 | 0.85989 | 724.895 |
| Exoc5 | 3459.45 | 0.2055 | 0.0968 | 2.122896 | 0.03376 | 0.12818 | 62.564 |
| Exoc6 | 1536.9 | 0.038557 | 0.07652 | 0.503864 | 0.61436 | 0.78043 | 163.181 |
| Exoc6b | 4497.23 | -0.0814 | 0.06714 | -1.21249 | 0.22532 | 0.42931 | 451.027 |
| Exoc7 | 2902.7 | 0.030139 | 0.07786 | 0.387081 | 0.6987 | 0.83678 | 19.233 |
| Exoc8 | 1512.28 | 0.082153 | 0.08885 | 0.92465 | 0.35515 | 0.56548 | 4.598 |
| Exog | 1220.37 | 0.034845 | 0.07879 | 0.442229 | 0.65832 | 0.81039 | 20.575 |
| Exosc1 | 519.743 | 0.040536 | 0.11352 | 0.357081 | 0.72103 | 0.85104 | 11.132 |
| Exosc10 | 2416.04 | 0.109083 | 0.07679 | 1.420553 | 0.15545 | 0.34204 | 23.973 |
| Exosc2 | 765.697 | 0.092883 | 0.08798 | 1.055701 | 0.2911 | 0.50115 | 10.635 |
| Exosc3 | 408.508 | 0.052919 | 0.11448 | 0.462233 | 0.64391 | 0.80136 | 26.12 |
| Exosc4 | 437.438 | 0.033169 | 0.11675 | 0.284108 | 0.77633 | 0.8836 | 3.281 |
| Exosc5 | 706.509 | 0.055082 | 0.10925 | 0.504199 | 0.61412 | 0.78029 | 12.205 |
| Exosc7 | 935.674 | 0.150902 | 0.08902 | 1.69524 | 0.09003 | 0.24339 | 22.915 |

| | | | | | | | |
|---|---|---|---|---|---|---|---|
| Exosc8 | 915.752 | 0.226126 | 0.08165 | 2.769426 | 0.00562 | 0.03939 | 6.716 |
| Exosc9 | 1028.76 | 0.09543 | 0.0832 | 1.147038 | 0.25137 | 0.4588 | 13.122 |
| Exph5 | 158.203 | 0.030153 | 0.17213 | 0.175175 | 0.86094 | 0.92962 | 75.845 |
| Ext1 | 2026.8 | -0.39698 | 0.18555 | -2.1395 | 0.0324 | 0.12455 | 282.122 |
| Ext2 | 1745.72 | 0.127571 | 0.07408 | 1.722103 | 0.08505 | 0.23388 | 161.541 |
| Extl1 | 2253.35 | -0.30202 | 0.12177 | -2.48022 | 0.01313 | 0.06863 | 27.479 |
| Extl2 | 3663.82 | 0.037047 | 0.09174 | 0.403822 | 0.68634 | 0.82888 | 21.556 |
| Extl3 | 7222.83 | -0.1959 | 0.07121 | -2.75126 | 0.00594 | 0.04094 | 97.796 |
| Eya1 | 543.058 | -0.30478 | 0.18463 | -1.65079 | 0.09878 | 0.25825 | 141.282 |
| Eya2 | 125.744 | -0.92674 | 0.35039 | -2.64491 | 0.00817 | 0.05023 | 176.696 |
| Eya3 | 1666.14 | 0.13911 | 0.06928 | 2.00794 | 0.04465 | 0.1535 | 85.779 |
| Eya4 | 203.528 | -0.67493 | 0.2409 | -2.80175 | 0.00508 | 0.03666 | 247.823 |
| Ezh1 | 3536.42 | 0.035618 | 0.0776 | 0.459006 | 0.64623 | 0.80297 | 35.349 |
| Ezh2 | 642.493 | 0.307776 | 0.13385 | 2.299417 | 0.02148 | 0.09394 | 65.203 |
| Ezr | 4707.82 | -0.54915 | 0.10041 | -5.4691 | 4.52E-08 | 1.05E-05 | 44.744 |
| F11r | 286.587 | -0.45918 | 0.1619 | -2.83624 | 0.00456 | 0.03426 | 27.069 |
| F12 | 32.6233 | 0.858362 | 0.33072 | 2.595452 | 0.00945 | 0.05528 | 8.836 |
| F13a1 | 177.384 | -0.69692 | 0.27307 | -2.55219 | 0.0107 | 0.06008 | 183.069 |
| F2r | 203.702 | -0.13799 | 0.13864 | -0.99533 | 0.31958 | 0.53026 | 16.685 |
| F2rl1 | 16.9939 | -1.44127 | 0.49845 | -2.89149 | 0.00383 | 0.03057 | 13.496 |
| F2rl3 | 32.3787 | -0.54976 | 0.34157 | -1.6095 | 0.10751 | 0.27228 | 1.995 |
| F3 | 2401.15 | -0.15857 | 0.10935 | -1.45013 | 0.14702 | 0.3302 | 11.512 |
| F420014N23Rik | 224.794 | -0.10181 | 0.13624 | -0.74728 | 0.45489 | 0.65702 | 7.395 |
| F5 | 28.4069 | -1.34276 | 0.5407 | -2.48339 | 0.01301 | 0.06826 | 68.44 |
| F530104D19Rik | 25.0558 | 0.161927 | 0.34933 | 0.463532 | 0.64298 | 0.80084 | 2.166 |
| F630040K05Rik | 63.6443 | -1.06028 | 0.23543 | -4.50355 | 6.68E-06 | 0.00041 | 54.089 |
| F630042J09Rik | 28.4504 | 0.521288 | 0.38079 | 1.368979 | 0.17101 | 0.36141 | 4.785 |
| F730043M19Rik | 1846.75 | -0.6385 | 0.29186 | -2.18774 | 0.02869 | 0.11431 | 35.902 |
| F730311O21Rik | 15.8294 | 0.60764 | 0.54632 | 1.112246 | 0.26603 | 0.47518 | 9.369 |
| F8 | 149.385 | 0.468399 | 0.24581 | 1.905502 | 0.05671 | 0.18033 | 209.901 |
| F830016B08Rik | 13.2007 | -1.03665 | 0.6285 | -1.6494 | 0.09907 | 0.25857 | 9.637 |
| F830212C03Rik | 19.659 | 0.941983 | 0.48361 | 1.947831 | 0.05144 | 0.16908 | 2.285 |
| F8a | 440.791 | -0.28075 | 0.1178 | -2.38326 | 0.01716 | 0.08148 | 2.505 |
| F9 | 12.9544 | -0.23798 | 0.63834 | -0.37282 | 0.70929 | 0.84343 | 31.296 |

| | | | | | | | |
|---|---|---|---|---|---|---|---|
| Fa2h | 1299.06 | -0.25656 | 0.12943 | -1.98215 | 0.04746 | 0.16012 | 48.69 |
| Faah | 2578.86 | -0.7063 | 0.2005 | -3.52273 | 0.00043 | 0.00721 | 50.782 |
| Faap100 | 657.785 | 0.122586 | 0.1223 | 1.002351 | 0.31617 | 0.52667 | 9.71 |
| Faap20 | 698.1 | 0.272673 | 0.08233 | 3.311952 | 0.00093 | 0.01217 | 6.886 |
| Faap24 | 165.277 | -0.13724 | 0.15331 | -0.89517 | 0.3707 | 0.58089 | 4.685 |
| Fabp3 | 1472.71 | -0.21682 | 0.09673 | -2.24158 | 0.02499 | 0.10395 | 6.869 |
| Fabp3-ps1 | 14.5604 | -0.57982 | 0.54581 | -1.06231 | 0.28809 | 0.49826 | 0.402 |
| Fabp4 | 27.4768 | 0.155481 | 0.3758 | 0.413729 | 0.67907 | 0.82468 | 4.489 |
| Fabp5 | 2583.15 | 0.00354 | 0.12467 | 0.028395 | 0.97735 | 0.98799 | 4.06 |
| Fabp7 | 578.909 | -0.28834 | 0.19711 | -1.46283 | 0.14351 | 0.32566 | 3.57 |
| Fadd | 62.5611 | -0.87694 | 0.25823 | -3.39599 | 0.00068 | 0.00984 | 5.146 |
| Fads1 | 10790.6 | -0.19137 | 0.09123 | -2.09769 | 0.03593 | 0.13339 | 13.99 |
| Fads2 | 6143.26 | -0.19938 | 0.12539 | -1.59 | 0.11183 | 0.27929 | 38.982 |
| Fads3 | 730.832 | -0.33512 | 0.11944 | -2.80568 | 0.00502 | 0.0364 | 18.542 |
| Fads6 | 2200.87 | -0.07711 | 0.1109 | -0.6953 | 0.48686 | 0.6826 | 18.042 |
| Faf1 | 1491.32 | 0.013493 | 0.07178 | 0.187966 | 0.8509 | 0.9241 | 287.373 |
| Faf2 | 2479.57 | -0.0097 | 0.08469 | -0.11456 | 0.9088 | 0.95435 | 42.28 |
| Fah | 659.905 | -0.32706 | 0.16748 | -1.9528 | 0.05084 | 0.16776 | 21.564 |
| Fahd1 | 685.649 | -0.14555 | 0.09081 | -1.60285 | 0.10897 | 0.27449 | 1.473 |
| Fahd2a | 553.521 | 0.001724 | 0.10678 | 0.016141 | 0.98712 | 0.99307 | 8.351 |
| Faim | 728.016 | -0.1754 | 0.10786 | -1.6261 | 0.10393 | 0.26644 | 15.649 |
| Faim2 | 16178.3 | -0.10964 | 0.07428 | -1.47601 | 0.13994 | 0.3212 | 31.154 |
| Fam102a | 4326.11 | 0.095834 | 0.09815 | 0.976412 | 0.32886 | 0.54005 | 34.425 |
| Fam102b | 7124.22 | 0.661744 | 0.14504 | 4.56241 | 5.06E-06 | 0.00034 | 56.611 |
| Fam104a | 828.091 | -0.05563 | 0.08225 | -0.6763 | 0.49885 | 0.69216 | 22.837 |
| Fam107a | 28618.9 | -0.41506 | 0.0954 | -4.35069 | 1.36E-05 | 0.0007 | 21.75 |
| Fam107b | 357.549 | -0.25057 | 0.17709 | -1.41494 | 0.15708 | 0.34408 | 211.655 |
| Fam110a | 190.964 | -0.13281 | 0.16358 | -0.81193 | 0.41683 | 0.62434 | 10.822 |
| Fam110b | 2363.35 | 0.397249 | 0.16269 | 2.441789 | 0.01461 | 0.07303 | 464.134 |
| Fam110d | 73.0079 | -0.33019 | 0.28728 | -1.14936 | 0.25041 | 0.45778 | 2.997 |
| Fam111a | 93.2545 | -0.61505 | 0.27549 | -2.23254 | 0.02558 | 0.10553 | 44.029 |
| Fam114a1 | 217.186 | -0.52718 | 0.15734 | -3.35052 | 0.00081 | 0.01103 | 71.816 |
| Fam114a2 | 2042.79 | 0.079975 | 0.0684 | 1.169229 | 0.24231 | 0.44829 | 35.625 |
| Fam117a | 385.168 | 0.786722 | 0.24124 | 3.261154 | 0.00111 | 0.0137 | 44.855 |

| | | | | | | | |
|---|---|---|---|---|---|---|---|
| Fam117b | 3589.05 | 0.140276 | 0.07356 | 1.906835 | 0.05654 | 0.18 | 72.341 |
| Fam118a | 971.93 | 0.021247 | 0.08963 | 0.237055 | 0.81261 | 0.90362 | 33.883 |
| Fam118b | 880.581 | 0.059631 | 0.09661 | 0.617213 | 0.53709 | 0.72389 | 50.841 |
| Fam120a | 10113.9 | 0.188755 | 0.05767 | 3.272895 | 0.00106 | 0.01335 | 88.799 |
| Fam120b | 3707.9 | 0.195926 | 0.06817 | 2.874054 | 0.00405 | 0.03175 | 37.382 |
| Fam120c | 3007.34 | 0.198909 | 0.07524 | 2.643818 | 0.0082 | 0.05033 | 127.668 |
| Fam122a | 410.525 | -0.14089 | 0.10309 | -1.36665 | 0.17174 | 0.3622 | 0.855 |
| Fam122b | 443.246 | 0.366648 | 0.18981 | 1.931682 | 0.0534 | 0.1731 | 26.391 |
| Fam124a | 737.752 | 0.008692 | 0.11917 | 0.072942 | 0.94185 | 0.97096 | 52.75 |
| Fam126a | 2561.21 | 0.356793 | 0.08639 | 4.130073 | 3.63E-05 | 0.00131 | 115.415 |
| Fam126b | 7068.53 | 0.289728 | 0.1174 | 2.467816 | 0.01359 | 0.07019 | 63.518 |
| Fam131a | 5554.1 | -0.32815 | 0.13027 | -2.51891 | 0.01177 | 0.06382 | 9.808 |
| Fam131b | 5452.59 | -0.16546 | 0.07832 | -2.11259 | 0.03464 | 0.13033 | 9.332 |
| Fam131c | 92.3124 | 0.069965 | 0.27087 | 0.258295 | 0.79618 | 0.89487 | 15.956 |
| Fam133b | 1654.17 | 0.0823 | 0.0809 | 1.01728 | 0.30902 | 0.51884 | 26.406 |
| Fam135a | 2056.96 | -0.06037 | 0.11961 | -0.50473 | 0.61375 | 0.78 | 89.249 |
| Fam135b | 4257.99 | -0.11654 | 0.14321 | -0.81377 | 0.41578 | 0.62362 | 296.23 |
| Fam136a | 848.795 | 0.085976 | 0.11062 | 0.777209 | 0.43704 | 0.64261 | 4.413 |
| Fam136b-ps | 15.2051 | 0.547875 | 0.4618 | 1.186398 | 0.23547 | 0.44058 | 0.412 |
| Fam13a | 848.86 | 0.005172 | 0.10549 | 0.049028 | 0.9609 | 0.98096 | 92.46 |
| Fam13b | 2971.36 | 0.010162 | 0.07665 | 0.132579 | 0.89453 | 0.94736 | 64.481 |
| Fam13c | 2726.22 | -0.10227 | 0.11603 | -0.88144 | 0.37808 | 0.58812 | 118.256 |
| Fam149a | 728.932 | -0.16854 | 0.11817 | -1.42623 | 0.1538 | 0.3396 | 45.575 |
| Fam149b | 1826.39 | 0.009598 | 0.06325 | 0.151764 | 0.87937 | 0.93962 | 35.33 |
| Fam151b | 252.26 | 0.064513 | 0.14243 | 0.452932 | 0.6506 | 0.80552 | 34.391 |
| Fam161a | 185.881 | 0.176593 | 0.18793 | 0.939659 | 0.34739 | 0.55793 | 23.258 |
| Fam161b | 760.558 | 0.106004 | 0.10804 | 0.981184 | 0.3265 | 0.53777 | 16.525 |
| Fam162a | 854.458 | -0.15886 | 0.09964 | -1.5943 | 0.11087 | 0.27788 | 27.834 |
| Fam163a | 249.036 | -0.2929 | 0.14109 | -2.07603 | 0.03789 | 0.13802 | 80.885 |
| Fam163b | 4949.89 | -0.89088 | 0.17871 | -4.98515 | 6.19E-07 | 7.64E-05 | 32.112 |
| Fam166a | 14.8551 | -0.1141 | 0.53263 | -0.21422 | 0.83037 | 0.91311 | 3.536 |
| Fam166b | 29.3712 | -0.02604 | 0.35298 | -0.07377 | 0.94119 | 0.97087 | 2.116 |
| Fam167a | 203.455 | -0.40789 | 0.29609 | -1.3776 | 0.16833 | 0.35831 | 29.105 |
| Fam168a | 15526.5 | -0.07774 | 0.0476 | -1.63316 | 0.10244 | 0.26424 | 135.022 |

| | | | | | | | |
|---|---|---|---|---|---|---|---|
| Fam168b | 17089 | -0.02263 | 0.0508 | -0.44539 | 0.65604 | 0.80893 | 29.847 |
| Fam169a | 3007.67 | -0.04756 | 0.10178 | -0.46726 | 0.64031 | 0.79859 | 63.728 |
| Fam169b | 21.7145 | -0.42371 | 0.45947 | -0.92216 | 0.35645 | 0.56684 | 89.254 |
| Fam171a1 | 5776.03 | 0.140964 | 0.07004 | 2.012689 | 0.04415 | 0.15229 | 113.583 |
| Fam171a2 | 3198.69 | 0.092783 | 0.07532 | 1.231775 | 0.21803 | 0.42003 | 10.702 |
| Fam171b | 19913.1 | -0.12361 | 0.09672 | -1.27795 | 0.20127 | 0.39952 | 70.851 |
| Fam172a | 1933.29 | -0.0083 | 0.06599 | -0.12581 | 0.89988 | 0.94981 | 464.203 |
| Fam174a | 1948.95 | -0.03491 | 0.07627 | -0.45773 | 0.64715 | 0.80359 | 21.662 |
| Fam174b | 1045.27 | -0.1586 | 0.12145 | -1.30587 | 0.1916 | 0.38826 | 36.618 |
| Fam174c | 109.552 | 0.308852 | 0.17069 | 1.809448 | 0.07038 | 0.20729 | 2.203 |
| Fam180a | 98.936 | -0.99717 | 0.38253 | -2.60681 | 0.00914 | 0.05397 | 13.474 |
| Fam181a | 36.9368 | -0.34925 | 0.38051 | -0.91785 | 0.3587 | 0.56892 | 6.091 |
| Fam181b | 193.44 | -0.33906 | 0.15846 | -2.13979 | 0.03237 | 0.12448 | 2.002 |
| Fam183b | 17.2731 | -0.74877 | 0.42818 | -1.74871 | 0.08034 | 0.22551 | 9.164 |
| Fam184a | 2218.3 | 0.025459 | 0.12359 | 0.205992 | 0.8368 | 0.91619 | 117.979 |
| Fam184b | 1327.87 | 0.844521 | 0.17401 | 4.853335 | 1.21E-06 | 0.00012 | 109.91 |
| Fam185a | 779.822 | 0.110715 | 0.09695 | 1.14196 | 0.25347 | 0.46086 | 57.167 |
| Fam186b | 134.747 | -0.13175 | 0.21798 | -0.6044 | 0.54557 | 0.73038 | 24.871 |
| Fam189a1 | 2061.69 | -0.12015 | 0.07285 | -1.64936 | 0.09907 | 0.25857 | 400.48 |
| Fam189a2 | 589.604 | -0.39593 | 0.12892 | -3.07111 | 0.00213 | 0.02103 | 58.27 |
| Fam189b | 1899.76 | 0.073551 | 0.06434 | 1.143147 | 0.25298 | 0.46034 | 6.153 |
| Fam193a | 2684.51 | 0.078083 | 0.07048 | 1.10783 | 0.26794 | 0.47725 | 116.524 |
| Fam193b | 6305.28 | 0.102765 | 0.0918 | 1.119494 | 0.26293 | 0.47176 | 31.805 |
| Fam199x | 795.684 | 0.041413 | 0.13065 | 0.316982 | 0.75126 | 0.86954 | 32.91 |
| Fam204a | 810.28 | 0.165568 | 0.11886 | 1.392995 | 0.16362 | 0.35238 | 28.116 |
| Fam205a1 | 61.3174 | -0.76246 | 0.30353 | -2.51197 | 0.01201 | 0.06467 | 5.818 |
| Fam205a2 | 53.6967 | 0.010002 | 0.35983 | 0.027796 | 0.97782 | 0.9881 | 5.596 |
| Fam205c | 360.267 | -0.71465 | 0.18771 | -3.80724 | 0.00014 | 0.00337 | 6.231 |
| Fam207a | 1663.16 | -0.0261 | 0.06336 | -0.41189 | 0.68042 | 0.82503 | 29.125 |
| Fam20a | 631.316 | 0.185279 | 0.11387 | 1.627178 | 0.1037 | 0.26622 | 52.531 |
| Fam20b | 4113.51 | -0.07469 | 0.06522 | -1.14513 | 0.25216 | 0.45946 | 40.555 |
| Fam20c | 2062.14 | 0.201389 | 0.07854 | 2.56416 | 0.01034 | 0.05872 | 55.564 |
| Fam210a | 1682 | 0.101706 | 0.06326 | 1.607766 | 0.10789 | 0.27288 | 40.147 |
| Fam210b | 1628.97 | 0.203051 | 0.09542 | 2.127878 | 0.03335 | 0.12684 | 10.185 |

| | | | | | | | |
|---|---|---|---|---|---|---|---|
| Fam214a | 4021.34 | 0.262506 | 0.12798 | 2.05107 | 0.04026 | 0.14335 | 79.585 |
| Fam214b | 1469.97 | -0.18947 | 0.07878 | -2.40497 | 0.01617 | 0.07822 | 13.807 |
| Fam216a | 690.866 | 0.272247 | 0.09147 | 2.976294 | 0.00292 | 0.02573 | 7.785 |
| Fam217a | 44.2191 | 0.109663 | 0.36171 | 0.303176 | 0.76176 | 0.87514 | 14.351 |
| Fam217b | 2554.21 | -0.02989 | 0.07808 | -0.38278 | 0.70188 | 0.83913 | 9.905 |
| Fam219a | 4328.6 | -0.12632 | 0.05945 | -2.12499 | 0.03359 | 0.12761 | 51.848 |
| Fam219b | 974.324 | -0.04075 | 0.08188 | -0.49771 | 0.61869 | 0.78312 | 5.657 |
| Fam220a | 31.2956 | -0.45916 | 0.34817 | -1.31875 | 0.18725 | 0.3825 | 15.826 |
| Fam221a | 52.6205 | 0.002684 | 0.3174 | 0.008457 | 0.99325 | 0.99618 | 22.801 |
| Fam221b | 61.9742 | -0.19382 | 0.26141 | -0.74143 | 0.45843 | 0.65956 | 9.524 |
| Fam222a | 626.394 | 0.118814 | 0.14222 | 0.835417 | 0.40348 | 0.61213 | 45.205 |
| Fam222b | 1406.42 | -0.16997 | 0.0767 | -2.21588 | 0.0267 | 0.10894 | 62.041 |
| Fam227a | 1241.91 | 0.416395 | 0.09661 | 4.309956 | 1.63E-05 | 0.00078 | 49.381 |
| Fam228a | 315.976 | 0.140684 | 0.11859 | 1.186341 | 0.23549 | 0.44059 | 24.761 |
| Fam228b | 463.503 | 0.044497 | 0.10411 | 0.427392 | 0.66909 | 0.81761 | 29.38 |
| Fam229b | 710.769 | 0.458135 | 0.12342 | 3.712127 | 0.00021 | 0.00435 | 15.111 |
| Fam234a | 643.532 | -0.18182 | 0.10258 | -1.77235 | 0.07634 | 0.21839 | 32.421 |
| Fam234b | 13344.3 | -0.02048 | 0.07036 | -0.29106 | 0.77101 | 0.88041 | 46.979 |
| Fam241a | 194.544 | 0.154507 | 0.1432 | 1.078977 | 0.2806 | 0.49035 | 27.267 |
| Fam241b | 862.757 | -0.13787 | 0.10837 | -1.27217 | 0.20331 | 0.40179 | 36.266 |
| Fam24a | 15.4282 | 0.08867 | 0.52171 | 0.16996 | 0.86504 | 0.93163 | 3.064 |
| Fam32a | 3387.51 | -0.17822 | 0.06747 | -2.64158 | 0.00825 | 0.0506 | 4.689 |
| Fam3a | 694.631 | -0.03889 | 0.10588 | -0.36731 | 0.71339 | 0.84584 | 8.554 |
| Fam3c | 3927.36 | -0.35351 | 0.11614 | -3.04373 | 0.00234 | 0.0223 | 49.724 |
| Fam43a | 836.385 | -0.09431 | 0.13414 | -0.70303 | 0.48204 | 0.67867 | 3.075 |
| Fam43b | 432.767 | -0.15582 | 0.12326 | -1.26424 | 0.20614 | 0.40534 | 2.437 |
| Fam50a | 1005.6 | -0.0362 | 0.09559 | -0.37871 | 0.7049 | 0.84107 | 7.117 |
| Fam53a | 1341.65 | 0.090375 | 0.09818 | 0.920467 | 0.35733 | 0.56765 | 29.289 |
| Fam53b | 1375.12 | -0.33491 | 0.10342 | -3.23847 | 0.0012 | 0.01435 | 101.805 |
| Fam53c | 3626.69 | -0.10258 | 0.0644 | -1.59293 | 0.11118 | 0.27839 | 14.855 |
| Fam71d | 23.118 | -0.11183 | 0.34814 | -0.32122 | 0.74804 | 0.86795 | 42.982 |
| Fam71e1 | 235.268 | -0.04848 | 0.13641 | -0.35538 | 0.7223 | 0.85185 | 4.906 |
| Fam71f2 | 30.6188 | 0.624099 | 0.40321 | 1.547807 | 0.12167 | 0.29511 | 11.088 |
| Fam72a | 54.2204 | -0.13143 | 0.24017 | -0.54721 | 0.58423 | 0.75912 | 11.97 |

| | | | | | | | |
|---|---|---|---|---|---|---|---|
| Fam76a | 2995.53 | 0.145647 | 0.08188 | 1.778837 | 0.07527 | 0.21642 | 23.346 |
| Fam76b | 1755.56 | 0.075197 | 0.10629 | 0.707461 | 0.47928 | 0.67694 | 27.272 |
| Fam78a | 359.596 | 0.280515 | 0.17129 | 1.637682 | 0.10149 | 0.26283 | 34.803 |
| Fam78b | 1717.74 | 0.15476 | 0.07069 | 2.189152 | 0.02859 | 0.11411 | 89.886 |
| Fam81a | 7197.17 | -0.91226 | 0.30872 | -2.955 | 0.00313 | 0.02678 | 54.05 |
| Fam83d | 197.314 | -0.43464 | 0.16417 | -2.64757 | 0.00811 | 0.04997 | 18.545 |
| Fam83h | 57.5143 | -0.73514 | 0.29737 | -2.47216 | 0.01343 | 0.06966 | 13.244 |
| Fam89a | 70.7407 | -0.03868 | 0.21778 | -0.17761 | 0.85903 | 0.9288 | 11.576 |
| Fam8a1 | 7311.14 | 0.208858 | 0.06568 | 3.180162 | 0.00147 | 0.01634 | 8.535 |
| Fam90a1b | 32.5591 | 0.777598 | 0.3479 | 2.235148 | 0.02541 | 0.10513 | 11.725 |
| Fam91a1 | 4304.82 | 0.138229 | 0.06931 | 1.994336 | 0.04612 | 0.15675 | 42.273 |
| Fam98a | 2099.5 | -0.21316 | 0.0938 | -2.27247 | 0.02306 | 0.09829 | 14.861 |
| Fam98b | 2947.94 | 0.158542 | 0.08065 | 1.965845 | 0.04932 | 0.16445 | 21.802 |
| Fam98c | 276.227 | 0.071709 | 0.14352 | 0.499648 | 0.61732 | 0.78222 | 3.725 |
| Fan1 | 682.43 | 0.031306 | 0.08502 | 0.368231 | 0.7127 | 0.84523 | 27.338 |
| Fanca | 113.393 | -0.40086 | 0.18608 | -2.15425 | 0.03122 | 0.12134 | 50.277 |
| Fancb | 249.296 | 0.549606 | 0.18074 | 3.040946 | 0.00236 | 0.02243 | 16.681 |
| Fancc | 354.126 | -0.06117 | 0.12606 | -0.48521 | 0.62752 | 0.78925 | 212.236 |
| Fancd2 | 350.356 | 0.130213 | 0.13145 | 0.990552 | 0.3219 | 0.53286 | 65.336 |
| Fance | 876.525 | 0.112614 | 0.08233 | 1.367802 | 0.17137 | 0.3618 | 13.054 |
| Fancf | 78.5235 | -0.43021 | 0.20078 | -2.14267 | 0.03214 | 0.12384 | 1.691 |
| Fancg | 838.106 | 0.299295 | 0.11535 | 2.594747 | 0.00947 | 0.05532 | 8.164 |
| Fanci | 31.5973 | -0.64881 | 0.37195 | -1.74434 | 0.0811 | 0.22667 | 58.336 |
| Fancl | 163.917 | 0.320611 | 0.18179 | 1.763653 | 0.07779 | 0.22069 | 85.742 |
| Fancm | 749.261 | 0.214942 | 0.09427 | 2.280185 | 0.0226 | 0.09709 | 56.456 |
| Fank1 | 139.363 | 0.11456 | 0.19771 | 0.579421 | 0.5623 | 0.74313 | 104.672 |
| Fap | 25.8204 | -1.07079 | 0.33609 | -3.18607 | 0.00144 | 0.01608 | 73.133 |
| Far1 | 8314.11 | 0.202974 | 0.11302 | 1.795946 | 0.0725 | 0.21164 | 57.678 |
| Far1os | 217.99 | -0.15513 | 0.14678 | -1.05692 | 0.29055 | 0.50076 | 34.499 |
| Far2 | 934.516 | -0.12101 | 0.08328 | -1.45307 | 0.1462 | 0.3292 | 135.5 |
| Far2os1 | 13.5494 | 0.66853 | 0.61504 | 1.086963 | 0.27705 | 0.48656 | 7.019 |
| Farp1 | 3673.96 | -0.45482 | 0.1447 | -3.14323 | 0.00167 | 0.0179 | 248.545 |
| Farp2 | 540.615 | -0.07284 | 0.09524 | -0.76485 | 0.44436 | 0.64834 | 109.898 |
| Fars2 | 583.94 | -0.03301 | 0.12112 | -0.27259 | 0.78517 | 0.88852 | 608.86 |

| | | | | | | | |
|---|---|---|---|---|---|---|---|
| Farsa | 1471.03 | 0.037333 | 0.06625 | 0.563562 | 0.57305 | 0.75178 | 12.269 |
| Farsb | 2076.3 | 0.045301 | 0.0692 | 0.654672 | 0.51268 | 0.70354 | 70.923 |
| Fas | 138.045 | -0.27924 | 0.19581 | -1.42608 | 0.15385 | 0.33962 | 37.107 |
| Fasn | 12491.6 | -0.03596 | 0.07586 | -0.47397 | 0.63552 | 0.79462 | 18.702 |
| Fastk | 2731.63 | -0.0547 | 0.06429 | -0.85075 | 0.39491 | 0.6039 | 6.848 |
| Fastkd1 | 775.292 | 0.1637 | 0.09446 | 1.733001 | 0.0831 | 0.23011 | 26.702 |
| Fastkd2 | 985.52 | -0.0513 | 0.10523 | -0.48752 | 0.62589 | 0.78827 | 24.042 |
| Fastkd3 | 519.24 | 0.014227 | 0.09563 | 0.14877 | 0.88174 | 0.94106 | 10.105 |
| Fastkd5 | 155.834 | 0.009065 | 0.1532 | 0.059175 | 0.95281 | 0.97684 | 16.188 |
| Fat1 | 6204.99 | 0.087208 | 0.07081 | 1.231608 | 0.2181 | 0.42011 | 116.811 |
| Fat2 | 16.4736 | -1.49568 | 0.55313 | -2.70403 | 0.00685 | 0.04485 | 85.956 |
| Fat3 | 10199.5 | 0.153641 | 0.09334 | 1.646075 | 0.09975 | 0.25974 | 591.097 |
| Fat4 | 633.56 | -0.41092 | 0.1978 | -2.07741 | 0.03776 | 0.13771 | 125.046 |
| Fau | 4583.04 | -0.13405 | 0.09278 | -1.4447 | 0.14854 | 0.33233 | 1.681 |
| Faxc | 2462.09 | -0.05951 | 0.07469 | -0.79683 | 0.42555 | 0.63215 | 65.511 |
| Fbf1 | 5527.79 | 0.125368 | 0.07143 | 1.755164 | 0.07923 | 0.22344 | 25.882 |
| Fbh1 | 4725.44 | 0.090321 | 0.07582 | 1.191268 | 0.23355 | 0.43857 | 35.01 |
| Fbl | 1455.16 | -0.05335 | 0.08176 | -0.65257 | 0.51403 | 0.7047 | 9.56 |
| Fblim1 | 70.4135 | -0.66327 | 0.20703 | -3.20378 | 0.00136 | 0.01541 | 30.035 |
| Fbll1 | 1069.69 | 0.181039 | 0.10439 | 1.734275 | 0.08287 | 0.22962 | 1.503 |
| Fbln1 | 506.93 | -0.74588 | 0.24594 | -3.03283 | 0.00242 | 0.02283 | 80.587 |
| Fbln2 | 392.974 | -0.56677 | 0.21962 | -2.58072 | 0.00986 | 0.05694 | 60.086 |
| Fbln5 | 243.993 | -0.76017 | 0.25943 | -2.9301 | 0.00339 | 0.02816 | 72.491 |
| Fbln7 | 62.5271 | -1.14771 | 0.31939 | -3.59342 | 0.00033 | 0.00599 | 34.054 |
| Fbn1 | 328.539 | -0.52607 | 0.18222 | -2.88704 | 0.00389 | 0.03084 | 207.4 |
| Fbn2 | 209.977 | 0.272645 | 0.16824 | 1.620543 | 0.10512 | 0.26824 | 201.865 |
| Fbrs | 2105.62 | 0.07929 | 0.07662 | 1.034833 | 0.30075 | 0.51072 | 12.513 |
| Fbrsl1 | 2456.41 | -0.12123 | 0.09033 | -1.34198 | 0.1796 | 0.37262 | 86.849 |
| Fbxl12 | 729.566 | 0.209221 | 0.08921 | 2.345309 | 0.01901 | 0.0866 | 26.65 |
| Fbxl12os | 231.559 | 0.330497 | 0.15851 | 2.085065 | 0.03706 | 0.1361 | 9.814 |
| Fbxl13 | 15.0891 | 0.409978 | 0.57754 | 0.709873 | 0.47778 | 0.67553 | 161.788 |
| Fbxl14 | 1386.41 | -0.11208 | 0.08001 | -1.40093 | 0.16124 | 0.34949 | 2.963 |
| Fbxl15 | 394.986 | -0.07689 | 0.12951 | -0.59372 | 0.5527 | 0.73559 | 2.263 |
| Fbxl16 | 41957.6 | 0.240438 | 0.12167 | 1.976184 | 0.04813 | 0.16167 | 12.171 |

| | | | | | | | |
|---|---|---|---|---|---|---|---|
| Fbxl17 | 2807.25 | 0.06195 | 0.07463 | 0.830073 | 0.4065 | 0.6148 | 442.566 |
| Fbxl18 | 3270.91 | -0.00504 | 0.07113 | -0.07087 | 0.9435 | 0.97177 | 28.476 |
| Fbxl19 | 4494.5 | 0.035912 | 0.07523 | 0.47735 | 0.63311 | 0.79325 | 24.941 |
| Fbxl2 | 1756.28 | 0.118479 | 0.06511 | 1.819776 | 0.06879 | 0.20391 | 82.555 |
| Fbxl20 | 4315.74 | 0.153186 | 0.07001 | 2.188165 | 0.02866 | 0.11427 | 67.848 |
| Fbxl21 | 290.275 | 0.290408 | 0.15361 | 1.890499 | 0.05869 | 0.18428 | 15.427 |
| Fbxl22 | 69.7864 | 0.20003 | 0.27708 | 0.721909 | 0.47035 | 0.6692 | 6.235 |
| Fbxl3 | 5732.21 | 0.088164 | 0.06333 | 1.392157 | 0.16387 | 0.35265 | 19.328 |
| Fbxl4 | 641.675 | -0.03227 | 0.09137 | -0.35315 | 0.72398 | 0.85274 | 76.549 |
| Fbxl5 | 5230.45 | 0.143672 | 0.05979 | 2.403101 | 0.01626 | 0.07853 | 77.024 |
| Fbxl6 | 1084.75 | 0.153213 | 0.09468 | 1.618244 | 0.10561 | 0.26914 | 3.026 |
| Fbxl7 | 49.0437 | -0.73877 | 0.30428 | -2.42795 | 0.01518 | 0.07507 | 355.122 |
| Fbxl8 | 72.3126 | -0.03199 | 0.27174 | -0.11774 | 0.90628 | 0.95309 | 4.679 |
| Fbxo10 | 2395.06 | 0.41567 | 0.13845 | 3.002385 | 0.00268 | 0.02442 | 50.358 |
| Fbxo11 | 6583.9 | 0.090274 | 0.06183 | 1.459921 | 0.14431 | 0.32703 | 74.433 |
| Fbxo16 | 354.197 | 0.197635 | 0.1509 | 1.30968 | 0.1903 | 0.38649 | 57.356 |
| Fbxo17 | 83.5845 | -0.3548 | 0.20115 | -1.76388 | 0.07775 | 0.22065 | 21.341 |
| Fbxo2 | 3185.34 | -0.40158 | 0.07092 | -5.66231 | 1.49E-08 | 4.61E-06 | 5.804 |
| Fbxo21 | 7130.51 | 0.052323 | 0.07436 | 0.703646 | 0.48165 | 0.67858 | 33.472 |
| Fbxo22 | 1231.79 | 0.016624 | 0.07961 | 0.208814 | 0.83459 | 0.91508 | 15.509 |
| Fbxo24 | 18.032 | 0.63606 | 0.49562 | 1.283368 | 0.19936 | 0.39732 | 16.5 |
| Fbxo25 | 2882.02 | 0.062167 | 0.09189 | 0.676549 | 0.49869 | 0.69208 | 32.72 |
| Fbxo27 | 356.079 | -0.31445 | 0.15602 | -2.01545 | 0.04386 | 0.15158 | 6.49 |
| Fbxo28 | 2777.22 | 0.000493 | 0.07731 | 0.006379 | 0.99491 | 0.99709 | 28.528 |
| Fbxo3 | 4875.9 | 0.149811 | 0.07561 | 1.981351 | 0.04755 | 0.16029 | 35.52 |
| Fbxo30 | 526.033 | -0.11152 | 0.13041 | -0.85514 | 0.39247 | 0.60165 | 16.723 |
| Fbxo31 | 5515.48 | -0.14982 | 0.06705 | -2.23439 | 0.02546 | 0.10521 | 29.367 |
| Fbxo32 | 1588.27 | -0.07483 | 0.12087 | -0.6191 | 0.53585 | 0.72278 | 39.054 |
| Fbxo33 | 2033.55 | 0.065771 | 0.06862 | 0.958496 | 0.33781 | 0.54856 | 19.071 |
| Fbxo34 | 2168.66 | -0.27332 | 0.09083 | -3.00906 | 0.00262 | 0.0241 | 81.542 |
| Fbxo36 | 95.5299 | -0.39761 | 0.20242 | -1.96432 | 0.04949 | 0.16471 | 60.647 |
| Fbxo38 | 2760.66 | 0.168011 | 0.07156 | 2.347717 | 0.01889 | 0.08627 | 44.541 |
| Fbxo4 | 436.906 | 0.284123 | 0.12006 | 2.366502 | 0.01796 | 0.08349 | 14.129 |
| Fbxo41 | 5232.89 | -0.04742 | 0.06861 | -0.69108 | 0.48951 | 0.68461 | 33.421 |

| | | | | | | | |
|---|---|---|---|---|---|---|---|
| Fbxo42 | 2261.51 | 0.058114 | 0.06043 | 0.961668 | 0.33622 | 0.54701 | 56.15 |
| Fbxo44 | 3688.85 | -0.13099 | 0.0691 | -1.89571 | 0.058 | 0.18306 | 7.683 |
| Fbxo45 | 2964.42 | -0.12802 | 0.07304 | -1.75284 | 0.07963 | 0.2241 | 17.047 |
| Fbxo46 | 570.793 | 0.003775 | 0.08696 | 0.043414 | 0.96537 | 0.98278 | 20.314 |
| Fbxo47 | 24.0154 | 0.155908 | 0.38641 | 0.403474 | 0.6866 | 0.82898 | 30.329 |
| Fbxo48 | 47.2799 | -0.06659 | 0.28578 | -0.23303 | 0.81574 | 0.90517 | 3.398 |
| Fbxo5 | 68.7407 | 0.108819 | 0.22579 | 0.481948 | 0.62984 | 0.79072 | 6.441 |
| Fbxo6 | 610.888 | 0.154233 | 0.09713 | 1.587847 | 0.11232 | 0.28 | 6.425 |
| Fbxo7 | 2484.86 | 0.025331 | 0.07936 | 0.319197 | 0.74958 | 0.86854 | 29.902 |
| Fbxo8 | 1058.6 | -0.02188 | 0.08664 | -0.25252 | 0.80064 | 0.89692 | 42.85 |
| Fbxo9 | 3275.49 | -0.21669 | 0.06087 | -3.55989 | 0.00037 | 0.00649 | 27.567 |
| Fbxw10 | 21.4634 | -0.5301 | 0.38254 | -1.38575 | 0.16582 | 0.35524 | 30.397 |
| Fbxw11 | 6730.28 | -0.04057 | 0.06155 | -0.65916 | 0.50979 | 0.70119 | 104.093 |
| Fbxw15 | 132.256 | -0.22339 | 0.2545 | -0.87776 | 0.38008 | 0.58974 | 15.661 |
| Fbxw17 | 514.469 | -0.00881 | 0.0972 | -0.09063 | 0.92779 | 0.96367 | 15.917 |
| Fbxw2 | 3229.93 | 0.001755 | 0.06231 | 0.028172 | 0.97753 | 0.98803 | 21.798 |
| Fbxw23 | 178.732 | 0.451925 | 0.29627 | 1.525364 | 0.12717 | 0.30336 | 18.471 |
| Fbxw4 | 678.49 | 0.090409 | 0.10261 | 0.881095 | 0.37827 | 0.58818 | 82.059 |
| Fbxw5 | 3814.95 | 0.040432 | 0.05681 | 0.711641 | 0.47669 | 0.67459 | 4.722 |
| Fbxw7 | 7108.57 | -0.19998 | 0.09971 | -2.00565 | 0.04489 | 0.15396 | 163.931 |
| Fbxw8 | 1259.14 | -0.10118 | 0.07959 | -1.27138 | 0.20359 | 0.40214 | 90.5 |
| Fbxw9 | 527.516 | -0.138 | 0.09683 | -1.42514 | 0.15412 | 0.33999 | 7.07 |
| Fcer1g | 244.923 | -0.53143 | 0.18451 | -2.88013 | 0.00398 | 0.03133 | 4.794 |
| Fcer2a | 79.0174 | -0.47075 | 0.24424 | -1.92736 | 0.05393 | 0.17445 | 12.439 |
| Fcf1 | 558.561 | 0.04698 | 0.09557 | 0.491559 | 0.62303 | 0.78589 | 12.407 |
| Fcgr1 | 141.497 | -0.54468 | 0.21922 | -2.48464 | 0.01297 | 0.06808 | 11.061 |
| Fcgr2b | 170.167 | -0.71506 | 0.22914 | -3.12068 | 0.0018 | 0.01883 | 17.931 |
| Fcgr3 | 511.403 | -0.51268 | 0.19327 | -2.65264 | 0.00799 | 0.04956 | 13.762 |
| Fcgrt | 676.801 | -0.4704 | 0.14323 | -3.28429 | 0.00102 | 0.01298 | 10.862 |
| Fcho1 | 4499.26 | 0.292973 | 0.08061 | 3.6343 | 0.00028 | 0.00533 | 17.33 |
| Fcho2 | 3079.38 | 0.038648 | 0.11995 | 0.322207 | 0.7473 | 0.86757 | 92.047 |
| Fchsd1 | 247.161 | 0.177686 | 0.13563 | 1.310028 | 0.19019 | 0.38629 | 12.344 |
| Fchsd2 | 2945.46 | 0.322031 | 0.09889 | 3.256365 | 0.00113 | 0.01378 | 191.543 |
| Fcna | 16.1177 | 0.322597 | 0.45339 | 0.711524 | 0.47676 | 0.67462 | 3.365 |

| | | | | | | | |
|---|---|---|---|---|---|---|---|
| Fcnaos | 59.9881 | 0.283843 | 0.23961 | 1.184609 | 0.23617 | 0.44147 | 23.87 |
| Fcor | 79.475 | -0.09659 | 0.22049 | -0.43806 | 0.66134 | 0.81269 | 1.175 |
| Fcrl1 | 14.8337 | -0.5983 | 0.5045 | -1.18595 | 0.23564 | 0.44084 | 26.548 |
| Fcrlb | 32.4286 | 0.044446 | 0.39189 | 0.113416 | 0.9097 | 0.95456 | 5.669 |
| Fcrls | 697.62 | -0.09835 | 0.14003 | -0.70238 | 0.48244 | 0.67901 | 12.981 |
| Fcsk | 1168.26 | 0.026296 | 0.07263 | 0.36204 | 0.71732 | 0.84903 | 20.033 |
| Fdft1 | 3471.87 | 0.016897 | 0.08486 | 0.199109 | 0.84218 | 0.91879 | 34.429 |
| Fdps | 49.7373 | -0.42867 | 0.27747 | -1.5449 | 0.12237 | 0.29603 | 8.372 |
| Fdx1 | 501.564 | 0.051129 | 0.1016 | 0.503261 | 0.61478 | 0.78079 | 20.251 |
| Fdx2 | 37.8819 | 0.319486 | 0.31177 | 1.024763 | 0.30548 | 0.51546 | 6.095 |
| Fdxacb1 | 113.601 | -0.02238 | 0.18697 | -0.11971 | 0.90471 | 0.95186 | 5.113 |
| Fdxr | 815.604 | 0.003326 | 0.14195 | 0.023433 | 0.9813 | 0.99019 | 9.027 |
| Fech | 2554.58 | -0.0693 | 0.08978 | -0.7719 | 0.44017 | 0.64519 | 37.338 |
| Fem1a | 3270.81 | 0.067719 | 0.0739 | 0.916401 | 0.35946 | 0.56983 | 6.801 |
| Fem1al | 20.3195 | -0.45281 | 0.45353 | -0.9984 | 0.31808 | 0.5286 | 3.274 |
| Fem1b | 7373.92 | 0.064348 | 0.05196 | 1.238437 | 0.21555 | 0.41717 | 20.114 |
| Fem1c | 1633.18 | 0.055365 | 0.09088 | 0.609199 | 0.54239 | 0.72795 | 24.226 |
| Fen1 | 14.4148 | -0.71433 | 0.48032 | -1.48721 | 0.13696 | 0.31768 | 5.038 |
| Fendrr | 67.7355 | -0.02452 | 0.29287 | -0.08374 | 0.93326 | 0.96643 | 28.229 |
| Fer | 741.673 | 0.008921 | 0.09949 | 0.08966 | 0.92856 | 0.96403 | 276.435 |
| Fer1l5 | 665.939 | -0.02323 | 0.0948 | -0.24504 | 0.80643 | 0.90039 | 49.82 |
| Fer1l6 | 12.6055 | -0.70657 | 0.61847 | -1.14244 | 0.25327 | 0.46062 | 155.045 |
| Fermt2 | 3878.7 | -0.14487 | 0.07927 | -1.82739 | 0.06764 | 0.20161 | 71.324 |
| Fermt3 | 240.526 | -0.33005 | 0.18169 | -1.8166 | 0.06928 | 0.20496 | 20.512 |
| Fes | 175.034 | -0.11334 | 0.16206 | -0.69937 | 0.48432 | 0.68067 | 10.191 |
| Fez1 | 8363.67 | -0.01924 | 0.06595 | -0.29178 | 0.77046 | 0.87998 | 57.061 |
| Fez2 | 1623.15 | -0.2627 | 0.0939 | -2.79758 | 0.00515 | 0.03699 | 48.941 |
| Fezf2 | 128.246 | -1.32507 | 0.27332 | -4.84802 | 1.25E-06 | 0.00012 | 6.096 |
| Fgd1 | 501.663 | -0.0214 | 0.10757 | -0.19889 | 0.84235 | 0.91879 | 43.372 |
| Fgd2 | 288.228 | -0.17693 | 0.11908 | -1.48583 | 0.13732 | 0.31798 | 18.748 |
| Fgd3 | 416.802 | -0.69268 | 0.20143 | -3.43884 | 0.00058 | 0.00882 | 58.758 |
| Fgd4 | 2078.72 | 0.073019 | 0.14548 | 0.501916 | 0.61573 | 0.78132 | 183.633 |
| Fgd5 | 527.615 | 0.096137 | 0.11927 | 0.806013 | 0.42024 | 0.62755 | 97.127 |
| Fgd6 | 1428.4 | -0.21021 | 0.07778 | -2.70269 | 0.00688 | 0.04491 | 109.339 |

| | | | | | | | |
|---|---|---|---|---|---|---|---|
| Fgf1 | 3230.34 | -0.25582 | 0.07754 | -3.2991 | 0.00097 | 0.01254 | 90.8 |
| Fgf10 | 496.849 | 0.375865 | 0.14519 | 2.588702 | 0.00963 | 0.05596 | 122.325 |
| Fgf11 | 2527.07 | 0.114325 | 0.07901 | 1.446945 | 0.14791 | 0.33165 | 5.79 |
| Fgf12 | 6951.42 | -0.11562 | 0.12545 | -0.92166 | 0.35671 | 0.56705 | 592.971 |
| Fgf13 | 3651.57 | -0.32898 | 0.09961 | -3.30276 | 0.00096 | 0.01243 | 505.927 |
| Fgf14 | 1807.02 | 0.175211 | 0.12291 | 1.42554 | 0.154 | 0.33978 | 699.221 |
| Fgf16 | 57.6172 | -0.81789 | 0.29172 | -2.80368 | 0.00505 | 0.03655 | 10.661 |
| Fgf17 | 22.5636 | -0.37719 | 0.36071 | -1.0457 | 0.2957 | 0.50616 | 6.066 |
| Fgf18 | 67.706 | -0.33599 | 0.23056 | -1.45728 | 0.14504 | 0.32787 | 29.971 |
| Fgf2 | 674.415 | 0.223859 | 0.10329 | 2.167334 | 0.03021 | 0.11872 | 61.763 |
| Fgf22 | 94.3856 | -0.36977 | 0.22678 | -1.63047 | 0.103 | 0.2651 | 3.521 |
| Fgf23 | 15.4369 | -0.55192 | 0.46872 | -1.1775 | 0.239 | 0.44478 | 8.507 |
| Fgf2os | 49.0028 | 1.110117 | 0.24339 | 4.561016 | 5.09E-06 | 0.00034 | 14.409 |
| Fgf3 | 66.9298 | 0.982119 | 0.47389 | 2.072476 | 0.03822 | 0.139 | 6.354 |
| Fgf5 | 30.207 | -0.9217 | 0.39164 | -2.35341 | 0.0186 | 0.08547 | 22.847 |
| Fgf7 | 47.5161 | 0.028066 | 0.41726 | 0.067263 | 0.94637 | 0.97302 | 56.528 |
| Fgf9 | 889.294 | -0.26699 | 0.10116 | -2.63944 | 0.0083 | 0.05077 | 42.174 |
| Fgfbp1 | 113.162 | -0.65904 | 0.324 | -2.0341 | 0.04194 | 0.1472 | 2.922 |
| Fgfbp3 | 344.678 | 0.012736 | 0.13853 | 0.091941 | 0.92675 | 0.9631 | 2.066 |
| Fgfr1 | 3492.05 | -0.19902 | 0.07833 | -2.54081 | 0.01106 | 0.06132 | 62.065 |
| Fgfr1op2 | 5077.54 | 0.189244 | 0.0785 | 2.410809 | 0.01592 | 0.07743 | 21.996 |
| Fgfr2 | 2161.13 | -0.21173 | 0.11275 | -1.87783 | 0.0604 | 0.18755 | 2960.899 |
| Fgfr3 | 5960.03 | -0.15493 | 0.10057 | -1.54047 | 0.12345 | 0.29772 | 15.395 |
| Fgfrl1 | 486.608 | -0.39631 | 0.13548 | -2.92512 | 0.00344 | 0.02843 | 14.543 |
| Fggy | 321.879 | -0.14496 | 0.1057 | -1.37147 | 0.17023 | 0.36055 | 369.433 |
| Fgl1 | 16.89 | 0.608818 | 0.44068 | 1.381543 | 0.16711 | 0.35694 | 23.723 |
| Fgl2 | 195.143 | -0.66798 | 0.46698 | -1.43041 | 0.1526 | 0.33794 | 5.733 |
| Fh1 | 2873.23 | -0.04456 | 0.06178 | -0.72123 | 0.47077 | 0.66942 | 25.262 |
| Fhad1 | 135.758 | -0.63254 | 0.21346 | -2.96329 | 0.00304 | 0.02634 | 124.645 |
| Fhad1os2 | 14.4494 | -0.20408 | 0.53652 | -0.38037 | 0.70367 | 0.83999 | 11.629 |
| Fhdc1 | 323.288 | -0.34515 | 0.14531 | -2.37519 | 0.01754 | 0.0823 | 38.232 |
| Fhip1a | 72.1153 | -0.45389 | 0.3317 | -1.36837 | 0.1712 | 0.36166 | 157.231 |
| Fhip1b | 4484.48 | 0.128368 | 0.05901 | 2.175225 | 0.02961 | 0.11714 | 28.844 |
| Fhip2a | 2132.65 | 0.081704 | 0.07046 | 1.159586 | 0.24622 | 0.45299 | 28.907 |

| | | | | | | | |
|---|---|---|---|---|---|---|---|
| Fhip2b | 2979.9 | 0.213345 | 0.08742 | 2.440339 | 0.01467 | 0.0732 | 16.54 |
| Fhit | 58.8947 | 0.214067 | 0.28913 | 0.740388 | 0.45906 | 0.65977 | 1611.944 |
| Fhl1 | 5928.34 | 0.273981 | 0.07459 | 3.67294 | 0.00024 | 0.00482 | 61.56 |
| Fhl2 | 962.936 | -0.95721 | 0.39848 | -2.40212 | 0.0163 | 0.07862 | 73.911 |
| Fhl3 | 48.013 | -0.2692 | 0.26756 | -1.00612 | 0.31436 | 0.52485 | 7.911 |
| Fhl4 | 42.7836 | 0.195341 | 0.33822 | 0.577555 | 0.56356 | 0.74412 | 6.209 |
| Fhod1 | 675.747 | 0.13496 | 0.09736 | 1.386215 | 0.16568 | 0.35498 | 18.791 |
| Fhod3 | 1449.28 | -0.48659 | 0.16103 | -3.02177 | 0.00251 | 0.02344 | 424.879 |
| Fibcd1 | 635.711 | -0.36273 | 0.20073 | -1.80704 | 0.07076 | 0.20791 | 32.716 |
| Fibin | 169.705 | -0.46708 | 0.53733 | -0.86927 | 0.3847 | 0.59416 | 2.267 |
| Fibp | 1176.45 | 0.006384 | 0.08512 | 0.075004 | 0.94021 | 0.97025 | 4.437 |
| Ficd | 604.569 | -0.17332 | 0.08849 | -1.95865 | 0.05015 | 0.16642 | 4.798 |
| Fig4 | 2022.84 | 0.250534 | 0.11182 | 2.240413 | 0.02506 | 0.10414 | 115.089 |
| Fign | 499.349 | -0.39818 | 0.19314 | -2.06164 | 0.03924 | 0.14118 | 126.532 |
| Fignl1 | 40.0725 | 0.272248 | 0.29533 | 0.921848 | 0.35661 | 0.56705 | 21.532 |
| Fignl2 | 88.0128 | -0.21866 | 0.19729 | -1.10828 | 0.26774 | 0.47712 | 28.374 |
| Filip1 | 3028.9 | 0.458582 | 0.13712 | 3.344375 | 0.00082 | 0.0112 | 207.758 |
| Filip1l | 204.459 | -0.34723 | 0.18449 | -1.88214 | 0.05982 | 0.18633 | 220.034 |
| Fip1l1 | 3675.27 | 0.154019 | 0.06805 | 2.263231 | 0.02362 | 0.09988 | 63.352 |
| Firre | 5710.64 | 0.243467 | 0.11799 | 2.063455 | 0.03907 | 0.1408 | 79.578 |
| Fis1 | 3998.8 | 0.009353 | 0.0804 | 0.116332 | 0.90739 | 0.95354 | 12.96 |
| Fitm2 | 1354.6 | -0.19496 | 0.09258 | -2.10578 | 0.03522 | 0.13165 | 6.251 |
| Fiz1 | 776.675 | -0.12489 | 0.0937 | -1.33293 | 0.18255 | 0.37638 | 7.639 |
| Fjx1 | 2721.82 | -0.38049 | 0.13287 | -2.86361 | 0.00419 | 0.03245 | 3.134 |
| Fkbp10 | 192.006 | -0.64114 | 0.22042 | -2.90876 | 0.00363 | 0.02954 | 9.128 |
| Fkbp11 | 38.0353 | -0.13367 | 0.33491 | -0.39912 | 0.68981 | 0.83108 | 3.833 |
| Fkbp14 | 719.497 | -0.12314 | 0.09266 | -1.32893 | 0.18387 | 0.37809 | 19.705 |
| Fkbp15 | 2653.48 | 0.033455 | 0.07529 | 0.44433 | 0.6568 | 0.80939 | 60.207 |
| Fkbp1a | 28904.6 | 0.315809 | 0.13132 | 2.404843 | 0.01618 | 0.07823 | 19.21 |
| Fkbp1b | 436.642 | -0.74183 | 0.13417 | -5.52918 | 3.22E-08 | 8.41E-06 | 8.418 |
| Fkbp2 | 2756.21 | -0.10002 | 0.08044 | -1.24344 | 0.21371 | 0.41503 | 2.761 |
| Fkbp3 | 3921.37 | 0.044333 | 0.08567 | 0.517465 | 0.60483 | 0.7735 | 11.584 |
| Fkbp4 | 5247.3 | -0.14389 | 0.08416 | -1.70974 | 0.08731 | 0.23804 | 8.943 |
| Fkbp5 | 6357.84 | -0.16218 | 0.07929 | -2.0455 | 0.04081 | 0.14467 | 118.434 |

| | | | | | | | |
|---|---|---|---|---|---|---|---|
| Fkbp7 | 145.811 | -0.41617 | 0.16796 | -2.47779 | 0.01322 | 0.06887 | 10.073 |
| Fkbp8 | 9358.71 | 0.017003 | 0.0762 | 0.223134 | 0.82343 | 0.90852 | 7.605 |
| Fkbp9 | 2047.07 | -0.20925 | 0.09171 | -2.2817 | 0.02251 | 0.09685 | 47.3 |
| Fkbpl | 323.88 | -0.1854 | 0.12993 | -1.42687 | 0.15362 | 0.33944 | 1.561 |
| Fkrp | 1815.62 | -0.14103 | 0.0766 | -1.84105 | 0.06561 | 0.19767 | 7.487 |
| Fktn | 3957.66 | 0.141473 | 0.06097 | 2.320427 | 0.02032 | 0.09037 | 63.893 |
| Flacc1 | 22.6452 | 0.434752 | 0.42991 | 1.011263 | 0.31189 | 0.52231 | 49.087 |
| Flad1 | 1284.98 | -0.05564 | 0.07559 | -0.73609 | 0.46167 | 0.66223 | 10.867 |
| Flcn | 2685.12 | 0.052831 | 0.06766 | 0.780854 | 0.43489 | 0.64066 | 18.609 |
| Fli1 | 175.018 | -0.45274 | 0.1843 | -2.45651 | 0.01403 | 0.07132 | 120.658 |
| Flicr | 36.41 | 0.3171 | 0.28289 | 1.120948 | 0.26231 | 0.47114 | 7.428 |
| Flii | 2864.05 | 0.043288 | 0.07039 | 0.614975 | 0.53857 | 0.72526 | 13.141 |
| Flna | 5245.97 | -0.03394 | 0.09493 | -0.35747 | 0.72074 | 0.85095 | 26.36 |
| Flnb | 5476.83 | 0.205411 | 0.08332 | 2.465351 | 0.01369 | 0.07038 | 133.632 |
| Flnc | 164.067 | -0.98618 | 0.34152 | -2.88762 | 0.00388 | 0.0308 | 28.628 |
| Flot1 | 2687.91 | -0.23558 | 0.11727 | -2.00885 | 0.04455 | 0.15336 | 9.562 |
| Flot2 | 3188.42 | -0.19734 | 0.08531 | -2.31326 | 0.02071 | 0.09171 | 22.504 |
| Flrt1 | 2129.59 | 0.153107 | 0.08083 | 1.894297 | 0.05819 | 0.18329 | 13.729 |
| Flrt2 | 3300.34 | -0.04992 | 0.13225 | -0.3775 | 0.7058 | 0.84147 | 92.99 |
| Flrt3 | 2314.5 | 0.502932 | 0.15318 | 3.283183 | 0.00103 | 0.01303 | 20.556 |
| Flt1 | 3455.27 | -0.43147 | 0.08339 | -5.17444 | 2.29E-07 | 3.32E-05 | 164.408 |
| Flt3 | 508.685 | -0.94357 | 0.19498 | -4.83929 | 1.30E-06 | 0.00013 | 69.749 |
| Flt3l | 225.521 | 0.253176 | 0.16861 | 1.50152 | 0.13322 | 0.31278 | 6.645 |
| Flt4 | 186.846 | -0.31843 | 0.14731 | -2.16165 | 0.03065 | 0.11978 | 43.477 |
| Flvcr1 | 919.041 | -0.01617 | 0.10192 | -0.15867 | 0.87393 | 0.9368 | 20.312 |
| Flvcr2 | 128.286 | -0.15955 | 0.18994 | -0.84002 | 0.4009 | 0.60962 | 67.047 |
| Flywch1 | 8053.32 | 0.022508 | 0.06684 | 0.336745 | 0.73631 | 0.86061 | 18.81 |
| Flywch2 | 237.054 | -0.04858 | 0.1688 | -0.28778 | 0.77352 | 0.88191 | 9.166 |
| Fmc1 | 474.594 | -0.39836 | 0.12748 | -3.12491 | 0.00178 | 0.01865 | 5.948 |
| Fmn1 | 541.815 | -0.59498 | 0.15045 | -3.95476 | 7.66E-05 | 0.00222 | 389.032 |
| Fmn2 | 3815.46 | -0.05783 | 0.06796 | -0.85085 | 0.39485 | 0.60389 | 320.905 |
| Fmnl1 | 3036.09 | -0.44429 | 0.19169 | -2.31781 | 0.02046 | 0.09094 | 27.795 |
| Fmnl2 | 3228.33 | -0.17372 | 0.10128 | -1.7152 | 0.08631 | 0.23622 | 275.945 |
| Fmnl3 | 567.253 | -0.14836 | 0.0898 | -1.65217 | 0.0985 | 0.25768 | 53.258 |

| | | | | | | | |
|---|---|---|---|---|---|---|---|
| Fmo1 | 474.359 | 0.152368 | 0.15576 | 0.978229 | 0.32796 | 0.5393 | 37.05 |
| Fmo2 | 675.075 | -0.80722 | 0.21482 | -3.7576 | 0.00017 | 0.00389 | 24.41 |
| Fmo5 | 196.774 | -0.53295 | 0.15676 | -3.39974 | 0.00067 | 0.00972 | 26.479 |
| Fmod | 1137.82 | -1.09686 | 0.44405 | -2.47013 | 0.01351 | 0.06993 | 11.024 |
| Fmr1 | 3926.26 | 0.03883 | 0.10673 | 0.363821 | 0.71599 | 0.8481 | 39.423 |
| Fn1 | 3866.24 | -0.63822 | 0.17574 | -3.63164 | 0.00028 | 0.00537 | 67.681 |
| Fn3k | 1938.08 | -0.01506 | 0.06448 | -0.2335 | 0.81537 | 0.90512 | 15.579 |
| Fn3krp | 1663.19 | -0.07396 | 0.07357 | -1.00537 | 0.31472 | 0.52524 | 9.888 |
| Fnbp1 | 6247.76 | -0.08319 | 0.099 | -0.84034 | 0.40072 | 0.60946 | 115.803 |
| Fnbp1l | 9392.5 | 0.434314 | 0.11708 | 3.709487 | 0.00021 | 0.00438 | 80.997 |
| Fnbp4 | 3678.88 | 0.175227 | 0.08294 | 2.112581 | 0.03464 | 0.13033 | 35.652 |
| Fndc1 | 587.326 | 0.471788 | 0.17578 | 2.684038 | 0.00727 | 0.04671 | 88.734 |
| Fndc10 | 820.952 | -0.10046 | 0.0971 | -1.03459 | 0.30086 | 0.51075 | 2.14 |
| Fndc3a | 4295.5 | 0.332032 | 0.06842 | 4.852795 | 1.22E-06 | 0.00012 | 172.058 |
| Fndc3b | 2247.02 | 0.349491 | 0.08538 | 4.093369 | 4.25E-05 | 0.00146 | 295.146 |
| Fndc4 | 2689.9 | -0.31707 | 0.10036 | -3.15922 | 0.00158 | 0.01725 | 3.839 |
| Fndc5 | 3005.93 | -0.24776 | 0.08475 | -2.92336 | 0.00346 | 0.02853 | 7.595 |
| Fndc8 | 214.368 | 0.155926 | 0.15178 | 1.027323 | 0.30427 | 0.51411 | 8.678 |
| Fndc9 | 644.083 | 0.693249 | 0.15591 | 4.446611 | 8.72E-06 | 0.0005 | 4.342 |
| Fnip1 | 4684.86 | 0.272319 | 0.11182 | 2.435314 | 0.01488 | 0.07397 | 80.037 |
| Fnip2 | 1263.44 | -0.00446 | 0.09391 | -0.04749 | 0.96212 | 0.98163 | 111.823 |
| Fnta | 2084.44 | -0.08397 | 0.09007 | -0.93232 | 0.35117 | 0.56165 | 16.929 |
| Fntb | 1178.47 | 0.134038 | 0.07316 | 1.832211 | 0.06692 | 0.20024 | 155.824 |
| Focad | 3030.12 | 0.312474 | 0.09004 | 3.470579 | 0.00052 | 0.00814 | 316.383 |
| Folh1 | 456.165 | 0.056692 | 0.11393 | 0.49759 | 0.61877 | 0.78313 | 56.967 |
| Fos | 562.989 | -0.49557 | 0.22524 | -2.20019 | 0.02779 | 0.11205 | 3.384 |
| Fosb | 847.282 | 0.09968 | 0.15285 | 0.652126 | 0.51432 | 0.70494 | 7.356 |
| Fosl1 | 14.9973 | 0.313077 | 0.53752 | 0.582447 | 0.56027 | 0.74164 | 8.4 |
| Fosl2 | 2793.11 | -0.24 | 0.17254 | -1.39098 | 0.16423 | 0.3531 | 22.042 |
| Foxc1 | 304.706 | -0.98389 | 0.19433 | -5.06295 | 4.13E-07 | 5.47E-05 | 5.844 |
| Foxc2 | 74.7599 | -1.48095 | 0.81024 | -1.82778 | 0.06758 | 0.20161 | 2.725 |
| Foxd1 | 86.2308 | -1.2503 | 0.37614 | -3.32406 | 0.00089 | 0.01177 | 5.064 |
| Foxd2 | 19.9065 | -1.48237 | 0.59509 | -2.49101 | 0.01274 | 0.0673 | 2.592 |
| Foxd2os | 20.5904 | -0.95497 | 0.56469 | -1.69116 | 0.09081 | 0.24468 | 12.741 |

| | | | | | | | |
|---|---|---|---|---|---|---|---|
| Foxf1 | 33.4934 | -0.40516 | 0.37356 | -1.08459 | 0.2781 | 0.48804 | 3.759 |
| Foxf2 | 141.684 | -0.49489 | 0.19348 | -2.55776 | 0.01053 | 0.05957 | 5.588 |
| Foxg1 | 4425.32 | 0.103772 | 0.07529 | 1.378241 | 0.16813 | 0.3581 | 4.202 |
| Foxj1 | 198.113 | -0.46946 | 0.16568 | -2.83352 | 0.0046 | 0.03446 | 4.696 |
| Foxj2 | 1821.15 | 0.023677 | 0.07915 | 0.299156 | 0.76482 | 0.87703 | 25.453 |
| Foxj3 | 4038.19 | 0.154975 | 0.05588 | 2.773404 | 0.00555 | 0.03899 | 92.116 |
| Foxk1 | 4114.63 | 0.096284 | 0.07627 | 1.262411 | 0.2068 | 0.40623 | 60.515 |
| Foxk2 | 5444.58 | 0.07655 | 0.07893 | 0.96983 | 0.33213 | 0.5431 | 49.907 |
| Foxl2 | 35.9556 | -0.42295 | 0.30194 | -1.40078 | 0.16128 | 0.34955 | 3.256 |
| Foxl2os | 30.5711 | -0.22633 | 0.3433 | -0.65928 | 0.50972 | 0.70118 | 5.519 |
| Foxm1 | 76.5846 | -0.30841 | 0.2409 | -1.28027 | 0.20045 | 0.39853 | 13.18 |
| Foxn2 | 1118.61 | 0.095129 | 0.09374 | 1.01481 | 0.3102 | 0.52029 | 49.823 |
| Foxn3 | 2711.57 | -0.29486 | 0.08936 | -3.29982 | 0.00097 | 0.01252 | 373.505 |
| Foxo1 | 3039.38 | 0.488516 | 0.16438 | 2.971792 | 0.00296 | 0.02597 | 84.886 |
| Foxo3 | 4751.48 | 0.040006 | 0.08992 | 0.44492 | 0.65638 | 0.8092 | 94.915 |
| Foxo4 | 1408.8 | -0.17161 | 0.12144 | -1.41311 | 0.15762 | 0.34467 | 6.346 |
| Foxo6 | 617.595 | 0.292885 | 0.08057 | 3.635122 | 0.00028 | 0.00533 | 20.271 |
| Foxp1 | 7708.1 | 0.46266 | 0.17109 | 2.704161 | 0.00685 | 0.04485 | 597.384 |
| Foxp2 | 3125.45 | 0.95407 | 0.33582 | 2.841036 | 0.0045 | 0.03392 | 540.629 |
| Foxp3 | 55.6244 | 0.235145 | 0.27467 | 0.85611 | 0.39194 | 0.60106 | 21.602 |
| Foxp4 | 1379.32 | -0.04413 | 0.07668 | -0.5755 | 0.56496 | 0.74523 | 57.513 |
| Foxq1 | 73.5394 | -0.45286 | 0.20695 | -2.18827 | 0.02865 | 0.11426 | 4.843 |
| Foxr2 | 50.7764 | 0.512575 | 0.30137 | 1.700794 | 0.08898 | 0.24129 | 14.076 |
| Foxred1 | 1937.14 | 0.048912 | 0.06041 | 0.809725 | 0.4181 | 0.62541 | 6.85 |
| Foxred2 | 869.494 | -0.24387 | 0.11341 | -2.15035 | 0.03153 | 0.12223 | 16.201 |
| Fpgs | 285.038 | 0.013172 | 0.11958 | 0.110148 | 0.91229 | 0.95576 | 21.537 |
| Fpgt | 1302.05 | 0.166625 | 0.09273 | 1.796875 | 0.07236 | 0.21133 | 8.486 |
| Fpr1 | 63.3082 | 0.71525 | 0.25416 | 2.814195 | 0.00489 | 0.03571 | 7.47 |
| Fpr2 | 13.8869 | 0.288658 | 0.61209 | 0.471592 | 0.63722 | 0.7959 | 6.129 |
| Fra10ac1 | 1019.89 | 0.272848 | 0.07814 | 3.491985 | 0.00048 | 0.00771 | 35.676 |
| Fras1 | 4895.23 | 0.358157 | 0.10251 | 3.493738 | 0.00048 | 0.00768 | 410.774 |
| Frat1 | 603.718 | 0.33389 | 0.12872 | 2.593874 | 0.00949 | 0.05541 | 2.614 |
| Frat2 | 289.187 | 0.415838 | 0.15614 | 2.663224 | 0.00774 | 0.04863 | 2.161 |
| Frem1 | 105.579 | -0.11957 | 0.28078 | -0.42586 | 0.67021 | 0.81828 | 154.42 |

| | | | | | | | |
|---|---|---|---|---|---|---|---|
| Frem2 | 375.655 | 0.146748 | 0.12647 | 1.160365 | 0.2459 | 0.45255 | 143.418 |
| Frem3 | 40.8609 | 1.629227 | 0.40697 | 4.003327 | 6.25E-05 | 0.00194 | 84.277 |
| Frg1 | 938.613 | 0.010028 | 0.0712 | 0.140847 | 0.88799 | 0.94406 | 19.616 |
| Frg2f1 | 391.463 | -0.05478 | 0.13287 | -0.41225 | 0.68015 | 0.82502 | 9.222 |
| Frk | 16.694 | -0.93191 | 0.58606 | -1.59012 | 0.11181 | 0.27929 | 127.88 |
| Frmd3 | 267.409 | 0.035373 | 0.17745 | 0.199336 | 0.842 | 0.91879 | 188.773 |
| Frmd4a | 2875.69 | 0.076103 | 0.06413 | 1.186754 | 0.23532 | 0.44049 | 596.327 |
| Frmd4b | 1040.46 | -0.33791 | 0.08143 | -4.14973 | 3.33E-05 | 0.00123 | 330.675 |
| Frmd5 | 2312.91 | -0.20892 | 0.11251 | -1.8569 | 0.06333 | 0.19338 | 261.559 |
| Frmd6 | 1976.14 | 0.093072 | 0.08602 | 1.081959 | 0.27927 | 0.48919 | 76.721 |
| Frmd7 | 156.102 | -1.78992 | 0.64134 | -2.79092 | 0.00526 | 0.03745 | 47.531 |
| Frmd8 | 796.505 | -0.31103 | 0.13423 | -2.31719 | 0.02049 | 0.09101 | 25.573 |
| Frmd8os | 14.1558 | 0.123647 | 0.43211 | 0.286148 | 0.77476 | 0.88253 | 5.766 |
| Frmpd1 | 823.055 | 0.09262 | 0.11577 | 0.800071 | 0.42367 | 0.63046 | 101.062 |
| Frmpd2 | 26.3348 | -0.28638 | 0.37589 | -0.76187 | 0.44614 | 0.64983 | 103.574 |
| Frmpd3 | 1716.58 | -0.4877 | 0.1582 | -3.08283 | 0.00205 | 0.02041 | 93.205 |
| Frmpd4 | 3663.73 | 0.349025 | 0.103 | 3.388639 | 0.0007 | 0.01 | 1105.923 |
| Frrs1 | 173.443 | -0.09442 | 0.20206 | -0.46727 | 0.64031 | 0.79859 | 48.714 |
| Frrs1l | 13645.5 | -0.04871 | 0.10452 | -0.46607 | 0.64117 | 0.79912 | 33.219 |
| Frs2 | 3440.23 | 0.073708 | 0.05861 | 1.257709 | 0.2085 | 0.40838 | 79.108 |
| Frs3 | 548.009 | 0.025767 | 0.10808 | 0.238414 | 0.81156 | 0.90289 | 15.257 |
| Frs3os | 13.4235 | -0.4322 | 0.48972 | -0.88255 | 0.37748 | 0.58759 | 2.026 |
| Fry | 11801.1 | 0.149234 | 0.06363 | 2.345166 | 0.01902 | 0.0866 | 379.109 |
| Fryl | 5856.81 | -0.14139 | 0.05856 | -2.41444 | 0.01576 | 0.07698 | 236.633 |
| Frzb | 658.405 | -1.09946 | 0.24141 | -4.55424 | 5.26E-06 | 0.00035 | 35.656 |
| Fsbp | 89.0982 | 0.646184 | 0.27084 | 2.385832 | 0.01704 | 0.08101 | 7.927 |
| Fscn1 | 7091.6 | -0.09053 | 0.07184 | -1.26001 | 0.20766 | 0.40722 | 12.843 |
| Fscn2 | 32.0778 | -0.09583 | 0.28927 | -0.33127 | 0.74044 | 0.86301 | 6.635 |
| Fsd1 | 1548.78 | -0.0212 | 0.06723 | -0.31537 | 0.75248 | 0.86997 | 10.373 |
| Fsd1l | 2523.7 | 0.122796 | 0.09546 | 1.286324 | 0.19833 | 0.39604 | 75.539 |
| Fsd2 | 20.1348 | -0.06222 | 0.37414 | -0.1663 | 0.86792 | 0.93316 | 33.637 |
| Fsip1 | 18.7267 | 0.433296 | 0.47907 | 0.904447 | 0.36576 | 0.57626 | 126.543 |
| Fst | 75.6867 | 0.645457 | 0.25954 | 2.486913 | 0.01289 | 0.0678 | 6.662 |
| Fstl1 | 2525.65 | -0.25608 | 0.07698 | -3.32666 | 0.00088 | 0.0117 | 59.642 |

| | | | | | | | |
|---|---|---|---|---|---|---|---|
| Fstl3 | 37.219 | 0.214935 | 0.30913 | 0.695298 | 0.48687 | 0.6826 | 5.359 |
| Fstl4 | 1705.74 | 0.321931 | 0.11219 | 2.869516 | 0.00411 | 0.03204 | 423.905 |
| Fstl5 | 1190.19 | 0.09777 | 0.13442 | 0.727337 | 0.46702 | 0.66622 | 635.75 |
| Fth1 | 52139.5 | -0.61715 | 0.10557 | -5.84578 | 5.04E-09 | 1.82E-06 | 4.501 |
| Ftl1 | 1042.17 | -0.33512 | 0.15195 | -2.20544 | 0.02742 | 0.11083 | 1.941 |
| Fto | 5107.69 | -0.24413 | 0.18555 | -1.31567 | 0.18829 | 0.38384 | 354.915 |
| Ftsj1 | 802.275 | -0.08055 | 0.0868 | -0.92792 | 0.35345 | 0.56371 | 13.739 |
| Ftsj3 | 1584.05 | 0.052794 | 0.06526 | 0.808984 | 0.41852 | 0.62568 | 6.938 |
| Ftx | 4531.66 | 0.402472 | 0.10489 | 3.837091 | 0.00012 | 0.00311 | 62.884 |
| Fubp1 | 10014.8 | 0.240711 | 0.08979 | 2.680852 | 0.00734 | 0.04707 | 26.405 |
| Fubp3 | 2516.65 | 0.162618 | 0.07001 | 2.322908 | 0.02018 | 0.09009 | 44.876 |
| Fuca1 | 1306.6 | -0.26662 | 0.0947 | -2.81552 | 0.00487 | 0.03561 | 19.577 |
| Fuca2 | 1807.45 | -0.07638 | 0.07852 | -0.97267 | 0.33072 | 0.54195 | 19.496 |
| Fundc1 | 2126.92 | 0.113928 | 0.08174 | 1.393802 | 0.16338 | 0.35208 | 15.762 |
| Fundc2 | 2090.74 | -0.0423 | 0.06153 | -0.68746 | 0.49179 | 0.68654 | 14.061 |
| Fuom | 828.763 | -0.008 | 0.08999 | -0.08887 | 0.92918 | 0.96438 | 5.672 |
| Furin | 1718.49 | -0.01697 | 0.08005 | -0.21194 | 0.83215 | 0.91398 | 16.852 |
| Fus | 22567.3 | 0.253989 | 0.11142 | 2.27956 | 0.02263 | 0.09721 | 18.245 |
| Fut10 | 587.315 | -0.23717 | 0.10327 | -2.29652 | 0.02165 | 0.09443 | 74.408 |
| Fut11 | 1551.79 | 0.069088 | 0.0832 | 0.830399 | 0.40631 | 0.61473 | 6.049 |
| Fut2 | 58.2163 | 0.120975 | 0.272 | 0.444756 | 0.6565 | 0.80921 | 17.804 |
| Fut4 | 13.2672 | -0.42072 | 0.50154 | -0.83887 | 0.40154 | 0.6101 | 4.074 |
| Fut7 | 15.4001 | -0.03513 | 0.61783 | -0.05686 | 0.95465 | 0.97773 | 3.108 |
| Fut8 | 2261.49 | 0.104551 | 0.08737 | 1.196644 | 0.23145 | 0.43602 | 238.214 |
| Fut9 | 5944.37 | 0.341164 | 0.35693 | 0.955828 | 0.33916 | 0.55007 | 190.913 |
| Fuz | 501.329 | 0.394062 | 0.13505 | 2.917909 | 0.00352 | 0.02888 | 6.553 |
| Fv1 | 318.982 | -0.02743 | 0.15366 | -0.17848 | 0.85835 | 0.92856 | 1.38 |
| Fxn | 199.519 | -0.00124 | 0.16238 | -0.00761 | 0.99393 | 0.99634 | 19.153 |
| Fxr1 | 3531.75 | 0.180222 | 0.08927 | 2.018792 | 0.04351 | 0.15088 | 50.38 |
| Fxr2 | 6566.01 | 0.014139 | 0.06144 | 0.230132 | 0.81799 | 0.90638 | 20.308 |
| Fxyd1 | 783.209 | -0.51444 | 0.13829 | -3.71996 | 0.0002 | 0.00426 | 5.522 |
| Fxyd2 | 122.527 | 0.479749 | 0.25926 | 1.850443 | 0.06425 | 0.19507 | 10.61 |
| Fxyd3 | 25.0125 | 0.053938 | 0.34982 | 0.154186 | 0.87746 | 0.93876 | 8.533 |
| Fxyd4 | 54.1191 | -0.00539 | 0.28034 | -0.01924 | 0.98465 | 0.99161 | 4.134 |

| | | | | | | | |
|---|---|---|---|---|---|---|---|
| Fxyd5 | 315.529 | -0.18068 | 0.19356 | -0.93343 | 0.3506 | 0.56126 | 9.76 |
| Fxyd6 | 2564.83 | -0.08073 | 0.12263 | -0.65837 | 0.5103 | 0.70169 | 25.975 |
| Fxyd7 | 517.131 | -0.32952 | 0.14997 | -2.19715 | 0.02801 | 0.11264 | 8.955 |
| Fyb | 211.554 | -0.35386 | 0.20038 | -1.76592 | 0.07741 | 0.2201 | 140.461 |
| Fyb2 | 75.0512 | -1.09085 | 0.364 | -2.99687 | 0.00273 | 0.02471 | 103.408 |
| Fyco1 | 4091.2 | 0.188348 | 0.11456 | 1.644103 | 0.10015 | 0.26034 | 62.4 |
| Fyn | 4684.7 | -0.0608 | 0.07076 | -0.8593 | 0.39018 | 0.59917 | 196.527 |
| Fyttd1 | 4652.32 | 0.191484 | 0.0819 | 2.337914 | 0.01939 | 0.08768 | 31.376 |
| Fzd1 | 427.552 | -0.41999 | 0.18035 | -2.32875 | 0.01987 | 0.08911 | 4.197 |
| Fzd10 | 24.2026 | -0.66686 | 0.39505 | -1.68802 | 0.09141 | 0.24568 | 3.25 |
| Fzd10os | 40.551 | -0.26734 | 0.27586 | -0.96913 | 0.33248 | 0.5434 | 21.644 |
| Fzd2 | 1905.26 | -0.08894 | 0.11659 | -0.76286 | 0.44555 | 0.64941 | 3.663 |
| Fzd3 | 8077.53 | 0.13184 | 0.07592 | 1.73648 | 0.08248 | 0.22902 | 70.015 |
| Fzd4 | 1155.09 | -0.0865 | 0.11841 | -0.73049 | 0.46509 | 0.66474 | 8.78 |
| Fzd5 | 355.343 | 0.199852 | 0.15641 | 1.27776 | 0.20133 | 0.39961 | 7.194 |
| Fzd6 | 478.071 | -0.3647 | 0.12778 | -2.8541 | 0.00432 | 0.03314 | 32.155 |
| Fzd7 | 558.486 | -0.50836 | 0.10948 | -4.64345 | 3.43E-06 | 0.00026 | 4.532 |
| Fzd8 | 230.489 | -0.48837 | 0.19793 | -2.46737 | 0.01361 | 0.07022 | 5.974 |
| Fzd9 | 112.151 | -0.2861 | 0.16185 | -1.76773 | 0.07711 | 0.21952 | 2.293 |
| Fzr1 | 2307.46 | 0.041571 | 0.08107 | 0.512783 | 0.6081 | 0.77608 | 12.295 |
| G0s2 | 119.495 | -0.31589 | 0.25283 | -1.24942 | 0.21151 | 0.41203 | 1.057 |
| G2e3 | 1286.69 | 0.147203 | 0.10302 | 1.428895 | 0.15303 | 0.33859 | 28.926 |
| G3bp1 | 3776.72 | -0.0811 | 0.06807 | -1.19149 | 0.23346 | 0.43851 | 35.154 |
| G3bp2 | 17134.6 | 0.123385 | 0.07759 | 1.590182 | 0.11179 | 0.27929 | 31.574 |
| G530011O06Rik | 152.668 | 0.264905 | 0.23568 | 1.124017 | 0.26101 | 0.46958 | 3.875 |
| G630016G05Rik | 1068.7 | 0.521579 | 0.11032 | 4.727747 | 2.27E-06 | 0.00019 | 18.698 |
| G630018N14Rik | 21.9633 | 0.639948 | 0.36707 | 1.743402 | 0.08126 | 0.22691 | 59.022 |
| G630022F23Rik | 37.4338 | -0.16548 | 0.35765 | -0.46268 | 0.64359 | 0.80116 | 3.627 |
| G630030J09Rik | 32.6622 | 0.09757 | 0.31032 | 0.314422 | 0.7532 | 0.87029 | 3.18 |
| G630093K05Rik | 19.8991 | 1.026444 | 0.45305 | 2.265607 | 0.02348 | 0.09941 | 23.867 |
| G6pc3 | 1083.92 | -0.04651 | 0.08135 | -0.57171 | 0.56752 | 0.7475 | 4.462 |
| G6pdx | 1310.53 | 0.02292 | 0.10305 | 0.222411 | 0.82399 | 0.90875 | 19.712 |
| G730003C15Rik | 18.2846 | 0.840984 | 0.41219 | 2.040272 | 0.04132 | 0.14588 | 2.313 |
| G730013B05Rik | 12.7255 | 0.623685 | 0.59408 | 1.049829 | 0.2938 | 0.50381 | 33.215 |

| | | | | | | | |
|---|---|---|---|---|---|---|---|
| Gaa | 16046.3 | 0.392035 | 0.09649 | 4.063023 | 4.84E-05 | 0.0016 | 17.568 |
| Gab1 | 1252.98 | -0.31724 | 0.09599 | -3.30501 | 0.00095 | 0.01235 | 116.082 |
| Gab2 | 1385.53 | -0.16766 | 0.10138 | -1.65375 | 0.09818 | 0.25724 | 227.361 |
| Gab3 | 89.7013 | 0.355785 | 0.19838 | 1.793482 | 0.0729 | 0.21236 | 118.616 |
| Gabarap | 4027.98 | -0.20848 | 0.08002 | -2.60527 | 0.00918 | 0.05412 | 3.809 |
| Gabarapl1 | 16635.1 | -0.09274 | 0.06447 | -1.43851 | 0.15029 | 0.33494 | 9.187 |
| Gabarapl2 | 5235.47 | -0.0068 | 0.08907 | -0.07639 | 0.93911 | 0.96951 | 12.91 |
| Gabbr1 | 40074.7 | 0.133213 | 0.06414 | 2.076981 | 0.0378 | 0.13782 | 29.102 |
| Gabbr2 | 6592.76 | -0.37739 | 0.14081 | -2.68025 | 0.00736 | 0.04714 | 329.569 |
| Gabpa | 2118.31 | 0.256063 | 0.08559 | 2.991847 | 0.00277 | 0.02495 | 28.855 |
| Gabpb1 | 1196.07 | 0.054328 | 0.07623 | 0.712718 | 0.47602 | 0.67416 | 48.896 |
| Gabpb2 | 2814.44 | 0.261557 | 0.09865 | 2.651403 | 0.00802 | 0.04964 | 36.151 |
| Gabra1 | 12998.7 | -0.00589 | 0.10931 | -0.05391 | 0.957 | 0.979 | 51.992 |
| Gabra2 | 4355.42 | 0.426547 | 0.16541 | 2.578728 | 0.00992 | 0.05714 | 138.253 |
| Gabra3 | 4873.33 | -0.1593 | 0.08392 | -1.89822 | 0.05767 | 0.1823 | 224.168 |
| Gabra4 | 6702.47 | 0.547945 | 0.14834 | 3.693828 | 0.00022 | 0.00457 | 88.56 |
| Gabra5 | 1142.93 | -0.00625 | 0.1522 | -0.04109 | 0.96723 | 0.98353 | 102.388 |
| Gabrb1 | 7552.55 | 0.255125 | 0.08916 | 2.861306 | 0.00422 | 0.03259 | 490.925 |
| Gabrb2 | 5584.08 | 0.278239 | 0.13499 | 2.061139 | 0.03929 | 0.14125 | 209.272 |
| Gabrb3 | 14696.8 | 0.327216 | 0.08724 | 3.750619 | 0.00018 | 0.00396 | 409.111 |
| Gabrd | 3784.67 | 0.469185 | 0.16392 | 2.862367 | 0.0042 | 0.03254 | 13.133 |
| Gabre | 19.8689 | 0.050057 | 0.53125 | 0.094225 | 0.92493 | 0.96217 | 18.805 |
| Gabrg1 | 2393.75 | 0.246647 | 0.10406 | 2.370241 | 0.01778 | 0.08294 | 91.571 |
| Gabrg2 | 6221.96 | -0.00743 | 0.11669 | -0.06371 | 0.9492 | 0.9749 | 90.655 |
| Gabrg3 | 3379.3 | 0.479895 | 0.139 | 3.452381 | 0.00056 | 0.00852 | 670.724 |
| Gabrq | 68.9136 | 1.158514 | 0.46004 | 2.518275 | 0.01179 | 0.06387 | 17.425 |
| Gabrr2 | 92.0022 | -0.6867 | 0.21602 | -3.17888 | 0.00148 | 0.01639 | 32.867 |
| Gad1 | 28344.7 | 0.16409 | 0.11248 | 1.458868 | 0.1446 | 0.32724 | 48.943 |
| Gad1os | 33.9775 | 0.68006 | 0.38973 | 1.744957 | 0.08099 | 0.22645 | 73.418 |
| Gad2 | 32599.1 | 0.571064 | 0.1645 | 3.471417 | 0.00052 | 0.00812 | 71.67 |
| Gadd45a | 100.453 | -0.21062 | 0.18393 | -1.14511 | 0.25216 | 0.45946 | 2.362 |
| Gadd45b | 542.554 | -0.33493 | 0.13667 | -2.45059 | 0.01426 | 0.07203 | 2.132 |
| Gadd45g | 548.363 | -0.07574 | 0.16442 | -0.46066 | 0.64504 | 0.80203 | 1.791 |
| Gadd45gip1 | 148.949 | -0.25084 | 0.15874 | -1.5802 | 0.11406 | 0.28278 | 3.961 |

| | | | | | | | |
|---|---|---|---|---|---|---|---|
| Gadl1 | 15.6012 | -0.07893 | 0.51638 | -0.15285 | 0.87851 | 0.93924 | 191.297 |
| Gak | 4044.16 | -0.06231 | 0.06556 | -0.95047 | 0.34187 | 0.55274 | 60.345 |
| Gal3st1 | 288.241 | -0.57282 | 0.20744 | -2.76141 | 0.00576 | 0.04009 | 15.691 |
| Gal3st3 | 739.107 | -0.272 | 0.09883 | -2.75222 | 0.00592 | 0.04089 | 10.408 |
| Galc | 2626.57 | 0.148436 | 0.09967 | 1.489329 | 0.1364 | 0.31682 | 57.166 |
| Gale | 396.505 | -0.15731 | 0.11742 | -1.33966 | 0.18036 | 0.37369 | 4.452 |
| Galk1 | 303.908 | -0.2111 | 0.18031 | -1.17076 | 0.2417 | 0.4477 | 4.263 |
| Galk2 | 765.613 | 0.115292 | 0.09155 | 1.259304 | 0.20792 | 0.40756 | 125.191 |
| Galm | 80.6669 | -0.43558 | 0.22491 | -1.93667 | 0.05279 | 0.17188 | 57.631 |
| Galns | 292.051 | -0.42611 | 0.13979 | -3.04816 | 0.0023 | 0.02206 | 33.222 |
| Galnt1 | 2898.04 | -0.12812 | 0.07427 | -1.72498 | 0.08453 | 0.23304 | 81.475 |
| Galnt10 | 693.717 | -0.14728 | 0.08863 | -1.66176 | 0.09656 | 0.25453 | 142.073 |
| Galnt11 | 2188.53 | 0.091548 | 0.0745 | 1.228761 | 0.21916 | 0.42137 | 44.015 |
| Galnt12 | 137.493 | 0.111924 | 0.20043 | 0.558424 | 0.57655 | 0.75404 | 31.162 |
| Galnt13 | 3540.33 | 0.612568 | 0.16061 | 3.814099 | 0.00014 | 0.00332 | 681.993 |
| Galnt14 | 719.125 | -0.45703 | 0.16657 | -2.74382 | 0.00607 | 0.04164 | 217.226 |
| Galnt15 | 544.202 | -0.58918 | 0.16905 | -3.4852 | 0.00049 | 0.00783 | 33.209 |
| Galnt16 | 5985.65 | 0.113347 | 0.07971 | 1.421992 | 0.15503 | 0.34134 | 112.696 |
| Galnt17 | 1737.82 | -0.16146 | 0.0961 | -1.68004 | 0.09295 | 0.24829 | 436.413 |
| Galnt18 | 2116.26 | 0.172999 | 0.11901 | 1.453623 | 0.14605 | 0.329 | 308.317 |
| Galnt2 | 3190.08 | -0.05048 | 0.07065 | -0.71443 | 0.47496 | 0.67299 | 114.334 |
| Galnt3 | 43.6338 | -0.41231 | 0.32872 | -1.25429 | 0.20974 | 0.40978 | 42.229 |
| Galnt4 | 220.002 | -0.13121 | 0.14946 | -0.87789 | 0.38 | 0.58972 | 5.089 |
| Galnt6 | 462.832 | -0.65963 | 0.1768 | -3.73087 | 0.00019 | 0.00417 | 37.564 |
| Galnt7 | 794.56 | -0.05157 | 0.0987 | -0.5225 | 0.60132 | 0.77152 | 129.205 |
| Galnt9 | 1311.31 | -0.77366 | 0.38409 | -2.01427 | 0.04398 | 0.15182 | 77.026 |
| Galntl6 | 902.222 | -0.11135 | 0.1283 | -0.8679 | 0.38545 | 0.59487 | 1138.589 |
| Galr1 | 58.2716 | 0.857518 | 0.37448 | 2.289864 | 0.02203 | 0.09565 | 14.398 |
| Galr2 | 68.3971 | -0.22889 | 0.22339 | -1.02463 | 0.30554 | 0.51553 | 3 |
| Galt | 881.824 | -0.06156 | 0.1026 | -0.60008 | 0.54846 | 0.73252 | 3.468 |
| Gamt | 352.907 | -0.22129 | 0.13428 | -1.64794 | 0.09937 | 0.25904 | 2.862 |
| Gan | 2485.46 | -0.02332 | 0.08738 | -0.26681 | 0.78961 | 0.89082 | 57.863 |
| Ganab | 6054.16 | 0.058397 | 0.05882 | 0.992818 | 0.3208 | 0.53155 | 18.602 |
| Ganc | 489.988 | -0.28863 | 0.09786 | -2.94954 | 0.00318 | 0.02714 | 57.805 |

| | | | | | | | |
|---|---|---|---|---|---|---|---|
| Gap43 | 11003.8 | -0.11028 | 0.08231 | -1.33985 | 0.18029 | 0.3736 | 92.21 |
| Gapdh | 4524.15 | -0.687 | 0.40751 | -1.68585 | 0.09183 | 0.24647 | 4.753 |
| Gapdhs | 391.965 | 0.500963 | 0.12565 | 3.986913 | 6.69E-05 | 0.00204 | 13.907 |
| Gapvd1 | 5697.6 | 0.040528 | 0.057 | 0.711074 | 0.47704 | 0.6749 | 80.639 |
| Gar1 | 726.767 | -0.00109 | 0.08843 | -0.01232 | 0.99017 | 0.99471 | 6.485 |
| Garem1 | 1760.14 | 0.086245 | 0.11452 | 0.753107 | 0.45139 | 0.65382 | 172.938 |
| Garem2 | 585.093 | -0.12982 | 0.13357 | -0.97188 | 0.33111 | 0.54208 | 13.218 |
| Garnl3 | 3576.41 | -0.52759 | 0.28986 | -1.82015 | 0.06874 | 0.20383 | 145.431 |
| Garre1 | 1143.36 | -0.12371 | 0.08428 | -1.46785 | 0.14214 | 0.32403 | 76.845 |
| Gars | 3974.52 | -0.08215 | 0.06678 | -1.2301 | 0.21866 | 0.42072 | 41.494 |
| Gart | 1415.26 | 0.008234 | 0.09483 | 0.086833 | 0.9308 | 0.96513 | 25.767 |
| Gas1 | 318.596 | -0.60356 | 0.15928 | -3.78939 | 0.00015 | 0.00355 | 2.961 |
| Gas2 | 563.225 | 0.101667 | 0.10168 | 0.999843 | 0.31739 | 0.5277 | 132.961 |
| Gas2l1 | 2898.13 | -0.11996 | 0.09181 | -1.30659 | 0.19135 | 0.38784 | 11.196 |
| Gas2l2 | 145.028 | 0.344849 | 0.16365 | 2.107235 | 0.0351 | 0.13139 | 7.554 |
| Gas2l3 | 169.144 | -0.31887 | 0.14381 | -2.21732 | 0.0266 | 0.10864 | 35.145 |
| Gas5 | 2727.86 | 0.086463 | 0.13312 | 0.649525 | 0.516 | 0.70609 | 4.125 |
| Gas6 | 2905.41 | -0.22262 | 0.06756 | -3.29521 | 0.00098 | 0.01266 | 29.117 |
| Gas7 | 9317.42 | 0.223208 | 0.08715 | 2.561132 | 0.01043 | 0.05907 | 229.884 |
| Gas8 | 1318.81 | 0.034896 | 0.06773 | 0.515236 | 0.60639 | 0.77468 | 17.817 |
| Gask1a | 373.103 | -0.01386 | 0.15443 | -0.08977 | 0.92847 | 0.96398 | 29.222 |
| Gask1b | 386.008 | -0.63009 | 0.27055 | -2.32888 | 0.01987 | 0.08909 | 61.748 |
| Gata2 | 273.259 | -0.22263 | 0.16164 | -1.37736 | 0.1684 | 0.35839 | 13.142 |
| Gatad1 | 2993.74 | -0.23251 | 0.06888 | -3.37585 | 0.00074 | 0.01036 | 24.254 |
| Gatad2a | 1111.85 | -0.30679 | 0.1069 | -2.86978 | 0.00411 | 0.03204 | 89.309 |
| Gatad2b | 6436.38 | 0.08668 | 0.0654 | 1.325298 | 0.18507 | 0.37981 | 70.23 |
| Gatb | 901.874 | 0.072782 | 0.09205 | 0.790708 | 0.42911 | 0.63573 | 81.504 |
| Gatc | 1704.35 | 0.125723 | 0.07484 | 1.679878 | 0.09298 | 0.24834 | 7.94 |
| Gatd1 | 895.897 | -0.047 | 0.09807 | -0.47926 | 0.63175 | 0.79208 | 6.343 |
| Gatd3a | 1953.07 | -0.16732 | 0.13672 | -1.22377 | 0.22104 | 0.42394 | 7.717 |
| Gatm | 4568.25 | -0.44403 | 0.1306 | -3.39984 | 0.00067 | 0.00972 | 16.837 |
| Gba | 1225.46 | -0.23227 | 0.08469 | -2.74239 | 0.0061 | 0.04175 | 6.039 |
| Gba2 | 4621.02 | 0.175405 | 0.08543 | 2.053175 | 0.04006 | 0.14298 | 11.946 |
| Gbe1 | 586.91 | -0.11408 | 0.10215 | -1.11683 | 0.26407 | 0.47294 | 255.768 |

| | | | | | | | |
|---|---|---|---|---|---|---|---|
| Gbf1 | 4363.42 | 0.043249 | 0.06188 | 0.698913 | 0.48461 | 0.68089 | 134.002 |
| Gbgt1 | 48.443 | -0.72381 | 0.25225 | -2.86943 | 0.00411 | 0.03204 | 8.525 |
| Gbp11 | 40.2142 | 1.248877 | 0.30073 | 4.152869 | 3.28E-05 | 0.00123 | 23.431 |
| Gbp2 | 119.145 | -1.11372 | 0.27284 | -4.08202 | 4.46E-05 | 0.00151 | 17.407 |
| Gbp3 | 203.218 | -0.57335 | 0.23123 | -2.47962 | 0.01315 | 0.06867 | 13.184 |
| Gbp4 | 49.1935 | -0.89577 | 0.273 | -3.28125 | 0.00103 | 0.01308 | 23.82 |
| Gbp5 | 154.704 | -0.69089 | 0.22969 | -3.00787 | 0.00263 | 0.02415 | 28.367 |
| Gbp6 | 22.441 | -1.20448 | 0.42613 | -2.82659 | 0.0047 | 0.03496 | 22.997 |
| Gbp7 | 240.704 | -0.33079 | 0.15659 | -2.11248 | 0.03465 | 0.13033 | 19.808 |
| Gbp9 | 147.095 | -0.445 | 0.18002 | -2.47189 | 0.01344 | 0.06967 | 61.91 |
| Gbx1 | 211.28 | 0.979448 | 0.47506 | 2.061739 | 0.03923 | 0.14117 | 24.446 |
| Gbx2 | 48.8346 | 1.335022 | 0.33469 | 3.988858 | 6.64E-05 | 0.00203 | 3.224 |
| Gc | 15.6204 | 1.242286 | 0.43438 | 2.859901 | 0.00424 | 0.0327 | 40.377 |
| Gca | 607.98 | -0.3382 | 0.12019 | -2.8139 | 0.00489 | 0.03572 | 29.825 |
| Gcat | 21.1065 | -0.47504 | 0.38975 | -1.21885 | 0.2229 | 0.4261 | 11.631 |
| Gcc1 | 1388.11 | -0.03237 | 0.07843 | -0.41277 | 0.67978 | 0.82495 | 12.3 |
| Gcc2 | 3792.61 | 0.244142 | 0.32275 | 0.756447 | 0.44938 | 0.65257 | 50.103 |
| Gcdh | 1185.72 | -0.02758 | 0.07457 | -0.36992 | 0.71144 | 0.84471 | 7.529 |
| Gcfc2 | 428.178 | 0.148291 | 0.11116 | 1.334079 | 0.18218 | 0.37594 | 36.247 |
| Gch1 | 77.9327 | -0.30238 | 0.23765 | -1.27238 | 0.20324 | 0.40179 | 35.519 |
| Gck | 47.7759 | 0.323695 | 0.2863 | 1.130629 | 0.25821 | 0.46624 | 49.262 |
| Gclc | 4088.4 | 0.202596 | 0.08228 | 2.462133 | 0.01381 | 0.07064 | 39.951 |
| Gclm | 3075.5 | -0.09351 | 0.06911 | -1.35303 | 0.17604 | 0.36794 | 25.176 |
| Gcn1 | 4468.42 | 0.07753 | 0.06756 | 1.147643 | 0.25112 | 0.45847 | 57.401 |
| Gcnt1 | 170.175 | -0.89769 | 0.30406 | -2.95231 | 0.00315 | 0.02696 | 46.704 |
| Gcnt2 | 1078.03 | 0.300911 | 0.12224 | 2.46159 | 0.01383 | 0.0707 | 101.139 |
| Gcnt4 | 328.188 | -0.37345 | 0.15933 | -2.34384 | 0.01909 | 0.08673 | 26.218 |
| Gcnt7 | 59.5317 | 0.695189 | 0.30189 | 2.302758 | 0.02129 | 0.09338 | 8.284 |
| Gcsh | 1601.12 | 0.009701 | 0.09605 | 0.100993 | 0.91956 | 0.95927 | 11.728 |
| Gda | 14467.8 | 0.255537 | 0.06656 | 3.839274 | 0.00012 | 0.00309 | 82.139 |
| Gdap1 | 6419.86 | 0.154185 | 0.07845 | 1.96538 | 0.04937 | 0.16457 | 18.91 |
| Gdap1l1 | 2196.95 | -0.14643 | 0.07811 | -1.87464 | 0.06084 | 0.18834 | 16.849 |
| Gdap2 | 1363.09 | 0.127449 | 0.08242 | 1.546401 | 0.12201 | 0.29544 | 44.601 |
| Gde1 | 5167.77 | -0.20441 | 0.08225 | -2.48517 | 0.01295 | 0.06803 | 17.234 |

| | | | | | | | |
|---|---|---|---|---|---|---|---|
| Gdf10 | 273.383 | -0.42851 | 0.18057 | -2.37305 | 0.01764 | 0.08252 | 14.397 |
| Gdf11 | 1031.2 | -0.35515 | 0.10798 | -3.28911 | 0.00101 | 0.01286 | 9.424 |
| Gdf7 | 34.6579 | 0.066451 | 0.37956 | 0.175076 | 0.86102 | 0.92965 | 4.037 |
| Gdf9 | 31.8391 | -0.01197 | 0.31861 | -0.03758 | 0.97002 | 0.98478 | 6.882 |
| Gdi1 | 25706.2 | -0.09068 | 0.05603 | -1.61822 | 0.10562 | 0.26914 | 6.865 |
| Gdi2 | 9853 | -0.08876 | 0.05784 | -1.53451 | 0.1249 | 0.29992 | 29.809 |
| Gdnf | 170.902 | 0.953172 | 0.29492 | 3.231974 | 0.00123 | 0.01456 | 26.73 |
| Gdpd1 | 2008.25 | -0.08399 | 0.06884 | -1.22014 | 0.22241 | 0.42566 | 40.196 |
| Gdpd2 | 94.6942 | 0.123893 | 0.30126 | 0.411252 | 0.68089 | 0.82531 | 9.023 |
| Gdpd3 | 80.6263 | 0.157755 | 0.19645 | 0.803024 | 0.42196 | 0.62879 | 9.316 |
| Gdpd5 | 1653.63 | 0.236583 | 0.12222 | 1.9357 | 0.0529 | 0.1721 | 80.464 |
| Gdpgp1 | 368.411 | -0.12477 | 0.10552 | -1.18245 | 0.23703 | 0.44215 | 9.197 |
| Gem | 163.541 | -0.11456 | 0.20827 | -0.55004 | 0.58229 | 0.75787 | 10.296 |
| Gemin2 | 422.075 | 0.242473 | 0.1052 | 2.304943 | 0.02117 | 0.09302 | 15.078 |
| Gemin4 | 18.5635 | -0.04636 | 0.39481 | -0.11743 | 0.90652 | 0.95319 | 7.094 |
| Gemin5 | 1473.28 | 0.17039 | 0.09819 | 1.735288 | 0.08269 | 0.22942 | 48.538 |
| Gemin6 | 164.353 | -0.01851 | 0.16994 | -0.10893 | 0.91326 | 0.95613 | 4.057 |
| Gemin7 | 567.821 | -0.34723 | 0.09916 | -3.50185 | 0.00046 | 0.00756 | 12.65 |
| Gemin8 | 495.509 | 0.196788 | 0.10607 | 1.855261 | 0.06356 | 0.1938 | 20.059 |
| Gen1 | 76.2634 | -0.05328 | 0.22317 | -0.23875 | 0.8113 | 0.9028 | 26.882 |
| Get1 | 2399.39 | 0.253072 | 0.08165 | 3.099448 | 0.00194 | 0.0197 | 12.446 |
| Get3 | 3082.58 | -0.17145 | 0.07899 | -2.17068 | 0.02996 | 0.11816 | 7.351 |
| Get4 | 1790.09 | 0.109117 | 0.07795 | 1.399804 | 0.16157 | 0.34979 | 17.728 |
| Gfap | 7210.81 | -1.34278 | 0.46296 | -2.90042 | 0.00373 | 0.03002 | 13.577 |
| Gfer | 645.819 | 0.002162 | 0.09887 | 0.021866 | 0.98255 | 0.99061 | 2.97 |
| Gfm1 | 2421.8 | 0.023252 | 0.06632 | 0.35061 | 0.72588 | 0.85388 | 46.434 |
| Gfm2 | 2017.32 | -0.01164 | 0.08147 | -0.14285 | 0.88641 | 0.94338 | 43.259 |
| Gfod1 | 5569.97 | -0.29004 | 0.10725 | -2.70438 | 0.00684 | 0.04485 | 108.928 |
| Gfod2 | 873.003 | -0.11176 | 0.10053 | -1.11172 | 0.26626 | 0.47555 | 44.811 |
| Gfpt1 | 4642.05 | 0.066087 | 0.05871 | 1.125563 | 0.26035 | 0.46882 | 49.352 |
| Gfpt2 | 594.212 | 0.146934 | 0.09621 | 1.527154 | 0.12672 | 0.30261 | 44.436 |
| Gfra1 | 3959.6 | 0.206023 | 0.14041 | 1.467262 | 0.1423 | 0.32422 | 220.306 |
| Gfra2 | 1655 | -0.75583 | 0.2917 | -2.59113 | 0.00957 | 0.05567 | 89.719 |
| Gfra4 | 1042.98 | -0.04466 | 0.07925 | -0.56356 | 0.57306 | 0.75178 | 3.457 |

| | | | | | | | |
|---|---|---|---|---|---|---|---|
| Gfus | 923.748 | -0.13081 | 0.08516 | -1.53609 | 0.12452 | 0.29945 | 5.157 |
| Gfy | 12.3875 | -0.73404 | 0.51385 | -1.42853 | 0.15314 | 0.33865 | 3.221 |
| Gga1 | 2721.73 | 0.103093 | 0.0594 | 1.73547 | 0.08266 | 0.22939 | 17.396 |
| Gga2 | 2044.34 | 0.309378 | 0.11012 | 2.809473 | 0.00496 | 0.03605 | 34.501 |
| Gga3 | 4384.14 | 0.124982 | 0.06628 | 1.885623 | 0.05935 | 0.1854 | 19.797 |
| Ggact | 320.28 | -0.37004 | 0.12955 | -2.85631 | 0.00429 | 0.03299 | 97.535 |
| Ggct | 809.924 | -0.03915 | 0.0813 | -0.4816 | 0.63009 | 0.79087 | 10.371 |
| Ggcx | 1179.33 | 0.109187 | 0.11207 | 0.974277 | 0.32992 | 0.54112 | 16.405 |
| Ggh | 468.722 | -0.16604 | 0.10798 | -1.53766 | 0.12413 | 0.29882 | 24.699 |
| Ggn | 49.0093 | -0.42661 | 0.3035 | -1.40563 | 0.15984 | 0.34778 | 3.767 |
| Ggnbp1 | 71.6729 | 0.424252 | 0.21127 | 2.008112 | 0.04463 | 0.1535 | 19.638 |
| Ggnbp2 | 4818.76 | 0.096713 | 0.09175 | 1.054057 | 0.29186 | 0.50175 | 38.457 |
| Ggps1 | 2615.64 | 0.094647 | 0.08187 | 1.156096 | 0.24764 | 0.45461 | 12.83 |
| Ggt1 | 86.4309 | -0.35741 | 0.22144 | -1.61404 | 0.10652 | 0.27074 | 24.597 |
| Ggt5 | 144.373 | -0.32263 | 0.20304 | -1.58897 | 0.11207 | 0.27957 | 27.861 |
| Ggt7 | 4435.85 | -0.16454 | 0.07364 | -2.23454 | 0.02545 | 0.1052 | 27.859 |
| Ggta1 | 273.899 | -0.62942 | 0.2181 | -2.8859 | 0.0039 | 0.03094 | 63.053 |
| Ghdc | 288.252 | -0.16274 | 0.15162 | -1.07338 | 0.2831 | 0.49331 | 4.926 |
| Ghitm | 21098.4 | 0.005167 | 0.06074 | 0.085079 | 0.9322 | 0.96589 | 15.181 |
| Ghr | 715.233 | 0.336055 | 0.11524 | 2.91619 | 0.00354 | 0.02898 | 265.733 |
| Ghrl | 191.03 | -0.08241 | 0.18468 | -0.44621 | 0.65544 | 0.80876 | 3.762 |
| Gid4 | 3820.37 | -0.07996 | 0.08165 | -0.97931 | 0.32743 | 0.53877 | 33.783 |
| Gid8 | 1868.42 | -0.01804 | 0.083 | -0.21732 | 0.82796 | 0.91174 | 8.617 |
| Gigyf1 | 5220.8 | 0.159078 | 0.07597 | 2.093871 | 0.03627 | 0.13417 | 9.388 |
| Gigyf2 | 4998.19 | 0.094203 | 0.07232 | 1.302512 | 0.19274 | 0.38957 | 123.799 |
| Gimap1 | 54.9861 | -0.51868 | 0.3076 | -1.68622 | 0.09175 | 0.24635 | 4.742 |
| Gimap3 | 14.0929 | -2 | 0.76642 | -2.94582 | 0.00322 | 0.02735 | 6.388 |
| Gimap4 | 15.7126 | -0.80461 | 0.62229 | -1.29298 | 0.19602 | 0.39362 | 7.512 |
| Gimap5 | 21.9653 | -1.02932 | 0.40745 | -2.52625 | 0.01153 | 0.06295 | 8.014 |
| Gimap6 | 178.187 | -0.55654 | 0.18789 | -2.96208 | 0.00306 | 0.02638 | 6.644 |
| Gimap8 | 279.696 | 0.557801 | 0.18707 | 2.981738 | 0.00287 | 0.02548 | 13.642 |
| Gimap9 | 18.4925 | -0.16513 | 0.63207 | -0.26125 | 0.7939 | 0.89334 | 2.986 |
| Gin1 | 508.874 | 0.228985 | 0.11854 | 1.931707 | 0.0534 | 0.1731 | 23.538 |
| Ginm1 | 617.097 | -0.08098 | 0.10953 | -0.73933 | 0.45971 | 0.66021 | 24.878 |

| | | | | | | | |
|---|---|---|---|---|---|---|---|
| Gins1 | 36.5915 | -0.50357 | 0.28829 | -1.74673 | 0.08068 | 0.22605 | 25.881 |
| Gins2 | 50.2238 | -0.30889 | 0.31268 | -0.98787 | 0.32322 | 0.53421 | 10.672 |
| Gins3 | 191.299 | 0.16178 | 0.14355 | 1.126957 | 0.25976 | 0.46825 | 11.608 |
| Gins4 | 591.818 | -0.20038 | 0.11747 | -1.70579 | 0.08805 | 0.23952 | 11.044 |
| Gipc1 | 1879.5 | -0.31364 | 0.10861 | -2.88766 | 0.00388 | 0.0308 | 12.018 |
| Gipc2 | 128.387 | 0.320978 | 0.27841 | 1.152876 | 0.24896 | 0.45629 | 72.698 |
| Gipc3 | 30.397 | -0.33895 | 0.31261 | -1.08424 | 0.27826 | 0.48808 | 8.002 |
| Gipr | 220.417 | 0.181484 | 0.16658 | 1.089458 | 0.27595 | 0.48512 | 10.067 |
| Git1 | 13999.9 | -0.37889 | 0.10829 | -3.49899 | 0.00047 | 0.00759 | 14.225 |
| Git2 | 2226.25 | 0.048992 | 0.06173 | 0.793611 | 0.42742 | 0.63401 | 48.111 |
| Gja1 | 10402.2 | -0.3612 | 0.10478 | -3.44709 | 0.00057 | 0.00863 | 25.184 |
| Gja4 | 63.7265 | -0.01752 | 0.28909 | -0.0606 | 0.95167 | 0.97626 | 2.619 |
| Gja5 | 90.6297 | 0.150071 | 0.27931 | 0.537286 | 0.59107 | 0.76376 | 172.724 |
| Gja6 | 12.5065 | 0.684658 | 0.6503 | 1.052838 | 0.29242 | 0.50224 | 4.937 |
| Gjb1 | 328.415 | -1.11701 | 0.20921 | -5.33911 | 9.34E-08 | 1.66E-05 | 9.252 |
| Gjb2 | 355.937 | -0.88364 | 0.39934 | -2.21275 | 0.02691 | 0.10951 | 6.103 |
| Gjb6 | 3680.16 | -0.38139 | 0.12807 | -2.97788 | 0.0029 | 0.02569 | 10.309 |
| Gjc1 | 212.97 | -0.23588 | 0.14184 | -1.66301 | 0.09631 | 0.25407 | 20.122 |
| Gjc2 | 455.9 | -0.48825 | 0.21247 | -2.29797 | 0.02156 | 0.09424 | 7.646 |
| Gjc3 | 3118.28 | -0.39742 | 0.16147 | -2.46128 | 0.01384 | 0.07073 | 9.638 |
| Gjd2 | 230.513 | -0.15875 | 0.1587 | -1.00027 | 0.31718 | 0.52753 | 4.019 |
| Gjd3 | 21.9158 | 0.217529 | 0.43056 | 0.505221 | 0.6134 | 0.77991 | 0.837 |
| Gk | 1668.02 | -0.1524 | 0.0983 | -1.55031 | 0.12107 | 0.29421 | 74.883 |
| Gk5 | 203.14 | -0.324 | 0.15582 | -2.07923 | 0.0376 | 0.13731 | 65.247 |
| Gkap1 | 613.951 | 0.175145 | 0.09003 | 1.945487 | 0.05172 | 0.16965 | 42.354 |
| Gkn3 | 53.6238 | -0.47028 | 0.33368 | -1.40939 | 0.15872 | 0.34621 | 5.68 |
| Gla | 493.681 | -0.40605 | 0.10947 | -3.70908 | 0.00021 | 0.00438 | 12.977 |
| Glb1 | 568.144 | -0.18186 | 0.13666 | -1.33073 | 0.18328 | 0.3774 | 73.823 |
| Glb1l | 496.355 | -0.05837 | 0.11084 | -0.52662 | 0.59846 | 0.76969 | 12.578 |
| Glb1l2 | 21.0575 | -0.50514 | 0.43168 | -1.17017 | 0.24193 | 0.44787 | 43.425 |
| Glcci1 | 245.95 | 0.145546 | 0.16876 | 0.862467 | 0.38843 | 0.59775 | 87.949 |
| Glce | 2504.54 | 0.638506 | 0.16559 | 3.856036 | 0.00012 | 0.00295 | 65.408 |
| Gldc | 1818.87 | -0.356 | 0.12233 | -2.91004 | 0.00361 | 0.02943 | 76.983 |
| Gldn | 59.0281 | 0.133802 | 0.28205 | 0.474392 | 0.63522 | 0.79438 | 55.301 |

| | | | | | | | |
|---|---|---|---|---|---|---|---|
| Gle1 | 2418.8 | -0.02455 | 0.06446 | -0.38082 | 0.70334 | 0.83991 | 24.946 |
| Glg1 | 14530.8 | 0.06869 | 0.06461 | 1.063178 | 0.2877 | 0.49783 | 104.796 |
| Gli1 | 66.813 | -0.49202 | 0.35196 | -1.39793 | 0.16213 | 0.35051 | 12.093 |
| Gli2 | 350.798 | -0.59157 | 0.14714 | -4.02048 | 5.81E-05 | 0.00184 | 219.488 |
| Gli3 | 575.157 | -0.11578 | 0.10525 | -1.10006 | 0.27131 | 0.48066 | 266.792 |
| Glipr2 | 236.669 | -0.40832 | 0.14498 | -2.81632 | 0.00486 | 0.03557 | 21.718 |
| Glis1 | 29.1743 | 0.178441 | 0.32332 | 0.551901 | 0.58102 | 0.75714 | 200.471 |
| Glis2 | 631.411 | -0.04114 | 0.10749 | -0.38271 | 0.70194 | 0.83914 | 30.212 |
| Glis3 | 370.444 | -0.30235 | 0.12871 | -2.34901 | 0.01882 | 0.08608 | 421.227 |
| Glmn | 364.419 | 0.52949 | 0.10909 | 4.853583 | 1.21E-06 | 0.00012 | 48.922 |
| Glmp | 1197.6 | -0.14398 | 0.07825 | -1.84 | 0.06577 | 0.19805 | 6.291 |
| Glns-ps1 | 40.0346 | -0.96478 | 0.29532 | -3.26687 | 0.00109 | 0.0135 | 1.118 |
| Glo1 | 2843.18 | -0.1265 | 0.08166 | -1.54915 | 0.12135 | 0.29465 | 41.936 |
| Glod4 | 2296 | -0.00938 | 0.08928 | -0.10503 | 0.91635 | 0.95722 | 122.739 |
| Glp1r | 274.861 | 0.723205 | 0.29227 | 2.474432 | 0.01334 | 0.0693 | 38.975 |
| Glp2r | 131.095 | 0.484788 | 0.18812 | 2.577013 | 0.00997 | 0.05737 | 109.652 |
| Glra1 | 17.8312 | -0.74888 | 0.52755 | -1.41955 | 0.15574 | 0.34238 | 93.961 |
| Glra2 | 740.347 | 0.026982 | 0.10918 | 0.24713 | 0.80481 | 0.89916 | 198.377 |
| Glra3 | 351.954 | 0.030235 | 0.25716 | 0.117574 | 0.9064 | 0.95317 | 189.611 |
| Glra4 | 19.6665 | 0.91317 | 0.39169 | 2.331387 | 0.01973 | 0.08874 | 22.468 |
| Glrb | 9431.17 | -0.16208 | 0.09319 | -1.73934 | 0.08198 | 0.22821 | 70.062 |
| Glrx | 776.994 | -0.37337 | 0.08744 | -4.26991 | 1.96E-05 | 0.00087 | 10.287 |
| Glrx2 | 4390.48 | 0.077671 | 0.09907 | 0.78397 | 0.43306 | 0.63916 | 33.339 |
| Glrx3 | 2730.76 | -0.06239 | 0.06263 | -0.99621 | 0.31915 | 0.52976 | 30.981 |
| Glrx5 | 1378.68 | -0.18064 | 0.07867 | -2.29621 | 0.02166 | 0.09443 | 10.219 |
| Gls | 16525.7 | -0.09081 | 0.09012 | -1.00758 | 0.31366 | 0.52419 | 69.785 |
| Gls2 | 757.36 | -0.19054 | 0.13962 | -1.3647 | 0.17235 | 0.36319 | 15.548 |
| Glt1d1 | 1704.08 | 0.038787 | 0.0932 | 0.416191 | 0.67727 | 0.82317 | 77.113 |
| Glt28d2 | 13.2557 | -0.61122 | 0.51743 | -1.18126 | 0.2375 | 0.44288 | 17.668 |
| Glt8d1 | 1864.66 | 0.119848 | 0.10856 | 1.103947 | 0.26962 | 0.47919 | 10.592 |
| Glt8d2 | 194.323 | 0.350735 | 0.22128 | 1.585045 | 0.11296 | 0.28124 | 40.218 |
| Gltp | 1227.41 | -0.50169 | 0.09135 | -5.49216 | 3.97E-08 | 9.54E-06 | 21.587 |
| Glud-ps | 21.0222 | 0.025318 | 0.41794 | 0.060578 | 0.9517 | 0.97626 | 0.657 |
| Glud1 | 12912 | -0.07467 | 0.08027 | -0.93024 | 0.35225 | 0.56254 | 34.539 |

| | | | | | | | |
|---|---|---|---|---|---|---|---|
| Glul | 90784 | -0.42575 | 0.07222 | -5.89518 | 3.74E-09 | 1.53E-06 | 9.78 |
| Glyctk | 127.611 | 0.316949 | 0.16488 | 1.922313 | 0.05457 | 0.17569 | 5.282 |
| Glyr1 | 8676.56 | 0.010109 | 0.05486 | 0.184278 | 0.8538 | 0.92588 | 35.955 |
| Gm10010 | 250.298 | -0.04187 | 0.17237 | -0.24293 | 0.80806 | 0.90131 | 12.828 |
| Gm10033 | 625.68 | 0.338756 | 0.36808 | 0.92033 | 0.3574 | 0.56765 | 29.03 |
| Gm10036 | 27.9521 | -0.44811 | 0.38147 | -1.17468 | 0.24012 | 0.44615 | 0.534 |
| Gm10037 | 366.184 | 0.269062 | 0.13949 | 1.928949 | 0.05374 | 0.17398 | 12.93 |
| Gm10038 | 21.9039 | 0.17371 | 0.43229 | 0.401837 | 0.6878 | 0.82989 | 0.447 |
| Gm10039 | 892.884 | -0.21614 | 0.08139 | -2.65555 | 0.00792 | 0.04931 | 0.411 |
| Gm10048 | 36.063 | -0.69543 | 0.28355 | -2.45258 | 0.01418 | 0.07176 | 3.787 |
| Gm10052 | 919.938 | 0.091337 | 0.10835 | 0.843002 | 0.39923 | 0.60812 | 0.96 |
| Gm10053 | 318.967 | -0.2005 | 0.13886 | -1.44389 | 0.14877 | 0.33266 | 0.946 |
| Gm10060 | 30.4309 | 0.292347 | 0.44004 | 0.66437 | 0.50645 | 0.69789 | 0.111 |
| Gm10069 | 115.991 | 0.154933 | 0.1924 | 0.805266 | 0.42067 | 0.6278 | 64.543 |
| Gm10073 | 38.0392 | -0.56363 | 0.3409 | -1.65338 | 0.09825 | 0.2573 | 0.342 |
| Gm10074 | 19.2279 | -0.19114 | 0.442 | -0.43244 | 0.66542 | 0.81526 | 0.249 |
| Gm10076 | 97.6377 | 0.063521 | 0.32898 | 0.193087 | 0.84689 | 0.92202 | 0.407 |
| Gm10080 | 14.3912 | 0.351082 | 0.49281 | 0.712415 | 0.47621 | 0.67419 | 0.438 |
| Gm10097 | 24.3147 | -0.33846 | 0.49725 | -0.68066 | 0.49609 | 0.68983 | 2.784 |
| Gm10109 | 28.8977 | -0.09578 | 0.37613 | -0.25466 | 0.79899 | 0.89629 | 6.405 |
| Gm10110 | 61.5879 | -0.11566 | 0.24894 | -0.46463 | 0.6422 | 0.80012 | 3.22 |
| Gm10118 | 277.162 | 0.686474 | 0.2067 | 3.321074 | 0.0009 | 0.01186 | 2.569 |
| Gm10123 | 1399.94 | 0.573117 | 0.35478 | 1.615422 | 0.10622 | 0.27017 | 0.49 |
| Gm10125 | 532.614 | 0.322235 | 0.13058 | 2.467787 | 0.0136 | 0.07019 | 102.003 |
| Gm10130 | 167.984 | 0.191241 | 0.18645 | 1.025691 | 0.30504 | 0.51494 | 51.641 |
| Gm10131 | 238.083 | 0.126259 | 0.13327 | 0.947416 | 0.34343 | 0.55414 | 0.736 |
| Gm10132 | 165.585 | -0.1518 | 0.16779 | -0.9047 | 0.36562 | 0.57622 | 0.468 |
| Gm10134 | 47.7654 | -0.23663 | 0.31462 | -0.7521 | 0.45199 | 0.65441 | 1.679 |
| Gm10136 | 42.4068 | -0.21464 | 0.25521 | -0.84104 | 0.40032 | 0.60915 | 0.435 |
| Gm10138 | 96.868 | 0.17002 | 0.2096 | 0.811181 | 0.41726 | 0.62471 | 46.757 |
| Gm10145 | 26.4412 | 0.463751 | 0.35956 | 1.289789 | 0.19712 | 0.39495 | 0.462 |
| Gm10146 | 133.645 | -0.2988 | 0.15727 | -1.8999 | 0.05745 | 0.18194 | 0.309 |
| Gm10154 | 64.0288 | -0.29646 | 0.24042 | -1.23312 | 0.21753 | 0.41947 | 0.354 |
| Gm10156 | 37.3861 | -0.31749 | 0.35859 | -0.88539 | 0.37595 | 0.5861 | 0.462 |

| | | | | | | | |
|---|---|---|---|---|---|---|---|
| Gm10157 | 14.2088 | -0.29283 | 0.43393 | -0.67483 | 0.49979 | 0.69285 | 0.343 |
| Gm10163 | 55.6113 | -0.41384 | 0.29079 | -1.42313 | 0.1547 | 0.34078 | 0.481 |
| Gm10165 | 17.4036 | 0.079676 | 0.42051 | 0.189475 | 0.84972 | 0.92352 | 33.651 |
| Gm10167 | 106.253 | 0.300735 | 0.22378 | 1.343888 | 0.17898 | 0.37164 | 1.479 |
| Gm10171 | 13.5217 | 0.52011 | 0.55128 | 0.943465 | 0.34544 | 0.55616 | 0.628 |
| Gm10175 | 21.1473 | -0.83423 | 0.437 | -1.909 | 0.05626 | 0.17945 | 0.653 |
| Gm10177 | 184.493 | -0.18721 | 0.17065 | -1.09706 | 0.27261 | 0.48182 | 0.444 |
| Gm10179 | 33.7355 | -0.44554 | 0.40155 | -1.10955 | 0.26719 | 0.47653 | 0.688 |
| Gm10184 | 18.5242 | -0.46393 | 0.46859 | -0.99007 | 0.32214 | 0.53314 | 1.988 |
| Gm10221 | 502.626 | -0.06364 | 0.17622 | -0.36112 | 0.71801 | 0.84944 | 0.309 |
| Gm10222 | 197.372 | 0.25817 | 0.3129 | 0.82509 | 0.40932 | 0.61713 | 0.294 |
| Gm10226 | 63.7982 | 0.204166 | 0.3024 | 0.675161 | 0.49957 | 0.69269 | 0.255 |
| Gm10232 | 189.92 | -0.11162 | 0.18499 | -0.60337 | 0.54626 | 0.73071 | 0.72 |
| Gm10240 | 65.2125 | -0.16374 | 0.29388 | -0.55717 | 0.57741 | 0.75448 | 0.483 |
| Gm10241 | 593.482 | -1.22701 | 0.74703 | -1.64252 | 0.10048 | 0.26096 | 0.286 |
| Gm10243 | 39.8302 | 0.077975 | 0.4276 | 0.182355 | 0.8553 | 0.92683 | 0.333 |
| Gm10250 | 461.559 | -0.21989 | 0.11969 | -1.83721 | 0.06618 | 0.19882 | 0.482 |
| Gm10253 | 12.7489 | 0.674537 | 0.50102 | 1.346326 | 0.1782 | 0.3705 | 0.94 |
| Gm10254 | 53.4549 | 0.025901 | 0.26273 | 0.098582 | 0.92147 | 0.96034 | 0.867 |
| Gm10263 | 22.6397 | -0.30387 | 0.36556 | -0.83126 | 0.40582 | 0.6144 | 0.3 |
| Gm10268 | 29.5593 | 0.615017 | 0.34498 | 1.782768 | 0.07462 | 0.21531 | 0.555 |
| Gm10269 | 46.9707 | 0.006748 | 0.29758 | 0.022677 | 0.98191 | 0.99056 | 0.37 |
| Gm10275 | 981.708 | -0.0721 | 0.1281 | -0.56286 | 0.57353 | 0.75223 | 0.659 |
| Gm10277 | 68.5559 | -0.08685 | 0.21698 | -0.40028 | 0.68895 | 0.83048 | 2.295 |
| Gm10288 | 25.7552 | 0.039954 | 0.34389 | 0.116183 | 0.90751 | 0.95354 | 0.537 |
| Gm10297 | 217.197 | 0.12273 | 0.16561 | 0.741055 | 0.45866 | 0.65962 | 0.651 |
| Gm10300 | 14.0103 | 0.47582 | 0.46726 | 1.018311 | 0.30853 | 0.51837 | 2.296 |
| Gm10309 | 31.7988 | 0.239762 | 0.3945 | 0.607758 | 0.54335 | 0.7288 | 7.501 |
| Gm10335 | 165.271 | 0.023813 | 0.14385 | 0.165543 | 0.86852 | 0.9334 | 0.468 |
| Gm10355 | 33.1838 | -0.42561 | 0.34732 | -1.22541 | 0.22042 | 0.4231 | 0.517 |
| Gm10356 | 13.0253 | -0.1281 | 0.50509 | -0.25363 | 0.79979 | 0.89655 | 1.258 |
| Gm10359 | 36.7782 | -0.23809 | 0.31428 | -0.75757 | 0.44871 | 0.65211 | 1.002 |
| Gm10369 | 16.7551 | -0.6556 | 0.46154 | -1.42046 | 0.15547 | 0.34204 | 5.296 |
| Gm10385 | 14.4439 | 0.216602 | 0.43857 | 0.493879 | 0.62139 | 0.78474 | 4.847 |

| | | | | | | | |
|---|---|---|---|---|---|---|---|
| Gm10390 | 19.9834 | 0.447364 | 0.49423 | 0.905181 | 0.36537 | 0.57607 | 18.168 |
| Gm10403 | 21.4502 | 0.581754 | 0.41041 | 1.417481 | 0.15634 | 0.34312 | 1.711 |
| Gm10406 | 20.7733 | 0.499892 | 0.45043 | 1.109804 | 0.26708 | 0.47644 | 21.335 |
| Gm10416 | 27.9317 | 0.581216 | 0.41067 | 1.415299 | 0.15698 | 0.34396 | 0.645 |
| Gm10419 | 6436.5 | 0.132364 | 0.10528 | 1.257211 | 0.20868 | 0.4086 | 10.474 |
| Gm10421 | 370.099 | 0.896234 | 0.21827 | 4.106131 | 4.02E-05 | 0.0014 | 7.132 |
| Gm1043 | 4104.6 | 0.51545 | 0.1439 | 3.582061 | 0.00034 | 0.00614 | 144.178 |
| Gm10432 | 200.611 | 0.507878 | 0.19165 | 2.650094 | 0.00805 | 0.04978 | 25.707 |
| Gm10433 | 12.8923 | -0.14384 | 0.54208 | -0.26536 | 0.79073 | 0.8913 | 5.573 |
| Gm10435 | 14.1475 | -0.02713 | 0.48625 | -0.05578 | 0.95551 | 0.97811 | 4.384 |
| Gm10443 | 224.284 | -0.37997 | 0.26275 | -1.44614 | 0.14814 | 0.33179 | 0.21 |
| Gm10451 | 100.39 | -0.36305 | 0.23895 | -1.51933 | 0.12868 | 0.30583 | 6.84 |
| Gm10461 | 72.8977 | 0.284474 | 0.2432 | 1.169724 | 0.24211 | 0.4481 | 4.786 |
| Gm10472 | 12.7337 | 0.456316 | 0.49201 | 0.927462 | 0.35369 | 0.56399 | 2.102 |
| Gm10478 | 36.7948 | 0.069663 | 0.29951 | 0.232593 | 0.81608 | 0.90539 | 0.4 |
| Gm10501 | 97.3531 | -0.20347 | 0.20036 | -1.01555 | 0.30984 | 0.51986 | 4.89 |
| Gm10509 | 98.2883 | 0.215413 | 0.23223 | 0.927583 | 0.35362 | 0.56394 | 0.842 |
| Gm10516 | 392.661 | 0.003519 | 0.12325 | 0.028555 | 0.97722 | 0.98797 | 14.128 |
| Gm10524 | 412.144 | -0.02392 | 0.1044 | -0.22908 | 0.81881 | 0.9068 | 1.893 |
| Gm10530 | 14.1099 | -0.61304 | 0.44039 | -1.39204 | 0.16391 | 0.35267 | 70.128 |
| Gm10535 | 84.8134 | 0.731841 | 0.28719 | 2.548299 | 0.01082 | 0.06059 | 4.418 |
| Gm10538 | 31.8367 | -0.36936 | 0.38913 | -0.9492 | 0.34252 | 0.55328 | 3.472 |
| Gm10544 | 97.9768 | 0.433117 | 0.21175 | 2.045455 | 0.04081 | 0.14467 | 18.865 |
| Gm10548 | 48.8894 | -0.56073 | 0.26478 | -2.11769 | 0.0342 | 0.12921 | 13.998 |
| Gm10557 | 406.065 | -0.01082 | 0.10589 | -0.10221 | 0.91859 | 0.95871 | 0.619 |
| Gm10558 | 23.0068 | 0.127131 | 0.44015 | 0.288835 | 0.77271 | 0.88166 | 6.521 |
| Gm10561 | 147.775 | -0.05427 | 0.15598 | -0.34793 | 0.72789 | 0.85483 | 9.374 |
| Gm10563 | 17.0033 | 0.43165 | 0.46367 | 0.930934 | 0.35189 | 0.56223 | 30.996 |
| Gm10564 | 13.3492 | 0.068117 | 0.49838 | 0.136678 | 0.89129 | 0.94585 | 3.377 |
| Gm10568 | 18.3174 | -0.00147 | 0.4876 | -0.00301 | 0.9976 | 0.99873 | 1.634 |
| Gm10570 | 45.2491 | 0.108916 | 0.31503 | 0.345734 | 0.72954 | 0.85623 | 2.119 |
| Gm10571 | 40.28 | 0.23136 | 0.26363 | 0.877584 | 0.38017 | 0.58975 | 1.64 |
| Gm10575 | 15.1009 | -0.78475 | 0.43752 | -1.79362 | 0.07287 | 0.21236 | 6.291 |
| Gm10584 | 13.4672 | -0.05248 | 0.48989 | -0.10713 | 0.91469 | 0.95656 | 2.628 |

| | | | | | | | |
|---|---|---|---|---|---|---|---|
| Gm10591 | 31.5285 | -0.04531 | 1.16244 | -0.03898 | 0.9689 | 0.98421 | 1.133 |
| Gm10599 | 59.5943 | -0.43158 | 0.27922 | -1.54563 | 0.12219 | 0.29575 | 2.284 |
| Gm10602 | 19.8482 | 0.17074 | 0.43865 | 0.38924 | 0.6971 | 0.8358 | 6.269 |
| Gm10603 | 191.206 | 0.54497 | 0.16688 | 3.265657 | 0.00109 | 0.01353 | 28.11 |
| Gm10605 | 197.055 | -0.41339 | 0.17995 | -2.29725 | 0.0216 | 0.09434 | 5.776 |
| Gm10608 | 15.3045 | -0.68996 | 0.44062 | -1.56588 | 0.11738 | 0.28827 | 0.676 |
| Gm10614 | 14.0104 | -0.70517 | 0.61533 | -1.14601 | 0.25179 | 0.45912 | 4.271 |
| Gm10629 | 46.3743 | 0.33934 | 0.3069 | 1.105696 | 0.26886 | 0.47813 | 16.711 |
| Gm10638 | 34.696 | -0.17505 | 0.29461 | -0.59419 | 0.55239 | 0.73536 | 3.203 |
| Gm10643 | 92.88 | -0.17614 | 0.25603 | -0.68799 | 0.49146 | 0.68643 | 1.577 |
| Gm10644 | 45.8033 | 0.074159 | 0.27642 | 0.26829 | 0.78848 | 0.89009 | 23.015 |
| Gm10647 | 17.4062 | -0.39665 | 0.49754 | -0.79723 | 0.42532 | 0.63195 | 2.036 |
| Gm10649 | 104.042 | 0.156122 | 0.20131 | 0.775524 | 0.43803 | 0.64338 | 295.396 |
| Gm10654 | 120.448 | -0.01206 | 0.17953 | -0.06719 | 0.94643 | 0.97302 | 1.633 |
| Gm10658 | 70.6986 | 0.120305 | 0.22529 | 0.53399 | 0.59335 | 0.76559 | 16.783 |
| Gm10676 | 26.4414 | -0.26638 | 0.42461 | -0.62734 | 0.53043 | 0.718 | 3.221 |
| Gm10684 | 18.6739 | 0.060302 | 0.44516 | 0.135461 | 0.89225 | 0.94617 | 28.431 |
| Gm10686 | 33.0697 | -0.67039 | 0.35545 | -1.88603 | 0.05929 | 0.18534 | 3.247 |
| Gm10687 | 25.8897 | -0.11287 | 0.43676 | -0.25843 | 0.79608 | 0.89487 | 12.354 |
| Gm10699 | 477.49 | 0.099349 | 0.09837 | 1.009933 | 0.31253 | 0.52299 | 2.786 |
| Gm10701 | 13.2177 | -0.16668 | 0.46099 | -0.36156 | 0.71768 | 0.8492 | 77.053 |
| Gm10705 | 106.958 | 0.665273 | 0.86483 | 0.76925 | 0.44175 | 0.64634 | 0.461 |
| Gm10709 | 300.976 | -0.37155 | 0.24874 | -1.4937 | 0.13525 | 0.31507 | 0.48 |
| Gm10710 | 92.8316 | 0.890568 | 0.25049 | 3.555335 | 0.00038 | 0.00657 | 3.971 |
| Gm10736 | 17.4099 | 0.208526 | 0.55362 | 0.376658 | 0.70643 | 0.8417 | 0.078 |
| Gm10754 | 255.272 | 0.748096 | 0.24779 | 3.019083 | 0.00254 | 0.02357 | 285.677 |
| Gm10762 | 90.6799 | -0.33101 | 0.21584 | -1.53358 | 0.12513 | 0.30014 | 2.54 |
| Gm10766 | 16.2971 | 0.517131 | 0.43621 | 1.185511 | 0.23582 | 0.44108 | 0.606 |
| Gm10767 | 192.629 | 0.270507 | 0.14407 | 1.87767 | 0.06043 | 0.18755 | 4.323 |
| Gm10773 | 14.7496 | 0.395575 | 0.48108 | 0.822264 | 0.41093 | 0.61891 | 3.935 |
| Gm10775 | 14.9877 | -0.27723 | 0.57801 | -0.47963 | 0.63149 | 0.792 | 1.288 |
| Gm10778 | 28.0095 | 0.922279 | 0.40436 | 2.280814 | 0.02256 | 0.09699 | 18.303 |
| Gm10785 | 113.491 | 0.365057 | 0.21045 | 1.734639 | 0.0828 | 0.22962 | 26.745 |
| Gm10790 | 69.4548 | -0.28352 | 0.22859 | -1.24034 | 0.21485 | 0.41624 | 47.291 |

| | | | | | | | |
|---|---|---|---|---|---|---|---|
| Gm10791 | 229.267 | 0.23772 | 0.15651 | 1.518925 | 0.12878 | 0.30595 | 7.238 |
| Gm10805 | 26.4174 | 0.165722 | 0.3608 | 0.459325 | 0.646 | 0.80278 | 0.401 |
| Gm10814 | 67.3927 | 0.41324 | 0.24428 | 1.691643 | 0.09071 | 0.24468 | 5.843 |
| Gm10817 | 29.7622 | -0.27567 | 0.34609 | -0.79654 | 0.42572 | 0.63229 | 1.292 |
| Gm10819 | 90.5295 | -0.4814 | 0.25339 | -1.8998 | 0.05746 | 0.18196 | 0.561 |
| Gm10827 | 65.2566 | -0.24197 | 0.2517 | -0.96134 | 0.33638 | 0.54715 | 10.967 |
| Gm10829 | 22.4257 | 0.008503 | 0.43232 | 0.019669 | 0.98431 | 0.99141 | 2.988 |
| Gm10855 | 35.1101 | -0.33191 | 0.354 | -0.93761 | 0.34845 | 0.55896 | 2.541 |
| Gm10856 | 14.2616 | 0.487304 | 0.45001 | 1.082873 | 0.27886 | 0.48881 | 3.601 |
| Gm10862 | 43.183 | 0.227194 | 0.3351 | 0.677984 | 0.49778 | 0.6911 | 3.35 |
| Gm10863 | 42.0034 | -0.58182 | 0.33802 | -1.72126 | 0.0852 | 0.23406 | 61.485 |
| Gm10912 | 13.7256 | 0.475333 | 0.52871 | 0.89905 | 0.36863 | 0.57929 | 1.69 |
| Gm10916 | 18.3411 | 0.487981 | 0.42308 | 1.153391 | 0.24875 | 0.45606 | 0.483 |
| Gm10925 | 4522.14 | 0.151345 | 0.12435 | 1.217103 | 0.22357 | 0.42677 | 0.678 |
| Gm10941 | 14.0304 | -1.00949 | 0.44711 | -2.25782 | 0.02396 | 0.10083 | 1.448 |
| Gm10997 | 23.0684 | -0.46118 | 0.41645 | -1.1074 | 0.26812 | 0.47741 | 1.5 |
| Gm11042 | 14.6016 | 0.485033 | 0.58442 | 0.829941 | 0.40657 | 0.6148 | 1.285 |
| Gm11084 | 20.316 | 0.330279 | 0.36517 | 0.904447 | 0.36576 | 0.57626 | 15.169 |
| Gm11100 | 28.7516 | -0.25593 | 0.38522 | -0.66438 | 0.50645 | 0.69789 | 32.799 |
| Gm11110 | 31.4193 | 0.625516 | 0.31726 | 1.971627 | 0.04865 | 0.16293 | 14.628 |
| Gm11149 | 59.2893 | 0.272192 | 0.29875 | 0.911118 | 0.36223 | 0.57278 | 54.304 |
| Gm11186 | 19.8243 | 0.446528 | 0.49269 | 0.906313 | 0.36477 | 0.57541 | 8.606 |
| Gm11190 | 69.8716 | 1.208102 | 0.24239 | 4.984098 | 6.23E-07 | 7.64E-05 | 17.162 |
| Gm11192 | 16.6984 | -0.91797 | 0.47898 | -1.91653 | 0.0553 | 0.17727 | 0.329 |
| Gm11201 | 38.3033 | -0.28648 | 0.31655 | -0.905 | 0.36546 | 0.57607 | 7.461 |
| Gm11221 | 46.1627 | -0.33751 | 0.27396 | -1.23199 | 0.21795 | 0.41996 | 0.597 |
| Gm11223 | 158.294 | -0.87048 | 0.2591 | -3.35968 | 0.00078 | 0.01076 | 0.45 |
| Gm11263 | 27.1032 | -0.98882 | 0.43471 | -2.27468 | 0.02293 | 0.09801 | 0.749 |
| Gm11266 | 92.2893 | -0.2997 | 0.18113 | -1.65462 | 0.098 | 0.25687 | 39.372 |
| Gm11274 | 24.9477 | -0.43737 | 0.35621 | -1.22784 | 0.21951 | 0.42192 | 1.352 |
| Gm11346 | 36.0901 | 1.123581 | 0.45105 | 2.491022 | 0.01274 | 0.0673 | 8.579 |
| Gm11353 | 22.9565 | -0.58049 | 0.35462 | -1.63692 | 0.10165 | 0.26311 | 0.627 |
| Gm11400 | 58.0377 | 0.234469 | 0.34226 | 0.685053 | 0.49331 | 0.6878 | 1.214 |
| Gm11401 | 15.3247 | 0.420709 | 0.53692 | 0.783561 | 0.4333 | 0.63924 | 0.344 |

| | | | | | | | |
|---|---|---|---|---|---|---|---|
| Gm1141 | 71.4042 | 0.403961 | 0.28462 | 1.419314 | 0.15581 | 0.34239 | 19.924 |
| Gm11423 | 25.0196 | -0.0184 | 0.34821 | -0.05284 | 0.95786 | 0.97944 | 1.503 |
| Gm11451 | 39.5858 | -0.87887 | 0.31051 | -2.8304 | 0.00465 | 0.03467 | 0.63 |
| Gm11457 | 45.5883 | 0.443233 | 0.34762 | 1.275034 | 0.2023 | 0.40074 | 15.425 |
| Gm11465 | 32.0251 | 0.274465 | 0.3013 | 0.910938 | 0.36233 | 0.57285 | 2.032 |
| Gm11508 | 447.043 | 0.257228 | 0.12827 | 2.005354 | 0.04493 | 0.15404 | 36.357 |
| Gm11513 | 16.8103 | 0.342816 | 0.50763 | 0.675332 | 0.49946 | 0.69266 | 5.525 |
| Gm11517 | 13.4551 | 0.049329 | 0.48966 | 0.100742 | 0.91975 | 0.95943 | 0.387 |
| Gm11518 | 67.7909 | -0.35932 | 0.23929 | -1.50162 | 0.13319 | 0.31278 | 0.411 |
| Gm11525 | 22.6998 | 0.058703 | 0.4678 | 0.125488 | 0.90014 | 0.94992 | 3.553 |
| Gm11537 | 46.6857 | 0.278698 | 0.26908 | 1.035726 | 0.30033 | 0.51042 | 1.468 |
| Gm11560 | 74.4977 | 0.016697 | 0.20002 | 0.083476 | 0.93347 | 0.9665 | 0.948 |
| Gm11585 | 138.566 | -0.09439 | 0.18363 | -0.514 | 0.60725 | 0.77542 | 0.569 |
| Gm11592 | 17.0373 | 0.623004 | 0.41556 | 1.499204 | 0.13382 | 0.31347 | 2.77 |
| Gm11611 | 374.293 | -0.03028 | 0.13873 | -0.21826 | 0.82723 | 0.91108 | 2.284 |
| Gm11613 | 481.379 | 0.053911 | 0.12147 | 0.443824 | 0.65717 | 0.80961 | 16.184 |
| Gm11614 | 37.7621 | -0.26412 | 0.30084 | -0.87796 | 0.37997 | 0.58971 | 2.061 |
| Gm11615 | 28.1512 | 1.155869 | 0.47268 | 2.445332 | 0.01447 | 0.07271 | 1.542 |
| Gm11620 | 50.3647 | 0.247822 | 0.25956 | 0.954786 | 0.33969 | 0.55054 | 1.845 |
| Gm11627 | 284.076 | -0.01679 | 0.20803 | -0.08069 | 0.93569 | 0.96778 | 3.064 |
| Gm11628 | 30.5819 | 0.539687 | 0.36753 | 1.468406 | 0.14199 | 0.32383 | 0.171 |
| Gm11638 | 23.9575 | 0.055631 | 0.47351 | 0.117485 | 0.90648 | 0.95319 | 27.391 |
| Gm11642 | 46.4092 | 0.385978 | 0.28845 | 1.338133 | 0.18085 | 0.37423 | 0.583 |
| Gm11646 | 18.3971 | 0.617095 | 0.44388 | 1.390226 | 0.16446 | 0.3534 | 0.702 |
| Gm11650 | 66.7347 | -0.45701 | 0.25993 | -1.75824 | 0.07871 | 0.22236 | 2.551 |
| Gm11652 | 18.7903 | -0.30569 | 0.39028 | -0.78327 | 0.43347 | 0.63935 | 1.267 |
| Gm11659 | 15.4781 | 0.101052 | 0.42335 | 0.238698 | 0.81134 | 0.9028 | 8.46 |
| Gm11670 | 19.975 | 0.417427 | 0.41708 | 1.000825 | 0.31691 | 0.52743 | 2.706 |
| Gm11681 | 14.1402 | -0.39961 | 0.6313 | -0.633 | 0.52673 | 0.7148 | 6.278 |
| Gm11696 | 164.824 | -0.04115 | 0.1618 | -0.25432 | 0.79925 | 0.89647 | 10.82 |
| Gm11703 | 15.92 | -0.25192 | 0.49814 | -0.50572 | 0.61306 | 0.77967 | 0.483 |
| Gm11706 | 22.7444 | 0.159838 | 0.38269 | 0.417675 | 0.67618 | 0.8224 | 1.339 |
| Gm11725 | 24.1271 | 0.48463 | 0.34225 | 1.415996 | 0.15678 | 0.34363 | 16.167 |
| Gm11729 | 13.9921 | 0.479027 | 0.4713 | 1.016399 | 0.30944 | 0.5194 | 0.98 |

| | | | | | | | |
|---|---|---|---|---|---|---|---|
| Gm11730 | 16.3396 | 1.07725 | 0.55491 | 1.941293 | 0.05222 | 0.1707 | 2.12 |
| Gm11732 | 20.0734 | -0.42091 | 0.49332 | -0.85322 | 0.39354 | 0.60261 | 12.698 |
| Gm11733 | 13.6189 | 0.215214 | 0.46451 | 0.463312 | 0.64314 | 0.80089 | 4.647 |
| Gm11734 | 22.6579 | -0.98081 | 0.39467 | -2.48514 | 0.01295 | 0.06803 | 8.694 |
| Gm11739 | 155.289 | 0.246513 | 0.15178 | 1.624156 | 0.10434 | 0.26727 | 7.956 |
| Gm11743 | 19.574 | 0.202467 | 0.41284 | 0.490427 | 0.62383 | 0.78645 | 2.254 |
| Gm11745 | 13.7471 | 1.184403 | 0.47856 | 2.47495 | 0.01333 | 0.06927 | 0.448 |
| Gm11748 | 20.7845 | -0.21499 | 0.43012 | -0.49983 | 0.6172 | 0.78211 | 2.989 |
| Gm11750 | 68.1008 | 0.425382 | 0.22618 | 1.880729 | 0.06001 | 0.18675 | 2.457 |
| Gm11762 | 67.2602 | -0.6618 | 0.29568 | -2.23825 | 0.0252 | 0.10449 | 20.417 |
| Gm11766 | 564.759 | 0.510508 | 0.10321 | 4.946326 | 7.56E-07 | 8.82E-05 | 3.564 |
| Gm11767 | 828.926 | 0.542281 | 0.11033 | 4.91522 | 8.87E-07 | 9.72E-05 | 10.625 |
| Gm11772 | 19.7207 | 0.609657 | 0.417 | 1.462001 | 0.14374 | 0.32606 | 0.889 |
| Gm11773 | 13.5634 | 0.942085 | 0.56323 | 1.672635 | 0.0944 | 0.25068 | 3.216 |
| Gm11808 | 2879.5 | -0.17864 | 0.11341 | -1.57511 | 0.11523 | 0.28461 | 0.504 |
| Gm11816 | 19.2099 | -0.60096 | 0.41537 | -1.44681 | 0.14795 | 0.33166 | 4.686 |
| Gm11817 | 30.008 | 0.562518 | 0.34519 | 1.629571 | 0.10319 | 0.26546 | 15.239 |
| Gm11827 | 42.6979 | -0.20892 | 0.3403 | -0.61393 | 0.53926 | 0.72582 | 4.194 |
| Gm11837 | 25.3962 | 0.030018 | 0.36738 | 0.081706 | 0.93488 | 0.96724 | 23.181 |
| Gm11839 | 74.7311 | 0.382501 | 0.29894 | 1.279511 | 0.20072 | 0.39886 | 1.953 |
| Gm11841 | 18.7088 | 0.31166 | 0.42888 | 0.726685 | 0.46742 | 0.66644 | 1.963 |
| Gm11847 | 988.285 | 0.088134 | 0.09495 | 0.928229 | 0.35329 | 0.56354 | 1.119 |
| Gm11870 | 72.6712 | -0.382 | 0.27821 | -1.37308 | 0.16973 | 0.36001 | 1.322 |
| Gm11872 | 18.0121 | 1.007009 | 0.49603 | 2.030127 | 0.04234 | 0.14806 | 4.113 |
| Gm11891 | 13.322 | 0.435025 | 0.53983 | 0.805849 | 0.42033 | 0.6276 | 1.716 |
| Gm11914 | 22.1177 | 0.944354 | 0.42488 | 2.222653 | 0.02624 | 0.10755 | 0.507 |
| Gm11915 | 14.5927 | 0.377502 | 0.47303 | 0.798057 | 0.42484 | 0.6314 | 0.78 |
| Gm11942 | 26.2891 | -0.24867 | 0.37283 | -0.66698 | 0.50479 | 0.6967 | 0.345 |
| Gm11944 | 107.462 | -0.18531 | 0.18495 | -1.00193 | 0.31638 | 0.52692 | 11.727 |
| Gm11948 | 60.592 | 0.120157 | 0.25698 | 0.467574 | 0.64009 | 0.79846 | 1.339 |
| Gm11955 | 28.3702 | -0.24821 | 0.39379 | -0.63032 | 0.52849 | 0.71632 | 0.47 |
| Gm11960 | 23.4433 | 0.166112 | 0.42051 | 0.395021 | 0.69283 | 0.83344 | 2.858 |
| Gm11963 | 16.0357 | -0.12476 | 0.42304 | -0.2949 | 0.76807 | 0.87883 | 8.099 |
| Gm11992 | 19.2307 | -0.32833 | 0.46839 | -0.70097 | 0.48332 | 0.67964 | 20.763 |

| | | | | | | | |
|---|---|---|---|---|---|---|---|
| Gm11993 | 105.068 | -0.10939 | 0.18095 | -0.60451 | 0.54551 | 0.73035 | 0.82 |
| Gm12002 | 13.022 | -1.00826 | 0.62849 | -1.60425 | 0.10866 | 0.27398 | 75.045 |
| Gm12033 | 134.429 | -0.23299 | 0.18617 | -1.25148 | 0.21076 | 0.4108 | 1.002 |
| Gm12057 | 14.2952 | -1.00329 | 0.50979 | -1.96803 | 0.04906 | 0.16382 | 1.427 |
| Gm12059 | 35.7218 | -0.31861 | 0.36334 | -0.87691 | 0.38054 | 0.59002 | 7.577 |
| Gm12063 | 170.722 | -0.2269 | 0.19405 | -1.16927 | 0.24229 | 0.44829 | 0.833 |
| Gm12064 | 31.0764 | 0.14309 | 0.37237 | 0.384264 | 0.70078 | 0.83841 | 15.718 |
| Gm12070 | 113.256 | -0.27674 | 0.15874 | -1.74335 | 0.08127 | 0.22691 | 1.002 |
| Gm12089 | 16.4806 | -0.32035 | 0.49511 | -0.64703 | 0.51761 | 0.70746 | 2.27 |
| Gm12097 | 13.5086 | 0.911281 | 0.60374 | 1.509393 | 0.1312 | 0.30967 | 1.664 |
| Gm12104 | 14.9462 | 0.15518 | 0.47286 | 0.328174 | 0.74278 | 0.86459 | 0.979 |
| Gm12108 | 21.474 | -0.25498 | 0.51165 | -0.49835 | 0.61824 | 0.78308 | 40.921 |
| Gm12111 | 20.4589 | 0.832438 | 0.4158 | 2.001996 | 0.04529 | 0.15478 | 23.024 |
| Gm12112 | 40.9429 | 0.165781 | 0.32429 | 0.511212 | 0.6092 | 0.77661 | 0.231 |
| Gm12116 | 55.724 | 0.723453 | 0.35856 | 2.01765 | 0.04363 | 0.15106 | 17.423 |
| Gm12119 | 20.4109 | 0.064887 | 0.39655 | 0.163629 | 0.87002 | 0.93428 | 2.015 |
| Gm12120 | 28.414 | 0.058579 | 0.37569 | 0.155923 | 0.87609 | 0.93819 | 5.886 |
| Gm12166 | 44.2542 | -0.44433 | 0.26663 | -1.66649 | 0.09562 | 0.25289 | 0.968 |
| Gm12174 | 70.5226 | 0.372552 | 0.24713 | 1.507528 | 0.13168 | 0.31044 | 0.813 |
| Gm12183 | 359.977 | -0.13431 | 0.18288 | -0.73443 | 0.46268 | 0.66283 | 1.848 |
| Gm12184 | 49.4417 | -0.58462 | 0.26506 | -2.20563 | 0.02741 | 0.11082 | 1.31 |
| Gm12191 | 340.79 | 0.101114 | 0.41569 | 0.243243 | 0.80782 | 0.90114 | 0.348 |
| Gm12195 | 15.9882 | 0.005207 | 0.49155 | 0.010594 | 0.99155 | 0.99531 | 0.923 |
| Gm12198 | 14.2742 | -0.25639 | 0.55651 | -0.46072 | 0.645 | 0.80203 | 3.151 |
| Gm12205 | 14.7017 | 1.076917 | 0.48748 | 2.209148 | 0.02716 | 0.1101 | 0.622 |
| Gm12207 | 55.1405 | 0.673556 | 0.24876 | 2.707667 | 0.00678 | 0.04451 | 2.896 |
| Gm12208 | 24.9505 | -0.30983 | 0.35944 | -0.86199 | 0.3887 | 0.59793 | 0.163 |
| Gm12212 | 19.8696 | -0.23275 | 0.41528 | -0.56045 | 0.57517 | 0.75324 | 1.235 |
| Gm12216 | 38.1881 | -0.96711 | 0.30592 | -3.1613 | 0.00157 | 0.01718 | 75.839 |
| Gm12224 | 49.423 | -0.02485 | 0.26408 | -0.09411 | 0.92502 | 0.96217 | 13.166 |
| Gm12227 | 26.1573 | 0.262245 | 0.4152 | 0.631614 | 0.52764 | 0.71577 | 5.647 |
| Gm12239 | 12.8978 | 0.361328 | 0.60532 | 0.596923 | 0.55056 | 0.73408 | 298.761 |
| Gm12251 | 130.009 | -0.37704 | 0.16691 | -2.25894 | 0.02389 | 0.10062 | 0.792 |
| Gm12254 | 15.1733 | -0.0664 | 0.55482 | -0.11969 | 0.90473 | 0.95186 | 0.471 |

| | | | | | | | |
|---|---|---|---|---|---|---|---|
| Gm12258 | 1179.95 | 0.213376 | 0.08066 | 2.645389 | 0.00816 | 0.05018 | 14.799 |
| Gm12263 | 24.786 | -0.10394 | 0.40735 | -0.25516 | 0.7986 | 0.89601 | 1.482 |
| Gm12264 | 149.649 | 0.2075 | 0.1797 | 1.154684 | 0.24822 | 0.45532 | 5.036 |
| Gm12272 | 17.4849 | 0.00036 | 0.44132 | 0.000817 | 0.99935 | 0.99967 | 0.823 |
| Gm12276 | 159.317 | 0.209365 | 0.1733 | 1.208103 | 0.22701 | 0.43117 | 0.301 |
| Gm12279 | 183.737 | 0.542624 | 0.16521 | 3.284426 | 0.00102 | 0.01298 | 6.805 |
| Gm12294 | 40.2925 | -0.05317 | 0.30415 | -0.17481 | 0.86123 | 0.92973 | 0.49 |
| Gm12295 | 15.0493 | -0.51395 | 0.57195 | -0.89859 | 0.36887 | 0.5795 | 92.33 |
| Gm12319 | 112.853 | -0.34049 | 0.21967 | -1.54998 | 0.12115 | 0.29433 | 1.759 |
| Gm12338 | 1065.34 | -0.31595 | 0.12262 | -2.57663 | 0.00998 | 0.0574 | 0.192 |
| Gm12339 | 114.728 | 0.894669 | 0.2039 | 4.3877 | 1.15E-05 | 0.00061 | 30.474 |
| Gm12346 | 358.636 | 0.128792 | 0.17822 | 0.72267 | 0.46988 | 0.66875 | 2.455 |
| Gm12353 | 20.7053 | 0.629885 | 0.49563 | 1.270881 | 0.20377 | 0.40241 | 45.68 |
| Gm12359 | 224.896 | 0.322909 | 0.14382 | 2.245237 | 0.02475 | 0.1033 | 12.381 |
| Gm12367 | 35.4072 | 0.25383 | 0.28459 | 0.891923 | 0.37243 | 0.58277 | 36.326 |
| Gm12371 | 89.2471 | -0.707 | 0.29735 | -2.37769 | 0.01742 | 0.08192 | 35.166 |
| Gm12391 | 19.6351 | 0.459879 | 0.41581 | 1.105979 | 0.26874 | 0.47804 | 0.447 |
| Gm12394 | 76.679 | -0.50372 | 0.28227 | -1.78456 | 0.07433 | 0.21492 | 74.844 |
| Gm12395 | 12.8082 | -0.79193 | 0.53631 | -1.47661 | 0.13978 | 0.3211 | 1.502 |
| Gm12397 | 19.5024 | -0.14749 | 0.42299 | -0.34869 | 0.72732 | 0.85443 | 0.615 |
| Gm12404 | 195.348 | 0.03073 | 0.1759 | 0.174702 | 0.86131 | 0.92974 | 5.13 |
| Gm12408 | 19.0735 | 0.536725 | 0.43572 | 1.231811 | 0.21802 | 0.42003 | 6.368 |
| Gm12411 | 131.557 | -0.60833 | 0.28304 | -2.14929 | 0.03161 | 0.12249 | 0.483 |
| Gm12418 | 28.6992 | -0.20698 | 0.36669 | -0.56446 | 0.57244 | 0.75117 | 0.942 |
| Gm12430 | 52.8889 | -0.72936 | 0.30246 | -2.41139 | 0.01589 | 0.07735 | 1.322 |
| Gm12435 | 14.8888 | -0.22848 | 0.45676 | -0.50022 | 0.61692 | 0.78199 | 1.016 |
| Gm12439 | 17.9693 | -0.18483 | 0.42088 | -0.43915 | 0.66056 | 0.81233 | 0.365 |
| Gm12446 | 27.9478 | -0.54691 | 0.38109 | -1.43513 | 0.15125 | 0.33614 | 45.509 |
| Gm12447 | 14.829 | -0.25115 | 0.49069 | -0.51183 | 0.60877 | 0.77635 | 0.47 |
| Gm12454 | 35.9237 | 0.216932 | 0.29153 | 0.744113 | 0.45681 | 0.65825 | 4.336 |
| Gm12472 | 17.5305 | -0.16039 | 0.4137 | -0.3877 | 0.69824 | 0.83649 | 20.684 |
| Gm12473 | 43.6571 | -0.25973 | 0.28991 | -0.89592 | 0.3703 | 0.58058 | 9.995 |
| Gm12500 | 24.3138 | -0.85355 | 0.39456 | -2.1633 | 0.03052 | 0.11945 | 2.428 |
| Gm12522 | 33.6003 | -0.12248 | 0.34098 | -0.3592 | 0.71944 | 0.85015 | 4.465 |

| | | | | | | | |
|---|---|---|---|---|---|---|---|
| Gm12525 | 20.7839 | 0.098799 | 0.4461 | 0.221474 | 0.82472 | 0.90935 | 0.706 |
| Gm12537 | 26.8741 | -0.3799 | 0.33666 | -1.12844 | 0.25914 | 0.46734 | 1.002 |
| Gm12540 | 27.388 | 0.920813 | 0.41226 | 2.233567 | 0.02551 | 0.10538 | 0.319 |
| Gm12542 | 15.4004 | 0.436689 | 0.45165 | 0.966885 | 0.3336 | 0.54445 | 4.939 |
| Gm12568 | 14.0474 | -0.62978 | 0.53358 | -1.18028 | 0.23789 | 0.44325 | 0.624 |
| Gm12571 | 32.4581 | 0.69801 | 0.37841 | 1.844577 | 0.0651 | 0.19674 | 0.243 |
| Gm12576 | 52.2385 | -0.32388 | 0.27991 | -1.15709 | 0.24724 | 0.45409 | 31.812 |
| Gm12592 | 111.657 | 0.165429 | 0.19003 | 0.870561 | 0.38399 | 0.59343 | 25.753 |
| Gm12655 | 83.549 | -0.33874 | 0.18291 | -1.85191 | 0.06404 | 0.19477 | 16.966 |
| Gm12657 | 13.9228 | -0.55395 | 0.52697 | -1.05118 | 0.29317 | 0.50308 | 0.411 |
| Gm12663 | 105.277 | -0.30921 | 0.25073 | -1.23327 | 0.21748 | 0.41946 | 0.958 |
| Gm12669 | 16.4381 | 0.104009 | 0.46454 | 0.223896 | 0.82284 | 0.90817 | 1.009 |
| Gm12671 | 15.0504 | -0.30851 | 0.48288 | -0.63888 | 0.5229 | 0.71143 | 1.002 |
| Gm12689 | 26.9479 | -0.11296 | 0.39158 | -0.28848 | 0.77298 | 0.88183 | 1.359 |
| Gm12691 | 13.8674 | -0.6548 | 0.45862 | -1.42775 | 0.15336 | 0.33903 | 4.197 |
| Gm12693 | 52.0738 | -0.065 | 0.30168 | -0.21546 | 0.82941 | 0.91247 | 0.483 |
| Gm12696 | 40.0322 | -0.42422 | 0.29955 | -1.41622 | 0.15671 | 0.34363 | 0.905 |
| Gm12709 | 120.029 | -0.18908 | 0.16427 | -1.15097 | 0.24975 | 0.45704 | 22.49 |
| Gm12715 | 313.388 | -0.12937 | 0.14753 | -0.87691 | 0.38054 | 0.59002 | 1.128 |
| Gm12735 | 15.1892 | 0.285709 | 0.58104 | 0.491718 | 0.62292 | 0.78589 | 0.748 |
| Gm12743 | 38.654 | -0.22483 | 0.38443 | -0.58486 | 0.55864 | 0.73997 | 27.425 |
| Gm12749 | 73.5625 | -0.32381 | 0.34829 | -0.92971 | 0.35252 | 0.56289 | 1.159 |
| Gm12751 | 264.326 | -0.29261 | 0.41406 | -0.70669 | 0.47976 | 0.67732 | 0.381 |
| Gm12758 | 41.2194 | 0.49907 | 0.39406 | 1.266491 | 0.20534 | 0.40437 | 5.831 |
| Gm12783 | 30.5392 | -0.05467 | 0.31967 | -0.17102 | 0.8642 | 0.93125 | 3.247 |
| Gm12786 | 20.9211 | 0.506506 | 0.38929 | 1.301088 | 0.19323 | 0.39017 | 4.565 |
| Gm12788 | 57.8709 | 0.0503 | 0.29175 | 0.172411 | 0.86311 | 0.93081 | 0.308 |
| Gm12795 | 23.3433 | -0.06297 | 0.45476 | -0.13846 | 0.88988 | 0.94485 | 0.714 |
| Gm12829 | 16.4015 | -0.46907 | 0.45025 | -1.0418 | 0.2975 | 0.5076 | 6.89 |
| Gm12854 | 65.2886 | -0.41435 | 0.2768 | -1.49694 | 0.13441 | 0.31425 | 0.297 |
| Gm12860 | 22.3577 | -0.09686 | 0.41895 | -0.2312 | 0.81716 | 0.90593 | 22.423 |
| Gm12867 | 28.3909 | -0.09449 | 0.34121 | -0.27693 | 0.78184 | 0.88646 | 0.71 |
| Gm12868 | 41.1584 | -0.27265 | 0.32823 | -0.83067 | 0.40616 | 0.61464 | 5.053 |
| Gm12869 | 13.8135 | 0.7358 | 0.47041 | 1.564181 | 0.11777 | 0.2889 | 2.315 |

| | | | | | | | |
|---|---|---|---|---|---|---|---|
| Gm12892 | 24.5917 | 0.791893 | 0.37417 | 2.116375 | 0.03431 | 0.12943 | 1.146 |
| Gm12902 | 19.1938 | -0.36736 | 0.43997 | -0.83496 | 0.40374 | 0.61241 | 0.322 |
| Gm12905 | 149.712 | 0.035496 | 0.17226 | 0.206063 | 0.83674 | 0.91619 | 12.962 |
| Gm12917 | 22.976 | 0.055164 | 0.36167 | 0.152524 | 0.87877 | 0.93928 | 0.639 |
| Gm12918 | 68.3781 | -0.25282 | 0.22903 | -1.10388 | 0.26965 | 0.47921 | 0.636 |
| Gm12925 | 47.28 | -0.18126 | 0.29613 | -0.61209 | 0.54048 | 0.72624 | 5.337 |
| Gm12928 | 33.5274 | 0.302858 | 0.28738 | 1.053857 | 0.29195 | 0.5018 | 0.573 |
| Gm12930 | 21.1191 | 0.780493 | 0.53229 | 1.466281 | 0.14257 | 0.32446 | 29.363 |
| Gm12932 | 37.478 | 0.647584 | 0.35227 | 1.838319 | 0.06602 | 0.19847 | 2.091 |
| Gm12940 | 2003.97 | 0.446084 | 0.14099 | 3.164012 | 0.00156 | 0.01704 | 15.668 |
| Gm12941 | 159.493 | 0.582796 | 0.20145 | 2.893068 | 0.00381 | 0.0305 | 0.22 |
| Gm12944 | 15.1666 | 0.683689 | 0.53062 | 1.288475 | 0.19758 | 0.39543 | 0.463 |
| Gm12960 | 125.05 | -0.23584 | 0.3131 | -0.75322 | 0.45132 | 0.65382 | 1.066 |
| Gm12963 | 81.701 | 0.39455 | 0.1998 | 1.974753 | 0.0483 | 0.16203 | 23.519 |
| Gm12966 | 325.79 | -0.0798 | 0.16977 | -0.47005 | 0.63832 | 0.79688 | 0.81 |
| Gm12971 | 17.7006 | 1.01796 | 0.4644 | 2.191991 | 0.02838 | 0.11359 | 3.817 |
| Gm12989 | 124.094 | -0.22792 | 0.16436 | -1.38676 | 0.16552 | 0.35477 | 0.731 |
| Gm12990 | 24.4795 | 0.59788 | 0.38834 | 1.539572 | 0.12366 | 0.29804 | 0.468 |
| Gm12992 | 391.154 | 0.604372 | 0.23914 | 2.527238 | 0.0115 | 0.06285 | 20.552 |
| Gm12993 | 45.1048 | 0.385466 | 0.28516 | 1.351749 | 0.17646 | 0.36846 | 0.878 |
| Gm13016 | 29.4066 | -0.13549 | 0.3756 | -0.36073 | 0.7183 | 0.84954 | 4.945 |
| Gm13029 | 12.8335 | -0.06485 | 0.47694 | -0.13598 | 0.89184 | 0.94605 | 1.761 |
| Gm13031 | 34.964 | -0.3249 | 0.37079 | -0.87623 | 0.38091 | 0.59049 | 10.237 |
| Gm13033 | 28.0634 | -0.13005 | 0.45118 | -0.28825 | 0.77315 | 0.88189 | 1.864 |
| Gm13034 | 17.5896 | 0.440011 | 0.39936 | 1.101792 | 0.27055 | 0.48006 | 3.177 |
| Gm13052 | 76.2313 | 0.925357 | 0.29563 | 3.130135 | 0.00175 | 0.01847 | 38.711 |
| Gm13054 | 13.0467 | 0.717812 | 0.52271 | 1.373255 | 0.16967 | 0.36001 | 2.924 |
| Gm13062 | 24.3107 | 1.290165 | 0.47815 | 2.69827 | 0.00697 | 0.0453 | 3.538 |
| Gm13067 | 20.3519 | 0.018718 | 0.38951 | 0.048055 | 0.96167 | 0.98141 | 9.243 |
| Gm13068 | 32.5591 | 0.547961 | 0.35907 | 1.526061 | 0.12699 | 0.30318 | 1.09 |
| Gm13070 | 15.8764 | 0.187858 | 0.71409 | 0.263073 | 0.79249 | 0.89235 | 4.144 |
| Gm13071 | 44.8108 | -0.11597 | 0.26926 | -0.4307 | 0.66668 | 0.8158 | 0.376 |
| Gm13090 | 13.072 | 0.554057 | 0.50773 | 1.091241 | 0.27517 | 0.48462 | 1.88 |
| Gm13091 | 327.895 | 0.628718 | 0.18154 | 3.463296 | 0.00053 | 0.0083 | 1.853 |

| | | | | | | | |
|---|---|---|---|---|---|---|---|
| Gm13092 | 17.0114 | -0.11606 | 0.51972 | -0.22332 | 0.82329 | 0.90846 | 0.26 |
| Gm13093 | 38.3565 | 0.10213 | 0.31982 | 0.319341 | 0.74947 | 0.86849 | 5.385 |
| Gm13110 | 86.0484 | -0.65374 | 0.23196 | -2.81826 | 0.00483 | 0.03547 | 9.565 |
| Gm13112 | 103.189 | 0.892913 | 0.23374 | 3.820152 | 0.00013 | 0.00328 | 7.498 |
| Gm13123 | 25.5689 | 0.482912 | 0.37517 | 1.287199 | 0.19802 | 0.39574 | 0.492 |
| Gm13127 | 47.295 | -0.0371 | 0.29214 | -0.127 | 0.89894 | 0.94937 | 1.389 |
| Gm13135 | 59.1153 | -0.14909 | 0.25247 | -0.59052 | 0.55484 | 0.73733 | 3.121 |
| Gm13136 | 38.9204 | 0.106587 | 0.31672 | 0.33653 | 0.73647 | 0.86069 | 2.95 |
| Gm13140 | 16.828 | -0.29697 | 0.54582 | -0.54408 | 0.58639 | 0.76036 | 3.35 |
| Gm13187 | 25.2987 | -0.06281 | 0.35691 | -0.17598 | 0.86031 | 0.92937 | 13.573 |
| Gm13189 | 16.8182 | 0.206 | 0.44382 | 0.464157 | 0.64253 | 0.80038 | 6.329 |
| Gm13192 | 46.5172 | -0.52154 | 0.28656 | -1.82002 | 0.06876 | 0.20383 | 0.208 |
| Gm13194 | 20.3971 | -0.91785 | 0.40213 | -2.28249 | 0.02246 | 0.09677 | 0.311 |
| Gm13199 | 57.5248 | 0.143035 | 0.32046 | 0.446348 | 0.65535 | 0.80876 | 0.855 |
| Gm13204 | 152.772 | 0.128954 | 0.19246 | 0.670045 | 0.50283 | 0.69537 | 0.821 |
| Gm13205 | 26.4106 | 0.56434 | 0.39479 | 1.429462 | 0.15287 | 0.33835 | 1.423 |
| Gm13207 | 12.6584 | -0.5119 | 0.53317 | -0.96011 | 0.337 | 0.54758 | 0.773 |
| Gm13217 | 18.6437 | 0.676369 | 0.45638 | 1.482022 | 0.13833 | 0.3195 | 1.361 |
| Gm13268 | 21.7999 | 0.912739 | 0.3826 | 2.385609 | 0.01705 | 0.08104 | 0.433 |
| Gm13270 | 20.796 | 0.116113 | 0.36311 | 0.319778 | 0.74914 | 0.86842 | 54.247 |
| Gm13292 | 18.9051 | 0.615825 | 0.62402 | 0.986862 | 0.32371 | 0.53477 | 1 |
| Gm13293 | 73.9673 | -0.40869 | 0.33374 | -1.22458 | 0.22073 | 0.42348 | 4.681 |
| Gm13301 | 14.0509 | -0.90623 | 0.58184 | -1.55752 | 0.11935 | 0.29146 | 3.241 |
| Gm13322 | 28.1295 | -0.16738 | 0.40625 | -0.41201 | 0.68033 | 0.82502 | 3.745 |
| Gm13325 | 19.718 | -0.15626 | 0.48876 | -0.31971 | 0.74919 | 0.86842 | 0.806 |
| Gm13340 | 63.1454 | -0.53826 | 0.28536 | -1.8863 | 0.05926 | 0.18525 | 1.545 |
| Gm13341 | 21.0966 | -0.51591 | 0.53471 | -0.96484 | 0.33463 | 0.54538 | 0.672 |
| Gm13351 | 14.17 | 0.22474 | 0.46815 | 0.480058 | 0.63119 | 0.79177 | 0.414 |
| Gm13369 | 17.3389 | -0.0821 | 0.50249 | -0.16339 | 0.87021 | 0.93443 | 0.987 |
| Gm13370 | 65.6135 | -0.22896 | 0.24634 | -0.92945 | 0.35266 | 0.56297 | 0.483 |
| Gm13375 | 95.1031 | -0.22004 | 0.22235 | -0.98965 | 0.32235 | 0.53334 | 1.336 |
| Gm13381 | 30.3513 | 0.144353 | 0.35703 | 0.404319 | 0.68598 | 0.82887 | 14.502 |
| Gm13387 | 13.031 | -0.47622 | 0.57739 | -0.82478 | 0.40949 | 0.6173 | 22.794 |
| Gm13389 | 13.5008 | 0.179029 | 0.63485 | 0.282004 | 0.77794 | 0.88423 | 2.655 |

| | | | | | | | |
|---|---|---|---|---|---|---|---|
| Gm13394 | 192.524 | -0.32324 | 0.13173 | -2.45389 | 0.01413 | 0.07164 | 1 |
| Gm13404 | 16.1582 | 0.071416 | 0.52474 | 0.136098 | 0.89174 | 0.94605 | 2.079 |
| Gm13408 | 13.8535 | -0.59794 | 0.46481 | -1.28641 | 0.1983 | 0.39603 | 0.798 |
| Gm13414 | 57.9176 | 0.011678 | 0.24517 | 0.047635 | 0.96201 | 0.98159 | 0.727 |
| Gm13416 | 166.307 | 0.670037 | 0.16594 | 4.037854 | 5.39E-05 | 0.00175 | 1.612 |
| Gm13421 | 147.558 | 0.260036 | 0.19828 | 1.311449 | 0.18971 | 0.38573 | 5.676 |
| Gm13423 | 25.0278 | 0.213625 | 0.42996 | 0.496846 | 0.6193 | 0.78335 | 1.651 |
| Gm13425 | 46.9618 | 0.095752 | 0.28017 | 0.341761 | 0.73253 | 0.85819 | 7.137 |
| Gm13436 | 18.4146 | -0.29043 | 0.43449 | -0.66844 | 0.50385 | 0.69618 | 0.567 |
| Gm13446 | 63.3017 | -0.92758 | 0.29798 | -3.1129 | 0.00185 | 0.01911 | 9.165 |
| Gm13456 | 109.296 | -0.17343 | 0.18972 | -0.9141 | 0.36066 | 0.57101 | 1.365 |
| Gm13461 | 27.6 | -0.32922 | 0.32052 | -1.02712 | 0.30437 | 0.51417 | 1.357 |
| Gm13479 | 12.5977 | -0.5612 | 0.64736 | -0.8669 | 0.386 | 0.59548 | 46.771 |
| Gm13480 | 12.8084 | 1.526407 | 0.82087 | 1.859495 | 0.06296 | 0.19263 | 6.973 |
| Gm13483 | 18.1452 | -0.10981 | 0.46121 | -0.23809 | 0.81181 | 0.90308 | 137.158 |
| Gm13493 | 104.953 | -0.61757 | 0.18735 | -3.29631 | 0.00098 | 0.01263 | 0.493 |
| Gm13523 | 15.4173 | -1.00929 | 0.45597 | -2.21349 | 0.02686 | 0.10939 | 2.758 |
| Gm13524 | 38.098 | -0.28197 | 0.31125 | -0.90591 | 0.36498 | 0.57561 | 1.983 |
| Gm13530 | 21.6459 | -0.24784 | 0.40287 | -0.61517 | 0.53844 | 0.72517 | 13.851 |
| Gm13541 | 15.64 | 1.134578 | 0.50618 | 2.241468 | 0.025 | 0.10396 | 0.283 |
| Gm13542 | 16.0782 | 0.393642 | 0.4506 | 0.873589 | 0.38234 | 0.59186 | 0.637 |
| Gm13544 | 22.1768 | 0.861445 | 0.39844 | 2.16206 | 0.03061 | 0.11973 | 10.124 |
| Gm13547 | 43.2304 | -0.04482 | 0.33599 | -0.13339 | 0.89388 | 0.94689 | 4.778 |
| Gm13552 | 18.3986 | 0.047289 | 0.41585 | 0.113716 | 0.90946 | 0.95456 | 0.417 |
| Gm13558 | 30.3782 | -0.84825 | 0.37387 | -2.26886 | 0.02328 | 0.09891 | 29.628 |
| Gm13562 | 141.317 | 0.161209 | 0.20187 | 0.798595 | 0.42453 | 0.63123 | 19.308 |
| Gm13563 | 128.359 | 0.024634 | 0.1938 | 0.127108 | 0.89885 | 0.94934 | 7.493 |
| Gm13577 | 125.772 | -0.21589 | 0.19094 | -1.13063 | 0.25821 | 0.46624 | 0.205 |
| Gm13584 | 19.6644 | -0.38957 | 0.39745 | -0.98018 | 0.327 | 0.53828 | 4.401 |
| Gm13597 | 26.7493 | -0.13452 | 0.35552 | -0.37839 | 0.70514 | 0.84115 | 0.375 |
| Gm136 | 28.6651 | 1.478928 | 0.384 | 3.851355 | 0.00012 | 0.003 | 12.48 |
| Gm13618 | 46.1819 | 0.110148 | 0.33588 | 0.327936 | 0.74296 | 0.86466 | 4.978 |
| Gm13629 | 163.529 | -0.3934 | 0.26313 | -1.49504 | 0.1349 | 0.31481 | 106.186 |
| Gm13630 | 79.1079 | 0.428525 | 0.2536 | 1.689772 | 0.09107 | 0.24511 | 12.095 |

| | | | | | | | |
|---|---|---|---|---|---|---|---|
| Gm13645 | 24.9445 | -0.20151 | 0.41231 | -0.48875 | 0.62502 | 0.78752 | 1.444 |
| Gm13661 | 20.3779 | 1.33053 | 0.47574 | 2.796782 | 0.00516 | 0.03703 | 2.321 |
| Gm13680 | 52.192 | 0.164082 | 0.3353 | 0.48936 | 0.62459 | 0.78719 | 0.793 |
| Gm13687 | 18.0407 | 0.36754 | 0.43486 | 0.845186 | 0.39801 | 0.60699 | 0.282 |
| Gm13688 | 74.1841 | 0.35667 | 0.19729 | 1.807819 | 0.07063 | 0.20776 | 0.608 |
| Gm13689 | 23.4301 | 0.095567 | 0.3814 | 0.250569 | 0.80215 | 0.89791 | 0.685 |
| Gm13690 | 26.1497 | 0.255312 | 0.3634 | 0.70256 | 0.48233 | 0.67895 | 0.285 |
| Gm13703 | 13.8731 | 0.304362 | 0.51728 | 0.588388 | 0.55627 | 0.73841 | 7.066 |
| Gm13707 | 17.0859 | 0.186557 | 0.46385 | 0.402191 | 0.68754 | 0.82964 | 7.234 |
| Gm13716 | 48.8911 | 0.075307 | 0.28707 | 0.262327 | 0.79307 | 0.8928 | 1.744 |
| Gm13736 | 213.423 | -0.09766 | 0.13381 | -0.72979 | 0.46552 | 0.66497 | 0.253 |
| Gm13748 | 13.0307 | -0.7066 | 0.56137 | -1.2587 | 0.20814 | 0.40775 | 12.129 |
| Gm13754 | 15.7233 | -1.80117 | 0.6499 | -2.77147 | 0.00558 | 0.03918 | 18.085 |
| Gm13778 | 14.1122 | -0.3493 | 0.48565 | -0.71925 | 0.47199 | 0.67045 | 9.842 |
| Gm13783 | 16.187 | -1.16008 | 0.61894 | -1.8743 | 0.06089 | 0.18836 | 6.873 |
| Gm13803 | 64.0842 | 0.150214 | 0.41787 | 0.359477 | 0.71924 | 0.85006 | 0.489 |
| Gm13814 | 25.4099 | 0.49126 | 0.33733 | 1.456312 | 0.14531 | 0.32812 | 46.001 |
| Gm13821 | 12.6668 | -0.0436 | 0.57481 | -0.07585 | 0.93954 | 0.96977 | 3.201 |
| Gm13830 | 110.301 | -0.24874 | 0.24713 | -1.00653 | 0.31416 | 0.52465 | 18.11 |
| Gm13833 | 59.5542 | -0.23659 | 0.32674 | -0.72411 | 0.469 | 0.66788 | 1.801 |
| Gm13835 | 890.717 | 0.010951 | 0.0988 | 0.110836 | 0.91175 | 0.95543 | 0.685 |
| Gm13841 | 235.301 | 0.081268 | 0.1687 | 0.481741 | 0.62999 | 0.79082 | 0.483 |
| Gm13844 | 102.687 | 0.416178 | 0.17698 | 2.351584 | 0.01869 | 0.08571 | 10.983 |
| Gm13848 | 19.267 | 0.956772 | 0.47771 | 2.002816 | 0.0452 | 0.15458 | 7.959 |
| Gm13856 | 47.414 | -0.1581 | 0.30918 | -0.51135 | 0.60911 | 0.77661 | 22.307 |
| Gm13861 | 23.7789 | -0.17213 | 0.44988 | -0.38262 | 0.702 | 0.83917 | 2.541 |
| Gm13872 | 17.1854 | -0.15099 | 0.52446 | -0.2879 | 0.77343 | 0.88189 | 8.854 |
| Gm13886 | 14.9717 | -0.33336 | 0.47927 | -0.69556 | 0.4867 | 0.68256 | 1.183 |
| Gm13889 | 764.521 | 0.168507 | 0.10966 | 1.536662 | 0.12438 | 0.29922 | 1.392 |
| Gm13974 | 29.3342 | 0.640744 | 0.34075 | 1.880378 | 0.06006 | 0.18684 | 189.88 |
| Gm13991 | 126.406 | 0.076798 | 0.17476 | 0.439443 | 0.66034 | 0.81225 | 0.648 |
| Gm14000 | 15.5186 | -0.03156 | 0.4638 | -0.06805 | 0.94575 | 0.973 | 0.233 |
| Gm14001 | 22.8034 | 0.096737 | 0.40862 | 0.236739 | 0.81286 | 0.90371 | 0.247 |
| Gm14004 | 142.713 | 0.589848 | 0.19308 | 3.055005 | 0.00225 | 0.02176 | 9.972 |

| | | | | | | | |
|---|---|---|---|---|---|---|---|
| Gm14018 | 74.0281 | 0.112449 | 0.19109 | 0.58846 | 0.55622 | 0.73841 | 6.081 |
| Gm14026 | 25.5959 | 0.333518 | 0.45724 | 0.729421 | 0.46574 | 0.665 | 1.789 |
| Gm14027 | 30.6783 | -0.35145 | 0.35749 | -0.9831 | 0.32556 | 0.53686 | 4.805 |
| Gm14033 | 33.34 | 0.633585 | 0.3153 | 2.009464 | 0.04449 | 0.15318 | 21.646 |
| Gm14036 | 32.5246 | -0.25508 | 0.37089 | -0.68775 | 0.49161 | 0.68651 | 0.48 |
| Gm14038 | 81.7755 | 0.165188 | 0.23964 | 0.689326 | 0.49062 | 0.68569 | 0.807 |
| Gm14042 | 174.686 | 0.553401 | 0.14761 | 3.749044 | 0.00018 | 0.00398 | 0.833 |
| Gm14057 | 15.448 | -0.13574 | 0.52064 | -0.26071 | 0.79431 | 0.89351 | 0.482 |
| Gm14066 | 229.626 | -0.17249 | 0.15862 | -1.08743 | 0.27685 | 0.48628 | 420.839 |
| Gm14113 | 20.3754 | 0.088937 | 0.4258 | 0.20887 | 0.83455 | 0.91508 | 0.976 |
| Gm14117 | 630.285 | -0.17573 | 0.13172 | -1.33414 | 0.18216 | 0.37594 | 0.843 |
| Gm14121 | 95.9662 | 0.091662 | 0.18184 | 0.504072 | 0.61421 | 0.78036 | 1.38 |
| Gm14122 | 47.2124 | 0.378375 | 0.35215 | 1.074467 | 0.28261 | 0.4928 | 1.144 |
| Gm14125 | 14.9927 | 0.242939 | 0.49914 | 0.486717 | 0.62646 | 0.78864 | 0.989 |
| Gm14137 | 280.121 | 0.389801 | 0.13078 | 2.980599 | 0.00288 | 0.02554 | 3.067 |
| Gm14150 | 41.4293 | -0.69673 | 0.28934 | -2.40802 | 0.01604 | 0.07783 | 1.354 |
| Gm14164 | 448.128 | 0.946236 | 0.19183 | 4.932603 | 8.11E-07 | 9.20E-05 | 29.656 |
| Gm14167 | 30.9783 | 0.028979 | 0.31854 | 0.090977 | 0.92751 | 0.96345 | 30.857 |
| Gm14168 | 88.7526 | 0.006047 | 0.19921 | 0.030356 | 0.97578 | 0.98762 | 5.666 |
| Gm14169 | 378.3 | 0.218712 | 0.14559 | 1.502215 | 0.13304 | 0.31265 | 4.217 |
| Gm14172 | 15.86 | 0.162385 | 0.46001 | 0.353003 | 0.72409 | 0.85282 | 0.644 |
| Gm14174 | 23.2095 | 0.019414 | 0.38848 | 0.049974 | 0.96014 | 0.98055 | 1.002 |
| Gm14176 | 38.8421 | 0.378417 | 0.26404 | 1.433156 | 0.15181 | 0.33699 | 0.477 |
| Gm14204 | 1701.72 | 0.701113 | 0.12083 | 5.802265 | 6.54E-09 | 2.16E-06 | 15.256 |
| Gm14207 | 15.4848 | 0.010354 | 0.47954 | 0.021591 | 0.98277 | 0.9907 | 5.02 |
| Gm14224 | 13.545 | 1.082988 | 0.4795 | 2.258572 | 0.02391 | 0.10068 | 3.746 |
| Gm14232 | 14.2049 | 0.60957 | 0.44125 | 1.381467 | 0.16714 | 0.35694 | 11.806 |
| Gm14239 | 46.8948 | 0.437727 | 0.31715 | 1.380207 | 0.16752 | 0.35731 | 1.392 |
| Gm14246 | 19.421 | -0.43217 | 0.45388 | -0.95217 | 0.34101 | 0.55206 | 1.075 |
| Gm14253 | 146.383 | 0.202947 | 0.19831 | 1.023362 | 0.30614 | 0.51628 | 0.639 |
| Gm14257 | 57.7215 | -0.11506 | 0.32187 | -0.35747 | 0.72074 | 0.85095 | 0.911 |
| Gm14269 | 88.2994 | -0.22591 | 0.2386 | -0.94682 | 0.34373 | 0.55441 | 0.306 |
| Gm14276 | 32.8037 | 0.053685 | 0.30843 | 0.174057 | 0.86182 | 0.92991 | 0.72 |
| Gm14286 | 294.766 | 0.48547 | 0.19758 | 2.457042 | 0.01401 | 0.07132 | 1.204 |

| | | | | | | | |
|---|---|---|---|---|---|---|---|
| Gm14287 | 203.664 | 0.236915 | 0.13481 | 1.757462 | 0.07884 | 0.22266 | 0.117 |
| Gm14288 | 103.354 | 0.648464 | 0.17441 | 3.717993 | 0.0002 | 0.00428 | 11.737 |
| Gm14291 | 69.2209 | -0.0284 | 0.32599 | -0.08713 | 0.93057 | 0.96505 | 1.224 |
| Gm14292 | 51.3989 | -0.26862 | 0.3365 | -0.79827 | 0.42471 | 0.63137 | 0.456 |
| Gm14295 | 537.866 | -0.0194 | 0.22854 | -0.08491 | 0.93234 | 0.96589 | 12.612 |
| Gm14296 | 178.004 | 0.247687 | 0.24299 | 1.019336 | 0.30804 | 0.51787 | 35.301 |
| Gm14302 | 17.3807 | 1.014817 | 0.51521 | 1.969725 | 0.04887 | 0.16341 | 1.06 |
| Gm14303 | 134.432 | -0.07088 | 0.16146 | -0.43897 | 0.66069 | 0.81233 | 0.171 |
| Gm14305 | 719.719 | 0.185933 | 0.20943 | 0.887801 | 0.37465 | 0.58497 | 13.486 |
| Gm14320 | 37.6159 | 0.856185 | 0.33849 | 2.529426 | 0.01142 | 0.0626 | 5.589 |
| Gm14321 | 15.0458 | 0.603518 | 0.49867 | 1.210266 | 0.22618 | 0.43019 | 5.103 |
| Gm14322 | 244.926 | 0.264335 | 0.19297 | 1.369805 | 0.17075 | 0.36109 | 11.185 |
| Gm14325 | 724.797 | 0.320551 | 0.19586 | 1.636653 | 0.1017 | 0.26315 | 8.546 |
| Gm14326 | 605.147 | 0.168376 | 0.24238 | 0.694668 | 0.48726 | 0.68271 | 21.305 |
| Gm14327 | 66.7032 | 0.580771 | 0.23781 | 2.442182 | 0.0146 | 0.07302 | 8.689 |
| Gm14330 | 21.2366 | -0.52136 | 0.42668 | -1.22188 | 0.22175 | 0.42471 | 9.288 |
| Gm14342 | 17.4064 | 0.030139 | 0.43093 | 0.069939 | 0.94424 | 0.97202 | 60.171 |
| Gm14372 | 40.4674 | 0.149247 | 0.30898 | 0.483025 | 0.62908 | 0.79023 | 38.886 |
| Gm14376 | 19.7707 | 0.105351 | 0.44184 | 0.238435 | 0.81154 | 0.90289 | 0.461 |
| Gm14377 | 69.2693 | 0.931711 | 0.23373 | 3.986219 | 6.71E-05 | 0.00204 | 15.14 |
| Gm14379 | 20.0636 | 0.408927 | 0.39593 | 1.032815 | 0.30169 | 0.51136 | 2.118 |
| Gm14387 | 36.6961 | 0.757462 | 0.32005 | 2.366701 | 0.01795 | 0.08348 | 3.418 |
| Gm14388 | 39.9677 | 0.077956 | 0.37572 | 0.207483 | 0.83563 | 0.91572 | 0.477 |
| Gm14391 | 56.9424 | 0.366337 | 0.81299 | 0.450603 | 0.65228 | 0.8066 | 15.853 |
| Gm14393 | 17.2434 | 0.610899 | 0.44532 | 1.371811 | 0.17012 | 0.36051 | 6.233 |
| Gm14397 | 36.367 | 0.575158 | 0.32627 | 1.762816 | 0.07793 | 0.221 | 4.558 |
| Gm14399 | 591.928 | 0.505341 | 0.18615 | 2.714711 | 0.00663 | 0.04392 | 12.52 |
| Gm14401 | 12.4942 | 0.087266 | 0.47075 | 0.185375 | 0.85293 | 0.92534 | 13.028 |
| Gm14403 | 249.633 | -0.17883 | 0.25265 | -0.70779 | 0.47907 | 0.6767 | 11.549 |
| Gm14410 | 661.538 | -0.04582 | 0.46068 | -0.09947 | 0.92077 | 0.95978 | 22.969 |
| Gm14411 | 39.8011 | 0.630193 | 0.30234 | 2.084359 | 0.03713 | 0.13624 | 2.247 |
| Gm14412 | 125.614 | 0.839933 | 0.64749 | 1.297211 | 0.19456 | 0.39189 | 9.788 |
| Gm14415 | 24.612 | 0.04125 | 0.49507 | 0.083321 | 0.9336 | 0.9665 | 2.925 |
| Gm14416 | 37.3792 | 0.252268 | 0.37079 | 0.680355 | 0.49628 | 0.68996 | 3.672 |

| | | | | | | | |
|---|---|---|---|---|---|---|---|
| Gm14418 | 158.441 | 0.413289 | 0.24243 | 1.704781 | 0.08824 | 0.23994 | 11.197 |
| Gm14419 | 26.7166 | 1.178276 | 1.17049 | 1.00665 | 0.3141 | 0.52459 | 11.832 |
| Gm14421 | 48.5904 | 0.781433 | 0.27628 | 2.828389 | 0.00468 | 0.03482 | 3.323 |
| Gm14443 | 160.883 | 0.096117 | 0.2445 | 0.39312 | 0.69423 | 0.83414 | 9.617 |
| Gm14448 | 12.6359 | 1.408623 | 0.6829 | 2.062705 | 0.03914 | 0.14099 | 0.21 |
| Gm14455 | 28.662 | 0.502402 | 0.37574 | 1.33711 | 0.18119 | 0.37458 | 14.757 |
| Gm14472 | 20.6383 | 0.835906 | 0.46677 | 1.790822 | 0.07332 | 0.2132 | 0.282 |
| Gm14489 | 21.4333 | 0.529643 | 0.4619 | 1.146653 | 0.25153 | 0.45892 | 0.901 |
| Gm14494 | 430.979 | 0.217591 | 0.16953 | 1.283518 | 0.19931 | 0.39726 | 0.435 |
| Gm14508 | 38.0071 | 0.097234 | 0.31058 | 0.313076 | 0.75422 | 0.87077 | 6.79 |
| Gm14532 | 36.2688 | 0.408954 | 0.29024 | 1.40902 | 0.15883 | 0.34634 | 15.627 |
| Gm14539 | 166.541 | -0.39471 | 0.18148 | -2.175 | 0.02963 | 0.11715 | 0.174 |
| Gm14549 | 15.9492 | 0.318667 | 0.45786 | 0.695996 | 0.48643 | 0.68236 | 2.43 |
| Gm14569 | 30.2932 | 0.619259 | 0.37267 | 1.661699 | 0.09657 | 0.25453 | 68.23 |
| Gm14584 | 122.374 | -0.04085 | 0.19189 | -0.2129 | 0.8314 | 0.91345 | 0.637 |
| Gm14586 | 30.0031 | -0.33894 | 0.31072 | -1.09084 | 0.27534 | 0.48468 | 0.408 |
| Gm14634 | 35.3798 | 0.25002 | 0.32817 | 0.761854 | 0.44615 | 0.64983 | 59.215 |
| Gm14636 | 258.661 | 0.333211 | 0.15459 | 2.155418 | 0.03113 | 0.12111 | 22.891 |
| Gm14648 | 43.0293 | -0.36262 | 0.3134 | -1.15706 | 0.24725 | 0.45409 | 0.483 |
| Gm14681 | 190.204 | 0.055593 | 0.12761 | 0.435634 | 0.6631 | 0.81387 | 0.411 |
| Gm14703 | 86.837 | -0.03785 | 0.23388 | -0.16184 | 0.87143 | 0.93501 | 6.153 |
| Gm14719 | 13.5042 | -0.44168 | 0.5397 | -0.81837 | 0.41315 | 0.62134 | 0.34 |
| Gm14760 | 21.8659 | -0.25479 | 0.34907 | -0.72992 | 0.46544 | 0.66495 | 0.999 |
| Gm14817 | 91.4048 | 0.253213 | 0.20906 | 1.211222 | 0.22581 | 0.4299 | 2.402 |
| Gm14820 | 12.6273 | -0.10719 | 0.55321 | -0.19376 | 0.84636 | 0.92184 | 31.931 |
| Gm14824 | 69.9033 | 0.333614 | 0.24372 | 1.368821 | 0.17106 | 0.36145 | 0.737 |
| Gm14827 | 639.866 | 0.27285 | 0.21807 | 1.251185 | 0.21087 | 0.4109 | 4.996 |
| Gm14830 | 60.6341 | 0.698409 | 0.28195 | 2.477091 | 0.01325 | 0.06895 | 0.7 |
| Gm14858 | 31.271 | -0.03374 | 0.36223 | -0.09315 | 0.92578 | 0.96257 | 112.977 |
| Gm14964 | 158.102 | -0.25058 | 0.16831 | -1.48883 | 0.13653 | 0.31704 | 1.301 |
| Gm14966 | 55.8113 | 0.16724 | 0.2637 | 0.634213 | 0.52594 | 0.71415 | 8.667 |
| Gm14978 | 28.7393 | -0.06221 | 0.34845 | -0.17854 | 0.8583 | 0.92856 | 2.306 |
| Gm15028 | 16.84 | 0.676337 | 0.45754 | 1.478194 | 0.13936 | 0.32067 | 0.37 |
| Gm15046 | 29.0747 | -0.44385 | 0.37471 | -1.18449 | 0.23622 | 0.44147 | 99.165 |

| | | | | | | | |
|---|---|---|---|---|---|---|---|
| Gm15082 | 18.886 | 0.027451 | 0.39862 | 0.068866 | 0.9451 | 0.97268 | 2.468 |
| Gm15122 | 186.99 | 0.563845 | 0.1573 | 3.58452 | 0.00034 | 0.00612 | 0.563 |
| Gm15155 | 19.68 | -0.48443 | 0.38653 | -1.25328 | 0.2101 | 0.41015 | 721.147 |
| Gm15163 | 48.3184 | -0.44158 | 0.28444 | -1.55243 | 0.12056 | 0.2934 | 1.302 |
| Gm15169 | 28.6074 | 0.480906 | 1.1152 | 0.431229 | 0.6663 | 0.8158 | 40.593 |
| Gm15178 | 43.1244 | 1.071525 | 0.2775 | 3.86135 | 0.00011 | 0.00291 | 4.233 |
| Gm15186 | 36.085 | -0.16779 | 0.3225 | -0.52029 | 0.60286 | 0.77254 | 0.563 |
| Gm15190 | 25.7597 | 0.105556 | 0.37351 | 0.282607 | 0.77748 | 0.88413 | 0.276 |
| Gm15201 | 43.3193 | -0.54408 | 0.27905 | -1.94978 | 0.0512 | 0.1685 | 1.536 |
| Gm15246 | 101.02 | 0.053772 | 0.26152 | 0.205617 | 0.83709 | 0.91629 | 47.569 |
| Gm15247 | 16.7427 | 0.01344 | 0.40905 | 0.032858 | 0.97379 | 0.98681 | 1.139 |
| Gm15265 | 127.521 | -0.19466 | 0.18592 | -1.04699 | 0.2951 | 0.5055 | 6.07 |
| Gm15270 | 13.8927 | -0.41407 | 0.60881 | -0.68012 | 0.49643 | 0.69002 | 47.944 |
| Gm15283 | 54.175 | 0.331818 | 0.33553 | 0.988926 | 0.3227 | 0.53366 | 106.776 |
| Gm15328 | 605.123 | 0.982323 | 0.13492 | 7.281046 | 3.31E-13 | 1.35E-09 | 7.063 |
| Gm15333 | 27.886 | -0.11919 | 0.52963 | -0.22505 | 0.82194 | 0.90781 | 0.364 |
| Gm15337 | 104.194 | 0.825104 | 0.19191 | 4.299465 | 1.71E-05 | 0.0008 | 22.388 |
| Gm15342 | 12.4979 | -0.73562 | 0.6715 | -1.09549 | 0.2733 | 0.48263 | 2.654 |
| Gm15344 | 124.059 | 0.148613 | 0.1794 | 0.828391 | 0.40745 | 0.61538 | 0.977 |
| Gm15347 | 135.997 | 0.555224 | 0.16886 | 3.287997 | 0.00101 | 0.0129 | 16.987 |
| Gm15351 | 13.8133 | 0.005529 | 0.54755 | 0.010098 | 0.99194 | 0.99537 | 16.613 |
| Gm15379 | 35.0338 | 0.694034 | 0.60538 | 1.146444 | 0.25161 | 0.45896 | 0.648 |
| Gm15382 | 45.6884 | 0.647201 | 0.38998 | 1.659596 | 0.097 | 0.25518 | 0.537 |
| Gm15387 | 349.877 | 0.184795 | 0.24399 | 0.757379 | 0.44882 | 0.65214 | 0.648 |
| Gm15408 | 63.3108 | -0.38891 | 0.28328 | -1.37289 | 0.16979 | 0.3601 | 6.52 |
| Gm15411 | 92.0617 | -0.5384 | 0.2622 | -2.05337 | 0.04004 | 0.14298 | 20.218 |
| Gm15413 | 26.6233 | 0.332933 | 0.43943 | 0.757653 | 0.44866 | 0.65208 | 10.117 |
| Gm15414 | 13.4825 | 0.450419 | 0.58852 | 0.765337 | 0.44407 | 0.6481 | 32.418 |
| Gm15417 | 115.384 | -0.33937 | 0.20494 | -1.65594 | 0.09773 | 0.2564 | 9.458 |
| Gm15418 | 43.6492 | -0.48849 | 0.33292 | -1.46732 | 0.14229 | 0.32422 | 16.749 |
| Gm15421 | 28.1869 | 0.415674 | 0.36419 | 1.141364 | 0.25372 | 0.46107 | 0.924 |
| Gm15422 | 16.3929 | 0.276509 | 0.50567 | 0.546812 | 0.58451 | 0.75928 | 0.541 |
| Gm15425 | 21.1866 | -0.6856 | 0.5774 | -1.18739 | 0.23508 | 0.44032 | 4.603 |
| Gm15427 | 845.162 | -0.21842 | 0.16839 | -1.2971 | 0.1946 | 0.39193 | 0.531 |

| | | | | | | | |
|---|---|---|---|---|---|---|---|
| Gm15429 | 37.9687 | -0.75752 | 0.3228 | -2.34668 | 0.01894 | 0.08642 | 1.248 |
| Gm15445 | 50.753 | -0.07012 | 0.29515 | -0.23759 | 0.8122 | 0.90334 | 0.844 |
| Gm15446 | 1701.39 | 0.2609 | 0.19626 | 1.329346 | 0.18373 | 0.37789 | 20.537 |
| Gm15459 | 1009 | -0.04266 | 0.1131 | -0.37722 | 0.70601 | 0.8415 | 1.941 |
| Gm15477 | 23.5257 | -0.10058 | 0.34027 | -0.2956 | 0.76753 | 0.87843 | 61.949 |
| Gm15478 | 59.3379 | -0.32961 | 0.35115 | -0.93866 | 0.34791 | 0.55841 | 4.376 |
| Gm15483 | 13.2344 | 0.98185 | 0.63072 | 1.556702 | 0.11954 | 0.29186 | 0.456 |
| Gm15489 | 72.1255 | -0.08321 | 0.22124 | -0.37611 | 0.70683 | 0.84193 | 20.394 |
| Gm15494 | 31.3301 | -0.53255 | 0.29478 | -1.80664 | 0.07082 | 0.20804 | 25.589 |
| Gm15496 | 46.3241 | 0.113785 | 0.28631 | 0.397424 | 0.69105 | 0.83227 | 31.305 |
| Gm15497 | 38.0581 | 0.225511 | 0.3156 | 0.714557 | 0.47488 | 0.67292 | 0.551 |
| Gm15500 | 628.196 | 0.144306 | 0.09385 | 1.537636 | 0.12414 | 0.29882 | 1.02 |
| Gm15501 | 87.9183 | -0.41927 | 0.62228 | -0.67376 | 0.50047 | 0.69333 | 5.194 |
| Gm15506 | 145.881 | 0.124707 | 0.17399 | 0.716751 | 0.47353 | 0.67184 | 20.653 |
| Gm15510 | 62.6406 | -0.05334 | 0.23407 | -0.22789 | 0.81973 | 0.90733 | 10.168 |
| Gm15513 | 446.119 | 0.292062 | 0.14151 | 2.063899 | 0.03903 | 0.14077 | 1.993 |
| Gm15518 | 28.1922 | -0.00737 | 0.36929 | -0.01995 | 0.98409 | 0.99128 | 40.295 |
| Gm15519 | 92.7082 | 0.111457 | 0.18217 | 0.611816 | 0.54066 | 0.72644 | 1.566 |
| Gm15527 | 29.298 | 0.967901 | 0.43536 | 2.22322 | 0.0262 | 0.10748 | 33.335 |
| Gm15533 | 31.7094 | 0.789117 | 0.35959 | 2.194514 | 0.0282 | 0.11309 | 3.292 |
| Gm15535 | 163.978 | 0.604175 | 0.21377 | 2.826348 | 0.00471 | 0.03497 | 17.9 |
| Gm15538 | 87.8394 | 0.106698 | 0.21389 | 0.498849 | 0.61789 | 0.78278 | 0.313 |
| Gm15539 | 25.7616 | 0.304701 | 0.35129 | 0.86738 | 0.38573 | 0.59517 | 0.566 |
| Gm15542 | 456.494 | 0.191424 | 0.13955 | 1.371686 | 0.17016 | 0.36052 | 1.194 |
| Gm15545 | 46.1396 | -0.52024 | 0.27887 | -1.8655 | 0.06211 | 0.19095 | 7.71 |
| Gm15558 | 16.8056 | -0.24163 | 0.44707 | -0.54047 | 0.58887 | 0.76203 | 14.292 |
| Gm15559 | 71.0172 | -0.59054 | 0.21932 | -2.69258 | 0.00709 | 0.04585 | 8.312 |
| Gm15560 | 20.3115 | 0.575359 | 0.39639 | 1.451505 | 0.14664 | 0.32981 | 0.643 |
| Gm15561 | 44.3264 | -0.03541 | 0.27896 | -0.12695 | 0.89898 | 0.94937 | 5.662 |
| Gm15563 | 30.1067 | 0.212144 | 0.37525 | 0.565343 | 0.57184 | 0.75062 | 10.059 |
| Gm15564 | 16.3533 | -0.19811 | 0.50775 | -0.39018 | 0.6964 | 0.83541 | 16.496 |
| Gm15567 | 102.577 | 0.183123 | 0.19148 | 0.95635 | 0.3389 | 0.54974 | 8.67 |
| Gm15570 | 12.7024 | -0.23824 | 0.55064 | -0.43266 | 0.66526 | 0.81526 | 0.625 |
| Gm15580 | 15.7596 | 0.168247 | 0.46246 | 0.363811 | 0.716 | 0.8481 | 0.556 |

| | | | | | | | |
|---|---|---|---|---|---|---|---|
| Gm15587 | 40.276 | -0.03467 | 0.34627 | -0.10011 | 0.92026 | 0.95961 | 17.423 |
| Gm1559 | 21.5155 | 1.725376 | 0.49544 | 3.482516 | 0.0005 | 0.00789 | 4.675 |
| Gm15591 | 24.4629 | 0.375247 | 0.3833 | 0.979001 | 0.32758 | 0.53893 | 0.816 |
| Gm15594 | 36.31 | -0.25406 | 0.34256 | -0.74166 | 0.45829 | 0.6595 | 3.202 |
| Gm15601 | 115.849 | 0.199886 | 0.16248 | 1.230234 | 0.21861 | 0.42068 | 34.679 |
| Gm15608 | 192.432 | 0.304925 | 0.17303 | 1.762298 | 0.07802 | 0.22104 | 23.254 |
| Gm15609 | 160.342 | 0.518372 | 0.16268 | 3.186408 | 0.00144 | 0.01608 | 3.582 |
| Gm15610 | 14.2222 | -0.02338 | 0.45542 | -0.05133 | 0.95906 | 0.98007 | 0.195 |
| Gm15612 | 66.096 | -0.03083 | 0.25256 | -0.12207 | 0.90284 | 0.95118 | 4.421 |
| Gm15624 | 380.378 | 0.411358 | 0.16765 | 2.453729 | 0.01414 | 0.07164 | 6.604 |
| Gm15627 | 25.8093 | 0.53 | 0.37636 | 1.408223 | 0.15906 | 0.34652 | 0.797 |
| Gm15631 | 393.287 | -0.18536 | 0.13347 | -1.38873 | 0.16492 | 0.35397 | 5.031 |
| Gm15635 | 58.6249 | -0.0786 | 0.31229 | -0.25168 | 0.80129 | 0.8972 | 7.371 |
| Gm15651 | 18.662 | -0.06691 | 0.50906 | -0.13144 | 0.89543 | 0.94744 | 2.597 |
| Gm15663 | 275.909 | -0.06474 | 0.19389 | -0.33388 | 0.73847 | 0.86209 | 56.353 |
| Gm15672 | 42.3523 | 0.389833 | 0.32885 | 1.185454 | 0.23584 | 0.44108 | 3.118 |
| Gm15675 | 30.6066 | 0.027436 | 0.32619 | 0.084111 | 0.93297 | 0.96629 | 7.411 |
| Gm15688 | 27.5571 | -0.17745 | 0.35088 | -0.50572 | 0.61305 | 0.77967 | 7.358 |
| Gm15691 | 20.3948 | 0.610233 | 0.37949 | 1.608029 | 0.10783 | 0.27277 | 2.457 |
| Gm15696 | 47.715 | 0.345145 | 0.27779 | 1.242464 | 0.21407 | 0.41549 | 0.445 |
| Gm15697 | 34.4452 | 0.980861 | 0.3804 | 2.578515 | 0.00992 | 0.05715 | 3.772 |
| Gm15704 | 22.1572 | 0.320116 | 0.52092 | 0.614522 | 0.53887 | 0.72547 | 7.581 |
| Gm15706 | 173.39 | -0.32815 | 0.14071 | -2.33217 | 0.01969 | 0.08869 | 1.321 |
| Gm15708 | 37.5989 | 0.012736 | 0.34324 | 0.037107 | 0.9704 | 0.9849 | 3.453 |
| Gm15710 | 157.243 | -0.23863 | 0.15739 | -1.5162 | 0.12947 | 0.30701 | 0.636 |
| Gm15718 | 19.8991 | 0.589662 | 0.65377 | 0.901945 | 0.36709 | 0.57746 | 3.971 |
| Gm15721 | 168.524 | 0.197687 | 0.14891 | 1.327537 | 0.18433 | 0.37877 | 7.285 |
| Gm15726 | 54.4936 | 0.095496 | 0.28858 | 0.330913 | 0.74071 | 0.86308 | 6.186 |
| Gm15728 | 152.928 | -0.21516 | 0.18044 | -1.19237 | 0.23312 | 0.43798 | 4.366 |
| Gm15738 | 102.053 | 0.660931 | 0.27357 | 2.415971 | 0.01569 | 0.07671 | 114.382 |
| Gm15743 | 54.2663 | 0.210928 | 0.26982 | 0.781751 | 0.43436 | 0.6402 | 1.901 |
| Gm15751 | 225.274 | 0.167656 | 0.1257 | 1.333784 | 0.18227 | 0.37605 | 5.617 |
| Gm15753 | 84.7691 | 0.403517 | 0.23389 | 1.725255 | 0.08448 | 0.233 | 1.387 |
| Gm15758 | 27.1238 | 0.65266 | 0.35654 | 1.830514 | 0.06717 | 0.20068 | 1.395 |

| | | | | | | | |
|---|---|---|---|---|---|---|---|
| Gm15760 | 102.148 | -0.35309 | 0.23579 | -1.49747 | 0.13427 | 0.31412 | 3.245 |
| Gm15764 | 185.519 | 0.479171 | 0.18848 | 2.542332 | 0.01101 | 0.06114 | 5.316 |
| Gm15772 | 242.71 | 0.160433 | 0.21722 | 0.738555 | 0.46018 | 0.66077 | 0.438 |
| Gm15774 | 86.5006 | 0.376504 | 0.17917 | 2.101391 | 0.03561 | 0.13257 | 9.79 |
| Gm15775 | 63.8873 | 0.112139 | 0.28361 | 0.395398 | 0.69255 | 0.8334 | 7.018 |
| Gm15777 | 278.502 | 0.247849 | 0.13452 | 1.842406 | 0.06542 | 0.19731 | 7.247 |
| Gm15787 | 150.197 | 0.13803 | 0.15115 | 0.913219 | 0.36113 | 0.57139 | 22.105 |
| Gm15788 | 38.3977 | 0.331337 | 0.29048 | 1.140653 | 0.25401 | 0.46137 | 15.107 |
| Gm15789 | 20.9585 | 0.411195 | 0.48352 | 0.850425 | 0.39509 | 0.60407 | 2.533 |
| Gm15798 | 18.4205 | 0.246056 | 0.45837 | 0.536807 | 0.5914 | 0.764 | 0.408 |
| Gm15809 | 26.6546 | 0.349023 | 0.50392 | 0.69261 | 0.48855 | 0.68367 | 7.557 |
| Gm15810 | 158.05 | 0.437884 | 0.31477 | 1.391132 | 0.16419 | 0.35308 | 5.567 |
| Gm15812 | 49.2045 | 0.558245 | 0.38977 | 1.43223 | 0.15208 | 0.33717 | 1.265 |
| Gm15813 | 417.871 | 0.783882 | 0.19306 | 4.060216 | 4.90E-05 | 0.00162 | 8.045 |
| Gm15816 | 82.5584 | 0.181861 | 0.20836 | 0.872833 | 0.38275 | 0.59205 | 10.792 |
| Gm15821 | 13.5224 | -0.1947 | 0.55732 | -0.34935 | 0.72683 | 0.85431 | 3.148 |
| Gm15829 | 33.3692 | -0.91847 | 0.31596 | -2.90695 | 0.00365 | 0.02963 | 1.074 |
| Gm15832 | 193.33 | 0.165509 | 0.20145 | 0.821594 | 0.41131 | 0.61936 | 4.433 |
| Gm15834 | 24.918 | -0.06102 | 0.48111 | -0.12682 | 0.89908 | 0.94943 | 18.083 |
| Gm15835 | 35.4614 | 0.020376 | 0.34082 | 0.059785 | 0.95233 | 0.97658 | 1.712 |
| Gm15842 | 18.6214 | 0.657004 | 0.39446 | 1.665586 | 0.0958 | 0.25322 | 0.618 |
| Gm15846 | 15.5565 | 0.237069 | 0.47333 | 0.500852 | 0.61647 | 0.78168 | 0.32 |
| Gm15850 | 33.5035 | 0.14494 | 0.38459 | 0.376869 | 0.70627 | 0.84156 | 5.057 |
| Gm15853 | 68.387 | -0.53638 | 0.24737 | -2.1683 | 0.03014 | 0.11861 | 2.616 |
| Gm15860 | 36.918 | -0.01124 | 0.28109 | -0.03997 | 0.96812 | 0.984 | 11.712 |
| Gm15867 | 42.1297 | 0.368461 | 0.35575 | 1.035733 | 0.30033 | 0.51042 | 2.029 |
| Gm15868 | 19.0675 | 0.362926 | 0.40197 | 0.902862 | 0.3666 | 0.57697 | 2.174 |
| Gm15878 | 12.8688 | 1.270586 | 0.66442 | 1.912323 | 0.05583 | 0.17857 | 0.306 |
| Gm15879 | 98.9681 | 0.160604 | 0.19542 | 0.821838 | 0.41117 | 0.61923 | 5.009 |
| Gm15880 | 20.1271 | 0.281785 | 0.38108 | 0.73944 | 0.45964 | 0.66018 | 8.368 |
| Gm15892 | 28.205 | 0.015391 | 0.3583 | 0.042957 | 0.96574 | 0.9828 | 8.411 |
| Gm15893 | 482.515 | 0.075114 | 0.12531 | 0.599435 | 0.54888 | 0.73283 | 7.657 |
| Gm15894 | 14.7196 | 0.045734 | 0.47231 | 0.096831 | 0.92286 | 0.96125 | 3.979 |
| Gm15898 | 22.54 | -0.27565 | 0.38874 | -0.70909 | 0.47827 | 0.67599 | 2.677 |

| | | | | | | | |
|---|---|---|---|---|---|---|---|
| Gm15899 | 31.0585 | -0.3934 | 0.44699 | -0.88012 | 0.37879 | 0.58851 | 0.238 |
| Gm15903 | 37.5341 | -0.20114 | 0.34201 | -0.58811 | 0.55646 | 0.73861 | 4.644 |
| Gm15904 | 53.9999 | 0.605463 | 0.25368 | 2.386744 | 0.017 | 0.08086 | 4.191 |
| Gm15906 | 17.4637 | 0.238439 | 0.4528 | 0.526582 | 0.59848 | 0.76969 | 1.82 |
| Gm15910 | 28.157 | -0.20418 | 0.35665 | -0.57249 | 0.56699 | 0.74704 | 5.698 |
| Gm15911 | 61.1252 | 0.126591 | 0.26263 | 0.482009 | 0.6298 | 0.79072 | 2.602 |
| Gm15912 | 106.124 | -0.46983 | 0.20014 | -2.34751 | 0.0189 | 0.08627 | 1.364 |
| Gm15927 | 78.7807 | -0.10847 | 0.22066 | -0.49156 | 0.62303 | 0.78589 | 12.774 |
| Gm15932 | 21.9489 | -0.02295 | 0.39153 | -0.05861 | 0.95327 | 0.97694 | 9.833 |
| Gm15933 | 49.4589 | 1.135106 | 0.31934 | 3.554528 | 0.00038 | 0.00658 | 1.645 |
| Gm15943 | 16.5843 | -0.27725 | 0.43688 | -0.63461 | 0.52568 | 0.71399 | 109.579 |
| Gm15945 | 15.0164 | 0.185742 | 0.50041 | 0.371177 | 0.71051 | 0.84419 | 2.123 |
| Gm15948 | 47.5053 | 0.031349 | 0.39035 | 0.08031 | 0.93599 | 0.96785 | 2.392 |
| Gm15950 | 126.541 | 0.433184 | 0.17875 | 2.423362 | 0.01538 | 0.0757 | 1.973 |
| Gm15952 | 14.9357 | 0.054161 | 0.59162 | 0.091548 | 0.92706 | 0.96325 | 57.796 |
| Gm15953 | 39.7997 | 0.26898 | 0.26636 | 1.009853 | 0.31257 | 0.52301 | 1.732 |
| Gm15956 | 14.5976 | 0.465697 | 0.49696 | 0.937096 | 0.34871 | 0.55921 | 4.908 |
| Gm15964 | 15.1316 | -0.05264 | 0.51694 | -0.10183 | 0.91889 | 0.95892 | 12.394 |
| Gm15965 | 13.8716 | -0.61518 | 0.57029 | -1.07871 | 0.28072 | 0.49045 | 3.331 |
| Gm15983 | 18.2102 | -0.76306 | 0.46228 | -1.65065 | 0.09881 | 0.25828 | 34.836 |
| Gm15990 | 22.4831 | -0.42736 | 0.43716 | -0.97758 | 0.32828 | 0.53961 | 4.028 |
| Gm15991 | 35.8743 | -0.44285 | 0.41913 | -1.0566 | 0.2907 | 0.50089 | 21.283 |
| Gm15994 | 30.2188 | 0.364223 | 0.34168 | 1.065986 | 0.28643 | 0.49639 | 0.402 |
| Gm15996 | 40.2457 | 0.455967 | 0.30522 | 1.493891 | 0.1352 | 0.31507 | 1.738 |
| Gm15997 | 36.091 | -0.53722 | 0.30144 | -1.78219 | 0.07472 | 0.21543 | 48.975 |
| Gm15998 | 47.2801 | 0.573765 | 0.26774 | 2.142998 | 0.03211 | 0.12379 | 3.469 |
| Gm16001 | 366.193 | 0.058017 | 0.09823 | 0.590616 | 0.55478 | 0.73729 | 7.716 |
| Gm16008 | 426.252 | 0.150545 | 0.13431 | 1.120863 | 0.26235 | 0.47116 | 9.656 |
| Gm16023 | 161.872 | 0.428594 | 0.15806 | 2.711604 | 0.0067 | 0.04418 | 16.458 |
| Gm16041 | 13.8737 | 0.301637 | 0.47268 | 0.638144 | 0.52338 | 0.71181 | 5.964 |
| Gm16044 | 62.612 | 1.154705 | 0.27913 | 4.136742 | 3.52E-05 | 0.00128 | 0.45 |
| Gm16046 | 33.6872 | -0.42292 | 0.33246 | -1.27209 | 0.20334 | 0.40179 | 7.855 |
| Gm1604a | 193.81 | 0.410321 | 0.23615 | 1.737521 | 0.0823 | 0.22875 | 118.72 |
| Gm16050 | 29.8729 | -0.29037 | 0.32935 | -0.88163 | 0.37798 | 0.58806 | 10.24 |

| | | | | | | | |
|---|---|---|---|---|---|---|---|
| Gm16053 | 24.676 | 0.444194 | 0.38259 | 1.161008 | 0.24564 | 0.45223 | 0.188 |
| Gm16054 | 148.041 | 0.463795 | 0.2231 | 2.078833 | 0.03763 | 0.13733 | 6.518 |
| Gm16057 | 13.4882 | -0.35756 | 0.69319 | -0.51582 | 0.60598 | 0.77458 | 1.682 |
| Gm16059 | 53.9318 | 0.187637 | 0.22861 | 0.820759 | 0.41178 | 0.61976 | 4.41 |
| Gm16070 | 27.8552 | 0.333761 | 0.32871 | 1.015358 | 0.30994 | 0.51992 | 51.62 |
| Gm16073 | 20.4456 | 0.415395 | 0.40098 | 1.035952 | 0.30022 | 0.51033 | 0.517 |
| Gm16075 | 46.1396 | 0.881973 | 0.32931 | 2.678238 | 0.0074 | 0.04732 | 0.526 |
| Gm16083 | 20.1377 | -0.22325 | 0.51306 | -0.43513 | 0.66346 | 0.81402 | 94.5 |
| Gm16089 | 44.3702 | 0.29354 | 0.30004 | 0.978331 | 0.32791 | 0.53926 | 0.336 |
| Gm16091 | 51.3276 | 0.044102 | 0.29538 | 0.149308 | 0.88131 | 0.94085 | 13.255 |
| Gm16096 | 47.8117 | 0.195561 | 0.30869 | 0.633526 | 0.52639 | 0.71457 | 3.44 |
| Gm16099 | 21.2106 | 0.491384 | 0.38822 | 1.265722 | 0.20561 | 0.40479 | 1.757 |
| Gm16104 | 15.8047 | 0.137403 | 0.55848 | 0.246028 | 0.80566 | 0.89982 | 2.08 |
| Gm16105 | 132.904 | 0.601041 | 0.19461 | 3.088432 | 0.00201 | 0.02016 | 1.814 |
| Gm16111 | 30.2047 | 1.354287 | 0.50084 | 2.704024 | 0.00685 | 0.04485 | 0.285 |
| Gm16124 | 22.933 | 0.260241 | 0.43832 | 0.59372 | 0.5527 | 0.73559 | 14.221 |
| Gm16127 | 15.1191 | -0.41482 | 0.57653 | -0.71951 | 0.47183 | 0.67032 | 1.294 |
| Gm16150 | 13.8341 | -0.06057 | 0.5064 | -0.11962 | 0.90479 | 0.95186 | 11.247 |
| Gm16153 | 177.879 | 0.249524 | 0.20599 | 1.211343 | 0.22576 | 0.42985 | 14.337 |
| Gm16159 | 38.7155 | -0.18358 | 0.26652 | -0.68878 | 0.49096 | 0.68593 | 7.423 |
| Gm16161 | 42.5023 | 0.562182 | 0.28111 | 1.999868 | 0.04551 | 0.1553 | 1.094 |
| Gm16166 | 18.5093 | 0.727407 | 0.47956 | 1.516816 | 0.12931 | 0.30684 | 0.787 |
| Gm16168 | 49.7083 | -0.77555 | 0.29069 | -2.66796 | 0.00763 | 0.04827 | 72.951 |
| Gm16170 | 33.4601 | -0.68299 | 0.35163 | -1.94237 | 0.05209 | 0.17044 | 9.846 |
| Gm16172 | 81.6001 | 0.20871 | 0.20587 | 1.013787 | 0.31068 | 0.52089 | 2.691 |
| Gm16178 | 24.9173 | 0.462204 | 0.39491 | 1.170416 | 0.24183 | 0.44781 | 77.044 |
| Gm16181 | 23.9419 | 0.525874 | 0.38604 | 1.362224 | 0.17313 | 0.3644 | 3.301 |
| Gm16183 | 236.133 | 0.598295 | 0.15417 | 3.880856 | 0.0001 | 0.00277 | 5.983 |
| Gm16184 | 45.0726 | 0.072435 | 0.26567 | 0.272653 | 0.78512 | 0.88852 | 0.573 |
| Gm16185 | 743.714 | 0.160507 | 0.14573 | 1.101388 | 0.27073 | 0.48026 | 3.285 |
| Gm16200 | 22.344 | 0.47972 | 0.50655 | 0.947033 | 0.34362 | 0.55429 | 0.331 |
| Gm16206 | 40.3705 | -0.02727 | 0.33071 | -0.08246 | 0.93428 | 0.96702 | 5.179 |
| Gm16209 | 84.9018 | 0.309396 | 0.19181 | 1.613029 | 0.10674 | 0.27102 | 0.627 |
| Gm16217 | 22.9794 | 0.035914 | 0.37848 | 0.09489 | 0.9244 | 0.96202 | 1.323 |

| | | | | | | | |
|---|---|---|---|---|---|---|---|
| Gm16230 | 15.3711 | 0.647407 | 0.45186 | 1.432762 | 0.15193 | 0.33702 | 5.121 |
| Gm16234 | 18.0446 | 0.486132 | 0.46551 | 1.044303 | 0.29635 | 0.50667 | 4.099 |
| Gm16235 | 28.1491 | -0.0043 | 0.35439 | -0.01214 | 0.99031 | 0.99473 | 5.652 |
| Gm16239 | 18.4869 | 0.204787 | 0.42727 | 0.47929 | 0.63173 | 0.79208 | 5.427 |
| Gm16244 | 24.636 | 0.380884 | 0.34564 | 1.101976 | 0.27047 | 0.47998 | 2.423 |
| Gm16253 | 20.6862 | -0.46092 | 0.438 | -1.05233 | 0.29265 | 0.50252 | 3.68 |
| Gm16254 | 29.723 | -0.13065 | 0.31954 | -0.40887 | 0.68263 | 0.82659 | 0.73 |
| Gm16267 | 41.9814 | -0.77217 | 0.37789 | -2.04338 | 0.04101 | 0.14503 | 26.984 |
| Gm16268 | 21.1231 | -0.25308 | 0.42472 | -0.59587 | 0.55126 | 0.73445 | 35.861 |
| Gm16275 | 23.9337 | -0.22142 | 0.38838 | -0.5701 | 0.56861 | 0.74839 | 5.923 |
| Gm16276 | 53.4034 | 0.323675 | 0.23547 | 1.374585 | 0.16926 | 0.35947 | 0.262 |
| Gm16278 | 52.6667 | 0.352526 | 0.29803 | 1.182845 | 0.23687 | 0.442 | 16.657 |
| Gm16279 | 42.3079 | 0.2994 | 0.30003 | 0.997894 | 0.31833 | 0.52892 | 15.27 |
| Gm16287 | 45.521 | -0.05803 | 0.2835 | -0.2047 | 0.8378 | 0.91655 | 5.251 |
| Gm16299 | 64.9792 | -0.2416 | 0.31143 | -0.77576 | 0.43789 | 0.64331 | 5.775 |
| Gm16302 | 15.2456 | 0.292943 | 0.45794 | 0.6397 | 0.52237 | 0.71129 | 2.223 |
| Gm16310 | 96.1536 | 0.648855 | 0.24984 | 2.597072 | 0.0094 | 0.05512 | 1.923 |
| Gm16315 | 74.2283 | 0.347595 | 0.23735 | 1.464515 | 0.14305 | 0.32512 | 1.232 |
| Gm16316 | 46.5744 | -0.33449 | 0.28673 | -1.16658 | 0.24338 | 0.44968 | 7.276 |
| Gm16333 | 78.4888 | 0.020685 | 0.2235 | 0.092547 | 0.92626 | 0.96294 | 0.463 |
| Gm16339 | 108.72 | 0.338723 | 0.23025 | 1.471097 | 0.14126 | 0.323 | 2.101 |
| Gm16342 | 12.8366 | -1.51763 | 0.5059 | -2.99986 | 0.0027 | 0.02455 | 20.457 |
| Gm16350 | 22.5511 | 0.077938 | 0.35427 | 0.219994 | 0.82588 | 0.91013 | 0.521 |
| Gm16353 | 14.5206 | -0.02802 | 0.56255 | -0.04981 | 0.96027 | 0.98057 | 1.223 |
| Gm16355 | 12.6756 | -0.23185 | 0.54947 | -0.42194 | 0.67307 | 0.82024 | 0.231 |
| Gm16364 | 209.949 | 0.216518 | 0.16292 | 1.328973 | 0.18386 | 0.37809 | 55.187 |
| Gm16378 | 58.7665 | 0.243144 | 0.27776 | 0.875363 | 0.38138 | 0.59084 | 0.342 |
| Gm16385 | 42.4368 | -0.05743 | 0.398 | -0.14431 | 0.88526 | 0.94299 | 1.102 |
| Gm16386 | 125.715 | 0.417079 | 0.29653 | 1.406548 | 0.15956 | 0.34742 | 30.52 |
| Gm16418 | 154.187 | -0.68652 | 0.24442 | -2.80881 | 0.00497 | 0.0361 | 0.171 |
| Gm16433 | 22.3235 | -0.20061 | 0.39918 | -0.50255 | 0.61528 | 0.78109 | 1.984 |
| Gm16437 | 50.8447 | -0.39509 | 0.23798 | -1.66017 | 0.09688 | 0.25499 | 0.504 |
| Gm16477 | 33.1145 | -0.193 | 0.33903 | -0.56926 | 0.56918 | 0.74881 | 0.801 |
| Gm16485 | 708.986 | 0.324723 | 0.15862 | 2.047154 | 0.04064 | 0.1442 | 3.983 |

| | | | | | | | |
|---|---|---|---|---|---|---|---|
| Gm16499 | 41.1407 | -0.15019 | 0.32247 | -0.46576 | 0.64139 | 0.79925 | 49.441 |
| Gm16505 | 25.1242 | -1.69086 | 0.53145 | -3.18158 | 0.00146 | 0.01628 | 5.59 |
| Gm16518 | 56.4709 | 0.069706 | 0.2747 | 0.253752 | 0.79969 | 0.89651 | 0.897 |
| Gm16523 | 14.2447 | 0.352095 | 0.58024 | 0.606811 | 0.54398 | 0.72924 | 4.75 |
| Gm16536 | 523.788 | 0.094019 | 0.12592 | 0.746666 | 0.45527 | 0.65728 | 1.589 |
| Gm16537 | 54.2296 | 0.351188 | 0.28629 | 1.2267 | 0.21994 | 0.42245 | 8.932 |
| Gm16540 | 24.558 | -0.21003 | 0.34078 | -0.61633 | 0.53768 | 0.72444 | 1.122 |
| Gm16564 | 17.5551 | 0.429899 | 0.45274 | 0.949547 | 0.34234 | 0.55307 | 19.033 |
| Gm16572 | 15.8768 | 0.437971 | 0.44951 | 0.974329 | 0.32989 | 0.54112 | 1.842 |
| Gm16574 | 17.7646 | 0.096436 | 0.45922 | 0.210001 | 0.83367 | 0.91471 | 0.745 |
| Gm16575 | 20.5713 | 0.45377 | 0.45513 | 0.997003 | 0.31876 | 0.5293 | 2.225 |
| Gm16576 | 107.221 | -0.25808 | 0.24302 | -1.06196 | 0.28825 | 0.49841 | 14.697 |
| Gm16577 | 43.7778 | 0.572693 | 0.27154 | 2.109049 | 0.03494 | 0.13107 | 0.7 |
| Gm16589 | 93.9223 | 0.161061 | 0.23239 | 0.693062 | 0.48827 | 0.68339 | 0.918 |
| Gm16596 | 17.4424 | -0.06385 | 0.42665 | -0.14964 | 0.88105 | 0.94067 | 18.376 |
| Gm16618 | 54.3879 | -0.08114 | 0.29131 | -0.27853 | 0.7806 | 0.88566 | 6.634 |
| Gm16630 | 28.908 | 0.496767 | 0.40784 | 1.218036 | 0.22321 | 0.42649 | 28.375 |
| Gm16638 | 417.694 | 0.771582 | 0.18936 | 4.074785 | 4.61E-05 | 0.00155 | 34.92 |
| Gm16675 | 35.4033 | 0.086596 | 0.33932 | 0.255205 | 0.79856 | 0.89601 | 70.708 |
| Gm16701 | 430.216 | 0.446457 | 0.14835 | 3.009417 | 0.00262 | 0.0241 | 25.706 |
| Gm1673 | 405.937 | -0.24194 | 0.13334 | -1.81441 | 0.06961 | 0.20569 | 1.581 |
| Gm16740 | 89.2793 | 0.072819 | 0.21915 | 0.332284 | 0.73968 | 0.86273 | 9.238 |
| Gm16754 | 92.5166 | 0.087036 | 0.23136 | 0.376193 | 0.70677 | 0.84193 | 7.793 |
| Gm16758 | 79.3832 | 0.382009 | 0.19756 | 1.93368 | 0.05315 | 0.1725 | 6.984 |
| Gm16759 | 22.7129 | 0.013896 | 0.41555 | 0.03344 | 0.97332 | 0.98671 | 170.091 |
| Gm16793 | 31.5844 | 0.583245 | 0.39029 | 1.494386 | 0.13507 | 0.31502 | 6.774 |
| Gm16794 | 98.6013 | 0.071714 | 0.20875 | 0.343542 | 0.73119 | 0.85734 | 17.349 |
| Gm16835 | 439.73 | 0.494837 | 0.19146 | 2.584502 | 0.00975 | 0.05647 | 16.11 |
| Gm16845 | 30.2794 | 0.057227 | 0.36586 | 0.156419 | 0.8757 | 0.93796 | 15.121 |
| Gm16853 | 55.6002 | 0.655854 | 0.24032 | 2.729125 | 0.00635 | 0.04276 | 6.479 |
| Gm16861 | 131.145 | 0.394196 | 0.15219 | 2.590218 | 0.00959 | 0.05577 | 4.585 |
| Gm16892 | 130.527 | 0.490208 | 0.15372 | 3.188943 | 0.00143 | 0.01601 | 7.013 |
| Gm16894 | 108.719 | 0.174154 | 0.20976 | 0.83026 | 0.40639 | 0.6148 | 9.624 |
| Gm16897 | 29.7511 | 0.044044 | 0.34587 | 0.127342 | 0.89867 | 0.94924 | 21.202 |

| | | | | | | | |
|---|---|---|---|---|---|---|---|
| Gm16938 | 13.906 | 0.107255 | 0.53897 | 0.199002 | 0.84226 | 0.91879 | 2.208 |
| Gm16958 | 29.0967 | 0.045025 | 0.35649 | 0.1263 | 0.89949 | 0.94963 | 6.985 |
| Gm16973 | 259.625 | 0.265228 | 0.12085 | 2.194705 | 0.02818 | 0.11309 | 4.989 |
| Gm16982 | 122.811 | 0.36881 | 0.21444 | 1.719849 | 0.08546 | 0.23458 | 3.682 |
| Gm16984 | 68.6191 | 0.269506 | 0.22997 | 1.171915 | 0.24123 | 0.44735 | 26.471 |
| Gm16998 | 13.5536 | 0.363908 | 0.50753 | 0.717022 | 0.47336 | 0.67166 | 5.16 |
| Gm17022 | 77.1565 | 0.296749 | 0.27037 | 1.097576 | 0.27239 | 0.48159 | 0.922 |
| Gm17023 | 56.9462 | 0.259634 | 0.23118 | 1.123067 | 0.26141 | 0.4701 | 6.664 |
| Gm17024 | 90.2647 | -0.08174 | 0.18027 | -0.4534 | 0.65026 | 0.80549 | 1.587 |
| Gm17029 | 58.588 | 0.037812 | 0.25111 | 0.150583 | 0.8803 | 0.94022 | 24.375 |
| Gm17058 | 14.149 | -0.51011 | 0.51175 | -0.99679 | 0.31887 | 0.52938 | 16.342 |
| Gm17066 | 331.345 | 0.207228 | 0.13932 | 1.487389 | 0.13691 | 0.31765 | 5.287 |
| Gm17080 | 29.8977 | 1.163475 | 0.33654 | 3.457209 | 0.00055 | 0.00841 | 2.258 |
| Gm17091 | 24.7767 | 0.165509 | 0.55733 | 0.296969 | 0.76649 | 0.87791 | 2.248 |
| Gm17092 | 48.1999 | 0.630897 | 0.25905 | 2.435421 | 0.01487 | 0.07397 | 2.486 |
| Gm17096 | 64.8848 | -0.24628 | 0.21499 | -1.14556 | 0.25198 | 0.45933 | 0.77 |
| Gm17098 | 15.2037 | -0.40219 | 0.53405 | -0.75309 | 0.45139 | 0.65382 | 0.836 |
| Gm17108 | 27.4804 | 0.781506 | 0.34523 | 2.263731 | 0.02359 | 0.09981 | 4.404 |
| Gm17111 | 18.7933 | -0.87029 | 0.46563 | -1.86906 | 0.06161 | 0.18981 | 0.517 |
| Gm17112 | 275.448 | 0.39349 | 0.13748 | 2.862155 | 0.00421 | 0.03254 | 8.296 |
| Gm17115 | 20.9897 | -0.52924 | 0.51379 | -1.03007 | 0.30298 | 0.51248 | 1.198 |
| Gm17116 | 98.255 | 0.03359 | 0.20666 | 0.162534 | 0.87089 | 0.93475 | 3.442 |
| Gm17120 | 12.6302 | -1.10188 | 0.82583 | -1.33427 | 0.18212 | 0.37592 | 1.938 |
| Gm17133 | 20.8 | 0.55013 | 0.37994 | 1.447936 | 0.14763 | 0.33114 | 5.247 |
| Gm17141 | 23.6045 | 1.158892 | 0.43971 | 2.635588 | 0.0084 | 0.05114 | 6.711 |
| Gm17146 | 38.0725 | 0.114664 | 0.3055 | 0.375331 | 0.70741 | 0.84212 | 0.581 |
| Gm17147 | 55.8236 | 0.282459 | 0.27364 | 1.032224 | 0.30197 | 0.51171 | 0.432 |
| Gm17167 | 32.4165 | 0.14197 | 0.47355 | 0.2998 | 0.76433 | 0.87657 | 28.316 |
| Gm17168 | 37.5132 | 0.399332 | 0.32433 | 1.231243 | 0.21823 | 0.4203 | 5.369 |
| Gm17180 | 47.9357 | 0.594316 | 0.31376 | 1.894174 | 0.0582 | 0.18329 | 1.59 |
| Gm17193 | 374.563 | 0.666979 | 0.16013 | 4.165254 | 3.11E-05 | 0.00119 | 0.971 |
| Gm17194 | 33.302 | -0.18022 | 0.37149 | -0.48514 | 0.62758 | 0.78926 | 0.348 |
| Gm17195 | 43.8444 | 0.118399 | 0.24388 | 0.485482 | 0.62733 | 0.78916 | 0.428 |
| Gm17200 | 68.0329 | 0.468009 | 0.24071 | 1.944255 | 0.05186 | 0.17002 | 22.951 |

| | | | | | | | |
|---|---|---|---|---|---|---|---|
| Gm17201 | 251.532 | 0.056755 | 0.13157 | 0.431359 | 0.66621 | 0.81576 | 3.722 |
| Gm17203 | 16.8555 | -0.35083 | 0.48538 | -0.7228 | 0.4698 | 0.6687 | 4.822 |
| Gm17205 | 21.2042 | -0.31653 | 0.41058 | -0.77094 | 0.44074 | 0.64559 | 1.207 |
| Gm17210 | 68.4015 | 0.218084 | 0.27125 | 0.803994 | 0.4214 | 0.62837 | 2.051 |
| Gm17227 | 26.6904 | 0.686089 | 0.41793 | 1.641645 | 0.10066 | 0.26126 | 6.871 |
| Gm17229 | 44.6612 | 0.498124 | 0.42312 | 1.177264 | 0.23909 | 0.44492 | 4.442 |
| Gm17235 | 35.0534 | 0.1151 | 0.32884 | 0.350021 | 0.72632 | 0.8541 | 1.119 |
| Gm17241 | 38.3626 | -0.0297 | 0.28322 | -0.10487 | 0.91648 | 0.95729 | 0.252 |
| Gm17244 | 30.0897 | 0.181106 | 0.37285 | 0.485733 | 0.62716 | 0.78903 | 3.416 |
| Gm17249 | 73.2745 | 0.349599 | 0.21274 | 1.643344 | 0.10031 | 0.26058 | 4.238 |
| Gm17251 | 60.2754 | -0.12436 | 0.27723 | -0.44858 | 0.65374 | 0.80782 | 1.9 |
| Gm17259 | 34.3275 | 0.199653 | 0.32934 | 0.60622 | 0.54437 | 0.72945 | 4.977 |
| Gm17276 | 935.173 | 0.779983 | 0.11091 | 7.032437 | 2.03E-12 | 4.60E-09 | 28.433 |
| Gm17315 | 16.5002 | 0.340247 | 0.50412 | 0.674938 | 0.49972 | 0.6928 | 3.324 |
| Gm17322 | 162.61 | -0.57989 | 0.19524 | -2.97004 | 0.00298 | 0.02605 | 9.411 |
| Gm17344 | 47.2628 | 0.280104 | 0.28332 | 0.988641 | 0.32284 | 0.53385 | 15.922 |
| Gm17354 | 277.08 | 0.095231 | 0.13477 | 0.706601 | 0.47981 | 0.67732 | 3.599 |
| Gm17382 | 23.491 | -0.38549 | 0.34455 | -1.1188 | 0.26323 | 0.47208 | 25.932 |
| Gm17396 | 165.244 | 0.356187 | 0.23169 | 1.537326 | 0.12421 | 0.29893 | 2.182 |
| Gm17399 | 24.8844 | -0.06299 | 0.4522 | -0.1393 | 0.88922 | 0.94457 | 20.146 |
| Gm17415 | 20.2772 | 0.331319 | 0.45561 | 0.727195 | 0.46711 | 0.66622 | 0.789 |
| Gm17428 | 32.9025 | 0.703762 | 0.47648 | 1.477014 | 0.13967 | 0.32098 | 0.093 |
| Gm17435 | 66.7012 | 0.113269 | 0.22755 | 0.497767 | 0.61865 | 0.78312 | 12.598 |
| Gm17477 | 73.6653 | 0.230616 | 0.25685 | 0.897854 | 0.36926 | 0.57974 | 3.043 |
| Gm17484 | 91.5852 | -0.01251 | 0.23609 | -0.05299 | 0.95774 | 0.97936 | 15.238 |
| Gm17491 | 265.747 | 0.280448 | 0.12592 | 2.227154 | 0.02594 | 0.10668 | 3.042 |
| Gm17494 | 474.417 | 0.532227 | 0.16273 | 3.2707 | 0.00107 | 0.01341 | 37.763 |
| Gm17501 | 61.9687 | -0.26365 | 0.24981 | -1.05538 | 0.29125 | 0.50117 | 27.698 |
| Gm17509 | 35.9453 | 0.059679 | 0.269 | 0.221857 | 0.82443 | 0.90912 | 2.375 |
| Gm17529 | 39.7312 | 0.036589 | 0.32202 | 0.113624 | 0.90954 | 0.95456 | 14.923 |
| Gm17530 | 109.319 | 0.198168 | 0.20343 | 0.974156 | 0.32998 | 0.54118 | 0.472 |
| Gm17545 | 15.3907 | -0.24114 | 0.52398 | -0.4602 | 0.64537 | 0.8023 | 7.773 |
| Gm17552 | 46.3787 | -0.2082 | 0.29799 | -0.69871 | 0.48474 | 0.68092 | 4.719 |
| Gm17597 | 49.0833 | 0.394594 | 0.25865 | 1.525566 | 0.12712 | 0.30327 | 1.823 |

| | | | | | | | |
|---|---|---|---|---|---|---|---|
| Gm17622 | 16.0494 | -0.52498 | 0.51211 | -1.02513 | 0.3053 | 0.51531 | 0.388 |
| Gm17638 | 20.9552 | 0.190335 | 0.41424 | 0.459476 | 0.64589 | 0.80278 | 5.453 |
| Gm17641 | 31.8848 | -0.47535 | 0.42989 | -1.10575 | 0.26884 | 0.47813 | 2.35 |
| Gm17655 | 71.5013 | 0.146612 | 0.26894 | 0.545149 | 0.58565 | 0.75999 | 26.375 |
| Gm17690 | 30.2132 | -0.22554 | 0.39521 | -0.57068 | 0.56822 | 0.74813 | 4.968 |
| Gm17705 | 69.672 | 0.14294 | 0.21895 | 0.652856 | 0.51385 | 0.70458 | 2.068 |
| Gm17709 | 147.346 | 0.620085 | 0.18289 | 3.3905 | 0.0007 | 0.00997 | 1.096 |
| Gm17720 | 17.819 | 0.074779 | 0.41488 | 0.180244 | 0.85696 | 0.92788 | 19.951 |
| Gm17733 | 17.9159 | -0.29946 | 0.46063 | -0.65011 | 0.51562 | 0.70584 | 7.236 |
| Gm17750 | 70.9361 | -0.62766 | 0.30058 | -2.08815 | 0.03678 | 0.13527 | 39.507 |
| Gm17753 | 95.9515 | -1.07353 | 0.26836 | -4.00033 | 6.33E-05 | 0.00196 | 10.648 |
| Gm17809 | 20.7484 | 0.054685 | 0.51838 | 0.105491 | 0.91599 | 0.95707 | 1.42 |
| Gm17816 | 17.0145 | -0.41326 | 0.51166 | -0.80768 | 0.41927 | 0.62648 | 2.617 |
| Gm17825 | 28.1917 | 0.290801 | 0.34288 | 0.848109 | 0.39638 | 0.60541 | 0.794 |
| Gm17828 | 25.6597 | 0.203934 | 0.53986 | 0.377753 | 0.70561 | 0.84142 | 0.799 |
| Gm17833 | 38.0927 | 0.408511 | 0.32182 | 1.269365 | 0.20431 | 0.40307 | 1.065 |
| Gm17853 | 29.5038 | -0.05121 | 0.35797 | -0.14305 | 0.88625 | 0.94336 | 1.349 |
| Gm17875 | 87.2984 | 0.437759 | 0.19312 | 2.266776 | 0.0234 | 0.09924 | 1.06 |
| Gm17891 | 15.3057 | -0.57523 | 0.59903 | -0.96027 | 0.33692 | 0.54752 | 0.779 |
| Gm17907 | 45.292 | 0.187132 | 0.35821 | 0.522408 | 0.60139 | 0.77152 | 0.71 |
| Gm17910 | 20.7379 | 0.442728 | 0.42262 | 1.047568 | 0.29484 | 0.50521 | 0.838 |
| Gm17916 | 70.4326 | 0.794703 | 0.24503 | 3.24328 | 0.00118 | 0.01415 | 15.637 |
| Gm17922 | 43.9872 | 0.446296 | 0.40529 | 1.101169 | 0.27082 | 0.48026 | 0.802 |
| Gm17931 | 50.6504 | 1.194184 | 0.28619 | 4.172684 | 3.01E-05 | 0.00116 | 0.634 |
| Gm17954 | 13.1158 | 0.519599 | 0.47189 | 1.101096 | 0.27085 | 0.48026 | 0.607 |
| Gm17969 | 20.1927 | 0.926537 | 0.53961 | 1.717036 | 0.08597 | 0.23558 | 3.671 |
| Gm17971 | 48.7044 | 0.472758 | 0.24599 | 1.921848 | 0.05462 | 0.17577 | 0.806 |
| Gm17981 | 13.1451 | -0.62642 | 0.55008 | -1.13877 | 0.2548 | 0.46229 | 0.739 |
| Gm18014 | 13.599 | 0.948036 | 0.46537 | 2.037185 | 0.04163 | 0.14663 | 0.817 |
| Gm18030 | 84.6127 | 0.075168 | 0.20813 | 0.361155 | 0.71798 | 0.84944 | 0.731 |
| Gm18095 | 33.0962 | 0.923456 | 0.31864 | 2.898134 | 0.00375 | 0.03016 | 1.018 |
| Gm18126 | 15.3115 | -0.43662 | 0.49866 | -0.8756 | 0.38125 | 0.59071 | 1.321 |
| Gm1818 | 91.022 | -0.21259 | 0.25573 | -0.83133 | 0.40579 | 0.6144 | 4.578 |
| Gm18180 | 13.8115 | 0.247913 | 0.45765 | 0.541704 | 0.58802 | 0.76147 | 1.484 |

| | | | | | | | |
|---|---|---|---|---|---|---|---|
| Gm18190 | 29.0992 | -0.35342 | 0.37541 | -0.94143 | 0.34649 | 0.55705 | 12.77 |
| Gm18194 | 180.233 | 0.194271 | 0.19771 | 0.982606 | 0.3258 | 0.53722 | 3.442 |
| Gm18225 | 12.9728 | 0.120676 | 0.70838 | 0.170354 | 0.86473 | 0.93154 | 0.742 |
| Gm18258 | 24.8409 | 1.01246 | 0.40255 | 2.515137 | 0.0119 | 0.06427 | 0.852 |
| Gm18271 | 25.7507 | 0.242184 | 0.36578 | 0.662096 | 0.50791 | 0.69921 | 0.945 |
| Gm18284 | 50.2563 | 0.042087 | 0.25897 | 0.162518 | 0.8709 | 0.93475 | 0.619 |
| Gm18290 | 58.0945 | -0.56924 | 0.29546 | -1.92663 | 0.05403 | 0.17469 | 0.844 |
| Gm18329 | 15.2376 | -0.36757 | 0.51346 | -0.71587 | 0.47407 | 0.67238 | 0.589 |
| Gm18336 | 14.0906 | 0.20643 | 0.47715 | 0.432629 | 0.66528 | 0.81526 | 2.298 |
| Gm18362 | 13.1385 | 0.039535 | 0.54456 | 0.072599 | 0.94213 | 0.97113 | 0.529 |
| Gm18406 | 104.389 | 0.789621 | 0.29827 | 2.647378 | 0.00811 | 0.04999 | 0.669 |
| Gm18499 | 19.0196 | 0.715563 | 0.5231 | 1.367925 | 0.17134 | 0.3618 | 1.021 |
| Gm18517 | 28.419 | -0.98407 | 0.33808 | -2.91079 | 0.00361 | 0.02937 | 2.174 |
| Gm18537 | 15.318 | 0.630512 | 0.44296 | 1.423404 | 0.15462 | 0.34066 | 1.523 |
| Gm18541 | 54.5554 | -0.13194 | 0.30816 | -0.42815 | 0.66854 | 0.81715 | 0.538 |
| Gm18588 | 164.225 | 0.018452 | 0.21183 | 0.08711 | 0.93058 | 0.96505 | 0.492 |
| Gm18609 | 647.578 | -0.12231 | 0.11804 | -1.03618 | 0.30012 | 0.51023 | 3.601 |
| Gm18637 | 31.6523 | 0.634009 | 0.35853 | 1.768367 | 0.077 | 0.2194 | 1.9 |
| Gm18649 | 176.378 | 0.091715 | 0.17604 | 0.521001 | 0.60237 | 0.77229 | 0.461 |
| Gm18670 | 18.1135 | -0.26471 | 0.47263 | -0.56007 | 0.57543 | 0.75343 | 0.696 |
| Gm18706 | 63.4128 | 0.086927 | 0.28321 | 0.306933 | 0.75889 | 0.87333 | 1.066 |
| Gm18755 | 29.5895 | 0.208153 | 0.35437 | 0.587397 | 0.55694 | 0.73891 | 1.099 |
| Gm18782 | 51.8374 | 0.368364 | 0.30435 | 1.210334 | 0.22615 | 0.43018 | 1.399 |
| Gm18856 | 25.6297 | 0.552259 | 0.40642 | 1.358833 | 0.1742 | 0.36573 | 6.851 |
| Gm18860 | 25.2085 | 0.458098 | 0.39064 | 1.172694 | 0.24092 | 0.4471 | 0.764 |
| Gm18890 | 17.535 | 0.072792 | 0.38975 | 0.186764 | 0.85185 | 0.92468 | 0.805 |
| Gm18917 | 17.0712 | 0.049724 | 0.53354 | 0.093196 | 0.92575 | 0.96257 | 0.55 |
| Gm18935 | 23.5096 | 0.658245 | 0.39596 | 1.662405 | 0.09643 | 0.25432 | 0.664 |
| Gm18949 | 14.5965 | -0.29914 | 0.53008 | -0.56433 | 0.57253 | 0.75123 | 1.303 |
| Gm18959 | 22.7756 | -0.08214 | 0.43728 | -0.18783 | 0.85101 | 0.92413 | 0.655 |
| Gm18960 | 32.8209 | -0.05976 | 0.3259 | -0.18338 | 0.8545 | 0.92645 | 0.713 |
| Gm18974 | 20.974 | 1.122405 | 0.44902 | 2.499664 | 0.01243 | 0.06618 | 2.229 |
| Gm18981 | 20.5457 | 0.750241 | 0.39697 | 1.889902 | 0.05877 | 0.18443 | 1.599 |
| Gm19026 | 50.4858 | -0.04997 | 0.27317 | -0.18292 | 0.85486 | 0.92662 | 11.642 |

| | | | | | | | |
|---|---|---|---|---|---|---|---|
| Gm19031 | 12.6958 | -0.06261 | 0.51554 | -0.12144 | 0.90334 | 0.95132 | 1.365 |
| Gm19040 | 16.7229 | -0.60395 | 0.44676 | -1.35184 | 0.17643 | 0.36846 | 4.726 |
| Gm19046 | 25.7683 | 0.46566 | 0.37746 | 1.233659 | 0.21733 | 0.41935 | 1.298 |
| Gm19144 | 63.0046 | 0.018492 | 0.23593 | 0.078381 | 0.93752 | 0.9688 | 2.41 |
| Gm19178 | 24.329 | -0.30948 | 0.34754 | -0.89048 | 0.37321 | 0.58357 | 1.819 |
| Gm19187 | 22.2739 | 0.086659 | 0.44682 | 0.193946 | 0.84622 | 0.92183 | 0.892 |
| Gm19196 | 144.816 | -0.1402 | 0.1846 | -0.75948 | 0.44757 | 0.65119 | 11.435 |
| Gm19221 | 121.203 | -0.37903 | 0.19338 | -1.95996 | 0.05 | 0.16599 | 1.583 |
| Gm19236 | 39.6387 | 0.311892 | 0.29638 | 1.052326 | 0.29265 | 0.50252 | 2.698 |
| Gm19265 | 65.8247 | 0.248882 | 0.24984 | 0.996155 | 0.31917 | 0.52977 | 40.304 |
| Gm19327 | 25.5774 | 0.242615 | 0.38403 | 0.631759 | 0.52754 | 0.71571 | 3.271 |
| Gm19391 | 29.8506 | -0.63886 | 0.32527 | -1.96408 | 0.04952 | 0.16478 | 2.363 |
| Gm19395 | 22.21 | -0.09022 | 0.45402 | -0.19871 | 0.84249 | 0.9189 | 3.406 |
| Gm19409 | 83.692 | -0.01447 | 0.24944 | -0.05801 | 0.95374 | 0.97729 | 2.858 |
| Gm19410 | 249.322 | -0.02794 | 0.18582 | -0.15039 | 0.88046 | 0.94029 | 52.258 |
| Gm19426 | 28.5588 | -0.62854 | 0.35973 | -1.74728 | 0.08059 | 0.22593 | 1.487 |
| Gm1943 | 141.378 | -0.1261 | 0.15998 | -0.78822 | 0.43057 | 0.63686 | 0.174 |
| Gm19434 | 25.4029 | 0.20271 | 0.45975 | 0.44091 | 0.65928 | 0.81122 | 6.481 |
| Gm19439 | 1201.16 | -0.47039 | 0.15606 | -3.01408 | 0.00258 | 0.02386 | 2.289 |
| Gm19445 | 19.7492 | -0.98358 | 0.44503 | -2.21014 | 0.0271 | 0.10994 | 2.438 |
| Gm19461 | 48.7813 | 0.398347 | 0.28979 | 1.374597 | 0.16926 | 0.35947 | 19.411 |
| Gm19466 | 15.8678 | 0.395762 | 0.51586 | 0.767187 | 0.44297 | 0.64746 | 66.315 |
| Gm19500 | 435.58 | -0.41013 | 0.21599 | -1.89884 | 0.05759 | 0.18222 | 10.814 |
| Gm19522 | 113.024 | 0.081076 | 0.18781 | 0.431684 | 0.66597 | 0.81557 | 27.696 |
| Gm19531 | 145.636 | 0.171019 | 0.1537 | 1.112671 | 0.26585 | 0.47503 | 4.537 |
| Gm19554 | 30.4551 | -0.5904 | 0.41172 | -1.43398 | 0.15158 | 0.33655 | 2.623 |
| Gm19557 | 25.5606 | 0.575775 | 0.38893 | 1.480399 | 0.13877 | 0.32002 | 3.597 |
| Gm19569 | 25.794 | 0.096463 | 0.44118 | 0.218649 | 0.82692 | 0.91102 | 1.509 |
| Gm19605 | 103.579 | -0.08367 | 0.20661 | -0.40494 | 0.68552 | 0.82866 | 11.705 |
| Gm19667 | 15.0658 | -1.16017 | 0.47561 | -2.43935 | 0.01471 | 0.07331 | 9.675 |
| Gm19680 | 17.8664 | -0.25187 | 0.45672 | -0.55147 | 0.58131 | 0.75725 | 1.513 |
| Gm19705 | 63.9585 | -0.17736 | 0.27056 | -0.65553 | 0.51213 | 0.70326 | 20.038 |
| Gm19710 | 78.7947 | -0.12652 | 0.24302 | -0.5206 | 0.60264 | 0.77235 | 22.686 |
| Gm19719 | 24.4593 | -0.52584 | 0.41854 | -1.25638 | 0.20898 | 0.40901 | 5.117 |

| | | | | | | | |
|---|---|---|---|---|---|---|---|
| Gm19721 | 78.5657 | 0.163096 | 0.20124 | 0.810468 | 0.41767 | 0.6251 | 1.764 |
| Gm19744 | 334.445 | -0.77605 | 0.27531 | -2.81878 | 0.00482 | 0.03545 | 27.062 |
| Gm1976 | 993.973 | 0.141715 | 0.10105 | 1.402429 | 0.16079 | 0.34897 | 74.884 |
| Gm19810 | 52.053 | -0.57619 | 0.32973 | -1.74743 | 0.08056 | 0.22592 | 0.251 |
| Gm19817 | 34.4416 | -0.0805 | 0.33367 | -0.24126 | 0.80936 | 0.90175 | 2.132 |
| Gm19863 | 26.6149 | -0.05055 | 0.40022 | -0.12629 | 0.8995 | 0.94963 | 2.218 |
| Gm19935 | 49.1036 | -0.20096 | 0.32805 | -0.61257 | 0.54016 | 0.72606 | 1.249 |
| Gm19938 | 53.8678 | -0.21293 | 0.26631 | -0.79955 | 0.42397 | 0.63068 | 11.596 |
| Gm19963 | 76.0609 | -0.98919 | 0.35029 | -2.82394 | 0.00474 | 0.03514 | 1.255 |
| Gm2000 | 146.474 | -0.33028 | 0.19667 | -1.6793 | 0.09309 | 0.2484 | 0.445 |
| Gm20008 | 553.374 | 0.565408 | 0.47785 | 1.183243 | 0.23671 | 0.44183 | 3.156 |
| Gm20033 | 129.348 | -0.96586 | 0.25987 | -3.7167 | 0.0002 | 0.00429 | 95.967 |
| Gm20036 | 16.7006 | 0.026082 | 0.46467 | 0.05613 | 0.95524 | 0.97801 | 2.096 |
| Gm2004 | 96.3331 | 1.253584 | 1.09845 | 1.141233 | 0.25377 | 0.46112 | 8.235 |
| Gm20045 | 697.618 | 0.509449 | 0.17758 | 2.868895 | 0.00412 | 0.03206 | 5.577 |
| Gm20063 | 72.7047 | 0.656492 | 0.30523 | 2.150813 | 0.03149 | 0.12213 | 32.727 |
| Gm20083 | 35.7354 | -0.13597 | 0.31484 | -0.43187 | 0.66584 | 0.81557 | 3.724 |
| Gm2011 | 821.417 | 0.267778 | 0.09904 | 2.703641 | 0.00686 | 0.04485 | 28.569 |
| Gm20149 | 15.4762 | -0.16206 | 0.52244 | -0.31019 | 0.75641 | 0.87186 | 14.596 |
| Gm20163 | 34.7424 | -0.4087 | 0.38067 | -1.07363 | 0.28299 | 0.49316 | 1.808 |
| Gm20164 | 25.4621 | -0.29799 | 0.42395 | -0.70289 | 0.48212 | 0.6787 | 0.316 |
| Gm20186 | 787.183 | -0.21121 | 0.10563 | -1.99946 | 0.04556 | 0.15537 | 5.6 |
| Gm20208 | 46.8409 | -0.4043 | 0.27811 | -1.45375 | 0.14602 | 0.32899 | 7.986 |
| Gm20242 | 43.3525 | 0.065183 | 0.34779 | 0.18742 | 0.85133 | 0.92439 | 1.855 |
| Gm20257 | 84.3579 | -0.0284 | 0.19263 | -0.14743 | 0.8828 | 0.9416 | 3.242 |
| Gm20274 | 60.099 | 0.662278 | 0.27253 | 2.430073 | 0.0151 | 0.07476 | 0.584 |
| Gm20300 | 2115.43 | 0.070834 | 0.10891 | 0.650392 | 0.51544 | 0.70576 | 3.44 |
| Gm20319 | 15.5888 | -1.04403 | 0.50434 | -2.07008 | 0.03844 | 0.13959 | 82.064 |
| Gm20324 | 32.4312 | 0.032049 | 0.31638 | 0.101298 | 0.91931 | 0.95917 | 6.739 |
| Gm20337 | 14.8565 | -0.75877 | 0.52353 | -1.44932 | 0.14725 | 0.33046 | 3.851 |
| Gm20342 | 600.483 | 0.341923 | 0.10239 | 3.339419 | 0.00084 | 0.01136 | 9.72 |
| Gm20371 | 21.7643 | -1.07943 | 0.38465 | -2.80623 | 0.00501 | 0.03636 | 1.756 |
| Gm20383 | 28.5032 | -0.67955 | 0.35136 | -1.93406 | 0.05311 | 0.17243 | 8.093 |
| Gm20387 | 134.025 | 0.477909 | 0.18217 | 2.623411 | 0.00871 | 0.05236 | 6.164 |

| | | | | | | | |
|---|---|---|---|---|---|---|---|
| Gm20394 | 138.882 | 1.133634 | 0.20347 | 5.571567 | 2.52E-08 | 7.18E-06 | 1.194 |
| Gm20405 | 15.0254 | -0.75247 | 0.61059 | -1.23237 | 0.21781 | 0.4199 | 0.488 |
| Gm20449 | 21.5052 | 0.441576 | 0.34859 | 1.266761 | 0.20524 | 0.40426 | 100.04 |
| Gm20457 | 152.781 | -0.00945 | 0.17786 | -0.05312 | 0.95763 | 0.97935 | 8.415 |
| Gm20468 | 18.5147 | -0.02308 | 0.45849 | -0.05033 | 0.95986 | 0.98049 | 47.849 |
| Gm20476 | 46.0397 | 0.651629 | 0.3286 | 1.983022 | 0.04737 | 0.15985 | 8.508 |
| Gm20492 | 49.7692 | 0.09457 | 0.28334 | 0.333771 | 0.73855 | 0.86214 | 1.337 |
| Gm20493 | 14.4449 | 0.422389 | 0.44459 | 0.950065 | 0.34208 | 0.55293 | 0.637 |
| Gm20501 | 101.596 | -0.0398 | 0.19282 | -0.20641 | 0.83647 | 0.91608 | 5.836 |
| Gm20511 | 16.4519 | -0.457 | 0.43765 | -1.04421 | 0.29639 | 0.50667 | 2.187 |
| Gm20515 | 122.112 | 0.017224 | 0.19307 | 0.089207 | 0.92892 | 0.96429 | 7.725 |
| Gm20517 | 28.8968 | 0.233889 | 0.32946 | 0.709913 | 0.47776 | 0.67553 | 72.273 |
| Gm20518 | 15.7356 | 0.244657 | 0.49695 | 0.492313 | 0.6225 | 0.78561 | 56.395 |
| Gm20535 | 42.5658 | 0.333736 | 0.25149 | 1.32705 | 0.18449 | 0.37899 | 3.835 |
| Gm20544 | 147.108 | -0.56139 | 0.21332 | -2.63168 | 0.0085 | 0.05147 | 8.45 |
| Gm20548 | 170.3 | 0.455618 | 0.17583 | 2.591286 | 0.00956 | 0.05566 | 8.995 |
| Gm20555 | 14.1133 | 0.077058 | 0.45855 | 0.168047 | 0.86655 | 0.93232 | 3.433 |
| Gm20559 | 160.619 | -0.88182 | 0.52355 | -1.68429 | 0.09212 | 0.24709 | 12.938 |
| Gm2058 | 78.5921 | -0.37874 | 0.25259 | -1.49945 | 0.13376 | 0.31339 | 0.787 |
| Gm20597 | 15.6248 | 0.085982 | 0.48933 | 0.175714 | 0.86052 | 0.92938 | 20.547 |
| Gm20604 | 225.108 | -0.33885 | 0.13379 | -2.53266 | 0.01132 | 0.06219 | 16.941 |
| Gm2061 | 147.041 | 0.361878 | 0.19882 | 1.820165 | 0.06873 | 0.20383 | 8.453 |
| Gm20627 | 55.8884 | -0.03356 | 0.30692 | -0.10936 | 0.91292 | 0.95597 | 1.659 |
| Gm20628 | 45.0786 | -0.33563 | 0.32298 | -1.03916 | 0.29873 | 0.50876 | 3.215 |
| Gm20632 | 41.4095 | -0.51049 | 0.34558 | -1.4772 | 0.13962 | 0.3209 | 2.427 |
| Gm20633 | 46.5154 | -0.41746 | 0.30385 | -1.3739 | 0.16947 | 0.35966 | 2.831 |
| Gm20636 | 76.55 | 0.350649 | 0.2472 | 1.418486 | 0.15605 | 0.3427 | 3.271 |
| Gm20646 | 102.542 | 0.177174 | 0.20354 | 0.870471 | 0.38404 | 0.59346 | 49.867 |
| Gm2065 | 77.6599 | 0.106725 | 0.21524 | 0.495839 | 0.62001 | 0.78385 | 2.757 |
| Gm20655 | 32.5133 | 0.069464 | 0.33696 | 0.206151 | 0.83667 | 0.91619 | 9.4 |
| Gm20681 | 55.7222 | 0.340512 | 0.2508 | 1.357717 | 0.17455 | 0.36617 | 2.408 |
| Gm20687 | 13.0137 | 0.449478 | 0.53292 | 0.843429 | 0.39899 | 0.60798 | 44.042 |
| Gm20689 | 24.4423 | -0.33195 | 0.36283 | -0.91491 | 0.36024 | 0.5706 | 0.279 |
| Gm20691 | 14.9887 | 0.249175 | 0.49004 | 0.508475 | 0.61112 | 0.77825 | 0.333 |

| | | | | | | | |
|---|---|---|---|---|---|---|---|
| Gm20692 | 85.5161 | 0.400411 | 0.21624 | 1.851697 | 0.06407 | 0.19484 | 0.784 |
| Gm20696 | 30.3852 | 0.565339 | 0.35963 | 1.572015 | 0.11595 | 0.28578 | 409.314 |
| Gm20700 | 23.8479 | 0.158832 | 0.35267 | 0.450369 | 0.65244 | 0.80666 | 9.106 |
| Gm20705 | 16.2652 | 0.357878 | 0.53164 | 0.67316 | 0.50085 | 0.69366 | 3.893 |
| Gm20712 | 127.251 | 0.124412 | 0.18024 | 0.690261 | 0.49003 | 0.68501 | 1.491 |
| Gm20717 | 24.2746 | 0.128252 | 0.4127 | 0.310766 | 0.75598 | 0.87157 | 3.081 |
| Gm20731 | 23.0973 | 0.097281 | 0.39881 | 0.243926 | 0.80729 | 0.90075 | 2.288 |
| Gm20743 | 61.7725 | -0.31772 | 0.27558 | -1.15294 | 0.24894 | 0.45629 | 73.643 |
| Gm20751 | 35.0297 | -0.39228 | 0.33114 | -1.18462 | 0.23617 | 0.44147 | 2.248 |
| Gm20753 | 18.0679 | 0.256481 | 0.39918 | 0.642525 | 0.52053 | 0.70978 | 18.9 |
| Gm20754 | 105.88 | 0.319813 | 0.22476 | 1.422903 | 0.15476 | 0.34087 | 528.541 |
| Gm20768 | 12.9288 | -0.32425 | 0.49748 | -0.65179 | 0.51454 | 0.70505 | 2.336 |
| Gm20784 | 17.5355 | -0.63375 | 0.491 | -1.29072 | 0.1968 | 0.39463 | 1.246 |
| Gm20939 | 99.0754 | 0.079785 | 0.32539 | 0.245198 | 0.8063 | 0.90039 | 13.433 |
| Gm21057 | 12.8285 | 0.284175 | 0.51464 | 0.552185 | 0.58082 | 0.75705 | 12.558 |
| Gm2106 | 16.8377 | 0.820501 | 0.48199 | 1.702333 | 0.08869 | 0.24064 | 1.55 |
| Gm21092 | 110.226 | 0.095696 | 0.17491 | 0.547117 | 0.5843 | 0.75915 | 15.175 |
| Gm21112 | 46.3967 | 0.449532 | 0.34612 | 1.298787 | 0.19402 | 0.39108 | 0.578 |
| Gm2115 | 903.209 | -0.22374 | 0.11086 | -2.01819 | 0.04357 | 0.151 | 54.578 |
| Gm21168 | 14.0672 | -0.93856 | 0.50485 | -1.85909 | 0.06302 | 0.19274 | 1.096 |
| Gm2136 | 205.178 | 0.143639 | 0.18199 | 0.789258 | 0.42996 | 0.63625 | 2.577 |
| Gm21378 | 63.8278 | -0.08199 | 0.25749 | -0.31842 | 0.75017 | 0.86885 | 1.246 |
| Gm21399 | 20.8653 | -0.022 | 0.49903 | -0.04408 | 0.96484 | 0.98273 | 0.597 |
| Gm2155 | 32.3353 | 0.538536 | 0.36142 | 1.490051 | 0.13621 | 0.31648 | 3.171 |
| Gm2164 | 49.457 | 1.972612 | 0.35919 | 5.491812 | 3.98E-08 | 9.54E-06 | 128.246 |
| Gm21691 | 29.0557 | 0.316299 | 0.37053 | 0.853643 | 0.3933 | 0.60236 | 23.905 |
| Gm21769 | 223.667 | 0.342147 | 0.17214 | 1.987649 | 0.04685 | 0.15876 | 0.111 |
| Gm21781 | 1578.41 | 0.218237 | 0.09017 | 2.420375 | 0.0155 | 0.0761 | 4.838 |
| Gm21811 | 777.072 | 0.24778 | 0.095 | 2.608277 | 0.0091 | 0.05383 | 15.164 |
| Gm21814 | 69.3002 | 0.727932 | 0.30738 | 2.368168 | 0.01788 | 0.08327 | 19.565 |
| Gm21844 | 100.661 | 0.090509 | 0.18305 | 0.494449 | 0.62099 | 0.78447 | 4.273 |
| Gm21846 | 16.8998 | -0.37592 | 0.51699 | -0.72713 | 0.46714 | 0.66622 | 3.073 |
| Gm21926 | 807.769 | 0.009674 | 0.09212 | 0.105017 | 0.91636 | 0.95722 | 15.099 |
| Gm21954 | 22.5179 | 0.033719 | 0.39922 | 0.084464 | 0.93269 | 0.96605 | 10.226 |

| | | | | | | | |
|---|---|---|---|---|---|---|---|
| Gm21955 | 720.621 | 0.159055 | 0.15401 | 1.032786 | 0.3017 | 0.51136 | 4.968 |
| Gm21984 | 77.4652 | -0.0405 | 0.20287 | -0.19964 | 0.84176 | 0.91879 | 10.73 |
| Gm21986 | 23.8948 | -0.08555 | 0.3941 | -0.21707 | 0.82815 | 0.91185 | 25.51 |
| Gm21992 | 141.134 | 0.124903 | 0.1797 | 0.695065 | 0.48701 | 0.68268 | 68.7 |
| Gm22009 | 462.721 | 0.183144 | 0.53739 | 0.340804 | 0.73325 | 0.85857 | 0.343 |
| Gm22027 | 16.0768 | 0.149244 | 0.50956 | 0.292889 | 0.76961 | 0.87977 | 0.127 |
| Gm22106 | 26.7662 | -0.01188 | 0.38534 | -0.03082 | 0.97541 | 0.9876 | 0.127 |
| Gm22192 | 15.7135 | 1.059146 | 0.64217 | 1.649332 | 0.09908 | 0.25857 | 0.085 |
| Gm22200 | 25.1042 | 0.510658 | 0.41172 | 1.24031 | 0.21486 | 0.41624 | 0.067 |
| Gm22215 | 17.093 | 0.105451 | 0.41152 | 0.256247 | 0.79776 | 0.89566 | 0.096 |
| Gm2223 | 122.446 | -0.23268 | 0.17152 | -1.35658 | 0.17492 | 0.36656 | 1.35 |
| Gm22270 | 43.9348 | 0.023615 | 0.26271 | 0.08989 | 0.92837 | 0.96398 | 0.132 |
| Gm22281 | 37.1152 | -0.01537 | 0.35882 | -0.04284 | 0.96583 | 0.9828 | 0.103 |
| Gm22358 | 22.1854 | 0.111154 | 0.36614 | 0.303585 | 0.76144 | 0.87503 | 0.122 |
| Gm2237 | 45.059 | -0.11346 | 0.27626 | -0.41071 | 0.68129 | 0.82569 | 21.336 |
| Gm2238 | 96.6446 | 0.285126 | 0.32386 | 0.880385 | 0.37865 | 0.5884 | 3.669 |
| Gm22442 | 47.8298 | 0.547956 | 0.33297 | 1.645663 | 0.09983 | 0.25983 | 0.144 |
| Gm22572 | 21.1571 | 0.700211 | 0.36303 | 1.928789 | 0.05376 | 0.17402 | 0.105 |
| Gm22573 | 19.1836 | 0.416526 | 0.4481 | 0.929531 | 0.35261 | 0.56294 | 0.102 |
| Gm22661 | 26.7652 | 0.090664 | 0.39958 | 0.226901 | 0.8205 | 0.90734 | 0.132 |
| Gm22716 | 24.309 | 0.180353 | 0.41891 | 0.430527 | 0.66681 | 0.81586 | 0.107 |
| Gm22720 | 21.3192 | 0.431324 | 0.40813 | 1.05683 | 0.29059 | 0.50076 | 0.087 |
| Gm22748 | 14.2295 | -0.03833 | 0.45764 | -0.08375 | 0.93326 | 0.96643 | 0.136 |
| Gm22753 | 30.2181 | -0.42451 | 0.30777 | -1.37931 | 0.1678 | 0.35771 | 0.107 |
| Gm22767 | 47.157 | 0.20243 | 0.29946 | 0.675988 | 0.49905 | 0.69225 | 0.165 |
| Gm22771 | 61.3457 | 0.155631 | 0.30814 | 0.505067 | 0.61351 | 0.77991 | 0.091 |
| Gm22772 | 13.7621 | 0.268861 | 0.56833 | 0.473075 | 0.63616 | 0.79507 | 0.121 |
| Gm22858 | 15.6031 | 0.580098 | 0.4528 | 1.281121 | 0.20015 | 0.39817 | 0.076 |
| Gm22888 | 13.3596 | 1.052992 | 0.5083 | 2.071577 | 0.0383 | 0.13922 | 0.077 |
| Gm2296 | 35.2677 | 0.382314 | 0.33278 | 1.148857 | 0.25061 | 0.45792 | 0.988 |
| Gm23034 | 13.7813 | 0.246212 | 0.47863 | 0.514414 | 0.60696 | 0.77526 | 0.147 |
| Gm23136 | 45.8923 | -0.02389 | 0.30316 | -0.07879 | 0.9372 | 0.96863 | 0.099 |
| Gm23205 | 43.5976 | 0.005977 | 0.28018 | 0.021335 | 0.98298 | 0.99076 | 0.103 |
| Gm23264 | 13.9223 | -0.245 | 0.50069 | -0.48933 | 0.62461 | 0.78719 | 0.107 |

| | | | | | | | |
|---|---|---|---|---|---|---|---|
| Gm23318 | 42.6599 | 0.304828 | 0.28486 | 1.070108 | 0.28457 | 0.4946 | 0.108 |
| Gm23442 | 16.3875 | 0.281503 | 0.49034 | 0.574098 | 0.5659 | 0.74585 | 0.134 |
| Gm23458 | 43.3249 | 0.010322 | 0.34892 | 0.029584 | 0.9764 | 0.98767 | 0.169 |
| Gm23573 | 28.444 | -0.33327 | 0.32162 | -1.03623 | 0.3001 | 0.51023 | 0.107 |
| Gm23608 | 16.4378 | 0.525457 | 0.58432 | 0.899268 | 0.36851 | 0.57916 | 0.128 |
| Gm23615 | 18.71 | 0.26979 | 0.46402 | 0.581425 | 0.56095 | 0.74209 | 0.065 |
| Gm23645 | 18.5284 | -0.55643 | 0.41138 | -1.35259 | 0.17619 | 0.3682 | 0.108 |
| Gm2367 | 60.0428 | 0.239744 | 0.26132 | 0.91742 | 0.35892 | 0.56923 | 0.472 |
| Gm23706 | 15.6408 | 1.082349 | 0.47272 | 2.289614 | 0.02204 | 0.09567 | 0.082 |
| Gm23747 | 13.0358 | 1.258488 | 0.57005 | 2.207672 | 0.02727 | 0.1103 | 0.131 |
| Gm23851 | 12.9268 | -0.07233 | 0.4708 | -0.15364 | 0.8779 | 0.93893 | 0.103 |
| Gm23887 | 17.5375 | -0.18874 | 0.57831 | -0.32636 | 0.74415 | 0.86539 | 0.172 |
| Gm23908 | 13.0752 | -0.21413 | 0.51312 | -0.41732 | 0.67645 | 0.82252 | 0.132 |
| Gm23966 | 52.8687 | 0.066478 | 0.27543 | 0.241365 | 0.80927 | 0.90175 | 0.107 |
| Gm24091 | 16.9494 | 0.295614 | 0.48898 | 0.604554 | 0.54548 | 0.73035 | 0.083 |
| Gm24119 | 28.7267 | 0.599037 | 0.43478 | 1.3778 | 0.16827 | 0.35827 | 0.246 |
| Gm24144 | 67.3467 | 0.160928 | 0.23171 | 0.694538 | 0.48735 | 0.68275 | 0.111 |
| Gm2415 | 927.954 | 0.281493 | 0.12453 | 2.260358 | 0.0238 | 0.10042 | 36.381 |
| Gm24166 | 13.0153 | -0.62503 | 0.53181 | -1.17529 | 0.23988 | 0.44581 | 0.135 |
| Gm2420 | 121.693 | 0.501709 | 0.26516 | 1.892118 | 0.05848 | 0.18391 | 15.865 |
| Gm2423 | 71.6305 | 0.224298 | 0.23812 | 0.941973 | 0.34621 | 0.55684 | 0.733 |
| Gm24379 | 20.8473 | 0.301849 | 0.44475 | 0.678687 | 0.49734 | 0.69077 | 0.086 |
| Gm24400 | 15.1375 | 0.760231 | 0.48717 | 1.560517 | 0.11864 | 0.29055 | 0.107 |
| Gm24409 | 27.5351 | -0.14502 | 0.32945 | -0.44018 | 0.65981 | 0.81183 | 0.152 |
| Gm24430 | 22.4826 | 0.181307 | 0.35172 | 0.515482 | 0.60622 | 0.77465 | 0.104 |
| Gm2446 | 27.1703 | -1.4827 | 1.25913 | -1.17756 | 0.23897 | 0.44478 | 12.785 |
| Gm2449 | 30.4447 | -0.34488 | 0.38875 | -0.88713 | 0.37501 | 0.58517 | 7.415 |
| Gm24494 | 29.8804 | 0.222951 | 0.36476 | 0.611223 | 0.54105 | 0.72668 | 0.081 |
| Gm24497 | 33.7397 | 0.62389 | 0.40425 | 1.54334 | 0.12275 | 0.29667 | 0.191 |
| Gm2451 | 12.4164 | -0.37055 | 0.45416 | -0.8159 | 0.41456 | 0.62254 | 0.999 |
| Gm24564 | 97.0021 | 0.04658 | 0.20655 | 0.225519 | 0.82158 | 0.90766 | 0.075 |
| Gm24608 | 12.7035 | 0.332729 | 0.51672 | 0.643929 | 0.51962 | 0.70925 | 0.136 |
| Gm24610 | 36.589 | 0.560728 | 0.32518 | 1.72438 | 0.08464 | 0.23318 | 0.109 |
| Gm2464 | 38.3467 | 0.127597 | 0.31986 | 0.398919 | 0.68995 | 0.83121 | 1.391 |

| | | | | | | | |
|---|---|---|---|---|---|---|---|
| Gm24644 | 30.5068 | 0.225883 | 0.37955 | 0.595138 | 0.55175 | 0.73471 | 0.139 |
| Gm24671 | 14.2013 | 1.048562 | 0.54885 | 1.910485 | 0.05607 | 0.17901 | 0.158 |
| Gm24727 | 12.5324 | 0.297792 | 0.50026 | 0.595274 | 0.55166 | 0.73468 | 0.144 |
| Gm24966 | 131 | 0.301207 | 0.22181 | 1.35798 | 0.17447 | 0.36611 | 0.079 |
| Gm25005 | 15.6351 | 0.331194 | 0.47055 | 0.703846 | 0.48153 | 0.67857 | 0.156 |
| Gm25014 | 29.8762 | 0.408486 | 0.41443 | 0.985667 | 0.3243 | 0.53539 | 0.103 |
| Gm25047 | 146.273 | 0.127749 | 0.1873 | 0.682062 | 0.4952 | 0.68921 | 0.104 |
| Gm25076 | 13.5789 | 0.088969 | 0.60298 | 0.147549 | 0.8827 | 0.9416 | 0.132 |
| Gm25107 | 28.7512 | -0.00165 | 0.43323 | -0.00382 | 0.99696 | 0.99828 | 0.129 |
| Gm25117 | 44.432 | -0.07445 | 0.28153 | -0.26443 | 0.79145 | 0.89159 | 0.144 |
| Gm25129 | 237.488 | -0.15091 | 0.13973 | -1.07999 | 0.28015 | 0.49017 | 0.133 |
| Gm25131 | 24.5062 | 0.821943 | 0.37771 | 2.176131 | 0.02955 | 0.11694 | 0.151 |
| Gm25134 | 14.3929 | 0.47446 | 0.54761 | 0.866417 | 0.38626 | 0.59571 | 0.16 |
| Gm25259 | 32.0157 | 0.309091 | 0.41974 | 0.73639 | 0.46149 | 0.66207 | 0.193 |
| Gm25321 | 13.4152 | 0.725566 | 0.54895 | 1.321731 | 0.18626 | 0.38115 | 0.102 |
| Gm25328 | 47.1079 | 0.220226 | 0.27598 | 0.797983 | 0.42488 | 0.6314 | 0.107 |
| Gm25364 | 21.2159 | 0.68878 | 0.44688 | 1.541292 | 0.12325 | 0.29741 | 0.107 |
| Gm2541 | 45.3283 | -0.32372 | 0.30045 | -1.07744 | 0.28128 | 0.49115 | 0.711 |
| Gm25411 | 31.4267 | 0.38396 | 0.39566 | 0.970429 | 0.33183 | 0.54278 | 0.151 |
| Gm25432 | 27.4612 | -0.20524 | 0.38756 | -0.52957 | 0.59641 | 0.76828 | 0.728 |
| Gm25492 | 42.1849 | -0.06052 | 0.33961 | -0.1782 | 0.85856 | 0.92859 | 0.129 |
| Gm25588 | 15.9763 | -0.25403 | 0.46139 | -0.55058 | 0.58192 | 0.75769 | 0.14 |
| Gm25596 | 46.8798 | -0.15903 | 0.26011 | -0.61138 | 0.54095 | 0.72658 | 0.107 |
| Gm25632 | 36.8245 | 0.108692 | 0.35505 | 0.306132 | 0.7595 | 0.87373 | 0.085 |
| Gm25658 | 16.9897 | 1.16224 | 0.52849 | 2.199154 | 0.02787 | 0.11224 | 0.128 |
| Gm2573 | 28.0516 | 0.237951 | 0.4036 | 0.589565 | 0.55548 | 0.73789 | 0.318 |
| Gm25777 | 27.8238 | -0.26945 | 0.32952 | -0.81771 | 0.41352 | 0.6216 | 0.126 |
| Gm25821 | 27.7213 | 0.460997 | 0.38461 | 1.198599 | 0.23068 | 0.43503 | 0.133 |
| Gm25835 | 27.506 | -0.05271 | 0.39156 | -0.1346 | 0.89293 | 0.94632 | 0.144 |
| Gm25857 | 16.7212 | 0.187467 | 0.42218 | 0.444046 | 0.65701 | 0.80951 | 0.129 |
| Gm25939 | 22.5931 | 0.421816 | 0.47061 | 0.896316 | 0.37008 | 0.58043 | 0.191 |
| Gm25965 | 17.8931 | 0.663411 | 0.40702 | 1.629929 | 0.10312 | 0.26535 | 0.093 |
| Gm25982 | 35.5031 | 0.260836 | 0.28498 | 0.915291 | 0.36004 | 0.57047 | 0.101 |
| Gm2606 | 45.6587 | 0.649089 | 0.32567 | 1.993096 | 0.04625 | 0.15713 | 1.002 |

| | | | | | | | |
|---|---|---|---|---|---|---|---|
| Gm26109 | 26.811 | -0.36387 | 0.40076 | -0.90793 | 0.36391 | 0.57434 | 0.07 |
| Gm26203 | 21.3841 | -0.15567 | 0.39736 | -0.39176 | 0.69523 | 0.83485 | 0.114 |
| Gm26230 | 12.4738 | -0.34479 | 0.58347 | -0.59093 | 0.55457 | 0.73711 | 0.079 |
| Gm26249 | 14.3009 | 0.000461 | 0.44722 | 0.001031 | 0.99918 | 0.99967 | 0.118 |
| Gm26254 | 19.9783 | 0.060775 | 0.39587 | 0.153523 | 0.87799 | 0.93893 | 0.118 |
| Gm2629 | 236.674 | -0.1811 | 0.13629 | -1.32879 | 0.18392 | 0.37815 | 81.514 |
| Gm26347 | 23.2739 | -0.0718 | 0.36363 | -0.19746 | 0.84347 | 0.91961 | 0.131 |
| Gm26387 | 15.8773 | -0.32832 | 0.51398 | -0.63878 | 0.52297 | 0.71143 | 0.132 |
| Gm26397 | 62.2026 | 0.65101 | 0.2923 | 2.227213 | 0.02593 | 0.10668 | 0.134 |
| Gm26446 | 28.7474 | 0.816755 | 0.38952 | 2.096806 | 0.03601 | 0.13358 | 0.103 |
| Gm26461 | 20.9337 | 0.990705 | 0.46451 | 2.132781 | 0.03294 | 0.1261 | 0.29 |
| Gm26493 | 19.8655 | -0.16881 | 0.39076 | -0.43201 | 0.66573 | 0.81552 | 0.136 |
| Gm26498 | 17.4106 | 0.18826 | 0.49941 | 0.376967 | 0.7062 | 0.84153 | 0.079 |
| Gm26512 | 82.8539 | -0.20156 | 0.19529 | -1.0321 | 0.30203 | 0.51176 | 131.035 |
| Gm26513 | 21.9381 | 0.39987 | 0.42636 | 0.937881 | 0.34831 | 0.55878 | 2.489 |
| Gm26520 | 14.4653 | 0.794067 | 0.51456 | 1.54319 | 0.12278 | 0.29669 | 13.338 |
| Gm26532 | 17.0647 | -0.7234 | 0.45333 | -1.59574 | 0.11055 | 0.27727 | 0.69 |
| Gm26533 | 12.5008 | 0.434988 | 0.46179 | 0.941963 | 0.34621 | 0.55684 | 7.82 |
| Gm26537 | 12.5509 | 0.485291 | 0.49888 | 0.972759 | 0.33067 | 0.54192 | 9.738 |
| Gm26542 | 30.4658 | 0.679164 | 0.35285 | 1.924799 | 0.05425 | 0.17515 | 51.887 |
| Gm26549 | 139.93 | 0.281997 | 0.14525 | 1.941409 | 0.05221 | 0.17068 | 13.903 |
| Gm26555 | 43.1352 | 1.121224 | 0.26928 | 4.163738 | 3.13E-05 | 0.00119 | 17.963 |
| Gm26559 | 46.1212 | -0.25801 | 0.37844 | -0.68178 | 0.49538 | 0.68937 | 5.132 |
| Gm26575 | 82.3022 | 0.414867 | 0.22098 | 1.877411 | 0.06046 | 0.18759 | 2.232 |
| Gm26577 | 49.3581 | 0.478483 | 0.29283 | 1.633969 | 0.10227 | 0.26399 | 9.281 |
| Gm26586 | 14.4848 | 0.301737 | 0.50432 | 0.598304 | 0.54964 | 0.73326 | 3.432 |
| Gm26588 | 119.938 | -0.18312 | 0.19635 | -0.93259 | 0.35103 | 0.56158 | 9.439 |
| Gm26592 | 81.2715 | 0.341488 | 0.24672 | 1.384086 | 0.16633 | 0.356 | 3.493 |
| Gm26594 | 77.2578 | -0.02306 | 0.21606 | -0.10674 | 0.91499 | 0.95656 | 5.52 |
| Gm266 | 158.848 | 0.367492 | 0.21061 | 1.744925 | 0.081 | 0.22645 | 1.215 |
| Gm26604 | 52.0353 | 1.094604 | 0.28762 | 3.805724 | 0.00014 | 0.00337 | 9.54 |
| Gm26614 | 47.6172 | 0.571472 | 0.28419 | 2.010905 | 0.04434 | 0.15276 | 8.059 |
| Gm26616 | 13.2024 | 0.189609 | 0.5087 | 0.372734 | 0.70935 | 0.84345 | 3.107 |
| Gm26618 | 34.191 | 0.567665 | 0.32579 | 1.742434 | 0.08143 | 0.22723 | 65.214 |

| | | | | | | | |
|---|---|---|---|---|---|---|---|
| Gm26621 | 90.0208 | -0.0339 | 0.23092 | -0.14678 | 0.8833 | 0.94179 | 2.716 |
| Gm26628 | 122.312 | 0.465657 | 0.21021 | 2.21522 | 0.02674 | 0.10907 | 2.631 |
| Gm26631 | 46.5308 | 0.030651 | 0.28862 | 0.106199 | 0.91542 | 0.95671 | 6.343 |
| Gm26633 | 15.0454 | -0.71261 | 0.53542 | -1.33094 | 0.18321 | 0.3773 | 35.271 |
| Gm26638 | 15.9119 | -0.71912 | 0.51284 | -1.40223 | 0.16085 | 0.34901 | 10.699 |
| Gm26642 | 123.4 | -0.01539 | 0.16878 | -0.09117 | 0.92736 | 0.96344 | 17.883 |
| Gm26645 | 80.4171 | -0.02115 | 0.24558 | -0.08614 | 0.93135 | 0.96532 | 1.316 |
| Gm26647 | 46.7552 | 1.076626 | 0.32047 | 3.359548 | 0.00078 | 0.01076 | 3.483 |
| Gm26648 | 227.841 | 0.500755 | 0.13411 | 3.733845 | 0.00019 | 0.00415 | 7.789 |
| Gm26652 | 26.6429 | 0.221391 | 0.3706 | 0.597392 | 0.55025 | 0.7338 | 35.494 |
| Gm26653 | 22.0056 | -0.01713 | 0.34482 | -0.04968 | 0.96038 | 0.98063 | 2.604 |
| Gm26658 | 24.8588 | 0.551367 | 0.46438 | 1.187322 | 0.2351 | 0.44032 | 8.782 |
| Gm26660 | 260.988 | 0.094314 | 0.13536 | 0.696772 | 0.48595 | 0.68206 | 63.737 |
| Gm26666 | 31.768 | -0.45419 | 0.31421 | -1.44547 | 0.14833 | 0.33196 | 2.872 |
| Gm26671 | 23.6736 | 0.121348 | 0.50645 | 0.239604 | 0.81064 | 0.90256 | 13.128 |
| Gm26672 | 35.8071 | 0.17049 | 0.29743 | 0.573216 | 0.5665 | 0.74654 | 21.684 |
| Gm26673 | 48.246 | -0.10755 | 0.26134 | -0.41151 | 0.6807 | 0.82518 | 22.918 |
| Gm26676 | 84.6928 | 0.28083 | 0.2397 | 1.171591 | 0.24136 | 0.44743 | 3.271 |
| Gm26679 | 21.8178 | -0.81691 | 0.35812 | -2.28111 | 0.02254 | 0.09695 | 4.127 |
| Gm26682 | 16.5215 | -0.796 | 0.46944 | -1.69564 | 0.08995 | 0.24325 | 2.893 |
| Gm26684 | 27.173 | -1.29737 | 0.44353 | -2.92507 | 0.00344 | 0.02843 | 19.023 |
| Gm26691 | 112.58 | 0.43116 | 0.24387 | 1.767994 | 0.07706 | 0.21943 | 409.974 |
| Gm26696 | 14.8815 | 2 | 0.79124 | 3.063621 | 0.00219 | 0.02131 | 2.848 |
| Gm2670 | 27.1427 | 0.102274 | 0.35042 | 0.291859 | 0.77039 | 0.87998 | 3.319 |
| Gm26703 | 263.894 | -0.08072 | 0.15405 | -0.524 | 0.60028 | 0.77088 | 10.471 |
| Gm26708 | 14.6051 | 0.22281 | 0.545 | 0.408825 | 0.68267 | 0.82659 | 42.976 |
| Gm26724 | 334.377 | 0.134525 | 0.1151 | 1.168743 | 0.24251 | 0.44857 | 2.04 |
| Gm26736 | 26.2444 | 0.572748 | 0.33129 | 1.728841 | 0.08384 | 0.23172 | 1.236 |
| Gm26737 | 22.4725 | 0.223078 | 0.41021 | 0.543808 | 0.58657 | 0.76038 | 9.038 |
| Gm26740 | 12.7794 | -1.18537 | 0.48856 | -2.42628 | 0.01525 | 0.07527 | 116.617 |
| Gm26744 | 15.4906 | 0.54524 | 0.47529 | 1.147181 | 0.25131 | 0.45873 | 2.326 |
| Gm26747 | 87.5074 | 0.022685 | 0.22575 | 0.100489 | 0.91996 | 0.95948 | 6.806 |
| Gm26748 | 16.5071 | 0.067997 | 0.51655 | 0.131638 | 0.89527 | 0.94744 | 6.783 |
| Gm26749 | 12.3984 | -0.10678 | 0.63031 | -0.16942 | 0.86547 | 0.9318 | 41.739 |

| | | | | | | | |
|---|---|---|---|---|---|---|---|
| Gm26756 | 32.0241 | 0.123524 | 0.29694 | 0.415996 | 0.67741 | 0.8232 | 10.311 |
| Gm26760 | 82.6948 | 0.390469 | 0.2236 | 1.746283 | 0.08076 | 0.22616 | 33.777 |
| Gm26761 | 18.5633 | -1.91401 | 0.58448 | -3.27474 | 0.00106 | 0.01329 | 3.061 |
| Gm26766 | 15.8531 | 0.551066 | 0.59495 | 0.926233 | 0.35433 | 0.56466 | 36.578 |
| Gm26770 | 29.8298 | -0.12105 | 0.33688 | -0.35933 | 0.71935 | 0.85009 | 2.834 |
| Gm26777 | 121.322 | -0.13153 | 0.18896 | -0.69604 | 0.4864 | 0.68236 | 8.966 |
| Gm26779 | 108.049 | 0.194471 | 0.21067 | 0.923125 | 0.35594 | 0.56636 | 4.907 |
| Gm26781 | 120.36 | -0.24615 | 0.17887 | -1.37613 | 0.16878 | 0.35886 | 15 |
| Gm26782 | 1100.55 | -0.31745 | 0.12479 | -2.5439 | 0.01096 | 0.06094 | 14.488 |
| Gm26788 | 30.5875 | -0.57203 | 0.35779 | -1.59881 | 0.10986 | 0.27606 | 12.591 |
| Gm26789 | 137.002 | -0.32823 | 0.19058 | -1.72229 | 0.08502 | 0.23388 | 3.508 |
| Gm26797 | 33.6109 | 0.284914 | 0.374 | 0.761807 | 0.44618 | 0.64983 | 15.06 |
| Gm26805 | 163.105 | 0.347954 | 0.1839 | 1.892056 | 0.05848 | 0.18391 | 6.09 |
| Gm26811 | 74.4183 | 0.585329 | 0.27401 | 2.136189 | 0.03266 | 0.12537 | 22.992 |
| Gm26814 | 46.3428 | 0.744179 | 0.31334 | 2.37497 | 0.01755 | 0.0823 | 4.149 |
| Gm26826 | 44.1046 | 0.456802 | 0.27171 | 1.681225 | 0.09272 | 0.24803 | 7.961 |
| Gm26839 | 13.7639 | -0.67693 | 0.56518 | -1.19773 | 0.23102 | 0.43554 | 56.243 |
| Gm26843 | 48.2206 | -0.49459 | 0.28035 | -1.76415 | 0.07771 | 0.22065 | 41.573 |
| Gm26852 | 107.245 | 0.265079 | 0.23179 | 1.143616 | 0.25278 | 0.46015 | 10.499 |
| Gm26868 | 17.9064 | -0.58628 | 0.56232 | -1.0426 | 0.29713 | 0.50731 | 7.781 |
| Gm26871 | 327.501 | 0.287235 | 0.13962 | 2.057238 | 0.03966 | 0.14223 | 99.699 |
| Gm26873 | 160.501 | 0.204469 | 0.16518 | 1.237826 | 0.21578 | 0.41743 | 22.051 |
| Gm26875 | 12.9373 | 0.008269 | 0.52339 | 0.015799 | 0.98739 | 0.99316 | 0.95 |
| Gm26876 | 68.029 | 0.067839 | 0.24922 | 0.272206 | 0.78546 | 0.88861 | 3.928 |
| Gm26879 | 30.8021 | -0.07122 | 0.36378 | -0.19579 | 0.84477 | 0.9206 | 7.274 |
| Gm26881 | 18.8202 | -0.22136 | 0.40431 | -0.54749 | 0.58404 | 0.75898 | 7.294 |
| Gm26901 | 109.006 | 0.435934 | 0.20361 | 2.141008 | 0.03227 | 0.1242 | 48.464 |
| Gm26902 | 23.6613 | -0.93692 | 0.38941 | -2.40598 | 0.01613 | 0.07809 | 6.739 |
| Gm26910 | 105.822 | 0.247569 | 0.17445 | 1.419122 | 0.15586 | 0.3424 | 1.167 |
| Gm26925 | 14.0934 | -0.19644 | 0.49779 | -0.39464 | 0.69311 | 0.83358 | 0.957 |
| Gm2693 | 16.0157 | 0.70037 | 0.4449 | 1.574205 | 0.11544 | 0.28498 | 8.44 |
| Gm26931 | 70.5949 | 0.567961 | 0.20815 | 2.728547 | 0.00636 | 0.0428 | 0.609 |
| Gm26935 | 112.283 | 0.33552 | 0.20206 | 1.660472 | 0.09682 | 0.25492 | 10.785 |
| Gm26937 | 25.692 | -0.45131 | 0.39754 | -1.13525 | 0.25627 | 0.46422 | 1.008 |

| | | | | | | | |
|---|---|---|---|---|---|---|---|
| Gm2694 | 47.855 | 0.106383 | 0.2731 | 0.389539 | 0.69688 | 0.83579 | 65.099 |
| Gm26944 | 124.003 | 0.437015 | 0.23759 | 1.839358 | 0.06586 | 0.19825 | 7.704 |
| Gm26947 | 35.4038 | -0.0196 | 0.37503 | -0.05227 | 0.95831 | 0.9796 | 19.657 |
| Gm26952 | 19.5043 | -0.14189 | 0.38379 | -0.3697 | 0.71161 | 0.84481 | 1.213 |
| Gm26953 | 17.8511 | 0.351177 | 0.4584 | 0.766099 | 0.44362 | 0.64794 | 18.26 |
| Gm26964 | 108.921 | -0.14885 | 0.1751 | -0.85007 | 0.39529 | 0.6043 | 2.165 |
| Gm26971 | 18.5517 | 0.47997 | 0.4506 | 1.06517 | 0.2868 | 0.4967 | 1.072 |
| Gm26981 | 72.3405 | 0.273624 | 0.27313 | 1.00181 | 0.31644 | 0.52698 | 3.909 |
| Gm26982 | 314.429 | -0.11762 | 0.1272 | -0.92468 | 0.35513 | 0.56548 | 1.151 |
| Gm26984 | 12.4127 | -0.9325 | 0.75491 | -1.23524 | 0.21674 | 0.41862 | 3.51 |
| Gm27000 | 143.548 | 0.30775 | 0.16915 | 1.819395 | 0.06885 | 0.20404 | 0.842 |
| Gm27002 | 34.9937 | -0.15532 | 0.29981 | -0.51807 | 0.60441 | 0.7733 | 3.478 |
| Gm27003 | 118.059 | 0.460671 | 0.17225 | 2.674504 | 0.00748 | 0.04763 | 7.902 |
| Gm27004 | 12.8624 | -0.93477 | 0.6266 | -1.49181 | 0.13575 | 0.31581 | 0.727 |
| Gm27008 | 29.3296 | 0.422682 | 0.31232 | 1.353369 | 0.17594 | 0.36783 | 159.513 |
| Gm27010 | 78.1152 | -0.13134 | 0.24397 | -0.53836 | 0.59033 | 0.76324 | 1.412 |
| Gm27028 | 41.4954 | -0.01236 | 0.27689 | -0.04466 | 0.96438 | 0.98268 | 0.399 |
| Gm27030 | 25.7473 | -0.17555 | 0.35793 | -0.49046 | 0.62381 | 0.78645 | 0.464 |
| Gm27032 | 686.369 | 0.019065 | 0.12961 | 0.147102 | 0.88305 | 0.94166 | 1.116 |
| Gm27042 | 311.588 | 0.143336 | 0.13317 | 1.076314 | 0.28179 | 0.49182 | 5.428 |
| Gm27151 | 14.5091 | -0.05889 | 0.44657 | -0.13186 | 0.89509 | 0.94743 | 10.346 |
| Gm27184 | 16.1492 | 0.085692 | 0.49889 | 0.171765 | 0.86362 | 0.93094 | 6.27 |
| Gm27188 | 17.4319 | 0.53804 | 0.50484 | 1.06577 | 0.28653 | 0.49648 | 22.601 |
| Gm27200 | 45.4367 | 0.361395 | 0.30139 | 1.199101 | 0.23049 | 0.43486 | 0.783 |
| Gm27202 | 120.583 | -0.39704 | 0.17009 | -2.33435 | 0.01958 | 0.08827 | 0.619 |
| Gm27206 | 63.8154 | 0.353725 | 0.2568 | 1.37741 | 0.16839 | 0.35839 | 7.983 |
| Gm27209 | 119.206 | 0.27258 | 0.20676 | 1.318338 | 0.18739 | 0.38267 | 1.878 |
| Gm2721 | 27.0067 | 0.398537 | 0.33547 | 1.187998 | 0.23483 | 0.44015 | 9.759 |
| Gm27219 | 19.9942 | -0.37504 | 0.42987 | -0.87246 | 0.38296 | 0.59222 | 0.742 |
| Gm27239 | 33.8469 | 0.08903 | 0.3504 | 0.254078 | 0.79944 | 0.89649 | 1.12 |
| Gm27243 | 33.4076 | 0.560825 | 0.37617 | 1.490874 | 0.13599 | 0.31623 | 0.521 |
| Gm27252 | 98.3884 | -0.138 | 0.20964 | -0.65826 | 0.51037 | 0.7017 | 3.77 |
| Gm27343 | 20.344 | -0.13607 | 0.40807 | -0.33345 | 0.7388 | 0.86228 | 0.063 |
| Gm27616 | 46.794 | 0.355656 | 0.28171 | 1.262476 | 0.20678 | 0.40622 | 0.063 |

| | | | | | | | |
|---|---|---|---|---|---|---|---|
| Gm2762 | 100.331 | -0.67845 | 0.21963 | -3.08912 | 0.00201 | 0.02016 | 62.013 |
| Gm28064 | 31.2182 | -0.59109 | 0.32632 | -1.8114 | 0.07008 | 0.2067 | 1.204 |
| Gm28085 | 24.6642 | -0.29183 | 0.37746 | -0.77313 | 0.43944 | 0.64481 | 13.573 |
| Gm2810 | 63.8293 | 0.064613 | 0.25878 | 0.249688 | 0.80283 | 0.89815 | 0.99 |
| Gm28100 | 28.2709 | 0.537308 | 0.50256 | 1.06915 | 0.285 | 0.49485 | 14.553 |
| Gm28119 | 93.633 | -0.01025 | 0.27496 | -0.03727 | 0.97027 | 0.98489 | 0.376 |
| Gm28151 | 243.735 | 0.122965 | 0.15511 | 0.792776 | 0.42791 | 0.6345 | 0.951 |
| Gm28153 | 14.0144 | -0.06489 | 0.54888 | -0.11823 | 0.90589 | 0.95277 | 63.503 |
| Gm28154 | 62.1353 | 0.02933 | 0.32188 | 0.091121 | 0.9274 | 0.96344 | 25.998 |
| Gm28175 | 50.9693 | -0.21399 | 0.32168 | -0.66522 | 0.50591 | 0.6974 | 44.664 |
| Gm28192 | 211.99 | 0.110256 | 0.16169 | 0.681901 | 0.4953 | 0.6893 | 0.921 |
| Gm28196 | 19.2468 | -0.00318 | 0.39632 | -0.00802 | 0.9936 | 0.99633 | 0.64 |
| Gm28198 | 25.2427 | 0.303689 | 0.34792 | 0.872859 | 0.38274 | 0.59205 | 13.975 |
| Gm28209 | 20.5329 | -0.58718 | 0.38236 | -1.53568 | 0.12462 | 0.29958 | 3.768 |
| Gm28229 | 77.6026 | 0.082543 | 0.27783 | 0.297094 | 0.76639 | 0.8779 | 2.153 |
| Gm2824 | 319 | -0.71803 | 0.50564 | -1.42004 | 0.1556 | 0.34222 | 45.739 |
| Gm28286 | 16.878 | -1.07102 | 0.47229 | -2.2677 | 0.02335 | 0.09911 | 6.939 |
| Gm28294 | 56.7117 | 0.172999 | 0.30755 | 0.562507 | 0.57377 | 0.75228 | 5.106 |
| Gm28307 | 136.068 | -0.19881 | 0.22315 | -0.89091 | 0.37298 | 0.58339 | 9.243 |
| Gm28320 | 14.0989 | -0.60115 | 0.66736 | -0.9008 | 0.3677 | 0.57815 | 2.273 |
| Gm28322 | 17.0509 | 0.029783 | 0.61558 | 0.048382 | 0.96141 | 0.98121 | 1.475 |
| Gm28370 | 1069.22 | 0.112348 | 0.12891 | 0.871522 | 0.38347 | 0.5928 | 4.817 |
| Gm28372 | 26.7101 | 0.04845 | 0.42085 | 0.115125 | 0.90835 | 0.95403 | 3.018 |
| Gm28373 | 17.2593 | -0.0788 | 0.43878 | -0.17959 | 0.85747 | 0.92804 | 54.317 |
| Gm28386 | 13.9663 | 0.185199 | 0.47185 | 0.392498 | 0.69469 | 0.8344 | 1.849 |
| Gm28424 | 120.628 | 0.283837 | 0.19079 | 1.487669 | 0.13684 | 0.31754 | 2.196 |
| Gm28437 | 2984.61 | -0.17323 | 0.09711 | -1.78384 | 0.07445 | 0.21505 | 0.783 |
| Gm28438 | 402.872 | -0.37539 | 0.18327 | -2.04829 | 0.04053 | 0.14386 | 0.345 |
| Gm28439 | 1533.1 | 0.17556 | 0.18599 | 0.943906 | 0.34522 | 0.55615 | 0.876 |
| Gm28455 | 47.9793 | 0.54219 | 0.25467 | 2.129022 | 0.03325 | 0.12662 | 9.364 |
| Gm28496 | 12.7269 | -1.07168 | 0.58952 | -1.81788 | 0.06908 | 0.20453 | 15.995 |
| Gm28512 | 41.1014 | 0.272762 | 0.26028 | 1.047941 | 0.29467 | 0.505 | 0.866 |
| Gm28514 | 15.1556 | -0.04814 | 0.43009 | -0.11192 | 0.91088 | 0.95502 | 1.311 |
| Gm28523 | 16.2571 | 0.677299 | 0.50901 | 1.330619 | 0.18331 | 0.3774 | 3.09 |

| | | | | | | | |
|---|---|---|---|---|---|---|---|
| Gm28528 | 33.2968 | 0.942835 | 0.392 | 2.405166 | 0.01617 | 0.07819 | 1.676 |
| Gm28529 | 15.3816 | -0.03982 | 0.46813 | -0.08505 | 0.93222 | 0.96589 | 6.82 |
| Gm28535 | 17.6791 | 0.131039 | 0.41293 | 0.317337 | 0.75099 | 0.86939 | 11.362 |
| Gm28551 | 33.9722 | 0.211868 | 0.44629 | 0.474727 | 0.63498 | 0.79438 | 27.161 |
| Gm28557 | 16.9781 | 0.243531 | 0.49877 | 0.488266 | 0.62536 | 0.7877 | 39.206 |
| Gm28578 | 404.778 | 0.508694 | 0.13272 | 3.832919 | 0.00013 | 0.00314 | 37.981 |
| Gm28609 | 81.3316 | 0.481894 | 0.23789 | 2.025717 | 0.04279 | 0.14911 | 21.71 |
| Gm28661 | 3655.4 | 0.22017 | 0.15244 | 1.444264 | 0.14866 | 0.33248 | 0.681 |
| Gm28722 | 159.605 | -0.80287 | 0.57635 | -1.39301 | 0.16362 | 0.35238 | 1.108 |
| Gm28756 | 149.208 | 0.46553 | 0.22035 | 2.112652 | 0.03463 | 0.13033 | 64.595 |
| Gm28784 | 20.3781 | 0.230035 | 0.44191 | 0.520548 | 0.60268 | 0.77235 | 4.701 |
| Gm28800 | 66.1002 | 0.137818 | 0.23275 | 0.59212 | 0.55377 | 0.73648 | 7.937 |
| Gm28836 | 13.8219 | -0.20037 | 0.64334 | -0.31145 | 0.75546 | 0.8714 | 21.235 |
| Gm2885 | 92.6209 | 0.3846 | 0.24132 | 1.593706 | 0.111 | 0.27814 | 29.637 |
| Gm28875 | 59.5078 | -0.30857 | 0.22216 | -1.38896 | 0.16485 | 0.35389 | 1.189 |
| Gm28876 | 24.4057 | -0.93179 | 0.40765 | -2.28574 | 0.02227 | 0.0962 | 6.987 |
| Gm28884 | 228.987 | 1.131239 | 0.17811 | 6.351321 | 2.13E-10 | 1.68E-07 | 4.086 |
| Gm28902 | 16.7611 | 0.865409 | 0.51354 | 1.68518 | 0.09195 | 0.24676 | 3.048 |
| Gm28905 | 33.2877 | 0.824382 | 0.32505 | 2.536156 | 0.01121 | 0.06178 | 231.355 |
| Gm28941 | 77.5061 | -0.12403 | 0.26899 | -0.46109 | 0.64473 | 0.80187 | 1.017 |
| Gm2897 | 38.803 | 0.223475 | 0.37356 | 0.598233 | 0.54968 | 0.73326 | 27.556 |
| Gm28988 | 92.8958 | 0.418382 | 0.18894 | 2.214371 | 0.0268 | 0.10921 | 1.947 |
| Gm28989 | 18.75 | 0.570195 | 0.54603 | 1.044258 | 0.29637 | 0.50667 | 8.727 |
| Gm29017 | 47.3594 | 0.063201 | 0.27199 | 0.232364 | 0.81626 | 0.9054 | 18.274 |
| Gm29019 | 19.9829 | 0.69156 | 0.42562 | 1.624817 | 0.1042 | 0.26697 | 5.106 |
| Gm29050 | 38.0923 | 0.10354 | 0.35846 | 0.288848 | 0.7727 | 0.88166 | 30.89 |
| Gm29054 | 14.0619 | -0.41524 | 0.60504 | -0.68631 | 0.49252 | 0.68714 | 1.45 |
| Gm29083 | 36.0086 | -0.60432 | 0.40429 | -1.49478 | 0.13497 | 0.31486 | 3.351 |
| Gm29093 | 122.44 | 0.166907 | 0.22366 | 0.74624 | 0.45552 | 0.65746 | 1.826 |
| Gm29123 | 20.3162 | -0.00756 | 0.4812 | -0.01571 | 0.98747 | 0.99316 | 25.89 |
| Gm29155 | 13.7277 | 0.422847 | 0.45723 | 0.924812 | 0.35506 | 0.56548 | 1.935 |
| Gm29156 | 27.3102 | -0.54391 | 0.39831 | -1.36555 | 0.17208 | 0.36278 | 0.499 |
| Gm29163 | 12.6597 | 0.270651 | 0.52197 | 0.518518 | 0.6041 | 0.7733 | 0.107 |
| Gm29170 | 393.009 | 0.241181 | 0.10169 | 2.371837 | 0.0177 | 0.08269 | 3.666 |

| | | | | | | | |
|---|---|---|---|---|---|---|---|
| Gm29216 | 1888.19 | -0.46096 | 0.35657 | -1.29278 | 0.19609 | 0.39368 | 0.492 |
| Gm29237 | 267.22 | 0.479025 | 0.11423 | 4.193408 | 2.75E-05 | 0.00109 | 1.727 |
| Gm29325 | 20.743 | -0.10578 | 0.43862 | -0.24117 | 0.80943 | 0.90175 | 2.528 |
| Gm29340 | 12.435 | -0.66954 | 0.61706 | -1.08505 | 0.2779 | 0.48787 | 3.854 |
| Gm29367 | 78.8256 | -0.25733 | 0.22886 | -1.12439 | 0.26085 | 0.46941 | 1.327 |
| Gm29374 | 33.994 | 0.877503 | 0.41212 | 2.129266 | 0.03323 | 0.12661 | 4.11 |
| Gm29394 | 19.9801 | 0.39157 | 0.50683 | 0.77259 | 0.43976 | 0.64503 | 48.183 |
| Gm29427 | 17.4014 | -0.54032 | 0.51364 | -1.05193 | 0.29283 | 0.50266 | 45.591 |
| Gm29438 | 89.3208 | -0.09619 | 0.26065 | -0.36904 | 0.7121 | 0.845 | 2.613 |
| Gm29443 | 28.8082 | -0.31496 | 0.34353 | -0.91683 | 0.35923 | 0.56963 | 18.82 |
| Gm29477 | 14.9294 | 1.379126 | 0.57625 | 2.393287 | 0.0167 | 0.07998 | 8.983 |
| Gm29480 | 87.9071 | 0.412638 | 0.1989 | 2.074547 | 0.03803 | 0.13837 | 6.595 |
| Gm29502 | 13.1416 | 0.636149 | 0.76631 | 0.830145 | 0.40646 | 0.6148 | 130.483 |
| Gm29508 | 27.0397 | -0.36512 | 0.37499 | -0.97368 | 0.33022 | 0.54139 | 2.403 |
| Gm29514 | 73.4046 | -0.41183 | 0.24886 | -1.65487 | 0.09795 | 0.2568 | 75.625 |
| Gm29529 | 24.4219 | 1.246369 | 0.44345 | 2.81061 | 0.00494 | 0.03597 | 2.266 |
| Gm29543 | 44.8959 | 0.229607 | 0.27535 | 0.83388 | 0.40435 | 0.61317 | 0.967 |
| Gm29587 | 20.0566 | -0.19082 | 0.36618 | -0.52112 | 0.60228 | 0.77228 | 12.643 |
| Gm29594 | 19.2767 | 0.335727 | 0.37972 | 0.884137 | 0.37662 | 0.58668 | 0.537 |
| Gm29609 | 70.2545 | 0.194573 | 0.28458 | 0.683711 | 0.49416 | 0.68845 | 35.173 |
| Gm2962 | 80.4743 | -0.59094 | 0.36805 | -1.60561 | 0.10836 | 0.27367 | 0.336 |
| Gm29642 | 64.5236 | 0.438407 | 0.23195 | 1.890076 | 0.05875 | 0.1844 | 23.576 |
| Gm29650 | 284.101 | 0.346143 | 0.1634 | 2.118364 | 0.03414 | 0.12905 | 0.742 |
| Gm29666 | 267.658 | 0.473907 | 0.13146 | 3.60492 | 0.00031 | 0.0058 | 9.601 |
| Gm29674 | 22.3994 | -0.64171 | 0.49027 | -1.3089 | 0.19057 | 0.38671 | 5.864 |
| Gm29675 | 116.927 | 0.012196 | 0.22469 | 0.054279 | 0.95671 | 0.9788 | 10.794 |
| Gm29676 | 40.6566 | 0.004778 | 0.27805 | 0.017183 | 0.98629 | 0.99257 | 30.99 |
| Gm29683 | 19.2704 | -0.92449 | 0.71333 | -1.29603 | 0.19497 | 0.39236 | 159.006 |
| Gm29695 | 321.284 | -0.45673 | 0.16964 | -2.69235 | 0.0071 | 0.04587 | 18.09 |
| Gm29707 | 12.4478 | 0.657228 | 0.65165 | 1.008557 | 0.31319 | 0.52364 | 3.734 |
| Gm29724 | 19.4157 | 1.171717 | 0.42465 | 2.759274 | 0.00579 | 0.04027 | 77.119 |
| Gm2974 | 22.762 | -0.34223 | 0.41457 | -0.8255 | 0.40909 | 0.61691 | 22.32 |
| Gm29773 | 35.5901 | 0.098552 | 0.36817 | 0.267682 | 0.78894 | 0.89044 | 4.348 |
| Gm29797 | 34.834 | -0.11669 | 0.33979 | -0.34342 | 0.73129 | 0.85734 | 2.434 |

| | | | | | | | |
|---|---|---|---|---|---|---|---|
| Gm2981 | 117.527 | 0.166129 | 0.19427 | 0.855148 | 0.39247 | 0.60165 | 0.694 |
| Gm2986 | 25.7427 | 0.804517 | 0.3898 | 2.063937 | 0.03902 | 0.14077 | 0.382 |
| Gm2990 | 20.4311 | 0.709082 | 0.52047 | 1.362382 | 0.17308 | 0.3644 | 18.507 |
| Gm2999 | 724.72 | -0.2647 | 0.1538 | -1.72109 | 0.08523 | 0.23406 | 0.538 |
| Gm29994 | 82.5433 | 0.450628 | 0.21341 | 2.111608 | 0.03472 | 0.13049 | 1.193 |
| Gm2a | 2088.57 | -0.26605 | 0.1021 | -2.60581 | 0.00917 | 0.05405 | 14.915 |
| Gm30015 | 21.763 | -0.73934 | 0.47846 | -1.54524 | 0.12229 | 0.29591 | 29.26 |
| Gm30018 | 16.2151 | 1.036861 | 0.56242 | 1.843583 | 0.06524 | 0.19707 | 5.649 |
| Gm30025 | 61.3843 | -0.00145 | 0.29196 | -0.00497 | 0.99603 | 0.99782 | 0.766 |
| Gm30042 | 85.0814 | 0.593883 | 0.22286 | 2.664829 | 0.0077 | 0.04849 | 2.496 |
| Gm30054 | 28.1696 | 0.401674 | 0.37367 | 1.074955 | 0.28239 | 0.49254 | 14.31 |
| Gm30055 | 13.5703 | -0.46792 | 0.44474 | -1.0521 | 0.29275 | 0.50265 | 0.803 |
| Gm30075 | 42.1916 | 0.34258 | 0.28947 | 1.183489 | 0.23662 | 0.44183 | 35.935 |
| Gm30082 | 38.1815 | -0.15243 | 0.29932 | -0.50924 | 0.61059 | 0.77784 | 1.494 |
| Gm30085 | 34.0843 | 0.039319 | 0.32137 | 0.122348 | 0.90262 | 0.95118 | 75.701 |
| Gm30108 | 12.7868 | -0.19838 | 0.46522 | -0.42642 | 0.6698 | 0.81795 | 2.391 |
| Gm30122 | 96.9317 | -0.21184 | 0.23365 | -0.90667 | 0.36458 | 0.57521 | 15.26 |
| Gm30124 | 13.5007 | 0.149769 | 0.51915 | 0.288491 | 0.77297 | 0.88183 | 49.893 |
| Gm30177 | 86.2196 | -1.16626 | 0.42961 | -2.71467 | 0.00663 | 0.04392 | 2.002 |
| Gm30238 | 185.87 | 0.135165 | 0.15291 | 0.883974 | 0.37671 | 0.58676 | 2.603 |
| Gm30329 | 13.7061 | -0.29126 | 0.50649 | -0.57507 | 0.56525 | 0.74547 | 7.295 |
| Gm30340 | 27.6017 | 0.017032 | 0.35732 | 0.047667 | 0.96198 | 0.98159 | 5.732 |
| Gm30371 | 39.9575 | 0.407447 | 0.29067 | 1.401756 | 0.16099 | 0.34918 | 24.988 |
| Gm30382 | 23.0426 | 0.311011 | 0.4128 | 0.753413 | 0.4512 | 0.65377 | 161.058 |
| Gm30400 | 50.8984 | 0.148002 | 0.26492 | 0.558666 | 0.57639 | 0.75396 | 20.456 |
| Gm30504 | 16.8923 | -0.64303 | 0.42527 | -1.51205 | 0.13052 | 0.30883 | 86.061 |
| Gm3052 | 115.635 | 0.448261 | 0.26459 | 1.694154 | 0.09024 | 0.24372 | 2.937 |
| Gm30531 | 18.9607 | 0.597134 | 0.41216 | 1.448798 | 0.14739 | 0.33067 | 40.027 |
| Gm30539 | 25.5184 | -0.23111 | 0.43412 | -0.53237 | 0.59447 | 0.7666 | 2.975 |
| Gm30541 | 60.9953 | -0.42814 | 0.23193 | -1.84602 | 0.06489 | 0.19621 | 5.146 |
| Gm3055 | 56.816 | 0.446508 | 0.27072 | 1.649321 | 0.09908 | 0.25857 | 28.341 |
| Gm30571 | 65.4461 | -0.60142 | 0.24416 | -2.46324 | 0.01377 | 0.07057 | 1.149 |
| Gm30648 | 21.987 | 0.389257 | 0.4355 | 0.893825 | 0.37142 | 0.58171 | 13.291 |
| Gm30655 | 80.0613 | 0.871625 | 0.27247 | 3.198941 | 0.00138 | 0.0156 | 7.512 |

| | | | | | | | |
|---|---|---|---|---|---|---|---|
| Gm30684 | 15.7439 | -1.85026 | 0.56873 | -3.25332 | 0.00114 | 0.01384 | 11.554 |
| Gm30694 | 13.8519 | -1.15904 | 0.56401 | -2.05502 | 0.03988 | 0.14266 | 15.452 |
| Gm30698 | 12.3781 | 0.914538 | 0.53407 | 1.712393 | 0.08682 | 0.23724 | 3.198 |
| Gm30731 | 90.1749 | 0.057003 | 0.24087 | 0.236659 | 0.81292 | 0.90371 | 2.579 |
| Gm30806 | 15.636 | 0.923361 | 0.51924 | 1.7783 | 0.07535 | 0.21653 | 39.674 |
| Gm3086 | 54.9804 | -0.03033 | 0.26635 | -0.11387 | 0.90934 | 0.95456 | 6.328 |
| Gm30873 | 30.4831 | 0.312004 | 0.34316 | 0.909215 | 0.36324 | 0.57357 | 9.775 |
| Gm30881 | 34.0725 | 0.69814 | 0.30659 | 2.277087 | 0.02278 | 0.09764 | 24.48 |
| Gm30929 | 14.1939 | 1.133533 | 0.68263 | 1.660541 | 0.09681 | 0.25492 | 5.582 |
| Gm30934 | 51.2989 | 0.242161 | 0.3305 | 0.732719 | 0.46373 | 0.66349 | 1.641 |
| Gm30938 | 82.1169 | 0.553052 | 0.23473 | 2.356113 | 0.01847 | 0.08514 | 10.436 |
| Gm30970 | 18.5593 | 0.238852 | 0.42802 | 0.558045 | 0.57681 | 0.75414 | 0.363 |
| Gm30984 | 19.3727 | -0.30826 | 0.43761 | -0.70443 | 0.48117 | 0.67822 | 1.242 |
| Gm31019 | 13.8706 | -0.06229 | 0.50008 | -0.12455 | 0.90088 | 0.95027 | 2.927 |
| Gm31024 | 68.4402 | 0.287605 | 0.26117 | 1.101213 | 0.2708 | 0.48026 | 2.353 |
| Gm31105 | 12.5202 | -0.21189 | 0.50471 | -0.41982 | 0.67461 | 0.82139 | 1.23 |
| Gm31152 | 200.562 | 0.579296 | 0.16964 | 3.414843 | 0.00064 | 0.00933 | 7.7 |
| Gm31166 | 330.084 | -0.35962 | 0.18054 | -1.99193 | 0.04638 | 0.15741 | 39.61 |
| Gm31218 | 40.7349 | -0.23084 | 0.257 | -0.89821 | 0.36907 | 0.57964 | 59.175 |
| Gm31224 | 18.9952 | 0.337068 | 0.39201 | 0.859838 | 0.38988 | 0.59898 | 16.025 |
| Gm31227 | 13.7517 | 0.313521 | 0.73396 | 0.427163 | 0.66926 | 0.81761 | 34.787 |
| Gm31251 | 92.4033 | 0.972976 | 0.37378 | 2.603048 | 0.00924 | 0.05436 | 12.209 |
| Gm31282 | 53.7747 | 0.340582 | 0.28442 | 1.197448 | 0.23113 | 0.43564 | 18.225 |
| Gm31305 | 242.199 | 0.158818 | 0.14114 | 1.125268 | 0.26048 | 0.46896 | 5.828 |
| Gm31323 | 35.7343 | 0.073159 | 0.4069 | 0.179796 | 0.85731 | 0.92794 | 33.341 |
| Gm31326 | 13.9869 | 1.406216 | 0.53613 | 2.622913 | 0.00872 | 0.05239 | 1.048 |
| Gm31333 | 20.7802 | 0.261824 | 0.42096 | 0.621966 | 0.53396 | 0.72115 | 60.602 |
| Gm31356 | 205.577 | 0.471337 | 0.23815 | 1.979144 | 0.0478 | 0.16084 | 18.314 |
| Gm31373 | 17.4361 | -0.19492 | 0.52572 | -0.37077 | 0.71081 | 0.84441 | 1.986 |
| Gm31462 | 77.2177 | 0.041632 | 0.23336 | 0.178397 | 0.85841 | 0.92856 | 1.004 |
| Gm31479 | 19.9751 | 0.487753 | 0.46661 | 1.045301 | 0.29588 | 0.50624 | 2.43 |
| Gm31508 | 19.8046 | 0.965662 | 0.40349 | 2.393302 | 0.0167 | 0.07998 | 131.646 |
| Gm31513 | 25.3307 | 0.18491 | 0.45147 | 0.409571 | 0.68212 | 0.82631 | 3.612 |
| Gm31518 | 49.4036 | -0.49036 | 0.28061 | -1.74747 | 0.08056 | 0.22592 | 28.088 |

| | | | | | | | |
|---|---|---|---|---|---|---|---|
| Gm31614 | 113.439 | -0.31712 | 0.21571 | -1.47012 | 0.14153 | 0.32333 | 7.278 |
| Gm31645 | 33.0826 | 0.360413 | 0.31673 | 1.137906 | 0.25516 | 0.46286 | 8.462 |
| Gm31728 | 29.8213 | 0.127936 | 0.33209 | 0.385251 | 0.70005 | 0.83797 | 19.853 |
| Gm31749 | 41.3048 | 0.029526 | 0.29642 | 0.099609 | 0.92066 | 0.95978 | 3.824 |
| Gm31763 | 113.764 | 0.124577 | 0.17326 | 0.719017 | 0.47213 | 0.67052 | 36.307 |
| Gm31793 | 42.953 | 0.916497 | 0.36104 | 2.538469 | 0.01113 | 0.0615 | 30.411 |
| Gm31805 | 20.5015 | 0.590094 | 0.55158 | 1.069823 | 0.2847 | 0.49463 | 5.078 |
| Gm31812 | 18.1302 | -0.82045 | 0.414 | -1.98175 | 0.04751 | 0.16022 | 5.631 |
| Gm31816 | 13.9184 | -1.19846 | 0.57411 | -2.0875 | 0.03684 | 0.13544 | 126.324 |
| Gm31848 | 15.9109 | 0.66974 | 0.44261 | 1.513162 | 0.13024 | 0.30834 | 3.109 |
| Gm31907 | 23.8612 | -0.13487 | 0.45509 | -0.29637 | 0.76695 | 0.87819 | 58.541 |
| Gm31938 | 19.3543 | 0.275236 | 0.45132 | 0.60985 | 0.54196 | 0.72752 | 5.01 |
| Gm32014 | 26.5015 | 0.65284 | 0.56467 | 1.156154 | 0.24762 | 0.4546 | 96.916 |
| Gm32031 | 42.7737 | -0.02029 | 0.30872 | -0.06571 | 0.94761 | 0.97364 | 1.497 |
| Gm32151 | 16.901 | 0.336722 | 0.43406 | 0.775751 | 0.4379 | 0.64331 | 4.245 |
| Gm32200 | 84.2486 | -0.48182 | 0.2112 | -2.28134 | 0.02253 | 0.09692 | 28.709 |
| Gm32219 | 13.7554 | -0.15831 | 0.47204 | -0.33538 | 0.73734 | 0.86132 | 21.99 |
| Gm32224 | 22.9694 | -0.16248 | 0.41233 | -0.39404 | 0.69355 | 0.83381 | 8.065 |
| Gm3226 | 30.807 | -0.76105 | 0.40532 | -1.87765 | 0.06043 | 0.18755 | 1.342 |
| Gm32261 | 214.339 | -0.19271 | 0.18051 | -1.06756 | 0.28572 | 0.49571 | 2.651 |
| Gm3227 | 44.5744 | -0.46964 | 0.38402 | -1.22295 | 0.22135 | 0.42428 | 6.344 |
| Gm32281 | 91.5252 | 0.605378 | 0.26066 | 2.322443 | 0.02021 | 0.09014 | 4.417 |
| Gm32282 | 44.6299 | -0.10288 | 0.36365 | -0.2829 | 0.77725 | 0.88407 | 0.77 |
| Gm32284 | 17.4679 | 2 | 0.56861 | 4.327486 | 1.51E-05 | 0.00074 | 26.427 |
| Gm3235 | 30.8041 | 0.072596 | 0.42894 | 0.169243 | 0.86561 | 0.93189 | 9.167 |
| Gm32391 | 269.277 | -0.19016 | 0.17892 | -1.06285 | 0.28785 | 0.49792 | 6.018 |
| Gm32442 | 30.8048 | -0.10432 | 0.32542 | -0.32056 | 0.74854 | 0.86814 | 176.764 |
| Gm32444 | 412.311 | 0.546 | 0.2307 | 2.366685 | 0.01795 | 0.08348 | 3.511 |
| Gm3248 | 30.6855 | -0.06466 | 0.36687 | -0.17625 | 0.8601 | 0.92937 | 22.521 |
| Gm32514 | 23.3138 | 0.438048 | 0.37926 | 1.154996 | 0.24809 | 0.45524 | 1.426 |
| Gm3252 | 19.4161 | -0.12883 | 0.39653 | -0.3249 | 0.74526 | 0.866 | 20.719 |
| Gm32568 | 28.2295 | 0.455646 | 0.54037 | 0.843211 | 0.39911 | 0.60807 | 1.888 |
| Gm3257 | 105.069 | 0.355767 | 0.27501 | 1.293662 | 0.19578 | 0.39346 | 0.675 |
| Gm32592 | 34.863 | 0.74441 | 0.3513 | 2.118991 | 0.03409 | 0.12893 | 126.816 |

| | | | | | | | |
|---|---|---|---|---|---|---|---|
| Gm32618 | 291.604 | 0.534297 | 0.12166 | 4.391577 | 1.13E-05 | 0.00061 | 17.154 |
| Gm32633 | 29.8198 | -0.41372 | 0.41605 | -0.9944 | 0.32003 | 0.53067 | 19.524 |
| Gm32635 | 20.1999 | 0.722706 | 0.41893 | 1.725118 | 0.08451 | 0.233 | 4.149 |
| Gm3264 | 14.5288 | -0.3186 | 0.50806 | -0.62709 | 0.5306 | 0.71813 | 20.276 |
| Gm32647 | 21.7852 | 0.746325 | 0.47172 | 1.58212 | 0.11362 | 0.28217 | 666.481 |
| Gm32679 | 27.7839 | -0.28916 | 0.33505 | -0.86305 | 0.38811 | 0.59748 | 6.232 |
| Gm32687 | 61.1314 | 0.531162 | 0.33546 | 1.583374 | 0.11334 | 0.28184 | 17.148 |
| Gm32699 | 204.744 | 0.663019 | 0.16555 | 4.004894 | 6.20E-05 | 0.00194 | 2.024 |
| Gm32786 | 15.1422 | -0.42042 | 0.47449 | -0.88606 | 0.37559 | 0.58577 | 29.432 |
| Gm32815 | 28.3453 | 0.967508 | 0.35378 | 2.734743 | 0.00624 | 0.04229 | 35.591 |
| Gm32834 | 17.5046 | 0.061767 | 0.38261 | 0.161436 | 0.87175 | 0.9352 | 40.551 |
| Gm32856 | 159.901 | -0.02971 | 0.19136 | -0.15527 | 0.87661 | 0.93831 | 19.265 |
| Gm32926 | 29.6461 | 0.599341 | 0.35161 | 1.704574 | 0.08827 | 0.23998 | 94.511 |
| Gm3294 | 330.297 | -0.2421 | 0.14257 | -1.69812 | 0.08949 | 0.24237 | 47.471 |
| Gm33023 | 22.7832 | 0.55414 | 0.40439 | 1.370309 | 0.17059 | 0.361 | 5.661 |
| Gm33027 | 56.4378 | -0.14572 | 0.32072 | -0.45433 | 0.64959 | 0.80496 | 22.051 |
| Gm33045 | 55.9244 | 0.311012 | 0.23086 | 1.347217 | 0.17791 | 0.3702 | 15.443 |
| Gm33195 | 105.68 | 0.155581 | 0.16851 | 0.923251 | 0.35588 | 0.56636 | 3.921 |
| Gm33228 | 45.209 | 0.119775 | 0.46628 | 0.256873 | 0.79728 | 0.89545 | 208.093 |
| Gm3325 | 67.7628 | 0.194267 | 0.32323 | 0.601014 | 0.54783 | 0.73193 | 35.382 |
| Gm33280 | 35.8858 | 0.015836 | 0.26773 | 0.059148 | 0.95283 | 0.97684 | 6.729 |
| Gm33320 | 66.2611 | 0.190841 | 0.22364 | 0.853338 | 0.39347 | 0.60255 | 0.53 |
| Gm33366 | 24.2368 | -0.13866 | 0.35223 | -0.39365 | 0.69384 | 0.83397 | 3.827 |
| Gm33373 | 15.5241 | -0.12553 | 0.47955 | -0.26176 | 0.79351 | 0.89304 | 10.479 |
| Gm33378 | 46.6541 | -0.20561 | 0.30867 | -0.66612 | 0.50534 | 0.69699 | 2.758 |
| Gm33467 | 15.2754 | 0.230582 | 0.53159 | 0.433759 | 0.66446 | 0.81456 | 0.961 |
| Gm33508 | 17.9463 | 0.136945 | 0.4351 | 0.314743 | 0.75296 | 0.87023 | 12.67 |
| Gm33524 | 35.3897 | 0.110137 | 0.30644 | 0.359405 | 0.71929 | 0.85007 | 39.137 |
| Gm33533 | 69.7735 | 0.269865 | 0.23929 | 1.127779 | 0.25941 | 0.46775 | 5.745 |
| Gm33594 | 19.4253 | -0.24052 | 0.45668 | -0.52668 | 0.59842 | 0.76969 | 3.826 |
| Gm336 | 25.9233 | -0.08705 | 0.3527 | -0.2468 | 0.80506 | 0.8993 | 1.48 |
| Gm3362 | 418.143 | 0.104577 | 0.42723 | 0.244779 | 0.80663 | 0.90042 | 0.466 |
| Gm3365 | 121.619 | -0.05845 | 0.19334 | -0.30233 | 0.7624 | 0.87544 | 30.361 |
| Gm33651 | 160.777 | 0.772885 | 0.29267 | 2.640834 | 0.00827 | 0.05066 | 12.557 |

| | | | | | | | |
|---|---|---|---|---|---|---|---|
| Gm33680 | 35.0628 | 0.939216 | 0.40411 | 2.324149 | 0.02012 | 0.08985 | 80.405 |
| Gm33699 | 21.1603 | -0.04284 | 0.48263 | -0.08876 | 0.92927 | 0.9644 | 26.905 |
| Gm3375 | 75.1654 | -0.51444 | 0.23138 | -2.22329 | 0.0262 | 0.10748 | 0.795 |
| Gm33869 | 428.479 | 0.738956 | 0.22523 | 3.280915 | 0.00103 | 0.01309 | 3.142 |
| Gm33948 | 28.3956 | 0.664632 | 0.35701 | 1.861688 | 0.06265 | 0.19209 | 135.327 |
| Gm33973 | 15.8687 | 0.92166 | 0.44971 | 2.049461 | 0.04042 | 0.14368 | 0.534 |
| Gm33989 | 1851.79 | -0.00043 | 0.14145 | -0.00306 | 0.99756 | 0.99873 | 36.188 |
| Gm34004 | 27.1632 | 0.880181 | 0.38763 | 2.270668 | 0.02317 | 0.09863 | 35.889 |
| Gm34006 | 38.3845 | -0.36083 | 0.29186 | -1.2363 | 0.21635 | 0.41825 | 3.318 |
| Gm34081 | 67.8617 | 1.446354 | 0.37053 | 3.903465 | 9.48E-05 | 0.00257 | 23.212 |
| Gm34121 | 48.7359 | 0.3908 | 0.24687 | 1.583012 | 0.11342 | 0.28194 | 11.109 |
| Gm34220 | 274.935 | 0.173013 | 0.15177 | 1.139942 | 0.25431 | 0.46169 | 2.709 |
| Gm34245 | 21.4874 | 0.070871 | 0.45954 | 0.154222 | 0.87743 | 0.93876 | 2.857 |
| Gm34256 | 26.2632 | 0.109318 | 0.37395 | 0.29233 | 0.77003 | 0.87989 | 3.025 |
| Gm34267 | 152.858 | -0.04814 | 0.16249 | -0.29624 | 0.76705 | 0.8782 | 4.284 |
| Gm34280 | 13.4268 | 0.36944 | 0.49913 | 0.740173 | 0.4592 | 0.65987 | 18.481 |
| Gm34294 | 25.5683 | 0.786383 | 0.3748 | 2.098147 | 0.03589 | 0.13334 | 4.263 |
| Gm34299 | 28.7816 | 0.270744 | 0.37115 | 0.729466 | 0.46572 | 0.665 | 3.853 |
| Gm34304 | 87.9348 | 0.454176 | 0.22186 | 2.047105 | 0.04065 | 0.1442 | 13.16 |
| Gm34333 | 22.0589 | 1.023572 | 0.51796 | 1.976162 | 0.04814 | 0.16167 | 13.458 |
| Gm3435 | 191.72 | 0.617472 | 0.16138 | 3.826208 | 0.00013 | 0.00322 | 12.274 |
| Gm34354 | 100.807 | -0.20472 | 0.30425 | -0.67286 | 0.50103 | 0.69382 | 58.455 |
| Gm34418 | 21.3178 | -0.25477 | 0.44253 | -0.57571 | 0.56481 | 0.74514 | 2.723 |
| Gm34425 | 20.6592 | 0.024171 | 0.39297 | 0.061509 | 0.95095 | 0.9759 | 4.701 |
| Gm34510 | 17.4403 | 0.165122 | 0.43022 | 0.38381 | 0.70112 | 0.83856 | 12.202 |
| Gm34552 | 29.0943 | 0.526389 | 0.35941 | 1.464583 | 0.14303 | 0.32511 | 2.962 |
| Gm34574 | 27.6437 | 0.379128 | 0.33946 | 1.116847 | 0.26406 | 0.47294 | 12.465 |
| Gm34583 | 991.351 | -1.09481 | 0.38903 | -2.81421 | 0.00489 | 0.03571 | 12.269 |
| Gm34589 | 19.1266 | -0.51249 | 0.41331 | -1.23997 | 0.21499 | 0.41637 | 16.595 |
| Gm34590 | 14.6401 | -0.06792 | 0.55581 | -0.1222 | 0.90274 | 0.95118 | 10.742 |
| Gm34655 | 68.0595 | 0.103683 | 0.22447 | 0.461902 | 0.64415 | 0.80146 | 8.226 |
| Gm34667 | 19.2512 | 1.22647 | 0.43372 | 2.827813 | 0.00469 | 0.03485 | 46.505 |
| Gm34678 | 14.6433 | 0.101481 | 0.44405 | 0.228538 | 0.81923 | 0.90697 | 16.669 |
| Gm3468 | 49.4409 | -0.087 | 0.30853 | -0.28198 | 0.77796 | 0.88423 | 21.371 |

| | | | | | | | |
|---|---|---|---|---|---|---|---|
| Gm34721 | 93.9795 | 0.994761 | 0.26062 | 3.81692 | 0.00014 | 0.0033 | 3.816 |
| Gm34730 | 12.5476 | -0.21023 | 0.6724 | -0.31266 | 0.75454 | 0.87093 | 16.316 |
| Gm34759 | 28.5733 | 0.128562 | 0.37559 | 0.342295 | 0.73213 | 0.85798 | 16.95 |
| Gm34776 | 54.7854 | -0.45513 | 0.3357 | -1.35577 | 0.17517 | 0.36685 | 9.555 |
| Gm34788 | 124.714 | 1.080551 | 0.24029 | 4.496846 | 6.90E-06 | 0.00042 | 7.637 |
| Gm34868 | 12.7154 | 0.736319 | 0.58401 | 1.260805 | 0.20738 | 0.40685 | 19.09 |
| Gm34883 | 44.9769 | 0.496097 | 0.29555 | 1.678538 | 0.09324 | 0.24855 | 1.724 |
| Gm34907 | 25.8067 | 0.556895 | 0.3923 | 1.419577 | 0.15573 | 0.34238 | 48.701 |
| Gm34934 | 37.7687 | -0.33405 | 0.31238 | -1.06936 | 0.28491 | 0.49485 | 16.182 |
| Gm35021 | 23.0472 | -0.22196 | 0.39516 | -0.56169 | 0.57433 | 0.75253 | 33.868 |
| Gm35025 | 14.019 | -0.39636 | 0.45172 | -0.87744 | 0.38025 | 0.58983 | 1.979 |
| Gm35040 | 2086.49 | -0.36307 | 0.07415 | -4.89666 | 9.75E-07 | 0.0001 | 24.022 |
| Gm3510 | 185.431 | 0.257878 | 0.23054 | 1.118577 | 0.26332 | 0.47208 | 9.652 |
| Gm35161 | 54.4176 | -1.15812 | 0.34063 | -3.39994 | 0.00067 | 0.00972 | 32.758 |
| Gm3519 | 15.5521 | 0.888156 | 0.48496 | 1.831408 | 0.06704 | 0.20046 | 2.094 |
| Gm35281 | 15.6615 | 0.679101 | 0.56438 | 1.203266 | 0.22887 | 0.43294 | 65.581 |
| Gm35290 | 57.9259 | 0.428066 | 0.21943 | 1.950795 | 0.05108 | 0.16824 | 1.298 |
| Gm35315 | 51.4422 | 0.455874 | 0.40066 | 1.137815 | 0.2552 | 0.46289 | 18.285 |
| Gm35339 | 683.619 | 0.1993 | 0.12953 | 1.538594 | 0.1239 | 0.2984 | 10.77 |
| Gm35394 | 64.9606 | -0.04985 | 0.41088 | -0.12133 | 0.90343 | 0.95134 | 5.19 |
| Gm35405 | 18.639 | 0.255346 | 0.42193 | 0.605188 | 0.54505 | 0.73012 | 0.4 |
| Gm35438 | 15.9868 | 0.020751 | 0.43808 | 0.047368 | 0.96222 | 0.98168 | 46.694 |
| Gm35501 | 69.3075 | -0.21053 | 0.28665 | -0.73444 | 0.46268 | 0.66283 | 6.844 |
| Gm3551 | 18.6876 | 0.117414 | 0.4163 | 0.28204 | 0.77791 | 0.88423 | 0.664 |
| Gm35549 | 59.2444 | 0.017253 | 0.24739 | 0.069739 | 0.9444 | 0.97211 | 21.26 |
| Gm3555 | 19.1512 | -0.55278 | 0.52149 | -1.05999 | 0.28915 | 0.49936 | 1.395 |
| Gm35558 | 171.753 | -0.16379 | 0.15788 | -1.03744 | 0.29953 | 0.5097 | 2.219 |
| Gm35569 | 24.9866 | -0.68263 | 0.37258 | -1.83216 | 0.06693 | 0.20024 | 6.713 |
| Gm35572 | 19.0602 | -0.32957 | 0.46539 | -0.70816 | 0.47884 | 0.67642 | 29.297 |
| Gm3558 | 21.6262 | 0.157693 | 0.49447 | 0.318912 | 0.74979 | 0.86869 | 23.416 |
| Gm35608 | 54.6291 | 1.670359 | 0.39293 | 4.25105 | 2.13E-05 | 0.00092 | 2.37 |
| Gm35638 | 18.3318 | -0.05054 | 0.46138 | -0.10955 | 0.91277 | 0.95597 | 0.652 |
| Gm3571 | 82.005 | -0.1737 | 0.24276 | -0.7155 | 0.4743 | 0.67238 | 1.06 |
| Gm35725 | 21.0285 | -0.09964 | 0.45243 | -0.22024 | 0.82569 | 0.91002 | 19.858 |

| | | | | | | | |
|---|---|---|---|---|---|---|---|
| Gm35755 | 12.9972 | 1.023538 | 0.64407 | 1.589177 | 0.11202 | 0.27952 | 2.562 |
| Gm35823 | 18.4676 | -1.17115 | 0.4519 | -2.5916 | 0.00955 | 0.05563 | 53.031 |
| Gm35835 | 32.7256 | 0.476256 | 0.40225 | 1.183975 | 0.23642 | 0.44163 | 57.252 |
| Gm35867 | 28.989 | 0.117467 | 0.37067 | 0.316907 | 0.75131 | 0.86954 | 51.618 |
| Gm35883 | 16.4565 | 0.225112 | 0.43648 | 0.515742 | 0.60603 | 0.77461 | 0.615 |
| Gm36012 | 20.6093 | -0.99584 | 0.44335 | -2.24618 | 0.02469 | 0.1031 | 7.909 |
| Gm36037 | 15.2269 | -0.24345 | 0.49113 | -0.49569 | 0.62011 | 0.78385 | 126.524 |
| Gm3604 | 36.3985 | 0.60925 | 0.36392 | 1.674128 | 0.09411 | 0.25029 | 14.85 |
| Gm36070 | 62.0886 | 0.280866 | 0.29901 | 0.939308 | 0.34757 | 0.55804 | 4.336 |
| Gm36120 | 25.5948 | 1.382132 | 0.43679 | 3.164272 | 0.00155 | 0.01704 | 9.445 |
| Gm3617 | 25.9628 | 0.592418 | 0.4085 | 1.450211 | 0.147 | 0.33019 | 0.555 |
| Gm36210 | 50.8901 | -0.51867 | 0.25147 | -2.06254 | 0.03916 | 0.14099 | 8.067 |
| Gm3625 | 33.2712 | -0.21519 | 0.31589 | -0.68122 | 0.49573 | 0.68962 | 3.233 |
| Gm36251 | 26.4092 | -1.73273 | 0.48624 | -3.5635 | 0.00037 | 0.00643 | 52.047 |
| Gm36266 | 39.8746 | 0.378421 | 0.31438 | 1.2037 | 0.22871 | 0.43274 | 2.396 |
| Gm36279 | 14.2196 | -0.37132 | 0.56088 | -0.66204 | 0.50795 | 0.69922 | 31.094 |
| Gm36283 | 23.0057 | -0.11743 | 0.51507 | -0.228 | 0.81965 | 0.90729 | 36.047 |
| Gm36298 | 147.554 | 0.422026 | 0.285 | 1.4808 | 0.13866 | 0.31984 | 3.367 |
| Gm36356 | 28.5635 | -0.0924 | 0.40781 | -0.22659 | 0.82074 | 0.90742 | 5.287 |
| Gm3636 | 96.8011 | -0.01358 | 0.20343 | -0.06674 | 0.94679 | 0.97324 | 6.648 |
| Gm36368 | 13.0645 | 1.27966 | 0.53575 | 2.388526 | 0.01692 | 0.08058 | 12.743 |
| Gm36371 | 70.3916 | -0.21207 | 0.24914 | -0.85123 | 0.39464 | 0.60371 | 1.103 |
| Gm36401 | 27.73 | 0.552675 | 0.48597 | 1.13726 | 0.25543 | 0.4631 | 2.028 |
| Gm36423 | 13.6778 | 0.592594 | 0.46408 | 1.276915 | 0.20163 | 0.40013 | 5.319 |
| Gm36445 | 113.38 | 0.442936 | 0.23523 | 1.882997 | 0.0597 | 0.18611 | 9.243 |
| Gm36447 | 33.5922 | 0.800736 | 0.33422 | 2.395858 | 0.01658 | 0.07964 | 43.743 |
| Gm36457 | 45.0071 | 0.287134 | 0.32549 | 0.882166 | 0.37769 | 0.58778 | 1.83 |
| Gm36495 | 16.351 | 0.07509 | 0.48506 | 0.154807 | 0.87697 | 0.93848 | 6.001 |
| Gm36529 | 15.9813 | 0.364268 | 0.64364 | 0.565948 | 0.57143 | 0.75033 | 19.159 |
| Gm36638 | 20.6211 | -0.42196 | 0.36889 | -1.14387 | 0.25268 | 0.46 | 2.169 |
| Gm3667 | 25.32 | -0.18551 | 0.42845 | -0.43299 | 0.66502 | 0.81509 | 21.461 |
| Gm36693 | 24.4782 | 0.632635 | 0.45339 | 1.395354 | 0.16291 | 0.3514 | 3.792 |
| Gm36737 | 167.062 | 1.122911 | 0.29648 | 3.787528 | 0.00015 | 0.00357 | 6.211 |
| Gm36742 | 30.7425 | -0.71095 | 0.37229 | -1.90967 | 0.05618 | 0.17923 | 1.444 |

| | | | | | | | |
|---|---|---|---|---|---|---|---|
| Gm36756 | 14.533 | 0.699104 | 0.46854 | 1.492074 | 0.13568 | 0.31568 | 28.2 |
| Gm36757 | 33.6288 | -0.5154 | 0.34251 | -1.50477 | 0.13238 | 0.31143 | 3.636 |
| Gm36787 | 60.2639 | -0.21083 | 0.2605 | -0.80934 | 0.41832 | 0.62561 | 6.53 |
| Gm36823 | 34.2868 | -0.10231 | 0.36335 | -0.28159 | 0.77826 | 0.88423 | 17.323 |
| Gm36839 | 36.7123 | 0.229676 | 0.33958 | 0.676356 | 0.49881 | 0.69216 | 2.921 |
| Gm36855 | 44.5836 | -0.45099 | 0.3874 | -1.16416 | 0.24436 | 0.45085 | 14.454 |
| Gm36864 | 22.1125 | 0.012335 | 0.42227 | 0.029212 | 0.9767 | 0.98784 | 7.015 |
| Gm36899 | 140.459 | 0.125877 | 0.18884 | 0.66658 | 0.50504 | 0.69681 | 3.615 |
| Gm36908 | 45.09 | -0.28896 | 0.30627 | -0.94351 | 0.34542 | 0.55616 | 5.963 |
| Gm3693 | 117.58 | 0.546271 | 0.1962 | 2.784311 | 0.00536 | 0.03798 | 14.975 |
| Gm36936 | 286.905 | 0.159599 | 0.15171 | 1.052036 | 0.29278 | 0.50266 | 2.749 |
| Gm36937 | 92.0316 | -0.6274 | 0.2224 | -2.82102 | 0.00479 | 0.0353 | 0.668 |
| Gm36944 | 27.6718 | -0.72009 | 0.37249 | -1.93318 | 0.05321 | 0.17267 | 2.759 |
| Gm36948 | 21.9463 | 0.345731 | 0.43966 | 0.786364 | 0.43165 | 0.63769 | 3.247 |
| Gm3695 | 34.3131 | -0.39497 | 0.3619 | -1.09137 | 0.27511 | 0.48456 | 1.009 |
| Gm36952 | 22.3686 | -0.02374 | 0.60694 | -0.03911 | 0.9688 | 0.98419 | 3.625 |
| Gm3696 | 53.1259 | -0.90639 | 0.24278 | -3.73336 | 0.00019 | 0.00415 | 17.414 |
| Gm36988 | 23.7846 | -0.22449 | 0.36184 | -0.62039 | 0.535 | 0.72197 | 2.408 |
| Gm36989 | 536.861 | 0.605166 | 0.18873 | 3.206494 | 0.00134 | 0.01534 | 2.16 |
| Gm36992 | 18.7184 | 0.204316 | 0.44878 | 0.455275 | 0.64891 | 0.80429 | 4.36 |
| Gm36993 | 12.5664 | 2 | 0.95091 | 2.470257 | 0.0135 | 0.06992 | 2.98 |
| Gm37003 | 31.17 | 0.016966 | 0.37725 | 0.044974 | 0.96413 | 0.98259 | 2.827 |
| Gm37006 | 17.2662 | 0.203286 | 0.59126 | 0.343818 | 0.73098 | 0.85734 | 2.674 |
| Gm37008 | 71.345 | 0.021941 | 0.22943 | 0.095633 | 0.92381 | 0.96165 | 1.079 |
| Gm37018 | 21.9572 | -0.38131 | 0.54799 | -0.69583 | 0.48653 | 0.68239 | 3.869 |
| Gm37019 | 170.604 | -0.04827 | 0.22549 | -0.21408 | 0.83048 | 0.91311 | 2.134 |
| Gm37027 | 101.266 | 0.250647 | 0.20735 | 1.208786 | 0.22674 | 0.43095 | 0.653 |
| Gm37033 | 13.2435 | -0.16232 | 0.51805 | -0.31334 | 0.75403 | 0.87067 | 7.35 |
| Gm37056 | 139.486 | -0.32204 | 0.1941 | -1.65912 | 0.09709 | 0.2554 | 1.865 |
| Gm37062 | 73.2177 | 0.208984 | 0.27688 | 0.75479 | 0.45038 | 0.65308 | 1.011 |
| Gm37067 | 38.3378 | -0.16723 | 0.28036 | -0.59648 | 0.55085 | 0.73431 | 1.628 |
| Gm37069 | 334.129 | -0.73494 | 0.18681 | -3.93426 | 8.35E-05 | 0.00237 | 4.142 |
| Gm37074 | 67.3209 | 0.121525 | 0.25408 | 0.478296 | 0.63244 | 0.79255 | 1.843 |
| Gm37077 | 42.1267 | -0.04149 | 0.32363 | -0.1282 | 0.89799 | 0.94882 | 3.691 |

| | | | | | | | |
|---|---|---|---|---|---|---|---|
| Gm37078 | 32.2038 | 0.262607 | 0.39206 | 0.669806 | 0.50298 | 0.69545 | 2.009 |
| Gm37084 | 20.3015 | -0.24612 | 0.40439 | -0.60863 | 0.54277 | 0.72841 | 5.898 |
| Gm37088 | 18.8077 | 0.263924 | 0.41848 | 0.630674 | 0.52825 | 0.71614 | 0.406 |
| Gm37090 | 536.734 | 0.130567 | 0.11903 | 1.09695 | 0.27266 | 0.48183 | 4.105 |
| Gm37095 | 52.6634 | -0.19463 | 0.24338 | -0.79969 | 0.42389 | 0.63068 | 0.205 |
| Gm37105 | 14.1478 | -0.89884 | 0.57305 | -1.56853 | 0.11676 | 0.28716 | 2.085 |
| Gm37106 | 18.1515 | -0.09905 | 0.47444 | -0.20876 | 0.83463 | 0.91508 | 1.532 |
| Gm37107 | 35.1497 | -0.33323 | 0.34878 | -0.95542 | 0.33937 | 0.5502 | 2.552 |
| Gm37111 | 103.219 | -0.90456 | 0.18355 | -4.92823 | 8.30E-07 | 9.34E-05 | 0.634 |
| Gm37116 | 33.0225 | 0.330346 | 0.324 | 1.019591 | 0.30792 | 0.51771 | 0.147 |
| Gm37120 | 46.659 | -0.79456 | 0.41459 | -1.91649 | 0.0553 | 0.17727 | 5.959 |
| Gm37121 | 102.723 | 0.139468 | 0.24172 | 0.576975 | 0.56396 | 0.74426 | 2.556 |
| Gm37123 | 74.9549 | 0.105744 | 0.22138 | 0.477666 | 0.63289 | 0.79306 | 1.171 |
| Gm37127 | 151.371 | 0.205224 | 0.19145 | 1.07194 | 0.28375 | 0.49384 | 3.036 |
| Gm37132 | 38.6365 | -0.84093 | 0.3574 | -2.35289 | 0.01863 | 0.08554 | 2.238 |
| Gm37139 | 35.035 | 0.49446 | 0.3298 | 1.499272 | 0.1338 | 0.31346 | 4.419 |
| Gm37144 | 29.6142 | 0.300934 | 0.34276 | 0.877967 | 0.37996 | 0.58971 | 1.15 |
| Gm37154 | 77.432 | 0.350136 | 0.19694 | 1.777841 | 0.07543 | 0.21668 | 1.29 |
| Gm37160 | 13.0272 | -0.57088 | 0.53826 | -1.06062 | 0.28886 | 0.49907 | 1.689 |
| Gm37166 | 47.8746 | 0.105791 | 0.30979 | 0.341495 | 0.73273 | 0.85824 | 2.474 |
| Gm37168 | 46.2498 | -0.8685 | 0.45465 | -1.91025 | 0.0561 | 0.17907 | 26.588 |
| Gm37174 | 82.053 | 0.190572 | 0.19582 | 0.973181 | 0.33046 | 0.54167 | 3.058 |
| Gm37176 | 46.8945 | -0.44371 | 0.2866 | -1.54816 | 0.12158 | 0.29498 | 4.051 |
| Gm37180 | 15.5373 | -0.53603 | 0.49412 | -1.08482 | 0.278 | 0.48797 | 2.819 |
| Gm37181 | 205.609 | 0.103868 | 0.17849 | 0.581924 | 0.56062 | 0.74193 | 1.727 |
| Gm37183 | 15.2847 | -0.18072 | 0.63384 | -0.28513 | 0.77555 | 0.88307 | 3.405 |
| Gm37186 | 18.1532 | -0.64076 | 0.56908 | -1.12596 | 0.26018 | 0.46875 | 2.861 |
| Gm37195 | 13.0596 | -0.47613 | 0.53126 | -0.89623 | 0.37013 | 0.58045 | 3.018 |
| Gm37204 | 21.1123 | -0.2496 | 0.55227 | -0.45194 | 0.65131 | 0.80589 | 3.961 |
| Gm37211 | 38.1773 | -0.14571 | 0.36666 | -0.39739 | 0.69108 | 0.83227 | 3.433 |
| Gm37212 | 149.989 | 0.660253 | 0.23646 | 2.792254 | 0.00523 | 0.03736 | 3.65 |
| Gm37226 | 336.806 | 0.455739 | 0.20798 | 2.191231 | 0.02844 | 0.1137 | 3.255 |
| Gm37233 | 333.595 | 0.232362 | 0.14334 | 1.621095 | 0.105 | 0.26814 | 6.339 |
| Gm37238 | 24.5788 | -0.8651 | 0.40408 | -2.1409 | 0.03228 | 0.1242 | 2.39 |

| | | | | | | | |
|---|---|---|---|---|---|---|---|
| Gm3724 | 41.8686 | -0.02857 | 0.26696 | -0.10701 | 0.91478 | 0.95656 | 0.349 |
| Gm37242 | 21.0494 | -0.71895 | 0.57738 | -1.24519 | 0.21306 | 0.41418 | 2.085 |
| Gm37247 | 16.8669 | -0.66384 | 0.49615 | -1.33797 | 0.18091 | 0.37426 | 0.73 |
| Gm37249 | 270.666 | 0.402715 | 0.17051 | 2.361822 | 0.01819 | 0.08428 | 2.869 |
| Gm37254 | 145.631 | 0.050376 | 0.15682 | 0.321234 | 0.74803 | 0.86795 | 3.901 |
| Gm37264 | 44.5513 | 0.334397 | 0.28558 | 1.170954 | 0.24162 | 0.44766 | 2.192 |
| Gm37268 | 28.2553 | 0.387127 | 0.40515 | 0.955517 | 0.33932 | 0.55016 | 2.614 |
| Gm37274 | 101.861 | 0.194663 | 0.2144 | 0.907956 | 0.3639 | 0.57434 | 1.682 |
| Gm37277 | 144.33 | 0.158439 | 0.19299 | 0.820984 | 0.41166 | 0.61973 | 1.111 |
| Gm37284 | 13.9976 | -0.84069 | 0.44748 | -1.87873 | 0.06028 | 0.18732 | 0.539 |
| Gm37285 | 35.9951 | 0.296025 | 0.32235 | 0.918349 | 0.35844 | 0.56863 | 1.139 |
| Gm37289 | 22.1429 | -0.06525 | 0.44194 | -0.14765 | 0.88262 | 0.9416 | 1.402 |
| Gm37292 | 30.5352 | 0.228955 | 0.37661 | 0.607932 | 0.54323 | 0.72869 | 0.758 |
| Gm37296 | 43.1791 | 0.178725 | 0.38808 | 0.460541 | 0.64513 | 0.80209 | 2.922 |
| Gm37297 | 15.5279 | 0.511865 | 0.43699 | 1.171347 | 0.24146 | 0.44749 | 2.384 |
| Gm37303 | 200.643 | 0.289481 | 0.1437 | 2.014535 | 0.04395 | 0.15175 | 1.559 |
| Gm37305 | 431.556 | 0.254017 | 0.12162 | 2.088567 | 0.03675 | 0.13521 | 4.279 |
| Gm37309 | 194.293 | 0.23179 | 0.18771 | 1.234801 | 0.2169 | 0.41877 | 1.639 |
| Gm37310 | 128.471 | -0.43468 | 0.22489 | -1.93288 | 0.05325 | 0.17276 | 2.968 |
| Gm37313 | 81.0783 | 0.185723 | 0.1984 | 0.936093 | 0.34923 | 0.55964 | 9.078 |
| Gm37314 | 32.118 | -0.05023 | 0.31613 | -0.15888 | 0.87376 | 0.93672 | 2.685 |
| Gm37320 | 24.7116 | 0.501729 | 0.37218 | 1.348073 | 0.17763 | 0.36986 | 4.953 |
| Gm37328 | 15.2272 | -0.58321 | 0.4285 | -1.36106 | 0.17349 | 0.3648 | 0.64 |
| Gm37331 | 18.4248 | -0.48524 | 0.51513 | -0.94198 | 0.3462 | 0.55684 | 3.375 |
| Gm37333 | 18.508 | -0.65098 | 0.54477 | -1.19496 | 0.2321 | 0.43702 | 2.251 |
| Gm37334 | 19.5359 | -0.95011 | 0.74256 | -1.2795 | 0.20072 | 0.39886 | 2.938 |
| Gm37342 | 20.4255 | 0.662089 | 0.45065 | 1.469173 | 0.14179 | 0.32357 | 4.965 |
| Gm37345 | 87.4048 | 0.2029 | 0.3228 | 0.628568 | 0.52963 | 0.71744 | 3.138 |
| Gm37352 | 22.7317 | -1.25912 | 0.47336 | -2.65996 | 0.00781 | 0.04892 | 2.181 |
| Gm37353 | 22.1807 | 0.352668 | 0.41379 | 0.85228 | 0.39406 | 0.60318 | 0.313 |
| Gm37357 | 14.3101 | 0.074752 | 0.60264 | 0.124041 | 0.90128 | 0.95046 | 3.123 |
| Gm37359 | 16.7851 | 0.078486 | 0.50349 | 0.155884 | 0.87612 | 0.93819 | 43.166 |
| Gm37366 | 100.175 | 0.298801 | 0.3126 | 0.955851 | 0.33915 | 0.55007 | 2.297 |
| Gm37367 | 20.3083 | -0.41476 | 0.44051 | -0.94154 | 0.34643 | 0.557 | 2.602 |

| | | | | | | | |
|---|---|---|---|---|---|---|---|
| Gm3739 | 60.8877 | 0.289955 | 0.25044 | 1.157766 | 0.24696 | 0.45383 | 21.402 |
| Gm37397 | 77.0927 | 0.036401 | 0.22314 | 0.163132 | 0.87041 | 0.93456 | 2.842 |
| Gm37403 | 20.0517 | -0.44449 | 0.41155 | -1.08005 | 0.28012 | 0.49017 | 3.521 |
| Gm37407 | 20.6317 | -0.19778 | 0.48824 | -0.40508 | 0.68542 | 0.8286 | 1.316 |
| Gm37411 | 40.0275 | 0.047807 | 0.30719 | 0.155627 | 0.87633 | 0.93824 | 2.282 |
| Gm37412 | 18.1675 | -0.75157 | 0.48302 | -1.55599 | 0.11971 | 0.29217 | 4.321 |
| Gm37422 | 156.848 | -0.24342 | 0.17546 | -1.38732 | 0.16534 | 0.35448 | 4.718 |
| Gm37426 | 19.5758 | 1.006268 | 0.49201 | 2.045206 | 0.04083 | 0.14468 | 2.47 |
| Gm37436 | 25.6778 | 0.051404 | 0.37598 | 0.136719 | 0.89125 | 0.94585 | 1.085 |
| Gm37437 | 15.2915 | 0.178213 | 0.57922 | 0.307679 | 0.75833 | 0.87287 | 1.937 |
| Gm37443 | 21.4555 | -0.77911 | 0.40761 | -1.91141 | 0.05595 | 0.17876 | 2.827 |
| Gm37447 | 63.4285 | 0.578835 | 0.27678 | 2.09132 | 0.0365 | 0.13459 | 2.928 |
| Gm37459 | 841.815 | -0.55424 | 0.23838 | -2.32505 | 0.02007 | 0.08969 | 43.988 |
| Gm37460 | 15.8335 | 0.695271 | 0.44592 | 1.559187 | 0.11895 | 0.29105 | 2.249 |
| Gm37464 | 35.8462 | 0.122332 | 0.3415 | 0.358222 | 0.72018 | 0.85067 | 2.22 |
| Gm37472 | 18.2668 | -0.71163 | 0.4058 | -1.75366 | 0.07949 | 0.22395 | 4.052 |
| Gm37475 | 22.078 | 0.149828 | 0.47059 | 0.318386 | 0.75019 | 0.86885 | 1.015 |
| Gm37478 | 47.7281 | -0.06484 | 0.36679 | -0.17679 | 0.85967 | 0.92904 | 3.495 |
| Gm37499 | 214.488 | 0.352225 | 0.16945 | 2.078677 | 0.03765 | 0.13734 | 3.478 |
| Gm37500 | 21.372 | 0.643817 | 0.43034 | 1.496081 | 0.13463 | 0.31449 | 1.443 |
| Gm37509 | 36.3862 | 0.198051 | 0.35371 | 0.559921 | 0.57553 | 0.75343 | 3.169 |
| Gm37519 | 12.8075 | 0.118143 | 0.5911 | 0.19987 | 0.84158 | 0.91879 | 2.21 |
| Gm37521 | 17.6334 | 0.126342 | 0.49947 | 0.252953 | 0.8003 | 0.89682 | 2.045 |
| Gm37524 | 39.8981 | -0.4912 | 0.38857 | -1.26413 | 0.20618 | 0.40534 | 1.692 |
| Gm37529 | 25.5301 | -0.15125 | 0.47488 | -0.31851 | 0.7501 | 0.86885 | 5.393 |
| Gm37530 | 26.7925 | -0.23055 | 0.41817 | -0.55133 | 0.58141 | 0.75733 | 4.932 |
| Gm37534 | 12.6013 | -1.13645 | 0.69517 | -1.63477 | 0.1021 | 0.26371 | 5.222 |
| Gm37537 | 14.5582 | 0.347633 | 0.50183 | 0.692726 | 0.48848 | 0.68364 | 1.738 |
| Gm37551 | 18.4702 | -0.59489 | 0.40976 | -1.45181 | 0.14655 | 0.3297 | 2.861 |
| Gm37559 | 17.928 | 0.166242 | 0.44004 | 0.377791 | 0.70559 | 0.84142 | 0.832 |
| Gm3756 | 182.645 | -0.51134 | 0.16047 | -3.18648 | 0.00144 | 0.01608 | 1.355 |
| Gm37570 | 145.78 | 0.114932 | 0.18199 | 0.631523 | 0.5277 | 0.71577 | 3.106 |
| Gm37573 | 32.4118 | -0.15573 | 0.34568 | -0.45049 | 0.65236 | 0.8066 | 5.109 |
| Gm37581 | 178.879 | 0.667385 | 0.21142 | 3.156683 | 0.0016 | 0.01734 | 0.988 |

| | | | | | | | |
|---|---|---|---|---|---|---|---|
| Gm37583 | 25.3799 | 1.014863 | 0.40925 | 2.479794 | 0.01315 | 0.06866 | 2.487 |
| Gm37584 | 64.615 | 0.303504 | 0.23983 | 1.265495 | 0.20569 | 0.40486 | 1.29 |
| Gm37589 | 31.3134 | 0.537938 | 0.43875 | 1.22606 | 0.22018 | 0.42272 | 3.994 |
| Gm37592 | 84.6996 | 0.139116 | 0.21753 | 0.639529 | 0.52248 | 0.71129 | 0.542 |
| Gm37602 | 25.2343 | -0.1383 | 0.36369 | -0.38028 | 0.70374 | 0.84002 | 3.31 |
| Gm37607 | 302.43 | 0.138572 | 0.21999 | 0.629909 | 0.52875 | 0.71654 | 2.623 |
| Gm37608 | 398.938 | 0.139165 | 0.13311 | 1.045506 | 0.29579 | 0.50616 | 1.88 |
| Gm37612 | 15.4557 | -0.01276 | 0.44986 | -0.02836 | 0.97737 | 0.98799 | 0.828 |
| Gm37621 | 126.001 | -0.04275 | 0.17336 | -0.24662 | 0.8052 | 0.89941 | 5.531 |
| Gm37628 | 28.2743 | -0.25163 | 0.41751 | -0.60269 | 0.54672 | 0.73092 | 5.44 |
| Gm37633 | 13.0107 | -0.48365 | 0.55675 | -0.8687 | 0.38501 | 0.59437 | 2.7 |
| Gm37637 | 151.656 | 0.404095 | 0.16959 | 2.382783 | 0.01718 | 0.08153 | 1.256 |
| Gm37639 | 51.3803 | 0.074114 | 0.2988 | 0.248043 | 0.8041 | 0.89887 | 2.785 |
| Gm3764 | 6645.84 | -0.25624 | 0.14694 | -1.74389 | 0.08118 | 0.22677 | 23.432 |
| Gm37653 | 12.6761 | -0.57137 | 0.48288 | -1.18324 | 0.23671 | 0.44183 | 3.578 |
| Gm37669 | 72.509 | 0.246106 | 0.22937 | 1.072972 | 0.28328 | 0.4934 | 0.82 |
| Gm37671 | 136.369 | 0.4509 | 0.23756 | 1.898073 | 0.05769 | 0.18234 | 2.096 |
| Gm37672 | 22.1013 | -0.52739 | 0.47019 | -1.12167 | 0.262 | 0.47075 | 4.014 |
| Gm37673 | 20.9279 | 0.082289 | 0.47824 | 0.172067 | 0.86339 | 0.93092 | 2.451 |
| Gm37674 | 17.528 | 0.646901 | 0.41668 | 1.552511 | 0.12054 | 0.2934 | 1.834 |
| Gm37677 | 88.7338 | -0.40937 | 0.35137 | -1.16507 | 0.24399 | 0.45038 | 7.634 |
| Gm37691 | 62.3786 | -0.29023 | 0.27087 | -1.07145 | 0.28397 | 0.49414 | 2.573 |
| Gm37694 | 60.0705 | 0.066734 | 0.23653 | 0.282142 | 0.77783 | 0.88423 | 2.133 |
| Gm37696 | 63.8523 | 0.253668 | 0.23225 | 1.092236 | 0.27473 | 0.48435 | 3.349 |
| Gm37702 | 14.4056 | -0.012 | 0.59788 | -0.02007 | 0.98399 | 0.99128 | 1.872 |
| Gm37711 | 850.162 | 0.30459 | 0.16477 | 1.848623 | 0.06451 | 0.19551 | 4.534 |
| Gm37717 | 13.3079 | -1.15996 | 0.54908 | -2.11255 | 0.03464 | 0.13033 | 3.027 |
| Gm37718 | 14.2557 | 0.4292 | 0.55312 | 0.775957 | 0.43777 | 0.64323 | 2.631 |
| Gm37728 | 41.5822 | 0.413316 | 0.31432 | 1.314959 | 0.18852 | 0.38421 | 2.292 |
| Gm37738 | 16.3901 | -0.52002 | 0.49833 | -1.04353 | 0.2967 | 0.50689 | 4.716 |
| Gm37768 | 74.1802 | 0.125024 | 0.23437 | 0.533438 | 0.59373 | 0.76599 | 12.627 |
| Gm37773 | 398.458 | 0.408105 | 0.17012 | 2.398956 | 0.01644 | 0.07912 | 2.12 |
| Gm37776 | 145.536 | 0.302867 | 0.14986 | 2.020998 | 0.04328 | 0.15032 | 2.497 |
| Gm37783 | 21.1452 | -0.11991 | 0.40175 | -0.29847 | 0.76534 | 0.87738 | 1.856 |

| | | | | | | | |
|---|---|---|---|---|---|---|---|
| Gm37785 | 60.6877 | -0.02291 | 0.27982 | -0.08186 | 0.93476 | 0.96717 | 2.408 |
| Gm37786 | 20.5687 | 0.218658 | 0.37216 | 0.587536 | 0.55684 | 0.73888 | 3.225 |
| Gm37790 | 14.903 | 0.355612 | 0.43305 | 0.821184 | 0.41154 | 0.61965 | 3.346 |
| Gm37792 | 22.2842 | 0.845587 | 0.45544 | 1.856639 | 0.06336 | 0.19338 | 3.413 |
| Gm37795 | 85.5727 | -0.53999 | 0.28919 | -1.86724 | 0.06187 | 0.19038 | 2.459 |
| Gm37805 | 38.6516 | -0.21691 | 0.27717 | -0.78261 | 0.43386 | 0.6396 | 1.165 |
| Gm37811 | 13.079 | -0.90144 | 0.585 | -1.54093 | 0.12333 | 0.29752 | 2.601 |
| Gm37814 | 51.8691 | -0.26136 | 0.26752 | -0.97701 | 0.32857 | 0.53979 | 3.644 |
| Gm37818 | 18.0062 | -0.14334 | 0.45973 | -0.31179 | 0.7552 | 0.87125 | 3.169 |
| Gm37824 | 90.4458 | -1.0367 | 0.80176 | -1.29302 | 0.196 | 0.39362 | 3.236 |
| Gm37844 | 66.6436 | 0.503919 | 0.27413 | 1.83823 | 0.06603 | 0.19848 | 4.097 |
| Gm37848 | 17.4518 | -0.98817 | 0.54565 | -1.811 | 0.07014 | 0.20676 | 3.103 |
| Gm37849 | 12.5305 | -0.01514 | 0.54018 | -0.02803 | 0.97764 | 0.98803 | 1.881 |
| Gm37850 | 170.119 | 0.122242 | 0.19451 | 0.628466 | 0.5297 | 0.71748 | 0.78 |
| Gm37859 | 74.8708 | 0.58929 | 0.26247 | 2.2452 | 0.02476 | 0.1033 | 2.173 |
| Gm37860 | 17.9727 | -0.10191 | 0.39561 | -0.2576 | 0.79671 | 0.89517 | 1.034 |
| Gm37861 | 15.1083 | 0.108969 | 0.42895 | 0.254034 | 0.79947 | 0.89649 | 2.869 |
| Gm37862 | 21.039 | -0.16303 | 0.37854 | -0.43067 | 0.66671 | 0.8158 | 2.533 |
| Gm37868 | 13.5256 | -0.49165 | 0.53608 | -0.91711 | 0.35908 | 0.56944 | 2.735 |
| Gm37876 | 38.3285 | 0.470941 | 0.32017 | 1.470903 | 0.14132 | 0.32308 | 10.583 |
| Gm3788 | 51.8257 | 0.025206 | 0.27296 | 0.092344 | 0.92642 | 0.96301 | 0.136 |
| Gm37880 | 19.7145 | -0.42328 | 0.62315 | -0.67926 | 0.49698 | 0.69055 | 3.257 |
| Gm37885 | 32.0995 | -0.39014 | 0.30383 | -1.28406 | 0.19912 | 0.39704 | 8.295 |
| Gm37891 | 94.8985 | -0.0101 | 0.22826 | -0.04424 | 0.96472 | 0.98273 | 1.326 |
| Gm37893 | 429.989 | 0.278372 | 0.1387 | 2.007078 | 0.04474 | 0.15361 | 2.437 |
| Gm37894 | 36.611 | 0.167068 | 0.42321 | 0.394764 | 0.69302 | 0.83352 | 2.599 |
| Gm37895 | 38.9479 | 0.289603 | 0.32273 | 0.897347 | 0.36953 | 0.57996 | 4.55 |
| Gm37899 | 143.603 | -0.19078 | 0.28105 | -0.67879 | 0.49727 | 0.69077 | 8.717 |
| Gm37902 | 47.9316 | -0.01432 | 0.26655 | -0.05373 | 0.95715 | 0.979 | 1.683 |
| Gm37906 | 12.5959 | -0.54251 | 0.48426 | -1.12028 | 0.26259 | 0.47144 | 3.035 |
| Gm37914 | 65.3378 | 0.02619 | 0.24449 | 0.107122 | 0.91469 | 0.95656 | 2.679 |
| Gm37928 | 595.951 | 0.605778 | 0.13465 | 4.498873 | 6.83E-06 | 0.00042 | 4.131 |
| Gm37931 | 33.2117 | 0.173086 | 0.33235 | 0.520798 | 0.60251 | 0.77234 | 2.866 |
| Gm37940 | 85.5333 | 0.221748 | 0.18184 | 1.21944 | 0.22268 | 0.42588 | 3.251 |

| | | | | | | | |
|---|---|---|---|---|---|---|---|
| Gm37941 | 77.0058 | -0.066 | 0.19955 | -0.33074 | 0.74084 | 0.86313 | 2.309 |
| Gm37942 | 12.8981 | 1.335905 | 0.5431 | 2.459791 | 0.0139 | 0.07096 | 0.628 |
| Gm37945 | 20.264 | 1.005334 | 0.4116 | 2.44251 | 0.01459 | 0.07302 | 4.367 |
| Gm37949 | 20.5441 | -0.4473 | 0.4797 | -0.93246 | 0.3511 | 0.56158 | 2.091 |
| Gm37953 | 62.8902 | 0.024284 | 0.21551 | 0.11268 | 0.91028 | 0.95475 | 1.884 |
| Gm37959 | 22.4712 | -0.35214 | 0.52331 | -0.67292 | 0.501 | 0.69382 | 2.897 |
| Gm37960 | 82.1007 | 0.345401 | 0.23702 | 1.457271 | 0.14504 | 0.32787 | 1.897 |
| Gm37962 | 46.5614 | 0.169064 | 0.25454 | 0.664203 | 0.50656 | 0.69789 | 2.034 |
| Gm37963 | 634.042 | 0.34083 | 0.10595 | 3.21679 | 0.0013 | 0.01495 | 1.124 |
| Gm37964 | 16.4426 | -0.13963 | 0.40748 | -0.34266 | 0.73185 | 0.85771 | 4.905 |
| Gm37965 | 13.3323 | 0.709974 | 0.57424 | 1.236382 | 0.21632 | 0.41823 | 1.362 |
| Gm37968 | 36.9258 | 0.083282 | 0.35735 | 0.233057 | 0.81572 | 0.90517 | 3.201 |
| Gm37974 | 32.8174 | 0.859683 | 0.32692 | 2.629651 | 0.00855 | 0.05167 | 1.318 |
| Gm37979 | 32.6699 | 0.065295 | 0.31715 | 0.205877 | 0.83689 | 0.91619 | 0.712 |
| Gm37983 | 19.9004 | 0.521847 | 0.39437 | 1.323237 | 0.18576 | 0.38054 | 2.006 |
| Gm37984 | 50.8283 | -0.34775 | 0.35286 | -0.98552 | 0.32437 | 0.53542 | 3.733 |
| Gm37986 | 38.4766 | 0.089589 | 0.29215 | 0.306659 | 0.7591 | 0.87347 | 0.383 |
| Gm37995 | 22.2081 | -0.20593 | 0.41774 | -0.49297 | 0.62203 | 0.7853 | 1.714 |
| Gm37999 | 21.6525 | 0.596228 | 0.38106 | 1.56466 | 0.11766 | 0.28879 | 1.809 |
| Gm38014 | 21.4111 | -0.53955 | 0.49939 | -1.08042 | 0.27996 | 0.49005 | 2.612 |
| Gm38020 | 1778.32 | 0.16985 | 0.13637 | 1.245533 | 0.21294 | 0.41405 | 4.676 |
| Gm38021 | 14.0936 | 0.269271 | 0.47321 | 0.56903 | 0.56934 | 0.74892 | 1.56 |
| Gm38024 | 24.5245 | 0.649082 | 0.41446 | 1.566095 | 0.11733 | 0.28827 | 2.532 |
| Gm38036 | 18.9092 | 0.289996 | 0.52362 | 0.553829 | 0.5797 | 0.75628 | 3.564 |
| Gm38043 | 61.72 | 0.205875 | 0.22563 | 0.912462 | 0.36153 | 0.57184 | 5.141 |
| Gm38056 | 33.7834 | -0.37624 | 0.33654 | -1.11797 | 0.26358 | 0.47241 | 5.027 |
| Gm38057 | 160.594 | 0.082322 | 0.1877 | 0.43859 | 0.66096 | 0.81246 | 3.203 |
| Gm38058 | 17.2801 | 0.274923 | 0.56193 | 0.489253 | 0.62466 | 0.78721 | 4.368 |
| Gm38059 | 14.1028 | 1.289923 | 0.47882 | 2.693973 | 0.00706 | 0.04574 | 1.999 |
| Gm38067 | 23.159 | 0.242472 | 0.38879 | 0.623653 | 0.53286 | 0.72018 | 1.911 |
| Gm38071 | 19.286 | -0.44572 | 0.46476 | -0.95903 | 0.33754 | 0.54833 | 3.378 |
| Gm38073 | 17.4692 | -1.79138 | 0.61865 | -2.89562 | 0.00378 | 0.03031 | 3.665 |
| Gm38077 | 877.539 | 0.396674 | 0.15305 | 2.591799 | 0.00955 | 0.05562 | 3.245 |
| Gm38080 | 34.076 | -0.33734 | 0.2868 | -1.17622 | 0.23951 | 0.44545 | 0.268 |

| | | | | | | | |
|---|---|---|---|---|---|---|---|
| Gm38082 | 30.8936 | 0.079698 | 0.31092 | 0.256332 | 0.79769 | 0.89566 | 6.882 |
| Gm38085 | 31.9853 | -0.24843 | 0.45714 | -0.54344 | 0.58683 | 0.7605 | 3.062 |
| Gm38091 | 30.8199 | -0.07495 | 0.48669 | -0.154 | 0.87761 | 0.93882 | 3.871 |
| Gm38099 | 38.7172 | 0.244391 | 0.28439 | 0.85935 | 0.39015 | 0.59917 | 3.17 |
| Gm38101 | 69.6328 | 0.663858 | 0.25069 | 2.64815 | 0.00809 | 0.04993 | 3.557 |
| Gm38102 | 252.389 | 0.249708 | 0.13846 | 1.803437 | 0.07132 | 0.20903 | 3.757 |
| Gm38104 | 152.995 | 0.273586 | 0.20525 | 1.332954 | 0.18255 | 0.37638 | 1.895 |
| Gm38109 | 15.1441 | -0.17907 | 0.60384 | -0.29655 | 0.76681 | 0.87812 | 0.181 |
| Gm38120 | 48.3661 | -0.34112 | 0.3529 | -0.96663 | 0.33373 | 0.54452 | 4.642 |
| Gm38124 | 109.307 | 0.409303 | 0.28881 | 1.417227 | 0.15642 | 0.34317 | 2.581 |
| Gm38139 | 17.4482 | 0.689814 | 0.6004 | 1.148931 | 0.25058 | 0.4579 | 23.569 |
| Gm38148 | 24.7777 | -0.47995 | 0.40325 | -1.19021 | 0.23396 | 0.43902 | 3.012 |
| Gm3815 | 50.2502 | 0.049701 | 0.29656 | 0.16759 | 0.86691 | 0.93246 | 0.502 |
| Gm38155 | 188.828 | 0.281684 | 0.14185 | 1.985807 | 0.04705 | 0.15917 | 32.5 |
| Gm38157 | 13.9111 | 0.167927 | 0.62998 | 0.26656 | 0.78981 | 0.89088 | 1.698 |
| Gm38162 | 80.5289 | 0.066013 | 0.31578 | 0.20905 | 0.83441 | 0.91508 | 4.597 |
| Gm38163 | 21.4836 | -0.26403 | 0.39721 | -0.66471 | 0.50624 | 0.69776 | 0.606 |
| Gm38190 | 440.372 | 0.279984 | 0.12337 | 2.269547 | 0.02324 | 0.09878 | 4.755 |
| Gm38192 | 205.982 | 0.157378 | 0.16379 | 0.96087 | 0.33662 | 0.54718 | 1.847 |
| Gm38198 | 53.9688 | -0.16305 | 0.27563 | -0.59154 | 0.55416 | 0.73681 | 3.581 |
| Gm38200 | 13.2762 | 0.092703 | 0.51029 | 0.181668 | 0.85584 | 0.92711 | 2.111 |
| Gm38214 | 176.494 | 0.545116 | 0.25902 | 2.104513 | 0.03533 | 0.13199 | 0.988 |
| Gm38217 | 17.797 | 0.543414 | 0.46532 | 1.16784 | 0.24287 | 0.44905 | 0.91 |
| Gm38227 | 20.8952 | -0.39976 | 0.37733 | -1.05945 | 0.2894 | 0.49971 | 1.602 |
| Gm38228 | 36.5239 | -0.20063 | 0.3449 | -0.58171 | 0.56077 | 0.74201 | 1.293 |
| Gm38235 | 18.6268 | -0.90309 | 0.44273 | -2.03981 | 0.04137 | 0.14593 | 4.952 |
| Gm38243 | 21.6767 | -0.20468 | 0.64462 | -0.31752 | 0.75085 | 0.86928 | 1.567 |
| Gm38245 | 32.9981 | 0.254642 | 0.2982 | 0.853944 | 0.39314 | 0.60231 | 2.621 |
| Gm38248 | 72.2053 | 0.552009 | 0.21843 | 2.527211 | 0.0115 | 0.06285 | 5.783 |
| Gm38250 | 107.305 | 0.088145 | 0.16622 | 0.530287 | 0.59591 | 0.76769 | 12.466 |
| Gm38251 | 16.7447 | -0.03124 | 0.46293 | -0.06748 | 0.9462 | 0.97302 | 19.365 |
| Gm38253 | 104.259 | 0.726382 | 0.30219 | 2.403687 | 0.01623 | 0.07844 | 4.678 |
| Gm38255 | 50.8897 | -0.05855 | 0.30376 | -0.19276 | 0.84714 | 0.92206 | 2.996 |
| Gm38260 | 19.3445 | -0.41671 | 0.63329 | -0.65801 | 0.51053 | 0.70183 | 3.995 |

| | | | | | | | |
|---|---|---|---|---|---|---|---|
| Gm38273 | 34.6213 | -0.11458 | 0.42072 | -0.27234 | 0.78536 | 0.88859 | 2.382 |
| Gm3828 | 266.611 | 1.200112 | 0.19855 | 6.044511 | 1.50E-09 | 7.45E-07 | 0.257 |
| Gm38297 | 89.3465 | 0.617103 | 0.22637 | 2.726037 | 0.00641 | 0.04297 | 1.057 |
| Gm38299 | 99.8361 | -0.47831 | 0.31508 | -1.51805 | 0.129 | 0.30621 | 3.789 |
| Gm38300 | 95.4464 | -0.11983 | 0.22252 | -0.5385 | 0.59023 | 0.76321 | 1.87 |
| Gm38308 | 13.1896 | -1.22792 | 0.49997 | -2.45599 | 0.01405 | 0.07136 | 1.809 |
| Gm38309 | 57.7698 | 0.194856 | 0.23574 | 0.826556 | 0.40849 | 0.61624 | 1.518 |
| Gm38317 | 20.0615 | -0.95634 | 0.47487 | -2.01387 | 0.04402 | 0.15194 | 0.343 |
| Gm38329 | 23.7622 | -0.06797 | 0.41198 | -0.16499 | 0.86895 | 0.93348 | 0.427 |
| Gm38331 | 26.1302 | 0.160882 | 0.37345 | 0.430799 | 0.66661 | 0.8158 | 2.671 |
| Gm38336 | 89.7801 | 0.21946 | 0.2266 | 0.968482 | 0.3328 | 0.54367 | 2.549 |
| Gm38340 | 21.1377 | -0.3871 | 0.41261 | -0.93819 | 0.34815 | 0.55865 | 4.008 |
| Gm38342 | 15.5321 | -0.09727 | 0.43147 | -0.22543 | 0.82165 | 0.90766 | 2.98 |
| Gm38344 | 14.8924 | 0.12645 | 0.46371 | 0.272689 | 0.78509 | 0.88852 | 1.353 |
| Gm38348 | 24.0746 | -0.03778 | 0.4686 | -0.08061 | 0.93575 | 0.96779 | 2.273 |
| Gm38355 | 17.6291 | -0.26353 | 0.49546 | -0.53188 | 0.59481 | 0.76686 | 1.57 |
| Gm38357 | 138.737 | -0.35507 | 0.22301 | -1.59216 | 0.11135 | 0.27873 | 2.464 |
| Gm38359 | 21.6087 | 0.421436 | 0.41149 | 1.024172 | 0.30575 | 0.51572 | 2.225 |
| Gm38364 | 13.0733 | 0.009152 | 0.52196 | 0.017534 | 0.98601 | 0.99244 | 0.179 |
| Gm38365 | 13.3329 | -0.68874 | 0.5154 | -1.33632 | 0.18145 | 0.37492 | 2.993 |
| Gm38366 | 51.9946 | -0.03306 | 0.29109 | -0.11356 | 0.90958 | 0.95456 | 1.934 |
| Gm38372 | 330.1 | -0.13165 | 0.20792 | -0.63317 | 0.52662 | 0.71469 | 4.028 |
| Gm38375 | 49.2949 | -0.33667 | 0.3665 | -0.91859 | 0.35831 | 0.56847 | 2.227 |
| Gm38379 | 14.5127 | 0.897612 | 0.48932 | 1.834413 | 0.06659 | 0.19977 | 1.525 |
| Gm38380 | 21.5957 | -0.01658 | 0.45948 | -0.03607 | 0.97122 | 0.98549 | 2.248 |
| Gm38381 | 42.2305 | 0.2999 | 0.35406 | 0.847035 | 0.39698 | 0.60609 | 2.275 |
| Gm38383 | 83.7752 | 0.414462 | 0.21023 | 1.971443 | 0.04867 | 0.16295 | 2.66 |
| Gm38384 | 294.427 | 0.427047 | 0.13613 | 3.137091 | 0.00171 | 0.01818 | 1.122 |
| Gm38387 | 100.761 | -0.08677 | 0.21049 | -0.41223 | 0.68017 | 0.82502 | 2.577 |
| Gm38393 | 787.508 | 0.102596 | 0.13025 | 0.787681 | 0.43088 | 0.63706 | 30.916 |
| Gm38413 | 758.834 | 0.605534 | 0.33432 | 1.811264 | 0.0701 | 0.2067 | 23.12 |
| Gm38431 | 21.0914 | 0.081158 | 0.41031 | 0.197798 | 0.8432 | 0.91948 | 24.484 |
| Gm38451 | 111.618 | 0.5094 | 0.23422 | 2.174878 | 0.02964 | 0.11715 | 2.646 |
| Gm3854 | 340.176 | 0.102235 | 0.10872 | 0.940326 | 0.34705 | 0.55765 | 12.198 |

| | | | | | | | |
|---|---|---|---|---|---|---|---|
| Gm38560 | 14.071 | -0.25973 | 0.63644 | -0.4081 | 0.6832 | 0.82693 | 13.276 |
| Gm38562 | 63.0694 | -0.53008 | 0.24939 | -2.12548 | 0.03355 | 0.12748 | 19.557 |
| Gm38575 | 75.4951 | 1.080871 | 0.34048 | 3.174588 | 0.0015 | 0.01657 | 45.451 |
| Gm38576 | 28.343 | -0.22932 | 0.35949 | -0.63791 | 0.52353 | 0.71197 | 10.185 |
| Gm38593 | 22.849 | -0.21055 | 0.46888 | -0.44904 | 0.6534 | 0.8075 | 7.386 |
| Gm38642 | 52.4146 | 0.017626 | 0.26746 | 0.065904 | 0.94745 | 0.97358 | 100.134 |
| Gm38661 | 15.0815 | -0.98548 | 0.65787 | -1.49798 | 0.13414 | 0.31389 | 7.231 |
| Gm38708 | 14.736 | -0.5063 | 0.48654 | -1.04061 | 0.29806 | 0.50816 | 2.285 |
| Gm3883 | 27.9093 | -0.1355 | 0.39995 | -0.33881 | 0.73476 | 0.85968 | 4.903 |
| Gm3888 | 13.5661 | 0.204072 | 0.48175 | 0.423603 | 0.67186 | 0.81948 | 1.24 |
| Gm38910 | 128.799 | -0.71419 | 0.18241 | -3.91533 | 9.03E-05 | 0.00251 | 2.448 |
| Gm38941 | 24.1925 | 0.165503 | 0.40349 | 0.410173 | 0.68168 | 0.82602 | 0.529 |
| Gm38948 | 37.9706 | 0.149242 | 0.34985 | 0.426589 | 0.66968 | 0.81794 | 12.772 |
| Gm39038 | 13.9592 | 0.141364 | 0.53968 | 0.261942 | 0.79337 | 0.89294 | 14.598 |
| Gm39041 | 44.1791 | 0.180083 | 0.32847 | 0.548241 | 0.58353 | 0.75854 | 3.902 |
| Gm39043 | 102.416 | 0.890976 | 0.29989 | 2.971037 | 0.00297 | 0.026 | 23.639 |
| Gm39090 | 33.0202 | -0.2771 | 0.35757 | -0.77497 | 0.43836 | 0.64363 | 9.713 |
| Gm39213 | 47.1233 | -0.57092 | 0.2799 | -2.03977 | 0.04137 | 0.14593 | 6.776 |
| Gm39244 | 74.6364 | 0.368402 | 0.26676 | 1.381038 | 0.16727 | 0.35697 | 10.922 |
| Gm39323 | 19.4882 | 0.11777 | 0.42161 | 0.279336 | 0.77999 | 0.88541 | 29.325 |
| Gm39326 | 142.26 | 0.345529 | 0.18236 | 1.89481 | 0.05812 | 0.18327 | 6.507 |
| Gm39456 | 81.0478 | 1.2364 | 0.28769 | 4.29764 | 1.73E-05 | 0.0008 | 8.646 |
| Gm39463 | 45.5226 | 0.899006 | 0.37263 | 2.412603 | 0.01584 | 0.07722 | 18.842 |
| Gm39468 | 44.4611 | -0.10534 | 0.30381 | -0.34672 | 0.7288 | 0.8556 | 1.531 |
| Gm39469 | 167.463 | 0.348058 | 0.22538 | 1.544322 | 0.12251 | 0.29627 | 11.984 |
| Gm39526 | 12.3576 | 1.223145 | 0.61146 | 2.000373 | 0.04546 | 0.15517 | 13.961 |
| Gm39822 | 41.1487 | -1.25666 | 0.44528 | -2.82219 | 0.00477 | 0.03524 | 10.659 |
| Gm3985 | 162.2 | 0.665979 | 0.1774 | 3.754154 | 0.00017 | 0.00392 | 61.522 |
| Gm40055 | 87.5546 | 0.749205 | 0.37448 | 2.000629 | 0.04543 | 0.15512 | 4.571 |
| Gm4013 | 17.8653 | -0.1898 | 0.47171 | -0.40238 | 0.68741 | 0.82961 | 0.85 |
| Gm4017 | 12.4238 | 0.206954 | 0.58228 | 0.355423 | 0.72227 | 0.85185 | 0.338 |
| Gm40309 | 13.4344 | -0.50531 | 0.51177 | -0.98738 | 0.32345 | 0.53448 | 1.752 |
| Gm40466 | 45.7368 | 0.648405 | 0.29495 | 2.198387 | 0.02792 | 0.11241 | 7.638 |
| Gm40493 | 19.4918 | 0.626244 | 0.38583 | 1.623115 | 0.10456 | 0.26745 | 11.304 |

| | | | | | | | |
|---|---|---|---|---|---|---|---|
| Gm40518 | 104.343 | -1.09092 | 0.31282 | -3.48737 | 0.00049 | 0.00779 | 26.462 |
| Gm40557 | 31.168 | 0.042615 | 0.3235 | 0.131733 | 0.8952 | 0.94744 | 4.691 |
| Gm40576 | 49.524 | 0.029713 | 0.34353 | 0.086495 | 0.93107 | 0.96526 | 10.848 |
| Gm40578 | 365.543 | -0.079 | 0.16714 | -0.47267 | 0.63645 | 0.79523 | 50.448 |
| Gm40603 | 13.2893 | 0.204485 | 0.64759 | 0.315762 | 0.75218 | 0.86997 | 27.836 |
| Gm40617 | 28.6274 | 0.523505 | 0.35 | 1.495722 | 0.13473 | 0.31451 | 7.824 |
| Gm40663 | 61.385 | 1.338747 | 0.31601 | 4.236419 | 2.27E-05 | 0.00096 | 36.799 |
| Gm40841 | 1370.76 | -0.19203 | 0.16838 | -1.14045 | 0.2541 | 0.46147 | 25.614 |
| Gm41077 | 126.104 | 0.33163 | 0.1683 | 1.970435 | 0.04879 | 0.16325 | 14.132 |
| Gm41115 | 32.64 | -0.19701 | 0.34298 | -0.57439 | 0.5657 | 0.74579 | 0.308 |
| Gm4117 | 31.7939 | 0.130279 | 0.37928 | 0.343494 | 0.73123 | 0.85734 | 147.707 |
| Gm4130 | 58.5504 | -0.2392 | 0.38617 | -0.61943 | 0.53564 | 0.72264 | 0.97 |
| Gm4131 | 21.7927 | -0.08183 | 0.41815 | -0.19569 | 0.84485 | 0.92064 | 103.508 |
| Gm4134 | 157.244 | 0.421605 | 0.16117 | 2.615907 | 0.0089 | 0.05309 | 2.927 |
| Gm41361 | 84.2393 | -0.09449 | 0.29507 | -0.32023 | 0.74879 | 0.86828 | 30.127 |
| Gm41381 | 53.0379 | 0.861418 | 0.27698 | 3.110074 | 0.00187 | 0.0192 | 1.521 |
| Gm41396 | 58.1969 | 0.978391 | 0.25005 | 3.912776 | 9.12E-05 | 0.00253 | 6.229 |
| Gm41414 | 62.1327 | -0.15711 | 0.29073 | -0.54039 | 0.58892 | 0.76205 | 145.458 |
| Gm41442 | 421.215 | 0.603571 | 0.14114 | 4.2764 | 1.90E-05 | 0.00086 | 14.724 |
| Gm41505 | 440.857 | -0.71636 | 0.38791 | -1.8467 | 0.06479 | 0.1961 | 31.605 |
| Gm41609 | 76.6473 | -0.94165 | 0.30964 | -3.04111 | 0.00236 | 0.02243 | 40.221 |
| Gm41662 | 38.4908 | 0.170971 | 0.35895 | 0.47631 | 0.63385 | 0.79385 | 0.768 |
| Gm41760 | 67.4308 | 0.440454 | 0.24271 | 1.814706 | 0.06957 | 0.20558 | 2.393 |
| Gm41804 | 20.3392 | -1.07731 | 0.41079 | -2.62256 | 0.00873 | 0.0524 | 65.212 |
| Gm41836 | 13.0873 | -0.05681 | 0.50579 | -0.11231 | 0.91058 | 0.9549 | 65.779 |
| Gm4189 | 43.5391 | 0.46835 | 0.2868 | 1.633024 | 0.10246 | 0.26428 | 1.094 |
| Gm4202 | 836.121 | 0.303925 | 0.12217 | 2.48782 | 0.01285 | 0.06768 | 0.733 |
| Gm4204 | 2048.92 | 0.231642 | 0.09499 | 2.438524 | 0.01475 | 0.07343 | 1.231 |
| Gm42047 | 779.346 | 0.066695 | 0.23843 | 0.279731 | 0.77968 | 0.88519 | 7.932 |
| Gm42064 | 25.1294 | 0.171685 | 0.32952 | 0.521013 | 0.60236 | 0.77229 | 1.642 |
| Gm42067 | 68.9524 | 0.006672 | 0.23026 | 0.028974 | 0.97688 | 0.98795 | 2.02 |
| Gm4208 | 15.7157 | 1.62729 | 0.54898 | 2.96423 | 0.00303 | 0.02632 | 11.02 |
| Gm4211 | 26.7166 | -0.70614 | 0.36496 | -1.9348 | 0.05301 | 0.17223 | 6.066 |
| Gm4219 | 17.7841 | -0.69639 | 0.42298 | -1.64639 | 0.09968 | 0.25965 | 4.677 |

| | | | | | | | |
|---|---|---|---|---|---|---|---|
| Gm4221 | 92.9592 | -0.19017 | 0.23628 | -0.80484 | 0.42091 | 0.62796 | 89.153 |
| Gm42303 | 47.96 | 0.025896 | 0.30658 | 0.084469 | 0.93268 | 0.96605 | 39.798 |
| Gm4240 | 13.7024 | 0.181108 | 0.53135 | 0.340846 | 0.73322 | 0.85857 | 0.733 |
| Gm42423 | 27.4967 | 1.226587 | 0.37447 | 3.275558 | 0.00105 | 0.01328 | 6.926 |
| Gm42432 | 19.1883 | 0.070577 | 0.53386 | 0.132201 | 0.89483 | 0.94736 | 1.617 |
| Gm42433 | 281.892 | -0.11939 | 0.15714 | -0.75977 | 0.44739 | 0.65112 | 0.94 |
| Gm42444 | 40.524 | 0.095443 | 0.30415 | 0.313807 | 0.75367 | 0.87065 | 4.558 |
| Gm42445 | 14.4043 | 0.128148 | 0.54636 | 0.234551 | 0.81456 | 0.90481 | 2.893 |
| Gm42446 | 14.8992 | -0.84773 | 0.61689 | -1.3742 | 0.16938 | 0.35964 | 3.284 |
| Gm42456 | 128.655 | 0.596754 | 0.2441 | 2.444717 | 0.0145 | 0.07276 | 14.57 |
| Gm42460 | 21.0047 | -0.12513 | 0.35762 | -0.34988 | 0.72643 | 0.8541 | 2.567 |
| Gm42462 | 21.3086 | -0.1685 | 0.43702 | -0.38557 | 0.69982 | 0.83779 | 1.363 |
| Gm42463 | 96.3876 | 0.111819 | 0.1995 | 0.560483 | 0.57515 | 0.75324 | 4.886 |
| Gm42464 | 23.449 | -0.63634 | 0.34837 | -1.82664 | 0.06775 | 0.20179 | 0.774 |
| Gm42467 | 26.6204 | 0.090415 | 0.33893 | 0.266763 | 0.78965 | 0.89082 | 1.447 |
| Gm4247 | 139.331 | 0.533652 | 0.1809 | 2.950048 | 0.00318 | 0.02711 | 1.042 |
| Gm42471 | 18.2174 | -0.01429 | 0.49678 | -0.02876 | 0.97706 | 0.98796 | 1.865 |
| Gm42477 | 13.0726 | 0.702802 | 0.56248 | 1.249475 | 0.21149 | 0.41203 | 5.049 |
| Gm42488 | 13.2234 | 0.499702 | 0.51783 | 0.964999 | 0.33455 | 0.54533 | 4.432 |
| Gm42489 | 42.9481 | 0.008718 | 0.26412 | 0.033008 | 0.97367 | 0.98681 | 0.669 |
| Gm42492 | 29.205 | 0.797811 | 0.32668 | 2.442199 | 0.0146 | 0.07302 | 2.915 |
| Gm42500 | 15.3876 | 0.74452 | 0.47005 | 1.583916 | 0.11321 | 0.28167 | 0.879 |
| Gm42502 | 51.8288 | -0.11479 | 0.36491 | -0.31458 | 0.75308 | 0.87029 | 1.089 |
| Gm42503 | 27.573 | 0.468404 | 0.41285 | 1.134562 | 0.25656 | 0.46453 | 3.706 |
| Gm42506 | 17.6977 | 1.117272 | 0.43902 | 2.544905 | 0.01093 | 0.06093 | 0.15 |
| Gm42507 | 24.5515 | -0.1763 | 0.3556 | -0.49578 | 0.62005 | 0.78385 | 2.555 |
| Gm42509 | 17.0585 | -0.53367 | 0.45908 | -1.1625 | 0.24503 | 0.45165 | 0.781 |
| Gm42517 | 1873.46 | 0.117683 | 0.08246 | 1.42707 | 0.15356 | 0.33935 | 1.85 |
| Gm42518 | 46.1463 | 0.741825 | 0.30583 | 2.425644 | 0.01528 | 0.07539 | 0.938 |
| Gm42521 | 16.8345 | 0.116511 | 0.42856 | 0.271867 | 0.78572 | 0.88863 | 2.04 |
| Gm42527 | 93.6526 | 0.255941 | 0.21256 | 1.204072 | 0.22856 | 0.43266 | 1.48 |
| Gm42528 | 233.907 | -1.08916 | 0.31692 | -3.43667 | 0.00059 | 0.00883 | 3.418 |
| Gm42531 | 18.4454 | 0.586435 | 0.44933 | 1.305137 | 0.19185 | 0.38861 | 4.048 |
| Gm42544 | 39.9573 | -0.74765 | 0.37685 | -1.98394 | 0.04726 | 0.15964 | 3.242 |

| | | | | | | | |
|---|---|---|---|---|---|---|---|
| Gm42547 | 560.769 | 0.036753 | 0.11892 | 0.309061 | 0.75727 | 0.87221 | 3.798 |
| Gm42548 | 22.5542 | -0.62445 | 0.49801 | -1.2539 | 0.20988 | 0.40994 | 3.933 |
| Gm42549 | 152.724 | 0.301511 | 0.17552 | 1.717831 | 0.08583 | 0.2354 | 7.004 |
| Gm42559 | 36.4044 | -0.04366 | 0.40039 | -0.10905 | 0.91316 | 0.95612 | 3.532 |
| Gm42568 | 77.0853 | -0.10153 | 0.22427 | -0.45272 | 0.65075 | 0.80552 | 1.841 |
| Gm42572 | 60.2561 | -0.48839 | 0.21618 | -2.25919 | 0.02387 | 0.10058 | 3.378 |
| Gm42576 | 26.3834 | -0.27039 | 0.42661 | -0.63382 | 0.5262 | 0.71436 | 2.846 |
| Gm42577 | 40.89 | 0.253622 | 0.32653 | 0.776716 | 0.43733 | 0.64276 | 3.211 |
| Gm42578 | 13.3028 | 0.523552 | 0.54477 | 0.961052 | 0.33653 | 0.54716 | 0.194 |
| Gm4258 | 18.249 | -1.34924 | 0.46776 | -2.88445 | 0.00392 | 0.03101 | 111.19 |
| Gm42583 | 189.044 | 0.081514 | 0.26699 | 0.305311 | 0.76013 | 0.87418 | 6.7 |
| Gm42585 | 20.4186 | -0.53331 | 0.46497 | -1.14697 | 0.25139 | 0.4588 | 4.02 |
| Gm42589 | 20.0789 | 0.439108 | 0.50749 | 0.865252 | 0.3869 | 0.59616 | 1.751 |
| Gm42595 | 120.596 | 0.116819 | 0.1845 | 0.633173 | 0.52662 | 0.71469 | 1.607 |
| Gm42600 | 45.8474 | 0.466888 | 0.28074 | 1.663053 | 0.0963 | 0.25407 | 3.588 |
| Gm42607 | 62.4559 | -0.11535 | 0.22037 | -0.52342 | 0.60068 | 0.77104 | 3.393 |
| Gm42608 | 446.763 | 0.38754 | 0.18987 | 2.04113 | 0.04124 | 0.14561 | 2.018 |
| Gm42611 | 61.0262 | 0.043696 | 0.23784 | 0.183722 | 0.85423 | 0.9262 | 3.407 |
| Gm42615 | 14.7908 | -0.02086 | 0.4971 | -0.04196 | 0.96653 | 0.98327 | 1.82 |
| Gm42616 | 22.6919 | -0.29572 | 0.39263 | -0.75317 | 0.45135 | 0.65382 | 4.184 |
| Gm42617 | 19.8482 | -0.46399 | 0.42917 | -1.08113 | 0.27964 | 0.48966 | 1.229 |
| Gm42632 | 39.5724 | 0.239146 | 0.32777 | 0.729607 | 0.46563 | 0.665 | 2.411 |
| Gm42633 | 12.4392 | -0.0927 | 0.54217 | -0.17099 | 0.86423 | 0.93125 | 2.764 |
| Gm42636 | 24.1565 | 0.272384 | 0.35423 | 0.768949 | 0.44192 | 0.64653 | 2.239 |
| Gm42639 | 12.7283 | -0.07257 | 0.46616 | -0.15568 | 0.87629 | 0.93824 | 1.871 |
| Gm42648 | 22.942 | -0.12864 | 0.4487 | -0.2867 | 0.77434 | 0.88244 | 2.505 |
| Gm42651 | 20.2038 | 0.255473 | 0.3798 | 0.672655 | 0.50117 | 0.69395 | 1.339 |
| Gm42656 | 101.645 | -0.41013 | 0.22342 | -1.83566 | 0.06641 | 0.19936 | 2.305 |
| Gm42658 | 47.243 | 0.4117 | 0.26916 | 1.529567 | 0.12612 | 0.3018 | 1.575 |
| Gm42659 | 67.2072 | -0.07228 | 0.26724 | -0.27046 | 0.78681 | 0.88907 | 1.754 |
| Gm42664 | 315.1 | 0.149878 | 0.11769 | 1.273496 | 0.20284 | 0.40148 | 3.239 |
| Gm42668 | 15.4442 | 1.465802 | 0.50624 | 2.895483 | 0.00379 | 0.03031 | 2.487 |
| Gm42672 | 137.957 | 0.126037 | 0.18235 | 0.69117 | 0.48946 | 0.68458 | 0.352 |
| Gm42686 | 12.9349 | -0.13965 | 0.4876 | -0.28641 | 0.77457 | 0.88245 | 4.108 |

| | | | | | | | |
|---|---|---|---|---|---|---|---|
| Gm42688 | 26.4994 | -0.28249 | 0.35363 | -0.79881 | 0.4244 | 0.63109 | 8.323 |
| Gm42690 | 18.2458 | 0.441478 | 0.44197 | 0.998898 | 0.31784 | 0.52829 | 1.901 |
| Gm42692 | 59.3831 | 0.326405 | 0.33184 | 0.983633 | 0.3253 | 0.53652 | 2.833 |
| Gm42693 | 14.0179 | -0.37198 | 0.43499 | -0.85515 | 0.39247 | 0.60165 | 0.631 |
| Gm42715 | 97.3025 | -0.1872 | 0.22475 | -0.83293 | 0.40489 | 0.61384 | 47.471 |
| Gm42717 | 47.0719 | 0.665426 | 0.32851 | 2.025583 | 0.04281 | 0.14911 | 3.463 |
| Gm42722 | 30.0824 | 1.019257 | 0.40796 | 2.49841 | 0.01248 | 0.06631 | 15.979 |
| Gm42723 | 62.5897 | -0.17728 | 0.31676 | -0.55968 | 0.5757 | 0.7536 | 15.188 |
| Gm42725 | 89.6156 | 0.46203 | 0.25685 | 1.798834 | 0.07204 | 0.21073 | 4.193 |
| Gm42726 | 31.8968 | 0.298304 | 0.35979 | 0.829096 | 0.40705 | 0.61503 | 3.725 |
| Gm42727 | 36.5686 | 0.133389 | 0.27157 | 0.491187 | 0.62329 | 0.78601 | 2.823 |
| Gm42731 | 14.5526 | 0.247261 | 0.65587 | 0.377 | 0.70617 | 0.84153 | 1.641 |
| Gm42732 | 1310.56 | 0.010182 | 0.08725 | 0.116698 | 0.9071 | 0.95354 | 1.197 |
| Gm42735 | 125.088 | -0.04725 | 0.19545 | -0.24176 | 0.80896 | 0.90168 | 1.646 |
| Gm42737 | 16.2982 | -0.16466 | 0.50641 | -0.32515 | 0.74507 | 0.866 | 1.755 |
| Gm42738 | 14.7457 | -0.88887 | 0.59 | -1.50656 | 0.13192 | 0.31074 | 1.364 |
| Gm42741 | 337.187 | -0.08978 | 0.15295 | -0.58697 | 0.55722 | 0.73902 | 2.278 |
| Gm42748 | 301.282 | 0.598803 | 0.15918 | 3.761914 | 0.00017 | 0.00386 | 2.341 |
| Gm42756 | 337.464 | -0.01069 | 0.26366 | -0.04053 | 0.96767 | 0.98368 | 17.343 |
| Gm42764 | 15.3425 | 1.596749 | 0.48876 | 3.266927 | 0.00109 | 0.0135 | 1.446 |
| Gm42767 | 15.827 | -0.5412 | 0.53702 | -1.00778 | 0.31356 | 0.52413 | 0.632 |
| Gm42768 | 27.5525 | -0.40299 | 0.43131 | -0.93434 | 0.35013 | 0.56078 | 2.897 |
| Gm42769 | 50.9748 | 0.647618 | 0.29272 | 2.212432 | 0.02694 | 0.10955 | 1.659 |
| Gm42770 | 54.5596 | -0.26878 | 0.33245 | -0.80849 | 0.41881 | 0.62588 | 4.155 |
| Gm42772 | 38.6479 | -0.60749 | 0.28551 | -2.12774 | 0.03336 | 0.12686 | 3.041 |
| Gm42775 | 17.5864 | 0.388871 | 0.50873 | 0.764399 | 0.44463 | 0.64863 | 4.986 |
| Gm42778 | 27.6164 | -0.67676 | 0.40445 | -1.67329 | 0.09427 | 0.25054 | 0.553 |
| Gm42782 | 16.638 | 0.298687 | 0.39541 | 0.755391 | 0.45001 | 0.65294 | 0.522 |
| Gm42783 | 37.3488 | -0.01191 | 0.28845 | -0.04128 | 0.96707 | 0.98348 | 0.795 |
| Gm42788 | 12.6109 | -1.04313 | 0.53026 | -1.9672 | 0.04916 | 0.16406 | 9.278 |
| Gm42790 | 17.6768 | -0.76955 | 0.52525 | -1.4651 | 0.14289 | 0.32494 | 0.969 |
| Gm42791 | 14.5492 | -0.42741 | 0.44269 | -0.96547 | 0.33431 | 0.54499 | 101.373 |
| Gm42797 | 16.4484 | 0.385016 | 0.47785 | 0.805726 | 0.4204 | 0.62766 | 1.58 |
| Gm42798 | 52.4226 | 0.119001 | 0.24429 | 0.487124 | 0.62617 | 0.78844 | 3.972 |

| | | | | | | | |
|---|---|---|---|---|---|---|---|
| Gm42808 | 16.4055 | -0.43919 | 0.47303 | -0.92847 | 0.35316 | 0.56347 | 0.907 |
| Gm42811 | 27.1565 | 0.202553 | 0.3941 | 0.51396 | 0.60728 | 0.77542 | 2.768 |
| Gm42812 | 13.6573 | 0.480791 | 0.57836 | 0.831305 | 0.4058 | 0.6144 | 9.074 |
| Gm42822 | 13.4764 | 0.591636 | 0.6294 | 0.939993 | 0.34722 | 0.55778 | 1.529 |
| Gm42829 | 362.392 | -0.11745 | 0.15886 | -0.7393 | 0.45973 | 0.66021 | 2.585 |
| Gm42837 | 34.5564 | 0.925923 | 0.3515 | 2.634172 | 0.00843 | 0.05118 | 2.829 |
| Gm42840 | 30.6775 | -0.53906 | 0.31624 | -1.70457 | 0.08827 | 0.23998 | 2.251 |
| Gm42841 | 13.4049 | 0.300893 | 0.58466 | 0.514645 | 0.6068 | 0.7751 | 1.932 |
| Gm42842 | 13.0434 | -0.55336 | 0.46453 | -1.1912 | 0.23357 | 0.43857 | 3.004 |
| Gm42843 | 19.3821 | -0.05781 | 0.46465 | -0.12442 | 0.90098 | 0.9503 | 3.616 |
| Gm42847 | 251.604 | -0.47055 | 0.1563 | -3.01055 | 0.00261 | 0.02405 | 2.553 |
| Gm42848 | 68.1067 | 0.238412 | 0.30927 | 0.770895 | 0.44077 | 0.64559 | 4.823 |
| Gm42849 | 45.6551 | -0.06315 | 0.28011 | -0.22544 | 0.82164 | 0.90766 | 1.807 |
| Gm4285 | 110.975 | -0.06191 | 0.1923 | -0.32196 | 0.74748 | 0.86763 | 1.6 |
| Gm42851 | 25.0135 | -0.08243 | 0.35729 | -0.23071 | 0.81754 | 0.90613 | 1.411 |
| Gm42853 | 122.063 | -0.39848 | 0.30955 | -1.28728 | 0.198 | 0.39572 | 1.387 |
| Gm42858 | 37.539 | 0.30618 | 0.32084 | 0.954314 | 0.33992 | 0.55075 | 1.704 |
| Gm42864 | 18.9436 | -1.14509 | 0.86264 | -1.32742 | 0.18437 | 0.37881 | 2.746 |
| Gm42869 | 27.8133 | 0.537213 | 0.36035 | 1.490789 | 0.13602 | 0.31625 | 2.121 |
| Gm42873 | 13.8572 | 0.554066 | 0.44638 | 1.241229 | 0.21452 | 0.4159 | 1.726 |
| Gm42875 | 57.2785 | 0.151313 | 0.25823 | 0.585965 | 0.5579 | 0.73951 | 8.314 |
| Gm42878 | 246.213 | 0.233307 | 0.15509 | 1.504294 | 0.13251 | 0.31164 | 26.87 |
| Gm42881 | 33.6669 | -0.2322 | 0.37001 | -0.62755 | 0.5303 | 0.71791 | 3.344 |
| Gm42890 | 24.5168 | 0.326833 | 0.3882 | 0.841925 | 0.39983 | 0.60858 | 25.098 |
| Gm42892 | 38.1617 | -0.10281 | 0.34532 | -0.29774 | 0.7659 | 0.8779 | 3.973 |
| Gm42894 | 12.6226 | -0.47063 | 0.57364 | -0.82042 | 0.41198 | 0.61994 | 3.39 |
| Gm42901 | 15.9434 | -0.11004 | 0.41366 | -0.26603 | 0.79022 | 0.89117 | 3.171 |
| Gm42902 | 143.244 | 0.052649 | 0.19681 | 0.267519 | 0.78907 | 0.89046 | 10.75 |
| Gm42913 | 93.7757 | 0.314218 | 0.23289 | 1.349223 | 0.17727 | 0.36947 | 1.853 |
| Gm42917 | 17.4634 | -0.51535 | 0.50453 | -1.02144 | 0.30705 | 0.51683 | 9.456 |
| Gm42918 | 37.8973 | -0.41028 | 0.32209 | -1.2738 | 0.20273 | 0.40133 | 1.583 |
| Gm42923 | 12.9035 | 0.185053 | 0.54616 | 0.338826 | 0.73474 | 0.85968 | 2.896 |
| Gm42929 | 15.6009 | -0.26443 | 0.48534 | -0.54482 | 0.58588 | 0.76012 | 1.286 |
| Gm42930 | 49.1195 | 0.180672 | 0.26414 | 0.684013 | 0.49397 | 0.68838 | 3.26 |

| | | | | | | | |
|---|---|---|---|---|---|---|---|
| Gm42937 | 46.9204 | -0.23719 | 0.30987 | -0.76546 | 0.444 | 0.6481 | 0.752 |
| Gm4294 | 21.1105 | -0.36111 | 0.4136 | -0.8731 | 0.38261 | 0.592 | 0.614 |
| Gm42940 | 36.844 | -0.10629 | 0.46499 | -0.22859 | 0.81919 | 0.90697 | 2.362 |
| Gm42941 | 24.4599 | 0.244961 | 0.44019 | 0.556483 | 0.57788 | 0.75477 | 1.321 |
| Gm42943 | 13.8179 | -0.71766 | 0.57317 | -1.25208 | 0.21054 | 0.41056 | 3.242 |
| Gm42944 | 62.4482 | -0.52781 | 0.25051 | -2.10698 | 0.03512 | 0.13139 | 3.98 |
| Gm42946 | 18.1978 | -1.75165 | 0.57022 | -3.07189 | 0.00213 | 0.02101 | 2.57 |
| Gm42967 | 76.4449 | 0.176003 | 0.21837 | 0.80599 | 0.42025 | 0.62755 | 2.312 |
| Gm42969 | 24.2548 | -0.54008 | 0.38114 | -1.41702 | 0.15648 | 0.34327 | 3.631 |
| Gm42970 | 36.941 | -0.27873 | 0.32644 | -0.85383 | 0.3932 | 0.60231 | 1.352 |
| Gm42971 | 22.6994 | -0.93799 | 0.49 | -1.91429 | 0.05558 | 0.1779 | 2.079 |
| Gm42972 | 35.6858 | -0.759 | 0.35345 | -2.14741 | 0.03176 | 0.12278 | 5.617 |
| Gm42973 | 26.3818 | -1.00184 | 0.43907 | -2.28172 | 0.02251 | 0.09685 | 3.2 |
| Gm42979 | 889.247 | 0.210772 | 0.11251 | 1.873403 | 0.06101 | 0.1886 | 2.228 |
| Gm42984 | 83.7161 | 0.398881 | 0.1906 | 2.092775 | 0.03637 | 0.13438 | 16.962 |
| Gm42986 | 17.8813 | -0.16846 | 0.54183 | -0.31091 | 0.75587 | 0.87157 | 3.385 |
| Gm42987 | 12.7085 | 1.112802 | 0.60076 | 1.85233 | 0.06398 | 0.19465 | 2.35 |
| Gm42989 | 17.7608 | 0.543218 | 0.47548 | 1.142454 | 0.25327 | 0.46062 | 2.306 |
| Gm42991 | 16.4395 | -0.25595 | 0.64987 | -0.39385 | 0.69369 | 0.83384 | 3.837 |
| Gm42992 | 32.1695 | -0.11814 | 0.44448 | -0.26579 | 0.7904 | 0.89129 | 6.836 |
| Gm42993 | 27.5635 | -0.63672 | 0.47784 | -1.33252 | 0.18269 | 0.37661 | 7.325 |
| Gm43011 | 15.8873 | -1.08479 | 0.52775 | -2.05551 | 0.03983 | 0.14259 | 4.433 |
| Gm43012 | 216.482 | 0.382548 | 0.13374 | 2.860309 | 0.00423 | 0.03267 | 0.937 |
| Gm43029 | 87.9611 | 0.22123 | 0.19912 | 1.111056 | 0.26654 | 0.47588 | 1.535 |
| Gm43031 | 240.744 | 0.449484 | 0.19805 | 2.269598 | 0.02323 | 0.09878 | 1.887 |
| Gm43032 | 134.252 | 0.147592 | 0.21275 | 0.693731 | 0.48785 | 0.68299 | 3.123 |
| Gm43034 | 26.5867 | -0.72629 | 0.37008 | -1.96255 | 0.0497 | 0.16518 | 1.364 |
| Gm43039 | 81.0914 | 0.289476 | 0.24973 | 1.159168 | 0.24639 | 0.45316 | 0.963 |
| Gm43046 | 15.6045 | -0.52042 | 0.45465 | -1.14466 | 0.25235 | 0.4596 | 2.451 |
| Gm43047 | 20.6776 | -0.66198 | 0.71824 | -0.92167 | 0.3567 | 0.56705 | 3.46 |
| Gm43051 | 14.4069 | -0.70755 | 0.55431 | -1.27645 | 0.2018 | 0.40022 | 2.678 |
| Gm43052 | 12.8788 | -1.5639 | 0.63764 | -2.45264 | 0.01418 | 0.07176 | 1.628 |
| Gm43054 | 25.6117 | -0.438 | 0.37672 | -1.16266 | 0.24497 | 0.45156 | 0.884 |
| Gm43059 | 124.646 | 0.116468 | 0.16782 | 0.694008 | 0.48768 | 0.68296 | 1.388 |

| | | | | | | | |
|---|---|---|---|---|---|---|---|
| Gm43061 | 33.0797 | -0.22014 | 0.29404 | -0.74867 | 0.45406 | 0.65623 | 2.241 |
| Gm43062 | 45.4302 | -0.00731 | 0.25526 | -0.02862 | 0.97717 | 0.98797 | 1.319 |
| Gm43066 | 64.412 | 0.403878 | 0.28165 | 1.433995 | 0.15157 | 0.33655 | 1.26 |
| Gm43071 | 151.66 | 0.385041 | 0.22604 | 1.703437 | 0.08849 | 0.24033 | 2.032 |
| Gm43080 | 359.278 | 0.426668 | 0.12659 | 3.370525 | 0.00075 | 0.01051 | 2.643 |
| Gm43081 | 24.035 | 0.344572 | 0.39649 | 0.869059 | 0.38482 | 0.59425 | 1.38 |
| Gm43088 | 74.2797 | 0.37794 | 0.24563 | 1.53867 | 0.12388 | 0.29839 | 2.347 |
| Gm43089 | 71.9747 | 0.376363 | 0.3108 | 1.210963 | 0.22591 | 0.42994 | 1.213 |
| Gm43099 | 39.907 | -0.70412 | 0.33613 | -2.09475 | 0.03619 | 0.134 | 5.241 |
| Gm43119 | 27.2662 | -0.34056 | 0.41037 | -0.82988 | 0.40661 | 0.6148 | 2.122 |
| Gm43128 | 40.209 | -0.7492 | 0.31055 | -2.4125 | 0.01584 | 0.07722 | 1.621 |
| Gm43133 | 30.66 | -0.43441 | 0.36956 | -1.17549 | 0.2398 | 0.44574 | 0.463 |
| Gm43136 | 15.0069 | -0.97155 | 0.52309 | -1.85734 | 0.06326 | 0.19325 | 0.636 |
| Gm43138 | 29.9123 | 0.147803 | 0.34165 | 0.432615 | 0.66529 | 0.81526 | 0.708 |
| Gm43140 | 106.525 | 0.576095 | 0.21203 | 2.717094 | 0.00659 | 0.04374 | 0.691 |
| Gm43143 | 27.7381 | -0.41624 | 0.40881 | -1.01817 | 0.3086 | 0.51837 | 2.023 |
| Gm43154 | 38.1906 | 0.29 | 0.32765 | 0.88508 | 0.37611 | 0.58618 | 192.864 |
| Gm4316 | 12.3882 | -0.11344 | 0.49748 | -0.22803 | 0.81963 | 0.90729 | 13.511 |
| Gm43162 | 30.2699 | 0.104125 | 0.55328 | 0.188194 | 0.85072 | 0.92407 | 1.77 |
| Gm43169 | 28.175 | -0.52391 | 0.4437 | -1.18078 | 0.23769 | 0.44312 | 4.734 |
| Gm43174 | 32.7384 | 0.058439 | 0.41216 | 0.141787 | 0.88725 | 0.94393 | 2.445 |
| Gm43175 | 28.5999 | -0.70999 | 0.34262 | -2.07223 | 0.03824 | 0.13906 | 3.81 |
| Gm43176 | 18.9227 | -0.0282 | 0.45995 | -0.06131 | 0.95111 | 0.97595 | 2.989 |
| Gm43185 | 96.7688 | 0.758028 | 0.60206 | 1.259049 | 0.20801 | 0.40767 | 2.775 |
| Gm43190 | 15.1736 | -0.83113 | 0.49997 | -1.66235 | 0.09644 | 0.25432 | 1.779 |
| Gm43195 | 15.3034 | 0.216346 | 0.40612 | 0.532718 | 0.59423 | 0.76639 | 2.332 |
| Gm43196 | 25.1258 | -0.79558 | 0.56808 | -1.40047 | 0.16137 | 0.3496 | 2.114 |
| Gm43197 | 34.0248 | -0.56695 | 0.43824 | -1.29368 | 0.19578 | 0.39346 | 2.583 |
| Gm43203 | 26.1592 | 0.089437 | 0.37063 | 0.241309 | 0.80932 | 0.90175 | 0.558 |
| Gm43205 | 63.3375 | -0.00142 | 0.22344 | -0.00635 | 0.99493 | 0.99709 | 1.968 |
| Gm43210 | 17.44 | 0.435676 | 0.58891 | 0.739798 | 0.45942 | 0.65996 | 6.152 |
| Gm43216 | 48.1644 | 0.809304 | 0.27339 | 2.960275 | 0.00307 | 0.02648 | 1.333 |
| Gm43221 | 62.8136 | 1.121239 | 0.27174 | 4.12619 | 3.69E-05 | 0.00132 | 2.071 |
| Gm43231 | 31.5717 | 0.046995 | 0.41847 | 0.112301 | 0.91059 | 0.9549 | 2.258 |

| | | | | | | | |
|---|---|---|---|---|---|---|---|
| Gm43237 | 13.0651 | -1.10815 | 0.53776 | -2.06066 | 0.03934 | 0.14138 | 1.044 |
| Gm43254 | 153.095 | 0.309148 | 0.18752 | 1.648608 | 0.09923 | 0.25882 | 2.491 |
| Gm43255 | 33.4026 | -0.19973 | 0.33972 | -0.58792 | 0.55659 | 0.73867 | 2.335 |
| Gm43266 | 23.8474 | 0.175214 | 0.38324 | 0.457197 | 0.64753 | 0.80377 | 1.771 |
| Gm43267 | 79.3841 | -0.1107 | 0.20383 | -0.54313 | 0.58704 | 0.76058 | 1.827 |
| Gm43268 | 666.33 | 0.261198 | 0.13858 | 1.88478 | 0.05946 | 0.1855 | 4.055 |
| Gm43272 | 276.341 | -0.05821 | 0.16978 | -0.34287 | 0.7317 | 0.85759 | 1.584 |
| Gm43273 | 16.8331 | -0.69778 | 0.45522 | -1.53282 | 0.12532 | 0.30036 | 2.594 |
| Gm43274 | 14.9153 | -0.40337 | 0.53333 | -0.75632 | 0.44946 | 0.65264 | 5 |
| Gm43279 | 60.9745 | 0.269619 | 0.23329 | 1.155703 | 0.2478 | 0.45486 | 2.512 |
| Gm43290 | 27.8438 | 0.08971 | 0.32328 | 0.277498 | 0.7814 | 0.88615 | 1.557 |
| Gm43292 | 13.0667 | 0.262886 | 0.53487 | 0.491492 | 0.62308 | 0.78589 | 2.449 |
| Gm43294 | 41.491 | 0.091639 | 0.3219 | 0.28468 | 0.77589 | 0.88331 | 2.49 |
| Gm43300 | 257.578 | 0.434649 | 0.21763 | 1.997175 | 0.04581 | 0.15602 | 2.043 |
| Gm43305 | 291.114 | -0.48786 | 0.25609 | -1.90504 | 0.05677 | 0.18035 | 5.014 |
| Gm43311 | 22.5611 | 0.166325 | 0.4477 | 0.371505 | 0.71026 | 0.8441 | 2.377 |
| Gm43313 | 13.7368 | -0.87255 | 0.56409 | -1.54683 | 0.12191 | 0.29538 | 2.235 |
| Gm43314 | 21.783 | 0.253997 | 0.41024 | 0.619135 | 0.53583 | 0.72278 | 2.231 |
| Gm43316 | 15.9197 | -1.0979 | 0.65809 | -1.66832 | 0.09525 | 0.25219 | 1.295 |
| Gm43317 | 28.64 | -0.35529 | 0.51887 | -0.68473 | 0.49352 | 0.68795 | 2.544 |
| Gm43318 | 26.6333 | 0.000813 | 0.3698 | 0.002199 | 0.99825 | 0.99908 | 1.949 |
| Gm43319 | 23.1808 | -0.84888 | 0.46453 | -1.82739 | 0.06764 | 0.20161 | 2.27 |
| Gm4332 | 172.97 | -0.29602 | 0.17185 | -1.7226 | 0.08496 | 0.23382 | 0.346 |
| Gm43320 | 22.9788 | -0.42534 | 0.44585 | -0.95399 | 0.34009 | 0.55085 | 2.916 |
| Gm43321 | 24.2352 | -0.07238 | 0.41352 | -0.17502 | 0.86106 | 0.92965 | 2.409 |
| Gm43322 | 60.9756 | 0.016256 | 0.3465 | 0.046914 | 0.96258 | 0.98185 | 1.612 |
| Gm43327 | 17.4236 | -0.39534 | 0.67947 | -0.58184 | 0.56067 | 0.74193 | 4.055 |
| Gm43328 | 105.437 | 0.158348 | 0.22249 | 0.711701 | 0.47665 | 0.67458 | 1.434 |
| Gm43335 | 13.9351 | 0.582556 | 0.76118 | 0.765334 | 0.44407 | 0.6481 | 3.276 |
| Gm43336 | 38.1205 | -0.29787 | 0.27933 | -1.06638 | 0.28625 | 0.49628 | 4.504 |
| Gm43338 | 36.0821 | 0.548215 | 0.31568 | 1.73663 | 0.08245 | 0.22898 | 2.223 |
| Gm43339 | 172.091 | -0.00143 | 0.2337 | -0.0061 | 0.99513 | 0.99724 | 1.606 |
| Gm43359 | 42.5229 | -0.66689 | 0.35353 | -1.88636 | 0.05925 | 0.18525 | 2.2 |
| Gm43360 | 27.3827 | -0.52867 | 0.50665 | -1.04346 | 0.29674 | 0.50689 | 2.887 |

| | | | | | | | |
|---|---|---|---|---|---|---|---|
| Gm43361 | 42.9886 | -0.67197 | 0.30601 | -2.19593 | 0.0281 | 0.11285 | 4.69 |
| Gm43362 | 28.2169 | 0.489694 | 0.38475 | 1.272746 | 0.20311 | 0.40172 | 2.47 |
| Gm43364 | 28.372 | -0.35327 | 0.36407 | -0.97034 | 0.33188 | 0.5428 | 3.274 |
| Gm43373 | 14.3878 | 0.155942 | 0.67017 | 0.232691 | 0.816 | 0.90537 | 2.87 |
| Gm43375 | 56.0509 | 0.065552 | 0.23089 | 0.283906 | 0.77648 | 0.88361 | 2.48 |
| Gm43379 | 305.841 | 0.099116 | 0.1689 | 0.586839 | 0.55731 | 0.73902 | 7.726 |
| Gm43387 | 13.3243 | 0.387457 | 0.52564 | 0.73712 | 0.46105 | 0.66169 | 1.066 |
| Gm43389 | 354.398 | 0.666706 | 0.14301 | 4.662049 | 3.13E-06 | 0.00024 | 2.595 |
| Gm43398 | 59.4582 | -0.72014 | 0.33145 | -2.1727 | 0.0298 | 0.11771 | 2.742 |
| Gm43413 | 142.261 | 0.357149 | 0.14633 | 2.44072 | 0.01466 | 0.07316 | 1.566 |
| Gm43414 | 38.9856 | 0.435783 | 0.31136 | 1.399616 | 0.16163 | 0.34982 | 3.188 |
| Gm43417 | 18.178 | -0.67711 | 0.52126 | -1.29899 | 0.19395 | 0.391 | 5.207 |
| Gm43420 | 24.1442 | 0.157261 | 0.38068 | 0.413105 | 0.67953 | 0.82491 | 1.857 |
| Gm43421 | 14.3259 | -0.31052 | 0.49343 | -0.62931 | 0.52914 | 0.71697 | 2.905 |
| Gm43422 | 14.3136 | 0.291868 | 0.53191 | 0.548719 | 0.5832 | 0.75835 | 3.571 |
| Gm43430 | 26.5064 | 0.219252 | 0.35078 | 0.625048 | 0.53194 | 0.71956 | 3.897 |
| Gm43437 | 254.05 | -0.06198 | 0.1833 | -0.33816 | 0.73525 | 0.85989 | 3.955 |
| Gm43442 | 18.5312 | -0.1963 | 0.60624 | -0.3238 | 0.74609 | 0.86672 | 0.414 |
| Gm43445 | 189.731 | 0.134107 | 0.17882 | 0.749949 | 0.45329 | 0.65558 | 2.091 |
| Gm43457 | 150.672 | 0.050271 | 0.24649 | 0.203944 | 0.8384 | 0.91681 | 2.409 |
| Gm43460 | 85.3247 | 0.121845 | 0.23268 | 0.523659 | 0.60052 | 0.77095 | 1.057 |
| Gm43461 | 89.6167 | 0.422946 | 0.23221 | 1.821416 | 0.06854 | 0.20352 | 0.996 |
| Gm43462 | 23.2814 | 0.20239 | 0.36042 | 0.561538 | 0.57443 | 0.75262 | 3.648 |
| Gm43481 | 551.082 | 0.236445 | 0.11302 | 2.091998 | 0.03644 | 0.13444 | 3.672 |
| Gm43482 | 25.5369 | 0.274005 | 0.42597 | 0.643253 | 0.52006 | 0.70963 | 1.861 |
| Gm43483 | 14.1469 | 0.158164 | 0.54153 | 0.292068 | 0.77023 | 0.87996 | 2.86 |
| Gm43485 | 77.6593 | -0.02577 | 0.2847 | -0.09053 | 0.92787 | 0.9637 | 1.71 |
| Gm43486 | 24.7577 | -0.19987 | 0.46298 | -0.4317 | 0.66596 | 0.81557 | 4.339 |
| Gm43487 | 16.2054 | 0.196717 | 0.47641 | 0.412918 | 0.67967 | 0.82491 | 1.031 |
| Gm43488 | 63.574 | 0.972177 | 0.34608 | 2.80907 | 0.00497 | 0.03608 | 2.504 |
| Gm4349 | 38.0196 | -0.76197 | 0.32006 | -2.38072 | 0.01728 | 0.08154 | 3.516 |
| Gm43502 | 19.549 | 0.489056 | 0.52906 | 0.924395 | 0.35528 | 0.56556 | 2.596 |
| Gm43511 | 59.8857 | 0.637966 | 0.25101 | 2.541616 | 0.01103 | 0.06125 | 2.297 |
| Gm43513 | 34.026 | 0.313831 | 0.35757 | 0.877673 | 0.38012 | 0.58974 | 1.076 |

| | | | | | | | |
|---|---|---|---|---|---|---|---|
| Gm43517 | 214.698 | 0.148959 | 0.16358 | 0.910627 | 0.36249 | 0.57294 | 8.954 |
| Gm43545 | 14.6911 | -0.30929 | 0.45077 | -0.68613 | 0.49263 | 0.68719 | 2.779 |
| Gm43555 | 41.6355 | 0.265895 | 0.27956 | 0.951123 | 0.34154 | 0.55245 | 0.31 |
| Gm4356 | 37.7119 | 0.371249 | 0.30035 | 1.236064 | 0.21643 | 0.41834 | 0.618 |
| Gm43566 | 23.1222 | -0.19309 | 0.3692 | -0.52299 | 0.60098 | 0.77129 | 0.808 |
| Gm43570 | 53.3937 | 0.05405 | 0.32996 | 0.163805 | 0.86988 | 0.93418 | 3.688 |
| Gm43571 | 14.6758 | 0.20128 | 0.54147 | 0.371727 | 0.7101 | 0.84395 | 3.098 |
| Gm43582 | 21.2392 | 0.767461 | 0.39352 | 1.950251 | 0.05115 | 0.16835 | 0.343 |
| Gm43587 | 18.8803 | -0.46018 | 0.42882 | -1.07313 | 0.28321 | 0.49338 | 0.204 |
| Gm43588 | 32.1241 | -0.22176 | 0.36793 | -0.60273 | 0.54669 | 0.73092 | 2.298 |
| Gm43593 | 81.1239 | -0.27578 | 0.32731 | -0.84257 | 0.39947 | 0.60831 | 1.346 |
| Gm43597 | 1684.9 | 0.111742 | 0.09801 | 1.140095 | 0.25425 | 0.46166 | 7.072 |
| Gm43598 | 91.0604 | -0.05927 | 0.24684 | -0.2401 | 0.81025 | 0.90233 | 2.479 |
| Gm43604 | 61.6788 | 0.005797 | 0.28974 | 0.020006 | 0.98404 | 0.99128 | 0.705 |
| Gm43606 | 34.7706 | -1.06318 | 0.32438 | -3.27761 | 0.00105 | 0.01321 | 3.556 |
| Gm43609 | 91.6901 | -0.08176 | 0.24665 | -0.33149 | 0.74028 | 0.863 | 2.712 |
| Gm43612 | 21.0279 | 1.132517 | 0.43428 | 2.607833 | 0.00911 | 0.05385 | 3.207 |
| Gm43622 | 147.728 | 0.084385 | 0.21262 | 0.396878 | 0.69146 | 0.83252 | 2.019 |
| Gm43625 | 21.6264 | -0.35023 | 0.38453 | -0.9108 | 0.3624 | 0.57292 | 1.539 |
| Gm43628 | 12.4752 | 0.103698 | 0.46989 | 0.220684 | 0.82534 | 0.90983 | 1.187 |
| Gm43633 | 16.522 | 0.122779 | 0.40653 | 0.302019 | 0.76264 | 0.87556 | 2.33 |
| Gm43647 | 27.9264 | 0.510748 | 0.41196 | 1.239787 | 0.21505 | 0.41642 | 68.102 |
| Gm43654 | 69.833 | -0.06291 | 0.23168 | -0.27156 | 0.78596 | 0.88868 | 3.417 |
| Gm4366 | 318.436 | -0.26094 | 0.11073 | -2.35648 | 0.01845 | 0.08512 | 1.342 |
| Gm43660 | 414.373 | 0.169959 | 0.15395 | 1.103968 | 0.26961 | 0.47919 | 8.261 |
| Gm43662 | 37.9601 | 0.093339 | 0.33764 | 0.276448 | 0.7822 | 0.88669 | 2.271 |
| Gm43665 | 23.3582 | -0.04326 | 0.50555 | -0.08557 | 0.9318 | 0.96568 | 1.659 |
| Gm43667 | 97.6067 | -0.01251 | 0.29858 | -0.0419 | 0.96658 | 0.98327 | 1.095 |
| Gm43668 | 293.636 | 0.146356 | 0.14222 | 1.029105 | 0.30343 | 0.51314 | 5.342 |
| Gm43672 | 244.207 | -0.31357 | 0.16211 | -1.93434 | 0.05307 | 0.17235 | 1.733 |
| Gm43674 | 47.4412 | -0.53406 | 0.34872 | -1.5315 | 0.12565 | 0.30089 | 0.791 |
| Gm43679 | 73.0631 | -0.12394 | 0.2878 | -0.43065 | 0.66672 | 0.8158 | 3.677 |
| Gm43681 | 432.896 | 0.249498 | 0.11886 | 2.099118 | 0.03581 | 0.13304 | 1.903 |
| Gm43682 | 630.381 | 0.322896 | 0.12674 | 2.547615 | 0.01085 | 0.06059 | 2.09 |

| | | | | | | | |
|---|---|---|---|---|---|---|---|
| Gm43684 | 50.6974 | 0.278401 | 0.33649 | 0.827363 | 0.40803 | 0.61578 | 6.573 |
| Gm43685 | 13.4558 | 0.494912 | 0.5664 | 0.87378 | 0.38224 | 0.59182 | 1.249 |
| Gm43686 | 40.3396 | 0.016174 | 0.32767 | 0.04936 | 0.96063 | 0.98079 | 2.439 |
| Gm43690 | 61.9746 | 0.272566 | 0.22598 | 1.206151 | 0.22776 | 0.43174 | 0.266 |
| Gm43692 | 65.2446 | 0.137664 | 0.25065 | 0.549228 | 0.58285 | 0.75814 | 1.062 |
| Gm43693 | 101.899 | 0.497141 | 0.21319 | 2.331901 | 0.01971 | 0.08869 | 1.249 |
| Gm43696 | 40.3268 | -0.35326 | 0.43404 | -0.81389 | 0.41571 | 0.62362 | 6.071 |
| Gm43702 | 18.7022 | -0.07051 | 0.39326 | -0.17931 | 0.8577 | 0.92823 | 2.546 |
| Gm43719 | 65.0448 | 0.063636 | 0.24564 | 0.259062 | 0.79559 | 0.8946 | 2.199 |
| Gm43737 | 400.185 | 0.00949 | 0.12643 | 0.07506 | 0.94017 | 0.97025 | 2.74 |
| Gm43743 | 25.0117 | -0.2548 | 0.3775 | -0.67498 | 0.49969 | 0.6928 | 0.939 |
| Gm43745 | 22.4396 | 0.464753 | 0.43703 | 1.063431 | 0.28759 | 0.49771 | 1.883 |
| Gm43747 | 15.9961 | 0.275574 | 0.42313 | 0.651279 | 0.51487 | 0.70526 | 1.33 |
| Gm43750 | 15.4673 | -0.13268 | 0.47056 | -0.28195 | 0.77798 | 0.88423 | 2.493 |
| Gm43751 | 36.3378 | -0.18568 | 0.33494 | -0.55437 | 0.57933 | 0.75592 | 3.071 |
| Gm43755 | 17.4947 | 0.164593 | 0.43165 | 0.38131 | 0.70297 | 0.8397 | 0.445 |
| Gm43758 | 30.2404 | 0.467037 | 0.30196 | 1.546676 | 0.12194 | 0.29538 | 0.367 |
| Gm43759 | 396.775 | 0.366035 | 0.11352 | 3.224305 | 0.00126 | 0.01476 | 1.261 |
| Gm43760 | 72.5387 | 1.138102 | 0.26614 | 4.27627 | 1.90E-05 | 0.00086 | 3.015 |
| Gm43766 | 51.5114 | 0.003132 | 0.33113 | 0.00946 | 0.99245 | 0.99567 | 5.417 |
| Gm43776 | 29.6086 | 0.461185 | 0.40647 | 1.13462 | 0.25653 | 0.46453 | 2.914 |
| Gm43787 | 13.5165 | 0.105269 | 0.55361 | 0.190149 | 0.84919 | 0.9233 | 1.037 |
| Gm43789 | 39.2825 | -0.29096 | 0.34071 | -0.85398 | 0.39312 | 0.60231 | 2.472 |
| Gm43790 | 17.4279 | 0.40537 | 0.4242 | 0.955615 | 0.33927 | 0.55014 | 2.013 |
| Gm43793 | 21.9898 | 0.052976 | 0.39395 | 0.134474 | 0.89303 | 0.94632 | 1.333 |
| Gm43800 | 56.5051 | 0.459038 | 0.24475 | 1.875544 | 0.06072 | 0.1881 | 2.731 |
| Gm43808 | 69.0066 | 0.067127 | 0.27737 | 0.242016 | 0.80877 | 0.90165 | 2.065 |
| Gm43813 | 22.0039 | 0.263304 | 0.55423 | 0.475083 | 0.63473 | 0.79438 | 5.378 |
| Gm43823 | 21.4877 | -0.42887 | 0.40618 | -1.05587 | 0.29103 | 0.50115 | 1.847 |
| Gm43829 | 26.7275 | -0.15937 | 0.35677 | -0.44669 | 0.6551 | 0.80862 | 3.802 |
| Gm43843 | 1146.5 | 0.010878 | 0.12786 | 0.085079 | 0.9322 | 0.96589 | 1.587 |
| Gm43844 | 58.4282 | 0.0835 | 0.2528 | 0.330306 | 0.74117 | 0.86331 | 5.605 |
| Gm43847 | 15.9689 | 0.503311 | 0.54294 | 0.927005 | 0.35392 | 0.5642 | 1.94 |
| Gm43848 | 68.4563 | 0.159352 | 0.22934 | 0.694819 | 0.48717 | 0.68271 | 1.247 |

| | | | | | | | |
|---|---|---|---|---|---|---|---|
| Gm43850 | 62.0409 | 0.193868 | 0.23875 | 0.812015 | 0.41678 | 0.62432 | 1.045 |
| Gm43852 | 30.3321 | 0.206518 | 0.33076 | 0.624374 | 0.53238 | 0.71981 | 0.917 |
| Gm43857 | 125.346 | 0.300702 | 0.25793 | 1.16582 | 0.24369 | 0.4501 | 3.079 |
| Gm43858 | 468.289 | 0.372459 | 0.15253 | 2.441914 | 0.01461 | 0.07303 | 1.778 |
| Gm43859 | 191.825 | 0.462796 | 0.1703 | 2.717555 | 0.00658 | 0.04371 | 1.412 |
| Gm43860 | 120.056 | 0.092822 | 0.18757 | 0.494873 | 0.62069 | 0.78424 | 2.705 |
| Gm43863 | 15.6542 | -0.10528 | 0.5027 | -0.20943 | 0.83411 | 0.9149 | 1.192 |
| Gm43864 | 15.0124 | -0.91271 | 0.54363 | -1.67893 | 0.09317 | 0.24849 | 4.028 |
| Gm43869 | 14.9892 | -0.15762 | 0.49852 | -0.31618 | 0.75186 | 0.86994 | 1.405 |
| Gm43872 | 31.0414 | 0.046525 | 0.41952 | 0.1109 | 0.9117 | 0.95543 | 2.213 |
| Gm43874 | 22.2252 | -0.09255 | 0.36461 | -0.25383 | 0.79963 | 0.89651 | 2.798 |
| Gm43879 | 36.9831 | 0.287304 | 0.50382 | 0.570252 | 0.56851 | 0.74835 | 3.76 |
| Gm43889 | 27.9837 | 0.023102 | 0.44161 | 0.052314 | 0.95828 | 0.9796 | 3.681 |
| Gm43890 | 57.3784 | 0.308789 | 0.23601 | 1.308394 | 0.19074 | 0.38701 | 2.799 |
| Gm43909 | 24.923 | -0.04615 | 0.3318 | -0.13908 | 0.88939 | 0.94464 | 1.876 |
| Gm43912 | 36.8908 | -0.31158 | 0.30461 | -1.0229 | 0.30635 | 0.51639 | 0.538 |
| Gm43915 | 25.6231 | 0.055352 | 0.42108 | 0.131454 | 0.89542 | 0.94744 | 38.655 |
| Gm43917 | 38.8032 | 0.629436 | 0.33025 | 1.905912 | 0.05666 | 0.18021 | 2.36 |
| Gm43937 | 36.2386 | 0.436899 | 0.3632 | 1.202912 | 0.22901 | 0.43312 | 3.122 |
| Gm43963 | 27.0432 | 0.280226 | 0.34873 | 0.803564 | 0.42165 | 0.62865 | 2.488 |
| Gm43980 | 1546.45 | 0.274917 | 0.10412 | 2.640339 | 0.00828 | 0.0507 | 1.612 |
| Gm43999 | 16.3127 | 0.826474 | 0.47147 | 1.752961 | 0.07961 | 0.22407 | 2.889 |
| Gm44001 | 30.0908 | 0.768083 | 0.32386 | 2.371679 | 0.01771 | 0.08269 | 3.773 |
| Gm44005 | 40.5385 | 0.297057 | 0.2619 | 1.134252 | 0.25669 | 0.46473 | 3.173 |
| Gm44008 | 18.7624 | 0.022832 | 0.46871 | 0.048713 | 0.96115 | 0.98112 | 0.412 |
| Gm44014 | 19.8893 | 1.069371 | 0.44983 | 2.377293 | 0.01744 | 0.08197 | 1.509 |
| Gm44017 | 15.3266 | 0.340979 | 0.52641 | 0.647742 | 0.51715 | 0.70706 | 1.444 |
| Gm44020 | 21.9312 | 0.761853 | 0.41268 | 1.846116 | 0.06488 | 0.1962 | 3.319 |
| Gm44021 | 51.94 | 0.388458 | 0.37539 | 1.034805 | 0.30076 | 0.51072 | 3.516 |
| Gm44022 | 25.4989 | -0.06755 | 0.48643 | -0.13887 | 0.88956 | 0.94471 | 5.429 |
| Gm44026 | 23.7445 | 0.232328 | 0.35621 | 0.652223 | 0.51426 | 0.7049 | 9.486 |
| Gm44033 | 76.7013 | -0.17162 | 0.23833 | -0.72011 | 0.47146 | 0.67022 | 0.695 |
| Gm44037 | 99.617 | 0.469594 | 0.21072 | 2.228557 | 0.02584 | 0.10644 | 1.891 |
| Gm44040 | 17.5292 | -0.00027 | 0.49044 | -0.00056 | 0.99955 | 0.9998 | 6.659 |

| | | | | | | | |
|---|---|---|---|---|---|---|---|
| Gm44043 | 31.1726 | -0.19386 | 0.35862 | -0.54057 | 0.58881 | 0.762 | 3.253 |
| Gm44053 | 207.943 | 0.182306 | 0.16248 | 1.122047 | 0.26184 | 0.47055 | 0.929 |
| Gm44063 | 15.9663 | 0.006419 | 0.41743 | 0.015378 | 0.98773 | 0.99316 | 0.376 |
| Gm44065 | 69.637 | -0.02189 | 0.23357 | -0.09374 | 0.92532 | 0.96233 | 3.527 |
| Gm44073 | 13.4175 | 0.215347 | 0.75825 | 0.284006 | 0.77641 | 0.8836 | 11.625 |
| Gm44089 | 13.757 | -0.11394 | 0.47188 | -0.24146 | 0.8092 | 0.90175 | 2.715 |
| Gm44090 | 54.8832 | 0.290214 | 0.24063 | 1.206047 | 0.2278 | 0.43174 | 1.834 |
| Gm44091 | 16.8883 | -0.17048 | 0.57322 | -0.29741 | 0.76615 | 0.8779 | 0.649 |
| Gm44093 | 12.6083 | -0.10856 | 0.47788 | -0.22717 | 0.82029 | 0.90734 | 1.802 |
| Gm44094 | 28.632 | 0.599572 | 0.36698 | 1.633809 | 0.1023 | 0.26399 | 2.03 |
| Gm44103 | 159.136 | 0.172337 | 0.14347 | 1.201238 | 0.22966 | 0.4339 | 3.662 |
| Gm44104 | 55.9522 | 0.151776 | 0.24733 | 0.613667 | 0.53944 | 0.72589 | 3.777 |
| Gm44117 | 32.0074 | 0.55833 | 0.5018 | 1.112647 | 0.26586 | 0.47503 | 1.758 |
| Gm44126 | 88.0561 | 0.181781 | 0.78682 | 0.231033 | 0.81729 | 0.90594 | 0.514 |
| Gm44136 | 173.115 | 0.147217 | 0.15181 | 0.969776 | 0.33216 | 0.5431 | 1.106 |
| Gm44151 | 41.2985 | 0.474461 | 0.26541 | 1.787664 | 0.07383 | 0.21412 | 9.331 |
| Gm44164 | 117.52 | -0.2822 | 0.21682 | -1.30155 | 0.19307 | 0.38993 | 2.561 |
| Gm44167 | 145.669 | 0.280309 | 0.17187 | 1.630891 | 0.10291 | 0.26494 | 1.878 |
| Gm44168 | 824.784 | 0.528637 | 0.11194 | 4.722644 | 2.33E-06 | 0.0002 | 3.539 |
| Gm44187 | 20.8626 | -0.74001 | 0.4501 | -1.64412 | 0.10015 | 0.26034 | 1.929 |
| Gm44189 | 24.603 | -0.0657 | 0.42237 | -0.15556 | 0.87638 | 0.93824 | 3.083 |
| Gm4419 | 176.624 | -0.03291 | 0.1733 | -0.1899 | 0.84939 | 0.92346 | 3.742 |
| Gm44190 | 33.5664 | 0.653069 | 0.29873 | 2.186117 | 0.02881 | 0.11468 | 1.401 |
| Gm44199 | 337.233 | 0.335927 | 0.13921 | 2.413092 | 0.01582 | 0.07717 | 2.128 |
| Gm44200 | 250.882 | 0.464868 | 0.15375 | 3.023468 | 0.0025 | 0.02336 | 4.053 |
| Gm44202 | 16.9949 | 0.816451 | 0.63188 | 1.292106 | 0.19632 | 0.39395 | 0.996 |
| Gm44207 | 30.4939 | 0.545189 | 0.3282 | 1.661141 | 0.09669 | 0.25473 | 1.424 |
| Gm44210 | 546.975 | 0.546316 | 0.18741 | 2.915074 | 0.00356 | 0.02907 | 2.497 |
| Gm44214 | 35.0656 | 0.21907 | 0.27523 | 0.795945 | 0.42606 | 0.63255 | 0.606 |
| Gm44220 | 59.2099 | -0.61114 | 0.27844 | -2.19486 | 0.02817 | 0.11309 | 2.487 |
| Gm44228 | 63.2103 | -0.01113 | 0.2531 | -0.044 | 0.96491 | 0.98273 | 2.026 |
| Gm44229 | 13.3755 | -0.58084 | 0.57694 | -1.00676 | 0.31405 | 0.52455 | 2.487 |
| Gm44230 | 47.7384 | -0.46712 | 0.25877 | -1.8052 | 0.07104 | 0.20855 | 3.55 |
| Gm44231 | 12.4059 | -0.92737 | 0.56349 | -1.64577 | 0.09981 | 0.25983 | 4.075 |

| | | | | | | | |
|---|---|---|---|---|---|---|---|
| Gm44234 | 20.1857 | 0.616719 | 0.45949 | 1.342187 | 0.17954 | 0.37256 | 0.9 |
| Gm44235 | 16.8881 | -1.24472 | 0.54685 | -2.27617 | 0.02284 | 0.09779 | 3.058 |
| Gm44240 | 55.1868 | 0.336355 | 0.28793 | 1.168191 | 0.24273 | 0.44886 | 2.654 |
| Gm44241 | 23.1171 | -1.77297 | 0.46727 | -3.79433 | 0.00015 | 0.0035 | 1.662 |
| Gm44242 | 60.7131 | 0.202303 | 0.34892 | 0.579796 | 0.56205 | 0.74291 | 2.721 |
| Gm44243 | 121.894 | 0.249946 | 0.19455 | 1.284719 | 0.19889 | 0.39678 | 0.692 |
| Gm44249 | 40.7031 | 0.641589 | 0.28914 | 2.218922 | 0.02649 | 0.10828 | 1.805 |
| Gm44250 | 138.982 | -0.27147 | 0.18119 | -1.49827 | 0.13406 | 0.31381 | 4.229 |
| Gm44257 | 281.684 | -0.14923 | 0.20874 | -0.71491 | 0.47466 | 0.6728 | 11.96 |
| Gm44260 | 14.0318 | 0.118628 | 0.50786 | 0.233583 | 0.81531 | 0.90512 | 2.094 |
| Gm44270 | 18.5495 | -0.33005 | 0.47216 | -0.69902 | 0.48454 | 0.68089 | 1.234 |
| Gm44284 | 77.9178 | -0.22045 | 0.26387 | -0.83543 | 0.40347 | 0.61213 | 22.15 |
| Gm44286 | 19.8647 | 0.044174 | 0.44058 | 0.100264 | 0.92013 | 0.95953 | 1.487 |
| Gm44292 | 44.6967 | 0.850987 | 0.3118 | 2.729314 | 0.00635 | 0.04275 | 2.364 |
| Gm4430 | 38.5267 | -0.19951 | 0.27026 | -0.7382 | 0.46039 | 0.6609 | 1.041 |
| Gm44317 | 14.3023 | 0.032931 | 0.46706 | 0.070508 | 0.94379 | 0.97192 | 7.117 |
| Gm44321 | 64.6214 | 0.233248 | 0.288 | 0.809882 | 0.41801 | 0.62541 | 2.855 |
| Gm44414 | 16.7873 | -0.34473 | 0.50984 | -0.67615 | 0.49895 | 0.6922 | 2.109 |
| Gm44415 | 13.8058 | -0.14926 | 0.5823 | -0.25633 | 0.7977 | 0.89566 | 2.112 |
| Gm44416 | 17.2655 | -1.07154 | 0.56145 | -1.90853 | 0.05632 | 0.17956 | 2.066 |
| Gm44418 | 58.4637 | -0.25315 | 0.33235 | -0.76169 | 0.44625 | 0.64987 | 2.006 |
| Gm44421 | 213.058 | -0.19016 | 0.15198 | -1.25117 | 0.21087 | 0.4109 | 1.221 |
| Gm44423 | 276.637 | 0.5234 | 0.1245 | 4.204025 | 2.62E-05 | 0.00106 | 3.158 |
| Gm44430 | 25.5993 | 0.562703 | 0.42328 | 1.329389 | 0.18372 | 0.37789 | 0.995 |
| Gm44432 | 42.1096 | 0.31025 | 0.31294 | 0.991389 | 0.3215 | 0.53233 | 0.938 |
| Gm44433 | 47.9063 | 0.148051 | 0.24612 | 0.601534 | 0.54748 | 0.73161 | 4.832 |
| Gm44434 | 17.3554 | -0.2586 | 0.52244 | -0.49498 | 0.62061 | 0.78424 | 3.745 |
| Gm44437 | 13.7756 | -1.01359 | 0.56297 | -1.80041 | 0.0718 | 0.21021 | 4.038 |
| Gm44438 | 15.9535 | -0.22947 | 0.50134 | -0.45771 | 0.64716 | 0.80359 | 3.236 |
| Gm44445 | 23.126 | 0.735356 | 0.43284 | 1.698891 | 0.08934 | 0.24214 | 2.683 |
| Gm44508 | 24.8069 | -0.07709 | 0.43187 | -0.17849 | 0.85834 | 0.92856 | 1.157 |
| Gm44509 | 1373.11 | -0.06601 | 0.09182 | -0.71888 | 0.47222 | 0.67054 | 1.324 |
| Gm44510 | 54.1848 | -0.27429 | 0.32228 | -0.85108 | 0.39473 | 0.60379 | 4.532 |
| Gm44518 | 34.3065 | -0.47229 | 0.35095 | -1.34573 | 0.17839 | 0.37086 | 4.37 |

| | | | | | | | |
|---|---|---|---|---|---|---|---|
| Gm44524 | 113.82 | 0.433182 | 0.21355 | 2.028476 | 0.04251 | 0.14846 | 0.271 |
| Gm44542 | 32.2801 | 0.031537 | 0.42544 | 0.074129 | 0.94091 | 0.97072 | 4.314 |
| Gm44544 | 21.7179 | 0.529436 | 0.3554 | 1.489676 | 0.13631 | 0.31668 | 3.174 |
| Gm44550 | 66.7133 | 0.115266 | 0.25212 | 0.45718 | 0.64754 | 0.80377 | 2.058 |
| Gm44553 | 17.6076 | 0.44362 | 0.42856 | 1.03515 | 0.3006 | 0.51066 | 2.037 |
| Gm44559 | 202.48 | 0.755676 | 0.50571 | 1.494279 | 0.1351 | 0.31502 | 1.012 |
| Gm44560 | 48.5598 | 0.066455 | 0.42027 | 0.158125 | 0.87436 | 0.93716 | 3.115 |
| Gm44562 | 302.811 | 0.721734 | 0.56581 | 1.275579 | 0.2021 | 0.40058 | 1.347 |
| Gm44567 | 15.7891 | -0.31333 | 0.44221 | -0.70855 | 0.47861 | 0.67618 | 2.631 |
| Gm44570 | 12.814 | -1.18213 | 0.66514 | -1.77725 | 0.07553 | 0.2169 | 4.058 |
| Gm44573 | 17.7374 | -0.11673 | 0.40223 | -0.29021 | 0.77165 | 0.88095 | 2.37 |
| Gm44584 | 13.7302 | 0.19101 | 0.49668 | 0.38457 | 0.70056 | 0.83833 | 2.88 |
| Gm44586 | 21.6977 | 0.901892 | 0.46805 | 1.926927 | 0.05399 | 0.1746 | 2.737 |
| Gm4459 | 189.405 | 0.160344 | 0.16245 | 0.987068 | 0.32361 | 0.53465 | 0.468 |
| Gm44593 | 110.417 | 1.044176 | 0.34341 | 3.040622 | 0.00236 | 0.02243 | 178.238 |
| Gm44597 | 57.1219 | 0.420871 | 0.30702 | 1.370818 | 0.17043 | 0.36076 | 11.727 |
| Gm44616 | 92.377 | 0.178621 | 0.25339 | 0.704921 | 0.48086 | 0.67805 | 1.732 |
| Gm44617 | 46.3854 | -1.48712 | 0.52929 | -2.80966 | 0.00496 | 0.03605 | 5.229 |
| Gm44618 | 17.6085 | -1.67693 | 0.64702 | -2.59178 | 0.00955 | 0.05562 | 52.689 |
| Gm44623 | 102.724 | -0.05764 | 0.20541 | -0.2806 | 0.77901 | 0.88484 | 0.434 |
| Gm44624 | 31.5439 | 0.311256 | 0.33366 | 0.932847 | 0.3509 | 0.56153 | 2.278 |
| Gm44625 | 27.0423 | 0.191819 | 0.32181 | 0.596071 | 0.55113 | 0.73445 | 1.561 |
| Gm44632 | 28.2513 | 0.600842 | 0.3888 | 1.54539 | 0.12225 | 0.29585 | 2.633 |
| Gm44636 | 43.818 | 0.012963 | 0.28505 | 0.045475 | 0.96373 | 0.98236 | 0.755 |
| Gm44639 | 34.4869 | 0.368448 | 0.38796 | 0.949702 | 0.34226 | 0.553 | 3.009 |
| Gm44643 | 23.3988 | 0.428604 | 0.37012 | 1.158007 | 0.24686 | 0.45374 | 38.464 |
| Gm44644 | 141.784 | 0.471453 | 0.19501 | 2.417565 | 0.01562 | 0.07651 | 7.361 |
| Gm44645 | 116.853 | -0.16696 | 0.19898 | -0.83907 | 0.40143 | 0.60999 | 1.653 |
| Gm44646 | 12.7734 | 0.125626 | 0.48973 | 0.256521 | 0.79755 | 0.89562 | 2.578 |
| Gm44649 | 15.3496 | -0.00361 | 0.45882 | -0.00786 | 0.99373 | 0.99633 | 6.791 |
| Gm44663 | 28.4002 | 0.040085 | 0.42438 | 0.094457 | 0.92475 | 0.96208 | 3.15 |
| Gm44664 | 22.7546 | 0.404508 | 0.40155 | 1.007365 | 0.31376 | 0.52419 | 1.786 |
| Gm44667 | 134.678 | -0.30033 | 0.18756 | -1.60128 | 0.10932 | 0.27513 | 3.337 |
| Gm44675 | 65.6984 | 0.453089 | 0.3059 | 1.481185 | 0.13856 | 0.31964 | 2.927 |

| | | | | | | | |
|---|---|---|---|---|---|---|---|
| Gm44676 | 20.4749 | 0.197504 | 0.55262 | 0.357398 | 0.72079 | 0.85096 | 3.055 |
| Gm44677 | 131.443 | 0.85288 | 0.17273 | 4.937736 | 7.90E-07 | 9.05E-05 | 3.739 |
| Gm44678 | 80.892 | 0.49505 | 0.23483 | 2.108091 | 0.03502 | 0.13124 | 2.775 |
| Gm44679 | 24.1451 | 0.439257 | 0.55276 | 0.794664 | 0.42681 | 0.63329 | 2.908 |
| Gm44680 | 24.0303 | -0.14714 | 0.47557 | -0.30939 | 0.75702 | 0.87207 | 2.91 |
| Gm44684 | 124.024 | 0.223919 | 0.18671 | 1.199301 | 0.23041 | 0.43483 | 3.186 |
| Gm44686 | 73.3126 | 0.216454 | 0.22945 | 0.943344 | 0.34551 | 0.55621 | 16.789 |
| Gm44698 | 202.232 | 0.156101 | 0.18931 | 0.824557 | 0.40962 | 0.61736 | 5.343 |
| Gm44699 | 20.8547 | 0.181188 | 0.41303 | 0.438684 | 0.66089 | 0.81242 | 22.578 |
| Gm44706 | 33.4975 | 0.397673 | 0.35333 | 1.1255 | 0.26038 | 0.46882 | 1.339 |
| Gm44718 | 26.095 | -0.77928 | 0.37075 | -2.10192 | 0.03556 | 0.13252 | 0.359 |
| Gm44735 | 61.4748 | 0.161076 | 0.25075 | 0.642368 | 0.52063 | 0.70983 | 1.752 |
| Gm44741 | 13.06 | -0.16488 | 0.57524 | -0.28663 | 0.77439 | 0.88245 | 2.127 |
| Gm44742 | 265.012 | 0.097644 | 0.22624 | 0.431598 | 0.66603 | 0.8156 | 1.412 |
| Gm44745 | 37.5591 | 0.538313 | 0.37662 | 1.429338 | 0.15291 | 0.33835 | 2.723 |
| Gm44746 | 62.5919 | 0.627402 | 0.32972 | 1.902818 | 0.05706 | 0.18096 | 1.349 |
| Gm44749 | 19.6375 | 0.369607 | 0.4127 | 0.895573 | 0.37048 | 0.58072 | 1.628 |
| Gm44758 | 34.9413 | -0.03441 | 0.32653 | -0.10538 | 0.91608 | 0.95711 | 2.068 |
| Gm44763 | 81.8969 | -0.32386 | 0.27893 | -1.16108 | 0.24561 | 0.45222 | 2.019 |
| Gm44769 | 310.126 | 0.507117 | 0.15432 | 3.286106 | 0.00102 | 0.01296 | 2.809 |
| Gm44770 | 90.112 | 0.257999 | 0.19398 | 1.330055 | 0.1835 | 0.3776 | 1.264 |
| Gm44776 | 16.3848 | 0.625223 | 0.41107 | 1.520949 | 0.12827 | 0.30524 | 0.381 |
| Gm44777 | 94.0089 | 0.345149 | 0.23107 | 1.493669 | 0.13526 | 0.31507 | 4.379 |
| Gm44797 | 24.0419 | -1.27349 | 0.44823 | -2.84117 | 0.00449 | 0.03392 | 1.837 |
| Gm44798 | 34.6284 | -0.73396 | 0.35239 | -2.08279 | 0.03727 | 0.13651 | 2.41 |
| Gm44799 | 18.2029 | -1.38973 | 0.4733 | -2.93623 | 0.00332 | 0.02779 | 1.202 |
| Gm44800 | 27.3378 | -0.65378 | 0.42095 | -1.55311 | 0.1204 | 0.29319 | 1.855 |
| Gm44801 | 30.2053 | -1.13003 | 0.45557 | -2.48045 | 0.01312 | 0.0686 | 2.06 |
| Gm44802 | 31.0397 | -1.15862 | 0.38562 | -3.00458 | 0.00266 | 0.02432 | 1.916 |
| Gm4482 | 45.2798 | -0.05692 | 0.2883 | -0.19742 | 0.8435 | 0.91961 | 0.742 |
| Gm44829 | 213.617 | 0.482524 | 0.24102 | 2.002004 | 0.04528 | 0.15478 | 3.193 |
| Gm44830 | 60.4736 | -0.26205 | 0.32277 | -0.81187 | 0.41687 | 0.62435 | 0.584 |
| Gm44831 | 153.633 | 0.273875 | 0.23647 | 1.158158 | 0.2468 | 0.45367 | 2.989 |
| Gm44832 | 70.448 | 0.221673 | 0.30081 | 0.736924 | 0.46117 | 0.66177 | 2.303 |

| | | | | | | | |
|---|---|---|---|---|---|---|---|
| Gm44836 | 24.5268 | 0.907515 | 0.38607 | 2.35067 | 0.01874 | 0.08588 | 1.14 |
| Gm44846 | 231.532 | 0.48595 | 0.25744 | 1.887643 | 0.05907 | 0.18503 | 3.243 |
| Gm44847 | 16.6535 | -0.53013 | 0.70778 | -0.749 | 0.45386 | 0.65609 | 1.991 |
| Gm44850 | 65.7031 | 0.079736 | 0.29613 | 0.269264 | 0.78773 | 0.88973 | 5.173 |
| Gm44860 | 52.9976 | 0.101185 | 0.25467 | 0.397316 | 0.69113 | 0.83229 | 2.346 |
| Gm44861 | 66.925 | 0.20692 | 0.22241 | 0.930356 | 0.35219 | 0.56253 | 1.22 |
| Gm44863 | 13.1018 | 0.345966 | 0.65144 | 0.531081 | 0.59536 | 0.76727 | 1.422 |
| Gm44866 | 191.835 | -0.77124 | 0.25626 | -3.00962 | 0.00262 | 0.0241 | 14.95 |
| Gm44874 | 15.9866 | 0.42287 | 0.39832 | 1.061631 | 0.2884 | 0.49847 | 1.542 |
| Gm44877 | 54.5169 | 0.090314 | 0.27451 | 0.328996 | 0.74216 | 0.86417 | 3.567 |
| Gm44878 | 17.4282 | 0.005864 | 0.51747 | 0.011331 | 0.99096 | 0.99495 | 1.31 |
| Gm44891 | 141.184 | 0.71485 | 0.18507 | 3.862487 | 0.00011 | 0.00291 | 2.223 |
| Gm44898 | 161.197 | -0.12347 | 0.20948 | -0.58939 | 0.5556 | 0.73792 | 2.223 |
| Gm44899 | 49.1433 | 0.075552 | 0.31744 | 0.238003 | 0.81188 | 0.9031 | 12.57 |
| Gm44901 | 36.468 | 0.319635 | 0.41848 | 0.763807 | 0.44498 | 0.64896 | 3.977 |
| Gm44907 | 40.126 | 0.040845 | 0.38104 | 0.107193 | 0.91464 | 0.95656 | 1.633 |
| Gm4491 | 47.9957 | 0.196445 | 0.30651 | 0.640909 | 0.52158 | 0.71075 | 3.378 |
| Gm44911 | 69.3597 | -0.15864 | 0.26985 | -0.58788 | 0.55661 | 0.73867 | 1.062 |
| Gm44913 | 22.7669 | -0.25147 | 0.4576 | -0.54955 | 0.58263 | 0.758 | 2.99 |
| Gm44914 | 30.0582 | 0.058506 | 0.39673 | 0.147472 | 0.88276 | 0.9416 | 2.72 |
| Gm44931 | 30.6674 | -0.09696 | 0.30194 | -0.32113 | 0.74811 | 0.86798 | 0.545 |
| Gm44941 | 41.7802 | 0.259637 | 0.31097 | 0.83493 | 0.40376 | 0.61241 | 2.748 |
| Gm44949 | 105.866 | 0.098326 | 0.18132 | 0.542291 | 0.58762 | 0.76109 | 1.45 |
| Gm44950 | 548.007 | 0.090934 | 0.14122 | 0.643919 | 0.51963 | 0.70925 | 2.296 |
| Gm44951 | 20.3847 | 0.128035 | 0.37891 | 0.337899 | 0.73544 | 0.85989 | 2.024 |
| Gm44954 | 62.3166 | 0.214414 | 0.27352 | 0.783909 | 0.43309 | 0.63917 | 0.227 |
| Gm44981 | 24.8914 | 0.250593 | 0.34819 | 0.719693 | 0.47171 | 0.67032 | 1.943 |
| Gm44982 | 30.5383 | 0.405512 | 0.35226 | 1.151167 | 0.24966 | 0.45704 | 0.471 |
| Gm44986 | 77.7461 | 0.157894 | 0.22836 | 0.691437 | 0.48929 | 0.68444 | 2.829 |
| Gm44997 | 61.1184 | 0.013971 | 0.31223 | 0.044747 | 0.96431 | 0.98268 | 16.822 |
| Gm45001 | 48.805 | 0.057957 | 0.25439 | 0.227826 | 0.81978 | 0.90734 | 2.515 |
| Gm45041 | 25.382 | 0.182417 | 0.36773 | 0.496058 | 0.61985 | 0.78377 | 3.108 |
| Gm45047 | 33.894 | 0.199364 | 0.30876 | 0.645687 | 0.51848 | 0.70819 | 0.309 |
| Gm45053 | 39.7597 | -0.24914 | 0.28536 | -0.87306 | 0.38263 | 0.592 | 2.499 |

| | | | | | | | |
|---|---|---|---|---|---|---|---|
| Gm45059 | 41.6819 | 0.849689 | 0.3713 | 2.288427 | 0.02211 | 0.09584 | 1.077 |
| Gm45073 | 81.9307 | 0.065438 | 0.2278 | 0.287266 | 0.77391 | 0.88219 | 4.247 |
| Gm45074 | 25.4987 | 0.399386 | 0.36833 | 1.084313 | 0.27823 | 0.48807 | 1.24 |
| Gm45094 | 42.6014 | 0.523292 | 0.25954 | 2.016229 | 0.04378 | 0.15142 | 0.617 |
| Gm45104 | 45.8246 | -0.08393 | 0.31267 | -0.26843 | 0.78837 | 0.89002 | 4.35 |
| Gm45110 | 269.83 | 0.28939 | 0.12763 | 2.267479 | 0.02336 | 0.09912 | 1.402 |
| Gm45112 | 16.6761 | 0.262951 | 0.42939 | 0.61239 | 0.54028 | 0.72614 | 5.125 |
| Gm45120 | 16.9519 | -0.03029 | 0.40594 | -0.07463 | 0.94051 | 0.97036 | 0.408 |
| Gm45124 | 163.944 | 0.304671 | 0.18105 | 1.682773 | 0.09242 | 0.24756 | 2.755 |
| Gm45129 | 13.4611 | 0.430116 | 0.46183 | 0.931321 | 0.35169 | 0.56204 | 2.224 |
| Gm45130 | 16.5988 | 0.379994 | 0.44932 | 0.845707 | 0.39772 | 0.60672 | 3.388 |
| Gm45133 | 1110.3 | 0.137015 | 0.12775 | 1.072548 | 0.28347 | 0.49354 | 2.823 |
| Gm45137 | 17.302 | -0.64162 | 0.54056 | -1.18696 | 0.23524 | 0.44045 | 4.415 |
| Gm45148 | 49.0722 | 0.638881 | 0.28096 | 2.273903 | 0.02297 | 0.09807 | 3.587 |
| Gm45151 | 96.6113 | 0.611413 | 0.19321 | 3.164522 | 0.00155 | 0.01704 | 2.1 |
| Gm45153 | 29.9281 | -0.52918 | 0.37784 | -1.40054 | 0.16135 | 0.3496 | 1.901 |
| Gm45159 | 116.84 | -0.10901 | 0.32267 | -0.33783 | 0.73549 | 0.85989 | 52.981 |
| Gm45163 | 15.9998 | 0.613532 | 0.48763 | 1.258195 | 0.20832 | 0.40807 | 9.637 |
| Gm45169 | 29.2145 | -0.09278 | 0.41593 | -0.22306 | 0.82349 | 0.90853 | 5.966 |
| Gm45176 | 46.7203 | -0.18759 | 0.32557 | -0.5762 | 0.56448 | 0.7448 | 4.096 |
| Gm45177 | 31.4739 | 0.043697 | 0.34703 | 0.125917 | 0.8998 | 0.94981 | 2.307 |
| Gm45178 | 39.8453 | 0.617785 | 0.38642 | 1.598758 | 0.10987 | 0.27606 | 4.11 |
| Gm45179 | 20.2727 | -0.12491 | 0.53862 | -0.2319 | 0.81661 | 0.90555 | 4.337 |
| Gm45182 | 145.588 | 0.518 | 0.25642 | 2.020137 | 0.04337 | 0.15051 | 2.404 |
| Gm45188 | 35.9122 | -0.24748 | 0.3261 | -0.7589 | 0.44791 | 0.65151 | 3.616 |
| Gm45191 | 103.311 | 0.301976 | 0.2343 | 1.288817 | 0.19746 | 0.39531 | 5.439 |
| Gm45201 | 104.363 | 0.722958 | 0.37982 | 1.903419 | 0.05699 | 0.18082 | 2.679 |
| Gm45203 | 53.8362 | 0.672206 | 0.37367 | 1.798915 | 0.07203 | 0.21072 | 1.831 |
| Gm45204 | 79.4089 | 0.22978 | 0.23135 | 0.993209 | 0.32061 | 0.53137 | 1.354 |
| Gm45205 | 24.8616 | 0.658131 | 0.40186 | 1.637721 | 0.10148 | 0.26283 | 3.33 |
| Gm45206 | 99.8143 | 0.265072 | 0.21653 | 1.22416 | 0.22089 | 0.42374 | 2.285 |
| Gm45216 | 65.0054 | 0.215844 | 0.25416 | 0.849246 | 0.39574 | 0.6048 | 9.971 |
| Gm45220 | 115.941 | 0.17683 | 0.16709 | 1.058292 | 0.28992 | 0.50018 | 2.509 |
| Gm45221 | 62.8026 | -0.19631 | 0.27205 | -0.7216 | 0.47054 | 0.6693 | 2.272 |

| | | | | | | | |
|---|---|---|---|---|---|---|---|
| Gm45222 | 21.4497 | -0.39127 | 0.46541 | -0.84071 | 0.40051 | 0.60926 | 4.307 |
| Gm45223 | 17.2624 | -0.48667 | 0.47102 | -1.03322 | 0.3015 | 0.51121 | 5.138 |
| Gm45226 | 12.5408 | -0.04191 | 0.58209 | -0.072 | 0.9426 | 0.97134 | 0.288 |
| Gm45234 | 30.8842 | 0.15729 | 0.35384 | 0.444519 | 0.65667 | 0.80928 | 16.833 |
| Gm45235 | 88.3334 | -0.31202 | 0.18549 | -1.68212 | 0.09255 | 0.24776 | 4.465 |
| Gm4524 | 29.0246 | -0.53399 | 0.39035 | -1.36798 | 0.17132 | 0.3618 | 3.868 |
| Gm45250 | 404.146 | 0.24483 | 0.13521 | 1.810748 | 0.07018 | 0.20685 | 3.592 |
| Gm45251 | 49.9735 | -0.18356 | 0.29967 | -0.61256 | 0.54017 | 0.72606 | 1.465 |
| Gm45263 | 17.2637 | -0.16038 | 0.55377 | -0.28961 | 0.77212 | 0.88128 | 1.375 |
| Gm45264 | 39.6896 | 0.014934 | 0.34835 | 0.042871 | 0.9658 | 0.9828 | 3.995 |
| Gm45265 | 27.6025 | -0.38637 | 0.35424 | -1.09071 | 0.2754 | 0.4847 | 2.497 |
| Gm45286 | 37.8021 | 0.07021 | 0.3563 | 0.197053 | 0.84379 | 0.91972 | 1.35 |
| Gm45290 | 24.0332 | -0.47819 | 0.4039 | -1.18394 | 0.23644 | 0.44163 | 3.89 |
| Gm45309 | 42.5835 | 0.103093 | 0.26483 | 0.389274 | 0.69707 | 0.8358 | 0.801 |
| Gm45311 | 44.2293 | 0.063649 | 0.26288 | 0.242127 | 0.80868 | 0.90165 | 1.33 |
| Gm45338 | 29.9575 | -0.26677 | 0.37825 | -0.70529 | 0.48063 | 0.67791 | 39.872 |
| Gm45342 | 95.028 | 0.151184 | 0.18402 | 0.821584 | 0.41131 | 0.61936 | 1.363 |
| Gm45343 | 40.346 | 0.247083 | 0.27681 | 0.892609 | 0.37207 | 0.58241 | 0.828 |
| Gm45344 | 35.6144 | 0.606607 | 0.33934 | 1.787627 | 0.07384 | 0.21412 | 0.341 |
| Gm45352 | 22.4781 | 0.130419 | 0.41242 | 0.316232 | 0.75183 | 0.86994 | 33.715 |
| Gm45353 | 33.2727 | 0.444725 | 0.34662 | 1.283046 | 0.19948 | 0.39743 | 3.93 |
| Gm45378 | 22.8557 | 0.296397 | 0.38263 | 0.774626 | 0.43856 | 0.64383 | 2.711 |
| Gm45380 | 135.411 | -0.06601 | 0.16662 | -0.3962 | 0.69195 | 0.83278 | 0.315 |
| Gm45407 | 44.6795 | 0.466852 | 0.37956 | 1.229975 | 0.21871 | 0.42078 | 3.676 |
| Gm45416 | 28.6361 | -0.23145 | 0.42258 | -0.54771 | 0.58389 | 0.75891 | 2.201 |
| Gm45426 | 83.0683 | -0.30995 | 0.33963 | -0.91262 | 0.36144 | 0.57177 | 3.672 |
| Gm45428 | 70.4866 | 0.466973 | 0.19962 | 2.339316 | 0.01932 | 0.08745 | 1.789 |
| Gm45434 | 14.337 | 0.744121 | 0.53366 | 1.394373 | 0.1632 | 0.35174 | 0.293 |
| Gm45437 | 17.5564 | -0.61836 | 0.63986 | -0.96641 | 0.33384 | 0.54457 | 2.414 |
| Gm4544 | 13.1943 | 0.385133 | 0.67768 | 0.568309 | 0.56983 | 0.74923 | 3.361 |
| Gm45441 | 33.5133 | -0.49311 | 0.34474 | -1.43037 | 0.15261 | 0.33794 | 4.355 |
| Gm45447 | 91.5871 | 0.150867 | 0.18278 | 0.825398 | 0.40915 | 0.61691 | 5.448 |
| Gm45457 | 58.7315 | -0.24865 | 0.28567 | -0.87041 | 0.38408 | 0.59346 | 3.089 |
| Gm45464 | 29.958 | 0.541732 | 0.3577 | 1.514494 | 0.1299 | 0.30775 | 2.91 |

| | | | | | | | |
|---|---|---|---|---|---|---|---|
| Gm45470 | 38.3104 | -0.61181 | 0.35039 | -1.74608 | 0.0808 | 0.22623 | 11.647 |
| Gm45472 | 13.4872 | 0.232269 | 0.51892 | 0.447601 | 0.65444 | 0.8083 | 4.385 |
| Gm45477 | 25.7677 | -0.1153 | 0.41965 | -0.27476 | 0.7835 | 0.88753 | 5.498 |
| Gm45481 | 16.8649 | -0.49604 | 0.4144 | -1.19699 | 0.23131 | 0.43586 | 0.877 |
| Gm45486 | 28.9789 | 0.596658 | 0.36205 | 1.64801 | 0.09935 | 0.25904 | 7.217 |
| Gm45487 | 14.7512 | 0.067202 | 0.6028 | 0.111483 | 0.91123 | 0.95519 | 0.545 |
| Gm45493 | 89.014 | 0.249471 | 0.23341 | 1.068825 | 0.28515 | 0.49493 | 3.948 |
| Gm45495 | 557.99 | 0.024439 | 0.09033 | 0.270539 | 0.78675 | 0.88907 | 1.7 |
| Gm45501 | 182.703 | 0.284599 | 0.14576 | 1.95246 | 0.05088 | 0.16778 | 2.526 |
| Gm45509 | 19.4372 | 0.608085 | 0.44121 | 1.378209 | 0.16814 | 0.3581 | 1.229 |
| Gm45510 | 14.1805 | 0.739725 | 0.50348 | 1.469228 | 0.14177 | 0.32357 | 2.714 |
| Gm45515 | 14.8343 | 0.048379 | 0.63169 | 0.076588 | 0.93895 | 0.96951 | 3.746 |
| Gm45516 | 31.2504 | 0.224285 | 0.35378 | 0.633965 | 0.5261 | 0.71427 | 4.182 |
| Gm45527 | 23.1463 | 0.457731 | 0.40986 | 1.116801 | 0.26408 | 0.47294 | 0.774 |
| Gm45532 | 22.8488 | 0.155493 | 0.42758 | 0.363662 | 0.71611 | 0.84818 | 3.62 |
| Gm45533 | 84.7296 | 0.501599 | 0.22248 | 2.254553 | 0.02416 | 0.10151 | 0.385 |
| Gm45540 | 16.7193 | -0.71684 | 0.65834 | -1.08886 | 0.27622 | 0.48542 | 3.606 |
| Gm45548 | 21.8548 | 0.478472 | 0.38436 | 1.244858 | 0.21318 | 0.41433 | 3.175 |
| Gm45552 | 77.4768 | -0.17798 | 0.20782 | -0.85642 | 0.39177 | 0.60089 | 2.228 |
| Gm45560 | 58.3885 | 0.772639 | 0.28984 | 2.665697 | 0.00768 | 0.04844 | 1.472 |
| Gm45570 | 32.5216 | 0.010982 | 0.36253 | 0.030292 | 0.97583 | 0.98762 | 0.653 |
| Gm45579 | 26.0114 | -0.23708 | 0.38068 | -0.62278 | 0.53343 | 0.72067 | 3.919 |
| Gm45591 | 383.651 | 0.344507 | 0.1354 | 2.544395 | 0.01095 | 0.06094 | 1.256 |
| Gm45592 | 38.0022 | 0.005493 | 0.27406 | 0.020042 | 0.98401 | 0.99128 | 1.043 |
| Gm45597 | 16.601 | 0.136952 | 0.57798 | 0.236948 | 0.8127 | 0.90367 | 1.994 |
| Gm45601 | 22.2238 | -0.09554 | 0.36442 | -0.26218 | 0.79318 | 0.89287 | 7.069 |
| Gm45605 | 377.852 | 0.32374 | 0.15382 | 2.104683 | 0.03532 | 0.13195 | 4.269 |
| Gm45606 | 162.778 | -0.55402 | 0.14567 | -3.80333 | 0.00014 | 0.0034 | 2.796 |
| Gm45609 | 18.6279 | -0.18199 | 0.50239 | -0.36225 | 0.71717 | 0.84899 | 25.33 |
| Gm45619 | 108.355 | 0.269061 | 0.19754 | 1.362079 | 0.17317 | 0.3644 | 0.528 |
| Gm45623 | 198.014 | -0.30796 | 0.16151 | -1.90673 | 0.05656 | 0.18002 | 1.457 |
| Gm45629 | 96.6396 | -0.10076 | 0.222 | -0.45386 | 0.64993 | 0.80518 | 2.546 |
| Gm45630 | 22.0153 | -0.84569 | 0.37603 | -2.24901 | 0.02451 | 0.10251 | 1.911 |
| Gm45631 | 24.1167 | 0.008948 | 0.39612 | 0.02259 | 0.98198 | 0.99058 | 1.372 |

| | | | | | | | |
|---|---|---|---|---|---|---|---|
| Gm45632 | 167.162 | 0.150917 | 0.16658 | 0.905957 | 0.36496 | 0.57561 | 1.888 |
| Gm45634 | 15.3808 | 0.235655 | 0.58667 | 0.401685 | 0.68792 | 0.82998 | 1.99 |
| Gm45640 | 12.7525 | 0.602935 | 0.49658 | 1.214177 | 0.22468 | 0.42839 | 1.896 |
| Gm45645 | 188.668 | 1.235065 | 0.27897 | 4.427192 | 9.55E-06 | 0.00054 | 38.053 |
| Gm45650 | 30.9295 | -0.69294 | 0.33145 | -2.09067 | 0.03656 | 0.13475 | 6.516 |
| Gm45670 | 32.0251 | 0.796695 | 0.32883 | 2.422829 | 0.0154 | 0.07575 | 6.464 |
| Gm45694 | 176.722 | 0.515767 | 0.20706 | 2.490926 | 0.01274 | 0.0673 | 3.251 |
| Gm45698 | 36.5141 | -0.55023 | 0.34884 | -1.5773 | 0.11473 | 0.28371 | 1.135 |
| Gm45702 | 92.5635 | 0.664131 | 0.24697 | 2.689125 | 0.00716 | 0.04623 | 1.219 |
| Gm45708 | 25.6268 | -0.14795 | 0.33905 | -0.43635 | 0.66258 | 0.81357 | 62.648 |
| Gm45709 | 41.4953 | -0.01494 | 0.28469 | -0.05247 | 0.95816 | 0.97954 | 2.237 |
| Gm45718 | 233.036 | 0.052038 | 0.14846 | 0.350524 | 0.72595 | 0.85388 | 3.794 |
| Gm45737 | 25.2155 | -0.10227 | 0.39925 | -0.25616 | 0.79783 | 0.89568 | 1.612 |
| Gm45743 | 354.723 | 0.444226 | 0.187 | 2.375602 | 0.01752 | 0.08225 | 2.795 |
| Gm45745 | 33.4553 | -0.65952 | 0.4504 | -1.46429 | 0.14312 | 0.32515 | 1.313 |
| Gm45758 | 34.2437 | 0.218137 | 0.36275 | 0.601348 | 0.54761 | 0.73172 | 1.006 |
| Gm45767 | 104.692 | -0.30997 | 0.19568 | -1.58406 | 0.11318 | 0.28162 | 4.561 |
| Gm45769 | 83.9689 | -0.14618 | 0.23217 | -0.62964 | 0.52893 | 0.71673 | 1.828 |
| Gm45779 | 41.2257 | 0.111944 | 0.30197 | 0.370708 | 0.71086 | 0.84441 | 4.064 |
| Gm45786 | 22.1725 | 0.372698 | 0.40936 | 0.910433 | 0.36259 | 0.57294 | 2.278 |
| Gm45790 | 15.6608 | -0.41963 | 0.46798 | -0.89669 | 0.36988 | 0.58027 | 30.942 |
| Gm45793 | 367.478 | -0.43651 | 0.12826 | -3.4033 | 0.00067 | 0.00964 | 1.342 |
| Gm45804 | 36.7704 | -0.12634 | 0.39469 | -0.3201 | 0.74889 | 0.86834 | 3.435 |
| Gm45809 | 91.6035 | 0.335215 | 0.2268 | 1.477992 | 0.13941 | 0.3207 | 3.894 |
| Gm45819 | 183.814 | 0.476505 | 0.18163 | 2.623498 | 0.0087 | 0.05236 | 1.751 |
| Gm45820 | 28.0869 | -0.11765 | 0.32627 | -0.36059 | 0.71841 | 0.84957 | 7.813 |
| Gm45828 | 26.7909 | 0.391978 | 0.33271 | 1.178144 | 0.23874 | 0.44446 | 1.836 |
| Gm45833 | 13.9681 | -0.24743 | 0.45718 | -0.5412 | 0.58837 | 0.76172 | 7.275 |
| Gm45838 | 328.584 | 1.004269 | 0.28595 | 3.512076 | 0.00044 | 0.00734 | 2.447 |
| Gm45844 | 241.869 | 0.138481 | 0.13387 | 1.034407 | 0.30095 | 0.51075 | 65.273 |
| Gm45847 | 222.531 | 0.370781 | 0.17511 | 2.117472 | 0.03422 | 0.12921 | 36.891 |
| Gm45853 | 65.3723 | -0.4201 | 0.27928 | -1.50424 | 0.13252 | 0.31164 | 3.171 |
| Gm45854 | 151.418 | 0.026657 | 0.18539 | 0.143784 | 0.88567 | 0.9432 | 3.488 |
| Gm45855 | 17.635 | 0.081256 | 0.39756 | 0.204384 | 0.83805 | 0.91666 | 0.88 |

| | | | | | | | |
|---|---|---|---|---|---|---|---|
| Gm45867 | 134.487 | -0.41996 | 0.23549 | -1.78335 | 0.07453 | 0.21518 | 2.317 |
| Gm45869 | 765.458 | 0.039348 | 0.14881 | 0.26442 | 0.79146 | 0.89159 | 6.301 |
| Gm45871 | 400.886 | 0.307296 | 0.18889 | 1.626839 | 0.10377 | 0.26624 | 12.752 |
| Gm45874 | 113.092 | 0.819018 | 0.26467 | 3.094482 | 0.00197 | 0.01994 | 0.346 |
| Gm45875 | 24.9041 | -0.2987 | 0.45791 | -0.65231 | 0.5142 | 0.70488 | 17.561 |
| Gm45885 | 24.9901 | 0.374381 | 0.35231 | 1.06266 | 0.28794 | 0.49803 | 1.579 |
| Gm45887 | 16.8971 | -0.16519 | 0.44087 | -0.37468 | 0.7079 | 0.84235 | 2.998 |
| Gm45890 | 16.3586 | -0.25077 | 0.42835 | -0.58544 | 0.55825 | 0.73975 | 10.146 |
| Gm45893 | 66.4896 | 0.613459 | 0.21773 | 2.817514 | 0.00484 | 0.03552 | 0.247 |
| Gm45902 | 417.927 | 0.083458 | 0.12694 | 0.657444 | 0.5109 | 0.70199 | 8.264 |
| Gm45904 | 43.4026 | 0.75063 | 0.28246 | 2.657427 | 0.00787 | 0.0492 | 124.538 |
| Gm45925 | 13.7609 | 0.995248 | 0.55673 | 1.78766 | 0.07383 | 0.21412 | 1.313 |
| Gm45949 | 15.2016 | -0.19135 | 0.55731 | -0.34334 | 0.73134 | 0.85734 | 8.169 |
| Gm4598 | 13.3775 | 1.523594 | 0.52513 | 2.90134 | 0.00372 | 0.02998 | 1.816 |
| Gm46123 | 146.026 | 0.583256 | 0.21728 | 2.684319 | 0.00727 | 0.04669 | 7.82 |
| Gm46136 | 25.9884 | -0.78964 | 0.41891 | -1.885 | 0.05943 | 0.18548 | 7.008 |
| Gm4617 | 28.3274 | -0.3111 | 0.30963 | -1.00473 | 0.31502 | 0.52562 | 0.331 |
| Gm46210 | 30.0443 | 0.063677 | 0.35551 | 0.179115 | 0.85785 | 0.9283 | 17.531 |
| Gm46218 | 121.125 | 0.407621 | 0.22141 | 1.841006 | 0.06562 | 0.19767 | 16.855 |
| Gm4631 | 102.043 | -0.41616 | 1.98094 | -0.21008 | 0.83361 | 0.91471 | 16.243 |
| Gm46339 | 16.8578 | -0.82195 | 0.54215 | -1.51608 | 0.1295 | 0.30701 | 3.539 |
| Gm46367 | 30.3387 | -0.03654 | 0.34306 | -0.10652 | 0.91517 | 0.95656 | 18.633 |
| Gm46376 | 24.4987 | 1.212933 | 0.39585 | 3.064088 | 0.00218 | 0.02131 | 7.987 |
| Gm46378 | 169.102 | 0.407703 | 0.18103 | 2.252165 | 0.02431 | 0.10193 | 4.288 |
| Gm46404 | 109.485 | 0.528756 | 0.2075 | 2.548259 | 0.01083 | 0.06059 | 4.114 |
| Gm46409 | 29.9384 | -0.40314 | 0.42457 | -0.94952 | 0.34236 | 0.55307 | 40.258 |
| Gm46415 | 29.6638 | 0.244498 | 0.30995 | 0.788825 | 0.43021 | 0.63658 | 0.345 |
| Gm46430 | 887.225 | 0.166384 | 0.16207 | 1.026593 | 0.30461 | 0.51448 | 38.598 |
| Gm46432 | 126.791 | -0.16418 | 0.17707 | -0.92718 | 0.35383 | 0.5641 | 0.638 |
| Gm46447 | 94.2644 | 0.206329 | 0.24625 | 0.837893 | 0.40209 | 0.6107 | 6.972 |
| Gm4651 | 16.1389 | 0.279713 | 0.44722 | 0.625442 | 0.53168 | 0.71934 | 5.491 |
| Gm4654 | 29.3102 | 0.059045 | 0.35831 | 0.164788 | 0.86911 | 0.93357 | 0.982 |
| Gm46555 | 16.8008 | 0.324254 | 0.41986 | 0.772286 | 0.43995 | 0.64503 | 5.014 |
| Gm46563 | 15.2803 | 0.096963 | 0.53833 | 0.180118 | 0.85706 | 0.9279 | 14.784 |

| | | | | | | | |
|---|---|---|---|---|---|---|---|
| Gm46565 | 12.3686 | 0.455044 | 0.51007 | 0.892126 | 0.37233 | 0.58269 | 22.44 |
| Gm4660 | 14.689 | 0.5716 | 0.49589 | 1.152674 | 0.24904 | 0.45629 | 0.403 |
| Gm46603 | 54.9847 | 1.373354 | 0.40515 | 3.389766 | 0.0007 | 0.00998 | 2.569 |
| Gm46620 | 483.423 | -0.07693 | 0.16216 | -0.4744 | 0.63522 | 0.79438 | 7.05 |
| Gm46633 | 27.0713 | -0.79753 | 0.42548 | -1.8744 | 0.06088 | 0.18836 | 1.443 |
| Gm46637 | 43.5032 | -0.4526 | 0.32182 | -1.40639 | 0.15961 | 0.34744 | 4.455 |
| Gm46658 | 40.7612 | -0.81117 | 0.28458 | -2.85044 | 0.00437 | 0.03339 | 3.029 |
| Gm4673 | 428.084 | -0.52986 | 0.12468 | -4.24977 | 2.14E-05 | 0.00092 | 4.711 |
| Gm4675 | 14.5274 | 0.052319 | 0.48332 | 0.10825 | 0.9138 | 0.95639 | 24.968 |
| Gm4681 | 28.5505 | 1.018441 | 0.38934 | 2.615788 | 0.0089 | 0.05309 | 105.51 |
| Gm4691 | 13.7527 | -0.12274 | 0.47789 | -0.25685 | 0.7973 | 0.89545 | 1.127 |
| Gm47003 | 35.0837 | 0.669727 | 0.30499 | 2.195865 | 0.0281 | 0.11285 | 6.963 |
| Gm47004 | 14.4329 | -0.99041 | 0.50693 | -1.95373 | 0.05073 | 0.16753 | 2.261 |
| Gm47006 | 15.0777 | 0.248337 | 0.62593 | 0.396751 | 0.69155 | 0.83252 | 1.726 |
| Gm47022 | 23.2632 | -0.32322 | 0.3958 | -0.8166 | 0.41415 | 0.62213 | 2.472 |
| Gm47024 | 17.2973 | 0.82361 | 0.72142 | 1.141652 | 0.2536 | 0.46097 | 2.571 |
| Gm47033 | 210.76 | 0.309482 | 0.26061 | 1.187549 | 0.23501 | 0.44032 | 2.132 |
| Gm4705 | 21.023 | -0.68939 | 0.46932 | -1.46893 | 0.14185 | 0.32366 | 0.354 |
| Gm47071 | 112.117 | 0.17968 | 0.18306 | 0.981558 | 0.32632 | 0.53755 | 258.6 |
| Gm47076 | 13.1835 | 1.066802 | 0.5869 | 1.817691 | 0.06911 | 0.20458 | 2.765 |
| Gm47079 | 31.626 | -0.08768 | 0.37162 | -0.23594 | 0.81348 | 0.9041 | 4.051 |
| Gm47096 | 21.905 | -0.3469 | 0.35506 | -0.97703 | 0.32855 | 0.53979 | 48.732 |
| Gm47097 | 26.8277 | -0.81791 | 0.47422 | -1.72475 | 0.08457 | 0.23312 | 1.48 |
| Gm47108 | 17.6763 | -1.20093 | 0.51118 | -2.34935 | 0.01881 | 0.08608 | 3.123 |
| Gm47113 | 115.484 | 0.38567 | 0.26473 | 1.456857 | 0.14516 | 0.32787 | 10.441 |
| Gm47119 | 26.3434 | 0.068929 | 0.40615 | 0.169712 | 0.86524 | 0.93172 | 1.153 |
| Gm47134 | 22.1696 | -0.02793 | 0.41991 | -0.06652 | 0.94696 | 0.97332 | 7.662 |
| Gm47135 | 248.129 | 0.393492 | 0.17269 | 2.278537 | 0.02269 | 0.09735 | 0.9 |
| Gm47136 | 26.3471 | 0.81712 | 0.41744 | 1.957445 | 0.0503 | 0.16665 | 27.266 |
| Gm47140 | 98.5411 | 0.202651 | 0.26785 | 0.756592 | 0.44929 | 0.65253 | 1.4 |
| Gm47155 | 1716.86 | 0.17776 | 0.10766 | 1.651179 | 0.0987 | 0.25807 | 2.175 |
| Gm47163 | 621.788 | -0.36016 | 0.11994 | -3.00275 | 0.00268 | 0.02441 | 1.665 |
| Gm47171 | 72.2163 | -0.74547 | 0.41743 | -1.78582 | 0.07413 | 0.21457 | 2.794 |
| Gm47173 | 61.3969 | -0.98965 | 0.38763 | -2.55308 | 0.01068 | 0.06001 | 2.324 |

| | | | | | | | |
|---|---|---|---|---|---|---|---|
| Gm47202 | 34.1232 | -0.9847 | 0.5074 | -1.94066 | 0.0523 | 0.17088 | 6.587 |
| Gm47204 | 44.191 | -0.25449 | 0.25443 | -1.00022 | 0.31721 | 0.52753 | 3.086 |
| Gm47205 | 33.1802 | 0.02298 | 0.32813 | 0.070035 | 0.94417 | 0.97202 | 2.29 |
| Gm47214 | 25.8653 | 1.801465 | 0.51287 | 3.51252 | 0.00044 | 0.00733 | 28.121 |
| Gm47218 | 89.8883 | 0.625049 | 0.22437 | 2.785753 | 0.00534 | 0.03789 | 1.319 |
| Gm47232 | 31.6322 | -0.08366 | 0.30561 | -0.27375 | 0.78427 | 0.888 | 0.866 |
| Gm47234 | 18.1324 | 0.008943 | 0.40889 | 0.021872 | 0.98255 | 0.99061 | 0.909 |
| Gm47239 | 27.7698 | -0.66849 | 0.40502 | -1.65051 | 0.09884 | 0.25828 | 0.704 |
| Gm47246 | 45.5072 | -0.15177 | 0.34853 | -0.43547 | 0.66322 | 0.81392 | 2.354 |
| Gm47251 | 75.3761 | 0.235533 | 0.22632 | 1.040687 | 0.29802 | 0.50814 | 1 |
| Gm47254 | 12.3626 | 0.408384 | 0.46767 | 0.873238 | 0.38253 | 0.59199 | 0.202 |
| Gm47258 | 30.5167 | -0.34201 | 0.50137 | -0.68215 | 0.49514 | 0.68919 | 1.342 |
| Gm47260 | 588.09 | 0.214458 | 0.11817 | 1.81477 | 0.06956 | 0.20558 | 3.105 |
| Gm47270 | 46.7941 | 0.19459 | 0.29323 | 0.663616 | 0.50694 | 0.69814 | 6.785 |
| Gm47283 | 148.82 | -0.07498 | 0.67727 | -0.11071 | 0.91184 | 0.95548 | 31.728 |
| Gm47322 | 27.361 | 1.018149 | 0.34338 | 2.965067 | 0.00303 | 0.02631 | 1.824 |
| Gm47339 | 12.4321 | -0.90314 | 0.48493 | -1.86241 | 0.06254 | 0.19192 | 3.086 |
| Gm47341 | 14.8194 | 0.343584 | 0.52897 | 0.64953 | 0.516 | 0.70609 | 2.172 |
| Gm4735 | 121.785 | -0.33931 | 0.16814 | -2.01803 | 0.04359 | 0.15103 | 1.302 |
| Gm47358 | 58.8476 | 1.52589 | 0.39815 | 3.832409 | 0.00013 | 0.00314 | 4.689 |
| Gm47359 | 16.0443 | -0.52665 | 0.61615 | -0.85475 | 0.39269 | 0.60193 | 12.184 |
| Gm47370 | 38.2021 | -0.21984 | 0.29207 | -0.75269 | 0.45164 | 0.65398 | 1.829 |
| Gm47372 | 102.988 | 0.117653 | 0.26155 | 0.449833 | 0.65283 | 0.80691 | 3.544 |
| Gm4739 | 20.58 | -0.19391 | 0.43514 | -0.44562 | 0.65587 | 0.80893 | 0.633 |
| Gm47391 | 15.4382 | -0.05291 | 0.45438 | -0.11644 | 0.9073 | 0.95354 | 4.556 |
| Gm47399 | 18.183 | 0.022341 | 0.51748 | 0.043173 | 0.96556 | 0.9828 | 0.474 |
| Gm47403 | 13.8952 | 0.311994 | 0.46564 | 0.670037 | 0.50283 | 0.69537 | 9.687 |
| Gm47414 | 38.99 | 0.258009 | 0.27145 | 0.950485 | 0.34187 | 0.55274 | 3.311 |
| Gm47423 | 16.3977 | 0.022551 | 0.57253 | 0.039389 | 0.96858 | 0.9841 | 2.777 |
| Gm47435 | 30.6976 | 0.311687 | 0.5104 | 0.610667 | 0.54142 | 0.72706 | 0.367 |
| Gm47446 | 20.9163 | 0.249156 | 0.39968 | 0.623388 | 0.53303 | 0.72034 | 2.89 |
| Gm47467 | 77.5176 | 0.07594 | 0.22722 | 0.334208 | 0.73822 | 0.86185 | 15.68 |
| Gm47469 | 475.224 | 0.222945 | 0.1665 | 1.339025 | 0.18056 | 0.37394 | 12.933 |
| Gm47472 | 70.8178 | -0.15369 | 0.29763 | -0.51637 | 0.6056 | 0.77424 | 4.563 |

| | | | | | | | |
|---|---|---|---|---|---|---|---|
| Gm47477 | 19.1986 | -0.21983 | 0.43842 | -0.50143 | 0.61607 | 0.78132 | 3.132 |
| Gm47483 | 283.691 | 0.041334 | 0.18054 | 0.228943 | 0.81891 | 0.90687 | 2.015 |
| Gm47484 | 40.131 | -0.1373 | 0.32377 | -0.42407 | 0.67152 | 0.81926 | 3.223 |
| Gm47493 | 21.8755 | 0.384345 | 0.42603 | 0.902152 | 0.36698 | 0.57733 | 4.714 |
| Gm47508 | 28.2886 | 0.32776 | 0.38725 | 0.846383 | 0.39734 | 0.60651 | 3.016 |
| Gm47512 | 16.7391 | 0.047208 | 0.50761 | 0.093 | 0.9259 | 0.96264 | 2.996 |
| Gm47550 | 59.5757 | 0.324768 | 0.26631 | 1.219505 | 0.22265 | 0.42587 | 1.705 |
| Gm47558 | 16.3741 | 0.161626 | 0.51969 | 0.311002 | 0.7558 | 0.87157 | 2.714 |
| Gm47572 | 142.048 | 0.247363 | 0.23432 | 1.055676 | 0.29112 | 0.50115 | 3.361 |
| Gm47573 | 111.279 | 0.311729 | 0.28145 | 1.107578 | 0.26804 | 0.47731 | 2.538 |
| Gm47580 | 19.3823 | -0.00443 | 0.471 | -0.0094 | 0.9925 | 0.99567 | 2.031 |
| Gm47583 | 28.4284 | 0.60397 | 0.36993 | 1.63264 | 0.10254 | 0.26442 | 3.425 |
| Gm47586 | 14.4937 | -0.96051 | 0.46526 | -2.06448 | 0.03897 | 0.14064 | 2.028 |
| Gm47588 | 17.1563 | 0.816233 | 0.56025 | 1.456906 | 0.14514 | 0.32787 | 3.85 |
| Gm47590 | 42.3938 | 0.516818 | 0.27575 | 1.874243 | 0.0609 | 0.18836 | 2.976 |
| Gm47591 | 93.4619 | 0.028006 | 0.31176 | 0.089832 | 0.92842 | 0.96398 | 7.371 |
| Gm47592 | 16.5852 | -0.07539 | 0.46419 | -0.16241 | 0.87098 | 0.93477 | 1.307 |
| Gm47594 | 27.1763 | -0.06646 | 0.36498 | -0.18209 | 0.85551 | 0.92695 | 4.154 |
| Gm47621 | 13.5448 | -0.58765 | 0.50583 | -1.16177 | 0.24533 | 0.45199 | 2.538 |
| Gm47624 | 49.8265 | 0.149998 | 0.28783 | 0.521139 | 0.60227 | 0.77228 | 3.836 |
| Gm47640 | 97.3825 | -0.07552 | 0.20374 | -0.37069 | 0.71087 | 0.84441 | 2.882 |
| Gm47657 | 24.9699 | 0.512138 | 0.46813 | 1.094016 | 0.27395 | 0.48339 | 1.804 |
| Gm47664 | 30.9427 | -0.11009 | 0.42866 | -0.25683 | 0.79731 | 0.89545 | 1.34 |
| Gm47666 | 28.1833 | 0.232518 | 0.30993 | 0.750233 | 0.45311 | 0.65552 | 1.396 |
| Gm47690 | 35.555 | -0.02226 | 0.33714 | -0.06603 | 0.94735 | 0.97353 | 2.946 |
| Gm47694 | 560.897 | 0.3227 | 0.08664 | 3.724615 | 0.0002 | 0.00421 | 5.446 |
| Gm47698 | 16.3654 | -1.03435 | 0.56104 | -1.84364 | 0.06524 | 0.19707 | 2.58 |
| Gm47700 | 19.1607 | -0.22056 | 0.42938 | -0.51367 | 0.60748 | 0.77558 | 2.51 |
| Gm47701 | 118.418 | 0.020406 | 0.19033 | 0.107216 | 0.91462 | 0.95656 | 2.787 |
| Gm47710 | 16.6777 | -0.05813 | 0.41516 | -0.14002 | 0.88864 | 0.9444 | 92.712 |
| Gm47765 | 40.418 | 1.120683 | 0.31255 | 3.585607 | 0.00034 | 0.00611 | 1.269 |
| Gm4779 | 18.8668 | 1.146137 | 0.44487 | 2.576321 | 0.00999 | 0.05742 | 4.371 |
| Gm47794 | 63.9693 | 0.333942 | 0.28292 | 1.180323 | 0.23787 | 0.44325 | 1.741 |
| Gm47797 | 16.6562 | 0.295255 | 0.46309 | 0.637578 | 0.52375 | 0.71217 | 0.327 |

| | | | | | | | |
|---|---|---|---|---|---|---|---|
| Gm47798 | 219.664 | 0.184061 | 0.1738 | 1.059031 | 0.28959 | 0.49978 | 3.815 |
| Gm47802 | 20.1931 | 0.260193 | 0.41631 | 0.624993 | 0.53198 | 0.71956 | 4.338 |
| Gm47806 | 32.42 | -0.01424 | 0.35703 | -0.03988 | 0.96819 | 0.98402 | 10.37 |
| Gm47814 | 199.706 | 0.365873 | 0.1959 | 1.867683 | 0.06181 | 0.19034 | 7.417 |
| Gm47817 | 24.7071 | -0.43498 | 0.40744 | -1.06759 | 0.2857 | 0.49571 | 3.153 |
| Gm47826 | 54.454 | -0.3048 | 0.39936 | -0.76321 | 0.44534 | 0.64922 | 3.139 |
| Gm47827 | 32.6165 | 0.491215 | 0.3473 | 1.414387 | 0.15725 | 0.3444 | 16.558 |
| Gm47854 | 17.8467 | 0.160107 | 0.47114 | 0.33983 | 0.73398 | 0.85925 | 33.703 |
| Gm47856 | 22.5101 | -0.27871 | 0.42754 | -0.65188 | 0.51448 | 0.70501 | 1.232 |
| Gm4786 | 38.771 | 0.286357 | 0.43835 | 0.653255 | 0.51359 | 0.70443 | 0.798 |
| Gm47863 | 147.71 | -0.30967 | 0.1873 | -1.65337 | 0.09826 | 0.2573 | 2.173 |
| Gm4787 | 67.5007 | 0.024019 | 0.28266 | 0.084977 | 0.93228 | 0.96589 | 2.474 |
| Gm47879 | 15.676 | -0.05251 | 0.45941 | -0.1143 | 0.909 | 0.95444 | 1.38 |
| Gm47882 | 17.0489 | -1.23727 | 0.53025 | -2.33338 | 0.01963 | 0.08846 | 31.446 |
| Gm47898 | 42.3627 | 0.098316 | 0.31357 | 0.313532 | 0.75388 | 0.87065 | 6.512 |
| Gm47920 | 18.5561 | -0.29195 | 0.3992 | -0.73133 | 0.46458 | 0.66431 | 2.408 |
| Gm4793 | 61.815 | -0.01562 | 0.25136 | -0.06216 | 0.95044 | 0.97552 | 15.176 |
| Gm47950 | 130.809 | 0.493778 | 0.16017 | 3.082835 | 0.00205 | 0.02041 | 1.015 |
| Gm47956 | 32.1974 | 0.225756 | 0.37575 | 0.600813 | 0.54796 | 0.73206 | 6.286 |
| Gm47962 | 38.0416 | -0.28032 | 0.32659 | -0.8583 | 0.39073 | 0.59983 | 26.231 |
| Gm47963 | 37.6817 | -0.72913 | 0.43866 | -1.66217 | 0.09648 | 0.25438 | 2.144 |
| Gm47976 | 106.388 | -0.39265 | 0.16889 | -2.32494 | 0.02008 | 0.0897 | 1.545 |
| Gm47980 | 23.0235 | 0.421256 | 0.37016 | 1.138034 | 0.25511 | 0.46281 | 2.921 |
| Gm47982 | 314.469 | 0.355767 | 0.27773 | 1.280984 | 0.2002 | 0.39817 | 2.34 |
| Gm47987 | 28.8343 | -0.32331 | 0.34696 | -0.93184 | 0.35142 | 0.56196 | 2.795 |
| Gm47992 | 13.7582 | 0.692254 | 0.69355 | 0.998127 | 0.31822 | 0.52878 | 1.987 |
| Gm47994 | 20.8014 | 0.473206 | 0.3675 | 1.287647 | 0.19787 | 0.39566 | 0.78 |
| Gm48001 | 17.7708 | 0.197049 | 0.56287 | 0.350077 | 0.72628 | 0.8541 | 0.758 |
| Gm48003 | 17.7354 | 0.116649 | 0.41661 | 0.279996 | 0.77948 | 0.88513 | 18.608 |
| Gm48006 | 20.4969 | 0.498338 | 0.56421 | 0.883255 | 0.3771 | 0.58718 | 22.295 |
| Gm48014 | 20.6712 | -0.48182 | 0.3904 | -1.23416 | 0.21714 | 0.41911 | 1.3 |
| Gm48027 | 41.2655 | 0.246331 | 0.26388 | 0.933497 | 0.35056 | 0.56125 | 3.436 |
| Gm48045 | 32.2809 | -0.05046 | 0.29215 | -0.17272 | 0.86287 | 0.93062 | 1.948 |
| Gm48057 | 26.5977 | 1.770261 | 0.47088 | 3.759475 | 0.00017 | 0.00387 | 2.02 |

| | | | | | | | |
|---|---|---|---|---|---|---|---|
| Gm48062 | 1164.05 | 0.301554 | 0.15895 | 1.897137 | 0.05781 | 0.18267 | 2.622 |
| Gm48063 | 53.228 | 0.128574 | 0.29629 | 0.433944 | 0.66433 | 0.8145 | 2.517 |
| Gm48069 | 34.0216 | 0.539436 | 0.36977 | 1.458826 | 0.14461 | 0.32724 | 1.484 |
| Gm48071 | 20.0024 | 0.427036 | 0.39894 | 1.070424 | 0.28443 | 0.49452 | 1.258 |
| Gm48079 | 15.7156 | 0.153297 | 0.4594 | 0.333687 | 0.73862 | 0.86216 | 0.55 |
| Gm48086 | 19.1953 | 0.031155 | 0.41584 | 0.07492 | 0.94028 | 0.97027 | 3.816 |
| Gm48102 | 30.3095 | 1.180971 | 0.33035 | 3.574952 | 0.00035 | 0.00623 | 7.234 |
| Gm48114 | 35.8047 | -0.51521 | 0.27294 | -1.88765 | 0.05907 | 0.18503 | 3.686 |
| Gm48123 | 13.0005 | 0.788097 | 0.48301 | 1.631627 | 0.10276 | 0.26467 | 1.341 |
| Gm48137 | 32.7125 | 0.229151 | 0.30025 | 0.76321 | 0.44534 | 0.64922 | 2.239 |
| Gm48139 | 12.3853 | 0.036941 | 0.66175 | 0.055824 | 0.95548 | 0.97811 | 3.915 |
| Gm48181 | 12.8334 | -0.06351 | 0.45383 | -0.13994 | 0.88871 | 0.9444 | 1.066 |
| Gm48194 | 70.7098 | 0.121207 | 0.20514 | 0.590859 | 0.55461 | 0.73712 | 0.517 |
| Gm48225 | 18.2605 | 0.388437 | 0.52193 | 0.744231 | 0.45674 | 0.65819 | 5.083 |
| Gm48226 | 393.105 | 0.068238 | 0.09902 | 0.689134 | 0.49074 | 0.68576 | 4.24 |
| Gm48239 | 1075.62 | -0.03195 | 0.1696 | -0.18841 | 0.85055 | 0.92407 | 10.145 |
| Gm48249 | 74.8468 | 0.251971 | 0.26493 | 0.951068 | 0.34157 | 0.55245 | 19.599 |
| Gm48261 | 22.985 | -0.49044 | 0.4808 | -1.02005 | 0.3077 | 0.51754 | 3.856 |
| Gm48267 | 96.0698 | -0.77071 | 0.24278 | -3.17451 | 0.0015 | 0.01657 | 1.533 |
| Gm48284 | 19.4136 | -0.00948 | 0.50285 | -0.01886 | 0.98495 | 0.99174 | 9.971 |
| Gm4829 | 43.0221 | 0.004155 | 0.27136 | 0.015312 | 0.98778 | 0.99316 | 0.594 |
| Gm48295 | 42.372 | -0.1735 | 0.32036 | -0.54158 | 0.58811 | 0.76153 | 2.435 |
| Gm48314 | 46.8628 | 0.035291 | 0.30883 | 0.114273 | 0.90902 | 0.95444 | 2.739 |
| Gm48327 | 12.617 | 0.048873 | 0.4979 | 0.098159 | 0.92181 | 0.96059 | 1.101 |
| Gm48342 | 22.8533 | -0.57434 | 0.38806 | -1.48003 | 0.13887 | 0.3201 | 6.19 |
| Gm48349 | 25.5417 | 1.376258 | 0.39159 | 3.514496 | 0.00044 | 0.00731 | 0.467 |
| Gm48350 | 68.7768 | 0.012303 | 0.23371 | 0.052644 | 0.95802 | 0.9795 | 11.11 |
| Gm48357 | 44.5188 | 0.319456 | 0.25447 | 1.255384 | 0.20934 | 0.40935 | 6.423 |
| Gm48375 | 25.6439 | -0.11107 | 0.38719 | -0.28687 | 0.77421 | 0.88239 | 1.274 |
| Gm48381 | 21.1209 | -0.58562 | 0.60412 | -0.96938 | 0.33235 | 0.54326 | 3.964 |
| Gm48382 | 24.0415 | 0.217297 | 0.47446 | 0.457991 | 0.64696 | 0.80353 | 3.675 |
| Gm48393 | 51.6435 | -0.46231 | 0.34932 | -1.32347 | 0.18568 | 0.38052 | 12.024 |
| Gm48405 | 35.4071 | 0.113564 | 0.3361 | 0.337885 | 0.73545 | 0.85989 | 2.223 |
| Gm48408 | 340.989 | 0.327596 | 0.12064 | 2.715465 | 0.00662 | 0.04386 | 1.131 |

| | | | | | | | |
|---|---|---|---|---|---|---|---|
| Gm48420 | 22.2861 | -0.21851 | 0.38377 | -0.56936 | 0.56911 | 0.74877 | 0.6 |
| Gm48452 | 25.5578 | 0.138539 | 0.35079 | 0.394934 | 0.69289 | 0.83344 | 0.256 |
| Gm48478 | 14.7558 | 0.580459 | 0.48637 | 1.193461 | 0.23269 | 0.43752 | 2.325 |
| Gm48482 | 280.309 | 0.364134 | 0.23303 | 1.562632 | 0.11814 | 0.28962 | 2.533 |
| Gm48485 | 161.973 | 2 | 0.66079 | 3.045298 | 0.00232 | 0.0222 | 13.614 |
| Gm48488 | 53.5836 | -0.43193 | 0.38964 | -1.10853 | 0.26763 | 0.477 | 6.181 |
| Gm48495 | 70.3915 | 0.484879 | 0.2104 | 2.304607 | 0.02119 | 0.09306 | 0.876 |
| Gm48496 | 14.2089 | -0.05129 | 0.44514 | -0.11522 | 0.90827 | 0.95403 | 0.256 |
| Gm48508 | 37.8776 | -0.62133 | 0.41801 | -1.4864 | 0.13717 | 0.31781 | 68.052 |
| Gm48510 | 20.4866 | 1.661827 | 0.50821 | 3.269954 | 0.00108 | 0.01343 | 2.143 |
| Gm48536 | 22.374 | 0.725431 | 0.51113 | 1.419263 | 0.15582 | 0.34239 | 3.33 |
| Gm48548 | 13.1638 | 1.755529 | 0.57551 | 3.050382 | 0.00229 | 0.02196 | 2.347 |
| Gm48552 | 40.1968 | 0.363591 | 0.28431 | 1.27886 | 0.20095 | 0.39907 | 2.263 |
| Gm48562 | 26.9352 | -0.02192 | 0.42652 | -0.0514 | 0.95901 | 0.98007 | 0.748 |
| Gm48582 | 66.33 | 0.157085 | 0.29865 | 0.52599 | 0.5989 | 0.76992 | 6.429 |
| Gm48583 | 126.076 | -0.21393 | 0.24291 | -0.88067 | 0.3785 | 0.58828 | 0.22 |
| Gm48600 | 27.9146 | 0.161962 | 0.32279 | 0.50176 | 0.61584 | 0.78132 | 4.48 |
| Gm48601 | 14.1366 | -0.169 | 0.60562 | -0.27905 | 0.7802 | 0.8855 | 3.032 |
| Gm48602 | 16.3053 | 0.342826 | 0.46735 | 0.733548 | 0.46322 | 0.66314 | 3.369 |
| Gm48603 | 136.647 | -0.02304 | 0.17514 | -0.13154 | 0.89535 | 0.94744 | 1.138 |
| Gm48604 | 431.059 | 0.10323 | 0.13551 | 0.761795 | 0.44618 | 0.64983 | 2.42 |
| Gm48611 | 16.6878 | -0.42581 | 0.52529 | -0.81062 | 0.41758 | 0.62501 | 2.555 |
| Gm48621 | 28.6589 | 0.063384 | 0.33275 | 0.190483 | 0.84893 | 0.92318 | 1.262 |
| Gm48622 | 30.1889 | 0.50174 | 0.35167 | 1.426739 | 0.15366 | 0.33948 | 17.976 |
| Gm48623 | 197.849 | 0.051481 | 0.18242 | 0.282215 | 0.77778 | 0.88423 | 10.846 |
| Gm48624 | 33.5943 | -0.05757 | 0.30344 | -0.18971 | 0.84953 | 0.92352 | 2.81 |
| Gm48627 | 22.2796 | 0.615604 | 0.38272 | 1.608495 | 0.10773 | 0.27261 | 1.351 |
| Gm48639 | 26.8201 | 0.222152 | 0.39228 | 0.566309 | 0.57118 | 0.75015 | 1.876 |
| Gm48653 | 107.81 | 0.316973 | 0.20879 | 1.518115 | 0.12899 | 0.30621 | 6.917 |
| Gm48655 | 17.75 | 0.203243 | 0.4234 | 0.480028 | 0.63121 | 0.79177 | 1.344 |
| Gm48677 | 211.611 | 0.57447 | 0.17262 | 3.328039 | 0.00087 | 0.01166 | 25.771 |
| Gm48678 | 82.8612 | -0.08322 | 0.30852 | -0.26974 | 0.78736 | 0.88947 | 60.549 |
| Gm48684 | 35.817 | 0.270006 | 0.34739 | 0.777247 | 0.43701 | 0.64261 | 0.978 |
| Gm48689 | 35.6025 | 0.29721 | 0.48171 | 0.61699 | 0.53724 | 0.72399 | 5.188 |

| | | | | | | | |
|---|---|---|---|---|---|---|---|
| Gm48691 | 32.5111 | 0.001197 | 0.33236 | 0.003601 | 0.99713 | 0.99835 | 14.551 |
| Gm48692 | 64.1933 | -0.10187 | 0.32864 | -0.30999 | 0.75657 | 0.87187 | 2.379 |
| Gm48699 | 17.4963 | 0.420781 | 0.51746 | 0.81317 | 0.41612 | 0.62369 | 1.525 |
| Gm48700 | 75.0771 | 0.368875 | 0.34476 | 1.069934 | 0.28465 | 0.49463 | 4.275 |
| Gm48701 | 72.0025 | 0.037442 | 0.25859 | 0.144793 | 0.88487 | 0.94273 | 2.293 |
| Gm48713 | 40.258 | 0.064577 | 0.41851 | 0.154301 | 0.87737 | 0.93876 | 2.846 |
| Gm48714 | 80.4342 | 0.30988 | 0.2194 | 1.412373 | 0.15784 | 0.34507 | 1.441 |
| Gm48716 | 21.4791 | 0.849079 | 0.40784 | 2.081886 | 0.03735 | 0.1367 | 1.409 |
| Gm48717 | 28.573 | 0.17856 | 0.33955 | 0.525872 | 0.59898 | 0.76993 | 1.072 |
| Gm48735 | 28.9986 | -0.60442 | 0.35192 | -1.7175 | 0.08589 | 0.23546 | 1.31 |
| Gm48737 | 18.0229 | -0.34129 | 0.40991 | -0.8326 | 0.40507 | 0.61387 | 2.312 |
| Gm48740 | 29.3954 | -0.37695 | 0.3604 | -1.04593 | 0.29559 | 0.50608 | 7.754 |
| Gm48747 | 206.446 | 0.058429 | 0.13785 | 0.423849 | 0.67168 | 0.81937 | 24.433 |
| Gm4875 | 29.0234 | 0.318533 | 0.34665 | 0.918878 | 0.35816 | 0.56828 | 0.786 |
| Gm4876 | 70.988 | 0.007878 | 0.21013 | 0.037493 | 0.97009 | 0.98478 | 106.824 |
| Gm48768 | 79.2379 | 0.402346 | 0.19833 | 2.028694 | 0.04249 | 0.14843 | 2.29 |
| Gm48780 | 30.0659 | 0.23221 | 0.3284 | 0.707093 | 0.47951 | 0.67717 | 0.997 |
| Gm48784 | 34.3678 | 0.502956 | 0.32845 | 1.53129 | 0.1257 | 0.30095 | 15.891 |
| Gm48796 | 331.204 | 0.159191 | 0.1538 | 1.035066 | 0.30064 | 0.51068 | 3.793 |
| Gm48798 | 23.4093 | 0.162638 | 0.4787 | 0.33975 | 0.73404 | 0.85925 | 0.793 |
| Gm48804 | 98.7903 | 0.595272 | 0.23468 | 2.536533 | 0.0112 | 0.06176 | 3.041 |
| Gm48822 | 48.4185 | 0.247709 | 0.31167 | 0.794788 | 0.42674 | 0.63327 | 2.371 |
| Gm48837 | 24.6583 | 1.242894 | 0.45402 | 2.737519 | 0.00619 | 0.04208 | 10.921 |
| Gm48838 | 48.0359 | 0.518061 | 0.27634 | 1.874745 | 0.06083 | 0.18832 | 1.234 |
| Gm48865 | 44.7136 | 0.015147 | 0.25803 | 0.058702 | 0.95319 | 0.97691 | 2.605 |
| Gm48870 | 26.9377 | -0.72676 | 0.42205 | -1.72197 | 0.08507 | 0.23389 | 4.024 |
| Gm48887 | 13.0204 | -0.95243 | 0.5692 | -1.67329 | 0.09427 | 0.25054 | 1.553 |
| Gm48893 | 49.2921 | 0.523884 | 0.3186 | 1.644329 | 0.10011 | 0.26033 | 47.561 |
| Gm48898 | 66.9568 | 0.180115 | 0.26378 | 0.682829 | 0.49472 | 0.68882 | 2.671 |
| Gm4890 | 59.4639 | 0.546795 | 0.36256 | 1.508155 | 0.13151 | 0.31024 | 37.411 |
| Gm48914 | 31.4095 | -0.01285 | 0.38416 | -0.03346 | 0.97331 | 0.98671 | 14.324 |
| Gm48928 | 20.7257 | 1.2747 | 0.4356 | 2.92631 | 0.00343 | 0.02836 | 0.547 |
| Gm48932 | 185.979 | 0.51591 | 0.20413 | 2.52737 | 0.01149 | 0.06285 | 3.25 |
| Gm48933 | 13.3365 | -0.03201 | 0.51144 | -0.0626 | 0.95009 | 0.97525 | 0.898 |

| | | | | | | | |
|---|---|---|---|---|---|---|---|
| Gm48949 | 41.5199 | 0.119702 | 0.28693 | 0.417175 | 0.67655 | 0.82256 | 0.668 |
| Gm4895 | 12.6156 | -0.05503 | 0.483 | -0.11394 | 0.90929 | 0.95456 | 1.137 |
| Gm48964 | 22.0343 | -0.10176 | 0.47531 | -0.21409 | 0.83048 | 0.91311 | 2.02 |
| Gm48967 | 34.1192 | -0.41183 | 0.53418 | -0.77095 | 0.44074 | 0.64559 | 4.184 |
| Gm48978 | 17.9002 | 0.022001 | 0.45313 | 0.048553 | 0.96128 | 0.98116 | 4.992 |
| Gm48992 | 44.0321 | -0.86467 | 0.32963 | -2.62314 | 0.00871 | 0.05239 | 1.101 |
| Gm48998 | 13.0852 | 0.414285 | 0.5825 | 0.711216 | 0.47695 | 0.67482 | 2.707 |
| Gm49012 | 19.7301 | -0.33398 | 0.5194 | -0.64302 | 0.52021 | 0.70963 | 14.609 |
| Gm49027 | 139.658 | 0.216622 | 0.16982 | 1.275591 | 0.2021 | 0.40058 | 13.407 |
| Gm49064 | 69.2334 | 0.260481 | 0.27322 | 0.953373 | 0.3404 | 0.55126 | 2.973 |
| Gm49066 | 40.6369 | 0.330764 | 0.29084 | 1.137277 | 0.25542 | 0.4631 | 0.903 |
| Gm49069 | 133.837 | -0.12685 | 0.32936 | -0.38514 | 0.70013 | 0.83799 | 4.011 |
| Gm49083 | 13.8895 | -1.6358 | 0.73534 | -2.22455 | 0.02611 | 0.10722 | 7.688 |
| Gm49085 | 134.945 | 0.366984 | 0.19932 | 1.841161 | 0.0656 | 0.19767 | 5.748 |
| Gm49089 | 22.3365 | 0.241066 | 0.42547 | 0.566587 | 0.57099 | 0.75014 | 2.906 |
| Gm49092 | 52.7469 | -0.23082 | 0.26384 | -0.87485 | 0.38166 | 0.59107 | 0.936 |
| Gm49096 | 28.9663 | -0.28921 | 0.39381 | -0.73437 | 0.46272 | 0.66283 | 2.308 |
| Gm49097 | 31.9651 | -0.33306 | 0.4254 | -0.78293 | 0.43367 | 0.63955 | 5.552 |
| Gm49125 | 182.34 | -0.07378 | 0.17072 | -0.43219 | 0.66561 | 0.81542 | 1.295 |
| Gm49130 | 62.6242 | -0.18736 | 0.24735 | -0.75748 | 0.44876 | 0.65214 | 2.498 |
| Gm49163 | 339.528 | 0.116725 | 0.11208 | 1.04141 | 0.29769 | 0.50774 | 1.858 |
| Gm49172 | 194.152 | 0.161707 | 0.14072 | 1.149109 | 0.25051 | 0.45785 | 21.391 |
| Gm49179 | 19.0983 | 0.107277 | 0.45668 | 0.234905 | 0.81428 | 0.90464 | 4.979 |
| Gm49182 | 15.5481 | -0.8133 | 0.43393 | -1.87425 | 0.0609 | 0.18836 | 1.973 |
| Gm49188 | 25.9122 | 0.263629 | 0.37684 | 0.699577 | 0.48419 | 0.68059 | 2.123 |
| Gm49194 | 68.0323 | 0.291761 | 0.24198 | 1.205716 | 0.22793 | 0.43183 | 1.474 |
| Gm49201 | 96.1712 | 0.215424 | 0.26013 | 0.828141 | 0.40759 | 0.61541 | 1.11 |
| Gm49204 | 367.276 | 0.288594 | 0.14992 | 1.925002 | 0.05423 | 0.1751 | 4.64 |
| Gm49209 | 52.9319 | 0.272124 | 0.30068 | 0.905026 | 0.36545 | 0.57607 | 2.494 |
| Gm49223 | 62.7505 | -0.1305 | 0.24201 | -0.53926 | 0.58971 | 0.76282 | 5.974 |
| Gm4924 | 320.969 | 0.259017 | 0.17531 | 1.477492 | 0.13954 | 0.32076 | 68.628 |
| Gm49244 | 16.6183 | 0.636123 | 0.52719 | 1.206621 | 0.22758 | 0.43174 | 0.442 |
| Gm4926 | 24.3008 | -0.61027 | 0.43529 | -1.402 | 0.16091 | 0.34906 | 8.491 |
| Gm49284 | 571.143 | 0.106954 | 0.09715 | 1.100934 | 0.27093 | 0.48035 | 2.064 |

| | | | | | | | |
|---|---|---|---|---|---|---|---|
| Gm49300 | 26.0729 | -0.54271 | 0.55114 | -0.98472 | 0.32476 | 0.53594 | 1.944 |
| Gm49308 | 20.8572 | -0.079 | 0.39289 | -0.20108 | 0.84064 | 0.91835 | 3.612 |
| Gm49312 | 27.0636 | 0.639358 | 0.38205 | 1.673498 | 0.09423 | 0.25054 | 0.527 |
| Gm49317 | 19.7974 | 0.002608 | 0.44397 | 0.005873 | 0.99531 | 0.99737 | 180.629 |
| Gm49322 | 87.1511 | 0.159511 | 0.22914 | 0.696137 | 0.48634 | 0.68233 | 9.764 |
| Gm49326 | 117.267 | 0.293519 | 0.19789 | 1.48326 | 0.13801 | 0.31902 | 0.126 |
| Gm49327 | 420.458 | 0.165066 | 0.11812 | 1.397387 | 0.1623 | 0.35067 | 6.465 |
| Gm49329 | 65.8583 | 0.151134 | 0.24845 | 0.608315 | 0.54298 | 0.72855 | 164.877 |
| Gm49331 | 76.5912 | 0.195664 | 0.20052 | 0.975769 | 0.32918 | 0.54039 | 2.287 |
| Gm49336 | 3095.53 | 0.096602 | 0.07501 | 1.287835 | 0.1978 | 0.39558 | 46.226 |
| Gm49338 | 194.221 | -0.18333 | 0.13248 | -1.38382 | 0.16641 | 0.356 | 2.079 |
| Gm49376 | 57.9437 | -0.25629 | 0.23926 | -1.07117 | 0.28409 | 0.49424 | 3.334 |
| Gm49396 | 166.226 | -0.06445 | 0.20106 | -0.32056 | 0.74855 | 0.86814 | 4.617 |
| Gm49405 | 126.841 | -0.00223 | 0.20024 | -0.01113 | 0.99112 | 0.99503 | 5.191 |
| Gm49411 | 12.6239 | 0.231871 | 0.55533 | 0.417538 | 0.67629 | 0.82247 | 2.476 |
| Gm49413 | 30.4603 | 0.45647 | 0.38645 | 1.181196 | 0.23752 | 0.44289 | 3.039 |
| Gm49417 | 89.9697 | -0.03071 | 0.18314 | -0.16768 | 0.86684 | 0.93243 | 49.964 |
| Gm49437 | 24.3182 | 0.328467 | 0.37209 | 0.88276 | 0.37737 | 0.58749 | 0.285 |
| Gm49445 | 31.3708 | -0.10613 | 0.30404 | -0.34908 | 0.72703 | 0.85431 | 21.701 |
| Gm49449 | 51.8372 | 0.855856 | 0.30036 | 2.849402 | 0.00438 | 0.03344 | 1.116 |
| Gm49484 | 123.683 | 0.390978 | 0.19152 | 2.041409 | 0.04121 | 0.14553 | 7.395 |
| Gm49486 | 14.5116 | 0.743478 | 0.59711 | 1.245124 | 0.21309 | 0.41418 | 27.284 |
| Gm49492 | 294.798 | -0.16257 | 0.12452 | -1.30564 | 0.19168 | 0.3883 | 2.563 |
| Gm49493 | 90.9366 | 0.454128 | 0.19612 | 2.3156 | 0.02058 | 0.09128 | 4.136 |
| Gm49494 | 44.5587 | 0.806924 | 0.2857 | 2.824419 | 0.00474 | 0.03512 | 2.063 |
| Gm4950 | 19.8547 | 0.247568 | 0.40924 | 0.604951 | 0.54521 | 0.73012 | 0.615 |
| Gm49502 | 183.054 | 0.176665 | 0.17694 | 0.998443 | 0.31806 | 0.5286 | 2.11 |
| Gm49503 | 16.6514 | -0.70639 | 0.6101 | -1.15782 | 0.24694 | 0.45383 | 4.199 |
| Gm4951 | 27.1008 | -1.33602 | 0.59677 | -2.23875 | 0.02517 | 0.10441 | 35.741 |
| Gm49519 | 49.355 | -0.0939 | 0.26354 | -0.35629 | 0.72163 | 0.8514 | 42.052 |
| Gm49520 | 17.9837 | 0.498503 | 0.45233 | 1.102073 | 0.27043 | 0.47998 | 0.221 |
| Gm49528 | 20.9427 | 0.017321 | 0.46594 | 0.037175 | 0.97035 | 0.98489 | 3.849 |
| Gm49542 | 15.8778 | 0.617092 | 0.45887 | 1.344822 | 0.17868 | 0.37128 | 8.764 |
| Gm49550 | 110.062 | 0.430003 | 0.1972 | 2.180545 | 0.02922 | 0.11598 | 0.252 |

| | | | | | | | |
|---|---|---|---|---|---|---|---|
| Gm49552 | 169.566 | -0.08058 | 0.15941 | -0.5055 | 0.61321 | 0.77981 | 2.193 |
| Gm49555 | 20.9643 | 0.281913 | 0.36939 | 0.763189 | 0.44535 | 0.64922 | 2.795 |
| Gm49565 | 54.2365 | -0.13822 | 0.46495 | -0.29727 | 0.76626 | 0.8779 | 3.414 |
| Gm49584 | 93.1661 | 0.29902 | 0.19885 | 1.503768 | 0.13264 | 0.31182 | 3.536 |
| Gm49592 | 29.9952 | 0.888027 | 0.32828 | 2.705072 | 0.00683 | 0.04479 | 0.228 |
| Gm49593 | 1279.84 | -0.38573 | 0.09369 | -4.11707 | 3.84E-05 | 0.00136 | 2.448 |
| Gm49594 | 49.8131 | -0.01768 | 0.24723 | -0.07153 | 0.94298 | 0.97157 | 0.615 |
| Gm49595 | 25.5224 | 0.406238 | 0.38733 | 1.048826 | 0.29426 | 0.50456 | 2.286 |
| Gm49599 | 105.929 | 0.188608 | 0.28727 | 0.656561 | 0.51146 | 0.70254 | 5.095 |
| Gm49600 | 43.3231 | 0.198815 | 0.35692 | 0.55703 | 0.57751 | 0.75451 | 1.29 |
| Gm49601 | 375.715 | 0.222978 | 0.13738 | 1.623089 | 0.10457 | 0.26745 | 5.962 |
| Gm49602 | 412.443 | 0.101339 | 0.12813 | 0.790918 | 0.42899 | 0.63565 | 2.97 |
| Gm49614 | 13.7403 | -0.95656 | 0.50498 | -1.89426 | 0.05819 | 0.18329 | 0.827 |
| Gm49654 | 879.955 | -0.35343 | 0.15441 | -2.28892 | 0.02208 | 0.09575 | 2.223 |
| Gm49668 | 492.528 | 0.167292 | 0.09374 | 1.784719 | 0.07431 | 0.21491 | 13.164 |
| Gm49678 | 18.6647 | -0.66272 | 0.52409 | -1.26451 | 0.20605 | 0.40531 | 64.585 |
| Gm49699 | 32.5266 | 0.330763 | 0.38222 | 0.865378 | 0.38683 | 0.5961 | 5.678 |
| Gm49700 | 39.722 | 0.504151 | 0.37503 | 1.344302 | 0.17885 | 0.37148 | 22.606 |
| Gm49702 | 1416.18 | 0.860357 | 0.22206 | 3.874448 | 0.00011 | 0.00281 | 1.797 |
| Gm49703 | 59.4448 | 0.55891 | 0.25285 | 2.210436 | 0.02707 | 0.10988 | 2.069 |
| Gm49705 | 14.9042 | 0.210722 | 0.62595 | 0.336641 | 0.73639 | 0.86065 | 5.219 |
| Gm49706 | 146.86 | 0.51289 | 0.20138 | 2.546849 | 0.01087 | 0.06069 | 2.355 |
| Gm49708 | 45.958 | -0.26657 | 0.33443 | -0.79708 | 0.4254 | 0.63203 | 8.048 |
| Gm4972 | 18.2111 | -0.66818 | 0.52584 | -1.27069 | 0.20384 | 0.4025 | 1.254 |
| Gm49727 | 33.2686 | -0.10966 | 0.34736 | -0.31568 | 0.75224 | 0.86997 | 2.579 |
| Gm49731 | 32.8006 | -0.61777 | 0.34579 | -1.78654 | 0.07401 | 0.2144 | 1.734 |
| Gm49741 | 17.5116 | -0.15402 | 0.50988 | -0.30207 | 0.7626 | 0.87556 | 0.769 |
| Gm49745 | 37.0954 | 0.329122 | 0.29332 | 1.122049 | 0.26184 | 0.47055 | 2.151 |
| Gm49747 | 137.395 | 0.365506 | 0.1735 | 2.106722 | 0.03514 | 0.13139 | 2.749 |
| Gm49759 | 1237.05 | 0.075051 | 0.09297 | 0.8073 | 0.41949 | 0.62676 | 3.887 |
| Gm49760 | 18.144 | -0.01222 | 0.47108 | -0.02594 | 0.97931 | 0.9889 | 1.785 |
| Gm49767 | 43.1352 | -0.32214 | 0.31518 | -1.0221 | 0.30673 | 0.51671 | 7.473 |
| Gm49768 | 295.467 | 0.174809 | 0.18095 | 0.966038 | 0.33403 | 0.54475 | 0.767 |
| Gm49769 | 19.0709 | 0.214739 | 0.40022 | 0.536552 | 0.59158 | 0.76416 | 0.633 |

| | | | | | | | |
|---|---|---|---|---|---|---|---|
| Gm49785 | 15.3722 | 0.336473 | 0.42766 | 0.786781 | 0.43141 | 0.63747 | 0.374 |
| Gm49789 | 83.7202 | 0.75388 | 0.269 | 2.802556 | 0.00507 | 0.0366 | 0.499 |
| Gm49794 | 63.5881 | 1.565893 | 0.43722 | 3.581444 | 0.00034 | 0.00615 | 13.382 |
| Gm49797 | 338.666 | 0.092187 | 0.14912 | 0.618209 | 0.53644 | 0.72344 | 113.011 |
| Gm49838 | 20.1599 | 0.354791 | 0.47498 | 0.74696 | 0.45509 | 0.65721 | 1.708 |
| Gm49863 | 198.096 | 0.157965 | 0.20098 | 0.785955 | 0.43189 | 0.6379 | 0.502 |
| Gm49864 | 20.3138 | -0.11355 | 0.47084 | -0.24117 | 0.80942 | 0.90175 | 2.13 |
| Gm49871 | 25.4928 | -0.29611 | 0.3443 | -0.86002 | 0.38978 | 0.59892 | 2.616 |
| Gm49876 | 18.5052 | -0.30433 | 0.46683 | -0.65191 | 0.51446 | 0.70501 | 26.892 |
| Gm49883 | 608.191 | 0.256335 | 0.17275 | 1.483878 | 0.13784 | 0.31867 | 3.732 |
| Gm49886 | 44.1048 | -0.05238 | 0.30487 | -0.1718 | 0.86359 | 0.93094 | 1.417 |
| Gm49894 | 34.3802 | 0.123927 | 0.29153 | 0.425085 | 0.67077 | 0.8186 | 2.005 |
| Gm49906 | 70.4236 | 1.005811 | 0.33703 | 2.984329 | 0.00284 | 0.02532 | 59.224 |
| Gm49911 | 224.181 | 0.522864 | 0.14936 | 3.500713 | 0.00046 | 0.00757 | 0.287 |
| Gm49915 | 24.0201 | -0.04851 | 0.39796 | -0.1219 | 0.90298 | 0.95118 | 4.866 |
| Gm49924 | 41.2266 | 0.4464 | 0.37137 | 1.202047 | 0.22935 | 0.43347 | 1.799 |
| Gm49927 | 53.9778 | 0.222904 | 0.25054 | 0.889696 | 0.37363 | 0.58383 | 0.238 |
| Gm49928 | 30.2583 | 0.502703 | 0.30724 | 1.636177 | 0.1018 | 0.26328 | 2.109 |
| Gm49932 | 27.9541 | -0.719 | 0.61492 | -1.16926 | 0.2423 | 0.44829 | 4.421 |
| Gm49937 | 80.9107 | -0.23189 | 0.20611 | -1.12508 | 0.26056 | 0.46902 | 2.746 |
| Gm49949 | 98.426 | -0.92909 | 0.29138 | -3.18859 | 0.00143 | 0.01601 | 2.503 |
| Gm49961 | 42.0298 | 0.237315 | 0.30519 | 0.77761 | 0.4368 | 0.6425 | 6.36 |
| Gm49968 | 27.3655 | -0.05422 | 0.3544 | -0.15299 | 0.87841 | 0.93918 | 2.773 |
| Gm49969 | 17.6783 | 0.271808 | 0.3984 | 0.682246 | 0.49508 | 0.68919 | 50.519 |
| Gm49980 | 288.292 | -0.34387 | 0.28946 | -1.18797 | 0.23484 | 0.44015 | 4.885 |
| Gm5 | 27.3923 | 1.219818 | 0.37465 | 3.255913 | 0.00113 | 0.01378 | 2.558 |
| Gm50009 | 36.5067 | 0.374336 | 0.33268 | 1.125199 | 0.2605 | 0.46897 | 1.312 |
| Gm50010 | 419.2 | 0.437851 | 0.1441 | 3.038604 | 0.00238 | 0.02253 | 1.323 |
| Gm50012 | 52.2217 | 0.160048 | 0.25007 | 0.640007 | 0.52217 | 0.71122 | 4.917 |
| Gm50021 | 31.982 | 0.228278 | 0.39482 | 0.578185 | 0.56314 | 0.74375 | 7.937 |
| Gm50023 | 29.3314 | 0.292677 | 0.33191 | 0.881795 | 0.37789 | 0.588 | 1.583 |
| Gm50024 | 12.4095 | 0.8166 | 0.4858 | 1.680954 | 0.09277 | 0.24807 | 25.302 |
| Gm50035 | 40.2047 | -0.05382 | 0.29924 | -0.17986 | 0.85726 | 0.92794 | 6.68 |
| Gm50048 | 40.3513 | -1.1191 | 0.29143 | -3.84 | 0.00012 | 0.00308 | 27.704 |

| | | | | | | | |
|---|---|---|---|---|---|---|---|
| Gm50050 | 14.4821 | 0.024478 | 0.43791 | 0.055897 | 0.95542 | 0.97811 | 14.281 |
| Gm50069 | 65.4124 | 0.060109 | 0.26129 | 0.230048 | 0.81805 | 0.90638 | 2.365 |
| Gm50070 | 41.1887 | 0.526867 | 0.2566 | 2.053244 | 0.04005 | 0.14298 | 1.448 |
| Gm50071 | 21.7705 | 0.098738 | 0.36388 | 0.271344 | 0.78613 | 0.88876 | 3.016 |
| Gm50087 | 62.2281 | 0.413218 | 0.27782 | 1.487385 | 0.13691 | 0.31765 | 3.584 |
| Gm50092 | 70.8802 | 0.292209 | 0.24848 | 1.176011 | 0.23959 | 0.44556 | 1.973 |
| Gm50100 | 27.7535 | -0.11725 | 0.36571 | -0.32061 | 0.7485 | 0.86814 | 6.301 |
| Gm50103 | 76.5884 | -0.00929 | 0.26318 | -0.0353 | 0.97184 | 0.98582 | 2.144 |
| Gm50105 | 44.4438 | 0.864121 | 0.34711 | 2.489488 | 0.01279 | 0.0675 | 2.711 |
| Gm50107 | 556.056 | 0.225341 | 0.11349 | 1.985613 | 0.04708 | 0.1592 | 1.692 |
| Gm50115 | 24.9141 | -0.14505 | 0.35715 | -0.40614 | 0.68464 | 0.82819 | 5.267 |
| Gm50138 | 72.5835 | 0.808466 | 0.27578 | 2.931515 | 0.00337 | 0.02811 | 15.346 |
| Gm50141 | 13.7313 | 0.112479 | 0.53437 | 0.210491 | 0.83328 | 0.91468 | 0.274 |
| Gm50144 | 24.5613 | 2 | 0.44875 | 4.522357 | 6.12E-06 | 0.00038 | 0.58 |
| Gm50146 | 13.8378 | 0.479617 | 0.58976 | 0.813246 | 0.41608 | 0.62368 | 8.428 |
| Gm50151 | 62.2084 | -0.06955 | 0.23115 | -0.30088 | 0.7635 | 0.87611 | 2.689 |
| Gm50152 | 173.988 | 0.236842 | 0.16357 | 1.44795 | 0.14763 | 0.33114 | 1.657 |
| Gm50169 | 151.754 | 0.35237 | 0.27161 | 1.297348 | 0.19451 | 0.39183 | 2.636 |
| Gm50170 | 39.6858 | 1.095855 | 0.31646 | 3.462834 | 0.00053 | 0.00831 | 2.594 |
| Gm50196 | 22.7536 | 0.545902 | 0.44946 | 1.214572 | 0.22453 | 0.42822 | 4.68 |
| Gm50203 | 15.4832 | 0.104747 | 0.41853 | 0.250273 | 0.80238 | 0.89807 | 0.833 |
| Gm50205 | 44.9668 | 0.22862 | 0.30663 | 0.745593 | 0.45591 | 0.65758 | 3.034 |
| Gm50230 | 27.0368 | -0.49453 | 0.39288 | -1.25874 | 0.20812 | 0.40775 | 22.306 |
| Gm50232 | 129.627 | 0.424206 | 0.1723 | 2.462059 | 0.01381 | 0.07064 | 5.726 |
| Gm50240 | 253.949 | 0.129647 | 0.15731 | 0.82414 | 0.40986 | 0.61767 | 6.295 |
| Gm50254 | 49.1568 | 0.120623 | 0.25113 | 0.48032 | 0.631 | 0.79169 | 4.117 |
| Gm50268 | 188.153 | 0.268114 | 0.17013 | 1.575926 | 0.11504 | 0.28432 | 2.756 |
| Gm50269 | 18.2972 | -0.17162 | 0.48443 | -0.35428 | 0.72313 | 0.85223 | 14.77 |
| Gm50270 | 44.7103 | 0.094476 | 0.34728 | 0.272048 | 0.78559 | 0.88863 | 7.623 |
| Gm50301 | 37.9294 | 0.223164 | 0.36734 | 0.607514 | 0.54351 | 0.72892 | 11.77 |
| Gm50313 | 14.2476 | -0.14325 | 0.45508 | -0.31478 | 0.75293 | 0.87023 | 0.427 |
| Gm50321 | 25.8494 | -0.2034 | 0.33712 | -0.60334 | 0.54628 | 0.73071 | 7.999 |
| Gm50322 | 66.1644 | 0.159705 | 0.2796 | 0.571199 | 0.56786 | 0.74779 | 4.072 |
| Gm50323 | 43.892 | -0.32991 | 0.31278 | -1.05476 | 0.29154 | 0.50128 | 20.775 |

| | | | | | | | |
|---|---|---|---|---|---|---|---|
| Gm50335 | 105.252 | 0.165865 | 0.22155 | 0.748674 | 0.45405 | 0.65623 | 1.495 |
| Gm50370 | 13.8296 | 0.391351 | 0.5666 | 0.6907 | 0.48975 | 0.68476 | 11.755 |
| Gm50387 | 106.381 | 0.090639 | 0.20168 | 0.44942 | 0.65313 | 0.80721 | 3.626 |
| Gm50403 | 55.067 | -0.18466 | 0.36355 | -0.50793 | 0.6115 | 0.77851 | 5.583 |
| Gm50425 | 13.7111 | -0.16071 | 0.50592 | -0.31766 | 0.75074 | 0.86925 | 2.653 |
| Gm50431 | 526.678 | -0.08818 | 0.09354 | -0.94268 | 0.34584 | 0.55658 | 1.945 |
| Gm50433 | 51.8513 | 0.04548 | 0.25893 | 0.175641 | 0.86058 | 0.92938 | 14.283 |
| Gm50455 | 144.997 | 0.14801 | 0.18159 | 0.815053 | 0.41504 | 0.6229 | 2.911 |
| Gm50462 | 163.048 | 0.000692 | 0.14412 | 0.004803 | 0.99617 | 0.99788 | 2.678 |
| Gm50467 | 13.1494 | 0.080358 | 0.494 | 0.162668 | 0.87078 | 0.93475 | 6.796 |
| Gm50475 | 63.6594 | 0.158438 | 0.36766 | 0.430942 | 0.66651 | 0.8158 | 0.713 |
| Gm50481 | 13.5276 | 0.045041 | 0.51525 | 0.087415 | 0.93034 | 0.9649 | 5.856 |
| Gm50490 | 183.521 | 0.507702 | 0.31567 | 1.608315 | 0.10777 | 0.27268 | 0.724 |
| Gm5067 | 41.5804 | -0.70951 | 0.2996 | -2.36822 | 0.01787 | 0.08327 | 1.009 |
| Gm5069 | 78.3745 | -0.51924 | 0.20135 | -2.57879 | 0.00991 | 0.05714 | 3.581 |
| Gm5083 | 36.3844 | -0.20188 | 0.29039 | -0.69518 | 0.48694 | 0.68266 | 4.013 |
| Gm5086 | 36.7296 | -0.53708 | 0.35148 | -1.52805 | 0.1265 | 0.3023 | 24.1 |
| Gm5089 | 110.437 | -0.21187 | 0.18101 | -1.17049 | 0.2418 | 0.4478 | 41.505 |
| Gm5093 | 29.6253 | -0.00504 | 0.4651 | -0.01084 | 0.99135 | 0.9952 | 0.52 |
| Gm5112 | 35.8587 | 0.286889 | 0.2974 | 0.964659 | 0.33472 | 0.54548 | 2.368 |
| Gm5113 | 1525.26 | -0.11859 | 0.08136 | -1.45757 | 0.14496 | 0.32781 | 12.45 |
| Gm5124 | 1980.76 | 0.145005 | 0.09181 | 1.57936 | 0.11425 | 0.28305 | 3.797 |
| Gm5141 | 194.252 | 0.604182 | 0.27591 | 2.189813 | 0.02854 | 0.11395 | 13.609 |
| Gm51425 | 124.093 | 0.700977 | 0.37873 | 1.850862 | 0.06419 | 0.19501 | 36.491 |
| Gm5148 | 360.168 | 0.580147 | 0.12953 | 4.478817 | 7.51E-06 | 0.00045 | 10.301 |
| Gm5165 | 194.518 | 0.611358 | 0.30799 | 1.985013 | 0.04714 | 0.15934 | 31.497 |
| Gm5176 | 79.4919 | 0.868813 | 0.29465 | 2.948625 | 0.00319 | 0.02718 | 1.227 |
| Gm5182 | 439.336 | -0.23189 | 0.13147 | -1.76389 | 0.07775 | 0.22065 | 6.092 |
| Gm5242 | 98.0782 | 0.09679 | 0.24568 | 0.393973 | 0.6936 | 0.83383 | 1.233 |
| Gm527 | 219.323 | 0.167538 | 0.13743 | 1.219064 | 0.22282 | 0.42605 | 6.687 |
| Gm52948 | 29.162 | -0.20815 | 0.35138 | -0.59236 | 0.55361 | 0.73632 | 16.29 |
| Gm52960 | 15.6847 | -0.59337 | 0.46164 | -1.28536 | 0.19867 | 0.39644 | 57.788 |
| Gm52964 | 136.195 | 0.212201 | 0.16429 | 1.291625 | 0.19649 | 0.3941 | 2.683 |
| Gm52965 | 110.265 | -0.15037 | 0.19494 | -0.77134 | 0.4405 | 0.64543 | 1.524 |

| | | | | | | | |
|---|---|---|---|---|---|---|---|
| Gm52969 | 18.0508 | -0.24175 | 0.47192 | -0.51227 | 0.60846 | 0.77621 | 259.692 |
| Gm52974 | 102.098 | 0.131939 | 0.23775 | 0.554939 | 0.57894 | 0.75565 | 22.742 |
| Gm52988 | 30.1289 | 0.573787 | 0.62893 | 0.912317 | 0.3616 | 0.57187 | 66.953 |
| Gm53012 | 190.214 | 0.339467 | 0.15929 | 2.131182 | 0.03307 | 0.1263 | 13.166 |
| Gm53028 | 37.516 | 0.376641 | 0.31568 | 1.193095 | 0.23283 | 0.43763 | 33.387 |
| Gm53029 | 27.1453 | 0.267896 | 0.33913 | 0.789958 | 0.42955 | 0.63601 | 39.69 |
| Gm53033 | 16.9313 | 0.794714 | 0.52741 | 1.506815 | 0.13186 | 0.31073 | 3.237 |
| Gm53041 | 35.2403 | 0.598794 | 0.32488 | 1.843122 | 0.06531 | 0.19714 | 1.773 |
| Gm53048 | 21.014 | 0.40338 | 0.45024 | 0.895918 | 0.3703 | 0.58058 | 1.516 |
| Gm53058 | 251.556 | 0.678873 | 0.2149 | 3.158956 | 0.00158 | 0.01726 | 19.344 |
| Gm5345 | 28.7688 | -0.09529 | 0.3178 | -0.29985 | 0.76429 | 0.87657 | 1.2 |
| Gm5353 | 21.656 | 0.56848 | 0.56315 | 1.009467 | 0.31275 | 0.52328 | 0.793 |
| Gm536 | 13.5797 | -1.48108 | 0.56534 | -2.6198 | 0.0088 | 0.05264 | 21.245 |
| Gm5417 | 219.727 | 0.506124 | 0.6869 | 0.736823 | 0.46123 | 0.66181 | 0.438 |
| Gm5421 | 23.3943 | 0.096655 | 0.42471 | 0.227578 | 0.81997 | 0.90734 | 2.354 |
| Gm5422 | 63.6501 | -0.20296 | 0.23496 | -0.86377 | 0.38771 | 0.59705 | 3.021 |
| Gm5424 | 479.751 | -0.04532 | 0.18912 | -0.23963 | 0.81062 | 0.90256 | 1.24 |
| Gm5425 | 14.2384 | 1.122167 | 0.4595 | 2.442142 | 0.0146 | 0.07302 | 0.459 |
| Gm5436 | 59.0309 | -0.1472 | 0.21188 | -0.69474 | 0.48722 | 0.68271 | 0.643 |
| Gm5441 | 275.084 | 0.439298 | 0.15071 | 2.914936 | 0.00356 | 0.02907 | 102.502 |
| Gm5451 | 23.1354 | -0.18969 | 0.41572 | -0.45629 | 0.64818 | 0.80406 | 0.579 |
| Gm5454 | 167.611 | 0.011727 | 0.27466 | 0.042698 | 0.96594 | 0.98287 | 1.453 |
| Gm5464 | 22.1266 | 0.60753 | 0.50372 | 1.206093 | 0.22778 | 0.43174 | 2.874 |
| Gm5466 | 38.033 | -0.08015 | 0.37424 | -0.21417 | 0.83041 | 0.91311 | 1.89 |
| Gm5468 | 96.4771 | -0.67112 | 0.28098 | -2.38846 | 0.01692 | 0.08058 | 49.341 |
| Gm5469 | 12.7681 | 2 | 0.69169 | 3.157934 | 0.00159 | 0.0173 | 1.076 |
| Gm5475 | 19.9278 | 0.267417 | 0.41256 | 0.648186 | 0.51686 | 0.70681 | 6.004 |
| Gm5493 | 17.231 | 0.308383 | 0.46252 | 0.666746 | 0.50493 | 0.69681 | 11.565 |
| Gm5499 | 24.261 | -0.313 | 0.35075 | -0.89237 | 0.37219 | 0.58257 | 0.885 |
| Gm5512 | 12.6115 | 0.585068 | 0.49434 | 1.183544 | 0.23659 | 0.44183 | 1.522 |
| Gm5518 | 314.98 | -0.22343 | 0.14661 | -1.52397 | 0.12752 | 0.30398 | 0.645 |
| Gm5527 | 70.0728 | -0.07087 | 0.21333 | -0.33219 | 0.73974 | 0.86273 | 0.642 |
| Gm5529 | 136.264 | -0.00019 | 0.18438 | -0.00103 | 0.99917 | 0.99967 | 0.891 |
| Gm5532 | 32.6651 | -0.69698 | 0.46477 | -1.49963 | 0.13371 | 0.31339 | 7.048 |

| | | | | | | | |
|---|---|---|---|---|---|---|---|
| Gm5535 | 41.4733 | -0.52086 | 0.30709 | -1.69613 | 0.08986 | 0.24306 | 109.176 |
| Gm5555 | 65.3881 | -0.13612 | 0.24036 | -0.56633 | 0.57117 | 0.75015 | 1.111 |
| Gm5559 | 540.594 | -0.05141 | 0.10642 | -0.48306 | 0.62905 | 0.79023 | 1.002 |
| Gm5577 | 375.818 | 0.55726 | 0.11951 | 4.662996 | 3.12E-06 | 0.00024 | 24.589 |
| Gm5602 | 53.2532 | -0.25533 | 0.23018 | -1.10927 | 0.26731 | 0.47664 | 5.046 |
| Gm5617 | 54.5462 | -0.14862 | 0.26882 | -0.55285 | 0.58037 | 0.75658 | 0.531 |
| Gm5620 | 23.0518 | -0.58617 | 0.43404 | -1.35052 | 0.17685 | 0.36898 | 1.348 |
| Gm5628 | 109.453 | 0.550178 | 0.20088 | 2.738816 | 0.00617 | 0.04202 | 1.049 |
| Gm5637 | 13.1988 | -0.479 | 0.56732 | -0.84432 | 0.39849 | 0.6075 | 1.119 |
| Gm5641 | 243.338 | 0.64042 | 0.29152 | 2.196843 | 0.02803 | 0.1127 | 1.488 |
| Gm5655 | 13.4556 | 1.006874 | 0.46469 | 2.166751 | 0.03025 | 0.11875 | 0.901 |
| Gm5668 | 25.7244 | 0.923253 | 0.49384 | 1.869526 | 0.06155 | 0.18978 | 0.436 |
| Gm5679 | 18.2575 | 0.169565 | 0.45554 | 0.372232 | 0.70972 | 0.8437 | 1.148 |
| Gm5741 | 24.0839 | 0.060395 | 0.38545 | 0.156687 | 0.87549 | 0.93793 | 0.415 |
| Gm5751 | 45.4442 | 0.637055 | 0.4165 | 1.529547 | 0.12613 | 0.3018 | 6.498 |
| Gm5763 | 13.6383 | -0.70042 | 0.48751 | -1.43675 | 0.15079 | 0.33567 | 2.533 |
| Gm5776 | 198.905 | -0.305 | 0.17219 | -1.77132 | 0.07651 | 0.21863 | 0.321 |
| Gm5784 | 61.4108 | -0.04225 | 0.21918 | -0.19279 | 0.84713 | 0.92206 | 45.273 |
| Gm5805 | 24.6699 | 0.212334 | 0.48494 | 0.437858 | 0.66149 | 0.81272 | 0.456 |
| Gm5815 | 17.0448 | -0.57291 | 0.54978 | -1.04208 | 0.29738 | 0.50755 | 1.227 |
| Gm5820 | 30.6247 | -0.55826 | 0.33926 | -1.64551 | 0.09986 | 0.25988 | 78.819 |
| Gm5844 | 17.6821 | -0.12461 | 0.43505 | -0.28643 | 0.77455 | 0.88245 | 2.191 |
| Gm5846 | 58.1339 | -0.06586 | 0.27168 | -0.2424 | 0.80847 | 0.90156 | 1.414 |
| Gm5854 | 180.691 | -0.44711 | 0.29181 | -1.5322 | 0.12547 | 0.30061 | 0.773 |
| Gm5857 | 55.1023 | -0.09077 | 0.25523 | -0.35563 | 0.72212 | 0.85173 | 0.493 |
| Gm5879 | 38.4466 | 0.045743 | 0.32617 | 0.140245 | 0.88847 | 0.94434 | 1.209 |
| Gm5881 | 23.3271 | -0.24909 | 0.5279 | -0.47185 | 0.63703 | 0.79576 | 0.367 |
| Gm5884 | 360.544 | -0.20452 | 0.16673 | -1.22666 | 0.21995 | 0.42245 | 1.479 |
| Gm5898 | 28.7585 | -0.25505 | 0.36128 | -0.70596 | 0.48021 | 0.67761 | 2.071 |
| Gm5903 | 51.4554 | 0.294031 | 0.24212 | 1.214395 | 0.2246 | 0.42827 | 2.325 |
| Gm5905 | 21.6921 | -0.50417 | 0.47586 | -1.05951 | 0.28937 | 0.4997 | 0.58 |
| Gm5914 | 34.9044 | 0.912251 | 0.2985 | 3.056127 | 0.00224 | 0.0217 | 13.15 |
| Gm5977 | 68.057 | 0.242198 | 0.23561 | 1.027951 | 0.30397 | 0.51382 | 5.369 |
| Gm6013 | 104.271 | 0.183577 | 0.20087 | 0.913925 | 0.36076 | 0.57103 | 0.699 |

| | | | | | | | |
|---|---|---|---|---|---|---|---|
| Gm6030 | 158.143 | 0.275332 | 0.18173 | 1.515021 | 0.12977 | 0.30754 | 0.39 |
| Gm6039 | 13.3022 | -0.30157 | 0.43675 | -0.69048 | 0.48989 | 0.68491 | 1.25 |
| Gm6055 | 685.594 | -0.31196 | 0.1132 | -2.75574 | 0.00586 | 0.04057 | 0.378 |
| Gm6087 | 14.799 | -1.44184 | 0.63378 | -2.27497 | 0.02291 | 0.09795 | 0.663 |
| Gm6089 | 26.4133 | 0.026551 | 0.42988 | 0.061763 | 0.95075 | 0.97574 | 0.938 |
| Gm609 | 21.8891 | -0.50053 | 0.40196 | -1.2452 | 0.21306 | 0.41418 | 81.202 |
| Gm6104 | 41.2352 | 0.29823 | 0.27802 | 1.072685 | 0.28341 | 0.49347 | 0.603 |
| Gm6123 | 35.6479 | -0.31849 | 0.30284 | -1.05168 | 0.29294 | 0.50281 | 1.069 |
| Gm6136 | 1176.35 | 0.008463 | 0.0773 | 0.109485 | 0.91282 | 0.95597 | 0.891 |
| Gm6142 | 29.2838 | 1.017812 | 0.34583 | 2.943063 | 0.00325 | 0.02752 | 0.325 |
| Gm6145 | 1026.79 | -0.01088 | 0.15365 | -0.0708 | 0.94356 | 0.97178 | 22.711 |
| Gm6161 | 35.9897 | -0.35072 | 0.28239 | -1.24197 | 0.21425 | 0.41558 | 1.102 |
| Gm6166 | 13.0396 | 0.381907 | 0.57018 | 0.669797 | 0.50299 | 0.69545 | 0.409 |
| Gm6180 | 57.0157 | -0.68428 | 0.2667 | -2.56572 | 0.0103 | 0.05859 | 0.498 |
| Gm6185 | 29.5786 | 0.135008 | 0.351 | 0.384644 | 0.7005 | 0.83831 | 60.372 |
| Gm6204 | 20.8472 | -0.97849 | 0.39508 | -2.47668 | 0.01326 | 0.06901 | 0.457 |
| Gm6206 | 377.461 | -0.40721 | 0.23106 | -1.76237 | 0.07801 | 0.22104 | 2.298 |
| Gm6209 | 15.3679 | -0.79912 | 0.44415 | -1.7992 | 0.07199 | 0.21065 | 90.828 |
| Gm6210 | 14.3771 | -0.00157 | 0.43294 | -0.00363 | 0.99711 | 0.99835 | 1.942 |
| Gm6225 | 173.136 | 0.426956 | 0.1441 | 2.962954 | 0.00305 | 0.02635 | 30.184 |
| Gm6252 | 28.7849 | -0.27271 | 0.51044 | -0.53426 | 0.59316 | 0.7655 | 0.58 |
| Gm6254 | 21.0771 | -0.00664 | 0.39098 | -0.01697 | 0.98646 | 0.99262 | 1.072 |
| Gm6263 | 14.2311 | -0.08352 | 0.54748 | -0.15255 | 0.87875 | 0.93928 | 1.342 |
| Gm6277 | 293.697 | 0.094404 | 0.16363 | 0.576927 | 0.56399 | 0.74426 | 20.199 |
| Gm6305 | 13.9862 | -0.00518 | 0.48173 | -0.01076 | 0.99142 | 0.99522 | 1.012 |
| Gm6345 | 15.3827 | -0.06749 | 0.51453 | -0.13116 | 0.89565 | 0.94747 | 1.691 |
| Gm6382 | 38.5616 | 0.118692 | 0.33974 | 0.349357 | 0.72682 | 0.85431 | 0.942 |
| Gm6419 | 22.9669 | -0.7518 | 0.41053 | -1.83129 | 0.06706 | 0.20048 | 0.429 |
| Gm6421 | 70.8949 | -0.15066 | 0.24273 | -0.62071 | 0.53479 | 0.72177 | 0.409 |
| Gm6439 | 25.5476 | -0.04898 | 0.35564 | -0.13773 | 0.89046 | 0.94517 | 0.862 |
| Gm6457 | 87.667 | -0.15434 | 0.19796 | -0.77966 | 0.43559 | 0.64118 | 0.492 |
| Gm6467 | 30.5398 | 0.092999 | 0.30578 | 0.304135 | 0.76103 | 0.8748 | 0.611 |
| Gm6475 | 20.6575 | -0.34675 | 0.48831 | -0.71011 | 0.47763 | 0.67546 | 1.936 |
| Gm6477 | 16.0458 | 0.382178 | 0.44571 | 0.857467 | 0.39119 | 0.60033 | 0.764 |

| | | | | | | | |
|---|---|---|---|---|---|---|---|
| Gm6483 | 64.1576 | 0.107237 | 0.28406 | 0.377513 | 0.70579 | 0.84147 | 15.288 |
| Gm6485 | 23.8069 | -0.13981 | 0.45197 | -0.30933 | 0.75707 | 0.87207 | 0.439 |
| Gm6493 | 51.1356 | -0.18943 | 0.29265 | -0.6473 | 0.51744 | 0.70733 | 2.831 |
| Gm6505 | 27.2113 | 0.031734 | 0.3338 | 0.095068 | 0.92426 | 0.96197 | 0.648 |
| Gm6525 | 17.6713 | -0.00548 | 0.45979 | -0.01192 | 0.99049 | 0.99473 | 1.029 |
| Gm6542 | 76.2383 | 0.136398 | 0.30121 | 0.452829 | 0.65067 | 0.80552 | 0.772 |
| Gm6545 | 19.5341 | -1.02978 | 0.50824 | -2.02615 | 0.04275 | 0.14903 | 1.786 |
| Gm6548 | 113.597 | -0.31997 | 0.20886 | -1.53197 | 0.12553 | 0.30071 | 1.165 |
| Gm6552 | 16.0573 | -0.1427 | 0.56144 | -0.25418 | 0.79936 | 0.89647 | 1.061 |
| Gm6556 | 72.191 | 0.281732 | 0.2373 | 1.18722 | 0.23514 | 0.44034 | 23.435 |
| Gm6560 | 296.817 | -0.14526 | 0.11136 | -1.3044 | 0.1921 | 0.38879 | 1.591 |
| Gm6563 | 630.657 | -0.15679 | 0.12358 | -1.26869 | 0.20455 | 0.40342 | 0.633 |
| Gm6565 | 230.185 | 0.020475 | 0.21696 | 0.094368 | 0.92482 | 0.9621 | 0.438 |
| Gm6566 | 147.777 | 0.08693 | 0.17938 | 0.484604 | 0.62796 | 0.78952 | 6.012 |
| Gm6568 | 19.3472 | -0.37104 | 0.46366 | -0.80025 | 0.42356 | 0.6304 | 0.697 |
| Gm6569 | 55.2876 | -0.12198 | 0.37069 | -0.32907 | 0.74211 | 0.86416 | 5.348 |
| Gm6583 | 48.7783 | 0.447677 | 0.27324 | 1.638428 | 0.10133 | 0.26262 | 2.233 |
| Gm6588 | 31.5968 | 0.042013 | 0.49484 | 0.084901 | 0.93234 | 0.96589 | 2.313 |
| Gm6592 | 37.1916 | 0.474846 | 0.39427 | 1.204352 | 0.22845 | 0.43251 | 5.614 |
| Gm6598 | 34.7823 | 0.111721 | 0.35176 | 0.317601 | 0.75079 | 0.86925 | 2.615 |
| Gm6606 | 18.6526 | 0.794984 | 0.42744 | 1.859879 | 0.0629 | 0.19257 | 9.045 |
| Gm6627 | 49.6085 | 0.055612 | 0.32491 | 0.171163 | 0.8641 | 0.93125 | 2.386 |
| Gm6658 | 61.1453 | 0.123454 | 0.23522 | 0.524842 | 0.59969 | 0.77046 | 1.885 |
| Gm6663 | 18.9822 | -0.42405 | 0.42377 | -1.00067 | 0.31699 | 0.52743 | 0.973 |
| Gm6682 | 16.0782 | -0.26122 | 0.439 | -0.59504 | 0.55182 | 0.73475 | 1.346 |
| Gm6685 | 24.7573 | 0.15814 | 0.43023 | 0.367574 | 0.71319 | 0.84571 | 0.687 |
| Gm6710 | 60.5986 | -0.33302 | 0.80542 | -0.41347 | 0.67926 | 0.82486 | 12.03 |
| Gm6712 | 213.735 | 0.430185 | 0.17983 | 2.392172 | 0.01675 | 0.08009 | 25.382 |
| Gm6723 | 22.0244 | -0.31421 | 0.43552 | -0.72147 | 0.47062 | 0.6693 | 0.441 |
| Gm6733 | 16.7648 | -0.82744 | 0.45791 | -1.80698 | 0.07076 | 0.20791 | 1.92 |
| Gm6741 | 60.1259 | 0.090195 | 0.44769 | 0.201466 | 0.84033 | 0.91812 | 0.747 |
| Gm6756 | 27.0604 | 0.164243 | 0.40972 | 0.400866 | 0.68852 | 0.83012 | 1.62 |
| Gm6781 | 17.0994 | 1.129952 | 0.44047 | 2.565317 | 0.01031 | 0.05861 | 1.242 |
| Gm6793 | 272.347 | -0.43544 | 0.59173 | -0.73588 | 0.4618 | 0.66226 | 1.056 |

| | | | | | | | |
|---|---|---|---|---|---|---|---|
| Gm6807 | 24.1545 | 0.871862 | 0.47221 | 1.846347 | 0.06484 | 0.19618 | 0.888 |
| Gm6855 | 19.7695 | 0.657366 | 0.39141 | 1.67949 | 0.09306 | 0.24838 | 0.79 |
| Gm6863 | 635.724 | 0.159996 | 0.10516 | 1.52149 | 0.12814 | 0.30506 | 0.636 |
| Gm6877 | 76.2663 | -0.20506 | 0.22249 | -0.92164 | 0.35672 | 0.56705 | 1.101 |
| Gm6939 | 150.436 | 0.327376 | 0.14607 | 2.241228 | 0.02501 | 0.10396 | 2.576 |
| Gm6961 | 12.9576 | -1.05131 | 0.65077 | -1.61548 | 0.10621 | 0.27017 | 26.509 |
| Gm6969 | 16.0125 | -0.25392 | 0.50064 | -0.50719 | 0.61202 | 0.77898 | 0.342 |
| Gm6980 | 15.4634 | 0.431161 | 0.50757 | 0.849467 | 0.39562 | 0.60466 | 18.185 |
| Gm6981 | 19.3511 | -0.09174 | 0.41719 | -0.2199 | 0.82595 | 0.91016 | 1.008 |
| Gm6999 | 14.6466 | 0.777356 | 0.52047 | 1.493556 | 0.13529 | 0.31507 | 169.615 |
| Gm7008 | 22.413 | -0.04605 | 0.38471 | -0.1197 | 0.90472 | 0.95186 | 6.427 |
| Gm7045 | 15.6427 | 0.304727 | 0.47288 | 0.644409 | 0.51931 | 0.70902 | 0.605 |
| Gm7061 | 140.08 | 0.189808 | 0.23273 | 0.815576 | 0.41474 | 0.62272 | 1.416 |
| Gm7067 | 15.319 | 0.831957 | 0.46346 | 1.795098 | 0.07264 | 0.21189 | 1.249 |
| Gm7072 | 707.849 | 0.372494 | 0.49639 | 0.750401 | 0.45301 | 0.65542 | 28.967 |
| Gm7099 | 17.5516 | -0.22089 | 0.42428 | -0.52063 | 0.60262 | 0.77235 | 0.554 |
| Gm7107 | 27.2637 | 0.335902 | 0.34765 | 0.966211 | 0.33394 | 0.54469 | 1.362 |
| Gm715 | 109.215 | -0.31046 | 0.17816 | -1.7426 | 0.0814 | 0.22718 | 1.21 |
| Gm7160 | 52.3408 | -0.43779 | 0.29675 | -1.47528 | 0.14014 | 0.32151 | 16.751 |
| Gm7162 | 27.73 | -1.19955 | 0.49989 | -2.39961 | 0.01641 | 0.07902 | 1.024 |
| Gm7180 | 17.769 | 0.572383 | 0.45869 | 1.247858 | 0.21208 | 0.41271 | 0.513 |
| Gm7206 | 14.5405 | 0.471447 | 0.51588 | 0.91387 | 0.36078 | 0.57103 | 9.188 |
| Gm7240 | 35.0356 | 0.607892 | 0.37564 | 1.618271 | 0.1056 | 0.26914 | 57.423 |
| Gm7244 | 18.337 | 1.399785 | 0.5399 | 2.592663 | 0.00952 | 0.05554 | 0.455 |
| Gm7265 | 23.3745 | -0.03622 | 0.44407 | -0.08157 | 0.93499 | 0.96725 | 3.502 |
| Gm7276 | 56.0103 | -0.46243 | 0.26521 | -1.7436 | 0.08123 | 0.22685 | 2.095 |
| Gm7278 | 14.2405 | -0.15237 | 0.4764 | -0.31984 | 0.74909 | 0.86842 | 1.401 |
| Gm7285 | 37.0097 | -0.02025 | 0.33298 | -0.06081 | 0.95151 | 0.97617 | 0.58 |
| Gm7292 | 175.475 | -0.05218 | 0.17845 | -0.29243 | 0.76996 | 0.87989 | 2.527 |
| Gm7308 | 128.25 | 0.030057 | 0.20357 | 0.147653 | 0.88262 | 0.9416 | 0.447 |
| Gm7324 | 15.1943 | -0.22664 | 0.44694 | -0.5071 | 0.61208 | 0.77898 | 17.551 |
| Gm7326 | 12.991 | -0.39068 | 0.5247 | -0.74456 | 0.45653 | 0.65813 | 0.255 |
| Gm7327 | 51.2269 | 0.706848 | 0.43943 | 1.608568 | 0.10771 | 0.27261 | 0.652 |
| Gm7328 | 14.5773 | 0.371201 | 0.49498 | 0.74993 | 0.4533 | 0.65558 | 2.225 |

| | | | | | | | |
|---|---|---|---|---|---|---|---|
| Gm7331 | 57.8273 | -0.24796 | 0.21366 | -1.16051 | 0.24584 | 0.45252 | 0.387 |
| Gm7332 | 15.6118 | -0.61482 | 0.45949 | -1.33805 | 0.18088 | 0.37425 | 0.454 |
| Gm7334 | 23.9429 | 0.567302 | 0.34519 | 1.643459 | 0.10029 | 0.26056 | 0.756 |
| Gm7335 | 125.628 | 0.167885 | 0.19216 | 0.873692 | 0.38229 | 0.59182 | 0.373 |
| Gm7336 | 91.9934 | -0.15731 | 0.28602 | -0.54998 | 0.58234 | 0.75787 | 1.232 |
| Gm7353 | 179.511 | -0.19521 | 0.17744 | -1.10013 | 0.27128 | 0.48066 | 2.944 |
| Gm7361 | 33.7644 | 0.509237 | 0.40678 | 1.251867 | 0.21062 | 0.41064 | 6.618 |
| Gm7367 | 67.4409 | -0.41354 | 0.25516 | -1.62074 | 0.10507 | 0.26817 | 0.492 |
| Gm7389 | 14.5211 | 0.639821 | 0.4502 | 1.421178 | 0.15527 | 0.34177 | 2.884 |
| Gm7390 | 28.9362 | 0.06598 | 0.40674 | 0.162217 | 0.87113 | 0.93489 | 1.216 |
| Gm7420 | 80.9246 | 0.56203 | 0.24618 | 2.283049 | 0.02243 | 0.09666 | 0.842 |
| Gm7435 | 49.874 | -0.5746 | 0.27338 | -2.10182 | 0.03557 | 0.13252 | 14.319 |
| Gm7452 | 21.8686 | -0.22288 | 0.57828 | -0.38543 | 0.69992 | 0.83787 | 3.087 |
| Gm7457 | 23.3204 | -0.09964 | 0.37679 | -0.26444 | 0.79144 | 0.89159 | 19.264 |
| Gm7467 | 20.0054 | -0.01012 | 0.46256 | -0.02187 | 0.98255 | 0.99061 | 0.652 |
| Gm7488 | 14.6763 | -0.25973 | 0.42631 | -0.60926 | 0.54235 | 0.72794 | 0.684 |
| Gm7517 | 28.2149 | -0.31483 | 0.39456 | -0.79792 | 0.42492 | 0.6314 | 1.672 |
| Gm7535 | 26.5437 | -1.33725 | 0.46593 | -2.87006 | 0.0041 | 0.03204 | 1.777 |
| Gm7536 | 384.466 | 0.207579 | 0.24343 | 0.852711 | 0.39382 | 0.60297 | 0.444 |
| Gm7591 | 12.7514 | 0.20856 | 0.51015 | 0.408821 | 0.68267 | 0.82659 | 0.853 |
| Gm7598 | 39.7315 | 0.265404 | 0.31598 | 0.839935 | 0.40094 | 0.60964 | 6.026 |
| Gm7599 | 23.1956 | -0.03022 | 0.33453 | -0.09034 | 0.92802 | 0.96381 | 0.796 |
| Gm7623 | 13.8906 | -0.42419 | 0.48806 | -0.86913 | 0.38478 | 0.59423 | 0.556 |
| Gm7628 | 20.5887 | 0.456255 | 0.36857 | 1.237892 | 0.21576 | 0.41742 | 3.953 |
| Gm7645 | 25.2229 | 0.955298 | 0.33023 | 2.892829 | 0.00382 | 0.0305 | 0.788 |
| Gm7658 | 47.0598 | 0.15443 | 0.26242 | 0.588485 | 0.55621 | 0.73841 | 1.124 |
| Gm7664 | 34.9573 | -0.04258 | 0.28136 | -0.15135 | 0.8797 | 0.93982 | 0.745 |
| Gm7666 | 268.52 | -0.48442 | 0.15178 | -3.19158 | 0.00141 | 0.01592 | 0.819 |
| Gm7672 | 50.6745 | 0.482313 | 0.25055 | 1.925045 | 0.05422 | 0.1751 | 0.872 |
| Gm7684 | 35.6562 | 0.646075 | 0.32816 | 1.968805 | 0.04898 | 0.1636 | 0.829 |
| Gm7694 | 275.96 | -0.03525 | 0.144 | -0.24478 | 0.80663 | 0.90042 | 8.134 |
| Gm7729 | 23.3968 | -0.05226 | 0.37507 | -0.13935 | 0.88918 | 0.94457 | 1.113 |
| Gm7730 | 38.2957 | -0.21691 | 0.28864 | -0.75149 | 0.45236 | 0.65477 | 0.454 |
| Gm7774 | 16.6916 | -0.34355 | 0.4184 | -0.82111 | 0.41159 | 0.61967 | 0.533 |

| | | | | | | | |
|---|---|---|---|---|---|---|---|
| Gm7785 | 26.6537 | -0.22021 | 0.31307 | -0.70341 | 0.4818 | 0.67858 | 0.892 |
| Gm7807 | 73.4538 | 0.276816 | 0.25341 | 1.092356 | 0.27468 | 0.4843 | 0.578 |
| Gm7816 | 33.423 | 0.052197 | 0.29465 | 0.177151 | 0.85939 | 0.92898 | 1.194 |
| Gm7818 | 321.527 | 0.42394 | 0.21414 | 1.979737 | 0.04773 | 0.16067 | 2.625 |
| Gm7846 | 26.3469 | -0.71157 | 0.35554 | -2.00137 | 0.04535 | 0.15493 | 0.274 |
| Gm7854 | 23.4926 | 0.161517 | 0.38337 | 0.421313 | 0.67353 | 0.82053 | 83.67 |
| Gm7879 | 46.4032 | 0.483405 | 0.29243 | 1.653059 | 0.09832 | 0.25734 | 0.774 |
| Gm7887 | 267.206 | 0.167856 | 0.13733 | 1.2223 | 0.22159 | 0.42449 | 0.537 |
| Gm7890 | 17.2342 | -0.38504 | 0.45503 | -0.84618 | 0.39745 | 0.60658 | 0.601 |
| Gm7895 | 17.9694 | -0.41624 | 0.49312 | -0.84409 | 0.39862 | 0.6076 | 1.173 |
| Gm7901 | 33.9958 | -0.77649 | 0.41106 | -1.88898 | 0.05889 | 0.18475 | 1.597 |
| Gm7902 | 29.1537 | -0.84595 | 0.32613 | -2.5939 | 0.00949 | 0.05541 | 0.947 |
| Gm7932 | 21.3831 | 0.816318 | 0.4218 | 1.935302 | 0.05295 | 0.17218 | 5.787 |
| Gm7962 | 15.6835 | -0.71353 | 0.63189 | -1.12921 | 0.25881 | 0.46703 | 2.308 |
| Gm7964 | 1381.32 | -0.0646 | 0.08552 | -0.75536 | 0.45003 | 0.65294 | 1.378 |
| Gm7967 | 38.5336 | -0.51344 | 0.28384 | -1.80894 | 0.07046 | 0.2074 | 17.868 |
| Gm7972 | 90.9584 | 0.535447 | 0.25151 | 2.128967 | 0.03326 | 0.12662 | 18.109 |
| Gm7984 | 17.7423 | -0.62238 | 0.51619 | -1.20573 | 0.22792 | 0.43183 | 0.538 |
| Gm7997 | 16.2786 | 0.554894 | 0.48182 | 1.151669 | 0.24946 | 0.45683 | 0.98 |
| Gm8010 | 23.5901 | 0.703715 | 0.39933 | 1.762231 | 0.07803 | 0.22104 | 0.972 |
| Gm8034 | 38.3964 | 0.245933 | 0.28378 | 0.866631 | 0.38614 | 0.59559 | 0.132 |
| Gm8066 | 291.28 | 0.263002 | 0.15451 | 1.702188 | 0.08872 | 0.24068 | 7.249 |
| Gm807 | 35.4116 | -0.12136 | 0.30037 | -0.40402 | 0.6862 | 0.82887 | 7.128 |
| Gm8098 | 138.322 | 0.685255 | 0.20288 | 3.377675 | 0.00073 | 0.01031 | 13.028 |
| Gm8113 | 12.9189 | -0.10808 | 0.63927 | -0.16906 | 0.86575 | 0.932 | 8.022 |
| Gm8116 | 177.746 | 0.266938 | 0.20838 | 1.280996 | 0.2002 | 0.39817 | 1.201 |
| Gm8121 | 211.559 | 0.285531 | 0.19777 | 1.443736 | 0.14881 | 0.33268 | 0.499 |
| Gm8129 | 32.1594 | -0.32248 | 0.47174 | -0.68361 | 0.49422 | 0.68845 | 0.213 |
| Gm8152 | 39.9633 | 0.474235 | 0.34166 | 1.388034 | 0.16513 | 0.3542 | 0.637 |
| Gm8170 | 25.7743 | 0.246575 | 0.34423 | 0.716313 | 0.4738 | 0.67213 | 0.945 |
| Gm8173 | 15.9457 | -0.54759 | 0.50251 | -1.08972 | 0.27584 | 0.48501 | 2.842 |
| Gm8186 | 64.5366 | -0.06164 | 0.27459 | -0.22449 | 0.82238 | 0.90789 | 0.231 |
| Gm8188 | 41.8036 | 0.396072 | 0.31395 | 1.261582 | 0.2071 | 0.4066 | 1.69 |
| Gm8189 | 20.2671 | -0.66118 | 0.52989 | -1.24779 | 0.21211 | 0.41272 | 68.486 |

| | | | | | | | |
|---|---|---|---|---|---|---|---|
| Gm8206 | 13.5027 | -0.28808 | 0.67538 | -0.42654 | 0.66971 | 0.81794 | 21.451 |
| Gm8234 | 25.4894 | -0.10627 | 0.3641 | -0.29187 | 0.77039 | 0.87998 | 1.107 |
| Gm8281 | 29.7847 | -0.23898 | 0.35791 | -0.66772 | 0.50431 | 0.69657 | 22.279 |
| Gm8319 | 30.8735 | 0.615091 | 0.33828 | 1.818269 | 0.06902 | 0.20444 | 0.641 |
| Gm8325 | 152.546 | -0.52174 | 0.15226 | -3.42667 | 0.00061 | 0.00909 | 1.86 |
| Gm8355 | 120.805 | -0.33611 | 0.17247 | -1.94878 | 0.05132 | 0.16879 | 1.941 |
| Gm8399 | 22.9525 | -0.4092 | 0.35477 | -1.15344 | 0.24873 | 0.45606 | 1.117 |
| Gm8407 | 59.4473 | -0.15476 | 0.25837 | -0.59897 | 0.54919 | 0.73299 | 0.427 |
| Gm8437 | 25.0952 | 0.374117 | 0.34406 | 1.087368 | 0.27687 | 0.48628 | 3.569 |
| Gm8451 | 17.3586 | -0.75142 | 0.38199 | -1.96711 | 0.04917 | 0.16407 | 0.447 |
| Gm8493 | 83.0924 | 0.786289 | 0.1991 | 3.949305 | 7.84E-05 | 0.00227 | 2.265 |
| Gm8494 | 68.9594 | 0.31428 | 0.28595 | 1.099064 | 0.27174 | 0.48106 | 1.233 |
| Gm8508 | 31.2354 | -0.12669 | 0.38821 | -0.32635 | 0.74416 | 0.86539 | 0.363 |
| Gm8540 | 53.4086 | 0.097048 | 0.31113 | 0.311919 | 0.7551 | 0.87123 | 7.812 |
| Gm8546 | 23.0646 | 0.132327 | 0.47659 | 0.277651 | 0.78128 | 0.88614 | 1.264 |
| Gm8566 | 58.0819 | 0.004936 | 0.22238 | 0.022195 | 0.98229 | 0.99059 | 0.465 |
| Gm8615 | 27.0657 | 0.83096 | 0.35804 | 2.320871 | 0.02029 | 0.09035 | 1.102 |
| Gm8642 | 20.259 | 0.146916 | 0.3769 | 0.389801 | 0.69668 | 0.83561 | 1.141 |
| Gm867 | 33.8872 | 0.414457 | 0.36576 | 1.133122 | 0.25716 | 0.46513 | 2.95 |
| Gm8717 | 76.3063 | -0.31507 | 0.33013 | -0.95438 | 0.33989 | 0.55075 | 0.814 |
| Gm8730 | 47.0389 | -0.71738 | 0.39141 | -1.83282 | 0.06683 | 0.20016 | 0.954 |
| Gm8738 | 20.6739 | 0.1577 | 0.44786 | 0.352115 | 0.72475 | 0.85331 | 0.329 |
| Gm8775 | 47.6213 | 0.011491 | 0.34396 | 0.033407 | 0.97335 | 0.98671 | 1.327 |
| Gm8783 | 22.0867 | 1.197892 | 0.6419 | 1.866152 | 0.06202 | 0.19074 | 0.522 |
| Gm8850 | 50.2247 | -0.33943 | 0.34932 | -0.97167 | 0.33122 | 0.54208 | 0.588 |
| Gm8899 | 28.133 | -0.45948 | 0.44251 | -1.03835 | 0.29911 | 0.50923 | 1.332 |
| Gm8991 | 30.7088 | -0.69227 | 0.49879 | -1.3879 | 0.16517 | 0.35425 | 1.075 |
| Gm9001 | 24.8883 | 0.226506 | 0.42142 | 0.537482 | 0.59093 | 0.7637 | 0.399 |
| Gm9003 | 29.6336 | -0.41704 | 0.47108 | -0.88529 | 0.376 | 0.5861 | 0.618 |
| Gm9017 | 24.7731 | 0.276157 | 0.35414 | 0.779793 | 0.43551 | 0.64111 | 6.471 |
| Gm9030 | 28.2037 | 0.148549 | 0.38988 | 0.381016 | 0.70319 | 0.83986 | 2.41 |
| Gm9057 | 13.1828 | 0.847918 | 0.62727 | 1.35176 | 0.17645 | 0.36846 | 0.296 |
| Gm9108 | 24.9573 | -0.44933 | 0.36906 | -1.2175 | 0.22341 | 0.42664 | 1.425 |
| Gm9118 | 55.218 | 0.229852 | 0.26121 | 0.879959 | 0.37888 | 0.58861 | 0.754 |

| | | | | | | | |
|---|---|---|---|---|---|---|---|
| Gm9234 | 52.9895 | 1.289742 | 0.28385 | 4.543823 | 5.52E-06 | 0.00036 | 0.492 |
| Gm9299 | 13.4581 | 0.079867 | 0.46063 | 0.173387 | 0.86235 | 0.93021 | 1.14 |
| Gm9311 | 33.6733 | 0.306947 | 0.35958 | 0.853618 | 0.39332 | 0.60236 | 1.428 |
| Gm9378 | 13.7116 | -0.27396 | 0.49711 | -0.55111 | 0.58156 | 0.75743 | 0.471 |
| Gm9392 | 131.217 | 0.037103 | 0.18085 | 0.205157 | 0.83745 | 0.91642 | 0.554 |
| Gm9403 | 24.5129 | -0.34915 | 0.32378 | -1.07837 | 0.28087 | 0.4906 | 2.181 |
| Gm9442 | 39.8042 | 0.125503 | 0.30135 | 0.416465 | 0.67707 | 0.82303 | 0.319 |
| Gm9493 | 162.392 | -0.22932 | 0.17122 | -1.33936 | 0.18045 | 0.37378 | 0.571 |
| Gm9517 | 37.9217 | 0.31293 | 0.32559 | 0.961106 | 0.3365 | 0.54716 | 0.874 |
| Gm9522 | 15.136 | -1.55704 | 0.5179 | -3.00642 | 0.00264 | 0.02422 | 0.926 |
| Gm9531 | 279.342 | 0.134857 | 0.13656 | 0.987526 | 0.32338 | 0.53445 | 1.08 |
| Gm9570 | 29.7048 | 0.062072 | 0.38289 | 0.162115 | 0.87122 | 0.93492 | 2.686 |
| Gm9574 | 16.1472 | 0.448254 | 0.40652 | 1.102665 | 0.27017 | 0.47976 | 1.918 |
| Gm9575 | 24.1224 | 0.052424 | 0.51848 | 0.10111 | 0.91946 | 0.95923 | 1.339 |
| Gm9586 | 15.8459 | -0.50881 | 0.54188 | -0.93897 | 0.34774 | 0.55823 | 1.048 |
| Gm960 | 234.547 | 0.100832 | 0.15615 | 0.645751 | 0.51844 | 0.70819 | 72.935 |
| Gm9616 | 327.011 | -0.18397 | 0.1419 | -1.29644 | 0.19482 | 0.39211 | 0.781 |
| Gm9625 | 13.0086 | -1.32521 | 0.51157 | -2.59049 | 0.00958 | 0.05574 | 0.955 |
| Gm9670 | 23.6144 | 0.464325 | 0.35354 | 1.313369 | 0.18906 | 0.38492 | 0.778 |
| Gm9725 | 20.812 | 0.188349 | 0.36502 | 0.51599 | 0.60586 | 0.77449 | 47.62 |
| Gm9727 | 12.8695 | -1.10661 | 0.57988 | -1.90833 | 0.05635 | 0.17956 | 0.555 |
| Gm973 | 361.804 | -0.32745 | 0.17944 | -1.82488 | 0.06802 | 0.20237 | 120.154 |
| Gm9732 | 84.8957 | 0.56204 | 0.28205 | 1.992702 | 0.04629 | 0.15723 | 30.976 |
| Gm9750 | 16.6812 | -0.00487 | 0.43001 | -0.01133 | 0.99096 | 0.99495 | 144.019 |
| Gm9752 | 97.2318 | -0.11648 | 0.2909 | -0.40041 | 0.68886 | 0.83047 | 2.958 |
| Gm9754 | 125.73 | 0.563937 | 0.31434 | 1.794046 | 0.07281 | 0.21225 | 4.351 |
| Gm9755 | 22.2958 | -0.38988 | 0.4044 | -0.9641 | 0.33499 | 0.54572 | 1.356 |
| Gm9761 | 187.682 | -0.00492 | 0.16613 | -0.02958 | 0.9764 | 0.98767 | 0.492 |
| Gm9762 | 53.583 | 0.209756 | 0.26562 | 0.789687 | 0.42971 | 0.63603 | 0.444 |
| Gm9768 | 26.7039 | 0.587832 | 0.3862 | 1.522105 | 0.12798 | 0.30479 | 2.861 |
| Gm9772 | 24.5392 | -0.2301 | 0.45924 | -0.50104 | 0.61634 | 0.78156 | 1.606 |
| Gm9774 | 32.9698 | -0.03467 | 0.30669 | -0.11305 | 0.90999 | 0.95466 | 1.392 |
| Gm9776 | 143.485 | -0.03557 | 0.17048 | -0.20863 | 0.83474 | 0.91511 | 3.093 |
| Gm9791 | 143.6 | -0.23921 | 0.52224 | -0.45806 | 0.64691 | 0.80352 | 0.447 |

| | | | | | | | |
|---|---|---|---|---|---|---|---|
| Gm9794 | 1765.38 | -0.03862 | 0.13338 | -0.28953 | 0.77217 | 0.8813 | 0.423 |
| Gm9795 | 31.966 | 0.765146 | 0.31678 | 2.415405 | 0.01572 | 0.07681 | 0.693 |
| Gm9796 | 37.4997 | -0.44783 | 0.30135 | -1.48611 | 0.13725 | 0.31789 | 2.25 |
| Gm9800 | 74.9665 | -0.53617 | 0.20461 | -2.62042 | 0.00878 | 0.05257 | 0.336 |
| Gm9801 | 246.547 | 0.015196 | 0.1648 | 0.09221 | 0.92653 | 0.96305 | 114.974 |
| Gm9816 | 871.956 | -0.00166 | 0.11845 | -0.01398 | 0.98884 | 0.99386 | 1.414 |
| Gm9821 | 310.426 | -0.00603 | 0.1371 | -0.04395 | 0.96494 | 0.98273 | 2.986 |
| Gm9824 | 713.265 | -0.10599 | 0.22067 | -0.4803 | 0.63101 | 0.79169 | 0.561 |
| Gm9826 | 29.1305 | 0.119635 | 0.40923 | 0.292338 | 0.77003 | 0.87989 | 1.299 |
| Gm9828 | 335.966 | 0.210398 | 0.1182 | 1.779975 | 0.07508 | 0.21608 | 10.808 |
| Gm9831 | 18.5722 | -1.22531 | 0.44901 | -2.72888 | 0.00636 | 0.04278 | 25.666 |
| Gm9839 | 18.8849 | 1.712378 | 0.46781 | 3.66041 | 0.00025 | 0.00497 | 1.437 |
| Gm9843 | 95.1854 | -0.37518 | 0.20669 | -1.81518 | 0.0695 | 0.20546 | 0.402 |
| Gm9844 | 140.574 | -0.01176 | 0.19925 | -0.05901 | 0.95294 | 0.97686 | 0.485 |
| Gm9856 | 222.294 | 0.510671 | 0.19168 | 2.664153 | 0.00772 | 0.04854 | 2.578 |
| Gm9866 | 1406.61 | -0.04266 | 0.07125 | -0.59869 | 0.54938 | 0.73299 | 109.684 |
| Gm9870 | 19.8927 | -0.07065 | 0.48294 | -0.14629 | 0.88369 | 0.94201 | 2.356 |
| Gm9873 | 15.2236 | 0.750205 | 0.44907 | 1.670588 | 0.0948 | 0.25135 | 1.361 |
| Gm9884 | 28.7602 | -0.44438 | 0.37424 | -1.1874 | 0.23507 | 0.44032 | 1.222 |
| Gm9885 | 69.0418 | -0.10883 | 0.25047 | -0.43451 | 0.66392 | 0.81428 | 2.439 |
| Gm9887 | 19.7921 | -0.26938 | 0.40919 | -0.65833 | 0.51033 | 0.70169 | 1.985 |
| Gm9892 | 232.165 | -0.08767 | 0.16444 | -0.53318 | 0.59391 | 0.76614 | 0.992 |
| Gm9895 | 57.4931 | -0.78546 | 0.29543 | -2.65872 | 0.00784 | 0.04906 | 31.283 |
| Gm9899 | 778.493 | 0.002979 | 0.08458 | 0.035217 | 0.97191 | 0.98584 | 14.633 |
| Gm9902 | 74.1863 | 1.342278 | 0.24147 | 5.558684 | 2.72E-08 | 7.49E-06 | 2.027 |
| Gm9903 | 16.0924 | -0.71955 | 0.50985 | -1.41129 | 0.15816 | 0.3455 | 2.168 |
| Gm9905 | 34.1771 | 0.318371 | 0.30157 | 1.0557 | 0.29111 | 0.50115 | 1.845 |
| Gm9908 | 27.0333 | 0.579491 | 0.37 | 1.566177 | 0.11731 | 0.28827 | 0.507 |
| Gm9910 | 45.0806 | -0.13732 | 0.28473 | -0.48227 | 0.62961 | 0.79064 | 1.55 |
| Gm9916 | 24.567 | -0.50729 | 0.35512 | -1.42852 | 0.15314 | 0.33865 | 1.228 |
| Gm9917 | 96.4908 | 0.052331 | 0.21684 | 0.241329 | 0.8093 | 0.90175 | 9.607 |
| Gm9923 | 155.641 | -0.23943 | 0.13813 | -1.73342 | 0.08302 | 0.22996 | 1.836 |
| Gm9924 | 12.5948 | -2 | 0.82028 | -2.53429 | 0.01127 | 0.06199 | 2.311 |
| Gm9925 | 59.5659 | -0.03439 | 0.23072 | -0.14907 | 0.8815 | 0.94099 | 0.483 |

| | | | | | | | |
|---|---|---|---|---|---|---|---|
| Gm9929 | 319.015 | 0.404384 | 0.1389 | 2.911415 | 0.0036 | 0.02933 | 13.703 |
| Gm9936 | 19.0235 | -0.62717 | 0.44909 | -1.39653 | 0.16256 | 0.35093 | 4.414 |
| Gm9939 | 13.911 | -1.17291 | 0.57784 | -2.02982 | 0.04237 | 0.14813 | 2.676 |
| Gm9945 | 21.6008 | -0.1437 | 0.47706 | -0.30123 | 0.76324 | 0.876 | 0.932 |
| Gm9947 | 16.6791 | -0.77393 | 0.49455 | -1.56492 | 0.1176 | 0.28868 | 23.995 |
| Gm9954 | 181.095 | -0.90366 | 0.34675 | -2.60607 | 0.00916 | 0.05404 | 4.725 |
| Gm9958 | 84.4665 | 0.310757 | 0.21009 | 1.479189 | 0.13909 | 0.32033 | 1.874 |
| Gm9962 | 98.4906 | 0.353546 | 0.20374 | 1.735279 | 0.08269 | 0.22942 | 22.67 |
| Gm9967 | 68.4038 | 0.192558 | 0.23932 | 0.804601 | 0.42105 | 0.62805 | 8.787 |
| Gm9970 | 13.5098 | -0.16187 | 0.49602 | -0.32634 | 0.74417 | 0.86539 | 0.767 |
| Gm9987 | 170.594 | -0.08894 | 0.21192 | -0.4197 | 0.67471 | 0.82139 | 0.569 |
| Gm9989 | 264.661 | 1.206957 | 0.24391 | 4.948368 | 7.48E-07 | 8.82E-05 | 6.456 |
| Gm9993 | 38.6132 | -0.08312 | 0.3499 | -0.23756 | 0.81222 | 0.90334 | 1.096 |
| Gm9996 | 18.4001 | 0.209078 | 0.47 | 0.444846 | 0.65643 | 0.8092 | 0.913 |
| Gmcl1 | 2657.92 | 0.233538 | 0.08535 | 2.736164 | 0.00622 | 0.04216 | 41.616 |
| Gmds | 206.176 | -0.07793 | 0.19103 | -0.40793 | 0.68333 | 0.82704 | 519.162 |
| Gmeb1 | 1234.67 | 0.105242 | 0.08229 | 1.278876 | 0.20094 | 0.39907 | 40.578 |
| Gmeb2 | 1165 | 0.051717 | 0.08778 | 0.589161 | 0.55575 | 0.73801 | 36.587 |
| Gmfb | 10772.7 | 0.0302 | 0.06556 | 0.460677 | 0.64503 | 0.80203 | 14.094 |
| Gmfg | 35.6508 | -0.78928 | 0.29115 | -2.71086 | 0.00671 | 0.04421 | 10.787 |
| Gmip | 632.636 | -0.01623 | 0.12043 | -0.1348 | 0.89277 | 0.94632 | 13.192 |
| Gmnc | 23.1745 | 0.282568 | 0.4239 | 0.666583 | 0.50504 | 0.69681 | 78.1 |
| Gmnn | 143.283 | 0.0013 | 0.16558 | 0.00785 | 0.99374 | 0.99633 | 10.079 |
| Gmppa | 1514.11 | 0.054583 | 0.06968 | 0.783283 | 0.43346 | 0.63935 | 7.25 |
| Gmppb | 401.493 | -0.02517 | 0.11079 | -0.22714 | 0.82031 | 0.90734 | 3.56 |
| Gmpr | 1630.12 | 0.254249 | 0.13361 | 1.902971 | 0.05704 | 0.18095 | 46.357 |
| Gmpr2 | 327.796 | 0.11741 | 0.12145 | 0.966707 | 0.33369 | 0.5445 | 7.26 |
| Gmps | 3981.32 | 0.154227 | 0.08081 | 1.908435 | 0.05633 | 0.17956 | 46.474 |
| Gna11 | 4966.63 | -0.03616 | 0.08771 | -0.41227 | 0.68014 | 0.82502 | 16.467 |
| Gna12 | 2462.22 | -0.28154 | 0.1077 | -2.61405 | 0.00895 | 0.05328 | 72.024 |
| Gna13 | 3290.21 | -0.10405 | 0.09677 | -1.07527 | 0.28226 | 0.49241 | 38.539 |
| Gna14 | 25.4602 | 1.266261 | 0.44183 | 2.865972 | 0.00416 | 0.03227 | 175.059 |
| Gna15 | 60.792 | -0.34338 | 0.27586 | -1.24475 | 0.21322 | 0.41433 | 21.92 |
| Gnai1 | 8260.79 | -0.09446 | 0.08567 | -1.10251 | 0.27024 | 0.47984 | 95.221 |

| | | | | | | | |
|---|---|---|---|---|---|---|---|
| Gnai2 | 6750.38 | -0.14062 | 0.09879 | -1.42349 | 0.15459 | 0.34066 | 21.243 |
| Gnai3 | 1785.96 | -0.11454 | 0.07887 | -1.45231 | 0.14642 | 0.32946 | 38.867 |
| Gnal | 26759.1 | 0.503539 | 0.14495 | 3.473836 | 0.00051 | 0.00806 | 138.577 |
| Gnao1 | 33368.9 | -0.07059 | 0.05822 | -1.21247 | 0.22533 | 0.42931 | 159.423 |
| Gnaq | 9729.34 | -0.03281 | 0.06714 | -0.48864 | 0.6251 | 0.78752 | 254.633 |
| Gnas | 1702.32 | -0.36125 | 0.14464 | -2.4975 | 0.01251 | 0.06641 | 62.425 |
| Gnat2 | 14.4463 | 0.044986 | 0.46096 | 0.09759 | 0.92226 | 0.96086 | 8.644 |
| Gnaz | 2649.44 | -0.16681 | 0.07116 | -2.34406 | 0.01908 | 0.08673 | 49.731 |
| Gnb1 | 23977.7 | -0.20164 | 0.06765 | -2.98067 | 0.00288 | 0.02554 | 67.909 |
| Gnb1l | 119.872 | -0.00035 | 0.16739 | -0.0021 | 0.99832 | 0.99911 | 67.967 |
| Gnb2 | 7155 | 0.108636 | 0.07852 | 1.383535 | 0.1665 | 0.356 | 5.384 |
| Gnb4 | 2653.94 | 0.365295 | 0.10352 | 3.528675 | 0.00042 | 0.00709 | 36.254 |
| Gnb5 | 7166.23 | 0.318986 | 0.11965 | 2.665968 | 0.00768 | 0.04843 | 39.589 |
| Gne | 1769.31 | 0.040602 | 0.07034 | 0.577189 | 0.56381 | 0.74426 | 50.103 |
| Gng10 | 57.6895 | 0.327849 | 0.25714 | 1.274973 | 0.20232 | 0.40074 | 6.816 |
| Gng11 | 248.958 | -0.626 | 0.12585 | -4.97431 | 6.55E-07 | 7.94E-05 | 4.542 |
| Gng12 | 3420.93 | -0.1259 | 0.07977 | -1.57814 | 0.11453 | 0.28343 | 124.954 |
| Gng13 | 15.3488 | -0.72226 | 0.43397 | -1.6643 | 0.09605 | 0.25372 | 2.236 |
| Gng2 | 3340.42 | -0.3183 | 0.15063 | -2.11311 | 0.03459 | 0.13028 | 105.069 |
| Gng3 | 4425.57 | -0.25094 | 0.10034 | -2.50083 | 0.01239 | 0.06605 | 2.262 |
| Gng4 | 2484.39 | -1.03238 | 0.29275 | -3.52644 | 0.00042 | 0.00714 | 43.838 |
| Gng5 | 657.197 | -0.32913 | 0.09795 | -3.36036 | 0.00078 | 0.01076 | 5.766 |
| Gng7 | 35389.4 | 0.618021 | 0.18476 | 3.344984 | 0.00082 | 0.01119 | 66.322 |
| Gng8 | 52.6312 | -0.74277 | 0.35013 | -2.12138 | 0.03389 | 0.12855 | 3.651 |
| Gngt2 | 46.9459 | -0.55847 | 0.30111 | -1.85469 | 0.06364 | 0.19397 | 8.519 |
| Gnl1 | 6889.15 | -0.14035 | 0.06469 | -2.16939 | 0.03005 | 0.1184 | 9.612 |
| Gnl2 | 2350.7 | 0.140202 | 0.10099 | 1.388295 | 0.16505 | 0.35406 | 38.796 |
| Gnl3 | 1276.98 | 0.164387 | 0.09718 | 1.691496 | 0.09074 | 0.24468 | 6.72 |
| Gnl3l | 11489.2 | 0.005563 | 0.06954 | 0.079996 | 0.93624 | 0.96801 | 34.182 |
| Gnmt | 55.0454 | 0.061934 | 0.22581 | 0.274282 | 0.78387 | 0.88762 | 3.505 |
| Gnpat | 2970.12 | 0.105852 | 0.06553 | 1.61534 | 0.10624 | 0.27019 | 27.025 |
| Gnpda1 | 1115.87 | 0.00737 | 0.08208 | 0.089787 | 0.92846 | 0.96398 | 11.469 |
| Gnpda2 | 811.819 | 0.199888 | 0.13184 | 1.516112 | 0.12949 | 0.30701 | 19.233 |
| Gnpnat1 | 45.2819 | 0.19435 | 0.29128 | 0.66722 | 0.50463 | 0.6967 | 28.58 |

| | | | | | | | |
|---|---|---|---|---|---|---|---|
| Gnptab | 3923.54 | -0.07705 | 0.07224 | -1.06663 | 0.28614 | 0.49619 | 68.198 |
| Gnptg | 3428.22 | 0.117361 | 0.07264 | 1.615711 | 0.10616 | 0.27012 | 6.794 |
| Gnrh1 | 56.2644 | -0.08863 | 0.31327 | -0.28292 | 0.77724 | 0.88407 | 4.259 |
| Gns | 4646.49 | -0.14833 | 0.08014 | -1.85093 | 0.06418 | 0.19501 | 32.16 |
| Golga1 | 2291.56 | 0.191701 | 0.08375 | 2.288917 | 0.02208 | 0.09575 | 49.387 |
| Golga2 | 4775.31 | 0.020938 | 0.05533 | 0.378388 | 0.70514 | 0.84115 | 20.538 |
| Golga3 | 3496.19 | 0.020608 | 0.05583 | 0.369091 | 0.71206 | 0.845 | 49.77 |
| Golga4 | 8835.47 | 0.269012 | 0.1154 | 2.331093 | 0.01975 | 0.08876 | 76.253 |
| Golga5 | 1440.78 | 0.067663 | 0.07964 | 0.849602 | 0.39555 | 0.60459 | 28.773 |
| Golga7 | 3228.74 | -0.27997 | 0.07746 | -3.61462 | 0.0003 | 0.00565 | 15.729 |
| Golga7b | 2470.93 | -0.99541 | 0.19476 | -5.11107 | 3.20E-07 | 4.38E-05 | 22.775 |
| Golgb1 | 7682.7 | 0.240263 | 0.11722 | 2.049713 | 0.04039 | 0.14367 | 57.946 |
| Golim4 | 1618.59 | -0.13874 | 0.09245 | -1.50075 | 0.13342 | 0.31307 | 81.866 |
| Golm1 | 1079.17 | -0.44803 | 0.16378 | -2.73546 | 0.00623 | 0.04221 | 41.186 |
| Golm2 | 5352.15 | 0.088885 | 0.08096 | 1.097832 | 0.27228 | 0.48159 | 69.251 |
| Golph3 | 3520.62 | -0.10972 | 0.06258 | -1.75318 | 0.07957 | 0.22403 | 30.142 |
| Golph3l | 801.031 | -0.0775 | 0.08729 | -0.88774 | 0.37468 | 0.58497 | 30.314 |
| Golt1b | 1493.84 | -0.04992 | 0.07471 | -0.66821 | 0.504 | 0.69627 | 16.638 |
| Gon4l | 2483.54 | 0.1084 | 0.06224 | 1.741757 | 0.08155 | 0.22753 | 74.876 |
| Gon7 | 190.573 | 0.093377 | 0.15237 | 0.612849 | 0.53998 | 0.726 | 4.529 |
| Gopc | 2721.03 | 0.081333 | 0.08585 | 0.94735 | 0.34346 | 0.55415 | 46.275 |
| Gorab | 457.834 | 0.114134 | 0.09678 | 1.179344 | 0.23826 | 0.44374 | 18.762 |
| Gorasp1 | 1250.5 | 0.003432 | 0.10302 | 0.033311 | 0.97343 | 0.98674 | 12.309 |
| Gorasp2 | 2911.08 | -0.009 | 0.06599 | -0.13644 | 0.89147 | 0.94595 | 51.061 |
| Gosr1 | 2783.35 | 0.050604 | 0.06899 | 0.733515 | 0.46324 | 0.66314 | 36.978 |
| Gosr2 | 3901.54 | -0.04111 | 0.07942 | -0.51756 | 0.60477 | 0.7735 | 21.05 |
| Got1 | 13857.7 | -0.12824 | 0.05896 | -2.17495 | 0.02963 | 0.11715 | 24.854 |
| Got2 | 7527.19 | -0.15829 | 0.0555 | -2.85217 | 0.00434 | 0.03326 | 24.414 |
| Gp1ba | 62.9941 | 0.263835 | 0.27712 | 0.952069 | 0.34106 | 0.55206 | 2.915 |
| Gp5 | 21.639 | -0.66206 | 0.36521 | -1.81282 | 0.06986 | 0.20626 | 3.095 |
| Gpaa1 | 1203.54 | -0.02423 | 0.0881 | -0.27504 | 0.78328 | 0.88742 | 3.677 |
| Gpalpp1 | 2337.23 | 0.13394 | 0.07534 | 1.777771 | 0.07544 | 0.21668 | 24.525 |
| Gpam | 4860.04 | -0.11002 | 0.1017 | -1.08186 | 0.27932 | 0.48922 | 59.971 |
| Gpank1 | 285.166 | 0.219266 | 0.11789 | 1.859936 | 0.06289 | 0.19257 | 3.36 |

| | | | | | | | |
|---|---|---|---|---|---|---|---|
| Gpat2 | 85.9107 | 0.504527 | 0.19667 | 2.565301 | 0.01031 | 0.05861 | 10.894 |
| Gpat3 | 290.337 | 0.222517 | 0.17408 | 1.278243 | 0.20116 | 0.39943 | 53.39 |
| Gpat4 | 4469.03 | -0.02393 | 0.06149 | -0.3891 | 0.6972 | 0.8358 | 37.082 |
| Gpatch1 | 1369.3 | 0.099539 | 0.08801 | 1.130936 | 0.25808 | 0.46613 | 41.905 |
| Gpatch11 | 1759.34 | 0.072248 | 0.0871 | 0.829535 | 0.4068 | 0.61497 | 12.742 |
| Gpatch2 | 1171.85 | 0.061737 | 0.08337 | 0.740568 | 0.45896 | 0.65977 | 136.197 |
| Gpatch2l | 1383.48 | -0.04187 | 0.0851 | -0.49193 | 0.62277 | 0.78585 | 49.927 |
| Gpatch3 | 274.393 | -0.17776 | 0.11626 | -1.52894 | 0.12628 | 0.30205 | 9.499 |
| Gpatch4 | 875.053 | -0.01118 | 0.10768 | -0.10378 | 0.91734 | 0.95781 | 12.886 |
| Gpatch8 | 7540.21 | 0.236905 | 0.09543 | 2.482511 | 0.01305 | 0.06836 | 80.478 |
| Gpbp1 | 3454.65 | 0.118924 | 0.0975 | 1.219731 | 0.22257 | 0.42583 | 64.432 |
| Gpbp1l1 | 1645.85 | 0.006796 | 0.06061 | 0.112131 | 0.91072 | 0.95499 | 36.225 |
| Gpc1 | 3105.45 | 0.084153 | 0.07046 | 1.194345 | 0.23234 | 0.43727 | 29.135 |
| Gpc2 | 35.4131 | -0.08257 | 0.34889 | -0.23666 | 0.81292 | 0.90371 | 6.351 |
| Gpc3 | 236.759 | 0.111523 | 0.26053 | 0.428063 | 0.6686 | 0.81717 | 341.525 |
| Gpc4 | 408.092 | -0.88348 | 0.23365 | -3.78124 | 0.00016 | 0.00365 | 112.232 |
| Gpc5 | 1529.85 | -0.03819 | 0.10236 | -0.37312 | 0.70906 | 0.84326 | 1432.945 |
| Gpc6 | 341.629 | 0.237714 | 0.11216 | 2.119413 | 0.03406 | 0.12885 | 1051.23 |
| Gpcpd1 | 5785.91 | -0.14893 | 0.10301 | -1.44574 | 0.14825 | 0.33188 | 58.648 |
| Gpd1 | 3016.83 | -0.26122 | 0.08493 | -3.07578 | 0.0021 | 0.0208 | 7.491 |
| Gpd1l | 5261.38 | -0.12163 | 0.08175 | -1.48795 | 0.13676 | 0.31745 | 34.647 |
| Gpd2 | 4684.47 | -0.22185 | 0.106 | -2.09287 | 0.03636 | 0.13438 | 133.085 |
| Gper1 | 106.844 | -0.1969 | 0.18982 | -1.0373 | 0.2996 | 0.50972 | 4.65 |
| Gpha2 | 20.0374 | 1.037708 | 0.41845 | 2.479903 | 0.01314 | 0.06865 | 1.389 |
| Gphn | 3526.42 | 0.156629 | 0.09983 | 1.56903 | 0.11664 | 0.28701 | 458.394 |
| Gpi1 | 28763.2 | 0.100033 | 0.06211 | 1.610592 | 0.10727 | 0.27183 | 29.007 |
| Gpkow | 2128.49 | 0.030144 | 0.08354 | 0.360854 | 0.71821 | 0.84954 | 13.132 |
| Gpld1 | 1888.54 | 0.168531 | 0.11695 | 1.441101 | 0.14956 | 0.3339 | 49.35 |
| Gpm6a | 52798 | -0.08749 | 0.08042 | -1.0879 | 0.27664 | 0.48595 | 106.029 |
| Gpm6b | 50963.4 | -0.06202 | 0.07058 | -0.87876 | 0.37953 | 0.5893 | 150.078 |
| Gpn1 | 975.464 | -0.01258 | 0.07516 | -0.16738 | 0.86707 | 0.93254 | 18.164 |
| Gpn2 | 283.598 | -0.02604 | 0.12422 | -0.20966 | 0.83393 | 0.9148 | 7.374 |
| Gpn3 | 814.433 | 0.161754 | 0.11712 | 1.381041 | 0.16727 | 0.35697 | 11.027 |
| Gpnmb | 325.771 | -2 | 0.63438 | -3.74552 | 0.00018 | 0.00401 | 34.384 |

| | | | | | | | |
|---|---|---|---|---|---|---|---|
| Gpr1 | 27.8804 | -0.00901 | 0.43712 | -0.0206 | 0.98356 | 0.991 | 31.853 |
| Gpr101 | 587.31 | 1.678191 | 0.46454 | 3.612622 | 0.0003 | 0.00567 | 7.09 |
| Gpr107 | 3510.94 | -0.04959 | 0.07257 | -0.68323 | 0.49446 | 0.68865 | 66.46 |
| Gpr108 | 1091.18 | -0.00301 | 0.07859 | -0.03828 | 0.96946 | 0.98458 | 13.812 |
| Gpr12 | 1181.57 | 0.465864 | 0.2046 | 2.276939 | 0.02279 | 0.09764 | 2.842 |
| Gpr135 | 586.637 | 0.114226 | 0.10072 | 1.134062 | 0.25677 | 0.46474 | 3.15 |
| Gpr137 | 2551.06 | -0.30983 | 0.09925 | -3.12189 | 0.0018 | 0.01879 | 4.424 |
| Gpr137b | 598.735 | -0.09847 | 0.10319 | -0.95434 | 0.33991 | 0.55075 | 36.395 |
| Gpr137b-ps | 625.488 | -0.2361 | 0.10052 | -2.34886 | 0.01883 | 0.08608 | 35.339 |
| Gpr137c | 2063.01 | 0.177964 | 0.10936 | 1.627292 | 0.10368 | 0.26622 | 63.009 |
| Gpr139 | 157.15 | 0.070174 | 0.28122 | 0.249532 | 0.80295 | 0.89822 | 43.857 |
| Gpr146 | 1071.66 | 0.045881 | 0.08926 | 0.514005 | 0.60725 | 0.77542 | 18.719 |
| Gpr149 | 1830.56 | 0.811488 | 0.33945 | 2.390624 | 0.01682 | 0.08027 | 76.064 |
| Gpr150 | 132.615 | 0.310611 | 0.18855 | 1.647351 | 0.09949 | 0.25923 | 2.146 |
| Gpr151 | 41.0761 | 0.483887 | 0.30158 | 1.604524 | 0.1086 | 0.2739 | 1.772 |
| Gpr153 | 1074.67 | -0.06804 | 0.0818 | -0.83182 | 0.40551 | 0.61425 | 11.106 |
| Gpr155 | 4013.51 | 0.228493 | 0.09189 | 2.486684 | 0.01289 | 0.06781 | 45.067 |
| Gpr156 | 80.9761 | -0.40554 | 0.22691 | -1.78727 | 0.07389 | 0.21422 | 91.035 |
| Gpr157 | 137.65 | -0.2794 | 0.18373 | -1.52068 | 0.12834 | 0.30535 | 18.563 |
| Gpr158 | 11107 | 0.140403 | 0.11303 | 1.242215 | 0.21416 | 0.41558 | 463.006 |
| Gpr160 | 40.7489 | -0.17346 | 0.31574 | -0.54937 | 0.58275 | 0.75806 | 41.243 |
| Gpr161 | 703.398 | 0.36094 | 0.13824 | 2.610958 | 0.00903 | 0.05353 | 30.957 |
| Gpr162 | 7553.18 | -0.2785 | 0.09579 | -2.9073 | 0.00365 | 0.02963 | 5.54 |
| Gpr165 | 435.462 | 0.31638 | 0.13398 | 2.361462 | 0.0182 | 0.08431 | 5.948 |
| Gpr17 | 2246.02 | -0.1666 | 0.10552 | -1.57888 | 0.11436 | 0.28327 | 6.64 |
| Gpr173 | 832.546 | -0.09897 | 0.0862 | -1.14815 | 0.25091 | 0.45825 | 25.107 |
| Gpr176 | 462.69 | -0.06941 | 0.1073 | -0.64689 | 0.51771 | 0.70754 | 96.31 |
| Gpr179 | 64.8874 | -0.08529 | 0.29094 | -0.29315 | 0.76941 | 0.87968 | 19.969 |
| Gpr18 | 13.0656 | 0.233637 | 0.54288 | 0.430366 | 0.66693 | 0.81591 | 4.529 |
| Gpr180 | 910.638 | 0.289486 | 0.09875 | 2.93139 | 0.00337 | 0.02811 | 26.104 |
| Gpr182 | 160.453 | -0.85681 | 0.27834 | -3.07829 | 0.00208 | 0.02064 | 4.457 |
| Gpr183 | 71.1517 | -0.37904 | 0.25935 | -1.46151 | 0.14387 | 0.32626 | 12.645 |
| Gpr19 | 591.125 | -0.07726 | 0.11135 | -0.69384 | 0.48778 | 0.68299 | 29.486 |
| Gpr21 | 174.179 | -0.2786 | 0.18656 | -1.49338 | 0.13534 | 0.31509 | 4.272 |

| | | | | | | | |
|---|---|---|---|---|---|---|---|
| Gpr22 | 1540.73 | -0.12922 | 0.20311 | -0.63619 | 0.52465 | 0.71306 | 7.081 |
| Gpr25 | 12.6982 | 0.812613 | 0.46424 | 1.750428 | 0.08004 | 0.22499 | 1.077 |
| Gpr26 | 16361.9 | -0.03242 | 0.07251 | -0.44702 | 0.65486 | 0.80847 | 27.946 |
| Gpr27 | 2722.92 | 0.19857 | 0.08761 | 2.266433 | 0.02342 | 0.09929 | 2.388 |
| Gpr3 | 251.71 | -0.50219 | 0.15475 | -3.24525 | 0.00117 | 0.01411 | 3.197 |
| Gpr34 | 568.706 | -0.14957 | 0.1221 | -1.22495 | 0.22059 | 0.42328 | 8.77 |
| Gpr35 | 37.3906 | -0.40654 | 0.29743 | -1.36687 | 0.17167 | 0.36213 | 35.527 |
| Gpr37 | 2239.35 | -0.44 | 0.16128 | -2.72817 | 0.00637 | 0.04283 | 24.852 |
| Gpr37l1 | 6211.83 | -0.34578 | 0.10626 | -3.25405 | 0.00114 | 0.01382 | 7.448 |
| Gpr4 | 189.283 | -0.04882 | 0.20281 | -0.24069 | 0.80979 | 0.90197 | 11.637 |
| Gpr45 | 489.573 | -0.07904 | 0.1074 | -0.7359 | 0.46179 | 0.66226 | 82.585 |
| Gpr50 | 58.8966 | -0.70887 | 0.25531 | -2.7765 | 0.00549 | 0.03875 | 5.591 |
| Gpr52 | 1886.59 | 0.790657 | 0.21179 | 3.733242 | 0.00019 | 0.00415 | 1.637 |
| Gpr55 | 295.645 | 0.895544 | 0.16076 | 5.570658 | 2.54E-08 | 7.18E-06 | 22.691 |
| Gpr6 | 1812.54 | 0.96138 | 0.38308 | 2.509595 | 0.01209 | 0.0649 | 2.309 |
| Gpr61 | 861.648 | 0.083531 | 0.08958 | 0.932498 | 0.35108 | 0.56158 | 6.562 |
| Gpr62 | 387.06 | -0.58951 | 0.15464 | -3.81229 | 0.00014 | 0.00333 | 1.977 |
| Gpr63 | 388.08 | 0.366979 | 0.14553 | 2.521712 | 0.01168 | 0.06353 | 42.827 |
| Gpr68 | 222.201 | -0.18761 | 0.16109 | -1.16458 | 0.24419 | 0.45062 | 31.517 |
| Gpr75 | 622.89 | 0.436696 | 0.13281 | 3.288116 | 0.00101 | 0.0129 | 8.372 |
| Gpr83 | 5906.56 | 0.963413 | 0.35203 | 2.736729 | 0.00621 | 0.04214 | 10.58 |
| Gpr85 | 1789.87 | 0.008838 | 0.09338 | 0.094645 | 0.9246 | 0.96202 | 5.485 |
| Gpr88 | 23740.2 | 0.700605 | 0.3109 | 2.253494 | 0.02423 | 0.10168 | 3.85 |
| Gpr89 | 1059.12 | 0.064952 | 0.08764 | 0.741095 | 0.45864 | 0.65962 | 37.066 |
| Gprasp1 | 29192.6 | 0.360318 | 0.09082 | 3.967257 | 7.27E-05 | 0.00215 | 60.742 |
| Gprasp2 | 6326.44 | 0.391829 | 0.07448 | 5.261076 | 1.43E-07 | 2.37E-05 | 5.697 |
| Gprc5b | 6805.71 | -0.43227 | 0.08689 | -4.97477 | 6.53E-07 | 7.94E-05 | 23.165 |
| Gprc5c | 122.371 | -0.93634 | 0.20026 | -4.67571 | 2.93E-06 | 0.00023 | 21.466 |
| Gprin1 | 3740.33 | -0.52665 | 0.17265 | -3.05047 | 0.00228 | 0.02196 | 13.204 |
| Gprin2 | 236.136 | 0.38413 | 0.18558 | 2.069899 | 0.03846 | 0.13963 | 15.966 |
| Gprin3 | 4474.09 | 0.621527 | 0.41205 | 1.508379 | 0.13146 | 0.31014 | 79.069 |
| Gps1 | 3527.92 | -0.0024 | 0.07569 | -0.03165 | 0.97475 | 0.98735 | 4.831 |
| Gps2 | 1010.9 | 0.082663 | 0.07716 | 1.071265 | 0.28405 | 0.4942 | 2.704 |
| Gpsm1 | 2943.1 | 0.216388 | 0.12413 | 1.743288 | 0.08128 | 0.22691 | 32.723 |

| | | | | | | | |
|---|---|---|---|---|---|---|---|
| Gpsm2 | 769.875 | -0.1157 | 0.10107 | -1.14471 | 0.25233 | 0.4596 | 43.672 |
| Gpsm3 | 91.9934 | -0.06248 | 0.21222 | -0.29441 | 0.76844 | 0.87901 | 1.949 |
| Gpt | 630.331 | -0.32737 | 0.18217 | -1.79705 | 0.07233 | 0.21128 | 3.971 |
| Gpt2 | 3849.35 | -0.10904 | 0.08388 | -1.30003 | 0.19359 | 0.39053 | 34.985 |
| Gpx1 | 18.68 | -1.13779 | 0.59718 | -1.90528 | 0.05674 | 0.18035 | 1.441 |
| Gpx2 | 12.4945 | 1.058673 | 0.59162 | 1.789442 | 0.07354 | 0.21355 | 3.167 |
| Gpx3 | 274.678 | -0.84079 | 0.21511 | -3.90872 | 9.28E-05 | 0.00255 | 7.925 |
| Gpx4 | 5715.73 | -0.28995 | 0.08939 | -3.24352 | 0.00118 | 0.01415 | 9.274 |
| Gpx4-ps2 | 808.777 | -0.32494 | 0.12712 | -2.55612 | 0.01058 | 0.05972 | 0.808 |
| Gpx6 | 34.1947 | 0.680715 | 0.41964 | 1.622147 | 0.10477 | 0.26777 | 7.5 |
| Gpx7 | 110.433 | -0.07527 | 0.18951 | -0.39719 | 0.69122 | 0.83234 | 6.572 |
| Gpx8 | 126.997 | -0.675 | 0.22031 | -3.06383 | 0.00219 | 0.02131 | 3.658 |
| Gramd1a | 4813.31 | 0.175608 | 0.07725 | 2.273373 | 0.023 | 0.09814 | 25.77 |
| Gramd1b | 12259.7 | -0.12738 | 0.07533 | -1.69089 | 0.09086 | 0.24474 | 238.151 |
| Gramd1c | 130.345 | 0.034021 | 0.25091 | 0.135593 | 0.89214 | 0.94617 | 82.996 |
| Gramd2 | 851.814 | 0.059631 | 0.08761 | 0.680655 | 0.49609 | 0.68983 | 38.731 |
| Gramd3 | 820.652 | -0.29649 | 0.09181 | -3.22941 | 0.00124 | 0.01462 | 103.456 |
| Gramd4 | 2558.79 | 0.135972 | 0.06524 | 2.084076 | 0.03715 | 0.13625 | 79.94 |
| Grap | 138.271 | -0.56191 | 0.23144 | -2.42786 | 0.01519 | 0.07507 | 19.52 |
| Grap2 | 25.6517 | -0.77252 | 0.46533 | -1.66015 | 0.09688 | 0.25499 | 80.261 |
| Grb10 | 3292.74 | 0.632238 | 0.12998 | 4.86423 | 1.15E-06 | 0.00012 | 108.176 |
| Grb14 | 806.618 | -0.29975 | 0.0894 | -3.35294 | 0.0008 | 0.01096 | 112.512 |
| Grb2 | 7790.58 | -0.07301 | 0.07991 | -0.91362 | 0.36092 | 0.57115 | 64.553 |
| Grb7 | 198.405 | 0.423801 | 0.22911 | 1.849746 | 0.06435 | 0.19522 | 8.98 |
| Grcc10 | 129.229 | 0.06419 | 0.17662 | 0.363427 | 0.71629 | 0.8483 | 2.192 |
| Greb1 | 67.4064 | -0.41634 | 0.27525 | -1.51259 | 0.13038 | 0.30861 | 130.272 |
| Greb1l | 872.163 | 0.4435 | 0.1528 | 2.90255 | 0.0037 | 0.0299 | 237.764 |
| Grem1 | 165.678 | 0.67274 | 0.21541 | 3.123011 | 0.00179 | 0.01874 | 12.483 |
| Grem2 | 1481.94 | -0.1697 | 0.10732 | -1.58119 | 0.11383 | 0.28246 | 88.035 |
| Grhl1 | 363.337 | -0.55434 | 0.15212 | -3.64406 | 0.00027 | 0.00519 | 45.109 |
| Grhl2 | 24.3276 | -0.84936 | 0.45161 | -1.88075 | 0.06001 | 0.18675 | 130.534 |
| Grhl3 | 68.5618 | -0.51585 | 0.28626 | -1.80207 | 0.07153 | 0.20947 | 31.743 |
| Grhpr | 888.679 | -0.18933 | 0.08978 | -2.10886 | 0.03496 | 0.13109 | 9.34 |
| Gria1 | 28295.9 | 0.238359 | 0.07688 | 3.100284 | 0.00193 | 0.01969 | 318.858 |

| | | | | | | | |
|---|---|---|---|---|---|---|---|
| Gria2 | 45922.8 | 0.261978 | 0.14762 | 1.774652 | 0.07596 | 0.21767 | 121.386 |
| Gria3 | 13106.4 | -0.06136 | 0.14842 | -0.41341 | 0.67931 | 0.82486 | 277.748 |
| Gria4 | 3629.09 | 0.121482 | 0.12919 | 0.940323 | 0.34705 | 0.55765 | 378.339 |
| Grid1 | 3661.18 | 0.089545 | 0.09646 | 0.928317 | 0.35324 | 0.56351 | 763.272 |
| Grid2 | 961.49 | 0.367753 | 0.11089 | 3.316372 | 0.00091 | 0.01201 | 1448.448 |
| Grid2ip | 135.835 | -0.59181 | 0.22891 | -2.58528 | 0.00973 | 0.05636 | 34.815 |
| Grik1 | 472.201 | -0.21612 | 0.15397 | -1.40364 | 0.16043 | 0.3486 | 394.366 |
| Grik2 | 7506.4 | 0.643252 | 0.18442 | 3.48806 | 0.00049 | 0.00778 | 693.934 |
| Grik3 | 7691.72 | 0.065273 | 0.10958 | 0.595646 | 0.55141 | 0.73445 | 223.474 |
| Grik4 | 883.032 | 0.244389 | 0.103 | 2.372704 | 0.01766 | 0.08258 | 426.359 |
| Grik5 | 11207.9 | 0.104187 | 0.07662 | 1.359848 | 0.17388 | 0.36528 | 62.498 |
| Grin1 | 12746.1 | 0.053694 | 0.0664 | 0.808605 | 0.41874 | 0.62584 | 28.007 |
| Grin1os | 1193.59 | 0.279698 | 0.14005 | 1.997076 | 0.04582 | 0.15602 | 7.706 |
| Grin2a | 10255.9 | -0.13271 | 0.09854 | -1.34678 | 0.17805 | 0.3703 | 427.663 |
| Grin2b | 34094.2 | 0.155341 | 0.09036 | 1.71914 | 0.08559 | 0.23485 | 460.279 |
| Grin2c | 3326.6 | -0.32319 | 0.1143 | -2.82761 | 0.00469 | 0.03486 | 18.075 |
| Grin2d | 621.628 | -0.02211 | 0.1449 | -0.15261 | 0.87871 | 0.93928 | 46.496 |
| Grin3a | 2153.97 | 0.162538 | 0.09677 | 1.679584 | 0.09304 | 0.24836 | 184.134 |
| Grin3b | 33.2376 | -0.06108 | 0.3579 | -0.17068 | 0.86448 | 0.93142 | 6.476 |
| Grina | 11355.9 | -0.06876 | 0.06067 | -1.13335 | 0.25707 | 0.46513 | 3.141 |
| Grip1 | 1135.54 | -0.10651 | 0.13678 | -0.7787 | 0.43615 | 0.64183 | 633.432 |
| Grip1os1 | 13.6038 | 0.107832 | 0.50038 | 0.215499 | 0.82938 | 0.91247 | 27.187 |
| Grip2 | 1674.66 | -0.19151 | 0.13006 | -1.47251 | 0.14088 | 0.32265 | 65.742 |
| Gripap1 | 7436.9 | 0.299252 | 0.09095 | 3.290329 | 0.001 | 0.01282 | 30.803 |
| Grk2 | 7056.78 | -0.25912 | 0.09519 | -2.7221 | 0.00649 | 0.04334 | 20.222 |
| Grk3 | 6563.32 | -0.34648 | 0.14151 | -2.44843 | 0.01435 | 0.07227 | 105.31 |
| Grk4 | 190.239 | -0.11514 | 0.15005 | -0.76734 | 0.44288 | 0.64738 | 94.927 |
| Grk5 | 1398.23 | 0.124003 | 0.09248 | 1.340809 | 0.17998 | 0.37318 | 206.22 |
| Grk6 | 3161.48 | -0.00216 | 0.06927 | -0.03122 | 0.97509 | 0.98745 | 15.856 |
| Grm1 | 3194.08 | 0.281047 | 0.10395 | 2.703591 | 0.00686 | 0.04485 | 396.298 |
| Grm2 | 397.69 | -0.80274 | 0.3703 | -2.16782 | 0.03017 | 0.11868 | 12.988 |
| Grm3 | 3093.33 | 0.23157 | 0.12844 | 1.802948 | 0.0714 | 0.20919 | 239.63 |
| Grm4 | 3321.26 | 0.518994 | 0.14737 | 3.521709 | 0.00043 | 0.00722 | 99.017 |
| Grm5 | 16189.4 | 0.464267 | 0.11479 | 4.044429 | 5.25E-05 | 0.00171 | 550.74 |

| | | | | | | | |
|---|---|---|---|---|---|---|---|
| Grm7 | 3430.71 | 0.478175 | 0.14901 | 3.209102 | 0.00133 | 0.01522 | 921.65 |
| Grm8 | 332.718 | -0.67882 | 0.22637 | -2.9987 | 0.00271 | 0.02459 | 860.06 |
| Grn | 2304.49 | -0.3144 | 0.08517 | -3.69171 | 0.00022 | 0.00458 | 6.734 |
| Grpel1 | 1286.37 | -0.09016 | 0.0911 | -0.9897 | 0.32232 | 0.53334 | 9.083 |
| Grpel2 | 1140.36 | 0.129763 | 0.06989 | 1.856743 | 0.06335 | 0.19338 | 13.902 |
| Grpr | 19.6541 | -0.66526 | 0.51492 | -1.29196 | 0.19637 | 0.39398 | 35.774 |
| Grsf1 | 4018.53 | 0.085835 | 0.07186 | 1.194517 | 0.23228 | 0.43719 | 16.724 |
| Grtp1 | 152.872 | -0.00151 | 0.15462 | -0.00974 | 0.99223 | 0.9955 | 28.599 |
| Grwd1 | 504.557 | -0.08601 | 0.09181 | -0.93686 | 0.34883 | 0.55931 | 5.722 |
| Gsap | 322.12 | -0.10069 | 0.15604 | -0.64527 | 0.51875 | 0.70849 | 128.878 |
| Gsdmd | 118.475 | -0.28103 | 0.2148 | -1.30836 | 0.19075 | 0.38701 | 5.082 |
| Gsdme | 1282.57 | 0.403449 | 0.11792 | 3.421465 | 0.00062 | 0.00921 | 74.975 |
| Gse1 | 2737.22 | -0.22923 | 0.13647 | -1.67968 | 0.09302 | 0.24834 | 352.935 |
| Gsg1 | 28.2573 | -0.25703 | 0.38757 | -0.6632 | 0.50721 | 0.69843 | 17.026 |
| Gsg1l | 5208.68 | 0.273077 | 0.138 | 1.978869 | 0.04783 | 0.16089 | 203.992 |
| Gsk3a | 4842.22 | -0.14103 | 0.09188 | -1.5349 | 0.12481 | 0.2998 | 9.594 |
| Gsk3b | 12898.7 | 0.060131 | 0.06415 | 0.93731 | 0.3486 | 0.55914 | 157.084 |
| Gskip | 1545.8 | 0.128672 | 0.09133 | 1.408846 | 0.15888 | 0.34639 | 18.195 |
| Gsn | 2481.02 | -0.65048 | 0.11741 | -5.54016 | 3.02E-08 | 8.00E-06 | 51.513 |
| Gspt1 | 3389.8 | 0.1348 | 0.06883 | 1.958369 | 0.05019 | 0.16648 | 35.034 |
| Gspt2 | 1842.92 | 0.194669 | 0.07331 | 2.655587 | 0.00792 | 0.04931 | 7.176 |
| Gsr | 1300.04 | -0.05143 | 0.10059 | -0.51126 | 0.60917 | 0.77661 | 45.641 |
| Gss | 989.188 | -0.11884 | 0.10138 | -1.17219 | 0.24112 | 0.44728 | 29.63 |
| Gsta3 | 125.346 | 0.818677 | 0.22908 | 3.573836 | 0.00035 | 0.00625 | 25.073 |
| Gsta4 | 884.458 | -0.33297 | 0.13214 | -2.51978 | 0.01174 | 0.06375 | 26.576 |
| Gstcd | 344.541 | 0.100333 | 0.12226 | 0.820684 | 0.41183 | 0.61976 | 110.282 |
| Gstk1 | 304.539 | -0.30896 | 0.13948 | -2.21509 | 0.02675 | 0.10907 | 4.513 |
| Gstm1 | 5621.24 | -0.23918 | 0.10708 | -2.23359 | 0.02551 | 0.10538 | 5.719 |
| Gstm2 | 71.4319 | -0.8689 | 0.43348 | -2.00449 | 0.04502 | 0.15421 | 4.752 |
| Gstm2-ps1 | 35.9223 | -0.20581 | 0.29191 | -0.70505 | 0.48078 | 0.67798 | 0.657 |
| Gstm4 | 290.783 | -0.18962 | 0.13724 | -1.38172 | 0.16706 | 0.35691 | 4.487 |
| Gstm5 | 2571.58 | -0.27163 | 0.09178 | -2.9596 | 0.00308 | 0.02652 | 2.866 |
| Gstm6 | 182.399 | 0.049278 | 0.1995 | 0.247006 | 0.8049 | 0.89917 | 4.903 |
| Gstm7 | 559.102 | -0.20543 | 0.11552 | -1.77837 | 0.07534 | 0.21652 | 5.484 |

| | | | | | | | |
|---|---|---|---|---|---|---|---|
| Gsto1 | 894.705 | -0.24992 | 0.11805 | -2.11714 | 0.03425 | 0.12928 | 9.821 |
| Gsto2 | 22.2923 | -0.63533 | 0.36181 | -1.75598 | 0.07909 | 0.22311 | 20.791 |
| Gstp-ps | 143.456 | -0.00863 | 0.22078 | -0.03907 | 0.96884 | 0.98419 | 0.631 |
| Gstp1 | 3407.98 | -0.13515 | 0.10502 | -1.28697 | 0.19811 | 0.39586 | 2.579 |
| Gstt1 | 236.387 | -0.06509 | 0.20956 | -0.31061 | 0.75609 | 0.87158 | 14.772 |
| Gstt2 | 84.5483 | -0.31741 | 0.20818 | -1.5247 | 0.12733 | 0.30358 | 6.311 |
| Gstt3 | 77.0085 | -0.64105 | 0.24601 | -2.60584 | 0.00916 | 0.05405 | 7.3 |
| Gstz1 | 1383.21 | -0.01219 | 0.09948 | -0.12256 | 0.90245 | 0.95112 | 17.559 |
| Gsx2 | 72.0534 | 0.327143 | 0.2928 | 1.117305 | 0.26386 | 0.47272 | 2.293 |
| Gt(ROSA)26Sor | 791.477 | 0.184929 | 0.11634 | 1.589519 | 0.11194 | 0.27943 | 10.452 |
| Gtdc1 | 1303.25 | -0.35958 | 0.13613 | -2.64144 | 0.00826 | 0.0506 | 363.246 |
| Gtf2a1 | 3012.4 | 0.034414 | 0.08053 | 0.427331 | 0.66914 | 0.81761 | 35.226 |
| Gtf2a2 | 1799.79 | 0.141552 | 0.09888 | 1.431624 | 0.15225 | 0.33745 | 10.317 |
| Gtf2b | 947.001 | -0.17984 | 0.07898 | -2.27698 | 0.02279 | 0.09764 | 18.562 |
| Gtf2e1 | 1013.79 | 0.023585 | 0.10684 | 0.220751 | 0.82529 | 0.90983 | 30 |
| Gtf2e2 | 693.993 | 0.163358 | 0.09229 | 1.770082 | 0.07671 | 0.21907 | 45.341 |
| Gtf2f1 | 2992.4 | -0.13751 | 0.06724 | -2.04499 | 0.04086 | 0.14474 | 7.884 |
| Gtf2f2 | 408.321 | -0.02209 | 0.12637 | -0.17484 | 0.86121 | 0.92973 | 113.929 |
| Gtf2h1 | 1932.55 | 0.152558 | 0.06849 | 2.227451 | 0.02592 | 0.10666 | 27.698 |
| Gtf2h2 | 1649.94 | 0.209046 | 0.10477 | 1.995295 | 0.04601 | 0.1565 | 32.362 |
| Gtf2h3 | 1066.91 | 0.03563 | 0.09231 | 0.385983 | 0.69951 | 0.83757 | 18.541 |
| Gtf2h4 | 505.568 | 0.057566 | 0.1072 | 0.536976 | 0.59128 | 0.76394 | 6.01 |
| Gtf2h5 | 2468.95 | -0.10155 | 0.06324 | -1.6058 | 0.10832 | 0.27367 | 6.732 |
| Gtf2i | 12031.7 | -0.02525 | 0.05804 | -0.43501 | 0.66356 | 0.81408 | 76.927 |
| Gtf2ird1 | 1649.51 | 0.144286 | 0.07262 | 1.986969 | 0.04693 | 0.15889 | 99.061 |
| Gtf2ird2 | 819.22 | -0.00404 | 0.09293 | -0.0435 | 0.9653 | 0.98278 | 40.352 |
| Gtf3a | 223.256 | -0.00555 | 0.13984 | -0.03966 | 0.96836 | 0.98409 | 6.958 |
| Gtf3c1 | 10924.8 | 0.135189 | 0.07335 | 1.843106 | 0.06531 | 0.19714 | 66.827 |
| Gtf3c2 | 1478.24 | 0.093848 | 0.0845 | 1.110599 | 0.26674 | 0.47595 | 24.14 |
| Gtf3c3 | 1945.12 | 0.098277 | 0.08291 | 1.185332 | 0.23589 | 0.44113 | 42.968 |
| Gtf3c4 | 2713.73 | 0.028712 | 0.06825 | 0.420692 | 0.67398 | 0.82099 | 18.062 |
| Gtf3c5 | 1144.21 | 0.334459 | 0.09765 | 3.425177 | 0.00061 | 0.00913 | 17.441 |
| Gtf3c6 | 955.119 | -0.10652 | 0.09972 | -1.06817 | 0.28545 | 0.49536 | 10.406 |
| Gtpbp1 | 2916.92 | 0.072627 | 0.07835 | 0.926906 | 0.35398 | 0.56422 | 30.635 |

| | | | | | | | |
|---|---|---|---|---|---|---|---|
| Gtpbp10 | 537.491 | 0.256989 | 0.10255 | 2.50601 | 0.01221 | 0.06535 | 22.085 |
| Gtpbp2 | 4442.5 | 0.077803 | 0.08344 | 0.932463 | 0.3511 | 0.56158 | 8.339 |
| Gtpbp3 | 1051.38 | -0.03107 | 0.07131 | -0.43577 | 0.663 | 0.81385 | 11.481 |
| Gtpbp4 | 1649.12 | 0.045047 | 0.06426 | 0.701058 | 0.48327 | 0.67961 | 29.753 |
| Gtpbp6 | 530.949 | -0.1035 | 0.11043 | -0.93725 | 0.34863 | 0.55914 | 4.223 |
| Gtpbp8 | 1258.22 | 0.037375 | 0.07223 | 0.517472 | 0.60483 | 0.7735 | 9.596 |
| Gtse1 | 29.2557 | -0.45923 | 0.45695 | -1.00499 | 0.3149 | 0.52545 | 16.829 |
| Guca1b | 60.8829 | 0.493136 | 0.26022 | 1.895098 | 0.05808 | 0.18321 | 7.575 |
| Gucd1 | 664.933 | 0.007273 | 0.09413 | 0.077268 | 0.93841 | 0.96937 | 166.862 |
| Gucy1a1 | 9692.5 | 0.696764 | 0.1971 | 3.535053 | 0.00041 | 0.00695 | 53.363 |
| Gucy1a2 | 549.576 | 0.130701 | 0.17583 | 0.743321 | 0.45729 | 0.65857 | 361.959 |
| Gucy1b1 | 12404.9 | 0.354118 | 0.11768 | 3.009178 | 0.00262 | 0.0241 | 42.684 |
| Gucy2c | 25.8981 | -0.16186 | 0.36301 | -0.4459 | 0.65567 | 0.80879 | 84.482 |
| Gucy2e | 181.04 | -0.14054 | 0.15566 | -0.90284 | 0.36661 | 0.57697 | 18.92 |
| Gucy2f | 67.8409 | 0.235642 | 0.288 | 0.818199 | 0.41324 | 0.62139 | 115.772 |
| Gucy2g | 266.387 | -1.47771 | 0.42799 | -3.45272 | 0.00055 | 0.00851 | 43.138 |
| Guf1 | 2731.92 | 0.227958 | 0.10099 | 2.257253 | 0.02399 | 0.10094 | 19.051 |
| Guk1 | 2309.8 | -0.14874 | 0.08383 | -1.77445 | 0.07599 | 0.21768 | 8.369 |
| Gulp1 | 596.932 | 0.941895 | 0.17368 | 5.423155 | 5.86E-08 | 1.22E-05 | 245.328 |
| Gusb | 740.043 | -0.26541 | 0.10445 | -2.54099 | 0.01105 | 0.06131 | 14.039 |
| Gvin-ps7 | 156.605 | 0.17744 | 0.29692 | 0.597612 | 0.5501 | 0.7337 | 7.443 |
| Gvin3 | 12.737 | -0.97429 | 0.8014 | -1.21574 | 0.22409 | 0.42749 | 7.293 |
| Gxylt1 | 1517.65 | 0.14522 | 0.09591 | 1.514142 | 0.12999 | 0.30786 | 35.438 |
| Gxylt2 | 138.034 | -0.37167 | 0.23399 | -1.58842 | 0.11219 | 0.27978 | 106.18 |
| Gyg | 866.305 | -0.16784 | 0.09188 | -1.82672 | 0.06774 | 0.20179 | 33.234 |
| Gypc | 64.8282 | -0.43875 | 0.40658 | -1.07914 | 0.28052 | 0.49031 | 31.713 |
| Gys1 | 839.577 | -0.22051 | 0.08831 | -2.49708 | 0.01252 | 0.06645 | 21.776 |
| Gzf1 | 2381.61 | 0.101509 | 0.08813 | 1.151766 | 0.24942 | 0.45679 | 11.927 |
| Gzmk | 30.4269 | 0.887562 | 0.34279 | 2.589237 | 0.00962 | 0.05589 | 54.301 |
| Gzmm | 88.9867 | 0.18979 | 0.21137 | 0.897898 | 0.36924 | 0.57974 | 6.242 |
| H13 | 3099.29 | 0.065592 | 0.06978 | 0.93995 | 0.34724 | 0.55778 | 39.21 |
| H1f0 | 2869.21 | -0.38426 | 0.09811 | -3.91658 | 8.98E-05 | 0.00251 | 2.055 |
| H1f10 | 343.188 | -0.05201 | 0.17132 | -0.30359 | 0.76144 | 0.87503 | 1.217 |
| H1f11-ps | 29.6937 | -0.06983 | 0.30733 | -0.22723 | 0.82024 | 0.90734 | 0.666 |

| | | | | | | | |
|---|---|---|---|---|---|---|---|
| H1f2 | 500.965 | -0.70841 | 0.17349 | -4.08319 | 4.44E-05 | 0.0015 | 1.56 |
| H1f4 | 37.9923 | -0.74727 | 0.3196 | -2.33815 | 0.01938 | 0.08767 | 1.93 |
| H2-Aa | 121.995 | -1.52615 | 0.38126 | -4.00289 | 6.26E-05 | 0.00194 | 5.08 |
| H2-Ab1 | 117.358 | -1.48874 | 0.58756 | -2.53375 | 0.01128 | 0.06204 | 11.731 |
| H2-Bl | 26.6502 | 0.396593 | 0.36636 | 1.082535 | 0.27901 | 0.48891 | 4.109 |
| H2-D1 | 2094.53 | -0.80453 | 0.16495 | -4.87756 | 1.07E-06 | 0.00011 | 4.77 |
| H2-DMa | 423.373 | 0.239205 | 0.12447 | 1.921727 | 0.05464 | 0.17579 | 19.561 |
| H2-DMb1 | 67.2578 | -0.06314 | 0.25126 | -0.25132 | 0.80157 | 0.89741 | 7.159 |
| H2-DMb2 | 47.9227 | 0.32009 | 0.32119 | 0.996586 | 0.31897 | 0.5295 | 8.249 |
| H2-Eb1 | 134.669 | -1.33359 | 0.38001 | -3.5094 | 0.00045 | 0.00739 | 10.808 |
| H2-K1 | 1283.34 | -0.9 | 0.18551 | -4.85142 | 1.23E-06 | 0.00012 | 4.331 |
| H2-K2 | 101.519 | -0.44695 | 0.23666 | -1.88855 | 0.05895 | 0.1848 | 4.204 |
| H2-M10.2 | 72.9075 | 0.028736 | 0.21593 | 0.13308 | 0.89413 | 0.9471 | 2.264 |
| H2-M3 | 100.304 | -0.20176 | 0.18413 | -1.09575 | 0.27319 | 0.48254 | 4.265 |
| H2-M5 | 48.9648 | -0.56009 | 0.3167 | -1.76851 | 0.07697 | 0.21936 | 5.477 |
| H2-M6-ps | 39.9975 | -0.0098 | 0.29816 | -0.03286 | 0.97379 | 0.98681 | 2.202 |
| H2-Ob | 17.0322 | -0.16915 | 0.63676 | -0.26565 | 0.79051 | 0.8913 | 15.512 |
| H2-Q1 | 62.0319 | -1.31858 | 0.49275 | -2.67598 | 0.00745 | 0.04751 | 4.695 |
| H2-Q2 | 72.6412 | 1.385045 | 0.30745 | 4.504912 | 6.64E-06 | 0.00041 | 4.521 |
| H2-Q4 | 118.858 | -0.96107 | 0.29277 | -3.28266 | 0.00103 | 0.01304 | 5.674 |
| H2-Q5 | 16.5593 | -0.92496 | 0.65933 | -1.40289 | 0.16065 | 0.34889 | 3.675 |
| H2-T10 | 138.869 | -0.20038 | 0.15533 | -1.29001 | 0.19705 | 0.39484 | 5.59 |
| H2-T22 | 832.222 | -0.2643 | 0.11162 | -2.36796 | 0.01789 | 0.0833 | 5.62 |
| H2-T23 | 586.559 | -0.34919 | 0.14255 | -2.44951 | 0.0143 | 0.07211 | 3.083 |
| H2-T24 | 697.395 | 0.035678 | 0.08575 | 0.416092 | 0.67734 | 0.82317 | 14.866 |
| H2ac6 | 41.7864 | -0.67871 | 0.28419 | -2.38825 | 0.01693 | 0.08059 | 2.482 |
| H2aj | 459.813 | -0.17328 | 0.12841 | -1.34949 | 0.17718 | 0.36946 | 1.831 |
| H2aw | 672.838 | 0.115083 | 0.1175 | 0.979469 | 0.32735 | 0.53868 | 2.146 |
| H2ax | 561.821 | 0.009689 | 0.10922 | 0.088713 | 0.92931 | 0.9644 | 1.384 |
| H2az1 | 3547.22 | -0.12069 | 0.06894 | -1.75077 | 0.07998 | 0.22488 | 2.436 |
| H2az2 | 1251.2 | 0.018649 | 0.10825 | 0.172276 | 0.86322 | 0.93085 | 17.215 |
| H2bc21 | 496.316 | -0.13269 | 0.11147 | -1.19031 | 0.23393 | 0.43899 | 2.62 |
| H2bc4 | 769.916 | -0.33573 | 0.11009 | -3.04962 | 0.00229 | 0.02199 | 8.71 |
| H2bc6 | 237.227 | -0.25144 | 0.19341 | -1.30001 | 0.1936 | 0.39053 | 37.455 |

| | | | | | | | |
|---|---|---|---|---|---|---|---|
| H2bc8 | 18.9225 | -0.82168 | 0.45387 | -1.81037 | 0.07024 | 0.20699 | 0.618 |
| H2bu2 | 358.124 | 0.113506 | 0.13438 | 0.844681 | 0.39829 | 0.60733 | 0.614 |
| H3c15 | 65.3313 | -0.09825 | 0.26224 | -0.37466 | 0.70792 | 0.84235 | 1.02 |
| H3c4 | 30.007 | -1.14814 | 0.39066 | -2.93896 | 0.00329 | 0.0277 | 1.249 |
| H3f3a | 6436.81 | 0.069036 | 0.08352 | 0.826554 | 0.40849 | 0.61624 | 13.112 |
| H3f3a-ps1 | 28.5666 | -0.26605 | 0.30867 | -0.86193 | 0.38873 | 0.59793 | 0.41 |
| H3f3aos | 60.2964 | 0.874646 | 0.27162 | 3.220123 | 0.00128 | 0.01486 | 3.084 |
| H3f3b | 7800.89 | -0.1032 | 0.07809 | -1.32156 | 0.18632 | 0.38123 | 6.051 |
| H3f3c | 33.7907 | -1.65568 | 0.41965 | -3.94537 | 7.97E-05 | 0.0023 | 0.411 |
| H3f4 | 40.3568 | -0.25454 | 0.33589 | -0.7578 | 0.44857 | 0.652 | 0.766 |
| H4c11 | 20.5489 | -0.42834 | 0.50741 | -0.84417 | 0.39857 | 0.60758 | 0.774 |
| H4c12 | 110.209 | 0.108234 | 0.24931 | 0.43414 | 0.66419 | 0.8145 | 0.312 |
| H4c14 | 121.864 | -0.05591 | 0.2424 | -0.23064 | 0.8176 | 0.90615 | 1.63 |
| H4c3 | 13.5766 | -0.34226 | 0.47054 | -0.72738 | 0.46699 | 0.66622 | 0.82 |
| H4c8 | 22.2865 | -1.04128 | 0.43903 | -2.37176 | 0.0177 | 0.08269 | 0.47 |
| H4c9 | 50.7037 | -0.47426 | 0.31363 | -1.51214 | 0.1305 | 0.30883 | 0.727 |
| H60b | 22.3834 | -0.41619 | 0.48561 | -0.85706 | 0.39141 | 0.60048 | 130.264 |
| H6pd | 442.085 | -0.25488 | 0.09698 | -2.62825 | 0.00858 | 0.05181 | 29.549 |
| Habp4 | 9355.31 | -0.007 | 0.08393 | -0.0834 | 0.93353 | 0.9665 | 24.714 |
| Hacd1 | 269.698 | 0.350102 | 0.18749 | 1.867295 | 0.06186 | 0.19038 | 205.854 |
| Hacd2 | 1449.23 | -0.21306 | 0.0908 | -2.34647 | 0.01895 | 0.08644 | 86.75 |
| Hacd3 | 7870.89 | -0.0556 | 0.06934 | -0.8019 | 0.42261 | 0.62939 | 34.711 |
| Hacd4 | 264.253 | -0.2626 | 0.1749 | -1.50142 | 0.13325 | 0.31281 | 42.785 |
| Hace1 | 3357.26 | 0.075702 | 0.08608 | 0.879483 | 0.37914 | 0.5888 | 134.517 |
| Hadh | 980.953 | -0.22789 | 0.12024 | -1.89526 | 0.05806 | 0.18317 | 38.683 |
| Hadha | 3553.58 | -0.06457 | 0.07492 | -0.8618 | 0.3888 | 0.59796 | 36.859 |
| Hadhb | 2831 | -0.13393 | 0.08507 | -1.57435 | 0.11541 | 0.28494 | 29.346 |
| Hadhb-ps | 670.966 | -0.03146 | 0.1244 | -0.25292 | 0.80033 | 0.89682 | 1.428 |
| Hagh | 2266.31 | -0.02386 | 0.06767 | -0.35261 | 0.72438 | 0.85307 | 24.308 |
| Haghl | 1847.04 | -0.15322 | 0.10173 | -1.5061 | 0.13204 | 0.31082 | 5.831 |
| Hap1 | 13538.9 | 0.850027 | 0.14383 | 5.909842 | 3.42E-09 | 1.45E-06 | 8.802 |
| Hapln1 | 1157.57 | 0.043117 | 0.09681 | 0.445367 | 0.65605 | 0.80893 | 71.857 |
| Hapln2 | 1460.35 | -1.01369 | 0.19439 | -5.2148 | 1.84E-07 | 2.89E-05 | 5.834 |
| Hapln3 | 75.3914 | 0.186658 | 0.22664 | 0.823598 | 0.41017 | 0.61804 | 15.917 |

| | | | | | | | |
|---|---|---|---|---|---|---|---|
| Hapln4 | 2271.36 | -0.65698 | 0.10676 | -6.15356 | 7.58E-10 | 4.41E-07 | 7.406 |
| Harbi1 | 290.49 | 0.02149 | 0.14282 | 0.150466 | 0.8804 | 0.94027 | 10.694 |
| Hars | 2743.87 | -0.11034 | 0.05909 | -1.8674 | 0.06185 | 0.19038 | 16.678 |
| Hars2 | 1950.49 | 0.183465 | 0.07138 | 2.570417 | 0.01016 | 0.0581 | 9.555 |
| Has1 | 206.457 | 0.252375 | 0.18253 | 1.382632 | 0.16678 | 0.35646 | 11.883 |
| Has3 | 509.685 | 0.214368 | 0.10387 | 2.063825 | 0.03903 | 0.14077 | 14.325 |
| Hat1 | 815.293 | 0.014764 | 0.10908 | 0.135357 | 0.89233 | 0.94617 | 52.665 |
| Haus1 | 279.892 | -0.02207 | 0.12641 | -0.17462 | 0.86138 | 0.92975 | 16.32 |
| Haus2 | 1065.64 | 0.028866 | 0.09134 | 0.316031 | 0.75198 | 0.86994 | 12.178 |
| Haus3 | 278.087 | -0.02118 | 0.1949 | -0.10866 | 0.91348 | 0.95625 | 15.648 |
| Haus4 | 78.2152 | -0.21563 | 0.2575 | -0.83738 | 0.40238 | 0.61109 | 12.832 |
| Haus5 | 31.7477 | 0.165661 | 0.33357 | 0.496627 | 0.61945 | 0.78345 | 11.284 |
| Haus6 | 629.639 | 0.185715 | 0.11037 | 1.682723 | 0.09243 | 0.24756 | 33.201 |
| Haus7 | 249.476 | -0.17408 | 0.13522 | -1.28737 | 0.19797 | 0.3957 | 21.73 |
| Haus8 | 249.828 | -0.25918 | 0.14332 | -1.80842 | 0.07054 | 0.20755 | 24.374 |
| Havcr2 | 227.421 | -0.4235 | 0.20102 | -2.10673 | 0.03514 | 0.13139 | 26.321 |
| Hax1 | 1735.99 | -0.00324 | 0.06807 | -0.04759 | 0.96204 | 0.98159 | 3.335 |
| Hba-a1 | 947.434 | -0.19003 | 0.31614 | -0.60107 | 0.54779 | 0.73192 | 0.955 |
| Hba-a2 | 91.3919 | -0.78306 | 0.3612 | -2.16793 | 0.03016 | 0.11868 | 0.81 |
| Hba-ps4 | 21.1123 | -0.24423 | 0.40708 | -0.59996 | 0.54854 | 0.73258 | 0.699 |
| Hbb-bs | 1786.47 | -0.27074 | 0.32645 | -0.82934 | 0.40691 | 0.615 | 1.563 |
| Hbb-bt | 426.478 | -0.40893 | 0.21865 | -1.87028 | 0.06145 | 0.18957 | 1.473 |
| Hbegf | 737.322 | 0.165509 | 0.14948 | 1.10722 | 0.2682 | 0.47751 | 10.879 |
| Hbp1 | 1150.01 | 0.122431 | 0.09344 | 1.310319 | 0.19009 | 0.38616 | 24.282 |
| Hbs1l | 2220.16 | 0.102017 | 0.06448 | 1.58218 | 0.11361 | 0.28217 | 72.92 |
| Hcar1 | 70.1509 | -1.68822 | 0.84052 | -2.00855 | 0.04458 | 0.15339 | 3.285 |
| Hccs | 1334.67 | -0.06688 | 0.08657 | -0.7726 | 0.43976 | 0.64503 | 70.18 |
| Hcfc1 | 6382.44 | 0.066788 | 0.06081 | 1.098245 | 0.2721 | 0.48148 | 23.566 |
| Hcfc1r1 | 1095.39 | -0.23341 | 0.08034 | -2.9052 | 0.00367 | 0.02974 | 1.632 |
| Hcfc2 | 1621.37 | 0.197348 | 0.08077 | 2.443422 | 0.01455 | 0.0729 | 46.269 |
| Hck | 112.188 | -0.23928 | 0.30273 | -0.7904 | 0.4293 | 0.63586 | 42.974 |
| Hcls1 | 207.402 | -0.34661 | 0.22249 | -1.55789 | 0.11926 | 0.29142 | 28.23 |
| Hcn1 | 3398.86 | -0.3194 | 0.14522 | -2.19944 | 0.02785 | 0.1122 | 385.099 |
| Hcn2 | 5716.97 | -0.30234 | 0.09954 | -3.03755 | 0.00239 | 0.02257 | 19.475 |

| | | | | | | | |
|---|---|---|---|---|---|---|---|
| Hcn3 | 901.023 | -0.03515 | 0.10783 | -0.32597 | 0.74445 | 0.8655 | 14.123 |
| Hcn4 | 506.003 | -0.47178 | 0.13438 | -3.51083 | 0.00045 | 0.00736 | 39.764 |
| Hcrtr1 | 43.0312 | -0.36892 | 0.28209 | -1.30783 | 0.19093 | 0.38718 | 9.143 |
| Hcrtr2 | 193.111 | 0.080724 | 0.16853 | 0.478997 | 0.63194 | 0.79217 | 97.977 |
| Hdac1 | 1246.61 | -0.09784 | 0.07793 | -1.25558 | 0.20927 | 0.4093 | 26.61 |
| Hdac1-ps | 50.7898 | -0.4179 | 0.26018 | -1.60621 | 0.10823 | 0.27351 | 1.444 |
| Hdac10 | 547.555 | -0.13258 | 0.12288 | -1.07895 | 0.28061 | 0.49035 | 5.394 |
| Hdac11 | 4352.06 | 0.079833 | 0.07025 | 1.13641 | 0.25579 | 0.46353 | 18.028 |
| Hdac2 | 4476.78 | 0.162576 | 0.06138 | 2.648688 | 0.00808 | 0.04991 | 27.346 |
| Hdac3 | 3244.84 | 0.027394 | 0.08485 | 0.32287 | 0.74679 | 0.86719 | 19.173 |
| Hdac4 | 2993.6 | 0.093809 | 0.12093 | 0.775704 | 0.43792 | 0.64331 | 266.921 |
| Hdac5 | 7558.44 | -0.23043 | 0.10749 | -2.14379 | 0.03205 | 0.12361 | 35.735 |
| Hdac6 | 2473.44 | 0.254963 | 0.08067 | 3.16072 | 0.00157 | 0.01719 | 17.77 |
| Hdac7 | 1754.76 | -0.14908 | 0.11103 | -1.34271 | 0.17936 | 0.37228 | 51.839 |
| Hdac8 | 616.134 | 0.018282 | 0.10606 | 0.172381 | 0.86314 | 0.93081 | 220.721 |
| Hdac9 | 1331.22 | -0.33972 | 0.17193 | -1.97601 | 0.04815 | 0.16168 | 869.516 |
| Hdc | 34.1743 | -0.15546 | 0.34756 | -0.44728 | 0.65468 | 0.80841 | 25.633 |
| Hddc2 | 390.43 | 0.173038 | 0.13639 | 1.268685 | 0.20455 | 0.40342 | 14.822 |
| Hddc3 | 154.883 | -0.23386 | 0.16179 | -1.44546 | 0.14833 | 0.33196 | 7.444 |
| Hdgf | 6441.56 | -0.02684 | 0.06498 | -0.41298 | 0.67962 | 0.82491 | 9.812 |
| Hdgfl2 | 4031.46 | 0.057396 | 0.05366 | 1.069638 | 0.28478 | 0.49472 | 20.974 |
| Hdgfl3 | 5424.91 | 0.123963 | 0.06696 | 1.851294 | 0.06413 | 0.19495 | 53.223 |
| Hdhd2 | 1142.2 | -0.10147 | 0.11145 | -0.9104 | 0.36261 | 0.57294 | 31.216 |
| Hdhd3 | 145.466 | -0.07051 | 0.18738 | -0.37626 | 0.70672 | 0.84193 | 4.46 |
| Hdhd5 | 2386.95 | 0.172023 | 0.07521 | 2.287219 | 0.02218 | 0.096 | 21.826 |
| Hdlbp | 10108.2 | 0.090633 | 0.05879 | 1.541568 | 0.12318 | 0.29739 | 72.876 |
| Hdnr | 13.3379 | 0.06014 | 0.47107 | 0.127665 | 0.89841 | 0.94902 | 47.224 |
| Hdx | 566.318 | 0.085712 | 0.2163 | 0.396262 | 0.69191 | 0.83278 | 127.149 |
| Heatr1 | 1896.83 | 0.117919 | 0.07729 | 1.525567 | 0.12712 | 0.30327 | 45.263 |
| Heatr3 | 1673.11 | 0.135303 | 0.10535 | 1.28437 | 0.19901 | 0.3969 | 34.173 |
| Heatr5a | 799.565 | -0.22002 | 0.08421 | -2.61278 | 0.00898 | 0.05334 | 95.451 |
| Heatr5b | 3234.5 | 0.178744 | 0.08723 | 2.049119 | 0.04045 | 0.14371 | 82.467 |
| Heatr6 | 1640.49 | 0.305483 | 0.08495 | 3.59611 | 0.00032 | 0.00595 | 30.059 |
| Hebp1 | 761.024 | 0.043945 | 0.10398 | 0.422626 | 0.67257 | 0.8199 | 60.501 |

| | | | | | | | |
|---|---|---|---|---|---|---|---|
| Hebp2 | 96.4075 | -0.58167 | 0.26554 | -2.19049 | 0.02849 | 0.11384 | 6.579 |
| Heca | 1719.76 | -0.08277 | 0.08275 | -1.00014 | 0.31724 | 0.52755 | 79.456 |
| Hectd1 | 7602.6 | 0.142259 | 0.07141 | 1.992033 | 0.04637 | 0.15739 | 85.815 |
| Hectd2 | 671.709 | 0.110436 | 0.15068 | 0.732904 | 0.46362 | 0.66342 | 66.497 |
| Hectd2os | 50.1884 | 0.011139 | 0.29595 | 0.037639 | 0.96998 | 0.98478 | 138.153 |
| Hectd3 | 2004.28 | 0.131449 | 0.08167 | 1.609455 | 0.10752 | 0.27228 | 9.961 |
| Hectd4 | 18986.1 | 0.076897 | 0.0598 | 1.285934 | 0.19847 | 0.39615 | 148.359 |
| Hecw1 | 2554.29 | 0.016815 | 0.09506 | 0.176889 | 0.8596 | 0.92904 | 296.791 |
| Hecw2 | 3309.86 | 0.20095 | 0.08456 | 2.376356 | 0.01748 | 0.08212 | 388.293 |
| Heg1 | 1812.65 | -0.11058 | 0.08342 | -1.32568 | 0.18495 | 0.37969 | 87.207 |
| Helb | 433.076 | -0.15732 | 0.11329 | -1.38864 | 0.16494 | 0.35399 | 29.38 |
| Hells | 222.754 | -0.20416 | 0.20886 | -0.97752 | 0.32831 | 0.53961 | 40.137 |
| Helq | 1112.18 | 0.215255 | 0.10267 | 2.096539 | 0.03603 | 0.13364 | 36.454 |
| Helz | 5203.99 | 0.207381 | 0.06779 | 3.058951 | 0.00222 | 0.02155 | 145.897 |
| Helz2 | 526.648 | -0.14463 | 0.10871 | -1.33043 | 0.18338 | 0.37745 | 14.413 |
| Hemk1 | 1158.01 | 0.218853 | 0.09971 | 2.194976 | 0.02817 | 0.11308 | 11.267 |
| Henmt1 | 56.5085 | -0.17391 | 0.30499 | -0.57021 | 0.56853 | 0.74835 | 20.716 |
| Hepacam | 4415.39 | -0.30061 | 0.0965 | -3.11506 | 0.00184 | 0.01906 | 19.225 |
| Hepacam2 | 17.8057 | -0.9188 | 0.65623 | -1.40014 | 0.16147 | 0.34975 | 41.203 |
| Heph | 487.045 | -0.2469 | 0.16958 | -1.45594 | 0.14541 | 0.32825 | 119.127 |
| Herc1 | 13960.6 | -0.0023 | 0.06269 | -0.03669 | 0.97074 | 0.98519 | 158.326 |
| Herc2 | 18748.2 | -0.03603 | 0.04959 | -0.72666 | 0.46744 | 0.66644 | 181.605 |
| Herc3 | 6470.23 | -0.24902 | 0.1207 | -2.06324 | 0.03909 | 0.14085 | 88.934 |
| Herc4 | 1772.23 | -0.05894 | 0.07912 | -0.74486 | 0.45636 | 0.65798 | 74.069 |
| Herc6 | 295.696 | -0.2842 | 0.20144 | -1.41085 | 0.15829 | 0.34558 | 83.633 |
| Herpud1 | 2018.75 | -0.07433 | 0.09355 | -0.79456 | 0.42687 | 0.63333 | 8.94 |
| Herpud2 | 2103.66 | 0.062623 | 0.06665 | 0.939579 | 0.34743 | 0.55795 | 43.689 |
| Hes1 | 313.942 | -0.61474 | 0.19904 | -3.08851 | 0.00201 | 0.02016 | 3.413 |
| Hes5 | 160.708 | 0.140074 | 0.26604 | 0.526516 | 0.59853 | 0.76969 | 1.449 |
| Hes6 | 305.268 | -0.29801 | 0.13024 | -2.2881 | 0.02213 | 0.09588 | 2.556 |
| Hexa | 2368.38 | -0.01116 | 0.07669 | -0.14558 | 0.88426 | 0.94244 | 25.57 |
| Hexb | 2351.19 | -0.28433 | 0.13786 | -2.06245 | 0.03917 | 0.141 | 22.027 |
| Hexdc | 1158.18 | 0.195267 | 0.12588 | 1.551278 | 0.12084 | 0.29387 | 18.223 |
| Hexim1 | 2989.58 | -0.12546 | 0.06075 | -2.06506 | 0.03892 | 0.14059 | 3.495 |

| | | | | | | | |
|---|---|---|---|---|---|---|---|
| Hexim2 | 261.958 | -0.09635 | 0.14059 | -0.68533 | 0.49314 | 0.68772 | 7.448 |
| Hey1 | 2412.88 | -0.03125 | 0.08868 | -0.35237 | 0.72456 | 0.85319 | 3.898 |
| Hey2 | 289.346 | -0.04513 | 0.14611 | -0.30888 | 0.75741 | 0.87222 | 10.443 |
| Heyl | 200.353 | 0.562219 | 0.21218 | 2.649692 | 0.00806 | 0.04983 | 16.32 |
| Hfe | 264.631 | -0.61943 | 0.14552 | -4.25651 | 2.08E-05 | 0.0009 | 8.821 |
| Hfm1 | 241.674 | 0.517021 | 0.18782 | 2.752726 | 0.00591 | 0.04085 | 86.13 |
| Hgf | 221.323 | -0.32652 | 0.21866 | -1.49329 | 0.13536 | 0.31509 | 66.658 |
| Hgfac | 13.953 | 0.115488 | 0.46693 | 0.247333 | 0.80465 | 0.89908 | 6.953 |
| Hgh1 | 409.513 | 0.003335 | 0.12914 | 0.025827 | 0.9794 | 0.9889 | 2.54 |
| Hgs | 2699.7 | -0.12892 | 0.06229 | -2.06972 | 0.03848 | 0.13966 | 16.345 |
| Hgsnat | 2323.12 | -0.12868 | 0.0739 | -1.74131 | 0.08163 | 0.22769 | 32.301 |
| Hhat | 70.9979 | 0.37052 | 0.27296 | 1.357425 | 0.17465 | 0.36622 | 274.513 |
| Hhatl | 432.761 | -0.59496 | 0.12543 | -4.74343 | 2.10E-06 | 0.00018 | 8.492 |
| Hhex | 78.4922 | -0.95594 | 0.26069 | -3.66699 | 0.00025 | 0.0049 | 5.922 |
| Hhip | 660.216 | -0.00975 | 0.13646 | -0.07145 | 0.94304 | 0.97157 | 92.156 |
| Hhipl1 | 985.069 | 0.04917 | 0.11108 | 0.442643 | 0.65802 | 0.81018 | 24.6 |
| Hhipl2 | 45.4929 | 0.011114 | 0.39691 | 0.028001 | 0.97766 | 0.98803 | 18.571 |
| Hibadh | 2699.08 | -0.01651 | 0.07798 | -0.21167 | 0.83237 | 0.91402 | 94.162 |
| Hibch | 844.023 | 0.095673 | 0.09161 | 1.04431 | 0.29634 | 0.50667 | 76.058 |
| Hic1 | 118.999 | -0.6123 | 0.23492 | -2.60645 | 0.00915 | 0.05401 | 4.955 |
| Hic2 | 426.125 | 0.075246 | 0.13936 | 0.53996 | 0.58922 | 0.76226 | 29.859 |
| Hid1 | 3751.71 | -0.10898 | 0.06767 | -1.61048 | 0.10729 | 0.27185 | 20.05 |
| Hif1a | 3228.83 | 0.085902 | 0.06706 | 1.280991 | 0.2002 | 0.39817 | 46.156 |
| Hif1an | 3539.52 | 0.026892 | 0.0625 | 0.430257 | 0.66701 | 0.81596 | 13.425 |
| Hif3a | 1322.84 | -0.07702 | 0.07985 | -0.96457 | 0.33476 | 0.54551 | 30.921 |
| Higd1a | 2985.72 | -0.14598 | 0.06344 | -2.30123 | 0.02138 | 0.09364 | 9.8 |
| Higd1b | 98.7798 | -0.33122 | 0.21988 | -1.50638 | 0.13197 | 0.31078 | 2.192 |
| Higd2a | 1889.06 | -0.30408 | 0.09556 | -3.18202 | 0.00146 | 0.01626 | 0.952 |
| Hikeshi | 1555.23 | 0.010846 | 0.0737 | 0.147161 | 0.88301 | 0.94166 | 23.676 |
| Hilpda | 120.019 | -0.05813 | 0.15532 | -0.37428 | 0.7082 | 0.84248 | 2.959 |
| Hinfp | 1846.72 | 0.162662 | 0.07661 | 2.123237 | 0.03373 | 0.1281 | 13.232 |
| Hint1 | 2855.54 | -0.30823 | 0.07358 | -4.1889 | 2.80E-05 | 0.0011 | 4.119 |
| Hint2 | 419.588 | -0.21138 | 0.12386 | -1.70665 | 0.08789 | 0.23915 | 2.24 |
| Hint3 | 533.106 | -0.00546 | 0.09735 | -0.0561 | 0.95526 | 0.97801 | 21.492 |

| | | | | | | | |
|---|---|---|---|---|---|---|---|
| Hip1 | 4660.16 | -0.29598 | 0.08085 | -3.66083 | 0.00025 | 0.00497 | 138.59 |
| Hip1r | 4700.15 | 0.009565 | 0.07557 | 0.126564 | 0.89929 | 0.9495 | 31.931 |
| Hipk1 | 8406.78 | -0.04311 | 0.0655 | -0.65814 | 0.51045 | 0.70176 | 51.749 |
| Hipk2 | 5847.35 | -0.35128 | 0.10019 | -3.50615 | 0.00045 | 0.00746 | 181.775 |
| Hipk3 | 5859.3 | 0.109966 | 0.05776 | 1.903804 | 0.05694 | 0.18072 | 67.966 |
| Hipk4 | 4179.84 | 0.056816 | 0.1305 | 0.435364 | 0.6633 | 0.81392 | 7.909 |
| Hira | 520.83 | -0.08579 | 0.11862 | -0.72317 | 0.46957 | 0.66851 | 93.273 |
| Hirip3 | 2057.51 | 0.236227 | 0.12286 | 1.922808 | 0.0545 | 0.17557 | 3.406 |
| Hivep1 | 2257.06 | -0.30038 | 0.11421 | -2.63007 | 0.00854 | 0.05162 | 140.517 |
| Hivep2 | 12508.9 | -0.05817 | 0.10408 | -0.55886 | 0.57626 | 0.75395 | 185.3 |
| Hivep3 | 2857.43 | -0.37703 | 0.16103 | -2.34138 | 0.01921 | 0.08706 | 404.262 |
| Hjurp | 2976.22 | 0.020431 | 0.07988 | 0.255782 | 0.79812 | 0.8958 | 15.163 |
| Hk1 | 14214.3 | -0.36301 | 0.09406 | -3.8593 | 0.00011 | 0.00293 | 111.054 |
| Hk1os | 23.8532 | 0.639193 | 0.41203 | 1.551334 | 0.12082 | 0.29387 | 24.293 |
| Hk2 | 220.599 | -0.58515 | 0.19975 | -2.9294 | 0.0034 | 0.02816 | 49.43 |
| Hk3 | 75.6766 | -0.59121 | 0.19694 | -3.002 | 0.00268 | 0.02444 | 15.401 |
| Hkdc1 | 20.0817 | -0.34827 | 0.45946 | -0.758 | 0.44845 | 0.65192 | 39.355 |
| Hlcs | 939.495 | -0.07011 | 0.12388 | -0.56594 | 0.57143 | 0.75033 | 184.69 |
| Hlf | 10228.4 | 0.24113 | 0.11571 | 2.083905 | 0.03717 | 0.13625 | 54.36 |
| Hltf | 1832.65 | 0.20557 | 0.1201 | 1.711639 | 0.08696 | 0.2374 | 60.68 |
| Hlx | 25.5787 | -0.63029 | 0.34561 | -1.8237 | 0.0682 | 0.2027 | 5.48 |
| Hmbox1 | 5186.87 | -0.15216 | 0.07879 | -1.93112 | 0.05347 | 0.17327 | 138.272 |
| Hmbs | 765.081 | 0.202796 | 0.12163 | 1.66729 | 0.09546 | 0.25257 | 7.89 |
| Hmces | 753.356 | 0.145523 | 0.11117 | 1.309026 | 0.19053 | 0.3867 | 22.695 |
| Hmcn1 | 282.798 | 0.079094 | 0.1383 | 0.5719 | 0.56739 | 0.74747 | 430.912 |
| Hmcn2 | 92.4002 | 1.375895 | 0.1939 | 7.095957 | 1.28E-12 | 3.72E-09 | 146.324 |
| Hmg20a | 5017.63 | 0.10902 | 0.06891 | 1.582015 | 0.11365 | 0.2822 | 78.328 |
| Hmg20b | 730.859 | -0.1881 | 0.10594 | -1.7756 | 0.0758 | 0.21741 | 4.433 |
| Hmga1 | 1250.54 | -0.05316 | 0.10663 | -0.49857 | 0.61808 | 0.78293 | 7.178 |
| Hmga1b | 166.071 | -0.01402 | 0.18059 | -0.07765 | 0.93811 | 0.96915 | 1.626 |
| Hmga2-ps1 | 23.3328 | -0.45072 | 0.40723 | -1.10679 | 0.26838 | 0.47771 | 1.552 |
| Hmgb1 | 2874.57 | -0.15428 | 0.08336 | -1.85065 | 0.06422 | 0.19501 | 137.788 |
| Hmgb1-ps2 | 76.8844 | 0.206693 | 0.36737 | 0.562634 | 0.57368 | 0.75227 | 0.636 |
| Hmgb1-rs16 | 15.8105 | 0.988015 | 0.55101 | 1.793102 | 0.07296 | 0.21243 | 0.613 |

| | | | | | | | |
|---|---|---|---|---|---|---|---|
| Hmgb2 | 367.336 | -0.47669 | 0.16243 | -2.93473 | 0.00334 | 0.0279 | 4.093 |
| Hmgb3 | 1460.51 | 0.243055 | 0.10295 | 2.360914 | 0.01823 | 0.08438 | 4.759 |
| Hmgcl | 1323.86 | 0.131833 | 0.08284 | 1.591431 | 0.11151 | 0.27897 | 16.17 |
| Hmgcll1 | 1112.44 | 0.278506 | 0.09869 | 2.821895 | 0.00477 | 0.03526 | 121.496 |
| Hmgcr | 5095.69 | -0.22123 | 0.11734 | -1.88543 | 0.05937 | 0.18544 | 21.97 |
| Hmgcs1 | 6438.71 | -0.42572 | 0.10655 | -3.99537 | 6.46E-05 | 0.00199 | 17.882 |
| Hmgcs2 | 806.192 | -0.06175 | 0.19043 | -0.32429 | 0.74572 | 0.86638 | 30.304 |
| Hmgn1 | 3310.26 | 0.155631 | 0.08658 | 1.797602 | 0.07224 | 0.21112 | 7.112 |
| Hmgn2 | 2915.63 | 0.159399 | 0.11744 | 1.357313 | 0.17468 | 0.36622 | 3.913 |
| Hmgn2-ps | 536.659 | -0.41519 | 0.35311 | -1.17581 | 0.23967 | 0.44563 | 1.151 |
| Hmgn2-ps1 | 60.0118 | 0.16752 | 0.31183 | 0.537225 | 0.59111 | 0.76377 | 0.279 |
| Hmgn3 | 1011.11 | -0.15052 | 0.10348 | -1.45464 | 0.14577 | 0.32869 | 36.738 |
| Hmgn5 | 446.534 | -0.07328 | 0.3938 | -0.18608 | 0.85238 | 0.92484 | 8.871 |
| Hmgxb3 | 1983.14 | 0.027425 | 0.08164 | 0.335932 | 0.73692 | 0.86107 | 45.774 |
| Hmgxb4 | 871.472 | 0.083576 | 0.10488 | 0.796885 | 0.42552 | 0.63215 | 38.623 |
| Hmmr | 36.2851 | -0.64305 | 0.45862 | -1.40213 | 0.16088 | 0.34901 | 32.028 |
| Hmox1 | 189.102 | -0.34695 | 0.18813 | -1.8442 | 0.06515 | 0.19687 | 6.969 |
| Hmox2 | 2175.47 | -0.03901 | 0.09732 | -0.40087 | 0.68851 | 0.83012 | 40.382 |
| Hnf1a | 12.6784 | 0.875895 | 0.50085 | 1.748807 | 0.08032 | 0.2255 | 22.115 |
| Hnf1b | 20.8967 | -0.39173 | 0.40131 | -0.97612 | 0.32901 | 0.54015 | 55.757 |
| Hnmt | 906.584 | 0.061887 | 0.10388 | 0.59574 | 0.55135 | 0.73445 | 46.485 |
| Hnrnpa0 | 8060.14 | -0.02639 | 0.07243 | -0.36442 | 0.71555 | 0.84781 | 2.678 |
| Hnrnpa1 | 15743.4 | 0.129953 | 0.09598 | 1.35397 | 0.17575 | 0.36754 | 6.261 |
| Hnrnpa2b1 | 29088 | 0.137501 | 0.09178 | 1.498226 | 0.13407 | 0.31381 | 8.963 |
| Hnrnpa3 | 5134.03 | 0.222197 | 0.09917 | 2.240635 | 0.02505 | 0.1041 | 10.147 |
| Hnrnpab | 2601.97 | -0.04584 | 0.07061 | -0.6492 | 0.51621 | 0.70625 | 6.748 |
| Hnrnpc | 9918.04 | 0.030603 | 0.05963 | 0.513178 | 0.60783 | 0.77583 | 30.652 |
| Hnrnpd | 8698.97 | 0.166056 | 0.10142 | 1.637347 | 0.10156 | 0.26294 | 23.004 |
| Hnrnpdl | 10942.5 | 0.186919 | 0.10203 | 1.831981 | 0.06695 | 0.20029 | 6.088 |
| Hnrnpf | 1679.88 | -0.11744 | 0.08202 | -1.43182 | 0.15219 | 0.33739 | 25.299 |
| Hnrnph1 | 21826.7 | 0.23461 | 0.09823 | 2.388337 | 0.01692 | 0.08059 | 9.539 |
| Hnrnph2 | 7476.74 | 0.007681 | 0.08374 | 0.091724 | 0.92692 | 0.96321 | 5.882 |
| Hnrnph3 | 3132.13 | 0.201171 | 0.10713 | 1.877796 | 0.06041 | 0.18755 | 9.554 |
| Hnrnpk | 14934.4 | 0.039864 | 0.06614 | 0.602684 | 0.54672 | 0.73092 | 12.202 |

| | | | | | | | |
|---|---|---|---|---|---|---|---|
| Hnrnpl | 12932.7 | 0.093514 | 0.0779 | 1.200476 | 0.22995 | 0.43425 | 13.726 |
| Hnrnpll | 4257.17 | 0.086967 | 0.07367 | 1.180514 | 0.2378 | 0.4432 | 32.848 |
| Hnrnpm | 7632.72 | 0.183758 | 0.0746 | 2.463191 | 0.01377 | 0.07057 | 40.628 |
| Hnrnpr | 7472.26 | 0.24831 | 0.07655 | 3.243612 | 0.00118 | 0.01415 | 48.506 |
| Hnrnpu | 29725.4 | 0.158886 | 0.10777 | 1.474314 | 0.1404 | 0.3219 | 16.69 |
| Hnrnpul1 | 5214.58 | 0.016616 | 0.06648 | 0.249922 | 0.80265 | 0.89815 | 33.593 |
| Hnrnpul2 | 14760.4 | 0.060168 | 0.05253 | 1.14544 | 0.25203 | 0.45939 | 14.134 |
| Hoga1 | 40.3103 | -0.18186 | 0.35108 | -0.51799 | 0.60446 | 0.7733 | 25.344 |
| Homer1 | 10730.5 | 0.198735 | 0.12199 | 1.629125 | 0.10329 | 0.26556 | 105.495 |
| Homer2 | 3266.94 | -0.98692 | 0.38242 | -2.58069 | 0.00986 | 0.05694 | 107.047 |
| Homer3 | 229.472 | -0.22739 | 0.18439 | -1.23322 | 0.21749 | 0.41946 | 11.535 |
| Homez | 736.208 | -0.04043 | 0.12473 | -0.32411 | 0.74585 | 0.86649 | 18.226 |
| Hook1 | 5449.19 | 0.208663 | 0.10926 | 1.9097 | 0.05617 | 0.17923 | 58.174 |
| Hook2 | 574.974 | -0.12771 | 0.13501 | -0.94594 | 0.34418 | 0.55487 | 12.747 |
| Hook3 | 5444.44 | 0.099243 | 0.10082 | 0.984392 | 0.32492 | 0.53614 | 97.804 |
| Hopx | 300.923 | -0.84065 | 0.18706 | -4.49407 | 6.99E-06 | 0.00043 | 28.134 |
| Hp | 12.8772 | -0.54441 | 0.58116 | -0.93676 | 0.34888 | 0.55931 | 4.045 |
| Hp1bp3 | 14438.6 | 0.173253 | 0.07096 | 2.441714 | 0.01462 | 0.07303 | 28.388 |
| Hpca | 36174.6 | 0.277775 | 0.16611 | 1.672286 | 0.09447 | 0.25083 | 10.467 |
| Hpcal1 | 1638.23 | -0.45857 | 0.15823 | -2.89817 | 0.00375 | 0.03016 | 101.078 |
| Hpcal4 | 38628.4 | -0.12104 | 0.13527 | -0.89473 | 0.37093 | 0.58108 | 11.475 |
| Hpdl | 40.6724 | -0.38724 | 0.303 | -1.27801 | 0.20125 | 0.39952 | 1.804 |
| Hpf1 | 1976.74 | -0.0285 | 0.10942 | -0.26049 | 0.79449 | 0.89365 | 18.254 |
| Hpgd | 213.756 | -0.28221 | 0.20491 | -1.37726 | 0.16843 | 0.3584 | 26.459 |
| Hpgds | 491.027 | -0.14526 | 0.1171 | -1.24049 | 0.21479 | 0.41624 | 27.616 |
| Hpn | 93.7093 | -0.83318 | 0.22988 | -3.62436 | 0.00029 | 0.00549 | 16.566 |
| Hprt | 4913.07 | -0.05058 | 0.07669 | -0.65951 | 0.50957 | 0.70111 | 33.523 |
| Hps1 | 874.215 | 0.269454 | 0.13814 | 1.950547 | 0.05111 | 0.1683 | 24.874 |
| Hps3 | 897.955 | 0.02201 | 0.07935 | 0.277388 | 0.78148 | 0.88617 | 39.371 |
| Hps4 | 1233.58 | 0.186503 | 0.07733 | 2.411779 | 0.01587 | 0.07732 | 35.332 |
| Hps5 | 1308.94 | 0.053556 | 0.08379 | 0.639143 | 0.52273 | 0.71143 | 35.599 |
| Hps6 | 218.49 | -0.22577 | 0.12334 | -1.83044 | 0.06718 | 0.20068 | 2.666 |
| Hpse | 83.8462 | 0.779677 | 0.38005 | 2.051499 | 0.04022 | 0.1433 | 40.233 |
| Hpse2 | 13.0631 | 0.487535 | 0.52482 | 0.928963 | 0.35291 | 0.56319 | 601.817 |

| | | | | | | | |
|---|---|---|---|---|---|---|---|
| Hpx | 22.7808 | -0.02237 | 0.3972 | -0.05632 | 0.95509 | 0.97798 | 8.525 |
| Hr | 966.594 | -0.75646 | 0.18026 | -4.19647 | 2.71E-05 | 0.00108 | 21.337 |
| Hras | 3676 | -0.09719 | 0.10151 | -0.95749 | 0.33832 | 0.54905 | 4.901 |
| Hrc | 33.7852 | -0.33101 | 0.30242 | -1.09454 | 0.27372 | 0.48313 | 3.685 |
| Hrct1 | 35.3789 | -0.83118 | 0.35481 | -2.34262 | 0.01915 | 0.08695 | 1.452 |
| Hrh1 | 236.817 | 0.124627 | 0.15052 | 0.827957 | 0.4077 | 0.61541 | 85.361 |
| Hrh2 | 434.31 | 0.090404 | 0.12386 | 0.72987 | 0.46547 | 0.66495 | 44.052 |
| Hrh3 | 4097.44 | 0.2404 | 0.1179 | 2.039008 | 0.04145 | 0.14608 | 5.024 |
| Hrk | 3244.59 | -0.06273 | 0.0933 | -0.6723 | 0.50139 | 0.69395 | 24.831 |
| Hrob | 25.0772 | -0.40225 | 0.33439 | -1.20293 | 0.229 | 0.43312 | 16.306 |
| Hs1bp3 | 636.959 | 0.072668 | 0.08571 | 0.847812 | 0.39654 | 0.60557 | 30.393 |
| Hs2st1 | 2853.47 | 0.100169 | 0.06874 | 1.457136 | 0.14508 | 0.32787 | 140.476 |
| Hs3st1 | 712.251 | -1.15134 | 0.19581 | -5.87977 | 4.11E-09 | 1.64E-06 | 141.541 |
| Hs3st2 | 926.646 | -0.9882 | 0.33641 | -2.93752 | 0.00331 | 0.02776 | 109.912 |
| Hs3st3a1 | 241.401 | -0.54446 | 0.18425 | -2.95504 | 0.00313 | 0.02678 | 87.51 |
| Hs3st3b1 | 171.69 | -0.97402 | 0.21809 | -4.46617 | 7.96E-06 | 0.00047 | 36.499 |
| Hs3st4 | 1420.33 | -0.18107 | 0.14132 | -1.28125 | 0.20011 | 0.39817 | 416.121 |
| Hs3st5 | 359.832 | 0.533081 | 0.17362 | 3.070374 | 0.00214 | 0.02106 | 327.584 |
| Hs3st6 | 14.1959 | 0.122022 | 0.46768 | 0.260908 | 0.79416 | 0.89349 | 5.697 |
| Hs6st1 | 2674.36 | -0.16145 | 0.0905 | -1.78402 | 0.07442 | 0.21503 | 38.047 |
| Hs6st2 | 3469.5 | 0.283845 | 0.12033 | 2.358945 | 0.01833 | 0.08467 | 294.645 |
| Hs6st3 | 690.388 | -0.22129 | 0.10761 | -2.05642 | 0.03974 | 0.14237 | 731.475 |
| Hsbp1 | 6845.27 | -0.1388 | 0.07472 | -1.85749 | 0.06324 | 0.19321 | 4.435 |
| Hsbp1l1 | 20.1464 | -0.61945 | 0.54692 | -1.13261 | 0.25738 | 0.46531 | 17.345 |
| Hscb | 241.655 | -0.06339 | 0.1444 | -0.43899 | 0.66067 | 0.81233 | 10.708 |
| Hsd11b1 | 229.606 | -0.55533 | 0.25495 | -2.17817 | 0.02939 | 0.11648 | 42.442 |
| Hsd11b2 | 22.9434 | 0.084104 | 0.45361 | 0.185412 | 0.85291 | 0.92534 | 5.234 |
| Hsd17b1 | 32.6956 | 0.115324 | 0.4265 | 0.270396 | 0.78686 | 0.88907 | 2.117 |
| Hsd17b10 | 1098.26 | 0.037896 | 0.10806 | 0.3507 | 0.72581 | 0.85388 | 2.598 |
| Hsd17b11 | 1848.37 | -0.28381 | 0.09664 | -2.93685 | 0.00332 | 0.02778 | 32.158 |
| Hsd17b12 | 2507.08 | -0.27976 | 0.0839 | -3.33453 | 0.00085 | 0.01148 | 125.276 |
| Hsd17b4 | 3095.18 | -0.10314 | 0.07671 | -1.34456 | 0.17877 | 0.37138 | 68.069 |
| Hsd17b7 | 1038.37 | -0.19628 | 0.07305 | -2.68689 | 0.00721 | 0.04645 | 19.707 |
| Hsdl1 | 2319.57 | 0.129754 | 0.09259 | 1.401436 | 0.16108 | 0.34924 | 13.195 |

| | | | | | | | |
|---|---|---|---|---|---|---|---|
| Hsdl2 | 1795.38 | -0.14881 | 0.0763 | -1.95039 | 0.05113 | 0.16832 | 37.127 |
| Hsf1 | 2039.53 | 0.174996 | 0.07091 | 2.46778 | 0.0136 | 0.07019 | 24.492 |
| Hsf2 | 1773.51 | 0.149739 | 0.07427 | 2.016088 | 0.04379 | 0.15143 | 26.751 |
| Hsf2bp | 13.0737 | 0.180159 | 0.49211 | 0.366095 | 0.71429 | 0.84667 | 91.581 |
| Hsf4 | 433.157 | -0.02574 | 0.12533 | -0.20536 | 0.83729 | 0.91639 | 6.045 |
| Hsf5 | 34.2698 | 0.048945 | 0.29758 | 0.164478 | 0.86935 | 0.93376 | 42.379 |
| Hsp90aa1 | 35471 | 0.087983 | 0.13053 | 0.674037 | 0.50029 | 0.69321 | 12.124 |
| Hsp90ab1 | 66541.5 | -0.04829 | 0.05824 | -0.82913 | 0.40703 | 0.61503 | 5.497 |
| Hsp90b1 | 8981.69 | 0.117852 | 0.08345 | 1.412267 | 0.15787 | 0.3451 | 15.301 |
| Hspa12a | 9688.11 | -0.20986 | 0.07963 | -2.63536 | 0.0084 | 0.05114 | 147.905 |
| Hspa12b | 582.453 | -0.012 | 0.11195 | -0.10718 | 0.91464 | 0.95656 | 19.042 |
| Hspa13 | 3054.39 | 0.242535 | 0.06934 | 3.49765 | 0.00047 | 0.00761 | 21.674 |
| Hspa14 | 1105.88 | 0.145086 | 0.08778 | 1.652847 | 0.09836 | 0.25742 | 23.965 |
| Hspa1a | 187.984 | -0.80829 | 0.22436 | -3.60268 | 0.00031 | 0.00584 | 2.967 |
| Hspa1b | 533.212 | -0.60953 | 0.39875 | -1.5286 | 0.12636 | 0.30211 | 2.803 |
| Hspa1l | 104.707 | 0.229331 | 0.20539 | 1.116542 | 0.26419 | 0.47298 | 6.583 |
| Hspa2 | 1826.33 | -0.31768 | 0.12577 | -2.52583 | 0.01154 | 0.06299 | 2.763 |
| Hspa4 | 12991.8 | 0.045643 | 0.07503 | 0.608318 | 0.54298 | 0.72855 | 40.644 |
| Hspa4l | 9910.34 | 0.272542 | 0.13108 | 2.079175 | 0.0376 | 0.13731 | 50.725 |
| Hspa5 | 15825.9 | 0.080985 | 0.12318 | 0.657448 | 0.51089 | 0.70199 | 5.578 |
| Hspa8 | 61966.9 | -0.07998 | 0.06281 | -1.27347 | 0.20285 | 0.40148 | 9.104 |
| Hspa9 | 10198.5 | 0.001792 | 0.05704 | 0.03142 | 0.97493 | 0.98738 | 16.944 |
| Hspb1 | 304.702 | -0.43443 | 0.17446 | -2.49018 | 0.01277 | 0.06741 | 1.645 |
| Hspb11 | 234.928 | 0.051927 | 0.15182 | 0.342036 | 0.73232 | 0.85811 | 26.346 |
| Hspb3 | 43.3173 | -1.01112 | 0.39322 | -2.5714 | 0.01013 | 0.05795 | 0.781 |
| Hspb6 | 112.303 | -0.01947 | 0.1955 | -0.09961 | 0.92065 | 0.95978 | 3.266 |
| Hspb8 | 689.366 | -0.37765 | 0.16517 | -2.28646 | 0.02223 | 0.0961 | 14.374 |
| Hspbap1 | 294.672 | 0.101718 | 0.1374 | 0.740303 | 0.45912 | 0.6598 | 58.103 |
| Hspbp1 | 2067.22 | -0.15747 | 0.08457 | -1.86204 | 0.0626 | 0.19199 | 24.548 |
| Hspd1 | 7898.47 | 0.125445 | 0.05074 | 2.472506 | 0.01342 | 0.06961 | 10.409 |
| Hspd1-ps3 | 133.041 | -0.4802 | 0.35571 | -1.34997 | 0.17702 | 0.3693 | 1.722 |
| Hspe1 | 2594.79 | 0.051998 | 0.08798 | 0.591018 | 0.55451 | 0.73708 | 3.176 |
| Hspe1-rs1 | 82.7845 | -0.05711 | 0.25142 | -0.22713 | 0.82032 | 0.90734 | 0.515 |
| Hspg2 | 579.644 | -0.18746 | 0.14856 | -1.26187 | 0.20699 | 0.40645 | 101.862 |

| | | | | | | | |
|---|---|---|---|---|---|---|---|
| Hsph1 | 19200.5 | 0.245418 | 0.06166 | 3.979952 | 6.89E-05 | 0.00207 | 22.09 |
| Htatip2 | 262.446 | -0.37421 | 0.13222 | -2.83031 | 0.00465 | 0.03467 | 14.861 |
| Htatsf1 | 4266.31 | 0.155308 | 0.09499 | 1.635005 | 0.10205 | 0.26371 | 13.601 |
| Htr1a | 726.313 | -0.22183 | 0.13277 | -1.67079 | 0.09476 | 0.25135 | 4.484 |
| Htr1b | 2366.5 | 0.736036 | 0.21092 | 3.489647 | 0.00048 | 0.00775 | 5.538 |
| Htr1d | 506.735 | 0.439622 | 0.17095 | 2.57159 | 0.01012 | 0.05794 | 20.875 |
| Htr1f | 180.768 | 0.301466 | 0.17823 | 1.691478 | 0.09075 | 0.24468 | 181.126 |
| Htr2a | 3422.53 | -0.01337 | 0.09499 | -0.14078 | 0.88804 | 0.94406 | 68.655 |
| Htr2b | 24.3283 | -0.41105 | 0.54641 | -0.75227 | 0.45189 | 0.6543 | 12.945 |
| Htr2c | 5772.01 | 0.172024 | 0.07982 | 2.155236 | 0.03114 | 0.12113 | 234.765 |
| Htr3a | 128.8 | -0.92761 | 0.28765 | -3.22475 | 0.00126 | 0.01476 | 11.887 |
| Htr4 | 727.579 | 0.93358 | 0.39245 | 2.378866 | 0.01737 | 0.08175 | 172.374 |
| Htr5a | 795.014 | -0.37092 | 0.13486 | -2.75037 | 0.00595 | 0.04101 | 12.94 |
| Htr5b | 39.1044 | -1.05369 | 0.49378 | -2.13391 | 0.03285 | 0.12589 | 18.624 |
| Htr6 | 492.872 | 0.465087 | 0.15489 | 3.002631 | 0.00268 | 0.02441 | 14.463 |
| Htr7 | 296.48 | -0.17668 | 0.19275 | -0.91664 | 0.35933 | 0.5697 | 98.774 |
| Htra1 | 8425.71 | -0.41513 | 0.1015 | -4.09011 | 4.31E-05 | 0.00147 | 49.55 |
| Htra2 | 803.481 | -0.01152 | 0.09444 | -0.122 | 0.9029 | 0.95118 | 4.008 |
| Htra3 | 176.857 | -0.43405 | 0.20092 | -2.16034 | 0.03075 | 0.12006 | 27.76 |
| Htra4 | 51.9605 | -0.41042 | 0.32392 | -1.26705 | 0.20514 | 0.40421 | 14.034 |
| Htt | 10172.4 | 0.078878 | 0.06157 | 1.281196 | 0.20012 | 0.39817 | 150.795 |
| Hunk | 655.368 | -0.07585 | 0.13856 | -0.54742 | 0.58409 | 0.75898 | 113.541 |
| Hus1 | 859.473 | -0.13777 | 0.08472 | -1.62611 | 0.10393 | 0.26644 | 18.055 |
| Huwe1 | 25193.1 | 0.093616 | 0.05681 | 1.647821 | 0.09939 | 0.25907 | 134.611 |
| Hvcn1 | 49.3061 | 0.053669 | 0.32109 | 0.167148 | 0.86725 | 0.93259 | 35.494 |
| Hyal1 | 171.731 | -0.455 | 0.14901 | -3.05349 | 0.00226 | 0.02182 | 4.784 |
| Hyal2 | 344.542 | -0.02891 | 0.11389 | -0.25388 | 0.79959 | 0.89651 | 4.835 |
| Hyal3 | 13.9322 | -0.43744 | 0.43362 | -1.00881 | 0.31306 | 0.52351 | 6.542 |
| Hydin | 65.8412 | -0.68372 | 0.38376 | -1.78165 | 0.07481 | 0.21551 | 343.277 |
| Hyi | 510.209 | 0.193 | 0.13402 | 1.440055 | 0.14985 | 0.33427 | 2.755 |
| Hykk | 212.537 | -0.22299 | 0.15709 | -1.41951 | 0.15575 | 0.34238 | 32.642 |
| Hyls1 | 318.22 | 0.209953 | 0.12229 | 1.716816 | 0.08601 | 0.23561 | 9.579 |
| Hyou1 | 6253.77 | 0.07768 | 0.09365 | 0.829464 | 0.40684 | 0.61498 | 12.88 |
| Hypk | 80.6317 | -0.01806 | 0.26794 | -0.06739 | 0.94627 | 0.97302 | 5.383 |

| | | | | | | | |
|---|---|---|---|---|---|---|---|
| I830077J02Rik | 58.012 | -0.15002 | 0.26951 | -0.55665 | 0.57776 | 0.75477 | 8.307 |
| I830134H01Rik | 12.5164 | 0.153708 | 0.64323 | 0.238964 | 0.81113 | 0.90277 | 2.755 |
| Iah1 | 422.85 | 0.106791 | 0.13559 | 0.7876 | 0.43093 | 0.63708 | 7.431 |
| Iars | 2803.08 | -0.06028 | 0.0885 | -0.68107 | 0.49583 | 0.68971 | 52.168 |
| Iars2 | 2148.53 | -0.08894 | 0.06894 | -1.29002 | 0.19704 | 0.39484 | 44.671 |
| Iba57 | 418.173 | 0.073305 | 0.119 | 0.61599 | 0.5379 | 0.72464 | 8.371 |
| Ibtk | 1595.14 | 0.108728 | 0.07287 | 1.492157 | 0.13566 | 0.31567 | 61.975 |
| Ica1 | 2242.11 | 0.186983 | 0.06826 | 2.739184 | 0.00616 | 0.042 | 147.962 |
| Ica1l | 1900.41 | 0.358747 | 0.0772 | 4.646952 | 3.37E-06 | 0.00026 | 60.695 |
| Icam1 | 111.541 | -1.32648 | 0.34867 | -3.80442 | 0.00014 | 0.00338 | 12.833 |
| Icam2 | 61.9669 | -0.03277 | 0.28151 | -0.1164 | 0.90733 | 0.95354 | 10.42 |
| Icam5 | 4735.1 | 0.056054 | 0.07505 | 0.746842 | 0.45516 | 0.65722 | 6.964 |
| Ice1 | 8800.27 | 0.258135 | 0.07211 | 3.57964 | 0.00034 | 0.00616 | 85.928 |
| Ice2 | 690.883 | 0.188288 | 0.10677 | 1.763477 | 0.07782 | 0.22075 | 35.217 |
| Icmt | 4228.86 | -0.02044 | 0.0691 | -0.29576 | 0.76742 | 0.87838 | 9.895 |
| Icosl | 210.349 | -0.59787 | 0.21648 | -2.76176 | 0.00575 | 0.04006 | 14.612 |
| Ict1os | 168.118 | 0.207917 | 0.17946 | 1.158576 | 0.24663 | 0.45346 | 1.729 |
| Id1 | 542.152 | -0.36291 | 0.10423 | -3.48171 | 0.0005 | 0.0079 | 1.16 |
| Id2 | 3380.97 | -0.5587 | 0.09559 | -5.84464 | 5.08E-09 | 1.82E-06 | 3.342 |
| Id3 | 1147.24 | -0.25821 | 0.10498 | -2.45964 | 0.01391 | 0.07096 | 2.259 |
| Id4 | 8668.55 | 0.18015 | 0.11562 | 1.558107 | 0.11921 | 0.2914 | 4.799 |
| Ide | 2548.68 | 0.020204 | 0.07359 | 0.274538 | 0.78367 | 0.88762 | 93.869 |
| Idh1 | 1712.5 | -0.276 | 0.10278 | -2.68537 | 0.00724 | 0.04657 | 27.885 |
| Idh2 | 1765.27 | -0.03214 | 0.08878 | -0.36198 | 0.71737 | 0.84903 | 20.547 |
| Idh3a | 9325.97 | -0.12942 | 0.06966 | -1.85807 | 0.06316 | 0.19305 | 18.328 |
| Idh3b | 7674.2 | -0.06547 | 0.05993 | -1.09236 | 0.27467 | 0.4843 | 5.239 |
| Idh3g | 3621.33 | -0.05065 | 0.06881 | -0.73608 | 0.46168 | 0.66223 | 7.935 |
| Idi1 | 785.802 | -0.35595 | 0.10201 | -3.48942 | 0.00048 | 0.00775 | 6.951 |
| Idnk | 838.712 | -0.20857 | 0.0952 | -2.19089 | 0.02846 | 0.11375 | 7.045 |
| Ido1 | 195.41 | 1.215132 | 0.33364 | 3.642062 | 0.00027 | 0.00521 | 12.874 |
| Ido2 | 80.6945 | -0.11289 | 0.20085 | -0.56205 | 0.57408 | 0.7525 | 44.442 |
| Ids | 27048 | -0.04031 | 0.0743 | -0.54257 | 0.58743 | 0.76091 | 22.016 |
| Idua | 990.585 | 0.122862 | 0.11098 | 1.107105 | 0.26825 | 0.47751 | 24.227 |
| Ier2 | 136.097 | -0.71799 | 0.24848 | -2.88958 | 0.00386 | 0.03072 | 1.524 |

| | | | | | | | |
|---|---|---|---|---|---|---|---|
| Ier3 | 247.976 | -0.09815 | 0.14352 | -0.68386 | 0.49407 | 0.68843 | 1.24 |
| Ier3ip1 | 1113.42 | 0.103342 | 0.08926 | 1.157701 | 0.24699 | 0.45383 | 11.596 |
| Ier5 | 1875.07 | -0.26503 | 0.11076 | -2.39283 | 0.01672 | 0.07998 | 3.276 |
| Ier5l | 702.252 | 0.071734 | 0.14012 | 0.511961 | 0.60868 | 0.77635 | 2.683 |
| Iffo1 | 2751.76 | -0.06177 | 0.05785 | -1.06768 | 0.28567 | 0.4957 | 16.542 |
| Iffo2 | 1190.1 | 0.093894 | 0.09109 | 1.030767 | 0.30265 | 0.51222 | 89.835 |
| Ifi203 | 57.2043 | -0.78746 | 0.36718 | -2.14461 | 0.03198 | 0.12345 | 22.266 |
| Ifi204 | 40.0139 | -1.66885 | 0.45538 | -3.66478 | 0.00025 | 0.00494 | 19.651 |
| Ifi207 | 19.8772 | -1.28838 | 0.63722 | -2.02188 | 0.04319 | 0.1501 | 18.321 |
| Ifi209 | 13.3051 | -1.95527 | 0.74568 | -2.62212 | 0.00874 | 0.05244 | 17.012 |
| Ifi27 | 1266.54 | -0.56569 | 0.14154 | -3.99668 | 6.42E-05 | 0.00198 | 6.029 |
| Ifi27l2a | 30.5295 | -1.10692 | 0.51854 | -2.13471 | 0.03278 | 0.12569 | 1.514 |
| Ifi30 | 77.6701 | -0.52693 | 0.24676 | -2.13542 | 0.03273 | 0.12556 | 3.889 |
| Ifi35 | 68.9838 | -0.39173 | 0.30564 | -1.28166 | 0.19996 | 0.39805 | 10.292 |
| Ifi44 | 24.3039 | -0.75321 | 0.5527 | -1.36278 | 0.17295 | 0.36423 | 19.039 |
| Ifi47 | 34.0455 | -1.16838 | 0.51046 | -2.28888 | 0.02209 | 0.09575 | 20.388 |
| Ifih1 | 187.421 | -0.57573 | 0.16416 | -3.5072 | 0.00045 | 0.00743 | 50.458 |
| Ifit1 | 109.418 | -0.85436 | 0.25174 | -3.39378 | 0.00069 | 0.00988 | 9.139 |
| Ifit1bl1 | 58.5982 | -1.056 | 0.29057 | -3.63428 | 0.00028 | 0.00533 | 9.066 |
| Ifit1bl2 | 20.537 | -0.98014 | 0.46523 | -2.10676 | 0.03514 | 0.13139 | 23.695 |
| Ifit2 | 440.897 | -0.63858 | 0.16689 | -3.82644 | 0.00013 | 0.00322 | 25.726 |
| Ifit3 | 387.862 | -0.61645 | 0.18408 | -3.34887 | 0.00081 | 0.01109 | 5.201 |
| Ifit3b | 196.465 | -0.44737 | 0.16577 | -2.69868 | 0.00696 | 0.04526 | 5.432 |
| Ifitm1 | 104.262 | -0.79253 | 0.23179 | -3.41919 | 0.00063 | 0.00925 | 2.605 |
| Ifitm10 | 261.722 | -0.53137 | 0.1753 | -3.03118 | 0.00244 | 0.02293 | 47.917 |
| Ifitm2 | 32.2092 | -0.81305 | 0.32943 | -2.46805 | 0.01358 | 0.07019 | 1.151 |
| Ifitm3 | 274.403 | -0.92213 | 0.20526 | -4.49241 | 7.04E-06 | 0.00043 | 1.185 |
| Ifitm7 | 35.4023 | -0.30539 | 0.34711 | -0.87982 | 0.37896 | 0.58864 | 5.19 |
| Ifnar1 | 2240.17 | -0.0229 | 0.06411 | -0.35716 | 0.72097 | 0.85102 | 22.204 |
| Ifnar2 | 466.271 | -0.17675 | 0.12013 | -1.47127 | 0.14122 | 0.32297 | 32.807 |
| Ifngr1 | 964.94 | -0.07374 | 0.11497 | -0.64137 | 0.52128 | 0.71042 | 18.281 |
| Ifngr2 | 1347.42 | -0.1301 | 0.08148 | -1.59678 | 0.11031 | 0.27686 | 18.552 |
| Ifnlr1 | 21.0176 | -0.1354 | 0.39367 | -0.34394 | 0.73089 | 0.85734 | 21.895 |
| Ifrd1 | 1287.74 | -0.19782 | 0.0762 | -2.59616 | 0.00943 | 0.05522 | 46.938 |

| | | | | | | | |
|---|---|---|---|---|---|---|---|
| Ifrd2 | 294.952 | -0.06389 | 0.11939 | -0.53508 | 0.59259 | 0.7651 | 5.744 |
| Ift122 | 1602.11 | 0.041968 | 0.07751 | 0.541473 | 0.58818 | 0.76158 | 73.23 |
| Ift140 | 1630.21 | 0.027047 | 0.12031 | 0.224806 | 0.82213 | 0.90785 | 83.411 |
| Ift172 | 4126.45 | -0.01213 | 0.06635 | -0.18289 | 0.85488 | 0.92662 | 37.84 |
| Ift20 | 1399.3 | -0.0801 | 0.07116 | -1.12565 | 0.26031 | 0.46882 | 5.377 |
| Ift22 | 1859.73 | 0.029246 | 0.07096 | 0.412131 | 0.68024 | 0.82502 | 7.255 |
| Ift27 | 809.139 | 0.029762 | 0.089 | 0.334405 | 0.73807 | 0.86177 | 14.645 |
| Ift43 | 352.52 | -0.2745 | 0.11997 | -2.28797 | 0.02214 | 0.09589 | 79.919 |
| Ift46 | 1301.49 | -0.01635 | 0.09205 | -0.1776 | 0.85904 | 0.9288 | 25.54 |
| Ift52 | 1480.24 | 0.037989 | 0.09657 | 0.393376 | 0.69404 | 0.83411 | 28.788 |
| Ift57 | 1305.61 | 0.021452 | 0.07299 | 0.293905 | 0.76883 | 0.87935 | 65.894 |
| Ift74 | 815.405 | 0.264263 | 0.15243 | 1.733669 | 0.08298 | 0.22987 | 78.739 |
| Ift80 | 1787.29 | -0.03994 | 0.09884 | -0.40413 | 0.68612 | 0.82887 | 112.072 |
| Ift81 | 1243.16 | 0.036488 | 0.10955 | 0.33307 | 0.73908 | 0.86241 | 64.315 |
| Ift88 | 900.336 | 0.155369 | 0.10583 | 1.468123 | 0.14207 | 0.32393 | 93.875 |
| Ift88os | 48.3074 | 0.250455 | 0.25385 | 0.986638 | 0.32382 | 0.53482 | 4.092 |
| Iftap | 807.329 | 0.13402 | 0.10074 | 1.330357 | 0.1834 | 0.37747 | 88.752 |
| Igbp1 | 1592.5 | 0.008713 | 0.07219 | 0.120686 | 0.90394 | 0.95146 | 21.835 |
| Igbp1b | 19.1682 | 0.633237 | 0.40688 | 1.55631 | 0.11963 | 0.29202 | 1.456 |
| Igdcc3 | 108.394 | -0.13372 | 0.18958 | -0.70536 | 0.48059 | 0.6779 | 44.684 |
| Igdcc4 | 2258.18 | -0.20605 | 0.12378 | -1.66466 | 0.09598 | 0.25359 | 36.455 |
| Igf1 | 355.04 | -0.45644 | 0.17843 | -2.55809 | 0.01052 | 0.05954 | 78.778 |
| Igf1r | 5400.62 | -0.12252 | 0.07684 | -1.59447 | 0.11083 | 0.27785 | 280.842 |
| Igf2 | 44.875 | -1.14518 | 0.47608 | -2.40543 | 0.01615 | 0.07816 | 16.051 |
| Igf2bp2 | 160.576 | -0.51809 | 0.16654 | -3.11086 | 0.00187 | 0.01917 | 104.291 |
| Igf2bp3 | 548.301 | 0.193976 | 0.1149 | 1.688226 | 0.09137 | 0.24564 | 129.735 |
| Igf2r | 2841.22 | 0.205836 | 0.06586 | 3.125369 | 0.00178 | 0.01864 | 87.259 |
| Igfbp2 | 1050.17 | -1.02019 | 0.22401 | -4.55418 | 5.26E-06 | 0.00035 | 27.972 |
| Igfbp3 | 580.268 | -0.27955 | 0.12691 | -2.20275 | 0.02761 | 0.11146 | 7.838 |
| Igfbp4 | 2090.95 | -0.23046 | 0.13381 | -1.72223 | 0.08503 | 0.23388 | 13.149 |
| Igfbp5 | 6638.14 | -0.66158 | 0.27782 | -2.38131 | 0.01725 | 0.08154 | 16.953 |
| Igfbp6 | 556.349 | -1.01778 | 0.14317 | -7.10889 | 1.17E-12 | 3.72E-09 | 5.15 |
| Igfbp7 | 522.893 | -0.65927 | 0.11853 | -5.56208 | 2.67E-08 | 7.44E-06 | 58.802 |
| Igfbpl1 | 138.62 | -0.36303 | 0.3059 | -1.18674 | 0.23533 | 0.44049 | 17.456 |

| | | | | | | | |
|---|---|---|---|---|---|---|---|
| Igfn1 | 441.903 | -2 | 0.57876 | -4.23081 | 2.33E-05 | 0.00097 | 52.765 |
| Igha | 451.118 | -2 | 1.23258 | -2.53483 | 0.01125 | 0.06195 | 5.407 |
| Ighg2c | 695.85 | -2 | 1.31414 | -3.11024 | 0.00187 | 0.0192 | 3.608 |
| Ighg3 | 45.4993 | -2 | 0.60704 | -6.83838 | 8.01E-12 | 1.17E-08 | 5.009 |
| Ighm | 924.798 | -1.18446 | 0.51332 | -2.30745 | 0.02103 | 0.09263 | 4.173 |
| Ighmbp2 | 586.257 | -0.0119 | 0.09356 | -0.12724 | 0.89875 | 0.94928 | 23.942 |
| Ighv1-26 | 99.4121 | -2 | 1.8782 | -1.38423 | 0.16629 | 0.35598 | 0.434 |
| Ighv1-53 | 23.9249 | -2 | 1.66636 | -2.29646 | 0.02165 | 0.09443 | 0.433 |
| Ighv1-55 | 13.1658 | -2 | 1.17315 | -4.10586 | 4.03E-05 | 0.0014 | 0.497 |
| Ighv1-61 | 24.5929 | -2 | 2.34118 | -1.03685 | 0.29981 | 0.50995 | 0.433 |
| Ighv1-78 | 12.8004 | -2 | 2.64031 | -1.62803 | 0.10352 | 0.26606 | 0.455 |
| Ighv1-82 | 15.7361 | -2 | 1.74337 | -4.33875 | 1.43E-05 | 0.00072 | 0.468 |
| Ighv1-85 | 111.561 | -2 | 2.10956 | -2.6516 | 0.00801 | 0.04964 | 0.434 |
| Ighv2-9-1 | 221.054 | -2 | 2.89364 | -1.4934 | 0.13533 | 0.31509 | 0.453 |
| Ighv5-6 | 91.1905 | -2 | 2.55108 | -1.25746 | 0.20859 | 0.40848 | 0.451 |
| Igip | 6087.11 | 0.189534 | 0.14931 | 1.269404 | 0.2043 | 0.40307 | 2.839 |
| Igkc | 1648.6 | -1.54609 | 1.35229 | -1.14331 | 0.25291 | 0.46033 | 0.532 |
| Igkv1-117 | 32.1288 | -2 | 2.01798 | -2.98705 | 0.00282 | 0.02517 | 0.782 |
| Igkv13-84 | 34.0369 | -2 | 2.22809 | -1.50415 | 0.13254 | 0.31166 | 0.466 |
| Igkv13-85 | 24.2819 | -2 | 1.72977 | -1.80015 | 0.07184 | 0.2103 | 0.466 |
| Igkv17-121 | 280.635 | -2 | 1.92942 | -2.0345 | 0.0419 | 0.1472 | 0.507 |
| Igkv3-2 | 400.898 | -2 | 2.48824 | -1.63693 | 0.10165 | 0.26311 | 0.619 |
| Igkv5-39 | 514.813 | -1.59076 | 1.5322 | -1.03823 | 0.29916 | 0.50926 | 0.554 |
| Igkv5-43 | 49.4549 | -2 | 1.85323 | -3.22577 | 0.00126 | 0.01474 | 0.589 |
| Igkv6-25 | 45.5154 | 0.515158 | 1.8476 | 0.278826 | 0.78038 | 0.8856 | 0.558 |
| Igkv8-27 | 74.6566 | -2 | 1.96116 | -4.26327 | 2.01E-05 | 0.00089 | 0.462 |
| Igkv9-120 | 24.74 | -2 | 1.68506 | -4.44378 | 8.84E-06 | 0.00051 | 0.474 |
| Iglon5 | 1216.68 | -0.29886 | 0.14376 | -2.07888 | 0.03763 | 0.13733 | 17.172 |
| Iglv1 | 16.544 | -2 | 2.75384 | -0.73254 | 0.46384 | 0.66359 | 0.484 |
| Igsf1 | 157.205 | 1.141227 | 0.35437 | 3.220435 | 0.00128 | 0.01485 | 15.214 |
| Igsf10 | 162.002 | -0.04533 | 0.16258 | -0.27883 | 0.78037 | 0.8856 | 27.66 |
| Igsf11 | 2524.78 | -0.34524 | 0.08659 | -3.98697 | 6.69E-05 | 0.00204 | 134.489 |
| Igsf21 | 1680.67 | -0.68499 | 0.169 | -4.05317 | 5.05E-05 | 0.00165 | 219.939 |
| Igsf3 | 1237.94 | -0.07909 | 0.11957 | -0.66144 | 0.50833 | 0.69969 | 85.977 |

| | | | | | | | |
|---|---|---|---|---|---|---|---|
| Igsf6 | 167.834 | 0.380102 | 0.17217 | 2.207677 | 0.02727 | 0.1103 | 10.506 |
| Igsf8 | 6315.07 | 0.064697 | 0.07386 | 0.875965 | 0.38105 | 0.59053 | 58.201 |
| Igsf9 | 508.544 | 0.455622 | 0.12948 | 3.518943 | 0.00043 | 0.00726 | 17.091 |
| Igsf9b | 8638.31 | -0.08435 | 0.09153 | -0.92152 | 0.35678 | 0.56706 | 58.343 |
| Igtp | 130.969 | -0.74767 | 0.25564 | -2.92466 | 0.00345 | 0.02844 | 8.036 |
| Ihh | 17.3814 | -0.64354 | 0.54778 | -1.17483 | 0.24006 | 0.44612 | 6.358 |
| Iho1 | 20.8055 | -0.77274 | 0.45375 | -1.70301 | 0.08857 | 0.24049 | 24.874 |
| Iigp1 | 72.6174 | -0.86614 | 0.4616 | -1.87638 | 0.0606 | 0.18787 | 16.608 |
| Ik | 5134.11 | 0.088551 | 0.06851 | 1.292478 | 0.19619 | 0.39377 | 12.984 |
| Ikbip | 685.486 | 0.072404 | 0.11241 | 0.64413 | 0.51949 | 0.70922 | 19.668 |
| Ikbkb | 3047.47 | 0.028156 | 0.06313 | 0.446032 | 0.65557 | 0.80877 | 47.378 |
| Ikbke | 116.118 | -0.25928 | 0.17503 | -1.48134 | 0.13852 | 0.31964 | 25.265 |
| Ikbkg | 2209.11 | 0.005599 | 0.07205 | 0.077708 | 0.93806 | 0.96915 | 60.565 |
| Ikzf1 | 292.976 | -0.35204 | 0.17465 | -2.01565 | 0.04384 | 0.15156 | 87.947 |
| Ikzf2 | 325.663 | -0.17304 | 0.1344 | -1.28748 | 0.19793 | 0.3957 | 156.032 |
| Ikzf3 | 85.3029 | 0.138624 | 0.2705 | 0.512477 | 0.60832 | 0.77621 | 81.13 |
| Ikzf4 | 952.059 | 0.230469 | 0.11687 | 1.972008 | 0.04861 | 0.16286 | 38.516 |
| Ikzf5 | 135.709 | 0.04953 | 0.16224 | 0.305282 | 0.76015 | 0.87418 | 21.87 |
| Il10ra | 308.809 | -0.19695 | 0.14259 | -1.38125 | 0.1672 | 0.35697 | 15.313 |
| Il10rb | 126.791 | -0.62586 | 0.17091 | -3.66203 | 0.00025 | 0.00496 | 19.671 |
| Il11ra1 | 1286.14 | -0.05166 | 0.07235 | -0.71405 | 0.4752 | 0.67328 | 69.486 |
| Il12a | 32.9947 | -1.06012 | 0.43369 | -2.44443 | 0.01451 | 0.07279 | 7.904 |
| Il12rb1 | 87.1876 | -0.79306 | 0.20223 | -3.92156 | 8.80E-05 | 0.00247 | 12.976 |
| Il12rb2 | 43.0247 | -0.4743 | 0.28174 | -1.68347 | 0.09229 | 0.24739 | 84.871 |
| Il13ra1 | 474.084 | -0.03218 | 0.13735 | -0.23431 | 0.81474 | 0.90481 | 59.15 |
| Il13ra2 | 38.6288 | -0.7968 | 0.51013 | -1.56195 | 0.1183 | 0.28987 | 45.717 |
| Il15 | 12.5522 | -1.19771 | 0.51364 | -2.3318 | 0.01971 | 0.0887 | 71.591 |
| Il15ra | 277.556 | -0.00913 | 0.13477 | -0.06774 | 0.94599 | 0.973 | 29.028 |
| Il16 | 143.81 | 0.376937 | 0.18816 | 2.003298 | 0.04515 | 0.15451 | 102.902 |
| Il17d | 646.443 | -0.18585 | 0.11591 | -1.60344 | 0.10884 | 0.27431 | 18.39 |
| Il17ra | 655.301 | -0.55933 | 0.15974 | -3.50142 | 0.00046 | 0.00757 | 24.313 |
| Il17rb | 115.803 | -0.54412 | 0.21157 | -2.57185 | 0.01012 | 0.05794 | 12.762 |
| Il17rc | 103.162 | -0.36195 | 0.18742 | -1.93117 | 0.05346 | 0.17327 | 11.714 |
| Il17rd | 291.855 | -0.32842 | 0.17722 | -1.85321 | 0.06385 | 0.19435 | 68.346 |

| | | | | | | | |
|---|---|---|---|---|---|---|---|
| Il17re | 12.9713 | 0.15635 | 0.51668 | 0.302606 | 0.76219 | 0.8753 | 12.275 |
| Il18 | 1019.51 | 0.111743 | 0.10083 | 1.108193 | 0.26778 | 0.47712 | 27.014 |
| Il18bp | 662.707 | 0.304092 | 0.08252 | 3.684859 | 0.00023 | 0.00466 | 3.806 |
| Il1a | 23.1121 | -0.80215 | 0.36629 | -2.18994 | 0.02853 | 0.11395 | 10.363 |
| Il1r1 | 482.423 | -0.55087 | 0.15613 | -3.52826 | 0.00042 | 0.00709 | 92.178 |
| Il1r2 | 31.3663 | 0.18241 | 0.30454 | 0.59897 | 0.54919 | 0.73299 | 51.153 |
| Il1rap | 1241.63 | -0.35961 | 0.11009 | -3.26643 | 0.00109 | 0.01351 | 148.414 |
| Il1rapl1 | 1136.58 | 0.15958 | 0.14748 | 1.082073 | 0.27922 | 0.48914 | 1374.776 |
| Il1rapl2 | 93.7917 | 0.594791 | 0.2697 | 2.205407 | 0.02743 | 0.11083 | 1276.339 |
| Il1rl2 | 78.4471 | -0.35557 | 0.26196 | -1.35734 | 0.17467 | 0.36622 | 42.953 |
| Il20ra | 178.474 | 0.69609 | 0.33907 | 2.052948 | 0.04008 | 0.14302 | 47.484 |
| Il20rb | 23.2361 | 0.317856 | 0.43707 | 0.727243 | 0.46708 | 0.66622 | 29.07 |
| Il21r | 57.7697 | -0.70379 | 0.28707 | -2.45159 | 0.01422 | 0.07191 | 30.142 |
| Il27ra | 27.826 | 0.052618 | 0.33435 | 0.157372 | 0.87495 | 0.9377 | 12.296 |
| Il2ra | 15.3924 | -0.64478 | 0.46134 | -1.39763 | 0.16222 | 0.35062 | 50.387 |
| Il2rb | 14.1331 | -0.9164 | 0.53643 | -1.70835 | 0.08757 | 0.23845 | 16.016 |
| Il2rg | 16.1643 | -0.65882 | 0.56184 | -1.17262 | 0.24095 | 0.4471 | 3.878 |
| Il31ra | 12.7681 | 0.132585 | 0.75456 | 0.175713 | 0.86052 | 0.92938 | 74.463 |
| Il33 | 2828.21 | -0.17405 | 0.13591 | -1.28057 | 0.20034 | 0.39838 | 35.605 |
| Il34 | 2092.42 | -0.06371 | 0.08081 | -0.78839 | 0.43047 | 0.63681 | 64.096 |
| Il3ra | 43.306 | -0.22053 | 0.31671 | -0.69631 | 0.48624 | 0.68228 | 9.737 |
| Il4 | 622.232 | 0.069954 | 0.12303 | 0.568598 | 0.56963 | 0.74912 | 15.688 |
| Il4ra | 157.703 | -0.48209 | 0.19636 | -2.45513 | 0.01408 | 0.07147 | 27.355 |
| Il6ra | 414.651 | -0.38802 | 0.14159 | -2.74045 | 0.00614 | 0.0419 | 49.138 |
| Il6st | 5370.71 | -0.17131 | 0.06831 | -2.50769 | 0.01215 | 0.06511 | 46.017 |
| Il7r | 89.5606 | -0.0711 | 0.21993 | -0.32328 | 0.74648 | 0.86693 | 24.389 |
| Ildr2 | 6445.69 | -0.25845 | 0.12096 | -2.13666 | 0.03263 | 0.12524 | 62.685 |
| Ilf2 | 3917.14 | 0.025662 | 0.06686 | 0.383794 | 0.70113 | 0.83856 | 12.254 |
| Ilf3 | 5100.02 | 0.188859 | 0.07396 | 2.553663 | 0.01066 | 0.05992 | 37.491 |
| Ilkap | 1900.6 | 0.081904 | 0.07515 | 1.089826 | 0.27579 | 0.48496 | 24.955 |
| Ilrun | 11760.8 | -0.25225 | 0.06424 | -3.9266 | 8.62E-05 | 0.00244 | 69.414 |
| Ilvbl | 895.268 | -0.16125 | 0.08207 | -1.96487 | 0.04943 | 0.16463 | 10.157 |
| Immp1l | 687.015 | -0.06962 | 0.13364 | -0.52094 | 0.60241 | 0.7723 | 60.921 |
| Immp2l | 92.824 | 0.012281 | 0.23157 | 0.053032 | 0.95771 | 0.97936 | 928.283 |

| | | | | | | | |
|---|---|---|---|---|---|---|---|
| Immt | 4699.34 | -0.01507 | 0.06367 | -0.23662 | 0.81295 | 0.90371 | 46.058 |
| Imp3 | 771.143 | 0.042005 | 0.09492 | 0.442514 | 0.65812 | 0.81023 | 0.924 |
| Imp4 | 1361.6 | -0.03069 | 0.07865 | -0.39027 | 0.69634 | 0.83541 | 9.506 |
| Impa1 | 2803.42 | 0.05175 | 0.07515 | 0.68861 | 0.49107 | 0.68604 | 19.484 |
| Impa2 | 456.315 | 0.085139 | 0.15375 | 0.553738 | 0.57976 | 0.75629 | 33.287 |
| Impact | 14145.9 | 0.203788 | 0.08952 | 2.276398 | 0.02282 | 0.09775 | 37.099 |
| Impdh1 | 2210.27 | 0.11723 | 0.08607 | 1.362075 | 0.17317 | 0.3644 | 15.931 |
| Impdh2 | 1422.91 | 0.054031 | 0.09343 | 0.578319 | 0.56305 | 0.74368 | 5.299 |
| Impdh2-ps | 458.36 | -0.14831 | 0.09766 | -1.51867 | 0.12885 | 0.30601 | 1.545 |
| Impg1 | 401.899 | 0.372199 | 0.38417 | 0.968838 | 0.33263 | 0.54346 | 126.923 |
| Impg2 | 122.773 | 0.065573 | 0.26032 | 0.251893 | 0.80112 | 0.8972 | 69.444 |
| Ina | 7245.02 | -0.26423 | 0.07639 | -3.45912 | 0.00054 | 0.00838 | 10.63 |
| Inafm1 | 1877.37 | -0.00774 | 0.08825 | -0.08774 | 0.93008 | 0.96478 | 1.605 |
| Inafm2 | 2305.55 | -0.10216 | 0.07409 | -1.37891 | 0.16792 | 0.35782 | 3.053 |
| Inava | 95.8397 | -0.66514 | 0.25354 | -2.62343 | 0.00871 | 0.05236 | 20.734 |
| Inca1 | 110.526 | 0.064813 | 0.24497 | 0.26458 | 0.79133 | 0.89159 | 11.795 |
| Incenp | 823.247 | -0.34607 | 0.12698 | -2.72539 | 0.00642 | 0.04301 | 27.256 |
| Inf2 | 19036.8 | 0.406751 | 0.15918 | 2.555289 | 0.01061 | 0.05981 | 26.774 |
| Ing1 | 1441.1 | -0.28128 | 0.0876 | -3.21093 | 0.00132 | 0.01516 | 7.681 |
| Ing2 | 967.827 | -0.02632 | 0.07664 | -0.34342 | 0.73128 | 0.85734 | 8.379 |
| Ing3 | 936.631 | -0.03227 | 0.09023 | -0.35763 | 0.72062 | 0.85095 | 26.468 |
| Ing4 | 116.519 | 0.170641 | 0.18385 | 0.928176 | 0.35332 | 0.56354 | 9.506 |
| Ing5 | 930.613 | 0.062448 | 0.10181 | 0.613388 | 0.53962 | 0.72589 | 18.137 |
| Inha | 2006.68 | 0.281253 | 0.06533 | 4.304832 | 1.67E-05 | 0.00079 | 3.29 |
| Inhba | 1509.91 | -0.05213 | 0.19685 | -0.26484 | 0.79113 | 0.89159 | 19.771 |
| Inhbb | 175.135 | -0.5271 | 0.24349 | -2.16475 | 0.03041 | 0.1191 | 6.784 |
| Inhca | 17.3475 | -0.20595 | 0.39677 | -0.51907 | 0.60371 | 0.77319 | 54.562 |
| Inip | 1222.91 | -0.09918 | 0.07181 | -1.38117 | 0.16723 | 0.35697 | 31.953 |
| Inka1 | 38.6969 | -0.44198 | 0.31581 | -1.39949 | 0.16167 | 0.34987 | 1.657 |
| Inka2 | 5348.27 | -0.14346 | 0.10447 | -1.3732 | 0.16969 | 0.36001 | 16.244 |
| Inmt | 55.0016 | -0.07895 | 0.32376 | -0.24386 | 0.80734 | 0.90076 | 4.418 |
| Ino80 | 1952.53 | 0.211181 | 0.06585 | 3.207114 | 0.00134 | 0.01532 | 104.646 |
| Ino80b | 284.356 | -0.08817 | 0.12294 | -0.71719 | 0.47326 | 0.6716 | 3.667 |
| Ino80c | 831.86 | -0.03466 | 0.10973 | -0.31591 | 0.75207 | 0.86995 | 17.197 |

| | | | | | | | |
|---|---|---|---|---|---|---|---|
| Ino80d | 3741.71 | 0.192947 | 0.07322 | 2.635313 | 0.00841 | 0.05114 | 156.273 |
| Ino80dos | 295.521 | 0.368393 | 0.13259 | 2.778453 | 0.00546 | 0.03854 | 45.216 |
| Ino80e | 1234.74 | 0.136638 | 0.07833 | 1.74429 | 0.08111 | 0.22667 | 11.418 |
| Inpp1 | 1740.09 | 0.004219 | 0.07932 | 0.053182 | 0.95759 | 0.97935 | 32.262 |
| Inpp4a | 8075.9 | 0.11068 | 0.05662 | 1.954722 | 0.05062 | 0.16736 | 110.872 |
| Inpp4b | 703.21 | 0.174229 | 0.14445 | 1.206128 | 0.22777 | 0.43174 | 785.359 |
| Inpp5a | 2465.92 | -0.06322 | 0.0939 | -0.67334 | 0.50073 | 0.69355 | 190.544 |
| Inpp5b | 1964.55 | -0.01019 | 0.09181 | -0.11104 | 0.91158 | 0.95536 | 59.662 |
| Inpp5d | 807.774 | -0.12652 | 0.101 | -1.25266 | 0.21033 | 0.41035 | 100.196 |
| Inpp5e | 1298.57 | 0.130742 | 0.0716 | 1.825919 | 0.06786 | 0.20204 | 12.955 |
| Inpp5f | 6191.44 | 0.307537 | 0.10342 | 2.973802 | 0.00294 | 0.02585 | 85.098 |
| Inpp5j | 3680.5 | -0.17744 | 0.15067 | -1.17766 | 0.23893 | 0.44474 | 10.447 |
| Inpp5k | 2163.85 | -0.08328 | 0.06977 | -1.19353 | 0.23266 | 0.43751 | 17.884 |
| Inppl1 | 1598.46 | -0.08458 | 0.08125 | -1.04101 | 0.29787 | 0.50794 | 15.598 |
| Insc | 90.5202 | -0.67354 | 0.24109 | -2.79376 | 0.00521 | 0.03725 | 106.69 |
| Insig1 | 2626.71 | -0.50147 | 0.11346 | -4.41995 | 9.87E-06 | 0.00055 | 7.3 |
| Insig2 | 2213.28 | -0.01405 | 0.07282 | -0.19293 | 0.84702 | 0.92206 | 28.237 |
| Insl5 | 60.853 | 0.20254 | 0.2248 | 0.901 | 0.36759 | 0.57807 | 8.971 |
| Insl6 | 46.3187 | 0.33317 | 0.3755 | 0.88727 | 0.37493 | 0.58517 | 4.013 |
| Insm1 | 556.214 | 0.634263 | 0.18189 | 3.487147 | 0.00049 | 0.00779 | 3.1 |
| Insr | 4704.2 | 0.064495 | 0.07892 | 0.817254 | 0.41378 | 0.62188 | 157.557 |
| Insrr | 88.8709 | 0.453449 | 0.30548 | 1.484361 | 0.13771 | 0.31856 | 19.151 |
| Insyn1 | 5846.13 | -0.24558 | 0.08754 | -2.80529 | 0.00503 | 0.03641 | 11.178 |
| Insyn2a | 528.876 | 0.459251 | 0.18747 | 2.44969 | 0.0143 | 0.07211 | 56.505 |
| Insyn2b | 413.822 | 0.533551 | 0.14216 | 3.753042 | 0.00017 | 0.00393 | 107.883 |
| Ints1 | 5047.11 | 0.032199 | 0.06571 | 0.489979 | 0.62415 | 0.78681 | 24.393 |
| Ints10 | 4279.1 | 0.211011 | 0.08634 | 2.444028 | 0.01452 | 0.07283 | 37.739 |
| Ints11 | 787.671 | 0.010346 | 0.10115 | 0.102282 | 0.91853 | 0.9587 | 19.558 |
| Ints12 | 458.774 | 0.008264 | 0.10053 | 0.082202 | 0.93449 | 0.96713 | 19.149 |
| Ints13 | 2217.5 | 0.24247 | 0.07725 | 3.138637 | 0.0017 | 0.01811 | 28.204 |
| Ints14 | 1021.98 | 0.174962 | 0.0758 | 2.308156 | 0.02099 | 0.09251 | 26.074 |
| Ints2 | 1637.57 | 0.260021 | 0.07604 | 3.419699 | 0.00063 | 0.00924 | 46.895 |
| Ints3 | 3575.04 | 0.068444 | 0.06593 | 1.038152 | 0.2992 | 0.50926 | 42.235 |
| Ints4 | 1622 | 0.086068 | 0.06777 | 1.270002 | 0.20408 | 0.40279 | 60.44 |

| | | | | | | | |
|---|---|---|---|---|---|---|---|
| Ints5 | 704.862 | 0.053224 | 0.1 | 0.532217 | 0.59458 | 0.76667 | 4.884 |
| Ints6 | 1740.74 | 0.092604 | 0.09992 | 0.926793 | 0.35403 | 0.56424 | 84.84 |
| Ints6l | 5386.73 | 0.083421 | 0.15212 | 0.548396 | 0.58342 | 0.75845 | 52.987 |
| Ints7 | 3105.09 | 0.133235 | 0.07559 | 1.762655 | 0.07796 | 0.22102 | 48.053 |
| Ints8 | 1477.41 | 0.143838 | 0.08915 | 1.613446 | 0.10665 | 0.27089 | 55.101 |
| Ints9 | 836.908 | 0.161035 | 0.0984 | 1.636494 | 0.10174 | 0.2632 | 89.791 |
| Intu | 824.572 | -0.06546 | 0.10777 | -0.60739 | 0.54359 | 0.72894 | 73.77 |
| Invs | 995.674 | 0.330176 | 0.11564 | 2.855311 | 0.0043 | 0.03307 | 152.195 |
| Ip6k1 | 9889.59 | 0.115157 | 0.0608 | 1.894156 | 0.0582 | 0.18329 | 46.282 |
| Ip6k2 | 4365.11 | 0.148048 | 0.0793 | 1.866869 | 0.06192 | 0.19052 | 22.542 |
| Ip6k3 | 28.2248 | 0.021893 | 0.36783 | 0.059519 | 0.95254 | 0.97669 | 23.831 |
| Ipcef1 | 3026.12 | -0.70709 | 0.39918 | -1.77136 | 0.0765 | 0.21863 | 166.683 |
| Ipmk | 1906.47 | -0.0234 | 0.07898 | -0.29631 | 0.767 | 0.87819 | 85.565 |
| Ipo11 | 2221.42 | -0.01442 | 0.06743 | -0.21381 | 0.83069 | 0.91321 | 142.52 |
| Ipo13 | 4362.49 | -0.01037 | 0.06996 | -0.14817 | 0.88221 | 0.94146 | 20.514 |
| Ipo4 | 1323.41 | 0.054843 | 0.08385 | 0.654065 | 0.51307 | 0.70394 | 10.558 |
| Ipo5 | 5790.76 | -0.12027 | 0.1016 | -1.18376 | 0.23651 | 0.44173 | 36.776 |
| Ipo7 | 8034.72 | 0.148363 | 0.09299 | 1.595551 | 0.11059 | 0.27734 | 38.336 |
| Ipo8 | 3607.91 | 0.090367 | 0.06531 | 1.383614 | 0.16648 | 0.356 | 60.785 |
| Ipo9 | 8686.41 | 0.005008 | 0.07123 | 0.0703 | 0.94396 | 0.97194 | 48.188 |
| Ipp | 667.98 | -0.01445 | 0.12157 | -0.11884 | 0.9054 | 0.95246 | 30.695 |
| Ippk | 1357.24 | 0.313459 | 0.12786 | 2.451548 | 0.01422 | 0.07191 | 43.325 |
| Iqank1 | 58.8461 | -0.65574 | 0.24681 | -2.65681 | 0.00789 | 0.04922 | 37.094 |
| Iqcb1 | 1378.32 | 0.257359 | 0.08073 | 3.187996 | 0.00143 | 0.01602 | 44.337 |
| Iqcc | 585.566 | -0.1684 | 0.16169 | -1.04151 | 0.29764 | 0.50774 | 3.974 |
| Iqcd | 83.6491 | 0.053069 | 0.23437 | 0.226436 | 0.82086 | 0.90745 | 18.103 |
| Iqce | 2837.53 | 0.171141 | 0.07686 | 2.226671 | 0.02597 | 0.10679 | 40.552 |
| Iqcg | 80.5259 | 0.091021 | 0.20979 | 0.433878 | 0.66438 | 0.8145 | 43.536 |
| Iqch | 19.9334 | 0.15439 | 0.39154 | 0.394311 | 0.69335 | 0.83377 | 181.039 |
| Iqck | 224.983 | 0.168763 | 0.16615 | 1.015704 | 0.30977 | 0.51979 | 116.901 |
| Iqcn | 16.05 | 0.151403 | 0.51221 | 0.295585 | 0.76755 | 0.87843 | 14.194 |
| Iqgap1 | 1604.36 | -0.32391 | 0.1158 | -2.79718 | 0.00516 | 0.03701 | 114.392 |
| Iqgap2 | 1031.7 | 0.339614 | 0.11966 | 2.838262 | 0.00454 | 0.0341 | 265.31 |
| Iqgap3 | 42.7242 | -0.21334 | 0.34264 | -0.62262 | 0.53354 | 0.72072 | 39.078 |

| | | | | | | | |
|---|---|---|---|---|---|---|---|
| Iqsec1 | 8448.85 | -0.11557 | 0.05935 | -1.94714 | 0.05152 | 0.16924 | 332.598 |
| Iqsec2 | 7446 | -0.04884 | 0.07418 | -0.65843 | 0.51026 | 0.70169 | 81.04 |
| Iqsec3 | 9264.28 | -0.19994 | 0.08188 | -2.44193 | 0.01461 | 0.07303 | 100.746 |
| Iqub | 145.625 | 0.360169 | 0.2094 | 1.720014 | 0.08543 | 0.23453 | 70.203 |
| Irag1 | 127.539 | -0.3645 | 0.23574 | -1.54616 | 0.12207 | 0.29547 | 114.196 |
| Irag2 | 154.961 | -1.08502 | 0.26552 | -4.0864 | 4.38E-05 | 0.00149 | 59.282 |
| Irak1 | 2560.27 | 0.108683 | 0.08302 | 1.309173 | 0.19048 | 0.3867 | 10.005 |
| Irak1bp1 | 866.487 | 0.275322 | 0.11514 | 2.39125 | 0.01679 | 0.08018 | 18.148 |
| Irak2 | 1183.8 | -0.06731 | 0.10278 | -0.65495 | 0.5125 | 0.70349 | 56.56 |
| Irak3 | 107.927 | 0.196419 | 0.22323 | 0.879908 | 0.37891 | 0.58861 | 60.483 |
| Irak4 | 124.996 | -0.52356 | 0.19731 | -2.6535 | 0.00797 | 0.04945 | 38.173 |
| Ireb2 | 4266.52 | 0.180112 | 0.06784 | 2.655032 | 0.00793 | 0.04932 | 48.742 |
| Irf1 | 147.79 | -0.46073 | 0.17631 | -2.61321 | 0.00897 | 0.05332 | 8.36 |
| Irf2 | 1133.68 | -0.07788 | 0.09203 | -0.84623 | 0.39742 | 0.60658 | 107.727 |
| Irf2bp1 | 2067.45 | 0.081798 | 0.08213 | 0.995954 | 0.31927 | 0.52984 | 2.734 |
| Irf2bp2 | 1390.66 | -0.03144 | 0.08682 | -0.36217 | 0.71723 | 0.84901 | 5.691 |
| Irf2bpl | 3588.07 | -0.06072 | 0.07108 | -0.85417 | 0.39301 | 0.60225 | 4.098 |
| Irf3 | 992.149 | -0.02105 | 0.09627 | -0.21861 | 0.82695 | 0.91102 | 5.201 |
| Irf4 | 40.7813 | -0.44579 | 0.3735 | -1.19356 | 0.23265 | 0.43751 | 17.751 |
| Irf5 | 156.674 | -0.36575 | 0.17626 | -2.07504 | 0.03798 | 0.13828 | 15.247 |
| Irf6 | 53.4955 | -0.56565 | 0.32501 | -1.74038 | 0.08179 | 0.22795 | 18.913 |
| Irf7 | 82.1559 | -0.54961 | 0.22376 | -2.45624 | 0.01404 | 0.07134 | 3.776 |
| Irf8 | 256.412 | -0.61511 | 0.20182 | -3.04784 | 0.0023 | 0.02207 | 20.337 |
| Irf9 | 890.387 | -0.09312 | 0.0866 | -1.07537 | 0.28221 | 0.49239 | 6.46 |
| Irgm1 | 375.594 | -0.27621 | 0.14655 | -1.88477 | 0.05946 | 0.1855 | 9.716 |
| Irgm2 | 199.966 | -0.59743 | 0.2095 | -2.85173 | 0.00435 | 0.0333 | 23.165 |
| Irgq | 4596.32 | -0.18521 | 0.06942 | -2.66791 | 0.00763 | 0.04827 | 7.912 |
| Irs1 | 424.214 | -0.32231 | 0.13176 | -2.44626 | 0.01443 | 0.07263 | 58.316 |
| Irs2 | 3713.5 | 0.069649 | 0.11179 | 0.623016 | 0.53327 | 0.7205 | 23.778 |
| Irs4 | 36.4472 | 0.841145 | 0.59902 | 1.404211 | 0.16026 | 0.34837 | 14.266 |
| Irx1 | 16.8335 | -0.48171 | 0.46796 | -1.0294 | 0.30329 | 0.51297 | 5.803 |
| Irx6 | 28.314 | 0.890546 | 0.55997 | 1.590343 | 0.11176 | 0.27929 | 6.669 |
| Isca1 | 5422.46 | 0.071697 | 0.07442 | 0.963358 | 0.33537 | 0.54619 | 14.402 |
| Isca2 | 1153.33 | 0.014405 | 0.08545 | 0.168569 | 0.86614 | 0.93222 | 1.838 |

| | | | | | | | |
|---|---|---|---|---|---|---|---|
| Iscu | 2268.14 | -0.13872 | 0.07562 | -1.83431 | 0.06661 | 0.19978 | 5.541 |
| Isg15 | 107.215 | -0.27312 | 0.23863 | -1.14454 | 0.2524 | 0.45965 | 1.342 |
| Isg20 | 31.9666 | -0.71802 | 0.37559 | -1.91171 | 0.05591 | 0.1787 | 6.973 |
| Isg20l2 | 706.635 | 0.041722 | 0.09009 | 0.46311 | 0.64329 | 0.80102 | 10.373 |
| Isl1 | 1049.03 | 0.757725 | 0.21281 | 3.560488 | 0.00037 | 0.00648 | 11.409 |
| Islr | 652.931 | -0.9905 | 0.37235 | -2.66014 | 0.00781 | 0.04892 | 48.039 |
| Islr2 | 4434.78 | 0.130705 | 0.11901 | 1.098304 | 0.27207 | 0.48148 | 8.023 |
| Ism1 | 531.446 | -0.35583 | 0.1476 | -2.41078 | 0.01592 | 0.07743 | 80.404 |
| Isoc1 | 2048.79 | 0.412681 | 0.13606 | 3.033107 | 0.00242 | 0.02282 | 20.903 |
| Isoc2a | 898.418 | -0.43948 | 0.10359 | -4.24249 | 2.21E-05 | 0.00094 | 19.821 |
| Isoc2b | 292.237 | 0.071665 | 0.11218 | 0.63881 | 0.52295 | 0.71143 | 25.3 |
| Ist1 | 5017.29 | 0.080594 | 0.06278 | 1.283698 | 0.19925 | 0.39717 | 21.936 |
| Isy1 | 861.007 | 0.073947 | 0.09413 | 0.785545 | 0.43213 | 0.63821 | 24.53 |
| Isyna1 | 1997.89 | 0.268187 | 0.11609 | 2.310175 | 0.02088 | 0.09216 | 2.918 |
| Itch | 3789.13 | 0.039337 | 0.064 | 0.614607 | 0.53881 | 0.72544 | 93.347 |
| Itfg1 | 7994.57 | 0.096781 | 0.08983 | 1.077378 | 0.28131 | 0.49116 | 123.344 |
| Itfg2 | 963.566 | 0.030331 | 0.09426 | 0.321785 | 0.74762 | 0.86765 | 15.488 |
| Itga1 | 483.313 | -0.32792 | 0.13983 | -2.34513 | 0.01902 | 0.0866 | 148.869 |
| Itga10 | 300.986 | -0.03324 | 0.1849 | -0.17978 | 0.85732 | 0.92794 | 18.936 |
| Itga11 | 359.467 | -0.07956 | 0.13734 | -0.57933 | 0.56237 | 0.74316 | 106.157 |
| Itga2 | 17.367 | -0.24588 | 0.49143 | -0.50034 | 0.61684 | 0.78194 | 99.02 |
| Itga2b | 175.378 | 0.477089 | 0.16821 | 2.836211 | 0.00457 | 0.03426 | 16.826 |
| Itga3 | 823.878 | -0.00818 | 0.09424 | -0.08685 | 0.93079 | 0.96513 | 32.328 |
| Itga4 | 305.005 | -0.63502 | 0.42502 | -1.49411 | 0.13515 | 0.31502 | 77.698 |
| Itga5 | 399.163 | 0.081204 | 0.21227 | 0.382548 | 0.70205 | 0.83919 | 22.478 |
| Itga6 | 1158.38 | -0.28574 | 0.13257 | -2.15537 | 0.03113 | 0.12111 | 112.801 |
| Itga7 | 502.734 | -0.44267 | 0.15773 | -2.80644 | 0.00501 | 0.03635 | 24.511 |
| Itga8 | 270.051 | -0.74143 | 0.29803 | -2.48773 | 0.01286 | 0.06768 | 195.291 |
| Itga9 | 723.15 | 0.302726 | 0.14088 | 2.148778 | 0.03165 | 0.12253 | 294.314 |
| Itgad | 60.7779 | -0.54273 | 0.27945 | -1.94217 | 0.05212 | 0.17047 | 69.441 |
| Itgae | 17.8357 | -0.54772 | 0.45594 | -1.20129 | 0.22964 | 0.4339 | 56.864 |
| Itgal | 33.1618 | -0.85484 | 0.56736 | -1.50669 | 0.13189 | 0.31073 | 38.879 |
| Itgam | 329.437 | -0.36522 | 0.16417 | -2.22465 | 0.0261 | 0.10722 | 55.852 |
| Itgav | 6549.19 | -0.15112 | 0.08922 | -1.69386 | 0.09029 | 0.24381 | 82.52 |

| | | | | | | | |
|---|---|---|---|---|---|---|---|
| Itgax | 19.4943 | -1.70559 | 0.70319 | -2.42551 | 0.01529 | 0.0754 | 21.111 |
| Itgb1 | 3431.19 | -0.1979 | 0.08068 | -2.45277 | 0.01418 | 0.07176 | 47.547 |
| Itgb1bp1 | 1081.05 | 0.461205 | 0.16908 | 2.727686 | 0.00638 | 0.04285 | 45.46 |
| Itgb1bp2 | 45.944 | 0.574461 | 0.27855 | 2.062345 | 0.03917 | 0.14101 | 4.454 |
| Itgb2 | 233.548 | -0.5416 | 0.20587 | -2.63086 | 0.00852 | 0.05156 | 35.457 |
| Itgb3 | 159.985 | -0.29991 | 0.17 | -1.7642 | 0.0777 | 0.22065 | 62.477 |
| Itgb3bp | 174.013 | -0.2471 | 0.21086 | -1.17182 | 0.24127 | 0.44737 | 133.372 |
| Itgb4 | 528.662 | -0.71302 | 0.13284 | -5.3677 | 7.97E-08 | 1.48E-05 | 33.704 |
| Itgb5 | 2106.27 | -0.45878 | 0.13683 | -3.35299 | 0.0008 | 0.01096 | 119.674 |
| Itgb7 | 45.4652 | -0.11733 | 0.28006 | -0.41893 | 0.67527 | 0.82187 | 15.95 |
| Itgb8 | 4601.94 | 0.110819 | 0.10596 | 1.045894 | 0.29561 | 0.50608 | 80.781 |
| Itgbl1 | 846.526 | 0.323804 | 0.15556 | 2.081538 | 0.03738 | 0.13674 | 315.648 |
| Itih2 | 287.967 | -1.26882 | 0.53357 | -2.378 | 0.01741 | 0.08189 | 36.804 |
| Itih3 | 1005.7 | 0.448192 | 0.18228 | 2.458803 | 0.01394 | 0.07106 | 15.189 |
| Itih5 | 2741.77 | -0.31303 | 0.10713 | -2.92184 | 0.00348 | 0.02864 | 102.959 |
| Itk | 14.5592 | 0.121691 | 0.5779 | 0.210573 | 0.83322 | 0.91466 | 64.366 |
| Itm2a | 2348.77 | -0.28165 | 0.09649 | -2.91891 | 0.00351 | 0.02881 | 6.278 |
| Itm2b | 23113.6 | -0.22386 | 0.06928 | -3.23118 | 0.00123 | 0.01456 | 23.064 |
| Itm2c | 41236.8 | 0.285591 | 0.12856 | 2.221381 | 0.02633 | 0.10784 | 14.395 |
| Itpa | 1716.01 | 0.16215 | 0.09444 | 1.716899 | 0.086 | 0.23561 | 14.005 |
| Itpa-ps1 | 49.1548 | 0.30649 | 0.32027 | 0.956963 | 0.33859 | 0.54937 | 0.597 |
| Itpk1 | 2705.63 | -0.53956 | 0.09829 | -5.48953 | 4.03E-08 | 9.55E-06 | 136.349 |
| Itpka | 4275.45 | 0.087678 | 0.09569 | 0.916258 | 0.35953 | 0.56984 | 8.927 |
| Itpkb | 1276.19 | -0.26471 | 0.13764 | -1.92324 | 0.05445 | 0.17542 | 94.318 |
| Itpkc | 206.372 | -0.07138 | 0.15482 | -0.46107 | 0.64475 | 0.80187 | 21.49 |
| Itpr1 | 41325.1 | 0.137343 | 0.09246 | 1.485399 | 0.13744 | 0.3181 | 338.014 |
| Itpr2 | 1693.6 | -0.34178 | 0.08481 | -4.02983 | 5.58E-05 | 0.00179 | 393.925 |
| Itpr3 | 378.932 | -0.20605 | 0.13389 | -1.53892 | 0.12382 | 0.29835 | 64.92 |
| Itprid2 | 2799.65 | -0.18144 | 0.09758 | -1.8594 | 0.06297 | 0.19265 | 37.615 |
| Itprip | 39.2176 | -0.8834 | 0.33807 | -2.61308 | 0.00897 | 0.05332 | 24.698 |
| Itpripl1 | 108.625 | -0.42382 | 0.23346 | -1.81537 | 0.06947 | 0.2054 | 4.671 |
| Itpripl2 | 552.314 | -0.42774 | 0.11961 | -3.57614 | 0.00035 | 0.00622 | 6.865 |
| Itsn1 | 10702.4 | -0.03995 | 0.0568 | -0.70343 | 0.48179 | 0.67858 | 191.317 |
| Itsn2 | 4649.82 | 0.133321 | 0.08103 | 1.64525 | 0.09992 | 0.25998 | 121.325 |

| | | | | | | | |
|---|---|---|---|---|---|---|---|
| Ivd | 5396.99 | 0.023323 | 0.09788 | 0.238282 | 0.81166 | 0.90296 | 20.956 |
| Ivns1abp | 15992.1 | 0.339849 | 0.1059 | 3.20913 | 0.00133 | 0.01522 | 19.946 |
| Iws1 | 3397.21 | 0.190213 | 0.06163 | 3.086271 | 0.00203 | 0.02025 | 36.588 |
| Izumo4 | 1296.15 | 0.234268 | 0.10598 | 2.2104 | 0.02708 | 0.10988 | 3.19 |
| Jade1 | 3082.18 | 0.260973 | 0.08869 | 2.94262 | 0.00325 | 0.02753 | 61.134 |
| Jade2 | 5406.63 | 0.135295 | 0.069 | 1.960703 | 0.04991 | 0.16576 | 44.199 |
| Jade3 | 834.205 | 0.54629 | 0.10162 | 5.376033 | 7.61E-08 | 1.44E-05 | 94.252 |
| Jag1 | 516.483 | -0.15833 | 0.13923 | -1.13719 | 0.25546 | 0.46311 | 35.189 |
| Jag2 | 2299.66 | 0.093213 | 0.07738 | 1.204675 | 0.22833 | 0.43232 | 21.958 |
| Jagn1 | 982.199 | 0.064495 | 0.08321 | 0.775131 | 0.43826 | 0.64363 | 5.661 |
| Jak1 | 6960.11 | -0.02484 | 0.07712 | -0.32207 | 0.7474 | 0.8676 | 112.916 |
| Jak2 | 4228.8 | 0.436356 | 0.07971 | 5.474316 | 4.39E-08 | 1.03E-05 | 61.253 |
| Jak3 | 325.646 | -0.38877 | 0.18039 | -2.15513 | 0.03115 | 0.12114 | 14.28 |
| Jakmip1 | 665.878 | 0.196595 | 0.13225 | 1.486591 | 0.13712 | 0.31781 | 122.273 |
| Jakmip2 | 2458.22 | 0.031751 | 0.10675 | 0.297438 | 0.76613 | 0.8779 | 156.367 |
| Jakmip3 | 2877.3 | 0.317315 | 0.0878 | 3.614206 | 0.0003 | 0.00565 | 121.434 |
| Jam2 | 2126.64 | -0.03034 | 0.07507 | -0.4042 | 0.68607 | 0.82887 | 51.806 |
| Jam3 | 761.441 | -0.06465 | 0.09506 | -0.68003 | 0.49648 | 0.69005 | 58.038 |
| Jaml | 27.873 | -0.68919 | 0.34586 | -1.99269 | 0.0463 | 0.15723 | 29.352 |
| Jarid2 | 2439.63 | 0.126946 | 0.09201 | 1.379691 | 0.16768 | 0.35761 | 192.17 |
| Jazf1 | 1054.38 | 0.179036 | 0.0883 | 2.027657 | 0.0426 | 0.14867 | 299.835 |
| Jcad | 10128.4 | -0.00511 | 0.08552 | -0.05974 | 0.95236 | 0.97658 | 47.992 |
| Jchain | 585.723 | -1.39185 | 1.07055 | -1.30013 | 0.19356 | 0.39052 | 8.083 |
| Jdp2 | 1379.92 | -0.53847 | 0.14465 | -3.72266 | 0.0002 | 0.00422 | 40.852 |
| Jkamp | 1656.47 | -0.02846 | 0.07222 | -0.39406 | 0.69354 | 0.83381 | 15.873 |
| Jmjd1c | 5067.49 | 0.155116 | 0.09764 | 1.588669 | 0.11214 | 0.27968 | 160.202 |
| Jmjd4 | 1810.57 | -0.02374 | 0.08709 | -0.27261 | 0.78515 | 0.88852 | 8.523 |
| Jmjd6 | 1266.36 | 0.172846 | 0.09933 | 1.740199 | 0.08182 | 0.22798 | 6.018 |
| Jmjd7 | 22.4512 | 0.356112 | 0.42827 | 0.831512 | 0.40568 | 0.61433 | 5.112 |
| Jmjd8 | 687.935 | -0.023 | 0.11224 | -0.20493 | 0.83763 | 0.91644 | 2.977 |
| Jmy | 3678.25 | 0.015482 | 0.06074 | 0.254888 | 0.79881 | 0.89614 | 69.708 |
| Josd1 | 4247.95 | 0.127217 | 0.09142 | 1.391505 | 0.16407 | 0.3529 | 13.669 |
| Josd1-ps | 86.5439 | 0.198773 | 0.2042 | 0.973446 | 0.33033 | 0.54149 | 0.609 |
| Josd2 | 2262 | -0.12261 | 0.12491 | -0.98163 | 0.32628 | 0.53755 | 3.683 |

| | | | | | | | |
|---|---|---|---|---|---|---|---|
| Jph1 | 1066.87 | -0.03999 | 0.10587 | -0.3777 | 0.70566 | 0.84142 | 133.33 |
| Jph2 | 67.2567 | -0.02478 | 0.2747 | -0.0902 | 0.92813 | 0.96388 | 61.752 |
| Jph3 | 10441.5 | 0.015254 | 0.08178 | 0.186534 | 0.85203 | 0.92477 | 64.654 |
| Jph4 | 19196.8 | 0.267869 | 0.08804 | 3.042543 | 0.00235 | 0.02234 | 10.11 |
| Jpt1 | 2098.48 | -0.06984 | 0.07993 | -0.87373 | 0.38227 | 0.59182 | 17.035 |
| Jpt2 | 294.197 | -0.12967 | 0.17762 | -0.73007 | 0.46535 | 0.66495 | 23.271 |
| Jpx | 556.231 | 0.223915 | 0.10673 | 2.097921 | 0.03591 | 0.13334 | 72.145 |
| Jrk | 393.457 | 0.064047 | 0.12854 | 0.498277 | 0.61829 | 0.7831 | 7.235 |
| Jrkl | 556.444 | 0.055094 | 0.16617 | 0.331544 | 0.74023 | 0.863 | 3.223 |
| Jsrp1 | 329.196 | 0.336462 | 0.22785 | 1.476697 | 0.13976 | 0.3211 | 5.003 |
| Jtb | 984.682 | -0.13043 | 0.08225 | -1.58582 | 0.11278 | 0.28083 | 4.242 |
| Jun | 1441.47 | -0.36063 | 0.10242 | -3.52092 | 0.00043 | 0.00723 | 3.189 |
| Junb | 1505.19 | -0.2143 | 0.12607 | -1.69981 | 0.08917 | 0.24176 | 4.235 |
| Jund | 522.643 | -0.30963 | 0.13496 | -2.29413 | 0.02178 | 0.09486 | 1.667 |
| Junos | 68.3766 | 0.241188 | 0.24376 | 0.989448 | 0.32244 | 0.53337 | 115.785 |
| Jup | 1918.35 | -0.08394 | 0.08384 | -1.00125 | 0.3167 | 0.52722 | 28.806 |
| K230015D01Rik | 36.888 | -0.56567 | 0.30479 | -1.85597 | 0.06346 | 0.19358 | 1.516 |
| Kalrn | 24528.7 | -0.17236 | 0.08674 | -1.9871 | 0.04691 | 0.15887 | 604.46 |
| Kank1 | 976.35 | -0.21102 | 0.09374 | -2.25123 | 0.02437 | 0.10211 | 197.522 |
| Kank2 | 665.921 | -0.32191 | 0.09706 | -3.31679 | 0.00091 | 0.01201 | 31.961 |
| Kank3 | 969.71 | -0.29845 | 0.12998 | -2.29607 | 0.02167 | 0.09443 | 12.4 |
| Kank4 | 235.952 | -0.31293 | 0.16233 | -1.92773 | 0.05389 | 0.17433 | 62.64 |
| Kansl1 | 4927.7 | 0.081119 | 0.06101 | 1.329679 | 0.18362 | 0.37774 | 135.633 |
| Kansl1l | 834.703 | 0.182612 | 0.13276 | 1.37555 | 0.16896 | 0.35907 | 98.315 |
| Kansl2 | 2403.01 | 0.054254 | 0.06038 | 0.898469 | 0.36894 | 0.57956 | 16.607 |
| Kansl2-ps | 332.883 | 0.105112 | 0.13473 | 0.780174 | 0.43529 | 0.64101 | 1.461 |
| Kansl3 | 4692.68 | 0.11478 | 0.06503 | 1.765042 | 0.07756 | 0.22037 | 33.452 |
| Kantr | 2170.95 | 0.341812 | 0.08255 | 4.140711 | 3.46E-05 | 0.00126 | 32.668 |
| Kars | 3208.89 | 0.072595 | 0.06553 | 1.107739 | 0.26797 | 0.47725 | 17.881 |
| Kash5 | 161.016 | -0.69138 | 0.16672 | -4.14692 | 3.37E-05 | 0.00124 | 21.269 |
| Kat14 | 1127.74 | 0.015166 | 0.07859 | 0.192986 | 0.84697 | 0.92206 | 38.694 |
| Kat2a | 2794.02 | 0.110041 | 0.0641 | 1.716792 | 0.08602 | 0.23561 | 7.72 |
| Kat2b | 1614.96 | -0.18277 | 0.08876 | -2.05917 | 0.03948 | 0.14175 | 105.86 |
| Kat5 | 1725.66 | 0.019103 | 0.07194 | 0.265561 | 0.79058 | 0.8913 | 7.224 |

| | | | | | | | |
|---|---|---|---|---|---|---|---|
| Kat6a | 5487.63 | 0.125693 | 0.05727 | 2.194608 | 0.02819 | 0.11309 | 83.725 |
| Kat6b | 4301.71 | 0.111763 | 0.07665 | 1.458102 | 0.14481 | 0.32758 | 191.045 |
| Kat7 | 3848.7 | 0.106513 | 0.07984 | 1.334111 | 0.18217 | 0.37594 | 35.988 |
| Kat8 | 1025.22 | 0.038683 | 0.07565 | 0.511346 | 0.60911 | 0.77661 | 13.322 |
| Katna1 | 563.787 | 0.162704 | 0.08934 | 1.821228 | 0.06857 | 0.20357 | 37.151 |
| Katnal1 | 3495.77 | -0.14403 | 0.08251 | -1.74556 | 0.08089 | 0.22639 | 57.737 |
| Katnal2 | 372.342 | 0.21573 | 0.14285 | 1.5102 | 0.13099 | 0.30945 | 70.161 |
| Katnb1 | 1418.58 | 0.042821 | 0.07382 | 0.58004 | 0.56189 | 0.74286 | 21.964 |
| Katnbl1 | 665.381 | 0.027028 | 0.10058 | 0.268719 | 0.78815 | 0.89 | 35.03 |
| Katnip | 1802.37 | 0.06112 | 0.09511 | 0.642616 | 0.52047 | 0.70975 | 166.906 |
| Kazald1 | 105.816 | -0.63815 | 0.21614 | -2.95245 | 0.00315 | 0.02696 | 4.049 |
| Kazn | 12444.5 | 0.266942 | 0.14344 | 1.861001 | 0.06274 | 0.19225 | 375.356 |
| Kbtbd11 | 8541.71 | 0.112839 | 0.09292 | 1.214393 | 0.2246 | 0.42827 | 22.309 |
| Kbtbd12 | 21.9248 | 0.736509 | 0.45456 | 1.620253 | 0.10518 | 0.26836 | 92.837 |
| Kbtbd2 | 2754.58 | 0.055009 | 0.07086 | 0.776256 | 0.4376 | 0.64306 | 20.29 |
| Kbtbd3 | 406.766 | 0.076046 | 0.15875 | 0.479026 | 0.63192 | 0.79217 | 21.9 |
| Kbtbd4 | 300.057 | -0.00079 | 0.11318 | -0.00697 | 0.99444 | 0.99669 | 6.887 |
| Kbtbd6 | 40.5275 | 0.055553 | 0.32595 | 0.170434 | 0.86467 | 0.93153 | 2.984 |
| Kbtbd7 | 2818.44 | 0.112626 | 0.07039 | 1.599954 | 0.10961 | 0.27556 | 4.526 |
| Kbtbd8 | 451.053 | 0.315457 | 0.09801 | 3.218782 | 0.00129 | 0.0149 | 12.551 |
| Kcmf1 | 2650.79 | -0.18122 | 0.06259 | -2.89548 | 0.00379 | 0.03031 | 58.866 |
| Kcna1 | 7756.22 | -0.00964 | 0.08475 | -0.1138 | 0.90939 | 0.95456 | 5.988 |
| Kcna2 | 14521.5 | -0.26169 | 0.07719 | -3.39018 | 0.0007 | 0.00997 | 13.86 |
| Kcna3 | 201.666 | -0.2441 | 0.16604 | -1.47011 | 0.14153 | 0.32333 | 1.902 |
| Kcna4 | 2431.71 | 0.529105 | 0.16156 | 3.275065 | 0.00106 | 0.01329 | 7.864 |
| Kcna5 | 2246.73 | 0.55575 | 0.19378 | 2.867952 | 0.00413 | 0.03212 | 2.862 |
| Kcna6 | 2495.67 | -0.412 | 0.15539 | -2.65136 | 0.00802 | 0.04964 | 32.346 |
| Kcnab1 | 9057.95 | 0.611334 | 0.18008 | 3.39477 | 0.00069 | 0.00985 | 429.028 |
| Kcnab2 | 20636.5 | -0.48656 | 0.09968 | -4.88112 | 1.05E-06 | 0.00011 | 87.169 |
| Kcnab3 | 1195.63 | -0.73469 | 0.16246 | -4.52227 | 6.12E-06 | 0.00038 | 6.785 |
| Kcnb1 | 15139.8 | -0.10095 | 0.05811 | -1.73734 | 0.08233 | 0.22879 | 94.187 |
| Kcnb2 | 931.428 | -0.03539 | 0.22544 | -0.15698 | 0.87526 | 0.93783 | 436.497 |
| Kcnc1 | 9439.57 | -0.1278 | 0.09052 | -1.41187 | 0.15799 | 0.34524 | 42.208 |
| Kcnc2 | 2623.2 | -0.49486 | 0.14617 | -3.38554 | 0.00071 | 0.01008 | 195.904 |

| | | | | | | | |
|---|---|---|---|---|---|---|---|
| Kcnc3 | 4095.69 | -0.13952 | 0.0743 | -1.87787 | 0.0604 | 0.18755 | 14.091 |
| Kcnc4 | 1620.02 | -0.88853 | 0.31863 | -2.78855 | 0.00529 | 0.03766 | 21.25 |
| Kcnd1 | 739.508 | 0.193435 | 0.10173 | 1.901401 | 0.05725 | 0.18143 | 15.992 |
| Kcnd2 | 6802.27 | 0.285938 | 0.10257 | 2.787619 | 0.00531 | 0.0377 | 514.303 |
| Kcnd3 | 4426.05 | -0.05278 | 0.12163 | -0.43393 | 0.66434 | 0.8145 | 221.673 |
| Kcnd3os | 111.428 | -0.08019 | 0.20839 | -0.3848 | 0.70039 | 0.83823 | 11.361 |
| Kcne1l | 216.868 | -0.25978 | 0.16847 | -1.54194 | 0.12309 | 0.29724 | 1.543 |
| Kcne4 | 96.9222 | -0.96997 | 0.25443 | -3.81226 | 0.00014 | 0.00333 | 3.271 |
| Kcnf1 | 6123.58 | -0.46824 | 0.15378 | -3.04491 | 0.00233 | 0.02222 | 4.789 |
| Kcng1 | 1424.93 | 0.118975 | 0.17474 | 0.680855 | 0.49596 | 0.6898 | 21.62 |
| Kcng2 | 205.753 | -0.01644 | 0.15857 | -0.10366 | 0.91744 | 0.95781 | 69.709 |
| Kcng3 | 452.334 | 0.706653 | 0.2465 | 2.866741 | 0.00415 | 0.03223 | 45.939 |
| Kcng4 | 84.1523 | -0.7389 | 0.25519 | -2.89548 | 0.00379 | 0.03031 | 11.827 |
| Kcnh1 | 6828.66 | 0.098027 | 0.08671 | 1.130503 | 0.25826 | 0.46625 | 319.386 |
| Kcnh2 | 1794.81 | 0.053945 | 0.10653 | 0.506363 | 0.6126 | 0.77934 | 32.016 |
| Kcnh3 | 6933.36 | 0.136027 | 0.07921 | 1.717385 | 0.08591 | 0.23546 | 17.957 |
| Kcnh4 | 1656.49 | 0.683014 | 0.20563 | 3.321506 | 0.0009 | 0.01185 | 19.567 |
| Kcnh5 | 811.903 | -0.11586 | 0.13482 | -0.85937 | 0.39014 | 0.59917 | 280.113 |
| Kcnh6 | 194.499 | 0.116215 | 0.15577 | 0.746053 | 0.45564 | 0.65749 | 26.426 |
| Kcnh7 | 3031.81 | -0.58145 | 0.38247 | -1.52024 | 0.12845 | 0.30549 | 490.874 |
| Kcnh8 | 58.1601 | -0.06969 | 0.32729 | -0.21292 | 0.83139 | 0.91345 | 376.486 |
| Kcnip1 | 1328.07 | 0.032286 | 0.09935 | 0.32496 | 0.74521 | 0.866 | 363.814 |
| Kcnip2 | 11602.1 | 0.701824 | 0.27588 | 2.543921 | 0.01096 | 0.06094 | 24.223 |
| Kcnip3 | 5322.38 | -0.31218 | 0.08555 | -3.64907 | 0.00026 | 0.00513 | 65.597 |
| Kcnip4 | 1632.65 | 0.267989 | 0.0978 | 2.740156 | 0.00614 | 0.04191 | 1135.406 |
| Kcnj10 | 8327.67 | -0.40966 | 0.11808 | -3.46939 | 0.00052 | 0.00817 | 32.876 |
| Kcnj11 | 1995.98 | 0.100601 | 0.11631 | 0.864918 | 0.38708 | 0.5964 | 6.812 |
| Kcnj12 | 452.228 | 0.287741 | 0.16783 | 1.714487 | 0.08644 | 0.23648 | 48.568 |
| Kcnj14 | 134.043 | -0.22964 | 0.20783 | -1.10494 | 0.26918 | 0.47859 | 8.323 |
| Kcnj16 | 905.628 | -0.42255 | 0.12763 | -3.31086 | 0.00093 | 0.01219 | 59.936 |
| Kcnj2 | 2567.01 | 0.569294 | 0.19424 | 2.930813 | 0.00338 | 0.02813 | 10.658 |
| Kcnj3 | 4741.08 | -0.73217 | 0.17066 | -4.29028 | 1.78E-05 | 0.00082 | 162.176 |
| Kcnj4 | 7392.66 | -0.08232 | 0.07147 | -1.15186 | 0.24938 | 0.45679 | 21.528 |
| Kcnj5 | 49.8973 | 1.061024 | 0.41246 | 2.572433 | 0.0101 | 0.05786 | 29.644 |

| | | | | | | | |
|---|---|---|---|---|---|---|---|
| Kcnj6 | 824.961 | -0.73212 | 0.37172 | -1.96958 | 0.04889 | 0.16344 | 249.066 |
| Kcnj8 | 133.343 | -0.18616 | 0.22019 | -0.84548 | 0.39784 | 0.60678 | 6.778 |
| Kcnj9 | 3264.42 | -0.52111 | 0.16513 | -3.15567 | 0.0016 | 0.01735 | 8.818 |
| Kcnk1 | 3342.46 | -0.39568 | 0.07733 | -5.11712 | 3.10E-07 | 4.36E-05 | 35.516 |
| Kcnk10 | 244.692 | -0.64285 | 0.15646 | -4.10885 | 3.98E-05 | 0.0014 | 148.874 |
| Kcnk12 | 226.815 | -0.5788 | 0.18152 | -3.18871 | 0.00143 | 0.01601 | 52.194 |
| Kcnk13 | 393.514 | -0.19224 | 0.14075 | -1.36581 | 0.172 | 0.36268 | 98.184 |
| Kcnk2 | 4061.36 | 0.207891 | 0.11711 | 1.775152 | 0.07587 | 0.21752 | 194.344 |
| Kcnk3 | 655.364 | -0.24655 | 0.12255 | -2.01193 | 0.04423 | 0.15247 | 37.102 |
| Kcnk4 | 189.311 | -0.62265 | 0.26474 | -2.35193 | 0.01868 | 0.08566 | 10.55 |
| Kcnk5 | 38.2676 | -1.36935 | 0.49343 | -2.77518 | 0.00552 | 0.03886 | 41.753 |
| Kcnk6 | 313.118 | -0.081 | 0.13542 | -0.59812 | 0.54976 | 0.7333 | 10.59 |
| Kcnk7 | 17.3688 | 0.650689 | 0.47082 | 1.382047 | 0.16696 | 0.35677 | 2.743 |
| Kcnk9 | 5507.55 | -0.38922 | 0.12557 | -3.0997 | 0.00194 | 0.0197 | 45.252 |
| Kcnma1 | 12509.8 | -0.34339 | 0.13107 | -2.61989 | 0.0088 | 0.05264 | 725.061 |
| Kcnmb1 | 22.1816 | -0.38785 | 0.47153 | -0.82254 | 0.41077 | 0.61876 | 10.629 |
| Kcnmb2 | 81.9205 | -0.27109 | 0.25396 | -1.06745 | 0.28577 | 0.49576 | 297.674 |
| Kcnmb4 | 63.7075 | -0.59754 | 0.29627 | -2.01685 | 0.04371 | 0.15125 | 56.018 |
| Kcnmb4os1 | 16.8886 | -0.30034 | 0.52828 | -0.56852 | 0.56968 | 0.74914 | 3.186 |
| Kcnmb4os2 | 57.426 | -0.19768 | 0.23644 | -0.8361 | 0.4031 | 0.6118 | 2.157 |
| Kcnn1 | 648.156 | -0.53009 | 0.12302 | -4.30891 | 1.64E-05 | 0.00078 | 21.21 |
| Kcnn2 | 1200.26 | -0.36218 | 0.14439 | -2.50843 | 0.01213 | 0.06503 | 417.165 |
| Kcnn3 | 3865.58 | 0.433818 | 0.11594 | 3.741595 | 0.00018 | 0.00406 | 154.969 |
| Kcnn4 | 17.4416 | -0.2193 | 0.52916 | -0.41443 | 0.67856 | 0.8241 | 16.428 |
| Kcnq1 | 61.3223 | -0.11024 | 0.43858 | -0.25136 | 0.80154 | 0.89741 | 320.681 |
| Kcnq1ot1 | 2193.27 | 0.397391 | 0.17244 | 2.304545 | 0.02119 | 0.09306 | 93.092 |
| Kcnq2 | 28867.2 | 0.176489 | 0.0612 | 2.883597 | 0.00393 | 0.03109 | 59.722 |
| Kcnq3 | 11119.4 | 0.14927 | 0.10173 | 1.467248 | 0.14231 | 0.32422 | 300.256 |
| Kcnq4 | 264.008 | -0.61964 | 0.17301 | -3.58158 | 0.00034 | 0.00615 | 52.475 |
| Kcnq5 | 6384.82 | 0.46745 | 0.1278 | 3.657803 | 0.00025 | 0.005 | 563.54 |
| Kcnrg | 62.7641 | 0.502794 | 0.25077 | 2.004987 | 0.04496 | 0.15415 | 5.37 |
| Kcns1 | 250.452 | -0.64634 | 0.13128 | -4.92323 | 8.51E-07 | 9.48E-05 | 7.495 |
| Kcns2 | 2156.24 | 0.236739 | 0.13162 | 1.798676 | 0.07207 | 0.21074 | 6.053 |
| Kcns3 | 145.043 | -0.40421 | 0.20723 | -1.95051 | 0.05112 | 0.1683 | 60.855 |

| | | | | | | | |
|---|---|---|---|---|---|---|---|
| Kcnt1 | 14005.6 | 0.228597 | 0.09912 | 2.306238 | 0.0211 | 0.09285 | 54.54 |
| Kcnt2 | 907.713 | 0.043857 | 0.15676 | 0.279774 | 0.77965 | 0.88519 | 365.91 |
| Kcnu1 | 102.321 | -0.01511 | 0.19322 | -0.07819 | 0.93768 | 0.96886 | 88.317 |
| Kcnv1 | 2983.21 | -0.01899 | 0.10599 | -0.17916 | 0.85781 | 0.9283 | 8.637 |
| Kcp | 115.887 | -0.01594 | 0.20987 | -0.07595 | 0.93946 | 0.96977 | 34.791 |
| Kctd1 | 4640.67 | 0.30265 | 0.1084 | 2.791845 | 0.00524 | 0.03739 | 182.762 |
| Kctd10 | 960.897 | 0.008069 | 0.11359 | 0.071037 | 0.94337 | 0.97176 | 16.942 |
| Kctd11 | 79.1049 | -0.16807 | 0.25638 | -0.65554 | 0.51212 | 0.70326 | 3.146 |
| Kctd12 | 7108.47 | 0.209091 | 0.06449 | 3.242094 | 0.00119 | 0.01419 | 6.057 |
| Kctd12b | 234.487 | 0.195904 | 0.14928 | 1.312301 | 0.18942 | 0.38536 | 11.238 |
| Kctd13 | 3052.54 | -0.08457 | 0.0791 | -1.06915 | 0.285 | 0.49485 | 16.753 |
| Kctd14 | 127.582 | -0.56824 | 0.17496 | -3.24781 | 0.00116 | 0.01403 | 8.235 |
| Kctd15 | 434.187 | -0.41081 | 0.15974 | -2.57169 | 0.01012 | 0.05794 | 17.719 |
| Kctd16 | 926.701 | 0.037698 | 0.16592 | 0.227211 | 0.82026 | 0.90734 | 274.669 |
| Kctd17 | 5881.91 | -0.12751 | 0.08186 | -1.55763 | 0.11932 | 0.29146 | 10.74 |
| Kctd18 | 1243.51 | 0.30055 | 0.10473 | 2.869848 | 0.00411 | 0.03204 | 63.856 |
| Kctd2 | 2287.72 | -0.28257 | 0.0762 | -3.70831 | 0.00021 | 0.00438 | 11.147 |
| Kctd20 | 3873.19 | 0.082974 | 0.06502 | 1.276085 | 0.20193 | 0.40043 | 17.6 |
| Kctd21 | 697.606 | -0.15629 | 0.10157 | -1.53878 | 0.12386 | 0.29838 | 17.887 |
| Kctd3 | 4095.08 | -0.15662 | 0.06981 | -2.24352 | 0.02486 | 0.10356 | 36.747 |
| Kctd4 | 609.41 | 0.235965 | 0.16805 | 1.40411 | 0.16029 | 0.3484 | 11.198 |
| Kctd5 | 1372.8 | -0.00983 | 0.10226 | -0.09614 | 0.92341 | 0.9614 | 25.752 |
| Kctd6 | 1371.87 | -0.13287 | 0.08916 | -1.49026 | 0.13616 | 0.31645 | 9.669 |
| Kctd7 | 646.3 | -0.0135 | 0.09274 | -0.14555 | 0.88428 | 0.94244 | 10.946 |
| Kctd8 | 684.063 | 0.750315 | 0.24153 | 3.106469 | 0.00189 | 0.01936 | 232.395 |
| Kctd9 | 844.866 | -0.04476 | 0.11548 | -0.38762 | 0.6983 | 0.83649 | 26.374 |
| Kdelr1 | 1851.46 | -0.05415 | 0.07865 | -0.68845 | 0.49117 | 0.68613 | 10.961 |
| Kdelr2 | 745.09 | -0.15596 | 0.09935 | -1.56977 | 0.11647 | 0.28669 | 18.064 |
| Kdelr3 | 20.8794 | -0.39223 | 0.46365 | -0.84596 | 0.39758 | 0.6066 | 11.329 |
| Kdm1a | 2778.87 | 0.110744 | 0.06009 | 1.84306 | 0.06532 | 0.19714 | 52.184 |
| Kdm1b | 677.684 | -0.03821 | 0.10918 | -0.34999 | 0.72634 | 0.8541 | 41.781 |
| Kdm2a | 5948.47 | 0.10567 | 0.05913 | 1.787137 | 0.07392 | 0.21422 | 83.867 |
| Kdm2b | 903.96 | -0.05993 | 0.07508 | -0.79821 | 0.42475 | 0.63137 | 119.159 |
| Kdm3a | 3470.93 | 0.296851 | 0.07662 | 3.874228 | 0.00011 | 0.00281 | 44.019 |

| | | | | | | | |
|---|---|---|---|---|---|---|---|
| Kdm3b | 3687.72 | 0.094708 | 0.07157 | 1.323212 | 0.18576 | 0.38054 | 61.614 |
| Kdm4a | 3281.81 | -0.02497 | 0.06067 | -0.41154 | 0.68068 | 0.82518 | 43.087 |
| Kdm4b | 3972.77 | 0.192349 | 0.08035 | 2.393935 | 0.01667 | 0.0799 | 76.809 |
| Kdm4c | 1686.16 | 0.25741 | 0.08914 | 2.887786 | 0.00388 | 0.0308 | 163.364 |
| Kdm4d | 73.8168 | 0.290453 | 0.29988 | 0.968563 | 0.33276 | 0.54364 | 37.935 |
| Kdm4dl | 27.8606 | 0.125253 | 0.35252 | 0.35531 | 0.72236 | 0.85187 | 1.187 |
| Kdm5a | 3508.98 | 0.08019 | 0.09653 | 0.830742 | 0.40612 | 0.61462 | 80.451 |
| Kdm5b | 5007.26 | 0.245038 | 0.05896 | 4.155873 | 3.24E-05 | 0.00122 | 75.115 |
| Kdm5c | 4208.36 | 0.097803 | 0.06145 | 1.591574 | 0.11148 | 0.27897 | 41.516 |
| Kdm5d | 2688.85 | 0.150653 | 0.09586 | 1.571633 | 0.11604 | 0.2859 | 58.999 |
| Kdm6a | 1848.67 | 0.183962 | 0.10068 | 1.827185 | 0.06767 | 0.20164 | 117.362 |
| Kdm6b | 1993.95 | -0.04709 | 0.11992 | -0.39272 | 0.69453 | 0.8343 | 15.168 |
| Kdm6bos | 52.0018 | 0.021177 | 0.27906 | 0.075886 | 0.93951 | 0.96977 | 1.188 |
| Kdm7a | 4845.71 | 0.102763 | 0.10342 | 0.993605 | 0.32042 | 0.53118 | 70.167 |
| Kdm8 | 275.175 | 0.157861 | 0.13265 | 1.190055 | 0.23402 | 0.4391 | 17.594 |
| Kdr | 445.019 | -0.48047 | 0.15366 | -3.12691 | 0.00177 | 0.01859 | 45.632 |
| Kdsr | 2438.31 | 0.047463 | 0.05998 | 0.791274 | 0.42878 | 0.63543 | 39.269 |
| Keap1 | 2378.56 | 0.08023 | 0.06983 | 1.148977 | 0.25057 | 0.4579 | 9.632 |
| Khdc3 | 71.7849 | 0.111511 | 0.26107 | 0.427134 | 0.66928 | 0.81761 | 2.603 |
| Khdc4 | 3983.98 | 0.12576 | 0.06793 | 1.851206 | 0.06414 | 0.19496 | 27.122 |
| Khdrbs1 | 6990.02 | 0.009462 | 0.07197 | 0.131461 | 0.89541 | 0.94744 | 39.14 |
| Khdrbs2 | 607.633 | 0.566673 | 0.18097 | 3.131326 | 0.00174 | 0.01841 | 485.855 |
| Khdrbs3 | 3551.42 | -0.36466 | 0.13063 | -2.7915 | 0.00525 | 0.03741 | 172.792 |
| Khk | 311.374 | -0.25096 | 0.12153 | -2.06504 | 0.03892 | 0.14059 | 9.818 |
| Khnyn | 949.501 | -0.11047 | 0.08527 | -1.29542 | 0.19517 | 0.39273 | 13.829 |
| Khsrp | 4901.53 | 0.054116 | 0.07039 | 0.768812 | 0.442 | 0.64653 | 10.472 |
| Kics2 | 544.991 | -3.20E-05 | 0.09647 | -0.00033 | 0.99974 | 0.99993 | 12.942 |
| Kidins220 | 23454.7 | 0.154245 | 0.04931 | 3.128184 | 0.00176 | 0.01854 | 88.228 |
| Kif11 | 120.444 | -0.08288 | 0.21968 | -0.37727 | 0.70598 | 0.8415 | 45.457 |
| Kif13a | 1946 | -0.20541 | 0.07505 | -2.73715 | 0.0062 | 0.04211 | 180.781 |
| Kif13b | 3431.66 | -0.19825 | 0.097 | -2.04378 | 0.04098 | 0.14493 | 162.353 |
| Kif14 | 35.4538 | 0.08844 | 0.27905 | 0.31693 | 0.7513 | 0.86954 | 65.169 |
| Kif15 | 185.39 | 0.558153 | 0.19616 | 2.845398 | 0.00444 | 0.03369 | 67.688 |
| Kif16b | 698.616 | 0.001027 | 0.08523 | 0.012051 | 0.99039 | 0.99473 | 284.058 |

| | | | | | | | |
|---|---|---|---|---|---|---|---|
| Kif17 | 3352.04 | 0.500551 | 0.13755 | 3.638976 | 0.00027 | 0.00526 | 51.533 |
| Kif18a | 48.8949 | -0.28471 | 0.25672 | -1.10902 | 0.26742 | 0.47675 | 61.01 |
| Kif18b | 12.3587 | -1.66334 | 0.76805 | -2.16567 | 0.03034 | 0.11892 | 19.596 |
| Kif19a | 162.676 | -0.20467 | 0.17824 | -1.14831 | 0.25084 | 0.45821 | 25.352 |
| Kif1a | 55426.3 | -0.15497 | 0.07598 | -2.03972 | 0.04138 | 0.14593 | 86.488 |
| Kif1b | 48112.3 | -0.13878 | 0.07916 | -1.75324 | 0.07956 | 0.22403 | 131.375 |
| Kif1c | 5841.22 | -0.37062 | 0.09045 | -4.09755 | 4.18E-05 | 0.00144 | 31.417 |
| Kif20a | 102.829 | 0.015439 | 0.20156 | 0.076598 | 0.93894 | 0.96951 | 8.665 |
| Kif20b | 46.5187 | -0.56828 | 0.38415 | -1.4793 | 0.13906 | 0.3203 | 53.385 |
| Kif21a | 10613.7 | 0.085068 | 0.08755 | 0.97167 | 0.33121 | 0.54208 | 116.673 |
| Kif21b | 5736.09 | -0.03002 | 0.09007 | -0.33333 | 0.73889 | 0.86228 | 46.61 |
| Kif22 | 271.646 | 0.395108 | 0.15899 | 2.485072 | 0.01295 | 0.06803 | 14.743 |
| Kif23 | 130.529 | 0.414171 | 0.20975 | 1.974554 | 0.04832 | 0.16205 | 30.87 |
| Kif24 | 111.15 | -0.0404 | 0.20811 | -0.19414 | 0.84607 | 0.92172 | 74.143 |
| Kif26a | 284.982 | -0.18119 | 0.16565 | -1.09381 | 0.27404 | 0.48344 | 35.54 |
| Kif26b | 989.377 | 0.412881 | 0.08765 | 4.710396 | 2.47E-06 | 0.00021 | 410.076 |
| Kif27 | 449.784 | 0.55204 | 0.16839 | 3.27828 | 0.00104 | 0.01319 | 71.621 |
| Kif2a | 4010.76 | -0.04381 | 0.11226 | -0.39022 | 0.69637 | 0.83541 | 63.131 |
| Kif3a | 6302.38 | 0.141177 | 0.07969 | 1.771493 | 0.07648 | 0.21863 | 34.589 |
| Kif3b | 4888.37 | -0.0038 | 0.06016 | -0.06313 | 0.94966 | 0.97495 | 41.978 |
| Kif3c | 6906.72 | -0.02795 | 0.05869 | -0.47622 | 0.63392 | 0.79385 | 41.363 |
| Kif4 | 43.3201 | -0.74216 | 0.33667 | -2.20441 | 0.0275 | 0.11105 | 101.478 |
| Kif5a | 115162 | -0.66139 | 0.14933 | -4.42912 | 9.46E-06 | 0.00054 | 37.653 |
| Kif5b | 13286.7 | -0.06863 | 0.13969 | -0.49128 | 0.62323 | 0.78598 | 41.173 |
| Kif5c | 39937.7 | -0.25606 | 0.09562 | -2.67794 | 0.00741 | 0.04733 | 155.481 |
| Kif6 | 115.492 | 0.259207 | 0.20904 | 1.239971 | 0.21499 | 0.41637 | 294.712 |
| Kif7 | 242.728 | 0.482713 | 0.16551 | 2.916469 | 0.00354 | 0.02897 | 17.623 |
| Kif9 | 329.1 | 0.309242 | 0.12106 | 2.55442 | 0.01064 | 0.05988 | 48.222 |
| Kifap3 | 12704.8 | 0.051459 | 0.07304 | 0.704539 | 0.4811 | 0.6782 | 137.527 |
| Kifbp | 4575.97 | -0.13482 | 0.05618 | -2.39981 | 0.0164 | 0.07899 | 39.832 |
| Kifc1 | 76.1922 | -0.31464 | 0.20957 | -1.50136 | 0.13326 | 0.31281 | 15.003 |
| Kifc2 | 18039.3 | -0.28349 | 0.08787 | -3.22607 | 0.00126 | 0.01474 | 8.339 |
| Kifc3 | 1002.46 | -0.01097 | 0.10133 | -0.10826 | 0.91379 | 0.95639 | 102.985 |
| Kifc5b | 194.362 | 0.086196 | 0.15876 | 0.54295 | 0.58716 | 0.76069 | 15.493 |

| | | | | | | | |
|---|---|---|---|---|---|---|---|
| Kin | 519.807 | 0.070277 | 0.11863 | 0.592412 | 0.55357 | 0.73632 | 12.214 |
| Kirrel | 975.892 | 0.586514 | 0.14189 | 4.133634 | 3.57E-05 | 0.0013 | 96.155 |
| Kirrel2 | 908.08 | 0.244177 | 0.12224 | 1.997606 | 0.04576 | 0.1559 | 10.157 |
| Kirrel3 | 2736.52 | 0.214547 | 0.08521 | 2.517894 | 0.01181 | 0.06392 | 550.823 |
| Kirrel3os | 29.2793 | 0.34534 | 0.31573 | 1.093788 | 0.27405 | 0.48344 | 11.277 |
| Kiss1r | 298.23 | 0.065106 | 0.12568 | 0.51804 | 0.60443 | 0.7733 | 5.303 |
| Kit | 1727.42 | -0.0584 | 0.09179 | -0.63627 | 0.5246 | 0.71304 | 81.807 |
| Kitl | 2950.65 | 0.197746 | 0.1566 | 1.262782 | 0.20667 | 0.40605 | 84.787 |
| Kiz | 599.293 | 0.072357 | 0.14718 | 0.491635 | 0.62298 | 0.78589 | 114.234 |
| Kl | 390.868 | 0.808504 | 0.22742 | 3.555102 | 0.00038 | 0.00657 | 41.211 |
| Klb | 38.0815 | 0.151455 | 0.32687 | 0.463346 | 0.64312 | 0.80089 | 35.694 |
| Klc1 | 28296.1 | -0.17136 | 0.0635 | -2.69876 | 0.00696 | 0.04526 | 48.996 |
| Klc2 | 7291.81 | -0.20699 | 0.07606 | -2.7213 | 0.0065 | 0.0434 | 10.815 |
| Klc3 | 109.26 | 0.187065 | 0.20337 | 0.919836 | 0.35766 | 0.56794 | 9.668 |
| Klc4 | 1151.28 | -0.02501 | 0.07163 | -0.3492 | 0.72694 | 0.85431 | 15.399 |
| Klf10 | 976.047 | -0.39745 | 0.15589 | -2.54952 | 0.01079 | 0.06047 | 9.244 |
| Klf11 | 314.092 | -0.47865 | 0.11024 | -4.342 | 1.41E-05 | 0.00071 | 11.516 |
| Klf12 | 1718.15 | -0.09343 | 0.06729 | -1.38852 | 0.16498 | 0.35399 | 414.048 |
| Klf13 | 9603.13 | -0.30424 | 0.10765 | -2.82625 | 0.00471 | 0.03497 | 52.565 |
| Klf14 | 20.7434 | -0.80733 | 0.41313 | -1.95418 | 0.05068 | 0.16741 | 3.056 |
| Klf15 | 1774.77 | -0.06331 | 0.11909 | -0.53163 | 0.59498 | 0.76702 | 12.663 |
| Klf16 | 2302.78 | 0.280389 | 0.13901 | 2.017025 | 0.04369 | 0.15121 | 10.198 |
| Klf2 | 496.686 | -0.34307 | 0.23201 | -1.4787 | 0.13922 | 0.32045 | 2.624 |
| Klf3 | 1731.9 | -0.54309 | 0.10319 | -5.26301 | 1.42E-07 | 2.37E-05 | 29.514 |
| Klf4 | 449.333 | -0.60818 | 0.21523 | -2.82567 | 0.00472 | 0.03502 | 5.324 |
| Klf5 | 748.903 | 0.337957 | 0.14245 | 2.372516 | 0.01767 | 0.0826 | 16.346 |
| Klf6 | 2142.81 | -0.55693 | 0.14814 | -3.75943 | 0.00017 | 0.00387 | 8.913 |
| Klf7 | 2896.05 | -0.2028 | 0.09617 | -2.10883 | 0.03496 | 0.13109 | 92.836 |
| Klf8 | 296.328 | 0.315833 | 0.13778 | 2.292278 | 0.02189 | 0.09522 | 158.667 |
| Klf9 | 16092.8 | 0.06452 | 0.07611 | 0.847674 | 0.39662 | 0.6056 | 26.909 |
| Klhdc1 | 1173.7 | 0.024771 | 0.08002 | 0.309544 | 0.75691 | 0.87207 | 43.457 |
| Klhdc10 | 6193.46 | 0.026879 | 0.06258 | 0.429533 | 0.66754 | 0.81626 | 53.312 |
| Klhdc2 | 4301.45 | -0.16199 | 0.0874 | -1.85356 | 0.0638 | 0.19426 | 14.007 |
| Klhdc3 | 4469.92 | -0.1916 | 0.08059 | -2.37739 | 0.01744 | 0.08197 | 6.365 |

| | | | | | | | |
|---|---|---|---|---|---|---|---|
| Klhdc4 | 1176.55 | -0.02781 | 0.06795 | -0.40923 | 0.68237 | 0.82651 | 33.257 |
| Klhdc7a | 1595.98 | -0.40789 | 0.10141 | -4.02206 | 5.77E-05 | 0.00183 | 7.807 |
| Klhdc7b | 19.2203 | 0.022948 | 0.48984 | 0.046847 | 0.96263 | 0.98185 | 3.951 |
| Klhdc8a | 686.737 | -0.61388 | 0.15971 | -3.84369 | 0.00012 | 0.00305 | 8.732 |
| Klhdc8b | 917.599 | -0.07852 | 0.10567 | -0.74309 | 0.45743 | 0.65867 | 13.946 |
| Klhdc9 | 135.448 | -0.07007 | 0.20711 | -0.33832 | 0.73512 | 0.85989 | 2.353 |
| Klhl1 | 244.153 | 0.119005 | 0.22075 | 0.5391 | 0.58982 | 0.76287 | 416.367 |
| Klhl10 | 48.3315 | 0.363699 | 0.30367 | 1.197686 | 0.23104 | 0.43554 | 15.106 |
| Klhl11 | 1383.41 | 0.010825 | 0.07156 | 0.151271 | 0.87976 | 0.93984 | 10.128 |
| Klhl12 | 2496.31 | 0.187253 | 0.07514 | 2.492124 | 0.0127 | 0.0672 | 35.488 |
| Klhl13 | 6173.87 | 0.421306 | 0.11543 | 3.649798 | 0.00026 | 0.00512 | 145.812 |
| Klhl14 | 266.913 | -0.95944 | 0.6195 | -1.54873 | 0.12145 | 0.29471 | 104.342 |
| Klhl15 | 1135.2 | 0.201289 | 0.08943 | 2.250911 | 0.02439 | 0.10216 | 43.126 |
| Klhl17 | 2064.33 | 0.178703 | 0.08319 | 2.14807 | 0.03171 | 0.12267 | 5.523 |
| Klhl18 | 2024.03 | -0.01735 | 0.06878 | -0.25228 | 0.80083 | 0.89707 | 50.769 |
| Klhl2 | 14739.1 | 0.16255 | 0.09686 | 1.678255 | 0.0933 | 0.2486 | 109.961 |
| Klhl20 | 1300.98 | 0.039062 | 0.08824 | 0.442682 | 0.658 | 0.81018 | 43.137 |
| Klhl21 | 918.777 | -0.07286 | 0.09173 | -0.79432 | 0.42701 | 0.63344 | 8.878 |
| Klhl22 | 3375.05 | -0.00342 | 0.06723 | -0.05083 | 0.95946 | 0.98032 | 33.765 |
| Klhl23 | 3521.92 | -0.0439 | 0.06864 | -0.63954 | 0.52247 | 0.71129 | 14.708 |
| Klhl24 | 6629.65 | 0.029417 | 0.07013 | 0.419465 | 0.67488 | 0.82154 | 31.68 |
| Klhl25 | 362.498 | -0.17359 | 0.11321 | -1.53332 | 0.1252 | 0.3002 | 25.822 |
| Klhl26 | 2114.94 | -0.04609 | 0.10089 | -0.45681 | 0.64781 | 0.80388 | 26.75 |
| Klhl28 | 1750.48 | 0.056902 | 0.10986 | 0.517966 | 0.60448 | 0.7733 | 26.702 |
| Klhl29 | 4767.82 | 0.244492 | 0.0673 | 3.632901 | 0.00028 | 0.00535 | 298.211 |
| Klhl3 | 4547.41 | 0.050432 | 0.0751 | 0.671532 | 0.50188 | 0.69441 | 113.365 |
| Klhl32 | 562.335 | 0.119074 | 0.16071 | 0.740913 | 0.45875 | 0.65969 | 238.571 |
| Klhl33 | 635.535 | 0.029914 | 0.14755 | 0.202738 | 0.83934 | 0.91737 | 8.926 |
| Klhl34 | 2790.94 | 0.268083 | 0.0752 | 3.565145 | 0.00036 | 0.0064 | 1.935 |
| Klhl35 | 97.5163 | 0.422099 | 0.21298 | 1.981894 | 0.04749 | 0.1602 | 8.019 |
| Klhl36 | 379.017 | 0.285718 | 0.13275 | 2.152338 | 0.03137 | 0.12183 | 14.73 |
| Klhl4 | 903.694 | -0.11008 | 0.13075 | -0.84196 | 0.39981 | 0.60858 | 86.497 |
| Klhl40 | 90.2411 | -0.84963 | 0.33682 | -2.52253 | 0.01165 | 0.06344 | 6.212 |
| Klhl41 | 19.0766 | 0.367541 | 0.51302 | 0.71642 | 0.47373 | 0.67209 | 14.111 |

| | | | | | | | |
|---|---|---|---|---|---|---|---|
| Klhl42 | 1410.39 | 0.080755 | 0.08129 | 0.993375 | 0.32053 | 0.53128 | 21.4 |
| Klhl5 | 2554.45 | 0.033416 | 0.07405 | 0.451239 | 0.65182 | 0.80618 | 60.65 |
| Klhl6 | 69.505 | -0.36092 | 0.21339 | -1.69133 | 0.09077 | 0.24468 | 36.542 |
| Klhl7 | 7865.84 | 0.581169 | 0.08969 | 6.479714 | 9.19E-11 | 9.37E-08 | 60.188 |
| Klhl8 | 1436.99 | 0.08597 | 0.07051 | 1.219237 | 0.22275 | 0.42599 | 49.287 |
| Klhl9 | 5276.02 | 0.206449 | 0.0636 | 3.245976 | 0.00117 | 0.0141 | 4.174 |
| Klk1 | 14.3623 | 0.049368 | 0.58214 | 0.084805 | 0.93242 | 0.96592 | 4.259 |
| Klk6 | 109.185 | -1.29827 | 0.25379 | -5.11552 | 3.13E-07 | 4.36E-05 | 7.532 |
| Klk8 | 35.5291 | -0.75053 | 0.34422 | -2.18039 | 0.02923 | 0.116 | 6.25 |
| Klra2 | 22.8908 | -0.14396 | 0.42556 | -0.33829 | 0.73514 | 0.85989 | 28.14 |
| Klrg2 | 23.1365 | -0.31365 | 0.46309 | -0.67729 | 0.49822 | 0.69157 | 11.55 |
| Kmo | 19.8662 | 0.24916 | 0.44763 | 0.556623 | 0.57779 | 0.75477 | 41.736 |
| Kmt2a | 16384 | 0.346694 | 0.08457 | 4.099562 | 4.14E-05 | 0.00143 | 77.943 |
| Kmt2b | 4529.04 | 0.163588 | 0.07473 | 2.1891 | 0.02859 | 0.11411 | 19.869 |
| Kmt2c | 7495.9 | 0.150008 | 0.07472 | 2.007622 | 0.04468 | 0.15353 | 226.986 |
| Kmt2d | 14108.8 | 0.21136 | 0.07803 | 2.70863 | 0.00676 | 0.04445 | 39.536 |
| Kmt2e | 7478.63 | 0.231475 | 0.08257 | 2.803335 | 0.00506 | 0.03656 | 69.795 |
| Kmt5a | 2075.49 | -0.15817 | 0.07794 | -2.02946 | 0.04241 | 0.14821 | 22.379 |
| Kmt5b | 4093.48 | 0.195947 | 0.06221 | 3.149676 | 0.00163 | 0.01761 | 50.883 |
| Kmt5c | 1058.44 | -0.08065 | 0.1177 | -0.68525 | 0.49319 | 0.68772 | 7.4 |
| Kncn | 16.6988 | 0.808237 | 0.4971 | 1.625894 | 0.10397 | 0.26648 | 5.127 |
| Kndc1 | 5023.33 | -0.42782 | 0.11059 | -3.86844 | 0.00011 | 0.00285 | 46.839 |
| Knl1 | 18.3046 | -0.24667 | 0.44728 | -0.55149 | 0.5813 | 0.75725 | 58.383 |
| Knop1 | 1246.24 | -0.09363 | 0.08158 | -1.14764 | 0.25112 | 0.45847 | 14.033 |
| Knstrn | 110.019 | -0.57102 | 0.2584 | -2.2098 | 0.02712 | 0.10996 | 39.955 |
| Kntc1 | 21.7703 | -0.55737 | 0.56396 | -0.98832 | 0.323 | 0.53402 | 71.878 |
| Kpna1 | 7482.26 | 0.1114 | 0.07377 | 1.510069 | 0.13103 | 0.30948 | 58.382 |
| Kpna2 | 814.55 | -0.02 | 0.09155 | -0.21842 | 0.8271 | 0.91108 | 10.913 |
| Kpna3 | 4317.17 | 0.097743 | 0.07484 | 1.306046 | 0.19154 | 0.38818 | 74.664 |
| Kpna4 | 5524.4 | 0.156907 | 0.05786 | 2.711666 | 0.00669 | 0.04418 | 59.965 |
| Kpna6 | 5052.61 | 0.024471 | 0.05867 | 0.417123 | 0.67659 | 0.82256 | 28.788 |
| Kpnb1 | 8163.35 | 0.012029 | 0.07275 | 0.165355 | 0.86866 | 0.93341 | 28.168 |
| Kptn | 1167.23 | 0.271141 | 0.10592 | 2.559827 | 0.01047 | 0.05926 | 7.622 |
| Kras | 5025.29 | -0.15933 | 0.11627 | -1.37028 | 0.1706 | 0.361 | 33.541 |

| | | | | | | | |
|---|---|---|---|---|---|---|---|
| Krba1 | 4254.98 | 0.242266 | 0.07408 | 3.270458 | 0.00107 | 0.01341 | 24.196 |
| Krcc1 | 631.239 | -0.13233 | 0.10336 | -1.28025 | 0.20046 | 0.39853 | 13.643 |
| Kremen1 | 1440.93 | -0.25857 | 0.07834 | -3.30068 | 0.00096 | 0.0125 | 70.007 |
| Kremen2 | 95.6855 | 0.541669 | 0.19755 | 2.74189 | 0.00611 | 0.04179 | 4.637 |
| Kri1 | 841.193 | 0.056629 | 0.07545 | 0.750596 | 0.4529 | 0.65531 | 14.513 |
| Krit1 | 4489.14 | 0.243292 | 0.09234 | 2.634806 | 0.00842 | 0.05116 | 42.381 |
| Krr1 | 1482.81 | 0.192742 | 0.08993 | 2.143211 | 0.0321 | 0.12375 | 15.769 |
| Krt1 | 145.052 | 0.462733 | 0.15378 | 3.00905 | 0.00262 | 0.0241 | 5.369 |
| Krt10 | 679.847 | 0.211125 | 0.15616 | 1.351963 | 0.17639 | 0.36846 | 4.111 |
| Krt12 | 114.447 | -0.02615 | 0.19772 | -0.13226 | 0.89478 | 0.94736 | 6.594 |
| Krt20 | 477.26 | 0.240252 | 0.16407 | 1.464345 | 0.1431 | 0.32515 | 9.748 |
| Krt222 | 2181.15 | 0.023247 | 0.07392 | 0.314497 | 0.75314 | 0.87029 | 11.325 |
| Krt23 | 20.6085 | 0.009747 | 0.44794 | 0.021759 | 0.98264 | 0.99061 | 15.164 |
| Krt25 | 35.0038 | 0.404621 | 0.31717 | 1.275727 | 0.20205 | 0.40058 | 7.436 |
| Krt26 | 36.5004 | 0.338514 | 0.41097 | 0.823687 | 0.41012 | 0.61801 | 9.417 |
| Krt73 | 57.9898 | 0.29941 | 0.2482 | 1.20635 | 0.22768 | 0.43174 | 9.039 |
| Krt77 | 485.97 | 0.720946 | 0.18512 | 3.894557 | 9.84E-05 | 0.00266 | 10.975 |
| Krt80 | 18.2339 | -0.96136 | 0.53602 | -1.79351 | 0.07289 | 0.21236 | 22.719 |
| Krt9 | 2350.15 | 0.83824 | 0.20128 | 4.164611 | 3.12E-05 | 0.00119 | 6.466 |
| Krtcap2 | 714.232 | -0.12068 | 0.11248 | -1.07283 | 0.28335 | 0.49341 | 3.941 |
| Krtcap3 | 210.42 | -0.00918 | 0.14559 | -0.06303 | 0.94974 | 0.97495 | 1.512 |
| Ksr1 | 4907.12 | -0.07096 | 0.07354 | -0.96493 | 0.33458 | 0.54535 | 132.968 |
| Ksr2 | 4862.53 | 0.15039 | 0.09458 | 1.590143 | 0.1118 | 0.27929 | 361.033 |
| Kti12 | 391.889 | 0.034013 | 0.12078 | 0.281626 | 0.77823 | 0.88423 | 1.629 |
| Ktn1 | 5932.8 | 0.327171 | 0.13969 | 2.342116 | 0.01917 | 0.087 | 91.447 |
| Kxd1 | 866.508 | 0.049617 | 0.08983 | 0.55233 | 0.58072 | 0.75697 | 19.685 |
| Ky | 187.793 | -0.05921 | 0.16826 | -0.35191 | 0.72491 | 0.85334 | 40.49 |
| Kyat1 | 728.716 | -0.00688 | 0.08733 | -0.07877 | 0.93722 | 0.96863 | 20.724 |
| Kyat3 | 307.059 | -0.07928 | 0.14751 | -0.53749 | 0.59093 | 0.7637 | 45.82 |
| L1cam | 7511.99 | 0.252304 | 0.05234 | 4.820362 | 1.43E-06 | 0.00014 | 42.328 |
| L2hgdh | 1199.9 | 0.195049 | 0.08124 | 2.40077 | 0.01636 | 0.07882 | 34.441 |
| L3hypdh | 165.837 | 0.078905 | 0.1672 | 0.471928 | 0.63698 | 0.79574 | 12.012 |
| L3mbtl1 | 928.63 | 0.234478 | 0.12975 | 1.807107 | 0.07075 | 0.20791 | 31.051 |
| L3mbtl2 | 1433.55 | 0.026332 | 0.08069 | 0.326353 | 0.74416 | 0.86539 | 24.427 |

| | | | | | | | |
|---|---|---|---|---|---|---|---|
| L3mbtl3 | 1397.91 | 0.0051 | 0.08883 | 0.05742 | 0.95421 | 0.97748 | 101.602 |
| L3mbtl4 | 77.6218 | -0.54314 | 0.24354 | -2.23021 | 0.02573 | 0.1061 | 506.29 |
| LTO1 | 778.802 | -0.04854 | 0.09483 | -0.51188 | 0.60874 | 0.77635 | 16.602 |
| Lacc1 | 256.767 | -0.40293 | 0.15362 | -2.62288 | 0.00872 | 0.05239 | 12.705 |
| Lactb | 822.959 | 0.23744 | 0.1167 | 2.034623 | 0.04189 | 0.1472 | 20.097 |
| Lactb2 | 1208.07 | -0.12356 | 0.09539 | -1.29538 | 0.19519 | 0.39273 | 37.217 |
| Lactbl1 | 13.6966 | 0.563639 | 0.56857 | 0.991319 | 0.32153 | 0.53233 | 16.49 |
| Lad1 | 81.0105 | -0.06278 | 0.26616 | -0.23586 | 0.81354 | 0.90411 | 14.745 |
| Lag3 | 219.727 | -0.58846 | 0.19958 | -2.94843 | 0.00319 | 0.02718 | 7.345 |
| Lage3 | 393.7 | -0.15634 | 0.10672 | -1.4649 | 0.14295 | 0.32502 | 2.343 |
| Lair1 | 461.093 | -0.30479 | 0.12885 | -2.36537 | 0.01801 | 0.08366 | 59.803 |
| Lama1 | 219.106 | -0.45354 | 0.20903 | -2.16981 | 0.03002 | 0.11832 | 125.389 |
| Lama2 | 843.213 | -0.61047 | 0.12554 | -4.8627 | 1.16E-06 | 0.00012 | 639.723 |
| Lama3 | 339.922 | -0.02578 | 0.17166 | -0.1502 | 0.88061 | 0.9404 | 249.195 |
| Lama4 | 272.591 | -0.47839 | 0.18833 | -2.54021 | 0.01108 | 0.06133 | 144.674 |
| Lama5 | 701.982 | -0.20086 | 0.09849 | -2.03943 | 0.04141 | 0.14598 | 49.487 |
| Lamb1 | 1716.68 | 0.539627 | 0.1833 | 2.944011 | 0.00324 | 0.02748 | 64.411 |
| Lamb2 | 1565.08 | -0.32074 | 0.10003 | -3.20644 | 0.00134 | 0.01534 | 10.795 |
| Lamb3 | 334.163 | 0.79717 | 0.19186 | 4.154889 | 3.25E-05 | 0.00122 | 136.18 |
| Lamc1 | 2335.86 | -0.17801 | 0.06811 | -2.61347 | 0.00896 | 0.05331 | 113.865 |
| Lamc2 | 492.808 | -0.33201 | 0.13249 | -2.50589 | 0.01221 | 0.06535 | 63.692 |
| Lamc3 | 395.196 | -0.48406 | 0.1575 | -3.07338 | 0.00212 | 0.02092 | 59.249 |
| Lamp1 | 8491.68 | -0.32516 | 0.08016 | -4.05626 | 4.99E-05 | 0.00163 | 16.178 |
| Lamp2 | 4341.19 | -0.23529 | 0.064 | -3.6765 | 0.00024 | 0.00478 | 55.098 |
| Lamp5 | 3304.96 | 0.487351 | 0.17977 | 2.710994 | 0.00671 | 0.04421 | 17.679 |
| Lamtor1 | 1953.89 | -0.1129 | 0.06325 | -1.78489 | 0.07428 | 0.21491 | 20.772 |
| Lamtor2 | 1301.64 | -0.16214 | 0.08651 | -1.87427 | 0.06089 | 0.18836 | 3.256 |
| Lamtor3 | 1933.78 | -0.10346 | 0.06989 | -1.48032 | 0.13879 | 0.32003 | 10.395 |
| Lamtor4 | 888.891 | -0.04125 | 0.09445 | -0.43677 | 0.66228 | 0.8134 | 3.917 |
| Lamtor5 | 1313.6 | -0.08352 | 0.06937 | -1.204 | 0.22859 | 0.43266 | 5.225 |
| Lancl1 | 6755.04 | 0.056651 | 0.08147 | 0.695323 | 0.48685 | 0.6826 | 38.356 |
| Lancl2 | 2673.74 | -0.18294 | 0.10868 | -1.6833 | 0.09232 | 0.2474 | 36.9 |
| Lancl3 | 393.253 | -0.04335 | 0.13836 | -0.31329 | 0.75406 | 0.87067 | 68.184 |
| Lao1 | 28.5119 | -0.33764 | 0.32737 | -1.03137 | 0.30237 | 0.51199 | 7.335 |

| | | | | | | | |
|---|---|---|---|---|---|---|---|
| Lap3 | 2230.93 | -0.28264 | 0.09827 | -2.87603 | 0.00403 | 0.03165 | 19.318 |
| Laptm4a | 7602.34 | -0.25208 | 0.08907 | -2.83023 | 0.00465 | 0.03467 | 17.145 |
| Laptm4b | 2684.49 | -0.45095 | 0.11099 | -4.06285 | 4.85E-05 | 0.0016 | 46.275 |
| Laptm5 | 1195.38 | -0.32309 | 0.12377 | -2.61041 | 0.00904 | 0.0536 | 23.017 |
| Large1 | 5668.82 | -0.31023 | 0.11806 | -2.62767 | 0.0086 | 0.05186 | 538.942 |
| Large2 | 13.2518 | -0.47242 | 0.49628 | -0.95191 | 0.34114 | 0.55206 | 9.824 |
| Larp1 | 7029.53 | -0.0368 | 0.05926 | -0.62092 | 0.53465 | 0.7217 | 52.971 |
| Larp1b | 595.738 | -0.34394 | 0.11664 | -2.94876 | 0.00319 | 0.02718 | 90.407 |
| Larp4 | 3064.65 | 0.118399 | 0.07599 | 1.558044 | 0.11922 | 0.2914 | 46.294 |
| Larp4b | 4470.9 | 0.080112 | 0.06684 | 1.198543 | 0.23071 | 0.43503 | 80.571 |
| Larp6 | 1312.86 | -0.06205 | 0.07367 | -0.84235 | 0.39959 | 0.6084 | 25.813 |
| Larp7 | 873.622 | 0.124698 | 0.09498 | 1.312914 | 0.18921 | 0.38511 | 16.636 |
| Lars | 2086.81 | 0.013799 | 0.082 | 0.168272 | 0.86637 | 0.93232 | 59.897 |
| Lars2 | 34098.6 | -0.08415 | 0.16309 | -0.51598 | 0.60587 | 0.77449 | 95.74 |
| Las1l | 3735.02 | 0.211166 | 0.05116 | 4.127427 | 3.67E-05 | 0.00131 | 21.628 |
| Lasp1 | 4853 | -0.23042 | 0.10414 | -2.21252 | 0.02693 | 0.10955 | 39.765 |
| Lat | 58.212 | 0.155077 | 0.21674 | 0.715515 | 0.47429 | 0.67238 | 5.747 |
| Lat2 | 62.1767 | -0.15141 | 0.29376 | -0.51542 | 0.60626 | 0.77465 | 15.004 |
| Lats1 | 3122.31 | 0.142358 | 0.06033 | 2.359707 | 0.01829 | 0.08456 | 35.247 |
| Lats2 | 968.803 | -0.3003 | 0.10154 | -2.95754 | 0.0031 | 0.02664 | 68.727 |
| Lax1 | 32.2943 | -0.44608 | 0.41288 | -1.08042 | 0.27995 | 0.49005 | 11.018 |
| Layn | 47.5382 | -0.99041 | 0.28893 | -3.4279 | 0.00061 | 0.00906 | 22.455 |
| Lbh | 2018.85 | -0.49683 | 0.10704 | -4.64157 | 3.46E-06 | 0.00026 | 23.643 |
| Lbhd1 | 169.258 | -0.05966 | 0.17041 | -0.35008 | 0.72628 | 0.8541 | 8.896 |
| Lbhd2 | 29.6641 | -0.39405 | 0.3856 | -1.02191 | 0.30682 | 0.51675 | 5.282 |
| Lbp | 70.0146 | -0.90247 | 0.43176 | -2.0902 | 0.0366 | 0.13482 | 26.36 |
| Lbr | 731.359 | -0.30548 | 0.12642 | -2.41632 | 0.01568 | 0.07666 | 27.712 |
| Lca5 | 934.409 | 0.334862 | 0.10632 | 3.149614 | 0.00163 | 0.01761 | 50.835 |
| Lca5l | 355.977 | 0.554336 | 0.11559 | 4.795685 | 1.62E-06 | 0.00015 | 33.865 |
| Lcat | 487.326 | -0.32565 | 0.10019 | -3.25011 | 0.00115 | 0.01396 | 3.832 |
| Lck | 84.1713 | 0.647075 | 0.21641 | 2.990066 | 0.00279 | 0.02505 | 25.298 |
| Lclat1 | 2186.03 | 0.04431 | 0.05946 | 0.745206 | 0.45615 | 0.65776 | 135.384 |
| Lcmt1 | 2807.38 | 0.0976 | 0.07007 | 1.392828 | 0.16367 | 0.35241 | 60.575 |
| Lcmt2 | 265.084 | -0.24387 | 0.14369 | -1.69719 | 0.08966 | 0.24268 | 12.347 |

| | | | | | | | |
|---|---|---|---|---|---|---|---|
| Lcn2 | 212.51 | -2 | 0.91855 | -2.91826 | 0.00352 | 0.02886 | 3.62 |
| Lcor | 1262.48 | 0.161844 | 0.09568 | 1.691493 | 0.09074 | 0.24468 | 103.9 |
| Lcorl | 849.19 | 0.371052 | 0.19387 | 1.913961 | 0.05563 | 0.17798 | 160.435 |
| Lcp1 | 356.452 | -0.55103 | 0.21402 | -2.57458 | 0.01004 | 0.05759 | 99.742 |
| Lcp2 | 114.044 | -0.29509 | 0.18996 | -1.55343 | 0.12032 | 0.29306 | 45.376 |
| Lctl | 14.9366 | -0.21373 | 0.54596 | -0.39148 | 0.69544 | 0.83491 | 20.972 |
| Ldah | 2160.7 | 0.1063 | 0.09966 | 1.066604 | 0.28615 | 0.49619 | 77.653 |
| Ldb1 | 5066.06 | 0.092221 | 0.06224 | 1.481801 | 0.13839 | 0.31955 | 13.645 |
| Ldb2 | 689.456 | 0.765074 | 0.21833 | 3.504288 | 0.00046 | 0.0075 | 327.549 |
| Ldb3 | 547.051 | 0.238521 | 0.09507 | 2.509 | 0.01211 | 0.06496 | 62.08 |
| Ldha | 8627.42 | -0.22604 | 0.08699 | -2.5984 | 0.00937 | 0.05497 | 14.153 |
| Ldha-ps2 | 58.7924 | -0.09958 | 0.24651 | -0.40396 | 0.68624 | 0.82887 | 1.057 |
| Ldhb | 19156.1 | -0.12456 | 0.08305 | -1.49995 | 0.13363 | 0.31339 | 17.709 |
| Ldhd | 404.267 | -0.22714 | 0.12035 | -1.88726 | 0.05913 | 0.18511 | 6.59 |
| Ldlr | 952.492 | -0.45799 | 0.15413 | -2.97146 | 0.00296 | 0.02597 | 26.437 |
| Ldlrad3 | 444.884 | -0.40764 | 0.14712 | -2.77078 | 0.00559 | 0.03925 | 236.183 |
| Ldlrad4 | 1233 | -0.54099 | 0.13538 | -3.99619 | 6.44E-05 | 0.00198 | 335.374 |
| Ldlrap1 | 209.9 | -0.88593 | 0.15311 | -5.78614 | 7.20E-09 | 2.33E-06 | 26.471 |
| Ldoc1 | 34.185 | -0.33581 | 0.37105 | -0.90504 | 0.36544 | 0.57607 | 1.459 |
| Lef1 | 125.965 | -0.42441 | 0.23286 | -1.82257 | 0.06837 | 0.20312 | 113.886 |
| Lefty1 | 23.0697 | -0.39009 | 0.41416 | -0.94189 | 0.34625 | 0.55684 | 3.379 |
| Lefty2 | 20.5623 | -1.60498 | 0.6745 | -2.37952 | 0.01734 | 0.08167 | 5.996 |
| Lekr1 | 356.372 | 0.189229 | 0.11985 | 1.578859 | 0.11437 | 0.28327 | 165.128 |
| Lemd1 | 31.1044 | -0.13679 | 0.39011 | -0.35065 | 0.72585 | 0.85388 | 67.797 |
| Lemd2 | 2001.74 | 0.114484 | 0.06864 | 1.667906 | 0.09533 | 0.25234 | 14.87 |
| Lemd3 | 2641.54 | -0.06137 | 0.06554 | -0.93637 | 0.34908 | 0.55954 | 55.92 |
| Leng1 | 374.189 | 0.111489 | 0.13434 | 0.829923 | 0.40658 | 0.6148 | 5.743 |
| Leng8 | 17165.8 | 0.206439 | 0.10665 | 1.93575 | 0.0529 | 0.1721 | 11.139 |
| Leng9 | 198.034 | -0.0339 | 0.17524 | -0.19344 | 0.84661 | 0.92187 | 2.46 |
| Leo1 | 983.15 | 0.081485 | 0.11273 | 0.722819 | 0.46979 | 0.6687 | 24.909 |
| Lepr | 148.07 | -0.86453 | 0.29657 | -2.91506 | 0.00356 | 0.02907 | 97.949 |
| Leprot | 1364.95 | -0.12081 | 0.06883 | -1.75504 | 0.07925 | 0.22347 | 11.647 |
| Leprotl1 | 3249.16 | -0.00672 | 0.06395 | -0.10514 | 0.91627 | 0.95722 | 11.462 |
| Letm1 | 3999.01 | -0.05239 | 0.06633 | -0.78989 | 0.42959 | 0.63602 | 43.145 |

| | | | | | | | |
|---|---|---|---|---|---|---|---|
| Letm2 | 1164.12 | 0.226579 | 0.09037 | 2.507244 | 0.01217 | 0.06517 | 19.093 |
| Letmd1 | 2296.64 | 0.015608 | 0.06824 | 0.228727 | 0.81908 | 0.90696 | 10.142 |
| Lexis1 | 18.3914 | -0.31548 | 0.3978 | -0.79305 | 0.42775 | 0.6344 | 12.792 |
| Lfng | 1111.22 | 0.133943 | 0.13728 | 0.975675 | 0.32923 | 0.54042 | 8.226 |
| Lgals1 | 300.792 | -1.03674 | 0.18465 | -5.61454 | 1.97E-08 | 5.82E-06 | 3.741 |
| Lgals12 | 40.885 | 0.545881 | 0.35649 | 1.531268 | 0.1257 | 0.30095 | 10.534 |
| Lgals3 | 28.6117 | -2 | 0.74241 | -2.73422 | 0.00625 | 0.04232 | 18.41 |
| Lgals3bp | 830.93 | -0.30731 | 0.167 | -1.84025 | 0.06573 | 0.19797 | 9.342 |
| Lgals4 | 179.983 | 0.304869 | 0.17169 | 1.775668 | 0.07579 | 0.21741 | 7.855 |
| Lgals8 | 1207.18 | -0.06416 | 0.0878 | -0.73085 | 0.46487 | 0.66451 | 25.53 |
| Lgals9 | 228.611 | -0.40774 | 0.15882 | -2.56736 | 0.01025 | 0.05844 | 21.973 |
| Lgalsl | 4776.62 | 0.09887 | 0.06449 | 1.533018 | 0.12527 | 0.30027 | 7.481 |
| Lgi1 | 4378 | 0.06704 | 0.08375 | 0.800511 | 0.42341 | 0.63022 | 47.679 |
| Lgi2 | 8106.75 | -0.43453 | 0.10104 | -4.30056 | 1.70E-05 | 0.0008 | 32.946 |
| Lgi3 | 4822.05 | -0.32504 | 0.09243 | -3.51656 | 0.00044 | 0.00729 | 7.64 |
| Lgi4 | 727.314 | -0.35084 | 0.14585 | -2.40553 | 0.01615 | 0.07815 | 11.594 |
| Lgmn | 3228.57 | -0.18805 | 0.10118 | -1.8586 | 0.06308 | 0.19285 | 45.73 |
| Lgr4 | 1509.88 | 0.076423 | 0.08213 | 0.930502 | 0.35211 | 0.5625 | 96.611 |
| Lgr5 | 395.926 | -0.73293 | 0.4237 | -1.72983 | 0.08366 | 0.23136 | 137.47 |
| Lgr6 | 155.722 | -1.98578 | 0.72186 | -2.7509 | 0.00594 | 0.04096 | 121.976 |
| Lhcgr | 129.982 | 1.009701 | 0.18588 | 5.431991 | 5.57E-08 | 1.20E-05 | 75.51 |
| Lhfp | 1170.54 | -0.52928 | 0.16899 | -3.1321 | 0.00174 | 0.01838 | 220.152 |
| Lhfpl2 | 577.836 | -0.6191 | 0.20852 | -2.96911 | 0.00299 | 0.02607 | 137.614 |
| Lhfpl3 | 1597.16 | 0.225431 | 0.12766 | 1.765838 | 0.07742 | 0.22011 | 529.539 |
| Lhfpl4 | 9061.51 | 0.037073 | 0.07376 | 0.502601 | 0.61524 | 0.78109 | 27.295 |
| Lhfpl5 | 162.995 | 0.191555 | 0.16188 | 1.183336 | 0.23668 | 0.44183 | 10.039 |
| Lhpp | 464.362 | 0.009609 | 0.12556 | 0.07653 | 0.939 | 0.96951 | 95.783 |
| Lhx2 | 1443.84 | -0.31143 | 0.12737 | -2.44511 | 0.01448 | 0.07271 | 30.453 |
| Lhx6 | 1070.96 | 0.45688 | 0.20024 | 2.281686 | 0.02251 | 0.09685 | 23.456 |
| Lhx8 | 302.713 | 1.355699 | 0.49695 | 2.728067 | 0.00637 | 0.04283 | 24.366 |
| Lhx9 | 20.814 | -0.48144 | 0.47148 | -1.02113 | 0.30719 | 0.51695 | 23.392 |
| Lias | 1242.35 | -0.0965 | 0.0737 | -1.30931 | 0.19043 | 0.38666 | 19.197 |
| Lif | 24.3622 | -0.15282 | 0.39979 | -0.38224 | 0.70228 | 0.83936 | 14.958 |
| Lifr | 2098.58 | -0.15424 | 0.08683 | -1.77628 | 0.07569 | 0.21723 | 106.876 |

| | | | | | | | |
|---|---|---|---|---|---|---|---|
| Lig1 | 368.388 | -0.34585 | 0.11697 | -2.95683 | 0.00311 | 0.02669 | 34.112 |
| Lig3 | 1541.11 | 0.151973 | 0.08407 | 1.807702 | 0.07065 | 0.20776 | 23.167 |
| Lig4 | 670.01 | 0.034114 | 0.1337 | 0.255161 | 0.7986 | 0.89601 | 8.638 |
| Lilrb4a | 17.3515 | -1.84157 | 0.5502 | -3.34706 | 0.00082 | 0.01114 | 5.656 |
| Lima1 | 989.737 | -0.06007 | 0.1098 | -0.54706 | 0.58434 | 0.75916 | 96.987 |
| Limch1 | 8249.67 | 0.016286 | 0.05524 | 0.294808 | 0.76814 | 0.87883 | 311.332 |
| Limd1 | 608.036 | -0.01894 | 0.09544 | -0.19842 | 0.84272 | 0.91905 | 42.847 |
| Limd2 | 2568.85 | -0.06664 | 0.09129 | -0.72994 | 0.46543 | 0.66495 | 4.605 |
| Limk1 | 1935.74 | -0.25745 | 0.08573 | -3.00301 | 0.00267 | 0.0244 | 32.56 |
| Limk2 | 5204.41 | 0.051247 | 0.07067 | 0.72512 | 0.46838 | 0.66728 | 64.934 |
| Lims1 | 1481.13 | 0.022988 | 0.07414 | 0.310064 | 0.75651 | 0.87187 | 101.226 |
| Lims2 | 381.213 | -0.40068 | 0.12429 | -3.22383 | 0.00126 | 0.01476 | 36.33 |
| Lin28b | 805.892 | 0.256779 | 0.12164 | 2.110967 | 0.03478 | 0.13064 | 109.776 |
| Lin37 | 645.623 | -0.00754 | 0.0982 | -0.0768 | 0.93878 | 0.96951 | 4.398 |
| Lin52 | 1285.65 | -0.1983 | 0.07501 | -2.64365 | 0.0082 | 0.05034 | 94.724 |
| Lin54 | 708.026 | -0.12992 | 0.08675 | -1.49765 | 0.13422 | 0.31405 | 58.594 |
| Lin7a | 1617.24 | 0.274312 | 0.12535 | 2.18839 | 0.02864 | 0.11426 | 149.789 |
| Lin7b | 827.637 | 0.004753 | 0.10047 | 0.047302 | 0.96227 | 0.98168 | 2.693 |
| Lin7c | 5752.84 | 0.29517 | 0.07688 | 3.839343 | 0.00012 | 0.00309 | 10.151 |
| Lin9 | 264.535 | 0.097874 | 0.12965 | 0.75493 | 0.45029 | 0.65308 | 49.545 |
| Lingo1 | 4795.92 | -0.68661 | 0.17445 | -3.93583 | 8.29E-05 | 0.00236 | 177.994 |
| Lingo2 | 660.026 | -0.18394 | 0.11597 | -1.5861 | 0.11272 | 0.28071 | 1245.101 |
| Lingo3 | 4121.45 | 0.615052 | 0.17976 | 3.421433 | 0.00062 | 0.00921 | 11.239 |
| Lingo4 | 32.039 | -0.64501 | 0.36927 | -1.74672 | 0.08069 | 0.22605 | 5.985 |
| Lins1 | 513.676 | 0.007429 | 0.09939 | 0.074751 | 0.94041 | 0.97035 | 27.368 |
| Lipa | 849.892 | -0.56457 | 0.15751 | -3.58441 | 0.00034 | 0.00612 | 35.157 |
| Lipe | 544.578 | 0.026185 | 0.10515 | 0.249018 | 0.80335 | 0.89848 | 19.184 |
| Lipg | 174.13 | -0.94068 | 0.22956 | -4.0978 | 4.17E-05 | 0.00144 | 21.942 |
| Liph | 50.17 | 0.103289 | 0.25732 | 0.401395 | 0.68813 | 0.83002 | 41.847 |
| Lipo3 | 533.064 | 0.181795 | 0.13294 | 1.367526 | 0.17146 | 0.36184 | 206.792 |
| Lipt1 | 164.852 | 0.238083 | 0.17441 | 1.365081 | 0.17223 | 0.36301 | 4.491 |
| Lipt2 | 120.826 | 0.063094 | 0.16469 | 0.38311 | 0.70164 | 0.83894 | 2.089 |
| Litaf | 641.678 | -0.59864 | 0.1591 | -3.76254 | 0.00017 | 0.00386 | 106.883 |
| Lix1 | 1864.58 | -0.26666 | 0.11568 | -2.30509 | 0.02116 | 0.093 | 56.716 |

| | | | | | | | |
|---|---|---|---|---|---|---|---|
| Lix1l | 1445.25 | -0.18268 | 0.09256 | -1.97373 | 0.04841 | 0.16228 | 25.023 |
| Llgl1 | 2670.6 | -0.00473 | 0.08573 | -0.05519 | 0.95599 | 0.9784 | 14.464 |
| Llgl2 | 70.0448 | -0.57394 | 0.23539 | -2.4383 | 0.01476 | 0.07344 | 31.732 |
| Llph | 1580.52 | -0.10232 | 0.08671 | -1.18005 | 0.23798 | 0.44334 | 5.513 |
| Llph-ps2 | 44.5986 | -0.22217 | 0.30005 | -0.74044 | 0.45903 | 0.65977 | 0.393 |
| Lman1 | 2893.26 | 0.319394 | 0.09906 | 3.224306 | 0.00126 | 0.01476 | 41.843 |
| Lman1l | 23.0265 | -0.47648 | 0.38639 | -1.23318 | 0.21751 | 0.41946 | 13.69 |
| Lman2 | 3876.74 | -0.16434 | 0.07748 | -2.12102 | 0.03392 | 0.12861 | 18.951 |
| Lman2l | 3028.39 | 0.166836 | 0.07961 | 2.095803 | 0.0361 | 0.13377 | 25.401 |
| Lmbr1 | 1804.66 | 0.086531 | 0.07065 | 1.224734 | 0.22068 | 0.4234 | 148.589 |
| Lmbr1l | 710.619 | 0.060209 | 0.10808 | 0.557086 | 0.57747 | 0.75451 | 14.315 |
| Lmbrd1 | 7979.76 | -0.05149 | 0.04912 | -1.04807 | 0.29461 | 0.50498 | 87.672 |
| Lmbrd2 | 6111.27 | 0.113929 | 0.09766 | 1.16655 | 0.24339 | 0.44968 | 61.933 |
| Lmcd1 | 200.701 | -0.41499 | 0.22892 | -1.81284 | 0.06986 | 0.20626 | 56.668 |
| Lmf1 | 971.706 | 0.291168 | 0.09418 | 3.091552 | 0.00199 | 0.02005 | 83.742 |
| Lmf2 | 703.08 | -0.28059 | 0.10542 | -2.66164 | 0.00778 | 0.04879 | 4.656 |
| Lmln | 1002.39 | 0.226091 | 0.10288 | 2.197637 | 0.02798 | 0.11256 | 65.175 |
| Lmna | 1038.94 | -0.2898 | 0.09725 | -2.98 | 0.00288 | 0.02558 | 29.81 |
| Lmnb1 | 506.718 | -0.18393 | 0.13027 | -1.41192 | 0.15797 | 0.34524 | 45.612 |
| Lmnb2 | 1374.32 | 0.028041 | 0.08382 | 0.334543 | 0.73797 | 0.86171 | 17.043 |
| Lmntd1 | 41.4938 | -0.49248 | 0.28397 | -1.73425 | 0.08287 | 0.22962 | 249.186 |
| Lmo1 | 169.019 | -0.01875 | 0.17265 | -0.10861 | 0.91351 | 0.95625 | 36.636 |
| Lmo2 | 1337.56 | 0.39482 | 0.1533 | 2.575496 | 0.01001 | 0.05751 | 23.893 |
| Lmo3 | 12093.6 | -0.54378 | 0.15032 | -3.61747 | 0.0003 | 0.0056 | 219.051 |
| Lmo4 | 5042.4 | 0.125653 | 0.09113 | 1.378834 | 0.16795 | 0.35782 | 16.691 |
| Lmo7 | 12555.1 | 0.315649 | 0.08023 | 3.934287 | 8.34E-05 | 0.00237 | 204.754 |
| Lmod1 | 139.826 | -0.92843 | 0.2182 | -4.25488 | 2.09E-05 | 0.00091 | 43.259 |
| Lmtk2 | 9096.71 | 0.081715 | 0.06375 | 1.281779 | 0.19992 | 0.39801 | 87.769 |
| Lmtk3 | 12913.9 | 0.219473 | 0.07967 | 2.754674 | 0.00588 | 0.04068 | 20.407 |
| Lncenc1 | 20.096 | 0.105014 | 0.54246 | 0.193591 | 0.8465 | 0.92184 | 73.95 |
| Lncpint | 409.922 | 0.09988 | 0.18989 | 0.525987 | 0.5989 | 0.76992 | 137.924 |
| Lncppara | 302.52 | -0.17731 | 0.14448 | -1.22721 | 0.21975 | 0.4222 | 53.972 |
| Lnp1 | 51.8655 | -1.16898 | 0.36786 | -3.1778 | 0.00148 | 0.01643 | 13.718 |
| Lnpep | 8186.61 | -0.06985 | 0.0657 | -1.06313 | 0.28772 | 0.49783 | 103.641 |

| | | | | | | | |
|---|---|---|---|---|---|---|---|
| Lnpk | 2319.06 | 0.04936 | 0.08815 | 0.559925 | 0.57553 | 0.75343 | 59.145 |
| Lnx1 | 614.712 | -0.45697 | 0.16332 | -2.79806 | 0.00514 | 0.03696 | 110.466 |
| Lnx2 | 333.616 | -0.24504 | 0.16989 | -1.44236 | 0.1492 | 0.33329 | 59.932 |
| Lonp1 | 4011.82 | 0.007122 | 0.05556 | 0.128201 | 0.89799 | 0.94882 | 12.591 |
| Lonp2 | 3437.34 | -0.08885 | 0.07366 | -1.20622 | 0.22773 | 0.43174 | 99.83 |
| Lonrf1 | 1488.47 | 0.292187 | 0.07811 | 3.740874 | 0.00018 | 0.00406 | 33.453 |
| Lonrf2 | 10838.4 | 0.197475 | 0.06347 | 3.111386 | 0.00186 | 0.01915 | 43.067 |
| Lonrf3 | 786.048 | 0.085872 | 0.12933 | 0.663985 | 0.5067 | 0.69797 | 33.989 |
| Lor | 399.838 | -0.05527 | 0.16909 | -0.32687 | 0.74377 | 0.86539 | 2.872 |
| Lox | 39.4909 | -1.0589 | 0.4169 | -2.53992 | 0.01109 | 0.06133 | 13.801 |
| Loxhd1 | 13.0816 | -1.02909 | 0.7131 | -1.44311 | 0.14899 | 0.33293 | 160.973 |
| Loxl1 | 109.84 | -0.82265 | 0.20016 | -4.10993 | 3.96E-05 | 0.00139 | 25.449 |
| Loxl2 | 23.5202 | -0.5814 | 0.44958 | -1.2932 | 0.19594 | 0.39362 | 86.767 |
| Loxl3 | 207.755 | -0.38757 | 0.13075 | -2.96412 | 0.00304 | 0.02632 | 18.39 |
| Loxl4 | 16.2271 | -0.56977 | 0.42794 | -1.33141 | 0.18305 | 0.37709 | 18.832 |
| Lpar1 | 1143.8 | -0.47717 | 0.15201 | -3.13915 | 0.00169 | 0.01808 | 118.644 |
| Lpar2 | 86.2427 | 0.340486 | 0.22908 | 1.486346 | 0.13719 | 0.31781 | 8.674 |
| Lpar3 | 23.2486 | -0.68321 | 0.48167 | -1.41843 | 0.15607 | 0.3427 | 65.224 |
| Lpar4 | 73.7531 | -0.25943 | 0.24394 | -1.06349 | 0.28756 | 0.49771 | 13.276 |
| Lpar5 | 21.6953 | -0.13231 | 0.41517 | -0.31868 | 0.74997 | 0.86875 | 14.553 |
| Lpar6 | 125.594 | -0.2336 | 0.16953 | -1.37794 | 0.16822 | 0.35824 | 5.4 |
| Lpcat1 | 1912.61 | -0.15284 | 0.07439 | -2.05462 | 0.03992 | 0.14277 | 49.226 |
| Lpcat2 | 271.822 | -0.10354 | 0.1652 | -0.62674 | 0.53083 | 0.71839 | 63.941 |
| Lpcat3 | 813.302 | -0.10535 | 0.08018 | -1.31398 | 0.18885 | 0.38461 | 41.392 |
| Lpcat4 | 3466.77 | -0.06274 | 0.08456 | -0.74198 | 0.4581 | 0.65937 | 7.644 |
| Lpgat1 | 17440.6 | -0.18759 | 0.10815 | -1.73455 | 0.08282 | 0.22962 | 66.422 |
| Lpin1 | 1891.19 | 0.059397 | 0.07589 | 0.782725 | 0.43379 | 0.63959 | 111.298 |
| Lpin2 | 3679.07 | 0.039501 | 0.07619 | 0.518433 | 0.60416 | 0.7733 | 67.258 |
| Lpin3 | 90.4032 | -0.45485 | 0.30731 | -1.48012 | 0.13884 | 0.32009 | 25.333 |
| Lpl | 2675.12 | 0.809593 | 0.29859 | 2.711354 | 0.0067 | 0.04418 | 26.958 |
| Lpo | 23.7876 | -0.32841 | 0.40536 | -0.81015 | 0.41785 | 0.62523 | 21.862 |
| Lpp | 811.714 | -0.15586 | 0.09353 | -1.66635 | 0.09564 | 0.25293 | 599.072 |
| Lpxn | 64.85 | -0.3652 | 0.27688 | -1.31896 | 0.18718 | 0.38244 | 37.615 |
| Lratd1 | 4048.19 | -0.04872 | 0.07943 | -0.61329 | 0.53968 | 0.72589 | 5.225 |

| | | | | | | | |
|---|---|---|---|---|---|---|---|
| Lratd2 | 251.731 | -0.03318 | 0.17611 | -0.18841 | 0.85056 | 0.92407 | 34.785 |
| Lrba | 3221.01 | 0.2387 | 0.10247 | 2.329394 | 0.01984 | 0.08901 | 558.013 |
| Lrch1 | 1135.99 | 0.080353 | 0.07743 | 1.037687 | 0.29942 | 0.50955 | 193.202 |
| Lrch2 | 2046.23 | 0.446598 | 0.12638 | 3.533751 | 0.00041 | 0.00698 | 83.707 |
| Lrch3 | 1502.54 | 0.148965 | 0.08924 | 1.669281 | 0.09506 | 0.25181 | 101.548 |
| Lrfn1 | 1202.53 | -0.32855 | 0.13256 | -2.47847 | 0.01319 | 0.06877 | 16.263 |
| Lrfn2 | 694.596 | -0.27736 | 0.11833 | -2.34397 | 0.01908 | 0.08673 | 165.21 |
| Lrfn3 | 773.869 | -0.01157 | 0.10758 | -0.10753 | 0.91437 | 0.95656 | 7.284 |
| Lrfn4 | 1658.67 | 0.086263 | 0.08365 | 1.031286 | 0.30241 | 0.51199 | 3.883 |
| Lrfn5 | 2128.84 | 0.306218 | 0.10541 | 2.905137 | 0.00367 | 0.02974 | 335.193 |
| Lrg1 | 26.0643 | -1.35881 | 0.45788 | -2.96764 | 0.003 | 0.02618 | 2.324 |
| Lrguk | 304.956 | 0.660786 | 0.18797 | 3.515416 | 0.00044 | 0.0073 | 104.591 |
| Lrif1 | 943.694 | 0.338166 | 0.18834 | 1.795551 | 0.07257 | 0.21177 | 51.591 |
| Lrig1 | 1512.39 | 0.117601 | 0.1218 | 0.965529 | 0.33428 | 0.54499 | 95.63 |
| Lrig2 | 2321 | 0.1254 | 0.06331 | 1.980767 | 0.04762 | 0.16041 | 115.501 |
| Lrig3 | 134.606 | -0.57992 | 0.23599 | -2.45744 | 0.01399 | 0.07126 | 49.192 |
| Lrmda | 26.4787 | 0.128636 | 0.44672 | 0.287957 | 0.77338 | 0.88189 | 1036.374 |
| Lrp1 | 31917.9 | 0.08201 | 0.05318 | 1.542146 | 0.12304 | 0.29719 | 82.988 |
| Lrp10 | 993.558 | -0.05176 | 0.0816 | -0.63434 | 0.52586 | 0.71408 | 7.361 |
| Lrp11 | 7120.54 | 0.059339 | 0.0808 | 0.73438 | 0.46272 | 0.66283 | 35.684 |
| Lrp12 | 2821.17 | 0.065194 | 0.06718 | 0.970451 | 0.33182 | 0.54278 | 73.406 |
| Lrp1b | 3487.48 | 0.235621 | 0.13322 | 1.768706 | 0.07694 | 0.21936 | 2058.379 |
| Lrp2bp | 57.5177 | 0.693314 | 0.22879 | 3.030339 | 0.00244 | 0.02296 | 30.175 |
| Lrp3 | 2992.01 | -0.04786 | 0.06135 | -0.78007 | 0.43535 | 0.64101 | 15.472 |
| Lrp4 | 2360.85 | -0.41439 | 0.12055 | -3.43745 | 0.00059 | 0.00882 | 56.269 |
| Lrp5 | 589.936 | -0.56578 | 0.10877 | -5.20173 | 1.97E-07 | 3.01E-05 | 101.737 |
| Lrp6 | 4285.01 | 0.128815 | 0.06583 | 1.956756 | 0.05038 | 0.16683 | 120.49 |
| Lrp8 | 4787.61 | -0.25135 | 0.1548 | -1.62374 | 0.10443 | 0.2674 | 74.972 |
| Lrp8os2 | 348.252 | 0.529786 | 0.22324 | 2.373216 | 0.01763 | 0.0825 | 16.99 |
| Lrp8os3 | 45.3652 | 0.171006 | 0.28306 | 0.604137 | 0.54575 | 0.73054 | 8.904 |
| Lrpap1 | 8741.69 | 0.015144 | 0.06995 | 0.216501 | 0.8286 | 0.91212 | 14.266 |
| Lrpprc | 3987.59 | 0.160295 | 0.08168 | 1.962583 | 0.04969 | 0.16518 | 85.543 |
| Lrrc1 | 1145.82 | 0.358304 | 0.15944 | 2.247292 | 0.02462 | 0.10289 | 114.048 |
| Lrrc10b | 6070.63 | 0.401663 | 0.31245 | 1.285511 | 0.19861 | 0.39637 | 2.103 |

| | | | | | | | |
|---|---|---|---|---|---|---|---|
| Lrrc14 | 959.251 | -0.07808 | 0.08146 | -0.95849 | 0.33782 | 0.54856 | 7.077 |
| Lrrc18 | 32.8557 | 0.781938 | 0.35373 | 2.210533 | 0.02707 | 0.10988 | 23.91 |
| Lrrc2 | 81.7924 | -0.34193 | 0.20467 | -1.6706 | 0.0948 | 0.25135 | 32.522 |
| Lrrc20 | 1135.71 | -0.11407 | 0.08564 | -1.33199 | 0.18286 | 0.37686 | 106.991 |
| Lrrc23 | 43.8467 | -0.00431 | 0.33973 | -0.0127 | 0.98987 | 0.99445 | 9.865 |
| Lrrc24 | 255.661 | -0.14209 | 0.11657 | -1.21899 | 0.22285 | 0.42605 | 6.898 |
| Lrrc25 | 18.4304 | -0.64226 | 0.40769 | -1.57537 | 0.11517 | 0.28453 | 5.329 |
| Lrrc26 | 162.219 | 0.287203 | 0.17663 | 1.626 | 0.10395 | 0.26646 | 1.266 |
| Lrrc27 | 239.844 | -0.23546 | 0.13054 | -1.80371 | 0.07128 | 0.20897 | 29.992 |
| Lrrc28 | 873.837 | -0.0368 | 0.08149 | -0.45151 | 0.65162 | 0.80613 | 131.859 |
| Lrrc29 | 123.103 | -0.00694 | 0.16778 | -0.04137 | 0.967 | 0.98345 | 14.014 |
| Lrrc3 | 535.572 | -0.05619 | 0.10195 | -0.55114 | 0.58153 | 0.75743 | 4.962 |
| Lrrc32 | 132.864 | -0.19294 | 0.22424 | -0.86041 | 0.38956 | 0.59881 | 12.899 |
| Lrrc34 | 14.5221 | 0.13768 | 0.53547 | 0.257118 | 0.79709 | 0.89539 | 23.603 |
| Lrrc38 | 116.693 | 0.629174 | 0.22913 | 2.745977 | 0.00603 | 0.04143 | 21.276 |
| Lrrc39 | 51.6034 | 0.082734 | 0.28107 | 0.294356 | 0.76849 | 0.87901 | 20.162 |
| Lrrc3b | 204.93 | 0.249425 | 0.16708 | 1.492806 | 0.13549 | 0.31534 | 81.473 |
| Lrrc4 | 2720.63 | -0.1235 | 0.06572 | -1.87918 | 0.06022 | 0.18722 | 169.917 |
| Lrrc40 | 1297.32 | 0.229205 | 0.08879 | 2.581327 | 0.00984 | 0.05688 | 31.826 |
| Lrrc41 | 955.422 | 0.035245 | 0.09951 | 0.354177 | 0.72321 | 0.85223 | 21.775 |
| Lrrc42 | 936.893 | 0.321064 | 0.09039 | 3.551838 | 0.00038 | 0.00663 | 20.019 |
| Lrrc45 | 3566.91 | 0.240025 | 0.10138 | 2.367623 | 0.0179 | 0.08334 | 7.204 |
| Lrrc46 | 21.3723 | 0.455454 | 0.36308 | 1.254431 | 0.20969 | 0.40977 | 6.806 |
| Lrrc47 | 1753.49 | -0.00987 | 0.07273 | -0.13576 | 0.89201 | 0.94613 | 9.783 |
| Lrrc49 | 2954.18 | 0.16787 | 0.09219 | 1.820968 | 0.06861 | 0.20362 | 119.3 |
| Lrrc4b | 7035.94 | -0.03885 | 0.07213 | -0.53864 | 0.59014 | 0.76314 | 34.334 |
| Lrrc4c | 4421.13 | 0.311514 | 0.09705 | 3.209892 | 0.00133 | 0.01521 | 1313.498 |
| Lrrc51 | 88.5037 | -0.0765 | 0.19115 | -0.40024 | 0.68898 | 0.83048 | 20.912 |
| Lrrc55 | 1127.16 | -0.10289 | 0.1424 | -0.72258 | 0.46994 | 0.66875 | 34.864 |
| Lrrc56 | 142.263 | -0.01757 | 0.17785 | -0.09876 | 0.92133 | 0.96024 | 15.899 |
| Lrrc57 | 1076.5 | -0.33464 | 0.0891 | -3.75561 | 0.00017 | 0.0039 | 5.283 |
| Lrrc58 | 7885.97 | -0.17295 | 0.05949 | -2.9069 | 0.00365 | 0.02963 | 20.47 |
| Lrrc59 | 3935.14 | -0.17666 | 0.08541 | -2.06836 | 0.03861 | 0.13992 | 15.45 |
| Lrrc6 | 179.33 | 0.256351 | 0.18697 | 1.371097 | 0.17034 | 0.36065 | 121.053 |

| | | | | | | | |
|---|---|---|---|---|---|---|---|
| Lrrc61 | 544.322 | 0.128085 | 0.1682 | 0.761522 | 0.44635 | 0.64993 | 15.927 |
| Lrrc7 | 5258.12 | 0.196418 | 0.13294 | 1.477542 | 0.13953 | 0.32076 | 479.331 |
| Lrrc70 | 23.1151 | 0.005775 | 0.42165 | 0.013697 | 0.98907 | 0.994 | 13.554 |
| Lrrc71 | 32.521 | -0.28194 | 0.33487 | -0.84193 | 0.39983 | 0.60858 | 11.703 |
| Lrrc73 | 1036.44 | 0.015277 | 0.08205 | 0.186202 | 0.85229 | 0.92484 | 3.785 |
| Lrrc74b | 34.4703 | 0.373148 | 0.30371 | 1.228631 | 0.21921 | 0.42143 | 16.783 |
| Lrrc75a | 269.738 | -0.63819 | 0.18305 | -3.48635 | 0.00049 | 0.0078 | 43.64 |
| Lrrc75b | 769.241 | -0.1845 | 0.1134 | -1.62699 | 0.10374 | 0.26624 | 10.208 |
| Lrrc8a | 572.353 | -0.31378 | 0.12633 | -2.48382 | 0.013 | 0.0682 | 26.076 |
| Lrrc8b | 4759.4 | -0.0788 | 0.09293 | -0.84798 | 0.39645 | 0.60547 | 74.3 |
| Lrrc8c | 2054.25 | -0.15086 | 0.10428 | -1.44672 | 0.14798 | 0.33168 | 93.631 |
| Lrrc8d | 1706.77 | 0.052011 | 0.08546 | 0.608573 | 0.54281 | 0.72841 | 132.468 |
| Lrrc9 | 323.866 | 0.240023 | 0.15189 | 1.580292 | 0.11404 | 0.28276 | 88.885 |
| Lrrcc1 | 1124.44 | 0.454926 | 0.34801 | 1.307209 | 0.19114 | 0.38745 | 38.871 |
| Lrrd1 | 27.4733 | -0.08229 | 0.32332 | -0.2545 | 0.79911 | 0.89638 | 21.424 |
| Lrrfip1 | 4147.41 | 0.261017 | 0.06215 | 4.199742 | 2.67E-05 | 0.00107 | 130.208 |
| Lrrfip2 | 2145.75 | 0.066147 | 0.08447 | 0.783067 | 0.43359 | 0.63948 | 108.077 |
| Lrriq1 | 75.448 | 0.224133 | 0.33593 | 0.667207 | 0.50464 | 0.6967 | 190.292 |
| Lrriq3 | 20.3573 | 0.188686 | 0.41663 | 0.452888 | 0.65063 | 0.80552 | 100.847 |
| Lrrk1 | 342.924 | -0.5305 | 0.12785 | -4.14951 | 3.33E-05 | 0.00123 | 161.439 |
| Lrrk2 | 6287.49 | 0.376014 | 0.14668 | 2.563477 | 0.01036 | 0.05879 | 142.946 |
| Lrrn1 | 3211.6 | -0.43407 | 0.12614 | -3.44126 | 0.00058 | 0.00877 | 40.447 |
| Lrrn2 | 4435.76 | -0.02266 | 0.1027 | -0.22061 | 0.82539 | 0.90984 | 59.733 |
| Lrrn3 | 3824.33 | 0.377551 | 0.11058 | 3.414314 | 0.00064 | 0.00934 | 34.764 |
| Lrrn4cl | 65.7308 | -1.0094 | 0.34332 | -2.9401 | 0.00328 | 0.02765 | 3.142 |
| Lrrtm1 | 1526.06 | 0.104697 | 0.07293 | 1.435618 | 0.15111 | 0.33587 | 15.103 |
| Lrrtm2 | 3383.72 | 0.026245 | 0.13792 | 0.190289 | 0.84908 | 0.92322 | 6.022 |
| Lrrtm3 | 3268.03 | 0.636402 | 0.17148 | 3.711293 | 0.00021 | 0.00436 | 161.806 |
| Lrrtm4 | 1059.78 | 0.168022 | 0.07848 | 2.140857 | 0.03229 | 0.1242 | 791.265 |
| Lrsam1 | 1392.78 | -0.08085 | 0.07466 | -1.0829 | 0.27885 | 0.48881 | 36.399 |
| Lrtm1 | 88.6278 | 0.276546 | 0.39465 | 0.700743 | 0.48346 | 0.67979 | 15.438 |
| Lrtm2 | 3178.43 | 0.227866 | 0.12183 | 1.870288 | 0.06144 | 0.18957 | 15.634 |
| Lrwd1 | 919.514 | 0.298293 | 0.08011 | 3.723745 | 0.0002 | 0.00421 | 13.303 |
| Lsamp | 12549.6 | 0.345528 | 0.0944 | 3.660191 | 0.00025 | 0.00497 | 2197.32 |

| | | | | | | | |
|---|---|---|---|---|---|---|---|
| Lsg1 | 1008.95 | 0.154308 | 0.08062 | 1.913952 | 0.05563 | 0.17798 | 27.099 |
| Lsm1 | 608.466 | 0.077852 | 0.0872 | 0.892777 | 0.37198 | 0.58232 | 18.688 |
| Lsm10 | 186.387 | 0.029046 | 0.14633 | 0.198496 | 0.84266 | 0.91903 | 1.962 |
| Lsm11 | 874.722 | -0.2946 | 0.10153 | -2.90167 | 0.00371 | 0.02996 | 16.667 |
| Lsm12 | 1123.48 | -0.05632 | 0.08234 | -0.68397 | 0.494 | 0.68838 | 22.8 |
| Lsm14a | 3978.93 | 0.047677 | 0.0663 | 0.719127 | 0.47206 | 0.67047 | 48.67 |
| Lsm14b | 6121.76 | 0.053961 | 0.0811 | 0.66533 | 0.50584 | 0.69735 | 10.479 |
| Lsm2 | 289.726 | -0.17379 | 0.12839 | -1.35356 | 0.17588 | 0.36778 | 4.03 |
| Lsm3 | 324.249 | -0.3468 | 0.11421 | -3.03664 | 0.00239 | 0.02263 | 6.698 |
| Lsm4 | 1029.13 | 0.018834 | 0.10463 | 0.180004 | 0.85715 | 0.92794 | 5.505 |
| Lsm5 | 154.243 | 0.055455 | 0.14671 | 0.377996 | 0.70543 | 0.8414 | 3.648 |
| Lsm6 | 1778.04 | -0.00111 | 0.06299 | -0.01768 | 0.98589 | 0.99237 | 16.276 |
| Lsm7 | 1003.37 | 0.068105 | 0.08621 | 0.790006 | 0.42952 | 0.63601 | 2.389 |
| Lsm8 | 846.607 | -0.05699 | 0.10466 | -0.54451 | 0.58609 | 0.76022 | 7.173 |
| Lsmem1 | 25.5912 | -0.58953 | 0.40134 | -1.46891 | 0.14186 | 0.32366 | 23.998 |
| Lsmem2 | 21.877 | 0.09576 | 0.39232 | 0.244089 | 0.80716 | 0.90071 | 4.446 |
| Lsp1 | 128.425 | -0.42518 | 0.26965 | -1.57678 | 0.11485 | 0.28393 | 34.059 |
| Lsr | 131.285 | -0.55521 | 0.19489 | -2.84887 | 0.00439 | 0.03347 | 15.695 |
| Lss | 2072.95 | -0.06913 | 0.06887 | -1.00375 | 0.3155 | 0.52606 | 25.551 |
| Lst1 | 46.1111 | 0.101087 | 0.25201 | 0.401129 | 0.68833 | 0.83003 | 3.345 |
| Lta4h | 2114.93 | -0.01194 | 0.0735 | -0.16249 | 0.87092 | 0.93475 | 31.465 |
| Ltb | 13.4536 | -0.69592 | 0.50052 | -1.39041 | 0.16441 | 0.35332 | 1.882 |
| Ltb4r2 | 17.1763 | 0.009271 | 0.49411 | 0.018762 | 0.98503 | 0.99174 | 3.711 |
| Ltbp1 | 343.248 | -0.63851 | 0.17109 | -3.732 | 0.00019 | 0.00416 | 386.945 |
| Ltbp2 | 79.9586 | 0.089424 | 0.32725 | 0.273256 | 0.78466 | 0.88833 | 93.321 |
| Ltbp3 | 2852.93 | 0.071045 | 0.08041 | 0.883525 | 0.37695 | 0.58704 | 17.629 |
| Ltbp4 | 2447.67 | -0.32412 | 0.12748 | -2.54247 | 0.01101 | 0.06114 | 32.557 |
| Ltbr | 271.586 | -0.38461 | 0.18626 | -2.06497 | 0.03893 | 0.14059 | 7.315 |
| Ltc4s | 21.6875 | -0.15946 | 0.39239 | -0.40637 | 0.68447 | 0.82807 | 2.156 |
| Ltk | 697.426 | -0.20186 | 0.09581 | -2.10679 | 0.03514 | 0.13139 | 9.112 |
| Ltn1 | 4702.62 | 0.214729 | 0.06847 | 3.136151 | 0.00171 | 0.01822 | 55.962 |
| Ltv1 | 660.972 | -0.01512 | 0.09356 | -0.16164 | 0.87159 | 0.93512 | 15.029 |
| Luc7l | 7586.92 | 0.118881 | 0.06681 | 1.779299 | 0.07519 | 0.21627 | 32.609 |
| Luc7l2 | 16794.4 | 0.14816 | 0.11363 | 1.303839 | 0.19229 | 0.38895 | 58.137 |

| | | | | | | | |
|---|---|---|---|---|---|---|---|
| Luc7l3 | 11180.6 | 0.19619 | 0.13751 | 1.426686 | 0.15367 | 0.33948 | 34.099 |
| Lum | 71.6608 | -1.11809 | 0.46221 | -2.41899 | 0.01556 | 0.07631 | 7.576 |
| Lurap1 | 994.872 | -0.03148 | 0.0882 | -0.35686 | 0.7212 | 0.85114 | 18.933 |
| Lurap1l | 420.772 | -0.87974 | 0.17614 | -4.99451 | 5.90E-07 | 7.38E-05 | 44.983 |
| Luzp1 | 4802.99 | -0.19014 | 0.06067 | -3.13385 | 0.00173 | 0.01828 | 85.02 |
| Luzp2 | 10897 | -0.25177 | 0.08178 | -3.07862 | 0.00208 | 0.02063 | 433.388 |
| Lxn | 635.304 | -0.04505 | 0.14446 | -0.31186 | 0.75514 | 0.87123 | 5.929 |
| Ly6a | 963.926 | -0.4788 | 0.14892 | -3.21521 | 0.0013 | 0.01501 | 3.155 |
| Ly6c1 | 1782.06 | -0.46406 | 0.07786 | -5.96041 | 2.52E-09 | 1.14E-06 | 4.813 |
| Ly6e | 2683.16 | -0.62305 | 0.1413 | -4.40951 | 1.04E-05 | 0.00057 | 4.853 |
| Ly6g5b | 14.7541 | 0.158695 | 0.56656 | 0.280101 | 0.7794 | 0.88509 | 1.481 |
| Ly6g6e | 78.7486 | -1.15501 | 0.46173 | -2.50145 | 0.01237 | 0.06597 | 1.903 |
| Ly6g6f | 18.1389 | -0.53398 | 0.40412 | -1.32135 | 0.18639 | 0.3813 | 5.131 |
| Ly6h | 5773.74 | 0.112083 | 0.10975 | 1.021292 | 0.30712 | 0.51686 | 15.988 |
| Ly75 | 99.4442 | -0.02535 | 0.23566 | -0.10759 | 0.91432 | 0.95656 | 91.201 |
| Ly86 | 285.455 | -0.25087 | 0.16451 | -1.52499 | 0.12726 | 0.30347 | 73.829 |
| Ly9 | 24.6928 | -1.63353 | 0.64017 | -2.55171 | 0.01072 | 0.06013 | 18.787 |
| Ly96 | 33.4567 | -0.56026 | 0.44664 | -1.25438 | 0.20971 | 0.40977 | 21.561 |
| Lyar | 432.412 | 0.048202 | 0.11346 | 0.42485 | 0.67095 | 0.81871 | 13.837 |
| Lyl1 | 77.9943 | -0.45835 | 0.2199 | -2.08436 | 0.03713 | 0.13624 | 3.492 |
| Lyn | 368.228 | -0.49537 | 0.17719 | -2.79566 | 0.00518 | 0.03711 | 135.008 |
| Lynx1 | 12400.4 | -0.55331 | 0.09039 | -6.12138 | 9.28E-10 | 5.11E-07 | 5.195 |
| Lypd1 | 1714.57 | 0.666336 | 0.15047 | 4.428318 | 9.50E-06 | 0.00054 | 45.48 |
| Lypd6 | 964.205 | 0.009053 | 0.16626 | 0.054451 | 0.95658 | 0.97876 | 127.141 |
| Lypd6b | 308.16 | -0.23318 | 0.18488 | -1.26124 | 0.20722 | 0.4067 | 161.162 |
| Lypla1 | 935.77 | -0.18792 | 0.09872 | -1.90362 | 0.05696 | 0.18077 | 40.623 |
| Lypla2 | 2304.03 | -0.13298 | 0.07837 | -1.6968 | 0.08973 | 0.24282 | 4.403 |
| Lyplal1 | 140.131 | -0.10111 | 0.19775 | -0.51129 | 0.60915 | 0.77661 | 29.58 |
| Lyrm1 | 409.01 | -0.05103 | 0.10295 | -0.49569 | 0.62011 | 0.78385 | 61.103 |
| Lyrm2 | 643.069 | 0.049533 | 0.10085 | 0.491146 | 0.62332 | 0.78601 | 1.307 |
| Lyrm4 | 645.524 | 0.149415 | 0.09779 | 1.527891 | 0.12654 | 0.30235 | 139.132 |
| Lyrm7 | 581.576 | -0.20952 | 0.11053 | -1.89558 | 0.05802 | 0.18309 | 34.051 |
| Lyrm9 | 3938.84 | 0.271495 | 0.06528 | 4.158731 | 3.20E-05 | 0.00121 | 20.287 |
| Lysmd1 | 370.729 | -0.04169 | 0.11271 | -0.36984 | 0.7115 | 0.84473 | 5.431 |

| | | | | | | | |
|---|---|---|---|---|---|---|---|
| Lysmd2 | 1270.1 | -0.0237 | 0.07726 | -0.30676 | 0.75902 | 0.87343 | 12.031 |
| Lysmd3 | 824.508 | 0.2293 | 0.12584 | 1.822154 | 0.06843 | 0.20328 | 15.209 |
| Lysmd4 | 1468.69 | 0.141894 | 0.06896 | 2.05755 | 0.03963 | 0.14221 | 84.787 |
| Lyst | 5990.25 | 0.207849 | 0.08202 | 2.533974 | 0.01128 | 0.06201 | 188.407 |
| Lyve1 | 102.24 | -0.97875 | 0.24359 | -4.01802 | 5.87E-05 | 0.00185 | 12.633 |
| Lyz2 | 878.651 | -2 | 0.67413 | -3.43777 | 0.00059 | 0.00882 | 4.991 |
| Lyzl4 | 32.6408 | -0.44851 | 0.35701 | -1.25628 | 0.20901 | 0.40904 | 64.254 |
| Lzic | 579.962 | -0.10946 | 0.09318 | -1.17472 | 0.2401 | 0.44615 | 11.44 |
| Lztfl1 | 2660.98 | 0.23681 | 0.13036 | 1.81653 | 0.06929 | 0.20496 | 23.282 |
| Lztr1 | 6588.27 | -0.08047 | 0.07743 | -1.03924 | 0.29869 | 0.50876 | 17.646 |
| Lzts1 | 7165.56 | 0.243373 | 0.09528 | 2.554216 | 0.01064 | 0.05988 | 51.557 |
| Lzts2 | 2128.25 | -0.40672 | 0.10901 | -3.73101 | 0.00019 | 0.00417 | 12.698 |
| Lzts3 | 19363 | 0.647181 | 0.1669 | 3.877604 | 0.00011 | 0.00279 | 9.965 |
| M5C1000I18Rik | 52.9753 | -0.75739 | 0.34431 | -2.19975 | 0.02782 | 0.11213 | 1.519 |
| M6pr | 4588.58 | -0.02598 | 0.06475 | -0.40131 | 0.68819 | 0.83002 | 8.961 |
| MIR628 | 163.932 | -0.27463 | 0.213 | -1.28938 | 0.19727 | 0.39516 | 0.122 |
| Mab21l1 | 21.5401 | -0.35324 | 0.36458 | -0.96891 | 0.33259 | 0.54345 | 2.492 |
| Macf1 | 27332.7 | 0.129277 | 0.07425 | 1.741046 | 0.08168 | 0.22777 | 334.728 |
| Macir | 1849.33 | -0.2777 | 0.11234 | -2.47184 | 0.01344 | 0.06967 | 55.757 |
| Maco1 | 3761.04 | 0.250995 | 0.07001 | 3.585382 | 0.00034 | 0.00611 | 50.587 |
| Macrod1 | 273.842 | -0.55759 | 0.16919 | -3.2957 | 0.00098 | 0.01264 | 141.245 |
| Macrod2 | 2960.32 | 0.313444 | 0.07053 | 4.444078 | 8.83E-06 | 0.00051 | 1997.658 |
| Macrod2os1 | 16.5014 | -0.51607 | 0.47651 | -1.08303 | 0.27879 | 0.48881 | 56.561 |
| Macrod2os2 | 18.2131 | 0.012097 | 0.40742 | 0.029691 | 0.97631 | 0.98767 | 5.757 |
| Macroh2a1 | 2574.54 | 0.059757 | 0.06926 | 0.862781 | 0.38826 | 0.59753 | 62.743 |
| Macroh2a2 | 972.467 | 0.010988 | 0.08048 | 0.136524 | 0.89141 | 0.94593 | 45.483 |
| Mad1l1 | 525.918 | -0.18281 | 0.10184 | -1.79519 | 0.07262 | 0.21188 | 312.864 |
| Mad2l1 | 440.687 | -0.12247 | 0.09915 | -1.23518 | 0.21676 | 0.41862 | 11.831 |
| Mad2l1bp | 274.945 | -0.25641 | 0.11728 | -2.18641 | 0.02879 | 0.11465 | 6.167 |
| Mad2l2 | 335.707 | 0.014305 | 0.13749 | 0.10405 | 0.91713 | 0.95777 | 15.316 |
| Madd | 10002.6 | 0.063655 | 0.04977 | 1.279018 | 0.20089 | 0.39907 | 46.478 |
| Maea | 2929.64 | -0.03412 | 0.07093 | -0.48105 | 0.63048 | 0.79112 | 37.788 |
| Maf | 1024.45 | -0.59438 | 0.10981 | -5.41255 | 6.21E-08 | 1.26E-05 | 24.853 |
| Maf1 | 3491.35 | -0.00996 | 0.07911 | -0.12589 | 0.89982 | 0.94981 | 3.087 |

| | | | | | | | |
|---|---|---|---|---|---|---|---|
| Mafa | 266.399 | -0.11399 | 0.16861 | -0.67608 | 0.49899 | 0.69222 | 2.739 |
| Mafb | 679.063 | -0.31664 | 0.11212 | -2.82417 | 0.00474 | 0.03514 | 3.363 |
| Maff | 106.037 | -0.58086 | 0.25405 | -2.28637 | 0.02223 | 0.0961 | 12.456 |
| Mafg | 5280.56 | -0.0412 | 0.06361 | -0.64779 | 0.51712 | 0.70706 | 8.484 |
| Mafk | 550.087 | -0.29391 | 0.11082 | -2.6521 | 0.008 | 0.04961 | 11.141 |
| Mag | 5525.92 | -0.63173 | 0.16947 | -3.72778 | 0.00019 | 0.0042 | 15.698 |
| Maged1 | 22423.3 | 0.174181 | 0.06178 | 2.819168 | 0.00481 | 0.03542 | 6.67 |
| Maged2 | 1409.95 | -0.05022 | 0.10382 | -0.48373 | 0.62858 | 0.79 | 7.976 |
| Magee1 | 6183.48 | 0.084061 | 0.072 | 1.167592 | 0.24297 | 0.44915 | 3.55 |
| Magee2 | 329.381 | 0.374822 | 0.12606 | 2.973301 | 0.00295 | 0.02586 | 2.316 |
| Magef1 | 364.384 | 0.14971 | 0.10589 | 1.413806 | 0.15742 | 0.34463 | 1.497 |
| Mageh1 | 1556.38 | -0.00429 | 0.09451 | -0.04537 | 0.96381 | 0.98236 | 1.41 |
| Magel2 | 148.158 | 0.909418 | 0.15532 | 5.855186 | 4.76E-09 | 1.80E-06 | 4.631 |
| Magi1 | 4767.71 | -0.19526 | 0.09858 | -1.98076 | 0.04762 | 0.16041 | 608.463 |
| Magi2 | 5952.86 | 0.044033 | 0.06347 | 0.693733 | 0.48785 | 0.68299 | 1477.757 |
| Magi3 | 2965.25 | 0.128155 | 0.09163 | 1.398647 | 0.16192 | 0.3502 | 207.116 |
| Magix | 17.6186 | -0.41106 | 0.4166 | -0.98669 | 0.32379 | 0.53482 | 8.088 |
| Magoh | 476.741 | -0.0015 | 0.11523 | -0.01305 | 0.98958 | 0.99424 | 7.67 |
| Magohb | 184.151 | 0.05631 | 0.13557 | 0.415357 | 0.67788 | 0.82352 | 8.857 |
| Magt1 | 1078.93 | -0.15946 | 0.08231 | -1.93738 | 0.0527 | 0.17178 | 43.823 |
| Maip1 | 678.071 | 0.068618 | 0.08535 | 0.80399 | 0.4214 | 0.62837 | 11.626 |
| Majin | 28.3894 | 1.113195 | 0.37294 | 2.984931 | 0.00284 | 0.02528 | 39.838 |
| Mak | 272.3 | 0.501304 | 0.11244 | 4.458597 | 8.25E-06 | 0.00048 | 54.699 |
| Mak16 | 340.022 | 0.310706 | 0.14478 | 2.146034 | 0.03187 | 0.12311 | 9.302 |
| Mal | 11428 | -0.43921 | 0.09228 | -4.75954 | 1.94E-06 | 0.00017 | 23.47 |
| Mal2 | 4412.53 | -0.26989 | 0.13373 | -2.01819 | 0.04357 | 0.151 | 31.656 |
| Malat1 | 156625 | -0.14801 | 0.09502 | -1.55771 | 0.1193 | 0.29146 | 6.988 |
| Mall | 55.4503 | -0.24994 | 0.31008 | -0.80603 | 0.42023 | 0.62755 | 25.547 |
| Malsu1 | 353.801 | -0.13224 | 0.14161 | -0.93387 | 0.35037 | 0.56099 | 12.957 |
| Malt1 | 773.052 | 0.270731 | 0.15088 | 1.794307 | 0.07276 | 0.21219 | 48.13 |
| Mamdc2 | 60.4854 | -0.41546 | 0.28793 | -1.44293 | 0.14904 | 0.33301 | 145.834 |
| Mamdc4 | 206.641 | 0.100589 | 0.17815 | 0.564642 | 0.57232 | 0.75105 | 11.731 |
| Maml1 | 859.461 | 0.029382 | 0.08986 | 0.326972 | 0.74369 | 0.86536 | 36.678 |
| Maml2 | 479.587 | -0.13543 | 0.12134 | -1.11609 | 0.26438 | 0.47315 | 322.379 |

| | | | | | | | |
|---|---|---|---|---|---|---|---|
| Maml3 | 701.939 | -0.0018 | 0.08482 | -0.02123 | 0.98306 | 0.99078 | 419.17 |
| Mamld1 | 1431.73 | -0.07878 | 0.08106 | -0.97193 | 0.33109 | 0.54208 | 105.817 |
| Mamstr | 108.877 | 0.067664 | 0.23488 | 0.288084 | 0.77328 | 0.88189 | 6.545 |
| Man1a | 1906.2 | 0.455492 | 0.10589 | 4.301658 | 1.70E-05 | 0.0008 | 171.825 |
| Man1a2 | 5936.08 | 0.253503 | 0.09284 | 2.730479 | 0.00632 | 0.04263 | 123.296 |
| Man1b1 | 3838.32 | -0.02262 | 0.07196 | -0.3144 | 0.75322 | 0.87029 | 19.875 |
| Man1c1 | 1511.61 | -0.33121 | 0.11837 | -2.79811 | 0.00514 | 0.03696 | 142.601 |
| Man2a1 | 1419.56 | 0.309688 | 0.13226 | 2.341579 | 0.0192 | 0.08705 | 154.375 |
| Man2a2 | 14001 | 0.11433 | 0.07737 | 1.477769 | 0.13947 | 0.32071 | 22.279 |
| Man2b1 | 2392.41 | -0.01992 | 0.07835 | -0.25418 | 0.79936 | 0.89647 | 15.013 |
| Man2b2 | 989.679 | -0.26213 | 0.09503 | -2.75831 | 0.00581 | 0.04034 | 23.733 |
| Man2c1 | 3419.58 | 0.080827 | 0.07785 | 1.038177 | 0.29919 | 0.50926 | 12.033 |
| Man2c1os | 57.8128 | 0.371413 | 0.25871 | 1.435641 | 0.1511 | 0.33587 | 31.353 |
| Manba | 410.976 | -0.30233 | 0.10148 | -2.97936 | 0.00289 | 0.02561 | 85.794 |
| Manbal | 1293.48 | -0.22801 | 0.09897 | -2.30376 | 0.02124 | 0.09321 | 29.17 |
| Manea | 2007.17 | 0.041128 | 0.09479 | 0.433899 | 0.66436 | 0.8145 | 22.386 |
| Maneal | 2147.7 | 0.02721 | 0.08078 | 0.336862 | 0.73622 | 0.8606 | 6.933 |
| Manf | 1528.41 | -0.02539 | 0.09208 | -0.27575 | 0.78274 | 0.88707 | 53.668 |
| Mansc1 | 311.049 | -0.19472 | 0.13857 | -1.40526 | 0.15994 | 0.34784 | 23.282 |
| Mansc4 | 46.7477 | -0.04594 | 0.28646 | -0.16038 | 0.87258 | 0.93573 | 11.973 |
| Maoa | 1600.78 | -0.11887 | 0.07654 | -1.55308 | 0.1204 | 0.29319 | 68.121 |
| Maob | 1250.58 | -0.23952 | 0.09715 | -2.46549 | 0.01368 | 0.07038 | 108.085 |
| Map10 | 204.937 | -0.12253 | 0.14467 | -0.84695 | 0.39702 | 0.60612 | 3.542 |
| Map1a | 75464.2 | -0.19297 | 0.07474 | -2.58167 | 0.00983 | 0.05687 | 21.233 |
| Map1b | 62312.2 | -0.0075 | 0.05665 | -0.13231 | 0.89474 | 0.94736 | 95.095 |
| Map1lc3a | 3579.5 | -0.35816 | 0.09911 | -3.61381 | 0.0003 | 0.00565 | 1.777 |
| Map1lc3b | 7403.56 | -0.19904 | 0.06487 | -3.06841 | 0.00215 | 0.02115 | 8.4 |
| Map1s | 2429.89 | 0.192351 | 0.06828 | 2.817291 | 0.00484 | 0.03553 | 11.598 |
| Map2 | 69088.3 | -0.04637 | 0.08433 | -0.54981 | 0.58245 | 0.75787 | 267.311 |
| Map2k1 | 15611.6 | 0.398493 | 0.1228 | 3.245087 | 0.00117 | 0.01411 | 67.862 |
| Map2k2 | 3038.81 | 0.032388 | 0.06802 | 0.476121 | 0.63399 | 0.79386 | 28.061 |
| Map2k3 | 585.739 | -0.14588 | 0.10564 | -1.38093 | 0.1673 | 0.35699 | 20.779 |
| Map2k3os | 36.1548 | 0.088277 | 0.2971 | 0.297127 | 0.76637 | 0.8779 | 11.037 |
| Map2k4 | 6966.53 | -0.12818 | 0.07413 | -1.72908 | 0.08379 | 0.23163 | 100.055 |

| | | | | | | | |
|---|---|---|---|---|---|---|---|
| Map2k5 | 1073.39 | -0.15575 | 0.10032 | -1.55258 | 0.12052 | 0.2934 | 214.135 |
| Map2k6 | 693.85 | 0.041283 | 0.10173 | 0.405798 | 0.68489 | 0.82838 | 126.401 |
| Map2k7 | 2642.97 | -0.01855 | 0.08324 | -0.22284 | 0.82366 | 0.90857 | 9.158 |
| Map3k1 | 1018.38 | -0.09258 | 0.10462 | -0.88498 | 0.37617 | 0.58622 | 62.566 |
| Map3k10 | 7644.38 | -0.23627 | 0.07514 | -3.14448 | 0.00166 | 0.01785 | 18.224 |
| Map3k11 | 907.293 | 0.040553 | 0.08694 | 0.466436 | 0.6409 | 0.79893 | 14.124 |
| Map3k12 | 4043.91 | 0.067872 | 0.06474 | 1.048418 | 0.29445 | 0.50482 | 19.419 |
| Map3k13 | 2487.11 | 0.268776 | 0.10495 | 2.561002 | 0.01044 | 0.05908 | 139.094 |
| Map3k14 | 326.271 | 0.207024 | 0.14979 | 1.382142 | 0.16693 | 0.35675 | 47.711 |
| Map3k15 | 110.073 | 0.521391 | 0.18864 | 2.763879 | 0.00571 | 0.0399 | 134.919 |
| Map3k19 | 524.222 | 0.064448 | 0.20471 | 0.314826 | 0.75289 | 0.87023 | 40.241 |
| Map3k2 | 3528.55 | 0.183593 | 0.06645 | 2.762962 | 0.00573 | 0.03996 | 73.663 |
| Map3k20 | 505.531 | -0.21555 | 0.12138 | -1.77575 | 0.07577 | 0.21741 | 156.974 |
| Map3k21 | 68.328 | -0.67609 | 0.27534 | -2.45544 | 0.01407 | 0.07142 | 36.991 |
| Map3k3 | 1748.33 | 0.169711 | 0.08341 | 2.03461 | 0.04189 | 0.1472 | 70.834 |
| Map3k4 | 3088.22 | 0.051829 | 0.07694 | 0.673608 | 0.50056 | 0.6934 | 91.176 |
| Map3k5 | 2068.72 | 0.402077 | 0.1325 | 3.034532 | 0.00241 | 0.02276 | 208.282 |
| Map3k6 | 452.12 | -0.23163 | 0.15485 | -1.49581 | 0.1347 | 0.31449 | 12.112 |
| Map3k7 | 3560.42 | 0.108775 | 0.05859 | 1.856641 | 0.06336 | 0.19338 | 59.371 |
| Map3k7cl | 36.8479 | -0.87841 | 0.3663 | -2.3981 | 0.01648 | 0.07926 | 42.007 |
| Map3k8 | 119.611 | -0.12432 | 0.19785 | -0.62836 | 0.52977 | 0.71748 | 21.689 |
| Map3k9 | 1717.49 | -0.37889 | 0.15379 | -2.46369 | 0.01375 | 0.07056 | 60.166 |
| Map4 | 21869.5 | -0.22582 | 0.07108 | -3.177 | 0.00149 | 0.01646 | 152.496 |
| Map4k1 | 118.882 | 0.26564 | 0.17172 | 1.546932 | 0.12188 | 0.29537 | 21.23 |
| Map4k2 | 4119.48 | 0.091505 | 0.06735 | 1.358675 | 0.17425 | 0.3658 | 14.481 |
| Map4k3 | 2738.54 | -0.12159 | 0.10597 | -1.14737 | 0.25123 | 0.45863 | 147.974 |
| Map4k4 | 7596.88 | -0.09411 | 0.09317 | -1.01016 | 0.31242 | 0.52285 | 125.398 |
| Map4k5 | 3092.34 | 0.187208 | 0.06868 | 2.725791 | 0.00641 | 0.04298 | 89.451 |
| Map6 | 5778.05 | 0.059685 | 0.06494 | 0.919071 | 0.35806 | 0.56817 | 69.691 |
| Map6d1 | 1616.37 | -0.12485 | 0.10084 | -1.23811 | 0.21568 | 0.41731 | 8.056 |
| Map7 | 6089.97 | -0.00075 | 0.07455 | -0.01009 | 0.99195 | 0.99537 | 133.12 |
| Map7d1 | 10819.4 | -0.08935 | 0.06165 | -1.44939 | 0.14723 | 0.33046 | 24.177 |
| Map7d2 | 8507.65 | -0.10333 | 0.07844 | -1.31734 | 0.18773 | 0.38316 | 84.186 |
| Map9 | 5964.53 | 0.161411 | 0.08468 | 1.906217 | 0.05662 | 0.18017 | 37.225 |

| | | | | | | | |
|---|---|---|---|---|---|---|---|
| Mapk1 | 32639.8 | 0.264574 | 0.09699 | 2.727717 | 0.00638 | 0.04285 | 64.072 |
| Mapk10 | 12880.1 | -0.15565 | 0.06722 | -2.31552 | 0.02058 | 0.09128 | 303.387 |
| Mapk11 | 988.192 | -0.82602 | 0.22579 | -3.65841 | 0.00025 | 0.00499 | 7.143 |
| Mapk12 | 468.627 | -0.89288 | 0.21295 | -4.19295 | 2.75E-05 | 0.00109 | 10.12 |
| Mapk14 | 4947.59 | 0.038109 | 0.06812 | 0.559414 | 0.57588 | 0.75379 | 57.078 |
| Mapk15 | 23.7138 | 0.67336 | 0.66512 | 1.012386 | 0.31135 | 0.52167 | 5.386 |
| Mapk1ip1 | 1928.86 | 0.458863 | 0.10376 | 4.422389 | 9.76E-06 | 0.00055 | 10.451 |
| Mapk1ip1l | 1833.99 | 0.140883 | 0.07118 | 1.979269 | 0.04779 | 0.16082 | 24.939 |
| Mapk3 | 4640.02 | -0.09584 | 0.10059 | -0.95275 | 0.34072 | 0.55173 | 6.219 |
| Mapk4 | 4175.15 | 0.034489 | 0.0875 | 0.394183 | 0.69345 | 0.83381 | 136.874 |
| Mapk6 | 3191.03 | 0.304137 | 0.11029 | 2.757625 | 0.00582 | 0.04041 | 40.944 |
| Mapk7 | 680.232 | -0.22449 | 0.12878 | -1.74318 | 0.0813 | 0.22693 | 5.595 |
| Mapk8 | 4938.97 | -0.11263 | 0.0958 | -1.17564 | 0.23974 | 0.44568 | 69.261 |
| Mapk8ip1 | 15604.9 | -0.13575 | 0.06917 | -1.96256 | 0.0497 | 0.16518 | 17.588 |
| Mapk8ip2 | 14971.7 | -0.18766 | 0.06824 | -2.75017 | 0.00596 | 0.04102 | 10.556 |
| Mapk8ip3 | 21357.7 | -0.04415 | 0.06036 | -0.73147 | 0.46449 | 0.66425 | 44.825 |
| Mapk9 | 11470.7 | -0.06484 | 0.05361 | -1.20945 | 0.22649 | 0.43063 | 39.671 |
| Mapkap1 | 2589.56 | 0.130563 | 0.0936 | 1.39484 | 0.16306 | 0.35163 | 218.18 |
| Mapkapk2 | 950.001 | -0.55116 | 0.14232 | -3.87268 | 0.00011 | 0.00282 | 44.127 |
| Mapkapk3 | 144.678 | -0.18042 | 0.19505 | -0.92499 | 0.35497 | 0.56542 | 34.951 |
| Mapkapk5 | 102.241 | 0.174653 | 0.2002 | 0.872408 | 0.38299 | 0.59222 | 20.868 |
| Mapkbp1 | 4356.66 | 0.260946 | 0.0782 | 3.336965 | 0.00085 | 0.01142 | 54.71 |
| Mapre1 | 4680.73 | -0.10161 | 0.06906 | -1.47146 | 0.14117 | 0.32297 | 32.037 |
| Mapre2 | 13533.3 | 0.053025 | 0.05915 | 0.896452 | 0.37001 | 0.58036 | 141.529 |
| Mapre3 | 7308.74 | -0.01676 | 0.06871 | -0.24392 | 0.80729 | 0.90075 | 51.466 |
| Mapt | 19808.1 | -0.14184 | 0.07 | -2.02624 | 0.04274 | 0.14903 | 100.701 |
| Marchf1 | 1087.35 | 0.452689 | 0.09659 | 4.686815 | 2.77E-06 | 0.00023 | 853.738 |
| Marchf10 | 56.637 | 0.264056 | 0.25395 | 1.039778 | 0.29844 | 0.50856 | 95.938 |
| Marchf11 | 116.505 | -0.03806 | 0.17578 | -0.21653 | 0.82858 | 0.91212 | 100.529 |
| Marchf2 | 2510.21 | -0.21465 | 0.06363 | -3.37347 | 0.00074 | 0.01043 | 32.979 |
| Marchf3 | 110.982 | 0.37952 | 0.22232 | 1.707094 | 0.0878 | 0.23896 | 163.833 |
| Marchf4 | 1326.44 | 0.138957 | 0.08334 | 1.667359 | 0.09544 | 0.25257 | 109.819 |
| Marchf5 | 2805.35 | -0.03096 | 0.06403 | -0.48357 | 0.62869 | 0.79004 | 14.609 |
| Marchf6 | 14913 | 0.042787 | 0.0639 | 0.669623 | 0.5031 | 0.69545 | 75.163 |

| | | | | | | | |
|---|---|---|---|---|---|---|---|
| Marchf7 | 4080.46 | 0.201109 | 0.15558 | 1.292638 | 0.19614 | 0.3937 | 40.79 |
| Marchf8 | 3357.01 | -0.25709 | 0.10454 | -2.45925 | 0.01392 | 0.071 | 71.517 |
| Marchf9 | 987.87 | 0.065208 | 0.08768 | 0.743741 | 0.45703 | 0.65829 | 5.174 |
| Marcks | 9732.59 | 0.173055 | 0.07146 | 2.421619 | 0.01545 | 0.07591 | 5.546 |
| Marcksl1 | 1975.06 | -0.65161 | 0.20856 | -3.12436 | 0.00178 | 0.01867 | 2.405 |
| Marf1 | 7816.59 | -0.02354 | 0.05747 | -0.40961 | 0.68209 | 0.82631 | 54.179 |
| Mark1 | 6084.84 | 0.124752 | 0.05131 | 2.431307 | 0.01504 | 0.07456 | 102.782 |
| Mark2 | 6017.54 | 0.020264 | 0.07258 | 0.279213 | 0.78008 | 0.88542 | 66.465 |
| Mark3 | 4235.51 | 0.0322 | 0.05677 | 0.567221 | 0.57056 | 0.74977 | 81.699 |
| Mark4 | 3050.64 | -0.12902 | 0.06382 | -2.02164 | 0.04321 | 0.15016 | 34.047 |
| Mars1 | 2380.51 | -0.10143 | 0.06266 | -1.61873 | 0.10551 | 0.269 | 15.566 |
| Mars2 | 1720.82 | -0.11755 | 0.10986 | -1.06997 | 0.28463 | 0.49463 | 11.294 |
| Marveld1 | 483.165 | -0.32116 | 0.13698 | -2.34448 | 0.01905 | 0.08668 | 4.304 |
| Marveld2 | 29.0689 | -0.63994 | 0.33231 | -1.92576 | 0.05413 | 0.17499 | 21.015 |
| Marveld3 | 45.3788 | 0.030582 | 0.372 | 0.08221 | 0.93448 | 0.96713 | 14.29 |
| Mas1 | 805.837 | -0.9053 | 0.21564 | -4.19831 | 2.69E-05 | 0.00107 | 27.065 |
| Maskbp3 | 54.9043 | 0.474359 | 0.24108 | 1.967613 | 0.04911 | 0.16396 | 18.058 |
| Masp1 | 465.987 | -0.13766 | 0.12088 | -1.13883 | 0.25478 | 0.46229 | 71.399 |
| Masp2 | 54.6499 | 0.748594 | 0.2634 | 2.842062 | 0.00448 | 0.03388 | 12.946 |
| Mast1 | 1434.11 | -0.56904 | 0.179 | -3.17904 | 0.00148 | 0.01639 | 25.457 |
| Mast2 | 5091.9 | -0.19544 | 0.09 | -2.17154 | 0.02989 | 0.11801 | 157.422 |
| Mast3 | 12606.5 | 0.069044 | 0.05312 | 1.299667 | 0.19371 | 0.39071 | 26.938 |
| Mast4 | 2720.48 | -0.39859 | 0.13454 | -2.96251 | 0.00305 | 0.02637 | 602.012 |
| Mastl | 50.6459 | -0.55851 | 0.26721 | -2.09018 | 0.0366 | 0.13482 | 40.419 |
| Mat2a | 14183.1 | -0.0313 | 0.06438 | -0.48612 | 0.62688 | 0.78888 | 6.76 |
| Mat2b | 4908.45 | -0.08681 | 0.06846 | -1.26808 | 0.20477 | 0.40364 | 15.89 |
| Matk | 8057.18 | 0.212982 | 0.09073 | 2.347474 | 0.0189 | 0.08627 | 10.431 |
| Matn1 | 25.2407 | 0.219463 | 0.43332 | 0.506463 | 0.61253 | 0.77934 | 11.091 |
| Matn2 | 1691.92 | -0.05856 | 0.19004 | -0.30813 | 0.75798 | 0.8726 | 129.597 |
| Matn4 | 462.897 | -0.14972 | 0.10453 | -1.43235 | 0.15204 | 0.33717 | 15.768 |
| Matr3 | 10484.2 | 0.276183 | 0.11377 | 2.427637 | 0.0152 | 0.0751 | 31.698 |
| Mau2 | 5146.42 | 0.104055 | 0.06691 | 1.555103 | 0.11992 | 0.29246 | 26.612 |
| Mavs | 365.258 | -0.28448 | 0.13364 | -2.12878 | 0.03327 | 0.12665 | 13.963 |
| Max | 3139.55 | -0.05826 | 0.06577 | -0.88577 | 0.37574 | 0.5859 | 24.933 |

| | | | | | | | |
|---|---|---|---|---|---|---|---|
| Maz | 6621.29 | -0.04301 | 0.07579 | -0.56743 | 0.57042 | 0.74969 | 4.908 |
| Mb | 14.9998 | -0.15513 | 0.52228 | -0.29703 | 0.76644 | 0.8779 | 35.184 |
| Mb21d2 | 1279.11 | 0.251276 | 0.08783 | 2.86078 | 0.00423 | 0.03263 | 103.498 |
| Mbd1 | 3860.63 | 0.115791 | 0.08194 | 1.413199 | 0.1576 | 0.34467 | 15.128 |
| Mbd2 | 2396.06 | 0.063105 | 0.09678 | 0.652037 | 0.51438 | 0.70497 | 57.943 |
| Mbd3 | 3948.88 | -0.05447 | 0.07975 | -0.68303 | 0.49459 | 0.68873 | 7.012 |
| Mbd4 | 600.869 | 0.207128 | 0.09544 | 2.170308 | 0.02998 | 0.11822 | 12.675 |
| Mbd5 | 3660.49 | 0.192892 | 0.0783 | 2.463649 | 0.01375 | 0.07056 | 375.898 |
| Mbd6 | 2877.22 | 0.141993 | 0.11525 | 1.232031 | 0.21794 | 0.41996 | 7.063 |
| Mbip | 1097.69 | 0.228269 | 0.11692 | 1.952352 | 0.0509 | 0.16779 | 17.592 |
| Mblac1 | 99.6435 | -0.55895 | 0.19586 | -2.85385 | 0.00432 | 0.03315 | 1.308 |
| Mblac2 | 1511.4 | 0.364234 | 0.08846 | 4.117341 | 3.83E-05 | 0.00136 | 41.935 |
| Mbnl1 | 6999.49 | 0.384742 | 0.09525 | 4.039317 | 5.36E-05 | 0.00174 | 156.921 |
| Mbnl2 | 14200.7 | 0.319992 | 0.11538 | 2.773481 | 0.00555 | 0.03899 | 156.029 |
| Mbnl3 | 12.7225 | 0.563802 | 0.62295 | 0.905055 | 0.36544 | 0.57607 | 89.264 |
| Mboat1 | 105.775 | -0.41088 | 0.21567 | -1.90511 | 0.05677 | 0.18035 | 110.229 |
| Mboat2 | 3058.49 | -0.17685 | 0.06618 | -2.67225 | 0.00753 | 0.04785 | 133.519 |
| Mboat7 | 5585.6 | 0.140217 | 0.05913 | 2.371359 | 0.01772 | 0.08274 | 15.735 |
| Mbp | 84644.9 | -0.74497 | 0.16324 | -4.56371 | 5.03E-06 | 0.00034 | 110.492 |
| Mbtd1 | 2248.29 | 0.310805 | 0.11724 | 2.650926 | 0.00803 | 0.04967 | 61.134 |
| Mbtps1 | 5765.21 | 0.270377 | 0.1038 | 2.604905 | 0.00919 | 0.05416 | 50.58 |
| Mbtps2 | 1344.3 | -0.03942 | 0.09463 | -0.41655 | 0.67701 | 0.82301 | 63.345 |
| Mc3r | 32.46 | 1.384853 | 0.37475 | 3.695426 | 0.00022 | 0.00454 | 2.623 |
| Mc4r | 158.08 | 0.659414 | 0.26384 | 2.499276 | 0.01244 | 0.06622 | 2.803 |
| Mc5r | 79.5278 | -0.92868 | 0.22302 | -4.16422 | 3.12E-05 | 0.00119 | 4.851 |
| Mcam | 479.409 | -0.24739 | 0.17245 | -1.43453 | 0.15142 | 0.33637 | 8.259 |
| Mcat | 671.407 | 0.019079 | 0.1156 | 0.16504 | 0.86891 | 0.93348 | 16.991 |
| Mcc | 924.038 | -0.23483 | 0.0986 | -2.38166 | 0.01723 | 0.08153 | 387.123 |
| Mccc1 | 1956.62 | 0.316268 | 0.08828 | 3.582744 | 0.00034 | 0.00614 | 41.367 |
| Mccc1os | 111.901 | 0.057825 | 0.24579 | 0.235263 | 0.814 | 0.90453 | 25.274 |
| Mccc2 | 1055.21 | 0.284722 | 0.11721 | 2.429098 | 0.01514 | 0.07491 | 67.11 |
| Mcee | 506.893 | -0.16899 | 0.11277 | -1.49854 | 0.13399 | 0.31369 | 19.519 |
| Mcemp1 | 49.5963 | 0.256777 | 0.25019 | 1.026329 | 0.30474 | 0.51453 | 3.506 |
| Mcf2 | 228.638 | 0.163584 | 0.20319 | 0.805074 | 0.42078 | 0.62785 | 123.134 |

| | | | | | | | |
|---|---|---|---|---|---|---|---|
| Mcf2l | 12829.1 | 0.094122 | 0.07932 | 1.186608 | 0.23538 | 0.44055 | 147.1 |
| Mcfd2 | 3727.47 | 0.067706 | 0.0908 | 0.745674 | 0.45586 | 0.65758 | 11.493 |
| Mchr1 | 1962.96 | 0.517281 | 0.18399 | 2.811493 | 0.00493 | 0.0359 | 3.466 |
| Mcidas | 57.2913 | 0.476457 | 0.25122 | 1.896606 | 0.05788 | 0.18283 | 6.55 |
| Mcl1 | 3614.43 | -0.27282 | 0.07099 | -3.84311 | 0.00012 | 0.00305 | 4.389 |
| Mcm10 | 39.2091 | -0.61712 | 0.34656 | -1.7807 | 0.07496 | 0.21585 | 23.078 |
| Mcm2 | 185.127 | -0.31329 | 0.19 | -1.64892 | 0.09916 | 0.25872 | 15.307 |
| Mcm3 | 111.314 | -0.52777 | 0.19578 | -2.69568 | 0.00702 | 0.04556 | 17.345 |
| Mcm3ap | 4007.26 | 0.241331 | 0.07133 | 3.383444 | 0.00072 | 0.01013 | 46.931 |
| Mcm4 | 709.718 | 0.094947 | 0.1007 | 0.942903 | 0.34573 | 0.55644 | 13.504 |
| Mcm5 | 315.453 | 0.025586 | 0.13606 | 0.188044 | 0.85084 | 0.9241 | 18.871 |
| Mcm6 | 241.889 | -0.33896 | 0.20896 | -1.62214 | 0.10477 | 0.26777 | 28.075 |
| Mcm7 | 829.833 | 0.085577 | 0.0886 | 0.965902 | 0.33409 | 0.54479 | 7.84 |
| Mcm8 | 301.894 | 0.343015 | 0.12962 | 2.646317 | 0.00814 | 0.05009 | 28.057 |
| Mcm9 | 576.15 | 0.173292 | 0.08893 | 1.948546 | 0.05135 | 0.16882 | 94.125 |
| Mcmbp | 1613.9 | 0.152473 | 0.08006 | 1.904597 | 0.05683 | 0.18045 | 44.055 |
| Mcmdc2 | 262.045 | 0.206223 | 0.14528 | 1.419505 | 0.15575 | 0.34238 | 33.448 |
| Mcoln1 | 3554.49 | -0.04855 | 0.07689 | -0.63137 | 0.5278 | 0.71577 | 14.776 |
| Mcph1 | 594.958 | 0.115784 | 0.09449 | 1.22537 | 0.22044 | 0.4231 | 208.059 |
| Mcrip1 | 1565.22 | -0.31247 | 0.08661 | -3.60781 | 0.00031 | 0.00575 | 6.84 |
| Mcrip2 | 341.811 | -0.1187 | 0.1247 | -0.95189 | 0.34115 | 0.55206 | 5.067 |
| Mcrs1 | 1727.62 | -0.04157 | 0.08359 | -0.49729 | 0.61898 | 0.78314 | 9.145 |
| Mctp1 | 9379.16 | 0.657084 | 0.17264 | 3.806141 | 0.00014 | 0.00337 | 647.671 |
| Mctp2 | 135.694 | -0.89332 | 0.29891 | -2.9886 | 0.0028 | 0.02513 | 228.779 |
| Mcts1 | 1823.24 | 0.01772 | 0.07804 | 0.22708 | 0.82036 | 0.90734 | 12.849 |
| Mcts2 | 270.901 | 0.220457 | 0.15625 | 1.410969 | 0.15825 | 0.34558 | 0.912 |
| Mcu | 1757.17 | -0.07509 | 0.08943 | -0.83966 | 0.4011 | 0.60978 | 169.709 |
| Mcub | 70.2603 | 0.033328 | 0.29546 | 0.112801 | 0.91019 | 0.95475 | 55.247 |
| Mcur1 | 1214.63 | -0.27284 | 0.08164 | -3.34213 | 0.00083 | 0.01127 | 21.799 |
| Mdc1 | 1732.44 | 0.222915 | 0.07847 | 2.84073 | 0.0045 | 0.03394 | 18.156 |
| Mdfi | 18.2875 | -0.95657 | 0.44246 | -2.16193 | 0.03062 | 0.11975 | 19.364 |
| Mdfic | 232.765 | -0.16249 | 0.1553 | -1.04628 | 0.29543 | 0.50589 | 81.509 |
| Mdga1 | 2523.05 | 0.27843 | 0.08756 | 3.179802 | 0.00147 | 0.01635 | 142.139 |
| Mdga2 | 3248.14 | 0.234818 | 0.12395 | 1.894486 | 0.05816 | 0.18329 | 756.49 |

| | | | | | | | |
|---|---|---|---|---|---|---|---|
| Mdh1 | 23625.9 | -0.10597 | 0.05133 | -2.06444 | 0.03898 | 0.14064 | 15.676 |
| Mdh1b | 35.8031 | 0.455392 | 0.38876 | 1.171397 | 0.24144 | 0.44749 | 31.5 |
| Mdh2 | 8434.23 | -0.06192 | 0.06659 | -0.92983 | 0.35246 | 0.56283 | 11.919 |
| Mdk | 665.536 | 0.32665 | 0.1173 | 2.784754 | 0.00536 | 0.03794 | 2.493 |
| Mdm1 | 535.011 | 0.344184 | 0.1111 | 3.097845 | 0.00195 | 0.01976 | 27.187 |
| Mdm2 | 3148.35 | 0.252853 | 0.07683 | 3.29128 | 0.001 | 0.0128 | 21.884 |
| Mdm4 | 7622.65 | 0.174806 | 0.06731 | 2.596865 | 0.00941 | 0.05514 | 71.078 |
| Mdn1 | 6157.02 | 0.234898 | 0.08844 | 2.656108 | 0.0079 | 0.04926 | 118.099 |
| Mdp1 | 1180.23 | -0.23089 | 0.09075 | -2.54433 | 0.01095 | 0.06094 | 2.63 |
| Me1 | 1603.49 | -0.43907 | 0.13442 | -3.26633 | 0.00109 | 0.01351 | 114.583 |
| Me2 | 999.087 | -0.19107 | 0.10588 | -1.80458 | 0.07114 | 0.20877 | 45.547 |
| Me3 | 3506.83 | -0.23063 | 0.08844 | -2.6079 | 0.00911 | 0.05385 | 221.968 |
| Mea1 | 1305.33 | -0.27701 | 0.09271 | -2.98789 | 0.00281 | 0.02514 | 4.319 |
| Meaf6 | 6605.59 | 0.235679 | 0.09962 | 2.365739 | 0.01799 | 0.08363 | 28.105 |
| Meak7 | 725.497 | 0.064411 | 0.09302 | 0.692474 | 0.48864 | 0.68372 | 18.341 |
| Mecom | 119.07 | -0.37304 | 0.19167 | -1.94624 | 0.05163 | 0.16941 | 596.713 |
| Mecp2 | 10355.1 | 0.085127 | 0.06488 | 1.311975 | 0.18953 | 0.38549 | 108.772 |
| Mecr | 832.065 | -0.05688 | 0.086 | -0.66139 | 0.50836 | 0.69969 | 24.317 |
| Med1 | 3536.78 | 0.055476 | 0.06774 | 0.818989 | 0.41279 | 0.62094 | 41.14 |
| Med10 | 1023.17 | -0.13805 | 0.11885 | -1.16148 | 0.24545 | 0.45204 | 13.71 |
| Med11 | 208.408 | -0.1984 | 0.13486 | -1.47115 | 0.14125 | 0.323 | 1.809 |
| Med12 | 2460.96 | 0.016007 | 0.0969 | 0.165187 | 0.8688 | 0.93346 | 23.436 |
| Med12l | 2285.36 | 0.220583 | 0.08059 | 2.73701 | 0.0062 | 0.04211 | 312.858 |
| Med13 | 5502.05 | 0.113875 | 0.09791 | 1.163048 | 0.24481 | 0.45139 | 90.57 |
| Med13l | 8132.75 | 0.000891 | 0.08459 | 0.010532 | 0.9916 | 0.99531 | 204.76 |
| Med14 | 3692.33 | 0.070089 | 0.06427 | 1.090475 | 0.2755 | 0.4847 | 86.705 |
| Med15 | 2866.86 | 0.009424 | 0.06676 | 0.141164 | 0.88774 | 0.94402 | 81.684 |
| Med16 | 2331.39 | 0.179332 | 0.09153 | 1.95923 | 0.05009 | 0.16622 | 14.216 |
| Med17 | 1413.8 | 0.13972 | 0.09352 | 1.493993 | 0.13518 | 0.31505 | 19.581 |
| Med18 | 156.804 | -0.18487 | 0.1768 | -1.04563 | 0.29573 | 0.50616 | 5.191 |
| Med19 | 1009.01 | -0.07101 | 0.08062 | -0.88082 | 0.37841 | 0.58827 | 9.814 |
| Med20 | 1062.06 | 0.156173 | 0.08225 | 1.89872 | 0.0576 | 0.18222 | 12.837 |
| Med21 | 516.291 | 0.204106 | 0.10361 | 1.969935 | 0.04885 | 0.16337 | 8.186 |
| Med22 | 1485.88 | 0.018718 | 0.09668 | 0.193609 | 0.84648 | 0.92184 | 5.416 |

| | | | | | | | |
|---|---|---|---|---|---|---|---|
| Med23 | 2360.26 | 0.164494 | 0.08692 | 1.892505 | 0.05842 | 0.18384 | 43.696 |
| Med24 | 5116.76 | 0.018785 | 0.05822 | 0.322645 | 0.74696 | 0.86734 | 24.845 |
| Med25 | 3319.6 | 0.02652 | 0.07098 | 0.373639 | 0.70867 | 0.8429 | 15.948 |
| Med26 | 299.181 | 0.15392 | 0.13607 | 1.131156 | 0.25799 | 0.46605 | 53.71 |
| Med27 | 480.556 | 0.165587 | 0.14211 | 1.165191 | 0.24394 | 0.45033 | 177.975 |
| Med28 | 2171.93 | 0.069211 | 0.06559 | 1.055265 | 0.2913 | 0.50121 | 9.048 |
| Med29 | 208.178 | -0.05272 | 0.13672 | -0.38559 | 0.6998 | 0.83779 | 6.563 |
| Med30 | 578.082 | -0.46017 | 0.11197 | -4.10981 | 3.96E-05 | 0.00139 | 18.157 |
| Med31 | 428.67 | 0.067738 | 0.11683 | 0.57978 | 0.56206 | 0.74291 | 3.869 |
| Med4 | 1311.09 | 0.114778 | 0.08852 | 1.296659 | 0.19475 | 0.39207 | 8.825 |
| Med6 | 805.536 | 0.166072 | 0.1016 | 1.634503 | 0.10215 | 0.26377 | 21.452 |
| Med7 | 756.46 | -0.08191 | 0.08833 | -0.92736 | 0.35374 | 0.56399 | 5.797 |
| Med8 | 1280.64 | 0.060925 | 0.0728 | 0.836932 | 0.40263 | 0.61129 | 6.446 |
| Med9 | 1742.09 | -0.09938 | 0.07655 | -1.29817 | 0.19423 | 0.39143 | 14 |
| Med9os | 15.8145 | -0.06358 | 0.45817 | -0.13876 | 0.88964 | 0.94474 | 1.291 |
| Medag | 173.733 | -0.87175 | 0.28697 | -3.03772 | 0.00238 | 0.02257 | 19.975 |
| Mef2a | 9775.97 | 0.213906 | 0.07915 | 2.702413 | 0.00688 | 0.04491 | 141.696 |
| Mef2c | 14509.4 | -0.26731 | 0.11973 | -2.23262 | 0.02557 | 0.10553 | 163.047 |
| Mef2d | 11670.6 | 0.012198 | 0.06684 | 0.182491 | 0.8552 | 0.92683 | 29.715 |
| Meg3 | 320102 | 0.306281 | 0.1337 | 2.290872 | 0.02197 | 0.09543 | 31.287 |
| Megf10 | 1977.08 | -0.28247 | 0.09418 | -2.99926 | 0.00271 | 0.02457 | 164.378 |
| Megf11 | 1204.75 | 0.154613 | 0.08258 | 1.87226 | 0.06117 | 0.18898 | 323.58 |
| Megf6 | 711.593 | -0.05636 | 0.14807 | -0.3806 | 0.7035 | 0.83993 | 104.984 |
| Megf8 | 11672.7 | 0.112651 | 0.0639 | 1.762893 | 0.07792 | 0.221 | 48.754 |
| Megf9 | 6811.23 | -0.20604 | 0.1058 | -1.94751 | 0.05147 | 0.16918 | 107.931 |
| Mei1 | 198.179 | 0.628856 | 0.18003 | 3.493158 | 0.00048 | 0.00769 | 56.819 |
| Mei4 | 130.715 | 1.377605 | 0.26175 | 5.263149 | 1.42E-07 | 2.37E-05 | 342.338 |
| Meig1 | 44.4616 | 0.25146 | 0.36712 | 0.684952 | 0.49337 | 0.6878 | 13.606 |
| Meioc | 47.8079 | -0.19006 | 0.32391 | -0.58676 | 0.55737 | 0.73904 | 18.522 |
| Meiosin | 21.9087 | -0.37284 | 0.37394 | -0.99705 | 0.31874 | 0.5293 | 19.323 |
| Meis1 | 355.917 | -0.18618 | 0.18188 | -1.02359 | 0.30603 | 0.51614 | 139.169 |
| Meis2 | 16452.4 | 0.412093 | 0.0986 | 4.179324 | 2.92E-05 | 0.00114 | 202.776 |
| Meis3 | 1703.68 | 0.296444 | 0.09106 | 3.25542 | 0.00113 | 0.01379 | 11.415 |
| Melk | 33.3747 | 0.049705 | 0.32804 | 0.151524 | 0.87956 | 0.93973 | 63.8 |

| | | | | | | | |
|---|---|---|---|---|---|---|---|
| Meltf | 17.8053 | -0.39243 | 0.44154 | -0.88876 | 0.37413 | 0.58439 | 20.211 |
| Memo1 | 1290.87 | 0.08086 | 0.09077 | 0.890843 | 0.37301 | 0.5834 | 96.486 |
| Men1 | 2704.46 | 0.025508 | 0.08218 | 0.31038 | 0.75627 | 0.87174 | 5.913 |
| Mepce | 2202.04 | 0.074541 | 0.0765 | 0.974397 | 0.32986 | 0.54112 | 5.75 |
| Mertk | 3696.13 | -0.19684 | 0.07721 | -2.54939 | 0.01079 | 0.06048 | 103.939 |
| Mesd | 1416.28 | 0.00405 | 0.08912 | 0.045441 | 0.96376 | 0.98236 | 17.067 |
| Mesp2 | 21.295 | -0.34103 | 0.44698 | -0.76296 | 0.44549 | 0.64938 | 2.713 |
| Mest | 1421.11 | 0.380679 | 0.0953 | 3.994512 | 6.48E-05 | 0.00199 | 24.919 |
| Met | 45.7727 | -1.62148 | 0.42686 | -3.79863 | 0.00015 | 0.00346 | 110.181 |
| Metap1 | 2272.46 | 0.113751 | 0.07156 | 1.589636 | 0.11192 | 0.2794 | 30.56 |
| Metap1d | 1240.45 | 0.121079 | 0.14567 | 0.831188 | 0.40587 | 0.61442 | 71.919 |
| Metap2 | 4119.56 | 0.127698 | 0.09279 | 1.376171 | 0.16877 | 0.35886 | 38.605 |
| Metazoa_SRP | 24.9816 | 0.141434 | 0.37488 | 0.377274 | 0.70597 | 0.8415 | 0.317 |
| Metrn | 1352.57 | -0.25371 | 0.10475 | -2.4222 | 0.01543 | 0.07584 | 3.916 |
| Metrnl | 86.925 | -0.20856 | 0.21588 | -0.9661 | 0.33399 | 0.54474 | 14.763 |
| Mettl1 | 460.128 | -0.01456 | 0.11399 | -0.12777 | 0.89833 | 0.94899 | 4.952 |
| Mettl11b | 143.878 | 0.272709 | 0.25774 | 1.058095 | 0.29001 | 0.50018 | 22.977 |
| Mettl14 | 1345.03 | 0.097072 | 0.09114 | 1.065083 | 0.28684 | 0.49672 | 17.812 |
| Mettl15 | 157.005 | -0.3459 | 0.15867 | -2.17995 | 0.02926 | 0.11606 | 188.452 |
| Mettl16 | 2267.93 | 0.032553 | 0.0675 | 0.482241 | 0.62963 | 0.79064 | 57.696 |
| Mettl17 | 1771.38 | 0.199691 | 0.08812 | 2.266249 | 0.02344 | 0.09932 | 8.198 |
| Mettl18 | 183.278 | 0.221046 | 0.14362 | 1.539082 | 0.12378 | 0.29829 | 2.355 |
| Mettl2 | 773.103 | 0.089478 | 0.09407 | 0.951189 | 0.34151 | 0.55244 | 13.97 |
| Mettl21a | 566.487 | 0.098127 | 0.10372 | 0.946081 | 0.34411 | 0.5548 | 10.77 |
| Mettl22 | 527.915 | -0.05483 | 0.12041 | -0.45537 | 0.64884 | 0.80429 | 19.897 |
| Mettl23 | 564.359 | 0.035345 | 0.09129 | 0.387166 | 0.69863 | 0.83677 | 10.986 |
| Mettl24 | 23.2328 | -0.12704 | 0.37353 | -0.3401 | 0.73378 | 0.85907 | 127.802 |
| Mettl25 | 267.951 | 0.121 | 0.11699 | 1.034316 | 0.30099 | 0.51075 | 78.192 |
| Mettl26 | 574.682 | -0.18713 | 0.10813 | -1.73064 | 0.08352 | 0.23102 | 2.096 |
| Mettl27 | 196.483 | -0.47641 | 0.17987 | -2.64861 | 0.00808 | 0.04991 | 10.27 |
| Mettl3 | 1099.5 | 0.176665 | 0.08447 | 2.091533 | 0.03648 | 0.13457 | 10.288 |
| Mettl4 | 1241.73 | 0.104883 | 0.07499 | 1.398633 | 0.16192 | 0.3502 | 27.43 |
| Mettl5 | 406.901 | 0.003988 | 0.14833 | 0.026889 | 0.97855 | 0.98863 | 14.418 |
| Mettl5os | 19.524 | 0.52492 | 0.51935 | 1.010721 | 0.31215 | 0.52253 | 2.428 |

| | | | | | | | |
|---|---|---|---|---|---|---|---|
| Mettl6 | 968.545 | -0.00142 | 0.08004 | -0.01776 | 0.98583 | 0.99235 | 21.463 |
| Mettl7a1 | 1602.5 | -0.4568 | 0.09658 | -4.72991 | 2.25E-06 | 0.00019 | 24.523 |
| Mettl8 | 626.961 | 0.061337 | 0.11282 | 0.543683 | 0.58666 | 0.76043 | 91.023 |
| Mettl9 | 2091.99 | 0.10103 | 0.10344 | 0.976681 | 0.32873 | 0.53991 | 43.885 |
| Mex3a | 243.654 | -0.20833 | 0.22275 | -0.93527 | 0.34965 | 0.56023 | 9.002 |
| Mex3b | 808.276 | -0.19644 | 0.12205 | -1.60944 | 0.10752 | 0.27228 | 4.183 |
| Mex3c | 2385.09 | 0.089417 | 0.06848 | 1.305711 | 0.19165 | 0.3883 | 19.532 |
| Mex3d | 570.512 | -0.12733 | 0.14582 | -0.87321 | 0.38255 | 0.59199 | 7.305 |
| Mfap1a | 1011.17 | -0.22865 | 0.17684 | -1.29299 | 0.19601 | 0.39362 | 14.755 |
| Mfap1b | 918.295 | 0.077786 | 0.21571 | 0.360602 | 0.7184 | 0.84957 | 13.833 |
| Mfap2 | 177.555 | 0.201646 | 0.17961 | 1.122666 | 0.26158 | 0.47032 | 5.567 |
| Mfap3 | 1853.54 | 0.124803 | 0.07795 | 1.600975 | 0.10938 | 0.2752 | 15.152 |
| Mfap3l | 4208.59 | -0.04742 | 0.11567 | -0.40997 | 0.68183 | 0.82616 | 43.903 |
| Mfap4 | 34.2093 | -1.20049 | 0.37336 | -3.21534 | 0.0013 | 0.01501 | 3.47 |
| Mff | 6565 | -0.05364 | 0.05299 | -1.01224 | 0.31142 | 0.52174 | 27.505 |
| Mfge8 | 7535.93 | -0.06619 | 0.12464 | -0.53108 | 0.59536 | 0.76727 | 15.293 |
| Mfhas1 | 2013.12 | -0.04161 | 0.09012 | -0.4617 | 0.6443 | 0.8016 | 91.652 |
| Mfn1 | 4645.08 | -0.09216 | 0.07791 | -1.18299 | 0.23681 | 0.44193 | 49.775 |
| Mfn2 | 10327.2 | -0.031 | 0.05628 | -0.55087 | 0.58172 | 0.75754 | 31.106 |
| Mfng | 148.802 | -0.24836 | 0.16329 | -1.52099 | 0.12826 | 0.30524 | 17.594 |
| Mfrp | 40.3462 | 0.260995 | 0.3387 | 0.770587 | 0.44095 | 0.64569 | 7.459 |
| Mfsd1 | 1483.73 | -0.2603 | 0.07007 | -3.71494 | 0.0002 | 0.00432 | 21.497 |
| Mfsd10 | 711.773 | 0.025242 | 0.08395 | 0.300689 | 0.76365 | 0.87623 | 3.571 |
| Mfsd11 | 1635.47 | 0.134195 | 0.0993 | 1.35138 | 0.17657 | 0.36865 | 23.196 |
| Mfsd12 | 724.716 | -0.34678 | 0.10719 | -3.23517 | 0.00122 | 0.01446 | 8.735 |
| Mfsd13a | 1312.85 | 0.22533 | 0.10138 | 2.22273 | 0.02623 | 0.10755 | 34.132 |
| Mfsd14a | 1369.02 | 0.085048 | 0.08931 | 0.952316 | 0.34094 | 0.552 | 31.514 |
| Mfsd14b | 2489.94 | 0.072612 | 0.08411 | 0.863341 | 0.38795 | 0.59728 | 48.313 |
| Mfsd2a | 1626 | -0.25018 | 0.10259 | -2.43855 | 0.01475 | 0.07343 | 14.339 |
| Mfsd2b | 212.32 | -0.07503 | 0.18618 | -0.40298 | 0.68697 | 0.82926 | 11.92 |
| Mfsd3 | 198.175 | -0.1209 | 0.14988 | -0.80666 | 0.41986 | 0.62714 | 2.775 |
| Mfsd4a | 2456.81 | -0.94025 | 0.31097 | -3.02358 | 0.0025 | 0.02336 | 45.257 |
| Mfsd4b1 | 46.5073 | 0.34186 | 0.25532 | 1.33895 | 0.18059 | 0.37394 | 23.694 |
| Mfsd4b4 | 865.15 | 0.187845 | 0.09773 | 1.922033 | 0.0546 | 0.17572 | 9.226 |

| | | | | | | | |
|---|---|---|---|---|---|---|---|
| Mfsd5 | 1116.6 | -0.20964 | 0.09226 | -2.27231 | 0.02307 | 0.09831 | 2.301 |
| Mfsd6 | 4453.14 | -0.23455 | 0.09268 | -2.53069 | 0.01138 | 0.06243 | 71.177 |
| Mfsd7a | 82.6117 | -0.66655 | 0.20114 | -3.31386 | 0.00092 | 0.0121 | 8.047 |
| Mfsd8 | 1491.6 | 0.20723 | 0.1037 | 1.998365 | 0.04568 | 0.1557 | 28.784 |
| Mfsd9 | 50.7204 | -0.1487 | 0.25255 | -0.58878 | 0.55601 | 0.73825 | 18.65 |
| Mga | 4997.7 | 0.232301 | 0.11464 | 2.026354 | 0.04273 | 0.14901 | 72.354 |
| Mgat1 | 2935.72 | 0.017377 | 0.05928 | 0.293135 | 0.76942 | 0.87968 | 18.84 |
| Mgat2 | 968.847 | -0.06238 | 0.08554 | -0.72928 | 0.46583 | 0.665 | 2.614 |
| Mgat3 | 7678.2 | -0.12764 | 0.05332 | -2.39403 | 0.01666 | 0.0799 | 41.799 |
| Mgat4a | 2731.04 | 0.230594 | 0.0663 | 3.478082 | 0.00051 | 0.00798 | 101.677 |
| Mgat4b | 2373.82 | 0.139528 | 0.10575 | 1.319428 | 0.18703 | 0.38229 | 24.214 |
| Mgat4c | 583.872 | 0.133621 | 0.12063 | 1.107713 | 0.26799 | 0.47725 | 709.983 |
| Mgat5 | 4699.58 | -0.30271 | 0.10322 | -2.93275 | 0.00336 | 0.02801 | 283.322 |
| Mgat5b | 3416.06 | 0.233621 | 0.11296 | 2.06814 | 0.03863 | 0.13993 | 68.086 |
| Mgl2 | 15.7843 | -0.20825 | 0.41721 | -0.49916 | 0.61767 | 0.78255 | 7.222 |
| Mgll | 9088.08 | 0.051758 | 0.09224 | 0.561109 | 0.57472 | 0.75281 | 103.949 |
| Mgme1 | 304.369 | -0.3269 | 0.12794 | -2.55511 | 0.01062 | 0.05981 | 10.565 |
| Mgmt | 41.6734 | -0.12055 | 0.35931 | -0.33551 | 0.73724 | 0.86132 | 235.653 |
| Mgp | 465.896 | -1.3963 | 0.48622 | -2.87177 | 0.00408 | 0.03194 | 3.389 |
| Mgrn1 | 8373.07 | 0.037761 | 0.05551 | 0.6803 | 0.49631 | 0.68996 | 52.048 |
| Mgst1 | 363.747 | -0.46721 | 0.22412 | -2.08468 | 0.0371 | 0.13621 | 16.44 |
| Mgst3 | 1759.15 | -0.79509 | 0.14607 | -5.44312 | 5.24E-08 | 1.16E-05 | 21.876 |
| Mhrt | 123.345 | 1.00189 | 0.3228 | 3.103745 | 0.00191 | 0.01951 | 5.817 |
| Mia2 | 2763.02 | 0.066573 | 0.08371 | 0.795296 | 0.42644 | 0.63297 | 95.785 |
| Mia3 | 6304.62 | 0.040245 | 0.08629 | 0.466374 | 0.64095 | 0.79894 | 43.213 |
| Miat | 12557 | -0.59335 | 0.27751 | -2.13815 | 0.0325 | 0.12487 | 16.168 |
| Mib1 | 13185.7 | 0.064015 | 0.05049 | 1.267919 | 0.20483 | 0.40371 | 93.157 |
| Mib2 | 4922.98 | 0.035878 | 0.06169 | 0.581566 | 0.56086 | 0.74205 | 14.522 |
| Mical1 | 351.716 | -0.20332 | 0.15787 | -1.2879 | 0.19778 | 0.39558 | 10.719 |
| Mical2 | 6981.15 | -0.64107 | 0.31981 | -2.0045 | 0.04502 | 0.15421 | 187.251 |
| Mical3 | 8879.27 | 0.17068 | 0.06887 | 2.478126 | 0.01321 | 0.06882 | 199.292 |
| Micall1 | 884.237 | -0.31384 | 0.12914 | -2.43032 | 0.01509 | 0.07475 | 28.003 |
| Micall2 | 62.7839 | -0.12314 | 0.28317 | -0.43488 | 0.66365 | 0.81414 | 29.641 |
| Micos10 | 3121.1 | -0.10671 | 0.0605 | -1.76388 | 0.07775 | 0.22065 | 29.3 |

| | | | | | | | |
|---|---|---|---|---|---|---|---|
| Micos13 | 1383.26 | -0.23158 | 0.0878 | -2.63757 | 0.00835 | 0.05095 | 2.32 |
| Micu1 | 3903.24 | -0.06488 | 0.0639 | -1.01533 | 0.30995 | 0.51992 | 161.656 |
| Micu2 | 1298.07 | 0.034715 | 0.06903 | 0.502902 | 0.61503 | 0.78097 | 83.002 |
| Micu3 | 4601.68 | 0.205849 | 0.08738 | 2.355784 | 0.01848 | 0.08517 | 78.851 |
| Mid1 | 411.776 | 0.296503 | 0.17997 | 1.64747 | 0.09946 | 0.25919 | 320.538 |
| Mid1-ps1 | 41.0295 | -0.22884 | 0.41352 | -0.55338 | 0.58 | 0.75648 | 10.429 |
| Mid1ip1 | 1816.23 | -0.36693 | 0.10611 | -3.45789 | 0.00054 | 0.0084 | 4.678 |
| Mid2 | 2372.18 | 0.284277 | 0.10969 | 2.591568 | 0.00955 | 0.05563 | 103.117 |
| Mideas | 1803.33 | 0.077802 | 0.11366 | 0.684498 | 0.49366 | 0.6881 | 69.706 |
| Midn | 1913.14 | -0.443 | 0.15275 | -2.90013 | 0.00373 | 0.03004 | 10.097 |
| Mief1 | 1354.94 | -0.13692 | 0.07308 | -1.87349 | 0.061 | 0.1886 | 19.354 |
| Mief2 | 571.136 | -0.12717 | 0.09866 | -1.28903 | 0.19739 | 0.39524 | 4.554 |
| Mien1 | 108.424 | -0.07831 | 0.25978 | -0.30144 | 0.76308 | 0.87597 | 1.284 |
| Mier1 | 2625.9 | 0.089641 | 0.08047 | 1.113942 | 0.2653 | 0.4745 | 51.365 |
| Mier2 | 1588.91 | 0.107067 | 0.08866 | 1.207582 | 0.22721 | 0.43135 | 14.955 |
| Mier3 | 1585.23 | 0.189835 | 0.08391 | 2.26225 | 0.02368 | 0.10007 | 37.618 |
| Mif | 3334.82 | -0.12632 | 0.08309 | -1.52032 | 0.12843 | 0.30549 | 0.888 |
| Mif4gd | 476.298 | -0.165 | 0.11083 | -1.48879 | 0.13654 | 0.31704 | 5.052 |
| Miga1 | 6888.11 | 0.152764 | 0.06541 | 2.33543 | 0.01952 | 0.08813 | 66.559 |
| Miga2 | 3487.32 | -0.12823 | 0.05891 | -2.17665 | 0.02951 | 0.11683 | 21.289 |
| Miip | 1086.22 | 0.125306 | 0.08325 | 1.505097 | 0.1323 | 0.3113 | 8.039 |
| Mill2 | 228.715 | -0.24331 | 0.21074 | -1.15456 | 0.24827 | 0.45535 | 25.437 |
| Minar1 | 221.475 | -0.0441 | 0.2236 | -0.19724 | 0.84364 | 0.91962 | 35.91 |
| Minar2 | 1418.42 | 0.627375 | 0.14394 | 4.358689 | 1.31E-05 | 0.00067 | 14.715 |
| Mindy1 | 846.89 | -0.24747 | 0.09537 | -2.5948 | 0.00946 | 0.05532 | 14.822 |
| Mindy2 | 4093.67 | -0.14364 | 0.08659 | -1.65888 | 0.09714 | 0.2554 | 58.161 |
| Mindy3 | 1564.33 | -0.13781 | 0.08354 | -1.64952 | 0.09904 | 0.25857 | 72.208 |
| Mindy4 | 395.544 | 0.048752 | 0.13616 | 0.358048 | 0.72031 | 0.85073 | 116.84 |
| Mindy4b-ps | 99.1739 | 0.148615 | 0.18454 | 0.805311 | 0.42064 | 0.6278 | 33.905 |
| Mink1 | 17752.6 | 0.12745 | 0.0707 | 1.802758 | 0.07143 | 0.20924 | 51.603 |
| Minpp1 | 1830.07 | 0.01583 | 0.10224 | 0.154824 | 0.87696 | 0.93848 | 29.596 |
| Minpp1-ps | 15.2834 | 0.197094 | 0.43584 | 0.452212 | 0.65112 | 0.8058 | 1.427 |
| Mios | 1336.07 | 0.17301 | 0.09667 | 1.78976 | 0.07349 | 0.21349 | 27.053 |
| Mip | 14.3294 | 0.259885 | 0.52575 | 0.494313 | 0.62108 | 0.7845 | 5.975 |

| | | | | | | | |
|---|---|---|---|---|---|---|---|
| Mipep | 1146.03 | 0.088805 | 0.09334 | 0.951458 | 0.34137 | 0.55226 | 120.906 |
| Mipol1 | 154.118 | 0.180739 | 0.14783 | 1.222588 | 0.22149 | 0.42444 | 266.773 |
| Mir100hg | 3862.98 | 0.175595 | 0.15974 | 1.099264 | 0.27165 | 0.48101 | 389.918 |
| Mir103-2 | 27.3554 | -0.01635 | 0.37928 | -0.04311 | 0.96561 | 0.9828 | 0.086 |
| Mir1188 | 159.64 | 0.402344 | 0.1526 | 2.636554 | 0.00838 | 0.05104 | 0.12 |
| Mir1192 | 17.039 | -0.03958 | 0.53749 | -0.07364 | 0.94129 | 0.97087 | 0.121 |
| Mir1198 | 45.6556 | 0.781954 | 0.33546 | 2.331015 | 0.01975 | 0.08876 | 0.121 |
| Mir124-2hg | 4357.45 | -0.14355 | 0.1101 | -1.30381 | 0.1923 | 0.38895 | 19.243 |
| Mir124a-1hg | 9489.53 | 0.061252 | 0.11657 | 0.525452 | 0.59927 | 0.77006 | 6.667 |
| Mir1258 | 23.5865 | 0.082281 | 0.41163 | 0.199888 | 0.84157 | 0.91879 | 0.06 |
| Mir128-1 | 24.1317 | -0.55308 | 0.38487 | -1.43707 | 0.1507 | 0.33561 | 0.07 |
| Mir132 | 22.9454 | -0.44636 | 0.44505 | -1.00294 | 0.31589 | 0.5265 | 0.066 |
| Mir135a-2 | 25.3251 | -0.43402 | 0.46026 | -0.94299 | 0.34569 | 0.55643 | 0.1 |
| Mir149 | 14.3462 | -0.05308 | 0.59717 | -0.08888 | 0.92917 | 0.96438 | 0.066 |
| Mir154 | 26.9786 | 0.175706 | 0.43988 | 0.39944 | 0.68957 | 0.83089 | 0.066 |
| Mir17hg | 49.5124 | -0.16752 | 0.31663 | -0.52908 | 0.59675 | 0.76862 | 3.849 |
| Mir181a-1hg | 43.7631 | -0.48544 | 0.3498 | -1.38776 | 0.16521 | 0.3543 | 3.388 |
| Mir186 | 200.911 | 0.099148 | 0.161 | 0.615833 | 0.538 | 0.72473 | 0.071 |
| Mir1898 | 66.3947 | 0.289194 | 0.28543 | 1.013183 | 0.31097 | 0.5212 | 0.083 |
| Mir1943 | 32.0676 | 0.343681 | 0.30458 | 1.128367 | 0.25917 | 0.46734 | 0.073 |
| Mir1969 | 13.4815 | 0.322697 | 0.49674 | 0.649628 | 0.51593 | 0.70609 | 0.094 |
| Mir1970 | 24.7525 | 0.586209 | 0.40752 | 1.438478 | 0.1503 | 0.33494 | 0.104 |
| Mir199a-1 | 19.9111 | 0.464068 | 0.40462 | 1.146914 | 0.25142 | 0.45881 | 0.07 |
| Mir199b | 19.0954 | 0.116531 | 0.39968 | 0.291558 | 0.77062 | 0.88002 | 0.11 |
| Mir208b | 74.3218 | 0.97137 | 0.38327 | 2.534431 | 0.01126 | 0.06199 | 0.077 |
| Mir212 | 43.8309 | 0.231011 | 0.32748 | 0.705425 | 0.48055 | 0.67789 | 0.091 |
| Mir22hg | 748.931 | -0.14825 | 0.10115 | -1.46566 | 0.14274 | 0.3247 | 5.138 |
| Mir23b | 16.0672 | -0.47715 | 0.62564 | -0.76266 | 0.44567 | 0.64947 | 0.074 |
| Mir26a-1 | 13.0894 | 0.300842 | 0.55979 | 0.537417 | 0.59098 | 0.7637 | 0.09 |
| Mir300 | 27.4586 | -0.01893 | 0.36041 | -0.05254 | 0.9581 | 0.97953 | 0.079 |
| Mir301 | 14.4643 | -0.15319 | 0.56924 | -0.26911 | 0.78784 | 0.88981 | 0.086 |
| Mir3060 | 79.7791 | 0.393949 | 0.21674 | 1.817638 | 0.06912 | 0.20458 | 0.083 |
| Mir3070a | 46.5849 | 0.481937 | 0.25282 | 1.906252 | 0.05662 | 0.18017 | 0.089 |
| Mir3070b | 66.1048 | 0.918056 | 0.272 | 3.375216 | 0.00074 | 0.01037 | 0.089 |

| | | | | | | | |
|---|---|---|---|---|---|---|---|
| Mir3072 | 17.0657 | 0.33091 | 0.44426 | 0.744853 | 0.45636 | 0.65798 | 0.083 |
| Mir341 | 100.754 | 0.583561 | 0.1784 | 3.27104 | 0.00107 | 0.01341 | 0.096 |
| Mir343 | 22.8783 | -0.05104 | 0.39137 | -0.13043 | 0.89623 | 0.94785 | 0.075 |
| Mir344-2 | 31.4721 | 0.117421 | 0.35411 | 0.331596 | 0.74019 | 0.863 | 0.08 |
| Mir3473g | 15.5589 | 0.234258 | 0.4896 | 0.478472 | 0.63231 | 0.79244 | 0.136 |
| Mir365-1 | 16.8309 | -0.15669 | 0.53138 | -0.29488 | 0.76809 | 0.87883 | 0.087 |
| Mir374b | 23.2216 | 0.369994 | 0.43933 | 0.842178 | 0.39969 | 0.6085 | 0.095 |
| Mir376c | 22.7152 | 0.344378 | 0.35633 | 0.966453 | 0.33382 | 0.54457 | 0.086 |
| Mir378b | 51.5387 | 0.210527 | 0.28427 | 0.740584 | 0.45895 | 0.65977 | 0.092 |
| Mir381 | 23.1487 | 0.948787 | 0.44155 | 2.148784 | 0.03165 | 0.12253 | 0.075 |
| Mir384 | 18.731 | 0.254195 | 0.68258 | 0.372403 | 0.70959 | 0.8436 | 0.088 |
| Mir421 | 121.177 | 0.582059 | 0.20386 | 2.855133 | 0.0043 | 0.03308 | 0.076 |
| Mir453 | 42.5372 | 0.939385 | 0.41083 | 2.286549 | 0.02222 | 0.0961 | 0.082 |
| Mir466i | 27.6224 | 0.561849 | 0.37837 | 1.484932 | 0.13756 | 0.31832 | 0.121 |
| Mir467f | 15.1939 | 0.018243 | 0.45054 | 0.040491 | 0.9677 | 0.98368 | 0.12 |
| Mir487b | 25.3372 | 0.456508 | 0.3952 | 1.155138 | 0.24803 | 0.4552 | 0.082 |
| Mir491 | 21.1605 | -0.02007 | 0.43665 | -0.04597 | 0.96334 | 0.98208 | 0.086 |
| Mir493 | 35.9051 | 0.233999 | 0.36883 | 0.634438 | 0.5258 | 0.71407 | 0.083 |
| Mir496a | 32.4133 | 0.479108 | 0.36184 | 1.324103 | 0.18547 | 0.38025 | 0.079 |
| Mir5119 | 23.5538 | 0.131889 | 0.39331 | 0.33533 | 0.73738 | 0.86132 | 0.057 |
| Mir5125 | 539.287 | 0.376188 | 0.156 | 2.411515 | 0.01589 | 0.07735 | 0.079 |
| Mir540 | 17.7302 | 0.458329 | 0.48118 | 0.952514 | 0.34084 | 0.55188 | 0.067 |
| Mir541 | 47.3503 | 0.471828 | 0.32449 | 1.454071 | 0.14593 | 0.32883 | 0.09 |
| Mir544 | 26.3697 | 0.723249 | 0.39041 | 1.852548 | 0.06395 | 0.19461 | 0.078 |
| Mir6392 | 12.8427 | 0.2122 | 0.67657 | 0.313642 | 0.75379 | 0.87065 | 0.108 |
| Mir6395 | 14.9664 | 0.475446 | 0.43177 | 1.101159 | 0.27083 | 0.48026 | 0.091 |
| Mir654 | 26.6143 | 0.492689 | 0.32762 | 1.503849 | 0.13262 | 0.3118 | 0.084 |
| Mir665 | 17.3489 | -0.04247 | 0.6076 | -0.0699 | 0.94427 | 0.97202 | 0.094 |
| Mir667 | 18.8513 | 0.476246 | 0.39534 | 1.204664 | 0.22833 | 0.43232 | 0.092 |
| Mir670hg | 326.174 | 0.114345 | 0.14855 | 0.769734 | 0.44146 | 0.64608 | 104.395 |
| Mir684-1 | 16.5595 | -0.19773 | 0.41074 | -0.48139 | 0.63024 | 0.79101 | 0.086 |
| Mir6941 | 19.9907 | -0.42468 | 0.41174 | -1.03143 | 0.30234 | 0.51199 | 0.059 |
| Mir6955 | 17.6744 | 0.201986 | 0.50311 | 0.401473 | 0.68807 | 0.83002 | 0.074 |
| Mir697 | 15.2259 | 0.155716 | 0.46752 | 0.333069 | 0.73908 | 0.86241 | 0.109 |

| | | | | | | | |
|---|---|---|---|---|---|---|---|
| Mir6982 | 32.5385 | 0.036516 | 0.39961 | 0.091379 | 0.92719 | 0.96334 | 0.068 |
| Mir7-1 | 40.526 | -0.11917 | 0.32729 | -0.36412 | 0.71577 | 0.84802 | 0.108 |
| Mir7027 | 13.6392 | -0.21439 | 0.48046 | -0.44623 | 0.65543 | 0.80876 | 0.072 |
| Mir7056 | 14.2923 | 0.659674 | 0.45078 | 1.463399 | 0.14336 | 0.32549 | 0.059 |
| Mir7220 | 119.945 | 0.328949 | 0.17886 | 1.839187 | 0.06589 | 0.1983 | 0.06 |
| Mir7240 | 73.8284 | -0.66615 | 0.22938 | -2.90407 | 0.00368 | 0.02981 | 0.058 |
| Mir7651 | 47.2503 | 0.353444 | 0.28195 | 1.253579 | 0.21 | 0.41005 | 0.064 |
| Mir7664 | 13.0572 | -0.28796 | 0.67292 | -0.42793 | 0.6687 | 0.81724 | 0.063 |
| Mir7669 | 23.2546 | 0.315171 | 0.32822 | 0.960233 | 0.33694 | 0.54752 | 0.058 |
| Mir8091 | 54.0122 | 0.1597 | 0.26032 | 0.613487 | 0.53955 | 0.72589 | 0.139 |
| Mir8103 | 24.5816 | 0.310653 | 0.3718 | 0.835545 | 0.40341 | 0.61211 | 0.107 |
| Mir874 | 19.4738 | 0.81401 | 0.39308 | 2.070839 | 0.03837 | 0.13938 | 0.076 |
| Mir9-3hg | 1607.69 | -0.09757 | 0.0921 | -1.05939 | 0.28942 | 0.49971 | 35.296 |
| Mir99ahg | 215.414 | 0.09121 | 0.21732 | 0.419711 | 0.6747 | 0.82139 | 537.132 |
| Mir99b | 25.0351 | 0.145259 | 0.41547 | 0.349624 | 0.72662 | 0.85423 | 0.07 |
| Mirg | 1696.07 | 0.29699 | 0.13389 | 2.218125 | 0.02655 | 0.10846 | 19.697 |
| Mirt1 | 37.713 | -0.6247 | 0.40052 | -1.55972 | 0.11883 | 0.29088 | 21.567 |
| Mis12 | 1097.22 | 0.081797 | 0.08161 | 1.002351 | 0.31617 | 0.52667 | 7.761 |
| Mis18a | 343.306 | 0.120242 | 0.1612 | 0.745934 | 0.45571 | 0.65754 | 8.093 |
| Mis18bp1 | 69.4826 | 0.001546 | 0.35571 | 0.004345 | 0.99653 | 0.9981 | 39.871 |
| Misp | 87.4344 | 0.675047 | 0.25799 | 2.616551 | 0.00888 | 0.05306 | 9.502 |
| Misp3 | 17.5841 | -1.14393 | 0.61848 | -1.84956 | 0.06438 | 0.19522 | 1.95 |
| Mitd1 | 344.579 | -0.07205 | 0.16269 | -0.44289 | 0.65785 | 0.81018 | 15.611 |
| Mitf | 779.061 | 0.041537 | 0.10689 | 0.388603 | 0.69757 | 0.83613 | 214.298 |
| Mki67 | 264.893 | 0.098964 | 0.183 | 0.540801 | 0.58865 | 0.76198 | 26.578 |
| Mkks | 740.626 | 0.101026 | 0.08212 | 1.230222 | 0.21861 | 0.42068 | 17.61 |
| Mkln1 | 6413.69 | 0.006164 | 0.07157 | 0.086124 | 0.93137 | 0.96532 | 118.077 |
| Mkln1os | 437.034 | 0.199139 | 0.12854 | 1.549237 | 0.12132 | 0.29465 | 32.831 |
| Mknk1 | 1716.43 | 0.051444 | 0.08426 | 0.610522 | 0.54152 | 0.72706 | 40.053 |
| Mknk2 | 2113.73 | 0.040733 | 0.09418 | 0.432518 | 0.66537 | 0.81526 | 12.786 |
| Mkrn1 | 6039.96 | 0.073162 | 0.07854 | 0.931579 | 0.35155 | 0.56204 | 22.659 |
| Mkrn2 | 1105.15 | 0.089415 | 0.10248 | 0.872477 | 0.38295 | 0.59222 | 20.723 |
| Mkrn2os | 97.0223 | -0.15431 | 0.18966 | -0.81361 | 0.41587 | 0.62362 | 9.033 |
| Mkrn3 | 49.8949 | -0.73619 | 0.27817 | -2.64651 | 0.00813 | 0.05008 | 2.547 |

| | | | | | | | |
|---|---|---|---|---|---|---|---|
| Mks1 | 343.036 | 0.109198 | 0.12334 | 0.885364 | 0.37596 | 0.5861 | 10.589 |
| Mkx | 337.331 | -0.43142 | 0.17791 | -2.42487 | 0.01531 | 0.07543 | 70.263 |
| Mlc1 | 8068.69 | -0.27316 | 0.12138 | -2.25047 | 0.02442 | 0.10223 | 23.124 |
| Mlec | 7757.36 | 0.038036 | 0.08232 | 0.462035 | 0.64406 | 0.80143 | 15.199 |
| Mlf1 | 14.167 | -1.04693 | 0.50083 | -2.09039 | 0.03658 | 0.1348 | 25.907 |
| Mlf2 | 17054.9 | -0.13145 | 0.06362 | -2.06622 | 0.03881 | 0.14041 | 5.853 |
| Mlh1 | 786.675 | 0.150193 | 0.09323 | 1.610928 | 0.1072 | 0.27174 | 43.564 |
| Mlh3 | 1796.92 | 0.157927 | 0.09569 | 1.650417 | 0.09886 | 0.25828 | 36.08 |
| Mlip | 678.203 | -0.42478 | 0.15744 | -2.69812 | 0.00697 | 0.0453 | 250.889 |
| Mlkl | 17.8244 | -0.09885 | 0.48768 | -0.20269 | 0.83938 | 0.91737 | 26.381 |
| Mllt1 | 3243.79 | -0.01771 | 0.07533 | -0.23517 | 0.81408 | 0.90456 | 42.805 |
| Mllt10 | 2436.42 | 0.234058 | 0.08121 | 2.882142 | 0.00395 | 0.03118 | 157.152 |
| Mllt11 | 5332.52 | -0.05996 | 0.07783 | -0.77033 | 0.4411 | 0.64575 | 13.037 |
| Mllt3 | 1904.37 | 0.176954 | 0.07587 | 2.332334 | 0.01968 | 0.08867 | 263.44 |
| Mllt6 | 15011.5 | 0.179141 | 0.07064 | 2.535941 | 0.01121 | 0.0618 | 22.05 |
| Mlph | 292.513 | -0.2767 | 0.16615 | -1.66533 | 0.09585 | 0.25327 | 36.058 |
| Mlst8 | 1002.36 | 0.138267 | 0.09653 | 1.432324 | 0.15205 | 0.33717 | 5.528 |
| Mlx | 983.259 | -0.04812 | 0.07575 | -0.63532 | 0.52522 | 0.71355 | 4.931 |
| Mlxip | 2103.51 | 0.030155 | 0.08355 | 0.360912 | 0.71816 | 0.84954 | 63.135 |
| Mlxipl | 152.469 | -0.18243 | 0.18229 | -1.0008 | 0.31692 | 0.52743 | 48.493 |
| Mlycd | 978.873 | 0.003389 | 0.0774 | 0.043789 | 0.96507 | 0.98277 | 16.225 |
| Mmaa | 1591.59 | 0.283787 | 0.07881 | 3.600747 | 0.00032 | 0.00587 | 31.34 |
| Mmab | 1594.07 | 0.090134 | 0.08258 | 1.091528 | 0.27504 | 0.48452 | 13.027 |
| Mmachc | 717.099 | -0.09182 | 0.12561 | -0.73095 | 0.46481 | 0.66447 | 6.128 |
| Mmadhc | 1455.04 | 0.024684 | 0.07113 | 0.34703 | 0.72857 | 0.85548 | 16.921 |
| Mmd | 7688.37 | 0.133155 | 0.07667 | 1.736751 | 0.08243 | 0.22898 | 29.134 |
| Mmd2 | 6911.67 | -0.20078 | 0.10569 | -1.89972 | 0.05747 | 0.18196 | 46.443 |
| Mme | 2732.65 | 0.527292 | 0.21714 | 2.428356 | 0.01517 | 0.07502 | 144.494 |
| Mmel1 | 53.4577 | 0.621255 | 0.26716 | 2.325384 | 0.02005 | 0.08969 | 25.944 |
| Mmgt1 | 2177.89 | -0.0939 | 0.06761 | -1.3889 | 0.16486 | 0.35389 | 12.558 |
| Mmgt2 | 454.067 | -0.35831 | 0.13474 | -2.65927 | 0.00783 | 0.04899 | 17.691 |
| Mmp11 | 79.2782 | 0.007119 | 0.23704 | 0.030032 | 0.97604 | 0.98762 | 13.275 |
| Mmp14 | 502.553 | -0.27136 | 0.10708 | -2.5341 | 0.01127 | 0.06201 | 13.753 |
| Mmp15 | 826.023 | -0.10242 | 0.13385 | -0.76517 | 0.44417 | 0.64819 | 22.813 |

| | | | | | | | |
|---|---|---|---|---|---|---|---|
| Mmp16 | 947.161 | 0.068403 | 0.14906 | 0.4589 | 0.64631 | 0.80301 | 266.253 |
| Mmp17 | 8469.84 | -0.22183 | 0.15617 | -1.42042 | 0.15548 | 0.34204 | 26.931 |
| Mmp19 | 48.6125 | -0.31075 | 0.26105 | -1.19036 | 0.23391 | 0.43899 | 9.915 |
| Mmp2 | 50.0983 | -0.56202 | 0.33091 | -1.69842 | 0.08943 | 0.24227 | 26.13 |
| Mmp21 | 17.8933 | -0.046 | 0.48287 | -0.09526 | 0.92411 | 0.96191 | 5.792 |
| Mmp23 | 105.982 | 0.16778 | 0.22422 | 0.748278 | 0.45429 | 0.65643 | 2.73 |
| Mmp24 | 1631 | -0.19943 | 0.08982 | -2.22043 | 0.02639 | 0.10797 | 43.025 |
| Mmp25 | 14.6959 | -0.26291 | 0.48529 | -0.54176 | 0.58798 | 0.76146 | 16.967 |
| Mmp28 | 108.468 | -0.10896 | 0.18737 | -0.58156 | 0.56087 | 0.74205 | 22.304 |
| Mmp9 | 48.5968 | -0.88101 | 0.38784 | -2.27156 | 0.02311 | 0.09843 | 15.071 |
| Mmrn2 | 161.676 | -0.32279 | 0.15632 | -2.06499 | 0.03892 | 0.14059 | 28.823 |
| Mms19 | 2196.47 | 0.211008 | 0.06804 | 3.101187 | 0.00193 | 0.01965 | 40.072 |
| Mms22l | 78.7901 | -0.2951 | 0.20711 | -1.42482 | 0.15421 | 0.34008 | 106.5 |
| Mmut | 2149.03 | 0.063219 | 0.08379 | 0.754526 | 0.45053 | 0.65319 | 27.304 |
| Mn1 | 2416.65 | -0.00426 | 0.06686 | -0.06369 | 0.94922 | 0.9749 | 39.672 |
| Mnat1 | 598.415 | -0.11999 | 0.09315 | -1.28812 | 0.1977 | 0.39553 | 150.272 |
| Mndal | 46.2694 | -0.32545 | 0.33807 | -0.96266 | 0.33572 | 0.54643 | 31.756 |
| Mns1 | 174.571 | 0.05429 | 0.15835 | 0.342848 | 0.73171 | 0.85759 | 24.015 |
| Mnt | 1518.63 | -0.16433 | 0.07323 | -2.244 | 0.02483 | 0.10351 | 14.806 |
| Mob1a | 1636.25 | 0.057909 | 0.08248 | 0.702059 | 0.48264 | 0.67906 | 17.761 |
| Mob1b | 2421.06 | 0.211974 | 0.11368 | 1.864648 | 0.06223 | 0.19119 | 43.366 |
| Mob2 | 1284.5 | -0.03499 | 0.06887 | -0.5081 | 0.61138 | 0.77842 | 52.531 |
| Mob3a | 364.093 | -0.33937 | 0.1406 | -2.41363 | 0.01579 | 0.0771 | 16.725 |
| Mob3b | 161.911 | -0.61149 | 0.15646 | -3.90838 | 9.29E-05 | 0.00255 | 208.411 |
| Mob3c | 323.02 | -0.10848 | 0.12014 | -0.90299 | 0.36653 | 0.57697 | 8.094 |
| Mob4 | 2869.73 | 0.022765 | 0.08103 | 0.280948 | 0.77875 | 0.88464 | 23.669 |
| Mobp | 38937 | -0.73187 | 0.12157 | -6.02023 | 1.74E-09 | 8.26E-07 | 31.778 |
| Mocos | 160.668 | -0.18177 | 0.17251 | -1.05367 | 0.29203 | 0.50186 | 47.885 |
| Mocs1 | 512.94 | -0.17938 | 0.1074 | -1.67013 | 0.09489 | 0.25147 | 27.074 |
| Mocs2 | 2036.59 | -0.07913 | 0.07027 | -1.12604 | 0.26015 | 0.46874 | 14.04 |
| Mocs3 | 309.556 | 0.095167 | 0.13047 | 0.729426 | 0.46574 | 0.665 | 1.973 |
| Mog | 1843.13 | -0.67806 | 0.14313 | -4.73739 | 2.16E-06 | 0.00019 | 12.656 |
| Mogs | 920.648 | -0.03872 | 0.09544 | -0.40575 | 0.68493 | 0.82838 | 3.403 |
| Mok | 787.146 | 0.459101 | 0.1026 | 4.474861 | 7.65E-06 | 0.00045 | 33.142 |

| | | | | | | | |
|---|---|---|---|---|---|---|---|
| Mon1a | 999.416 | 0.043212 | 0.09103 | 0.474701 | 0.635 | 0.79438 | 14.989 |
| Mon1b | 1828.22 | -0.20729 | 0.0809 | -2.56236 | 0.0104 | 0.05893 | 9.593 |
| Mon2 | 5694.07 | 0.086887 | 0.06477 | 1.341468 | 0.17977 | 0.37289 | 84.446 |
| Morc2a | 3119.6 | 0.068275 | 0.076 | 0.898319 | 0.36902 | 0.57964 | 41.287 |
| Morc2b | 314.158 | 0.249085 | 0.16658 | 1.495241 | 0.13485 | 0.31476 | 26.333 |
| Morc3 | 1499.16 | 0.224229 | 0.06447 | 3.477894 | 0.00051 | 0.00798 | 43.953 |
| Morc4 | 330.216 | 0.127355 | 0.13063 | 0.974947 | 0.32959 | 0.5409 | 50.046 |
| Morf4l1 | 14062.1 | 0.001169 | 0.06057 | 0.019297 | 0.9846 | 0.99161 | 23.11 |
| Morf4l1-ps1 | 1000.55 | 0.103489 | 0.09555 | 1.083056 | 0.27878 | 0.48881 | 0.969 |
| Morf4l2 | 10098.4 | 0.058559 | 0.054 | 1.084526 | 0.27813 | 0.48804 | 10.749 |
| Morn1 | 379.618 | 0.181987 | 0.11584 | 1.571087 | 0.11616 | 0.28615 | 58.929 |
| Morn2 | 236.551 | 0.508491 | 0.11862 | 4.28686 | 1.81E-05 | 0.00083 | 7.274 |
| Morn4 | 2292.09 | -0.05814 | 0.08257 | -0.70411 | 0.48136 | 0.67838 | 11.419 |
| Morrbid | 19.5479 | -0.51866 | 0.46205 | -1.12252 | 0.26164 | 0.4704 | 324.447 |
| Mosmo | 3788.36 | 0.103787 | 0.06599 | 1.572706 | 0.11579 | 0.28563 | 57.237 |
| Mospd1 | 590.883 | -0.11266 | 0.10637 | -1.05905 | 0.28958 | 0.49978 | 25.905 |
| Mospd2 | 1092.08 | 0.263866 | 0.10644 | 2.478997 | 0.01318 | 0.06872 | 44.207 |
| Mospd3 | 524.108 | -0.03401 | 0.10475 | -0.32467 | 0.74543 | 0.86615 | 4.414 |
| Mov10 | 1107.94 | 0.355524 | 0.11582 | 3.069652 | 0.00214 | 0.02109 | 23.728 |
| Moxd1 | 371.389 | -0.23539 | 0.15151 | -1.55363 | 0.12027 | 0.29298 | 79.274 |
| Mpc1 | 838.037 | 0.515921 | 0.13314 | 3.875103 | 0.00011 | 0.00281 | 14.758 |
| Mpc1-ps | 1090.03 | -0.04442 | 0.12138 | -0.36596 | 0.71439 | 0.84674 | 0.33 |
| Mpc2 | 2364.52 | -0.15549 | 0.06954 | -2.23591 | 0.02536 | 0.10497 | 20.578 |
| Mpdu1 | 1635.41 | -0.19193 | 0.0872 | -2.20094 | 0.02774 | 0.11188 | 5.946 |
| Mpdz | 3321.03 | -0.42538 | 0.18736 | -2.27043 | 0.02318 | 0.09867 | 164.316 |
| Mpeg1 | 1601.85 | -0.50982 | 0.1988 | -2.56449 | 0.01033 | 0.0587 | 4.505 |
| Mpg | 370.086 | -0.01005 | 0.11035 | -0.09108 | 0.92743 | 0.96344 | 6.196 |
| Mphosph10 | 674.934 | 0.25495 | 0.12152 | 2.098002 | 0.0359 | 0.13334 | 15.742 |
| Mphosph6 | 721.3 | -0.06077 | 0.10321 | -0.58883 | 0.55598 | 0.73825 | 10.299 |
| Mphosph8 | 2521.3 | 0.203629 | 0.14601 | 1.394628 | 0.16313 | 0.35169 | 29.183 |
| Mphosph9 | 2379.52 | 0.098197 | 0.08555 | 1.147772 | 0.25106 | 0.45847 | 77.014 |
| Mpi | 2244.91 | 0.091072 | 0.09579 | 0.950769 | 0.34172 | 0.55261 | 8.508 |
| Mplkip | 356.411 | -0.05704 | 0.13395 | -0.42583 | 0.67023 | 0.81828 | 4.557 |
| Mpnd | 2729.91 | 0.004663 | 0.06441 | 0.072391 | 0.94229 | 0.97116 | 11.08 |

| | | | | | | | |
|---|---|---|---|---|---|---|---|
| Mpp1 | 1609.82 | 0.115413 | 0.10188 | 1.132788 | 0.2573 | 0.46522 | 21.284 |
| Mpp2 | 6346.73 | -0.11969 | 0.10679 | -1.12079 | 0.26238 | 0.47117 | 31.501 |
| Mpp3 | 2703.27 | -0.07965 | 0.09211 | -0.86466 | 0.38722 | 0.59657 | 28.81 |
| Mpp4 | 46.7118 | 0.007847 | 0.2753 | 0.028505 | 0.97726 | 0.98797 | 42.455 |
| Mpp7 | 876.951 | 0.113358 | 0.1915 | 0.591936 | 0.55389 | 0.7366 | 278.908 |
| Mppe1 | 901.528 | 0.116452 | 0.10595 | 1.099123 | 0.27171 | 0.48106 | 20.788 |
| Mpped1 | 5822.09 | -0.77897 | 0.41349 | -1.8839 | 0.05958 | 0.18581 | 79.028 |
| Mpped2 | 2029.46 | 0.48461 | 0.17476 | 2.773046 | 0.00555 | 0.03901 | 175.088 |
| Mprip | 21699.8 | 0.251981 | 0.07695 | 3.27443 | 0.00106 | 0.0133 | 119.556 |
| Mpst | 590.46 | -0.35743 | 0.10258 | -3.48442 | 0.00049 | 0.00785 | 7.598 |
| Mpv17 | 787.745 | -0.36653 | 0.10207 | -3.59089 | 0.00033 | 0.00602 | 13.598 |
| Mpv17l | 3592.75 | 0.129621 | 0.0664 | 1.952221 | 0.05091 | 0.1678 | 46.459 |
| Mpv17l2 | 476.673 | -0.29867 | 0.12794 | -2.3344 | 0.01957 | 0.08827 | 2.298 |
| Mpzl1 | 953.337 | -0.09871 | 0.09115 | -1.08292 | 0.27884 | 0.48881 | 42.299 |
| Mpzl2 | 38.5635 | -1.1747 | 0.53534 | -2.19433 | 0.02821 | 0.11309 | 14.217 |
| Mpzl3 | 294.929 | 0.225374 | 0.16341 | 1.379198 | 0.16783 | 0.35774 | 22.251 |
| Mr1 | 230.609 | -0.14712 | 0.13739 | -1.07087 | 0.28423 | 0.49434 | 19.538 |
| Mrap2 | 478.652 | 0.279153 | 0.14182 | 1.96839 | 0.04902 | 0.16374 | 39.74 |
| Mras | 8474.07 | 0.00642 | 0.07175 | 0.089466 | 0.92871 | 0.96414 | 51.962 |
| Mrc1 | 295.813 | -0.89623 | 0.2501 | -3.58343 | 0.00034 | 0.00613 | 102.666 |
| Mrc2 | 469.322 | -0.75325 | 0.22034 | -3.41866 | 0.00063 | 0.00926 | 58.497 |
| Mre11a | 958.791 | 0.447943 | 0.13032 | 3.437242 | 0.00059 | 0.00882 | 52.47 |
| Mreg | 135.849 | -0.50123 | 0.21406 | -2.34156 | 0.0192 | 0.08705 | 52.866 |
| Mrfap1 | 11535.2 | -0.07975 | 0.07589 | -1.05093 | 0.29329 | 0.50323 | 1.906 |
| Mrgbp | 982.689 | 0.199621 | 0.07809 | 2.556403 | 0.01058 | 0.05969 | 5.001 |
| Mrgpre | 327.323 | 0.071974 | 0.15131 | 0.47567 | 0.63431 | 0.79416 | 6.138 |
| Mrgprf | 36.8835 | -1.03365 | 0.3565 | -2.89944 | 0.00374 | 0.03007 | 8.73 |
| Mri1 | 318.978 | 0.033701 | 0.12948 | 0.26028 | 0.79465 | 0.89379 | 7.421 |
| Mrip-ps | 20.5772 | 0.147725 | 0.38154 | 0.387187 | 0.69862 | 0.83677 | 0.94 |
| Mrm1 | 303.943 | -0.38507 | 0.12296 | -3.1316 | 0.00174 | 0.0184 | 6.455 |
| Mrm2 | 366.732 | 0.037925 | 0.10562 | 0.359076 | 0.71954 | 0.85017 | 4.225 |
| Mrm3 | 393.241 | 0.075415 | 0.12158 | 0.620279 | 0.53507 | 0.72203 | 6.905 |
| Mrnip | 156.202 | -0.24751 | 0.19459 | -1.27195 | 0.20339 | 0.40181 | 25.672 |
| Mro | 1149.78 | -0.14735 | 0.07801 | -1.88884 | 0.05891 | 0.18475 | 21.949 |

| | | | | | | | |
|---|---|---|---|---|---|---|---|
| Mroh1 | 3301.32 | -0.35835 | 0.13171 | -2.72079 | 0.00651 | 0.04345 | 72.778 |
| Mroh2a | 158.699 | -0.22241 | 0.15191 | -1.46413 | 0.14316 | 0.32518 | 35.304 |
| Mroh3 | 24.1981 | 0.124304 | 0.40788 | 0.30476 | 0.76055 | 0.87444 | 31.177 |
| Mroh5 | 80.9129 | -0.32479 | 0.27134 | -1.19697 | 0.23132 | 0.43586 | 79.633 |
| Mroh6 | 13.9193 | 0.485163 | 0.51815 | 0.936337 | 0.3491 | 0.55954 | 4.634 |
| Mroh7 | 483.35 | 0.547496 | 0.20571 | 2.661489 | 0.00778 | 0.04879 | 50.509 |
| Mroh8 | 62.8317 | 0.036662 | 0.29591 | 0.123894 | 0.9014 | 0.95046 | 71 |
| Mrpl1 | 705.097 | 0.094512 | 0.11814 | 0.799994 | 0.42371 | 0.63048 | 57.235 |
| Mrpl10 | 1180.47 | 0.15507 | 0.08391 | 1.848073 | 0.06459 | 0.19561 | 7.654 |
| Mrpl11 | 1250.27 | 0.000855 | 0.11128 | 0.007687 | 0.99387 | 0.99634 | 4.853 |
| Mrpl12 | 1479.84 | -0.15226 | 0.08812 | -1.72783 | 0.08402 | 0.23207 | 4.453 |
| Mrpl13 | 935.69 | -0.28506 | 0.07957 | -3.58264 | 0.00034 | 0.00614 | 23.655 |
| Mrpl14 | 270.377 | -0.05592 | 0.13051 | -0.42848 | 0.6683 | 0.81705 | 12.174 |
| Mrpl15 | 1077.28 | -0.08319 | 0.09263 | -0.89813 | 0.36912 | 0.57967 | 12.534 |
| Mrpl16 | 876.578 | -0.09496 | 0.10389 | -0.91405 | 0.36069 | 0.57101 | 4.57 |
| Mrpl17 | 1127.97 | 0.035326 | 0.0832 | 0.424619 | 0.67111 | 0.81882 | 7.894 |
| Mrpl18 | 665.684 | -0.10155 | 0.10111 | -1.00434 | 0.31522 | 0.52568 | 4.997 |
| Mrpl19 | 1175.81 | 0.033913 | 0.06959 | 0.487301 | 0.62604 | 0.78842 | 8.108 |
| Mrpl2 | 938.599 | -0.06903 | 0.0797 | -0.86615 | 0.38641 | 0.59577 | 3.911 |
| Mrpl20 | 1356.23 | -0.15824 | 0.07974 | -1.98441 | 0.04721 | 0.15951 | 7.098 |
| Mrpl21 | 722.013 | -0.10805 | 0.1111 | -0.97256 | 0.33077 | 0.54197 | 9.937 |
| Mrpl22 | 459.028 | -0.04859 | 0.11684 | -0.41586 | 0.67751 | 0.82322 | 7.905 |
| Mrpl23 | 849.57 | 0.077784 | 0.10627 | 0.731982 | 0.46418 | 0.6639 | 8.062 |
| Mrpl23-ps1 | 18.4762 | -1.15489 | 0.43447 | -2.65813 | 0.00786 | 0.04912 | 0.58 |
| Mrpl24 | 1422.06 | 0.02783 | 0.07166 | 0.388336 | 0.69777 | 0.83624 | 4.167 |
| Mrpl27 | 671.841 | -0.05945 | 0.08951 | -0.66414 | 0.5066 | 0.69789 | 6.323 |
| Mrpl27-ps | 28.9211 | 0.037588 | 0.40716 | 0.092316 | 0.92645 | 0.96301 | 0.447 |
| Mrpl28 | 1083.69 | -0.15128 | 0.10095 | -1.49865 | 0.13396 | 0.31366 | 3.114 |
| Mrpl3 | 3514.72 | 0.14981 | 0.06862 | 2.183089 | 0.02903 | 0.11537 | 26.65 |
| Mrpl30 | 1403.75 | -0.05967 | 0.07469 | -0.79889 | 0.42436 | 0.63107 | 8.059 |
| Mrpl32 | 798.337 | 0.027612 | 0.08082 | 0.341661 | 0.73261 | 0.85819 | 4.982 |
| Mrpl33 | 371.929 | 0.10242 | 0.13367 | 0.766222 | 0.44354 | 0.64792 | 67.45 |
| Mrpl34 | 475.081 | -0.14019 | 0.10852 | -1.29179 | 0.19643 | 0.39402 | 0.789 |
| Mrpl35 | 1344.45 | 0.048617 | 0.07373 | 0.659416 | 0.50963 | 0.70111 | 13.973 |

| | | | | | | | |
|---|---|---|---|---|---|---|---|
| Mrpl36 | 627.949 | -0.22723 | 0.09536 | -2.38274 | 0.01718 | 0.08153 | 3.666 |
| Mrpl37 | 2168.1 | 0.084943 | 0.06866 | 1.237168 | 0.21602 | 0.41777 | 10.995 |
| Mrpl38 | 1511.27 | -0.00973 | 0.08345 | -0.11655 | 0.90721 | 0.95354 | 7.052 |
| Mrpl39 | 980.387 | 0.003593 | 0.12016 | 0.029898 | 0.97615 | 0.98767 | 18.167 |
| Mrpl4 | 1486.22 | -0.05069 | 0.07397 | -0.68526 | 0.49318 | 0.68772 | 6.102 |
| Mrpl40 | 746.752 | -0.1256 | 0.10031 | -1.25215 | 0.21051 | 0.41056 | 4.845 |
| Mrpl41 | 1525.83 | -0.04148 | 0.09914 | -0.41844 | 0.67563 | 0.82192 | 0.982 |
| Mrpl42 | 1242.74 | 0.034836 | 0.08455 | 0.412015 | 0.68033 | 0.82502 | 21.136 |
| Mrpl43 | 238.604 | -0.25267 | 0.15601 | -1.61955 | 0.10533 | 0.26862 | 1.614 |
| Mrpl44 | 759.132 | 0.069184 | 0.09741 | 0.710226 | 0.47756 | 0.67546 | 5.428 |
| Mrpl45 | 1232.05 | -0.01145 | 0.08092 | -0.14146 | 0.88751 | 0.94401 | 14.205 |
| Mrpl46 | 772.948 | -0.05972 | 0.08502 | -0.70248 | 0.48238 | 0.67897 | 8.296 |
| Mrpl47 | 644.015 | 0.180195 | 0.1052 | 1.712822 | 0.08675 | 0.23709 | 12.209 |
| Mrpl48 | 2374.69 | -0.13039 | 0.07728 | -1.68727 | 0.09155 | 0.24597 | 62.551 |
| Mrpl48-ps | 23.2606 | 0.075808 | 0.435 | 0.17427 | 0.86165 | 0.9299 | 0.636 |
| Mrpl49 | 1497.09 | -0.03015 | 0.08049 | -0.37453 | 0.70801 | 0.84235 | 4.164 |
| Mrpl50 | 1640.65 | -0.03498 | 0.07717 | -0.45323 | 0.65038 | 0.80552 | 8.498 |
| Mrpl51 | 1541.69 | -0.04299 | 0.07527 | -0.57116 | 0.56789 | 0.74779 | 4.469 |
| Mrpl52 | 516.478 | 0.046296 | 0.1213 | 0.381665 | 0.70271 | 0.8396 | 2.848 |
| Mrpl54 | 399.111 | -0.16309 | 0.11319 | -1.44083 | 0.14963 | 0.33397 | 2.222 |
| Mrpl55 | 815.056 | -0.00949 | 0.08255 | -0.11501 | 0.90844 | 0.95408 | 3.649 |
| Mrpl57 | 783.668 | 0.098328 | 0.09221 | 1.066353 | 0.28626 | 0.49628 | 3.063 |
| Mrpl58 | 842.425 | -0.1729 | 0.08241 | -2.09802 | 0.0359 | 0.13334 | 8.923 |
| Mrpl9 | 1263.19 | -0.05561 | 0.09286 | -0.59888 | 0.54926 | 0.73299 | 7.813 |
| Mrps10 | 1140.85 | 0.098833 | 0.08272 | 1.19473 | 0.23219 | 0.43715 | 12.531 |
| Mrps11 | 597.433 | 0.023667 | 0.08734 | 0.270972 | 0.78641 | 0.88899 | 9.871 |
| Mrps12 | 897.771 | -0.25617 | 0.09591 | -2.67105 | 0.00756 | 0.04797 | 2.18 |
| Mrps14 | 1310.99 | 0.015283 | 0.07405 | 0.206392 | 0.83649 | 0.91608 | 6.956 |
| Mrps15 | 1308.74 | -0.00086 | 0.0841 | -0.01021 | 0.99186 | 0.99537 | 8.608 |
| Mrps16 | 661.446 | -0.04186 | 0.09921 | -0.42197 | 0.67305 | 0.82024 | 4.331 |
| Mrps17 | 3127.24 | 0.108247 | 0.07014 | 1.54326 | 0.12277 | 0.29668 | 7.06 |
| Mrps18a | 1223.87 | -0.04598 | 0.07596 | -0.60531 | 0.54497 | 0.73012 | 17.925 |
| Mrps18b | 302.504 | 0.18951 | 0.13584 | 1.395084 | 0.16299 | 0.35154 | 6.011 |
| Mrps18c | 128.442 | 0.10073 | 0.22741 | 0.442943 | 0.65781 | 0.81018 | 5.845 |

| | | | | | | | |
|---|---|---|---|---|---|---|---|
| Mrps2 | 1089.13 | 0.025873 | 0.08374 | 0.308959 | 0.75735 | 0.87222 | 3.113 |
| Mrps21 | 994.925 | -0.09994 | 0.10638 | -0.93946 | 0.34749 | 0.55799 | 8.886 |
| Mrps22 | 748.48 | -0.02992 | 0.09193 | -0.32542 | 0.74486 | 0.86583 | 12.931 |
| Mrps23 | 1432.26 | -0.23311 | 0.08486 | -2.74696 | 0.00602 | 0.04135 | 7.12 |
| Mrps24 | 661.743 | -0.15623 | 0.08729 | -1.78993 | 0.07347 | 0.21344 | 11.698 |
| Mrps25 | 1123.87 | -0.01314 | 0.10007 | -0.1313 | 0.89554 | 0.94746 | 14.509 |
| Mrps26 | 1383.48 | -0.08605 | 0.07384 | -1.1653 | 0.2439 | 0.45033 | 4.954 |
| Mrps27 | 642.015 | -0.02348 | 0.10297 | -0.22803 | 0.81962 | 0.90729 | 70.777 |
| Mrps28 | 294.542 | -0.08439 | 0.15739 | -0.5362 | 0.59182 | 0.76439 | 121.773 |
| Mrps30 | 621.448 | 0.174839 | 0.08812 | 1.984189 | 0.04723 | 0.15957 | 8.872 |
| Mrps31 | 845.323 | 0.014341 | 0.09417 | 0.152278 | 0.87897 | 0.93943 | 18.305 |
| Mrps33 | 1449.56 | 0.072239 | 0.1004 | 0.719521 | 0.47182 | 0.67032 | 9.185 |
| Mrps34 | 378.8 | -0.19601 | 0.126 | -1.55566 | 0.11979 | 0.29232 | 2.387 |
| Mrps35 | 880.096 | -0.09512 | 0.08285 | -1.14817 | 0.2509 | 0.45825 | 31.228 |
| Mrps36 | 774.407 | -0.14688 | 0.08402 | -1.74809 | 0.08045 | 0.22572 | 8.705 |
| Mrps5 | 2488.41 | -0.04321 | 0.06352 | -0.6803 | 0.49632 | 0.68996 | 19.608 |
| Mrps6 | 357.663 | 0.16547 | 0.12263 | 1.349291 | 0.17724 | 0.36947 | 53.958 |
| Mrps7 | 1379.75 | -0.08549 | 0.07192 | -1.18864 | 0.23458 | 0.43974 | 4.112 |
| Mrps9 | 1619.26 | 0.083966 | 0.06393 | 1.313364 | 0.18906 | 0.38492 | 54.451 |
| Mrrf | 652.377 | 0.086751 | 0.10335 | 0.839416 | 0.40124 | 0.6099 | 54.259 |
| Mrs2 | 3870.48 | 0.075456 | 0.05479 | 1.377232 | 0.16844 | 0.3584 | 32.897 |
| Mrtfa | 4977.49 | 0.46186 | 0.13102 | 3.525157 | 0.00042 | 0.00716 | 178.477 |
| Mrtfb | 15782.4 | 0.209864 | 0.0647 | 3.243444 | 0.00118 | 0.01415 | 161.049 |
| Mrto4 | 780.675 | -0.0351 | 0.09414 | -0.37283 | 0.70928 | 0.84343 | 5.142 |
| Mrto4-ps1 | 36.8203 | -0.33842 | 0.31036 | -1.09043 | 0.27552 | 0.4847 | 0.721 |
| Ms4a15 | 18.0748 | -2 | 0.84482 | -5.3386 | 9.37E-08 | 1.66E-05 | 14.944 |
| Ms4a4a | 37.1311 | -1.3602 | 0.32802 | -4.1467 | 3.37E-05 | 0.00124 | 17.268 |
| Ms4a6b | 108.881 | -0.76175 | 0.27322 | -2.78806 | 0.0053 | 0.03768 | 14.745 |
| Ms4a6c | 41.221 | -0.9485 | 0.4628 | -2.04949 | 0.04041 | 0.14368 | 12.827 |
| Ms4a6d | 77.7137 | -1.0299 | 0.38183 | -2.69731 | 0.00699 | 0.04538 | 18.246 |
| Ms4a7 | 21.5727 | -0.66351 | 0.42909 | -1.54632 | 0.12203 | 0.29545 | 15.108 |
| Msantd1 | 235.866 | 0.459215 | 0.1482 | 3.098515 | 0.00194 | 0.01973 | 15.828 |
| Msantd2 | 933.759 | 0.115676 | 0.13728 | 0.842647 | 0.39943 | 0.60828 | 35.605 |
| Msantd3 | 511.865 | -0.21787 | 0.12029 | -1.81119 | 0.07011 | 0.20671 | 21.985 |

| | | | | | | | |
|---|---|---|---|---|---|---|---|
| Msantd4 | 3327.8 | -0.03615 | 0.07801 | -0.46333 | 0.64312 | 0.80089 | 10.309 |
| Msc | 31.609 | -0.50327 | 0.33828 | -1.48774 | 0.13682 | 0.31754 | 2.647 |
| Msh2 | 1408.11 | 0.067258 | 0.07549 | 0.89095 | 0.37296 | 0.58339 | 51.384 |
| Msh3 | 1163.02 | 0.232967 | 0.10539 | 2.210468 | 0.02707 | 0.10988 | 143.132 |
| Msh5 | 32.0684 | -0.32067 | 0.32347 | -0.99135 | 0.32151 | 0.53233 | 18.141 |
| Msh6 | 1181.59 | 0.197639 | 0.07814 | 2.529215 | 0.01143 | 0.0626 | 15.831 |
| Msi1 | 1402.59 | -0.2124 | 0.09573 | -2.21884 | 0.0265 | 0.10829 | 26.1 |
| Msi2 | 9328.32 | -0.07533 | 0.06946 | -1.08443 | 0.27817 | 0.48806 | 379.132 |
| Msl1 | 8339.59 | -0.06932 | 0.06306 | -1.09939 | 0.2716 | 0.48096 | 12.344 |
| Msl2 | 3149.73 | -0.00648 | 0.08428 | -0.07687 | 0.93872 | 0.9695 | 25.846 |
| Msl3 | 987.14 | 0.009528 | 0.08712 | 0.109361 | 0.91292 | 0.95597 | 19.782 |
| Msl3l2 | 550.085 | 0.109671 | 0.08831 | 1.241952 | 0.21425 | 0.41558 | 9.964 |
| Msln | 21.1262 | -1.75874 | 0.42604 | -4.12809 | 3.66E-05 | 0.00131 | 5.766 |
| Msmo1 | 3116.99 | -0.34374 | 0.09103 | -3.77604 | 0.00016 | 0.00369 | 15.654 |
| Msn | 1347.92 | -0.51799 | 0.13046 | -3.97052 | 7.17E-05 | 0.00213 | 72.511 |
| Msra | 880.121 | 0.192716 | 0.08147 | 2.365364 | 0.01801 | 0.08366 | 333.279 |
| Msrb1 | 615.472 | -0.28481 | 0.09075 | -3.13829 | 0.0017 | 0.01812 | 6.137 |
| Msrb2 | 727.2 | -0.01012 | 0.08843 | -0.11446 | 0.90887 | 0.95438 | 23.537 |
| Msrb3 | 372.471 | -0.33515 | 0.13506 | -2.48151 | 0.01308 | 0.06847 | 118.006 |
| Mss51 | 55.1105 | 0.359029 | 0.29437 | 1.219639 | 0.2226 | 0.42586 | 14.035 |
| Mst1 | 41.3305 | -0.65792 | 0.36925 | -1.78177 | 0.07479 | 0.21551 | 4.568 |
| Mst1r | 80.0443 | -0.32885 | 0.23607 | -1.39301 | 0.16362 | 0.35238 | 13.511 |
| Mstn | 15.7052 | -0.49077 | 0.66325 | -0.73994 | 0.45934 | 0.65991 | 6.44 |
| Msto1 | 567.977 | 0.000388 | 0.09409 | 0.004128 | 0.99671 | 0.99821 | 8.893 |
| Msx1 | 67.1957 | -0.66227 | 0.3197 | -2.07153 | 0.03831 | 0.13922 | 4.099 |
| Msx2 | 19.1235 | -0.59548 | 0.412 | -1.44533 | 0.14837 | 0.33201 | 6.191 |
| Msx3 | 27.8063 | 0.610351 | 0.42491 | 1.436428 | 0.15088 | 0.33576 | 2.933 |
| Mt1 | 6734.79 | -0.4044 | 0.1106 | -3.65628 | 0.00026 | 0.00503 | 1.246 |
| Mt3 | 6379.8 | -0.42466 | 0.13176 | -3.22285 | 0.00127 | 0.01478 | 1.54 |
| Mta1 | 2721.35 | -0.03465 | 0.06301 | -0.54991 | 0.58238 | 0.75787 | 38.929 |
| Mta2 | 2636.49 | 0.022104 | 0.06604 | 0.334691 | 0.73786 | 0.86163 | 10.429 |
| Mta3 | 2782.45 | -0.29238 | 0.109 | -2.68245 | 0.00731 | 0.04692 | 115.354 |
| Mtag2 | 18.2148 | -1.05575 | 0.45197 | -2.33589 | 0.0195 | 0.08804 | 1.786 |
| Mtap | 437.894 | -0.15386 | 0.09802 | -1.5697 | 0.11649 | 0.2867 | 43.96 |

| | | | | | | | |
|---|---|---|---|---|---|---|---|
| Mtarc2 | 1556.66 | -0.2778 | 0.10162 | -2.73373 | 0.00626 | 0.04235 | 33.384 |
| Mtbp | 143.047 | 0.181226 | 0.17587 | 1.030443 | 0.3028 | 0.51232 | 69.016 |
| Mtch1 | 14699.4 | 0.034489 | 0.06909 | 0.499197 | 0.61764 | 0.78255 | 15.863 |
| Mtch2 | 3389.24 | -0.15591 | 0.05344 | -2.91774 | 0.00353 | 0.02888 | 19.656 |
| Mtcl1 | 6042.12 | 0.123183 | 0.08818 | 1.396894 | 0.16245 | 0.35091 | 112.769 |
| Mtcp1 | 595.652 | 0.25333 | 0.12325 | 2.05543 | 0.03984 | 0.14259 | 6.145 |
| Mtdh | 5602.04 | 0.106259 | 0.07615 | 1.395476 | 0.16287 | 0.35138 | 62.931 |
| Mterf1a | 207.725 | 0.366316 | 0.14468 | 2.531836 | 0.01135 | 0.06226 | 3.353 |
| Mterf1b | 135.217 | 0.042828 | 0.20591 | 0.207991 | 0.83524 | 0.91549 | 5.285 |
| Mterf2 | 491.581 | 0.287575 | 0.15326 | 1.876342 | 0.06061 | 0.18787 | 8.595 |
| Mterf3 | 1271.15 | 0.279078 | 0.0813 | 3.432664 | 0.0006 | 0.00894 | 26.121 |
| Mterf4 | 553.309 | -0.02538 | 0.10387 | -0.24438 | 0.80693 | 0.90065 | 6.705 |
| Mtf1 | 1838.08 | 0.089599 | 0.07688 | 1.165515 | 0.24381 | 0.45025 | 47.697 |
| Mtf2 | 3076.79 | 0.316334 | 0.09663 | 3.273726 | 0.00106 | 0.01332 | 43.331 |
| Mtfmt | 751.852 | 0.209785 | 0.09827 | 2.134872 | 0.03277 | 0.12566 | 17.273 |
| Mtfp1 | 1876.31 | -0.73616 | 0.20561 | -3.58033 | 0.00034 | 0.00616 | 3.966 |
| Mtfr1 | 351.066 | -0.3397 | 0.11812 | -2.8758 | 0.00403 | 0.03166 | 33.491 |
| Mtfr1l | 2697.05 | 0.008345 | 0.07166 | 0.116439 | 0.9073 | 0.95354 | 9.838 |
| Mtfr2 | 63.8473 | 0.303843 | 0.23918 | 1.270342 | 0.20396 | 0.40263 | 13.535 |
| Mtg1 | 850.78 | 0.062556 | 0.08054 | 0.776727 | 0.43732 | 0.64276 | 13.223 |
| Mtg2 | 946.236 | 0.287856 | 0.0883 | 3.259992 | 0.00111 | 0.0137 | 15.315 |
| Mthfd1 | 2311.57 | 0.188091 | 0.07941 | 2.368598 | 0.01786 | 0.08321 | 64.506 |
| Mthfd1l | 649.822 | 0.020779 | 0.12101 | 0.171709 | 0.86367 | 0.93094 | 193.964 |
| Mthfd2 | 408.438 | -0.11653 | 0.12741 | -0.91454 | 0.36043 | 0.57074 | 20.218 |
| Mthfd2l | 502.637 | 0.28836 | 0.10053 | 2.868305 | 0.00413 | 0.03209 | 90.252 |
| Mthfr | 1634.27 | -0.07849 | 0.097 | -0.80914 | 0.41844 | 0.62566 | 20.475 |
| Mthfs | 77.7285 | 0.093789 | 0.26865 | 0.349109 | 0.72701 | 0.85431 | 167.038 |
| Mthfsd | 1936.82 | -0.02257 | 0.08505 | -0.26537 | 0.79073 | 0.8913 | 16.765 |
| Mthfsl | 257.72 | 0.135817 | 0.13734 | 0.98893 | 0.3227 | 0.53366 | 31.695 |
| Mtif2 | 1728.39 | 0.171354 | 0.10008 | 1.71212 | 0.08687 | 0.23726 | 18.872 |
| Mtif3 | 432.255 | -0.06002 | 0.0994 | -0.60388 | 0.54593 | 0.73067 | 12.228 |
| Mtln | 822.51 | 0.011065 | 0.10605 | 0.104336 | 0.9169 | 0.95763 | 1.101 |
| Mtm1 | 552.511 | 0.067924 | 0.11203 | 0.606324 | 0.5443 | 0.72945 | 104.925 |
| Mtmr1 | 2868.06 | 0.068598 | 0.06363 | 1.07803 | 0.28102 | 0.49082 | 54.437 |

| | | | | | | | |
|---|---|---|---|---|---|---|---|
| Mtmr10 | 974.084 | -0.17654 | 0.09364 | -1.88525 | 0.0594 | 0.18544 | 53.154 |
| Mtmr11 | 182.198 | -0.49206 | 0.18462 | -2.66522 | 0.00769 | 0.04845 | 9.714 |
| Mtmr12 | 1851.4 | -0.12265 | 0.09949 | -1.23275 | 0.21767 | 0.41969 | 69.469 |
| Mtmr14 | 770.37 | -0.01263 | 0.09285 | -0.13599 | 0.89183 | 0.94605 | 43.55 |
| Mtmr2 | 2864.2 | -0.0594 | 0.07789 | -0.76263 | 0.44568 | 0.64947 | 58.072 |
| Mtmr3 | 5018.02 | 0.109535 | 0.05883 | 1.861792 | 0.06263 | 0.19207 | 113.996 |
| Mtmr4 | 5883.34 | 0.114458 | 0.07245 | 1.579736 | 0.11417 | 0.28298 | 24.141 |
| Mtmr6 | 7510.51 | 0.119258 | 0.06293 | 1.895028 | 0.05809 | 0.18321 | 37.143 |
| Mtmr7 | 3416.92 | 0.211656 | 0.07445 | 2.842927 | 0.00447 | 0.03384 | 83.705 |
| Mtmr9 | 4097.68 | 0.154952 | 0.0837 | 1.851376 | 0.06412 | 0.19495 | 24.332 |
| Mto1 | 893.79 | 0.00945 | 0.07598 | 0.124366 | 0.90103 | 0.9503 | 27.141 |
| Mtor | 7282.62 | 0.065256 | 0.06178 | 1.056204 | 0.29088 | 0.50107 | 109.073 |
| Mtpap | 893.594 | 0.08836 | 0.08412 | 1.050405 | 0.29353 | 0.50348 | 21.739 |
| Mtpn | 16683.3 | 0.1399 | 0.0637 | 2.196075 | 0.02809 | 0.11283 | 30.983 |
| Mtr | 2067.57 | 0.140389 | 0.10777 | 1.302724 | 0.19267 | 0.38947 | 75.402 |
| Mtres1 | 789.123 | -0.10358 | 0.11538 | -0.89768 | 0.36936 | 0.57978 | 15.862 |
| Mtrex | 3633.42 | 0.195984 | 0.07948 | 2.46583 | 0.01367 | 0.07038 | 59.981 |
| Mtrf1 | 310.45 | 0.257352 | 0.1225 | 2.100773 | 0.03566 | 0.13272 | 25.816 |
| Mtrf1l | 523.349 | 0.155384 | 0.09642 | 1.611458 | 0.10708 | 0.27158 | 12.024 |
| Mtrfr | 514.813 | -0.09431 | 0.12526 | -0.75287 | 0.45153 | 0.65392 | 13.756 |
| Mtrr | 692.492 | 0.05543 | 0.09947 | 0.557238 | 0.57736 | 0.75448 | 21.37 |
| Mtss1 | 1845.84 | 0.029731 | 0.08583 | 0.346402 | 0.72904 | 0.85579 | 140.772 |
| Mtss2 | 7320.52 | -0.22911 | 0.10298 | -2.2248 | 0.02609 | 0.10722 | 19.925 |
| Mttp | 960.078 | 0.62967 | 0.16765 | 3.755793 | 0.00017 | 0.0039 | 55.115 |
| Mturn | 4151.31 | -0.38271 | 0.12268 | -3.11959 | 0.00181 | 0.01885 | 22.228 |
| Mtus1 | 2868.55 | 0.215829 | 0.11398 | 1.893504 | 0.05829 | 0.18353 | 142.813 |
| Mtus2 | 2369.89 | 0.134951 | 0.07694 | 1.754088 | 0.07942 | 0.22378 | 358.746 |
| Mtx1 | 612.892 | 0.032314 | 0.09408 | 0.343477 | 0.73124 | 0.85734 | 18.008 |
| Mtx2 | 1503.25 | 0.172897 | 0.08779 | 1.969336 | 0.04891 | 0.16347 | 52.629 |
| Mtx3 | 4694.88 | 0.219061 | 0.06797 | 3.22296 | 0.00127 | 0.01478 | 13.471 |
| Muc1 | 50.8147 | 0.141418 | 0.28034 | 0.504445 | 0.61395 | 0.78017 | 4.325 |
| Muc15 | 30.7766 | -0.04364 | 0.34827 | -0.12531 | 0.90027 | 0.94992 | 18.188 |
| Muc2 | 36.9617 | -0.16955 | 0.36734 | -0.46158 | 0.64439 | 0.80166 | 31.848 |
| Muc3a | 857.583 | 0.62669 | 0.34132 | 1.836089 | 0.06634 | 0.1992 | 3.583 |

| | | | | | | | |
|---|---|---|---|---|---|---|---|
| Muc6 | 136.367 | 0.182676 | 0.19301 | 0.946447 | 0.34392 | 0.55458 | 28.269 |
| Mul1 | 2089.79 | -0.09681 | 0.06948 | -1.39334 | 0.16352 | 0.3523 | 7.595 |
| Mus81 | 964.296 | 0.202088 | 0.09534 | 2.119549 | 0.03404 | 0.12885 | 6.058 |
| Musk | 235.332 | 0.995626 | 0.25354 | 3.926834 | 8.61E-05 | 0.00244 | 88.344 |
| Mustn1 | 33.7615 | -0.64873 | 0.44249 | -1.46611 | 0.14262 | 0.32449 | 2.411 |
| Mutyh | 115.424 | 0.27473 | 0.19671 | 1.396628 | 0.16253 | 0.35091 | 11.718 |
| Mvb12a | 611.894 | -0.15396 | 0.08123 | -1.89536 | 0.05804 | 0.18315 | 5.164 |
| Mvb12b | 5117.55 | -0.12827 | 0.07711 | -1.66361 | 0.09619 | 0.25395 | 157.994 |
| Mvd | 522.446 | -0.39038 | 0.09806 | -3.98115 | 6.86E-05 | 0.00207 | 9.822 |
| Mvk | 580.723 | -0.31888 | 0.08962 | -3.55827 | 0.00037 | 0.00652 | 16.323 |
| Mvp | 476.319 | -0.25946 | 0.09255 | -2.80335 | 0.00506 | 0.03656 | 27.762 |
| Mx1 | 16.831 | -0.78352 | 0.77767 | -1.00752 | 0.31368 | 0.52419 | 15.873 |
| Mx2 | 66.1338 | 0.088899 | 0.28687 | 0.309891 | 0.75664 | 0.87187 | 25.593 |
| Mxd1 | 2022.22 | 0.306124 | 0.11794 | 2.59565 | 0.00944 | 0.05527 | 22.12 |
| Mxd3 | 54.1891 | -0.32844 | 0.39649 | -0.82837 | 0.40746 | 0.61538 | 4.656 |
| Mxd4 | 5565.65 | -0.12874 | 0.10489 | -1.22729 | 0.21971 | 0.42219 | 13.838 |
| Mxi1 | 2986.15 | -0.0161 | 0.07595 | -0.212 | 0.8321 | 0.91398 | 65.293 |
| Mxra7 | 453.722 | -0.09843 | 0.14138 | -0.69621 | 0.4863 | 0.68232 | 24.985 |
| Mxra8 | 238.802 | -0.36688 | 0.15218 | -2.41073 | 0.01592 | 0.07743 | 4.409 |
| Myadm | 1644.59 | -0.34788 | 0.11902 | -2.92283 | 0.00347 | 0.02856 | 11.354 |
| Myadml2 | 251.65 | -0.09534 | 0.12271 | -0.77695 | 0.43719 | 0.64268 | 2.128 |
| Myb | 84.1852 | 0.711944 | 0.24544 | 2.900629 | 0.00372 | 0.03002 | 36.05 |
| Mybbp1a | 3211.56 | 0.084852 | 0.06433 | 1.319074 | 0.18714 | 0.38243 | 10.414 |
| Mybl1 | 198.506 | 0.026134 | 0.17826 | 0.146604 | 0.88344 | 0.94185 | 32.795 |
| Mybl2 | 97.162 | -0.07778 | 0.25626 | -0.30352 | 0.76149 | 0.87504 | 30.002 |
| Mybpc1 | 53.1816 | -1.7056 | 0.32779 | -5.20342 | 1.96E-07 | 3.01E-05 | 86.874 |
| Mybpc2 | 24.3788 | -0.02371 | 0.37558 | -0.06313 | 0.94966 | 0.97495 | 22.958 |
| Mybpc3 | 24.5133 | 0.857021 | 0.43846 | 1.954622 | 0.05063 | 0.16737 | 18.373 |
| Mybph | 16.3298 | 0.329343 | 0.52579 | 0.626371 | 0.53107 | 0.71862 | 7.785 |
| Myc | 180.104 | -0.00977 | 0.15388 | -0.06349 | 0.94938 | 0.97495 | 4.984 |
| Mycbp | 246.588 | 0.052574 | 0.16879 | 0.311483 | 0.75543 | 0.8714 | 7.438 |
| Mycbp2 | 22918.8 | 0.130061 | 0.06792 | 1.914878 | 0.05551 | 0.17779 | 233.404 |
| Mycbpap | 43.0584 | -0.10448 | 0.335 | -0.31189 | 0.75513 | 0.87123 | 20.396 |
| Mycl | 717.794 | -0.30388 | 0.1554 | -1.95545 | 0.05053 | 0.16718 | 6.834 |

| | | | | | | | |
|---|---|---|---|---|---|---|---|
| Mycn | 192.243 | -0.52466 | 0.21595 | -2.42949 | 0.01512 | 0.07486 | 5.822 |
| Myct1 | 13.5713 | -0.71243 | 0.57388 | -1.24144 | 0.21444 | 0.4158 | 13.13 |
| Myd88 | 180.212 | -0.46336 | 0.22323 | -2.07569 | 0.03792 | 0.13811 | 5.478 |
| Mydgf | 1064.81 | -0.1295 | 0.09842 | -1.31581 | 0.18824 | 0.38384 | 8.177 |
| Myef2 | 4311.33 | 0.07831 | 0.0965 | 0.811519 | 0.41707 | 0.62456 | 39.034 |
| Myef2l | 569.1 | 0.079648 | 0.12463 | 0.639053 | 0.52279 | 0.71143 | 4.456 |
| Myg1 | 992.092 | 0.141901 | 0.07623 | 1.861366 | 0.06269 | 0.19214 | 6.431 |
| Myh10 | 22178.1 | 0.185139 | 0.07394 | 2.50403 | 0.01228 | 0.06564 | 125.074 |
| Myh11 | 428.931 | -0.56784 | 0.23687 | -2.39723 | 0.01652 | 0.07941 | 96.838 |
| Myh14 | 2514.64 | -0.1136 | 0.09271 | -1.22525 | 0.22048 | 0.42315 | 65.041 |
| Myh15 | 72.2644 | -0.28745 | 0.24371 | -1.17946 | 0.23821 | 0.44369 | 141.619 |
| Myh2 | 16.1658 | 0.017421 | 0.42071 | 0.041407 | 0.96697 | 0.98345 | 26.491 |
| Myh3 | 14.008 | -0.15227 | 0.4794 | -0.31762 | 0.75077 | 0.86925 | 23.992 |
| Myh6 | 285.885 | -0.40627 | 0.16688 | -2.4345 | 0.01491 | 0.07409 | 25.007 |
| Myh7 | 1365.94 | 1.102582 | 0.34564 | 3.190007 | 0.00142 | 0.01598 | 23.943 |
| Myh7b | 425.606 | -0.03694 | 0.17038 | -0.21684 | 0.82833 | 0.91195 | 23.096 |
| Myh8 | 16.3115 | -0.79851 | 0.56291 | -1.41854 | 0.15603 | 0.3427 | 31.511 |
| Myh9 | 6316.86 | -0.12991 | 0.07766 | -1.67268 | 0.09439 | 0.25068 | 81.589 |
| Myl1 | 19.8797 | -0.40345 | 0.42289 | -0.95402 | 0.34007 | 0.55085 | 21.11 |
| Myl12a | 314.822 | -0.34672 | 0.10375 | -3.34187 | 0.00083 | 0.01127 | 9.223 |
| Myl12b | 245.8 | -0.04801 | 0.14582 | -0.3292 | 0.742 | 0.86409 | 17.758 |
| Myl4 | 53.9108 | -0.8552 | 0.38977 | -2.19408 | 0.02823 | 0.11312 | 45.091 |
| Myl6 | 8443.76 | -0.13938 | 0.06307 | -2.21 | 0.02711 | 0.10995 | 3.286 |
| Myl6b | 426.188 | -0.19941 | 0.12978 | -1.53651 | 0.12441 | 0.29928 | 4.529 |
| Myl9 | 381.131 | -0.39201 | 0.152 | -2.57897 | 0.00991 | 0.05714 | 6.239 |
| Mylip | 635.707 | -0.30614 | 0.12447 | -2.45966 | 0.01391 | 0.07096 | 22.281 |
| Mylk | 875.744 | -0.76012 | 0.16238 | -4.68116 | 2.85E-06 | 0.00023 | 257.211 |
| Mylpf | 26.2413 | -0.30035 | 0.41304 | -0.72716 | 0.46713 | 0.66622 | 5.409 |
| Mynn | 1003.19 | 0.159774 | 0.07584 | 2.106847 | 0.03513 | 0.13139 | 17.809 |
| Myo10 | 3047.63 | -0.12374 | 0.07322 | -1.69005 | 0.09102 | 0.24507 | 191.149 |
| Myo15b | 156.998 | -0.2925 | 0.33199 | -0.88106 | 0.37829 | 0.58818 | 34.198 |
| Myo16 | 2845.06 | 0.232599 | 0.14134 | 1.645725 | 0.09982 | 0.25983 | 480.832 |
| Myo18a | 13607.9 | 0.122038 | 0.06924 | 1.762583 | 0.07797 | 0.22102 | 102.735 |
| Myo18b | 125.885 | 0.059767 | 0.21157 | 0.282487 | 0.77757 | 0.88414 | 207.487 |

| | | | | | | | |
|---|---|---|---|---|---|---|---|
| Myo19 | 726.44 | 0.032653 | 0.13349 | 0.244602 | 0.80676 | 0.90051 | 31.079 |
| Myo1a | 111.589 | 0.361668 | 0.21504 | 1.681901 | 0.09259 | 0.24781 | 15.771 |
| Myo1b | 3003.43 | 0.399141 | 0.18856 | 2.116791 | 0.03428 | 0.12937 | 166.307 |
| Myo1c | 825.924 | -0.3249 | 0.12519 | -2.59523 | 0.00945 | 0.05529 | 23.407 |
| Myo1d | 667.55 | -0.27506 | 0.13195 | -2.08453 | 0.03711 | 0.13623 | 297.9 |
| Myo1e | 852.271 | -0.79696 | 0.16959 | -4.69946 | 2.61E-06 | 0.00022 | 192.417 |
| Myo1f | 136.078 | -0.48235 | 0.23191 | -2.07994 | 0.03753 | 0.13715 | 52.058 |
| Myo1g | 30.0841 | -0.61475 | 0.46451 | -1.32345 | 0.18568 | 0.38052 | 14.418 |
| Myo1h | 28.5926 | -0.25034 | 0.39524 | -0.63338 | 0.52649 | 0.71461 | 76.192 |
| Myo3b | 192.131 | 0.394136 | 0.16568 | 2.378879 | 0.01737 | 0.08175 | 390.073 |
| Myo5a | 29514 | 0.106563 | 0.0711 | 1.498825 | 0.13392 | 0.3136 | 152.674 |
| Myo5b | 2962.16 | 0.421186 | 0.13437 | 3.134597 | 0.00172 | 0.01828 | 330.558 |
| Myo5c | 357.383 | -0.19247 | 0.16597 | -1.15962 | 0.2462 | 0.45299 | 73.432 |
| Myo6 | 7385.93 | -0.33335 | 0.08898 | -3.74618 | 0.00018 | 0.00401 | 146.699 |
| Myo7a | 1001.07 | 0.107869 | 0.09185 | 1.174388 | 0.24024 | 0.44624 | 68.465 |
| Myo9a | 7045.89 | 0.304043 | 0.12833 | 2.369216 | 0.01783 | 0.08313 | 177.971 |
| Myo9b | 5011.57 | 0.164796 | 0.06923 | 2.380559 | 0.01729 | 0.08154 | 88 |
| Myoc | 989.867 | -0.48511 | 0.18115 | -2.67798 | 0.00741 | 0.04733 | 10.539 |
| Myocd | 22.9852 | -1.68718 | 0.60533 | -2.78722 | 0.00532 | 0.03773 | 93.429 |
| Myof | 327.408 | -0.7872 | 0.17987 | -4.3766 | 1.21E-05 | 0.00063 | 144.542 |
| Myom1 | 131.634 | -0.21548 | 0.2152 | -1.00129 | 0.31669 | 0.52722 | 124.224 |
| Myom3 | 129.969 | 0.051192 | 0.22787 | 0.224656 | 0.82225 | 0.90785 | 55.85 |
| Myorg | 2092.32 | -0.21602 | 0.10088 | -2.14133 | 0.03225 | 0.12412 | 7.473 |
| Myot | 19.1061 | 1.713934 | 0.63903 | 2.682108 | 0.00732 | 0.04693 | 21.651 |
| Mypn | 78.9594 | -0.94803 | 0.20334 | -4.66233 | 3.13E-06 | 0.00024 | 88.158 |
| Mypop | 413.933 | -0.02731 | 0.12031 | -0.22696 | 0.82045 | 0.90734 | 10.522 |
| Mypopos | 270.79 | 0.044624 | 0.11978 | 0.372557 | 0.70948 | 0.84351 | 7.999 |
| Myrf | 2838.49 | -0.53298 | 0.14447 | -3.68924 | 0.00022 | 0.0046 | 32.477 |
| Myrip | 2720.67 | -0.19353 | 0.08262 | -2.34238 | 0.01916 | 0.08697 | 173.329 |
| Mysm1 | 2296.64 | 0.21979 | 0.12966 | 1.695097 | 0.09006 | 0.24343 | 37.061 |
| Myt1 | 1054.95 | 0.029595 | 0.07699 | 0.384417 | 0.70067 | 0.83841 | 64.466 |
| Myt1l | 17197.8 | 0.236731 | 0.09519 | 2.486931 | 0.01289 | 0.0678 | 394.83 |
| Myzap | 40.8975 | -0.86756 | 0.42954 | -2.01976 | 0.04341 | 0.15061 | 88.014 |
| Mzf1 | 1157.44 | 0.385221 | 0.10285 | 3.745357 | 0.00018 | 0.00401 | 12.462 |

| | | | | | | | |
|---|---|---|---|---|---|---|---|
| Mzt1 | 4224.03 | 0.178024 | 0.0665 | 2.677126 | 0.00743 | 0.04738 | 11.591 |
| Mzt2 | 695.778 | -0.28393 | 0.10307 | -2.75483 | 0.00587 | 0.04067 | 14.929 |
| N4bp1 | 3656.38 | 0.179784 | 0.05527 | 3.253047 | 0.00114 | 0.01384 | 77.099 |
| N4bp2 | 2015.05 | 0.834576 | 0.19644 | 4.248482 | 2.15E-05 | 0.00092 | 66.588 |
| N4bp2l1 | 2140.06 | 0.258237 | 0.07935 | 3.254444 | 0.00114 | 0.01382 | 25.545 |
| N4bp2l2 | 3294.24 | 0.201857 | 0.07935 | 2.544024 | 0.01096 | 0.06094 | 58.177 |
| N4bp2os | 69.6196 | 0.28458 | 0.21851 | 1.302352 | 0.1928 | 0.38957 | 3.389 |
| N4bp3 | 646.333 | -0.37214 | 0.16187 | -2.29904 | 0.0215 | 0.09402 | 7.78 |
| N6amt1 | 1036.11 | 0.095162 | 0.08554 | 1.112518 | 0.26592 | 0.47506 | 14.558 |
| Naa10 | 438.363 | 0.125886 | 0.10819 | 1.163517 | 0.24462 | 0.45129 | 5.072 |
| Naa12 | 2424.38 | -1.71E-05 | 0.06494 | -0.00026 | 0.99979 | 0.99994 | 5.977 |
| Naa15 | 3772.23 | 0.233321 | 0.10418 | 2.239546 | 0.02512 | 0.10427 | 61.36 |
| Naa16 | 921.047 | 0.316588 | 0.08712 | 3.634117 | 0.00028 | 0.00533 | 65.51 |
| Naa20 | 2001.95 | 0.05285 | 0.06676 | 0.791604 | 0.42859 | 0.63524 | 14.327 |
| Naa25 | 1775.27 | 0.082936 | 0.07237 | 1.146061 | 0.25177 | 0.45912 | 46.443 |
| Naa30 | 2928.55 | 0.110479 | 0.0803 | 1.375779 | 0.16889 | 0.35902 | 18.806 |
| Naa35 | 2076.66 | 0.04829 | 0.07546 | 0.639985 | 0.52218 | 0.71122 | 50.664 |
| Naa38 | 372.05 | -0.21994 | 0.16635 | -1.32214 | 0.18612 | 0.38103 | 1.194 |
| Naa40 | 2853.15 | 0.190457 | 0.06602 | 2.884947 | 0.00391 | 0.03099 | 15.516 |
| Naa50 | 4604.14 | 0.059795 | 0.07607 | 0.786054 | 0.43184 | 0.63787 | 23.537 |
| Naa60 | 2147.33 | 0.045127 | 0.07493 | 0.602253 | 0.54701 | 0.73125 | 32.396 |
| Naa80 | 239.144 | -0.12693 | 0.1219 | -1.04121 | 0.29778 | 0.50786 | 5.164 |
| Naaa | 1982.47 | -0.36345 | 0.10568 | -3.43922 | 0.00058 | 0.00882 | 20.512 |
| Naalad2 | 162.032 | -0.06715 | 0.15886 | -0.42269 | 0.67252 | 0.8199 | 81.045 |
| Naaladl1 | 33.6484 | -0.36145 | 0.31077 | -1.1631 | 0.24479 | 0.45139 | 10.011 |
| Naaladl2 | 128.291 | 0.295234 | 0.18294 | 1.61387 | 0.10656 | 0.27076 | 1346.16 |
| Nab1 | 1570.7 | -0.42316 | 0.13074 | -3.2367 | 0.00121 | 0.01441 | 43.386 |
| Nab2 | 922.842 | 0.213979 | 0.09668 | 2.213184 | 0.02688 | 0.10945 | 7.651 |
| Nabp1 | 551.77 | -0.18635 | 0.11669 | -1.59701 | 0.11026 | 0.27686 | 12.564 |
| Nabp2 | 1101.17 | -0.12967 | 0.07995 | -1.62182 | 0.10484 | 0.26787 | 10.098 |
| Naca | 5066.14 | 0.105234 | 0.06369 | 1.652159 | 0.0985 | 0.25768 | 13.063 |
| Nacad | 1759.81 | -0.02485 | 0.07481 | -0.33217 | 0.73976 | 0.86273 | 8.231 |
| Nacc1 | 5002.91 | -0.15713 | 0.066 | -2.38091 | 0.01727 | 0.08154 | 17.424 |
| Nacc2 | 7131.41 | -0.47044 | 0.10266 | -4.58252 | 4.59E-06 | 0.00032 | 67.686 |

| | | | | | | | |
|---|---|---|---|---|---|---|---|
| Nadk | 3443.88 | -0.0142 | 0.07068 | -0.20089 | 0.84079 | 0.91835 | 28.624 |
| Nadk2 | 2882.28 | -0.12006 | 0.07854 | -1.52876 | 0.12632 | 0.30205 | 39.64 |
| Nadsyn1 | 259.714 | 0.025566 | 0.12962 | 0.19723 | 0.84365 | 0.91962 | 27.266 |
| Nae1 | 2455.62 | 0.202447 | 0.07106 | 2.848946 | 0.00439 | 0.03347 | 23.61 |
| Naf1 | 884.35 | 0.022262 | 0.08986 | 0.247743 | 0.80433 | 0.89895 | 30.348 |
| Naga | 910.634 | 0.000797 | 0.07852 | 0.010145 | 0.99191 | 0.99537 | 9.394 |
| Nagk | 1300.7 | -0.0741 | 0.09778 | -0.75785 | 0.44854 | 0.652 | 9.037 |
| Naglu | 387.912 | -0.0266 | 0.1384 | -0.19217 | 0.84761 | 0.92246 | 7.661 |
| Nagpa | 400.246 | -0.50451 | 0.12338 | -4.08898 | 4.33E-05 | 0.00147 | 8.724 |
| Naif1 | 237.476 | -0.05243 | 0.12433 | -0.42174 | 0.67322 | 0.8203 | 6.497 |
| Naip1 | 22.8938 | -0.14336 | 0.48632 | -0.29479 | 0.76815 | 0.87883 | 45.106 |
| Naip2 | 67.4546 | -0.40651 | 0.28416 | -1.43055 | 0.15256 | 0.33794 | 58.03 |
| Naip5 | 76.9724 | -0.34513 | 0.28514 | -1.2104 | 0.22613 | 0.43018 | 34.585 |
| Naip6 | 45.597 | -0.57252 | 0.27415 | -2.08831 | 0.03677 | 0.13524 | 36.554 |
| Nalcn | 6916.22 | 0.254831 | 0.06909 | 3.688389 | 0.00023 | 0.00461 | 350.511 |
| Nalf1 | 2224.2 | -0.67552 | 0.1821 | -3.70962 | 0.00021 | 0.00438 | 565.26 |
| Nalf2 | 1178.9 | 0.057366 | 0.07897 | 0.72639 | 0.4676 | 0.66659 | 27.77 |
| Nampt | 2535.95 | 0.062867 | 0.08657 | 0.72622 | 0.4677 | 0.66669 | 33.805 |
| Nanos1 | 1342.19 | 0.161811 | 0.08331 | 1.94227 | 0.0521 | 0.17045 | 3.928 |
| Nanos2 | 21.9362 | -0.2625 | 0.47825 | -0.54888 | 0.58309 | 0.75831 | 1.563 |
| Nanos3 | 23.4118 | -0.35587 | 0.39719 | -0.89596 | 0.37027 | 0.58058 | 11.487 |
| Nanp | 24.1275 | 0.562002 | 0.36594 | 1.53577 | 0.12459 | 0.29957 | 9.714 |
| Nans | 504.318 | -0.26225 | 0.12828 | -2.04438 | 0.04092 | 0.14485 | 14.385 |
| Nap1l1 | 12978.6 | 0.272544 | 0.06882 | 3.960293 | 7.49E-05 | 0.00219 | 24.928 |
| Nap1l2 | 5658.69 | 0.04238 | 0.08933 | 0.474412 | 0.63521 | 0.79438 | 2.465 |
| Nap1l3 | 2331.23 | 0.13674 | 0.10765 | 1.270204 | 0.20401 | 0.40269 | 2.837 |
| Nap1l4 | 4691.28 | -0.07557 | 0.06591 | -1.14651 | 0.25158 | 0.45895 | 35.528 |
| Nap1l5 | 11987.1 | 0.143481 | 0.12844 | 1.117116 | 0.26394 | 0.47282 | 1.844 |
| Napa | 6880.82 | -0.03219 | 0.05868 | -0.54852 | 0.58334 | 0.75845 | 19.518 |
| Napb | 21398.5 | -0.28056 | 0.08863 | -3.16553 | 0.00155 | 0.01702 | 38.483 |
| Napepld | 521.604 | -0.60383 | 0.18261 | -3.30664 | 0.00094 | 0.01229 | 38.496 |
| Napg | 7955.86 | -0.08308 | 0.07218 | -1.15113 | 0.24968 | 0.45704 | 21.62 |
| Naprt | 201.726 | -0.22445 | 0.13119 | -1.71089 | 0.0871 | 0.23762 | 3.526 |
| Napsa | 77.6177 | 0.021217 | 0.19234 | 0.110307 | 0.91217 | 0.9557 | 14.483 |

| | | | | | | | |
|---|---|---|---|---|---|---|---|
| Narf | 2901.88 | -0.10651 | 0.05986 | -1.77922 | 0.0752 | 0.21627 | 18.604 |
| Nars | 8836.34 | -0.1648 | 0.06632 | -2.48494 | 0.01296 | 0.06804 | 17.006 |
| Nars2 | 706.905 | 0.128417 | 0.11864 | 1.082366 | 0.27909 | 0.48899 | 113.254 |
| Nasp | 1097.78 | -0.10584 | 0.11012 | -0.96117 | 0.33646 | 0.54716 | 26.89 |
| Nat1 | 119.108 | -0.30665 | 0.18831 | -1.62844 | 0.10343 | 0.26587 | 11.184 |
| Nat10 | 2412.16 | 0.243485 | 0.07012 | 3.472193 | 0.00052 | 0.00811 | 40.015 |
| Nat14 | 2850.28 | -0.02508 | 0.06713 | -0.37367 | 0.70865 | 0.8429 | 2.979 |
| Nat2 | 43.2438 | -0.48437 | 0.29114 | -1.66367 | 0.09618 | 0.25395 | 7.727 |
| Nat8 | 106.577 | 0.26009 | 0.25114 | 1.035626 | 0.30038 | 0.51045 | 1.695 |
| Nat8f1 | 171.967 | -0.03476 | 0.14051 | -0.24738 | 0.80462 | 0.89908 | 5.534 |
| Nat8f2 | 21.9366 | 0.05279 | 0.43948 | 0.120118 | 0.90439 | 0.95168 | 3.737 |
| Nat8f3 | 147.469 | 0.211892 | 0.20378 | 1.039806 | 0.29843 | 0.50856 | 33.242 |
| Nat8f4 | 693.219 | -0.27792 | 0.15827 | -1.75602 | 0.07909 | 0.22311 | 5.834 |
| Nat8f5 | 71.8242 | 0.070204 | 0.2366 | 0.296724 | 0.76668 | 0.87802 | 3.755 |
| Nat8f6 | 37.1177 | 0.519356 | 0.27925 | 1.859842 | 0.06291 | 0.19257 | 12.931 |
| Nat8f7 | 16.5895 | 0.716817 | 0.42449 | 1.688675 | 0.09128 | 0.24554 | 6.479 |
| Nat8l | 15093 | -0.41377 | 0.07646 | -5.41128 | 6.26E-08 | 1.26E-05 | 9.933 |
| Nat9 | 584.616 | -0.14432 | 0.09262 | -1.5581 | 0.11921 | 0.2914 | 5.028 |
| Natd1 | 522.716 | -0.19841 | 0.09357 | -2.12052 | 0.03396 | 0.12867 | 12.505 |
| Nav1 | 7391.85 | -0.27024 | 0.11157 | -2.42229 | 0.01542 | 0.07584 | 253.526 |
| Nav2 | 5443.88 | -0.19126 | 0.10368 | -1.84462 | 0.06509 | 0.19674 | 701.375 |
| Nav3 | 3800.73 | -0.2297 | 0.1379 | -1.66575 | 0.09576 | 0.25318 | 774.946 |
| Naxd | 1897.17 | -0.06322 | 0.06057 | -1.04381 | 0.29657 | 0.50689 | 17.455 |
| Naxe | 1383.74 | -0.09184 | 0.07258 | -1.26546 | 0.20571 | 0.40486 | 1.976 |
| Nbas | 2303.81 | -0.04977 | 0.06929 | -0.71818 | 0.47265 | 0.6709 | 314.679 |
| Nbdy | 391.07 | 0.333006 | 0.11178 | 2.979088 | 0.00289 | 0.02562 | 18.331 |
| Nbea | 15889.1 | 0.207014 | 0.078 | 2.654179 | 0.00795 | 0.04941 | 558.507 |
| Nbeal1 | 2009.28 | 0.170613 | 0.16874 | 1.01108 | 0.31198 | 0.52237 | 157.73 |
| Nbeal2 | 1377.76 | 0.220217 | 0.09509 | 2.315927 | 0.02056 | 0.09126 | 29.373 |
| Nbl1 | 1400.16 | -0.71955 | 0.16964 | -4.2416 | 2.22E-05 | 0.00094 | 10.71 |
| Nbn | 711.536 | -0.01498 | 0.11079 | -0.13518 | 0.89247 | 0.94617 | 34.665 |
| Nbr1 | 10135.7 | 0.036049 | 0.05306 | 0.679431 | 0.49686 | 0.69044 | 29.803 |
| Ncald | 8752.19 | -0.83776 | 0.36088 | -2.32143 | 0.02026 | 0.0903 | 426.396 |
| Ncam1 | 8802.79 | 0.112537 | 0.06432 | 1.749745 | 0.08016 | 0.22519 | 296.79 |

| | | | | | | | |
|---|---|---|---|---|---|---|---|
| Ncam2 | 3441.51 | 0.449674 | 0.17584 | 2.557345 | 0.01055 | 0.0596 | 426.132 |
| Ncan | 11162.4 | 0.008321 | 0.08367 | 0.09944 | 0.92079 | 0.95978 | 27.789 |
| Ncapd2 | 1647.27 | -0.03826 | 0.07332 | -0.5218 | 0.60181 | 0.77182 | 23.695 |
| Ncapd3 | 1688.66 | 0.127447 | 0.09025 | 1.412138 | 0.15791 | 0.34515 | 65.141 |
| Ncapg | 21.6826 | -0.33374 | 0.42896 | -0.77801 | 0.43656 | 0.64224 | 30.628 |
| Ncapg2 | 350.713 | -0.03218 | 0.12905 | -0.24933 | 0.8031 | 0.89834 | 58.33 |
| Ncaph | 66.0415 | -0.23262 | 0.28971 | -0.80292 | 0.42202 | 0.62883 | 30.146 |
| Ncaph2 | 3252.2 | -0.02594 | 0.06541 | -0.39651 | 0.69173 | 0.83261 | 17.108 |
| Ncbp1 | 2812.93 | 0.150784 | 0.06625 | 2.276068 | 0.02284 | 0.0978 | 33.791 |
| Ncbp2 | 2672.69 | 0.033239 | 0.06104 | 0.544561 | 0.58606 | 0.76022 | 13.269 |
| Ncbp3 | 1633.77 | -0.02851 | 0.06903 | -0.41304 | 0.67958 | 0.82491 | 41.535 |
| Ncdn | 51832.8 | 0.127973 | 0.09536 | 1.342029 | 0.17959 | 0.37262 | 9.689 |
| Nceh1 | 6324.44 | -0.05759 | 0.05035 | -1.14389 | 0.25267 | 0.46 | 101.644 |
| Ncf1 | 385.096 | -0.2843 | 0.15106 | -1.88201 | 0.05983 | 0.18635 | 9.573 |
| Ncf2 | 207.968 | -0.32681 | 0.1936 | -1.68813 | 0.09139 | 0.24566 | 36.798 |
| Ncf4 | 28.0939 | -1.18714 | 0.43052 | -2.75747 | 0.00583 | 0.04041 | 17.78 |
| Nck1 | 900.096 | 0.062246 | 0.10381 | 0.599609 | 0.54877 | 0.73274 | 53.842 |
| Nck2 | 1137.64 | -0.1898 | 0.10986 | -1.72773 | 0.08404 | 0.23208 | 125.937 |
| Nckap1 | 23622.5 | 0.041466 | 0.06635 | 0.624939 | 0.53201 | 0.71956 | 80.869 |
| Nckap1l | 678.022 | -0.17402 | 0.11945 | -1.45686 | 0.14515 | 0.32787 | 45.017 |
| Nckap5 | 153.42 | -0.27202 | 0.20806 | -1.30743 | 0.19107 | 0.38734 | 917.18 |
| Nckap5l | 851.613 | 0.0104 | 0.10995 | 0.094593 | 0.92464 | 0.96202 | 35.714 |
| Nckap5los | 106.571 | 0.09098 | 0.19872 | 0.457839 | 0.64707 | 0.80359 | 7.613 |
| Nckipsd | 2398.43 | 0.003074 | 0.06683 | 0.045998 | 0.96331 | 0.98208 | 10.477 |
| Ncl | 15614.4 | 0.204347 | 0.08857 | 2.307176 | 0.02104 | 0.09265 | 14.682 |
| Ncln | 2319.02 | 0.080363 | 0.06989 | 1.149905 | 0.25018 | 0.45761 | 10.144 |
| Ncmap | 53.2801 | 0.04124 | 0.2887 | 0.142849 | 0.88641 | 0.94338 | 28.652 |
| Ncoa1 | 6981.21 | 0.078942 | 0.06584 | 1.19899 | 0.23053 | 0.43487 | 229.821 |
| Ncoa2 | 6750.53 | 0.159768 | 0.06585 | 2.426276 | 0.01525 | 0.07527 | 234.979 |
| Ncoa3 | 3529.5 | 0.017594 | 0.08005 | 0.219781 | 0.82604 | 0.91021 | 80.607 |
| Ncoa4 | 2138.22 | 0.094895 | 0.08258 | 1.149138 | 0.2505 | 0.45785 | 19.991 |
| Ncoa4-ps | 338.797 | -0.02805 | 0.12022 | -0.23335 | 0.81549 | 0.90514 | 1.878 |
| Ncoa5 | 3235.98 | 0.087554 | 0.08071 | 1.084829 | 0.278 | 0.48797 | 34.511 |
| Ncoa6 | 3909.29 | 0.085039 | 0.06519 | 1.304428 | 0.19209 | 0.38879 | 83.239 |

| | | | | | | | |
|---|---|---|---|---|---|---|---|
| Ncoa7 | 3484.69 | 0.041605 | 0.09555 | 0.435439 | 0.66324 | 0.81392 | 174.328 |
| Ncor1 | 13218.1 | 0.019215 | 0.0665 | 0.288953 | 0.77262 | 0.88166 | 142.116 |
| Ncor2 | 8498.35 | -0.14007 | 0.08582 | -1.63215 | 0.10265 | 0.26456 | 162.067 |
| Ncr3-ps | 25.4404 | 0.797394 | 0.39813 | 2.002828 | 0.0452 | 0.15458 | 4.128 |
| Ncs1 | 15855.6 | 0.09533 | 0.0702 | 1.357925 | 0.17449 | 0.36611 | 50.167 |
| Ncstn | 2733.84 | -0.07013 | 0.06815 | -1.02907 | 0.30344 | 0.51314 | 16.783 |
| Ndc1 | 1123.99 | 0.018954 | 0.08919 | 0.212511 | 0.83171 | 0.91374 | 48.563 |
| Nde1 | 1293.67 | -0.55915 | 0.11079 | -5.04691 | 4.49E-07 | 5.83E-05 | 29.654 |
| Ndel1 | 4086.48 | 0.051769 | 0.06385 | 0.810836 | 0.41746 | 0.62487 | 50.425 |
| Ndfip1 | 22056.8 | -0.01982 | 0.07573 | -0.26171 | 0.79354 | 0.89304 | 54.908 |
| Ndfip2 | 5421.59 | -0.06983 | 0.06396 | -1.09171 | 0.27496 | 0.48452 | 50.725 |
| Ndn | 3493.1 | 0.074003 | 0.10144 | 0.729545 | 0.46567 | 0.665 | 3.694 |
| Ndnf | 1230.36 | -0.82711 | 0.36619 | -2.25868 | 0.0239 | 0.10067 | 40.737 |
| Ndor1 | 343.003 | -0.27985 | 0.12922 | -2.16566 | 0.03034 | 0.11892 | 11.202 |
| Ndp | 270.075 | -0.20307 | 0.1373 | -1.47905 | 0.13913 | 0.32034 | 26.254 |
| Ndrg1 | 2996.97 | -0.62914 | 0.1292 | -4.8697 | 1.12E-06 | 0.00012 | 40.323 |
| Ndrg2 | 28877.8 | -0.20622 | 0.10734 | -1.92115 | 0.05471 | 0.17597 | 8.888 |
| Ndrg3 | 19491.6 | -0.40119 | 0.09891 | -4.05622 | 4.99E-05 | 0.00163 | 64.712 |
| Ndrg4 | 127664 | 0.196521 | 0.09932 | 1.978655 | 0.04785 | 0.16092 | 38.14 |
| Ndst1 | 6275.02 | -0.04482 | 0.06936 | -0.64617 | 0.51817 | 0.70789 | 64.157 |
| Ndst2 | 1003.7 | 0.198468 | 0.08696 | 2.282214 | 0.02248 | 0.09681 | 10.833 |
| Ndst3 | 1392.56 | -0.75393 | 0.40153 | -1.87764 | 0.06043 | 0.18755 | 164.688 |
| Ndst4 | 3641.97 | -1.06275 | 0.47354 | -2.24429 | 0.02481 | 0.10348 | 324.824 |
| Ndufa1 | 1736.05 | 0.208685 | 0.2136 | 0.977 | 0.32857 | 0.53979 | 3.576 |
| Ndufa10 | 5002.57 | -0.2006 | 0.06401 | -3.13407 | 0.00172 | 0.01828 | 34.851 |
| Ndufa11 | 398.536 | -0.13073 | 0.15633 | -0.83625 | 0.40301 | 0.61174 | 4.375 |
| Ndufa12 | 1983.34 | -0.28006 | 0.2862 | -0.97857 | 0.32779 | 0.53919 | 22.724 |
| Ndufa12-ps | 462.987 | -0.725 | 0.2139 | -3.38945 | 0.0007 | 0.00998 | 0.402 |
| Ndufa13 | 3110.24 | -0.24614 | 0.0875 | -2.81305 | 0.00491 | 0.03576 | 8.379 |
| Ndufa2 | 1726.37 | -0.1618 | 0.11775 | -1.37416 | 0.16939 | 0.35964 | 2.226 |
| Ndufa3 | 2142.94 | -0.29005 | 0.09256 | -3.13378 | 0.00173 | 0.01828 | 2.955 |
| Ndufa4 | 6675.8 | -0.23082 | 0.06959 | -3.31672 | 0.00091 | 0.01201 | 7.206 |
| Ndufa4l2 | 58.5656 | -0.39554 | 0.24392 | -1.62157 | 0.1049 | 0.26798 | 2.188 |
| Ndufa5 | 2034.31 | -0.20681 | 0.088 | -2.35011 | 0.01877 | 0.08597 | 9.348 |

| | | | | | | | |
|---|---|---|---|---|---|---|---|
| Ndufa6 | 2685.44 | 0.007733 | 0.08004 | 0.096626 | 0.92302 | 0.96131 | 4.183 |
| Ndufa7 | 1942.35 | -0.14079 | 0.08107 | -1.73664 | 0.08245 | 0.22898 | 13.746 |
| Ndufa8 | 3364.86 | -0.20652 | 0.06203 | -3.32916 | 0.00087 | 0.01163 | 13.081 |
| Ndufa9 | 3840.69 | -0.14302 | 0.07412 | -1.92953 | 0.05367 | 0.17378 | 27.416 |
| Ndufab1 | 1160.71 | 0.177615 | 0.1009 | 1.760369 | 0.07835 | 0.22159 | 16.484 |
| Ndufab1-ps | 62.4447 | -0.25047 | 0.2637 | -0.94982 | 0.3422 | 0.553 | 0.468 |
| Ndufaf1 | 1000.35 | -0.13978 | 0.10194 | -1.37111 | 0.17034 | 0.36065 | 7.382 |
| Ndufaf2 | 513.222 | -0.17518 | 0.10072 | -1.73938 | 0.08197 | 0.22821 | 155.909 |
| Ndufaf3 | 879.332 | -0.03292 | 0.0922 | -0.35701 | 0.72108 | 0.85106 | 1.898 |
| Ndufaf4 | 1963.05 | 0.20023 | 0.09399 | 2.130383 | 0.03314 | 0.12648 | 6.919 |
| Ndufaf5 | 1179.95 | 0.067856 | 0.08167 | 0.830832 | 0.40607 | 0.61461 | 33.041 |
| Ndufaf6 | 335.676 | 0.395783 | 0.10537 | 3.756023 | 0.00017 | 0.0039 | 25.161 |
| Ndufaf7 | 2115.13 | 0.110817 | 0.07552 | 1.467428 | 0.14226 | 0.32422 | 10.917 |
| Ndufaf8 | 332.852 | -0.15641 | 0.13791 | -1.1341 | 0.25675 | 0.46474 | 1.778 |
| Ndufb1 | 981.531 | -0.18785 | 0.16739 | -1.12224 | 0.26176 | 0.47052 | 8.979 |
| Ndufb10 | 5087.24 | -0.20267 | 0.08612 | -2.35337 | 0.0186 | 0.08547 | 2.419 |
| Ndufb11 | 2125.23 | -0.26014 | 0.08809 | -2.95308 | 0.00315 | 0.02691 | 2.294 |
| Ndufb2 | 958.178 | 0.053722 | 0.10317 | 0.52073 | 0.60255 | 0.77234 | 10.809 |
| Ndufb3 | 2391.26 | -0.1295 | 0.07416 | -1.74634 | 0.08075 | 0.22616 | 9.581 |
| Ndufb4 | 2194.21 | -0.07235 | 0.08527 | -0.84853 | 0.39614 | 0.60532 | 7.284 |
| Ndufb4c | 38.1895 | 0.042636 | 0.3331 | 0.127998 | 0.89815 | 0.94885 | 0.39 |
| Ndufb5 | 2251.87 | -0.08496 | 0.06547 | -1.2976 | 0.19442 | 0.39169 | 14.577 |
| Ndufb6 | 1271.84 | -0.29232 | 0.08741 | -3.34403 | 0.00083 | 0.01121 | 8.831 |
| Ndufb7 | 1969.21 | -0.14404 | 0.07787 | -1.84964 | 0.06437 | 0.19522 | 4.956 |
| Ndufb8 | 1931.02 | -0.2959 | 0.0958 | -3.08881 | 0.00201 | 0.02016 | 6.869 |
| Ndufb9 | 595.09 | -0.49888 | 0.17007 | -2.9334 | 0.00335 | 0.02799 | 5.733 |
| Ndufc1 | 1420.51 | -0.20362 | 0.0827 | -2.46209 | 0.01381 | 0.07064 | 4.312 |
| Ndufc2 | 2979.58 | -0.20994 | 0.07317 | -2.86912 | 0.00412 | 0.03205 | 7.8 |
| Ndufs1 | 5667.79 | 0.043235 | 0.07876 | 0.548963 | 0.58303 | 0.75828 | 33.238 |
| Ndufs2 | 7691.52 | -0.12343 | 0.05524 | -2.23465 | 0.02544 | 0.1052 | 16.536 |
| Ndufs3 | 3734.78 | -0.03738 | 0.06385 | -0.58541 | 0.55828 | 0.73975 | 10.194 |
| Ndufs4 | 2126.07 | -0.01445 | 0.07822 | -0.1847 | 0.85346 | 0.92567 | 100.464 |
| Ndufs5 | 2717.51 | -0.17894 | 0.06689 | -2.67533 | 0.00747 | 0.04758 | 5.493 |
| Ndufs5-ps | 17.2073 | -0.24247 | 0.44617 | -0.54344 | 0.58683 | 0.7605 | 0.319 |

| | | | | | | | |
|---|---|---|---|---|---|---|---|
| Ndufs6 | 1482.38 | -0.21261 | 0.07938 | -2.67841 | 0.0074 | 0.04731 | 8.705 |
| Ndufs7 | 2224.61 | -0.15529 | 0.06869 | -2.26062 | 0.02378 | 0.10039 | 7.674 |
| Ndufs8 | 2096.61 | -0.13467 | 0.08475 | -1.58901 | 0.11206 | 0.27957 | 3.912 |
| Ndufv1 | 6251.34 | -0.09067 | 0.06414 | -1.4137 | 0.15745 | 0.34464 | 5.423 |
| Ndufv2 | 4640.2 | -0.06437 | 0.06322 | -1.01821 | 0.30858 | 0.51837 | 22.922 |
| Ndufv3 | 1882.07 | -0.02402 | 0.06392 | -0.37576 | 0.7071 | 0.84196 | 11.217 |
| Neat1 | 5234.34 | -0.73948 | 0.20057 | -3.68696 | 0.00023 | 0.00463 | 21.163 |
| Neb | 145.847 | 0.696294 | 0.23067 | 3.018634 | 0.00254 | 0.02359 | 241.828 |
| Nebl | 5050.24 | 0.030983 | 0.09374 | 0.330515 | 0.74101 | 0.86328 | 387.556 |
| Necab1 | 5833.1 | 0.221928 | 0.18736 | 1.184503 | 0.23621 | 0.44147 | 197.55 |
| Necab2 | 4924.94 | 0.657316 | 0.19638 | 3.347125 | 0.00082 | 0.01114 | 25.922 |
| Necab3 | 1482.02 | -0.98454 | 0.2058 | -4.78403 | 1.72E-06 | 0.00016 | 14.492 |
| Necap1 | 7421.32 | 0.055453 | 0.07012 | 0.790841 | 0.42904 | 0.63567 | 14.468 |
| Necap2 | 1256.92 | -0.28858 | 0.10494 | -2.74991 | 0.00596 | 0.04104 | 11.846 |
| Nectin1 | 2009.09 | -1.02084 | 0.40837 | -2.49977 | 0.01243 | 0.06618 | 88.675 |
| Nectin2 | 157.373 | -0.28114 | 0.14173 | -1.98362 | 0.0473 | 0.15968 | 33.84 |
| Nectin3 | 966.923 | -0.14378 | 0.11712 | -1.22756 | 0.21961 | 0.42204 | 110.82 |
| Nectin4 | 175.948 | -0.81926 | 0.21128 | -3.87751 | 0.00011 | 0.00279 | 18.5 |
| Nedd1 | 206.125 | -0.14627 | 0.14059 | -1.04043 | 0.29814 | 0.50826 | 37.675 |
| Nedd4 | 25032 | 0.210447 | 0.05974 | 3.522763 | 0.00043 | 0.00721 | 87.507 |
| Nedd4l | 13237.6 | 0.299502 | 0.10634 | 2.816426 | 0.00486 | 0.03557 | 330.124 |
| Nedd8 | 2028.71 | -0.17378 | 0.0795 | -2.18579 | 0.02883 | 0.11472 | 9.862 |
| Nedd9 | 487.043 | -0.01794 | 0.10766 | -0.16662 | 0.86767 | 0.93294 | 177.782 |
| Nefh | 3271 | -0.44437 | 0.11993 | -3.70536 | 0.00021 | 0.00442 | 9.311 |
| Nefl | 11870.2 | -0.23055 | 0.08596 | -2.68209 | 0.00732 | 0.04693 | 5.233 |
| Nefm | 14016.6 | -0.33987 | 0.13617 | -2.49589 | 0.01256 | 0.06664 | 42.257 |
| Negr1 | 11336.4 | 0.010958 | 0.08793 | 0.12462 | 0.90082 | 0.95027 | 754.652 |
| Neil1 | 448.917 | -0.00543 | 0.1279 | -0.04245 | 0.96614 | 0.98302 | 5.506 |
| Neil2 | 117.605 | -0.2939 | 0.22659 | -1.29702 | 0.19462 | 0.39194 | 11.414 |
| Nek1 | 2448.66 | -0.00869 | 0.15217 | -0.0571 | 0.95446 | 0.97761 | 138.152 |
| Nek10 | 185.783 | 1.195014 | 0.1904 | 6.276249 | 3.47E-10 | 2.44E-07 | 208.645 |
| Nek11 | 37.9052 | 0.28501 | 0.28377 | 1.004375 | 0.3152 | 0.52568 | 233.369 |
| Nek2 | 109.665 | -0.01768 | 0.20517 | -0.08616 | 0.93134 | 0.96532 | 11.607 |
| Nek3 | 92.1199 | -0.46155 | 0.2299 | -2.00758 | 0.04469 | 0.15353 | 38.153 |

| | | | | | | | |
|---|---|---|---|---|---|---|---|
| Nek4 | 1426.16 | 0.117664 | 0.08499 | 1.384461 | 0.16622 | 0.3559 | 37.445 |
| Nek6 | 1008.74 | -0.12766 | 0.08892 | -1.43567 | 0.1511 | 0.33587 | 82.964 |
| Nek7 | 2375.11 | -0.39539 | 0.09716 | -4.06928 | 4.72E-05 | 0.00158 | 137.267 |
| Nek8 | 172.993 | -0.40674 | 0.13829 | -2.94121 | 0.00327 | 0.02757 | 10.57 |
| Nek9 | 4381.09 | -0.09154 | 0.05776 | -1.58467 | 0.11304 | 0.28135 | 39.849 |
| Nelfa | 950.094 | 0.203789 | 0.08818 | 2.311164 | 0.02082 | 0.09204 | 38.498 |
| Nelfb | 1767.17 | -0.18955 | 0.08997 | -2.10676 | 0.03514 | 0.13139 | 11.778 |
| Nelfcd | 1879.74 | -0.15836 | 0.09011 | -1.75742 | 0.07885 | 0.22266 | 11.699 |
| Nelfe | 642.637 | 0.024744 | 0.0977 | 0.253265 | 0.80006 | 0.89676 | 5.982 |
| Nell1 | 2176.83 | -0.58498 | 0.1958 | -2.98761 | 0.00281 | 0.02514 | 891.745 |
| Nell2 | 17618.9 | -0.27148 | 0.14342 | -1.89292 | 0.05837 | 0.18369 | 453.33 |
| Nemf | 4155.38 | 0.131531 | 0.13101 | 1.003952 | 0.3154 | 0.52594 | 46.644 |
| Nemp1 | 499.268 | 0.266496 | 0.10468 | 2.545928 | 0.0109 | 0.06082 | 34.149 |
| Nemp2 | 402.137 | -0.5355 | 0.16732 | -3.20053 | 0.00137 | 0.01554 | 21.217 |
| Nenf | 887.484 | -0.33803 | 0.08048 | -4.20015 | 2.67E-05 | 0.00107 | 11.406 |
| Neo1 | 5677.48 | -0.01623 | 0.0601 | -0.27002 | 0.78714 | 0.88931 | 161.755 |
| Nepro | 401.668 | 0.132778 | 0.10299 | 1.289231 | 0.19732 | 0.3952 | 12.984 |
| Nes | 643.35 | 0.080195 | 0.11611 | 0.690707 | 0.48975 | 0.68476 | 9.374 |
| Net1 | 1868.59 | -0.17482 | 0.1297 | -1.34782 | 0.17772 | 0.36999 | 36.203 |
| Neto1 | 10009.1 | 0.399743 | 0.13373 | 2.989113 | 0.0028 | 0.02511 | 111.767 |
| Neto2 | 4161.55 | 0.320423 | 0.10288 | 3.114593 | 0.00184 | 0.01906 | 64.337 |
| Neu1 | 1996.1 | -0.1336 | 0.06871 | -1.94442 | 0.05184 | 0.17001 | 4.701 |
| Neu2 | 157.34 | 0.003787 | 0.2184 | 0.017338 | 0.98617 | 0.9925 | 87.957 |
| Neu3 | 169.516 | -0.20095 | 0.17345 | -1.15855 | 0.24664 | 0.45346 | 16.979 |
| Neu4 | 587.447 | -0.28536 | 0.10616 | -2.68808 | 0.00719 | 0.04631 | 7.884 |
| Neurl1a | 7482.51 | -0.11779 | 0.0948 | -1.24247 | 0.21406 | 0.41549 | 80.622 |
| Neurl1b | 2778.26 | -0.02554 | 0.10676 | -0.23929 | 0.81088 | 0.90259 | 31.521 |
| Neurl2 | 50.8483 | -0.12978 | 0.27229 | -0.47661 | 0.63364 | 0.79376 | 2.725 |
| Neurl3 | 31.9065 | -0.87086 | 0.40519 | -2.14924 | 0.03162 | 0.12249 | 10.083 |
| Neurl4 | 6404.79 | 0.017831 | 0.06483 | 0.275062 | 0.78327 | 0.88742 | 12.749 |
| Neurod1 | 144.816 | -1.39673 | 0.63506 | -2.19936 | 0.02785 | 0.1122 | 4.231 |
| Neurod2 | 1661.88 | -1.2329 | 0.49302 | -2.50072 | 0.01239 | 0.06605 | 4.234 |
| Neurod6 | 565.741 | -0.77804 | 0.47585 | -1.63506 | 0.10204 | 0.26371 | 3.442 |
| Nexmif | 2310.68 | 0.120963 | 0.13143 | 0.920388 | 0.35737 | 0.56765 | 123.752 |

| | | | | | | | |
|---|---|---|---|---|---|---|---|
| Nexn | 1084.71 | 0.704246 | 0.53668 | 1.312223 | 0.18944 | 0.38536 | 29.369 |
| Nf1 | 6631.22 | 0.053058 | 0.06886 | 0.770505 | 0.441 | 0.64569 | 241.92 |
| Nf2 | 6720.26 | 0.033017 | 0.07067 | 0.467209 | 0.64035 | 0.79859 | 83.692 |
| Nfam1 | 241.355 | 0.105134 | 0.1547 | 0.679588 | 0.49677 | 0.69035 | 40.334 |
| Nfasc | 18479.8 | -0.13197 | 0.08002 | -1.64928 | 0.09909 | 0.25857 | 177.108 |
| Nfat5 | 7809.49 | 0.14374 | 0.08581 | 1.675095 | 0.09392 | 0.24995 | 86.048 |
| Nfatc1 | 279.675 | -0.43379 | 0.12581 | -3.44786 | 0.00057 | 0.00861 | 106.867 |
| Nfatc2 | 1878.61 | 0.501546 | 0.19207 | 2.611214 | 0.00902 | 0.05351 | 125.248 |
| Nfatc2ip | 335.252 | -0.12936 | 0.12762 | -1.01367 | 0.31074 | 0.5209 | 13.884 |
| Nfatc3 | 711.816 | -0.17334 | 0.09746 | -1.77853 | 0.07532 | 0.21648 | 71.698 |
| Nfatc4 | 84.6649 | -0.7564 | 0.26507 | -2.85355 | 0.00432 | 0.03317 | 10.8 |
| Nfe2l1 | 16336.7 | -0.14994 | 0.0591 | -2.53702 | 0.01118 | 0.06169 | 12.555 |
| Nfe2l2 | 1010.08 | -0.38416 | 0.14031 | -2.738 | 0.00618 | 0.04207 | 29.129 |
| Nfe2l3 | 241.703 | -0.14252 | 0.17366 | -0.82073 | 0.4118 | 0.61976 | 26.099 |
| Nfia | 2214.52 | -0.5704 | 0.28519 | -2.00006 | 0.04549 | 0.15526 | 346.141 |
| Nfib | 3904.27 | -0.49339 | 0.28773 | -1.71476 | 0.08639 | 0.2364 | 415.578 |
| Nfic | 3277.37 | -0.42204 | 0.08658 | -4.87441 | 1.09E-06 | 0.00011 | 59.45 |
| Nfil3 | 302.583 | -0.3639 | 0.16069 | -2.26454 | 0.02354 | 0.09962 | 13.865 |
| Nfix | 3384.19 | -0.66426 | 0.16779 | -3.959 | 7.53E-05 | 0.0022 | 100.469 |
| Nfkb1 | 1154.98 | 0.123804 | 0.09455 | 1.309453 | 0.19038 | 0.3866 | 106.893 |
| Nfkb2 | 133.561 | -0.49175 | 0.17371 | -2.83087 | 0.00464 | 0.03467 | 8.066 |
| Nfkbia | 1098.09 | -0.16103 | 0.1492 | -1.07929 | 0.28046 | 0.4903 | 3.238 |
| Nfkbib | 602.3 | -0.22092 | 0.10225 | -2.16066 | 0.03072 | 0.11999 | 9.262 |
| Nfkbid | 171.168 | 0.169227 | 0.16809 | 1.006765 | 0.31405 | 0.52455 | 7.015 |
| Nfkbie | 59.984 | -0.33497 | 0.23513 | -1.42463 | 0.15427 | 0.34017 | 7.467 |
| Nfkbil1 | 129.955 | -0.09155 | 0.17385 | -0.52664 | 0.59844 | 0.76969 | 15.618 |
| Nfkbiz | 454.563 | -0.37896 | 0.17526 | -2.16225 | 0.0306 | 0.11972 | 27.525 |
| Nfrkb | 2380.56 | 0.216696 | 0.07969 | 2.719103 | 0.00655 | 0.04362 | 35.142 |
| Nfs1 | 1152.21 | 0.07628 | 0.08429 | 0.904967 | 0.36548 | 0.57607 | 20.548 |
| Nfu1 | 1435.83 | -0.03005 | 0.09697 | -0.3099 | 0.75664 | 0.87187 | 19.226 |
| Nfx1 | 6646.46 | 0.33494 | 0.10128 | 3.307192 | 0.00094 | 0.01228 | 55.088 |
| Nfxl1 | 823.333 | 0.14316 | 0.11984 | 1.194585 | 0.23225 | 0.43719 | 46.384 |
| Nfya | 1526.51 | -0.00687 | 0.10706 | -0.06415 | 0.94885 | 0.97462 | 23.022 |
| Nfyb | 2567.4 | 0.097626 | 0.10723 | 0.910439 | 0.36259 | 0.57294 | 15.444 |

| | | | | | | | |
|---|---|---|---|---|---|---|---|
| Nfyc | 1703.02 | 0.018467 | 0.08559 | 0.215768 | 0.82917 | 0.91246 | 74.135 |
| Ngb | 112.537 | 0.722698 | 0.31607 | 2.286539 | 0.02222 | 0.0961 | 5.009 |
| Ngdn | 650.714 | -0.05232 | 0.09304 | -0.56228 | 0.57392 | 0.75244 | 8.684 |
| Ngef | 13322.2 | 0.384056 | 0.13518 | 2.841043 | 0.0045 | 0.03392 | 97.037 |
| Ngf | 88.6866 | -1.03755 | 0.22442 | -4.62322 | 3.78E-06 | 0.00028 | 51.095 |
| Ngfr | 153.331 | 0.485558 | 0.71229 | 0.681684 | 0.49544 | 0.6894 | 18.918 |
| Ngly1 | 1107.86 | 0.044674 | 0.09361 | 0.477251 | 0.63318 | 0.79329 | 62.647 |
| Ngrn | 1370.99 | 0.040669 | 0.09089 | 0.447463 | 0.65454 | 0.80832 | 4.16 |
| Nhej1 | 116.541 | -0.26842 | 0.16383 | -1.63838 | 0.10134 | 0.26262 | 95.547 |
| Nhlh2 | 27.2956 | -0.30829 | 0.36886 | -0.8358 | 0.40327 | 0.61194 | 5.417 |
| Nhlrc1 | 715.396 | 0.06189 | 0.08815 | 0.702129 | 0.4826 | 0.67906 | 2.294 |
| Nhlrc2 | 1804.09 | 0.050163 | 0.08377 | 0.598816 | 0.5493 | 0.73299 | 55.243 |
| Nhlrc3 | 786.547 | -0.01917 | 0.08111 | -0.23637 | 0.81314 | 0.90382 | 14.75 |
| Nhp2 | 1063.1 | -0.20186 | 0.08711 | -2.31737 | 0.02048 | 0.09101 | 3.98 |
| Nhs | 801.83 | 0.615446 | 0.16986 | 3.62315 | 0.00029 | 0.0055 | 326.435 |
| Nhsl1 | 1167.94 | -0.57181 | 0.13492 | -4.23816 | 2.25E-05 | 0.00095 | 214.908 |
| Nhsl2 | 6805.98 | 0.439016 | 0.12326 | 3.561829 | 0.00037 | 0.00646 | 242.671 |
| Niban1 | 102.369 | -1.06779 | 0.2501 | -4.26945 | 1.96E-05 | 0.00087 | 150.754 |
| Niban2 | 1277.64 | -0.54224 | 0.10425 | -5.20144 | 1.98E-07 | 3.01E-05 | 49.141 |
| Niban3 | 242.96 | 0.593165 | 0.17639 | 3.362749 | 0.00077 | 0.01071 | 10.289 |
| Nicn1 | 3620.64 | -0.03219 | 0.06118 | -0.52612 | 0.5988 | 0.76992 | 6.07 |
| Nid1 | 409.487 | -0.51744 | 0.20833 | -2.48382 | 0.013 | 0.0682 | 74.719 |
| Nid2 | 113.814 | -0.21884 | 0.21133 | -1.03553 | 0.30042 | 0.51048 | 60.523 |
| Nif3l1 | 875.7 | 0.08295 | 0.09995 | 0.829869 | 0.40661 | 0.6148 | 36.666 |
| Nifk | 779.795 | 0.082826 | 0.10794 | 0.767358 | 0.44287 | 0.64738 | 11.984 |
| Nim1k | 1021.47 | -0.14599 | 0.09638 | -1.51476 | 0.12983 | 0.30763 | 45.789 |
| Nin | 5506.39 | 0.325554 | 0.10987 | 2.963188 | 0.00304 | 0.02634 | 102.283 |
| Ninj1 | 560.335 | -0.03517 | 0.12704 | -0.27688 | 0.78187 | 0.88646 | 8.76 |
| Ninj2 | 43.3372 | -0.60884 | 0.34041 | -1.78853 | 0.07369 | 0.21385 | 106.99 |
| Ninl | 328.215 | -0.21876 | 0.16493 | -1.3264 | 0.18471 | 0.37935 | 104.864 |
| Nip7 | 1229.62 | -0.02118 | 0.09858 | -0.21484 | 0.82989 | 0.9128 | 4.055 |
| Nipa1 | 2618.91 | -0.07836 | 0.06466 | -1.21197 | 0.22552 | 0.42955 | 42.388 |
| Nipa2 | 1534.29 | -0.00282 | 0.07515 | -0.0375 | 0.97009 | 0.98478 | 31.19 |
| Nipal2 | 525.205 | 0.564178 | 0.1226 | 4.60183 | 4.19E-06 | 0.0003 | 106.414 |

| | | | | | | | |
|---|---|---|---|---|---|---|---|
| Nipal3 | 5952.33 | -0.04626 | 0.05457 | -0.84773 | 0.39659 | 0.60559 | 49.619 |
| Nipal4 | 144.215 | -0.92026 | 0.28607 | -3.21684 | 0.0013 | 0.01495 | 18.354 |
| Nipbl | 3998.84 | 0.249174 | 0.18554 | 1.342979 | 0.17928 | 0.37214 | 153.847 |
| Nipsnap1 | 2585.36 | 0.041424 | 0.07971 | 0.519708 | 0.60327 | 0.77281 | 20.25 |
| Nipsnap2 | 4138.46 | 0.056956 | 0.07196 | 0.791532 | 0.42863 | 0.63525 | 33.265 |
| Nipsnap3b | 328.19 | -0.18376 | 0.11103 | -1.65504 | 0.09792 | 0.25674 | 10.181 |
| Nisch | 23471.9 | 0.113991 | 0.06523 | 1.747421 | 0.08056 | 0.22592 | 46.017 |
| Nit1 | 1267.93 | 0.206527 | 0.0764 | 2.703396 | 0.00686 | 0.04486 | 7.639 |
| Nit2 | 467.34 | 0.32537 | 0.14869 | 2.188271 | 0.02865 | 0.11426 | 10.677 |
| Nkain1 | 881.355 | -0.5761 | 0.14355 | -4.01316 | 5.99E-05 | 0.00188 | 42.448 |
| Nkain2 | 3972.16 | 0.088516 | 0.06805 | 1.300739 | 0.19335 | 0.39022 | 1201.153 |
| Nkain3 | 339.663 | 0.325703 | 0.129 | 2.52492 | 0.01157 | 0.06307 | 659.993 |
| Nkain4 | 721.869 | -0.41693 | 0.10798 | -3.86124 | 0.00011 | 0.00291 | 19.928 |
| Nkap | 1095.46 | 0.494648 | 0.20122 | 2.458275 | 0.01396 | 0.07114 | 23.952 |
| Nkapd1 | 1091.35 | -0.03531 | 0.07579 | -0.46589 | 0.64129 | 0.79917 | 12.225 |
| Nkd1 | 361.663 | -0.44453 | 0.13377 | -3.32313 | 0.00089 | 0.0118 | 73.531 |
| Nkd2 | 124.587 | -0.3845 | 0.20909 | -1.83892 | 0.06593 | 0.19835 | 29.098 |
| Nkiras1 | 2400.92 | -0.1695 | 0.08271 | -2.04943 | 0.04042 | 0.14368 | 12.868 |
| Nkiras2 | 381.042 | -0.0496 | 0.14015 | -0.35395 | 0.72338 | 0.85232 | 8.364 |
| Nkpd1 | 37.6458 | 0.380661 | 0.37085 | 1.026449 | 0.30468 | 0.51453 | 7.219 |
| Nkrf | 1395.69 | -0.02111 | 0.11677 | -0.1808 | 0.85652 | 0.92765 | 15.974 |
| Nktr | 9707.64 | 0.060051 | 0.0705 | 0.851804 | 0.39432 | 0.60344 | 37.675 |
| Nkx2-1 | 134.458 | 1.167725 | 0.34115 | 3.422921 | 0.00062 | 0.00918 | 4.951 |
| Nkx2-2 | 112.377 | -0.49377 | 0.20062 | -2.46117 | 0.01385 | 0.07073 | 16.698 |
| Nkx2-2os | 22.0711 | -0.50307 | 0.34942 | -1.43971 | 0.14995 | 0.33445 | 147.599 |
| Nkx3-1 | 26.6315 | -0.47056 | 0.50582 | -0.93029 | 0.35222 | 0.56254 | 4.025 |
| Nkx6-2 | 269.032 | -0.54289 | 0.21368 | -2.5406 | 0.01107 | 0.06133 | 3.422 |
| Nle1 | 577.002 | 0.093448 | 0.09539 | 0.979675 | 0.32725 | 0.5386 | 7.644 |
| Nlgn1 | 2995.12 | 0.298637 | 0.13392 | 2.230044 | 0.02574 | 0.10612 | 906.231 |
| Nlgn2 | 12530.2 | 0.00694 | 0.05745 | 0.120803 | 0.90385 | 0.95146 | 14.663 |
| Nlgn3 | 2599.1 | 0.03001 | 0.07293 | 0.411503 | 0.6807 | 0.82518 | 26.796 |
| Nlk | 5093.94 | -0.18061 | 0.07152 | -2.52516 | 0.01156 | 0.06306 | 130.206 |
| Nln | 1627.45 | 0.003695 | 0.07161 | 0.051597 | 0.95885 | 0.97995 | 86.558 |
| Nlrc3 | 147.901 | 0.024351 | 0.19948 | 0.12207 | 0.90284 | 0.95118 | 31.626 |

| | | | | | | | |
|---|---|---|---|---|---|---|---|
| Nlrc4 | 38.6372 | -0.01694 | 0.29687 | -0.05707 | 0.95449 | 0.97761 | 33.705 |
| Nlrc5 | 83.679 | -0.97678 | 0.34485 | -2.83245 | 0.00462 | 0.03454 | 92.917 |
| Nlrp10 | 120.331 | 0.794976 | 0.37955 | 2.094535 | 0.03621 | 0.13401 | 8.327 |
| Nlrp1b | 26.3109 | -0.21769 | 0.38552 | -0.56467 | 0.5723 | 0.75105 | 77.632 |
| Nlrp1c-ps | 17.0618 | -0.69414 | 0.39567 | -1.75437 | 0.07937 | 0.22367 | 42.89 |
| Nlrp3 | 47.9264 | -0.01371 | 0.27457 | -0.04995 | 0.96017 | 0.98055 | 25.389 |
| Nlrp5-ps | 28.4099 | -0.45894 | 0.44062 | -1.04158 | 0.29761 | 0.50773 | 91.828 |
| Nlrp6 | 62.1765 | 0.631166 | 0.27494 | 2.295669 | 0.02169 | 0.09451 | 8.291 |
| Nlrx1 | 302.672 | -0.12994 | 0.18724 | -0.69398 | 0.4877 | 0.68296 | 15.883 |
| Nmb | 79.7011 | 0.082425 | 0.19246 | 0.428265 | 0.66846 | 0.81714 | 2.85 |
| Nmbr | 816.826 | -1.08654 | 0.50313 | -2.15957 | 0.03081 | 0.12025 | 65.26 |
| Nmd3 | 1255.76 | 0.15991 | 0.09772 | 1.636356 | 0.10177 | 0.26325 | 34.389 |
| Nme1 | 1069.03 | -0.3729 | 0.08734 | -4.26966 | 1.96E-05 | 0.00087 | 11.543 |
| Nme2 | 27.8625 | -0.26614 | 0.35282 | -0.75433 | 0.45065 | 0.65326 | 6.446 |
| Nme4 | 96.1458 | 0.181259 | 0.22994 | 0.788288 | 0.43053 | 0.63686 | 3.869 |
| Nme5 | 292.395 | 0.085967 | 0.13621 | 0.63113 | 0.52796 | 0.71583 | 16.482 |
| Nme6 | 195.882 | 0.23665 | 0.15471 | 1.529642 | 0.12611 | 0.3018 | 10.368 |
| Nme7 | 681.807 | 0.11564 | 0.10506 | 1.100704 | 0.27103 | 0.48038 | 133.605 |
| Nmi | 123.678 | -0.30847 | 0.16811 | -1.83491 | 0.06652 | 0.19961 | 25.008 |
| Nmnat1 | 319.247 | -0.12586 | 0.1104 | -1.14002 | 0.25428 | 0.46167 | 17.631 |
| Nmnat2 | 6646.9 | -0.06679 | 0.05502 | -1.21385 | 0.22481 | 0.42855 | 164.264 |
| Nmnat3 | 239.67 | 0.215253 | 0.24417 | 0.88157 | 0.37801 | 0.58806 | 133.004 |
| Nmral1 | 684.808 | -0.09689 | 0.11381 | -0.85137 | 0.39456 | 0.60364 | 9.298 |
| Nmrk1 | 227.983 | -0.57259 | 0.15515 | -3.69051 | 0.00022 | 0.00459 | 20.479 |
| Nmrk2 | 17.5091 | 1.296346 | 0.46499 | 2.78791 | 0.0053 | 0.03768 | 3.882 |
| Nmt1 | 8262.89 | -0.05562 | 0.07706 | -0.72182 | 0.4704 | 0.66923 | 40.723 |
| Nmt2 | 5425 | -0.22763 | 0.11642 | -1.95523 | 0.05056 | 0.16723 | 44.666 |
| Nmur2 | 12.8977 | -0.15 | 0.4981 | -0.30115 | 0.7633 | 0.876 | 16.024 |
| Nnat | 5742.78 | 0.156867 | 0.1498 | 1.047174 | 0.29502 | 0.50544 | 2.445 |
| Nnmt | 23.02 | -1.09247 | 0.64592 | -1.69135 | 0.09077 | 0.24468 | 13.277 |
| Nnt | 35.7559 | 0.236193 | 0.33003 | 0.715668 | 0.4742 | 0.67238 | 73.55 |
| Noa1 | 723.655 | 0.13186 | 0.09755 | 1.351762 | 0.17645 | 0.36846 | 15.903 |
| Nob1 | 1373.71 | 0.090724 | 0.08161 | 1.111614 | 0.2663 | 0.47559 | 12.566 |
| Noc2l | 3573.5 | 0.230527 | 0.07569 | 3.045487 | 0.00232 | 0.0222 | 11.698 |

| | | | | | | | |
|---|---|---|---|---|---|---|---|
| Noc3l | 1008.12 | 0.089808 | 0.07988 | 1.124345 | 0.26087 | 0.46941 | 31.11 |
| Noc4l | 663.586 | -0.03982 | 0.0972 | -0.40968 | 0.68204 | 0.82631 | 5 |
| Noct | 2906.88 | 0.000741 | 0.07276 | 0.010184 | 0.99187 | 0.99537 | 27.198 |
| Nod1 | 330.808 | -0.5278 | 0.14693 | -3.59228 | 0.00033 | 0.006 | 48.664 |
| Nod2 | 188.218 | 0.81027 | 0.24069 | 3.366453 | 0.00076 | 0.01062 | 41.16 |
| Nog | 233.45 | -0.67811 | 0.20836 | -3.25449 | 0.00114 | 0.01382 | 1.695 |
| Nol10 | 1001.72 | 0.061412 | 0.08604 | 0.713736 | 0.47539 | 0.6735 | 81.638 |
| Nol11 | 1338.38 | 0.17648 | 0.07372 | 2.39396 | 0.01667 | 0.0799 | 22.719 |
| Nol12 | 756.499 | -0.07339 | 0.08025 | -0.91456 | 0.36042 | 0.57074 | 8.706 |
| Nol3 | 584.29 | -0.3061 | 0.1179 | -2.59635 | 0.00942 | 0.05521 | 5.8 |
| Nol4 | 3227.88 | 0.019132 | 0.1076 | 0.1778 | 0.85888 | 0.92877 | 348.473 |
| Nol4l | 2189.36 | -0.04406 | 0.07245 | -0.60815 | 0.54309 | 0.72864 | 122.51 |
| Nol6 | 5562.9 | 0.136477 | 0.07325 | 1.863271 | 0.06242 | 0.19158 | 10.029 |
| Nol7 | 1696.46 | 0.177208 | 0.09115 | 1.944155 | 0.05188 | 0.17004 | 8.708 |
| Nol8 | 1431.4 | 0.34048 | 0.19305 | 1.763727 | 0.07778 | 0.22069 | 25.939 |
| Nol9 | 1133.81 | -0.00546 | 0.07946 | -0.06875 | 0.94519 | 0.97272 | 22.174 |
| Nolc1 | 3243.84 | 0.066716 | 0.05889 | 1.132877 | 0.25727 | 0.46522 | 9.668 |
| Nom1 | 1534.18 | 0.038539 | 0.07178 | 0.536926 | 0.59132 | 0.76394 | 23.18 |
| Nomo1 | 6189.09 | 0.036658 | 0.06245 | 0.587033 | 0.55718 | 0.73902 | 50.515 |
| Nono | 6780.06 | 0.047285 | 0.06101 | 0.775072 | 0.4383 | 0.64363 | 19.274 |
| Nop10 | 958.499 | 0.014962 | 0.08658 | 0.172805 | 0.8628 | 0.9306 | 1.344 |
| Nop14 | 863.199 | 0.06835 | 0.08855 | 0.771889 | 0.44018 | 0.64519 | 21.613 |
| Nop16 | 676.609 | 0.031691 | 0.13057 | 0.242702 | 0.80824 | 0.90141 | 5.906 |
| Nop2 | 1154.95 | 0.047307 | 0.08405 | 0.562841 | 0.57354 | 0.75223 | 12.845 |
| Nop53 | 3058.79 | -0.06979 | 0.07855 | -0.88849 | 0.37427 | 0.58452 | 9.892 |
| Nop56 | 3489.71 | 0.098755 | 0.08292 | 1.191015 | 0.23365 | 0.43864 | 4.884 |
| Nop58 | 1981.12 | 0.138259 | 0.08613 | 1.605315 | 0.10842 | 0.27367 | 34.074 |
| Nop9 | 530.127 | -0.04053 | 0.1155 | -0.35091 | 0.72566 | 0.85388 | 9.808 |
| Nopchap1 | 3277.61 | -0.02021 | 0.08681 | -0.23277 | 0.81594 | 0.90534 | 128.285 |
| Norad | 76560.7 | -0.045 | 0.05766 | -0.78039 | 0.43516 | 0.64096 | 4.942 |
| Nos1 | 4047.3 | 0.215277 | 0.08105 | 2.656104 | 0.0079 | 0.04926 | 177.809 |
| Nos1ap | 4183.1 | 0.077128 | 0.09033 | 0.853888 | 0.39317 | 0.60231 | 287.194 |
| Nos2 | 17.7229 | -0.81733 | 0.52475 | -1.55756 | 0.11934 | 0.29146 | 39.468 |
| Nos3 | 211.589 | -0.00498 | 0.14269 | -0.03489 | 0.97216 | 0.98602 | 19.665 |

| | | | | | | | |
|---|---|---|---|---|---|---|---|
| Nosip | 1212.18 | -0.04747 | 0.07928 | -0.59872 | 0.54936 | 0.73299 | 15.782 |
| Nostrin | 278.002 | -0.21618 | 0.18057 | -1.19716 | 0.23124 | 0.4358 | 53.531 |
| Notch1 | 1490.83 | -0.29505 | 0.10753 | -2.74388 | 0.00607 | 0.04164 | 58.761 |
| Notch2 | 1393.56 | -0.36217 | 0.11901 | -3.04324 | 0.00234 | 0.02231 | 136.835 |
| Notch3 | 1621.77 | 0.506683 | 0.08647 | 5.859529 | 4.64E-09 | 1.80E-06 | 46.033 |
| Notch4 | 747.63 | -0.17836 | 0.11561 | -1.54273 | 0.1229 | 0.29692 | 24.236 |
| Notum | 100.406 | -0.22837 | 0.2209 | -1.03386 | 0.3012 | 0.51092 | 7.388 |
| Notumos | 22.6812 | -0.22613 | 0.39314 | -0.57519 | 0.56516 | 0.74541 | 2.659 |
| Nova1 | 3399.16 | 0.406204 | 0.11588 | 3.505302 | 0.00046 | 0.00747 | 124.413 |
| Nova2 | 5461.16 | 0.003989 | 0.06169 | 0.064661 | 0.94844 | 0.97435 | 39.432 |
| Nox1 | 20.9457 | 0.237847 | 0.37508 | 0.63413 | 0.526 | 0.71417 | 135.536 |
| Noxo1 | 40.8602 | -0.25307 | 0.34053 | -0.74316 | 0.45738 | 0.65866 | 4.296 |
| Noxred1 | 171.644 | 0.184643 | 0.17669 | 1.045014 | 0.29602 | 0.50634 | 17.693 |
| Npas1 | 248.897 | -0.56113 | 0.18496 | -3.03379 | 0.00242 | 0.02279 | 22.008 |
| Npas2 | 3181.83 | 0.55558 | 0.15377 | 3.613074 | 0.0003 | 0.00566 | 169.506 |
| Npas3 | 451.134 | 0.080599 | 0.11603 | 0.694638 | 0.48728 | 0.68271 | 824.019 |
| Npas4 | 896.397 | -0.10756 | 0.20615 | -0.52175 | 0.60185 | 0.77182 | 5.962 |
| Npat | 1631.01 | 0.173352 | 0.11103 | 1.561337 | 0.11844 | 0.29019 | 37.296 |
| Npbwr1 | 71.9004 | -0.14133 | 0.28173 | -0.50165 | 0.61591 | 0.78132 | 3.692 |
| Npc1 | 2830.35 | -0.18923 | 0.09581 | -1.97516 | 0.04825 | 0.16192 | 46.709 |
| Npc1l1 | 76.0835 | 0.30902 | 0.26046 | 1.18646 | 0.23544 | 0.44058 | 19.131 |
| Npc2 | 2694.98 | -0.35574 | 0.09494 | -3.74697 | 0.00018 | 0.004 | 18.591 |
| Npdc1 | 5056.72 | -0.13953 | 0.07853 | -1.77665 | 0.07563 | 0.21709 | 10.144 |
| Npepl1 | 593.706 | -0.18731 | 0.10831 | -1.72942 | 0.08373 | 0.23153 | 12.722 |
| Npepps | 8630.04 | -0.00913 | 0.05678 | -0.16083 | 0.87223 | 0.93547 | 74.797 |
| Npff | 60.7162 | -0.04683 | 0.24705 | -0.18956 | 0.84966 | 0.92352 | 1.104 |
| Npffr1 | 53.6873 | 0.029047 | 0.30222 | 0.096112 | 0.92343 | 0.9614 | 33.074 |
| Npffr2 | 30.1096 | 0.241772 | 0.41125 | 0.587891 | 0.55661 | 0.73867 | 56.312 |
| Nphp1 | 840.327 | 0.162855 | 0.10346 | 1.574051 | 0.11548 | 0.28501 | 48.166 |
| Nphp3 | 452.482 | -0.16042 | 0.11732 | -1.36742 | 0.17149 | 0.36185 | 41.275 |
| Nphp4 | 1534.96 | 0.243315 | 0.0922 | 2.638941 | 0.00832 | 0.05082 | 86.478 |
| Nphs1 | 141.805 | 0.396977 | 0.19364 | 2.050076 | 0.04036 | 0.14359 | 28.909 |
| Nphs2 | 36.3302 | -0.75921 | 0.50829 | -1.49364 | 0.13527 | 0.31507 | 17.309 |
| Npl | 239.14 | 0.264123 | 0.16241 | 1.626321 | 0.10388 | 0.26639 | 47.031 |

| | | | | | | | |
|---|---|---|---|---|---|---|---|
| Nploc4 | 3112.68 | -0.13306 | 0.06769 | -1.96565 | 0.04934 | 0.16449 | 57.339 |
| Npm1 | 5365.79 | 0.207312 | 0.09385 | 2.208998 | 0.02717 | 0.11013 | 10.92 |
| Npm2 | 32.1222 | -0.03967 | 0.30744 | -0.12903 | 0.89733 | 0.94857 | 11.943 |
| Npm3 | 175.811 | -0.02191 | 0.14422 | -0.1519 | 0.87927 | 0.93962 | 1.858 |
| Npnt | 563.14 | -0.99515 | 0.22928 | -4.3403 | 1.42E-05 | 0.00071 | 68.547 |
| Nppc | 157.449 | -1.07459 | 0.22369 | -4.80404 | 1.55E-06 | 0.00014 | 4.281 |
| Npr1 | 94.451 | -0.37884 | 0.22025 | -1.72008 | 0.08542 | 0.23453 | 15.276 |
| Npr2 | 2193.15 | -0.02064 | 0.09105 | -0.22669 | 0.82066 | 0.90738 | 19.31 |
| Npr3 | 356.548 | -0.64968 | 0.52071 | -1.24769 | 0.21214 | 0.41275 | 67.392 |
| Nprl2 | 1103.3 | 0.172017 | 0.078 | 2.205312 | 0.02743 | 0.11084 | 3.481 |
| Nprl3 | 1255.38 | 0.040647 | 0.0934 | 0.435194 | 0.66342 | 0.81401 | 42.08 |
| Npsr1 | 134.049 | -0.71607 | 0.28107 | -2.54766 | 0.01084 | 0.06059 | 218.403 |
| Nptn | 24568.4 | 0.057453 | 0.06482 | 0.886364 | 0.37542 | 0.5856 | 75.716 |
| Nptx1 | 8932.48 | -0.79869 | 0.3051 | -2.61775 | 0.00885 | 0.05291 | 9.035 |
| Nptx2 | 1145.84 | -0.63415 | 0.17703 | -3.58207 | 0.00034 | 0.00614 | 11.577 |
| Nptxr | 14336.1 | -1.08086 | 0.45338 | -2.38401 | 0.01713 | 0.08135 | 18.416 |
| Npy | 1651.44 | -0.43772 | 0.13154 | -3.32775 | 0.00088 | 0.01166 | 6.798 |
| Npy1r | 1577.99 | 0.097147 | 0.07601 | 1.278052 | 0.20123 | 0.39952 | 9.601 |
| Npy2r | 433.113 | 0.719489 | 0.21441 | 3.355619 | 0.00079 | 0.0109 | 9.702 |
| Npy5r | 493.316 | 0.459259 | 0.16336 | 2.811368 | 0.00493 | 0.0359 | 8.164 |
| Npy6r | 14.7893 | -0.24882 | 0.47976 | -0.51862 | 0.60402 | 0.7733 | 7.574 |
| Nqo1 | 188.809 | -0.0411 | 0.17 | -0.24177 | 0.80896 | 0.90168 | 14.982 |
| Nqo2 | 307.328 | -0.24588 | 0.14608 | -1.6832 | 0.09234 | 0.2474 | 23.757 |
| Nr1d1 | 2529.85 | 0.114447 | 0.08374 | 1.366709 | 0.17172 | 0.3622 | 7.402 |
| Nr1d2 | 8729.5 | 0.305156 | 0.07286 | 4.188117 | 2.81E-05 | 0.0011 | 35.074 |
| Nr1h2 | 1807.27 | -0.17813 | 0.0716 | -2.48794 | 0.01285 | 0.06768 | 4.336 |
| Nr1h3 | 203.781 | -0.20897 | 0.13979 | -1.49492 | 0.13493 | 0.31481 | 18.774 |
| Nr2c1 | 732.209 | 0.107894 | 0.11432 | 0.943771 | 0.34529 | 0.55616 | 49.189 |
| Nr2c2 | 3597.17 | 0.07846 | 0.06301 | 1.245259 | 0.21304 | 0.41418 | 82.905 |
| Nr2c2ap | 33.6677 | -0.65224 | 0.39815 | -1.63819 | 0.10138 | 0.26262 | 2.417 |
| Nr2e1 | 754.055 | -0.14482 | 0.13002 | -1.11387 | 0.26534 | 0.4745 | 21.67 |
| Nr2f1 | 1048.1 | -0.98655 | 0.34383 | -2.8693 | 0.00411 | 0.03204 | 10.785 |
| Nr2f2 | 344.412 | -0.84997 | 0.18024 | -4.7159 | 2.41E-06 | 0.0002 | 14.792 |
| Nr2f6 | 336.528 | 0.019696 | 0.12975 | 0.151793 | 0.87935 | 0.93962 | 7.838 |

| | | | | | | | |
|---|---|---|---|---|---|---|---|
| Nr3c1 | 4080.76 | -0.07034 | 0.08879 | -0.79218 | 0.42825 | 0.63491 | 108.877 |
| Nr3c2 | 2350.62 | 0.22291 | 0.09833 | 2.266904 | 0.0234 | 0.09923 | 345.572 |
| Nr4a1 | 3047.05 | -0.40256 | 0.157 | -2.56407 | 0.01035 | 0.05872 | 20.527 |
| Nr4a2 | 932.239 | -1.01482 | 0.39729 | -2.55434 | 0.01064 | 0.05988 | 17.174 |
| Nr4a3 | 1471.17 | -0.25474 | 0.3384 | -0.75277 | 0.45159 | 0.65396 | 41.295 |
| Nr6a1 | 622.029 | 0.063211 | 0.10454 | 0.604661 | 0.5454 | 0.73031 | 204.319 |
| Nradd | 103.997 | 0.563082 | 0.20993 | 2.682245 | 0.00731 | 0.04693 | 3.318 |
| Nrap | 82.4095 | -0.18492 | 0.24987 | -0.74006 | 0.45926 | 0.65991 | 70.003 |
| Nrarp | 75.3652 | 0.219411 | 0.31252 | 0.702068 | 0.48264 | 0.67906 | 2.582 |
| Nras | 2669.19 | -0.02909 | 0.07088 | -0.41038 | 0.68152 | 0.82588 | 9.63 |
| Nrbf2 | 541.296 | -0.06205 | 0.1117 | -0.55548 | 0.57857 | 0.75537 | 18.617 |
| Nrbp1 | 2660.89 | -0.03 | 0.06032 | -0.49726 | 0.61901 | 0.78314 | 10.703 |
| Nrbp2 | 4031.37 | -0.40177 | 0.11343 | -3.54205 | 0.0004 | 0.00684 | 6.11 |
| Nrcam | 13508.3 | -0.07629 | 0.10232 | -0.74566 | 0.45587 | 0.65758 | 273.08 |
| Nrd1 | 7808.48 | 0.103829 | 0.06988 | 1.485739 | 0.13735 | 0.318 | 61.123 |
| Nrde2 | 979.957 | 0.169162 | 0.07173 | 2.358359 | 0.01836 | 0.08475 | 34.202 |
| Nrep | 1535.9 | -0.10182 | 0.13348 | -0.76279 | 0.44559 | 0.64942 | 27.011 |
| Nrf1 | 1111.4 | 0.108027 | 0.0742 | 1.455872 | 0.14543 | 0.32825 | 105.471 |
| Nrg1 | 740.896 | 0.266211 | 0.10756 | 2.475074 | 0.01332 | 0.06927 | 1070.247 |
| Nrg2 | 767.743 | 0.229457 | 0.10841 | 2.116649 | 0.03429 | 0.12939 | 179.729 |
| Nrg3 | 1275.15 | -0.41133 | 0.16507 | -2.49181 | 0.01271 | 0.06724 | 1104.137 |
| Nrg3os | 26.044 | -0.41972 | 0.37144 | -1.13 | 0.25848 | 0.46651 | 60.493 |
| Nrg4 | 170.881 | 0.28668 | 0.1559 | 1.838928 | 0.06593 | 0.19835 | 106.623 |
| Nrgn | 3035.05 | -0.66974 | 0.13094 | -5.11475 | 3.14E-07 | 4.36E-05 | 8.413 |
| Nrip1 | 2207.33 | -0.12001 | 0.12069 | -0.99434 | 0.32006 | 0.53067 | 86.428 |
| Nrip2 | 573.295 | 0.172191 | 0.11985 | 1.436737 | 0.15079 | 0.33567 | 17.132 |
| Nrip3 | 3694.62 | -0.80097 | 0.19006 | -4.21439 | 2.50E-05 | 0.00103 | 23.971 |
| Nrn1 | 901.685 | -1.44383 | 0.47375 | -3.04766 | 0.00231 | 0.02207 | 9.771 |
| Nron | 27.7829 | -0.08752 | 0.35548 | -0.2462 | 0.80553 | 0.89972 | 7.776 |
| Nrp1 | 632.17 | 0.21029 | 0.1372 | 1.532708 | 0.12535 | 0.30036 | 144.76 |
| Nrp2 | 4253.18 | 0.291209 | 0.10582 | 2.752059 | 0.00592 | 0.04089 | 115.411 |
| Nrros | 336.003 | -0.15917 | 0.13119 | -1.2133 | 0.22502 | 0.42883 | 22.81 |
| Nrsn1 | 14406.2 | -0.04479 | 0.07431 | -0.60276 | 0.54667 | 0.73092 | 18.463 |
| Nrsn2 | 2706.12 | 0.537062 | 0.15939 | 3.369512 | 0.00075 | 0.01054 | 7.884 |

| | | | | | | | |
|---|---|---|---|---|---|---|---|
| Nrtn | 80.7062 | -0.4544 | 0.1899 | -2.39283 | 0.01672 | 0.07998 | 6.206 |
| Nrxn1 | 13658.3 | -0.16128 | 0.11308 | -1.42621 | 0.15381 | 0.3396 | 1059.441 |
| Nrxn2 | 14107.5 | -0.39376 | 0.10341 | -3.80781 | 0.00014 | 0.00337 | 125.439 |
| Nrxn3 | 12035.2 | 0.309852 | 0.13608 | 2.276941 | 0.02279 | 0.09764 | 1612.064 |
| Nsa2 | 1706.95 | -0.07353 | 0.08147 | -0.90247 | 0.36681 | 0.57719 | 8.514 |
| Nsa2-ps1 | 76.7524 | 0.077854 | 0.21778 | 0.357495 | 0.72072 | 0.85095 | 0.78 |
| Nscme3l | 197.181 | 0.167703 | 0.14756 | 1.136533 | 0.25573 | 0.46349 | 1.021 |
| Nsd1 | 9955.99 | 0.116024 | 0.06111 | 1.898697 | 0.0576 | 0.18222 | 108.544 |
| Nsd2 | 5884.16 | 0.068627 | 0.06597 | 1.040319 | 0.29819 | 0.50831 | 77.251 |
| Nsd3 | 6252.46 | 0.015894 | 0.05796 | 0.274243 | 0.7839 | 0.88762 | 118.078 |
| Nsdhl | 728.576 | -0.14503 | 0.09826 | -1.476 | 0.13994 | 0.3212 | 39.958 |
| Nsf | 32620.9 | -0.20044 | 0.08013 | -2.50149 | 0.01237 | 0.06597 | 132.275 |
| Nsfl1c | 3495.74 | -0.01298 | 0.08066 | -0.16093 | 0.87215 | 0.93543 | 17.233 |
| Nsg1 | 15616.1 | 0.294761 | 0.12193 | 2.417415 | 0.01563 | 0.07651 | 22.716 |
| Nsg2 | 40971.8 | 0.336452 | 0.10799 | 3.115641 | 0.00184 | 0.01906 | 58.74 |
| Nsl1 | 63.2405 | 0.255273 | 0.26622 | 0.958896 | 0.33761 | 0.5484 | 23.463 |
| Nsmaf | 2644.76 | 0.191953 | 0.08719 | 2.2016 | 0.02769 | 0.11172 | 58.065 |
| Nsmce1 | 363.608 | 0.04837 | 0.12923 | 0.374299 | 0.70818 | 0.84248 | 23.957 |
| Nsmce2 | 974.36 | -0.03436 | 0.08505 | -0.40393 | 0.68626 | 0.82887 | 227.438 |
| Nsmce3 | 885.96 | -0.02568 | 0.11379 | -0.2257 | 0.82143 | 0.90764 | 5.946 |
| Nsmce4a | 1225.47 | -0.06275 | 0.08234 | -0.76205 | 0.44603 | 0.64983 | 40.593 |
| Nsmf | 14077 | -0.58793 | 0.1475 | -3.986 | 6.72E-05 | 0.00204 | 8.527 |
| Nsrp1 | 926.829 | 0.226442 | 0.17141 | 1.321082 | 0.18647 | 0.38141 | 34.144 |
| Nsun2 | 2721.85 | 0.014781 | 0.06362 | 0.232312 | 0.8163 | 0.9054 | 102.035 |
| Nsun3 | 476.211 | -0.15921 | 0.12531 | -1.27054 | 0.20389 | 0.40253 | 54.375 |
| Nsun4 | 569.055 | 0.115116 | 0.09177 | 1.254388 | 0.2097 | 0.40977 | 21.035 |
| Nsun5 | 676.762 | -0.04889 | 0.10598 | -0.46131 | 0.64458 | 0.80175 | 6.853 |
| Nsun6 | 417.151 | -0.00983 | 0.13461 | -0.07304 | 0.94178 | 0.97095 | 59.939 |
| Nsun7 | 1390.18 | 0.292408 | 0.11437 | 2.556594 | 0.01057 | 0.05968 | 38.13 |
| Nt5c | 700.192 | -0.14082 | 0.10049 | -1.40136 | 0.16111 | 0.34925 | 1.443 |
| Nt5c1a | 146.653 | -0.27241 | 0.24396 | -1.11661 | 0.26416 | 0.47298 | 14.772 |
| Nt5c2 | 4606.95 | 0.05654 | 0.05898 | 0.958556 | 0.33778 | 0.54856 | 131.837 |
| Nt5c3 | 1352.71 | -0.08581 | 0.07604 | -1.1286 | 0.25907 | 0.46729 | 41.533 |
| Nt5c3b | 1577.86 | 0.019078 | 0.0699 | 0.272953 | 0.78489 | 0.88852 | 19.488 |

| | | | | | | | |
|---|---|---|---|---|---|---|---|
| Nt5dc1 | 153.614 | 0.169676 | 0.15568 | 1.089909 | 0.27575 | 0.48496 | 130.265 |
| Nt5dc2 | 229.915 | -0.5083 | 0.14848 | -3.42342 | 0.00062 | 0.00917 | 8.072 |
| Nt5dc3 | 4234.79 | -0.21636 | 0.06575 | -3.2905 | 0.001 | 0.01282 | 59.385 |
| Nt5e | 1500.58 | 0.73728 | 0.21431 | 3.440316 | 0.00058 | 0.00879 | 44.896 |
| Nt5m | 1008.55 | -0.11158 | 0.08506 | -1.31179 | 0.18959 | 0.38558 | 41.522 |
| Ntan1 | 1226.44 | -0.07958 | 0.07436 | -1.07022 | 0.28452 | 0.49456 | 16.607 |
| Ntf3 | 12.5078 | -1.41346 | 0.60789 | -2.32519 | 0.02006 | 0.08969 | 65.499 |
| Nthl1 | 198.048 | 0.151105 | 0.14108 | 1.071026 | 0.28416 | 0.4943 | 6.158 |
| Ntm | 6984.7 | -0.39979 | 0.12027 | -3.32405 | 0.00089 | 0.01177 | 968.392 |
| Ntmt1 | 861.991 | -0.07716 | 0.09987 | -0.77255 | 0.43979 | 0.64503 | 15.208 |
| Ntn1 | 2552.65 | 0.531352 | 0.19272 | 2.757093 | 0.00583 | 0.04045 | 191.46 |
| Ntn3 | 445.439 | 0.011254 | 0.14238 | 0.079043 | 0.937 | 0.9685 | 5.684 |
| Ntn4 | 482.674 | -0.44779 | 0.10735 | -4.17144 | 3.03E-05 | 0.00117 | 106.527 |
| Ntn5 | 37.1457 | -1.0749 | 0.58824 | -1.82731 | 0.06765 | 0.20161 | 10.535 |
| Ntng1 | 865.049 | -0.41314 | 0.18643 | -2.21604 | 0.02669 | 0.10891 | 363.972 |
| Ntng2 | 1354.9 | -0.48683 | 0.16567 | -2.93852 | 0.0033 | 0.02772 | 58.465 |
| Ntpcr | 589.725 | -0.04258 | 0.10566 | -0.40298 | 0.68697 | 0.82926 | 18.273 |
| Ntrk1 | 324.337 | 0.913906 | 0.39324 | 2.324016 | 0.02012 | 0.08986 | 16.919 |
| Ntrk2 | 38449.5 | -0.2047 | 0.09647 | -2.12189 | 0.03385 | 0.12843 | 327.402 |
| Ntrk3 | 11540.1 | 0.368525 | 0.09315 | 3.956328 | 7.61E-05 | 0.00221 | 562.054 |
| Nts | 1082.2 | 0.36195 | 0.30605 | 1.182663 | 0.23694 | 0.44205 | 8.731 |
| Ntsr1 | 211.809 | -0.57275 | 0.22262 | -2.5728 | 0.01009 | 0.05783 | 45.005 |
| Ntsr2 | 4944.64 | -0.16812 | 0.07409 | -2.26898 | 0.02327 | 0.0989 | 6.846 |
| Nuak1 | 2587.71 | -0.33511 | 0.14775 | -2.2681 | 0.02332 | 0.09902 | 69.693 |
| Nuak2 | 73.8606 | -0.41227 | 0.23232 | -1.77454 | 0.07597 | 0.21768 | 17.363 |
| Nub1 | 2949.93 | 0.108521 | 0.06853 | 1.583518 | 0.1133 | 0.28179 | 24.847 |
| Nubp1 | 464.699 | 0.008204 | 0.11332 | 0.072394 | 0.94229 | 0.97116 | 12.481 |
| Nubp2 | 778.921 | -0.25661 | 0.10082 | -2.54532 | 0.01092 | 0.06087 | 3.739 |
| Nubpl | 231.018 | 0.193826 | 0.13212 | 1.467019 | 0.14237 | 0.32433 | 215.008 |
| Nucb1 | 2834.3 | -0.10969 | 0.08126 | -1.34993 | 0.17704 | 0.3693 | 19.949 |
| Nucb2 | 776.563 | 0.263153 | 0.11531 | 2.282168 | 0.02248 | 0.09681 | 36.216 |
| Nucks1 | 7538.11 | 0.008581 | 0.06377 | 0.134571 | 0.89295 | 0.94632 | 25.788 |
| Nudc | 4536.29 | -0.19463 | 0.07482 | -2.60127 | 0.00929 | 0.05459 | 13.455 |
| Nudc-ps1 | 299.169 | -0.1559 | 0.14438 | -1.07981 | 0.28023 | 0.49019 | 0.551 |

| | | | | | | | |
|---|---|---|---|---|---|---|---|
| Nudcd1 | 838.605 | 0.070871 | 0.10217 | 0.693642 | 0.48791 | 0.68302 | 55.145 |
| Nudcd2 | 600.648 | 0.092721 | 0.10425 | 0.889412 | 0.37378 | 0.58398 | 6.38 |
| Nudcd3 | 6334.07 | -0.04691 | 0.05766 | -0.81357 | 0.41589 | 0.62362 | 94.725 |
| Nudt1 | 113.103 | -0.22495 | 0.22397 | -1.00436 | 0.31521 | 0.52568 | 16.482 |
| Nudt10 | 1014.01 | 0.190024 | 0.08315 | 2.285365 | 0.02229 | 0.09623 | 4.32 |
| Nudt11 | 1635.38 | 0.13648 | 0.09486 | 1.438824 | 0.1502 | 0.33479 | 7.658 |
| Nudt12 | 262.961 | -0.35909 | 0.18478 | -1.94331 | 0.05198 | 0.17024 | 13.755 |
| Nudt13 | 896.261 | 0.082397 | 0.09381 | 0.878348 | 0.37975 | 0.58952 | 22.882 |
| Nudt14 | 158.124 | 0.034164 | 0.16748 | 0.203995 | 0.83836 | 0.91681 | 7.834 |
| Nudt15 | 262.832 | 0.316301 | 0.18194 | 1.738488 | 0.08212 | 0.22844 | 29.366 |
| Nudt16 | 510.836 | 0.029276 | 0.11994 | 0.244079 | 0.80717 | 0.90071 | 2.922 |
| Nudt16l1 | 828.694 | 0.154669 | 0.09572 | 1.615849 | 0.10613 | 0.27007 | 2.272 |
| Nudt17 | 137.687 | -0.14367 | 0.16451 | -0.87331 | 0.3825 | 0.59199 | 2.496 |
| Nudt18 | 1089.89 | -0.21422 | 0.08562 | -2.50185 | 0.01235 | 0.06594 | 5.052 |
| Nudt19 | 1798.84 | -0.17588 | 0.07512 | -2.34132 | 0.01922 | 0.08706 | 9.12 |
| Nudt2 | 459.981 | 0.013985 | 0.11461 | 0.122024 | 0.90288 | 0.95118 | 15.776 |
| Nudt21 | 3855.75 | 0.053038 | 0.06352 | 0.834994 | 0.40372 | 0.61241 | 21.536 |
| Nudt22 | 199.32 | -0.30824 | 0.14667 | -2.10165 | 0.03558 | 0.13252 | 3.1 |
| Nudt3 | 7275.33 | -0.29647 | 0.07846 | -3.77857 | 0.00016 | 0.00367 | 44.114 |
| Nudt4 | 4807.77 | -0.47103 | 0.08907 | -5.28836 | 1.23E-07 | 2.11E-05 | 17.14 |
| Nudt5 | 789.806 | 0.013932 | 0.09722 | 0.143299 | 0.88605 | 0.94329 | 26.877 |
| Nudt6 | 83.6891 | -0.2782 | 0.23383 | -1.18975 | 0.23414 | 0.43924 | 15.302 |
| Nudt7 | 271.133 | 0.054929 | 0.16076 | 0.341675 | 0.7326 | 0.85819 | 21.183 |
| Nudt9 | 1418.83 | -0.19219 | 0.07636 | -2.51682 | 0.01184 | 0.06404 | 19.074 |
| Nuf2 | 80.1345 | -0.33217 | 0.31423 | -1.0571 | 0.29047 | 0.50076 | 33.531 |
| Nufip1 | 657.1 | -0.06362 | 0.09111 | -0.69822 | 0.48504 | 0.68125 | 26.489 |
| Nufip2 | 2967.25 | 0.069127 | 0.09409 | 0.734703 | 0.46252 | 0.66283 | 55.767 |
| Numa1 | 7554.54 | 0.131499 | 0.06167 | 2.132406 | 0.03297 | 0.12614 | 80.854 |
| Numb | 2486.75 | -0.08419 | 0.09407 | -0.89492 | 0.37083 | 0.58097 | 127.901 |
| Numbl | 4220.61 | 0.029529 | 0.06607 | 0.44693 | 0.65493 | 0.8085 | 23.712 |
| Nup107 | 883.136 | 0.161403 | 0.10223 | 1.578753 | 0.11439 | 0.28327 | 42.085 |
| Nup133 | 1418.91 | 0.019554 | 0.08358 | 0.233943 | 0.81503 | 0.90503 | 52.143 |
| Nup153 | 3119.93 | 0.129647 | 0.0686 | 1.889804 | 0.05878 | 0.18443 | 48.036 |
| Nup155 | 1807.65 | 0.07431 | 0.06602 | 1.125498 | 0.26038 | 0.46882 | 51.975 |

| | | | | | | | |
|---|---|---|---|---|---|---|---|
| Nup160 | 971.736 | -0.00868 | 0.08696 | -0.09979 | 0.92051 | 0.95978 | 59.114 |
| Nup188 | 1023.22 | 0.121693 | 0.08208 | 1.482694 | 0.13816 | 0.31929 | 57.87 |
| Nup205 | 2054.11 | 0.218081 | 0.09448 | 2.308183 | 0.02099 | 0.09251 | 70.176 |
| Nup210 | 2281.21 | 0.194933 | 0.06907 | 2.822155 | 0.00477 | 0.03524 | 103.762 |
| Nup210l | 46.6912 | -0.27077 | 0.33793 | -0.80126 | 0.42298 | 0.62985 | 107.917 |
| Nup214 | 1799.82 | 0.13776 | 0.08307 | 1.658442 | 0.09723 | 0.25553 | 79.54 |
| Nup35 | 742.039 | 0.16369 | 0.10557 | 1.550527 | 0.12102 | 0.29412 | 41.671 |
| Nup37 | 212.789 | -0.02623 | 0.14355 | -0.18275 | 0.85499 | 0.92668 | 31.399 |
| Nup43 | 200.708 | 0.106171 | 0.13477 | 0.787778 | 0.43083 | 0.63702 | 11.379 |
| Nup50 | 2353.05 | 0.075286 | 0.06364 | 1.183014 | 0.2368 | 0.44193 | 19.553 |
| Nup54 | 865.443 | 0.065886 | 0.08377 | 0.786488 | 0.43158 | 0.63765 | 19.68 |
| Nup62 | 868.419 | -0.15 | 0.08376 | -1.79088 | 0.07331 | 0.2132 | 14.725 |
| Nup85 | 1512.94 | 0.146872 | 0.06549 | 2.242587 | 0.02492 | 0.1037 | 19.552 |
| Nup88 | 3113.89 | 0.26571 | 0.092 | 2.888308 | 0.00387 | 0.03079 | 26.916 |
| Nup93 | 2949.12 | 0.348918 | 0.13591 | 2.567241 | 0.01025 | 0.05844 | 102.664 |
| Nup98 | 2417.74 | 0.112856 | 0.06935 | 1.627375 | 0.10366 | 0.26622 | 90.779 |
| Nupl2 | 443.097 | 0.183245 | 0.12999 | 1.409728 | 0.15862 | 0.34603 | 19.051 |
| Nupr1 | 189.256 | -0.73656 | 0.17745 | -4.15077 | 3.31E-05 | 0.00123 | 7.613 |
| Nupr1l | 39.3004 | -0.38979 | 0.26524 | -1.46961 | 0.14167 | 0.3235 | 5.25 |
| Nus1 | 7613.94 | 0.284944 | 0.07937 | 3.589938 | 0.00033 | 0.00603 | 22.637 |
| Nusap1 | 278.503 | 0.171346 | 0.14352 | 1.193893 | 0.23252 | 0.43737 | 32.947 |
| Nutf2 | 979.018 | -0.09306 | 0.08695 | -1.07026 | 0.2845 | 0.49456 | 18.751 |
| Nutf2-ps1 | 83.1867 | -0.06126 | 0.27141 | -0.22572 | 0.82142 | 0.90764 | 0.901 |
| Nutm1 | 12.4461 | -0.98863 | 0.6701 | -1.47534 | 0.14012 | 0.3215 | 11.344 |
| Nvl | 4710.06 | 0.062939 | 0.10359 | 0.607553 | 0.54348 | 0.72892 | 57.067 |
| Nwd1 | 6738.09 | -0.19808 | 0.08414 | -2.35403 | 0.01857 | 0.08539 | 71.166 |
| Nwd2 | 643.471 | -1.022 | 0.39804 | -2.5676 | 0.01024 | 0.05844 | 161.445 |
| Nxf1 | 5286.21 | 0.113135 | 0.07859 | 1.439568 | 0.14999 | 0.33447 | 13.85 |
| Nxf7 | 47.3309 | 0.185008 | 0.40765 | 0.453845 | 0.64994 | 0.80518 | 19.223 |
| Nxn | 751.182 | -0.13604 | 0.09892 | -1.37521 | 0.16907 | 0.35922 | 141.943 |
| Nxpe2 | 28.4164 | -0.36768 | 0.33851 | -1.0862 | 0.27739 | 0.48702 | 35.449 |
| Nxpe3 | 1146.21 | -0.24088 | 0.09153 | -2.6317 | 0.0085 | 0.05147 | 55.333 |
| Nxpe4 | 593.218 | 0.055398 | 0.10823 | 0.511838 | 0.60876 | 0.77635 | 238.005 |
| Nxph1 | 1322.82 | -0.17251 | 0.08757 | -1.96991 | 0.04885 | 0.16337 | 300.602 |

| | | | | | | | |
|---|---|---|---|---|---|---|---|
| Nxph2 | 277.103 | -0.43088 | 0.21222 | -2.03037 | 0.04232 | 0.14806 | 80.728 |
| Nxph3 | 581.904 | -0.83512 | 0.38025 | -2.19627 | 0.02807 | 0.1128 | 4.726 |
| Nxph4 | 174.531 | -0.73357 | 0.24411 | -3.00503 | 0.00266 | 0.0243 | 9.087 |
| Nxt1 | 289.792 | 0.02441 | 0.13113 | 0.186148 | 0.85233 | 0.92484 | 3.427 |
| Nxt2 | 1563.38 | 0.044503 | 0.08178 | 0.544173 | 0.58632 | 0.76033 | 12.923 |
| Nyap1 | 3711.66 | 0.010427 | 0.07954 | 0.131084 | 0.89571 | 0.94749 | 11.709 |
| Nyap2 | 2007.21 | 0.536088 | 0.1449 | 3.699612 | 0.00022 | 0.00449 | 264.813 |
| Nynrin | 644.761 | 0.020228 | 0.13183 | 0.153434 | 0.87806 | 0.93895 | 20.727 |
| Nyx | 57.5562 | 0.475378 | 0.25252 | 1.882532 | 0.05976 | 0.18622 | 23.204 |
| Oacyl | 112.111 | 0.048136 | 0.23605 | 0.203918 | 0.83842 | 0.91681 | 53.334 |
| Oaf | 676.103 | -0.40123 | 0.1077 | -3.72546 | 0.00019 | 0.00421 | 18.677 |
| Oard1 | 483.308 | -0.15836 | 0.10964 | -1.44437 | 0.14864 | 0.33248 | 7.253 |
| Oas1a | 36.607 | -1.40844 | 0.45637 | -3.08617 | 0.00203 | 0.02025 | 11.266 |
| Oas1b | 34.1267 | -0.39182 | 0.39284 | -0.99741 | 0.31857 | 0.52914 | 11.529 |
| Oas1c | 36.7029 | -0.46428 | 0.33203 | -1.39831 | 0.16202 | 0.35036 | 12.321 |
| Oas2 | 69.686 | -0.13292 | 0.33648 | -0.39501 | 0.69283 | 0.83344 | 19.521 |
| Oasl2 | 186.706 | -1.02032 | 0.24522 | -4.16085 | 3.17E-05 | 0.0012 | 15.299 |
| Oat | 3893.28 | -0.00297 | 0.10083 | -0.02949 | 0.97647 | 0.98767 | 18.921 |
| Oaz1 | 9833.18 | 0.040914 | 0.07517 | 0.544255 | 0.58627 | 0.7603 | 2.635 |
| Oaz1-ps | 125.219 | 0.020808 | 0.16876 | 0.1233 | 0.90187 | 0.95075 | 0.685 |
| Oaz2 | 6857.75 | 0.012322 | 0.07906 | 0.155851 | 0.87615 | 0.93819 | 22.3 |
| Oaz2-ps | 75.6637 | 0.09894 | 0.31703 | 0.312079 | 0.75498 | 0.87123 | 0.569 |
| Oaz3 | 53.5542 | -0.16355 | 0.24626 | -0.66413 | 0.50661 | 0.69789 | 11.502 |
| Obi1 | 610.679 | 0.181878 | 0.08072 | 2.25324 | 0.02424 | 0.10171 | 45.11 |
| Obscn | 292.544 | 0.046673 | 0.13321 | 0.350378 | 0.72605 | 0.85396 | 144.915 |
| Obsl1 | 1628.86 | 0.093682 | 0.08535 | 1.097609 | 0.27238 | 0.48159 | 27.143 |
| Oca2 | 19.2432 | -0.61204 | 0.4182 | -1.4635 | 0.14333 | 0.32546 | 296.759 |
| Ocel1 | 876.316 | 0.049334 | 0.09836 | 0.501578 | 0.61596 | 0.78132 | 8.064 |
| Ociad1 | 11029.4 | 0.012392 | 0.05567 | 0.222589 | 0.82386 | 0.90869 | 21.286 |
| Ociad2 | 6264.51 | -0.43907 | 0.14926 | -2.9416 | 0.00327 | 0.02757 | 18.83 |
| Ocln | 451.807 | -0.45926 | 0.10763 | -4.267 | 1.98E-05 | 0.00087 | 56.212 |
| Ocrl | 4603.24 | 0.226946 | 0.06546 | 3.46689 | 0.00053 | 0.00822 | 53.482 |
| Odad1 | 127.011 | 0.042428 | 0.18478 | 0.229609 | 0.8184 | 0.90654 | 24.892 |
| Odad3 | 401.124 | 0.480082 | 0.11983 | 4.006233 | 6.17E-05 | 0.00193 | 12.764 |

| | | | | | | | |
|---|---|---|---|---|---|---|---|
| Odad4 | 82.0466 | 0.344785 | 0.21957 | 1.57024 | 0.11636 | 0.28653 | 26.962 |
| Odc1 | 2543.64 | -0.04372 | 0.09869 | -0.44293 | 0.65781 | 0.81018 | 6.712 |
| Odf2 | 5004.89 | 0.027839 | 0.07525 | 0.369934 | 0.71143 | 0.84471 | 42.526 |
| Odf2l | 986.902 | 0.296242 | 0.12796 | 2.315153 | 0.0206 | 0.09132 | 35.328 |
| Odf3 | 27.3098 | 0.617733 | 0.41013 | 1.506185 | 0.13202 | 0.31082 | 3.214 |
| Odf3b | 25.1711 | -0.33575 | 0.38611 | -0.86957 | 0.38454 | 0.594 | 2.139 |
| Odf4 | 74.0817 | 0.827172 | 0.34016 | 2.431736 | 0.01503 | 0.07451 | 5.247 |
| Odr4 | 2353.38 | 0.151296 | 0.11455 | 1.320781 | 0.18657 | 0.38153 | 31.776 |
| Ofd1 | 639.035 | 0.373311 | 0.14606 | 2.555833 | 0.01059 | 0.05973 | 50.672 |
| Oga | 12190.4 | 0.115055 | 0.09186 | 1.252445 | 0.21041 | 0.41047 | 33.579 |
| Ogdh | 11015.3 | -0.06086 | 0.05976 | -1.01828 | 0.30855 | 0.51837 | 65.01 |
| Ogdhl | 6297.28 | -0.05381 | 0.07126 | -0.75504 | 0.45022 | 0.65305 | 26.133 |
| Ogfod1 | 6315.97 | 0.115116 | 0.07077 | 1.626565 | 0.10383 | 0.26629 | 30.724 |
| Ogfod2 | 960.429 | -0.0429 | 0.07902 | -0.5429 | 0.5872 | 0.76069 | 3.187 |
| Ogfod3 | 160.82 | -0.1417 | 0.1851 | -0.76557 | 0.44393 | 0.6481 | 27.121 |
| Ogfr | 1698.66 | 0.052236 | 0.07857 | 0.664834 | 0.50616 | 0.69769 | 6.592 |
| Ogfrl1 | 5936.91 | 0.044309 | 0.09256 | 0.478731 | 0.63213 | 0.79231 | 31.349 |
| Ogg1 | 228.883 | -0.0578 | 0.14667 | -0.39407 | 0.69353 | 0.83381 | 8.097 |
| Ogn | 513.782 | -0.68687 | 0.65032 | -1.0562 | 0.29088 | 0.50107 | 16.456 |
| Ogt | 28027.9 | 0.239021 | 0.09049 | 2.641375 | 0.00826 | 0.0506 | 44.292 |
| Oip5 | 42.5875 | 0.137396 | 0.27831 | 0.493674 | 0.62154 | 0.78483 | 8.958 |
| Oip5os1 | 26558.5 | -0.47934 | 0.16369 | -2.92836 | 0.00341 | 0.02822 | 13.227 |
| Ola1 | 4159.16 | -0.03771 | 0.06666 | -0.56577 | 0.57155 | 0.75039 | 126.124 |
| Olfm1 | 16452.3 | -1.0155 | 0.36753 | -2.76308 | 0.00573 | 0.03996 | 37.745 |
| Olfm2 | 1917.12 | -0.55267 | 0.1502 | -3.6796 | 0.00023 | 0.00475 | 78.66 |
| Olfm3 | 3243.62 | 0.537075 | 0.13991 | 3.838757 | 0.00012 | 0.00309 | 221.645 |
| Olfm4 | 55.0195 | -0.01708 | 0.3637 | -0.04696 | 0.96255 | 0.98185 | 39.059 |
| Olfml1 | 382.178 | -0.73373 | 0.11314 | -6.48494 | 8.88E-11 | 9.37E-08 | 23.649 |
| Olfml2a | 280.457 | -0.34023 | 0.20428 | -1.66555 | 0.0958 | 0.25322 | 31.776 |
| Olfml2b | 182.493 | -0.65873 | 0.24794 | -2.65679 | 0.00789 | 0.04922 | 38.258 |
| Olfml3 | 576.846 | -0.43621 | 0.11169 | -3.90562 | 9.40E-05 | 0.00257 | 15.78 |
| Olfr1033 | 24.9545 | -2 | 0.76641 | -2.62123 | 0.00876 | 0.0525 | 24.539 |
| Olfr111 | 16.2615 | -0.73506 | 0.44199 | -1.66308 | 0.0963 | 0.25407 | 24.42 |
| Olfr1372-ps1 | 38.0034 | -0.8942 | 0.44116 | -2.02695 | 0.04267 | 0.14888 | 13.438 |

| | | | | | | | |
|---|---|---|---|---|---|---|---|
| Olfr1392 | 34.0717 | 0.459325 | 0.30824 | 1.490168 | 0.13618 | 0.31645 | 10.61 |
| Olfr1393 | 329.143 | 0.9381 | 0.20791 | 4.512069 | 6.42E-06 | 0.0004 | 4.466 |
| Olfr1417 | 40.4878 | 0.096659 | 0.48392 | 0.199744 | 0.84168 | 0.91879 | 12.417 |
| Olfr1420 | 12.764 | 0.412018 | 0.55639 | 0.740519 | 0.45899 | 0.65977 | 11.515 |
| Olfr1564 | 52.597 | -0.41942 | 0.28533 | -1.46997 | 0.14157 | 0.32333 | 10.485 |
| Olfr239 | 15.6622 | -0.1943 | 0.56459 | -0.34415 | 0.73073 | 0.85728 | 11.205 |
| Olfr277-ps1 | 14.7713 | 0.473463 | 0.58554 | 0.808595 | 0.41875 | 0.62584 | 0.285 |
| Olfr287 | 193.063 | -0.65334 | 0.16955 | -3.85334 | 0.00012 | 0.00298 | 14.086 |
| Olfr316 | 41.3944 | -2 | 0.59229 | -3.84683 | 0.00012 | 0.00303 | 6.938 |
| Olfr539 | 25.1833 | -1.1849 | 0.50869 | -2.32931 | 0.01984 | 0.08901 | 18.651 |
| Olfr543 | 31.4694 | 0.059401 | 0.28261 | 0.210189 | 0.83352 | 0.91471 | 5.101 |
| Olfr544 | 29.5942 | 0.338785 | 0.33651 | 1.00676 | 0.31405 | 0.52455 | 6.283 |
| Olfr545 | 47.2399 | 0.258097 | 0.38494 | 0.670488 | 0.50255 | 0.69511 | 8.108 |
| Olfr55 | 24.8239 | -0.3187 | 0.41369 | -0.77037 | 0.44108 | 0.64575 | 12.307 |
| Olfr689 | 20.272 | -0.72493 | 0.37306 | -1.94321 | 0.05199 | 0.17024 | 8.498 |
| Olfr690 | 24.474 | 0.024726 | 0.36163 | 0.068374 | 0.94549 | 0.97293 | 5.88 |
| Olfr691 | 14.4605 | -0.30232 | 0.47035 | -0.64276 | 0.52038 | 0.70971 | 6.762 |
| Olfr692 | 41.8275 | 0.077437 | 0.36177 | 0.214049 | 0.83051 | 0.91311 | 1.126 |
| Olfr718-ps1 | 16.9301 | 1.392926 | 0.57646 | 2.416356 | 0.01568 | 0.07666 | 13.568 |
| Olfr856-ps1 | 17.6427 | -0.26555 | 0.65274 | -0.40682 | 0.68414 | 0.82778 | 1.591 |
| Olfr920 | 15.5928 | -0.05368 | 0.47944 | -0.11196 | 0.91086 | 0.95502 | 7.819 |
| Olig1 | 2885.11 | -0.43016 | 0.13629 | -3.15616 | 0.0016 | 0.01735 | 2.162 |
| Olig2 | 1001.31 | -0.35087 | 0.1178 | -2.97844 | 0.0029 | 0.02565 | 3.221 |
| Oma1 | 307.468 | -0.06962 | 0.14335 | -0.48568 | 0.62719 | 0.78903 | 58.057 |
| Omd | 79.8923 | -1.05941 | 0.45324 | -2.33743 | 0.01942 | 0.08776 | 10.361 |
| Omg | 3897.9 | 0.009168 | 0.08187 | 0.111989 | 0.91083 | 0.95502 | 3.103 |
| Onecut2 | 1013.93 | 0.375799 | 0.37072 | 1.013693 | 0.31073 | 0.5209 | 60.968 |
| Opa1 | 9193.73 | 0.188457 | 0.07641 | 2.466288 | 0.01365 | 0.07034 | 75.551 |
| Opa3 | 2397.35 | -0.23907 | 0.07695 | -3.1067 | 0.00189 | 0.01936 | 28.21 |
| Opalin | 412.879 | -0.19518 | 0.24016 | -0.81271 | 0.41638 | 0.62395 | 15.404 |
| Opcml | 7344.88 | -0.34796 | 0.13656 | -2.54805 | 0.01083 | 0.06059 | 1134.636 |
| Ophn1 | 3343.87 | 0.252721 | 0.07501 | 3.369113 | 0.00075 | 0.01055 | 336.749 |
| Oplah | 1212.67 | -0.24695 | 0.07937 | -3.11146 | 0.00186 | 0.01915 | 31.415 |
| Opn1mw | 17.5429 | 0.726535 | 0.42674 | 1.702543 | 0.08865 | 0.2406 | 23.297 |

| | | | | | | | |
|---|---|---|---|---|---|---|---|
| Opn3 | 72.2085 | 0.237582 | 0.23006 | 1.032715 | 0.30174 | 0.51136 | 30.356 |
| Opn4 | 25.6229 | -0.28981 | 0.59301 | -0.4887 | 0.62505 | 0.78752 | 9.525 |
| Oprd1 | 1796.78 | 0.474206 | 0.1409 | 3.365515 | 0.00076 | 0.01064 | 33.761 |
| Oprk1 | 2572.98 | 0.956821 | 0.36181 | 2.644536 | 0.00818 | 0.05026 | 17.666 |
| Oprl1 | 1766.34 | 0.057591 | 0.07604 | 0.757411 | 0.4488 | 0.65214 | 5.97 |
| Oprm1 | 244.873 | 0.624492 | 0.18209 | 3.42965 | 0.0006 | 0.00901 | 279.693 |
| Optc | 29.3288 | -1.1658 | 0.43802 | -2.66155 | 0.00778 | 0.04879 | 10.801 |
| Optn | 616.757 | -0.34157 | 0.14036 | -2.43351 | 0.01495 | 0.0742 | 43.41 |
| Orai1 | 188.718 | -0.15684 | 0.16678 | -0.94041 | 0.34701 | 0.55765 | 15.383 |
| Orai2 | 4203.39 | -0.07973 | 0.07025 | -1.13496 | 0.25639 | 0.46435 | 23.255 |
| Orai3 | 470.965 | -0.37341 | 0.14166 | -2.63599 | 0.00839 | 0.05111 | 5.336 |
| Orc1 | 77.8647 | -0.5265 | 0.24318 | -2.16505 | 0.03038 | 0.11904 | 35.411 |
| Orc2 | 1009.75 | 0.151717 | 0.07837 | 1.9358 | 0.05289 | 0.1721 | 42.339 |
| Orc3 | 2327.37 | 0.215068 | 0.11069 | 1.943007 | 0.05202 | 0.17024 | 44.149 |
| Orc4 | 2872.36 | 0.186421 | 0.12605 | 1.478945 | 0.13916 | 0.32037 | 47.454 |
| Orc5 | 877.55 | 0.208726 | 0.1008 | 2.070781 | 0.03838 | 0.13938 | 63.945 |
| Orc6 | 985.898 | 0.124512 | 0.09486 | 1.312642 | 0.1893 | 0.38522 | 8.647 |
| Ormdl1 | 298.732 | 0.072223 | 0.12549 | 0.575522 | 0.56494 | 0.74523 | 13.215 |
| Ormdl2 | 99.3242 | -0.23937 | 0.23531 | -1.01726 | 0.30903 | 0.51884 | 3.744 |
| Ormdl3 | 2593.64 | 0.013841 | 0.08234 | 0.168101 | 0.8665 | 0.93232 | 6.113 |
| Os9 | 6296.36 | -0.17964 | 0.07072 | -2.54015 | 0.01108 | 0.06133 | 25.482 |
| Osbp | 4023.22 | -0.01395 | 0.06187 | -0.22541 | 0.82166 | 0.90766 | 28.172 |
| Osbp2 | 3704.92 | -0.09077 | 0.07894 | -1.14986 | 0.2502 | 0.45761 | 160.173 |
| Osbpl10 | 1359.4 | -0.15692 | 0.08669 | -1.8102 | 0.07027 | 0.20701 | 253.657 |
| Osbpl11 | 1193.71 | 0.099906 | 0.08515 | 1.173254 | 0.24069 | 0.44693 | 58.242 |
| Osbpl1a | 4970 | -0.31285 | 0.09158 | -3.41602 | 0.00064 | 0.00932 | 186.528 |
| Osbpl2 | 2990.36 | 0.101076 | 0.0732 | 1.380733 | 0.16736 | 0.35707 | 43.375 |
| Osbpl3 | 2103.16 | -0.06661 | 0.12502 | -0.53277 | 0.59419 | 0.76639 | 162.872 |
| Osbpl5 | 809.483 | 0.03248 | 0.08639 | 0.375957 | 0.70695 | 0.84193 | 68.224 |
| Osbpl6 | 3770.68 | 0.102909 | 0.07561 | 1.361031 | 0.1735 | 0.3648 | 194.14 |
| Osbpl7 | 845.988 | 0.112566 | 0.08773 | 1.283135 | 0.19944 | 0.39741 | 18.277 |
| Osbpl8 | 13739.2 | 0.567992 | 0.13795 | 4.117386 | 3.83E-05 | 0.00136 | 132.498 |
| Osbpl9 | 3471.9 | 0.226495 | 0.10269 | 2.205623 | 0.02741 | 0.11082 | 141.128 |
| Oscar | 248.822 | 0.230484 | 0.18462 | 1.248412 | 0.21188 | 0.41257 | 6.345 |

| | | | | | | | |
|---|---|---|---|---|---|---|---|
| Oscp1 | 1924.91 | 0.032833 | 0.07778 | 0.422139 | 0.67292 | 0.82019 | 33.631 |
| Oser1 | 524.178 | -0.01965 | 0.09003 | -0.2183 | 0.82719 | 0.91108 | 18.876 |
| Osgep | 1042.74 | 0.030431 | 0.07725 | 0.393914 | 0.69364 | 0.83383 | 18.416 |
| Osgepl1 | 539.926 | 0.162218 | 0.12569 | 1.290634 | 0.19683 | 0.39463 | 12.72 |
| Osgin1 | 30.3702 | -0.10424 | 0.41968 | -0.24837 | 0.80385 | 0.89875 | 12.133 |
| Osgin2 | 1076.12 | -0.06092 | 0.10101 | -0.60312 | 0.54643 | 0.73077 | 16.768 |
| Osmr | 350.688 | -0.5926 | 0.19111 | -3.1008 | 0.00193 | 0.01967 | 61.393 |
| Osr1 | 58.1602 | -1.07266 | 0.79493 | -1.34937 | 0.17722 | 0.36947 | 11.385 |
| Ost4 | 187.667 | -0.1502 | 0.18435 | -0.81473 | 0.41522 | 0.62304 | 1.904 |
| Ostc | 1347.39 | 0.02015 | 0.06991 | 0.288236 | 0.77317 | 0.88189 | 13.533 |
| Ostf1 | 393.172 | -0.50012 | 0.12558 | -3.98243 | 6.82E-05 | 0.00206 | 115.687 |
| Ostm1 | 2350.6 | -0.03281 | 0.08446 | -0.38851 | 0.69764 | 0.83617 | 118.638 |
| Otof | 4864.92 | 0.606722 | 0.33635 | 1.80385 | 0.07125 | 0.20896 | 94.871 |
| Otogl | 67.5977 | -0.37699 | 0.3215 | -1.17258 | 0.24096 | 0.4471 | 151.604 |
| Otub1 | 6460.9 | -0.06626 | 0.0749 | -0.8847 | 0.37632 | 0.58637 | 8.115 |
| Otub2 | 1156.81 | -0.36076 | 0.15532 | -2.32268 | 0.0202 | 0.09011 | 17.669 |
| Otud1 | 934.556 | -0.44253 | 0.14442 | -3.06427 | 0.00218 | 0.02131 | 2.832 |
| Otud3 | 627.615 | 0.009584 | 0.11616 | 0.082513 | 0.93424 | 0.96702 | 18.567 |
| Otud4 | 5068.82 | 0.289571 | 0.08577 | 3.376282 | 0.00073 | 0.01036 | 38.107 |
| Otud5 | 4251.39 | 0.112854 | 0.07242 | 1.558409 | 0.11914 | 0.29132 | 35.263 |
| Otud6b | 3500.14 | 0.109367 | 0.0745 | 1.468066 | 0.14209 | 0.32393 | 17.09 |
| Otud7a | 1893 | -0.29963 | 0.10616 | -2.82252 | 0.00476 | 0.03524 | 314.278 |
| Otud7b | 4660.69 | -0.33601 | 0.10877 | -3.08923 | 0.00201 | 0.02016 | 56.608 |
| Otulin | 653.005 | -0.04318 | 0.10016 | -0.43109 | 0.6664 | 0.8158 | 88.059 |
| Otulinl | 1082.84 | 0.254799 | 0.11357 | 2.243594 | 0.02486 | 0.10356 | 26.511 |
| Otx1 | 83.3048 | -1.37232 | 0.2764 | -4.96495 | 6.87E-07 | 8.29E-05 | 8.134 |
| Otx2 | 40.3948 | 0.156643 | 0.34567 | 0.453163 | 0.65043 | 0.80552 | 16.143 |
| Ovca2 | 343.208 | 0.04693 | 0.11489 | 0.408483 | 0.68292 | 0.8267 | 2.894 |
| Ovgp1 | 242.24 | 0.5459 | 0.196 | 2.785157 | 0.00535 | 0.03791 | 13.713 |
| Oxa1l | 1810.77 | 0.025658 | 0.07613 | 0.337046 | 0.73608 | 0.86049 | 8.84 |
| Oxct1 | 12500.6 | -0.01832 | 0.0576 | -0.31814 | 0.75038 | 0.86898 | 128.962 |
| Oxgr1 | 26.3352 | 1.195876 | 0.63712 | 1.877004 | 0.06052 | 0.18773 | 22.851 |
| Oxld1 | 102.857 | 0.036462 | 0.1734 | 0.210274 | 0.83345 | 0.91471 | 1.463 |
| Oxnad1 | 839.784 | 0.014821 | 0.09763 | 0.151805 | 0.87934 | 0.93962 | 17.829 |

| | | | | | | | |
|---|---|---|---|---|---|---|---|
| Oxr1 | 10339.8 | -0.42154 | 0.15134 | -2.78541 | 0.00535 | 0.0379 | 413.567 |
| Oxsm | 453 | -0.20631 | 0.12454 | -1.65655 | 0.09761 | 0.25633 | 11.157 |
| Oxsr1 | 3321.99 | -0.30461 | 0.09124 | -3.33856 | 0.00084 | 0.01137 | 83.996 |
| Oxtr | 677.98 | -0.88407 | 0.47045 | -1.87922 | 0.06021 | 0.18722 | 16.261 |
| P2rx3 | 58.9031 | -0.3322 | 0.32344 | -1.02709 | 0.30438 | 0.51417 | 38.88 |
| P2rx4 | 564.385 | -0.21455 | 0.10057 | -2.1333 | 0.0329 | 0.12604 | 22.195 |
| P2rx5 | 32.0418 | 0.056577 | 0.35017 | 0.161569 | 0.87164 | 0.93514 | 12.265 |
| P2rx6 | 350.103 | 0.146305 | 0.10823 | 1.351815 | 0.17643 | 0.36846 | 15.916 |
| P2rx7 | 433.875 | -0.28194 | 0.11854 | -2.37842 | 0.01739 | 0.08183 | 47.522 |
| P2ry1 | 677.413 | 0.906042 | 0.21621 | 4.190624 | 2.78E-05 | 0.0011 | 6.188 |
| P2ry10b | 20.6849 | -0.52714 | 0.38772 | -1.35958 | 0.17396 | 0.36537 | 24.735 |
| P2ry12 | 1375.36 | -0.12235 | 0.11998 | -1.01981 | 0.30782 | 0.51763 | 46.6 |
| P2ry13 | 416.992 | -0.38743 | 0.14781 | -2.62102 | 0.00877 | 0.0525 | 2.991 |
| P2ry14 | 165.121 | -0.49136 | 0.19147 | -2.56628 | 0.01028 | 0.05853 | 39.764 |
| P2ry2 | 21.4511 | -0.59717 | 0.439 | -1.36031 | 0.17373 | 0.36516 | 16.299 |
| P2ry6 | 74.5853 | -0.71013 | 0.22522 | -3.15298 | 0.00162 | 0.01748 | 37.02 |
| P3h1 | 665.549 | 0.468459 | 0.08642 | 5.420967 | 5.93E-08 | 1.22E-05 | 16.061 |
| P3h2 | 119.333 | -0.52322 | 0.25103 | -2.08429 | 0.03713 | 0.13624 | 146.497 |
| P3h3 | 1439.61 | 0.243493 | 0.10654 | 2.285505 | 0.02228 | 0.09621 | 16.664 |
| P3h4 | 364.989 | -0.40074 | 0.13468 | -2.97563 | 0.00292 | 0.02575 | 6.381 |
| P4ha1 | 2766.11 | -0.20331 | 0.08651 | -2.35019 | 0.01876 | 0.08597 | 50.009 |
| P4ha2 | 1494.17 | -0.06346 | 0.11953 | -0.53089 | 0.59549 | 0.76734 | 31.571 |
| P4ha3 | 207.436 | -0.28512 | 0.18366 | -1.55243 | 0.12056 | 0.2934 | 34.18 |
| P4hb | 4808.64 | -0.11894 | 0.0778 | -1.5288 | 0.12631 | 0.30205 | 12.956 |
| P4htm | 2616.76 | 0.056608 | 0.0859 | 0.658987 | 0.5099 | 0.70125 | 18.806 |
| Pa2g4 | 2416.82 | 0.009563 | 0.06245 | 0.15313 | 0.8783 | 0.93916 | 8.222 |
| Pabpc1 | 8127.84 | -0.02593 | 0.06464 | -0.40119 | 0.68828 | 0.83003 | 14.008 |
| Pabpc1l | 55.0933 | -0.34257 | 0.28011 | -1.22299 | 0.22133 | 0.42428 | 25.089 |
| Pabpc1l2a | 26.0412 | 0.282645 | 0.36723 | 0.769678 | 0.44149 | 0.64609 | 0.513 |
| Pabpc1l2b | 22.8812 | 0.469086 | 0.37163 | 1.262249 | 0.20686 | 0.4063 | 0.69 |
| Pabpc4 | 1625.08 | -0.00664 | 0.06835 | -0.0971 | 0.92265 | 0.96107 | 36.575 |
| Pabpc4l | 12.9255 | -0.25456 | 0.5686 | -0.4477 | 0.65437 | 0.80826 | 6.023 |
| Pabpc5 | 136.177 | 0.280969 | 0.18388 | 1.528039 | 0.1265 | 0.3023 | 2.97 |
| Pabpn1 | 1892.1 | 0.043682 | 0.11744 | 0.37195 | 0.70993 | 0.84385 | 5.67 |

| | | | | | | | |
|---|---|---|---|---|---|---|---|
| Pacc1 | 421.407 | 0.017326 | 0.11179 | 0.154989 | 0.87683 | 0.93843 | 25.003 |
| Pacrg | 405.3 | -0.16794 | 0.1305 | -1.2869 | 0.19813 | 0.39587 | 437.258 |
| Pacs1 | 3295.37 | 0.117299 | 0.0665 | 1.764012 | 0.07773 | 0.22065 | 139.962 |
| Pacs2 | 6591.34 | -0.12115 | 0.09078 | -1.3345 | 0.18204 | 0.37584 | 59.894 |
| Pacsin1 | 16651.5 | -0.15338 | 0.06251 | -2.45388 | 0.01413 | 0.07164 | 55.974 |
| Pacsin2 | 5018.39 | 0.0439 | 0.08045 | 0.545682 | 0.58528 | 0.75985 | 89 |
| Pacsin3 | 1038.39 | -0.58404 | 0.08366 | -6.98103 | 2.93E-12 | 5.43E-09 | 8.726 |
| Padi2 | 1256.03 | -0.71544 | 0.17889 | -3.99935 | 6.35E-05 | 0.00196 | 46.243 |
| Paf1 | 2681.24 | 0.072396 | 0.07088 | 1.021338 | 0.30709 | 0.51686 | 6.438 |
| Pafah1b1 | 27581 | -0.00366 | 0.08306 | -0.044 | 0.9649 | 0.98273 | 50.722 |
| Pafah1b2 | 10352.2 | -0.03601 | 0.06555 | -0.54942 | 0.58272 | 0.75806 | 49.832 |
| Pafah1b3 | 231.062 | -0.07841 | 0.18845 | -0.41608 | 0.67735 | 0.82317 | 2.938 |
| Pafah2 | 373.271 | -0.13589 | 0.11443 | -1.18753 | 0.23502 | 0.44032 | 31.094 |
| Pag1 | 2452.9 | 0.201676 | 0.06968 | 2.89418 | 0.0038 | 0.03041 | 146.201 |
| Paics | 4195.03 | -0.0468 | 0.07333 | -0.63814 | 0.52338 | 0.71181 | 16.203 |
| Paip1 | 2820.3 | -0.08626 | 0.0715 | -1.20654 | 0.22761 | 0.43174 | 29.618 |
| Paip2 | 6626.49 | 0.112256 | 0.07358 | 1.525653 | 0.1271 | 0.30327 | 18.571 |
| Paip2b | 4487.59 | 0.019453 | 0.07527 | 0.25845 | 0.79606 | 0.89487 | 26.341 |
| Pak1 | 10793.9 | -0.45388 | 0.10601 | -4.28163 | 1.86E-05 | 0.00084 | 123.841 |
| Pak1ip1 | 1597.45 | 0.064608 | 0.07425 | 0.870133 | 0.38423 | 0.59361 | 11.993 |
| Pak2 | 3051.5 | -0.0175 | 0.06951 | -0.25176 | 0.80122 | 0.8972 | 63.053 |
| Pak3 | 7171.6 | 0.37865 | 0.08837 | 4.284738 | 1.83E-05 | 0.00083 | 279.206 |
| Pak4 | 342.457 | -0.12251 | 0.12807 | -0.95659 | 0.33877 | 0.54959 | 39.367 |
| Pak5 | 1156.85 | 0.089909 | 0.09338 | 0.962848 | 0.33562 | 0.54639 | 306.864 |
| Pak6 | 3970.44 | -0.19215 | 0.09886 | -1.94377 | 0.05192 | 0.17016 | 34.718 |
| Pakap | 594.013 | -0.19672 | 0.13846 | -1.42077 | 0.15538 | 0.34197 | 328.804 |
| Palb2 | 84.6103 | -0.07208 | 0.21624 | -0.33335 | 0.73887 | 0.86228 | 25.724 |
| Pald1 | 516.796 | -0.21317 | 0.1314 | -1.62229 | 0.10474 | 0.26777 | 63.875 |
| Palld | 266.134 | -0.19452 | 0.14326 | -1.35776 | 0.17454 | 0.36617 | 391.258 |
| Palm | 10298.9 | -0.04122 | 0.08241 | -0.50013 | 0.61698 | 0.78203 | 27.325 |
| Palm3 | 117.476 | -0.74255 | 0.2448 | -3.03325 | 0.00242 | 0.02282 | 8.825 |
| Palmd | 510.719 | -0.10533 | 0.10966 | -0.96048 | 0.33681 | 0.54741 | 50.73 |
| Pals1 | 2605.45 | 0.239852 | 0.10374 | 2.312144 | 0.02077 | 0.09184 | 91.808 |
| Pals2 | 1871.46 | 0.39292 | 0.11164 | 3.519484 | 0.00043 | 0.00725 | 88.699 |

| | | | | | | | |
|---|---|---|---|---|---|---|---|
| Pam | 8178.04 | -0.25228 | 0.10014 | -2.51922 | 0.01176 | 0.06378 | 300.533 |
| Pam16 | 487.807 | -0.00065 | 0.11929 | -0.00547 | 0.99564 | 0.99764 | 8.525 |
| Pamr1 | 403.724 | -0.79182 | 0.1084 | -7.30439 | 2.79E-13 | 1.35E-09 | 93.03 |
| Pan2 | 1955.63 | 0.172759 | 0.1054 | 1.639061 | 0.1012 | 0.26235 | 18.024 |
| Pan3 | 2562.49 | 0.016803 | 0.08161 | 0.2059 | 0.83687 | 0.91619 | 118.342 |
| Panct2 | 195.307 | -0.00148 | 0.16076 | -0.00922 | 0.99264 | 0.99577 | 36.671 |
| Pank1 | 1867.62 | -0.16127 | 0.09552 | -1.68842 | 0.09133 | 0.2456 | 72.516 |
| Pank2 | 2257.26 | -0.04905 | 0.07187 | -0.68247 | 0.49494 | 0.68908 | 36.694 |
| Pank3 | 4088.46 | 0.012765 | 0.07688 | 0.166039 | 0.86813 | 0.93328 | 21.802 |
| Pank4 | 2165.72 | 0.173364 | 0.06485 | 2.67322 | 0.00751 | 0.04775 | 16.816 |
| Pantr1 | 1183.7 | -0.10954 | 0.08739 | -1.25345 | 0.21004 | 0.4101 | 104.831 |
| Pantr2 | 224.419 | -0.04739 | 0.13394 | -0.35385 | 0.72345 | 0.85232 | 6.735 |
| Panx1 | 997.822 | 0.297914 | 0.10191 | 2.923276 | 0.00346 | 0.02853 | 43.351 |
| Panx2 | 3534.02 | -0.53594 | 0.10575 | -5.06822 | 4.02E-07 | 5.42E-05 | 13.834 |
| Paox | 384.406 | -0.31384 | 0.10838 | -2.8957 | 0.00378 | 0.03031 | 21.426 |
| Papln | 908.474 | 0.40882 | 0.1991 | 2.05339 | 0.04003 | 0.14298 | 28.749 |
| Papola | 7466.54 | 0.166392 | 0.06598 | 2.521901 | 0.01167 | 0.06352 | 54.251 |
| Papolb | 42.0824 | 0.853242 | 0.29113 | 2.93084 | 0.00338 | 0.02813 | 4.239 |
| Papolg | 1211.36 | 0.140022 | 0.08116 | 1.725337 | 0.08447 | 0.23299 | 32.608 |
| Pappa | 58.0475 | -1.25788 | 0.38762 | -3.2451 | 0.00117 | 0.01411 | 233.336 |
| Pappa2 | 97.9846 | -1.33794 | 0.27245 | -4.91069 | 9.08E-07 | 9.89E-05 | 268.764 |
| Papss1 | 3489.87 | -0.07214 | 0.08142 | -0.88611 | 0.37556 | 0.58576 | 78.904 |
| Papss2 | 1136.6 | -0.14942 | 0.10994 | -1.35904 | 0.17413 | 0.36563 | 71.398 |
| Paqr3 | 706.701 | 0.113867 | 0.08034 | 1.417222 | 0.15642 | 0.34317 | 29.268 |
| Paqr4 | 1695.4 | -0.23806 | 0.07972 | -2.9861 | 0.00283 | 0.02521 | 4.682 |
| Paqr5 | 323.041 | -0.21712 | 0.11207 | -1.93734 | 0.0527 | 0.17178 | 73.376 |
| Paqr6 | 184.085 | -0.21343 | 0.14436 | -1.47852 | 0.13927 | 0.32052 | 3.958 |
| Paqr7 | 2413.31 | -0.16478 | 0.1152 | -1.43039 | 0.1526 | 0.33794 | 13.539 |
| Paqr8 | 10091.4 | -0.40084 | 0.09928 | -4.0376 | 5.40E-05 | 0.00175 | 49.045 |
| Paqr9 | 3673.19 | 0.233259 | 0.09882 | 2.360395 | 0.01826 | 0.08442 | 8.367 |
| Pard3 | 1834.47 | 0.513568 | 0.15415 | 3.33171 | 0.00086 | 0.01155 | 548.125 |
| Pard3b | 313.698 | 0.031302 | 0.16918 | 0.185025 | 0.85321 | 0.92555 | 1003.461 |
| Pard6a | 538.008 | -0.34342 | 0.09058 | -3.79148 | 0.00015 | 0.00352 | 2.349 |
| Pard6b | 341.146 | 0.314028 | 0.18144 | 1.730759 | 0.08349 | 0.23102 | 20.2 |

| | | | | | | | |
|---|---|---|---|---|---|---|---|
| Pard6g | 299.513 | 0.044616 | 0.13062 | 0.34158 | 0.73267 | 0.85822 | 72.75 |
| Parg | 1990.36 | 0.084338 | 0.06922 | 1.218438 | 0.22306 | 0.42629 | 95.602 |
| Park7 | 3838.69 | -0.22826 | 0.0716 | -3.18801 | 0.00143 | 0.01602 | 17.305 |
| Parl | 1208.3 | -0.18331 | 0.1201 | -1.52625 | 0.12695 | 0.30312 | 22.57 |
| Parm1 | 1907.25 | 0.007692 | 0.10505 | 0.073221 | 0.94163 | 0.97087 | 109.28 |
| Parn | 1029.36 | -0.07575 | 0.08744 | -0.86628 | 0.38634 | 0.59577 | 130.211 |
| Parp1 | 3988.78 | -0.14305 | 0.10695 | -1.33755 | 0.18104 | 0.37447 | 32.331 |
| Parp10 | 134.473 | -0.48305 | 0.1809 | -2.67025 | 0.00758 | 0.04806 | 12.268 |
| Parp11 | 1024.64 | 0.102166 | 0.09262 | 1.103079 | 0.26999 | 0.47957 | 47.422 |
| Parp12 | 655.483 | -0.57536 | 0.13083 | -4.39774 | 1.09E-05 | 0.0006 | 31.94 |
| Parp14 | 225.284 | -0.7999 | 0.24887 | -3.21406 | 0.00131 | 0.01505 | 38.671 |
| Parp16 | 150.347 | -0.15417 | 0.16017 | -0.96259 | 0.33575 | 0.54643 | 43.872 |
| Parp2 | 1075.47 | 0.140432 | 0.08913 | 1.575621 | 0.11511 | 0.28442 | 13.461 |
| Parp3 | 294.584 | -0.18502 | 0.14492 | -1.2767 | 0.20171 | 0.40016 | 6.628 |
| Parp4 | 464.14 | -0.20162 | 0.09675 | -2.08405 | 0.03716 | 0.13625 | 84.176 |
| Parp6 | 5265.08 | -0.02078 | 0.06393 | -0.32497 | 0.7452 | 0.866 | 33.002 |
| Parp8 | 930.74 | 0.045538 | 0.08517 | 0.534685 | 0.59287 | 0.76531 | 170.718 |
| Parp9 | 193.09 | -0.48925 | 0.15782 | -3.10004 | 0.00193 | 0.01969 | 34.136 |
| Pars2 | 326.091 | 0.127333 | 0.11842 | 1.075231 | 0.28227 | 0.49241 | 4.214 |
| Particl | 237.976 | -0.04092 | 0.12387 | -0.33035 | 0.74114 | 0.86331 | 1.56 |
| Parva | 1959.54 | -0.32884 | 0.07573 | -4.34208 | 1.41E-05 | 0.00071 | 164.188 |
| Parvb | 1054.05 | -0.13682 | 0.11779 | -1.16159 | 0.2454 | 0.45204 | 83.646 |
| Parvg | 157.883 | -0.72803 | 0.22149 | -3.2869 | 0.00101 | 0.01293 | 18.953 |
| Pask | 402.033 | 0.512269 | 0.1533 | 3.341586 | 0.00083 | 0.01127 | 34.713 |
| Pate2 | 15.3225 | 0.165569 | 0.54671 | 0.302845 | 0.76201 | 0.87518 | 120.035 |
| Patj | 1053.84 | 0.292302 | 0.16202 | 1.804096 | 0.07122 | 0.20889 | 323.819 |
| Patl1 | 1535.16 | 0.026847 | 0.07242 | 0.370693 | 0.71087 | 0.84441 | 32.693 |
| Patz1 | 1351.78 | 0.074786 | 0.08642 | 0.865385 | 0.38683 | 0.5961 | 20.21 |
| Pawr | 118.659 | -0.92072 | 0.37701 | -2.44215 | 0.0146 | 0.07302 | 82.12 |
| Pax6 | 762.951 | -0.71338 | 0.15831 | -4.5061 | 6.60E-06 | 0.00041 | 28.465 |
| Paxbp1 | 4081.03 | 0.279187 | 0.07776 | 3.590273 | 0.00033 | 0.00602 | 30.507 |
| Paxip1 | 2471.47 | -0.03831 | 0.07302 | -0.52473 | 0.59977 | 0.77047 | 51.612 |
| Paxx | 1022.66 | -0.00215 | 0.09698 | -0.02215 | 0.98233 | 0.99059 | 5.954 |
| Pbdc1 | 544.983 | 0.003321 | 0.09346 | 0.035531 | 0.97166 | 0.98578 | 37.335 |

| | | | | | | | |
|---|---|---|---|---|---|---|---|
| Pbk | 24.2879 | -0.70106 | 0.62905 | -1.11447 | 0.26508 | 0.47415 | 11.986 |
| Pbld1 | 13.4862 | 0.084909 | 0.48868 | 0.173751 | 0.86206 | 0.93002 | 17.801 |
| Pbld2 | 59.2989 | -0.41366 | 0.2439 | -1.69604 | 0.08988 | 0.24308 | 34.499 |
| Pbrm1 | 3474.21 | 0.237643 | 0.08583 | 2.76871 | 0.00563 | 0.03945 | 102.455 |
| Pbx1 | 10477.5 | -0.02864 | 0.08447 | -0.33905 | 0.73457 | 0.85961 | 312.907 |
| Pbx2 | 3068.13 | 0.127252 | 0.08241 | 1.544148 | 0.12255 | 0.29632 | 6.135 |
| Pbx3 | 1875.06 | 0.430543 | 0.15593 | 2.761213 | 0.00576 | 0.0401 | 201.686 |
| Pbx4 | 353.335 | 0.02667 | 0.12454 | 0.214153 | 0.83043 | 0.91311 | 39.802 |
| Pbxip1 | 3090.49 | -0.47232 | 0.09915 | -4.76358 | 1.90E-06 | 0.00017 | 14.247 |
| Pcare | 24.6293 | 0.165562 | 0.35964 | 0.460349 | 0.64527 | 0.80221 | 9.34 |
| Pcbd1 | 107.868 | 0.03208 | 0.23717 | 0.135262 | 0.8924 | 0.94617 | 4.994 |
| Pcbd2 | 103.675 | -0.13734 | 0.23398 | -0.58697 | 0.55722 | 0.73902 | 49.463 |
| Pcbp1 | 5192.07 | -0.06384 | 0.07245 | -0.88117 | 0.37823 | 0.58818 | 1.83 |
| Pcbp2 | 6381.1 | 0.082683 | 0.05317 | 1.554957 | 0.11996 | 0.29246 | 29.523 |
| Pcbp3 | 1891.71 | -0.34615 | 0.13352 | -2.59246 | 0.00953 | 0.05556 | 200.031 |
| Pcbp4 | 4218.18 | -0.18408 | 0.08977 | -2.05055 | 0.04031 | 0.14348 | 10.722 |
| Pcca | 1454.5 | 0.034011 | 0.09056 | 0.375557 | 0.70725 | 0.84204 | 356.777 |
| Pccb | 2316.55 | -0.15817 | 0.08943 | -1.76862 | 0.07696 | 0.21936 | 52.867 |
| Pcdh1 | 10640.2 | -0.14078 | 0.09515 | -1.4796 | 0.13898 | 0.3202 | 26.14 |
| Pcdh10 | 13399.7 | 0.314907 | 0.09546 | 3.298661 | 0.00097 | 0.01255 | 57.226 |
| Pcdh11x | 1363.25 | 0.266598 | 0.10165 | 2.622698 | 0.00872 | 0.05239 | 620.361 |
| Pcdh15 | 1261.19 | 0.203634 | 0.14486 | 1.405765 | 0.15979 | 0.34774 | 1550.396 |
| Pcdh17 | 7870.61 | 0.272266 | 0.09332 | 2.917652 | 0.00353 | 0.02888 | 95.44 |
| Pcdh18 | 276.572 | 0.026648 | 0.16987 | 0.156876 | 0.87534 | 0.93787 | 14.03 |
| Pcdh19 | 1796.95 | 0.100624 | 0.13165 | 0.764326 | 0.44467 | 0.64865 | 106.128 |
| Pcdh20 | 853.448 | -0.65556 | 0.17713 | -3.701 | 0.00021 | 0.00447 | 6.654 |
| Pcdh7 | 3952.98 | -0.199 | 0.12889 | -1.54393 | 0.12261 | 0.29639 | 415.264 |
| Pcdh8 | 1313.65 | 0.307939 | 0.13973 | 2.203795 | 0.02754 | 0.11118 | 4.538 |
| Pcdh9 | 4283.9 | -0.05171 | 0.11413 | -0.45304 | 0.65052 | 0.80552 | 877.27 |
| Pcdha1 | 66.6128 | -0.42671 | 0.26382 | -1.61742 | 0.10579 | 0.26938 | 257.478 |
| Pcdha10 | 108.16 | 0.00899 | 0.20623 | 0.043593 | 0.96523 | 0.98278 | 182.455 |
| Pcdha12 | 200.075 | 0.119113 | 0.17944 | 0.6638 | 0.50682 | 0.69804 | 167.509 |
| Pcdha2 | 125.719 | 0.086306 | 0.24384 | 0.353948 | 0.72338 | 0.85232 | 248.453 |
| Pcdha3 | 166.773 | -0.04617 | 0.1717 | -0.26888 | 0.78802 | 0.88992 | 241.553 |

| | | | | | | | |
|---|---|---|---|---|---|---|---|
| Pcdha5 | 220.288 | -0.10718 | 0.15177 | -0.7062 | 0.48006 | 0.67758 | 227.218 |
| Pcdha6 | 107.846 | 0.033619 | 0.19002 | 0.176921 | 0.85957 | 0.92904 | 220.027 |
| Pcdha7 | 101.922 | -0.33193 | 0.26806 | -1.23825 | 0.21562 | 0.41724 | 213.856 |
| Pcdha8 | 32.3636 | -0.20858 | 0.41947 | -0.49725 | 0.61902 | 0.78314 | 195.287 |
| Pcdha9 | 182.36 | -0.1558 | 0.20847 | -0.74734 | 0.45486 | 0.65702 | 189.778 |
| Pcdhac1 | 104.535 | -0.24614 | 0.17381 | -1.41612 | 0.15674 | 0.34363 | 97.645 |
| Pcdhac2 | 1168.13 | 0.120117 | 0.08463 | 1.419246 | 0.15583 | 0.34239 | 44.155 |
| Pcdhb1 | 16.4197 | 1.694956 | 0.77811 | 2.178298 | 0.02938 | 0.11646 | 2.588 |
| Pcdhb10 | 257.651 | 0.096983 | 0.15209 | 0.637648 | 0.5237 | 0.71215 | 2.851 |
| Pcdhb11 | 270.525 | 0.286217 | 0.14507 | 1.972961 | 0.0485 | 0.16252 | 4.419 |
| Pcdhb12 | 172.968 | 0.006767 | 0.15565 | 0.043478 | 0.96532 | 0.98278 | 3.034 |
| Pcdhb13 | 420.994 | 0.286027 | 0.12578 | 2.273992 | 0.02297 | 0.09807 | 3.71 |
| Pcdhb14 | 672.142 | 0.137962 | 0.13164 | 1.048001 | 0.29464 | 0.50499 | 8.695 |
| Pcdhb15 | 185.248 | 0.283964 | 0.17597 | 1.613722 | 0.10659 | 0.27077 | 2.801 |
| Pcdhb16 | 656.107 | 0.226256 | 0.14752 | 1.53378 | 0.12508 | 0.30014 | 5.225 |
| Pcdhb17 | 895.824 | 0.351577 | 0.12885 | 2.728671 | 0.00636 | 0.04279 | 4.66 |
| Pcdhb18 | 371.815 | 0.199278 | 0.12284 | 1.622275 | 0.10474 | 0.26777 | 5.041 |
| Pcdhb19 | 725.818 | 0.179657 | 0.11098 | 1.618751 | 0.1055 | 0.269 | 7.138 |
| Pcdhb2 | 162.415 | 0.002324 | 0.15491 | 0.015004 | 0.98803 | 0.99319 | 2.813 |
| Pcdhb20 | 782.619 | 0.203649 | 0.08572 | 2.375843 | 0.01751 | 0.08222 | 3.559 |
| Pcdhb21 | 224.911 | -0.00334 | 0.15542 | -0.02149 | 0.98285 | 0.9907 | 4.705 |
| Pcdhb22 | 715.733 | 0.066071 | 0.0954 | 0.692586 | 0.48857 | 0.68367 | 6.609 |
| Pcdhb3 | 277.555 | 0.034774 | 0.12583 | 0.27636 | 0.78227 | 0.88672 | 3.787 |
| Pcdhb4 | 118.342 | 0.00841 | 0.18835 | 0.044653 | 0.96438 | 0.98268 | 3.718 |
| Pcdhb5 | 286.733 | 0.097546 | 0.11263 | 0.866087 | 0.38644 | 0.59577 | 3.535 |
| Pcdhb6 | 321.947 | 0.288223 | 0.17208 | 1.674971 | 0.09394 | 0.24995 | 3.754 |
| Pcdhb7 | 532.782 | 0.253185 | 0.10869 | 2.329423 | 0.01984 | 0.08901 | 3.501 |
| Pcdhb8 | 213.186 | 0.194027 | 0.20549 | 0.944199 | 0.34507 | 0.55603 | 3.484 |
| Pcdhb9 | 898.106 | 0.130289 | 0.09126 | 1.427657 | 0.15339 | 0.33905 | 3.055 |
| Pcdhga1 | 161.834 | -0.10805 | 0.16003 | -0.67521 | 0.49954 | 0.69269 | 180.081 |
| Pcdhga11 | 388.332 | -0.15961 | 0.10974 | -1.45436 | 0.14585 | 0.32872 | 86.143 |
| Pcdhga12 | 507.018 | 0.039674 | 0.1154 | 0.343797 | 0.731 | 0.85734 | 76.294 |
| Pcdhga2 | 410.494 | 0.035848 | 0.12902 | 0.277854 | 0.78112 | 0.88606 | 172.918 |
| Pcdhga3 | 651.618 | 0.173798 | 0.15212 | 1.142486 | 0.25325 | 0.46062 | 167.567 |

| | | | | | | | |
|---|---|---|---|---|---|---|---|
| Pcdhga4 | 344.667 | 0.203455 | 0.1287 | 1.58082 | 0.11392 | 0.28258 | 156.664 |
| Pcdhga5 | 486.951 | -0.14775 | 0.14842 | -0.99555 | 0.31947 | 0.53013 | 147.494 |
| Pcdhga6 | 728.111 | 0.100777 | 0.11604 | 0.86843 | 0.38516 | 0.59451 | 134.835 |
| Pcdhga7 | 525.428 | 0.197239 | 0.13633 | 1.446826 | 0.14795 | 0.33166 | 127.11 |
| Pcdhga8 | 442.797 | -0.07814 | 0.14867 | -0.52558 | 0.59918 | 0.77005 | 116.168 |
| Pcdhga9 | 618.093 | 0.100027 | 0.13211 | 0.757175 | 0.44895 | 0.65222 | 104.935 |
| Pcdhgb1 | 375.897 | -0.03901 | 0.15184 | -0.25692 | 0.79724 | 0.89545 | 161.638 |
| Pcdhgb2 | 368.793 | 0.115912 | 0.12304 | 0.942061 | 0.34616 | 0.55684 | 152.046 |
| Pcdhgb4 | 257.668 | 0.185807 | 0.16979 | 1.09436 | 0.2738 | 0.48317 | 121.502 |
| Pcdhgb5 | 306.842 | 0.102536 | 0.18822 | 0.544782 | 0.5859 | 0.76012 | 110.865 |
| Pcdhgb6 | 878.564 | 0.27616 | 0.13227 | 2.087898 | 0.03681 | 0.13533 | 99.78 |
| Pcdhgb7 | 279.019 | 0.273116 | 0.16862 | 1.619682 | 0.1053 | 0.26859 | 90.274 |
| Pcdhgb8 | 144.733 | 0.036534 | 0.16314 | 0.223941 | 0.8228 | 0.90817 | 80.07 |
| Pcdhgc3 | 1038.19 | -0.08318 | 0.12107 | -0.68703 | 0.49207 | 0.68677 | 35.51 |
| Pcdhgc4 | 548.56 | 0.172765 | 0.11779 | 1.466757 | 0.14244 | 0.32435 | 26.795 |
| Pcdhgc5 | 4458.89 | 0.131545 | 0.10905 | 1.206336 | 0.22769 | 0.43174 | 22.361 |
| Pced1a | 1076.06 | 0.03568 | 0.10021 | 0.356049 | 0.7218 | 0.85151 | 7.455 |
| Pced1b | 166.222 | 0.144566 | 0.21065 | 0.686288 | 0.49253 | 0.68714 | 138.574 |
| Pcf11 | 2742.21 | 0.189329 | 0.11213 | 1.688515 | 0.09131 | 0.24559 | 26.392 |
| Pcgf1 | 509.405 | 0.114997 | 0.09258 | 1.242131 | 0.21419 | 0.41558 | 2.987 |
| Pcgf2 | 1386.05 | 0.222618 | 0.10159 | 2.191261 | 0.02843 | 0.1137 | 11.675 |
| Pcgf3 | 3474.52 | 0.173733 | 0.07047 | 2.465292 | 0.01369 | 0.07038 | 45.745 |
| Pcgf5 | 1523.24 | 0.199169 | 0.07439 | 2.677216 | 0.00742 | 0.04738 | 112.662 |
| Pcgf6 | 1563.78 | 0.152817 | 0.10845 | 1.409136 | 0.15879 | 0.3463 | 17.29 |
| Pcid2 | 1359.76 | 0.034047 | 0.08862 | 0.384178 | 0.70085 | 0.83843 | 28.271 |
| Pcif1 | 2388.87 | 0.03853 | 0.0765 | 0.503642 | 0.61451 | 0.78055 | 15.151 |
| Pck2 | 922.556 | -0.04231 | 0.0836 | -0.50606 | 0.61281 | 0.77951 | 10.977 |
| Pclaf | 59.4149 | -0.40924 | 0.27815 | -1.47127 | 0.14122 | 0.32297 | 13.03 |
| Pclo | 16748 | 0.180947 | 0.09824 | 1.841924 | 0.06549 | 0.19743 | 348.542 |
| Pcm1 | 5494.21 | 0.41328 | 0.19152 | 2.15787 | 0.03094 | 0.1206 | 92.593 |
| Pcmt1 | 6203.37 | -0.03688 | 0.06012 | -0.61341 | 0.5396 | 0.72589 | 51.764 |
| Pcmtd1 | 9863.15 | 0.20782 | 0.12439 | 1.67067 | 0.09479 | 0.25135 | 84.709 |
| Pcmtd2 | 6594.86 | 0.187404 | 0.09276 | 2.020319 | 0.04335 | 0.15049 | 19.608 |
| Pcna | 951.089 | -0.22003 | 0.08771 | -2.50864 | 0.01212 | 0.06501 | 4.153 |

| | | | | | | | |
|---|---|---|---|---|---|---|---|
| Pcnp | 6120.48 | 0.005142 | 0.0759 | 0.067741 | 0.94599 | 0.973 | 22.495 |
| Pcnt | 2001.42 | -0.06253 | 0.08116 | -0.77053 | 0.44098 | 0.64569 | 91.524 |
| Pcnx | 8279.43 | 0.165025 | 0.06094 | 2.708194 | 0.00677 | 0.04448 | 140.902 |
| Pcnx2 | 4497.62 | -0.03568 | 0.05695 | -0.62654 | 0.53096 | 0.71852 | 146.81 |
| Pcnx3 | 3465.54 | 0.151129 | 0.0595 | 2.539932 | 0.01109 | 0.06133 | 24.274 |
| Pcnx4 | 1937.01 | 0.037834 | 0.06844 | 0.552834 | 0.58038 | 0.75658 | 43.737 |
| Pcolce | 314.663 | -0.57029 | 0.24289 | -2.3479 | 0.01888 | 0.08625 | 8.682 |
| Pcolce2 | 392.912 | -0.81794 | 0.15024 | -5.44432 | 5.20E-08 | 1.16E-05 | 60.496 |
| Pcp2 | 14.7404 | -0.38724 | 0.47428 | -0.81647 | 0.41423 | 0.62214 | 2.175 |
| Pcp4 | 135.603 | -0.31947 | 0.28873 | -1.10644 | 0.26853 | 0.47785 | 58.188 |
| Pcp4l1 | 18224.2 | 0.169558 | 0.1413 | 1.199995 | 0.23014 | 0.43449 | 23.007 |
| Pcsk1 | 1749.35 | -0.28367 | 0.19291 | -1.4705 | 0.14143 | 0.32326 | 45.036 |
| Pcsk1n | 12644.4 | -0.13501 | 0.08903 | -1.51651 | 0.12939 | 0.3069 | 4.589 |
| Pcsk2 | 10930.8 | 0.162278 | 0.11655 | 1.39237 | 0.16381 | 0.3526 | 270.13 |
| Pcsk2os1 | 148.336 | 0.288403 | 0.19579 | 1.473024 | 0.14074 | 0.32246 | 12.362 |
| Pcsk2os2 | 153.759 | 0.476154 | 0.20729 | 2.297056 | 0.02162 | 0.09437 | 48.383 |
| Pcsk4 | 241.182 | 0.686305 | 0.20095 | 3.415268 | 0.00064 | 0.00933 | 8.216 |
| Pcsk5 | 682.857 | -0.67657 | 0.21986 | -3.07733 | 0.00209 | 0.0207 | 405.314 |
| Pcsk6 | 672.809 | -0.22444 | 0.1119 | -2.00576 | 0.04488 | 0.15394 | 188.653 |
| Pcsk7 | 1835.06 | 0.213979 | 0.07358 | 2.908002 | 0.00364 | 0.02958 | 23.23 |
| Pcsk9 | 256.638 | -0.59883 | 0.17157 | -3.49026 | 0.00048 | 0.00774 | 22.001 |
| Pctp | 130.649 | -0.22064 | 0.18928 | -1.16564 | 0.24376 | 0.4502 | 20.23 |
| Pcx | 3921.96 | -0.03225 | 0.07202 | -0.44784 | 0.65427 | 0.80821 | 111.281 |
| Pcyox1 | 5432.14 | 0.003358 | 0.06556 | 0.051217 | 0.95915 | 0.98012 | 11.149 |
| Pcyox1l | 830.324 | -0.10796 | 0.10147 | -1.0639 | 0.28737 | 0.49748 | 10.799 |
| Pcyt1a | 1842.63 | -0.11045 | 0.07107 | -1.55414 | 0.12015 | 0.29275 | 44.15 |
| Pcyt1b | 2331.2 | -0.23515 | 0.10182 | -2.30942 | 0.02092 | 0.09224 | 95.089 |
| Pcyt2 | 3094.51 | -0.10964 | 0.08492 | -1.29116 | 0.19665 | 0.39438 | 7.85 |
| Pdap1 | 3619.94 | -0.17736 | 0.06765 | -2.6219 | 0.00874 | 0.05244 | 11.47 |
| Pdcd10 | 1731.69 | 0.109329 | 0.07863 | 1.390496 | 0.16438 | 0.35332 | 40.367 |
| Pdcd11 | 1808.89 | -0.09054 | 0.07126 | -1.27058 | 0.20388 | 0.40253 | 41.098 |
| Pdcd2l | 483.702 | -0.04992 | 0.09455 | -0.52794 | 0.59754 | 0.76924 | 12.163 |
| Pdcd4 | 4286.2 | 0.082787 | 0.06328 | 1.308224 | 0.1908 | 0.38706 | 37.63 |
| Pdcd5 | 796.222 | 0.1131 | 0.09524 | 1.187582 | 0.235 | 0.44032 | 5.54 |

| | | | | | | | |
|---|---|---|---|---|---|---|---|
| Pdcd5-ps | 293.096 | -0.08987 | 0.15039 | -0.59756 | 0.55013 | 0.7337 | 0.381 |
| Pdcd6 | 2239.71 | -0.0179 | 0.07792 | -0.22968 | 0.81834 | 0.90653 | 14.072 |
| Pdcd6ip | 4336.9 | -0.0121 | 0.05705 | -0.21203 | 0.83208 | 0.91398 | 56.516 |
| Pdcd7 | 1077.07 | 0.004836 | 0.08337 | 0.058001 | 0.95375 | 0.97729 | 13.54 |
| Pdcl | 1442.9 | -0.14366 | 0.08338 | -1.72302 | 0.08488 | 0.2337 | 9.259 |
| Pdcl3 | 1085.05 | 0.069867 | 0.10902 | 0.64089 | 0.52159 | 0.71075 | 11.631 |
| Pde10a | 49134.6 | 0.640947 | 0.29959 | 2.139401 | 0.0324 | 0.12455 | 461.277 |
| Pde11a | 243.139 | 0.568776 | 0.19852 | 2.86515 | 0.00417 | 0.03233 | 349.634 |
| Pde12 | 745.475 | 0.043611 | 0.08772 | 0.497145 | 0.61909 | 0.78314 | 9.926 |
| Pde1a | 6179.8 | 0.096012 | 0.12731 | 0.754126 | 0.45077 | 0.65334 | 295.006 |
| Pde1b | 34869.4 | 0.658519 | 0.27738 | 2.374085 | 0.01759 | 0.08244 | 27.019 |
| Pde1c | 2853.52 | 0.527301 | 0.18867 | 2.794877 | 0.00519 | 0.03717 | 582.684 |
| Pde2a | 376.836 | 0.674859 | 0.21283 | 3.170898 | 0.00152 | 0.01675 | 91.137 |
| Pde3a | 478.458 | 0.504943 | 0.18042 | 2.798727 | 0.00513 | 0.03691 | 258.18 |
| Pde3b | 898.845 | 0.114467 | 0.08638 | 1.32514 | 0.18512 | 0.37985 | 123.971 |
| Pde4a | 6046.35 | -0.32156 | 0.10846 | -2.96468 | 0.00303 | 0.02631 | 47.535 |
| Pde4b | 9476.18 | 0.349981 | 0.12327 | 2.839112 | 0.00452 | 0.03407 | 519.717 |
| Pde4c | 284.185 | -0.27167 | 0.1692 | -1.60556 | 0.10837 | 0.27367 | 27.467 |
| Pde4d | 1892.08 | 0.016736 | 0.07276 | 0.230013 | 0.81808 | 0.90638 | 1503.514 |
| Pde4dip | 13400.4 | 0.115026 | 0.06934 | 1.658948 | 0.09713 | 0.2554 | 198.884 |
| Pde5a | 295.843 | -0.61747 | 0.17878 | -3.45377 | 0.00055 | 0.00849 | 130.428 |
| Pde6a | 22.0592 | 0.316984 | 0.37595 | 0.843161 | 0.39914 | 0.60807 | 69.609 |
| Pde6b | 28.9302 | 0.304561 | 0.375 | 0.812159 | 0.4167 | 0.62431 | 44.007 |
| Pde6d | 1678.09 | 0.000213 | 0.08696 | 0.002455 | 0.99804 | 0.99897 | 39.636 |
| Pde6g | 21.2795 | -0.62284 | 0.37998 | -1.63911 | 0.10119 | 0.26235 | 5.896 |
| Pde6h | 28.1126 | 0.555243 | 0.34503 | 1.60924 | 0.10756 | 0.27234 | 45.034 |
| Pde7a | 1972.41 | 0.209827 | 0.086 | 2.439757 | 0.0147 | 0.07327 | 88.215 |
| Pde7b | 7815.16 | 0.600579 | 0.17741 | 3.385335 | 0.00071 | 0.01008 | 327.075 |
| Pde8a | 1095.59 | -0.32742 | 0.13786 | -2.37501 | 0.01755 | 0.0823 | 120.938 |
| Pde8b | 6536.5 | 0.434165 | 0.15331 | 2.83202 | 0.00463 | 0.03457 | 225.883 |
| Pde9a | 1338.78 | 0.407587 | 0.1262 | 3.22969 | 0.00124 | 0.01462 | 90.101 |
| Pdf | 19.7457 | 0.310798 | 0.39975 | 0.777481 | 0.43688 | 0.64256 | 3.917 |
| Pdgfa | 880.72 | -0.29607 | 0.12087 | -2.44941 | 0.01431 | 0.07211 | 21.357 |
| Pdgfb | 1094.67 | -0.26208 | 0.12619 | -2.07692 | 0.03781 | 0.13782 | 19.104 |

| | | | | | | | |
|---|---|---|---|---|---|---|---|
| Pdgfc | 424.564 | -0.39015 | 0.18554 | -2.10279 | 0.03548 | 0.1323 | 177.625 |
| Pdgfd | 194.68 | -0.24687 | 0.19188 | -1.28657 | 0.19824 | 0.39602 | 210.267 |
| Pdgfra | 3113.01 | -0.19584 | 0.06249 | -3.13393 | 0.00172 | 0.01828 | 45.924 |
| Pdgfrb | 1037.4 | -0.34577 | 0.09417 | -3.67184 | 0.00024 | 0.00483 | 39.912 |
| Pdgfrl | 38.9085 | -0.47175 | 0.31009 | -1.52134 | 0.12817 | 0.30512 | 64.574 |
| Pdha1 | 6265.23 | -0.11964 | 0.07298 | -1.63935 | 0.10114 | 0.26232 | 16.205 |
| Pdhb | 6082.6 | 0.006384 | 0.06835 | 0.093397 | 0.92559 | 0.96248 | 7.03 |
| Pdhx | 1993.57 | -0.05109 | 0.06913 | -0.73906 | 0.45987 | 0.66038 | 52.439 |
| Pdia2 | 17.9904 | 0.087404 | 0.50223 | 0.174029 | 0.86184 | 0.92991 | 3.089 |
| Pdia3 | 7962.02 | -0.02655 | 0.09784 | -0.27141 | 0.78608 | 0.88876 | 24.913 |
| Pdia4 | 2205.92 | -0.01835 | 0.11986 | -0.15307 | 0.87834 | 0.93916 | 17.29 |
| Pdia5 | 416.041 | 0.411519 | 0.1526 | 2.696637 | 0.007 | 0.04544 | 93.562 |
| Pdia6 | 5059.86 | -0.02712 | 0.11858 | -0.22867 | 0.81913 | 0.90696 | 18.226 |
| Pdik1l | 717.582 | 0.129784 | 0.10473 | 1.239172 | 0.21528 | 0.41674 | 12.894 |
| Pdk1 | 2160.33 | -0.09135 | 0.08346 | -1.09452 | 0.27373 | 0.48313 | 30.635 |
| Pdk2 | 3192.25 | -0.24372 | 0.08414 | -2.89652 | 0.00377 | 0.03027 | 15.097 |
| Pdk3 | 1504.12 | -0.01768 | 0.07469 | -0.23665 | 0.81292 | 0.90371 | 67.595 |
| Pdk4 | 1245.7 | -0.07289 | 0.20776 | -0.35083 | 0.72572 | 0.85388 | 12.959 |
| Pdlim1 | 176.401 | -1.18323 | 0.17595 | -6.72485 | 1.76E-11 | 2.24E-08 | 50.67 |
| Pdlim2 | 304.602 | -0.45741 | 0.19849 | -2.30441 | 0.0212 | 0.09307 | 13.464 |
| Pdlim3 | 144.053 | -0.98045 | 0.2273 | -4.31355 | 1.61E-05 | 0.00077 | 34.088 |
| Pdlim4 | 325.743 | -0.09473 | 0.21236 | -0.44609 | 0.65553 | 0.80876 | 14.087 |
| Pdlim5 | 2168.83 | -0.21946 | 0.08903 | -2.46485 | 0.01371 | 0.07043 | 156.107 |
| Pdlim7 | 1886.38 | 0.120448 | 0.08923 | 1.349841 | 0.17707 | 0.36932 | 17.882 |
| Pdp1 | 6162.7 | -0.01105 | 0.07806 | -0.1415 | 0.88748 | 0.94401 | 8.269 |
| Pdp2 | 607.455 | -0.11534 | 0.08564 | -1.34682 | 0.17804 | 0.3703 | 7.576 |
| Pdpk1 | 9527.45 | 0.24466 | 0.07942 | 3.080534 | 0.00207 | 0.02052 | 77.245 |
| Pdpn | 491.821 | -0.53139 | 0.12626 | -4.20881 | 2.57E-05 | 0.00104 | 32.134 |
| Pdpr | 1920.56 | -0.02254 | 0.07003 | -0.32192 | 0.74751 | 0.86763 | 42.445 |
| Pdrg1 | 2319.7 | 0.138922 | 0.09638 | 1.441382 | 0.14948 | 0.33376 | 6.538 |
| Pds5a | 3716.51 | 0.116113 | 0.07686 | 1.510734 | 0.13086 | 0.3093 | 92.553 |
| Pds5b | 7997.65 | 0.360208 | 0.11371 | 3.1678 | 0.00154 | 0.0169 | 136.952 |
| Pdss1 | 620.115 | 0.270211 | 0.11144 | 2.424828 | 0.01532 | 0.07543 | 44.745 |
| Pdss2 | 407.349 | -0.05017 | 0.10766 | -0.466 | 0.64121 | 0.79912 | 243.397 |

| | | | | | | | |
|---|---|---|---|---|---|---|---|
| Pdxdc1 | 3613.29 | -0.01614 | 0.07259 | -0.2224 | 0.824 | 0.90875 | 69.984 |
| Pdxk | 7409.1 | 0.320973 | 0.11261 | 2.850341 | 0.00437 | 0.03339 | 28.232 |
| Pdxp | 1357.4 | -0.08445 | 0.09819 | -0.86005 | 0.38976 | 0.59892 | 5.599 |
| Pdyn | 8898.41 | 0.812357 | 0.34456 | 2.357695 | 0.01839 | 0.08488 | 13.28 |
| Pdzd11 | 695.261 | -0.09015 | 0.11745 | -0.76762 | 0.44271 | 0.64727 | 3.686 |
| Pdzd2 | 13821.9 | 0.551929 | 0.19168 | 2.879367 | 0.00398 | 0.0314 | 380.214 |
| Pdzd3 | 63.5935 | 0.332239 | 0.27177 | 1.222479 | 0.22153 | 0.42444 | 4.165 |
| Pdzd4 | 6076.08 | 0.105616 | 0.07709 | 1.370037 | 0.17068 | 0.36101 | 31.611 |
| Pdzd7 | 802.885 | 0.042275 | 0.10001 | 0.422692 | 0.67252 | 0.8199 | 18.812 |
| Pdzd8 | 4117.07 | 0.166099 | 0.07493 | 2.21681 | 0.02664 | 0.10874 | 48.603 |
| Pdzd9 | 25.4014 | 0.206933 | 0.37067 | 0.558263 | 0.57666 | 0.75408 | 11.613 |
| Pdzk1 | 14.9454 | -0.01355 | 0.46996 | -0.02883 | 0.977 | 0.98796 | 41.643 |
| Pdzk1ip1 | 13.7554 | -2 | 0.77798 | -2.76379 | 0.00571 | 0.0399 | 5.192 |
| Pdzph1 | 215.293 | 0.076546 | 0.15716 | 0.487057 | 0.62622 | 0.78844 | 112.568 |
| Pdzrn3 | 541.888 | -1.19589 | 0.25638 | -4.66454 | 3.09E-06 | 0.00024 | 228.289 |
| Pdzrn4 | 363.308 | 0.153975 | 0.131 | 1.175381 | 0.23984 | 0.44579 | 374.939 |
| Pea15a | 16853.9 | -0.15453 | 0.09612 | -1.60767 | 0.10791 | 0.2729 | 10.077 |
| Peak1 | 3133.3 | 0.083609 | 0.09705 | 0.861507 | 0.38896 | 0.59811 | 216.942 |
| Pear1 | 301.392 | -0.56016 | 0.2066 | -2.7113 | 0.0067 | 0.04418 | 76.417 |
| Pebp1 | 11452.7 | -0.16159 | 0.08482 | -1.90516 | 0.05676 | 0.18035 | 4.972 |
| Pecam1 | 970.703 | -0.45724 | 0.14703 | -3.10978 | 0.00187 | 0.01921 | 96.412 |
| Pecr | 238.019 | 0.254114 | 0.12159 | 2.089879 | 0.03663 | 0.13489 | 25.148 |
| Peds1 | 819.275 | -0.34278 | 0.10245 | -3.34587 | 0.00082 | 0.01117 | 47.215 |
| Pef1 | 1623.91 | -0.00534 | 0.08245 | -0.06483 | 0.94831 | 0.97427 | 25.578 |
| Peg10 | 3471.75 | 1.280924 | 0.31941 | 4.010248 | 6.07E-05 | 0.0019 | 13.212 |
| Peg13 | 8317.31 | -0.18854 | 0.0637 | -2.96007 | 0.00308 | 0.02649 | 4.725 |
| Peg3 | 27709 | 0.24392 | 0.10617 | 2.297341 | 0.0216 | 0.09434 | 26.54 |
| Peli1 | 3077.33 | 0.223671 | 0.07952 | 2.812896 | 0.00491 | 0.03577 | 59.033 |
| Peli2 | 2030.04 | 0.041346 | 0.09411 | 0.439325 | 0.66043 | 0.81229 | 160.753 |
| Peli3 | 1339.38 | 0.047435 | 0.10214 | 0.464392 | 0.64237 | 0.80022 | 13.411 |
| Pelp1 | 1738.56 | 0.02858 | 0.07094 | 0.402881 | 0.68704 | 0.82926 | 17.149 |
| Pemt | 50.5029 | -0.27855 | 0.29731 | -0.93692 | 0.3488 | 0.55931 | 75.876 |
| Penk | 25104.2 | 0.817297 | 0.3789 | 2.157022 | 0.031 | 0.12075 | 5.289 |
| Pepd | 962.374 | -0.22393 | 0.10555 | -2.12163 | 0.03387 | 0.12849 | 132.33 |

| | | | | | | | |
|---|---|---|---|---|---|---|---|
| Per1 | 5811.31 | 0.017736 | 0.0625 | 0.283769 | 0.77659 | 0.88361 | 14.744 |
| Per2 | 3280.23 | 0.211391 | 0.1173 | 1.802148 | 0.07152 | 0.20947 | 43.343 |
| Per3 | 4959.37 | 0.342301 | 0.10499 | 3.260316 | 0.00111 | 0.0137 | 41.014 |
| Perm1 | 384.443 | 0.298398 | 0.12919 | 2.309716 | 0.0209 | 0.09219 | 5.44 |
| Perp | 386.402 | 0.22652 | 0.1913 | 1.184087 | 0.23638 | 0.44163 | 12.054 |
| Pes1 | 1550.79 | -0.13215 | 0.06927 | -1.90772 | 0.05643 | 0.17969 | 16.03 |
| Pet100 | 449.764 | -0.48178 | 0.14362 | -3.35451 | 0.0008 | 0.01094 | 4.301 |
| Pex1 | 1399.08 | 0.114785 | 0.08064 | 1.423392 | 0.15462 | 0.34066 | 41.167 |
| Pex10 | 558.994 | 0.057692 | 0.08647 | 0.667185 | 0.50465 | 0.6967 | 5.418 |
| Pex11a | 561.378 | -0.08264 | 0.08797 | -0.93942 | 0.34751 | 0.55799 | 7.175 |
| Pex11b | 612.347 | -0.01236 | 0.08972 | -0.13773 | 0.89046 | 0.94517 | 9.991 |
| Pex11g | 123.073 | 0.585858 | 0.18964 | 3.089237 | 0.00201 | 0.02016 | 10.576 |
| Pex12 | 666.515 | 0.039635 | 0.09166 | 0.432419 | 0.66544 | 0.81526 | 7.945 |
| Pex13 | 1486.01 | 0.141399 | 0.08525 | 1.658548 | 0.09721 | 0.25551 | 19.481 |
| Pex14 | 354.931 | -0.09309 | 0.10616 | -0.87695 | 0.38051 | 0.59002 | 139.342 |
| Pex16 | 823.046 | -0.21556 | 0.0835 | -2.5816 | 0.00983 | 0.05687 | 6.542 |
| Pex19 | 3249.75 | -0.18968 | 0.05773 | -3.28558 | 0.00102 | 0.01296 | 9.739 |
| Pex2 | 1040.87 | -0.05906 | 0.08678 | -0.68061 | 0.49612 | 0.68983 | 16.052 |
| Pex26 | 1138.93 | 0.122986 | 0.09987 | 1.231486 | 0.21814 | 0.42016 | 15.171 |
| Pex3 | 984.145 | 0.118549 | 0.08633 | 1.373145 | 0.16971 | 0.36001 | 29.301 |
| Pex5 | 2614.26 | -0.02919 | 0.06837 | -0.42699 | 0.66938 | 0.81764 | 18.252 |
| Pex5l | 4865.02 | -0.55609 | 0.17573 | -3.16457 | 0.00155 | 0.01704 | 193.84 |
| Pex6 | 2368.54 | -0.0688 | 0.08833 | -0.77891 | 0.43603 | 0.64169 | 14.079 |
| Pex7 | 1425.62 | 0.304511 | 0.06906 | 4.409377 | 1.04E-05 | 0.00057 | 53.79 |
| Pf4 | 20.6684 | -0.49861 | 0.45727 | -1.0904 | 0.27554 | 0.4847 | 0.949 |
| Pfas | 2096.46 | -0.03679 | 0.06 | -0.61312 | 0.5398 | 0.72592 | 22.764 |
| Pfdn1 | 1357.45 | -0.07321 | 0.07708 | -0.94973 | 0.34225 | 0.553 | 50.849 |
| Pfdn2 | 2854.85 | -0.33722 | 0.07925 | -4.25516 | 2.09E-05 | 0.00091 | 13.585 |
| Pfdn4 | 470.758 | 0.035728 | 0.09539 | 0.374563 | 0.70799 | 0.84235 | 22.696 |
| Pfdn5 | 3503.32 | -0.0953 | 0.09359 | -1.01827 | 0.30855 | 0.51837 | 5.758 |
| Pfdn6 | 1254.84 | -0.1781 | 0.08829 | -2.01721 | 0.04367 | 0.15117 | 1.523 |
| Pfkfb1 | 26.1164 | 0.037983 | 0.33982 | 0.111775 | 0.911 | 0.95504 | 55.65 |
| Pfkfb2 | 2174.32 | 0.174893 | 0.08949 | 1.954299 | 0.05067 | 0.16741 | 40.072 |
| Pfkfb3 | 1944 | 0.002457 | 0.07954 | 0.030891 | 0.97536 | 0.9876 | 82.645 |

| | | | | | | | |
|---|---|---|---|---|---|---|---|
| Pfkfb4 | 568.268 | -0.12874 | 0.11155 | -1.15409 | 0.24846 | 0.45566 | 40.451 |
| Pfkl | 6352.85 | -0.10703 | 0.07713 | -1.38769 | 0.16523 | 0.35431 | 23.137 |
| Pfkm | 17598.2 | -0.07653 | 0.05635 | -1.35802 | 0.17446 | 0.36611 | 39.863 |
| Pfkp | 18412.1 | -0.16469 | 0.09207 | -1.78866 | 0.07367 | 0.21382 | 69.01 |
| Pfn1 | 3408.42 | -0.16834 | 0.07421 | -2.26857 | 0.02329 | 0.09892 | 2.795 |
| Pfn2 | 18212.8 | -0.07485 | 0.05835 | -1.28278 | 0.19957 | 0.39758 | 6.185 |
| Pfn4 | 371.446 | 0.016583 | 0.11721 | 0.141481 | 0.88749 | 0.94401 | 9.792 |
| Pgam1 | 10531.7 | -0.12723 | 0.08105 | -1.5699 | 0.11644 | 0.28668 | 6.738 |
| Pgam1-ps2 | 127.82 | -0.09456 | 0.17197 | -0.54987 | 0.58241 | 0.75787 | 0.765 |
| Pgam2 | 333.762 | -0.01985 | 0.1375 | -0.14434 | 0.88523 | 0.94299 | 2.094 |
| Pgam5 | 1991.32 | -0.08312 | 0.10545 | -0.78818 | 0.43059 | 0.63686 | 10.784 |
| Pgap1 | 2773.98 | 0.530529 | 0.09567 | 5.545148 | 2.94E-08 | 7.88E-06 | 84.691 |
| Pgap2 | 447.74 | -0.07936 | 0.10111 | -0.78484 | 0.43255 | 0.63868 | 28.36 |
| Pgap3 | 399.179 | 0.046808 | 0.12563 | 0.372583 | 0.70946 | 0.84351 | 11.814 |
| Pgap4 | 2043.44 | -0.21292 | 0.08827 | -2.41226 | 0.01585 | 0.07724 | 13.371 |
| Pgap6 | 746.522 | -0.17358 | 0.12986 | -1.33667 | 0.18133 | 0.3748 | 9.956 |
| Pgbd1 | 175.188 | 0.20633 | 0.2104 | 0.98067 | 0.32676 | 0.53806 | 19.784 |
| Pgbd5 | 7461.74 | -0.16685 | 0.09211 | -1.81154 | 0.07006 | 0.20667 | 70.61 |
| Pgd | 1717.38 | -0.15969 | 0.07434 | -2.14798 | 0.03172 | 0.12267 | 16.781 |
| Pgf | 76.0749 | -0.50342 | 0.23936 | -2.1032 | 0.03545 | 0.13227 | 11.156 |
| Pgghg | 229.592 | -0.28695 | 0.13243 | -2.16677 | 0.03025 | 0.11875 | 6.274 |
| Pggt1b | 1591.14 | 0.119105 | 0.13865 | 0.859068 | 0.3903 | 0.59927 | 45.643 |
| Pgk1 | 13368.5 | -0.20411 | 0.0541 | -3.77288 | 0.00016 | 0.00372 | 16.6 |
| Pgk1-rs7 | 63.5627 | -0.6017 | 0.26123 | -2.30333 | 0.02126 | 0.09326 | 1.255 |
| Pgls | 665.494 | 0.036379 | 0.09825 | 0.370268 | 0.71118 | 0.8446 | 8.917 |
| Pglyrp1 | 416.913 | -0.48078 | 0.13766 | -3.4924 | 0.00048 | 0.00771 | 19.129 |
| Pgm1 | 2290.86 | 0.101875 | 0.09824 | 1.037 | 0.29974 | 0.50987 | 57.881 |
| Pgm2 | 383.167 | -0.27175 | 0.09901 | -2.74475 | 0.00606 | 0.04156 | 35.402 |
| Pgm2l1 | 12748.9 | -0.11505 | 0.13321 | -0.86368 | 0.38776 | 0.59708 | 51.475 |
| Pgm3 | 478.281 | 0.246888 | 0.11285 | 2.187845 | 0.02868 | 0.1143 | 17.466 |
| Pgm5 | 243.66 | -0.85118 | 0.19596 | -4.34371 | 1.40E-05 | 0.00071 | 178.84 |
| Pgp | 2695.53 | -0.34062 | 0.0763 | -4.46432 | 8.03E-06 | 0.00047 | 2.719 |
| Pgpep1 | 508.225 | -0.40769 | 0.13094 | -3.11342 | 0.00185 | 0.0191 | 13.954 |
| Pgpep1l | 24.6672 | -0.94968 | 0.5369 | -1.76883 | 0.07692 | 0.21936 | 27.624 |

| | | | | | | | |
|---|---|---|---|---|---|---|---|
| Pgr | 1588.87 | 0.158944 | 0.07757 | 2.049011 | 0.04046 | 0.14371 | 68.779 |
| Pgrmc1 | 12376.7 | 0.072607 | 0.08499 | 0.854305 | 0.39294 | 0.6022 | 7.874 |
| Pgrmc2 | 3656.11 | 0.02659 | 0.06703 | 0.39666 | 0.69162 | 0.83252 | 16.721 |
| Pgs1 | 2405.82 | -0.08483 | 0.07097 | -1.1952 | 0.23201 | 0.43692 | 37.72 |
| Phactr1 | 41275.1 | 0.410104 | 0.14919 | 2.748876 | 0.00598 | 0.04115 | 457.904 |
| Phactr2 | 1852.48 | -0.84652 | 0.19177 | -4.41416 | 1.01E-05 | 0.00056 | 266.696 |
| Phactr3 | 9193.59 | -0.09505 | 0.07462 | -1.27386 | 0.20271 | 0.40133 | 219.518 |
| Phactr4 | 817.179 | -0.36842 | 0.12937 | -2.84785 | 0.0044 | 0.03354 | 66.567 |
| Phaf1 | 1606.57 | 0.107106 | 0.08308 | 1.289206 | 0.19733 | 0.3952 | 27.909 |
| Phax | 1314.52 | 0.052198 | 0.07677 | 0.679899 | 0.49657 | 0.69012 | 25.27 |
| Phb | 2735.76 | -0.08448 | 0.08226 | -1.02694 | 0.30445 | 0.51424 | 13.817 |
| Phb2 | 1671.16 | -0.05036 | 0.0783 | -0.64317 | 0.52011 | 0.70963 | 4.615 |
| Phc1 | 2529.53 | 0.050791 | 0.08169 | 0.621725 | 0.53412 | 0.72132 | 22.831 |
| Phc2 | 4007.56 | 0.148932 | 0.06425 | 2.317997 | 0.02045 | 0.09091 | 98.18 |
| Phc3 | 5767.39 | 0.070143 | 0.06924 | 1.013072 | 0.31103 | 0.52125 | 70.045 |
| Pheta1 | 572.652 | 0.153041 | 0.11824 | 1.294335 | 0.19555 | 0.39318 | 5.649 |
| Pheta2 | 27.4899 | -0.36537 | 0.39723 | -0.9198 | 0.35768 | 0.56794 | 6.802 |
| Phex | 151.8 | 1.707942 | 0.24381 | 7.0052 | 2.47E-12 | 5.03E-09 | 253.238 |
| Phf1 | 902.522 | 0.030719 | 0.1073 | 0.286304 | 0.77465 | 0.88249 | 4.857 |
| Phf10 | 2302.66 | 0.187801 | 0.07069 | 2.656548 | 0.00789 | 0.04923 | 16.265 |
| Phf11b | 13.3945 | -1.05711 | 0.57932 | -1.82473 | 0.06804 | 0.20237 | 20.388 |
| Phf11c | 66.6664 | 0.162297 | 0.26474 | 0.613043 | 0.53985 | 0.72592 | 14.549 |
| Phf11d | 34.7666 | -0.91766 | 0.40077 | -2.28976 | 0.02204 | 0.09565 | 18.064 |
| Phf12 | 2752.2 | 0.171473 | 0.06078 | 2.821153 | 0.00479 | 0.0353 | 47.786 |
| Phf13 | 556.077 | 0.00292 | 0.10899 | 0.026787 | 0.97863 | 0.98863 | 6.626 |
| Phf14 | 4148.29 | 0.231688 | 0.12094 | 1.915748 | 0.0554 | 0.17752 | 173.397 |
| Phf19 | 62.9193 | 0.189841 | 0.21724 | 0.87387 | 0.38219 | 0.59181 | 20.27 |
| Phf2 | 3541.5 | 0.015269 | 0.0708 | 0.215673 | 0.82924 | 0.91246 | 69.37 |
| Phf20 | 3991.82 | -0.03941 | 0.07405 | -0.53219 | 0.5946 | 0.76667 | 113.487 |
| Phf20l1 | 4766.14 | 0.180794 | 0.13821 | 1.308072 | 0.19085 | 0.3871 | 70.417 |
| Phf21a | 3522.69 | 0.127726 | 0.07611 | 1.678272 | 0.09329 | 0.2486 | 271.55 |
| Phf21b | 399.229 | 0.147798 | 0.11672 | 1.266238 | 0.20543 | 0.40451 | 70.669 |
| Phf23 | 1541.99 | -0.04953 | 0.07678 | -0.64506 | 0.51889 | 0.70858 | 4.233 |
| Phf24 | 15397.2 | 0.120732 | 0.04796 | 2.517508 | 0.01182 | 0.06397 | 28.093 |

| | | | | | | | |
|---|---|---|---|---|---|---|---|
| Phf2os1 | 124.207 | 0.071251 | 0.20109 | 0.354321 | 0.7231 | 0.85223 | 4.349 |
| Phf3 | 5854.93 | 0.174176 | 0.08928 | 1.950973 | 0.05106 | 0.1682 | 71.583 |
| Phf5a | 850.934 | 0.099967 | 0.12007 | 0.832569 | 0.40509 | 0.61387 | 7.392 |
| Phf6 | 1708.67 | 0.134974 | 0.08364 | 1.613785 | 0.10657 | 0.27077 | 44.678 |
| Phf7 | 203.638 | 0.20569 | 0.13847 | 1.48545 | 0.13742 | 0.3181 | 13.629 |
| Phf8 | 1851.98 | 0.093284 | 0.07358 | 1.26776 | 0.20488 | 0.40376 | 113.188 |
| Phgdh | 2713.55 | -0.31474 | 0.11853 | -2.65531 | 0.00792 | 0.04932 | 26.821 |
| Phip | 5110.62 | 0.50026 | 0.20177 | 2.479324 | 0.01316 | 0.06871 | 109.358 |
| Phka1 | 1558.72 | -0.14285 | 0.07356 | -1.94193 | 0.05215 | 0.17053 | 130.272 |
| Phka2 | 1724.51 | -0.20535 | 0.06513 | -3.15296 | 0.00162 | 0.01748 | 96.713 |
| Phkb | 2209.46 | -0.0582 | 0.06435 | -0.90437 | 0.3658 | 0.57628 | 220.418 |
| Phkg1 | 818.268 | -0.20942 | 0.10314 | -2.03041 | 0.04231 | 0.14806 | 35.129 |
| Phkg2 | 1380.51 | -0.02685 | 0.08661 | -0.31 | 0.75656 | 0.87187 | 9.968 |
| Phlda1 | 491.287 | 0.065307 | 0.14565 | 0.448393 | 0.65387 | 0.80788 | 2.36 |
| Phlda3 | 720.434 | -0.34917 | 0.1398 | -2.49758 | 0.0125 | 0.06641 | 3.018 |
| Phldb1 | 5302.8 | -0.31976 | 0.14247 | -2.24445 | 0.0248 | 0.10346 | 48.895 |
| Phldb2 | 396.27 | -0.37856 | 0.20342 | -1.86098 | 0.06275 | 0.19225 | 207.356 |
| Phlpp1 | 5337.34 | -0.17332 | 0.09195 | -1.88494 | 0.05944 | 0.18548 | 222.499 |
| Phlpp2 | 4193.12 | 0.158394 | 0.06401 | 2.474467 | 0.01334 | 0.0693 | 76.13 |
| Phospho1 | 21.7905 | -1.3399 | 0.55248 | -2.42527 | 0.0153 | 0.07541 | 7.642 |
| Phospho2 | 1190.22 | -0.05187 | 0.0677 | -0.76619 | 0.44356 | 0.64792 | 10.383 |
| Phpt1 | 233.017 | -0.21983 | 0.13984 | -1.57204 | 0.11594 | 0.28578 | 1.795 |
| Phrf1 | 2766.48 | 0.017326 | 0.061 | 0.284031 | 0.77639 | 0.8836 | 33.967 |
| Phtf1 | 2673.4 | 0.125471 | 0.07076 | 1.77313 | 0.07621 | 0.21808 | 56.489 |
| Phtf1os | 44.2692 | 0.60254 | 0.30599 | 1.96917 | 0.04893 | 0.16349 | 4.046 |
| Phtf2 | 1095.18 | 0.152545 | 0.09701 | 1.572404 | 0.11586 | 0.2857 | 123.462 |
| Phyh | 2539.37 | -0.24746 | 0.09399 | -2.63291 | 0.00847 | 0.05131 | 19.712 |
| Phyhd1 | 241.293 | -0.24669 | 0.12624 | -1.95404 | 0.0507 | 0.16743 | 15.947 |
| Phyhip | 18117.8 | 0.160995 | 0.1092 | 1.474279 | 0.14041 | 0.3219 | 11.357 |
| Phyhipl | 6556.2 | -0.23616 | 0.08386 | -2.81605 | 0.00486 | 0.03557 | 98.284 |
| Phykpl | 721.56 | 0.227828 | 0.08986 | 2.535307 | 0.01123 | 0.0619 | 19.733 |
| Pi15 | 26.6206 | -0.16746 | 0.41447 | -0.40403 | 0.68619 | 0.82887 | 29.039 |
| Pi16 | 166.169 | 0.980549 | 0.23101 | 4.244542 | 2.19E-05 | 0.00093 | 12.914 |
| Pi4k2a | 3710.52 | -0.03863 | 0.07126 | -0.54207 | 0.58777 | 0.76124 | 31.618 |

| | | | | | | | |
|---|---|---|---|---|---|---|---|
| Pi4k2b | 146.869 | 0.196513 | 0.18438 | 1.065819 | 0.28651 | 0.49648 | 27.767 |
| Pi4ka | 13144.8 | 0.062323 | 0.05319 | 1.171758 | 0.24129 | 0.44737 | 125.964 |
| Pi4kb | 1613.91 | 0.070737 | 0.07313 | 0.967212 | 0.33344 | 0.54427 | 32.113 |
| Pianp | 6522.67 | 0.205485 | 0.06508 | 3.157243 | 0.00159 | 0.01732 | 6.403 |
| Pias1 | 3549.79 | 0.236332 | 0.10962 | 2.155979 | 0.03109 | 0.12099 | 109.557 |
| Pias2 | 3489.36 | 0.108616 | 0.05672 | 1.914817 | 0.05552 | 0.17779 | 88.593 |
| Pias3 | 697.724 | -0.03288 | 0.07801 | -0.42147 | 0.67342 | 0.82048 | 9.687 |
| Pias4 | 1038.04 | -0.16216 | 0.07185 | -2.25707 | 0.024 | 0.10097 | 14.658 |
| Pibf1 | 567.638 | 0.285906 | 0.13129 | 2.17767 | 0.02943 | 0.1166 | 155.07 |
| Picalm | 8891.33 | 0.070379 | 0.06085 | 1.156516 | 0.24747 | 0.45441 | 83.253 |
| Pick1 | 829.378 | 0.038222 | 0.13421 | 0.284793 | 0.7758 | 0.88326 | 20.312 |
| Pid1 | 2257.53 | -0.34323 | 0.12812 | -2.67895 | 0.00739 | 0.04728 | 327.885 |
| Pidd1 | 75.7134 | -0.22181 | 0.23748 | -0.93403 | 0.35029 | 0.5609 | 5.913 |
| Piezo1 | 293.864 | -0.22404 | 0.17665 | -1.26826 | 0.2047 | 0.40355 | 69.632 |
| Piezo2 | 49.5555 | -0.45795 | 0.29192 | -1.56876 | 0.1167 | 0.28707 | 376.971 |
| Pif1 | 21.0092 | -0.24902 | 0.57949 | -0.42971 | 0.6674 | 0.8162 | 8.808 |
| Pifo | 19.8581 | 1.094848 | 0.45958 | 2.38227 | 0.01721 | 0.08153 | 17.69 |
| Piga | 498.188 | -0.16758 | 0.13772 | -1.21678 | 0.22369 | 0.4269 | 14.13 |
| Pigb | 1167.9 | -0.0476 | 0.09492 | -0.50148 | 0.61604 | 0.78132 | 32.96 |
| Pigbos1 | 73.6333 | 0.354151 | 0.25596 | 1.383617 | 0.16648 | 0.356 | 3.062 |
| Pigc | 731.118 | 0.314193 | 0.10618 | 2.959131 | 0.00309 | 0.02655 | 4.25 |
| Pigf | 313.858 | 0.188264 | 0.12686 | 1.484001 | 0.13781 | 0.31864 | 28.151 |
| Pigg | 1251.49 | 0.087699 | 0.09418 | 0.931223 | 0.35174 | 0.56208 | 36.747 |
| Pigh | 242.084 | 0.117214 | 0.15963 | 0.734277 | 0.46278 | 0.66287 | 8.988 |
| Pigk | 3233.89 | 0.066046 | 0.0794 | 0.831811 | 0.40552 | 0.61425 | 266.309 |
| Pigl | 408.889 | -0.00364 | 0.13655 | -0.02669 | 0.97871 | 0.98863 | 55.958 |
| Pigm | 1048.55 | -0.12811 | 0.07718 | -1.65975 | 0.09696 | 0.25513 | 7.554 |
| Pign | 1216.13 | 0.123853 | 0.09004 | 1.375527 | 0.16897 | 0.35907 | 145.256 |
| Pigo | 1414.26 | 0.128962 | 0.09111 | 1.415496 | 0.15692 | 0.34387 | 8.185 |
| Pigp | 783.272 | 0.082159 | 0.09952 | 0.825514 | 0.40908 | 0.61691 | 13.08 |
| Pigq | 3622.7 | -0.01312 | 0.07029 | -0.18673 | 0.85187 | 0.92468 | 18.516 |
| Pigs | 1989.61 | -0.13695 | 0.06947 | -1.97143 | 0.04867 | 0.16295 | 14.368 |
| Pigt | 3008.76 | -0.08113 | 0.06908 | -1.17434 | 0.24026 | 0.44624 | 10.782 |
| Pigu | 1088.03 | 0.121684 | 0.08392 | 1.449932 | 0.14708 | 0.33022 | 79.188 |

| | | | | | | | |
|---|---|---|---|---|---|---|---|
| Pigv | 515 | 0.0996 | 0.11018 | 0.903968 | 0.36601 | 0.57648 | 12.261 |
| Pigw | 214.878 | -0.03286 | 0.15254 | -0.21543 | 0.82944 | 0.91247 | 3.971 |
| Pigx | 516.431 | 0.069672 | 0.0925 | 0.753178 | 0.45134 | 0.65382 | 15.325 |
| Pigyl | 560.943 | -0.21478 | 0.10692 | -2.00882 | 0.04456 | 0.15336 | 1.513 |
| Pigz | 269.81 | -0.29114 | 0.18591 | -1.56605 | 0.11734 | 0.28827 | 12.196 |
| Pih1d1 | 608.644 | -0.26036 | 0.10813 | -2.40791 | 0.01604 | 0.07783 | 5.763 |
| Pih1d2 | 74.5837 | 0.333137 | 0.22955 | 1.451287 | 0.1467 | 0.32983 | 7.68 |
| Pik3ap1 | 181.497 | -0.70183 | 0.17088 | -4.10705 | 4.01E-05 | 0.0014 | 112.726 |
| Pik3c2a | 1689.65 | 0.134149 | 0.11972 | 1.120513 | 0.2625 | 0.4713 | 106.185 |
| Pik3c2b | 4473.02 | 0.433673 | 0.15035 | 2.884514 | 0.00392 | 0.03101 | 63.021 |
| Pik3c3 | 2708.57 | 0.151137 | 0.07086 | 2.132896 | 0.03293 | 0.12609 | 75.38 |
| Pik3ca | 3625.44 | 0.109446 | 0.08678 | 1.26126 | 0.20722 | 0.4067 | 70.816 |
| Pik3cb | 1543.43 | -0.00632 | 0.07022 | -0.09008 | 0.92823 | 0.96393 | 103.968 |
| Pik3cd | 1329.62 | -0.11426 | 0.08208 | -1.39212 | 0.16389 | 0.35265 | 53.404 |
| Pik3cg | 272.254 | -0.03061 | 0.13802 | -0.2218 | 0.82447 | 0.90912 | 35.187 |
| Pik3ip1 | 1605.5 | 0.013956 | 0.10316 | 0.135277 | 0.89239 | 0.94617 | 12.571 |
| Pik3r1 | 8013.86 | -0.01101 | 0.08155 | -0.13497 | 0.89264 | 0.94631 | 87.655 |
| Pik3r2 | 4640.54 | -0.10409 | 0.07045 | -1.47751 | 0.13954 | 0.32076 | 8.538 |
| Pik3r3 | 2116.35 | 0.091024 | 0.08322 | 1.093725 | 0.27408 | 0.48345 | 81.439 |
| Pik3r4 | 2457.93 | 0.157599 | 0.093 | 1.69454 | 0.09016 | 0.24359 | 44.68 |
| Pik3r5 | 209.564 | 0.16782 | 0.14429 | 1.163098 | 0.24479 | 0.45139 | 65.729 |
| Pik3r6 | 86.8204 | -0.23024 | 0.24401 | -0.94354 | 0.3454 | 0.55616 | 49.68 |
| Pikfyve | 3531.4 | 0.136096 | 0.08325 | 1.634756 | 0.1021 | 0.26371 | 92.013 |
| Pim1 | 137.168 | -0.46142 | 0.22789 | -2.02472 | 0.0429 | 0.14934 | 5.36 |
| Pim2 | 1796.34 | 0.0697 | 0.10934 | 0.637439 | 0.52384 | 0.7122 | 5.171 |
| Pim3 | 1565.82 | -0.25309 | 0.09993 | -2.53254 | 0.01132 | 0.06219 | 3.541 |
| Pimreg | 20.1453 | -0.84783 | 0.53566 | -1.58277 | 0.11347 | 0.282 | 5.339 |
| Pin1 | 2512.88 | -0.21727 | 0.07872 | -2.76013 | 0.00578 | 0.04021 | 14.49 |
| Pin4 | 513.758 | -0.03809 | 0.15562 | -0.24476 | 0.80664 | 0.90042 | 8.223 |
| Pin4-ps | 43.0561 | -0.53853 | 0.31887 | -1.68887 | 0.09124 | 0.2455 | 0.395 |
| Pink1 | 10991.4 | -0.16961 | 0.09061 | -1.87189 | 0.06122 | 0.18908 | 12.899 |
| Pinx1 | 243.554 | 0.23695 | 0.15197 | 1.559235 | 0.11894 | 0.29105 | 59.496 |
| Pip4k2a | 4690.1 | -0.28172 | 0.08532 | -3.30174 | 0.00096 | 0.01246 | 155.872 |
| Pip4k2b | 8337.75 | -0.06512 | 0.06382 | -1.02026 | 0.3076 | 0.51751 | 29.548 |

| | | | | | | | |
|---|---|---|---|---|---|---|---|
| Pip4k2c | 4194.1 | -0.41479 | 0.11591 | -3.57864 | 0.00035 | 0.00618 | 14.519 |
| Pip4p1 | 2526.25 | -0.07321 | 0.06987 | -1.04771 | 0.29477 | 0.50514 | 4.789 |
| Pip4p2 | 2227.67 | 0.197981 | 0.06544 | 3.025586 | 0.00248 | 0.02325 | 51.101 |
| Pip5k1a | 3424.08 | 0.161303 | 0.09908 | 1.627962 | 0.10353 | 0.26606 | 48.401 |
| Pip5k1b | 696.108 | -0.64592 | 0.2033 | -3.17721 | 0.00149 | 0.01646 | 261.079 |
| Pip5k1c | 12580.6 | -0.24975 | 0.09052 | -2.75916 | 0.0058 | 0.04027 | 27.011 |
| Pip5kl1 | 214.653 | -0.1323 | 0.15911 | -0.83153 | 0.40568 | 0.61433 | 8.986 |
| Pipox | 120.457 | 0.204405 | 0.19532 | 1.046537 | 0.29531 | 0.50577 | 13.482 |
| Pir | 161.664 | -0.45247 | 0.18285 | -2.47457 | 0.01334 | 0.0693 | 103.641 |
| Pirb | 32.3722 | -0.54908 | 0.37724 | -1.4555 | 0.14553 | 0.32841 | 8.983 |
| Pisd | 730.451 | -0.04925 | 0.08901 | -0.55331 | 0.58005 | 0.75648 | 49.346 |
| Pisd-ps1 | 1170.15 | -0.07372 | 0.11809 | -0.62433 | 0.53241 | 0.71981 | 7.957 |
| Pisd-ps2 | 193.506 | 0.153323 | 0.13803 | 1.11077 | 0.26667 | 0.47594 | 7.606 |
| Pithd1 | 2592.45 | -0.00125 | 0.08057 | -0.01545 | 0.98767 | 0.99316 | 11.805 |
| Pitpna | 12782.5 | -0.3612 | 0.08344 | -4.32869 | 1.50E-05 | 0.00074 | 40.708 |
| Pitpnb | 2903.92 | -0.12806 | 0.09662 | -1.32532 | 0.18507 | 0.37981 | 57.597 |
| Pitpnc1 | 4294.99 | 0.04742 | 0.10159 | 0.466761 | 0.64067 | 0.79889 | 262.808 |
| Pitpnm1 | 7821.63 | -0.02366 | 0.07807 | -0.30301 | 0.76189 | 0.87518 | 13.968 |
| Pitpnm2 | 25064.9 | 0.288428 | 0.09667 | 2.983702 | 0.00285 | 0.02535 | 131.071 |
| Pitpnm2os1 | 20.083 | 1.128327 | 0.52562 | 2.146669 | 0.03182 | 0.12296 | 7.413 |
| Pitpnm2os2 | 81.0068 | 0.710849 | 0.21156 | 3.360046 | 0.00078 | 0.01076 | 5.44 |
| Pitpnm3 | 3253.32 | -0.37592 | 0.14486 | -2.59502 | 0.00946 | 0.05531 | 88.251 |
| Pitrm1 | 1743.87 | -0.01277 | 0.07084 | -0.18028 | 0.85694 | 0.92788 | 32.367 |
| Piwil2 | 50.3955 | 0.005537 | 0.24577 | 0.02253 | 0.98203 | 0.99058 | 56.907 |
| Pja1 | 9274.5 | 0.069405 | 0.06918 | 1.003281 | 0.31573 | 0.52631 | 5.54 |
| Pja2 | 30176.9 | 0.172189 | 0.08573 | 2.008591 | 0.04458 | 0.15339 | 50.912 |
| Pkd1 | 11365.1 | 0.210626 | 0.06361 | 3.311027 | 0.00093 | 0.01219 | 46.675 |
| Pkd1l2 | 17.4096 | -0.44412 | 0.39569 | -1.12239 | 0.2617 | 0.47045 | 86.771 |
| Pkd1l3 | 156.682 | 0.13201 | 0.16167 | 0.816536 | 0.41419 | 0.62213 | 59.87 |
| Pkd2 | 4159.42 | 0.106103 | 0.12154 | 0.872982 | 0.38267 | 0.59202 | 46.37 |
| Pkd2l1 | 79.6566 | -0.99598 | 0.21599 | -4.61119 | 4.00E-06 | 0.00029 | 44.806 |
| Pkd2l2 | 779.294 | 0.121285 | 0.11544 | 1.050598 | 0.29344 | 0.50337 | 35.628 |
| Pkdcc | 227.861 | -0.56384 | 0.15374 | -3.66741 | 0.00025 | 0.0049 | 9.779 |
| Pkdrej | 78.992 | -0.23503 | 0.22875 | -1.02745 | 0.30421 | 0.51405 | 7.065 |

| | | | | | | | |
|---|---|---|---|---|---|---|---|
| Pkhd1l1 | 46.0513 | 0.639133 | 0.28188 | 2.267403 | 0.02337 | 0.09912 | 143.876 |
| Pkia | 11153.8 | 0.356338 | 0.09393 | 3.793605 | 0.00015 | 0.0035 | 78.698 |
| Pkib | 330.91 | 0.033847 | 0.12333 | 0.274449 | 0.78374 | 0.88762 | 109.132 |
| Pkig | 2595.17 | -0.03446 | 0.09621 | -0.35814 | 0.72024 | 0.85069 | 67.773 |
| Pkm | 49339 | -0.06515 | 0.06478 | -1.00565 | 0.31458 | 0.5251 | 23.008 |
| Pkmyt1 | 224.936 | 0.01825 | 0.13172 | 0.138555 | 0.8898 | 0.94482 | 10.4 |
| Pkn1 | 1684.62 | 0.198283 | 0.06595 | 3.006379 | 0.00264 | 0.02422 | 32.644 |
| Pkn2 | 1955.26 | 0.123938 | 0.08161 | 1.518645 | 0.12885 | 0.30601 | 91.103 |
| Pkn3 | 74.6205 | 0.05069 | 0.23957 | 0.211588 | 0.83243 | 0.91404 | 13.339 |
| Pknox1 | 860.579 | 0.097056 | 0.09292 | 1.044486 | 0.29626 | 0.50667 | 42.96 |
| Pknox2 | 3039.15 | 0.214302 | 0.07873 | 2.721948 | 0.00649 | 0.04334 | 256.426 |
| Pkp2 | 866.878 | 0.793251 | 0.21367 | 3.712532 | 0.00021 | 0.00435 | 59.395 |
| Pkp4 | 10488.5 | -0.3509 | 0.09562 | -3.66963 | 0.00024 | 0.00487 | 194.359 |
| Pla1a | 109.433 | -0.39443 | 0.27024 | -1.45959 | 0.1444 | 0.32717 | 37.029 |
| Pla2g12a | 903.888 | 0.107917 | 0.09902 | 1.089826 | 0.27579 | 0.48496 | 17.22 |
| Pla2g15 | 1194.31 | -0.10559 | 0.08693 | -1.21474 | 0.22447 | 0.42814 | 14.317 |
| Pla2g3 | 627.161 | -0.29202 | 0.18451 | -1.58273 | 0.11348 | 0.282 | 11.067 |
| Pla2g4a | 128.195 | -0.63075 | 0.18723 | -3.36883 | 0.00075 | 0.01055 | 131.673 |
| Pla2g4b | 100.388 | 0.071801 | 0.21528 | 0.33353 | 0.73873 | 0.86225 | 9.614 |
| Pla2g4c | 24.9191 | -0.15011 | 0.38228 | -0.39266 | 0.69457 | 0.8343 | 36.018 |
| Pla2g4e | 192.247 | -0.47877 | 0.16293 | -2.93857 | 0.0033 | 0.02772 | 78.924 |
| Pla2g5 | 83.4819 | -0.50212 | 0.24728 | -2.03061 | 0.04229 | 0.14806 | 64.239 |
| Pla2g6 | 712.819 | -0.06621 | 0.09218 | -0.71825 | 0.4726 | 0.6709 | 42.163 |
| Pla2g7 | 3279.85 | -0.3806 | 0.09778 | -3.89249 | 9.92E-05 | 0.00268 | 44.104 |
| Pla2r1 | 35.6782 | -0.24491 | 0.37931 | -0.64566 | 0.5185 | 0.70819 | 135.766 |
| Plaa | 2064.45 | 0.070122 | 0.07474 | 0.938215 | 0.34813 | 0.55865 | 35.731 |
| Plaat1 | 1130.55 | 0.570951 | 0.18457 | 3.093436 | 0.00198 | 0.02 | 20.837 |
| Plaat3 | 2213.7 | -0.2002 | 0.10594 | -1.88985 | 0.05878 | 0.18443 | 31.087 |
| Plac9 | 169.463 | -0.74305 | 0.22443 | -3.31084 | 0.00093 | 0.01219 | 15.595 |
| Plag1 | 335.363 | 0.142405 | 0.12596 | 1.130573 | 0.25823 | 0.46624 | 37.428 |
| Plagl1 | 1164.99 | -0.02117 | 0.10488 | -0.20185 | 0.84003 | 0.91792 | 71.191 |
| Plagl2 | 652.096 | 0.342211 | 0.11198 | 3.055987 | 0.00224 | 0.0217 | 13.668 |
| Plat | 2188.38 | -0.34317 | 0.11644 | -2.94722 | 0.00321 | 0.02727 | 25.118 |
| Platr25 | 450.761 | 0.544487 | 0.20214 | 2.693638 | 0.00707 | 0.04576 | 88.926 |

| | | | | | | | |
|---|---|---|---|---|---|---|---|
| Platr31 | 18.2118 | -0.01171 | 0.45516 | -0.02572 | 0.97948 | 0.98894 | 10.171 |
| Plau | 52.4669 | -0.67772 | 0.3128 | -2.16665 | 0.03026 | 0.11875 | 6.726 |
| Plaur | 40.172 | 0.203107 | 0.27114 | 0.749098 | 0.4538 | 0.65609 | 13.485 |
| Plb1 | 22.3475 | -0.15044 | 0.46126 | -0.32616 | 0.7443 | 0.86539 | 133.813 |
| Plbd2 | 2575.12 | -0.07212 | 0.06445 | -1.11901 | 0.26314 | 0.472 | 20.344 |
| Plcb1 | 12817.2 | 0.126461 | 0.11576 | 1.092404 | 0.27466 | 0.4843 | 689.192 |
| Plcb2 | 287.768 | -0.01298 | 0.18297 | -0.07092 | 0.94346 | 0.97177 | 20.922 |
| Plcb3 | 961.971 | -0.32812 | 0.11328 | -2.89654 | 0.00377 | 0.03027 | 24.146 |
| Plcb4 | 1695.09 | -0.07207 | 0.1515 | -0.47573 | 0.63426 | 0.79416 | 355.583 |
| Plcd1 | 349.438 | -0.53164 | 0.1052 | -5.05345 | 4.34E-07 | 5.71E-05 | 21.976 |
| Plcd3 | 437.098 | -0.403 | 0.12365 | -3.25907 | 0.00112 | 0.01373 | 31.355 |
| Plcd4 | 1078.07 | -0.29797 | 0.10226 | -2.91384 | 0.00357 | 0.02915 | 24.907 |
| Plce1 | 1333.89 | 0.225215 | 0.15589 | 1.44472 | 0.14854 | 0.33233 | 303.918 |
| Plcg1 | 5098.81 | 0.085333 | 0.08967 | 0.951659 | 0.34127 | 0.55219 | 44.461 |
| Plcg2 | 235.961 | -0.38665 | 0.13165 | -2.93692 | 0.00331 | 0.02778 | 136.852 |
| Plch1 | 360.369 | -0.01959 | 0.14619 | -0.13399 | 0.89341 | 0.94662 | 203.239 |
| Plch2 | 4624.7 | -0.71969 | 0.28838 | -2.49562 | 0.01257 | 0.06666 | 73.67 |
| Plcl1 | 2333.3 | -0.49906 | 0.11927 | -4.18415 | 2.86E-05 | 0.00112 | 348.365 |
| Plcl2 | 3857.22 | 0.289982 | 0.1004 | 2.888175 | 0.00387 | 0.03079 | 179.082 |
| Plcxd1 | 395.044 | 0.290876 | 0.16173 | 1.798556 | 0.07209 | 0.21077 | 5.985 |
| Plcxd2 | 2537.2 | -0.737 | 0.19501 | -3.77934 | 0.00016 | 0.00367 | 50.956 |
| Plcxd3 | 626.227 | -0.08678 | 0.19014 | -0.45637 | 0.64812 | 0.80406 | 200.063 |
| Plcz1 | 46.5045 | 0.27804 | 0.36415 | 0.763529 | 0.44515 | 0.64913 | 51.785 |
| Pld1 | 780.19 | -0.41708 | 0.09601 | -4.34413 | 1.40E-05 | 0.00071 | 194.668 |
| Pld2 | 1309.52 | -0.02659 | 0.09578 | -0.27758 | 0.78133 | 0.88614 | 18.047 |
| Pld3 | 15597.6 | -0.23463 | 0.08067 | -2.90846 | 0.00363 | 0.02954 | 21.219 |
| Pld4 | 308.896 | -0.56949 | 0.15475 | -3.68007 | 0.00023 | 0.00475 | 8.336 |
| Pld5 | 396.171 | 0.19974 | 0.14718 | 1.357121 | 0.17474 | 0.36631 | 313.007 |
| Pld6 | 110.16 | 0.999912 | 0.2433 | 4.109837 | 3.96E-05 | 0.00139 | 3.749 |
| Plec | 13296.4 | -0.06273 | 0.05714 | -1.0978 | 0.27229 | 0.48159 | 61.601 |
| Plek | 915.366 | -0.04511 | 0.08988 | -0.5019 | 0.61574 | 0.78132 | 81.176 |
| Plek2 | 14.8391 | -0.33952 | 0.54695 | -0.62075 | 0.53476 | 0.72177 | 18.274 |
| Plekha1 | 6616.07 | -0.19028 | 0.09152 | -2.07904 | 0.03761 | 0.13733 | 47.557 |
| Plekha2 | 1073.78 | -0.6386 | 0.20516 | -3.11268 | 0.00185 | 0.01911 | 63.233 |

| | | | | | | | |
|---|---|---|---|---|---|---|---|
| Plekha3 | 2143.47 | 0.048225 | 0.07229 | 0.667067 | 0.50473 | 0.6967 | 21.548 |
| Plekha4 | 87.989 | -0.31109 | 0.21265 | -1.46292 | 0.14349 | 0.32566 | 27.9 |
| Plekha5 | 7471.19 | 0.442219 | 0.09504 | 4.652893 | 3.27E-06 | 0.00025 | 173.057 |
| Plekha6 | 6554.23 | 0.29015 | 0.08404 | 3.452693 | 0.00056 | 0.00851 | 139.226 |
| Plekha7 | 1520.53 | -0.41529 | 0.14305 | -2.90301 | 0.0037 | 0.02987 | 184.892 |
| Plekha8 | 1125.41 | 0.081985 | 0.09084 | 0.902535 | 0.36677 | 0.57719 | 50.729 |
| Plekhb1 | 12005.9 | -0.58073 | 0.13886 | -4.18213 | 2.89E-05 | 0.00113 | 19.523 |
| Plekhb2 | 13412.6 | -0.02071 | 0.0762 | -0.27178 | 0.78579 | 0.88863 | 29.605 |
| Plekhd1 | 242.503 | -0.7609 | 0.1246 | -6.10654 | 1.02E-09 | 5.32E-07 | 31.624 |
| Plekhf1 | 303.658 | -0.3994 | 0.15858 | -2.5185 | 0.01179 | 0.06387 | 11.045 |
| Plekhf2 | 397.734 | -0.04408 | 0.11012 | -0.40028 | 0.68895 | 0.83048 | 19.266 |
| Plekhg1 | 954.16 | -0.36547 | 0.1116 | -3.27472 | 0.00106 | 0.01329 | 226.94 |
| Plekhg2 | 328.742 | -0.22129 | 0.12841 | -1.72331 | 0.08483 | 0.23363 | 12.996 |
| Plekhg3 | 1134.95 | -0.48743 | 0.11999 | -4.06209 | 4.86E-05 | 0.00161 | 49.598 |
| Plekhg4 | 707.568 | 0.660659 | 0.1746 | 3.78386 | 0.00015 | 0.00361 | 9.589 |
| Plekhg5 | 3539.66 | -0.20881 | 0.11264 | -1.85388 | 0.06376 | 0.19418 | 42.903 |
| Plekhg6 | 17.5553 | -1.08374 | 0.5314 | -2.03941 | 0.04141 | 0.14598 | 18.134 |
| Plekhh1 | 1688.68 | -0.47599 | 0.14654 | -3.24819 | 0.00116 | 0.01403 | 52.493 |
| Plekhh2 | 853.471 | -0.00954 | 0.1215 | -0.0785 | 0.93743 | 0.96877 | 110.244 |
| Plekhh3 | 347.245 | -0.2915 | 0.14341 | -2.03265 | 0.04209 | 0.14754 | 8.673 |
| Plekhj1 | 846.098 | -0.03917 | 0.08911 | -0.43957 | 0.66025 | 0.81222 | 2.528 |
| Plekhm1 | 1661.44 | -0.04682 | 0.0693 | -0.67561 | 0.49929 | 0.69253 | 48.413 |
| Plekhm2 | 4168.56 | -0.22817 | 0.09605 | -2.37554 | 0.01752 | 0.08225 | 39.166 |
| Plekhm3 | 3203.79 | 0.108217 | 0.06804 | 1.59042 | 0.11174 | 0.27929 | 170.842 |
| Plekhn1 | 844.488 | 0.358501 | 0.13463 | 2.662933 | 0.00775 | 0.04864 | 8.262 |
| Plekho1 | 777.841 | -0.75599 | 0.15656 | -4.82889 | 1.37E-06 | 0.00013 | 7.573 |
| Plekho2 | 491.009 | -0.40891 | 0.11115 | -3.67874 | 0.00023 | 0.00476 | 27.343 |
| Plekhs1 | 45.5653 | 1.162053 | 0.29244 | 3.973619 | 7.08E-05 | 0.00212 | 25.113 |
| Plgrkt | 809.756 | 0.115417 | 0.11658 | 0.990038 | 0.32216 | 0.53314 | 18.792 |
| Plin2 | 272.293 | -0.6166 | 0.1479 | -4.16908 | 3.06E-05 | 0.00117 | 21.675 |
| Plin3 | 678.355 | -0.66131 | 0.11751 | -5.6279 | 1.82E-08 | 5.47E-06 | 15.398 |
| Plin4 | 3389.77 | -0.55504 | 0.12929 | -4.29288 | 1.76E-05 | 0.00081 | 9.213 |
| Plin5 | 76.9397 | -0.27244 | 0.24713 | -1.10242 | 0.27028 | 0.47985 | 5.996 |
| Plk-ps1 | 92.8719 | -0.22012 | 0.20181 | -1.09076 | 0.27538 | 0.4847 | 1.787 |

| | | | | | | | |
|---|---|---|---|---|---|---|---|
| Plk1 | 94.1859 | -0.17335 | 0.22663 | -0.76488 | 0.44434 | 0.64834 | 10.435 |
| Plk2 | 9323.05 | 0.182416 | 0.09085 | 2.00788 | 0.04466 | 0.1535 | 5.799 |
| Plk3 | 346.292 | -0.45238 | 0.19579 | -2.31048 | 0.02086 | 0.09212 | 5.309 |
| Plk4 | 401.409 | 0.292115 | 0.12261 | 2.382563 | 0.01719 | 0.08153 | 16.865 |
| Plk5 | 1089.64 | 0.122675 | 0.12689 | 0.966773 | 0.33366 | 0.54449 | 9.031 |
| Pllp | 123.407 | -0.11361 | 0.20704 | -0.54874 | 0.58319 | 0.75835 | 21.385 |
| Pln | 15.6002 | -0.31415 | 0.49064 | -0.64029 | 0.52199 | 0.71105 | 8.321 |
| Plod1 | 919.915 | -0.36636 | 0.12376 | -2.96024 | 0.00307 | 0.02648 | 27.015 |
| Plod2 | 269.873 | -0.49882 | 0.17545 | -2.84307 | 0.00447 | 0.03383 | 66.206 |
| Plod3 | 1158.61 | -0.01276 | 0.08471 | -0.15064 | 0.88026 | 0.94022 | 9.63 |
| Plp1 | 56713.6 | -0.43851 | 0.18343 | -2.39057 | 0.01682 | 0.08027 | 17.063 |
| Plp2 | 68.1146 | -0.53388 | 0.25283 | -2.11159 | 0.03472 | 0.13049 | 3.45 |
| Plpbp | 3528.83 | 0.063954 | 0.06729 | 0.950354 | 0.34193 | 0.55274 | 13.578 |
| Plpp1 | 366.028 | -0.08575 | 0.11171 | -0.76757 | 0.44274 | 0.64727 | 67.103 |
| Plpp2 | 77.7297 | -0.47978 | 0.20623 | -2.32643 | 0.02 | 0.08956 | 7.367 |
| Plpp3 | 11580.8 | -0.27195 | 0.07926 | -3.43125 | 0.0006 | 0.00897 | 75.418 |
| Plpp4 | 162.667 | -0.00847 | 0.16779 | -0.0505 | 0.95972 | 0.98045 | 134.382 |
| Plpp5 | 1101.88 | 0.045369 | 0.09178 | 0.494328 | 0.62107 | 0.7845 | 4.851 |
| Plpp6 | 2218.64 | -0.25191 | 0.11523 | -2.18606 | 0.02881 | 0.11468 | 2.859 |
| Plpp7 | 781.15 | 0.018281 | 0.08698 | 0.210177 | 0.83353 | 0.91471 | 15.293 |
| Plppr1 | 2114.86 | 0.791098 | 0.29552 | 2.677002 | 0.00743 | 0.04738 | 280.987 |
| Plppr2 | 5309.03 | -0.11594 | 0.09511 | -1.21905 | 0.22283 | 0.42605 | 11.875 |
| Plppr3 | 2839.78 | 0.387476 | 0.12272 | 3.157468 | 0.00159 | 0.01732 | 14.16 |
| Plppr4 | 15723.3 | 0.294295 | 0.08065 | 3.648854 | 0.00026 | 0.00513 | 41.738 |
| Plppr5 | 2780.99 | 0.12698 | 0.09475 | 1.340134 | 0.1802 | 0.37349 | 204.424 |
| Plrg1 | 1469.67 | 0.069314 | 0.09713 | 0.713602 | 0.47547 | 0.67357 | 16.834 |
| Pls1 | 793.761 | 0.043695 | 0.14404 | 0.303361 | 0.76161 | 0.87511 | 92.67 |
| Pls3 | 4718.25 | -0.34838 | 0.14688 | -2.3718 | 0.0177 | 0.08269 | 89.529 |
| Plscr1 | 66.7379 | -0.49608 | 0.26754 | -1.85422 | 0.06371 | 0.19405 | 22.529 |
| Plscr2 | 66.1457 | -0.80972 | 0.2726 | -2.9704 | 0.00297 | 0.02603 | 22.151 |
| Plscr3 | 435.761 | -0.0607 | 0.11831 | -0.51311 | 0.60788 | 0.77584 | 5.683 |
| Plscr4 | 177.314 | -0.02556 | 0.20449 | -0.12497 | 0.90055 | 0.95009 | 35.084 |
| Pltp | 1472.86 | -0.16703 | 0.07916 | -2.11006 | 0.03485 | 0.13081 | 18.194 |
| Plvap | 212.5 | -0.06292 | 0.17493 | -0.35967 | 0.71909 | 0.84995 | 13.988 |

| | | | | | | | |
|---|---|---|---|---|---|---|---|
| Plxdc1 | 587.415 | -0.31057 | 0.10886 | -2.85309 | 0.00433 | 0.03319 | 63.207 |
| Plxdc2 | 1602.87 | -0.17981 | 0.09549 | -1.883 | 0.0597 | 0.18611 | 399.536 |
| Plxna1 | 2952.97 | 0.112982 | 0.09802 | 1.152667 | 0.24905 | 0.45629 | 46.307 |
| Plxna2 | 11057.5 | 0.263439 | 0.08872 | 2.969193 | 0.00299 | 0.02607 | 198.652 |
| Plxna3 | 1522.91 | 0.012787 | 0.13619 | 0.093891 | 0.9252 | 0.96225 | 15.624 |
| Plxna4 | 7148.87 | -0.02202 | 0.07552 | -0.29156 | 0.77062 | 0.88002 | 443.925 |
| Plxna4os1 | 13.6309 | -1.2177 | 0.52451 | -2.3216 | 0.02025 | 0.09028 | 4.504 |
| Plxna4os2 | 28.893 | 0.185625 | 0.37028 | 0.501304 | 0.61616 | 0.78137 | 3.581 |
| Plxnb1 | 6137.2 | -0.08038 | 0.08756 | -0.91801 | 0.35861 | 0.56887 | 24.529 |
| Plxnb2 | 3465.67 | -0.05096 | 0.08989 | -0.56688 | 0.57079 | 0.75002 | 25.24 |
| Plxnb3 | 1550.83 | -0.353 | 0.16958 | -2.08157 | 0.03738 | 0.13674 | 15.425 |
| Plxnc1 | 2251.15 | 0.150327 | 0.09709 | 1.548404 | 0.12153 | 0.29487 | 153.97 |
| Plxnd1 | 3775.19 | 0.025516 | 0.07986 | 0.319521 | 0.74933 | 0.86849 | 40.195 |
| Pm20d1 | 151.395 | -0.16937 | 0.28063 | -0.60354 | 0.54615 | 0.73071 | 24.093 |
| Pm20d2 | 197.201 | 0.128036 | 0.14993 | 0.853981 | 0.39312 | 0.60231 | 15.508 |
| Pmaip1 | 100.334 | -0.36304 | 0.20956 | -1.73243 | 0.0832 | 0.23033 | 7.033 |
| Pmel | 24.1636 | -1.12607 | 0.42384 | -2.65683 | 0.00789 | 0.04922 | 16.044 |
| Pmepa1 | 3748.65 | 0.243904 | 0.10871 | 2.243539 | 0.02486 | 0.10356 | 52.076 |
| Pmf1 | 135.95 | 0.075857 | 0.15632 | 0.485269 | 0.62749 | 0.78925 | 16.189 |
| Pmfbp1 | 21.6528 | -0.60681 | 0.47288 | -1.28322 | 0.19941 | 0.39739 | 48.614 |
| Pml | 776.535 | 0.176752 | 0.09368 | 1.886804 | 0.05919 | 0.18518 | 31.711 |
| Pmm1 | 4423.33 | -0.21129 | 0.07927 | -2.66546 | 0.00769 | 0.04845 | 9.823 |
| Pmm2 | 1485.85 | 0.243373 | 0.1268 | 1.919271 | 0.05495 | 0.17653 | 24.794 |
| Pmp22 | 790.527 | -0.65783 | 0.10723 | -6.13468 | 8.53E-10 | 4.83E-07 | 30.566 |
| Pmpca | 3241.48 | -0.02684 | 0.0643 | -0.41749 | 0.67632 | 0.82247 | 7.784 |
| Pmpcb | 2006.36 | -0.01136 | 0.07256 | -0.15662 | 0.87555 | 0.93793 | 20.012 |
| Pms1 | 220.175 | -0.1371 | 0.18389 | -0.74557 | 0.45593 | 0.65758 | 107.832 |
| Pms2 | 760.078 | 0.018553 | 0.09113 | 0.20359 | 0.83867 | 0.9169 | 24.005 |
| Pmvk | 1190.78 | -0.18436 | 0.10103 | -1.82483 | 0.06803 | 0.20237 | 14.473 |
| Pnck | 3196.75 | 0.600588 | 0.09422 | 6.374554 | 1.83E-10 | 1.64E-07 | 4.124 |
| Pnisr | 19299.6 | 0.25505 | 0.15158 | 1.682624 | 0.09245 | 0.24756 | 28.893 |
| Pnkd | 6287.03 | -0.11103 | 0.05872 | -1.89067 | 0.05867 | 0.18424 | 68.765 |
| Pnkp | 996.659 | -0.17843 | 0.0855 | -2.08689 | 0.0369 | 0.13557 | 5.854 |
| Pnldc1 | 134.13 | 0.173849 | 0.19743 | 0.880584 | 0.37854 | 0.58831 | 21.502 |

| | | | | | | | |
|---|---|---|---|---|---|---|---|
| Pnlip | 27.984 | -0.3971 | 0.39074 | -1.01628 | 0.3095 | 0.51945 | 29.291 |
| Pnma1 | 116.286 | -0.2311 | 0.18424 | -1.25436 | 0.20971 | 0.40977 | 2.359 |
| Pnma2 | 3322.89 | 0.155468 | 0.08389 | 1.853211 | 0.06385 | 0.19435 | 9.854 |
| Pnma3 | 959.985 | -0.06182 | 0.12453 | -0.49646 | 0.61957 | 0.78351 | 3.405 |
| Pnma8a | 8360.5 | 0.134166 | 0.06915 | 1.940192 | 0.05236 | 0.17103 | 4.929 |
| Pnma8b | 22184.4 | 0.401796 | 0.10324 | 3.891893 | 9.95E-05 | 0.00268 | 4.147 |
| Pnma8c | 6971.3 | 0.108312 | 0.07035 | 1.539655 | 0.12364 | 0.29802 | 5.067 |
| Pnmt | 43.3981 | -1.46303 | 0.61436 | -2.3814 | 0.01725 | 0.08154 | 1.732 |
| Pnn | 6882.37 | 0.152433 | 0.13695 | 1.113021 | 0.2657 | 0.4749 | 7.115 |
| Pno1 | 713.205 | 0.101472 | 0.08936 | 1.135555 | 0.25614 | 0.46403 | 8.371 |
| Pnoc | 165.052 | -0.81533 | 0.20771 | -3.92535 | 8.66E-05 | 0.00245 | 24.8 |
| Pnp | 94.5285 | -0.07041 | 0.23218 | -0.30327 | 0.76169 | 0.87511 | 34.156 |
| Pnp2 | 63.1284 | 0.536631 | 0.31243 | 1.717601 | 0.08587 | 0.23546 | 8.758 |
| Pnpla1 | 22.4145 | 0.669355 | 0.38438 | 1.741401 | 0.08161 | 0.22767 | 31.899 |
| Pnpla2 | 1101.26 | -0.53632 | 0.09318 | -5.7555 | 8.64E-09 | 2.75E-06 | 5.546 |
| Pnpla3 | 258.069 | -0.22153 | 0.15458 | -1.43306 | 0.15184 | 0.33701 | 19.4 |
| Pnpla6 | 3622.23 | 0.125372 | 0.08083 | 1.550959 | 0.12091 | 0.29397 | 28.884 |
| Pnpla7 | 1093.75 | -0.53158 | 0.09137 | -5.81811 | 5.95E-09 | 2.02E-06 | 78.025 |
| Pnpla8 | 5695.37 | 0.082485 | 0.09339 | 0.883276 | 0.37709 | 0.58718 | 101.163 |
| Pnpo | 1122.64 | 0.087798 | 0.10077 | 0.871303 | 0.38359 | 0.59294 | 6.162 |
| Pnpt1 | 1327.84 | 0.14713 | 0.13236 | 1.111553 | 0.26633 | 0.47559 | 31.085 |
| Pnrc1 | 1499.64 | -0.10526 | 0.07429 | -1.41693 | 0.1565 | 0.34329 | 44.741 |
| Pnrc2 | 2243.41 | -0.01857 | 0.09077 | -0.20462 | 0.83787 | 0.91656 | 2.933 |
| Poc1a | 388.342 | -0.38992 | 0.15606 | -2.49847 | 0.01247 | 0.06631 | 69.461 |
| Poc1b | 277.527 | -0.03024 | 0.11436 | -0.2644 | 0.79147 | 0.89159 | 91.039 |
| Poc5 | 2143.15 | 0.097979 | 0.07163 | 1.367772 | 0.17138 | 0.3618 | 29.444 |
| Podn | 161.795 | -0.52599 | 0.23133 | -2.27377 | 0.02298 | 0.09808 | 81.655 |
| Podnl1 | 16.243 | 0.615006 | 0.48851 | 1.258955 | 0.20805 | 0.40769 | 6.539 |
| Podxl | 1655.74 | -0.09366 | 0.08822 | -1.06174 | 0.28835 | 0.49846 | 44.494 |
| Podxl2 | 2262.28 | -0.39204 | 0.10435 | -3.75678 | 0.00017 | 0.0039 | 32.487 |
| Pof1b | 19.4914 | -0.13517 | 0.54323 | -0.24883 | 0.80349 | 0.89853 | 60.221 |
| Pofut1 | 1750.23 | 0.165906 | 0.06465 | 2.56617 | 0.01028 | 0.05853 | 28.715 |
| Pofut2 | 1145.83 | -0.04188 | 0.07576 | -0.55283 | 0.58038 | 0.75658 | 10.358 |
| Pogk | 3753.61 | -0.10696 | 0.08316 | -1.28615 | 0.19839 | 0.3961 | 25.242 |

| | | | | | | | |
|---|---|---|---|---|---|---|---|
| Poglut1 | 2397.85 | 0.076656 | 0.06294 | 1.217943 | 0.22325 | 0.42649 | 25.122 |
| Poglut2 | 416.62 | 0.071708 | 0.10665 | 0.672394 | 0.50133 | 0.69395 | 12.263 |
| Poglut3 | 368.158 | -0.05219 | 0.11901 | -0.4385 | 0.66102 | 0.81248 | 17.843 |
| Pogz | 4773.6 | 0.291306 | 0.07249 | 4.018508 | 5.86E-05 | 0.00185 | 44.76 |
| Pola1 | 271.444 | -0.16145 | 0.18197 | -0.88721 | 0.37497 | 0.58517 | 327.389 |
| Pola2 | 284.415 | -0.16883 | 0.11544 | -1.46258 | 0.14358 | 0.32574 | 23.661 |
| Polb | 2475.96 | 0.076033 | 0.0807 | 0.942124 | 0.34613 | 0.55684 | 25.31 |
| Pold1 | 458.108 | -0.03606 | 0.11519 | -0.31301 | 0.75427 | 0.87077 | 16.104 |
| Pold2 | 474.178 | 0.035209 | 0.11775 | 0.299003 | 0.76494 | 0.87707 | 6.113 |
| Pold3 | 1448.94 | 0.21115 | 0.06756 | 3.125407 | 0.00178 | 0.01864 | 39.455 |
| Pold4 | 82.0318 | -0.06247 | 0.21345 | -0.29268 | 0.76977 | 0.87983 | 1.736 |
| Poldip2 | 3314.95 | -0.13423 | 0.07153 | -1.8766 | 0.06057 | 0.18785 | 10.544 |
| Poldip3 | 3865.57 | 0.16148 | 0.08639 | 1.869263 | 0.06159 | 0.18978 | 23.409 |
| Pole | 74.5535 | -0.46696 | 0.25134 | -1.85785 | 0.06319 | 0.19309 | 51.169 |
| Pole2 | 39.4114 | 0.49812 | 0.28464 | 1.749995 | 0.08012 | 0.22514 | 26.423 |
| Pole3 | 1018.19 | -0.03883 | 0.0955 | -0.40662 | 0.68429 | 0.8279 | 2.42 |
| Pole4 | 1246.28 | -0.10831 | 0.07878 | -1.37492 | 0.16916 | 0.35933 | 86.374 |
| Polg | 3927.15 | 0.096685 | 0.06463 | 1.49591 | 0.13468 | 0.31449 | 20.132 |
| Polg2 | 177.692 | 0.14756 | 0.13945 | 1.058188 | 0.28997 | 0.50018 | 11.285 |
| Polh | 269.565 | -0.15912 | 0.12076 | -1.31765 | 0.18762 | 0.38302 | 31.291 |
| Poli | 541.545 | 0.17317 | 0.09912 | 1.747149 | 0.08061 | 0.22596 | 21.941 |
| Polk | 805.824 | -0.04628 | 0.10972 | -0.42182 | 0.67316 | 0.82028 | 61.89 |
| Poll | 346.587 | 0.018431 | 0.12717 | 0.144934 | 0.88476 | 0.94271 | 8.257 |
| Polm | 436.411 | 0.075617 | 0.10833 | 0.698022 | 0.48516 | 0.68138 | 10.157 |
| Poln | 23.692 | 0.38401 | 0.40489 | 0.948423 | 0.34291 | 0.55368 | 162.27 |
| Polq | 569.917 | 0.261908 | 0.13257 | 1.975581 | 0.0482 | 0.16182 | 83.632 |
| Polr1a | 2496.36 | 0.203484 | 0.07437 | 2.736078 | 0.00622 | 0.04216 | 75.883 |
| Polr1b | 1137.47 | 0.022791 | 0.09814 | 0.232225 | 0.81636 | 0.90542 | 25.6 |
| Polr1c | 974.291 | -0.09199 | 0.0756 | -1.21677 | 0.22369 | 0.4269 | 4.135 |
| Polr1d | 1045.11 | -0.14143 | 0.12471 | -1.13402 | 0.25679 | 0.46474 | 34.548 |
| Polr1e | 367.729 | -0.02249 | 0.10374 | -0.21684 | 0.82833 | 0.91195 | 17.983 |
| Polr1f | 562.584 | 0.148684 | 0.12614 | 1.17871 | 0.23851 | 0.44413 | 11.758 |
| Polr1h | 438.037 | 0.023447 | 0.09785 | 0.239611 | 0.81063 | 0.90256 | 4.204 |
| Polr1has | 98.1575 | 0.216017 | 0.20303 | 1.063939 | 0.28736 | 0.49748 | 7.034 |

| | | | | | | | |
|---|---|---|---|---|---|---|---|
| Polr2a | 4408.29 | -0.0762 | 0.07458 | -1.02171 | 0.30692 | 0.5168 | 24.641 |
| Polr2b | 3055.88 | 0.086535 | 0.06899 | 1.254392 | 0.2097 | 0.40977 | 39.178 |
| Polr2c | 1287.93 | -0.04233 | 0.08053 | -0.52568 | 0.59911 | 0.77005 | 16.086 |
| Polr2d | 586.935 | -0.01318 | 0.12002 | -0.10978 | 0.91259 | 0.95592 | 7.543 |
| Polr2e | 1445.07 | -0.00891 | 0.09113 | -0.09781 | 0.92208 | 0.96078 | 3.847 |
| Polr2f | 894.412 | -0.14575 | 0.11294 | -1.29051 | 0.19687 | 0.39467 | 10.766 |
| Polr2g | 1791.38 | -0.02544 | 0.07197 | -0.35351 | 0.72371 | 0.85252 | 5.451 |
| Polr2h | 413.711 | -0.09766 | 0.10556 | -0.92511 | 0.35491 | 0.56542 | 4.603 |
| Polr2i | 357.189 | -0.11253 | 0.13376 | -0.8413 | 0.40018 | 0.60898 | 1.443 |
| Polr2j | 613.515 | -0.18541 | 0.08764 | -2.11563 | 0.03438 | 0.1296 | 6.317 |
| Polr2k | 601.593 | -0.02858 | 0.08503 | -0.33618 | 0.73674 | 0.86096 | 3.001 |
| Polr2l | 778.309 | -0.16194 | 0.08319 | -1.94668 | 0.05157 | 0.16934 | 3.273 |
| Polr2m | 7113.53 | -0.15107 | 0.06378 | -2.36865 | 0.01785 | 0.08321 | 7.499 |
| Polr3a | 1891.22 | 0.146272 | 0.06721 | 2.176179 | 0.02954 | 0.11694 | 38.363 |
| Polr3b | 915.762 | -0.05332 | 0.11735 | -0.45438 | 0.64955 | 0.80496 | 104.887 |
| Polr3c | 884.341 | -0.1476 | 0.10828 | -1.36311 | 0.17285 | 0.36406 | 16.139 |
| Polr3d | 751.232 | -0.0216 | 0.08276 | -0.26101 | 0.79409 | 0.89345 | 4.691 |
| Polr3e | 1910.19 | 0.145764 | 0.0718 | 2.030249 | 0.04233 | 0.14806 | 29.689 |
| Polr3f | 1552.34 | 0.088664 | 0.06544 | 1.354814 | 0.17548 | 0.36728 | 14.278 |
| Polr3g | 409.037 | 0.537095 | 0.14113 | 3.805792 | 0.00014 | 0.00337 | 37.171 |
| Polr3gl | 790.485 | -0.43429 | 0.14784 | -2.93755 | 0.00331 | 0.02776 | 16.31 |
| Polr3h | 1562.45 | -0.15375 | 0.07112 | -2.1617 | 0.03064 | 0.11978 | 38.013 |
| Polr3k | 2055.98 | -0.13489 | 0.07887 | -1.71029 | 0.08721 | 0.23786 | 6.494 |
| Polrmt | 1303.19 | 0.008444 | 0.06965 | 0.121244 | 0.9035 | 0.95134 | 10.459 |
| Pom121 | 2797.21 | 0.007654 | 0.05913 | 0.129434 | 0.89701 | 0.94835 | 18.406 |
| Pomc | 50.3335 | 0.345857 | 0.26213 | 1.319386 | 0.18704 | 0.38229 | 5.692 |
| Pomgnt1 | 2329.19 | 0.024289 | 0.06923 | 0.350843 | 0.72571 | 0.85388 | 36.01 |
| Pomgnt2 | 1282.89 | 0.276146 | 0.10564 | 2.613907 | 0.00895 | 0.05329 | 15.505 |
| Pomk | 1479.37 | -0.04281 | 0.09097 | -0.47058 | 0.63794 | 0.79651 | 13.53 |
| Pomp | 2826.23 | -0.26355 | 0.06654 | -3.96085 | 7.47E-05 | 0.00219 | 15.996 |
| Pomt1 | 906.217 | 0.017092 | 0.09372 | 0.182379 | 0.85529 | 0.92683 | 18.416 |
| Pomt2 | 1331.36 | 0.039935 | 0.09149 | 0.436481 | 0.66249 | 0.8135 | 41.108 |
| Pon2 | 2259.75 | -0.328 | 0.0958 | -3.42375 | 0.00062 | 0.00917 | 34.309 |
| Pon3 | 79.6614 | -0.52417 | 0.23937 | -2.18983 | 0.02854 | 0.11395 | 35.435 |

| | | | | | | | |
|---|---|---|---|---|---|---|---|
| Pop1 | 606.848 | 0.21429 | 0.09894 | 2.165958 | 0.03031 | 0.11888 | 35.345 |
| Pop4 | 749.831 | -0.03511 | 0.09527 | -0.36852 | 0.71248 | 0.84516 | 9.428 |
| Pop5 | 1397.25 | -0.16392 | 0.08877 | -1.8465 | 0.06482 | 0.19615 | 9.516 |
| Pop7 | 421.294 | 0.029271 | 0.10338 | 0.283156 | 0.77706 | 0.88395 | 1.081 |
| Popdc2 | 13.9502 | -0.11521 | 0.61622 | -0.18696 | 0.85169 | 0.92458 | 16.008 |
| Popdc3 | 22.5582 | 0.000909 | 0.39719 | 0.00229 | 0.99817 | 0.99906 | 140.355 |
| Por | 3834.32 | -0.0024 | 0.08874 | -0.02708 | 0.9784 | 0.98853 | 65.294 |
| Porcn | 2149.35 | -0.11002 | 0.08412 | -1.30787 | 0.19092 | 0.38718 | 12.678 |
| Postn | 282.34 | -0.25621 | 0.27007 | -0.94871 | 0.34277 | 0.55356 | 29.929 |
| Pot1a | 976.35 | -0.10416 | 0.07985 | -1.30446 | 0.19208 | 0.38879 | 65.51 |
| Pot1b | 649.627 | -0.01269 | 0.1192 | -0.10646 | 0.91522 | 0.95656 | 60.678 |
| Potefam3c | 25.0527 | 0.568513 | 0.49658 | 1.144868 | 0.25226 | 0.45957 | 25.586 |
| Potefam3d | 19.8801 | 0.138133 | 0.4324 | 0.319456 | 0.74938 | 0.86849 | 41.631 |
| Potegl | 15.4626 | -0.18777 | 0.51128 | -0.36726 | 0.71343 | 0.84584 | 59.654 |
| Pou2af1 | 14.6384 | -1.71062 | 0.95048 | -1.79975 | 0.0719 | 0.21043 | 26.373 |
| Pou2f1 | 1968.65 | 0.144564 | 0.09597 | 1.506395 | 0.13197 | 0.31078 | 137.525 |
| Pou2f2 | 674.167 | -0.00081 | 0.1635 | -0.00494 | 0.99606 | 0.99782 | 92.383 |
| Pou3f1 | 721.918 | 0.585875 | 0.19915 | 2.941905 | 0.00326 | 0.02757 | 3.849 |
| Pou3f2 | 1343.92 | -0.36948 | 0.17347 | -2.12985 | 0.03318 | 0.12657 | 5.587 |
| Pou3f3 | 6132.69 | -0.13656 | 0.07845 | -1.74067 | 0.08174 | 0.22788 | 8.252 |
| Pou3f4 | 1595.45 | 0.614207 | 0.1415 | 4.340818 | 1.42E-05 | 0.00071 | 2.928 |
| Pou6f1 | 5582.54 | -0.14278 | 0.08792 | -1.62397 | 0.10438 | 0.26734 | 24.667 |
| Pou6f2 | 61.4701 | -0.34296 | 0.37912 | -0.9046 | 0.36568 | 0.57626 | 276.589 |
| Ppa1 | 2676.15 | -0.10985 | 0.06536 | -1.68072 | 0.09282 | 0.24813 | 25.617 |
| Ppa2 | 1146.55 | -0.10933 | 0.08194 | -1.33427 | 0.18211 | 0.37592 | 68.126 |
| Ppan | 600.002 | 0.071492 | 0.10692 | 0.668629 | 0.50373 | 0.69618 | 4.004 |
| Ppara | 262.584 | -0.3035 | 0.17258 | -1.75862 | 0.07864 | 0.22224 | 67.837 |
| Ppard | 1001.01 | -0.35228 | 0.1007 | -3.49818 | 0.00047 | 0.0076 | 68.775 |
| Pparg | 46.5507 | -1.19136 | 0.44083 | -2.70255 | 0.00688 | 0.04491 | 129.449 |
| Ppargc1a | 2884.78 | -0.22122 | 0.12092 | -1.82948 | 0.06733 | 0.20106 | 113.477 |
| Ppargc1b | 1065.47 | 0.349487 | 0.10604 | 3.29592 | 0.00098 | 0.01264 | 102.296 |
| Ppat | 992.751 | 0.026528 | 0.08176 | 0.324478 | 0.74558 | 0.86627 | 38.33 |
| Ppcdc | 658.665 | -0.03925 | 0.08919 | -0.44009 | 0.65987 | 0.81186 | 55.03 |
| Ppcs | 176.739 | 0.1907 | 0.15657 | 1.218019 | 0.22322 | 0.42649 | 3.891 |

| | | | | | | | |
|---|---|---|---|---|---|---|---|
| Ppdpf | 1400.82 | -0.07137 | 0.09856 | -0.72418 | 0.46896 | 0.66787 | 1.525 |
| Ppef1 | 26.5994 | 0.369268 | 0.35385 | 1.043565 | 0.29669 | 0.50689 | 112.672 |
| Ppef2 | 200.699 | 0.181464 | 0.16401 | 1.106444 | 0.26853 | 0.47785 | 29.6 |
| Ppfia1 | 3013.31 | 0.208268 | 0.07606 | 2.738371 | 0.00617 | 0.04206 | 76.972 |
| Ppfia2 | 4467.81 | -0.01579 | 0.12254 | -0.12888 | 0.89745 | 0.94865 | 465.614 |
| Ppfia3 | 7127.17 | -0.28026 | 0.08889 | -3.15286 | 0.00162 | 0.01748 | 27.898 |
| Ppfia4 | 5583.79 | 0.376933 | 0.15313 | 2.461501 | 0.01384 | 0.0707 | 36.146 |
| Ppfibp1 | 2112.36 | -0.78054 | 0.17744 | -4.3988 | 1.09E-05 | 0.0006 | 143.539 |
| Ppfibp2 | 569.864 | -0.56628 | 0.19093 | -2.96597 | 0.00302 | 0.02624 | 153.377 |
| Pphln1 | 2193.09 | 0.044871 | 0.06913 | 0.649067 | 0.5163 | 0.70632 | 93.161 |
| Ppia | 20079.8 | -0.29338 | 0.07838 | -3.74323 | 0.00018 | 0.00404 | 4.375 |
| Ppib | 2630.72 | -0.22294 | 0.09322 | -2.3917 | 0.01677 | 0.08013 | 6.402 |
| Ppic | 49.5557 | -0.61154 | 0.26349 | -2.32093 | 0.02029 | 0.09035 | 11.784 |
| Ppid | 4151.22 | 0.078571 | 0.06939 | 1.132271 | 0.25752 | 0.46553 | 12.309 |
| Ppie | 351.649 | -0.19472 | 0.136 | -1.43171 | 0.15223 | 0.33743 | 12.837 |
| Ppif | 816.161 | -0.04294 | 0.09113 | -0.47122 | 0.63749 | 0.79609 | 6.315 |
| Ppig | 5155.71 | 0.234239 | 0.15622 | 1.49946 | 0.13375 | 0.31339 | 31.468 |
| Ppih | 173.403 | 0.119391 | 0.2524 | 0.473017 | 0.6362 | 0.79507 | 20.537 |
| Ppil1 | 689.66 | 0.044418 | 0.08051 | 0.551726 | 0.58114 | 0.75715 | 13.384 |
| Ppil2 | 2904.41 | 0.134745 | 0.06616 | 2.036631 | 0.04169 | 0.14679 | 24.703 |
| Ppil3 | 418.282 | 0.078591 | 0.10437 | 0.753 | 0.45145 | 0.65385 | 14.493 |
| Ppil4 | 2882.03 | 0.25856 | 0.10499 | 2.462622 | 0.01379 | 0.07062 | 30.245 |
| Ppil6 | 240.334 | -0.07681 | 0.1258 | -0.61056 | 0.54149 | 0.72706 | 23.848 |
| Ppip5k1 | 7917.1 | 0.154524 | 0.06838 | 2.259624 | 0.02384 | 0.10049 | 44.836 |
| Ppip5k2 | 2215.07 | 0.300469 | 0.10388 | 2.892444 | 0.00382 | 0.03052 | 64.364 |
| Ppl | 155.747 | 0.062502 | 0.16377 | 0.381637 | 0.70273 | 0.8396 | 46.131 |
| Ppm1a | 12956.2 | -0.03014 | 0.05829 | -0.51712 | 0.60508 | 0.77377 | 42.365 |
| Ppm1b | 6975.14 | 0.048463 | 0.06304 | 0.768789 | 0.44202 | 0.64653 | 67.251 |
| Ppm1d | 868.464 | 0.133661 | 0.07835 | 1.705877 | 0.08803 | 0.23951 | 35.823 |
| Ppm1e | 10931.4 | -0.74682 | 0.35078 | -2.12901 | 0.03325 | 0.12662 | 132.118 |
| Ppm1f | 4812.4 | 0.173498 | 0.08961 | 1.936222 | 0.05284 | 0.17201 | 30.896 |
| Ppm1g | 2717.49 | 0.013236 | 0.07068 | 0.187253 | 0.85146 | 0.92445 | 18.024 |
| Ppm1h | 3497.78 | -0.19656 | 0.08694 | -2.2609 | 0.02377 | 0.10038 | 267.034 |
| Ppm1j | 22.9849 | -1.06191 | 0.461 | -2.30348 | 0.02125 | 0.09324 | 4.963 |

| | | | | | | | |
|---|---|---|---|---|---|---|---|
| Ppm1k | 3962.1 | -0.05391 | 0.06986 | -0.77165 | 0.44032 | 0.64526 | 28.967 |
| Ppm1l | 9075.2 | -0.46167 | 0.12667 | -3.64456 | 0.00027 | 0.00518 | 243.942 |
| Ppm1m | 527.332 | -0.44607 | 0.09759 | -4.57102 | 4.85E-06 | 0.00033 | 4.8 |
| Ppme1 | 6880.38 | -0.1166 | 0.06709 | -1.73785 | 0.08224 | 0.22863 | 45.571 |
| Ppox | 1148.26 | 0.071642 | 0.09346 | 0.766555 | 0.44335 | 0.64783 | 5.198 |
| Ppp1ca | 5138.8 | -0.02015 | 0.07863 | -0.25629 | 0.79773 | 0.89566 | 3.356 |
| Ppp1cb | 16589.9 | 0.11558 | 0.07355 | 1.571348 | 0.1161 | 0.28603 | 58.591 |
| Ppp1cc | 9968.32 | 0.191487 | 0.07342 | 2.608098 | 0.0091 | 0.05384 | 16.996 |
| Ppp1ccb | 634.758 | 0.019656 | 0.10913 | 0.180108 | 0.85707 | 0.9279 | 2.301 |
| Ppp1r10 | 2535.46 | -0.02707 | 0.08446 | -0.32056 | 0.74855 | 0.86814 | 15.85 |
| Ppp1r11 | 1946.76 | -0.1715 | 0.08054 | -2.12943 | 0.03322 | 0.12659 | 3.386 |
| Ppp1r12a | 7060.17 | 0.155102 | 0.08341 | 1.859597 | 0.06294 | 0.19263 | 122.283 |
| Ppp1r12b | 8300.37 | 0.00309 | 0.06969 | 0.044332 | 0.96464 | 0.98273 | 201.285 |
| Ppp1r12c | 5217.62 | 0.020523 | 0.06177 | 0.33222 | 0.73972 | 0.86273 | 20.161 |
| Ppp1r13b | 4077.43 | 0.09486 | 0.0736 | 1.288822 | 0.19746 | 0.39531 | 79.654 |
| Ppp1r13l | 207.292 | 0.090031 | 0.14435 | 0.623683 | 0.53284 | 0.72018 | 18.785 |
| Ppp1r14a | 408.631 | -0.69447 | 0.23118 | -3.00396 | 0.00266 | 0.02435 | 4.494 |
| Ppp1r14b | 573.083 | -0.27467 | 0.12698 | -2.16308 | 0.03053 | 0.11949 | 2.357 |
| Ppp1r14c | 342.023 | -0.17904 | 0.10237 | -1.74888 | 0.08031 | 0.22549 | 98.919 |
| Ppp1r15a | 504.032 | -0.22745 | 0.09004 | -2.52595 | 0.01154 | 0.06299 | 3.353 |
| Ppp1r15b | 3757.55 | 0.02279 | 0.06907 | 0.329956 | 0.74143 | 0.86347 | 8.641 |
| Ppp1r16a | 2541.6 | -0.07562 | 0.09579 | -0.78944 | 0.42985 | 0.63616 | 23.305 |
| Ppp1r16b | 12596.2 | 0.111563 | 0.10417 | 1.070978 | 0.28418 | 0.4943 | 100.937 |
| Ppp1r18 | 353.402 | -0.33741 | 0.11491 | -2.93624 | 0.00332 | 0.02779 | 10.004 |
| Ppp1r1a | 2630.21 | -0.05392 | 0.09268 | -0.58185 | 0.56067 | 0.74193 | 7.725 |
| Ppp1r1b | 39259.6 | 0.532315 | 0.30681 | 1.735027 | 0.08274 | 0.22952 | 9.393 |
| Ppp1r1c | 18.7835 | -1.4248 | 0.50465 | -2.82332 | 0.00475 | 0.03518 | 110.717 |
| Ppp1r2 | 25588.1 | 0.52335 | 0.15927 | 3.285927 | 0.00102 | 0.01296 | 23.741 |
| Ppp1r2-ps4 | 29.7739 | -0.317 | 0.35342 | -0.89693 | 0.36975 | 0.58022 | 0.633 |
| Ppp1r21 | 3263.2 | -0.00461 | 0.07286 | -0.06328 | 0.94955 | 0.97495 | 58.25 |
| Ppp1r26 | 902.071 | -0.14218 | 0.10861 | -1.3091 | 0.1905 | 0.3867 | 8.709 |
| Ppp1r32 | 57.9556 | 0.603746 | 0.30658 | 1.969293 | 0.04892 | 0.16347 | 8.694 |
| Ppp1r35 | 363.974 | -0.19678 | 0.11064 | -1.77859 | 0.07531 | 0.21648 | 1.262 |
| Ppp1r36 | 24.9051 | -0.73909 | 0.40934 | -1.80556 | 0.07099 | 0.20844 | 21.955 |

| | | | | | | | |
|---|---|---|---|---|---|---|---|
| Ppp1r37 | 5579.53 | -0.00968 | 0.05953 | -0.16258 | 0.87085 | 0.93475 | 32.277 |
| Ppp1r3a | 24.7817 | 1.296994 | 0.48011 | 2.701475 | 0.0069 | 0.04499 | 41.298 |
| Ppp1r3b | 68.048 | -0.37791 | 0.25895 | -1.45943 | 0.14445 | 0.32717 | 12.401 |
| Ppp1r3c | 2285.71 | -0.27528 | 0.12878 | -2.1376 | 0.03255 | 0.12502 | 4.923 |
| Ppp1r3d | 404.469 | -0.0064 | 0.13191 | -0.04855 | 0.96128 | 0.98116 | 3.267 |
| Ppp1r3e | 500.48 | -0.07454 | 0.10858 | -0.68653 | 0.49238 | 0.68711 | 4.882 |
| Ppp1r3f | 142.246 | -0.06093 | 0.17178 | -0.35471 | 0.7228 | 0.85222 | 16.988 |
| Ppp1r3g | 415.876 | -0.37786 | 0.21179 | -1.78417 | 0.0744 | 0.21502 | 11.55 |
| Ppp1r42 | 37.4303 | -0.32968 | 0.30817 | -1.06982 | 0.2847 | 0.49463 | 40.513 |
| Ppp1r7 | 8604.13 | 0.373296 | 0.1579 | 2.364155 | 0.01807 | 0.08383 | 30.636 |
| Ppp1r8 | 2141.5 | 0.032102 | 0.07848 | 0.409049 | 0.6825 | 0.82653 | 16.241 |
| Ppp1r9a | 26998.6 | 0.387407 | 0.12272 | 3.156947 | 0.00159 | 0.01733 | 262.745 |
| Ppp1r9b | 59055.5 | -0.04394 | 0.07984 | -0.55039 | 0.58205 | 0.75769 | 15.865 |
| Ppp2ca | 12293.8 | -0.04909 | 0.05228 | -0.93899 | 0.34774 | 0.55823 | 29.098 |
| Ppp2cb | 4142.57 | -0.10669 | 0.06981 | -1.52835 | 0.12643 | 0.30219 | 19.817 |
| Ppp2r1a | 17656.6 | -0.07948 | 0.07887 | -1.00775 | 0.31358 | 0.52413 | 20.606 |
| Ppp2r1b | 1118.64 | 0.002989 | 0.08692 | 0.034381 | 0.97257 | 0.98622 | 48.929 |
| Ppp2r2a | 5890.26 | 0.293133 | 0.09768 | 3.000849 | 0.00269 | 0.02451 | 58.389 |
| Ppp2r2b | 4728.83 | -0.10159 | 0.08167 | -1.24397 | 0.21351 | 0.41479 | 445.696 |
| Ppp2r2c | 26745.5 | -0.07777 | 0.06797 | -1.14421 | 0.25254 | 0.45986 | 86.566 |
| Ppp2r2cos | 151.394 | 0.719305 | 0.16311 | 4.409886 | 1.03E-05 | 0.00057 | 3.285 |
| Ppp2r2d | 2582.7 | 0.011509 | 0.09571 | 0.120249 | 0.90429 | 0.95167 | 36.979 |
| Ppp2r3a | 2816.9 | -0.11849 | 0.09124 | -1.29877 | 0.19402 | 0.39108 | 146.712 |
| Ppp2r3c | 745.037 | -0.05805 | 0.11019 | -0.52683 | 0.59831 | 0.76969 | 24.008 |
| Ppp2r3d | 1414.2 | 0.106005 | 0.07919 | 1.338613 | 0.1807 | 0.37405 | 8.933 |
| Ppp2r5a | 4550.54 | -0.08315 | 0.0633 | -1.31371 | 0.18894 | 0.38476 | 51.298 |
| Ppp2r5b | 8230.19 | 0.347828 | 0.13489 | 2.578687 | 0.00992 | 0.05714 | 8.108 |
| Ppp2r5c | 9770.28 | 0.039169 | 0.05181 | 0.756028 | 0.44963 | 0.65275 | 135.943 |
| Ppp2r5d | 5994.22 | 0.244013 | 0.12057 | 2.023792 | 0.04299 | 0.14957 | 22.121 |
| Ppp2r5e | 5153.49 | 0.099122 | 0.05491 | 1.805061 | 0.07107 | 0.20858 | 145.365 |
| Ppp3ca | 57559.8 | 0.365502 | 0.12252 | 2.983105 | 0.00285 | 0.02539 | 267.604 |
| Ppp3cb | 16029.5 | -0.03882 | 0.06722 | -0.5776 | 0.56354 | 0.74412 | 47.21 |
| Ppp3cc | 731.543 | -0.14344 | 0.10174 | -1.40992 | 0.15856 | 0.34594 | 71.607 |
| Ppp3r1 | 29782.2 | 0.070119 | 0.07378 | 0.950373 | 0.34192 | 0.55274 | 41.113 |

| | | | | | | | |
|---|---|---|---|---|---|---|---|
| Ppp3r2 | 17.2632 | 0.287867 | 0.46496 | 0.619118 | 0.53584 | 0.72278 | 3.278 |
| Ppp4c | 1611.07 | 0.057214 | 0.06867 | 0.83313 | 0.40477 | 0.61372 | 6.631 |
| Ppp4r1 | 2077.79 | 0.170041 | 0.1273 | 1.33573 | 0.18164 | 0.37528 | 59.354 |
| Ppp4r1l-ps | 1506.07 | -0.00873 | 0.08973 | -0.09732 | 0.92248 | 0.96099 | 80.336 |
| Ppp4r2 | 4822.54 | 0.172959 | 0.07591 | 2.278569 | 0.02269 | 0.09735 | 36.313 |
| Ppp4r3a | 2710.27 | 0.194798 | 0.10258 | 1.898919 | 0.05758 | 0.18222 | 44.294 |
| Ppp4r3b | 3167.93 | 0.280302 | 0.12999 | 2.156325 | 0.03106 | 0.12093 | 47.908 |
| Ppp4r4 | 2588.56 | 0.503756 | 0.11755 | 4.2853 | 1.82E-05 | 0.00083 | 81.549 |
| Ppp5c | 3552.37 | -0.23527 | 0.07653 | -3.07436 | 0.00211 | 0.02087 | 23.285 |
| Ppp6c | 2580.4 | 0.129442 | 0.06936 | 1.866268 | 0.062 | 0.19072 | 32.098 |
| Ppp6r1 | 7317.96 | 0.099682 | 0.065 | 1.533683 | 0.12511 | 0.30014 | 27.525 |
| Ppp6r2 | 5882.9 | 0.049351 | 0.0564 | 0.875094 | 0.38152 | 0.59096 | 75.458 |
| Ppp6r3 | 5246.96 | 0.064149 | 0.0562 | 1.141515 | 0.25366 | 0.461 | 120.822 |
| Pprc1 | 1224.95 | 0.185083 | 0.08333 | 2.22108 | 0.02635 | 0.10784 | 28.03 |
| Ppt1 | 4529.46 | 0.077802 | 0.06812 | 1.142153 | 0.25339 | 0.46076 | 22.934 |
| Ppt2 | 318.776 | -0.06037 | 0.13782 | -0.43804 | 0.66136 | 0.81269 | 11.849 |
| Pptc7 | 5315.65 | 0.299998 | 0.10454 | 2.869646 | 0.00411 | 0.03204 | 39.917 |
| Ppwd1 | 577.75 | 0.151668 | 0.10013 | 1.514758 | 0.12983 | 0.30763 | 23.72 |
| Pqbp1 | 2004.1 | -0.01619 | 0.08886 | -0.18215 | 0.85546 | 0.92695 | 4.751 |
| Pqlc3 | 233.016 | -0.25498 | 0.20169 | -1.26422 | 0.20615 | 0.40534 | 11.761 |
| Pradc1 | 420.139 | -0.06748 | 0.11773 | -0.57314 | 0.56655 | 0.74656 | 5.161 |
| Praf2 | 556.929 | 0.041328 | 0.11684 | 0.353716 | 0.72355 | 0.85239 | 2.626 |
| Prag1 | 750.705 | 0.347095 | 0.14165 | 2.450366 | 0.01427 | 0.07203 | 52.96 |
| Pram1 | 32.6319 | 0.143274 | 0.34517 | 0.415085 | 0.67808 | 0.82367 | 7.651 |
| Pramel12 | 816.296 | -0.02135 | 0.08871 | -0.2407 | 0.80979 | 0.90197 | 8.666 |
| Prap1 | 21.7792 | -0.12391 | 0.41624 | -0.2977 | 0.76593 | 0.8779 | 3.806 |
| Prc1 | 272.588 | 0.218225 | 0.14625 | 1.492166 | 0.13566 | 0.31567 | 21.81 |
| Prcc | 1933.87 | 0.015466 | 0.09036 | 0.171155 | 0.8641 | 0.93125 | 26.706 |
| Prcd | 78.3018 | -0.35712 | 0.2395 | -1.49112 | 0.13593 | 0.31614 | 14.856 |
| Prcp | 418.701 | -0.23729 | 0.09882 | -2.40124 | 0.01634 | 0.07875 | 60.114 |
| Prdm1 | 64.6946 | -0.93204 | 0.26527 | -3.51362 | 0.00044 | 0.00731 | 91.325 |
| Prdm10 | 1579.95 | 0.107756 | 0.09286 | 1.160455 | 0.24586 | 0.45252 | 101.186 |
| Prdm11 | 752.059 | -0.0172 | 0.12233 | -0.14064 | 0.88816 | 0.94406 | 74.15 |
| Prdm12 | 81.211 | -0.61283 | 0.24418 | -2.50976 | 0.01208 | 0.06489 | 15.759 |

| | | | | | | | |
|---|---|---|---|---|---|---|---|
| Prdm15 | 1375.65 | 0.170555 | 0.06575 | 2.593873 | 0.00949 | 0.05541 | 60.384 |
| Prdm16 | 690.029 | -0.14726 | 0.10643 | -1.38362 | 0.16647 | 0.356 | 320.749 |
| Prdm16os | 1221.41 | -0.53911 | 0.13683 | -3.93995 | 8.15E-05 | 0.00233 | 7.149 |
| Prdm2 | 4341.17 | -0.06296 | 0.07097 | -0.88715 | 0.375 | 0.58517 | 105.605 |
| Prdm4 | 1472.99 | 0.096057 | 0.06719 | 1.429564 | 0.15284 | 0.33835 | 25.177 |
| Prdm5 | 180.335 | 0.103146 | 0.13663 | 0.754906 | 0.45031 | 0.65308 | 158.023 |
| Prdm6 | 30.6285 | -1.59547 | 0.53106 | -3.00428 | 0.00266 | 0.02433 | 112.325 |
| Prdm8 | 295.594 | -0.78303 | 0.45735 | -1.7121 | 0.08688 | 0.23726 | 21.794 |
| Prdm9 | 185.196 | -0.16664 | 0.18813 | -0.88577 | 0.37574 | 0.5859 | 21.276 |
| Prdx1 | 4211.51 | -0.14879 | 0.10009 | -1.48651 | 0.13714 | 0.31781 | 15.279 |
| Prdx2 | 4759.78 | -0.09247 | 0.06863 | -1.34745 | 0.17783 | 0.37009 | 5.248 |
| Prdx3 | 2480.86 | -0.02472 | 0.0787 | -0.31406 | 0.75348 | 0.87054 | 10.506 |
| Prdx4 | 334.414 | -0.45777 | 0.13142 | -3.48321 | 0.0005 | 0.00788 | 16.837 |
| Prdx5 | 4758.29 | -0.2334 | 0.06922 | -3.37179 | 0.00075 | 0.01048 | 3.41 |
| Prdx6 | 9054.44 | -0.30291 | 0.11924 | -2.54035 | 0.01107 | 0.06133 | 11.108 |
| Preb | 3107.25 | 0.023057 | 0.06022 | 0.382907 | 0.70179 | 0.83907 | 9.509 |
| Prelid1 | 2859.42 | -0.2101 | 0.06943 | -3.02607 | 0.00248 | 0.02322 | 4.773 |
| Prelid3a | 1850.91 | 0.197997 | 0.09464 | 2.092099 | 0.03643 | 0.13444 | 14.97 |
| Prelid3b | 1826.94 | -0.19125 | 0.09343 | -2.04702 | 0.04066 | 0.1442 | 8.015 |
| Prelp | 567.74 | -0.87316 | 0.18853 | -4.63146 | 3.63E-06 | 0.00027 | 11.111 |
| Prep | 1302.5 | 0.0111 | 0.08278 | 0.134087 | 0.89333 | 0.9466 | 99.996 |
| Prepl | 13507.8 | 0.00472 | 0.06499 | 0.072622 | 0.94211 | 0.97113 | 26.772 |
| Prex1 | 8558.1 | -0.16196 | 0.07625 | -2.12404 | 0.03367 | 0.12789 | 147.491 |
| Prex2 | 4853.19 | -0.09371 | 0.11006 | -0.85145 | 0.39452 | 0.60364 | 310.217 |
| Prg4 | 381.534 | -1.47715 | 0.64918 | -2.27542 | 0.02288 | 0.09786 | 16.754 |
| Prickle1 | 2662.74 | -0.35606 | 0.10736 | -3.31636 | 0.00091 | 0.01201 | 96.778 |
| Prickle2 | 14228.4 | -0.05697 | 0.07301 | -0.78034 | 0.43519 | 0.64096 | 335.248 |
| Prickle3 | 61.6023 | -0.08471 | 0.30113 | -0.28131 | 0.77847 | 0.88438 | 10.927 |
| Prim1 | 319.009 | 0.042638 | 0.11552 | 0.369092 | 0.71206 | 0.845 | 14.87 |
| Prim2 | 311.191 | 0.028912 | 0.14623 | 0.197716 | 0.84327 | 0.9195 | 215.986 |
| Prima1 | 395.078 | 0.468484 | 0.26718 | 1.753466 | 0.07952 | 0.22402 | 45.243 |
| Primpol | 161.471 | 0.070801 | 0.16187 | 0.437399 | 0.66182 | 0.81293 | 41.619 |
| Prkaa1 | 1890.1 | 0.078073 | 0.08035 | 0.971629 | 0.33123 | 0.54208 | 38.038 |
| Prkaa2 | 2052.86 | -0.16643 | 0.08308 | -2.00335 | 0.04514 | 0.15451 | 80.017 |

| | | | | | | | |
|---|---|---|---|---|---|---|---|
| Prkab1 | 590.724 | -0.18548 | 0.11025 | -1.68246 | 0.09248 | 0.24762 | 10.923 |
| Prkab2 | 3963.14 | 0.071144 | 0.09939 | 0.715815 | 0.47411 | 0.67238 | 15.62 |
| Prkaca | 7511.5 | -0.13057 | 0.07073 | -1.84597 | 0.0649 | 0.19621 | 23.451 |
| Prkacb | 24145.8 | 0.338877 | 0.09934 | 3.411208 | 0.00065 | 0.00942 | 83.417 |
| Prkag1 | 322.692 | 0.023091 | 0.13061 | 0.176791 | 0.85967 | 0.92904 | 18.725 |
| Prkag2 | 3531.98 | 0.077367 | 0.07498 | 1.031905 | 0.30212 | 0.51187 | 237.899 |
| Prkag2os1 | 37.304 | -0.33572 | 0.32454 | -1.03444 | 0.30093 | 0.51075 | 4.282 |
| Prkag2os2 | 186.848 | 0.390834 | 0.13808 | 2.830517 | 0.00465 | 0.03467 | 2.391 |
| Prkar1a | 35756.2 | -0.03346 | 0.06606 | -0.50644 | 0.61255 | 0.77934 | 20.252 |
| Prkar1b | 20390.3 | -0.31174 | 0.10136 | -3.07546 | 0.0021 | 0.02081 | 132.696 |
| Prkar2a | 3396.74 | 0.197585 | 0.06283 | 3.144697 | 0.00166 | 0.01785 | 61.123 |
| Prkar2b | 8151.13 | 0.72887 | 0.16758 | 4.349384 | 1.37E-05 | 0.0007 | 102.821 |
| Prkca | 12083.4 | -0.46247 | 0.16491 | -2.80438 | 0.00504 | 0.03649 | 410.542 |
| Prkcb | 23591.3 | 0.127168 | 0.10657 | 1.193304 | 0.23275 | 0.43754 | 345.652 |
| Prkcd | 726.301 | -0.22679 | 0.11427 | -1.98464 | 0.04718 | 0.15945 | 30.857 |
| Prkce | 21573.6 | -0.22904 | 0.09111 | -2.51405 | 0.01194 | 0.06441 | 490.135 |
| Prkcg | 9659.63 | -0.26589 | 0.12323 | -2.1577 | 0.03095 | 0.12063 | 41.912 |
| Prkch | 2014.61 | 0.904552 | 0.32116 | 2.816546 | 0.00485 | 0.03557 | 193.39 |
| Prkci | 3447.26 | -0.01521 | 0.06267 | -0.24263 | 0.80829 | 0.90142 | 57.213 |
| Prkcq | 638.518 | -0.06588 | 0.13621 | -0.48367 | 0.62862 | 0.79 | 129.115 |
| Prkcsh | 3972.47 | 0.03416 | 0.06766 | 0.504897 | 0.61363 | 0.77996 | 11.417 |
| Prkcz | 12982.2 | 0.11839 | 0.07463 | 1.586326 | 0.11267 | 0.28066 | 101.233 |
| Prkcz2 | 374.112 | 0.002833 | 0.1058 | 0.026778 | 0.97864 | 0.98863 | 1.756 |
| Prkd1 | 736.801 | 0.107305 | 0.11029 | 0.972928 | 0.33059 | 0.54183 | 307.868 |
| Prkd2 | 383.427 | -0.20737 | 0.11271 | -1.83989 | 0.06578 | 0.19807 | 27.563 |
| Prkd3 | 1147.46 | -0.29191 | 0.08826 | -3.30716 | 0.00094 | 0.01228 | 71.412 |
| Prkdc | 3481.22 | 0.552483 | 0.13095 | 4.218918 | 2.45E-05 | 0.00101 | 204.37 |
| Prkg1 | 618.54 | 0.413813 | 0.13879 | 2.981549 | 0.00287 | 0.02548 | 1200.545 |
| Prkg2 | 452.433 | 0.302108 | 0.15718 | 1.922048 | 0.0546 | 0.17572 | 107.579 |
| Prkn | 728.864 | 0.304394 | 0.1782 | 1.708189 | 0.0876 | 0.23847 | 1222.978 |
| Prkra | 546.491 | -0.07591 | 0.11867 | -0.63964 | 0.52241 | 0.71129 | 18.118 |
| Prkrip1 | 951.67 | 0.012715 | 0.09399 | 0.135287 | 0.89239 | 0.94617 | 20.837 |
| Prkx | 1061.85 | 0.089547 | 0.09213 | 0.971969 | 0.33107 | 0.54208 | 34.868 |
| Prlhr | 47 | 1.274379 | 0.34455 | 3.698636 | 0.00022 | 0.0045 | 1.573 |

| | | | | | | | |
|---|---|---|---|---|---|---|---|
| Prlr | 44.0008 | 0.349594 | 0.41775 | 0.836839 | 0.40268 | 0.61133 | 171.943 |
| Prmt1 | 3062.93 | 0.148133 | 0.09227 | 1.605379 | 0.10841 | 0.27367 | 10.58 |
| Prmt2 | 2311.73 | 0.156585 | 0.09412 | 1.663695 | 0.09617 | 0.25395 | 30.644 |
| Prmt3 | 1011.38 | -0.06544 | 0.0854 | -0.76623 | 0.44354 | 0.64792 | 79.92 |
| Prmt5 | 2352.44 | 0.073406 | 0.08492 | 0.864405 | 0.38737 | 0.59665 | 10.339 |
| Prmt6 | 354.763 | -0.02547 | 0.10086 | -0.25255 | 0.80061 | 0.89692 | 4.89 |
| Prmt7 | 1823.19 | 0.036043 | 0.07479 | 0.481939 | 0.62985 | 0.79072 | 41.859 |
| Prmt8 | 2780.33 | -0.10366 | 0.07305 | -1.41912 | 0.15586 | 0.3424 | 80.462 |
| Prmt9 | 1075.32 | 0.165709 | 0.09502 | 1.743941 | 0.08117 | 0.22677 | 31.942 |
| Prnp | 21469.7 | -0.1602 | 0.09658 | -1.65868 | 0.09718 | 0.25547 | 28.502 |
| Prob1 | 304.677 | -0.39932 | 0.17465 | -2.28634 | 0.02223 | 0.0961 | 4.888 |
| Proca1 | 149.599 | 0.170192 | 0.18744 | 0.908007 | 0.36387 | 0.57434 | 12.372 |
| Procr | 28.0211 | -0.8925 | 0.42608 | -2.09466 | 0.0362 | 0.134 | 4.355 |
| Prodh | 2293.62 | -0.27406 | 0.06983 | -3.92448 | 8.69E-05 | 0.00245 | 29.847 |
| Prok2 | 226.263 | 1.064686 | 0.43563 | 2.444021 | 0.01452 | 0.07283 | 15.094 |
| Prokr2 | 130.526 | -0.86444 | 0.31615 | -2.73432 | 0.00625 | 0.04232 | 47.715 |
| Prom1 | 651.399 | -0.41496 | 0.1151 | -3.60534 | 0.00031 | 0.00579 | 108.413 |
| Prom2 | 26.9797 | -0.3137 | 0.3494 | -0.89783 | 0.36928 | 0.57974 | 14.995 |
| Prorp | 238.056 | -0.26006 | 0.15092 | -1.72319 | 0.08485 | 0.23364 | 82.957 |
| Prorsd1 | 210.615 | -0.0632 | 0.16109 | -0.3923 | 0.69483 | 0.83452 | 3.277 |
| Pros1 | 732.635 | -0.28478 | 0.14274 | -1.99512 | 0.04603 | 0.15654 | 75.04 |
| Proser1 | 1493.52 | 0.104021 | 0.09954 | 1.045019 | 0.29601 | 0.50634 | 18.09 |
| Proser2 | 116.9 | -0.24181 | 0.18549 | -1.30362 | 0.19236 | 0.38897 | 32.605 |
| Proser3 | 165.195 | 0.081448 | 0.16929 | 0.481099 | 0.63045 | 0.79112 | 13.166 |
| Prox1 | 561.546 | -0.61397 | 0.18066 | -3.3985 | 0.00068 | 0.00976 | 52.68 |
| Prox1os | 13.6472 | -0.45758 | 0.47204 | -0.96936 | 0.33237 | 0.54326 | 50.103 |
| Prox2 | 315.565 | 0.338539 | 0.12878 | 2.628725 | 0.00857 | 0.05176 | 24.375 |
| Prox2os | 55.8563 | 0.682731 | 0.3831 | 1.782114 | 0.07473 | 0.21543 | 1.855 |
| Proz | 899.044 | 0.160947 | 0.07948 | 2.024938 | 0.04287 | 0.14929 | 15.113 |
| Prpf18 | 3197.32 | 0.155884 | 0.06636 | 2.34912 | 0.01882 | 0.08608 | 30.056 |
| Prpf19 | 6155.04 | 0.012211 | 0.06888 | 0.177273 | 0.85929 | 0.92893 | 14.329 |
| Prpf3 | 1639.74 | 0.188458 | 0.07358 | 2.561187 | 0.01043 | 0.05907 | 25.762 |
| Prpf31 | 1340.37 | -0.17837 | 0.0695 | -2.56644 | 0.01027 | 0.05852 | 12.502 |
| Prpf38a | 1057.39 | 0.148597 | 0.08698 | 1.708434 | 0.08756 | 0.23845 | 16.165 |

| | | | | | | | |
|---|---|---|---|---|---|---|---|
| Prpf38b | 6090.68 | 0.274944 | 0.09277 | 2.963636 | 0.00304 | 0.02634 | 8.923 |
| Prpf39 | 3393.35 | 0.143727 | 0.13976 | 1.028384 | 0.30377 | 0.51361 | 27.054 |
| Prpf4 | 1189.64 | 0.023515 | 0.07456 | 0.315393 | 0.75246 | 0.86997 | 18.194 |
| Prpf40a | 2837.39 | 0.186272 | 0.11237 | 1.657704 | 0.09738 | 0.25589 | 56.999 |
| Prpf40b | 3655.78 | 0.083627 | 0.07844 | 1.066057 | 0.2864 | 0.49639 | 21.932 |
| Prpf4b | 7964.66 | 0.243629 | 0.12056 | 2.02086 | 0.04329 | 0.15032 | 30.763 |
| Prpf6 | 2982.39 | 0.095292 | 0.06283 | 1.516654 | 0.12935 | 0.30688 | 63.793 |
| Prpf8 | 12020.9 | 0.127186 | 0.0718 | 1.771435 | 0.07649 | 0.21863 | 22.634 |
| Prph | 15.0825 | -0.6178 | 0.62665 | -0.98587 | 0.3242 | 0.53531 | 3.805 |
| Prps1 | 2890.65 | -0.01759 | 0.07293 | -0.24125 | 0.80936 | 0.90175 | 19.528 |
| Prps1l3 | 437.519 | 0.087605 | 0.11474 | 0.763504 | 0.44516 | 0.64913 | 9.471 |
| Prps2 | 1218.53 | -0.16647 | 0.10371 | -1.60514 | 0.10846 | 0.27373 | 36.428 |
| Prpsap1 | 1925.43 | 0.103362 | 0.09307 | 1.110615 | 0.26673 | 0.47595 | 23.791 |
| Prpsap2 | 1724.87 | 0.234782 | 0.10458 | 2.245087 | 0.02476 | 0.10331 | 32.439 |
| Prr11 | 44.5692 | -0.5648 | 0.36461 | -1.54903 | 0.12138 | 0.29466 | 19.556 |
| Prr12 | 4217.55 | 0.148549 | 0.06604 | 2.249464 | 0.02448 | 0.10241 | 25.319 |
| Prr13 | 768.333 | -0.66883 | 0.14107 | -4.74108 | 2.13E-06 | 0.00019 | 3.779 |
| Prr14 | 1106.23 | -0.10068 | 0.07343 | -1.37112 | 0.17034 | 0.36065 | 17.149 |
| Prr14l | 3647.69 | 0.110626 | 0.06958 | 1.58985 | 0.11187 | 0.27932 | 64.437 |
| Prr15 | 15.6334 | -0.43545 | 0.44864 | -0.9706 | 0.33175 | 0.54275 | 4.367 |
| Prr15l | 28.7067 | 0.163782 | 0.34701 | 0.471983 | 0.63694 | 0.79574 | 9.727 |
| Prr16 | 419.312 | -0.35022 | 0.16587 | -2.11137 | 0.03474 | 0.13053 | 302.312 |
| Prr18 | 1804.54 | -0.66037 | 0.16136 | -4.09244 | 4.27E-05 | 0.00146 | 6.655 |
| Prr22 | 43.3366 | 0.205951 | 0.29464 | 0.698985 | 0.48456 | 0.68089 | 1.959 |
| Prr3 | 1112.7 | 0.03473 | 0.07616 | 0.456012 | 0.64838 | 0.80408 | 7.696 |
| Prr33 | 15.3823 | -0.63311 | 0.49976 | -1.26683 | 0.20522 | 0.40426 | 15.698 |
| Prr36 | 679.894 | -0.12614 | 0.12848 | -0.98179 | 0.3262 | 0.53752 | 7.917 |
| Prr5 | 388.984 | 0.046228 | 0.12109 | 0.381763 | 0.70264 | 0.8396 | 34.054 |
| Prr5l | 401.222 | -0.70513 | 0.17315 | -4.07243 | 4.65E-05 | 0.00156 | 168.743 |
| Prr7 | 1075.98 | 0.243107 | 0.10174 | 2.389404 | 0.01688 | 0.08047 | 8.888 |
| Prrc1 | 1077.57 | 0.085078 | 0.08884 | 0.95766 | 0.33823 | 0.54905 | 38.216 |
| Prrc2a | 22226.8 | 0.083168 | 0.05996 | 1.387136 | 0.1654 | 0.35456 | 15.822 |
| Prrc2b | 28993 | 0.086623 | 0.06247 | 1.3867 | 0.16553 | 0.35477 | 83.456 |
| Prrc2c | 15350.9 | 0.184774 | 0.07476 | 2.471653 | 0.01345 | 0.06969 | 69.832 |

| | | | | | | | |
|---|---|---|---|---|---|---|---|
| Prrg1 | 287.815 | -0.4014 | 0.16501 | -2.43256 | 0.01499 | 0.07436 | 134.284 |
| Prrg2 | 172.298 | -0.23845 | 0.15468 | -1.54153 | 0.12319 | 0.29739 | 9.094 |
| Prrg3 | 1772.46 | -0.12393 | 0.10415 | -1.18995 | 0.23407 | 0.43913 | 10.099 |
| Prrg4 | 35.7266 | 0.082005 | 0.28992 | 0.282856 | 0.77729 | 0.88407 | 19.136 |
| Prrt1 | 5226.43 | 0.053127 | 0.07762 | 0.684432 | 0.4937 | 0.68811 | 3.594 |
| Prrt2 | 48.8672 | 0.244887 | 0.28651 | 0.854709 | 0.39271 | 0.60193 | 3.681 |
| Prrt3 | 2266.44 | -0.12896 | 0.09059 | -1.42364 | 0.15455 | 0.34062 | 8.293 |
| Prrt4 | 283.014 | 0.877528 | 0.2248 | 3.903662 | 9.47E-05 | 0.00257 | 10.353 |
| Prrx1 | 491.563 | 0.019055 | 0.10908 | 0.174687 | 0.86133 | 0.92974 | 68.592 |
| Prrx2 | 12.6891 | -1.36831 | 0.55849 | -2.44999 | 0.01429 | 0.07208 | 46.28 |
| Prss12 | 557.513 | -0.98606 | 0.40417 | -2.43974 | 0.0147 | 0.07327 | 59.685 |
| Prss22 | 18.9885 | -0.85692 | 0.57847 | -1.48135 | 0.13851 | 0.31964 | 4.567 |
| Prss23 | 638.819 | -1.19639 | 0.44073 | -2.71455 | 0.00664 | 0.04392 | 19.405 |
| Prss23os | 71.1111 | -0.6169 | 0.47035 | -1.31156 | 0.18967 | 0.38569 | 1.867 |
| Prss30 | 70.5062 | -0.93629 | 0.42603 | -2.1977 | 0.02797 | 0.11256 | 8.441 |
| Prss35 | 65.8862 | -0.89371 | 0.44737 | -1.99771 | 0.04575 | 0.15589 | 14.795 |
| Prss36 | 242.914 | -0.15131 | 0.15176 | -0.99702 | 0.31876 | 0.5293 | 14.088 |
| Prss50 | 37.1181 | 0.160447 | 0.30199 | 0.531307 | 0.59521 | 0.76726 | 6.663 |
| Prss52 | 24.0123 | -0.68903 | 0.40331 | -1.70842 | 0.08756 | 0.23845 | 9.47 |
| Prss53 | 27.029 | -0.14517 | 0.35126 | -0.41328 | 0.6794 | 0.82491 | 5.13 |
| Prss56 | 24.6191 | 0.125604 | 0.4189 | 0.299842 | 0.7643 | 0.87657 | 5.093 |
| Prss57 | 26.4046 | 0.048474 | 0.40999 | 0.118231 | 0.90588 | 0.95277 | 9.488 |
| Prtg | 214.61 | -0.03296 | 0.13228 | -0.2492 | 0.80321 | 0.89841 | 110.418 |
| Prtn3 | 60.1821 | 0.60048 | 0.2434 | 2.467066 | 0.01362 | 0.07023 | 8.699 |
| Prune1 | 1038.86 | -0.0841 | 0.09354 | -0.89901 | 0.36865 | 0.57929 | 28.403 |
| Prune2 | 4630.92 | 0.047605 | 0.08171 | 0.582629 | 0.56014 | 0.74154 | 267.815 |
| Prx | 86.2977 | 0.048046 | 0.25397 | 0.189179 | 0.84995 | 0.92371 | 22.827 |
| Prxl2a | 5173.77 | -0.18138 | 0.09078 | -1.99791 | 0.04573 | 0.15584 | 20.049 |
| Prxl2b | 1596.88 | -0.12856 | 0.10351 | -1.24202 | 0.21423 | 0.41558 | 3.632 |
| Prxl2c | 576.736 | 0.132365 | 0.14466 | 0.914978 | 0.3602 | 0.5706 | 37.431 |
| Psap | 67454.4 | -0.16565 | 0.06536 | -2.53428 | 0.01127 | 0.06199 | 24.928 |
| Psat1 | 6101.68 | -0.28571 | 0.09172 | -3.11508 | 0.00184 | 0.01906 | 42.66 |
| Psd | 17618.6 | 0.188085 | 0.09744 | 1.93021 | 0.05358 | 0.17353 | 15.07 |
| Psd2 | 2887.03 | -0.10581 | 0.1169 | -0.90509 | 0.36542 | 0.57607 | 49.886 |

| | | | | | | | |
|---|---|---|---|---|---|---|---|
| Psd3 | 26159.2 | 0.073346 | 0.06795 | 1.079484 | 0.28037 | 0.49023 | 522.946 |
| Psd4 | 86.8096 | -0.09769 | 0.20939 | -0.46654 | 0.64083 | 0.79893 | 47.375 |
| Psen1 | 2911.25 | 0.02283 | 0.0623 | 0.366451 | 0.71403 | 0.8464 | 47.048 |
| Psen2 | 539.488 | -0.1779 | 0.11482 | -1.54942 | 0.12128 | 0.29459 | 36.435 |
| Psenen | 1107.2 | -0.08388 | 0.10656 | -0.7871 | 0.43122 | 0.63728 | 1.765 |
| Psg16 | 55.0438 | -0.24815 | 0.27391 | -0.90596 | 0.36496 | 0.57561 | 59.411 |
| Psip1 | 8326.79 | 0.186138 | 0.12318 | 1.51116 | 0.13075 | 0.30908 | 30.78 |
| Pskh1 | 490.496 | -0.02384 | 0.12106 | -0.19695 | 0.84387 | 0.91976 | 31.338 |
| Psma1 | 2985.26 | -0.00185 | 0.06122 | -0.03028 | 0.97584 | 0.98762 | 11.513 |
| Psma2 | 2874.53 | -0.08351 | 0.06988 | -1.19503 | 0.23207 | 0.43701 | 60.997 |
| Psma3 | 2958.98 | 0.028858 | 0.05669 | 0.509037 | 0.61073 | 0.77794 | 21.727 |
| Psma4 | 2420.41 | -0.03383 | 0.0637 | -0.53102 | 0.59541 | 0.76727 | 7.241 |
| Psma5 | 2294.1 | -0.09208 | 0.08576 | -1.07373 | 0.28294 | 0.49316 | 23.049 |
| Psma5-ps | 121.381 | 0.201989 | 0.1805 | 1.119076 | 0.26311 | 0.47199 | 0.952 |
| Psma6 | 3188.91 | -0.11406 | 0.07963 | -1.43228 | 0.15206 | 0.33717 | 34.233 |
| Psma7 | 2766.48 | -0.02475 | 0.06688 | -0.37004 | 0.71135 | 0.84471 | 6.06 |
| Psmb1 | 4041.59 | -0.02312 | 0.07639 | -0.30267 | 0.76214 | 0.87529 | 24.731 |
| Psmb10 | 696.384 | -0.46612 | 0.12799 | -3.64191 | 0.00027 | 0.00521 | 2.71 |
| Psmb2 | 1419.64 | -0.03563 | 0.09018 | -0.39509 | 0.69278 | 0.83344 | 32.085 |
| Psmb3 | 1771.83 | -0.02557 | 0.09571 | -0.26717 | 0.78934 | 0.89071 | 10.102 |
| Psmb4 | 3899.57 | -0.09426 | 0.07068 | -1.33359 | 0.18234 | 0.37612 | 2.869 |
| Psmb5 | 1087.53 | -0.34551 | 0.10478 | -3.29765 | 0.00097 | 0.01259 | 3.904 |
| Psmb6 | 2444.09 | -0.01348 | 0.06725 | -0.20038 | 0.84119 | 0.91861 | 2.492 |
| Psmb7 | 2323.44 | -0.12757 | 0.08511 | -1.49883 | 0.13392 | 0.3136 | 56.052 |
| Psmb8 | 43.1961 | -0.88622 | 0.32194 | -2.75276 | 0.00591 | 0.04085 | 3.734 |
| Psmb9 | 44.9257 | -0.62017 | 0.30438 | -2.03747 | 0.0416 | 0.14657 | 5.778 |
| Psmc1 | 6315.46 | -0.20254 | 0.06491 | -3.12025 | 0.00181 | 0.01883 | 13.252 |
| Psmc2 | 3185.03 | -0.01396 | 0.07557 | -0.18472 | 0.85345 | 0.92567 | 18.505 |
| Psmc3 | 5000.49 | -0.25757 | 0.07996 | -3.22103 | 0.00128 | 0.01485 | 12.361 |
| Psmc3ip | 161.386 | 0.20086 | 0.18107 | 1.109313 | 0.2673 | 0.47664 | 3.614 |
| Psmc4 | 2871.33 | -0.07564 | 0.06693 | -1.13003 | 0.25846 | 0.46651 | 8.395 |
| Psmc5 | 5458.22 | -0.22758 | 0.06213 | -3.66319 | 0.00025 | 0.00496 | 6.967 |
| Psmc6 | 3764.71 | 0.089407 | 0.08738 | 1.023183 | 0.30622 | 0.51631 | 19.918 |
| Psmd1 | 4603.36 | 0.10964 | 0.08006 | 1.369446 | 0.17086 | 0.36125 | 74.765 |

| | | | | | | | |
|---|---|---|---|---|---|---|---|
| Psmd10 | 697.52 | -0.11775 | 0.08637 | -1.36329 | 0.17279 | 0.36397 | 8.3 |
| Psmd11 | 2914.39 | 0.011788 | 0.07673 | 0.153643 | 0.87789 | 0.93893 | 44.634 |
| Psmd12 | 2732.09 | -0.01847 | 0.05891 | -0.31356 | 0.75385 | 0.87065 | 24.879 |
| Psmd13 | 2645.25 | 0.047003 | 0.07149 | 0.657462 | 0.51088 | 0.70199 | 16.676 |
| Psmd14 | 3129.57 | 0.01975 | 0.06925 | 0.285212 | 0.77548 | 0.88307 | 88.683 |
| Psmd2 | 9837.6 | -0.04741 | 0.08066 | -0.58778 | 0.55668 | 0.73871 | 11.763 |
| Psmd3 | 4262.73 | -0.01675 | 0.05804 | -0.28853 | 0.77294 | 0.88183 | 13.426 |
| Psmd4 | 3460.67 | -0.12869 | 0.06611 | -1.94663 | 0.05158 | 0.16934 | 9.921 |
| Psmd5 | 2294.36 | 0.087913 | 0.09184 | 0.957226 | 0.33845 | 0.5492 | 25.235 |
| Psmd6 | 2789.67 | 0.068809 | 0.07688 | 0.895042 | 0.37076 | 0.58094 | 8.811 |
| Psmd7 | 4407.89 | -0.03005 | 0.06188 | -0.48568 | 0.62719 | 0.78903 | 8.084 |
| Psmd8 | 2988.7 | -0.01467 | 0.08088 | -0.1814 | 0.85605 | 0.92729 | 6.514 |
| Psmd9 | 1304.7 | 0.163758 | 0.0796 | 2.057204 | 0.03967 | 0.14223 | 80.719 |
| Psme1 | 945.344 | 0.026742 | 0.13024 | 0.205334 | 0.83731 | 0.91639 | 3.407 |
| Psme2 | 782.426 | 0.05575 | 0.09814 | 0.568045 | 0.57 | 0.74937 | 3.673 |
| Psme2b | 251.807 | -0.1686 | 0.12961 | -1.30079 | 0.19333 | 0.39022 | 0.837 |
| Psme3 | 4831.2 | -0.11253 | 0.06068 | -1.85452 | 0.06367 | 0.19397 | 7.325 |
| Psme3ip1 | 1752.77 | 0.009708 | 0.07578 | 0.12811 | 0.89806 | 0.94885 | 27.86 |
| Psme4 | 3505.52 | 0.104121 | 0.11885 | 0.876095 | 0.38098 | 0.59053 | 108.636 |
| Psmf1 | 1222.17 | -0.00678 | 0.08799 | -0.07709 | 0.93855 | 0.96942 | 28.375 |
| Psmg1 | 527.102 | 0.07835 | 0.11213 | 0.698761 | 0.4847 | 0.68092 | 11.028 |
| Psmg2 | 621.387 | 0.130344 | 0.09731 | 1.339512 | 0.1804 | 0.37372 | 12.564 |
| Psmg3 | 137.185 | 0.051251 | 0.16189 | 0.316579 | 0.75156 | 0.86975 | 3.294 |
| Psmg4 | 211.443 | -0.17237 | 0.1362 | -1.26562 | 0.20565 | 0.40482 | 15.21 |
| Pspc1 | 2143.55 | 0.104527 | 0.08692 | 1.202613 | 0.22913 | 0.43326 | 57.607 |
| Psph | 825.924 | -0.18459 | 0.09614 | -1.91989 | 0.05487 | 0.17636 | 21.892 |
| Pspn | 14.7622 | 0.637156 | 0.46121 | 1.38148 | 0.16713 | 0.35694 | 0.562 |
| Psrc1 | 100.471 | -1.05836 | 0.33237 | -3.18426 | 0.00145 | 0.01617 | 4.393 |
| Pstk | 824.863 | 0.002692 | 0.08552 | 0.031471 | 0.97489 | 0.98738 | 23.342 |
| Pstpip1 | 47.9236 | -0.52823 | 0.36633 | -1.44195 | 0.14932 | 0.33352 | 38.927 |
| Pstpip2 | 423.139 | -0.49079 | 0.14169 | -3.46378 | 0.00053 | 0.00829 | 92.966 |
| Ptafr | 60.9461 | -0.75928 | 0.29471 | -2.57637 | 0.00998 | 0.05742 | 18.617 |
| Ptar1 | 3147.9 | -0.22132 | 0.12741 | -1.73702 | 0.08238 | 0.22892 | 44.24 |
| Ptbp1 | 976.582 | 0.184664 | 0.0893 | 2.067875 | 0.03865 | 0.13997 | 10.345 |

| | | | | | | | |
|---|---|---|---|---|---|---|---|
| Ptbp2 | 5723.12 | 0.126959 | 0.06614 | 1.919533 | 0.05492 | 0.17648 | 65.725 |
| Ptbp3 | 1397.17 | -0.10688 | 0.10145 | -1.05351 | 0.29211 | 0.50188 | 77.497 |
| Ptcd1 | 1558.49 | 0.325606 | 0.15442 | 2.108544 | 0.03498 | 0.13116 | 19.595 |
| Ptcd2 | 1080.72 | -0.12651 | 0.07129 | -1.77473 | 0.07594 | 0.21766 | 25.059 |
| Ptcd3 | 2620.77 | 0.100589 | 0.06183 | 1.626783 | 0.10378 | 0.26624 | 28.113 |
| Ptch1 | 4935.23 | 0.352704 | 0.09254 | 3.811209 | 0.00014 | 0.00334 | 65.271 |
| Ptch2 | 411.058 | 0.471221 | 0.21101 | 2.233168 | 0.02554 | 0.10544 | 20.027 |
| Ptchd1 | 326.767 | 0.154762 | 0.12591 | 1.229111 | 0.21903 | 0.42125 | 50.36 |
| Ptchd4 | 137.809 | 0.104585 | 0.22588 | 0.463003 | 0.64336 | 0.80102 | 189.131 |
| Ptdss1 | 2255.62 | -0.17811 | 0.09637 | -1.84823 | 0.06457 | 0.1956 | 65.572 |
| Ptdss2 | 1929.21 | -0.03303 | 0.07203 | -0.45863 | 0.6465 | 0.80316 | 35.225 |
| Pten | 9177.82 | 0.130763 | 0.06518 | 2.006212 | 0.04483 | 0.15385 | 68.664 |
| Pter | 1612.17 | -0.95389 | 0.66435 | -1.43582 | 0.15105 | 0.33587 | 79.415 |
| Ptgdr | 35.3628 | -0.71596 | 0.46379 | -1.54372 | 0.12266 | 0.29648 | 8.141 |
| Ptgds | 30276.9 | -1.04249 | 0.30223 | -3.44938 | 0.00056 | 0.00859 | 3.338 |
| Ptger1 | 70.6755 | 0.317948 | 0.22847 | 1.391642 | 0.16403 | 0.35285 | 6.076 |
| Ptger2 | 12.4261 | -0.12201 | 0.52054 | -0.23439 | 0.81468 | 0.90481 | 15.626 |
| Ptger3 | 127.705 | -0.08132 | 0.23372 | -0.34794 | 0.72789 | 0.85483 | 78.997 |
| Ptger4 | 48.8375 | -0.15285 | 0.27692 | -0.55196 | 0.58098 | 0.75714 | 37.527 |
| Ptges | 160.367 | -0.59807 | 0.19755 | -3.02744 | 0.00247 | 0.02314 | 40.393 |
| Ptges2 | 1152.99 | -0.24362 | 0.07535 | -3.23298 | 0.00123 | 0.01453 | 9.877 |
| Ptges3 | 3876.44 | -0.0557 | 0.10659 | -0.52255 | 0.60129 | 0.77152 | 18.319 |
| Ptges3-ps | 1041.65 | -0.04226 | 0.10901 | -0.38767 | 0.69826 | 0.83649 | 0.48 |
| Ptges3l | 57.8487 | -0.74095 | 0.30631 | -2.41899 | 0.01556 | 0.07631 | 6.522 |
| Ptgfr | 52.2203 | -0.78713 | 0.36602 | -2.15054 | 0.03151 | 0.12219 | 41.129 |
| Ptgfrn | 1497.64 | 0.073853 | 0.0977 | 0.755886 | 0.44972 | 0.65278 | 70.047 |
| Ptgis | 151.3 | -0.51243 | 0.27401 | -1.87014 | 0.06146 | 0.18957 | 48.8 |
| Ptgr1 | 32.1503 | -0.55362 | 0.3644 | -1.51928 | 0.12869 | 0.30583 | 21.681 |
| Ptgr2 | 1230.34 | -0.16463 | 0.11181 | -1.47248 | 0.14089 | 0.32265 | 30.601 |
| Ptgs1 | 531.157 | -0.26433 | 0.10191 | -2.59381 | 0.00949 | 0.05541 | 21.847 |
| Ptgs2 | 300.282 | -0.73645 | 0.24736 | -2.97721 | 0.00291 | 0.02572 | 8.197 |
| Ptgs2os | 15.5559 | -0.59183 | 0.47395 | -1.24872 | 0.21177 | 0.4124 | 25.091 |
| Pth1r | 214.131 | -0.67267 | 0.18734 | -3.59065 | 0.00033 | 0.00602 | 25.061 |
| Pth2r | 49.3046 | -0.33683 | 0.40807 | -0.82542 | 0.40913 | 0.61691 | 107.189 |

| | | | | | | | |
|---|---|---|---|---|---|---|---|
| Pthlh | 123.428 | -0.04419 | 0.24142 | -0.18305 | 0.85476 | 0.92658 | 12.083 |
| Ptk2 | 5467.95 | -0.02189 | 0.06612 | -0.33103 | 0.74062 | 0.86302 | 218.179 |
| Ptk2b | 27835.2 | 0.071551 | 0.07205 | 0.993106 | 0.32066 | 0.53141 | 127.796 |
| Ptk6 | 12.9762 | 0.803943 | 0.55082 | 1.459532 | 0.14442 | 0.32717 | 9.069 |
| Ptk7 | 706.871 | 0.278987 | 0.12168 | 2.292778 | 0.02186 | 0.09513 | 65.034 |
| Ptma | 12219.9 | -0.16222 | 0.07487 | -2.16667 | 0.03026 | 0.11875 | 3.987 |
| Ptma-ps2 | 22.1872 | -0.10387 | 0.37839 | -0.27451 | 0.78369 | 0.88762 | 2.231 |
| Ptms | 15179.5 | -0.02474 | 0.08967 | -0.27589 | 0.78264 | 0.88707 | 6.423 |
| Ptn | 14452.2 | -0.41396 | 0.08979 | -4.61033 | 4.02E-06 | 0.00029 | 95.292 |
| Ptov1 | 4686.06 | 0.037961 | 0.08198 | 0.463042 | 0.64333 | 0.80102 | 6.722 |
| Ptp4a1 | 58.6207 | -0.11205 | 0.28685 | -0.39061 | 0.69609 | 0.83541 | 9.469 |
| Ptp4a2 | 7357.81 | -0.04451 | 0.0603 | -0.73819 | 0.4604 | 0.6609 | 38.785 |
| Ptp4a3 | 3389.11 | -0.23326 | 0.09709 | -2.40242 | 0.01629 | 0.0786 | 35.622 |
| Ptpa | 7575.67 | -0.04286 | 0.06419 | -0.66773 | 0.5043 | 0.69657 | 31.768 |
| Ptpdc1 | 2126.15 | -0.12097 | 0.11468 | -1.0548 | 0.29152 | 0.50128 | 47.793 |
| Ptpmt1 | 281.004 | -0.17827 | 0.13222 | -1.34824 | 0.17758 | 0.36981 | 9.33 |
| Ptpn1 | 2873.19 | -0.08147 | 0.07624 | -1.06868 | 0.28521 | 0.495 | 47.329 |
| Ptpn11 | 13564.5 | -0.11422 | 0.06384 | -1.78923 | 0.07358 | 0.2136 | 60.865 |
| Ptpn12 | 2103.51 | 0.105287 | 0.07512 | 1.40166 | 0.16102 | 0.34918 | 69.267 |
| Ptpn13 | 1866.18 | 0.13432 | 0.11125 | 1.207317 | 0.22731 | 0.4315 | 173.112 |
| Ptpn14 | 858.785 | -0.24899 | 0.17818 | -1.39742 | 0.16229 | 0.35067 | 148.428 |
| Ptpn18 | 16.3705 | -0.01392 | 0.4579 | -0.0304 | 0.97574 | 0.98762 | 15.972 |
| Ptpn2 | 1291.56 | 0.12444 | 0.08695 | 1.431107 | 0.1524 | 0.33764 | 59.085 |
| Ptpn20 | 19.6493 | -0.64559 | 0.4131 | -1.56281 | 0.1181 | 0.28958 | 51.548 |
| Ptpn21 | 480.434 | -0.13246 | 0.0914 | -1.44931 | 0.14725 | 0.33046 | 60.665 |
| Ptpn22 | 52.9239 | -0.16512 | 0.26179 | -0.63074 | 0.52821 | 0.71613 | 52.453 |
| Ptpn23 | 3451.52 | 0.211561 | 0.07499 | 2.821184 | 0.00478 | 0.0353 | 23.132 |
| Ptpn3 | 1694.78 | -1.16858 | 0.40771 | -2.86618 | 0.00415 | 0.03226 | 110.997 |
| Ptpn4 | 5925 | -0.01241 | 0.14332 | -0.08662 | 0.93097 | 0.96526 | 185.147 |
| Ptpn5 | 18718.6 | 0.691272 | 0.17699 | 3.905734 | 9.39E-05 | 0.00257 | 55.89 |
| Ptpn6 | 131.651 | -0.41294 | 0.19131 | -2.15848 | 0.03089 | 0.12052 | 18.008 |
| Ptpn7 | 210.338 | 0.497687 | 0.24663 | 2.017947 | 0.0436 | 0.15103 | 12.618 |
| Ptpn9 | 1729.58 | 0.112567 | 0.07241 | 1.554573 | 0.12005 | 0.29261 | 67.885 |
| Ptpra | 5535.48 | 0.023532 | 0.06019 | 0.390957 | 0.69583 | 0.83522 | 105.847 |

| | | | | | | | |
|---|---|---|---|---|---|---|---|
| Ptprb | 2030.16 | -0.52147 | 0.1024 | -5.09254 | 3.53E-07 | 4.80E-05 | 114.013 |
| Ptprc | 201.19 | -0.39453 | 0.19389 | -2.03484 | 0.04187 | 0.1472 | 112.848 |
| Ptprd | 8837.29 | -0.36272 | 0.13181 | -2.75189 | 0.00593 | 0.04089 | 2270.724 |
| Ptpre | 2138.27 | 0.465361 | 0.11064 | 4.206088 | 2.60E-05 | 0.00105 | 148.813 |
| Ptprf | 5433.9 | 0.040161 | 0.07606 | 0.528004 | 0.5975 | 0.76924 | 83.193 |
| Ptprh | 135.178 | 0.19542 | 0.21692 | 0.900892 | 0.36765 | 0.57811 | 55.43 |
| Ptprj | 6663.5 | 0.021033 | 0.05698 | 0.369143 | 0.71202 | 0.845 | 150.842 |
| Ptprk | 2280.74 | -0.76917 | 0.36599 | -2.10163 | 0.03559 | 0.13252 | 522.578 |
| Ptprm | 2054.09 | 0.168125 | 0.08687 | 1.935379 | 0.05294 | 0.17218 | 687.511 |
| Ptprn | 22281.9 | 0.055247 | 0.06921 | 0.798223 | 0.42474 | 0.63137 | 17.476 |
| Ptprn2 | 10810.7 | 0.091793 | 0.06901 | 1.330121 | 0.18348 | 0.37759 | 791.13 |
| Ptpro | 3120.08 | 0.096417 | 0.10833 | 0.890064 | 0.37343 | 0.58357 | 210.915 |
| Ptprr | 961.073 | 0.200462 | 0.10123 | 1.980184 | 0.04768 | 0.1606 | 256.72 |
| Ptprs | 20890.8 | -0.08636 | 0.06836 | -1.26337 | 0.20645 | 0.40578 | 64.058 |
| Ptprt | 8780.84 | -0.25611 | 0.08719 | -2.93741 | 0.00331 | 0.02776 | 1139.158 |
| Ptpru | 2216.25 | -0.37537 | 0.19216 | -1.95344 | 0.05077 | 0.16761 | 69.832 |
| Ptprv | 2135.51 | 0.560461 | 0.38282 | 1.464037 | 0.14318 | 0.3252 | 24.098 |
| Ptprz1 | 13723.7 | -0.077 | 0.09462 | -0.81381 | 0.41575 | 0.62362 | 177.415 |
| Ptrh1 | 106.147 | -0.28731 | 0.20734 | -1.38567 | 0.16585 | 0.35526 | 1.812 |
| Ptrh2 | 999.106 | 0.19828 | 0.11558 | 1.715569 | 0.08624 | 0.23607 | 8.473 |
| Ptrhd1 | 214.626 | 0.00163 | 0.18105 | 0.009005 | 0.99282 | 0.99589 | 5.924 |
| Pts | 1252.32 | -0.01465 | 0.07142 | -0.20512 | 0.83748 | 0.91642 | 7.108 |
| Pttg1 | 760.247 | -0.27404 | 0.09099 | -3.01181 | 0.0026 | 0.02399 | 6.002 |
| Pttg1ip | 7205.91 | 0.414169 | 0.11837 | 3.498858 | 0.00047 | 0.00759 | 17.013 |
| Puf60 | 5054.25 | -0.08505 | 0.06697 | -1.26994 | 0.20411 | 0.4028 | 10.743 |
| Pum1 | 6611.59 | 0.114988 | 0.07405 | 1.552763 | 0.12048 | 0.29334 | 118.244 |
| Pum2 | 11886.2 | 0.132253 | 0.06403 | 2.065446 | 0.03888 | 0.14058 | 78.448 |
| Pum3 | 1347.11 | 0.043465 | 0.07335 | 0.592545 | 0.55349 | 0.7363 | 41.128 |
| Pura | 4517.33 | -0.0019 | 0.09337 | -0.0204 | 0.98373 | 0.99112 | 11.439 |
| Purb | 30639.7 | -0.02585 | 0.06425 | -0.40228 | 0.68748 | 0.82964 | 8.319 |
| Purg | 1016.43 | 0.16105 | 0.12775 | 1.260634 | 0.20744 | 0.40694 | 31.143 |
| Pus1 | 442.376 | -0.16335 | 0.09482 | -1.72274 | 0.08494 | 0.23381 | 6.993 |
| Pus10 | 878.164 | 0.121375 | 0.0982 | 1.236054 | 0.21644 | 0.41834 | 67.203 |
| Pus7 | 1056.58 | 0.099794 | 0.08641 | 1.154927 | 0.24812 | 0.45524 | 43.064 |

| | | | | | | | |
|---|---|---|---|---|---|---|---|
| Pus7l | 144.194 | -0.07749 | 0.18087 | -0.42843 | 0.66834 | 0.81705 | 20.86 |
| Pusl1 | 427.496 | -0.11631 | 0.10246 | -1.13517 | 0.2563 | 0.46423 | 3.903 |
| Pvalb | 572.233 | -0.46714 | 0.1497 | -3.1204 | 0.00181 | 0.01883 | 15.287 |
| Pvr | 572.1 | 0.069011 | 0.10202 | 0.676474 | 0.49874 | 0.6921 | 17.583 |
| Pvt1 | 507.214 | -0.19588 | 0.11163 | -1.75473 | 0.07931 | 0.22353 | 340.129 |
| Pwp1 | 964.091 | 0.090438 | 0.08377 | 1.079588 | 0.28033 | 0.49019 | 59.603 |
| Pwp2 | 1538.25 | 0.15276 | 0.07038 | 2.170645 | 0.02996 | 0.11816 | 14.241 |
| Pwwp2a | 1546.9 | 0.139796 | 0.09087 | 1.538418 | 0.12395 | 0.29847 | 39.494 |
| Pwwp2b | 899.531 | 0.293205 | 0.13524 | 2.168007 | 0.03016 | 0.11868 | 21.422 |
| Pwwp3a | 2156.5 | 0.001719 | 0.06986 | 0.02461 | 0.98037 | 0.98964 | 17.47 |
| Pwwp3b | 411.488 | 0.152601 | 0.15098 | 1.010742 | 0.31214 | 0.52253 | 28.294 |
| Pxdc1 | 157.41 | -0.22514 | 0.14897 | -1.51129 | 0.13071 | 0.30907 | 25.653 |
| Pxdn | 4262.51 | -0.0486 | 0.06163 | -0.78855 | 0.43037 | 0.63672 | 80.051 |
| Pxk | 1753.27 | 0.087627 | 0.074 | 1.184204 | 0.23633 | 0.44163 | 66.907 |
| Pxmp2 | 101.863 | -0.31613 | 0.18551 | -1.70408 | 0.08837 | 0.24017 | 11.905 |
| Pxmp4 | 617.549 | -0.27577 | 0.10591 | -2.60383 | 0.00922 | 0.0543 | 17.951 |
| Pxn | 1702.16 | -0.11294 | 0.08086 | -1.39673 | 0.16249 | 0.35091 | 49.312 |
| Pxylp1 | 818.834 | 0.343914 | 0.11144 | 3.086192 | 0.00203 | 0.02025 | 69.334 |
| Pycard | 107.741 | 0.12306 | 0.18139 | 0.678433 | 0.4975 | 0.69094 | 4.16 |
| Pycr1 | 146.31 | -0.09456 | 0.16366 | -0.57777 | 0.56342 | 0.74402 | 8.058 |
| Pycr2 | 985.732 | -0.02618 | 0.10295 | -0.25428 | 0.79928 | 0.89647 | 3.796 |
| Pycrl | 499.171 | -0.11216 | 0.10512 | -1.06692 | 0.28601 | 0.49604 | 5.084 |
| Pygb | 10104.3 | -0.26974 | 0.09172 | -2.94092 | 0.00327 | 0.02759 | 45.024 |
| Pygl | 237.753 | 0.082434 | 0.15534 | 0.530655 | 0.59566 | 0.76745 | 40.678 |
| Pygm | 1050.9 | -0.34471 | 0.12771 | -2.69909 | 0.00695 | 0.04524 | 14.061 |
| Pygo1 | 2750.14 | -0.1414 | 0.08022 | -1.7626 | 0.07797 | 0.22102 | 26.537 |
| Pygo2 | 2075.54 | -0.04877 | 0.08942 | -0.5454 | 0.58548 | 0.75994 | 4.915 |
| Pym1 | 546.187 | -0.1396 | 0.08848 | -1.57787 | 0.1146 | 0.28352 | 18.676 |
| Pyroxd1 | 326.636 | 0.313022 | 0.12247 | 2.555973 | 0.01059 | 0.05973 | 17.604 |
| Pyroxd2 | 62.6581 | 0.031824 | 0.27429 | 0.116023 | 0.90763 | 0.95362 | 26.918 |
| Pyurf | 550.478 | -0.12796 | 0.09763 | -1.31062 | 0.18999 | 0.38603 | 5.503 |
| Pzp | 21.9192 | -0.41628 | 0.46823 | -0.88905 | 0.37398 | 0.58424 | 43.154 |
| Qars | 2185.1 | -0.03204 | 0.07054 | -0.45428 | 0.64963 | 0.80496 | 8.236 |
| Qdpr | 2501.57 | -0.56686 | 0.15178 | -3.73472 | 0.00019 | 0.00414 | 16.216 |

| | | | | | | | |
|---|---|---|---|---|---|---|---|
| Qk | 19372.9 | -0.13819 | 0.11002 | -1.25597 | 0.20913 | 0.40918 | 117.254 |
| Qpct | 614.311 | 0.583342 | 0.13592 | 4.291951 | 1.77E-05 | 0.00081 | 38.473 |
| Qpctl | 759.114 | 0.105274 | 0.08217 | 1.281169 | 0.20013 | 0.39817 | 9.59 |
| Qprt | 16.5274 | -1.45919 | 0.55281 | -2.63957 | 0.0083 | 0.05077 | 15.113 |
| Qrfpr | 72.5812 | -0.69953 | 0.30774 | -2.27315 | 0.02302 | 0.09815 | 42.89 |
| Qrfprl | 180.742 | 1.113504 | 0.26821 | 4.151648 | 3.30E-05 | 0.00123 | 77.046 |
| Qrich1 | 3569.1 | 0.094669 | 0.06369 | 1.486358 | 0.13718 | 0.31781 | 43.358 |
| Qrsl1 | 756.905 | -0.00204 | 0.08478 | -0.0241 | 0.98078 | 0.98991 | 27.558 |
| Qser1 | 1255.71 | -0.00242 | 0.09507 | -0.02541 | 0.97973 | 0.98909 | 61.966 |
| Qsox1 | 1093.06 | -0.02733 | 0.09263 | -0.29504 | 0.76796 | 0.87881 | 36.861 |
| Qsox2 | 1786.53 | 0.144056 | 0.08353 | 1.724518 | 0.08461 | 0.23317 | 28.889 |
| Qtrt1 | 802.056 | 0.212239 | 0.08081 | 2.626243 | 0.00863 | 0.05203 | 8.438 |
| Qtrt2 | 1730.67 | 0.370303 | 0.06824 | 5.426384 | 5.75E-08 | 1.21E-05 | 65.403 |
| R3hcc1 | 852.904 | -0.25889 | 0.0918 | -2.82027 | 0.0048 | 0.03534 | 10.278 |
| R3hcc1l | 552.775 | 0.025099 | 0.08819 | 0.284606 | 0.77595 | 0.88333 | 73.585 |
| R3hdm1 | 17259.5 | -0.0775 | 0.11616 | -0.66714 | 0.50468 | 0.6967 | 134.436 |
| R3hdm2 | 8813.75 | -0.23825 | 0.1039 | -2.29299 | 0.02185 | 0.0951 | 119.058 |
| R3hdm4 | 7598.28 | 0.047725 | 0.07441 | 0.641418 | 0.52125 | 0.71042 | 11.157 |
| R74862 | 154.145 | 0.114862 | 0.14344 | 0.800752 | 0.42328 | 0.63011 | 31.903 |
| Rab10 | 6492.8 | -0.01875 | 0.05455 | -0.34379 | 0.731 | 0.85734 | 62.542 |
| Rab10os | 3020.33 | 0.202372 | 0.1054 | 1.920059 | 0.05485 | 0.17632 | 14.942 |
| Rab11a | 3315.93 | -0.04764 | 0.06151 | -0.77453 | 0.43861 | 0.64387 | 22.46 |
| Rab11b | 5055.41 | -0.09644 | 0.06608 | -1.45929 | 0.14448 | 0.32717 | 18.047 |
| Rab11b-ps2 | 474.788 | -0.15041 | 0.10123 | -1.48587 | 0.13731 | 0.31798 | 0.654 |
| Rab11fip1 | 193.678 | -0.31622 | 0.13519 | -2.33901 | 0.01933 | 0.0875 | 35.874 |
| Rab11fip2 | 3838.67 | 0.17484 | 0.08218 | 2.127466 | 0.03338 | 0.12693 | 40.771 |
| Rab11fip3 | 5785.01 | 0.120229 | 0.06205 | 1.937694 | 0.05266 | 0.17169 | 80.52 |
| Rab11fip4 | 18377.3 | -0.08191 | 0.0539 | -1.51969 | 0.12859 | 0.30578 | 106.812 |
| Rab11fip4os1 | 36.3286 | 0.233479 | 0.32913 | 0.709374 | 0.47809 | 0.67583 | 16.085 |
| Rab11fip4os2 | 28.9235 | 0.637583 | 0.38659 | 1.649258 | 0.09909 | 0.25857 | 4.757 |
| Rab11fip5 | 3255.12 | -0.10581 | 0.06895 | -1.5346 | 0.12488 | 0.29992 | 39.673 |
| Rab12 | 4161.83 | 0.004488 | 0.06738 | 0.066612 | 0.94689 | 0.9733 | 25.218 |
| Rab13 | 223.726 | 0.109601 | 0.12209 | 0.897728 | 0.36933 | 0.57978 | 12.691 |
| Rab14 | 13188.9 | 0.029572 | 0.05599 | 0.528184 | 0.59737 | 0.7692 | 20.916 |

| | | | | | | | |
|---|---|---|---|---|---|---|---|
| Rab15 | 13977.6 | 0.182939 | 0.10515 | 1.739779 | 0.0819 | 0.22809 | 24.949 |
| Rab18 | 6235.8 | -0.092 | 0.05416 | -1.69864 | 0.08939 | 0.2422 | 60.525 |
| Rab1a | 8585.29 | -0.07809 | 0.07389 | -1.05682 | 0.29059 | 0.50076 | 25.425 |
| Rab1b | 4643.49 | -0.10448 | 0.08582 | -1.21743 | 0.22344 | 0.42664 | 7.868 |
| Rab21 | 4817.39 | -0.13326 | 0.0595 | -2.23975 | 0.02511 | 0.10427 | 28.511 |
| Rab22a | 3474.26 | -0.03205 | 0.05963 | -0.53748 | 0.59094 | 0.7637 | 47.584 |
| Rab23 | 1604.94 | 0.068056 | 0.07285 | 0.934225 | 0.35019 | 0.56083 | 22.678 |
| Rab24 | 275.346 | -0.05309 | 0.12551 | -0.423 | 0.6723 | 0.81977 | 2.204 |
| Rab26 | 1695.02 | -0.20828 | 0.08666 | -2.40344 | 0.01624 | 0.07847 | 6.97 |
| Rab26os | 77.0122 | -0.11633 | 0.23039 | -0.50491 | 0.61362 | 0.77996 | 0.519 |
| Rab27a | 696.763 | 0.091148 | 0.11181 | 0.815185 | 0.41497 | 0.6229 | 52.776 |
| Rab27b | 2352.44 | 0.476432 | 0.10916 | 4.364577 | 1.27E-05 | 0.00066 | 162.475 |
| Rab28 | 3357.63 | 0.01249 | 0.07014 | 0.17808 | 0.85866 | 0.92864 | 83.182 |
| Rab29 | 124.124 | -0.7762 | 0.17314 | -4.48305 | 7.36E-06 | 0.00044 | 5.664 |
| Rab2a | 10150.9 | 0.071105 | 0.05924 | 1.200237 | 0.23005 | 0.43435 | 72.135 |
| Rab2b | 231.612 | -0.05888 | 0.16109 | -0.3655 | 0.71474 | 0.84705 | 17.787 |
| Rab30 | 3199.54 | 0.252463 | 0.15897 | 1.588138 | 0.11226 | 0.2799 | 102.933 |
| Rab31 | 4101.26 | -0.45231 | 0.12145 | -3.72436 | 0.0002 | 0.00421 | 121.024 |
| Rab32 | 104.223 | -0.46292 | 0.20262 | -2.28462 | 0.02233 | 0.09639 | 13.264 |
| Rab33a | 495.665 | -0.27997 | 0.09415 | -2.97356 | 0.00294 | 0.02585 | 10.948 |
| Rab33b | 1169.1 | -0.19845 | 0.09646 | -2.05732 | 0.03966 | 0.14223 | 12.313 |
| Rab34 | 1016.75 | 0.234543 | 0.12381 | 1.894331 | 0.05818 | 0.18329 | 3.764 |
| Rab35 | 2836.91 | -0.09815 | 0.06725 | -1.45936 | 0.14447 | 0.32717 | 15.829 |
| Rab36 | 1938.53 | 0.485723 | 0.09279 | 5.23442 | 1.66E-07 | 2.66E-05 | 17.691 |
| Rab37 | 96.9294 | -1.02146 | 0.22903 | -4.46001 | 8.20E-06 | 0.00047 | 70.806 |
| Rab39 | 119.211 | -0.36651 | 0.22781 | -1.60886 | 0.10765 | 0.27251 | 22.123 |
| Rab39b | 1527.63 | 0.223096 | 0.10873 | 2.05177 | 0.04019 | 0.14325 | 6.186 |
| Rab3a | 14711.5 | -0.23605 | 0.06936 | -3.40323 | 0.00067 | 0.00964 | 3.996 |
| Rab3b | 2211.73 | -0.10369 | 0.09003 | -1.15177 | 0.24942 | 0.45679 | 64.262 |
| Rab3c | 23866.3 | -0.0948 | 0.10637 | -0.89124 | 0.3728 | 0.58329 | 226.81 |
| Rab3d | 367.964 | -0.67688 | 0.18815 | -3.59751 | 0.00032 | 0.00593 | 10.702 |
| Rab3gap1 | 6151.67 | 0.154053 | 0.06398 | 2.407993 | 0.01604 | 0.07783 | 75.096 |
| Rab3gap2 | 4567.59 | 0.067062 | 0.07147 | 0.938316 | 0.34808 | 0.55864 | 82.643 |
| Rab3il1 | 367.723 | -0.54628 | 0.13465 | -4.05701 | 4.97E-05 | 0.00163 | 36.712 |

| | | | | | | | |
|---|---|---|---|---|---|---|---|
| Rab3ip | 910.951 | 0.008968 | 0.08149 | 0.110046 | 0.91237 | 0.95579 | 44.986 |
| Rab40b | 3246.93 | 0.487682 | 0.15326 | 3.182122 | 0.00146 | 0.01626 | 32.127 |
| Rab40c | 2169.5 | 0.309983 | 0.10687 | 2.9007 | 0.00372 | 0.03002 | 37.614 |
| Rab42 | 40.1666 | 0.344609 | 0.25508 | 1.350969 | 0.17671 | 0.36877 | 1.189 |
| Rab43 | 115.648 | 0.178368 | 0.24516 | 0.727567 | 0.46688 | 0.66621 | 23.312 |
| Rab4a | 2441.22 | 0.063287 | 0.07495 | 0.844391 | 0.39845 | 0.6075 | 29.303 |
| Rab4b | 62.0008 | -0.21502 | 0.2149 | -1.00055 | 0.31704 | 0.52743 | 10.473 |
| Rab5a | 4919.8 | 0.053784 | 0.06417 | 0.838102 | 0.40197 | 0.6106 | 28.447 |
| Rab5b | 7122.63 | -0.07315 | 0.07852 | -0.93158 | 0.35155 | 0.56204 | 19.09 |
| Rab5c | 5733.27 | -0.11146 | 0.08711 | -1.2795 | 0.20072 | 0.39886 | 23.207 |
| Rab5if | 633.049 | -0.08759 | 0.09166 | -0.95559 | 0.33928 | 0.55014 | 10.436 |
| Rab6a | 17764.6 | -0.23336 | 0.0837 | -2.78812 | 0.0053 | 0.03768 | 33.859 |
| Rab6b | 41223.7 | -0.1946 | 0.05212 | -3.73393 | 0.00019 | 0.00415 | 73.49 |
| Rab7 | 9607.46 | -0.14976 | 0.06612 | -2.26498 | 0.02351 | 0.09952 | 46.165 |
| Rab7-ps1 | 195.844 | -0.0723 | 0.15178 | -0.47633 | 0.63384 | 0.79385 | 0.622 |
| Rab7b | 70.9257 | -0.68588 | 0.24827 | -2.76261 | 0.00573 | 0.03997 | 26.745 |
| Rab8a | 1709.76 | -0.12295 | 0.076 | -1.61787 | 0.10569 | 0.26923 | 22.705 |
| Rab8b | 2338.11 | 0.156847 | 0.06935 | 2.261562 | 0.02372 | 0.10023 | 76.024 |
| Rab9 | 1010.46 | -0.00849 | 0.08011 | -0.10595 | 0.91562 | 0.95673 | 22.615 |
| Rab9b | 1940.08 | 0.679035 | 0.15528 | 4.372985 | 1.23E-05 | 0.00064 | 10.609 |
| Rabac1 | 3804.12 | -0.11879 | 0.0807 | -1.47203 | 0.14101 | 0.32275 | 3.003 |
| Rabep1 | 6957.31 | 0.193085 | 0.08303 | 2.325499 | 0.02005 | 0.08969 | 98.328 |
| Rabep2 | 384.122 | 0.154354 | 0.10052 | 1.535533 | 0.12465 | 0.2996 | 20.487 |
| Rabepk | 1131.89 | -0.0814 | 0.0907 | -0.89745 | 0.36948 | 0.57993 | 22.357 |
| Rabgap1 | 4359.71 | 0.069978 | 0.06464 | 1.082544 | 0.27901 | 0.48891 | 123.176 |
| Rabgap1l | 6083.59 | 0.004963 | 0.10753 | 0.04615 | 0.96319 | 0.98208 | 574.038 |
| Rabgef1 | 1599.6 | -0.13969 | 0.07866 | -1.77587 | 0.07576 | 0.2174 | 42.545 |
| Rabggta | 1528.14 | 0.109674 | 0.07922 | 1.384341 | 0.16625 | 0.35594 | 6.848 |
| Rabggtb | 1844.39 | 0.046098 | 0.063 | 0.731709 | 0.46435 | 0.66409 | 5.723 |
| Rabif | 1391.91 | -0.00739 | 0.06767 | -0.10923 | 0.91302 | 0.95603 | 14.127 |
| Rabl2 | 1017.53 | 0.050684 | 0.08762 | 0.578449 | 0.56296 | 0.74365 | 9.391 |
| Rabl3 | 492.302 | 0.135305 | 0.10215 | 1.324594 | 0.18531 | 0.38008 | 32.501 |
| Rabl6 | 8174.12 | 0.148761 | 0.06007 | 2.476308 | 0.01327 | 0.06906 | 25.504 |
| Rac1 | 10490 | -0.06252 | 0.05725 | -1.09204 | 0.27482 | 0.48444 | 24.403 |

| | | | | | | | |
|---|---|---|---|---|---|---|---|
| Rac2 | 71.4077 | -0.33632 | 0.24895 | -1.35095 | 0.17671 | 0.36877 | 13.617 |
| Rac3 | 481.543 | 0.07618 | 0.11079 | 0.687614 | 0.4917 | 0.68654 | 2.5 |
| Racgap1 | 910.536 | 0.53193 | 0.11116 | 4.785169 | 1.71E-06 | 0.00016 | 31.161 |
| Rack1 | 5372.96 | -0.05833 | 0.09024 | -0.64646 | 0.51798 | 0.70768 | 6.103 |
| Rad1 | 423.946 | -0.06461 | 0.10868 | -0.59451 | 0.55217 | 0.73517 | 13.046 |
| Rad17 | 842.626 | 0.203986 | 0.0822 | 2.481611 | 0.01308 | 0.06846 | 33.888 |
| Rad18 | 342.588 | 0.021616 | 0.10638 | 0.203206 | 0.83897 | 0.91715 | 76.837 |
| Rad21 | 6311.64 | -0.09228 | 0.07319 | -1.26081 | 0.20738 | 0.40685 | 29.508 |
| Rad21l | 32.5669 | 0.163557 | 0.3068 | 0.533108 | 0.59396 | 0.76614 | 23.13 |
| Rad23a | 1502.17 | -0.15573 | 0.06849 | -2.27388 | 0.02297 | 0.09807 | 6.647 |
| Rad23b | 8922.51 | -0.05061 | 0.06843 | -0.73959 | 0.45955 | 0.6601 | 42.195 |
| Rad50 | 887.804 | 0.226469 | 0.12319 | 1.838344 | 0.06601 | 0.19847 | 57.801 |
| Rad51 | 44.7136 | 0.108548 | 0.32876 | 0.330171 | 0.74127 | 0.86333 | 34.653 |
| Rad51ap1 | 91.2138 | -0.23293 | 0.21252 | -1.09605 | 0.27306 | 0.48241 | 16.538 |
| Rad51ap2 | 27.4577 | -0.69976 | 0.4401 | -1.59001 | 0.11183 | 0.27929 | 6.85 |
| Rad51c | 188.962 | -0.41982 | 0.16513 | -2.54233 | 0.01101 | 0.06114 | 28.31 |
| Rad51d | 828.147 | 0.073433 | 0.10684 | 0.687345 | 0.49187 | 0.68654 | 14.181 |
| Rad52 | 893.73 | 0.324714 | 0.09369 | 3.465792 | 0.00053 | 0.00825 | 20.131 |
| Rad54b | 88.8769 | -0.50789 | 0.22247 | -2.28302 | 0.02243 | 0.09666 | 56.884 |
| Rad54l | 61.0835 | -0.12293 | 0.22494 | -0.54648 | 0.58474 | 0.75938 | 29.427 |
| Rad54l2 | 2652.94 | 0.018327 | 0.06 | 0.305473 | 0.76001 | 0.87418 | 101.113 |
| Rad9a | 431.605 | -0.00558 | 0.12832 | -0.04345 | 0.96534 | 0.98278 | 6.467 |
| Rad9b | 434.931 | 0.529645 | 0.15675 | 3.378833 | 0.00073 | 0.01029 | 31.011 |
| Radil | 1264.69 | 0.278762 | 0.14242 | 1.95732 | 0.05031 | 0.16667 | 66.26 |
| Rae1 | 1518.68 | 0.099183 | 0.07229 | 1.3721 | 0.17003 | 0.3604 | 15.623 |
| Raf1 | 1758.93 | 0.139577 | 0.07256 | 1.923529 | 0.05441 | 0.17539 | 58.569 |
| Rai1 | 5712.33 | 0.180065 | 0.0765 | 2.353674 | 0.01859 | 0.08546 | 94.185 |
| Rai14 | 410.906 | -0.6325 | 0.183 | -3.45628 | 0.00055 | 0.00843 | 145.656 |
| Rai2 | 300.371 | -0.30303 | 0.17883 | -1.69453 | 0.09016 | 0.24359 | 62.428 |
| Rala | 3063.91 | 0.003286 | 0.07364 | 0.044618 | 0.96441 | 0.98268 | 63.669 |
| Ralb | 799.061 | -0.32551 | 0.09611 | -3.38683 | 0.00071 | 0.01004 | 34.49 |
| Ralbp1 | 4547.8 | 0.008603 | 0.08009 | 0.107426 | 0.91445 | 0.95656 | 37.381 |
| Ralgapa1 | 7778.69 | 0.005517 | 0.10251 | 0.053819 | 0.95708 | 0.979 | 218.272 |
| Ralgapa2 | 1963.65 | 0.203209 | 0.07102 | 2.86145 | 0.00422 | 0.03259 | 272.466 |

| | | | | | | | |
|---|---|---|---|---|---|---|---|
| Ralgapb | 9118.4 | 0.120136 | 0.05688 | 2.112266 | 0.03466 | 0.13034 | 89.406 |
| Ralgds | 4040.04 | 0.072609 | 0.07783 | 0.932953 | 0.35084 | 0.56148 | 39.957 |
| Ralgps1 | 4581.62 | 0.07396 | 0.06501 | 1.137598 | 0.25529 | 0.46297 | 238.07 |
| Ralgps2 | 2832.16 | 0.118521 | 0.08431 | 1.40581 | 0.15978 | 0.34774 | 135.461 |
| Raly | 2499.21 | 0.042265 | 0.08974 | 0.470989 | 0.63765 | 0.7962 | 76.166 |
| Ralyl | 3462.8 | 0.168106 | 0.09372 | 1.793752 | 0.07285 | 0.21233 | 710.633 |
| Ramac | 3288.21 | -0.05828 | 0.07852 | -0.74224 | 0.45794 | 0.65918 | 6.567 |
| Ramp1 | 2586.76 | 0.148304 | 0.129 | 1.149652 | 0.25029 | 0.45769 | 45.375 |
| Ramp2 | 508.718 | -0.36228 | 0.10851 | -3.33866 | 0.00084 | 0.01137 | 13.519 |
| Ramp3 | 124.1 | 0.568559 | 0.22938 | 2.478678 | 0.01319 | 0.06875 | 18.955 |
| Ran | 6939.22 | 0.009923 | 0.07428 | 0.133587 | 0.89373 | 0.9468 | 4.255 |
| Ranbp1 | 740.029 | -0.19633 | 0.0842 | -2.33158 | 0.01972 | 0.08871 | 8.949 |
| Ranbp10 | 2286.42 | 0.043266 | 0.07214 | 0.599798 | 0.54864 | 0.73267 | 59.043 |
| Ranbp17 | 486.927 | 0.377711 | 0.1156 | 3.267504 | 0.00109 | 0.01349 | 301.952 |
| Ranbp2 | 9120 | 0.292928 | 0.15588 | 1.87915 | 0.06022 | 0.18722 | 47.437 |
| Ranbp3 | 3446.04 | 0.004535 | 0.06386 | 0.071007 | 0.94339 | 0.97176 | 38.471 |
| Ranbp3l | 412.714 | -0.7421 | 0.21208 | -3.49914 | 0.00047 | 0.00759 | 69.985 |
| Ranbp6 | 2206.4 | 0.086458 | 0.10737 | 0.805204 | 0.4207 | 0.6278 | 4.575 |
| Ranbp9 | 3976.99 | -0.0893 | 0.068 | -1.31309 | 0.18915 | 0.38507 | 78.608 |
| Rangap1 | 8585.53 | -0.10833 | 0.06607 | -1.63979 | 0.10105 | 0.26212 | 41.283 |
| Rangrf | 16.8204 | 0.44494 | 0.42099 | 1.056893 | 0.29056 | 0.50076 | 2.702 |
| Rap1a | 1960.05 | 0.030218 | 0.09125 | 0.331158 | 0.74052 | 0.86301 | 74.07 |
| Rap1b | 2828.5 | -0.10261 | 0.08049 | -1.27485 | 0.20236 | 0.40075 | 32.165 |
| Rap1gap | 21025.3 | 0.652658 | 0.15303 | 4.264947 | 2.00E-05 | 0.00088 | 65.136 |
| Rap1gap2 | 8090.31 | -0.29326 | 0.1177 | -2.49166 | 0.01271 | 0.06724 | 227.56 |
| Rap1gapos | 69.5723 | 1.055303 | 0.24684 | 4.275208 | 1.91E-05 | 0.00086 | 9.628 |
| Rap1gds1 | 11342.8 | -0.02725 | 0.05479 | -0.49738 | 0.61892 | 0.78314 | 149.3 |
| Rap2a | 4073.84 | -0.25034 | 0.14136 | -1.7709 | 0.07658 | 0.21877 | 28.751 |
| Rap2b | 2322.61 | 0.22817 | 0.11076 | 2.06003 | 0.0394 | 0.14148 | 6.793 |
| Rap2c | 1919.9 | 0.055441 | 0.07854 | 0.705898 | 0.48025 | 0.67761 | 14.107 |
| Rapgef1 | 6024.64 | 0.027036 | 0.07339 | 0.368377 | 0.71259 | 0.8452 | 121.259 |
| Rapgef2 | 10113.3 | 0.201522 | 0.09424 | 2.138468 | 0.03248 | 0.1248 | 224.002 |
| Rapgef3 | 1922.32 | -0.07538 | 0.10374 | -0.72661 | 0.46746 | 0.66644 | 23.203 |
| Rapgef3os2 | 16.0035 | -0.55067 | 0.45273 | -1.21632 | 0.22386 | 0.42711 | 2.165 |

| | | | | | | | |
|---|---|---|---|---|---|---|---|
| Rapgef4 | 16414.6 | -0.44253 | 0.12801 | -3.45704 | 0.00055 | 0.00841 | 276.235 |
| Rapgef4os1 | 20.5331 | -0.01698 | 0.45657 | -0.03719 | 0.97033 | 0.98489 | 51.194 |
| Rapgef4os2 | 31.7582 | 0.136577 | 0.3369 | 0.405391 | 0.68519 | 0.82845 | 7.615 |
| Rapgef4os3 | 241.791 | 0.212215 | 0.14398 | 1.473869 | 0.14052 | 0.32203 | 4.011 |
| Rapgef5 | 7409.38 | 0.041484 | 0.10331 | 0.401537 | 0.68802 | 0.83002 | 243.374 |
| Rapgef6 | 3617.15 | 0.284687 | 0.10505 | 2.710081 | 0.00673 | 0.04429 | 176.439 |
| Rapgefl1 | 6720.29 | -0.33236 | 0.10052 | -3.30657 | 0.00094 | 0.01229 | 16.913 |
| Raph1 | 2170.11 | -0.11961 | 0.13005 | -0.91967 | 0.35775 | 0.56794 | 84.813 |
| Rapsn | 42.6026 | 0.129732 | 0.27326 | 0.474757 | 0.63496 | 0.79438 | 10.11 |
| Rara | 342.33 | -0.41395 | 0.15452 | -2.67897 | 0.00738 | 0.04728 | 47.125 |
| Rarb | 1299.23 | 0.709123 | 0.33236 | 2.133615 | 0.03287 | 0.12596 | 388.385 |
| Rarg | 717.703 | 0.283392 | 0.12652 | 2.239957 | 0.02509 | 0.10424 | 22.58 |
| Rarres1 | 24.1708 | -0.4165 | 0.44657 | -0.93266 | 0.351 | 0.56158 | 36.552 |
| Rarres2 | 102.255 | -0.12365 | 0.24413 | -0.50647 | 0.61253 | 0.77934 | 3.094 |
| Rars | 1765.21 | 0.027504 | 0.08748 | 0.314411 | 0.75321 | 0.87029 | 26.126 |
| Rars2 | 995.37 | 0.195842 | 0.09378 | 2.088362 | 0.03677 | 0.13524 | 45.211 |
| Rasa1 | 2772.79 | 0.175881 | 0.13789 | 1.27548 | 0.20214 | 0.40058 | 74.351 |
| Rasa2 | 1992.4 | 0.032634 | 0.09743 | 0.334959 | 0.73766 | 0.86154 | 92.318 |
| Rasa3 | 2973.24 | -0.36316 | 0.11201 | -3.24211 | 0.00119 | 0.01419 | 110.656 |
| Rasa4 | 242.66 | -0.51449 | 0.17355 | -2.96445 | 0.00303 | 0.02631 | 27.945 |
| Rasal1 | 2892.92 | 0.520941 | 0.10871 | 4.792094 | 1.65E-06 | 0.00015 | 30.786 |
| Rasal2 | 6134.26 | -0.31003 | 0.12965 | -2.39126 | 0.01679 | 0.08018 | 277.414 |
| Rasal3 | 127.124 | -0.08225 | 0.22798 | -0.36079 | 0.71826 | 0.84954 | 12.925 |
| Rasd1 | 577.59 | 0.773329 | 0.23261 | 3.324624 | 0.00089 | 0.01177 | 1.764 |
| Rasd2 | 14987.5 | 0.507656 | 0.33921 | 1.496577 | 0.1345 | 0.31436 | 10.17 |
| Rasgef1a | 18200.5 | 0.136122 | 0.07918 | 1.719133 | 0.08559 | 0.23485 | 80.109 |
| Rasgef1b | 2291.69 | -0.08288 | 0.09244 | -0.8966 | 0.36993 | 0.58027 | 511.64 |
| Rasgef1c | 1273.55 | 0.033246 | 0.09437 | 0.352279 | 0.72463 | 0.85321 | 79.16 |
| Rasgrf1 | 16515.5 | 0.175766 | 0.0647 | 2.716688 | 0.00659 | 0.04378 | 117.07 |
| Rasgrf2 | 5835.58 | -0.12062 | 0.08683 | -1.38905 | 0.16482 | 0.35389 | 239.646 |
| Rasgrp1 | 24540.9 | 0.248601 | 0.12922 | 1.923807 | 0.05438 | 0.17534 | 63.009 |
| Rasgrp2 | 6764.16 | 0.571472 | 0.18739 | 3.049716 | 0.00229 | 0.02199 | 15.877 |
| Rasgrp3 | 693.799 | -0.3729 | 0.1749 | -2.13205 | 0.033 | 0.12614 | 93.159 |
| Rasgrp4 | 57.1778 | 1.474287 | 0.34788 | 4.237965 | 2.26E-05 | 0.00095 | 19.111 |

| | | | | | | | |
|---|---|---|---|---|---|---|---|
| Rasip1 | 93.9234 | -0.29764 | 0.24367 | -1.22152 | 0.22189 | 0.42493 | 11.606 |
| Rasl10a | 924.645 | -0.70123 | 0.17804 | -3.93861 | 8.20E-05 | 0.00234 | 2.258 |
| Rasl10b | 22793.5 | 0.101601 | 0.09983 | 1.01772 | 0.30881 | 0.51865 | 11.903 |
| Rasl11a | 78.4179 | -0.31293 | 0.23376 | -1.33865 | 0.18068 | 0.37405 | 2.656 |
| Rasl11b | 1258.08 | -0.44883 | 0.12601 | -3.56173 | 0.00037 | 0.00646 | 4.196 |
| Rasl12 | 115.439 | -0.64802 | 0.21939 | -2.95372 | 0.00314 | 0.02688 | 16.348 |
| Rassf1 | 316.532 | -0.20238 | 0.10954 | -1.84753 | 0.06467 | 0.19576 | 10.713 |
| Rassf10 | 37.1934 | 0.172016 | 0.46937 | 0.366485 | 0.714 | 0.8464 | 3.496 |
| Rassf2 | 3215.19 | -0.44658 | 0.08619 | -5.18152 | 2.20E-07 | 3.27E-05 | 40.844 |
| Rassf3 | 869.783 | -0.21404 | 0.09053 | -2.36423 | 0.01807 | 0.08383 | 65.998 |
| Rassf4 | 450.646 | -0.29639 | 0.15967 | -1.85624 | 0.06342 | 0.19352 | 40.945 |
| Rassf5 | 769 | -0.29189 | 0.14159 | -2.0615 | 0.03926 | 0.1412 | 68.849 |
| Rassf6 | 13.3117 | -0.23875 | 0.53737 | -0.44429 | 0.65683 | 0.80939 | 37.582 |
| Rassf7 | 50.8843 | 0.319702 | 0.33785 | 0.946276 | 0.34401 | 0.55468 | 2.799 |
| Rassf8 | 1509.65 | -0.19852 | 0.09595 | -2.06899 | 0.03855 | 0.13979 | 74.332 |
| Rassf9 | 50.2339 | -0.10891 | 0.36086 | -0.30181 | 0.76279 | 0.87569 | 37.515 |
| Raver2 | 1173.72 | 0.099126 | 0.11083 | 0.894405 | 0.37111 | 0.58131 | 83.388 |
| Rb1 | 1973.01 | -0.03886 | 0.06078 | -0.63932 | 0.52262 | 0.71139 | 142.15 |
| Rb1cc1 | 4535.87 | 0.179711 | 0.18485 | 0.972206 | 0.33095 | 0.54208 | 70.452 |
| Rbak | 671.015 | 0.099048 | 0.08447 | 1.172632 | 0.24094 | 0.4471 | 8.59 |
| Rbbp4 | 5472.16 | 0.085896 | 0.07124 | 1.205721 | 0.22793 | 0.43183 | 28.271 |
| Rbbp5 | 3561.54 | 0.073711 | 0.07543 | 0.977249 | 0.32845 | 0.53979 | 28.295 |
| Rbbp6 | 6122.34 | 0.264122 | 0.07183 | 3.676988 | 0.00024 | 0.00478 | 36.872 |
| Rbbp7 | 5987.65 | 0.104906 | 0.07362 | 1.424973 | 0.15417 | 0.34002 | 18.691 |
| Rbbp8 | 150.419 | -0.01449 | 0.21447 | -0.06757 | 0.94613 | 0.97302 | 111.946 |
| Rbbp9 | 1580.64 | -0.09219 | 0.08652 | -1.06553 | 0.28664 | 0.4965 | 8.605 |
| Rbck1 | 5407.33 | 0.336292 | 0.11255 | 2.987828 | 0.00281 | 0.02514 | 16.32 |
| Rbfa | 373.952 | -0.09785 | 0.11963 | -0.81797 | 0.41337 | 0.62154 | 8.394 |
| Rbfox1 | 14283.4 | 0.24727 | 0.10502 | 2.35449 | 0.01855 | 0.08535 | 1526.172 |
| Rbfox2 | 19081.9 | 0.032479 | 0.07634 | 0.425462 | 0.6705 | 0.81841 | 228.015 |
| Rbfox3 | 11566.9 | -0.20119 | 0.0643 | -3.12885 | 0.00175 | 0.01853 | 421.836 |
| Rbis | 745.109 | 0.032353 | 0.09755 | 0.331642 | 0.74016 | 0.863 | 5.002 |
| Rbks | 145.583 | -0.27667 | 0.15583 | -1.77546 | 0.07582 | 0.21743 | 73.189 |
| Rbl1 | 147.816 | -0.0487 | 0.19035 | -0.25584 | 0.79807 | 0.8958 | 58.642 |

| | | | | | | | |
|---|---|---|---|---|---|---|---|
| Rbl2 | 2152.52 | -0.0547 | 0.07292 | -0.7501 | 0.4532 | 0.65555 | 53.788 |
| Rbm10 | 3328.33 | 0.094286 | 0.06438 | 1.464424 | 0.14308 | 0.32514 | 33.399 |
| Rbm11 | 202.542 | 0.038736 | 0.17212 | 0.225054 | 0.82194 | 0.90781 | 9.986 |
| Rbm12 | 665.357 | 0.146691 | 0.10876 | 1.348798 | 0.1774 | 0.36951 | 20.021 |
| Rbm12b1 | 647.662 | 0.377883 | 0.10755 | 3.513467 | 0.00044 | 0.00731 | 6.468 |
| Rbm12b2 | 777.289 | 0.236689 | 0.11177 | 2.117604 | 0.03421 | 0.12921 | 6.805 |
| Rbm14 | 1029.54 | 0.189244 | 0.11447 | 1.653174 | 0.0983 | 0.25734 | 11.066 |
| Rbm15 | 1038.42 | 0.11071 | 0.08908 | 1.24283 | 0.21393 | 0.41531 | 8.253 |
| Rbm15b | 2748.88 | -0.0942 | 0.06962 | -1.35303 | 0.17605 | 0.36794 | 6.511 |
| Rbm17 | 2361.3 | 0.003458 | 0.06927 | 0.049925 | 0.96018 | 0.98055 | 18.717 |
| Rbm18 | 2505.73 | 0.05585 | 0.065 | 0.85919 | 0.39024 | 0.59921 | 20.695 |
| Rbm19 | 481.788 | -0.07265 | 0.12404 | -0.58566 | 0.5581 | 0.73969 | 82.517 |
| Rbm20 | 578.018 | -0.06913 | 0.13885 | -0.49787 | 0.61857 | 0.78312 | 189.775 |
| Rbm22 | 2115.79 | 0.064554 | 0.07097 | 0.909551 | 0.36306 | 0.57343 | 12.075 |
| Rbm24 | 1679.65 | 0.488273 | 0.10528 | 4.637888 | 3.52E-06 | 0.00026 | 12.662 |
| Rbm25 | 9906.26 | 0.194726 | 0.13216 | 1.473421 | 0.14064 | 0.32225 | 51.888 |
| Rbm26 | 4212.96 | 0.33108 | 0.12166 | 2.72138 | 0.0065 | 0.0434 | 70.577 |
| Rbm27 | 2520.28 | 0.152381 | 0.09562 | 1.593609 | 0.11102 | 0.27814 | 66.19 |
| Rbm28 | 3708.48 | 0.225377 | 0.12259 | 1.838514 | 0.06599 | 0.19847 | 41.431 |
| Rbm3 | 3465.85 | -0.06432 | 0.09279 | -0.69315 | 0.48822 | 0.68339 | 6.906 |
| Rbm3-ps | 45.9968 | -0.35872 | 0.30885 | -1.16149 | 0.24544 | 0.45204 | 0.465 |
| Rbm33 | 7944.39 | 0.189515 | 0.10889 | 1.740501 | 0.08177 | 0.22793 | 102.119 |
| Rbm34 | 1352.78 | 0.281408 | 0.13527 | 2.080303 | 0.0375 | 0.13705 | 23.9 |
| Rbm38 | 81.465 | 0.194211 | 0.2895 | 0.670848 | 0.50232 | 0.69489 | 14.237 |
| Rbm39 | 14448.6 | 0.224249 | 0.09657 | 2.322138 | 0.02023 | 0.09019 | 33 |
| Rbm4 | 290.435 | 0.043108 | 0.13748 | 0.313564 | 0.75385 | 0.87065 | 9.609 |
| Rbm41 | 637.305 | 0.300934 | 0.16339 | 1.841847 | 0.0655 | 0.19743 | 109.085 |
| Rbm43 | 511.938 | 0.015453 | 0.09834 | 0.157132 | 0.87514 | 0.93783 | 10.716 |
| Rbm45 | 1297.07 | 0.132776 | 0.07494 | 1.771679 | 0.07645 | 0.21863 | 13.785 |
| Rbm46 | 18.774 | -1.04901 | 0.45788 | -2.29102 | 0.02196 | 0.09543 | 39.256 |
| Rbm47 | 44.8646 | -0.3367 | 0.37289 | -0.90295 | 0.36655 | 0.57697 | 156.57 |
| Rbm48 | 370.533 | 0.044227 | 0.12317 | 0.359084 | 0.71953 | 0.85017 | 12.608 |
| Rbm4b | 1610.75 | 0.259744 | 0.09688 | 2.681061 | 0.00734 | 0.04706 | 10.569 |
| Rbm5 | 10313.2 | 0.186094 | 0.07536 | 2.469551 | 0.01353 | 0.06999 | 30.626 |

| | | | | | | | |
|---|---|---|---|---|---|---|---|
| Rbm6 | 3925.37 | 0.229923 | 0.09408 | 2.443902 | 0.01453 | 0.07284 | 99.679 |
| Rbm7 | 1626.46 | 0.21209 | 0.06652 | 3.188221 | 0.00143 | 0.01602 | 6.599 |
| Rbm8a | 2309.47 | -0.11975 | 0.0791 | -1.51399 | 0.13003 | 0.30791 | 3.859 |
| Rbms1 | 1973.92 | 0.361159 | 0.15348 | 2.353091 | 0.01862 | 0.08551 | 213 |
| Rbms2 | 493.328 | -0.37597 | 0.10648 | -3.53079 | 0.00041 | 0.00704 | 48.74 |
| Rbms3 | 1877.88 | 0.45908 | 0.17091 | 2.686124 | 0.00723 | 0.04653 | 1299.936 |
| Rbmx | 4285.76 | 0.258825 | 0.09526 | 2.717084 | 0.00659 | 0.04374 | 9.507 |
| Rbmx2 | 240.081 | -0.03721 | 0.13282 | -0.28014 | 0.77937 | 0.88509 | 15.72 |
| Rbmxl1 | 1425.97 | 0.00743 | 0.08168 | 0.09096 | 0.92752 | 0.96345 | 3.63 |
| Rbp1 | 867.741 | -0.02882 | 0.16319 | -0.17659 | 0.85983 | 0.92916 | 23.615 |
| Rbp4 | 315.784 | -0.19407 | 0.11078 | -1.75183 | 0.0798 | 0.2245 | 8.653 |
| Rbpj | 1482.28 | -0.13088 | 0.07307 | -1.79117 | 0.07327 | 0.21317 | 191.211 |
| Rbpj-ps3 | 24.4398 | -0.40334 | 0.3638 | -1.10866 | 0.26758 | 0.47694 | 1.441 |
| Rbpjl | 34.803 | -0.02306 | 0.30455 | -0.07571 | 0.93965 | 0.96981 | 12.308 |
| Rbpms | 183.088 | -0.4345 | 0.16374 | -2.65358 | 0.00796 | 0.04945 | 147.221 |
| Rbpms2 | 101.212 | -0.31603 | 0.20252 | -1.56047 | 0.11865 | 0.29055 | 30.881 |
| Rbsn | 3168.31 | 0.085686 | 0.06631 | 1.292256 | 0.19627 | 0.39389 | 28.214 |
| Rbx1 | 4060.1 | -0.01076 | 0.06166 | -0.17448 | 0.86149 | 0.92982 | 10.074 |
| Rc3h1 | 3363.86 | 0.017775 | 0.07489 | 0.237339 | 0.81239 | 0.90348 | 68.561 |
| Rc3h2 | 5476.88 | 0.297996 | 0.07406 | 4.023495 | 5.73E-05 | 0.00182 | 52.835 |
| Rcan1 | 2494.02 | 0.156902 | 0.10068 | 1.558352 | 0.11915 | 0.29132 | 78.915 |
| Rcan2 | 14112.3 | -0.06293 | 0.07678 | -0.81959 | 0.41245 | 0.62047 | 238.166 |
| Rcan3 | 2834.63 | 0.17273 | 0.09836 | 1.756121 | 0.07907 | 0.22311 | 21.546 |
| Rcbtb1 | 3084.65 | -0.12646 | 0.08826 | -1.4328 | 0.15191 | 0.33702 | 36.057 |
| Rcbtb2 | 934.989 | -0.02453 | 0.09744 | -0.25176 | 0.80123 | 0.8972 | 84.807 |
| Rcc1 | 638.073 | 0.163539 | 0.08811 | 1.85607 | 0.06344 | 0.19357 | 21.687 |
| Rcc1l | 570.045 | -0.04254 | 0.08324 | -0.51112 | 0.60927 | 0.77665 | 28.721 |
| Rcc2 | 4349.37 | 0.082627 | 0.06619 | 1.248352 | 0.2119 | 0.41257 | 22.68 |
| Rccd1 | 771.662 | 0.263504 | 0.08783 | 3.000247 | 0.0027 | 0.02453 | 31.226 |
| Rce1 | 693.95 | 0.182014 | 0.11149 | 1.63254 | 0.10257 | 0.26445 | 3.051 |
| Rchy1 | 1209.43 | -0.14615 | 0.08582 | -1.70292 | 0.08858 | 0.2405 | 14.165 |
| Rcl1 | 695.94 | 0.040568 | 0.09243 | 0.438923 | 0.66072 | 0.81233 | 42.555 |
| Rcn1 | 1880.29 | 0.632067 | 0.17126 | 3.69059 | 0.00022 | 0.00459 | 13.029 |
| Rcn2 | 5352.1 | -0.14322 | 0.07049 | -2.03184 | 0.04217 | 0.14777 | 20.038 |

| | | | | | | | |
|---|---|---|---|---|---|---|---|
| Rcn3 | 109.065 | -0.97574 | 0.35781 | -2.727 | 0.00639 | 0.04288 | 9.309 |
| Rcor1 | 1569.08 | 0.201284 | 0.08114 | 2.48057 | 0.01312 | 0.06859 | 76.551 |
| Rcor2 | 459.485 | -0.55557 | 0.1742 | -3.18918 | 0.00143 | 0.01601 | 7.901 |
| Rcor3 | 998.631 | -0.08235 | 0.09852 | -0.83595 | 0.40318 | 0.61186 | 39.514 |
| Rcsd1 | 375.728 | -0.15439 | 0.11597 | -1.33126 | 0.1831 | 0.37714 | 63.242 |
| Rcvrn | 49.6583 | 0.806531 | 0.31108 | 2.59269 | 0.00952 | 0.05554 | 8.008 |
| Rdh10 | 813.036 | 0.517703 | 0.12049 | 4.296576 | 1.73E-05 | 0.0008 | 27.961 |
| Rdh11 | 1113.24 | -0.0937 | 0.09357 | -1.00143 | 0.31662 | 0.5272 | 17.957 |
| Rdh12 | 118.218 | 0.025092 | 0.21436 | 0.117057 | 0.90682 | 0.9534 | 13.752 |
| Rdh13 | 1176.34 | 0.065144 | 0.07289 | 0.893687 | 0.37149 | 0.58178 | 20.88 |
| Rdh14 | 951.036 | 0.106183 | 0.10055 | 1.056015 | 0.29096 | 0.50114 | 4.791 |
| Rdh5 | 75.3523 | -0.25713 | 0.22498 | -1.14293 | 0.25307 | 0.46046 | 9.296 |
| Rdm1 | 204.217 | -0.28726 | 0.14508 | -1.97997 | 0.04771 | 0.16063 | 8.906 |
| Rdx | 4446.52 | -0.14612 | 0.07522 | -1.94258 | 0.05207 | 0.17039 | 53.291 |
| Rec114 | 117.653 | 0.528292 | 0.17364 | 3.042432 | 0.00235 | 0.02234 | 91.115 |
| Rec8 | 478.437 | 0.432034 | 0.2088 | 2.069129 | 0.03853 | 0.13979 | 7.359 |
| Reck | 794.216 | -0.11115 | 0.11842 | -0.93866 | 0.34791 | 0.55841 | 69.277 |
| Recql | 852.777 | 0.062351 | 0.08486 | 0.734786 | 0.46247 | 0.66283 | 36.746 |
| Recql4 | 115.243 | -0.19082 | 0.21104 | -0.90415 | 0.36591 | 0.57637 | 6.996 |
| Recql5 | 1426.85 | 0.125409 | 0.09153 | 1.370163 | 0.17064 | 0.361 | 40.883 |
| Reep1 | 6286.5 | 0.065106 | 0.04965 | 1.311234 | 0.18978 | 0.38584 | 103.15 |
| Reep2 | 3936.13 | -0.01146 | 0.05991 | -0.19133 | 0.84827 | 0.92288 | 6.875 |
| Reep3 | 2358.81 | -0.21937 | 0.0789 | -2.78018 | 0.00543 | 0.03839 | 87.757 |
| Reep4 | 109.662 | -0.38845 | 0.22691 | -1.71192 | 0.08691 | 0.23732 | 3.683 |
| Reep5 | 15961.3 | -0.13371 | 0.06129 | -2.18151 | 0.02915 | 0.11574 | 29.186 |
| Reep6 | 322.261 | 0.016882 | 0.13159 | 0.128297 | 0.89791 | 0.94882 | 6.489 |
| Rel | 343.084 | -0.19569 | 0.10538 | -1.85704 | 0.0633 | 0.19335 | 34.124 |
| Rela | 920.481 | -0.18181 | 0.09254 | -1.96475 | 0.04944 | 0.16463 | 10.648 |
| Relb | 172.172 | -0.29729 | 0.19992 | -1.48703 | 0.13701 | 0.31772 | 23.222 |
| Relch | 3849.27 | 0.123389 | 0.063 | 1.958526 | 0.05017 | 0.16644 | 91.331 |
| Rell1 | 256.119 | -0.54029 | 0.15346 | -3.52063 | 0.00043 | 0.00723 | 60.001 |
| Rell2 | 1866.06 | -0.247 | 0.08604 | -2.87081 | 0.00409 | 0.032 | 4.101 |
| Reln | 3090.91 | -0.14327 | 0.09311 | -1.53875 | 0.12387 | 0.29838 | 460.249 |
| Relt | 527.765 | -0.10332 | 0.13485 | -0.76617 | 0.44357 | 0.64792 | 17.6 |

| | | | | | | | |
|---|---|---|---|---|---|---|---|
| Rem1 | 21.0345 | -0.39675 | 0.42996 | -0.92277 | 0.35613 | 0.56647 | 8.248 |
| Rem2 | 1364.91 | 0.748762 | 0.33679 | 2.223253 | 0.0262 | 0.10748 | 4.332 |
| Renbp | 134.169 | -0.53106 | 0.16465 | -3.22545 | 0.00126 | 0.01475 | 8.73 |
| Rep15 | 41.4105 | -0.80618 | 0.4909 | -1.64227 | 0.10053 | 0.26106 | 1.662 |
| Repin1 | 1766.56 | 0.208751 | 0.10534 | 1.981604 | 0.04752 | 0.16023 | 5.2 |
| Reps1 | 1679.6 | 0.015102 | 0.06776 | 0.222871 | 0.82364 | 0.90857 | 69.295 |
| Reps2 | 13296.5 | 0.263339 | 0.08179 | 3.219808 | 0.00128 | 0.01487 | 231.696 |
| Rer1 | 2327.67 | -0.06471 | 0.08278 | -0.78168 | 0.4344 | 0.64022 | 12.273 |
| Rere | 9271.14 | 0.092545 | 0.05955 | 1.554149 | 0.12015 | 0.29275 | 340.321 |
| Rerg | 1011.45 | 0.52943 | 0.17157 | 3.085792 | 0.00203 | 0.02026 | 115.673 |
| Resf1 | 851.663 | 0.176676 | 0.17179 | 1.028411 | 0.30376 | 0.51361 | 26.25 |
| Resp18 | 478.653 | -0.33538 | 0.17249 | -1.94434 | 0.05185 | 0.17002 | 6.217 |
| Rest | 339.699 | -0.16807 | 0.12068 | -1.39264 | 0.16373 | 0.35247 | 20.942 |
| Ret | 230.201 | 0.189331 | 0.14976 | 1.264196 | 0.20616 | 0.40534 | 45.974 |
| Retn | 52.9842 | 0.655349 | 0.32571 | 2.012065 | 0.04421 | 0.15247 | 4.341 |
| Retreg1 | 2194.57 | -0.01701 | 0.06784 | -0.25083 | 0.80195 | 0.89774 | 130.508 |
| Retreg2 | 6364.14 | 0.006103 | 0.06337 | 0.096312 | 0.92327 | 0.9614 | 5.136 |
| Retreg3 | 2973.19 | -0.08994 | 0.07067 | -1.27267 | 0.20313 | 0.40174 | 23.572 |
| Retsat | 539.909 | -0.20068 | 0.11502 | -1.74484 | 0.08101 | 0.22646 | 9.951 |
| Rev1 | 1377.4 | 0.162818 | 0.07912 | 2.057951 | 0.03959 | 0.1421 | 77.016 |
| Rev3l | 2375.35 | 0.164534 | 0.17711 | 0.929 | 0.35289 | 0.56319 | 143.094 |
| Rex1bd | 966.099 | -0.15798 | 0.09231 | -1.71146 | 0.087 | 0.23745 | 2.461 |
| Rexo1 | 2609.23 | 0.034502 | 0.06952 | 0.496282 | 0.6197 | 0.78361 | 20.639 |
| Rexo2 | 2242.9 | -0.07074 | 0.05851 | -1.20905 | 0.22665 | 0.43083 | 12.112 |
| Rexo4 | 2962.65 | 0.054225 | 0.07514 | 0.721688 | 0.47049 | 0.6693 | 10.824 |
| Rexo5 | 80.1396 | -0.43051 | 0.22897 | -1.88019 | 0.06008 | 0.18687 | 54.938 |
| Rfc1 | 2162.6 | -0.01596 | 0.08373 | -0.19057 | 0.84886 | 0.92318 | 73.821 |
| Rfc2 | 1376.11 | -0.15442 | 0.08677 | -1.77961 | 0.07514 | 0.21621 | 20.44 |
| Rfc3 | 229.23 | 0.274572 | 0.16582 | 1.655815 | 0.09776 | 0.25643 | 8.487 |
| Rfc4 | 106.185 | 0.051971 | 0.17313 | 0.300181 | 0.76404 | 0.8765 | 13.795 |
| Rfc5 | 657.916 | 0.020978 | 0.09037 | 0.232141 | 0.81643 | 0.90544 | 10.945 |
| Rfesd | 450.535 | 0.19689 | 0.10311 | 1.909474 | 0.0562 | 0.17928 | 17.185 |
| Rffl | 720.701 | -0.07844 | 0.12241 | -0.64078 | 0.52167 | 0.71081 | 68.762 |
| Rfk | 2520.88 | -0.0528 | 0.07506 | -0.7034 | 0.48181 | 0.67858 | 7.307 |

| | | | | | | | |
|---|---|---|---|---|---|---|---|
| Rflna | 49.3034 | 0.040499 | 0.27044 | 0.149753 | 0.88096 | 0.94067 | 9.327 |
| Rflnb | 633.531 | 0.050008 | 0.16548 | 0.3022 | 0.7625 | 0.8755 | 8.589 |
| Rfng | 2236.72 | -0.06363 | 0.06922 | -0.91923 | 0.35797 | 0.56813 | 3.462 |
| Rft1 | 543.45 | -0.08616 | 0.10266 | -0.83926 | 0.40132 | 0.60994 | 36.958 |
| Rftn1 | 540.669 | -0.45226 | 0.35227 | -1.28385 | 0.19919 | 0.39711 | 198.418 |
| Rftn2 | 1127.61 | -0.21328 | 0.09551 | -2.23305 | 0.02555 | 0.10545 | 56.624 |
| Rfwd3 | 1875.83 | 0.144577 | 0.06883 | 2.100636 | 0.03567 | 0.13274 | 29.279 |
| Rfx1 | 2103.39 | 0.391634 | 0.08565 | 4.572324 | 4.82E-06 | 0.00033 | 30.159 |
| Rfx2 | 113.061 | 0.069147 | 0.23355 | 0.296074 | 0.76717 | 0.8783 | 55.117 |
| Rfx3 | 4968.13 | 0.143386 | 0.07892 | 1.816766 | 0.06925 | 0.20491 | 249.446 |
| Rfx4 | 846.152 | 0.387377 | 0.10833 | 3.57598 | 0.00035 | 0.00622 | 150.477 |
| Rfx5 | 647.385 | -0.02085 | 0.11203 | -0.18607 | 0.85239 | 0.92484 | 7.487 |
| Rfx7 | 2200.16 | 0.223846 | 0.11583 | 1.932606 | 0.05328 | 0.17284 | 90.698 |
| Rfxank | 296.152 | -0.10238 | 0.12102 | -0.84597 | 0.39757 | 0.6066 | 8.361 |
| Rfxap | 1179.17 | 0.081341 | 0.07684 | 1.058575 | 0.28979 | 0.50005 | 4.677 |
| Rgcc | 764.822 | -0.23303 | 0.14445 | -1.6132 | 0.1067 | 0.27096 | 12.89 |
| Rgl1 | 6417.15 | 0.090288 | 0.06814 | 1.325035 | 0.18516 | 0.37986 | 249.592 |
| Rgl2 | 2970.37 | 0.051399 | 0.10082 | 0.509812 | 0.61018 | 0.77757 | 8.145 |
| Rgl3 | 226.874 | 0.132757 | 0.12546 | 1.058136 | 0.28999 | 0.50018 | 20.736 |
| Rgma | 2087.25 | -0.16018 | 0.09351 | -1.71295 | 0.08672 | 0.23706 | 44.391 |
| Rgmb | 3341.28 | -0.04832 | 0.07683 | -0.62888 | 0.52943 | 0.71727 | 27.852 |
| Rgp1 | 1774.84 | -0.06508 | 0.07667 | -0.84882 | 0.39598 | 0.60512 | 8.773 |
| Rgs10 | 666.818 | 0.051967 | 0.14338 | 0.362436 | 0.71703 | 0.84892 | 45.138 |
| Rgs11 | 1768.82 | 0.445966 | 0.10429 | 4.276133 | 1.90E-05 | 0.00086 | 8.374 |
| Rgs12 | 2365.69 | 0.325987 | 0.12047 | 2.706063 | 0.00681 | 0.04467 | 90.2 |
| Rgs13 | 13.1251 | -1.05699 | 0.59574 | -1.77425 | 0.07602 | 0.21771 | 38.719 |
| Rgs14 | 3560.09 | 0.097486 | 0.09405 | 1.03655 | 0.29995 | 0.51006 | 14.956 |
| Rgs16 | 411.982 | 0.469367 | 0.17838 | 2.631342 | 0.0085 | 0.0515 | 5.12 |
| Rgs17 | 5110.48 | 0.530992 | 0.15006 | 3.538459 | 0.0004 | 0.00691 | 96.738 |
| Rgs19 | 598.914 | -0.01897 | 0.10551 | -0.17977 | 0.85733 | 0.92794 | 5.559 |
| Rgs2 | 3263.74 | 0.123523 | 0.10223 | 1.208226 | 0.22696 | 0.43116 | 4.824 |
| Rgs20 | 1485.84 | 0.267982 | 0.13155 | 2.037136 | 0.04164 | 0.14663 | 160.71 |
| Rgs22 | 26.306 | 0.005776 | 0.38341 | 0.015064 | 0.98798 | 0.99319 | 130.922 |
| Rgs3 | 1056.34 | -0.51181 | 0.11436 | -4.47537 | 7.63E-06 | 0.00045 | 144.155 |

| | | | | | | | |
|---|---|---|---|---|---|---|---|
| Rgs4 | 14081.4 | -0.08205 | 0.09727 | -0.84348 | 0.39896 | 0.60798 | 6.166 |
| Rgs5 | 4224.85 | -0.31913 | 0.15168 | -2.10397 | 0.03538 | 0.13213 | 40.313 |
| Rgs6 | 1187.51 | -0.54348 | 0.11365 | -4.78209 | 1.73E-06 | 0.00016 | 573.765 |
| Rgs7 | 2406.66 | -0.05903 | 0.07828 | -0.75413 | 0.45077 | 0.65334 | 433.414 |
| Rgs7bp | 19829.6 | 0.141203 | 0.11429 | 1.23549 | 0.21665 | 0.41857 | 109.027 |
| Rgs8 | 9488.41 | 0.587311 | 0.19521 | 3.008541 | 0.00263 | 0.02412 | 47.299 |
| Rgs9 | 21061.9 | 0.798189 | 0.27926 | 2.858214 | 0.00426 | 0.03285 | 72.775 |
| Rgs9bp | 21.3875 | -0.28812 | 0.48847 | -0.58984 | 0.5553 | 0.73781 | 6.59 |
| Rgsl1 | 199.814 | 0.303262 | 0.46611 | 0.650619 | 0.51529 | 0.70568 | 64.762 |
| Rhbdd1 | 390.003 | -0.20256 | 0.10054 | -2.01476 | 0.04393 | 0.15174 | 128.915 |
| Rhbdd2 | 2717.51 | 0.006482 | 0.06261 | 0.103533 | 0.91754 | 0.95781 | 13.831 |
| Rhbdd3 | 712.374 | 0.1021 | 0.13728 | 0.743744 | 0.45703 | 0.65829 | 7.168 |
| Rhbdf1 | 504.811 | -0.30189 | 0.10811 | -2.7925 | 0.00523 | 0.03734 | 12.716 |
| Rhbdf2 | 74.8968 | -0.30119 | 0.20333 | -1.4813 | 0.13853 | 0.31964 | 28.855 |
| Rhbdl1 | 41.2842 | 0.634137 | 0.35601 | 1.781216 | 0.07488 | 0.21567 | 2.831 |
| Rhbdl2 | 20.8959 | -0.24801 | 0.45594 | -0.54396 | 0.58647 | 0.76037 | 42.031 |
| Rhbdl3 | 2423.11 | 0.337719 | 0.12066 | 2.798901 | 0.00513 | 0.03691 | 55.044 |
| Rhcg | 45.3697 | 0.550034 | 0.26902 | 2.044579 | 0.0409 | 0.14483 | 24.295 |
| Rheb | 4994.66 | -0.10905 | 0.08678 | -1.25662 | 0.20889 | 0.40895 | 39.802 |
| Rhebl1 | 290.388 | 0.04271 | 0.14184 | 0.301121 | 0.76332 | 0.876 | 16.763 |
| Rhno1 | 406.409 | 0.104434 | 0.10712 | 0.974904 | 0.32961 | 0.5409 | 5.912 |
| Rho | 41.1007 | 0.037226 | 0.26229 | 0.14193 | 0.88714 | 0.94391 | 8.289 |
| Rhoa | 4853.83 | -0.31231 | 0.07085 | -4.40781 | 1.04E-05 | 0.00057 | 31.806 |
| Rhob | 16153.4 | -0.00178 | 0.11684 | -0.01522 | 0.98786 | 0.99317 | 2.349 |
| Rhobtb1 | 246.79 | -0.36133 | 0.13537 | -2.66913 | 0.0076 | 0.04816 | 140.358 |
| Rhobtb2 | 6609.84 | 0.094985 | 0.06863 | 1.383956 | 0.16637 | 0.356 | 20.647 |
| Rhobtb3 | 1592.93 | -0.10175 | 0.09991 | -1.01845 | 0.30846 | 0.51837 | 74.389 |
| Rhoc | 437.886 | -0.30024 | 0.12216 | -2.45772 | 0.01398 | 0.07122 | 6.085 |
| Rhod | 15.3874 | -0.96431 | 0.45272 | -2.13002 | 0.03317 | 0.12657 | 13.974 |
| Rhof | 1215.12 | -0.14421 | 0.09791 | -1.47285 | 0.14079 | 0.32253 | 29.649 |
| Rhog | 739.981 | -0.54987 | 0.14683 | -3.74493 | 0.00018 | 0.00402 | 16.932 |
| Rhoh | 42.6459 | -1.03692 | 0.28426 | -3.64777 | 0.00026 | 0.00514 | 35.488 |
| Rhoj | 208.61 | -0.47935 | 0.14819 | -3.2346 | 0.00122 | 0.01448 | 93.135 |
| Rhoq | 1736.52 | 0.251932 | 0.08848 | 2.847199 | 0.00441 | 0.03356 | 36.988 |

| | | | | | | | |
|---|---|---|---|---|---|---|---|
| Rhot1 | 3988.03 | 0.038219 | 0.06722 | 0.568599 | 0.56963 | 0.74912 | 58.889 |
| Rhot2 | 1479.6 | -0.05914 | 0.07816 | -0.7567 | 0.44923 | 0.65249 | 6.395 |
| Rhou | 2708.94 | -0.46552 | 0.14247 | -3.2675 | 0.00109 | 0.01349 | 9.956 |
| Rhov | 209.084 | 0.471337 | 0.15011 | 3.139881 | 0.00169 | 0.01805 | 2.072 |
| Rhpn1 | 443.352 | -0.16979 | 0.11661 | -1.45602 | 0.14539 | 0.32825 | 11.206 |
| Rhpn2 | 569.58 | -0.46433 | 0.12636 | -3.67468 | 0.00024 | 0.00479 | 58.12 |
| Rian | 40332.5 | 0.280145 | 0.10975 | 2.552551 | 0.01069 | 0.06007 | 57.909 |
| Ribc1 | 49.087 | -0.06445 | 0.288 | -0.22377 | 0.82293 | 0.90818 | 11.714 |
| Ric1 | 4264.15 | 0.108472 | 0.07758 | 1.398271 | 0.16203 | 0.35036 | 84.548 |
| Ric3 | 3775.2 | -0.0114 | 0.07946 | -0.14341 | 0.88597 | 0.94329 | 49.02 |
| Ric8a | 1605.25 | 0.072639 | 0.0741 | 0.980318 | 0.32693 | 0.53825 | 7.416 |
| Ric8b | 5000.15 | 0.515803 | 0.14838 | 3.476253 | 0.00051 | 0.008 | 100.722 |
| Rictor | 4173.26 | 0.137574 | 0.1028 | 1.338211 | 0.18083 | 0.37423 | 92.023 |
| Rida | 1452.81 | -0.23049 | 0.09363 | -2.46177 | 0.01383 | 0.07068 | 11.235 |
| Rif1 | 1933.42 | 0.378729 | 0.18325 | 2.066714 | 0.03876 | 0.14026 | 49.552 |
| Riiad1 | 70.6948 | -0.06805 | 0.23975 | -0.28382 | 0.77655 | 0.88361 | 15.359 |
| Rilp | 61.9407 | 0.140703 | 0.26139 | 0.538284 | 0.59038 | 0.76326 | 3.075 |
| Rilpl1 | 1625.33 | -0.1906 | 0.07298 | -2.6118 | 0.00901 | 0.05345 | 38.312 |
| Rilpl2 | 430.879 | 0.098944 | 0.13246 | 0.746991 | 0.45507 | 0.65721 | 15.102 |
| Rimbp2 | 7983.29 | 0.124599 | 0.09554 | 1.304211 | 0.19216 | 0.38882 | 195.696 |
| Rimbp3 | 249.093 | 0.268406 | 0.13451 | 1.995396 | 0.046 | 0.15648 | 5.38 |
| Rimkla | 857.319 | 0.399142 | 0.09738 | 4.098725 | 4.15E-05 | 0.00144 | 27.346 |
| Rimklb | 2152.25 | 0.217835 | 0.12017 | 1.812651 | 0.06989 | 0.20631 | 51.466 |
| Rims1 | 8119.86 | -0.4044 | 0.12827 | -3.15277 | 0.00162 | 0.01748 | 488.729 |
| Rims2 | 5940.01 | 0.307113 | 0.08007 | 3.835446 | 0.00013 | 0.00312 | 486.113 |
| Rims3 | 4328.54 | 0.238665 | 0.14184 | 1.682623 | 0.09245 | 0.24756 | 41.764 |
| Rims4 | 4193.34 | -0.48183 | 0.13385 | -3.5997 | 0.00032 | 0.00589 | 58.933 |
| Rin1 | 4565.56 | 0.308396 | 0.11035 | 2.794592 | 0.0052 | 0.03718 | 6.591 |
| Rin2 | 2309.37 | -0.19177 | 0.1149 | -1.66911 | 0.0951 | 0.25184 | 212.402 |
| Rin3 | 219.994 | -0.66097 | 0.18214 | -3.62889 | 0.00028 | 0.00541 | 107.808 |
| Ring1 | 2197.03 | 0.237403 | 0.08831 | 2.688166 | 0.00718 | 0.04631 | 3.889 |
| Rinl | 114.34 | -0.5151 | 0.21277 | -2.4209 | 0.01548 | 0.07605 | 9.995 |
| Rint1 | 967.643 | 0.080047 | 0.08027 | 0.997238 | 0.31865 | 0.52924 | 32.659 |
| Riok1 | 987.637 | 0.191577 | 0.09951 | 1.925122 | 0.05421 | 0.1751 | 24.439 |

| | | | | | | | |
|---|---|---|---|---|---|---|---|
| Riok2 | 954.104 | 0.111153 | 0.11781 | 0.943469 | 0.34544 | 0.55616 | 20.591 |
| Riok3 | 4433.03 | 0.154093 | 0.08781 | 1.754918 | 0.07927 | 0.2235 | 28.647 |
| Riox1 | 319.227 | -0.17187 | 0.1227 | -1.40073 | 0.16129 | 0.34955 | 2.344 |
| Riox2 | 419.504 | 0.4265 | 0.15413 | 2.767228 | 0.00565 | 0.03958 | 20.687 |
| Ripk1 | 341.597 | -0.28639 | 0.14856 | -1.92774 | 0.05389 | 0.17433 | 34.785 |
| Ripk2 | 339.357 | 0.125193 | 0.14134 | 0.885733 | 0.37576 | 0.5859 | 40.915 |
| Ripk3 | 12.5243 | -0.36189 | 0.55857 | -0.64789 | 0.51705 | 0.70702 | 3.871 |
| Ripk4 | 42.3662 | -0.17239 | 0.30344 | -0.56811 | 0.56996 | 0.74936 | 21.855 |
| Ripor1 | 4107.77 | 0.080286 | 0.06223 | 1.290222 | 0.19697 | 0.3948 | 16.965 |
| Ripor2 | 4730.18 | 0.47333 | 0.09975 | 4.745114 | 2.08E-06 | 0.00018 | 232.292 |
| Ripor3 | 36.582 | -0.46672 | 0.45143 | -1.03386 | 0.3012 | 0.51092 | 30.455 |
| Ripply2 | 95.2811 | -0.03256 | 0.21714 | -0.14994 | 0.88081 | 0.94057 | 4.381 |
| Rit1 | 1136.12 | -0.16923 | 0.10242 | -1.65225 | 0.09848 | 0.25768 | 14.212 |
| Rit2 | 3393.05 | 0.184916 | 0.11745 | 1.574409 | 0.11539 | 0.28494 | 343.802 |
| Rita1 | 591.065 | -0.07971 | 0.09485 | -0.84039 | 0.40069 | 0.60946 | 18.285 |
| Rlbp1 | 676.366 | -0.94425 | 0.18247 | -5.17489 | 2.28E-07 | 3.32E-05 | 12.179 |
| Rlf | 1835.5 | 0.162001 | 0.07893 | 2.052535 | 0.04012 | 0.14312 | 69.712 |
| Rlim | 3218.22 | 0.051619 | 0.07419 | 0.695808 | 0.48655 | 0.68239 | 24.122 |
| Rln1 | 43.8756 | -0.1571 | 0.29229 | -0.53747 | 0.59095 | 0.7637 | 3.501 |
| Rmc1 | 1462.66 | 0.047531 | 0.06795 | 0.699513 | 0.48423 | 0.68059 | 21.942 |
| Rmdn1 | 647.093 | 0.076095 | 0.16849 | 0.451626 | 0.65154 | 0.80608 | 31.771 |
| Rmdn2 | 349.424 | 0.005132 | 0.12269 | 0.041825 | 0.96664 | 0.98328 | 81.33 |
| Rmdn3 | 1707.59 | -0.01391 | 0.08394 | -0.16566 | 0.86843 | 0.9334 | 20.034 |
| Rmi1 | 626.28 | 0.097074 | 0.09683 | 1.002564 | 0.31607 | 0.52659 | 8.901 |
| Rmnd1 | 1176.61 | -0.1842 | 0.10424 | -1.76709 | 0.07721 | 0.21967 | 30.474 |
| Rmnd5a | 9197.27 | 0.213129 | 0.06969 | 3.058302 | 0.00223 | 0.02157 | 52.004 |
| Rmnd5b | 1810.86 | 0.114857 | 0.09413 | 1.220255 | 0.22237 | 0.42565 | 12.226 |
| Rmst | 182.516 | -0.64873 | 0.16335 | -3.97133 | 7.15E-05 | 0.00213 | 94.234 |
| Rn18s-rs5 | 801590 | -0.0017 | 0.11939 | -0.01421 | 0.98866 | 0.99378 | 1.849 |
| Rn7sk | 551.338 | -0.02001 | 0.16548 | -0.12092 | 0.90375 | 0.95146 | 0.331 |
| Rnase4 | 542.978 | -0.2879 | 0.14332 | -2.00877 | 0.04456 | 0.15336 | 15.075 |
| Rnase6 | 12.6047 | -0.1772 | 0.45074 | -0.39314 | 0.69422 | 0.83414 | 8.28 |
| Rnaseh1 | 712.236 | 0.026285 | 0.08372 | 0.313956 | 0.75355 | 0.87058 | 9.988 |
| Rnaseh2a | 612.939 | 0.011098 | 0.10379 | 0.106929 | 0.91485 | 0.95656 | 13.158 |

| | | | | | | | |
|---|---|---|---|---|---|---|---|
| Rnaseh2b | 264.781 | 0.06686 | 0.14114 | 0.473701 | 0.63571 | 0.79475 | 80.404 |
| Rnaseh2c | 481.194 | -0.21314 | 0.15406 | -1.38351 | 0.16651 | 0.356 | 5.147 |
| Rnasek | 19.8869 | -0.45585 | 0.39758 | -1.14655 | 0.25157 | 0.45895 | 1.719 |
| Rnasel | 551.937 | 0.312151 | 0.12266 | 2.54477 | 0.01093 | 0.06093 | 14.796 |
| Rnaset2b | 35.7829 | 0.471407 | 0.35525 | 1.326983 | 0.18451 | 0.37899 | 40.666 |
| Rnd1 | 376.616 | -0.46907 | 0.16375 | -2.86451 | 0.00418 | 0.03238 | 14.041 |
| Rnd2 | 4938.14 | 0.041606 | 0.12804 | 0.324938 | 0.74523 | 0.866 | 6.855 |
| Rnd3 | 600.66 | -0.54676 | 0.18589 | -2.94127 | 0.00327 | 0.02757 | 18.674 |
| Rnf10 | 9915.32 | -0.15312 | 0.06978 | -2.19435 | 0.02821 | 0.11309 | 31.487 |
| Rnf103 | 3165.67 | 0.180955 | 0.08748 | 2.068549 | 0.03859 | 0.13989 | 16.988 |
| Rnf11 | 9915.6 | -0.09912 | 0.0703 | -1.41006 | 0.15852 | 0.34589 | 33.631 |
| Rnf111 | 2339.11 | 0.082264 | 0.06125 | 1.343026 | 0.17926 | 0.37214 | 78.302 |
| Rnf112 | 14680.9 | 0.242842 | 0.09855 | 2.464123 | 0.01373 | 0.0705 | 5.69 |
| Rnf113a1 | 244.191 | 0.472473 | 0.28415 | 1.662753 | 0.09636 | 0.25417 | 1.223 |
| Rnf113a2 | 505.247 | -0.03153 | 0.12618 | -0.24991 | 0.80266 | 0.89815 | 1.383 |
| Rnf114 | 2670.56 | -0.0283 | 0.0811 | -0.34896 | 0.72712 | 0.85437 | 23.529 |
| Rnf115 | 2203.57 | -0.15478 | 0.06922 | -2.23617 | 0.02534 | 0.10493 | 63.975 |
| Rnf121 | 1799.14 | 0.013014 | 0.07509 | 0.173314 | 0.8624 | 0.93022 | 46.333 |
| Rnf122 | 251.574 | 0.33628 | 0.12754 | 2.636603 | 0.00837 | 0.05104 | 19.663 |
| Rnf123 | 2889.56 | 0.099077 | 0.0731 | 1.355419 | 0.17528 | 0.36699 | 31.813 |
| Rnf125 | 61.8037 | -0.5242 | 0.25048 | -2.09278 | 0.03637 | 0.13438 | 39.238 |
| Rnf126 | 1921.19 | -0.00298 | 0.07547 | -0.03955 | 0.96845 | 0.98409 | 8.438 |
| Rnf128 | 942.446 | -0.04169 | 0.10235 | -0.40736 | 0.68375 | 0.82745 | 109.83 |
| Rnf13 | 4715.34 | -0.14869 | 0.08044 | -1.84839 | 0.06455 | 0.19559 | 99.172 |
| Rnf130 | 8367.49 | -0.27806 | 0.12885 | -2.15797 | 0.03093 | 0.1206 | 100.374 |
| Rnf135 | 169.586 | -0.29245 | 0.19473 | -1.50182 | 0.13314 | 0.31274 | 15.907 |
| Rnf138 | 1192.16 | 0.200666 | 0.10013 | 2.00397 | 0.04507 | 0.15434 | 26.884 |
| Rnf138rt1 | 36.1395 | 0.429094 | 0.33149 | 1.294442 | 0.19551 | 0.39318 | 1.191 |
| Rnf139 | 1027.46 | 0.042631 | 0.07824 | 0.544887 | 0.58583 | 0.76012 | 17.829 |
| Rnf14 | 13602.7 | -0.0454 | 0.07644 | -0.59402 | 0.5525 | 0.73541 | 33.313 |
| Rnf141 | 3424.16 | -0.20059 | 0.07499 | -2.67477 | 0.00748 | 0.04763 | 44.026 |
| Rnf144a | 1561.76 | 0.145203 | 0.08649 | 1.678825 | 0.09319 | 0.2485 | 114.291 |
| Rnf144b | 1259.44 | -0.16574 | 0.0929 | -1.78401 | 0.07442 | 0.21503 | 125.336 |
| Rnf145 | 4947.44 | 0.119323 | 0.08492 | 1.405169 | 0.15997 | 0.34786 | 46.557 |

| | | | | | | | |
|---|---|---|---|---|---|---|---|
| Rnf146 | 690.776 | -0.04858 | 0.10201 | -0.47619 | 0.63394 | 0.79385 | 18.267 |
| Rnf149 | 1141.71 | 0.135839 | 0.07973 | 1.703745 | 0.08843 | 0.24021 | 26.11 |
| Rnf150 | 7319.64 | 0.425268 | 0.12178 | 3.491962 | 0.00048 | 0.00771 | 227.913 |
| Rnf152 | 555.623 | -0.31113 | 0.2268 | -1.37181 | 0.17012 | 0.36051 | 79.797 |
| Rnf157 | 11899.9 | -0.24824 | 0.0846 | -2.93438 | 0.00334 | 0.02791 | 76.68 |
| Rnf165 | 3972.61 | -0.18254 | 0.1296 | -1.40845 | 0.159 | 0.34641 | 109.052 |
| Rnf166 | 1195.68 | 0.310366 | 0.10543 | 2.943923 | 0.00324 | 0.02748 | 9.918 |
| Rnf167 | 2114.57 | 0.015695 | 0.06572 | 0.238794 | 0.81127 | 0.9028 | 4.187 |
| Rnf168 | 2077.38 | 0.084287 | 0.08241 | 1.022782 | 0.30641 | 0.51644 | 23.976 |
| Rnf169 | 3934.25 | 0.363165 | 0.09754 | 3.723073 | 0.0002 | 0.00422 | 60.195 |
| Rnf17 | 49.8831 | 0.599851 | 0.41674 | 1.439384 | 0.15004 | 0.33455 | 122.452 |
| Rnf170 | 1427.19 | 0.187572 | 0.11295 | 1.660712 | 0.09677 | 0.25492 | 32.423 |
| Rnf170-ps | 35.3515 | -0.23437 | 0.35855 | -0.65367 | 0.51333 | 0.70424 | 0.859 |
| Rnf180 | 703.67 | 0.121998 | 0.10388 | 1.17445 | 0.24021 | 0.44624 | 164.815 |
| Rnf181 | 2330.49 | -0.1274 | 0.06766 | -1.8828 | 0.05973 | 0.18616 | 7.243 |
| Rnf182 | 490.958 | -0.56204 | 0.13033 | -4.31241 | 1.61E-05 | 0.00077 | 55.236 |
| Rnf185 | 1483.2 | -0.0546 | 0.07784 | -0.70151 | 0.48298 | 0.6794 | 36.382 |
| Rnf187 | 12882 | -0.03453 | 0.06665 | -0.51808 | 0.6044 | 0.7733 | 6.629 |
| Rnf19a | 2046.81 | -0.05961 | 0.07261 | -0.82093 | 0.41168 | 0.61973 | 43.215 |
| Rnf19b | 1443.03 | 0.011772 | 0.08368 | 0.140677 | 0.88812 | 0.94406 | 27.616 |
| Rnf2 | 1490.64 | 0.150579 | 0.07713 | 1.952209 | 0.05091 | 0.1678 | 42.952 |
| Rnf20 | 5263.12 | 0.129168 | 0.05965 | 2.165603 | 0.03034 | 0.11892 | 24.882 |
| Rnf207 | 1192.96 | 0.344753 | 0.12242 | 2.816053 | 0.00486 | 0.03557 | 11.975 |
| Rnf208 | 2238.88 | -0.07283 | 0.08284 | -0.87918 | 0.3793 | 0.589 | 1.334 |
| Rnf213 | 990.371 | -0.37134 | 0.15474 | -2.39984 | 0.0164 | 0.07899 | 94.319 |
| Rnf214 | 2329.29 | 0.055965 | 0.07069 | 0.791662 | 0.42856 | 0.63523 | 43.487 |
| Rnf215 | 1125.04 | 0.047822 | 0.07218 | 0.662492 | 0.50766 | 0.69896 | 5.971 |
| Rnf216 | 2504.4 | 0.01183 | 0.06947 | 0.170304 | 0.86477 | 0.93154 | 122.102 |
| Rnf217 | 1401.8 | -0.15461 | 0.12857 | -1.20255 | 0.22915 | 0.43326 | 115.992 |
| Rnf220 | 5836.07 | -0.1658 | 0.06963 | -2.38103 | 0.01726 | 0.08154 | 225.59 |
| Rnf223 | 94.4836 | 0.199957 | 0.17065 | 1.171725 | 0.24131 | 0.44737 | 2.908 |
| Rnf225 | 27.011 | -0.04912 | 0.31395 | -0.15646 | 0.87567 | 0.93796 | 3.657 |
| Rnf227 | 5092.69 | 0.028726 | 0.06132 | 0.46844 | 0.63947 | 0.79788 | 2.181 |
| Rnf24 | 1544.33 | -0.04573 | 0.07361 | -0.62124 | 0.53444 | 0.72156 | 54.829 |

| | | | | | | | |
|---|---|---|---|---|---|---|---|
| Rnf25 | 1963.76 | 0.202114 | 0.07991 | 2.529191 | 0.01143 | 0.0626 | 7.65 |
| Rnf26 | 28.3364 | -0.44081 | 0.37463 | -1.17665 | 0.23933 | 0.44533 | 17.887 |
| Rnf31 | 1621.99 | 0.164141 | 0.07201 | 2.279492 | 0.02264 | 0.09721 | 11.986 |
| Rnf32 | 880.333 | 0.257896 | 0.08066 | 3.197458 | 0.00139 | 0.01566 | 32.466 |
| Rnf34 | 2672.94 | 0.061339 | 0.08608 | 0.712586 | 0.4761 | 0.67418 | 21.244 |
| Rnf38 | 2829.63 | 0.141027 | 0.05989 | 2.354678 | 0.01854 | 0.08534 | 107.58 |
| Rnf39 | 100.526 | -0.22911 | 0.22811 | -1.00441 | 0.31518 | 0.52568 | 5.069 |
| Rnf4 | 3515.2 | 0.041827 | 0.07487 | 0.558621 | 0.57642 | 0.75396 | 19.341 |
| Rnf40 | 2666.47 | 0.038314 | 0.064 | 0.598685 | 0.54938 | 0.73299 | 16.033 |
| Rnf41 | 2970.93 | -0.11928 | 0.07635 | -1.56236 | 0.1182 | 0.2897 | 29.785 |
| Rnf43 | 168.339 | -0.02963 | 0.18296 | -0.16195 | 0.87135 | 0.93501 | 72.818 |
| Rnf44 | 8761.83 | 0.009996 | 0.0651 | 0.153541 | 0.87797 | 0.93893 | 14.509 |
| Rnf5 | 1805.32 | -0.21896 | 0.11505 | -1.90322 | 0.05701 | 0.18088 | 2.6 |
| Rnf6 | 3922.35 | 0.138961 | 0.08933 | 1.555591 | 0.11981 | 0.29233 | 12.364 |
| Rnf7 | 2522.6 | -0.09087 | 0.06967 | -1.30418 | 0.19217 | 0.38882 | 7.739 |
| Rnf8 | 468.457 | 0.201031 | 0.13401 | 1.50017 | 0.13357 | 0.31333 | 30.202 |
| Rnft1 | 1407.67 | 0.183068 | 0.08627 | 2.12201 | 0.03384 | 0.12842 | 14.369 |
| Rnft2 | 4418.93 | 0.02735 | 0.06437 | 0.424914 | 0.6709 | 0.8187 | 54.381 |
| Rngtt | 1705.11 | 0.217348 | 0.06964 | 3.121102 | 0.0018 | 0.01882 | 192.304 |
| Rnh1 | 1235.1 | -0.00947 | 0.1 | -0.09467 | 0.92457 | 0.96202 | 12.532 |
| Rnls | 26.2811 | -0.38582 | 0.37011 | -1.04242 | 0.29722 | 0.50736 | 254.519 |
| Rnmt | 3332.13 | 0.038105 | 0.07129 | 0.534471 | 0.59302 | 0.76545 | 24.498 |
| Rnpc3 | 1757.51 | 0.754777 | 0.13956 | 5.408102 | 6.37E-08 | 1.26E-05 | 25.083 |
| Rnpep | 1830.25 | 0.034929 | 0.07477 | 0.467144 | 0.6404 | 0.7986 | 21.373 |
| Rnpepl1 | 1564.52 | 0.059606 | 0.0662 | 0.900356 | 0.36793 | 0.57836 | 13.602 |
| Rnps1 | 6136.52 | -0.00495 | 0.0621 | -0.07974 | 0.93644 | 0.96817 | 11.337 |
| Ro60 | 3793.67 | 0.140926 | 0.11297 | 1.247431 | 0.21224 | 0.41289 | 26.279 |
| Robo1 | 4000.94 | 0.653195 | 0.17542 | 3.723566 | 0.0002 | 0.00421 | 737.79 |
| Robo2 | 5912.86 | -0.34492 | 0.12992 | -2.65497 | 0.00793 | 0.04932 | 519.987 |
| Robo3 | 216.872 | -0.4606 | 0.24288 | -1.89642 | 0.0579 | 0.18288 | 17.578 |
| Robo4 | 371.754 | -0.32184 | 0.1689 | -1.90553 | 0.05671 | 0.18033 | 13.219 |
| Rock1 | 2268.35 | 0.203057 | 0.18507 | 1.09716 | 0.27257 | 0.48182 | 117.645 |
| Rock2 | 11586.9 | 0.136565 | 0.14663 | 0.931355 | 0.35167 | 0.56204 | 92.929 |
| Rogdi | 3410.36 | -0.00042 | 0.05503 | -0.00759 | 0.99395 | 0.99634 | 4.824 |

| | | | | | | | |
|---|---|---|---|---|---|---|---|
| Rom1 | 180.408 | -0.2751 | 0.18132 | -1.51723 | 0.12921 | 0.30665 | 1.966 |
| Romo1 | 953.716 | -0.15602 | 0.08648 | -1.80407 | 0.07122 | 0.20889 | 1.759 |
| Ror1 | 106.014 | 0.644826 | 0.23683 | 2.722741 | 0.00647 | 0.04328 | 348.975 |
| Ror2 | 19.5528 | -0.88919 | 0.42146 | -2.10977 | 0.03488 | 0.13086 | 176.813 |
| Rora | 2563.91 | -0.26469 | 0.13182 | -2.00792 | 0.04465 | 0.1535 | 734.461 |
| Rorb | 1381.03 | -0.33813 | 0.11157 | -3.03055 | 0.00244 | 0.02295 | 180.592 |
| Rorc | 106.361 | 0.114672 | 0.21829 | 0.525322 | 0.59936 | 0.77013 | 25.483 |
| Rp2 | 731.836 | 0.071633 | 0.10726 | 0.667855 | 0.50423 | 0.69654 | 41.173 |
| Rp9 | 972.374 | -0.10714 | 0.08545 | -1.25374 | 0.20994 | 0.40997 | 58.697 |
| Rpa1 | 2170.74 | -0.06831 | 0.08232 | -0.82979 | 0.40666 | 0.6148 | 50.159 |
| Rpa2 | 542.753 | 0.14764 | 0.12126 | 1.217536 | 0.2234 | 0.42664 | 10.421 |
| Rpa3 | 361.955 | -0.00858 | 0.12661 | -0.06774 | 0.94599 | 0.973 | 3.238 |
| Rpain | 804.698 | 0.121997 | 0.11207 | 1.088618 | 0.27632 | 0.48548 | 7.621 |
| Rpap1 | 1407.59 | 0.158492 | 0.07879 | 2.011558 | 0.04427 | 0.15256 | 24.234 |
| Rpap2 | 643.782 | 0.023814 | 0.10211 | 0.233216 | 0.81559 | 0.90517 | 64.466 |
| Rpap3 | 1151.32 | 0.0391 | 0.09672 | 0.404239 | 0.68604 | 0.82887 | 30.729 |
| Rpe | 1190.36 | 0.143422 | 0.0803 | 1.786017 | 0.0741 | 0.21451 | 18.975 |
| Rpe65 | 157.727 | 0.960248 | 0.23773 | 4.039234 | 5.36E-05 | 0.00174 | 26.147 |
| Rpf1 | 562.078 | -0.0071 | 0.09902 | -0.07168 | 0.94286 | 0.9715 | 15.474 |
| Rpf2 | 600.871 | -0.12533 | 0.10497 | -1.19397 | 0.23249 | 0.43737 | 23.791 |
| Rpgr | 1713.55 | 0.39438 | 0.16453 | 2.397076 | 0.01653 | 0.07941 | 143.3 |
| Rpgrip1 | 35.4751 | -0.58849 | 0.30597 | -1.92334 | 0.05444 | 0.17541 | 52.843 |
| Rpgrip1l | 2142.02 | 0.230605 | 0.08754 | 2.634406 | 0.00843 | 0.05116 | 96.233 |
| Rph3a | 11238.1 | -0.65066 | 0.14077 | -4.62204 | 3.80E-06 | 0.00028 | 69.594 |
| Rph3al | 163.657 | 1.011328 | 0.16293 | 6.207299 | 5.39E-10 | 3.33E-07 | 107.431 |
| Rpia | 613.55 | 0.202551 | 0.09105 | 2.224636 | 0.02611 | 0.10722 | 26.513 |
| Rpl10 | 7084.68 | -0.00864 | 0.08583 | -0.10067 | 0.91981 | 0.95944 | 2.324 |
| Rpl10-ps5 | 14.2934 | -0.69242 | 0.51162 | -1.35338 | 0.17593 | 0.36783 | 0.642 |
| Rpl10-ps6 | 18.9065 | 0.232702 | 0.37803 | 0.615572 | 0.53818 | 0.72492 | 0.642 |
| Rpl10a | 3959.37 | -0.06471 | 0.07804 | -0.82918 | 0.407 | 0.61503 | 2.563 |
| Rpl10a-ps1 | 215.815 | -0.00861 | 0.14924 | -0.05766 | 0.95402 | 0.97747 | 0.654 |
| Rpl11 | 5553.32 | -0.0272 | 0.0915 | -0.29729 | 0.76625 | 0.8779 | 25.164 |
| Rpl12 | 4271.07 | -0.11542 | 0.10127 | -1.1398 | 0.25437 | 0.46172 | 3.787 |
| Rpl13 | 7725.63 | -0.07634 | 0.09174 | -0.83209 | 0.40536 | 0.61415 | 2.895 |

| | | | | | | | |
|---|---|---|---|---|---|---|---|
| Rpl13-ps5 | 22.2657 | 0.782281 | 0.35425 | 2.208247 | 0.02723 | 0.11025 | 0.63 |
| Rpl13a | 236.924 | 0.113414 | 0.14705 | 0.771261 | 0.44055 | 0.64545 | 3.204 |
| Rpl14 | 6315.79 | -0.01046 | 0.09826 | -0.10647 | 0.91521 | 0.95656 | 3.211 |
| Rpl14-ps1 | 1898.23 | -0.0674 | 0.11978 | -0.56271 | 0.57363 | 0.75227 | 0.653 |
| Rpl15 | 5868.63 | 0.003156 | 0.0802 | 0.039357 | 0.96861 | 0.9841 | 3.569 |
| Rpl15-ps3 | 57.4447 | 0.163559 | 0.27063 | 0.604373 | 0.5456 | 0.73038 | 0.615 |
| Rpl15-ps6 | 15.8867 | -0.48177 | 0.49354 | -0.97615 | 0.32899 | 0.54015 | 0.612 |
| Rpl17 | 3332.95 | -0.31883 | 0.51313 | -0.62134 | 0.53437 | 0.72154 | 4.824 |
| Rpl17-ps10 | 2351.24 | 0.319375 | 0.60902 | 0.524405 | 0.6 | 0.77061 | 0.555 |
| Rpl17-ps3 | 1084.46 | 0.209211 | 0.13793 | 1.516784 | 0.12932 | 0.30684 | 0.556 |
| Rpl17-ps5 | 113.781 | 0.041947 | 0.18145 | 0.23118 | 0.81718 | 0.90593 | 0.555 |
| Rpl17-ps9 | 13.5073 | -0.0196 | 0.6051 | -0.0324 | 0.97416 | 0.98704 | 0.555 |
| Rpl18 | 3375.06 | -0.14143 | 0.08425 | -1.67863 | 0.09322 | 0.24855 | 5.38 |
| Rpl18a | 5885.4 | -0.11601 | 0.08134 | -1.42622 | 0.1538 | 0.3396 | 2.691 |
| Rpl19 | 2034.57 | -0.22948 | 0.16504 | -1.3904 | 0.16441 | 0.35332 | 3.783 |
| Rpl19-ps11 | 33.5305 | -0.3792 | 0.37814 | -1.00279 | 0.31596 | 0.52651 | 0.591 |
| Rpl21 | 944.923 | 0.462338 | 0.37272 | 1.240432 | 0.21482 | 0.41624 | 4.143 |
| Rpl21-ps15 | 601.414 | 0.194851 | 0.18949 | 1.028304 | 0.30381 | 0.51363 | 0.483 |
| Rpl21-ps8 | 32.8299 | 0.173278 | 0.46326 | 0.374043 | 0.70837 | 0.84264 | 0.483 |
| Rpl22 | 3333.3 | -0.24423 | 0.09512 | -2.56759 | 0.01024 | 0.05844 | 8.33 |
| Rpl22l1 | 564.626 | -0.12627 | 0.22716 | -0.55584 | 0.57832 | 0.75514 | 1.989 |
| Rpl23 | 4230.26 | -0.04437 | 0.07066 | -0.62799 | 0.53001 | 0.71762 | 4.911 |
| Rpl23a | 207.805 | -0.37424 | 0.25608 | -1.46145 | 0.14389 | 0.32626 | 2.653 |
| Rpl23a-ps3 | 113.789 | -0.16153 | 0.194 | -0.83262 | 0.40506 | 0.61387 | 0.468 |
| Rpl24 | 28.7054 | 0.38601 | 0.34567 | 1.116692 | 0.26413 | 0.47298 | 5.161 |
| Rpl26 | 6339.32 | -0.03146 | 0.0839 | -0.37492 | 0.70772 | 0.8423 | 5.407 |
| Rpl26-ps6 | 46.6415 | -0.23896 | 0.3385 | -0.70593 | 0.48023 | 0.67761 | 0.434 |
| Rpl27 | 2900.64 | -0.06742 | 0.10978 | -0.61407 | 0.53917 | 0.72582 | 3.232 |
| Rpl27a | 6278.88 | -0.05675 | 0.08669 | -0.65468 | 0.51268 | 0.70354 | 3.221 |
| Rpl28 | 3567.08 | -0.16116 | 0.08292 | -1.94364 | 0.05194 | 0.17019 | 2.193 |
| Rpl28-ps1 | 224.566 | -0.10608 | 0.17498 | -0.60625 | 0.54435 | 0.72945 | 0.414 |
| Rpl28-ps3 | 19.498 | -0.349 | 0.39537 | -0.88272 | 0.37739 | 0.58749 | 0.393 |
| Rpl29 | 2204.9 | -0.02674 | 0.09498 | -0.28149 | 0.77834 | 0.88427 | 2.115 |
| Rpl29-ps2 | 14.803 | -0.32745 | 0.53042 | -0.61733 | 0.53701 | 0.72388 | 5.692 |

| | | | | | | | |
|---|---|---|---|---|---|---|---|
| Rpl3 | 19794.6 | 0.092956 | 0.10189 | 0.912336 | 0.36159 | 0.57187 | 14.078 |
| Rpl3-ps1 | 882.451 | 0.085041 | 0.09616 | 0.884406 | 0.37648 | 0.58653 | 1.21 |
| Rpl30 | 658.482 | 0.203879 | 0.20937 | 0.973771 | 0.33017 | 0.54139 | 3.136 |
| Rpl30-ps1 | 29.7614 | 0.34006 | 0.49647 | 0.684953 | 0.49337 | 0.6878 | 0.348 |
| Rpl30-ps10 | 249.541 | -0.19922 | 0.22062 | -0.90299 | 0.36653 | 0.57697 | 0.347 |
| Rpl31 | 6346.74 | 0.016861 | 0.09202 | 0.183231 | 0.85462 | 0.92652 | 4.07 |
| Rpl31-ps8 | 644.288 | -0.03777 | 0.1444 | -0.26158 | 0.79364 | 0.8931 | 0.378 |
| Rpl32 | 3199.03 | -0.1894 | 0.09893 | -1.91444 | 0.05556 | 0.17789 | 3.243 |
| Rpl34 | 2347.71 | -0.09074 | 0.09868 | -0.91953 | 0.35782 | 0.56801 | 3.568 |
| Rpl34-ps1 | 372.14 | -0.03266 | 0.17113 | -0.19086 | 0.84864 | 0.92307 | 0.354 |
| Rpl35 | 328.653 | -0.43365 | 0.26166 | -1.6573 | 0.09746 | 0.25604 | 4.045 |
| Rpl35a | 5467.67 | 0.000244 | 0.09677 | 0.002525 | 0.99799 | 0.99897 | 3.737 |
| Rpl35a-ps2 | 13.4304 | 0.346914 | 0.46853 | 0.740427 | 0.45904 | 0.65977 | 0.333 |
| Rpl36 | 2273.2 | -0.18036 | 0.09985 | -1.80634 | 0.07086 | 0.20814 | 0.828 |
| Rpl36-ps12 | 36.7396 | -0.57271 | 0.29189 | -1.96211 | 0.04975 | 0.1653 | 0.39 |
| Rpl36a | 2271.89 | -0.02031 | 0.10247 | -0.19823 | 0.84286 | 0.91916 | 2.409 |
| Rpl36a-ps2 | 224.241 | -0.1621 | 0.15999 | -1.01321 | 0.31096 | 0.5212 | 0.321 |
| Rpl36al | 14.5686 | -0.3634 | 0.47911 | -0.75849 | 0.44816 | 0.65172 | 1.353 |
| Rpl37 | 4669.69 | -0.06298 | 0.08602 | -0.73217 | 0.46407 | 0.66379 | 2.496 |
| Rpl37a | 3842.28 | -0.14688 | 0.09723 | -1.51058 | 0.1309 | 0.30933 | 2.524 |
| Rpl37rt | 653.023 | -0.08207 | 0.16956 | -0.48399 | 0.62839 | 0.78986 | 7.346 |
| Rpl38 | 2410.74 | -0.27919 | 0.13173 | -2.11943 | 0.03405 | 0.12885 | 3.808 |
| Rpl38-ps2 | 427.699 | -0.30495 | 0.20607 | -1.47983 | 0.13892 | 0.32015 | 0.213 |
| Rpl39 | 1850.71 | -0.19595 | 0.11505 | -1.70311 | 0.08855 | 0.24047 | 2.883 |
| Rpl39-ps | 15.6539 | -0.66305 | 0.43243 | -1.53329 | 0.1252 | 0.3002 | 0.151 |
| Rpl3l | 34.9517 | 0.325276 | 0.44291 | 0.734405 | 0.4627 | 0.66283 | 8.324 |
| Rpl4 | 16311.1 | -0.01084 | 0.08058 | -0.13451 | 0.893 | 0.94632 | 5.292 |
| Rpl41 | 8305.12 | -0.12045 | 0.10521 | -1.14479 | 0.2523 | 0.45959 | 1.192 |
| Rpl5 | 7488.39 | 0.005068 | 0.07187 | 0.070523 | 0.94378 | 0.97192 | 8.504 |
| Rpl6 | 7476.56 | -0.00157 | 0.10005 | -0.01568 | 0.98749 | 0.99316 | 4.761 |
| Rpl6l | 291.541 | -0.11799 | 0.1711 | -0.68958 | 0.49046 | 0.68551 | 0.871 |
| Rpl7 | 7858.84 | 0.104683 | 0.08017 | 1.305702 | 0.19165 | 0.3883 | 3.368 |
| Rpl7-ps9 | 18.9259 | -0.12168 | 0.69159 | -0.17594 | 0.86034 | 0.92937 | 0.797 |
| Rpl7a | 3478.69 | 0.055144 | 0.30237 | 0.182374 | 0.85529 | 0.92683 | 2.555 |

| | | | | | | | |
|---|---|---|---|---|---|---|---|
| Rpl7a-ps11 | 18.7244 | -0.54711 | 0.39178 | -1.39647 | 0.16257 | 0.35093 | 0.801 |
| Rpl7a-ps3 | 21.005 | -0.74444 | 0.47098 | -1.58061 | 0.11397 | 0.28262 | 0.796 |
| Rpl7a-ps5 | 132.521 | 0.235848 | 0.29323 | 0.804299 | 0.42122 | 0.6282 | 0.792 |
| Rpl7l1 | 1861.64 | -0.132 | 0.07203 | -1.83248 | 0.06688 | 0.20022 | 8.766 |
| Rpl8 | 7181.3 | -0.01194 | 0.08541 | -0.1398 | 0.88882 | 0.94447 | 2.237 |
| Rpl9 | 7509.5 | -0.01611 | 0.10291 | -0.15657 | 0.87559 | 0.93793 | 3.081 |
| Rpl9-ps3 | 16.0716 | 0.563159 | 0.45768 | 1.230463 | 0.21852 | 0.42062 | 0.576 |
| Rpl9-ps4 | 294.573 | 0.142433 | 0.19109 | 0.745372 | 0.45605 | 0.65766 | 0.579 |
| Rpl9-ps6 | 765.75 | 0.009257 | 0.14013 | 0.066057 | 0.94733 | 0.97353 | 0.714 |
| Rpl9-ps7 | 79.179 | 0.139225 | 0.21793 | 0.638844 | 0.52292 | 0.71143 | 0.579 |
| Rplp0 | 6039.68 | -0.18535 | 0.08453 | -2.19267 | 0.02833 | 0.11346 | 4.261 |
| Rplp1 | 11235.8 | -0.043 | 0.09979 | -0.43087 | 0.66656 | 0.8158 | 1.259 |
| Rplp2 | 2907.16 | -0.06056 | 0.10702 | -0.56591 | 0.57146 | 0.75033 | 4.216 |
| Rpn1 | 5023.99 | -0.02991 | 0.07805 | -0.38319 | 0.70158 | 0.83894 | 20.823 |
| Rpn2 | 4752.73 | 0.111731 | 0.07624 | 1.465518 | 0.14278 | 0.32475 | 47.303 |
| Rpp14 | 1320.52 | 0.063675 | 0.08892 | 0.716092 | 0.47393 | 0.67223 | 11.468 |
| Rpp21 | 415.884 | -0.1069 | 0.10751 | -0.99439 | 0.32003 | 0.53067 | 2.425 |
| Rpp25 | 482.238 | 0.009903 | 0.123 | 0.080506 | 0.93583 | 0.96779 | 1.423 |
| Rpp25l | 394.962 | -0.20663 | 0.10155 | -2.03475 | 0.04188 | 0.1472 | 1.502 |
| Rpp30 | 480.671 | 0.078379 | 0.10686 | 0.733489 | 0.46326 | 0.66314 | 21.062 |
| Rpp38 | 193.05 | 0.086859 | 0.18095 | 0.480021 | 0.63121 | 0.79177 | 3.695 |
| Rpp40 | 263.609 | 0.292299 | 0.12206 | 2.394681 | 0.01663 | 0.07984 | 12.888 |
| Rprd1a | 5134.21 | 0.161376 | 0.06404 | 2.519857 | 0.01174 | 0.06375 | 45.245 |
| Rprd1b | 1485.71 | 0.123116 | 0.08827 | 1.394836 | 0.16307 | 0.35163 | 103.801 |
| Rprd2 | 3243.62 | 0.090966 | 0.06152 | 1.478697 | 0.13922 | 0.32045 | 58.523 |
| Rprm | 1110.82 | -0.58986 | 0.14655 | -4.02504 | 5.70E-05 | 0.00181 | 1.46 |
| Rprml | 1325.7 | -0.37762 | 0.14279 | -2.6447 | 0.00818 | 0.05025 | 1.01 |
| Rps10 | 143.319 | 0.008634 | 0.17435 | 0.049519 | 0.96051 | 0.98071 | 6.252 |
| Rps10-ps1 | 35.8252 | -0.09482 | 0.28905 | -0.32804 | 0.74288 | 0.86462 | 0.495 |
| Rps10-ps2 | 1404.35 | -0.11884 | 0.12738 | -0.93298 | 0.35083 | 0.56148 | 0.496 |
| Rps11 | 696.002 | -0.32116 | 0.13656 | -2.35182 | 0.01868 | 0.08567 | 2.061 |
| Rps11-ps2 | 18.8546 | 0.273764 | 0.41117 | 0.665822 | 0.50552 | 0.69706 | 0.456 |
| Rps12 | 1884.11 | -0.16323 | 0.1098 | -1.48653 | 0.13714 | 0.31781 | 2.093 |
| Rps12-ps26 | 12.6837 | 0.918453 | 0.49107 | 1.87031 | 0.06144 | 0.18957 | 0.333 |

| | | | | | | | |
|---|---|---|---|---|---|---|---|
| Rps12-ps3 | 609.958 | -0.00545 | 0.18134 | -0.03005 | 0.97602 | 0.98762 | 0.396 |
| Rps12-ps9 | 27.9282 | 0.094197 | 0.40303 | 0.233723 | 0.8152 | 0.90512 | 0.399 |
| Rps13 | 2907.52 | -0.02452 | 0.07977 | -0.30736 | 0.75857 | 0.8731 | 2.691 |
| Rps13-ps1 | 45.075 | 0.029537 | 0.24691 | 0.119628 | 0.90478 | 0.95186 | 0.538 |
| Rps13-ps2 | 17.0746 | 0.102215 | 0.40912 | 0.24984 | 0.80271 | 0.89815 | 0.453 |
| Rps13-ps4 | 26.0979 | 0.192662 | 0.34098 | 0.565023 | 0.57206 | 0.75086 | 0.456 |
| Rps14 | 7875.76 | -0.17277 | 0.08432 | -2.04893 | 0.04047 | 0.14371 | 31.449 |
| Rps15 | 8837.99 | -0.08252 | 0.09692 | -0.85137 | 0.39456 | 0.60364 | 1.662 |
| Rps15-ps3 | 48.0212 | 0.667414 | 0.40316 | 1.655463 | 0.09783 | 0.25657 | 3.686 |
| Rps15a | 5314.41 | -0.0894 | 0.0823 | -1.08629 | 0.27735 | 0.48702 | 11.817 |
| Rps16 | 4014.65 | -0.08836 | 0.0883 | -1.00058 | 0.31703 | 0.52743 | 2.504 |
| Rps16-ps2 | 655.511 | -0.10643 | 0.12939 | -0.82258 | 0.41075 | 0.61876 | 0.441 |
| Rps17 | 3954.41 | -0.11183 | 0.08013 | -1.39559 | 0.16284 | 0.35136 | 2.523 |
| Rps18 | 4023.48 | -0.12119 | 0.08371 | -1.44774 | 0.14769 | 0.33123 | 4.003 |
| Rps18-ps5 | 28.7412 | 0.504959 | 0.33563 | 1.50452 | 0.13245 | 0.31154 | 0.607 |
| Rps19 | 4954.66 | 0.005871 | 0.09995 | 0.058744 | 0.95316 | 0.97691 | 5.436 |
| Rps19-ps3 | 67.0618 | 0.19207 | 0.23488 | 0.817739 | 0.41351 | 0.6216 | 0.426 |
| Rps19-ps6 | 53.8735 | -0.30047 | 0.26988 | -1.11335 | 0.26556 | 0.47475 | 0.438 |
| Rps19bp1 | 618.895 | -0.0374 | 0.11711 | -0.31937 | 0.74945 | 0.86849 | 3.701 |
| Rps2 | 12274.9 | 0.02979 | 0.10026 | 0.297114 | 0.76638 | 0.8779 | 3.814 |
| Rps2-ps10 | 69.0594 | -0.21199 | 0.28582 | -0.74171 | 0.45827 | 0.6595 | 0.981 |
| Rps2-ps11 | 35.8318 | 1.085075 | 0.29735 | 3.64909 | 0.00026 | 0.00513 | 0.868 |
| Rps2-ps13 | 14.913 | 0.333835 | 0.5529 | 0.603785 | 0.54599 | 0.73068 | 0.882 |
| Rps2-ps6 | 32.0844 | 0.188986 | 0.37976 | 0.497645 | 0.61873 | 0.78313 | 0.855 |
| Rps20 | 3647.69 | -0.15606 | 0.09039 | -1.72654 | 0.08425 | 0.23242 | 4.332 |
| Rps21 | 5090.89 | -0.15225 | 0.10562 | -1.44145 | 0.14946 | 0.33376 | 1.069 |
| Rps23 | 7702.75 | -0.00996 | 0.09341 | -0.10663 | 0.91508 | 0.95656 | 1.993 |
| Rps24 | 7936.75 | -0.14095 | 0.08634 | -1.63236 | 0.1026 | 0.26448 | 9.835 |
| Rps25 | 5349.9 | -0.01859 | 0.0869 | -0.21395 | 0.83058 | 0.91314 | 3.289 |
| Rps26 | 4268.17 | -0.02013 | 0.10238 | -0.19663 | 0.84412 | 0.91999 | 2.214 |
| Rps26-ps1 | 79.043 | 0.878438 | 0.90411 | 0.971608 | 0.33125 | 0.54208 | 0.449 |
| Rps27 | 2009.27 | -0.02359 | 0.11408 | -0.2068 | 0.83616 | 0.91592 | 1.13 |
| Rps27a | 3109.02 | -0.91261 | 0.18831 | -4.84627 | 1.26E-06 | 0.00012 | 2.264 |
| Rps27a-ps2 | 70.6354 | -0.11412 | 0.30258 | -0.37714 | 0.70607 | 0.8415 | 0.467 |

| | | | | | | | |
|---|---|---|---|---|---|---|---|
| Rps27a-ps3 | 140.328 | 1.167022 | 0.36062 | 3.236194 | 0.00121 | 0.01442 | 0.471 |
| Rps27l | 342.69 | -0.35963 | 0.1356 | -2.65211 | 0.008 | 0.04961 | 3.431 |
| Rps27rt | 660.812 | 0.246611 | 0.21394 | 1.152722 | 0.24902 | 0.45629 | 0.374 |
| Rps28 | 1607.33 | -0.20012 | 0.11172 | -1.79128 | 0.07325 | 0.21314 | 5.536 |
| Rps29 | 4579.18 | -0.21875 | 0.19333 | -1.13145 | 0.25787 | 0.46595 | 1.465 |
| Rps3 | 11211.1 | -0.10801 | 0.07717 | -1.39964 | 0.16162 | 0.34982 | 5.843 |
| Rps3a1 | 10722.9 | -0.02242 | 0.0764 | -0.29343 | 0.76919 | 0.87962 | 4.763 |
| Rps3a2 | 26.3492 | -0.12735 | 0.42014 | -0.30311 | 0.7618 | 0.87515 | 0.792 |
| Rps3a3 | 32.6681 | 0.070812 | 0.29592 | 0.239293 | 0.81088 | 0.90259 | 0.795 |
| Rps4l | 411.637 | -0.31915 | 0.11387 | -2.80267 | 0.00507 | 0.0366 | 1.334 |
| Rps4x | 7474.59 | -0.05296 | 0.08298 | -0.63825 | 0.52331 | 0.71181 | 4.454 |
| Rps5 | 4940.04 | -0.01455 | 0.09368 | -0.15528 | 0.8766 | 0.93831 | 4.397 |
| Rps6 | 8654.88 | -0.03067 | 0.08068 | -0.38012 | 0.70386 | 0.84011 | 2.753 |
| Rps6-ps1 | 12.7585 | -0.14641 | 0.57976 | -0.25253 | 0.80063 | 0.89692 | 0.747 |
| Rps6-ps4 | 379.904 | -0.02813 | 0.15831 | -0.17771 | 0.85895 | 0.9288 | 0.75 |
| Rps6ka1 | 478.219 | -0.24985 | 0.15368 | -1.62585 | 0.10398 | 0.26648 | 40.508 |
| Rps6ka2 | 4647.42 | 0.279639 | 0.09198 | 3.040324 | 0.00236 | 0.02243 | 133.201 |
| Rps6ka3 | 5056.66 | 0.221163 | 0.10558 | 2.094707 | 0.0362 | 0.134 | 157.938 |
| Rps6ka4 | 1365.06 | 0.129653 | 0.09683 | 1.338969 | 0.18058 | 0.37394 | 11.427 |
| Rps6ka5 | 2712.22 | 0.466572 | 0.1399 | 3.335138 | 0.00085 | 0.01147 | 178.545 |
| Rps6ka6 | 384.979 | 0.665732 | 0.17466 | 3.811483 | 0.00014 | 0.00334 | 149.768 |
| Rps6kb1 | 3064.43 | 0.081469 | 0.09181 | 0.887376 | 0.37488 | 0.58517 | 45.935 |
| Rps6kb2 | 1594.62 | 0.138961 | 0.10035 | 1.384766 | 0.16612 | 0.35578 | 9.364 |
| Rps6kc1 | 2149.45 | 0.057614 | 0.09168 | 0.628403 | 0.52974 | 0.71748 | 211.569 |
| Rps6kl1 | 1427.27 | 0.142171 | 0.08078 | 1.759988 | 0.07841 | 0.22174 | 16.016 |
| Rps7 | 6305.85 | -0.05201 | 0.08946 | -0.58135 | 0.561 | 0.74209 | 5.1 |
| Rps7-ps2 | 24.5761 | 0.217297 | 0.48146 | 0.45133 | 0.65175 | 0.80618 | 1.347 |
| Rps7-ps3 | 1625.26 | 0.013377 | 0.16311 | 0.08201 | 0.93464 | 0.96717 | 0.651 |
| Rps8 | 6658.23 | -0.05297 | 0.08694 | -0.60928 | 0.54234 | 0.72794 | 2.417 |
| Rps9 | 6771.44 | -0.08387 | 0.08861 | -0.94653 | 0.34388 | 0.55456 | 2.905 |
| Rpsa | 12623.6 | 0.079219 | 0.08566 | 0.9248 | 0.35507 | 0.56548 | 4.681 |
| Rpsa-ps10 | 210.92 | -0.0095 | 0.13834 | -0.06868 | 0.94524 | 0.97273 | 0.885 |
| Rpsa-ps2 | 21.8815 | -0.33288 | 0.37796 | -0.88074 | 0.37846 | 0.58827 | 0.88 |
| Rpsa-ps7 | 275.35 | -0.25415 | 0.24352 | -1.04366 | 0.29664 | 0.50689 | 0.885 |

| | | | | | | | |
|---|---|---|---|---|---|---|---|
| Rptor | 4043.86 | -0.05689 | 0.07195 | -0.79061 | 0.42917 | 0.63577 | 296.672 |
| Rptoros | 16.8209 | 0.543556 | 0.41836 | 1.29924 | 0.19386 | 0.39087 | 13.703 |
| Rpusd1 | 1554.71 | 0.013779 | 0.07205 | 0.191242 | 0.84834 | 0.92291 | 3.795 |
| Rpusd2 | 279.906 | 0.034827 | 0.12966 | 0.268591 | 0.78824 | 0.89 | 4.98 |
| Rpusd3 | 63.3234 | -0.17307 | 0.22814 | -0.75861 | 0.44809 | 0.65167 | 4.022 |
| Rpusd4 | 307.596 | 0.055596 | 0.11753 | 0.473049 | 0.63618 | 0.79507 | 9.867 |
| Rrad | 26.1317 | -0.11612 | 0.37303 | -0.3113 | 0.75557 | 0.87148 | 3.258 |
| Rraga | 3609.64 | 0.049468 | 0.08804 | 0.561879 | 0.5742 | 0.75253 | 1.618 |
| Rragb | 3475.52 | 0.091556 | 0.07437 | 1.231079 | 0.21829 | 0.42038 | 31.963 |
| Rragc | 2680.44 | 0.01392 | 0.08427 | 0.165191 | 0.86879 | 0.93346 | 19.552 |
| Rragd | 11012.4 | 0.32293 | 0.14904 | 2.166736 | 0.03025 | 0.11875 | 39.144 |
| Rras | 307.363 | -0.48514 | 0.11538 | -4.20481 | 2.61E-05 | 0.00106 | 3.687 |
| Rras2 | 381.774 | -0.69971 | 0.18171 | -3.85062 | 0.00012 | 0.003 | 71 |
| Rrbp1 | 2818.41 | -0.24031 | 0.08161 | -2.94474 | 0.00323 | 0.02743 | 63.869 |
| Rreb1 | 6033.84 | 1.260884 | 0.32523 | 3.876902 | 0.00011 | 0.00279 | 173.606 |
| Rrh | 19.809 | 0.088621 | 0.43088 | 0.205675 | 0.83705 | 0.91629 | 18.18 |
| Rrm1 | 925.36 | -0.17607 | 0.09655 | -1.8237 | 0.0682 | 0.2027 | 28.077 |
| Rrm2 | 76.1219 | -0.89052 | 0.27667 | -3.21872 | 0.00129 | 0.0149 | 5.906 |
| Rrm2b | 2327.98 | 0.170937 | 0.09664 | 1.768803 | 0.07693 | 0.21936 | 37.367 |
| Rrn3 | 3249.52 | 0.085223 | 0.08925 | 0.954919 | 0.33962 | 0.55048 | 34.132 |
| Rrnad1 | 2244.23 | 0.156173 | 0.07174 | 2.176845 | 0.02949 | 0.1168 | 8.117 |
| Rrp1 | 12847.2 | -0.03249 | 0.05909 | -0.5498 | 0.58246 | 0.75787 | 12.66 |
| Rrp12 | 2064.63 | 0.012787 | 0.06615 | 0.193316 | 0.84671 | 0.92193 | 33.323 |
| Rrp15 | 428.274 | 0.069296 | 0.11622 | 0.596247 | 0.55101 | 0.73439 | 28.381 |
| Rrp1b | 1981.01 | 0.400002 | 0.07901 | 5.062683 | 4.13E-07 | 5.47E-05 | 26.766 |
| Rrp36 | 685.132 | -0.06034 | 0.10051 | -0.60036 | 0.54827 | 0.73231 | 8.029 |
| Rrp7a | 2530.77 | -0.02781 | 0.06577 | -0.42278 | 0.67246 | 0.8199 | 9.369 |
| Rrp8 | 767.673 | 0.082435 | 0.1266 | 0.651149 | 0.51495 | 0.70533 | 5.656 |
| Rrp9 | 985.266 | 0.115757 | 0.09306 | 1.243938 | 0.21352 | 0.41479 | 9.462 |
| Rrs1 | 1000.2 | -0.00955 | 0.07231 | -0.13206 | 0.89493 | 0.94736 | 2.048 |
| Rs1 | 16.731 | -0.03181 | 0.51353 | -0.06194 | 0.95061 | 0.97565 | 31.651 |
| Rsad1 | 674.819 | -0.03664 | 0.09418 | -0.38909 | 0.69721 | 0.8358 | 9.458 |
| Rsad2 | 79.3231 | -0.71316 | 0.27798 | -2.56547 | 0.0103 | 0.05861 | 13.707 |
| Rsbn1 | 1377.17 | 0.301545 | 0.09341 | 3.228049 | 0.00125 | 0.01466 | 52.517 |

| | | | | | | | |
|---|---|---|---|---|---|---|---|
| Rsbn1l | 1921.17 | 0.191595 | 0.09142 | 2.09588 | 0.03609 | 0.13377 | 63.371 |
| Rsf1 | 3743.96 | 0.206939 | 0.11438 | 1.809284 | 0.07041 | 0.20732 | 112.89 |
| Rskr | 138.669 | 0.134012 | 0.17665 | 0.758635 | 0.44807 | 0.65167 | 5.965 |
| Rsl1 | 222.438 | -0.14672 | 0.13947 | -1.05197 | 0.29281 | 0.50266 | 10.863 |
| Rsl1d1 | 2043.65 | 0.094891 | 0.0634 | 1.496733 | 0.13446 | 0.31432 | 10.362 |
| Rsl24d1 | 1081.89 | -0.07832 | 0.09264 | -0.84549 | 0.39784 | 0.60678 | 9.908 |
| Rsph1 | 113.255 | 0.140602 | 0.18335 | 0.766859 | 0.44317 | 0.6477 | 22.365 |
| Rsph10b | 254.929 | -0.11346 | 0.13518 | -0.83933 | 0.40128 | 0.60993 | 52.685 |
| Rsph3a | 284.623 | -0.19678 | 0.19398 | -1.01442 | 0.31038 | 0.52052 | 34.211 |
| Rsph3b | 286.716 | 0.380846 | 0.13814 | 2.756952 | 0.00583 | 0.04045 | 44.27 |
| Rsph4a | 113.244 | 0.256591 | 0.23115 | 1.11007 | 0.26697 | 0.47627 | 10.911 |
| Rsph6a | 20.1648 | -0.40622 | 0.46522 | -0.87317 | 0.38257 | 0.59199 | 19.758 |
| Rsph9 | 328.023 | 0.019782 | 0.13845 | 0.142883 | 0.88638 | 0.94338 | 22.182 |
| Rspo1 | 146.355 | 0.291136 | 0.31993 | 0.910009 | 0.36282 | 0.57322 | 22.67 |
| Rspo2 | 80.9701 | -1.05944 | 0.31521 | -3.36106 | 0.00078 | 0.01076 | 150.033 |
| Rspo3 | 274.151 | -1.04696 | 0.40874 | -2.56141 | 0.01042 | 0.05907 | 83.452 |
| Rspry1 | 1443.29 | -0.03594 | 0.07876 | -0.45626 | 0.64821 | 0.80406 | 58.339 |
| Rsrc1 | 2096.42 | -0.07947 | 0.07283 | -1.0911 | 0.27523 | 0.48468 | 377.007 |
| Rsrc2 | 4752.25 | 0.113273 | 0.12036 | 0.941105 | 0.34665 | 0.55727 | 20.989 |
| Rsrp1 | 16112.5 | -0.17203 | 0.14941 | -1.15142 | 0.24956 | 0.45697 | 4.08 |
| Rsu1 | 676.766 | -0.23217 | 0.11285 | -2.05735 | 0.03965 | 0.14223 | 194.595 |
| Rtbdn | 101.388 | -0.13175 | 0.25135 | -0.52416 | 0.60017 | 0.77078 | 9.613 |
| Rtca | 1725.92 | -0.08569 | 0.06765 | -1.26652 | 0.20533 | 0.40437 | 19.246 |
| Rtcb | 3849.36 | 0.032058 | 0.06947 | 0.461463 | 0.64447 | 0.80171 | 19.187 |
| Rtel1 | 1329.88 | 0.240362 | 0.12298 | 1.954408 | 0.05065 | 0.1674 | 36.878 |
| Rtf1 | 4300.88 | 0.039318 | 0.06198 | 0.634404 | 0.52582 | 0.71407 | 60.34 |
| Rtf2 | 2522.03 | -0.1062 | 0.06526 | -1.62736 | 0.10366 | 0.26622 | 29.353 |
| Rtkn | 811.68 | -0.18047 | 0.09606 | -1.87874 | 0.06028 | 0.18732 | 17.117 |
| Rtkn2 | 229.773 | -0.39885 | 0.15647 | -2.5491 | 0.0108 | 0.06051 | 80.171 |
| Rtl1 | 834.701 | 0.676666 | 0.10618 | 6.372763 | 1.86E-10 | 1.64E-07 | 11.138 |
| Rtl10 | 47.9454 | 0.214577 | 0.30019 | 0.714814 | 0.47472 | 0.67282 | 2.758 |
| Rtl4 | 72.8167 | 0.112965 | 0.26996 | 0.418446 | 0.67562 | 0.82192 | 433.505 |
| Rtl5 | 1760.88 | 0.514657 | 0.15773 | 3.262821 | 0.0011 | 0.01364 | 4.761 |
| Rtl6 | 3243.39 | -0.092 | 0.0752 | -1.22335 | 0.2212 | 0.42412 | 4.426 |

| | | | | | | | |
|---|---|---|---|---|---|---|---|
| Rtl8a | 1023.18 | 0.021186 | 0.09616 | 0.220318 | 0.82562 | 0.91 | 1.195 |
| Rtl8b | 1106.12 | 0.00802 | 0.08101 | 0.099008 | 0.92113 | 0.96008 | 1.232 |
| Rtl8c | 3179.94 | 0.172108 | 0.1036 | 1.66129 | 0.09666 | 0.25469 | 0.401 |
| Rtl9 | 126.707 | 1.309194 | 0.305 | 4.292505 | 1.77E-05 | 0.00081 | 4.704 |
| Rtn1 | 38699.3 | -0.10921 | 0.07059 | -1.54715 | 0.12183 | 0.2953 | 197.303 |
| Rtn2 | 3231.92 | -0.30124 | 0.09555 | -3.15265 | 0.00162 | 0.01748 | 13.537 |
| Rtn3 | 41018.4 | -0.1853 | 0.07347 | -2.52195 | 0.01167 | 0.06352 | 57.381 |
| Rtn4 | 32843.5 | -0.18827 | 0.10013 | -1.88029 | 0.06007 | 0.18685 | 51.385 |
| Rtn4ip1 | 712.291 | -6.45E-06 | 0.08489 | -7.60E-05 | 0.99994 | 0.99999 | 55.395 |
| Rtn4r | 536.354 | -1.0543 | 0.39227 | -2.68772 | 0.00719 | 0.04635 | 24.767 |
| Rtn4rl1 | 2049.42 | -0.21396 | 0.10817 | -1.97798 | 0.04793 | 0.16115 | 73.987 |
| Rtn4rl2 | 558.809 | -1.19247 | 0.51834 | -2.30055 | 0.02142 | 0.09374 | 14.787 |
| Rtp1 | 59.5127 | -0.23881 | 0.41984 | -0.56881 | 0.56948 | 0.74902 | 4.828 |
| Rtp4 | 90.6848 | -0.41368 | 0.27083 | -1.52742 | 0.12666 | 0.30249 | 93.932 |
| Rtraf | 1794.67 | 0.014288 | 0.08381 | 0.170471 | 0.86464 | 0.93153 | 12.474 |
| Rttn | 367.036 | 0.149095 | 0.12317 | 1.210448 | 0.22611 | 0.43018 | 159.227 |
| Rubcn | 5343.66 | 0.153644 | 0.06413 | 2.395711 | 0.01659 | 0.07965 | 56.064 |
| Rubcnl | 51.7889 | -0.33652 | 0.28619 | -1.17584 | 0.23966 | 0.44563 | 36.506 |
| Rufy1 | 1735.1 | -0.23316 | 0.09314 | -2.50328 | 0.0123 | 0.06575 | 41.84 |
| Rufy2 | 4890.19 | -0.02137 | 0.0909 | -0.23507 | 0.81416 | 0.90456 | 36.988 |
| Rufy3 | 11999.5 | 0.041733 | 0.07744 | 0.538941 | 0.58993 | 0.76292 | 86.353 |
| Rundc1 | 2518.29 | 0.140749 | 0.0805 | 1.748334 | 0.08041 | 0.22564 | 10.589 |
| Rundc3a | 9748.74 | -0.15101 | 0.07465 | -2.02298 | 0.04308 | 0.14979 | 9.153 |
| Rundc3b | 2370.2 | 0.407944 | 0.11688 | 3.490196 | 0.00048 | 0.00774 | 132.619 |
| Runx1 | 86.6082 | -0.31856 | 0.28884 | -1.1029 | 0.27007 | 0.47962 | 224.684 |
| Runx1t1 | 2241.79 | 0.451576 | 0.1458 | 3.097274 | 0.00195 | 0.01979 | 150.214 |
| Runx2 | 399.612 | -0.83249 | 0.41082 | -2.0264 | 0.04272 | 0.14901 | 318.811 |
| Rusc1 | 2053.04 | -0.18966 | 0.10662 | -1.77878 | 0.07528 | 0.21642 | 9.331 |
| Rusc2 | 5548.86 | -0.16883 | 0.07572 | -2.22964 | 0.02577 | 0.10621 | 45.11 |
| Rusf1 | 582.053 | -0.04585 | 0.08814 | -0.52017 | 0.60295 | 0.7726 | 26.792 |
| Ruvbl1 | 1495.98 | 0.141092 | 0.10066 | 1.401615 | 0.16103 | 0.34918 | 32.164 |
| Ruvbl2 | 963.053 | 0.052936 | 0.1019 | 0.519515 | 0.6034 | 0.77292 | 16.337 |
| Rwdd1 | 879.111 | 0.061484 | 0.07958 | 0.772628 | 0.43974 | 0.64503 | 23.07 |
| Rwdd2a | 450.597 | -0.13479 | 0.1342 | -1.00437 | 0.3152 | 0.52568 | 2.909 |

| | | | | | | | |
|---|---|---|---|---|---|---|---|
| Rwdd2b | 490.297 | 0.073852 | 0.1147 | 0.643877 | 0.51966 | 0.70925 | 7.167 |
| Rwdd3 | 310.724 | 0.223475 | 0.13682 | 1.633345 | 0.1024 | 0.26421 | 16.298 |
| Rwdd4a | 1748.63 | 0.046 | 0.07028 | 0.654544 | 0.51276 | 0.70361 | 19.292 |
| Rxfp1 | 528.315 | -1.33863 | 0.47167 | -2.83809 | 0.00454 | 0.0341 | 96.27 |
| Rxfp2 | 18.8009 | 0.321209 | 0.58623 | 0.54792 | 0.58375 | 0.75877 | 63.51 |
| Rxfp3 | 89.8114 | -0.66691 | 0.25555 | -2.6097 | 0.00906 | 0.0537 | 4.275 |
| Rxra | 2139.41 | -0.13846 | 0.09503 | -1.45704 | 0.14511 | 0.32787 | 86.518 |
| Rxrb | 2905.12 | -0.03093 | 0.0783 | -0.39503 | 0.69282 | 0.83344 | 6.588 |
| Rxrg | 2296.34 | 0.581051 | 0.19848 | 2.927456 | 0.00342 | 0.02827 | 41.24 |
| Rxylt1 | 833.12 | 6.81E-05 | 0.0828 | 0.000822 | 0.99934 | 0.99967 | 19.258 |
| Rybp | 1707.47 | 0.210712 | 0.09617 | 2.190947 | 0.02846 | 0.11375 | 58.921 |
| Ryk | 1447.54 | -0.05622 | 0.07141 | -0.78719 | 0.43117 | 0.63725 | 73.389 |
| Ryr1 | 3616.78 | 1.01322 | 0.20655 | 4.905431 | 9.32E-07 | 0.0001 | 121.836 |
| Ryr2 | 9622.82 | -0.11515 | 0.07675 | -1.50035 | 0.13352 | 0.31328 | 553.844 |
| Ryr3 | 9456.22 | 0.637992 | 0.26782 | 2.382188 | 0.01721 | 0.08153 | 585.742 |
| S100a1 | 1669.46 | -0.4363 | 0.11012 | -3.96205 | 7.43E-05 | 0.00218 | 3.359 |
| S100a10 | 371.803 | -0.95689 | 0.20209 | -4.73497 | 2.19E-06 | 0.00019 | 9.564 |
| S100a11 | 76.2604 | -0.81946 | 0.32518 | -2.52002 | 0.01173 | 0.06375 | 5.8 |
| S100a13 | 465.876 | -0.4155 | 0.14018 | -2.96399 | 0.00304 | 0.02632 | 10.147 |
| S100a16 | 1746.21 | -0.29077 | 0.10718 | -2.71291 | 0.00667 | 0.04408 | 5.898 |
| S100a6 | 239.96 | -0.71136 | 0.26213 | -2.71381 | 0.00665 | 0.04399 | 11.3 |
| S100a7a | 13.2337 | 0.445979 | 0.46548 | 0.958105 | 0.33801 | 0.54883 | 4.653 |
| S100b | 3670.79 | -0.51451 | 0.11185 | -4.60018 | 4.22E-06 | 0.0003 | 7.307 |
| S100pbp | 4806.46 | 0.419961 | 0.11774 | 3.566746 | 0.00036 | 0.00637 | 41.723 |
| S1pr1 | 7319.64 | -0.28828 | 0.09002 | -3.20256 | 0.00136 | 0.01546 | 4.64 |
| S1pr2 | 85.8401 | -0.09937 | 0.20363 | -0.48798 | 0.62556 | 0.7879 | 14.421 |
| S1pr3 | 243.17 | -0.64057 | 0.19697 | -3.2522 | 0.00115 | 0.01387 | 14.159 |
| S1pr4 | 22.4796 | -0.32804 | 0.36947 | -0.88789 | 0.3746 | 0.58497 | 2.386 |
| S1pr5 | 453.729 | -0.78273 | 0.23117 | -3.38591 | 0.00071 | 0.01007 | 5.532 |
| Saal1 | 603.872 | -0.10282 | 0.08346 | -1.23202 | 0.21794 | 0.41996 | 24.573 |
| Sac3d1 | 330.812 | 0.098893 | 0.12064 | 0.819744 | 0.41236 | 0.62038 | 2.651 |
| Sacm1l | 1981.48 | 0.149893 | 0.09854 | 1.521101 | 0.12823 | 0.30522 | 62.842 |
| Sacs | 4976.87 | 0.240836 | 0.18746 | 1.284702 | 0.1989 | 0.39678 | 102.239 |
| Sae1 | 3384.37 | 0.020848 | 0.06928 | 0.300915 | 0.76348 | 0.87611 | 67.573 |

| | | | | | | | |
|---|---|---|---|---|---|---|---|
| Safb | 9432.03 | 0.178341 | 0.05971 | 2.986851 | 0.00282 | 0.02517 | 21.47 |
| Safb2 | 8055.58 | 0.182581 | 0.07323 | 2.493144 | 0.01266 | 0.06707 | 23.621 |
| Sag | 104.292 | -0.06653 | 0.189 | -0.35201 | 0.72483 | 0.85334 | 41.479 |
| Sall1 | 1200.99 | -0.2946 | 0.13227 | -2.22719 | 0.02593 | 0.10668 | 16.928 |
| Sall2 | 3315.63 | 0.193189 | 0.12146 | 1.590529 | 0.11172 | 0.27929 | 17.591 |
| Sall3 | 488.446 | -0.4236 | 0.15484 | -2.73567 | 0.00623 | 0.0422 | 20.396 |
| Samd1 | 962.172 | -0.09894 | 0.09401 | -1.05249 | 0.29258 | 0.50248 | 2.909 |
| Samd10 | 586.562 | 0.099147 | 0.09805 | 1.011141 | 0.31195 | 0.52237 | 3.995 |
| Samd11 | 124.271 | 0.633383 | 0.23952 | 2.644346 | 0.00818 | 0.05027 | 9.295 |
| Samd12 | 4291.44 | 0.423548 | 0.13745 | 3.081433 | 0.00206 | 0.02049 | 448.728 |
| Samd14 | 2216.7 | 0.262279 | 0.11189 | 2.344006 | 0.01908 | 0.08673 | 16.209 |
| Samd15 | 189.949 | 0.396346 | 0.20704 | 1.914352 | 0.05558 | 0.1779 | 13.819 |
| Samd3 | 48.0192 | -1.3195 | 0.44638 | -2.95603 | 0.00312 | 0.02673 | 42.466 |
| Samd4 | 1698.86 | 0.110779 | 0.1049 | 1.056058 | 0.29094 | 0.50114 | 222.962 |
| Samd4b | 3272.99 | -0.22962 | 0.09054 | -2.53618 | 0.01121 | 0.06178 | 198.623 |
| Samd5 | 677.948 | 0.272814 | 0.19301 | 1.413463 | 0.15752 | 0.34464 | 52.273 |
| Samd8 | 4672.71 | 0.032068 | 0.06632 | 0.483498 | 0.62874 | 0.79006 | 48.196 |
| Samd9l | 414.003 | -0.21942 | 0.49693 | -0.44156 | 0.65881 | 0.81094 | 27.316 |
| Samhd1 | 1372.54 | 0.080251 | 0.07447 | 1.077649 | 0.28119 | 0.49107 | 37.733 |
| Samm50 | 4498.48 | 0.040296 | 0.05477 | 0.735762 | 0.46188 | 0.66232 | 25.027 |
| Samsn1 | 51.6241 | -0.11652 | 0.3336 | -0.3493 | 0.72687 | 0.85431 | 163.489 |
| Sanbr | 2109.42 | 0.042393 | 0.08309 | 0.510211 | 0.6099 | 0.77731 | 68.679 |
| Sap130 | 2271.49 | 0.135935 | 0.07891 | 1.722584 | 0.08496 | 0.23382 | 88.691 |
| Sap18 | 3143.66 | -0.05788 | 0.08426 | -0.68695 | 0.49211 | 0.68679 | 8.459 |
| Sap18b | 392.571 | 0.084706 | 0.1256 | 0.674431 | 0.50004 | 0.69306 | 0.789 |
| Sap30 | 141.579 | 0.018562 | 0.16897 | 0.109858 | 0.91252 | 0.9559 | 5.154 |
| Sap30bp | 2606.53 | -0.04082 | 0.08202 | -0.49776 | 0.61866 | 0.78312 | 33.444 |
| Sap30bpos | 102.769 | 0.179598 | 0.22609 | 0.79438 | 0.42697 | 0.63344 | 5.656 |
| Sap30l | 925.256 | -0.1103 | 0.09895 | -1.11464 | 0.265 | 0.47405 | 8.581 |
| Sapcd2 | 56.2042 | 0.145394 | 0.32011 | 0.454196 | 0.64969 | 0.80496 | 5.893 |
| Sar1a | 5768.7 | -0.02615 | 0.06335 | -0.41276 | 0.67978 | 0.82495 | 12.999 |
| Sar1b | 2353.46 | -0.03118 | 0.0769 | -0.4054 | 0.68518 | 0.82845 | 28.239 |
| Saraf | 8670.36 | -0.14627 | 0.07367 | -1.98558 | 0.04708 | 0.1592 | 16.283 |
| Sardh | 559.569 | -0.2248 | 0.11008 | -2.04215 | 0.04114 | 0.14535 | 59.945 |

| | | | | | | | |
|---|---|---|---|---|---|---|---|
| Sarm1 | 1233.92 | -0.14235 | 0.08426 | -1.68954 | 0.09112 | 0.24519 | 25.425 |
| Sarnp | 698.754 | 0.051445 | 0.18315 | 0.280889 | 0.7788 | 0.88465 | 60.297 |
| Sars | 5624.89 | -0.19211 | 0.05995 | -3.20473 | 0.00135 | 0.01537 | 20.345 |
| Sars2 | 473.113 | -0.04276 | 0.10998 | -0.38882 | 0.69741 | 0.83599 | 11.88 |
| Sart1 | 3235.03 | -0.04725 | 0.0724 | -0.65266 | 0.51397 | 0.7047 | 11.181 |
| Sart3 | 2711.08 | 0.110323 | 0.06234 | 1.769802 | 0.07676 | 0.21911 | 30.065 |
| Sash1 | 3513.1 | -0.12828 | 0.10626 | -1.20725 | 0.22734 | 0.43151 | 163.832 |
| Sash3 | 64.0858 | -0.21977 | 0.23993 | -0.91598 | 0.35968 | 0.56997 | 15.13 |
| Sass6 | 188.102 | -0.14365 | 0.1595 | -0.90064 | 0.36778 | 0.57819 | 36.023 |
| Sat1 | 1985.68 | -0.11791 | 0.083 | -1.42059 | 0.15544 | 0.34204 | 3.318 |
| Sat2 | 242.379 | -0.32402 | 0.15797 | -2.05113 | 0.04025 | 0.14335 | 1.847 |
| Satb1 | 1589.98 | -0.61985 | 0.1845 | -3.35956 | 0.00078 | 0.01076 | 97.104 |
| Satb2 | 171.323 | -1.14727 | 0.51139 | -2.24344 | 0.02487 | 0.10356 | 184.67 |
| Sav1 | 646.938 | -0.28749 | 0.12633 | -2.27567 | 0.02287 | 0.09783 | 21.991 |
| Saxo2 | 122.31 | 0.377862 | 0.20019 | 1.887486 | 0.05909 | 0.18507 | 15.569 |
| Saysd1 | 249.131 | -0.23774 | 0.15661 | -1.51805 | 0.129 | 0.30621 | 62.899 |
| Sbds | 2530.83 | -0.1166 | 0.07309 | -1.59532 | 0.11064 | 0.27744 | 9.8 |
| Sbf1 | 8577.3 | 0.007706 | 0.06385 | 0.120681 | 0.90394 | 0.95146 | 27.076 |
| Sbf2 | 4231.17 | 0.10162 | 0.06592 | 1.541509 | 0.12319 | 0.29739 | 306.91 |
| Sbk1 | 4968.23 | -0.16687 | 0.09561 | -1.74531 | 0.08093 | 0.22645 | 46.155 |
| Sbk2 | 14.0619 | -0.70426 | 0.5605 | -1.25648 | 0.20894 | 0.40901 | 23.895 |
| Sbk3 | 139.394 | 0.532827 | 0.22852 | 2.331662 | 0.01972 | 0.08871 | 5.909 |
| Sbno1 | 9852.85 | 0.088947 | 0.07218 | 1.23235 | 0.21782 | 0.4199 | 57.3 |
| Sbno2 | 1350.17 | -0.13967 | 0.0739 | -1.89011 | 0.05874 | 0.1844 | 48.58 |
| Sbsn | 608.314 | 0.28691 | 0.13032 | 2.201651 | 0.02769 | 0.11172 | 4.664 |
| Sbspon | 32.3895 | -0.46112 | 0.39201 | -1.1763 | 0.23948 | 0.44543 | 38.861 |
| Sc5d | 3810.55 | -0.17583 | 0.08833 | -1.99055 | 0.04653 | 0.1579 | 12.662 |
| Scaf1 | 7595.74 | -0.095 | 0.07303 | -1.30082 | 0.19332 | 0.39022 | 13.416 |
| Scaf11 | 3789.71 | 0.033244 | 0.07484 | 0.444181 | 0.65691 | 0.80944 | 49.145 |
| Scaf4 | 2814.9 | 0.212049 | 0.0829 | 2.557927 | 0.01053 | 0.05956 | 58.824 |
| Scaf8 | 2897.27 | 0.117351 | 0.06441 | 1.821802 | 0.06849 | 0.20338 | 83.888 |
| Scai | 4137.26 | 0.175619 | 0.09619 | 1.82582 | 0.06788 | 0.20204 | 124.521 |
| Scamp1 | 14730.4 | 0.076201 | 0.09159 | 0.831974 | 0.40542 | 0.61421 | 84.548 |
| Scamp2 | 1107.57 | -0.00996 | 0.10815 | -0.09205 | 0.92666 | 0.9631 | 27.853 |

| | | | | | | | |
|---|---|---|---|---|---|---|---|
| Scamp3 | 1099.39 | -0.00039 | 0.08622 | -0.00447 | 0.99643 | 0.99805 | 5.293 |
| Scamp5 | 14930.2 | -0.26614 | 0.0692 | -3.84618 | 0.00012 | 0.00303 | 26.734 |
| Scand1 | 167.94 | -0.24276 | 0.14621 | -1.66033 | 0.09685 | 0.25496 | 0.902 |
| Scap | 3180.94 | 0.019085 | 0.06635 | 0.287642 | 0.77362 | 0.88191 | 51.663 |
| Scaper | 1714.85 | 0.278118 | 0.10451 | 2.661188 | 0.00779 | 0.04882 | 388.241 |
| Scara3 | 1377.55 | -0.442 | 0.13676 | -3.232 | 0.00123 | 0.01456 | 34.542 |
| Scara5 | 45.2639 | -0.97362 | 0.39077 | -2.49157 | 0.01272 | 0.06724 | 98.424 |
| Scarb1 | 2275.06 | 0.248516 | 0.1014 | 2.450939 | 0.01425 | 0.072 | 64.008 |
| Scarb2 | 7682.97 | -0.20524 | 0.07515 | -2.73098 | 0.00631 | 0.04258 | 65.52 |
| Scarf1 | 113.727 | -0.48714 | 0.15909 | -3.06203 | 0.0022 | 0.02139 | 13.043 |
| Scarf2 | 207.762 | 0.147614 | 0.15284 | 0.965819 | 0.33413 | 0.54479 | 11.012 |
| Scarna17 | 13.1992 | 0.442129 | 0.46448 | 0.951872 | 0.34116 | 0.55206 | 0.079 |
| Scarna2 | 14.2629 | -1.06147 | 0.53408 | -1.98747 | 0.04687 | 0.15876 | 0.414 |
| Scarna6 | 16.8635 | 0.474512 | 0.44974 | 1.055088 | 0.29139 | 0.50126 | 0.265 |
| Sccpdh | 2808.66 | -0.43018 | 0.14545 | -2.95759 | 0.0031 | 0.02664 | 18.98 |
| Scd1 | 8946.1 | -0.37685 | 0.13556 | -2.7799 | 0.00544 | 0.03841 | 13.425 |
| Scd2 | 72086 | -0.24546 | 0.08459 | -2.90191 | 0.00371 | 0.02995 | 13.191 |
| Scd3 | 172.371 | -0.85426 | 0.22144 | -3.85773 | 0.00011 | 0.00294 | 40.748 |
| Scel | 100.689 | -0.16406 | 0.26664 | -0.61527 | 0.53837 | 0.72514 | 99.456 |
| Scfd1 | 1343.71 | 0.202811 | 0.08809 | 2.302379 | 0.02131 | 0.09343 | 72.592 |
| Scfd2 | 787.708 | 0.111464 | 0.10313 | 1.080785 | 0.27979 | 0.48989 | 326.944 |
| Scg2 | 20410.7 | 0.134582 | 0.07733 | 1.740274 | 0.08181 | 0.22797 | 5.452 |
| Scg3 | 9160.59 | -0.00958 | 0.08315 | -0.11516 | 0.90832 | 0.95403 | 40.868 |
| Scg5 | 5642.55 | 0.082169 | 0.07543 | 1.089406 | 0.27598 | 0.48512 | 52.76 |
| Scgb1c1 | 12.6856 | -0.59617 | 0.53099 | -1.12275 | 0.26154 | 0.4703 | 1.223 |
| Scgn | 35.2738 | -2 | 0.57608 | -3.62903 | 0.00028 | 0.00541 | 37.759 |
| Schip1 | 202.54 | -0.62972 | 0.22112 | -2.84784 | 0.0044 | 0.03354 | 561.68 |
| Scin | 14.2899 | -1.06201 | 0.51348 | -2.06826 | 0.03862 | 0.13992 | 74.46 |
| Sclt1 | 752.64 | 0.23272 | 0.1258 | 1.849907 | 0.06433 | 0.19522 | 115.795 |
| Scly | 529.988 | -0.3156 | 0.12965 | -2.43424 | 0.01492 | 0.07412 | 22.738 |
| Scmh1 | 2354.13 | 0.262211 | 0.08553 | 3.065762 | 0.00217 | 0.02127 | 124.906 |
| Scml2 | 68.0525 | -0.1256 | 0.31008 | -0.40506 | 0.68544 | 0.8286 | 175.689 |
| Scml4 | 88.531 | -0.16665 | 0.22595 | -0.73754 | 0.46079 | 0.66142 | 100.411 |
| Scn10a | 36.4082 | -0.82028 | 0.30355 | -2.70232 | 0.00689 | 0.04491 | 110.867 |

| | | | | | | | |
|---|---|---|---|---|---|---|---|
| Scn11a | 12.8671 | 1.707883 | 0.56414 | 3.027416 | 0.00247 | 0.02314 | 71.698 |
| Scn1a | 9327.19 | -0.09671 | 0.12479 | -0.77499 | 0.43835 | 0.64363 | 170.063 |
| Scn1b | 8073.89 | -0.66765 | 0.09994 | -6.68074 | 2.38E-11 | 2.85E-08 | 10.48 |
| Scn2a | 16277.3 | 0.10043 | 0.08386 | 1.197549 | 0.23109 | 0.4356 | 146.677 |
| Scn2b | 19977 | 0.209894 | 0.08331 | 2.519298 | 0.01176 | 0.06378 | 12.289 |
| Scn3a | 4026.23 | 0.395932 | 0.10108 | 3.916865 | 8.97E-05 | 0.00251 | 110.51 |
| Scn3b | 7433.63 | 0.302481 | 0.08653 | 3.495574 | 0.00047 | 0.00765 | 22.402 |
| Scn4b | 29776.4 | 0.454614 | 0.32488 | 1.399319 | 0.16172 | 0.3499 | 15.758 |
| Scn5a | 801.317 | 0.51964 | 0.18532 | 2.804024 | 0.00505 | 0.03652 | 95.609 |
| Scn7a | 68.9581 | -0.47917 | 0.33263 | -1.44055 | 0.14971 | 0.33411 | 111.49 |
| Scn8a | 17679.1 | -0.1505 | 0.06908 | -2.17874 | 0.02935 | 0.11635 | 176.081 |
| Scn9a | 463.63 | 0.725428 | 0.21792 | 3.328848 | 0.00087 | 0.01163 | 154.883 |
| Scnm1 | 1049.57 | -0.08393 | 0.08216 | -1.02157 | 0.30699 | 0.5168 | 4.476 |
| Scnn1a | 78.6143 | -0.62666 | 0.29916 | -2.09473 | 0.0362 | 0.134 | 24.285 |
| Sco1 | 396.383 | 0.050967 | 0.10244 | 0.497508 | 0.61883 | 0.78314 | 14.401 |
| Scoc | 3692.32 | 0.088293 | 0.09707 | 0.909602 | 0.36303 | 0.57343 | 28.308 |
| Scp2 | 4565.62 | -0.04991 | 0.06416 | -0.77791 | 0.43662 | 0.64228 | 101.16 |
| Scp2-ps2 | 60.0979 | -0.04035 | 0.33174 | -0.12163 | 0.90319 | 0.95127 | 11.399 |
| Scpep1 | 535.021 | -0.34213 | 0.09516 | -3.5953 | 0.00032 | 0.00597 | 31.446 |
| Scpep1os | 19.2406 | -0.09919 | 0.37515 | -0.26441 | 0.79147 | 0.89159 | 30.134 |
| Scrg1 | 847.888 | -0.18573 | 0.11529 | -1.61097 | 0.10719 | 0.27174 | 21.663 |
| Scrib | 2526.35 | 0.127132 | 0.07415 | 1.71441 | 0.08645 | 0.23648 | 22.627 |
| Scrn1 | 13437.8 | 0.053675 | 0.07532 | 0.712635 | 0.47607 | 0.67418 | 65.317 |
| Scrn2 | 154.625 | -0.25062 | 0.13666 | -1.83387 | 0.06667 | 0.19992 | 4.021 |
| Scrn3 | 1632.41 | -0.01076 | 0.08682 | -0.12399 | 0.90132 | 0.95046 | 25.218 |
| Scrt1 | 6576.78 | -0.39622 | 0.09648 | -4.10671 | 4.01E-05 | 0.0014 | 6.297 |
| Scrt2 | 2095.84 | 0.118297 | 0.10838 | 1.09147 | 0.27507 | 0.48453 | 14.274 |
| Sctr | 15.4741 | -0.21195 | 0.53286 | -0.39776 | 0.69081 | 0.83213 | 56.643 |
| Scube1 | 1162.68 | -1.25207 | 0.38872 | -3.22101 | 0.00128 | 0.01485 | 120.023 |
| Scube2 | 78.6936 | -0.49599 | 0.24707 | -2.00746 | 0.0447 | 0.15355 | 67.004 |
| Scube3 | 1561.96 | 0.63352 | 0.19651 | 3.223839 | 0.00126 | 0.01476 | 32.714 |
| Scx | 128.007 | 0.228985 | 0.1821 | 1.257451 | 0.20859 | 0.40848 | 2.007 |
| Scyl1 | 2534.82 | 0.023611 | 0.06459 | 0.365525 | 0.71472 | 0.84705 | 13.069 |
| Scyl2 | 1888.37 | 0.136786 | 0.06369 | 2.147521 | 0.03175 | 0.12278 | 47.565 |

| | | | | | | | |
|---|---|---|---|---|---|---|---|
| Scyl3 | 1343.09 | 0.193922 | 0.08884 | 2.182928 | 0.02904 | 0.11539 | 26.027 |
| Sdad1 | 1266.61 | 0.073077 | 0.08937 | 0.817683 | 0.41354 | 0.6216 | 26.47 |
| Sdc1 | 39.9119 | -1.07343 | 0.36688 | -2.92584 | 0.00344 | 0.0284 | 22.393 |
| Sdc2 | 2960.03 | 0.372021 | 0.12534 | 2.968008 | 0.003 | 0.02616 | 114.213 |
| Sdc3 | 7339.82 | -0.15276 | 0.08874 | -1.72131 | 0.08519 | 0.23406 | 33.783 |
| Sdc4 | 4813.54 | -0.38753 | 0.11383 | -3.40448 | 0.00066 | 0.00962 | 19.641 |
| Sdcbp | 7419.55 | 0.121903 | 0.06331 | 1.925557 | 0.05416 | 0.17504 | 42.774 |
| Sdcbp2 | 162.832 | 0.109773 | 0.15795 | 0.694996 | 0.48706 | 0.68268 | 17.384 |
| Sdccag8 | 1367.08 | 0.086284 | 0.08605 | 1.002767 | 0.31597 | 0.52651 | 205.778 |
| Sde2 | 1465 | 0.086857 | 0.07229 | 1.201447 | 0.22958 | 0.43383 | 16.987 |
| Sdf2 | 1834.22 | -0.09694 | 0.07047 | -1.37572 | 0.16891 | 0.35902 | 9.751 |
| Sdf2l1 | 517.533 | -0.23287 | 0.1593 | -1.4618 | 0.1438 | 0.32616 | 2.246 |
| Sdf4 | 10163.3 | -0.012 | 0.06768 | -0.17728 | 0.85929 | 0.92893 | 20.739 |
| Sdha | 20733.7 | -0.03032 | 0.05837 | -0.51942 | 0.60347 | 0.77292 | 28.027 |
| Sdhaf1 | 22.3982 | -0.89058 | 0.51715 | -1.72209 | 0.08505 | 0.23388 | 2.49 |
| Sdhaf2 | 1777.21 | 0.018461 | 0.06434 | 0.286918 | 0.77417 | 0.88239 | 25.609 |
| Sdhaf3 | 551.547 | 0.157987 | 0.14291 | 1.105529 | 0.26893 | 0.47818 | 86.976 |
| Sdhaf4 | 310.589 | -0.18241 | 0.1261 | -1.44659 | 0.14801 | 0.33173 | 9.718 |
| Sdhb | 3735.47 | -0.09223 | 0.07128 | -1.294 | 0.19567 | 0.39335 | 17.991 |
| Sdhc | 3237.71 | -0.14023 | 0.0864 | -1.62308 | 0.10457 | 0.26745 | 23.439 |
| Sdhd | 2954.57 | -0.00905 | 0.06725 | -0.13459 | 0.89294 | 0.94632 | 7.456 |
| Sdk1 | 397.614 | 0.161167 | 0.14362 | 1.122153 | 0.2618 | 0.47055 | 974.097 |
| Sdk2 | 1743.22 | -0.04414 | 0.11299 | -0.39066 | 0.69605 | 0.83541 | 290.673 |
| Sdr39u1 | 976.6 | -0.04258 | 0.09794 | -0.43474 | 0.66375 | 0.81417 | 3.011 |
| Sdr42e1 | 361.378 | -0.13683 | 0.12106 | -1.13032 | 0.25834 | 0.46635 | 12.291 |
| Sds | 36.7867 | 0.520562 | 0.40295 | 1.291889 | 0.1964 | 0.39399 | 7.407 |
| Sdsl | 110.449 | 0.111718 | 0.19742 | 0.565882 | 0.57147 | 0.75033 | 14.625 |
| Sebox | 58.7606 | -0.07793 | 0.37823 | -0.20605 | 0.83675 | 0.91619 | 1.635 |
| Sec1 | 28.8535 | -1.16397 | 0.46645 | -2.49539 | 0.01258 | 0.06668 | 16.717 |
| Sec11a | 1107.84 | -0.10289 | 0.08567 | -1.2009 | 0.22979 | 0.43407 | 42.892 |
| Sec11c | 1869.89 | -0.25186 | 0.08906 | -2.82795 | 0.00468 | 0.03485 | 17.17 |
| Sec13 | 1751.59 | 0.019911 | 0.07503 | 0.265389 | 0.79071 | 0.8913 | 12.682 |
| Sec14l1 | 11748.2 | 0.064136 | 0.06699 | 0.957458 | 0.33834 | 0.54905 | 44.101 |
| Sec14l2 | 1833.69 | -0.34199 | 0.08735 | -3.91532 | 9.03E-05 | 0.00251 | 26.377 |

| | | | | | | | |
|---|---|---|---|---|---|---|---|
| Sec14l3 | 26.5534 | 0.755484 | 0.46703 | 1.617619 | 0.10574 | 0.26933 | 12.896 |
| Sec14l5 | 2204.4 | -0.21955 | 0.1116 | -1.96728 | 0.04915 | 0.16405 | 40.163 |
| Sec16a | 4587.36 | 0.123633 | 0.05836 | 2.118303 | 0.03415 | 0.12905 | 35.786 |
| Sec16b | 76.3133 | -0.82457 | 0.37159 | -2.21902 | 0.02649 | 0.10828 | 61.698 |
| Sec22a | 761.8 | 0.130471 | 0.08267 | 1.578241 | 0.11451 | 0.28343 | 52.788 |
| Sec22b | 2832.54 | 0.010501 | 0.06952 | 0.151044 | 0.87994 | 0.93998 | 22.087 |
| Sec22c | 2799.77 | 0.22671 | 0.06088 | 3.723689 | 0.0002 | 0.00421 | 25.446 |
| Sec23a | 4426.41 | -0.03485 | 0.07369 | -0.4729 | 0.63629 | 0.79512 | 53.635 |
| Sec23b | 1978.44 | -0.06529 | 0.0807 | -0.80907 | 0.41847 | 0.62566 | 34.521 |
| Sec23ip | 2163.89 | 0.032865 | 0.07107 | 0.462423 | 0.64378 | 0.80129 | 39.894 |
| Sec24a | 1740.39 | 0.146334 | 0.07654 | 1.911923 | 0.05589 | 0.17867 | 71.371 |
| Sec24b | 3372.87 | 0.223776 | 0.06983 | 3.20469 | 0.00135 | 0.01537 | 78.795 |
| Sec24c | 4626.18 | 0.03808 | 0.05682 | 0.670132 | 0.50277 | 0.69537 | 20.545 |
| Sec24d | 710.366 | -0.36098 | 0.19813 | -1.82198 | 0.06846 | 0.20333 | 98.187 |
| Sec31a | 4525.81 | 0.021895 | 0.06662 | 0.328662 | 0.74241 | 0.86432 | 54.586 |
| Sec31b | 17.2246 | -0.19973 | 0.44356 | -0.4503 | 0.6525 | 0.80668 | 28.908 |
| Sec61a1 | 2308.24 | -0.01602 | 0.07405 | -0.21641 | 0.82867 | 0.91212 | 15.327 |
| Sec61a2 | 3119.58 | 0.120673 | 0.06316 | 1.910618 | 0.05605 | 0.17898 | 24.446 |
| Sec61b | 521.906 | -0.19271 | 0.11371 | -1.69475 | 0.09012 | 0.24355 | 8.585 |
| Sec61g | 819.385 | -0.10501 | 0.10153 | -1.03429 | 0.301 | 0.51075 | 7.955 |
| Sec62 | 7206.12 | -0.04837 | 0.12007 | -0.4029 | 0.68703 | 0.82926 | 28.389 |
| Sec63 | 4718.21 | -0.03082 | 0.08113 | -0.37988 | 0.70404 | 0.84028 | 71.019 |
| Secisbp2 | 1253.75 | 0.055886 | 0.08767 | 0.637438 | 0.52384 | 0.7122 | 32.348 |
| Secisbp2l | 8109.55 | -0.04592 | 0.06467 | -0.71015 | 0.47761 | 0.67546 | 45.885 |
| Seh1l | 1607.09 | 0.086923 | 0.08166 | 1.064445 | 0.28713 | 0.49718 | 20.612 |
| Sel1l | 10151.5 | 0.001064 | 0.0693 | 0.015353 | 0.98775 | 0.99316 | 43.115 |
| Sel1l3 | 1590.62 | -0.84046 | 0.18576 | -4.52434 | 6.06E-06 | 0.00038 | 106.845 |
| Selenbp1 | 405.072 | -0.21038 | 0.11886 | -1.76996 | 0.07673 | 0.2191 | 11.703 |
| Selenof | 6421.65 | -0.12363 | 0.08669 | -1.42607 | 0.15385 | 0.33962 | 27.377 |
| Selenoh | 316.381 | -0.36809 | 0.15827 | -2.3257 | 0.02003 | 0.08968 | 1.556 |
| Selenoi | 5095.85 | 0.268627 | 0.09384 | 2.862697 | 0.0042 | 0.03252 | 39.847 |
| Selenok | 3851.31 | -0.17865 | 0.07117 | -2.51027 | 0.01206 | 0.06482 | 7.355 |
| Selenok-ps1 | 32.2082 | 0.456971 | 0.35643 | 1.282066 | 0.19982 | 0.39799 | 0.785 |
| Selenom | 1833 | -0.2662 | 0.10194 | -2.61143 | 0.00902 | 0.05349 | 2.668 |

| | | | | | | | |
|---|---|---|---|---|---|---|---|
| Selenon | 1206.79 | -0.11615 | 0.09783 | -1.1873 | 0.23511 | 0.44032 | 14.275 |
| Selenoo | 534.275 | -0.14257 | 0.10485 | -1.35968 | 0.17393 | 0.36536 | 11.257 |
| Selenop | 9548.52 | -0.36271 | 0.07343 | -4.93967 | 7.83E-07 | 9.01E-05 | 11.962 |
| Selenos | 1244.64 | -0.13961 | 0.09242 | -1.51056 | 0.1309 | 0.30933 | 9.757 |
| Selenot | 10873.3 | -0.00151 | 0.06748 | -0.02234 | 0.98218 | 0.99059 | 16.498 |
| Selenov | 94.7864 | 0.778908 | 0.28999 | 2.685996 | 0.00723 | 0.04653 | 6.535 |
| Selenow | 17079.2 | -0.34021 | 0.08917 | -3.81519 | 0.00014 | 0.00331 | 5.195 |
| Selplg | 1208.37 | -0.20632 | 0.13225 | -1.56003 | 0.11875 | 0.29077 | 14.109 |
| Sem1 | 1111.87 | 0.00824 | 0.08408 | 0.098002 | 0.92193 | 0.96067 | 21.37 |
| Sema3a | 1340.85 | -0.12155 | 0.15144 | -0.80263 | 0.42219 | 0.62898 | 477.153 |
| Sema3b | 164.37 | -0.68947 | 0.2209 | -3.12117 | 0.0018 | 0.01882 | 11.556 |
| Sema3c | 1048.83 | -0.28967 | 0.18517 | -1.56434 | 0.11774 | 0.28887 | 155.988 |
| Sema3d | 410.64 | -0.45271 | 0.21614 | -2.09446 | 0.03622 | 0.13402 | 205.564 |
| Sema3e | 2171.3 | 0.228441 | 0.11787 | 1.938027 | 0.05262 | 0.17162 | 231.414 |
| Sema3f | 774.401 | 0.89046 | 0.20858 | 4.269233 | 1.96E-05 | 0.00087 | 28.976 |
| Sema3g | 413.079 | -0.606 | 0.12248 | -4.94786 | 7.50E-07 | 8.82E-05 | 12.493 |
| Sema4a | 2273.74 | 0.051013 | 0.09242 | 0.55198 | 0.58096 | 0.75714 | 25.224 |
| Sema4b | 862.421 | 0.012678 | 0.08731 | 0.145217 | 0.88454 | 0.94257 | 39.687 |
| Sema4c | 580.562 | 0.116724 | 0.12011 | 0.971785 | 0.33116 | 0.54208 | 9.711 |
| Sema4d | 2724.18 | -0.34698 | 0.10976 | -3.16125 | 0.00157 | 0.01718 | 108.219 |
| Sema4f | 1953.34 | 0.059234 | 0.07403 | 0.800128 | 0.42364 | 0.63046 | 27.885 |
| Sema4g | 1080.48 | 0.034775 | 0.11908 | 0.29203 | 0.77026 | 0.87996 | 14.297 |
| Sema5a | 1574.24 | -0.23177 | 0.09133 | -2.53765 | 0.01116 | 0.0616 | 451.532 |
| Sema5b | 2433.58 | 0.265024 | 0.14602 | 1.815035 | 0.06952 | 0.20549 | 123.588 |
| Sema6a | 1040.31 | -0.20466 | 0.12406 | -1.64968 | 0.09901 | 0.25857 | 133.273 |
| Sema6b | 5002.2 | -0.03967 | 0.06171 | -0.64291 | 0.52028 | 0.70963 | 17.259 |
| Sema6c | 841.624 | -0.10474 | 0.11788 | -0.88854 | 0.37425 | 0.58452 | 13.568 |
| Sema6d | 3957.43 | -0.31219 | 0.10858 | -2.87524 | 0.00404 | 0.03169 | 577.802 |
| Sema7a | 4389.21 | -0.54274 | 0.10242 | -5.29898 | 1.16E-07 | 2.01E-05 | 22.754 |
| Semp2l1 | 30.8641 | 1.052478 | 0.4499 | 2.339383 | 0.01932 | 0.08745 | 3.609 |
| Senp1 | 1083.71 | 0.093556 | 0.08865 | 1.055386 | 0.29125 | 0.50117 | 55.001 |
| Senp2 | 6210.42 | 0.131337 | 0.06014 | 2.183807 | 0.02898 | 0.11518 | 39.786 |
| Senp3 | 3828.84 | 0.031537 | 0.06157 | 0.512205 | 0.60851 | 0.77621 | 8.97 |
| Senp5 | 3608.48 | 0.088675 | 0.06094 | 1.455046 | 0.14566 | 0.32855 | 43.616 |

| | | | | | | | |
|---|---|---|---|---|---|---|---|
| Senp6 | 3408.47 | 0.178402 | 0.1063 | 1.678269 | 0.09329 | 0.2486 | 78.051 |
| Senp7 | 2895.66 | 0.325205 | 0.09723 | 3.344764 | 0.00082 | 0.01119 | 141.675 |
| Senp8 | 478.889 | 0.062487 | 0.11118 | 0.56205 | 0.57408 | 0.7525 | 16.452 |
| Sephs1 | 1476.79 | 0.050956 | 0.08544 | 0.596396 | 0.55091 | 0.73431 | 28.994 |
| Sephs2 | 780.99 | -0.04512 | 0.09123 | -0.49455 | 0.62092 | 0.78443 | 2.177 |
| Sepsecs | 1073.36 | 0.093601 | 0.07945 | 1.178098 | 0.23876 | 0.44446 | 29.622 |
| Septin1 | 112.929 | 0.15215 | 0.18669 | 0.814993 | 0.41508 | 0.6229 | 18.684 |
| Septin10 | 242.099 | -0.14264 | 0.16276 | -0.87638 | 0.38082 | 0.59041 | 80.221 |
| Septin11 | 8592.23 | 0.097644 | 0.06501 | 1.50201 | 0.13309 | 0.3127 | 83.011 |
| Septin2 | 132.643 | -0.03489 | 0.15421 | -0.22624 | 0.82102 | 0.90748 | 31.297 |
| Septin3 | 23070 | -0.22775 | 0.07376 | -3.08776 | 0.00202 | 0.02018 | 25.773 |
| Septin4 | 4834.77 | -0.34727 | 0.13675 | -2.53949 | 0.0111 | 0.06139 | 23.887 |
| Septin5 | 1388.35 | -0.36814 | 0.12603 | -2.92098 | 0.00349 | 0.02869 | 9.453 |
| Septin6 | 2157.85 | -0.24438 | 0.07478 | -3.26818 | 0.00108 | 0.01349 | 80.469 |
| Septin7 | 13555.1 | -0.08139 | 0.13808 | -0.58939 | 0.5556 | 0.73792 | 56.133 |
| Septin8 | 12458 | -0.32779 | 0.10628 | -3.0843 | 0.00204 | 0.02034 | 30.309 |
| Septin9 | 1999.92 | -0.6315 | 0.14464 | -4.36614 | 1.26E-05 | 0.00066 | 162.665 |
| Serac1 | 1499.56 | -0.00689 | 0.07163 | -0.09616 | 0.9234 | 0.9614 | 37.546 |
| Serbp1 | 11274 | 0.032736 | 0.06838 | 0.478769 | 0.6321 | 0.79231 | 59.561 |
| Serf1 | 50.3624 | 0.078423 | 0.25139 | 0.311959 | 0.75507 | 0.87123 | 6.56 |
| Serf2 | 330.528 | -0.17182 | 0.11592 | -1.48229 | 0.13826 | 0.31942 | 9.119 |
| Sergef | 891.485 | 0.512056 | 0.09438 | 5.425589 | 5.78E-08 | 1.21E-05 | 196.654 |
| Serhl | 119.276 | -0.11471 | 0.17878 | -0.64159 | 0.52114 | 0.71037 | 27.169 |
| Serinc1 | 38862.8 | -0.06036 | 0.07454 | -0.80973 | 0.4181 | 0.62541 | 16.757 |
| Serinc2 | 169.102 | -0.58284 | 0.33974 | -1.71555 | 0.08624 | 0.23607 | 25.711 |
| Serinc3 | 12345 | -0.21889 | 0.07475 | -2.92823 | 0.00341 | 0.02822 | 21.86 |
| Serinc5 | 5389.53 | -0.06938 | 0.09788 | -0.70886 | 0.47841 | 0.67609 | 100.857 |
| Serp1 | 2264.19 | -0.16706 | 0.08582 | -1.94661 | 0.05158 | 0.16934 | 6.076 |
| Serp2 | 2403.76 | 0.148149 | 0.08218 | 1.802714 | 0.07143 | 0.20924 | 24.076 |
| Serpina10 | 14.3159 | 0.360893 | 0.60577 | 0.595761 | 0.55133 | 0.73445 | 16.659 |
| Serpina3h | 23.3332 | -1.41428 | 0.50146 | -2.82031 | 0.0048 | 0.03534 | 6.551 |
| Serpina3n | 817.737 | -1.69869 | 0.62871 | -2.70187 | 0.0069 | 0.04496 | 7.601 |
| Serpina9 | 1617.38 | 0.881484 | 0.29182 | 3.020623 | 0.00252 | 0.02351 | 19.013 |
| Serpinb1a | 278.112 | -0.81389 | 0.17455 | -4.66275 | 3.12E-06 | 0.00024 | 9.094 |

| | | | | | | | |
|---|---|---|---|---|---|---|---|
| Serpinb1b | 78.3465 | 0.493456 | 0.31022 | 1.590652 | 0.11169 | 0.27927 | 15.806 |
| Serpinb5 | 21.2718 | -1.04279 | 0.44766 | -2.32944 | 0.01984 | 0.08901 | 22.176 |
| Serpinb6a | 872.098 | -0.25316 | 0.09565 | -2.64664 | 0.00813 | 0.05008 | 84.877 |
| Serpinb6b | 50.3809 | -0.20869 | 0.3131 | -0.66652 | 0.50508 | 0.69681 | 13.859 |
| Serpinb8 | 197.777 | -0.97256 | 0.51433 | -1.89094 | 0.05863 | 0.18415 | 20.479 |
| Serpinb9 | 274.823 | -0.57063 | 0.17527 | -3.25567 | 0.00113 | 0.01378 | 14.708 |
| Serpind1 | 170.916 | -0.7957 | 0.31336 | -2.53926 | 0.01111 | 0.06142 | 12.205 |
| Serpine1 | 142.376 | 0.392468 | 0.19409 | 2.022125 | 0.04316 | 0.15004 | 10.765 |
| Serpine2 | 8686.56 | 0.015307 | 0.10283 | 0.148851 | 0.88167 | 0.94104 | 66.984 |
| Serpinf1 | 331.021 | -0.39993 | 0.14578 | -2.74328 | 0.00608 | 0.04167 | 12.933 |
| Serping1 | 382.273 | -1.25478 | 0.41151 | -3.04921 | 0.00229 | 0.02201 | 10.058 |
| Serpinh1 | 725.53 | -0.45135 | 0.12066 | -3.74064 | 0.00018 | 0.00406 | 7.864 |
| Serpini1 | 10909.5 | -0.69493 | 0.28561 | -2.43315 | 0.01497 | 0.07425 | 85.949 |
| Sertad1 | 208.963 | -0.14079 | 0.15318 | -0.9191 | 0.35805 | 0.56817 | 3.407 |
| Sertad2 | 1624.94 | 0.076851 | 0.08342 | 0.921212 | 0.35694 | 0.56723 | 109.769 |
| Sertad3 | 92.3642 | -0.57073 | 0.17649 | -3.23373 | 0.00122 | 0.01451 | 3.597 |
| Sertad4 | 1853.23 | 0.565492 | 0.16512 | 3.424669 | 0.00062 | 0.00915 | 11.269 |
| Sertm1 | 403.724 | -0.271 | 0.18917 | -1.43258 | 0.15198 | 0.3371 | 18.82 |
| Sesn1 | 4398.41 | 0.155077 | 0.09636 | 1.609395 | 0.10753 | 0.27228 | 98.49 |
| Sesn2 | 589.291 | -0.19767 | 0.14027 | -1.40921 | 0.15877 | 0.34629 | 18.47 |
| Sesn3 | 18064.8 | 0.383681 | 0.12721 | 3.016124 | 0.00256 | 0.02375 | 58.035 |
| Sestd1 | 3585.53 | -0.00116 | 0.09581 | -0.01207 | 0.99037 | 0.99473 | 100.253 |
| Set | 8798.47 | -0.09188 | 0.05764 | -1.59418 | 0.1109 | 0.27791 | 15.2 |
| Setbp1 | 1729.88 | -0.03295 | 0.12134 | -0.27155 | 0.78596 | 0.88868 | 359.012 |
| Setd1a | 1470.71 | 0.09152 | 0.08645 | 1.05869 | 0.28974 | 0.50001 | 23.453 |
| Setd1b | 3276.15 | 0.013851 | 0.05863 | 0.236233 | 0.81325 | 0.90389 | 26.437 |
| Setd2 | 4355.44 | 0.277361 | 0.10283 | 2.697344 | 0.00699 | 0.04538 | 86.037 |
| Setd3 | 7605.49 | 0.025075 | 0.04854 | 0.516618 | 0.60542 | 0.77411 | 72.884 |
| Setd4 | 883.068 | 0.177297 | 0.09643 | 1.838539 | 0.06598 | 0.19847 | 20.607 |
| Setd5 | 6457.28 | -0.01375 | 0.06437 | -0.21367 | 0.83081 | 0.91321 | 76.071 |
| Setd6 | 837.091 | 0.190001 | 0.09819 | 1.934974 | 0.05299 | 0.17223 | 3.13 |
| Setd7 | 10530.1 | -0.20772 | 0.06716 | -3.09283 | 0.00198 | 0.02003 | 45.561 |
| Setdb1 | 3065.32 | 0.17918 | 0.0626 | 2.862275 | 0.00421 | 0.03254 | 33.678 |
| Setdb2 | 348.394 | -0.1257 | 0.15012 | -0.83728 | 0.40243 | 0.61113 | 38.876 |

| | | | | | | | |
|---|---|---|---|---|---|---|---|
| Setmar | 274.469 | 0.041173 | 0.13239 | 0.310997 | 0.7558 | 0.87157 | 12.078 |
| Setx | 5161.37 | 0.215034 | 0.09245 | 2.326042 | 0.02002 | 0.08963 | 58.291 |
| Sez6 | 19832.9 | 0.410014 | 0.12719 | 3.223677 | 0.00127 | 0.01476 | 48.249 |
| Sez6l | 7288.15 | -0.44643 | 0.14572 | -3.06352 | 0.00219 | 0.02131 | 158.035 |
| Sez6l2 | 10597.6 | -0.24772 | 0.08452 | -2.93079 | 0.00338 | 0.02813 | 20.044 |
| Sf1 | 10867.6 | 0.185959 | 0.06292 | 2.955369 | 0.00312 | 0.02677 | 14.341 |
| Sf3a1 | 4422.42 | 0.085577 | 0.07548 | 1.133819 | 0.25687 | 0.46485 | 22.192 |
| Sf3a2 | 1128.79 | 0.048003 | 0.08077 | 0.594345 | 0.55228 | 0.73527 | 6.727 |
| Sf3a3 | 1421.61 | -0.01386 | 0.06867 | -0.20182 | 0.84006 | 0.91792 | 17.685 |
| Sf3b1 | 20429.3 | 0.225784 | 0.07477 | 3.019708 | 0.00253 | 0.02353 | 42.313 |
| Sf3b2 | 9342.47 | 0.039149 | 0.06242 | 0.627214 | 0.53052 | 0.71807 | 21.524 |
| Sf3b3 | 4250.99 | 0.290095 | 0.09238 | 3.140393 | 0.00169 | 0.01803 | 36.549 |
| Sf3b4 | 1280.34 | 0.07192 | 0.08707 | 0.826006 | 0.4088 | 0.61662 | 5.233 |
| Sf3b5 | 891.852 | 0.1218 | 0.08194 | 1.486523 | 0.13714 | 0.31781 | 3.841 |
| Sf3b6 | 1514.63 | 0.00822 | 0.10023 | 0.082018 | 0.93463 | 0.96717 | 10.116 |
| Sfi1 | 1796.83 | 0.326063 | 0.10004 | 3.259404 | 0.00112 | 0.01372 | 61.614 |
| Sfmbt1 | 1705.45 | -0.07886 | 0.07599 | -1.03767 | 0.29942 | 0.50955 | 107.873 |
| Sfmbt2 | 223.858 | 0.177234 | 0.16101 | 1.100741 | 0.27101 | 0.48038 | 224.744 |
| Sfn | 136.973 | 0.424984 | 0.19857 | 2.140169 | 0.03234 | 0.12439 | 1.613 |
| Sfpq | 13673.8 | 0.172065 | 0.06902 | 2.493061 | 0.01266 | 0.06707 | 15.69 |
| Sfr1 | 3560.73 | -0.01627 | 0.08055 | -0.20197 | 0.83994 | 0.91789 | 3.907 |
| Sfrp1 | 688.941 | -0.23727 | 0.12562 | -1.88877 | 0.05892 | 0.18475 | 38.131 |
| Sfrp2 | 40.5422 | 0.221882 | 0.27067 | 0.819764 | 0.41235 | 0.62038 | 7.996 |
| Sfrp4 | 58.905 | -0.9522 | 0.29064 | -3.27616 | 0.00105 | 0.01327 | 9.891 |
| Sfrp5 | 51.3339 | 0.166524 | 0.3745 | 0.444655 | 0.65657 | 0.80921 | 4.282 |
| Sfswap | 5244.77 | -0.08054 | 0.06586 | -1.22292 | 0.22136 | 0.42428 | 70.164 |
| Sft2d1 | 1229.48 | 0.0666 | 0.09186 | 0.725046 | 0.46842 | 0.6673 | 16.341 |
| Sft2d2 | 3323.5 | -0.13629 | 0.08843 | -1.54124 | 0.12326 | 0.29741 | 20.102 |
| Sft2d3 | 438.427 | 0.081967 | 0.11307 | 0.724913 | 0.46851 | 0.66737 | 2.727 |
| Sfta3-ps | 92.2535 | 1.468863 | 0.3416 | 4.299941 | 1.71E-05 | 0.0008 | 41.268 |
| Sftpc | 18.2805 | -1.12013 | 0.41311 | -2.71147 | 0.0067 | 0.04418 | 3.133 |
| Sfxn1 | 3058.71 | 0.104468 | 0.08444 | 1.237263 | 0.21599 | 0.41776 | 36.474 |
| Sfxn2 | 974.715 | 0.196917 | 0.10964 | 1.796017 | 0.07249 | 0.21164 | 24.702 |
| Sfxn3 | 6079.71 | 0.011361 | 0.06881 | 0.165122 | 0.86885 | 0.93346 | 8.881 |

| | | | | | | | |
|---|---|---|---|---|---|---|---|
| Sfxn4 | 893.169 | 0.236451 | 0.12471 | 1.896047 | 0.05795 | 0.18301 | 24.203 |
| Sfxn5 | 7125.36 | -0.20416 | 0.07849 | -2.6012 | 0.00929 | 0.05459 | 120.374 |
| Sgcb | 3779.59 | 0.003297 | 0.06524 | 0.050539 | 0.95969 | 0.98045 | 15.042 |
| Sgcd | 125.082 | 0.244145 | 0.2593 | 0.941555 | 0.34642 | 0.557 | 1092.717 |
| Sgce | 1399.41 | 0.146208 | 0.10315 | 1.417373 | 0.15637 | 0.34315 | 72.858 |
| Sgcz | 165.974 | 0.440479 | 0.24712 | 1.782439 | 0.07468 | 0.21534 | 1139.211 |
| Sgf29 | 363.894 | 0.044399 | 0.12338 | 0.359842 | 0.71897 | 0.8499 | 23.617 |
| Sgip1 | 8817.22 | 0.120326 | 0.06272 | 1.918529 | 0.05504 | 0.17678 | 232.332 |
| Sgk1 | 8883.99 | -0.13918 | 0.14178 | -0.98161 | 0.32629 | 0.53755 | 117.72 |
| Sgk2 | 52.9998 | -0.60691 | 0.27364 | -2.21787 | 0.02656 | 0.10851 | 26.798 |
| Sgk3 | 2677.07 | 0.204397 | 0.10562 | 1.935243 | 0.05296 | 0.17218 | 102.739 |
| Sgms1 | 2285.51 | -0.18185 | 0.07137 | -2.54803 | 0.01083 | 0.06059 | 266.988 |
| Sgms2 | 184.481 | -0.1807 | 0.15818 | -1.14239 | 0.25329 | 0.46062 | 84.539 |
| Sgo1 | 15.2765 | -0.11443 | 0.53539 | -0.21372 | 0.83076 | 0.91321 | 14.548 |
| Sgo2a | 18.1002 | 0.267771 | 0.43848 | 0.610686 | 0.54141 | 0.72706 | 29.929 |
| Sgpl1 | 1863.74 | -0.1353 | 0.09687 | -1.39669 | 0.16251 | 0.35091 | 49.062 |
| Sgpp1 | 2308.37 | 0.194402 | 0.09243 | 2.103346 | 0.03544 | 0.13224 | 21.481 |
| Sgpp2 | 713.338 | -0.12933 | 0.14932 | -0.86611 | 0.38643 | 0.59577 | 109.945 |
| Sgsh | 170.927 | -0.53845 | 0.19332 | -2.7853 | 0.00535 | 0.0379 | 12.112 |
| Sgsm1 | 5368.16 | 0.245402 | 0.07556 | 3.247686 | 0.00116 | 0.01403 | 67.567 |
| Sgsm2 | 5594.6 | 0.130016 | 0.0689 | 1.887017 | 0.05916 | 0.18515 | 47.8 |
| Sgsm3 | 2381.77 | 0.150523 | 0.05993 | 2.511675 | 0.01202 | 0.06468 | 34.526 |
| Sgta | 4075.51 | -0.11011 | 0.06352 | -1.73332 | 0.08304 | 0.22998 | 16.107 |
| Sgtb | 5575.99 | -0.16719 | 0.06851 | -2.44041 | 0.01467 | 0.0732 | 31.995 |
| Sh2b1 | 3559.35 | -0.01359 | 0.05916 | -0.22975 | 0.81828 | 0.90651 | 8.431 |
| Sh2b2 | 370.535 | -0.38523 | 0.18488 | -2.08365 | 0.03719 | 0.13628 | 28.41 |
| Sh2b3 | 933.759 | 0.074659 | 0.09975 | 0.748495 | 0.45416 | 0.65629 | 22.159 |
| Sh2d3c | 397.564 | -0.27274 | 0.14178 | -1.92366 | 0.0544 | 0.17536 | 34.458 |
| Sh2d4b | 49.2157 | -1.00988 | 0.26781 | -3.7708 | 0.00016 | 0.00375 | 79.478 |
| Sh2d5 | 9060.96 | 0.440082 | 0.13909 | 3.164089 | 0.00156 | 0.01704 | 10.93 |
| Sh2d6 | 63.8723 | 0.737109 | 0.23357 | 3.155859 | 0.0016 | 0.01735 | 8.067 |
| Sh3bgr | 32.5499 | 0.027032 | 0.38372 | 0.070448 | 0.94384 | 0.97192 | 28.486 |
| Sh3bgrl | 4657.32 | 0.112971 | 0.12104 | 0.9333 | 0.35066 | 0.56133 | 102.509 |
| Sh3bgrl2 | 2536.25 | -0.32847 | 0.09874 | -3.32653 | 0.00088 | 0.0117 | 90.475 |

| | | | | | | | |
|---|---|---|---|---|---|---|---|
| Sh3bgrl3 | 1235.58 | -0.07902 | 0.09351 | -0.84502 | 0.3981 | 0.60708 | 1.384 |
| Sh3bp1 | 133.886 | 0.272544 | 0.17871 | 1.525101 | 0.12723 | 0.30346 | 12.284 |
| Sh3bp2 | 251.981 | -0.10152 | 0.21061 | -0.48204 | 0.62978 | 0.79072 | 37.804 |
| Sh3bp4 | 1028.7 | -0.19267 | 0.1195 | -1.61235 | 0.10688 | 0.27129 | 84.654 |
| Sh3bp5 | 2147.57 | -0.03456 | 0.09707 | -0.35608 | 0.72178 | 0.85151 | 76.199 |
| Sh3bp5l | 2202.86 | -0.10084 | 0.07313 | -1.3788 | 0.16796 | 0.35782 | 17.005 |
| Sh3d19 | 667.892 | -0.38977 | 0.14795 | -2.63452 | 0.00843 | 0.05116 | 159.418 |
| Sh3d21 | 61.6941 | -0.05625 | 0.2743 | -0.20506 | 0.83752 | 0.91642 | 12.89 |
| Sh3gl1 | 1389.47 | -0.09974 | 0.08449 | -1.18051 | 0.2378 | 0.4432 | 19.886 |
| Sh3gl2 | 8040.35 | -0.19511 | 0.09161 | -2.12983 | 0.03319 | 0.12657 | 434.07 |
| Sh3gl3 | 1970.49 | 0.157448 | 0.10858 | 1.450017 | 0.14705 | 0.3302 | 133.04 |
| Sh3glb1 | 5044.26 | -0.02477 | 0.0617 | -0.40149 | 0.68806 | 0.83002 | 36.658 |
| Sh3glb2 | 8081.57 | 0.104273 | 0.07904 | 1.319219 | 0.1871 | 0.38237 | 14.529 |
| Sh3kbp1 | 2829.1 | 0.241272 | 0.0955 | 2.526364 | 0.01152 | 0.06295 | 350.798 |
| Sh3pxd2a | 5624.45 | 0.108511 | 0.0801 | 1.354757 | 0.1755 | 0.36728 | 204.228 |
| Sh3pxd2b | 1231.44 | -0.29634 | 0.17178 | -1.72514 | 0.0845 | 0.233 | 80.354 |
| Sh3rf1 | 1535.88 | -0.49295 | 0.17451 | -2.82487 | 0.00473 | 0.03509 | 172.2 |
| Sh3rf2 | 1169.7 | 0.537365 | 0.3694 | 1.454678 | 0.14576 | 0.32869 | 105.294 |
| Sh3rf3 | 1858.93 | 0.167739 | 0.12318 | 1.361721 | 0.17329 | 0.36452 | 325.558 |
| Sh3tc1 | 99.6366 | -0.10127 | 0.1916 | -0.52853 | 0.59713 | 0.76906 | 42.808 |
| Sh3tc2 | 140.509 | -0.3156 | 0.19502 | -1.61827 | 0.1056 | 0.26914 | 71.328 |
| Sh3yl1 | 320.513 | 0.058236 | 0.12291 | 0.473812 | 0.63563 | 0.7947 | 48.495 |
| Shank1 | 18802 | -0.65629 | 0.18973 | -3.45911 | 0.00054 | 0.00838 | 50.32 |
| Shank2 | 10959.2 | 0.000985 | 0.0976 | 0.010097 | 0.99194 | 0.99537 | 422.567 |
| Shank3 | 13955.9 | 0.284924 | 0.09742 | 2.924666 | 0.00345 | 0.02844 | 60.639 |
| Sharpin | 1299.44 | 0.056879 | 0.08432 | 0.674595 | 0.49993 | 0.693 | 4.072 |
| Shb | 733.713 | 0.336894 | 0.14298 | 2.356197 | 0.01846 | 0.08514 | 109.193 |
| Shc1 | 733.267 | -0.36491 | 0.11116 | -3.28271 | 0.00103 | 0.01304 | 11.585 |
| Shc2 | 2717.12 | 0.174731 | 0.06494 | 2.690819 | 0.00713 | 0.04604 | 19.868 |
| Shc3 | 2175.74 | 0.028698 | 0.10724 | 0.267601 | 0.78901 | 0.89044 | 138.447 |
| Shc4 | 283.327 | -0.21193 | 0.13598 | -1.55851 | 0.11911 | 0.29131 | 96.702 |
| Shcbp1 | 36.7316 | 0.059355 | 0.33308 | 0.178204 | 0.85856 | 0.92859 | 43.592 |
| Shcbp1l | 349.191 | 0.193608 | 0.13803 | 1.402658 | 0.16072 | 0.34889 | 27.413 |
| Shd | 528.111 | -0.1009 | 0.13358 | -0.75533 | 0.45005 | 0.65294 | 6.159 |

| | | | | | | | |
|---|---|---|---|---|---|---|---|
| She | 237.364 | -0.07621 | 0.15347 | -0.49658 | 0.61948 | 0.78345 | 27.465 |
| Shf | 883.76 | 0.0704 | 0.08102 | 0.868901 | 0.3849 | 0.59434 | 20.271 |
| Shfl | 3376.33 | 0.282889 | 0.09209 | 3.07173 | 0.00213 | 0.02101 | 5.666 |
| Shh | 142.225 | -0.0357 | 0.27349 | -0.13053 | 0.89615 | 0.94785 | 10.442 |
| Shisa2 | 294.286 | 0.12099 | 0.17121 | 0.706674 | 0.47977 | 0.67732 | 6.353 |
| Shisa3 | 44.7248 | -0.99413 | 0.46392 | -2.14288 | 0.03212 | 0.1238 | 15.671 |
| Shisa4 | 2195.48 | -0.67036 | 0.12518 | -5.35509 | 8.55E-08 | 1.56E-05 | 4.181 |
| Shisa5 | 1605.37 | -0.52218 | 0.11624 | -4.49236 | 7.04E-06 | 0.00043 | 19.213 |
| Shisa6 | 691.406 | -0.29365 | 0.15562 | -1.88692 | 0.05917 | 0.18516 | 314.24 |
| Shisa7 | 13509.3 | 0.389537 | 0.06918 | 5.630532 | 1.80E-08 | 5.47E-06 | 19.145 |
| Shisa8 | 343.684 | -0.56308 | 0.18058 | -3.11815 | 0.00182 | 0.01894 | 6.648 |
| Shisa9 | 5187.25 | 0.452705 | 0.0839 | 5.39597 | 6.82E-08 | 1.31E-05 | 286.79 |
| Shisal1 | 3573.29 | -0.64726 | 0.19614 | -3.29999 | 0.00097 | 0.01252 | 67.895 |
| Shkbp1 | 1564.75 | 0.426967 | 0.11174 | 3.821181 | 0.00013 | 0.00327 | 13.887 |
| Shld1 | 306.023 | 0.016497 | 0.1416 | 0.116509 | 0.90725 | 0.95354 | 64.125 |
| Shld2 | 339.594 | 0.071863 | 0.11302 | 0.635832 | 0.52489 | 0.71319 | 73.461 |
| Shld3 | 52.4766 | -0.10452 | 0.25989 | -0.40218 | 0.68755 | 0.82964 | 4.718 |
| Shmt1 | 187.924 | -0.07925 | 0.15106 | -0.52461 | 0.59986 | 0.77052 | 23.615 |
| Shmt2 | 915.169 | -0.17823 | 0.10318 | -1.72734 | 0.08411 | 0.23225 | 5.322 |
| Shoc2 | 2789.7 | 0.082685 | 0.06592 | 1.254371 | 0.20971 | 0.40977 | 88.828 |
| Shpk | 243.559 | -0.03715 | 0.13304 | -0.27926 | 0.78004 | 0.88542 | 25.052 |
| Shprh | 3232.15 | 0.257372 | 0.09755 | 2.638399 | 0.00833 | 0.05087 | 68.169 |
| Shq1 | 274.753 | -0.08015 | 0.12228 | -0.65545 | 0.51218 | 0.70328 | 102.902 |
| Shroom1 | 344.062 | -0.31073 | 0.10386 | -2.99189 | 0.00277 | 0.02495 | 10.562 |
| Shroom2 | 4963.21 | -0.03936 | 0.06931 | -0.56792 | 0.57009 | 0.74938 | 159.957 |
| Shroom3 | 139.801 | -1.00675 | 0.23997 | -4.19524 | 2.73E-05 | 0.00108 | 281.884 |
| Shroom4 | 421.094 | -0.2343 | 0.1202 | -1.94928 | 0.05126 | 0.16863 | 237.595 |
| Shtn1 | 2083.5 | -0.01208 | 0.1005 | -0.12019 | 0.90434 | 0.95168 | 102.745 |
| Siae | 845.162 | -0.12062 | 0.08507 | -1.41788 | 0.15623 | 0.34301 | 93.958 |
| Siah1a | 1017.01 | -0.1282 | 0.07728 | -1.65893 | 0.09713 | 0.2554 | 21.928 |
| Siah1b | 236.01 | 0.272733 | 0.16356 | 1.667469 | 0.09542 | 0.25254 | 5.789 |
| Siah2 | 501.83 | -0.2323 | 0.08905 | -2.60852 | 0.00909 | 0.05382 | 17.463 |
| Siah3 | 335.943 | -0.28403 | 0.13771 | -2.0626 | 0.03915 | 0.14099 | 73.727 |
| Sidt1 | 2013.07 | -0.74825 | 0.40432 | -1.85064 | 0.06422 | 0.19501 | 93.017 |

| | | | | | | | |
|---|---|---|---|---|---|---|---|
| Sidt2 | 3312.4 | 0.064258 | 0.05743 | 1.118842 | 0.26321 | 0.47208 | 17.402 |
| Sigirr | 53.9476 | -0.3867 | 0.29205 | -1.32406 | 0.18548 | 0.38025 | 9.398 |
| Siglec1 | 41.5804 | -0.85224 | 0.41259 | -2.06561 | 0.03887 | 0.14057 | 17.546 |
| Siglece | 42.8803 | -0.84269 | 0.28867 | -2.91924 | 0.00351 | 0.02879 | 9.092 |
| Siglech | 645.962 | -0.04055 | 0.13304 | -0.30481 | 0.76051 | 0.87444 | 10.748 |
| Sigmar1 | 959.009 | -0.13079 | 0.10821 | -1.20869 | 0.22678 | 0.43097 | 17.665 |
| Sik1 | 1164.28 | -0.1604 | 0.11051 | -1.45144 | 0.14666 | 0.32981 | 11.555 |
| Sik2 | 1594.94 | 0.104676 | 0.09814 | 1.066576 | 0.28616 | 0.49619 | 116.273 |
| Sik3 | 10298.3 | 0.193377 | 0.07748 | 2.495767 | 0.01257 | 0.06665 | 211.375 |
| Sike1 | 3191.25 | 0.014375 | 0.07791 | 0.184505 | 0.85362 | 0.92574 | 12.752 |
| Sil1 | 764.061 | -0.14544 | 0.10724 | -1.35622 | 0.17503 | 0.36668 | 233.385 |
| Simc1 | 999.434 | 0.274604 | 0.08357 | 3.285779 | 0.00102 | 0.01296 | 47.512 |
| Sin3a | 1982.2 | 0.11545 | 0.08963 | 1.288118 | 0.1977 | 0.39553 | 56.327 |
| Sin3b | 4507.83 | 0.035862 | 0.05639 | 0.635957 | 0.5248 | 0.71317 | 34.917 |
| Sinhcaf | 48.804 | -0.92142 | 0.29651 | -3.10756 | 0.00189 | 0.01933 | 25.433 |
| Sipa1 | 714.985 | -0.20149 | 0.10449 | -1.9283 | 0.05382 | 0.17416 | 12.523 |
| Sipa1l1 | 27529.1 | 0.210304 | 0.07405 | 2.839942 | 0.00451 | 0.034 | 282.467 |
| Sipa1l2 | 2964.62 | -0.46619 | 0.10532 | -4.4263 | 9.59E-06 | 0.00054 | 151.746 |
| Sipa1l3 | 3373.51 | 0.152459 | 0.06715 | 2.27033 | 0.02319 | 0.09868 | 198.27 |
| Sirpa | 15236.8 | -0.11657 | 0.08676 | -1.34356 | 0.17909 | 0.37183 | 39.394 |
| Sirt1 | 1351.1 | 0.237073 | 0.08727 | 2.71639 | 0.0066 | 0.04379 | 62.7 |
| Sirt2 | 4878.39 | -0.23616 | 0.10547 | -2.23905 | 0.02515 | 0.10438 | 21.927 |
| Sirt3 | 2164.61 | -0.19293 | 0.07161 | -2.69396 | 0.00706 | 0.04574 | 18.644 |
| Sirt4 | 1136.71 | 0.0629 | 0.11198 | 0.561712 | 0.57431 | 0.75253 | 6.716 |
| Sirt5 | 632.365 | 0.051644 | 0.09888 | 0.522314 | 0.60145 | 0.77155 | 29.708 |
| Sirt6 | 632.532 | -0.13465 | 0.08742 | -1.54021 | 0.12351 | 0.2978 | 6.013 |
| Sirt7 | 2057.85 | 0.039635 | 0.08066 | 0.491407 | 0.62314 | 0.78592 | 6.869 |
| Siva1 | 331.868 | 0.007633 | 0.13949 | 0.05472 | 0.95636 | 0.97864 | 4.471 |
| Six2 | 24.0425 | -1.18874 | 0.50265 | -2.36492 | 0.01803 | 0.08372 | 3.998 |
| Six3 | 1783.03 | 0.894465 | 0.35344 | 2.530727 | 0.01138 | 0.06243 | 15.695 |
| Six3os1 | 1035.28 | 1.048526 | 0.22227 | 4.717432 | 2.39E-06 | 0.0002 | 17.988 |
| Six4 | 34.9415 | -0.80047 | 0.33845 | -2.36507 | 0.01803 | 0.0837 | 13.848 |
| Six5 | 103.895 | -0.25526 | 0.17647 | -1.44647 | 0.14804 | 0.33174 | 3.956 |
| Ska2 | 645.289 | 0.146379 | 0.11073 | 1.321888 | 0.18621 | 0.38114 | 15.402 |

| | | | | | | | |
|---|---|---|---|---|---|---|---|
| Ska3 | 68.9169 | 0.074119 | 0.22104 | 0.335311 | 0.73739 | 0.86132 | 19.625 |
| Skap1 | 64.6774 | -0.67015 | 0.29977 | -2.23552 | 0.02538 | 0.10506 | 294.538 |
| Skap2 | 1003.16 | -0.21085 | 0.11803 | -1.78637 | 0.07404 | 0.21441 | 155.128 |
| Ski | 9184.9 | 0.139785 | 0.08906 | 1.569587 | 0.11651 | 0.28673 | 68.518 |
| Skida1 | 423.558 | -0.26862 | 0.09539 | -2.81588 | 0.00486 | 0.03558 | 8.376 |
| Skil | 5004.48 | 0.017266 | 0.0715 | 0.241489 | 0.80918 | 0.90175 | 27.52 |
| Skiv2l | 3296.71 | -0.01448 | 0.07479 | -0.19357 | 0.84651 | 0.92184 | 10.983 |
| Skor1 | 16.856 | 0.907639 | 0.44051 | 2.06043 | 0.03936 | 0.14142 | 10.792 |
| Skp1 | 12171.2 | -0.06248 | 0.06184 | -1.01045 | 0.31228 | 0.5227 | 14.864 |
| Skp2 | 629.012 | 0.229424 | 0.11966 | 1.917327 | 0.0552 | 0.17713 | 43.44 |
| Sla | 535.627 | -1.08677 | 0.47421 | -2.29173 | 0.02192 | 0.09532 | 51.011 |
| Sla2 | 36.6601 | -0.61965 | 0.34132 | -1.81546 | 0.06945 | 0.20539 | 14.736 |
| Slain1 | 1683.33 | -0.31334 | 0.09749 | -3.21416 | 0.00131 | 0.01505 | 54.68 |
| Slain1os | 195.692 | 0.188169 | 0.1853 | 1.015503 | 0.30987 | 0.51986 | 6.46 |
| Slain2 | 1318.09 | 0.034497 | 0.07817 | 0.441287 | 0.65901 | 0.81103 | 64.526 |
| Slamf6 | 13.3485 | -0.43349 | 0.57653 | -0.75188 | 0.45212 | 0.65455 | 35.656 |
| Slamf9 | 21.0453 | -0.59371 | 0.48628 | -1.22093 | 0.22211 | 0.42532 | 3.218 |
| Slbp | 617.725 | -0.0577 | 0.12631 | -0.45682 | 0.6478 | 0.80388 | 17.623 |
| Slc10a1 | 39.1129 | 0.115965 | 0.37887 | 0.306084 | 0.75954 | 0.87373 | 15.523 |
| Slc10a3 | 78.2366 | -0.11684 | 0.20512 | -0.56964 | 0.56892 | 0.74862 | 4.133 |
| Slc10a4 | 628.725 | 0.891387 | 0.37649 | 2.367627 | 0.0179 | 0.08334 | 6.073 |
| Slc10a7 | 376.818 | -0.13224 | 0.10388 | -1.27304 | 0.203 | 0.40164 | 224.658 |
| Slc11a1 | 98.1652 | -0.28119 | 0.20764 | -1.35426 | 0.17565 | 0.36746 | 10.868 |
| Slc11a2 | 2800.53 | 0.047747 | 0.06564 | 0.727358 | 0.46701 | 0.66622 | 37.175 |
| Slc12a1 | 12.8491 | 0.745198 | 0.58821 | 1.266898 | 0.20519 | 0.40426 | 77.498 |
| Slc12a2 | 5805.6 | -0.27357 | 0.12888 | -2.1226 | 0.03379 | 0.12825 | 68.144 |
| Slc12a3 | 80.8642 | 0.604805 | 0.28686 | 2.108354 | 0.035 | 0.1312 | 37.014 |
| Slc12a4 | 740.333 | -0.26777 | 0.10066 | -2.66015 | 0.00781 | 0.04892 | 22.508 |
| Slc12a5 | 28479.8 | 0.126421 | 0.05834 | 2.166886 | 0.03024 | 0.11875 | 38.93 |
| Slc12a6 | 4213.52 | 0.068619 | 0.07411 | 0.92587 | 0.35451 | 0.56492 | 97.339 |
| Slc12a7 | 257.764 | -0.63709 | 0.22031 | -2.89181 | 0.00383 | 0.03055 | 83.661 |
| Slc12a8 | 48.6873 | 0.025759 | 0.23711 | 0.108634 | 0.91349 | 0.95625 | 146.808 |
| Slc12a9 | 1951.95 | 0.147112 | 0.08736 | 1.683944 | 0.09219 | 0.2472 | 19.04 |
| Slc13a3 | 893.291 | -1.00089 | 0.20718 | -4.83095 | 1.36E-06 | 0.00013 | 68.203 |

| | | | | | | | |
|---|---|---|---|---|---|---|---|
| Slc13a4 | 650.611 | -1.13585 | 0.42582 | -2.66744 | 0.00764 | 0.04831 | 40.175 |
| Slc13a5 | 468.876 | -0.00766 | 0.13787 | -0.05556 | 0.9557 | 0.9782 | 25.234 |
| Slc14a1 | 292.836 | -0.38672 | 0.17628 | -2.19381 | 0.02825 | 0.11317 | 42.029 |
| Slc15a2 | 1581.37 | -0.20296 | 0.15652 | -1.29667 | 0.19474 | 0.39207 | 34.786 |
| Slc15a3 | 56.3647 | -0.16006 | 0.30559 | -0.52378 | 0.60043 | 0.77095 | 19.636 |
| Slc15a4 | 993.851 | -0.09795 | 0.13062 | -0.74987 | 0.45333 | 0.65558 | 37.234 |
| Slc16a1 | 1224.4 | -0.28488 | 0.1002 | -2.8432 | 0.00447 | 0.03383 | 19.795 |
| Slc16a10 | 89.4309 | -0.33878 | 0.21987 | -1.54083 | 0.12336 | 0.29754 | 108.727 |
| Slc16a11 | 911.114 | 0.32287 | 0.1149 | 2.80989 | 0.00496 | 0.03604 | 3.666 |
| Slc16a12 | 204.336 | -0.50622 | 0.18527 | -2.73231 | 0.00629 | 0.04245 | 78.887 |
| Slc16a13 | 206.207 | 0.011175 | 0.1391 | 0.08034 | 0.93597 | 0.96785 | 4.276 |
| Slc16a14 | 377.354 | -0.42559 | 0.18284 | -2.32769 | 0.01993 | 0.08932 | 29.237 |
| Slc16a2 | 2024.88 | 0.23852 | 0.11541 | 2.066736 | 0.03876 | 0.14026 | 124.57 |
| Slc16a3 | 225.145 | 0.181855 | 0.16577 | 1.097023 | 0.27263 | 0.48182 | 12.389 |
| Slc16a4 | 406.285 | 0.404322 | 0.13805 | 2.928773 | 0.0034 | 0.02821 | 20.886 |
| Slc16a5 | 43.8036 | -0.14802 | 0.41749 | -0.35456 | 0.72292 | 0.85223 | 11.925 |
| Slc16a6 | 723.458 | 0.015932 | 0.10694 | 0.148985 | 0.88157 | 0.94099 | 22.744 |
| Slc16a7 | 1764.75 | -0.24884 | 0.14068 | -1.7688 | 0.07693 | 0.21936 | 170.196 |
| Slc16a8 | 97.2973 | 0.099382 | 0.21226 | 0.468211 | 0.63963 | 0.79795 | 3.748 |
| Slc16a9 | 106.931 | -0.21629 | 0.24331 | -0.88893 | 0.37404 | 0.58429 | 40.869 |
| Slc17a5 | 1109.31 | 0.010176 | 0.10538 | 0.09657 | 0.92307 | 0.96131 | 51.554 |
| Slc17a6 | 556.369 | -0.62818 | 0.24903 | -2.52253 | 0.01165 | 0.06344 | 49.114 |
| Slc17a7 | 24909.3 | -1.3963 | 0.50042 | -2.79026 | 0.00527 | 0.0375 | 12.194 |
| Slc17a8 | 872.873 | 0.718661 | 0.22884 | 3.140432 | 0.00169 | 0.01803 | 47.234 |
| Slc17a9 | 36.2591 | -0.5847 | 0.29905 | -1.95517 | 0.05056 | 0.16723 | 17.018 |
| Slc18a2 | 325.13 | 0.224566 | 0.13979 | 1.606502 | 0.10816 | 0.27338 | 35.117 |
| Slc18a3 | 279.668 | 1.040699 | 0.41535 | 2.505602 | 0.01222 | 0.06538 | 2.413 |
| Slc18b1 | 1041.94 | 0.224478 | 0.06894 | 3.256065 | 0.00113 | 0.01378 | 30.983 |
| Slc19a1 | 642.866 | 0.107243 | 0.09638 | 1.112737 | 0.26582 | 0.47503 | 28.762 |
| Slc19a2 | 838.064 | 0.052641 | 0.08004 | 0.657686 | 0.51074 | 0.70199 | 16.34 |
| Slc19a3 | 200.087 | -0.07931 | 0.14657 | -0.54115 | 0.5884 | 0.76172 | 25.926 |
| Slc1a1 | 3787.7 | -0.21367 | 0.10877 | -1.96438 | 0.04949 | 0.16471 | 78.912 |
| Slc1a2 | 157532 | -0.18019 | 0.07374 | -2.44374 | 0.01454 | 0.07286 | 132.126 |
| Slc1a3 | 30755.2 | -0.29505 | 0.08378 | -3.52184 | 0.00043 | 0.00722 | 76.641 |

| | | | | | | | |
|---|---|---|---|---|---|---|---|
| Slc1a4 | 2610.53 | -0.09471 | 0.08259 | -1.14678 | 0.25147 | 0.45887 | 30.534 |
| Slc1a5 | 268.478 | -0.64512 | 0.18865 | -3.41965 | 0.00063 | 0.00924 | 16.935 |
| Slc1a6 | 285.732 | -0.14821 | 0.15097 | -0.98176 | 0.32622 | 0.53752 | 34.27 |
| Slc20a1 | 5161.13 | 0.048861 | 0.06944 | 0.70363 | 0.48166 | 0.67858 | 12.853 |
| Slc20a2 | 2394.63 | -0.12846 | 0.07108 | -1.80719 | 0.07073 | 0.20791 | 92.825 |
| Slc22a12 | 44.7675 | 0.101981 | 0.32744 | 0.311445 | 0.75546 | 0.8714 | 7.188 |
| Slc22a13b | 69.9356 | -0.23915 | 0.24531 | -0.97487 | 0.32963 | 0.5409 | 11.205 |
| Slc22a15 | 577.97 | 0.138107 | 0.09941 | 1.389239 | 0.16476 | 0.35385 | 68.678 |
| Slc22a17 | 26858.3 | 0.122607 | 0.06458 | 1.898445 | 0.05764 | 0.18229 | 6.829 |
| Slc22a18 | 29.0419 | -0.40923 | 0.43373 | -0.94351 | 0.34542 | 0.55616 | 25.599 |
| Slc22a2 | 35.586 | -2 | 0.95553 | -2.11228 | 0.03466 | 0.13034 | 44.357 |
| Slc22a21 | 65.8651 | 0.175774 | 0.26459 | 0.664337 | 0.50647 | 0.69789 | 30.368 |
| Slc22a23 | 4626.78 | 0.160738 | 0.08008 | 2.00719 | 0.04473 | 0.15361 | 166.025 |
| Slc22a3 | 305.868 | -0.1223 | 0.22023 | -0.55532 | 0.57868 | 0.75546 | 88.745 |
| Slc22a4 | 315.582 | 0.028135 | 0.17446 | 0.161267 | 0.87188 | 0.93524 | 44.968 |
| Slc22a5 | 707.639 | -0.06867 | 0.08891 | -0.77234 | 0.43991 | 0.64503 | 27.119 |
| Slc22a6 | 357.927 | -0.89922 | 0.45444 | -1.97874 | 0.04785 | 0.16091 | 10.261 |
| Slc22a7 | 19.5514 | -0.35055 | 0.53663 | -0.65324 | 0.5136 | 0.70443 | 6.275 |
| Slc22a8 | 774.643 | -0.7417 | 0.22046 | -3.36427 | 0.00077 | 0.01068 | 20.582 |
| Slc23a1 | 70.1387 | 0.006763 | 0.23566 | 0.028698 | 0.97711 | 0.98796 | 12.641 |
| Slc23a2 | 7807.43 | -0.23016 | 0.07086 | -3.24828 | 0.00116 | 0.01403 | 92.613 |
| Slc23a4 | 29.5952 | 0.064361 | 0.31614 | 0.203585 | 0.83868 | 0.9169 | 43.127 |
| Slc24a1 | 24.2351 | -0.12536 | 0.35558 | -0.35255 | 0.72442 | 0.85307 | 28.747 |
| Slc24a2 | 28630.9 | -0.43383 | 0.13187 | -3.28988 | 0.001 | 0.01283 | 247.354 |
| Slc24a3 | 5256.87 | -0.57349 | 0.17891 | -3.20543 | 0.00135 | 0.01537 | 474.413 |
| Slc24a4 | 5730.72 | 0.604394 | 0.19284 | 3.134101 | 0.00172 | 0.01828 | 138.359 |
| Slc24a5 | 195.318 | -0.03756 | 0.16714 | -0.22472 | 0.82219 | 0.90785 | 20.554 |
| Slc25a1 | 1232.6 | 0.21899 | 0.09503 | 2.304538 | 0.02119 | 0.09306 | 2.997 |
| Slc25a10 | 461.623 | -0.1097 | 0.12868 | -0.85251 | 0.39393 | 0.60303 | 7.348 |
| Slc25a11 | 3280.48 | 0.032642 | 0.08988 | 0.363184 | 0.71647 | 0.84846 | 3.284 |
| Slc25a12 | 7073.68 | 0.052693 | 0.06142 | 0.857987 | 0.3909 | 0.60005 | 96.687 |
| Slc25a13 | 383.244 | -0.3153 | 0.14078 | -2.2397 | 0.02511 | 0.10427 | 175.956 |
| Slc25a14 | 2404.56 | 0.160387 | 0.09027 | 1.776768 | 0.07561 | 0.21706 | 38.882 |
| Slc25a15 | 749.269 | -0.14198 | 0.09567 | -1.48405 | 0.1378 | 0.31864 | 23.047 |

| | | | | | | | |
|---|---|---|---|---|---|---|---|
| Slc25a16 | 1885.09 | -0.07742 | 0.0684 | -1.13185 | 0.2577 | 0.46576 | 25.866 |
| Slc25a17 | 1876.35 | 0.026744 | 0.07013 | 0.381378 | 0.70292 | 0.8397 | 41.902 |
| Slc25a18 | 1247.09 | -0.37349 | 0.1324 | -2.82095 | 0.00479 | 0.0353 | 20.759 |
| Slc25a19 | 2653.72 | -0.00945 | 0.07355 | -0.12848 | 0.89777 | 0.94882 | 14.118 |
| Slc25a20 | 696.767 | -0.04366 | 0.09642 | -0.45278 | 0.65071 | 0.80552 | 22.554 |
| Slc25a21 | 35.7454 | 0.281996 | 0.32622 | 0.86443 | 0.38735 | 0.59665 | 484.839 |
| Slc25a22 | 6872.35 | -0.22006 | 0.06984 | -3.15086 | 0.00163 | 0.01757 | 8.149 |
| Slc25a23 | 14402.1 | -0.20441 | 0.06794 | -3.00841 | 0.00263 | 0.02412 | 16.153 |
| Slc25a24 | 150.592 | -0.19603 | 0.15188 | -1.29063 | 0.19683 | 0.39463 | 45.309 |
| Slc25a25 | 2877.93 | 0.035749 | 0.08665 | 0.412585 | 0.67991 | 0.82501 | 36.959 |
| Slc25a26 | 805.728 | 0.065326 | 0.09171 | 0.712273 | 0.4763 | 0.67427 | 104.342 |
| Slc25a27 | 3499.57 | 0.049843 | 0.10023 | 0.497304 | 0.61897 | 0.78314 | 25.324 |
| Slc25a28 | 2034.85 | 0.070987 | 0.08095 | 0.876919 | 0.38053 | 0.59002 | 11.081 |
| Slc25a29 | 585.474 | -0.01556 | 0.10588 | -0.147 | 0.88313 | 0.94166 | 10.011 |
| Slc25a3 | 13596.6 | -0.08081 | 0.05967 | -1.35441 | 0.17561 | 0.36743 | 7.486 |
| Slc25a30 | 635.413 | 0.306777 | 0.13709 | 2.237784 | 0.02524 | 0.10457 | 26.921 |
| Slc25a32 | 1094.57 | 0.043331 | 0.10517 | 0.412018 | 0.68033 | 0.82502 | 21.486 |
| Slc25a33 | 1031.83 | -0.01817 | 0.10418 | -0.17437 | 0.86158 | 0.92987 | 30.242 |
| Slc25a34 | 715.146 | -0.05073 | 0.14552 | -0.34861 | 0.72738 | 0.85443 | 4.998 |
| Slc25a35 | 257.587 | -0.15161 | 0.12792 | -1.18519 | 0.23594 | 0.44119 | 6.235 |
| Slc25a36 | 3836.31 | 0.077672 | 0.10435 | 0.744318 | 0.45668 | 0.65819 | 36.197 |
| Slc25a37 | 2855.54 | 0.256799 | 0.08245 | 3.114488 | 0.00184 | 0.01906 | 43.265 |
| Slc25a38 | 948.824 | -0.13961 | 0.08727 | -1.59973 | 0.10966 | 0.27562 | 14.131 |
| Slc25a39 | 1559.07 | -0.11673 | 0.08273 | -1.41095 | 0.15826 | 0.34558 | 4.962 |
| Slc25a4 | 20100.1 | -0.1514 | 0.04871 | -3.10817 | 0.00188 | 0.0193 | 4.488 |
| Slc25a40 | 916.245 | 0.113866 | 0.09893 | 1.150954 | 0.24975 | 0.45704 | 31.948 |
| Slc25a42 | 2579.84 | 0.244679 | 0.07295 | 3.354003 | 0.0008 | 0.01095 | 27.966 |
| Slc25a44 | 4428.92 | -0.05122 | 0.06723 | -0.76186 | 0.44615 | 0.64983 | 14.642 |
| Slc25a45 | 171.937 | -0.50391 | 0.17816 | -2.82843 | 0.00468 | 0.03482 | 8.071 |
| Slc25a46 | 3850.22 | 0.05979 | 0.07965 | 0.750661 | 0.45286 | 0.65531 | 53.801 |
| Slc25a47 | 175.454 | -0.10504 | 0.14606 | -0.71917 | 0.47204 | 0.67047 | 20.77 |
| Slc25a5 | 9004.12 | -0.02602 | 0.07841 | -0.33178 | 0.74006 | 0.86295 | 3.157 |
| Slc25a51 | 5264.85 | 0.130464 | 0.07031 | 1.855642 | 0.0635 | 0.1937 | 12.844 |
| Slc25a53 | 872.522 | 0.137177 | 0.0801 | 1.712608 | 0.08678 | 0.23717 | 57.187 |

| | | | | | | | |
|---|---|---|---|---|---|---|---|
| Slc26a1 | 166.588 | 0.499023 | 0.15264 | 3.269367 | 0.00108 | 0.01345 | 5.692 |
| Slc26a10 | 816.991 | 0.64649 | 0.21163 | 3.054809 | 0.00225 | 0.02176 | 9.253 |
| Slc26a11 | 190.56 | -0.45616 | 0.14588 | -3.12704 | 0.00177 | 0.01859 | 25.523 |
| Slc26a2 | 625.98 | -0.31288 | 0.13322 | -2.34856 | 0.01885 | 0.08613 | 18.694 |
| Slc26a4 | 106.038 | -0.78066 | 0.38882 | -2.00777 | 0.04467 | 0.15352 | 40.143 |
| Slc26a5 | 25.6752 | 0.527332 | 0.36868 | 1.430329 | 0.15262 | 0.33794 | 54.95 |
| Slc26a6 | 455.321 | 0.204456 | 0.11286 | 1.81152 | 0.07006 | 0.20667 | 11.636 |
| Slc26a7 | 89.4762 | -1.16652 | 0.92904 | -1.25562 | 0.20925 | 0.4093 | 119.376 |
| Slc26a8 | 153.785 | -0.25868 | 0.17892 | -1.44579 | 0.14824 | 0.33188 | 52.451 |
| Slc27a1 | 5571.68 | -0.03377 | 0.08916 | -0.3787 | 0.70491 | 0.84107 | 18.421 |
| Slc27a2 | 404.734 | 0.556064 | 0.17073 | 3.256999 | 0.00113 | 0.01378 | 35.837 |
| Slc27a3 | 261.695 | 0.292896 | 0.14535 | 2.01509 | 0.0439 | 0.15165 | 4.7 |
| Slc27a4 | 3236.8 | -0.19098 | 0.05982 | -3.19281 | 0.00141 | 0.01586 | 14.889 |
| Slc27a5 | 15.2427 | -0.6506 | 0.42744 | -1.52207 | 0.12799 | 0.30479 | 9.847 |
| Slc29a1 | 1330.44 | 0.047773 | 0.08528 | 0.560214 | 0.57533 | 0.75336 | 14.407 |
| Slc29a2 | 537.319 | 0.011445 | 0.09252 | 0.123696 | 0.90156 | 0.95057 | 8.113 |
| Slc29a3 | 870.568 | -0.47963 | 0.11052 | -4.33979 | 1.43E-05 | 0.00071 | 40.723 |
| Slc29a4 | 277.099 | -0.14106 | 0.15506 | -0.9097 | 0.36298 | 0.57343 | 29.979 |
| Slc2a1 | 7255.65 | -0.26804 | 0.1155 | -2.32075 | 0.0203 | 0.09035 | 29.273 |
| Slc2a10 | 77.6794 | -0.48578 | 0.24668 | -1.9693 | 0.04892 | 0.16347 | 16.131 |
| Slc2a12 | 134.164 | 0.10712 | 0.22165 | 0.483287 | 0.62889 | 0.79019 | 59.275 |
| Slc2a13 | 10638.8 | 0.466338 | 0.13253 | 3.518687 | 0.00043 | 0.00726 | 305.566 |
| Slc2a2 | 24.9824 | -0.15773 | 0.35943 | -0.43884 | 0.66078 | 0.81233 | 33.457 |
| Slc2a3 | 9746.33 | 0.142337 | 0.08242 | 1.727015 | 0.08416 | 0.23231 | 73.832 |
| Slc2a4 | 65.0842 | 0.150413 | 0.36183 | 0.415699 | 0.67763 | 0.82332 | 5.65 |
| Slc2a4rg-ps | 870.863 | 0.191116 | 0.16289 | 1.173248 | 0.2407 | 0.44693 | 3.348 |
| Slc2a5 | 204.144 | -0.19577 | 0.15017 | -1.30366 | 0.19235 | 0.38897 | 24.887 |
| Slc2a6 | 672.708 | -0.33682 | 0.15884 | -2.12047 | 0.03397 | 0.12867 | 6.636 |
| Slc2a8 | 807.378 | -0.22875 | 0.08114 | -2.81924 | 0.00481 | 0.03542 | 9.094 |
| Slc2a9 | 48.0456 | -0.32007 | 0.24859 | -1.28755 | 0.1979 | 0.39569 | 153.871 |
| Slc30a1 | 2413.58 | 0.095024 | 0.10167 | 0.934653 | 0.34997 | 0.56056 | 6.481 |
| Slc30a10 | 1099.67 | 0.276522 | 0.11842 | 2.335018 | 0.01954 | 0.08818 | 13.915 |
| Slc30a2 | 184.995 | -0.44092 | 0.16628 | -2.65171 | 0.00801 | 0.04963 | 11.304 |
| Slc30a3 | 1506.39 | -1.7762 | 0.55036 | -3.22733 | 0.00125 | 0.01469 | 22.132 |

| | | | | | | | |
|---|---|---|---|---|---|---|---|
| Slc30a4 | 2940.02 | 0.190758 | 0.07375 | 2.58647 | 0.0097 | 0.0562 | 21.431 |
| Slc30a5 | 2299.52 | 0.132589 | 0.08485 | 1.562552 | 0.11816 | 0.28963 | 30.78 |
| Slc30a6 | 534.697 | 0.035858 | 0.10522 | 0.340787 | 0.73326 | 0.85857 | 28.614 |
| Slc30a7 | 1207.67 | 0.132407 | 0.09369 | 1.413241 | 0.15758 | 0.34467 | 68.434 |
| Slc30a9 | 8095.62 | 0.042956 | 0.05748 | 0.747307 | 0.45488 | 0.65702 | 51.489 |
| Slc31a1 | 1153.67 | -0.10034 | 0.07692 | -1.30447 | 0.19207 | 0.38879 | 31.043 |
| Slc31a2 | 594.957 | 0.026921 | 0.09345 | 0.288079 | 0.77329 | 0.88189 | 35.85 |
| Slc32a1 | 6984.13 | 0.353553 | 0.16668 | 2.12111 | 0.03391 | 0.12861 | 4.982 |
| Slc33a1 | 1185.44 | -0.03128 | 0.08111 | -0.38561 | 0.69979 | 0.83779 | 31.262 |
| Slc35a1 | 1013.4 | 0.15468 | 0.09596 | 1.611984 | 0.10697 | 0.27139 | 24.182 |
| Slc35a2 | 1333.4 | -0.10029 | 0.07184 | -1.39606 | 0.1627 | 0.35113 | 10.465 |
| Slc35a3 | 1566.02 | 0.214989 | 0.08061 | 2.667006 | 0.00765 | 0.04834 | 43.362 |
| Slc35a4 | 2087.14 | -0.23172 | 0.07211 | -3.21341 | 0.00131 | 0.01508 | 4.647 |
| Slc35a5 | 2310.47 | 0.014431 | 0.06909 | 0.208877 | 0.83454 | 0.91508 | 19.134 |
| Slc35b1 | 1487.88 | 0.021251 | 0.06915 | 0.307307 | 0.75861 | 0.8731 | 7.085 |
| Slc35b2 | 781.485 | -0.14888 | 0.11992 | -1.24146 | 0.21444 | 0.4158 | 3.798 |
| Slc35b3 | 605.895 | 0.073771 | 0.10792 | 0.683595 | 0.49423 | 0.68845 | 28.74 |
| Slc35b4 | 3178.23 | 0.125275 | 0.07544 | 1.660502 | 0.09681 | 0.25492 | 23.732 |
| Slc35c1 | 643.275 | -0.04973 | 0.10105 | -0.49214 | 0.62262 | 0.7857 | 7.775 |
| Slc35c2 | 1803.29 | -0.04095 | 0.06471 | -0.63277 | 0.52689 | 0.71496 | 11.316 |
| Slc35d1 | 955.888 | 0.026824 | 0.06947 | 0.386103 | 0.69942 | 0.83751 | 44.516 |
| Slc35d2 | 27.3177 | -0.77331 | 0.37786 | -2.04655 | 0.0407 | 0.14434 | 33.061 |
| Slc35d3 | 1565.63 | 0.716491 | 0.31291 | 2.289794 | 0.02203 | 0.09565 | 3.544 |
| Slc35e1 | 3536.26 | -0.02035 | 0.0913 | -0.22284 | 0.82366 | 0.90857 | 11.974 |
| Slc35e2 | 2747.24 | -0.03607 | 0.06311 | -0.57159 | 0.5676 | 0.74756 | 21.925 |
| Slc35e3 | 1425.18 | -0.06135 | 0.08939 | -0.6864 | 0.49246 | 0.68714 | 12.687 |
| Slc35e4 | 967.573 | -0.11633 | 0.12944 | -0.8987 | 0.36881 | 0.5795 | 7.644 |
| Slc35f1 | 6155.43 | 0.313669 | 0.07648 | 4.101415 | 4.11E-05 | 0.00143 | 421.09 |
| Slc35f2 | 180.319 | 0.126382 | 0.16201 | 0.780103 | 0.43533 | 0.64101 | 46.617 |
| Slc35f3 | 2743.08 | 0.196256 | 0.09666 | 2.030403 | 0.04232 | 0.14806 | 256.984 |
| Slc35f4 | 502.908 | 0.123919 | 0.12125 | 1.022018 | 0.30677 | 0.51671 | 227.528 |
| Slc35f5 | 711.714 | -0.23301 | 0.0832 | -2.80072 | 0.0051 | 0.03675 | 35.226 |
| Slc35f6 | 1147.52 | -0.112 | 0.10668 | -1.04989 | 0.29377 | 0.5038 | 11.797 |
| Slc35g1 | 140.09 | -0.49455 | 0.18527 | -2.66938 | 0.0076 | 0.04816 | 9.631 |

| | | | | | | | |
|---|---|---|---|---|---|---|---|
| Slc35g2 | 627.337 | 0.060811 | 0.1091 | 0.557367 | 0.57728 | 0.75445 | 18.903 |
| Slc36a1 | 3750.91 | -0.49068 | 0.15867 | -3.09257 | 0.00198 | 0.02003 | 31.981 |
| Slc36a2 | 16.747 | -0.41555 | 0.69672 | -0.59644 | 0.55088 | 0.73431 | 26.608 |
| Slc36a4 | 3625.37 | -0.01301 | 0.05727 | -0.22711 | 0.82034 | 0.90734 | 33.624 |
| Slc37a1 | 481.957 | 0.413261 | 0.16531 | 2.49988 | 0.01242 | 0.06617 | 63.948 |
| Slc37a2 | 203.266 | -0.164 | 0.14227 | -1.15274 | 0.24902 | 0.45629 | 28.154 |
| Slc37a3 | 2723.45 | -0.14644 | 0.10801 | -1.3559 | 0.17513 | 0.36682 | 42.903 |
| Slc37a4 | 776.823 | -0.29775 | 0.08464 | -3.51785 | 0.00044 | 0.00728 | 6.117 |
| Slc38a1 | 9238.21 | -0.15412 | 0.08164 | -1.8877 | 0.05907 | 0.18503 | 71.496 |
| Slc38a10 | 3555.86 | 0.097347 | 0.09338 | 1.042455 | 0.2972 | 0.50736 | 47.387 |
| Slc38a11 | 107.581 | -0.09171 | 0.17172 | -0.53407 | 0.59329 | 0.76557 | 47.605 |
| Slc38a2 | 8807.32 | -0.49533 | 0.12917 | -3.83476 | 0.00013 | 0.00313 | 12.339 |
| Slc38a3 | 3337.48 | -0.37673 | 0.06972 | -5.40344 | 6.54E-08 | 1.28E-05 | 18.897 |
| Slc38a4 | 38.9183 | -0.05925 | 0.31128 | -0.19034 | 0.84904 | 0.92322 | 61.137 |
| Slc38a5 | 59.6085 | -0.54938 | 0.36057 | -1.52364 | 0.1276 | 0.30407 | 9.047 |
| Slc38a6 | 925.674 | 0.197007 | 0.09384 | 2.099303 | 0.03579 | 0.13303 | 67.271 |
| Slc38a7 | 1317.02 | -0.22348 | 0.0807 | -2.76939 | 0.00562 | 0.03939 | 17.62 |
| Slc38a8 | 15.9116 | 0.731774 | 0.52986 | 1.381073 | 0.16726 | 0.35697 | 22.097 |
| Slc38a9 | 1944.26 | 0.097372 | 0.06597 | 1.475999 | 0.13994 | 0.3212 | 77.999 |
| Slc39a1 | 1506.87 | -0.23334 | 0.07833 | -2.97889 | 0.00289 | 0.02563 | 5.441 |
| Slc39a10 | 7027.98 | -0.05682 | 0.085 | -0.6685 | 0.50381 | 0.69618 | 85.309 |
| Slc39a11 | 644.493 | -0.05426 | 0.11941 | -0.45439 | 0.64955 | 0.80496 | 405.227 |
| Slc39a12 | 2052.06 | -0.20802 | 0.09326 | -2.2305 | 0.02571 | 0.10604 | 106.662 |
| Slc39a13 | 869.854 | -0.01809 | 0.08368 | -0.21617 | 0.82885 | 0.91223 | 8.627 |
| Slc39a14 | 1942.32 | 0.083694 | 0.07266 | 1.151932 | 0.24935 | 0.45679 | 47.957 |
| Slc39a2 | 1112.61 | 0.33507 | 0.12793 | 2.619104 | 0.00882 | 0.05273 | 3.857 |
| Slc39a3 | 3804.69 | 0.092991 | 0.06891 | 1.349494 | 0.17718 | 0.36946 | 8.889 |
| Slc39a5 | 46.6922 | 0.011214 | 0.27515 | 0.040758 | 0.96749 | 0.98365 | 5.299 |
| Slc39a6 | 1934.5 | -0.3013 | 0.14469 | -2.08245 | 0.0373 | 0.13656 | 23.937 |
| Slc39a8 | 149.83 | -0.39206 | 0.14162 | -2.76847 | 0.00563 | 0.03946 | 63.294 |
| Slc39a9 | 2625.85 | 0.005708 | 0.07177 | 0.079537 | 0.93661 | 0.96829 | 39.46 |
| Slc3a2 | 4238.74 | -0.27282 | 0.08932 | -3.05441 | 0.00226 | 0.02177 | 16.488 |
| Slc40a1 | 389.355 | -0.42899 | 0.12438 | -3.44909 | 0.00056 | 0.00859 | 18.456 |
| Slc41a1 | 4552.76 | 0.009684 | 0.11616 | 0.083365 | 0.93356 | 0.9665 | 21.373 |

| | | | | | | | |
|---|---|---|---|---|---|---|---|
| Slc41a2 | 876.674 | -0.07731 | 0.08382 | -0.92235 | 0.35635 | 0.56677 | 107.035 |
| Slc41a3 | 114.552 | -0.22016 | 0.17388 | -1.26613 | 0.20547 | 0.40454 | 41.688 |
| Slc43a1 | 26.086 | -0.47968 | 0.40227 | -1.19245 | 0.23309 | 0.43798 | 24.745 |
| Slc43a2 | 3280.08 | -0.27577 | 0.06417 | -4.29766 | 1.73E-05 | 0.0008 | 45.882 |
| Slc43a3 | 35.017 | -0.74325 | 0.49029 | -1.51595 | 0.12953 | 0.30705 | 21.931 |
| Slc44a1 | 3626.46 | -0.43702 | 0.10266 | -4.25695 | 2.07E-05 | 0.0009 | 182.066 |
| Slc44a2 | 2152.8 | -0.12171 | 0.07575 | -1.60671 | 0.10812 | 0.2733 | 34.331 |
| Slc44a5 | 779.635 | 0.149345 | 0.08538 | 1.749076 | 0.08028 | 0.22543 | 298.287 |
| Slc45a1 | 1781.1 | -0.06976 | 0.07273 | -0.95916 | 0.33748 | 0.54829 | 23.603 |
| Slc45a3 | 35.1039 | -0.58842 | 0.36402 | -1.61646 | 0.10599 | 0.26977 | 20.003 |
| Slc45a4 | 2523.94 | -0.13712 | 0.07732 | -1.77333 | 0.07617 | 0.21802 | 68.339 |
| Slc46a1 | 727.149 | -0.20113 | 0.088 | -2.2855 | 0.02228 | 0.09621 | 6.363 |
| Slc46a3 | 398.328 | 0.027528 | 0.11328 | 0.243015 | 0.80799 | 0.90129 | 16.379 |
| Slc47a1 | 80.1851 | -1.45562 | 0.94116 | -1.54662 | 0.12195 | 0.29538 | 34.945 |
| Slc48a1 | 4750.61 | -0.24564 | 0.08337 | -2.94657 | 0.00321 | 0.0273 | 14.173 |
| Slc49a4 | 3266.49 | -0.16975 | 0.07509 | -2.26072 | 0.02378 | 0.10039 | 75.295 |
| Slc4a10 | 11298.7 | 0.136566 | 0.08608 | 1.586439 | 0.11264 | 0.28066 | 280.269 |
| Slc4a11 | 568.074 | 1.162247 | 0.40405 | 2.87651 | 0.00402 | 0.03161 | 13.407 |
| Slc4a1ap | 1509.08 | 0.093684 | 0.07283 | 1.286397 | 0.1983 | 0.39603 | 29.938 |
| Slc4a2 | 1398.58 | -0.31461 | 0.09097 | -3.45822 | 0.00054 | 0.0084 | 17.114 |
| Slc4a3 | 10573 | 0.160325 | 0.07215 | 2.222023 | 0.02628 | 0.10771 | 15.907 |
| Slc4a4 | 13569.4 | 0.049291 | 0.09563 | 0.515445 | 0.60624 | 0.77465 | 353.168 |
| Slc4a7 | 2406.38 | 0.258178 | 0.10022 | 2.576137 | 0.00999 | 0.05743 | 97.662 |
| Slc4a8 | 6642.63 | -0.33577 | 0.11326 | -2.9645 | 0.00303 | 0.02631 | 62.222 |
| Slc4a9 | 18.8322 | 0.767593 | 0.46853 | 1.6383 | 0.10136 | 0.26262 | 28.127 |
| Slc50a1 | 980.511 | 0.080631 | 0.09874 | 0.816631 | 0.41414 | 0.62213 | 2.325 |
| Slc52a2 | 582.855 | 0.115683 | 0.09037 | 1.280159 | 0.20049 | 0.39855 | 5.777 |
| Slc52a3 | 109.053 | -0.10075 | 0.19345 | -0.52079 | 0.60252 | 0.77234 | 12.748 |
| Slc5a11 | 52.1498 | 0.059486 | 0.37223 | 0.159811 | 0.87303 | 0.93613 | 58.474 |
| Slc5a12 | 13.4725 | -0.06715 | 0.5157 | -0.13021 | 0.8964 | 0.94793 | 50.482 |
| Slc5a3 | 3006.12 | 0.39724 | 0.09381 | 4.234325 | 2.29E-05 | 0.00096 | 29.152 |
| Slc5a4b | 13.2795 | 0.9261 | 0.48725 | 1.900653 | 0.05735 | 0.18169 | 53.468 |
| Slc5a5 | 521.39 | -0.4774 | 0.12101 | -3.94511 | 7.98E-05 | 0.0023 | 9.869 |
| Slc5a6 | 1006.03 | -0.10208 | 0.0896 | -1.13924 | 0.25461 | 0.46206 | 12.889 |

| | | | | | | | |
|---|---|---|---|---|---|---|---|
| Slc5a7 | 1737.84 | 1.123598 | 0.39421 | 2.850262 | 0.00437 | 0.03339 | 25.445 |
| Slc66a1 | 599.188 | -0.02179 | 0.10698 | -0.20372 | 0.83857 | 0.91688 | 16.68 |
| Slc66a2 | 739.701 | 0.071835 | 0.08678 | 0.827799 | 0.40778 | 0.6155 | 39.434 |
| Slc6a1 | 25936.6 | 0.084344 | 0.08573 | 0.983876 | 0.32518 | 0.53645 | 34.898 |
| Slc6a11 | 6869.45 | -0.24712 | 0.12266 | -2.01472 | 0.04393 | 0.15174 | 118.712 |
| Slc6a12 | 86.2055 | -1.01815 | 0.38946 | -2.61424 | 0.00894 | 0.05327 | 22.7 |
| Slc6a13 | 634.53 | -1.31971 | 0.43829 | -3.01102 | 0.0026 | 0.02403 | 37.507 |
| Slc6a15 | 3013.41 | 0.261185 | 0.06327 | 4.12811 | 3.66E-05 | 0.00131 | 51.595 |
| Slc6a16 | 46.6645 | 0.15555 | 0.27095 | 0.574096 | 0.5659 | 0.74585 | 32.283 |
| Slc6a17 | 25323.7 | -0.30205 | 0.09636 | -3.1345 | 0.00172 | 0.01828 | 50.476 |
| Slc6a20a | 632.533 | -1.02211 | 0.42783 | -2.38907 | 0.01689 | 0.08052 | 44.116 |
| Slc6a3 | 27.3984 | -0.59827 | 0.4225 | -1.41603 | 0.15677 | 0.34363 | 41.926 |
| Slc6a6 | 9539.38 | 0.148453 | 0.07343 | 2.021581 | 0.04322 | 0.15016 | 75.014 |
| Slc6a7 | 2713.11 | -0.63155 | 0.18604 | -3.39478 | 0.00069 | 0.00985 | 18.843 |
| Slc6a8 | 5167.44 | -0.18068 | 0.08333 | -2.16829 | 0.03014 | 0.11861 | 9.353 |
| Slc6a9 | 1158.15 | -0.37642 | 0.12191 | -3.08776 | 0.00202 | 0.02018 | 40.693 |
| Slc7a1 | 2546.67 | -0.13871 | 0.0798 | -1.73822 | 0.08217 | 0.22854 | 72.495 |
| Slc7a10 | 980.527 | -0.16455 | 0.10657 | -1.54411 | 0.12256 | 0.29632 | 14.73 |
| Slc7a11 | 2993.28 | -0.19719 | 0.12649 | -1.55893 | 0.11901 | 0.29113 | 551.089 |
| Slc7a14 | 7304.34 | 0.329836 | 0.09837 | 3.353155 | 0.0008 | 0.01096 | 107.521 |
| Slc7a2 | 3507.65 | -0.1791 | 0.08615 | -2.07889 | 0.03763 | 0.13733 | 59.913 |
| Slc7a3 | 342.79 | -0.38834 | 0.16063 | -2.41755 | 0.01563 | 0.07651 | 6.811 |
| Slc7a4 | 3152.5 | 0.194101 | 0.09546 | 2.033377 | 0.04201 | 0.14738 | 11.197 |
| Slc7a5 | 2634.09 | -0.30962 | 0.08222 | -3.76597 | 0.00017 | 0.00381 | 26.545 |
| Slc7a6 | 1670.73 | 0.222573 | 0.08102 | 2.747217 | 0.00601 | 0.04133 | 29.85 |
| Slc7a6os | 1109.71 | 0.091616 | 0.07295 | 1.255914 | 0.20915 | 0.40918 | 10.219 |
| Slc7a7 | 139.709 | 0.006276 | 0.16275 | 0.038561 | 0.96924 | 0.98441 | 48.339 |
| Slc7a8 | 2611.93 | -0.05371 | 0.08869 | -0.6056 | 0.54478 | 0.7299 | 59.738 |
| Slc8a1 | 8824.35 | -0.14794 | 0.17012 | -0.86961 | 0.38451 | 0.594 | 365.273 |
| Slc8a2 | 12338 | 0.065601 | 0.08303 | 0.79012 | 0.42946 | 0.63601 | 31.238 |
| Slc8a3 | 1972.48 | -0.02063 | 0.10342 | -0.19947 | 0.8419 | 0.91879 | 135.266 |
| Slc8b1 | 255.658 | -0.41275 | 0.12647 | -3.26363 | 0.0011 | 0.01361 | 22.857 |
| Slc9a1 | 3025.59 | -0.09437 | 0.07056 | -1.33741 | 0.18109 | 0.37449 | 53.997 |
| Slc9a2 | 536.59 | -0.13926 | 0.14636 | -0.95148 | 0.34136 | 0.55226 | 88.7 |

| | | | | | | | |
|---|---|---|---|---|---|---|---|
| Slc9a3 | 37.3604 | -0.76698 | 0.28388 | -2.70174 | 0.0069 | 0.04496 | 47.986 |
| Slc9a3r1 | 1972.78 | -0.23765 | 0.11821 | -2.01041 | 0.04439 | 0.15289 | 17.841 |
| Slc9a3r2 | 1578.91 | -0.26453 | 0.11328 | -2.33527 | 0.01953 | 0.08815 | 11.042 |
| Slc9a4 | 13.4455 | -1.84818 | 0.68139 | -2.71236 | 0.00668 | 0.04414 | 50.645 |
| Slc9a5 | 2934.48 | 0.48349 | 0.13064 | 3.700857 | 0.00021 | 0.00447 | 21.039 |
| Slc9a6 | 6705.52 | 0.100002 | 0.06073 | 1.646752 | 0.09961 | 0.25951 | 54.474 |
| Slc9a7 | 1014.85 | 0.079025 | 0.10794 | 0.732151 | 0.46408 | 0.66379 | 186.054 |
| Slc9a8 | 1914.28 | -0.01462 | 0.06844 | -0.21364 | 0.83082 | 0.91321 | 55.289 |
| Slc9a9 | 641.57 | -0.56884 | 0.15571 | -3.65329 | 0.00026 | 0.00506 | 560.537 |
| Slc9b1 | 43.5745 | -0.02506 | 0.31661 | -0.07916 | 0.93691 | 0.9685 | 49.799 |
| Slc9b2 | 157.456 | 0.471956 | 0.18524 | 2.547822 | 0.01084 | 0.06059 | 37.688 |
| Slco1a4 | 2688.14 | -0.25358 | 0.10903 | -2.32591 | 0.02002 | 0.08964 | 50.76 |
| Slco1b2 | 17.7944 | -0.38186 | 0.57089 | -0.66888 | 0.50357 | 0.69601 | 57.129 |
| Slco1c1 | 2963.05 | -0.08492 | 0.13277 | -0.63956 | 0.52246 | 0.71129 | 45.81 |
| Slco2a1 | 89.8269 | -0.68247 | 0.26584 | -2.56721 | 0.01025 | 0.05844 | 107.291 |
| Slco2b1 | 755.376 | -0.36317 | 0.11139 | -3.26031 | 0.00111 | 0.0137 | 53.537 |
| Slco3a1 | 1758.42 | -0.22869 | 0.09296 | -2.46021 | 0.01389 | 0.0709 | 279.362 |
| Slco4a1 | 149.035 | -0.44699 | 0.223 | -2.00447 | 0.04502 | 0.15421 | 18.623 |
| Slco4c1 | 234.74 | 0.048664 | 0.18256 | 0.266558 | 0.78981 | 0.89088 | 55.902 |
| Slco5a1 | 1461.61 | 0.392305 | 0.16044 | 2.445208 | 0.01448 | 0.07271 | 126.102 |
| Slf1 | 1162.1 | 0.333108 | 0.1378 | 2.417298 | 0.01564 | 0.07651 | 92.386 |
| Slf2 | 4330.59 | 0.246885 | 0.12208 | 2.022308 | 0.04314 | 0.15 | 52.149 |
| Slfn2 | 81.4888 | -0.34786 | 0.23904 | -1.45525 | 0.1456 | 0.32852 | 5.567 |
| Slfn3 | 20.5897 | -0.28549 | 0.42632 | -0.66967 | 0.50307 | 0.69545 | 23.825 |
| Slfn5 | 410.508 | -0.39333 | 0.11071 | -3.55299 | 0.00038 | 0.00661 | 13.492 |
| Slfn8 | 61.01 | -0.40128 | 0.29919 | -1.34124 | 0.17984 | 0.37297 | 18.653 |
| Slfn9 | 29.8003 | -0.70888 | 0.4331 | -1.63676 | 0.10168 | 0.26313 | 13.441 |
| Slit1 | 3224.99 | -0.45524 | 0.30576 | -1.48886 | 0.13652 | 0.31704 | 143.409 |
| Slit2 | 1894.94 | 0.030753 | 0.15764 | 0.195089 | 0.84532 | 0.92096 | 324.596 |
| Slit3 | 2751.39 | 0.294394 | 0.13561 | 2.170818 | 0.02994 | 0.11816 | 587.284 |
| Slitrk1 | 2881.59 | -0.48373 | 0.16126 | -2.99965 | 0.0027 | 0.02455 | 5.359 |
| Slitrk2 | 1687.83 | -0.03634 | 0.11382 | -0.31931 | 0.74949 | 0.86849 | 12.076 |
| Slitrk3 | 3768.9 | 0.024742 | 0.08314 | 0.297578 | 0.76602 | 0.8779 | 10.539 |
| Slitrk4 | 2672.88 | 0.021837 | 0.17458 | 0.125082 | 0.90046 | 0.95005 | 12.112 |

| | | | | | | | |
|---|---|---|---|---|---|---|---|
| Slitrk5 | 2681.22 | 0.088444 | 0.06258 | 1.413246 | 0.15758 | 0.34467 | 8.045 |
| Slitrk6 | 213.255 | 1.348551 | 0.22159 | 6.085658 | 1.16E-09 | 5.91E-07 | 6.57 |
| Slk | 8552.31 | 0.182042 | 0.07343 | 2.479262 | 0.01317 | 0.06871 | 65.569 |
| Slmap | 12002.7 | 0.505904 | 0.13753 | 3.678592 | 0.00023 | 0.00476 | 121.764 |
| Slpi | 25.1172 | -1.22827 | 0.54557 | -2.25134 | 0.02436 | 0.1021 | 35.026 |
| Sltm | 4490.55 | 0.207173 | 0.13116 | 1.579498 | 0.11422 | 0.28303 | 49.481 |
| Slu7 | 2743.19 | -0.05805 | 0.08274 | -0.70154 | 0.48297 | 0.6794 | 14.238 |
| Slx1b | 536.125 | -0.02025 | 0.10612 | -0.19082 | 0.84866 | 0.92307 | 6.317 |
| Slx4 | 1556.02 | 0.109636 | 0.08363 | 1.3109 | 0.18989 | 0.38596 | 24.666 |
| Slx4ip | 704.603 | 0.112424 | 0.15248 | 0.737308 | 0.46094 | 0.66158 | 180.733 |
| Smad1 | 1127.42 | -0.1724 | 0.09488 | -1.81712 | 0.0692 | 0.20478 | 61.124 |
| Smad2 | 1705.02 | 0.04702 | 0.07663 | 0.613558 | 0.53951 | 0.72589 | 69.384 |
| Smad3 | 4415.98 | 0.379852 | 0.11772 | 3.226805 | 0.00125 | 0.01471 | 111.228 |
| Smad4 | 2857.07 | 0.141568 | 0.09571 | 1.479091 | 0.13912 | 0.32034 | 64.772 |
| Smad5 | 3042.48 | 0.136617 | 0.06843 | 1.996509 | 0.04588 | 0.15618 | 39.368 |
| Smad6 | 119.013 | -0.57813 | 0.16896 | -3.42179 | 0.00062 | 0.00921 | 68.984 |
| Smad7 | 1445.65 | -0.52062 | 0.12704 | -4.09793 | 4.17E-05 | 0.00144 | 28.407 |
| Smad9 | 769.333 | -0.29695 | 0.13571 | -2.18802 | 0.02867 | 0.11429 | 45.676 |
| Smagp | 36.5118 | -0.32264 | 0.39766 | -0.81136 | 0.41716 | 0.62461 | 15.602 |
| Smap1 | 1291.31 | -0.22442 | 0.10813 | -2.07553 | 0.03794 | 0.13814 | 77.472 |
| Smap2 | 8176.01 | -0.01595 | 0.07297 | -0.21852 | 0.82703 | 0.91105 | 48.931 |
| Smarca1 | 2639.96 | 0.651261 | 0.12319 | 5.286786 | 1.24E-07 | 2.11E-05 | 83.607 |
| Smarca2 | 11261.8 | 0.074335 | 0.06773 | 1.097558 | 0.2724 | 0.48159 | 173.273 |
| Smarca4 | 7175.88 | 0.010347 | 0.05159 | 0.200552 | 0.84105 | 0.91851 | 88.062 |
| Smarca5 | 3818.2 | 0.245704 | 0.09669 | 2.541253 | 0.01105 | 0.0613 | 40.991 |
| Smarcad1 | 1693.64 | 0.166432 | 0.08593 | 1.936935 | 0.05275 | 0.17186 | 73.479 |
| Smarcal1 | 2505.34 | 0.139838 | 0.07172 | 1.949713 | 0.05121 | 0.1685 | 49.884 |
| Smarcb1 | 2679.12 | 0.108127 | 0.08481 | 1.274867 | 0.20236 | 0.40075 | 24.849 |
| Smarcc1 | 3684.41 | 0.192073 | 0.06387 | 3.007462 | 0.00263 | 0.02416 | 122.471 |
| Smarcc2 | 15213 | -0.18796 | 0.07054 | -2.66459 | 0.00771 | 0.04851 | 31.235 |
| Smarcd1 | 7406.85 | -0.04747 | 0.06382 | -0.74368 | 0.45707 | 0.6583 | 11.863 |
| Smarcd2 | 853.299 | -0.17556 | 0.10755 | -1.6324 | 0.1026 | 0.26448 | 9.794 |
| Smarcd3 | 2951.58 | 0.160162 | 0.10043 | 1.594752 | 0.11077 | 0.27772 | 33.594 |
| Smarce1 | 2585.93 | 0.111786 | 0.07473 | 1.495959 | 0.13466 | 0.31449 | 21.971 |

| | | | | | | | |
|---|---|---|---|---|---|---|---|
| Smarce1-ps1 | 64.8525 | 0.128281 | 0.25383 | 0.505374 | 0.6133 | 0.77988 | 1.077 |
| Smc1a | 4601.15 | 0.047023 | 0.10735 | 0.438043 | 0.66136 | 0.81269 | 46.267 |
| Smc2 | 605.845 | 0.280089 | 0.16955 | 1.651956 | 0.09854 | 0.25776 | 49.018 |
| Smc2os | 15.3274 | 0.053451 | 0.44392 | 0.120407 | 0.90416 | 0.95164 | 8.681 |
| Smc3 | 3474.27 | 0.242112 | 0.16337 | 1.481998 | 0.13834 | 0.3195 | 45.436 |
| Smc4 | 441.769 | 0.159392 | 0.22582 | 0.70585 | 0.48028 | 0.67761 | 29.886 |
| Smc5 | 2429.27 | 0.381262 | 0.13571 | 2.809414 | 0.00496 | 0.03605 | 67.447 |
| Smc6 | 2183.95 | 0.251625 | 0.16966 | 1.48308 | 0.13805 | 0.31909 | 53.9 |
| Smchd1 | 2681.2 | 0.222648 | 0.08328 | 2.673469 | 0.00751 | 0.04775 | 130.855 |
| Smco4 | 86.7109 | -0.33261 | 0.23563 | -1.41161 | 0.15806 | 0.34534 | 51.829 |
| Smcr8 | 2508.09 | 0.136013 | 0.0752 | 1.808585 | 0.07052 | 0.20751 | 10.764 |
| Smdt1 | 2994.6 | -0.24103 | 0.08558 | -2.81645 | 0.00486 | 0.03557 | 10.133 |
| Smg1 | 13795.4 | 0.20892 | 0.08975 | 2.32782 | 0.01992 | 0.08931 | 112.363 |
| Smg5 | 7435.79 | 0.282665 | 0.12826 | 2.203925 | 0.02753 | 0.11117 | 26.079 |
| Smg6 | 2406.16 | 0.124913 | 0.06969 | 1.792427 | 0.07306 | 0.21267 | 238.626 |
| Smg7 | 5503.63 | -0.00214 | 0.06624 | -0.03224 | 0.97428 | 0.98712 | 65.652 |
| Smg8 | 692.078 | -0.07324 | 0.11106 | -0.65945 | 0.50961 | 0.70111 | 9.043 |
| Smg9 | 1434.1 | 0.104152 | 0.07431 | 1.401575 | 0.16104 | 0.34918 | 23.16 |
| Smim1 | 304.76 | -0.21785 | 0.13289 | -1.63924 | 0.10116 | 0.26232 | 5.761 |
| Smim10l1 | 6130.83 | 0.069685 | 0.10224 | 0.681576 | 0.49551 | 0.68945 | 5.991 |
| Smim10l2a | 824.967 | 0.074727 | 0.10441 | 0.715678 | 0.47419 | 0.67238 | 3.214 |
| Smim11 | 353.6 | 0.007999 | 0.103 | 0.077658 | 0.9381 | 0.96915 | 11.756 |
| Smim12 | 814.856 | -0.15036 | 0.09972 | -1.50785 | 0.13159 | 0.31032 | 4.026 |
| Smim13 | 9984.88 | -0.12663 | 0.06852 | -1.84806 | 0.06459 | 0.19561 | 26.734 |
| Smim14 | 5400.06 | 0.043 | 0.05858 | 0.734086 | 0.4629 | 0.66297 | 90.341 |
| Smim15 | 1773.9 | -0.00189 | 0.10001 | -0.01885 | 0.98496 | 0.99174 | 4.727 |
| Smim17 | 920.854 | 0.319528 | 0.11497 | 2.779147 | 0.00545 | 0.03847 | 33.379 |
| Smim18 | 152.853 | -0.10136 | 0.14554 | -0.69641 | 0.48617 | 0.68228 | 5.659 |
| Smim19 | 654.787 | -0.02649 | 0.12309 | -0.21519 | 0.82962 | 0.91263 | 14.269 |
| Smim20 | 992.664 | 0.102754 | 0.09599 | 1.070519 | 0.28439 | 0.49449 | 11.458 |
| Smim24 | 52.4255 | 0.199791 | 0.30988 | 0.644738 | 0.5191 | 0.70877 | 2.342 |
| Smim26 | 442.025 | -0.07275 | 0.11133 | -0.65341 | 0.51349 | 0.70439 | 1.313 |
| Smim3 | 702.485 | -0.27511 | 0.15421 | -1.78394 | 0.07443 | 0.21503 | 27.797 |
| Smim30 | 996.111 | -0.37407 | 0.1036 | -3.61086 | 0.00031 | 0.00569 | 3.784 |

| | | | | | | | |
|---|---|---|---|---|---|---|---|
| Smim36 | 112.781 | -0.10316 | 0.24478 | -0.42143 | 0.67344 | 0.82048 | 63.573 |
| Smim38 | 34.0828 | -0.34512 | 0.29561 | -1.16746 | 0.24302 | 0.4492 | 2.295 |
| Smim4 | 243.455 | -0.0412 | 0.12526 | -0.32892 | 0.74222 | 0.86419 | 40.825 |
| Smim41 | 18.477 | 0.477229 | 0.51591 | 0.925023 | 0.35495 | 0.56542 | 4.215 |
| Smim43 | 801.287 | -0.93737 | 0.39711 | -2.36051 | 0.01825 | 0.08442 | 17.328 |
| Smim45 | 3891.07 | -0.06168 | 0.07895 | -0.78122 | 0.43467 | 0.64042 | 8.357 |
| Smim5 | 48.7292 | 0.369297 | 0.23877 | 1.546683 | 0.12194 | 0.29538 | 6.304 |
| Smim6 | 56.7109 | 0.463407 | 0.29886 | 1.550572 | 0.121 | 0.29412 | 1.904 |
| Smim7 | 3284.12 | -0.10185 | 0.08775 | -1.16068 | 0.24577 | 0.45244 | 5.901 |
| Smim8 | 404.047 | -0.07558 | 0.15478 | -0.48831 | 0.62533 | 0.7877 | 9.76 |
| Smn1 | 708.457 | 0.05335 | 0.12752 | 0.418378 | 0.67567 | 0.82192 | 12.839 |
| Smndc1 | 1477 | -0.05908 | 0.07971 | -0.74117 | 0.45859 | 0.65962 | 11.189 |
| Smo | 639.89 | -0.17758 | 0.11277 | -1.57463 | 0.11534 | 0.28485 | 25.863 |
| Smoc1 | 657.779 | -0.90597 | 0.16258 | -5.57252 | 2.51E-08 | 7.18E-06 | 159.607 |
| Smoc2 | 87.7649 | -1.10434 | 0.33563 | -3.29035 | 0.001 | 0.01282 | 125.285 |
| Smok4a | 18.3325 | -0.25684 | 0.57394 | -0.4475 | 0.65451 | 0.80832 | 6.978 |
| Smox | 2688.44 | -0.36932 | 0.09538 | -3.8721 | 0.00011 | 0.00282 | 34.427 |
| Smpd1 | 3444.14 | -0.23604 | 0.08851 | -2.6668 | 0.00766 | 0.04834 | 4.03 |
| Smpd2 | 620.647 | -0.01742 | 0.10491 | -0.16607 | 0.86811 | 0.93328 | 4.728 |
| Smpd3 | 7239.68 | 0.43835 | 0.14609 | 3.000465 | 0.0027 | 0.02452 | 85.441 |
| Smpd4 | 3759.19 | 0.060908 | 0.09375 | 0.649669 | 0.51591 | 0.70609 | 25.475 |
| Smpd5 | 14.4032 | -0.4894 | 0.4644 | -1.05383 | 0.29196 | 0.5018 | 2.549 |
| Smpdl3a | 1104.59 | -0.31389 | 0.11774 | -2.66598 | 0.00768 | 0.04843 | 17.496 |
| Smpdl3b | 503.492 | 0.649666 | 0.19597 | 3.315146 | 0.00092 | 0.01205 | 24.287 |
| Sms | 4673.71 | 0.170269 | 0.07135 | 2.386404 | 0.01701 | 0.0809 | 48.433 |
| Sms-ps | 1069.91 | 0.20717 | 0.13146 | 1.575974 | 0.11503 | 0.28432 | 1.101 |
| Smtn | 289.985 | -0.20976 | 0.1227 | -1.7095 | 0.08736 | 0.23805 | 23.09 |
| Smtnl2 | 34.3594 | -0.30134 | 0.51968 | -0.57985 | 0.56201 | 0.74291 | 22.55 |
| Smu1 | 2009.5 | 0.048174 | 0.08774 | 0.54905 | 0.58297 | 0.75825 | 21.382 |
| Smug1 | 1751.76 | 0.30156 | 0.16302 | 1.849801 | 0.06434 | 0.19522 | 13.803 |
| Smurf1 | 3563.47 | 0.277605 | 0.10526 | 2.637227 | 0.00836 | 0.05098 | 89.353 |
| Smurf2 | 2416.69 | 0.315569 | 0.10612 | 2.973638 | 0.00294 | 0.02585 | 100.65 |
| Smyd1 | 86.5642 | 0.487822 | 0.33977 | 1.435753 | 0.15107 | 0.33587 | 108.294 |
| Smyd2 | 717.587 | -0.53463 | 0.14355 | -3.72442 | 0.0002 | 0.00421 | 41.872 |

| | | | | | | | |
|---|---|---|---|---|---|---|---|
| Smyd3 | 1693.06 | 0.277475 | 0.08321 | 3.334559 | 0.00085 | 0.01148 | 566.082 |
| Smyd4 | 556.508 | -0.19638 | 0.1222 | -1.60701 | 0.10805 | 0.27317 | 57.273 |
| Smyd5 | 2157.06 | 0.17666 | 0.07686 | 2.298587 | 0.02153 | 0.09411 | 14.447 |
| Snai1 | 19.2243 | -1.11034 | 0.47256 | -2.34963 | 0.01879 | 0.08604 | 4.62 |
| Snai2 | 23.6829 | -0.25012 | 0.38053 | -0.6573 | 0.51099 | 0.70208 | 3.534 |
| Snai3 | 26.6631 | 0.486297 | 0.36366 | 1.337234 | 0.18115 | 0.37453 | 6.526 |
| Snap23 | 1269.22 | -0.00767 | 0.09693 | -0.07911 | 0.93695 | 0.9685 | 33.585 |
| Snap25 | 70873.1 | -0.30971 | 0.1094 | -2.83103 | 0.00464 | 0.03467 | 68.976 |
| Snap29 | 1262.58 | -0.14298 | 0.07739 | -1.84759 | 0.06466 | 0.19576 | 24.842 |
| Snap47 | 9376.96 | -0.1171 | 0.07423 | -1.57748 | 0.11469 | 0.28365 | 44.053 |
| Snap91 | 13417.1 | -0.12158 | 0.06947 | -1.75 | 0.08012 | 0.22514 | 114.732 |
| Snapc1 | 2423.95 | 0.009943 | 0.07528 | 0.132073 | 0.89493 | 0.94736 | 24.486 |
| Snapc2 | 725.008 | 0.125413 | 0.11234 | 1.116352 | 0.26427 | 0.47305 | 3.141 |
| Snapc3 | 1630.3 | 0.273885 | 0.09202 | 2.976437 | 0.00292 | 0.02573 | 49.953 |
| Snapc4 | 2164.26 | 0.165036 | 0.08126 | 2.030963 | 0.04226 | 0.14806 | 17.889 |
| Snapc5 | 813.684 | -0.14877 | 0.12286 | -1.21095 | 0.22592 | 0.42994 | 3.859 |
| Snapin | 1802.25 | -0.13587 | 0.08815 | -1.54132 | 0.12324 | 0.29741 | 3.008 |
| Snca | 9262.86 | -0.12564 | 0.10105 | -1.24333 | 0.21375 | 0.41506 | 98.281 |
| Sncaip | 350.09 | 0.320717 | 0.12767 | 2.512088 | 0.012 | 0.06467 | 148.227 |
| Sncb | 9055.78 | -0.66478 | 0.12995 | -5.11547 | 3.13E-07 | 4.36E-05 | 7.736 |
| Sncg | 54.2256 | -0.22001 | 0.43879 | -0.50141 | 0.61608 | 0.78132 | 4.516 |
| Snd1 | 3436.08 | 0.106584 | 0.0692 | 1.540287 | 0.12349 | 0.29779 | 460.024 |
| Sned1 | 704.248 | -0.33621 | 0.17342 | -1.9387 | 0.05254 | 0.1715 | 65.225 |
| Snf8 | 2111.44 | -0.18959 | 0.06886 | -2.75322 | 0.0059 | 0.04083 | 12.546 |
| Snhg1 | 254.463 | 0.143465 | 0.11551 | 1.241987 | 0.21424 | 0.41558 | 3.011 |
| Snhg10 | 92.3674 | -0.14916 | 0.25567 | -0.58341 | 0.55962 | 0.74102 | 1.693 |
| Snhg11 | 125304 | 0.109293 | 0.12323 | 0.886918 | 0.37512 | 0.58522 | 10.514 |
| Snhg12 | 571.894 | 0.029304 | 0.10383 | 0.282235 | 0.77776 | 0.88423 | 2.831 |
| Snhg14 | 4796.48 | 0.169322 | 0.12915 | 1.311073 | 0.18983 | 0.38591 | 667.836 |
| Snhg15 | 102.069 | -0.18672 | 0.19332 | -0.96585 | 0.33412 | 0.54479 | 3.547 |
| Snhg16 | 107.849 | 0.005702 | 0.20741 | 0.027492 | 0.97807 | 0.98827 | 13.469 |
| Snhg17 | 803.529 | 0.110293 | 0.08309 | 1.327347 | 0.18439 | 0.37882 | 7.882 |
| Snhg18 | 66.2512 | -0.19681 | 0.26984 | -0.72938 | 0.46577 | 0.665 | 4.138 |
| Snhg20 | 570.935 | 0.077598 | 0.09352 | 0.829781 | 0.40666 | 0.6148 | 2.957 |

| | | | | | | | |
|---|---|---|---|---|---|---|---|
| Snhg4 | 195.888 | -0.00654 | 0.20588 | -0.03175 | 0.97467 | 0.98734 | 34.305 |
| Snhg5 | 413.195 | 0.117984 | 0.10304 | 1.145067 | 0.25218 | 0.45946 | 28.295 |
| Snhg6 | 366.859 | -0.19353 | 0.14879 | -1.30073 | 0.19335 | 0.39022 | 87.59 |
| Snhg8 | 678.472 | -0.20363 | 0.13884 | -1.46663 | 0.14248 | 0.32436 | 0.86 |
| Snip1 | 608.132 | 0.071158 | 0.08829 | 0.805957 | 0.42027 | 0.62755 | 7.371 |
| Snn | 7696.97 | 0.002943 | 0.06421 | 0.04584 | 0.96344 | 0.98213 | 14.041 |
| Snora30 | 37.3185 | -0.15627 | 0.32214 | -0.48509 | 0.62761 | 0.78926 | 0.13 |
| Snora31 | 40.2712 | -0.07164 | 0.28324 | -0.25293 | 0.80032 | 0.89682 | 0.13 |
| Snora47 | 22.4057 | 0.113214 | 0.34186 | 0.33117 | 0.74052 | 0.86301 | 0.138 |
| Snorc | 25.0755 | 0.6329 | 0.46053 | 1.374279 | 0.16936 | 0.35963 | 5.212 |
| Snord3b-ps2 | 43.6142 | 0.216457 | 0.28453 | 0.760759 | 0.4468 | 0.65034 | 0.214 |
| Snord58b | 26.2728 | -0.26019 | 0.36401 | -0.71479 | 0.47474 | 0.67282 | 0.066 |
| Snord7 | 29.8689 | 0.52383 | 0.36354 | 1.440932 | 0.1496 | 0.33394 | 0.097 |
| Snord71 | 24.503 | -0.10378 | 0.37342 | -0.27792 | 0.78108 | 0.88605 | 0.085 |
| Snord72 | 18.0941 | -0.14554 | 0.58485 | -0.24886 | 0.80347 | 0.89853 | 0.081 |
| Snord73b | 23.9272 | 0.611197 | 0.3946 | 1.54891 | 0.1214 | 0.29466 | 0.071 |
| Snord88c | 41.3717 | 0.509118 | 0.30374 | 1.676179 | 0.0937 | 0.24952 | 0.09 |
| Snord89 | 23.8669 | 0.473051 | 0.33539 | 1.410469 | 0.1584 | 0.34574 | 0.11 |
| Snord91a | 193.843 | 0.057196 | 0.13669 | 0.418429 | 0.67563 | 0.82192 | 0.093 |
| Snord98 | 14.5779 | -0.46304 | 0.60116 | -0.77025 | 0.44115 | 0.64576 | 0.065 |
| Snph | 14024.6 | -0.13796 | 0.07146 | -1.93053 | 0.05354 | 0.17346 | 42.045 |
| Snrk | 4262.54 | -0.03703 | 0.06555 | -0.56487 | 0.57217 | 0.75095 | 52.437 |
| Snrnp200 | 7793.31 | 0.146777 | 0.08835 | 1.661272 | 0.09666 | 0.25469 | 32.066 |
| Snrnp25 | 417.423 | -0.31762 | 0.12914 | -2.45947 | 0.01391 | 0.07098 | 3.57 |
| Snrnp27 | 1396.99 | -0.03082 | 0.08282 | -0.37208 | 0.70983 | 0.84378 | 9.372 |
| Snrnp35 | 457.669 | -0.31217 | 0.1145 | -2.72631 | 0.0064 | 0.04294 | 7.991 |
| Snrnp40 | 914.042 | 0.037366 | 0.1017 | 0.367425 | 0.7133 | 0.84579 | 29.895 |
| Snrnp48 | 2408.98 | 0.087091 | 0.07565 | 1.151179 | 0.24966 | 0.45704 | 22.728 |
| Snrnp70 | 22611.6 | 0.110548 | 0.10705 | 1.032709 | 0.30174 | 0.51136 | 19.29 |
| Snrpa | 1225.63 | 0.089597 | 0.08688 | 1.031307 | 0.3024 | 0.51199 | 9.267 |
| Snrpa1 | 492.37 | -0.00866 | 0.10386 | -0.0834 | 0.93353 | 0.9665 | 15.585 |
| Snrpb | 2083.6 | -0.05429 | 0.09759 | -0.55629 | 0.57801 | 0.75483 | 7.769 |
| Snrpb2 | 1035.49 | -0.03779 | 0.08694 | -0.43469 | 0.66379 | 0.81417 | 9.815 |
| Snrpc | 1115.15 | -0.11048 | 0.10231 | -1.07988 | 0.28019 | 0.49019 | 13.819 |

| | | | | | | | |
|---|---|---|---|---|---|---|---|
| Snrpd1 | 1307.06 | 0.030384 | 0.07441 | 0.408358 | 0.68301 | 0.82675 | 24.305 |
| Snrpd2 | 1518.32 | -0.01317 | 0.07847 | -0.16785 | 0.8667 | 0.9324 | 3.821 |
| Snrpd3 | 2718.87 | -0.10831 | 0.0865 | -1.25216 | 0.21051 | 0.41056 | 19.831 |
| Snrpe | 492.276 | 0.265122 | 0.18078 | 1.466539 | 0.1425 | 0.32436 | 6.421 |
| Snrpert | 66.3216 | -0.47053 | 0.31433 | -1.49691 | 0.13442 | 0.31425 | 0.279 |
| Snrpf | 32.1654 | 0.465877 | 0.3261 | 1.428613 | 0.15312 | 0.33865 | 6.678 |
| Snrpg | 604.544 | -0.01012 | 0.11514 | -0.08787 | 0.92998 | 0.96471 | 8.025 |
| Snrpn | 6147.92 | -0.36416 | 0.08898 | -4.09259 | 4.27E-05 | 0.00146 | 157.725 |
| Snta1 | 1345.79 | -0.30053 | 0.11097 | -2.70831 | 0.00676 | 0.04448 | 31.787 |
| Sntb1 | 336.216 | 1.016632 | 0.25223 | 4.030577 | 5.56E-05 | 0.00179 | 270.562 |
| Sntb2 | 2515.01 | 0.160123 | 0.09423 | 1.699293 | 0.08926 | 0.24196 | 83.965 |
| Sntg1 | 1031.69 | 0.065836 | 0.10083 | 0.652935 | 0.5138 | 0.70456 | 938.405 |
| Sntg2 | 177.584 | 0.12465 | 0.14998 | 0.831137 | 0.4059 | 0.61442 | 198.894 |
| Snu13 | 3252.29 | -0.01118 | 0.07633 | -0.14644 | 0.88357 | 0.94193 | 7.074 |
| Snupn | 570.461 | -0.06752 | 0.11529 | -0.58567 | 0.5581 | 0.73969 | 32.329 |
| Snw1 | 1766.58 | 0.12982 | 0.08406 | 1.544384 | 0.1225 | 0.29627 | 23.2 |
| Snx1 | 3944.31 | 0.058375 | 0.09292 | 0.62825 | 0.52984 | 0.71748 | 38.455 |
| Snx10 | 5776.4 | -0.04725 | 0.08055 | -0.58655 | 0.55751 | 0.73919 | 66.779 |
| Snx11 | 1818.33 | -0.01117 | 0.06751 | -0.16543 | 0.86861 | 0.9334 | 10.005 |
| Snx12 | 2461.64 | -0.0855 | 0.07428 | -1.15101 | 0.24973 | 0.45704 | 124.778 |
| Snx13 | 4137.47 | 0.091756 | 0.10431 | 0.879664 | 0.37904 | 0.58872 | 100.294 |
| Snx14 | 2334.68 | 0.154461 | 0.07595 | 2.033827 | 0.04197 | 0.14726 | 62.212 |
| Snx15 | 1532.86 | 0.098847 | 0.1069 | 0.924708 | 0.35512 | 0.56548 | 8.906 |
| Snx16 | 973.868 | 0.203118 | 0.09145 | 2.22115 | 0.02634 | 0.10784 | 22.286 |
| Snx17 | 2448.34 | -0.03182 | 0.07043 | -0.45186 | 0.65137 | 0.80592 | 5.917 |
| Snx18 | 1350.45 | -0.37356 | 0.07984 | -4.67855 | 2.89E-06 | 0.00023 | 26.386 |
| Snx19 | 4184.4 | 0.018483 | 0.0799 | 0.231326 | 0.81706 | 0.90593 | 39.626 |
| Snx2 | 2525.71 | 0.199748 | 0.08664 | 2.305402 | 0.02114 | 0.09295 | 44.55 |
| Snx20 | 44.4155 | -0.31422 | 0.26201 | -1.19926 | 0.23043 | 0.43483 | 9.566 |
| Snx21 | 1141.01 | -0.02012 | 0.0836 | -0.24067 | 0.80981 | 0.90197 | 7.994 |
| Snx22 | 192.501 | -0.18934 | 0.14289 | -1.3251 | 0.18514 | 0.37985 | 4.544 |
| Snx24 | 672.98 | 0.113325 | 0.11324 | 1.000764 | 0.31694 | 0.52743 | 145.259 |
| Snx25 | 3874.64 | 0.407906 | 0.10953 | 3.724039 | 0.0002 | 0.00421 | 118.899 |
| Snx27 | 9322.9 | 0.248099 | 0.0816 | 3.040558 | 0.00236 | 0.02243 | 85.173 |

| | | | | | | | |
|---|---|---|---|---|---|---|---|
| Snx29 | 695.533 | -0.06258 | 0.09784 | -0.63965 | 0.5224 | 0.71129 | 432.565 |
| Snx3 | 4576.43 | 0.013696 | 0.07664 | 0.178715 | 0.85816 | 0.92854 | 33.352 |
| Snx30 | 2354.26 | -0.13458 | 0.08158 | -1.64956 | 0.09903 | 0.25857 | 98.898 |
| Snx31 | 37.6258 | -0.61591 | 0.29526 | -2.08597 | 0.03698 | 0.13585 | 51.511 |
| Snx32 | 3582.7 | 0.078805 | 0.07563 | 1.041977 | 0.29742 | 0.50755 | 18.304 |
| Snx33 | 527.911 | -0.51651 | 0.12797 | -4.03624 | 5.43E-05 | 0.00176 | 11.179 |
| Snx4 | 1505.55 | -0.04717 | 0.07293 | -0.64674 | 0.5178 | 0.70757 | 48.828 |
| Snx5 | 3723.22 | -0.06241 | 0.06789 | -0.91922 | 0.35798 | 0.56813 | 20.784 |
| Snx6 | 1681.43 | 0.036624 | 0.07293 | 0.502181 | 0.61554 | 0.78122 | 49.355 |
| Snx7 | 795.235 | 0.101952 | 0.11218 | 0.908785 | 0.36346 | 0.57384 | 87.29 |
| Snx8 | 412.833 | -0.40385 | 0.136 | -2.9695 | 0.00298 | 0.02607 | 48.964 |
| Snx9 | 799.156 | -0.13592 | 0.07605 | -1.78724 | 0.0739 | 0.21422 | 90.626 |
| Soat1 | 2790.05 | -0.00848 | 0.07143 | -0.11866 | 0.90555 | 0.95256 | 49.807 |
| Sobp | 2301.69 | -0.53732 | 0.15177 | -3.54038 | 0.0004 | 0.00687 | 172.031 |
| Socs1 | 14.4084 | -0.52136 | 0.61457 | -0.84834 | 0.39625 | 0.60539 | 3.297 |
| Socs2 | 704.359 | -0.30139 | 0.18446 | -1.63393 | 0.10227 | 0.26399 | 31.819 |
| Socs3 | 112.085 | -1.31109 | 0.31772 | -4.12653 | 3.68E-05 | 0.00132 | 3.969 |
| Socs4 | 1366.48 | 0.093953 | 0.09533 | 0.985524 | 0.32437 | 0.53542 | 19.172 |
| Socs5 | 7154.82 | 0.031057 | 0.06162 | 0.503995 | 0.61427 | 0.78038 | 30.161 |
| Socs6 | 1310.22 | 0.226638 | 0.09466 | 2.394183 | 0.01666 | 0.0799 | 262.258 |
| Socs7 | 8346.54 | 0.093615 | 0.0595 | 1.573353 | 0.11564 | 0.28537 | 36.108 |
| Sod1 | 5972.05 | 0.064364 | 0.07308 | 0.880782 | 0.37844 | 0.58827 | 5.576 |
| Sod2 | 9970.84 | -0.00119 | 0.05705 | -0.02088 | 0.98334 | 0.99092 | 33.218 |
| Sod3 | 470.171 | -0.60394 | 0.12491 | -4.83502 | 1.33E-06 | 0.00013 | 7.628 |
| Soga1 | 2597.4 | -0.01526 | 0.08665 | -0.1761 | 0.86022 | 0.92937 | 63.456 |
| Soga3 | 2606.92 | -0.04462 | 0.07141 | -0.62475 | 0.53213 | 0.71963 | 55.792 |
| Son | 27053 | 0.160384 | 0.08365 | 1.917246 | 0.05521 | 0.17713 | 31.716 |
| Sorbs1 | 5443.5 | 0.205237 | 0.10001 | 2.052116 | 0.04016 | 0.14321 | 219.027 |
| Sorbs2 | 8712.63 | 0.082496 | 0.09726 | 0.848235 | 0.39631 | 0.60539 | 320.119 |
| Sorbs2os | 340.936 | 0.228866 | 0.20313 | 1.126698 | 0.25987 | 0.46839 | 95.973 |
| Sorbs3 | 1444.19 | -0.24977 | 0.07932 | -3.14881 | 0.00164 | 0.01763 | 31.522 |
| Sorcs1 | 2224.81 | 0.389655 | 0.15629 | 2.493146 | 0.01266 | 0.06707 | 535.348 |
| Sorcs2 | 3795.33 | -0.17827 | 0.11423 | -1.56067 | 0.1186 | 0.29054 | 380.96 |
| Sorcs3 | 2566.62 | 0.004304 | 0.08625 | 0.049902 | 0.9602 | 0.98055 | 599.481 |

| | | | | | | | |
|---|---|---|---|---|---|---|---|
| Sord | 799.084 | -0.07474 | 0.09536 | -0.78379 | 0.43317 | 0.63921 | 30.592 |
| Sorl1 | 16571.3 | 0.005937 | 0.06729 | 0.088225 | 0.9297 | 0.96459 | 159.578 |
| Sort1 | 23180.5 | -0.14986 | 0.06394 | -2.34367 | 0.0191 | 0.08675 | 77.43 |
| Sos1 | 3882.56 | 0.149227 | 0.08475 | 1.760687 | 0.07829 | 0.2215 | 86.703 |
| Sos2 | 2505.4 | 0.165058 | 0.06029 | 2.737721 | 0.00619 | 0.04207 | 98.091 |
| Sostdc1 | 12.5853 | -0.57854 | 0.51649 | -1.12013 | 0.26266 | 0.47151 | 4.314 |
| Sowaha | 8418.66 | 0.480852 | 0.14402 | 3.338853 | 0.00084 | 0.01137 | 3.702 |
| Sowahb | 403.757 | -0.67503 | 0.2273 | -2.96984 | 0.00298 | 0.02606 | 3.9 |
| Sowahc | 505.822 | -0.25174 | 0.164 | -1.53501 | 0.12478 | 0.2998 | 4.482 |
| Sox1 | 2175.85 | 0.430806 | 0.15156 | 2.842467 | 0.00448 | 0.03387 | 4.832 |
| Sox10 | 1225.73 | -0.53078 | 0.15962 | -3.32536 | 0.00088 | 0.01174 | 10.333 |
| Sox11 | 2842.11 | 0.419837 | 0.09186 | 4.570431 | 4.87E-06 | 0.00033 | 8.311 |
| Sox12 | 1122.44 | 0.192603 | 0.07212 | 2.670597 | 0.00757 | 0.04802 | 4.453 |
| Sox13 | 386.127 | 0.042402 | 0.13401 | 0.316416 | 0.75169 | 0.86985 | 42.075 |
| Sox17 | 358.096 | -0.1887 | 0.15214 | -1.24035 | 0.21485 | 0.41624 | 6.424 |
| Sox18 | 210.143 | -0.5804 | 0.16371 | -3.54539 | 0.00039 | 0.00676 | 1.805 |
| Sox1ot | 4390.63 | 0.77016 | 0.18634 | 4.133084 | 3.58E-05 | 0.0013 | 50.998 |
| Sox2 | 1647.28 | -0.27037 | 0.08101 | -3.33739 | 0.00085 | 0.01141 | 2.057 |
| Sox21 | 330.05 | -0.54333 | 0.14242 | -3.81514 | 0.00014 | 0.00331 | 3.799 |
| Sox2ot | 1327.82 | 0.026265 | 0.1474 | 0.178194 | 0.85857 | 0.92859 | 578.35 |
| Sox3 | 44.682 | -0.24652 | 0.28606 | -0.86179 | 0.38881 | 0.59796 | 2.065 |
| Sox4 | 1643.02 | -0.60944 | 0.15455 | -3.94341 | 8.03E-05 | 0.00231 | 4.795 |
| Sox5 | 1659.53 | -0.40876 | 0.20095 | -2.03413 | 0.04194 | 0.1472 | 953.553 |
| Sox6 | 1326.63 | -0.10103 | 0.1057 | -0.95579 | 0.33918 | 0.55007 | 567.925 |
| Sox7 | 78.1468 | -0.14526 | 0.28348 | -0.51241 | 0.60836 | 0.77621 | 7.06 |
| Sox8 | 2019.02 | 0.145793 | 0.09504 | 1.533942 | 0.12504 | 0.30014 | 4.795 |
| Sox9 | 1213.45 | -0.55981 | 0.12384 | -4.5204 | 6.17E-06 | 0.00039 | 5.537 |
| Sp1 | 1897.97 | -0.06867 | 0.0701 | -0.97968 | 0.32724 | 0.5386 | 30.262 |
| Sp100 | 170.611 | -0.49279 | 0.15061 | -3.272 | 0.00107 | 0.01339 | 60.011 |
| Sp110 | 39.5026 | -0.41252 | 0.30529 | -1.35123 | 0.17662 | 0.36865 | 21.919 |
| Sp2 | 706.785 | -0.06207 | 0.13147 | -0.47213 | 0.63683 | 0.79566 | 29.619 |
| Sp3 | 2911.38 | 0.192783 | 0.08374 | 2.302281 | 0.02132 | 0.09344 | 44.02 |
| Sp3os | 260.824 | 0.067331 | 0.1333 | 0.505095 | 0.61349 | 0.77991 | 9.829 |
| Sp4 | 1624.66 | 0.19309 | 0.09289 | 2.078655 | 0.03765 | 0.13734 | 66.508 |

| | | | | | | | |
|---|---|---|---|---|---|---|---|
| Sp5 | 16.2569 | -0.79914 | 0.5121 | -1.56051 | 0.11864 | 0.29055 | 2.807 |
| Sp7 | 92.0493 | -0.95834 | 0.3831 | -2.50152 | 0.01237 | 0.06597 | 10.577 |
| Sp8 | 207.646 | -1.67408 | 0.66002 | -2.53641 | 0.0112 | 0.06177 | 6.248 |
| Sp9 | 988.864 | 0.487008 | 0.16148 | 3.015958 | 0.00256 | 0.02375 | 12.782 |
| Spa17 | 47.6654 | -0.12344 | 0.26872 | -0.45937 | 0.64597 | 0.80278 | 10.433 |
| Spaar | 71.7631 | -0.36833 | 0.30552 | -1.20557 | 0.22798 | 0.43189 | 29.429 |
| Spaca6 | 3766.54 | 0.183033 | 0.12296 | 1.488561 | 0.1366 | 0.31714 | 15.852 |
| Spag1 | 1147.79 | 0.151948 | 0.08769 | 1.73273 | 0.08314 | 0.23021 | 57.523 |
| Spag4 | 154.47 | -0.02985 | 0.17586 | -0.16971 | 0.86524 | 0.93172 | 5.443 |
| Spag5 | 215.469 | -0.36605 | 0.186 | -1.96804 | 0.04906 | 0.16382 | 20.929 |
| Spag6 | 78.7073 | -0.41689 | 0.32969 | -1.26449 | 0.20606 | 0.40531 | 56.382 |
| Spag6l | 242.231 | 0.357656 | 0.17181 | 2.08169 | 0.03737 | 0.13674 | 76.441 |
| Spag7 | 1458.59 | 0.098921 | 0.07295 | 1.356001 | 0.1751 | 0.36679 | 5.646 |
| Spag8 | 58.7148 | 0.008657 | 0.27612 | 0.031351 | 0.97499 | 0.98739 | 2.26 |
| Spag9 | 22163.5 | 0.092154 | 0.07149 | 1.289067 | 0.19737 | 0.39524 | 129.995 |
| Sparc | 8113.46 | -0.00857 | 0.15395 | -0.05569 | 0.95559 | 0.97814 | 28.684 |
| Sparcl1 | 75090.6 | -0.19752 | 0.07665 | -2.57689 | 0.00997 | 0.05737 | 34.623 |
| Spast | 2965.33 | -0.07369 | 0.08914 | -0.82672 | 0.4084 | 0.61619 | 52.129 |
| Spata1 | 150.079 | 0.175587 | 0.17185 | 1.021746 | 0.3069 | 0.5168 | 42.558 |
| Spata13 | 3143.85 | 0.974218 | 0.28333 | 3.438513 | 0.00058 | 0.00882 | 130.556 |
| Spata17 | 20.9553 | -0.40204 | 0.42697 | -0.94161 | 0.34639 | 0.557 | 170.818 |
| Spata2 | 2314.54 | -0.11747 | 0.08847 | -1.32771 | 0.18427 | 0.37873 | 11.755 |
| Spata21 | 20.1009 | -0.4021 | 0.46965 | -0.85615 | 0.39191 | 0.60106 | 24.428 |
| Spata24 | 33.789 | -0.08257 | 0.2904 | -0.28434 | 0.77615 | 0.88346 | 5.498 |
| Spata2l | 2008.17 | 0.038316 | 0.10448 | 0.366739 | 0.71381 | 0.84625 | 6.433 |
| Spata3 | 30.397 | -0.87882 | 0.37414 | -2.34889 | 0.01883 | 0.08608 | 13.143 |
| Spata32 | 37.0345 | -0.17211 | 0.35893 | -0.4795 | 0.63158 | 0.79201 | 10.306 |
| Spata33 | 397.905 | 0.111468 | 0.10388 | 1.073063 | 0.28324 | 0.49339 | 9.215 |
| Spata4 | 64.7764 | 0.845032 | 0.24512 | 3.44747 | 0.00057 | 0.00862 | 9.318 |
| Spata45 | 39.4967 | 0.042624 | 0.30594 | 0.139322 | 0.8892 | 0.94457 | 6.304 |
| Spata46 | 51.4115 | 0.063148 | 0.2823 | 0.223691 | 0.823 | 0.90818 | 3.323 |
| Spata5 | 531.201 | -0.04543 | 0.13588 | -0.33435 | 0.73811 | 0.86177 | 159.201 |
| Spata5l1 | 497.133 | 0.203348 | 0.11643 | 1.746538 | 0.08072 | 0.2261 | 13.631 |
| Spata6 | 145.02 | -0.47538 | 0.16985 | -2.79885 | 0.00513 | 0.03691 | 109.201 |

| | | | | | | | |
|---|---|---|---|---|---|---|---|
| Spata6l | 128.486 | -0.40928 | 0.22331 | -1.83283 | 0.06683 | 0.20016 | 74.759 |
| Spata7 | 816.683 | 0.0873 | 0.09696 | 0.90033 | 0.36794 | 0.57836 | 41.659 |
| Spata9 | 86.2714 | -0.1291 | 0.22371 | -0.57711 | 0.56386 | 0.74426 | 31.823 |
| Spats1 | 133.37 | 0.268543 | 0.19528 | 1.375197 | 0.16907 | 0.35922 | 34.838 |
| Spats2 | 1368.27 | -0.03343 | 0.07068 | -0.47304 | 0.63619 | 0.79507 | 87.3 |
| Spats2l | 715.507 | 0.327739 | 0.09969 | 3.287551 | 0.00101 | 0.01291 | 174.233 |
| Spc24 | 103.155 | 0.201339 | 0.19444 | 1.035462 | 0.30045 | 0.5105 | 4.862 |
| Spc25 | 28.3884 | -0.59082 | 0.40814 | -1.44757 | 0.14774 | 0.33129 | 12.3 |
| Spcs1 | 2272.5 | 0.037942 | 0.0853 | 0.444827 | 0.65644 | 0.8092 | 1.847 |
| Spcs2 | 4152.84 | -0.12392 | 0.08994 | -1.37778 | 0.16827 | 0.35827 | 25.894 |
| Spcs3 | 3778.05 | 0.118688 | 0.0635 | 1.869071 | 0.06161 | 0.18981 | 9.566 |
| Spdl1 | 27.8401 | 0.175784 | 0.46101 | 0.381302 | 0.70298 | 0.8397 | 24.452 |
| Spdya | 52.2566 | -0.06267 | 0.23013 | -0.27234 | 0.78536 | 0.88859 | 37.473 |
| Specc1 | 5305.69 | -0.07299 | 0.06632 | -1.10061 | 0.27107 | 0.48039 | 266.251 |
| Specc1l | 3896.83 | 0.152726 | 0.073 | 2.092077 | 0.03643 | 0.13444 | 100.673 |
| Speer4b | 15.5785 | -0.11689 | 0.62287 | -0.18766 | 0.85114 | 0.92423 | 5.8 |
| Spef1 | 386.401 | -0.15145 | 0.127 | -1.19249 | 0.23307 | 0.43798 | 17.022 |
| Spef2 | 56.3303 | -0.66807 | 0.24683 | -2.70655 | 0.0068 | 0.04463 | 170.676 |
| Speg | 11063.5 | 0.139982 | 0.07296 | 1.918671 | 0.05503 | 0.17675 | 57.024 |
| Spem3 | 16.9787 | 0.682356 | 0.46801 | 1.457999 | 0.14484 | 0.32761 | 4.334 |
| Spen | 6055.25 | 0.097438 | 0.05501 | 1.77119 | 0.07653 | 0.21867 | 70.708 |
| Spg11 | 1706.42 | 0.13471 | 0.08364 | 1.610687 | 0.10725 | 0.27181 | 64.867 |
| Spg20 | 2120.13 | -0.07452 | 0.096 | -0.77621 | 0.43763 | 0.64306 | 25.215 |
| Spg21 | 1269.85 | 0.198008 | 0.10954 | 1.807604 | 0.07067 | 0.20777 | 27.524 |
| Spg7 | 2642.81 | -0.23493 | 0.07186 | -3.26909 | 0.00108 | 0.01345 | 34.819 |
| Sphk1 | 147.475 | -0.99371 | 0.27587 | -3.60216 | 0.00032 | 0.00585 | 5.751 |
| Sphk2 | 993.827 | -0.09038 | 0.08479 | -1.06601 | 0.28642 | 0.49639 | 8.536 |
| Sphkap | 6594.53 | -0.67168 | 0.29544 | -2.27351 | 0.023 | 0.09812 | 154.062 |
| Spi1 | 101.457 | -0.49774 | 0.19498 | -2.55281 | 0.01069 | 0.06004 | 33.367 |
| Spice1 | 770.489 | -0.02228 | 0.08641 | -0.25787 | 0.79651 | 0.89514 | 41.377 |
| Spidr | 130.109 | -0.44105 | 0.1789 | -2.46538 | 0.01369 | 0.07038 | 257.628 |
| Spin1 | 10676.4 | 0.086087 | 0.06181 | 1.39283 | 0.16367 | 0.35241 | 51.667 |
| Spin2c | 151.006 | -0.16595 | 0.20312 | -0.81699 | 0.41394 | 0.622 | 2.019 |
| Spin4 | 46.8456 | 0.095976 | 0.26747 | 0.358824 | 0.71973 | 0.85033 | 4.173 |

| | | | | | | | |
|---|---|---|---|---|---|---|---|
| Spindoc | 873.831 | 0.080893 | 0.07782 | 1.039476 | 0.29858 | 0.50864 | 28.917 |
| Spink10 | 315.06 | 0.036408 | 0.27752 | 0.131193 | 0.89562 | 0.94747 | 119.16 |
| Spink8 | 173.06 | 0.890475 | 0.17213 | 5.173342 | 2.30E-07 | 3.32E-05 | 14.354 |
| Spint1 | 387.881 | 0.695282 | 0.20341 | 3.418113 | 0.00063 | 0.00927 | 12.166 |
| Spint2 | 645.556 | 0.331168 | 0.13325 | 2.485301 | 0.01294 | 0.06803 | 25.59 |
| Spire1 | 8745.4 | -0.20458 | 0.06959 | -2.93977 | 0.00328 | 0.02767 | 122.582 |
| Spire2 | 701.189 | 0.272942 | 0.10167 | 2.684597 | 0.00726 | 0.04667 | 36.803 |
| Spns1 | 2719.58 | 0.058552 | 0.0646 | 0.906398 | 0.36473 | 0.57539 | 7.391 |
| Spns2 | 2001.23 | -0.44085 | 0.12813 | -3.44065 | 0.00058 | 0.00878 | 38.267 |
| Spns3 | 19.739 | 0.747622 | 0.39528 | 1.891385 | 0.05857 | 0.18403 | 55.588 |
| Spo11 | 35.4597 | -0.82708 | 0.57908 | -1.42825 | 0.15322 | 0.33878 | 15.877 |
| Spocd1 | 48.5254 | 0.486682 | 0.25519 | 1.907113 | 0.05651 | 0.17992 | 27.867 |
| Spock1 | 8641.22 | -0.10267 | 0.07758 | -1.32347 | 0.18568 | 0.38052 | 487.139 |
| Spock2 | 48986.6 | 0.060291 | 0.06833 | 0.882294 | 0.37762 | 0.58773 | 28.98 |
| Spock3 | 8679.28 | 0.612583 | 0.17616 | 3.477346 | 0.00051 | 0.00798 | 406.095 |
| Spon1 | 3596.62 | -0.39447 | 0.13848 | -2.84851 | 0.00439 | 0.0335 | 277.371 |
| Spop | 12731.3 | 0.035131 | 0.062 | 0.566645 | 0.57096 | 0.75014 | 79.327 |
| Spopl | 1234.4 | 0.29875 | 0.10683 | 2.79654 | 0.00517 | 0.03705 | 65.887 |
| Spout1 | 1082.16 | -0.01353 | 0.08251 | -0.16393 | 0.86979 | 0.93413 | 5.007 |
| Spp1 | 395.594 | -0.74787 | 0.20858 | -3.58563 | 0.00034 | 0.00611 | 5.933 |
| Sppl2a | 3434.37 | 0.206045 | 0.08023 | 2.568153 | 0.01022 | 0.05839 | 42.845 |
| Sppl2b | 2135.17 | 0.160132 | 0.08223 | 1.947305 | 0.0515 | 0.16923 | 13.434 |
| Sppl3 | 1882.14 | -0.00062 | 0.0705 | -0.00882 | 0.99297 | 0.99595 | 87.654 |
| Spr | 782.263 | -0.23515 | 0.09877 | -2.38092 | 0.01727 | 0.08154 | 7.591 |
| Spred1 | 11126.1 | 0.191046 | 0.06421 | 2.975418 | 0.00293 | 0.02575 | 60.906 |
| Spred2 | 2922.03 | -0.10349 | 0.10007 | -1.03413 | 0.30107 | 0.51079 | 99.652 |
| Spred3 | 2498.64 | -0.18208 | 0.11157 | -1.63196 | 0.10269 | 0.26462 | 11.583 |
| Spring1 | 2363.84 | 0.112665 | 0.0937 | 1.202432 | 0.2292 | 0.43327 | 20.867 |
| Sprn | 3771.1 | -0.25971 | 0.13276 | -1.95629 | 0.05043 | 0.16691 | 4.25 |
| Sprtn | 1202.48 | 0.122874 | 0.08558 | 1.435789 | 0.15106 | 0.33587 | 8.276 |
| Spry1 | 270.547 | -0.01952 | 0.13851 | -0.14091 | 0.88794 | 0.94406 | 4.652 |
| Spry2 | 2644.89 | -0.05737 | 0.08921 | -0.64314 | 0.52014 | 0.70963 | 4.873 |
| Spry4 | 563.862 | -0.38104 | 0.19148 | -1.98995 | 0.0466 | 0.15807 | 15.169 |
| Spryd3 | 6504.17 | -0.00947 | 0.08644 | -0.10954 | 0.91278 | 0.95597 | 19.707 |

| | | | | | | | |
|---|---|---|---|---|---|---|---|
| Spryd4 | 237.325 | -0.12543 | 0.13775 | -0.91058 | 0.36251 | 0.57294 | 1.9 |
| Spryd7 | 2267.72 | 0.067465 | 0.09214 | 0.732209 | 0.46404 | 0.66379 | 24.894 |
| Spsb1 | 795.929 | -0.51006 | 0.15527 | -3.28493 | 0.00102 | 0.01298 | 58.761 |
| Spsb2 | 131.333 | -0.07304 | 0.19105 | -0.38232 | 0.70222 | 0.83934 | 1.959 |
| Spsb3 | 231.921 | -0.18539 | 0.12078 | -1.53495 | 0.1248 | 0.2998 | 5.51 |
| Spsb4 | 99.8741 | -0.60952 | 0.17446 | -3.49386 | 0.00048 | 0.00768 | 75.362 |
| Sptan1 | 46123 | -0.00453 | 0.05622 | -0.08057 | 0.93579 | 0.96779 | 65.892 |
| Sptb | 16407.1 | 0.289642 | 0.10545 | 2.746844 | 0.00602 | 0.04135 | 130.06 |
| Sptbn1 | 52447.2 | -0.01784 | 0.04902 | -0.36389 | 0.71594 | 0.8481 | 168.781 |
| Sptbn2 | 41766.1 | -0.10708 | 0.09641 | -1.11068 | 0.26671 | 0.47595 | 41.194 |
| Sptbn4 | 19048.1 | 0.278591 | 0.07759 | 3.590736 | 0.00033 | 0.00602 | 91.304 |
| Sptbn5 | 165.94 | 0.136346 | 0.16933 | 0.80519 | 0.42071 | 0.6278 | 44.186 |
| Sptlc1 | 1872.31 | -0.05585 | 0.06643 | -0.84071 | 0.40051 | 0.60926 | 44.65 |
| Sptlc2 | 1708.02 | -0.08953 | 0.08281 | -1.08114 | 0.27964 | 0.48966 | 83.298 |
| Sptssa | 965.699 | -0.00557 | 0.08429 | -0.06603 | 0.94735 | 0.97353 | 11.182 |
| Sptssb | 73.4976 | -0.54564 | 0.33302 | -1.63847 | 0.10132 | 0.26262 | 40.399 |
| Spty2d1 | 1082.69 | 0.120554 | 0.07938 | 1.518749 | 0.12883 | 0.30601 | 18.016 |
| Spx | 153.131 | 0.010853 | 0.22438 | 0.04837 | 0.96142 | 0.98121 | 6.29 |
| Sqle | 2011.78 | -0.29298 | 0.12379 | -2.36666 | 0.01795 | 0.08348 | 16.116 |
| Sqor | 198.528 | -0.31005 | 0.15311 | -2.02501 | 0.04287 | 0.14929 | 44.333 |
| Sqstm1 | 15780.4 | -0.09341 | 0.07256 | -1.28736 | 0.19797 | 0.3957 | 11.462 |
| Sra1 | 1519.35 | -0.00264 | 0.07641 | -0.03456 | 0.97243 | 0.98616 | 4.135 |
| Srarp | 106.832 | -0.17897 | 0.19262 | -0.92917 | 0.3528 | 0.56311 | 3.434 |
| Srbd1 | 632.593 | 0.075408 | 0.09709 | 0.776651 | 0.43736 | 0.64277 | 160.511 |
| Src | 4283.79 | -0.15987 | 0.06975 | -2.29206 | 0.0219 | 0.09525 | 53.419 |
| Srcap | 310.192 | 0.104513 | 0.15163 | 0.689245 | 0.49067 | 0.68571 | 49.237 |
| Srcin1 | 19112 | -0.28874 | 0.09399 | -3.07193 | 0.00213 | 0.02101 | 66.847 |
| Srd5a1 | 269.193 | -0.38462 | 0.12542 | -3.06654 | 0.00217 | 0.02125 | 37.994 |
| Srd5a3 | 504.304 | -0.0221 | 0.09612 | -0.22996 | 0.81812 | 0.90638 | 15.234 |
| Srebf1 | 2576.15 | 0.003546 | 0.09924 | 0.03573 | 0.9715 | 0.98567 | 23.493 |
| Srebf2 | 10040 | -0.15862 | 0.06442 | -2.46212 | 0.01381 | 0.07064 | 58.199 |
| Srek1 | 6589.55 | 0.25944 | 0.12296 | 2.109939 | 0.03486 | 0.13083 | 35.261 |
| Srek1ip1 | 974.083 | 0.160283 | 0.08061 | 1.988323 | 0.04678 | 0.1586 | 46.791 |
| Srf | 2357.32 | 0.159895 | 0.0892 | 1.792479 | 0.07306 | 0.21267 | 7.847 |

| | | | | | | | |
|---|---|---|---|---|---|---|---|
| Srfbp1 | 818.668 | 0.040058 | 0.12531 | 0.31967 | 0.74922 | 0.86842 | 26.239 |
| Srgap1 | 2375.58 | 0.524916 | 0.13113 | 4.00293 | 6.26E-05 | 0.00194 | 266.325 |
| Srgap2 | 5391.27 | -0.01731 | 0.06144 | -0.28168 | 0.77819 | 0.88423 | 242.102 |
| Srgap3 | 19770.1 | -0.08877 | 0.07631 | -1.16326 | 0.24472 | 0.45139 | 229.296 |
| Srgn | 354.828 | -0.39055 | 0.13983 | -2.79297 | 0.00522 | 0.03732 | 33.619 |
| Sri | 2166.06 | -0.25802 | 0.07655 | -3.37077 | 0.00075 | 0.01051 | 23.302 |
| Srl | 317.173 | -0.29901 | 0.1557 | -1.92039 | 0.05481 | 0.17623 | 61.601 |
| Srm | 1679.31 | 0.051258 | 0.08779 | 0.583848 | 0.55932 | 0.74077 | 3.491 |
| Srms | 22.0186 | 0.001719 | 0.3631 | 0.004735 | 0.99622 | 0.99789 | 7.624 |
| Sros1 | 15.8723 | -0.5108 | 0.46199 | -1.10565 | 0.26888 | 0.47813 | 4.869 |
| Srp14 | 2500.4 | -0.01377 | 0.09701 | -0.14194 | 0.88713 | 0.94391 | 3.862 |
| Srp19 | 1486.79 | -0.051 | 0.07387 | -0.69037 | 0.48996 | 0.68496 | 5.753 |
| Srp54a | 285.451 | -0.06197 | 0.19437 | -0.31881 | 0.74987 | 0.8687 | 35.106 |
| Srp54b | 304.932 | 0.424954 | 1.4878 | 0.285625 | 0.77517 | 0.88288 | 34.47 |
| Srp54c | 789.241 | -0.1198 | 0.11283 | -1.06175 | 0.28835 | 0.49846 | 33.669 |
| Srp68 | 2990.59 | 0.0239 | 0.06762 | 0.353455 | 0.72375 | 0.85252 | 29.052 |
| Srp72 | 5282.64 | 0.102799 | 0.0618 | 1.66342 | 0.09623 | 0.25398 | 25.255 |
| Srp9 | 2497.91 | 0.01666 | 0.07119 | 0.234029 | 0.81496 | 0.905 | 7.671 |
| Srpk1 | 4112.26 | 0.16498 | 0.06803 | 2.425178 | 0.0153 | 0.07541 | 35.062 |
| Srpk2 | 11110.2 | 0.027612 | 0.07091 | 0.389398 | 0.69698 | 0.8358 | 181.354 |
| Srpk3 | 270.532 | -0.4309 | 0.17622 | -2.44517 | 0.01448 | 0.07271 | 4.521 |
| Srpr | 3579.24 | -0.00147 | 0.08454 | -0.01742 | 0.9861 | 0.99248 | 47.799 |
| Srprb | 1709.46 | -0.02349 | 0.09117 | -0.25771 | 0.79663 | 0.89517 | 14.106 |
| Srpx | 19.1797 | 0.013788 | 0.39956 | 0.034508 | 0.97247 | 0.98616 | 79.733 |
| Srpx2 | 31.5096 | 0.103042 | 0.33319 | 0.309257 | 0.75713 | 0.87208 | 24.021 |
| Srr | 2014 | 0.004183 | 0.06574 | 0.063634 | 0.94926 | 0.9749 | 19.59 |
| Srrd | 316.343 | 0.199969 | 0.11581 | 1.726662 | 0.08423 | 0.23239 | 5.649 |
| Srrm1 | 10398.2 | 0.3692 | 0.07374 | 5.006704 | 5.54E-07 | 6.97E-05 | 32.838 |
| Srrm2 | 43610.6 | 0.241337 | 0.10141 | 2.379828 | 0.01732 | 0.08165 | 34.21 |
| Srrm3 | 3868.27 | 0.07151 | 0.07678 | 0.931352 | 0.35167 | 0.56204 | 67.883 |
| Srrm3os | 68.2531 | 0.203647 | 0.27359 | 0.744354 | 0.45666 | 0.65819 | 1.89 |
| Srrm4 | 1944.31 | -0.35013 | 0.15223 | -2.29995 | 0.02145 | 0.09385 | 152.543 |
| Srrm4os | 268.068 | -0.21164 | 0.17926 | -1.18065 | 0.23774 | 0.44318 | 26.776 |
| Srrt | 7209.98 | 0.157501 | 0.07967 | 1.976824 | 0.04806 | 0.16148 | 11.971 |

| | | | | | | | |
|---|---|---|---|---|---|---|---|
| Srsf1 | 10872.7 | 0.231797 | 0.08762 | 2.645341 | 0.00816 | 0.05018 | 6.383 |
| Srsf10 | 4730.24 | 0.119578 | 0.07449 | 1.605381 | 0.10841 | 0.27367 | 14.162 |
| Srsf11 | 11741.9 | 0.252527 | 0.0962 | 2.624994 | 0.00867 | 0.0522 | 26.167 |
| Srsf12 | 916.878 | 0.119005 | 0.09554 | 1.245654 | 0.21289 | 0.414 | 24.35 |
| Srsf2 | 4358.32 | 0.073795 | 0.09113 | 0.809755 | 0.41808 | 0.62541 | 3.194 |
| Srsf3 | 9030.29 | 0.112152 | 0.06352 | 1.765625 | 0.07746 | 0.22018 | 10.746 |
| Srsf4 | 1128.33 | 0.138276 | 0.09636 | 1.435001 | 0.15129 | 0.33615 | 28.09 |
| Srsf5 | 21742.5 | 0.15424 | 0.09571 | 1.611524 | 0.10707 | 0.27158 | 5.004 |
| Srsf6 | 9395.35 | 0.127844 | 0.06339 | 2.016667 | 0.04373 | 0.15129 | 5.594 |
| Srsf7 | 7073.82 | 0.252635 | 0.07187 | 3.514985 | 0.00044 | 0.0073 | 7.228 |
| Srsf9 | 2762.29 | 0.112937 | 0.08161 | 1.383846 | 0.16641 | 0.356 | 5.904 |
| Srxn1 | 5080.22 | 0.13399 | 0.09181 | 1.459424 | 0.14445 | 0.32717 | 5.861 |
| Ss18 | 1679.7 | -0.08791 | 0.09152 | -0.96056 | 0.33677 | 0.54739 | 58.717 |
| Ss18l1 | 2718.28 | 0.100661 | 0.06711 | 1.499875 | 0.13365 | 0.31339 | 27.693 |
| Ss18l2 | 1452.26 | -0.11926 | 0.081 | -1.47236 | 0.14092 | 0.32265 | 3.06 |
| Ssb | 3810.64 | 0.221374 | 0.14618 | 1.514374 | 0.12993 | 0.30778 | 10.285 |
| Ssbp1 | 951.283 | 0.098172 | 0.08782 | 1.117824 | 0.26364 | 0.47241 | 13.349 |
| Ssbp2 | 3203.44 | 0.266849 | 0.08754 | 3.048228 | 0.0023 | 0.02206 | 243.147 |
| Ssbp3 | 4713.33 | -0.12097 | 0.07491 | -1.61494 | 0.10632 | 0.27034 | 138.994 |
| Ssbp4 | 3915.05 | 0.159054 | 0.07587 | 2.096473 | 0.03604 | 0.13364 | 11.383 |
| Ssc4d | 45.6343 | -0.08601 | 0.26296 | -0.3271 | 0.74359 | 0.8653 | 14.321 |
| Ssc5d | 303.794 | 0.14116 | 0.14219 | 0.99273 | 0.32084 | 0.53158 | 19.042 |
| Ssh1 | 1235.96 | -0.10475 | 0.10356 | -1.0115 | 0.31178 | 0.52216 | 56.801 |
| Ssh2 | 2990.23 | -0.03691 | 0.08177 | -0.45134 | 0.65174 | 0.80618 | 243.934 |
| Ssh3 | 2610.1 | 0.136735 | 0.09997 | 1.367798 | 0.17138 | 0.3618 | 7.513 |
| Ssna1 | 893.147 | -0.13682 | 0.10214 | -1.33953 | 0.1804 | 0.37372 | 1.4 |
| Sspn | 677.161 | -0.36423 | 0.11868 | -3.06903 | 0.00215 | 0.02112 | 33.583 |
| Sspo | 357.032 | -0.34798 | 0.14395 | -2.41733 | 0.01564 | 0.07651 | 53.022 |
| Ssr1 | 7943.64 | 0.102674 | 0.05406 | 1.899401 | 0.05751 | 0.18207 | 27.613 |
| Ssr2 | 1640.59 | -0.10628 | 0.07274 | -1.46109 | 0.14399 | 0.32645 | 12.544 |
| Ssr3 | 7441.72 | 0.076308 | 0.0641 | 1.190388 | 0.23389 | 0.43899 | 12.969 |
| Ssr4 | 1292.31 | 0.002056 | 0.09256 | 0.022212 | 0.98228 | 0.99059 | 3.803 |
| Ssrp1 | 4436.94 | 0.047057 | 0.05912 | 0.795892 | 0.42609 | 0.63255 | 9.876 |
| Sst | 7010.39 | -0.34604 | 0.13267 | -2.60825 | 0.0091 | 0.05383 | 1.386 |

| | | | | | | | |
|---|---|---|---|---|---|---|---|
| Sstr1 | 339.637 | -0.65635 | 0.21176 | -3.09954 | 0.00194 | 0.0197 | 2.673 |
| Sstr2 | 298.362 | -0.5143 | 0.16964 | -3.03175 | 0.00243 | 0.02289 | 6.687 |
| Sstr3 | 792.858 | -1.17589 | 0.40006 | -2.93932 | 0.00329 | 0.02768 | 7.678 |
| Sstr4 | 422.669 | -0.61564 | 0.22653 | -2.71776 | 0.00657 | 0.04371 | 1.424 |
| Sstr5 | 22.7893 | 1.064397 | 0.54514 | 1.952508 | 0.05088 | 0.16778 | 7.414 |
| Ssu72 | 1454.68 | -0.1481 | 0.08099 | -1.82855 | 0.06747 | 0.20144 | 29.08 |
| Ssx2ip | 3702.97 | -0.05381 | 0.07066 | -0.76158 | 0.44631 | 0.64992 | 35.503 |
| St13 | 8612.13 | 0.080409 | 0.05365 | 1.498805 | 0.13392 | 0.3136 | 36.409 |
| St14 | 46.8476 | -0.85042 | 0.29762 | -2.85743 | 0.00427 | 0.03289 | 42.452 |
| St18 | 363.443 | 0.15752 | 0.1571 | 1.002678 | 0.31602 | 0.52654 | 373.71 |
| St3gal1 | 2404.92 | -0.12229 | 0.14355 | -0.85194 | 0.39425 | 0.60338 | 73.956 |
| St3gal2 | 5258.76 | -0.14967 | 0.07253 | -2.06353 | 0.03906 | 0.1408 | 52.559 |
| St3gal3 | 1756.24 | -0.25569 | 0.06923 | -3.6931 | 0.00022 | 0.00457 | 202.761 |
| St3gal4 | 973.389 | -0.26658 | 0.10985 | -2.42678 | 0.01523 | 0.07524 | 72.693 |
| St3gal5 | 5196.96 | -0.18168 | 0.09634 | -1.88585 | 0.05932 | 0.18536 | 56.98 |
| St3gal6 | 594.362 | -0.40466 | 0.10765 | -3.75897 | 0.00017 | 0.00388 | 56.119 |
| St6gal1 | 1168.9 | -0.56924 | 0.10451 | -5.44694 | 5.12E-08 | 1.16E-05 | 135.611 |
| St6gal2 | 1797.57 | 0.070694 | 0.10893 | 0.648974 | 0.51636 | 0.70635 | 69.2 |
| St6galnac2 | 249.918 | 0.123791 | 0.1631 | 0.759006 | 0.44785 | 0.65146 | 19.566 |
| St6galnac3 | 589.458 | -0.18619 | 0.1445 | -1.28851 | 0.19757 | 0.39543 | 526.909 |
| St6galnac4 | 1205.79 | 0.022419 | 0.08376 | 0.267642 | 0.78897 | 0.89044 | 12.604 |
| St6galnac5 | 2762.89 | 0.227424 | 0.09546 | 2.382424 | 0.0172 | 0.08153 | 165.166 |
| St6galnac6 | 4951.65 | -0.15312 | 0.08868 | -1.72672 | 0.08422 | 0.23239 | 21.098 |
| St7 | 689.974 | 0.162612 | 0.11677 | 1.392632 | 0.16373 | 0.35247 | 250.093 |
| St7l | 1295.49 | -0.0251 | 0.07647 | -0.32828 | 0.7427 | 0.86456 | 66.06 |
| St8sia1 | 3112.82 | -0.20905 | 0.10272 | -2.03507 | 0.04184 | 0.14718 | 142.908 |
| St8sia2 | 349.621 | 0.845283 | 0.18591 | 4.546832 | 5.45E-06 | 0.00035 | 74.572 |
| St8sia3 | 8586.66 | 0.549577 | 0.13102 | 4.194558 | 2.73E-05 | 0.00108 | 25.253 |
| St8sia3os | 25.5686 | 1.373319 | 0.41502 | 3.309043 | 0.00094 | 0.01223 | 175.986 |
| St8sia4 | 794.044 | 0.216281 | 0.14712 | 1.470098 | 0.14154 | 0.32333 | 79.89 |
| St8sia5 | 1584.41 | -1.01889 | 0.35966 | -2.83292 | 0.00461 | 0.0345 | 72.933 |
| St8sia6 | 187.513 | 0.086628 | 0.18989 | 0.456205 | 0.64824 | 0.80406 | 143.044 |
| Stab1 | 1043.78 | -0.30377 | 0.10325 | -2.94214 | 0.00326 | 0.02756 | 29.629 |
| Stab2 | 78.1557 | -0.22318 | 0.24737 | -0.9022 | 0.36695 | 0.57733 | 166.828 |

| | | | | | | | |
|---|---|---|---|---|---|---|---|
| Stac | 155.647 | 0.515239 | 0.30287 | 1.701203 | 0.0889 | 0.24112 | 128.912 |
| Stac2 | 3603.78 | -0.70233 | 0.15434 | -4.55049 | 5.35E-06 | 0.00035 | 16.84 |
| Stac3 | 346.472 | 0.472226 | 0.18119 | 2.606284 | 0.00915 | 0.05402 | 7.138 |
| Stag1 | 3038.77 | 0.413772 | 0.09746 | 4.245394 | 2.18E-05 | 0.00093 | 361.578 |
| Stag2 | 3990.67 | 0.222878 | 0.13275 | 1.678891 | 0.09317 | 0.24849 | 127.869 |
| Stag3 | 53.2666 | -0.21189 | 0.29331 | -0.72242 | 0.47004 | 0.66885 | 32.154 |
| Stam | 3017.81 | -0.06601 | 0.06946 | -0.95035 | 0.34194 | 0.55274 | 75.537 |
| Stam2 | 1602.4 | 0.200819 | 0.10175 | 1.973599 | 0.04843 | 0.16231 | 50.618 |
| Stambp | 1179.57 | -0.011 | 0.09094 | -0.12097 | 0.90372 | 0.95146 | 29.519 |
| Stambp-ps1 | 71.4552 | 0.382184 | 0.21356 | 1.789623 | 0.07351 | 0.21353 | 1.315 |
| Stambpl1 | 562.604 | -0.07914 | 0.09071 | -0.87243 | 0.38297 | 0.59222 | 48.105 |
| Stap1 | 91.9675 | 0.14413 | 0.26686 | 0.540093 | 0.58913 | 0.76226 | 34.38 |
| Stap2 | 78.9258 | -0.19475 | 0.23175 | -0.84031 | 0.40073 | 0.60946 | 8.508 |
| Star | 553.141 | 0.029877 | 0.11539 | 0.258922 | 0.7957 | 0.89467 | 9.428 |
| Stard10 | 2888.68 | -0.06134 | 0.10831 | -0.56635 | 0.57116 | 0.75015 | 29.541 |
| Stard13 | 771.645 | -0.0557 | 0.10984 | -0.50711 | 0.61208 | 0.77898 | 196.327 |
| Stard3 | 1403.95 | 0.085149 | 0.08082 | 1.053583 | 0.29207 | 0.50187 | 22.745 |
| Stard3nl | 1988.48 | -0.00322 | 0.06952 | -0.04626 | 0.9631 | 0.98208 | 38.12 |
| Stard4 | 1522.92 | -0.04207 | 0.07991 | -0.52652 | 0.59853 | 0.76969 | 14.508 |
| Stard5 | 1732.72 | 0.862329 | 0.32495 | 2.653766 | 0.00796 | 0.04944 | 21.169 |
| Stard6 | 65.2218 | -0.05545 | 0.23844 | -0.23255 | 0.81611 | 0.90539 | 28.613 |
| Stard7 | 4751.19 | 0.006397 | 0.07728 | 0.082773 | 0.93403 | 0.9669 | 28.715 |
| Stard8 | 917.044 | -0.56498 | 0.13068 | -4.32322 | 1.54E-05 | 0.00075 | 71.481 |
| Stard9 | 1584.37 | -0.39205 | 0.10856 | -3.6114 | 0.0003 | 0.00568 | 102.775 |
| Stat1 | 482.024 | -0.46644 | 0.11622 | -4.01351 | 5.98E-05 | 0.00188 | 42.426 |
| Stat2 | 1627.35 | -0.14003 | 0.07963 | -1.75848 | 0.07867 | 0.22228 | 22.291 |
| Stat3 | 3090.41 | -0.14165 | 0.06524 | -2.17104 | 0.02993 | 0.11811 | 54.443 |
| Stat4 | 14.8266 | -1.40446 | 0.604 | -2.32528 | 0.02006 | 0.08969 | 120.042 |
| Stat5a | 119.505 | -0.65904 | 0.16592 | -3.97198 | 7.13E-05 | 0.00212 | 25.819 |
| Stat5b | 2082.28 | 0.091496 | 0.09219 | 0.992445 | 0.32098 | 0.53177 | 69.994 |
| Stat6 | 570.541 | -0.44468 | 0.16249 | -2.73657 | 0.00621 | 0.04214 | 17.972 |
| Stau1 | 3492.48 | -0.01676 | 0.06777 | -0.24727 | 0.8047 | 0.89909 | 48.751 |
| Stau2 | 5113.53 | -0.15085 | 0.0771 | -1.95651 | 0.05041 | 0.16686 | 291.439 |
| Stbd1 | 37.1359 | -0.11657 | 0.39104 | -0.29811 | 0.76562 | 0.87765 | 3.539 |

| | | | | | | | |
|---|---|---|---|---|---|---|---|
| Stc1 | 186.259 | 0.201185 | 0.17656 | 1.139453 | 0.25451 | 0.46194 | 12.567 |
| Stc2 | 297.578 | 0.284318 | 0.1771 | 1.605421 | 0.1084 | 0.27367 | 12.768 |
| Steap2 | 833.686 | -0.12716 | 0.16278 | -0.78119 | 0.43469 | 0.64042 | 29.75 |
| Steap3 | 202.425 | -0.30563 | 0.19785 | -1.54476 | 0.1224 | 0.29608 | 81.949 |
| Steep1 | 2553.8 | -0.0484 | 0.08615 | -0.5618 | 0.57426 | 0.75253 | 50.375 |
| Stil | 38.1945 | -0.3404 | 0.45079 | -0.75512 | 0.45018 | 0.65304 | 43.038 |
| Stim1 | 5084.61 | 0.020147 | 0.05577 | 0.361273 | 0.7179 | 0.84941 | 169.514 |
| Stim2 | 3013.91 | 0.197885 | 0.08613 | 2.297488 | 0.02159 | 0.09434 | 122.559 |
| Stimate | 1363.1 | -0.05625 | 0.08989 | -0.6257 | 0.53151 | 0.71917 | 51.621 |
| Sting1 | 108.184 | -0.71426 | 0.2459 | -2.90464 | 0.00368 | 0.02977 | 6.876 |
| Stip1 | 8075.42 | -0.0264 | 0.06717 | -0.39304 | 0.69429 | 0.83417 | 19.266 |
| Stk10 | 349.25 | -0.25913 | 0.11418 | -2.26951 | 0.02324 | 0.09878 | 91.283 |
| Stk11 | 3252.77 | -0.14985 | 0.07506 | -1.99643 | 0.04589 | 0.15618 | 14.88 |
| Stk11ip | 1990.21 | 0.000994 | 0.07425 | 0.013387 | 0.98932 | 0.99415 | 15.807 |
| Stk16 | 1836.08 | 0.036573 | 0.0625 | 0.585167 | 0.55844 | 0.73984 | 4.769 |
| Stk17b | 458.207 | -0.26799 | 0.1131 | -2.36952 | 0.01781 | 0.08308 | 29.719 |
| Stk19 | 1476.91 | 0.10038 | 0.06538 | 1.535355 | 0.1247 | 0.29967 | 12.953 |
| Stk24 | 2493.55 | -0.08606 | 0.06127 | -1.4046 | 0.16014 | 0.34819 | 93.669 |
| Stk25 | 7928.38 | -0.06377 | 0.06039 | -1.0559 | 0.29101 | 0.50115 | 38.909 |
| Stk26 | 1342.92 | 0.930464 | 0.3282 | 2.835015 | 0.00458 | 0.03435 | 52.051 |
| Stk3 | 490.884 | -0.1869 | 0.15128 | -1.23547 | 0.21666 | 0.41857 | 303.426 |
| Stk32a | 837.865 | 0.84612 | 0.19121 | 4.425169 | 9.64E-06 | 0.00054 | 111.1 |
| Stk32b | 128.471 | 0.474274 | 0.30698 | 1.544991 | 0.12235 | 0.29602 | 270.347 |
| Stk32c | 6161.28 | 0.19554 | 0.10672 | 1.832243 | 0.06692 | 0.20024 | 109.67 |
| Stk33 | 61.6195 | -0.09541 | 0.27789 | -0.34335 | 0.73134 | 0.85734 | 159.859 |
| Stk35 | 1646.73 | -0.1101 | 0.06918 | -1.59147 | 0.1115 | 0.27897 | 31.771 |
| Stk36 | 366.384 | 0.084162 | 0.12768 | 0.659183 | 0.50978 | 0.70119 | 35.45 |
| Stk38 | 3339.92 | 0.069695 | 0.06704 | 1.03965 | 0.2985 | 0.50856 | 37.116 |
| Stk38l | 1829.18 | -0.14551 | 0.07626 | -1.90814 | 0.05637 | 0.17956 | 53.818 |
| Stk39 | 2678.15 | -0.14061 | 0.08418 | -1.67043 | 0.09484 | 0.25135 | 261.824 |
| Stk4 | 2411.03 | -0.00868 | 0.07567 | -0.11477 | 0.90863 | 0.95422 | 85.203 |
| Stk40 | 1496.23 | -0.3221 | 0.10419 | -3.09161 | 0.00199 | 0.02005 | 37.073 |
| Stkld1 | 24.1608 | -0.73153 | 0.3565 | -2.05198 | 0.04017 | 0.14323 | 19.45 |
| Stmn1 | 10668.7 | -0.79713 | 0.18973 | -4.20132 | 2.65E-05 | 0.00107 | 5.524 |

| | | | | | | | |
|---|---|---|---|---|---|---|---|
| Stmn2 | 7929.04 | -0.59967 | 0.15638 | -3.83459 | 0.00013 | 0.00313 | 52.247 |
| Stmn3 | 9900.14 | -0.38371 | 0.08373 | -4.5825 | 4.59E-06 | 0.00032 | 8.042 |
| Stmn4 | 6103.47 | -0.25068 | 0.07094 | -3.5337 | 0.00041 | 0.00698 | 17.385 |
| Stn1 | 534.394 | 0.179279 | 0.09895 | 1.811796 | 0.07002 | 0.20664 | 36.475 |
| Stom | 589.125 | -0.36384 | 0.14336 | -2.53793 | 0.01115 | 0.06157 | 22.991 |
| Stoml1 | 1807.26 | -0.19356 | 0.09903 | -1.95454 | 0.05064 | 0.16737 | 9.357 |
| Stoml2 | 1380.97 | -0.28174 | 0.09079 | -3.10308 | 0.00192 | 0.01954 | 4.021 |
| Stoml3 | 14.6463 | 0.503302 | 0.47069 | 1.069285 | 0.28494 | 0.49485 | 19.85 |
| Ston1 | 259.459 | -0.53937 | 0.16905 | -3.19061 | 0.00142 | 0.01595 | 64.903 |
| Ston2 | 1786.65 | -0.14618 | 0.09401 | -1.55495 | 0.11996 | 0.29246 | 155.379 |
| Stox1 | 130.776 | 0.10044 | 0.19484 | 0.515508 | 0.6062 | 0.77465 | 67.086 |
| Stox2 | 9436.2 | 0.103923 | 0.06562 | 1.583662 | 0.11327 | 0.28175 | 266.315 |
| Stpg1 | 55.2499 | -0.97097 | 0.52944 | -1.83394 | 0.06666 | 0.19992 | 43.014 |
| Stra6 | 959.367 | 0.295466 | 0.20341 | 1.452527 | 0.14636 | 0.32939 | 90.209 |
| Strada | 1767.23 | -0.0722 | 0.1074 | -0.67224 | 0.50143 | 0.69395 | 38.839 |
| Stradb | 1499.39 | 0.020109 | 0.06518 | 0.3085 | 0.7577 | 0.87241 | 22.194 |
| Strap | 5590.71 | 0.035032 | 0.06146 | 0.570027 | 0.56866 | 0.74839 | 16.855 |
| Strbp | 10439.5 | -0.18307 | 0.11877 | -1.54136 | 0.12323 | 0.29741 | 220.632 |
| Strc | 39.8147 | -0.37426 | 0.34379 | -1.08863 | 0.27632 | 0.48548 | 23.441 |
| Strip1 | 2054.83 | -0.43804 | 0.13637 | -3.21216 | 0.00132 | 0.01512 | 19.163 |
| Strip2 | 13842.6 | 0.529744 | 0.1637 | 3.2361 | 0.00121 | 0.01442 | 42.67 |
| Strn | 13728.3 | 0.43745 | 0.16519 | 2.648185 | 0.00809 | 0.04993 | 87.284 |
| Strn3 | 3952.6 | 0.179801 | 0.08741 | 2.056978 | 0.03969 | 0.14226 | 82.266 |
| Strn4 | 8940.77 | 0.019856 | 0.05769 | 0.344189 | 0.7307 | 0.85728 | 25.043 |
| Stt3a | 2101.42 | -0.05312 | 0.07552 | -0.70332 | 0.48186 | 0.67858 | 38.336 |
| Stt3b | 5132.49 | 0.021869 | 0.07009 | 0.312038 | 0.75501 | 0.87123 | 67.841 |
| Stub1 | 1323.74 | -0.03878 | 0.06777 | -0.57229 | 0.56712 | 0.74717 | 2.9 |
| Stum | 13047.7 | -0.36463 | 0.19067 | -1.9123 | 0.05584 | 0.17857 | 51.118 |
| Stx11 | 15.943 | -0.42875 | 0.74344 | -0.57672 | 0.56413 | 0.74438 | 26.09 |
| Stx12 | 5606.1 | 0.029226 | 0.06107 | 0.478594 | 0.63223 | 0.79238 | 31.009 |
| Stx16 | 6098.01 | 0.222676 | 0.07124 | 3.125518 | 0.00177 | 0.01864 | 23.464 |
| Stx17 | 1473.55 | -0.11756 | 0.11308 | -1.03962 | 0.29851 | 0.50856 | 61.593 |
| Stx18 | 819.14 | -0.0216 | 0.0799 | -0.27041 | 0.78684 | 0.88907 | 99.024 |
| Stx19 | 23.4122 | -0.72622 | 0.41327 | -1.75725 | 0.07887 | 0.22271 | 9.671 |

| | | | | | | | |
|---|---|---|---|---|---|---|---|
| Stx1a | 3567.19 | -0.76439 | 0.32266 | -2.36905 | 0.01783 | 0.08315 | 27.619 |
| Stx1b | 23694.4 | -0.13443 | 0.05866 | -2.29162 | 0.02193 | 0.09532 | 20.65 |
| Stx2 | 1203.91 | 0.322021 | 0.09007 | 3.57508 | 0.00035 | 0.00623 | 24.018 |
| Stx3 | 1994.09 | 0.199587 | 0.09998 | 1.996319 | 0.0459 | 0.1562 | 44.286 |
| Stx4a | 1907.1 | -0.2352 | 0.09146 | -2.57152 | 0.01013 | 0.05794 | 24.726 |
| Stx5a | 2764.57 | 0.120754 | 0.07627 | 1.58324 | 0.11337 | 0.28188 | 14.657 |
| Stx6 | 4300.54 | 0.041867 | 0.06783 | 0.617191 | 0.53711 | 0.72389 | 49.542 |
| Stx7 | 5150.11 | -0.17298 | 0.07324 | -2.36178 | 0.01819 | 0.08428 | 40.921 |
| Stx8 | 740.146 | -0.05705 | 0.08874 | -0.64294 | 0.52027 | 0.70963 | 240.956 |
| Stxbp1 | 35951.7 | -0.24557 | 0.07222 | -3.40032 | 0.00067 | 0.00972 | 59.644 |
| Stxbp2 | 3449.94 | 0.174942 | 0.10832 | 1.615015 | 0.10631 | 0.27033 | 12.69 |
| Stxbp3 | 1371.73 | -0.28287 | 0.11836 | -2.38993 | 0.01685 | 0.08037 | 47.351 |
| Stxbp3-ps | 40.3139 | -0.24859 | 0.41205 | -0.6033 | 0.54631 | 0.73071 | 1.392 |
| Stxbp4 | 1163.58 | 0.108483 | 0.08988 | 1.20691 | 0.22747 | 0.43164 | 161.593 |
| Stxbp5 | 7541.54 | -0.03016 | 0.11251 | -0.26809 | 0.78863 | 0.89021 | 145.533 |
| Stxbp5l | 2647.55 | 0.090904 | 0.10994 | 0.82685 | 0.40832 | 0.61613 | 270.021 |
| Stxbp6 | 2458.69 | 0.06567 | 0.15693 | 0.418467 | 0.67561 | 0.82192 | 222.226 |
| Styx | 775.608 | 0.179925 | 0.11611 | 1.54963 | 0.12123 | 0.2945 | 71.728 |
| Styx-ps | 72.9699 | 0.301364 | 0.29406 | 1.024842 | 0.30544 | 0.51544 | 4.226 |
| Sub1 | 9115.31 | 0.15524 | 0.08151 | 1.904607 | 0.05683 | 0.18045 | 15.645 |
| Sucla2 | 6001.92 | 0.073949 | 0.0821 | 0.900692 | 0.36775 | 0.57819 | 70.824 |
| Suclg1 | 3646.35 | -0.10859 | 0.05661 | -1.91831 | 0.05507 | 0.17684 | 28.53 |
| Suclg2 | 1087.22 | -0.10648 | 0.12234 | -0.87037 | 0.3841 | 0.59346 | 245.792 |
| Suco | 2461.87 | 0.246398 | 0.08812 | 2.796143 | 0.00517 | 0.03708 | 60.569 |
| Suds3 | 5730.44 | -0.14145 | 0.07296 | -1.93867 | 0.05254 | 0.1715 | 24.434 |
| Sufu | 1097.19 | 0.14881 | 0.08692 | 1.71212 | 0.08687 | 0.23726 | 91.909 |
| Sugct | 214.517 | 0.013151 | 0.16112 | 0.081623 | 0.93495 | 0.96725 | 838.082 |
| Sugp1 | 2498.6 | -0.02222 | 0.06878 | -0.323 | 0.7467 | 0.86713 | 29.535 |
| Sugp2 | 5695.65 | 0.106892 | 0.06067 | 1.761796 | 0.0781 | 0.22118 | 45.69 |
| Sugt1 | 5132.53 | 0.053531 | 0.07095 | 0.754511 | 0.45054 | 0.65319 | 47.506 |
| Sulf1 | 2489.25 | 0.194797 | 0.11149 | 1.747269 | 0.08059 | 0.22593 | 168.916 |
| Sulf2 | 4374.97 | -0.37671 | 0.11951 | -3.15219 | 0.00162 | 0.0175 | 82.575 |
| Sult1a1 | 898.528 | -0.3658 | 0.13356 | -2.73892 | 0.00616 | 0.04202 | 3.568 |
| Sult2b1 | 425.053 | 0.295354 | 0.14423 | 2.04783 | 0.04058 | 0.144 | 54.687 |

| | | | | | | | |
|---|---|---|---|---|---|---|---|
| Sult4a1 | 14728.3 | -0.32018 | 0.08106 | -3.95021 | 7.81E-05 | 0.00226 | 29.658 |
| Sult5a1 | 23.9907 | -1.14938 | 0.46111 | -2.49265 | 0.01268 | 0.06713 | 18.124 |
| Sumf1 | 1443.92 | -0.10625 | 0.09499 | -1.11862 | 0.2633 | 0.47208 | 78.555 |
| Sumf2 | 450.536 | -0.04746 | 0.113 | -0.42 | 0.67449 | 0.82131 | 17.065 |
| Sumo1 | 3394.43 | -0.00223 | 0.07228 | -0.03082 | 0.97541 | 0.9876 | 84.277 |
| Sumo2 | 2357.1 | 0.953633 | 0.53466 | 1.783615 | 0.07449 | 0.21509 | 13.175 |
| Sumo3 | 4721.37 | 0.171752 | 0.09397 | 1.827729 | 0.06759 | 0.20161 | 12.235 |
| Sun1 | 5552.13 | 0.07354 | 0.0688 | 1.068963 | 0.28509 | 0.49488 | 49.204 |
| Sun2 | 4748.1 | 0.115833 | 0.10376 | 1.116302 | 0.26429 | 0.47305 | 18.467 |
| Suox | 723.168 | -0.03855 | 0.11641 | -0.33116 | 0.74052 | 0.86301 | 4.18 |
| Supt16 | 2652.41 | 0.107986 | 0.07773 | 1.389194 | 0.16477 | 0.35385 | 37.003 |
| Supt20 | 2677.57 | 0.169855 | 0.09319 | 1.822754 | 0.06834 | 0.20307 | 35.96 |
| Supt3 | 261.395 | -0.06658 | 0.13149 | -0.50635 | 0.61261 | 0.77934 | 342.139 |
| Supt4a | 1300.56 | -0.12284 | 0.06934 | -1.77161 | 0.07646 | 0.21863 | 6.072 |
| Supt5 | 6090.83 | 0.055273 | 0.0684 | 0.808131 | 0.41902 | 0.62614 | 23.856 |
| Supt6 | 11239.2 | 0.095546 | 0.0726 | 1.316061 | 0.18815 | 0.38372 | 39.242 |
| Supt7l | 803.716 | -0.05791 | 0.10441 | -0.55463 | 0.57915 | 0.75584 | 13.244 |
| Supv3l1 | 1043.73 | 0.032492 | 0.07759 | 0.418755 | 0.6754 | 0.82188 | 20.53 |
| Surf1 | 1646.77 | 0.090385 | 0.09135 | 0.989479 | 0.32243 | 0.53337 | 3.15 |
| Surf2 | 940.166 | 0.025543 | 0.09516 | 0.268426 | 0.78837 | 0.89002 | 3.817 |
| Surf4 | 2987.2 | -0.04694 | 0.09029 | -0.51988 | 0.60315 | 0.77277 | 13.889 |
| Surf6 | 926.512 | -0.02013 | 0.09727 | -0.20694 | 0.83606 | 0.9159 | 14.252 |
| Susd1 | 626.047 | -0.05785 | 0.1202 | -0.4813 | 0.6303 | 0.79104 | 123.951 |
| Susd2 | 321.74 | 0.208219 | 0.14024 | 1.48474 | 0.13761 | 0.3184 | 7.303 |
| Susd3 | 40.0047 | -0.55291 | 0.40297 | -1.37209 | 0.17003 | 0.3604 | 18.017 |
| Susd4 | 4994.3 | 0.037444 | 0.07708 | 0.485796 | 0.62711 | 0.78903 | 132.732 |
| Susd5 | 243.434 | -0.25048 | 0.15901 | -1.57525 | 0.1152 | 0.28456 | 41.589 |
| Susd6 | 2867.83 | -0.22485 | 0.10157 | -2.21364 | 0.02685 | 0.10939 | 90.326 |
| Suv39h1 | 784.529 | -0.04765 | 0.09434 | -0.50507 | 0.61351 | 0.77991 | 13.59 |
| Suv39h2 | 258.52 | -0.07171 | 0.14739 | -0.48654 | 0.62659 | 0.78871 | 19.217 |
| Suz12 | 2771.23 | 0.124038 | 0.11295 | 1.098166 | 0.27213 | 0.48148 | 41.018 |
| Sv2a | 21988.8 | -0.14117 | 0.06664 | -2.11847 | 0.03414 | 0.12905 | 14.371 |
| Sv2b | 14513.8 | -0.68974 | 0.29913 | -2.30582 | 0.02112 | 0.0929 | 194.369 |
| Sv2c | 14502.7 | 0.559343 | 0.17174 | 3.256863 | 0.00113 | 0.01378 | 177.984 |

| | | | | | | | |
|---|---|---|---|---|---|---|---|
| Svbp | 390.291 | -0.02033 | 0.15741 | -0.12917 | 0.89723 | 0.94851 | 5.993 |
| Svep1 | 82.6274 | -0.70966 | 0.31352 | -2.26352 | 0.0236 | 0.09984 | 164.418 |
| Svet1 | 13.5494 | -0.87413 | 0.52465 | -1.66612 | 0.09569 | 0.25302 | 3.94 |
| Svil | 1195.85 | 0.144453 | 0.10622 | 1.359928 | 0.17385 | 0.36527 | 198.76 |
| Svip | 3179.47 | -0.09206 | 0.08254 | -1.11525 | 0.26474 | 0.47371 | 8.848 |
| Svop | 6514.42 | 0.09438 | 0.06724 | 1.403667 | 0.16042 | 0.3486 | 64.661 |
| Swap70 | 463.402 | -0.20247 | 0.12298 | -1.64636 | 0.09969 | 0.25965 | 61.796 |
| Swi5 | 3920.13 | 0.073152 | 0.07078 | 1.033453 | 0.30139 | 0.5111 | 9.26 |
| Swsap1 | 491.061 | -0.25886 | 0.11583 | -2.23485 | 0.02543 | 0.10518 | 2.51 |
| Swt1 | 762.005 | 0.304978 | 0.13186 | 2.312858 | 0.02073 | 0.09173 | 60.757 |
| Syap1 | 1460.71 | 0.116902 | 0.08013 | 1.458896 | 0.14459 | 0.32724 | 31.391 |
| Sybu | 3272.65 | -0.013 | 0.05791 | -0.22456 | 0.82232 | 0.90788 | 116.208 |
| Syce1 | 12.451 | -0.20123 | 0.48735 | -0.41292 | 0.67967 | 0.82491 | 10.624 |
| Syce2 | 115.123 | -0.17492 | 0.21374 | -0.8184 | 0.41313 | 0.62134 | 16.111 |
| Sycp2 | 241.31 | 0.438416 | 0.2716 | 1.614215 | 0.10648 | 0.27067 | 62.393 |
| Sycp2l | 51.9071 | 1.904669 | 0.35681 | 5.338062 | 9.39E-08 | 1.66E-05 | 60.045 |
| Sycp3 | 49.902 | -0.05433 | 0.26899 | -0.20196 | 0.83995 | 0.91789 | 13.668 |
| Syde1 | 274.348 | -0.45226 | 0.12686 | -3.56489 | 0.00036 | 0.0064 | 7.462 |
| Syde2 | 339.922 | 0.050126 | 0.10565 | 0.474442 | 0.63518 | 0.79438 | 33.851 |
| Syf2 | 1978.57 | -0.07837 | 0.0849 | -0.92311 | 0.35595 | 0.56636 | 6.651 |
| Syk | 201.753 | -0.56128 | 0.19755 | -2.84128 | 0.00449 | 0.03392 | 65.62 |
| Sympk | 6810.38 | 0.040419 | 0.06593 | 0.613094 | 0.53981 | 0.72592 | 30.242 |
| Syn1 | 12942.5 | -0.37601 | 0.10596 | -3.54876 | 0.00039 | 0.00668 | 60.494 |
| Syn2 | 17915.7 | 0.057394 | 0.06054 | 0.948069 | 0.34309 | 0.55382 | 147.105 |
| Syn3 | 897.035 | 0.24886 | 0.11671 | 2.132312 | 0.03298 | 0.12614 | 443.772 |
| Syna | 187.85 | -0.62371 | 0.18794 | -3.31865 | 0.0009 | 0.01195 | 2.026 |
| Sync | 22.9097 | 0.089621 | 0.43973 | 0.203811 | 0.8385 | 0.91686 | 20.943 |
| Syncrip | 4780.24 | 0.093151 | 0.09087 | 1.025057 | 0.30534 | 0.51531 | 35.566 |
| Syndig1 | 2188.26 | 0.623402 | 0.2009 | 3.103036 | 0.00192 | 0.01954 | 175.182 |
| Syndig1l | 3331.44 | 0.734383 | 0.30689 | 2.392957 | 0.01671 | 0.07998 | 21.555 |
| Syne1 | 46968.5 | 0.21192 | 0.08539 | 2.481912 | 0.01307 | 0.06843 | 530.566 |
| Syne2 | 1693.94 | -0.12861 | 0.08081 | -1.59145 | 0.11151 | 0.27897 | 292.793 |
| Syne3 | 226.386 | -0.47013 | 0.1352 | -3.47734 | 0.00051 | 0.00798 | 79.877 |
| Syne4 | 310.75 | 0.280471 | 0.1289 | 2.175822 | 0.02957 | 0.11701 | 4.24 |

| | | | | | | | |
|---|---|---|---|---|---|---|---|
| Syngap1 | 19196.2 | 0.10568 | 0.06166 | 1.713797 | 0.08657 | 0.23673 | 31.182 |
| Syngr1 | 20197.1 | -0.38807 | 0.09618 | -4.03478 | 5.47E-05 | 0.00176 | 28.168 |
| Syngr2 | 244.578 | -0.55529 | 0.17606 | -3.15403 | 0.00161 | 0.01744 | 4.616 |
| Syngr3 | 5089.55 | -0.1215 | 0.0673 | -1.80533 | 0.07102 | 0.20852 | 4.864 |
| Syngr4 | 23.3993 | 0.42088 | 0.41262 | 1.020021 | 0.30772 | 0.51754 | 11.493 |
| Synj1 | 21129 | -0.01002 | 0.06792 | -0.1476 | 0.88266 | 0.9416 | 75.217 |
| Synj2 | 2395.82 | -0.72901 | 0.3691 | -1.9751 | 0.04826 | 0.16192 | 103.011 |
| Synj2bp | 1875.75 | 0.078491 | 0.09706 | 0.80869 | 0.41869 | 0.62584 | 34.971 |
| Synm | 2184.69 | -0.43796 | 0.13556 | -3.23065 | 0.00124 | 0.01458 | 29.583 |
| Synpo | 7983.45 | -0.07447 | 0.08943 | -0.83266 | 0.40504 | 0.61387 | 66.17 |
| Synpo2 | 376.781 | -0.51807 | 0.15381 | -3.36824 | 0.00076 | 0.01056 | 159.631 |
| Synpo2l | 12.8689 | 0.748784 | 0.52819 | 1.417651 | 0.15629 | 0.34308 | 9.409 |
| Synpr | 5801.66 | 0.506782 | 0.13262 | 3.82141 | 0.00013 | 0.00327 | 331.561 |
| Synrg | 4638.77 | 0.065467 | 0.06326 | 1.034844 | 0.30074 | 0.51072 | 80.151 |
| Syp | 23458.7 | -0.38774 | 0.08983 | -4.31617 | 1.59E-05 | 0.00077 | 14.786 |
| Sypl | 2373.69 | -0.05275 | 0.06892 | -0.76548 | 0.44398 | 0.6481 | 25.97 |
| Sypl2 | 91.9174 | -0.28676 | 0.26791 | -1.07037 | 0.28445 | 0.49452 | 15.177 |
| Sys1 | 819.295 | 0.118893 | 0.09662 | 1.230562 | 0.21849 | 0.42062 | 22.675 |
| Syt1 | 57345.1 | -0.17873 | 0.07576 | -2.3592 | 0.01831 | 0.08465 | 513.333 |
| Syt10 | 494.823 | 0.762209 | 0.18925 | 4.027525 | 5.64E-05 | 0.00181 | 59.468 |
| Syt11 | 33329.9 | -0.17761 | 0.06075 | -2.92363 | 0.00346 | 0.02852 | 30.465 |
| Syt12 | 1314.5 | 0.052317 | 0.10817 | 0.483675 | 0.62862 | 0.79 | 31.54 |
| Syt13 | 5263.1 | -0.81544 | 0.31748 | -2.56844 | 0.01022 | 0.05838 | 40.961 |
| Syt14 | 1721.83 | 0.17911 | 0.08724 | 2.053156 | 0.04006 | 0.14298 | 144.543 |
| Syt15 | 38.4683 | -0.02753 | 0.29955 | -0.09192 | 0.92676 | 0.9631 | 10.826 |
| Syt16 | 2528.6 | -0.16663 | 0.11705 | -1.42363 | 0.15455 | 0.34062 | 275.607 |
| Syt17 | 1648.18 | -1.3125 | 0.45958 | -2.85589 | 0.00429 | 0.03302 | 67.506 |
| Syt2 | 3413.18 | -0.0132 | 0.1785 | -0.07396 | 0.94104 | 0.97076 | 115.917 |
| Syt3 | 2042.16 | -0.06981 | 0.06814 | -1.02458 | 0.30556 | 0.51553 | 16.086 |
| Syt4 | 6481.87 | 0.138012 | 0.07346 | 1.878753 | 0.06028 | 0.18732 | 9.599 |
| Syt5 | 6716.42 | 0.129877 | 0.08007 | 1.621974 | 0.10481 | 0.26782 | 7.777 |
| Syt6 | 5433.8 | 0.775379 | 0.17988 | 4.310653 | 1.63E-05 | 0.00078 | 70.339 |
| Syt7 | 15872.4 | -0.08616 | 0.06272 | -1.3737 | 0.16953 | 0.35975 | 64.091 |
| Syt9 | 406.25 | -0.39336 | 0.18198 | -2.16152 | 0.03066 | 0.11979 | 177.929 |

| | | | | | | | |
|---|---|---|---|---|---|---|---|
| Sytl1 | 52.2998 | 0.212355 | 0.34132 | 0.622164 | 0.53383 | 0.72102 | 10.024 |
| Sytl2 | 1085.27 | -0.308 | 0.10349 | -2.97599 | 0.00292 | 0.02574 | 108.468 |
| Sytl4 | 32.5396 | -0.51359 | 0.33978 | -1.51154 | 0.13065 | 0.30899 | 45.428 |
| Sytl5 | 357.209 | 0.814533 | 0.23514 | 3.464052 | 0.00053 | 0.00829 | 108.922 |
| Syvn1 | 2169.84 | -0.04862 | 0.08833 | -0.55039 | 0.58205 | 0.75769 | 7.137 |
| Szrd1 | 1806.86 | -0.22385 | 0.08235 | -2.7182 | 0.00656 | 0.04367 | 26.775 |
| Szt2 | 4587.07 | 0.23858 | 0.08858 | 2.693403 | 0.00707 | 0.04577 | 46.531 |
| T2 | 21.4143 | -0.34519 | 0.46906 | -0.73592 | 0.46178 | 0.66226 | 67.522 |
| Tab1 | 1013.82 | 0.043286 | 0.07673 | 0.564129 | 0.57267 | 0.75137 | 28.581 |
| Tab2 | 3568.7 | 0.026959 | 0.06323 | 0.426355 | 0.66985 | 0.81796 | 50.578 |
| Tab3 | 1235.65 | 0.202872 | 0.12028 | 1.686696 | 0.09166 | 0.2462 | 60.448 |
| Tac1 | 13392.8 | 0.573816 | 0.18745 | 3.061168 | 0.0022 | 0.02144 | 10.956 |
| Tac2 | 179.444 | 0.343379 | 0.16355 | 2.099566 | 0.03577 | 0.13297 | 7.291 |
| Tacc1 | 14408.5 | -0.04676 | 0.06181 | -0.75655 | 0.44932 | 0.65253 | 102.037 |
| Tacc2 | 3196.36 | 0.051886 | 0.06949 | 0.746699 | 0.45525 | 0.65728 | 187.348 |
| Tacc3 | 199.709 | -0.31385 | 0.17526 | -1.79077 | 0.07333 | 0.2132 | 20.868 |
| Taco1 | 415.7 | -0.2494 | 0.10633 | -2.34562 | 0.019 | 0.08658 | 7.552 |
| Tacr1 | 959.493 | 0.486225 | 0.18195 | 2.672346 | 0.00753 | 0.04785 | 157.771 |
| Tacr3 | 297.545 | 0.883202 | 0.39966 | 2.209908 | 0.02711 | 0.10996 | 105.573 |
| Tada1 | 774.492 | 0.244141 | 0.076 | 3.212246 | 0.00132 | 0.01512 | 14.525 |
| Tada2a | 1468.98 | 0.072583 | 0.07876 | 0.921518 | 0.35678 | 0.56706 | 50.681 |
| Tada2b | 1620.87 | -0.11373 | 0.07675 | -1.48193 | 0.13836 | 0.31951 | 10.616 |
| Tada3 | 1240.42 | -0.07394 | 0.07637 | -0.96818 | 0.33295 | 0.54382 | 11.859 |
| Taf1 | 5051.28 | 0.09986 | 0.08127 | 1.228785 | 0.21915 | 0.42137 | 69.056 |
| Taf10 | 865.375 | -0.0659 | 0.08098 | -0.8138 | 0.41576 | 0.62362 | 4.969 |
| Taf11 | 1462.61 | 0.039946 | 0.07988 | 0.500069 | 0.61703 | 0.78204 | 8.723 |
| Taf12 | 760.366 | -0.0433 | 0.09481 | -0.45673 | 0.64786 | 0.80388 | 21.392 |
| Taf13 | 1461.44 | -0.35214 | 0.07893 | -4.46116 | 8.15E-06 | 0.00047 | 11.525 |
| Taf15 | 4388.24 | 0.092268 | 0.12456 | 0.740747 | 0.45885 | 0.65977 | 33.658 |
| Taf1a | 980.978 | 0.121149 | 0.08942 | 1.354782 | 0.17549 | 0.36728 | 20.696 |
| Taf1b | 995.223 | 0.128524 | 0.09289 | 1.383684 | 0.16646 | 0.356 | 60.181 |
| Taf1c | 525.126 | 0.032994 | 0.13289 | 0.24828 | 0.80392 | 0.89875 | 7.352 |
| Taf1d | 1026.56 | 0.390326 | 0.1072 | 3.641214 | 0.00027 | 0.00522 | 10.778 |
| Taf2 | 2435.23 | 0.091917 | 0.06489 | 1.416485 | 0.15663 | 0.3435 | 57.022 |

| | | | | | | | |
|---|---|---|---|---|---|---|---|
| Taf3 | 1004.95 | -0.00565 | 0.09335 | -0.06048 | 0.95177 | 0.9763 | 134.045 |
| Taf4 | 1785.15 | 0.263948 | 0.10017 | 2.634924 | 0.00842 | 0.05116 | 64.495 |
| Taf4b | 235.084 | 0.065402 | 0.15459 | 0.423065 | 0.67225 | 0.81976 | 117.115 |
| Taf5 | 342.906 | -0.03309 | 0.13683 | -0.24183 | 0.80891 | 0.90168 | 15.734 |
| Taf5l | 799.542 | 0.073984 | 0.09578 | 0.772473 | 0.43983 | 0.64503 | 25.08 |
| Taf6 | 986.389 | -0.08394 | 0.08008 | -1.0482 | 0.29454 | 0.50492 | 8.835 |
| Taf6l | 274.937 | 0.094342 | 0.14266 | 0.661313 | 0.50841 | 0.69971 | 11.931 |
| Taf7 | 325.545 | -0.07188 | 0.17864 | -0.40238 | 0.6874 | 0.82961 | 3.714 |
| Taf7l | 45.8891 | -0.0986 | 0.32477 | -0.30361 | 0.76143 | 0.87503 | 16.373 |
| Taf8 | 1032.38 | 0.079624 | 0.0807 | 0.986651 | 0.32381 | 0.53482 | 18.929 |
| Taf9 | 14.1469 | 0.287345 | 0.53144 | 0.54069 | 0.58872 | 0.762 | 4.482 |
| Taf9b | 1640.29 | 0.095504 | 0.07528 | 1.268659 | 0.20456 | 0.40342 | 14.276 |
| Tafa1 | 691.514 | -1.41845 | 0.69108 | -2.05252 | 0.04012 | 0.14312 | 544.025 |
| Tafa2 | 2675.91 | -0.25008 | 0.08607 | -2.90554 | 0.00367 | 0.02974 | 477.209 |
| Tafa4 | 53.7368 | 0.802133 | 0.3019 | 2.656932 | 0.00789 | 0.04922 | 229.211 |
| Tafa5 | 3831.39 | -0.0678 | 0.07924 | -0.85568 | 0.39218 | 0.60133 | 215.066 |
| Tafazzin | 2929.72 | 0.296245 | 0.06866 | 4.314503 | 1.60E-05 | 0.00077 | 8.24 |
| Tagap | 97.3727 | 0.553082 | 0.21564 | 2.564833 | 0.01032 | 0.05866 | 8.898 |
| Tagap1 | 949.571 | -0.20247 | 0.07528 | -2.68968 | 0.00715 | 0.04616 | 6.146 |
| Tagln2 | 251.004 | -0.55143 | 0.16398 | -3.36281 | 0.00077 | 0.01071 | 7.334 |
| Tagln3 | 4998.25 | -0.29816 | 0.1024 | -2.91189 | 0.00359 | 0.0293 | 13.379 |
| Tal1 | 85.6777 | -0.40904 | 0.21841 | -1.87282 | 0.06109 | 0.1888 | 15.33 |
| Taldo1 | 3115.78 | -0.14522 | 0.07884 | -1.8418 | 0.0655 | 0.19743 | 10.77 |
| Tamalin | 1962.22 | -0.12535 | 0.09803 | -1.27862 | 0.20103 | 0.3992 | 8.569 |
| Tamm41 | 421.568 | 0.034777 | 0.10496 | 0.331341 | 0.74039 | 0.86301 | 33.496 |
| Tanc1 | 2412.96 | -0.64495 | 0.2042 | -3.15844 | 0.00159 | 0.01728 | 234.108 |
| Tanc2 | 11673.4 | 0.19507 | 0.05666 | 3.442835 | 0.00058 | 0.00873 | 339.319 |
| Tango2 | 2601.32 | -0.27977 | 0.07612 | -3.67559 | 0.00024 | 0.00478 | 47.279 |
| Tango6 | 380.502 | 0.025733 | 0.11387 | 0.22598 | 0.82122 | 0.90759 | 168.372 |
| Tank | 1873.09 | 0.005364 | 0.08974 | 0.059773 | 0.95234 | 0.97658 | 75.587 |
| Taok1 | 9814.39 | 0.191176 | 0.11091 | 1.723646 | 0.08477 | 0.23351 | 78.654 |
| Taok2 | 8629.53 | -0.03167 | 0.06047 | -0.52366 | 0.60052 | 0.77095 | 19.026 |
| Taok3 | 2988.21 | 0.120561 | 0.07118 | 1.693748 | 0.09031 | 0.24383 | 155.091 |
| Tap1 | 138.283 | -0.33974 | 0.21827 | -1.55649 | 0.11959 | 0.29195 | 9.673 |

| | | | | | | | |
|---|---|---|---|---|---|---|---|
| Tap2 | 221.252 | -0.38555 | 0.15173 | -2.54113 | 0.01105 | 0.0613 | 12.795 |
| Tapbp | 944.925 | -0.10911 | 0.10902 | -1.0008 | 0.31692 | 0.52743 | 13.394 |
| Tapbpl | 379.895 | -0.17453 | 0.1291 | -1.35189 | 0.17641 | 0.36846 | 7.928 |
| Tapt1 | 2376.82 | 0.02871 | 0.07663 | 0.374639 | 0.70793 | 0.84235 | 51.473 |
| Tarbp1 | 894.807 | 0.147837 | 0.08064 | 1.833262 | 0.06676 | 0.20013 | 49.737 |
| Tarbp2 | 945.058 | 0.061981 | 0.12031 | 0.515168 | 0.60644 | 0.77468 | 5.485 |
| Tardbp | 12143.3 | 0.167843 | 0.08021 | 2.092592 | 0.03639 | 0.13441 | 14.638 |
| Tars | 1525.35 | -0.03979 | 0.07027 | -0.5663 | 0.57119 | 0.75015 | 17.365 |
| Tars2 | 2130.17 | 0.205599 | 0.0709 | 2.899784 | 0.00373 | 0.03006 | 20.231 |
| Tarsl2 | 1931.16 | 0.100058 | 0.07865 | 1.272192 | 0.2033 | 0.40179 | 47.194 |
| Tas1r1 | 220.609 | 0.341273 | 0.13832 | 2.467285 | 0.01361 | 0.07022 | 10.655 |
| Tas1r3 | 110.476 | 0.463908 | 0.25307 | 1.833094 | 0.06679 | 0.20013 | 4.095 |
| Tasor | 2419.27 | 0.181581 | 0.12503 | 1.452315 | 0.14641 | 0.32946 | 54.722 |
| Tasor2 | 2012.66 | 0.301374 | 0.15735 | 1.915288 | 0.05546 | 0.17765 | 45.074 |
| Tasp1 | 1109.61 | 0.195127 | 0.08529 | 2.287868 | 0.02215 | 0.0959 | 233.326 |
| Tatdn1 | 749.875 | 0.075662 | 0.10375 | 0.729271 | 0.46584 | 0.665 | 43.584 |
| Tatdn2 | 2068.86 | -0.05577 | 0.1051 | -0.53067 | 0.59565 | 0.76745 | 14.02 |
| Tatdn3 | 1641.96 | 0.189919 | 0.06436 | 2.951031 | 0.00317 | 0.02706 | 17.107 |
| Tax1bp1 | 6409.97 | 0.145746 | 0.09506 | 1.533265 | 0.12521 | 0.3002 | 53.052 |
| Tax1bp3 | 148.491 | -0.40308 | 0.22356 | -1.80298 | 0.07139 | 0.20919 | 6.08 |
| Tbata | 73.9817 | -0.45395 | 0.69395 | -0.65415 | 0.51302 | 0.70391 | 16.888 |
| Tbc1d1 | 744.771 | -0.34282 | 0.18272 | -1.8762 | 0.06063 | 0.1879 | 195.182 |
| Tbc1d10a | 1056.09 | -0.10935 | 0.0813 | -1.34506 | 0.17861 | 0.37116 | 28.717 |
| Tbc1d10b | 3458.26 | -0.06595 | 0.06239 | -1.05706 | 0.29049 | 0.50076 | 11.01 |
| Tbc1d10c | 22.077 | 0.911225 | 0.38638 | 2.358362 | 0.01836 | 0.08475 | 7.874 |
| Tbc1d12 | 921.522 | 0.052948 | 0.09543 | 0.55484 | 0.579 | 0.75569 | 83.345 |
| Tbc1d13 | 2176.13 | -0.10116 | 0.07616 | -1.32815 | 0.18413 | 0.37854 | 18.268 |
| Tbc1d14 | 4290.36 | -0.04358 | 0.06803 | -0.64058 | 0.52179 | 0.71093 | 102.673 |
| Tbc1d15 | 2343.51 | 0.274346 | 0.07845 | 3.497279 | 0.00047 | 0.00761 | 53.596 |
| Tbc1d16 | 7882.42 | 0.222416 | 0.11362 | 1.95753 | 0.05029 | 0.16664 | 85.455 |
| Tbc1d17 | 1929.77 | 0.085084 | 0.0584 | 1.456821 | 0.14517 | 0.32787 | 14.457 |
| Tbc1d19 | 1772.09 | 0.080289 | 0.08536 | 0.940611 | 0.3469 | 0.55759 | 94.36 |
| Tbc1d2 | 209.174 | -0.30725 | 0.18101 | -1.69744 | 0.08961 | 0.24259 | 45.82 |
| Tbc1d20 | 3613.12 | 0.191865 | 0.10905 | 1.759409 | 0.07851 | 0.22199 | 20.169 |

| | | | | | | | |
|---|---|---|---|---|---|---|---|
| Tbc1d22a | 1323.25 | -0.09548 | 0.08896 | -1.07332 | 0.28313 | 0.49331 | 284.045 |
| Tbc1d22b | 1577.11 | -0.11423 | 0.08016 | -1.42499 | 0.15416 | 0.34002 | 57.114 |
| Tbc1d22bos | 31.8383 | 0.550333 | 0.37122 | 1.482496 | 0.13821 | 0.31934 | 17.49 |
| Tbc1d23 | 1911.79 | 0.117202 | 0.09127 | 1.284087 | 0.19911 | 0.39704 | 62.643 |
| Tbc1d24 | 4978.79 | -0.10906 | 0.06584 | -1.65649 | 0.09762 | 0.25633 | 30.132 |
| Tbc1d25 | 388.506 | -0.03034 | 0.11144 | -0.27225 | 0.78543 | 0.88861 | 21.71 |
| Tbc1d2b | 1449.27 | 0.269509 | 0.08787 | 3.067175 | 0.00216 | 0.02122 | 68.778 |
| Tbc1d30 | 4395.39 | -0.12718 | 0.07913 | -1.60714 | 0.10802 | 0.27316 | 87.459 |
| Tbc1d31 | 506.472 | 0.116833 | 0.09521 | 1.227171 | 0.21976 | 0.4222 | 57.869 |
| Tbc1d32 | 829.28 | 0.240534 | 0.14866 | 1.618043 | 0.10565 | 0.26917 | 214.397 |
| Tbc1d4 | 961.316 | -0.05294 | 0.11148 | -0.47488 | 0.63487 | 0.79438 | 166.832 |
| Tbc1d5 | 2065.24 | -0.01039 | 0.07477 | -0.13901 | 0.88944 | 0.94464 | 446.229 |
| Tbc1d7 | 537.355 | 0.005591 | 0.10151 | 0.055082 | 0.95607 | 0.97844 | 19.762 |
| Tbc1d8 | 4028.21 | 0.669757 | 0.18642 | 3.592827 | 0.00033 | 0.006 | 107.264 |
| Tbc1d8b | 528.07 | 0.067349 | 0.17842 | 0.37748 | 0.70582 | 0.84147 | 68.41 |
| Tbc1d9 | 3542.13 | -0.19641 | 0.09493 | -2.06908 | 0.03854 | 0.13979 | 107.583 |
| Tbc1d9b | 9209.32 | 0.007866 | 0.07864 | 0.100016 | 0.92033 | 0.95964 | 41.39 |
| Tbca | 1147.37 | 0.08046 | 0.10939 | 0.735509 | 0.46203 | 0.66235 | 54.013 |
| Tbcb | 1595.29 | -0.2313 | 0.09198 | -2.51465 | 0.01191 | 0.06432 | 8.142 |
| Tbcc | 767.731 | -0.1894 | 0.09065 | -2.08942 | 0.03667 | 0.135 | 1.164 |
| Tbccd1 | 720.778 | -0.01483 | 0.08363 | -0.17736 | 0.85922 | 0.92893 | 44.456 |
| Tbcd | 2239.61 | -0.00469 | 0.07842 | -0.05977 | 0.95234 | 0.97658 | 165.216 |
| Tbce | 2663.97 | 0.209731 | 0.08788 | 2.386429 | 0.01701 | 0.0809 | 41.69 |
| Tbcel | 3089.51 | -0.04277 | 0.0718 | -0.59564 | 0.55142 | 0.73445 | 95.494 |
| Tbck | 1689.32 | 0.150731 | 0.08232 | 1.831035 | 0.0671 | 0.20057 | 157.545 |
| Tbk1 | 2879.54 | 0.142244 | 0.07419 | 1.917399 | 0.05519 | 0.17713 | 40.333 |
| Tbkbp1 | 3876.75 | -0.15905 | 0.09241 | -1.72125 | 0.08521 | 0.23406 | 15.325 |
| Tbl1x | 4718.38 | 0.245813 | 0.06329 | 3.883717 | 0.0001 | 0.00274 | 151.971 |
| Tbl1xr1 | 3465.35 | -0.20556 | 0.08749 | -2.3497 | 0.01879 | 0.08604 | 139.943 |
| Tbl2 | 1531.5 | -0.07889 | 0.09519 | -0.82875 | 0.40725 | 0.61523 | 16.104 |
| Tbl3 | 978.552 | 0.000357 | 0.08974 | 0.003979 | 0.99683 | 0.99821 | 9.712 |
| Tbp | 1707.31 | 0.076762 | 0.08339 | 0.920508 | 0.35731 | 0.56765 | 28.492 |
| Tbpl1 | 2282.59 | 0.200993 | 0.08619 | 2.331957 | 0.0197 | 0.08869 | 28.06 |
| Tbr1 | 3175.16 | -1.17951 | 0.45146 | -2.61264 | 0.00898 | 0.05335 | 11.185 |

| | | | | | | | |
|---|---|---|---|---|---|---|---|
| Tbrg1 | 1097.43 | -0.13718 | 0.09862 | -1.39101 | 0.16422 | 0.3531 | 8.55 |
| Tbrg4 | 1471.47 | -0.01118 | 0.07002 | -0.1596 | 0.8732 | 0.93626 | 10.47 |
| Tbx1 | 54.6307 | -0.65637 | 0.27859 | -2.35606 | 0.01847 | 0.08514 | 9.693 |
| Tbx15 | 26.4151 | -1.03815 | 0.48841 | -2.12556 | 0.03354 | 0.12748 | 113.879 |
| Tbx18 | 104.494 | -0.93538 | 0.27502 | -3.40115 | 0.00067 | 0.00971 | 28.461 |
| Tbx2 | 88.3149 | -0.14274 | 0.25372 | -0.56259 | 0.57371 | 0.75227 | 9.398 |
| Tbx3 | 245.8 | -0.50294 | 0.12822 | -3.92247 | 8.76E-05 | 0.00246 | 14.056 |
| Tbx3os1 | 77.4044 | 0.127555 | 0.2841 | 0.448981 | 0.65345 | 0.80751 | 101.379 |
| Tbx6 | 62.8398 | 0.245578 | 0.24378 | 1.007362 | 0.31376 | 0.52419 | 4.078 |
| Tbxa2r | 20.4129 | 0.205557 | 0.52274 | 0.393233 | 0.69415 | 0.83414 | 6.442 |
| Tbxas1 | 98.8341 | -0.36411 | 0.19963 | -1.82398 | 0.06816 | 0.20267 | 209.182 |
| Tcaf1 | 17100.1 | 0.330903 | 0.08523 | 3.882391 | 0.0001 | 0.00276 | 42.087 |
| Tcaf2 | 35.6269 | -0.50826 | 0.31699 | -1.60342 | 0.10884 | 0.27431 | 22.239 |
| Tcaim | 1079.66 | 0.068553 | 0.1136 | 0.603456 | 0.54621 | 0.73071 | 30.796 |
| Tcap | 20.1868 | -0.36308 | 0.48476 | -0.74899 | 0.45386 | 0.65609 | 1.143 |
| Tcea1 | 2651.03 | 0.249542 | 0.06731 | 3.70748 | 0.00021 | 0.00439 | 40.096 |
| Tcea1-ps1 | 176.976 | 0.205072 | 0.17091 | 1.199862 | 0.23019 | 0.43455 | 0.904 |
| Tcea2 | 1247.94 | -0.19183 | 0.11362 | -1.68837 | 0.09134 | 0.2456 | 7.762 |
| Tcea3 | 12.4876 | -1.07521 | 0.48057 | -2.23737 | 0.02526 | 0.10462 | 27.17 |
| Tceal1 | 855.511 | 0.051315 | 0.0973 | 0.527408 | 0.59791 | 0.76947 | 3.497 |
| Tceal3 | 3581.05 | 0.006563 | 0.07585 | 0.086521 | 0.93105 | 0.96526 | 77.537 |
| Tceal5 | 3922.4 | -0.07674 | 0.07328 | -1.04729 | 0.29497 | 0.50539 | 2.929 |
| Tceal6 | 2780.73 | 0.142184 | 0.10606 | 1.340634 | 0.18004 | 0.37326 | 2.001 |
| Tceal7 | 25.3537 | -0.17836 | 0.34874 | -0.51143 | 0.60905 | 0.77661 | 11.322 |
| Tceal8 | 1825.71 | 5.85E-05 | 0.0694 | 0.000843 | 0.99933 | 0.99967 | 3.359 |
| Tceal9 | 2455.32 | 0.083738 | 0.11481 | 0.72936 | 0.46578 | 0.665 | 2.061 |
| Tceanc | 97.3714 | -0.17911 | 0.23401 | -0.76538 | 0.44405 | 0.6481 | 18.756 |
| Tceanc2 | 750.606 | -0.02333 | 0.08901 | -0.26213 | 0.79322 | 0.89287 | 44.962 |
| Tcerg1 | 3836.45 | -0.01142 | 0.0937 | -0.1219 | 0.90298 | 0.95118 | 64.307 |
| Tcerg1l | 892.071 | 0.679877 | 0.21372 | 3.18109 | 0.00147 | 0.0163 | 188.757 |
| Tcf12 | 2813.91 | 0.110446 | 0.06395 | 1.72706 | 0.08416 | 0.23231 | 269.184 |
| Tcf15 | 18.4955 | 0.036482 | 0.38508 | 0.09474 | 0.92452 | 0.96202 | 5.537 |
| Tcf19 | 97.6097 | -0.76941 | 0.18531 | -4.15191 | 3.30E-05 | 0.00123 | 4.091 |
| Tcf20 | 8056.43 | 0.245044 | 0.08029 | 3.052098 | 0.00227 | 0.02188 | 179.437 |

| | | | | | | | |
|---|---|---|---|---|---|---|---|
| Tcf23 | 45.6434 | -1.1943 | 0.3079 | -3.8788 | 0.0001 | 0.00279 | 8.357 |
| Tcf24 | 26.4112 | -0.19432 | 0.37462 | -0.51872 | 0.60396 | 0.7733 | 7.77 |
| Tcf25 | 19313.9 | -0.21776 | 0.05814 | -3.74573 | 0.00018 | 0.00401 | 30.083 |
| Tcf3 | 1251.09 | -0.00339 | 0.08238 | -0.04115 | 0.96718 | 0.98353 | 24.134 |
| Tcf4 | 10988.2 | -0.4589 | 0.16098 | -2.85066 | 0.00436 | 0.03338 | 345.724 |
| Tcf7 | 462.945 | 0.568469 | 0.20322 | 2.797354 | 0.00515 | 0.03701 | 30.961 |
| Tcf7l1 | 101.26 | -0.54473 | 0.21412 | -2.54404 | 0.01096 | 0.06094 | 162.877 |
| Tcf7l2 | 438.705 | 0.1046 | 0.17257 | 0.606148 | 0.54442 | 0.72947 | 191.835 |
| Tcfl5 | 35.3428 | 1.692859 | 0.48064 | 3.522085 | 0.00043 | 0.00722 | 20.753 |
| Tchh | 498.306 | -0.0158 | 0.11199 | -0.14109 | 0.8878 | 0.94402 | 6.748 |
| Tchp | 732.18 | 0.301069 | 0.11281 | 2.668831 | 0.00761 | 0.04818 | 14.568 |
| Tcim | 142.483 | -0.26887 | 0.20302 | -1.32436 | 0.18538 | 0.38017 | 1.805 |
| Tcirg1 | 710.429 | -0.15694 | 0.11403 | -1.37635 | 0.16871 | 0.35879 | 11.084 |
| Tcn2 | 875.382 | -0.48285 | 0.10661 | -4.52927 | 5.92E-06 | 0.00038 | 14.968 |
| Tcof1 | 1227.95 | 0.159172 | 0.07879 | 2.020114 | 0.04337 | 0.15051 | 35.217 |
| Tcp1 | 6190.57 | 0.180823 | 0.05653 | 3.198456 | 0.00138 | 0.01562 | 9.367 |
| Tcp10b | 21.8555 | -0.06029 | 0.41967 | -0.14366 | 0.88577 | 0.9432 | 21.378 |
| Tcp11l1 | 1619.51 | -0.08169 | 0.09427 | -0.86662 | 0.38615 | 0.59559 | 54.882 |
| Tcp11l2 | 945.295 | -0.318 | 0.11464 | -2.77398 | 0.00554 | 0.03896 | 37.734 |
| Tcta | 1031.2 | -0.13343 | 0.07722 | -1.72793 | 0.084 | 0.23204 | 3.203 |
| Tcte1 | 29.2358 | -1.04831 | 0.36042 | -2.90856 | 0.00363 | 0.02954 | 19.247 |
| Tcte2 | 281.502 | 0.365983 | 0.12064 | 3.033804 | 0.00241 | 0.02279 | 49.382 |
| Tctn1 | 1539.42 | 0.244112 | 0.09238 | 2.64251 | 0.00823 | 0.05049 | 26.613 |
| Tctn2 | 1047.75 | 0.0327 | 0.07929 | 0.412406 | 0.68004 | 0.82502 | 28.992 |
| Tctn3 | 1581.48 | 0.055185 | 0.09969 | 0.553568 | 0.57987 | 0.75639 | 15.788 |
| Tdg | 1595.53 | 0.195601 | 0.07941 | 2.463162 | 0.01377 | 0.07057 | 20.972 |
| Tdg-ps2 | 38.8415 | -0.41329 | 0.37719 | -1.09571 | 0.27321 | 0.48254 | 1.194 |
| Tdgf1 | 23.3816 | -0.11235 | 0.57429 | -0.19562 | 0.8449 | 0.92065 | 6.556 |
| Tdo2 | 39.4594 | 0.093532 | 0.28103 | 0.332816 | 0.73927 | 0.86253 | 19.113 |
| Tdp1 | 852.034 | -0.08155 | 0.137 | -0.59526 | 0.55167 | 0.73468 | 70.703 |
| Tdp2 | 900.776 | 0.016899 | 0.11473 | 0.147296 | 0.8829 | 0.94161 | 10.475 |
| Tdrd1 | 39.5995 | 0.112078 | 0.31369 | 0.357291 | 0.72087 | 0.85098 | 43.804 |
| Tdrd12 | 49.1173 | -0.38898 | 0.2872 | -1.35442 | 0.1756 | 0.36743 | 68.648 |
| Tdrd3 | 1059.74 | -0.00833 | 0.08896 | -0.09361 | 0.92542 | 0.96239 | 128.866 |

| | | | | | | | |
|---|---|---|---|---|---|---|---|
| Tdrd5 | 137.756 | -0.06664 | 0.19046 | -0.34988 | 0.72643 | 0.8541 | 48.369 |
| Tdrd6 | 75.9973 | 0.07256 | 0.27632 | 0.262597 | 0.79286 | 0.89266 | 14.965 |
| Tdrd7 | 1955.98 | 0.228715 | 0.07656 | 2.987547 | 0.00281 | 0.02514 | 69.428 |
| Tdrd9 | 60.7855 | -0.21117 | 0.26256 | -0.8043 | 0.42122 | 0.6282 | 97.296 |
| Tdrkh | 1572.85 | 0.02808 | 0.06931 | 0.405113 | 0.68539 | 0.8286 | 21.396 |
| Tdrp | 1269.37 | 0.23023 | 0.15667 | 1.469481 | 0.1417 | 0.3235 | 23.025 |
| Tead1 | 1368.9 | -0.13757 | 0.08304 | -1.65665 | 0.09759 | 0.25632 | 227.49 |
| Tead2 | 118.045 | -0.625 | 0.18885 | -3.30953 | 0.00093 | 0.01223 | 17.892 |
| Tead3 | 118.405 | -0.03521 | 0.16944 | -0.20779 | 0.83539 | 0.91561 | 19.147 |
| Tead4 | 39.2199 | 0.043233 | 0.32674 | 0.132318 | 0.89473 | 0.94736 | 76.536 |
| Tec | 133.464 | -0.18833 | 0.17925 | -1.05066 | 0.29341 | 0.50336 | 112.768 |
| Tecpr1 | 4619.83 | -0.031 | 0.05521 | -0.56139 | 0.57453 | 0.75265 | 29.174 |
| Tecpr2 | 4272.26 | 0.012462 | 0.05955 | 0.209277 | 0.83423 | 0.91498 | 83.131 |
| Tecr | 10525.8 | -0.22745 | 0.06843 | -3.32399 | 0.00089 | 0.01177 | 36.274 |
| Tecrl | 24.3209 | -0.04569 | 0.33584 | -0.13603 | 0.8918 | 0.94605 | 77.051 |
| Tecta | 15.7549 | 0.699453 | 0.44462 | 1.573135 | 0.11569 | 0.28546 | 70.311 |
| Tectb | 23.2086 | -1.13157 | 0.44783 | -2.52677 | 0.01151 | 0.06289 | 15.581 |
| Tedc1 | 87.2744 | -0.41636 | 0.1833 | -2.27153 | 0.02311 | 0.09843 | 9.628 |
| Tedc2 | 279.97 | -0.07829 | 0.15344 | -0.51027 | 0.60987 | 0.77731 | 5.798 |
| Teddm1b | 13.3775 | 0.827098 | 0.53448 | 1.547473 | 0.12175 | 0.2952 | 2.527 |
| Teddm2 | 126.388 | 0.029257 | 0.23336 | 0.125369 | 0.90023 | 0.94992 | 50.687 |
| Tef | 26270.1 | 0.286145 | 0.11523 | 2.483152 | 0.01302 | 0.06827 | 24.443 |
| Tefm | 267.64 | 0.15494 | 0.18575 | 0.834145 | 0.4042 | 0.61299 | 5.498 |
| Tek | 531.659 | -0.33834 | 0.10535 | -3.21172 | 0.00132 | 0.01513 | 135.688 |
| Tekt1 | 38.0654 | -0.48309 | 0.32203 | -1.50014 | 0.13358 | 0.31333 | 17.721 |
| Tekt2 | 82.5103 | 0.565248 | 0.2118 | 2.668748 | 0.00761 | 0.04818 | 3.568 |
| Tekt4 | 100.614 | -0.52054 | 0.21601 | -2.40977 | 0.01596 | 0.07759 | 21.988 |
| Tekt5 | 490.713 | -0.36229 | 0.11195 | -3.23621 | 0.00121 | 0.01442 | 37.543 |
| Telo2 | 1300.03 | 0.097349 | 0.08189 | 1.188756 | 0.23454 | 0.43973 | 16.398 |
| Ten1 | 492.258 | -0.09627 | 0.12748 | -0.75518 | 0.45014 | 0.65303 | 16.464 |
| Tenm1 | 3278.14 | 0.333601 | 0.11491 | 2.903079 | 0.0037 | 0.02987 | 901.261 |
| Tenm2 | 3058.56 | 0.352655 | 0.09234 | 3.819058 | 0.00013 | 0.00329 | 1229.309 |
| Tenm3 | 8162.35 | -0.2561 | 0.18605 | -1.3765 | 0.16867 | 0.35873 | 616.27 |
| Tenm4 | 11828.5 | 0.275168 | 0.09382 | 2.932801 | 0.00336 | 0.02801 | 739.848 |

| | | | | | | | |
|---|---|---|---|---|---|---|---|
| Tent2 | 1899.32 | 0.371195 | 0.08441 | 4.397592 | 1.09E-05 | 0.0006 | 46.104 |
| Tent4a | 2202.81 | -0.01941 | 0.06995 | -0.27747 | 0.78142 | 0.88615 | 36.659 |
| Tent4b | 3168.99 | 0.096987 | 0.06906 | 1.404348 | 0.16022 | 0.34832 | 60.51 |
| Tent5a | 417.992 | -0.45375 | 0.12503 | -3.62926 | 0.00028 | 0.00541 | 6.91 |
| Tent5b | 21.5618 | -0.26897 | 0.41416 | -0.64944 | 0.51606 | 0.70609 | 7.807 |
| Tent5c | 183.892 | -0.65494 | 0.27811 | -2.35494 | 0.01853 | 0.0853 | 37.697 |
| Tep1 | 324.532 | -0.59995 | 0.13684 | -4.38427 | 1.16E-05 | 0.00062 | 46.502 |
| Tepsin | 728.262 | -0.04171 | 0.0807 | -0.51686 | 0.60526 | 0.77395 | 8.201 |
| Terb1 | 74.3802 | -0.07227 | 0.2409 | -0.30001 | 0.76417 | 0.87653 | 63.192 |
| Terf1 | 489.325 | 0.170957 | 0.10745 | 1.591099 | 0.11159 | 0.27913 | 38.407 |
| Terf2 | 1794.79 | 0.027261 | 0.074 | 0.368371 | 0.7126 | 0.8452 | 27.148 |
| Terf2ip | 2523.69 | 0.158813 | 0.07091 | 2.239631 | 0.02511 | 0.10427 | 9.131 |
| Tert | 120.504 | 0.270452 | 0.18807 | 1.438022 | 0.15043 | 0.33519 | 22.933 |
| Tes | 116.357 | -0.26521 | 0.16225 | -1.63463 | 0.10213 | 0.26375 | 40.68 |
| Tes3-ps | 36.2449 | -0.17635 | 0.3156 | -0.5588 | 0.5763 | 0.75395 | 3.272 |
| Tesc | 2727.15 | 0.239933 | 0.14996 | 1.599994 | 0.1096 | 0.27556 | 34.136 |
| Tesk1 | 1787.63 | -0.16191 | 0.0754 | -2.14748 | 0.03175 | 0.12278 | 6.126 |
| Tesk2 | 196.942 | -0.0536 | 0.18489 | -0.28992 | 0.77188 | 0.88116 | 85.009 |
| Tet1 | 1125.7 | 0.050733 | 0.09957 | 0.509534 | 0.61038 | 0.77775 | 104.427 |
| Tet2 | 4106.34 | 0.191571 | 0.1168 | 1.64021 | 0.10096 | 0.26197 | 81.461 |
| Tet3 | 5225.77 | 0.120845 | 0.07234 | 1.670454 | 0.09483 | 0.25135 | 96.712 |
| Tex10 | 966.115 | 0.15918 | 0.07686 | 2.07114 | 0.03835 | 0.1393 | 42.602 |
| Tex101 | 13.5383 | 1.807242 | 0.62023 | 2.913818 | 0.00357 | 0.02915 | 8.669 |
| Tex14 | 29.1788 | -0.68237 | 0.40324 | -1.69222 | 0.0906 | 0.24452 | 150.759 |
| Tex15 | 70.4697 | -0.1769 | 0.39649 | -0.44616 | 0.65548 | 0.80876 | 68.845 |
| Tex2 | 4356.16 | -0.00192 | 0.06069 | -0.03171 | 0.9747 | 0.98734 | 111.277 |
| Tex261 | 1936.02 | -0.14892 | 0.09282 | -1.60439 | 0.10863 | 0.27394 | 6.929 |
| Tex264 | 1714.3 | -0.08184 | 0.07914 | -1.03413 | 0.30108 | 0.51079 | 27.182 |
| Tex29 | 16.4321 | -0.19067 | 0.55601 | -0.34292 | 0.73166 | 0.85759 | 15.288 |
| Tex30 | 91.2848 | -0.20708 | 0.23639 | -0.87601 | 0.38102 | 0.59053 | 15.829 |
| Tex45 | 16.4039 | 0.181976 | 0.47825 | 0.380504 | 0.70357 | 0.83997 | 16.32 |
| Tex49 | 15.9251 | -0.6565 | 0.59945 | -1.09518 | 0.27344 | 0.48283 | 18.577 |
| Tex50 | 156.06 | 0.18745 | 0.16113 | 1.163356 | 0.24468 | 0.45137 | 4.757 |
| Tex52 | 56.9412 | 0.163077 | 0.33667 | 0.484378 | 0.62812 | 0.78956 | 9.689 |

| | | | | | | | |
|---|---|---|---|---|---|---|---|
| Tex9 | 520.876 | 0.08953 | 0.12152 | 0.736755 | 0.46127 | 0.66183 | 41.819 |
| Tfam | 1479.27 | -0.03577 | 0.07048 | -0.50751 | 0.61179 | 0.77879 | 12.817 |
| Tfap2b | 98.4938 | -1.53751 | 0.39386 | -3.90369 | 9.47E-05 | 0.00257 | 29.663 |
| Tfap2c | 13.6943 | -1.42066 | 0.54252 | -2.61863 | 0.00883 | 0.05279 | 9.03 |
| Tfap2d | 93.4824 | -2 | 0.96811 | -2.08365 | 0.03719 | 0.13628 | 63.325 |
| Tfap4 | 181.068 | 0.245831 | 0.13282 | 1.850845 | 0.06419 | 0.19501 | 15.194 |
| Tfb1m | 173.729 | 0.038498 | 0.13068 | 0.294604 | 0.7683 | 0.87894 | 38.526 |
| Tfb2m | 984.011 | 0.080716 | 0.07528 | 1.072164 | 0.28365 | 0.49375 | 18.213 |
| Tfcp2 | 1336.71 | -0.10049 | 0.08731 | -1.15096 | 0.24975 | 0.45704 | 53.997 |
| Tfcp2l1 | 1493.33 | -0.37288 | 0.14223 | -2.6217 | 0.00875 | 0.05244 | 57.224 |
| Tfdp1 | 3023.15 | 0.170825 | 0.06158 | 2.774069 | 0.00554 | 0.03896 | 39.698 |
| Tfdp2 | 3104.13 | 0.145036 | 0.08936 | 1.623099 | 0.10457 | 0.26745 | 127.372 |
| Tfe3 | 1740.79 | 0.064821 | 0.07218 | 0.898001 | 0.36918 | 0.57973 | 12.643 |
| Tfeb | 318.411 | -0.42423 | 0.12457 | -3.40562 | 0.00066 | 0.00958 | 55.39 |
| Tfg | 2362.17 | -0.08815 | 0.07126 | -1.23714 | 0.21604 | 0.41777 | 27.119 |
| Tfip11 | 1810.06 | 0.038356 | 0.07484 | 0.512504 | 0.6083 | 0.77621 | 11.716 |
| Tfpi | 218.837 | -0.65557 | 0.27001 | -2.42799 | 0.01518 | 0.07507 | 43.921 |
| Tfpt | 866.601 | -0.08654 | 0.09576 | -0.90373 | 0.36614 | 0.57663 | 9.606 |
| Tfr2 | 202.851 | -0.19416 | 0.17787 | -1.09158 | 0.27502 | 0.48452 | 17.642 |
| Tfrc | 4006.02 | -0.31708 | 0.09952 | -3.18615 | 0.00144 | 0.01608 | 23.875 |
| Tgds | 426.043 | 0.181865 | 0.1022 | 1.779437 | 0.07517 | 0.21626 | 20.845 |
| Tgfa | 3165.19 | 0.334783 | 0.14937 | 2.241287 | 0.02501 | 0.10396 | 80.497 |
| Tgfb1 | 202.307 | -0.19844 | 0.14182 | -1.39925 | 0.16174 | 0.34991 | 18.076 |
| Tgfb1i1 | 266.557 | -0.68905 | 0.17332 | -3.97555 | 7.02E-05 | 0.00211 | 8.888 |
| Tgfb2 | 611.986 | 0.138184 | 0.09071 | 1.523349 | 0.12767 | 0.30417 | 83.198 |
| Tgfb3 | 596.809 | -0.32405 | 0.09313 | -3.4796 | 0.0005 | 0.00794 | 22.297 |
| Tgfbi | 174.621 | -0.57785 | 0.18568 | -3.11212 | 0.00186 | 0.01912 | 30.04 |
| Tgfbr1 | 1810.59 | -0.21716 | 0.11507 | -1.88725 | 0.05913 | 0.18511 | 61.71 |
| Tgfbr2 | 812.752 | -0.64569 | 0.09122 | -7.0784 | 1.46E-12 | 3.72E-09 | 91.068 |
| Tgfbr3 | 1018.63 | 0.157023 | 0.1565 | 1.00335 | 0.31569 | 0.5263 | 183.06 |
| Tgfbr3l | 207.365 | 0.288572 | 0.229 | 1.260149 | 0.20762 | 0.40716 | 3.21 |
| Tgfbrap1 | 2854.84 | -0.02555 | 0.06318 | -0.40436 | 0.68595 | 0.82887 | 51.438 |
| Tgif1 | 77.9866 | -0.33152 | 0.28928 | -1.14603 | 0.25178 | 0.45912 | 9.342 |
| Tgm1 | 46.32 | -0.66885 | 0.53383 | -1.25292 | 0.21023 | 0.4102 | 13.918 |

| | | | | | | | |
|---|---|---|---|---|---|---|---|
| Tgm2 | 604.71 | -0.40268 | 0.13319 | -3.02327 | 0.0025 | 0.02336 | 30.035 |
| Tgm3 | 18.3455 | -0.3843 | 0.40081 | -0.95881 | 0.33765 | 0.54842 | 38.051 |
| Tgm4 | 57.7987 | -0.3798 | 0.22359 | -1.69866 | 0.08938 | 0.2422 | 32.836 |
| Tgm7 | 34.0052 | 0.068293 | 0.35274 | 0.193608 | 0.84648 | 0.92184 | 22.832 |
| Tgoln1 | 7031.59 | -0.08882 | 0.05273 | -1.68449 | 0.09209 | 0.24705 | 8.569 |
| Tgs1 | 1754.29 | 0.095379 | 0.09369 | 1.018036 | 0.30866 | 0.51844 | 41.745 |
| Tgtp2 | 16.8663 | -1.18245 | 0.46539 | -2.54076 | 0.01106 | 0.06132 | 7.013 |
| Th | 245.172 | -0.25766 | 0.17707 | -1.45516 | 0.14562 | 0.32854 | 38.377 |
| Tha1 | 192.586 | -0.19914 | 0.16702 | -1.19228 | 0.23315 | 0.43798 | 5.53 |
| Thada | 938.657 | 0.154762 | 0.11954 | 1.294603 | 0.19546 | 0.39311 | 276.13 |
| Thap1 | 546.228 | 0.124238 | 0.10778 | 1.15265 | 0.24905 | 0.45629 | 6.011 |
| Thap11 | 1031.94 | -0.01657 | 0.10185 | -0.16267 | 0.87078 | 0.93475 | 1.819 |
| Thap12 | 3612.81 | 0.049958 | 0.06385 | 0.782467 | 0.43394 | 0.63968 | 14.96 |
| Thap2 | 1564.42 | 0.214078 | 0.08695 | 2.462126 | 0.01381 | 0.07064 | 16.04 |
| Thap3 | 614.903 | 0.095424 | 0.09913 | 0.962614 | 0.33574 | 0.54643 | 6.359 |
| Thap4 | 435.212 | 0.196718 | 0.12114 | 1.623845 | 0.10441 | 0.26737 | 52.93 |
| Thap6 | 138.848 | 0.034039 | 0.17071 | 0.199391 | 0.84196 | 0.91879 | 9.678 |
| Thap7 | 670.474 | 0.016383 | 0.10875 | 0.150639 | 0.88026 | 0.94022 | 3.155 |
| Thbd | 756.799 | -0.89168 | 0.34867 | -2.55737 | 0.01055 | 0.0596 | 3.723 |
| Thbs1 | 89.3624 | -0.4128 | 0.26063 | -1.58384 | 0.11323 | 0.28168 | 15.258 |
| Thbs2 | 225.679 | -1.02001 | 0.22402 | -4.55327 | 5.28E-06 | 0.00035 | 28.736 |
| Thbs3 | 1306.03 | 0.705862 | 0.08822 | 8.001086 | 1.23E-15 | 2.51E-11 | 11.658 |
| Thbs4 | 203.438 | -1.54681 | 0.6315 | -2.44943 | 0.01431 | 0.07211 | 43.229 |
| Theg | 23.9802 | 1.325705 | 0.40694 | 3.257711 | 0.00112 | 0.01378 | 10.953 |
| Them4 | 738.116 | 0.038876 | 0.08917 | 0.435975 | 0.66285 | 0.81376 | 22.452 |
| Them6 | 2006.82 | -0.00523 | 0.09308 | -0.05616 | 0.95521 | 0.97801 | 3.436 |
| Themis2 | 53.3897 | -0.58534 | 0.33437 | -1.75061 | 0.08001 | 0.22493 | 14.545 |
| Thg1l | 379.931 | 0.101283 | 0.0958 | 1.057264 | 0.29039 | 0.50071 | 8.652 |
| Thnsl1 | 945.276 | 0.044868 | 0.10657 | 0.421029 | 0.67373 | 0.82074 | 9.286 |
| Thnsl2 | 122.56 | -0.20971 | 0.18533 | -1.13156 | 0.25782 | 0.46595 | 16.274 |
| Thoc1 | 1795.99 | 0.129767 | 0.12526 | 1.035942 | 0.30023 | 0.51033 | 37.584 |
| Thoc2 | 3705.34 | 0.331294 | 0.15352 | 2.157947 | 0.03093 | 0.1206 | 125.684 |
| Thoc2l | 2005.77 | 0.048303 | 0.18245 | 0.264743 | 0.79121 | 0.89159 | 45.858 |
| Thoc3 | 958.724 | -0.13548 | 0.09541 | -1.41999 | 0.15561 | 0.34222 | 10.013 |

| | | | | | | | |
|---|---|---|---|---|---|---|---|
| Thoc5 | 1348.14 | -0.02152 | 0.07821 | -0.27518 | 0.78318 | 0.88742 | 33.548 |
| Thoc6 | 130.057 | -0.22547 | 0.21355 | -1.05584 | 0.29104 | 0.50115 | 5.269 |
| Thoc7 | 1339.52 | 0.073477 | 0.10083 | 0.728699 | 0.46619 | 0.66546 | 12.841 |
| Thop1 | 1571.34 | 0.072034 | 0.08092 | 0.89016 | 0.37338 | 0.58357 | 12.525 |
| Thpo | 346.997 | 0.170482 | 0.19385 | 0.879464 | 0.37915 | 0.5888 | 10.058 |
| Thra | 25029 | -0.02067 | 0.08463 | -0.24419 | 0.80708 | 0.90071 | 28.369 |
| Thrap3 | 8597.2 | 0.070334 | 0.06219 | 1.13099 | 0.25806 | 0.46613 | 38.679 |
| Thrb | 1916.27 | 0.298962 | 0.06893 | 4.337347 | 1.44E-05 | 0.00072 | 377.825 |
| Thrsp | 405.976 | -1.11829 | 0.14361 | -7.78673 | 6.88E-15 | 7.01E-11 | 4.793 |
| Thsd1 | 188.529 | -0.10671 | 0.14608 | -0.73048 | 0.4651 | 0.66474 | 34.022 |
| Thsd4 | 635.94 | 0.071143 | 0.17787 | 0.399963 | 0.68918 | 0.83062 | 555.116 |
| Thsd7a | 3778.28 | 0.415653 | 0.14475 | 2.871591 | 0.00408 | 0.03195 | 437.801 |
| Thsd7b | 327.47 | 0.443564 | 0.14017 | 3.164542 | 0.00155 | 0.01704 | 945.977 |
| Thtpa | 380.423 | -0.11962 | 0.10624 | -1.1259 | 0.26021 | 0.46875 | 4.203 |
| Thumpd1 | 2699.58 | 0.121408 | 0.08394 | 1.446307 | 0.14809 | 0.33176 | 5.706 |
| Thumpd2 | 261.041 | 0.136483 | 0.12841 | 1.062873 | 0.28784 | 0.49792 | 38.751 |
| Thumpd3 | 875.222 | 0.187272 | 0.10235 | 1.829653 | 0.0673 | 0.20101 | 22.049 |
| Thy1 | 20895.9 | -0.27401 | 0.07047 | -3.88816 | 0.0001 | 0.00271 | 5.196 |
| Thyn1 | 1678.1 | 0.049252 | 0.06463 | 0.762035 | 0.44604 | 0.64983 | 7.627 |
| Tia1 | 8672.5 | 0.115075 | 0.08293 | 1.387631 | 0.16525 | 0.35431 | 29.187 |
| Tiaf2 | 24.9872 | -0.06451 | 0.41744 | -0.15455 | 0.87718 | 0.93864 | 1.897 |
| Tial1 | 7015.13 | 0.306571 | 0.09917 | 3.091384 | 0.00199 | 0.02005 | 21.941 |
| Tiam1 | 11484.9 | 0.188419 | 0.08002 | 2.354537 | 0.01855 | 0.08535 | 356.659 |
| Tiam2 | 6067.75 | 0.220301 | 0.0808 | 2.726581 | 0.0064 | 0.04292 | 204.772 |
| Ticam1 | 276.934 | 0.053793 | 0.12713 | 0.423133 | 0.6722 | 0.81975 | 7.468 |
| Ticam2 | 24.9734 | -0.71641 | 0.37757 | -1.89743 | 0.05777 | 0.18257 | 17.243 |
| Tie1 | 457.757 | -0.28566 | 0.09997 | -2.85759 | 0.00427 | 0.03288 | 18.871 |
| Tifa | 270.307 | -0.39719 | 0.15551 | -2.55411 | 0.01065 | 0.05988 | 42.36 |
| Tifab | 131.592 | -0.56361 | 0.16253 | -3.46767 | 0.00052 | 0.00821 | 5.182 |
| Tigar | 1297.08 | -0.32349 | 0.11015 | -2.93688 | 0.00332 | 0.02778 | 24.442 |
| Tigd2 | 602.444 | 0.129412 | 0.09658 | 1.339901 | 0.18028 | 0.3736 | 3.164 |
| Tigd3 | 141.26 | -0.12737 | 0.15856 | -0.80329 | 0.4218 | 0.6287 | 3.012 |
| Tigd5 | 217.568 | -0.22279 | 0.15311 | -1.45511 | 0.14564 | 0.32854 | 4.801 |
| Timeless | 222.069 | -0.06656 | 0.15333 | -0.43407 | 0.66424 | 0.8145 | 20.877 |

| | | | | | | | |
|---|---|---|---|---|---|---|---|
| Timm10 | 545.235 | -0.12517 | 0.11197 | -1.11782 | 0.26364 | 0.47241 | 3.217 |
| Timm10b | 306.39 | 0.089241 | 0.13894 | 0.642302 | 0.52068 | 0.70984 | 3.582 |
| Timm13 | 1716.91 | -0.14109 | 0.08937 | -1.57873 | 0.1144 | 0.28327 | 1.52 |
| Timm17a | 3513.24 | -0.08744 | 0.06644 | -1.31607 | 0.18815 | 0.38372 | 18.566 |
| Timm17b | 484.6 | -0.17025 | 0.11401 | -1.49332 | 0.13535 | 0.31509 | 8.995 |
| Timm21 | 684.146 | 0.065947 | 0.09111 | 0.723784 | 0.4692 | 0.66807 | 5.334 |
| Timm22 | 1225.16 | 0.109983 | 0.09739 | 1.129356 | 0.25875 | 0.46696 | 9.341 |
| Timm23 | 1234.4 | -0.12726 | 0.06942 | -1.83307 | 0.06679 | 0.20013 | 21.737 |
| Timm29 | 4008.96 | 0.113114 | 0.09099 | 1.24317 | 0.21381 | 0.41511 | 3.249 |
| Timm44 | 2887.4 | 0.104434 | 0.08577 | 1.217666 | 0.22335 | 0.42664 | 16.183 |
| Timm50 | 942.435 | -0.16828 | 0.08269 | -2.03514 | 0.04184 | 0.14718 | 6.557 |
| Timm8a1 | 345.749 | -0.09538 | 0.14935 | -0.63865 | 0.52305 | 0.7115 | 4.61 |
| Timm8b | 1594.78 | -0.07693 | 0.07005 | -1.09821 | 0.27211 | 0.48148 | 1.411 |
| Timm9 | 1865.64 | 0.232697 | 0.07733 | 3.009252 | 0.00262 | 0.0241 | 13.512 |
| Timmdc1 | 822.38 | -0.09345 | 0.0874 | -1.06915 | 0.285 | 0.49485 | 24.317 |
| Timp1 | 79.1632 | 0.113569 | 0.22435 | 0.506208 | 0.61271 | 0.77942 | 4.57 |
| Timp2 | 16242.1 | 0.03608 | 0.08241 | 0.437805 | 0.66153 | 0.81272 | 54.672 |
| Timp3 | 5758.01 | -0.33519 | 0.14467 | -2.31699 | 0.0205 | 0.09104 | 49.135 |
| Timp4 | 1394.7 | -0.13172 | 0.10739 | -1.2265 | 0.22001 | 0.42246 | 10.314 |
| Tinagl1 | 597.843 | -0.2388 | 0.16289 | -1.46598 | 0.14265 | 0.32454 | 10.669 |
| Tincr | 200.988 | -0.13554 | 0.17288 | -0.784 | 0.43304 | 0.63916 | 2.974 |
| Tinf2 | 515.116 | -0.16083 | 0.1192 | -1.34921 | 0.17727 | 0.36947 | 7.132 |
| Tiparp | 1123.14 | 0.11737 | 0.09258 | 1.267735 | 0.20489 | 0.40376 | 27.109 |
| Tipin | 376.19 | -0.16534 | 0.17312 | -0.95505 | 0.33955 | 0.55041 | 23.844 |
| Tiprl | 2151.42 | 0.101379 | 0.0734 | 1.381117 | 0.16724 | 0.35697 | 24.711 |
| Tirap | 117.108 | -0.40213 | 0.21034 | -1.91177 | 0.05591 | 0.1787 | 15.741 |
| Tjap1 | 1361.74 | 0.04994 | 0.07248 | 0.68904 | 0.4908 | 0.6858 | 25.176 |
| Tjp1 | 6358.49 | -0.00775 | 0.05455 | -0.14216 | 0.88696 | 0.94383 | 231.617 |
| Tjp2 | 1565.74 | -0.24497 | 0.10882 | -2.25112 | 0.02438 | 0.10212 | 130.526 |
| Tjp3 | 13.3497 | 0.169356 | 0.47795 | 0.354343 | 0.72308 | 0.85223 | 18.375 |
| Tk1 | 103.961 | -0.09837 | 0.22881 | -0.42992 | 0.66725 | 0.8162 | 10.567 |
| Tk2 | 1412.75 | -0.15197 | 0.08692 | -1.74833 | 0.08041 | 0.22564 | 21.874 |
| Tkfc | 408.386 | -0.00228 | 0.10477 | -0.02179 | 0.98261 | 0.99061 | 17.864 |
| Tkt | 3842.51 | -0.14495 | 0.07807 | -1.85674 | 0.06335 | 0.19338 | 26.362 |

| | | | | | | | |
|---|---|---|---|---|---|---|---|
| Tlcd1 | 480.125 | -0.21332 | 0.12069 | -1.76746 | 0.07715 | 0.21958 | 5.199 |
| Tlcd3a | 844.465 | 0.130409 | 0.08612 | 1.514323 | 0.12994 | 0.30778 | 35.157 |
| Tlcd3b | 2062.78 | -0.27673 | 0.09893 | -2.7971 | 0.00516 | 0.03701 | 32.552 |
| Tlcd4 | 2780.49 | 0.006852 | 0.07874 | 0.087021 | 0.93065 | 0.96507 | 81.338 |
| Tlcd5 | 1293.5 | 0.010791 | 0.07012 | 0.153893 | 0.87769 | 0.93886 | 7.921 |
| Tle1 | 2344.08 | 0.371053 | 0.11491 | 3.229122 | 0.00124 | 0.01463 | 83.778 |
| Tle2 | 1143.85 | -0.32051 | 0.15823 | -2.02558 | 0.04281 | 0.14911 | 16.285 |
| Tle3 | 2417.69 | -0.65231 | 0.13568 | -4.80778 | 1.53E-06 | 0.00014 | 46.132 |
| Tle4 | 3317.64 | 0.613966 | 0.13242 | 4.636556 | 3.54E-06 | 0.00026 | 149.902 |
| Tle5 | 13928.2 | -0.19457 | 0.08086 | -2.40621 | 0.01612 | 0.07806 | 6.875 |
| Tle6 | 90.899 | -0.36794 | 0.25363 | -1.45069 | 0.14687 | 0.32996 | 10.17 |
| Tlk1 | 4234.16 | -0.02109 | 0.06522 | -0.32338 | 0.74641 | 0.86693 | 113.322 |
| Tlk2 | 2299.33 | 0.019607 | 0.07998 | 0.245139 | 0.80635 | 0.90039 | 105.153 |
| Tll1 | 235.055 | -0.60586 | 0.22728 | -2.66575 | 0.00768 | 0.04844 | 191.341 |
| Tll2 | 33.6137 | 0.060609 | 0.38063 | 0.159236 | 0.87348 | 0.93647 | 124.083 |
| Tln1 | 2925.99 | -0.24898 | 0.07646 | -3.25649 | 0.00113 | 0.01378 | 31.173 |
| Tln2 | 9990.25 | 0.358136 | 0.103 | 3.476927 | 0.00051 | 0.00799 | 342.617 |
| Tlnrd1 | 1815.86 | 0.50905 | 0.15593 | 3.264512 | 0.0011 | 0.01358 | 4.433 |
| Tlr1 | 28.6373 | -0.46879 | 0.41005 | -1.14325 | 0.25293 | 0.46033 | 8.885 |
| Tlr12 | 28.7543 | -0.10395 | 0.37558 | -0.27678 | 0.78195 | 0.8865 | 3.177 |
| Tlr13 | 141.737 | -0.30051 | 0.21236 | -1.41512 | 0.15703 | 0.34404 | 17.29 |
| Tlr2 | 70.8069 | -0.97942 | 0.33911 | -2.88825 | 0.00387 | 0.03079 | 5.496 |
| Tlr3 | 605.335 | -0.31121 | 0.12628 | -2.46442 | 0.01372 | 0.07048 | 15.416 |
| Tlr4 | 95.9084 | -0.49344 | 0.249 | -1.98168 | 0.04751 | 0.16022 | 102.701 |
| Tlr5 | 32.6994 | -0.1916 | 0.33644 | -0.56949 | 0.56902 | 0.7487 | 21.658 |
| Tlr6 | 34.8466 | -0.28551 | 0.31227 | -0.91431 | 0.36056 | 0.57089 | 8.067 |
| Tlr7 | 326.421 | -0.24526 | 0.16037 | -1.52933 | 0.12618 | 0.30189 | 25.63 |
| Tlr8 | 12.805 | -1.26141 | 0.57001 | -2.21294 | 0.0269 | 0.10948 | 21.634 |
| Tlr9 | 57.4148 | -0.54875 | 0.31337 | -1.75114 | 0.07992 | 0.22477 | 4.286 |
| Tm2d1 | 1101.5 | -0.08684 | 0.08321 | -1.04363 | 0.29666 | 0.50689 | 27.937 |
| Tm2d2 | 1738.71 | -0.08278 | 0.07981 | -1.03725 | 0.29962 | 0.50972 | 6.05 |
| Tm2d3 | 827.623 | -0.12626 | 0.08932 | -1.41346 | 0.15752 | 0.34464 | 10.745 |
| Tm4sf1 | 380.154 | -0.42727 | 0.12181 | -3.50778 | 0.00045 | 0.00742 | 16.378 |
| Tm6sf1 | 595.876 | -0.34456 | 0.15318 | -2.24947 | 0.02448 | 0.10241 | 25.434 |

| | | | | | | | |
|---|---|---|---|---|---|---|---|
| Tm6sf2 | 276.326 | 0.327139 | 0.1378 | 2.373981 | 0.0176 | 0.08244 | 7.143 |
| Tm7sf2 | 880.73 | -0.05758 | 0.08566 | -0.67211 | 0.50151 | 0.69397 | 5.553 |
| Tm7sf3 | 2479.54 | -0.04733 | 0.07281 | -0.65009 | 0.51564 | 0.70584 | 40.473 |
| Tm9sf1 | 1360.52 | -0.0559 | 0.08246 | -0.67786 | 0.49786 | 0.69116 | 7.842 |
| Tm9sf2 | 5834.32 | 0.016116 | 0.06814 | 0.236526 | 0.81302 | 0.90373 | 52.567 |
| Tm9sf3 | 9617.75 | 0.013573 | 0.05592 | 0.242724 | 0.80822 | 0.90141 | 53.154 |
| Tm9sf4 | 4012.81 | 0.048359 | 0.07101 | 0.680991 | 0.49588 | 0.68973 | 49.164 |
| Tma16 | 1170.11 | 0.144242 | 0.1335 | 1.080491 | 0.27992 | 0.49005 | 13.413 |
| Tma7 | 2657.11 | -0.13417 | 0.07517 | -1.78483 | 0.07429 | 0.21491 | 8.02 |
| Tma7-ps | 138.022 | -0.29469 | 0.1699 | -1.73453 | 0.08282 | 0.22962 | 0.195 |
| Tmbim1 | 1527.91 | -0.33724 | 0.10541 | -3.19932 | 0.00138 | 0.01559 | 17.376 |
| Tmbim4 | 1088.84 | -0.06289 | 0.07851 | -0.80099 | 0.42314 | 0.62995 | 23.328 |
| Tmbim6 | 14519.3 | -0.13549 | 0.07867 | -1.72227 | 0.08502 | 0.23388 | 17.168 |
| Tmc1 | 27.5048 | -1.89477 | 0.55047 | -3.44209 | 0.00058 | 0.00875 | 170.745 |
| Tmc3 | 74.6515 | 0.805133 | 0.30059 | 2.678498 | 0.0074 | 0.04731 | 40.688 |
| Tmc4 | 488.389 | 0.385695 | 0.15279 | 2.524321 | 0.01159 | 0.06316 | 11.764 |
| Tmc6 | 166.407 | -0.51395 | 0.18389 | -2.79495 | 0.00519 | 0.03717 | 16.211 |
| Tmc7 | 1070.19 | -0.1813 | 0.08928 | -2.03071 | 0.04228 | 0.14806 | 48.896 |
| Tmc8 | 17.4269 | 0.107647 | 0.51675 | 0.208314 | 0.83498 | 0.91532 | 11.035 |
| Tmcc1 | 4358.61 | 0.101271 | 0.07794 | 1.299288 | 0.19385 | 0.39087 | 174.876 |
| Tmcc2 | 10367.5 | -0.21349 | 0.08117 | -2.63011 | 0.00854 | 0.05162 | 34.967 |
| Tmcc3 | 2707.17 | -0.10815 | 0.07324 | -1.47663 | 0.13978 | 0.3211 | 279.008 |
| Tmcc3os | 32.422 | -0.26169 | 0.33427 | -0.78286 | 0.43371 | 0.63957 | 44.257 |
| Tmco1 | 1507.22 | -0.07618 | 0.09346 | -0.81508 | 0.41503 | 0.6229 | 25.601 |
| Tmco3 | 2040.65 | -0.00462 | 0.07308 | -0.06322 | 0.94959 | 0.97495 | 34.735 |
| Tmco4 | 220.172 | -0.14003 | 0.14842 | -0.94346 | 0.34545 | 0.55616 | 86.284 |
| Tmco5 | 68.9264 | 0.493618 | 0.26506 | 1.862262 | 0.06257 | 0.19195 | 13.804 |
| Tmco6 | 560.29 | 0.156149 | 0.09294 | 1.680095 | 0.09294 | 0.24829 | 7.359 |
| Tmed1 | 327.278 | 0.037898 | 0.11087 | 0.341817 | 0.73249 | 0.85819 | 2.7 |
| Tmed10 | 5948.85 | -0.07368 | 0.05963 | -1.23573 | 0.21656 | 0.4185 | 34.234 |
| Tmed10-ps | 134.892 | -0.16961 | 0.18629 | -0.91046 | 0.36258 | 0.57294 | 0.66 |
| Tmed2 | 3153.15 | -0.06663 | 0.08559 | -0.77849 | 0.43628 | 0.64192 | 9.812 |
| Tmed2b | 269.204 | -0.30264 | 0.1334 | -2.26863 | 0.02329 | 0.09892 | 0.604 |
| Tmed3 | 716.772 | -0.28174 | 0.13396 | -2.10307 | 0.03546 | 0.13228 | 5.863 |

| | | | | | | | |
|---|---|---|---|---|---|---|---|
| Tmed4 | 2892.32 | -0.03847 | 0.06687 | -0.57522 | 0.56515 | 0.74541 | 4.502 |
| Tmed5 | 1723.07 | 0.227145 | 0.15597 | 1.456349 | 0.1453 | 0.32812 | 26.255 |
| Tmed7 | 5993.53 | 0.076522 | 0.05814 | 1.316213 | 0.1881 | 0.38372 | 37.301 |
| Tmed8 | 3959.86 | 0.071943 | 0.05612 | 1.281869 | 0.19989 | 0.39801 | 34.218 |
| Tmed9 | 3777.31 | -0.23908 | 0.07279 | -3.28456 | 0.00102 | 0.01298 | 23.966 |
| Tmeff1 | 4191.46 | -0.28716 | 0.10853 | -2.64587 | 0.00815 | 0.05013 | 77.958 |
| Tmeff2 | 2217.06 | -0.02481 | 0.10696 | -0.23191 | 0.81661 | 0.90555 | 286.624 |
| Tmem100 | 531.557 | -0.21595 | 0.09778 | -2.20854 | 0.02721 | 0.11019 | 6.161 |
| Tmem101 | 840.637 | -0.11476 | 0.08617 | -1.33181 | 0.18292 | 0.3769 | 3.859 |
| Tmem102 | 50.0152 | -0.02833 | 0.27332 | -0.10365 | 0.91744 | 0.95781 | 2.022 |
| Tmem104 | 867.292 | -0.13776 | 0.07674 | -1.79519 | 0.07262 | 0.21188 | 59.537 |
| Tmem106a | 63.1002 | -0.30076 | 0.24471 | -1.22906 | 0.21905 | 0.42125 | 9.547 |
| Tmem106b | 10385 | 0.206979 | 0.08719 | 2.373965 | 0.0176 | 0.08244 | 19.511 |
| Tmem106c | 1001.82 | -0.04611 | 0.08358 | -0.55169 | 0.58116 | 0.75715 | 6.076 |
| Tmem107 | 332.9 | 0.115718 | 0.11845 | 0.976917 | 0.32861 | 0.53979 | 2.488 |
| Tmem108 | 264.459 | -0.45001 | 0.19456 | -2.31298 | 0.02072 | 0.09173 | 278.891 |
| Tmem109 | 1268.35 | -0.07401 | 0.09227 | -0.80203 | 0.42253 | 0.62935 | 11.342 |
| Tmem11 | 959.124 | -0.11328 | 0.08143 | -1.39111 | 0.16419 | 0.35308 | 14.821 |
| Tmem114 | 31.064 | 0.254057 | 0.35856 | 0.708551 | 0.4786 | 0.67618 | 15.862 |
| Tmem115 | 939.266 | -0.01864 | 0.08507 | -0.2191 | 0.82657 | 0.91075 | 4.729 |
| Tmem116 | 248.622 | 0.158553 | 0.15324 | 1.034686 | 0.30082 | 0.51075 | 72.291 |
| Tmem117 | 378.416 | -0.36764 | 0.1424 | -2.58168 | 0.00983 | 0.05687 | 466.867 |
| Tmem119 | 1074.74 | -0.32565 | 0.14261 | -2.28355 | 0.0224 | 0.09661 | 6.788 |
| Tmem120a | 399.679 | -0.19213 | 0.10774 | -1.7833 | 0.07454 | 0.21518 | 8.964 |
| Tmem120b | 395.494 | 0.27377 | 0.09645 | 2.838502 | 0.00453 | 0.0341 | 49.335 |
| Tmem121 | 449.653 | -0.0109 | 0.10687 | -0.10202 | 0.91874 | 0.95882 | 3.594 |
| Tmem121b | 3088.97 | -0.11661 | 0.07517 | -1.55125 | 0.12084 | 0.29387 | 4.869 |
| Tmem123 | 830.472 | -0.06622 | 0.1105 | -0.59928 | 0.54899 | 0.73289 | 30.293 |
| Tmem125 | 182.13 | -0.82689 | 0.23901 | -3.4597 | 0.00054 | 0.00838 | 3.104 |
| Tmem126a | 894.672 | -0.04744 | 0.07265 | -0.65295 | 0.51379 | 0.70456 | 6.53 |
| Tmem126b | 1888.39 | 0.138499 | 0.09821 | 1.410297 | 0.15845 | 0.34581 | 8.564 |
| Tmem127 | 5384.19 | -0.08308 | 0.05768 | -1.44048 | 0.14973 | 0.33411 | 13.2 |
| Tmem128 | 993.946 | -0.12748 | 0.09124 | -1.39722 | 0.16235 | 0.35074 | 9.45 |
| Tmem129 | 942.251 | -0.21406 | 0.09186 | -2.33026 | 0.01979 | 0.08891 | 5.201 |

| | | | | | | | |
|---|---|---|---|---|---|---|---|
| Tmem130 | 8404.13 | 0.137465 | 0.08574 | 1.603317 | 0.10886 | 0.27434 | 25.904 |
| Tmem131 | 5464.75 | 0.16001 | 0.06889 | 2.322652 | 0.0202 | 0.09011 | 151.476 |
| Tmem131l | 1440.4 | 0.524695 | 0.08819 | 5.94951 | 2.69E-09 | 1.19E-06 | 142.521 |
| Tmem132a | 4303.81 | 0.175505 | 0.09601 | 1.82803 | 0.06755 | 0.20161 | 12.119 |
| Tmem132b | 10348.8 | 0.0359 | 0.0777 | 0.462027 | 0.64406 | 0.80143 | 260.81 |
| Tmem132c | 263.214 | -0.96159 | 0.1734 | -5.54541 | 2.93E-08 | 7.88E-06 | 323.986 |
| Tmem132d | 1336.3 | -0.48448 | 0.10712 | -4.52283 | 6.10E-06 | 0.00038 | 651.448 |
| Tmem132e | 951.572 | 0.291221 | 0.10235 | 2.845444 | 0.00443 | 0.03369 | 57.433 |
| Tmem134 | 738.05 | -0.15484 | 0.08678 | -1.78423 | 0.07439 | 0.21502 | 6.373 |
| Tmem135 | 2768.12 | 0.066074 | 0.06528 | 1.0121 | 0.31149 | 0.52177 | 264.509 |
| Tmem138 | 162.776 | 0.05108 | 0.19718 | 0.259056 | 0.79559 | 0.8946 | 6.885 |
| Tmem140 | 56.9319 | -0.08496 | 0.28593 | -0.29713 | 0.76637 | 0.8779 | 11.801 |
| Tmem141 | 245.38 | -0.40209 | 0.17603 | -2.28414 | 0.02236 | 0.0965 | 1.939 |
| Tmem143 | 837.168 | -0.03313 | 0.09567 | -0.3463 | 0.72912 | 0.85583 | 20.473 |
| Tmem144 | 440.756 | -0.71004 | 0.1818 | -3.9057 | 9.40E-05 | 0.00257 | 40.21 |
| Tmem145 | 646.376 | -0.33569 | 0.16257 | -2.06487 | 0.03894 | 0.14059 | 10.09 |
| Tmem147 | 1603.58 | -0.07568 | 0.07343 | -1.0306 | 0.30273 | 0.51231 | 1.84 |
| Tmem147os | 275.03 | 0.527363 | 0.14526 | 3.63044 | 0.00028 | 0.00539 | 21.388 |
| Tmem14a | 1023.82 | 0.047804 | 0.09827 | 0.486449 | 0.62665 | 0.78874 | 11.593 |
| Tmem14c | 1201.57 | -0.14541 | 0.07046 | -2.06362 | 0.03905 | 0.1408 | 6.295 |
| Tmem150a | 438.021 | -1.64E-05 | 0.1158 | -0.00014 | 0.99989 | 0.99999 | 4.316 |
| Tmem150b | 40.5264 | 0.829041 | 0.2896 | 2.862743 | 0.0042 | 0.03252 | 18.418 |
| Tmem150c | 1752.78 | -0.22896 | 0.10956 | -2.0898 | 0.03664 | 0.13489 | 81.937 |
| Tmem150cos | 23.6639 | 0.114491 | 0.40907 | 0.279883 | 0.77957 | 0.88518 | 17.312 |
| Tmem151a | 5887.09 | -0.38191 | 0.08982 | -4.25176 | 2.12E-05 | 0.00092 | 14.673 |
| Tmem151b | 3940.58 | -0.76667 | 0.17606 | -4.35458 | 1.33E-05 | 0.00068 | 7.726 |
| Tmem154 | 28.4645 | -0.61587 | 0.45784 | -1.34517 | 0.17857 | 0.37112 | 38.384 |
| Tmem158 | 6138.52 | 0.743125 | 0.29566 | 2.513414 | 0.01196 | 0.0645 | 1.712 |
| Tmem159 | 252.416 | -0.39123 | 0.16794 | -2.32955 | 0.01983 | 0.08901 | 18.64 |
| Tmem160 | 719.227 | -0.58926 | 0.11332 | -5.19981 | 1.99E-07 | 3.01E-05 | 2.716 |
| Tmem161a | 1358.06 | -0.05426 | 0.08542 | -0.63526 | 0.52526 | 0.71356 | 11.326 |
| Tmem161b | 1372.78 | 0.129768 | 0.08755 | 1.482251 | 0.13827 | 0.31942 | 73.846 |
| Tmem163 | 471.071 | -0.59948 | 0.168 | -3.56839 | 0.00036 | 0.00634 | 193.003 |
| Tmem164 | 2136.4 | 0.399649 | 0.14626 | 2.732417 | 0.00629 | 0.04245 | 162.089 |

| | | | | | | | |
|---|---|---|---|---|---|---|---|
| Tmem165 | 1195.48 | 0.122196 | 0.0724 | 1.687764 | 0.09146 | 0.24577 | 25.366 |
| Tmem167 | 3083.32 | 0.145845 | 0.09075 | 1.60707 | 0.10804 | 0.27317 | 54.785 |
| Tmem167b | 2539.23 | 0.036302 | 0.08745 | 0.415101 | 0.67807 | 0.82367 | 6.042 |
| Tmem168 | 978.203 | 0.014865 | 0.08169 | 0.181974 | 0.8556 | 0.927 | 27.414 |
| Tmem169 | 810.761 | -0.01347 | 0.08999 | -0.14965 | 0.88104 | 0.94067 | 18.736 |
| Tmem17 | 219.606 | 0.056894 | 0.16689 | 0.340915 | 0.73317 | 0.85857 | 7.147 |
| Tmem170 | 373.894 | 0.061605 | 0.1064 | 0.579021 | 0.56257 | 0.74339 | 13.713 |
| Tmem170b | 4557.8 | -0.00274 | 0.07305 | -0.03753 | 0.97006 | 0.98478 | 35.449 |
| Tmem175 | 1376.81 | 0.087024 | 0.10101 | 0.861581 | 0.38892 | 0.59809 | 19.013 |
| Tmem176a | 484.944 | -0.06043 | 0.14041 | -0.43038 | 0.66692 | 0.81591 | 6.153 |
| Tmem176b | 734.007 | -0.27446 | 0.11649 | -2.35604 | 0.01847 | 0.08514 | 7.679 |
| Tmem177 | 459.679 | -0.31831 | 0.13747 | -2.3154 | 0.02059 | 0.09128 | 5.319 |
| Tmem178 | 982.446 | -0.8479 | 0.21411 | -3.96005 | 7.49E-05 | 0.00219 | 57.185 |
| Tmem178b | 8865.86 | 0.182946 | 0.06756 | 2.707866 | 0.00677 | 0.0445 | 383.69 |
| Tmem179 | 5191.13 | 0.035974 | 0.06943 | 0.518172 | 0.60434 | 0.7733 | 16.559 |
| Tmem179b | 187.329 | -0.11751 | 0.1686 | -0.69694 | 0.48584 | 0.682 | 1.943 |
| Tmem18 | 2214.95 | -0.05165 | 0.09839 | -0.52496 | 0.59961 | 0.7704 | 6.79 |
| Tmem181a | 3058.19 | 0.268732 | 0.07271 | 3.696192 | 0.00022 | 0.00453 | 51.455 |
| Tmem181b-ps | 10820 | 0.437508 | 0.11203 | 3.905238 | 9.41E-05 | 0.00257 | 28.499 |
| Tmem181c-ps | 24.0146 | 0.048545 | 0.39858 | 0.121793 | 0.90306 | 0.95122 | 26.038 |
| Tmem183a | 2853.18 | -0.09423 | 0.06699 | -1.40657 | 0.15955 | 0.34742 | 15.883 |
| Tmem184b | 3579.58 | -0.00747 | 0.06537 | -0.11421 | 0.90907 | 0.95444 | 42.886 |
| Tmem184c | 2884.74 | 0.066558 | 0.12193 | 0.545884 | 0.58515 | 0.75977 | 14.717 |
| Tmem185a | 1988 | 0.194819 | 0.1029 | 1.893223 | 0.05833 | 0.1836 | 17.428 |
| Tmem185b | 513.692 | 0.085959 | 0.10337 | 0.831556 | 0.40566 | 0.61433 | 2.824 |
| Tmem186 | 1145.7 | 0.122531 | 0.09294 | 1.31842 | 0.18736 | 0.38265 | 4.484 |
| Tmem19 | 1731.44 | -0.16106 | 0.07423 | -2.16986 | 0.03002 | 0.11832 | 22.828 |
| Tmem191c | 3829.04 | 0.294094 | 0.09826 | 2.992944 | 0.00276 | 0.02493 | 7.153 |
| Tmem192 | 380.671 | -0.19278 | 0.12103 | -1.5929 | 0.11118 | 0.27839 | 21.849 |
| Tmem196 | 984.162 | -0.03508 | 0.1547 | -0.22675 | 0.82061 | 0.90737 | 75.425 |
| Tmem198 | 1628.49 | -0.10559 | 0.08411 | -1.25535 | 0.20935 | 0.40935 | 6.174 |
| Tmem198b | 244.061 | 0.147171 | 0.14237 | 1.033737 | 0.30126 | 0.51097 | 4.335 |
| Tmem199 | 1436.32 | 0.029335 | 0.0811 | 0.361697 | 0.71758 | 0.84915 | 5.118 |
| Tmem200a | 254.952 | -0.27463 | 0.18599 | -1.47656 | 0.13979 | 0.3211 | 166.722 |

| | | | | | | | |
|---|---|---|---|---|---|---|---|
| Tmem200b | 340.706 | 0.396792 | 0.21676 | 1.830555 | 0.06717 | 0.20068 | 1.941 |
| Tmem200c | 578.803 | -0.57451 | 0.17324 | -3.31624 | 0.00091 | 0.01201 | 16.517 |
| Tmem201 | 4730.5 | 0.221205 | 0.08179 | 2.704643 | 0.00684 | 0.04483 | 22.67 |
| Tmem202 | 45.0032 | -0.00426 | 0.30383 | -0.01401 | 0.98882 | 0.99386 | 21.163 |
| Tmem203 | 84.1957 | 0.072679 | 0.24135 | 0.301138 | 0.76331 | 0.876 | 0.854 |
| Tmem204 | 363.847 | -0.30268 | 0.12577 | -2.40668 | 0.0161 | 0.07804 | 25.89 |
| Tmem205 | 624.391 | -0.07623 | 0.08866 | -0.85973 | 0.38994 | 0.59902 | 6.549 |
| Tmem208 | 687.783 | -0.01567 | 0.10266 | -0.15262 | 0.8787 | 0.93928 | 8.866 |
| Tmem209 | 1745.89 | 0.133265 | 0.0727 | 1.8332 | 0.06677 | 0.20013 | 30.731 |
| Tmem210 | 104.225 | -0.01472 | 0.17804 | -0.08268 | 0.93411 | 0.96693 | 1.045 |
| Tmem214 | 2439.05 | 0.115249 | 0.06366 | 1.810246 | 0.07026 | 0.20701 | 11.169 |
| Tmem215 | 80.8238 | -0.52384 | 0.39252 | -1.33453 | 0.18203 | 0.37584 | 4.989 |
| Tmem216 | 276.869 | 0.198847 | 0.11465 | 1.734339 | 0.08286 | 0.22962 | 22.374 |
| Tmem218 | 546.163 | -0.31266 | 0.0937 | -3.33677 | 0.00085 | 0.01142 | 16.478 |
| Tmem219 | 168.781 | -0.22558 | 0.13852 | -1.62851 | 0.10342 | 0.26586 | 36.747 |
| Tmem220 | 45.9544 | -0.30742 | 0.27448 | -1.12003 | 0.2627 | 0.47155 | 10.159 |
| Tmem222 | 2052.12 | -0.13149 | 0.07259 | -1.81126 | 0.0701 | 0.2067 | 11.748 |
| Tmem223 | 761.286 | -0.00434 | 0.0995 | -0.04366 | 0.96518 | 0.98278 | 1.48 |
| Tmem229a | 3457.53 | -0.29308 | 0.11776 | -2.48877 | 0.01282 | 0.06759 | 5.157 |
| Tmem229b | 1419.05 | -0.10995 | 0.07217 | -1.52336 | 0.12767 | 0.30417 | 45.833 |
| Tmem230 | 1178.5 | 0.070857 | 0.08902 | 0.795935 | 0.42607 | 0.63255 | 8.316 |
| Tmem231 | 862.601 | 0.134342 | 0.10219 | 1.314686 | 0.18862 | 0.38428 | 21.871 |
| Tmem234 | 2831.16 | -0.04205 | 0.06422 | -0.65475 | 0.51263 | 0.70354 | 23.392 |
| Tmem236 | 13.8668 | 0.000502 | 0.64212 | 0.000782 | 0.99938 | 0.99967 | 47.471 |
| Tmem237 | 702.753 | -0.03008 | 0.089 | -0.33802 | 0.73535 | 0.85989 | 19.819 |
| Tmem238 | 54.5682 | -0.23493 | 0.27427 | -0.85655 | 0.39169 | 0.60082 | 5.1 |
| Tmem239 | 15.1797 | 0.016574 | 0.46171 | 0.035898 | 0.97136 | 0.98558 | 1.425 |
| Tmem240 | 1820.26 | -0.06636 | 0.06986 | -0.94987 | 0.34218 | 0.553 | 5.765 |
| Tmem241 | 301.022 | -0.08147 | 0.15462 | -0.52689 | 0.59827 | 0.76969 | 157.088 |
| Tmem242 | 1557.49 | -0.09883 | 0.10248 | -0.96438 | 0.33486 | 0.54562 | 29.39 |
| Tmem243 | 698.019 | -0.06371 | 0.08386 | -0.7597 | 0.44743 | 0.65114 | 60.319 |
| Tmem245 | 6115.82 | -0.11378 | 0.08185 | -1.39009 | 0.1645 | 0.35345 | 80.515 |
| Tmem248 | 2129.7 | -0.0745 | 0.07582 | -0.98256 | 0.32582 | 0.53722 | 26.685 |
| Tmem249 | 21.1033 | -0.03036 | 0.41396 | -0.07333 | 0.94154 | 0.97087 | 2.154 |

| | | | | | | | |
|---|---|---|---|---|---|---|---|
| Tmem25 | 1823.89 | -0.46786 | 0.12727 | -3.67628 | 0.00024 | 0.00478 | 5.539 |
| Tmem250-ps | 2015.07 | 0.108639 | 0.09241 | 1.17567 | 0.23973 | 0.44568 | 3.708 |
| Tmem251 | 457.78 | 0.062457 | 0.09941 | 0.628302 | 0.52981 | 0.71748 | 3.8 |
| Tmem252 | 209.123 | -0.76612 | 0.20132 | -3.8054 | 0.00014 | 0.00338 | 8.226 |
| Tmem253 | 136.908 | 0.550217 | 0.18195 | 3.023967 | 0.00249 | 0.02334 | 4.137 |
| Tmem254 | 810.023 | -0.41855 | 0.14769 | -2.83397 | 0.0046 | 0.03443 | 4.373 |
| Tmem255a | 1703 | 0.532182 | 0.14711 | 3.617592 | 0.0003 | 0.0056 | 55.867 |
| Tmem255b | 54.2632 | -0.19366 | 0.31201 | -0.62067 | 0.53481 | 0.72177 | 33.285 |
| Tmem256 | 782.228 | -0.20845 | 0.11979 | -1.74004 | 0.08185 | 0.22803 | 1.105 |
| Tmem258 | 812.704 | -0.11825 | 0.10953 | -1.07962 | 0.28031 | 0.49019 | 3.811 |
| Tmem259 | 3619.75 | -0.02497 | 0.06053 | -0.41248 | 0.67998 | 0.82501 | 9.383 |
| Tmem26 | 23.3749 | 1.092542 | 0.54206 | 2.015553 | 0.04385 | 0.15156 | 59.005 |
| Tmem260 | 1925.97 | 0.123719 | 0.06459 | 1.915583 | 0.05542 | 0.17756 | 78.123 |
| Tmem263 | 2803.33 | 0.363881 | 0.09763 | 3.727171 | 0.00019 | 0.0042 | 17.992 |
| Tmem266 | 376.05 | -0.06346 | 0.1212 | -0.52364 | 0.60053 | 0.77095 | 111.433 |
| Tmem267 | 411.361 | -0.04305 | 0.14701 | -0.29286 | 0.76963 | 0.87977 | 448.021 |
| Tmem268 | 837.325 | -0.04435 | 0.08484 | -0.52276 | 0.60114 | 0.7714 | 27.577 |
| Tmem269 | 92.2184 | 0.193081 | 0.21534 | 0.896635 | 0.36991 | 0.58027 | 13.163 |
| Tmem273 | 39.6483 | -0.16964 | 0.32283 | -0.52548 | 0.59925 | 0.77006 | 32.022 |
| Tmem29 | 772.342 | 0.024447 | 0.0819 | 0.298485 | 0.76533 | 0.87738 | 61.382 |
| Tmem30a | 18627 | 0.10545 | 0.09896 | 1.065538 | 0.28663 | 0.4965 | 24.565 |
| Tmem30c | 41.3166 | 0.149955 | 0.27505 | 0.545192 | 0.58562 | 0.75999 | 26.727 |
| Tmem33 | 5594.56 | 0.023978 | 0.06195 | 0.387043 | 0.69872 | 0.83678 | 30.897 |
| Tmem35a | 3135.82 | -0.10344 | 0.06905 | -1.49812 | 0.1341 | 0.31384 | 10.745 |
| Tmem35b | 146.38 | -0.18445 | 0.23653 | -0.77984 | 0.43548 | 0.64111 | 5.954 |
| Tmem37 | 29.5046 | -0.77935 | 0.34131 | -2.28344 | 0.0224 | 0.09661 | 6.698 |
| Tmem38a | 4204.75 | -0.04731 | 0.0811 | -0.58335 | 0.55966 | 0.74103 | 15.228 |
| Tmem38b | 645.646 | -0.09299 | 0.09921 | -0.93724 | 0.34864 | 0.55914 | 35.975 |
| Tmem39a | 511.108 | -0.22025 | 0.10153 | -2.16939 | 0.03005 | 0.1184 | 33.465 |
| Tmem39b | 382.403 | 0.038216 | 0.11714 | 0.32625 | 0.74424 | 0.86539 | 20.484 |
| Tmem40 | 86.0105 | -0.75458 | 0.37166 | -2.0303 | 0.04233 | 0.14806 | 33.28 |
| Tmem41a | 679.951 | 0.280767 | 0.13037 | 2.153564 | 0.03127 | 0.12152 | 13.227 |
| Tmem41b | 1700.3 | 0.069372 | 0.07799 | 0.889463 | 0.37375 | 0.58398 | 14.743 |
| Tmem42 | 179.369 | -0.19276 | 0.15605 | -1.23522 | 0.21675 | 0.41862 | 2.38 |

| | | | | | | | |
|---|---|---|---|---|---|---|---|
| Tmem43 | 1525.19 | -0.01482 | 0.10295 | -0.14395 | 0.88554 | 0.94315 | 14.761 |
| Tmem44 | 4476.38 | -0.04932 | 0.10799 | -0.45673 | 0.64786 | 0.80388 | 38.988 |
| Tmem45a | 33.4784 | -0.28263 | 0.41632 | -0.67887 | 0.49722 | 0.69076 | 81.006 |
| Tmem47 | 6556.33 | 0.167552 | 0.06792 | 2.46684 | 0.01363 | 0.07025 | 27.175 |
| Tmem50a | 3874.4 | -0.10953 | 0.0794 | -1.37946 | 0.16775 | 0.35765 | 17.176 |
| Tmem50b | 3653.32 | -0.1837 | 0.06758 | -2.71827 | 0.00656 | 0.04367 | 23.298 |
| Tmem51 | 134.052 | -0.03663 | 0.20526 | -0.17846 | 0.85836 | 0.92856 | 53.313 |
| Tmem52 | 43.8343 | -0.33768 | 0.30124 | -1.12099 | 0.26229 | 0.47114 | 1.745 |
| Tmem53 | 77.7069 | 0.274214 | 0.24676 | 1.111263 | 0.26646 | 0.47577 | 16.632 |
| Tmem59 | 5700.61 | -0.02253 | 0.07402 | -0.30433 | 0.76087 | 0.8747 | 22.598 |
| Tmem59l | 4949.1 | -0.20311 | 0.10064 | -2.01828 | 0.04356 | 0.151 | 3.492 |
| Tmem60 | 796.918 | -0.09016 | 0.07789 | -1.15746 | 0.24708 | 0.45391 | 4.68 |
| Tmem62 | 1381.77 | 0.082976 | 0.077 | 1.07756 | 0.28123 | 0.4911 | 30.836 |
| Tmem63a | 1027.06 | -0.4923 | 0.15556 | -3.16474 | 0.00155 | 0.01704 | 32.769 |
| Tmem63b | 4603.21 | -0.06453 | 0.07324 | -0.88096 | 0.37834 | 0.58821 | 26.735 |
| Tmem63c | 1962.41 | 0.100994 | 0.07856 | 1.285569 | 0.19859 | 0.39637 | 68.704 |
| Tmem64 | 2847.3 | -0.00138 | 0.09026 | -0.0153 | 0.9878 | 0.99316 | 20.923 |
| Tmem65 | 3663.76 | -0.21163 | 0.10836 | -1.95304 | 0.05081 | 0.1677 | 41.37 |
| Tmem67 | 794.02 | 0.104199 | 0.0952 | 1.094519 | 0.27373 | 0.48313 | 50.666 |
| Tmem68 | 2187.1 | 0.108018 | 0.09842 | 1.097532 | 0.27241 | 0.48159 | 25.813 |
| Tmem69 | 382.341 | -0.0163 | 0.11196 | -0.14556 | 0.88427 | 0.94244 | 4.304 |
| Tmem70 | 1753.03 | -0.0154 | 0.07249 | -0.21244 | 0.83176 | 0.91375 | 13.069 |
| Tmem71 | 16.7704 | -0.12901 | 0.48346 | -0.26684 | 0.78959 | 0.89082 | 34.892 |
| Tmem72 | 20.3981 | 0.009736 | 0.40814 | 0.023855 | 0.98097 | 0.99 | 37.851 |
| Tmem74 | 372.843 | -0.24574 | 0.20104 | -1.22234 | 0.22158 | 0.42449 | 5.391 |
| Tmem74b | 165.488 | 0.542061 | 0.14624 | 3.706643 | 0.00021 | 0.0044 | 10.15 |
| Tmem74bos | 77.9681 | 0.653278 | 0.32826 | 1.990154 | 0.04657 | 0.15802 | 6.979 |
| Tmem79 | 37.1364 | -0.49594 | 0.28434 | -1.74419 | 0.08113 | 0.22669 | 7.494 |
| Tmem80 | 1280.03 | 0.004549 | 0.08457 | 0.053795 | 0.9571 | 0.979 | 9.418 |
| Tmem81 | 305.978 | 0.190488 | 0.11512 | 1.654672 | 0.09799 | 0.25687 | 2.41 |
| Tmem82 | 66.2016 | 0.28964 | 0.29034 | 0.997586 | 0.31848 | 0.52909 | 5.258 |
| Tmem86a | 470.126 | -0.30594 | 0.12009 | -2.5477 | 0.01084 | 0.06059 | 4.177 |
| Tmem86b | 600.798 | 0.366148 | 0.13994 | 2.616408 | 0.00889 | 0.05306 | 3.383 |
| Tmem87a | 2011.13 | 0.115013 | 0.07134 | 1.612211 | 0.10692 | 0.2713 | 48.802 |

| | | | | | | | |
|---|---|---|---|---|---|---|---|
| Tmem87b | 2514.51 | 0.070449 | 0.0972 | 0.72477 | 0.46859 | 0.66745 | 36.144 |
| Tmem88 | 33.6938 | -0.37216 | 0.32621 | -1.14084 | 0.25394 | 0.4613 | 1.716 |
| Tmem88b | 2521.68 | -0.74324 | 0.17692 | -4.20097 | 2.66E-05 | 0.00107 | 4.284 |
| Tmem8b | 3941.6 | -0.09649 | 0.08589 | -1.12345 | 0.26124 | 0.46984 | 23.698 |
| Tmem9 | 2009.23 | -0.0737 | 0.0807 | -0.91323 | 0.36112 | 0.57139 | 27.193 |
| Tmem91 | 869.864 | 0.364666 | 0.15682 | 2.325407 | 0.02005 | 0.08969 | 6.028 |
| Tmem94 | 4585.74 | 0.083539 | 0.06112 | 1.366863 | 0.17167 | 0.36213 | 33.934 |
| Tmem97 | 359.359 | -0.28638 | 0.13614 | -2.10366 | 0.03541 | 0.13219 | 8.961 |
| Tmem98 | 326.87 | -0.65903 | 0.19009 | -3.46691 | 0.00053 | 0.00822 | 11.859 |
| Tmem9b | 3372.23 | -0.16938 | 0.07509 | -2.25574 | 0.02409 | 0.10128 | 17.042 |
| Tmf1 | 2151.31 | 0.091513 | 0.10548 | 0.867611 | 0.38561 | 0.59502 | 26.126 |
| Tmie | 219.512 | 0.157897 | 0.1774 | 0.890082 | 0.37342 | 0.58357 | 14.403 |
| Tmod1 | 2816 | 0.244463 | 0.11573 | 2.112446 | 0.03465 | 0.13033 | 77.098 |
| Tmod2 | 27395.3 | -0.06788 | 0.05785 | -1.17327 | 0.24069 | 0.44693 | 45.705 |
| Tmod3 | 2216.48 | 0.064822 | 0.07716 | 0.840053 | 0.40088 | 0.60962 | 61.951 |
| Tmod4 | 24.2872 | 0.040641 | 0.36489 | 0.111377 | 0.91132 | 0.95523 | 4.734 |
| Tmpo | 2224 | -0.18823 | 0.09011 | -2.0888 | 0.03673 | 0.13515 | 33.745 |
| Tmppe | 355.637 | -0.03992 | 0.14361 | -0.27797 | 0.78104 | 0.88605 | 6.172 |
| Tmprss11a | 16.9605 | 1.279121 | 0.52662 | 2.428937 | 0.01514 | 0.07492 | 58.581 |
| Tmprss4 | 35.1956 | 1.066729 | 0.34794 | 3.065817 | 0.00217 | 0.02127 | 31.367 |
| Tmprss5 | 52.7793 | -0.18521 | 0.29756 | -0.62243 | 0.53366 | 0.72083 | 36.328 |
| Tmprss6 | 77.3022 | 0.13482 | 0.23141 | 0.582605 | 0.56016 | 0.74154 | 28.968 |
| Tmprss7 | 45.4639 | 0.250746 | 0.27089 | 0.925623 | 0.35464 | 0.56503 | 37.344 |
| Tmprss9 | 190.835 | 1.028869 | 0.19778 | 5.202077 | 1.97E-07 | 3.01E-05 | 27.645 |
| Tmsb10 | 5994.72 | -0.13433 | 0.08919 | -1.50607 | 0.13205 | 0.31082 | 1.402 |
| Tmsb4x | 21668.6 | -0.23773 | 0.10152 | -2.34179 | 0.01919 | 0.08705 | 2.223 |
| Tmtc1 | 7015.86 | 0.328611 | 0.1345 | 2.443239 | 0.01456 | 0.07292 | 211.96 |
| Tmtc2 | 732.075 | 0.10661 | 0.106 | 1.00578 | 0.31452 | 0.52507 | 386.789 |
| Tmtc3 | 1308.14 | 0.272097 | 0.11147 | 2.441061 | 0.01464 | 0.07311 | 43.449 |
| Tmtc4 | 450.149 | -0.15238 | 0.12576 | -1.21166 | 0.22564 | 0.42969 | 65.065 |
| Tmub1 | 509.262 | -0.03129 | 0.09408 | -0.33258 | 0.73945 | 0.86264 | 2.397 |
| Tmub2 | 2348.89 | -0.06623 | 0.06139 | -1.07875 | 0.2807 | 0.49045 | 4.307 |
| Tmx1 | 2591.28 | 0.0991 | 0.06677 | 1.48413 | 0.13777 | 0.31863 | 14.946 |
| Tmx2 | 6284.23 | -0.07237 | 0.0706 | -1.02507 | 0.30533 | 0.51531 | 7.825 |

| | | | | | | | |
|---|---|---|---|---|---|---|---|
| Tmx3 | 4908.46 | 0.17867 | 0.08467 | 2.110228 | 0.03484 | 0.13081 | 33.114 |
| Tmx4 | 13977.3 | 0.048667 | 0.06185 | 0.786837 | 0.43138 | 0.63746 | 49.961 |
| Tnc | 91.6432 | -0.08565 | 0.23671 | -0.36184 | 0.71747 | 0.8491 | 87.231 |
| Tnfaip1 | 1939.4 | -0.10464 | 0.07485 | -1.39802 | 0.16211 | 0.35049 | 13.483 |
| Tnfaip2 | 87.036 | -0.75711 | 0.32733 | -2.31296 | 0.02072 | 0.09173 | 12.55 |
| Tnfaip3 | 316.639 | 0.115507 | 0.15086 | 0.765673 | 0.44387 | 0.6481 | 14.748 |
| Tnfaip6 | 383.527 | -0.6743 | 0.19964 | -3.37764 | 0.00073 | 0.01031 | 18.678 |
| Tnfaip8 | 311.072 | 0.212473 | 0.14817 | 1.433965 | 0.15158 | 0.33655 | 127.747 |
| Tnfaip8l1 | 179.486 | -0.2326 | 0.16751 | -1.38854 | 0.16497 | 0.35399 | 11.479 |
| Tnfaip8l2 | 72.9264 | -0.40698 | 0.2229 | -1.82585 | 0.06787 | 0.20204 | 2.84 |
| Tnfaip8l3 | 1232.76 | -0.42137 | 0.17305 | -2.43491 | 0.0149 | 0.07404 | 44.984 |
| Tnfrsf10b | 59.8736 | -0.19444 | 0.22874 | -0.85005 | 0.3953 | 0.6043 | 16.94 |
| Tnfrsf11a | 251.813 | 0.050814 | 0.16667 | 0.304873 | 0.76046 | 0.87444 | 67.267 |
| Tnfrsf11b | 71.1062 | -0.70579 | 0.37316 | -1.89141 | 0.05857 | 0.18403 | 27.866 |
| Tnfrsf12a | 147.167 | -0.19575 | 0.17231 | -1.13604 | 0.25594 | 0.4637 | 2.024 |
| Tnfrsf13b | 52.0395 | 0.023997 | 0.27832 | 0.086223 | 0.93129 | 0.96532 | 22.618 |
| Tnfrsf13c | 22.0036 | -0.21293 | 0.36711 | -0.58002 | 0.5619 | 0.74286 | 2.627 |
| Tnfrsf14 | 42.6186 | -0.26107 | 0.32171 | -0.81152 | 0.41707 | 0.62456 | 6.631 |
| Tnfrsf18 | 217.295 | 0.058577 | 0.15524 | 0.377341 | 0.70592 | 0.8415 | 2.732 |
| Tnfrsf19 | 2390.07 | -0.0253 | 0.09446 | -0.26786 | 0.7888 | 0.89036 | 82.616 |
| Tnfrsf1a | 386.758 | -0.676 | 0.1627 | -4.15479 | 3.26E-05 | 0.00122 | 13.123 |
| Tnfrsf1b | 202.896 | -0.45988 | 0.1876 | -2.45146 | 0.01423 | 0.07191 | 33.408 |
| Tnfrsf21 | 3502.03 | 0.027465 | 0.07886 | 0.34828 | 0.72763 | 0.85462 | 72.635 |
| Tnfrsf22 | 76.1074 | -0.21097 | 0.20983 | -1.00544 | 0.31469 | 0.52522 | 14.856 |
| Tnfrsf23 | 61.8294 | -0.59208 | 0.2829 | -2.09292 | 0.03636 | 0.13438 | 20.064 |
| Tnfrsf25 | 842.441 | -0.07031 | 0.14117 | -0.49806 | 0.61844 | 0.78312 | 4.186 |
| Tnfrsf4 | 89.4508 | 0.373372 | 0.20134 | 1.854473 | 0.06367 | 0.19397 | 2.77 |
| Tnfrsf8 | 16.7567 | -1.02116 | 0.49853 | -2.04832 | 0.04053 | 0.14386 | 48.028 |
| Tnfrsf9 | 44.732 | 0.262766 | 0.24753 | 1.06156 | 0.28844 | 0.49847 | 31.541 |
| Tnfsf10 | 87.3292 | -0.97135 | 0.32434 | -2.99485 | 0.00275 | 0.02485 | 25.4 |
| Tnfsf12 | 69.2464 | 0.158398 | 0.27019 | 0.586246 | 0.55771 | 0.73936 | 9.6 |
| Tnfsf13 | 34.9691 | -0.42957 | 0.3569 | -1.20361 | 0.22874 | 0.43277 | 2.939 |
| Tnfsf13b | 91.1588 | -0.04577 | 0.19697 | -0.23239 | 0.81623 | 0.9054 | 32.606 |
| Tnfsf13os | 29.6753 | -0.5134 | 0.31677 | -1.62074 | 0.10507 | 0.26817 | 17.251 |

| | | | | | | | |
|---|---|---|---|---|---|---|---|
| Tnfsf9 | 14.7501 | -0.05075 | 0.48525 | -0.10459 | 0.9167 | 0.95748 | 2.433 |
| Tnik | 14838.9 | 0.110315 | 0.07451 | 1.480462 | 0.13875 | 0.32001 | 412.645 |
| Tnip1 | 1144.98 | -0.0278 | 0.08814 | -0.31546 | 0.75241 | 0.86997 | 52.133 |
| Tnip2 | 984.849 | 0.047139 | 0.09417 | 0.500579 | 0.61667 | 0.78182 | 17.905 |
| Tnk2 | 12162.8 | 0.065946 | 0.08264 | 0.797951 | 0.4249 | 0.6314 | 39.62 |
| Tnk2os | 92.448 | 0.235578 | 0.19309 | 1.220034 | 0.22245 | 0.42568 | 2.975 |
| Tnks | 7339.2 | 0.12671 | 0.06026 | 2.102586 | 0.0355 | 0.13235 | 139.231 |
| Tnks1bp1 | 3075.34 | -0.40492 | 0.15152 | -2.67238 | 0.00753 | 0.04785 | 25.027 |
| Tnks2 | 9339.04 | 0.172234 | 0.05943 | 2.898136 | 0.00375 | 0.03016 | 59.246 |
| Tnnc1 | 42.9012 | 0.224662 | 0.32216 | 0.697362 | 0.48558 | 0.68176 | 3.418 |
| Tnni1 | 112.09 | -0.35126 | 0.21359 | -1.64451 | 0.10007 | 0.26028 | 31.556 |
| Tnni3 | 75.1453 | -0.55134 | 0.23259 | -2.3705 | 0.01776 | 0.08291 | 5.925 |
| Tnnt1 | 315.254 | -0.82244 | 0.25227 | -3.26014 | 0.00111 | 0.0137 | 11.813 |
| Tnnt2 | 224.26 | 1.026303 | 0.20348 | 5.043807 | 4.56E-07 | 5.85E-05 | 15.907 |
| Tnpo1 | 7007.95 | -0.07083 | 0.06533 | -1.08415 | 0.2783 | 0.48811 | 87.366 |
| Tnpo2 | 7375.9 | -0.15855 | 0.07522 | -2.10774 | 0.03505 | 0.13131 | 20.669 |
| Tnpo3 | 2888.57 | 0.104994 | 0.07935 | 1.323177 | 0.18578 | 0.38054 | 69.061 |
| Tnr | 11353.6 | -0.0638 | 0.07827 | -0.81514 | 0.41499 | 0.6229 | 407.961 |
| Tnrc18 | 8736.96 | 0.004064 | 0.0618 | 0.065764 | 0.94757 | 0.97364 | 93.002 |
| Tnrc6a | 7336.6 | 0.208509 | 0.08488 | 2.456631 | 0.01402 | 0.07132 | 71.412 |
| Tnrc6b | 9260.83 | 0.019638 | 0.06539 | 0.300326 | 0.76393 | 0.87645 | 229.773 |
| Tnrc6c | 7839.97 | 0.007482 | 0.06063 | 0.123395 | 0.90179 | 0.95072 | 109.151 |
| Tns1 | 3751.62 | -0.28565 | 0.07996 | -3.57257 | 0.00035 | 0.00627 | 214.219 |
| Tns2 | 1340.03 | -0.22982 | 0.10154 | -2.26337 | 0.02361 | 0.09986 | 15.989 |
| Tns3 | 5701.75 | -0.35909 | 0.0981 | -3.66048 | 0.00025 | 0.00497 | 233.03 |
| Tnxb | 296.402 | -0.7766 | 0.24923 | -3.11602 | 0.00183 | 0.01905 | 59.359 |
| Tob1 | 2089.58 | 0.017043 | 0.08193 | 0.208015 | 0.83522 | 0.91549 | 4.042 |
| Tob2 | 1632.54 | -0.14794 | 0.10076 | -1.46818 | 0.14205 | 0.32393 | 10.525 |
| Toe1 | 194.294 | 0.098136 | 0.14219 | 0.690163 | 0.49009 | 0.68505 | 13.331 |
| Togaram1 | 3243.74 | 0.240548 | 0.08095 | 2.971588 | 0.00296 | 0.02597 | 56.77 |
| Togaram2 | 696.4 | 0.591362 | 0.14248 | 4.150514 | 3.32E-05 | 0.00123 | 56.409 |
| Tollip | 7183.9 | -0.15212 | 0.06925 | -2.19664 | 0.02805 | 0.11271 | 43.695 |
| Tom1 | 561.281 | -0.21326 | 0.10141 | -2.10297 | 0.03547 | 0.13229 | 36.417 |
| Tom1l1 | 1120.81 | 0.127009 | 0.0719 | 1.766582 | 0.0773 | 0.21988 | 44.902 |

| | | | | | | | |
|---|---|---|---|---|---|---|---|
| Tom1l2 | 11151.7 | -0.11785 | 0.07034 | -1.6753 | 0.09388 | 0.24988 | 126.192 |
| Tomm20 | 19668.6 | 0.315231 | 0.11851 | 2.65985 | 0.00782 | 0.04893 | 15.178 |
| Tomm22 | 1871.17 | -0.04175 | 0.07488 | -0.55764 | 0.57709 | 0.75441 | 2.54 |
| Tomm34 | 2638.07 | 0.063075 | 0.0829 | 0.760831 | 0.44676 | 0.65034 | 17.63 |
| Tomm40 | 1724.7 | -0.05166 | 0.07929 | -0.65153 | 0.5147 | 0.70514 | 14.126 |
| Tomm40l | 934.83 | -0.00648 | 0.07974 | -0.0813 | 0.93521 | 0.96743 | 6.504 |
| Tomm5 | 852.193 | -0.19867 | 0.09165 | -2.16762 | 0.03019 | 0.11869 | 2.907 |
| Tomm6os | 21.3618 | -0.38681 | 0.41362 | -0.93519 | 0.34969 | 0.56025 | 3.474 |
| Tomm7 | 1498.49 | -0.14904 | 0.07689 | -1.93839 | 0.05258 | 0.17154 | 5.218 |
| Tomm70a | 16140.2 | 0.447745 | 0.12544 | 3.569329 | 0.00036 | 0.00633 | 35.003 |
| Tomt | 13.0001 | 0.143646 | 0.49267 | 0.291566 | 0.77062 | 0.88002 | 7.99 |
| Tonsl | 294.689 | -0.30683 | 0.19321 | -1.58808 | 0.11227 | 0.2799 | 13.957 |
| Top1 | 6229.03 | 0.264295 | 0.11271 | 2.344998 | 0.01903 | 0.0866 | 76.877 |
| Top1mt | 581.37 | -0.0618 | 0.10054 | -0.61464 | 0.53879 | 0.72544 | 21.766 |
| Top2a | 122.87 | -1.09037 | 0.70947 | -1.53688 | 0.12432 | 0.29916 | 31.247 |
| Top2b | 6294.44 | 0.158908 | 0.12781 | 1.243295 | 0.21376 | 0.41506 | 70.284 |
| Top3a | 344.937 | 0.189147 | 0.11666 | 1.621404 | 0.10493 | 0.26801 | 37.308 |
| Top3b | 1935.26 | 0.137093 | 0.07348 | 1.865664 | 0.06209 | 0.19092 | 22.255 |
| Topbp1 | 1082.32 | -0.10224 | 0.08745 | -1.16915 | 0.24234 | 0.44831 | 45.214 |
| Topors | 1184.28 | 0.134021 | 0.08923 | 1.502015 | 0.13309 | 0.3127 | 10.25 |
| Tor1a | 1051.24 | -0.07932 | 0.08199 | -0.96734 | 0.33337 | 0.54424 | 7.307 |
| Tor1aip1 | 1673.16 | -0.19351 | 0.08001 | -2.41857 | 0.01558 | 0.07637 | 31.882 |
| Tor1aip2 | 3013.87 | 0.018509 | 0.062 | 0.298557 | 0.76528 | 0.87738 | 33.459 |
| Tor1b | 2225.06 | 0.051284 | 0.07125 | 0.719793 | 0.47165 | 0.67032 | 6.057 |
| Tor2a | 653.144 | 0.045359 | 0.11655 | 0.389198 | 0.69713 | 0.8358 | 5.011 |
| Tor3a | 420.436 | -0.35264 | 0.14642 | -2.40851 | 0.01602 | 0.07776 | 20.74 |
| Tor4a | 132.275 | -0.91306 | 0.19606 | -4.65714 | 3.21E-06 | 0.00025 | 3.921 |
| Tox | 460.973 | -0.12277 | 0.11815 | -1.03918 | 0.29872 | 0.50876 | 305.205 |
| Tox2 | 921.844 | -0.07222 | 0.10828 | -0.66697 | 0.50479 | 0.6967 | 121.046 |
| Tox3 | 532.654 | -0.64846 | 0.16359 | -3.96393 | 7.37E-05 | 0.00217 | 101.304 |
| Tox4 | 3149.01 | 0.010277 | 0.06304 | 0.163019 | 0.8705 | 0.9346 | 17.256 |
| Tpbg | 1075.27 | -0.27648 | 0.13714 | -2.01607 | 0.04379 | 0.15143 | 4.661 |
| Tpbgl | 273.418 | 0.241296 | 0.14832 | 1.62683 | 0.10377 | 0.26624 | 3.022 |
| Tpcn1 | 2471.72 | -0.09045 | 0.0884 | -1.02316 | 0.30623 | 0.51631 | 54.521 |

| | | | | | | | |
|---|---|---|---|---|---|---|---|
| Tpcn2 | 408.927 | -0.20201 | 0.13513 | -1.49496 | 0.13493 | 0.31481 | 97.488 |
| Tpd52 | 2443.11 | 0.075689 | 0.0722 | 1.04839 | 0.29446 | 0.50482 | 79.131 |
| Tpd52l1 | 785.778 | -0.10592 | 0.12339 | -0.8584 | 0.39067 | 0.59979 | 113.583 |
| Tpd52l2 | 2208.6 | -0.0407 | 0.07389 | -0.5508 | 0.58177 | 0.75756 | 20.825 |
| Tpgs1 | 1195.67 | -0.1168 | 0.09392 | -1.24353 | 0.21367 | 0.41501 | 6.759 |
| Tpgs2 | 2143.25 | -0.11131 | 0.06243 | -1.78296 | 0.07459 | 0.21525 | 41.785 |
| Tph2 | 18.2754 | -0.93444 | 0.46633 | -2.00379 | 0.04509 | 0.15438 | 106.382 |
| Tpi-rs11 | 71.1284 | -0.12495 | 0.25906 | -0.48233 | 0.62957 | 0.79064 | 0.31 |
| Tpi1 | 18297.2 | -0.1968 | 0.06992 | -2.8147 | 0.00488 | 0.03568 | 3.711 |
| Tpk1 | 80.0291 | -0.29305 | 0.23339 | -1.25563 | 0.20925 | 0.4093 | 321.278 |
| Tpm1 | 8954.58 | 0.026753 | 0.07779 | 0.343905 | 0.73092 | 0.85734 | 26.817 |
| Tpm2 | 460.719 | -0.44085 | 0.10427 | -4.22784 | 2.36E-05 | 0.00098 | 9.055 |
| Tpm3 | 6250.82 | -0.05221 | 0.06742 | -0.77444 | 0.43867 | 0.64389 | 28.254 |
| Tpm3-rs7 | 194.849 | -0.07216 | 0.19487 | -0.37029 | 0.71117 | 0.8446 | 2.147 |
| Tpm4 | 1213.06 | -0.3522 | 0.09326 | -3.77654 | 0.00016 | 0.00369 | 22.969 |
| Tpmt | 335.927 | -0.21096 | 0.11765 | -1.79308 | 0.07296 | 0.21243 | 22.256 |
| Tpo | 42.3232 | 0.366267 | 0.37689 | 0.971811 | 0.33114 | 0.54208 | 77.966 |
| Tpp1 | 3871.07 | 0.035981 | 0.0649 | 0.554389 | 0.57931 | 0.75592 | 7.425 |
| Tpp2 | 3037.14 | 0.165023 | 0.08896 | 1.854969 | 0.0636 | 0.1939 | 69.354 |
| Tppp | 17235.6 | -0.20107 | 0.08185 | -2.45662 | 0.01402 | 0.07132 | 26.347 |
| Tppp3 | 1674.45 | -0.38703 | 0.0881 | -4.39297 | 1.12E-05 | 0.0006 | 4.034 |
| Tpr | 5544.63 | 0.380183 | 0.28365 | 1.340305 | 0.18015 | 0.37341 | 57.098 |
| Tpra1 | 1219.68 | 0.269959 | 0.08692 | 3.105832 | 0.0019 | 0.0194 | 9.988 |
| Tprgl | 5109.35 | -0.02028 | 0.08865 | -0.22873 | 0.81908 | 0.90696 | 3.182 |
| Tprkb | 3206.15 | -0.19804 | 0.08006 | -2.47352 | 0.01338 | 0.06943 | 18.42 |
| Tprn | 766.328 | -0.41853 | 0.12696 | -3.29655 | 0.00098 | 0.01263 | 7.268 |
| Tpst1 | 1510.25 | 0.230347 | 0.122 | 1.888014 | 0.05902 | 0.18496 | 62.404 |
| Tpst2 | 904.287 | -0.12644 | 0.08593 | -1.47133 | 0.1412 | 0.32297 | 38.671 |
| Tpt1 | 12747.3 | -0.23339 | 0.0744 | -3.13689 | 0.00171 | 0.01819 | 3.433 |
| Tpt1-ps3 | 568.211 | -0.32375 | 0.11791 | -2.74569 | 0.00604 | 0.04146 | 0.519 |
| Tpx2 | 123.655 | -0.40336 | 0.22096 | -1.8255 | 0.06793 | 0.20213 | 47.358 |
| Tra2a | 8920.59 | 0.228617 | 0.08148 | 2.805639 | 0.00502 | 0.0364 | 20.11 |
| Tra2b | 4487.75 | 0.160713 | 0.06616 | 2.429225 | 0.01513 | 0.0749 | 21.457 |
| Trabd | 845.562 | 0.065577 | 0.09894 | 0.662775 | 0.50747 | 0.69875 | 11.954 |

| | | | | | | | |
|---|---|---|---|---|---|---|---|
| Trabd2b | 80.9973 | -1.28944 | 0.34915 | -3.69311 | 0.00022 | 0.00457 | 208.375 |
| Tradd | 256.175 | -0.18306 | 0.13324 | -1.37395 | 0.16946 | 0.35966 | 6.324 |
| Traf1 | 38.4231 | -0.92324 | 0.32131 | -2.87333 | 0.00406 | 0.03181 | 20.023 |
| Traf2 | 520.179 | -0.13342 | 0.12151 | -1.09798 | 0.27221 | 0.48159 | 28.959 |
| Traf3 | 2921.93 | -0.05982 | 0.09583 | -0.62428 | 0.53244 | 0.71981 | 100.784 |
| Traf3ip1 | 1239.74 | 0.085801 | 0.08685 | 0.987901 | 0.3232 | 0.53421 | 34.661 |
| Traf3ip2 | 623.967 | 0.390413 | 0.09804 | 3.982223 | 6.83E-05 | 0.00206 | 42.374 |
| Traf3ip3 | 33.6146 | -0.07641 | 0.32586 | -0.2345 | 0.8146 | 0.90481 | 26.251 |
| Traf4 | 341.941 | -0.1935 | 0.14149 | -1.36763 | 0.17143 | 0.36181 | 7.091 |
| Traf5 | 80.9664 | -0.12297 | 0.21565 | -0.57021 | 0.56853 | 0.74835 | 47.703 |
| Traf6 | 1261.19 | 0.042159 | 0.08207 | 0.513668 | 0.60748 | 0.77558 | 23.241 |
| Traf7 | 2379.4 | 0.108879 | 0.07783 | 1.398858 | 0.16186 | 0.35013 | 19.377 |
| Trafd1 | 2896.91 | -0.17204 | 0.0862 | -1.99572 | 0.04596 | 0.15639 | 13.908 |
| Traip | 2843 | 0.856075 | 0.17664 | 4.846325 | 1.26E-06 | 0.00012 | 22.135 |
| Trak1 | 9712.63 | -0.17795 | 0.08673 | -2.05183 | 0.04019 | 0.14325 | 177.417 |
| Trak2 | 11438.4 | -0.08746 | 0.08421 | -1.03852 | 0.29903 | 0.50914 | 72.982 |
| Tram1 | 1131.29 | -0.23117 | 0.08717 | -2.65186 | 0.008 | 0.04963 | 25.213 |
| Tram1l1 | 1423.48 | 0.02994 | 0.0878 | 0.341003 | 0.7331 | 0.85857 | 3.889 |
| Tram2 | 91.5573 | -0.63296 | 0.27753 | -2.28068 | 0.02257 | 0.09701 | 82.931 |
| Trank1 | 42625.3 | 0.358278 | 0.12251 | 2.924549 | 0.00345 | 0.02844 | 84.037 |
| Trap1 | 2823.04 | -0.00499 | 0.07424 | -0.06724 | 0.94639 | 0.97302 | 37.857 |
| Trappc1 | 839.502 | -0.09895 | 0.10022 | -0.98738 | 0.32346 | 0.53448 | 1.814 |
| Trappc10 | 5883.91 | -0.0689 | 0.07222 | -0.95404 | 0.34006 | 0.55085 | 57.917 |
| Trappc11 | 4023.77 | 0.166118 | 0.06476 | 2.564989 | 0.01032 | 0.05865 | 43.356 |
| Trappc12 | 3364.37 | 0.167912 | 0.088 | 1.908101 | 0.05638 | 0.17956 | 59.845 |
| Trappc13 | 2916.07 | 0.080106 | 0.06056 | 1.322788 | 0.18591 | 0.38074 | 36.321 |
| Trappc14 | 2197.68 | 0.017872 | 0.07003 | 0.255192 | 0.79858 | 0.89601 | 4.378 |
| Trappc2 | 1588.95 | -0.11061 | 0.08301 | -1.33245 | 0.18271 | 0.37662 | 12.567 |
| Trappc2l | 1097.31 | 0.028903 | 0.09369 | 0.30849 | 0.75771 | 0.87241 | 5.021 |
| Trappc3 | 2565.26 | 0.017571 | 0.06705 | 0.262081 | 0.79326 | 0.89287 | 13.559 |
| Trappc4 | 1002.09 | -0.02268 | 0.10424 | -0.21761 | 0.82774 | 0.91159 | 3.903 |
| Trappc5 | 1422.57 | -0.05627 | 0.07326 | -0.76805 | 0.44246 | 0.64699 | 4.957 |
| Trappc6a | 260.331 | 0.22896 | 0.15016 | 1.524808 | 0.12731 | 0.30355 | 7.751 |
| Trappc6b | 3055.31 | -0.02772 | 0.07936 | -0.34926 | 0.72689 | 0.85431 | 18.381 |

| | | | | | | | |
|---|---|---|---|---|---|---|---|
| Trappc8 | 3543.87 | 0.047105 | 0.06129 | 0.768571 | 0.44215 | 0.64668 | 78.871 |
| Trappc9 | 2599.01 | 0.080083 | 0.08444 | 0.948343 | 0.34295 | 0.55368 | 471.585 |
| Trarg1 | 16.2477 | -0.65774 | 0.55216 | -1.19122 | 0.23357 | 0.43857 | 18.857 |
| Trat1 | 54.6243 | 0.420756 | 0.41763 | 1.007475 | 0.31371 | 0.52419 | 41.248 |
| Trdmt1 | 326.282 | -0.03846 | 0.11031 | -0.34866 | 0.72734 | 0.85443 | 35.655 |
| Trdn | 27.1054 | 0.132741 | 0.66297 | 0.200222 | 0.84131 | 0.91864 | 396.156 |
| Trem2 | 413.46 | -0.40211 | 0.20311 | -1.97979 | 0.04773 | 0.16067 | 7.747 |
| Treml2 | 20.5168 | -0.82806 | 0.50145 | -1.65132 | 0.09867 | 0.25803 | 13.036 |
| Trerf1 | 5016.01 | 0.426769 | 0.12133 | 3.517443 | 0.00044 | 0.00729 | 221.083 |
| Trf | 8859.38 | -0.87397 | 0.13944 | -6.26775 | 3.66E-10 | 2.49E-07 | 26.444 |
| Trh | 145.624 | -0.1598 | 0.222 | -0.71984 | 0.47162 | 0.67032 | 2.59 |
| Trhde | 604.128 | -0.29033 | 0.16264 | -1.78511 | 0.07424 | 0.21485 | 403.487 |
| Trhr | 363.181 | 1.208481 | 0.25834 | 4.677807 | 2.90E-06 | 0.00023 | 39.778 |
| Trhr2 | 286.821 | 0.093272 | 0.14519 | 0.642431 | 0.52059 | 0.70982 | 3.78 |
| Triap1 | 520.559 | -0.18113 | 0.09989 | -1.81328 | 0.06979 | 0.20614 | 2.345 |
| Trib1 | 755.65 | -0.06444 | 0.12659 | -0.50901 | 0.61074 | 0.77794 | 8.75 |
| Trib2 | 1686.01 | 0.068038 | 0.08445 | 0.805654 | 0.42044 | 0.62768 | 25.196 |
| Trib3 | 93.7048 | 0.573545 | 0.3001 | 1.911188 | 0.05598 | 0.1788 | 6.611 |
| Tril | 1648.73 | -0.09447 | 0.12969 | -0.72842 | 0.46635 | 0.66565 | 5.363 |
| Trim11 | 1647.21 | 0.196515 | 0.09685 | 2.029023 | 0.04246 | 0.14834 | 13.366 |
| Trim12a | 192.987 | -0.52512 | 0.21401 | -2.45375 | 0.01414 | 0.07164 | 15.573 |
| Trim12c | 148.216 | -0.209 | 0.19786 | -1.05634 | 0.29081 | 0.50105 | 14.609 |
| Trim13 | 339.216 | -0.01883 | 0.16195 | -0.11627 | 0.90744 | 0.95354 | 7.7 |
| Trim14 | 34.5902 | -0.30439 | 0.31735 | -0.95913 | 0.33749 | 0.54829 | 42.361 |
| Trim15 | 32.2877 | 0.493529 | 0.30439 | 1.621381 | 0.10494 | 0.26801 | 6.52 |
| Trim16 | 94.8697 | -0.11374 | 0.21936 | -0.51849 | 0.60412 | 0.7733 | 30.578 |
| Trim17 | 368.486 | -0.06724 | 0.1723 | -0.39025 | 0.69635 | 0.83541 | 18.414 |
| Trim2 | 17791.8 | -0.07386 | 0.07198 | -1.02604 | 0.30487 | 0.5147 | 146.439 |
| Trim21 | 137.376 | -0.34711 | 0.21675 | -1.60148 | 0.10927 | 0.27509 | 7.566 |
| Trim23 | 3885.49 | 0.232919 | 0.08649 | 2.692951 | 0.00708 | 0.04582 | 24.576 |
| Trim24 | 1521.22 | 0.154425 | 0.07524 | 2.052355 | 0.04014 | 0.14315 | 95.486 |
| Trim25 | 568.014 | -0.38951 | 0.14154 | -2.75189 | 0.00593 | 0.04089 | 20.918 |
| Trim26 | 1254.61 | -0.18962 | 0.10841 | -1.74914 | 0.08027 | 0.22543 | 22.265 |
| Trim27 | 936.084 | 0.044007 | 0.08392 | 0.524405 | 0.6 | 0.77061 | 15.28 |

| | | | | | | | |
|---|---|---|---|---|---|---|---|
| Trim28 | 4890.32 | 0.096729 | 0.07352 | 1.315685 | 0.18828 | 0.38384 | 31.922 |
| Trim3 | 4400.7 | -0.17106 | 0.06808 | -2.51269 | 0.01198 | 0.06461 | 29.109 |
| Trim30a | 155.276 | -0.55177 | 0.23887 | -2.30995 | 0.02089 | 0.09217 | 56.169 |
| Trim30d | 94.5283 | -0.56711 | 0.27218 | -2.08356 | 0.0372 | 0.13629 | 37.836 |
| Trim32 | 7230.92 | -0.03387 | 0.07717 | -0.43888 | 0.66075 | 0.81233 | 11.253 |
| Trim33 | 4035.71 | 0.097582 | 0.09552 | 1.021531 | 0.307 | 0.5168 | 79.483 |
| Trim34a | 61.1701 | -0.78717 | 0.28613 | -2.75109 | 0.00594 | 0.04095 | 17.778 |
| Trim35 | 9075.29 | 0.112928 | 0.07531 | 1.499516 | 0.13374 | 0.31339 | 14.394 |
| Trim36 | 267.858 | -0.93938 | 0.23036 | -4.07785 | 4.55E-05 | 0.00153 | 47.308 |
| Trim37 | 14916.9 | -0.02336 | 0.05983 | -0.39037 | 0.69626 | 0.83541 | 93.607 |
| Trim39 | 2472.3 | 0.060864 | 0.10939 | 0.556414 | 0.57793 | 0.75477 | 13.375 |
| Trim41 | 3316.35 | 0.007339 | 0.05734 | 0.127993 | 0.89815 | 0.94885 | 10.95 |
| Trim43a | 114.086 | 0.292598 | 0.19437 | 1.505373 | 0.13223 | 0.31117 | 7.929 |
| Trim43c | 15.8945 | -0.22246 | 0.48215 | -0.4614 | 0.64451 | 0.80172 | 9.027 |
| Trim44 | 17696.2 | -0.02941 | 0.06187 | -0.47539 | 0.63451 | 0.79427 | 107.71 |
| Trim45 | 501.032 | 0.121824 | 0.15125 | 0.805443 | 0.42056 | 0.62777 | 14.719 |
| Trim46 | 5111.83 | 0.36087 | 0.08612 | 4.190455 | 2.78E-05 | 0.0011 | 12.133 |
| Trim47 | 137.57 | -0.5032 | 0.17745 | -2.83571 | 0.00457 | 0.0343 | 21.459 |
| Trim5 | 36.5575 | -0.44012 | 0.27827 | -1.58161 | 0.11374 | 0.28233 | 24.709 |
| Trim56 | 747.479 | -0.47071 | 0.12048 | -3.90689 | 9.35E-05 | 0.00256 | 10.566 |
| Trim59 | 710.126 | -0.57674 | 0.19423 | -2.96943 | 0.00298 | 0.02607 | 9.468 |
| Trim62 | 1628.09 | 0.33754 | 0.08645 | 3.904608 | 9.44E-05 | 0.00257 | 27.749 |
| Trim63 | 15.7966 | -1.09798 | 0.54615 | -2.0104 | 0.04439 | 0.15289 | 14.51 |
| Trim65 | 340.677 | -0.21706 | 0.13587 | -1.5975 | 0.11015 | 0.27663 | 9.283 |
| Trim66 | 1946.07 | -0.1129 | 0.11036 | -1.02301 | 0.3063 | 0.51635 | 59.129 |
| Trim67 | 240.329 | -0.30405 | 0.19826 | -1.53354 | 0.12514 | 0.30014 | 41.622 |
| Trim68 | 304.868 | 0.106784 | 0.15165 | 0.704128 | 0.48135 | 0.67838 | 9.746 |
| Trim7 | 117.912 | -0.36745 | 0.20806 | -1.7661 | 0.07738 | 0.22005 | 26.07 |
| Trim71 | 83.8189 | -0.59466 | 0.24749 | -2.40274 | 0.01627 | 0.07857 | 57.237 |
| Trim72 | 40.6233 | 0.696105 | 0.29491 | 2.36042 | 0.01825 | 0.08442 | 7.085 |
| Trim8 | 2716.52 | -0.11872 | 0.06754 | -1.7579 | 0.07876 | 0.2225 | 15.147 |
| Trim9 | 14899.5 | -0.12708 | 0.08827 | -1.43959 | 0.14998 | 0.33447 | 103.082 |
| Trio | 14536.8 | 0.018869 | 0.06797 | 0.277604 | 0.78132 | 0.88614 | 295.198 |
| Triobp | 367.283 | -0.43779 | 0.12546 | -3.48939 | 0.00048 | 0.00775 | 58.146 |

| | | | | | | | |
|---|---|---|---|---|---|---|---|
| Trip10 | 282.278 | -0.6035 | 0.18636 | -3.23831 | 0.0012 | 0.01435 | 14.211 |
| Trip11 | 3234.25 | 0.108559 | 0.15991 | 0.678856 | 0.49723 | 0.69076 | 79.225 |
| Trip12 | 11567.6 | 0.1203 | 0.06322 | 1.90284 | 0.05706 | 0.18096 | 119.328 |
| Trip13 | 13.1596 | -0.53547 | 0.52777 | -1.0146 | 0.3103 | 0.52042 | 26.438 |
| Trip4 | 2684.34 | 0.229237 | 0.07575 | 3.026235 | 0.00248 | 0.02322 | 79.865 |
| Trip6 | 419.094 | 0.05677 | 0.12934 | 0.438902 | 0.66073 | 0.81233 | 4.751 |
| Triqk | 951.004 | 0.373024 | 0.09697 | 3.846687 | 0.00012 | 0.00303 | 76.716 |
| Trir | 1945.28 | -0.18338 | 0.07425 | -2.46966 | 0.01352 | 0.06999 | 3.464 |
| Trit1 | 857.629 | 0.162665 | 0.08286 | 1.963091 | 0.04964 | 0.16511 | 38.353 |
| Trmo | 143.83 | 0.167552 | 0.16773 | 0.99895 | 0.31782 | 0.52829 | 12.933 |
| Trmt1 | 2885.16 | 0.034811 | 0.07627 | 0.456427 | 0.64808 | 0.80406 | 13.502 |
| Trmt10a | 312.663 | 0.137929 | 0.13939 | 0.989507 | 0.32242 | 0.53337 | 16.374 |
| Trmt10b | 793.199 | 0.248117 | 0.08969 | 2.766354 | 0.00567 | 0.03965 | 19.005 |
| Trmt10c | 575.535 | 0.089606 | 0.12092 | 0.741049 | 0.45866 | 0.65962 | 5.211 |
| Trmt11 | 326.01 | 0.013165 | 0.11218 | 0.117364 | 0.90657 | 0.95319 | 66.525 |
| Trmt112 | 418.545 | 0.031152 | 0.10384 | 0.300011 | 0.76417 | 0.87653 | 1.352 |
| Trmt112-ps2 | 116.27 | 0.030922 | 0.19106 | 0.161845 | 0.87143 | 0.93501 | 0.378 |
| Trmt12 | 392.644 | -0.11997 | 0.10948 | -1.09581 | 0.27316 | 0.48254 | 4.454 |
| Trmt13 | 401.954 | -0.02446 | 0.12812 | -0.19093 | 0.84858 | 0.92307 | 33.495 |
| Trmt1l | 1794.31 | 0.200237 | 0.08497 | 2.356589 | 0.01844 | 0.08512 | 29.62 |
| Trmt2a | 1657.91 | 0.091505 | 0.08501 | 1.076415 | 0.28174 | 0.49178 | 6.094 |
| Trmt2b | 726.298 | -0.26256 | 0.12695 | -2.06823 | 0.03862 | 0.13992 | 54.646 |
| Trmt44 | 423.086 | 0.190538 | 0.12249 | 1.555477 | 0.11983 | 0.29236 | 18.868 |
| Trmt5 | 398.211 | -0.01202 | 0.15171 | -0.07923 | 0.93685 | 0.9685 | 6.7 |
| Trmt6 | 1594.8 | 0.117422 | 0.08678 | 1.353153 | 0.17601 | 0.36793 | 11.849 |
| Trmt61a | 752.131 | 0.02584 | 0.09818 | 0.263196 | 0.7924 | 0.89229 | 5.798 |
| Trmt61b | 367.098 | 0.142095 | 0.09999 | 1.421146 | 0.15527 | 0.34177 | 41.772 |
| Trmt9b | 760.752 | -0.33411 | 0.1231 | -2.71407 | 0.00665 | 0.04397 | 59.023 |
| Trmu | 992.666 | 0.118076 | 0.07822 | 1.509563 | 0.13115 | 0.30963 | 18.083 |
| Trnau1ap | 636.182 | 0.048962 | 0.12543 | 0.390347 | 0.69628 | 0.83541 | 17.776 |
| Trnp1 | 5630.95 | -0.20896 | 0.07764 | -2.69161 | 0.00711 | 0.04596 | 7.451 |
| Trnt1 | 2106.89 | 0.110837 | 0.10787 | 1.027545 | 0.30416 | 0.51405 | 13.355 |
| Tro | 12740.1 | 0.408888 | 0.08826 | 4.6329 | 3.61E-06 | 0.00027 | 12.28 |
| Trp53 | 1008.41 | 0.125273 | 0.08773 | 1.427875 | 0.15333 | 0.33898 | 11.515 |

| | | | | | | | |
|---|---|---|---|---|---|---|---|
| Trp53bp1 | 6509.93 | 0.222822 | 0.06812 | 3.271045 | 0.00107 | 0.01341 | 78.127 |
| Trp53bp2 | 2220.8 | -0.1615 | 0.07278 | -2.21917 | 0.02647 | 0.10826 | 53.261 |
| Trp53cor1 | 19.0208 | -0.08177 | 0.40685 | -0.20098 | 0.84072 | 0.91835 | 21.75 |
| Trp53i11 | 2684.62 | -0.18786 | 0.11767 | -1.5965 | 0.11038 | 0.27695 | 14.212 |
| Trp53i13 | 269.544 | -0.26502 | 0.12555 | -2.11082 | 0.03479 | 0.13066 | 7.882 |
| Trp53inp1 | 1778.3 | -0.11447 | 0.11903 | -0.9617 | 0.3362 | 0.54701 | 17.949 |
| Trp53inp2 | 8052.91 | -0.53754 | 0.09224 | -5.82735 | 5.63E-09 | 1.98E-06 | 8.792 |
| Trp53rka | 605.969 | -0.12236 | 0.11092 | -1.10313 | 0.26997 | 0.47957 | 3.215 |
| Trp53rkb | 217.142 | -0.08441 | 0.13456 | -0.62734 | 0.53043 | 0.718 | 6.975 |
| Trp53tg5 | 62.8822 | 0.469305 | 0.25604 | 1.832923 | 0.06681 | 0.20016 | 3.424 |
| Trp63 | 37.8165 | -0.97565 | 0.39849 | -2.44835 | 0.01435 | 0.07227 | 208.34 |
| Trp73 | 59.5252 | 0.033599 | 0.32323 | 0.103945 | 0.91721 | 0.95781 | 83.96 |
| Trpc1 | 2842.54 | 0.370099 | 0.09985 | 3.706563 | 0.00021 | 0.0044 | 45.294 |
| Trpc3 | 826.383 | 0.454121 | 0.17758 | 2.557311 | 0.01055 | 0.0596 | 69.686 |
| Trpc4 | 699.581 | -0.41748 | 0.2352 | -1.775 | 0.0759 | 0.21756 | 162.437 |
| Trpc4ap | 6623.26 | 0.008329 | 0.06668 | 0.124918 | 0.90059 | 0.95009 | 58.114 |
| Trpc5 | 312.738 | 0.198386 | 0.16636 | 1.192548 | 0.23305 | 0.43798 | 306.51 |
| Trpc6 | 124.179 | -0.07812 | 0.21217 | -0.36818 | 0.71274 | 0.84523 | 136.6 |
| Trpc7 | 575.163 | 0.654865 | 0.12042 | 5.438055 | 5.39E-08 | 1.17E-05 | 122.881 |
| Trpm1 | 16.0355 | 0.219162 | 0.55047 | 0.398139 | 0.69053 | 0.83185 | 115.941 |
| Trpm2 | 2711.38 | -0.02248 | 0.07935 | -0.2833 | 0.77695 | 0.88393 | 62.842 |
| Trpm3 | 3711.51 | -0.20166 | 0.10777 | -1.87111 | 0.06133 | 0.18933 | 856.365 |
| Trpm4 | 511.785 | -0.30972 | 0.11585 | -2.67334 | 0.00751 | 0.04775 | 31.149 |
| Trpm6 | 40.1679 | -0.21592 | 0.28817 | -0.74928 | 0.45369 | 0.65598 | 142.529 |
| Trpm7 | 3672.73 | 0.163188 | 0.10749 | 1.518235 | 0.12896 | 0.30621 | 84.666 |
| Trps1 | 2602.01 | 0.020648 | 0.11426 | 0.180707 | 0.8566 | 0.92769 | 235.712 |
| Trpt1 | 276.815 | -0.11703 | 0.12907 | -0.90675 | 0.36454 | 0.57518 | 2.916 |
| Trpv2 | 504.6 | -0.06759 | 0.12846 | -0.52611 | 0.59881 | 0.76992 | 26.03 |
| Trpv3 | 109.993 | 0.101933 | 0.18287 | 0.557392 | 0.57726 | 0.75445 | 32.976 |
| Trpv4 | 144.566 | -0.27656 | 0.20205 | -1.36875 | 0.17108 | 0.36145 | 36.27 |
| Trpv6 | 112.35 | 0.227052 | 0.1909 | 1.189352 | 0.2343 | 0.43949 | 15.782 |
| Trrap | 13545.2 | 0.249388 | 0.07441 | 3.351518 | 0.0008 | 0.01101 | 92.047 |
| Trub1 | 1614.77 | 0.147864 | 0.08363 | 1.768095 | 0.07704 | 0.21943 | 38.094 |
| Trub2 | 1128.12 | -0.11635 | 0.09203 | -1.26435 | 0.2061 | 0.40534 | 27.279 |

| | | | | | | | |
|---|---|---|---|---|---|---|---|
| Tsacc | 37.2962 | 0.111486 | 0.28514 | 0.390991 | 0.6958 | 0.83522 | 14.243 |
| Tsc1 | 6222.69 | 0.326292 | 0.0594 | 5.493207 | 3.95E-08 | 9.54E-06 | 49.94 |
| Tsc2 | 6659.48 | 0.084169 | 0.06118 | 1.375831 | 0.16887 | 0.35902 | 36.815 |
| Tsc22d1 | 25783.6 | 0.28037 | 0.12436 | 2.254472 | 0.02417 | 0.10151 | 92.805 |
| Tsc22d2 | 2156.31 | 0.021 | 0.11103 | 0.189144 | 0.84998 | 0.92371 | 52.069 |
| Tsc22d3 | 6998.8 | -0.1999 | 0.13359 | -1.49636 | 0.13456 | 0.31443 | 61.132 |
| Tsc22d4 | 3163.31 | -0.42161 | 0.0775 | -5.44048 | 5.31E-08 | 1.16E-05 | 22.721 |
| Tsen15 | 911.358 | 0.085191 | 0.07804 | 1.091631 | 0.275 | 0.48452 | 15.954 |
| Tsen2 | 846.502 | -0.0178 | 0.08601 | -0.20696 | 0.83604 | 0.9159 | 33.965 |
| Tsen34 | 1989.06 | -0.05028 | 0.09696 | -0.51856 | 0.60407 | 0.7733 | 8.162 |
| Tsen54 | 390.213 | -0.01467 | 0.12868 | -0.11402 | 0.90922 | 0.95455 | 8.371 |
| Tsfm | 606.95 | 0.028725 | 0.0918 | 0.312922 | 0.75434 | 0.87077 | 19.269 |
| Tsg101 | 2403.87 | 0.01262 | 0.07474 | 0.168844 | 0.86592 | 0.93209 | 31.021 |
| Tsga10 | 1361.57 | 0.023208 | 0.12674 | 0.183113 | 0.85471 | 0.92658 | 111.654 |
| Tshb | 50.1878 | -0.13469 | 0.3042 | -0.44279 | 0.65792 | 0.81018 | 7.254 |
| Tshr | 78.9167 | 0.423924 | 0.1908 | 2.221805 | 0.0263 | 0.10774 | 165.246 |
| Tshz1 | 3679.8 | 0.241773 | 0.12815 | 1.886567 | 0.05922 | 0.18523 | 76.08 |
| Tshz2 | 1306.85 | 0.377286 | 0.19376 | 1.947173 | 0.05151 | 0.16924 | 438.804 |
| Tshz3 | 345.978 | -0.27634 | 0.10708 | -2.58074 | 0.00986 | 0.05694 | 75.436 |
| Tsks | 44.1965 | 0.705943 | 0.33362 | 2.115985 | 0.03435 | 0.12953 | 14.832 |
| Tsku | 73.8914 | -0.69692 | 0.26942 | -2.58672 | 0.00969 | 0.05617 | 10.661 |
| Tsn | 4326.45 | -0.0175 | 0.07066 | -0.24771 | 0.80436 | 0.89895 | 16.962 |
| Tsnax | 15078.2 | 0.229183 | 0.14943 | 1.53369 | 0.12511 | 0.30014 | 21.196 |
| Tsnaxip1 | 51.0674 | 0.956391 | 0.29934 | 3.194966 | 0.0014 | 0.01578 | 16.95 |
| Tspan1 | 12.7485 | 0.975356 | 0.71388 | 1.366278 | 0.17185 | 0.36241 | 5.733 |
| Tspan11 | 59.904 | -1.12498 | 0.51733 | -2.1746 | 0.02966 | 0.11722 | 66.389 |
| Tspan12 | 1154.6 | -0.16594 | 0.10171 | -1.63152 | 0.10278 | 0.2647 | 81.121 |
| Tspan13 | 7902.64 | -0.24784 | 0.10411 | -2.38052 | 0.01729 | 0.08154 | 27.944 |
| Tspan14 | 1050.15 | -0.46903 | 0.13744 | -3.41268 | 0.00064 | 0.00939 | 60.363 |
| Tspan15 | 647.806 | -0.41626 | 0.12204 | -3.41072 | 0.00065 | 0.00943 | 45.856 |
| Tspan17 | 637.798 | -0.55967 | 0.18796 | -2.97762 | 0.0029 | 0.0257 | 7.4 |
| Tspan18 | 85.3935 | -0.46448 | 0.26244 | -1.76985 | 0.07675 | 0.21911 | 132.746 |
| Tspan2 | 6296.9 | -0.34195 | 0.12648 | -2.70363 | 0.00686 | 0.04485 | 66.985 |
| Tspan3 | 10630.3 | -0.09766 | 0.07316 | -1.335 | 0.18188 | 0.37561 | 33.753 |

| | | | | | | | |
|---|---|---|---|---|---|---|---|
| Tspan31 | 1405.17 | -0.12343 | 0.08342 | -1.47967 | 0.13896 | 0.3202 | 2.985 |
| Tspan32 | 17.3718 | -0.56904 | 0.42251 | -1.3468 | 0.17804 | 0.3703 | 14.599 |
| Tspan32os | 27.7817 | -0.42998 | 0.34943 | -1.23051 | 0.21851 | 0.42062 | 19.065 |
| Tspan33 | 852.368 | -0.40005 | 0.15893 | -2.51709 | 0.01183 | 0.06401 | 24.338 |
| Tspan4 | 621.38 | -0.09471 | 0.11981 | -0.79053 | 0.42922 | 0.6358 | 18.188 |
| Tspan5 | 8399.84 | -0.32989 | 0.08299 | -3.97515 | 7.03E-05 | 0.00211 | 165.711 |
| Tspan6 | 450.059 | -0.13126 | 0.12586 | -1.04287 | 0.29701 | 0.50722 | 7.362 |
| Tspan7 | 42815.5 | -0.17504 | 0.08223 | -2.12864 | 0.03328 | 0.12667 | 111.448 |
| Tspan8 | 15.9427 | -1.7156 | 0.60835 | -2.82008 | 0.0048 | 0.03534 | 33.062 |
| Tspan9 | 1550.62 | -0.0369 | 0.10301 | -0.35824 | 0.72016 | 0.85067 | 182.199 |
| Tspear | 59.9736 | -0.01077 | 0.25551 | -0.04217 | 0.96637 | 0.9832 | 200.453 |
| Tspo | 77.4341 | -0.90046 | 0.26555 | -3.391 | 0.0007 | 0.00996 | 10.612 |
| Tspoap1 | 20613.3 | 0.288758 | 0.09139 | 3.159652 | 0.00158 | 0.01724 | 25.388 |
| Tspyl1 | 9756.47 | -0.01231 | 0.072 | -0.17103 | 0.8642 | 0.93125 | 3.086 |
| Tspyl2 | 6259.69 | 0.258616 | 0.12546 | 2.061303 | 0.03927 | 0.14122 | 5.574 |
| Tspyl3 | 1498.72 | 0.175368 | 0.09303 | 1.885028 | 0.05943 | 0.18548 | 3.072 |
| Tspyl4 | 14911 | -0.05175 | 0.0614 | -0.84294 | 0.39926 | 0.60812 | 3.879 |
| Tspyl5 | 2019.93 | 0.042771 | 0.07935 | 0.538988 | 0.5899 | 0.76292 | 4.01 |
| Tsr1 | 1560.54 | 0.084156 | 0.06533 | 1.288112 | 0.19771 | 0.39553 | 11.272 |
| Tsr2 | 2304.46 | -0.0632 | 0.082 | -0.77077 | 0.44084 | 0.64565 | 9.45 |
| Tsr3 | 796.708 | 0.151914 | 0.08523 | 1.782462 | 0.07467 | 0.21534 | 2.629 |
| Tssc4 | 844.274 | -0.0532 | 0.08102 | -0.65666 | 0.5114 | 0.70252 | 1.845 |
| Tssk1 | 16.9325 | -0.18758 | 0.49876 | -0.37609 | 0.70685 | 0.84193 | 3.7 |
| Tssk4 | 25.3896 | -0.14007 | 0.37322 | -0.37528 | 0.70745 | 0.84212 | 2.356 |
| Tssk6 | 32.3714 | -0.19268 | 0.31899 | -0.60403 | 0.54583 | 0.73059 | 1.335 |
| Tst | 678.906 | -0.24297 | 0.1781 | -1.36424 | 0.17249 | 0.36342 | 6.352 |
| Tstd1 | 40.7347 | 0.097543 | 0.30326 | 0.321647 | 0.74772 | 0.86772 | 1.481 |
| Tstd2 | 536.785 | 0.096635 | 0.09942 | 0.971946 | 0.33108 | 0.54208 | 23.949 |
| Tstd3 | 403.088 | -0.11392 | 0.14382 | -0.79214 | 0.42828 | 0.63491 | 9.831 |
| Ttbk1 | 9948.55 | -0.11986 | 0.07469 | -1.60485 | 0.10853 | 0.27381 | 45.228 |
| Ttbk2 | 5573.85 | 0.045406 | 0.07942 | 0.571743 | 0.5675 | 0.7475 | 117.789 |
| Ttc1 | 1924.72 | -0.04665 | 0.06191 | -0.7535 | 0.45115 | 0.65374 | 18.497 |
| Ttc12 | 176.441 | 0.263996 | 0.19603 | 1.346736 | 0.17807 | 0.3703 | 49.263 |
| Ttc13 | 2121.72 | 0.196384 | 0.06529 | 3.007812 | 0.00263 | 0.02415 | 50.652 |

| | | | | | | | |
|---|---|---|---|---|---|---|---|
| Ttc14 | 10382.1 | 0.296956 | 0.16308 | 1.820942 | 0.06862 | 0.20362 | 15.029 |
| Ttc16 | 44.2049 | -0.20303 | 0.3032 | -0.66962 | 0.5031 | 0.69545 | 18.608 |
| Ttc17 | 1996.2 | 0.155771 | 0.10554 | 1.475928 | 0.13996 | 0.32121 | 105.923 |
| Ttc19 | 4770.15 | 0.007406 | 0.07295 | 0.101529 | 0.91913 | 0.95913 | 46.979 |
| Ttc21a | 58.9534 | 0.526653 | 0.25972 | 2.02778 | 0.04258 | 0.14866 | 30.188 |
| Ttc21b | 942.761 | 0.166403 | 0.07869 | 2.114731 | 0.03445 | 0.12984 | 72.291 |
| Ttc22 | 83.55 | 0.096199 | 0.19772 | 0.486538 | 0.62659 | 0.78871 | 17.758 |
| Ttc23 | 155.982 | -0.25264 | 0.1801 | -1.40279 | 0.16068 | 0.34889 | 115.503 |
| Ttc26 | 642.213 | 0.104844 | 0.10162 | 1.031742 | 0.30219 | 0.51192 | 46.179 |
| Ttc27 | 740.572 | -0.02401 | 0.10388 | -0.2311 | 0.81724 | 0.90594 | 145.839 |
| Ttc28 | 723.428 | -0.43553 | 0.12646 | -3.44393 | 0.00057 | 0.00871 | 409.978 |
| Ttc3 | 41944.9 | 0.132954 | 0.07499 | 1.773035 | 0.07622 | 0.2181 | 98.726 |
| Ttc30a1 | 200.927 | -0.04729 | 0.15089 | -0.31337 | 0.754 | 0.87067 | 3.721 |
| Ttc30b | 546.262 | 0.081776 | 0.1056 | 0.7744 | 0.43869 | 0.64389 | 2.737 |
| Ttc32 | 183.259 | 0.102875 | 0.18936 | 0.543273 | 0.58694 | 0.76058 | 11.992 |
| Ttc33 | 1454.42 | -0.01994 | 0.0625 | -0.31899 | 0.74974 | 0.86868 | 32.812 |
| Ttc34 | 148.874 | 0.369342 | 0.2086 | 1.770602 | 0.07663 | 0.21888 | 29.669 |
| Ttc37 | 1490.07 | 0.198472 | 0.09399 | 2.111738 | 0.03471 | 0.13049 | 91.583 |
| Ttc38 | 475.653 | -0.19359 | 0.12229 | -1.58295 | 0.11343 | 0.28194 | 26.517 |
| Ttc39a | 146.576 | 0.099594 | 0.1887 | 0.527791 | 0.59764 | 0.76924 | 38.123 |
| Ttc39aos1 | 195.382 | 0.199545 | 0.22783 | 0.875833 | 0.38112 | 0.59057 | 3.955 |
| Ttc39b | 4323.51 | -0.20154 | 0.08182 | -2.46331 | 0.01377 | 0.07057 | 103.956 |
| Ttc39c | 2406.89 | 0.330934 | 0.09933 | 3.331676 | 0.00086 | 0.01155 | 138.968 |
| Ttc4 | 2362.05 | 0.101903 | 0.11381 | 0.895344 | 0.3706 | 0.58084 | 16.689 |
| Ttc41 | 122.386 | 0.133393 | 0.19753 | 0.675305 | 0.49948 | 0.69266 | 71.034 |
| Ttc5 | 2480.57 | 0.071887 | 0.06889 | 1.043557 | 0.29669 | 0.50689 | 20.105 |
| Ttc7 | 769.553 | 0.192251 | 0.10301 | 1.866359 | 0.06199 | 0.19071 | 98.884 |
| Ttc7b | 5732.21 | -0.09912 | 0.0941 | -1.05343 | 0.29215 | 0.5019 | 220.057 |
| Ttc8 | 1601.84 | 0.297582 | 0.08381 | 3.550875 | 0.00038 | 0.00664 | 62.665 |
| Ttc9 | 2228.13 | 0.078325 | 0.09276 | 0.844371 | 0.39846 | 0.6075 | 36.309 |
| Ttc9b | 2218.15 | -0.75305 | 0.16963 | -4.43942 | 9.02E-06 | 0.00052 | 3.432 |
| Ttc9c | 1922.74 | 0.071937 | 0.0839 | 0.857449 | 0.3912 | 0.60033 | 10.259 |
| Ttf1 | 1053.23 | 0.232726 | 0.09497 | 2.450424 | 0.01427 | 0.07203 | 27.395 |
| Ttf2 | 101.881 | -0.46367 | 0.23159 | -2.0021 | 0.04527 | 0.15478 | 30.804 |

| | | | | | | | |
|---|---|---|---|---|---|---|---|
| Tti1 | 1015.56 | -0.02455 | 0.08671 | -0.28316 | 0.77706 | 0.88395 | 46.631 |
| Tti2 | 955.293 | 0.000486 | 0.09093 | 0.00534 | 0.99574 | 0.99765 | 14.388 |
| Ttk | 17.1028 | -0.83653 | 0.56538 | -1.47958 | 0.13899 | 0.3202 | 37.701 |
| Ttl | 3734.65 | -0.01018 | 0.06529 | -0.15585 | 0.87615 | 0.93819 | 30.342 |
| Ttll1 | 1881.51 | 0.103749 | 0.082 | 1.26524 | 0.20579 | 0.40497 | 27.125 |
| Ttll11 | 1437.36 | -0.06154 | 0.08304 | -0.74106 | 0.45866 | 0.65962 | 228.673 |
| Ttll12 | 577.278 | -0.1645 | 0.12466 | -1.31963 | 0.18696 | 0.3822 | 20.068 |
| Ttll13 | 62.6671 | 0.097933 | 0.22603 | 0.433281 | 0.66481 | 0.81493 | 14.446 |
| Ttll3 | 1068.3 | 0.144399 | 0.0845 | 1.708882 | 0.08747 | 0.23831 | 25.328 |
| Ttll4 | 2198.34 | 0.104997 | 0.06739 | 1.55797 | 0.11924 | 0.2914 | 41.986 |
| Ttll5 | 1554.55 | -0.24517 | 0.10453 | -2.34545 | 0.019 | 0.0866 | 237.235 |
| Ttll7 | 14449.4 | 0.095849 | 0.08607 | 1.113571 | 0.26546 | 0.47463 | 131.643 |
| Ttll8 | 72.4721 | -0.50681 | 0.27192 | -1.86379 | 0.06235 | 0.1915 | 63.786 |
| Ttll9 | 57.5296 | -0.1115 | 0.27579 | -0.40428 | 0.68601 | 0.82887 | 45.998 |
| Ttn | 120.923 | 0.006768 | 0.22857 | 0.02961 | 0.97638 | 0.98767 | 278.568 |
| Ttpa | 266.855 | 0.030135 | 0.14115 | 0.213496 | 0.83094 | 0.91329 | 22.848 |
| Ttpal | 3276.24 | -0.05475 | 0.07756 | -0.70585 | 0.48028 | 0.67761 | 16.7 |
| Ttyh1 | 22843.8 | 0.052295 | 0.06414 | 0.815349 | 0.41487 | 0.62287 | 17.301 |
| Ttyh2 | 1237.53 | -0.51071 | 0.17058 | -2.9939 | 0.00275 | 0.0249 | 45.547 |
| Ttyh3 | 13697.4 | 0.000687 | 0.05751 | 0.011942 | 0.99047 | 0.99473 | 28.454 |
| Tub | 9395.49 | 0.070998 | 0.08606 | 0.82496 | 0.40939 | 0.6172 | 84.123 |
| Tuba1a | 12567.2 | -0.34914 | 0.10731 | -3.25369 | 0.00114 | 0.01383 | 3.867 |
| Tuba1b | 3667.46 | -0.46439 | 0.12343 | -3.76231 | 0.00017 | 0.00386 | 3.141 |
| Tuba1c | 335.717 | -0.3151 | 0.14908 | -2.1137 | 0.03454 | 0.13012 | 8.22 |
| Tuba3a | 38.3075 | 0.409023 | 0.33154 | 1.23371 | 0.21731 | 0.41935 | 7.769 |
| Tuba4a | 9043.23 | -0.36384 | 0.09038 | -4.02561 | 5.68E-05 | 0.00181 | 5.638 |
| Tuba8 | 267.043 | -1.31403 | 0.29458 | -4.46061 | 8.17E-06 | 0.00047 | 16.159 |
| Tubb2a | 10118 | -0.34763 | 0.12067 | -2.88075 | 0.00397 | 0.0313 | 3.734 |
| Tubb2b | 3201.53 | -0.28154 | 0.09434 | -2.98421 | 0.00284 | 0.02532 | 3.719 |
| Tubb3 | 9223.82 | -0.34289 | 0.07466 | -4.593 | 4.37E-06 | 0.00031 | 10.592 |
| Tubb4a | 15419.8 | -0.54291 | 0.14948 | -3.632 | 0.00028 | 0.00537 | 7.717 |
| Tubb4b | 5335.14 | -0.43075 | 0.11559 | -3.7267 | 0.00019 | 0.0042 | 2.543 |
| Tubb4b-ps1 | 53.8138 | -0.68927 | 0.24434 | -2.82093 | 0.00479 | 0.0353 | 1.336 |
| Tubb5 | 13163.4 | -0.32386 | 0.09774 | -3.31353 | 0.00092 | 0.01211 | 4.386 |

| | | | | | | | |
|---|---|---|---|---|---|---|---|
| Tubb6 | 258.912 | 0.034712 | 0.12618 | 0.275096 | 0.78324 | 0.88742 | 12.033 |
| Tubd1 | 458.521 | 0.060205 | 0.12555 | 0.479543 | 0.63155 | 0.79201 | 22.37 |
| Tube1 | 169.616 | -0.23901 | 0.17461 | -1.36879 | 0.17107 | 0.36145 | 18.567 |
| Tubg1 | 1097.16 | -0.11907 | 0.08287 | -1.43685 | 0.15076 | 0.33567 | 6.482 |
| Tubg2 | 1351.91 | -0.16782 | 0.09914 | -1.69278 | 0.0905 | 0.2443 | 5.881 |
| Tubgcp2 | 1576.43 | -0.10457 | 0.0768 | -1.36167 | 0.1733 | 0.36452 | 40.715 |
| Tubgcp3 | 2518.1 | 0.037496 | 0.05892 | 0.636392 | 0.52452 | 0.71298 | 57.972 |
| Tubgcp4 | 1330.85 | 0.139624 | 0.07815 | 1.786524 | 0.07401 | 0.2144 | 28.117 |
| Tubgcp5 | 2008.66 | 0.126555 | 0.09751 | 1.297837 | 0.19434 | 0.39157 | 37.524 |
| Tubgcp6 | 2235.96 | -0.0206 | 0.06814 | -0.30236 | 0.76238 | 0.87544 | 24.756 |
| Tufm | 2078.86 | -0.14928 | 0.0648 | -2.30361 | 0.02124 | 0.09323 | 3.371 |
| Tuft1 | 63.9643 | -0.39981 | 0.23778 | -1.68141 | 0.09268 | 0.24799 | 46.116 |
| Tug1 | 5059.59 | 0.231686 | 0.08674 | 2.671141 | 0.00756 | 0.04797 | 9.889 |
| Tulp3 | 1404.79 | -0.02237 | 0.07104 | -0.31493 | 0.75281 | 0.87023 | 34.691 |
| Tulp4 | 8886.06 | 0.041357 | 0.07097 | 0.582705 | 0.56009 | 0.74154 | 144.692 |
| Tunar | 1668.94 | -0.31705 | 0.13393 | -2.36736 | 0.01792 | 0.08338 | 50.063 |
| Tusc1 | 169.809 | -0.32247 | 0.18001 | -1.79139 | 0.07323 | 0.21312 | 1.374 |
| Tusc2 | 1401.71 | -0.20786 | 0.09119 | -2.27948 | 0.02264 | 0.09721 | 2.858 |
| Tusc3 | 6463.15 | 0.118063 | 0.08437 | 1.399424 | 0.16169 | 0.34987 | 146.369 |
| Tut1 | 1260.05 | 0.178215 | 0.10075 | 1.768928 | 0.07691 | 0.21936 | 12.364 |
| Tut4 | 2093.76 | 0.308498 | 0.15831 | 1.948675 | 0.05133 | 0.1688 | 99.996 |
| Tut7 | 3069.58 | -0.01687 | 0.0635 | -0.26562 | 0.79053 | 0.8913 | 51.587 |
| Tvp23a | 715.26 | 0.060309 | 0.11016 | 0.547468 | 0.58406 | 0.75898 | 26.806 |
| Tvp23b | 1420.4 | -0.06492 | 0.08104 | -0.80107 | 0.42309 | 0.62995 | 37.326 |
| Twf1 | 2143.48 | -0.15213 | 0.06214 | -2.4484 | 0.01435 | 0.07227 | 11.939 |
| Twf2 | 1096.89 | 0.009072 | 0.09901 | 0.091628 | 0.92699 | 0.96324 | 12.282 |
| Twist1 | 19.0676 | -0.92895 | 0.64322 | -1.44423 | 0.14868 | 0.33248 | 2.159 |
| Twnk | 973.597 | -0.01199 | 0.09347 | -0.12829 | 0.89792 | 0.94882 | 7.1 |
| Twsg1 | 2027.1 | 0.145262 | 0.0942 | 1.542007 | 0.12307 | 0.29724 | 29.255 |
| Txlna | 2981.56 | 0.06452 | 0.07113 | 0.907115 | 0.36435 | 0.57497 | 14.988 |
| Txlnb | 111.014 | -0.27565 | 0.26721 | -1.03158 | 0.30227 | 0.51199 | 49.44 |
| Txlng | 2099.67 | 0.484077 | 0.09958 | 4.861061 | 1.17E-06 | 0.00012 | 50.536 |
| Txn1 | 1274.02 | -0.15845 | 0.0887 | -1.78638 | 0.07404 | 0.21441 | 13.039 |
| Txn2 | 3314.65 | -0.1346 | 0.07062 | -1.90604 | 0.05665 | 0.18019 | 13.96 |

| | | | | | | | |
|---|---|---|---|---|---|---|---|
| Txndc11 | 1747.01 | -0.02382 | 0.06784 | -0.35109 | 0.72552 | 0.85387 | 59.74 |
| Txndc12 | 1026.08 | -0.06163 | 0.08655 | -0.71201 | 0.47646 | 0.67441 | 27.527 |
| Txndc15 | 2465.16 | -0.01808 | 0.06498 | -0.27822 | 0.78084 | 0.88589 | 11.578 |
| Txndc16 | 5629.47 | 0.117772 | 0.07009 | 1.680274 | 0.0929 | 0.24829 | 86.864 |
| Txndc17 | 661.363 | -0.01503 | 0.09465 | -0.15875 | 0.87386 | 0.93678 | 2.469 |
| Txndc5 | 2375.11 | 0.115938 | 0.10202 | 1.136373 | 0.2558 | 0.46353 | 28.746 |
| Txndc9 | 2232.31 | 0.14611 | 0.07063 | 2.068556 | 0.03859 | 0.13989 | 12.705 |
| Txnip | 1134.83 | -0.49516 | 0.1994 | -2.48325 | 0.01302 | 0.06827 | 3.927 |
| Txnl1 | 3489.32 | 0.071394 | 0.06419 | 1.112273 | 0.26602 | 0.47518 | 47.708 |
| Txnl4a | 609.086 | -0.20125 | 0.10049 | -2.00265 | 0.04521 | 0.15461 | 19.061 |
| Txnl4b | 519.454 | -0.2016 | 0.10466 | -1.92619 | 0.05408 | 0.17484 | 8.16 |
| Txnrd1 | 4029.03 | 0.281553 | 0.1023 | 2.752101 | 0.00592 | 0.04089 | 63.762 |
| Txnrd2 | 384.434 | -0.21942 | 0.11884 | -1.84629 | 0.06485 | 0.19618 | 52.69 |
| Txnrd3 | 153.726 | -0.25031 | 0.19665 | -1.2729 | 0.20305 | 0.40165 | 31.542 |
| Tyk2 | 1341.43 | -0.10117 | 0.08277 | -1.22224 | 0.22162 | 0.42449 | 27.176 |
| Tyms | 98.4799 | -0.16859 | 0.22767 | -0.74052 | 0.45898 | 0.65977 | 15.416 |
| Tyro3 | 12244.3 | 0.167351 | 0.06844 | 2.445334 | 0.01447 | 0.07271 | 20.372 |
| Tyrobp | 280.192 | -0.27808 | 0.18774 | -1.4812 | 0.13855 | 0.31964 | 3.826 |
| Tysnd1 | 692.726 | -0.09949 | 0.0981 | -1.01417 | 0.3105 | 0.52067 | 7.26 |
| Tyw1 | 846.459 | 0.024736 | 0.09099 | 0.271842 | 0.78574 | 0.88863 | 85.945 |
| Tyw3 | 501.37 | 0.133253 | 0.10215 | 1.304505 | 0.19206 | 0.38879 | 20.674 |
| Tyw5 | 474.724 | 0.034498 | 0.15605 | 0.221074 | 0.82503 | 0.90964 | 18.865 |
| U1 | 30.4923 | -0.28235 | 0.3613 | -0.7815 | 0.43451 | 0.64028 | 0.184 |
| U2af1 | 2082.97 | 0.033828 | 0.08829 | 0.383133 | 0.70162 | 0.83894 | 11.812 |
| U2af2 | 7381.59 | 0.136414 | 0.06054 | 2.253154 | 0.02425 | 0.10171 | 17.796 |
| U2surp | 4590.87 | 0.255654 | 0.13065 | 1.956792 | 0.05037 | 0.16683 | 55.099 |
| U6 | 370.193 | 0.213164 | 0.1251 | 1.703992 | 0.08838 | 0.24018 | 0.118 |
| Uaca | 547.517 | -0.53675 | 0.18279 | -2.93642 | 0.00332 | 0.02779 | 85.829 |
| Uap1 | 1723.41 | -0.17545 | 0.07283 | -2.40908 | 0.01599 | 0.07767 | 33.02 |
| Uap1l1 | 576.503 | -0.08689 | 0.11587 | -0.74984 | 0.45335 | 0.65558 | 5.794 |
| Uba1 | 15188.4 | -0.07033 | 0.06356 | -1.10636 | 0.26857 | 0.47788 | 24.854 |
| Uba1y | 23.7484 | 0.036208 | 0.4108 | 0.08814 | 0.92977 | 0.96459 | 25.576 |
| Uba2 | 4380.19 | 0.034299 | 0.06201 | 0.553102 | 0.58019 | 0.75652 | 28.912 |
| Uba3 | 3786.31 | 0.023124 | 0.08756 | 0.264109 | 0.7917 | 0.89165 | 21.978 |

| | | | | | | | |
|---|---|---|---|---|---|---|---|
| Uba5 | 2640.94 | 0.079594 | 0.07227 | 1.101333 | 0.27075 | 0.48026 | 16.536 |
| Uba52 | 163.162 | -0.3283 | 0.16649 | -1.9719 | 0.04862 | 0.16287 | 2.539 |
| Uba6 | 1586.88 | 0.271047 | 0.10998 | 2.464561 | 0.01372 | 0.07047 | 63.517 |
| Uba7 | 149.772 | -0.00068 | 0.17134 | -0.00397 | 0.99683 | 0.99821 | 8.556 |
| Ubac1 | 1883.81 | -0.0092 | 0.07413 | -0.12411 | 0.90122 | 0.95046 | 23.205 |
| Ubac2 | 1320.4 | -0.12313 | 0.09003 | -1.36771 | 0.1714 | 0.3618 | 142.415 |
| Ubald1 | 3491.65 | 0.039476 | 0.06839 | 0.577256 | 0.56377 | 0.74426 | 5.538 |
| Ubald2 | 522.107 | -0.29625 | 0.10113 | -2.92942 | 0.0034 | 0.02816 | 4.984 |
| Ubap1 | 2360.07 | -0.03944 | 0.06813 | -0.57881 | 0.56272 | 0.74348 | 41.53 |
| Ubap1l | 344.852 | 0.177836 | 0.16167 | 1.099983 | 0.27134 | 0.48066 | 19.337 |
| Ubap2 | 2416.51 | -0.0816 | 0.0657 | -1.24205 | 0.21422 | 0.41558 | 80.832 |
| Ubap2l | 5594.61 | 0.095039 | 0.05819 | 1.633192 | 0.10243 | 0.26424 | 52.489 |
| Ubash3b | 1505.13 | -0.61557 | 0.15747 | -3.90902 | 9.27E-05 | 0.00255 | 150.6 |
| Ubb | 23653.9 | -0.15218 | 0.07475 | -2.03583 | 0.04177 | 0.14702 | 2.043 |
| Ubb-ps | 301.343 | 0.082189 | 0.12455 | 0.659893 | 0.50932 | 0.70083 | 0.975 |
| Ubc | 2284.53 | -0.34466 | 0.15726 | -2.19159 | 0.02841 | 0.11366 | 4.238 |
| Ube2a | 1651.49 | -0.04067 | 0.07531 | -0.53999 | 0.5892 | 0.76226 | 83.048 |
| Ube2b | 7808.39 | 0.10493 | 0.06838 | 1.534579 | 0.12489 | 0.29992 | 15.266 |
| Ube2c | 19.8278 | -1.0095 | 0.47918 | -2.10672 | 0.03514 | 0.13139 | 8.925 |
| Ube2cbp | 116.845 | 0.422874 | 0.18191 | 2.32458 | 0.02009 | 0.08977 | 157.645 |
| Ube2d-ps | 636.888 | -0.01767 | 0.09803 | -0.18026 | 0.85695 | 0.92788 | 22.898 |
| Ube2d1 | 2201.56 | 0.024669 | 0.08007 | 0.30811 | 0.758 | 0.8726 | 30.283 |
| Ube2d2a | 3478.47 | -0.07916 | 0.0635 | -1.24672 | 0.2125 | 0.41332 | 37.937 |
| Ube2d3 | 13157.4 | 0.117878 | 0.07056 | 1.670648 | 0.09479 | 0.25135 | 30.05 |
| Ube2e1 | 834.702 | -0.2315 | 0.0861 | -2.68858 | 0.00718 | 0.04629 | 49.138 |
| Ube2e2 | 1563.18 | -0.22248 | 0.09169 | -2.42654 | 0.01524 | 0.07527 | 320.693 |
| Ube2e3 | 2795.96 | -0.18406 | 0.07369 | -2.49789 | 0.01249 | 0.06637 | 53.17 |
| Ube2f | 1802.96 | -0.04753 | 0.07898 | -0.60181 | 0.5473 | 0.73143 | 40.034 |
| Ube2g1 | 2796.12 | -0.01208 | 0.06693 | -0.18041 | 0.85683 | 0.92788 | 79.199 |
| Ube2g2 | 2454.86 | -0.19335 | 0.08052 | -2.4012 | 0.01634 | 0.07875 | 23.719 |
| Ube2h | 6107.01 | -0.0483 | 0.05899 | -0.81877 | 0.41292 | 0.62109 | 93.251 |
| Ube2i | 5980.96 | 0.142354 | 0.07122 | 1.998755 | 0.04563 | 0.15558 | 12.707 |
| Ube2j1 | 4690.4 | -0.06205 | 0.05646 | -1.09903 | 0.27175 | 0.48106 | 20.948 |
| Ube2j2 | 1589.14 | -0.04112 | 0.08321 | -0.49421 | 0.62116 | 0.78454 | 15.774 |

| | | | | | | | |
|---|---|---|---|---|---|---|---|
| Ube2k | 5887.61 | -0.09405 | 0.06878 | -1.3674 | 0.1715 | 0.36185 | 61.756 |
| Ube2l3 | 4248.17 | -0.10268 | 0.07076 | -1.45107 | 0.14676 | 0.32983 | 50.637 |
| Ube2l6 | 143.869 | -0.48545 | 0.24022 | -2.0209 | 0.04329 | 0.15032 | 11.508 |
| Ube2m | 3820.37 | -0.20289 | 0.07336 | -2.7659 | 0.00568 | 0.03968 | 3.156 |
| Ube2n | 7608.88 | -0.01143 | 0.06453 | -0.17707 | 0.85945 | 0.929 | 30.513 |
| Ube2n-ps1 | 79.1147 | -0.20535 | 0.21669 | -0.94769 | 0.34329 | 0.55397 | 0.459 |
| Ube2o | 6632.11 | -0.00395 | 0.06962 | -0.05668 | 0.9548 | 0.97783 | 43.708 |
| Ube2q1 | 4139.77 | -0.09023 | 0.06643 | -1.35841 | 0.17433 | 0.36594 | 10.385 |
| Ube2q2 | 3399.07 | 0.087039 | 0.0768 | 1.133262 | 0.2571 | 0.46513 | 58.676 |
| Ube2ql1 | 4679.83 | -0.68084 | 0.18878 | -3.60652 | 0.00031 | 0.00578 | 37.058 |
| Ube2r2 | 2930.4 | -0.00661 | 0.09116 | -0.07255 | 0.94216 | 0.97113 | 57.638 |
| Ube2s | 1458.82 | -0.08473 | 0.07271 | -1.16524 | 0.24392 | 0.45033 | 18.045 |
| Ube2t | 54.3388 | 0.061488 | 0.27366 | 0.224685 | 0.82222 | 0.90785 | 11.598 |
| Ube2v1 | 25.3897 | -0.01492 | 0.33913 | -0.044 | 0.9649 | 0.98273 | 24.458 |
| Ube2v2 | 1928.13 | 0.203589 | 0.17087 | 1.191451 | 0.23348 | 0.43851 | 43.697 |
| Ube2w | 2625.03 | 0.013806 | 0.07284 | 0.189527 | 0.84968 | 0.92352 | 78.7 |
| Ube2z | 4280.21 | -0.06017 | 0.07512 | -0.80103 | 0.42311 | 0.62995 | 17.957 |
| Ube3a | 5333.51 | 0.230589 | 0.11609 | 1.986271 | 0.047 | 0.15907 | 82.787 |
| Ube3b | 4876.42 | -0.22204 | 0.07373 | -3.01128 | 0.0026 | 0.02402 | 40.563 |
| Ube3c | 4561.08 | 0.122164 | 0.08828 | 1.38383 | 0.16641 | 0.356 | 106.851 |
| Ube4a | 4532.67 | 0.057507 | 0.06481 | 0.887311 | 0.37491 | 0.58517 | 42.474 |
| Ube4b | 6180.4 | 0.295334 | 0.0962 | 3.07003 | 0.00214 | 0.02108 | 98.334 |
| Ube4bos1 | 65.3288 | 0.379339 | 0.27726 | 1.368182 | 0.17126 | 0.36175 | 17.393 |
| Ube4bos2 | 19.2465 | 0.262794 | 0.42933 | 0.612102 | 0.54047 | 0.72624 | 0.445 |
| Ube4bos3 | 64.7228 | 0.698793 | 0.27866 | 2.507645 | 0.01215 | 0.06511 | 1.259 |
| Ubfd1 | 3416.45 | 0.092647 | 0.07303 | 1.268557 | 0.2046 | 0.40342 | 15.02 |
| Ubiad1 | 429.938 | -0.07139 | 0.10223 | -0.6984 | 0.48493 | 0.68115 | 10.277 |
| Ubl3 | 9219.11 | 0.004382 | 0.05651 | 0.077549 | 0.93819 | 0.96919 | 48.155 |
| Ubl4a | 2581.8 | -0.08306 | 0.06588 | -1.26089 | 0.20735 | 0.40685 | 7.501 |
| Ubl5 | 1932.76 | -0.21452 | 0.08358 | -2.56655 | 0.01027 | 0.05852 | 4.263 |
| Ubl7 | 2013.2 | -0.07921 | 0.06593 | -1.20147 | 0.22957 | 0.43383 | 18.99 |
| Ublcp1 | 1722.15 | -0.09162 | 0.07857 | -1.16619 | 0.24354 | 0.44987 | 15.928 |
| Ubn1 | 3851.48 | 0.016848 | 0.06377 | 0.264195 | 0.79163 | 0.89165 | 36.233 |
| Ubn2 | 7042.13 | 0.178391 | 0.09409 | 1.895978 | 0.05796 | 0.18301 | 90.876 |

| | | | | | | | |
|---|---|---|---|---|---|---|---|
| Ubox5 | 524.593 | -0.06396 | 0.09727 | -0.65752 | 0.51085 | 0.70199 | 40.037 |
| Ubp1 | 5397.3 | 0.150281 | 0.08183 | 1.836558 | 0.06628 | 0.19905 | 46.269 |
| Ubqln1 | 9941.29 | -0.01331 | 0.05599 | -0.23768 | 0.81213 | 0.90333 | 39.498 |
| Ubqln2 | 10769.4 | 0.142617 | 0.05402 | 2.640073 | 0.00829 | 0.05072 | 3.344 |
| Ubqln4 | 5926.15 | -0.14409 | 0.07733 | -1.8633 | 0.06242 | 0.19158 | 15.968 |
| Ubr1 | 3507.22 | 0.149302 | 0.0946 | 1.578186 | 0.11452 | 0.28343 | 110.447 |
| Ubr2 | 6735.81 | 0.137134 | 0.06491 | 2.112601 | 0.03463 | 0.13033 | 82.262 |
| Ubr3 | 14827.4 | -0.04258 | 0.08678 | -0.49061 | 0.6237 | 0.78639 | 126.768 |
| Ubr4 | 15180.1 | 0.129576 | 0.06817 | 1.900729 | 0.05734 | 0.18168 | 136.98 |
| Ubr5 | 18274 | 0.373648 | 0.10962 | 3.408599 | 0.00065 | 0.00949 | 111.527 |
| Ubr7 | 1999.14 | 0.093089 | 0.07858 | 1.184672 | 0.23615 | 0.44147 | 19.741 |
| Ubtd1 | 321.077 | -0.24042 | 0.13369 | -1.79829 | 0.07213 | 0.21084 | 52.881 |
| Ubtd2 | 481.182 | 0.037738 | 0.1099 | 0.34338 | 0.73131 | 0.85734 | 61.318 |
| Ubtf | 7204.28 | 0.002596 | 0.0713 | 0.036414 | 0.97095 | 0.98532 | 15.183 |
| Ubxn1 | 2175.24 | -0.2164 | 0.0823 | -2.62938 | 0.00855 | 0.05169 | 4.368 |
| Ubxn10 | 55.3393 | -0.08957 | 0.23354 | -0.38354 | 0.70132 | 0.8387 | 27.331 |
| Ubxn11 | 785.806 | 0.378726 | 0.10779 | 3.513434 | 0.00044 | 0.00731 | 25.024 |
| Ubxn2a | 2319.19 | -0.13596 | 0.10388 | -1.30891 | 0.19057 | 0.38671 | 28.674 |
| Ubxn2b | 4100.75 | 0.318822 | 0.09866 | 3.231366 | 0.00123 | 0.01456 | 30.591 |
| Ubxn4 | 3767.9 | 0.062366 | 0.06271 | 0.994452 | 0.32 | 0.53067 | 35.415 |
| Ubxn6 | 546.081 | 0.011205 | 0.09875 | 0.113471 | 0.90966 | 0.95456 | 7.984 |
| Ubxn7 | 4025.16 | 0.197959 | 0.07792 | 2.540518 | 0.01107 | 0.06133 | 61.491 |
| Ubxn8 | 1168.91 | -0.10728 | 0.09067 | -1.18323 | 0.23672 | 0.44183 | 22.382 |
| Uchl1 | 10748.8 | -0.07158 | 0.06601 | -1.08432 | 0.27822 | 0.48807 | 11.144 |
| Uchl3 | 684.853 | 0.14777 | 0.12488 | 1.183262 | 0.23671 | 0.44183 | 42.159 |
| Uchl4 | 17.7041 | -0.01458 | 0.47723 | -0.03055 | 0.97563 | 0.98762 | 1.162 |
| Uchl5 | 1513.71 | -0.01934 | 0.11367 | -0.17011 | 0.86492 | 0.93163 | 30.195 |
| Uck1 | 1636.24 | 0.091921 | 0.08004 | 1.14837 | 0.25082 | 0.45821 | 5.158 |
| Uck2 | 1297.45 | -0.14011 | 0.07626 | -1.8373 | 0.06617 | 0.19881 | 62.438 |
| Uckl1 | 1110.72 | 0.186433 | 0.0855 | 2.180575 | 0.02921 | 0.11598 | 15.744 |
| Uckl1os | 177.935 | 0.288194 | 0.14675 | 1.963884 | 0.04954 | 0.16483 | 6.691 |
| Ucma | 13.6404 | -0.21857 | 0.75973 | -0.2877 | 0.77358 | 0.88191 | 9.627 |
| Ucp2 | 1526.65 | -0.53182 | 0.14889 | -3.57175 | 0.00035 | 0.00628 | 8.684 |
| Ucp3 | 152.855 | -0.14483 | 0.19404 | -0.74638 | 0.45544 | 0.65743 | 13.443 |

| | | | | | | | |
|---|---|---|---|---|---|---|---|
| Uevld | 830.05 | 0.290745 | 0.0893 | 3.255742 | 0.00113 | 0.01378 | 35.312 |
| Ufc1 | 1874.02 | -0.24701 | 0.08674 | -2.84756 | 0.00441 | 0.03356 | 6.462 |
| Ufd1 | 3611.66 | 0.131979 | 0.06966 | 1.894563 | 0.05815 | 0.18329 | 23.483 |
| Ufl1 | 1941.74 | 0.277273 | 0.10397 | 2.666885 | 0.00766 | 0.04834 | 33.222 |
| Ufm1 | 2993.25 | -0.14433 | 0.07094 | -2.0345 | 0.0419 | 0.1472 | 10.455 |
| Ufsp1 | 353.91 | 0.33341 | 0.18551 | 1.797245 | 0.0723 | 0.21123 | 1.037 |
| Ufsp2 | 1606.2 | -0.07855 | 0.11357 | -0.69158 | 0.4892 | 0.68436 | 21.431 |
| Ugcg | 3470.53 | -0.1647 | 0.07603 | -2.16638 | 0.03028 | 0.11878 | 33.577 |
| Ugdh | 1186.39 | 0.051102 | 0.10306 | 0.495854 | 0.62 | 0.78385 | 22.729 |
| Uggt1 | 4026.79 | 0.068694 | 0.08146 | 0.843279 | 0.39907 | 0.60807 | 104.694 |
| Uggt2 | 1997.94 | 0.164976 | 0.11204 | 1.472461 | 0.1409 | 0.32265 | 114.392 |
| Ugp2 | 3326.76 | -0.13919 | 0.07738 | -1.7987 | 0.07207 | 0.21074 | 50.064 |
| Ugt1a7c | 20.7415 | -0.47421 | 0.39946 | -1.18713 | 0.23518 | 0.44037 | 124.941 |
| Ugt8a | 1782.92 | -0.51694 | 0.20969 | -2.46521 | 0.01369 | 0.07038 | 73.349 |
| Uhmk1 | 16439.1 | 0.01537 | 0.05404 | 0.284395 | 0.77611 | 0.88346 | 21.978 |
| Uhrf1 | 58.1634 | -0.75765 | 0.34266 | -2.21109 | 0.02703 | 0.10985 | 20.166 |
| Uhrf1bp1 | 1665.49 | 0.105701 | 0.07476 | 1.41385 | 0.15741 | 0.34463 | 43.601 |
| Uhrf1bp1l | 5253.29 | 0.016116 | 0.08157 | 0.197582 | 0.84337 | 0.91957 | 74.881 |
| Uhrf2 | 3451.13 | 0.170449 | 0.05693 | 2.994277 | 0.00275 | 0.02488 | 63.21 |
| Uimc1 | 890.578 | 0.087629 | 0.07785 | 1.12556 | 0.26035 | 0.46882 | 72.421 |
| Ulk1 | 3608.44 | -0.04811 | 0.0659 | -0.73006 | 0.46535 | 0.66495 | 25.61 |
| Ulk2 | 5752.33 | 0.011566 | 0.06569 | 0.176065 | 0.86024 | 0.92937 | 79.425 |
| Ulk3 | 1382.86 | 0.165789 | 0.13675 | 1.212315 | 0.22539 | 0.42934 | 6.782 |
| Ulk4 | 402.233 | 0.219562 | 0.13795 | 1.591657 | 0.11146 | 0.27897 | 321.847 |
| Umad1 | 1307.51 | 0.142098 | 0.07666 | 1.853688 | 0.06378 | 0.19423 | 200.183 |
| Umps | 745.455 | -0.02706 | 0.08296 | -0.32623 | 0.74425 | 0.86539 | 12.257 |
| Unc119 | 565.098 | -0.21111 | 0.10971 | -1.92434 | 0.05431 | 0.17523 | 5.683 |
| Unc119b | 1680.06 | -0.01008 | 0.0784 | -0.12854 | 0.89772 | 0.94882 | 12.426 |
| Unc13a | 18594.6 | -0.05146 | 0.10254 | -0.5018 | 0.61581 | 0.78132 | 47.341 |
| Unc13b | 2579.66 | -0.3778 | 0.14621 | -2.58398 | 0.00977 | 0.05654 | 205.919 |
| Unc13c | 4586.85 | 1.117344 | 0.37726 | 2.961759 | 0.00306 | 0.02639 | 489.545 |
| Unc13d | 57.721 | 0.197427 | 0.23786 | 0.830009 | 0.40653 | 0.6148 | 15.867 |
| Unc45a | 1775.99 | 0.049602 | 0.07206 | 0.688367 | 0.49122 | 0.68615 | 22.702 |
| Unc45b | 68.8165 | -0.17147 | 0.23632 | -0.72557 | 0.4681 | 0.66717 | 32.854 |

| | | | | | | | |
|---|---|---|---|---|---|---|---|
| Unc50 | 1601.61 | -0.07057 | 0.08207 | -0.85995 | 0.38981 | 0.59893 | 8.986 |
| Unc5a | 4274.09 | -0.25157 | 0.09425 | -2.66915 | 0.0076 | 0.04816 | 56.608 |
| Unc5b | 973.925 | -0.51723 | 0.15049 | -3.43693 | 0.00059 | 0.00882 | 68.989 |
| Unc5c | 2209.01 | -0.06561 | 0.08169 | -0.8032 | 0.42186 | 0.6287 | 369.709 |
| Unc5d | 1661.38 | -0.15579 | 0.14039 | -1.10969 | 0.26713 | 0.47648 | 572.92 |
| Unc79 | 5593.55 | 0.493622 | 0.09054 | 5.451857 | 4.98E-08 | 1.14E-05 | 235.207 |
| Unc80 | 18879 | 0.045052 | 0.08862 | 0.50835 | 0.61121 | 0.77829 | 230.782 |
| Unc93b1 | 509.646 | -0.42822 | 0.13782 | -3.10708 | 0.00189 | 0.01935 | 14.155 |
| Ung | 151.598 | -0.08417 | 0.16255 | -0.5178 | 0.6046 | 0.7734 | 8.938 |
| Unk | 1183.61 | 0.127358 | 0.07816 | 1.629456 | 0.10322 | 0.26548 | 30.893 |
| Unkl | 1318.9 | 0.319882 | 0.08905 | 3.592264 | 0.00033 | 0.006 | 46.047 |
| Upb1 | 350.864 | 0.929282 | 0.63933 | 1.453518 | 0.14608 | 0.32903 | 40.565 |
| Upf1 | 4146.37 | 0.182024 | 0.06907 | 2.635236 | 0.00841 | 0.05114 | 21.754 |
| Upf2 | 3579.17 | 0.322549 | 0.1607 | 2.007164 | 0.04473 | 0.15361 | 105.235 |
| Upf3a | 1898.87 | -0.05629 | 0.07966 | -0.7066 | 0.47981 | 0.67732 | 13.579 |
| Upf3b | 2239.7 | 0.556315 | 0.13397 | 4.152571 | 3.29E-05 | 0.00123 | 18.645 |
| Upk1a | 61.2414 | 0.210767 | 0.32795 | 0.642671 | 0.52044 | 0.70975 | 9.756 |
| Upk1b | 44.6763 | 0.53229 | 0.27646 | 1.925408 | 0.05418 | 0.17506 | 27.145 |
| Upp1 | 61.658 | 0.017198 | 0.2521 | 0.068219 | 0.94561 | 0.97296 | 18.068 |
| Upp2 | 1045.08 | -0.22895 | 0.07627 | -3.00177 | 0.00268 | 0.02445 | 225.674 |
| Uprt | 401.365 | 0.2136 | 0.12058 | 1.771482 | 0.07648 | 0.21863 | 24.673 |
| Uqcc1 | 2446.61 | 0.027666 | 0.07601 | 0.363975 | 0.71588 | 0.8481 | 83.417 |
| Uqcc2 | 1759.37 | -0.16932 | 0.10447 | -1.62073 | 0.10508 | 0.26817 | 11.254 |
| Uqcc3 | 613.672 | -0.13583 | 0.10649 | -1.27544 | 0.20215 | 0.40058 | 0.91 |
| Uqcr10 | 2496 | -0.27692 | 0.09354 | -2.96059 | 0.00307 | 0.02648 | 2.37 |
| Uqcr11 | 2721.32 | -0.29951 | 0.09199 | -3.25607 | 0.00113 | 0.01378 | 3.834 |
| Uqcrb | 1248.33 | -0.11384 | 0.09979 | -1.14084 | 0.25394 | 0.4613 | 4.762 |
| Uqcrc1 | 6206.62 | -0.14944 | 0.05979 | -2.4995 | 0.01244 | 0.06619 | 12.991 |
| Uqcrc2 | 5805.03 | -0.03082 | 0.06193 | -0.49771 | 0.61869 | 0.78312 | 24.349 |
| Uqcrfs1 | 3807 | -0.09821 | 0.06983 | -1.40642 | 0.1596 | 0.34744 | 5.055 |
| Uqcrh | 4324.28 | -0.17277 | 0.06947 | -2.48681 | 0.01289 | 0.0678 | 8.107 |
| Uqcrq | 3134.95 | -0.17833 | 0.07436 | -2.39808 | 0.01648 | 0.07926 | 2.91 |
| Urb1 | 948.135 | 0.095026 | 0.0921 | 1.031823 | 0.30215 | 0.5119 | 58.887 |
| Urb2 | 736.598 | 0.235269 | 0.10225 | 2.300899 | 0.0214 | 0.09368 | 26.998 |

| | | | | | | | |
|---|---|---|---|---|---|---|---|
| Urgcp | 2607.19 | 0.024124 | 0.06795 | 0.355012 | 0.72258 | 0.85208 | 48.96 |
| Uri1 | 2279.02 | -0.00048 | 0.0894 | -0.00537 | 0.99572 | 0.99765 | 63.56 |
| Urm1 | 1473.43 | -0.25868 | 0.09376 | -2.75907 | 0.0058 | 0.04027 | 24.928 |
| Urod | 1301.47 | 0.08421 | 0.0946 | 0.890138 | 0.37339 | 0.58357 | 4.449 |
| Uros | 1078.91 | -0.08863 | 0.10983 | -0.80697 | 0.41969 | 0.62696 | 23.83 |
| Usb1 | 315.587 | -0.24199 | 0.15415 | -1.56985 | 0.11645 | 0.28668 | 15.224 |
| Use1 | 789.161 | -0.09986 | 0.09722 | -1.02709 | 0.30438 | 0.51417 | 2.885 |
| Usf1 | 2091.49 | -0.05034 | 0.08458 | -0.59522 | 0.5517 | 0.73468 | 7.83 |
| Usf2 | 1510.24 | -0.12681 | 0.07561 | -1.67718 | 0.09351 | 0.24909 | 11.556 |
| Usf3 | 4765.6 | 0.049515 | 0.08168 | 0.606244 | 0.54435 | 0.72945 | 54.22 |
| Ush1c | 14.7468 | 1.144903 | 0.50204 | 2.280502 | 0.02258 | 0.09703 | 43.154 |
| Ush1g | 14.9611 | -1.30801 | 0.58699 | -2.22832 | 0.02586 | 0.10649 | 6.85 |
| Ushbp1 | 255.11 | -0.18233 | 0.1735 | -1.05088 | 0.29331 | 0.50323 | 11.531 |
| Uso1 | 4774.53 | 0.088094 | 0.05866 | 1.501671 | 0.13318 | 0.31278 | 64.861 |
| Usp1 | 1459.04 | 0.107797 | 0.07365 | 1.463667 | 0.14329 | 0.32539 | 11.734 |
| Usp10 | 2935.43 | 0.202188 | 0.09814 | 2.060126 | 0.03939 | 0.14147 | 47.201 |
| Usp11 | 8594.6 | 0.390123 | 0.07629 | 5.113519 | 3.16E-07 | 4.36E-05 | 16.634 |
| Usp12 | 3246.68 | -0.08212 | 0.07011 | -1.17137 | 0.24145 | 0.44749 | 60.198 |
| Usp13 | 1749.12 | -0.21557 | 0.09587 | -2.24856 | 0.02454 | 0.10261 | 120.526 |
| Usp14 | 4922.24 | 0.041142 | 0.06106 | 0.673767 | 0.50046 | 0.69333 | 52.054 |
| Usp15 | 4535.33 | 0.246128 | 0.08229 | 2.99101 | 0.00278 | 0.02499 | 91.99 |
| Usp16 | 1944.09 | 0.076317 | 0.08761 | 0.871062 | 0.38372 | 0.5931 | 28.815 |
| Usp18 | 58.0719 | -0.85412 | 0.35322 | -2.41807 | 0.0156 | 0.07646 | 25.012 |
| Usp19 | 8560.84 | 0.130648 | 0.06409 | 2.038426 | 0.04151 | 0.14626 | 11.736 |
| Usp2 | 4048.28 | 0.099641 | 0.07982 | 1.248254 | 0.21194 | 0.41258 | 28.607 |
| Usp20 | 3791.18 | 0.112094 | 0.05688 | 1.970688 | 0.04876 | 0.16318 | 41.308 |
| Usp21 | 1963.71 | 0.146302 | 0.0677 | 2.161113 | 0.03069 | 0.11988 | 6.046 |
| Usp22 | 9583.15 | 0.096525 | 0.06589 | 1.464849 | 0.14296 | 0.32502 | 23.271 |
| Usp24 | 6839.12 | 0.198177 | 0.07444 | 2.662187 | 0.00776 | 0.04874 | 125.11 |
| Usp25 | 1513.9 | -0.11891 | 0.09845 | -1.20782 | 0.22712 | 0.43132 | 103.075 |
| Usp27x | 815.865 | 0.332197 | 0.11701 | 2.838946 | 0.00453 | 0.03407 | 4.557 |
| Usp28 | 2836.58 | 0.486159 | 0.13275 | 3.662094 | 0.00025 | 0.00496 | 57.143 |
| Usp29 | 2441.01 | 0.505361 | 0.12743 | 3.965928 | 7.31E-05 | 0.00216 | 236.642 |
| Usp3 | 2501.39 | 0.409542 | 0.10436 | 3.924412 | 8.69E-05 | 0.00245 | 78.506 |

| | | | | | | | |
|---|---|---|---|---|---|---|---|
| Usp30 | 1974.72 | 0.151775 | 0.07675 | 1.97749 | 0.04799 | 0.1613 | 59.26 |
| Usp31 | 6825.62 | -0.07383 | 0.09403 | -0.78526 | 0.4323 | 0.63842 | 65.233 |
| Usp32 | 7048.64 | -0.06212 | 0.08609 | -0.72151 | 0.47059 | 0.6693 | 155.72 |
| Usp33 | 3507.35 | 0.093591 | 0.07805 | 1.199038 | 0.23051 | 0.43487 | 47.14 |
| Usp34 | 6784.4 | 0.160009 | 0.10713 | 1.493563 | 0.13529 | 0.31507 | 183.666 |
| Usp35 | 1539.1 | -0.09062 | 0.08545 | -1.0605 | 0.28892 | 0.49909 | 22.641 |
| Usp36 | 2674.68 | 0.155403 | 0.0596 | 2.607478 | 0.00912 | 0.05389 | 30.594 |
| Usp37 | 1708.94 | 0.112522 | 0.07872 | 1.429404 | 0.15289 | 0.33835 | 108.774 |
| Usp38 | 1360.62 | 0.138556 | 0.07863 | 1.762158 | 0.07804 | 0.22104 | 34.196 |
| Usp39 | 1196.51 | -0.07303 | 0.08611 | -0.84818 | 0.39634 | 0.60539 | 26.507 |
| Usp4 | 5746.63 | 0.192543 | 0.0723 | 2.663117 | 0.00774 | 0.04863 | 44.693 |
| Usp40 | 1893.52 | 0.116849 | 0.07529 | 1.551911 | 0.12068 | 0.29359 | 63.433 |
| Usp42 | 1320.77 | 0.067633 | 0.07675 | 0.881226 | 0.3782 | 0.58818 | 21.956 |
| Usp43 | 35.9908 | -1.26742 | 0.31681 | -4.0006 | 6.32E-05 | 0.00196 | 67.631 |
| Usp45 | 3591.76 | 0.228435 | 0.06634 | 3.443587 | 0.00057 | 0.00871 | 70.717 |
| Usp46 | 5510.9 | -0.0449 | 0.07196 | -0.624 | 0.53263 | 0.71997 | 69.979 |
| Usp47 | 5002.42 | 0.145394 | 0.0597 | 2.435577 | 0.01487 | 0.07396 | 88.158 |
| Usp48 | 9220.43 | 0.259719 | 0.06684 | 3.885476 | 0.0001 | 0.00273 | 64.783 |
| Usp49 | 602.275 | 0.180892 | 0.10582 | 1.709461 | 0.08737 | 0.23805 | 56.049 |
| Usp5 | 8976.47 | 0.088488 | 0.06451 | 1.371637 | 0.17018 | 0.36052 | 14.466 |
| Usp50 | 31.5319 | 0.19732 | 0.36843 | 0.535568 | 0.59226 | 0.76487 | 74.375 |
| Usp51 | 90.3803 | 0.135418 | 0.25744 | 0.526012 | 0.59888 | 0.76992 | 2.948 |
| Usp53 | 1985.85 | 0.167116 | 0.10046 | 1.663516 | 0.09621 | 0.25397 | 53.018 |
| Usp54 | 4781.8 | -0.20776 | 0.07483 | -2.7763 | 0.0055 | 0.03875 | 92.152 |
| Usp6nl | 1227.17 | -0.3803 | 0.12404 | -3.066 | 0.00217 | 0.02127 | 123.724 |
| Usp7 | 9548.06 | 0.11477 | 0.05585 | 2.05504 | 0.03988 | 0.14266 | 102.714 |
| Usp8 | 4007.73 | -0.00913 | 0.08107 | -0.1126 | 0.91035 | 0.95475 | 51.97 |
| Usp9x | 9425.34 | 0.191716 | 0.11053 | 1.734579 | 0.08282 | 0.22962 | 101.831 |
| Uspl1 | 1929.62 | -0.05742 | 0.08264 | -0.69486 | 0.48714 | 0.68271 | 31.085 |
| Ust | 2363.84 | -0.34602 | 0.07996 | -4.32735 | 1.51E-05 | 0.00074 | 314.07 |
| Utp11 | 1219.92 | 0.254568 | 0.08844 | 2.878486 | 0.004 | 0.03147 | 15.441 |
| Utp14a | 1001.53 | 0.156402 | 0.09458 | 1.653635 | 0.0982 | 0.25724 | 25.592 |
| Utp14b | 1481.49 | -0.25574 | 0.15285 | -1.67314 | 0.0943 | 0.25058 | 13.475 |
| Utp15 | 1750.13 | 0.098949 | 0.06509 | 1.520263 | 0.12844 | 0.30549 | 16.197 |

| | | | | | | | |
|---|---|---|---|---|---|---|---|
| Utp18 | 1149.02 | 0.240294 | 0.08028 | 2.993027 | 0.00276 | 0.02493 | 26.524 |
| Utp20 | 1968.03 | 0.162627 | 0.08182 | 1.987581 | 0.04686 | 0.15876 | 80.198 |
| Utp23 | 366.79 | -0.14764 | 0.11324 | -1.30374 | 0.19232 | 0.38895 | 7.186 |
| Utp25 | 1134.51 | 0.125911 | 0.08823 | 1.427097 | 0.15355 | 0.33935 | 39.169 |
| Utp3 | 1282.09 | -0.04203 | 0.09161 | -0.45885 | 0.64634 | 0.80301 | 1.629 |
| Utp4 | 1334.16 | -0.00546 | 0.09564 | -0.05709 | 0.95448 | 0.97761 | 29.453 |
| Utp6 | 2313.15 | 0.000502 | 0.06828 | 0.007347 | 0.99414 | 0.99648 | 30.07 |
| Utrn | 4914.11 | 0.111703 | 0.09933 | 1.124527 | 0.26079 | 0.46936 | 487.178 |
| Uts2 | 53.6294 | 0.981544 | 0.28092 | 3.49407 | 0.00048 | 0.00768 | 4.714 |
| Uty | 1053.96 | 0.236263 | 0.10816 | 2.184475 | 0.02893 | 0.11505 | 148.899 |
| Uvrag | 1605.72 | -0.22458 | 0.08826 | -2.54442 | 0.01095 | 0.06094 | 256.121 |
| Uvssa | 820.285 | 0.117925 | 0.16375 | 0.720157 | 0.47143 | 0.67022 | 41.206 |
| Uxs1 | 2500.83 | 0.095353 | 0.06372 | 1.496327 | 0.13457 | 0.31443 | 80.835 |
| Uxt | 88.5498 | -0.13147 | 0.21499 | -0.61152 | 0.54086 | 0.72656 | 20.296 |
| Vac14 | 1537.36 | 0.101287 | 0.07649 | 1.324103 | 0.18547 | 0.38025 | 101.814 |
| Vamp1 | 5700.28 | -0.25603 | 0.10209 | -2.50777 | 0.01215 | 0.06511 | 30.414 |
| Vamp2 | 36993.1 | -0.32269 | 0.07549 | -4.27455 | 1.92E-05 | 0.00086 | 3.895 |
| Vamp3 | 1633.82 | -0.10003 | 0.08882 | -1.12616 | 0.2601 | 0.46869 | 10.664 |
| Vamp4 | 4835.73 | 0.141416 | 0.0681 | 2.076484 | 0.03785 | 0.13792 | 28.57 |
| Vamp5 | 101.8 | -0.57593 | 0.17604 | -3.27164 | 0.00107 | 0.0134 | 11.675 |
| Vamp8 | 300.659 | -0.68155 | 0.14946 | -4.55998 | 5.12E-06 | 0.00034 | 5.481 |
| Vamp9 | 34.1585 | 0.735826 | 0.34113 | 2.157034 | 0.031 | 0.12075 | 44.839 |
| Vangl1 | 270.957 | 0.035696 | 0.15703 | 0.227323 | 0.82017 | 0.90734 | 51.111 |
| Vangl2 | 601.699 | -0.37963 | 0.16816 | -2.25751 | 0.02398 | 0.10089 | 27.485 |
| Vapa | 5705.55 | -0.01354 | 0.06443 | -0.21022 | 0.8335 | 0.91471 | 35.229 |
| Vapb | 7877.33 | -0.09934 | 0.05712 | -1.73934 | 0.08198 | 0.22821 | 46.829 |
| Vars | 2065.29 | 0.101186 | 0.07926 | 1.276561 | 0.20176 | 0.40021 | 15.336 |
| Vars2 | 1175.85 | 0.139685 | 0.08333 | 1.676207 | 0.0937 | 0.24952 | 11.959 |
| Vash1 | 851.763 | -0.67829 | 0.19727 | -3.43835 | 0.00059 | 0.00882 | 13.392 |
| Vash2 | 106.633 | -0.51159 | 0.21499 | -2.37956 | 0.01733 | 0.08167 | 31.651 |
| Vasn | 775.661 | 0.089381 | 0.09467 | 0.944151 | 0.34509 | 0.55603 | 10.862 |
| Vasp | 578.943 | -0.09167 | 0.11528 | -0.79523 | 0.42648 | 0.63298 | 14.889 |
| Vat1 | 1670.03 | 0.065248 | 0.11676 | 0.558797 | 0.5763 | 0.75395 | 7.486 |
| Vat1l | 4118.45 | 0.961974 | 0.37515 | 2.56424 | 0.01034 | 0.05872 | 168.46 |

| | | | | | | | |
|---|---|---|---|---|---|---|---|
| Vav1 | 208.297 | -0.40396 | 0.1776 | -2.27451 | 0.02294 | 0.09803 | 48.932 |
| Vav2 | 651.434 | -0.11569 | 0.12364 | -0.93573 | 0.34941 | 0.5599 | 164.93 |
| Vav3 | 553.581 | 0.272195 | 0.13528 | 2.012085 | 0.04421 | 0.15247 | 345.046 |
| Vax1 | 80.7315 | 0.358353 | 0.21995 | 1.629276 | 0.10325 | 0.26552 | 3.87 |
| Vbp1 | 2567.51 | 0.123757 | 0.10227 | 1.210065 | 0.22625 | 0.43026 | 20.644 |
| Vcam1 | 1608.82 | -0.43339 | 0.11137 | -3.89149 | 9.96E-05 | 0.00268 | 19.74 |
| Vcan | 1850.03 | 0.090029 | 0.11494 | 0.783267 | 0.43347 | 0.63935 | 87.198 |
| Vcl | 1956.6 | 0.398864 | 0.13551 | 2.943476 | 0.00325 | 0.02751 | 104.279 |
| Vcp | 9202.22 | -0.01883 | 0.07525 | -0.25029 | 0.80236 | 0.89807 | 20.545 |
| Vcp-rs | 1714.78 | -0.1031 | 0.10343 | -0.9968 | 0.31886 | 0.52938 | 2.406 |
| Vcpip1 | 4131.1 | 0.080611 | 0.09038 | 0.891925 | 0.37243 | 0.58277 | 29.761 |
| Vcpkmt | 663.904 | -0.0059 | 0.10008 | -0.05894 | 0.953 | 0.97687 | 5.483 |
| Vdac1 | 19139.1 | -0.0845 | 0.05775 | -1.46314 | 0.14343 | 0.32561 | 28.538 |
| Vdac2 | 7131.17 | -0.04177 | 0.05597 | -0.74619 | 0.45555 | 0.65746 | 20.642 |
| Vdac3 | 3975.88 | -0.04798 | 0.08331 | -0.57596 | 0.56464 | 0.74496 | 16.739 |
| Vdac3-ps1 | 596.008 | -0.0446 | 0.09433 | -0.47283 | 0.63633 | 0.79513 | 0.853 |
| Vegfa | 4075.63 | -0.24511 | 0.14559 | -1.68355 | 0.09227 | 0.24738 | 15.377 |
| Vegfb | 1646.56 | -0.33116 | 0.09566 | -3.46174 | 0.00054 | 0.00834 | 5.179 |
| Vegfc | 102.129 | -0.96454 | 0.24497 | -3.93736 | 8.24E-05 | 0.00235 | 109.491 |
| Vegfd | 50.6695 | -0.8821 | 0.3171 | -2.78177 | 0.00541 | 0.03825 | 29.273 |
| Vezf1 | 2213.34 | -0.04414 | 0.07556 | -0.58417 | 0.55911 | 0.74054 | 16.451 |
| Vezt | 4087.03 | 0.096223 | 0.06378 | 1.508642 | 0.13139 | 0.31002 | 96.653 |
| Vgf | 10312.4 | -0.1288 | 0.09811 | -1.31275 | 0.18927 | 0.38518 | 6.96 |
| Vgll3 | 40.5946 | 0.078979 | 0.34719 | 0.22748 | 0.82005 | 0.90734 | 51.112 |
| Vgll4 | 1168.15 | -0.2853 | 0.07729 | -3.6911 | 0.00022 | 0.00459 | 109.367 |
| Vhl | 2793.38 | -0.07506 | 0.09927 | -0.75604 | 0.44963 | 0.65275 | 7.675 |
| Vil1 | 35.9322 | -0.72057 | 0.30727 | -2.34503 | 0.01903 | 0.0866 | 26.184 |
| Vill | 252.044 | 0.130621 | 0.17811 | 0.733368 | 0.46333 | 0.66317 | 18.748 |
| Vim | 1828.75 | -0.91657 | 0.34906 | -2.62581 | 0.00864 | 0.05209 | 8.9 |
| Vinac1 | 84.9783 | -0.02471 | 0.24843 | -0.09945 | 0.92078 | 0.95978 | 23.1 |
| Vip | 708.288 | -1.11681 | 0.38964 | -2.86628 | 0.00415 | 0.03226 | 8.4 |
| Vipas39 | 2377.78 | -0.03533 | 0.05669 | -0.62327 | 0.5331 | 0.72037 | 27.389 |
| Vipr1 | 1453.81 | 0.018303 | 0.09008 | 0.203178 | 0.839 | 0.91715 | 30.239 |
| Virma | 2882.07 | 0.171612 | 0.06695 | 2.5634 | 0.01037 | 0.05879 | 64.727 |

| | | | | | | | |
|---|---|---|---|---|---|---|---|
| Vit | 107.116 | -0.19044 | 0.22972 | -0.82899 | 0.40711 | 0.61507 | 119.345 |
| Vkorc1 | 20.8041 | -0.14556 | 0.40742 | -0.35727 | 0.72089 | 0.85098 | 2.555 |
| Vkorc1l1 | 3542.76 | -0.00976 | 0.06598 | -0.14799 | 0.88235 | 0.94157 | 44.723 |
| Vldlr | 6308.38 | 0.219082 | 0.10771 | 2.034086 | 0.04194 | 0.1472 | 37.748 |
| Vma21 | 3195.75 | 0.044368 | 0.06901 | 0.64291 | 0.52028 | 0.70963 | 23.834 |
| Vmac | 436.808 | 0.125893 | 0.11402 | 1.104095 | 0.26955 | 0.47916 | 4.087 |
| Vmn1r-ps109 | 32.4505 | 1.090928 | 0.52824 | 2.065204 | 0.0389 | 0.14059 | 0.725 |
| Vmn1r-ps110 | 58.6649 | 0.998987 | 0.36103 | 2.767054 | 0.00566 | 0.03958 | 1.094 |
| Vmn1r206 | 209.509 | 1.091892 | 0.25204 | 4.332213 | 1.48E-05 | 0.00073 | 6.796 |
| Vmn1r90 | 15.2387 | 0.577874 | 0.42669 | 1.354319 | 0.17563 | 0.36746 | 8.033 |
| Vmn2r1 | 64.2161 | -0.62135 | 0.28501 | -2.18009 | 0.02925 | 0.11605 | 27.938 |
| Vmn2r27 | 13.5951 | 0.97633 | 0.58529 | 1.668104 | 0.0953 | 0.25227 | 42.101 |
| Vmn2r29 | 37.9066 | 0.144043 | 0.3426 | 0.420438 | 0.67417 | 0.82111 | 16.002 |
| Vmn2r57 | 101.173 | 0.466455 | 0.18667 | 2.49882 | 0.01246 | 0.06628 | 52.752 |
| Vmn2r84 | 70.8159 | -0.17126 | 0.30794 | -0.55613 | 0.57812 | 0.75492 | 16.408 |
| Vmn2r85 | 30.5731 | -0.39888 | 0.37176 | -1.07294 | 0.2833 | 0.4934 | 15.64 |
| Vmn2r86 | 16.1704 | -0.43924 | 0.41768 | -1.05163 | 0.29297 | 0.50281 | 14.489 |
| Vmn2r87 | 64.7021 | 0.180532 | 0.2843 | 0.635006 | 0.52542 | 0.71368 | 26.345 |
| Vmp1 | 4280.6 | -0.21916 | 0.06434 | -3.40624 | 0.00066 | 0.00957 | 99.972 |
| Vopp1 | 4721.91 | -0.0829 | 0.08665 | -0.95672 | 0.33871 | 0.54953 | 97.353 |
| Vps11 | 3470.85 | 0.074553 | 0.06959 | 1.071288 | 0.28404 | 0.4942 | 13.922 |
| Vps13a | 3448.03 | 0.315778 | 0.16414 | 1.923778 | 0.05438 | 0.17534 | 165.568 |
| Vps13b | 3881.23 | 0.176963 | 0.07231 | 2.447338 | 0.01439 | 0.07245 | 560.07 |
| Vps13c | 6829.6 | 0.001693 | 0.08128 | 0.020829 | 0.98338 | 0.99092 | 155.243 |
| Vps13d | 8132.24 | 0.002502 | 0.07483 | 0.033438 | 0.97333 | 0.98671 | 222.384 |
| Vps16 | 2075.89 | -0.03519 | 0.08037 | -0.43791 | 0.66145 | 0.81272 | 19.931 |
| Vps18 | 2436.87 | 0.004741 | 0.06595 | 0.071886 | 0.94269 | 0.97138 | 9.714 |
| Vps25 | 24.8776 | 0.151066 | 0.37339 | 0.404582 | 0.68578 | 0.82885 | 5.843 |
| Vps26a | 2478.98 | -0.13507 | 0.07801 | -1.73158 | 0.08335 | 0.23071 | 31.571 |
| Vps26b | 5479.85 | -0.01031 | 0.07722 | -0.13356 | 0.89375 | 0.9468 | 22.295 |
| Vps26c | 1194.23 | -0.21355 | 0.08088 | -2.64044 | 0.00828 | 0.0507 | 29.048 |
| Vps28 | 1770.11 | -0.09386 | 0.0708 | -1.32578 | 0.18491 | 0.37966 | 4 |
| Vps29 | 4885.46 | 0.005229 | 0.05898 | 0.088668 | 0.92935 | 0.9644 | 10.616 |
| Vps33a | 5673.45 | -0.01018 | 0.06548 | -0.15551 | 0.87642 | 0.93824 | 44.38 |

| | | | | | | | |
|---|---|---|---|---|---|---|---|
| Vps33b | 969.674 | 0.086175 | 0.08843 | 0.974466 | 0.32983 | 0.54112 | 22.106 |
| Vps35 | 7522.07 | 0.073378 | 0.069 | 1.063394 | 0.2876 | 0.49771 | 39.411 |
| Vps35l | 2160.89 | -0.05461 | 0.0713 | -0.76593 | 0.44372 | 0.64799 | 102.741 |
| Vps36 | 1304.66 | -0.17567 | 0.0855 | -2.05453 | 0.03992 | 0.14278 | 28.035 |
| Vps37a | 5051.92 | 0.254982 | 0.10884 | 2.342804 | 0.01914 | 0.08693 | 38.975 |
| Vps37b | 640.008 | -0.2923 | 0.118 | -2.47724 | 0.01324 | 0.06894 | 27.63 |
| Vps37c | 820.422 | -0.06656 | 0.08484 | -0.78453 | 0.43273 | 0.63891 | 25.893 |
| Vps37d | 476.187 | -0.03941 | 0.11315 | -0.34829 | 0.72763 | 0.85462 | 5.367 |
| Vps39 | 4390.29 | 0.126341 | 0.06363 | 1.985433 | 0.0471 | 0.15923 | 36.677 |
| Vps41 | 6745.69 | 0.037251 | 0.06632 | 0.561687 | 0.57433 | 0.75253 | 149.526 |
| Vps45 | 1693.13 | 0.010414 | 0.08986 | 0.115893 | 0.90774 | 0.95368 | 58.635 |
| Vps4a | 3358.98 | -0.04812 | 0.07187 | -0.66948 | 0.50319 | 0.69554 | 14.534 |
| Vps4b | 2798.17 | 0.140851 | 0.0632 | 2.228826 | 0.02583 | 0.10639 | 32.658 |
| Vps50 | 3508.04 | 0.146399 | 0.08556 | 1.711151 | 0.08705 | 0.23758 | 105.15 |
| Vps51 | 1526.76 | 0.130084 | 0.08264 | 1.574155 | 0.11545 | 0.28498 | 9.39 |
| Vps52 | 3156.04 | 0.008553 | 0.05511 | 0.155187 | 0.87667 | 0.93831 | 11.224 |
| Vps53 | 2832.51 | 0.012683 | 0.07575 | 0.167436 | 0.86703 | 0.93254 | 133.422 |
| Vps54 | 2380.39 | 0.09278 | 0.09431 | 0.98376 | 0.32523 | 0.5365 | 81.856 |
| Vps72 | 1141.74 | 0.036261 | 0.07209 | 0.502981 | 0.61498 | 0.78094 | 12.03 |
| Vps8 | 2019.85 | -0.28791 | 0.09247 | -3.11352 | 0.00185 | 0.0191 | 221.563 |
| Vps9d1 | 2246.4 | 0.132577 | 0.08558 | 1.54913 | 0.12135 | 0.29465 | 11.993 |
| Vrk1 | 1684.22 | 0.630935 | 0.14776 | 4.270089 | 1.95E-05 | 0.00087 | 67.199 |
| Vrk2 | 45.7435 | 0.084963 | 0.31369 | 0.270852 | 0.7865 | 0.88899 | 122.678 |
| Vrk3 | 852.653 | -0.02084 | 0.12296 | -0.16951 | 0.8654 | 0.93177 | 29.103 |
| Vsig10 | 92.066 | -0.80113 | 0.17833 | -4.4923 | 7.05E-06 | 0.00043 | 35.923 |
| Vsig10l | 125.394 | 0.127202 | 0.19619 | 0.648377 | 0.51674 | 0.70669 | 8.869 |
| Vsig2 | 479.964 | -0.299 | 0.18563 | -1.61067 | 0.10725 | 0.27181 | 4.951 |
| Vsig8 | 20.1287 | 0.513748 | 0.37001 | 1.388457 | 0.165 | 0.354 | 7.78 |
| Vsir | 1436.14 | -0.32288 | 0.1439 | -2.2438 | 0.02485 | 0.10355 | 25.834 |
| Vsnl1 | 13511.2 | -0.57614 | 0.14411 | -3.99806 | 6.39E-05 | 0.00197 | 111.372 |
| Vstm2a | 4910.45 | 0.233928 | 0.09249 | 2.52914 | 0.01143 | 0.0626 | 169.587 |
| Vstm2b | 1869.86 | -0.53173 | 0.16626 | -3.19824 | 0.00138 | 0.01562 | 32.267 |
| Vstm2l | 2654.13 | 0.565223 | 0.17255 | 3.275636 | 0.00105 | 0.01328 | 30.067 |
| Vstm4 | 184.763 | -0.76383 | 0.23783 | -3.2117 | 0.00132 | 0.01513 | 82.736 |

| | | | | | | | |
|---|---|---|---|---|---|---|---|
| Vstm5 | 951.022 | 0.511503 | 0.16238 | 3.149949 | 0.00163 | 0.01761 | 20.372 |
| Vta1 | 1005.31 | -0.02677 | 0.09659 | -0.27718 | 0.78164 | 0.8863 | 50.806 |
| Vtcn1 | 17.0608 | 1.194606 | 0.46175 | 2.587126 | 0.00968 | 0.05614 | 71.464 |
| Vti1a | 2501.76 | -0.09629 | 0.06574 | -1.46473 | 0.14299 | 0.32506 | 311.015 |
| Vti1b | 4649.67 | 0.028395 | 0.07965 | 0.356508 | 0.72146 | 0.85135 | 16.651 |
| Vtn | 2207.84 | -0.72484 | 0.16751 | -4.32727 | 1.51E-05 | 0.00074 | 3.234 |
| Vwa1 | 1096.87 | -0.35567 | 0.11398 | -3.12053 | 0.00181 | 0.01883 | 6.55 |
| Vwa3a | 196.734 | -0.62682 | 0.24518 | -2.55652 | 0.01057 | 0.05968 | 66.425 |
| Vwa3b | 31.1102 | 0.046336 | 0.34593 | 0.133947 | 0.89344 | 0.94662 | 161.018 |
| Vwa5a | 801.274 | -0.51157 | 0.11229 | -4.55557 | 5.22E-06 | 0.00035 | 25.07 |
| Vwa5b1 | 68.6669 | 0.20151 | 0.36102 | 0.558164 | 0.57673 | 0.75408 | 70.525 |
| Vwa5b2 | 1832.39 | -0.44656 | 0.17078 | -2.61485 | 0.00893 | 0.05321 | 15.907 |
| Vwa7 | 63.0879 | 0.35471 | 0.29124 | 1.217939 | 0.22325 | 0.42649 | 10.163 |
| Vwa8 | 2458.45 | 0.020471 | 0.06488 | 0.315505 | 0.75238 | 0.86997 | 353.259 |
| Vwc2 | 282.999 | -0.17386 | 0.21855 | -0.79553 | 0.4263 | 0.63281 | 154.709 |
| Vwc2l | 403.713 | 0.598436 | 0.19915 | 3.004928 | 0.00266 | 0.0243 | 159.683 |
| Vwce | 51.6575 | 0.273137 | 0.32295 | 0.845751 | 0.39769 | 0.60672 | 32.001 |
| Vwf | 1219.78 | -0.74515 | 0.17652 | -4.22131 | 2.43E-05 | 0.001 | 139.906 |
| Vxn | 7306.31 | -0.90899 | 0.47798 | -1.90173 | 0.05721 | 0.18132 | 25.945 |
| Wac | 4994.84 | 0.062214 | 0.06085 | 1.022502 | 0.30654 | 0.5165 | 60.197 |
| Wapl | 3326.17 | 0.144147 | 0.08171 | 1.76408 | 0.07772 | 0.22065 | 74.056 |
| Wars | 3846.75 | -0.08873 | 0.06748 | -1.31482 | 0.18857 | 0.38427 | 34.145 |
| Wars2 | 502.228 | 0.139144 | 0.09416 | 1.477741 | 0.13948 | 0.32071 | 98.119 |
| Was | 75.9998 | -0.8358 | 0.27932 | -2.99221 | 0.00277 | 0.02495 | 9.046 |
| Wasf1 | 10076.9 | 0.041846 | 0.06854 | 0.610531 | 0.54151 | 0.72706 | 55.096 |
| Wasf2 | 1892.57 | -0.12317 | 0.10766 | -1.14405 | 0.2526 | 0.45994 | 69.252 |
| Wasf3 | 9481.04 | 0.010022 | 0.07504 | 0.133553 | 0.89376 | 0.9468 | 88.631 |
| Washc1 | 1403.67 | 0.077084 | 0.07933 | 0.971644 | 0.33123 | 0.54208 | 8.959 |
| Washc2 | 4875.3 | -0.14324 | 0.06182 | -2.31692 | 0.02051 | 0.09104 | 54.649 |
| Washc3 | 437.821 | -0.04983 | 0.10287 | -0.48442 | 0.62809 | 0.78956 | 45.066 |
| Washc4 | 2560.75 | 0.061145 | 0.12173 | 0.502305 | 0.61545 | 0.78121 | 52.722 |
| Washc5 | 2091.47 | 0.134476 | 0.07202 | 1.867276 | 0.06186 | 0.19038 | 42.171 |
| Wasl | 6834.03 | 0.008293 | 0.07123 | 0.116429 | 0.90731 | 0.95354 | 51.205 |
| Wbp1 | 1106.66 | -0.0441 | 0.07538 | -0.58503 | 0.55852 | 0.73991 | 2.516 |

| | | | | | | | |
|---|---|---|---|---|---|---|---|
| Wbp11 | 3608.07 | 0.027483 | 0.07193 | 0.382079 | 0.7024 | 0.83946 | 14.58 |
| Wbp1l | 2591.24 | -0.29162 | 0.07874 | -3.70354 | 0.00021 | 0.00444 | 58.306 |
| Wbp2 | 8933.42 | -0.08099 | 0.0681 | -1.1893 | 0.23432 | 0.43949 | 8.423 |
| Wbp4 | 2909.17 | 0.089162 | 0.08145 | 1.094684 | 0.27365 | 0.48313 | 21.584 |
| Wdcp | 955.173 | -0.04141 | 0.07615 | -0.54381 | 0.58657 | 0.76038 | 16.741 |
| Wdfy1 | 1244.23 | 0.08602 | 0.08745 | 0.983694 | 0.32527 | 0.53651 | 73.882 |
| Wdfy2 | 1672.25 | -0.12751 | 0.09682 | -1.31693 | 0.18786 | 0.38336 | 123.832 |
| Wdfy3 | 16067.3 | 0.207819 | 0.07448 | 2.790273 | 0.00527 | 0.0375 | 236.966 |
| Wdfy4 | 122.812 | -0.13804 | 0.20654 | -0.66833 | 0.50392 | 0.69622 | 225.962 |
| Wdhd1 | 398.156 | 0.170514 | 0.15534 | 1.097658 | 0.27235 | 0.48159 | 35.914 |
| Wdpcp | 222.013 | -0.0738 | 0.14589 | -0.50583 | 0.61298 | 0.77967 | 326.755 |
| Wdr1 | 4588.72 | -0.12566 | 0.08449 | -1.48715 | 0.13698 | 0.31768 | 36.409 |
| Wdr11 | 2624.06 | 0.056686 | 0.06842 | 0.828547 | 0.40736 | 0.61536 | 43.876 |
| Wdr12 | 1878.83 | 0.05685 | 0.06756 | 0.841466 | 0.40009 | 0.60888 | 28.861 |
| Wdr13 | 13213.7 | 0.038362 | 0.05679 | 0.675564 | 0.49932 | 0.69253 | 9.592 |
| Wdr17 | 2679.24 | 0.515558 | 0.13833 | 3.726949 | 0.00019 | 0.0042 | 258.13 |
| Wdr18 | 2368.04 | -0.01563 | 0.0744 | -0.2101 | 0.83359 | 0.91471 | 10.052 |
| Wdr19 | 1511.05 | 0.043076 | 0.08618 | 0.499827 | 0.6172 | 0.78211 | 60.72 |
| Wdr20 | 1376.41 | -0.07336 | 0.06932 | -1.05824 | 0.28995 | 0.50018 | 66.295 |
| Wdr24 | 1053.55 | -0.0836 | 0.08911 | -0.93812 | 0.34818 | 0.55865 | 5.104 |
| Wdr25 | 407.948 | 0.081397 | 0.11092 | 0.733863 | 0.46303 | 0.66305 | 134.822 |
| Wdr26 | 8119.34 | 0.0162 | 0.06433 | 0.251816 | 0.80118 | 0.8972 | 38.774 |
| Wdr27 | 148.241 | 0.674499 | 0.18521 | 3.641886 | 0.00027 | 0.00521 | 124.64 |
| Wdr3 | 906.734 | 0.124264 | 0.08608 | 1.443614 | 0.14885 | 0.33272 | 24.228 |
| Wdr31 | 383.713 | 0.187507 | 0.12288 | 1.525939 | 0.12703 | 0.30319 | 22.244 |
| Wdr33 | 3076.77 | 0.03095 | 0.05958 | 0.519475 | 0.60343 | 0.77292 | 103.308 |
| Wdr35 | 2577.25 | 0.240921 | 0.08034 | 2.998729 | 0.00271 | 0.02459 | 54.956 |
| Wdr36 | 1343.67 | 0.098902 | 0.07831 | 1.262906 | 0.20662 | 0.40604 | 30.37 |
| Wdr37 | 3879.16 | 0.040006 | 0.07365 | 0.543185 | 0.587 | 0.76058 | 68.942 |
| Wdr38 | 16.7294 | -0.59681 | 0.39899 | -1.4958 | 0.13471 | 0.31449 | 8.693 |
| Wdr4 | 1142.93 | 0.073179 | 0.08212 | 0.891118 | 0.37287 | 0.58335 | 25.659 |
| Wdr41 | 2034.95 | 0.115658 | 0.07692 | 1.503589 | 0.13269 | 0.31189 | 46.971 |
| Wdr43 | 1513.79 | 0.187817 | 0.07835 | 2.397172 | 0.01652 | 0.07941 | 43.137 |
| Wdr44 | 1288.4 | 0.297094 | 0.11335 | 2.621002 | 0.00877 | 0.0525 | 112.975 |

| | | | | | | | |
|---|---|---|---|---|---|---|---|
| Wdr45 | 456.636 | 0.040403 | 0.10958 | 0.36871 | 0.71234 | 0.84509 | 13.874 |
| Wdr45b | 2621.27 | 0.151592 | 0.09059 | 1.673378 | 0.09425 | 0.25054 | 27.222 |
| Wdr46 | 512.789 | 0.034613 | 0.1033 | 0.335086 | 0.73756 | 0.86147 | 9.038 |
| Wdr47 | 6462.6 | 0.051948 | 0.07078 | 0.733913 | 0.463 | 0.66305 | 54.441 |
| Wdr48 | 3615.79 | 0.18359 | 0.0535 | 3.431582 | 0.0006 | 0.00896 | 31.71 |
| Wdr5 | 1371.58 | 0.050725 | 0.06594 | 0.76923 | 0.44176 | 0.64634 | 21.379 |
| Wdr53 | 343.319 | -0.0903 | 0.13298 | -0.67904 | 0.49711 | 0.69069 | 9.857 |
| Wdr54 | 650.307 | -0.03305 | 0.12385 | -0.26687 | 0.78957 | 0.89082 | 7.038 |
| Wdr55 | 402.444 | -0.04488 | 0.12654 | -0.35469 | 0.72282 | 0.85222 | 3.591 |
| Wdr59 | 1685.8 | 0.17725 | 0.07973 | 2.223162 | 0.0262 | 0.10748 | 73.296 |
| Wdr5b | 284.301 | 0.012955 | 0.1072 | 0.120843 | 0.90382 | 0.95146 | 2.739 |
| Wdr6 | 5459.73 | 0.34743 | 0.14154 | 2.454598 | 0.0141 | 0.07154 | 6.429 |
| Wdr61 | 1433.59 | -0.02232 | 0.06935 | -0.32186 | 0.74756 | 0.86763 | 19.785 |
| Wdr62 | 71.4125 | -0.34787 | 0.21869 | -1.59069 | 0.11168 | 0.27927 | 40.282 |
| Wdr66 | 256.988 | 0.286396 | 0.20971 | 1.365704 | 0.17203 | 0.36271 | 75.383 |
| Wdr7 | 8484.26 | 0.000592 | 0.06027 | 0.009816 | 0.99217 | 0.99549 | 281.092 |
| Wdr70 | 966.352 | 0.033359 | 0.08246 | 0.404563 | 0.6858 | 0.82885 | 226.158 |
| Wdr73 | 1136.79 | 0.028754 | 0.0698 | 0.411928 | 0.68039 | 0.82503 | 10.547 |
| Wdr74 | 917.169 | -0.0539 | 0.0773 | -0.69722 | 0.48567 | 0.6818 | 4.798 |
| Wdr75 | 1606.55 | 0.198533 | 0.08224 | 2.414115 | 0.01577 | 0.07703 | 28.454 |
| Wdr76 | 256.276 | -0.32068 | 0.17228 | -1.86137 | 0.06269 | 0.19214 | 38.138 |
| Wdr77 | 1362.92 | 0.113581 | 0.07339 | 1.547604 | 0.12172 | 0.29516 | 10.669 |
| Wdr81 | 1642.01 | -0.14381 | 0.079 | -1.82048 | 0.06869 | 0.2038 | 13.774 |
| Wdr82 | 7299.68 | 0.043213 | 0.07207 | 0.599636 | 0.54875 | 0.73274 | 20.212 |
| Wdr83 | 918.577 | -0.11009 | 0.10262 | -1.07282 | 0.28335 | 0.49341 | 6.272 |
| Wdr83os | 184.646 | -0.00784 | 0.18125 | -0.04327 | 0.96549 | 0.9828 | 1.6 |
| Wdr86 | 28.5405 | 0.02514 | 0.47739 | 0.052661 | 0.958 | 0.9795 | 18.99 |
| Wdr89 | 118.326 | 0.321908 | 0.17062 | 1.886705 | 0.0592 | 0.1852 | 38.942 |
| Wdr90 | 1461.1 | 0.036151 | 0.13566 | 0.266472 | 0.78988 | 0.89088 | 16.731 |
| Wdr91 | 1042.65 | -0.09817 | 0.09623 | -1.02021 | 0.30763 | 0.51751 | 30.451 |
| Wdr95 | 89.3589 | -0.10044 | 0.20201 | -0.4972 | 0.61905 | 0.78314 | 83.216 |
| Wdsub1 | 434.62 | 0.269426 | 0.09519 | 2.830487 | 0.00465 | 0.03467 | 30.228 |
| Wdtc1 | 3682.8 | -0.05402 | 0.07216 | -0.74862 | 0.45409 | 0.65623 | 61.023 |
| Wdyhv1 | 586.71 | 0.013685 | 0.10654 | 0.128447 | 0.8978 | 0.94882 | 17.289 |

| | | | | | | | |
|---|---|---|---|---|---|---|---|
| Wee1 | 1374.13 | 0.130813 | 0.10809 | 1.210211 | 0.2262 | 0.43019 | 21.241 |
| Wfdc1 | 299.882 | -0.29798 | 0.16228 | -1.83616 | 0.06633 | 0.1992 | 21.858 |
| Wfdc18 | 19.4824 | -0.82171 | 0.41343 | -1.98752 | 0.04687 | 0.15876 | 2.334 |
| Wfdc3 | 57.4221 | 0.193172 | 0.29098 | 0.663879 | 0.50677 | 0.69802 | 14.752 |
| Wfikkn1 | 57.6781 | 0.383549 | 0.25774 | 1.488098 | 0.13673 | 0.31739 | 3.231 |
| Wfikkn2 | 89.4421 | -1.45077 | 0.33619 | -4.31533 | 1.59E-05 | 0.00077 | 10.05 |
| Wfs1 | 14500.9 | 1.017597 | 0.38813 | 2.621814 | 0.00875 | 0.05244 | 23.102 |
| Whamm | 617.434 | -0.27421 | 0.11094 | -2.4718 | 0.01344 | 0.06967 | 25.571 |
| Whrn | 1186.1 | -0.00208 | 0.10587 | -0.01969 | 0.98429 | 0.99141 | 81.082 |
| Wif1 | 39.7675 | -0.98737 | 0.2872 | -3.43789 | 0.00059 | 0.00882 | 66.691 |
| Wipf1 | 845.929 | -0.45205 | 0.1052 | -4.29691 | 1.73E-05 | 0.0008 | 100.125 |
| Wipf2 | 4753.47 | -0.07857 | 0.08314 | -0.94502 | 0.34465 | 0.55549 | 41.403 |
| Wipf3 | 12124.7 | -0.11184 | 0.06385 | -1.75151 | 0.07986 | 0.22462 | 74.166 |
| Wipi1 | 948.5 | -0.50885 | 0.08688 | -5.857 | 4.71E-09 | 1.80E-06 | 38.637 |
| Wipi2 | 6022.97 | -0.04753 | 0.07311 | -0.65013 | 0.51561 | 0.70584 | 42.891 |
| Wiz | 1987.6 | -0.05175 | 0.08435 | -0.61354 | 0.53952 | 0.72589 | 35.385 |
| Wls | 1687.71 | -0.52787 | 0.08198 | -6.43897 | 1.20E-10 | 1.17E-07 | 98.992 |
| Wnk1 | 15792.7 | -0.02514 | 0.07428 | -0.33847 | 0.73501 | 0.85987 | 114.704 |
| Wnk2 | 7547.61 | 0.020871 | 0.07285 | 0.286486 | 0.77451 | 0.88245 | 111.712 |
| Wnk3 | 2034.81 | 0.131421 | 0.07822 | 1.680059 | 0.09295 | 0.24829 | 122.075 |
| Wnk4 | 804.458 | 0.182465 | 0.23473 | 0.777332 | 0.43696 | 0.6426 | 16.843 |
| Wnt1 | 93.911 | 0.027645 | 0.21148 | 0.130722 | 0.896 | 0.94774 | 3.981 |
| Wnt10a | 85.5896 | -0.21553 | 0.22738 | -0.94788 | 0.34319 | 0.55393 | 12.664 |
| Wnt10b | 98.4062 | 0.27513 | 0.19147 | 1.43697 | 0.15073 | 0.33564 | 7.439 |
| Wnt11 | 21.7251 | -0.61252 | 0.47 | -1.30323 | 0.19249 | 0.38919 | 20.084 |
| Wnt16 | 46.4466 | 0.014967 | 0.39307 | 0.038078 | 0.96963 | 0.98465 | 10.296 |
| Wnt2 | 490.092 | 0.691746 | 0.18334 | 3.77297 | 0.00016 | 0.00372 | 41.646 |
| Wnt2b | 237.682 | 0.05807 | 0.13701 | 0.423836 | 0.67169 | 0.81937 | 16.65 |
| Wnt3 | 145.545 | -0.86282 | 0.16427 | -5.25243 | 1.50E-07 | 2.47E-05 | 43.808 |
| Wnt4 | 292.307 | -0.93593 | 0.2421 | -3.86589 | 0.00011 | 0.00287 | 22.238 |
| Wnt5a | 660.75 | -0.62175 | 0.15175 | -4.09717 | 4.18E-05 | 0.00144 | 22.699 |
| Wnt5b | 65.4823 | -0.725 | 0.24638 | -2.94264 | 0.00325 | 0.02753 | 111.817 |
| Wnt6 | 77.8067 | -1.08888 | 0.44597 | -2.44156 | 0.01462 | 0.07304 | 13.431 |
| Wnt7a | 770.725 | 0.177471 | 0.14068 | 1.261487 | 0.20713 | 0.40661 | 47.383 |

| | | | | | | | |
|---|---|---|---|---|---|---|---|
| Wnt7b | 1725.3 | -0.53244 | 0.3163 | -1.68335 | 0.09231 | 0.2474 | 47.037 |
| Wnt9a | 207.532 | -0.95347 | 0.49279 | -1.93486 | 0.05301 | 0.17223 | 26.625 |
| Wrap53 | 422.72 | -0.15164 | 0.13887 | -1.09196 | 0.27485 | 0.48444 | 18.498 |
| Wrap73 | 931.642 | 0.084594 | 0.07309 | 1.157387 | 0.24711 | 0.45392 | 25.049 |
| Wrn | 687.895 | 0.014325 | 0.12422 | 0.115319 | 0.90819 | 0.95401 | 151.144 |
| Wrnip1 | 1871.67 | 0.07525 | 0.06652 | 1.131194 | 0.25797 | 0.46605 | 20.572 |
| Wsb1 | 5677.06 | 0.052187 | 0.13705 | 0.380796 | 0.70335 | 0.83991 | 15.3 |
| Wsb2 | 7613.15 | -0.18871 | 0.07452 | -2.53217 | 0.01134 | 0.06222 | 21.298 |
| Wscd1 | 2314.08 | -0.47926 | 0.10937 | -4.38191 | 1.18E-05 | 0.00062 | 39.728 |
| Wscd2 | 4576.29 | 0.105543 | 0.07871 | 1.340828 | 0.17998 | 0.37318 | 89.588 |
| Wt1 | 14.9512 | 0.443259 | 0.61101 | 0.725459 | 0.46817 | 0.66717 | 47.088 |
| Wtap | 2624.82 | 0.08554 | 0.06596 | 1.296908 | 0.19466 | 0.39198 | 27.374 |
| Wtip | 273.527 | -0.05887 | 0.13953 | -0.42191 | 0.67309 | 0.82024 | 23.726 |
| Wwc1 | 4794.15 | -0.21868 | 0.09259 | -2.36193 | 0.01818 | 0.08428 | 142.128 |
| Wwc2 | 2429.57 | 0.04297 | 0.06344 | 0.677332 | 0.4982 | 0.69157 | 166.966 |
| Wwox | 733.176 | -0.19817 | 0.10508 | -1.88585 | 0.05932 | 0.18536 | 913.053 |
| Wwp1 | 2647.57 | -0.03926 | 0.12176 | -0.3224 | 0.74715 | 0.8675 | 100.691 |
| Wwp2 | 1575.85 | 0.064278 | 0.07863 | 0.81751 | 0.41364 | 0.62171 | 122.231 |
| Wwtr1 | 1094.97 | -0.1696 | 0.1024 | -1.65623 | 0.09767 | 0.25637 | 120.262 |
| Xab2 | 1684.47 | 0.059331 | 0.07659 | 0.774701 | 0.43852 | 0.64381 | 12.896 |
| Xaf1 | 247.934 | -0.35424 | 0.16523 | -2.14394 | 0.03204 | 0.12359 | 12.105 |
| Xbp1 | 3740.13 | 0.027973 | 0.10556 | 0.265011 | 0.791 | 0.89151 | 5.235 |
| Xdh | 1570.35 | -0.56165 | 0.13216 | -4.24987 | 2.14E-05 | 0.00092 | 66.302 |
| Xiap | 7043.55 | 0.159941 | 0.06711 | 2.383403 | 0.01715 | 0.08147 | 49.978 |
| Xist | 19.0953 | 0.16935 | 0.59068 | 0.286702 | 0.77434 | 0.88244 | 22.889 |
| Xk | 1111.01 | 0.0989 | 0.08002 | 1.236003 | 0.21646 | 0.41834 | 40.495 |
| Xkr4 | 1742.32 | -0.07198 | 0.13458 | -0.53486 | 0.59275 | 0.76521 | 465.598 |
| Xkr6 | 670.094 | 0.390281 | 0.20203 | 1.931771 | 0.05339 | 0.1731 | 214.307 |
| Xkr7 | 218.581 | -0.20022 | 0.15137 | -1.32271 | 0.18593 | 0.38075 | 23.924 |
| Xkr8 | 473.39 | 0.537078 | 0.20386 | 2.634515 | 0.00843 | 0.05116 | 7.65 |
| Xkrx | 166.921 | 0.315345 | 0.1865 | 1.690846 | 0.09087 | 0.24474 | 13.034 |
| Xlr | 31.114 | -0.62344 | 0.30772 | -2.026 | 0.04276 | 0.14905 | 20.589 |
| Xlr3a | 64.2753 | -0.18232 | 0.39562 | -0.46086 | 0.6449 | 0.802 | 10.803 |
| Xlr3b | 265.062 | -0.53425 | 0.4809 | -1.11092 | 0.2666 | 0.47589 | 10.724 |

| | | | | | | | |
|---|---|---|---|---|---|---|---|
| Xlr4a | 25.5771 | 0.397926 | 0.64511 | 0.616838 | 0.53734 | 0.72408 | 8.134 |
| Xlr4b | 49.8377 | -0.42997 | 0.56397 | -0.7624 | 0.44582 | 0.64963 | 8.121 |
| Xndc1 | 389.008 | 0.354369 | 0.10284 | 3.445751 | 0.00057 | 0.00866 | 18.255 |
| Xntrpc | 13.6301 | 0.471563 | 0.48346 | 0.9754 | 0.32936 | 0.5406 | 31.152 |
| Xpa | 456.313 | 0.035988 | 0.11502 | 0.312892 | 0.75436 | 0.87077 | 40.965 |
| Xpc | 1377.4 | 0.289897 | 0.11361 | 2.551782 | 0.01072 | 0.06013 | 26.584 |
| Xpnpep1 | 1721.33 | -0.13183 | 0.07801 | -1.68981 | 0.09106 | 0.24511 | 108.289 |
| Xpnpep3 | 1096.44 | 0.052352 | 0.07763 | 0.674342 | 0.50009 | 0.69308 | 57.345 |
| Xpo1 | 5000.94 | 0.13842 | 0.08507 | 1.627216 | 0.10369 | 0.26622 | 42.209 |
| Xpo4 | 1326.02 | -0.02782 | 0.08016 | -0.34705 | 0.72856 | 0.85548 | 87.91 |
| Xpo5 | 3219.22 | 0.107093 | 0.0689 | 1.554333 | 0.12011 | 0.29271 | 40.817 |
| Xpo6 | 5069.37 | 0.055097 | 0.05659 | 0.97361 | 0.33025 | 0.5414 | 98.787 |
| Xpo7 | 5291.84 | 0.144624 | 0.08734 | 1.655957 | 0.09773 | 0.2564 | 112.383 |
| Xpot | 5176.17 | 0.124275 | 0.05774 | 2.152336 | 0.03137 | 0.12183 | 38.953 |
| Xpr1 | 10562.7 | 0.18177 | 0.0664 | 2.737684 | 0.00619 | 0.04207 | 141.715 |
| Xrcc1 | 1205.24 | 0.091302 | 0.08563 | 1.06622 | 0.28632 | 0.49634 | 27.152 |
| Xrcc2 | 371.67 | 0.001094 | 0.13402 | 0.008165 | 0.99349 | 0.99627 | 16.014 |
| Xrcc3 | 760.915 | 0.180677 | 0.1139 | 1.586267 | 0.11268 | 0.28066 | 10.682 |
| Xrcc4 | 331.181 | 0.229327 | 0.17619 | 1.301564 | 0.19307 | 0.38993 | 315.582 |
| Xrcc5 | 921.039 | 0.105568 | 0.08769 | 1.203923 | 0.22862 | 0.43266 | 87.526 |
| Xrcc6 | 942.706 | 0.155481 | 0.09213 | 1.687718 | 0.09147 | 0.24577 | 52.251 |
| Xrn1 | 2838.26 | 0.198793 | 0.06105 | 3.255996 | 0.00113 | 0.01378 | 103.044 |
| Xrn2 | 2574.16 | 0.165876 | 0.10493 | 1.5808 | 0.11392 | 0.28258 | 65.005 |
| Xrra1 | 64.03 | 0.027189 | 0.24108 | 0.11278 | 0.91021 | 0.95475 | 58.707 |
| Xxylt1 | 1465.54 | 0.250478 | 0.11415 | 2.194251 | 0.02822 | 0.11309 | 126.289 |
| Xylb | 393.393 | -0.05182 | 0.10695 | -0.48453 | 0.62801 | 0.78952 | 36.417 |
| Xylt1 | 3856.45 | -0.56413 | 0.19622 | -2.87499 | 0.00404 | 0.0317 | 292.59 |
| Xylt2 | 1035.36 | -0.11206 | 0.08126 | -1.37915 | 0.16785 | 0.35774 | 13.665 |
| Yae1d1 | 1391.45 | -0.09614 | 0.11468 | -0.83829 | 0.40187 | 0.6105 | 12.02 |
| Yaf2 | 3244.26 | -0.17176 | 0.06658 | -2.57963 | 0.00989 | 0.0571 | 52.658 |
| Yap1 | 1709.98 | -0.11202 | 0.11591 | -0.96647 | 0.33381 | 0.54457 | 72.598 |
| Yars | 4501.34 | -0.05836 | 0.05191 | -1.12416 | 0.26094 | 0.46951 | 29.848 |
| Yars2 | 621.342 | 0.085749 | 0.085 | 1.008809 | 0.31307 | 0.52351 | 6.676 |
| Ybey | 241.473 | -0.13871 | 0.15161 | -0.91496 | 0.36021 | 0.5706 | 9.548 |

| | | | | | | | |
|---|---|---|---|---|---|---|---|
| Ybx1 | 8486.1 | 0.089678 | 0.08385 | 1.069487 | 0.28485 | 0.49479 | 16.624 |
| Ybx2 | 259.845 | -0.39932 | 0.1804 | -2.21351 | 0.02686 | 0.10939 | 5.81 |
| Ybx3 | 1329.34 | -0.22957 | 0.09711 | -2.36405 | 0.01808 | 0.08384 | 23.622 |
| Ydjc | 505.161 | 0.079499 | 0.10689 | 0.743761 | 0.45702 | 0.65829 | 16.081 |
| Yeats2 | 2270.09 | -0.01636 | 0.10143 | -0.16132 | 0.87184 | 0.93524 | 91.511 |
| Yeats4 | 1166.21 | -0.12183 | 0.06926 | -1.759 | 0.07858 | 0.22215 | 9.287 |
| Yes1 | 982.726 | 0.182074 | 0.12547 | 1.451149 | 0.14674 | 0.32983 | 75.887 |
| Yif1a | 359.849 | -0.22635 | 0.1373 | -1.64865 | 0.09922 | 0.25882 | 4.344 |
| Yif1b | 535.575 | -0.07143 | 0.10623 | -0.6724 | 0.50133 | 0.69395 | 9.269 |
| Yipf1 | 1436.16 | 0.169748 | 0.10701 | 1.586247 | 0.11268 | 0.28066 | 45.461 |
| Yipf2 | 293.884 | -0.26149 | 0.11262 | -2.32184 | 0.02024 | 0.09025 | 4.147 |
| Yipf3 | 1804.67 | -0.09835 | 0.06895 | -1.42637 | 0.15376 | 0.3396 | 4.458 |
| Yipf4 | 2335.95 | 0.089739 | 0.07985 | 1.123887 | 0.26106 | 0.46964 | 10.785 |
| Yipf5 | 2109.99 | -0.03846 | 0.07907 | -0.48634 | 0.62672 | 0.78878 | 15.534 |
| Yipf6 | 2990.31 | 0.10179 | 0.06972 | 1.460015 | 0.14429 | 0.32701 | 12.702 |
| Yjefn3 | 549.194 | -0.12948 | 0.13321 | -0.97199 | 0.33105 | 0.54208 | 14.925 |
| Yju2 | 329.947 | -0.07801 | 0.12657 | -0.61634 | 0.53767 | 0.72444 | 9.187 |
| Ykt6 | 4167.64 | -0.12166 | 0.07236 | -1.68136 | 0.09269 | 0.24799 | 12.088 |
| Ylpm1 | 8497.13 | 0.222546 | 0.07714 | 2.885074 | 0.00391 | 0.03099 | 74.195 |
| Yme1l1 | 5219.91 | 0.203576 | 0.10015 | 2.032625 | 0.04209 | 0.14754 | 42.892 |
| Yod1 | 1271.7 | 0.086285 | 0.08918 | 0.967529 | 0.33328 | 0.54421 | 7.032 |
| Ypel1 | 916.94 | -0.22832 | 0.18098 | -1.26155 | 0.20711 | 0.4066 | 17.35 |
| Ypel2 | 6337.61 | 0.335705 | 0.12793 | 2.62412 | 0.00869 | 0.0523 | 57.283 |
| Ypel3 | 4189.04 | -0.1737 | 0.05812 | -2.98836 | 0.0028 | 0.02513 | 3.56 |
| Ypel4 | 2450.46 | -0.05981 | 0.11535 | -0.5185 | 0.60411 | 0.7733 | 4.598 |
| Ypel5 | 5634.58 | -0.14672 | 0.06339 | -2.31463 | 0.02063 | 0.09143 | 14.743 |
| Yrdc | 633.722 | 0.003385 | 0.11235 | 0.03013 | 0.97596 | 0.98762 | 4.564 |
| Ythdc1 | 4687.26 | 0.194503 | 0.07724 | 2.518293 | 0.01179 | 0.06387 | 32.439 |
| Ythdc2 | 2396.48 | 0.181071 | 0.08689 | 2.083881 | 0.03717 | 0.13625 | 61.979 |
| Ythdf1 | 2661.74 | -0.08958 | 0.06067 | -1.47653 | 0.1398 | 0.3211 | 16.573 |
| Ythdf2 | 1646.37 | -0.05017 | 0.07721 | -0.64985 | 0.51579 | 0.70601 | 27.392 |
| Ythdf3 | 5191.79 | 0.193712 | 0.08132 | 2.381995 | 0.01722 | 0.08153 | 33.826 |
| Ywhab | 22582.3 | -0.04255 | 0.06337 | -0.6715 | 0.5019 | 0.69441 | 23.629 |
| Ywhae | 31975.4 | -0.07217 | 0.0558 | -1.29339 | 0.19588 | 0.39361 | 32.977 |

| | | | | | | | |
|---|---|---|---|---|---|---|---|
| Ywhag | 37616.9 | -0.28333 | 0.09055 | -3.12898 | 0.00175 | 0.01853 | 26.208 |
| Ywhah | 24710.8 | -0.27621 | 0.08572 | -3.22212 | 0.00127 | 0.01481 | 9.151 |
| Ywhaq | 9592.8 | -0.00638 | 0.08928 | -0.07142 | 0.94307 | 0.97157 | 27.567 |
| Ywhaq-ps3 | 111.9 | -0.35312 | 0.16434 | -2.14874 | 0.03165 | 0.12253 | 0.738 |
| Ywhaz | 46379.9 | 0.080088 | 0.07768 | 1.030945 | 0.30257 | 0.51217 | 26.16 |
| Yy1 | 4752.05 | 0.026397 | 0.05559 | 0.474851 | 0.63489 | 0.79438 | 27.176 |
| Zadh2 | 2097.65 | 0.250728 | 0.12118 | 2.069066 | 0.03854 | 0.13979 | 9.466 |
| Zan | 20.4317 | 0.08172 | 0.38322 | 0.213244 | 0.83114 | 0.91334 | 98.428 |
| Zap70 | 65.9237 | -0.33465 | 0.22203 | -1.50724 | 0.13175 | 0.31054 | 21.021 |
| Zar1 | 15.4019 | 0.085827 | 0.60382 | 0.142141 | 0.88697 | 0.94383 | 4.449 |
| Zar1l | 88.4228 | 0.908276 | 0.30649 | 2.96348 | 0.00304 | 0.02634 | 11.11 |
| Zbbx | 27.9528 | -0.4561 | 0.41892 | -1.08875 | 0.27626 | 0.48546 | 127.128 |
| Zbed3 | 673.667 | -0.08271 | 0.11508 | -0.71869 | 0.47233 | 0.67066 | 14.23 |
| Zbed4 | 320.135 | 0.049226 | 0.11323 | 0.434745 | 0.66375 | 0.81417 | 32.857 |
| Zbed5 | 329.054 | 0.232212 | 0.1064 | 2.182504 | 0.02907 | 0.1155 | 7.901 |
| Zbed6 | 3285.15 | 0.20345 | 0.09976 | 2.039294 | 0.04142 | 0.146 | 41.379 |
| Zbp1 | 74.3133 | 0.724454 | 0.26738 | 2.709478 | 0.00674 | 0.04435 | 12.312 |
| Zbtb1 | 1085.37 | 0.108072 | 0.10242 | 1.055221 | 0.29132 | 0.50121 | 26.685 |
| Zbtb10 | 488.569 | 0.099677 | 0.12065 | 0.826196 | 0.40869 | 0.6165 | 34.732 |
| Zbtb11 | 2723.23 | 0.112135 | 0.0839 | 1.336577 | 0.18136 | 0.37482 | 35.031 |
| Zbtb11os1 | 54.5314 | -0.18257 | 0.25305 | -0.72149 | 0.47061 | 0.6693 | 1.35 |
| Zbtb12 | 281.926 | 0.145537 | 0.1138 | 1.278922 | 0.20092 | 0.39907 | 17.385 |
| Zbtb14 | 1195.35 | 0.019393 | 0.09002 | 0.215435 | 0.82943 | 0.91247 | 8.15 |
| Zbtb16 | 3216.54 | -0.40574 | 0.15235 | -2.66317 | 0.00774 | 0.04863 | 181.926 |
| Zbtb17 | 1649.39 | 0.045049 | 0.08535 | 0.5278 | 0.59764 | 0.76924 | 23.277 |
| Zbtb18 | 5980.46 | -0.25372 | 0.10379 | -2.44466 | 0.0145 | 0.07276 | 8.414 |
| Zbtb2 | 668.794 | 0.100907 | 0.08906 | 1.132966 | 0.25723 | 0.46521 | 21.034 |
| Zbtb20 | 18523.3 | 0.310283 | 0.11781 | 2.633694 | 0.00845 | 0.05121 | 766.722 |
| Zbtb21 | 1177.97 | 0.072088 | 0.0837 | 0.861245 | 0.3891 | 0.59824 | 19.266 |
| Zbtb22 | 925.034 | -0.1953 | 0.08217 | -2.37672 | 0.01747 | 0.08208 | 3.422 |
| Zbtb24 | 819.789 | 0.037523 | 0.08214 | 0.456798 | 0.64782 | 0.80388 | 15.192 |
| Zbtb25 | 518.109 | 0.158169 | 0.09565 | 1.653605 | 0.09821 | 0.25724 | 21.821 |
| Zbtb26 | 851.209 | 0.209414 | 0.10921 | 1.917588 | 0.05516 | 0.17708 | 10.968 |
| Zbtb3 | 203.675 | 0.189423 | 0.18013 | 1.051573 | 0.293 | 0.50281 | 2.292 |

| | | | | | | | |
|---|---|---|---|---|---|---|---|
| Zbtb32 | 84.7454 | 0.120973 | 0.21126 | 0.572617 | 0.5669 | 0.74698 | 9.229 |
| Zbtb33 | 1717.28 | 0.298383 | 0.07665 | 3.893022 | 9.90E-05 | 0.00267 | 7.254 |
| Zbtb34 | 1252.78 | -0.07542 | 0.08768 | -0.86025 | 0.38965 | 0.59887 | 25.217 |
| Zbtb37 | 1739.43 | -0.05331 | 0.09284 | -0.57427 | 0.56578 | 0.74583 | 31.928 |
| Zbtb38 | 3524.48 | 0.072652 | 0.07302 | 0.994987 | 0.31974 | 0.53041 | 70.062 |
| Zbtb39 | 653.884 | -0.03858 | 0.10707 | -0.36035 | 0.71859 | 0.84962 | 7.812 |
| Zbtb4 | 7941.23 | -0.05386 | 0.06118 | -0.88024 | 0.37873 | 0.58846 | 18.112 |
| Zbtb40 | 1249.84 | 0.115152 | 0.10664 | 1.079787 | 0.28024 | 0.49019 | 69.07 |
| Zbtb41 | 1701.08 | 0.164318 | 0.13517 | 1.215639 | 0.22412 | 0.42752 | 30.718 |
| Zbtb42 | 61.948 | -0.21522 | 0.2192 | -0.98183 | 0.32618 | 0.53752 | 3.92 |
| Zbtb43 | 1977.65 | 0.014115 | 0.0622 | 0.226935 | 0.82047 | 0.90734 | 18.273 |
| Zbtb44 | 1818.23 | 0.076411 | 0.07491 | 1.020085 | 0.30769 | 0.51754 | 45.242 |
| Zbtb45 | 934.854 | -0.06816 | 0.09823 | -0.69382 | 0.48779 | 0.68299 | 6.137 |
| Zbtb46 | 330.043 | -0.19218 | 0.11857 | -1.6208 | 0.10506 | 0.26817 | 71.665 |
| Zbtb48 | 608.509 | 0.106974 | 0.11378 | 0.940161 | 0.34714 | 0.55774 | 7.898 |
| Zbtb49 | 711.63 | 0.27879 | 0.10113 | 2.75683 | 0.00584 | 0.04045 | 30.723 |
| Zbtb5 | 667.176 | 0.227156 | 0.08029 | 2.829095 | 0.00467 | 0.03477 | 21.171 |
| Zbtb6 | 1399.62 | 0.141187 | 0.08834 | 1.598211 | 0.11 | 0.27633 | 17.672 |
| Zbtb7a | 6512.4 | -0.0465 | 0.06868 | -0.67708 | 0.49836 | 0.69171 | 17.776 |
| Zbtb7b | 946.143 | -0.54246 | 0.1234 | -4.39583 | 1.10E-05 | 0.0006 | 17.133 |
| Zbtb7c | 492.389 | 1.005545 | 0.2596 | 3.873482 | 0.00011 | 0.00281 | 328.387 |
| Zbtb8a | 188.709 | 0.115206 | 0.14529 | 0.792952 | 0.42781 | 0.6344 | 24.489 |
| Zbtb8b | 133.186 | -0.21104 | 0.24618 | -0.85727 | 0.39129 | 0.60039 | 15.089 |
| Zbtb8os | 769.091 | 0.146102 | 0.09798 | 1.491086 | 0.13594 | 0.31614 | 18.853 |
| Zbtb9 | 35.5443 | -0.06343 | 0.31664 | -0.20031 | 0.84124 | 0.91862 | 35.21 |
| Zc2hc1a | 2887.55 | 0.205023 | 0.07986 | 2.567372 | 0.01025 | 0.05844 | 50.354 |
| Zc2hc1c | 171.217 | -0.13572 | 0.17831 | -0.76116 | 0.44656 | 0.65015 | 25.073 |
| Zc3h10 | 291.363 | -0.15032 | 0.14582 | -1.03089 | 0.30259 | 0.51217 | 5.812 |
| Zc3h11a | 681.813 | 0.25187 | 0.10942 | 2.301842 | 0.02134 | 0.09352 | 41.519 |
| Zc3h12a | 58.6021 | -0.5546 | 0.30409 | -1.82379 | 0.06818 | 0.2027 | 9.418 |
| Zc3h12b | 1439.09 | 0.099006 | 0.08446 | 1.172173 | 0.24113 | 0.44728 | 220.96 |
| Zc3h12c | 1531.24 | 0.286855 | 0.08829 | 3.248846 | 0.00116 | 0.01402 | 57.229 |
| Zc3h13 | 5554.03 | 0.143606 | 0.08025 | 1.789461 | 0.07354 | 0.21355 | 60.054 |
| Zc3h14 | 2699.32 | 0.146485 | 0.08048 | 1.820209 | 0.06873 | 0.20383 | 40.79 |

| | | | | | | | |
|---|---|---|---|---|---|---|---|
| Zc3h15 | 4531.64 | 0.09514 | 0.09499 | 1.001592 | 0.31654 | 0.52711 | 20.188 |
| Zc3h18 | 2784.5 | 0.098823 | 0.06179 | 1.599395 | 0.10973 | 0.27577 | 40.752 |
| Zc3h3 | 725.539 | 0.137374 | 0.09042 | 1.519328 | 0.12868 | 0.30583 | 87.486 |
| Zc3h4 | 3660.89 | 0.039807 | 0.06254 | 0.636487 | 0.52446 | 0.71294 | 36.787 |
| Zc3h6 | 795.324 | 0.28658 | 0.13438 | 2.132553 | 0.03296 | 0.12614 | 51.162 |
| Zc3h7a | 3418.93 | 0.16471 | 0.07738 | 2.128547 | 0.03329 | 0.12668 | 39.802 |
| Zc3h7b | 9689.51 | -0.01948 | 0.07687 | -0.25346 | 0.79992 | 0.89665 | 51.204 |
| Zc3h8 | 387.947 | 0.022985 | 0.11529 | 0.199367 | 0.84198 | 0.91879 | 17.81 |
| Zc3hav1 | 320.898 | -0.45987 | 0.13236 | -3.47443 | 0.00051 | 0.00805 | 49.318 |
| Zc3hav1l | 515.748 | 0.288839 | 0.12066 | 2.393872 | 0.01667 | 0.0799 | 11.864 |
| Zc3hc1 | 764.276 | 0.083453 | 0.08522 | 0.979248 | 0.32746 | 0.53878 | 24.649 |
| Zc4h2 | 793.677 | -0.07827 | 0.09911 | -0.78967 | 0.42972 | 0.63603 | 19.317 |
| Zcchc10 | 356.122 | 0.1766 | 0.1116 | 1.582445 | 0.11355 | 0.28213 | 8.623 |
| Zcchc12 | 6398.57 | 0.593868 | 0.14226 | 4.174474 | 2.99E-05 | 0.00116 | 3.255 |
| Zcchc14 | 7279.94 | 0.31695 | 0.07711 | 4.110554 | 3.95E-05 | 0.00139 | 54.199 |
| Zcchc17 | 2238.16 | -0.16876 | 0.07915 | -2.13221 | 0.03299 | 0.12614 | 44.561 |
| Zcchc18 | 12970.7 | 0.226878 | 0.07749 | 2.927936 | 0.00341 | 0.02824 | 5.318 |
| Zcchc2 | 1516.29 | 0.061388 | 0.06958 | 0.882281 | 0.37762 | 0.58773 | 43.669 |
| Zcchc24 | 5051.6 | -0.46063 | 0.07383 | -6.23899 | 4.40E-10 | 2.81E-07 | 57.398 |
| Zcchc3 | 534.478 | 0.048055 | 0.08811 | 0.545387 | 0.58549 | 0.75994 | 3.073 |
| Zcchc4 | 750.508 | 0.189393 | 0.07951 | 2.382085 | 0.01721 | 0.08153 | 49.257 |
| Zcchc7 | 4662.66 | 0.343251 | 0.11641 | 2.94864 | 0.00319 | 0.02718 | 176.339 |
| Zcchc8 | 2406.8 | 0.14099 | 0.07156 | 1.970233 | 0.04881 | 0.1633 | 22.807 |
| Zcchc9 | 700.493 | 0.179645 | 0.08838 | 2.03273 | 0.04208 | 0.14754 | 11.173 |
| Zcrb1 | 2611.98 | 0.010152 | 0.07844 | 0.12941 | 0.89703 | 0.94835 | 12.238 |
| Zcwpw1 | 259.95 | 0.113284 | 0.15398 | 0.735705 | 0.46191 | 0.66232 | 34.824 |
| Zcwpw2 | 81.6863 | 0.099129 | 0.25792 | 0.384341 | 0.70073 | 0.83841 | 164.26 |
| Zdbf2 | 3429.17 | 0.74602 | 0.13899 | 5.367571 | 7.98E-08 | 1.48E-05 | 41.312 |
| Zdhhc1 | 2657.62 | 0.192683 | 0.07802 | 2.469541 | 0.01353 | 0.06999 | 24.48 |
| Zdhhc12 | 164.847 | -0.07785 | 0.17099 | -0.45528 | 0.64891 | 0.80429 | 2.705 |
| Zdhhc13 | 875.91 | 0.229964 | 0.08128 | 2.829379 | 0.00466 | 0.03475 | 38.438 |
| Zdhhc14 | 2353.94 | 0.301998 | 0.12239 | 2.467407 | 0.01361 | 0.07022 | 261.255 |
| Zdhhc15 | 612.99 | 0.340201 | 0.11288 | 3.013894 | 0.00258 | 0.02386 | 134.096 |
| Zdhhc16 | 767.61 | 0.048624 | 0.08663 | 0.561293 | 0.5746 | 0.75269 | 10.625 |

| | | | | | | | |
|---|---|---|---|---|---|---|---|
| Zdhhc17 | 8435.07 | 0.300623 | 0.07104 | 4.231824 | 2.32E-05 | 0.00097 | 68.361 |
| Zdhhc18 | 1915.54 | 0.211707 | 0.11351 | 1.865095 | 0.06217 | 0.19102 | 44.856 |
| Zdhhc2 | 2543.78 | 0.27453 | 0.07192 | 3.817359 | 0.00013 | 0.0033 | 86.454 |
| Zdhhc20 | 4132.3 | 0.189592 | 0.10567 | 1.794146 | 0.07279 | 0.21224 | 57.574 |
| Zdhhc21 | 3209.32 | 0.040935 | 0.08066 | 0.507524 | 0.61179 | 0.77879 | 61.221 |
| Zdhhc22 | 1194.78 | -0.27621 | 0.11823 | -2.33629 | 0.01948 | 0.08797 | 9.668 |
| Zdhhc23 | 2089.75 | 0.225881 | 0.10113 | 2.233658 | 0.02551 | 0.10538 | 14.759 |
| Zdhhc24 | 1084.35 | -0.17729 | 0.08111 | -2.18582 | 0.02883 | 0.11472 | 6.73 |
| Zdhhc3 | 3307.88 | -0.17242 | 0.06563 | -2.62727 | 0.00861 | 0.05191 | 47.046 |
| Zdhhc4 | 979.267 | 0.12336 | 0.07978 | 1.546179 | 0.12206 | 0.29547 | 12.768 |
| Zdhhc5 | 2521.44 | -0.15415 | 0.09428 | -1.63513 | 0.10202 | 0.26371 | 27.211 |
| Zdhhc6 | 1217.2 | 0.018824 | 0.08872 | 0.212174 | 0.83197 | 0.91393 | 44.742 |
| Zdhhc7 | 1370.8 | -0.05634 | 0.0801 | -0.70335 | 0.48184 | 0.67858 | 20.593 |
| Zdhhc8 | 3137.55 | -0.14797 | 0.07566 | -1.95566 | 0.0505 | 0.16713 | 14.384 |
| Zdhhc9 | 1683.99 | -0.14301 | 0.0998 | -1.43292 | 0.15188 | 0.33702 | 36.91 |
| Zeb1 | 5198.18 | 0.337825 | 0.10817 | 3.122957 | 0.00179 | 0.01874 | 183.608 |
| Zeb2 | 4571.17 | -0.72762 | 0.3146 | -2.31281 | 0.02073 | 0.09173 | 133.764 |
| Zeb2os | 21.8341 | -0.87736 | 0.36189 | -2.42439 | 0.01533 | 0.0755 | 3.381 |
| Zer1 | 9564.53 | 0.072228 | 0.04951 | 1.458992 | 0.14457 | 0.32724 | 27.303 |
| Zfand1 | 1244.24 | 0.141723 | 0.09567 | 1.481381 | 0.13851 | 0.31964 | 11.365 |
| Zfand2a | 2603.9 | -0.21854 | 0.07881 | -2.77306 | 0.00555 | 0.03901 | 13.339 |
| Zfand2b | 443.138 | -0.12372 | 0.09841 | -1.25718 | 0.20869 | 0.4086 | 2.984 |
| Zfand3 | 3776.71 | -0.13068 | 0.07721 | -1.69249 | 0.09055 | 0.24441 | 206.096 |
| Zfand4 | 383.959 | -0.3152 | 0.15351 | -2.05331 | 0.04004 | 0.14298 | 66.081 |
| Zfand5 | 7499.39 | 0.034216 | 0.05421 | 0.631131 | 0.52795 | 0.71583 | 14.563 |
| Zfand6 | 2317.4 | 0.027062 | 0.08913 | 0.303633 | 0.76141 | 0.87503 | 76.194 |
| Zfas1 | 707.707 | -0.18084 | 0.09214 | -1.96257 | 0.0497 | 0.16518 | 2.93 |
| Zfat | 455.294 | 0.341433 | 0.11408 | 2.992816 | 0.00276 | 0.02493 | 175.093 |
| Zfc3h1 | 4611.31 | 0.268767 | 0.14003 | 1.91939 | 0.05494 | 0.17651 | 47.814 |
| Zfhx2 | 5266.05 | 0.103017 | 0.08799 | 1.170732 | 0.24171 | 0.4477 | 32.063 |
| Zfhx3 | 910.499 | 0.339223 | 0.12381 | 2.739938 | 0.00615 | 0.04192 | 1018.987 |
| Zfhx4 | 1467.9 | -0.04194 | 0.12881 | -0.32558 | 0.74474 | 0.86574 | 197.332 |
| Zfp1 | 900.315 | 0.172399 | 0.0824 | 2.092322 | 0.03641 | 0.13444 | 27.611 |
| Zfp101 | 473.097 | -0.16215 | 0.159 | -1.0198 | 0.30782 | 0.51763 | 34.135 |

| | | | | | | | |
|---|---|---|---|---|---|---|---|
| Zfp105 | 141.129 | -0.29279 | 0.16852 | -1.73748 | 0.0823 | 0.22875 | 7.957 |
| Zfp106 | 14827.5 | -0.06897 | 0.06959 | -0.99102 | 0.32168 | 0.53253 | 57.024 |
| Zfp108 | 366.675 | 0.140523 | 0.10938 | 1.284777 | 0.19887 | 0.39678 | 7.652 |
| Zfp109 | 470.029 | 0.253097 | 0.11046 | 2.291377 | 0.02194 | 0.09536 | 9.989 |
| Zfp11 | 386.538 | 0.02713 | 0.12011 | 0.225876 | 0.8213 | 0.90759 | 15.507 |
| Zfp110 | 814.321 | -0.00277 | 0.08024 | -0.03451 | 0.97247 | 0.98616 | 15.817 |
| Zfp111 | 645.972 | 0.079597 | 0.09559 | 0.832656 | 0.40504 | 0.61387 | 17.497 |
| Zfp112 | 570.29 | 0.094747 | 0.09766 | 0.970193 | 0.33195 | 0.54288 | 15.639 |
| Zfp113 | 598.285 | 0.048758 | 0.11146 | 0.437451 | 0.66178 | 0.81293 | 16.043 |
| Zfp114 | 227.15 | 0.036693 | 0.12746 | 0.28789 | 0.77343 | 0.88189 | 8.129 |
| Zfp119a | 181.056 | 0.205928 | 0.22081 | 0.932618 | 0.35102 | 0.56158 | 14.039 |
| Zfp119b | 174.783 | 0.405423 | 0.17941 | 2.259757 | 0.02384 | 0.10049 | 11.12 |
| Zfp12 | 1935.75 | 0.02517 | 0.07355 | 0.342208 | 0.73219 | 0.85801 | 13.672 |
| Zfp120 | 878.257 | 0.471205 | 0.19092 | 2.468119 | 0.01358 | 0.07019 | 22.303 |
| Zfp125 | 227.877 | 0.315747 | 0.15448 | 2.043913 | 0.04096 | 0.14491 | 44.179 |
| Zfp128 | 335.066 | -0.00245 | 0.1266 | -0.01936 | 0.98455 | 0.9916 | 12.246 |
| Zfp13 | 667.073 | -0.04012 | 0.10049 | -0.39923 | 0.68973 | 0.83103 | 23.644 |
| Zfp131 | 2326.05 | 0.077023 | 0.07934 | 0.970835 | 0.33163 | 0.54262 | 25.703 |
| Zfp133-ps | 158.792 | 0.241057 | 0.1548 | 1.557203 | 0.11942 | 0.2916 | 40.988 |
| Zfp14 | 620.072 | 0.178822 | 0.09291 | 1.924644 | 0.05427 | 0.17519 | 15.022 |
| Zfp141 | 692.438 | 0.241138 | 0.10641 | 2.266083 | 0.02345 | 0.09933 | 32.355 |
| Zfp142 | 2068.01 | 0.100038 | 0.06855 | 1.459332 | 0.14447 | 0.32717 | 23.12 |
| Zfp143 | 568.42 | 0.036143 | 0.09474 | 0.381483 | 0.70284 | 0.83969 | 34.178 |
| Zfp146 | 1960.59 | 0.153153 | 0.07039 | 2.175732 | 0.02958 | 0.11702 | 8.482 |
| Zfp148 | 5664.46 | 0.140602 | 0.0831 | 1.691861 | 0.09067 | 0.24467 | 123.528 |
| Zfp157 | 1387.18 | 0.350072 | 0.18748 | 1.867268 | 0.06186 | 0.19038 | 19.227 |
| Zfp160 | 1176 | 0.246001 | 0.16779 | 1.466129 | 0.14261 | 0.32449 | 34.168 |
| Zfp169 | 1073.3 | 0.006847 | 0.0933 | 0.073386 | 0.9415 | 0.97087 | 25.805 |
| Zfp174 | 434.687 | 0.032494 | 0.15126 | 0.214817 | 0.82991 | 0.9128 | 11.613 |
| Zfp180 | 3568.29 | 0.086178 | 0.07903 | 1.090382 | 0.27554 | 0.4847 | 25.79 |
| Zfp182 | 800.325 | 0.314629 | 0.15861 | 1.983636 | 0.0473 | 0.15968 | 33.378 |
| Zfp184 | 101.872 | 0.036795 | 0.17554 | 0.209609 | 0.83397 | 0.9148 | 15.686 |
| Zfp185 | 50.1067 | -0.8518 | 0.40487 | -2.10391 | 0.03539 | 0.13213 | 44.205 |
| Zfp189 | 693.31 | 0.271943 | 0.14594 | 1.863391 | 0.06241 | 0.19158 | 10.342 |

| | | | | | | | |
|---|---|---|---|---|---|---|---|
| Zfp2 | 1319.68 | 0.33307 | 0.08856 | 3.760844 | 0.00017 | 0.00387 | 17.454 |
| Zfp202 | 201.826 | -0.00598 | 0.13584 | -0.04404 | 0.96487 | 0.98273 | 21.289 |
| Zfp207 | 8814.98 | 0.237239 | 0.06103 | 3.887467 | 0.0001 | 0.00271 | 22.455 |
| Zfp212 | 680.13 | -0.15255 | 0.10412 | -1.46522 | 0.14286 | 0.3249 | 12.164 |
| Zfp213 | 324.908 | 0.007405 | 0.12856 | 0.057602 | 0.95407 | 0.97747 | 7.472 |
| Zfp217 | 265.301 | -0.27584 | 0.16924 | -1.62988 | 0.10313 | 0.26535 | 39.461 |
| Zfp219 | 1808.25 | 0.050326 | 0.08674 | 0.580214 | 0.56177 | 0.74286 | 14.657 |
| Zfp229 | 197.092 | -0.07666 | 0.17986 | -0.4262 | 0.66996 | 0.81804 | 38.548 |
| Zfp235 | 970.14 | -0.00945 | 0.07625 | -0.12388 | 0.90141 | 0.95046 | 9.073 |
| Zfp236 | 4088.86 | 0.183621 | 0.07619 | 2.409908 | 0.01596 | 0.07759 | 99.291 |
| Zfp239 | 1884.75 | 0.174475 | 0.08879 | 1.965116 | 0.0494 | 0.1646 | 11.192 |
| Zfp24 | 2858.42 | 0.038281 | 0.06702 | 0.571224 | 0.56785 | 0.74779 | 11.109 |
| Zfp248 | 510.24 | 0.169634 | 0.10388 | 1.632959 | 0.10248 | 0.26428 | 28.21 |
| Zfp251 | 2387.4 | -0.10233 | 0.08281 | -1.23563 | 0.2166 | 0.41853 | 19.827 |
| Zfp26 | 1775.08 | 0.141114 | 0.08105 | 1.741006 | 0.08168 | 0.22777 | 31.714 |
| Zfp260 | 2802.39 | -0.02166 | 0.09899 | -0.21878 | 0.82682 | 0.91097 | 12.846 |
| Zfp263 | 1343.75 | 0.066098 | 0.0698 | 0.947035 | 0.34362 | 0.55429 | 6.698 |
| Zfp266 | 3024.94 | 0.104463 | 0.07403 | 1.411112 | 0.15821 | 0.34558 | 26.35 |
| Zfp267 | 1153.51 | 0.009515 | 0.12455 | 0.076393 | 0.93911 | 0.96951 | 19.326 |
| Zfp27 | 755.696 | -0.04908 | 0.08477 | -0.57895 | 0.56263 | 0.74341 | 13.24 |
| Zfp273 | 289.087 | 0.358231 | 0.18912 | 1.894157 | 0.0582 | 0.18329 | 13.263 |
| Zfp275 | 1922.09 | 0.016832 | 0.07287 | 0.230993 | 0.81732 | 0.90594 | 16.46 |
| Zfp276 | 898.56 | -0.08573 | 0.09211 | -0.93077 | 0.35197 | 0.56232 | 15.551 |
| Zfp277 | 835.627 | 0.307771 | 0.08665 | 3.551802 | 0.00038 | 0.00663 | 130.857 |
| Zfp28 | 592.01 | 0.198045 | 0.11667 | 1.697482 | 0.08961 | 0.24259 | 13.621 |
| Zfp280b | 1311.61 | 0.172764 | 0.09789 | 1.764898 | 0.07758 | 0.22041 | 10.585 |
| Zfp280c | 1612.82 | 0.093361 | 0.14425 | 0.647231 | 0.51748 | 0.70733 | 52.88 |
| Zfp280d | 1928.71 | 0.404319 | 0.12255 | 3.299153 | 0.00097 | 0.01254 | 88.918 |
| Zfp281 | 2662.11 | 0.184239 | 0.07723 | 2.38549 | 0.01706 | 0.08104 | 5.153 |
| Zfp282 | 868.999 | -0.09812 | 0.10903 | -0.89997 | 0.36814 | 0.57862 | 31.282 |
| Zfp286 | 820.914 | 0.160483 | 0.10054 | 1.596179 | 0.11045 | 0.27709 | 36.886 |
| Zfp287 | 1014.41 | 0.184215 | 0.08173 | 2.253872 | 0.0242 | 0.10162 | 31.55 |
| Zfp292 | 2637.38 | 0.421511 | 0.19379 | 2.175089 | 0.02962 | 0.11715 | 79.848 |
| Zfp296 | 22.2601 | -0.22369 | 0.55143 | -0.40566 | 0.68499 | 0.82841 | 3.37 |

| | | | | | | | |
|---|---|---|---|---|---|---|---|
| Zfp3 | 790.105 | 0.277774 | 0.09786 | 2.838352 | 0.00453 | 0.0341 | 8.72 |
| Zfp30 | 643.355 | 0.111695 | 0.08563 | 1.304374 | 0.19211 | 0.38879 | 10.705 |
| Zfp300 | 139.919 | -0.01807 | 0.24998 | -0.07228 | 0.94238 | 0.9712 | 10.354 |
| Zfp316 | 1404 | -0.15314 | 0.09318 | -1.64351 | 0.10028 | 0.26056 | 20.326 |
| Zfp317 | 1741.49 | 0.127743 | 0.08905 | 1.434481 | 0.15143 | 0.33637 | 27.63 |
| Zfp318 | 3650.02 | 0.203711 | 0.07432 | 2.741148 | 0.00612 | 0.04184 | 37.19 |
| Zfp319 | 1216.85 | 0.039193 | 0.07651 | 0.512278 | 0.60846 | 0.77621 | 8.94 |
| Zfp322a | 1708.95 | 0.141461 | 0.12988 | 1.08918 | 0.27607 | 0.48521 | 16.106 |
| Zfp324 | 566.115 | -0.04369 | 0.0877 | -0.49817 | 0.61836 | 0.7831 | 8.399 |
| Zfp326 | 2449.15 | 0.192592 | 0.07273 | 2.647969 | 0.0081 | 0.04993 | 39.254 |
| Zfp329 | 2697.42 | 0.308085 | 0.12535 | 2.457719 | 0.01398 | 0.07122 | 13.882 |
| Zfp330 | 1099.96 | -0.0208 | 0.08022 | -0.25934 | 0.79537 | 0.89446 | 10.554 |
| Zfp334 | 1977.72 | 0.156957 | 0.06861 | 2.287708 | 0.02215 | 0.09592 | 13.996 |
| Zfp335 | 1354.91 | 0.128425 | 0.06884 | 1.865433 | 0.06212 | 0.19095 | 19.876 |
| Zfp335os | 110.113 | 0.226694 | 0.20722 | 1.093971 | 0.27397 | 0.48339 | 8.605 |
| Zfp341 | 352.699 | 0.098549 | 0.11383 | 0.865728 | 0.38664 | 0.59598 | 33.525 |
| Zfp346 | 1712.11 | 0.038134 | 0.07302 | 0.522222 | 0.60152 | 0.77159 | 29.515 |
| Zfp35 | 938.633 | 0.056122 | 0.11942 | 0.469949 | 0.63839 | 0.79692 | 15.745 |
| Zfp354a | 474.926 | 0.201842 | 0.09777 | 2.064548 | 0.03897 | 0.14064 | 13.543 |
| Zfp354b | 195.14 | 0.448215 | 0.16759 | 2.674521 | 0.00748 | 0.04763 | 9.811 |
| Zfp354c | 2594.93 | 0.076998 | 0.08125 | 0.947678 | 0.34329 | 0.55397 | 16.639 |
| Zfp358 | 836.687 | 0.149161 | 0.11367 | 1.312263 | 0.18943 | 0.38536 | 4.071 |
| Zfp36 | 199.25 | -0.44923 | 0.2 | -2.24618 | 0.02469 | 0.1031 | 3.47 |
| Zfp362 | 1144.3 | 0.059514 | 0.07314 | 0.813707 | 0.41581 | 0.62362 | 32.958 |
| Zfp365 | 22701.5 | -0.25055 | 0.09354 | -2.67844 | 0.0074 | 0.04731 | 26.56 |
| Zfp366 | 480.744 | -0.04163 | 0.13399 | -0.31072 | 0.75601 | 0.87157 | 65.834 |
| Zfp367 | 739.372 | -0.39768 | 0.13002 | -3.05856 | 0.00222 | 0.02156 | 20.185 |
| Zfp369 | 2275.7 | 0.143249 | 0.08239 | 1.738673 | 0.08209 | 0.22838 | 25.408 |
| Zfp36l1 | 1381.39 | -0.41692 | 0.10843 | -3.84505 | 0.00012 | 0.00304 | 5.26 |
| Zfp36l2 | 1321.13 | -0.27063 | 0.10874 | -2.48881 | 0.01282 | 0.06759 | 4.017 |
| Zfp37 | 785.548 | 0.151064 | 0.16256 | 0.929268 | 0.35275 | 0.56307 | 41.508 |
| Zfp382 | 704.432 | 0.097693 | 0.09664 | 1.01092 | 0.31205 | 0.52246 | 13.009 |
| Zfp383 | 798.845 | 0.035732 | 0.08943 | 0.399531 | 0.6895 | 0.83086 | 8.591 |
| Zfp384 | 1975.76 | 0.085273 | 0.09423 | 0.90494 | 0.3655 | 0.57607 | 28.726 |

| | | | | | | | |
|---|---|---|---|---|---|---|---|
| Zfp385a | 2376.94 | -0.19494 | 0.08558 | -2.27798 | 0.02273 | 0.09747 | 26.199 |
| Zfp385b | 2272.64 | 0.394627 | 0.1386 | 2.847273 | 0.00441 | 0.03356 | 409.006 |
| Zfp385c | 35.0438 | -0.22611 | 0.31755 | -0.71205 | 0.47643 | 0.67441 | 64.913 |
| Zfp386 | 830.73 | 0.285998 | 0.14812 | 1.930845 | 0.0535 | 0.17336 | 15.637 |
| Zfp389 | 38.4546 | -0.36989 | 0.27178 | -1.36102 | 0.17351 | 0.3648 | 2.221 |
| Zfp39 | 330.052 | 0.180962 | 0.11187 | 1.617566 | 0.10576 | 0.26933 | 16.073 |
| Zfp395 | 426.774 | -0.11205 | 0.12547 | -0.8931 | 0.3718 | 0.58214 | 40.542 |
| Zfp397 | 2787.52 | 0.192954 | 0.10905 | 1.769465 | 0.07682 | 0.21921 | 9.982 |
| Zfp398 | 2127.63 | 0.079632 | 0.06858 | 1.161156 | 0.24558 | 0.4522 | 37.877 |
| Zfp40 | 990.252 | 0.509583 | 0.32407 | 1.572444 | 0.11585 | 0.2857 | 19.384 |
| Zfp407 | 868.628 | 0.140883 | 0.11575 | 1.217163 | 0.22354 | 0.42677 | 461.699 |
| Zfp408 | 664.482 | 0.114097 | 0.09559 | 1.193649 | 0.23262 | 0.4375 | 6.123 |
| Zfp41 | 382.29 | 0.005934 | 0.12707 | 0.0467 | 0.96275 | 0.98189 | 13.149 |
| Zfp410 | 1402.17 | -0.09928 | 0.09244 | -1.07399 | 0.28283 | 0.49308 | 26.872 |
| Zfp414 | 548.392 | 0.115932 | 0.12926 | 0.896871 | 0.36979 | 0.58023 | 2.69 |
| Zfp418 | 209.928 | 0.107766 | 0.15065 | 0.715324 | 0.47441 | 0.67249 | 12.233 |
| Zfp420 | 1104.16 | 0.190715 | 0.11921 | 1.599819 | 0.10964 | 0.2756 | 17.314 |
| Zfp422 | 1048.99 | 0.002399 | 0.07523 | 0.031892 | 0.97456 | 0.9873 | 4.984 |
| Zfp423 | 680.269 | -0.58436 | 0.12697 | -4.6025 | 4.17E-06 | 0.0003 | 297.786 |
| Zfp426 | 1301.86 | -0.12573 | 0.0854 | -1.47224 | 0.14096 | 0.32269 | 24.198 |
| Zfp428 | 523.535 | 0.07545 | 0.10111 | 0.746227 | 0.45553 | 0.65746 | 8.677 |
| Zfp429 | 78.7624 | -0.1744 | 0.28305 | -0.61615 | 0.5378 | 0.72455 | 11.915 |
| Zfp433 | 38.7306 | 0.363987 | 0.32538 | 1.118641 | 0.26329 | 0.47208 | 21.862 |
| Zfp438 | 230.254 | -0.18129 | 0.13464 | -1.34648 | 0.17815 | 0.37043 | 124.779 |
| Zfp442 | 166.46 | 0.455699 | 0.28537 | 1.59687 | 0.11029 | 0.27686 | 44.346 |
| Zfp444 | 1490.06 | 0.049212 | 0.06567 | 0.749341 | 0.45365 | 0.65597 | 21.586 |
| Zfp445 | 13398 | 0.260683 | 0.13127 | 1.985805 | 0.04706 | 0.15917 | 21.478 |
| Zfp446 | 966.728 | 0.352502 | 0.11683 | 3.01726 | 0.00255 | 0.02368 | 7.867 |
| Zfp449 | 364.128 | 0.304629 | 0.14387 | 2.117421 | 0.03422 | 0.12921 | 19.275 |
| Zfp451 | 3680.84 | 0.1159 | 0.06238 | 1.857912 | 0.06318 | 0.19309 | 53.051 |
| Zfp454 | 246.522 | 0.05673 | 0.13865 | 0.409163 | 0.68242 | 0.82653 | 14.932 |
| Zfp455 | 178.709 | 0.099052 | 0.16795 | 0.589777 | 0.55534 | 0.73781 | 14.793 |
| Zfp456 | 20.2137 | 0.355477 | 0.47917 | 0.741861 | 0.45817 | 0.65942 | 13.698 |
| Zfp457 | 38.54 | -0.46968 | 0.29987 | -1.56627 | 0.11729 | 0.28825 | 18.348 |

| | | | | | | | |
|---|---|---|---|---|---|---|---|
| Zfp458 | 278.379 | 0.157533 | 0.16126 | 0.976918 | 0.32861 | 0.53979 | 23.549 |
| Zfp459 | 45.8309 | 0.144624 | 0.25909 | 0.558201 | 0.57671 | 0.75408 | 15.71 |
| Zfp46 | 2363.88 | -0.23576 | 0.07556 | -3.12005 | 0.00181 | 0.01883 | 9.26 |
| Zfp462 | 5456.46 | 0.277293 | 0.08653 | 3.20473 | 0.00135 | 0.01537 | 138.516 |
| Zfp467 | 1502.63 | 0.271994 | 0.11349 | 2.396695 | 0.01654 | 0.07948 | 18.129 |
| Zfp469 | 326.638 | -0.42238 | 0.16765 | -2.51946 | 0.01175 | 0.06378 | 14.031 |
| Zfp472 | 229.271 | 0.092889 | 0.14094 | 0.659067 | 0.50985 | 0.70122 | 13.42 |
| Zfp473 | 50.4753 | -0.19936 | 0.28054 | -0.71064 | 0.47731 | 0.67519 | 19.571 |
| Zfp488 | 244.29 | -0.34455 | 0.17118 | -2.01278 | 0.04414 | 0.15229 | 11.695 |
| Zfp493 | 154.07 | 0.300432 | 0.2007 | 1.496925 | 0.13441 | 0.31425 | 12.82 |
| Zfp503 | 1298.99 | 0.646319 | 0.22461 | 2.877538 | 0.00401 | 0.03156 | 5.643 |
| Zfp507 | 1744.01 | 0.177655 | 0.07948 | 2.235089 | 0.02541 | 0.10513 | 30.661 |
| Zfp51 | 212.353 | 0.308205 | 0.28001 | 1.100677 | 0.27104 | 0.48038 | 15.214 |
| Zfp511 | 606.741 | 0.11244 | 0.10577 | 1.063034 | 0.28777 | 0.49786 | 4.212 |
| Zfp512 | 3039.41 | -0.01567 | 0.06065 | -0.25844 | 0.79607 | 0.89487 | 29.324 |
| Zfp512b | 4160.68 | 0.237594 | 0.08667 | 2.741224 | 0.00612 | 0.04184 | 10.701 |
| Zfp513 | 992.041 | 0.072487 | 0.08374 | 0.865662 | 0.38668 | 0.59599 | 3.323 |
| Zfp516 | 1803.7 | 0.457881 | 0.12562 | 3.644905 | 0.00027 | 0.00518 | 94.652 |
| Zfp518a | 562.586 | 0.126747 | 0.35044 | 0.361678 | 0.71759 | 0.84915 | 23.243 |
| Zfp518b | 889.544 | -0.04481 | 0.10686 | -0.41939 | 0.67493 | 0.82156 | 16.343 |
| Zfp52 | 259.703 | 0.084153 | 0.22704 | 0.370649 | 0.7109 | 0.84441 | 27.063 |
| Zfp521 | 579.439 | 0.249209 | 0.14461 | 1.723283 | 0.08484 | 0.23363 | 285.775 |
| Zfp523 | 4025.18 | -0.01565 | 0.09266 | -0.16895 | 0.86584 | 0.93204 | 28.758 |
| Zfp524 | 42.2292 | -0.30592 | 0.28318 | -1.08033 | 0.28 | 0.49005 | 4.074 |
| Zfp526 | 579.636 | 0.081289 | 0.11466 | 0.708965 | 0.47835 | 0.67605 | 6.083 |
| Zfp53 | 235.322 | 0.312493 | 0.18947 | 1.649266 | 0.09909 | 0.25857 | 21.476 |
| Zfp532 | 2044.87 | -0.01044 | 0.06611 | -0.15795 | 0.8745 | 0.93726 | 109.532 |
| Zfp536 | 2067.88 | -0.68341 | 0.16843 | -4.05755 | 4.96E-05 | 0.00163 | 455.618 |
| Zfp54 | 42.0313 | 0.145098 | 0.37404 | 0.387925 | 0.69807 | 0.83639 | 12.414 |
| Zfp541 | 24.9712 | -0.46987 | 0.35578 | -1.32065 | 0.18662 | 0.38158 | 34.521 |
| Zfp551 | 282.928 | 0.077893 | 0.13939 | 0.558814 | 0.57629 | 0.75395 | 7.599 |
| Zfp553 | 925.955 | -0.12478 | 0.08351 | -1.49415 | 0.13514 | 0.31502 | 5.119 |
| Zfp558 | 92.3239 | 0.246165 | 0.21356 | 1.152674 | 0.24904 | 0.45629 | 22.694 |
| Zfp560 | 489.813 | 0.18288 | 0.11702 | 1.562815 | 0.1181 | 0.28958 | 40.042 |

| | | | | | | | |
|---|---|---|---|---|---|---|---|
| Zfp563 | 1292.21 | 0.158401 | 0.10968 | 1.444242 | 0.14867 | 0.33248 | 21.396 |
| Zfp566 | 207.36 | 0.696486 | 0.21726 | 3.205837 | 0.00135 | 0.01536 | 13.191 |
| Zfp568 | 268.888 | 0.048945 | 0.13733 | 0.356408 | 0.72154 | 0.85138 | 44.328 |
| Zfp57 | 267.312 | -0.52438 | 0.22595 | -2.32077 | 0.0203 | 0.09035 | 9.473 |
| Zfp574 | 1705.45 | 0.049123 | 0.07615 | 0.645077 | 0.51888 | 0.70858 | 10.926 |
| Zfp575 | 199.103 | 0.03985 | 0.15153 | 0.262986 | 0.79256 | 0.89238 | 3.804 |
| Zfp579 | 921.962 | 0.151938 | 0.10985 | 1.383107 | 0.16663 | 0.35623 | 12.676 |
| Zfp58 | 168.224 | 0.005505 | 0.14651 | 0.037574 | 0.97003 | 0.98478 | 10.389 |
| Zfp580 | 465.197 | -0.19203 | 0.10993 | -1.7468 | 0.08067 | 0.22605 | 2.186 |
| Zfp583 | 318.147 | 0.172976 | 0.11606 | 1.490419 | 0.13611 | 0.31644 | 15.626 |
| Zfp59 | 360.413 | 0.266965 | 0.10562 | 2.527704 | 0.01148 | 0.06281 | 17.832 |
| Zfp592 | 1865.11 | -0.0489 | 0.06355 | -0.76942 | 0.44164 | 0.64627 | 51.484 |
| Zfp593 | 14.5338 | -0.62301 | 0.50508 | -1.23349 | 0.21739 | 0.4194 | 2.304 |
| Zfp595 | 174.03 | -0.03461 | 0.16715 | -0.20707 | 0.83596 | 0.91589 | 19.573 |
| Zfp597 | 1352.59 | 0.066633 | 0.08053 | 0.827431 | 0.40799 | 0.61577 | 26.241 |
| Zfp598 | 2795.42 | 0.129306 | 0.08253 | 1.566828 | 0.11715 | 0.28807 | 12.33 |
| Zfp599 | 244.218 | 0.458602 | 0.16493 | 2.780556 | 0.00543 | 0.03835 | 12.466 |
| Zfp60 | 2583.73 | 0.360794 | 0.09158 | 3.939857 | 8.15E-05 | 0.00233 | 22.314 |
| Zfp605 | 697.149 | 0.337358 | 0.1452 | 2.32335 | 0.02016 | 0.09 | 19.703 |
| Zfp606 | 744.666 | 0.239437 | 0.13029 | 1.837741 | 0.0661 | 0.19864 | 17.943 |
| Zfp607a | 85.5654 | -0.09116 | 0.18358 | -0.49659 | 0.61948 | 0.78345 | 23.299 |
| Zfp607b | 855.855 | 0.200389 | 0.07978 | 2.511816 | 0.01201 | 0.06468 | 17.145 |
| Zfp608 | 1135.67 | -0.30788 | 0.1509 | -2.04021 | 0.04133 | 0.14588 | 104.508 |
| Zfp609 | 3999.19 | 0.130469 | 0.06722 | 1.941054 | 0.05225 | 0.17077 | 135.174 |
| Zfp61 | 749.753 | 0.095398 | 0.10826 | 0.881188 | 0.37822 | 0.58818 | 10.194 |
| Zfp612 | 2558.4 | 0.177556 | 0.08644 | 2.053974 | 0.03998 | 0.14292 | 12.996 |
| Zfp617 | 1205.93 | 0.144923 | 0.09725 | 1.490284 | 0.13615 | 0.31645 | 25.995 |
| Zfp618 | 666.017 | -0.35115 | 0.13546 | -2.59225 | 0.00954 | 0.05558 | 174.136 |
| Zfp619 | 283.014 | 0.026384 | 0.13924 | 0.189485 | 0.84971 | 0.92352 | 22.655 |
| Zfp62 | 1912.16 | 0.248691 | 0.18575 | 1.338869 | 0.18061 | 0.37396 | 15.525 |
| Zfp622 | 1838.01 | -0.04467 | 0.07499 | -0.5957 | 0.55138 | 0.73445 | 14.116 |
| Zfp623 | 543.104 | -0.11821 | 0.09808 | -1.20525 | 0.22811 | 0.43202 | 8.449 |
| Zfp626 | 860.931 | 0.076319 | 0.09095 | 0.839084 | 0.40142 | 0.60999 | 15.721 |
| Zfp628 | 546.644 | 0.016002 | 0.12103 | 0.132217 | 0.89481 | 0.94736 | 6.786 |

| | | | | | | | |
|---|---|---|---|---|---|---|---|
| Zfp629 | 1965.36 | 0.033972 | 0.07008 | 0.484736 | 0.62786 | 0.78949 | 8.767 |
| Zfp637 | 800.879 | 0.047916 | 0.09996 | 0.479337 | 0.6317 | 0.79208 | 4.777 |
| Zfp638 | 6064.05 | 0.424585 | 0.18138 | 2.340891 | 0.01924 | 0.08714 | 122.442 |
| Zfp639 | 1605.16 | 0.224304 | 0.08502 | 2.638299 | 0.00833 | 0.05087 | 10.575 |
| Zfp64 | 643.685 | 0.06718 | 0.09227 | 0.728078 | 0.46657 | 0.66581 | 62.257 |
| Zfp641 | 642.878 | 0.131915 | 0.09755 | 1.352218 | 0.17631 | 0.36841 | 10.577 |
| Zfp644 | 3023.94 | 0.338222 | 0.11133 | 3.038003 | 0.00238 | 0.02256 | 80.549 |
| Zfp646 | 1835.55 | 0.045844 | 0.07362 | 0.622707 | 0.53348 | 0.72068 | 9.776 |
| Zfp647 | 94.235 | -0.01019 | 0.22039 | -0.04622 | 0.96314 | 0.98208 | 15.078 |
| Zfp648 | 18.7704 | 0.289529 | 0.50143 | 0.577411 | 0.56366 | 0.7442 | 4.503 |
| Zfp65 | 521.011 | 0.107435 | 0.16962 | 0.633388 | 0.52648 | 0.71461 | 23.865 |
| Zfp651 | 3878.67 | -0.1838 | 0.09442 | -1.94671 | 0.05157 | 0.16934 | 12.413 |
| Zfp652 | 4251.37 | 0.094094 | 0.06811 | 1.381444 | 0.16714 | 0.35694 | 122.443 |
| Zfp652os | 22.6359 | 0.214692 | 0.45009 | 0.477001 | 0.63336 | 0.79346 | 12.844 |
| Zfp653 | 702.568 | 0.132992 | 0.10002 | 1.329688 | 0.18362 | 0.37774 | 15.966 |
| Zfp654 | 1654.47 | 0.31711 | 0.15193 | 2.087201 | 0.03687 | 0.13549 | 71.306 |
| Zfp655 | 1433.1 | 0.059923 | 0.1124 | 0.53313 | 0.59394 | 0.76614 | 15.588 |
| Zfp658 | 440.393 | 0.12511 | 0.09618 | 1.300757 | 0.19334 | 0.39022 | 13.206 |
| Zfp660 | 139.954 | -0.10071 | 0.21207 | -0.47492 | 0.63485 | 0.79438 | 10.909 |
| Zfp661 | 454.222 | 0.030291 | 0.10722 | 0.282513 | 0.77755 | 0.88414 | 12.433 |
| Zfp664 | 7830.6 | 0.037853 | 0.0526 | 0.719648 | 0.47174 | 0.67032 | 40.003 |
| Zfp667 | 1134.58 | 0.39628 | 0.08847 | 4.479026 | 7.50E-06 | 0.00045 | 21.305 |
| Zfp668 | 1028.27 | 0.102964 | 0.08966 | 1.148342 | 0.25083 | 0.45821 | 13.782 |
| Zfp672 | 1698.64 | 0.122297 | 0.06762 | 1.808578 | 0.07052 | 0.20751 | 15.226 |
| Zfp677 | 98.2968 | 0.189308 | 0.32564 | 0.581347 | 0.56101 | 0.74209 | 15.518 |
| Zfp68 | 1612.66 | 0.276873 | 0.13359 | 2.072513 | 0.03822 | 0.139 | 16.11 |
| Zfp687 | 1560.04 | 0.126744 | 0.07268 | 1.743754 | 0.0812 | 0.2268 | 8.859 |
| Zfp688 | 302.924 | -0.1095 | 0.11612 | -0.94297 | 0.34569 | 0.55643 | 3.102 |
| Zfp689 | 202.196 | -0.12102 | 0.1315 | -0.92034 | 0.35739 | 0.56765 | 7.023 |
| Zfp69 | 70.0897 | -0.2954 | 0.23202 | -1.27318 | 0.20295 | 0.40161 | 21.563 |
| Zfp691 | 203.002 | 0.113303 | 0.12652 | 0.895544 | 0.3705 | 0.58072 | 4.683 |
| Zfp692 | 1987.2 | 0.093236 | 0.11918 | 0.782304 | 0.43404 | 0.63977 | 7.559 |
| Zfp697 | 903.699 | -0.30948 | 0.13724 | -2.25505 | 0.02413 | 0.10143 | 371.352 |
| Zfp7 | 279.731 | -0.1791 | 0.13813 | -1.29653 | 0.19479 | 0.39211 | 13.137 |

| | | | | | | | |
|---|---|---|---|---|---|---|---|
| Zfp703 | 1277.4 | -0.77378 | 0.17039 | -4.54123 | 5.59E-06 | 0.00036 | 4.137 |
| Zfp704 | 3262.8 | -0.08006 | 0.07971 | -1.00436 | 0.31521 | 0.52568 | 183.066 |
| Zfp706 | 9832.36 | 0.173637 | 0.08257 | 2.1028 | 0.03548 | 0.1323 | 10.747 |
| Zfp707 | 417.703 | -0.01353 | 0.12574 | -0.10764 | 0.91428 | 0.95656 | 6.746 |
| Zfp709 | 633.63 | 0.163685 | 0.12073 | 1.355745 | 0.17518 | 0.36685 | 13.709 |
| Zfp710 | 1324.87 | 0.095632 | 0.08153 | 1.172913 | 0.24083 | 0.44705 | 69.36 |
| Zfp711 | 629.341 | 0.180129 | 0.21752 | 0.828097 | 0.40762 | 0.61541 | 34.545 |
| Zfp712 | 299.124 | 0.124669 | 0.18044 | 0.690898 | 0.48963 | 0.68473 | 22.575 |
| Zfp715 | 1471.53 | 0.103718 | 0.11889 | 0.872367 | 0.38301 | 0.59222 | 17.098 |
| Zfp719 | 700.304 | -0.08676 | 0.13874 | -0.62539 | 0.53172 | 0.71934 | 13.626 |
| Zfp72 | 370.493 | 0.213881 | 0.13989 | 1.528912 | 0.12629 | 0.30205 | 21.511 |
| Zfp729a | 727.019 | 0.05721 | 0.1723 | 0.332028 | 0.73987 | 0.86278 | 25.029 |
| Zfp729b | 334.683 | 0.182708 | 0.24397 | 0.748891 | 0.45392 | 0.65613 | 20.269 |
| Zfp738 | 2028.17 | 0.174724 | 0.13358 | 1.308052 | 0.19086 | 0.3871 | 28.387 |
| Zfp74 | 789.887 | 0.183278 | 0.12566 | 1.458527 | 0.1447 | 0.32739 | 23.275 |
| Zfp740 | 5197.98 | 0.084308 | 0.07034 | 1.198544 | 0.23071 | 0.43503 | 12.358 |
| Zfp746 | 1550.98 | 0.055606 | 0.07986 | 0.696316 | 0.48623 | 0.68228 | 24.199 |
| Zfp747 | 234.377 | 0.039879 | 0.14463 | 0.275726 | 0.78276 | 0.88707 | 3.514 |
| Zfp748 | 475.764 | 0.134532 | 0.16223 | 0.829271 | 0.40695 | 0.61501 | 15.19 |
| Zfp750 | 20.5014 | 0.626237 | 0.55054 | 1.137506 | 0.25533 | 0.463 | 8.356 |
| Zfp758 | 508.282 | 0.624723 | 0.38928 | 1.604808 | 0.10854 | 0.27381 | 15.829 |
| Zfp759 | 324.779 | 0.331748 | 0.24659 | 1.345328 | 0.17852 | 0.37106 | 20.128 |
| Zfp760 | 654.072 | -0.03872 | 0.21287 | -0.18189 | 0.85567 | 0.92702 | 17.895 |
| Zfp763 | 951.547 | 0.095962 | 0.11792 | 0.81381 | 0.41575 | 0.62362 | 16.54 |
| Zfp764 | 165.902 | -0.00754 | 0.16152 | -0.04668 | 0.96277 | 0.98189 | 3.155 |
| Zfp768 | 321.356 | -0.21543 | 0.10962 | -1.96526 | 0.04938 | 0.16459 | 2.795 |
| Zfp770 | 1390.91 | 0.246112 | 0.10342 | 2.379774 | 0.01732 | 0.08165 | 8.009 |
| Zfp771 | 1183.28 | -0.13762 | 0.07905 | -1.74085 | 0.08171 | 0.22782 | 10.53 |
| Zfp772 | 798.949 | 0.06591 | 0.08961 | 0.735511 | 0.46203 | 0.66235 | 7.882 |
| Zfp773 | 176.293 | -0.25102 | 0.21685 | -1.15756 | 0.24704 | 0.45387 | 9.683 |
| Zfp775 | 301.604 | 0.057736 | 0.13747 | 0.419994 | 0.67449 | 0.82131 | 21.165 |
| Zfp777 | 1133.76 | 0.005634 | 0.08255 | 0.068255 | 0.94558 | 0.97296 | 24.724 |
| Zfp78 | 217.688 | 0.27962 | 0.12047 | 2.321067 | 0.02028 | 0.09035 | 19.326 |
| Zfp780b | 901.316 | 0.126429 | 0.12491 | 1.012121 | 0.31148 | 0.52177 | 20.037 |

| | | | | | | | |
|---|---|---|---|---|---|---|---|
| Zfp781 | 49.922 | 0.664628 | 0.2529 | 2.628076 | 0.00859 | 0.05182 | 28.57 |
| Zfp783 | 561.66 | -0.10903 | 0.11344 | -0.96115 | 0.33648 | 0.54716 | 12.781 |
| Zfp786 | 250.379 | 0.434831 | 0.15284 | 2.845008 | 0.00444 | 0.0337 | 11.602 |
| Zfp787 | 648.456 | -0.09668 | 0.10037 | -0.96324 | 0.33542 | 0.54624 | 24.507 |
| Zfp788 | 2255.54 | 0.371896 | 0.14685 | 2.532529 | 0.01132 | 0.06219 | 18.328 |
| Zfp790 | 870.838 | 0.103124 | 0.07425 | 1.388905 | 0.16486 | 0.35389 | 14.892 |
| Zfp791 | 187.519 | -0.09477 | 0.15659 | -0.60523 | 0.54502 | 0.73012 | 14.543 |
| Zfp799 | 1763.34 | 0.184774 | 0.07625 | 2.423183 | 0.01539 | 0.0757 | 14.813 |
| Zfp800 | 584.832 | 0.210982 | 0.12091 | 1.744963 | 0.08099 | 0.22645 | 158.079 |
| Zfp804a | 1745.05 | 0.230755 | 0.12482 | 1.848775 | 0.06449 | 0.19551 | 206.658 |
| Zfp804b | 29.896 | -0.07017 | 0.56798 | -0.12354 | 0.90168 | 0.95065 | 575.747 |
| Zfp808 | 76.7539 | 0.314695 | 0.23325 | 1.349158 | 0.17729 | 0.36947 | 106.259 |
| Zfp809 | 669.819 | 0.096746 | 0.10539 | 0.91795 | 0.35865 | 0.56888 | 17.641 |
| Zfp81 | 1641.17 | 0.248477 | 0.14835 | 1.674996 | 0.09394 | 0.24995 | 29.537 |
| Zfp810 | 960.584 | 0.277997 | 0.10223 | 2.719317 | 0.00654 | 0.04361 | 30.901 |
| Zfp811 | 699.796 | 0.016128 | 0.13274 | 0.121505 | 0.90329 | 0.95131 | 14.256 |
| Zfp82 | 54.8691 | 0.56433 | 0.29123 | 1.937758 | 0.05265 | 0.17169 | 18.412 |
| Zfp820 | 35.5854 | 0.438729 | 0.31039 | 1.413497 | 0.15751 | 0.34464 | 28.888 |
| Zfp821 | 1533.45 | 0.067587 | 0.06699 | 1.008974 | 0.31299 | 0.5235 | 19.387 |
| Zfp825 | 298.181 | 0.350476 | 0.15845 | 2.211871 | 0.02698 | 0.10969 | 13.939 |
| Zfp827 | 3470.51 | 0.333721 | 0.09976 | 3.345287 | 0.00082 | 0.01118 | 165.33 |
| Zfp830 | 485.521 | 0.027826 | 0.11248 | 0.247395 | 0.8046 | 0.89908 | 1.591 |
| Zfp831 | 2306.64 | 0.241406 | 0.07057 | 3.420845 | 0.00062 | 0.00922 | 67.299 |
| Zfp839 | 739.679 | 0.062006 | 0.11091 | 0.559066 | 0.57612 | 0.75395 | 19.744 |
| Zfp84 | 979.047 | 0.093022 | 0.09113 | 1.020806 | 0.30735 | 0.51717 | 11.27 |
| Zfp846 | 1664 | 0.127432 | 0.10277 | 1.24001 | 0.21497 | 0.41637 | 24.119 |
| Zfp85 | 148.376 | 0.282953 | 0.22075 | 1.281788 | 0.19992 | 0.39801 | 7.429 |
| Zfp850 | 398.87 | 0.272845 | 0.12814 | 2.12922 | 0.03324 | 0.12661 | 29.262 |
| Zfp853 | 132.15 | -0.26115 | 0.2002 | -1.30446 | 0.19208 | 0.38879 | 6.586 |
| Zfp85os | 124.097 | 0.202303 | 0.19167 | 1.055456 | 0.29122 | 0.50117 | 27.633 |
| Zfp862-ps | 891.727 | 0.118884 | 0.1018 | 1.167792 | 0.24289 | 0.44905 | 30.496 |
| Zfp865 | 1713.8 | 0.058336 | 0.07409 | 0.787384 | 0.43106 | 0.63717 | 16.393 |
| Zfp866 | 1615.53 | 0.142117 | 0.07513 | 1.891511 | 0.05856 | 0.18403 | 29.853 |
| Zfp867 | 645.477 | 0.003554 | 0.10925 | 0.032527 | 0.97405 | 0.98698 | 11.278 |

| | | | | | | | |
|---|---|---|---|---|---|---|---|
| Zfp868 | 683.974 | 0.087303 | 0.09374 | 0.931364 | 0.35167 | 0.56204 | 14.692 |
| Zfp869 | 1752.1 | 0.028362 | 0.07238 | 0.391829 | 0.69518 | 0.83485 | 14.328 |
| Zfp87 | 485.216 | -0.03937 | 0.09544 | -0.41248 | 0.67999 | 0.82501 | 10.397 |
| Zfp870 | 501.792 | -0.07237 | 0.10232 | -0.70726 | 0.4794 | 0.67707 | 12.424 |
| Zfp871 | 9459.82 | 0.08985 | 0.07862 | 1.142822 | 0.25311 | 0.4605 | 22.792 |
| Zfp873 | 379.503 | 0.50935 | 0.16352 | 3.114824 | 0.00184 | 0.01906 | 16.623 |
| Zfp874a | 643.113 | 0.018264 | 0.11974 | 0.152534 | 0.87877 | 0.93928 | 27.076 |
| Zfp874b | 434.303 | 0.023325 | 0.11181 | 0.208613 | 0.83475 | 0.91511 | 12.746 |
| Zfp879 | 175.394 | 0.344572 | 0.15397 | 2.2379 | 0.02523 | 0.10456 | 9.522 |
| Zfp882 | 487.45 | 0.382104 | 0.1107 | 3.451752 | 0.00056 | 0.00853 | 7.747 |
| Zfp9 | 1070.94 | 0.001689 | 0.10688 | 0.015807 | 0.98739 | 0.99316 | 17.371 |
| Zfp90 | 750.715 | 0.265492 | 0.08309 | 3.195093 | 0.0014 | 0.01578 | 11.272 |
| Zfp91 | 5543.25 | -0.05541 | 0.07672 | -0.72226 | 0.47014 | 0.66894 | 29.107 |
| Zfp92 | 337.513 | 0.430825 | 0.12819 | 3.360861 | 0.00078 | 0.01076 | 17.29 |
| Zfp93 | 482.153 | 0.025863 | 0.12517 | 0.206618 | 0.83631 | 0.91603 | 7.375 |
| Zfp930 | 622.186 | 0.520418 | 0.34706 | 1.499507 | 0.13374 | 0.31339 | 21.499 |
| Zfp931 | 572.319 | 0.158425 | 0.12233 | 1.295091 | 0.19529 | 0.39285 | 10.782 |
| Zfp932 | 2016.4 | 0.468361 | 0.18885 | 2.480084 | 0.01314 | 0.06864 | 13.891 |
| Zfp933 | 1434.07 | 0.326411 | 0.13252 | 2.463058 | 0.01378 | 0.07057 | 25.381 |
| Zfp934 | 204.593 | 0.392697 | 0.17465 | 2.248437 | 0.02455 | 0.10262 | 91.373 |
| Zfp935 | 137.073 | 0.479563 | 0.15652 | 3.063875 | 0.00218 | 0.02131 | 13.817 |
| Zfp937 | 201.61 | 0.588059 | 0.19627 | 2.996129 | 0.00273 | 0.02476 | 22.653 |
| Zfp938 | 917.994 | 0.216062 | 0.14634 | 1.476437 | 0.13983 | 0.32112 | 16.431 |
| Zfp939 | 408.256 | 0.224241 | 0.1047 | 2.141735 | 0.03221 | 0.12406 | 28.236 |
| Zfp94 | 423.977 | 0.024615 | 0.10075 | 0.244325 | 0.80698 | 0.90065 | 14.963 |
| Zfp940 | 844.871 | 0.104018 | 0.0945 | 1.100699 | 0.27103 | 0.48038 | 20.05 |
| Zfp941 | 1324.39 | 0.260394 | 0.1055 | 2.468094 | 0.01358 | 0.07019 | 14.73 |
| Zfp942 | 954.204 | 0.374168 | 0.35448 | 1.055527 | 0.29118 | 0.50117 | 35.505 |
| Zfp943 | 1081.08 | 0.280226 | 0.19316 | 1.45073 | 0.14686 | 0.32996 | 108.617 |
| Zfp944 | 721.044 | 0.433308 | 0.21283 | 2.035919 | 0.04176 | 0.14701 | 23.412 |
| Zfp945 | 829.641 | 0.17587 | 0.20049 | 0.877204 | 0.38038 | 0.58999 | 20.438 |
| Zfp946 | 922.38 | 0.425274 | 0.37111 | 1.145948 | 0.25182 | 0.45913 | 32.468 |
| Zfp947 | 64.3537 | 1.102354 | 0.32257 | 3.417411 | 0.00063 | 0.00928 | 21.832 |
| Zfp948 | 567.143 | 0.231627 | 0.23587 | 0.982007 | 0.3261 | 0.53752 | 21.692 |

| | | | | | | | |
|---|---|---|---|---|---|---|---|
| Zfp949 | 1314.52 | 0.274402 | 0.10339 | 2.653926 | 0.00796 | 0.04943 | 23.042 |
| Zfp950 | 4725.66 | 0.601471 | 0.34067 | 1.765549 | 0.07747 | 0.22019 | 87.001 |
| Zfp951 | 97.2704 | 0.183372 | 0.32219 | 0.569147 | 0.56926 | 0.74887 | 46.902 |
| Zfp952 | 1046.42 | 0.290454 | 0.09499 | 3.057845 | 0.00223 | 0.02159 | 12.329 |
| Zfp953 | 52.5799 | -0.05372 | 0.32677 | -0.1644 | 0.86942 | 0.93378 | 21.355 |
| Zfp954 | 390.37 | 0.067567 | 0.12126 | 0.557196 | 0.57739 | 0.75448 | 6.799 |
| Zfp955a | 482.357 | 0.148048 | 0.10468 | 1.414304 | 0.15727 | 0.34442 | 15.428 |
| Zfp955b | 1549.45 | 0.223568 | 0.13583 | 1.645913 | 0.09978 | 0.2598 | 17.698 |
| Zfp956 | 795.345 | 0.066804 | 0.08252 | 0.80952 | 0.41822 | 0.6255 | 11.911 |
| Zfp958 | 368.309 | 0.25524 | 0.17787 | 1.435008 | 0.15128 | 0.33615 | 17.065 |
| Zfp959 | 387.26 | 0.524506 | 0.17078 | 3.071162 | 0.00213 | 0.02103 | 6.836 |
| Zfp960 | 68.1596 | 0.598093 | 0.36401 | 1.643059 | 0.10037 | 0.2607 | 25.516 |
| Zfp961 | 650.766 | 0.147376 | 0.12351 | 1.193263 | 0.23277 | 0.43754 | 42.755 |
| Zfp963 | 116.142 | -0.22379 | 0.20353 | -1.09952 | 0.27154 | 0.4809 | 8.332 |
| Zfp964 | 291.656 | 0.262115 | 0.13065 | 2.006216 | 0.04483 | 0.15385 | 12.504 |
| Zfp969 | 48.4892 | 1.369056 | 0.80403 | 1.702749 | 0.08861 | 0.24052 | 11.424 |
| Zfp97 | 105.548 | 0.780668 | 0.27425 | 2.846577 | 0.00442 | 0.03361 | 25.417 |
| Zfp970 | 585.512 | 0.650347 | 0.22443 | 2.897764 | 0.00376 | 0.03017 | 13.397 |
| Zfp971 | 503.981 | 0.317123 | 0.19339 | 1.639838 | 0.10104 | 0.26212 | 10.739 |
| Zfp972 | 22.4768 | 1.006945 | 0.44248 | 2.27566 | 0.02287 | 0.09783 | 22.274 |
| Zfp973 | 20.3308 | 0.141529 | 0.4582 | 0.30888 | 0.75741 | 0.87222 | 48.564 |
| Zfp974 | 533.204 | 0.23343 | 0.15651 | 1.491518 | 0.13583 | 0.31595 | 22.069 |
| Zfp975 | 302.547 | 0.79362 | 0.2893 | 2.743202 | 0.00608 | 0.04167 | 32.638 |
| Zfp976 | 187.622 | 0.521048 | 0.31433 | 1.657642 | 0.09739 | 0.25589 | 33.063 |
| Zfp977 | 44.3083 | 0.701217 | 0.38865 | 1.80426 | 0.07119 | 0.20889 | 12.765 |
| Zfp979 | 31.1659 | 0.580526 | 0.48031 | 1.208648 | 0.2268 | 0.43097 | 30.577 |
| Zfp982 | 46.837 | 0.606104 | 0.57498 | 1.054126 | 0.29183 | 0.50173 | 21.07 |
| Zfp983 | 731.049 | 0.079188 | 0.13211 | 0.599405 | 0.5489 | 0.73283 | 14.338 |
| Zfp984 | 103.794 | 0.091188 | 0.33172 | 0.274892 | 0.7834 | 0.8875 | 55.815 |
| Zfp991 | 43.9879 | 0.269722 | 0.43423 | 0.621143 | 0.53451 | 0.72159 | 48.542 |
| Zfp992 | 14.9374 | 0.070678 | 0.50505 | 0.139942 | 0.88871 | 0.9444 | 21.27 |
| Zfp994 | 371.675 | 0.334177 | 0.26541 | 1.259093 | 0.208 | 0.40767 | 28.352 |
| Zfp995 | 300.58 | 0.253374 | 0.21217 | 1.194186 | 0.23241 | 0.43733 | 30.357 |
| Zfpl1 | 539.217 | 0.074747 | 0.10061 | 0.742947 | 0.45751 | 0.65871 | 4.195 |

| | | | | | | | |
|---|---|---|---|---|---|---|---|
| Zfpm1 | 853.08 | -0.03621 | 0.10174 | -0.35592 | 0.7219 | 0.85157 | 55.111 |
| Zfpm2 | 244.516 | -0.00932 | 0.1521 | -0.0613 | 0.95112 | 0.97595 | 449.558 |
| Zfr | 15511.8 | 0.139837 | 0.04898 | 2.854708 | 0.00431 | 0.03311 | 67.853 |
| Zfr2 | 3247.49 | 0.136262 | 0.08635 | 1.577937 | 0.11458 | 0.28352 | 18.969 |
| Zfta | 2372.8 | -0.25197 | 0.10128 | -2.48782 | 0.01285 | 0.06768 | 8.28 |
| Zfx | 1561.64 | 0.187122 | 0.11137 | 1.680111 | 0.09294 | 0.24829 | 49.072 |
| Zfyve1 | 3146.05 | 0.104648 | 0.08151 | 1.283837 | 0.1992 | 0.39711 | 50.665 |
| Zfyve16 | 1288.68 | 0.039875 | 0.1081 | 0.368876 | 0.71222 | 0.84507 | 43.761 |
| Zfyve19 | 698.993 | 0.21169 | 0.10295 | 2.056246 | 0.03976 | 0.14241 | 8.433 |
| Zfyve21 | 725.697 | 0.076119 | 0.10865 | 0.70059 | 0.48356 | 0.67988 | 18.023 |
| Zfyve26 | 1821.71 | 0.12413 | 0.06942 | 1.788192 | 0.07375 | 0.21398 | 63.959 |
| Zfyve27 | 4304.2 | -0.03541 | 0.05504 | -0.6433 | 0.52003 | 0.70963 | 30.64 |
| Zfyve28 | 3086.63 | 0.237789 | 0.11473 | 2.072651 | 0.0382 | 0.13899 | 93.557 |
| Zfyve9 | 3965.27 | -0.01806 | 0.06566 | -0.27505 | 0.78328 | 0.88742 | 143.333 |
| Zgpat | 2988.23 | 0.098481 | 0.07886 | 1.248872 | 0.21171 | 0.41234 | 18.701 |
| Zgrf1 | 522.922 | 0.34548 | 0.11833 | 2.919596 | 0.0035 | 0.02878 | 64.535 |
| Zhx1 | 3404.04 | 0.145359 | 0.11033 | 1.317444 | 0.18769 | 0.38312 | 29.539 |
| Zhx2 | 875.641 | -0.1488 | 0.09858 | -1.50952 | 0.13117 | 0.30963 | 145.168 |
| Zhx3 | 5760.07 | -0.01533 | 0.06182 | -0.24795 | 0.80418 | 0.8989 | 124.291 |
| Zic1 | 2019.92 | -0.25949 | 0.14095 | -1.84109 | 0.06561 | 0.19767 | 7.753 |
| Zic2 | 516.481 | -0.73381 | 0.23191 | -3.16424 | 0.00155 | 0.01704 | 4.418 |
| Zic3 | 420.229 | -0.08653 | 0.12867 | -0.67254 | 0.50124 | 0.69395 | 19.037 |
| Zic4 | 291.226 | -0.46546 | 0.20955 | -2.22118 | 0.02634 | 0.10784 | 26.936 |
| Zic5 | 158.964 | -0.72431 | 0.25043 | -2.89222 | 0.00383 | 0.03053 | 8.883 |
| Zik1 | 1096.88 | 0.343927 | 0.0787 | 4.369907 | 1.24E-05 | 0.00065 | 8.166 |
| Zim1 | 673.971 | 0.256264 | 0.15124 | 1.694423 | 0.09018 | 0.24362 | 25.182 |
| Zkscan1 | 2458.86 | 0.025745 | 0.06849 | 0.375911 | 0.70698 | 0.84193 | 22.739 |
| Zkscan14 | 426.781 | -0.018 | 0.10611 | -0.16963 | 0.8653 | 0.93172 | 6.923 |
| Zkscan16 | 2184.6 | 0.559518 | 0.12328 | 4.538611 | 5.66E-06 | 0.00036 | 14.728 |
| Zkscan17 | 363.771 | -0.26245 | 0.12529 | -2.09471 | 0.0362 | 0.134 | 41.232 |
| Zkscan2 | 2358.52 | 0.270887 | 0.12776 | 2.120247 | 0.03399 | 0.12872 | 25.066 |
| Zkscan3 | 3611.51 | 0.130447 | 0.07521 | 1.734398 | 0.08285 | 0.22962 | 15.753 |
| Zkscan4 | 98.7513 | -0.00542 | 0.23255 | -0.02332 | 0.98139 | 0.99023 | 6.601 |
| Zkscan5 | 753.761 | -0.03977 | 0.09534 | -0.4171 | 0.6766 | 0.82256 | 17.189 |

| | | | | | | | |
|---|---|---|---|---|---|---|---|
| Zkscan6 | 605.801 | -0.05978 | 0.10213 | -0.58532 | 0.55833 | 0.73975 | 22.065 |
| Zkscan7 | 476.778 | 0.101648 | 0.11381 | 0.893133 | 0.37179 | 0.58214 | 12.934 |
| Zkscan8 | 2017.99 | 0.22425 | 0.08983 | 2.496311 | 0.01255 | 0.06658 | 17.899 |
| Zmat1 | 1162.91 | 0.481722 | 0.16848 | 2.859145 | 0.00425 | 0.03276 | 37.838 |
| Zmat2 | 6164.23 | -0.18804 | 0.06074 | -3.09578 | 0.00196 | 0.01988 | 5.791 |
| Zmat3 | 5496.7 | -0.03964 | 0.05483 | -0.72296 | 0.4697 | 0.66865 | 31.223 |
| Zmat4 | 1447.02 | -0.16537 | 0.128 | -1.29197 | 0.19637 | 0.39398 | 520.567 |
| Zmat5 | 240.829 | -0.00531 | 0.12128 | -0.0438 | 0.96506 | 0.98277 | 32.992 |
| Zmiz1 | 5462.79 | -0.36526 | 0.12668 | -2.88337 | 0.00393 | 0.0311 | 207.559 |
| Zmiz1os1 | 90.1157 | -0.27164 | 0.28171 | -0.96427 | 0.33491 | 0.54565 | 203.606 |
| Zmiz2 | 10729.3 | -0.19649 | 0.08211 | -2.39303 | 0.01671 | 0.07998 | 17.085 |
| Zmpste24 | 2066.76 | 0.017384 | 0.09066 | 0.191741 | 0.84795 | 0.92278 | 39.005 |
| Zmym1 | 1322.79 | 0.545541 | 0.10188 | 5.354506 | 8.58E-08 | 1.56E-05 | 14.059 |
| Zmym2 | 5074.09 | 0.152075 | 0.11327 | 1.342583 | 0.17941 | 0.37233 | 76.049 |
| Zmym3 | 9239.36 | 0.365542 | 0.12188 | 2.999146 | 0.00271 | 0.02457 | 16.466 |
| Zmym4 | 3408.87 | 0.117746 | 0.07602 | 1.548878 | 0.12141 | 0.29466 | 106.197 |
| Zmym5 | 2792.02 | 0.265031 | 0.10598 | 2.500784 | 0.01239 | 0.06605 | 21.132 |
| Zmym6 | 2190.62 | 0.221626 | 0.10984 | 2.017729 | 0.04362 | 0.15106 | 46.99 |
| Zmynd10 | 107.979 | 0.879405 | 0.21411 | 4.107291 | 4.00E-05 | 0.0014 | 4.022 |
| Zmynd11 | 11138.9 | 0.173859 | 0.07183 | 2.420573 | 0.0155 | 0.07608 | 80.498 |
| Zmynd12 | 21.2689 | 0.546213 | 0.4813 | 1.134862 | 0.25643 | 0.46439 | 44.631 |
| Zmynd15 | 45.1431 | -0.48883 | 0.35451 | -1.37887 | 0.16793 | 0.35782 | 6.77 |
| Zmynd19 | 1150.28 | -0.01527 | 0.08189 | -0.18648 | 0.85207 | 0.92477 | 12.284 |
| Zmynd8 | 9207.8 | 0.119253 | 0.07397 | 1.612272 | 0.1069 | 0.2713 | 114.865 |
| Znf41-ps | 35.3396 | 0.051314 | 0.34771 | 0.147577 | 0.88268 | 0.9416 | 4.706 |
| Znfx1 | 5116.89 | 0.073261 | 0.09348 | 0.783687 | 0.43322 | 0.63922 | 27.223 |
| Znhit1 | 926.623 | -0.03959 | 0.09417 | -0.42039 | 0.6742 | 0.82111 | 5.858 |
| Znhit2 | 48.0839 | 0.005207 | 0.31224 | 0.016677 | 0.98669 | 0.99277 | 1.281 |
| Znhit3 | 584.701 | 0.051128 | 0.11367 | 0.449816 | 0.65284 | 0.80691 | 5.417 |
| Znhit6 | 805.934 | 0.211287 | 0.0907 | 2.329474 | 0.01983 | 0.08901 | 28.591 |
| Znrf1 | 6419.53 | 0.066749 | 0.08132 | 0.820789 | 0.41177 | 0.61976 | 89.934 |
| Znrf2 | 1173.18 | -0.23228 | 0.09359 | -2.48187 | 0.01307 | 0.06843 | 76.585 |
| Znrf3 | 990.595 | 0.131212 | 0.09336 | 1.405484 | 0.15988 | 0.34781 | 168.524 |
| Zp1 | 12.5694 | 0.734255 | 0.62275 | 1.179052 | 0.23838 | 0.44391 | 6.346 |

| | | | | | | | |
|---|---|---|---|---|---|---|---|
| Zp2 | 13.899 | 1.785145 | 0.54435 | 3.279382 | 0.00104 | 0.01316 | 18.52 |
| Zpbp | 103.708 | 0.38071 | 0.24324 | 1.56517 | 0.11754 | 0.28858 | 182.783 |
| Zpr1 | 1363.95 | -0.06653 | 0.07291 | -0.9126 | 0.36145 | 0.57177 | 9.58 |
| Zranb1 | 2895.32 | 0.121502 | 0.07451 | 1.63077 | 0.10294 | 0.26497 | 55.25 |
| Zranb2 | 10985.6 | 0.085948 | 0.12884 | 0.667079 | 0.50472 | 0.6967 | 14.251 |
| Zranb3 | 396.433 | 0.060527 | 0.14993 | 0.403703 | 0.68643 | 0.82888 | 148.864 |
| Zrsr1 | 2298.76 | -0.01328 | 0.07729 | -0.17186 | 0.86355 | 0.93094 | 4.492 |
| Zrsr2 | 2557.31 | -0.02874 | 0.07269 | -0.39542 | 0.69253 | 0.8334 | 23.219 |
| Zscan12 | 393.696 | -0.04124 | 0.10526 | -0.39177 | 0.69523 | 0.83485 | 9.47 |
| Zscan18 | 1377.13 | 0.367526 | 0.09886 | 3.71764 | 0.0002 | 0.00428 | 35.546 |
| Zscan2 | 329.485 | 0.14501 | 0.12477 | 1.16226 | 0.24513 | 0.45174 | 15.618 |
| Zscan20 | 514.385 | 0.087195 | 0.10345 | 0.842891 | 0.39929 | 0.60812 | 26.56 |
| Zscan21 | 735.421 | 0.167903 | 0.08873 | 1.892305 | 0.05845 | 0.18389 | 17.363 |
| Zscan22 | 583.879 | 0.238294 | 0.09417 | 2.530545 | 0.01139 | 0.06243 | 11.269 |
| Zscan25 | 535.935 | 0.111877 | 0.09317 | 1.200835 | 0.22982 | 0.43407 | 10.779 |
| Zscan26 | 3951.46 | 0.243217 | 0.09687 | 2.510714 | 0.01205 | 0.0648 | 11.551 |
| Zscan29 | 614.943 | -0.12371 | 0.11832 | -1.04548 | 0.2958 | 0.50616 | 12.853 |
| Zscan30 | 376.047 | 0.275134 | 0.14671 | 1.875344 | 0.06075 | 0.18815 | 53.317 |
| Zswim1 | 593.203 | -0.12587 | 0.10396 | -1.21073 | 0.226 | 0.43006 | 4.186 |
| Zswim3 | 415.702 | 0.102733 | 0.10595 | 0.969662 | 0.33221 | 0.5431 | 17.033 |
| Zswim4 | 837.724 | -0.05936 | 0.08164 | -0.72706 | 0.46719 | 0.66624 | 26.378 |
| Zswim5 | 1709.64 | 0.326397 | 0.15021 | 2.172953 | 0.02978 | 0.11766 | 111.889 |
| Zswim6 | 3361.96 | 0.338758 | 0.13024 | 2.60105 | 0.00929 | 0.05459 | 165.447 |
| Zswim7 | 264.774 | -0.12297 | 0.16194 | -0.75938 | 0.44763 | 0.65123 | 14.152 |
| Zswim8 | 8728.06 | 0.128425 | 0.05296 | 2.424856 | 0.01531 | 0.07543 | 16.068 |
| Zswim9 | 1086.34 | 0.047887 | 0.08172 | 0.585994 | 0.55788 | 0.73951 | 19.754 |
| Zup1 | 698.33 | 0.254834 | 0.22599 | 1.127649 | 0.25947 | 0.46776 | 32.128 |
| Zw10 | 1216.78 | 0.07041 | 0.07839 | 0.898244 | 0.36906 | 0.57964 | 23.163 |
| Zwilch | 150.996 | 0.108188 | 0.18345 | 0.589746 | 0.55536 | 0.73781 | 35.961 |
| Zwint | 29163.7 | 0.056072 | 0.07625 | 0.735409 | 0.46209 | 0.66239 | 20.12 |
| Zxda | 30.7109 | 0.203363 | 0.36683 | 0.554373 | 0.57932 | 0.75592 | 2.621 |
| Zxdb | 772.721 | 0.161182 | 0.09569 | 1.68441 | 0.0921 | 0.24706 | 5.619 |
| Zxdc | 1343.13 | 0.000707 | 0.0859 | 0.00823 | 0.99343 | 0.99627 | 33.999 |
| Zyg11b | 12499.8 | 0.08569 | 0.05392 | 1.58924 | 0.11201 | 0.27952 | 71.373 |

| | | | | | | | |
|---|---|---|---|---|---|---|---|
| Zyx | 785.787 | -0.57566 | 0.10075 | -5.71351 | 1.11E-08 | 3.47E-06 | 10.584 |
| Zzef1 | 5170.68 | 0.166729 | 0.06934 | 2.404356 | 0.0162 | 0.07831 | 130.895 |
| Zzz3 | 3694.76 | 0.13718 | 0.08821 | 1.555207 | 0.1199 | 0.29246 | 67.354 |
| mt-Atp6 | 1667.65 | 0.198812 | 0.18209 | 1.09185 | 0.2749 | 0.48448 | 0.681 |
| mt-Co1 | 921069 | -0.04188 | 0.07374 | -0.56794 | 0.57008 | 0.74938 | 1.545 |
| mt-Co2 | 4187.47 | -0.02151 | 0.27517 | -0.07818 | 0.93768 | 0.96886 | 0.684 |
| mt-Co3 | 3477.16 | -0.05192 | 0.14324 | -0.36245 | 0.71702 | 0.84892 | 0.784 |
| mt-Cytb | 325732 | 0.128008 | 0.132 | 0.969735 | 0.33218 | 0.5431 | 1.144 |
| mt-Nd1 | 231549 | 0.089239 | 0.11061 | 0.806818 | 0.41977 | 0.62704 | 0.957 |
| mt-Nd2 | 160317 | 0.10236 | 0.13364 | 0.765933 | 0.44372 | 0.64799 | 1.038 |
| mt-Nd3 | 472.567 | -0.04376 | 0.11783 | -0.37137 | 0.71036 | 0.84412 | 0.348 |
| mt-Nd4 | 150508 | 0.25034 | 0.1893 | 1.322472 | 0.18601 | 0.38084 | 1.378 |
| mt-Nd4l | 167.626 | -0.05391 | 0.55298 | -0.09749 | 0.92234 | 0.9609 | 0.297 |
| mt-Nd5 | 201241 | 0.195483 | 0.18175 | 1.075575 | 0.28212 | 0.49231 | 1.824 |
| mt-Nd6 | 30052.4 | 0.049925 | 0.19118 | 0.261143 | 0.79398 | 0.89338 | 0.519 |
| mt-Rnr1 | 140714 | -0.05327 | 0.11531 | -0.462 | 0.64408 | 0.80143 | 0.955 |
| mt-Rnr2 | 195612 | 0.062582 | 0.10369 | 0.603576 | 0.54613 | 0.73071 | 1.582 |
| mt-Tp | 2160.53 | -0.03639 | 0.13267 | -0.27429 | 0.78386 | 0.88762 | 0.067 |
| n-R5s197 | 16.2371 | 0.123632 | 0.46438 | 0.26623 | 0.79006 | 0.89104 | 0.119 |
