## Supplementary material for "Reversible m^6^Am methylation of snRNA by FTO controls morphine reward and tolerance without altering analgesia": Table S5

### Table S5. GSEA_KEGG_Fto_cKO

| | ID | Description | setSize | enrichme | NES | pvalue | p.adjust | qvalue | rank | leading_e | core_enrichment |
|---|---|---|---|---|---|---|---|---|---|---|---|
| mmu05032 | mmu05032 | Morphine addiction | 76 | 0.3713 | 1.44568 | 0.01639 | 0.0585 | 0.03687 | 1159 | tags=29% | 13488/207728/18574/23984/18390/14708/29863/224129/241489/14397/18575/54611/14407/14403/218461/14395/14696/22348/18578/18749/14402/14697 |
| mmu04145 | mmu04145 | Phagocytosis | 175 | -0.5393 | -2.022 | 6.71E-08 | 2.28E-05 | 1.43E-05 | 3046 | tags=41% | [illegible] |
| mmu05152 | mmu05152 | Tuberculosis | 116 | -0.5688 | -2.0644 | 4.73E-07 | 8.02E-05 | 5.06E-05 | 1829 | tags=30% | [illegible] |
| mmu05140 | mmu05140 | Leishmaniasis | 49 | -0.7021 | -2.1909 | 1.10E-06 | 8.09E-05 | 5.10E-05 | 1732 | tags=45% | 21809/17970/16476/16409/13057/15170/17874/20846/14281/14131/266632/16414/14129/16401/17357/19225/19094/29857/14969/14961/14960/12266 |
| mmu04932 | mmu04932 | Non-alcoholic fatty liv | 130 | -0.557 | -2.0368 | 1.19E-06 | 8.09E-05 | 5.10E-05 | 3916 | tags=57% | [illegible] |
| mmu05169 | mmu05169 | Epstein-Barr virus infe | 151 | -0.525 | -1.9448 | 1.09E-06 | 8.09E-05 | 5.10E-05 | 3947 | tags=49% | [illegible] |
| mmu04518 | mmu04518 | Integrin signaling | 120 | -0.5529 | -2.0159 | 3.80E-06 | 0.00021 | 0.00014 | 2395 | tags=47% | [illegible] |
| mmu05171 | mmu05171 | Coronavirus disease | 162 | -0.5074 | -1.887 | 6.68E-06 | 0.00032 | 0.0002 | 3786 | tags=48% | [illegible] |
| mmu05133 | mmu05133 | Pertussis | 46 | -0.696 | -2.1512 | 8.16E-06 | 0.00034 | 0.00022 | 1508 | tags=48% | 12631/11848/12315/16476/16409/117149/16362/17874/12475/14281/266632/107607/16414/12262/12260/15900/12259/12268/19094/29857/12258/12266 |
| mmu04610 | mmu04610 | Complement and coa | 32 | -0.7473 | -2.1462 | 9.99E-06 | 0.00034 | 0.00022 | 1631 | tags=62% | 12759/18791/16409/12509/12628/16414/12262/12260/12259/21788/74145/22370/22371/15160/12268/12267/21824/12258/232345/12266 |
| mmu05168 | mmu05168 | Herpes simplex virus 1 | 126 | -0.5332 | -1.9434 | 1.01E-05 | 0.00034 | 0.00022 | 2258 | tags=26% | [illegible] |
| mmu04640 | mmu04640 | Hematopoietic cell lin | 44 | -0.6736 | -2.0596 | 2.16E-05 | 0.00067 | 0.00042 | 2068 | tags=61% | [illegible] |
| mmu05020 | mmu05020 | Prion disease | 206 | -0.4567 | -1.7482 | 2.53E-05 | 0.00071 | 0.00045 | 4225 | tags=52% | [illegible] |
| mmu05142 | mmu05142 | Chagas disease | 71 | -0.5758 | -1.9394 | 3.07E-05 | 0.0008 | 0.0005 | 1732 | tags=25% | 21809/18797/16476/14683/11421/12633/17874/14281/266632/12262/12260/21813/12259/432530/21937/19094/29857/12266 |
| mmu05322 | mmu05322 | Systemic lupus erythe | 37 | -0.7022 | -2.1002 | 3.55E-05 | 0.00086 | 0.00054 | 831 | tags=30% | 14131/14129/12262/12260/12259/12524/12268/14969/14961/14960/12266 |
| mmu05167 | mmu05167 | Kaposi sarcoma-asso | 146 | -0.4993 | -1.8389 | 4.29E-05 | 0.00093 | 0.00059 | 3792 | tags=50% | [illegible] |
| mmu05415 | mmu05415 | Diabetic cardiomyopa | 150 | -0.4884 | -1.8029 | 4.41E-05 | 0.00093 | 0.00059 | 4225 | tags=55% | [illegible] |
| mmu04514 | mmu04514 | Cell adhesion molecul | 94 | -0.5368 | -1.8911 | 6.17E-05 | 0.00123 | 0.00078 | 1744 | tags=37% | [illegible] |
| mmu04936 | mmu04936 | Alcoholic liver disease | 96 | -0.5286 | -1.865 | 7.12E-05 | 0.00134 | 0.00084 | 2672 | tags=34% | [illegible] |
| mmu04670 | mmu04670 | Leukocyte transendot | 79 | -0.5607 | -1.9277 | 8.87E-05 | 0.0015 | 0.00095 | 2206 | tags=39% | [illegible] |
| mmu05205 | mmu05205 | Proteoglycans in canc | 162 | -0.4744 | -1.7646 | 8.54E-05 | 0.0015 | 0.00095 | 2230 | tags=31% | [illegible] |
| mmu05150 | mmu05150 | Staphylococcus aureu | 25 | -0.7531 | -2.0937 | 0.00012 | 0.00184 | 0.00116 | 926 | tags=60% | 12628/14131/16414/14129/12262/12260/12259/14130/12268/12267/15894/14969/14961/14960/12266 |
| mmu04082 | mmu04082 | Neuroactive ligand sig | 146 | 0.39843 | 1.74043 | 0.00012 | 0.00184 | 0.00116 | 1159 | tags=29% | [illegible] |
| mmu04510 | mmu04510 | Focal adhesion | 174 | -0.4465 | -1.672 | 0.00015 | 0.00223 | 0.0014 | 2395 | tags=36% | [illegible] |
| mmu04612 | mmu04612 | Antigen processing an | 47 | -0.6314 | -1.9643 | 0.00018 | 0.00242 | 0.00152 | 1780 | tags=34% | 15512/13039/21354/15040/21355/13040/15511/12010/14964/193740/14972/15015/14969/14961/14960/16149 |
| mmu05203 | mmu05203 | Viral carcinogenesis | 151 | -0.4626 | -1.7139 | 0.00018 | 0.00242 | 0.00152 | 3924 | tags=45% | [illegible] |
| mmu04613 | mmu04613 | Neutrophil extracellul | 86 | -0.5172 | -1.8042 | 0.0004 | 0.00527 | 0.00332 | 2536 | tags=28% | 15184/170743/319179/74244/18752/69146/17969/16416/17970/18797/68024/79221/16409/234779/13057/18750/14131/16414/14129/20963/22371/19094/29857/12266 |
| mmu04666 | mmu04666 | Fc gamma R-mediated | 84 | -0.5292 | -1.8405 | 0.00042 | 0.00531 | 0.00335 | 2738 | tags=39% | [illegible] |
| mmu05166 | mmu05166 | Human T-cell leukemi | 173 | -0.4424 | -1.6602 | 0.00045 | 0.00546 | 0.00344 | 2219 | tags=27% | [illegible] |
| mmu04380 | mmu04380 | Osteoclast differentia | 99 | -0.5096 | -1.8113 | 0.00051 | 0.00592 | 0.00373 | 2113 | tags=33% | [illegible] |
| mmu05332 | mmu05332 | Graft-versus-host dise | 14 | -0.8136 | -1.9819 | 0.00054 | 0.00605 | 0.00381 | 268 | tags=86% | 15024/14991/15039/14102/15040/14964/12524/14972/15015/14969/14961/14960 |
| mmu05320 | mmu05320 | Autoimmune thyroid d | 15 | -0.8074 | -2.016 | 0.00064 | 0.00641 | 0.00404 | 268 | tags=67% | 15039/14102/15040/14964/12524/14972/15015/14969/14961/14960 |
| mmu05330 | mmu05330 | Allograft rejection | 15 | -0.8074 | -2.016 | 0.00064 | 0.00641 | 0.00404 | 268 | tags=67% | [illegible] |
| mmu05206 | mmu05206 | MicroRNAs in cancer | 138 | -0.4488 | -1.6467 | 0.0006 | 0.00641 | 0.00404 | 3041 | tags=39% | [illegible] |
| mmu04151 | mmu04151 | PI3K-Akt signaling pat | 251 | -0.3988 | -1.5579 | 0.00064 | 0.00641 | 0.00404 | 2434 | tags=33% | [illegible] |
| mmu04933 | mmu04933 | AGE-RAGE signaling pa | 82 | -0.5185 | -1.7975 | 0.00072 | 0.00675 | 0.00426 | 1732 | tags=34% | [illegible] |
| mmu04926 | mmu04926 | Relaxin signaling path | 101 | -0.4945 | -1.7649 | 0.0007 | 0.00675 | 0.00426 | 1895 | tags=32% | [illegible] |
| mmu03013 | mmu03013 | Nucleocytoplasmic tra | 99 | 0.40986 | 1.67944 | 0.00079 | 0.0072 | 0.00454 | 4974 | tags=51% | [illegible] |
| mmu04064 | mmu04064 | NF-kappa B signaling p | 64 | -0.535 | -1.7627 | 0.00101 | 0.00833 | 0.00525 | 2038 | tags=36% | [illegible] |
| mmu05418 | mmu05418 | Fluid shear stress and | 113 | -0.4814 | -1.7409 | 0.00099 | 0.00833 | 0.00525 | 2492 | tags=38% | [illegible] |
| mmu05208 | mmu05208 | Chemical carcinogene | 168 | -0.433 | -1.6175 | 0.00094 | 0.00833 | 0.00525 | 4225 | tags=52% | [illegible] |
| mmu04820 | mmu04820 | Cytoskeleton in muscl | 162 | -0.4296 | -1.5978 | 0.001 | 0.00833 | 0.00525 | 2043 | tags=34% | [illegible] |
| mmu05163 | mmu05163 | Human cytomegalovir | 171 | -0.4244 | -1.586 | 0.00114 | 0.00923 | 0.00582 | 3736 | tags=45% | [illegible] |
| mmu05010 | mmu05010 | Alzheimer disease | 301 | -0.3755 | -1.49 | 0.00123 | 0.00968 | 0.0061 | 3958 | tags=40% | [illegible] |
| mmu01040 | mmu01040 | Biosynthesis of unsat | 26 | -0.6538 | -1.8333 | 0.00132 | 0.0102 | 0.00643 | 3107 | tags=65% | [illegible] |
| mmu04672 | mmu04672 | Intestinal immune ne | 15 | -0.7875 | -1.9663 | 0.00144 | 0.01083 | 0.00683 | 1355 | tags=53% | [illegible] |
| mmu05200 | mmu05200 | Pathways in cancer | 398 | -0.3525 | -1.417 | 0.00159 | 0.01173 | 0.0074 | 2345 | tags=29% | [illegible] |
| mmu05012 | mmu05012 | Parkinson disease | 216 | -0.4035 | -1.5527 | 0.00164 | 0.01185 | 0.00747 | 4225 | tags=52% | [illegible] |
| mmu05134 | mmu05134 | Legionellosis | 40 | -0.5991 | -1.8193 | 0.00178 | 0.01257 | 0.00792 | 987 | tags=20% | [illegible] |
| mmu05145 | mmu05145 | Toxoplasmosis | 85 | -0.4966 | -1.7276 | 0.00203 | 0.01404 | 0.00885 | 3808 | tags=53% | [illegible] |
| mmu05016 | mmu05016 | Huntington disease | 247 | -0.3845 | -1.4975 | 0.0021 | 0.01422 | 0.00897 | 4228 | tags=44% | [illegible] |
| mmu04060 | mmu04060 | Cytokine-cytokine rec | 87 | -0.4764 | -1.6629 | 0.00219 | 0.01453 | 0.00916 | 3292 | tags=52% | [illegible] |
| mmu05022 | mmu05022 | Pathways of neurodeg | 380 | -0.355 | -1.4226 | 0.00233 | 0.01517 | 0.00956 | 3932 | tags=39% | [illegible] |
| mmu04935 | mmu04935 | Growth hormone synt | 101 | -0.4644 | -1.6574 | 0.00263 | 0.01681 | 0.0106 | 2206 | tags=30% | [illegible] |
| mmu04010 | mmu04010 | MAPK signaling pathw | 246 | -0.3799 | -1.479 | 0.00273 | 0.01714 | 0.0108 | 2763 | tags=39% | [illegible] |
| mmu04940 | mmu04940 | Type I diabetes mellit | 21 | -0.6972 | -1.8672 | 0.00282 | 0.01736 | 0.01095 | 268 | tags=52% | [illegible] |
| mmu04911 | mmu04911 | Insulin secretion | 65 | -0.5174 | -1.7108 | 0.00411 | 0.02473 | 0.01559 | 2206 | tags=51% | [illegible] |
| mmu04512 | mmu04512 | ECM-receptor interact | 65 | -0.5173 | -1.7106 | 0.00416 | 0.02473 | 0.01559 | 1758 | tags=49% | [illegible] |
| mmu04390 | mmu04390 | Hippo signaling pathw | 127 | -0.4362 | -1.593 | 0.00453 | 0.02647 | 0.01668 | 2297 | tags=30% | [illegible] |
| mmu04913 | mmu04913 | Ovarian steroidogene | 33 | -0.6056 | -1.7557 | 0.00499 | 0.02733 | 0.01723 | 1902 | tags=55% | [illegible] |
| mmu05416 | mmu05416 | Viral myocarditis | 50 | -0.5447 | -1.7048 | 0.005 | 0.02733 | 0.01723 | 1561 | tags=32% | [illegible] |
| mmu05162 | mmu05162 | Measles | 90 | -0.4579 | -1.6108 | 0.0049 | 0.02733 | 0.01723 | 3808 | tags=48% | [illegible] |
| mmu03015 | mmu03015 | mRNA surveillance pa | 85 | 0.3845 | 1.53048 | 0.00484 | 0.02733 | 0.01723 | 4335 | tags=40% | [illegible] |
| mmu05165 | mmu05165 | Human papillomaviru | 267 | -0.3588 | -1.4023 | 0.00519 | 0.02793 | 0.0176 | 2289 | tags=27% | [illegible] |
| mmu04216 | mmu04216 | Ferroptosis | 36 | -0.5719 | -1.7029 | 0.00529 | 0.02805 | 0.01768 | 3425 | tags=72% | [illegible] |
| mmu04210 | mmu04210 | Apoptosis | 111 | -0.4412 | -1.5853 | 0.00599 | 0.03125 | 0.0197 | 3903 | tags=51% | [illegible] |
| mmu04659 | mmu04659 | Th17 cell differentiati | 65 | -0.502 | -1.6601 | 0.00626 | 0.03213 | 0.02025 | 1502 | tags=28% | [illegible] |
| mmu03272 | mmu03272 | Virion - Hepatitis virus | 26 | -0.6004 | -1.6837 | 0.00695 | 0.03363 | 0.0212 | 1863 | tags=38% | [illegible] |
| mmu04918 | mmu04918 | Thyroid hormone synt | 50 | -0.5354 | -1.6755 | 0.00686 | 0.03363 | 0.0212 | 1895 | tags=40% | [illegible] |
| mmu04971 | mmu04971 | Gastric acid secretion | 57 | -0.5176 | -1.669 | 0.00667 | 0.03363 | 0.0212 | 2206 | tags=46% | [illegible] |
| mmu05132 | mmu05132 | Salmonella infection | 205 | -0.3844 | -1.4711 | 0.00681 | 0.03363 | 0.0212 | 3321 | tags=31% | [illegible] |
| mmu05323 | mmu05323 | Rheumatoid arthritis | 41 | -0.5548 | -1.6962 | 0.00717 | 0.03374 | 0.02127 | 1732 | tags=61% | [illegible] |
| mmu03083 | mmu03083 | Polycomb repressive c | 75 | 0.38604 | 1.51233 | 0.00714 | 0.03374 | 0.02127 | 4192 | tags=55% | [illegible] |
| mmu04916 | mmu04916 | Melanogenesis | 76 | -0.4669 | -1.6021 | 0.00739 | 0.03382 | 0.02132 | 2206 | tags=36% | [illegible] |
| mmu05164 | mmu05164 | Influenza A | 98 | -0.4405 | -1.5619 | 0.00748 | 0.03382 | 0.02132 | 3906 | tags=40% | [illegible] |
| mmu04310 | mmu04310 | Wnt signaling pathwa | 139 | -0.413 | -1.5167 | 0.00743 | 0.03382 | 0.02132 | 2630 | tags=34% | [illegible] |
| mmu04668 | mmu04668 | TNF signaling pathway | 85 | -0.4623 | -1.6082 | 0.0079 | 0.03524 | 0.02221 | 2174 | tags=32% | [illegible] |
| mmu00590 | mmu00590 | Arachidonic acid meta | 31 | -0.5719 | -1.6325 | 0.00813 | 0.0358 | 0.02257 | 4000 | tags=68% | [illegible] |
| mmu04625 | mmu04625 | C-type lectin receptor | 77 | -0.4604 | -1.5848 | 0.00853 | 0.03706 | 0.02336 | 1949 | tags=29% | [illegible] |
| mmu04517 | mmu04517 | IgSF CAM signaling | 247 | -0.3594 | -1.3999 | 0.00864 | 0.03709 | 0.02338 | 2432 | tags=28% | [illegible] |
| mmu05170 | mmu05170 | Human immunodefici | 160 | -0.3889 | -1.4434 | 0.0089 | 0.03772 | 0.02377 | 3853 | tags=40% | [illegible] |
| mmu00190 | mmu00190 | Oxidative phosphoryla | 92 | -0.4618 | -1.6283 | 0.00923 | 0.03863 | 0.02435 | 4775 | tags=73% | [illegible] |
| mmu04630 | mmu04630 | JAK-STAT signaling pat | 88 | -0.4476 | -1.5652 | 0.00975 | 0.04033 | 0.02542 | 2381 | tags=30% | [illegible] |
| mmu01200 | mmu01200 | Carbon metabolism | 96 | -0.4361 | -1.5386 | 0.01062 | 0.04336 | 0.02733 | 5385 | tags=66% | [illegible] |
| mmu04658 | mmu04658 | Th1 and Th2 cell differ | 55 | -0.5102 | -1.644 | 0.01094 | 0.04365 | 0.02751 | 1497 | tags=27% | [illegible] |
| mmu00620 | mmu00620 | Pyruvate metabolism | 35 | -0.555 | -1.6237 | 0.01091 | 0.04365 | 0.02751 | 4339 | tags=63% | [illegible] |
| mmu04979 | mmu04979 | Cholesterol metabolis | 30 | -0.5732 | -1.6247 | 0.01164 | 0.04588 | 0.02892 | 1874 | tags=33% | [illegible] |
| mmu05146 | mmu05146 | Amoebiasis | 62 | -0.4768 | -1.5717 | 0.01287 | 0.05014 | 0.03161 | 1758 | tags=37% | [illegible] |
| mmu04742 | mmu04742 | Taste transduction | 40 | 0.4631 | 1.5847 | 0.01371 | 0.0528 | 0.03328 | 1678 | tags=40% | [illegible] |
| mmu04066 | mmu04066 | HIF-1 signaling pathwa | 89 | -0.4347 | -1.5264 | 0.01442 | 0.05494 | 0.03463 | 3365 | tags=36% | [illegible] |
| mmu05144 | mmu05144 | Malaria | 21 | -0.6292 | -1.6852 | 0.01471 | 0.05542 | 0.03493 | 2177 | tags=57% | [illegible] |
| mmu04620 | mmu04620 | Toll-like receptor sign | 63 | -0.4814 | -1.5902 | 0.01496 | 0.05573 | 0.03513 | 4566 | tags=52% | [illegible] |
| mmu04144 | mmu04144 | Endocytosis | 223 | -0.3583 | -1.3773 | 0.01514 | 0.05577 | 0.03516 | 4345 | tags=40% | [illegible] |
| mmu00565 | mmu00565 | Ether lipid metabolism | 33 | -0.5643 | -1.6363 | 0.01616 | 0.05829 | 0.03675 | 2486 | tags=42% | [illegible] |
| mmu04142 | mmu04142 | Lysosome biogenesis | 216 | -0.3598 | -1.3848 | 0.01613 | 0.05829 | 0.03675 | 3940 | tags=37% | [illegible] |
| mmu04650 | mmu04650 | Natural killer cell med | 61 | -0.4639 | -1.5226 | 0.01722 | 0.06023 | 0.03796 | 2206 | tags=31% | [illegible] |
| mmu05161 | mmu05161 | Hepatitis B | 126 | -0.3992 | -1.4549 | 0.01723 | 0.06023 | 0.03796 | 3808 | tags=40% | [illegible] |
| mmu05202 | mmu05202 | Transcriptional misre | 125 | -0.395 | -1.4357 | 0.01814 | 0.06276 | 0.03956 | 2195 | tags=33% | [illegible] |
| mmu04148 | mmu04148 | Efferocytosis | 124 | -0.39 | -1.4191 | 0.01948 | 0.06669 | 0.04204 | 2938 | tags=36% | [illegible] |
| mmu04976 | mmu04976 | Bile secretion | 41 | -0.514 | -1.5713 | 0.02021 | 0.06851 | 0.04318 | 2819 | tags=59% | [illegible] |
| mmu04270 | mmu04270 | Vascular smooth mus | 94 | -0.4175 | -1.4709 | 0.02102 | 0.07055 | 0.04447 | 2206 | tags=34% | [illegible] |
| mmu04740 | mmu04740 | Olfactory transductio | 31 | 0.47634 | 1.55092 | 0.02214 | 0.07186 | 0.0453 | 1268 | tags=32% | [illegible] |
| mmu03320 | mmu03320 | PPAR signaling pathwa | 48 | -0.4851 | -1.5199 | 0.02264 | 0.07186 | 0.0453 | 3544 | tags=60% | [illegible] |
| mmu04370 | mmu04370 | VEGF signaling pathwa | 51 | -0.4814 | -1.5159 | 0.0221 | 0.07186 | 0.0453 | 1347 | tags=22% | [illegible] |
| mmu01212 | mmu01212 | Fatty acid metabolism | 54 | -0.4678 | -1.5002 | 0.02242 | 0.07186 | 0.0453 | 3502 | tags=50% | [illegible] |
| mmu04350 | mmu04350 | TGF-beta signaling pat | 85 | -0.4305 | -1.4976 | 0.02268 | 0.07186 | 0.0453 | 1950 | tags=27% | [illegible] |
| mmu04072 | mmu04072 | Phospholipase D signa | 123 | -0.3822 | -1.3947 | 0.02211 | 0.07186 | 0.0453 | 2378 | tags=31% | [illegible] |
| mmu04928 | mmu04928 | Parathyroid hormone | 91 | -0.4209 | -1.4825 | 0.02426 | 0.07614 | 0.04799 | 2206 | tags=34% | [illegible] |
| mmu05160 | mmu05160 | Hepatitis C | 107 | -0.3971 | -1.4206 | 0.02466 | 0.0767 | 0.04835 | 3853 | tags=40% | [illegible] |
| mmu04925 | mmu04925 | Aldosterone synthesis | 77 | -0.4273 | -1.4706 | 0.02887 | 0.08899 | 0.05609 | 2819 | tags=44% | [illegible] |
| mmu04934 | mmu04934 | Cushing syndrome | 122 | -0.3961 | -1.4484 | 0.02919 | 0.08915 | 0.0562 | 2041 | tags=30% | [illegible] |
| mmu04920 | mmu04920 | Adipocytokine signalin | 55 | -0.4659 | -1.5013 | 0.03139 | 0.09375 | 0.05909 | 3904 | tags=51% | [illegible] |
| mmu05417 | mmu05417 | Lipid and atheroscler | 145 | -0.3763 | -1.3856 | 0.03153 | 0.09375 | 0.05909 | 2258 | tags=37% | [illegible] |
| mmu04014 | mmu04014 | Ras signaling pathway | 177 | -0.3659 | -1.3743 | 0.03152 | 0.09375 | 0.05909 | 3504 | tags=40% | [illegible] |
| mmu05321 | mmu05321 | Inflammatory bowel d | 25 | -0.5837 | -1.6228 | 0.03317 | 0.09756 | 0.06149 | 1732 | tags=40% | [illegible] |
| mmu00100 | mmu00100 | Steroid biosynthesis | 15 | -0.6437 | -1.6073 | 0.03338 | 0.09756 | 0.06149 | 3661 | tags=67% | [illegible] |
| mmu04015 | mmu04015 | Rap1 signaling pathwa | 168 | -0.3676 | -1.3731 | 0.0344 | 0.09967 | 0.06282 | 2969 | tags=38% | [illegible] |
| mmu05204 | mmu05204 | Chemical carcinogene | 22 | -0.5936 | -1.608 | 0.03609 | 0.09988 | 0.06296 | 3053 | tags=64% | [illegible] |
| mmu05221 | mmu05221 | Acute myeloid leukem | 59 | -0.4563 | -1.4875 | 0.03541 | 0.09988 | 0.06296 | 1546 | tags=25% | [illegible] |
| mmu04912 | mmu04912 | GnRH signaling pathw | 74 | -0.4236 | -1.4438 | 0.03654 | 0.09988 | 0.06296 | 2598 | tags=32% | [illegible] |
| mmu04611 | mmu04611 | Platelet activation | 98 | -0.401 | -1.4219 | 0.03548 | 0.09988 | 0.06296 | 1968 | tags=28% | [illegible] |
| mmu04217 | mmu04217 | Necroptosis | 95 | -0.4025 | -1.4197 | 0.03586 | 0.09988 | 0.06296 | 3941 | tags=45% | [illegible] |
| mmu04621 | mmu04621 | NOD-like receptor sig | 115 | -0.3847 | -1.3987 | 0.03581 | 0.09988 | 0.06296 | 2258 | tags=36% | [illegible] |
| mmu04714 | mmu04714 | Thermogenesis | 173 | -0.3627 | -1.3611 | 0.03636 | 0.09988 | 0.06296 | 3954 | tags=49% | [illegible] |

### Table S5. GSEA_KEGG_FB23

[illegible]
